## Supplementary Fig S1 for "Whole-Genome Sequence of African Swine Fever Virus isolate from India provides insights into diversity and evolution"

```
FR682468.2_Georgia_2007/1        -------------------------------------------------------------------------------- [    80]
ASF_IND_20_CAD543                nnnaagtatattatatattttataattttatatatat-aaagcaggt-----------acaattttattatannnnnnag [    80]
LR899193.1_Germany_2020/1        -------------------------------------------------------------------------------- [    80]
MN393477.1_Wuhan_2019            --------------------------------------aaagcaggt-----------acaattttattatatagtgcag [    80]
MW396979.1_Timor-Leste/2019/1    --------------------------------------aaagcaggt-----------acaattttattatatagtgcag [    80]
MK940252.1_CN/2019/InnerMongolia -------------------------------------------------------------------------------- [    80]
MK128995.1_China/2018/AnhuiXCGQ  -------------------------------------------------------------------------------- [    80]
MH681419.1_ASFV/POL/2015         -------------------------------------------------------------------------------- [    80]
LS478113.1_Estonia_2014          -------------------------ttatatacttataaaagcaggt-----------acaattttattatatagtgcag [    80]
KP843857.1_Odintsovo_02/14       --------------------------------------a----------------------------------------- [    80]
FN557520.1_strain_E75            -------------------------------------------------------------------------------- [    80]
AY261366.1_Warthog               -------------------------------------------------------------------------------- [    80]
AY261365.1_Warmbaths             --------------------------------------------------------------tcttatt----------- [    80]
AY261364.1_Tengani_62            --------------------------------------------------------ctacaattttattatatagtgcag [    80]
AY261363.1_Pretoriuskop/96/4     -------------------------------ggatatcggactcggtc-------cccacaattttattatatagtgcag [    80]
AY261362.1_Mkuzi_1979            -------------------------------------------------------------------ttatatagtgcag [    80]
AY261361.1_Malawi_Lil-20/1_1983  -----------------------tatcggttggtggagagaccgggtctgtgcaaaagaaagtttta-----------aa [    80]
AY261360.1_Kenya_1950            ----------------------------------------------------------acaattttatattatagtgcag [    80]
AM712240.1_OURT_88/3             -------------------------------------------------------------------------------- [    80]
AM712239.1_Benin_97/1            -------------------------------------------------------------------------------- [    80]
KM111295.1_Ken06.Bus             -------------------------------------------------------------------------------- [    80]
KM111294.1_Ken05/Tk1             -------------------------------------------------------------------------------- [    80]
KM262844.1_strain_L60            -------------------------------------------------------------------------------- [    80]
KM102979.1_26544/OG10_Italy      -------------------------------------------------------------------------------- [    80]
KM262845.1_strain_NHV            ------c------------------------------------------------------------------------- [    80]
KX354450.1_47/Ss/2008            ------tatattatatattt-ataattttatatatat-aaagcaggt-----------acaattttattatatagtgcag [    80]
KP055815.1_strain_BA71           -------------------------------------------------------------------------------- [    80]
MH025920.1_strain_R35            --------------------------------------aaagcagg-----------tacaattttatattatagtgcag [    80]
MH025919.1_strain_N10            --------------------------------------aaagcagg-----------tacaattttatattatagtgcag [    80]
MH025918.1_strain_R25            --------------------------------------aaagcagg-----------tacaattttatattatagtgcag [    80]
MH025917.1_strain_R7             --------------------------------------aaagcagg-----------tacaattttatattatagtgcag [    80]
MH025916.1_strain_R8             --------------------------------------aaagcagg-----------tacaattttatattatagtgcag [    80]

FR682468.2_Georgia_2007/1        -------------------------------------------------------------------------------- [   160]
ASF_IND_20_CAD543                atcttttttgtgaaaaaaaaataattatttttttgcgcgcc--ggaataattattttttttnnnnnnnnnnnnnnnnnnn [   160]
LR899193.1_Germany_2020/1        -------------------------------------------------------------------------------- [   160]
MN393477.1_Wuhan_2019            atcttttttgtgaaaaaaaaataattatt--tttgcgcgcc--ggaataattatttttttttcgcgccggaaagt----- [   160]
MW396979.1_Timor-Leste/2019/1    atcttttttgtgaaaaaaaaataattatt--tttgcgcgcc--ggaataattatttttttttcgcgccggaaagt----- [   160]
MK940252.1_CN/2019/InnerMongolia -------------------------------------------------------------------------------- [   160]
MK128995.1_China/2018/AnhuiXCGQ  -------------------------------------------------------------------------------- [   160]
MH681419.1_ASFV/POL/2015         -------------------------------------------------------------------------------- [   160]
LS478113.1_Estonia_2014          atcttttttgtgaaaaaaaaataattatt--tttgcgcgcc--ggaataattatttttttttcgcgccggaaagt----- [   160]
KP843857.1_Odintsovo_02/14       -------------------------------------------------------------------------------- [   160]
```

```
FN557520.1_strain_E75             -------------------------------------------------------------------------------- [   160]
AY261366.1_Warthog                -------------------------------------------------------------------------------- [   160]
AY261365.1_Warmbaths              -------------------------------------------------------------------------------- [   160]
AY261364.1_Tengani_62             atcttttttgtgaaaaaaaataa-ttatttttttgcgcgac--ggaataattatttttttgcgtggccggaatat----- [   160]
AY261363.1_Pretoriuskop/96/4      atcttttttgtgaaaaaaaataa--ttattttttttgcgcc--ggaataattattttttacggggcg------------ [   160]
AY261362.1_Mkuzi_1979             atcttttttgtgaaaaaaaaaaa---ttattttttcgcggc--ggaataattatttttttgggccggaaatatttttgta [   160]
AY261361.1_Malawi_Lil-20/1_1983   gacttaggagcgcagataaatta-ttaaaacatcgggcg-c--agaataatgggtattttttgcggggcgaaaaa----- [   160]
AY261360.1_Kenya_1950             atcttttttgtgaaaaaaaaataattatttttttttatggggtggaataattatttttttacggggagtaattat----- [   160]
AM712240.1_OURT_88/3              -------------------------------------------------------------------------------- [   160]
AM712239.1_Benin_97/1             -------------------------------------------------------------------------------- [   160]
KM111295.1_Ken06.Bus              -------------------------------------------------------------------------------- [   160]
KM111294.1_Ken05/Tk1              -------------------------------------------------------------------------------- [   160]
KM262844.1_strain_L60             -------------------------------------------------------------------------------- [   160]
KM102979.1_26544/OG10_Italy       -------------------------------------------------------------------------------- [   160]
KM262845.1_strain_NHV             -------------------------------------------------------------------------------- [   160]
KX354450.1_47/Ss/2008             atcttttttgtgaaaaaaaaataattatttttttgcgcgcc--ggaataattattttttttgcgcggaaatatttttgca [   160]
KP055815.1_strain_BA71            --------------------------atttttttgcgcgcc--ggaataattattttttttgcgcggaaatatttttgca [   160]
MH025920.1_strain_R35             atcttttttgtgaaaaaaaaataattatttttttttacggg--ggaataattatttttgc-------------gg----- [   160]
MH025919.1_strain_N10             atcttttttgtgaaaaaaaaataattatttttttttacggg--ggaataattatttttgc-------------gg----- [   160]
MH025918.1_strain_R25             atcttttttgtgaaaaaaaaataattatttttttttacggg--ggaataattatttttgc-------------gg----- [   160]
MH025917.1_strain_R7              atcttttttgtgaaaaaaaaataattatttttttttacggg--ggaataattatttttgc-------------gg----- [   160]
MH025916.1_strain_R8              atcttttttgtgaaaaaaaaataattatttttttttacggg--ggaataattatttttgc-------------gg----- [   160]

FR682468.2_Georgia_2007/1         -------------------------------------------------------------------------------- [   240]
ASF_IND_20_CAD543                 nnnnnnattgatttttttttcagttgtnnnnnnnnnnnnnnnnnnnnnnnnnnnnnnnnnnnnnnnnnnnn------nnn [   240]
LR899193.1_Germany_2020/1         -------------------------------------------------------------------------------- [   240]
MN393477.1_Wuhan_2019             ----------attttgtgtaccgtag------------------------------------------------------ [   240]
MW396979.1_Timor-Leste/2019/1     ----------attttgtgtaccgtag------------------------------------------------------ [   240]
MK940252.1_CN/2019/InnerMongolia  -------------------------------------------------------------------------------- [   240]
MK128995.1_China/2018/AnhuiXCGQ   -------------------------------------------------------------------------------- [   240]
MH681419.1_ASFV/POL/2015          -------------------------------------------------------------------------------- [   240]
LS478113.1_Estonia_2014           ----------attttgtgtaccgtag------------------------------------------------------ [   240]
KP843857.1_Odintsovo_02/14        -------------------------------------------------------------------------------- [   240]
FN557520.1_strain_E75             -------------------------------------------------------------------------------- [   240]
AY261366.1_Warthog                -------------------------------------------------------------------------------- [   240]
AY261365.1_Warmbaths              -------------------------------------------------------------------------------- [   240]
AY261364.1_Tengani_62             ---------------------------------------------------tttcgccgcgccgtattttggtcccgtag [   240]
AY261363.1_Pretoriuskop/96/4      -------------------------------------------------------gccggaaagcattttg------tag [   240]
AY261362.1_Mkuzi_1979             gtggcgattgatttttttttcggttgtctgcccg-ggcggaataa------tttcgccggaaagcattttg------tag [   240]
AY261361.1_Malawi_Lil-20/1_1983   ---------------------------------------------------------------------------gaaag [   240]
AY261360.1_Kenya_1950             ---------------------------------------------------ttttgcggtgcggaa--------aggtag [   240]
AM712240.1_OURT_88/3              -------------------------------------------------------------------------------- [   240]
AM712239.1_Benin_97/1             -------------------------------------------------------------------------------- [   240]
KM111295.1_Ken06.Bus              -------------------------------------------------------------------------------- [   240]
```

```
KM111294.1_Ken05/Tk1             -------------------------------------------------------------------------------- [   240]
KM262844.1_strain_L60            -------------------------------------------------------------------------------- [   240]
KM102979.1_26544/OG10_Italy      -------------------------------------------------------------------------------- [   240]
KM262845.1_strain_NHV            -------------------------------------------------------------------------------- [   240]
KX354450.1_47/Ss/2008            gtggcgattgatttttttttcagttgtctgcccgtggcggaataattattttttcgccggaaagcattttg------tag [   240]
KP055815.1_strain_BA71           gtggcgattgatttttttttcagttgtctgcccgtggcggaataattattttttcgccggaaagcattttg------tag [   240]
MH025920.1_strain_R35            ---------------------------------------------------ttttgcggtgcggaa--------aggtag [   240]
MH025919.1_strain_N10            ---------------------------------------------------ttttgcggtgcggaa--------aggtag [   240]
MH025918.1_strain_R25            ---------------------------------------------------ttttgcggtgcggaa--------aggtag [   240]
MH025917.1_strain_R7             ---------------------------------------------------ttttgcggtgcggaa--------aggtag [   240]
MH025916.1_strain_R8             ---------------------------------------------------ttttgcggtgcggaa--------aggtag [   240]

FR682468.2_Georgia_2007/1        -------------------------------------------------------------------------------- [   320]
ASF_IND_20_CAD543                tttcacaagat--------------------------------------------------------------gcagaat [   320]
LR899193.1_Germany_2020/1        -------------------------------------------------------------------------------- [   320]
MN393477.1_Wuhan_2019            tttcactagat-----tt---------------------------------------------------tgcaacatagt [   320]
MW396979.1_Timor-Leste/2019/1    tttcactagat-----tt---------------------------------------------------tgcaacatagt [   320]
MK940252.1_CN/2019/InnerMongolia -------------------------------------------------------------------------------- [   320]
MK128995.1_China/2018/AnhuiXCGQ  -------------------------------------------------------------------------------- [   320]
MH681419.1_ASFV/POL/2015         -------------------------------------------------------------------------------- [   320]
LS478113.1_Estonia_2014          tttcactagat-----tt---------------------------------------------------tgcaacatagt [   320]
KP843857.1_Odintsovo_02/14       -------------------------------------------------------------------------------- [   320]
FN557520.1_strain_E75            -------------------------------------------------------------------------------- [   320]
AY261366.1_Warthog               -------------------------------------------------------------------------------- [   320]
AY261365.1_Warmbaths             -------------------------------------------------------------------------------- [   320]
AY261364.1_Tengani_62            tttcactagatttcgcgcggaattttgcga-------------------------------------------tcatagt [   320]
AY261363.1_Pretoriuskop/96/4     tttcacaagat-----gcggaattttgcga-------------------------------------------tcatagt [   320]
AY261362.1_Mkuzi_1979            tttcacaagat-----gcagaatttggcggtacgtag---------tttcacaagatgcggaatttttgcggcgcgtagt [   320]
AY261361.1_Malawi_Lil-20/1_1983  ctacccacgtt-----gtaaagtggggggg--------------------------------------gggcgccggaaa [   320]
AY261360.1_Kenya_1950            ttacacaagat-----gcagaatttc------------------------------------------------------ [   320]
AM712240.1_OURT_88/3             -------------------------------------------------------------------------------- [   320]
AM712239.1_Benin_97/1            -------------------------------------------------------------------------------- [   320]
KM111295.1_Ken06.Bus             -------------------------------------------------------------------------------- [   320]
KM111294.1_Ken05/Tk1             --------gaa-----acagatcgcc------------------------------------------------------ [   320]
KM262844.1_strain_L60            -------------------------------------------------------------------------------- [   320]
KM102979.1_26544/OG10_Italy      -------------------------------------------------------------------------------- [   320]
KM262845.1_strain_NHV            -------------------------------------------------------------------------------- [   320]
KX354450.1_47/Ss/2008            tttcacaagat-----gcagaatttggcgggccggaataattattttttcgccgga---aagtattttgtgtaccgtagt [   320]
KP055815.1_strain_BA71           tttcacaagat-----gcagaatttggcgggccggaataattattttttcgccgga---aagtattttgtgtaccgtagt [   320]
MH025920.1_strain_R35            ttacacaagat-----gcagaatttcgcgg-------------------------------------------------- [   320]
MH025919.1_strain_N10            ttacacaagat-----gcagaatttcgcgg-------------------------------------------------- [   320]
MH025918.1_strain_R25            ttacacaagat-----gcagaatttcgcgg-------------------------------------------------- [   320]
MH025917.1_strain_R7             ttacacaagat-----gcagaatttcgcgg-------------------------------------------------- [   320]
MH025916.1_strain_R8             ttacacaagat-----gcagaatttcgcgg-------------------------------------------------- [   320]
```

```
FR682468.2_Georgia_2007/1        -------------------------------------------------------------------------------- [   400]
ASF_IND_20_CAD543                ttnnnnnn-------nnnnnnnnnnnnnnnn--------nnnnnnnnnnnnnnnnnnnnnnnnnnnnnnnnnnnnnnnnn [   400]
LR899193.1_Germany_2020/1        -------------------------------------------------------------------------------- [   400]
MN393477.1_Wuhan_2019            ttcactag-------atttcgcggtgcatac--------cggtgggc--------------------------------- [   400]
MW396979.1_Timor-Leste/2019/1    ttcactag-------atttcgcggtgcatac--------cggtgggc--------------------------------- [   400]
MK940252.1_CN/2019/InnerMongolia -------------------------------------------------------------------------------- [   400]
MK128995.1_China/2018/AnhuiXCGQ  -------------------------------------------------------------------------------- [   400]
MH681419.1_ASFV/POL/2015         -------------------------------------------------------------------------------- [   400]
LS478113.1_Estonia_2014          ttcactag-------atttcgcggtgcatac--------cggtgggc--------------------------------- [   400]
KP843857.1_Odintsovo_02/14       -------------------------------------------------------------------------------- [   400]
FN557520.1_strain_E75            -------------------------------------------------------------------------------- [   400]
AY261366.1_Warthog               -------------------------------------------------------------------------------- [   400]
AY261365.1_Warmbaths             -------------------------------------------------------------------------------- [   400]
AY261364.1_Tengani_62            ttcactag-------atttcgcggtgcatac--------cggtgggc--------------------------------- [   400]
AY261363.1_Pretoriuskop/96/4     ttcactag-------atttcgcggtgcatac--------cggtgggc--------------------------------- [   400]
AY261362.1_Mkuzi_1979            ttcacaagatgcggaatttcgcggtgcataccgttttgacgataggt--------------------------------- [   400]
AY261361.1_Malawi_Lil-20/1_1983  gtaataggtggattaattgcagggcggcgac--atagagctagagac--------------------------------- [   400]
AY261360.1_Kenya_1950            ---------------------------------------------gcggcg----------------------------- [   400]
AM712240.1_OURT_88/3             -------------------------------------------------------------------------------- [   400]
AM712239.1_Benin_97/1            -------------------------------------------------------------------------------- [   400]
KM111295.1_Ken06.Bus             -------------------------------------------------------------------------------- [   400]
KM111294.1_Ken05/Tk1             --------------------------------------------ggctaaa----------------------------- [   400]
KM262844.1_strain_L60            -------------------------------------------------------------------------------- [   400]
KM102979.1_26544/OG10_Italy      -------------------------------------------------------------------------------- [   400]
KM262845.1_strain_NHV            -------------------------------------------------------------------------------- [   400]
KX354450.1_47/Ss/2008            ttcactag-------atttcgcggtgcatac--------cggtgggc--------------------------------- [   400]
KP055815.1_strain_BA71           ttcactag-------atttcgcggtgcatac--------cggtgggc--------------------------------- [   400]
MH025920.1_strain_R35            -------------------------------------------cggcggcg----------------------------- [   400]
MH025919.1_strain_N10            -------------------------------------------cggcggcg----------------------------- [   400]
MH025918.1_strain_R25            -------------------------------------------cggcggcg----------------------------- [   400]
MH025917.1_strain_R7             -------------------------------------------cggcggcg----------------------------- [   400]
MH025916.1_strain_R8             -------------------------------------------cggcggcg----------------------------- [   400]

FR682468.2_Georgia_2007/1        -------------------------------------------------------------------------------- [   480]
ASF_IND_20_CAD543                nnnnnnnnnnnnnnnnnnnnnnnnnnnnnnnnnnnnnnnnnnnnnnnnnnnnnnnnnnnnnnnnta-----aaattatta [   480]
LR899193.1_Germany_2020/1        -------------------------------------------------------------------------------- [   480]
MN393477.1_Wuhan_2019            ------------------------------------------------------------------------gataggca [   480]
MW396979.1_Timor-Leste/2019/1    ------------------------------------------------------------------------gataggca [   480]
MK940252.1_CN/2019/InnerMongolia -------------------------------------------------------------------------------- [   480]
MK128995.1_China/2018/AnhuiXCGQ  -------------------------------------------------------------------------------- [   480]
MH681419.1_ASFV/POL/2015         -------------------------------------------------------------------------------- [   480]
LS478113.1_Estonia_2014          ------------------------------------------------------------------------gataggca [   480]
KP843857.1_Odintsovo_02/14       -------------------------------------------------------------------------------- [   480]
```

```
FN557520.1_strain_E75             -------------------------------------------------------------------------------- [   480]
AY261366.1_Warthog                -------------------------------------------------------------------------------- [   480]
AY261365.1_Warmbaths              -------------------------------------------------------------------------------a [   480]
AY261364.1_Tengani_62             -------------------------------------------------------gataggtatcaataggcgttaaaga [   480]
AY261363.1_Pretoriuskop/96/4      ------------------------------------------------------------------------gttaggca [   480]
AY261362.1_Mkuzi_1979             -------------------------------------------------------gtcaaacatta-----aaat----- [   480]
AY261361.1_Malawi_Lil-20/1_1983   -----------------------------------------------------------agcataaaaaggaga------ [   480]
AY261360.1_Kenya_1950             -------------------------------------------------------------------------------- [   480]
AM712240.1_OURT_88/3              -------------------------------------------------------------------------------- [   480]
AM712239.1_Benin_97/1             -------------------------------------------------------------------------------- [   480]
KM111295.1_Ken06.Bus              -------------------------------------------------------------------------------- [   480]
KM111294.1_Ken05/Tk1              -------------------------------------------------------------------------------- [   480]
KM262844.1_strain_L60             -------------------------------------------------------------------------------- [   480]
KM102979.1_26544/OG10_Italy       -------------------------------------------------------------------------------- [   480]
KM262845.1_strain_NHV             -------------------------------------------------------------------------------- [   480]
KX354450.1_47/Ss/2008             -------------------------------------------------------gataggtatta-----aaataattt [   480]
KP055815.1_strain_BA71            -------------------------------------------------------gataggtatta-----aaataattt [   480]
MH025920.1_strain_R35             -------------------------------------------------------------------------------- [   480]
MH025919.1_strain_N10             -------------------------------------------------------------------------------- [   480]
MH025918.1_strain_R25             -------------------------------------------------------------------------------- [   480]
MH025917.1_strain_R7              -------------------------------------------------------------------------------- [   480]
MH025916.1_strain_R8              -------------------------------------------------------------------------------- [   480]

FR682468.2_Georgia_2007/1         -------------------------------------------------------------------------------- [   560]
ASF_IND_20_CAD543                 ttactgctnnnnnnnnnnnnnnnnnnnnnnnnnnnnnnnnnnnnnnnnnnnnnnnnnnnnnnnnnnnnnnnnnnntgtag [   560]
LR899193.1_Germany_2020/1         -------------------------------------------------------------------------------- [   560]
MN393477.1_Wuhan_2019             ttattgc--------------------------------------------------------------------tgtag [   560]
MW396979.1_Timor-Leste/2019/1     ttattgc--------------------------------------------------------------------tgtag [   560]
MK940252.1_CN/2019/InnerMongolia  -------------------------------------------------------------------------------- [   560]
MK128995.1_China/2018/AnhuiXCGQ   -------------------------------------------------------------------------------- [   560]
MH681419.1_ASFV/POL/2015          -------------------------------------------------------------------------------- [   560]
LS478113.1_Estonia_2014           ttattgc--------------------------------------------------------------------tgtag [   560]
KP843857.1_Odintsovo_02/14        -------------------------------------------------------------------------------- [   560]
FN557520.1_strain_E75             -------------------------------------------------------------------------------- [   560]
AY261366.1_Warthog                ----tgc--------------------------------------------------------------------tgtag [   560]
AY261365.1_Warmbaths              ctactgc--------------------------------------------------------------------tgtag [   560]
AY261364.1_Tengani_62             ttactgc--------------------------------------------------------------------tgtag [   560]
AY261363.1_Pretoriuskop/96/4      ttactgc--------------------------------------------------------------------tgtag [   560]
AY261362.1_Mkuzi_1979             -tattac--------------------------------------------------------------------tgtag [   560]
AY261361.1_Malawi_Lil-20/1_1983   ---------------------------------------------------------------------------tgcag [   560]
AY261360.1_Kenya_1950             ---------------------------------------------------------------------------gaaag [   560]
AM712240.1_OURT_88/3              -------------------------------------------------------------------------------- [   560]
AM712239.1_Benin_97/1             -------------------------------------------------------------------------------- [   560]
KM111295.1_Ken06.Bus              -------------------------------------------------------------------------------- [   560]
```

```
KM111294.1_Ken05/Tk1               ---------------------------------------------------------------------------aaaag [   560]
KM262844.1_strain_L60              -------------------------------------------------------------------------------- [   560]
KM102979.1_26544/OG10_Italy        -------------------------------------------------------------------------------- [   560]
KM262845.1_strain_NHV              -------------------------------------------------------------------------------- [   560]
KX354450.1_47/Ss/2008              gtactgc--------------------------------------------------------------------tgtag [   560]
KP055815.1_strain_BA71             gtactgc--------------------------------------------------------------------tgtag [   560]
MH025920.1_strain_R35              ---------------------------------------------------------------------------gagag [   560]
MH025919.1_strain_N10              ---------------------------------------------------------------------------gagag [   560]
MH025918.1_strain_R25              ---------------------------------------------------------------------------gagag [   560]
MH025917.1_strain_R7               ---------------------------------------------------------------------------gagag [   560]
MH025916.1_strain_R8               ---------------------------------------------------------------------------gagag [   560]

FR682468.2_Georgia_2007/1          ----------------------------actg-----ctgtaggcgtcaaagattaaaattat----------------- [   640]
ASF_IND_20_CAD543                  gc-gttaaacattaaaatt---attact....-----..........................----------------- [   640]
LR899193.1_Germany_2020/1          ----------------------------....-----..........................----------------- [   640]
MN393477.1_Wuhan_2019              gc-gttaaacattaaaatt---attact....-----......................a...----------------- [   640]
MW396979.1_Timor-Leste/2019/1      gc-gttaaacattaaaatt---attact....-----......................a...----------------- [   640]
MK940252.1_CN/2019/InnerMongolia   -------------------------------------------------------------------------------- [   640]
MK128995.1_China/2018/AnhuiXCGQ    -------------------------------------------------------------------------------- [   640]
MH681419.1_ASFV/POL/2015           -------------------------------------------------------------------------------- [   640]
LS478113.1_Estonia_2014            gc-gttaaacattaaaatt---attact....-----..........................----------------- [   640]
KP843857.1_Odintsovo_02/14         -------------------------------------------------------------------------------- [   640]
FN557520.1_strain_E75              -------------------------------------------------------------------------------- [   640]
AY261366.1_Warthog                 gc-gttaaacattaaaatt------------------------------------------------------------- [   640]
AY261365.1_Warmbaths               gc-gttaaacattaaaattattatttgt....-----t.........t...c...........tcgccggaaagtatttt [   640]
AY261364.1_Tengani_62              gc-gttaaacattaaaatt---attact....-----......a...t...............----------------- [   640]
AY261363.1_Pretoriuskop/96/4       gc-gttaaagattaaaatt---atttgt....-----........a.....c....g......----------------- [   640]
AY261362.1_Mkuzi_1979              gt-gtcaaagattaaaatt---attact....-----........a.t...c......c....----------------- [   640]
AY261361.1_Malawi_Lil-20/1_1983    gcggatggacatgcggggtaaaatgcct.gc.gccgt...a.a.tact.tgaggagg...c.c----------------- [   640]
AY261360.1_Kenya_1950              gtagttacaca----------agatgc----------------------------.g....tc----------------- [   640]
AM712240.1_OURT_88/3               -------------------------------------------------------------------------------- [   640]
AM712239.1_Benin_97/1              -------------------------------------------------------------------------------- [   640]
KM111295.1_Ken06.Bus               -------------------------------------------------------------------------------- [   640]
KM111294.1_Ken05/Tk1               gccg---------------------------------------------------------------------------- [   640]
KM262844.1_strain_L60              -------------------------------------------------------------------------------- [   640]
KM102979.1_26544/OG10_Italy        -------------------------------------------------------------------------------- [   640]
KM262845.1_strain_NHV              -------------------------------------------------------------------------------- [   640]
KX354450.1_47/Ss/2008              gc-gttaaagattaaaatt---attatt....-----..........t...c......c....----------------- [   640]
KP055815.1_strain_BA71             gc-gttaaagattaaaatt---attatt....-----..........t...c......c....----------------- [   640]
MH025920.1_strain_R35              gtagttacaca----------agttgcgg...gccg----------------gaagg....tc----------------- [   640]
MH025919.1_strain_N10              gtagttacaca----------agttgcgg...gccg----------------gaagg....tc----------------- [   640]
MH025918.1_strain_R25              gtagttacaca----------agttgcgg...gccg----------------gaagg....tc----------------- [   640]
MH025917.1_strain_R7               gtagttacaca----------agttgcgg...gccg----------------gaagg....tc----------------- [   640]
MH025916.1_strain_R8               gtagttacaca----------agttgcgg...gccg----------------gaagg....tc----------------- [   640]
```

```
FR682468.2_Georgia_2007/1        ---tactactgct-------------------------gtaggcgtta------aacattaaaattattactactgctgt [   720]
ASF_IND_20_CAD543                ---..........-------------------------..........------.........cnnnnnnnnnnnnnnn. [   720]
LR899193.1_Germany_2020/1        ---..........-------------------------..........------.......................... [   720]
MN393477.1_Wuhan_2019            ---..g.......-------------------------.c........------.......................... [   720]
MW396979.1_Timor-Leste/2019/1    ---..g.......-------------------------.c........------.......................... [   720]
MK940252.1_CN/2019/InnerMongolia -------------------------------------------------------------------------------- [   720]
MK128995.1_China/2018/AnhuiXCGQ  -------------------------------------------------------------------------------- [   720]
MH681419.1_ASFV/POL/2015         -------------------------------------------------------------------------------- [   720]
LS478113.1_Estonia_2014          ---..........-------------------------..........------.......................... [   720]
KP843857.1_Odintsovo_02/14       -------------------------------------------------------------------------------- [   720]
FN557520.1_strain_E75            -------------------------------------------------------------------------------- [   720]
AY261366.1_Warthog               ------------------------------------------------------------------...tg......... [   720]
AY261365.1_Warmbaths             gtg...cgta.t.tcactagatttcgcggtgcataccg..g....a..------ggt....g...a...tg......... [   720]
AY261364.1_Tengani_62            ---..........-------------------------....a.....------..g............tg......... [   720]
AY261363.1_Pretoriuskop/96/4     ---....g.....-------------------------..........------.......................... [   720]
AY261362.1_Mkuzi_1979            ---..........-------------------------..........------..------.................. [   720]
AY261361.1_Malawi_Lil-20/1_1983  acgg.gcgta.t.tcaca--------------------ag.....ga.tatcgcggtgcc.....gca...cgt.ttga. [   720]
AY261360.1_Kenya_1950            ----------------------------------------------------gcggtgcc.....gca...cgt.ttg.. [   720]
AM712240.1_OURT_88/3             -------------------------------------------------------------------------------- [   720]
AM712239.1_Benin_97/1            -------------------------------------------------------------------------------- [   720]
KM111295.1_Ken06.Bus             -------------------------------------------------------------------------------- [   720]
KM111294.1_Ken05/Tk1             ---------------------------------------------------------gcc....aa-------------- [   720]
KM262844.1_strain_L60            -------------------------------------------------------------------------------- [   720]
KM102979.1_26544/OG10_Italy      -------------------------------------------------------------------------------- [   720]
KM262845.1_strain_NHV            -------------------------------------------------------------------------------- [   720]
KX354450.1_47/Ss/2008            ---..........-------------------------a.........------......................t... [   720]
KP055815.1_strain_BA71           ---..........-------------------------a.........------......................t... [   720]
MH025920.1_strain_R35            ----------------------------------------------------gcggtgcc.....gca...cgt.ttg.. [   720]
MH025919.1_strain_N10            ----------------------------------------------------gcggtgcc.....gca...cgt.ttg.. [   720]
MH025918.1_strain_R25            ----------------------------------------------------gcggtgcc.....gca...cgt.ttg.. [   720]
MH025917.1_strain_R7             ----------------------------------------------------gcggtgcc.....gca...cgt.ttg.. [   720]
MH025916.1_strain_R8             ----------------------------------------------------gcggtgcc.....gca...cgt.ttg.. [   720]

FR682468.2_Georgia_2007/1        aggcgttaaagattaaaaatat--tagtactgctg--------------------------caggcgttaaagattaaaa [   800]
ASF_IND_20_CAD543                ..........c.......t...--..c.....t..--------------------------t......c........... [   800]
LR899193.1_Germany_2020/1        ......................--...........--------------------------................... [   800]
MN393477.1_Wuhan_2019            ......................--...........--------------------------...........c....... [   800]
MW396979.1_Timor-Leste/2019/1    ......................--...........--------------------------...........c....... [   800]
MK940252.1_CN/2019/InnerMongolia -------------------------------------------------------------------------------- [   800]
MK128995.1_China/2018/AnhuiXCGQ  -------------------------------------------------------------------------------- [   800]
MH681419.1_ASFV/POL/2015         -------------------------------------------------------------------------------- [   800]
LS478113.1_Estonia_2014          ......................--...........--------------------------................... [   800]
KP843857.1_Odintsovo_02/14       -------------------------------------------------------------------------------- [   800]
```

```
FN557520.1_strain_E75            -------------------------------------------------------------------------------- [   800]
AY261366.1_Warthog               ..........c.......t...--.t.........--------------------------t.................. [   800]
AY261365.1_Warmbaths             ..........c.......t...--..c.gt..ta.--------------------------t..........c....... [   800]
AY261364.1_Tengani_62            ..........c.......t...--..c........--------------------------t..........c....... [   800]
AY261363.1_Pretoriuskop/96/4     ..................t...--.t.........--------------------------t..........c....... [   800]
AY261362.1_Mkuzi_1979            ..........c......ct...--..t........--------------------------t..........c....--- [   800]
AY261361.1_Malawi_Lil-20/1_1983  g..a.ac....gc.....ca..--c.t.g...a..-------------------------c.g...a.a...c....g.. [   800]
AY261360.1_Kenya_1950            g.....c....t......t...tc.gt.g...tg.-----cggacgagttttgcggtgctgtg..............--- [   800]
AM712240.1_OURT_88/3             -------------------------------------------------------------------------------- [   800]
AM712239.1_Benin_97/1            -------------------------------------------------------------------------------- [   800]
KM111295.1_Ken06.Bus             -------------------------------------------------------------------------------- [   800]
KM111294.1_Ken05/Tk1             -------....cccg...---------------------------------------------------.t....c.--- [   800]
KM262844.1_strain_L60            -------------------------------------------------------------------------------- [   800]
KM102979.1_26544/OG10_Italy      -------------------------------------------------------------------------------- [   800]
KM262845.1_strain_NHV            -------------------------------------------------------------------------------- [   800]
KX354450.1_47/Ss/2008            ..........c.......t...--..c........--------------------------t......c........... [   800]
KP055815.1_strain_BA71           ..........c.......t...--..c........--------------------------t......c........... [   800]
MH025920.1_strain_R35            g.....c....t......t...tc.gt.g...tg.cgggccggacgaattttgcgg----gtg................. [   800]
MH025919.1_strain_N10            g.....c....t......t...tc.gt.g...tg.cgggccggacgaattttgcgg----gtg................. [   800]
MH025918.1_strain_R25            g.....c....t......t...tc.gt.g...tg.cgggccggacgaattttgcgg----gtg................. [   800]
MH025917.1_strain_R7             g.....c....t......t...tc.gt.g...tg.cgggccggacgaattttgcgg----gtg................. [   800]
MH025916.1_strain_R8             g.....c....t......t...tc.gt.g...tg.cgggccggacgaattttgcgg----gtg................. [   800]

FR682468.2_Georgia_2007/1        atattagtactgctgcaggcgttaaacattaaaattattg----tactgctgtaggcgttaaaca-ttaaaattattac- [   880]
ASF_IND_20_CAD543                t.....c........t.......................ac---.....................-.............- [   880]
LR899193.1_Germany_2020/1        ........................................----.....................-.............- [   880]
MN393477.1_Wuhan_2019            t.......................................----.....................-.............- [   880]
MW396979.1_Timor-Leste/2019/1    t.......................................----.....................-.............- [   880]
MK940252.1_CN/2019/InnerMongolia -------------------------------------------------------------------------------- [   880]
MK128995.1_China/2018/AnhuiXCGQ  -------------------------------------------------------------------------------- [   880]
MH681419.1_ASFV/POL/2015         -------------------------------------------------------------------------------- [   880]
LS478113.1_Estonia_2014          ........................................----.....................-.............- [   880]
KP843857.1_Odintsovo_02/14       -------------------------------------------------------------------------------- [   880]
FN557520.1_strain_E75            -------------------------------------------------------------------------------- [   880]
AY261366.1_Warthog               t....t.........t.......................tg--------------------------------------- [   880]
AY261365.1_Warmbaths             t....t........at.......................tgtac.g.................g.-............tt [   880]
AY261364.1_Tengani_62            t....t.........t......c...g............tg---...............c...g.-.............- [   880]
AY261363.1_Pretoriuskop/96/4     t....t.........t.......................tg---.....................-.............- [   880]
AY261362.1_Mkuzi_1979            ---...t........t.....-------.....c.....ac---....................-------.......t- [   880]
AY261361.1_Malawi_Lil-20/1_1983  t.....c........gg...cga......g.........ac---........gg.........ggt.............- [   880]
AY261360.1_Kenya_1950            t.....c.g...ta.t...............g......------.c...t...t.........g.-...tt........- [   880]
AM712240.1_OURT_88/3             -------------------------------------------------------------------------------- [   880]
AM712239.1_Benin_97/1            -------------------------------------------------------------------------------- [   880]
KM111295.1_Ken06.Bus             -------------------------------------------------------------------------------- [   880]
```

```
KM111294.1_Ken05/Tk1             -------------------------------------------------------------------------------- [   880]
KM262844.1_strain_L60            -------------------------------------------------------------------------------- [   880]
KM102979.1_26544/OG10_Italy      -------------------------------------------------------------------------------- [   880]
KM262845.1_strain_NHV            -------------------------------------------------------------------------------- [   880]
KX354450.1_47/Ss/2008            t.....c........t.................c.....ac---.......a.............-.............- [   880]
KP055815.1_strain_BA71           t.....c........t.................c.....ac---.......a.............-.............- [   880]
MH025920.1_strain_R35            t....cc.gt..t..tt......................ac---.g.......t...........-...........c.- [   880]
MH025919.1_strain_N10            t....cc.gt..t..tt......................ac---.g.......t...........-...........c.- [   880]
MH025918.1_strain_R25            t....cc.gt..t..tt......................ac---.g.......t...........-...........c.- [   880]
MH025917.1_strain_R7             t....cc.gt..t..tt......................ac---.g.......t...........-...........c.- [   880]
MH025916.1_strain_R8             t....cc.gt..t..tt......................ac---.g.......t...........-...........c.- [   880]

FR682468.2_Georgia_2007/1        --tactg------------------------------------------------------------------------- [   960]
ASF_IND_20_CAD543                --.....ttgtaggcgttaaacattaaaattattactactgtt---gtaggcgtcaaagattaaaattattactactg-- [   960]
LR899193.1_Germany_2020/1        --.....------------------------------------------------------------------------- [   960]
MN393477.1_Wuhan_2019            --.....------------------------------------------------------------------------- [   960]
MW396979.1_Timor-Leste/2019/1    --.....ctgtaggcgtcaaagattaaaattattactactgct---gtaggcgttaaacattaaaattattactactgct [   960]
MK940252.1_CN/2019/InnerMongolia -------------------------------------------------------------------------------- [   960]
MK128995.1_China/2018/AnhuiXCGQ  -------------------------------------------------------------------------------- [   960]
MH681419.1_ASFV/POL/2015         -------------------------------------------------------------------------------- [   960]
LS478113.1_Estonia_2014          --.....ctgtaggcgtcaaagattaaaattattactactgct---gtaggcgttaaacattaaaattattactactg-- [   960]
KP843857.1_Odintsovo_02/14       -------------------------------------------------------------------------------- [   960]
FN557520.1_strain_E75            -------------------------------------------------------------------------------- [   960]
AY261366.1_Warthog               -------------------------------------------------------------------------tactg-- [   960]
AY261365.1_Warmbaths             tg.....ttgtaggcgttaaacattaaaattattatttgtact---gctggcgtcaaacattaaaattattactactg-- [   960]
AY261364.1_Tengani_62            --.....ctgtaggcgttaaacattaaaattattactactgct---gtaggcgttaaacattaaaattatttgtactg-- [   960]
AY261363.1_Pretoriuskop/96/4     --.....ctgtaggcgttaaagattaaaattatttgtactgct---gtaggcgttaaacattaaaattatttgtactg-- [   960]
AY261362.1_Mkuzi_1979            --.....ctgtaggcgttaaac------attattattactgct---gtaggcgttaaac-------ttattactactg-- [   960]
AY261361.1_Malawi_Lil-20/1_1983  --.....ctgtgggcggtaaagattaaaattattactactggt---gtggacgttaaacattaaaattattattactg-- [   960]
AY261360.1_Kenya_1950            --.g...tggtaggcgttaaacatt---attattattactgctgtggtaggcgttaaacatt---attattattactgct [   960]
AM712240.1_OURT_88/3             -------------------------------------------------------------------------------- [   960]
AM712239.1_Benin_97/1            -------------------------------------------------------------------------------- [   960]
KM111295.1_Ken06.Bus             -------------------------------------------------------------------------------- [   960]
KM111294.1_Ken05/Tk1             ---------------------cactga--------------------------ttaaacttt------------------ [   960]
KM262844.1_strain_L60            -------------------------------------------------------------------------------- [   960]
KM102979.1_26544/OG10_Italy      -------------------------------------------------------------------------------- [   960]
KM262845.1_strain_NHV            -------------------------------------------------------------------------------- [   960]
KX354450.1_47/Ss/2008            --.....ttgtaggcgtcaaag---------------------------------------------------------- [   960]
KP055815.1_strain_BA71           --.....ttgtaggcgtcaaagattaaaattattactactgct---gtaggcgttaaacattaaaattattactactg-- [   960]
MH025920.1_strain_R35            --.gt..ttgttggcgttaaacattaaaattattactgctgctgtggtaggcgttaaacattaaaattattaatactgct [   960]
MH025919.1_strain_N10            --.gt..ttgttggcgttaaacattaaaattattactgctgctgtggtaggcgttaaacattaaaattattaatactgct [   960]
MH025918.1_strain_R25            --.gt..ttgttggcgttaaacattaaaattattactgctgctgtggtaggcgttaaacattaaaattattaatactgct [   960]
MH025917.1_strain_R7             --.gt..ttgttggcgttaaacattaaaattattactgctgctgtggtaggcgttaaacattaaaattattaatactgct [   960]
MH025916.1_strain_R8             --.gt..ttgttggcgttaaacattaaaattattactgctgctgtggtaggcgttaaacattaaaattattaatactgct [   960]
```

```
FR682468.2_Georgia_2007/1        -------------------------------------------------------------------------------- [  1040]
ASF_IND_20_CAD543                -------------------------------------------------------------------------------- [  1040]
LR899193.1_Germany_2020/1        -------------------------------------------------------------------------------- [  1040]
MN393477.1_Wuhan_2019            -------------------------------------------------------------------------------- [  1040]
MW396979.1_Timor-Leste/2019/1    gcaggcgttaaacattaaaattattgtactgctgtaggcgttaaacattaaaattattactactgctgtaggcgtcaaag [  1040]
MK940252.1_CN/2019/InnerMongolia -------------------------------------------------------------------------------- [  1040]
MK128995.1_China/2018/AnhuiXCGQ  -------------------------------------------------------------------------------- [  1040]
MH681419.1_ASFV/POL/2015         -------------------------------------------------------------------------------- [  1040]
LS478113.1_Estonia_2014          -------------------------------------------------------------------------------- [  1040]
KP843857.1_Odintsovo_02/14       -------------------------------------------------------------------------------- [  1040]
FN557520.1_strain_E75            -------------------------------------------------------------------------------- [  1040]
AY261366.1_Warthog               -------------------------------------------------------------------------------- [  1040]
AY261365.1_Warmbaths             -------------------------------------------------------------------------------- [  1040]
AY261364.1_Tengani_62            -------------------------------------------------------------------------------- [  1040]
AY261363.1_Pretoriuskop/96/4     -------------------------------------------------------------------------------- [  1040]
AY261362.1_Mkuzi_1979            -------------------------------------------------------------------------------- [  1040]
AY261361.1_Malawi_Lil-20/1_1983  -------------------------------------------------------------------------------- [  1040]
AY261360.1_Kenya_1950            g------------------------------------------------------------------------------- [  1040]
AM712240.1_OURT_88/3             -------------------------------------------------------------------------------- [  1040]
AM712239.1_Benin_97/1            -------------------------------------------------------------------------------- [  1040]
KM111295.1_Ken06.Bus             -------------------------------------------------------------------------------- [  1040]
KM111294.1_Ken05/Tk1             -------------------------------------------------------------------------------- [  1040]
KM262844.1_strain_L60            -------------------------------------------------------------------------------- [  1040]
KM102979.1_26544/OG10_Italy      -------------------------------------------------------------------------------- [  1040]
KM262845.1_strain_NHV            -------------------------------------------------------------------------------- [  1040]
KX354450.1_47/Ss/2008            -------------------------------------------------------------------------------- [  1040]
KP055815.1_strain_BA71           -------------------------------------------------------------------------------- [  1040]
MH025920.1_strain_R35            g------------------------------------------------------------------------------- [  1040]
MH025919.1_strain_N10            g------------------------------------------------------------------------------- [  1040]
MH025918.1_strain_R25            g------------------------------------------------------------------------------- [  1040]
MH025917.1_strain_R7             g------------------------------------------------------------------------------- [  1040]
MH025916.1_strain_R8             g------------------------------------------------------------------------------- [  1040]

FR682468.2_Georgia_2007/1        -------------------------------------------------------------------------------- [  1120]
ASF_IND_20_CAD543                -------------------------------------------------------------------------------- [  1120]
LR899193.1_Germany_2020/1        -------------------------------------------------------------------------------- [  1120]
MN393477.1_Wuhan_2019            -------------------------------------------------------------------------------- [  1120]
MW396979.1_Timor-Leste/2019/1    attaaaattattactactgctgtaggcgttaaacattaaaattattactactgctgtaggcgtcaaagattaaaattatt [  1120]
MK940252.1_CN/2019/InnerMongolia -------------------------------------------------------------------------------- [  1120]
MK128995.1_China/2018/AnhuiXCGQ  -------------------------------------------------------------------------------- [  1120]
MH681419.1_ASFV/POL/2015         -------------------------------------------------------------------------------- [  1120]
LS478113.1_Estonia_2014          -------------------------------------------------------------------------------- [  1120]
KP843857.1_Odintsovo_02/14       -------------------------------------------------------------------------------- [  1120]
```

```
FN557520.1_strain_E75            -------------------------------------------------------------------------------- [  1120]
AY261366.1_Warthog               -------------------------------------------------------------------------------- [  1120]
AY261365.1_Warmbaths             -------------------------------------------------------------------------------- [  1120]
AY261364.1_Tengani_62            -------------------------------------------------------------------------------- [  1120]
AY261363.1_Pretoriuskop/96/4     -------------------------------------------------------------------------------- [  1120]
AY261362.1_Mkuzi_1979            -------------------------------------------------------------------------------- [  1120]
AY261361.1_Malawi_Lil-20/1_1983  -------------------------------------------------------------------------------- [  1120]
AY261360.1_Kenya_1950            -------------------------------------------------------------------------------- [  1120]
AM712240.1_OURT_88/3             -------------------------------------------------------------------------------- [  1120]
AM712239.1_Benin_97/1            -------------------------------------------------------------------------------- [  1120]
KM111295.1_Ken06.Bus             -------------------------------------------------------------------------------- [  1120]
KM111294.1_Ken05/Tk1             -------------------------------------------------------------------------------- [  1120]
KM262844.1_strain_L60            -------------------------------------------------------------------------------- [  1120]
KM102979.1_26544/OG10_Italy      -------------------------------------------------------------------------------- [  1120]
KM262845.1_strain_NHV            -------------------------------------------------------------------------------- [  1120]
KX354450.1_47/Ss/2008            -------------------------------------------------------------------------------- [  1120]
KP055815.1_strain_BA71           -------------------------------------------------------------------------------- [  1120]
MH025920.1_strain_R35            -------------------------------------------------------------------------------- [  1120]
MH025919.1_strain_N10            -------------------------------------------------------------------------------- [  1120]
MH025918.1_strain_R25            -------------------------------------------------------------------------------- [  1120]
MH025917.1_strain_R7             -------------------------------------------------------------------------------- [  1120]
MH025916.1_strain_R8             -------------------------------------------------------------------------------- [  1120]

FR682468.2_Georgia_2007/1        -------------------------------------------------------------------------------- [  1200]
ASF_IND_20_CAD543                -------------------------------------------------------------------------------- [  1200]
LR899193.1_Germany_2020/1        -------------------------------------------------------------------------------- [  1200]
MN393477.1_Wuhan_2019            -------------------------------------------------------------------------------- [  1200]
MW396979.1_Timor-Leste/2019/1    actactgctgtaggcgttaaacattaaaattattactactgctgtaggcgttaaagattaaaaatattagtactgctgca [  1200]
MK940252.1_CN/2019/InnerMongolia -------------------------------------------------------------------------------- [  1200]
MK128995.1_China/2018/AnhuiXCGQ  -------------------------------------------------------------------------------- [  1200]
MH681419.1_ASFV/POL/2015         -------------------------------------------------------------------------------- [  1200]
LS478113.1_Estonia_2014          -------------------------------------------------------------------------------- [  1200]
KP843857.1_Odintsovo_02/14       -------------------------------------------------------------------------------- [  1200]
FN557520.1_strain_E75            -------------------------------------------------------------------------------- [  1200]
AY261366.1_Warthog               -------------------------------------------------------------------------------- [  1200]
AY261365.1_Warmbaths             -------------------------------------------------------------------------------- [  1200]
AY261364.1_Tengani_62            -------------------------------------------------------------------------------- [  1200]
AY261363.1_Pretoriuskop/96/4     -------------------------------------------------------------------------------- [  1200]
AY261362.1_Mkuzi_1979            -------------------------------------------------------------------------------- [  1200]
AY261361.1_Malawi_Lil-20/1_1983  -------------------------------------------------------------------------------- [  1200]
AY261360.1_Kenya_1950            -------------------------------------------------------------------------------- [  1200]
AM712240.1_OURT_88/3             -------------------------------------------------------------------------------- [  1200]
AM712239.1_Benin_97/1            -------------------------------------------------------------------------------- [  1200]
KM111295.1_Ken06.Bus             -------------------------------------------------------------------------------- [  1200]
```

```
KM111294.1_Ken05/Tk1             -------------------------------------------------------------------------------- [  1200]
KM262844.1_strain_L60            -------------------------------------------------------------------------------- [  1200]
KM102979.1_26544/OG10_Italy      -------------------------------------------------------------------------------- [  1200]
KM262845.1_strain_NHV            -------------------------------------------------------------------------------- [  1200]
KX354450.1_47/Ss/2008            -------------------------------------------------------------------------------- [  1200]
KP055815.1_strain_BA71           -------------------------------------------------------------------------------- [  1200]
MH025920.1_strain_R35            -------------------------------------------------------------------------------- [  1200]
MH025919.1_strain_N10            -------------------------------------------------------------------------------- [  1200]
MH025918.1_strain_R25            -------------------------------------------------------------------------------- [  1200]
MH025917.1_strain_R7             -------------------------------------------------------------------------------- [  1200]
MH025916.1_strain_R8             -------------------------------------------------------------------------------- [  1200]

FR682468.2_Georgia_2007/1        -------------------------------------------------------------------------------- [  1280]
ASF_IND_20_CAD543                -------------------------------------------------------------------------------- [  1280]
LR899193.1_Germany_2020/1        -------------------------------------------------------------------------------- [  1280]
MN393477.1_Wuhan_2019            -------------------------------------------------------------------------------- [  1280]
MW396979.1_Timor-Leste/2019/1    ggcgttaaagattaaaaatattagtactgctgcaggcgttaaacattaaaattattgtactgctgtaggcgttaaacatt [  1280]
MK940252.1_CN/2019/InnerMongolia -------------------------------------------------------------------------------- [  1280]
MK128995.1_China/2018/AnhuiXCGQ  -------------------------------------------------------------------------------- [  1280]
MH681419.1_ASFV/POL/2015         -------------------------------------------------------------------------------- [  1280]
LS478113.1_Estonia_2014          -------------------------------------------------------------------------------- [  1280]
KP843857.1_Odintsovo_02/14       -------------------------------------------------------------------------------- [  1280]
FN557520.1_strain_E75            -------------------------------------------------------------------------------- [  1280]
AY261366.1_Warthog               -------------------------------------------------------------------------------- [  1280]
AY261365.1_Warmbaths             -------------------------------------------------------------------------------- [  1280]
AY261364.1_Tengani_62            -------------------------------------------------------------------------------- [  1280]
AY261363.1_Pretoriuskop/96/4     -------------------------------------------------------------------------------- [  1280]
AY261362.1_Mkuzi_1979            -------------------------------------------------------------------------------- [  1280]
AY261361.1_Malawi_Lil-20/1_1983  -------------------------------------------------------------------------------- [  1280]
AY261360.1_Kenya_1950            -------------------------------------------------------------------------------- [  1280]
AM712240.1_OURT_88/3             -------------------------------------------------------------------------------- [  1280]
AM712239.1_Benin_97/1            -------------------------------------------------------------------------------- [  1280]
KM111295.1_Ken06.Bus             -------------------------------------------------------------------------------- [  1280]
KM111294.1_Ken05/Tk1             -------------------------------------------------------------------------------- [  1280]
KM262844.1_strain_L60            -------------------------------------------------------------------------------- [  1280]
KM102979.1_26544/OG10_Italy      -------------------------------------------------------------------------------- [  1280]
KM262845.1_strain_NHV            -------------------------------------------------------------------------------- [  1280]
KX354450.1_47/Ss/2008            -------------------------------------------------------------------------------- [  1280]
KP055815.1_strain_BA71           -------------------------------------------------------------------------------- [  1280]
MH025920.1_strain_R35            -------------------------------------------------------------------------------- [  1280]
MH025919.1_strain_N10            -------------------------------------------------------------------------------- [  1280]
MH025918.1_strain_R25            -------------------------------------------------------------------------------- [  1280]
MH025917.1_strain_R7             -------------------------------------------------------------------------------- [  1280]
MH025916.1_strain_R8             -------------------------------------------------------------------------------- [  1280]
```

```
FR682468.2_Georgia_2007/1        -------------------------------------------------------------------------------- [  1360]
ASF_IND_20_CAD543                -------------------------------------------------------------------------------- [  1360]
LR899193.1_Germany_2020/1        -------------------------------------------------------------------------------- [  1360]
MN393477.1_Wuhan_2019            -------------------------------------------------------------------------------- [  1360]
MW396979.1_Timor-Leste/2019/1    aaaattattactactgctgtaggcgtcaaagattaaaattattactactgctgtaggcgttaaacattaaaattattact [  1360]
MK940252.1_CN/2019/InnerMongolia -------------------------------------------------------------------------------- [  1360]
MK128995.1_China/2018/AnhuiXCGQ  -------------------------------------------------------------------------------- [  1360]
MH681419.1_ASFV/POL/2015         -------------------------------------------------------------------------------- [  1360]
LS478113.1_Estonia_2014          -------------------------------------------------------------------------------- [  1360]
KP843857.1_Odintsovo_02/14       -------------------------------------------------------------------------------- [  1360]
FN557520.1_strain_E75            -------------------------------------------------------------------------------- [  1360]
AY261366.1_Warthog               -------------------------------------------------------------------------------- [  1360]
AY261365.1_Warmbaths             -------------------------------------------------------------------------------- [  1360]
AY261364.1_Tengani_62            -------------------------------------------------------------------------------- [  1360]
AY261363.1_Pretoriuskop/96/4     -------------------------------------------------------------------------------- [  1360]
AY261362.1_Mkuzi_1979            -------------------------------------------------------------------------------- [  1360]
AY261361.1_Malawi_Lil-20/1_1983  -------------------------------------------------------------------------------- [  1360]
AY261360.1_Kenya_1950            -------------------------------------------------------------------------------- [  1360]
AM712240.1_OURT_88/3             -------------------------------------------------------------------------------- [  1360]
AM712239.1_Benin_97/1            -------------------------------------------------------------------------------- [  1360]
KM111295.1_Ken06.Bus             -------------------------------------------------------------------------------- [  1360]
KM111294.1_Ken05/Tk1             -------------------------------------------------------------------------------- [  1360]
KM262844.1_strain_L60            -------------------------------------------------------------------------------- [  1360]
KM102979.1_26544/OG10_Italy      -------------------------------------------------------------------------------- [  1360]
KM262845.1_strain_NHV            -------------------------------------------------------------------------------- [  1360]
KX354450.1_47/Ss/2008            -------------------------------------------------------------------------------- [  1360]
KP055815.1_strain_BA71           -------------------------------------------------------------------------------- [  1360]
MH025920.1_strain_R35            -------------------------------------------------------------------------------- [  1360]
MH025919.1_strain_N10            -------------------------------------------------------------------------------- [  1360]
MH025918.1_strain_R25            -------------------------------------------------------------------------------- [  1360]
MH025917.1_strain_R7             -------------------------------------------------------------------------------- [  1360]
MH025916.1_strain_R8             -------------------------------------------------------------------------------- [  1360]

FR682468.2_Georgia_2007/1        -------------------------------------------------------------------------------- [  1440]
ASF_IND_20_CAD543                -------------------------------------------------------------------------------- [  1440]
LR899193.1_Germany_2020/1        -------------------------------------------------------------------------------- [  1440]
MN393477.1_Wuhan_2019            -------------------------------------------------------------------------------- [  1440]
MW396979.1_Timor-Leste/2019/1    actgctgtaggcgttaaagattaaaaatattagtactgctgcaggcgttaaagattaaaaatattagtactgctgcaggc [  1440]
MK940252.1_CN/2019/InnerMongolia -------------------------------------------------------------------------------- [  1440]
MK128995.1_China/2018/AnhuiXCGQ  -------------------------------------------------------------------------------- [  1440]
MH681419.1_ASFV/POL/2015         -------------------------------------------------------------------------------- [  1440]
LS478113.1_Estonia_2014          -------------------------------------------------------------------------------- [  1440]
KP843857.1_Odintsovo_02/14       -------------------------------------------------------------------------------- [  1440]
```

```
FN557520.1_strain_E75          -------------------------------------------------------------------------------- [  1440]
AY261366.1_Warthog             -------------------------------------------------------------------------------- [  1440]
AY261365.1_Warmbaths           -------------------------------------------------------------------------------- [  1440]
AY261364.1_Tengani_62          -------------------------------------------------------------------------------- [  1440]
AY261363.1_Pretoriuskop/96/4   -------------------------------------------------------------------------------- [  1440]
AY261362.1_Mkuzi_1979          -------------------------------------------------------------------------------- [  1440]
AY261361.1_Malawi_Lil-20/1_1983 -------------------------------------------------------------------------------- [  1440]
AY261360.1_Kenya_1950          -------------------------------------------------------------------------------- [  1440]
AM712240.1_OURT_88/3           -------------------------------------------------------------------------------- [  1440]
AM712239.1_Benin_97/1          -------------------------------------------------------------------------------- [  1440]
KM111295.1_Ken06.Bus           -------------------------------------------------------------------------------- [  1440]
KM111294.1_Ken05/Tk1           -------------------------------------------------------------------------------- [  1440]
KM262844.1_strain_L60          -------------------------------------------------------------------------------- [  1440]
KM102979.1_26544/OG10_Italy    -------------------------------------------------------------------------------- [  1440]
KM262845.1_strain_NHV          -------------------------------------------------------------------------------- [  1440]
KX354450.1_47/Ss/2008          -------------------------------------------------------------------------------- [  1440]
KP055815.1_strain_BA71         -------------------------------------------------------------------------------- [  1440]
MH025920.1_strain_R35          -------------------------------------------------------------------------------- [  1440]
MH025919.1_strain_N10          -------------------------------------------------------------------------------- [  1440]
MH025918.1_strain_R25          -------------------------------------------------------------------------------- [  1440]
MH025917.1_strain_R7           -------------------------------------------------------------------------------- [  1440]
MH025916.1_strain_R8           -------------------------------------------------------------------------------- [  1440]

FR682468.2_Georgia_2007/1      -----------------------------------------------------------ctgtaggcgtcaaagattaaa [  1520]
ASF_IND_20_CAD543              -----------------------------------------------------------..........t...c...... [  1520]
LR899193.1_Germany_2020/1      -----------------------------------------------------------..................... [  1520]
MN393477.1_Wuhan_2019          -----------------------------------------------------------..................... [  1520]
MW396979.1_Timor-Leste/2019/1  gttaaacattaaaattattgtactgctgtaggcgttaaacattaaaattattactactg..................... [  1520]
MK940252.1_CN/2019/InnerMongolia -------------------------------------------------------------------------------- [  1520]
MK128995.1_China/2018/AnhuiXCGQ -------------------------------------------------------------------------------- [  1520]
MH681419.1_ASFV/POL/2015       -------------------------------------------------------------------------------- [  1520]
LS478113.1_Estonia_2014        -----------------------------------------------------------..........t.......... [  1520]
KP843857.1_Odintsovo_02/14     -------------------------------------------------------------------------------- [  1520]
FN557520.1_strain_E75          -------------------------------------------------------------------------------- [  1520]
AY261366.1_Warthog             -----------------------------------------------------------t.........t.......... [  1520]
AY261365.1_Warmbaths           -----------------------------------------------------------..........t...c...... [  1520]
AY261364.1_Tengani_62          -----------------------------------------------------------..................... [  1520]
AY261363.1_Pretoriuskop/96/4   -----------------------------------------------------------..........t...c...... [  1520]
AY261362.1_Mkuzi_1979          -----------------------------------------------------------..........t.......... [  1520]
AY261361.1_Malawi_Lil-20/1_1983 -----------------------------------------------------------t.........t...c...... [  1520]
AY261360.1_Kenya_1950          -----------------------------------------------------------tg........t...c....tt [  1520]
AM712240.1_OURT_88/3           -------------------------------------------------------------------------------- [  1520]
AM712239.1_Benin_97/1          -------------------------------------------------------------------------------- [  1520]
KM111295.1_Ken06.Bus           -------------------------------------------------------------------------------- [  1520]
```

```
KM111294.1_Ken05/Tk1            -------------------------------------------------------------------------------- [  1520]
KM262844.1_strain_L60           -------------------------------------------------------------------------------- [  1520]
KM102979.1_26544/OG10_Italy     -------------------------------------------------------------------------------- [  1520]
KM262845.1_strain_NHV           -------------------------------------------------------------------------------- [  1520]
KX354450.1_47/Ss/2008           -------------------------------------------------------------------------------- [  1520]
KP055815.1_strain_BA71          -----------------------------------------------------------..........t...c...... [  1520]
MH025920.1_strain_R35           -----------------------------------------------------------tg........t...c...... [  1520]
MH025919.1_strain_N10           -----------------------------------------------------------tg........t...c...... [  1520]
MH025918.1_strain_R25           -----------------------------------------------------------tg........t...c...... [  1520]
MH025917.1_strain_R7            -----------------------------------------------------------tg........t...c...... [  1520]
MH025916.1_strain_R8            -----------------------------------------------------------tg........t...c...... [  1520]

FR682468.2_Georgia_2007/1       attattactactgctgtaggcgttaaacattaaaattattactactgctgta--ggcgttaaagattaaaaatattagta [  1600]
ASF_IND_20_CAD543               ...............................................t....--.........c.......t.....c.. [  1600]
LR899193.1_Germany_2020/1       ....................................................--.......................... [  1600]
MN393477.1_Wuhan_2019           ....................................................--.......................... [  1600]
MW396979.1_Timor-Leste/2019/1   ....................................................--.......................... [  1600]
MK940252.1_CN/2019/InnerMongolia -------------------------------------------------------------------------------- [  1600]
MK128995.1_China/2018/AnhuiXCGQ -------------------------------------------------------------------------------- [  1600]
MH681419.1_ASFV/POL/2015        -------------------------------------------------------------------------------- [  1600]
LS478113.1_Estonia_2014         .a.....g........c..........g........................--.................t........ [  1600]
KP843857.1_Odintsovo_02/14      -------------------------------------------------------------------------------- [  1600]
FN557520.1_strain_E75           -------------------------------------------------------------------------------- [  1600]
AY261366.1_Warthog              .............t.........c...g....g.......tg..........gg.....c...c.......t.....c.. [  1600]
AY261365.1_Warmbaths            .............t..........................tg.......a..--.........c.......t....t... [  1600]
AY261364.1_Tengani_62           ......tg................................tg..........--.....c...........t....t... [  1600]
AY261363.1_Pretoriuskop/96/4    ........................................tg.....t....--.....c...........t....t... [  1600]
AY261362.1_Mkuzi_1979           c...........................------.......t..........--....-------.....ct.....c.. [  1600]
AY261361.1_Malawi_Lil-20/1_1983 .................g.....c...gt.......................-g.........c.......t.....c.. [  1600]
AY261360.1_Kenya_1950           .........g...ta.................tt.........g...tg...--.........c...---.t.....c.g [  1600]
AM712240.1_OURT_88/3            -------------------------------------------------------------------------------- [  1600]
AM712239.1_Benin_97/1           -------------------------------------------------------------------------------- [  1600]
KM111295.1_Ken06.Bus            -------------------------------------------------------------------------------- [  1600]
KM111294.1_Ken05/Tk1            -------------------------------------------------------------------------------- [  1600]
KM262844.1_strain_L60           -------------------------------------------------------------------------------- [  1600]
KM102979.1_26544/OG10_Italy     -------------------------------------------------------------------------------- [  1600]
KM262845.1_strain_NHV           -------------------------------------------------------------------------------- [  1600]
KX354450.1_47/Ss/2008           ----------------------------...................t....--.....c...........t.....c.. [  1600]
KP055815.1_strain_BA71          .............t.................................t....--.....c...........t.....c.. [  1600]
MH025920.1_strain_R35           .......a.....t...t.........g...............g...t...t--...a.............t....cc.g [  1600]
MH025919.1_strain_N10           .......a.....t...t.........g...............g...t...t--...a.............t....cc.g [  1600]
MH025918.1_strain_R25           .......a.....t...t.........g...............g...t...t--...a.............t....cc.g [  1600]
MH025917.1_strain_R7            .......a.....t...t.........g...............g...t...t--...a.............t....cc.g [  1600]
MH025916.1_strain_R8            .......a.....t...t.........g...............g...t...t--...a.............t....cc.g [  1600]
```

```
FR682468.2_Georgia_2007/1        ctgctgcaggcgttaaagatta-aaattattactactgctgtaggtgtcaaagattaaaattattactgt---------- [  1680]
ASF_IND_20_CAD543                ......t......c........-......................c..t...c......c........acnnnnnnnnnn [  1680]
LR899193.1_Germany_2020/1        ......................-...............................................---------- [  1680]
MN393477.1_Wuhan_2019            ......................-...............................................---------- [  1680]
MW396979.1_Timor-Leste/2019/1    ......................-...............................................---------- [  1680]
MK940252.1_CN/2019/InnerMongolia -------------------------------------------------------------------------------- [  1680]
MK128995.1_China/2018/AnhuiXCGQ  -------------------------------------------------------------------------------- [  1680]
MH681419.1_ASFV/POL/2015         -------------------------------------------------------------------------------- [  1680]
LS478113.1_Estonia_2014          ......................-...............................................---------- [  1680]
KP843857.1_Odintsovo_02/14       -------------------------------------------------------------------------------- [  1680]
FN557520.1_strain_E75            -------------------------------------------------------------------------------- [  1680]
AY261366.1_Warthog               ......t......c.g.t-----------.atagg.....t.gatcaag...at..c.cc.atac.tca.---------- [  1680]
AY261365.1_Warmbaths             .....at..........c....-........tg.......a....c........................---------- [  1680]
AY261364.1_Tengani_62            ......t......c........-......................c..t...c............tg.ac---------- [  1680]
AY261363.1_Pretoriuskop/96/4     ......t..a.......c....-........tg.....t......c...........g.....c.g....---------- [  1680]
AY261362.1_Mkuzi_1979            ......t..........c-------.......t............c..t...c------.......t.ac---------- [  1680]
AY261361.1_Malawi_Lil-20/1_1983  ...t.atg.........c....g...............t..g..c..t...c....g.....c....ag---------- [  1680]
AY261360.1_Kenya_1950            ...ta.t...-....------.-.g......c..-------------------------g.....g.a..---------- [  1680]
AM712240.1_OURT_88/3             -------------------------------------------------------------------------------- [  1680]
AM712239.1_Benin_97/1            -------------------------------------------------------------------------------- [  1680]
KM111295.1_Ken06.Bus             -------------------------------------------------------------------------------- [  1680]
KM111294.1_Ken05/Tk1             -------------------------------------------------------------------------------- [  1680]
KM262844.1_strain_L60            -------------------------------------------------------------------------------- [  1680]
KM102979.1_26544/OG10_Italy      -------------------------------------------------------------------------------- [  1680]
KM262845.1_strain_NHV            -------------------------------------------------------------------------------- [  1680]
KX354450.1_47/Ss/2008            ......t..........c....-......................c..t...c...............ac---------- [  1680]
KP055815.1_strain_BA71           ......t..........c....-......................c..t...c...............ac---------- [  1680]
MH025920.1_strain_R35            ...t..tt.........c....-........c..g.......g.----------.gggcg.....g.a..---------- [  1680]
MH025919.1_strain_N10            ..at..tt.........c....-........c..g.......g.----------.gggcg.....g.a..---------- [  1680]
MH025918.1_strain_R25            ...t..tt.........c....-........c..g.......g.----------.gggcg.....g.a..---------- [  1680]
MH025917.1_strain_R7             ...t..tt.........c....-........c..g.......g.----------.gggcg.....g.a..---------- [  1680]
MH025916.1_strain_R8             ...t..tt.........c....-........c..g.......g.----------.gggcg.....g.a..---------- [  1680]

FR682468.2_Georgia_2007/1        ------------------------------aggcg--------------------------------------------- [  1760]
ASF_IND_20_CAD543                nnnnnnnnnnnnnnnnnnnnnnnnnnnnnn.....--------------------------------------------- [  1760]
LR899193.1_Germany_2020/1        ------------------------------.....--------------------------------------------- [  1760]
MN393477.1_Wuhan_2019            ------------------------------.....--------------------------------------------- [  1760]
MW396979.1_Timor-Leste/2019/1    ------------------------------.....--------------------------------------------- [  1760]
MK940252.1_CN/2019/InnerMongolia -------------------------------------------------------------------------------- [  1760]
MK128995.1_China/2018/AnhuiXCGQ  -------------------------------------------------------------------------------- [  1760]
MH681419.1_ASFV/POL/2015         -------------------------------------------------------------------------------- [  1760]
LS478113.1_Estonia_2014          ------------------------------.....--------------------------------------------- [  1760]
KP843857.1_Odintsovo_02/14       -------------------------------------------------------------------------------- [  1760]
```

```
FN557520.1_strain_E75            -------------------------------------------------------------------------------- [  1760]
AY261366.1_Warthog               ------------------------------....---------------------------------------------- [  1760]
AY261365.1_Warmbaths             ------------------------------.....--------------------------------------------- [  1760]
AY261364.1_Tengani_62            ------------------------tgctgt.....tcaaaga-------------------------------------- [  1760]
AY261363.1_Pretoriuskop/96/4     ------------------------------.....tcaaagattattgctttgatcaagaaaatttcaccatatacatca [  1760]
AY261362.1_Mkuzi_1979            ------------------------tgctgt.....--------------------------------------------- [  1760]
AY261361.1_Malawi_Lil-20/1_1983  -------------------------------------------------------------------------------- [  1760]
AY261360.1_Kenya_1950            ------------------------------....atcaaaaa-------------------------------------- [  1760]
AM712240.1_OURT_88/3             -------------------------------------------------------------------------------- [  1760]
AM712239.1_Benin_97/1            -------------------------------------------------------------------------------- [  1760]
KM111295.1_Ken06.Bus             -------------------------------------------------------------------------------- [  1760]
KM111294.1_Ken05/Tk1             -------------------------------.c.attatgaa-------------------------------------- [  1760]
KM262844.1_strain_L60            -------------------------------------------------------------------------------- [  1760]
KM102979.1_26544/OG10_Italy      -------------------------------------------------------------------------------- [  1760]
KM262845.1_strain_NHV            -------------------------------------------------------------------------------- [  1760]
KX354450.1_47/Ss/2008            ------------------------tgttgt.....--------------------------------------------- [  1760]
KP055815.1_strain_BA71           ------------------------tgttgt.....--------------------------------------------- [  1760]
MH025920.1_strain_R35            ------------------------------....atcaaaaa-------------------------------------- [  1760]
MH025919.1_strain_N10            ------------------------------....atcaaaaa-------------------------------------- [  1760]
MH025918.1_strain_R25            ------------------------------....atcaaaaa-------------------------------------- [  1760]
MH025917.1_strain_R7             ------------------------------....atcaaaaa-------------------------------------- [  1760]
MH025916.1_strain_R8             ------------------------------....atcaaaaa-------------------------------------- [  1760]

FR682468.2_Georgia_2007/1        ----ttcatttcacaagatgcggaattatttcgcaaagattatttttgaaacgccgcggccggaaatattttttttttgc [  1840]
ASF_IND_20_CAD543                ----..--------..acattaa......---------.c..c.g..gt.gg..t.aaa.atta....ta..ac.ac... [  1840]
LR899193.1_Germany_2020/1        ----............................................................................ [  1840]
MN393477.1_Wuhan_2019            ----............................................................................ [  1840]
MW396979.1_Timor-Leste/2019/1    ----............................................................................ [  1840]
MK940252.1_CN/2019/InnerMongolia -------------------------------------------------------------------------------- [  1840]
MK128995.1_China/2018/AnhuiXCGQ  -------------------------------------------------------------------------------- [  1840]
MH681419.1_ASFV/POL/2015         -------------------------------------------------------------------------------- [  1840]
LS478113.1_Estonia_2014          ----............................................................................ [  1840]
KP843857.1_Odintsovo_02/14       -------------------------------------------------------------------------------- [  1840]
FN557520.1_strain_E75            -------------------------------------------------------------------------------- [  1840]
AY261366.1_Warthog               -----cagccgg............--------------...............t......................c... [  1840]
AY261365.1_Warmbaths             ----............................................................................ [  1840]
AY261364.1_Tengani_62            -------------------------.....a.tagcg................t.......................... [  1840]
AY261363.1_Pretoriuskop/96/4     taggccagccgg............--------------..............................------------ [  1840]
AY261362.1_Mkuzi_1979            ----..--------..acatta------.---------....c.gc.gt.gg..tta------a.c..ta..ac.ac... [  1840]
AY261361.1_Malawi_Lil-20/1_1983  -------------------------------------------------------------------------------- [  1840]
AY261360.1_Kenya_1950            ----a.gcc-ggct..a.cttaat--------------g..t...cg.c.g.aat...t...a.g.tg.a.a...a.c-- [  1840]
AM712240.1_OURT_88/3             -------------------------------------------------------------------------------- [  1840]
AM712239.1_Benin_97/1            -------------------------------------------------------------------------------- [  1840]
KM111295.1_Ken06.Bus             -------------------------------------------------------------------------------- [  1840]
```

```
KM111294.1_Ken05/Tk1              ----a.g..----------------------------------------------------------------------- [  1840]
KM262844.1_strain_L60             -------------------------------------------------------------------------------- [  1840]
KM102979.1_26544/OG10_Italy       -------------------------------------------------------------------------------- [  1840]
KM262845.1_strain_NHV             -------------------------------------------------------------------------------- [  1840]
KX354450.1_47/Ss/2008             ----..--------..acattaa.......--------.c..c.gc.gt.gg..t.aaa.atta....ta..ac.ac... [  1840]
KP055815.1_strain_BA71            ----..--------..acattaa.......--------.c..c.gc.gt.gg..t.aaa.atta....ta..ac.ac... [  1840]
MH025920.1_strain_R35             ----a.gccgggct..a..ttaat--------------g..t...cg.c...aaa...c...a.g.tg.a.a...a.c-- [  1840]
MH025919.1_strain_N10             ----a.gccgggct..a..ttaat--------------g..t...cg.c...aaa...c...a.g.tg.a.a...a.c-- [  1840]
MH025918.1_strain_R25             ----a.gccgggct..a..ttaat--------------g..t...cg.c...aaa...c...a.g.tg.a.a...a.c-- [  1840]
MH025917.1_strain_R7              ----a.gccgggct..a..ttaat--------------g..t...cg.c...aaa...c...a.g.tg.a.a...a.c-- [  1840]
MH025916.1_strain_R8              ----a.gccgggct..a..ttaat--------------g..t...cg.c...aaa...c...a.g.tg.a.a...a.c-- [  1840]

FR682468.2_Georgia_2007/1         ggttgtaattga-tttttttt--tgcggctgggcggcgggccagacaaaattgaccataactggtgttacgccgccgg-- [  1920]
ASF_IND_20_CAD543                 t..a.gcg..a.------------------------------.c.tt...a.t.tt.ct...---------.t.ta..-- [  1920]
LR899193.1_Germany_2020/1         ............-........--.......................................................-- [  1920]
MN393477.1_Wuhan_2019             ............t........--.......................................................-- [  1920]
MW396979.1_Timor-Leste/2019/1     ............t........--.......................................................-- [  1920]
MK940252.1_CN/2019/InnerMongolia -------------------------------------------------------------------------------- [  1920]
MK128995.1_China/2018/AnhuiXCGQ   -------------------------------------------------------------------------------- [  1920]
MH681419.1_ASFV/POL/2015          -------------------------------------------------------------------------------- [  1920]
LS478113.1_Estonia_2014           ............t........--.......................................................-- [  1920]
KP843857.1_Odintsovo_02/14        -------------------------------------------------------------------------------- [  1920]
FN557520.1_strain_E75             -------------------------------------------------------------------------------- [  1920]
AY261366.1_Warthog                ..c.........t......c.gcg..t.................................t.....ca..........ta [  1920]
AY261365.1_Warmbaths              ..c.........t........gcg..t...................................................-- [  1920]
AY261364.1_Tengani_62             ..c------...t........------------t..........................t.....ca........t.t- [  1920]
AY261363.1_Pretoriuskop/96/4      -----------.t......c.gcg..t.................................t.....ca..........-- [  1920]
AY261362.1_Mkuzi_1979             t..a.gcg..a.------------------------------.c.------.t.tt.ct...---------.t.ta..-- [  1920]
AY261361.1_Malawi_Lil-20/1_1983   -------------------------------------------------------------------------------- [  1920]
AY261360.1_Kenya_1950             --------...ca.gg...c.----.ta.atacta.t....a.cctg.t...ag.a.----------------------- [  1920]
AM712240.1_OURT_88/3              -------------------------------------------------------------------------------- [  1920]
AM712239.1_Benin_97/1             -------------------------------------------------------------------------------- [  1920]
KM111295.1_Ken06.Bus              -------------------------------------------------------------------------------- [  1920]
KM111294.1_Ken05/Tk1              ------------------------------------------------g.a.a.ta.----------------------- [  1920]
KM262844.1_strain_L60             -------------------------------------------------------------------------------- [  1920]
KM102979.1_26544/OG10_Italy       -------------------------------------------------------------------------------- [  1920]
KM262845.1_strain_NHV             -------------------------------------------------------------------------------- [  1920]
KX354450.1_47/Ss/2008             t..a.gcg..a.------------------------------.c.tt...c.t.tt.ct...---------.tata..-- [  1920]
KP055815.1_strain_BA71            t..a.gcg..a.------------------------------.c.tt...c.t.tt.ct...---------.tata..-- [  1920]
MH025920.1_strain_R35             --------..acg.gg...c.----.t..atacta.t....a.cctg.t...ag.a.----------------------- [  1920]
MH025919.1_strain_N10             --------..acg.gg...c.----.t..atacta.t....a.cctg.t...ag.a.----------------------- [  1920]
MH025918.1_strain_R25             --------..acg.gg...c.----.t..atacta.t....a.cctg.t...ag.a.----------------------- [  1920]
MH025917.1_strain_R7              --------..acg.gg...c.----.t..atacta.t....a.cctg.t...ag.a.----------------------- [  1920]
MH025916.1_strain_R8              --------..acg.gg...c.----.t..atacta.t....a.cctg.t...ag.a.----------------------- [  1920]
```

```
FR682468.2_Georgia_2007/1        -taataaaccctaccgtaaatacttttttttggggcggccaga---gagacattatcgccgtaggtatcaattactgctg [  2000]
ASF_IND_20_CAD543                -cgt.....at..aaa.t.t....ac.g..gta....tt--------.a......-------.aa.tatt.c.....t.. [  2000]
LR899193.1_Germany_2020/1        -..........................................---.................................. [  2000]
MN393477.1_Wuhan_2019            -..........................................---.................................. [  2000]
MW396979.1_Timor-Leste/2019/1    -..........................................---.................................. [  2000]
MK940252.1_CN/2019/InnerMongolia -------------------------------------------------------------------------------- [  2000]
MK128995.1_China/2018/AnhuiXCGQ  -------------------------------------------------------------------------------- [  2000]
MH681419.1_ASFV/POL/2015         -------------------------------------------------------------------------------- [  2000]
LS478113.1_Estonia_2014          -..........................................---.................................. [  2000]
KP843857.1_Odintsovo_02/14       -------------------------------------------------------------------------------- [  2000]
FN557520.1_strain_E75            -------------------------------------------------------------------------------- [  2000]
AY261366.1_Warthog               a...........g......c.......................---tg.g...----------------------c.... [  2000]
AY261365.1_Warmbaths             -..........g.......c.......................---tg......----------------.c...c.... [  2000]
AY261364.1_Tengani_62            --------...........cc......................tgat......cg-------ct..tggt........g. [  2000]
AY261363.1_Pretoriuskop/96/4     -..........g.......c.......................---tg.g...cgct.tt.gtttc.att.c...c.... [  2000]
AY261362.1_Mkuzi_1979            -cgt....gat..aac.t.t....ac.gc.gta....t.--------.a.g....-------.aa.tatt.c.......a [  2000]
AY261361.1_Malawi_Lil-20/1_1983  -------------------------------------------------------------------------------- [  2000]
AY261360.1_Kenya_1950            -.cg..t.ttgc.tga..g..gagg.g.g.gataatat..gt.taat.aga.a.g.t--------------gcc.c..c. [  2000]
AM712240.1_OURT_88/3             -------------------------------------------------------------------------------- [  2000]
AM712239.1_Benin_97/1            -------------------------------------------------------------------------------- [  2000]
KM111295.1_Ken06.Bus             -------------------------------------------------------------------------------- [  2000]
KM111294.1_Ken05/Tk1             -.t.a.t.t...gtta.....g---------------------------------------------------------- [  2000]
KM262844.1_strain_L60            -------------------------------------------------------------------------------- [  2000]
KM102979.1_26544/OG10_Italy      -------------------------------------------------------------------------------- [  2000]
KM262845.1_strain_NHV            -------------------------------------------------------------------------------- [  2000]
KX354450.1_47/Ss/2008            -cgt.....at..aaa.t.t....ac.g.-------------------------------------------------.. [  2000]
KP055815.1_strain_BA71           -cgt.....at..aaa.t.t....ac.g..gta....t.--------.a.g....-------.aa.tatt.c........ [  2000]
MH025920.1_strain_R35            -.cg..t.ttgc.tga..g..gagg.g.g.gataatat...t.taat.aga.a.g.t..-----------cgcc.c..c. [  2000]
MH025919.1_strain_N10            -.cg..t.ttgc.tga..g..gagg.g.g.gataatat...t.taat.aga.a.g.t..-----------cgcc.c..c. [  2000]
MH025918.1_strain_R25            -.cg..t.ttgc.tga..g..gagg.g.g.gataatat...t.taat.aga.a.g.t..-----------cgcc.c..c. [  2000]
MH025917.1_strain_R7             -.cg..t.ttgc.tga..g..gagg.g.g.gataatat...t.taat.aga.a.g.t..-----------cgcc.c..c. [  2000]
MH025916.1_strain_R8             -.cg..t.ttgc.tga..g..gagg.g.g.gataatat...t.taat.aga.a.g.t..-----------cgcc.c..c. [  2000]

FR682468.2_Georgia_2007/1        taggtatcaattatta-------------tactacaggcgttaaacattattagta-cacaggcgtcaa----------- [  2080]
ASF_IND_20_CAD543                ....cg....ag....aaattattactac.g..gt...............aa.t..tt..tact.ctgt----------- [  2080]
LR899193.1_Germany_2020/1        ................-------------...........................-............----------- [  2080]
MN393477.1_Wuhan_2019            ................-------------...........................-............----------- [  2080]
MW396979.1_Timor-Leste/2019/1    ................-------------...........................-............----------- [  2080]
MK940252.1_CN/2019/InnerMongolia -------------------------------------------------------------------------------- [  2080]
MK128995.1_China/2018/AnhuiXCGQ  -------------------------------------------------------------------------------- [  2080]
MH681419.1_ASFV/POL/2015         -------------------------------------------------------------------------------- [  2080]
LS478113.1_Estonia_2014          ................-------------...........................-............----------- [  2080]
KP843857.1_Odintsovo_02/14       -------------------------------------------------------------------------------- [  2080]
```

```
FN557520.1_strain_E75            ----------------------------------------------------.t..ctg.t.ta.g.gt----------- [  2080]
AY261366.1_Warthog               .t...t....---------------------------------------------------------------------- [  2080]
AY261365.1_Warmbaths             .....t....---------------------------------------------------------------------- [  2080]
AY261364.1_Tengani_62            .t.t.t....------------------t..t.gt......c..------------------------------------ [  2080]
AY261363.1_Pretoriuskop/96/4     .....t..........------------c..t..t......--------------------------------------- [  2080]
AY261362.1_Mkuzi_1979            ....cg.t..ac.------ttattattac.g..gt...............aa.t..tt..t.ta.g.gttcatttcacaa [  2080]
AY261361.1_Malawi_Lil-20/1_1983  -------------------------------------------------------------------------------- [  2080]
AY261360.1_Kenya_1950            cc.cctgag.-----------------------------------------------tg.c.c..c.gc----------- [  2080]
AM712240.1_OURT_88/3             -------------------------------------------------------------------------------- [  2080]
AM712239.1_Benin_97/1            -------------------------------------------------------------------------------- [  2080]
KM111295.1_Ken06.Bus             -------------------------------------------------------------------------------- [  2080]
KM111294.1_Ken05/Tk1             -------------------------------------------------------------------------------- [  2080]
KM262844.1_strain_L60            -------------------------------------------------------------------------------- [  2080]
KM102979.1_26544/OG10_Italy      -------------------------------------------------------------------------------- [  2080]
KM262845.1_strain_NHV            -------------------------------------------------------------------------------- [  2080]
KX354450.1_47/Ss/2008            ....cg.t..ac....aaattattactac.g..gt..............---.t..ctg.t.ta.g.gt----------- [  2080]
KP055815.1_strain_BA71           ....cg.t..ac....aaattattactac.g..gt..............---.t..ctg.t.ta.g.gt----------- [  2080]
MH025920.1_strain_R35            cc.cctgag.-----------------------------------------------t..c.c..c.gc----------- [  2080]
MH025919.1_strain_N10            cc.cctgag.-----------------------------------------------t..c.c..c.gc----------- [  2080]
MH025918.1_strain_R25            cc.cctgag.-----------------------------------------------t..c.c..c.gc----------- [  2080]
MH025917.1_strain_R7             cc.cctgag.-----------------------------------------------t..c.c..c.gc----------- [  2080]
MH025916.1_strain_R8             cc.cctgag.-----------------------------------------------t..c.c..c.gc----------- [  2080]

FR682468.2_Georgia_2007/1        -------------------------------------------------------------------------------- [  2160]
ASF_IND_20_CAD543                -------------------------------------------------------------------------------- [  2160]
LR899193.1_Germany_2020/1        -------------------------------------------------------------------------------- [  2160]
MN393477.1_Wuhan_2019            -------------------------------------------------------------------------------- [  2160]
MW396979.1_Timor-Leste/2019/1    -------------------------------------------------------------------------------- [  2160]
MK940252.1_CN/2019/InnerMongolia -------------------------------------------------------------------------------- [  2160]
MK128995.1_China/2018/AnhuiXCGQ  -------------------------------------------------------------------------------- [  2160]
MH681419.1_ASFV/POL/2015         -------------------------------------------------------------------------------- [  2160]
LS478113.1_Estonia_2014          -------------------------------------------------------------------------------- [  2160]
KP843857.1_Odintsovo_02/14       -------------------------------------------------------------------------------- [  2160]
FN557520.1_strain_E75            -------------------------------------------------------------------------------- [  2160]
AY261366.1_Warthog               -------------------------------------------------------------------------------- [  2160]
AY261365.1_Warmbaths             -------------------------------------------------------------------------------- [  2160]
AY261364.1_Tengani_62            -------------------------------------------------------------------------------- [  2160]
AY261363.1_Pretoriuskop/96/4     -------------------------------------------------------------------------------- [  2160]
AY261362.1_Mkuzi_1979            gatgtggaattatttcgcaaagattattttttgaaacgccgcggccggaaatattttttttgcggctgggccagacaaaa [  2160]
AY261361.1_Malawi_Lil-20/1_1983  -------------------------------------------------------------------------------- [  2160]
AY261360.1_Kenya_1950            -------------------------------------------------------------------------------- [  2160]
AM712240.1_OURT_88/3             -------------------------------------------------------------------------------- [  2160]
AM712239.1_Benin_97/1            -------------------------------------------------------------------------------- [  2160]
KM111295.1_Ken06.Bus             -------------------------------------------------------------------------------- [  2160]
```

```
KM111294.1_Ken05/Tk1           -------------------------------------------------------------------------------- [  2160]
KM262844.1_strain_L60          -------------------------------------------------------------------------------- [  2160]
KM102979.1_26544/OG10_Italy    -------------------------------------------------------------------------------- [  2160]
KM262845.1_strain_NHV          -------------------------------------------------------------------------------- [  2160]
KX354450.1_47/Ss/2008          -------------------------------------------------------------------------------- [  2160]
KP055815.1_strain_BA71         -------------------------------------------------------------------------------- [  2160]
MH025920.1_strain_R35          -------------------------------------------------------------------------------- [  2160]
MH025919.1_strain_N10          -------------------------------------------------------------------------------- [  2160]
MH025918.1_strain_R25          -------------------------------------------------------------------------------- [  2160]
MH025917.1_strain_R7           -------------------------------------------------------------------------------- [  2160]
MH025916.1_strain_R8           -------------------------------------------------------------------------------- [  2160]

FR682468.2_Georgia_2007/1      -------------------------------------------------------------------------------- [  2240]
ASF_IND_20_CAD543              -------------------------------------------------------------------------------- [  2240]
LR899193.1_Germany_2020/1      -------------------------------------------------------------------------------- [  2240]
MN393477.1_Wuhan_2019          -------------------------------------------------------------------------------- [  2240]
MW396979.1_Timor-Leste/2019/1  -------------------------------------------------------------------------------- [  2240]
MK940252.1_CN/2019/InnerMongolia -------------------------------------------------------------------------------- [  2240]
MK128995.1_China/2018/AnhuiXCGQ -------------------------------------------------------------------------------- [  2240]
MH681419.1_ASFV/POL/2015       -------------------------------------------------------------------------------- [  2240]
LS478113.1_Estonia_2014        -------------------------------------------------------------------------------- [  2240]
KP843857.1_Odintsovo_02/14     -------------------------------------------------------------------------------- [  2240]
FN557520.1_strain_E75          -------------------------------------------------------------------------------- [  2240]
AY261366.1_Warthog             -------------------------------------------------------------------------------- [  2240]
AY261365.1_Warmbaths           -------------------------------------------------------------------------------- [  2240]
AY261364.1_Tengani_62          -------------------------------------------------------------------------------- [  2240]
AY261363.1_Pretoriuskop/96/4   ---------actgg------------------------------------------------------------------ [  2240]
AY261362.1_Mkuzi_1979          ttgaccataactggtgttacgccgccggtaataaaccccaccgtaaatacttttttttggggcggccagatggacattac [  2240]
AY261361.1_Malawi_Lil-20/1_1983 -------------------------------------------------------------------------------- [  2240]
AY261360.1_Kenya_1950          -------------------------------------------------------------------------------- [  2240]
AM712240.1_OURT_88/3           -------------------------------------------------------------------------------- [  2240]
AM712239.1_Benin_97/1          -------------------------------------------------------------------------------- [  2240]
KM111295.1_Ken06.Bus           -------------------------------------------------------------------------------- [  2240]
KM111294.1_Ken05/Tk1           -------------------------------------------------------------------------------- [  2240]
KM262844.1_strain_L60          -------------------------------------------------------------------------------- [  2240]
KM102979.1_26544/OG10_Italy    -------------------------------------------------------------------------------- [  2240]
KM262845.1_strain_NHV          -------------------------------------------------------------------------------- [  2240]
KX354450.1_47/Ss/2008          -------------------------------------------------------------------------------- [  2240]
KP055815.1_strain_BA71         -------------------------------------------------------------------------------- [  2240]
MH025920.1_strain_R35          -------------------------------------------------------------------------------- [  2240]
MH025919.1_strain_N10          -------------------------------------------------------------------------------- [  2240]
MH025918.1_strain_R25          -------------------------------------------------------------------------------- [  2240]
MH025917.1_strain_R7           -------------------------------------------------------------------------------- [  2240]
MH025916.1_strain_R8           -------------------------------------------------------------------------------- [  2240]
```

```
FR682468.2_Georgia_2007/1        -----------------------------------------------agaagctaaaacttaatgttttttcgtcaaaaa [  2320]
ASF_IND_20_CAD543                -----------------------------------------------..gc.t....ca....aa..a..a.-------- [  2320]
LR899193.1_Germany_2020/1        -----------------------------------------------................................. [  2320]
MN393477.1_Wuhan_2019            -----------------------------------------------................................. [  2320]
MW396979.1_Timor-Leste/2019/1    -----------------------------------------------................................. [  2320]
MK940252.1_CN/2019/InnerMongolia -------------------------------------------------------------------------------- [  2320]
MK128995.1_China/2018/AnhuiXCGQ  -------------------------------------------------------------------------------- [  2320]
MH681419.1_ASFV/POL/2015         -------------------------------------------------------------------------------- [  2320]
LS478113.1_Estonia_2014          -----------------------------------------------................................. [  2320]
KP843857.1_Odintsovo_02/14       -------------------------------------------------------------------------------- [  2320]
FN557520.1_strain_E75            --------------------------------------------------.......tt...............a..... [  2320]
AY261366.1_Warthog               -------------------------------------------------------...aga.gct...cg...------- [  2320]
AY261365.1_Warmbaths             -------------------------------------------------------...aga.gct...cg...------- [  2320]
AY261364.1_Tengani_62            ------------------------------------------------------.g..gc.....c-------------g [  2320]
AY261363.1_Pretoriuskop/96/4     ------------------------------------------------------cg..g..g...........------- [  2320]
AY261362.1_Mkuzi_1979            cgctgtgggtatcaattattattaggcgttaaacattattcgcgtca.ag.a............................ [  2320]
AY261361.1_Malawi_Lil-20/1_1983  ----------------------------------------------------.....tt..................... [  2320]
AY261360.1_Kenya_1950            ------------------------------------------------.gta........................c... [  2320]
AM712240.1_OURT_88/3             -------------------------------------------------------------------------------- [  2320]
AM712239.1_Benin_97/1            -------------------------------------------------------------------------------- [  2320]
KM111295.1_Ken06.Bus             -------------------------------------------------------------------------------- [  2320]
KM111294.1_Ken05/Tk1             -----------------------------------------------------....ca...c.a......tt.t..... [  2320]
KM262844.1_strain_L60            -------------------------------------------------------------------------------- [  2320]
KM102979.1_26544/OG10_Italy      -------------------------------------------------------------------------------- [  2320]
KM262845.1_strain_NHV            -------------------------------------------------------------------------------- [  2320]
KX354450.1_47/Ss/2008            --------------------------------------------------.......tt...............a..... [  2320]
KP055815.1_strain_BA71           --------------------------------------------------.......tt...............a..... [  2320]
MH025920.1_strain_R35            ------------------------------------------------.gta...................t....c... [  2320]
MH025919.1_strain_N10            ------------------------------------------------.gta...................t....c... [  2320]
MH025918.1_strain_R25            ------------------------------------------------.gta...................t....c... [  2320]
MH025917.1_strain_R7             ------------------------------------------------.gta...................t....c... [  2320]
MH025916.1_strain_R8             ------------------------------------------------.gta...................t....c... [  2320]

FR682468.2_Georgia_2007/1        tc-------gccatgaa--tatctatcttgtgtggtttctctacatactattggggaacctgatattagcagtaatatat [  2400]
ASF_IND_20_CAD543                ---------------------------------------------....g...-------------...gc..t.a.c.. [  2400]
LR899193.1_Germany_2020/1        ..-------........--............................................................. [  2400]
MN393477.1_Wuhan_2019            ..-------........--............................................................. [  2400]
MW396979.1_Timor-Leste/2019/1    ..-------........--............................................................. [  2400]
MK940252.1_CN/2019/InnerMongolia -------------------------------------------------------------------------------- [  2400]
MK128995.1_China/2018/AnhuiXCGQ  -------------------------------------------------------------------------------- [  2400]
MH681419.1_ASFV/POL/2015         -------------------------------------------------------------------------------- [  2400]
LS478113.1_Estonia_2014          ..-------........--............................................................. [  2400]
KP843857.1_Odintsovo_02/14       -------------------------------------------------------------------------------- [  2400]
```

FN557520.1_strain_E75 ..-------....------------------------------------------------------------------- [ 2400]
AY261366.1_Warthog c.-------.....ag.--------------------------------------------------------------- [ 2400]
AY261365.1_Warmbaths c.-------.....ag.--------------------------------------------------------------- [ 2400]
AY261364.1_Tengani_62 c.-------.....ag.--------------------------------------------------------------- [ 2400]
AY261363.1_Pretoriuskop/96/4 ..-------.t......at...t..---------------------------------------..c.ag.aa..a...- [ 2400]
AY261362.1_Mkuzi_1979 ..-------a.......--............................................................. [ 2400]
AY261361.1_Malawi_Lil-20/1_1983 .t-------........--...........c.........t..............................a.cg..... [ 2400]
AY261360.1_Kenya_1950 ..-------........--c.g......g...............g.g..ggc..tc......c.gg.gctgt.c....gc [ 2400]
AM712240.1_OURT_88/3 -------------------------------------------------------------------------------- [ 2400]
AM712239.1_Benin_97/1 -------------------------------------------------------------------------------- [ 2400]
KM111295.1_Ken06.Bus -------------------------------------------------------------------------------- [ 2400]
KM111294.1_Ken05/Tk1 aa------------------------------------------------------------------------------ [ 2400]
KM262844.1_strain_L60 -------------------------------------------------------------------------------- [ 2400]
KM102979.1_26544/OG10_Italy -------------------------------------------------------------------------------- [ 2400]
KM262845.1_strain_NHV -------------------------------------------------------------------------------- [ 2400]
KX354450.1_47/Ss/2008 ..-------....------------------------------------------------------------------- [ 2400]
KP055815.1_strain_BA71 ..-------....------------------------------------------------------------------- [ 2400]
MH025920.1_strain_R35 ..gccgcca........--cgg......g........c......g.g..ggc..tc......c.gg.gctgt.c....gc [ 2400]
MH025919.1_strain_N10 ..gccgcca........--cgg......g........c......g.g..ggc..tc......c.gg.gctgt.c....gc [ 2400]
MH025918.1_strain_R25 ..gccgcca........--cgg......g........c......g.g..ggc..tc......c.gg.gctgt.c....gc [ 2400]
MH025917.1_strain_R7 ..gccgcca........--cgg......g........c......g.g..ggc..tc......c.gg.gctgt.c....gc [ 2400]
MH025916.1_strain_R8 ..gccgcca........--cgg......g........c......g.g..ggc..tc......c.gg.gctgt.c....gc [ 2400]

FR682468.2_Georgia_2007/1 tgcgtcatagatgaggtggtgtgtgacaatatccatataaaaaaaaatgttgccgcccctgagatgccgcggcggttcta [ 2480]
ASF_IND_20_CAD543 .-----------------------------------------....t.a..a.ta.tg...------------------- [ 2480]
LR899193.1_Germany_2020/1 ................................................................................ [ 2480]
MN393477.1_Wuhan_2019 ................................................................................ [ 2480]
MW396979.1_Timor-Leste/2019/1 ......................................-......................................... [ 2480]
MK940252.1_CN/2019/InnerMongolia -------------------------------------------------------------------------------- [ 2480]
MK128995.1_China/2018/AnhuiXCGQ -------------------------------------------------------------------------------- [ 2480]
MH681419.1_ASFV/POL/2015 -------------------------------------------------------------------------------- [ 2480]
LS478113.1_Estonia_2014 ................................................................................ [ 2480]
KP843857.1_Odintsovo_02/14 -------------------------------------------------------------------------------- [ 2480]
FN557520.1_strain_E75 -------------------------------------------------------------------------------- [ 2480]
AY261366.1_Warthog -------------------------------------------------------------------------------- [ 2480]
AY261365.1_Warmbaths -------------------------------------------------------------------------------- [ 2480]
AY261364.1_Tengani_62 -------------------------------------------------------------------------------- [ 2480]
AY261363.1_Pretoriuskop/96/4 ------------a.a....ca.taat...c..at....------------------------------------------ [ 2480]
AY261362.1_Mkuzi_1979 ...a.........g....a......................------------t.ttt.ca..g......c......... [ 2480]
AY261361.1_Malawi_Lil-20/1_1983 ..ta.gg...........-------------------------------------------------------------- [ 2480]
AY261360.1_Kenya_1950 .a..g...c..g.gcc..c.------...gc..tgcgccgccc....aaagc.g...gag...gc.....---------- [ 2480]
AM712240.1_OURT_88/3 -------------------------------------------------------------------------------- [ 2480]
AM712239.1_Benin_97/1 -------------------------------------------------------------------------------- [ 2480]
KM111295.1_Ken06.Bus -------------------------------------------------------------------------------- [ 2480]

```
KM111294.1_Ken05/Tk1             -------------------------------------------------------------------------------- [  2480]
KM262844.1_strain_L60            -------------------------------------------------------------------------------- [  2480]
KM102979.1_26544/OG10_Italy      -------------------------------------------------------------------------------- [  2480]
KM262845.1_strain_NHV            -------------------------------------------------------------------------------- [  2480]
KX354450.1_47/Ss/2008            -------------------------------------------------------------------------------- [  2480]
KP055815.1_strain_BA71           -------------------------------------------------------------------------------- [  2480]
MH025920.1_strain_R35            .a..g...c..g.gcc..c.------...gc..tgcgcggcgc....aaagc.g...gag.c.gc.....---------- [  2480]
MH025919.1_strain_N10            .a..g...c..g.gcc..c.------...gc..tgcgcggcgc....aaagc.g...gag.c.gc.....---------- [  2480]
MH025918.1_strain_R25            .a..g...c..g.gcc..c.------...gc..tgcgcggcgc....aaagc.g...gag.c.gc.....---------- [  2480]
MH025917.1_strain_R7             .a..g...c..g.gcc..c.------...gc..tgcgcggcgc....aaagc.g...gag.c.gc.....---------- [  2480]
MH025916.1_strain_R8             .a..g...c..g.gcc..c.------...gc..tgcgcggcgc....aaagc.g...gag.c.gc.....---------- [  2480]

FR682468.2_Georgia_2007/1        aatttt------aatgtttttttcggcgaacat----------------------------------------------- [  2560]
ASF_IND_20_CAD543                -------------------------------------------------------------------------------- [  2560]
LR899193.1_Germany_2020/1        ......------.....................----------------------------------------------- [  2560]
MN393477.1_Wuhan_2019            ......------.....................----------------------------------------------- [  2560]
MW396979.1_Timor-Leste/2019/1    ......------.....................----------------------------------------------- [  2560]
MK940252.1_CN/2019/InnerMongolia -------------------------------------------------------------------------------- [  2560]
MK128995.1_China/2018/AnhuiXCGQ  -------------------------------------------------------------------------------- [  2560]
MH681419.1_ASFV/POL/2015         -------------------------------------------------------------------------------- [  2560]
LS478113.1_Estonia_2014          ......------.....................----------------------------------------------- [  2560]
KP843857.1_Odintsovo_02/14       -------------------------------------------------------------------------------- [  2560]
FN557520.1_strain_E75            ---------------------g..acaa..a.a----------------------------------------------- [  2560]
AY261366.1_Warthog               -------------------------------------------------------------------------------- [  2560]
AY261365.1_Warmbaths             -------------------------------------------------------------------------------- [  2560]
AY261364.1_Tengani_62            -------------------------------------------------------------------------------- [  2560]
AY261363.1_Pretoriuskop/96/4     ------------tg.a..a...caa.taggt..-------------taatcat--ccctattatg-----------atgc [  2560]
AY261362.1_Mkuzi_1979            ...g..tttcgcg.aa..a..ac..caa..a.atcttaagtggcattaatcatgcccctattacgccaatcgacatatat [  2560]
AY261361.1_Malawi_Lil-20/1_1983  -------------------------------------------------------------------------------- [  2560]
AY261360.1_Kenya_1950            -------------------------------------------------------ccccgccgtac-------------- [  2560]
AM712240.1_OURT_88/3             -------------------------------------------------------------------------------- [  2560]
AM712239.1_Benin_97/1            -------------------------------------------------------------------------------- [  2560]
KM111295.1_Ken06.Bus             -------------------------------------------------------------------------------- [  2560]
KM111294.1_Ken05/Tk1             -------------------------------------------------------------------------------- [  2560]
KM262844.1_strain_L60            -------------------------------------------------------------------------------- [  2560]
KM102979.1_26544/OG10_Italy      -------------------------------------------------------------------------------- [  2560]
KM262845.1_strain_NHV            -------------------------------------------------------------------------------- [  2560]
KX354450.1_47/Ss/2008            ---------------------g..acaa..a.a----------------------------------------------- [  2560]
KP055815.1_strain_BA71           ---------------------g..acaa..a.a----------------------------------------------- [  2560]
MH025920.1_strain_R35            -------------------------------------------------------ccccgccgtacg-----------gc [  2560]
MH025919.1_strain_N10            -------------------------------------------------------ccccgccgtacg-----------gc [  2560]
MH025918.1_strain_R25            -------------------------------------------------------ccccgccgtacg-----------gc [  2560]
MH025917.1_strain_R7             -------------------------------------------------------ccccgccgtacg-----------gc [  2560]
MH025916.1_strain_R8             -------------------------------------------------------ccccgccgtacg-----------gc [  2560]
```

```
FR682468.2_Georgia_2007/1        -------------ttttcacatatgcgatattggcgctaaagcgagcgtattaccgcttgtaacaacattttttttcgat [  2640]
ASF_IND_20_CAD543                ------------------------------.a....tc....-------....--------------------------- [  2640]
LR899193.1_Germany_2020/1        -------------................................................................... [  2640]
MN393477.1_Wuhan_2019            -------------................................................................... [  2640]
MW396979.1_Timor-Leste/2019/1    -------------.......................................................-........... [  2640]
MK940252.1_CN/2019/InnerMongolia -------------------------------------------------------------------------------- [  2640]
MK128995.1_China/2018/AnhuiXCGQ  -------------------------------------------------------------------------------- [  2640]
MH681419.1_ASFV/POL/2015         -------------------------------------------------------------------------------- [  2640]
LS478113.1_Estonia_2014          -------------................................................................... [  2640]
KP843857.1_Odintsovo_02/14       -------------------------------------------------------------------------------- [  2640]
FN557520.1_strain_E75            -------------------------------------------------------------------------------- [  2640]
AY261366.1_Warthog               ----------------------------------------------------------...g....t..----------- [  2640]
AY261365.1_Warmbaths             ----------------------------------------------------------...g....t..----------- [  2640]
AY261364.1_Tengani_62            ----------------------------------------------------------...g....t..----------- [  2640]
AY261363.1_Pretoriuskop/96/4     gtcaaccgcagca......t.....a........a.t......t.a.......tt.t....................... [  2640]
AY261362.1_Mkuzi_1979            tgttactacagca............a............................t...c..................... [  2640]
AY261361.1_Malawi_Lil-20/1_1983  ----------------------------------------------------------...g.t..t..----------- [  2640]
AY261360.1_Kenya_1950            ----------------------c.at...t...--------.ga.t.ag..c.....c..cg.agtt..----------- [  2640]
AM712240.1_OURT_88/3             -------------------------------------------------------------------------------- [  2640]
AM712239.1_Benin_97/1            -------------------------------------------------------------------------------- [  2640]
KM111295.1_Ken06.Bus             -------------------------------------------------------------------------------- [  2640]
KM111294.1_Ken05/Tk1             -------------------------------------------------------------------------------- [  2640]
KM262844.1_strain_L60            -------------------------------------------------------------------------------- [  2640]
KM102979.1_26544/OG10_Italy      -------------------------------------------------------------------------------- [  2640]
KM262845.1_strain_NHV            -------------------------------------------------------------------------------- [  2640]
KX354450.1_47/Ss/2008            -------------------------------------------------------------------------------- [  2640]
KP055815.1_strain_BA71           -------------------------------------------------------------------------------- [  2640]
MH025920.1_strain_R35            agcgacgaca-cgccgcggt.cgcat...t....gatgtg..ga.t.ag..cg..a.c..cgga.tt..----------- [  2640]
MH025919.1_strain_N10            agcgacgaca-cgccgcggt.cgcat...t....gatgtg..ga.t.ag..cg..a.c..cgga.tt..----------- [  2640]
MH025918.1_strain_R25            agcgacgaca-cgccgcggt.cgcat...t....gatgtg..ga.t.ag..cg..a.c..cgga.tt..----------- [  2640]
MH025917.1_strain_R7             agcgacgaca-cgccgcggt.cgcat...t....gatgtg..ga.t.ag..cg..a.c..cgga.tt..----------- [  2640]
MH025916.1_strain_R8             agcgacgaca-cgccgcggt.cgcat...t....gatgtg..ga.t.ag..cg..a.c..cgga.tt..----------- [  2640]

FR682468.2_Georgia_2007/1        cggcaatagataagtagaatataccatattattgctattgccatcaatgagaatgccacgtaggcataggtcat----cc [  2720]
ASF_IND_20_CAD543                -------------------------.a......a...c...-----------------t......g.taaa...taaa.t [  2720]
LR899193.1_Germany_2020/1        ..........................................................................----.. [  2720]
MN393477.1_Wuhan_2019            ..........................................................................----.. [  2720]
MW396979.1_Timor-Leste/2019/1    ..........................................................................----.. [  2720]
MK940252.1_CN/2019/InnerMongolia ------------------........................................................----.. [  2720]
MK128995.1_China/2018/AnhuiXCGQ  ------------------........................................................----.. [  2720]
MH681419.1_ASFV/POL/2015         ----------------..........................................................----.. [  2720]
LS478113.1_Estonia_2014          ..........................................................................----.. [  2720]
KP843857.1_Odintsovo_02/14       ------------------........................................................----.. [  2720]
```

```
FN557520.1_strain_E75             -------------------------------------------------------------------------------- [  2720]
AY261366.1_Warthog                -----------------------......--------------------------------------------------- [  2720]
AY261365.1_Warmbaths              -----------------------......--------------------------------------------------- [  2720]
AY261364.1_Tengani_62             -----------------------......--------------------------------------------------- [  2720]
AY261363.1_Pretoriuskop/96/4      ................t......................a.........t.....at.t...............----.. [  2720]
AY261362.1_Mkuzi_1979             ..........................................................................----.. [  2720]
AY261361.1_Malawi_Lil-20/1_1983   -----------------------..g...a..------------------------------------------------ [  2720]
AY261360.1_Kenya_1950             -----------------..a.ctta....-----------------------------------------------gttt [  2720]
AM712240.1_OURT_88/3              -------------------------------------------------------------------------------- [  2720]
AM712239.1_Benin_97/1             -------------------------------------------------------------------------------- [  2720]
KM111295.1_Ken06.Bus              -------------------------------------------------------------------------------- [  2720]
KM111294.1_Ken05/Tk1              -----------------.....tta.------------------------------------------------------ [  2720]
KM262844.1_strain_L60             -------------------------------------------------------------------------------- [  2720]
KM102979.1_26544/OG10_Italy       -------------------------------------------------------------------------------- [  2720]
KM262845.1_strain_NHV             -------------------------------------------------------------------------------- [  2720]
KX354450.1_47/Ss/2008             -------------------------------------------------------------------------------- [  2720]
KP055815.1_strain_BA71            -------------------------------------------------------------------------------- [  2720]
MH025920.1_strain_R35             -----------------..a.ctta....-----------------------------------------------attt [  2720]
MH025919.1_strain_N10             -----------------..a.ctta....-----------------------------------------------attt [  2720]
MH025918.1_strain_R25             -----------------..a.ctta....-----------------------------------------------attt [  2720]
MH025917.1_strain_R7              -----------------..a.ctta....-----------------------------------------------attt [  2720]
MH025916.1_strain_R8              -----------------..a.ctta....-----------------------------------------------attt [  2720]

FR682468.2_Georgia_2007/1         tatggccggaccaatccatggctgcacttaaaaatatcaaaa-------------------------------------- [  2800]
ASF_IND_20_CAD543                 ...ta.tnnnnnnnnnnnnnntaaaca......t...t.ct.nnnnnnnnnnnnnnnnnnnnnnnnnnnnnnnnnnnnnn [  2800]
LR899193.1_Germany_2020/1         ..........................................-------------------------------------- [  2800]
MN393477.1_Wuhan_2019             ..........................................-------------------------------------- [  2800]
MW396979.1_Timor-Leste/2019/1     ..........................................-------------------------------------- [  2800]
MK940252.1_CN/2019/InnerMongolia  ..........................................-------------------------------------- [  2800]
MK128995.1_China/2018/AnhuiXCGQ   ......................................-...-------------------------------------- [  2800]
MH681419.1_ASFV/POL/2015          ..........................................-------------------------------------- [  2800]
LS478113.1_Estonia_2014           ..........................................-------------------------------------- [  2800]
KP843857.1_Odintsovo_02/14        ..........................................-------------------------------------- [  2800]
FN557520.1_strain_E75             -------------------------------------------------------------------------------- [  2800]
AY261366.1_Warthog                ---------------------------------------...-------------------------------------- [  2800]
AY261365.1_Warmbaths              ---------------------------------------...-------------------------------------- [  2800]
AY261364.1_Tengani_62             ---------------------------------------...-------------------------------------- [  2800]
AY261363.1_Pretoriuskop/96/4      ......t..---......................cc......-------------------------------------- [  2800]
AY261362.1_Mkuzi_1979             ..........................................-------------------------------------- [  2800]
AY261361.1_Malawi_Lil-20/1_1983   -------------------------------------------------------------------------------- [  2800]
AY261360.1_Kenya_1950             .cc.aggaa.a..gattgcc..c.tg.ac.ctg..t.gc-g.-------------------------------------- [  2800]
AM712240.1_OURT_88/3              -------------------------------------------------------------------------------- [  2800]
AM712239.1_Benin_97/1             -------------------------------------------------------------------------------- [  2800]
KM111295.1_Ken06.Bus              -------------------------------------------------------------------------------- [  2800]
```

```
KM111294.1_Ken05/Tk1              ----------------------------------------c.-------------------------------------- [  2800]
KM262844.1_strain_L60             -------------------------------------------------------------------------------- [  2800]
KM102979.1_26544/OG10_Italy       -------------------------------------------------------------------------------- [  2800]
KM262845.1_strain_NHV             -------------------------------------------------------------------------------- [  2800]
KX354450.1_47/Ss/2008             -------------------------------------------------------------------------------- [  2800]
KP055815.1_strain_BA71            -------------------------------------------------------------------------------- [  2800]
MH025920.1_strain_R35             .cc.aagaa.at..attgcc..c.tg.a..ctgg.t.gcgg.-------------------------------------- [  2800]
MH025919.1_strain_N10             .cc.aagaa.at..attgcc..c.tg.a..ctgg.t.gcgg.-------------------------------------- [  2800]
MH025918.1_strain_R25             .cc.aagaa.at..attgcc..c.tg.a..ctgg.t.gcgg.-------------------------------------- [  2800]
MH025917.1_strain_R7              .cc.aagaa.at..attgcc..c.tg.a..ctgg.t.gcgg.-------------------------------------- [  2800]
MH025916.1_strain_R8              .cc.aagaa.at..attgcc..c.tg.a..ctgg.t.gcgg.-------------------------------------- [  2800]

FR682468.2_Georgia_2007/1         ----------------------------------------------aaagtttaagttttggg-------------ccgg [  2880]
ASF_IND_20_CAD543                 nnnnnnnnnnnnnnnnnnnnnnnnnnnnnnnnnnnnnnnnnnnnnnnnnnnnnnnnnnnnnnn-------------nnnn [  2880]
LR899193.1_Germany_2020/1         ----------------------------------------------.................-------------.... [  2880]
MN393477.1_Wuhan_2019             ----------------------------------------------.................-------------.... [  2880]
MW396979.1_Timor-Leste/2019/1     ----------------------------------------------.................-------------.... [  2880]
MK940252.1_CN/2019/InnerMongolia  ----------------------------------------------.................-------------.... [  2880]
MK128995.1_China/2018/AnhuiXCGQ   ----------------------------------------------.................-------------.... [  2880]
MH681419.1_ASFV/POL/2015          -----------------------------------------------................-------------.... [  2880]
LS478113.1_Estonia_2014           ----------------------------------------------.................-------------.... [  2880]
KP843857.1_Odintsovo_02/14        -----------------------------------------------................-------------.... [  2880]
FN557520.1_strain_E75             -------------------------------------------------------------------------------- [  2880]
AY261366.1_Warthog                ---------------------------------------------a...------------------------------- [  2880]
AY261365.1_Warmbaths              ---------------------------------------------a...------------------------------- [  2880]
AY261364.1_Tengani_62             ---------------------------------------------a...------------------------------- [  2880]
AY261363.1_Pretoriuskop/96/4      ---------------------------------------------a...ca...........aagagggtaatgacg.a. [  2880]
AY261362.1_Mkuzi_1979             ----------------------------------------------.................-------------..c. [  2880]
AY261361.1_Malawi_Lil-20/1_1983   -------------------------------------------------------------------------------- [  2880]
AY261360.1_Kenya_1950             ---------------------------------------------g...------------------------------- [  2880]
AM712240.1_OURT_88/3              -------------------------------------------------------------------------------- [  2880]
AM712239.1_Benin_97/1             -------------------------------------------------------------------------------- [  2880]
KM111295.1_Ken06.Bus              -------------------------------------------------------------------------------- [  2880]
KM111294.1_Ken05/Tk1              ---------------------------------------------a.c.------------------------------- [  2880]
KM262844.1_strain_L60             -------------------------------------------------------------------------------- [  2880]
KM102979.1_26544/OG10_Italy       -------------------------------------------------------------------------------- [  2880]
KM262845.1_strain_NHV             -------------------------------------------------------------------------------- [  2880]
KX354450.1_47/Ss/2008             -------------------------------------------------------------------------------- [  2880]
KP055815.1_strain_BA71            -------------------------------------------------------------------------------- [  2880]
MH025920.1_strain_R35             ---------------------------------------------g...------------------------------- [  2880]
MH025919.1_strain_N10             ---------------------------------------------g...------------------------------- [  2880]
MH025918.1_strain_R25             ---------------------------------------------g...------------------------------- [  2880]
MH025917.1_strain_R7              ---------------------------------------------g...------------------------------- [  2880]
MH025916.1_strain_R8              ---------------------------------------------g...------------------------------- [  2880]
```

```
FR682468.2_Georgia_2007/1         cgttaaaatttaaacc-ttttctggttga-------------------tctttagccatgtatagctgcgatgtttggtg [  2960]
ASF_IND_20_CAD543                 nnnct......t..tg-....t.c..aa.aaatcgccannnnnnnnnnnnnnn...........t...t........... [  2960]
LR899193.1_Germany_2020/1         ................-............-------------------................................ [  2960]
MN393477.1_Wuhan_2019             ................-............-------------------................................ [  2960]
MW396979.1_Timor-Leste/2019/1     ................-............-------------------................................ [  2960]
MK940252.1_CN/2019/InnerMongolia  ................-............-------------------................................ [  2960]
MK128995.1_China/2018/AnhuiXCGQ   ................-............-------------------................................ [  2960]
MH681419.1_ASFV/POL/2015          ................-............-------------------................................ [  2960]
LS478113.1_Estonia_2014           ................-............-------------------................................ [  2960]
KP843857.1_Odintsovo_02/14        ................-............-------------------................................ [  2960]
FN557520.1_strain_E75             --------------------------------------------------agc...........t...t........... [  2960]
AY261366.1_Warthog                -------------------------------------------------------------------------------- [  2960]
AY261365.1_Warmbaths              -------------------------------------------------------------------------------- [  2960]
AY261364.1_Tengani_62             -------------------------------------------------------------------------------- [  2960]
AY261363.1_Pretoriuskop/96/4      tac...........a.-.......a....-------------------............a...t...t........... [  2960]
AY261362.1_Mkuzi_1979             tac.............t............-------------------................t............... [  2960]
AY261361.1_Malawi_Lil-20/1_1983   -------------------------------------------------------------------------------- [  2960]
AY261360.1_Kenya_1950             ------------------------------------------------........t....t..tt..t.....c..... [  2960]
AM712240.1_OURT_88/3              -------------------------------------------------------------------------------- [  2960]
AM712239.1_Benin_97/1             -------------------------------------------------------------------------------- [  2960]
KM111295.1_Ken06.Bus              -------------------------------------------------------------------------------- [  2960]
KM111294.1_Ken05/Tk1              ------------------------------------------------.t....aat....t.----------------- [  2960]
KM262844.1_strain_L60             -------------------------------------------------------------------------------- [  2960]
KM102979.1_26544/OG10_Italy       -------------------------------------------------------------------------------- [  2960]
KM262845.1_strain_NHV             -------------------------------------------------------------------------------- [  2960]
KX354450.1_47/Ss/2008             --------------------------------------------------aac...........t...t........... [  2960]
KP055815.1_strain_BA71            --------------------------------------------------agc...........t...t........... [  2960]
MH025920.1_strain_R35             ------------------------------------------------........t....t..tt..t.....c..... [  2960]
MH025919.1_strain_N10             ------------------------------------------------........t....t..tt..t.....c..... [  2960]
MH025918.1_strain_R25             ------------------------------------------------........t....t..tt..t.....c..... [  2960]
MH025917.1_strain_R7              ------------------------------------------------........t....t..tt..t.....c..... [  2960]
MH025916.1_strain_R8              ------------------------------------------------........t....t..tt..t.....c..... [  2960]

FR682468.2_Georgia_2007/1         ccttatctacatgctattggcattcctgatattcgcactaaagtgctatgttacaaccgtcttatgcgtgatttttatcc [  3040]
ASF_IND_20_CAD543                 tt................................nnnnnnnnnnnnnnnnnnnnn------------------------- [  3040]
LR899193.1_Germany_2020/1         ................................................................................ [  3040]
MN393477.1_Wuhan_2019             ................................................................................ [  3040]
MW396979.1_Timor-Leste/2019/1     ................................................................................ [  3040]
MK940252.1_CN/2019/InnerMongolia  ................................................................................ [  3040]
MK128995.1_China/2018/AnhuiXCGQ   ................................................................................ [  3040]
MH681419.1_ASFV/POL/2015          ................................................................................ [  3040]
LS478113.1_Estonia_2014           ................................................................................ [  3040]
KP843857.1_Odintsovo_02/14        ................................................................................ [  3040]
```

FN557520.1_strain_E75 tt................................atgt...........ac..t....a.......t.g.c.c.....a. [ 3040]
AY261366.1_Warthog -------------------------------------------------------------------------------- [ 3040]
AY261365.1_Warmbaths -------------------------------------------------------------------------------- [ 3040]
AY261364.1_Tengani_62 -------------------------------------------------------------------------------- [ 3040]
AY261363.1_Pretoriuskop/96/4 tt...........t...................t...t...........a...t....a.......t...........a. [ 3040]
AY261362.1_Mkuzi_1979 t..................a.............................a.............................. [ 3040]
AY261361.1_Malawi_Lil-20/1_1983 -------------------------------------------------------------------------------- [ 3040]
AY261360.1_Kenya_1950 tt.g.........g....a...c.....t.....at.t....a..g...a...t.g.aa.....att.a.c.c.c..aa. [ 3040]
AM712240.1_OURT_88/3 -------------------------------------------------------------------------------- [ 3040]
AM712239.1_Benin_97/1 -------------------------------------------------------------------------------- [ 3040]
KM111295.1_Ken06.Bus -------------------------------------------------------------------------------- [ 3040]
KM111294.1_Ken05/Tk1 -------------------------------------t.g..a.tt..ca.----------------------------- [ 3040]
KM262844.1_strain_L60 ----------------------............atgt...........ac..t....a.......t.g.c.c.....a. [ 3040]
KM102979.1_26544/OG10_Italy -------------------------------------------------------------------------------- [ 3040]
KM262845.1_strain_NHV -------------------------.........atgt...........ac..t....a.......t.g.c.c.....a. [ 3040]
KX354450.1_47/Ss/2008 tt................................atgt...........ac..t....a.......t.g.c.c.....a. [ 3040]
KP055815.1_strain_BA71 tt................................atgt...........ac..t....a.......t.g.c.c.....a. [ 3040]
MH025920.1_strain_R35 tt.gg......c.t.g..................at.t.......tc.aa...tgg.aa.......ttatt.c.c...aa [ 3040]
MH025919.1_strain_N10 tt.gg......c.t.g..................at.t.......tc.aa...tgg.aa.......ttatt.c.c...aa [ 3040]
MH025918.1_strain_R25 tt.gg......c.t.g..................at.t.......tc.aa...tgg.aa.......ttatt.c.c...aa [ 3040]
MH025917.1_strain_R7 tt.gg......c.t.g..................at.t.......tc.aa...tgg.aa.......ttatt.c.c...aa [ 3040]
MH025916.1_strain_R8 tt.gg......c.t.g..................at.t.......tc.aa...tgg.aa.......ttatt.c.c...aa [ 3040]

FR682468.2_Georgia_2007/1 accttattggccgaagggccgccttgtatttcctgttaggtggtttggccgtattctactggtggcaag-cagctatcaa [ 3120]
ASF_IND_20_CAD543 ------------------------------------nnnnnnnnnnnnnnnnnnnnnnnnnnnnnnnnn-nnnnnnnnnn [ 3120]
LR899193.1_Germany_2020/1 .....................................................................-.......... [ 3120]
MN393477.1_Wuhan_2019 .....................................................................-.......... [ 3120]
MW396979.1_Timor-Leste/2019/1 .....................................................................-.......... [ 3120]
MK940252.1_CN/2019/InnerMongolia .....................................................................-.......... [ 3120]
MK128995.1_China/2018/AnhuiXCGQ .....................................................................-.......... [ 3120]
MH681419.1_ASFV/POL/2015 .....................................................................-.......... [ 3120]
LS478113.1_Estonia_2014 .....................................................................-.......... [ 3120]
KP843857.1_Odintsovo_02/14 .....................................................................-.......... [ 3120]
FN557520.1_strain_E75 .t.........a.g...c..a.ta.a...a........a..a....a..t....a..----------------------- [ 3120]
AY261366.1_Warthog -----------------------------------------------------------------------g.t.gc... [ 3120]
AY261365.1_Warmbaths -----------------------------------------------------------------------g.t.gc... [ 3120]
AY261364.1_Tengani_62 -----------------------------------------------------------------------g.t.gc... [ 3120]
AY261363.1_Pretoriuskop/96/4 .t....g....a.....a..a.t..a...a........a.......a..--...aa....at......at.......... [ 3120]
AY261362.1_Mkuzi_1979 ......................................................a.............at.......... [ 3120]
AY261361.1_Malawi_Lil-20/1_1983 -------------------------------------------------------------------------------- [ 3120]
AY261360.1_Kenya_1950 .tt.......tat.gattt.a.t..a...a.ga..c.....at...a..t....a---------------ttat.gg... [ 3120]
AM712240.1_OURT_88/3 -------------------------------------------------------------------------------- [ 3120]
AM712239.1_Benin_97/1 -------------------------------------------------------------------------------- [ 3120]
KM111295.1_Ken06.Bus -------------------------------------------------------------------------------- [ 3120]

```
KM111294.1_Ken05/Tk1           ----------------------------------------------------------------------------gt.. [  3120]
KM262844.1_strain_L60          .t.........a.g...c..a.ta.a...a........a..a....a..t....a..----------------------- [  3120]
KM102979.1_26544/OG10_Italy    -------------------------------------------------------------------------------- [  3120]
KM262845.1_strain_NHV          .t.........a.g...c..a.ta.a...a........a..a....a..t....a..----------------------- [  3120]
KX354450.1_47/Ss/2008          .t.........a.g...c..a.ta.a...a........a..a....a..t....a..----------------------- [  3120]
KP055815.1_strain_BA71         .t.........a.g...c..a.ta.a...a........a..a....a..t....a..----------------------- [  3120]
MH025920.1_strain_R35          .at.......ta..gaatt.a.t..a...a.t...c..a..at......t....a---------------.tat.gg... [  3120]
MH025919.1_strain_N10          .at.......ta..gaatt.a.t..a...a.t...c..a..at......t....a---------------.tat.gg... [  3120]
MH025918.1_strain_R25          .at.......ta..gaatt.a.t..a...a.t...c..a..at......t....a---------------.tat.gg... [  3120]
MH025917.1_strain_R7           .at.......ta..gaatt.a.t..a...a.t...c..a..at......t....a---------------.tat.gg... [  3120]
MH025916.1_strain_R8           .at.......ta..gaatt.a.t..a...a.t...c..a..at......t....a---------------.tat.gg... [  3120]

FR682468.2_Georgia_2007/1      taaaatttaatggctctcacttaagatccttgctgtaagcgggcgtttacatactttgatcaagaaa-----aaaaatta [  3200]
ASF_IND_20_CAD543              nnnnnnnnnnnnnnnnnnnnnnnnnnnnnnnnnnnnnnnnnnnnnnnnnnnnnnnnnnn........tatac......nn [  3200]
LR899193.1_Germany_2020/1      ...................................................................-----........ [  3200]
MN393477.1_Wuhan_2019          ...................................................................-----........ [  3200]
MW396979.1_Timor-Leste/2019/1  ...................................................................-----........ [  3200]
MK940252.1_CN/2019/InnerMongolia ...................................................................-----........ [  3200]
MK128995.1_China/2018/AnhuiXCGQ ...................................................................-----........ [  3200]
MH681419.1_ASFV/POL/2015       ...................................................................-----........ [  3200]
LS478113.1_Estonia_2014        ...................................................................-----........ [  3200]
KP843857.1_Odintsovo_02/14     ...................................................................-----.-...... [  3200]
FN557520.1_strain_E75          ----........tt.t..g...-----------------------------------..........tatac......c- [  3200]
AY261366.1_Warthog             a.tt.....c.----------------------------------------------------....-----.....a.. [  3200]
AY261365.1_Warmbaths           a.tt.....c.----------------------------------------------------....-----.....a.. [  3200]
AY261364.1_Tengani_62          a.tt.....c.----------------------------------------------------....-----.....a.. [  3200]
AY261363.1_Pretoriuskop/96/4   .............t..a..........t.........----...a......................-----.....a.. [  3200]
AY261362.1_Mkuzi_1979          ...........................t........t-.ta..........................---aa...t.a.- [  3200]
AY261361.1_Malawi_Lil-20/1_1983 ------------------------------------------------------------------------.....a.g [  3200]
AY261360.1_Kenya_1950          acctg....gat.t.------------------------------------------------....gtggct......t [  3200]
AM712240.1_OURT_88/3           -------------------------------------------------------------------------------- [  3200]
AM712239.1_Benin_97/1          -------------------------------------------------------------------------------- [  3200]
KM111295.1_Ken06.Bus           -------------------------------------------------------------------------------- [  3200]
KM111294.1_Ken05/Tk1           acc..c...----------------------------------------------------------------------- [  3200]
KM262844.1_strain_L60          ----........tt.t..g...-----------------------------------..........tatac......c- [  3200]
KM102979.1_26544/OG10_Italy    -------------------------------------------------------------------------------- [  3200]
KM262845.1_strain_NHV          ----........tt.t..g...-----------------------------------..........tatac......c- [  3200]
KX354450.1_47/Ss/2008          ----........tt.t..g...-----------------------------------..........tatac......c- [  3200]
KP055815.1_strain_BA71         ----........tt.t..g...-----------------------------------..........tatac......c- [  3200]
MH025920.1_strain_R35          acct.c...gacat.------------------------------------------------....gcgact......t [  3200]
MH025919.1_strain_N10          acct.c...gacat.------------------------------------------------....gcgact......t [  3200]
MH025918.1_strain_R25          acct.c...gacat.------------------------------------------------....gcgact......t [  3200]
MH025917.1_strain_R7           acct.c...gacat.------------------------------------------------....gcgact......t [  3200]
MH025916.1_strain_R8           acct.c...gacat.------------------------------------------------....gcgact......t [  3200]
```

```
FR682468.2_Georgia_2007/1        tttttgga------------------------------------cccccccccatgttttatacaaaaatcatataataa [  3280]
ASF_IND_20_CAD543                nnnnnnnnnnnnnnnnnnnnnnnnnnnnnnnnnnnnnnnnnnnnnnnnnnnnnnnnnnnnnnnnnnnnnnnnnnnnnnnn [  3280]
LR899193.1_Germany_2020/1        ........-----------------------------------c.................................... [  3280]
MN393477.1_Wuhan_2019            ........-----------------------------------c.................................... [  3280]
MW396979.1_Timor-Leste/2019/1    ........-----------------------------------c.................................... [  3280]
MK940252.1_CN/2019/InnerMongolia ........c-------------------------------cccc.................................... [  3280]
MK128995.1_China/2018/AnhuiXCGQ  ........------------------------------------.................................... [  3280]
MH681419.1_ASFV/POL/2015         ........----------------------------------cc.................................... [  3280]
LS478113.1_Estonia_2014          .....--------------------------------------------------------------------------- [  3280]
KP843857.1_Odintsovo_02/14       ........-----------------------------------c.................................... [  3280]
FN557520.1_strain_E75            -----------------------------------------------------------------------------... [  3280]
AY261366.1_Warthog               a..a.--------------------------------------------------------------------------. [  3280]
AY261365.1_Warmbaths             a..a.--------------------------------------------------------------------------. [  3280]
AY261364.1_Tengani_62            a..a.--------------------------------------------------------------------------. [  3280]
AY261363.1_Pretoriuskop/96/4     a......----------------------------------------------------------------------.c. [  3280]
AY261362.1_Mkuzi_1979            -----------------------------------------------------------------------------.tt [  3280]
AY261361.1_Malawi_Lil-20/1_1983  ..------------------------------------------------------------------------------ [  3280]
AY261360.1_Kenya_1950            aa.g.--------------------------------------------------------------------------- [  3280]
AM712240.1_OURT_88/3             -------------------------------------------------------------------------------- [  3280]
AM712239.1_Benin_97/1            -------------------------------------------------------------------------------- [  3280]
KM111295.1_Ken06.Bus             -------------------------------------------------------------------------------- [  3280]
KM111294.1_Ken05/Tk1             -------------------------------------------------------------------------------- [  3280]
KM262844.1_strain_L60            -----------------------------------------------------------------------------... [  3280]
KM102979.1_26544/OG10_Italy      -------------------------------------------------------------------------------- [  3280]
KM262845.1_strain_NHV            -----------------------------------------------------------------------------... [  3280]
KX354450.1_47/Ss/2008            -----------------------------------------------------------------------------... [  3280]
KP055815.1_strain_BA71           -----------------------------------------------------------------------------... [  3280]
MH025920.1_strain_R35            aa.g.--------------------------------------------------------------------------- [  3280]
MH025919.1_strain_N10            aa.g.--------------------------------------------------------------------------- [  3280]
MH025918.1_strain_R25            aa.g.--------------------------------------------------------------------------- [  3280]
MH025917.1_strain_R7             aa.g.--------------------------------------------------------------------------- [  3280]
MH025916.1_strain_R8             aa.g.--------------------------------------------------------------------------- [  3280]

FR682468.2_Georgia_2007/1        agtggcgacaatcaacatattaatcaaccacagcattttatga---------tgtgttaatcaacatataccatattaat [  3360]
ASF_IND_20_CAD543                nnnnnnnnnnnnnnnnnnnnnnnnnnnnnnnnnnnnnnnnnnn---------nnnnnnnnnnnnnnnnnnnnnnnnnnnn [  3360]
LR899193.1_Germany_2020/1        ...........................................---------............................ [  3360]
MN393477.1_Wuhan_2019            ...........................................---------............................ [  3360]
MW396979.1_Timor-Leste/2019/1    ...........................................---------............................ [  3360]
MK940252.1_CN/2019/InnerMongolia ...........................................---------............................ [  3360]
MK128995.1_China/2018/AnhuiXCGQ  ...........................................---------............................ [  3360]
MH681419.1_ASFV/POL/2015         ...........................................---------............................ [  3360]
LS478113.1_Estonia_2014          -------------------------------------------------------------------------------- [  3360]
KP843857.1_Odintsovo_02/14       ...........................................---------............................ [  3360]
```

```
FN557520.1_strain_E75          gtgca.c.ttt.atgggc..c.....t........c...ggcgtga--a.caa.c.ca.c.tt.tatgggcg.c... [  3360]
AY261366.1_Warthog             caacc.c.tgt.tt.t.c.aa...t.caac.g.....a-..c.actctatgcgtc..c.....cagc..ttt..ga.--- [  3360]
AY261365.1_Warmbaths           caacc.c.tgt.tt.t.c.aa...t.caac.g.....a-..c.actctatgcgtc..c.....----------------- [  3360]
AY261364.1_Tengani_62          caacc.c.tgt.tt.t.c.aa...t.caac.g.....a-..c.actctatgcatc..c.....cagc..ttt..ga.--- [  3360]
AY261363.1_Pretoriuskop/96/4   caacc.c.tgt.tt.t.c.aa.....ttactta...aa..ac.aataaagtag.c.ac...............a.... [  3360]
AY261362.1_Mkuzi_1979          g..ca.a..cccatgtt...ac.aa..t..----------------------.tac...tat..t.a.gtaggcgac... [  3360]
AY261361.1_Malawi_Lil-20/1_1983 -------------------------------------------------------------------------------- [  3360]
AY261360.1_Kenya_1950          ----------------------------------------------------.t.t.c..aa..g..c.-tt...aa--- [  3360]
AM712240.1_OURT_88/3           -------------------------------------------------------------------------------- [  3360]
AM712239.1_Benin_97/1          -------------------------------------------------------------------------------- [  3360]
KM111295.1_Ken06.Bus           -------------------------------------------------------------------------------- [  3360]
KM111294.1_Ken05/Tk1           -----------------------------------------------------c.c.c.g...gg.ac..gt...----- [  3360]
KM262844.1_strain_L60          gtgca.c.ttt.atgggc..c.....t........c...ggcgtga--a.caa.c.ca.c.tt.tatgggcg.c... [  3360]
KM102979.1_26544/OG10_Italy    -------------------------------------------------------------------------------- [  3360]
KM262845.1_strain_NHV          gtgca.c.ttt.atgggc..c.....t........c...ggcgtga--a.caa.c.ca.c.tt.tatgggcg.c... [  3360]
KX354450.1_47/Ss/2008          gtgca.c.ttt.atgggc..c.....t........c...ggcgtga--a.caa.c.ca.c.tt.tatgggcg.c... [  3360]
KP055815.1_strain_BA71         gtgca.c.ttt.atgggc..c.....t........c...ggcgtga--a.caa.c.ca.c.tt.tatgggcg.c... [  3360]
MH025920.1_strain_R35          ----------------------------------------------------.t.t.c..aa..g..c.ttt...aa--- [  3360]
MH025919.1_strain_N10          ----------------------------------------------------.t.t.c..aa..g..c.ttt...aa--- [  3360]
MH025918.1_strain_R25          ----------------------------------------------------.t.t.c..aa..g..c.ttt...aa--- [  3360]
MH025917.1_strain_R7           ----------------------------------------------------.t.t.c..aa..g..c.ttt...aa--- [  3360]
MH025916.1_strain_R8           ----------------------------------------------------.t.t.c..aa..g..c.ttt...aa--- [  3360]

FR682468.2_Georgia_2007/1      caaccacagcattttatgatgcgtcaatcaacat--attattacggaga-----gcg-----tcaatcaa-tataatatt [  3440]
ASF_IND_20_CAD543              nnnnnnnnnnnnnnnnnnnnnnnnnnnnnnnnnn--nnnnnnnnnnnnn-----nnn-----nnnnnnnn-nnnnnnnnn [  3440]
LR899193.1_Germany_2020/1      ..................................--.............-----...-----........-......... [  3440]
MN393477.1_Wuhan_2019          ..................................--.............-----...-----........-......... [  3440]
MW396979.1_Timor-Leste/2019/1  ..................................--.............-----...-----........-......... [  3440]
MK940252.1_CN/2019/InnerMongolia ..................................--.............-----...-----........-......... [  3440]
MK128995.1_China/2018/AnhuiXCGQ ..................................--.............-----...-----........-......... [  3440]
MH681419.1_ASFV/POL/2015       ..................................--.............-----...-----........-......... [  3440]
LS478113.1_Estonia_2014        -----------------------.tgg......c--c..t..gt-------------------------.-cg..tag.. [  3440]
KP843857.1_Odintsovo_02/14     ..................................--.............-----...-----........-......... [  3440]
FN557520.1_strain_E75          ...t....c........ggcatca.tca.t..c--.gc...ctatg.------...-----.g......-c.c.t.... [  3440]
AY261366.1_Warthog             -t...t..................c..------.c...attatgat-----a..-----.......cc...t.... [  3440]
AY261365.1_Warmbaths           ------..................c..------.c...attatgat-----a..-----.......cc...t.... [  3440]
AY261364.1_Tengani_62          -t...t...............a....c..------.c...attatgat-----a..-----.......cc...t.... [  3440]
AY261363.1_Pretoriuskop/96/4   ......................t.......cat.cc..attattacggaga...-----.......-......... [  3440]
AY261362.1_Mkuzi_1979          ....---------------atat.a.tca.c..c--.gc..attatgat-----...-----.......-......... [  3440]
AY261361.1_Malawi_Lil-20/1_1983 ------------------------------------gcc.ccc..cc.cc----...-----gt.c.a..t.t....g-- [  3440]
AY261360.1_Kenya_1950          ---------------------------ct.tggaaggcggcgttatgcttttcg...catac.t...a..-.g.gg.ga. [  3440]
AM712240.1_OURT_88/3           -------------------------------------------------------------------------------- [  3440]
AM712239.1_Benin_97/1          -------------------------------------------------------------------------------- [  3440]
KM111295.1_Ken06.Bus           -------------------------------------------------------------------------------- [  3440]
```

```
KM111294.1_Ken05/Tk1             --------------------------------------------------------------.t.t.ag.-c-------- [  3440]
KM262844.1_strain_L60            ...t....c.........ggcatca.tca.t..c--.gc...ctatg.------...-----.g......-c.c.t.... [  3440]
KM102979.1_26544/OG10_Italy      -----------.......ggcatca.tca.t..c--.gc...ctatg.------...-----.g......-c.c.t.... [  3440]
KM262845.1_strain_NHV            ...t....c.........ggcatca.tca.t..c--.gc...ctatg.------...-----.g......-c.c.t.... [  3440]
KX354450.1_47/Ss/2008            ...t....c.........ggcatca.tca.t..c--.gc...ctatg.------...-----.g......-c.c.t.... [  3440]
KP055815.1_strain_BA71           ...t....c.........ggcatca.tca.t..c--.gc...ctatg.------...-----.g......-c.c.t.... [  3440]
MH025920.1_strain_R35            ---------------------------c..tggaagccgg.gttatgcttttct...tatac.t...a..-.g.g..ga. [  3440]
MH025919.1_strain_N10            ---------------------------c..tggaagccgg.gttatgcttttct...tatac.t...a..-.g.g..ga. [  3440]
MH025918.1_strain_R25            ---------------------------c..tggaagccgg.gttatgcttttct...tatac.t...a..-.g.g..ga. [  3440]
MH025917.1_strain_R7             ---------------------------c..tggaagccgg.gttatgcttttct...tatac.t...a..-.g.g..ga. [  3440]
MH025916.1_strain_R8             ---------------------------c..tggaagccgg.gttatgcttttct...tatac.t...a..-.g.g..ga. [  3440]

FR682468.2_Georgia_2007/1        gagaacagcgacttgataccgtgtatggtggtggc------------------------------ggcggcatgttgttt [  3520]
ASF_IND_20_CAD543                nnnnnnnnnnnnnnnnnnnnnnnnnnnnnnnnnnn------------------------------nnnnnnnnnnnnnnn [  3520]
LR899193.1_Germany_2020/1        ...................................------------------------------............... [  3520]
MN393477.1_Wuhan_2019            ...................................------------------------------............... [  3520]
MW396979.1_Timor-Leste/2019/1    ...................................------------------------------............... [  3520]
MK940252.1_CN/2019/InnerMongolia ...................................------------------------------............... [  3520]
MK128995.1_China/2018/AnhuiXCGQ  ...................................------------------------------............... [  3520]
MH681419.1_ASFV/POL/2015         ...................................------------------------------............... [  3520]
LS478113.1_Estonia_2014          ..t..a..----------------------------------------------------------------..c..gca [  3520]
KP843857.1_Odintsovo_02/14       ...................................------------------------------............... [  3520]
FN557520.1_strain_E75            .g.g.....t..c.....t................accgtgttgtttgaaggcgtgtatggtatta...c.......... [  3520]
AY261366.1_Warthog               a....t...t......c.....a........c..t---------------------------gat..t............ [  3520]
AY261365.1_Warmbaths             a....t...t......c.....a........c..t---------------------------ggt..t............ [  3520]
AY261364.1_Tengani_62            a....t...t......c.....a........c..t---------------------------ggt..t............ [  3520]
AY261363.1_Pretoriuskop/96/4     ................c..................---------------------------ggt............... [  3520]
AY261362.1_Mkuzi_1979            ...................................------------------------------............... [  3520]
AY261361.1_Malawi_Lil-20/1_1983  -------------------------------------------------------------------------------- [  3520]
AY261360.1_Kenya_1950            a.c....a.ata.at.acttt.t...atctaa.at----------------------ccgagtacaaaaaa.....aa.. [  3520]
AM712240.1_OURT_88/3             -------------------------------------------gtttgaaggcgtgtatggtatta...c.......... [  3520]
AM712239.1_Benin_97/1            -------------------------------------------gtttgaaggcgtgtatggtatta...c.......... [  3520]
KM111295.1_Ken06.Bus             -------------------------------------------------------------------------------- [  3520]
KM111294.1_Ken05/Tk1             ----------------ac.aa.......ct.a------------------------------------------------ [  3520]
KM262844.1_strain_L60            .g.g.....t..c.....t................accgtgttgtttgaaggcgtgtatggtatta...c.......... [  3520]
KM102979.1_26544/OG10_Italy      .g.g.....t..c.....t................accgtgttgtttgaaggcgtgtatggtatta...c.......... [  3520]
KM262845.1_strain_NHV            .g.g.....t..c.....t................accgtgttgtttgaaggcgtgtatggtatta...c.......... [  3520]
KX354450.1_47/Ss/2008            .g.g.....t..c.....t................accgtgttgtttgaaggcgtgtatggtatta...c.......... [  3520]
KP055815.1_strain_BA71           .g.g.....t..c.....t................accgtgttgtttgaaggcgtgtatggtatta...c.......... [  3520]
MH025920.1_strain_R35            a.c....a.ata.atcac.tt.c...atctaa.at----------------------ccgagtacaaaaaa.....aa.. [  3520]
MH025919.1_strain_N10            a.c....a.ata.atcac.tt.c...atctaa.at----------------------ccgagtacaaaaaa.....aa.. [  3520]
MH025918.1_strain_R25            a.c....a.ata.atcac.tt.c...atctaa.at----------------------ccgagtacaaaaaa.....aa.. [  3520]
MH025917.1_strain_R7             a.c....a.ata.atcac.tt.c...atctaa.at----------------------ccgagtacaaaaaa.....aa.. [  3520]
MH025916.1_strain_R8             a.c....a.ata.atcac.tt.c...atctaa.at----------------------ccgagtacaaaaaa.....aa.. [  3520]
```

```
FR682468.2_Georgia_2007/1        ---------------gtaacagcatttttcatcattcgaagcttacaaaagatatgtataagatagcatattaatgttat [  3600]
ASF_IND_20_CAD543                ---------------nnnnnnnnnnnnnnnnnnnnnnnnnnnnnnnnnn...................a.......c... [  3600]
LR899193.1_Germany_2020/1        ---------------................................................................. [  3600]
MN393477.1_Wuhan_2019            ---------------................................................................. [  3600]
MW396979.1_Timor-Leste/2019/1    ---------------................................................................. [  3600]
MK940252.1_CN/2019/InnerMongolia ---------------................................................................. [  3600]
MK128995.1_China/2018/AnhuiXCGQ  ---------------................................................................. [  3600]
MH681419.1_ASFV/POL/2015         ---------------................................................................. [  3600]
LS478113.1_Estonia_2014          ---------------a...t.a.....a.a..tt..ttct.ggcga....a...at...gtt.cgt.....--------- [  3600]
KP843857.1_Odintsovo_02/14       ---------------................................................................. [  3600]
FN557520.1_strain_E75            gaaggcagcgcgtta......a.g.....tt..g.at.------....c...................a.......c... [  3600]
AY261366.1_Warthog               ---------------a....ga.......tt..ga...------....c.....a..........c.........a.g.. [  3600]
AY261365.1_Warmbaths             ---------------a.g...a.......tt..ga...------....c.....a..........c.........a.g.. [  3600]
AY261364.1_Tengani_62            ---------------.....ga.......tt..ga...------....c.....a..........c.........a.... [  3600]
AY261363.1_Pretoriuskop/96/4     ---------------......ct.......t................................................. [  3600]
AY261362.1_Mkuzi_1979            ---------------.....t........................................................... [  3600]
AY261361.1_Malawi_Lil-20/1_1983  -------------------------....tt..g.c..------------------------------------------ [  3600]
AY261360.1_Kenya_1950            ----atgacgcg---a.c..ttt.a....t-.tgg..t------.gg...a----------------------------- [  3600]
AM712240.1_OURT_88/3             gaaggcagcgcgtta......a.g.....tt..g.at.------....c...................a.......c... [  3600]
AM712239.1_Benin_97/1            gaaggcagcgcgtta......a.g.....tt..g.at.------....c...................a......ac... [  3600]
KM111295.1_Ken06.Bus             -------------------------------------------------------------------------------- [  3600]
KM111294.1_Ken05/Tk1             ---------------------tt..c...t-------------------------------------------------- [  3600]
KM262844.1_strain_L60            gaaggcagcgcgtta......a.g.....tt..g.at.------....c...................a.......c... [  3600]
KM102979.1_26544/OG10_Italy      gaaggcagcgcgtta......a.g.....tt..g.at.------....c...................a......ac... [  3600]
KM262845.1_strain_NHV            gaaggcagcgcgtta......a.g.....tt..g.at.------....c...................a.......c... [  3600]
KX354450.1_47/Ss/2008            gaaggcagcgcgtta......a.g.....tt..g.at.------....c...................a......ac... [  3600]
KP055815.1_strain_BA71           gaaggcagcgcgtta......a.g.....tt..g.at.------....c...................a.......c... [  3600]
MH025920.1_strain_R35            ----ataacacg---a.c..ttt......tt.tgg..t------.gg...a.a.aa..----.a.tt...g.c....--- [  3600]
MH025919.1_strain_N10            ----ataacacg---a.c..ttt......tt.tgg..t------.gg...a.a.aa..----.a.tt...g.c....--- [  3600]
MH025918.1_strain_R25            ----ataacacg---a.c..ttt......tt.tgg..t------.gg...a.a.aa..----.a.tt...g.c....--- [  3600]
MH025917.1_strain_R7             ----ataacacg---a.c..ttt......tt.tgg..t------.gg...a.a.aa..----.a.tt...g.c....--- [  3600]
MH025916.1_strain_R8             ----ataacacg---a.c..ttt......tt.tgg..t------.gg...a.a.aa..----.a.tt...g.c....--- [  3600]

FR682468.2_Georgia_2007/1        taacagtaatatcaataaggcgtagctatagatcttcacttt---ggtagaccaataatccatggttgcgcttaaaaata [  3680]
ASF_IND_20_CAD543                .nnnnnnnnnnnnnnnnnnnnnnnnnnnnnnnnnnnnnnn..---...........g......-.......c.......c [  3680]
LR899193.1_Germany_2020/1        ..........................................---................................... [  3680]
MN393477.1_Wuhan_2019            ..........................................---................................... [  3680]
MW396979.1_Timor-Leste/2019/1    ..........................................---................................... [  3680]
MK940252.1_CN/2019/InnerMongolia ..........................................---................................... [  3680]
MK128995.1_China/2018/AnhuiXCGQ  ..........................................---................................... [  3680]
MH681419.1_ASFV/POL/2015         ..........................................---................................... [  3680]
LS478113.1_Estonia_2014          -----------------------------------------------------------------------------..t [  3680]
KP843857.1_Odintsovo_02/14       ..........................................---................................... [  3680]
```

```
FN557520.1_strain_E75            .gt..tc...gag..ac.taa....a.......c..g.ctc..---.....t.....g.....-......a.c......c [  3680]
AY261366.1_Warthog               ..c.....t.ga.....c.at..........tc.g..gtcc.atg.c.g.t...g...ca..........a........c [  3680]
AY261365.1_Warmbaths             ..c.....t.ga.....c.at..........tc.g..gtcc.atg.c.g.t...g...ca..........a........c [  3680]
AY261364.1_Tengani_62            ..c.....t.g......c.at..........tc.g..gtcc.atg.c.g.t...g...ca....a.....a........c [  3680]
AY261363.1_Pretoriuskop/96/4     ..c.......g.........t......................---......................a.a........c [  3680]
AY261362.1_Mkuzi_1979            ..g........................................---.......................t.........c [  3680]
AY261361.1_Malawi_Lil-20/1_1983  --------ccgc...gcc.tgc..t.gg..a.c..g.tgc..gt----------t..ggcttg.-.c..ta.gc....ag [  3680]
AY261360.1_Kenya_1950            -------------------------------------------------------------------------------- [  3680]
AM712240.1_OURT_88/3             .gt..tc...gag..ac.taa....a.......c..g.ctc..---.....t.....g.....-......a.c......c [  3680]
AM712239.1_Benin_97/1            .gt..tc...gag..ac.taa....a.......c..a.ctc..---.....t.....g.....-......a.c......c [  3680]
KM111295.1_Ken06.Bus             -------------------------------------------------------------------------------- [  3680]
KM111294.1_Ken05/Tk1             -------------------------------------------------------------------------------- [  3680]
KM262844.1_strain_L60            .gt..tc...gag..ac.taa....a.......c..g.ctc..---.....t.....g.....-......a.c......c [  3680]
KM102979.1_26544/OG10_Italy      .gt..tc...gag..ac.taa....a.......c..a.ctc..---.....t.....g.....-......a.c......c [  3680]
KM262845.1_strain_NHV            .gt..tc...gag..ac.taa....a.......c..g.ctc..---.....t.....g.....-......a.c......c [  3680]
KX354450.1_47/Ss/2008            .gt..tc...gag..ac.taa....a.......c..a.ctc..---.....t.....g.....-......a.c......c [  3680]
KP055815.1_strain_BA71           .gt..tc...gag..ac.taa....a.......c..g.ctc..---.....t.....g.....-......a.c......c [  3680]
MH025920.1_strain_R35            ----------------------------------------------.a.cctat..a......--ca..a.ca...cc.c [  3680]
MH025919.1_strain_N10            ----------------------------------------------.a.cctat..a......--ca..a.ca...cc.c [  3680]
MH025918.1_strain_R25            ----------------------------------------------.a.cctat..a......--ca..a.ca...cc.c [  3680]
MH025917.1_strain_R7             ----------------------------------------------.a.cctat..a......--ca..a.ca...cc.c [  3680]
MH025916.1_strain_R8             ----------------------------------------------.a.cctat..a......--ca..a.ca...cc.c [  3680]

FR682468.2_Georgia_2007/1        cc--aaaaaaaa-cattaagtttt---ggagggt----aagattggtttttcaccattggtaaagattattattctaaat [  3760]
ASF_IND_20_CAD543                ..--........a...........---.......----.......................................... [  3760]
LR899193.1_Germany_2020/1        ..--........-...........---.......----.......................................... [  3760]
MN393477.1_Wuhan_2019            ..--........-...........---.......----.......................................... [  3760]
MW396979.1_Timor-Leste/2019/1    ..--........-...........---.......----.......................................... [  3760]
MK940252.1_CN/2019/InnerMongolia ..--........-...........---.......----.......................................... [  3760]
MK128995.1_China/2018/AnhuiXCGQ  ..--........-...........---.......----.......................................... [  3760]
MH681419.1_ASFV/POL/2015         ..--........-...........---.......----.......................................... [  3760]
LS478113.1_Estonia_2014          ..--t.tcgg.----------------------------------.a...g......---c.g...g.tc.g..a....a [  3760]
KP843857.1_Odintsovo_02/14       ..--.......--...........---.......----.......................................... [  3760]
FN557520.1_strain_E75            ..--........a...........---.......----.......................................... [  3760]
AY261366.1_Warthog               .a---.......a......a....---a..aa..aata..a...a........t...............-.......gg. [  3760]
AY261365.1_Warmbaths             .----.......a......a....---a..aa..aata..a...a........t...............-.......gg. [  3760]
AY261364.1_Tengani_62            .a---.......a......a....---a...a..aata..a...a........t...............-.......gg. [  3760]
AY261363.1_Pretoriuskop/96/4     ..---.......a...........ggaa......aata..a...a...............c................gg. [  3760]
AY261362.1_Mkuzi_1979            ..---.......a...........---.......----.......................................... [  3760]
AY261361.1_Malawi_Lil-20/1_1983  ..---ggccgccg..c...aac..-----------------.a..t......g.t.caaa.---------cgcca.gg.. [  3760]
AY261360.1_Kenya_1950            ----....ttt.gt..cg-----------------------------------------------..g.----------- [  3760]
AM712240.1_OURT_88/3             ..--........a...........---.......----.......................................... [  3760]
AM712239.1_Benin_97/1            ..--........a...........---.......----.......................................... [  3760]
KM111295.1_Ken06.Bus             -------------------------------------------------------------------------------- [  3760]
```

```
KM111294.1_Ken05/Tk1              ----..t...t.a..gc.-------------------------------------------------------------- [  3760]
KM262844.1_strain_L60             ..--........a...........---.......----.......................................... [  3760]
KM102979.1_26544/OG10_Italy       ..---.......a...........---.......----.......................................... [  3760]
KM262845.1_strain_NHV             ..--........a...........---.......----.......................................... [  3760]
KX354450.1_47/Ss/2008             ..---.......a...........---.......----.......................................... [  3760]
KP055815.1_strain_BA71            ..--........a...........---.......----.......................................... [  3760]
MH025920.1_strain_R35             tatc....t.t.t...cgcta..atg-----------a..a.a.aa....-------------.t....----------- [  3760]
MH025919.1_strain_N10             tatc....t.t.t...cgcta..atg-----------a..a.a.aa....-------------.t....----------- [  3760]
MH025918.1_strain_R25             tatc....t.t.t...cgcta..atg-----------a..a.a.aa....-------------.t....----------- [  3760]
MH025917.1_strain_R7              tatc....t.t.t...cgcta..atg-----------a..a.a.aa....-------------.t....----------- [  3760]
MH025916.1_strain_R8              tatc....t.t.t...cgcta..atg-----------a..a.a.aa....-------------.t....----------- [  3760]

FR682468.2_Georgia_2007/1         gtttaccccatagatgtgaaacaatgattcttcatatattaacatattttttgacttatacttttcttcatctagtaagg [  3840]
ASF_IND_20_CAD543                 ................................................................................ [  3840]
LR899193.1_Germany_2020/1         ................................................................................ [  3840]
MN393477.1_Wuhan_2019             ................................................................................ [  3840]
MW396979.1_Timor-Leste/2019/1     ................................................................................ [  3840]
MK940252.1_CN/2019/InnerMongolia  ................................................................................ [  3840]
MK128995.1_China/2018/AnhuiXCGQ   ................................................................................ [  3840]
MH681419.1_ASFV/POL/2015          ................................................................................ [  3840]
LS478113.1_Estonia_2014           aa.a.ttttg..tcaa...---g..a..........c...................c.................a..... [  3840]
KP843857.1_Odintsovo_02/14        ................................................................................ [  3840]
FN557520.1_strain_E75             ................................................................................ [  3840]
AY261366.1_Warthog                ....gt..............................c.....................................a....a [  3840]
AY261365.1_Warmbaths              ....gt..............................c.....................................a..... [  3840]
AY261364.1_Tengani_62             ....gt..............................c.....................................a..... [  3840]
AY261363.1_Pretoriuskop/96/4      a......t............................cc.....................................c.... [  3840]
AY261362.1_Mkuzi_1979             ................................................................................ [  3840]
AY261361.1_Malawi_Lil-20/1_1983   agc....tg..gtg-----------.t....ct.c..gc.gg.ggccaacc..----c.gg.gc.g.....a.gc..c.. [  3840]
AY261360.1_Kenya_1950             -------------------------..................tc...........c...tg....a.......a...aa [  3840]
AM712240.1_OURT_88/3              ................................................................................ [  3840]
AM712239.1_Benin_97/1             ................................................................................ [  3840]
KM111295.1_Ken06.Bus              -------------------------------------------------------------------------------- [  3840]
KM111294.1_Ken05/Tk1              --------------------------------------------..c....a.tt..g..g...------.....c...a [  3840]
KM262844.1_strain_L60             ................................................................................ [  3840]
KM102979.1_26544/OG10_Italy       ................................................................................ [  3840]
KM262845.1_strain_NHV             ................................................................................ [  3840]
KX354450.1_47/Ss/2008             ................................................................................ [  3840]
KP055815.1_strain_BA71            ................................................................................ [  3840]
MH025920.1_strain_R35             -------------------------..................cc...........c...a.....c.......a....a [  3840]
MH025919.1_strain_N10             -------------------------..................cc...........c...a.....c.......a....a [  3840]
MH025918.1_strain_R25             -------------------------..................cc...........c...a.....c.......a....a [  3840]
MH025917.1_strain_R7              -------------------------..................cc...........c...a.....c.......a....a [  3840]
MH025916.1_strain_R8              -------------------------..................cc...........c...a.....c.......a....a [  3840]
```

| | | |
|---|---|---|
| FR682468.2_Georgia_2007/1 | cgttaattttttccggatctgtc---gtttttattgataaaagagaagagtctggact---gtaatttttaaataataag | [ 3920] |
| ASF_IND_20_CAD543 | .......................---................................---................... | [ 3920] |
| LR899193.1_Germany_2020/1 | .......................---................................---................... | [ 3920] |
| MN393477.1_Wuhan_2019 | .......................---................................---................... | [ 3920] |
| MW396979.1_Timor-Leste/2019/1 | .......................---................................---................... | [ 3920] |
| MK940252.1_CN/2019/InnerMongolia | .......................---................................---................... | [ 3920] |
| MK128995.1_China/2018/AnhuiXCGQ | .......................---................................---................... | [ 3920] |
| MH681419.1_ASFV/POL/2015 | .......................---................................---................... | [ 3920] |
| LS478113.1_Estonia_2014 | .a.....................---.........--------------------------------------------- | [ 3920] |
| KP843857.1_Odintsovo_02/14 | .......................---................................---................... | [ 3920] |
| FN557520.1_strain_E75 | .......................---................................---................... | [ 3920] |
| AY261366.1_Warthog | .a.....................---.......a..................c.....---a.................. | [ 3920] |
| AY261365.1_Warmbaths | .a.....................---.......a..................c.....---a.................. | [ 3920] |
| AY261364.1_Tengani_62 | .a.....................---.......a..................c.....---a.................. | [ 3920] |
| AY261363.1_Pretoriuskop/96/4 | .a.....................---.......a.a...................t..---................... | [ 3920] |
| AY261362.1_Mkuzi_1979 | .......................---................................---................... | [ 3920] |
| AY261361.1_Malawi_Lil-20/1_1983 | .a.cg.--------..gc...cttaa.c.c.gcgc.gcgc..a.a...cccga...gc---------------------- | [ 3920] |
| AY261360.1_Kenya_1950 | ta........c...........t---a......a.....t...g..gcta........ataa.........t...t...t | [ 3920] |
| AM712240.1_OURT_88/3 | .......................---................................---................... | [ 3920] |
| AM712239.1_Benin_97/1 | .......................---................................---................... | [ 3920] |
| KM111295.1_Ken06.Bus | -------------------------------------------------------------------------------- | [ 3920] |
| KM111294.1_Ken05/Tk1 | .t........c..t.........---.......a............g..t........---a............g....t | [ 3920] |
| KM262844.1_strain_L60 | .......................---................................---................... | [ 3920] |
| KM102979.1_26544/OG10_Italy | .......................---................................---................... | [ 3920] |
| KM262845.1_strain_NHV | .......................---................................---................... | [ 3920] |
| KX354450.1_47/Ss/2008 | .......................---................................---................... | [ 3920] |
| KP055815.1_strain_BA71 | .......................---................................---................... | [ 3920] |
| MH025920.1_strain_R35 | ta........c...........t---a......a.....t...g..gcta........ataa........c...g....t | [ 3920] |
| MH025919.1_strain_N10 | ta........c...........t---a......a.....t...g..gcta........ataa........c...g....t | [ 3920] |
| MH025918.1_strain_R25 | ta........c...........t---a......a.....t...g..gcta........ataa........c...g....t | [ 3920] |
| MH025917.1_strain_R7 | ta........c...........t---a......a.....t...g..gcta........ataa........c...g....t | [ 3920] |
| MH025916.1_strain_R8 | ta........c...........t---a......a.....t...g..gcta........ataa........c...g....t | [ 3920] |

| | | |
|---|---|---|
| FR682468.2_Georgia_2007/1 | atatttattaatatccaattattcgtttggctcgctatttccatgctctcttcgaaagcatcagctcctaaatctataca | [ 4000] |
| ASF_IND_20_CAD543 | .....................................................a..c....................... | [ 4000] |
| LR899193.1_Germany_2020/1 | ................................................................................ | [ 4000] |
| MN393477.1_Wuhan_2019 | ................................................................................ | [ 4000] |
| MW396979.1_Timor-Leste/2019/1 | ................................................................................ | [ 4000] |
| MK940252.1_CN/2019/InnerMongolia | ................................................................................ | [ 4000] |
| MK128995.1_China/2018/AnhuiXCGQ | ................................................................................ | [ 4000] |
| MH681419.1_ASFV/POL/2015 | ................................................................................ | [ 4000] |
| LS478113.1_Estonia_2014 | -------------------------------------------------------------------------------- | [ 4000] |
| KP843857.1_Odintsovo_02/14 | ................................................................................ | [ 4000] |

```
FN557520.1_strain_E75            .............................a.......................a..c....................... [  4000]
AY261366.1_Warthog               ........c................c...............t..........ta..t....................... [  4000]
AY261365.1_Warmbaths             ........c................c...............t..........ta..t....................... [  4000]
AY261364.1_Tengani_62            .........................c...............t..........ta..t....................... [  4000]
AY261363.1_Pretoriuskop/96/4     .......................t.............................a..c....................... [  4000]
AY261362.1_Mkuzi_1979            .....................................................a..c....................... [  4000]
AY261361.1_Malawi_Lil-20/1_1983  -----------------------------c.c...cgccc..g.a....taat..----------.tt.cggg.cg...t [  4000]
AY261360.1_Kenya_1950            ..t.....cg..tcaat..c....tg...tt.t........t.aa.a.........g...gt...c.............. [  4000]
AM712240.1_OURT_88/3             .............................a.......................a..c....................... [  4000]
AM712239.1_Benin_97/1            .............................a.......................a..c....................... [  4000]
KM111295.1_Ken06.Bus             -------------------------------------------------------------------------------- [  4000]
KM111294.1_Ken05/Tk1             ..t....c......att..c....t..c.tt.t........t.aa.a.........g...gt.................. [  4000]
KM262844.1_strain_L60            .............................a.......................a..c....................... [  4000]
KM102979.1_26544/OG10_Italy      .............................a.......................a..c....................... [  4000]
KM262845.1_strain_NHV            .............................a.......................a..c....................... [  4000]
KX354450.1_47/Ss/2008            .............................a.......................a..c....................... [  4000]
KP055815.1_strain_BA71           .............................a.......................a..c....................... [  4000]
MH025920.1_strain_R35            ..t....c....tcatt.....c.tg...tt.t........t.aa.a.........g...gt.................. [  4000]
MH025919.1_strain_N10            ..t....c....tcatt.....c.tg...tt.t........t.aa.a.........g...gt.................. [  4000]
MH025918.1_strain_R25            ..t....c....tcatt.....c.tg...tt.t........t.aa.a.........g...gt.................. [  4000]
MH025917.1_strain_R7             ..t....c....tcatt.....c.tg...tt.t........t.aa.a.........g...gt.................. [  4000]
MH025916.1_strain_R8             ..t....c....tcatt.....c.tg...tt.t........t.aa.a.........g...gt.................. [  4000]

FR682468.2_Georgia_2007/1        aaggaata----agttaccttcacaaaaattcattaccgaggtaatcattgcccgattaatgtcagcccccaacataaaa [  4080]
ASF_IND_20_CAD543                ........----..............................................r..................... [  4080]
LR899193.1_Germany_2020/1        ........----.................................................................... [  4080]
MN393477.1_Wuhan_2019            ........----.................................................................... [  4080]
MW396979.1_Timor-Leste/2019/1    ........----.................................................................... [  4080]
MK940252.1_CN/2019/InnerMongolia ........----.................................................................... [  4080]
MK128995.1_China/2018/AnhuiXCGQ  ........----.................................................................... [  4080]
MH681419.1_ASFV/POL/2015         ........----.................................................................... [  4080]
LS478113.1_Estonia_2014          -------------------------------------------------------------------------------- [  4080]
KP843857.1_Odintsovo_02/14       ........----.................................................................... [  4080]
FN557520.1_strain_E75            ........----.......................................................t.........c.. [  4080]
AY261366.1_Warthog               ........----..............................................g........t.........c.. [  4080]
AY261365.1_Warmbaths             ........----.........................t....................g........t.........c.. [  4080]
AY261364.1_Tengani_62            ........----..............................................g........t............ [  4080]
AY261363.1_Pretoriuskop/96/4     ........----..............................................g..................... [  4080]
AY261362.1_Mkuzi_1979            ........----.......................................................t.........c.. [  4080]
AY261361.1_Malawi_Lil-20/1_1983  gc..gccgctgcg.gccg..g.g..gg--------g...g.c...---------a.c.t.aac.g.tttt.c.a.c.... [  4080]
AY261360.1_Kenya_1950            ..a...c.----......aaat.t..g.t..t.....t...............t.............t....t....... [  4080]
AM712240.1_OURT_88/3             ........----.......................................................t.........c.. [  4080]
AM712239.1_Benin_97/1            ........----.......................................................t.........c.. [  4080]
KM111295.1_Ken06.Bus             -------------------------------------------------------------------------------- [  4080]
```

```
KM111294.1_Ken05/Tk1             ..a...a.----t....gt.atttg.c.gagtt........a...........a.......a...a.......t.c.... [  4080]
KM262844.1_strain_L60            ........----.......................................................t.........c.. [  4080]
KM102979.1_26544/OG10_Italy      ........----.......................................................t.........c.. [  4080]
KM262845.1_strain_NHV            ........----.......................................................t.........c.. [  4080]
KX354450.1_47/Ss/2008            ........----.......................................................t.........c.. [  4080]
KP055815.1_strain_BA71           ........----.......................................................t.........c.. [  4080]
MH025920.1_strain_R35            ..a...c.----......aagt.t..g.t..tg.a................................t....tt.c.... [  4080]
MH025919.1_strain_N10            ..a...c.----......aagt.t..g.t..tg.a................................t....tt.c.... [  4080]
MH025918.1_strain_R25            ..a...c.----......aagt.t..g.t..tg.a................................t....tt.c.... [  4080]
MH025917.1_strain_R7             ..a...c.----......aagt.t..g.t..tg.a................................t....tt.c.... [  4080]
MH025916.1_strain_R8             ..a...c.----......aagt.t..g.t..tg.a................................t....tt.c.... [  4080]

FR682468.2_Georgia_2007/1        ca---ataatatatagttgtataattacaatcatacatacaggccaactgcatcatttcatcaatgtctatatttgtctt [  4160]
ASF_IND_20_CAD543                ..---......w.....y.w........m..............y.....k.......................yy..... [  4160]
LR899193.1_Germany_2020/1        ..---........................................................................... [  4160]
MN393477.1_Wuhan_2019            ..---........................................................................... [  4160]
MW396979.1_Timor-Leste/2019/1    ..---........................................................................... [  4160]
MK940252.1_CN/2019/InnerMongolia ..---........................................................................... [  4160]
MK128995.1_China/2018/AnhuiXCGQ  ..---........................................................................... [  4160]
MH681419.1_ASFV/POL/2015         ..---........................................................................... [  4160]
LS478113.1_Estonia_2014          -------------------------------------------------------------------------------- [  4160]
KP843857.1_Odintsovo_02/14       ..---........................................................................... [  4160]
FN557520.1_strain_E75            ..---.....................................................................c..... [  4160]
AY261366.1_Warthog               ..---......a................c.............................................c..... [  4160]
AY261365.1_Warmbaths             ..---......a................c.............................................c..... [  4160]
AY261364.1_Tengani_62            ..---......a......aa........c...............t.............................c..... [  4160]
AY261363.1_Pretoriuskop/96/4     ..---......a.......a........c............a................................c..... [  4160]
AY261362.1_Mkuzi_1979            ..---.....................................................................c..... [  4160]
AY261361.1_Malawi_Lil-20/1_1983  ..tat...t.t...t.g.ag.g....------------------------------------------------------ [  4160]
AY261360.1_Kenya_1950            ..---......a..g..g.a.......t.cct...t..........gt.c........gg.t...a..c...ccaac... [  4160]
AM712240.1_OURT_88/3             ..---.....................................................................c..... [  4160]
AM712239.1_Benin_97/1            ..---.....................................................................c..... [  4160]
KM111295.1_Ken06.Bus             -------------------------------------------------------------------------------- [  4160]
KM111294.1_Ken05/Tk1             ..---...g..a......aa.a....g.t...tct.g......a..ga.a......a....t....ctc....caa.... [  4160]
KM262844.1_strain_L60            ..---.....................................................................c..... [  4160]
KM102979.1_26544/OG10_Italy      ..---.....................................................................c..... [  4160]
KM262845.1_strain_NHV            ..---.....................................................................c..... [  4160]
KX354450.1_47/Ss/2008            ..---.....................................................................c..... [  4160]
KP055815.1_strain_BA71           ..---.....................................................................c..... [  4160]
MH025920.1_strain_R35            ..---...g..a.....g.a....c....tat...tgc.....tt.t..ct.......g..t...a..c...c.aac... [  4160]
MH025919.1_strain_N10            ..---...g..a.....g.a....c....tat...tgc.....tt.t..ct.......g..t...a..c...c.aac... [  4160]
MH025918.1_strain_R25            ..---...g..a.....g.a....c....tat...tgc.....tt.t..ct.......g..t...a..c...c.aac... [  4160]
MH025917.1_strain_R7             ..---...g..a.....g.a....c....tat...tgc.....tt.t..ct.......g..t...a..c...c.aac... [  4160]
MH025916.1_strain_R8             ..---...g..a.....g.a....c....tat...tgc.....tt.t..ct.......g..t...a..c...c.aac... [  4160]
```

```
FR682468.2_Georgia_2007/1        ctctttgttataaatttcatgaaggtcaaagacgttgttataagcaaccccacatattaaccgccaatctttaaaatgac [  4240]
ASF_IND_20_CAD543                .....n..................................w...m..s...........r..................m. [  4240]
LR899193.1_Germany_2020/1        ................................................................................ [  4240]
MN393477.1_Wuhan_2019            ................................................................................ [  4240]
MW396979.1_Timor-Leste/2019/1    ................................................................................ [  4240]
MK940252.1_CN/2019/InnerMongolia ................................................................................ [  4240]
MK128995.1_China/2018/AnhuiXCGQ  ................................................................................ [  4240]
MH681419.1_ASFV/POL/2015         ................................................................................ [  4240]
LS478113.1_Estonia_2014          -------------------------------------------------------------------------------- [  4240]
KP843857.1_Odintsovo_02/14       ................................................................................ [  4240]
FN557520.1_strain_E75            ..............................................................------------------ [  4240]
AY261366.1_Warthog               ................................................................................ [  4240]
AY261365.1_Warmbaths             ................................................................................ [  4240]
AY261364.1_Tengani_62            ....a............................a...........................................a.. [  4240]
AY261363.1_Pretoriuskop/96/4     ................................................................................ [  4240]
AY261362.1_Mkuzi_1979            ................................................................................ [  4240]
AY261361.1_Malawi_Lil-20/1_1983  ---.act.....t...ct..t...----------------------------t..t...g.t..---------------- [  4240]
AY261360.1_Kenya_1950            ............t.a.........a.....c..............g.g...........gt.................ct [  4240]
AM712240.1_OURT_88/3             ................................................................................ [  4240]
AM712239.1_Benin_97/1            ................................................................................ [  4240]
KM111295.1_Ken06.Bus             -------------------------------------------------------------------------------- [  4240]
KM111294.1_Ken05/Tk1             t.t.a.a....tc..........aa...t.c..a......a...at.g............t.................tt [  4240]
KM262844.1_strain_L60            ................................................................................ [  4240]
KM102979.1_26544/OG10_Italy      ................................................................................ [  4240]
KM262845.1_strain_NHV            ................................................................................ [  4240]
KX354450.1_47/Ss/2008            ................................................................................ [  4240]
KP055815.1_strain_BA71           ................................................................................ [  4240]
MH025920.1_strain_R35            ............t...........a.....c..a..a...a......g...g........t..............c..ct [  4240]
MH025919.1_strain_N10            ............t...........a.....c..a..a...a......g...g........t..............c..ct [  4240]
MH025918.1_strain_R25            ............t...........a.....c..a..a...a......g...g........t..............c..ct [  4240]
MH025917.1_strain_R7             ............t...........a.....c..a..a...a......g...g........t..............c..ct [  4240]
MH025916.1_strain_R8             ............t...........a.....c..a..a...a......g...g........t..............c..ct [  4240]

FR682468.2_Georgia_2007/1        tatatcgttgataaaaatattggatggcttcagtaagcttatatagtatcgccatactataccaatacctagttagcatt [  4320]
ASF_IND_20_CAD543                .....................................s...y...r......y..............y.......r.... [  4320]
LR899193.1_Germany_2020/1        ................................................................................ [  4320]
MN393477.1_Wuhan_2019            ................................................................................ [  4320]
MW396979.1_Timor-Leste/2019/1    ................................................................................ [  4320]
MK940252.1_CN/2019/InnerMongolia ................................................................................ [  4320]
MK128995.1_China/2018/AnhuiXCGQ  ................................................................................ [  4320]
MH681419.1_ASFV/POL/2015         ................................................................................ [  4320]
LS478113.1_Estonia_2014          -------------------------------------------------------------------------------- [  4320]
KP843857.1_Odintsovo_02/14       ................................................................................ [  4320]
```

```
FN557520.1_strain_E75            -------------------------------------------------------------------------------- [  4320]
AY261366.1_Warthog               .................................................t.............................. [  4320]
AY261365.1_Warmbaths             .................................................t.............................. [  4320]
AY261364.1_Tengani_62            ................................................................................ [  4320]
AY261363.1_Pretoriuskop/96/4     .................................................t.............................. [  4320]
AY261362.1_Mkuzi_1979            ................................................................................ [  4320]
AY261361.1_Malawi_Lil-20/1_1983  -...cta....c...cc....----------------------------------------------------------- [  4320]
AY261360.1_Kenya_1950            .....tt...............a..a...........g..........c....tg.........ga.t..g......t.. [  4320]
AM712240.1_OURT_88/3             ................................................................................ [  4320]
AM712239.1_Benin_97/1            ................................................................................ [  4320]
KM111295.1_Ken06.Bus             ----.tt......g...........------------------------------------------------------- [  4320]
KM111294.1_Ken05/Tk1             ......t.........g.....a..a.........taa......c...ct....atgcg......a.t.....c.a.t.. [  4320]
KM262844.1_strain_L60            ................................................................................ [  4320]
KM102979.1_26544/OG10_Italy      ......a......................................................................... [  4320]
KM262845.1_strain_NHV            ................................................................................ [  4320]
KX354450.1_47/Ss/2008            ......a......................................................................... [  4320]
KP055815.1_strain_BA71           ................................................................................ [  4320]
MH025920.1_strain_R35            .....tt......g...........------------------------------------------------------- [  4320]
MH025919.1_strain_N10            .....tt......g...........------------------------------------------------------- [  4320]
MH025918.1_strain_R25            .....tt......g...........------------------------------------------------------- [  4320]
MH025917.1_strain_R7             .....tt......g...........------------------------------------------------------- [  4320]
MH025916.1_strain_R8             .....tt......g...........------------------------------------------------------- [  4320]

FR682468.2_Georgia_2007/1        tcgttgaatgaaatattatccaatgtaaagttaattgataatgtatctagttcaccaaaaattcttaatttcagttgag- [  4400]
ASF_IND_20_CAD543                ...y.r..........y.......r.......r..............................................- [  4400]
LR899193.1_Germany_2020/1        ...............................................................................- [  4400]
MN393477.1_Wuhan_2019            ...............................................................................- [  4400]
MW396979.1_Timor-Leste/2019/1    ...............................................................................- [  4400]
MK940252.1_CN/2019/InnerMongolia ...............................................................................- [  4400]
MK128995.1_China/2018/AnhuiXCGQ  ...............................................................................- [  4400]
MH681419.1_ASFV/POL/2015         ...............................................................................- [  4400]
LS478113.1_Estonia_2014          -------------------------------------------------------------------------------- [  4400]
KP843857.1_Odintsovo_02/14       ...............................................................................- [  4400]
FN557520.1_strain_E75            -------------------------------------------------------------------------------- [  4400]
AY261366.1_Warthog               ....................t.......t...g..........c...................................- [  4400]
AY261365.1_Warmbaths             ....................t.......t...g..........c...................................- [  4400]
AY261364.1_Tengani_62            .................................................a.............................- [  4400]
AY261363.1_Pretoriuskop/96/4     ................................g..............................................- [  4400]
AY261362.1_Mkuzi_1979            ...............................................................................- [  4400]
AY261361.1_Malawi_Lil-20/1_1983  --------------------------------------------------------------------------..aga- [  4400]
AY261360.1_Kenya_1950            ..a..a........t.c.....g.a...............t.c.ccg...a.t.ta.tt....a..ct.c...a....ac [  4400]
AM712240.1_OURT_88/3             ................................g..............................................- [  4400]
AM712239.1_Benin_97/1            ................................g..............................................- [  4400]
KM111295.1_Ken06.Bus             ----------------------------------------------------------------..cc..gt.aa...a- [  4400]
```

```
KM111294.1_Ken05/Tk1           ...c.a.......c..c.......a.g.....g.......t.t.c..c.aa..c.a.t.....a..gt.....a.----- [  4400]
KM262844.1_strain_L60          ................................g..............................................- [  4400]
KM102979.1_26544/OG10_Italy    ................................g..............................................- [  4400]
KM262845.1_strain_NHV          ................................g..............................................- [  4400]
KX354450.1_47/Ss/2008          ................................g..............................................- [  4400]
KP055815.1_strain_BA71         ................................g..............................................- [  4400]
MH025920.1_strain_R35          ----------------------------------------------------------------..cc..gt.aa...a- [  4400]
MH025919.1_strain_N10          ----------------------------------------------------------------..cc..gt.aa...a- [  4400]
MH025918.1_strain_R25          ----------------------------------------------------------------..cc..gt.aa...a- [  4400]
MH025917.1_strain_R7           ----------------------------------------------------------------..cc..gt.aa...a- [  4400]
MH025916.1_strain_R8           ----------------------------------------------------------------..cc..gt.aa...a- [  4400]

FR682468.2_Georgia_2007/1      -----cattatttaggaaaaggggattatcagataataattcatggcatagaataatattactgctagttttaacatact [  4480]
ASF_IND_20_CAD543              -----........................................................................... [  4480]
LR899193.1_Germany_2020/1      -----........................................................................... [  4480]
MN393477.1_Wuhan_2019          -----........................................................................... [  4480]
MW396979.1_Timor-Leste/2019/1  -----........................................................................... [  4480]
MK940252.1_CN/2019/InnerMongolia -----........................................................................... [  4480]
MK128995.1_China/2018/AnhuiXCGQ -----........................................................................... [  4480]
MH681419.1_ASFV/POL/2015       -----........................................................................... [  4480]
LS478113.1_Estonia_2014        -------------------------------------------------------------------------------- [  4480]
KP843857.1_Odintsovo_02/14     -----........................................................................... [  4480]
FN557520.1_strain_E75          -------------------------------------------------------------------------------- [  4480]
AY261366.1_Warthog             -----..........a...........g.t.................................................. [  4480]
AY261365.1_Warmbaths           -----..........a...........g.t.................................................. [  4480]
AY261364.1_Tengani_62          -----..........a...........g.t....................c............................. [  4480]
AY261363.1_Pretoriuskop/96/4   -----..........a...........g.t.....................t......................t..... [  4480]
AY261362.1_Mkuzi_1979          -----........................................................................... [  4480]
AY261361.1_Malawi_Lil-20/1_1983 -----tg....aa.aa..c.taaa...--------------------------.......tt....g------------- [  4480]
AY261360.1_Kenya_1950          attat.t.c.c....a.....a.......t......c........a....a..........t.a...c......a.cgt. [  4480]
AM712240.1_OURT_88/3           -----.............t............................................................. [  4480]
AM712239.1_Benin_97/1          -----.............t............................................................. [  4480]
KM111295.1_Ken06.Bus           -------------------------------------------------------------------------------- [  4480]
KM111294.1_Ken05/Tk1           -atat.ta.......a......aa.....t........g...........t.................a.....t.cgg. [  4480]
KM262844.1_strain_L60          -----.............t............................................................. [  4480]
KM102979.1_26544/OG10_Italy    -----.............t............................................................. [  4480]
KM262845.1_strain_NHV          -----.............t............................................................. [  4480]
KX354450.1_47/Ss/2008          -----.............t............................................................. [  4480]
KP055815.1_strain_BA71         -----.............t............................................................. [  4480]
MH025920.1_strain_R35          -------------------------------------------------------------------------------- [  4480]
MH025919.1_strain_N10          -------------------------------------------------------------------------------- [  4480]
MH025918.1_strain_R25          -------------------------------------------------------------------------------- [  4480]
MH025917.1_strain_R7           -------------------------------------------------------------------------------- [  4480]
MH025916.1_strain_R8           -------------------------------------------------------------------------------- [  4480]
```

```
FR682468.2_Georgia_2007/1        gtacattataaaatatttctaaaattttattttcactcaaagctttcctcgcacctaacttttggcataggtcct-ggtg [  4560]
ASF_IND_20_CAD543                ...........................................................................g.... [  4560]
LR899193.1_Germany_2020/1        ...........................................................................-.... [  4560]
MN393477.1_Wuhan_2019            ...........................................................................-.... [  4560]
MW396979.1_Timor-Leste/2019/1    ...........................................................................-.... [  4560]
MK940252.1_CN/2019/InnerMongolia ...........................................................................-.... [  4560]
MK128995.1_China/2018/AnhuiXCGQ  ...........................................................................-.... [  4560]
MH681419.1_ASFV/POL/2015         ...........................................................................-.... [  4560]
LS478113.1_Estonia_2014          -------------------------------------------------------------------------------- [  4560]
KP843857.1_Odintsovo_02/14       ...........................................................................-.... [  4560]
FN557520.1_strain_E75            -------------------------------------------------------------------------------- [  4560]
AY261366.1_Warthog               t...........................g.................t............................g.... [  4560]
AY261365.1_Warmbaths             t...........................g..............................................g.... [  4560]
AY261364.1_Tengani_62            t..................................t.....a.c.c.....................c.......g.... [  4560]
AY261363.1_Pretoriuskop/96/4     t..t........................c......t.....a.c.c.....................c.......g.... [  4560]
AY261362.1_Mkuzi_1979            ........................................c..................................g.... [  4560]
AY261361.1_Malawi_Lil-20/1_1983  --------c....g..cgt..t..aca.g--------------------------------------------------- [  4560]
AY261360.1_Kenya_1950            t...tg.........ca.......c..ccca..t.t.t..g.a..c.t.g..........c.c........cttcg.a.. [  4560]
AM712240.1_OURT_88/3             ...........................................................................g.... [  4560]
AM712239.1_Benin_97/1            ...........................................................................g.... [  4560]
KM111295.1_Ken06.Bus             -..t...g.......cg.-------------------------------------------------------------- [  4560]
KM111294.1_Ken05/Tk1             a...t...c.......a.t..t.gca.ccg......c.....t....t.............cc.a..g.a..t.ca...a [  4560]
KM262844.1_strain_L60            ...........................................................................g.... [  4560]
KM102979.1_26544/OG10_Italy      ...........................................................................g.... [  4560]
KM262845.1_strain_NHV            ...........................................................................g.... [  4560]
KX354450.1_47/Ss/2008            ...........................................................................g.... [  4560]
KP055815.1_strain_BA71           ...........................................................................g.... [  4560]
MH025920.1_strain_R35            -..t...g.......cg.-------------------------------------------------------------- [  4560]
MH025919.1_strain_N10            -..t...g.......cg.-------------------------------------------------------------- [  4560]
MH025918.1_strain_R25            -..t...g.......cg.-------------------------------------------------------------- [  4560]
MH025917.1_strain_R7             -..t...g.......cg.-------------------------------------------------------------- [  4560]
MH025916.1_strain_R8             -..t...g.......cg.-------------------------------------------------------------- [  4560]

FR682468.2_Georgia_2007/1        cactccatattgacagtaaccaacccaaagctgatgtctgcaccccattcggtaaacagctctattaaaccatgattgtt [  4640]
ASF_IND_20_CAD543                ................................................................................ [  4640]
LR899193.1_Germany_2020/1        ................................................................................ [  4640]
MN393477.1_Wuhan_2019            ................................................................................ [  4640]
MW396979.1_Timor-Leste/2019/1    ................................................................................ [  4640]
MK940252.1_CN/2019/InnerMongolia ................................................................................ [  4640]
MK128995.1_China/2018/AnhuiXCGQ  ................................................................................ [  4640]
MH681419.1_ASFV/POL/2015         ................................................................................ [  4640]
LS478113.1_Estonia_2014          -------------------------------------------------------------------------------- [  4640]
KP843857.1_Odintsovo_02/14       ................................................................................ [  4640]
```

```
FN557520.1_strain_E75          -------------------------------------------------------------------------------- [  4640]
AY261366.1_Warthog             ..t..............g...................................................t.......... [  4640]
AY261365.1_Warmbaths           .....................................................................t.......... [  4640]
AY261364.1_Tengani_62          ..t................................a.................................t.......... [  4640]
AY261363.1_Pretoriuskop/96/4   .................g.....t.............................................t.......... [  4640]
AY261362.1_Mkuzi_1979          ................................................................................ [  4640]
AY261361.1_Malawi_Lil-20/1_1983 --------------------------g......g...ta-------------------------------------..c. [  4640]
AY261360.1_Kenya_1950          ttt...g.g........g.t...t.....at.a.....c...........aa...............gg...c....... [  4640]
AM712240.1_OURT_88/3           ................................................................................ [  4640]
AM712239.1_Benin_97/1          ................................................................................ [  4640]
KM111295.1_Ken06.Bus           -------------------------------------------------------------------------------- [  4640]
KM111294.1_Ken05/Tk1           ......g.....g....gg...........t.a..a.tc........c..a..........t...g.g.tt..ag..a.. [  4640]
KM262844.1_strain_L60          ................................................................................ [  4640]
KM102979.1_26544/OG10_Italy    ................................................................................ [  4640]
KM262845.1_strain_NHV          ................................................................................ [  4640]
KX354450.1_47/Ss/2008          ................................................................................ [  4640]
KP055815.1_strain_BA71         ................................................................................ [  4640]
MH025920.1_strain_R35          -------------------------------------------------------------------------------- [  4640]
MH025919.1_strain_N10          -------------------------------------------------------------------------------- [  4640]
MH025918.1_strain_R25          -------------------------------------------------------------------------------- [  4640]
MH025917.1_strain_R7           -------------------------------------------------------------------------------- [  4640]
MH025916.1_strain_R8           -------------------------------------------------------------------------------- [  4640]

FR682468.2_Georgia_2007/1      ttcctgtacagccttcattaatg------------caacatttaatgttaaaccatgtttaaaacttgctgtttttatta [  4720]
ASF_IND_20_CAD543              .......................------------............................................. [  4720]
LR899193.1_Germany_2020/1      .......................------------............................................. [  4720]
MN393477.1_Wuhan_2019          .......................------------............................................. [  4720]
MW396979.1_Timor-Leste/2019/1  .......................------------............................................. [  4720]
MK940252.1_CN/2019/InnerMongolia .......................------------............................................. [  4720]
MK128995.1_China/2018/AnhuiXCGQ .......................------------............................................. [  4720]
MH681419.1_ASFV/POL/2015       .......................------------............................................. [  4720]
LS478113.1_Estonia_2014        -------------------------------------------------------------------------------- [  4720]
KP843857.1_Odintsovo_02/14     .......................------------............................................. [  4720]
FN557520.1_strain_E75          -------------------------------------------------------------------------------- [  4720]
AY261366.1_Warthog             .......................------------..........................................c.. [  4720]
AY261365.1_Warmbaths           .......................------------..........................................c.. [  4720]
AY261364.1_Tengani_62          .......................------------............................................. [  4720]
AY261363.1_Pretoriuskop/96/4   .......................------------............................................. [  4720]
AY261362.1_Mkuzi_1979          .......................------------............................................. [  4720]
AY261361.1_Malawi_Lil-20/1_1983 ..tg...gt.ta...a..a..a.tggtggcaacaat........tc.c.tttttgcacc.....t.c.-----ag...a. [  4720]
AY261360.1_Kenya_1950          ...t..c..g..t..........------------.g.........tc...g....c......g.......a......c. [  4720]
AM712240.1_OURT_88/3           .......................------------............................................. [  4720]
AM712239.1_Benin_97/1          .......................------------............................................. [  4720]
KM111295.1_Ken06.Bus           -------------------------------------------------------------------------------- [  4720]
```

```
KM111294.1_Ken05/Tk1              ...t........t..........------------.g.g.....tctc......g.c......gg..a...a......c. [  4720]
KM262844.1_strain_L60             .......................------------............................................. [  4720]
KM102979.1_26544/OG10_Italy       .......................------------............................................. [  4720]
KM262845.1_strain_NHV             .......................------------............................................. [  4720]
KX354450.1_47/Ss/2008             .......................------------............................................. [  4720]
KP055815.1_strain_BA71            .......................------------............................................. [  4720]
MH025920.1_strain_R35             -------------------------------------------------------------------------------- [  4720]
MH025919.1_strain_N10             -------------------------------------------------------------------------------- [  4720]
MH025918.1_strain_R25             -------------------------------------------------------------------------------- [  4720]
MH025917.1_strain_R7              -------------------------------------------------------------------------------- [  4720]
MH025916.1_strain_R8              -------------------------------------------------------------------------------- [  4720]

FR682468.2_Georgia_2007/1         atatttgttcatctatacaagta--------------------------------------------------------- [  4800]
ASF_IND_20_CAD543                 .........y.............--------------------------------------------------------- [  4800]
LR899193.1_Germany_2020/1         .......................--------------------------------------------------------- [  4800]
MN393477.1_Wuhan_2019             .......................--------------------------------------------------------- [  4800]
MW396979.1_Timor-Leste/2019/1     .......................--------------------------------------------------------- [  4800]
MK940252.1_CN/2019/InnerMongolia  .......................--------------------------------------------------------- [  4800]
MK128995.1_China/2018/AnhuiXCGQ   .......................--------------------------------------------------------- [  4800]
MH681419.1_ASFV/POL/2015          .......................--------------------------------------------------------- [  4800]
LS478113.1_Estonia_2014           -------------------------------------------------------------------------------- [  4800]
KP843857.1_Odintsovo_02/14        .......................--------------------------------------------------------- [  4800]
FN557520.1_strain_E75             -------------------------------------------------------------------------------- [  4800]
AY261366.1_Warthog                .........t.............--------------------------------------------------------- [  4800]
AY261365.1_Warmbaths              .........t.............--------------------------------------------------------- [  4800]
AY261364.1_Tengani_62             ......c..t..t.........---------------------------------------------------------- [  4800]
AY261363.1_Pretoriuskop/96/4      .........t............---------------------------------------------------------- [  4800]
AY261362.1_Mkuzi_1979             .........t.............--------------------------------------------------------- [  4800]
AY261361.1_Malawi_Lil-20/1_1983   .a.aa....t..t.ta.t.g..---------------------------------------------------------- [  4800]
AY261360.1_Kenya_1950             tc.....a.t...ctc.ttc...gaaagcatgattggacctccatgccaccataaaccacaacattttaatatatgacaa [  4800]
AM712240.1_OURT_88/3              .........t.............--------------------------------------------------------- [  4800]
AM712239.1_Benin_97/1             .........t.............--------------------------------------------------------- [  4800]
KM111295.1_Ken06.Bus              -------------------------------------------------------------------------------- [  4800]
KM111294.1_Ken05/Tk1              t.tga..a..c..atc.tt....gaaagcataattgggcccccatgccaccataaaccacaacattttaatatatgacaa [  4800]
KM262844.1_strain_L60             .........t.............--------------------------------------------------------- [  4800]
KM102979.1_26544/OG10_Italy       .........t.............--------------------------------------------------------- [  4800]
KM262845.1_strain_NHV             .........t.............--------------------------------------------------------- [  4800]
KX354450.1_47/Ss/2008             .........t.............--------------------------------------------------------- [  4800]
KP055815.1_strain_BA71            .........t.............--------------------------------------------------------- [  4800]
MH025920.1_strain_R35             -------------------------------------------------------------------------------- [  4800]
MH025919.1_strain_N10             -------------------------------------------------------------------------------- [  4800]
MH025918.1_strain_R25             -------------------------------------------------------------------------------- [  4800]
MH025917.1_strain_R7              -------------------------------------------------------------------------------- [  4800]
MH025916.1_strain_R8              -------------------------------------------------------------------------------- [  4800]
```

```
FR682468.2_Georgia_2007/1         -------------------------------------------------------------------------------- [  4880]
ASF_IND_20_CAD543                 -------------------------------------------------------------------------------- [  4880]
LR899193.1_Germany_2020/1         -------------------------------------------------------------------------------- [  4880]
MN393477.1_Wuhan_2019             -------------------------------------------------------------------------------- [  4880]
MW396979.1_Timor-Leste/2019/1     -------------------------------------------------------------------------------- [  4880]
MK940252.1_CN/2019/InnerMongolia  -------------------------------------------------------------------------------- [  4880]
MK128995.1_China/2018/AnhuiXCGQ   -------------------------------------------------------------------------------- [  4880]
MH681419.1_ASFV/POL/2015          -------------------------------------------------------------------------------- [  4880]
LS478113.1_Estonia_2014           -------------------------------------------------------------------------------- [  4880]
KP843857.1_Odintsovo_02/14        -------------------------------------------------------------------------------- [  4880]
FN557520.1_strain_E75             -------------------------------------------------------------------------------- [  4880]
AY261366.1_Warthog                -------------------------------------------------------------------------------- [  4880]
AY261365.1_Warmbaths              -------------------------------------------------------------------------------- [  4880]
AY261364.1_Tengani_62             -------------------------------------------------------------------------------- [  4880]
AY261363.1_Pretoriuskop/96/4      -------------------------------------------------------------------------------- [  4880]
AY261362.1_Mkuzi_1979             -------------------------------------------------------------------------------- [  4880]
AY261361.1_Malawi_Lil-20/1_1983   -------------------------------------------------------------------------------- [  4880]
AY261360.1_Kenya_1950             tattccctagattgagaatattctttagatatatgctgtgtggccagtacttttttagtaagggcctgcagagaagatgg [  4880]
AM712240.1_OURT_88/3              -------------------------------------------------------------------------------- [  4880]
AM712239.1_Benin_97/1             -------------------------------------------------------------------------------- [  4880]
KM111295.1_Ken06.Bus              -------------------------------------------------------------------------------- [  4880]
KM111294.1_Ken05/Tk1              tattccctagattgagaatattctttagatatatgctgtgtggccagtacttttttagtaagggcctgcagagaagatgg [  4880]
KM262844.1_strain_L60             -------------------------------------------------------------------------------- [  4880]
KM102979.1_26544/OG10_Italy       -------------------------------------------------------------------------------- [  4880]
KM262845.1_strain_NHV             -------------------------------------------------------------------------------- [  4880]
KX354450.1_47/Ss/2008             -------------------------------------------------------------------------------- [  4880]
KP055815.1_strain_BA71            -------------------------------------------------------------------------------- [  4880]
MH025920.1_strain_R35             -------------------------------------------------------------------------------- [  4880]
MH025919.1_strain_N10             -------------------------------------------------------------------------------- [  4880]
MH025918.1_strain_R25             -------------------------------------------------------------------------------- [  4880]
MH025917.1_strain_R7              -------------------------------------------------------------------------------- [  4880]
MH025916.1_strain_R8              -------------------------------------------------------------------------------- [  4880]

FR682468.2_Georgia_2007/1         -------------------------------------------------------------------------------- [  4960]
ASF_IND_20_CAD543                 -------------------------------------------------------------------------------- [  4960]
LR899193.1_Germany_2020/1         -------------------------------------------------------------------------------- [  4960]
MN393477.1_Wuhan_2019             -------------------------------------------------------------------------------- [  4960]
MW396979.1_Timor-Leste/2019/1     -------------------------------------------------------------------------------- [  4960]
MK940252.1_CN/2019/InnerMongolia  -------------------------------------------------------------------------------- [  4960]
MK128995.1_China/2018/AnhuiXCGQ   -------------------------------------------------------------------------------- [  4960]
MH681419.1_ASFV/POL/2015          -------------------------------------------------------------------------------- [  4960]
LS478113.1_Estonia_2014           -------------------------------------------------------------------------------- [  4960]
KP843857.1_Odintsovo_02/14        -------------------------------------------------------------------------------- [  4960]
```

```
FN557520.1_strain_E75             -------------------------------------------------------------------------------- [  4960]
AY261366.1_Warthog                -------------------------------------------------------------------------------- [  4960]
AY261365.1_Warmbaths              -------------------------------------------------------------------------------- [  4960]
AY261364.1_Tengani_62             -------------------------------------------------------------------------------- [  4960]
AY261363.1_Pretoriuskop/96/4      -------------------------------------------------------------------------------- [  4960]
AY261362.1_Mkuzi_1979             -------------------------------------------------------------------------------- [  4960]
AY261361.1_Malawi_Lil-20/1_1983   -------------------------------------------------------------------------------- [  4960]
AY261360.1_Kenya_1950             tgtagacatgtttcttcatctacaaaataaatcaaattaaattatttatttctggcgctctatggccttgttgtataagc [  4960]
AM712240.1_OURT_88/3              -------------------------------------------------------------------------------- [  4960]
AM712239.1_Benin_97/1             -------------------------------------------------------------------------------- [  4960]
KM111295.1_Ken06.Bus              -------------------------------------------------------------------------------- [  4960]
KM111294.1_Ken05/Tk1              tgtaggcatgttccttcatctacaaaataaatcaaattaaattatttatttctggcgctctatggccttattatataagc [  4960]
KM262844.1_strain_L60             -------------------------------------------------------------------------------- [  4960]
KM102979.1_26544/OG10_Italy       -------------------------------------------------------------------------------- [  4960]
KM262845.1_strain_NHV             -------------------------------------------------------------------------------- [  4960]
KX354450.1_47/Ss/2008             -------------------------------------------------------------------------------- [  4960]
KP055815.1_strain_BA71            -------------------------------------------------------------------------------- [  4960]
MH025920.1_strain_R35             -------------------------------------------------------------------------------- [  4960]
MH025919.1_strain_N10             -------------------------------------------------------------------------------- [  4960]
MH025918.1_strain_R25             -------------------------------------------------------------------------------- [  4960]
MH025917.1_strain_R7              -------------------------------------------------------------------------------- [  4960]
MH025916.1_strain_R8              -------------------------------------------------------------------------------- [  4960]

FR682468.2_Georgia_2007/1         -------------------------------------------------------------------------------- [  5040]
ASF_IND_20_CAD543                 -------------------------------------------------------------------------------- [  5040]
LR899193.1_Germany_2020/1         -------------------------------------------------------------------------------- [  5040]
MN393477.1_Wuhan_2019             -------------------------------------------------------------------------------- [  5040]
MW396979.1_Timor-Leste/2019/1     -------------------------------------------------------------------------------- [  5040]
MK940252.1_CN/2019/InnerMongolia  -------------------------------------------------------------------------------- [  5040]
MK128995.1_China/2018/AnhuiXCGQ   -------------------------------------------------------------------------------- [  5040]
MH681419.1_ASFV/POL/2015          -------------------------------------------------------------------------------- [  5040]
LS478113.1_Estonia_2014           -------------------------------------------------------------------------------- [  5040]
KP843857.1_Odintsovo_02/14        -------------------------------------------------------------------------------- [  5040]
FN557520.1_strain_E75             -------------------------------------------------------------------------------- [  5040]
AY261366.1_Warthog                -------------------------------------------------------------------------------- [  5040]
AY261365.1_Warmbaths              -------------------------------------------------------------------------------- [  5040]
AY261364.1_Tengani_62             -------------------------------------------------------------------------------- [  5040]
AY261363.1_Pretoriuskop/96/4      -------------------------------------------------------------------------------- [  5040]
AY261362.1_Mkuzi_1979             -------------------------------------------------------------------------------- [  5040]
AY261361.1_Malawi_Lil-20/1_1983   -------------------------------------------------------------------------------- [  5040]
AY261360.1_Kenya_1950             tctatacaaaaaaaaagttgttacacccgctgtgacgatgcatacttttatttgtggctaaaatttaatattttcttatg [  5040]
AM712240.1_OURT_88/3              -------------------------------------------------------------------------------- [  5040]
AM712239.1_Benin_97/1             -------------------------------------------------------------------------------- [  5040]
KM111295.1_Ken06.Bus              -----------------------------------cgattcgtggtatcatttgggggcaaaa----------------- [  5040]
```

```
KM111294.1_Ken05/Tk1            tctatacacaaaaaaaaaggttacacccgctgtgacgatgtatacttttatttgtggctaaaatt---tattttcctatg [  5040]
KM262844.1_strain_L60           -------------------------------------------------------------------------------- [  5040]
KM102979.1_26544/OG10_Italy     -------------------------------------------------------------------------------- [  5040]
KM262845.1_strain_NHV           -------------------------------------------------------------------------------- [  5040]
KX354450.1_47/Ss/2008           -------------------------------------------------------------------------------- [  5040]
KP055815.1_strain_BA71          -------------------------------------------------------------------------------- [  5040]
MH025920.1_strain_R35           -----------------------------------cgattcgtggtatcatttgggggcaaaa----------------- [  5040]
MH025919.1_strain_N10           -----------------------------------cgattcgtggtatcatttgggggcaaaa----------------- [  5040]
MH025918.1_strain_R25           -----------------------------------cgattcgtggtatcatttgggggcaaaa----------------- [  5040]
MH025917.1_strain_R7            -----------------------------------cgattcgtggtatcatttgggggcaaaa----------------- [  5040]
MH025916.1_strain_R8            -----------------------------------cgattcgtggtatcatttgggggcaaaa----------------- [  5040]

FR682468.2_Georgia_2007/1       -------------------------------------------------------------------------------- [  5120]
ASF_IND_20_CAD543               -------------------------------------------------------------------------------- [  5120]
LR899193.1_Germany_2020/1       -------------------------------------------------------------------------------- [  5120]
MN393477.1_Wuhan_2019           -------------------------------------------------------------------------------- [  5120]
MW396979.1_Timor-Leste/2019/1   -------------------------------------------------------------------------------- [  5120]
MK940252.1_CN/2019/InnerMongolia -------------------------------------------------------------------------------- [  5120]
MK128995.1_China/2018/AnhuiXCGQ -------------------------------------------------------------------------------- [  5120]
MH681419.1_ASFV/POL/2015        -------------------------------------------------------------------------------- [  5120]
LS478113.1_Estonia_2014         -------------------------------------------------------------------------------- [  5120]
KP843857.1_Odintsovo_02/14      -------------------------------------------------------------------------------- [  5120]
FN557520.1_strain_E75           -------------------------------------------------------------------------------- [  5120]
AY261366.1_Warthog              -------------------------------------------------------------------------------- [  5120]
AY261365.1_Warmbaths            -------------------------------------------------------------------------------- [  5120]
AY261364.1_Tengani_62           -------------------------------------------------------------------------------- [  5120]
AY261363.1_Pretoriuskop/96/4    -------------------------------------------------------------------------------- [  5120]
AY261362.1_Mkuzi_1979           -------------------------------------------------------------------------------- [  5120]
AY261361.1_Malawi_Lil-20/1_1983 -------------------------------------------------------------------------------- [  5120]
AY261360.1_Kenya_1950           caacatcctattatagaaggtcatattaaatatactttt---gaaaaaaatcttgagttctattaatatatcctattctt [  5120]
AM712240.1_OURT_88/3            -------------------------------------------------------------------------------- [  5120]
AM712239.1_Benin_97/1           -------------------------------------------------------------------------------- [  5120]
KM111295.1_Ken06.Bus            -------------------------------------------------------------------------------- [  5120]
KM111294.1_Ken05/Tk1            caacatcctattatagaaggtcatattaaatatacccttttgaaaaaaaaatcttgagttctattaatatatcctattctt [  5120]
KM262844.1_strain_L60           -------------------------------------------------------------------------------- [  5120]
KM102979.1_26544/OG10_Italy     -------------------------------------------------------------------------------- [  5120]
KM262845.1_strain_NHV           -------------------------------------------------------------------------------- [  5120]
KX354450.1_47/Ss/2008           -------------------------------------------------------------------------------- [  5120]
KP055815.1_strain_BA71          -------------------------------------------------------------------------------- [  5120]
MH025920.1_strain_R35           -------------------------------------------------------------------------------- [  5120]
MH025919.1_strain_N10           -------------------------------------------------------------------------------- [  5120]
MH025918.1_strain_R25           -------------------------------------------------------------------------------- [  5120]
MH025917.1_strain_R7            -------------------------------------------------------------------------------- [  5120]
MH025916.1_strain_R8            -------------------------------------------------------------------------------- [  5120]
```

```
FR682468.2_Georgia_2007/1         -------------------------------------------------------------------------------- [  5200]
ASF_IND_20_CAD543                 -------------------------------------------------------------------------------- [  5200]
LR899193.1_Germany_2020/1         -------------------------------------------------------------------------------- [  5200]
MN393477.1_Wuhan_2019             -------------------------------------------------------------------------------- [  5200]
MW396979.1_Timor-Leste/2019/1     -------------------------------------------------------------------------------- [  5200]
MK940252.1_CN/2019/InnerMongolia  -------------------------------------------------------------------------------- [  5200]
MK128995.1_China/2018/AnhuiXCGQ   -------------------------------------------------------------------------------- [  5200]
MH681419.1_ASFV/POL/2015          -------------------------------------------------------------------------------- [  5200]
LS478113.1_Estonia_2014           -------------------------------------------------------------------------------- [  5200]
KP843857.1_Odintsovo_02/14        -------------------------------------------------------------------------------- [  5200]
FN557520.1_strain_E75             -------------------------------------------------------------------------------- [  5200]
AY261366.1_Warthog                -------------------------------------------------------------------------------- [  5200]
AY261365.1_Warmbaths              -------------------------------------------------------------------------------- [  5200]
AY261364.1_Tengani_62             -------------------------------------------------------------------------------- [  5200]
AY261363.1_Pretoriuskop/96/4      -------------------------------------------------------------------------------- [  5200]
AY261362.1_Mkuzi_1979             -------------------------------------------------------------------------------- [  5200]
AY261361.1_Malawi_Lil-20/1_1983   -------------------------------------------------------------------------------- [  5200]
AY261360.1_Kenya_1950             gacaagattaaaaatacccaatacacatatagtcaaaagttgacatattctttgagtaataggatttcaacatggtatta [  5200]
AM712240.1_OURT_88/3              -------------------------------------------------------------------------------- [  5200]
AM712239.1_Benin_97/1             -------------------------------------------------------------------------------- [  5200]
KM111295.1_Ken06.Bus              -------------------------------------------------------------------------------- [  5200]
KM111294.1_Ken05/Tk1              gacaagattaaaaatacccaatacacatatagtcaaaggttgacatattctttgagtaataggatttcaacatggtatta [  5200]
KM262844.1_strain_L60             -------------------------------------------------------------------------------- [  5200]
KM102979.1_26544/OG10_Italy       -------------------------------------------------------------------------------- [  5200]
KM262845.1_strain_NHV             -------------------------------------------------------------------------------- [  5200]
KX354450.1_47/Ss/2008             -------------------------------------------------------------------------------- [  5200]
KP055815.1_strain_BA71            -------------------------------------------------------------------------------- [  5200]
MH025920.1_strain_R35             -------------------------------------------------------------------------------- [  5200]
MH025919.1_strain_N10             -------------------------------------------------------------------------------- [  5200]
MH025918.1_strain_R25             -------------------------------------------------------------------------------- [  5200]
MH025917.1_strain_R7              -------------------------------------------------------------------------------- [  5200]
MH025916.1_strain_R8              -------------------------------------------------------------------------------- [  5200]

FR682468.2_Georgia_2007/1         -------------------------------------------------------------------------------- [  5280]
ASF_IND_20_CAD543                 -------------------------------------------------------------------------------- [  5280]
LR899193.1_Germany_2020/1         -------------------------------------------------------------------------------- [  5280]
MN393477.1_Wuhan_2019             -------------------------------------------------------------------------------- [  5280]
MW396979.1_Timor-Leste/2019/1     -------------------------------------------------------------------------------- [  5280]
MK940252.1_CN/2019/InnerMongolia  -------------------------------------------------------------------------------- [  5280]
MK128995.1_China/2018/AnhuiXCGQ   -------------------------------------------------------------------------------- [  5280]
MH681419.1_ASFV/POL/2015          -------------------------------------------------------------------------------- [  5280]
LS478113.1_Estonia_2014           -------------------------------------------------------------------------------- [  5280]
KP843857.1_Odintsovo_02/14        -------------------------------------------------------------------------------- [  5280]
```

```
FN557520.1_strain_E75         -------------------------------------------------------------------------------- [  5280]
AY261366.1_Warthog            -------------------------------------------------------------------------------- [  5280]
AY261365.1_Warmbaths          -------------------------------------------------------------------------------- [  5280]
AY261364.1_Tengani_62         -------------------------------------------------------------------------------- [  5280]
AY261363.1_Pretoriuskop/96/4  -------------------------------------------------------------------------------- [  5280]
AY261362.1_Mkuzi_1979         -------------------------------------------------------------------------------- [  5280]
AY261361.1_Malawi_Lil-20/1_1983 -------------------------------------------------------------------------------- [  5280]
AY261360.1_Kenya_1950         atttttttttgggtcaattatttttgagattaggaaaggatccggactataatttttaaacgataacatatttactattaa [  5280]
AM712240.1_OURT_88/3          -------------------------------------------------------------------------------- [  5280]
AM712239.1_Benin_97/1         -------------------------------------------------------------------------------- [  5280]
KM111295.1_Ken06.Bus          -------------------------------------------------------------------------------- [  5280]
KM111294.1_Ken05/Tk1          atttttttttgggtcaattatttttgggattaggaagggatccggactataatttttaaatgataacatatttactattaa [  5280]
KM262844.1_strain_L60         -------------------------------------------------------------------------------- [  5280]
KM102979.1_26544/OG10_Italy   -------------------------------------------------------------------------------- [  5280]
KM262845.1_strain_NHV         -------------------------------------------------------------------------------- [  5280]
KX354450.1_47/Ss/2008         -------------------------------------------------------------------------------- [  5280]
KP055815.1_strain_BA71        -------------------------------------------------------------------------------- [  5280]
MH025920.1_strain_R35         -------------------------------------------------------------------------------- [  5280]
MH025919.1_strain_N10         -------------------------------------------------------------------------------- [  5280]
MH025918.1_strain_R25         -------------------------------------------------------------------------------- [  5280]
MH025917.1_strain_R7          -------------------------------------------------------------------------------- [  5280]
MH025916.1_strain_R8          -------------------------------------------------------------------------------- [  5280]

FR682468.2_Georgia_2007/1     -------------------------------------------------------------------------------- [  5360]
ASF_IND_20_CAD543             -------------------------------------------------------------------------------- [  5360]
LR899193.1_Germany_2020/1     -------------------------------------------------------------------------------- [  5360]
MN393477.1_Wuhan_2019         -------------------------------------------------------------------------------- [  5360]
MW396979.1_Timor-Leste/2019/1 -------------------------------------------------------------------------------- [  5360]
MK940252.1_CN/2019/InnerMongolia -------------------------------------------------------------------------------- [  5360]
MK128995.1_China/2018/AnhuiXCGQ -------------------------------------------------------------------------------- [  5360]
MH681419.1_ASFV/POL/2015      -------------------------------------------------------------------------------- [  5360]
LS478113.1_Estonia_2014       -------------------------------------------------------------------------------- [  5360]
KP843857.1_Odintsovo_02/14    -------------------------------------------------------------------------------- [  5360]
FN557520.1_strain_E75         -------------------------------------------------------------------------------- [  5360]
AY261366.1_Warthog            -------------------------------------------------------------------------------- [  5360]
AY261365.1_Warmbaths          -------------------------------------------------------------------------------- [  5360]
AY261364.1_Tengani_62         -------------------------------------------------------------------------------- [  5360]
AY261363.1_Pretoriuskop/96/4  -------------------------------------------------------------------------------- [  5360]
AY261362.1_Mkuzi_1979         -------------------------------------------------------------------------------- [  5360]
AY261361.1_Malawi_Lil-20/1_1983 -------------------------------------------------------------------------------- [  5360]
AY261360.1_Kenya_1950         atgataatttttttgttctgccaatgccttactctcttcaatagcattagcccccaaatccatacaaaagaacatgttga [  5360]
AM712240.1_OURT_88/3          -------------------------------------------------------------------------------- [  5360]
AM712239.1_Benin_97/1         -------------------------------------------------------------------------------- [  5360]
KM111295.1_Ken06.Bus          -------------------------------------------------------------------------------- [  5360]
```

```
KM111294.1_Ken05/Tk1             atgataatttttttgttctgccaatgccttactctcttcaatagcattagcccccaagtccatacaaaagaacatgttaa [  5360]
KM262844.1_strain_L60            -------------------------------------------------------------------------------- [  5360]
KM102979.1_26544/OG10_Italy      -------------------------------------------------------------------------------- [  5360]
KM262845.1_strain_NHV            -------------------------------------------------------------------------------- [  5360]
KX354450.1_47/Ss/2008            -------------------------------------------------------------------------------- [  5360]
KP055815.1_strain_BA71           -------------------------------------------------------------------------------- [  5360]
MH025920.1_strain_R35            -------------------------------------------------------------------------------- [  5360]
MH025919.1_strain_N10            -------------------------------------------------------------------------------- [  5360]
MH025918.1_strain_R25            -------------------------------------------------------------------------------- [  5360]
MH025917.1_strain_R7             -------------------------------------------------------------------------------- [  5360]
MH025916.1_strain_R8             -------------------------------------------------------------------------------- [  5360]

FR682468.2_Georgia_2007/1        -------------------------------------------------------------------------------- [  5440]
ASF_IND_20_CAD543                -------------------------------------------------------------------------------- [  5440]
LR899193.1_Germany_2020/1        -------------------------------------------------------------------------------- [  5440]
MN393477.1_Wuhan_2019            -------------------------------------------------------------------------------- [  5440]
MW396979.1_Timor-Leste/2019/1    -------------------------------------------------------------------------------- [  5440]
MK940252.1_CN/2019/InnerMongolia -------------------------------------------------------------------------------- [  5440]
MK128995.1_China/2018/AnhuiXCGQ  -------------------------------------------------------------------------------- [  5440]
MH681419.1_ASFV/POL/2015         -------------------------------------------------------------------------------- [  5440]
LS478113.1_Estonia_2014          -------------------------------------------------------------------------------- [  5440]
KP843857.1_Odintsovo_02/14       -------------------------------------------------------------------------------- [  5440]
FN557520.1_strain_E75            -------------------------------------------------------------------------------- [  5440]
AY261366.1_Warthog               -------------------------------------------------------------------------------- [  5440]
AY261365.1_Warmbaths             -------------------------------------------------------------------------------- [  5440]
AY261364.1_Tengani_62            -------------------------------------------------------------------------------- [  5440]
AY261363.1_Pretoriuskop/96/4     -------------------------------------------------------------------------------- [  5440]
AY261362.1_Mkuzi_1979            -------------------------------------------------------------------------------- [  5440]
AY261361.1_Malawi_Lil-20/1_1983  -------------------------------------------------------------------------------- [  5440]
AY261360.1_Kenya_1950            aaacattatagtttaatattgaggtaaacattgcctgattgatgtcggcccccaatgcaaaacagtagtaaatagtcaaa [  5440]
AM712240.1_OURT_88/3             -------------------------------------------------------------------------------- [  5440]
AM712239.1_Benin_97/1            -------------------------------------------------------------------------------- [  5440]
KM111295.1_Ken06.Bus             -------------------------------------------------------------------------------- [  5440]
KM111294.1_Ken05/Tk1             aaacattatagtttaatattgaggtaaacattgcctgattcatgtcggcccccaatgcaaaacagtagtaaatagtcaaa [  5440]
KM262844.1_strain_L60            -------------------------------------------------------------------------------- [  5440]
KM102979.1_26544/OG10_Italy      -------------------------------------------------------------------------------- [  5440]
KM262845.1_strain_NHV            -------------------------------------------------------------------------------- [  5440]
KX354450.1_47/Ss/2008            -------------------------------------------------------------------------------- [  5440]
KP055815.1_strain_BA71           -------------------------------------------------------------------------------- [  5440]
MH025920.1_strain_R35            -------------------------------------------------------------------------------- [  5440]
MH025919.1_strain_N10            -------------------------------------------------------------------------------- [  5440]
MH025918.1_strain_R25            -------------------------------------------------------------------------------- [  5440]
MH025917.1_strain_R7             -------------------------------------------------------------------------------- [  5440]
MH025916.1_strain_R8             -------------------------------------------------------------------------------- [  5440]
```

```
FR682468.2_Georgia_2007/1        -------------------------------------------------------------------------------- [  5520]
ASF_IND_20_CAD543                -------------------------------------------------------------------------------- [  5520]
LR899193.1_Germany_2020/1        -------------------------------------------------------------------------------- [  5520]
MN393477.1_Wuhan_2019            -------------------------------------------------------------------------------- [  5520]
MW396979.1_Timor-Leste/2019/1    -------------------------------------------------------------------------------- [  5520]
MK940252.1_CN/2019/InnerMongolia -------------------------------------------------------------------------------- [  5520]
MK128995.1_China/2018/AnhuiXCGQ  -------------------------------------------------------------------------------- [  5520]
MH681419.1_ASFV/POL/2015         -------------------------------------------------------------------------------- [  5520]
LS478113.1_Estonia_2014          -------------------------------------------------------------------------------- [  5520]
KP843857.1_Odintsovo_02/14       -------------------------------------------------------------------------------- [  5520]
FN557520.1_strain_E75            -------------------------------------------------------------------------------- [  5520]
AY261366.1_Warthog               -------------------------------------------------------------------------------- [  5520]
AY261365.1_Warmbaths             -------------------------------------------------------------------------------- [  5520]
AY261364.1_Tengani_62            -------------------------------------------------------------------------------- [  5520]
AY261363.1_Pretoriuskop/96/4     -------------------------------------------------------------------------------- [  5520]
AY261362.1_Mkuzi_1979            -------------------------------------------------------------------------------- [  5520]
AY261361.1_Malawi_Lil-20/1_1983  -------------------------------------------------------------------------------- [  5520]
AY261360.1_Kenya_1950            aaattgttgtcacggatacaggccagccgcatcatttcatcaatatccatataaacttttcctgtattatgtatctcatg [  5520]
AM712240.1_OURT_88/3             -------------------------------------------------------------------------------- [  5520]
AM712239.1_Benin_97/1            -------------------------------------------------------------------------------- [  5520]
KM111295.1_Ken06.Bus             -------------------------------------------------------------------------------- [  5520]
KM111294.1_Ken05/Tk1             aaattgttgtcacggatacaggccagctgcatcatttcatcaatatcgatatgaacttttcctgtgatatatatctcatg [  5520]
KM262844.1_strain_L60            -------------------------------------------------------------------------------- [  5520]
KM102979.1_26544/OG10_Italy      -------------------------------------------------------------------------------- [  5520]
KM262845.1_strain_NHV            -------------------------------------------------------------------------------- [  5520]
KX354450.1_47/Ss/2008            -------------------------------------------------------------------------------- [  5520]
KP055815.1_strain_BA71           -------------------------------------------------------------------------------- [  5520]
MH025920.1_strain_R35            -------------------------------------------------------------------------------- [  5520]
MH025919.1_strain_N10            -------------------------------------------------------------------------------- [  5520]
MH025918.1_strain_R25            -------------------------------------------------------------------------------- [  5520]
MH025917.1_strain_R7             -------------------------------------------------------------------------------- [  5520]
MH025916.1_strain_R8             -------------------------------------------------------------------------------- [  5520]

FR682468.2_Georgia_2007/1        -------------------------------------------------------------------------------- [  5600]
ASF_IND_20_CAD543                -------------------------------------------------------------------------------- [  5600]
LR899193.1_Germany_2020/1        -------------------------------------------------------------------------------- [  5600]
MN393477.1_Wuhan_2019            -------------------------------------------------------------------------------- [  5600]
MW396979.1_Timor-Leste/2019/1    -------------------------------------------------------------------------------- [  5600]
MK940252.1_CN/2019/InnerMongolia -------------------------------------------------------------------------------- [  5600]
MK128995.1_China/2018/AnhuiXCGQ  -------------------------------------------------------------------------------- [  5600]
MH681419.1_ASFV/POL/2015         -------------------------------------------------------------------------------- [  5600]
LS478113.1_Estonia_2014          -------------------------------------------------------------------------------- [  5600]
KP843857.1_Odintsovo_02/14       -------------------------------------------------------------------------------- [  5600]
```

```
FN557520.1_strain_E75          -------------------------------------------------------------------------------- [  5600]
AY261366.1_Warthog             -------------------------------------------------------------------------------- [  5600]
AY261365.1_Warmbaths           -------------------------------------------------------------------------------- [  5600]
AY261364.1_Tengani_62          -------------------------------------------------------------------------------- [  5600]
AY261363.1_Pretoriuskop/96/4   -------------------------------------------------------------------------------- [  5600]
AY261362.1_Mkuzi_1979          -------------------------------------------------------------------------------- [  5600]
AY261361.1_Malawi_Lil-20/1_1983 -------------------------------------------------------------------------------- [  5600]
AY261360.1_Kenya_1950          aagatcaaacacattattaaaagaaagcgcacaatttaaacgccactcattgagctgctcgtattcttgatagaaatatt [  5600]
AM712240.1_OURT_88/3           -------------------------------------------------------------------------------- [  5600]
AM712239.1_Benin_97/1          -------------------------------------------------------------------------------- [  5600]
KM111295.1_Ken06.Bus           -------------------------------------------------------------------------------- [  5600]
KM111294.1_Ken05/Tk1           aagatcaaacacattgttaaaagaaagcgcacaatttaaacgccactcagtgagctgctcatattcttgatagaaatatt [  5600]
KM262844.1_strain_L60          -------------------------------------------------------------------------------- [  5600]
KM102979.1_26544/OG10_Italy    -------------------------------------------------------------------------------- [  5600]
KM262845.1_strain_NHV          -------------------------------------------------------------------------------- [  5600]
KX354450.1_47/Ss/2008          -------------------------------------------------------------------------------- [  5600]
KP055815.1_strain_BA71         -------------------------------------------------------------------------------- [  5600]
MH025920.1_strain_R35          -------------------------------------------------------------------------------- [  5600]
MH025919.1_strain_N10          -------------------------------------------------------------------------------- [  5600]
MH025918.1_strain_R25          -------------------------------------------------------------------------------- [  5600]
MH025917.1_strain_R7           -------------------------------------------------------------------------------- [  5600]
MH025916.1_strain_R8           -------------------------------------------------------------------------------- [  5600]

FR682468.2_Georgia_2007/1      -------------------------------------------------------------------------------- [  5680]
ASF_IND_20_CAD543              -------------------------------------------------------------------------------- [  5680]
LR899193.1_Germany_2020/1      -------------------------------------------------------------------------------- [  5680]
MN393477.1_Wuhan_2019          -------------------------------------------------------------------------------- [  5680]
MW396979.1_Timor-Leste/2019/1  -------------------------------------------------------------------------------- [  5680]
MK940252.1_CN/2019/InnerMongolia -------------------------------------------------------------------------------- [  5680]
MK128995.1_China/2018/AnhuiXCGQ -------------------------------------------------------------------------------- [  5680]
MH681419.1_ASFV/POL/2015       -------------------------------------------------------------------------------- [  5680]
LS478113.1_Estonia_2014        -------------------------------------------------------------------------------- [  5680]
KP843857.1_Odintsovo_02/14     -------------------------------------------------------------------------------- [  5680]
FN557520.1_strain_E75          -------------------------------------------------------------------------------- [  5680]
AY261366.1_Warthog             -------------------------------------------------------------------------------- [  5680]
AY261365.1_Warmbaths           -------------------------------------------------------------------------------- [  5680]
AY261364.1_Tengani_62          -------------------------------------------------------------------------------- [  5680]
AY261363.1_Pretoriuskop/96/4   -------------------------------------------------------------------------------- [  5680]
AY261362.1_Mkuzi_1979          -------------------------------------------------------------------------------- [  5680]
AY261361.1_Malawi_Lil-20/1_1983 -------------------------------------------------------------------------------- [  5680]
AY261360.1_Kenya_1950          gaatcgcttctttaagcttatatcttaccgctataccataccaatatttagttaacatctcactcaatgaagtattatca [  5680]
AM712240.1_OURT_88/3           -------------------------------------------------------------------------------- [  5680]
AM712239.1_Benin_97/1          -------------------------------------------------------------------------------- [  5680]
KM111295.1_Ken06.Bus           -------------------------------------------------------------------------------- [  5680]
```

```
KM111294.1_Ken05/Tk1            gaatcgcttctttaagcttatatcttaccgctataccataccaatatttagttaacatctcactcaatgaagtattatca [  5680]
KM262844.1_strain_L60           -------------------------------------------------------------------------------- [  5680]
KM102979.1_26544/OG10_Italy     -------------------------------------------------------------------------------- [  5680]
KM262845.1_strain_NHV           -------------------------------------------------------------------------------- [  5680]
KX354450.1_47/Ss/2008           -------------------------------------------------------------------------------- [  5680]
KP055815.1_strain_BA71          -------------------------------------------------------------------------------- [  5680]
MH025920.1_strain_R35           -------------------------------------------------------------------------------- [  5680]
MH025919.1_strain_N10           -------------------------------------------------------------------------------- [  5680]
MH025918.1_strain_R25           -------------------------------------------------------------------------------- [  5680]
MH025917.1_strain_R7            -------------------------------------------------------------------------------- [  5680]
MH025916.1_strain_R8            -------------------------------------------------------------------------------- [  5680]

FR682468.2_Georgia_2007/1       -------------------------------------------------------------------------------- [  5760]
ASF_IND_20_CAD543               -------------------------------------------------------------------------------- [  5760]
LR899193.1_Germany_2020/1       -------------------------------------------------------------------------------- [  5760]
MN393477.1_Wuhan_2019           -------------------------------------------------------------------------------- [  5760]
MW396979.1_Timor-Leste/2019/1   -------------------------------------------------------------------------------- [  5760]
MK940252.1_CN/2019/InnerMongolia -------------------------------------------------------------------------------- [  5760]
MK128995.1_China/2018/AnhuiXCGQ -------------------------------------------------------------------------------- [  5760]
MH681419.1_ASFV/POL/2015        -------------------------------------------------------------------------------- [  5760]
LS478113.1_Estonia_2014         -------------------------------------------------------------------------------- [  5760]
KP843857.1_Odintsovo_02/14      -------------------------------------------------------------------------------- [  5760]
FN557520.1_strain_E75           -------------------------------------------------------------------------------- [  5760]
AY261366.1_Warthog              -------------------------------------------------------------------------------- [  5760]
AY261365.1_Warmbaths            -------------------------------------------------------------------------------- [  5760]
AY261364.1_Tengani_62           -------------------------------------------------------------------------------- [  5760]
AY261363.1_Pretoriuskop/96/4    -------------------------------------------------------------------------------- [  5760]
AY261362.1_Mkuzi_1979           -------------------------------------------------------------------------------- [  5760]
AY261361.1_Malawi_Lil-20/1_1983 -------------------------------------------------------------------------------- [  5760]
AY261360.1_Kenya_1950           tccaatatgatattgtctgttagatcctttagatcccaataaattatcatctttaactcctccatatctacattctgtaa [  5760]
AM712240.1_OURT_88/3            -------------------------------------------------------------------------------- [  5760]
AM712239.1_Benin_97/1           -------------------------------------------------------------------------------- [  5760]
KM111295.1_Ken06.Bus            -------------------------------------------------------------------------------- [  5760]
KM111294.1_Ken05/Tk1            tccaatatgatattgtctgttatatcctttagatcccaataaattatcatctttaactcctccatatctacattctgtaa [  5760]
KM262844.1_strain_L60           -------------------------------------------------------------------------------- [  5760]
KM102979.1_26544/OG10_Italy     -------------------------------------------------------------------------------- [  5760]
KM262845.1_strain_NHV           -------------------------------------------------------------------------------- [  5760]
KX354450.1_47/Ss/2008           -------------------------------------------------------------------------------- [  5760]
KP055815.1_strain_BA71          -------------------------------------------------------------------------------- [  5760]
MH025920.1_strain_R35           -------------------------------------------------------------------------------- [  5760]
MH025919.1_strain_N10           -------------------------------------------------------------------------------- [  5760]
MH025918.1_strain_R25           -------------------------------------------------------------------------------- [  5760]
MH025917.1_strain_R7            -------------------------------------------------------------------------------- [  5760]
MH025916.1_strain_R8            -------------------------------------------------------------------------------- [  5760]
```

```
FR682468.2_Georgia_2007/1        -------------------------------------------------------------------------------- [  5840]
ASF_IND_20_CAD543                -------------------------------------------------------------------------------- [  5840]
LR899193.1_Germany_2020/1        -------------------------------------------------------------------------------- [  5840]
MN393477.1_Wuhan_2019            -------------------------------------------------------------------------------- [  5840]
MW396979.1_Timor-Leste/2019/1    -------------------------------------------------------------------------------- [  5840]
MK940252.1_CN/2019/InnerMongolia -------------------------------------------------------------------------------- [  5840]
MK128995.1_China/2018/AnhuiXCGQ  -------------------------------------------------------------------------------- [  5840]
MH681419.1_ASFV/POL/2015         -------------------------------------------------------------------------------- [  5840]
LS478113.1_Estonia_2014          -------------------------------------------------------------------------------- [  5840]
KP843857.1_Odintsovo_02/14       -------------------------------------------------------------------------------- [  5840]
FN557520.1_strain_E75            -------------------------------------------------------------------------------- [  5840]
AY261366.1_Warthog               -------------------------------------------------------------------------------- [  5840]
AY261365.1_Warmbaths             -------------------------------------------------------------------------------- [  5840]
AY261364.1_Tengani_62            -------------------------------------------------------------------------------- [  5840]
AY261363.1_Pretoriuskop/96/4     -------------------------------------------------------------------------------- [  5840]
AY261362.1_Mkuzi_1979            -------------------------------------------------------------------------------- [  5840]
AY261361.1_Malawi_Lil-20/1_1983  -------------------------------------------------------------------------------- [  5840]
AY261360.1_Kenya_1950            aaggagattattagaaaacaattcatggcatagaatgatattactactcgttttatgacgctttgttttaaagaaaaatt [  5840]
AM712240.1_OURT_88/3             -------------------------------------------------------------------------------- [  5840]
AM712239.1_Benin_97/1            -------------------------------------------------------------------------------- [  5840]
KM111295.1_Ken06.Bus             -------------------------------------------------------------------------------- [  5840]
KM111294.1_Ken05/Tk1             aaggagattattagaaaacaattcatggcatagaatgatattactactcgttttatgacgctttgttttaaagaaaaatc [  5840]
KM262844.1_strain_L60            -------------------------------------------------------------------------------- [  5840]
KM102979.1_26544/OG10_Italy      -------------------------------------------------------------------------------- [  5840]
KM262845.1_strain_NHV            -------------------------------------------------------------------------------- [  5840]
KX354450.1_47/Ss/2008            -------------------------------------------------------------------------------- [  5840]
KP055815.1_strain_BA71           -------------------------------------------------------------------------------- [  5840]
MH025920.1_strain_R35            -------------------------------------------------------------------------------- [  5840]
MH025919.1_strain_N10            -------------------------------------------------------------------------------- [  5840]
MH025918.1_strain_R25            -------------------------------------------------------------------------------- [  5840]
MH025917.1_strain_R7             -------------------------------------------------------------------------------- [  5840]
MH025916.1_strain_R8             -------------------------------------------------------------------------------- [  5840]

FR682468.2_Georgia_2007/1        -------------------------------------------------------------------------------- [  5920]
ASF_IND_20_CAD543                -------------------------------------------------------------------------------- [  5920]
LR899193.1_Germany_2020/1        -------------------------------------------------------------------------------- [  5920]
MN393477.1_Wuhan_2019            -------------------------------------------------------------------------------- [  5920]
MW396979.1_Timor-Leste/2019/1    -------------------------------------------------------------------------------- [  5920]
MK940252.1_CN/2019/InnerMongolia -------------------------------------------------------------------------------- [  5920]
MK128995.1_China/2018/AnhuiXCGQ  -------------------------------------------------------------------------------- [  5920]
MH681419.1_ASFV/POL/2015         -------------------------------------------------------------------------------- [  5920]
LS478113.1_Estonia_2014          -------------------------------------------------------------------------------- [  5920]
KP843857.1_Odintsovo_02/14       -------------------------------------------------------------------------------- [  5920]
```

```
FN557520.1_strain_E75             -------------------------------------------------------------------------------- [  5920]
AY261366.1_Warthog                -------------------------------------------------------------------------------- [  5920]
AY261365.1_Warmbaths              -------------------------------------------------------------------------------- [  5920]
AY261364.1_Tengani_62             -------------------------------------------------------------------------------- [  5920]
AY261363.1_Pretoriuskop/96/4      -------------------------------------------------------------------------------- [  5920]
AY261362.1_Mkuzi_1979             -------------------------------------------------------------------------------- [  5920]
AY261361.1_Malawi_Lil-20/1_1983   -------------------------------------------------------------------------------- [  5920]
AY261360.1_Kenya_1950             gtaaaatttcacttgcttcgagaccttctttggcacccaattttcggcataagtttcgagtatgctccatgttaacagaa [  5920]
AM712240.1_OURT_88/3              -------------------------------------------------------------------------------- [  5920]
AM712239.1_Benin_97/1             -------------------------------------------------------------------------------- [  5920]
KM111295.1_Ken06.Bus              -------------------------------------------------------------------------------- [  5920]
KM111294.1_Ken05/Tk1              gtaaaatttcactagcctcaagaccttctttggcacccaattttcggcataggtttcgggtatgctcctgattaacagaa [  5920]
KM262844.1_strain_L60             -------------------------------------------------------------------------------- [  5920]
KM102979.1_26544/OG10_Italy       -------------------------------------------------------------------------------- [  5920]
KM262845.1_strain_NHV             -------------------------------------------------------------------------------- [  5920]
KX354450.1_47/Ss/2008             -------------------------------------------------------------------------------- [  5920]
KP055815.1_strain_BA71            -------------------------------------------------------------------------------- [  5920]
MH025920.1_strain_R35             -------------------------------------------------------------------------------- [  5920]
MH025919.1_strain_N10             -------------------------------------------------------------------------------- [  5920]
MH025918.1_strain_R25             -------------------------------------------------------------------------------- [  5920]
MH025917.1_strain_R7              -------------------------------------------------------------------------------- [  5920]
MH025916.1_strain_R8              -------------------------------------------------------------------------------- [  5920]

FR682468.2_Georgia_2007/1         -------------------------------------------------------------------------------- [  6000]
ASF_IND_20_CAD543                 -------------------------------------------------------------------------------- [  6000]
LR899193.1_Germany_2020/1         -------------------------------------------------------------------------------- [  6000]
MN393477.1_Wuhan_2019             -------------------------------------------------------------------------------- [  6000]
MW396979.1_Timor-Leste/2019/1     -------------------------------------------------------------------------------- [  6000]
MK940252.1_CN/2019/InnerMongolia  -------------------------------------------------------------------------------- [  6000]
MK128995.1_China/2018/AnhuiXCGQ   -------------------------------------------------------------------------------- [  6000]
MH681419.1_ASFV/POL/2015          -------------------------------------------------------------------------------- [  6000]
LS478113.1_Estonia_2014           -------------------------------------------------------------------------------- [  6000]
KP843857.1_Odintsovo_02/14        -------------------------------------------------------------------------------- [  6000]
FN557520.1_strain_E75             -------------------------------------------------------------------------------- [  6000]
AY261366.1_Warthog                -------------------------------------------------------------------------------- [  6000]
AY261365.1_Warmbaths              -------------------------------------------------------------------------------- [  6000]
AY261364.1_Tengani_62             -------------------------------------------------------------------------------- [  6000]
AY261363.1_Pretoriuskop/96/4      -------------------------------------------------------------------------------- [  6000]
AY261362.1_Mkuzi_1979             -------------------------------------------------------------------------------- [  6000]
AY261361.1_Malawi_Lil-20/1_1983   -------------------------------------------------------------------------------- [  6000]
AY261360.1_Kenya_1950             agtaatccataattaatatttgcaccccattcagtaaacaacatgatcaattcatagttattttcctgtacagctttcat [  6000]
AM712240.1_OURT_88/3              -------------------------------------------------------------------------------- [  6000]
AM712239.1_Benin_97/1             -------------------------------------------------------------------------------- [  6000]
KM111295.1_Ken06.Bus              -------------------------------------------------------------------------------- [  6000]
```

```
KM111294.1_Ken05/Tk1              agtaacccataattaatatttgcaccccattcagtaaacaacatgatcaattcatagttattttcctgcacagctttcat [  6000]
KM262844.1_strain_L60             -------------------------------------------------------------------------------- [  6000]
KM102979.1_26544/OG10_Italy       -------------------------------------------------------------------------------- [  6000]
KM262845.1_strain_NHV             -------------------------------------------------------------------------------- [  6000]
KX354450.1_47/Ss/2008             -------------------------------------------------------------------------------- [  6000]
KP055815.1_strain_BA71            -------------------------------------------------------------------------------- [  6000]
MH025920.1_strain_R35             -------------------------------------------------------------------------------- [  6000]
MH025919.1_strain_N10             -------------------------------------------------------------------------------- [  6000]
MH025918.1_strain_R25             -------------------------------------------------------------------------------- [  6000]
MH025917.1_strain_R7              -------------------------------------------------------------------------------- [  6000]
MH025916.1_strain_R8              -------------------------------------------------------------------------------- [  6000]

FR682468.2_Georgia_2007/1         ----------------------------------------------------------------------tgataaatcg [  6080]
ASF_IND_20_CAD543                 ----------------------------------------------------------------------.......... [  6080]
LR899193.1_Germany_2020/1         ----------------------------------------------------------------------.......... [  6080]
MN393477.1_Wuhan_2019             ----------------------------------------------------------------------.......... [  6080]
MW396979.1_Timor-Leste/2019/1     ----------------------------------------------------------------------.......... [  6080]
MK940252.1_CN/2019/InnerMongolia  ----------------------------------------------------------------------.......... [  6080]
MK128995.1_China/2018/AnhuiXCGQ   ----------------------------------------------------------------------.......... [  6080]
MH681419.1_ASFV/POL/2015          ----------------------------------------------------------------------.......... [  6080]
LS478113.1_Estonia_2014           -------------------------------------------------------------------------------- [  6080]
KP843857.1_Odintsovo_02/14        ----------------------------------------------------------------------.......... [  6080]
FN557520.1_strain_E75             -------------------------------------------------------------------------------- [  6080]
AY261366.1_Warthog                ----------------------------------------------------------------------....g..... [  6080]
AY261365.1_Warmbaths              ----------------------------------------------------------------------....g..... [  6080]
AY261364.1_Tengani_62             ------------------------------------------------------------------------..g..... [  6080]
AY261363.1_Pretoriuskop/96/4      ------------------------------------------------------------------------..g..... [  6080]
AY261362.1_Mkuzi_1979             ----------------------------------------------------------------------....g..... [  6080]
AY261361.1_Malawi_Lil-20/1_1983   -------------------------------------------------------------------------------- [  6080]
AY261360.1_Kenya_1950             taatgcagtattaagtaacaagccctctttaagtattggtgtcttcactaacatttgatggttcttatca.a.c...g.a [  6080]
AM712240.1_OURT_88/3              ----------------------------------------------------------------------....g..... [  6080]
AM712239.1_Benin_97/1             ----------------------------------------------------------------------....g..... [  6080]
KM111295.1_Ken06.Bus              ------------------------------------tggcgtc------------------------------------- [  6080]
KM111294.1_Ken05/Tk1              taatgcagtattaagtaacaagccctctttaagtattggtgtcttcactaacatttgatggttcttatca.a.c...g.a [  6080]
KM262844.1_strain_L60             ----------------------------------------------------------------------....g..... [  6080]
KM102979.1_26544/OG10_Italy       ----------------------------------------------------------------------....g..... [  6080]
KM262845.1_strain_NHV             ----------------------------------------------------------------------....g..... [  6080]
KX354450.1_47/Ss/2008             ----------------------------------------------------------------------....g..... [  6080]
KP055815.1_strain_BA71            ----------------------------------------------------------------------....g..... [  6080]
MH025920.1_strain_R35             ------------------------------------tggcgtc------------------------------------- [  6080]
MH025919.1_strain_N10             ------------------------------------tggcgtc------------------------------------- [  6080]
MH025918.1_strain_R25             ------------------------------------tggcgtc------------------------------------- [  6080]
MH025917.1_strain_R7              ------------------------------------tggcgtc------------------------------------- [  6080]
MH025916.1_strain_R8              ------------------------------------tggcgtc------------------------------------- [  6080]
```

```
FR682468.2_Georgia_2007/1        taattggggcttcatgccaccacaaaccacaacgctctaaaatacaataatcatcttttaacacaggctgtgtagctagt [  6160]
ASF_IND_20_CAD543                ..........................................................k..................... [  6160]
LR899193.1_Germany_2020/1        ................................................................................ [  6160]
MN393477.1_Wuhan_2019            ................................................................................ [  6160]
MW396979.1_Timor-Leste/2019/1    ................................................................................ [  6160]
MK940252.1_CN/2019/InnerMongolia ................................................................................ [  6160]
MK128995.1_China/2018/AnhuiXCGQ  ................................................................................ [  6160]
MH681419.1_ASFV/POL/2015         ................................................................................ [  6160]
LS478113.1_Estonia_2014          -------------------------------------------------------------------------------- [  6160]
KP843857.1_Odintsovo_02/14       ................................................................................ [  6160]
FN557520.1_strain_E75            -------------------------------------------------------------------------------- [  6160]
AY261366.1_Warthog               ..........................................................g..t.t................ [  6160]
AY261365.1_Warmbaths             ..........c...............................................g..................... [  6160]
AY261364.1_Tengani_62            ............t...t...........................t.............g..t.t.........g...... [  6160]
AY261363.1_Pretoriuskop/96/4     ..........................................................g..t.t................ [  6160]
AY261362.1_Mkuzi_1979            ..........................................................g..................... [  6160]
AY261361.1_Malawi_Lil-20/1_1983  -----------------------------------------tc..tt..g.tt.t...gg-------------------- [  6160]
AY261360.1_Kenya_1950            ...g..................t.....g....a..t...tg..t............cgg.t.t.a....c..g..c.a. [  6160]
AM712240.1_OURT_88/3             ..........................................................g..................... [  6160]
AM712239.1_Benin_97/1            ..........................................................g..................... [  6160]
KM111295.1_Ken06.Bus             -------------------------------------------------------------------------------- [  6160]
KM111294.1_Ken05/Tk1             .t.g..................t..........at.t...t...t............cag.......t.....g...... [  6160]
KM262844.1_strain_L60            ..........................................................g..................... [  6160]
KM102979.1_26544/OG10_Italy      ..........................................................g..................... [  6160]
KM262845.1_strain_NHV            ..........................................................g..................... [  6160]
KX354450.1_47/Ss/2008            ..........................................................g..................... [  6160]
KP055815.1_strain_BA71           ..........................................................g..................... [  6160]
MH025920.1_strain_R35            -------------------------------------------------------------------------------- [  6160]
MH025919.1_strain_N10            -------------------------------------------------------------------------------- [  6160]
MH025918.1_strain_R25            -------------------------------------------------------------------------------- [  6160]
MH025917.1_strain_R7             -------------------------------------------------------------------------------- [  6160]
MH025916.1_strain_R8             -------------------------------------------------------------------------------- [  6160]

FR682468.2_Georgia_2007/1        acttttttagtaagtgcttgtaaagtagatggca-------tcttctatctgcaaaataattatttccgaaaaaaaaatc [  6240]
ASF_IND_20_CAD543                ..................................-------....................................... [  6240]
LR899193.1_Germany_2020/1        ..................................-------....................................... [  6240]
MN393477.1_Wuhan_2019            ..................................-------....................................... [  6240]
MW396979.1_Timor-Leste/2019/1    ..................................-------....................................... [  6240]
MK940252.1_CN/2019/InnerMongolia ..................................-------....................................... [  6240]
MK128995.1_China/2018/AnhuiXCGQ  ..................................-------....................................... [  6240]
MH681419.1_ASFV/POL/2015         ..................................-------....................................... [  6240]
LS478113.1_Estonia_2014          -------------------------------------------------------------------------------- [  6240]
KP843857.1_Odintsovo_02/14       ..................................-------....................................... [  6240]
```

```
FN557520.1_strain_E75           -------------------------------------------------------------------------------- [  6240]
AY261366.1_Warthog              ..................................-------.........................t............. [  6240]
AY261365.1_Warmbaths            ..................................-------....................................... [  6240]
AY261364.1_Tengani_62           ..................................-------............................-.......... [  6240]
AY261363.1_Pretoriuskop/96/4    ..................................-------....................................... [  6240]
AY261362.1_Mkuzi_1979           ..................................-------............................-.......... [  6240]
AY261361.1_Malawi_Lil-20/1_1983 ------------------------------------------------------------------..cg.........t [  6240]
AY261360.1_Kenya_1950           ................................t.gcatact..........a.g...........a..--.......... [  6240]
AM712240.1_OURT_88/3            ..................................-------....................................... [  6240]
AM712239.1_Benin_97/1           ..................................-------....................................... [  6240]
KM111295.1_Ken06.Bus            -------------------------------------------------------------------------------- [  6240]
KM111294.1_Ken05/Tk1            ........................c.......t.gcatact..........a.g.........c...t.g.......... [  6240]
KM262844.1_strain_L60           ..................................-------....................................... [  6240]
KM102979.1_26544/OG10_Italy     ..................................-------....................................... [  6240]
KM262845.1_strain_NHV           ..................................-------....................................... [  6240]
KX354450.1_47/Ss/2008           ..................................-------....................................... [  6240]
KP055815.1_strain_BA71          ..................................-------....................................... [  6240]
MH025920.1_strain_R35           -------------------------------------------------------------------------------- [  6240]
MH025919.1_strain_N10           -------------------------------------------------------------------------------- [  6240]
MH025918.1_strain_R25           -------------------------------------------------------------------------------- [  6240]
MH025917.1_strain_R7            -------------------------------------------------------------------------------- [  6240]
MH025916.1_strain_R8            -------------------------------------------------------------------------------- [  6240]

FR682468.2_Georgia_2007/1       aaattaaaatactaaattctatttttttttttaataaagcctgtaa------attatataataaatctc---gcccaccg [  6320]
ASF_IND_20_CAD543               ........................--....................------.................---........ [  6320]
LR899193.1_Germany_2020/1       ..............................................------.................---........ [  6320]
MN393477.1_Wuhan_2019           .........................-....................------.................---........ [  6320]
MW396979.1_Timor-Leste/2019/1   .........................-....................------.................---........ [  6320]
MK940252.1_CN/2019/InnerMongolia .........................-....................------.................---........ [  6320]
MK128995.1_China/2018/AnhuiXCGQ .........................-....................------.................---........ [  6320]
MH681419.1_ASFV/POL/2015        ..............................................------.................---........ [  6320]
LS478113.1_Estonia_2014         -------------------------------------------------------------------------------- [  6320]
KP843857.1_Odintsovo_02/14      ........................--....................------.................---........ [  6320]
FN557520.1_strain_E75           -------------------------------------------------------------------------------- [  6320]
AY261366.1_Warthog              ..........c........----..-.............t......attatt..........c......---.t..---- [  6320]
AY261365.1_Warmbaths            ....................t....-....................------....c............---........ [  6320]
AY261364.1_Tengani_62           ............c..c....--...-....................attact........t.c......---........ [  6320]
AY261363.1_Pretoriuskop/96/4    ...................----..-.................c..------....c............---........ [  6320]
AY261362.1_Mkuzi_1979           ............c........-..--....................------....c............---........ [  6320]
AY261361.1_Malawi_Lil-20/1_1983 t...---....a.c.g...c--..ca.a.a....c.g.ttt.t.g.---------------------------------- [  6320]
AY261360.1_Kenya_1950           .........atact.ga....g.a..................a...attact.........ac......ggg.aaa...a [  6320]
AM712240.1_OURT_88/3            ........................--....................------.................---........ [  6320]
AM712239.1_Benin_97/1           ........................--....................------.................---........ [  6320]
KM111295.1_Ken06.Bus            -------------------------------------------------------------------------------- [  6320]
```

```
KM111294.1_Ken05/Tk1             ..............g.a....a.a.a.....--.at..........attact.........at.t...t---c...g... [  6320]
KM262844.1_strain_L60            ........................--....................------.................---........ [  6320]
KM102979.1_26544/OG10_Italy      ........................--....................------.................---........ [  6320]
KM262845.1_strain_NHV            ........................--....................------.................---........ [  6320]
KX354450.1_47/Ss/2008            ........................--....................------.................---........ [  6320]
KP055815.1_strain_BA71           ........................--....................------.................---........ [  6320]
MH025920.1_strain_R35            -------------------------------------------------------------------------------- [  6320]
MH025919.1_strain_N10            -------------------------------------------------------------------------------- [  6320]
MH025918.1_strain_R25            -------------------------------------------------------------------------------- [  6320]
MH025917.1_strain_R7             -------------------------------------------------------------------------------- [  6320]
MH025916.1_strain_R8             -------------------------------------------------------------------------------- [  6320]

FR682468.2_Georgia_2007/1        tattatttccggacacaacttttttatacctcattatatttttagatctatagttttttaacaaggcattaatttttttctg [  6400]
ASF_IND_20_CAD543                ................................................................................ [  6400]
LR899193.1_Germany_2020/1        ................................................................................ [  6400]
MN393477.1_Wuhan_2019            ................................................................................ [  6400]
MW396979.1_Timor-Leste/2019/1    ................................................................................ [  6400]
MK940252.1_CN/2019/InnerMongolia ................................................................................ [  6400]
MK128995.1_China/2018/AnhuiXCGQ  ................................................................................ [  6400]
MH681419.1_ASFV/POL/2015         ................................................................................ [  6400]
LS478113.1_Estonia_2014          -------------------------------------------------------------------------------- [  6400]
KP843857.1_Odintsovo_02/14       .........................................................................-...... [  6400]
FN557520.1_strain_E75            -------------------------------------------------------------------------------- [  6400]
AY261366.1_Warthog               -------------------------------------------------------------------------------- [  6400]
AY261365.1_Warmbaths             .........t..g..........................................g........................ [  6400]
AY261364.1_Tengani_62            ....c..gt...gat..t..........ac..ac.............................................. [  6400]
AY261363.1_Pretoriuskop/96/4     ............g-................................t................................. [  6400]
AY261362.1_Mkuzi_1979            ........gt.............................................g........................ [  6400]
AY261361.1_Malawi_Lil-20/1_1983  -------------------c.ca...gtt....c..--------------------c..g....t.............c. [  6400]
AY261360.1_Kenya_1950            .t..t..g...aga..gt.---.....tg...ac..................a...........c..........c..c. [  6400]
AM712240.1_OURT_88/3             ........t....................................................................... [  6400]
AM712239.1_Benin_97/1            ........t....................................................................... [  6400]
KM111295.1_Ken06.Bus             -------------------------------------------------------------------------------- [  6400]
KM111294.1_Ken05/Tk1             ....c..a...aggg.gt.........tgc..ac..............................t..........c.... [  6400]
KM262844.1_strain_L60            ........t....................................................................... [  6400]
KM102979.1_26544/OG10_Italy      ........t....................................................................... [  6400]
KM262845.1_strain_NHV            ........t....................................................................... [  6400]
KX354450.1_47/Ss/2008            ........t....................................................................... [  6400]
KP055815.1_strain_BA71           ........t....................................................................... [  6400]
MH025920.1_strain_R35            -------------------------------------------------------------------------------- [  6400]
MH025919.1_strain_N10            -------------------------------------------------------------------------------- [  6400]
MH025918.1_strain_R25            -------------------------------------------------------------------------------- [  6400]
MH025917.1_strain_R7             -------------------------------------------------------------------------------- [  6400]
MH025916.1_strain_R8             -------------------------------------------------------------------------------- [  6400]
```

```
FR682468.2_Georgia_2007/1        gatctgtcgtttttaaagataaaagagagacgtttgaactataataatctttaaatgataatattt-ctactaatatatc [  6480]
ASF_IND_20_CAD543                ......................................y.....w.....................-............. [  6480]
LR899193.1_Germany_2020/1        ..................................................................-............. [  6480]
MN393477.1_Wuhan_2019            ..................................................................-............. [  6480]
MW396979.1_Timor-Leste/2019/1    ..................................................................-............. [  6480]
MK940252.1_CN/2019/InnerMongolia ..................................................................-............. [  6480]
MK128995.1_China/2018/AnhuiXCGQ  ..................................................................-............. [  6480]
MH681419.1_ASFV/POL/2015         ..................................................................-............. [  6480]
LS478113.1_Estonia_2014          -----------------............gta................t.................-............t [  6480]
KP843857.1_Odintsovo_02/14       ..................................................................-............. [  6480]
FN557520.1_strain_E75            -------------------------------------------------------------------------------- [  6480]
AY261366.1_Warthog               -------------------------------------------------------------------------------- [  6480]
AY261365.1_Warmbaths             ................t.............t.......t.....a.....................-............. [  6480]
AY261364.1_Tengani_62            ................t........g...g....................................-............. [  6480]
AY261363.1_Pretoriuskop/96/4     ................t........g...g.................................a..-..g.......... [  6480]
AY261362.1_Mkuzi_1979            ................t.............t.............a.....................-........t.... [  6480]
AY261361.1_Malawi_Lil-20/1_1983  ................t............ct..c..g...........t.................-t.......gc.aa [  6480]
AY261360.1_Kenya_1950            .......t........t....t...g...gta....gt.........ct.................-.........a.c. [  6480]
AM712240.1_OURT_88/3             ......................................t.....a.....................-............. [  6480]
AM712239.1_Benin_97/1            ......................................t.....a.....................-............. [  6480]
KM111295.1_Ken06.Bus             -------------------------------------------------------------------------------- [  6480]
KM111294.1_Ken05/Tk1             ................t........c...g.a........................ct........-...........c. [  6480]
KM262844.1_strain_L60            ......................................t.....a.....................-............. [  6480]
KM102979.1_26544/OG10_Italy      ......................................t.....a.....................c............. [  6480]
KM262845.1_strain_NHV            ......................................t.....a.....................-............. [  6480]
KX354450.1_47/Ss/2008            ......................................t.....a.....................-............. [  6480]
KP055815.1_strain_BA71           ......................................t.....a.....................-............. [  6480]
MH025920.1_strain_R35            -------------------------------------------------------------------------------- [  6480]
MH025919.1_strain_N10            -------------------------------------------------------------------------------- [  6480]
MH025918.1_strain_R25            -------------------------------------------------------------------------------- [  6480]
MH025917.1_strain_R7             -------------------------------------------------------------------------------- [  6480]
MH025916.1_strain_R8             -------------------------------------------------------------------------------- [  6480]

FR682468.2_Georgia_2007/1        atgattcttttgttttgctaattctaagctctcttcgaaagcattagctcctaaatctatacaaaagaacaagttattca [  6560]
ASF_IND_20_CAD543                ....................................k.........................................ag [  6560]
LR899193.1_Germany_2020/1        ................................................................................ [  6560]
MN393477.1_Wuhan_2019            ................................................................................ [  6560]
MW396979.1_Timor-Leste/2019/1    ................................................................................ [  6560]
MK940252.1_CN/2019/InnerMongolia ................................................................................ [  6560]
MK128995.1_China/2018/AnhuiXCGQ  ................................................................................ [  6560]
MH681419.1_ASFV/POL/2015         ................................................................................ [  6560]
LS478113.1_Estonia_2014          ..t.................g...c.t...g.............c....................g..........cca. [  6560]
KP843857.1_Odintsovo_02/14       ................................................................................ [  6560]
```

```
FN557520.1_strain_E75              -------------------------------------------------------------------------------- [  6560]
AY261366.1_Warthog                 -------------------------------------------------------------------------------- [  6560]
AY261365.1_Warmbaths               ..a.....................................................................t......ag [  6560]
AY261364.1_Tengani_62              ..t...t................................g........................................ [  6560]
AY261363.1_Pretoriuskop/96/4       ..t.................................a.........................................ag [  6560]
AY261362.1_Mkuzi_1979              .......................................................................t......ag [  6560]
AY261361.1_Malawi_Lil-20/1_1983    ..t....g......c.....t......a.a.........g..g.c...c.......................a....aa. [  6560]
AY261360.1_Kenya_1950              ..c.................t......a.a......a......g...................................a. [  6560]
AM712240.1_OURT_88/3               ..............................................................................ag [  6560]
AM712239.1_Benin_97/1              ..............................................................................ag [  6560]
KM111295.1_Ken06.Bus               -------------------------------------------------------------------------------- [  6560]
KM111294.1_Ken05/Tk1               ..t...---------........t...a.a.............g.......c...........................a. [  6560]
KM262844.1_strain_L60              ..............................................................................ag [  6560]
KM102979.1_26544/OG10_Italy        ..............................................................................ag [  6560]
KM262845.1_strain_NHV              ..............................................................................ag [  6560]
KX354450.1_47/Ss/2008              ..............................................................................ag [  6560]
KP055815.1_strain_BA71             ..............................................................................ag [  6560]
MH025920.1_strain_R35              -------------------------------------------------------------------------------- [  6560]
MH025919.1_strain_N10              -------------------------------------------------------------------------------- [  6560]
MH025918.1_strain_R25              -------------------------------------------------------------------------------- [  6560]
MH025917.1_strain_R7               -------------------------------------------------------------------------------- [  6560]
MH025916.1_strain_R8               -------------------------------------------------------------------------------- [  6560]

FR682468.2_Georgia_2007/1          tataaaagttttttaccgaggtaaccattgcccgattgatgtcagcccccaatacaaaacaatagtaaatggttaaaaaa [  6640]
ASF_IND_20_CAD543                  .................................................................w.............. [  6640]
LR899193.1_Germany_2020/1          ................................................................................ [  6640]
MN393477.1_Wuhan_2019              ................................................................................ [  6640]
MW396979.1_Timor-Leste/2019/1      ................................................................................ [  6640]
MK940252.1_CN/2019/InnerMongolia   ................................................................................ [  6640]
MK128995.1_China/2018/AnhuiXCGQ    ................................................................................ [  6640]
MH681419.1_ASFV/POL/2015           ................................................................................ [  6640]
LS478113.1_Estonia_2014            ...g.......a..........t.a...........................c.t.........a.....a..cg..t.. [  6640]
KP843857.1_Odintsovo_02/14         ................................................................................ [  6640]
FN557520.1_strain_E75              -------------------------------------------------------------------------------- [  6640]
AY261366.1_Warthog                 -------------------------------------------------------------------------------- [  6640]
AY261365.1_Warmbaths               ...................a.............................t...g...........a.............. [  6640]
AY261364.1_Tengani_62              ......................................................................a......... [  6640]
AY261363.1_Pretoriuskop/96/4       .......t........t.ca.............................t...............a....c......... [  6640]
AY261362.1_Mkuzi_1979              ...................a.............................t...g...........a.............. [  6640]
AY261361.1_Malawi_Lil-20/1_1983    .....g.t...a....t.......t............a........t....tc.t.........a..g...t.ggg.t.. [  6640]
AY261360.1_Kenya_1950              .......ca.......t..a...ca............................gt......g........a..c...... [  6640]
AM712240.1_OURT_88/3               ...................a......................t......t...............a.............. [  6640]
AM712239.1_Benin_97/1              ...................a......................t......t...............a.............. [  6640]
KM111295.1_Ken06.Bus               -------------------------------------------------------------------------------- [  6640]
```

```
KM111294.1_Ken05/Tk1            .......c...............ca.......t....................g................a......... [  6640]
KM262844.1_strain_L60           ...................a......................t......t...............a.............. [  6640]
KM102979.1_26544/OG10_Italy     ...................a......................t......t...............a.............. [  6640]
KM262845.1_strain_NHV           ...................a......................t......t...............a.............. [  6640]
KX354450.1_47/Ss/2008           ...................a......................t......t...............a.............. [  6640]
KP055815.1_strain_BA71          ...................a......................t......t...............a.............. [  6640]
MH025920.1_strain_R35           -------------------------------------------------------------------------------- [  6640]
MH025919.1_strain_N10           -------------------------------------------------------------------------------- [  6640]
MH025918.1_strain_R25           -------------------------------------------------------------------------------- [  6640]
MH025917.1_strain_R7            -------------------------------------------------------------------------------- [  6640]
MH025916.1_strain_R8            -------------------------------------------------------------------------------- [  6640]

FR682468.2_Georgia_2007/1       ttgctatctctca---tacaggccagatatatcatttcatcaatattcatatcaaccttttttatatgatacatttcatg [  6720]
ASF_IND_20_CAD543               .............---.......................................................y........ [  6720]
LR899193.1_Germany_2020/1       .............---................................................................ [  6720]
MN393477.1_Wuhan_2019           .............---................................................................ [  6720]
MW396979.1_Timor-Leste/2019/1   .............---................................................................ [  6720]
MK940252.1_CN/2019/InnerMongolia .............---................................................................ [  6720]
MK128995.1_China/2018/AnhuiXCGQ .............---................................................................ [  6720]
MH681419.1_ASFV/POL/2015        .............---................................................................ [  6720]
LS478113.1_Estonia_2014         ..a.c...ata.gtac......t..ac.tc..............g.ct.....cgt...c.c..ag.t...t.c...... [  6720]
KP843857.1_Odintsovo_02/14      .............---................................................................ [  6720]
FN557520.1_strain_E75           -------------------------------------------------------------------------------- [  6720]
AY261366.1_Warthog              -------------------------------------------------------------------------------- [  6720]
AY261365.1_Warmbaths            .............---.......t.....c.........................................t........ [  6720]
AY261364.1_Tengani_62           .............---.............c.................................................. [  6720]
AY261363.1_Pretoriuskop/96/4    ..a..........---.............c.......................................------..... [  6720]
AY261362.1_Mkuzi_1979           .............---....t........c.........................................t........ [  6720]
AY261361.1_Malawi_Lil-20/1_1983 ..atacctatatg---..........t.cc.......g..t.....ct.....ggt...c.c.t.g.t...t........ [  6720]
AY261360.1_Kenya_1950           ...t.g..a.ga.---..........ccgc................c.....g..tt.c---.t...t...t..c..... [  6720]
AM712240.1_OURT_88/3            .............---.............c.........................................t........ [  6720]
AM712239.1_Benin_97/1           .............---.............c.........................................t........ [  6720]
KM111295.1_Ken06.Bus            -------------------------------------------------------------------------------- [  6720]
KM111294.1_Ken05/Tk1            .......tat.t.---.........ac.gc..........tc....c.........t.c..c.t...t...t..c..... [  6720]
KM262844.1_strain_L60           .............---.............c.........................................t........ [  6720]
KM102979.1_26544/OG10_Italy     .............---.............c.........................................t........ [  6720]
KM262845.1_strain_NHV           .............---.............c.........................................t........ [  6720]
KX354450.1_47/Ss/2008           .............---.............c.........................................t........ [  6720]
KP055815.1_strain_BA71          .............---.............c.........................................t........ [  6720]
MH025920.1_strain_R35           -------------------------------------------------------------------------------- [  6720]
MH025919.1_strain_N10           -------------------------------------------------------------------------------- [  6720]
MH025918.1_strain_R25           -------------------------------------------------------------------------------- [  6720]
MH025917.1_strain_R7            -------------------------------------------------------------------------------- [  6720]
MH025916.1_strain_R8            -------------------------------------------------------------------------------- [  6720]
```

```
FR682468.2_Georgia_2007/1        aagatcagacacgttattaaaagaaagcccacatattagccgccaatctttaaaatgactatatcgttgataaaaatatt [  6800]
ASF_IND_20_CAD543                ................................................................................ [  6800]
LR899193.1_Germany_2020/1        ................................................................................ [  6800]
MN393477.1_Wuhan_2019            ................................................................................ [  6800]
MW396979.1_Timor-Leste/2019/1    ................................................................................ [  6800]
MK940252.1_CN/2019/InnerMongolia ................................................................................ [  6800]
MK128995.1_China/2018/AnhuiXCGQ  ................................................................................ [  6800]
MH681419.1_ASFV/POL/2015         ................................................................................ [  6800]
LS478113.1_Estonia_2014          ...g.......aa..g....................a....................c...................... [  6800]
KP843857.1_Odintsovo_02/14       ................................................................................ [  6800]
FN557520.1_strain_E75            -----------------------------------------................c.....a................ [  6800]
AY261366.1_Warthog               -------------------------------------------------------------------------------- [  6800]
AY261365.1_Warmbaths             ................................................................................ [  6800]
AY261364.1_Tengani_62            ................................................................................ [  6800]
AY261363.1_Pretoriuskop/96/4     .........t...................................................................... [  6800]
AY261362.1_Mkuzi_1979            .........................................................t.....a................ [  6800]
AY261361.1_Malawi_Lil-20/1_1983  ......ga...gt....---...c..c............t.................ct.....tt.............. [  6800]
AY261360.1_Kenya_1950            g...........a..g.......c..............at................ac......t............... [  6800]
AM712240.1_OURT_88/3             .........................................................c.....a................ [  6800]
AM712239.1_Benin_97/1            .........................................................c.....a................ [  6800]
KM111295.1_Ken06.Bus             -------------------------------------------------------------------------------- [  6800]
KM111294.1_Ken05/Tk1             .......a...........t...c..............at.................c......t............... [  6800]
KM262844.1_strain_L60            .........................................................c.....a................ [  6800]
KM102979.1_26544/OG10_Italy      .........................................................c.....a................ [  6800]
KM262845.1_strain_NHV            .........................................................c.....a................ [  6800]
KX354450.1_47/Ss/2008            .........................................................c.....a................ [  6800]
KP055815.1_strain_BA71           .........................................................c.....a................ [  6800]
MH025920.1_strain_R35            -------------------------------------------------------------------------------- [  6800]
MH025919.1_strain_N10            -------------------------------------------------------------------------------- [  6800]
MH025918.1_strain_R25            -------------------------------------------------------------------------------- [  6800]
MH025917.1_strain_R7             -------------------------------------------------------------------------------- [  6800]
MH025916.1_strain_R8             -------------------------------------------------------------------------------- [  6800]

FR682468.2_Georgia_2007/1        ggatggcttcagtaagcttacatagtatcgctatactataccaatatctagttagcatttcgttgaatgttatttcattc [  6880]
ASF_IND_20_CAD543                ................g.g.t..c...................ta.c................................. [  6880]
LR899193.1_Germany_2020/1        ................................................................................ [  6880]
MN393477.1_Wuhan_2019            ................................................................................ [  6880]
MW396979.1_Timor-Leste/2019/1    ................................................................................ [  6880]
MK940252.1_CN/2019/InnerMongolia ................................................................................ [  6880]
MK128995.1_China/2018/AnhuiXCGQ  ................................................................................ [  6880]
MH681419.1_ASFV/POL/2015         ................................................................................ [  6880]
LS478113.1_Estonia_2014          ................g...t...a......c...........g..........a.......c.a....aa..a....c. [  6880]
KP843857.1_Odintsovo_02/14       ................................................................................ [  6880]
```

```
FN557520.1_strain_E75            ....a........t..g.g.t..c...c...c...........ta.c....c............c............... [  6880]
AY261366.1_Warthog               -------------------------------------------------------------------------------- [  6880]
AY261365.1_Warmbaths             .............t..g...t..........c...................c.................aa..aa..... [  6880]
AY261364.1_Tengani_62            ....................t........................................................... [  6880]
AY261363.1_Pretoriuskop/96/4     ................a...t..t.......c...............................................t [  6880]
AY261362.1_Mkuzi_1979            ....a.....c..t..g.g.t..c...c...............ta......c............c............... [  6880]
AY261361.1_Malawi_Lil-20/1_1983  .a..a..........ag...t..tc..ct..c.atgcg......a....g......t....ac.a....a........c. [  6880]
AY261360.1_Kenya_1950            .............g..g...t..t...ct..............ta.c.......a......ac......a........c. [  6880]
AM712240.1_OURT_88/3             ....a........t..g.g.t..c...c...c...........ta.c....c............c............... [  6880]
AM712239.1_Benin_97/1            ....a........t..g.g.t..c...c...c...........ta.c....c............c............... [  6880]
KM111295.1_Ken06.Bus             -------------------------------------------------------------------------------- [  6880]
KM111294.1_Ken05/Tk1             ................g...tg.c...c...............ta.c...............c......a........c. [  6880]
KM262844.1_strain_L60            ....a........t..g.g.t..c...c...c...........ta.c....c............c............... [  6880]
KM102979.1_26544/OG10_Italy      ....a........t..g.g.t..c...c...c...........ta.c....c............c............... [  6880]
KM262845.1_strain_NHV            ....a........t..g.g.t..c...c...c...........ta.c....c............c............... [  6880]
KX354450.1_47/Ss/2008            ....a........t..g.g.t..c...c...c...........ta.c....c............c............... [  6880]
KP055815.1_strain_BA71           ....a........t..g.g.t..c...c...c...........ta.c....c............c............... [  6880]
MH025920.1_strain_R35            -------------------------------------------------------------------------------- [  6880]
MH025919.1_strain_N10            -------------------------------------------------------------------------------- [  6880]
MH025918.1_strain_R25            -------------------------------------------------------------------------------- [  6880]
MH025917.1_strain_R7             -------------------------------------------------------------------------------- [  6880]
MH025916.1_strain_R8             -------------------------------------------------------------------------------- [  6880]

FR682468.2_Georgia_2007/1        aatataaagttgatcgatatcttctctagaaaacaacaaattattacttttaat------tcctctatattctgg---aa [  6960]
ASF_IND_20_CAD543                ..............................................y.......------...............---.. [  6960]
LR899193.1_Germany_2020/1        ......................................................------...............---.. [  6960]
MN393477.1_Wuhan_2019            ......................................................------...............---.. [  6960]
MW396979.1_Timor-Leste/2019/1    ......................................................------...............---.. [  6960]
MK940252.1_CN/2019/InnerMongolia ......................................................------...............---.. [  6960]
MK128995.1_China/2018/AnhuiXCGQ  ......................................................------...............---.. [  6960]
MH681419.1_ASFV/POL/2015         ......................................................------...............---.. [  6960]
LS478113.1_Estonia_2014          ..............t....ag..t.tc.agccc.t.t...c.....aac.c...------ct.........t...---.. [  6960]
KP843857.1_Odintsovo_02/14       ......................................................------...............---.. [  6960]
FN557520.1_strain_E75            ..............................................t.......------...............---.. [  6960]
AY261366.1_Warthog               -------------------------------------------------------------------------------- [  6960]
AY261365.1_Warmbaths             ................g.............................t...c...------c..............---.. [  6960]
AY261364.1_Tengani_62            ................c.............................t...c...------c..............---.. [  6960]
AY261363.1_Pretoriuskop/96/4     ................g.............................t...c...------c..............---.. [  6960]
AY261362.1_Mkuzi_1979            ......................................g.......t.......------...............---.. [  6960]
AY261361.1_Malawi_Lil-20/1_1983  ...........a..t....aa....t..attt.t..tt.......ct.c.c...tgaaca.ta.t.tc.c.ta.a---.. [  6960]
AY261360.1_Kenya_1950            ..............t..........t.............g...a......c...------c......c...t.cattc.t [  6960]
AM712240.1_OURT_88/3             ..............................................t.......------...............---.. [  6960]
AM712239.1_Benin_97/1            ..............................................t.......------...............---.. [  6960]
KM111295.1_Ken06.Bus             -------------------------------------------------------------------------------- [  6960]
```

```
KM111294.1_Ken05/Tk1            ..............t...................................c...------c......c...t.cattc.t [  6960]
KM262844.1_strain_L60           ..............................................t.......------...............---.. [  6960]
KM102979.1_26544/OG10_Italy     ..............................................t.......------...............---.. [  6960]
KM262845.1_strain_NHV           ..............................................t.......------...............---.. [  6960]
KX354450.1_47/Ss/2008           ..............................................t.......------...............---.. [  6960]
KP055815.1_strain_BA71          ..............................................t.......------...............---.. [  6960]
MH025920.1_strain_R35           -------------------------------------------------------------------------------- [  6960]
MH025919.1_strain_N10           -------------------------------------------------------------------------------- [  6960]
MH025918.1_strain_R25           -------------------------------------------------------------------------------- [  6960]
MH025917.1_strain_R7            -------------------------------------------------------------------------------- [  6960]
MH025916.1_strain_R8            -------------------------------------------------------------------------------- [  6960]

FR682468.2_Georgia_2007/1       aaggggattattagataacaatttatggcataaaataatattactactagttttaatacgatgtattttataaaa----- [  7040]
ASF_IND_20_CAD543               ...........................................................................----- [  7040]
LR899193.1_Germany_2020/1       ...........................................................................----- [  7040]
MN393477.1_Wuhan_2019           ...........................................................................----- [  7040]
MW396979.1_Timor-Leste/2019/1   ...........................................................................----- [  7040]
MK940252.1_CN/2019/InnerMongolia ...........................................................................----- [  7040]
MK128995.1_China/2018/AnhuiXCGQ ...........................................................................----- [  7040]
MH681419.1_ASFV/POL/2015        ...........................................................................----- [  7040]
LS478113.1_Estonia_2014         ....a..........a.......c..tat................g..........g.t..c..g..........----- [  7040]
KP843857.1_Odintsovo_02/14      ...........................................................................----- [  7040]
FN557520.1_strain_E75           ................................g..........................................----- [  7040]
AY261366.1_Warthog              -------------------------------------------------------------------------------- [  7040]
AY261365.1_Warmbaths            ......c.............g......................................................----- [  7040]
AY261364.1_Tengani_62           ....a.c.............g...........g..........................a...............----- [  7040]
AY261363.1_Pretoriuskop/96/4    ......c.............g...........g..........................................----- [  7040]
AY261362.1_Mkuzi_1979           ................................g..........................................----- [  7040]
AY261361.1_Malawi_Lil-20/1_1983 ...a..............t....c...................t.....c......a...t.t..c.g.......----- [  7040]
AY261360.1_Kenya_1950           .........g.............cg..a...tg...c.....gt.g..c...cg.........a......c....aatgt [  7040]
AM712240.1_OURT_88/3            ................................g..........................................----- [  7040]
AM712239.1_Benin_97/1           ................................g..........................................----- [  7040]
KM111295.1_Ken06.Bus            -------------------------------------------------------------------------------- [  7040]
KM111294.1_Ken05/Tk1            .t....g..g.............c...a...............t.g..c...cg.........a......c....aatat [  7040]
KM262844.1_strain_L60           ................................g..........................................----- [  7040]
KM102979.1_26544/OG10_Italy     ................................g..........................................----- [  7040]
KM262845.1_strain_NHV           ................................g..........................................----- [  7040]
KX354450.1_47/Ss/2008           ................................g..........................................----- [  7040]
KP055815.1_strain_BA71          ................................g..........................................----- [  7040]
MH025920.1_strain_R35           -------------------------------------------------------------------------------- [  7040]
MH025919.1_strain_N10           -------------------------------------------------------------------------------- [  7040]
MH025918.1_strain_R25           -------------------------------------------------------------------------------- [  7040]
MH025917.1_strain_R7            -------------------------------------------------------------------------------- [  7040]
MH025916.1_strain_R8            -------------------------------------------------------------------------------- [  7040]
```

```
FR682468.2_Georgia_2007/1        -------tatttgtacaatatccatttcattcaaaatt---tttgcgcctaactcccggcagaaattccaagtatgctcc [  7120]
ASF_IND_20_CAD543                -------...............................---....................................... [  7120]
LR899193.1_Germany_2020/1        -------...............................---....................................... [  7120]
MN393477.1_Wuhan_2019            -------...............................---....................................... [  7120]
MW396979.1_Timor-Leste/2019/1    -------...............................---....................................... [  7120]
MK940252.1_CN/2019/InnerMongolia -------...............................---....................................... [  7120]
MK128995.1_China/2018/AnhuiXCGQ  -------...............................---....................................... [  7120]
MH681419.1_ASFV/POL/2015         -------...............................---....................................... [  7120]
LS478113.1_Estonia_2014          -------........a.......c...........gc.tcc...........t.t.t.a..a.gg....gg....a.... [  7120]
KP843857.1_Odintsovo_02/14       -------...............................---....................................... [  7120]
FN557520.1_strain_E75            -------...............................---....................................... [  7120]
AY261366.1_Warthog               -------------------------------------------------------------------------------- [  7120]
AY261365.1_Warmbaths             -------................g......c.......---.....a................................. [  7120]
AY261364.1_Tengani_62            -------................g............c.---.....a................................. [  7120]
AY261363.1_Pretoriuskop/96/4     -------...a...........................---....................................... [  7120]
AY261362.1_Mkuzi_1979            -------...............................---....................................... [  7120]
AY261361.1_Malawi_Lil-20/1_1983  -------..cg.t..a..gt...ca..t...t...gc.tct..g..a........t..a..t.ggc.t.gga.g.tt... [  7120]
AY261360.1_Kenya_1950            tacaaaa...g.c..a......tc.....c.....gc.tcc..c.......g...t..a..a.g..c..gg..g.a.... [  7120]
AM712240.1_OURT_88/3             -------...............................---....................................... [  7120]
AM712239.1_Benin_97/1            -------...............................---....................................... [  7120]
KM111295.1_Ken06.Bus             -------------------------------------------------------------------------------- [  7120]
KM111294.1_Ken05/Tk1             tacaaaa.....c..a.......c.....a......cgtcc..c..a....g...t..a..a.g...t.gg..g.a.... [  7120]
KM262844.1_strain_L60            -------...............................---....................................... [  7120]
KM102979.1_26544/OG10_Italy      -------...............................---....................................... [  7120]
KM262845.1_strain_NHV            -------...............................---....................................... [  7120]
KX354450.1_47/Ss/2008            -------...............................---....................................... [  7120]
KP055815.1_strain_BA71           -------...............................---....................................... [  7120]
MH025920.1_strain_R35            -------------------------------------------------------------------------------- [  7120]
MH025919.1_strain_N10            -------------------------------------------------------------------------------- [  7120]
MH025918.1_strain_R25            -------------------------------------------------------------------------------- [  7120]
MH025917.1_strain_R7             -------------------------------------------------------------------------------- [  7120]
MH025916.1_strain_R8             -------------------------------------------------------------------------------- [  7120]

FR682468.2_Georgia_2007/1        gtattgacagtgactaagctagagttgatgtctgcaccccattcagtaaacaactctattagatcatagttgttttcctg [  7200]
ASF_IND_20_CAD543                ............................................r................................... [  7200]
LR899193.1_Germany_2020/1        ................................................................................ [  7200]
MN393477.1_Wuhan_2019            ................................................................................ [  7200]
MW396979.1_Timor-Leste/2019/1    ................................................................................ [  7200]
MK940252.1_CN/2019/InnerMongolia ................................................................................ [  7200]
MK128995.1_China/2018/AnhuiXCGQ  ................................................................................ [  7200]
MH681419.1_ASFV/POL/2015         ................................................................................ [  7200]
LS478113.1_Estonia_2014          .....a.....a.................a...........c.tg................a.....g...a.....t.. [  7200]
KP843857.1_Odintsovo_02/14       ................................................................................ [  7200]
```

```
FN557520.1_strain_E75             ............................................g................................... [  7200]
AY261366.1_Warthog                -------------------------------------------------------------------------------- [  7200]
AY261365.1_Warmbaths              .............t............t.................g.......g........................... [  7200]
AY261364.1_Tengani_62             ............................................g.......g..................a........ [  7200]
AY261363.1_Pretoriuskop/96/4      .............t..........g...................g.......g........................... [  7200]
AY261362.1_Mkuzi_1979             ........g...................................g................................... [  7200]
AY261361.1_Malawi_Lil-20/1_1983   ..g..........tc..t.c.a.a..a..............c...a......g.........gc..cga........t.. [  7200]
AY261360.1_Kenya_1950             a....a...........c.c.aca..a.................g.......g..........c..cga........... [  7200]
AM712240.1_OURT_88/3              ............................................g................................... [  7200]
AM712239.1_Benin_97/1             ............................................g................................... [  7200]
KM111295.1_Ken06.Bus              -------------------------------------------------------------------------------- [  7200]
KM111294.1_Ken05/Tk1              .....a...........c.c.aca..a.................g.......g..g.....c.ct.cga........... [  7200]
KM262844.1_strain_L60             ............................................g................................... [  7200]
KM102979.1_26544/OG10_Italy       ............................................g................................... [  7200]
KM262845.1_strain_NHV             ............................................g................................... [  7200]
KX354450.1_47/Ss/2008             ............................................g................................... [  7200]
KP055815.1_strain_BA71            ............................................g................................... [  7200]
MH025920.1_strain_R35             -------------------------------------------------------------------------------- [  7200]
MH025919.1_strain_N10             -------------------------------------------------------------------------------- [  7200]
MH025918.1_strain_R25             -------------------------------------------------------------------------------- [  7200]
MH025917.1_strain_R7              -------------------------------------------------------------------------------- [  7200]
MH025916.1_strain_R8              -------------------------------------------------------------------------------- [  7200]

FR682468.2_Georgia_2007/1         cacagttttcattaatgcgagatttaactctaaaccatctttaaaaattgctgattttatcatcaattgattatcc---t [  7280]
ASF_IND_20_CAD543                 ............................................................................---. [  7280]
LR899193.1_Germany_2020/1         ............................................................................---. [  7280]
MN393477.1_Wuhan_2019             ............................................................................---. [  7280]
MW396979.1_Timor-Leste/2019/1     ............................................................................---. [  7280]
MK940252.1_CN/2019/InnerMongolia  ..............................................t.............................---. [  7280]
MK128995.1_China/2018/AnhuiXCGQ   ............................................................................---. [  7280]
MH681419.1_ASFV/POL/2015          ............................................................................---. [  7280]
LS478113.1_Estonia_2014           ...gac..............t..c...aga...g..t.....g..gc....c..........at.t...gc.....---. [  7280]
KP843857.1_Odintsovo_02/14        ............................................................................---. [  7280]
FN557520.1_strain_E75             .....c......................................................................---. [  7280]
AY261366.1_Warthog                -------------------------------------------------------------------------------- [  7280]
AY261365.1_Warmbaths              .....c.....................a..............................................t.---. [  7280]
AY261364.1_Tengani_62             .....c.....................t..............................................t.---. [  7280]
AY261363.1_Pretoriuskop/96/4      .....c.....................a..............................................t.---. [  7280]
AY261362.1_Mkuzi_1979             .....c.....................a..............................................t.---. [  7280]
AY261361.1_Malawi_Lil-20/1_1983   ...g.c...........................g...........gc.............................tca. [  7280]
AY261360.1_Kenya_1950             t..g.c............a.c.....tt......t..........gc.....a.......t..---..........---. [  7280]
AM712240.1_OURT_88/3              .....c......................................................................---. [  7280]
AM712239.1_Benin_97/1             .....c......................................................................---. [  7280]
KM111295.1_Ken06.Bus              --.g.c............a.c.....tt......t..........gc.....a.......t..---.....g....---. [  7280]
```

```
KM111294.1_Ken05/Tk1             ...g.c............a.c.....t..................gc................---.....g....tca. [  7280]
KM262844.1_strain_L60            .....c......................................................................---. [  7280]
KM102979.1_26544/OG10_Italy      .....c......................................................................---. [  7280]
KM262845.1_strain_NHV            .....c......................................................................---. [  7280]
KX354450.1_47/Ss/2008            .....c......................................................................---. [  7280]
KP055815.1_strain_BA71           .....c......................................................................---. [  7280]
MH025920.1_strain_R35            --.g.c............a.c.....tt......t..........gc.....a.......t..---.....g....---. [  7280]
MH025919.1_strain_N10            --.g.c............a.c.....tt......t..........gc.....a.......t..---.....g....---. [  7280]
MH025918.1_strain_R25            --.g.c............a.c.....tt......t..........gc.....a.......t..---.....g....---. [  7280]
MH025917.1_strain_R7             --.g.c............a.c.....tt......t..........gc.....a.......t..---.....g....---. [  7280]
MH025916.1_strain_R8             --.g.c............a.c.....tt......t..........gc.....a.......t..---.....g....---. [  7280]

FR682468.2_Georgia_2007/1        cattagtagaaagcataattggagctccatgccaccacaaaccacaatatttcaaaataaagtagtgttctttagatatg [  7360]
ASF_IND_20_CAD543                ...................................................................a.t.......... [  7360]
LR899193.1_Germany_2020/1        ................................................................................ [  7360]
MN393477.1_Wuhan_2019            ................................................................................ [  7360]
MW396979.1_Timor-Leste/2019/1    ................................................................................ [  7360]
MK940252.1_CN/2019/InnerMongolia ................................................................................ [  7360]
MK128995.1_China/2018/AnhuiXCGQ  ................................................................................ [  7360]
MH681419.1_ASFV/POL/2015         ................................................................................ [  7360]
LS478113.1_Estonia_2014          ...c.a.gc...t.g....c..c...t..........t.......................................... [  7360]
KP843857.1_Odintsovo_02/14       ................................................................................ [  7360]
FN557520.1_strain_E75            ............a......................................................a.t...g...... [  7360]
AY261366.1_Warthog               -------------------------------------------------------------------------------- [  7360]
AY261365.1_Warmbaths             ................g............................................................... [  7360]
AY261364.1_Tengani_62            ................................................................c............... [  7360]
AY261363.1_Pretoriuskop/96/4     ................g............................................................... [  7360]
AY261362.1_Mkuzi_1979            ................g............................................................... [  7360]
AY261361.1_Malawi_Lil-20/1_1983  .......g......g.g.....ga.......................cgc.ct......c.a..a.ca....c...c.ca [  7360]
AY261360.1_Kenya_1950            ....c...........g......a......c........g....t..cgc.ct......c.a..a.ca........c.ca [  7360]
AM712240.1_OURT_88/3             ............a......................................................a.t...g...... [  7360]
AM712239.1_Benin_97/1            ............a......................................................a.t...g...... [  7360]
KM111295.1_Ken06.Bus             ....c...........g......a..t...c........g....t..cgc.ct......c.a..a.ca..........ca [  7360]
KM111294.1_Ken05/Tk1             ................g......c................c......cgc.ct......c.a..a.ca........c.ca [  7360]
KM262844.1_strain_L60            ............a......................................................a.t...g...... [  7360]
KM102979.1_26544/OG10_Italy      ............a......................................................a.t...g...... [  7360]
KM262845.1_strain_NHV            ............a......................................................a.t...g...... [  7360]
KX354450.1_47/Ss/2008            ............a......................................................a.t...g...... [  7360]
KP055815.1_strain_BA71           ............a......................................................a.t...g...... [  7360]
MH025920.1_strain_R35            ....c...........g......a..t...c........g....t..cgc.ct......c.a..a.ca..........ca [  7360]
MH025919.1_strain_N10            ....c...........g......a..t...c........g....t..cgc.ct......c.a..a.ca..........ca [  7360]
MH025918.1_strain_R25            ....c...........g......a..t...c........g....t..cgc.ct......c.a..a.ca..........ca [  7360]
MH025917.1_strain_R7             ....c...........g......a..t...c........g....t..cgc.ct......c.a..a.ca..........ca [  7360]
MH025916.1_strain_R8             ....c...........g......a..t...c........g....t..cgc.ct......c.a..a.ca..........ca [  7360]
```

```
FR682468.2_Georgia_2007/1        tgctgtgtggccagtat-ttttttagcaagagcctgcagagaaattggagtagacata--tttttttttgcaaaatggtt [  7440]
ASF_IND_20_CAD543                .................-.....................................--.--.................... [  7440]
LR899193.1_Germany_2020/1        .................-........................................--.................... [  7440]
MN393477.1_Wuhan_2019            .................-........................................--.................... [  7440]
MW396979.1_Timor-Leste/2019/1    .................-........................................--.................... [  7440]
MK940252.1_CN/2019/InnerMongolia .................-........................................--.................... [  7440]
MK128995.1_China/2018/AnhuiXCGQ  .................-........................................--.................... [  7440]
MH681419.1_ASFV/POL/2015         .................-........................................--.................... [  7440]
LS478113.1_Estonia_2014          ..............c.c-.......a......t...t.......g..........--.--.........a......c... [  7440]
KP843857.1_Odintsovo_02/14       .................-........................................--.................... [  7440]
FN557520.1_strain_E75            .................-..........................g..........--.--.................... [  7440]
AY261366.1_Warthog               ---------------------------------------------------.a.....--.................... [  7440]
AY261365.1_Warmbaths             ..............c.c-.......a................cga.............--.................... [  7440]
AY261364.1_Tengani_62            ..t.............c-c.....ga................cga...t......--.-t.................... [  7440]
AY261363.1_Pretoriuskop/96/4     ................c-c.....ga................cga...t.........-t.................c.. [  7440]
AY261362.1_Mkuzi_1979            .................-..........................g..........--.---................... [  7440]
AY261361.1_Malawi_Lil-20/1_1983  .....gaa.t......cccc....ga....gatt.........ga...t.a.......--.................... [  7440]
AY261360.1_Kenya_1950            .....gaa.t......c-......ga....ga...........gg...t.........--................a... [  7440]
AM712240.1_OURT_88/3             .................-..........................g..........--.--.................... [  7440]
AM712239.1_Benin_97/1            .................-..........................g..........--.--.................... [  7440]
KM111295.1_Ken06.Bus             .....gaa.t......c-c.....ga....ga............g...t.........-t.........a......a... [  7440]
KM111294.1_Ken05/Tk1             .....gaa.t......c-cc....ga....gat...t......ga...t.........ttc...............a... [  7440]
KM262844.1_strain_L60            .................-..........................g..........--.--.................... [  7440]
KM102979.1_26544/OG10_Italy      .................-..........................g..........--.--.................... [  7440]
KM262845.1_strain_NHV            .................-..........................g..........--.--.................... [  7440]
KX354450.1_47/Ss/2008            .................-..........................g..........--.--.................... [  7440]
KP055815.1_strain_BA71           .................-..........................g..........--.--.................... [  7440]
MH025920.1_strain_R35            .....gaa.t......c-c.....ga....ga............g...t.........-t.........a......a... [  7440]
MH025919.1_strain_N10            .....gaa.t......c-c.....ga....ga............g...t.........-t.........a......a... [  7440]
MH025918.1_strain_R25            .....gaa.t......c-c.....ga....ga............g...t.........-t.........a......a... [  7440]
MH025917.1_strain_R7             .....gaa.t......c-c.....ga....ga............g...t.........-t.........a......a... [  7440]
MH025916.1_strain_R8             .....gaa.t......c-c.....ga....ga............g...t.........-t.........a......a... [  7440]

FR682468.2_Georgia_2007/1        taagttttt-------caagaatacagattggataaattaggttgttgac-ttagttac------aggaggtattaaata [  7520]
ASF_IND_20_CAD543                ........aatgtttt.............................aca---.....w..------..r.r.......... [  7520]
LR899193.1_Germany_2020/1        .........-------..................................-........------............... [  7520]
MN393477.1_Wuhan_2019            .........-------..................................-........------............... [  7520]
MW396979.1_Timor-Leste/2019/1    .........-------..................................-........------............... [  7520]
MK940252.1_CN/2019/InnerMongolia .........-------..................................-........------............... [  7520]
MK128995.1_China/2018/AnhuiXCGQ  .........-------..................................-........------............... [  7520]
MH681419.1_ASFV/POL/2015         .........-------..................................-........------............... [  7520]
LS478113.1_Estonia_2014          ........------------------------------------------------------------..tgg.c.ggcc [  7520]
KP843857.1_Odintsovo_02/14       .........-------..................................-........------............... [  7520]
```

```
FN557520.1_strain_E75          ........aatgtttt.............................aca---.....a..------..a.a.....c.... [  7520]
AY261366.1_Warthog             .g---------ttttt..................................-........------............... [  7520]
AY261365.1_Warmbaths           ........aatctttt.g................................-........------............... [  7520]
AY261364.1_Tengani_62          ........-------t..................................-........------............... [  7520]
AY261363.1_Pretoriuskop/96/4   ........aatctttt......-....g......................-........------............... [  7520]
AY261362.1_Mkuzi_1979          ........aatgtttt.............................aca---.....a..------....a.....c.... [  7520]
AY261361.1_Malawi_Lil-20/1_1983 ........aatctttt.......g......a.g..........a......a........------....a....a..... [  7520]
AY261360.1_Kenya_1950          ........aatctttt...a.---------atgc.g.....a.a......agat.ac.tcggtac........a.....g [  7520]
AM712240.1_OURT_88/3           ........aatgtttt.............................aca---.....a..------..a.a.....c.... [  7520]
AM712239.1_Benin_97/1          ........aatgtttt.............................aca---.....a..------..a.a.....c.... [  7520]
KM111295.1_Ken06.Bus           ........aatctttt...a.---------atgc.g.....a.a......agataac.ttggtac........a.....g [  7520]
KM111294.1_Ken05/Tk1           ...a....agtctgtaa.ga.t----....a......c..a..aa.....g.......t---------...c.......g [  7520]
KM262844.1_strain_L60          ........aatgtttt.............................aca---.....a..------..a.a.....c.... [  7520]
KM102979.1_26544/OG10_Italy    ........aatgtttt.............................aca---.....a..------..a.a.....c.... [  7520]
KM262845.1_strain_NHV          ........aatgtttt.............................aca---.....a..------..a.a.....c.... [  7520]
KX354450.1_47/Ss/2008          ........aatgtttt.............................aca---.....a..------..a.a.....c.... [  7520]
KP055815.1_strain_BA71         ........aatgtttt.............................aca---.....a..------..a.a.....c.... [  7520]
MH025920.1_strain_R35          ........aatctttt...a.---------atgc.g.....a.a......agataac.ttggtac........a.....g [  7520]
MH025919.1_strain_N10          ........aatctttt...a.---------atgc.g.....a.a......agataac.ttggtac........a.....g [  7520]
MH025918.1_strain_R25          ........aatctttt...a.---------atgc.g.....a.a......agataac.ttggtac........a.....g [  7520]
MH025917.1_strain_R7           ........aatctttt...a.---------atgc.g.....a.a......agataac.ttggtac........a.....g [  7520]
MH025916.1_strain_R8           ........aatctttt...a.---------atgc.g.....a.a......agataac.ttggtac........a.....g [  7520]

FR682468.2_Georgia_2007/1      ttatgtagacata---aaaatgagatcctccaaaaa--aataaacaaca-aaaaaaaatatgtttaatattaaaatgaca [  7600]
ASF_IND_20_CAD543              ..........c..taa.........y..........at...........c.............................. [  7600]
LR899193.1_Germany_2020/1      .............---....................--...........-.............................. [  7600]
MN393477.1_Wuhan_2019          .............---....................--...........-.............................. [  7600]
MW396979.1_Timor-Leste/2019/1  .............---....................--...........-.............................. [  7600]
MK940252.1_CN/2019/InnerMongolia .............---....................--...........-.............................. [  7600]
MK128995.1_China/2018/AnhuiXCGQ .............---....................--...........-.............................. [  7600]
MH681419.1_ASFV/POL/2015       .............---....................--...........-.............................. [  7600]
LS478113.1_Estonia_2014        ...at..a...-----------------------------------------------...tac.g.cc.......t..t [  7600]
KP843857.1_Odintsovo_02/14     .............---....................--...........-.....-........................ [  7600]
FN557520.1_strain_E75          ..........c..taa.........c..........at...........c.............................. [  7600]
AY261366.1_Warthog             ..........c..taa.........c..........at...........c.............................. [  7600]
AY261365.1_Warmbaths           ..........c..taa.........c..........--...........-.............................. [  7600]
AY261364.1_Tengani_62          .............---.........c..........--...........-.............................. [  7600]
AY261363.1_Pretoriuskop/96/4   ..........c..taa.........c..........--...........-.............................g [  7600]
AY261362.1_Mkuzi_1979          ..........c..taa....................at...........c.............................. [  7600]
AY261361.1_Malawi_Lil-20/1_1983 ..c.......c..taa........cctgat..g...a-..a.t....a.-c............................. [  7600]
AY261360.1_Kenya_1950          ..........c..taa...g....-.ggc...c...--.........a.t.............ac............... [  7600]
AM712240.1_OURT_88/3           ..........c..taa.........c..........at...........c.............................. [  7600]
AM712239.1_Benin_97/1          ..........c..taa.........c..........at...........c.............................. [  7600]
KM111295.1_Ken06.Bus           ..........c..taa...g....-.ggc...c...--.....t...a.t.............ac............... [  7600]
```

```
KM111294.1_Ken05/Tk1           .....c...tc..taa-----------------...--.......g...c.............................. [  7600]
KM262844.1_strain_L60          ..........c..taa.........c..........at...........c.............................. [  7600]
KM102979.1_26544/OG10_Italy    ..........c..taa.........c..........at...........c.............................. [  7600]
KM262845.1_strain_NHV          ..........c..taa.........c..........at...........c.............................. [  7600]
KX354450.1_47/Ss/2008          ..........c..taa.........c..........at...........c.............................. [  7600]
KP055815.1_strain_BA71         ..........c..taa.........c..........at...........c.............................. [  7600]
MH025920.1_strain_R35          ..........c..taa...g....-.ggc...c...--.....t...a.t.............ac............... [  7600]
MH025919.1_strain_N10          ..........c..taa...g....-.ggc...c...--.....t...a.t.............ac............... [  7600]
MH025918.1_strain_R25          ..........c..taa...g....-.ggc...c...--.....t...a.t.............ac............... [  7600]
MH025917.1_strain_R7           ..........c..taa...g....-.ggc...c...--.....t...a.t.............ac............... [  7600]
MH025916.1_strain_R8           ..........c..taa...g....-.ggc...c...--.....t...a.t.............ac............... [  7600]

FR682468.2_Georgia_2007/1      atttctacattgctta---------ttgctcttattatactacttattattattttagtagtgtttttatactataagaa [  7680]
ASF_IND_20_CAD543              ................---------.................s...................r........y.....r.. [  7680]
LR899193.1_Germany_2020/1      ................---------....................................................... [  7680]
MN393477.1_Wuhan_2019          ................---------....................................................... [  7680]
MW396979.1_Timor-Leste/2019/1  ................---------....................................................... [  7680]
MK940252.1_CN/2019/InnerMongolia ................---------....................................................... [  7680]
MK128995.1_China/2018/AnhuiXCGQ ................---------....................................................... [  7680]
MH681419.1_ASFV/POL/2015       ................---------....................................................... [  7680]
LS478113.1_Estonia_2014        g-..g..gg.gttaa.---------..a.....gc.g..ggc....-a...gc-------a..c...gt.cta....... [  7680]
KP843857.1_Odintsovo_02/14     ................---------....................................................... [  7680]
FN557520.1_strain_E75          ................---------.................g................c..a........t.....a.. [  7680]
AY261366.1_Warthog             .....gt.........---------...........g........................................... [  7680]
AY261365.1_Warmbaths           .....gt.........---------...........g........................................... [  7680]
AY261364.1_Tengani_62          ................---------....................................................... [  7680]
AY261363.1_Pretoriuskop/96/4   .....gt.........---------...........g........................................... [  7680]
AY261362.1_Mkuzi_1979          ................---------.................g................c..a........t.....a.. [  7680]
AY261361.1_Malawi_Lil-20/1_1983 ..g.............ttgctctta...........g......................c..a........t.....a.. [  7680]
AY261360.1_Kenya_1950          .........c......---------...........g......................g..a........tc....... [  7680]
AM712240.1_OURT_88/3           ................---------.................g................c..a........t.....a.. [  7680]
AM712239.1_Benin_97/1          ................---------.................g................c..a........t.....a.. [  7680]
KM111295.1_Ken06.Bus           .........c......---------...........g......................g..a........t........ [  7680]
KM111294.1_Ken05/Tk1           ..g.............---------...........g.t..............ga....c..a........t.....a.. [  7680]
KM262844.1_strain_L60          ................---------.................g................c..a........t.....a.. [  7680]
KM102979.1_26544/OG10_Italy    ................---------.................g................c..a........t.....a.. [  7680]
KM262845.1_strain_NHV          ................---------.................g................c..a........t.....a.. [  7680]
KX354450.1_47/Ss/2008          ................---------.................g................c..a........t.....a.. [  7680]
KP055815.1_strain_BA71         ................---------.................g................c..a........t.....a.. [  7680]
MH025920.1_strain_R35          .........c......---------...........g......................g..a........t........ [  7680]
MH025919.1_strain_N10          .........c......---------...........g......................g..a........t........ [  7680]
MH025918.1_strain_R25          .........c......---------...........g......................g..a........t........ [  7680]
MH025917.1_strain_R7           .........c......---------...........g......................g..a........t........ [  7680]
MH025916.1_strain_R8           .........c......---------...........g......................g..a........t........ [  7680]
```

```
FR682468.2_Georgia_2007/1         acaacaaccaccgaaaaaggtctgtaaagtagataaagattgtggtagtggagagcattgtgttcgtggatcatgtagct [  7760]
ASF_IND_20_CAD543                 ............r................................................................... [  7760]
LR899193.1_Germany_2020/1         ................................................................................ [  7760]
MN393477.1_Wuhan_2019             ................................................................................ [  7760]
MW396979.1_Timor-Leste/2019/1     ................................................................................ [  7760]
MK940252.1_CN/2019/InnerMongolia  ................................................................................ [  7760]
MK128995.1_China/2018/AnhuiXCGQ   ................................................................................ [  7760]
MH681419.1_ASFV/POL/2015          ................................................................................ [  7760]
LS478113.1_Estonia_2014           .a..t..-ag..ctc.c.aac.c...ttt..t.c...a..ca.------------------------------------- [  7760]
KP843857.1_Odintsovo_02/14        ................................................................................ [  7760]
FN557520.1_strain_E75             ............a.........................................................a....c...a [  7760]
AY261366.1_Warthog                ............a................................................................... [  7760]
AY261365.1_Warmbaths              ............a................................................................... [  7760]
AY261364.1_Tengani_62             ................................................................................ [  7760]
AY261363.1_Pretoriuskop/96/4      ............a................................................................... [  7760]
AY261362.1_Mkuzi_1979             ............a.........................................................a....c.... [  7760]
AY261361.1_Malawi_Lil-20/1_1983   ............a....................................a....................a......... [  7760]
AY261360.1_Kenya_1950             g...........a.........................................................a......... [  7760]
AM712240.1_OURT_88/3              ............a.........................................................a....c...a [  7760]
AM712239.1_Benin_97/1             ............a.........................................................a....c...a [  7760]
KM111295.1_Ken06.Bus              g...........a......................................g..................a......... [  7760]
KM111294.1_Ken05/Tk1              ......................................................................a......... [  7760]
KM262844.1_strain_L60             ............a.........................................................a....c...a [  7760]
KM102979.1_26544/OG10_Italy       ............a.........................................................a....c...a [  7760]
KM262845.1_strain_NHV             ............a.........................................................a....c...a [  7760]
KX354450.1_47/Ss/2008             ............a.........................................................a....c...a [  7760]
KP055815.1_strain_BA71            ............a.........................................................a....c...a [  7760]
MH025920.1_strain_R35             g...........a......................................g..................a......... [  7760]
MH025919.1_strain_N10             g...........a......................................g..................a......... [  7760]
MH025918.1_strain_R25             g...........a......................................g..................a......... [  7760]
MH025917.1_strain_R7              g...........a......................................g..................a......... [  7760]
MH025916.1_strain_R8              g...........a......................................g..................a......... [  7760]

FR682468.2_Georgia_2007/1         cattgagctgcttagatgccgtaaaaatggacaaacgaaatattaagatagattctaagatttcctcatgcgaattcact [  7840]
ASF_IND_20_CAD543                 ......................................................n......................... [  7840]
LR899193.1_Germany_2020/1         ................................................................................ [  7840]
MN393477.1_Wuhan_2019             ................................................................................ [  7840]
MW396979.1_Timor-Leste/2019/1     ................................................................................ [  7840]
MK940252.1_CN/2019/InnerMongolia  ................................................................................ [  7840]
MK128995.1_China/2018/AnhuiXCGQ   ................................................................................ [  7840]
MH681419.1_ASFV/POL/2015          ................................................................................ [  7840]
LS478113.1_Estonia_2014           --------.a....t..aataa.gt.ggc.....t.a.c......--..ca.ccac.gc....ta.g.....-------- [  7840]
KP843857.1_Odintsovo_02/14        ................................................................................ [  7840]
```

```
FN557520.1_strain_E75            ................c..t..................................................t......... [  7840]
AY261366.1_Warthog               ................c..t.......c..............................c..................... [  7840]
AY261365.1_Warmbaths             ................c..t.......c..............................c..................... [  7840]
AY261364.1_Tengani_62            ................c..t.......c.................t.................................. [  7840]
AY261363.1_Pretoriuskop/96/4     ................c..t.......c.................................................... [  7840]
AY261362.1_Mkuzi_1979            ................c..t..............................................c............. [  7840]
AY261361.1_Malawi_Lil-20/1_1983  ...................t..................g..g.......g.....................a....t... [  7840]
AY261360.1_Kenya_1950            ...................t..................g..g.......g.....................a....t... [  7840]
AM712240.1_OURT_88/3             ................c..t..................................................t......... [  7840]
AM712239.1_Benin_97/1            ................c..t..................................................t......... [  7840]
KM111295.1_Ken06.Bus             ...................t..................g..g.......g.....................a....t... [  7840]
KM111294.1_Ken05/Tk1             ................c..t.......c..................a.............................t... [  7840]
KM262844.1_strain_L60            ................c..t..................................................t......... [  7840]
KM102979.1_26544/OG10_Italy      ................c..t..................................................t......... [  7840]
KM262845.1_strain_NHV            ................c..t..................................................t......... [  7840]
KX354450.1_47/Ss/2008            ................c..t..................................................t......... [  7840]
KP055815.1_strain_BA71           ................c..t..................................................t......... [  7840]
MH025920.1_strain_R35            ..................tt..................g..g.......g.....................a....t... [  7840]
MH025919.1_strain_N10            ..................tt..................g..g.......g.....................a....t... [  7840]
MH025918.1_strain_R25            ..................tt..................g..g.......g.....................a....t... [  7840]
MH025917.1_strain_R7             ..................tt..................g..g.......g.....................a....t... [  7840]
MH025916.1_strain_R8             ..................tt..................g..g.......g.....................a....t... [  7840]

FR682468.2_Georgia_2007/1        cccaatttttaccgttttacggatactgctgctgatgagcagcaagaatttggaaaaacacggcatcctataaaaataac [  7920]
ASF_IND_20_CAD543                ................................................................................ [  7920]
LR899193.1_Germany_2020/1        ................................................................................ [  7920]
MN393477.1_Wuhan_2019            ................................................................................ [  7920]
MW396979.1_Timor-Leste/2019/1    ................................................................................ [  7920]
MK940252.1_CN/2019/InnerMongolia ................................................................................ [  7920]
MK128995.1_China/2018/AnhuiXCGQ  ................................................................................ [  7920]
MH681419.1_ASFV/POL/2015         ................................................................................ [  7920]
LS478113.1_Estonia_2014          -t....caac.tat.a.....a.g.gc.tc-------------------------------------------------- [  7920]
KP843857.1_Odintsovo_02/14       ................................................................................ [  7920]
FN557520.1_strain_E75            ................................c............................................... [  7920]
AY261366.1_Warthog               ...................t......a.......................................t............. [  7920]
AY261365.1_Warmbaths             ...................t......a.......................................t............. [  7920]
AY261364.1_Tengani_62            ..........................a.......................................t............. [  7920]
AY261363.1_Pretoriuskop/96/4     ....................a.....a.....................................g............... [  7920]
AY261362.1_Mkuzi_1979            ..................................................................t............. [  7920]
AY261361.1_Malawi_Lil-20/1_1983  .............a....................------....................t.....t............. [  7920]
AY261360.1_Kenya_1950            .............a...............c....------....................t.....t............. [  7920]
AM712240.1_OURT_88/3             ................................c............................................... [  7920]
AM712239.1_Benin_97/1            ................................c............................................... [  7920]
KM111295.1_Ken06.Bus             .............a....................------....................t.....t............. [  7920]
```

```
KM111294.1_Ken05/Tk1             ..............................................................c................. [  7920]
KM262844.1_strain_L60            ................................c............................................... [  7920]
KM102979.1_26544/OG10_Italy      ................................c............................................... [  7920]
KM262845.1_strain_NHV            ................................c............................................... [  7920]
KX354450.1_47/Ss/2008            ................................c............................................... [  7920]
KP055815.1_strain_BA71           ................................c............................................... [  7920]
MH025920.1_strain_R35            .............a....................------....................t.....t............. [  7920]
MH025919.1_strain_N10            .............a....................------....................t.....t............. [  7920]
MH025918.1_strain_R25            .............a....................------....................t.....t............. [  7920]
MH025917.1_strain_R7             .............a....................------....................t.....t............. [  7920]
MH025916.1_strain_R8             .............a....................------....................t.....t............. [  7920]

FR682468.2_Georgia_2007/1        tccatctccaagtgaatcccatagcccccaagaggtgtgtgaaaaatattgttcatggggaaccgatgactgtacaggtt [  8000]
ASF_IND_20_CAD543                ................................................................................ [  8000]
LR899193.1_Germany_2020/1        ................................................................................ [  8000]
MN393477.1_Wuhan_2019            ................................................................................ [  8000]
MW396979.1_Timor-Leste/2019/1    ................................................................................ [  8000]
MK940252.1_CN/2019/InnerMongolia ................................................................................ [  8000]
MK128995.1_China/2018/AnhuiXCGQ  ................................................................................ [  8000]
MH681419.1_ASFV/POL/2015         ................................................................................ [  8000]
LS478113.1_Estonia_2014          ----------.a.c..c..t..tattgag..c..c.act...-------------------------c..c..gt.t.g. [  8000]
KP843857.1_Odintsovo_02/14       ................................................................................ [  8000]
FN557520.1_strain_E75            ................................................................................ [  8000]
AY261366.1_Warthog               .........g..................................g..................a................ [  8000]
AY261365.1_Warmbaths             ............................................g..................a................ [  8000]
AY261364.1_Tengani_62            .........g..................................g..................a................ [  8000]
AY261363.1_Pretoriuskop/96/4     .........g..................................g..................a................ [  8000]
AY261362.1_Mkuzi_1979            ................................................................................ [  8000]
AY261361.1_Malawi_Lil-20/1_1983  c.........g............c.t.......aa.a.......g........t.........a................ [  8000]
AY261360.1_Kenya_1950            c.........g............c.t.......aa.a.......g........t.........a................ [  8000]
AM712240.1_OURT_88/3             ................................................................................ [  8000]
AM712239.1_Benin_97/1            ................................................................................ [  8000]
KM111295.1_Ken06.Bus             c.........g.........g..c.t.......aa.a.......g........t.........a................ [  8000]
KM111294.1_Ken05/Tk1             ................c........t.t...................................a................ [  8000]
KM262844.1_strain_L60            ................................................................................ [  8000]
KM102979.1_26544/OG10_Italy      ................................................................................ [  8000]
KM262845.1_strain_NHV            ................................................................................ [  8000]
KX354450.1_47/Ss/2008            ................................................................................ [  8000]
KP055815.1_strain_BA71           ................................................................................ [  8000]
MH025920.1_strain_R35            c.........g............c.t.......aa.a.......g........t.........a................ [  8000]
MH025919.1_strain_N10            c.........g.........g..c.t.......aa.a.......g........t.........a................ [  8000]
MH025918.1_strain_R25            c.........g............c.t.......aa.a.......g........t.........a................ [  8000]
MH025917.1_strain_R7             c.........g............c.t.......aa.a.......g........t.........a................ [  8000]
MH025916.1_strain_R8             c.........g............c.t.......aa.a.......g........t.........a................ [  8000]
```

```
FR682468.2_Georgia_2007/1        gggaatatgttggtgatgaaaaggagggaacatgttatgtatataataatccacatcacccggttcttaaatatggtaag [  8080]
ASF_IND_20_CAD543                ................................................................................ [  8080]
LR899193.1_Germany_2020/1        ................................................................................ [  8080]
MN393477.1_Wuhan_2019            ................................................................................ [  8080]
MW396979.1_Timor-Leste/2019/1    ................................................................................ [  8080]
MK940252.1_CN/2019/InnerMongolia ................................................................................ [  8080]
MK128995.1_China/2018/AnhuiXCGQ  ................................................................................ [  8080]
MH681419.1_ASFV/POL/2015         ................................................................................ [  8080]
LS478113.1_Estonia_2014          ..cggc......t.tg.a-------------------------------------------------------------- [  8080]
KP843857.1_Odintsovo_02/14       ................................................................................ [  8080]
FN557520.1_strain_E75            ................................................................................ [  8080]
AY261366.1_Warthog               .........a....a.......................a...................t..a.................. [  8080]
AY261365.1_Warmbaths             .........a....a.......................a...................t..a.................. [  8080]
AY261364.1_Tengani_62            .........a....a.......................a...................t..a.................. [  8080]
AY261363.1_Pretoriuskop/96/4     .........ag...a.......................a...................t..a.................. [  8080]
AY261362.1_Mkuzi_1979            .........a....a.......................a...................t..................... [  8080]
AY261361.1_Malawi_Lil-20/1_1983  ........t..............a.t.........a..a..............t..---.ta.c.........cac...a [  8080]
AY261360.1_Kenya_1950            ........t................t.........a.........c.......t..---.ta.c.........cac...a [  8080]
AM712240.1_OURT_88/3             ................................................................................ [  8080]
AM712239.1_Benin_97/1            ................................................................................ [  8080]
KM111295.1_Ken06.Bus             ........t................t.........a..a......t..........---.ta.c.........cac...a [  8080]
KM111294.1_Ken05/Tk1             ......................a...............a.................---...tc................ [  8080]
KM262844.1_strain_L60            ................................................................................ [  8080]
KM102979.1_26544/OG10_Italy      ................................................................................ [  8080]
KM262845.1_strain_NHV            ................................................................................ [  8080]
KX354450.1_47/Ss/2008            ................................................................................ [  8080]
KP055815.1_strain_BA71           ................................................................................ [  8080]
MH025920.1_strain_R35            ........t................t.........a..a......t..........---.ta.c.........cac...a [  8080]
MH025919.1_strain_N10            ........t................t.........a..a......t..........---.ta.c.........cac...a [  8080]
MH025918.1_strain_R25            ........t................t.........a..a......t..........---.ta.c.........cac...a [  8080]
MH025917.1_strain_R7             ........t................t.........a..a......t..........---.ta.c.........cac...a [  8080]
MH025916.1_strain_R8             ........t................t.........a..a......t..........---.ta.c.........cac...a [  8080]

FR682468.2_Georgia_2007/1        gatcacatcatagccttacctagaaatcataaacatgcataaataaatac-attaggctcatcgtatcttttta--aaat [  8160]
ASF_IND_20_CAD543                ..................................................-.................c....t-a.... [  8160]
LR899193.1_Germany_2020/1        ..................................................-.......................--.... [  8160]
MN393477.1_Wuhan_2019            ..................................................-.......................--.... [  8160]
MW396979.1_Timor-Leste/2019/1    ..................................................-.......................--.... [  8160]
MK940252.1_CN/2019/InnerMongolia ..................................................-.......................--.... [  8160]
MK128995.1_China/2018/AnhuiXCGQ  ..................................................-.......................--.... [  8160]
MH681419.1_ASFV/POL/2015         ..................................................-.......................--.... [  8160]
LS478113.1_Estonia_2014          ------------------------------------------------------------.ct.c..t.....------. [  8160]
KP843857.1_Odintsovo_02/14       ..................................................-.......................--.... [  8160]
```

```
FN557520.1_strain_E75            ..................................................-.................c....t-a.... [  8160]
AY261366.1_Warthog               a.....g.t...........c.............................a............a..........-a.... [  8160]
AY261365.1_Warmbaths             a.....g.t...........c.............................a............a..........-a.... [  8160]
AY261364.1_Tengani_62            ......g.t...........c.............................a............a....c....taa.... [  8160]
AY261363.1_Pretoriuskop/96/4     ......g.t...........c.............................a.........tcaac..a.c....-a.... [  8160]
AY261362.1_Mkuzi_1979            ....................c........................c.----................-c....t-a.... [  8160]
AY261361.1_Malawi_Lil-20/1_1983  ......g..t..ta......c............t........gc.t.a.ta.a..ca---........c.....-a.... [  8160]
AY261360.1_Kenya_1950            ......g..t..ta......c............t....g...........a..g.....a..t.....c....--a.c.. [  8160]
AM712240.1_OURT_88/3             ..................................................-.................c....t-a.... [  8160]
AM712239.1_Benin_97/1            ..................................................-.................c....t-a.... [  8160]
KM111295.1_Ken06.Bus             ......g..t..ta......c............t....g...........a..g.....a..t.....c....--a.c.. [  8160]
KM111294.1_Ken05/Tk1             ......g.............c.................g.........tta.....ttaatgta.cct....a--a.... [  8160]
KM262844.1_strain_L60            ..................................................-.................c....t-a.... [  8160]
KM102979.1_26544/OG10_Italy      ..................................................-.................c....t-a.... [  8160]
KM262845.1_strain_NHV            ..................................................-.................c....t-a.... [  8160]
KX354450.1_47/Ss/2008            ..................................................-.................c....t-a.... [  8160]
KP055815.1_strain_BA71           ..................................................-.................c....t-a.... [  8160]
MH025920.1_strain_R35            ......g..t..ta......c............t....g...........a..g.....a..t.....c....--a.c.. [  8160]
MH025919.1_strain_N10            ......g..t..ta......c............t....g...........a..g.....a..t.....c....--a.c.. [  8160]
MH025918.1_strain_R25            ......g..t..ta......c............t....g...........a..g.....a..t.....c....--a.c.. [  8160]
MH025917.1_strain_R7             ......g..t..ta......c............t....g...........a..g.....a..t.....c....--a.c.. [  8160]
MH025916.1_strain_R8             ......g..t..ta......c............t....g...........a..g.....a..t.....c....--a.c.. [  8160]

FR682468.2_Georgia_2007/1        ccata-------------------aatattcg--tttgatatatgctgaaatttt--tataaaaaaaaataactatttcc [  8240]
ASF_IND_20_CAD543                .....-------------------........--.....................--....................... [  8240]
LR899193.1_Germany_2020/1        .....-------------------........--.....................--....................... [  8240]
MN393477.1_Wuhan_2019            .....-------------------........--.....................--....................... [  8240]
MW396979.1_Timor-Leste/2019/1    .....-------------------........--.....................--....................... [  8240]
MK940252.1_CN/2019/InnerMongolia .....-------------------........--.....................--....................... [  8240]
MK128995.1_China/2018/AnhuiXCGQ  .....-------------------........--.....................--....................... [  8240]
MH681419.1_ASFV/POL/2015         .....-------------------........--.....................--....................... [  8240]
LS478113.1_Estonia_2014          .g..c-------------------g.c.a.a.--..aag....a.a.acc..a..--a..-----------.g....a.. [  8240]
KP843857.1_Odintsovo_02/14       .....-------------------........--.....................--...-................... [  8240]
FN557520.1_strain_E75            .....-------------------........--.....................--......................t [  8240]
AY261366.1_Warthog               .....-------------------......atac.............a.......--...-..................t [  8240]
AY261365.1_Warmbaths             .....-------------------......atac.............a.......--......................t [  8240]
AY261364.1_Tengani_62            .....-------------------......ac--.............a.......--......................t [  8240]
AY261363.1_Pretoriuskop/96/4     .....-------------------.......t--.....................--......................t [  8240]
AY261362.1_Mkuzi_1979            .....-------------------.......t--....t................--...-..................t [  8240]
AY261361.1_Malawi_Lil-20/1_1983  .t...-------------------......tt--..ca......a..........--...g..................t [  8240]
AY261360.1_Kenya_1950            .t...aatattctttaacatctat......-t--...a.......t.........t-......................t [  8240]
AM712240.1_OURT_88/3             .....-------------------........--.....................--......................t [  8240]
AM712239.1_Benin_97/1            .....-------------------........--.....................--......................t [  8240]
KM111295.1_Ken06.Bus             .t...-------------------......-t--...a.......t.........t-...-..................t [  8240]
```

```
KM111294.1_Ken05/Tk1           .t...-------------------.......t--c.c..................ta...............t.....at [  8240]
KM262844.1_strain_L60          .....-------------------........--.....................--......................t [  8240]
KM102979.1_26544/OG10_Italy    .....-------------------........--.....................--......................t [  8240]
KM262845.1_strain_NHV          .....-------------------........--.....................--......................t [  8240]
KX354450.1_47/Ss/2008          .....-------------------........--.....................--......................t [  8240]
KP055815.1_strain_BA71         .....-------------------........--.....................--......................t [  8240]
MH025920.1_strain_R35          .t...-------------------......-t--...a.......t.........t-...-..................t [  8240]
MH025919.1_strain_N10          .t...-------------------......-t--...a.......t.........t-...-..................t [  8240]
MH025918.1_strain_R25          .t...-------------------......-t--...a.......t.........t-...-..................t [  8240]
MH025917.1_strain_R7           .t...-------------------......-t--...a.......t.........t-...-..................t [  8240]
MH025916.1_strain_R8           .t...-------------------......-t--...a.......t.........t-...-..................t [  8240]

FR682468.2_Georgia_2007/1      tataaatcatctagaaatagt-cctcgtt----------ttgatcggtttatatcttataatattgtgcatcgatgcaca [  8320]
ASF_IND_20_CAD543              .....................-.......----------......................................... [  8320]
LR899193.1_Germany_2020/1      .....................-.......----------......................................... [  8320]
MN393477.1_Wuhan_2019          .....................-.......----------......................................... [  8320]
MW396979.1_Timor-Leste/2019/1  .....................-.......----------......................................... [  8320]
MK940252.1_CN/2019/InnerMongolia .....................-.......----------......................................... [  8320]
MK128995.1_China/2018/AnhuiXCGQ .....................-.......----------......................................... [  8320]
MH681419.1_ASFV/POL/2015       .....................-.......----------......................................... [  8320]
LS478113.1_Estonia_2014        ag..--------------------------------------------------t.g.c.....ga..t.---------- [  8320]
KP843857.1_Odintsovo_02/14     .....................-.......----------......................................... [  8320]
FN557520.1_strain_E75          .....................-.......----------......................................... [  8320]
AY261366.1_Warthog             ...................t.c...a...agaatcgct-..a..aa........-------................... [  8320]
AY261365.1_Warmbaths           ...................t.c...a...agaatcgct-..a..aa........-------................... [  8320]
AY261364.1_Tengani_62          ...................t.c...a...agaatcgct-..a..aa........-------................... [  8320]
AY261363.1_Pretoriuskop/96/4   .............a.....c.-...aa..agaatc-tt-.....aaa.......-------................... [  8320]
AY261362.1_Mkuzi_1979          ...................c.-...aa..agaatctttac....aa........a.....t................... [  8320]
AY261361.1_Malawi_Lil-20/1_1983 ........----.a...c.ta---------------------------......-------.........a........g [  8320]
AY261360.1_Kenya_1950          ..........t........cc-...aa..agaaactttac....at........-------..................g [  8320]
AM712240.1_OURT_88/3           .....................-.......----------......................................... [  8320]
AM712239.1_Benin_97/1          .....................-.......----------......................................... [  8320]
KM111295.1_Ken06.Bus           ..........t........cc-...aa..agaaactttac....at........-------..................g [  8320]
KM111294.1_Ken05/Tk1           .....t............-----------------tttaca...aaa.......-------c........c........g [  8320]
KM262844.1_strain_L60          .....................-.......----------......................................... [  8320]
KM102979.1_26544/OG10_Italy    .....................-.......----------......................................... [  8320]
KM262845.1_strain_NHV          .....................-.......----------......................................... [  8320]
KX354450.1_47/Ss/2008          .....................-.......----------......................................... [  8320]
KP055815.1_strain_BA71         .....................-.......----------......................................... [  8320]
MH025920.1_strain_R35          ..........t........cc-...aa..agaaactttac....at........-------..................g [  8320]
MH025919.1_strain_N10          ..........t........cc-...aa..agaaactttac....at........-------..................g [  8320]
MH025918.1_strain_R25          ..........t........cc-...aa..agaaactttac....at........-------..................g [  8320]
MH025917.1_strain_R7           ..........t........cc-...aa..agaaactttac....at........-------..................g [  8320]
MH025916.1_strain_R8           ..........t........cc-...aa..agaaactttac....at........-------..................g [  8320]
```

```
FR682468.2_Georgia_2007/1        actgcttttttt------ggtccttctggaacatcattatattttctttcattaatataccattcagatgtaaacgttga [  8400]
ASF_IND_20_CAD543                ............------........................................................y..... [  8400]
LR899193.1_Germany_2020/1        ............------.............................................................. [  8400]
MN393477.1_Wuhan_2019            ............------.............................................................. [  8400]
MW396979.1_Timor-Leste/2019/1    ............------.............................................................. [  8400]
MK940252.1_CN/2019/InnerMongolia ............------.............................................................. [  8400]
MK128995.1_China/2018/AnhuiXCGQ  ............------.............................................................. [  8400]
MH681419.1_ASFV/POL/2015         ............------.............................................................. [  8400]
LS478113.1_Estonia_2014          -----------------------------------gc....g..ct.c..cc.g...gt.-------------------- [  8400]
KP843857.1_Odintsovo_02/14       ............------.............................................................. [  8400]
FN557520.1_strain_E75            ............------........................................................t..... [  8400]
AY261366.1_Warthog               t...........------..a........................................................... [  8400]
AY261365.1_Warmbaths             t...........------..a........................................................... [  8400]
AY261364.1_Tengani_62            t...........------..a........................................................... [  8400]
AY261363.1_Pretoriuskop/96/4     t...........------..a........c................................c...........t..... [  8400]
AY261362.1_Mkuzi_1979            ....t.......------..a.....................................................t..... [  8400]
AY261361.1_Malawi_Lil-20/1_1983  ....a.......ggacta..a.......c..t.....c...............c................a...t..... [  8400]
AY261360.1_Kenya_1950            .t..a.......gcacta..a...............a....a........t....c..c.....g..gc.c...t..... [  8400]
AM712240.1_OURT_88/3             ............------........................................................t..... [  8400]
AM712239.1_Benin_97/1            ............------...........................t............................t..... [  8400]
KM111295.1_Ken06.Bus             .t..a.......gcacta..a..........t....a....a.............c..c.....g..gc.c...t..... [  8400]
KM111294.1_Ken05/Tk1             ....a......gggacca..a.......c..t......................................a...t..... [  8400]
KM262844.1_strain_L60            ............------........................................................t..... [  8400]
KM102979.1_26544/OG10_Italy      ............------........................................................t..... [  8400]
KM262845.1_strain_NHV            ............------........................................................t..... [  8400]
KX354450.1_47/Ss/2008            ............------........................................................t..... [  8400]
KP055815.1_strain_BA71           ............------........................................................t..... [  8400]
MH025920.1_strain_R35            .t..a.......gcacta..a..........t....a....a.............c..c.....g..gc.c...t..... [  8400]
MH025919.1_strain_N10            .t..a.......gcacta..a..........t....a....a.............c..c.....g..gc.c...t..... [  8400]
MH025918.1_strain_R25            .t..a.......gcacta..a..........t....a....a.............c..c.....g..gc.c...t..... [  8400]
MH025917.1_strain_R7             .t..a.......gcacta..a..........t....a....a.............c..c.....g..gc.c...t..... [  8400]
MH025916.1_strain_R8             .t..a.......gcacta..a..........t....a....a.............c..c.....g..gc.c...t..... [  8400]

FR682468.2_Georgia_2007/1        ataattttatggcaacaatctaccattgaattatatttagtaacatctaatacatcgtttgttttatcaggctcagctc [  8480]
ASF_IND_20_CAD543                ...............................................................................y [  8480]
LR899193.1_Germany_2020/1        ................................................................................ [  8480]
MN393477.1_Wuhan_2019            ................................................................................ [  8480]
MW396979.1_Timor-Leste/2019/1    ................................................................................ [  8480]
MK940252.1_CN/2019/InnerMongolia ................................................................................ [  8480]
MK128995.1_China/2018/AnhuiXCGQ  ................................................................................ [  8480]
MH681419.1_ASFV/POL/2015         ................................................................................ [  8480]
LS478113.1_Estonia_2014          -------------........c.t-----gg..gc.....aa..t..--------------------------------- [  8480]
KP843857.1_Odintsovo_02/14       ................................................................................ [  8480]
```

```
FN557520.1_strain_E75           ..........................c...............................c....................t [  8480]
AY261366.1_Warthog              ...................................g..............g.......c.............t...t..t [  8480]
AY261365.1_Warmbaths            ...................................g..............g.......c.............t...t..t [  8480]
AY261364.1_Tengani_62           ............a.....g......g......c.........................c..........g......t..t [  8480]
AY261363.1_Pretoriuskop/96/4    ..................g.......................................c.................t..t [  8480]
AY261362.1_Mkuzi_1979           ............a...........t..........................a......c.................t..t [  8480]
AY261361.1_Malawi_Lil-20/1_1983 ...................................g....a..g...t..t......a...a.g...c...a....t..t [  8480]
AY261360.1_Kenya_1950           ..................c........................t.............ac..a..............t..t [  8480]
AM712240.1_OURT_88/3            ..........................c...............................c....................t [  8480]
AM712239.1_Benin_97/1           ..........................c...............................c....................t [  8480]
KM111295.1_Ken06.Bus            ..................c.........a..............t.g...........ac..a..............t..t [  8480]
KM111294.1_Ken05/Tk1            ...................................t.......gtt....t......a...a.........a....t..t [  8480]
KM262844.1_strain_L60           ..........................c...............................c....................t [  8480]
KM102979.1_26544/OG10_Italy     ..........................c...............................c....................t [  8480]
KM262845.1_strain_NHV           ..........................c...............................c....................t [  8480]
KX354450.1_47/Ss/2008           ..........................c...............................c....................t [  8480]
KP055815.1_strain_BA71          ..........................c...............................c....................t [  8480]
MH025920.1_strain_R35           ..................c.........a..............t.g...........ac..a..............t..t [  8480]
MH025919.1_strain_N10           ..................c.........a..............t.g...........ac..a..............t..t [  8480]
MH025918.1_strain_R25           ..................c.........a..............t.g...........ac..a..............t..t [  8480]
MH025917.1_strain_R7            ..................c.........a..............t.g...........ac..a..............t..t [  8480]
MH025916.1_strain_R8            ..................c.........a..............t.g...........ac..a..............t..t [  8480]

FR682468.2_Georgia_2007/1       tataatcttgataa-tttttgttatcagcttctaaagctccatcattatttttcaaagaagtatccataattatgtttgg [  8560]
ASF_IND_20_CAD543               ..............-................................................................. [  8560]
LR899193.1_Germany_2020/1       ..............-................................................................. [  8560]
MN393477.1_Wuhan_2019           ..............-................................................................. [  8560]
MW396979.1_Timor-Leste/2019/1   ..............-................................................................. [  8560]
MK940252.1_CN/2019/InnerMongolia ..............-................................................................. [  8560]
MK128995.1_China/2018/AnhuiXCGQ ..............-................................................................. [  8560]
MH681419.1_ASFV/POL/2015        ..............-................................................................. [  8560]
LS478113.1_Estonia_2014         -------------------------------------------------------------------------------- [  8560]
KP843857.1_Odintsovo_02/14      ..............t................................................................. [  8560]
FN557520.1_strain_E75           ..............-...............................................g........c........ [  8560]
AY261366.1_Warthog              ..............-................................................................. [  8560]
AY261365.1_Warmbaths            ..............-................................................................. [  8560]
AY261364.1_Tengani_62           ..............-..............a........a......................................... [  8560]
AY261363.1_Pretoriuskop/96/4    ..............-..............a........a........................................a [  8560]
AY261362.1_Mkuzi_1979           .g..g.........-..............a.................................................a [  8560]
AY261361.1_Malawi_Lil-20/1_1983 ....g.........-.................cg.g..........................g.....g........a.a [  8560]
AY261360.1_Kenya_1950           ....g.t.......-.......................c....t...c....................g..g.......a [  8560]
AM712240.1_OURT_88/3            ..............-...............................................g........c........ [  8560]
AM712239.1_Benin_97/1           ..............-...............................................g........c........ [  8560]
KM111295.1_Ken06.Bus            ....g.t.......-.....................................................g..g.......a [  8560]
```

```
KM111294.1_Ken05/Tk1             ....g.........-........c...a........a.ata..................g........g..g..ac..a. [  8560]
KM262844.1_strain_L60            ..............-...............................................g........c........ [  8560]
KM102979.1_26544/OG10_Italy      ..............-...............................................g........c........ [  8560]
KM262845.1_strain_NHV            ..............-...............................................g........c........ [  8560]
KX354450.1_47/Ss/2008            ..............-...............................................g........c........ [  8560]
KP055815.1_strain_BA71           ..............-...............................................g........c........ [  8560]
MH025920.1_strain_R35            ....g.t.......-.....................................................g..g.......a [  8560]
MH025919.1_strain_N10            ....g.t.......-.....................................................g..g.......a [  8560]
MH025918.1_strain_R25            ....g.t.......-.....................................................g..g.......a [  8560]
MH025917.1_strain_R7             ....g.t.......-.....................................................g..g.......a [  8560]
MH025916.1_strain_R8             ....g.t.......-.....................................................g..g.......a [  8560]

FR682468.2_Georgia_2007/1        taaaaatactttaagttttaatgt--gatatttaaaatggttgttatataaatttaccgcttacaggt------------ [  8640]
ASF_IND_20_CAD543                ......y...............r.--..........................................------------ [  8640]
LR899193.1_Germany_2020/1        ........................--..........................................------------ [  8640]
MN393477.1_Wuhan_2019            ........................--..........................................------------ [  8640]
MW396979.1_Timor-Leste/2019/1    ........................--..........................................------------ [  8640]
MK940252.1_CN/2019/InnerMongolia ........................--..........................................------------ [  8640]
MK128995.1_China/2018/AnhuiXCGQ  ........................--..........................................------------ [  8640]
MH681419.1_ASFV/POL/2015         ........................--..........................................------------ [  8640]
LS478113.1_Estonia_2014          ......a..a..........-----------------------------------ga.aaca...aa.------------ [  8640]
KP843857.1_Odintsovo_02/14       ........................--..........................................------------ [  8640]
FN557520.1_strain_E75            ......c...............a.--..........................................------------ [  8640]
AY261366.1_Warthog               ........................aa..........................................------------ [  8640]
AY261365.1_Warmbaths             ........................aa............a---..c.....................a.---------aac [  8640]
AY261364.1_Tengani_62            ........................aa.g......................................a.---------aac [  8640]
AY261363.1_Pretoriuskop/96/4     c.......................aaa............---........................a.---------atc [  8640]
AY261362.1_Mkuzi_1979            ........................--a.g.c--------------------..............aa.aaaacttataac [  8640]
AY261361.1_Malawi_Lil-20/1_1983  .c...................ac.aa..c.a........a.....g............a..c...aa.aaaacttataac [  8640]
AY261360.1_Kenya_1950            .....................a.aaa..c......c..a.....a---.....g...ta..c...aa.aatacttatatc [  8640]
AM712240.1_OURT_88/3             ......c...............a.--..........................................------------ [  8640]
AM712239.1_Benin_97/1            ......c...............a.--..........................................------------ [  8640]
KM111295.1_Ken06.Bus             .....................a.aaa..c......c..aa.....---.....g...ta..c...aa.aatacctatatc [  8640]
KM111294.1_Ken05/Tk1             .....................gt.aa..c..........a.....g............at.c...aa.aaaacttataac [  8640]
KM262844.1_strain_L60            ......c...............a.--..........................................------------ [  8640]
KM102979.1_26544/OG10_Italy      ......c...............a.--..........................................------------ [  8640]
KM262845.1_strain_NHV            ......c...............a.--..........................................------------ [  8640]
KX354450.1_47/Ss/2008            ......c...............a.--..........................................------------ [  8640]
KP055815.1_strain_BA71           ......c...............a.--..........................................------------ [  8640]
MH025920.1_strain_R35            .....................a.aaa..c......c..aa.....---.....g...ta..c...aa.aatacctatatc [  8640]
MH025919.1_strain_N10            .....................a.aaa..c......c..aa.....---.....g...ta..c...aa.aatacctatatc [  8640]
MH025918.1_strain_R25            .....................a.aaa..c......c..aa.....---.....g...ta..c...aa.aatacctatatc [  8640]
MH025917.1_strain_R7             .....................a.aaa..c......c..aa.....---.....g...ta..c...aa.aatacctatatc [  8640]
MH025916.1_strain_R8             .....................a.aaa..c......c..aa.....---.....g...ta..c...aa.aatacctatatc [  8640]
```

```
FR682468.2_Georgia_2007/1        ----------------aatctttattcagtgtcataaactatacttttga------------------------------ [  8720]
ASF_IND_20_CAD543                ----------------..................................------------------------------ [  8720]
LR899193.1_Germany_2020/1        ----------------..................................------------------------------ [  8720]
MN393477.1_Wuhan_2019            ----------------..................................------------------------------ [  8720]
MW396979.1_Timor-Leste/2019/1    ----------------..................................------------------------------ [  8720]
MK940252.1_CN/2019/InnerMongolia ----------------..................................------------------------------ [  8720]
MK128995.1_China/2018/AnhuiXCGQ  ----------------..................................------------------------------ [  8720]
MH681419.1_ASFV/POL/2015         ----------------..................................------------------------------ [  8720]
LS478113.1_Estonia_2014          ----------------.g.t.gccgc.g--.c.gc.......tt..c--------------------------------- [  8720]
KP843857.1_Odintsovo_02/14       ----------------..................................------------------------------ [  8720]
FN557520.1_strain_E75            ----------------..............a...................tggttcagtattctgtaaatcggtatttt- [  8720]
AY261366.1_Warthog               ----------------..........t.......................tgattcagtattctg--------------- [  8720]
AY261365.1_Warmbaths             attacatat-------..........t.........g.............tgattcagtgttc----------------- [  8720]
AY261364.1_Tengani_62            attacatat-------..................a.g.............tgattcattgttctc--------------- [  8720]
AY261363.1_Pretoriuskop/96/4     attacatat-------....................g.............tgattcagtgttctc--------------- [  8720]
AY261362.1_Mkuzi_1979            attacatataattgcc....................g.a...........------------------------------ [  8720]
AY261361.1_Malawi_Lil-20/1_1983  attacatataattgcc.t..g.....t.a.c.....g.a.........tctgaatcagtattttctgaatcagtattttc [  8720]
AY261360.1_Kenya_1950            attacatataaatgcc............a..c...gg.a.......c...tgagtcagtattctgtgagtcagtattct- [  8720]
AM712240.1_OURT_88/3             ----------------..............a...................tggttcagtattctgtaaatcggtatttt- [  8720]
AM712239.1_Benin_97/1            ----------------..............a...................tggttcagtattctgtaaatcggtatttt- [  8720]
KM111295.1_Ken06.Bus             attacatatagatgcc............a..ct...g.a...........tgaatcagtattctgtgagtcagtatctt- [  8720]
KM111294.1_Ken05/Tk1             attacatataattgcc.t..g.......a.c.....g.a.......c...tggttcaacattctgtgaatcagtattc-- [  8720]
KM262844.1_strain_L60            ----------------..............a...................tggttcagtattctgtaaatcggtatttt- [  8720]
KM102979.1_26544/OG10_Italy      ----------------..............a...................tggttcagtattctgtaaatcggtatttt- [  8720]
KM262845.1_strain_NHV            ----------------..............a...................tggttcagtattctgtaaatcggtatttt- [  8720]
KX354450.1_47/Ss/2008            ----------------..............a...................tggttcagtattctgtaaatcggtatttt- [  8720]
KP055815.1_strain_BA71           ----------------..............a...................tggttcagtattctgtaaatcggtatttt- [  8720]
MH025920.1_strain_R35            attacatatagatgcc.......g....a..ct...g.a...........tgaatcagtattctgtgagtcagtatctt- [  8720]
MH025919.1_strain_N10            attacatatagatgcc............a..ct...g.a...........tgaatcagtattctgtgagtcagtatctt- [  8720]
MH025918.1_strain_R25            attacatatagatgcc.......g....a..ct...g.a...........tgaatcagtattctgtgagtcagtatctt- [  8720]
MH025917.1_strain_R7             attacatatagatgcc.......g....a..ct...g.a...........tgaatcagtattctgtgagtcagtatctt- [  8720]
MH025916.1_strain_R8             attacatatagatgcc.......g....a..ct...g.a...........tgaatcagtattctgtgagtcagtatctt- [  8720]

FR682468.2_Georgia_2007/1        ------------------------------tgat---------------tcagtattt---------------------- [  8800]
ASF_IND_20_CAD543                ------------------------------....tcagtattttgtaaa.........---------------------- [  8800]
LR899193.1_Germany_2020/1        ------------------------------....---------------.........---------------------- [  8800]
MN393477.1_Wuhan_2019            ------------------------------....---------------.........---------------------- [  8800]
MW396979.1_Timor-Leste/2019/1    ------------------------------....---------------.........---------------------- [  8800]
MK940252.1_CN/2019/InnerMongolia ------------------------------....---------------.........---------------------- [  8800]
MK128995.1_China/2018/AnhuiXCGQ  ------------------------------....---------------.........---------------------- [  8800]
MH681419.1_ASFV/POL/2015         ------------------------------....---------------.........---------------------- [  8800]
LS478113.1_Estonia_2014          ----------------------------------------------------..g...---------------------- [  8800]
KP843857.1_Odintsovo_02/14       ------------------------------....---------------.........---------------------- [  8800]
```

```
FN557520.1_strain_E75           -----------------------------g....tcagtattctgtaaa..g......---------------------- [  8800]
AY261366.1_Warthog              ------------------------------.a.atcggtattttgtgat..g......---------------------- [  8800]
AY261365.1_Warmbaths            -------------------------------------------tctgat..g......---------------------- [  8800]
AY261364.1_Tengani_62           ------------------------------....tcggtattttgtgaa.........---------------------- [  8800]
AY261363.1_Pretoriuskop/96/4    ------------------------------....tcggtattttgtgaa.........---------------------- [  8800]
AY261362.1_Mkuzi_1979           ------------------------------....tcagcattctgcgat....c...c---------------------- [  8800]
AY261361.1_Malawi_Lil-20/1_1983 tgaatcagtgttctgtgaatcagtattctgcagatcagtattctgcgga.....g..c---------------------- [  8800]
AY261360.1_Kenya_1950           -----------------------------g...atcggtatgctttgag........c---------------------- [  8800]
AM712240.1_OURT_88/3            -----------------------------g....tcagtattctgtaaa..g......---------------------- [  8800]
AM712239.1_Benin_97/1           -----------------------------g....tcagtattctgtaaa..g......---------------------- [  8800]
KM111295.1_Ken06.Bus            -----------------------------g...gtcagtattctgtgag........ctgtgagtcagtattctgtgaat [  8800]
KM111294.1_Ken05/Tk1            -------------------------------------------------------------------------------- [  8800]
KM262844.1_strain_L60           -----------------------------g....tcagtattctgtaaa..g......---------------------- [  8800]
KM102979.1_26544/OG10_Italy     -----------------------------g....tcagtattctgtaaa..g......---------------------- [  8800]
KM262845.1_strain_NHV           -----------------------------g....tcagtattctgtaaa..g......---------------------- [  8800]
KX354450.1_47/Ss/2008           -----------------------------g....tcagtattctgtaaa..g......---------------------- [  8800]
KP055815.1_strain_BA71          -----------------------------g....tcagtattctgtaaa..g......---------------------- [  8800]
MH025920.1_strain_R35           -----------------------------g...gtcagtattctgtgag........ctgcgagtcagtattctgtgaat [  8800]
MH025919.1_strain_N10           -----------------------------g...gtcagtattctgtgag........ctgcgagtcagtattctgtgaat [  8800]
MH025918.1_strain_R25           -----------------------------g...gtcagtattctgtgag........ctgcgagtcagtattctgtgaat [  8800]
MH025917.1_strain_R7            -----------------------------g...gtcagtattctgtgag........ctgcgagtcagtattctgtgaat [  8800]
MH025916.1_strain_R8            -----------------------------g...gtcagtattctgtgag........ctgcgagtcagtattctgtgaat [  8800]

FR682468.2_Georgia_2007/1       --------tgtgaatcagtacatttattatcattaatatttttaggctgtttttccaatgttttattgttgcaatgagcc [  8880]
ASF_IND_20_CAD543               --------........................................................................ [  8880]
LR899193.1_Germany_2020/1       --------........................................................................ [  8880]
MN393477.1_Wuhan_2019           --------........................................................................ [  8880]
MW396979.1_Timor-Leste/2019/1   --------........................................................................ [  8880]
MK940252.1_CN/2019/InnerMongolia --------........................................................................ [  8880]
MK128995.1_China/2018/AnhuiXCGQ --------........................................................................ [  8880]
MH681419.1_ASFV/POL/2015        --------........................................................................ [  8880]
LS478113.1_Estonia_2014         --------.t..t.ct.ta.a..----------------------a..t.....gg.g..ggc.gc..c.at.ggcga.t [  8880]
KP843857.1_Odintsovo_02/14      --------........................................................................ [  8880]
FN557520.1_strain_E75           --------.........................................a.............................. [  8880]
AY261366.1_Warthog              --------........................................................................ [  8880]
AY261365.1_Warmbaths            --------........................................................................ [  8880]
AY261364.1_Tengani_62           --------....t...g............................................................... [  8880]
AY261363.1_Pretoriuskop/96/4    --------........ga......................................................------.. [  8880]
AY261362.1_Mkuzi_1979           --------...............................................t.g...................... [  8880]
AY261361.1_Malawi_Lil-20/1_1983 --------....g....a................................................c............. [  8880]
AY261360.1_Kenya_1950           --------........g................c....a.......t..a......g..........a............ [  8880]
AM712240.1_OURT_88/3            --------.........................................a.............................. [  8880]
AM712239.1_Benin_97/1           --------.........................................a.............................. [  8880]
KM111295.1_Ken06.Bus            cggtattt.........................c....a.......t..a......g....................... [  8880]
```

```
KM111294.1_Ken05/Tk1            --------..ctc...........................................g.........c............. [  8880]
KM262844.1_strain_L60           --------.........................................a.............................. [  8880]
KM102979.1_26544/OG10_Italy     --------.........................................a.............................. [  8880]
KM262845.1_strain_NHV           --------.........................................a.............................. [  8880]
KX354450.1_47/Ss/2008           --------.........................................a.............................. [  8880]
KP055815.1_strain_BA71          --------.........................................a.............................. [  8880]
MH025920.1_strain_R35           cggtattt.........................c....a.......t..a......g....................... [  8880]
MH025919.1_strain_N10           cggtattt.........................c....a.......t..a......g....................... [  8880]
MH025918.1_strain_R25           cggtattt.........................c....a.......t..a......g....................... [  8880]
MH025917.1_strain_R7            cggtattt.........................c....a.......t..a......g....................... [  8880]
MH025916.1_strain_R8            cggtattt.........................c....a.......t..a......g....................... [  8880]

FR682468.2_Georgia_2007/1       tgctcctcctttgacgaggaag---tgtctgttggagtcatctgtttaggaagagtatcatccatatctattatgaagaa [  8960]
ASF_IND_20_CAD543               ......................---....................................................... [  8960]
LR899193.1_Germany_2020/1       ......................---....................................................... [  8960]
MN393477.1_Wuhan_2019           ......................---....................................................... [  8960]
MW396979.1_Timor-Leste/2019/1   ......................---....................................................... [  8960]
MK940252.1_CN/2019/InnerMongolia ......................---....................................................... [  8960]
MK128995.1_China/2018/AnhuiXCGQ ......................---....................................................... [  8960]
MH681419.1_ASFV/POL/2015        ......................---....................................................... [  8960]
LS478113.1_Estonia_2014         .------------------------------------.a....a-------------------...at....t..g..-- [  8960]
KP843857.1_Odintsovo_02/14      ......................---....................................................... [  8960]
FN557520.1_strain_E75           ............a...g.....---....................................................... [  8960]
AY261366.1_Warthog              ................g.....tat.a................................g.................... [  8960]
AY261365.1_Warmbaths            ................ga....tat.a..................................................... [  8960]
AY261364.1_Tengani_62           ......................tat.a..................................................... [  8960]
AY261363.1_Pretoriuskop/96/4    .............g..g.....---....................................................... [  8960]
AY261362.1_Mkuzi_1979           ..t.............g.a...---.........a..a.....c.................................... [  8960]
AY261361.1_Malawi_Lil-20/1_1983 ..t.........a.t.ga.g..tag.a.......a..a...t............a.g....................... [  8960]
AY261360.1_Kenya_1950           ..t........ga...g.ag..---.c.......a..a...........a..........c.........c....gt... [  8960]
AM712240.1_OURT_88/3            ............a...g.....---....................................................... [  8960]
AM712239.1_Benin_97/1           ............a...g.....---....................................................... [  8960]
KM111295.1_Ken06.Bus            ..t........ga.....ag..---.c.......a..a...........a..........c.........c....gt... [  8960]
KM111294.1_Ken05/Tk1            ............a.t.ga.g..---.a.......a..a..................g....................... [  8960]
KM262844.1_strain_L60           ............a...g.....---....................................................... [  8960]
KM102979.1_26544/OG10_Italy     ............a...g.....---....................................................... [  8960]
KM262845.1_strain_NHV           ............a...g.....---....................................................... [  8960]
KX354450.1_47/Ss/2008           ............a...g.....---....................................................... [  8960]
KP055815.1_strain_BA71          ............a...g.....---....................................................... [  8960]
MH025920.1_strain_R35           ..t........ga.....ag..---.c.......a..a...........a..........c.........c....gt... [  8960]
MH025919.1_strain_N10           ..t........ga.....ag..---.c.......a..a...........a..........c.........c....gt... [  8960]
MH025918.1_strain_R25           ..t........ga.....ag..---.c.......a..a...........a..........c.........c....gt... [  8960]
MH025917.1_strain_R7            ..t........ga.....ag..---.c.......a..a...........a..........c.........c....gt... [  8960]
MH025916.1_strain_R8            ..t........ga.....ag..---.c.......a..a...........a..........c.........c....gt... [  8960]
```

```
FR682468.2_Georgia_2007/1        aatata-taaatattgatatacaatcaaaaatatttttgatcacgtctttgttatctatcgatattgttgataacgtctt [  9040]
ASF_IND_20_CAD543                ......-.......................................................................... [  9040]
LR899193.1_Germany_2020/1        ......-.......................................................................... [  9040]
MN393477.1_Wuhan_2019            ......-.......................................................................... [  9040]
MW396979.1_Timor-Leste/2019/1    ......-.......................................................................... [  9040]
MK940252.1_CN/2019/InnerMongolia ......-.......................................................................... [  9040]
MK128995.1_China/2018/AnhuiXCGQ  ......-.......................................................................... [  9040]
MH681419.1_ASFV/POL/2015         ......-.......................................................................... [  9040]
LS478113.1_Estonia_2014          ----------------------------------c......--------------------------------------- [  9040]
KP843857.1_Odintsovo_02/14       ......-.......................................................................... [  9040]
FN557520.1_strain_E75            ......-.......................................................................... [  9040]
AY261366.1_Warthog               ......t...g...................................................c..........c...... [  9040]
AY261365.1_Warmbaths             ......t...g...................................................c................. [  9040]
AY261364.1_Tengani_62            ......t...g.............at....................................c......a.....a---- [  9040]
AY261363.1_Pretoriuskop/96/4     ......t...g.ta.......t...........--.......g...................c...a............. [  9040]
AY261362.1_Mkuzi_1979            ..c...t...g.t.....tgtat.......c............t......a...........c................. [  9040]
AY261361.1_Malawi_Lil-20/1_1983  ..c...t...g.t........--.....g........................t........c............a..c. [  9040]
AY261360.1_Kenya_1950            ..c...t...g.t..c.c.c.tc...........a....................t....a.c...a.....-..a..c. [  9040]
AM712240.1_OURT_88/3             ......-.......................................................................... [  9040]
AM712239.1_Benin_97/1            ......-.......................................................................... [  9040]
KM111295.1_Ken06.Bus             ..c...t...g.t..c.c...ac...........a....................t....a.c...a........a..c. [  9040]
KM111294.1_Ken05/Tk1             ..c...t...g.t..a...c.--.....g............t.t...--.a..t.t.............a..-------- [  9040]
KM262844.1_strain_L60            ......-.......................................................................... [  9040]
KM102979.1_26544/OG10_Italy      ......-.......................................................................... [  9040]
KM262845.1_strain_NHV            ......-.......................................................................... [  9040]
KX354450.1_47/Ss/2008            ......-.......................................................................... [  9040]
KP055815.1_strain_BA71           ......-.......................................................................... [  9040]
MH025920.1_strain_R35            ..c...t...g.t..c.c...ac...........a....................t....a.c...a........a..c. [  9040]
MH025919.1_strain_N10            ..c...t...g.t..c.c...ac...........a....................t....a.c...a........a..c. [  9040]
MH025918.1_strain_R25            ..c...t...g.t..c.c...ac...........a....................t....a.c...a........a..c. [  9040]
MH025917.1_strain_R7             ..c...t...g.t..c.c...ac...........a....................t....a.c...a........a..c. [  9040]
MH025916.1_strain_R8             ..c...t...g.t..c.c...ac...........a....................t....a.c...a........a..c. [  9040]

FR682468.2_Georgia_2007/1        gaataacctacatcatttttttacata-aaaaaatagatata--------atttttattatatctcaat----------- [  9120]
ASF_IND_20_CAD543                ...........................-..............--------...................tatcttaagat [  9120]
LR899193.1_Germany_2020/1        ...........................-..............--------...................----------- [  9120]
MN393477.1_Wuhan_2019            ...........................-..............--------...................----------- [  9120]
MW396979.1_Timor-Leste/2019/1    ...........................-..............--------...................----------- [  9120]
MK940252.1_CN/2019/InnerMongolia ...........................-..............--------...................----------- [  9120]
MK128995.1_China/2018/AnhuiXCGQ  ...........................-..............--------...................----------- [  9120]
MH681419.1_ASFV/POL/2015         ...........................-..............--------...................----------- [  9120]
LS478113.1_Estonia_2014          ----------------------------..g...a.a.a...--------......g.agcggtc.-------------- [  9120]
KP843857.1_Odintsovo_02/14       ...........................-..............--------...................----------- [  9120]
```

```
FN557520.1_strain_E75            ...........................-..............--------...................tatcttaagat [  9120]
AY261366.1_Warthog               ...........................-..............at--------....g............tattttaagat [  9120]
AY261365.1_Warmbaths             a....t................-t.cg-t.....ag......gtgaca-t.a............-....tatcttaaaat [  9120]
AY261364.1_Tengani_62            .....t................-t.cg-t........tg.c.-------t.a............-....tatcttaaaat [  9120]
AY261363.1_Pretoriuskop/96/4     ......................--.cg-t.....ag......gtgaca-t.a............-....tatcttaaaat [  9120]
AY261362.1_Mkuzi_1979            .............tt.....acgt.-.-..............gtaata-t...aaa.a......-....tatgttaagat [  9120]
AY261361.1_Malawi_Lil-20/1_1983  ......ta..a..t.........tg..t......a......ggtaacata..c......c.g..-....tatcttatgac [  9120]
AY261360.1_Kenya_1950            c....t.t..a..g.g....---.g..c........a....ggtaatatg..c......t.g..-....----------- [  9120]
AM712240.1_OURT_88/3             ...........................-..............--------...................tatcttaagat [  9120]
AM712239.1_Benin_97/1            ...........................-..............--------...................tatcttaagat [  9120]
KM111295.1_Ken06.Bus             c.........a..g.g....---.g..c........--------aatatg..c......t.g..-....tatcttaagta [  9120]
KM111294.1_Ken05/Tk1             -..c.t.ta.at.at.....---t..gt......a......ggtaaca--..c......c.g..-....tatcttatgat [  9120]
KM262844.1_strain_L60            ...........................-..............--------...................tatcttaagat [  9120]
KM102979.1_26544/OG10_Italy      ...........................-..............--------...................tatcttaagat [  9120]
KM262845.1_strain_NHV            ...........................-..............--------...................tatcttaagat [  9120]
KX354450.1_47/Ss/2008            ...........................-..............--------...................tatcttaagat [  9120]
KP055815.1_strain_BA71           ...........................-..............--------...................tatcttaagat [  9120]
MH025920.1_strain_R35            c.........a..g.g....---.g..c........--------aatatg..c......t.g..-....tatcttaagta [  9120]
MH025919.1_strain_N10            c.........a..g.g....---.g..c........--------aatatg..c......t.g..-....tatcttaagta [  9120]
MH025918.1_strain_R25            c.........a..g.g....---.g..c........--------aatatg..c......t.g..-....tatcttaagta [  9120]
MH025917.1_strain_R7             c.........a..g.g....---.g..c........--------aatatg..c......t.g..-....tatcttaagta [  9120]
MH025916.1_strain_R8             c.........a..g.g....---.g..c........--------aatatg..c......t.g..-....tatcttaagta [  9120]

FR682468.2_Georgia_2007/1        ------------------------------------------------tatttaaagataat---tatcaatacagcaaa [  9200]
ASF_IND_20_CAD543                aa--------------------------------ttatcaaagtaac...............---..........a.... [  9200]
LR899193.1_Germany_2020/1        -----------------------------------------------...............---............... [  9200]
MN393477.1_Wuhan_2019            -----------------------------------------------...............---............... [  9200]
MW396979.1_Timor-Leste/2019/1    -----------------------------------------------...............---............... [  9200]
MK940252.1_CN/2019/InnerMongolia -----------------------------------------------...............---............... [  9200]
MK128995.1_China/2018/AnhuiXCGQ  -----------------------------------------------...............---............... [  9200]
MH681419.1_ASFV/POL/2015         -----------------------------------------------...............---............... [  9200]
LS478113.1_Estonia_2014          --------------------------------------------------.c...ga.....ataa.c.ggggtt....c [  9200]
KP843857.1_Odintsovo_02/14       -----------------------------------------------...............---............... [  9200]
FN557520.1_strain_E75            aa--------------------------------ttatcaaagtaac...............---..........a.... [  9200]
AY261366.1_Warthog               aa--------------------------------ttatcaaagtaac...............---...........t... [  9200]
AY261365.1_Warmbaths             aa--------------------------------ttatcaaagtaac...............---............... [  9200]
AY261364.1_Tengani_62            aa--------------------------------ttatcaaagtaac...............---............... [  9200]
AY261363.1_Pretoriuskop/96/4     aa--------------------------------ttatcaaagtaac..-............---............... [  9200]
AY261362.1_Mkuzi_1979            aattatct-----------------------tacttatcaaagtaac...............---.......g.g..... [  9200]
AY261361.1_Malawi_Lil-20/1_1983  aattatcttacgacaattatattaagaaagataattatcaaaataac...............---.......g.g.t... [  9200]
AY261360.1_Kenya_1950            -t--------------------------------atcttaaagtaaa..g............---......c.tt..... [  9200]
AM712240.1_OURT_88/3             aa--------------------------------ttatcaaagtaac...............---..........a.... [  9200]
AM712239.1_Benin_97/1            aa--------------------------------ttatcaaagtaac...............---..........a.... [  9200]
KM111295.1_Ken06.Bus             at--------------------------------atcttaaagtaaa..g............---......c.tt..... [  9200]
```

```
KM111294.1_Ken05/Tk1            aa--------------------------------ttatcaaagt-ac............g..---.......g.g..... [  9200]
KM262844.1_strain_L60           aa--------------------------------ttatcaaagtaac...............---..........a.... [  9200]
KM102979.1_26544/OG10_Italy     aa--------------------------------ttatcaaagtaac...............---..........a.... [  9200]
KM262845.1_strain_NHV           aa--------------------------------ttatcaaagtaac...............---..........a.... [  9200]
KX354450.1_47/Ss/2008           aa--------------------------------ttatcaaagtaac...............---..........a.... [  9200]
KP055815.1_strain_BA71          aa--------------------------------ttatcaaagtaac...............---..........a.... [  9200]
MH025920.1_strain_R35           at--------------------------------atcttaaagtaaa..g............---......c.tt..... [  9200]
MH025919.1_strain_N10           at--------------------------------atcttaaagtaaa..g............---......c.tt..... [  9200]
MH025918.1_strain_R25           at--------------------------------atcttaaagtaaa..g............---......c.tt..... [  9200]
MH025917.1_strain_R7            at--------------------------------atcttaaagtaaa..g............---......c.tt..... [  9200]
MH025916.1_strain_R8            at--------------------------------atcttaaagtaaa..g............---......c.tt..... [  9200]

FR682468.2_Georgia_2007/1       tatcataagctaacatatttttcgaataatagtttttagtaaagtattaatcttttcaggattggtttcttttgataat [  9280]
ASF_IND_20_CAD543               ................................................................................ [  9280]
LR899193.1_Germany_2020/1       ................................................................................ [  9280]
MN393477.1_Wuhan_2019           ................................................................................ [  9280]
MW396979.1_Timor-Leste/2019/1   ................................................................................ [  9280]
MK940252.1_CN/2019/InnerMongolia ................................................................................ [  9280]
MK128995.1_China/2018/AnhuiXCGQ ................................................................................ [  9280]
MH681419.1_ASFV/POL/2015        ................................................................................ [  9280]
LS478113.1_Estonia_2014         g....a....g....g......t.g....a.........ac....a.c....a.g......t.---c.at.......g.g [  9280]
KP843857.1_Odintsovo_02/14      ................................................................................ [  9280]
FN557520.1_strain_E75           ................................................................a............... [  9280]
AY261366.1_Warthog              ................................................................................ [  9280]
AY261365.1_Warmbaths            ................................................................................ [  9280]
AY261364.1_Tengani_62           ................................................................................ [  9280]
AY261363.1_Pretoriuskop/96/4    ................................................................................ [  9280]
AY261362.1_Mkuzi_1979           ........................g.......................g............................... [  9280]
AY261361.1_Malawi_Lil-20/1_1983 ........................g..............a............t...........a..c.......c.... [  9280]
AY261360.1_Kenya_1950           ........................g...........................t..........ca..c............ [  9280]
AM712240.1_OURT_88/3            ................................................................a............... [  9280]
AM712239.1_Benin_97/1           ................................................................a............... [  9280]
KM111295.1_Ken06.Bus            ........................g...........................t..........ca..c.......c.... [  9280]
KM111294.1_Ken05/Tk1            ........................g...........................t...........a..c.......c.... [  9280]
KM262844.1_strain_L60           ................................................................a............... [  9280]
KM102979.1_26544/OG10_Italy     ................................................................a............... [  9280]
KM262845.1_strain_NHV           ................................................................a............... [  9280]
KX354450.1_47/Ss/2008           ................................................................a............... [  9280]
KP055815.1_strain_BA71          ................................................................a............... [  9280]
MH025920.1_strain_R35           ........................g...........................t..........ca..c.......c.... [  9280]
MH025919.1_strain_N10           ........................g...........................t..........ca..c.......c.... [  9280]
MH025918.1_strain_R25           ........................g...........................t..........ca..c.......c.... [  9280]
MH025917.1_strain_R7            ........................g...........................t..........ca..c.......c.... [  9280]
MH025916.1_strain_R8            ........................g...........................t..........ca..c.......c.... [  9280]
```

```
FR682468.2_Georgia_2007/1        aagataggattcgctttataa---atttttaaagataatatattcacaatgatagaataaccgtatatatctgctaatgt [  9360]
ASF_IND_20_CAD543                .....................---........................................................ [  9360]
LR899193.1_Germany_2020/1        .....................---........................................................ [  9360]
MN393477.1_Wuhan_2019            .....................---........................................................ [  9360]
MW396979.1_Timor-Leste/2019/1    .....................---........................................................ [  9360]
MK940252.1_CN/2019/InnerMongolia .....................---........................................................ [  9360]
MK128995.1_China/2018/AnhuiXCGQ  .....................---........................................................ [  9360]
MH681419.1_ASFV/POL/2015         .....................---........................................................ [  9360]
LS478113.1_Estonia_2014          ...tc..t...t.gac.....atg..a.c...t.....g..t.c...t..ttc.tt...c..c.t.tgt.tc..c.gg.. [  9360]
KP843857.1_Odintsovo_02/14       .....................---........................................................ [  9360]
FN557520.1_strain_E75            .....................---........................................................ [  9360]
AY261366.1_Warthog               .....................---...................................g.................... [  9360]
AY261365.1_Warmbaths             .....................---........................................................ [  9360]
AY261364.1_Tengani_62            ....c................---......................t...t........g..a................. [  9360]
AY261363.1_Pretoriuskop/96/4     .....................---..........................t........g.................... [  9360]
AY261362.1_Mkuzi_1979            .....................---........................................................ [  9360]
AY261361.1_Malawi_Lil-20/1_1983  .....................---...................ca..t..t.c...........g.cgt.........tc [  9360]
AY261360.1_Kenya_1950            .....................---...................ca..t..t.c...........g.cgt.........tc [  9360]
AM712240.1_OURT_88/3             .....................---........................................................ [  9360]
AM712239.1_Benin_97/1            .....................---........................................................ [  9360]
KM111295.1_Ken06.Bus             .....................---...................ca..t..t.c...........g..gt.........tc [  9360]
KM111294.1_Ken05/Tk1             .....................---...................ca.tt..t.c...........g.cgt.........tc [  9360]
KM262844.1_strain_L60            .....................---........................................................ [  9360]
KM102979.1_26544/OG10_Italy      .....................---........................................................ [  9360]
KM262845.1_strain_NHV            .....................---........................................................ [  9360]
KX354450.1_47/Ss/2008            .....................---........................................................ [  9360]
KP055815.1_strain_BA71           .....................---........................................................ [  9360]
MH025920.1_strain_R35            .....................---...................ca..t..t.c...........g..gt.........tc [  9360]
MH025919.1_strain_N10            .....................---...................ca..t..t.c.........a.g..gt.........tc [  9360]
MH025918.1_strain_R25            .....................---...................ca..t..t.c...........g..gt.........tc [  9360]
MH025917.1_strain_R7             .....................---...................ca..t..t.c...........g..gt.........tc [  9360]
MH025916.1_strain_R8             .....................---...................ca..t..t.c...........g..gt.........tc [  9360]

FR682468.2_Georgia_2007/1        cttactgtgttcaataacattagcccctaaatccatacaaaagaacatattttcaatacaaaagttttttaccgagatta [  9440]
ASF_IND_20_CAD543                ................................................................................ [  9440]
LR899193.1_Germany_2020/1        ................................................................................ [  9440]
MN393477.1_Wuhan_2019            ................................................................................ [  9440]
MW396979.1_Timor-Leste/2019/1    ................................................................................ [  9440]
MK940252.1_CN/2019/InnerMongolia ................................................................................ [  9440]
MK128995.1_China/2018/AnhuiXCGQ  ................................................................................ [  9440]
MH681419.1_ASFV/POL/2015         ................................................................................ [  9440]
LS478113.1_Estonia_2014          .....cc.c.....agg...c.........g..t...........t.ag..aaa....tt.t.a...aa...t...g... [  9440]
KP843857.1_Odintsovo_02/14       ................................................................................ [  9440]
```

```
FN557520.1_strain_E75            ................................................................................ [  9440]
AY261366.1_Warthog               ................................................................................ [  9440]
AY261365.1_Warmbaths             ................................................................................ [  9440]
AY261364.1_Tengani_62            ...........................................................g...............a.... [  9440]
AY261363.1_Pretoriuskop/96/4     ...........................................................g.................... [  9440]
AY261362.1_Mkuzi_1979            ........................................................................t....... [  9440]
AY261361.1_Malawi_Lil-20/1_1983  taa...c.c...g.agg...c....................................t.g.......ga...t...ccat [  9440]
AY261360.1_Kenya_1950            taa...c.c...g.ag....c....................................t.g.......ga...t...ccat [  9440]
AM712240.1_OURT_88/3             ................................................................................ [  9440]
AM712239.1_Benin_97/1            ................................................................................ [  9440]
KM111295.1_Ken06.Bus             taa...c.c....agg...c......................................g.......ga...t...tcat [  9440]
KM111294.1_Ken05/Tk1             taa...c.c...g.ag....c....................................t.g.......ga...t...ccat [  9440]
KM262844.1_strain_L60            ................................................................................ [  9440]
KM102979.1_26544/OG10_Italy      ................................................................................ [  9440]
KM262845.1_strain_NHV            ................................................................................ [  9440]
KX354450.1_47/Ss/2008            ................................................................................ [  9440]
KP055815.1_strain_BA71           ................................................................................ [  9440]
MH025920.1_strain_R35            taa...c.c....agg...c......................................g.......ga...t...tcat [  9440]
MH025919.1_strain_N10            taa...c.c....agg...c......................................g.......ga...t...tcat [  9440]
MH025918.1_strain_R25            taa...c.c....agg...c......................................g.......ga...t...tcat [  9440]
MH025917.1_strain_R7             taa...c.c....agg...c......................................g.......ga...t...tcat [  9440]
MH025916.1_strain_R8             taa...c.c....agg...c......................................g.......ga...t...tcat [  9440]

FR682468.2_Georgia_2007/1        acattgctcgattagcgttgg-ctcccaatgcaaaacagtagtaaatggtcaaaaaattattatcgcgca---tacaggc [  9520]
ASF_IND_20_CAD543                .....................-................................................---....... [  9520]
LR899193.1_Germany_2020/1        .....................-................................................---....... [  9520]
MN393477.1_Wuhan_2019            .....................-................................................---....... [  9520]
MW396979.1_Timor-Leste/2019/1    .....................-................................................---....... [  9520]
MK940252.1_CN/2019/InnerMongolia .....................-................................................---....... [  9520]
MK128995.1_China/2018/AnhuiXCGQ  .....................-................................................---....... [  9520]
MH681419.1_ASFV/POL/2015         .....................-................................................---....... [  9520]
LS478113.1_Estonia_2014          ........t....gat...a.c.c......ag......a...........tg.t.....tc.g..tt.a.tgc.g..... [  9520]
KP843857.1_Odintsovo_02/14       .....................-................................................---....... [  9520]
FN557520.1_strain_E75            .....................-........................................g.......---....... [  9520]
AY261366.1_Warthog               .....................-.c.......................a..............g.......---....... [  9520]
AY261365.1_Warmbaths             .....................-........................................g.......---....... [  9520]
AY261364.1_Tengani_62            ..............a.a....-.c.......................a..............g.......---....... [  9520]
AY261363.1_Pretoriuskop/96/4     ..............a......-.c.......................a..............g.......---....... [  9520]
AY261362.1_Mkuzi_1979            .....................-.c......................................g.......---....... [  9520]
AY261361.1_Malawi_Lil-20/1_1983  .......c.....gat..c..-................a..a.....a..t........g..g..a....---....... [  9520]
AY261360.1_Kenya_1950            .......c.....gat..c..-................a..a.....a..t........g..g..a....---....... [  9520]
AM712240.1_OURT_88/3             .....................-........................................g.......---....... [  9520]
AM712239.1_Benin_97/1            .....................-........................................g.......---....... [  9520]
KM111295.1_Ken06.Bus             .......g.a...gat..c..-.c..............a........a..t........g..g..a....---....... [  9520]
```

```
KM111294.1_Ken05/Tk1            .......c.....gat..c..-................a..a.....a..t........g..g..a....---....... [  9520]
KM262844.1_strain_L60           .....................-........................................g.......---....... [  9520]
KM102979.1_26544/OG10_Italy     .....................-........................................g.......---....... [  9520]
KM262845.1_strain_NHV           .....................-........................................g.......---....... [  9520]
KX354450.1_47/Ss/2008           .....................-........................................g.......---....... [  9520]
KP055815.1_strain_BA71          .....................-........................................g.......---....... [  9520]
MH025920.1_strain_R35           .......g.a...gat..c..-.c..............a........a..t........g..g..a....---....... [  9520]
MH025919.1_strain_N10           .......g.a...gat..c..-.c..............a........a..t........g..g..a....---....... [  9520]
MH025918.1_strain_R25           .......g.a...gat..c..-.c..............a........a..t........g..g..a....---....... [  9520]
MH025917.1_strain_R7            .......g.a...gat..c..-.c..............a........a..t........g..g..a....---....... [  9520]
MH025916.1_strain_R8            .......g.a...gat..c..-.c..............a........a..t........g..g..a....---....... [  9520]

FR682468.2_Georgia_2007/1       cagctccatcattttattaatactcatatgaattttcgttgtgttacatatttcatgaaggtcaaacacattgttgaaag [  9600]
ASF_IND_20_CAD543               ................................................................................ [  9600]
LR899193.1_Germany_2020/1       ................................................................................ [  9600]
MN393477.1_Wuhan_2019           ................................................................................ [  9600]
MW396979.1_Timor-Leste/2019/1   ................................................................................ [  9600]
MK940252.1_CN/2019/InnerMongolia ................................................................................ [  9600]
MK128995.1_China/2018/AnhuiXCGQ ................................................................................ [  9600]
MH681419.1_ASFV/POL/2015        ................................................................................ [  9600]
LS478113.1_Estonia_2014         t..a.t........cg.c.t..t.........cc...tc...ta..t.c..c.t......a...tt......a..a.... [  9600]
KP843857.1_Odintsovo_02/14      ................................................................................ [  9600]
FN557520.1_strain_E75           ................................................................................ [  9600]
AY261366.1_Warthog              ................................c..................c............................ [  9600]
AY261365.1_Warmbaths            ................................c..................c............................ [  9600]
AY261364.1_Tengani_62           ...................................................c............................ [  9600]
AY261363.1_Pretoriuskop/96/4    ................................c..................c..............t............. [  9600]
AY261362.1_Mkuzi_1979           .....................................................................g.......... [  9600]
AY261361.1_Malawi_Lil-20/1_1983 ....cg...............gt......c...a...a.............c........c.....t..g.....a.... [  9600]
AY261360.1_Kenya_1950           ....cg................t......c...a...a.............c.....t........t............. [  9600]
AM712240.1_OURT_88/3            ................................................................................ [  9600]
AM712239.1_Benin_97/1           ................................................................................ [  9600]
KM111295.1_Ken06.Bus            ...t.g................t...------.a...a.............c........c.....t..g..a..a.... [  9600]
KM111294.1_Ken05/Tk1            ....cg................t......c...a...a.............c.....t........t..g.....a.... [  9600]
KM262844.1_strain_L60           ................................................................................ [  9600]
KM102979.1_26544/OG10_Italy     ................................................................................ [  9600]
KM262845.1_strain_NHV           ................................................................................ [  9600]
KX354450.1_47/Ss/2008           ................................................................................ [  9600]
KP055815.1_strain_BA71          ................................................................................ [  9600]
MH025920.1_strain_R35           ...t.g................t...------.a...a.............c........c.....t..g..a..a.... [  9600]
MH025919.1_strain_N10           ...t.g................t...------.a...a.............c........c.....t..g..a..a.... [  9600]
MH025918.1_strain_R25           ...t.g................t...------.a...a.............c........c.....t..g..a..a.... [  9600]
MH025917.1_strain_R7            ...t.g................t...------.a...a.............c........c.....t..g..a..a.... [  9600]
MH025916.1_strain_R8            ...t.g................t...------.a...a.............c........c.....t..g..a..a.... [  9600]
```

```
FR682468.2_Georgia_2007/1        aaagtgcacaaattaatcgccattcatcaaaatgcctgtattcttgacaaaaatattgaatagcttctttaagattatat [  9680]
ASF_IND_20_CAD543                ................................................................................ [  9680]
LR899193.1_Germany_2020/1        ................................................................................ [  9680]
MN393477.1_Wuhan_2019            ................................................................................ [  9680]
MW396979.1_Timor-Leste/2019/1    ................................................................................ [  9680]
MK940252.1_CN/2019/InnerMongolia ................................................................................ [  9680]
MK128995.1_China/2018/AnhuiXCGQ  ................................................................................ [  9680]
MH681419.1_ASFV/POL/2015         ................................................................................ [  9680]
LS478113.1_Estonia_2014          ....a.....tg...c......c..g.tg..g.c.a..a..ctc..gt..........g..c..a..c............ [  9680]
KP843857.1_Odintsovo_02/14       ................................................................................ [  9680]
FN557520.1_strain_E75            .................................................g.............................. [  9680]
AY261366.1_Warthog               .................................................g.............................. [  9680]
AY261365.1_Warmbaths             .................................................g.............................. [  9680]
AY261364.1_Tengani_62            ............................g....................g........g..................... [  9680]
AY261363.1_Pretoriuskop/96/4     ............................g....................g.............................. [  9680]
AY261362.1_Mkuzi_1979            ............................g....................g.............................. [  9680]
AY261361.1_Malawi_Lil-20/1_1983  ................g..........tg.g....tc............g.............................. [  9680]
AY261360.1_Kenya_1950            ......................c....tg.g....tc............g.............................. [  9680]
AM712240.1_OURT_88/3             .................................................g.............................. [  9680]
AM712239.1_Benin_97/1            .................................................g.............................. [  9680]
KM111295.1_Ken06.Bus             ................g.....c....t..g...ttca...........g.............................. [  9680]
KM111294.1_Ken05/Tk1             ................g.....c....t..g...ttca...........g.............................. [  9680]
KM262844.1_strain_L60            .................................................g.............................. [  9680]
KM102979.1_26544/OG10_Italy      .................................................g.............................. [  9680]
KM262845.1_strain_NHV            .................................................g.............................. [  9680]
KX354450.1_47/Ss/2008            .................................................g.............................. [  9680]
KP055815.1_strain_BA71           .................................................g.............................. [  9680]
MH025920.1_strain_R35            ................g.....c....t..g...ttca...........g.............................. [  9680]
MH025919.1_strain_N10            ................g.....c....t..g...ttca...........g.............................. [  9680]
MH025918.1_strain_R25            ................g.....c....t..g...ttca...........g.............................. [  9680]
MH025917.1_strain_R7             ................g.....c....t..g...ttca...........g.............................. [  9680]
MH025916.1_strain_R8             ................g.....c....t..g...ttca...........g.............................. [  9680]

FR682468.2_Georgia_2007/1        tttaccgctatgccataccaatatttggttagcatctcac---------------taaatgagatc-------------- [  9760]
ASF_IND_20_CAD543                ........................................---------------...........-------------- [  9760]
LR899193.1_Germany_2020/1        ........................................---------------...........-------------- [  9760]
MN393477.1_Wuhan_2019            ........................................---------------...........-------------- [  9760]
MW396979.1_Timor-Leste/2019/1    ........................................---------------...........-------------- [  9760]
MK940252.1_CN/2019/InnerMongolia ........................................---------------...........-------------- [  9760]
MK128995.1_China/2018/AnhuiXCGQ  ........................................---------------...........-------------- [  9760]
MH681419.1_ASFV/POL/2015         ........................................---------------...........-------------- [  9760]
LS478113.1_Estonia_2014          .....t.....a........ga....a....at..t....---------------...ta...tca-------------- [  9760]
KP843857.1_Odintsovo_02/14       ........................................---------------...........-------------- [  9760]
```

```
FN557520.1_strain_E75           ........................................---------------...........-------------- [  9760]
AY261366.1_Warthog              ........................................---------------...........-------------- [  9760]
AY261365.1_Warmbaths            ........................................---------------...........-------------- [  9760]
AY261364.1_Tengani_62           ...............................a........taaatgaaatctcat...........-------------- [  9760]
AY261363.1_Pretoriuskop/96/4    ...............................a........---------------......g....-------------- [  9760]
AY261362.1_Mkuzi_1979           ........................................---------------.t.........-------------- [  9760]
AY261361.1_Malawi_Lil-20/1_1983 .....t..g..c..............a....a.......g---------------........t..-------------- [  9760]
AY261360.1_Kenya_1950           .....t..g.....g...........a....a.......g---------------........tc.-------------- [  9760]
AM712240.1_OURT_88/3            ........................................---------------...........-------------- [  9760]
AM712239.1_Benin_97/1           ........................................---------------...........-------------- [  9760]
KM111295.1_Ken06.Bus            .....t..a..c..............a....a.......g---------------........ga.-------------- [  9760]
KM111294.1_Ken05/Tk1            .....t..g.....g...........a....a........---------------.t.....a...tcacttaatgggat [  9760]
KM262844.1_strain_L60           ........................................---------------...........-------------- [  9760]
KM102979.1_26544/OG10_Italy     ........................................---------------...........-------------- [  9760]
KM262845.1_strain_NHV           ........................................---------------...........-------------- [  9760]
KX354450.1_47/Ss/2008           ........................................---------------...........-------------- [  9760]
KP055815.1_strain_BA71          ........................................---------------...........-------------- [  9760]
MH025920.1_strain_R35           .....t..a..c..............a....a.......g---------------........ga.-------------- [  9760]
MH025919.1_strain_N10           .....t..a..c..............a....a.......g---------------........ga.-------------- [  9760]
MH025918.1_strain_R25           .....t..a..c..............a....a.......g---------------........ga.-------------- [  9760]
MH025917.1_strain_R7            .....t..a..c..............a....a.......g---------------........ga.-------------- [  9760]
MH025916.1_strain_R8            .....t..a..c..............a....a.......g---------------........ga.-------------- [  9760]

FR682468.2_Georgia_2007/1       -tcatttaacatagaatttgttgttaaatccttcaactcccaataaatgatcatccttaaatccaccatgtttacatttt [  9840]
ASF_IND_20_CAD543               -............................................................................... [  9840]
LR899193.1_Germany_2020/1       -............................................................................... [  9840]
MN393477.1_Wuhan_2019           -............................................................................... [  9840]
MW396979.1_Timor-Leste/2019/1   -............................................................................... [  9840]
MK940252.1_CN/2019/InnerMongolia -............................................................................... [  9840]
MK128995.1_China/2018/AnhuiXCGQ -............................................................................... [  9840]
MH681419.1_ASFV/POL/2015        -............................................................................... [  9840]
LS478113.1_Estonia_2014         -.t......t.atagg.---.c.......tt....t.ct.........t........c.tt...c....a........c. [  9840]
KP843857.1_Odintsovo_02/14      -............................................................................... [  9840]
FN557520.1_strain_E75           -......................g........................a............................... [  9840]
AY261366.1_Warthog              -...............................................a............................... [  9840]
AY261365.1_Warmbaths            -...............................................a............................... [  9840]
AY261364.1_Tengani_62           -.............g.................................t............................... [  9840]
AY261363.1_Pretoriuskop/96/4    -..g............................................t...g........................... [  9840]
AY261362.1_Mkuzi_1979           -............ag.....................t...........t...........t.............tt.... [  9840]
AY261361.1_Malawi_Lil-20/1_1983 -.......ct...ag.......a.........................a...........t...........g....... [  9840]
AY261360.1_Kenya_1950           -.......ct...ag.................................t...........t...........g....... [  9840]
AM712240.1_OURT_88/3            -......................g........................a............................... [  9840]
AM712239.1_Benin_97/1           -......................g........................a............................... [  9840]
KM111295.1_Ken06.Bus            -.......ct...ag.................................t...........t...........g....... [  9840]
```

```
KM111294.1_Ken05/Tk1            a.......ct...ag...gc................t.......................t................... [  9840]
KM262844.1_strain_L60           -......................g........................a............................... [  9840]
KM102979.1_26544/OG10_Italy     -......................g........................a............................... [  9840]
KM262845.1_strain_NHV           -......................g........................a............................... [  9840]
KX354450.1_47/Ss/2008           -......................g........................a............................... [  9840]
KP055815.1_strain_BA71          -......................g........................a............................... [  9840]
MH025920.1_strain_R35           -.......ct...ag.................................t...........t...........g....... [  9840]
MH025919.1_strain_N10           -.......ct...ag.................................t...........t...........g....... [  9840]
MH025918.1_strain_R25           -.......ct...ag.................................t...........t...........g....... [  9840]
MH025917.1_strain_R7            -.......ct...ag.................................t...........t...........g....... [  9840]
MH025916.1_strain_R8            -.......ct...ag.................................t...........t...........g....... [  9840]

FR682468.2_Georgia_2007/1       gtaaaaaagggttattagaaaataattcatgacacaaaatgacattactacttgttattttacactttgtttcaaagaaa [  9920]
ASF_IND_20_CAD543               ................................................................................ [  9920]
LR899193.1_Germany_2020/1       ................................................................................ [  9920]
MN393477.1_Wuhan_2019           ................................................................................ [  9920]
MW396979.1_Timor-Leste/2019/1   ................................................................................ [  9920]
MK940252.1_CN/2019/InnerMongolia ................................................................................ [  9920]
MK128995.1_China/2018/AnhuiXCGQ ................................................................................ [  9920]
MH681419.1_ASFV/POL/2015        ................................................................................ [  9920]
LS478113.1_Estonia_2014         c.....g...a..g..g.tg..............t.....t.t.........g..gg.a.c.tctaa.....tg..t.t. [  9920]
KP843857.1_Odintsovo_02/14      ................................................................................ [  9920]
FN557520.1_strain_E75           ................................................................................ [  9920]
AY261366.1_Warthog              ................................................................................ [  9920]
AY261365.1_Warmbaths            ................................................................................ [  9920]
AY261364.1_Tengani_62           ...............................................................g................ [  9920]
AY261363.1_Pretoriuskop/96/4    ..................................t.......t...................tg................ [  9920]
AY261362.1_Mkuzi_1979           ..........................................g...............a....g.....a.......... [  9920]
AY261361.1_Malawi_Lil-20/1_1983 .......c..a.............................a.t.........c...t.a.g..g................ [  9920]
AY261360.1_Kenya_1950           .......c..a...............................t.........c...t.a.g..g................ [  9920]
AM712240.1_OURT_88/3            ................................................................................ [  9920]
AM712239.1_Benin_97/1           ................................................................................ [  9920]
KM111295.1_Ken06.Bus            .......c..a................t..............t.............t.a.g..g................ [  9920]
KM111294.1_Ken05/Tk1            t.........................................t.........c...t.a.g..g................ [  9920]
KM262844.1_strain_L60           ................................................................................ [  9920]
KM102979.1_26544/OG10_Italy     ................................................................................ [  9920]
KM262845.1_strain_NHV           ................................................................................ [  9920]
KX354450.1_47/Ss/2008           ................................................................................ [  9920]
KP055815.1_strain_BA71          ................................................................................ [  9920]
MH025920.1_strain_R35           .......c..a................t..............t.............t.a.g..g................ [  9920]
MH025919.1_strain_N10           .......c..a................t..............t.............t.a.g..g................ [  9920]
MH025918.1_strain_R25           .......c..a................t..............t.............t.a.g..g................ [  9920]
MH025917.1_strain_R7            .......c..a................t..............t.............t.a.g..g................ [  9920]
MH025916.1_strain_R8            .......c..a................t..............t.............t.a.g..g................ [  9920]
```

```
FR682468.2_Georgia_2007/1        aatcgtaaaatttcacttgtctcaagctcttctttagc-tcccaattttcggcataggtttcgagtatgctcgttattaa [ 10000]
ASF_IND_20_CAD543                ......................................-y........................................ [ 10000]
LR899193.1_Germany_2020/1        ......................................-......................................... [ 10000]
MN393477.1_Wuhan_2019            ......................................-......................................... [ 10000]
MW396979.1_Timor-Leste/2019/1    ......................................-......................................... [ 10000]
MK940252.1_CN/2019/InnerMongolia ......................................-......................................... [ 10000]
MK128995.1_China/2018/AnhuiXCGQ  ......................................-......................................... [ 10000]
MH681419.1_ASFV/POL/2015         ......................................-......................................... [ 10000]
LS478113.1_Estonia_2014          .t.t...t..a....t..cct......at.........-...t........a.....a.cc..g.c.......g...... [ 10000]
KP843857.1_Odintsovo_02/14       ......................................-......................................... [ 10000]
FN557520.1_strain_E75            ......................................cc........................................ [ 10000]
AY261366.1_Warthog               ......................................-c........................................ [ 10000]
AY261365.1_Warmbaths             ......................................-c........................................ [ 10000]
AY261364.1_Tengani_62            ...t..................................-......................................... [ 10000]
AY261363.1_Pretoriuskop/96/4     ...................................g..-..................a...................... [ 10000]
AY261362.1_Mkuzi_1979            ......................................-.................................a....... [ 10000]
AY261361.1_Malawi_Lil-20/1_1983  ..........c........ct.....ac.......g..-......a..........a...........a...c....... [ 10000]
AY261360.1_Kenya_1950            ..........c........ct.....ac.......g..-.................................c....... [ 10000]
AM712240.1_OURT_88/3             ......................................-c........................................ [ 10000]
AM712239.1_Benin_97/1            ......................................cc........................................ [ 10000]
KM111295.1_Ken06.Bus             ..........c........ct.....ac.......g..-.................................c....... [ 10000]
KM111294.1_Ken05/Tk1             ..........................ac.......g..-.................................c....... [ 10000]
KM262844.1_strain_L60            ......................................-c........................................ [ 10000]
KM102979.1_26544/OG10_Italy      ......................................cc........................................ [ 10000]
KM262845.1_strain_NHV            ......................................-c........................................ [ 10000]
KX354450.1_47/Ss/2008            ......................................cc........................................ [ 10000]
KP055815.1_strain_BA71           ......................................-c........................................ [ 10000]
MH025920.1_strain_R35            ..........c........ct.....ac.......g..-.................................c....... [ 10000]
MH025919.1_strain_N10            ..........c........ct.....ac.......g..-.................................c....... [ 10000]
MH025918.1_strain_R25            ..........c........ct.....ac.......g..-.................................c....... [ 10000]
MH025917.1_strain_R7             ..........c........ct.....ac.......g..-.................................c....... [ 10000]
MH025916.1_strain_R8             ..........c........ct.....ac.......g..-.................................c....... [ 10000]

FR682468.2_Georgia_2007/1        taaaaagtaacccataattaatatttgcaccccattcagtaaacaacatgattagatcatcattgttttccttaactgcc [ 10080]
ASF_IND_20_CAD543                ..........................................................w..................... [ 10080]
LR899193.1_Germany_2020/1        ................................................................................ [ 10080]
MN393477.1_Wuhan_2019            ................................................................................ [ 10080]
MW396979.1_Timor-Leste/2019/1    ................................................................................ [ 10080]
MK940252.1_CN/2019/InnerMongolia ................................................................................ [ 10080]
MK128995.1_China/2018/AnhuiXCGQ  ................................................................................ [ 10080]
MH681419.1_ASFV/POL/2015         ................................................................................ [ 10080]
LS478113.1_Estonia_2014          .gg...t...t........g..g.....t..................t.t....at....a...a..c....gt..a..t [ 10080]
KP843857.1_Odintsovo_02/14       ................................................................................ [ 10080]
```

```
FN557520.1_strain_E75          ............................g.............................t..................... [ 10080]
AY261366.1_Warthog             ............................g.............................t..................... [ 10080]
AY261365.1_Warmbaths           ............................g.............................t..................... [ 10080]
AY261364.1_Tengani_62          ................................................................................ [ 10080]
AY261363.1_Pretoriuskop/96/4   .............g.................................................................. [ 10080]
AY261362.1_Mkuzi_1979          c...............g....c........................a..a..c..c....a...a.....t..g..c... [ 10080]
AY261361.1_Malawi_Lil-20/1_1983 c.g........g.............c....................a.ca..c.......a.........t..g..c... [ 10080]
AY261360.1_Kenya_1950          c.g........g....................................ca.....g....a...a.....t..g..c... [ 10080]
AM712240.1_OURT_88/3           ............................g.............................t..................... [ 10080]
AM712239.1_Benin_97/1          ............................g.............................t..................... [ 10080]
KM111295.1_Ken06.Bus           c.g.............................................ca..c..g....a...a.....t..g..c... [ 10080]
KM111294.1_Ken05/Tk1           c.g........g....................................ca.....g....a...a.....t..g..c... [ 10080]
KM262844.1_strain_L60          ............................g.............................t..................... [ 10080]
KM102979.1_26544/OG10_Italy    ............................g.............................t..................... [ 10080]
KM262845.1_strain_NHV          ............................g.............................t..................... [ 10080]
KX354450.1_47/Ss/2008          ............................g.............................t..................... [ 10080]
KP055815.1_strain_BA71         ............................g.............................t...................t. [ 10080]
MH025920.1_strain_R35          c.g.............................................ca..c..g....a...a.....t..g..c... [ 10080]
MH025919.1_strain_N10          c.g.............................................ca..c..g....a...a.....t..g..c... [ 10080]
MH025918.1_strain_R25          c.g.............................................ca..c..g....a...a.....t..g..c... [ 10080]
MH025917.1_strain_R7           c.g.............................................ca..c..g....a...a.....t..g..c... [ 10080]
MH025916.1_strain_R8           c.g.............................................ca..c..g....a...a.....t..g..c... [ 10080]

FR682468.2_Georgia_2007/1      aataccaatgcagtattaagccttataccctctttaaagcataatgtccttatcattatttgattatcatcatctatata [ 10160]
ASF_IND_20_CAD543              ................................................................................ [ 10160]
LR899193.1_Germany_2020/1      ................................................................................ [ 10160]
MN393477.1_Wuhan_2019          ................................................................................ [ 10160]
MW396979.1_Timor-Leste/2019/1  ................................................................................ [ 10160]
MK940252.1_CN/2019/InnerMongolia ................................................................................ [ 10160]
MK128995.1_China/2018/AnhuiXCGQ ................................................................................ [ 10160]
MH681419.1_ASFV/POL/2015       ................................................................................ [ 10160]
LS478113.1_Estonia_2014        ttc.tt..............taa...g..t....c....at.gga...t.......c.....c.c.ctcc....ata.c. [ 10160]
KP843857.1_Odintsovo_02/14     ................................................................................ [ 10160]
FN557520.1_strain_E75          .....................t.......................................................... [ 10160]
AY261366.1_Warthog             .....................t................................................t......... [ 10160]
AY261365.1_Warmbaths           .....................t................................................t......... [ 10160]
AY261364.1_Tengani_62          .....................t..............t......gc..tt....t.......................... [ 10160]
AY261363.1_Pretoriuskop/96/4   .....t...............t.......t......t.......c..tt....t.......................... [ 10160]
AY261362.1_Mkuzi_1979          .....................t.......t......t.....ggc..tt....t........---t...c.....g...g [ 10160]
AY261361.1_Malawi_Lil-20/1_1983 .....t...............t.......t......t..t..ggc..tt....t.a......---t...c.....g.... [ 10160]
AY261360.1_Kenya_1950          ..c..t...............t.......t......t.....ggc..tt....t.a......---t...c.....g.... [ 10160]
AM712240.1_OURT_88/3           .....................t.......................................................... [ 10160]
AM712239.1_Benin_97/1          .....................t.......................................................... [ 10160]
KM111295.1_Ken06.Bus           ..c..t...............t.......t......t.....ggc..tt....t.a......---t...c.....g.... [ 10160]
```

```
KM111294.1_Ken05/Tk1              ..c..t...............t.......t......t.....ggc..tt....t.a......---t...ct....g.... [ 10160]
KM262844.1_strain_L60             .....................t.......................................................... [ 10160]
KM102979.1_26544/OG10_Italy       .....................t.......................................................... [ 10160]
KM262845.1_strain_NHV             .....................t.......................................................... [ 10160]
KX354450.1_47/Ss/2008             .....................t.......................................................... [ 10160]
KP055815.1_strain_BA71            .....................t.......................................................... [ 10160]
MH025920.1_strain_R35             ..c..t...............t.......t......t.....ggc..tt....t.a......---t...c.....g.... [ 10160]
MH025919.1_strain_N10             ..c..t...............t.......t......t.....ggc..tt....t.a......---t...c.....g.... [ 10160]
MH025918.1_strain_R25             ..c..t...............t.......t......t.....ggc..tt....t.a......---t...c.....g.... [ 10160]
MH025917.1_strain_R7              ..c..t...............t.......t......t.....ggc..tt....t.a......---t...c.....g.... [ 10160]
MH025916.1_strain_R8              ..c..t...............t.......t......t.....ggc..tt....t.a......---t...c.....g.... [ 10160]

FR682468.2_Georgia_2007/1         cattgagataggagcttcatgccaccataaaccataacgctctaaaatataataatca---tctttagatacgtgtt--- [ 10240]
ASF_IND_20_CAD543                 ..........................................................---................--- [ 10240]
LR899193.1_Germany_2020/1         ..........................................................---................--- [ 10240]
MN393477.1_Wuhan_2019             ..........................................................---................--- [ 10240]
MW396979.1_Timor-Leste/2019/1     ..........................................................---................--- [ 10240]
MK940252.1_CN/2019/InnerMongolia  ..........................................................---................--- [ 10240]
MK128995.1_China/2018/AnhuiXCGQ   ..........................................................---................--- [ 10240]
MH681419.1_ASFV/POL/2015          ..........................................................---................--- [ 10240]
LS478113.1_Estonia_2014           a.g.ata..c.......................gc...a..t...t....g.....a.tat...c......ta....cat [ 10240]
KP843857.1_Odintsovo_02/14        ..........................................................---................--- [ 10240]
FN557520.1_strain_E75             a.................................c.......................---................--- [ 10240]
AY261366.1_Warthog                a.................................c.......................---................--- [ 10240]
AY261365.1_Warmbaths              a.................................c.......................---................--- [ 10240]
AY261364.1_Tengani_62             ..........................................................---................--- [ 10240]
AY261363.1_Pretoriuskop/96/4      .........g........................c.......................---................--- [ 10240]
AY261362.1_Mkuzi_1979             a.g..t.........a...........c......c..............c......a.---........g.......--- [ 10240]
AY261361.1_Malawi_Lil-20/1_1983   a.g..t.........a..g...............c..............c........---..g.....g..a..c.--- [ 10240]
AY261360.1_Kenya_1950             a.g..t.........a..g..............ga......t.......c........---..a.....g.....c.--- [ 10240]
AM712240.1_OURT_88/3              a.................................c.......................---................--- [ 10240]
AM712239.1_Benin_97/1             a.................................c.......................---................--- [ 10240]
KM111295.1_Ken06.Bus              a.g..t.........a..................c......t.......c........---........g.....c.--- [ 10240]
KM111294.1_Ken05/Tk1              a.g..t.........a..g..............ga......t.......c........---..a.....g.....c.--- [ 10240]
KM262844.1_strain_L60             a.................................c.......................---................--- [ 10240]
KM102979.1_26544/OG10_Italy       a.................................c.......................---................--- [ 10240]
KM262845.1_strain_NHV             a.................................c.......................---................--- [ 10240]
KX354450.1_47/Ss/2008             a.................................c.......................---................--- [ 10240]
KP055815.1_strain_BA71            a.................................c.......................---................--- [ 10240]
MH025920.1_strain_R35             a.g..t.........a..................c......t.......c........---........g.....c.--- [ 10240]
MH025919.1_strain_N10             a.g..t.........a..................c......t.......c........---........g.....c.--- [ 10240]
MH025918.1_strain_R25             a.g..t.........a..................c......t.......c........---........g.....c.--- [ 10240]
MH025917.1_strain_R7              a.g..t.........a..................c......t.......c........---........g.....c.--- [ 10240]
MH025916.1_strain_R8              a.g..t.........a..................c......t.......c........---........g.....c.--- [ 10240]
```

```
FR682468.2_Georgia_2007/1          ---------------gcgtggccaatgcccttttagcaagtgcttgtaaagtcgatggctgcatgtttattctgtt--aa [ 10320]
ASF_IND_20_CAD543                  ---------------........................................................y....--.. [ 10320]
LR899193.1_Germany_2020/1          ---------------.............................................................--.. [ 10320]
MN393477.1_Wuhan_2019              ---------------.............................................................--.. [ 10320]
MW396979.1_Timor-Leste/2019/1      ---------------.............................................................--.. [ 10320]
MK940252.1_CN/2019/InnerMongolia   ---------------.............................................................--.. [ 10320]
MK128995.1_China/2018/AnhuiXCGQ    ---------------.............................................................--.. [ 10320]
MH681419.1_ASFV/POL/2015           ---------------.............................................................--.. [ 10320]
LS478113.1_Estonia_2014            tttctttatgctcccctaa.......a..t..........ca......g...a....---....a...t..a...----- [ 10320]
KP843857.1_Odintsovo_02/14         ---------------.............................................................--.. [ 10320]
FN557520.1_strain_E75              ---------------........................................................t....--.. [ 10320]
AY261366.1_Warthog                 ---------------.............................................................a-.. [ 10320]
AY261365.1_Warmbaths               ---------------.................................................c...........a-.. [ 10320]
AY261364.1_Tengani_62              ---------------.............................................................a-.. [ 10320]
AY261363.1_Pretoriuskop/96/4       ---------------..............t......t.....t.................................a-.. [ 10320]
AY261362.1_Mkuzi_1979              ---------------.............................................................a-.. [ 10320]
AY261361.1_Malawi_Lil-20/1_1983    ---------------.t.........a.........t.......................................t-.. [ 10320]
AY261360.1_Kenya_1950              ---------------.t.........a.......g.t..................................t....ta.. [ 10320]
AM712240.1_OURT_88/3               ---------------........................................................t....---. [ 10320]
AM712239.1_Benin_97/1              ---------------........................................................t....--.. [ 10320]
KM111295.1_Ken06.Bus               ---------------.t.........a.......g.t.....t.....................c...........t-.. [ 10320]
KM111294.1_Ken05/Tk1               ---------------.t.........a.......g.t.......................................t-.. [ 10320]
KM262844.1_strain_L60              ---------------........................................................t....--.. [ 10320]
KM102979.1_26544/OG10_Italy        ---------------........................................................t....--.. [ 10320]
KM262845.1_strain_NHV              ---------------........................................................t....---. [ 10320]
KX354450.1_47/Ss/2008              ---------------........................................................t....--.. [ 10320]
KP055815.1_strain_BA71             ---------------........................................................t....--.. [ 10320]
MH025920.1_strain_R35              ---------------.t.........a.......g.t.....t.....................c...........t-.. [ 10320]
MH025919.1_strain_N10              ---------------.t.........a.......g.t.....t.....................c...........t-.. [ 10320]
MH025918.1_strain_R25              ---------------.t.........a.......g.t.....t.....................c...........t-.. [ 10320]
MH025917.1_strain_R7               ---------------.t.........a.......g.t.....t.....................c...........t-.. [ 10320]
MH025916.1_strain_R8               ---------------.t.........a.......g.t.....t.....................c...........t-.. [ 10320]

FR682468.2_Georgia_2007/1          aaaaaaatcaaattatcgggtaaacataaggatcaacccgtagttaatatttgcagtagtattttttaacaatgaattat [ 10400]
ASF_IND_20_CAD543                  .........................................................gac.....-.............. [ 10400]
LR899193.1_Germany_2020/1          ................................................................................ [ 10400]
MN393477.1_Wuhan_2019              ................................................................................ [ 10400]
MW396979.1_Timor-Leste/2019/1      ................................................................................ [ 10400]
MK940252.1_CN/2019/InnerMongolia   ................................................................................ [ 10400]
MK128995.1_China/2018/AnhuiXCGQ    ................................................................................ [ 10400]
MH681419.1_ASFV/POL/2015           ................................................................................ [ 10400]
LS478113.1_Estonia_2014            ------------------------------------------------------.a.g....c..--------------. [ 10400]
KP843857.1_Odintsovo_02/14         ................................................................................ [ 10400]
```

```
FN557520.1_strain_E75            .........................................................gac.-.................. [ 10400]
AY261366.1_Warthog               ......................................t..................gac.................... [ 10400]
AY261365.1_Warmbaths             ......................................t.............a....gac.................... [ 10400]
AY261364.1_Tengani_62            ................ta......g...--.........a.......c.........g.c.-.................. [ 10400]
AY261363.1_Pretoriuskop/96/4     .........................................t...............g.c.................... [ 10400]
AY261362.1_Mkuzi_1979            .........................................................gac.................... [ 10400]
AY261361.1_Malawi_Lil-20/1_1983  ...........................g.a...........................ga..............t.....g [ 10400]
AY261360.1_Kenya_1950            ................taa..t.......a.............g..g..a.a.tt.c..c--ca..cggt..ac...ct. [ 10400]
AM712240.1_OURT_88/3             .........................................................gac.-.................. [ 10400]
AM712239.1_Benin_97/1            .........................................................gac.-.................. [ 10400]
KM111295.1_Ken06.Bus             ................taa..t.......a.............g..g..a.a.tt.c..c--ca..cggt..ac...ct. [ 10400]
KM111294.1_Ken05/Tk1             ................taa..t.......a.............g..g..a.a.tt.c..c--ca..cggt..ac...ct. [ 10400]
KM262844.1_strain_L60            .........................................................gac.-.................. [ 10400]
KM102979.1_26544/OG10_Italy      .........................................................gac.-.................. [ 10400]
KM262845.1_strain_NHV            .........................................................gac.-.................. [ 10400]
KX354450.1_47/Ss/2008            .........................................................gac.-.................. [ 10400]
KP055815.1_strain_BA71           .........................................................gac.-.................. [ 10400]
MH025920.1_strain_R35            ................taa..t.......a.............g..g..a.a.tt.c..c--ca..cggt..ac...ct. [ 10400]
MH025919.1_strain_N10            ................taa..t.......a.............g..g..a.a.tt.c..c--ca..cggt..ac...ct. [ 10400]
MH025918.1_strain_R25            ................taa..t.......a.............g..g..a.a.tt.c..c--ca..cggt..ac...ct. [ 10400]
MH025917.1_strain_R7             ................taa..t.......a.............g..g..a.a.tt.c..c--ca..cggt..ac...ct. [ 10400]
MH025916.1_strain_R8             ................taa..t.......a.............g..g..a.a.tt.c..c--ca..cggt..ac...ct. [ 10400]

FR682468.2_Georgia_2007/1        aataaaaaaataattcattactat---ctattataaaacccatcttt------------aactttaaagaaga-actaga [ 10480]
ASF_IND_20_CAD543                r.................k.....---....................------------...w........r.-..y... [ 10480]
LR899193.1_Germany_2020/1        ........................---....................------------..............-...... [ 10480]
MN393477.1_Wuhan_2019            ........................---....................------------..............-...... [ 10480]
MW396979.1_Timor-Leste/2019/1    ........................---....................------------..............-...... [ 10480]
MK940252.1_CN/2019/InnerMongolia ........................---....................------------..............-...... [ 10480]
MK128995.1_China/2018/AnhuiXCGQ  ........................---....................------------..............-...... [ 10480]
MH681419.1_ASFV/POL/2015         ........................---....................------------..............-...... [ 10480]
LS478113.1_Estonia_2014          ..ccg.........------------------------------------------------------------------ [ 10480]
KP843857.1_Odintsovo_02/14       ........................---....................------------..............-...... [ 10480]
FN557520.1_strain_E75            g.................g.....---....c..gg...........------------...a........a.-..c... [ 10480]
AY261366.1_Warthog               g.................g.....---....c...............------------..............-...... [ 10480]
AY261365.1_Warmbaths             g..........c......g.....---....c...............------------..............-...... [ 10480]
AY261364.1_Tengani_62            g..-..............a.....---....c...............------------..............-...... [ 10480]
AY261363.1_Pretoriuskop/96/4     g...............t.......---....c...............------------..............-...... [ 10480]
AY261362.1_Mkuzi_1979            g.................g.....---....c...............------------..............-...... [ 10480]
AY261361.1_Malawi_Lil-20/1_1983  g.......................---....c...............------------...a..........-...... [ 10480]
AY261360.1_Kenya_1950            .t..g.tc.ta.t.gtt..c..g.acg..t....g..tgtg..a...aattagta----gg.c.c.......gcc...-- [ 10480]
AM712240.1_OURT_88/3             g.................g.....---....c..gg...........------------...a........a.-..c... [ 10480]
AM712239.1_Benin_97/1            g.................g.....---....c..gg...........------------...a........a.-..c... [ 10480]
KM111295.1_Ken06.Bus             .t..g.tc.ta.t.gtt..c..g.atg..t.c..g..tgtg..a...aattagtaggccgg.c.c.......g-c...-- [ 10480]
```

```
KM111294.1_Ken05/Tk1           .t..g.tc.ta.t.gtt..c..g.acg..t.c..g..tgtg..a...aattagta----gg.c.c.......g-c...-- [ 10480]
KM262844.1_strain_L60          g.................g.....---....c..gg...........------------...a........a.-..c... [ 10480]
KM102979.1_26544/OG10_Italy    g.................g.....---....c..gg...........------------...a........a.-..c... [ 10480]
KM262845.1_strain_NHV          g.................g.....---....c..gg...........------------...a........a.-..c... [ 10480]
KX354450.1_47/Ss/2008          g.................g.....---....c..gg...........------------...a........a.-..c... [ 10480]
KP055815.1_strain_BA71         g.................g.....---....c..gg...........------------...a........a.-..c... [ 10480]
MH025920.1_strain_R35          .t..g.tc.ta.t.gtt..c..g.atg..t.c..g..tgtg..a...aattagtaggccgg.c.c.......g-c...-- [ 10480]
MH025919.1_strain_N10          .t..g.tc.ta.t.gtt..c..g.atg..t.c..g..tgtg..a...aattagtaggccgg.c.c.......g-c...-- [ 10480]
MH025918.1_strain_R25          .t..g.tc.ta.t.gtt..c..g.atg..t.c..g..tgtg..a...aattagtaggccgg.c.c.......g-c...-- [ 10480]
MH025917.1_strain_R7           .t..g.tc.ta.t.gtt..c..g.atg..t.c..g..tgtg..a...aattagtaggccgg.c.c.......g-c...-- [ 10480]
MH025916.1_strain_R8           .t..g.tc.ta.t.gtt..c..g.atg..t.c..g..tgtg..a...aattagtaggccgg.c.c.......g-c...-- [ 10480]

FR682468.2_Georgia_2007/1      tcatc-tttttttttgttgtgtcagaa---cttcttcaatttattacc----cacattttatct-aaaaaaaataaaaac [ 10560]
ASF_IND_20_CAD543              .....a..............r.y..r.---.........w........----............-.......--t..... [ 10560]
LR899193.1_Germany_2020/1      .....--....................---..................----............-............... [ 10560]
MN393477.1_Wuhan_2019          .....--....................---..................----............-............... [ 10560]
MW396979.1_Timor-Leste/2019/1  .....--....................---..................----............-............... [ 10560]
MK940252.1_CN/2019/InnerMongolia .....--....................---..................----............-............... [ 10560]
MK128995.1_China/2018/AnhuiXCGQ .....--....................---..................----............-............... [ 10560]
MH681419.1_ASFV/POL/2015       .....-.....................---..................----............-............... [ 10560]
LS478113.1_Estonia_2014        -------.a.....c.c-------------------------------------------------------------.. [ 10560]
KP843857.1_Odintsovo_02/14     .....--....................---..................----............-............... [ 10560]
FN557520.1_strain_E75          .....a..............a.t..g.---.........a........----............-.......--t..... [ 10560]
AY261366.1_Warthog             c....t-...........a...t..g.ctt.........a......ttacca............-.......--t..... [ 10560]
AY261365.1_Warmbaths           c....--....................---.g................----............a.......--t..... [ 10560]
AY261364.1_Tengani_62          .....--....................---.....a............----............a.......--t..... [ 10560]
AY261363.1_Pretoriuskop/96/4   .....a-...............t..g.---.........a........----............-.......--t..... [ 10560]
AY261362.1_Mkuzi_1979          c....--....................---.g...a............----............-.......--t..... [ 10560]
AY261361.1_Malawi_Lil-20/1_1983 .....a----.....t........ag.---..................----.t..........a.......--t..... [ 10560]
AY261360.1_Kenya_1950          ----------------------------------------------------.ta.........-.......t---.... [ 10560]
AM712240.1_OURT_88/3           .....a..............a.t..g.---.........a........----............-.......--t..... [ 10560]
AM712239.1_Benin_97/1          .....a..............a.t..g.---.........a........----............-.......--t..... [ 10560]
KM111295.1_Ken06.Bus           ----------------------------------------------------.ta.........-.......t---.... [ 10560]
KM111294.1_Ken05/Tk1           ----------------------------------------------------.ta.........-.......t---.... [ 10560]
KM262844.1_strain_L60          .....a..............a.t..g.---.........a........----............-.......--t..... [ 10560]
KM102979.1_26544/OG10_Italy    .....a..............a.t..g.---.........a........----............-.......--t..... [ 10560]
KM262845.1_strain_NHV          .....a..............a.t..g.---.........a........----............-.......--t..... [ 10560]
KX354450.1_47/Ss/2008          .....a..............a.t..g.---.........a........----............-.......--t..... [ 10560]
KP055815.1_strain_BA71         .....a..............a.t..g.---.........a........----............-.......--t..... [ 10560]
MH025920.1_strain_R35          ----------------------------------------------------.ta.........-.......t---.... [ 10560]
MH025919.1_strain_N10          ----------------------------------------------------.ta.........-.......t---.... [ 10560]
MH025918.1_strain_R25          ----------------------------------------------------.ta.........-.......t---.... [ 10560]
MH025917.1_strain_R7           ----------------------------------------------------.ta.........-.......t---.... [ 10560]
MH025916.1_strain_R8           ----------------------------------------------------.ta.........-.......t---.... [ 10560]
```

```
FR682468.2_Georgia_2007/1        tacatcatatcttgtttct--tcatcaaattatcataccatttatagggtgtaggttgggaacattccatcatgtggtaa [ 10640]
ASF_IND_20_CAD543                .....m.............--.................................................a..a...gt. [ 10640]
LR899193.1_Germany_2020/1        ...................--........................................................... [ 10640]
MN393477.1_Wuhan_2019            ...................--........................................................... [ 10640]
MW396979.1_Timor-Leste/2019/1    ...................--........................................................... [ 10640]
MK940252.1_CN/2019/InnerMongolia ...................--........................................................... [ 10640]
MK128995.1_China/2018/AnhuiXCGQ  ...................--........................................................... [ 10640]
MH681419.1_ASFV/POL/2015         ...................--........................................................... [ 10640]
LS478113.1_Estonia_2014          gg...t....ta..a..tcag.......---------------------------------------------------- [ 10640]
KP843857.1_Odintsovo_02/14       ...................--........................................................... [ 10640]
FN557520.1_strain_E75            .....a.............--.......................a..t......................a..a...gt. [ 10640]
AY261366.1_Warthog               .....a.............--.......................................................a... [ 10640]
AY261365.1_Warmbaths             .....a.............--.......................................................a... [ 10640]
AY261364.1_Tengani_62            .....ac............--.......................a..t..........a...........a..a...gt. [ 10640]
AY261363.1_Pretoriuskop/96/4     ...................--....t...................................................... [ 10640]
AY261362.1_Mkuzi_1979            ...................--.......................a..t......................a..a...gt. [ 10640]
AY261361.1_Malawi_Lil-20/1_1983  ...---.............--.......................a..t..........a...........a..a...gt. [ 10640]
AY261360.1_Kenya_1950            .....a.............--.......................a..t......................a..a...gt. [ 10640]
AM712240.1_OURT_88/3             .....a.............--.......................a..t......................a..a...gt. [ 10640]
AM712239.1_Benin_97/1            .....a.............--.......................a..t......................a..a...gt. [ 10640]
KM111295.1_Ken06.Bus             .....a.............--..........................t......................a......gt. [ 10640]
KM111294.1_Ken05/Tk1             .....a.............--.......................a..t......................a......gt. [ 10640]
KM262844.1_strain_L60            .....a.............--.......................a..t......................a..a...gt. [ 10640]
KM102979.1_26544/OG10_Italy      .....a.............--.......................a..t......................a..a...gt. [ 10640]
KM262845.1_strain_NHV            .....a.............--.......................a..t......................a..a...gt. [ 10640]
KX354450.1_47/Ss/2008            .....a.............--.......................a..t......................a..a...gt. [ 10640]
KP055815.1_strain_BA71           .....a.............--.......................a..t......................a..a...gt. [ 10640]
MH025920.1_strain_R35            .....a.............--..........................t......................a......gt. [ 10640]
MH025919.1_strain_N10            .....a.............--..........................t......................a......gt. [ 10640]
MH025918.1_strain_R25            .....a.............--..........................t......................a......gt. [ 10640]
MH025917.1_strain_R7             .....a.............--..........................t......................a......gt. [ 10640]
MH025916.1_strain_R8             .....a.............--..........................t......................a......gt. [ 10640]

FR682468.2_Georgia_2007/1        tcagggtatttatatatttttgatagtaacatctatttggcagatgtattgtccaacaatcatgtctaataaaatcatt [ 10720]
ASF_IND_20_CAD543                .t.aa..g..............s.........................r...............r............... [ 10720]
LR899193.1_Germany_2020/1        ................................................................................ [ 10720]
MN393477.1_Wuhan_2019            ................................................................................ [ 10720]
MW396979.1_Timor-Leste/2019/1    ................................................................................ [ 10720]
MK940252.1_CN/2019/InnerMongolia ................................................................................ [ 10720]
MK128995.1_China/2018/AnhuiXCGQ  ................................................................................ [ 10720]
MH681419.1_ASFV/POL/2015         ................................................................................ [ 10720]
LS478113.1_Estonia_2014          -----------------..a..a.a.t..gt..tac.g.c.t..g..--------------------------------- [ 10720]
KP843857.1_Odintsovo_02/14       ................................................................................ [ 10720]
```

```
FN557520.1_strain_E75            .t.aa..g..............c....g.....t..............g...............a............... [ 10720]
AY261366.1_Warthog               .......................g.........t......a....................................... [ 10720]
AY261365.1_Warmbaths             .......................g.........t......a....................................... [ 10720]
AY261364.1_Tengani_62            ...aa..g..............c..........t.....a.t......g...............a............... [ 10720]
AY261363.1_Pretoriuskop/96/4     .........................................t...................................... [ 10720]
AY261362.1_Mkuzi_1979            .t.aa..g..............cg...g.....t..............g...............a............... [ 10720]
AY261361.1_Malawi_Lil-20/1_1983  ...aa..g..............c.....t....t.....a.t......g...............a............... [ 10720]
AY261360.1_Kenya_1950            .t.aa..g............c.c..........t.....a.t......g...............a............... [ 10720]
AM712240.1_OURT_88/3             ...aa..g..............c....g.....t..............g...............a............... [ 10720]
AM712239.1_Benin_97/1            .t.aa..g..............c....g.....t..............g...............a............... [ 10720]
KM111295.1_Ken06.Bus             ....a...........c......g.................t...................t.................g [ 10720]
KM111294.1_Ken05/Tk1             ....................c..g...a.....t..............g..........t.t.................. [ 10720]
KM262844.1_strain_L60            ...aa..g..............c....g.....t..............g...............a............... [ 10720]
KM102979.1_26544/OG10_Italy      .t.aa..g..............c....g.....t..............g...............a............... [ 10720]
KM262845.1_strain_NHV            ...aa..g..............c....g.....t..............g...............a............... [ 10720]
KX354450.1_47/Ss/2008            .t.aa..g..............c....g.....t..............g...............a............... [ 10720]
KP055815.1_strain_BA71           .t.aa..g..............c....g.....t..............g...............a............... [ 10720]
MH025920.1_strain_R35            ....a...........c......g.................t.....................................g [ 10720]
MH025919.1_strain_N10            ....a...........c......g.................t......g............t.................g [ 10720]
MH025918.1_strain_R25            ....a...........c......g.................t.....................................g [ 10720]
MH025917.1_strain_R7             ....a...........c......g.................t.....................................g [ 10720]
MH025916.1_strain_R8             ....a...........c......g.................t.....................................g [ 10720]

FR682468.2_Georgia_2007/1        ttcacctatgggggaatca-tcttaaaaaccttattcctacagattccattttgacagtcccagcaaaagtcacaatatt [ 10800]
ASF_IND_20_CAD543                ..............g..g.t........r................................t.................. [ 10800]
LR899193.1_Germany_2020/1        ...................-.........................................t.................. [ 10800]
MN393477.1_Wuhan_2019            ...................-.........................................t.................. [ 10800]
MW396979.1_Timor-Leste/2019/1    ...................-.........................................t.................. [ 10800]
MK940252.1_CN/2019/InnerMongolia ...................-.........................................t.................. [ 10800]
MK128995.1_China/2018/AnhuiXCGQ  ...................-.........................................t.................. [ 10800]
MH681419.1_ASFV/POL/2015         ...................-............................................................ [ 10800]
LS478113.1_Estonia_2014          ---------------------------------------..c...cta.aa..t.ac..ttttt...g.t.ttt.ga.a. [ 10800]
KP843857.1_Odintsovo_02/14       ...................-.........................................t.................. [ 10800]
FN557520.1_strain_E75            ..............g..g.t........g........t.......g.........................t........ [ 10800]
AY261366.1_Warthog               ..............g..g.t.........................a.................................. [ 10800]
AY261365.1_Warmbaths             ..............g..g.t.........................a.................................. [ 10800]
AY261364.1_Tengani_62            ..............g..g.t.........................g..g............................... [ 10800]
AY261363.1_Pretoriuskop/96/4     ..............g..g.t......................a..g.................................. [ 10800]
AY261362.1_Mkuzi_1979            ..............g..g.t.................t.......g.........................t........ [ 10800]
AY261361.1_Malawi_Lil-20/1_1983  ..............g..g.t..........t......t.......g...........a.............t........ [ 10800]
AY261360.1_Kenya_1950            .........a....g..g.t.........................g.........................t........ [ 10800]
AM712240.1_OURT_88/3             ..............g..g.t........g........t.......g.........................t........ [ 10800]
AM712239.1_Benin_97/1            ..............g..g.t........g........t.......g.........................t........ [ 10800]
KM111295.1_Ken06.Bus             ..............g..g.t................t.....a..g.................................. [ 10800]
```

```
KM111294.1_Ken05/Tk1             ....t.........g..g.t.........................g...........a...................... [ 10800]
KM262844.1_strain_L60            ..............g..g.t........g........t.......g.........................t........ [ 10800]
KM102979.1_26544/OG10_Italy      ..............g..g.t........g........t.......g.........................t........ [ 10800]
KM262845.1_strain_NHV            ..............g..g.t........g........t.......g.........................t........ [ 10800]
KX354450.1_47/Ss/2008            ..............g..g.t........g........t.......g.........................t........ [ 10800]
KP055815.1_strain_BA71           ..............g..g.t........g........t.......g.........................t........ [ 10800]
MH025920.1_strain_R35            ..............g..g.t................t.....a..g.................................. [ 10800]
MH025919.1_strain_N10            ..............g..g.t................t.....a..g.................................. [ 10800]
MH025918.1_strain_R25            ..............g..g.t................t.....a..g.................................. [ 10800]
MH025917.1_strain_R7             ..............g..g.t................t.....a..g.................................. [ 10800]
MH025916.1_strain_R8             ..............g..g.t................t.....a..g.................................. [ 10800]

FR682468.2_Georgia_2007/1        ttccatgagtacaccaatgttcaagctctctttcgggaggaatgctgccaattttatgttttttagcttctaactctctg [ 10880]
ASF_IND_20_CAD543                .................................................................k............ca [ 10880]
LR899193.1_Germany_2020/1        ................................................................................ [ 10880]
MN393477.1_Wuhan_2019            ................................................................................ [ 10880]
MW396979.1_Timor-Leste/2019/1    ................................................................................ [ 10880]
MK940252.1_CN/2019/InnerMongolia ................................................................................ [ 10880]
MK128995.1_China/2018/AnhuiXCGQ  ................................................................................ [ 10880]
MH681419.1_ASFV/POL/2015         ................................................................................ [ 10880]
LS478113.1_Estonia_2014          aa.....ga..--------------------------------------------------...g.ta.t..-------- [ 10880]
KP843857.1_Odintsovo_02/14       ................................................................................ [ 10880]
FN557520.1_strain_E75            .................................t...............................t............ca [ 10880]
AY261366.1_Warthog               .................................t..................c............t.............. [ 10880]
AY261365.1_Warmbaths             .................................t..................c............t.............. [ 10880]
AY261364.1_Tengani_62            .................................................................tg............. [ 10880]
AY261363.1_Pretoriuskop/96/4     .................................t..................c............t.............. [ 10880]
AY261362.1_Mkuzi_1979            .................................t................g..............t............ca [ 10880]
AY261361.1_Malawi_Lil-20/1_1983  .................................................................t.............. [ 10880]
AY261360.1_Kenya_1950            ..............................................a...g..............t.............. [ 10880]
AM712240.1_OURT_88/3             .................................t...............................t............ca [ 10880]
AM712239.1_Benin_97/1            .................................t...............................t............ca [ 10880]
KM111295.1_Ken06.Bus             ......a..........................................................tg............. [ 10880]
KM111294.1_Ken05/Tk1             ......................g.............................c.......g....t..agc.....a.g. [ 10880]
KM262844.1_strain_L60            .................................t...............................t............ca [ 10880]
KM102979.1_26544/OG10_Italy      .................................t...............................t............ca [ 10880]
KM262845.1_strain_NHV            .................................t...............................t............ca [ 10880]
KX354450.1_47/Ss/2008            .................................t...............................t............ca [ 10880]
KP055815.1_strain_BA71           .................................t...............................t............ca [ 10880]
MH025920.1_strain_R35            ......a..........................................................tg............. [ 10880]
MH025919.1_strain_N10            ......a..........................................................tg............. [ 10880]
MH025918.1_strain_R25            ......a..........................................................tg............. [ 10880]
MH025917.1_strain_R7             ......a..........................................................tg............. [ 10880]
MH025916.1_strain_R8             ......a..........................................................tg............. [ 10880]
```

```
FR682468.2_Georgia_2007/1        tacaacatcagttgggaaagcagaaagaagattaccaggagaaccattaaatatataatagtctgcaaactacgtttgcg [ 10960]
ASF_IND_20_CAD543                ...........................r............ag...................................... [ 10960]
LR899193.1_Germany_2020/1        ................................................................................ [ 10960]
MN393477.1_Wuhan_2019            ................................................................................ [ 10960]
MW396979.1_Timor-Leste/2019/1    ................................................................................ [ 10960]
MK940252.1_CN/2019/InnerMongolia ................................................................................ [ 10960]
MK128995.1_China/2018/AnhuiXCGQ  ................................................................................ [ 10960]
MH681419.1_ASFV/POL/2015         ................................................................................ [ 10960]
LS478113.1_Estonia_2014          ------------------------------------------------..g...g.c..t.c..a..t....tta...ac [ 10960]
KP843857.1_Odintsovo_02/14       ................................................................................ [ 10960]
FN557520.1_strain_E75            ....g......................g............ag....gc.....c....c......t.............. [ 10960]
AY261366.1_Warthog               ....g........................a................gc.....c....c..................... [ 10960]
AY261365.1_Warmbaths             ....g........................a................gc.....c....c..................... [ 10960]
AY261364.1_Tengani_62            ..............................................gc.....c....c..................... [ 10960]
AY261363.1_Pretoriuskop/96/4     ....g.........................................gc.....c....c..................... [ 10960]
AY261362.1_Mkuzi_1979            ....g.........................................gc.....c....c..................... [ 10960]
AY261361.1_Malawi_Lil-20/1_1983  ....g..........................................c.....c.......................... [ 10960]
AY261360.1_Kenya_1950            ....g.......................................t.ac.......c........................ [ 10960]
AM712240.1_OURT_88/3             ....g......................g............ag....gc.....c....c......t.............. [ 10960]
AM712239.1_Benin_97/1            ....g......................g............ag....gc.....c....c......t.............. [ 10960]
KM111295.1_Ken06.Bus             ....g.........................................cc...............c.......g........ [ 10960]
KM111294.1_Ken05/Tk1             ....g.........a.......................a.....t.gc................................ [ 10960]
KM262844.1_strain_L60            ....g......................g............ag....gc.....c....c......t.............. [ 10960]
KM102979.1_26544/OG10_Italy      ....g......................g............ag....gc.....c....c......t.............. [ 10960]
KM262845.1_strain_NHV            ....g......................g............ag....gc.....c....c......t.............. [ 10960]
KX354450.1_47/Ss/2008            ....g......................g............ag....gc.....c....c......t.............. [ 10960]
KP055815.1_strain_BA71           ....g......................g............ag....gc.....c....c......t.............. [ 10960]
MH025920.1_strain_R35            ....g.........................................cc...............c.......g........ [ 10960]
MH025919.1_strain_N10            ....g.........................................cc...............c.......g........ [ 10960]
MH025918.1_strain_R25            ....g.........................................cc...............c.......g........ [ 10960]
MH025917.1_strain_R7             ....g.........................................cc...............c.......g........ [ 10960]
MH025916.1_strain_R8             ....g.........................................cc...............c.......g........ [ 10960]

FR682468.2_Georgia_2007/1        aatgtaatttgcaactaaaacacaacccacaaggtaaaatccataagttaataacttttgccattttcgtatgacagcct [ 11040]
ASF_IND_20_CAD543                .........ca...........................................t..c..............at...... [ 11040]
LR899193.1_Germany_2020/1        ................................................................................ [ 11040]
MN393477.1_Wuhan_2019            ................................................................................ [ 11040]
MW396979.1_Timor-Leste/2019/1    ................................................................................ [ 11040]
MK940252.1_CN/2019/InnerMongolia ................................................................................ [ 11040]
MK128995.1_China/2018/AnhuiXCGQ  ................................................................................ [ 11040]
MH681419.1_ASFV/POL/2015         ................................................................................ [ 11040]
LS478113.1_Estonia_2014          .....--------------------------------------------------------------------------- [ 11040]
KP843857.1_Odintsovo_02/14       ................................................................................ [ 11040]
```

```
FN557520.1_strain_E75           .........ca..g......t...t...g....a...........g.....c..t..c..............at...... [ 11040]
AY261366.1_Warthog              ..c..............................a.............................................. [ 11040]
AY261365.1_Warmbaths            ..c..............................a.............................................. [ 11040]
AY261364.1_Tengani_62           ..c..............................a...........g..........................at...... [ 11040]
AY261363.1_Pretoriuskop/96/4    ..c..............................a...........g.................................. [ 11040]
AY261362.1_Mkuzi_1979           ..c..............................a...........g........t..c...................... [ 11040]
AY261361.1_Malawi_Lil-20/1_1983 ..c..............................a...........g........t..c.................t.... [ 11040]
AY261360.1_Kenya_1950           ..........a......................a...........g..........................a..t.... [ 11040]
AM712240.1_OURT_88/3            .........ca..g......t...t...g....a...........g.....c..t..c..............at...... [ 11040]
AM712239.1_Benin_97/1           .........ca..g......t...t...g....a...........g.....c..t..c..............at...... [ 11040]
KM111295.1_Ken06.Bus            ...a......a......................a...........g........t..c.................g.... [ 11040]
KM111294.1_Ken05/Tk1            ..c..............................a...........g........t..c..............a..t.... [ 11040]
KM262844.1_strain_L60           .........ca..g......t...t...g....a...........g.....c..t..c..............at...... [ 11040]
KM102979.1_26544/OG10_Italy     .........ca..g......t...t...g....a...........g.....c..t..c..............at...... [ 11040]
KM262845.1_strain_NHV           .........ca..g......t...t...g....a...........g.....c..t..c..............at...... [ 11040]
KX354450.1_47/Ss/2008           .........ca..g......t...t...g....a...........g.....c..t..c..............at...... [ 11040]
KP055815.1_strain_BA71          .........ca..g......t...t...g....a...........g.....c..t..c..............at...... [ 11040]
MH025920.1_strain_R35           ...a......a......................a...........g........t..c.................g.... [ 11040]
MH025919.1_strain_N10           ...a......a......................a...........g........t..c.................g.... [ 11040]
MH025918.1_strain_R25           ...a......a......................a...........g........t..c.................g.... [ 11040]
MH025917.1_strain_R7            ...a......a......................a...........g........t..c.................g.... [ 11040]
MH025916.1_strain_R8            ...a......a......................a...........g........t..c.................g.... [ 11040]

FR682468.2_Georgia_2007/1       cgtgccattcatggt---tgtgttgtgggcattctgttcggtaaacttcatgaggctttatagaagttacatagtaggta [ 11120]
ASF_IND_20_CAD543               ...............tta.........................cgt..t....................tg......... [ 11120]
LR899193.1_Germany_2020/1       ...............---.............................................................. [ 11120]
MN393477.1_Wuhan_2019           ...............---.............................................................. [ 11120]
MW396979.1_Timor-Leste/2019/1   ...............---.............................................................. [ 11120]
MK940252.1_CN/2019/InnerMongolia ...............---.............................................................. [ 11120]
MK128995.1_China/2018/AnhuiXCGQ ...............---.............................................................. [ 11120]
MH681419.1_ASFV/POL/2015        ...............---.............................................................. [ 11120]
LS478113.1_Estonia_2014         -----------....---aa.agga.taat..a.c..atta..gttc....a------.....g...at.----c...cg [ 11120]
KP843857.1_Odintsovo_02/14      ...............---.............................................................. [ 11120]
FN557520.1_strain_E75           .a....t........tta..........a..............cgt..t....................tg......a.. [ 11120]
AY261366.1_Warthog              .............ac---.c..................................................g.....a... [ 11120]
AY261365.1_Warmbaths            .............ac---.c..................................................g.....a... [ 11120]
AY261364.1_Tengani_62           .a....g.......ctta..........a..............c.t..t....................tg......a.. [ 11120]
AY261363.1_Pretoriuskop/96/4    .............ac---.c..................................................g.....a... [ 11120]
AY261362.1_Mkuzi_1979           ...............tta..........a..............cgt..t....................tg......a.. [ 11120]
AY261361.1_Malawi_Lil-20/1_1983 .a............ctca...a......a..............c.t..t....................tg......a.. [ 11120]
AY261360.1_Kenya_1950           .a............cttaa.....c..aa................t..t....................tg..a...a.. [ 11120]
AM712240.1_OURT_88/3            .a....t........tta..........a..............cgt..t....................tg.g....a.. [ 11120]
AM712239.1_Benin_97/1           .a....t........tta..........a..............cgt..t....................tg......a.. [ 11120]
KM111295.1_Ken06.Bus            .a............ctta...........................t..t.....................g......... [ 11120]
```

```
KM111294.1_Ken05/Tk1           .a............ctta..........a................t..t....................tg......a.. [ 11120]
KM262844.1_strain_L60          .a....t........tta..........a..............cgt..t....................tg......a.. [ 11120]
KM102979.1_26544/OG10_Italy    .a....t........tta..........a..............cgt..t....................tg......a.. [ 11120]
KM262845.1_strain_NHV          .a....t........tta..........a..............cgt..t....................tg.g....a.. [ 11120]
KX354450.1_47/Ss/2008          .a....t........tta..........a..............cgt..t....................tg......a.. [ 11120]
KP055815.1_strain_BA71         .a....t........tta..........a..............cgt..t....................tg......a.. [ 11120]
MH025920.1_strain_R35          .a............ctta...........................t..t.....................g......... [ 11120]
MH025919.1_strain_N10          .a............ctta...........................t..t.....................g......... [ 11120]
MH025918.1_strain_R25          .a............ctta...........................t..t.....................g......... [ 11120]
MH025917.1_strain_R7           .a............ctta...........................t..t.....................g......... [ 11120]
MH025916.1_strain_R8           .a............ctta...........................t..t.....................g......... [ 11120]

FR682468.2_Georgia_2007/1      cagaattcattgtgacgaaaaacactgcagttagctatgtagtcattttcaagaatgggagaatggttttcaaagacctt [ 11200]
ASF_IND_20_CAD543              ......................................r....y..........................t...ag.... [ 11200]
LR899193.1_Germany_2020/1      ................................................................................ [ 11200]
MN393477.1_Wuhan_2019          ................................................................................ [ 11200]
MW396979.1_Timor-Leste/2019/1  ................................................................................ [ 11200]
MK940252.1_CN/2019/InnerMongolia ................................................................................ [ 11200]
MK128995.1_China/2018/AnhuiXCGQ ................................................................................ [ 11200]
MH681419.1_ASFV/POL/2015       ................................................................................ [ 11200]
LS478113.1_Estonia_2014        ..--------------a.....tgt..tttg..a.a...cgc..gc...------------------------------- [ 11200]
KP843857.1_Odintsovo_02/14     ................................................................................ [ 11200]
FN557520.1_strain_E75          .....a......c.....t...t...............a...............................t...ag.... [ 11200]
AY261366.1_Warthog             ............c................................................................... [ 11200]
AY261365.1_Warmbaths           ............c..............................................................g.... [ 11200]
AY261364.1_Tengani_62          .....a......c.....t...t...............a...............................t...ag..c. [ 11200]
AY261363.1_Pretoriuskop/96/4   ............c................................................................... [ 11200]
AY261362.1_Mkuzi_1979          .....a......c.....t...................a...............................t...ag.... [ 11200]
AY261361.1_Malawi_Lil-20/1_1983 .....a.t....ct....t...................a...............................t...ag.... [ 11200]
AY261360.1_Kenya_1950          .....a......c.....t..........a........................................t...ag.... [ 11200]
AM712240.1_OURT_88/3           .....a......c.....t...t...............a...............................t...ag.... [ 11200]
AM712239.1_Benin_97/1          .....a......c.....t...t...............a...............................t...ag.... [ 11200]
KM111295.1_Ken06.Bus           .....a.t......gt..t..............................................a....t....g.... [ 11200]
KM111294.1_Ken05/Tk1           .....a......c..t..t............................................................. [ 11200]
KM262844.1_strain_L60          .....a......c.....t...t...............a...............................t...ag.... [ 11200]
KM102979.1_26544/OG10_Italy    .....a......c.....t...t...............a...............................t...ag.... [ 11200]
KM262845.1_strain_NHV          .....a......c.....t...t...............a...............................t...ag.... [ 11200]
KX354450.1_47/Ss/2008          .....a......c.....t...t...............a...............................t...ag.... [ 11200]
KP055815.1_strain_BA71         .....a......c.....t...t...............a...............................t...ag.... [ 11200]
MH025920.1_strain_R35          .....a.t......gt..t..............................................a....t....g.... [ 11200]
MH025919.1_strain_N10          .....a........gt..t..............................................a....t....g.... [ 11200]
MH025918.1_strain_R25          .....a.t......gt..t..............................................a....t....g.... [ 11200]
MH025917.1_strain_R7           .....a.t......gt..t..............................................a....t....g.... [ 11200]
MH025916.1_strain_R8           .....a.t......gt..t..............................................a....t....g.... [ 11200]
```

```
FR682468.2_Georgia_2007/1         attcttacagatgccatcttgacagtcccaacagaacctacaatgatttgcataggtgcaccagtattcaagctcctttt [ 11280]
ASF_IND_20_CAD543                 ......................................................r............y............ [ 11280]
LR899193.1_Germany_2020/1         ................................................................................ [ 11280]
MN393477.1_Wuhan_2019             ................................................................................ [ 11280]
MW396979.1_Timor-Leste/2019/1     ................................................................................ [ 11280]
MK940252.1_CN/2019/InnerMongolia  ................................................................................ [ 11280]
MK128995.1_China/2018/AnhuiXCGQ   ................................................................................ [ 11280]
MH681419.1_ASFV/POL/2015          ................................................................................ [ 11280]
LS478113.1_Estonia_2014           -----------------------------------.tt.g.t...----------------t....a.t.g.tatt.... [ 11280]
KP843857.1_Odintsovo_02/14        ................................................................................ [ 11280]
FN557520.1_strain_E75             g...c.......a.......................t........g........a............c............ [ 11280]
AY261366.1_Warthog                ....c..........................................................a................ [ 11280]
AY261365.1_Warmbaths              ....c..........................................................a................ [ 11280]
AY261364.1_Tengani_62             g......................................g..............a..a.........c............ [ 11280]
AY261363.1_Pretoriuskop/96/4      ................................................................................ [ 11280]
AY261362.1_Mkuzi_1979             g...c.......a.......................t.....g..g.....................c............ [ 11280]
AY261361.1_Malawi_Lil-20/1_1983   g...c................g...........a..t.....g..g........a............c............ [ 11280]
AY260360.1_Kenya_1950             g...c....a..a.......................a........g........a............c............ [ 11280]
AM712240.1_OURT_88/3              g...c.......a.......................t........g........a............c...a........ [ 11280]
AM712239.1_Benin_97/1             g...c.......a.......................t........g........a............c............ [ 11280]
KM111295.1_Ken06.Bus              g...c................g..............t.....g..g.................a...c............ [ 11280]
KM111294.1_Ken05/Tk1              g...c....a..a.......................t.....g..g........a............c....a....... [ 11280]
KM262844.1_strain_L60             g...c.......a.......................t........g........a............c............ [ 11280]
KM102979.1_26544/OG10_Italy       g...c.......a.......................t........g........a............c............ [ 11280]
KM262845.1_strain_NHV             g...c.......a.......................t........g........a............c............ [ 11280]
KX354450.1_47/Ss/2008             g...c.......a.......................t........g........a............c............ [ 11280]
KP055815.1_strain_BA71            g...c.......a.......................t........g........a............c............ [ 11280]
MH025920.1_strain_R35             g...c................g..............t.....g..g.................a...c............ [ 11280]
MH025919.1_strain_N10             g...c................g..............t.....g..g.................a...c............ [ 11280]
MH025918.1_strain_R25             g...c................g..............t.....g..g.................a...c............ [ 11280]
MH025917.1_strain_R7              g...c................g..............t.....g..g.................a...c............ [ 11280]
MH025916.1_strain_R8              g...c................g..............t.....g..g.................a...c............ [ 11280]

FR682468.2_Georgia_2007/1         caggaggggttcttgttagat---ccaggagct---ctagctcatatgtataaagaagagttggaatggatagtaaagta [ 11360]
ASF_IND_20_CAD543                 ....w................---.........---.......................................g.... [ 11360]
LR899193.1_Germany_2020/1         .....................---.........---............................................ [ 11360]
MN393477.1_Wuhan_2019             .....................---.........---............................................ [ 11360]
MW396979.1_Timor-Leste/2019/1     .....................---.........---............................................ [ 11360]
MK940252.1_CN/2019/InnerMongolia  .....................---.........---............................................ [ 11360]
MK128995.1_China/2018/AnhuiXCGQ   .....................---.........---............................................ [ 11360]
MH681419.1_ASFV/POL/2015          .....................---.........---............................................ [ 11360]
LS478113.1_Estonia_2014           .-------------------------------------------------.....c.at....t..c------------- [ 11360]
KP843857.1_Odintsovo_02/14        .....................---.........---............................................ [ 11360]
```

```
FN557520.1_strain_E75           ....t.....a......c.t.---.....g...---...t...........................a.......gg..g [ 11360]
AY261366.1_Warthog              .....................---.........---..........c............................g.... [ 11360]
AY261365.1_Warmbaths            .....................---.........---..........c............................g.... [ 11360]
AY261364.1_Tengani_62           ....t..a..a......c.t.---.....g...---..........c............................g.... [ 11360]
AY261363.1_Pretoriuskop/96/4    .....................---.........---...............................a.......gg..g [ 11360]
AY261362.1_Mkuzi_1979           .................c.t.---.....g...---...t...........................a.......gg..g [ 11360]
AY261361.1_Malawi_Lil-20/1_1983 ....ta....a......c.t.---.....g...---..........c....................a.......gg..g [ 11360]
AY261360.1_Kenya_1950           .ta.c......tc....c.t.---.....g...---............c...................a......gg..g [ 11360]
AM712240.1_OURT_88/3            ....t.....a......c.t.---.....g...---...t...........................a.......gg..g [ 11360]
AM712239.1_Benin_97/1           ....t.....a......c.t.---.....g...---...t...........................a.......gg..g [ 11360]
KM111295.1_Ken06.Bus            ....t.............at.---...a.g...cta....t.............a....................gg... [ 11360]
KM111294.1_Ken05/Tk1            ....t.....a......ctt.cca...a.g...---...............................a....a..gg... [ 11360]
KM262844.1_strain_L60           ....t.....a......c.t.---.....g...---...t...........................a.......gg..g [ 11360]
KM102979.1_26544/OG10_Italy     ....t.....a......c.t.---.....g...---...t...........................a.......gg..g [ 11360]
KM262845.1_strain_NHV           ....t.....a......c.t.---.....g...---...t...........................a.......gg..g [ 11360]
KX354450.1_47/Ss/2008           ....t.....a......c.t.---.....g...---...t...........................a.......gg..g [ 11360]
KP055815.1_strain_BA71          ....t.....a......c.t.---.....g...---...t...........................a.......gg..g [ 11360]
MH025920.1_strain_R35           ....t.............at.---...a.g...cta....t.............a....................gg... [ 11360]
MH025919.1_strain_N10           ....t.............at.---...a.g...cta....t.............a....................gg... [ 11360]
MH025918.1_strain_R25           ....t.............at.---...a.g...cta....t.............a....................gg... [ 11360]
MH025917.1_strain_R7            ....t.............at.---...a.g...cta....t.............a....................gg... [ 11360]
MH025916.1_strain_R8            ....t.............at.---...a.g...cta....t.............a....................gg... [ 11360]

FR682468.2_Georgia_2007/1       aatatttgcagaccaagcatggctacttgtgaacaagtggctgctcgtcaacaaatagctgtttatcagcaaatagctgt [ 11440]
ASF_IND_20_CAD543               ..................................................r..........------------------- [ 11440]
LR899193.1_Germany_2020/1       ................................................................................ [ 11440]
MN393477.1_Wuhan_2019           ................................................................................ [ 11440]
MW396979.1_Timor-Leste/2019/1   ................................................................................ [ 11440]
MK940252.1_CN/2019/InnerMongolia ................................................................................ [ 11440]
MK128995.1_China/2018/AnhuiXCGQ ................................................................................ [ 11440]
MH681419.1_ASFV/POL/2015        ................................................................................ [ 11440]
LS478113.1_Estonia_2014         -------------------------------------------------------------------------------- [ 11440]
KP843857.1_Odintsovo_02/14      ................................................................................ [ 11440]
FN557520.1_strain_E75           ..................................................g..........c.cg.........g....c [ 11440]
AY261366.1_Warthog              ........................................t.........gt...........................c [ 11440]
AY261365.1_Warmbaths            ........................................t.........gt...........................c [ 11440]
AY261364.1_Tengani_62           .............................................-------------------g............... [ 11440]
AY261363.1_Pretoriuskop/96/4    ...........................................t......g...................-..c...... [ 11440]
AY261362.1_Mkuzi_1979           .....................................------------------------------------.g....c [ 11440]
AY261361.1_Malawi_Lil-20/1_1983 ..............................a.......t...a..t..g............c.cg.......g....... [ 11440]
AY261360.1_Kenya_1950           .t...................a..gt....tc....c........t..g.....g.g....c.cg............... [ 11440]
AM712240.1_OURT_88/3            ..................................................g..........c.cg.........g....c [ 11440]
AM712239.1_Benin_97/1           ..................................................g..........c.cg.........g....c [ 11440]
KM111295.1_Ken06.Bus            ............g...a....a------------------.....t..a.......g...ac.cg.........g..... [ 11440]
```

```
KM111294.1_Ken05/Tk1            ................a.................................g...c.g....c.cg............... [ 11440]
KM262844.1_strain_L60           ..................................................g..........c.cg.........g....c [ 11440]
KM102979.1_26544/OG10_Italy     ..................................................g..........c.cg.........g....c [ 11440]
KM262845.1_strain_NHV           ..................................................g..........c.cg.........g....c [ 11440]
KX354450.1_47/Ss/2008           ..................................................g..........c.cg.........g....c [ 11440]
KP055815.1_strain_BA71          ..................................................g..........c.cg.........g....c [ 11440]
MH025920.1_strain_R35           ............g...a....a------------------.....t..a.......g...ac.cg.........g..... [ 11440]
MH025919.1_strain_N10           ............g...a....a------------------.....t..a.......g...ac.cg.........g..... [ 11440]
MH025918.1_strain_R25           ............g...a....a------------------.....t..a.......g...ac.cg.........g..... [ 11440]
MH025917.1_strain_R7            ............g...a....a------------------.....t..a.......g...ac.cg.........g..... [ 11440]
MH025916.1_strain_R8            ............g...a....a------------------.....t..a.......g...ac.cg.........g..... [ 11440]

FR682468.2_Georgia_2007/1       ttatcagcaacaactaattatcagcaaatgctgcttgtgggtaagccaataaataggccatacccttgaaaggagaattc [ 11520]
ASF_IND_20_CAD543               ---------------------------------.......................................r....... [ 11520]
LR899193.1_Germany_2020/1       ................................................................................ [ 11520]
MN393477.1_Wuhan_2019           ................................................................................ [ 11520]
MW396979.1_Timor-Leste/2019/1   ................................................................................ [ 11520]
MK940252.1_CN/2019/InnerMongolia ................................................................................ [ 11520]
MK128995.1_China/2018/AnhuiXCGQ ................................................................................ [ 11520]
MH681419.1_ASFV/POL/2015        ................................................................................ [ 11520]
LS478113.1_Estonia_2014         -------------------------------------------------------------------------------- [ 11520]
KP843857.1_Odintsovo_02/14      ................................................................................ [ 11520]
FN557520.1_strain_E75           -----------------.cg..................................g......................... [ 11520]
AY261366.1_Warthog              -----------------.cg..................................c......................... [ 11520]
AY261365.1_Warmbaths            -----------------.cg..................................c......................... [ 11520]
AY261364.1_Tengani_62           -----------------.cg.......g.........................cc......................... [ 11520]
AY261363.1_Pretoriuskop/96/4    -----------------..........g.........................cc......................... [ 11520]
AY261362.1_Mkuzi_1979           -----------------.cg..................................g......................... [ 11520]
AY261361.1_Malawi_Lil-20/1_1983 -----------------..........g.........................cc............a............ [ 11520]
AY261360.1_Kenya_1950           -----------------..........g.........................cc............a............ [ 11520]
AM712240.1_OURT_88/3            -----------------.cg..................................g......................... [ 11520]
AM712239.1_Benin_97/1           -----------------.cg..................................g......................... [ 11520]
KM111295.1_Ken06.Bus            -----------------c..g............a.a.....-......c...g-ta.................g...... [ 11520]
KM111294.1_Ken05/Tk1            -----------------..........g.........................cc............a............ [ 11520]
KM262844.1_strain_L60           -----------------.cg..................................g......................... [ 11520]
KM102979.1_26544/OG10_Italy     -----------------.cg..................................g......................... [ 11520]
KM262845.1_strain_NHV           -----------------.cg..................................g......................... [ 11520]
KX354450.1_47/Ss/2008           -----------------.cg..................................g......................... [ 11520]
KP055815.1_strain_BA71          -----------------.cg..................................g......................... [ 11520]
MH025920.1_strain_R35           -----------------c..g............a.a.....-......c...g-ta.................g...... [ 11520]
MH025919.1_strain_N10           -----------------c..g............a.a.....-......c...g-ta.................g...... [ 11520]
MH025918.1_strain_R25           -----------------c..g............a.a.....-......c...g-ta.................g...... [ 11520]
MH025917.1_strain_R7            -----------------c..g............a.a.....-......c...g-ta.................g...... [ 11520]
MH025916.1_strain_R8            -----------------c..g............a.a.....-......c...g-ta.................g...... [ 11520]
```

```
FR682468.2_Georgia_2007/1        ag-tttgataaaaaaaataacgagttttctaataacccggtcaagcatttaataaatgaatagcatcacacgtctgcatc [ 11600]
ASF_IND_20_CAD543                ..-............................................................................. [ 11600]
LR899193.1_Germany_2020/1        ..-............................................................................. [ 11600]
MN393477.1_Wuhan_2019            ..-............................................................................. [ 11600]
MW396979.1_Timor-Leste/2019/1    ..-............................................................................. [ 11600]
MK940252.1_CN/2019/InnerMongolia ..-............................................................................. [ 11600]
MK128995.1_China/2018/AnhuiXCGQ  ..-............................................................................. [ 11600]
MH681419.1_ASFV/POL/2015         ..-............................................................................. [ 11600]
LS478113.1_Estonia_2014          --------.g..........tc.ac.-----------------.aa........-------------------------- [ 11600]
KP843857.1_Odintsovo_02/14       ..-............................................................................. [ 11600]
FN557520.1_strain_E75            ..-................................................t.......g.g......g.t......... [ 11600]
AY261366.1_Warthog               ..-................................................t.......g.g.................. [ 11600]
AY261365.1_Warmbaths             ..-................................................t.......g.g.................. [ 11600]
AY261364.1_Tengani_62            ..-........................c....c.......................c....g.................. [ 11600]
AY261363.1_Pretoriuskop/96/4     ..-.......................cct--.ac......................c....g.................. [ 11600]
AY261362.1_Mkuzi_1979            ..-....................................a...........t.......g.g.......tt......... [ 11600]
AY261361.1_Malawi_Lil-20/1_1983  ..-.............................c...........a...........c....g.................. [ 11600]
AY261360.1_Kenya_1950            ..-..................t.....c...gc..t.....t.........g.........g.................. [ 11600]
AM712240.1_OURT_88/3             ..t................................................t.......g.g......g.t.....t... [ 11600]
AM712239.1_Benin_97/1            ..-................................................t.......g.g......g.t......... [ 11600]
KM111295.1_Ken06.Bus             ..-..................t.....c...gc..t..a..t.........g.........ga.....t........... [ 11600]
KM111294.1_Ken05/Tk1             ..-..................t.....c...gc..t..a..t.........g.........g.................. [ 11600]
KM262844.1_strain_L60            ..-................................................t.......g.g......g.t......... [ 11600]
KM102979.1_26544/OG10_Italy      ..-................................................t.......g.g......g.t......... [ 11600]
KM262845.1_strain_NHV            ..t................................................t.......g.g......g.t.....t... [ 11600]
KX354450.1_47/Ss/2008            ..-................................................t.......g.g......g.t......... [ 11600]
KP055815.1_strain_BA71           ..-................................................t.......g.g......g.t......... [ 11600]
MH025920.1_strain_R35            ..-..................t.....c...gc..t..a..t.........g.........ga.....t........... [ 11600]
MH025919.1_strain_N10            ..-..................t.....c...gc..t..a..t.........g.........ga.....t........... [ 11600]
MH025918.1_strain_R25            ..-..................t.....c...gc..t..a..t.........g.........ga.....t........... [ 11600]
MH025917.1_strain_R7             ..-..................t.....c...gc..t..a..t.........g.........ga.....t........... [ 11600]
MH025916.1_strain_R8             ..-..................t.....c...gc..t..a..t.........g.........ga.....t........... [ 11600]

FR682468.2_Georgia_2007/1        gtgcattctgcctggaaaatgggcccatctctaatatatttacactgacggtgaatcatacagtgttccatgggatagct [ 11680]
ASF_IND_20_CAD543                ................................................................................ [ 11680]
LR899193.1_Germany_2020/1        ................................................................................ [ 11680]
MN393477.1_Wuhan_2019            ................................................................................ [ 11680]
MW396979.1_Timor-Leste/2019/1    ................................................................................ [ 11680]
MK940252.1_CN/2019/InnerMongolia ................................................................................ [ 11680]
MK128995.1_China/2018/AnhuiXCGQ  ................................................................................ [ 11680]
MH681419.1_ASFV/POL/2015         ................................................................................ [ 11680]
LS478113.1_Estonia_2014          -------------------------------------------------------------------------------- [ 11680]
KP843857.1_Odintsovo_02/14       ................................................................................ [ 11680]
```

```
FN557520.1_strain_E75          .........a.t.................................................................... [ 11680]
AY261366.1_Warthog             ................................................................................ [ 11680]
AY261365.1_Warmbaths           ................................................................................ [ 11680]
AY261364.1_Tengani_62          .........a.............................................g........................ [ 11680]
AY261363.1_Pretoriuskop/96/4   .........a....................g........................g........a............... [ 11680]
AY261362.1_Mkuzi_1979          .........a.t.................................................................... [ 11680]
AY261361.1_Malawi_Lil-20/1_1983 .........a...................c.........................g........................ [ 11680]
AY261360.1_Kenya_1950          a........a.t....................t.c..........................................a.. [ 11680]
AM712240.1_OURT_88/3           .........a.t.......................................................t............ [ 11680]
AM712239.1_Benin_97/1          .........a.t.................................................................... [ 11680]
KM111295.1_Ken06.Bus           a........a.t....................t.c............................................. [ 11680]
KM111294.1_Ken05/Tk1           a........a.t..a.................t.c............................................. [ 11680]
KM262844.1_strain_L60          .........a.t.................................................................... [ 11680]
KM102979.1_26544/OG10_Italy    .........a.t.................................................................... [ 11680]
KM262845.1_strain_NHV          .........a.t.......................................................t............ [ 11680]
KX354450.1_47/Ss/2008          .........a.t.................................................................... [ 11680]
KP055815.1_strain_BA71         .........a.t..............................t..................................... [ 11680]
MH025920.1_strain_R35          a........a.t....................t.c............................................. [ 11680]
MH025919.1_strain_N10          a........a.t....................t.c............................................. [ 11680]
MH025918.1_strain_R25          a........a.t....................t.c............................................. [ 11680]
MH025917.1_strain_R7           a........a.t....................t.c............................................. [ 11680]
MH025916.1_strain_R8           a........a.t....................t.c............................................. [ 11680]

FR682468.2_Georgia_2007/1      atgctcctgtacaggaggcatatcttttagaactttattcttacaaagaccatcttgacaagcccagcaaaaccgacaat [ 11760]
ASF_IND_20_CAD543              ...........................w.................................................... [ 11760]
LR899193.1_Germany_2020/1      ................................................................................ [ 11760]
MN393477.1_Wuhan_2019          ................................................................................ [ 11760]
MW396979.1_Timor-Leste/2019/1  ................................................................................ [ 11760]
MK940252.1_CN/2019/InnerMongolia ................................................................................ [ 11760]
MK128995.1_China/2018/AnhuiXCGQ ................................................................................ [ 11760]
MH681419.1_ASFV/POL/2015       ................................................................................ [ 11760]
LS478113.1_Estonia_2014        ----------------------------------..t..t.c.a...a.t.t.t-------------------------- [ 11760]
KP843857.1_Odintsovo_02/14     ................................................................................ [ 11760]
FN557520.1_strain_E75          ..............a..a......a.ccca.........t.......t.....tcct...tt.......g..atc...g. [ 11760]
AY261366.1_Warthog             ..............a............a.................................................... [ 11760]
AY261365.1_Warmbaths           ..............a............a.................................................... [ 11760]
AY261364.1_Tengani_62          .........c....a................................................................. [ 11760]
AY261363.1_Pretoriuskop/96/4   .........c...................................................................... [ 11760]
AY261362.1_Mkuzi_1979          ..............a..a.........a..................................t................. [ 11760]
AY261361.1_Malawi_Lil-20/1_1983 .........c....a........t..................g....ag.....c......................... [ 11760]
AY261360.1_Kenya_1950          ..............a........t..................g....ag.....c......................... [ 11760]
AM712240.1_OURT_88/3           ..............a..a.........a..................................t................. [ 11760]
AM712239.1_Benin_97/1          ..............a..a......a.ccca.........t.......t.....tcct...tt.......g..atc...g. [ 11760]
KM111295.1_Ken06.Bus           ..............a..............a............g..........t........................g. [ 11760]
```

```
KM111294.1_Ken05/Tk1              .........c...aa...........................g....ag.....c......................... [ 11760]
KM262844.1_strain_L60             ..............a..a......a.ccca.........t.......t.....tcct...tt.......g..atc...g. [ 11760]
KM102979.1_26544/OG10_Italy       ..............a..a......a.ccca.........t.......t.....tcct...tt.......g..atc...g. [ 11760]
KM262845.1_strain_NHV             ..............a..a.........a..................................t................. [ 11760]
KX354450.1_47/Ss/2008             ..............a..a......a.ccca.........t.......t.....tcct...tt.......g..atc...g. [ 11760]
KP055815.1_strain_BA71            ..............a..a.........a.....................------------------------------- [ 11760]
MH025920.1_strain_R35             ..............a....g.........a............g..........t........................g. [ 11760]
MH025919.1_strain_N10             ..............a....g.........a............g..........t........................g. [ 11760]
MH025918.1_strain_R25             ..............a....g.........a............g..........t........................g. [ 11760]
MH025917.1_strain_R7              ..............a....g.........a............g..........t........................g. [ 11760]
MH025916.1_strain_R8              ..............a....g.........a............g..........t........................g. [ 11760]

FR682468.2_Georgia_2007/1         ttttcacatattgacaccagtatctaagctcctcttccaggggattgtcggtcgaaaacccctgt--------------- [ 11840]
ASF_IND_20_CAD543                 .......r.........................................................--------------- [ 11840]
LR899193.1_Germany_2020/1         .................................................................--------------- [ 11840]
MN393477.1_Wuhan_2019             .................................................................--------------- [ 11840]
MW396979.1_Timor-Leste/2019/1     .................................................................--------------- [ 11840]
MK940252.1_CN/2019/InnerMongolia  .................................................................--------------- [ 11840]
MK128995.1_China/2018/AnhuiXCGQ   .................................................................--------------- [ 11840]
MH681419.1_ASFV/POL/2015          .................................................................--------------- [ 11840]
LS478113.1_Estonia_2014           -------------------------------------------------------------------------------- [ 11840]
KP843857.1_Odintsovo_02/14        .................................................................--------------- [ 11840]
FN557520.1_strain_E75             g...tc....cgtg.........tc......tct.atag.a..tg.a.aa..---------------------------- [ 11840]
AY261366.1_Warthog                ..........................................................a......--------------- [ 11840]
AY261365.1_Warmbaths              ..........................................................a......--------------- [ 11840]
AY261364.1_Tengani_62             ....................c.....................................t......--------------- [ 11840]
AY261363.1_Pretoriuskop/96/4      ...........a........c.....................................a......--------------- [ 11840]
AY261362.1_Mkuzi_1979             .........................t...............a................a......--------------- [ 11840]
AY261361.1_Malawi_Lil-20/1_1983   ....................c.....................................at.t...agattagctagacta [ 11840]
AY261360.1_Kenya_1950             ....................c................t....................at.....agattagctagacta [ 11840]
AM712240.1_OURT_88/3              .........................t...............a................a......--------------- [ 11840]
AM712239.1_Benin_97/1             g...tc....cgtg.........tc......tct.atag.a..tg.a.aa..---------------------------- [ 11840]
KM111295.1_Ken06.Bus              ...g..t...................................................at.....agattagctagacta [ 11840]
KM111294.1_Ken05/Tk1              ....................c................t....................at.t...agattagctggacta [ 11840]
KM262844.1_strain_L60             g...tc....cgtg.........tc......tct.atag.a..tg.a.aa..---------------------------- [ 11840]
KM102979.1_26544/OG10_Italy       g...tc....cgtg.........tc......tct.atag.a..tg.a.aa..---------------------------- [ 11840]
KM262845.1_strain_NHV             .........................t...............a................a......--------------- [ 11840]
KX354450.1_47/Ss/2008             g...tc....cgtg.........tc......tct.atag.a..tg.a.aa..---------------------------- [ 11840]
KP055815.1_strain_BA71            -------------------------------------------------------------------------------- [ 11840]
MH025920.1_strain_R35             ...g..t...................................................at.....agattagctagacta [ 11840]
MH025919.1_strain_N10             ...g..t...................................................at.....agattagctagacta [ 11840]
MH025918.1_strain_R25             ...g..t...................................................at.....agattagctagacta [ 11840]
MH025917.1_strain_R7              ...g..t...................................................at.....agattagctagacta [ 11840]
MH025916.1_strain_R8              ...g..t...................................................at.....agattagctagacta [ 11840]
```

```
FR682468.2_Georgia_2007/1        ---------------------agactagctaggccagctagcagcaagccgaggtaactaaagaacctcattgtagt--- [ 11920]
ASF_IND_20_CAD543                ---------------------........................................................--- [ 11920]
LR899193.1_Germany_2020/1        ---------------------........................................................--- [ 11920]
MN393477.1_Wuhan_2019            ---------------------........................................................--- [ 11920]
MW396979.1_Timor-Leste/2019/1    ---------------------........................................................--- [ 11920]
MK940252.1_CN/2019/InnerMongolia ---------------------........................................................--- [ 11920]
MK128995.1_China/2018/AnhuiXCGQ  ---------------------........................................................--- [ 11920]
MH681419.1_ASFV/POL/2015         ---------------------........................................................--- [ 11920]
LS478113.1_Estonia_2014          -------------------------------------------------------------------------------- [ 11920]
KP843857.1_Odintsovo_02/14       ---------------------........................................................--- [ 11920]
FN557520.1_strain_E75            -------------------------------------------------------------------------------- [ 11920]
AY261366.1_Warthog               ---------------------......a.................................................--- [ 11920]
AY261365.1_Warmbaths             ---------------------......a....t..g..............a................a.........--- [ 11920]
AY261364.1_Tengani_62            ---------------------........................................................--- [ 11920]
AY261363.1_Pretoriuskop/96/4     ---------------------..............g.........................................--- [ 11920]
AY261362.1_Mkuzi_1979            ---------------------........................................................--- [ 11920]
AY261361.1_Malawi_Lil-20/1_1983  actagaccagctagactagct...........a............................................--- [ 11920]
AY261360.1_Kenya_1950            gctaaactagtt---------...........a...a......c..g....................c.........--- [ 11920]
AM712240.1_OURT_88/3             ---------------------.......t...a............................................--- [ 11920]
AM712239.1_Benin_97/1            -------------------------------------------------------------------------------- [ 11920]
KM111295.1_Ken06.Bus             gctagactagtt---------...........a...a......c..g..t.................c.........--- [ 11920]
KM111294.1_Ken05/Tk1             actagaccagct---------...........a...a......ct.g....................c.........gtt [ 11920]
KM262844.1_strain_L60            -------------------------------------------------------------------------------- [ 11920]
KM102979.1_26544/OG10_Italy      -------------------------------------------------------------------------------- [ 11920]
KM262845.1_strain_NHV            ---------------------.......t...a............................................--- [ 11920]
KX354450.1_47/Ss/2008            -------------------------------------------------------------------------------- [ 11920]
KP055815.1_strain_BA71           -------------------------------------------------------------------------------- [ 11920]
MH025920.1_strain_R35            gctagactagtt---------...........a...a......c..g..t.................c.........--- [ 11920]
MH025919.1_strain_N10            gctagactagtt---------...........a...a......c..g..t.................c.........--- [ 11920]
MH025918.1_strain_R25            gctagactagtt---------...........a...a......c..g..t.................c.........--- [ 11920]
MH025917.1_strain_R7             gctagactagtt---------...........a...a......c..g..t.................c.........--- [ 11920]
MH025916.1_strain_R8             gctagactagtt---------...........a...a......c..g..t.................c.........--- [ 11920]

FR682468.2_Georgia_2007/1        -----------gttatattacgaaaaaacatgttaaaatttgg--aaaaaaaagccctttttatagatctggaaaaaaat [ 12000]
ASF_IND_20_CAD543                -----------.................y..............--................................... [ 12000]
LR899193.1_Germany_2020/1        -----------................................--................................... [ 12000]
MN393477.1_Wuhan_2019            -----------................................--................................... [ 12000]
MW396979.1_Timor-Leste/2019/1    -----------................................--................................... [ 12000]
MK940252.1_CN/2019/InnerMongolia -----------................................--................................... [ 12000]
MK128995.1_China/2018/AnhuiXCGQ  -----------................................--................................... [ 12000]
MH681419.1_ASFV/POL/2015         -----------................................--................................... [ 12000]
LS478113.1_Estonia_2014          ----------------------.g.t....c.c..-------------------------.....a...at..gg...g- [ 12000]
KP843857.1_Odintsovo_02/14       -----------................................--................................... [ 12000]
```

```
FN557520.1_strain_E75             -------------------------------------------------------------------------------- [ 12000]
AY261366.1_Warthog                -----------...........-....................--.................................t. [ 12000]
AY261365.1_Warmbaths              -----------................................--................................... [ 12000]
AY261364.1_Tengani_62             -----------..........................t.....--.c...................g.....g......a [ 12000]
AY261363.1_Pretoriuskop/96/4      -----------................g...............--................................... [ 12000]
AY261362.1_Mkuzi_1979             -----------................................--.....................g.....g......a [ 12000]
AY261361.1_Malawi_Lil-20/1_1983   -----------..........................t.....gc.......g...t...aat................. [ 12000]
AY261360.1_Kenya_1950             -----------...........g..............t.....a-c........a........................a [ 12000]
AM712240.1_OURT_88/3              -----------.................t..............--................................... [ 12000]
AM712239.1_Benin_97/1             -------------------------------------------------------------------------------- [ 12000]
KM111295.1_Ken06.Bus              -----------.......a..................t.....a-c........a.................-......a [ 12000]
KM111294.1_Ken05/Tk1              atataacgaaa.......a..................t.....a-c........a........................a [ 12000]
KM262844.1_strain_L60             -------------------------------------------------------------------------------- [ 12000]
KM102979.1_26544/OG10_Italy       -------------------------------------------------------------------------------- [ 12000]
KM262845.1_strain_NHV             -----------.................t..............--................................... [ 12000]
KX354450.1_47/Ss/2008             -------------------------------------------------------------------------------- [ 12000]
KP055815.1_strain_BA71            -------------------------------------------------------------------------------- [ 12000]
MH025920.1_strain_R35             -----------.......a..................t.....a-c........a........................a [ 12000]
MH025919.1_strain_N10             -----------.......a..................t.....a-c........a........................a [ 12000]
MH025918.1_strain_R25             -----------.......a..................t.....a-c........a........................a [ 12000]
MH025917.1_strain_R7              -----------.......a..................t.....a-c........a........................a [ 12000]
MH025916.1_strain_R8              -----------.......a..................t.....a-c........a........................a [ 12000]

FR682468.2_Georgia_2007/1         tt-tcacaaatctaattaaaa-gccttacagatcatccttttcataaattttcattaacaattggtgggggcggttgtga [ 12080]
ASF_IND_20_CAD543                 ..-..................-............w............................................. [ 12080]
LR899193.1_Germany_2020/1         ..-..................-.......................................................... [ 12080]
MN393477.1_Wuhan_2019             ..-..................-.......................................................... [ 12080]
MW396979.1_Timor-Leste/2019/1     ..-..................-.......................................................... [ 12080]
MK940252.1_CN/2019/InnerMongolia  ..-..................-.......................................................... [ 12080]
MK128995.1_China/2018/AnhuiXCGQ   ..-..................-.......................................................... [ 12080]
MH681419.1_ASFV/POL/2015          ..-..................-.......................................................... [ 12080]
LS478113.1_Estonia_2014           --------------------------------------g.a..g.gcg...ctg..tgtt....t.ta.t.t...----- [ 12080]
KP843857.1_Odintsovo_02/14        ..-..................-.......................................................... [ 12080]
FN557520.1_strain_E75             ---------------------------------------c..gg........gt..c-------a.ataaaa.a.g.aa. [ 12080]
AY261366.1_Warthog                ..-c-................-....c.......t..................t.............t..a..a...... [ 12080]
AY261365.1_Warmbaths              ..-.--...............-....c.......t................................t..a..a...... [ 12080]
AY261364.1_Tengani_62             a.----..g.......c....aat..c.......t..................c..cc....c....t..a..a...... [ 12080]
AY261363.1_Pretoriuskop/96/4      ..-..................-....c.......t..................c.............t..a..a...... [ 12080]
AY261362.1_Mkuzi_1979             a.----..g.......c....a....c.......t................................t..a..a...... [ 12080]
AY261361.1_Malawi_Lil-20/1_1983   ..-..................-t...c.......t.....................ct............a..a...... [ 12080]
AY261360.1_Kenya_1950             ---.....g....c.......a....c.......c..tc.................cc....c.a..a..t.t...a... [ 12080]
AM712240.1_OURT_88/3              ..-..................-t...c.......t..................c..cc....c....t..a..a...... [ 12080]
AM712239.1_Benin_97/1             ---------------------------------------c..gg........gt..c-------a.ataaaa.a.g.aa. [ 12080]
KM111295.1_Ken06.Bus              a.c.....g....c.......a....c.......c..tc.................cc....c....a..agta..a... [ 12080]
```

```
KM111294.1_Ken05/Tk1            a.c.....g....c.......ac...c.......c..tc.................cc....c.a..a..t.t...a... [ 12080]
KM262844.1_strain_L60           ---------------------------------------c..gg........gt..c-------a.ataaaa.a.g.aa. [ 12080]
KM102979.1_26544/OG10_Italy     ---------------------------------------c..gg........gt..c-------a.ataaaa.a.g.aa. [ 12080]
KM262845.1_strain_NHV           ..-..................-t...c.......t..................c..cc....c....t..a..a...... [ 12080]
KX354450.1_47/Ss/2008           ---------------------------------------c..gg........gt..c-------a.ataaaa.a.g.aa. [ 12080]
KP055815.1_strain_BA71          -------------------------------------------------------------------------------- [ 12080]
MH025920.1_strain_R35           a.c.....g....c.......a....c.......c..tc.................cc....c....a..agta..a... [ 12080]
MH025919.1_strain_N10           a.c.....g....c.......a....c.......c..tc.................cc....c....a..agta..a... [ 12080]
MH025918.1_strain_R25           a.c.....g....c.......a....c.......c..tc.................cc....c....a..agta..a... [ 12080]
MH025917.1_strain_R7            a.c.....g....c.......a....c.......c..tc.................cc....c....a..agta..a... [ 12080]
MH025916.1_strain_R8            a.c.....g....c.......a....c.......c..tc.................cc....c....a..agta..a... [ 12080]

FR682468.2_Georgia_2007/1       ggtactggatcagaacaatccataacatggtaatgtc---catttccttcaccatatgtacactggttataccagcgaga [ 12160]
ASF_IND_20_CAD543               ...................................y.---............y........................... [ 12160]
LR899193.1_Germany_2020/1       .....................................---........................................ [ 12160]
MN393477.1_Wuhan_2019           .....................................---........................................ [ 12160]
MW396979.1_Timor-Leste/2019/1   .....................................---........................................ [ 12160]
MK940252.1_CN/2019/InnerMongolia .....................................---........................................ [ 12160]
MK128995.1_China/2018/AnhuiXCGQ .....................................---........................................ [ 12160]
MH681419.1_ASFV/POL/2015        .....................................---........................................ [ 12160]
LS478113.1_Estonia_2014         -------------------------------------------------------------------------------- [ 12160]
KP843857.1_Odintsovo_02/14      .....................................---........................................ [ 12160]
FN557520.1_strain_E75           ..ggtc..t.t.a.c.---------------------------------------------------------------- [ 12160]
AY261366.1_Warthog              ...................................c.---............t........................... [ 12160]
AY261365.1_Warmbaths            ...................................c.---............t........................... [ 12160]
AY261364.1_Tengani_62           ...................................c.---...........t............................ [ 12160]
AY261363.1_Pretoriuskop/96/4    ...................................c.---............t........................... [ 12160]
AY261362.1_Mkuzi_1979           ..........t............t.............---........................................ [ 12160]
AY261361.1_Malawi_Lil-20/1_1983 ..........t..................a.......---........................................ [ 12160]
AY261360.1_Kenya_1950           ..a.......t.............gt...a......tttg....c.....tt....ca....t..a.c......a..g.. [ 12160]
AM712240.1_OURT_88/3            ..........t............t.............---...........t............................ [ 12160]
AM712239.1_Benin_97/1           ..ggtc..t.t.a.c.---------------------------------------------------------------- [ 12160]
KM111295.1_Ken06.Bus            ..a.....................gt...a..t....---...........t....a.....t...........a..... [ 12160]
KM111294.1_Ken05/Tk1            ..a.....................gt...a..t....---...........t....a.....t...........a..... [ 12160]
KM262844.1_strain_L60           ..ggtc..t.t.a.c.---------------------------------------------------------------- [ 12160]
KM102979.1_26544/OG10_Italy     ..ggtc..t.t.a.c.---------------------------------------------------------------- [ 12160]
KM262845.1_strain_NHV           ..........t............t.............---...........t............................ [ 12160]
KX354450.1_47/Ss/2008           ..ggtc..t.t.a.c.---------------------------------------------------------------- [ 12160]
KP055815.1_strain_BA71          -------------------------------------------------------------------------------- [ 12160]
MH025920.1_strain_R35           ..a.....................gt...a..t....---...........t....a.....t...........a..... [ 12160]
MH025919.1_strain_N10           ..a.....................gt...a..t....---...........t....a.....t...........a..... [ 12160]
MH025918.1_strain_R25           ..a.....................gt...a..t....---...........t....a.....t...........a..... [ 12160]
MH025917.1_strain_R7            ..a.....................gt...a..t....---...........t....a.....t...........a..... [ 12160]
MH025916.1_strain_R8            ..a.....................gt...a..t....---...........t....a.....t...........a..... [ 12160]
```

```
FR682468.2_Georgia_2007/1        aacctcacaagatgtcaaataactgttctcaacaatcaatggcatgctcttattcaccttgttcttgcaaattccatgtg [ 12240]
ASF_IND_20_CAD543                ................................y............................................... [ 12240]
LR899193.1_Germany_2020/1        ................................................................................ [ 12240]
MN393477.1_Wuhan_2019            ................................................................................ [ 12240]
MW396979.1_Timor-Leste/2019/1    ................................................................................ [ 12240]
MK940252.1_CN/2019/InnerMongolia ................................................................................ [ 12240]
MK128995.1_China/2018/AnhuiXCGQ  ................................................................................ [ 12240]
MH681419.1_ASFV/POL/2015         ................................................................................ [ 12240]
LS478113.1_Estonia_2014          -------------------------------------------------------------------------------- [ 12240]
KP843857.1_Odintsovo_02/14       ................................................................................ [ 12240]
FN557520.1_strain_E75            -------------------------------------------------------------------------------- [ 12240]
AY261366.1_Warthog               ................................t............................................... [ 12240]
AY261365.1_Warmbaths             ................................t................a.............................. [ 12240]
AY261364.1_Tengani_62            ................................t....c.......................................... [ 12240]
AY261363.1_Pretoriuskop/96/4     ................................t................g.............................. [ 12240]
AY261362.1_Mkuzi_1979            ..................g..g.....t....t................c.............................t [ 12240]
AY261361.1_Malawi_Lil-20/1_1983  ................................t.................c............................. [ 12240]
AY261360.1_Kenya_1950            ...tg....t.t......g..g..........t....c.......a....c.........a...........a......t [ 12240]
AM712240.1_OURT_88/3             ................................t....c...........c.............................t [ 12240]
AM712239.1_Benin_97/1            -------------------------------------------------------------------------------- [ 12240]
KM111295.1_Ken06.Bus             ....at...t.g.........g..........t....c.......a....c.........a...........a......t [ 12240]
KM111294.1_Ken05/Tk1             .....t...tag.........g.......t..t.................c.....................a......t [ 12240]
KM262844.1_strain_L60            -------------------------------------------------------------------------------- [ 12240]
KM102979.1_26544/OG10_Italy      -------------------------------------------------------------------------------- [ 12240]
KM262845.1_strain_NHV            ................................t....c...........c.............................t [ 12240]
KX354450.1_47/Ss/2008            -------------------------------------------------------------------------------- [ 12240]
KP055815.1_strain_BA71           -------------------------------------------------------------------------------- [ 12240]
MH025920.1_strain_R35            ....at...t.g.........g..........t....c.......a....c.........a...........a......t [ 12240]
MH025919.1_strain_N10            ....at...t.g.........g..........t....c.......a....c.........a...........a......t [ 12240]
MH025918.1_strain_R25            ....at...t.g.........g..........t....c.......a....c.........a...........a......t [ 12240]
MH025917.1_strain_R7             ....at...t.g.........g..........t....c.......a....c.........a...........a......t [ 12240]
MH025916.1_strain_R8             ....at...t.g.........g..........t....c.......a....c.........a...........a......t [ 12240]

FR682468.2_Georgia_2007/1        cacattcccagcaaaacttgcagttttccatgtaagtacaccagtatccaagttcttcttgtggaggattatccgttgaa [ 12320]
ASF_IND_20_CAD543                .....................................r.........y................................ [ 12320]
LR899193.1_Germany_2020/1        ................................................................................ [ 12320]
MN393477.1_Wuhan_2019            ................................................................................ [ 12320]
MW396979.1_Timor-Leste/2019/1    ................................................................................ [ 12320]
MK940252.1_CN/2019/InnerMongolia ................................................................................ [ 12320]
MK128995.1_China/2018/AnhuiXCGQ  ................................................................................ [ 12320]
MH681419.1_ASFV/POL/2015         ................................................................................ [ 12320]
LS478113.1_Estonia_2014          ---------.....c...at..t..a.tg..t...tggatatta.....g.-------..a.aa.aa.a...taa..--- [ 12320]
KP843857.1_Odintsovo_02/14       ................................................................................ [ 12320]
```

```
FN557520.1_strain_E75            -------------------------------------------------------------------------------- [ 12320]
AY261366.1_Warthog               .......................c.............g.......................................... [ 12320]
AY261365.1_Warmbaths             .......................c.............g.......................................... [ 12320]
AY261364.1_Tengani_62            .......................c.............g.......................................... [ 12320]
AY261363.1_Pretoriuskop/96/4     .......................c.............g.......................................... [ 12320]
AY261362.1_Mkuzi_1979            .......................c.............g.......................................... [ 12320]
AY261361.1_Malawi_Lil-20/1_1983  .......................c........................................................ [ 12320]
AY261360.1_Kenya_1950            ....a............c.....c..........g.........a.............c.c.........c..t...... [ 12320]
AM712240.1_OURT_88/3             .......................c.............g.......................................... [ 12320]
AM712239.1_Benin_97/1            -------------------------------------------------------------------------------- [ 12320]
KM111295.1_Ken06.Bus             ....g............c.....c..........g.........................t......c.cc..t...... [ 12320]
KM111294.1_Ken05/Tk1             ....a............c.....c..........g........aa.............c.c.........c..t...... [ 12320]
KM262844.1_strain_L60            -------------------------------------------------------------------------------- [ 12320]
KM102979.1_26544/OG10_Italy      -------------------------------------------------------------------------------- [ 12320]
KM262845.1_strain_NHV            .......................c.............g.......................................... [ 12320]
KX354450.1_47/Ss/2008            -------------------------------------------------------------------------------- [ 12320]
KP055815.1_strain_BA71           -------------------------------------------------------------------------------- [ 12320]
MH025920.1_strain_R35            ....g............c.....c..........g.........................t......c.cc..t...... [ 12320]
MH025919.1_strain_N10            ....g............c.....c..........g.........................t......c.cc..t...... [ 12320]
MH025918.1_strain_R25            ....g............c.....c..........g.........................t......c.cc..t...... [ 12320]
MH025917.1_strain_R7             ....g............c.....c..........g.........................t......c.cc..t...... [ 12320]
MH025916.1_strain_R8             ....g............c.....c..........g.........................t......c.cc..t...... [ 12320]

FR682468.2_Georgia_2007/1        cgaagatgccctcctgcctgagtaggtagtcctaagacctgattggccagcaggccaagaatttcc---aagaagatcac [ 12400]
ASF_IND_20_CAD543                ..................................................................---........... [ 12400]
LR899193.1_Germany_2020/1        ..................................................................---........... [ 12400]
MN393477.1_Wuhan_2019            ..................................................................---........... [ 12400]
MW396979.1_Timor-Leste/2019/1    ..................................................................---........... [ 12400]
MK940252.1_CN/2019/InnerMongolia ..................................................................---........... [ 12400]
MK128995.1_China/2018/AnhuiXCGQ  ..................................................................---........... [ 12400]
MH681419.1_ASFV/POL/2015         ..................................................................---........... [ 12400]
LS478113.1_Estonia_2014          -------------------------------------------------------------------------------- [ 12400]
KP843857.1_Odintsovo_02/14       ..................................................................---........... [ 12400]
FN557520.1_strain_E75            -------------------------------------------------------------------------------- [ 12400]
AY261366.1_Warthog               .....................................t.........................c..aag.......c... [ 12400]
AY261365.1_Warmbaths             .....................................t.........................c..---.......c... [ 12400]
AY261364.1_Tengani_62            ..........................c....................................c..---.......ct.. [ 12400]
AY261363.1_Pretoriuskop/96/4     ...............................................................c..---........... [ 12400]
AY261362.1_Mkuzi_1979            ............t...t..............................................c..---........... [ 12400]
AY261361.1_Malawi_Lil-20/1_1983  .a...................a.........................................c..---........... [ 12400]
AY261360.1_Kenya_1950            tag.....t..ag.att..tt.ct........a.....t........................c..---........... [ 12400]
AM712240.1_OURT_88/3             ............t..................................................c..---........... [ 12400]
AM712239.1_Benin_97/1            -------------------------------------------------------------------------------- [ 12400]
KM111295.1_Ken06.Bus             t....g..t..ag.att..tt.ct........a........g.....................c..---........... [ 12400]
```

```
KM111294.1_Ken05/Tk1              tag.....t..ag.att..tt.ct........a.....t........................c..---........... [ 12400]
KM262844.1_strain_L60             -------------------------------------------------------------------------------- [ 12400]
KM102979.1_26544/OG10_Italy       -------------------------------------------------------------------------------- [ 12400]
KM262845.1_strain_NHV             ............t..................................................c..---........... [ 12400]
KX354450.1_47/Ss/2008             -------------------------------------------------------------------------------- [ 12400]
KP055815.1_strain_BA71            -------------------------------------------------------------------------------- [ 12400]
MH025920.1_strain_R35             t....g..t..ag.att..tt.ct........a........g.....................c..---........... [ 12400]
MH025919.1_strain_N10             t....g..t..ag.att..tt.ct........a........g.....................c..---........... [ 12400]
MH025918.1_strain_R25             t....g..t..ag.att..tt.ct........a........g.....................c..---........... [ 12400]
MH025917.1_strain_R7              t....g..t..ag.att..tt.ct........a........g.....................c..---........... [ 12400]
MH025916.1_strain_R8              t....g..t..ag.att..tt.ct........a........g.....................c..---........... [ 12400]

FR682468.2_Georgia_2007/1         caacattgctacggctg-----------------------gctgaacagctgg-cagatagcta----------gctaat [ 12480]
ASF_IND_20_CAD543                 .................-----------------------.............c.ga.......----------...... [ 12480]
LR899193.1_Germany_2020/1         .................-----------------------.............-..........----------...... [ 12480]
MN393477.1_Wuhan_2019             .................-----------------------.............-..........----------...... [ 12480]
MW396979.1_Timor-Leste/2019/1     .................-----------------------.............-..........----------...... [ 12480]
MK940252.1_CN/2019/InnerMongolia  .................-----------------------.............-..........----------...... [ 12480]
MK128995.1_China/2018/AnhuiXCGQ   .................-----------------------.............-..........----------...... [ 12480]
MH681419.1_ASFV/POL/2015          .................-----------------------.............-..........----------...... [ 12480]
LS478113.1_Estonia_2014           -------------------------------------------------------------------------------- [ 12480]
KP843857.1_Odintsovo_02/14        .................-----------------------.............-..........----------...... [ 12480]
FN557520.1_strain_E75             -----------......--------------------------------------.......------------------ [ 12480]
AY261366.1_Warthog                .................-----------------------.............ct.........----------...... [ 12480]
AY261365.1_Warmbaths              .................-----------------------.............ct.........----------...... [ 12480]
AY261364.1_Tengani_62             .................-----------------------.............-..........----------...... [ 12480]
AY261363.1_Pretoriuskop/96/4      .................-----------------------.............ct.........----------...... [ 12480]
AY261362.1_Mkuzi_1979             ........t........-----------------------.........t...ct.a......gctaaatagct...g.a [ 12480]
AY261361.1_Malawi_Lil-20/1_1983   .................-----------------------............act.........----------...... [ 12480]
AY261360.1_Kenya_1950             .................gctaaacaacgagctagatagct............c-t........gctgaatagct...... [ 12480]
AM712240.1_OURT_88/3              ........t........-----------------------.............ct.a......-----------...g.a [ 12480]
AM712239.1_Benin_97/1             -----------......--------------------------------------.......------------------ [ 12480]
KM111295.1_Ken06.Bus              ...........--....-----------------------............c-t........gctgaatagct...... [ 12480]
KM111294.1_Ken05/Tk1              ..........g--....gctg-------------------............c-t........gctgaatagct...... [ 12480]
KM262844.1_strain_L60             -----------......--------------------------------------.......------------------ [ 12480]
KM102979.1_26544/OG10_Italy       -----------......--------------------------------------.......------------------ [ 12480]
KM262845.1_strain_NHV             ........t........-----------------------.............ct.a......-----------...g.a [ 12480]
KX354450.1_47/Ss/2008             -----------......--------------------------------------.......------------------ [ 12480]
KP055815.1_strain_BA71            -------------------------------------------------------------------------------- [ 12480]
MH025920.1_strain_R35             ...........--....-----------------------............c-t........gctgaatagct...... [ 12480]
MH025919.1_strain_N10             ...........--....-----------------------............c-t........gctgaatagct...... [ 12480]
MH025918.1_strain_R25             ...........--....-----------------------............c-t........gctgaatagct...... [ 12480]
MH025917.1_strain_R7              ...........--....-----------------------............c-t........gctgaatagct...... [ 12480]
MH025916.1_strain_R8              ...........--....-----------------------............c-t........gctgaatagct...... [ 12480]
```

```
FR682468.2_Georgia_2007/1        tagcaaaccaagtgactcgccctctctactcttaatatgagaatttaagattcggtccggcttttttcccatgttttaca [ 12560]
ASF_IND_20_CAD543                ......................-.............................nnnnnna..a.......tg......... [ 12560]
LR899193.1_Germany_2020/1        ................................................................................ [ 12560]
MN393477.1_Wuhan_2019            ................................................................................ [ 12560]
MW396979.1_Timor-Leste/2019/1    ................................................................................ [ 12560]
MK940252.1_CN/2019/InnerMongolia ................................................................................ [ 12560]
MK128995.1_China/2018/AnhuiXCGQ  ................................................................................ [ 12560]
MH681419.1_ASFV/POL/2015         ................................................................................ [ 12560]
LS478113.1_Estonia_2014          -----------c.a..a..tt.a..t..a.ta.t..g...-------------------at......a...ca...---- [ 12560]
KP843857.1_Odintsovo_02/14       ................................................................................ [ 12560]
FN557520.1_strain_E75            ---....t...--------------------------------------------------------------------- [ 12560]
AY261366.1_Warthog               .....................................................a....a..a.................. [ 12560]
AY261365.1_Warmbaths             .....................................................a....a..a.................. [ 12560]
AY261364.1_Tengani_62            ..................a...c............................c.................g..a....... [ 12560]
AY261363.1_Pretoriuskop/96/4     ..........................................................a..a.................. [ 12560]
AY261362.1_Mkuzi_1979            ..........................................................a..a.................. [ 12560]
AY261361.1_Malawi_Lil-20/1_1983  ........................................a.............a...a..a.......tg......... [ 12560]
AY261360.1_Kenya_1950            .................................c...a..............g....ta..a.................. [ 12560]
AM712240.1_OURT_88/3             ........................................a.................a..a.................. [ 12560]
AM712239.1_Benin_97/1            ---....t...--------------------------------------------------------------------- [ 12560]
KM111295.1_Ken06.Bus             .................................c.........a..............a..a.................. [ 12560]
KM111294.1_Ken05/Tk1             .................................c...at.............g.....a..a.................. [ 12560]
KM262844.1_strain_L60            ---....t...--------------------------------------------------------------------- [ 12560]
KM102979.1_26544/OG10_Italy      ---....t...--------------------------------------------------------------------- [ 12560]
KM262845.1_strain_NHV            ........................................a.................a..a.................. [ 12560]
KX354450.1_47/Ss/2008            ---....t...--------------------------------------------------------------------- [ 12560]
KP055815.1_strain_BA71           -------------------------------------------------------------------------------- [ 12560]
MH025920.1_strain_R35            .................................c........................a..a.................. [ 12560]
MH025919.1_strain_N10            .................................c........................a..a.................. [ 12560]
MH025918.1_strain_R25            .................................c........................a..a.................. [ 12560]
MH025917.1_strain_R7             .................................c........................a..a.................. [ 12560]
MH025916.1_strain_R8             .................................c........................a..a.................. [ 12560]

FR682468.2_Georgia_2007/1        gggaaaaggtatttttagcctatgaatgtacatggtt-ccgcacattaa-aaaaaaataaaagaaattatttaatattgg [ 12640]
ASF_IND_20_CAD543                .a......a.........t.....nnnnnnnnnnnnn-nn.......--------------------------------- [ 12640]
LR899193.1_Germany_2020/1        .....................................-...........-.............................. [ 12640]
MN393477.1_Wuhan_2019            .....................................-...........-.............................. [ 12640]
MW396979.1_Timor-Leste/2019/1    .....................................-...........-.............................. [ 12640]
MK940252.1_CN/2019/InnerMongolia .....................................-...........-.............................. [ 12640]
MK128995.1_China/2018/AnhuiXCGQ  .....................................-...........-.............................. [ 12640]
MH681419.1_ASFV/POL/2015         .....................................-...........-.............................. [ 12640]
LS478113.1_Estonia_2014          -------------------------------------------------------------------------------- [ 12640]
KP843857.1_Odintsovo_02/14       .....................................-.........-.-.............................. [ 12640]
```

```
FN557520.1_strain_E75             ------------------------...ac.----------------.--------..a.g.gc..g.ag...c...gc.. [ 12640]
AY261366.1_Warthog                ........a.........t.........c........-...t.t...---............t..........------- [ 12640]
AY261365.1_Warmbaths              ........a.........t.........c........-.....t...---............t..........------- [ 12640]
AY261364.1_Tengani_62             a.a....aa.........-...ca...ac..t------t.at.t..cc--ct.....a.c.ga..t........------ [ 12640]
AY261363.1_Pretoriuskop/96/4      ........a.........t......g..c........c.....t...t.ag...........t..........------- [ 12640]
AY261362.1_Mkuzi_1979             ........a.........t.c.......c........-.........--------..a...t............t.... [ 12640]
AY261361.1_Malawi_Lil-20/1_1983   .a......a.........t.c.......c...--...-.........-------...a...t...........------ [ 12640]
AY261360.1_Kenya_1950             ........a.........t.......c.c........-.....t...---t.g....a.t..a.c.......tc.t.cct [ 12640]
AM712240.1_OURT_88/3              ........a........tt.........c........-.........--------..a...t............t.... [ 12640]
AM712239.1_Benin_97/1             ------------------------...ac.----------------.--------..a.g.gc..g.ag...c...gc.. [ 12640]
KM111295.1_Ken06.Bus              ........a.......c.t.........c........-.....t.a.---t.g....a.t..a.c.......tc.t.cct [ 12640]
KM111294.1_Ken05/Tk1              ........a.........t.......c.c........-.........---t.g....a.t..a.c..--...tc.t.cct [ 12640]
KM262844.1_strain_L60             ------------------------...ac.----------------.--------..a.g.gc..g.ag...c...gc.. [ 12640]
KM102979.1_26544/OG10_Italy       ------------------------...ac.----------------.--------..a.g.gc..g.ag...c...gc.. [ 12640]
KM262845.1_strain_NHV             ........a........tt.........c........-.........--------..a...t............t.... [ 12640]
KX354450.1_47/Ss/2008             ------------------------...ac.----------------.--------..a.g.gc..g.ag...c...gc.. [ 12640]
KP055815.1_strain_BA71            -------------------------------------------------------------------------------- [ 12640]
MH025920.1_strain_R35             ........a.......c.t.........c........-.....t.a.---t.g....a.t..a.c.......tc.t.cct [ 12640]
MH025919.1_strain_N10             ........a.......c.t.........c........-.....t.a.---t.g....a.t..a.c.......tc.t.cct [ 12640]
MH025918.1_strain_R25             ........a.......c.t.........c........-.....t.a.---t.g....a.t..a.c.......tc.t.cct [ 12640]
MH025917.1_strain_R7              ........a.......c.t.........c........-.....t.a.---t.g....a.t..a.c.......tc.t.cct [ 12640]
MH025916.1_strain_R8              ........a.......c.t.........c........-.....t.a.---t.g....a.t..a.c.......tc.t.cct [ 12640]

FR682468.2_Georgia_2007/1         ctgttattttctttcaactagcaac-aagccaggtaactaaagaacttcattgtagttttatattacggaaa--ag---g [ 12720]
ASF_IND_20_CAD543                 -------------------------------------------------------------------------------- [ 12720]
LR899193.1_Germany_2020/1         .........................-..............................................--..---. [ 12720]
MN393477.1_Wuhan_2019             .........................-..............................................--..---. [ 12720]
MW396979.1_Timor-Leste/2019/1     .........................-..............................................--..---. [ 12720]
MK940252.1_CN/2019/InnerMongolia  .........................-..............................................--..---. [ 12720]
MK128995.1_China/2018/AnhuiXCGQ   .........................-..............................................--..---. [ 12720]
MH681419.1_ASFV/POL/2015          .........................-..............................................--..---. [ 12720]
LS478113.1_Estonia_2014           ---------------------------.ttt.aa..gt.t..a.....-------------------------------- [ 12720]
KP843857.1_Odintsovo_02/14        .........................-..............................................--..---. [ 12720]
FN557520.1_strain_E75             tat------------------------ttaggt....-----------------.t.......g....c...--tacaat [ 12720]
AY261366.1_Warthog                -------------------------------------------------------------------------------- [ 12720]
AY261365.1_Warmbaths              -------------------------------------------------------------------------------- [ 12720]
AY261364.1_Tengani_62             -------------------------------------------------------------------------------- [ 12720]
AY261363.1_Pretoriuskop/96/4      -------------------------------------------------------------------------------- [ 12720]
AY261362.1_Mkuzi_1979             ..a------------------------gc.gg..........a.--c..........g..........a...--.acat. [ 12720]
AY261361.1_Malawi_Lil-20/1_1983   -------------------------------------------------------------------------------- [ 12720]
AY261360.1_Kenya_1950             ..t------....c.........g.a.gc.a...............c..........g..............aa.acat. [ 12720]
AM712240.1_OURT_88/3              ..a------------------------gc.gg..........a.--c..........g..........a...--.acat. [ 12720]
AM712239.1_Benin_97/1             tat------------------------ttaggt....-----------------.t.......g....c...--tacaat [ 12720]
KM111295.1_Ken06.Bus              ..t------....c.........g.a.gc.a...............c..........g..............aa.acat. [ 12720]
```

```
KM111294.1_Ken05/Tk1             ..t------....c.........g.a.gc.a...............c..........g..............a-.acat. [ 12720]
KM262844.1_strain_L60            tat------------------------ttaggt....-----------------.t.......g....c...--tacaat [ 12720]
KM102979.1_26544/OG10_Italy      tat------------------------ttaggt....-----------------.t.......g....c...--tacaat [ 12720]
KM262845.1_strain_NHV            ..a------------------------gc.gg..........a.--c..........g..........a...--.acat. [ 12720]
KX354450.1_47/Ss/2008            tat------------------------ttaggt....-----------------.t.......g....c...--tacaat [ 12720]
KP055815.1_strain_BA71           -------------------------------------------------------------------------------- [ 12720]
MH025920.1_strain_R35            ..t------....c.........g.a.gc.a...............c..........g..............aa.acat. [ 12720]
MH025919.1_strain_N10            ..t------....c.........g.a.gc.a...............c..........g..............aa.acat. [ 12720]
MH025918.1_strain_R25            ..t------....c.........g.a.gc.a...............c..........g..............aa.acat. [ 12720]
MH025917.1_strain_R7             ..t------....c.........g.a.gc.a...............c..........g..............aa.acat. [ 12720]
MH025916.1_strain_R8             ..t------....c.........g.a.gc.a...............c..........g..............aa.acat. [ 12720]

FR682468.2_Georgia_2007/1        ttaaatttt---------------------------ggacaaaaaaaatc----atatctaattaaaaatcctcacagat [ 12800]
ASF_IND_20_CAD543                -------------------------------------a.a....t....------------------------------- [ 12800]
LR899193.1_Germany_2020/1        .........---------------------------..............----.......................... [ 12800]
MN393477.1_Wuhan_2019            .........---------------------------..............----.......................... [ 12800]
MW396979.1_Timor-Leste/2019/1    .........---------------------------..............----.......................... [ 12800]
MK940252.1_CN/2019/InnerMongolia .........---------------------------..............----.......................... [ 12800]
MK128995.1_China/2018/AnhuiXCGQ  .........---------------------------..............----.......................... [ 12800]
MH681419.1_ASFV/POL/2015         .........---------------------------..............----.......................... [ 12800]
LS478113.1_Estonia_2014          -------------------------------------.ct.........------------------------------- [ 12800]
KP843857.1_Odintsovo_02/14       .........---------------------------..............----.......................... [ 12800]
FN557520.1_strain_E75            g...---------------------------------------------------------------------------- [ 12800]
AY261366.1_Warthog               -------------------------------------------------------------------------------- [ 12800]
AY261365.1_Warmbaths             -------------------------------------------------------------------------------- [ 12800]
AY261364.1_Tengani_62            -------------------------------------------------------------------------------- [ 12800]
AY261363.1_Pretoriuskop/96/4     -------------------------------------------------------------------------------- [ 12800]
AY261362.1_Mkuzi_1979            .........ggacaaaaaagccc-tttttataggtct.gg..........----.g.......c.....g.......... [ 12800]
AY261361.1_Malawi_Lil-20/1_1983  -------------------------------------------------------------------------------- [ 12800]
AY261360.1_Kenya_1950            ........cggtcaaaaaagcctttttttatagttct..a.........atcac.g....c........g.......... [ 12800]
AM712240.1_OURT_88/3             .........ggacaaaaaagcccttttttataggtct.gg..........----.g.............-.......... [ 12800]
AM712239.1_Benin_97/1            g...---------------------------------------------------------------------------- [ 12800]
KM111295.1_Ken06.Bus             ........cggtcaaaaaaggccttttttatagatct.-a..........tcac.g....c.........g......a.. [ 12800]
KM111294.1_Ken05/Tk1             ........cggtcaaaaaag-ccttttttatggatct.--.........atcac.g....c........g.......... [ 12800]
KM262844.1_strain_L60            g...---------------------------------------------------------------------------- [ 12800]
KM102979.1_26544/OG10_Italy      g...---------------------------------------------------------------------------- [ 12800]
KM262845.1_strain_NHV            .........ggacaaaaaagcccttttttataggtct.gg..........----.g.............-.......... [ 12800]
KX354450.1_47/Ss/2008            g...---------------------------------------------------------------------------- [ 12800]
KP055815.1_strain_BA71           -------------------------------------------------------------------------------- [ 12800]
MH025920.1_strain_R35            ........cggtcaaaaaaggccttttttatagatct.-a..........tcac.g....c.........g......a.. [ 12800]
MH025919.1_strain_N10            ........cggtcaaaaaaggccttttttatagatct.-a..........tcac.g....c.........g......a.. [ 12800]
MH025918.1_strain_R25            ........cggtcaaaaaaggccttttttatagatct.-a..........tcac.g....c.........g......a.. [ 12800]
MH025917.1_strain_R7             ........cggtcaaaaaaggccttttttatagatct.-a..........tcac.g....c.........g......a.. [ 12800]
MH025916.1_strain_R8             ........cggtcaaaaaaggccttttttatagatct.-a..........tcac.g....c.........g......a.. [ 12800]
```

```
FR682468.2_Georgia_2007/1        ctttcttttcataaattttcattaacaattggtaggggcggttgtgaggtactggatcagaacaatccataacatggtaa [ 12880]
ASF_IND_20_CAD543                -------------------------------------------------------------------------------- [ 12880]
LR899193.1_Germany_2020/1        ................................................................................ [ 12880]
MN393477.1_Wuhan_2019            ................................................................................ [ 12880]
MW396979.1_Timor-Leste/2019/1    ..........................................c..................................... [ 12880]
MK940252.1_CN/2019/InnerMongolia ................................................................................ [ 12880]
MK128995.1_China/2018/AnhuiXCGQ  ................................................................................ [ 12880]
MH681419.1_ASFV/POL/2015         ................................................................................ [ 12880]
LS478113.1_Estonia_2014          --------------.....tt....tt...ct.ct..at..--------------------------------------- [ 12880]
KP843857.1_Odintsovo_02/14       ................................................................................ [ 12880]
FN557520.1_strain_E75            -------------------------------------------------------------------------------- [ 12880]
AY261366.1_Warthog               -------------------------------------------------------------------------------- [ 12880]
AY261365.1_Warmbaths             -------------------------------------------------------------------------------- [ 12880]
AY261364.1_Tengani_62            -------------------------------------------------------------------------------- [ 12880]
AY261363.1_Pretoriuskop/96/4     -------------------------------------------------------------------------------- [ 12880]
AY261362.1_Mkuzi_1979            ...c.....t.......................gt..a..a...................................a... [ 12880]
AY261361.1_Malawi_Lil-20/1_1983  -------------------------------------------------------------------------------- [ 12880]
AY261360.1_Kenya_1950            ...c.....t.............ct....c...ga..a..a........c.............................. [ 12880]
AM712240.1_OURT_88/3             ...c.........................c...gt..a..a....................................... [ 12880]
AM712239.1_Benin_97/1            -------------------------------------------------------------------------------- [ 12880]
KM111295.1_Ken06.Bus             ...c.....t.....a.......cc....c...ga..a..a........c.............................. [ 12880]
KM111294.1_Ken05/Tk1             ...c...................ct........ga..a..a....................................... [ 12880]
KM262844.1_strain_L60            -------------------------------------------------------------------------------- [ 12880]
KM102979.1_26544/OG10_Italy      -------------------------------------------------------------------------------- [ 12880]
KM262845.1_strain_NHV            ...c.........................c...gt..a..a....................................... [ 12880]
KX354450.1_47/Ss/2008            -------------------------------------------------------------------------------- [ 12880]
KP055815.1_strain_BA71           -------------------------------------------------------------------------------- [ 12880]
MH025920.1_strain_R35            ...c.....t.....a.......cc....c...ga..a..a........c.............................. [ 12880]
MH025919.1_strain_N10            ...c.....t.....a.......cc....c...ga..a..a........c.............................. [ 12880]
MH025918.1_strain_R25            ...c.....t.....a.......cc....c...ga..a..a........c.............................. [ 12880]
MH025917.1_strain_R7             ...c.....t.....a.......cc....c...ga..a..a........c.............................. [ 12880]
MH025916.1_strain_R8             ...c.....t.....a.......cc....c...ga..a..a........c.............................. [ 12880]

FR682468.2_Georgia_2007/1        tgcccatttccttcaccatatgtacactggttataccagcgagaaacctcacatgttgtcaagtagctgttttcaataat [ 12960]
ASF_IND_20_CAD543                -------------------------------------------------------------------------------- [ 12960]
LR899193.1_Germany_2020/1        ................................................................................ [ 12960]
MN393477.1_Wuhan_2019            ................................................................................ [ 12960]
MW396979.1_Timor-Leste/2019/1    ................................................................................ [ 12960]
MK940252.1_CN/2019/InnerMongolia ................................................................................ [ 12960]
MK128995.1_China/2018/AnhuiXCGQ  ................................................................................ [ 12960]
MH681419.1_ASFV/POL/2015         ................................................................................ [ 12960]
LS478113.1_Estonia_2014          -------------------------------------------------------------------------------- [ 12960]
KP843857.1_Odintsovo_02/14       ................................................................................ [ 12960]
```

| | | |
|---|---|---|
| FN557520.1_strain_E75 | -------------------------------------------------------------------------------- | [ 12960] |
| AY261366.1_Warthog | -------------------------------------------------------------------------------- | [ 12960] |
| AY261365.1_Warmbaths | -------------------------------------------------------------------------------- | [ 12960] |
| AY261364.1_Tengani_62 | -------------------------------------------------------------------------------- | [ 12960] |
| AY261363.1_Pretoriuskop/96/4 | -------------------------------------------------------------------------------- | [ 12960] |
| AY261362.1_Mkuzi_1979 | ..t..................................................a.a......a..a.....c........ | [ 12960] |
| AY261361.1_Malawi_Lil-20/1_1983 | -------------------------------------------------------------------------------- | [ 12960] |
| AY261360.1_Kenya_1950 | ...............t..........t............................a......a..a.....c........ | [ 12960] |
| AM712240.1_OURT_88/3 | ................t......g.............................a.a......a..a.....c........ | [ 12960] |
| AM712239.1_Benin_97/1 | -------------------------------------------------------------------------------- | [ 12960] |
| KM111295.1_Ken06.Bus | ...............t.......................................a......a..a.....c........ | [ 12960] |
| KM111294.1_Ken05/Tk1 | ...............t.....................................a.a......a..a.....c........ | [ 12960] |
| KM262844.1_strain_L60 | -------------------------------------------------------------------------------- | [ 12960] |
| KM102979.1_26544/OG10_Italy | -------------------------------------------------------------------------------- | [ 12960] |
| KM262845.1_strain_NHV | ................t......g.............................a.a......a..a.....c........ | [ 12960] |
| KX354450.1_47/Ss/2008 | -------------------------------------------------------------------------------- | [ 12960] |
| KP055815.1_strain_BA71 | -------------------------------------------------------------------------------- | [ 12960] |
| MH025920.1_strain_R35 | ...............t.......................................a......a..a.....c........ | [ 12960] |
| MH025919.1_strain_N10 | ...............t.......................................a......a..a.....c........ | [ 12960] |
| MH025918.1_strain_R25 | ...............t.......................................a......a..a.....c........ | [ 12960] |
| MH025917.1_strain_R7 | ...............t.......................................a......a..a.....c........ | [ 12960] |
| MH025916.1_strain_R8 | ...............t.......................................a......a..a.....c........ | [ 12960] |

| | | |
|---|---|---|
| FR682468.2_Georgia_2007/1 | caatggcatgctattattcaccttgttcttgcaaattccatgtgcacattcccagcaaaacttgcacctttccatgtaag | [ 13040] |
| ASF_IND_20_CAD543 | -------------------------------------------------------------------------------- | [ 13040] |
| LR899193.1_Germany_2020/1 | ................................................................................ | [ 13040] |
| MN393477.1_Wuhan_2019 | ................................................................................ | [ 13040] |
| MW396979.1_Timor-Leste/2019/1 | ................................................................................ | [ 13040] |
| MK940252.1_CN/2019/InnerMongolia | ................................................................................ | [ 13040] |
| MK128995.1_China/2018/AnhuiXCGQ | ................................................................................ | [ 13040] |
| MH681419.1_ASFV/POL/2015 | ................................................................................ | [ 13040] |
| LS478113.1_Estonia_2014 | ------------------------------------------------------------------------------.. | [ 13040] |
| KP843857.1_Odintsovo_02/14 | ................................................................................ | [ 13040] |
| FN557520.1_strain_E75 | ---------------------------.c.a.....g.a..ac---------------------------------...a | [ 13040] |
| AY261366.1_Warthog | -------------------------------------------------------------------------------- | [ 13040] |
| AY261365.1_Warmbaths | -------------------------------------------------------------------------------- | [ 13040] |
| AY261364.1_Tengani_62 | -------------------------------------------------------------------------------- | [ 13040] |
| AY261363.1_Pretoriuskop/96/4 | -------------------------------------------------------------------------------- | [ 13040] |
| AY261362.1_Mkuzi_1979 | ............c.c...................................................g............. | [ 13040] |
| AY261361.1_Malawi_Lil-20/1_1983 | -------------------------------------------------------------------------------- | [ 13040] |
| AY261360.1_Kenya_1950 | .gt.........c.c....................g........t.........a...........g............. | [ 13040] |
| AM712240.1_OURT_88/3 | ............c.c...................................................g............. | [ 13040] |
| AM712239.1_Benin_97/1 | ---------------------------.c.a.....g.a..ac---------------------------------...a | [ 13040] |
| KM111295.1_Ken06.Bus | .gt.........c.c....................g........t.........a...........g............. | [ 13040] |

```
KM111294.1_Ken05/Tk1             ............c.c...................................................g............. [ 13040]
KM262844.1_strain_L60            ---------------------------.c.a.....g.a..ac---------------------------------...a [ 13040]
KM102979.1_26544/OG10_Italy      ---------------------------.c.a.....g.a..ac---------------------------------...a [ 13040]
KM262845.1_strain_NHV            ............c.c...................................................g............. [ 13040]
KX354450.1_47/Ss/2008            ---------------------------.c.a.....g.a..ac---------------------------------...a [ 13040]
KP055815.1_strain_BA71           -------------------------------------------------------------------------------- [ 13040]
MH025920.1_strain_R35            .gt.........c.c....................g........t.........a...........g............. [ 13040]
MH025919.1_strain_N10            .gt.........c.c....................g........t.........a...........g............. [ 13040]
MH025918.1_strain_R25            .gt.........c.c....................g........t.........a...........g............. [ 13040]
MH025917.1_strain_R7             .gt.........c.c....................g........t.........a...........g............. [ 13040]
MH025916.1_strain_R8             .gt.........c.c....................g........t.........a...........g............. [ 13040]

FR682468.2_Georgia_2007/1        tgcaccagtatccaagttcttcttgtggaggattatccgttgaacgaagatgccctcctgcctgagtaggtagtcctacg [ 13120]
ASF_IND_20_CAD543                -------------------------------------------------------------------------------- [ 13120]
LR899193.1_Germany_2020/1        ................................................................................ [ 13120]
MN393477.1_Wuhan_2019            ................................................................................ [ 13120]
MW396979.1_Timor-Leste/2019/1    ................................................................................ [ 13120]
MK940252.1_CN/2019/InnerMongolia ................................................................................ [ 13120]
MK128995.1_China/2018/AnhuiXCGQ  ................................................................................ [ 13120]
MH681419.1_ASFV/POL/2015         ................................................................................ [ 13120]
LS478113.1_Estonia_2014          c...tt..---------------------....cccttacca..tact..aaa.t.tt..t...cact...ca.tt.t.- [ 13120]
KP843857.1_Odintsovo_02/14       ................................................................................ [ 13120]
FN557520.1_strain_E75            .a..---------------------------------------------------------------.....a------- [ 13120]
AY261366.1_Warthog               -------------------------------------------------------------------------------- [ 13120]
AY261365.1_Warmbaths             -------------------------------------------------------------------------------- [ 13120]
AY261364.1_Tengani_62            -------------------------------------------------------------------------------- [ 13120]
AY261363.1_Pretoriuskop/96/4     -------------------------------------------------------------------------------- [ 13120]
AY261362.1_Mkuzi_1979            .a............................................................................a. [ 13120]
AY261361.1_Malawi_Lil-20/1_1983  -------------------------------------------------------------------------------- [ 13120]
AY261360.1_Kenya_1950            .......a...tt...........c............t........................................a. [ 13120]
AM712240.1_OURT_88/3             .a............................................................................a. [ 13120]
AM712239.1_Benin_97/1            .a..---------------------------------------------------------------.....a------- [ 13120]
KM111295.1_Ken06.Bus             ...........t............c............t..............t.................c.tc...ga. [ 13120]
KM111294.1_Ken05/Tk1             .a....................................................................c.......a. [ 13120]
KM262844.1_strain_L60            .a..---------------------------------------------------------------.....a------- [ 13120]
KM102979.1_26544/OG10_Italy      .a..---------------------------------------------------------------.....a------- [ 13120]
KM262845.1_strain_NHV            .a............................................................................a. [ 13120]
KX354450.1_47/Ss/2008            .a..---------------------------------------------------------------.....a------- [ 13120]
KP055815.1_strain_BA71           -------------------------------------------------------------------------------- [ 13120]
MH025920.1_strain_R35            ...........t............c............t..............t.................c.tc...ga. [ 13120]
MH025919.1_strain_N10            ...........t............c............t..............t.................c.tc...ga. [ 13120]
MH025918.1_strain_R25            ...........t............c............t..............t.................c.tc...ga. [ 13120]
MH025917.1_strain_R7             ...........t............c............t..............t.................c.tc...ga. [ 13120]
MH025916.1_strain_R8             ...........t............c............t..............t.................c.tc...ga. [ 13120]
```

```
FR682468.2_Georgia_2007/1        acctgattggccagcaggccaagaattcccaagaagactaccaacattgctacggctggctgaacagct--------gg- [ 13200]
ASF_IND_20_CAD543                -------------------------------------------------------------------------------- [ 13200]
LR899193.1_Germany_2020/1        .....................................................................--------..- [ 13200]
MN393477.1_Wuhan_2019            .....................................................................--------..- [ 13200]
MW396979.1_Timor-Leste/2019/1    .....................................................................--------..- [ 13200]
MK940252.1_CN/2019/InnerMongolia .....................................................................--------..- [ 13200]
MK128995.1_China/2018/AnhuiXCGQ  .....................................................................--------..- [ 13200]
MH681419.1_ASFV/POL/2015         .....................................................................--------..- [ 13200]
LS478113.1_Estonia_2014          -------------------------------------------------------------------------------- [ 13200]
KP843857.1_Odintsovo_02/14       .....................................................................--------..- [ 13200]
FN557520.1_strain_E75            -------------------------------------------------------------------------------- [ 13200]
AY261366.1_Warthog               -------------------------------------------------------------------------------- [ 13200]
AY261365.1_Warmbaths             -------------------------------------------------------------------------------- [ 13200]
AY261364.1_Tengani_62            -------------------------------------------------------------------------------- [ 13200]
AY261363.1_Pretoriuskop/96/4     -------------------------------------------------------------------------------- [ 13200]
AY261362.1_Mkuzi_1979            .....................................tc..............................--------.ac [ 13200]
AY261361.1_Malawi_Lil-20/1_1983  -------------------------------------------------------------------------------- [ 13200]
AY261360.1_Kenya_1950            .....g...............................tc..............................--------..c [ 13200]
AM712240.1_OURT_88/3             .......------------------------------------------------------------------------- [ 13200]
AM712239.1_Benin_97/1            -------------------------------------------------------------------------------- [ 13200]
KM111295.1_Ken06.Bus             .g...g.............................a.tc......................ag.t....ctgaaagct.c [ 13200]
KM111294.1_Ken05/Tk1             .t.................................a.tc..............................---------.c [ 13200]
KM262844.1_strain_L60            -------------------------------------------------------------------------------- [ 13200]
KM102979.1_26544/OG10_Italy      -------------------------------------------------------------------------------- [ 13200]
KM262845.1_strain_NHV            .......------------------------------------------------------------------------- [ 13200]
KX354450.1_47/Ss/2008            -------------------------------------------------------------------------------- [ 13200]
KP055815.1_strain_BA71           -------------------------------------------------------------------------------- [ 13200]
MH025920.1_strain_R35            .g...g.............................a.tc......................ag.t....ctgaaagct.c [ 13200]
MH025919.1_strain_N10            .g...g.............................a.tc......................ag.t....ctgaaagct.c [ 13200]
MH025918.1_strain_R25            .g...g.............................a.tc......................ag.t....ctgaaagct.c [ 13200]
MH025917.1_strain_R7             .g...g.............................a.tc......................ag.t....ctgaaagct.c [ 13200]
MH025916.1_strain_R8             .g...g.............................a.tc......................ag.t....ctgaaagct.c [ 13200]

FR682468.2_Georgia_2007/1        cagatagctagctaattagcaaaccaagtgactcaccctctctactcttaatatgagaatttaagatccggtccgacatt [ 13280]
ASF_IND_20_CAD543                -------------------------------------------------------------------------------- [ 13280]
LR899193.1_Germany_2020/1        ................................................................................ [ 13280]
MN393477.1_Wuhan_2019            ................................................................................ [ 13280]
MW396979.1_Timor-Leste/2019/1    ................................................................................ [ 13280]
MK940252.1_CN/2019/InnerMongolia ................................................................................ [ 13280]
MK128995.1_China/2018/AnhuiXCGQ  ................................................................................ [ 13280]
MH681419.1_ASFV/POL/2015         ................................................................................ [ 13280]
LS478113.1_Estonia_2014          ---------------..g.ac...a.ca..g.a.gt.a.t..c..-------------------------..ttt.t.cc [ 13280]
KP843857.1_Odintsovo_02/14       ................................................................................ [ 13280]
```

```
FN557520.1_strain_E75              -------------------------------------------------------------------------------- [ 13280]
AY261366.1_Warthog                 -------------------------------------------------------------------------------- [ 13280]
AY261365.1_Warmbaths               -------------------------------------------------------------------------------- [ 13280]
AY261364.1_Tengani_62              -------------------------------------------------------------------------------- [ 13280]
AY261363.1_Pretoriuskop/96/4       -------------------------------------------------------------------------------- [ 13280]
AY261362.1_Mkuzi_1979              .ga.........c.....................g...-.................a..........t..a.t.ag.... [ 13280]
AY261361.1_Malawi_Lil-20/1_1983    -------------------------------------------------------------------------------- [ 13280]
AY261360.1_Kenya_1950              t........-...g------------------------------------------------------------------ [ 13280]
AM712240.1_OURT_88/3               -------------------------------------------------------------------------------- [ 13280]
AM712239.1_Benin_97/1              -------------------------------------------------------------------------------- [ 13280]
KM111295.1_Ken06.Bus               t.a......-...g...................................c...a.............t....t..g.... [ 13280]
KM111294.1_Ken05/Tk1               t........-........................g..............c...a.............t....t..g.... [ 13280]
KM262844.1_strain_L60              -------------------------------------------------------------------------------- [ 13280]
KM102979.1_26544/OG10_Italy        -------------------------------------------------------------------------------- [ 13280]
KM262845.1_strain_NHV              -------------------------------------------------------------------------------- [ 13280]
KX354450.1_47/Ss/2008              -------------------------------------------------------------------------------- [ 13280]
KP055815.1_strain_BA71             -------------------------------------------------------------------------------- [ 13280]
MH025920.1_strain_R35              t.a......-...g...................................c...a.............t....t..g.... [ 13280]
MH025919.1_strain_N10              t.a......-...g...................................c...a.............t....t..g.... [ 13280]
MH025918.1_strain_R25              t.a......-...g...................................c...a.............t....t..g.... [ 13280]
MH025917.1_strain_R7               t.a......-...g...................................c...a.............t....t..g.... [ 13280]
MH025916.1_strain_R8               t.a......-...g...................................c...a.............t....t..g.... [ 13280]

FR682468.2_Georgia_2007/1          tttccgatattttacaagaaaaagatatttttag-ctacaaatacac-------ttcatatatccctaaaaaaaacaaaa [ 13360]
ASF_IND_20_CAD543                  -------------------------------------------------------------------------------- [ 13360]
LR899193.1_Germany_2020/1          ..................................-............-------.......................... [ 13360]
MN393477.1_Wuhan_2019              ..................................-............-------.......................... [ 13360]
MW396979.1_Timor-Leste/2019/1      ..................................-............-------.......................... [ 13360]
MK940252.1_CN/2019/InnerMongolia   ..................................-............-------.......................... [ 13360]
MK128995.1_China/2018/AnhuiXCGQ    ..................................-............-------.......................... [ 13360]
MH681419.1_ASFV/POL/2015           ..................................-............-------.......................... [ 13360]
LS478113.1_Estonia_2014            .....a.....cag...--------------------------------------------------------------- [ 13360]
KP843857.1_Odintsovo_02/14         ..................................-............-------.......................... [ 13360]
FN557520.1_strain_E75              -------------------------------------------------------------------------------- [ 13360]
AY261366.1_Warthog                 -------------------------------------------------------------------------------- [ 13360]
AY261365.1_Warmbaths               -------------------------------------------------------------------------------- [ 13360]
AY261364.1_Tengani_62              -------------------------------------------------------------------------------- [ 13360]
AY261363.1_Pretoriuskop/96/4       -------------------------------------------------------------------------------- [ 13360]
AY261362.1_Mkuzi_1979              .....tg.g.......gag...............t.c.tg...g...a----gt.c.gc.c..t-----......a..t. [ 13360]
AY261361.1_Malawi_Lil-20/1_1983    -------------------------------------------------------------------------------- [ 13360]
AY261360.1_Kenya_1950              -------------------------------------------------------------------------------- [ 13360]
AM712240.1_OURT_88/3               -------------------------------------------------------------------------------- [ 13360]
AM712239.1_Benin_97/1              -------------------------------------------------------------------------------- [ 13360]
KM111295.1_Ken06.Bus               ...-t...........................g.-...t.......tttcaaata..g..c...tag..........g.. [ 13360]
```

```
KM111294.1_Ken05/Tk1           ...-t.............................-...t.......tttcaaata..g..c...tag..........g.. [ 13360]
KM262844.1_strain_L60          -------------------------------------------------------------------------------- [ 13360]
KM102979.1_26544/OG10_Italy    -------------------------------------------------------------------------------- [ 13360]
KM262845.1_strain_NHV          -------------------------------------------------------------------------------- [ 13360]
KX354450.1_47/Ss/2008          -------------------------------------------------------------------------------- [ 13360]
KP055815.1_strain_BA71         -------------------------------------------------------------------------------- [ 13360]
MH025920.1_strain_R35          ...-t...........................g.-...t.......tttcaaata..g..c...tag..........g.. [ 13360]
MH025919.1_strain_N10          ...-t...........................g.-...t.......tttcaaata..g..c...tag..........g.. [ 13360]
MH025918.1_strain_R25          ...-t...........................g.-...t.......tttcaaata..g..c...tag..........g.. [ 13360]
MH025917.1_strain_R7           ...-t...........................g.-...t.......tttcaaata..g..c...tag..........g.. [ 13360]
MH025916.1_strain_R8           ...-t...........................g.-...t.......tttcaaata..g..c...tag..........g.. [ 13360]

FR682468.2_Georgia_2007/1      atttatttaattttaactattattttctttccactctctctttaagattttgtaaggattccagggctttggttcagaac [ 13440]
ASF_IND_20_CAD543              -------------------------------------------------------------------------------- [ 13440]
LR899193.1_Germany_2020/1      ................................................................................ [ 13440]
MN393477.1_Wuhan_2019          ................................................................................ [ 13440]
MW396979.1_Timor-Leste/2019/1  ................................................................................ [ 13440]
MK940252.1_CN/2019/InnerMongolia ................................................................................ [ 13440]
MK128995.1_China/2018/AnhuiXCGQ ................................................................................ [ 13440]
MH681419.1_ASFV/POL/2015       ................................................................................ [ 13440]
LS478113.1_Estonia_2014        -------------------------------------------------------------------------------- [ 13440]
KP843857.1_Odintsovo_02/14     ................................................................................ [ 13440]
FN557520.1_strain_E75          -------------------------------------------------------------------------------- [ 13440]
AY261366.1_Warthog             ----------....gg..g..................t.............a............tat.......g..... [ 13440]
AY261365.1_Warmbaths           ----------....gg..g..................t.............a............tat.......g..... [ 13440]
AY261364.1_Tengani_62          ----------...........................t.............a............tat.......g..... [ 13440]
AY261363.1_Pretoriuskop/96/4   ----------....gg..g..................t.............a............tat.....c.g..... [ 13440]
AY261362.1_Mkuzi_1979          .a............g...g..................t.............a............ta........g..... [ 13440]
AY261361.1_Malawi_Lil-20/1_1983 ----------....gg..g..................t.............a............ta........g..... [ 13440]
AY261360.1_Kenya_1950          -------------------------------------------------------------------------------- [ 13440]
AM712240.1_OURT_88/3           -------------------------------------------------------------------------------- [ 13440]
AM712239.1_Benin_97/1          -------------------------------------------------------------------------------- [ 13440]
KM111295.1_Ken06.Bus           ..............gg..gac................t.......................................... [ 13440]
KM111294.1_Ken05/Tk1           ..............gg..gac.....t..................................................... [ 13440]
KM262844.1_strain_L60          -------------------------------------------------------------------------------- [ 13440]
KM102979.1_26544/OG10_Italy    -------------------------------------------------------------------------------- [ 13440]
KM262845.1_strain_NHV          -------------------------------------------------------------------------------- [ 13440]
KX354450.1_47/Ss/2008          -------------------------------------------------------------------------------- [ 13440]
KP055815.1_strain_BA71         -------------------------------------------------------------------------------- [ 13440]
MH025920.1_strain_R35          ..............gg..gac................t.......................................... [ 13440]
MH025919.1_strain_N10          ..............gg..gac................t.......................................... [ 13440]
MH025918.1_strain_R25          ..............gg..gac................t.......................................... [ 13440]
MH025917.1_strain_R7           ..............gg..gac................t.......................................... [ 13440]
MH025916.1_strain_R8           ..............gg..gac................t.......................................... [ 13440]
```

```
FR682468.2_Georgia_2007/1        aggccattacatggtgaatcccctgtcctagatcatacatacatttatttagccagcgggaaactatacatgattgcaca [ 13520]
ASF_IND_20_CAD543                -----------------------------------------t..............y...............ga...... [ 13520]
LR899193.1_Germany_2020/1        ................................................................................ [ 13520]
MN393477.1_Wuhan_2019            ................................................................................ [ 13520]
MW396979.1_Timor-Leste/2019/1    ................................................................................ [ 13520]
MK940252.1_CN/2019/InnerMongolia ................................................................................ [ 13520]
MK128995.1_China/2018/AnhuiXCGQ  ................................................................................ [ 13520]
MH681419.1_ASFV/POL/2015         ................................................................................ [ 13520]
LS478113.1_Estonia_2014          -------------------------------------------------------------------------------- [ 13520]
KP843857.1_Odintsovo_02/14       ................................................................................ [ 13520]
FN557520.1_strain_E75            -------------------------------------------------------------------------------- [ 13520]
AY261366.1_Warthog               ..a..........a.c...g..t..a..c.....g.....................................ga...... [ 13520]
AY261365.1_Warmbaths             ..a..........a.c...g..t..a..c.....g.....................................ga...... [ 13520]
AY261364.1_Tengani_62            ..a..........a.c...g..t..a..c...........................................ga...... [ 13520]
AY261363.1_Pretoriuskop/96/4     ..a..........a.c...g..t..a..c...........................................ga...... [ 13520]
AY261362.1_Mkuzi_1979            ..a..........a.c...g..t..a..c...........g...............................ga...... [ 13520]
AY261361.1_Malawi_Lil-20/1_1983  ..a..........a.c...g..t..a..c...........................................ga...... [ 13520]
AY261360.1_Kenya_1950            -------------------------------------------------------------------------------- [ 13520]
AM712240.1_OURT_88/3             -------------------------------------------------------------------------------- [ 13520]
AM712239.1_Benin_97/1            -------------------------------------------------------------------------------- [ 13520]
KM111295.1_Ken06.Bus             ...g............................................................................ [ 13520]
KM111294.1_Ken05/Tk1             ...............c................................................................ [ 13520]
KM262844.1_strain_L60            -------------------------------------------------------------------------------- [ 13520]
KM102979.1_26544/OG10_Italy      -------------------------------------------------------------------------------- [ 13520]
KM262845.1_strain_NHV            -------------------------------------------------------------------------------- [ 13520]
KX354450.1_47/Ss/2008            -------------------------------------------------------------------------------- [ 13520]
KP055815.1_strain_BA71           -------------------------------------------------------------------------------- [ 13520]
MH025920.1_strain_R35            ...g............................................................................ [ 13520]
MH025919.1_strain_N10            ...g............................................................................ [ 13520]
MH025918.1_strain_R25            ...g............................................................................ [ 13520]
MH025917.1_strain_R7             ...g............................................................................ [ 13520]
MH025916.1_strain_R8             ...g............................................................................ [ 13520]

FR682468.2_Georgia_2007/1        tactcattttcaagaattgttgtattctccaatttgccctcacaaaggccattttgacaattccagcaaaacttgcagtt [ 13600]
ASF_IND_20_CAD543                ..........................tc....c..att.a.................................g..nnnn [ 13600]
LR899193.1_Germany_2020/1        ................................................................................ [ 13600]
MN393477.1_Wuhan_2019            ................................................................................ [ 13600]
MW396979.1_Timor-Leste/2019/1    ................................................................................ [ 13600]
MK940252.1_CN/2019/InnerMongolia ................................................................................ [ 13600]
MK128995.1_China/2018/AnhuiXCGQ  ................................................................................ [ 13600]
MH681419.1_ASFV/POL/2015         ................................................................................ [ 13600]
LS478113.1_Estonia_2014          -----------------------------------------------t.....c...tg..gt..--------------- [ 13600]
KP843857.1_Odintsovo_02/14       ................................................................................ [ 13600]
```

```
FN557520.1_strain_E75          -------------------------------------------------------------------------------- [ 13600]
AY261366.1_Warthog             ..........................tc....c..att.a...................g.c...........g...aca [ 13600]
AY261365.1_Warmbaths           ..........................tc....c..att.a...................g.c...........g...aca [ 13600]
AY261364.1_Tengani_62          ..........................tc....c..att.a...................g.c...........g...aca [ 13600]
AY261363.1_Pretoriuskop/96/4   ..........................tc....c..att.a...................g.c...........g...aca [ 13600]
AY261362.1_Mkuzi_1979          ..........................tc....c..att.a............c......g.c...........g...aca [ 13600]
AY261361.1_Malawi_Lil-20/1_1983 ..........................tc....c..att.a............c......g.c...........g...aca [ 13600]
AY261360.1_Kenya_1950          -------------------------------------------------------------------------------- [ 13600]
AM712240.1_OURT_88/3           -------------------------------------------------------------------------------- [ 13600]
AM712239.1_Benin_97/1          -------------------------------------------------------------------------------- [ 13600]
KM111295.1_Ken06.Bus           ...........................................................g...........t..a...c. [ 13600]
KM111294.1_Ken05/Tk1           ...........................................................g...........t..a...c. [ 13600]
KM262844.1_strain_L60          -------------------------------------------------------------------------------- [ 13600]
KM102979.1_26544/OG10_Italy    -------------------------------------------------------------------------------- [ 13600]
KM262845.1_strain_NHV          -------------------------------------------------------------------------------- [ 13600]
KX354450.1_47/Ss/2008          -------------------------------------------------------------------------------- [ 13600]
KP055815.1_strain_BA71         -------------------------------------------------------------------------------- [ 13600]
MH025920.1_strain_R35          ...........................................................g...........t..a...c. [ 13600]
MH025919.1_strain_N10          ...........................................................g...........t..a...c. [ 13600]
MH025918.1_strain_R25          ...........................................................g...........t..a...c. [ 13600]
MH025917.1_strain_R7           ...........................................................g...........t..a...c. [ 13600]
MH025916.1_strain_R8           ...........................................................g...........t..a...c. [ 13600]

FR682468.2_Georgia_2007/1      ttctgtataagtgcaccagtattcaag----------------------------------------------------- [ 13680]
ASF_IND_20_CAD543              nnnnnnnn..........a........nnnnnnnnnnnnnnnnnnnnnnnnnnnnnnnnnnnnnnnnnnnnnnnnn---- [ 13680]
LR899193.1_Germany_2020/1      ...........................----------------------------------------------------- [ 13680]
MN393477.1_Wuhan_2019          ...........................----------------------------------------------------- [ 13680]
MW396979.1_Timor-Leste/2019/1  ...........................----------------------------------------------------- [ 13680]
MK940252.1_CN/2019/InnerMongolia ...........................----------------------------------------------------- [ 13680]
MK128995.1_China/2018/AnhuiXCGQ ...........................----------------------------------------------------- [ 13680]
MH681419.1_ASFV/POL/2015       ...........................----------------------------------------------------- [ 13680]
LS478113.1_Estonia_2014        -------------------------------------------------------------------------------- [ 13680]
KP843857.1_Odintsovo_02/14     ...........................----------------------------------------------------- [ 13680]
FN557520.1_strain_E75          -------------------------------------------------------------------------------- [ 13680]
AY261366.1_Warthog             ...ca.g....................ttcattctctgaaggattttccgttggatgaagttgtccaacgagctgg---- [ 13680]
AY261365.1_Warmbaths           ...ca.g....................ttcattctctgaaggattttccgttggatgaagttgtccaacgagctgg---- [ 13680]
AY261364.1_Tengani_62          ...ca.g....................ttcattctctgaaggattttccgttggatgaagttgtccaactggctgg---- [ 13680]
AY261363.1_Pretoriuskop/96/4   ...ca.g....................ttcattctctgaaggattttccgttggatgaagttgtccaacgagctgg---- [ 13680]
AY261362.1_Mkuzi_1979          ...ca.g....................ttcattctctgaaggattttccgttggatgaagttgtccaacgagctgg---- [ 13680]
AY261361.1_Malawi_Lil-20/1_1983 ...ca.g....................ttcattctctgaaggattttccgttggatgaagttgtccaacgagctgg---- [ 13680]
AY261360.1_Kenya_1950          -------------------------------------------------------------------------------- [ 13680]
AM712240.1_OURT_88/3           -------------------------------------------------------------------------------- [ 13680]
AM712239.1_Benin_97/1          -------------------------------------------------------------------------------- [ 13680]
KM111295.1_Ken06.Bus           ...c..g...............c....ttcttcttttggaggattatccgttggatgaagttgtccagctggttgattag [ 13680]
```

```
KM111294.1_Ken05/Tk1            ...c..g...............c....ttcttcttttggaggattatccgttggatgaagttgtccagctggttgattag [ 13680]
KM262844.1_strain_L60           -------------------------------------------------------------------------------- [ 13680]
KM102979.1_26544/OG10_Italy     -------------------------------------------------------------------------------- [ 13680]
KM262845.1_strain_NHV           -------------------------------------------------------------------------------- [ 13680]
KX354450.1_47/Ss/2008           -------------------------------------------------------------------------------- [ 13680]
KP055815.1_strain_BA71          -------------------------------------------------------------------------------- [ 13680]
MH025920.1_strain_R35           ...c..g...............c....ttcttcttttggaggattatccgttggatgaagttgtccagctggttgattag [ 13680]
MH025919.1_strain_N10           ...c..g...............c....ttcttcttttggaggattatccgttggatgaagttgtccagctggttgattag [ 13680]
MH025918.1_strain_R25           ...c..g...............c....ttcttcttttggaggattatccgttggatgaagttgtccagctggttgattag [ 13680]
MH025917.1_strain_R7            ...c..g...............c....ttcttcttttggaggattatccgttggatgaagttgtccagctggttgattag [ 13680]
MH025916.1_strain_R8            ...c..g...............c....ttcttcttttggaggattatccgttggatgaagttgtccagctggttgattag [ 13680]

FR682468.2_Georgia_2007/1       -------------------------------------------------------------------------------- [ 13760]
ASF_IND_20_CAD543               -----nnnnnnnnnnnnnnnnnnnnnnnnnnnnnnnnnnnnnnnnnnnnnnnnnnnnnnnnnnnnnnn----------- [ 13760]
LR899193.1_Germany_2020/1       -------------------------------------------------------------------------------- [ 13760]
MN393477.1_Wuhan_2019           -------------------------------------------------------------------------------- [ 13760]
MW396979.1_Timor-Leste/2019/1   -------------------------------------------------------------------------------- [ 13760]
MK940252.1_CN/2019/InnerMongolia -------------------------------------------------------------------------------- [ 13760]
MK128995.1_China/2018/AnhuiXCGQ -------------------------------------------------------------------------------- [ 13760]
MH681419.1_ASFV/POL/2015        -------------------------------------------------------------------------------- [ 13760]
LS478113.1_Estonia_2014         -------------------------------------------------------------------------------- [ 13760]
KP843857.1_Odintsovo_02/14      -------------------------------------------------------------------------------- [ 13760]
FN557520.1_strain_E75           -------------------------------------------------------------------------------- [ 13760]
AY261366.1_Warthog              -----cttgagacctggttggccagaaggccaagaattcccaagaagattactaacattgctacggctg----------- [ 13760]
AY261365.1_Warmbaths            -----cttgagacctggttggccagaaggccaagaattcccaaaaagattactaacattgctacggctg----------- [ 13760]
AY261364.1_Tengani_62           -----cttgagacctggttgaccagaaggccaagaattcccaagaagatgactaacattgctacggccg----------- [ 13760]
AY261363.1_Pretoriuskop/96/4    -----cttgagacctggttggccagaaggccaagaattcccaagaagattactaacattgctatggctg----------- [ 13760]
AY261362.1_Mkuzi_1979           -----cttgagacctggttggccagaaggccaagaattcccaagaagattactaacattgctacggccg----------- [ 13760]
AY261361.1_Malawi_Lil-20/1_1983 -----cttgagacctggttggccagaaggccaagaattcccaagaagataaccaacattgctacggctg----------- [ 13760]
AY261360.1_Kenya_1950           -------------------------------------------------------------------------------- [ 13760]
AM712240.1_OURT_88/3            -------------------------------------------------------------------------------- [ 13760]
AM712239.1_Benin_97/1           -------------------------------------------------------------------------------- [ 13760]
KM111295.1_Ken06.Bus            gtagccctaagacctggttggccagcaggccaaggagtcccaagaagatcaccaacattgctatggctg----------- [ 13760]
KM111294.1_Ken05/Tk1            gtagccctaagacctggttggccagcaggccaaggagtcccaagaagatcaccaacattgctatggctgactaaacagcc [ 13760]
KM262844.1_strain_L60           -------------------------------------------------------------------------------- [ 13760]
KM102979.1_26544/OG10_Italy     -------------------------------------------------------------------------------- [ 13760]
KM262845.1_strain_NHV           -------------------------------------------------------------------------------- [ 13760]
KX354450.1_47/Ss/2008           -------------------------------------------------------------------------------- [ 13760]
KP055815.1_strain_BA71          -------------------------------------------------------------------------------- [ 13760]
MH025920.1_strain_R35           gtagccctaagacctggttggccagcaggccaaggagtcccaagaagatcaccaacattgctatggctg----------- [ 13760]
MH025919.1_strain_N10           gtagccctaagacctggttggccagcaggccaaggagtcccaagaagatcaccaacattgctatggctg----------- [ 13760]
MH025918.1_strain_R25           gtagccctaagacctggttggccagcaggccaaggagtcccaagaagatcaccaacattgctatggctg----------- [ 13760]
MH025917.1_strain_R7            gtagccctaagacctggttggccagcaggccaaggagtcccaagaagatcaccaacattgctatggctg----------- [ 13760]
MH025916.1_strain_R8            gtagccctaagacctggttggccagcaggccaaggagtcccaagaagatcaccaacattgctatggctg----------- [ 13760]
```

```
FR682468.2_Georgia_2007/1        -------------------------------------------------------------------------------- [ 13840]
ASF_IND_20_CAD543                -nnnnnnnnnnnnnnnnnnn-n------------------------nnnnnnnnnnnnnnnnnnnnnnnn--nnnnnnnn [ 13840]
LR899193.1_Germany_2020/1        -------------------------------------------------------------------------------- [ 13840]
MN393477.1_Wuhan_2019            -------------------------------------------------------------------------------- [ 13840]
MW396979.1_Timor-Leste/2019/1    -------------------------------------------------------------------------------- [ 13840]
MK940252.1_CN/2019/InnerMongolia -------------------------------------------------------------------------------- [ 13840]
MK128995.1_China/2018/AnhuiXCGQ  -------------------------------------------------------------------------------- [ 13840]
MH681419.1_ASFV/POL/2015         -------------------------------------------------------------------------------- [ 13840]
LS478113.1_Estonia_2014          -------------------------------------------------------------------------------- [ 13840]
KP843857.1_Odintsovo_02/14       -------------------------------------------------------------------------------- [ 13840]
FN557520.1_strain_E75            -------------------------------------------------------------------------------- [ 13840]
AY261366.1_Warthog               -gctagatagctatttagc-----------------------------tagctattagcaaaccaagtgacttgccctct [ 13840]
AY261365.1_Warmbaths             -gctagatagctatttagc-----------------------------tagctattagcaaaccaagtgacttgccctct [ 13840]
AY261364.1_Tengani_62            -gctgaacagctggctaaatag------------------------ctagctaattagcaaaccaagtgacttgccctct [ 13840]
AY261363.1_Pretoriuskop/96/4     -gctagctagc---------------------------------------gctattagcaaaccaagtga--tgccctct [ 13840]
AY261362.1_Mkuzi_1979            -gctgaacagctgctaggta-g------------------------ctagctaattagcaaaccaagtga--cgccctct [ 13840]
AY261361.1_Malawi_Lil-20/1_1983  -gctgaacagctggctagatagctagctaattagctagctaattatctagctaattagcaaaccaagtga--cgccctct [ 13840]
AY261360.1_Kenya_1950            -----aatagctgctgaaatagc-------------------------tgctgaatagcaaaccaagtgattcgccctct [ 13840]
AM712240.1_OURT_88/3             -------------------------------------------------------------------------------- [ 13840]
AM712239.1_Benin_97/1            -------------------------------------------------------------------------------- [ 13840]
KM111295.1_Ken06.Bus             -gctaaacagctgactaaacagc-------------------------ggctaattagcaaaccaagcga----cccact [ 13840]
KM111294.1_Ken05/Tk1             gactaaacagctgactaaacagc-------------------------ggctaattagcaaaccaagcga----cccact [ 13840]
KM262844.1_strain_L60            -------------------------------------------------------------------------------- [ 13840]
KM102979.1_26544/OG10_Italy      -------------------------------------------------------------------------------- [ 13840]
KM262845.1_strain_NHV            -------------------------------------------------------------------------------- [ 13840]
KX354450.1_47/Ss/2008            -------------------------------------------------------------------------------- [ 13840]
KP055815.1_strain_BA71           -------------------------------------------------------------------------------- [ 13840]
MH025920.1_strain_R35            -gctaaacagctgactaaacagc-------------------------ggctaattagcaaaccaagcga----cccact [ 13840]
MH025919.1_strain_N10            -gctaaacagctgactaaacagc-------------------------ggctaattagcaaaccaagcga----cccact [ 13840]
MH025918.1_strain_R25            -gctaaacagctgactaaacagc-------------------------ggctaattagcaaaccaagcga----cccact [ 13840]
MH025917.1_strain_R7             -gctaaacagctgactaaacagc-------------------------ggctaattagcaaaccaagcga----cccact [ 13840]
MH025916.1_strain_R8             -gctaaacagctgactaaacagc-------------------------ggctaattagcaaaccaagcga----cccact [ 13840]

FR682468.2_Georgia_2007/1        -------------------------------------------------------------------------------- [ 13920]
ASF_IND_20_CAD543                nnnnnnnnnnnnnnnnnnnnnnnnnnnnnnnnnnnnnnnnnnnnnnnnnnttacaggaaaaaggtatttttaannnnnnn [ 13920]
LR899193.1_Germany_2020/1        -------------------------------------------------------------------------------- [ 13920]
MN393477.1_Wuhan_2019            -------------------------------------------------------------------------------- [ 13920]
MW396979.1_Timor-Leste/2019/1    -------------------------------------------------------------------------------- [ 13920]
MK940252.1_CN/2019/InnerMongolia -------------------------------------------------------------------------------- [ 13920]
MK128995.1_China/2018/AnhuiXCGQ  -------------------------------------------------------------------------------- [ 13920]
MH681419.1_ASFV/POL/2015         -------------------------------------------------------------------------------- [ 13920]
LS478113.1_Estonia_2014          -------------------------------------------------------------------------------- [ 13920]
KP843857.1_Odintsovo_02/14       -------------------------------------------------------------------------------- [ 13920]
```

```
FN557520.1_strain_E75          -------------------------------------------------------------------------------- [ 13920]
AY261366.1_Warthog             ctattctttatatgatgatttaagattgagtccaacatttttccaacgttttacaggaaaaaggta-ttttaactatgaa [ 13920]
AY261365.1_Warmbaths           ctattctttatatgatgatttaagattgagtccaacatttttccaacgttttacaggaaaaaggta-ttttaactatgaa [ 13920]
AY261364.1_Tengani_62          ctattctttatatgatgatttaagattgagtccaacatttttccaacgttttacaggaaaaaggtattttttaactatgaa [ 13920]
AY261363.1_Pretoriuskop/96/4   ctattctttatatgatgttttaagattgagtccaacatttttccaacgttttacaggaaaaaggtattttttaactatgaa [ 13920]
AY261362.1_Mkuzi_1979          ctattctttatatgatgatttaagattgagtccaacatttttccaacgttttacaggaaaaaggtattttttaactatgaa [ 13920]
AY261361.1_Malawi_Lil-20/1_1983 ctattctttatatgatgatttaagattcggtccaacatttttccaatgttttacaggaaaaaggtattttttaactatgaa [ 13920]
AY261360.1_Kenya_1950          ctacttttcataagatagtttagaattcggtccaacgtttttccaatgttttataggaaaaagatatttttagctatgaa [ 13920]
AM712240.1_OURT_88/3           -------------------------------------------------------------------------------- [ 13920]
AM712239.1_Benin_97/1          -------------------------------------------------------------------------------- [ 13920]
KM111295.1_Ken06.Bus           ctacttttcataagatagtttagaattcggtccaacgtttttccaatggtttacaggaaaaagatatttttagccatgaa [ 13920]
KM111294.1_Ken05/Tk1           ctacttttcataagatagtttagaattcggtctaacgtttttccaatgttttataggaaaaagatatttttagctatgaa [ 13920]
KM262844.1_strain_L60          -------------------------------------------------------------------------------- [ 13920]
KM102979.1_26544/OG10_Italy    -------------------------------------------------------------------------------- [ 13920]
KM262845.1_strain_NHV          -------------------------------------------------------------------------------- [ 13920]
KX354450.1_47/Ss/2008          -------------------------------------------------------------------------------- [ 13920]
KP055815.1_strain_BA71         -------------------------------------------------------------------------------- [ 13920]
MH025920.1_strain_R35          ctacttttcataagatagtttagaattcggtccaacgtttttccaatggtttacaggaaaaagatatttttagccatgaa [ 13920]
MH025919.1_strain_N10          ctacttttcataagatagtttagaattcggtccaacgtttttccaatggtttacaggaaaaagatatttttagccatgaa [ 13920]
MH025918.1_strain_R25          ctacttttcataagatagtttagaattcggtccaacgtttttccaatggtttacaggaaaaagatatttttagccatgaa [ 13920]
MH025917.1_strain_R7           ctacttttcataagatagtttagaattcggtccaacgtttttccaatggtttacaggaaaaagatatttttagccatgaa [ 13920]
MH025916.1_strain_R8           ctacttttcataagatagtttagaattcggtccaacgtttttccaatggtttacaggaaaaagatatttttagccatgaa [ 13920]

FR682468.2_Georgia_2007/1      ---------------ttcttcttgt-ggaggattatccgttggat----------------------------------- [ 14000]
ASF_IND_20_CAD543              nnnnnnnnnnn--nnnn..at..ag-aa.aa.a..aaa.acac..ttatttaattttannnnnnnnnnnnnnnnnnnnnn [ 14000]
LR899193.1_Germany_2020/1      ---------------..........-...................----------------------------------- [ 14000]
MN393477.1_Wuhan_2019          ---------------..........-...................----------------------------------- [ 14000]
MW396979.1_Timor-Leste/2019/1  ---------------..........-...................----------------------------------- [ 14000]
MK940252.1_CN/2019/InnerMongolia ---------------..........-...................----------------------------------- [ 14000]
MK128995.1_China/2018/AnhuiXCGQ ---------------..........-...................----------------------------------- [ 14000]
MH681419.1_ASFV/POL/2015       ---------------..........-...................----------------------------------- [ 14000]
LS478113.1_Estonia_2014        -----------------t...gc.g-.cc.ac..c..a.g.at..----------------------------------- [ 14000]
KP843857.1_Odintsovo_02/14     ---------------..........-...................----------------------------------- [ 14000]
FN557520.1_strain_E75          -------------------------------------------------------------------------------- [ 14000]
AY261366.1_Warthog             attaatcttgac-ccact.at..ag-aa.aa.a..aaa.acac..ttatttaattttagctgaggcaaatttgctgacaa [ 14000]
AY261365.1_Warmbaths           attaatcttgac-ccact.at..ag-aa.aa.a..aaa.acac..ttatttaattttagctgaggcaaatttgctgacaa [ 14000]
AY261364.1_Tengani_62          attaatcttga--ccact.at..ag-aa.aa.a..aaa.acac..ttatttaattttagctgaggcaaatttgctgacaa [ 14000]
AY261363.1_Pretoriuskop/96/4   attgatcttgac-ccact.at..aa-aa.aa.a------a.aa..ttatttaattttagctgaggcaaatttgctgacaa [ 14000]
AY261362.1_Mkuzi_1979          attaatcttga--ccca..at..ag-aa.aa.-..aaa.acac..ttatttaattttagctgaggcaaatttgctgacaa [ 14000]
AY261361.1_Malawi_Lil-20/1_1983 attaatcttga--ccact.at..ag-aa.aa.g..aaa.acaa..ttatttaattttagctgaggcaaatttgctgacaa [ 14000]
AY261360.1_Kenya_1950          tgcaattcacatatcc.acat.cag--a.aa.aa...aaaccat.ttattgaattttggctggaacaaatttgctgat-- [ 14000]
AM712240.1_OURT_88/3           -------------------------------------------------------------------------------- [ 14000]
AM712239.1_Benin_97/1          -------------------------------------------------------------------------------- [ 14000]
KM111295.1_Ken06.Bus           tgcacttcacgtatcc.aca.ccag-aa.aa.aa...aaacta..ttattgaattttggctggaacaaatttgctgat-- [ 14000]
```

```
KM111294.1_Ken05/Tk1            tgcacttcacatatcc.acat.cag--a.aa.aa...aaacca..ttattgaattttgactgtaacaaatttactgacaa [ 14000]
KM262844.1_strain_L60           -------------------------------------------------------------------------------- [ 14000]
KM102979.1_26544/OG10_Italy     -------------------------------------------------------------------------------- [ 14000]
KM262845.1_strain_NHV           -------------------------------------------------------------------------------- [ 14000]
KX354450.1_47/Ss/2008           -------------------------------------------------------------------------------- [ 14000]
KP055815.1_strain_BA71          -------------------------------------------------------------------------------- [ 14000]
MH025920.1_strain_R35           tgcacttcacgtatcc.aca.ccagaaa.aa.aa...aaacta..ttattgaattttggctggaacaaatttgctgat-- [ 14000]
MH025919.1_strain_N10           tgcacttcacgtatcc.aca.ccagaaa.aa.aa...aaacta..ttattgaattttggctggaacaaatttgctgat-- [ 14000]
MH025918.1_strain_R25           tgcacttcacgtatcc.aca.ccagaaa.aa.aa...aaacta..ttattgaattttggctggaacaaatttgctgat-- [ 14000]
MH025917.1_strain_R7            tgcacttcacgtatcc.aca.ccagaaa.aa.aa...aaacta..ttattgaattttggctggaacaaatttgctgat-- [ 14000]
MH025916.1_strain_R8            tgcacttcacgtatcc.aca.ccagaaa.aa.aa...aaacta..ttattgaattttggctggaacaaatttgctgat-- [ 14000]

FR682468.2_Georgia_2007/1       ------------gaagttgtccagctggttgattaggtag---------ccctaagac-----ctggttgcaattcatgg [ 14080]
ASF_IND_20_CAD543               nnnnnnnnnnnnnnnn..t.t.cat.cc..cc.g.annnnnnnnnnnnntt.c.....catgg...agaa.......... [ 14080]
LR899193.1_Germany_2020/1       ------------............................---------.........-----................. [ 14080]
MN393477.1_Wuhan_2019           ------------............................---------.........-----................. [ 14080]
MW396979.1_Timor-Leste/2019/1   ------------............................---------.........-----................. [ 14080]
MK940252.1_CN/2019/InnerMongolia ------------............................---------.........-----................. [ 14080]
MK128995.1_China/2018/AnhuiXCGQ ------------............................---------.........-----................. [ 14080]
MH681419.1_ASFV/POL/2015        ------------............................---------.........-----................. [ 14080]
LS478113.1_Estonia_2014         ------------..ttcgc...........tc.cg...g----------------------------------------- [ 14080]
KP843857.1_Odintsovo_02/14      ------------............................---------.........-----................. [ 14080]
FN557520.1_strain_E75           -------------------------------------------------------------------------------- [ 14080]
AY261366.1_Warthog              actctagagttcatct..t.t.cat.cc..cc.g.a...tttgtaaggatt.c.....catgg...agaa.......... [ 14080]
AY261365.1_Warmbaths            actctagagttcatct..t.t.cat.cc..cc.g.a...tttgtaaggatt.c.....catgg...agaa.......... [ 14080]
AY261364.1_Tengani_62           actctagagttcatct..t.t.cat.cc..cc.g.a...tttgtaaggatt.c.....catgg...agaa.......... [ 14080]
AY261363.1_Pretoriuskop/96/4    actctagagttcatct..t.t.cat.cc..cc.g.a...tttgtaaggatt.c.....catgg...agaa.......... [ 14080]
AY261362.1_Mkuzi_1979           actctagagttcatct..t.t.cat.cc..cc.g.a...tttgtaaggatt.c.....catgg...agaa.......... [ 14080]
AY261361.1_Malawi_Lil-20/1_1983 actctagagttcatct..t.t.cat.cc..cc.g.a...tttgtagggatt.c.g.c.tttggg.cagaa..gc.....c [ 14080]
AY261360.1_Kenya_1950           attctagagttcatct..t.t.ca..cc..cc.g.a...tttgtagggatt.c.....tgtgg...agaa........t. [ 14080]
AM712240.1_OURT_88/3            -------------------------------------------------------------------------------- [ 14080]
AM712239.1_Benin_97/1           -------------------------------------------------------------------------------- [ 14080]
KM111295.1_Ken06.Bus            attctagagttcatct..t.t.ca..cc..cc.g.aac.tttgtagggatt.c.....cgtgg...agaa..g.....t. [ 14080]
KM111294.1_Ken05/Tk1            actctagagttcatct..t.t.ca..cc..cc.g.a...tttgtagggatt.c.....tgtgg...agaa........t. [ 14080]
KM262844.1_strain_L60           -------------------------------------------------------------------------------- [ 14080]
KM102979.1_26544/OG10_Italy     -------------------------------------------------------------------------------- [ 14080]
KM262845.1_strain_NHV           -------------------------------------------------------------------------------- [ 14080]
KX354450.1_47/Ss/2008           -------------------------------------------------------------------------------- [ 14080]
KP055815.1_strain_BA71          -------------------------------------------------------------------------------- [ 14080]
MH025920.1_strain_R35           attctagagttcatct..t.t.ca..cc..cc.g.aac.tttgtagggatt.c.....cgtgg...agaa..g.....t. [ 14080]
MH025919.1_strain_N10           attctagagttcatct..t.t.ca..cc..cc.g.aac.tttgtagggatt.c.....cgtgg...agaa..g.....t. [ 14080]
MH025918.1_strain_R25           attctagagttcatct..t.t.ca..cc..cc.g.aac.tttgtagggatt.c.....cgtgg...agaa..g.....t. [ 14080]
MH025917.1_strain_R7            attctagagttcatct..t.t.ca..cc..cc.g.aac.tttgtagggatt.c.....cgtgg...agaa..g.....t. [ 14080]
MH025916.1_strain_R8            attctagagttcatct..t.t.ca..cc..cc.g.aac.tttgtagggatt.c.....cgtgg...agaa..g.....t. [ 14080]
```

```
FR682468.2_Georgia_2007/1        tatggtagatacccttatctaaatcatacatacatttatccagccaacgggaaaccagacatgatttcacatactcattc [ 14160]
ASF_IND_20_CAD543                .............y......................................m..........m................ [ 14160]
LR899193.1_Germany_2020/1        ................................................................................ [ 14160]
MN393477.1_Wuhan_2019            ................................................................................ [ 14160]
MW396979.1_Timor-Leste/2019/1    ................................................................................ [ 14160]
MK940252.1_CN/2019/InnerMongolia ................................................................................ [ 14160]
MK128995.1_China/2018/AnhuiXCGQ  ................................................................................ [ 14160]
MH681419.1_ASFV/POL/2015         ................................................................................ [ 14160]
LS478113.1_Estonia_2014          ------------.a.....c---------------------------------..g.c.ac.......t..g.tat..cg [ 14160]
KP843857.1_Odintsovo_02/14       ................................................................................ [ 14160]
FN557520.1_strain_E75            -------------------------------------------------------------------------------- [ 14160]
AY261366.1_Warthog               .............t.................................................................. [ 14160]
AY261365.1_Warmbaths             .............t.................................................................. [ 14160]
AY261364.1_Tengani_62            ................................................................................ [ 14160]
AY261363.1_Pretoriuskop/96/4     .............t.................................................................. [ 14160]
AY261362.1_Mkuzi_1979            .............t.................................................c................ [ 14160]
AY261361.1_Malawi_Lil-20/1_1983  ..................t.......................t.g..........tga.....g.cg............t [ 14160]
AY261360.1_Kenya_1950            ....a.............t.......................t...g........tga.....g.cg....a.......t [ 14160]
AM712240.1_OURT_88/3             -------------------------------------------------------------------------------- [ 14160]
AM712239.1_Benin_97/1            -------------------------------------------------------------------------------- [ 14160]
KM111295.1_Ken06.Bus             ....a..................................tt.....g........tga.....g..g............t [ 14160]
KM111294.1_Ken05/Tk1             ....a.............t....................t..t...g........tga.....g.cg....a.......t [ 14160]
KM262844.1_strain_L60            -------------------------------------------------------------------------------- [ 14160]
KM102979.1_26544/OG10_Italy      -------------------------------------------------------------------------------- [ 14160]
KM262845.1_strain_NHV            -------------------------------------------------------------------------------- [ 14160]
KX354450.1_47/Ss/2008            -------------------------------------------------------------------------------- [ 14160]
KP055815.1_strain_BA71           -------------------------------------------------------------------------------- [ 14160]
MH025920.1_strain_R35            ....a..................................tt.....g........tga.....g..g............t [ 14160]
MH025919.1_strain_N10            ....a..................................tt.....g........tga.....g..g............t [ 14160]
MH025918.1_strain_R25            ....a..................................tt.....g........tga.....g..g............t [ 14160]
MH025917.1_strain_R7             ....a..................................tt.....g........tga.....g..g............t [ 14160]
MH025916.1_strain_R8             ....a..................................tt.....g........tga.....g..g............t [ 14160]

FR682468.2_Georgia_2007/1        ttgtaaattactgacccatctattttgtttatacaagtgccgtcttggcagtcccagcaaaattggcaactttccatgta [ 14240]
ASF_IND_20_CAD543                ................................................................................ [ 14240]
LR899193.1_Germany_2020/1        ................................................................................ [ 14240]
MN393477.1_Wuhan_2019            ................................................................................ [ 14240]
MW396979.1_Timor-Leste/2019/1    ................................................................................ [ 14240]
MK940252.1_CN/2019/InnerMongolia ................................................................................ [ 14240]
MK128995.1_China/2018/AnhuiXCGQ  ................................................................................ [ 14240]
MH681419.1_ASFV/POL/2015         ................................................................................ [ 14240]
LS478113.1_Estonia_2014          ..cat.tca..c.gtttc...------....g.g..caat.a.--------------------------------g.... [ 14240]
KP843857.1_Odintsovo_02/14       ................................................................................ [ 14240]
```

```
FN557520.1_strain_E75            -------------------------------------------------------------------------------- [ 14240]
AY261366.1_Warthog               .........................................a...................................... [ 14240]
AY261365.1_Warmbaths             .........................................a...................................... [ 14240]
AY261364.1_Tengani_62            .........................................a...................................... [ 14240]
AY261363.1_Pretoriuskop/96/4     .........................................a...................................... [ 14240]
AY261362.1_Mkuzi_1979            .........................................a...................................... [ 14240]
AY261361.1_Malawi_Lil-20/1_1983  ..a.....g...........................t....a....................c................. [ 14240]
AY261360.1_Kenya_1950            ..a.....g.......................g...t....a....................c................. [ 14240]
AM712240.1_OURT_88/3             -------------------------------------------------------------------------------- [ 14240]
AM712239.1_Benin_97/1            -------------------------------------------------------------------------------- [ 14240]
KM111295.1_Ken06.Bus             ..a.....g.............c.............t....a.....a..............c................. [ 14240]
KM111294.1_Ken05/Tk1             ..a.....g.............c.............t....a..............a.....c................. [ 14240]
KM262844.1_strain_L60            -------------------------------------------------------------------------------- [ 14240]
KM102979.1_26544/OG10_Italy      -------------------------------------------------------------------------------- [ 14240]
KM262845.1_strain_NHV            -------------------------------------------------------------------------------- [ 14240]
KX354450.1_47/Ss/2008            -------------------------------------------------------------------------------- [ 14240]
KP055815.1_strain_BA71           -------------------------------------------------------------------------------- [ 14240]
MH025920.1_strain_R35            ..a.....g.............c.............t....a.....a..............c................. [ 14240]
MH025919.1_strain_N10            ..a.....g.............c.............t....a.....a..............c................. [ 14240]
MH025918.1_strain_R25            ..a.....g.............c.............t....a.....a..............c................. [ 14240]
MH025917.1_strain_R7             ..a.....g.............c.............t....a.....a..............c................. [ 14240]
MH025916.1_strain_R8             ..a.....g.............c.............t....a.....a..............c................. [ 14240]

FR682468.2_Georgia_2007/1        ggcacaccagtattcgagttcttcctctggaggctcctctgttggacgaagttgtccaacgagctga------------c [ 14320]
ASF_IND_20_CAD543                ...................................................................------------. [ 14320]
LR899193.1_Germany_2020/1        ...................................................................------------. [ 14320]
MN393477.1_Wuhan_2019            ...................................................................------------. [ 14320]
MW396979.1_Timor-Leste/2019/1    ...................................................................------------. [ 14320]
MK940252.1_CN/2019/InnerMongolia ...................................................................------------. [ 14320]
MK128995.1_China/2018/AnhuiXCGQ  ...................................................................------------. [ 14320]
MH681419.1_ASFV/POL/2015         ...................................................................------------. [ 14320]
LS478113.1_Estonia_2014          .a..tg.t.a..g.------------------------------------------------------------------ [ 14320]
KP843857.1_Odintsovo_02/14       ...................................................................------------. [ 14320]
FN557520.1_strain_E75            -------------------------------------------------------------------------------- [ 14320]
AY261366.1_Warthog               ...g.............................a...........t.....................------------. [ 14320]
AY261365.1_Warmbaths             ...g.............................a.................................------------. [ 14320]
AY261364.1_Tengani_62            .................................a.................................------------. [ 14320]
AY261363.1_Pretoriuskop/96/4     ...g.............................a.................................------------. [ 14320]
AY261362.1_Mkuzi_1979            ...g.............................a...........t.................t...------------. [ 14320]
AY261361.1_Malawi_Lil-20/1_1983  ...g.............................a.................................------------. [ 14320]
AY261360.1_Kenya_1950            .................................a........................gttg.t...ttagatggtagc. [ 14320]
AM712240.1_OURT_88/3             -------------------------------------------------------------------------------- [ 14320]
AM712239.1_Benin_97/1            -------------------------------------------------------------------------------- [ 14320]
KM111295.1_Ken06.Bus             ..tg.....a...........a.t.........g..........................tg.....------------. [ 14320]
```

```
KM111294.1_Ken05/Tk1              ..tg.....a....ta........t........a................................t.tg.t...------------. [ 14320]
KM262844.1_strain_L60             -------------------------------------------------------------------------------- [ 14320]
KM102979.1_26544/OG10_Italy       -------------------------------------------------------------------------------- [ 14320]
KM262845.1_strain_NHV             -------------------------------------------------------------------------------- [ 14320]
KX354450.1_47/Ss/2008             -------------------------------------------------------------------------------- [ 14320]
KP055815.1_strain_BA71            -------------------------------------------------------------------------------- [ 14320]
MH025920.1_strain_R35             ..tg.....a...........a.t.........g..........................tg.....------------. [ 14320]
MH025919.1_strain_N10             ..tg.....a...........a.t.........g..........................tg.....------------. [ 14320]
MH025918.1_strain_R25             ..tg.....a...........a.t.........g..........................tg.....------------. [ 14320]
MH025917.1_strain_R7              ..tg.....a...........a.t.........g..........................tg.....------------. [ 14320]
MH025916.1_strain_R8              ..tg.....a...........a.t.........g..........................tg.....------------. [ 14320]

FR682468.2_Georgia_2007/1         ttgaaacctggctggccagaaggccaagaattcccaagaagatcaccaacattgctacggctggctgaaca--------- [ 14400]
ASF_IND_20_CAD543                 .......................................................................--------- [ 14400]
LR899193.1_Germany_2020/1         .......................................................................--------- [ 14400]
MN393477.1_Wuhan_2019             .......................................................................--------- [ 14400]
MW396979.1_Timor-Leste/2019/1     .......................................................................--------- [ 14400]
MK940252.1_CN/2019/InnerMongolia  .......................................................................--------- [ 14400]
MK128995.1_China/2018/AnhuiXCGQ   .......................................................................--------- [ 14400]
MH681419.1_ASFV/POL/2015          .......................................................................--------- [ 14400]
LS478113.1_Estonia_2014           -------------------------------------------------------------------------------- [ 14400]
KP843857.1_Odintsovo_02/14        .......................................................................--------- [ 14400]
FN557520.1_strain_E75             -------------------------------------------------------------------------------- [ 14400]
AY261366.1_Warthog                .......................................................................--------- [ 14400]
AY261365.1_Warmbaths              .......................................................................--------- [ 14400]
AY261364.1_Tengani_62             ...........................a........................g..................--------- [ 14400]
AY261363.1_Pretoriuskop/96/4      .......................................................................--------- [ 14400]
AY261362.1_Mkuzi_1979             .....................................a.................................--------- [ 14400]
AY261361.1_Malawi_Lil-20/1_1983   ....g................................a...g.....................a.......--------- [ 14400]
AY261360.1_Kenya_1950             c.a.g............t...................a...................t............ggctgctgaa [ 14400]
AM712240.1_OURT_88/3              -------------------------------------------------------------------------------- [ 14400]
AM712239.1_Benin_97/1             -------------------------------------------------------------------------------- [ 14400]
KM111295.1_Ken06.Bus              ....g................................a........................a.......g--------- [ 14400]
KM111294.1_Ken05/Tk1              ....g................................a.....t..........................ggct------ [ 14400]
KM262844.1_strain_L60             -------------------------------------------------------------------------------- [ 14400]
KM102979.1_26544/OG10_Italy       -------------------------------------------------------------------------------- [ 14400]
KM262845.1_strain_NHV             -------------------------------------------------------------------------------- [ 14400]
KX354450.1_47/Ss/2008             -------------------------------------------------------------------------------- [ 14400]
KP055815.1_strain_BA71            -------------------------------------------------------------------------------- [ 14400]
MH025920.1_strain_R35             ....g................................a........................a.......g--------- [ 14400]
MH025919.1_strain_N10             ....g................................a........................a.......g--------- [ 14400]
MH025918.1_strain_R25             ....g................................a........................a.......g--------- [ 14400]
MH025917.1_strain_R7              ....g................................a........................a.......g--------- [ 14400]
MH025916.1_strain_R8              ....g................................a........................a.......g--------- [ 14400]
```

```
FR682468.2_Georgia_2007/1        --gctgactgaatagctagccaatt--------------------------------agcaatcca--ctgtacttttca [ 14480]
ASF_IND_20_CAD543                --.......................--------------------------------.........--............ [ 14480]
LR899193.1_Germany_2020/1        --.......................--------------------------------.........--............ [ 14480]
MN393477.1_Wuhan_2019            --.......................--------------------------------.........--............ [ 14480]
MW396979.1_Timor-Leste/2019/1    --.......................--------------------------------.........--............ [ 14480]
MK940252.1_CN/2019/InnerMongolia --.......................--------------------------------.........--............ [ 14480]
MK128995.1_China/2018/AnhuiXCGQ  --.......................--------------------------------.........--............ [ 14480]
MH681419.1_ASFV/POL/2015         --.......................--------------------------------.........--............ [ 14480]
LS478113.1_Estonia_2014          ----------------------------------------------------------------t.--..a...------ [ 14480]
KP843857.1_Odintsovo_02/14       --.......................--------------------------------.........--............ [ 14480]
FN557520.1_strain_E75            -------------------------------------------------------------------------------- [ 14480]
AY261366.1_Warthog               --.......................--------------------------------.........--............ [ 14480]
AY261365.1_Warmbaths             --.......................--------------------------------.........--............ [ 14480]
AY261364.1_Tengani_62            --.......................--------------------------------.........--............ [ 14480]
AY261363.1_Pretoriuskop/96/4     --....g..ag.........t....--------------------------------.........--............ [ 14480]
AY261362.1_Mkuzi_1979            --......c................--------------------------------.........--............ [ 14480]
AY261361.1_Malawi_Lil-20/1_1983  --...a........a......g...a-----------------------gcaaacca.........--............ [ 14480]
AY261360.1_Kenya_1950            ca....g..a.......g..t..a.agctgactgaatagctgctgattagcaaacca.........ct..c.gt...... [ 14480]
AM712240.1_OURT_88/3             -------------------------------------------------------------------------------- [ 14480]
AM712239.1_Benin_97/1            -------------------------------------------------------------------------------- [ 14480]
KM111295.1_Ken06.Bus             --....g.....c....g..t..a.agctgactgaatagctgctgattagcaaacca.......t.ct..c.g....... [ 14480]
KM111294.1_Ken05/Tk1             ------g.....c....g..t..a.agctgactgaatagctgctgattagcaaacca.........--..c......... [ 14480]
KM262844.1_strain_L60            -------------------------------------------------------------------------------- [ 14480]
KM102979.1_26544/OG10_Italy      -------------------------------------------------------------------------------- [ 14480]
KM262845.1_strain_NHV            -------------------------------------------------------------------------------- [ 14480]
KX354450.1_47/Ss/2008            -------------------------------------------------------------------------------- [ 14480]
KP055815.1_strain_BA71           -------------------------------------------------------------------------------- [ 14480]
MH025920.1_strain_R35            --....g.....c....g..t..a.agctgactgaatagctgctgattagcaaacca.......t.ct..c.g....... [ 14480]
MH025919.1_strain_N10            --....g.....c....g..t..a.agctgactgaatagctgctgattagcaaacca.......t.ct..c.g....... [ 14480]
MH025918.1_strain_R25            --....g.....c....g..t..a.agctgactgaatagctgctgattagcaaacca.......t.ct..c.g....... [ 14480]
MH025917.1_strain_R7             --....g.....c....g..t..a.agctgactgaatagctgctgattagcaaacca.......t.ct..c.g....... [ 14480]
MH025916.1_strain_R8             --....g.....c....g..t..a.agctgactgaatagctgctgattagcaaacca.......t.ct..c.g....... [ 14480]

FR682468.2_Georgia_2007/1        taagatcatttaagattcggtc-ggcattttttcaatagtttgctag-gaaaaaatttttaattttatagattcacacta [ 14560]
ASF_IND_20_CAD543                ......................-........................-...............y................ [ 14560]
LR899193.1_Germany_2020/1        ......................-........................-................................ [ 14560]
MN393477.1_Wuhan_2019            ......................-........................-................................ [ 14560]
MW396979.1_Timor-Leste/2019/1    ......................-........................-................................ [ 14560]
MK940252.1_CN/2019/InnerMongolia ......................-........................-................................ [ 14560]
MK128995.1_China/2018/AnhuiXCGQ  ......................-........................-................................ [ 14560]
MH681419.1_ASFV/POL/2015         ......................-........................-................................ [ 14560]
LS478113.1_Estonia_2014          a..a..ag......t..t...----------------------------------------------------------- [ 14560]
KP843857.1_Odintsovo_02/14       ......................-........................-................................ [ 14560]
```

```
FN557520.1_strain_E75              -------------------------------------------------------------------------------- [ 14560]
AY261366.1_Warthog                 ......................-........................-a..............c..............c. [ 14560]
AY261365.1_Warmbaths               ......................-........................--..............c..............c. [ 14560]
AY261364.1_Tengani_62              ......................-........................g...............c..............c. [ 14560]
AY261363.1_Pretoriuskop/96/4       ......................-....................t...--..............c.c............c. [ 14560]
AY261362.1_Mkuzi_1979              ......................-.a......................-...............c..............c. [ 14560]
AY261361.1_Malawi_Lil-20/1_1983    ......................-........................-...............c.........t....c- [ 14560]
AY261360.1_Kenya_1950              ......................t.a.......c............ga-...............c....c....t..cact [ 14560]
AM712240.1_OURT_88/3               -------------------------------------------------------------------------------- [ 14560]
AM712239.1_Benin_97/1              -------------------------------------------------------------------------------- [ 14560]
KM111295.1_Ken06.Bus               ....g.................ta........c....g.......g.-...............c....c....t..cact [ 14560]
KM111294.1_Ken05/Tk1               ......................t...............c......g.-...............c....c....t..cact [ 14560]
KM262844.1_strain_L60              -------------------------------------------------------------------------------- [ 14560]
KM102979.1_26544/OG10_Italy        -------------------------------------------------------------------------------- [ 14560]
KM262845.1_strain_NHV              -------------------------------------------------------------------------------- [ 14560]
KX354450.1_47/Ss/2008              -------------------------------------------------------------------------------- [ 14560]
KP055815.1_strain_BA71             -------------------------------------------------------------------------------- [ 14560]
MH025920.1_strain_R35              ....g.................ta........c....g.......g.-...............c....c....t..cact [ 14560]
MH025919.1_strain_N10              ....g.................ta........c....g.......g.-...............c....c....t..cact [ 14560]
MH025918.1_strain_R25              ....g.................ta........c....g.......g.-...............c....c....t..cact [ 14560]
MH025917.1_strain_R7               ....g.................ta........c....g.......g.-...............c....c....t..cact [ 14560]
MH025916.1_strain_R8               ....g.................ta........c....g.......g.-...............c....c....t..cact [ 14560]

FR682468.2_Georgia_2007/1          cttcattctcatgcttaggaaaaaaa-------caaactaaatcttacaatgtatctggatctaa-tgagaagctagaat [ 14640]
ASF_IND_20_CAD543                  .........................t--------...............................-......r....... [ 14640]
LR899193.1_Germany_2020/1          ..........................-------................................-.............. [ 14640]
MN393477.1_Wuhan_2019              ..........................-------................................-.............. [ 14640]
MW396979.1_Timor-Leste/2019/1      ..........................-------................................-.............. [ 14640]
MK940252.1_CN/2019/InnerMongolia   ..........................-------................................-.............. [ 14640]
MK128995.1_China/2018/AnhuiXCGQ    ..........................-------................................-.............. [ 14640]
MH681419.1_ASFV/POL/2015           ..........................-------................................-.............. [ 14640]
LS478113.1_Estonia_2014            ----------------------------------.t.t.........a..------------------------------ [ 14640]
KP843857.1_Odintsovo_02/14         ..........................-------................................-.............. [ 14640]
FN557520.1_strain_E75              -------------------------------------------------------------------------------- [ 14640]
AY261366.1_Warthog                 ..........................-------.....c..........................-.............. [ 14640]
AY261365.1_Warmbaths               ..........................-------.....c..........................-.............. [ 14640]
AY261364.1_Tengani_62              ...t.....t.c..cc..........-------.....c..........................-.............. [ 14640]
AY261363.1_Pretoriuskop/96/4       ..........................-------.....c..........................-.............. [ 14640]
AY261362.1_Mkuzi_1979              .........................t--------....c....t.........c...........-......a....... [ 14640]
AY261361.1_Malawi_Lil-20/1_1983    .........................t--------....c....t.........c...........-......a....... [ 14640]
AY261360.1_Kenya_1950              t.attg.ta..c...cg..g......acaaacctg..tc.......t...--.............t......a....... [ 14640]
AM712240.1_OURT_88/3               -------------------------------------------------------------------------------- [ 14640]
AM712239.1_Benin_97/1              -------------------------------------------------------------------------------- [ 14640]
KM111295.1_Ken06.Bus               t.attg.ta..c...cg..g......acaaacctg..tc...........--......a...c..t......a....... [ 14640]
```

```
KM111294.1_Ken05/Tk1            g.attg.ta..c...cg..g.....acaaacctg..tc........t...--.............t......a....... [ 14640]
KM262844.1_strain_L60           -------------------------------------------------------------------------------- [ 14640]
KM102979.1_26544/OG10_Italy     -------------------------------------------------------------------------------- [ 14640]
KM262845.1_strain_NHV           -------------------------------------------------------------------------------- [ 14640]
KX354450.1_47/Ss/2008           -------------------------------------------------------------------------------- [ 14640]
KP055815.1_strain_BA71          -------------------------------------------------------------------------------- [ 14640]
MH025920.1_strain_R35           t.attg.ta..c...cg..g.....acaaacctg..tc............--......a...c..t......a....... [ 14640]
MH025919.1_strain_N10           t.attg.ta..c...cg..g.....acaaacctg..tc............--......a...c..t......a....... [ 14640]
MH025918.1_strain_R25           t.attg.ta..c...cg..g.....acaaacctg..tc............--......a...c..t......a....... [ 14640]
MH025917.1_strain_R7            t.attg.ta..c...cg..g.....acaaacctg..tc............--......a...c..t......a....... [ 14640]
MH025916.1_strain_R8            t.attg.ta..c...cg..g.....acaaacctg..tc............--......a...c..t......a....... [ 14640]

FR682468.2_Georgia_2007/1       tcatcttttttcaaatcctttctgggatgttcattctttttccactccttcc-ttgcaattttataaggattccagggct [ 14720]
ASF_IND_20_CAD543               ..............m..........r.....y..y.................-..........r................ [ 14720]
LR899193.1_Germany_2020/1       ....................................................-........................... [ 14720]
MN393477.1_Wuhan_2019           ....................................................-........................... [ 14720]
MW396979.1_Timor-Leste/2019/1   ....................................................-........................... [ 14720]
MK940252.1_CN/2019/InnerMongolia ....................................................-........................... [ 14720]
MK128995.1_China/2018/AnhuiXCGQ ....................................................-........................... [ 14720]
MH681419.1_ASFV/POL/2015        ....................................................-........................... [ 14720]
LS478113.1_Estonia_2014         -------------------------------------------------------------------------------- [ 14720]
KP843857.1_Odintsovo_02/14      ....................................................-........................... [ 14720]
FN557520.1_strain_E75           -------------------------------------------------------------------------------- [ 14720]
AY261366.1_Warthog              ....................................................-........................... [ 14720]
AY261365.1_Warmbaths            ....................................................-........................... [ 14720]
AY261364.1_Tengani_62           ....................................................-........................... [ 14720]
AY261363.1_Pretoriuskop/96/4    ........................a...a.......................-........................... [ 14720]
AY261362.1_Mkuzi_1979           ................t........a........c.................-........................... [ 14720]
AY261361.1_Malawi_Lil-20/1_1983 ................t........a........c.................t........................... [ 14720]
AY261360.1_Kenya_1950           ........................aa..........................-........................... [ 14720]
AM712240.1_OURT_88/3            -------------------------------------------------------------------------------- [ 14720]
AM712239.1_Benin_97/1           -------------------------------------------------------------------------------- [ 14720]
KM111295.1_Ken06.Bus            .........................a......................ac..-...t..................a.... [ 14720]
KM111294.1_Ken05/Tk1            ........................aa......................ac..-...t.....................t. [ 14720]
KM262844.1_strain_L60           -------------------------------------------------------------------------------- [ 14720]
KM102979.1_26544/OG10_Italy     -------------------------------------------------------------------------------- [ 14720]
KM262845.1_strain_NHV           -------------------------------------------------------------------------------- [ 14720]
KX354450.1_47/Ss/2008           -------------------------------------------------------------------------------- [ 14720]
KP055815.1_strain_BA71          -------------------------------------------------------------------------------- [ 14720]
MH025920.1_strain_R35           .........................a......................ac..-...t..................a.... [ 14720]
MH025919.1_strain_N10           .........................a......................ac..-...t..................a.... [ 14720]
MH025918.1_strain_R25           .........................a......................ac..-...t..................a.... [ 14720]
MH025917.1_strain_R7            .........................a......................ac..-...t..................a.... [ 14720]
MH025916.1_strain_R8            .........................a......................ac..-...t..................a.... [ 14720]
```

```
FR682468.2_Georgia_2007/1        ttgggtcagaacagttcatgctatggtaaatgtgctcctccacatcatatctacataggtcaccccagcgggaaacctca [ 14800]
ASF_IND_20_CAD543                ......-------------------------------------------------------------------------- [ 14800]
LR899193.1_Germany_2020/1        ................................................................................ [ 14800]
MN393477.1_Wuhan_2019            ................................................................................ [ 14800]
MW396979.1_Timor-Leste/2019/1    ................................................................................ [ 14800]
MK940252.1_CN/2019/InnerMongolia ................................................................................ [ 14800]
MK128995.1_China/2018/AnhuiXCGQ  ................................................................................ [ 14800]
MH681419.1_ASFV/POL/2015         ................................................................................ [ 14800]
LS478113.1_Estonia_2014          ---------------.t...t...ta..gg.a.----------------------------------------------- [ 14800]
KP843857.1_Odintsovo_02/14       ................................................................................ [ 14800]
FN557520.1_strain_E75            -------------------------------------------------------------------------------- [ 14800]
AY261366.1_Warthog               ..........g.................................................t................... [ 14800]
AY261365.1_Warmbaths             ..........g.................................................t................... [ 14800]
AY261364.1_Tengani_62            ................................................................................ [ 14800]
AY261363.1_Pretoriuskop/96/4     ........................a...................................tt.................. [ 14800]
AY261362.1_Mkuzi_1979            ....................g...a...............t...............t.............a......... [ 14800]
AY261361.1_Malawi_Lil-20/1_1983  ....................g...a...............................t.............a......... [ 14800]
AY261360.1_Kenya_1950            ....................g...a..............................ct...................t... [ 14800]
AM712240.1_OURT_88/3             -------------------------------------------------------------------------------- [ 14800]
AM712239.1_Benin_97/1            -------------------------------------------------------------------------------- [ 14800]
KM111295.1_Ken06.Bus             ....................g...a...............................t..c..tt............t... [ 14800]
KM111294.1_Ken05/Tk1             ....................g...........c......................ct...................t... [ 14800]
KM262844.1_strain_L60            -------------------------------------------------------------------------------- [ 14800]
KM102979.1_26544/OG10_Italy      -------------------------------------------------------------------------------- [ 14800]
KM262845.1_strain_NHV            -------------------------------------------------------------------------------- [ 14800]
KX354450.1_47/Ss/2008            -------------------------------------------------------------------------------- [ 14800]
KP055815.1_strain_BA71           -------------------------------------------------------------------------------- [ 14800]
MH025920.1_strain_R35            ....................g...a...............................t..c..tt............t... [ 14800]
MH025919.1_strain_N10            ....................g...a...............................t..c..tt............t... [ 14800]
MH025918.1_strain_R25            ....................g...a...............................t..c..tt............t... [ 14800]
MH025917.1_strain_R7             ....................g...a...............................t..c..tt............t... [ 14800]
MH025916.1_strain_R8             ....................g...a...............................t..c..tt............t... [ 14800]

FR682468.2_Georgia_2007/1        caatattttacatagtcattctcaataatacttgtggagttgtttccccaaaccctgctggtacaaatcccatcttcaca [ 14880]
ASF_IND_20_CAD543                -------------------------------------------------------------------------------- [ 14880]
LR899193.1_Germany_2020/1        ................................................................................ [ 14880]
MN393477.1_Wuhan_2019            ................................................................................ [ 14880]
MW396979.1_Timor-Leste/2019/1    ................................................................................ [ 14880]
MK940252.1_CN/2019/InnerMongolia ................................................................................ [ 14880]
MK128995.1_China/2018/AnhuiXCGQ  ................................................................................ [ 14880]
MH681419.1_ASFV/POL/2015         ................................................................................ [ 14880]
LS478113.1_Estonia_2014          ------------------------------------------------t....a..------------------------ [ 14880]
KP843857.1_Odintsovo_02/14       ................................................................................ [ 14880]
```

```
FN557520.1_strain_E75              -------------------------------------------------------------------------------- [ 14880]
AY261366.1_Warthog                 .........................c...................................................... [ 14880]
AY261365.1_Warmbaths               .........................c...................................................... [ 14880]
AY261364.1_Tengani_62              ................................................................................ [ 14880]
AY261363.1_Pretoriuskop/96/4       .........................c...................................................... [ 14880]
AY261362.1_Mkuzi_1979              .................g.......c.................................a.................... [ 14880]
AY261361.1_Malawi_Lil-20/1_1983    .........................c.................................a.....g.c............ [ 14880]
AY261360.1_Kenya_1950              ..........t....ctg.......c..................c.....g.t......a.....g.c............ [ 14880]
AM712240.1_OURT_88/3               -------------------------------------------------------------------------------- [ 14880]
AM712239.1_Benin_97/1              -------------------------------------------------------------------------------- [ 14880]
KM111295.1_Ken06.Bus               ..........t....ctg..a....c..................c..............a.................... [ 14880]
KM111294.1_Ken05/Tk1               ...............ctg.......c..................c.....g.t......a.....g.c............ [ 14880]
KM262844.1_strain_L60              -------------------------------------------------------------------------------- [ 14880]
KM102979.1_26544/OG10_Italy        -------------------------------------------------------------------------------- [ 14880]
KM262845.1_strain_NHV              -------------------------------------------------------------------------------- [ 14880]
KX354450.1_47/Ss/2008              -------------------------------------------------------------------------------- [ 14880]
KP055815.1_strain_BA71             -------------------------------------------------------------------------------- [ 14880]
MH025920.1_strain_R35              ..........t....ctg..a....c..................c..............a.................... [ 14880]
MH025919.1_strain_N10              ..........t....ctg..a....c..................c..............a.................... [ 14880]
MH025918.1_strain_R25              ..........t....ctg..a....c..................c..............a.................... [ 14880]
MH025917.1_strain_R7               ..........t....ctg..a....c..................c..............a.................... [ 14880]
MH025916.1_strain_R8               ..........t....ctg..a....c..................c..............a.................... [ 14880]

FR682468.2_Georgia_2007/1          atcccagcagaaccgacagctttccacataagtgcaccagtatccaagttcattctct-gggggttcaaatgttagagga [ 14960]
ASF_IND_20_CAD543                  -------------------------------------------------------------------------------- [ 14960]
LR899193.1_Germany_2020/1          ..........................................................-..................... [ 14960]
MN393477.1_Wuhan_2019              ..........................................................-..................... [ 14960]
MW396979.1_Timor-Leste/2019/1      ..........................................................-..................... [ 14960]
MK940252.1_CN/2019/InnerMongolia   ..........................................................-..................... [ 14960]
MK128995.1_China/2018/AnhuiXCGQ    ..........................................................-..................... [ 14960]
MH681419.1_ASFV/POL/2015           ..........................................................-..................... [ 14960]
LS478113.1_Estonia_2014            --------------------------------------------------------------------...c...c.... [ 14960]
KP843857.1_Odintsovo_02/14         ..........................................................-..................... [ 14960]
FN557520.1_strain_E75              -------------------------------------------------------------------------------- [ 14960]
AY261366.1_Warthog                 .........................................................g-.....a............... [ 14960]
AY261365.1_Warmbaths               ..........................................................-.....a............... [ 14960]
AY261364.1_Tengani_62              ......a...................................................g.....a............... [ 14960]
AY261363.1_Pretoriuskop/96/4       ..........................................................-.....a............... [ 14960]
AY261362.1_Mkuzi_1979              ..........................................................-..a..a..ctc.......... [ 14960]
AY261361.1_Malawi_Lil-20/1_1983    ..................a...............t.......................-..a..a............... [ 14960]
AY261360.1_Kenya_1950              ..........................................................-..a..a............... [ 14960]
AM712240.1_OURT_88/3               -------------------------------------------------------------------------------- [ 14960]
AM712239.1_Benin_97/1              -------------------------------------------------------------------------------- [ 14960]
KM111295.1_Ken06.Bus               ..t..................................t....................-..a..a..ctc.....a.... [ 14960]
```

```
KM111294.1_Ken05/Tk1             ..........................................................-..a..a............... [ 14960]
KM262844.1_strain_L60            -------------------------------------------------------------------------------- [ 14960]
KM102979.1_26544/OG10_Italy      -------------------------------------------------------------------------------- [ 14960]
KM262845.1_strain_NHV            -------------------------------------------------------------------------------- [ 14960]
KX354450.1_47/Ss/2008            -------------------------------------------------------------------------------- [ 14960]
KP055815.1_strain_BA71           -------------------------------------------------------------------------------- [ 14960]
MH025920.1_strain_R35            ..t.......................................................-..a..a..ctc.....a.... [ 14960]
MH025919.1_strain_N10            ..t.......................................................-..a..a..ctc.....a.... [ 14960]
MH025918.1_strain_R25            ..t.......................................................-..a..a..ctc.....a.... [ 14960]
MH025917.1_strain_R7             ..t.......................................................-..a..a..ctc.....a.... [ 14960]
MH025916.1_strain_R8             ..t.......................................................-..a..a..ctc.....a.... [ 14960]

FR682468.2_Georgia_2007/1        agatgtccacctacccgagtagaagtggaggatgaaaccaggttgctactggccagcaggccaataattcccaggataat [ 15040]
ASF_IND_20_CAD543                -------------------------------------------------------------------------------- [ 15040]
LR899193.1_Germany_2020/1        .............................a.................................................. [ 15040]
MN393477.1_Wuhan_2019            ................................................................................ [ 15040]
MW396979.1_Timor-Leste/2019/1    ................................................................................ [ 15040]
MK940252.1_CN/2019/InnerMongolia ................................................................................ [ 15040]
MK128995.1_China/2018/AnhuiXCGQ  ................................................................................ [ 15040]
MH681419.1_ASFV/POL/2015         ................................................................................ [ 15040]
LS478113.1_Estonia_2014          ..------------------------------------------------------------------------------ [ 15040]
KP843857.1_Odintsovo_02/14       ................................................................................ [ 15040]
FN557520.1_strain_E75            -------------------------------------------------------------------------------- [ 15040]
AY261366.1_Warthog               ................................................................................ [ 15040]
AY261365.1_Warmbaths             ................................................................................ [ 15040]
AY261364.1_Tengani_62            .............................................t.................................. [ 15040]
AY261363.1_Pretoriuskop/96/4     ................................................................................ [ 15040]
AY261362.1_Mkuzi_1979            ........................................a..........................c............ [ 15040]
AY261361.1_Malawi_Lil-20/1_1983  ........................................a..........................c............ [ 15040]
AY261360.1_Kenya_1950            ........................................a..........................c............ [ 15040]
AM712240.1_OURT_88/3             -------------------------------------------------..................c......a..... [ 15040]
AM712239.1_Benin_97/1            -------------------------------------------------------------------------------- [ 15040]
KM111295.1_Ken06.Bus             .................................................c.................c............ [ 15040]
KM111294.1_Ken05/Tk1             .............t..........................a..........................c............ [ 15040]
KM262844.1_strain_L60            -------------------------------------------------------------------------------- [ 15040]
KM102979.1_26544/OG10_Italy      -------------------------------------------------------------------------------- [ 15040]
KM262845.1_strain_NHV            -------------------------------------------------..................c......a..... [ 15040]
KX354450.1_47/Ss/2008            -------------------------------------------------------------------------------- [ 15040]
KP055815.1_strain_BA71           -------------------------------------------------------------------------------- [ 15040]
MH025920.1_strain_R35            .................................................c.................c............ [ 15040]
MH025919.1_strain_N10            .................................................c.................c............ [ 15040]
MH025918.1_strain_R25            .................................................c.................c............ [ 15040]
MH025917.1_strain_R7             .................................................c.................c............ [ 15040]
MH025916.1_strain_R8             .................................................c.................c............ [ 15040]
```

```
FR682468.2_Georgia_2007/1        caccagcattgtgctcaacc-----------agcaac-ggctagcaacgactagcaactgactagcaatagctagaaatg [ 15120]
ASF_IND_20_CAD543                -------------------------------------------------------------------------------- [ 15120]
LR899193.1_Germany_2020/1        ....................-----------......-.......................................... [ 15120]
MN393477.1_Wuhan_2019            ....................-----------......-.......................................... [ 15120]
MW396979.1_Timor-Leste/2019/1    ....................-----------......-.......................................... [ 15120]
MK940252.1_CN/2019/InnerMongolia ....................-----------......-.......................................... [ 15120]
MK128995.1_China/2018/AnhuiXCGQ  ....................-----------......-.......................................... [ 15120]
MH681419.1_ASFV/POL/2015         ....................-----------......-.......................................... [ 15120]
LS478113.1_Estonia_2014          -------------------------------.at..t-..gg..gcggcg.a.aa...g...g.a.g.c----------- [ 15120]
KP843857.1_Odintsovo_02/14       ....................-----------......-.......................................... [ 15120]
FN557520.1_strain_E75            ----------------..a.-----------.a...t-.t..tat..t..t.g..c.a.a.tc.---------------- [ 15120]
AY261366.1_Warthog               ....................--------------------------------...........................a [ 15120]
AY261365.1_Warmbaths             ....................--------------------------------..t...................-..... [ 15120]
AY261364.1_Tengani_62            ....................-----------......-.......................................... [ 15120]
AY261363.1_Pretoriuskop/96/4     ....................agcaactgact......-.......................................... [ 15120]
AY261362.1_Mkuzi_1979            ....................-----------.a....-...............................c.......... [ 15120]
AY261361.1_Malawi_Lil-20/1_1983  .............t.t....-----------.a....t.a.........g.........................c...a [ 15120]
AY261360.1_Kenya_1950            ....................--------------------------------.......................c...a [ 15120]
AM712240.1_OURT_88/3             ....................-----------......-.......................................... [ 15120]
AM712239.1_Benin_97/1            ----------------..a.-----------.a...t-.t..tat..t..t.g..c.a.a.tc.---------------- [ 15120]
KM111295.1_Ken06.Bus             ....................--------------------------------..t..............g.....c...a [ 15120]
KM111294.1_Ken05/Tk1             ....................--------------------------------..................---------- [ 15120]
KM262844.1_strain_L60            ----------------..a.-----------.a...t-.t..tat..t..t.g..c.a.a.tc.c--------------- [ 15120]
KM102979.1_26544/OG10_Italy      ----------------..a.-----------.a...t-.t..tat..t..t.g..c.a.a.t------------------ [ 15120]
KM262845.1_strain_NHV            ....................-----------......-.......................................... [ 15120]
KX354450.1_47/Ss/2008            ----------------..a.-----------.a...t-.t..tat..t..t.g..c.a.a.t------------------ [ 15120]
KP055815.1_strain_BA71           -------------------------------------------------------------------------------- [ 15120]
MH025920.1_strain_R35            ....................--------------------------------..t..............g.....c...a [ 15120]
MH025919.1_strain_N10            ....................--------------------------------..t..............g.....c...a [ 15120]
MH025918.1_strain_R25            ....................--------------------------------..t..............g.....c...a [ 15120]
MH025917.1_strain_R7             ....................--------------------------------..t..............g.....c...a [ 15120]
MH025916.1_strain_R8             ....................--------------------------------..t..............g.....c...a [ 15120]

FR682468.2_Georgia_2007/1        g-------------------------------ctagcaatcagtagtagctaacgctctactctttataagaaatttaaa [ 15200]
ASF_IND_20_CAD543                -------------------------------------------------------------------------------- [ 15200]
LR899193.1_Germany_2020/1        .-------------------------------................................................ [ 15200]
MN393477.1_Wuhan_2019            .-------------------------------................................................ [ 15200]
MW396979.1_Timor-Leste/2019/1    .-------------------------------................................................ [ 15200]
MK940252.1_CN/2019/InnerMongolia .-------------------------------................................................ [ 15200]
MK128995.1_China/2018/AnhuiXCGQ  .-------------------------------................................................ [ 15200]
MH681419.1_ASFV/POL/2015         .-------------------------------................................................ [ 15200]
LS478113.1_Estonia_2014          -------------------------------------------------------------------------------- [ 15200]
KP843857.1_Odintsovo_02/14       .-------------------------------................................................ [ 15200]
```

```
FN557520.1_strain_E75             -------------------------------------------------------------------------------- [ 15200]
AY261366.1_Warthog                .------------------------------...............g......................a.......... [ 15200]
AY261365.1_Warmbaths              .------------------------------...............g................................. [ 15200]
AY261364.1_Tengani_62             .------------------------------.......c.......g................................. [ 15200]
AY261363.1_Pretoriuskop/96/4      .------------------------------...............g................................. [ 15200]
AY261362.1_Mkuzi_1979             .------------------------------.......c.......g................................. [ 15200]
AY261361.1_Malawi_Lil-20/1_1983   .ttagcaata--------------------a...............g..............t.................. [ 15200]
AY261360.1_Kenya_1950             .ctagaaatgactagcaatggctagaaatag...............g......a.......................... [ 15200]
AM712240.1_OURT_88/3              .------------------------------.......c.......g......a.......................... [ 15200]
AM712239.1_Benin_97/1             -------------------------------------------------------------------------------- [ 15200]
KM111295.1_Ken06.Bus              .ctaacaatg--------------------g...............g......a.......................... [ 15200]
KM111294.1_Ken05/Tk1              ------------------------------g...............g......a...............a...t...... [ 15200]
KM262844.1_strain_L60             -------------------------------------------------------------------------------- [ 15200]
KM102979.1_26544/OG10_Italy       -------------------------------------------------------------------------------- [ 15200]
KM262845.1_strain_NHV             .------------------------------.......c.......g......a.......................... [ 15200]
KX354450.1_47/Ss/2008             -------------------------------------------------------------------------------- [ 15200]
KP055815.1_strain_BA71            -------------------------------------------------------------------------------- [ 15200]
MH025920.1_strain_R35             .ctaacaatg--------------------g...............g......a.......................... [ 15200]
MH025919.1_strain_N10             .ctaacaatg--------------------g...............g......a.......................... [ 15200]
MH025918.1_strain_R25             .ctaacaatg--------------------g...............g......a.......................... [ 15200]
MH025917.1_strain_R7              .ctaacaatg--------------------g...............g......a.......................... [ 15200]
MH025916.1_strain_R8              .ctaacaatg--------------------g...............g......a.......................... [ 15200]

FR682468.2_Georgia_2007/1         attcgatcagatttttttagaattgagaatgagtaaaacgcttatattctttttctagctag-aaaaaataagctagttt [ 15280]
ASF_IND_20_CAD543                 -------------------------------------------------------------------------------- [ 15280]
LR899193.1_Germany_2020/1         ..............................................................-................. [ 15280]
MN393477.1_Wuhan_2019             ..............................................................-................. [ 15280]
MW396979.1_Timor-Leste/2019/1     ..............................................................-................. [ 15280]
MK940252.1_CN/2019/InnerMongolia  ..............................................................-................. [ 15280]
MK128995.1_China/2018/AnhuiXCGQ   ..............................................................-................. [ 15280]
MH681419.1_ASFV/POL/2015          ..............................................................-................. [ 15280]
LS478113.1_Estonia_2014           --------------------------------.cg..g.atg.cc...t.gcca.agc.--------------------- [ 15280]
KP843857.1_Odintsovo_02/14        ..............................................................-................. [ 15280]
FN557520.1_strain_E75             --------------------------------------------cccc.ccccc.c.----------------------- [ 15280]
AY261366.1_Warthog                ..................g..................................c........-................. [ 15280]
AY261365.1_Warmbaths              ..................g...................t..............c........-................. [ 15280]
AY261364.1_Tengani_62             ..................g....................a.............c........a................. [ 15280]
AY261363.1_Pretoriuskop/96/4      ..................g..................................c..g.....a................. [ 15280]
AY261362.1_Mkuzi_1979             .................g-..................................c........a................. [ 15280]
AY261361.1_Malawi_Lil-20/1_1983   ..............................a......................c........-................. [ 15280]
AY261360.1_Kenya_1950             ....................g.....a......c....ta...g.........ct.......-................. [ 15280]
AM712240.1_OURT_88/3              .......t..........-.g.....a..a..ac...................c.....g..-................. [ 15280]
AM712239.1_Benin_97/1             ------------------------------------------------.ccccc.c.----------------------- [ 15280]
KM111295.1_Ken06.Bus              ....................g.....a......c....ta...g.........ct.......-................. [ 15280]
```

```
KM111294.1_Ken05/Tk1           .......t............g.....a......c.....a...g.........c........--................ [ 15280]
KM262844.1_strain_L60          -------------------------------------------ccccc.ccccc.c.----------------------- [ 15280]
KM102979.1_26544/OG10_Italy    -----------------------------------------------c.ccccc.c.----------------------- [ 15280]
KM262845.1_strain_NHV          .......t..........-.g.....a..a..ac...................c.....g..-................. [ 15280]
KX354450.1_47/Ss/2008          ------------------------------------------------.ccccc.c.----------------------- [ 15280]
KP055815.1_strain_BA71         -------------------------------------------------------------------------------- [ 15280]
MH025920.1_strain_R35          ....................g.....a......c....ta...g.........ct.......-................. [ 15280]
MH025919.1_strain_N10          ....................g.....a......c....ta...g.........ct.......-................. [ 15280]
MH025918.1_strain_R25          ....................g.....a......c....ta...g.........ct.......-................. [ 15280]
MH025917.1_strain_R7           ....................g.....a......c....ta...g.........ct.......-................. [ 15280]
MH025916.1_strain_R8           ....................g.....a......c....ta...g.........ct.......-................. [ 15280]

FR682468.2_Georgia_2007/1      aagataggatttcccttac-taacggtttaa-tttttagcaaaggtataggtaaaa---tacacttgtacttagctgcaa [ 15360]
ASF_IND_20_CAD543              -------------------------------------------------------------------------------- [ 15360]
LR899193.1_Germany_2020/1      ...................-...........-........................---..................... [ 15360]
MN393477.1_Wuhan_2019          ...................-...........-........................---..................... [ 15360]
MW396979.1_Timor-Leste/2019/1  ...................-...........-........................---..................... [ 15360]
MK940252.1_CN/2019/InnerMongolia ...................-...........-........................---..................... [ 15360]
MK128995.1_China/2018/AnhuiXCGQ ...................-...........-........................---..................... [ 15360]
MH681419.1_ASFV/POL/2015       ...................-...........-........................---..................... [ 15360]
LS478113.1_Estonia_2014        --------------------------------------.tcg....g.g..----------------------------- [ 15360]
KP843857.1_Odintsovo_02/14     ...................-...........-........................---..................... [ 15360]
FN557520.1_strain_E75          ----------..tttcc.t-g..tat..c.t-..cc.gtat.g...c....atg-------------------------- [ 15360]
AY261366.1_Warthog             .........-.........-......g....-.....g.t...........gt...a--......g...........g.. [ 15360]
AY261365.1_Warmbaths           .........-.........-......g....-.....g.t...........gt-..a--......g...........g.. [ 15360]
AY261364.1_Tengani_62          .........-.........-...t.......-....cg.............gt...a--......g...........g.. [ 15360]
AY261363.1_Pretoriuskop/96/4   ...................-...t.......t.....g.....a....g..gt...a--.......a........c.g.. [ 15360]
AY261362.1_Mkuzi_1979          .........-.........-......g....-.....g............agt...aaa......g...........g.. [ 15360]
AY261361.1_Malawi_Lil-20/1_1983 ...................-....a......-.....g..................---..................g.. [ 15360]
AY261360.1_Kenya_1950          ..a.........t......a..t.a......-.....g..................t-----........c...cag-. [ 15360]
AM712240.1_OURT_88/3           ......t............-...........-.....g....g......a.gt...ata--------c.t......-g.. [ 15360]
AM712239.1_Benin_97/1          ----------..tttcc.t-g..tat..c.t-..cc.gtat.g...c....atg-------------------------- [ 15360]
KM111295.1_Ken06.Bus           ..a.........t......a..t.a......--....g..................t-----........c...cag.. [ 15360]
KM111294.1_Ken05/Tk1           ..a.........t......a..t.a......-.....g.g................t-----........c...cag.. [ 15360]
KM262844.1_strain_L60          ----------..tttcc.t-g..tat..c.t-..cc.gtat.g...c....atg-------------------------- [ 15360]
KM102979.1_26544/OG10_Italy    ----------..tttcc.t-g..tat..c.t-..cc.gtat.g...c....atg-------------------------- [ 15360]
KM262845.1_strain_NHV          ......t............-...........-.....g....g......a.gt...ata--------c.t......-g.. [ 15360]
KX354450.1_47/Ss/2008          ----------..tttcc.t-g..tat..c.t-..cc.gtat.g...c....atg-------------------------- [ 15360]
KP055815.1_strain_BA71         -------------------------------------------------------------------------------- [ 15360]
MH025920.1_strain_R35          ..a.........t......a..t.a......--....g..................t-----........c...cag.. [ 15360]
MH025919.1_strain_N10          ..a.........t......a..t.a......--....g..................t-----........c...cag.. [ 15360]
MH025918.1_strain_R25          ..a.........t......a..t.a......--....g..................t-----........c...cag.. [ 15360]
MH025917.1_strain_R7           ..a.........t......a..t.a......--....g..................t-----........c...cag.. [ 15360]
MH025916.1_strain_R8           ..a.........t......a..t.a......--....g..................t-----........c...cag.. [ 15360]
```

```
FR682468.2_Georgia_2007/1        aaaaataagcttatggcgtataagccgccataagtttatttaattaaaatgttaaactctgtgataagactggaatctta [ 15440]
ASF_IND_20_CAD543                -------------------------------------------------------------------------------- [ 15440]
LR899193.1_Germany_2020/1        ................................................................................ [ 15440]
MN393477.1_Wuhan_2019            ................................................................................ [ 15440]
MW396979.1_Timor-Leste/2019/1    ................................................................................ [ 15440]
MK940252.1_CN/2019/InnerMongolia ................................................................................ [ 15440]
MK128995.1_China/2018/AnhuiXCGQ  ................................................................................ [ 15440]
MH681419.1_ASFV/POL/2015         ................................................................................ [ 15440]
LS478113.1_Estonia_2014          ---------------------......----------------------------------------------------- [ 15440]
KP843857.1_Odintsovo_02/14       ................................................................................ [ 15440]
FN557520.1_strain_E75            ---------------------------------------------tg..ca..cc.tg..a....g.----------... [ 15440]
AY261366.1_Warthog               .................a........................--------a............................. [ 15440]
AY261365.1_Warmbaths             ..................................c.......--------a....t........................ [ 15440]
AY261364.1_Tengani_62            ............................................................a................... [ 15440]
AY261363.1_Pretoriuskop/96/4     .............c............................--------a............................. [ 15440]
AY261362.1_Mkuzi_1979            ..................gc.t.ta.................--------a............................. [ 15440]
AY261361.1_Malawi_Lil-20/1_1983  ................................................................................ [ 15440]
AY261360.1_Kenya_1950            ........a.c..c....ac.t.ta.a.....c...........--..t.t............................. [ 15440]
AM712240.1_OURT_88/3             ................t......a..a..................................................... [ 15440]
AM712239.1_Benin_97/1            ---------------------------------------------tg..ca..cc.tg..a....g.----------... [ 15440]
KM111295.1_Ken06.Bus             ........a.c..c....ac.t.ta.a.....c...........--..-.t............................. [ 15440]
KM111294.1_Ken05/Tk1             ........a.c..c....ac.t.ta.a.....c...........--..t.t............................. [ 15440]
KM262844.1_strain_L60            ---------------------------------------------tg..ca..cc.tg..a....g.----------... [ 15440]
KM102979.1_26544/OG10_Italy      ---------------------------------------------tg..ca..cc.tg..a....g.----------... [ 15440]
KM262845.1_strain_NHV            ................t......a..a..................................................... [ 15440]
KX354450.1_47/Ss/2008            ---------------------------------------------tg..ca..cc.tg..a....g.----------... [ 15440]
KP055815.1_strain_BA71           -------------------------------------------------------------------------------- [ 15440]
MH025920.1_strain_R35            ........a.c..c....ac.t.ta.a.....c...........--..-.t............................. [ 15440]
MH025919.1_strain_N10            ........a.c..c....ac.t.ta.a.....c...........--..-.t............................. [ 15440]
MH025918.1_strain_R25            ........a.c..c....ac.t.ta.a.....c...........--..-.t............................. [ 15440]
MH025917.1_strain_R7             ........a.c..c....ac.t.ta.a.....c...........--..-.t............................. [ 15440]
MH025916.1_strain_R8             ........a.c..c....ac.t.ta.a.....c...........--..-.t............................. [ 15440]

FR682468.2_Georgia_2007/1        ggcaggtttgatgtggagaacagcatgaaatacaagagtgcctgttacacgaataagttctctcaaaccggggatggtca [ 15520]
ASF_IND_20_CAD543                -------------------------------------------------------------------------------- [ 15520]
LR899193.1_Germany_2020/1        ................................................................................ [ 15520]
MN393477.1_Wuhan_2019            ................................................................................ [ 15520]
MW396979.1_Timor-Leste/2019/1    ................................................................................ [ 15520]
MK940252.1_CN/2019/InnerMongolia ................................................................................ [ 15520]
MK128995.1_China/2018/AnhuiXCGQ  ................................................................................ [ 15520]
MH681419.1_ASFV/POL/2015         ................................................................................ [ 15520]
LS478113.1_Estonia_2014          -------------------..ga...cg.gcg....g..c....-------------------gg.g.a..t.gc....g [ 15520]
KP843857.1_Odintsovo_02/14       ................................................................................ [ 15520]
```

```
FN557520.1_strain_E75          ....---------------------------------------------------------------------------- [ 15520]
AY261366.1_Warthog             ......c...............................................................a......... [ 15520]
AY261365.1_Warmbaths           ......c...............................................................a......... [ 15520]
AY261364.1_Tengani_62          .a....c........................g......................................a......... [ 15520]
AY261363.1_Pretoriuskop/96/4   .a....c...............................................................a......... [ 15520]
AY261362.1_Mkuzi_1979          ......c...............................................................a......... [ 15520]
AY261361.1_Malawi_Lil-20/1_1983 ......c.............................................................t.a......... [ 15520]
AY261360.1_Kenya_1950          .a....c...............a....g.........a...........ga...g.............t.a......... [ 15520]
AM712240.1_OURT_88/3           ......c.....................................c....t....................a......... [ 15520]
AM712239.1_Benin_97/1          ....---------------------------------------------------------------------------- [ 15520]
KM111295.1_Ken06.Bus           .a....c...............a....g.........a...........ga...g.............t.a......... [ 15520]
KM111294.1_Ken05/Tk1           .a....c...............a....g.........a...........ga...g.............t.a......... [ 15520]
KM262844.1_strain_L60          ....---------------------------------------------------------------------------- [ 15520]
KM102979.1_26544/OG10_Italy    ....---------------------------------------------------------------------------- [ 15520]
KM262845.1_strain_NHV          ......c.....................................c....t....................a......... [ 15520]
KX354450.1_47/Ss/2008          ....---------------------------------------------------------------------------- [ 15520]
KP055815.1_strain_BA71         -------------------------------------------------------------------------------- [ 15520]
MH025920.1_strain_R35          .a....c...............a....g.........a...........ga...g.............t.a......... [ 15520]
MH025919.1_strain_N10          .a....c...............a....g.........a...........ga...g.............t.a......... [ 15520]
MH025918.1_strain_R25          .a....c...............a....g.........a...........ga...g.............t.a......... [ 15520]
MH025917.1_strain_R7           .a....c...............a....g.........a...........ga...g.............t.a......... [ 15520]
MH025916.1_strain_R8           .a....c...............a....g.........a...........ga...g.............t.a......... [ 15520]

FR682468.2_Georgia_2007/1      tactcacatctatgaaatcctggtctaggagattcatttgatgcatgatggccgcacccacacttatgagacactgaaga [ 15600]
ASF_IND_20_CAD543              -------------------------------------------------------------------------------- [ 15600]
LR899193.1_Germany_2020/1      ................................................................................ [ 15600]
MN393477.1_Wuhan_2019          ................................................................................ [ 15600]
MW396979.1_Timor-Leste/2019/1  ................................................................................ [ 15600]
MK940252.1_CN/2019/InnerMongolia ................................................................................ [ 15600]
MK128995.1_China/2018/AnhuiXCGQ ................................................................................ [ 15600]
MH681419.1_ASFV/POL/2015       ................................................................................ [ 15600]
LS478113.1_Estonia_2014        -------------------------------------------------------------------------------- [ 15600]
KP843857.1_Odintsovo_02/14     ................................................................................ [ 15600]
FN557520.1_strain_E75          -------------------------------------------------------------------------------- [ 15600]
AY261366.1_Warthog             .............................t.....c..........t.......................g......... [ 15600]
AY261365.1_Warmbaths           ...................................c..........t...c...................g......... [ 15600]
AY261364.1_Tengani_62          ...........................................................g..........g......... [ 15600]
AY261363.1_Pretoriuskop/96/4   ........c..........................c..........t.......................g......... [ 15600]
AY261362.1_Mkuzi_1979          ......................................................................g......... [ 15600]
AY261361.1_Malawi_Lil-20/1_1983 ......................................................................g......... [ 15600]
AY261360.1_Kenya_1950          ..........c...................................................t.......g......... [ 15600]
AM712240.1_OURT_88/3           ......................................................................g......... [ 15600]
AM712239.1_Benin_97/1          -------------------------------------------------------------------------------- [ 15600]
KM111295.1_Ken06.Bus           ............................................c.t..........t....tc......g....---.. [ 15600]
```

```
KM111294.1_Ken05/Tk1            ..........c...................................................t.......g......... [ 15600]
KM262844.1_strain_L60           -------------------------------------------------------------------------------- [ 15600]
KM102979.1_26544/OG10_Italy     -------------------------------------------------------------------------------- [ 15600]
KM262845.1_strain_NHV           ......................................................................g......... [ 15600]
KX354450.1_47/Ss/2008           -------------------------------------------------------------------------------- [ 15600]
KP055815.1_strain_BA71          -------------------------------------------------------------------------------- [ 15600]
MH025920.1_strain_R35           ............................................c.t..........t....tc......g....---.. [ 15600]
MH025919.1_strain_N10           ............................................c.t..........t....tc......g....---.. [ 15600]
MH025918.1_strain_R25           ............................................c.t..........t....tc......g....---.. [ 15600]
MH025917.1_strain_R7            ............................................c.t..........t....tc......g....---.. [ 15600]
MH025916.1_strain_R8            ............................................c.t..........t....tc......g....---.. [ 15600]

FR682468.2_Georgia_2007/1       actaaagggtttaattttgatctgaatggtactatataggatgatggcaatccatatcaagattagagcaatcaaaatca [ 15680]
ASF_IND_20_CAD543               -------------------------------------------------------------------------------- [ 15680]
LR899193.1_Germany_2020/1       ................................................................................ [ 15680]
MN393477.1_Wuhan_2019           ................................................................................ [ 15680]
MW396979.1_Timor-Leste/2019/1   ................................................................................ [ 15680]
MK940252.1_CN/2019/InnerMongolia ................................................................................ [ 15680]
MK128995.1_China/2018/AnhuiXCGQ ................................................................................ [ 15680]
MH681419.1_ASFV/POL/2015        ................................................................................ [ 15680]
LS478113.1_Estonia_2014         -------------------------------------------------------------------------------- [ 15680]
KP843857.1_Odintsovo_02/14      ................................................................................ [ 15680]
FN557520.1_strain_E75           -------------------t.t.a.c..a..t.tc...aa.--------------------------------------. [ 15680]
AY261366.1_Warthog              ..c.............c.............g................................................. [ 15680]
AY261365.1_Warmbaths            ..c.............c.............g................................................. [ 15680]
AY261364.1_Tengani_62           ................c.............g................................g................ [ 15680]
AY261363.1_Pretoriuskop/96/4    ..c............gc.............g................................................. [ 15680]
AY261362.1_Mkuzi_1979           ................c.............g................................................. [ 15680]
AY261361.1_Malawi_Lil-20/1_1983 .tc.............c..........a..g.......ct........................................ [ 15680]
AY261360.1_Kenya_1950           .ga.cca........ccc........a.c..........t..a.......c........g.....a............tc [ 15680]
AM712240.1_OURT_88/3            ................c.............g................................c................ [ 15680]
AM712239.1_Benin_97/1           -------------------t.t.a.c..a..t.tc...aa.--------------------------------------. [ 15680]
KM111295.1_Ken06.Bus            .ga.cca.........c..........ac..........t...................gt................... [ 15680]
KM111294.1_Ken05/Tk1            ..c..ta........ccc........a............t..a.......c........g.....a............tc [ 15680]
KM262844.1_strain_L60           -------------------t.t.a.c..a..t.tc...aa.--------------------------------------. [ 15680]
KM102979.1_26544/OG10_Italy     -------------------t.t.a.c..a..t.tc...aa.--------------------------------------. [ 15680]
KM262845.1_strain_NHV           ................c.............g................................c................ [ 15680]
KX354450.1_47/Ss/2008           -------------------t.t.a.c..a..t.tc...aa.--------------------------------------. [ 15680]
KP055815.1_strain_BA71          -------------------------------------------------------------------------------- [ 15680]
MH025920.1_strain_R35           .ga.cca.........c..........ac..........t...................gt................... [ 15680]
MH025919.1_strain_N10           .ga.cca.........c..........ac..........t...................gt................... [ 15680]
MH025918.1_strain_R25           .ga.cca.........c..........ac..........t...................gt................... [ 15680]
MH025917.1_strain_R7            .ga.cca.........c..........ac..........t...................gt................... [ 15680]
MH025916.1_strain_R8            .ga.cca.........c..........ac..........t...................gt................... [ 15680]
```

```
FR682468.2_Georgia_2007/1        cctcctcaagaagcatgatgtagccttaaatct----------------------------------------------- [ 15760]
ASF_IND_20_CAD543                -------------------------------------------------------------------------------- [ 15760]
LR899193.1_Germany_2020/1        .................................----------------------------------------------- [ 15760]
MN393477.1_Wuhan_2019            .................................----------------------------------------------- [ 15760]
MW396979.1_Timor-Leste/2019/1    .................................----------------------------------------------- [ 15760]
MK940252.1_CN/2019/InnerMongolia .................................----------------------------------------------- [ 15760]
MK128995.1_China/2018/AnhuiXCGQ  .................................----------------------------------------------- [ 15760]
MH681419.1_ASFV/POL/2015         .................................----------------------------------------------- [ 15760]
LS478113.1_Estonia_2014          -------------------------------------------------------------------------------- [ 15760]
KP843857.1_Odintsovo_02/14       .................................----------------------------------------------- [ 15760]
FN557520.1_strain_E75            .ac..c..g...t.gc...ta.ct..ac.g.t.----------------------------------------------- [ 15760]
AY261366.1_Warthog               .................................taggccgcc----------------------------------ttaa [ 15760]
AY261365.1_Warmbaths             .................................taggccgcc----------------------------------ttaa [ 15760]
AY261364.1_Tengani_62            .................................taggccgct----------------------------------ttaa [ 15760]
AY261363.1_Pretoriuskop/96/4     .................................taggccgccttaaatcttaggccgccttaaatcttaggccgctttaa [ 15760]
AY261362.1_Mkuzi_1979            .................................----------------------------------------------- [ 15760]
AY261361.1_Malawi_Lil-20/1_1983  .........a....................c..caagccgcc----------------------------------ttaa [ 15760]
AY261360.1_Kenya_1950            ....................--.g.........caggctgcc----------------------------------ttaa [ 15760]
AM712240.1_OURT_88/3             .................................taggctgcc-----------------ttaaatcttagaccgctttaa [ 15760]
AM712239.1_Benin_97/1            .ac..c..g...t.gc...ta.ct..ac.g.t.----------------------------------------------- [ 15760]
KM111295.1_Ken06.Bus             ....................--.g.........caggctgcc----------------------------------ttaa [ 15760]
KM111294.1_Ken05/Tk1             ....................--.g.........caggctgcc----------------------------------ttaa [ 15760]
KM262844.1_strain_L60            .ac..c..g...t.gc...ta.ct..ac.g.t.----------------------------------------------- [ 15760]
KM102979.1_26544/OG10_Italy      .ac..c..g...t.gc...ta.ct..ac.g.t.----------------------------------------------- [ 15760]
KM262845.1_strain_NHV            .................................taggctgcc-----------------ttaaatcttagaccgctttaa [ 15760]
KX354450.1_47/Ss/2008            .ac..c..g...t.gc...ta.ct..ac.g.t.----------------------------------------------- [ 15760]
KP055815.1_strain_BA71           -------------------------------------------------------------------------------- [ 15760]
MH025920.1_strain_R35            ....................--.g.........caggctgcc----------------------------------ttaa [ 15760]
MH025919.1_strain_N10            ....................--.g.........caggctgcc----------------------------------ttaa [ 15760]
MH025918.1_strain_R25            ....................--.g.........caggctgcc----------------------------------ttaa [ 15760]
MH025917.1_strain_R7             ....................--.g.........caggctgcc----------------------------------ttaa [ 15760]
MH025916.1_strain_R8             ....................--.g.........caggctgcc----------------------------------ttaa [ 15760]

FR682468.2_Georgia_2007/1        ----tagactgctttaaaccttaggccctcactatctttaatgaaggagtttaaattttgat--ccctttttcaagaccc [ 15840]
ASF_IND_20_CAD543                -------------------------------------------------------------------------------- [ 15840]
LR899193.1_Germany_2020/1        ----..........................................................--................ [ 15840]
MN393477.1_Wuhan_2019            ----..........................................................--................ [ 15840]
MW396979.1_Timor-Leste/2019/1    ----..........................................................--................ [ 15840]
MK940252.1_CN/2019/InnerMongolia ----..........................................................--................ [ 15840]
MK128995.1_China/2018/AnhuiXCGQ  ----..........................................................--................ [ 15840]
MH681419.1_ASFV/POL/2015         ----..........................................................--................ [ 15840]
LS478113.1_Estonia_2014          --------------------------------------------------------a.c.g.--t..gc..c..gcg..g [ 15840]
KP843857.1_Odintsovo_02/14       ----..........................................................--................ [ 15840]
```

```
FN557520.1_strain_E75            -------------------------------------------------------------------------------- [ 15840]
AY261366.1_Warthog               acct..ag.c..c.................................................--................ [ 15840]
AY261365.1_Warmbaths             acct..ag.c..c.................................................--t............... [ 15840]
AY261364.1_Tengani_62            acct..ag.c..c.................................................--................ [ 15840]
AY261363.1_Pretoriuskop/96/4     acct..ag.c..c.................................................---............... [ 15840]
AY261362.1_Mkuzi_1979            ----...g.c.............a..................a...a............a..--................ [ 15840]
AY261361.1_Malawi_Lil-20/1_1983  aact..agtc..c...................................c.............---...........g... [ 15840]
AY261360.1_Kenya_1950            a--tc.ag.c..c.......-......t..................................--...........ag... [ 15840]
AM712240.1_OURT_88/3             atct..ag.c..c.....t...........................a...............cc............g..g [ 15840]
AM712239.1_Benin_97/1            -------------------------------------------------------------------------------- [ 15840]
KM111295.1_Ken06.Bus             a--tc.ag.c..c..g....-......t..................................--...........ag... [ 15840]
KM111294.1_Ken05/Tk1             a--tc.ag.c..c.......-......t..............a...................--..--....t...g... [ 15840]
KM262844.1_strain_L60            -------------------------------------------------------------------------------- [ 15840]
KM102979.1_26544/OG10_Italy      -------------------------------------------------------------------------------- [ 15840]
KM262845.1_strain_NHV            atct..ag.c..c.....t...........................a...............cc............g..g [ 15840]
KX354450.1_47/Ss/2008            -------------------------------------------------------------------------------- [ 15840]
KP055815.1_strain_BA71           -------------------------------------------------------------------------------- [ 15840]
MH025920.1_strain_R35            a--tc.ag.c..c..g....-......t..................................--...........ag... [ 15840]
MH025919.1_strain_N10            a--tc.ag.c..c..g....-......t..................................--...........ag... [ 15840]
MH025918.1_strain_R25            a--tc.ag.c..c..g....-......t..................................--...........ag... [ 15840]
MH025917.1_strain_R7             a--tc.ag.c..c..g....-......t..................................--...........ag... [ 15840]
MH025916.1_strain_R8             a--tc.ag.c..c..g....-......t..................................--...........ag... [ 15840]

FR682468.2_Georgia_2007/1        atttagaagaaaaaaataaagtttatatcaatctaattcataagtcatctctttcataaatcttcatgtattctctatgt [ 15920]
ASF_IND_20_CAD543                -------------------------------------------------------------------------------- [ 15920]
LR899193.1_Germany_2020/1        ................................................................................ [ 15920]
MN393477.1_Wuhan_2019            ................................................................................ [ 15920]
MW396979.1_Timor-Leste/2019/1    ................................................................................ [ 15920]
MK940252.1_CN/2019/InnerMongolia ................................................................................ [ 15920]
MK128995.1_China/2018/AnhuiXCGQ  ................................................................................ [ 15920]
MH681419.1_ASFV/POL/2015         ................................................................................ [ 15920]
LS478113.1_Estonia_2014          ga.cgcg..--------------------------------------------------------------------c.. [ 15920]
KP843857.1_Odintsovo_02/14       .........-...................................................................... [ 15920]
FN557520.1_strain_E75            ---------------------------a.....g......---------------------------------------- [ 15920]
AY261366.1_Warthog               ....................a......c..............................g...............g..... [ 15920]
AY261365.1_Warmbaths             ....................a......c..............................g..................... [ 15920]
AY261364.1_Tengani_62            ........................g..c..........t...................g..................... [ 15920]
AY261363.1_Pretoriuskop/96/4     .....................--....c..............................g..................... [ 15920]
AY261362.1_Mkuzi_1979            ....................a......c..............................g...............g..... [ 15920]
AY261361.1_Malawi_Lil-20/1_1983  ..a...........-.....a......a......g...t......................t.................. [ 15920]
AY261360.1_Kenya_1950            ..g............ta..g...ata.ca.t...gg......g..........a...................aa.g... [ 15920]
AM712240.1_OURT_88/3             ..a..a........-............a..............g...............g..................... [ 15920]
AM712239.1_Benin_97/1            ---------------------------a.....g......---------------------------------------- [ 15920]
KM111295.1_Ken06.Bus             ..g............ta..g...ata.ca.t...g.......g..........a...................aa..... [ 15920]
```

```
KM111294.1_Ken05/Tk1             ..g.......................................g...............g..............aa.g... [ 15920]
KM262844.1_strain_L60            ---------------------------a.....g......---------------------------------------- [ 15920]
KM102979.1_26544/OG10_Italy      ---------------------------a.....g......---------------------------------------- [ 15920]
KM262845.1_strain_NHV            ..a..a........-............a..............g...............g..................... [ 15920]
KX354450.1_47/Ss/2008            ---------------------------a.....g......---------------------------------------- [ 15920]
KP055815.1_strain_BA71           -------------------------------------------------------------------------------- [ 15920]
MH025920.1_strain_R35            ..g............ta..g...ata.ca.t...g.......g..........a...................aa..... [ 15920]
MH025919.1_strain_N10            ..g............ta..g...ata.ca.t...g.......g..........a...................aa..... [ 15920]
MH025918.1_strain_R25            ..g............ta..g...ata.ca.t...g.......g..........a...................aa..... [ 15920]
MH025917.1_strain_R7             ..g............ta..g...ata.ca.t...g.......g..........a...................aa..... [ 15920]
MH025916.1_strain_R8             ..g............ta..g...ata.ca.t...g.......g..........a...................aa..... [ 15920]

FR682468.2_Georgia_2007/1        ggataagtatgggatgttggatttgcgcagtccatttgatgatctgtatggtttttgggtccttcataataactacatat [ 16000]
ASF_IND_20_CAD543                -------------------------------------------------------------------------------- [ 16000]
LR899193.1_Germany_2020/1        ................................................................................ [ 16000]
MN393477.1_Wuhan_2019            ................................................................................ [ 16000]
MW396979.1_Timor-Leste/2019/1    ................................................................................ [ 16000]
MK940252.1_CN/2019/InnerMongolia ................................................................................ [ 16000]
MK128995.1_China/2018/AnhuiXCGQ  ................................................................................ [ 16000]
MH681419.1_ASFV/POL/2015         ................................................................................ [ 16000]
LS478113.1_Estonia_2014          ...gg..ctc--------------------------------------------------------------...tccgc [ 16000]
KP843857.1_Odintsovo_02/14       ................................................................................ [ 16000]
FN557520.1_strain_E75            ----------------------------------------------------------------...a...gact...t. [ 16000]
AY261366.1_Warthog               .....g..................ata..........c.....gcag..at.g..ga.aa............tg....tg [ 16000]
AY261365.1_Warmbaths             ..........................a...........................c......................... [ 16000]
AY261364.1_Tengani_62            ..........................a...........................c......................... [ 16000]
AY261363.1_Pretoriuskop/96/4     ..........................a...........................c......................... [ 16000]
AY261362.1_Mkuzi_1979            .....g...................ta..........c.....g.ag..at.g..ga.aa............tg....tg [ 16000]
AY261361.1_Malawi_Lil-20/1_1983  .....g....................a...........................c......................... [ 16000]
AY261360.1_Kenya_1950            ............t.............a..........c.......a.....cc.c................c........ [ 16000]
AM712240.1_OURT_88/3             ..........................a..........c......cag..at.g.c.a.aa...........t......tc [ 16000]
AM712239.1_Benin_97/1            ----------------------------------------------------------------...a...gact...t. [ 16000]
KM111295.1_Ken06.Bus             ..........................a..........c.......a.....cc.c................c........ [ 16000]
KM111294.1_Ken05/Tk1             ..........................a..........c.......a.....ac.c................c........ [ 16000]
KM262844.1_strain_L60            ----------------------------------------------------------------...a...gact...t. [ 16000]
KM102979.1_26544/OG10_Italy      ----------------------------------------------------------------...a...gact...t. [ 16000]
KM262845.1_strain_NHV            ..........................a..........c......cag..at.g.c.a.aa...........t......tc [ 16000]
KX354450.1_47/Ss/2008            ----------------------------------------------------------------...a...gact...t. [ 16000]
KP055815.1_strain_BA71           -------------------------------------------------------------------------------- [ 16000]
MH025920.1_strain_R35            ..........................a..........c.......a.....cc.c................c........ [ 16000]
MH025919.1_strain_N10            ..........................a..........c.......a.....cc.c................c........ [ 16000]
MH025918.1_strain_R25            ..........................a..........c.......a.....cc.c................c........ [ 16000]
MH025917.1_strain_R7             ..........................a..........c.......a.....cc.c................c........ [ 16000]
MH025916.1_strain_R8             ..........................a..........c.......a.....cc.c................c........ [ 16000]
```

```
FR682468.2_Georgia_2007/1        accattccagcgggaaaccgtgcaatttataatccagtcattttgatgaataactggccaatctgtttgaatcctgtttc [ 16080]
ASF_IND_20_CAD543                -------------------------------------------------------------------------------- [ 16080]
LR899193.1_Germany_2020/1        ................................................................................ [ 16080]
MN393477.1_Wuhan_2019            ................................................................................ [ 16080]
MW396979.1_Timor-Leste/2019/1    ................................................................................ [ 16080]
MK940252.1_CN/2019/InnerMongolia ................................................................................ [ 16080]
MK128995.1_China/2018/AnhuiXCGQ  ................................................................................ [ 16080]
MH681419.1_ASFV/POL/2015         ................................................................................ [ 16080]
LS478113.1_Estonia_2014          ggtgc...t....c...--------------------------------------------------------------- [ 16080]
KP843857.1_Odintsovo_02/14       ................................................................................ [ 16080]
FN557520.1_strain_E75            gtt-------------------------------------------------------------------...t.a...t [ 16080]
AY261366.1_Warthog               g................................................................c.............. [ 16080]
AY261365.1_Warmbaths             ............a.....t..........................g...................c.............. [ 16080]
AY261364.1_Tengani_62            .............................................g...................c.............. [ 16080]
AY261363.1_Pretoriuskop/96/4     .................................................................c.............. [ 16080]
AY261362.1_Mkuzi_1979            g.....................................................c..........c.............. [ 16080]
AY261361.1_Malawi_Lil-20/1_1983  .............................................g........c..........c.........a.... [ 16080]
AY261360.1_Kenya_1950            ..................g..........................g.....c..a..........c......t....... [ 16080]
AM712240.1_OURT_88/3             g................................................................c.............. [ 16080]
AM712239.1_Benin_97/1            gtt-------------------------------------------------------------------...t.a...t [ 16080]
KM111295.1_Ken06.Bus             ..................a..........................g.....c..g..........c......t....... [ 16080]
KM111294.1_Ken05/Tk1             ..................g..........................g.....c..a..........c......t....... [ 16080]
KM262844.1_strain_L60            gtt-------------------------------------------------------------------...t.a...t [ 16080]
KM102979.1_26544/OG10_Italy      gtt-------------------------------------------------------------------...t.a...t [ 16080]
KM262845.1_strain_NHV            g................................................................c.............. [ 16080]
KX354450.1_47/Ss/2008            gtt-------------------------------------------------------------------...t.a...t [ 16080]
KP055815.1_strain_BA71           -------------------------------------------------------------------------------- [ 16080]
MH025920.1_strain_R35            ..................a..........................g.....c..g..........c......t....... [ 16080]
MH025919.1_strain_N10            ..................a..........................g.....c..g..........c......t....... [ 16080]
MH025918.1_strain_R25            ..................a..........................g.....c..g..........c......t....... [ 16080]
MH025917.1_strain_R7             ..................a..........................g.....c..g..........c......t....... [ 16080]
MH025916.1_strain_R8             ..................a..........................g.....c..g..........c......t....... [ 16080]

FR682468.2_Georgia_2007/1        ggcagataccgtggacgcattcccagcaaaagtcacattggtttgcgtaagtgcaccaataaactagctcatgttcagga [ 16160]
ASF_IND_20_CAD543                -------------------------------------------------------------------------------- [ 16160]
LR899193.1_Germany_2020/1        ................................................................................ [ 16160]
MN393477.1_Wuhan_2019            ................................................................................ [ 16160]
MW396979.1_Timor-Leste/2019/1    ................................................................................ [ 16160]
MK940252.1_CN/2019/InnerMongolia ................................................................................ [ 16160]
MK128995.1_China/2018/AnhuiXCGQ  ................................................................................ [ 16160]
MH681419.1_ASFV/POL/2015         ................................................................................ [ 16160]
LS478113.1_Estonia_2014          ---------------------------.....ct..------------tg.a...g..g...-----------------g [ 16160]
KP843857.1_Odintsovo_02/14       ................................................................................ [ 16160]
```

```
FN557520.1_strain_E75           ta..a..g..a.tttg...a........g........a.tc...a.a..c..a........tgga.....c.cc.t.... [ 16160]
AY261366.1_Warthog              .......g.....a.t..............................a..g................at....t..gg... [ 16160]
AY261365.1_Warmbaths            ................................................................................ [ 16160]
AY261364.1_Tengani_62           .....................................c...........g.............................. [ 16160]
AY261363.1_Pretoriuskop/96/4    ................................................................................ [ 16160]
AY261362.1_Mkuzi_1979           ....a............................................g..............c............... [ 16160]
AY261361.1_Malawi_Lil-20/1_1983 .............t...................................g.......................g...... [ 16160]
AY261360.1_Kenya_1950           .......t.....t.ta..............a.................g................a.....t..gg... [ 16160]
AM712240.1_OURT_88/3            ........................................a........g..............c..........t.... [ 16160]
AM712239.1_Benin_97/1           ta..a..g..a.tttg...a........g........a.tc...a.a..c..a........tgga.....c.cc.t.... [ 16160]
KM111295.1_Ken06.Bus            .......t.....t..a................................g................a.....t..gg... [ 16160]
KM111294.1_Ken05/Tk1            .......t.....t..a................................g........g.............t..gg... [ 16160]
KM262844.1_strain_L60           ta..a..g..a.tttg...a........g........a.tc...a.a..c..a........tgga.....c.cc.t.... [ 16160]
KM102979.1_26544/OG10_Italy     ta..a..g..a.tttg...a........g........a.tc...a.a..c..a........tgga.....c.cc.t.... [ 16160]
KM262845.1_strain_NHV           ........................................a........g..............c..........t.... [ 16160]
KX354450.1_47/Ss/2008           ta..a..g..a.tttg...a........g........a.tc...a.a..c..a........tgga.....c.cc.t.... [ 16160]
KP055815.1_strain_BA71          --------..a.cttga..agt.........ccg...a.tt..ca.a..ttga.....g..tct......c.c...ca.g [ 16160]
MH025920.1_strain_R35           .......t.....t..a................................g................a.....t..gg... [ 16160]
MH025919.1_strain_N10           .......t.....t..a................................g................a.....t..gg... [ 16160]
MH025918.1_strain_R25           .......t.....t..a................................g................a.....t..gg... [ 16160]
MH025917.1_strain_R7            .......t.....t..a................................g................a.....t..gg... [ 16160]
MH025916.1_strain_R8            .......t.....t..a................................g................a.....t..gg... [ 16160]

FR682468.2_Georgia_2007/1       ggataacgggttggtagtaaatcttctaatttacg-tataggagcggcttgaaggacaacca------cc------ccca [ 16240]
ASF_IND_20_CAD543               -------------------------------------------------------------------------------- [ 16240]
LR899193.1_Germany_2020/1       ...................................-..........................------..------.... [ 16240]
MN393477.1_Wuhan_2019           ...................................-..........................------..------.... [ 16240]
MW396979.1_Timor-Leste/2019/1   ...................................-..........................------..------.... [ 16240]
MK940252.1_CN/2019/InnerMongolia ...................................-..........................------..------.... [ 16240]
MK128995.1_China/2018/AnhuiXCGQ ...................................-..........................------..------.... [ 16240]
MH681419.1_ASFV/POL/2015        ...................................-..........................------..------.... [ 16240]
LS478113.1_Estonia_2014         aag.cgt...g..aa..a.cg..c.tctt...g..c.gcga.c.ta.t...gc.cg..ga.g------ttt----a.g.. [ 16240]
KP843857.1_Odintsovo_02/14      ...................................-..........................------..------.... [ 16240]
FN557520.1_strain_E75           ....gct....gat.ga...g...gg.....c.t.-.g------------------------------------------ [ 16240]
AY261366.1_Warthog              ....t.........c....g..t.....c......-............c.....a.t.....------..------.... [ 16240]
AY261365.1_Warmbaths            ...................g...............-............c.....a.t.....------..------.... [ 16240]
AY261364.1_Tengani_62           ...................g...............-..........................------..c-----.... [ 16240]
AY261363.1_Pretoriuskop/96/4    ...................g...............-............c.......t.....------..------.... [ 16240]
AY261362.1_Mkuzi_1979           ...................g...............-............c.....a.t..tac------..cccccc.... [ 16240]
AY261361.1_Malawi_Lil-20/1_1983 ...................g...............-....................t.....------..------g... [ 16240]
AY261360.1_Kenya_1950           ....t.........c....gg..............-.........a..c.......t...g.------..------.... [ 16240]
AM712240.1_OURT_88/3            ...................g...............-......t.....c.......t..tac------..cccccc-... [ 16240]
AM712239.1_Benin_97/1           ....gct....gat.ga...g...gg.....c.t.-.g------------------------------------------ [ 16240]
KM111295.1_Ken06.Bus            ....t.........c....gg..............-.........a..c.......t...g-------..------.... [ 16240]
```

```
KM111294.1_Ken05/Tk1            ....t.........c....gg..............-.........a..c.......t...g.------..------.... [ 16240]
KM262844.1_strain_L60           ....gct....gat.ga...g...gg.....c.t.-.g------------------------------------------ [ 16240]
KM102979.1_26544/OG10_Italy     ....gct....gat.ga...g...gg.....c.t.-.g------------------------------------------ [ 16240]
KM262845.1_strain_NHV           ...................g...............-......t.....c.......t..tac------..cccccc-... [ 16240]
KX354450.1_47/Ss/2008           ....gct....gat.ga...g...gg.....c.t.-.g------------------------------------------ [ 16240]
KP055815.1_strain_BA71          a..---tt.tcg.tcgaa...c.c.g..---------------------------------------------------- [ 16240]
MH025920.1_strain_R35           ....t.........c....gg..............-...t.....a..c.......t...g-cccccc..------.... [ 16240]
MH025919.1_strain_N10           ....t.........c....gg..............-...t.....a..c.......t...g--ccccc..------.... [ 16240]
MH025918.1_strain_R25           ....t.........c....gg..............-...t.....a..c.......t...g-cccccc..------.... [ 16240]
MH025917.1_strain_R7            ....t.........c....gg..............-...t.....a..c.......t...gccccccc..------.... [ 16240]
MH025916.1_strain_R8            ....t.........c....gg..............-...t.....a..c.......t...g--ccccc..------.... [ 16240]

FR682468.2_Georgia_2007/1       gtagtactagaatcagtacctttatagtggccaccctа-cactagacctctaagttgaagacaaagaactaaaatttaga [ 16320]
ASF_IND_20_CAD543               -------------------------------------------------------------------------------- [ 16320]
LR899193.1_Germany_2020/1       ......................................-......................................... [ 16320]
MN393477.1_Wuhan_2019           ......................................-......................................... [ 16320]
MW396979.1_Timor-Leste/2019/1   ......................................-......................................... [ 16320]
MK940252.1_CN/2019/InnerMongolia ......................................-......................................... [ 16320]
MK128995.1_China/2018/AnhuiXCGQ ......................................-......................................... [ 16320]
MH681419.1_ASFV/POL/2015        ......................................-......................................... [ 16320]
LS478113.1_Estonia_2014         ..g.gg.ggag.ct...c....g.ggc..c.tc..a.c-a.g..ag.--------------------------------- [ 16320]
KP843857.1_Odintsovo_02/14      ......................................-......................................... [ 16320]
FN557520.1_strain_E75           -------------------------------------------------...gaa...g...tg..t.gc---------- [ 16320]
AY261366.1_Warthog              .............t..c.....c...............-......................................... [ 16320]
AY261365.1_Warmbaths            .............t..c.....c...............-......................................... [ 16320]
AY261364.1_Tengani_62           .............t..c.....c...............-t...................t.................... [ 16320]
AY261363.1_Pretoriuskop/96/4    .............t..c.....c...............-.............................t........... [ 16320]
AY261362.1_Mkuzi_1979           ................c.....c...........t...-t........................................ [ 16320]
AY261361.1_Malawi_Lil-20/1_1983 .c..............c.....c...............-t............c........-....g............. [ 16320]
AY261360.1_Kenya_1950           ................c.....c..c........ga..ct.g..............a....a.gg..t..........a. [ 16320]
AM712240.1_OURT_88/3            .............t..c.....c........t......-t........................................ [ 16320]
AM712239.1_Benin_97/1           -------------------------------------------------...gaa...g...tg..t.gc---------- [ 16320]
KM111295.1_Ken06.Bus            ................c.....c..g.....t..ga..ct.g..............ag...a.gg..t..........a. [ 16320]
KM111294.1_Ken05/Tk1            ................c.....c..g........ga..ct.g..............a....a.gg..t..........a. [ 16320]
KM262844.1_strain_L60           -------------------------------------------------...gaa...g...tg..t.gc---------- [ 16320]
KM102979.1_26544/OG10_Italy     -------------------------------------------------...gaa...g...tg..t.gc---------- [ 16320]
KM262845.1_strain_NHV           .............t..c.....c........t......-t........................................ [ 16320]
KX354450.1_47/Ss/2008           -------------------------------------------------...gaa...g...tg..t.gc---------- [ 16320]
KP055815.1_strain_BA71          -----------------------------------------------------------...t.gtt.ga---------- [ 16320]
MH025920.1_strain_R35           ................c.....c..g.....t..ga..ct.g..............ag...a.gg..t..........a. [ 16320]
MH025919.1_strain_N10           ................c.....c..g.....t..ga..ct.g..............ag...a.gg..t..........a. [ 16320]
MH025918.1_strain_R25           ................c.....c..g.....t..ga..ct.g..............ag...a.gg..t..........a. [ 16320]
MH025917.1_strain_R7            ................c.....c..g.....t..ga..ct.g..............ag...a.gg..t..........a. [ 16320]
MH025916.1_strain_R8            ................c.....c..g.....t..ga..ct.g..............ag...a.gg..t..........a. [ 16320]
```

```
FR682468.2_Georgia_2007/1        gccgtttaattactactaataattata-----------ttttttattgtctacaataggattctat-taaaaaataatga [ 16400]
ASF_IND_20_CAD543                -------------------------------------------------------------------------------- [ 16400]
LR899193.1_Germany_2020/1        ...........................-----------............................-............. [ 16400]
MN393477.1_Wuhan_2019            ...........................-----------............................-............. [ 16400]
MW396979.1_Timor-Leste/2019/1    ...........................-----------............................-............. [ 16400]
MK940252.1_CN/2019/InnerMongolia ...........................-----------............................-............. [ 16400]
MK128995.1_China/2018/AnhuiXCGQ  ...........................-----------............................-............. [ 16400]
MH681419.1_ASFV/POL/2015         ...........................-----------............................-............. [ 16400]
LS478113.1_Estonia_2014          ---------------------------------------------------.t.gc.aatcc...c-c------.gg.tg [ 16400]
KP843857.1_Odintsovo_02/14       ...........................-----------............................-............. [ 16400]
FN557520.1_strain_E75            -------------------------------------------------------------------------------- [ 16400]
AY261366.1_Warthog               .g........cg...............----------t........................-...-............. [ 16400]
AY261365.1_Warmbaths             .g........cg...............----------t........................-...-............. [ 16400]
AY261364.1_Tengani_62            ..........c................----------t..............t.........-...-............. [ 16400]
AY261363.1_Pretoriuskop/96/4     .g........cg........g......----------t..............t..c......-...-............. [ 16400]
AY261362.1_Mkuzi_1979            ..........a.t..t........ctaataattatat..........t......t..........c-.t.....g..... [ 16400]
AY261361.1_Malawi_Lil-20/1_1983  ..........a.t..t........ctaatatttatat..........t...............c..-.--.....c..t. [ 16400]
AY261360.1_Kenya_1950            ..........c.t....c..c...--------------------....t.t....g...g..a...a.t......c..tg [ 16400]
AM712240.1_OURT_88/3             ............t..........g.-------------.a............t..........c.a-.t.....g..... [ 16400]
AM712239.1_Benin_97/1            -------------------------------------------------------------------------------- [ 16400]
KM111295.1_Ken06.Bus             ..........c.t....c..c......-------------........t.t....g...g..a...a.t......c..t. [ 16400]
KM111294.1_Ken05/Tk1             ..........c.t....c..c..---------------------....t.t....a...g..a...a.t......c.gtg [ 16400]
KM262844.1_strain_L60            -------------------------------------------------------------------------------- [ 16400]
KM102979.1_26544/OG10_Italy      -------------------------------------------------------------------------------- [ 16400]
KM262845.1_strain_NHV            ............t..........g.-------------.a............t..........c.a-.t.....g..... [ 16400]
KX354450.1_47/Ss/2008            -------------------------------------------------------------------------------- [ 16400]
KP055815.1_strain_BA71           -------------------------------------------------------------------------------- [ 16400]
MH025920.1_strain_R35            ..........c.t....c..c......-------------........t.t....g...g..a...a.t......c..t. [ 16400]
MH025919.1_strain_N10            ..........c.t....c..c......-------------........t.t....g...g..a...a.t......c..t. [ 16400]
MH025918.1_strain_R25            ..........c.t....c..c......-------------........t.t....g...g..a...a.t......c..t. [ 16400]
MH025917.1_strain_R7             ..........c.t....c..c......-------------........t.t....g...g..a...a.t......c..t. [ 16400]
MH025916.1_strain_R8             ..........c.t....c..c......-------------........t.t....g...g..a...a.t......c..t. [ 16400]

FR682468.2_Georgia_2007/1        tttttaccaagaaat------atttttat-aaaaaattaatatattttgtaataaactttatttccaatgactgttaaaa [ 16480]
ASF_IND_20_CAD543                -------------------------------------------------------------------------------- [ 16480]
LR899193.1_Germany_2020/1        ...............------........-.................................................. [ 16480]
MN393477.1_Wuhan_2019            ...............------........-.................................................. [ 16480]
MW396979.1_Timor-Leste/2019/1    ...............------........-.................................................. [ 16480]
MK940252.1_CN/2019/InnerMongolia ...............------........-.................................................. [ 16480]
MK128995.1_China/2018/AnhuiXCGQ  ...............------........-.................................................. [ 16480]
MH681419.1_ASFV/POL/2015         ...............------........-.................................................. [ 16480]
LS478113.1_Estonia_2014          ..gc.ct.....ct.------..ccgg..-gg---------------------------c..ccgt..ga....cgcggg [ 16480]
KP843857.1_Odintsovo_02/14       ...............------........-.................................................. [ 16480]
```

```
FN557520.1_strain_E75            ---cc.a...-----------.ag.ccta-g..g..ccttc..g..g...c----------------------------- [ 16480]
AY261366.1_Warthog               ......a........------.a......-.................................................. [ 16480]
AY261365.1_Warmbaths             ......a........------.a......-.................................................. [ 16480]
AY261364.1_Tengani_62            ......a........------.a......-.................................................. [ 16480]
AY261363.1_Pretoriuskop/96/4     ......a........------.a......-.................................................. [ 16480]
AY261362.1_Mkuzi_1979            ..-...a........------.a......-.................................................. [ 16480]
AY261361.1_Malawi_Lil-20/1_1983  .c...ga........------.a......-.................c.................t.............. [ 16480]
AY261360.1_Kenya_1950            ......ag.......------.a......a...................................t......--..t... [ 16480]
AM712240.1_OURT_88/3             ......a........------.a......-...................................t.............. [ 16480]
AM712239.1_Benin_97/1            ---cc.a...-----------.ag.ccta-g..g..ccttc..g..g...c----------------------------- [ 16480]
KM111295.1_Ken06.Bus             ...c..a........------.a......-...................................t..c.......t... [ 16480]
KM111294.1_Ken05/Tk1             ...c..a........------.a......-...................................t..........t... [ 16480]
KM262844.1_strain_L60            ---cc.a...-----------.ag.ccta-g..g..ccttc..g..g...c----------------------------- [ 16480]
KM102979.1_26544/OG10_Italy      ---cc.a...-----------.ag.ccta-g..g..ccttc..g..g...c----------------------------- [ 16480]
KM262845.1_strain_NHV            ......a........------.a......-...................................t.............. [ 16480]
KX354450.1_47/Ss/2008            ---cc.a...-----------.ag.ccta-g..g..ccttc..g..g...c----------------------------- [ 16480]
KP055815.1_strain_BA71           ---cc.g.t..c.gcaagccg.gg.aac.-...g..cctcat.g.ag...------------------------------ [ 16480]
MH025920.1_strain_R35            ...c..a........------.a......-..................................tt..c.......t... [ 16480]
MH025919.1_strain_N10            ...c..a........------.a......-..................................tt..c.......t... [ 16480]
MH025918.1_strain_R25            ...c..a........------.a......-..................................tt..c.......t... [ 16480]
MH025917.1_strain_R7             ...c..a........------.a......-..................................tt..c.......t... [ 16480]
MH025916.1_strain_R8             ...c..a........------.a......-..................................tt..c.......t... [ 16480]

FR682468.2_Georgia_2007/1        taaggaaactatccttagttagtcgaggaagatggttaggttatttcgcaatccgataaaatgtttattttatcgtaggt [ 16560]
ASF_IND_20_CAD543                -------------------------------------------------------------------------------- [ 16560]
LR899193.1_Germany_2020/1        ................................................................................ [ 16560]
MN393477.1_Wuhan_2019            ................................................................................ [ 16560]
MW396979.1_Timor-Leste/2019/1    ................................................................................ [ 16560]
MK940252.1_CN/2019/InnerMongolia ................................................................................ [ 16560]
MK128995.1_China/2018/AnhuiXCGQ  ................................................................................ [ 16560]
MH681419.1_ASFV/POL/2015         ................................................................................ [ 16560]
LS478113.1_Estonia_2014          .c..aggca...at..---------------------------------------g..g.gcaaa..ca...c.tg.... [ 16560]
KP843857.1_Odintsovo_02/14       ................................................................................ [ 16560]
FN557520.1_strain_E75            --------------------------------------------------------c...tg.cacc.g.c...t..--- [ 16560]
AY261366.1_Warthog               ................................................................a............... [ 16560]
AY261365.1_Warmbaths             ................................................................a............... [ 16560]
AY261364.1_Tengani_62            ........................a....................................................... [ 16560]
AY261363.1_Pretoriuskop/96/4     ................................................................a............... [ 16560]
AY261362.1_Mkuzi_1979            ................................................................a............... [ 16560]
AY261361.1_Malawi_Lil-20/1_1983  .......................................a.......a............................g.a. [ 16560]
AY261360.1_Kenya_1950            .......tg..............................a......tat.............ac..............a. [ 16560]
AM712240.1_OURT_88/3             ................................................................a............... [ 16560]
AM712239.1_Benin_97/1            --------------------------------------------------------c...tg.cacc.g.c...t..--- [ 16560]
KM111295.1_Ken06.Bus             .......tg.....................a.......aa......tat..............cc.............a. [ 16560]
```

```
KM111294.1_Ken05/Tk1           ......tg...............................a......tat..............c..............a. [ 16560]
KM262844.1_strain_L60          ---------------------------------------------------------c..tg.cacc.g.c...t..--- [ 16560]
KM102979.1_26544/OG10_Italy    ---------------------------------------------------------c..tg.cacc.g.c...t..--- [ 16560]
KM262845.1_strain_NHV          ................................................................a............... [ 16560]
KX354450.1_47/Ss/2008          ---------------------------------------------------------c..tg.cacc.g.c...t..--- [ 16560]
KP055815.1_strain_BA71         -----------------------------------------------------------------...a...cgaa.--- [ 16560]
MH025920.1_strain_R35          ......tg......................a.......aa......tat.............cc..............a. [ 16560]
MH025919.1_strain_N10          ......tg......................a.......aa......tat.............cc..............a. [ 16560]
MH025918.1_strain_R25          ......tg......................a.......aa......tat.............cc..............a. [ 16560]
MH025917.1_strain_R7           ......tg......................a.......aa......tat.............cc..............a. [ 16560]
MH025916.1_strain_R8           ......tg......................a.......aa......tat.............cc..............a. [ 16560]

FR682468.2_Georgia_2007/1      ctcgtaaaatccaggaaaaaaaattacggaagagtttaaaaaagctaaatttttaccaccctccagaagattgttgtcaa [ 16640]
ASF_IND_20_CAD543              -------------------------------------------------------------------------------- [ 16640]
LR899193.1_Germany_2020/1      ................................................................................ [ 16640]
MN393477.1_Wuhan_2019          ................................................................................ [ 16640]
MW396979.1_Timor-Leste/2019/1  ................................................................................ [ 16640]
MK940252.1_CN/2019/InnerMongolia ................................................................................ [ 16640]
MK128995.1_China/2018/AnhuiXCGQ ................................................................................ [ 16640]
MH681419.1_ASFV/POL/2015       ................................................................................ [ 16640]
LS478113.1_Estonia_2014        tcacc...g.--.a.c.gc.c..a..aa.gcc....c.gc....tccgg..cc.-------------------------- [ 16640]
KP843857.1_Odintsovo_02/14     ................................................................................ [ 16640]
FN557520.1_strain_E75          -------------------------------------.....gat.t..a....gtt..--------------------- [ 16640]
AY261366.1_Warthog             ................................................................................ [ 16640]
AY261365.1_Warmbaths           ................................................................................ [ 16640]
AY261364.1_Tengani_62          .........................g...................................................... [ 16640]
AY261363.1_Pretoriuskop/96/4   ..........................................................t..................... [ 16640]
AY261362.1_Mkuzi_1979          ..................................c....................t..t.....g............... [ 16640]
AY261361.1_Malawi_Lil-20/1_1983 ....g....c..g.............................................t.......g............. [ 16640]
AY261360.1_Kenya_1950          ..t....---------.....g......a...gc.....................t..t......a....a......... [ 16640]
AM712240.1_OURT_88/3           ..................................c....................t..t...................t. [ 16640]
AM712239.1_Benin_97/1          -------------------------------------.....gat.t..a....gtt..--------------------- [ 16640]
KM111295.1_Ken06.Bus           ..t....---------............a...gc.....................t..t......a....a......... [ 16640]
KM111294.1_Ken05/Tk1           ..t....---------.....g......a...gc.....................t..t......a....a......... [ 16640]
KM262844.1_strain_L60          -------------------------------------.....gat.t..a....gtt..--------------------- [ 16640]
KM102979.1_26544/OG10_Italy    -------------------------------------.....gat.t..a....gtt..--------------------- [ 16640]
KM262845.1_strain_NHV          ..................................c....................t..t...................t. [ 16640]
KX354450.1_47/Ss/2008          -------------------------------------.....gat.t..a....gtt..--------------------- [ 16640]
KP055815.1_strain_BA71         -------------------------------------...t.t.t....a..---------------------------- [ 16640]
MH025920.1_strain_R35          ..t....---------............a...gc.....................t..t......a....a......... [ 16640]
MH025919.1_strain_N10          ..t....---------............a...gc.....................t..t......a....a......... [ 16640]
MH025918.1_strain_R25          ..t....---------............a...gc.....................t..t......a....a......... [ 16640]
MH025917.1_strain_R7           ..t....---------............a...gc.....................t..t......a....a......... [ 16640]
MH025916.1_strain_R8           ..t....---------............a...gc.....................t..t......a....a......... [ 16640]
```

```
FR682468.2_Georgia_2007/1        atatatcgtttgctagaaaatgttcctggaggaacttactttattacagaaaatatgacgaatgatttaattatggtcgt [ 16720]
ASF_IND_20_CAD543                -------------------------------------------------------------------------------- [ 16720]
LR899193.1_Germany_2020/1        ................................................................................ [ 16720]
MN393477.1_Wuhan_2019            ................................................................................ [ 16720]
MW396979.1_Timor-Leste/2019/1    ................................................................................ [ 16720]
MK940252.1_CN/2019/InnerMongolia ................................................................................ [ 16720]
MK128995.1_China/2018/AnhuiXCGQ  ................................................................................ [ 16720]
MH681419.1_ASFV/POL/2015         ................................................................................ [ 16720]
LS478113.1_Estonia_2014          -------------------------------------------------------------------------------- [ 16720]
KP843857.1_Odintsovo_02/14       ................................................................................ [ 16720]
FN557520.1_strain_E75            -------------------------------------------------------------------------------- [ 16720]
AY261366.1_Warthog               .............................g.................................................. [ 16720]
AY261365.1_Warmbaths             .............................g.................................................. [ 16720]
AY261364.1_Tengani_62            .............................g.................................................. [ 16720]
AY261363.1_Pretoriuskop/96/4     .............................g.........................................a........ [ 16720]
AY261362.1_Mkuzi_1979            ...........a............t.a..g.................................................. [ 16720]
AY261361.1_Malawi_Lil-20/1_1983  ...........a..............a..g...................................g.....a...a.... [ 16720]
AY261360.1_Kenya_1950            ......................c...a..g.............................c.....g.....a...a...c [ 16720]
AM712240.1_OURT_88/3             .............................g...................................g.....a........ [ 16720]
AM712239.1_Benin_97/1            -------------------------------------------------------------------------------- [ 16720]
KM111295.1_Ken06.Bus             ......................c...a..g.............................c.....g.....a...a.t.c [ 16720]
KM111294.1_Ken05/Tk1             ......................c...a..g.............................c.....g.....a...a...c [ 16720]
KM262844.1_strain_L60            -------------------------------------------------------------------------------- [ 16720]
KM102979.1_26544/OG10_Italy      -------------------------------------------------------------------------------- [ 16720]
KM262845.1_strain_NHV            .............................g...................................g.....a........ [ 16720]
KX354450.1_47/Ss/2008            -------------------------------------------------------------------------------- [ 16720]
KP055815.1_strain_BA71           -------------------------------------------------------------------------------- [ 16720]
MH025920.1_strain_R35            ......................c...a..g.............................c.....g.....a...a.t.c [ 16720]
MH025919.1_strain_N10            ......................c...a..g.............................c.....g.....a...a.t.c [ 16720]
MH025918.1_strain_R25            ......................c...a..g.............................c.....g.....a...a.t.c [ 16720]
MH025917.1_strain_R7             ......................c...a..g.............................c.....g.....a...a.t.c [ 16720]
MH025916.1_strain_R8             ......................c...a..g.............................c.....g.....a...a.t.c [ 16720]

FR682468.2_Georgia_2007/1        aaaggattcggtggataaaaaaattaaaagcattaaattatatcttcatggaagttatattaagattcatcagcactatt [ 16800]
ASF_IND_20_CAD543                -------------------------------------------------------------------------------- [ 16800]
LR899193.1_Germany_2020/1        ................................................................................ [ 16800]
MN393477.1_Wuhan_2019            ................................................................................ [ 16800]
MW396979.1_Timor-Leste/2019/1    ................................................................................ [ 16800]
MK940252.1_CN/2019/InnerMongolia ................................................................................ [ 16800]
MK128995.1_China/2018/AnhuiXCGQ  ................................................................................ [ 16800]
MH681419.1_ASFV/POL/2015         ................................................................................ [ 16800]
LS478113.1_Estonia_2014          -------------------------------------------------------------------------------- [ 16800]
KP843857.1_Odintsovo_02/14       ................................................................................ [ 16800]
```

```
FN557520.1_strain_E75            -----------c.c.a........cc.----------------------------------------------------- [ 16800]
AY261366.1_Warthog               ................................................................................ [ 16800]
AY261365.1_Warmbaths             ................................................................................ [ 16800]
AY261364.1_Tengani_62            ................................................................................ [ 16800]
AY261363.1_Pretoriuskop/96/4     ................................................................................ [ 16800]
AY261362.1_Mkuzi_1979            ................................................................................ [ 16800]
AY261361.1_Malawi_Lil-20/1_1983  .........t.....................g........a..t..t....t............................ [ 16800]
AY261360.1_Kenya_1950            c......c....a..c...............g.......g......ac...t.a.............a............ [ 16800]
AM712240.1_OURT_88/3             ...............................g..............t....t...............a.......t.... [ 16800]
AM712239.1_Benin_97/1            -----------c.c.a........cc.----------------------------------------------------- [ 16800]
KM111295.1_Ken06.Bus             .......c....a..c...............g.......g......ac...t.a.............a............ [ 16800]
KM111294.1_Ken05/Tk1             .......c....a..c...............g.......g......ac...t.a.............a............ [ 16800]
KM262844.1_strain_L60            -----------c.c.a........cc.----------------------------------------------------- [ 16800]
KM102979.1_26544/OG10_Italy      -----------c.c.a........cc.----------------------------------------------------- [ 16800]
KM262845.1_strain_NHV            ...............................g..............t....t...............a.......t.... [ 16800]
KX354450.1_47/Ss/2008            -----------c.c-a........cc.----------------------------------------------------- [ 16800]
KP055815.1_strain_BA71           ------------...a......gccc------------------------------------------------------ [ 16800]
MH025920.1_strain_R35            .......c....a..c...............g.......g......ac...t.a.............a............ [ 16800]
MH025919.1_strain_N10            .......c....a..c...............g.......g......ac...t.a.............a............ [ 16800]
MH025918.1_strain_R25            .......c....a..c...............g.......g......ac...t.a.............a............ [ 16800]
MH025917.1_strain_R7             .......c....a..c...............g.......g......ac...t.a.............a............ [ 16800]
MH025916.1_strain_R8             .......c....a..c...............g.......g......ac...t.a.............a............ [ 16800]

FR682468.2_Georgia_2007/1        atattaatatttatatgtatcttatgagatatacccaaatttataaatatcccttaatttgttttaacaa---------- [ 16880]
ASF_IND_20_CAD543                -------------------------------------------------------------------------------- [ 16880]
LR899193.1_Germany_2020/1        ......................................................................---------- [ 16880]
MN393477.1_Wuhan_2019            ......................................................................---------- [ 16880]
MW396979.1_Timor-Leste/2019/1    ......................................................................---------- [ 16880]
MK940252.1_CN/2019/InnerMongolia ......................................................................---------- [ 16880]
MK128995.1_China/2018/AnhuiXCGQ  ......................................................................---------- [ 16880]
MH681419.1_ASFV/POL/2015         ......................................................................---------- [ 16880]
LS478113.1_Estonia_2014          -------------------------------------------------------------------..t---------- [ 16880]
KP843857.1_Odintsovo_02/14       ......................................................................---------- [ 16880]
FN557520.1_strain_E75            -------------------------------------------------------------------------------- [ 16880]
AY261366.1_Warthog               ......................................................................---------- [ 16880]
AY261365.1_Warmbaths             .................................................c....................---------- [ 16880]
AY261364.1_Tengani_62            ......................................................................---------- [ 16880]
AY261363.1_Pretoriuskop/96/4     ......................................................................---------- [ 16880]
AY261362.1_Mkuzi_1979            ......................................................................---------- [ 16880]
AY261361.1_Malawi_Lil-20/1_1983  ...................................................................t..atatt----c [ 16880]
AY261360.1_Kenya_1950            .................................a......a...........t.............gt..atattcaaaa [ 16880]
AM712240.1_OURT_88/3             ...............................................................a...t..---------- [ 16880]
AM712239.1_Benin_97/1            -------------------------------------------------------------------------------- [ 16880]
KM111295.1_Ken06.Bus             .................................a......a...........t.............gt..atattcaaaa [ 16880]
```

```
KM111294.1_Ken05/Tk1           .................................a......a...........t.............gt..atattcaaaa [ 16880]
KM262844.1_strain_L60          -------------------------------------------------------------------------------- [ 16880]
KM102979.1_26544/OG10_Italy    -------------------------------------------------------------------------------- [ 16880]
KM262845.1_strain_NHV          ...............................................................a...t..---------- [ 16880]
KX354450.1_47/Ss/2008          -------------------------------------------------------------------------------- [ 16880]
KP055815.1_strain_BA71         -------------------------------------------------------------------------------- [ 16880]
MH025920.1_strain_R35          .................................a......a...........t.............gt..atattcaaaa [ 16880]
MH025919.1_strain_N10          .................................a......a...........t.............gt..atattcaaaa [ 16880]
MH025918.1_strain_R25          .................................a......a...........t.............gt..atattcaaaa [ 16880]
MH025917.1_strain_R7           .................................a......a...........t.............gt..atattcaaaa [ 16880]
MH025916.1_strain_R8           .................................a......a...........t.............gt..atattcaaaa [ 16880]

FR682468.2_Georgia_2007/1      atattataacatctaagtaaatattcttggaatggattttcttatagaatggttacaggatatgtcagcgacagg---ct [ 16960]
ASF_IND_20_CAD543              -------------------------------------------------------------------------------- [ 16960]
LR899193.1_Germany_2020/1      ...........................................................................---.. [ 16960]
MN393477.1_Wuhan_2019          ...........................................................................---.. [ 16960]
MW396979.1_Timor-Leste/2019/1  ...........................................................................---.. [ 16960]
MK940252.1_CN/2019/InnerMongolia ...........................................................................---.. [ 16960]
MK128995.1_China/2018/AnhuiXCGQ ...........................................................................---.. [ 16960]
MH681419.1_ASFV/POL/2015       ...........................................................................---.. [ 16960]
LS478113.1_Estonia_2014        ..cg..c..t......---------------------------------------------------------------- [ 16960]
KP843857.1_Odintsovo_02/14     ...........................................................................---.. [ 16960]
FN557520.1_strain_E75          ----------------...tg....t..tt...acg.a.aa....t...ctc...t.a....aa---------------- [ 16960]
AY261366.1_Warthog             ...................................................a.......................---t. [ 16960]
AY261365.1_Warmbaths           ...................................c...............a.............-.........---g. [ 16960]
AY261364.1_Tengani_62          ..................................................aa.......................---g. [ 16960]
AY261363.1_Pretoriuskop/96/4   ........................................-........c.a....g..........at......---t. [ 16960]
AY261362.1_Mkuzi_1979          ...................................................a....................-a.---g. [ 16960]
AY261361.1_Malawi_Lil-20/1_1983 .....g....tc..........................a.....c---..aa...t.a............g..a.cctg. [ 16960]
AY261360.1_Kenya_1950          ..........tc.cg......-----..atgt.a............t...aa.......t...a.tc.t..t.a.----- [ 16960]
AM712240.1_OURT_88/3           ...................................................a..........g..........a.---a. [ 16960]
AM712239.1_Benin_97/1          ----------------...tg....t..tt...acg.a.aa....t...ctc...t.a....aa---------------- [ 16960]
KM111295.1_Ken06.Bus           ..........tc.cg......-----..atgt.a............t...aa.......t...a.g..t..t.a.----- [ 16960]
KM111294.1_Ken05/Tk1           ..........tc.cg......-----..atgt.a............t...aa.......t...a.tc.t..t.a.----- [ 16960]
KM262844.1_strain_L60          ----------------...tg....t..tt...acg.a.aa....t...ctc...t.a....aa---------------- [ 16960]
KM102979.1_26544/OG10_Italy    ----------------...tg....t..tt...acg.a.aa....t...ctc...t.a....aa---------------- [ 16960]
KM262845.1_strain_NHV          ...................................................a..........g..........a.---a. [ 16960]
KX354450.1_47/Ss/2008          ----------------...tg....t..tt...acg.a.aa....t...ctc...t.a....aa---------------- [ 16960]
KP055815.1_strain_BA71         -------------------------------------------------------------------------------- [ 16960]
MH025920.1_strain_R35          ..........tc.cg......-----..atgt.a............t...aa.......t...a.g..t..t.a.----- [ 16960]
MH025919.1_strain_N10          ..........tc.cg......-----..atgt.a............t...aa.......t...a.g..t..t.a.----- [ 16960]
MH025918.1_strain_R25          ..........tc.cg......-----..atgt.a............t...aa.......t...a.g..t..t.a.----- [ 16960]
MH025917.1_strain_R7           ..........tc.cg......-----..atgt.a............t...aa.......t...a.g..t..t.a.----- [ 16960]
MH025916.1_strain_R8           ..........tc.cg......-----..atgt.a............t...aa.......t...a.g..t..t.a.----- [ 16960]
```

```
FR682468.2_Georgia_2007/1        taataacaaatttg-----ttaatatttttttgttaaataaatgaacaggccac-catttaatattacccgttgcaaaat [ 17040]
ASF_IND_20_CAD543                -------------------------------------------------------------------------------- [ 17040]
LR899193.1_Germany_2020/1        ..............-----...................................-......................... [ 17040]
MN393477.1_Wuhan_2019            ..............-----...................................-......................... [ 17040]
MW396979.1_Timor-Leste/2019/1    ..............-----...................................-......................... [ 17040]
MK940252.1_CN/2019/InnerMongolia ..............-----...................................-......................... [ 17040]
MK128995.1_China/2018/AnhuiXCGQ  ..............-----...................................-......................... [ 17040]
MH681419.1_ASFV/POL/2015         ..............-----...................................-......................... [ 17040]
LS478113.1_Estonia_2014          -----------------------------------..c....g..g..t.a..----------------------..t.. [ 17040]
KP843857.1_Odintsovo_02/14       ..............-----...................................-......................... [ 17040]
FN557520.1_strain_E75            -------------------------------------------------------------------------------- [ 17040]
AY261366.1_Warthog               ..............ttatt......-............................-........g............g... [ 17040]
AY261365.1_Warmbaths             ..............ttatt......-............................-........g............g... [ 17040]
AY261364.1_Tengani_62            ..............ttatt......-..................g.........-........g..t.........g... [ 17040]
AY261363.1_Pretoriuskop/96/4     ..............ttatt......-....................t.......-........g............g... [ 17040]
AY261362.1_Mkuzi_1979            ..............ttatt.................................ggc........g............g... [ 17040]
AY261361.1_Malawi_Lil-20/1_1983  ..t.......c...ctatt......---c...a......g...t......a..t-........g....t.......g... [ 17040]
AY261360.1_Kenya_1950            ---------...gctattt......--------------------------.gt-t.......g.....aa.....gc.g [ 17040]
AM712240.1_OURT_88/3             ..............ctatt......--....................g----------.....g...........----- [ 17040]
AM712239.1_Benin_97/1            -------------------------------------------------------------------------------- [ 17040]
KM111295.1_Ken06.Bus             ---------...gctattt......--------------------------.g.-t...........t.aa..a..g--- [ 17040]
KM111294.1_Ken05/Tk1             ---------...gctattt......--------------------------.gt-t.......g.....aa.....gc.g [ 17040]
KM262844.1_strain_L60            -------------------------------------------------------------------------------- [ 17040]
KM102979.1_26544/OG10_Italy      -------------------------------------------------------------------------------- [ 17040]
KM262845.1_strain_NHV            ..............ctatt......--....................g----------.....g...........----- [ 17040]
KX354450.1_47/Ss/2008            -------------------------------------------------------------------------------- [ 17040]
KP055815.1_strain_BA71           -------------------------------------------------------------------------------- [ 17040]
MH025920.1_strain_R35            ---------...gctattt......--------------------------.g.-t...........t.aa..a..g--- [ 17040]
MH025919.1_strain_N10            ---------...gctattt......--------------------------.g.-t...........t.aa..a..g--- [ 17040]
MH025918.1_strain_R25            ---------...gctattt......--------------------------.g.-t...........t.aa..a..g--- [ 17040]
MH025917.1_strain_R7             ---------...gctattt......--------------------------.g.-t...........t.aa..a..g--- [ 17040]
MH025916.1_strain_R8             ---------...gctattt......--------------------------.g.-t...........t.aa..a..g--- [ 17040]

FR682468.2_Georgia_2007/1        aagaaaaaaaaaacaaacttatagttacaaatcatcttgattaatcacatgtcgttttaactcaatgaaccattctaaat [ 17120]
ASF_IND_20_CAD543                -------------------------------------------------------------------------------- [ 17120]
LR899193.1_Germany_2020/1        ................................................................................ [ 17120]
MN393477.1_Wuhan_2019            ...-............................................................................ [ 17120]
MW396979.1_Timor-Leste/2019/1    ...-............................................................................ [ 17120]
MK940252.1_CN/2019/InnerMongolia ...-............................................................................ [ 17120]
MK128995.1_China/2018/AnhuiXCGQ  ...-............................................................................ [ 17120]
MH681419.1_ASFV/POL/2015         ...-............................................................................ [ 17120]
LS478113.1_Estonia_2014          gga------------------------------------------------------..g...ggaa....--------- [ 17120]
KP843857.1_Odintsovo_02/14       ...-............................................................................ [ 17120]
```

```
FN557520.1_strain_E75           -------------------------------------------------------------------------------- [ 17120]
AY261366.1_Warthog              ...--..........................................g...............t................ [ 17120]
AY261365.1_Warmbaths            ...---.........................................................t................ [ 17120]
AY261364.1_Tengani_62           ...---.......................................................................... [ 17120]
AY261363.1_Pretoriuskop/96/4    ...------....a........ca........................................................ [ 17120]
AY261362.1_Mkuzi_1979           ...--...........c............................t.................t................ [ 17120]
AY261361.1_Malawi_Lil-20/1_1983 ...--............t...........................t.................t................ [ 17120]
AY261360.1_Kenya_1950           .gc---...........tg...g...................g..t.................t................ [ 17120]
AM712240.1_OURT_88/3            --.--g...........t.....a.....c...------......t.................t................ [ 17120]
AM712239.1_Benin_97/1           -------------------------------------------------------------------------------- [ 17120]
KM111295.1_Ken06.Bus            -..---c..........tg...g......................t..............ag.tg.............t. [ 17120]
KM111294.1_Ken05/Tk1            .gc---...........tg...g...................g..t.................t................ [ 17120]
KM262844.1_strain_L60           -------------------------------------------------------------------------------- [ 17120]
KM102979.1_26544/OG10_Italy     -------------------------------------------------------------------------------- [ 17120]
KM262845.1_strain_NHV           --.--g...........t.....a.....c...------......t.................t................ [ 17120]
KX354450.1_47/Ss/2008           -------------------------------------------------------------------------------- [ 17120]
KP055815.1_strain_BA71          -------------------------------------------------------------------------------- [ 17120]
MH025920.1_strain_R35           -..---c..........tg...g......................t..............ag.tg.............t. [ 17120]
MH025919.1_strain_N10           -..---c..........tg...g......................t..............ag.tg.............t. [ 17120]
MH025918.1_strain_R25           -..---c..........tg...g......................t..............ag.tg.............t. [ 17120]
MH025917.1_strain_R7            -..---c..........tg...g......................t..............ag.tg.............t. [ 17120]
MH025916.1_strain_R8            -..---c..........tg...g......................t..............ag.tg.............t. [ 17120]

FR682468.2_Georgia_2007/1       ctttgggttgtgaacaattcatgttatgttgatagtgtatcctaaagtgagcttcatacatacaccggtcatgccaccgg [ 17200]
ASF_IND_20_CAD543               -------------------------------------------------------------------------------- [ 17200]
LR899193.1_Germany_2020/1       ................................................................................ [ 17200]
MN393477.1_Wuhan_2019           ................................................................................ [ 17200]
MW396979.1_Timor-Leste/2019/1   ................................................................................ [ 17200]
MK940252.1_CN/2019/InnerMongolia ................................................................................ [ 17200]
MK128995.1_China/2018/AnhuiXCGQ ................................................................................ [ 17200]
MH681419.1_ASFV/POL/2015        ................................................................................ [ 17200]
LS478113.1_Estonia_2014         --------------------------------c...a..g.tc.------------------------------------ [ 17200]
KP843857.1_Odintsovo_02/14      ................................................................................ [ 17200]
FN557520.1_strain_E75           ------.cc.a.....t.at..t..g.a.a...ga.....---------------------------------------- [ 17200]
AY261366.1_Warthog              ................................................................................ [ 17200]
AY261365.1_Warmbaths            ...............................................................................a [ 17200]
AY261364.1_Tengani_62           ................................................................................ [ 17200]
AY261363.1_Pretoriuskop/96/4    ................................................................................ [ 17200]
AY261362.1_Mkuzi_1979           ...........................................g..a..g.....g........................ [ 17200]
AY261361.1_Malawi_Lil-20/1_1983 ...........................................g..a..g.....g........................ [ 17200]
AY261360.1_Kenya_1950           ...........................................g..a..g..c..g........................ [ 17200]
AM712240.1_OURT_88/3            ...........................................g..a..g.....g........................ [ 17200]
AM712239.1_Benin_97/1           ------.cc.a.....t.at..t..g.a.a...ga.....---------------------------------------- [ 17200]
KM111295.1_Ken06.Bus            .ag....c....................c....................t.............................. [ 17200]
```

```
KM111294.1_Ken05/Tk1            ...........................................g..a..g.....g........................ [ 17200]
KM262844.1_strain_L60           ------.cc.a.....t.at..t..g.a.a...ga.....---------------------------------------- [ 17200]
KM102979.1_26544/OG10_Italy     ------.cc.a.....t.at..t..g.a.a...ga.....---------------------------------------- [ 17200]
KM262845.1_strain_NHV           ...........................................g..a..g.....g........................ [ 17200]
KX354450.1_47/Ss/2008           ------.cc.a.....t.at..t..g.a.a...ga.....---------------------------------------- [ 17200]
KP055815.1_strain_BA71          ----------------------t..t.a.a...----------------------------------------------- [ 17200]
MH025920.1_strain_R35           .ag....c....................c....................t.............................. [ 17200]
MH025919.1_strain_N10           .ag....c....................c....................t.............................. [ 17200]
MH025918.1_strain_R25           .ag....c....................c....................t.............................. [ 17200]
MH025917.1_strain_R7            .ag....c....................c....................t.............................. [ 17200]
MH025916.1_strain_R8            .ag....c....................c....................t.............................. [ 17200]

FR682468.2_Georgia_2007/1       gaaactgtacaattaacaatataatcattttgcgtaataatagggtggtcactaaacactttatttttacacattccatc [ 17280]
ASF_IND_20_CAD543               -------------------------------------------------------------------------------- [ 17280]
LR899193.1_Germany_2020/1       ................................................................................ [ 17280]
MN393477.1_Wuhan_2019           ................................................................................ [ 17280]
MW396979.1_Timor-Leste/2019/1   ................................................................................ [ 17280]
MK940252.1_CN/2019/InnerMongolia ................................................................................ [ 17280]
MK128995.1_China/2018/AnhuiXCGQ ................................................................................ [ 17280]
MH681419.1_ASFV/POL/2015        ................................................................................ [ 17280]
LS478113.1_Estonia_2014         --------------------...gc.t.g.....------------------------------------------a.ct [ 17280]
KP843857.1_Odintsovo_02/14      ................................................................................ [ 17280]
FN557520.1_strain_E75           -------------------------------------------------------------------------------- [ 17280]
AY261366.1_Warthog              ................................a...................................g........... [ 17280]
AY261365.1_Warmbaths            ................................a...................................g........... [ 17280]
AY261364.1_Tengani_62           ................................a...................................g........... [ 17280]
AY261363.1_Pretoriuskop/96/4    ................................a...................................g........... [ 17280]
AY261362.1_Mkuzi_1979           ....................................................................g........... [ 17280]
AY261361.1_Malawi_Lil-20/1_1983 ................................a...........a.....c.................g........... [ 17280]
AY261360.1_Kenya_1950           ................................a...................................g........... [ 17280]
AM712240.1_OURT_88/3            ................................a..................................-g........... [ 17280]
AM712239.1_Benin_97/1           -------------------------------------------------------------------------------- [ 17280]
KM111295.1_Ken06.Bus            ...............................caa.....g....a..a..ct....g...........g..........t [ 17280]
KM111294.1_Ken05/Tk1            ................................a...........a.......................g........... [ 17280]
KM262844.1_strain_L60           -------------------------------------------------------------------------------- [ 17280]
KM102979.1_26544/OG10_Italy     -------------------------------------------------------------------------------- [ 17280]
KM262845.1_strain_NHV           ................................a..................................-g........... [ 17280]
KX354450.1_47/Ss/2008           -------------------------------------------------------------------------------- [ 17280]
KP055815.1_strain_BA71          -------------------------------------------------------------------------------- [ 17280]
MH025920.1_strain_R35           .........................t.....caa.....g....a..a..ct....g...........g..........t [ 17280]
MH025919.1_strain_N10           .........................t.....caa.....g....a..a..ct....g...........g..........t [ 17280]
MH025918.1_strain_R25           .........................t.....caa.....g....a..a..ct....g...........g..........t [ 17280]
MH025917.1_strain_R7            .........................t.....caa.....g....a..a..ct....g...........g..........t [ 17280]
MH025916.1_strain_R8            .........................t.....caa.....g....a..a..ct....g...........g..........t [ 17280]
```

```
FR682468.2_Georgia_2007/1        tttacaggtccagcagaagtcacagtgttttgcataggtgcaccagaacttgagatccctttcaggaggcctacgcattt [ 17360]
ASF_IND_20_CAD543                -------------------------------------------------------------------------------- [ 17360]
LR899193.1_Germany_2020/1        ................................................................................ [ 17360]
MN393477.1_Wuhan_2019            ................................................................................ [ 17360]
MW396979.1_Timor-Leste/2019/1    ................................................................................ [ 17360]
MK940252.1_CN/2019/InnerMongolia ................................................................................ [ 17360]
MK128995.1_China/2018/AnhuiXCGQ  ................................................................................ [ 17360]
MH681419.1_ASFV/POL/2015         ................................................................................ [ 17360]
LS478113.1_Estonia_2014          ...t....c.--....cc.ag....a.a...----------------------------------------..aat...a [ 17360]
KP843857.1_Odintsovo_02/14       ................................................................................ [ 17360]
FN557520.1_strain_E75            ------------------------------------------....t.t.ca.ac..t...at.a.-------------- [ 17360]
AY261366.1_Warthog               ................................................................................ [ 17360]
AY261365.1_Warmbaths             ................................................................................ [ 17360]
AY261364.1_Tengani_62            .....................g.......................................................... [ 17360]
AY261363.1_Pretoriuskop/96/4     ................................................................................ [ 17360]
AY261362.1_Mkuzi_1979            .....................................................................t.......... [ 17360]
AY261361.1_Malawi_Lil-20/1_1983  ..............................................t.t.ca..c..t...a.g......---------- [ 17360]
AY261360.1_Kenya_1950            ......a...................................................tc..t................. [ 17360]
AM712240.1_OURT_88/3             ................................................................................ [ 17360]
AM712239.1_Benin_97/1            ------------------------------------------....t.t.ca.ac..t...at.a.-------------- [ 17360]
KM111295.1_Ken06.Bus             ..........................................................tc..t....a............ [ 17360]
KM111294.1_Ken05/Tk1             .............................................a............tc..t................. [ 17360]
KM262844.1_strain_L60            ------------------------------------------....t.t.ca.ac..t...at.a.-------------- [ 17360]
KM102979.1_26544/OG10_Italy      ------------------------------------------....t.t.ca.ac..t...at.a.-------------- [ 17360]
KM262845.1_strain_NHV            ................................................................................ [ 17360]
KX354450.1_47/Ss/2008            ------------------------------------------....t.t.ca.ac..t...at.a.-------------- [ 17360]
KP055815.1_strain_BA71           -------------------------------------------------------------------------------- [ 17360]
MH025920.1_strain_R35            ..........................................................tc..t....a............ [ 17360]
MH025919.1_strain_N10            ..........................................................tc..t....a............ [ 17360]
MH025918.1_strain_R25            ..........................................................tc..t....a............ [ 17360]
MH025917.1_strain_R7             ..........................................................tc..t....a............ [ 17360]
MH025916.1_strain_R8             ..........................................................tc..t....a............ [ 17360]

FR682468.2_Georgia_2007/1        gcatcggattatctgtggaaagaggtaggttcattattatgttcgtcatcaaaattcctaaaagaacatagaagccaaga [ 17440]
ASF_IND_20_CAD543                -------------------------------------------------------------------------------- [ 17440]
LR899193.1_Germany_2020/1        ................................................................................ [ 17440]
MN393477.1_Wuhan_2019            ................................................................................ [ 17440]
MW396979.1_Timor-Leste/2019/1    ................................................................................ [ 17440]
MK940252.1_CN/2019/InnerMongolia ................................................................................ [ 17440]
MK128995.1_China/2018/AnhuiXCGQ  ................................................................................ [ 17440]
MH681419.1_ASFV/POL/2015         ................................................................................ [ 17440]
LS478113.1_Estonia_2014          ......t..a.c.---------------....------------------------------------------------ [ 17440]
KP843857.1_Odintsovo_02/14       ................................................................................ [ 17440]
```

```
FN557520.1_strain_E75           ---------..ca...a.g..at...caa..---------a...aaa..tggctgaga...c.a...ca.a.tattc.a. [ 17440]
AY261366.1_Warthog              ...........................................a............................g....... [ 17440]
AY261365.1_Warmbaths            ........................................................................g....... [ 17440]
AY261364.1_Tengani_62           ...........................a............................................g....... [ 17440]
AY261363.1_Pretoriuskop/96/4    ...........................................a............................g....... [ 17440]
AY261362.1_Mkuzi_1979           .....................................g..................................tt...... [ 17440]
AY261361.1_Malawi_Lil-20/1_1983 -------------------------------------------------------------------------------- [ 17440]
AY261360.1_Kenya_1950           .........................c...........g..a..a........t.....c.......g.....tt.....g [ 17440]
AM712240.1_OURT_88/3            ..................t.....................................................g....... [ 17440]
AM712239.1_Benin_97/1           ---------..ca...a.g..at...caa..---------a...aaa..tggctgaga...c.a...ca.a.tattc.a. [ 17440]
KM111295.1_Ken06.Bus            .........................c...........g.....a........t.....c.......g.....tt.....g [ 17440]
KM111294.1_Ken05/Tk1            .........................c...........g..a..a........t.....c.......g.....tt.....g [ 17440]
KM262844.1_strain_L60           ---------..ca...a.g..at...caa..---------a...aaa..tggctgaga...c.a...ca.a.tattc.a. [ 17440]
KM102979.1_26544/OG10_Italy     ---------..ca...a.g..at...caa..---------a...aaa..tggctgaga...c.a...ca.a.tattc.a. [ 17440]
KM262845.1_strain_NHV           ..................t.....................................................g....... [ 17440]
KX354450.1_47/Ss/2008           ---------..ca...a.g..at...caa..---------a...aaa..tggctgaga...c.a...ca.a.tattc.a. [ 17440]
KP055815.1_strain_BA71          -------------------------------------------------------------ct.g.a.a.a.ttttc.c. [ 17440]
MH025920.1_strain_R35           .....................................g.....a........t.....c.......g.....tt.....g [ 17440]
MH025919.1_strain_N10           .....................................g.....a........t.....c.......g.....tt.....g [ 17440]
MH025918.1_strain_R25           .....................................g.....a........t.....c.......g.....tt.....g [ 17440]
MH025917.1_strain_R7            .....................................g.....a........t.....c.......g.....tt.....g [ 17440]
MH025916.1_strain_R8            .....................................g.....a........t.....c.......g.....tt.....g [ 17440]

FR682468.2_Georgia_2007/1       aagataagcagtcttgtagcggcttgcattc--gcattcgtgagtattgtttgcg-aacatagcttatgagagcaatggt [ 17520]
ASF_IND_20_CAD543               -------------------------------------------------------------------------------- [ 17520]
LR899193.1_Germany_2020/1       ...............................--......................-........................ [ 17520]
MN393477.1_Wuhan_2019           ...............................--......................-........................ [ 17520]
MW396979.1_Timor-Leste/2019/1   ...............................--......................-........................ [ 17520]
MK940252.1_CN/2019/InnerMongolia ...............................--......................-........................ [ 17520]
MK128995.1_China/2018/AnhuiXCGQ ...............................--......................-........................ [ 17520]
MH681419.1_ASFV/POL/2015        ...............................--......................-........................ [ 17520]
LS478113.1_Estonia_2014         -----------------...at....acc..--aaggc..c.caac.gacc...-------------------------- [ 17520]
KP843857.1_Odintsovo_02/14      ...............................--......................-........................ [ 17520]
FN557520.1_strain_E75           .gc.....t.a.t.ca.g.tt.taca..g..gtag...tt..cag..c.caaat.c...gc.a.cagca.at....a.c. [ 17520]
AY261366.1_Warthog              ............................g.t--..........a..c........-..t.....c......g........ [ 17520]
AY261365.1_Warmbaths            .................g.............--......................-........c......g........ [ 17520]
AY261364.1_Tengani_62           ...............................--......................-........c......g........ [ 17520]
AY261363.1_Pretoriuskop/96/4    ............................g.t--..........a..c........-..t.....c......g........ [ 17520]
AY261362.1_Mkuzi_1979           ...............................--..........a...........-..tg...acc.....g........ [ 17520]
AY261361.1_Malawi_Lil-20/1_1983 -------------------------------------------------------------------------------- [ 17520]
AY261360.1_Kenya_1950           .....g.........................--......................-..tg...acc.....g........ [ 17520]
AM712240.1_OURT_88/3            ...............................--..........a..c........-........c......g.....a.. [ 17520]
AM712239.1_Benin_97/1           .gc.....t.a.t.ca.g.tt.taca..g..gtag...tt..cag..c.caaat.c...gc.a.cagca.at....a.c. [ 17520]
KM111295.1_Ken06.Bus            ...............................--......................-..tg...acc.............. [ 17520]
```

```
KM111294.1_Ken05/Tk1             ...............................--......................-..tg...acc.............. [ 17520]
KM262844.1_strain_L60            .gc.....t.a.t.ca.g.tt.taca..g..gtag...tt..cag..c.caaat.c...gc.a.cagca.at....a.c. [ 17520]
KM102979.1_26544/OG10_Italy      .gc.....t.a.t.ca.g.tt.taca..g..gtag...tt..cag..c.caaat.c...gc.a.cagca.at....a.c. [ 17520]
KM262845.1_strain_NHV            ...............................--..........a..c........-........c......g.....a.. [ 17520]
KX354450.1_47/Ss/2008            .gc.....t.a.t.ca.g.tt.taca..g..gtag...tt..cag..c.caaat.c...gc.a.cagca.at....a.c. [ 17520]
KP055815.1_strain_BA71           ..tc...tt.-----------------------aa..c.tcacag..c-------------------------------- [ 17520]
MH025920.1_strain_R35            ...............................--......................-..tg...acc.............. [ 17520]
MH025919.1_strain_N10            ...............................--......................-..tg...acc.............. [ 17520]
MH025918.1_strain_R25            ...............................--......................-..tg...acc.............. [ 17520]
MH025917.1_strain_R7             ...............................--......................-..tg...acc.............. [ 17520]
MH025916.1_strain_R8             ...............................--......................-..tg...acc.............. [ 17520]

FR682468.2_Georgia_2007/1        agctatcatacaaagacaagtatgtttgatattctcagtgtcaatgaccctatcctcctttatttgcattaactcatcaa [ 17600]
ASF_IND_20_CAD543                -------------------------------------------------------------------------------- [ 17600]
LR899193.1_Germany_2020/1        ................................................................................ [ 17600]
MN393477.1_Wuhan_2019            ................................................................................ [ 17600]
MW396979.1_Timor-Leste/2019/1    ................................................................................ [ 17600]
MK940252.1_CN/2019/InnerMongolia ................................................................................ [ 17600]
MK128995.1_China/2018/AnhuiXCGQ  ................................................................................ [ 17600]
MH681419.1_ASFV/POL/2015         ................................................................................ [ 17600]
LS478113.1_Estonia_2014          ------------------------c...gg.....g.------------------------------------------- [ 17600]
KP843857.1_Odintsovo_02/14       ................................................................................ [ 17600]
FN557520.1_strain_E75            .aa..ca.ggt...a....ta..acc.t...a.---------g..tga..a..t..ta.--------------------- [ 17600]
AY261366.1_Warthog               .....................................ac.........t.............c................. [ 17600]
AY261365.1_Warmbaths             .....................................ac.........t.............c................. [ 17600]
AY261364.1_Tengani_62            ...............g................................................................ [ 17600]
AY261363.1_Pretoriuskop/96/4     ...............g...........a....................t.............c......c.......... [ 17600]
AY261362.1_Mkuzi_1979            ...........................t.........ac........tt.........c...c................. [ 17600]
AY261361.1_Malawi_Lil-20/1_1983  -------------------------------------------------------------------------------- [ 17600]
AY261360.1_Kenya_1950            .....c.....................t.........aca.......tt.............c................. [ 17600]
AM712240.1_OURT_88/3             ...............g.....................ac..............t........c................. [ 17600]
AM712239.1_Benin_97/1            .aa..ca.ggt...a....ta..acc.t...a.---------g..tga..a..t..ta.--------------------- [ 17600]
KM111295.1_Ken06.Bus             .....c.....................t.........ac........tt.........c...c................. [ 17600]
KM111294.1_Ken05/Tk1             .....c.....................t.........ac.........tt........c...c................. [ 17600]
KM262844.1_strain_L60            .aa..ca.ggt...a....ta..acc.t...a.---------g..tga..a..t..ta.--------------------- [ 17600]
KM102979.1_26544/OG10_Italy      .aa..ca.ggt...a....ta..acc.t...a.---------g..tga..a..t..ta.--------------------- [ 17600]
KM262845.1_strain_NHV            ...............g.....................ac..............t........c................. [ 17600]
KX354450.1_47/Ss/2008            .aa..ca.ggt...a....ta..acc.t...a.---------g..tga..a..t..ta.--------------------- [ 17600]
KP055815.1_strain_BA71           ----------------------------------------------------.t.ctt.--------------------- [ 17600]
MH025920.1_strain_R35            .....c.....................t.........ac........tt.........c...c................. [ 17600]
MH025919.1_strain_N10            .....c.....................t.........ac........tt.........c...c................. [ 17600]
MH025918.1_strain_R25            .....c.....................t.........ac........tt.........c...c................. [ 17600]
MH025917.1_strain_R7             .....c.....................t.........ac........tt.........c...c................. [ 17600]
MH025916.1_strain_R8             .....c.....................t.........ac........tt.........c...c................. [ 17600]
```

```
FR682468.2_Georgia_2007/1        accaatcataatatgtgggatttgtacagctcatgatgtgaaagcggcgtatcctagagtctgtaaagtagctacatctt [ 17680]
ASF_IND_20_CAD543                -------------------------------------------------------------------------------- [ 17680]
LR899193.1_Germany_2020/1        ................................................................................ [ 17680]
MN393477.1_Wuhan_2019            ................................................................................ [ 17680]
MW396979.1_Timor-Leste/2019/1    ................................................................................ [ 17680]
MK940252.1_CN/2019/InnerMongolia ................................................................................ [ 17680]
MK128995.1_China/2018/AnhuiXCGQ  ................................................................................ [ 17680]
MH681419.1_ASFV/POL/2015         ................................................................................ [ 17680]
LS478113.1_Estonia_2014          ------.ga....g...at..cc.-------------------------------------------------------g [ 17680]
KP843857.1_Odintsovo_02/14       ................................................................................ [ 17680]
FN557520.1_strain_E75            -------------------------------------------------------------------------------- [ 17680]
AY261366.1_Warthog               ...............................................................................a [ 17680]
AY261365.1_Warmbaths             ................................................................................ [ 17680]
AY261364.1_Tengani_62            ...................................................................a............ [ 17680]
AY261363.1_Pretoriuskop/96/4     .........................................................t...................... [ 17680]
AY261362.1_Mkuzi_1979            ..............................................t.t....t.......................t.. [ 17680]
AY261361.1_Malawi_Lil-20/1_1983  -------------------------------------------------------------------------------- [ 17680]
AY261360.1_Kenya_1950            ..............................................t.t....t.......................t.. [ 17680]
AM712240.1_OURT_88/3             ...............................................................................a [ 17680]
AM712239.1_Benin_97/1            -------------------------------------------------------------------------------- [ 17680]
KM111295.1_Ken06.Bus             ....g.........................................t.t....t................t......t.. [ 17680]
KM111294.1_Ken05/Tk1             .......................t......................t.t....t.......................t.. [ 17680]
KM262844.1_strain_L60            -------------------------------------------------------------------------------- [ 17680]
KM102979.1_26544/OG10_Italy      -------------------------------------------------------------------------------- [ 17680]
KM262845.1_strain_NHV            ...............................................................................a [ 17680]
KX354450.1_47/Ss/2008            -------------------------------------------------------------------------------- [ 17680]
KP055815.1_strain_BA71           -------------------------------------------------------------------------------- [ 17680]
MH025920.1_strain_R35            ....g.........................................t.t....t................t......t.. [ 17680]
MH025919.1_strain_N10            ....g.........................................t.t....t................t......t.. [ 17680]
MH025918.1_strain_R25            ....g.........................................t.t....t................t......t.. [ 17680]
MH025917.1_strain_R7             ....g.........................................t.t....t................t......t.. [ 17680]
MH025916.1_strain_R8             ....g.........................................t.t....t................t......t.. [ 17680]

FR682468.2_Georgia_2007/1        tcattatagcgagaaaccctacatatttgtatgtaatcattttttttgatgagagggtgtttttcaaaaaccttattttt [ 17760]
ASF_IND_20_CAD543                -------------------------------------------------------------------------------- [ 17760]
LR899193.1_Germany_2020/1        ................................................................................ [ 17760]
MN393477.1_Wuhan_2019            ................................................................................ [ 17760]
MW396979.1_Timor-Leste/2019/1    ................................................................................ [ 17760]
MK940252.1_CN/2019/InnerMongolia ................................................................................ [ 17760]
MK128995.1_China/2018/AnhuiXCGQ  ................................................................................ [ 17760]
MH681419.1_ASFV/POL/2015         ................................................................................ [ 17760]
LS478113.1_Estonia_2014          .t.aa...t.a...ggg...--------------------------------......---------------------- [ 17760]
KP843857.1_Odintsovo_02/14       ................................................................................ [ 17760]
```

```
FN557520.1_strain_E75            ----------------....c...t...cc...----------c.a......ttaa..a....----------------- [ 17760]
AY261366.1_Warthog               ...........g............................................a....................... [ 17760]
AY261365.1_Warmbaths             ...........g............................................a....................... [ 17760]
AY261364.1_Tengani_62            ...........g............................................a....................... [ 17760]
AY261363.1_Pretoriuskop/96/4     ...........g............................................a....................... [ 17760]
AY261362.1_Mkuzi_1979            ...cc......g..............c.......................a.....a.............t......... [ 17760]
AY261361.1_Malawi_Lil-20/1_1983  -------------------------------------------------------------------------------- [ 17760]
AY261360.1_Kenya_1950            ...cc......g..............c.............................a....................... [ 17760]
AM712240.1_OURT_88/3             .t.........g............................................a....................... [ 17760]
AM712239.1_Benin_97/1            ----------------....c...t...cc...----------c.a......ttaa..a....----------------- [ 17760]
KM111295.1_Ken06.Bus             ...cc......g..............c.............................a....................... [ 17760]
KM111294.1_Ken05/Tk1             ...cc......g..............c.............................a....................... [ 17760]
KM262844.1_strain_L60            ----------------....c...t...cc...----------c.a......ttaa..a....----------------- [ 17760]
KM102979.1_26544/OG10_Italy      ----------------....c...t...cc...----------c.a......ttaa..a....----------------- [ 17760]
KM262845.1_strain_NHV            .t.........g............................................a....................... [ 17760]
KX354450.1_47/Ss/2008            ----------------....c...t...cc...----------c.a......ttaa..a....----------------- [ 17760]
KP055815.1_strain_BA71           ----------------t.a..a..t..cc--------------------------------------------------- [ 17760]
MH025920.1_strain_R35            ...cc......g..............c.............................a....................... [ 17760]
MH025919.1_strain_N10            ...cc......g..............c.............................a....................... [ 17760]
MH025918.1_strain_R25            ...cc......g..............c.............................a....................... [ 17760]
MH025917.1_strain_R7             ...cc......g..............c.............................a....................... [ 17760]
MH025916.1_strain_R8             ...cc......g..............c.............................a....................... [ 17760]

FR682468.2_Georgia_2007/1        acaaaccccgtgtcgacaattccagcagaagtcacacgattttgcataggtgcaccaatactcaagctctctctttggag [ 17840]
ASF_IND_20_CAD543                -------------------------------------------------------------------------------- [ 17840]
LR899193.1_Germany_2020/1        ................................................................................ [ 17840]
MN393477.1_Wuhan_2019            ................................................................................ [ 17840]
MW396979.1_Timor-Leste/2019/1    ................................................................................ [ 17840]
MK940252.1_CN/2019/InnerMongolia ................................................................................ [ 17840]
MK128995.1_China/2018/AnhuiXCGQ  ................................................................................ [ 17840]
MH681419.1_ASFV/POL/2015         ................................................................................ [ 17840]
LS478113.1_Estonia_2014          -------t.ta..a...gg..aagaag.gg.c..g..ag.acc---------------.c....ga..gc.ca.c..ag- [ 17840]
KP843857.1_Odintsovo_02/14       ................................................................................ [ 17840]
FN557520.1_strain_E75            ---------------------------------.agt...a..t....aaaa.....----------------------- [ 17840]
AY261366.1_Warthog               g.......................a.............................................t......... [ 17840]
AY261365.1_Warmbaths             g........a............................................................t......... [ 17840]
AY261364.1_Tengani_62            .......................................................................c........ [ 17840]
AY261363.1_Pretoriuskop/96/4     ........................a.............................................t......... [ 17840]
AY261362.1_Mkuzi_1979            g....t...........................g.....................................c........ [ 17840]
AY261361.1_Malawi_Lil-20/1_1983  -------------------------------------------------------------------------------- [ 17840]
AY261360.1_Kenya_1950            g.................g.................a...c...............................t.....g. [ 17840]
AM712240.1_OURT_88/3             g....t...........................g.....................................c........ [ 17840]
AM712239.1_Benin_97/1            ---------------------------------.agt...a..t....aaaa.....----------------------- [ 17840]
KM111295.1_Ken06.Bus             g.................g.c...............a...c...............................t....... [ 17840]
```

```
KM111294.1_Ken05/Tk1          g.................g.c...............a...c...............................t.....g. [ 17840]
KM262844.1_strain_L60         ---------------------------------.agt...a..t....aaaa....------------------------ [ 17840]
KM102979.1_26544/OG10_Italy   ---------------------------------.agt...a..t....aaaa....------------------------ [ 17840]
KM262845.1_strain_NHV         g....t...........................g.....................................c........ [ 17840]
KX354450.1_47/Ss/2008         ---------------------------------.agt...a..t....aaaa....------------------------ [ 17840]
KP055815.1_strain_BA71        -------------------------------------------------------------------------------- [ 17840]
MH025920.1_strain_R35         g.................g.c...............a...c...............................t....... [ 17840]
MH025919.1_strain_N10         g.................g.c...............a...c...............................t....... [ 17840]
MH025918.1_strain_R25         g.................g.c...............a...c...............................t....... [ 17840]
MH025917.1_strain_R7          g.................g.c...............a...c...............................t....... [ 17840]
MH025916.1_strain_R8          g.................g.c...............a...c...............................t....... [ 17840]

FR682468.2_Georgia_2007/1     gtctccgggtcattggtaactctcctgttcctggaaaagattggctttgaatgaccggctgcatgaccgccagtaccaaa [ 17920]
ASF_IND_20_CAD543             -------------------------------------------------------------------------------- [ 17920]
LR899193.1_Germany_2020/1     ................................................................................ [ 17920]
MN393477.1_Wuhan_2019         ................................................................................ [ 17920]
MW396979.1_Timor-Leste/2019/1 ................................................................................ [ 17920]
MK940252.1_CN/2019/InnerMongolia ................................................................................ [ 17920]
MK128995.1_China/2018/AnhuiXCGQ ................................................................................ [ 17920]
MH681419.1_ASFV/POL/2015      ................................................................................ [ 17920]
LS478113.1_Estonia_2014       ----------...aa.a.gtc...ta.c....---t.gt.......c-------------------------------g. [ 17920]
KP843857.1_Odintsovo_02/14    ................................................................................ [ 17920]
FN557520.1_strain_E75         ----.t....tg.....g.t.gata...----aag..t..ggatgg......a...-----------------..gt... [ 17920]
AY261366.1_Warthog            .......................t....c..........g........................................ [ 17920]
AY261365.1_Warmbaths          .......................t....c........g.g........................................ [ 17920]
AY261364.1_Tengani_62         ............................c........g.g........................................ [ 17920]
AY261363.1_Pretoriuskop/96/4  ............................c..........g..............................t......... [ 17920]
AY261362.1_Mkuzi_1979         .......................t.............g.g................................a....... [ 17920]
AY261361.1_Malawi_Lil-20/1_1983 -------------------------------------------------------------------------------- [ 17920]
AY261360.1_Kenya_1950         .........................c..c........g.g....t...........c.......tt..ag.......... [ 17920]
AM712240.1_OURT_88/3          .......................t.............g.g................................a....... [ 17920]
AM712239.1_Benin_97/1         ----.t....tg.....g.t.gata...----aag..t..ggatgg......a...-----------------..gt... [ 17920]
KM111295.1_Ken06.Bus          .........................c..c........g.g................c......ctt..at.......... [ 17920]
KM111294.1_Ken05/Tk1          .........................c..c........g.g................c......ctt..a........... [ 17920]
KM262844.1_strain_L60         ----.t....tg.....g.t.gata...----aag..t..ggatgg......a...-----------------..gt... [ 17920]
KM102979.1_26544/OG10_Italy   ----.t....tg.....g.t.gata...----aag..t..ggatgg......a...-----------------..gt... [ 17920]
KM262845.1_strain_NHV         .......................t.............g.g................................a....... [ 17920]
KX354450.1_47/Ss/2008         ----.t....tg.....g.t.gata...----aag..t..ggatgg......a...-----------------..gt... [ 17920]
KP055815.1_strain_BA71        ------------..ccc..tcggtg..g----acg.tt.tga..tac..g..---------------------------- [ 17920]
MH025920.1_strain_R35         .........................c..c........g.g................c......ctt..at.......... [ 17920]
MH025919.1_strain_N10         .........................c..c........g.g................c......ctt..at.......... [ 17920]
MH025918.1_strain_R25         .........................c..c........g.g................c......ctt..at.......... [ 17920]
MH025917.1_strain_R7          .........................c..c........g.g................c......ctt..at.......... [ 17920]
MH025916.1_strain_R8          .........................c..c........g.g................c......ctt..at.......... [ 17920]
```

```
FR682468.2_Georgia_2007/1        aggaacacaatcaccttcatggctgcaacttataag----------------------------ttgcaacttatgggtt [ 18000]
ASF_IND_20_CAD543                -------------------------------------------------------------------------------- [ 18000]
LR899193.1_Germany_2020/1        ....................................----------------------------................ [ 18000]
MN393477.1_Wuhan_2019            ....................................----------------------------................ [ 18000]
MW396979.1_Timor-Leste/2019/1    ....................................----------------------------................ [ 18000]
MK940252.1_CN/2019/InnerMongolia ....................................----------------------------................ [ 18000]
MK128995.1_China/2018/AnhuiXCGQ  ....................................----------------------------................ [ 18000]
MH681419.1_ASFV/POL/2015         ....................................----------------------------................ [ 18000]
LS478113.1_Estonia_2014          .a...gg...ctg..--------------------------------------------------c..tg..g.g....a [ 18000]
KP843857.1_Odintsovo_02/14       ....................................----------------------------................ [ 18000]
FN557520.1_strain_E75            ..tcct.g..aa.........----------------------------------------------------------- [ 18000]
AY261366.1_Warthog               ....gt............................g.----------------------------...........a.... [ 18000]
AY261365.1_Warmbaths             ....gt.......................g....g.----------------------------................ [ 18000]
AY261364.1_Tengani_62            ....................................----------------------------................ [ 18000]
AY261363.1_Pretoriuskop/96/4     ....gt..............................----------------------------...............c [ 18000]
AY261362.1_Mkuzi_1979            ....g......................c......g.----------------------------....------------ [ 18000]
AY261361.1_Malawi_Lil-20/1_1983  -------------------------------------------------------------------------------- [ 18000]
AY261360.1_Kenya_1950            ..................................g.ttgcaacttataggttgcgacttataga....g......a.a.. [ 18000]
AM712240.1_OURT_88/3             ....g......................----------------------------------------------------- [ 18000]
AM712239.1_Benin_97/1            ..tcct.g..aa.........----------------------------------------------------------- [ 18000]
KM111295.1_Ken06.Bus             ......................------------------------------------------......tg...a.a.. [ 18000]
KM111294.1_Ken05/Tk1             ..................................g.tt--------------------------g..........a.... [ 18000]
KM262844.1_strain_L60            ..tcct.g..aa.........----------------------------------------------------------- [ 18000]
KM102979.1_26544/OG10_Italy      ..tcct.g..aa.........----------------------------------------------------------- [ 18000]
KM262845.1_strain_NHV            ....g......................----------------------------------------------------- [ 18000]
KX354450.1_47/Ss/2008            ..tcct.g..aa.........----------------------------------------------------------- [ 18000]
KP055815.1_strain_BA71           -----t.g..ca.--.c...a----------------------------------------------------------- [ 18000]
MH025920.1_strain_R35            ......................------------------------------------------......tg...a.a.. [ 18000]
MH025919.1_strain_N10            ......................------------------------------------------......tg...a.a.. [ 18000]
MH025918.1_strain_R25            ......................------------------------------------------......tg...a.a.. [ 18000]
MH025917.1_strain_R7             ......................------------------------------------------......tg...a.a.. [ 18000]
MH025916.1_strain_R8             ......................------------------------------------------......tg...a.a.. [ 18000]

FR682468.2_Georgia_2007/1        gcaat-------actgcaacgtataggt--tgcaccttatagatcgcgactcaaaaggtatgaaaaccttaccctcaata [ 18080]
ASF_IND_20_CAD543                -------------------------------------------------------------------------------- [ 18080]
LR899193.1_Germany_2020/1        .....-------................--.................................................. [ 18080]
MN393477.1_Wuhan_2019            .....-------................--.................................................. [ 18080]
MW396979.1_Timor-Leste/2019/1    .....-------................--.................................................. [ 18080]
MK940252.1_CN/2019/InnerMongolia .....-------................--.................................................. [ 18080]
MK128995.1_China/2018/AnhuiXCGQ  .....-------................--.................................................. [ 18080]
MH681419.1_ASFV/POL/2015         .....-------................--.................................................. [ 18080]
LS478113.1_Estonia_2014          ag..c-------c.g.agc..---------------c.gc..g...---.ct.gc..cgt..cc..tac..tttct..c. [ 18080]
KP843857.1_Odintsovo_02/14       .....-------................--.................................................. [ 18080]
```

```
FN557520.1_strain_E75           -------------------------------------------------------------------------------- [ 18080]
AY261366.1_Warthog              ....-------------------------------............................................. [ 18080]
AY261365.1_Warmbaths            .....-------g...............--.................................................. [ 18080]
AY261364.1_Tengani_62           .....-------................--..............t.......................g....------- [ 18080]
AY261363.1_Pretoriuskop/96/4    .....-------g...............--............g.t.................-................. [ 18080]
AY261362.1_Mkuzi_1979           -----------------.....g.....--...g.............a..............g.....a....t...... [ 18080]
AY261361.1_Malawi_Lil-20/1_1983 -------------------------------------------------------------------------------- [ 18080]
AY261360.1_Kenya_1950           ....cttataggg......t........--...ga......a....................g........t........ [ 18080]
AM712240.1_OURT_88/3            ----------------------------------..g........................................... [ 18080]
AM712239.1_Benin_97/1           -------------------------------------------------------------------------------- [ 18080]
KM111295.1_Ken06.Bus            ....cttataggg......t......a.--...ga......a....................g................. [ 18080]
KM111294.1_Ken05/Tk1            .-------------......t.......tg....a.........t...............c.g................. [ 18080]
KM262844.1_strain_L60           -------------------------------------------------------------------------------- [ 18080]
KM102979.1_26544/OG10_Italy     -------------------------------------------------------------------------------- [ 18080]
KM262845.1_strain_NHV           ----------------------------------..g........................................... [ 18080]
KX354450.1_47/Ss/2008           -------------------------------------------------------------------------------- [ 18080]
KP055815.1_strain_BA71          -------------------------------------------------------------------------------- [ 18080]
MH025920.1_strain_R35           ....cttataggg......t......a.--...ga......a....................g................. [ 18080]
MH025919.1_strain_N10           ....cttataggg......t......a.--...ga......a....................g................. [ 18080]
MH025918.1_strain_R25           ....cttataggg......t......a.--...ga......a....................g................. [ 18080]
MH025917.1_strain_R7            ....cttataggg......t......a.--...ga......a....................g................. [ 18080]
MH025916.1_strain_R8            ....cttataggg......t......a.--...ga......a....................g................. [ 18080]

FR682468.2_Georgia_2007/1       cagaatttaagttttaatcctgataatgtatc----tgtttatgaaaaaaaattttttttactcatgtatgaattcttat [ 18160]
ASF_IND_20_CAD543               -------------------------------------------------------------------------------- [ 18160]
LR899193.1_Germany_2020/1       ................................----............................................ [ 18160]
MN393477.1_Wuhan_2019           ................................----............................................ [ 18160]
MW396979.1_Timor-Leste/2019/1   ................................----............................................ [ 18160]
MK940252.1_CN/2019/InnerMongolia ................................----............................................ [ 18160]
MK128995.1_China/2018/AnhuiXCGQ ................................----............................................ [ 18160]
MH681419.1_ASFV/POL/2015        ................................----............................................ [ 18160]
LS478113.1_Estonia_2014         t-----------------------------c.----...a..ga........---------------------------- [ 18160]
KP843857.1_Odintsovo_02/14      ................................----............................................ [ 18160]
FN557520.1_strain_E75           -----------------.tgc.t.c..acca.caga....at.-.........a..-----------------------. [ 18160]
AY261366.1_Warthog              ..................t.............----............................................ [ 18160]
AY261365.1_Warmbaths            ..................t.............----............................................ [ 18160]
AY261364.1_Tengani_62           -------------------------------------------------------------------------------- [ 18160]
AY261363.1_Pretoriuskop/96/4    ....g.....a......a...a.....a....----.t..............a....----acgc.ac..a......... [ 18160]
AY261362.1_Mkuzi_1979           ....g...........................----.....................--......g.....g...t.... [ 18160]
AY261361.1_Malawi_Lil-20/1_1983 -------------------------------------------------------------------------------- [ 18160]
AY261360.1_Kenya_1950           ....g.....a...........g.........----................a-.......................... [ 18160]
AM712240.1_OURT_88/3            t...g...........................----.......a............c---.................... [ 18160]
AM712239.1_Benin_97/1           -----------------.tgc.t.c..acca.caga....at.a.........a..-----------------------. [ 18160]
KM111295.1_Ken06.Bus            ....g.....a........t..g.........----................a........................... [ 18160]
```

```
KM111294.1_Ken05/Tk1             ....g.....a....................t----................a........................... [ 18160]
KM262844.1_strain_L60            -----------------.tgc.t.c..acca.caga....at.-.........a..-----------------------. [ 18160]
KM102979.1_26544/OG10_Italy      -----------------.tgc.t.c..acca.caga....at.a.........a..-----------------------. [ 18160]
KM262845.1_strain_NHV            t...g...........................----.......a............c---.................... [ 18160]
KX354450.1_47/Ss/2008            -----------------.tgc.t.c..acca.caga....at.a.........a..-----------------------. [ 18160]
KP055815.1_strain_BA71           -----------------a.a..g.....cc------ca...cc------------------------------------- [ 18160]
MH025920.1_strain_R35            ....g.....a........t..g.........----................a........................... [ 18160]
MH025919.1_strain_N10            ....g.....a........t..g.........----................a........................... [ 18160]
MH025918.1_strain_R25            ....g.....a........t..g.........----................a........................... [ 18160]
MH025917.1_strain_R7             ....g.....a........t..g.........----................a........................... [ 18160]
MH025916.1_strain_R8             ....g.....a........t..g.........----................a........................... [ 18160]

FR682468.2_Georgia_2007/1        acgaatcataatatgtaggctgagaataata------attcatatacggtgttgcgggctcaataaaaattttgttacca [ 18240]
ASF_IND_20_CAD543                -------------------------------------------------------------------------------- [ 18240]
LR899193.1_Germany_2020/1        ...............................------........................................... [ 18240]
MN393477.1_Wuhan_2019            ...............................------........................................... [ 18240]
MW396979.1_Timor-Leste/2019/1    ...............................------........................................... [ 18240]
MK940252.1_CN/2019/InnerMongolia ...............................------........................................... [ 18240]
MK128995.1_China/2018/AnhuiXCGQ  ...............................------........................................... [ 18240]
MH681419.1_ASFV/POL/2015         ...............................------........................................... [ 18240]
LS478113.1_Estonia_2014          -------------------------------------------------------------------------------g [ 18240]
KP843857.1_Odintsovo_02/14       ...............................------........................................... [ 18240]
FN557520.1_strain_E75            .aatt.t.c.g.....gtaa.ac--------------------------------------c.c......a..c...tat [ 18240]
AY261366.1_Warthog               ..........................c...tc---------......aag.............................. [ 18240]
AY261365.1_Warmbaths             ..................a....a..c...tc---------........g.............................. [ 18240]
AY261364.1_Tengani_62            -------------------------------------------------------------------------------- [ 18240]
AY261363.1_Pretoriuskop/96/4     .tt..c.......------------------------------------------------------------------- [ 18240]
AY261362.1_Mkuzi_1979            ..........................c...tcatata..ga......-......t............c............ [ 18240]
AY261361.1_Malawi_Lil-20/1_1983  -------------------------------------------------------------------------------- [ 18240]
AY261360.1_Kenya_1950            ...---------------------------------------------......................g......... [ 18240]
AM712240.1_OURT_88/3             ..............................t----------..............a...-tc..t............... [ 18240]
AM712239.1_Benin_97/1            .aatt.t.c.g.....gtaa.ac--------------------------------------c.c......a..c...tat [ 18240]
KM111295.1_Ken06.Bus             ...---------------------------------------------......................g......... [ 18240]
KM111294.1_Ken05/Tk1             .........c................g...tcatata..gg.....-c......t............c..........at [ 18240]
KM262844.1_strain_L60            .aatt.t.c.g.....gtaa.ac--------------------------------------c.c......a..c...tat [ 18240]
KM102979.1_26544/OG10_Italy      .aatt.t.c.g.....gtaa.ac--------------------------------------c.c......a..c...tat [ 18240]
KM262845.1_strain_NHV            ..............................t----------..............a...-tc..t............... [ 18240]
KX354450.1_47/Ss/2008            .aatt.t.c.g.....gtaa.ac--------------------------------------c.c......a..c...tat [ 18240]
KP055815.1_strain_BA71           ----t...ct......gca...---------------------------------------------------....tac [ 18240]
MH025920.1_strain_R35            ...---------------------------------------------......................g......... [ 18240]
MH025919.1_strain_N10            ...---------------------------------------------......................g......... [ 18240]
MH025918.1_strain_R25            ...---------------------------------------------......................g......... [ 18240]
MH025917.1_strain_R7             ...---------------------------------------------......................g......... [ 18240]
MH025916.1_strain_R8             ...---------------------------------------------......................g......... [ 18240]
```

```
FR682468.2_Georgia_2007/1        caaaaaat---aaatgctgga--tttttaagat--atatatctattaatgactaaaccctttatacgctgtaggctgaaa [ 18320]
ASF_IND_20_CAD543                -------------------------------------------------------------------------------- [ 18320]
LR899193.1_Germany_2020/1        ........---..........--..........--............................................. [ 18320]
MN393477.1_Wuhan_2019            ........---..........--..........--............................................. [ 18320]
MW396979.1_Timor-Leste/2019/1    ........---..........--..........--............................................. [ 18320]
MK940252.1_CN/2019/InnerMongolia ........---..........--..........--............................................. [ 18320]
MK128995.1_China/2018/AnhuiXCGQ  ........---..........--..........--............................................. [ 18320]
MH681419.1_ASFV/POL/2015         ........---..........--..........--............................................. [ 18320]
LS478113.1_Estonia_2014          a...ggt.---...att...g--a.....ttt.--g..---------------------..c.c..a..----------- [ 18320]
KP843857.1_Odintsovo_02/14       ........---..........--..........--............................................. [ 18320]
FN557520.1_strain_E75            t..cg..a---.t...tg.------------------------------------------------------------- [ 18320]
AY261366.1_Warthog               ........---..........--.......----c..............................t........t...g. [ 18320]
AY261365.1_Warmbaths             ........---..........--.......----c..............................t........t...g. [ 18320]
AY261364.1_Tengani_62            -------------------------------------------------------------------------------- [ 18320]
AY261363.1_Pretoriuskop/96/4     --------------------------------------------------------------------...t..ta..g. [ 18320]
AY261362.1_Mkuzi_1979            .......a--t..........--.....t.ag---.c........g................................g. [ 18320]
AY261361.1_Malawi_Lil-20/1_1983  -------------------------------------------------------------------------------- [ 18320]
AY261360.1_Kenya_1950            .......aaat......ca..--.....t.ag---.c.g......g..........t....c...a............g. [ 18320]
AM712240.1_OURT_88/3             ........---..........--.c........at..............................t..------------ [ 18320]
AM712239.1_Benin_97/1            t..cg..a---.t...tg.------------------------------------------------------------- [ 18320]
KM111295.1_Ken06.Bus             .......a-at......ca..--.....t.ag---.c........g..........t....c...a............g. [ 18320]
KM111294.1_Ken05/Tk1             ..c....a-at.......a..tt.....t.ag---.c........g..........t....c...a............g. [ 18320]
KM262844.1_strain_L60            t..cg..a---.t...tg.------------------------------------------------------------- [ 18320]
KM102979.1_26544/OG10_Italy      t..cg..a---.t...tg.------------------------------------------------------------- [ 18320]
KM262845.1_strain_NHV            ........---..........--.c........at..............................t..------------ [ 18320]
KX354450.1_47/Ss/2008            t..cg..a---.t...tg.------------------------------------------------------------- [ 18320]
KP055815.1_strain_BA71           ..gcg.ga---.-------------------------------------------------------------------- [ 18320]
MH025920.1_strain_R35            .......aaat......ca..--.....t.ag---.c........g..........t....c...a............g. [ 18320]
MH025919.1_strain_N10            .......aaat......ca..--.....t.ag---.c........g..........t....c...a............g. [ 18320]
MH025918.1_strain_R25            .......aaat......ca..--.....t.ag---.c........g..........t....c...a............g. [ 18320]
MH025917.1_strain_R7             .......aaat......ca..--.....t.ag---.c........g..........t....c...a............g. [ 18320]
MH025916.1_strain_R8             .......aaat......ca..--.....t.ag---.c........g..........t....c...a............g. [ 18320]

FR682468.2_Georgia_2007/1        acaatccatataatgaatatacggtgatttgggtttaataaaatacatacaacggtcaaaatagcgggcaatactacatt [ 18400]
ASF_IND_20_CAD543                -------------------------------------------------------------------------------- [ 18400]
LR899193.1_Germany_2020/1        ................................................................................ [ 18400]
MN393477.1_Wuhan_2019            ................................................................................ [ 18400]
MW396979.1_Timor-Leste/2019/1    ................................................................................ [ 18400]
MK940252.1_CN/2019/InnerMongolia ................................................................................ [ 18400]
MK128995.1_China/2018/AnhuiXCGQ  ................................................................................ [ 18400]
MH681419.1_ASFV/POL/2015         ................................................................................ [ 18400]
LS478113.1_Estonia_2014          ----------------------------------------------......aaa.a....cca.a.a------------ [ 18400]
KP843857.1_Odintsovo_02/14       ................................................................................ [ 18400]
```

```
FN557520.1_strain_E75             -------------------------------------------------------------------------------- [ 18400]
AY261366.1_Warthog                ..........................t..................................................... [ 18400]
AY261365.1_Warmbaths              ..........................t..................................................... [ 18400]
AY261364.1_Tengani_62             -------------------------------------------------------------------------------- [ 18400]
AY261363.1_Pretoriuskop/96/4      .....t.........g.....t...at...a..c.c..............................a............. [ 18400]
AY261362.1_Mkuzi_1979             ..........................t..................................................... [ 18400]
AY261361.1_Malawi_Lil-20/1_1983   -------------------------------------------------------------------------------- [ 18400]
AY261360.1_Kenya_1950             ...............t..........t.......................................a............. [ 18400]
AM712240.1_OURT_88/3              -------------------------------------------------------------------------------- [ 18400]
AM712239.1_Benin_97/1             -------------------------------------------------------------------------------- [ 18400]
KM111295.1_Ken06.Bus              ..........................t.............................................g....... [ 18400]
KM111294.1_Ken05/Tk1              ..........ag...t..........t..................................................... [ 18400]
KM262844.1_strain_L60             -------------------------------------------------------------------------------- [ 18400]
KM102979.1_26544/OG10_Italy       -------------------------------------------------------------------------------- [ 18400]
KM262845.1_strain_NHV             -------------------------------------------------------------------------------- [ 18400]
KX354450.1_47/Ss/2008             -------------------------------------------------------------------------------- [ 18400]
KP055815.1_strain_BA71            -------------------------------------------------------------------------------- [ 18400]
MH025920.1_strain_R35             ..........................t.............................................g....... [ 18400]
MH025919.1_strain_N10             ..........................t.............................................g....... [ 18400]
MH025918.1_strain_R25             ..........................t.............................................g....... [ 18400]
MH025917.1_strain_R7              ..........................t.............................................g....... [ 18400]
MH025916.1_strain_R8              ..........................t.............................................g....... [ 18400]

FR682468.2_Georgia_2007/1         gactaatataatcattttgtttaataa---gaggcatatcatcccacactttatttttacaaataccgttcctacattcc [ 18480]
ASF_IND_20_CAD543                 -------------------------------------------------------------------------------- [ 18480]
LR899193.1_Germany_2020/1         ...........................---.................................................. [ 18480]
MN393477.1_Wuhan_2019             ...........................---.................................................. [ 18480]
MW396979.1_Timor-Leste/2019/1     ...........................---.................................................. [ 18480]
MK940252.1_CN/2019/InnerMongolia  ...........................---.................................................. [ 18480]
MK128995.1_China/2018/AnhuiXCGQ   ...........................---.................................................. [ 18480]
MH681419.1_ASFV/POL/2015          ...........................---.................................................. [ 18480]
LS478113.1_Estonia_2014           -------...tctt....c....t...---a.cct...ct..ttt..--------------------------------- [ 18480]
KP843857.1_Odintsovo_02/14        ...........................---.................................................. [ 18480]
FN557520.1_strain_E75             -------------------------------------------------------------------------------- [ 18480]
AY261366.1_Warthog                ...........................---................a....................a............ [ 18480]
AY261365.1_Warmbaths              ...........................---................a....................a............ [ 18480]
AY261364.1_Tengani_62             -------------------------------------------------------------------------------- [ 18480]
AY261363.1_Pretoriuskop/96/4      t.t................ga......agg.g...........ta.a..............g.....c............ [ 18480]
AY261362.1_Mkuzi_1979             ...........................---................a....................a............ [ 18480]
AY261361.1_Malawi_Lil-20/1_1983   -------------------------------------------------------------------------------- [ 18480]
AY261360.1_Kenya_1950             ...........................---a...............a.................g..a.g.......... [ 18480]
AM712240.1_OURT_88/3              -------------------------------------------------------------------------------- [ 18480]
AM712239.1_Benin_97/1             -------------------------------------------------------------------------------- [ 18480]
KM111295.1_Ken06.Bus              ...........................---................a.................g..a.g.......... [ 18480]
```

```
KM111294.1_Ken05/Tk1             ...........................---................a.................g..a.g.......... [ 18480]
KM262844.1_strain_L60            -------------------------------------------------------------------------------- [ 18480]
KM102979.1_26544/OG10_Italy      -------------------------------------------------------------------------------- [ 18480]
KM262845.1_strain_NHV            -------------------------------------------------------------------------------- [ 18480]
KX354450.1_47/Ss/2008            -------------------------------------------------------------------------------- [ 18480]
KP055815.1_strain_BA71           -------------------------------------------------------------------------------- [ 18480]
MH025920.1_strain_R35            ...........................---................a.................g..a.g.......... [ 18480]
MH025919.1_strain_N10            ...........................---................a.................g..a.g.......... [ 18480]
MH025918.1_strain_R25            ...........................---................a.................g..a.g.......... [ 18480]
MH025917.1_strain_R7             ...........................---................a.................g..a.g.......... [ 18480]
MH025916.1_strain_R8             ...........................---................a.................g..a.g.......... [ 18480]

FR682468.2_Georgia_2007/1        cagcagaaatcacagtgttttccatacgtgcaccagtattcaagctctcttataggaggcgtataagtccttggtaaatt [ 18560]
ASF_IND_20_CAD543                -------------------------------------------------------------------------------- [ 18560]
LR899193.1_Germany_2020/1        ................................................................................ [ 18560]
MN393477.1_Wuhan_2019            ................................................................................ [ 18560]
MW396979.1_Timor-Leste/2019/1    ................................................................................ [ 18560]
MK940252.1_CN/2019/InnerMongolia ................................................................................ [ 18560]
MK128995.1_China/2018/AnhuiXCGQ  ................................................................................ [ 18560]
MH681419.1_ASFV/POL/2015         ................................................................................ [ 18560]
LS478113.1_Estonia_2014          -----------------------------------------------------------......--------------- [ 18560]
KP843857.1_Odintsovo_02/14       ................................................................................ [ 18560]
FN557520.1_strain_E75            -------------------------------------------------------------------------------- [ 18560]
AY261366.1_Warthog               ................................................................................ [ 18560]
AY261365.1_Warmbaths             ...........................................................t.................... [ 18560]
AY261364.1_Tengani_62            -------------------------------------------------------------------------------- [ 18560]
AY261363.1_Pretoriuskop/96/4     .....a..g............g....g........a..a............g.g.....t.....g............aa [ 18560]
AY261362.1_Mkuzi_1979            ..........t.........................................cg..t..t......c............. [ 18560]
AY261361.1_Malawi_Lil-20/1_1983  ------------------------------------------------------------.................... [ 18560]
AY261360.1_Kenya_1950            ..........g...a.....................................cg.....t.................... [ 18560]
AM712240.1_OURT_88/3             -------------------------------------------------------------------------------- [ 18560]
AM712239.1_Benin_97/1            -------------------------------------------------------------------------------- [ 18560]
KM111295.1_Ken06.Bus             ..........g...a.....................................cg.....t.................... [ 18560]
KM111294.1_Ken05/Tk1             ..............a.....................................cg.....t..g................. [ 18560]
KM262844.1_strain_L60            -------------------------------------------------------------------------------- [ 18560]
KM102979.1_26544/OG10_Italy      -------------------------------------------------------------------------------- [ 18560]
KM262845.1_strain_NHV            -------------------------------------------------------------------------------- [ 18560]
KX354450.1_47/Ss/2008            -------------------------------------------------------------------------------- [ 18560]
KP055815.1_strain_BA71           -------------------------------------------------------------------------------- [ 18560]
MH025920.1_strain_R35            ..........g...a.....................................cg.....t.................... [ 18560]
MH025919.1_strain_N10            ..........g...a.....................................cg.....t.................... [ 18560]
MH025918.1_strain_R25            ..........g...a.....................................cg.....t.................... [ 18560]
MH025917.1_strain_R7             ..........g...a.....................................cg.....t.................... [ 18560]
MH025916.1_strain_R8             ..........g...a.....................................cg.....t.................... [ 18560]
```

```
FR682468.2_Georgia_2007/1        ttgtttcatataaaagatggaaaggggtcgatttaaacccggctgagatagccaaatcaaaatacataaaagagcaagta [ 18640]
ASF_IND_20_CAD543                -------------------------------------------------------------------------------- [ 18640]
LR899193.1_Germany_2020/1        ................................................................................ [ 18640]
MN393477.1_Wuhan_2019            ................................................................................ [ 18640]
MW396979.1_Timor-Leste/2019/1    ................................................................................ [ 18640]
MK940252.1_CN/2019/InnerMongolia ................................................................................ [ 18640]
MK128995.1_China/2018/AnhuiXCGQ  ................................................................................ [ 18640]
MH681419.1_ASFV/POL/2015         ................................................................................ [ 18640]
LS478113.1_Estonia_2014          ----------------------------------------..t..g...catttc..a..gg.ga...tt..----ga.g [ 18640]
KP843857.1_Odintsovo_02/14       ................................................................................ [ 18640]
FN557520.1_strain_E75            -------------------------------------------------------------------------------- [ 18640]
AY261366.1_Warthog               ..............................g..............................................t.. [ 18640]
AY261365.1_Warmbaths             ..............................................................................t.. [ 18640]
AY261364.1_Tengani_62            -------------------------------------------------------------------------------- [ 18640]
AY261363.1_Pretoriuskop/96/4     .......g................t.....g......a.t.....t.cc.agt...c........tc.......tc.a.. [ 18640]
AY261362.1_Mkuzi_1979            .......g......................g...........t..g...................t.......a...... [ 18640]
AY261361.1_Malawi_Lil-20/1_1983  ..............................g......g...........................t.............. [ 18640]
AY261360.1_Kenya_1950            ...........................a..g...........t......................t.............. [ 18640]
AM712240.1_OURT_88/3             -------------------------------------------------------------------------------- [ 18640]
AM712239.1_Benin_97/1            -------------------------------------------------------------------------------- [ 18640]
KM111295.1_Ken06.Bus             .......g......................g......a.....................g.....t.......a...... [ 18640]
KM111294.1_Ken05/Tk1             .......g......................g......a...........................t.......a...... [ 18640]
KM262844.1_strain_L60            -------------------------------------------------------------------------------- [ 18640]
KM102979.1_26544/OG10_Italy      -------------------------------------------------------------------------------- [ 18640]
KM262845.1_strain_NHV            -------------------------------------------------------------------------------- [ 18640]
KX354450.1_47/Ss/2008            -------------------------------------------------------------------------------- [ 18640]
KP055815.1_strain_BA71           -------------------------------------------------------------------------------- [ 18640]
MH025920.1_strain_R35            .......g......................g......a.....................g.....t.......a...... [ 18640]
MH025919.1_strain_N10            .......g......................g......a.....................g.....t.......a...... [ 18640]
MH025918.1_strain_R25            .......g......................g......a.....................g.....t.......a...... [ 18640]
MH025917.1_strain_R7             .......g......................g......a.....................g.....t.......a...... [ 18640]
MH025916.1_strain_R8             .......g......................g......a.....................g.....t.......a...... [ 18640]

FR682468.2_Georgia_2007/1        gtttcatagtggtatttagatgtaaatttttatagtatgcaaatacaatgtaacctacaaatacaatactaaatacaagg [ 18720]
ASF_IND_20_CAD543                -------------------------------------------------------------------------------- [ 18720]
LR899193.1_Germany_2020/1        ................................................................................ [ 18720]
MN393477.1_Wuhan_2019            ................................................................................ [ 18720]
MW396979.1_Timor-Leste/2019/1    ................................................................................ [ 18720]
MK940252.1_CN/2019/InnerMongolia ................................................................................ [ 18720]
MK128995.1_China/2018/AnhuiXCGQ  ................................................................................ [ 18720]
MH681419.1_ASFV/POL/2015         ................................................................................ [ 18720]
LS478113.1_Estonia_2014          ...gt...t.ac.....g.------------ta....ca.t...gtg....----------------------------- [ 18720]
KP843857.1_Odintsovo_02/14       ................................................................................ [ 18720]
```

```
FN557520.1_strain_E75          -------------------------------------------------------------------------------- [ 18720]
AY261366.1_Warthog             ...................g.................c........................g................. [ 18720]
AY261365.1_Warmbaths           ...................g..........................................g................. [ 18720]
AY261364.1_Tengani_62          -------------------------------------------------------------------------------- [ 18720]
AY261363.1_Pretoriuskop/96/4   ....t.....t..g.....tc...g.......cgtac....t.c..g.aac.....g.....g.t.......t..t..a. [ 18720]
AY261362.1_Mkuzi_1979          ...................g.............g............................g................. [ 18720]
AY261361.1_Malawi_Lil-20/1_1983 ...................g.............t...ca.......................g................. [ 18720]
AY261360.1_Kenya_1950          ...................g.............g...c........................g................. [ 18720]
AM712240.1_OURT_88/3           -------------------------------------------------------------------------------- [ 18720]
AM712239.1_Benin_97/1          -------------------------------------------------------------------------------- [ 18720]
KM111295.1_Ken06.Bus           ...................g.........c...g...c......g.g...c...........g................. [ 18720]
KM111294.1_Ken05/Tk1           ...................g.........c...g...c......g.....c...........g................. [ 18720]
KM262844.1_strain_L60          -------------------------------------------------------------------------------- [ 18720]
KM102979.1_26544/OG10_Italy    -------------------------------------------------------------------------------- [ 18720]
KM262845.1_strain_NHV          -------------------------------------------------------------------------------- [ 18720]
KX354450.1_47/Ss/2008          -------------------------------------------------------------------------------- [ 18720]
KP055815.1_strain_BA71         -------------------------------------------------------------------------------- [ 18720]
MH025920.1_strain_R35          ...................g.........c...g...c......g.g...c...........g................. [ 18720]
MH025919.1_strain_N10          ...................g.........c...g...c......g.g...c...........g................. [ 18720]
MH025918.1_strain_R25          ...................g.........c...g...c......g.g...c...........g................. [ 18720]
MH025917.1_strain_R7           ...................g.........c...g...c......g.g...c...........g................. [ 18720]
MH025916.1_strain_R8           ...................g.........c...g...c......g.g...c...........g................. [ 18720]

FR682468.2_Georgia_2007/1      taaaaacaacaatgtcttataatgattgg---ccaataa-tcacccccccccccccatttttccatgaatatttcatttc [ 18800]
ASF_IND_20_CAD543              -------------------------------------------------------------------------------- [ 18800]
LR899193.1_Germany_2020/1      .............................---.......-...----................................. [ 18800]
MN393477.1_Wuhan_2019          .............................---.......-...---.................................. [ 18800]
MW396979.1_Timor-Leste/2019/1  .............................---.......-...---.................................. [ 18800]
MK940252.1_CN/2019/InnerMongolia .............................---.......-...--................................... [ 18800]
MK128995.1_China/2018/AnhuiXCGQ .............................---.......-...--................................... [ 18800]
MH681419.1_ASFV/POL/2015       .............................---.......-...--................................... [ 18800]
LS478113.1_Estonia_2014        --c...t..tt....g.at....---------------------------------..cc.......g..t...tt..g. [ 18800]
KP843857.1_Odintsovo_02/14     .............................---.......-...--................................... [ 18800]
FN557520.1_strain_E75          -------------------------------------------------------------------------------- [ 18800]
AY261366.1_Warthog             ..............c..............---.......-...----................................. [ 18800]
AY261365.1_Warmbaths           ..............c........a.....---.......-...-----................................ [ 18800]
AY261364.1_Tengani_62          -------------------------------------------------------------------------------- [ 18800]
AY261363.1_Pretoriuskop/96/4   ..t.g....t.t...t...a.g.t.a.a.agt.at....-...------..t...ac.....t...a............. [ 18800]
AY261362.1_Mkuzi_1979          .............................---.......-...---.................................. [ 18800]
AY261361.1_Malawi_Lil-20/1_1983 .............................---.......-........................................ [ 18800]
AY261360.1_Kenya_1950          ........g.....c..............---.......-...---.................................. [ 18800]
AM712240.1_OURT_88/3           -------------------------------------------------------------------------------- [ 18800]
AM712239.1_Benin_97/1          -------------------------------------------------------------------------------- [ 18800]
KM111295.1_Ken06.Bus           ........g.....c..............---.....t.--------................................. [ 18800]
```

```
KM111294.1_Ken05/Tk1            ...g....g.....c..............---.....g.--tt---a................................. [ 18800]
KM262844.1_strain_L60           -------------------------------------------------------------------------------- [ 18800]
KM102979.1_26544/OG10_Italy     -------------------------------------------------------------------------------- [ 18800]
KM262845.1_strain_NHV           -------------------------------------------------------------------------------- [ 18800]
KX354450.1_47/Ss/2008           -------------------------------------------------------------------------------- [ 18800]
KP055815.1_strain_BA71          -------------------------------------------------------------------------------- [ 18800]
MH025920.1_strain_R35           ........g.....c..............---.....t.cc.c---.................................. [ 18800]
MH025919.1_strain_N10           ........g.....c..............---.....t.---------................................ [ 18800]
MH025918.1_strain_R25           ........g.....c..............---.....t.-c.c---.................................. [ 18800]
MH025917.1_strain_R7            ........g.....c..............---.....t.--.c---.................................. [ 18800]
MH025916.1_strain_R8            ........g.....c..............---.....t.--.c---.................................. [ 18800]

FR682468.2_Georgia_2007/1       ctgtatagggtctaggatgtgaacactccatgttatgatgattaggcattttaactgatatttcataaaaacacccccag [ 18880]
ASF_IND_20_CAD543               -------------------------------------------------------------------------------- [ 18880]
LR899193.1_Germany_2020/1       ................................................................................ [ 18880]
MN393477.1_Wuhan_2019           ................................................................................ [ 18880]
MW396979.1_Timor-Leste/2019/1   ................................................................................ [ 18880]
MK940252.1_CN/2019/InnerMongolia ................................................................................ [ 18880]
MK128995.1_China/2018/AnhuiXCGQ ................................................................................ [ 18880]
MH681419.1_ASFV/POL/2015        ................................................................................ [ 18880]
LS478113.1_Estonia_2014         --------------------------------------------------------------.a.a.....t.------- [ 18880]
KP843857.1_Odintsovo_02/14      ................................................................................ [ 18880]
FN557520.1_strain_E75           -------------------------------------------------------------------------------- [ 18880]
AY261366.1_Warthog              ...............................................................................- [ 18880]
AY261365.1_Warmbaths            ...............................................................................- [ 18880]
AY261364.1_Tengani_62           -------------------------------------------------------------------------------- [ 18880]
AY261363.1_Pretoriuskop/96/4    ..a....a...g.............aga....c..aa...............gga.........g........ggt.tg. [ 18880]
AY261362.1_Mkuzi_1979           .........................t.................................................a.tg. [ 18880]
AY261361.1_Malawi_Lil-20/1_1983 .......a.................t......c........................................gga.tg. [ 18880]
AY261360.1_Kenya_1950           .......a........ta.......t.............................c...................a.tg. [ 18880]
AM712240.1_OURT_88/3            -------------------------------------------------------------------------------- [ 18880]
AM712239.1_Benin_97/1           -------------------------------------------------------------------------------- [ 18880]
KM111295.1_Ken06.Bus            .......a........ta.......t.....a....................................t....t...... [ 18880]
KM111294.1_Ken05/Tk1            .......a.................t.............a.....a........g..................atgttg. [ 18880]
KM262844.1_strain_L60           -------------------------------------------------------------------------------- [ 18880]
KM102979.1_26544/OG10_Italy     -------------------------------------------------------------------------------- [ 18880]
KM262845.1_strain_NHV           -------------------------------------------------------------------------------- [ 18880]
KX354450.1_47/Ss/2008           -------------------------------------------------------------------------------- [ 18880]
KP055815.1_strain_BA71          -------------------------------------------------------------------------------- [ 18880]
MH025920.1_strain_R35           .......a........ta.......t.....a....................................t....t...... [ 18880]
MH025919.1_strain_N10           .......a........ta.......t.....a....................................t....t...... [ 18880]
MH025918.1_strain_R25           .......a........ta.......t.....a....................................t....t...... [ 18880]
MH025917.1_strain_R7            .......a........ta.......t.....a....................................t....t...... [ 18880]
MH025916.1_strain_R8            .......a........ta.......t.....a....................................t....t...... [ 18880]
```

```
FR682468.2_Georgia_2007/1        ------gaattgcgattaactatacagtttacaatcgaattcatcgaattagactcatt-------tgttatcttatttt [ 18960]
ASF_IND_20_CAD543                -------------------------------------------------------------------------------- [ 18960]
LR899193.1_Germany_2020/1        ------.....................................................-------.............. [ 18960]
MN393477.1_Wuhan_2019            ------.....................................................-------.............. [ 18960]
MW396979.1_Timor-Leste/2019/1    ------.....................................................-------.............. [ 18960]
MK940252.1_CN/2019/InnerMongolia ------.....................................................-------.............. [ 18960]
MK128995.1_China/2018/AnhuiXCGQ  ------.....................................................-------.............. [ 18960]
MH681419.1_ASFV/POL/2015         ------.....................................................-------.............. [ 18960]
LS478113.1_Estonia_2014          ------------------------------------------------------------------------t..t.... [ 18960]
KP843857.1_Odintsovo_02/14       ------.....................................................-------.............. [ 18960]
FN557520.1_strain_E75            -------------------------------------------------------------------------------- [ 18960]
AY261366.1_Warthog               -----g..........................................a..........-------.............. [ 18960]
AY261365.1_Warmbaths             -----g...........................t..............a..........-------.............. [ 18960]
AY261364.1_Tengani_62            -------------------------------------------------------------------------------- [ 18960]
AY261363.1_Pretoriuskop/96/4     gtctcg.....a..gga....c....a........a...-.g..t.g.a.c..t.a...gagtttcca....t....... [ 18960]
AY261362.1_Mkuzi_1979            tgtata....c....g.....c...................t......a..........-------.............. [ 18960]
AY261361.1_Malawi_Lil-20/1_1983  gatggg....c....g.....c...................t......a..........-------.............. [ 18960]
AY261360.1_Kenya_1950            tgtata....c....g.....c...................t......a..........-------.............. [ 18960]
AM712240.1_OURT_88/3             -------------------------------------------------------------------------------- [ 18960]
AM712239.1_Benin_97/1            -------------------------------------------------------------------------------- [ 18960]
KM111295.1_Ken06.Bus             ------.g..c....g.....c...................t......a..........-------.............. [ 18960]
KM111294.1_Ken05/Tk1             tctata..t.c....g.....c....a..............t......a..........-------.............. [ 18960]
KM262844.1_strain_L60            -------------------------------------------------------------------------------- [ 18960]
KM102979.1_26544/OG10_Italy      -------------------------------------------------------------------------------- [ 18960]
KM262845.1_strain_NHV            -------------------------------------------------------------------------------- [ 18960]
KX354450.1_47/Ss/2008            -------------------------------------------------------------------------------- [ 18960]
KP055815.1_strain_BA71           -------------------------------------------------------------------------------- [ 18960]
MH025920.1_strain_R35            ------.g..c....g.....c...................t......a..........-------.............. [ 18960]
MH025919.1_strain_N10            ------.g..c....g.....c...................t......a..........-------.............. [ 18960]
MH025918.1_strain_R25            ------.g..c....g.....c...................t......a..........-------.............. [ 18960]
MH025917.1_strain_R7             ------.g..c....g.....c...................t......a..........-------.............. [ 18960]
MH025916.1_strain_R8             ------.g..c....g.....c...................t......a..........-------.............. [ 18960]

FR682468.2_Georgia_2007/1        tacaaatgcca------ttttgacaatcccagcagaagtcacaattctttacatacgtacaccaatatggaagctcctcc [ 19040]
ASF_IND_20_CAD543                -------------------------------------------------------------------------------- [ 19040]
LR899193.1_Germany_2020/1        ...........------............................................................... [ 19040]
MN393477.1_Wuhan_2019            ...........------............................................................... [ 19040]
MW396979.1_Timor-Leste/2019/1    ...........------............................................................... [ 19040]
MK940252.1_CN/2019/InnerMongolia ...........------............................................................... [ 19040]
MK128995.1_China/2018/AnhuiXCGQ  ...........------............................................................... [ 19040]
MH681419.1_ASFV/POL/2015         ...........------............................................................... [ 19040]
LS478113.1_Estonia_2014          .gt.g.caa..ggatgt.gga.c....gtt..t.atg.c.c.t....c.cttg...tc.t.tttt...a----------- [ 19040]
KP843857.1_Odintsovo_02/14       ...........------............................................................... [ 19040]
```

```
FN557520.1_strain_E75          -------------------------------------------------------------------------------- [ 19040]
AY261366.1_Warthog             ...........------............................................................... [ 19040]
AY261365.1_Warmbaths           ........t..------............................................................... [ 19040]
AY261364.1_Tengani_62          -------------------------------------------------------------------------------- [ 19040]
AY261363.1_Pretoriuskop/96/4   .g.....a...------.....g..g.....a..a.....g..t.gtgg......a............tt.......ct. [ 19040]
AY261362.1_Mkuzi_1979          ........t..------.....g................................t..g........ctc......t... [ 19040]
AY261361.1_Malawi_Lil-20/1_1983 ........t..------.....g.................g...................................t... [ 19040]
AY261360.1_Kenya_1950          ........t..------.....g..............................................c......t... [ 19040]
AM712240.1_OURT_88/3           -------------------------------------------------------------------------------- [ 19040]
AM712239.1_Benin_97/1          -------------------------------------------------------------------------------- [ 19040]
KM111295.1_Ken06.Bus           ........t..------.....g..............................................c......t... [ 19040]
KM111294.1_Ken05/Tk1           ........t..------.....g..............a...............................c......t... [ 19040]
KM262844.1_strain_L60          -------------------------------------------------------------------------------- [ 19040]
KM102979.1_26544/OG10_Italy    -------------------------------------------------------------------------------- [ 19040]
KM262845.1_strain_NHV          -------------------------------------------------------------------------------- [ 19040]
KX354450.1_47/Ss/2008          -------------------------------------------------------------------------------- [ 19040]
KP055815.1_strain_BA71         -------------------------------------------------------------------------------- [ 19040]
MH025920.1_strain_R35          ........t..------.....g..............................................c......t... [ 19040]
MH025919.1_strain_N10          ........t..------.....g..............................................c......t... [ 19040]
MH025918.1_strain_R25          ........t..------.....g..............................................c......t... [ 19040]
MH025917.1_strain_R7           ........t..------.....g..............................................c......t... [ 19040]
MH025916.1_strain_R8           ........t..------.....g..............................................c......t... [ 19040]

FR682468.2_Georgia_2007/1      ttaggaggatgctgggttcttggtaattctggtaattcatgtgcaagaatgaggactgagtagcccaacaaaagtcctag [ 19120]
ASF_IND_20_CAD543              -------------------------------------------------------------------------------- [ 19120]
LR899193.1_Germany_2020/1      ................................................................................ [ 19120]
MN393477.1_Wuhan_2019          ................................................................................ [ 19120]
MW396979.1_Timor-Leste/2019/1  ................................................................................ [ 19120]
MK940252.1_CN/2019/InnerMongolia ................................................................................ [ 19120]
MK128995.1_China/2018/AnhuiXCGQ ................................................................................ [ 19120]
MH681419.1_ASFV/POL/2015       ................................................................................ [ 19120]
LS478113.1_Estonia_2014        -------------------------------------------------------------------------------- [ 19120]
KP843857.1_Odintsovo_02/14     ................................................................................ [ 19120]
FN557520.1_strain_E75          -------------------------------------------------------------------------------- [ 19120]
AY261366.1_Warthog             ............................................g................................... [ 19120]
AY261365.1_Warmbaths           ............................................g................................... [ 19120]
AY261364.1_Tengani_62          -------------------------------------------------------------------------------- [ 19120]
AY261363.1_Pretoriuskop/96/4   ..g.......................g..........gt...............ctga.ta............cg...-- [ 19120]
AY261362.1_Mkuzi_1979          ..................gc..a...g..a.............t..................a................. [ 19120]
AY261361.1_Malawi_Lil-20/1_1983 ............c.....ga......g.a..............................................t.c.. [ 19120]
AY261360.1_Kenya_1950          ..................gc..a...g.....c..........t.................................... [ 19120]
AM712240.1_OURT_88/3           -------------------------------------------------------------------------------- [ 19120]
AM712239.1_Benin_97/1          -------------------------------------------------------------------------------- [ 19120]
KM111295.1_Ken06.Bus           ............c.....ac......g................t......c............................. [ 19120]
```

```
KM111294.1_Ken05/Tk1           ............c.....gc......g.....c..........t...................................a [ 19120]
KM262844.1_strain_L60          -------------------------------------------------------------------------------- [ 19120]
KM102979.1_26544/OG10_Italy    -------------------------------------------------------------------------------- [ 19120]
KM262845.1_strain_NHV          -------------------------------------------------------------------------------- [ 19120]
KX354450.1_47/Ss/2008          -------------------------------------------------------------------------------- [ 19120]
KP055815.1_strain_BA71         -------------------------------------------------------------------------------- [ 19120]
MH025920.1_strain_R35          ............c.....ac......g................t......c............................. [ 19120]
MH025919.1_strain_N10          ............c.....ac......g................t......c............................. [ 19120]
MH025918.1_strain_R25          ............c.....ac......g................t......c............................. [ 19120]
MH025917.1_strain_R7           ............c.....ac......g................t......c............................. [ 19120]
MH025916.1_strain_R8           ............c.....ac......g................t......c............................. [ 19120]

FR682468.2_Georgia_2007/1      aa---ccttcatgt---------------------------------------tgtgtccaaatggcacctgtcatttta [ 19200]
ASF_IND_20_CAD543              -------------------------------------------------------------------------------- [ 19200]
LR899193.1_Germany_2020/1      ..---.........---------------------------------------........................... [ 19200]
MN393477.1_Wuhan_2019          ..---.........---------------------------------------........................... [ 19200]
MW396979.1_Timor-Leste/2019/1  ..---.........---------------------------------------........................... [ 19200]
MK940252.1_CN/2019/InnerMongolia ..---.........---------------------------------------........................... [ 19200]
MK128995.1_China/2018/AnhuiXCGQ ..---.........---------------------------------------........................... [ 19200]
MH681419.1_ASFV/POL/2015       ..---.........---------------------------------------........................... [ 19200]
LS478113.1_Estonia_2014        ---------------------------------------------------------------------t.c.t.c.... [ 19200]
KP843857.1_Odintsovo_02/14     ..---.........---------------------------------------........................... [ 19200]
FN557520.1_strain_E75          -------------------------------------------------------------------------------- [ 19200]
AY261366.1_Warthog             ..gaa.........---------------------------------------........................... [ 19200]
AY261365.1_Warmbaths           ..gaa.........---------------------------------------........................... [ 19200]
AY261364.1_Tengani_62          -------------------------------------------------------------------------------- [ 19200]
AY261363.1_Pretoriuskop/96/4   -.acc.........---------------------------------------..cac..............cat..... [ 19200]
AY261362.1_Mkuzi_1979          ..gag.........---------------------------------------.......................g... [ 19200]
AY261361.1_Malawi_Lil-20/1_1983 ..gaa.........---------------------------------------...a....................... [ 19200]
AY261360.1_Kenya_1950          ..gga.........--------------------------tgcgtttaaatgac.c.........a.............. [ 19200]
AM712240.1_OURT_88/3           -------------------------------------------------------------------------------- [ 19200]
AM712239.1_Benin_97/1          -------------------------------------------------------------------------------- [ 19200]
KM111295.1_Ken06.Bus           ..gga.........tgcgtctaaatgctgcgtctaaatgctgcgtttaaatgcc.c...t.....a........-c.... [ 19200]
KM111294.1_Ken05/Tk1           ..gga.........--------------------------tgcgtttaaatgac.c.........a.............. [ 19200]
KM262844.1_strain_L60          -------------------------------------------------------------------------------- [ 19200]
KM102979.1_26544/OG10_Italy    -------------------------------------------------------------------------------- [ 19200]
KM262845.1_strain_NHV          -------------------------------------------------------------------------------- [ 19200]
KX354450.1_47/Ss/2008          -------------------------------------------------------------------------------- [ 19200]
KP055815.1_strain_BA71         -------------------------------------------------------------------------------- [ 19200]
MH025920.1_strain_R35          ..gga.........tgcgtctaaatgctgcgtctaaatgctgcgtttaaatgcc.c...t.....a........-c.... [ 19200]
MH025919.1_strain_N10          ..gga.........tgcgtctaaatgctgcgtctaaatgctgcgtttaaatgcc.c...t.....a........-c.... [ 19200]
MH025918.1_strain_R25          ..gga.........tgcgtctaaatgctgcgtctaaatgctgcgtttaaatgcc.c...t.....a........-c.... [ 19200]
MH025917.1_strain_R7           ..gga.........tgcgtctaaatgctgcgtctaaatgctgcgtttaaatgcc.c...t.....a........-c.... [ 19200]
MH025916.1_strain_R8           ..gga.........tgcgtctaaatgctgcgtctaaatgctgcgtttaaatgcc.c...t.....a........-c.... [ 19200]
```

```
FR682468.2_Georgia_2007/1        aaaaagatttaaattttgctaccgc-aaaaaaaaatccagtatgtat----ttttttaatacatataattattgaag-tc [ 19280]
ASF_IND_20_CAD543                -------------------------------------------------------------------------------- [ 19280]
LR899193.1_Germany_2020/1        .........................-.....................----..........................-.. [ 19280]
MN393477.1_Wuhan_2019            .........................-.....................----..........................-.. [ 19280]
MW396979.1_Timor-Leste/2019/1    .........................-.....................----..........................-.. [ 19280]
MK940252.1_CN/2019/InnerMongolia .........................-.....................----..........................-.. [ 19280]
MK128995.1_China/2018/AnhuiXCGQ  .........................-.....................----..........................-.. [ 19280]
MH681419.1_ASFV/POL/2015         .........................-.....................----..........................-.. [ 19280]
LS478113.1_Estonia_2014          ...t.a.g...c..--------------------------------------------.a..t.....ca....c.c-.. [ 19280]
KP843857.1_Odintsovo_02/14       .........................--....................----..........................-.. [ 19280]
FN557520.1_strain_E75            -------------------------------------------------------------------------------- [ 19280]
AY261366.1_Warthog               ..................t....a.-.....................----.........................c-.. [ 19280]
AY261365.1_Warmbaths             ..................t....a.-....................-----.........................c-.. [ 19280]
AY261364.1_Tengani_62            -------------------------------------------------------------------------------- [ 19280]
AY261363.1_Pretoriuskop/96/4     .....a............a....a.-......g..------...c.---------------------------------- [ 19280]
AY261362.1_Mkuzi_1979            ..................t....a.-.....................----.........................---. [ 19280]
AY261361.1_Malawi_Lil-20/1_1983  ..................t......---...................----.........................c-c. [ 19280]
AY261360.1_Kenya_1950            .....a.a..........t.t..a.-.............ac......attt....aat--..............g.a-c. [ 19280]
AM712240.1_OURT_88/3             -------------------------------------------------------------------------------- [ 19280]
AM712239.1_Benin_97/1            -------------------------------------------------------------------------------- [ 19280]
KM111295.1_Ken06.Bus             .....a.a..........t.t..a.-..........-..ac......attt.......--.t.......c....g.acc. [ 19280]
KM111294.1_Ken05/Tk1             .....a.a..........t.t..a.g..............c......attt..........t.......c....g.acc. [ 19280]
KM262844.1_strain_L60            -------------------------------------------------------------------------------- [ 19280]
KM102979.1_26544/OG10_Italy      -------------------------------------------------------------------------------- [ 19280]
KM262845.1_strain_NHV            -------------------------------------------------------------------------------- [ 19280]
KX354450.1_47/Ss/2008            -------------------------------------------------------------------------------- [ 19280]
KP055815.1_strain_BA71           -------------------------------------------------------------------------------- [ 19280]
MH025920.1_strain_R35            .....a.a..........t.t..a.-..........-..ac......attt.......--.t.......c....g.acc. [ 19280]
MH025919.1_strain_N10            .....a.a..........t.t..a.-..........-..ac......attt.......--.t.......c....g.acc. [ 19280]
MH025918.1_strain_R25            .....a.a..........t.t..a.-..........-..ac......attt.......--.t.......c....g.acc. [ 19280]
MH025917.1_strain_R7             .....a.a..........t.t..a.-..........-..ac......attt.......--.t.......c....g.acc. [ 19280]
MH025916.1_strain_R8             .....a.a..........t.t..a.-..........-..ac......attt.......--.t.......c....g.acc. [ 19280]

FR682468.2_Georgia_2007/1        ttataagata-aagccgagaacactatattttgtatagatgatg--tatccggtattcaaactctcttataa-------g [ 19360]
ASF_IND_20_CAD543                -------------------------------------------------------------------------------- [ 19360]
LR899193.1_Germany_2020/1        ..........-.................................--..........................-------. [ 19360]
MN393477.1_Wuhan_2019            ..........-.................................--..........................-------. [ 19360]
MW396979.1_Timor-Leste/2019/1    ..........-.................................--..........................-------. [ 19360]
MK940252.1_CN/2019/InnerMongolia ..........-.................................--..........................-------. [ 19360]
MK128995.1_China/2018/AnhuiXCGQ  ..........-.................................--..........................-------. [ 19360]
MH681419.1_ASFV/POL/2015         ..........-.................................--..........................-------. [ 19360]
LS478113.1_Estonia_2014          ..t.t..-----------..tgtt...t...ca...-------------------------------------------- [ 19360]
KP843857.1_Odintsovo_02/14       ..........-.................................--..........................-------. [ 19360]
```

```
FN557520.1_strain_E75             -------------------------------------------------------------------------------- [ 19360]
AY261366.1_Warthog                ..........t..ag.............................--.....-....................-------. [ 19360]
AY261365.1_Warmbaths              ..........t..ag.............................--.....-....................-------. [ 19360]
AY261364.1_Tengani_62             -------------------------------------------------------------------------------- [ 19360]
AY261363.1_Pretoriuskop/96/4      ------------------------gg......t....------------------------------------------- [ 19360]
AY261362.1_Mkuzi_1979             .c........-.................................--..........................gtacttg. [ 19360]
AY261361.1_Malawi_Lil-20/1_1983   .......g.gt.................................--.....-....................gtacttg. [ 19360]
AY261360.1_Kenya_1950             .......g..t.................................--....t-a...................gtacttg. [ 19360]
AM712240.1_OURT_88/3              -------------------------------------------------------------------------------- [ 19360]
AM712239.1_Benin_97/1             -------------------------------------------------------------------------------- [ 19360]
KM111295.1_Ken06.Bus              .....t.gggt....t........c...................--.....-....................gtacttga [ 19360]
KM111294.1_Ken05/Tk1              .......gggt....t........c........c.....g....ta.....-........g...........gttcttg. [ 19360]
KM262844.1_strain_L60             -------------------------------------------------------------------------------- [ 19360]
KM102979.1_26544/OG10_Italy       -------------------------------------------------------------------------------- [ 19360]
KM262845.1_strain_NHV             -------------------------------------------------------------------------------- [ 19360]
KX354450.1_47/Ss/2008             -------------------------------------------------------------------------------- [ 19360]
KP055815.1_strain_BA71            -------------------------------------------------------------------------------- [ 19360]
MH025920.1_strain_R35             .....t.gggt....t........c...................--.....-....................gtacttga [ 19360]
MH025919.1_strain_N10             .....t.gggt....t........c...................--.....-....................gtacttga [ 19360]
MH025918.1_strain_R25             .....t.gggt....t........c...................--.....-....................gtacttga [ 19360]
MH025917.1_strain_R7              .....t.gggt....t........c...................--.....-....................gtacttga [ 19360]
MH025916.1_strain_R8              .....t.gggt....t........c...................--.....-....................gtacttga [ 19360]

FR682468.2_Georgia_2007/1         tacatgtaggaaatggtcaattattcaagattggctgagataacaacaaaaccaaaatactcaaaagcataagtaatttc [ 19440]
ASF_IND_20_CAD543                 -------------------------------------------------------------------------------- [ 19440]
LR899193.1_Germany_2020/1         ................................................................................ [ 19440]
MN393477.1_Wuhan_2019             ................................................................................ [ 19440]
MW396979.1_Timor-Leste/2019/1     ................................................................................ [ 19440]
MK940252.1_CN/2019/InnerMongolia  ................................................................................ [ 19440]
MK128995.1_China/2018/AnhuiXCGQ   ................................................................................ [ 19440]
MH681419.1_ASFV/POL/2015          ................................................................................ [ 19440]
LS478113.1_Estonia_2014           -------------------------------------------------------------------------------- [ 19440]
KP843857.1_Odintsovo_02/14        ................................................................................ [ 19440]
FN557520.1_strain_E75             -------------------------------------------------------------------------------- [ 19440]
AY261366.1_Warthog                .........................................---.................................... [ 19440]
AY261365.1_Warmbaths              .........................................---.................................... [ 19440]
AY261364.1_Tengani_62             -------------------------------------------------------------------------------- [ 19440]
AY261363.1_Pretoriuskop/96/4      -------------------------------------------------------------------------------- [ 19440]
AY261362.1_Mkuzi_1979             ...............a.........................---................c................... [ 19440]
AY261361.1_Malawi_Lil-20/1_1983   ..............aa.........................---.................................... [ 19440]
AY261360.1_Kenya_1950             ...............a.........................---...c...................a.......g...t [ 19440]
AM712240.1_OURT_88/3              -------------------------------------------------------------------------------- [ 19440]
AM712239.1_Benin_97/1             -------------------------------------------------------------------------------- [ 19440]
KM111295.1_Ken06.Bus              ...............a.........................---..gc................................ [ 19440]
```

```
KM111294.1_Ken05/Tk1          .........a.....a...---........c...t.....c---...c...................a.......g...t [ 19440]
KM262844.1_strain_L60         -------------------------------------------------------------------------------- [ 19440]
KM102979.1_26544/OG10_Italy   -------------------------------------------------------------------------------- [ 19440]
KM262845.1_strain_NHV         -------------------------------------------------------------------------------- [ 19440]
KX354450.1_47/Ss/2008         -------------------------------------------------------------------------------- [ 19440]
KP055815.1_strain_BA71        -------------------------------------------------------------------------------- [ 19440]
MH025920.1_strain_R35         .............a.a.........................---..gc................................ [ 19440]
MH025919.1_strain_N10         .............a.a.........................---..gc................................ [ 19440]
MH025918.1_strain_R25         .............a.a.........................---..gc................................ [ 19440]
MH025917.1_strain_R7          .............a.a.........................---..gc................................ [ 19440]
MH025916.1_strain_R8          .............a.a.........................---..gc................................ [ 19440]

FR682468.2_Georgia_2007/1     atggttgtactcagtcgtagatttttgcagatcgcaaatgcaacgcaaccagcaaatacaaagctaaatacaaggtaaaa [ 19520]
ASF_IND_20_CAD543             -------------------------------------------------------------------------------- [ 19520]
LR899193.1_Germany_2020/1     ................................................................................ [ 19520]
MN393477.1_Wuhan_2019         ................................................................................ [ 19520]
MW396979.1_Timor-Leste/2019/1 ................................................................................ [ 19520]
MK940252.1_CN/2019/InnerMongolia ................................................................................ [ 19520]
MK128995.1_China/2018/AnhuiXCGQ ................................................................................ [ 19520]
MH681419.1_ASFV/POL/2015      ................................................................................ [ 19520]
LS478113.1_Estonia_2014       --------------.acg.........tttcc.a.cc...tt.t.g.ct..at------------------------t.c [ 19520]
KP843857.1_Odintsovo_02/14    ................................................................................ [ 19520]
FN557520.1_strain_E75         -------------------------------------------------------------------------------- [ 19520]
AY261366.1_Warthog            ..a.......a................................t.............g....a................. [ 19520]
AY261365.1_Warmbaths          ..a.......a................................t.............g....a................. [ 19520]
AY261364.1_Tengani_62         -------------------------------------------------------------------------------- [ 19520]
AY261363.1_Pretoriuskop/96/4  -------------------------------------------------------------------------------- [ 19520]
AY261362.1_Mkuzi_1979         ..........a.......................................g......g..........c........... [ 19520]
AY261361.1_Malawi_Lil-20/1_1983 ..........g..............................................g...................... [ 19520]
AY261360.1_Kenya_1950         ..a.......g.........g....................................g...................... [ 19520]
AM712240.1_OURT_88/3          -------------------------------------------------------------------------------- [ 19520]
AM712239.1_Benin_97/1         -------------------------------------------------------------------------------- [ 19520]
KM111295.1_Ken06.Bus          ..a......................................................g...................... [ 19520]
KM111294.1_Ken05/Tk1          ..a.......g....................................t.........g...t.................. [ 19520]
KM262844.1_strain_L60         -------------------------------------------------------------------------------- [ 19520]
KM102979.1_26544/OG10_Italy   -------------------------------------------------------------------------------- [ 19520]
KM262845.1_strain_NHV         -------------------------------------------------------------------------------- [ 19520]
KX354450.1_47/Ss/2008         -------------------------------------------------------------------------------- [ 19520]
KP055815.1_strain_BA71        -------------------------------------------------------------------------------- [ 19520]
MH025920.1_strain_R35         ..a......................................................g...................... [ 19520]
MH025919.1_strain_N10         ..a......................................................g...................... [ 19520]
MH025918.1_strain_R25         ..a......................................................g...................... [ 19520]
MH025917.1_strain_R7          ..a......................................................g...................... [ 19520]
MH025916.1_strain_R8          ..a......................................................g...................... [ 19520]
```

```
FR682468.2_Georgia_2007/1        acaataataccttataatgattggccaattcttatccctccatttttccatgaacatttcatgttcataaagtctaggat [ 19600]
ASF_IND_20_CAD543                -------------------------------------------------------------------------------- [ 19600]
LR899193.1_Germany_2020/1        ................................................................................ [ 19600]
MN393477.1_Wuhan_2019            ................................................................................ [ 19600]
MW396979.1_Timor-Leste/2019/1    ................................................................................ [ 19600]
MK940252.1_CN/2019/InnerMongolia ................................................................................ [ 19600]
MK128995.1_China/2018/AnhuiXCGQ  ................................................................................ [ 19600]
MH681419.1_ASFV/POL/2015         ................................................................................ [ 19600]
LS478113.1_Estonia_2014          ......t...t.c.--------------------------------..t..t..t....t.------------------- [ 19600]
KP843857.1_Odintsovo_02/14       ................................................................................ [ 19600]
FN557520.1_strain_E75            -------------------------------------------------------------------------------- [ 19600]
AY261366.1_Warthog               .......................a.......................t.........gaa.....t.............. [ 19600]
AY261365.1_Warmbaths             .......................a..............................t..gaa.....t.............. [ 19600]
AY261364.1_Tengani_62            -------------------------------------------------------------------------------- [ 19600]
AY261363.1_Pretoriuskop/96/4     -------------------------------------------------------------------------------- [ 19600]
AY261362.1_Mkuzi_1979            .......................a...........................----------------------------- [ 19600]
AY261361.1_Malawi_Lil-20/1_1983  ........gt.......g..............c...................g....gaa.....t.............. [ 19600]
AY261360.1_Kenya_1950            ....c.c.gtt..................a..c..................a............................ [ 19600]
AM712240.1_OURT_88/3             -------------------------------------------------------------------------------- [ 19600]
AM712239.1_Benin_97/1            -------------------------------------------------------------------------------- [ 19600]
KM111295.1_Ken06.Bus             ........gtt............a........c.............................t......cga..c..... [ 19600]
KM111294.1_Ken05/Tk1             ....c.c.gtt..................a..c..................a............................ [ 19600]
KM262844.1_strain_L60            -------------------------------------------------------------------------------- [ 19600]
KM102979.1_26544/OG10_Italy      -------------------------------------------------------------------------------- [ 19600]
KM262845.1_strain_NHV            -------------------------------------------------------------------------------- [ 19600]
KX354450.1_47/Ss/2008            -------------------------------------------------------------------------------- [ 19600]
KP055815.1_strain_BA71           -------------------------------------------------------------------------------- [ 19600]
MH025920.1_strain_R35            ........gtt............a........c.............................t......cga..c..... [ 19600]
MH025919.1_strain_N10            ........gtt............a........c.............................t......cga..c..... [ 19600]
MH025918.1_strain_R25            ........gtt............a........c.............................t......cga..c..... [ 19600]
MH025917.1_strain_R7             ........gtt............a........c.............................t......cga..c..... [ 19600]
MH025916.1_strain_R8             ........gtt............a........c.............................t......cga..c..... [ 19600]

FR682468.2_Georgia_2007/1        acgaacaacatttcatgctatgatgattaggtattttaagtgatatttcataaaaaca---------------------- [ 19680]
ASF_IND_20_CAD543                -------------------------------------------------------------------------------- [ 19680]
LR899193.1_Germany_2020/1        ..........................................................---------------------- [ 19680]
MN393477.1_Wuhan_2019            ..........................................................---------------------- [ 19680]
MW396979.1_Timor-Leste/2019/1    ..........................................................---------------------- [ 19680]
MK940252.1_CN/2019/InnerMongolia ..........................................................---------------------- [ 19680]
MK128995.1_China/2018/AnhuiXCGQ  ..........................................................---------------------- [ 19680]
MH681419.1_ASFV/POL/2015         ..........................................................---------------------- [ 19680]
LS478113.1_Estonia_2014          ---------------------.g.a..a..tct...c....tcag.g.a.....t.------------------------ [ 19680]
KP843857.1_Odintsovo_02/14       ..........................................................---------------------- [ 19680]
```

```
FN557520.1_strain_E75            -------------------------------------------------------------------------------- [ 19680]
AY261366.1_Warthog               .t.---.g....c.............................................---------------------- [ 19680]
AY261365.1_Warmbaths             .t.---......c.............................................---------------------- [ 19680]
AY261364.1_Tengani_62            -------------------------------------------------------------------------------- [ 19680]
AY261363.1_Pretoriuskop/96/4     -------------------------------------------------------------------------------- [ 19680]
AY261362.1_Mkuzi_1979            ----------------.............a............................---------------------- [ 19680]
AY261361.1_Malawi_Lil-20/1_1983  .t.---......c...........a.................................ggactgggattgggaatcgcga [ 19680]
AY261360.1_Kenya_1950            .t.---......c..........................c......a...........ggac------tgggaatcgcga [ 19680]
AM712240.1_OURT_88/3             -------------------------------------------------------------------------------- [ 19680]
AM712239.1_Benin_97/1            -------------------------------------------------------------------------------- [ 19680]
KM111295.1_Ken06.Bus             .a.---......c............................a.............g..ggactgagattgggaatcgcaa [ 19680]
KM111294.1_Ken05/Tk1             .t.---......c..........................c..................ggat------tgggaatcgcga [ 19680]
KM262844.1_strain_L60            -------------------------------------------------------------------------------- [ 19680]
KM102979.1_26544/OG10_Italy      -------------------------------------------------------------------------------- [ 19680]
KM262845.1_strain_NHV            -------------------------------------------------------------------------------- [ 19680]
KX354450.1_47/Ss/2008            -------------------------------------------------------------------------------- [ 19680]
KP055815.1_strain_BA71           -------------------------------------------------------------------------------- [ 19680]
MH025920.1_strain_R35            .a.---......c............................a.............g..ggactgagattgggaatcgcaa [ 19680]
MH025919.1_strain_N10            .a.---......c............................a.............g..ggactgagattgggaatcgcaa [ 19680]
MH025918.1_strain_R25            .a.---......c............................a.............g..ggactgagattgggaatcgcaa [ 19680]
MH025917.1_strain_R7             .a.---......c............................a.............g..ggactgagattgggaatcgcaa [ 19680]
MH025916.1_strain_R8             .a.---......c............................a.............g..ggactgagattgggaatcgcaa [ 19680]

FR682468.2_Georgia_2007/1        -------------------------------------------------------------------------------- [ 19760]
ASF_IND_20_CAD543                -------------------------------------------------------------------------------- [ 19760]
LR899193.1_Germany_2020/1        -------------------------------------------------------------------------------- [ 19760]
MN393477.1_Wuhan_2019            -------------------------------------------------------------------------------- [ 19760]
MW396979.1_Timor-Leste/2019/1    -------------------------------------------------------------------------------- [ 19760]
MK940252.1_CN/2019/InnerMongolia -------------------------------------------------------------------------------- [ 19760]
MK128995.1_China/2018/AnhuiXCGQ  -------------------------------------------------------------------------------- [ 19760]
MH681419.1_ASFV/POL/2015         -------------------------------------------------------------------------------- [ 19760]
LS478113.1_Estonia_2014          -------------------------------------------------------------------------------- [ 19760]
KP843857.1_Odintsovo_02/14       -------------------------------------------------------------------------------- [ 19760]
FN557520.1_strain_E75            -------------------------------------------------------------------------------- [ 19760]
AY261366.1_Warthog               -------------------------------------------------------------------------------- [ 19760]
AY261365.1_Warmbaths             -------------------------------------------------------------------------------- [ 19760]
AY261364.1_Tengani_62            -------------------------------------------------------------------------------- [ 19760]
AY261363.1_Pretoriuskop/96/4     -------------------------------------------------------------------------------- [ 19760]
AY261362.1_Mkuzi_1979            -------------------------------------------------------------------------------- [ 19760]
AY261361.1_Malawi_Lil-20/1_1983  gtaactctacagtttacaatcgaattcatcgaaatagattcatttattaccttatttatacaagtgtcattttggcaatc [ 19760]
AY261360.1_Kenya_1950            gtaactctacagtttacaatcgaagttatcgaaatagactcatttattaccttatttatacaagtgtcattttggcaatc [ 19760]
AM712240.1_OURT_88/3             -------------------------------------------------------------------------------- [ 19760]
AM712239.1_Benin_97/1            -------------------------------------------------------------------------------- [ 19760]
KM111295.1_Ken06.Bus             gtaactctacagtttataatcgaatttatcgaaatagactcacttattaccttgttccaatagatgctatcttgacaatc [ 19760]
```

```
KM111294.1_Ken05/Tk1             gtaactctacagtttacaatcgaagttatcgaaatagactcatttattaccttatttatacaagtgtcattttggcaatc [ 19760]
KM262844.1_strain_L60            -------------------------------------------------------------------------------- [ 19760]
KM102979.1_26544/OG10_Italy      -------------------------------------------------------------------------------- [ 19760]
KM262845.1_strain_NHV            -------------------------------------------------------------------------------- [ 19760]
KX354450.1_47/Ss/2008            -------------------------------------------------------------------------------- [ 19760]
KP055815.1_strain_BA71           -------------------------------------------------------------------------------- [ 19760]
MH025920.1_strain_R35            gtaactctacagtttataatcgaatttatcgaaatagactcacttattaccttgttccaatagatgctatcttgacaatc [ 19760]
MH025919.1_strain_N10            gtaactctacagtttataatcgaatttatcgaaatagactcacttattaccttgttccaatagatgctatcttgacaatc [ 19760]
MH025918.1_strain_R25            gtaactctacagtttataatcgaatttatcgaaatagactcacttattaccttgttccaatagatgctatcttgacaatc [ 19760]
MH025917.1_strain_R7             gtaactctacagtttataatcgaatttatcgaaatagactcacttattaccttgttccaatagatgctatcttgacaatc [ 19760]
MH025916.1_strain_R8             gtaactctacagtttataatcgaatttatcgaaatagactcacttattaccttgttccaatagatgctatcttgacaatc [ 19760]

FR682468.2_Georgia_2007/1        ------------------------------------------------------------ccacggggttgttggtgatt [ 19840]
ASF_IND_20_CAD543                -------------------------------------------------------------------------------- [ 19840]
LR899193.1_Germany_2020/1        ------------------------------------------------------------.................... [ 19840]
MN393477.1_Wuhan_2019            ------------------------------------------------------------.................... [ 19840]
MW396979.1_Timor-Leste/2019/1    ------------------------------------------------------------.................... [ 19840]
MK940252.1_CN/2019/InnerMongolia ------------------------------------------------------------.................... [ 19840]
MK128995.1_China/2018/AnhuiXCGQ  ------------------------------------------------------------.................... [ 19840]
MH681419.1_ASFV/POL/2015         ------------------------------------------------------------.................... [ 19840]
LS478113.1_Estonia_2014          -------------------------------------------------------------------------------- [ 19840]
KP843857.1_Odintsovo_02/14       ------------------------------------------------------------.................... [ 19840]
FN557520.1_strain_E75            -------------------------------------------------------------------------------- [ 19840]
AY261366.1_Warthog               ------------------------------------------------------------....t......c........ [ 19840]
AY261365.1_Warmbaths             ------------------------------------------------------------....t......c........ [ 19840]
AY261364.1_Tengani_62            -------------------------------------------------------------------------------- [ 19840]
AY261363.1_Pretoriuskop/96/4     -------------------------------------------------------------------------------- [ 19840]
AY261362.1_Mkuzi_1979            ------------------------------------------------------------....t.......c....... [ 19840]
AY261361.1_Malawi_Lil-20/1_1983  ccagcagaagtcacaagtttttgcataagtgcaccaatactcaagctcttccttaggaggatg.t............... [ 19840]
AY261360.1_Kenya_1950            ccagcagaagtcacaagattttgcataagtgcaccagtacgcaagctcttccttaggaggatg.t.......c....... [ 19840]
AM712240.1_OURT_88/3             -------------------------------------------------------------------------------- [ 19840]
AM712239.1_Benin_97/1            -------------------------------------------------------------------------------- [ 19840]
KM111295.1_Ken06.Bus             ccaacagaggtcgcaattctttacatacgtacaccaatatggaagctcctccttaggaggatg.t............t.. [ 19840]
KM111294.1_Ken05/Tk1             ccagcagaagtcacaagattttgcataagtgcaccagtacgcaagctcttccttaggaggatg.t.a.....c....... [ 19840]
KM262844.1_strain_L60            -------------------------------------------------------------------------------- [ 19840]
KM102979.1_26544/OG10_Italy      -------------------------------------------------------------------------------- [ 19840]
KM262845.1_strain_NHV            -------------------------------------------------------------------------------- [ 19840]
KX354450.1_47/Ss/2008            -------------------------------------------------------------------------------- [ 19840]
KP055815.1_strain_BA71           -------------------------------------------------------------------------------- [ 19840]
MH025920.1_strain_R35            ccaacagaggtcgcaattctttacatacgtacaccaatatggaagctcctccttaggaggatg.t............t.. [ 19840]
MH025919.1_strain_N10            ccaacagaggtcgcaattctttacatacgtacaccaatatggaagctcctccttaggaggatg.t............t.. [ 19840]
MH025918.1_strain_R25            ccaacagaggtcgcaattctttacatacgtacaccaatatggaagctcctccttaggaggatg.t............t.. [ 19840]
MH025917.1_strain_R7             ccaacagaggtcgcaattctttacatacgtacaccaatatggaagctcctccttaggaggatg.t............t.. [ 19840]
MH025916.1_strain_R8             ccaacagaggtcgcaattctttacatacgtacaccaatatggaagctcctccttaggaggatg.t............t.. [ 19840]
```

```
FR682468.2_Georgia_2007/1        gataggtaagaataaggatggttgaataacctagtaaaagtcctagaaaaaccttcat-attgcgtt------------c [ 19920]
ASF_IND_20_CAD543                -------------------------------------------------------------------------------- [ 19920]
LR899193.1_Germany_2020/1        ..........................................................-........------------. [ 19920]
MN393477.1_Wuhan_2019            ..........................................................-........------------. [ 19920]
MW396979.1_Timor-Leste/2019/1    ..........................................................-........------------. [ 19920]
MK940252.1_CN/2019/InnerMongolia ..........................................................-........------------. [ 19920]
MK128995.1_China/2018/AnhuiXCGQ  ..........................................................-........------------. [ 19920]
MH681419.1_ASFV/POL/2015         ..........................................................-........------------. [ 19920]
LS478113.1_Estonia_2014          ---.a....ct...t..t.--..a...gt.ta.aa..t.t......---------------------------------- [ 19920]
KP843857.1_Odintsovo_02/14       ..........................................................-........------------. [ 19920]
FN557520.1_strain_E75            -------------------------------------------------------------------------------- [ 19920]
AY261366.1_Warthog               ....t........g............................................gg.......------------. [ 19920]
AY261365.1_Warmbaths             ....t........g............................................gg.......------------. [ 19920]
AY261364.1_Tengani_62            -------------------------------------------------------------------------------- [ 19920]
AY261363.1_Pretoriuskop/96/4     -------------------------------------------------------------------------------- [ 19920]
AY261362.1_Mkuzi_1979            ....t........g............................................gg.......------------. [ 19920]
AY261361.1_Malawi_Lil-20/1_1983  ....t........g............................................-.......-------------- [ 19920]
AY261360.1_Kenya_1950            ....t........g......a.....a...............................-g....ag.atgtaatataca. [ 19920]
AM712240.1_OURT_88/3             -------------------------------------------------------------------------------- [ 19920]
AM712239.1_Benin_97/1            -------------------------------------------------------------------------------- [ 19920]
KM111295.1_Ken06.Bus             ....t........g.....a...a.................g.c..............-g.......------------. [ 19920]
KM111294.1_Ken05/Tk1             ....t........g......a.....a...............................-g....ag.atgtaa------t [ 19920]
KM262844.1_strain_L60            -------------------------------------------------------------------------------- [ 19920]
KM102979.1_26544/OG10_Italy      -------------------------------------------------------------------------------- [ 19920]
KM262845.1_strain_NHV            -------------------------------------------------------------------------------- [ 19920]
KX354450.1_47/Ss/2008            -------------------------------------------------------------------------------- [ 19920]
KP055815.1_strain_BA71           -------------------------------------------------------------------------------- [ 19920]
MH025920.1_strain_R35            ....t........g.....a...a.................g.c..............-g.......------------. [ 19920]
MH025919.1_strain_N10            ....t........g.....a...a.................g.c..............-g.......------------. [ 19920]
MH025918.1_strain_R25            ....t........g.....a...a.................g.c..............-g.......------------. [ 19920]
MH025917.1_strain_R7             ....t........g.....a...a.................g.c..............-g.......------------. [ 19920]
MH025916.1_strain_R8             ....t........g.....a...a.................g.c..............-g.......------------. [ 19920]

FR682468.2_Georgia_2007/1        ataccacagatgttatt-taaaaaaaata--taaattttacagtatgtgatatacacataccacaaaaatgttcttatat [ 20000]
ASF_IND_20_CAD543                -------------------------------------------------------------------------------- [ 20000]
LR899193.1_Germany_2020/1        .................-...........--................................................. [ 20000]
MN393477.1_Wuhan_2019            .................-...........--................................................. [ 20000]
MW396979.1_Timor-Leste/2019/1    .................-...........--................................................. [ 20000]
MK940252.1_CN/2019/InnerMongolia .................-...........--................................................. [ 20000]
MK128995.1_China/2018/AnhuiXCGQ  .................-...........--................................................. [ 20000]
MH681419.1_ASFV/POL/2015         .................-...........--................................................. [ 20000]
LS478113.1_Estonia_2014          ---------------------------------------...tc..a..t..------------......a..t..cc.- [ 20000]
KP843857.1_Odintsovo_02/14       .................-...........--................................................. [ 20000]
```

```
FN557520.1_strain_E75           -------------------------------------------------------------------------------- [ 20000]
AY261366.1_Warthog              .................--t.........tt..................tat............................ [ 20000]
AY261365.1_Warmbaths            .................-a..........tt..................tat............................ [ 20000]
AY261364.1_Tengani_62           -------------------------------------------------------------------------------- [ 20000]
AY261363.1_Pretoriuskop/96/4    -------------------------------------------------------------------------------- [ 20000]
AY261362.1_Mkuzi_1979           ...t.............-a..........tt..................ta............................. [ 20000]
AY261361.1_Malawi_Lil-20/1_1983 ---..............ta..........tt..........g...................................... [ 20000]
AY261360.1_Kenya_1950           .................ta..........tt........g.c.......tatatt......tg................. [ 20000]
AM712240.1_OURT_88/3            -------------------------------------------------------------------------------- [ 20000]
AM712239.1_Benin_97/1           -------------------------------------------------------------------------------- [ 20000]
KM111295.1_Ken06.Bus            ....a............ta..........tt........g..c.g..-atata-........g.-.....ac........ [ 20000]
KM111294.1_Ken05/Tk1            .................ta..........tt........g.c.......tatatt......tg................. [ 20000]
KM262844.1_strain_L60           -------------------------------------------------------------------------------- [ 20000]
KM102979.1_26544/OG10_Italy     -------------------------------------------------------------------------------- [ 20000]
KM262845.1_strain_NHV           -------------------------------------------------------------------------------- [ 20000]
KX354450.1_47/Ss/2008           -------------------------------------------------------------------------------- [ 20000]
KP055815.1_strain_BA71          -------------------------------------------------------------------------------- [ 20000]
MH025920.1_strain_R35           ....a............ta..........tt........g..c.g..-atata-........g.-.....ac........ [ 20000]
MH025919.1_strain_N10           ....a............t-..........tt........g..c.g..-atata-........g.-.....ac........ [ 20000]
MH025918.1_strain_R25           ....a............ta..........tt........g..c.g..-atata-........g.-.....ac........ [ 20000]
MH025917.1_strain_R7            ....a............ta..........tt........g..c.g..-atata-........g.-.....ac........ [ 20000]
MH025916.1_strain_R8            ....a............ta..........tt........g..c.g..-atata-........g.-.....ac........ [ 20000]

FR682468.2_Georgia_2007/1       taactaaaatatgtgggcagagagcaattcatataatgaatatatggtattttaggctcaataaagtacatacaacgatc [ 20080]
ASF_IND_20_CAD543               -------------------------------------------------------------------------------- [ 20080]
LR899193.1_Germany_2020/1       ................................................................................ [ 20080]
MN393477.1_Wuhan_2019           ................................................................................ [ 20080]
MW396979.1_Timor-Leste/2019/1   ................................................................................ [ 20080]
MK940252.1_CN/2019/InnerMongolia ................................................................................ [ 20080]
MK128995.1_China/2018/AnhuiXCGQ ................................................................................ [ 20080]
MH681419.1_ASFV/POL/2015        ................................................................................ [ 20080]
LS478113.1_Estonia_2014         ---------------------------------------------aa..c....a------------------------- [ 20080]
KP843857.1_Odintsovo_02/14      ................................................................................ [ 20080]
FN557520.1_strain_E75           ---------------................................................................. [ 20080]
AY261366.1_Warthog              .................t.....a................................t....................... [ 20080]
AY261365.1_Warmbaths            .......................a........................................................ [ 20080]
AY261364.1_Tengani_62           -------------------------------------------------------------------------------- [ 20080]
AY261363.1_Pretoriuskop/96/4    --------------------------------------------------------------...a......t..t.g.. [ 20080]
AY261362.1_Mkuzi_1979           ................................................................................ [ 20080]
AY261361.1_Malawi_Lil-20/1_1983 ................................................................................ [ 20080]
AY261360.1_Kenya_1950           ............a.........ta........g...........c....a...............a........g..... [ 20080]
AM712240.1_OURT_88/3            -----------...a..tt....a....c...............g...ga...g..t.t......a..t........g.. [ 20080]
AM712239.1_Benin_97/1           ---------------................................................................. [ 20080]
KM111295.1_Ken06.Bus            ......................ta........g...........c................................... [ 20080]
```

```
KM111294.1_Ken05/Tk1          ............................................c....................a.............. [ 20080]
KM262844.1_strain_L60         ---------------................................................................. [ 20080]
KM102979.1_26544/OG10_Italy   ---------------................................................................. [ 20080]
KM262845.1_strain_NHV         -----------...a..tt....a....c...............g...ga...g..t.t......a..t........g.. [ 20080]
KX354450.1_47/Ss/2008         ---------------................................................................. [ 20080]
KP055815.1_strain_BA71        -------------------------.cc...c.ag...------------------------------------------ [ 20080]
MH025920.1_strain_R35         ......................ta........g...........c................................... [ 20080]
MH025919.1_strain_N10         ......................ta........g...........c................................... [ 20080]
MH025918.1_strain_R25         ......................ta........g...........c................................... [ 20080]
MH025917.1_strain_R7          ......................ta........g...........c................................... [ 20080]
MH025916.1_strain_R8          ......................ta........g...........c................................... [ 20080]

FR682468.2_Georgia_2007/1     aataaaacgggtaatactacatttactgatgtaatcattttgaacaataagaggcatatcatccaaaaccttatttttac [ 20160]
ASF_IND_20_CAD543             -------------------------------------------------------------------------------- [ 20160]
LR899193.1_Germany_2020/1     ................................................................................ [ 20160]
MN393477.1_Wuhan_2019         ................................................................................ [ 20160]
MW396979.1_Timor-Leste/2019/1 ................................................................................ [ 20160]
MK940252.1_CN/2019/InnerMongolia ................................................................................ [ 20160]
MK128995.1_China/2018/AnhuiXCGQ ................................................................................ [ 20160]
MH681419.1_ASFV/POL/2015      ................................................................................ [ 20160]
LS478113.1_Estonia_2014       ------------....t..a.......t.gcct.a..a.a.-----------att.c..t..tgt.c...........tt [ 20160]
KP843857.1_Odintsovo_02/14    ................................................................................ [ 20160]
FN557520.1_strain_E75         ..............................a............................t.................... [ 20160]
AY261366.1_Warthog            ................................................................................ [ 20160]
AY261365.1_Warmbaths          ................................................................................ [ 20160]
AY261364.1_Tengani_62         -------------------------------------------------------------------------------- [ 20160]
AY261363.1_Pretoriuskop/96/4  .------------------------------------------------------------------------------- [ 20160]
AY261362.1_Mkuzi_1979         ................................................................................ [ 20160]
AY261361.1_Malawi_Lil-20/1_1983 ....................................g........................................... [ 20160]
AY261360.1_Kenya_1950         .........................................................g.t.................... [ 20160]
AM712240.1_OURT_88/3          ..a.t.g....c...........g...a..a...........ttt...................c....t.......... [ 20160]
AM712239.1_Benin_97/1         ..............................a............................t.................... [ 20160]
KM111295.1_Ken06.Bus          ................................................................................ [ 20160]
KM111294.1_Ken05/Tk1          ......g......................................................................... [ 20160]
KM262844.1_strain_L60         ..............................a............................t.................... [ 20160]
KM102979.1_26544/OG10_Italy   ..............................a............................t.................... [ 20160]
KM262845.1_strain_NHV         ..a.t.g....c...........g...a..a...........ttt...................c....t.......... [ 20160]
KX354450.1_47/Ss/2008         ..............................a............................t.................... [ 20160]
KP055815.1_strain_BA71        --------------------------.c.aa...ctg..c.c..t...c.at.....gctc..attc.....g..c..g. [ 20160]
MH025920.1_strain_R35         ................................................................................ [ 20160]
MH025919.1_strain_N10         ................................................................................ [ 20160]
MH025918.1_strain_R25         ................................................................................ [ 20160]
MH025917.1_strain_R7          ................................................................................ [ 20160]
MH025916.1_strain_R8          ................................................................................ [ 20160]
```

```
FR682468.2_Georgia_2007/1        aaataccattcttacaatcccagcagaaatcacagtgttttccatacgtacaccaatattcaagttctctcataggaggc [ 20240]
ASF_IND_20_CAD543                -------------------------------------------------------------------------------- [ 20240]
LR899193.1_Germany_2020/1        ................................................................................ [ 20240]
MN393477.1_Wuhan_2019            ................................................................................ [ 20240]
MW396979.1_Timor-Leste/2019/1    ................................................................................ [ 20240]
MK940252.1_CN/2019/InnerMongolia ................................................................................ [ 20240]
MK128995.1_China/2018/AnhuiXCGQ  ................................................................................ [ 20240]
MH681419.1_ASFV/POL/2015         ................................................................................ [ 20240]
LS478113.1_Estonia_2014          tgg-.......a.------------------------..catt..tt...g.a..c....-------------------- [ 20240]
KP843857.1_Odintsovo_02/14       ................................................................................ [ 20240]
FN557520.1_strain_E75            ................................................................................ [ 20240]
AY261366.1_Warthog               ..................................a......................................g...... [ 20240]
AY261365.1_Warmbaths             ................................................................................ [ 20240]
AY261364.1_Tengani_62            -------------------------------------------------------------------------------- [ 20240]
AY261363.1_Pretoriuskop/96/4     --------------------------....ag..a....g.-t...taggtgt...........c...tat.cga.---- [ 20240]
AY261362.1_Mkuzi_1979            ................t............................................................... [ 20240]
AY261361.1_Malawi_Lil-20/1_1983  .........................................................................g...... [ 20240]
AY261360.1_Kenya_1950            ................tgt...........................t..........................g.....t [ 20240]
AM712240.1_OURT_88/3             ...........c....t................................g.....g........c.....t........t [ 20240]
AM712239.1_Benin_97/1            ................................................................................ [ 20240]
KM111295.1_Ken06.Bus             ................tgt......................................................g.....t [ 20240]
KM111294.1_Ken05/Tk1             ................tgt......................................................g...... [ 20240]
KM262844.1_strain_L60            ................................................................................ [ 20240]
KM102979.1_26544/OG10_Italy      ................................................................................ [ 20240]
KM262845.1_strain_NHV            ...........c....t................................g.....g........c.....t........t [ 20240]
KX354450.1_47/Ss/2008            ................................................................................ [ 20240]
KP055815.1_strain_BA71           ....t....gtgc...t........a..c.tg...ct..ccatg..a........g...c........tcttgt.....a [ 20240]
MH025920.1_strain_R35            ................tgt......................................................g.....t [ 20240]
MH025919.1_strain_N10            ................tgt......................................................g.....t [ 20240]
MH025918.1_strain_R25            ................tgt......................................................g.....t [ 20240]
MH025917.1_strain_R7             ................tgt......................................................g.....t [ 20240]
MH025916.1_strain_R8             ................tgt......................................................g.....t [ 20240]

FR682468.2_Georgia_2007/1        gtataggtccttggtaaaatttgtttcgtataaaagatggaaaggggtcgatttaaaactggctgtgctaaccaaaccaa [ 20320]
ASF_IND_20_CAD543                -------------------------------------------------------------------------------- [ 20320]
LR899193.1_Germany_2020/1        ................................................................................ [ 20320]
MN393477.1_Wuhan_2019            ................................................................................ [ 20320]
MW396979.1_Timor-Leste/2019/1    ................................................................................ [ 20320]
MK940252.1_CN/2019/InnerMongolia ................................................................................ [ 20320]
MK128995.1_China/2018/AnhuiXCGQ  ................................................................................ [ 20320]
MH681419.1_ASFV/POL/2015         ................................................................................ [ 20320]
LS478113.1_Estonia_2014          ----------------------------.c.....a..-------------.....t...act..gat.-----..atct [ 20320]
KP843857.1_Odintsovo_02/14       ................................................................................ [ 20320]
```

```
FN557520.1_strain_E75          ................................................................................ [ 20320]
AY261366.1_Warthog             ................................................................................ [ 20320]
AY261365.1_Warmbaths           ................................................................................ [ 20320]
AY261364.1_Tengani_62          -------------------------------------------------------------------------------- [ 20320]
AY261363.1_Pretoriuskop/96/4   .....aa.tt.c......g.gat..aac...-----g.a.g..at...t.......g........g.............. [ 20320]
AY261362.1_Mkuzi_1979          ................................................................................ [ 20320]
AY261361.1_Malawi_Lil-20/1_1983 .............................................................................t.. [ 20320]
AY261360.1_Kenya_1950          ..................................................g...........t..g.............. [ 20320]
AM712240.1_OURT_88/3           .....a............t........a......................g......c.c.....a.a..g.....t... [ 20320]
AM712239.1_Benin_97/1          ................................................................................ [ 20320]
KM111295.1_Ken06.Bus           ........t.........................................g...........t..g.............. [ 20320]
KM111294.1_Ken05/Tk1           ..............................................................t..g.............. [ 20320]
KM262844.1_strain_L60          ................................................................................ [ 20320]
KM102979.1_26544/OG10_Italy    ................................................................................ [ 20320]
KM262845.1_strain_NHV          .....a............t........a......................g......c.c.....a.a..g.....t... [ 20320]
KX354450.1_47/Ss/2008          ............a................................................................... [ 20320]
KP055815.1_strain_BA71         t...cc..tgaac.a.g.tgccc.cct.cc.---------g.gta...a.tcc...g..ct.a.tg..c.g.aggc.a.g [ 20320]
MH025920.1_strain_R35          ........t.........................................g...........t..g.............. [ 20320]
MH025919.1_strain_N10          ........t.........................................g...........t..g.............. [ 20320]
MH025918.1_strain_R25          ........t.........................................g...........t..g.............. [ 20320]
MH025917.1_strain_R7           ........t.........................................g...........t..g.............. [ 20320]
MH025916.1_strain_R8           ........t.........................................g...........t..g.............. [ 20320]

FR682468.2_Georgia_2007/1      aatactcaaaagaacgaaaagtttcatggttgtactcagacgcagattcttacaaagcgcacatacaaagcagcctgtat [ 20400]
ASF_IND_20_CAD543              -------------------------------------------------------------------------------- [ 20400]
LR899193.1_Germany_2020/1      ................................................................................ [ 20400]
MN393477.1_Wuhan_2019          ................................................................................ [ 20400]
MW396979.1_Timor-Leste/2019/1  ................................................................................ [ 20400]
MK940252.1_CN/2019/InnerMongolia ................................................................................ [ 20400]
MK128995.1_China/2018/AnhuiXCGQ ................................................................................ [ 20400]
MH681419.1_ASFV/POL/2015       ................................................................................ [ 20400]
LS478113.1_Estonia_2014        ................................................................................ [ 20400]
KP843857.1_Odintsovo_02/14     ................................................................................ [ 20400]
FN557520.1_strain_E75          ...........................a.................................................... [ 20400]
AY261366.1_Warthog             ..............t..................t.............................................. [ 20400]
AY261365.1_Warmbaths           ................................................................................ [ 20400]
AY261364.1_Tengani_62          -------------------------------------------------------------------------------- [ 20400]
AY261363.1_Pretoriuskop/96/4   .............c.a.gt..c.....a..a.........t..a.....c...gg.t....a..g...ca..a.tc.c.a [ 20400]
AY261362.1_Mkuzi_1979          .......................................t........................................ [ 20400]
AY261361.1_Malawi_Lil-20/1_1983 .................................................................t.......t...... [ 20400]
AY261360.1_Kenya_1950          ......t......g....t.c............................................g.............. [ 20400]
AM712240.1_OURT_88/3           .....at......g.a.gt........a.cg...t.t..gt.t.a...t...t.gta....a......t.t.a...ac.a [ 20400]
AM712239.1_Benin_97/1          ...........................a.................................................... [ 20400]
KM111295.1_Ken06.Bus           .............g....t.c........................................................... [ 20400]
```

```
KM111294.1_Ken05/Tk1              ......t......g....t.c.................a......................................... [ 20400]
KM262844.1_strain_L60             ...........................a.................................................... [ 20400]
KM102979.1_26544/OG10_Italy       ...........................a.................................................... [ 20400]
KM262845.1_strain_NHV             .....at......g.a.gt........a.cg...t.t..gt.t.a...t...t.gta....a......t.t.a...ac.a [ 20400]
KX354450.1_47/Ss/2008             ...........................a.................................................... [ 20400]
KP055815.1_strain_BA71            ...t.c...g.ag.tc.cc.ac-----------------------...gc...ggctg..---.ga.c...t.a.c.a.. [ 20400]
MH025920.1_strain_R35             .............g....t.c........................................................... [ 20400]
MH025919.1_strain_N10             .............g....t.c........................................................... [ 20400]
MH025918.1_strain_R25             .............g....t.c........................................................... [ 20400]
MH025917.1_strain_R7              .............g....t.c........................................................... [ 20400]
MH025916.1_strain_R8              .............g....t.c........................................................... [ 20400]

FR682468.2_Georgia_2007/1         atgcaataccaatgatgaaatagagacagtattgctttatagataattgttga-tggtca---ccccccccccccccccc [ 20480]
ASF_IND_20_CAD543                 -------------------------------------------------------------------------------- [ 20480]
LR899193.1_Germany_2020/1         .....................................................-......------.............. [ 20480]
MN393477.1_Wuhan_2019             .....................................................-......-------............. [ 20480]
MW396979.1_Timor-Leste/2019/1     .....................................................-......----................ [ 20480]
MK940252.1_CN/2019/InnerMongolia  .....................................................-......c-c................. [ 20480]
MK128995.1_China/2018/AnhuiXCGQ   .....................................................-......----................ [ 20480]
MH681419.1_ASFV/POL/2015          .....................................................-......----................ [ 20480]
LS478113.1_Estonia_2014           .....................................................-......-----............... [ 20480]
KP843857.1_Odintsovo_02/14        .....................................................-......----------.......... [ 20480]
FN557520.1_strain_E75             .a...................................................-......-----............... [ 20480]
AY261366.1_Warthog                ............................aa.......................-......----------.......... [ 20480]
AY261365.1_Warmbaths              .....................................................-......-----------......... [ 20480]
AY261364.1_Tengani_62             -------------------------------------------------------------------------------- [ 20480]
AY261363.1_Pretoriuskop/96/4      .........t..at.ca.g...a.a...ac.a..tc.....at....g.cca.-.a..tg----------t.at..tt.. [ 20480]
AY261362.1_Mkuzi_1979             ..................................t..................-......-----------......... [ 20480]
AY261361.1_Malawi_Lil-20/1_1983   .....................................................-......--------............ [ 20480]
AY261360.1_Kenya_1950             .......g................ag...........................-......c---------.......... [ 20480]
AM712240.1_OURT_88/3              .........t..at.ca.gg..a.a...ac.a..tc.....atggt.g.cca.-.aa...-----------......... [ 20480]
AM712239.1_Benin_97/1             .a...................................................-......-----------......... [ 20480]
KM111295.1_Ken06.Bus              .......g................ag...........................-.....gtca--------......... [ 20480]
KM111294.1_Ken05/Tk1              .......g................ag...............t...........-......cca-------.......... [ 20480]
KM262844.1_strain_L60             .a...................................................-......-----------......... [ 20480]
KM102979.1_26544/OG10_Italy       .a...................................................t......----------.......... [ 20480]
KM262845.1_strain_NHV             .........t..at.ca.gg..a.a...ac.a..tc.....atggt.g.cca.-.aa...----------.......... [ 20480]
KX354450.1_47/Ss/2008             .a...................................................t......----------.......... [ 20480]
KP055815.1_strain_BA71            .gct.--g.....t.gc...cca..-----------------------------..actc-----------g....t.ta [ 20480]
MH025920.1_strain_R35             .......g................ag.......a...................-.....gtca------........... [ 20480]
MH025919.1_strain_N10             .......g................ag.......a...................-.....gtca-------.......... [ 20480]
MH025918.1_strain_R25             .......g................ag.......a...................-.....gtca------........... [ 20480]
MH025917.1_strain_R7              .......g................ag.......a...................-.....gtca------........... [ 20480]
MH025916.1_strain_R8              .......g................ag.......a...................-.....gtca------........... [ 20480]
```

```
FR682468.2_Georgia_2007/1        atgtttgcatgaatatttcatttcctgtatagggtctaggatgtaaacattcca---tgctaaagtgattaggcatttt- [ 20560]
ASF_IND_20_CAD543                -------------------------------------------------------------------------------- [ 20560]
LR899193.1_Germany_2020/1        ......................................................---......................- [ 20560]
MN393477.1_Wuhan_2019            ......................................................---......................- [ 20560]
MW396979.1_Timor-Leste/2019/1    ......................................................---......................- [ 20560]
MK940252.1_CN/2019/InnerMongolia ......................................................---......................- [ 20560]
MK128995.1_China/2018/AnhuiXCGQ  ......................................................---......................- [ 20560]
MH681419.1_ASFV/POL/2015         ......................................................---......................- [ 20560]
LS478113.1_Estonia_2014          ......................................................---......................- [ 20560]
KP843857.1_Odintsovo_02/14       ......................................................---......................- [ 20560]
FN557520.1_strain_E75            .................................a....................---......................- [ 20560]
AY261366.1_Warthog               .....c.........................t......................---..t...................- [ 20560]
AY261365.1_Warmbaths             .....c.........................t............g.........---..t..tga.a............- [ 20560]
AY261364.1_Tengani_62            -------------------------------------------------------------------------------- [ 20560]
AY261363.1_Pretoriuskop/96/4     ..t....t..a..c.......a....a....a.a.a........g......g..tac.a...t...cg....at.....- [ 20560]
AY261362.1_Mkuzi_1979            .....c.................t.......t............g.........---..t..tga.a............- [ 20560]
AY261361.1_Malawi_Lil-20/1_1983  ...............................t......................---......................- [ 20560]
AY261360.1_Kenya_1950            ...g..t........................a............g.........---..t...................- [ 20560]
AM712240.1_OURT_88/3             ..t...c.....................................g.........---.....tga........t.....- [ 20560]
AM712239.1_Benin_97/1            .................................a....................---......................- [ 20560]
KM111295.1_Ken06.Bus             ......t........................a............g.........---.a....................- [ 20560]
KM111294.1_Ken05/Tk1             ...g..t........................a......................---......................- [ 20560]
KM262844.1_strain_L60            .................................a....................---......................- [ 20560]
KM102979.1_26544/OG10_Italy      .................................a....................---......................- [ 20560]
KM262845.1_strain_NHV            ..t...c.....................................g.........---.....tga........t.....- [ 20560]
KX354450.1_47/Ss/2008            .................................a....................---......................- [ 20560]
KP055815.1_strain_BA71           c.c..aat.....a....a.ga.t.ga.tc..ca.t.ttcc...---------g---.tt..c..a..aa..at.....t [ 20560]
MH025920.1_strain_R35            ......t........................a............g.........---.a....................- [ 20560]
MH025919.1_strain_N10            ......t........................a............g.........---.a....................- [ 20560]
MH025918.1_strain_R25            ......t........................a............g.........---.a....................- [ 20560]
MH025917.1_strain_R7             ......t........................a............g.........---.a....................- [ 20560]
MH025916.1_strain_R8             ......t........................a............g.........---.a....................- [ 20560]

FR682468.2_Georgia_2007/1        -------------agatgaaatttcatataaaca----ggattgagtcttggaatcacggaaaactctaca--------- [ 20640]
ASF_IND_20_CAD543                -------------------------------------------------------------------------------- [ 20640]
LR899193.1_Germany_2020/1        -------------.....................----.................................--------- [ 20640]
MN393477.1_Wuhan_2019            -------------.....................----.................................--------- [ 20640]
MW396979.1_Timor-Leste/2019/1    -------------.....................----.................................--------- [ 20640]
MK940252.1_CN/2019/InnerMongolia -------------.....................----....c............................--------- [ 20640]
MK128995.1_China/2018/AnhuiXCGQ  -------------.....................----.................................--------- [ 20640]
MH681419.1_ASFV/POL/2015         -------------.....................----.................................--------- [ 20640]
LS478113.1_Estonia_2014          -------------.....................----.................................--------- [ 20640]
KP843857.1_Odintsovo_02/14       -------------.....................----.................................--------- [ 20640]
```

```
FN557520.1_strain_E75           -------------.....................----.....................a...........--------- [ 20640]
AY261366.1_Warthog              -------------.....................----.................................--------- [ 20640]
AY261365.1_Warmbaths            -------------...c.................----...........c.....................--------- [ 20640]
AY261364.1_Tengani_62           -------------------------------------------------------------------------------- [ 20640]
AY261363.1_Pretoriuskop/96/4    -------------...c..t....tg..a.....----c.g------a.caa..atg.ca....t.a....--------- [ 20640]
AY261362.1_Mkuzi_1979           -------------.....................----.................t...............--------- [ 20640]
AY261361.1_Malawi_Lil-20/1_1983 -------------.....................----.........t.......................--------- [ 20640]
AY261360.1_Kenya_1950           -------------.....................----.................................--------- [ 20640]
AM712240.1_OURT_88/3            -------------.ac...t..............----.....................a...........--------- [ 20640]
AM712239.1_Benin_97/1           -------------.....................----.....................a...........--------- [ 20640]
KM111295.1_Ken06.Bus            -------------g.g...t..............----...c......c.a....................--------- [ 20640]
KM111294.1_Ken05/Tk1            -------------......t..............----...c...a..c.a....................--------- [ 20640]
KM262844.1_strain_L60           -------------.....................----.....................a...........--------- [ 20640]
KM102979.1_26544/OG10_Italy     -------------.t...................----.....................a...........--------- [ 20640]
KM262845.1_strain_NHV           -------------.ac...t..............----.....................a...........--------- [ 20640]
KX354450.1_47/Ss/2008           -------------.t...................----.....................a...........--------- [ 20640]
KP055815.1_strain_BA71          agtctatgaatgc.c...gttccg..c..t..a.aaataa...att.a..tagccag...g.....a....tggatgcac [ 20640]
MH025920.1_strain_R35           -------------g.g...t..............----...c......c.a....................--------- [ 20640]
MH025919.1_strain_N10           -------------g.g...t..............----...c......c.a....................--------- [ 20640]
MH025918.1_strain_R25           -------------g.g...t..............----...c......c.a....................--------- [ 20640]
MH025917.1_strain_R7            -------------g.g...t..............----...c......c.a....................--------- [ 20640]
MH025916.1_strain_R8            -------------g.g...t..............----...c......c.a....................--------- [ 20640]

FR682468.2_Georgia_2007/1       ------gtttacaatagaatgattggagtcaatgaaacgagattccgttatcttatttttgcaaatgccatcttgacagt [ 20720]
ASF_IND_20_CAD543               -------------------------------------------------------------------------------- [ 20720]
LR899193.1_Germany_2020/1       ------.......................................................................... [ 20720]
MN393477.1_Wuhan_2019           ------.......................................................................... [ 20720]
MW396979.1_Timor-Leste/2019/1   ------.......................................................................... [ 20720]
MK940252.1_CN/2019/InnerMongolia ------.......................................................................... [ 20720]
MK128995.1_China/2018/AnhuiXCGQ ------.......................................................................... [ 20720]
MH681419.1_ASFV/POL/2015        ------.......................................................................... [ 20720]
LS478113.1_Estonia_2014         ------.......................................................................... [ 20720]
KP843857.1_Odintsovo_02/14      ------.......................................................................... [ 20720]
FN557520.1_strain_E75           ------.......................................................................... [ 20720]
AY261366.1_Warthog              ------...............g.......................................................... [ 20720]
AY261365.1_Warmbaths            ------.........................................c................................ [ 20720]
AY261364.1_Tengani_62           -------------------------------------------------------------------------------- [ 20720]
AY261363.1_Pretoriuskop/96/4    ------a......t.g---------------------------------------------------------------- [ 20720]
AY261362.1_Mkuzi_1979           ------...............g.......................................................... [ 20720]
AY261361.1_Malawi_Lil-20/1_1983 ------...............................t.......................................... [ 20720]
AY261360.1_Kenya_1950           ------...............g.......t.............c......................a............. [ 20720]
AM712240.1_OURT_88/3            ------.......................................................................... [ 20720]
AM712239.1_Benin_97/1           ------.......................................................................... [ 20720]
KM111295.1_Ken06.Bus            ------...............g.......t.............c...................t................ [ 20720]
```

```
KM111294.1_Ken05/Tk1             ------.......................t........t......................................... [ 20720]
KM262844.1_strain_L60            ------.......................................................................... [ 20720]
KM102979.1_26544/OG10_Italy      ------.......................................................................... [ 20720]
KM262845.1_strain_NHV            ------.......................................................................... [ 20720]
KX354450.1_47/Ss/2008            ------.......................................................................... [ 20720]
KP055815.1_strain_BA71           atactca...t..--.....tg...t.----------------.ttccc.a......caca....g.....t........ [ 20720]
MH025920.1_strain_R35            ------...............g.......t.............c...................t................ [ 20720]
MH025919.1_strain_N10            ------...............g.......t.............c...................t................ [ 20720]
MH025918.1_strain_R25            ------...............g.......t.............c...................t................ [ 20720]
MH025917.1_strain_R7             ------...............g.......t.............c...................t................ [ 20720]
MH025916.1_strain_R8             ------...............g.......t.............c...................t................ [ 20720]

FR682468.2_Georgia_2007/1        cccaacagaaatcgcattgtggtacatacgtacaccaatatgaaagctcactcttgggaggatgctgggttcttggtaag [ 20800]
ASF_IND_20_CAD543                -------------------------------------------------------------------------------- [ 20800]
LR899193.1_Germany_2020/1        ................................................................................ [ 20800]
MN393477.1_Wuhan_2019            ................................................................................ [ 20800]
MW396979.1_Timor-Leste/2019/1    ................................................................................ [ 20800]
MK940252.1_CN/2019/InnerMongolia ................................................................................ [ 20800]
MK128995.1_China/2018/AnhuiXCGQ  ................................................................................ [ 20800]
MH681419.1_ASFV/POL/2015         ................................................................................ [ 20800]
LS478113.1_Estonia_2014          ................................................................................ [ 20800]
KP843857.1_Odintsovo_02/14       ................................................................................ [ 20800]
FN557520.1_strain_E75            ..........g..................................................................... [ 20800]
AY261366.1_Warthog               ..........g..................................................................... [ 20800]
AY261365.1_Warmbaths             ..........g..................................................................... [ 20800]
AY261364.1_Tengani_62            -------------------------------------------------------------------------------- [ 20800]
AY261363.1_Pretoriuskop/96/4     -------------------------------------------------------------------------------- [ 20800]
AY261362.1_Mkuzi_1979            ..........g..................................................................... [ 20800]
AY261361.1_Malawi_Lil-20/1_1983  ..........g..................................................................... [ 20800]
AY261360.1_Kenya_1950            ..........g..................................................................... [ 20800]
AM712240.1_OURT_88/3             ..........g..................................................................... [ 20800]
AM712239.1_Benin_97/1            ..........g..................................................................... [ 20800]
KM111295.1_Ken06.Bus             ..........g..............g...................................................... [ 20800]
KM111294.1_Ken05/Tk1             ..........g..................................................................... [ 20800]
KM262844.1_strain_L60            ..........g..................................................................... [ 20800]
KM102979.1_26544/OG10_Italy      ..........g..................................................................... [ 20800]
KM262845.1_strain_NHV            ..........g..................................................................... [ 20800]
KX354450.1_47/Ss/2008            ..........g..................................................................... [ 20800]
KP055815.1_strain_BA71           ....g..a..c.g...aca.tcc.tg..a..g.........tc...t...t...ct.a.....tt.cc...--------- [ 20800]
MH025920.1_strain_R35            ..........g..............g...................................................... [ 20800]
MH025919.1_strain_N10            ..........g..............g...................................................... [ 20800]
MH025918.1_strain_R25            ..........g..............g...................................................... [ 20800]
MH025917.1_strain_R7             ..........g..............g...................................................... [ 20800]
MH025916.1_strain_R8             ..........g..............g...................................................... [ 20800]
```

```
FR682468.2_Georgia_2007/1         tctggtaattcatgtgcgagaatgaggactgagtagcc---caac---aaaagtcccagaagaaccttc----------- [ 20880]
ASF_IND_20_CAD543                 -------------------------------------------------------------------------------- [ 20880]
LR899193.1_Germany_2020/1         ......................................---....---.....................----------- [ 20880]
MN393477.1_Wuhan_2019             ......................................---....---.....................----------- [ 20880]
MW396979.1_Timor-Leste/2019/1     ......................................---....---.....................----------- [ 20880]
MK940252.1_CN/2019/InnerMongolia  ......................................---....---.....................----------- [ 20880]
MK128995.1_China/2018/AnhuiXCGQ   ......................................---....---.....................----------- [ 20880]
MH681419.1_ASFV/POL/2015          ......................................---....---.....................----------- [ 20880]
LS478113.1_Estonia_2014           ......................................---....---.....................----------- [ 20880]
KP843857.1_Odintsovo_02/14        ......................................---....---.....................----------- [ 20880]
FN557520.1_strain_E75             ..................................g...---....---.....................----------- [ 20880]
AY261366.1_Warthog                ......................................---....---.....................----------- [ 20880]
AY261365.1_Warmbaths              ......................................---....---.....................----------- [ 20880]
AY261364.1_Tengani_62             -------------------------------------------------------------------------------- [ 20880]
AY261363.1_Pretoriuskop/96/4      ---------......a.........t.g.ca.a....tcag...t---....a......g.agg.....at--------- [ 20880]
AY261362.1_Mkuzi_1979             ......................................---....---.....................----------- [ 20880]
AY261361.1_Malawi_Lil-20/1_1983   ........................t.............---....---......t..............----------- [ 20880]
AY261360.1_Kenya_1950             ..c..........a....t...................---...t---.g...c...............atgttgcgtct [ 20880]
AM712240.1_OURT_88/3              ..................................g...---....---.....................----------- [ 20880]
AM712239.1_Benin_97/1             ..................................g...---....---.....................----------- [ 20880]
KM111295.1_Ken06.Bus              .............a....t........g..........---....aga.g...................atgttgcgtct [ 20880]
KM111294.1_Ken05/Tk1              .............a....t...................---....---........tg...........atgttgcaccc [ 20880]
KM262844.1_strain_L60             ..................................g...---....---.....................----------- [ 20880]
KM102979.1_26544/OG10_Italy       ..................................g...---....---.....................----------- [ 20880]
KM262845.1_strain_NHV             ..................................g...---....---.....................----------- [ 20880]
KX354450.1_47/Ss/2008             ..................................g...---....---.....................----------- [ 20880]
KP055815.1_strain_BA71            -------------------------...tga...t.t.---....---g.------------------------------ [ 20880]
MH025920.1_strain_R35             .............a....t........g..........---....aga.g...................atgttgcgtct [ 20880]
MH025919.1_strain_N10             .............a....t........g..........---....aga.g...................atgttgcgtct [ 20880]
MH025918.1_strain_R25             .............a....t........g..........---....aga.g...................atgttgcgtct [ 20880]
MH025917.1_strain_R7              .............a....t........g..........---....aga.g...................atgttgcgtct [ 20880]
MH025916.1_strain_R8              .............a....t........g..........---....aga.g...................atgttgcgtct [ 20880]

FR682468.2_Georgia_2007/1         -------------------------------------------------------------------------------- [ 20960]
ASF_IND_20_CAD543                 -------------------------------------------------------------------------------- [ 20960]
LR899193.1_Germany_2020/1         -------------------------------------------------------------------------------- [ 20960]
MN393477.1_Wuhan_2019             -------------------------------------------------------------------------------- [ 20960]
MW396979.1_Timor-Leste/2019/1     -------------------------------------------------------------------------------- [ 20960]
MK940252.1_CN/2019/InnerMongolia  -------------------------------------------------------------------------------- [ 20960]
MK128995.1_China/2018/AnhuiXCGQ   -------------------------------------------------------------------------------- [ 20960]
MH681419.1_ASFV/POL/2015          -------------------------------------------------------------------------------- [ 20960]
LS478113.1_Estonia_2014           -------------------------------------------------------------------------------- [ 20960]
KP843857.1_Odintsovo_02/14        -------------------------------------------------------------------------------- [ 20960]
```

```
FN557520.1_strain_E75          -------------------------------------------------------------------------------- [ 20960]
AY261366.1_Warthog             -------------------------------------------------------------------------------- [ 20960]
AY261365.1_Warmbaths           -------------------------------------------------------------------------------- [ 20960]
AY261364.1_Tengani_62          -------------------------------------------------------------------------------- [ 20960]
AY261363.1_Pretoriuskop/96/4   -------------------------------------------------------------------------------- [ 20960]
AY261362.1_Mkuzi_1979          -------------------------------------------------------------------------------- [ 20960]
AY261361.1_Malawi_Lil-20/1_1983 -------------------------------------------------------------------------------- [ 20960]
AY261360.1_Kenya_1950          aaaggccgc----------------------------------------------------------------------- [ 20960]
AM712240.1_OURT_88/3           -------------------------------------------------------------------------------- [ 20960]
AM712239.1_Benin_97/1          -------------------------------------------------------------------------------- [ 20960]
KM111295.1_Ken06.Bus           aaatgccgc----------------------------------------------------------------------- [ 20960]
KM111294.1_Ken05/Tk1           aaatggcacctgcatttttaaaaaagatttaaatttgataccacaaaaaagaaatactggtattttttataaaaataca [ 20960]
KM262844.1_strain_L60          -------------------------------------------------------------------------------- [ 20960]
KM102979.1_26544/OG10_Italy    -------------------------------------------------------------------------------- [ 20960]
KM262845.1_strain_NHV          -------------------------------------------------------------------------------- [ 20960]
KX354450.1_47/Ss/2008          -------------------------------------------------------------------------------- [ 20960]
KP055815.1_strain_BA71         -------------------------------------------------------------------------------- [ 20960]
MH025920.1_strain_R35          aaatgccgc----------------------------------------------------------------------- [ 20960]
MH025919.1_strain_N10          aaatgccgc----------------------------------------------------------------------- [ 20960]
MH025918.1_strain_R25          aaatgccgc----------------------------------------------------------------------- [ 20960]
MH025917.1_strain_R7           aaatgccgc----------------------------------------------------------------------- [ 20960]
MH025916.1_strain_R8           aaatgccgc----------------------------------------------------------------------- [ 20960]

FR682468.2_Georgia_2007/1      -------------------------------------------------------------------------------- [ 21040]
ASF_IND_20_CAD543              -------------------------------------------------------------------------------- [ 21040]
LR899193.1_Germany_2020/1      -------------------------------------------------------------------------------- [ 21040]
MN393477.1_Wuhan_2019          -------------------------------------------------------------------------------- [ 21040]
MW396979.1_Timor-Leste/2019/1  -------------------------------------------------------------------------------- [ 21040]
MK940252.1_CN/2019/InnerMongolia -------------------------------------------------------------------------------- [ 21040]
MK128995.1_China/2018/AnhuiXCGQ -------------------------------------------------------------------------------- [ 21040]
MH681419.1_ASFV/POL/2015       -------------------------------------------------------------------------------- [ 21040]
LS478113.1_Estonia_2014        -------------------------------------------------------------------------------- [ 21040]
KP843857.1_Odintsovo_02/14     -------------------------------------------------------------------------------- [ 21040]
FN557520.1_strain_E75          -------------------------------------------------------------------------------- [ 21040]
AY261366.1_Warthog             -------------------------------------------------------------------------------- [ 21040]
AY261365.1_Warmbaths           -------------------------------------------------------------------------------- [ 21040]
AY261364.1_Tengani_62          -------------------------------------------------------------------------------- [ 21040]
AY261363.1_Pretoriuskop/96/4   -------------------------------------------------------------------------------- [ 21040]
AY261362.1_Mkuzi_1979          -------------------------------------------------------------------------------- [ 21040]
AY261361.1_Malawi_Lil-20/1_1983 -------------------------------------------------------------------------------- [ 21040]
AY261360.1_Kenya_1950          -------------------------------------------------------------------------------- [ 21040]
AM712240.1_OURT_88/3           -------------------------------------------------------------------------------- [ 21040]
AM712239.1_Benin_97/1          -------------------------------------------------------------------------------- [ 21040]
KM111295.1_Ken06.Bus           -------------------------------------------------------------------------------- [ 21040]
```

```
KM111294.1_Ken05/Tk1            tatctatgaattattaaactctttataaagtatataggctgagaacattgcatataataaaaatagggtgtttagattta [ 21040]
KM262844.1_strain_L60           -------------------------------------------------------------------------------- [ 21040]
KM102979.1_26544/OG10_Italy     -------------------------------------------------------------------------------- [ 21040]
KM262845.1_strain_NHV           -------------------------------------------------------------------------------- [ 21040]
KX354450.1_47/Ss/2008           -------------------------------------------------------------------------------- [ 21040]
KP055815.1_strain_BA71          -------------------------------------------------------------------------------- [ 21040]
MH025920.1_strain_R35           -------------------------------------------------------------------------------- [ 21040]
MH025919.1_strain_N10           -------------------------------------------------------------------------------- [ 21040]
MH025918.1_strain_R25           -------------------------------------------------------------------------------- [ 21040]
MH025917.1_strain_R7            -------------------------------------------------------------------------------- [ 21040]
MH025916.1_strain_R8            -------------------------------------------------------------------------------- [ 21040]

FR682468.2_Georgia_2007/1       -------------------------------------------------------------------------------- [ 21120]
ASF_IND_20_CAD543               -------------------------------------------------------------------------------- [ 21120]
LR899193.1_Germany_2020/1       -------------------------------------------------------------------------------- [ 21120]
MN393477.1_Wuhan_2019           -------------------------------------------------------------------------------- [ 21120]
MW396979.1_Timor-Leste/2019/1   -------------------------------------------------------------------------------- [ 21120]
MK940252.1_CN/2019/InnerMongolia -------------------------------------------------------------------------------- [ 21120]
MK128995.1_China/2018/AnhuiXCGQ -------------------------------------------------------------------------------- [ 21120]
MH681419.1_ASFV/POL/2015        -------------------------------------------------------------------------------- [ 21120]
LS478113.1_Estonia_2014         -------------------------------------------------------------------------------- [ 21120]
KP843857.1_Odintsovo_02/14      -------------------------------------------------------------------------------- [ 21120]
FN557520.1_strain_E75           -------------------------------------------------------------------------------- [ 21120]
AY261366.1_Warthog              -------------------------------------------------------------------------------- [ 21120]
AY261365.1_Warmbaths            -------------------------------------------------------------------------------- [ 21120]
AY261364.1_Tengani_62           -------------------------------------------------------------------------------- [ 21120]
AY261363.1_Pretoriuskop/96/4    -------------------------------------------------------------------------------- [ 21120]
AY261362.1_Mkuzi_1979           -------------------------------------------------------------------------------- [ 21120]
AY261361.1_Malawi_Lil-20/1_1983 -------------------------------------------------------------------------------- [ 21120]
AY261360.1_Kenya_1950           -------------------------------------------------------------------------------- [ 21120]
AM712240.1_OURT_88/3            -------------------------------------------------------------------------------- [ 21120]
AM712239.1_Benin_97/1           -------------------------------------------------------------------------------- [ 21120]
KM111295.1_Ken06.Bus            -------------------------------------------------------------------------------- [ 21120]
KM111294.1_Ken05/Tk1            taaaatatatataatgatcaaaatagcaatgttgttataggtgtcaatattcaagctcttatgcgagatgtatatgtttt [ 21120]
KM262844.1_strain_L60           -------------------------------------------------------------------------------- [ 21120]
KM102979.1_26544/OG10_Italy     -------------------------------------------------------------------------------- [ 21120]
KM262845.1_strain_NHV           -------------------------------------------------------------------------------- [ 21120]
KX354450.1_47/Ss/2008           -------------------------------------------------------------------------------- [ 21120]
KP055815.1_strain_BA71          -------------------------------------------------------------------------------- [ 21120]
MH025920.1_strain_R35           -------------------------------------------------------------------------------- [ 21120]
MH025919.1_strain_N10           -------------------------------------------------------------------------------- [ 21120]
MH025918.1_strain_R25           -------------------------------------------------------------------------------- [ 21120]
MH025917.1_strain_R7            -------------------------------------------------------------------------------- [ 21120]
MH025916.1_strain_R8            -------------------------------------------------------------------------------- [ 21120]
```

```
FR682468.2_Georgia_2007/1        -------------------------------------------------------------------------------- [ 21200]
ASF_IND_20_CAD543                -------------------------------------------------------------------------------- [ 21200]
LR899193.1_Germany_2020/1        -------------------------------------------------------------------------------- [ 21200]
MN393477.1_Wuhan_2019            -------------------------------------------------------------------------------- [ 21200]
MW396979.1_Timor-Leste/2019/1    -------------------------------------------------------------------------------- [ 21200]
MK940252.1_CN/2019/InnerMongolia -------------------------------------------------------------------------------- [ 21200]
MK128995.1_China/2018/AnhuiXCGQ  -------------------------------------------------------------------------------- [ 21200]
MH681419.1_ASFV/POL/2015         -------------------------------------------------------------------------------- [ 21200]
LS478113.1_Estonia_2014          -------------------------------------------------------------------------------- [ 21200]
KP843857.1_Odintsovo_02/14       -------------------------------------------------------------------------------- [ 21200]
FN557520.1_strain_E75            -------------------------------------------------------------------------------- [ 21200]
AY261366.1_Warthog               -------------------------------------------------------------------------------- [ 21200]
AY261365.1_Warmbaths             -------------------------------------------------------------------------------- [ 21200]
AY261364.1_Tengani_62            -------------------------------------------------------------------------------- [ 21200]
AY261363.1_Pretoriuskop/96/4     -------------------------------------------------------------------------------- [ 21200]
AY261362.1_Mkuzi_1979            -------------------------------------------------------------------------------- [ 21200]
AY261361.1_Malawi_Lil-20/1_1983  -------------------------------------------------------------------------------- [ 21200]
AY261360.1_Kenya_1950            -------------------------------------------------------------------------------- [ 21200]
AM712240.1_OURT_88/3             -------------------------------------------------------------------------------- [ 21200]
AM712239.1_Benin_97/1            -------------------------------------------------------------------------------- [ 21200]
KM111295.1_Ken06.Bus             -------------------------------------------------------------------------------- [ 21200]
KM111294.1_Ken05/Tk1             tggtaaaatgatttaactatatgtaggaaatggttgatttaagactggctggactatccaaaccaaaatactcaaaagag [ 21200]
KM262844.1_strain_L60            -------------------------------------------------------------------------------- [ 21200]
KM102979.1_26544/OG10_Italy      -------------------------------------------------------------------------------- [ 21200]
KM262845.1_strain_NHV            -------------------------------------------------------------------------------- [ 21200]
KX354450.1_47/Ss/2008            -------------------------------------------------------------------------------- [ 21200]
KP055815.1_strain_BA71           -------------------------------------------------------------------------------- [ 21200]
MH025920.1_strain_R35            -------------------------------------------------------------------------------- [ 21200]
MH025919.1_strain_N10            -------------------------------------------------------------------------------- [ 21200]
MH025918.1_strain_R25            -------------------------------------------------------------------------------- [ 21200]
MH025917.1_strain_R7             -------------------------------------------------------------------------------- [ 21200]
MH025916.1_strain_R8             -------------------------------------------------------------------------------- [ 21200]

FR682468.2_Georgia_2007/1        -------------------------------------------------------------------------------- [ 21280]
ASF_IND_20_CAD543                -------------------------------------------------------------------------------- [ 21280]
LR899193.1_Germany_2020/1        -------------------------------------------------------------------------------- [ 21280]
MN393477.1_Wuhan_2019            -------------------------------------------------------------------------------- [ 21280]
MW396979.1_Timor-Leste/2019/1    -------------------------------------------------------------------------------- [ 21280]
MK940252.1_CN/2019/InnerMongolia -------------------------------------------------------------------------------- [ 21280]
MK128995.1_China/2018/AnhuiXCGQ  -------------------------------------------------------------------------------- [ 21280]
MH681419.1_ASFV/POL/2015         -------------------------------------------------------------------------------- [ 21280]
LS478113.1_Estonia_2014          -------------------------------------------------------------------------------- [ 21280]
KP843857.1_Odintsovo_02/14       -------------------------------------------------------------------------------- [ 21280]
```

```
FN557520.1_strain_E75            -------------------------------------------------------------------------------- [ 21280]
AY261366.1_Warthog               -------------------------------------------------------------------------------- [ 21280]
AY261365.1_Warmbaths             -------------------------------------------------------------------------------- [ 21280]
AY261364.1_Tengani_62            -------------------------------------------------------------------------------- [ 21280]
AY261363.1_Pretoriuskop/96/4     -------------------------------------------------------------------------------- [ 21280]
AY261362.1_Mkuzi_1979            -------------------------------------------------------------------------------- [ 21280]
AY261361.1_Malawi_Lil-20/1_1983  -------------------------------------------------------------------------------- [ 21280]
AY261360.1_Kenya_1950            -------------------------------------------------------------------------------- [ 21280]
AM712240.1_OURT_88/3             -------------------------------------------------------------------------------- [ 21280]
AM712239.1_Benin_97/1            -------------------------------------------------------------------------------- [ 21280]
KM111295.1_Ken06.Bus             -------------------------------------------------------------------------------- [ 21280]
KM111294.1_Ken05/Tk1             caagtagcttcatagtagtactcagatgcaaatttctatggatcgcaaatgcaacacaactcgcaaatgcaatactaaat [ 21280]
KM262844.1_strain_L60            -------------------------------------------------------------------------------- [ 21280]
KM102979.1_26544/OG10_Italy      -------------------------------------------------------------------------------- [ 21280]
KM262845.1_strain_NHV            -------------------------------------------------------------------------------- [ 21280]
KX354450.1_47/Ss/2008            -------------------------------------------------------------------------------- [ 21280]
KP055815.1_strain_BA71           -------------------------------------------------------------------------------- [ 21280]
MH025920.1_strain_R35            -------------------------------------------------------------------------------- [ 21280]
MH025919.1_strain_N10            -------------------------------------------------------------------------------- [ 21280]
MH025918.1_strain_R25            -------------------------------------------------------------------------------- [ 21280]
MH025917.1_strain_R7             -------------------------------------------------------------------------------- [ 21280]
MH025916.1_strain_R8             -------------------------------------------------------------------------------- [ 21280]

FR682468.2_Georgia_2007/1        -------------------------------------------------------------------------------- [ 21360]
ASF_IND_20_CAD543                -------------------------------------------------------------------------------- [ 21360]
LR899193.1_Germany_2020/1        -------------------------------------------------------------------------------- [ 21360]
MN393477.1_Wuhan_2019            -------------------------------------------------------------------------------- [ 21360]
MW396979.1_Timor-Leste/2019/1    -------------------------------------------------------------------------------- [ 21360]
MK940252.1_CN/2019/InnerMongolia -------------------------------------------------------------------------------- [ 21360]
MK128995.1_China/2018/AnhuiXCGQ  -------------------------------------------------------------------------------- [ 21360]
MH681419.1_ASFV/POL/2015         -------------------------------------------------------------------------------- [ 21360]
LS478113.1_Estonia_2014          -------------------------------------------------------------------------------- [ 21360]
KP843857.1_Odintsovo_02/14       -------------------------------------------------------------------------------- [ 21360]
FN557520.1_strain_E75            -------------------------------------------------------------------------------- [ 21360]
AY261366.1_Warthog               -------------------------------------------------------------------------------- [ 21360]
AY261365.1_Warmbaths             -------------------------------------------------------------------------------- [ 21360]
AY261364.1_Tengani_62            -------------------------------------------------------------------------------- [ 21360]
AY261363.1_Pretoriuskop/96/4     -------------------------------------------------------------------------------- [ 21360]
AY261362.1_Mkuzi_1979            -------------------------------------------------------------------------------- [ 21360]
AY261361.1_Malawi_Lil-20/1_1983  -------------------------------------------------------------------------------- [ 21360]
AY261360.1_Kenya_1950            -------------------------------------------------------------------------------- [ 21360]
AM712240.1_OURT_88/3             -------------------------------------------------------------------------------- [ 21360]
AM712239.1_Benin_97/1            -------------------------------------------------------------------------------- [ 21360]
KM111295.1_Ken06.Bus             -------------------------------------------------------------------------------- [ 21360]
```

```
KM111294.1_Ken05/Tk1             acaagataaaaacaataatgtcttatcattaatggccaatagttatcgtcctcccatttttgtataaacatttcacatcc [ 21360]
KM262844.1_strain_L60            -------------------------------------------------------------------------------- [ 21360]
KM102979.1_26544/OG10_Italy      -------------------------------------------------------------------------------- [ 21360]
KM262845.1_strain_NHV            -------------------------------------------------------------------------------- [ 21360]
KX354450.1_47/Ss/2008            -------------------------------------------------------------------------------- [ 21360]
KP055815.1_strain_BA71           -------------------------------------------------------------------------------- [ 21360]
MH025920.1_strain_R35            -------------------------------------------------------------------------------- [ 21360]
MH025919.1_strain_N10            -------------------------------------------------------------------------------- [ 21360]
MH025918.1_strain_R25            -------------------------------------------------------------------------------- [ 21360]
MH025917.1_strain_R7             -------------------------------------------------------------------------------- [ 21360]
MH025916.1_strain_R8             -------------------------------------------------------------------------------- [ 21360]

FR682468.2_Georgia_2007/1        -------------------------------------------------------------------------------- [ 21440]
ASF_IND_20_CAD543                -------------------------------------------------------------------------------- [ 21440]
LR899193.1_Germany_2020/1        -------------------------------------------------------------------------------- [ 21440]
MN393477.1_Wuhan_2019            -------------------------------------------------------------------------------- [ 21440]
MW396979.1_Timor-Leste/2019/1    -------------------------------------------------------------------------------- [ 21440]
MK940252.1_CN/2019/InnerMongolia -------------------------------------------------------------------------------- [ 21440]
MK128995.1_China/2018/AnhuiXCGQ  -------------------------------------------------------------------------------- [ 21440]
MH681419.1_ASFV/POL/2015         -------------------------------------------------------------------------------- [ 21440]
LS478113.1_Estonia_2014          -------------------------------------------------------------------------------- [ 21440]
KP843857.1_Odintsovo_02/14       -------------------------------------------------------------------------------- [ 21440]
FN557520.1_strain_E75            -------------------------------------------------------------------------------- [ 21440]
AY261366.1_Warthog               -------------------------------------------------------------------------------- [ 21440]
AY261365.1_Warmbaths             -------------------------------------------------------------------------------- [ 21440]
AY261364.1_Tengani_62            -------------------------------------------------------------------------------- [ 21440]
AY261363.1_Pretoriuskop/96/4     -------------------------------------------------------------------------------- [ 21440]
AY261362.1_Mkuzi_1979            -------------------------------------------------------------------------------- [ 21440]
AY261361.1_Malawi_Lil-20/1_1983  -------------------------------------------------------------------------------- [ 21440]
AY261360.1_Kenya_1950            -------------------------------------------------------------------------------- [ 21440]
AM712240.1_OURT_88/3             -------------------------------------------------------------------------------- [ 21440]
AM712239.1_Benin_97/1            -------------------------------------------------------------------------------- [ 21440]
KM111295.1_Ken06.Bus             -------------------------------------------------------------------------------- [ 21440]
KM111294.1_Ken05/Tk1             tatataaaatataggatgtgaacattgcatactatagtcattagatattttaggcgatatttcgtaaaaacacgaattaa [ 21440]
KM262844.1_strain_L60            -------------------------------------------------------------------------------- [ 21440]
KM102979.1_26544/OG10_Italy      -------------------------------------------------------------------------------- [ 21440]
KM262845.1_strain_NHV            -------------------------------------------------------------------------------- [ 21440]
KX354450.1_47/Ss/2008            -------------------------------------------------------------------------------- [ 21440]
KP055815.1_strain_BA71           -------------------------------------------------------------------------------- [ 21440]
MH025920.1_strain_R35            -------------------------------------------------------------------------------- [ 21440]
MH025919.1_strain_N10            -------------------------------------------------------------------------------- [ 21440]
MH025918.1_strain_R25            -------------------------------------------------------------------------------- [ 21440]
MH025917.1_strain_R7             -------------------------------------------------------------------------------- [ 21440]
MH025916.1_strain_R8             -------------------------------------------------------------------------------- [ 21440]
```

```
FR682468.2_Georgia_2007/1        -------------------------------------------------------------------------------- [ 21520]
ASF_IND_20_CAD543                -------------------------------------------------------------------------------- [ 21520]
LR899193.1_Germany_2020/1        -------------------------------------------------------------------------------- [ 21520]
MN393477.1_Wuhan_2019            -------------------------------------------------------------------------------- [ 21520]
MW396979.1_Timor-Leste/2019/1    -------------------------------------------------------------------------------- [ 21520]
MK940252.1_CN/2019/InnerMongolia -------------------------------------------------------------------------------- [ 21520]
MK128995.1_China/2018/AnhuiXCGQ  -------------------------------------------------------------------------------- [ 21520]
MH681419.1_ASFV/POL/2015         -------------------------------------------------------------------------------- [ 21520]
LS478113.1_Estonia_2014          -------------------------------------------------------------------------------- [ 21520]
KP843857.1_Odintsovo_02/14       -------------------------------------------------------------------------------- [ 21520]
FN557520.1_strain_E75            -------------------------------------------------------------------------------- [ 21520]
AY261366.1_Warthog               -------------------------------------------------------------------------------- [ 21520]
AY261365.1_Warmbaths             -------------------------------------------------------------------------------- [ 21520]
AY261364.1_Tengani_62            -------------------------------------------------------------------------------- [ 21520]
AY261363.1_Pretoriuskop/96/4     -------------------------------------------------------------------------------- [ 21520]
AY261362.1_Mkuzi_1979            -------------------------------------------------------------------------------- [ 21520]
AY261361.1_Malawi_Lil-20/1_1983  -------------------------------------------------------------------------------- [ 21520]
AY261360.1_Kenya_1950            -------------------------------------------------------------------------------- [ 21520]
AM712240.1_OURT_88/3             -------------------------------------------------------------------------------- [ 21520]
AM712239.1_Benin_97/1            -------------------------------------------------------------------------------- [ 21520]
KM111295.1_Ken06.Bus             -------------------------------------------------------------------------------- [ 21520]
KM111294.1_Ken05/Tk1             aaatgcgataaattatacaatttatattgtcatgtatgagaatgatagctaaataactcagcaataaaaatcccaggaag [ 21520]
KM262844.1_strain_L60            -------------------------------------------------------------------------------- [ 21520]
KM102979.1_26544/OG10_Italy      -------------------------------------------------------------------------------- [ 21520]
KM262845.1_strain_NHV            -------------------------------------------------------------------------------- [ 21520]
KX354450.1_47/Ss/2008            -------------------------------------------------------------------------------- [ 21520]
KP055815.1_strain_BA71           -------------------------------------------------------------------------------- [ 21520]
MH025920.1_strain_R35            -------------------------------------------------------------------------------- [ 21520]
MH025919.1_strain_N10            -------------------------------------------------------------------------------- [ 21520]
MH025918.1_strain_R25            -------------------------------------------------------------------------------- [ 21520]
MH025917.1_strain_R7             -------------------------------------------------------------------------------- [ 21520]
MH025916.1_strain_R8             -------------------------------------------------------------------------------- [ 21520]

FR682468.2_Georgia_2007/1        ------atgttgcgtctaaatgacacctgcacttac-aaaaaaaaatttaaattttgaatataacacaaa--aaaaccac [ 21600]
ASF_IND_20_CAD543                -------------------------------------------------------------------------------- [ 21600]
LR899193.1_Germany_2020/1        ------..............................-.................................--........ [ 21600]
MN393477.1_Wuhan_2019            ------..............................-.................................--........ [ 21600]
MW396979.1_Timor-Leste/2019/1    ------..............................-.................................--........ [ 21600]
MK940252.1_CN/2019/InnerMongolia ------..............................-.................................--........ [ 21600]
MK128995.1_China/2018/AnhuiXCGQ  ------..............................-.................................--........ [ 21600]
MH681419.1_ASFV/POL/2015         ------..............................-.................................--........ [ 21600]
LS478113.1_Estonia_2014          ------..............................-.................................--........ [ 21600]
KP843857.1_Odintsovo_02/14       ------..............................-.....-...........................--........ [ 21600]
```

```
FN557520.1_strain_E75            ------..............................-.................................---....... [ 21600]
AY261366.1_Warthog               ------..............................--......g..................g.g.-..-a........ [ 21600]
AY261365.1_Warmbaths             ------..............................--......g.............t....g.g....-a........ [ 21600]
AY261364.1_Tengani_62            -------------------------------------------------------------------------------- [ 21600]
AY261363.1_Pretoriuskop/96/4     -----g...a...a......g..t....a----..t--.c....g........g.........t.cgc..-a....t... [ 21600]
AY261362.1_Mkuzi_1979            ------..............................--......g..................g.g....aa........ [ 21600]
AY261361.1_Malawi_Lil-20/1_1983  ------..............................--.........................g.g.----a........ [ 21600]
AY261360.1_Kenya_1950            gtctaa.a.cc.........................--......g..............g...cgc---.aa....t... [ 21600]
AM712240.1_OURT_88/3             ------...............a..............a.................................---....... [ 21600]
AM712239.1_Benin_97/1            ------..............................-.................................---....... [ 21600]
KM111295.1_Ken06.Bus             gtctaa.g.cc.........................--.....................g...ggc--..aa....t... [ 21600]
KM111294.1_Ken05/Tk1             gccttc.......a.........t....at.t....---.....g....................t-gc.aa....t... [ 21600]
KM262844.1_strain_L60            ------..............................--................................---....... [ 21600]
KM102979.1_26544/OG10_Italy      ------..............................-.................................---....... [ 21600]
KM262845.1_strain_NHV            ------...............a..............-.................................---....... [ 21600]
KX354450.1_47/Ss/2008            ------..............................-.................................---....... [ 21600]
KP055815.1_strain_BA71           -------------------------------------------------------------------------------- [ 21600]
MH025920.1_strain_R35            gtctaa.g.cc.........................--.....................g...ggc-...aa....t... [ 21600]
MH025919.1_strain_N10            gtctaa.g.cc.........................--.....................g...ggca...aa....t... [ 21600]
MH025918.1_strain_R25            gtctaa.g.cc.........................--.....................g...ggc-...aa....t... [ 21600]
MH025917.1_strain_R7             gtctaa.g.cc.........................--.....................g...ggc-...aa....t... [ 21600]
MH025916.1_strain_R8             gtctaa.g.cc.........................--.....................g...ggc-...aa....t... [ 21600]

FR682468.2_Georgia_2007/1        cttaaaatttcttata-----------ttatttcttggatctgccccgacgtcatacaa-tg----tattaaaattata- [ 21680]
ASF_IND_20_CAD543                -------------------------------------------------------------------------------- [ 21680]
LR899193.1_Germany_2020/1        ................-----------................................-..----.............- [ 21680]
MN393477.1_Wuhan_2019            ................-----------................................-..----.............- [ 21680]
MW396979.1_Timor-Leste/2019/1    ................-----------................................-..----.............- [ 21680]
MK940252.1_CN/2019/InnerMongolia ................-----------................................-..----.............- [ 21680]
MK128995.1_China/2018/AnhuiXCGQ  ................-----------................................-..----.............- [ 21680]
MH681419.1_ASFV/POL/2015         ................-----------................................-..----.............- [ 21680]
LS478113.1_Estonia_2014          ................-----------................................-..----.............- [ 21680]
KP843857.1_Odintsovo_02/14       ................-----------................................-..----.............- [ 21680]
FN557520.1_strain_E75            ................-----------................-...............-..----.............- [ 21680]
AY261366.1_Warthog               ................-----------................-...t...........-.a----.............- [ 21680]
AY261365.1_Warmbaths             ................-----------................-...............-..----.............- [ 21680]
AY261364.1_Tengani_62            -------------------------------------------------------------------------------- [ 21680]
AY261363.1_Pretoriuskop/96/4     ...-.c......a...acaaacattat..t...t...a.....--a..........tg.cca----.......c..t..- [ 21680]
AY261362.1_Mkuzi_1979            ................-----------................-...............-..----.............- [ 21680]
AY261361.1_Malawi_Lil-20/1_1983  .......c........tcctttata--..g......a......-...............-..----.............- [ 21680]
AY261360.1_Kenya_1950            ...-...c...c....tcctttata--.........a...a..-.t-............-..----.............t [ 21680]
AM712240.1_OURT_88/3             ................-----------................-...............-..----.............- [ 21680]
AM712239.1_Benin_97/1            ................-----------................-...............-..----.............- [ 21680]
KM111295.1_Ken06.Bus             ...-...c...c....tcctttata--.........a...a..-.t.............-..----.............- [ 21680]
```

```
KM111294.1_Ken05/Tk1             ...-......t.a...ac--aaaca--......t..ttt.gaa-t.t.t..at...tg.-..tcca.......c.at..- [ 21680]
KM262844.1_strain_L60            ................-----------................-...............-..----.............- [ 21680]
KM102979.1_26544/OG10_Italy      ................-----------................-...............-..----.............- [ 21680]
KM262845.1_strain_NHV            ................-----------................-...............-..----.............- [ 21680]
KX354450.1_47/Ss/2008            ................-----------................-...............-..----.............- [ 21680]
KP055815.1_strain_BA71           -------------------------------------------------------------------------------- [ 21680]
MH025920.1_strain_R35            ...-...c...c....tcctttata--.........a...a..-.t.............-..----.............- [ 21680]
MH025919.1_strain_N10            ...-...c...c....tcctttata--.........a...a..-.t.............-..----.............- [ 21680]
MH025918.1_strain_R25            ...-...c...c....tcctttata--.........a...a..-.t.............-..----.............- [ 21680]
MH025917.1_strain_R7             ...-...c...c....tcctttata--.........a...a..-.t.............-..----.............- [ 21680]
MH025916.1_strain_R8             ...-...c...c....tcctttata--.........a...a..-.t.............-..----.............- [ 21680]

FR682468.2_Georgia_2007/1        gaccaatcatcttt-------------------------------------------ttgtatataggctaatcatcttt [ 21760]
ASF_IND_20_CAD543                -------------------------------------------------------------------------------- [ 21760]
LR899193.1_Germany_2020/1        ..............-------------------------------------------....................... [ 21760]
MN393477.1_Wuhan_2019            ..............-------------------------------------------....................... [ 21760]
MW396979.1_Timor-Leste/2019/1    ..............-------------------------------------------....................... [ 21760]
MK940252.1_CN/2019/InnerMongolia ..............-------------------------------------------....................... [ 21760]
MK128995.1_China/2018/AnhuiXCGQ  ..............-------------------------------------------....................... [ 21760]
MH681419.1_ASFV/POL/2015         ..............-------------------------------------------....................... [ 21760]
LS478113.1_Estonia_2014          ..............-------------------------------------------....................... [ 21760]
KP843857.1_Odintsovo_02/14       ..............-------------------------------------------....................... [ 21760]
FN557520.1_strain_E75            .........------------------------------------------------------------------..... [ 21760]
AY261366.1_Warthog               .........--.c.-------------------------------------------.c..................... [ 21760]
AY261365.1_Warmbaths             a...------------------------------------------------------------------.......... [ 21760]
AY261364.1_Tengani_62            -------------------------------------------------------------------------------- [ 21760]
AY261363.1_Pretoriuskop/96/4     ----------------------------------------------------------------................ [ 21760]
AY261362.1_Mkuzi_1979            .........------------------------------------------------------------------..... [ 21760]
AY261361.1_Malawi_Lil-20/1_1983  ..............gtatataaactaattatatataaactaattatatataaactaa..a.......a............ [ 21760]
AY261360.1_Kenya_1950            ac....a.......---------------------------------------------........a.........g.. [ 21760]
AM712240.1_OURT_88/3             .........------------------------------------------------------------------..... [ 21760]
AM712239.1_Benin_97/1            .........------------------------------------------------------------------..... [ 21760]
KM111295.1_Ken06.Bus             .g....a.......---------------------------------------------........a.........g.. [ 21760]
KM111294.1_Ken05/Tk1             ----------------------------------------------------------------................ [ 21760]
KM262844.1_strain_L60            .........------------------------------------------------------------------..... [ 21760]
KM102979.1_26544/OG10_Italy      .........------------------------------------------------------------------..... [ 21760]
KM262845.1_strain_NHV            .........------------------------------------------------------------------..... [ 21760]
KX354450.1_47/Ss/2008            .........------------------------------------------------------------------..... [ 21760]
KP055815.1_strain_BA71           -------------------------------------------------------------------------------- [ 21760]
MH025920.1_strain_R35            .g....a.......---------------------------------------------........a.........g.. [ 21760]
MH025919.1_strain_N10            .g....a.......---------------------------------------------........a.........g.. [ 21760]
MH025918.1_strain_R25            .g....a.......---------------------------------------------........a.........g.. [ 21760]
MH025917.1_strain_R7             .g....a.......---------------------------------------------........a.........g.. [ 21760]
MH025916.1_strain_R8             .g....a.......---------------------------------------------........a.........g.. [ 21760]
```

```
FR682468.2_Georgia_2007/1        atatatagattttagatg-tttgcttgttgtatcaacttaactgctagcgaagaaaatggataaaaactttctgtatttt [ 21840]
ASF_IND_20_CAD543                -------------------------------------------------------------------------------- [ 21840]
LR899193.1_Germany_2020/1        ..................-............................................................. [ 21840]
MN393477.1_Wuhan_2019            ..................-............................................................. [ 21840]
MW396979.1_Timor-Leste/2019/1    ..................-............................................................. [ 21840]
MK940252.1_CN/2019/InnerMongolia ..................-............................................................. [ 21840]
MK128995.1_China/2018/AnhuiXCGQ  ..................-............................................................. [ 21840]
MH681419.1_ASFV/POL/2015         ..................-............................................................. [ 21840]
LS478113.1_Estonia_2014          ..................-............................................................. [ 21840]
KP843857.1_Odintsovo_02/14       ..................-............................................................. [ 21840]
FN557520.1_strain_E75            g..............t.at.............................t............................... [ 21840]
AY261366.1_Warthog               g..............t.at.............................t............................... [ 21840]
AY261365.1_Warmbaths             g..............t.at.............................t............................... [ 21840]
AY261364.1_Tengani_62            -------------------------------------------------------------------------------- [ 21840]
AY261363.1_Pretoriuskop/96/4     g..............t.at............................................................. [ 21840]
AY261362.1_Mkuzi_1979            g..............t.at............................at............................... [ 21840]
AY261361.1_Malawi_Lil-20/1_1983  g..............t.at.............................g............................... [ 21840]
AY261360.1_Kenya_1950            g.....g...........-..................c.......c...tc.........g......t............ [ 21840]
AM712240.1_OURT_88/3             g..............t.at.............................t............................... [ 21840]
AM712239.1_Benin_97/1            g..............t.at.............................t............................... [ 21840]
KM111295.1_Ken06.Bus             g.....ga..........-..................c..g....c...tc.........g...t..t............ [ 21840]
KM111294.1_Ken05/Tk1             g................a-................................................g...t...gg... [ 21840]
KM262844.1_strain_L60            g..............t.at.............................t............................... [ 21840]
KM102979.1_26544/OG10_Italy      g..............t.at.............................t............................... [ 21840]
KM262845.1_strain_NHV            g..............t.at.............................t............................... [ 21840]
KX354450.1_47/Ss/2008            g..............t.at.............................t............................... [ 21840]
KP055815.1_strain_BA71           -------------------------------gctgg...g.gac..g.ttggcc.g.a..cc..g..t.cc.-------- [ 21840]
MH025920.1_strain_R35            g.....ga..........-..................c..g....c...tc.........g...t..t............ [ 21840]
MH025919.1_strain_N10            g.....ga..........-..................c..g....c...tc.........g...t..t............ [ 21840]
MH025918.1_strain_R25            g.....ga..........-..................c..g....c...tc.........g...t..t............ [ 21840]
MH025917.1_strain_R7             g.....ga..........-..................c..g....c...tc.........g...t..t............ [ 21840]
MH025916.1_strain_R8             g.....ga..........-..................c..g....c...tc.........g...t..t............ [ 21840]

FR682468.2_Georgia_2007/1        tataggttgaaatcattttatgcacatcgctaggatctaatattttattttgaagaaccgaatgtgggcttaaaattttt [ 21920]
ASF_IND_20_CAD543                -------------------------------------------------------------------------------- [ 21920]
LR899193.1_Germany_2020/1        ................................................................................ [ 21920]
MN393477.1_Wuhan_2019            ................................................................................ [ 21920]
MW396979.1_Timor-Leste/2019/1    ................................................................................ [ 21920]
MK940252.1_CN/2019/InnerMongolia ...............................................................................c [ 21920]
MK128995.1_China/2018/AnhuiXCGQ  ................................................................................ [ 21920]
MH681419.1_ASFV/POL/2015         ................................................................................ [ 21920]
LS478113.1_Estonia_2014          ................................................................................ [ 21920]
KP843857.1_Odintsovo_02/14       ................................................................................ [ 21920]
```

| Strain | Sequence | |
|---|---|---|
| FN557520.1_strain_E75 | ....................c.......a.............................t..................... | [ 21920] |
| AY261366.1_Warthog | c....................a......a.............................t..............t.g.... | [ 21920] |
| AY261365.1_Warmbaths | ....................c.....g.a.............................t..................... | [ 21920] |
| AY261364.1_Tengani_62 | -------------------------------------------------------------------------------- | [ 21920] |
| AY261363.1_Pretoriuskop/96/4 | c....................a......a.............................t..............t.g.... | [ 21920] |
| AY261362.1_Mkuzi_1979 | ....................c.......a.............................t..................... | [ 21920] |
| AY261361.1_Malawi_Lil-20/1_1983 | .....................a......a.............................t..................... | [ 21920] |
| AY261360.1_Kenya_1950 | c..ga...a...........c........g...............................................c.. | [ 21920] |
| AM712240.1_OURT_88/3 | ....................c.......a.............................t..................... | [ 21920] |
| AM712239.1_Benin_97/1 | ....................c.......a.............................t....a................ | [ 21920] |
| KM111295.1_Ken06.Bus | ...g....a...........c...t....g....g...........g..............................g.. | [ 21920] |
| KM111294.1_Ken05/Tk1 | c...........a.........t..t..a...............................................g... | [ 21920] |
| KM262844.1_strain_L60 | ....................c.......a.............................t..................... | [ 21920] |
| KM102979.1_26544/OG10_Italy | ....................c.......a.............................t..................... | [ 21920] |
| KM262845.1_strain_NHV | ....................c.......a.............................t..................... | [ 21920] |
| KX354450.1_47/Ss/2008 | ....................c.......a.............................t..................... | [ 21920] |
| KP055815.1_strain_BA71 | -------------------------------------------------------------------------------- | [ 21920] |
| MH025920.1_strain_R35 | ...g....a...........c...t....g....g...........g..............................g.. | [ 21920] |
| MH025919.1_strain_N10 | ...g....a...........c...t....g....g...........g..............................g.. | [ 21920] |
| MH025918.1_strain_R25 | ...g....a...........c...t....g....g...........g..............................g.. | [ 21920] |
| MH025917.1_strain_R7 | ...g....a...........c...t....g....g...........g..............................g.. | [ 21920] |
| MH025916.1_strain_R8 | ...g....a...........c...t....g....g...........g..............................g.. | [ 21920] |

| Strain | Sequence | |
|---|---|---|
| FR682468.2_Georgia_2007/1 | ttct---tagaaaaaagtagaatcataatattgctatgtttttgtttaatgatttcttgtatcttttttgtatacgggtt | [ 22000] |
| ASF_IND_20_CAD543 | -------------------------------------------------------------------------------- | [ 22000] |
| LR899193.1_Germany_2020/1 | ....---......................................................................... | [ 22000] |
| MN393477.1_Wuhan_2019 | ....---......................................................................... | [ 22000] |
| MW396979.1_Timor-Leste/2019/1 | ....---......................................................................... | [ 22000] |
| MK940252.1_CN/2019/InnerMongolia | ....---......................................................................... | [ 22000] |
| MK128995.1_China/2018/AnhuiXCGQ | ....---......................................................................... | [ 22000] |
| MH681419.1_ASFV/POL/2015 | ....---......................................................................... | [ 22000] |
| LS478113.1_Estonia_2014 | ....---......................................................................... | [ 22000] |
| KP843857.1_Odintsovo_02/14 | ....---......................................................................... | [ 22000] |
| FN557520.1_strain_E75 | ....---.......g....................................g............................ | [ 22000] |
| AY261366.1_Warthog | ....---...........................................ag.a...............a....a..a.. | [ 22000] |
| AY261365.1_Warmbaths | ....---......gg......................t.............g............................ | [ 22000] |
| AY261364.1_Tengani_62 | -------------------------------------------------------------------------------- | [ 22000] |
| AY261363.1_Pretoriuskop/96/4 | ....---...........................................ag.a...............a....a..a.. | [ 22000] |
| AY261362.1_Mkuzi_1979 | ....---............................................g.a........t......a....a..a.. | [ 22000] |
| AY261361.1_Malawi_Lil-20/1_1983 | ....---.....t.g......................t.............g............................ | [ 22000] |
| AY261360.1_Kenya_1950 | ....tcc..........g...................t.............g.a...............a....a..a.. | [ 22000] |
| AM712240.1_OURT_88/3 | ....---.......g....................................g............................ | [ 22000] |
| AM712239.1_Benin_97/1 | ....---.......g....................................g............................ | [ 22000] |
| KM111295.1_Ken06.Bus | ....tcc.........-g.....t.............t.............-------...........a....a..a.. | [ 22000] |

```
KM111294.1_Ken05/Tk1              ....---.....................g........t.............g.a....................a..a.. [ 22000]
KM262844.1_strain_L60             ....---.......g....................................g............................ [ 22000]
KM102979.1_26544/OG10_Italy       ....---.......g....................................g............................ [ 22000]
KM262845.1_strain_NHV             ....---.......g....................................g............................ [ 22000]
KX354450.1_47/Ss/2008             ....---.......g....................................g......................-..... [ 22000]
KP055815.1_strain_BA71            --------------...a...t.ac...c.......c.------------------------------------------ [ 22000]
MH025920.1_strain_R35             ....tcc.........-g.....t.............t.............-------...........a....a..a.. [ 22000]
MH025919.1_strain_N10             ....tcc.........-g.....t.............t.............-------...........a....a..a.. [ 22000]
MH025918.1_strain_R25             ....tcc.........-g.....t.............t.............-------...........a....a..a.. [ 22000]
MH025917.1_strain_R7              ....tcc.........-g.....t.............t.............-------...........a....a..a.. [ 22000]
MH025916.1_strain_R8              ....tcc.........-g.....t.............t.............-------...........a....a..a.. [ 22000]

FR682468.2_Georgia_2007/1         ggcacccaaacctatacaaaaatatacattactcaaataactaccttctatacataatcttttttccccacgtattttcc [ 22080]
ASF_IND_20_CAD543                 -------------------------------------------------------------------------------- [ 22080]
LR899193.1_Germany_2020/1         ................................................................................ [ 22080]
MN393477.1_Wuhan_2019             ................................................................................ [ 22080]
MW396979.1_Timor-Leste/2019/1     ................................................................................ [ 22080]
MK940252.1_CN/2019/InnerMongolia  ................................................................................ [ 22080]
MK128995.1_China/2018/AnhuiXCGQ   ................................................................................ [ 22080]
MH681419.1_ASFV/POL/2015          ................................................................................ [ 22080]
LS478113.1_Estonia_2014           ................................................................................ [ 22080]
KP843857.1_Odintsovo_02/14        ................................................................................ [ 22080]
FN557520.1_strain_E75             .............................................................g.........t........ [ 22080]
AY261366.1_Warthog                ...........t..............t....a.........c..g.......tta..aa.ga.......----------- [ 22080]
AY261365.1_Warmbaths              ..........t...............t..................................a......t.---------- [ 22080]
AY261364.1_Tengani_62             -------------------------------------------------------------------------------- [ 22080]
AY261363.1_Pretoriuskop/96/4      ...........t..............t....a.........c..g.......tta..aa.ga.......t.tgg...c.. [ 22080]
AY261362.1_Mkuzi_1979             ...........t...................a.........c.ta.......t....---ag.......c.cgg...c.. [ 22080]
AY261361.1_Malawi_Lil-20/1_1983   ..........................t..................................g...t.....t........ [ 22080]
AY261360.1_Kenya_1950             ...t..........................cgat.......cc.a.......t....a...a..c...t..cag...ct. [ 22080]
AM712240.1_OURT_88/3              .............................................................g.........t........ [ 22080]
AM712239.1_Benin_97/1             .............................................................g.........t........ [ 22080]
KM111295.1_Ken06.Bus              ...t.......t..............t...cg.t.......c.tg.......t....a...a.a.at....tgg...c.. [ 22080]
KM111294.1_Ken05/Tk1              ......t....t...................g.........cc.a.......t....a...a..c...t..cgg...ct. [ 22080]
KM262844.1_strain_L60             .............................................................g.........t........ [ 22080]
KM102979.1_26544/OG10_Italy       .............................................................g.........t........ [ 22080]
KM262845.1_strain_NHV             .............................................................g.........t........ [ 22080]
KX354450.1_47/Ss/2008             .............................................................g.........t........ [ 22080]
KP055815.1_strain_BA71            ---------------------------gc.ggctg..c.g..g.------------------------------------ [ 22080]
MH025920.1_strain_R35             ...t.......t..............t...cg.t.......c.ta.......t....a...a.a.at....tgg...c.. [ 22080]
MH025919.1_strain_N10             ...t.......t..............t...cg.t.......c.ta.......t....a...a.a.at....tgg...c.. [ 22080]
MH025918.1_strain_R25             ...t.......t..............t...cg.t.......c.ta.......t....a...a.a.at....tgg...c.. [ 22080]
MH025917.1_strain_R7              ...t.......t..............t...cg.t.......c.ta.......t....a...a.a.at....tgg...c.. [ 22080]
MH025916.1_strain_R8              ...t.......t..............t...cg.t.......c.ta.......t....a...a.a.at....tgg...c.. [ 22080]
```

```
FR682468.2_Georgia_2007/1        tatttatttccctatttatggaattaaaggatatcaatctctctaaggcacggtcaaggtctgcgcctaaggcaaaacaa [ 22160]
ASF_IND_20_CAD543                -------------------------------------------------------------------------------- [ 22160]
LR899193.1_Germany_2020/1        ................................................................................ [ 22160]
MN393477.1_Wuhan_2019            ................................................................................ [ 22160]
MW396979.1_Timor-Leste/2019/1    ................................................................................ [ 22160]
MK940252.1_CN/2019/InnerMongolia ................................................................................ [ 22160]
MK128995.1_China/2018/AnhuiXCGQ  ................................................................................ [ 22160]
MH681419.1_ASFV/POL/2015         ................................................................................ [ 22160]
LS478113.1_Estonia_2014          ................................................................................ [ 22160]
KP843857.1_Odintsovo_02/14       ................................................................................ [ 22160]
FN557520.1_strain_E75            ................................................................................ [ 22160]
AY261366.1_Warthog               -------------------------------------------------------------------------------- [ 22160]
AY261365.1_Warmbaths             --------------....................t............................................. [ 22160]
AY261364.1_Tengani_62            -------------------------------------------------------------------------------- [ 22160]
AY261363.1_Pretoriuskop/96/4     ...g.....g.t.....g.ttc..c...acg..g....a.a.....a...tt...t.ta...........a......... [ 22160]
AY261362.1_Mkuzi_1979            ...g.......a.....g.ttc..c...acg..g....a.a.....a...tta..t.ta..c........a......... [ 22160]
AY261361.1_Malawi_Lil-20/1_1983  ..................................................t............................. [ 22160]
AY261360.1_Kenya_1950            .g.........t.....g.ttc..c...acg..g....a.a.....a...tta..t.ta.....a.....a......... [ 22160]
AM712240.1_OURT_88/3             ................................................................................ [ 22160]
AM712239.1_Benin_97/1            ................................................................................ [ 22160]
KM111295.1_Ken06.Bus             ...a.......tc.gc.g.ttc..c...acg..a....a.a.t...a...tt..tt.ta..ca.a.....aa........ [ 22160]
KM111294.1_Ken05/Tk1             .g.........t...c.g.ttt..c...acg..g....a.a.....a...tta..t.ta..c........a......... [ 22160]
KM262844.1_strain_L60            ................................................................................ [ 22160]
KM102979.1_26544/OG10_Italy      ................................................................................ [ 22160]
KM262845.1_strain_NHV            ................................................................................ [ 22160]
KX354450.1_47/Ss/2008            ................................................................................ [ 22160]
KP055815.1_strain_BA71           -------------------------------------------------------------------------------- [ 22160]
MH025920.1_strain_R35            ...a.......tc.gc.g.ttc..c...acg..a....a.a.t...a...tt..tt.ta..ca.a.....aa........ [ 22160]
MH025919.1_strain_N10            ...a.......tc.gc.g.ttc..c...acg..a....a.a.t...a...tt..tt.ta..ca.a.....aa........ [ 22160]
MH025918.1_strain_R25            ...a.......tc.gc.g.ttc..c...acg..a....a.a.t...a...tt..tt.ta..ca.a.....aa........ [ 22160]
MH025917.1_strain_R7             ...a.......tc.gc.g.ttc..c...acg..a....a.a.t...a...tt..tt.ta..ca.a.....aa........ [ 22160]
MH025916.1_strain_R8             ...a.......tc.gc.g.ttc..c...acg..a....a.a.t...a...tt..tt.ta..ca.a.....aa........ [ 22160]

FR682468.2_Georgia_2007/1        taatatatacctaatttattcccagg--gcgtgcacaggcaagaaacatcatgacgtttagccctaaacgtatattttcc [ 22240]
ASF_IND_20_CAD543                -------------------------------------------------------------------------------- [ 22240]
LR899193.1_Germany_2020/1        ..........................--.................................................... [ 22240]
MN393477.1_Wuhan_2019            ..........................--.................................................... [ 22240]
MW396979.1_Timor-Leste/2019/1    ..........................--.................................................... [ 22240]
MK940252.1_CN/2019/InnerMongolia ..........................--.................................................... [ 22240]
MK128995.1_China/2018/AnhuiXCGQ  ..........................--.................................................... [ 22240]
MH681419.1_ASFV/POL/2015         ..........................--.................................................... [ 22240]
LS478113.1_Estonia_2014          ..........................--.................................................... [ 22240]
KP843857.1_Odintsovo_02/14       ..........................--.................................................... [ 22240]
```

```
FN557520.1_strain_E75            ............................................................aga................ [ 22240]
AY261366.1_Warthog               -------------------------------------------------------------------------------- [ 22240]
AY261365.1_Warmbaths             ..........................--.................................gga................ [ 22240]
AY261364.1_Tengani_62            -------------------------------------------------------------------------------- [ 22240]
AY261363.1_Pretoriuskop/96/4     .....................tgg..--atacatg........t.........t.a...g.at................. [ 22240]
AY261362.1_Mkuzi_1979            .....................ttg..--atacttg........t.........t.a...g.at................. [ 22240]
AY261361.1_Malawi_Lil-20/1_1983  ..........................--...........................a.......t................ [ 22240]
AY261360.1_Kenya_1950            ......................ag..--at..at........acg.......atga...g.at................. [ 22240]
AM712240.1_OURT_88/3             ..........................--.................................aga................ [ 22240]
AM712239.1_Benin_97/1            ..........................--.................................aga................ [ 22240]
KM111295.1_Ken06.Bus             ....................t.ag..atat..at........ttg.......atga...g.at................. [ 22240]
KM111294.1_Ken05/Tk1             .......................g..--at.cat.........cg........tga...g.at................. [ 22240]
KM262844.1_strain_L60            ..........................--.................................aga................ [ 22240]
KM102979.1_26544/OG10_Italy      ..........................--.................................aga................ [ 22240]
KM262845.1_strain_NHV            ..........................--.................................aga................ [ 22240]
KX354450.1_47/Ss/2008            ..........................--.................................aga................ [ 22240]
KP055815.1_strain_BA71           ---..g...g...gc.a..------------------------------------------------------------- [ 22240]
MH025920.1_strain_R35            ....................t.ag..atat..at...a....ttg.......atga...g.at................. [ 22240]
MH025919.1_strain_N10            ....................t.ag..atat..at...a....ttg.......atga...g.at................. [ 22240]
MH025918.1_strain_R25            ....................t.ag..atat..at...a....ttg.......atga...g.at................. [ 22240]
MH025917.1_strain_R7             ....................t.ag..atat..at...a....ttg.......atga...g.at................. [ 22240]
MH025916.1_strain_R8             ....................t.ag..atat..at...a....ttg.......atga...g.at................. [ 22240]

FR682468.2_Georgia_2007/1        tgaaaatacgcatgatgaacttcatcaatattacctaagtatatggccgtttgtaaacgccaaagatctaaatgaggaaa [ 22320]
ASF_IND_20_CAD543                -------------------------------------------------------------------------------- [ 22320]
LR899193.1_Germany_2020/1        ................................................................................ [ 22320]
MN393477.1_Wuhan_2019            ................................................................................ [ 22320]
MW396979.1_Timor-Leste/2019/1    ................................................................................ [ 22320]
MK940252.1_CN/2019/InnerMongolia ................................................................................ [ 22320]
MK128995.1_China/2018/AnhuiXCGQ  ................................................................................ [ 22320]
MH681419.1_ASFV/POL/2015         ................................................................................ [ 22320]
LS478113.1_Estonia_2014          ................................................................................ [ 22320]
KP843857.1_Odintsovo_02/14       ................................................................................ [ 22320]
FN557520.1_strain_E75            ................................................................................ [ 22320]
AY261366.1_Warthog               -------------------------------------------------------------------------------- [ 22320]
AY261365.1_Warmbaths             ................................................................................ [ 22320]
AY261364.1_Tengani_62            -------------------------------------------------------------------------------- [ 22320]
AY261363.1_Pretoriuskop/96/4     ............c...............................a....c..t...g.......t.........g..... [ 22320]
AY261362.1_Mkuzi_1979            ................................g...........a.......t..cg....................... [ 22320]
AY261361.1_Malawi_Lil-20/1_1983  ................................................................................ [ 22320]
AY261360.1_Kenya_1950            ..............g..g.t........c...c..c........a.......t..........tc.....gg........ [ 22320]
AM712240.1_OURT_88/3             ................................................................................ [ 22320]
AM712239.1_Benin_97/1            ................................................................................ [ 22320]
KM111295.1_Ken06.Bus             c............a..a.......g...c...c...........a.......t..........ta.....g......... [ 22320]
```

```
KM111294.1_Ken05/Tk1            ............................c...c...........a.......t..........t......gg........ [ 22320]
KM262844.1_strain_L60           ................................................................................ [ 22320]
KM102979.1_26544/OG10_Italy     ................................................................................ [ 22320]
KM262845.1_strain_NHV           ................................................................................ [ 22320]
KX354450.1_47/Ss/2008           ................................................................................ [ 22320]
KP055815.1_strain_BA71          -------------------------------------------------------------------------------- [ 22320]
MH025920.1_strain_R35           c............a..a.......a...c...c...........a.......t..........ta.....g......... [ 22320]
MH025919.1_strain_N10           c............a..a.......a...c...c...........a.......t..........ta.....g......... [ 22320]
MH025918.1_strain_R25           c............a..a.......a...c...c...........a.......t..........ta.....g......... [ 22320]
MH025917.1_strain_R7            c............a..a.......a...c...c...........a.......t..........ta.....g......... [ 22320]
MH025916.1_strain_R8            c............a..a.......a...c...c...........a.......t..........ta.....g......... [ 22320]

FR682468.2_Georgia_2007/1       ttttttactaagataatgaataggttttgtgagattaaaatctatggcgaacttataccaaaattttaatacaagtgtat [ 22400]
ASF_IND_20_CAD543               -------------------------------------------------------------------------------- [ 22400]
LR899193.1_Germany_2020/1       ................................................................................ [ 22400]
MN393477.1_Wuhan_2019           ................................................................................ [ 22400]
MW396979.1_Timor-Leste/2019/1   ................................................................................ [ 22400]
MK940252.1_CN/2019/InnerMongolia ................................a............................................... [ 22400]
MK128995.1_China/2018/AnhuiXCGQ ................................................................................ [ 22400]
MH681419.1_ASFV/POL/2015        ................................................................................ [ 22400]
LS478113.1_Estonia_2014         ................................................................................ [ 22400]
KP843857.1_Odintsovo_02/14      ................................................................................ [ 22400]
FN557520.1_strain_E75           ................c....g........a..t.............................................. [ 22400]
AY261366.1_Warthog              -------------------------------------------------------------------------------- [ 22400]
AY261365.1_Warmbaths            ................c....g........a..t.............................................. [ 22400]
AY261364.1_Tengani_62           -------------------------------------------------------------------------------- [ 22400]
AY261363.1_Pretoriuskop/96/4    ................c...........aca..t...........................................c.. [ 22400]
AY261362.1_Mkuzi_1979           ...............cc...........aca................................................. [ 22400]
AY261361.1_Malawi_Lil-20/1_1983 ................c.............a..t..............................c............... [ 22400]
AY261360.1_Kenya_1950           ................c.g.........a.a................................................. [ 22400]
AM712240.1_OURT_88/3            ................c....g........a..t.............................................. [ 22400]
AM712239.1_Benin_97/1           ................c....g........a..t.............................................. [ 22400]
KM111295.1_Ken06.Bus            ................c.g.........aca...................................c....t........ [ 22400]
KM111294.1_Ken05/Tk1            .............a..c.g.........a.a.................a......a........................ [ 22400]
KM262844.1_strain_L60           ................c....g........a..t.............................................. [ 22400]
KM102979.1_26544/OG10_Italy     ................c....g........a..t.............................................. [ 22400]
KM262845.1_strain_NHV           ................c....g........a..t.............................................. [ 22400]
KX354450.1_47/Ss/2008           ................c....g........a..t.............................................. [ 22400]
KP055815.1_strain_BA71          --------------------------------------------.a..a...-------------------......acg [ 22400]
MH025920.1_strain_R35           ................c.g.........aca...................................c....t........ [ 22400]
MH025919.1_strain_N10           ................c.g.........aca...................................c....t........ [ 22400]
MH025918.1_strain_R25           ................c.g.........aca...................................c....t........ [ 22400]
MH025917.1_strain_R7            ................c.g.........aca...................................c....t........ [ 22400]
MH025916.1_strain_R8            ................c.g.........aca...................................c....t........ [ 22400]
```

```
FR682468.2_Georgia_2007/1        ttctcgtcatttcttcttctttttcatctaaatataagataaaacgattgtaaacaaagtctatcaataggtgaaaatca [ 22480]
ASF_IND_20_CAD543                -------------------------------------------------------------------------------- [ 22480]
LR899193.1_Germany_2020/1        ................................................................................ [ 22480]
MN393477.1_Wuhan_2019            ................................................................................ [ 22480]
MW396979.1_Timor-Leste/2019/1    ................................................................................ [ 22480]
MK940252.1_CN/2019/InnerMongolia ................................................................................ [ 22480]
MK128995.1_China/2018/AnhuiXCGQ  ................................................................................ [ 22480]
MH681419.1_ASFV/POL/2015         ................................................................................ [ 22480]
LS478113.1_Estonia_2014          ................................................................................ [ 22480]
KP843857.1_Odintsovo_02/14       ................................................................................ [ 22480]
FN557520.1_strain_E75            ...................................................................c.c.......... [ 22480]
AY261366.1_Warthog               -------------------------------------------------------------------------------- [ 22480]
AY261365.1_Warmbaths             ...................................................................c.c.......... [ 22480]
AY261364.1_Tengani_62            -------------------------------------------------------------------------------- [ 22480]
AY261363.1_Pretoriuskop/96/4     ................g...................g...............c.t......................... [ 22480]
AY261362.1_Mkuzi_1979            .................................t...........t......c.t............c.t.......... [ 22480]
AY261361.1_Malawi_Lil-20/1_1983  ...............t...................................................c.c.......... [ 22480]
AY261360.1_Kenya_1950            ....t..........---.......g.......t..................c.t...a.....t..c.ta......... [ 22480]
AM712240.1_OURT_88/3             ...................................................................c.c.......... [ 22480]
AM712239.1_Benin_97/1            ...................................................................c.c.......... [ 22480]
KM111295.1_Ken06.Bus             ....t..........---.......g.......t..................c.t...a........c.t.......... [ 22480]
KM111294.1_Ken05/Tk1             ....t..........---...............t..................c.t............c.t.......... [ 22480]
KM262844.1_strain_L60            ...................................................................c.c.......... [ 22480]
KM102979.1_26544/OG10_Italy      ...................................................................c.c.......... [ 22480]
KM262845.1_strain_NHV            ...................................................................c.c.......... [ 22480]
KX354450.1_47/Ss/2008            ...................................................................c.c.......... [ 22480]
KP055815.1_strain_BA71           cc...tct...ct..-----------------------...tg.t....------------------------------- [ 22480]
MH025920.1_strain_R35            ....t..........---.......g.......t..................c.....a........c.t.......... [ 22480]
MH025919.1_strain_N10            ....t..........---.......g.......t..................c.....a........c.t.......... [ 22480]
MH025918.1_strain_R25            ....t..........---.......g.......t..................c.....a........c.t.......... [ 22480]
MH025917.1_strain_R7             ....t..........---.......g.......t..................c.....a........c.t.......... [ 22480]
MH025916.1_strain_R8             ....t..........---.......g.......t..................c.....a........c.t.......... [ 22480]

FR682468.2_Georgia_2007/1        ttgctattaaagctgtcgagaatcaaaatattgtcataataaatttcgatcgccagtaaaaccttttttcgtttgacgag [ 22560]
ASF_IND_20_CAD543                -------------------------------------------------------------------------------- [ 22560]
LR899193.1_Germany_2020/1        ................................................................................ [ 22560]
MN393477.1_Wuhan_2019            ................................................................................ [ 22560]
MW396979.1_Timor-Leste/2019/1    ................................................................................ [ 22560]
MK940252.1_CN/2019/InnerMongolia ................................................................................ [ 22560]
MK128995.1_China/2018/AnhuiXCGQ  ................................................................................ [ 22560]
MH681419.1_ASFV/POL/2015         ................................................................................ [ 22560]
LS478113.1_Estonia_2014          ................................................................................ [ 22560]
KP843857.1_Odintsovo_02/14       ................................................................................ [ 22560]
```

```
FN557520.1_strain_E75            .g...g.c...a............g...................a..a.....................t......ta.. [ 22560]
AY261366.1_Warthog               -------------------------------------------------------------------------------- [ 22560]
AY261365.1_Warmbaths             .g...g.c...a............g...................a..a.....................t......ta.. [ 22560]
AY261364.1_Tengani_62            -------------------------------------------------------------------------------- [ 22560]
AY261363.1_Pretoriuskop/96/4     .a.....c......a................................a.............................a.. [ 22560]
AY261362.1_Mkuzi_1979            .a.....c...a....................a...........a..a.............t.................. [ 22560]
AY261361.1_Malawi_Lil-20/1_1983  .g.....c......a.........g.......a..............a................................ [ 22560]
AY261360.1_Kenya_1950            .g.....c...............t........a...........a..a.....................t.......t.. [ 22560]
AM712240.1_OURT_88/3             .g...g.c...a............g...................a..a.....................t......ta.. [ 22560]
AM712239.1_Benin_97/1            .g...g.c...a............g...................a..a.....................t......ta.. [ 22560]
KM111295.1_Ken06.Bus             .g.....c........................a...........a..a.....................t.......t.. [ 22560]
KM111294.1_Ken05/Tk1             .g.....cg...............g.......a...........a..t..a..t.a...g.......g.t.......t.. [ 22560]
KM262844.1_strain_L60            .g...g.c...a............g...................a..a.....................t......ta.. [ 22560]
KM102979.1_26544/OG10_Italy      .g...g.c...a............g...................a..a.....................t......ta.. [ 22560]
KM262845.1_strain_NHV            .g...g.c...a............g...................a..a.....................t......ta.. [ 22560]
KX354450.1_47/Ss/2008            .g...g.c...a............g...................a..a.....................t......ta.. [ 22560]
KP055815.1_strain_BA71           -------------------------------------------------------------------------------- [ 22560]
MH025920.1_strain_R35            .g.....c........................a...........a..a.....................t.......t.. [ 22560]
MH025919.1_strain_N10            .g.....c........................a...........a..a.....................t.......t.. [ 22560]
MH025918.1_strain_R25            .g.....c........................a...........a..a.....................t.......t.. [ 22560]
MH025917.1_strain_R7             .g.....c........................a...........a..a.....................t.......t.. [ 22560]
MH025916.1_strain_R8             .g.....c........................a...........a..a.....................t.......t.. [ 22560]

FR682468.2_Georgia_2007/1        ataaacaaacatattatacaaccctacatctaaaaattctggattggctcctagttggatacacaggtctttagtctgct [ 22640]
ASF_IND_20_CAD543                -------------------------------------------------------------------------------- [ 22640]
LR899193.1_Germany_2020/1        ................................................................................ [ 22640]
MN393477.1_Wuhan_2019            ................................................................................ [ 22640]
MW396979.1_Timor-Leste/2019/1    ................................................................................ [ 22640]
MK940252.1_CN/2019/InnerMongolia ................................................................................ [ 22640]
MK128995.1_China/2018/AnhuiXCGQ  ................................................................................ [ 22640]
MH681419.1_ASFV/POL/2015         ................................................................................ [ 22640]
LS478113.1_Estonia_2014          ................................................................................ [ 22640]
KP843857.1_Odintsovo_02/14       ................................................................................ [ 22640]
FN557520.1_strain_E75            ............................................c................................... [ 22640]
AY261366.1_Warthog               -------------------------------------------------------------------------------- [ 22640]
AY261365.1_Warmbaths             ............................................c................................... [ 22640]
AY261364.1_Tengani_62            -------------------------------------------------------------------------------- [ 22640]
AY261363.1_Pretoriuskop/96/4     ......................................g.....c................................... [ 22640]
AY261362.1_Mkuzi_1979            ......................................t.....c...........................c....... [ 22640]
AY261361.1_Malawi_Lil-20/1_1983  ............................................c................................... [ 22640]
AY261360.1_Kenya_1950            ..................t..t...t............t.....c...........................ca...tt. [ 22640]
AM712240.1_OURT_88/3             ............................................c................................... [ 22640]
AM712239.1_Benin_97/1            ............................................c................................... [ 22640]
KM111295.1_Ken06.Bus             ..............c..........tt.................c...........................ca...tt. [ 22640]
```

```
KM111294.1_Ken05/Tk1            ............................................c...........................ca...... [ 22640]
KM262844.1_strain_L60           ............................................c................................... [ 22640]
KM102979.1_26544/OG10_Italy     ............................................c................................... [ 22640]
KM262845.1_strain_NHV           ............................................c................................... [ 22640]
KX354450.1_47/Ss/2008           ............................................c................................... [ 22640]
KP055815.1_strain_BA71          -------------------------------------------------------------------------------- [ 22640]
MH025920.1_strain_R35           ..............c..........tt.................c...........................ca...tt. [ 22640]
MH025919.1_strain_N10           ..............c..........tt.................c...........................ca...tt. [ 22640]
MH025918.1_strain_R25           ..............c..........tt.................c...........................ca...tt. [ 22640]
MH025917.1_strain_R7            ..............c..........tt.................c...........................ca...tt. [ 22640]
MH025916.1_strain_R8            ..............c..........tt.................c...........................ca...tt. [ 22640]

FR682468.2_Georgia_2007/1       tcgttttggcacacatgatgccaaaattaatatcagcaccccataaaacaaataacttgattagatcagtctggttttcc [ 22720]
ASF_IND_20_CAD543               -------------------------------------------------------------------------------- [ 22720]
LR899193.1_Germany_2020/1       ................................................................................ [ 22720]
MN393477.1_Wuhan_2019           ................................................................................ [ 22720]
MW396979.1_Timor-Leste/2019/1   ................................................................................ [ 22720]
MK940252.1_CN/2019/InnerMongolia ................................................................................ [ 22720]
MK128995.1_China/2018/AnhuiXCGQ ................................................................................ [ 22720]
MH681419.1_ASFV/POL/2015        ................................................................................ [ 22720]
LS478113.1_Estonia_2014         ................................................................................ [ 22720]
KP843857.1_Odintsovo_02/14      ................................................................................ [ 22720]
FN557520.1_strain_E75           ................a..............t............g...g.....t......................... [ 22720]
AY261366.1_Warthog              ---------------------------------------------------------------------........c.. [ 22720]
AY261365.1_Warmbaths            .......................t.......t..g.........g.............a..................... [ 22720]
AY261364.1_Tengani_62           -------------------------------------------------------------------------------- [ 22720]
AY261363.1_Pretoriuskop/96/4    ...............................t..g.......................a..................... [ 22720]
AY261362.1_Mkuzi_1979           .t..............a...........................g.........t......................... [ 22720]
AY261361.1_Malawi_Lil-20/1_1983 .............t....................c............................................. [ 22720]
AY261360.1_Kenya_1950           ....c..........c...............t..g..g.......................................... [ 22720]
AM712240.1_OURT_88/3            ................a..............t............g...g.....t......................... [ 22720]
AM712239.1_Benin_97/1           ................a..............t............g...g.....t......................... [ 22720]
KM111295.1_Ken06.Bus            c...c........t.........t.......t..g..g.......................................... [ 22720]
KM111294.1_Ken05/Tk1            ................a..a.t.........g..g..g..................................ta...... [ 22720]
KM262844.1_strain_L60           ................a..............t............g...g.....t......................... [ 22720]
KM102979.1_26544/OG10_Italy     ................a..............t............g...g.....t......................... [ 22720]
KM262845.1_strain_NHV           ................a..............t............g...g.....t......................... [ 22720]
KX354450.1_47/Ss/2008           ................a..............t............g...g.....t......................... [ 22720]
KP055815.1_strain_BA71          ---------------------t..g...g.gtc..a..tttt-------------------------------------- [ 22720]
MH025920.1_strain_R35           c...c........t.........t.......t..g..g.......................................... [ 22720]
MH025919.1_strain_N10           c...c........t.........t.......t..g..g.......................................... [ 22720]
MH025918.1_strain_R25           c...c........t.........t.......t..g..g.......................................... [ 22720]
MH025917.1_strain_R7            c...c........t.........t.......t..g..g.......................................... [ 22720]
MH025916.1_strain_R8            c...c........t.........t.......t..g..g.......................................... [ 22720]
```

```
FR682468.2_Georgia_2007/1        ttcacagcttttactaaggctctgtcaagctcatagctgtcgacatcagagcatgacatagagccaccggttaccatttt [ 22800]
ASF_IND_20_CAD543                -------------------------------------------------------------------------------- [ 22800]
LR899193.1_Germany_2020/1        ................................................................................ [ 22800]
MN393477.1_Wuhan_2019            ................................................................................ [ 22800]
MW396979.1_Timor-Leste/2019/1    ................................................................................ [ 22800]
MK940252.1_CN/2019/InnerMongolia ................................................................................ [ 22800]
MK128995.1_China/2018/AnhuiXCGQ  ................................................................................ [ 22800]
MH681419.1_ASFV/POL/2015         ................................................................................ [ 22800]
LS478113.1_Estonia_2014          ................................................................................ [ 22800]
KP843857.1_Odintsovo_02/14       ................................................................................ [ 22800]
FN557520.1_strain_E75            ...................................a....t.........................t............. [ 22800]
AY261366.1_Warthog               .atg---------------------------------------------------------------------------- [ 22800]
AY261365.1_Warmbaths             ........c................................a..g................................... [ 22800]
AY261364.1_Tengani_62            -------------------------------------------------------------------------------- [ 22800]
AY261363.1_Pretoriuskop/96/4     ...................................a............................................ [ 22800]
AY261362.1_Mkuzi_1979            ...................................a............................................ [ 22800]
AY261361.1_Malawi_Lil-20/1_1983  ...................................a..............a................t............ [ 22800]
AY261360.1_Kenya_1950            ..................................................a...a......................... [ 22800]
AM712240.1_OURT_88/3             ...................................a....t.........................t............. [ 22800]
AM712239.1_Benin_97/1            ...................................a....t.........................t............. [ 22800]
KM111295.1_Ken06.Bus             ........c..................................................t......at............ [ 22800]
KM111294.1_Ken05/Tk1             .............t..................g........a........a................t............ [ 22800]
KM262844.1_strain_L60            ...................................a....t.........................t............. [ 22800]
KM102979.1_26544/OG10_Italy      ...................................a....t.........................t............. [ 22800]
KM262845.1_strain_NHV            ...................................a....t.........................t............. [ 22800]
KX354450.1_47/Ss/2008            ...................................a....t.........................t............. [ 22800]
KP055815.1_strain_BA71           -------------------------------------------------------------------------------- [ 22800]
MH025920.1_strain_R35            ........c..................................................t......at............ [ 22800]
MH025919.1_strain_N10            ........c..................................................t......at............ [ 22800]
MH025918.1_strain_R25            ........c..................................................t......at............ [ 22800]
MH025917.1_strain_R7             ........c..................................................t......at............ [ 22800]
MH025916.1_strain_R8             ........c..................................................t......at............ [ 22800]

FR682468.2_Georgia_2007/1        acattgcttacaaaaacctatgggtccgttttcccaccatagtccaagctgttgtagaataaaaatatcatcctcatgat [ 22880]
ASF_IND_20_CAD543                -------------------------------------------------------------------------------- [ 22880]
LR899193.1_Germany_2020/1        ................................................................................ [ 22880]
MN393477.1_Wuhan_2019            ................................................................................ [ 22880]
MW396979.1_Timor-Leste/2019/1    ................................................................................ [ 22880]
MK940252.1_CN/2019/InnerMongolia ................................................................................ [ 22880]
MK128995.1_China/2018/AnhuiXCGQ  ................................................................................ [ 22880]
MH681419.1_ASFV/POL/2015         ................................................................................ [ 22880]
LS478113.1_Estonia_2014          ................................................................................ [ 22880]
KP843857.1_Odintsovo_02/14       ................................................................................ [ 22880]
```

```
FN557520.1_strain_E75            ................................................................................ [ 22880]
AY261366.1_Warthog               -------------------------------------------------------------------------------- [ 22880]
AY261365.1_Warmbaths             ............................a...............................gt.................. [ 22880]
AY261364.1_Tengani_62            -------------------------------------------------------------------------------- [ 22880]
AY261363.1_Pretoriuskop/96/4     ............................a...............................gt.................. [ 22880]
AY261362.1_Mkuzi_1979            ............................a...........................a.................t..... [ 22880]
AY261361.1_Malawi_Lil-20/1_1983  ......t..................................a...................................... [ 22880]
AY261360.1_Kenya_1950            ......t..............a.................c................a.......c.......t....... [ 22880]
AM712240.1_OURT_88/3             ................................................................................ [ 22880]
AM712239.1_Benin_97/1            ................................................................................ [ 22880]
KM111295.1_Ken06.Bus             ......t..............a.................c...........c....a....c.................. [ 22880]
KM111294.1_Ken05/Tk1             ......t...................................a.....................c.......t....... [ 22880]
KM262844.1_strain_L60            ................................................................................ [ 22880]
KM102979.1_26544/OG10_Italy      ................................................................................ [ 22880]
KM262845.1_strain_NHV            ................................................................................ [ 22880]
KX354450.1_47/Ss/2008            ................................................................................ [ 22880]
KP055815.1_strain_BA71           ------------------------------------------.....cg.t..ac..g.a...gg...tt.t-------- [ 22880]
MH025920.1_strain_R35            ......t..............a.................c...........c....a....c.................. [ 22880]
MH025919.1_strain_N10            ......t..............a.................c...........c....a....c.................. [ 22880]
MH025918.1_strain_R25            ......t..............a.................c...........c....a....c.................. [ 22880]
MH025917.1_strain_R7             ......t..............a.................c...........c....a....c.................. [ 22880]
MH025916.1_strain_R8             ......t..............a.................c...........c....a....c.................. [ 22880]

FR682468.2_Georgia_2007/1        aatttgaaaaagccttggtttctatcaagactttttttgtaagaacctgtaaagagttcatcgtattattatgaataaca [ 22960]
ASF_IND_20_CAD543                -------------------------------------------------------------------------------- [ 22960]
LR899193.1_Germany_2020/1        ................................................................................ [ 22960]
MN393477.1_Wuhan_2019            ................................................................................ [ 22960]
MW396979.1_Timor-Leste/2019/1    ................................................................................ [ 22960]
MK940252.1_CN/2019/InnerMongolia ................................................................................ [ 22960]
MK128995.1_China/2018/AnhuiXCGQ  ................................................................................ [ 22960]
MH681419.1_ASFV/POL/2015         ................................................................................ [ 22960]
LS478113.1_Estonia_2014          ................................................................................ [ 22960]
KP843857.1_Odintsovo_02/14       ................................................................................ [ 22960]
FN557520.1_strain_E75            .c...................................a.......................................... [ 22960]
AY261366.1_Warthog               -------------------------------------------------------------------------------- [ 22960]
AY261365.1_Warmbaths             ................................................................................ [ 22960]
AY261364.1_Tengani_62            -------------------------------------------------------------------------------- [ 22960]
AY261363.1_Pretoriuskop/96/4     ................................................................................ [ 22960]
AY261362.1_Mkuzi_1979            ..a........tt................................................................... [ 22960]
AY261361.1_Malawi_Lil-20/1_1983  ................................................................................ [ 22960]
AY261360.1_Kenya_1950            .................t.....................................a..........c-............ [ 22960]
AM712240.1_OURT_88/3             .c...................................a.......................................... [ 22960]
AM712239.1_Benin_97/1            .c...................................a.......................................... [ 22960]
KM111295.1_Ken06.Bus             .g.........a.....t............................................................t. [ 22960]
```

```
KM111294.1_Ken05/Tk1             .g...............t..........a.....................................c-............ [ 22960]
KM262844.1_strain_L60            .c...................................a.......................................... [ 22960]
KM102979.1_26544/OG10_Italy      .c...................................a.......................................... [ 22960]
KM262845.1_strain_NHV            .c...................................a.......................................... [ 22960]
KX354450.1_47/Ss/2008            .c...................................a.......................................... [ 22960]
KP055815.1_strain_BA71           -------------------------------------------------------------------------..ct.tg [ 22960]
MH025920.1_strain_R35            .g.........a.....t............................................................t. [ 22960]
MH025919.1_strain_N10            .g.........a.....t............................................................t. [ 22960]
MH025918.1_strain_R25            .g.........a.....t............................................................t. [ 22960]
MH025917.1_strain_R7             .g.........a.....t............................................................t. [ 22960]
MH025916.1_strain_R8             .g.........a.....t............................................................t. [ 22960]

FR682468.2_Georgia_2007/1        ggagtaaacgtaatcaattat-aaaagtgattttttcgaaaaaaactttagatggttgaaaatgataatgtacatgtt-- [ 23040]
ASF_IND_20_CAD543                -------------------------------------------------------------------------------- [ 23040]
LR899193.1_Germany_2020/1        .....................-........................................................-- [ 23040]
MN393477.1_Wuhan_2019            .....................-........................................................-- [ 23040]
MW396979.1_Timor-Leste/2019/1    .....................-........................................................-- [ 23040]
MK940252.1_CN/2019/InnerMongolia .....................-........................................................-- [ 23040]
MK128995.1_China/2018/AnhuiXCGQ  .....................-........................................................-- [ 23040]
MH681419.1_ASFV/POL/2015         .....................-........................................................-- [ 23040]
LS478113.1_Estonia_2014          .....................-........................................................-- [ 23040]
KP843857.1_Odintsovo_02/14       .....................-........................................................-- [ 23040]
FN557520.1_strain_E75            .....................a........c.....t......t-...c.a...........c...............-- [ 23040]
AY261366.1_Warthog               -------------------------------------------------------------------------------- [ 23040]
AY261365.1_Warmbaths             .....................-........c............t-...c.a...t..--...c...............-- [ 23040]
AY261364.1_Tengani_62            -------------------------------------------------------------------------------- [ 23040]
AY261363.1_Pretoriuskop/96/4     .....................-........c............t-...c.a...c.......c.........-------- [ 23040]
AY261362.1_Mkuzi_1979            .....................-........c.....t......t-...c.a...........c...............-- [ 23040]
AY261361.1_Malawi_Lil-20/1_1983  .....................-........c.............t...c.....t..............a........-- [ 23040]
AY261360.1_Kenya_1950            .a...................-........c.....t..g---------.a...c......g.........g..ca..-- [ 23040]
AM712240.1_OURT_88/3             .....................a........c.....t......t-...c.a...........c...............-- [ 23040]
AM712239.1_Benin_97/1            .....................a........c.....t......t-...c.a...........c...............-- [ 23040]
KM111295.1_Ken06.Bus             .a...................-......--c........g...t-...c.a...c......gca..g..a.....a..ca [ 23040]
KM111294.1_Ken05/Tk1             .a...................-........c.....t..g...t-...c.a...c......g.........g..ca..-- [ 23040]
KM262844.1_strain_L60            .....................a........c.....t......t-...c.a...........c...............-- [ 23040]
KM102979.1_26544/OG10_Italy      .....................a........c.....t......t-...c.a...........c...............-- [ 23040]
KM262845.1_strain_NHV            .....................a........c.....t......t-...c.a...........c...............-- [ 23040]
KX354450.1_47/Ss/2008            .....................a........c.....t......t-...c.a...........c...............-- [ 23040]
KP055815.1_strain_BA71           .a.t...t.t...c.c.c..--------------..tag....------------------------------------- [ 23040]
MH025920.1_strain_R35            .a...................-......--c........g...t-...c.a...c......gca..g..a.....a..ca [ 23040]
MH025919.1_strain_N10            .a...................-......--c........g...t-...c.a...c......gca..g..a.....a..ca [ 23040]
MH025918.1_strain_R25            .a...................-......--c........g...t-...c.a...c......gca..g..a.....a..ca [ 23040]
MH025917.1_strain_R7             .a...................-......--c........g...t-...c.a...c......gca..g..a.....a..ca [ 23040]
MH025916.1_strain_R8             .a...................-......--c........g...t-...c.a...c......gca..g..a.....a..ca [ 23040]
```

```
FR682468.2_Georgia_2007/1        catacaaaaaatagatgcagtgatgtctaaaatcaaaatttaattttctatgtaaaaagtacagac-ttacttatttggg [ 23120]
ASF_IND_20_CAD543                -------------------------------------------------------------------------------- [ 23120]
LR899193.1_Germany_2020/1        ..................................................................-............. [ 23120]
MN393477.1_Wuhan_2019            ..................................................................-............. [ 23120]
MW396979.1_Timor-Leste/2019/1    ..................................................................-............. [ 23120]
MK940252.1_CN/2019/InnerMongolia ..................................................................-............. [ 23120]
MK128995.1_China/2018/AnhuiXCGQ  ..................................................................-............. [ 23120]
MH681419.1_ASFV/POL/2015         ..................................................................-............. [ 23120]
LS478113.1_Estonia_2014          ..................................................................-............. [ 23120]
KP843857.1_Odintsovo_02/14       ..................................................................-............. [ 23120]
FN557520.1_strain_E75            .............t..t..........c......................................g..g.c....c... [ 23120]
AY261366.1_Warthog               -------------------------------------------------------------------------------- [ 23120]
AY261365.1_Warmbaths             .............t..t..........c......................................g............. [ 23120]
AY261364.1_Tengani_62            -------------------------------------------------------------------------------- [ 23120]
AY261363.1_Pretoriuskop/96/4     .............t...........c.c......................................g....c........ [ 23120]
AY261362.1_Mkuzi_1979            ...........................c....c..............t..................-............. [ 23120]
AY261361.1_Malawi_Lil-20/1_1983  ...........................c......................................-------------- [ 23120]
AY261360.1_Kenya_1950            atacg..g...c.c..a.....g.-..c.tg.............g............g........g....c.g...... [ 23120]
AM712240.1_OURT_88/3             .............t..t..........c......................................g..g.c....c... [ 23120]
AM712239.1_Benin_97/1            .............t..t..........c......................................g..g.c....c... [ 23120]
KM111295.1_Ken06.Bus             tgat.......c.c..a.....g....c..g.............g.....................g....c.g...... [ 23120]
KM111294.1_Ken05/Tk1             atacg..g...c.c........g....c.tg.............g.....................g....c.g...... [ 23120]
KM262844.1_strain_L60            .............t..t..........c......................................g..g.c....c... [ 23120]
KM102979.1_26544/OG10_Italy      .............t..t..........c......................................g..g.c....c... [ 23120]
KM262845.1_strain_NHV            .............t..t..........c......................................g..g.c....c... [ 23120]
KX354450.1_47/Ss/2008            .............t..t..........c......................................g..g.c....c... [ 23120]
KP055815.1_strain_BA71           -.a.t....g.c.c..tt-------------------..........--------------------------------- [ 23120]
MH025920.1_strain_R35            tgat.......c.c..a.....g....c..g.............g.....................g....c.g...... [ 23120]
MH025919.1_strain_N10            tgat.......c.c..a.....g....c..g.............g.....................g....c.g...... [ 23120]
MH025918.1_strain_R25            tgat.......c.c..a.....g....c..g.............g.....................g....c.g...... [ 23120]
MH025917.1_strain_R7             tgat.......c.c..a.....g....c..g.............g.....................g....c.g...... [ 23120]
MH025916.1_strain_R8             tgat.......c.c..a.....g....c..g.............g.....................g....c.g...... [ 23120]

FR682468.2_Georgia_2007/1        ttaaattgtttattttaaactttaattaaccgtttgagttagcgatgtttgatttatcttccatactcatcc---ggggg [ 23200]
ASF_IND_20_CAD543                -------------------------------------------------------------------------------- [ 23200]
LR899193.1_Germany_2020/1        ........................................................................----.... [ 23200]
MN393477.1_Wuhan_2019            ........................................................................------.. [ 23200]
MW396979.1_Timor-Leste/2019/1    ........................................................................------.. [ 23200]
MK940252.1_CN/2019/InnerMongolia ........................................................................----.... [ 23200]
MK128995.1_China/2018/AnhuiXCGQ  ........................................................................-----... [ 23200]
MH681419.1_ASFV/POL/2015         ........................................................................g-g..... [ 23200]
LS478113.1_Estonia_2014          ........................................................................ggg..... [ 23200]
KP843857.1_Odintsovo_02/14       ........................................................................--g..... [ 23200]
```

```
FN557520.1_strain_E75           ...............................a................................g.......ggg..... [ 23200]
AY261366.1_Warthog              -------------------------------------------------------------------------------- [ 23200]
AY261365.1_Warmbaths            ...g........-..................a....................ac.......t..g.......---..... [ 23200]
AY261364.1_Tengani_62           -------------------------------------------------------------------------------- [ 23200]
AY261363.1_Pretoriuskop/96/4    ...........g........................................ac..........g.......----.... [ 23200]
AY261362.1_Mkuzi_1979           ....................g...............................a...................--g..... [ 23200]
AY261361.1_Malawi_Lil-20/1_1983 -------------------------------------------------------------------------------- [ 23200]
AY261360.1_Kenya_1950           ...c................................................ac..........g.......--aa.... [ 23200]
AM712240.1_OURT_88/3            ...............................a................................g.......-------- [ 23200]
AM712239.1_Benin_97/1           ...............................a................................g.......---..... [ 23200]
KM111295.1_Ken06.Bus            ...c...........................a...a............c..gac..................------.. [ 23200]
KM111294.1_Ken05/Tk1            ...c................g..............a............c..gc........t..........---..... [ 23200]
KM262844.1_strain_L60           ...............................a................................g.......ggg..... [ 23200]
KM102979.1_26544/OG10_Italy     ...............................a................................g.......-----... [ 23200]
KM262845.1_strain_NHV           ...............................a................................g.......-------- [ 23200]
KX354450.1_47/Ss/2008           ...............................a................................g.......-----... [ 23200]
KP055815.1_strain_BA71          -------------------------------------------------------------------------------- [ 23200]
MH025920.1_strain_R35           ...c...........................a...a............c..gac..................----.... [ 23200]
MH025919.1_strain_N10           ...c...........................a...a............c..gac..................----.... [ 23200]
MH025918.1_strain_R25           ...c...........................a...a............c..gac..................----.... [ 23200]
MH025917.1_strain_R7            ...c...........................a...a............c..gac..................----.... [ 23200]
MH025916.1_strain_R8            ...c...........................a...a............c..gac..................----.... [ 23200]

FR682468.2_Georgia_2007/1       gggggtccttatagctctgacattattgtggattattgaatataatgaatacttcatagatg-ctaaacattttaat--- [ 23280]
ASF_IND_20_CAD543               -------------------------------------------------------------------------------- [ 23280]
LR899193.1_Germany_2020/1       ..............................................................-..............--- [ 23280]
MN393477.1_Wuhan_2019           ..............................................................-..............--- [ 23280]
MW396979.1_Timor-Leste/2019/1   ..............................................................-..............--- [ 23280]
MK940252.1_CN/2019/InnerMongolia ..............................................................-..............--- [ 23280]
MK128995.1_China/2018/AnhuiXCGQ ..............................................................-..............--- [ 23280]
MH681419.1_ASFV/POL/2015        ..............................................................-..............--- [ 23280]
LS478113.1_Estonia_2014         ..............................................................-..............--- [ 23280]
KP843857.1_Odintsovo_02/14      ....tc.g......................................................-..............--- [ 23280]
FN557520.1_strain_E75           .........a....................................................-...g.t..a...g.att [ 23280]
AY261366.1_Warthog              -------------------------------------------------------------------------------- [ 23280]
AY261365.1_Warmbaths            ..............................................................-............g.aat [ 23280]
AY261364.1_Tengani_62           -------------------------------------------------------------------------------- [ 23280]
AY261363.1_Pretoriuskop/96/4    ..................a...........................................-........a.....agt [ 23280]
AY261362.1_Mkuzi_1979           .........a............................................t.......-........gc..c.ttt [ 23280]
AY261361.1_Malawi_Lil-20/1_1983 -------------------------------------------------------------------------------- [ 23280]
AY261360.1_Kenya_1950           .........c..........t.c....a......g..............a..........c.c...t....g.g...--- [ 23280]
AM712240.1_OURT_88/3            -------------------------------------------------------------------------------- [ 23280]
AM712239.1_Benin_97/1           .........a....................................................-...g.t..a...g.att [ 23280]
KM111295.1_Ken06.Bus            .........c..........t.............g....t.........a............-.....t..a.....c-- [ 23280]
```

```
KM111294.1_Ken05/Tk1              ...........................................................g..-...c..g.ag...aagt [ 23280]
KM262844.1_strain_L60             .........a....................................................-...g.t..a...g.att [ 23280]
KM102979.1_26544/OG10_Italy       .........a....................................................-...g.t..a...g.att [ 23280]
KM262845.1_strain_NHV             -------------------------------------------------------------------------------- [ 23280]
KX354450.1_47/Ss/2008             .........a....................................................-...g.t..a...g.att [ 23280]
KP055815.1_strain_BA71            -------------------------------------------------------------------------------- [ 23280]
MH025920.1_strain_R35             .........c..........t.............g....t.........a............-.....t..a.....c-- [ 23280]
MH025919.1_strain_N10             .........c..........t.............g....t.........a............-.....t..a.....c-- [ 23280]
MH025918.1_strain_R25             .........c..........t.............g....t.........a............-.....t..a.....c-- [ 23280]
MH025917.1_strain_R7              .........c..........t.............g....t.........a............-.....t..a.....c-- [ 23280]
MH025916.1_strain_R8              .........c..........t.............g....t.........a............-.....t..a.....c-- [ 23280]

FR682468.2_Georgia_2007/1         agtagttctgaggcttaattgtactctataaatttataaaaactttttgatcaaaatttaatttcttat-aaaaagagta [ 23360]
ASF_IND_20_CAD543                 -------------------------------------------------------------------------------- [ 23360]
LR899193.1_Germany_2020/1         .....................................................................-.......... [ 23360]
MN393477.1_Wuhan_2019             .....................................................................-.......... [ 23360]
MW396979.1_Timor-Leste/2019/1     .....................................................................-.......... [ 23360]
MK940252.1_CN/2019/InnerMongolia  .....................................................................-.......... [ 23360]
MK128995.1_China/2018/AnhuiXCGQ   .....................................................................-.......... [ 23360]
MH681419.1_ASFV/POL/2015          .....................................................................-.......... [ 23360]
LS478113.1_Estonia_2014           .....................................................................-.......... [ 23360]
KP843857.1_Odintsovo_02/14        .....................................................................a.......... [ 23360]
FN557520.1_strain_E75             tt..a..a.act.........................................................-.......... [ 23360]
AY261366.1_Warthog                -------------------------------------------------------------------------------- [ 23360]
AY261365.1_Warmbaths              ....---..........................................g...................-.......... [ 23360]
AY261364.1_Tengani_62             -------------------------------------------------------------------------------- [ 23360]
AY261363.1_Pretoriuskop/96/4      ..........t..........................................................-.......... [ 23360]
AY261362.1_Mkuzi_1979             .a..a..t.ac.a....................................g...................-.......... [ 23360]
AY261361.1_Malawi_Lil-20/1_1983   -------------------------------------------------------------------------------- [ 23360]
AY261360.1_Kenya_1950             --------------.....aa..........g.c.t..t...--c....t...................-.g........ [ 23360]
AM712240.1_OURT_88/3              -------------------------------------------------------------------------------- [ 23360]
AM712239.1_Benin_97/1             tt..a..a.act.........................................................-.......... [ 23360]
KM111295.1_Ken06.Bus              -t.....g.agt.......g.....t.....g..........a..........................-.......... [ 23360]
KM111294.1_Ken05/Tk1              tt..a..g.tct...................g...........cc....g..............t....-.g........ [ 23360]
KM262844.1_strain_L60             tt..a..a.act.........................................................-.......... [ 23360]
KM102979.1_26544/OG10_Italy       tt..a..a.act.........................................................-.......... [ 23360]
KM262845.1_strain_NHV             -------------------------------------------------------------------------------- [ 23360]
KX354450.1_47/Ss/2008             tt..a..a.act.........................................................-.......... [ 23360]
KP055815.1_strain_BA71            ------------------------------.gc.g.ggc...t..gc...ca..ctc.ag.g.----------------- [ 23360]
MH025920.1_strain_R35             -t.....g.agt.......g.....t.....g..........a..........................-.......... [ 23360]
MH025919.1_strain_N10             -t.....g.agt.......g.....t.....g..........a..........................-.......... [ 23360]
MH025918.1_strain_R25             -t.....g.agt.......g.....t.....g..........a..........................-.......... [ 23360]
MH025917.1_strain_R7              -t.....g.agt.......g.....t.....g..........a..........................-.......... [ 23360]
MH025916.1_strain_R8              -t.....g.agt.......g.....t.....g..........a..........................-.......... [ 23360]
```

```
FR682468.2_Georgia_2007/1         cagacgtcgcttgtttaagcttcatcatgtttcattcattactttctacaattac--------gggggggggagtcccct [ 23440]
ASF_IND_20_CAD543                 -------------------------------------------------------------------------------- [ 23440]
LR899193.1_Germany_2020/1         .......................................................-------g................. [ 23440]
MN393477.1_Wuhan_2019             .......................................................--------................. [ 23440]
MW396979.1_Timor-Leste/2019/1     .......................................................---------................ [ 23440]
MK940252.1_CN/2019/InnerMongolia  .......................................................--------................. [ 23440]
MK128995.1_China/2018/AnhuiXCGQ   .......................................................--------................. [ 23440]
MH681419.1_ASFV/POL/2015          .......................................................g-----gg................. [ 23440]
LS478113.1_Estonia_2014           .......................................................--------................. [ 23440]
KP843857.1_Odintsovo_02/14        .......................................................cggggggg........agtc..... [ 23440]
FN557520.1_strain_E75             t........t.........t...................................--------t........g.gt.... [ 23440]
AY261366.1_Warthog                -------------------------------------------------------------------------------- [ 23440]
AY261365.1_Warmbaths              .......................................................---------t.......g..-.tt. [ 23440]
AY261364.1_Tengani_62             -------------------------------------------------------------------------------- [ 23440]
AY261363.1_Pretoriuskop/96/4      .......................................................---------t.......g....... [ 23440]
AY261362.1_Mkuzi_1979             ..........c............................................--------t........g.gt.t.. [ 23440]
AY261361.1_Malawi_Lil-20/1_1983   -----...........................a......................---------t.......g.gt.t.. [ 23440]
AY261360.1_Kenya_1950             ..-..a......ac...........a......a......................t-----------.....g.gt.t.. [ 23440]
AM712240.1_OURT_88/3              ----------------------------------------------------------------.....----------- [ 23440]
AM712239.1_Benin_97/1             t........t.........t...................................--------t.....-..g.gt.... [ 23440]
KM111295.1_Ken06.Bus              ............a.......a...........t.....a................t----------......g.gt.... [ 23440]
KM111294.1_Ken05/Tk1              ..-..a......a...................a......................t-----------.....g.gt--.. [ 23440]
KM262844.1_strain_L60             t........t.........t...................................--------t.....--.g.gt.... [ 23440]
KM102979.1_26544/OG10_Italy       t........t.........t...................................--------t....---.g.gt.... [ 23440]
KM262845.1_strain_NHV             ----------------------------------------------------------------.....----------- [ 23440]
KX354450.1_47/Ss/2008             t........t.........t...................................--------t.....--.g.gt.... [ 23440]
KP055815.1_strain_BA71            ------------------------....c...tt.c....c...c..ga.g.---------------------------- [ 23440]
MH025920.1_strain_R35             ............a.......a...........t.....a................t--------........g.gt.... [ 23440]
MH025919.1_strain_N10             ............a.......a...........t.....a................t-----------.....g.gt.... [ 23440]
MH025918.1_strain_R25             ............a.......a...........t.....a................tg-------........g.gt.... [ 23440]
MH025917.1_strain_R7              ............a.......a...........t.....a................t--------........g.gt.... [ 23440]
MH025916.1_strain_R8              ............a.......a...........t.....a................t--------........g.gt.... [ 23440]

FR682468.2_Georgia_2007/1         catagctttagtattgctatggtttactaattattatgtagaatttatagaagcatatgtacctgaaag----------- [ 23520]
ASF_IND_20_CAD543                 -------------------------------------------------------------------------------- [ 23520]
LR899193.1_Germany_2020/1         .....................................................................----------- [ 23520]
MN393477.1_Wuhan_2019             .....................................................................----------- [ 23520]
MW396979.1_Timor-Leste/2019/1     .....................................................................----------- [ 23520]
MK940252.1_CN/2019/InnerMongolia  .....................................................................----------- [ 23520]
MK128995.1_China/2018/AnhuiXCGQ   .....................................................................----------- [ 23520]
MH681419.1_ASFV/POL/2015          .....................................................................----------- [ 23520]
LS478113.1_Estonia_2014           .....................................................................----------- [ 23520]
KP843857.1_Odintsovo_02/14        .....................................................................----------- [ 23520]
```

```
FN557520.1_strain_E75            .....t..........t......................................g..a..ta...tg.----------- [ 23520]
AY261366.1_Warthog               -------------------------------------------------------------------------------- [ 23520]
AY261365.1_Warmbaths             ...c............t......................................g.aa..ta.a.ttataattataata [ 23520]
AY261364.1_Tengani_62            -------------------------------------------------------------------------------- [ 23520]
AY261363.1_Pretoriuskop/96/4     .......................................................g.aa..ta...t..tagtaattgt- [ 23520]
AY261362.1_Mkuzi_1979            ..........................................................ac.ta...tg.tg--------- [ 23520]
AY261361.1_Malawi_Lil-20/1_1983  ................t......................................g.aa..ta.a.t..ta--------- [ 23520]
AY261360.1_Kenya_1950            .....t...................g.............................g....gta.t.tg.----------- [ 23520]
AM712240.1_OURT_88/3             -------------------------------------------------------------------------------- [ 23520]
AM712239.1_Benin_97/1            .....t..........t......................................g..a..ta...tg.----------- [ 23520]
KM111295.1_Ken06.Bus             .........................g.............................gtaa.cta.a.tga----------- [ 23520]
KM111294.1_Ken05/Tk1             .........................g.............................g.aa..ta....g.----------- [ 23520]
KM262844.1_strain_L60            .....t..........t......................................g..a..ta...tg.----------- [ 23520]
KM102979.1_26544/OG10_Italy      .....t..........t......................................g..a..ta...tg.----------- [ 23520]
KM262845.1_strain_NHV            -------------------------------------------------------------------------------- [ 23520]
KX354450.1_47/Ss/2008            .....t..........t......................................g..a..ta...tg.----------- [ 23520]
KP055815.1_strain_BA71           -------------------------------------------------------------------------------- [ 23520]
MH025920.1_strain_R35            .........................g.............................gtaa.cta.a.tga----------- [ 23520]
MH025919.1_strain_N10            .........................g.............................gtaa.cta.a.tga----------- [ 23520]
MH025918.1_strain_R25            .........................g.............................gtaa.cta.a.tga----------- [ 23520]
MH025917.1_strain_R7             .........................g.............................gtaa.cta.a.tga----------- [ 23520]
MH025916.1_strain_R8             .........................g.............................gtaa.cta.a.tga----------- [ 23520]

FR682468.2_Georgia_2007/1        -------tatacctactctataaaatta-aataatttcagtata-ttttttttatgaatagaacggaaatgatataaaaa [ 23600]
ASF_IND_20_CAD543                -------------------------------------------------------------------------------- [ 23600]
LR899193.1_Germany_2020/1        -------.....................-...............-................................... [ 23600]
MN393477.1_Wuhan_2019            -------.....................-...............-................................... [ 23600]
MW396979.1_Timor-Leste/2019/1    -------.....................-...............-................................... [ 23600]
MK940252.1_CN/2019/InnerMongolia -------.....................-...............-................................... [ 23600]
MK128995.1_China/2018/AnhuiXCGQ  -------.....................-...............-................................... [ 23600]
MH681419.1_ASFV/POL/2015         -------.....................-...............-................................... [ 23600]
LS478113.1_Estonia_2014          -------.....................-...............-................................... [ 23600]
KP843857.1_Odintsovo_02/14       -------.....................-...............-................................... [ 23600]
FN557520.1_strain_E75            -------..attg..........gt...t..a............-..................-................ [ 23600]
AY261366.1_Warthog               -------------------------------------------------------------------------------- [ 23600]
AY261365.1_Warmbaths             atattacc.acag...............-...............-................................... [ 23600]
AY261364.1_Tengani_62            -------------------------------------------------------------------------------- [ 23600]
AY261363.1_Pretoriuskop/96/4     -----------a...g.......gt...t..a............-..................-................ [ 23600]
AY261362.1_Mkuzi_1979            ------a..attg....t.....gt...t..a............-..........a.......t...........-.... [ 23600]
AY261361.1_Malawi_Lil-20/1_1983  ------a..at.t....t..........-........t......t................................... [ 23600]
AY261360.1_Kenya_1950            -------..attg..........gt..gt..a............-..........a........................ [ 23600]
AM712240.1_OURT_88/3             -------------------------------------------------------------------------------- [ 23600]
AM712239.1_Benin_97/1            -------..attg..........gt...t..a............-..................-................ [ 23600]
KM111295.1_Ken06.Bus             -------..gttg....t.g......c.-...............-..................t................ [ 23600]
```

```
KM111294.1_Ken05/Tk1              -------...c..cgt............-...............--.................................. [ 23600]
KM262844.1_strain_L60             -------..attg..........gt...t..a............--.................-................ [ 23600]
KM102979.1_26544/OG10_Italy       -------..attg..........gt...t..a............-..................-................ [ 23600]
KM262845.1_strain_NHV             -------------------------------------------------------------------------------- [ 23600]
KX354450.1_47/Ss/2008             -------..attg..........gt...t..a............-..................-................ [ 23600]
KP055815.1_strain_BA71            -----------------------.t..gt..gg...c..ag.c------------------------------------- [ 23600]
MH025920.1_strain_R35             -------..gttg....t.g......c.-...............-..................t................ [ 23600]
MH025919.1_strain_N10             -------..gttg....t.g......c.-...............-..................t................ [ 23600]
MH025918.1_strain_R25             -------..gttg....t.g......c.-...............-..................t................ [ 23600]
MH025917.1_strain_R7              -------..gttg....t.g......c.-...............-..................t................ [ 23600]
MH025916.1_strain_R8              -------..gttg....t.g......c.-...............-..................t................ [ 23600]

FR682468.2_Georgia_2007/1         taatttaatattgc-aa-------aaaaa--attcataatgttggtatgtattataa--acataatagcatgtgtaattt [ 23680]
ASF_IND_20_CAD543                 -------------------------------------------------------------------------------- [ 23680]
LR899193.1_Germany_2020/1         ..............-..-------.....--..........................--..................... [ 23680]
MN393477.1_Wuhan_2019             ..............-..-------.....--..........................--..................... [ 23680]
MW396979.1_Timor-Leste/2019/1     ..............-..-------.....--..........................--..................... [ 23680]
MK940252.1_CN/2019/InnerMongolia  ..............-..-------.....--..........................--..................... [ 23680]
MK128995.1_China/2018/AnhuiXCGQ   ..............-..-------.....--..........................--..................... [ 23680]
MH681419.1_ASFV/POL/2015          ..............-..-------.....--..........................--..................... [ 23680]
LS478113.1_Estonia_2014           ..............-..-------.....--..........................--..................... [ 23680]
KP843857.1_Odintsovo_02/14        ..............-..-------.....--..........................--..................... [ 23680]
FN557520.1_strain_E75             ..............-..-------.....at..........................at.t......a.....a...... [ 23680]
AY261366.1_Warthog                -------------------------------------------------------------------------------- [ 23680]
AY261365.1_Warmbaths              ..............a..-------.....at..............c..a..---------------------------.. [ 23680]
AY261364.1_Tengani_62             -------------------------------------------------------------------------------- [ 23680]
AY261363.1_Pretoriuskop/96/4      ..............-..-------.....at........c.................at.t................... [ 23680]
AY261362.1_Mkuzi_1979             ..............-..-------.....at..........................--.t................... [ 23680]
AY261361.1_Malawi_Lil-20/1_1983   ..............--.-------.....at.........a....c..a....------.t........c....a..... [ 23680]
AY261360.1_Kenya_1950             ..............a..-------.....at.........a....c-------....at.t......a............ [ 23680]
AM712240.1_OURT_88/3              -------------------------------------------------------------------------------- [ 23680]
AM712239.1_Benin_97/1             ..............-..-------.....at..........................at.t......a.....a...... [ 23680]
KM111295.1_Ken06.Bus              ..............a..acattgc.....aa..............c..a........at.t...............t... [ 23680]
KM111294.1_Ken05/Tk1              ..............a..-----------.aa.a............c-------....at.t........-.......... [ 23680]
KM262844.1_strain_L60             ..............-..-------.....at..........................at.t......a.....a...... [ 23680]
KM102979.1_26544/OG10_Italy       ..............-..-------.....at..........................at.t......a.....a...... [ 23680]
KM262845.1_strain_NHV             -------------------------------------------------------------------------------- [ 23680]
KX354450.1_47/Ss/2008             ..............-..-------.....at..........................at.t......a.....a...... [ 23680]
KP055815.1_strain_BA71            --------c..g..-tg-------.g..---------------------------------------------------- [ 23680]
MH025920.1_strain_R35             ..............a..acattgc.....aa..............c..a........at.t...............t... [ 23680]
MH025919.1_strain_N10             ..............a..acattgc.....aa..............c..a........at.t...............t... [ 23680]
MH025918.1_strain_R25             ..............a..acattgc.....aa..............c..a........at.t...............t... [ 23680]
MH025917.1_strain_R7              ..............a..acattgc.....aa..............c..a........at.t...............t... [ 23680]
MH025916.1_strain_R8              ..............a..acattgc.....aa..............c..a........at.t...............t... [ 23680]
```

```
FR682468.2_Georgia_2007/1        ataaactgactcctctatataattattagatgaggtaccaacctacttatgatatgccgatgatagatattgtata-cta [ 23760]
ASF_IND_20_CAD543                -------------------------------------------------------------------------------- [ 23760]
LR899193.1_Germany_2020/1        ............................................................................-... [ 23760]
MN393477.1_Wuhan_2019            ............................................................................-... [ 23760]
MW396979.1_Timor-Leste/2019/1    ............................................................................-... [ 23760]
MK940252.1_CN/2019/InnerMongolia ............................................................................-... [ 23760]
MK128995.1_China/2018/AnhuiXCGQ  ............................................................................-... [ 23760]
MH681419.1_ASFV/POL/2015         ............................................................................-... [ 23760]
LS478113.1_Estonia_2014          ............................................................................-... [ 23760]
KP843857.1_Odintsovo_02/14       ............................................................................-... [ 23760]
FN557520.1_strain_E75            ...........t...c.a.a......----...a....t..t...t.....................c........-.c. [ 23760]
AY261366.1_Warthog               -------------------------------------------------------------------------------- [ 23760]
AY261365.1_Warmbaths             ......c....t...c.a.a..................t......t.....................g........c.c. [ 23760]
AY261364.1_Tengani_62            -------------------------------------------------------------------------------- [ 23760]
AY261363.1_Pretoriuskop/96/4     ......c....t...c.a....a............................................c........c.c. [ 23760]
AY261362.1_Mkuzi_1979            ....----...t...c.a......................g..........................c........-.c. [ 23760]
AY261361.1_Malawi_Lil-20/1_1983  ........t..t...c.a..........a...t....tt......t.....................c........c.c. [ 23760]
AY261360.1_Kenya_1950            .....a.........c.a...........t........t......t....a.......-........c.-......t.c. [ 23760]
AM712240.1_OURT_88/3             -------------------------------------------------------------------------------- [ 23760]
AM712239.1_Benin_97/1            ...........t...c.a.a......----...a....t..t...t.....................c........-.c. [ 23760]
KM111295.1_Ken06.Bus             ...............c.............................t.....g...............c........c.c. [ 23760]
KM111294.1_Ken05/Tk1             ...............c.a....................t-....gt............a........c........ttc. [ 23760]
KM262844.1_strain_L60            ...........t...c.a.a......----...a....t..t...t.....................c........-.c. [ 23760]
KM102979.1_26544/OG10_Italy      ...........t...c.a.a......----...a....t..t...t.....................c........-.c. [ 23760]
KM262845.1_strain_NHV            -------------------------------------------------------------------------------- [ 23760]
KX354450.1_47/Ss/2008            ...........t...c.a.a......----...a....t..t...t.....................c........-.c. [ 23760]
KP055815.1_strain_BA71           ------------------------------------------.a.t.c...g....------------------------ [ 23760]
MH025920.1_strain_R35            ...............c.............................t.....g...............c........c.c. [ 23760]
MH025919.1_strain_N10            ...............c.............................t.....g...............c........c.c. [ 23760]
MH025918.1_strain_R25            ...............c.............................t.....g...............c........c.c. [ 23760]
MH025917.1_strain_R7             ...............c.............................t.....g...............c........c.c. [ 23760]
MH025916.1_strain_R8             ...............c.............................t.....g...............c........c.c. [ 23760]

FR682468.2_Georgia_2007/1        taaaacaaaattattttaaatgtattcatggataca-ttataacat--------------------------------tt [ 23840]
ASF_IND_20_CAD543                -------------------------------------------------------------------------------- [ 23840]
LR899193.1_Germany_2020/1        ....................................-.........--------------------------------.. [ 23840]
MN393477.1_Wuhan_2019            ....................................-.........--------------------------------.. [ 23840]
MW396979.1_Timor-Leste/2019/1    ....................................-.........--------------------------------.. [ 23840]
MK940252.1_CN/2019/InnerMongolia ....................................-.........--------------------------------.. [ 23840]
MK128995.1_China/2018/AnhuiXCGQ  ....................................-.........--------------------------------.. [ 23840]
MH681419.1_ASFV/POL/2015         ....................................-.........--------------------------------.. [ 23840]
LS478113.1_Estonia_2014          ....................................-.........--------------------------------.. [ 23840]
KP843857.1_Odintsovo_02/14       ....................................-.........--------------------------------.. [ 23840]
```

```
FN557520.1_strain_E75            .....................a..............-catag.t..a--------------------catcataata-.. [ 23840]
AY261366.1_Warthog               -------------------------------------------------------------------------------- [ 23840]
AY261365.1_Warmbaths             ......---.....................t..ct.t.a.at....a--------------tggatacatcataaca-.. [ 23840]
AY261364.1_Tengani_62            -------------------------------------------------------------------------------- [ 23840]
AY261363.1_Pretoriuskop/96/4     ....................................-.atag.t..a--------------tggatacatcataacat.. [ 23840]
AY261362.1_Mkuzi_1979            .....t..............................-.c.......--------------------------------.. [ 23840]
AY261361.1_Malawi_Lil-20/1_1983  ..............................t..c..-.agatt...ggatacatagatccatagatacatcataacat.. [ 23840]
AY261360.1_Kenya_1950            ..............................a.....-.gtatt...agataca--------tggacacatcataacatg. [ 23840]
AM712240.1_OURT_88/3             -------------------------------------------------------------------------------- [ 23840]
AM712239.1_Benin_97/1            .....................a..............-catag.t..a--------------------catcataata-.. [ 23840]
KM111295.1_Ken06.Bus             .....t...............t...c...a......-.gtat--..ggataca--------tggacacatcataacat-. [ 23840]
KM111294.1_Ken05/Tk1             ...........................-----------------..gaataca--------tggatacatcataacat-. [ 23840]
KM262844.1_strain_L60            .....................a..............-catag.t..a--------------------catcataata-.. [ 23840]
KM102979.1_26544/OG10_Italy      .....................a..............-catag.t..a--------------------catcataata-.. [ 23840]
KM262845.1_strain_NHV            -------------------------------------------------------------------------------- [ 23840]
KX354450.1_47/Ss/2008            .....................a..............-catag.t..a--------------------catcataata-.. [ 23840]
KP055815.1_strain_BA71           ------gt.ga..cc..t.tcta.a....ac.....-..ta--------------------------------------- [ 23840]
MH025920.1_strain_R35            .....t...............t...c...a......-.gtat--..ggataca--------tggacacatcataacat-. [ 23840]
MH025919.1_strain_N10            .....t...............t...c...a......-.gtat--..ggataca--------tggacacatcataacat-. [ 23840]
MH025918.1_strain_R25            .....t...............t...c...a......-.gtat--..ggataca--------tggacacatcataacat-. [ 23840]
MH025917.1_strain_R7             .....t...............t...c...a......-.gtat--..ggataca--------tggacacatcataacat-. [ 23840]
MH025916.1_strain_R8             .....t...............t...c...a......-.gtat--..ggataca--------tggacacatcataacat-. [ 23840]

FR682468.2_Georgia_2007/1        ttaccgcaaattgtctctcagcgaagaaaatgaatgaaacgtttctgtatattcataggttgaaattattttacgcactt [ 23920]
ASF_IND_20_CAD543                -------------------------------------------------------------------------------- [ 23920]
LR899193.1_Germany_2020/1        ................................................................................ [ 23920]
MN393477.1_Wuhan_2019            ................................................................................ [ 23920]
MW396979.1_Timor-Leste/2019/1    ................................................................................ [ 23920]
MK940252.1_CN/2019/InnerMongolia ................................................................................ [ 23920]
MK128995.1_China/2018/AnhuiXCGQ  ................................................................................ [ 23920]
MH681419.1_ASFV/POL/2015         ................................................................................ [ 23920]
LS478113.1_Estonia_2014          ................................................................................ [ 23920]
KP843857.1_Odintsovo_02/14       ................................................................................ [ 23920]
FN557520.1_strain_E75            ......................a......................c.................................. [ 23920]
AY261366.1_Warthog               -------------------------------------------------------------------------------- [ 23920]
AY261365.1_Warmbaths             .....................ta......................................................... [ 23920]
AY261364.1_Tengani_62            -------------------------------------------------------------------------------- [ 23920]
AY261363.1_Pretoriuskop/96/4     ......................a......................c........g......................... [ 23920]
AY261362.1_Mkuzi_1979            ....t.................a......................................................... [ 23920]
AY261361.1_Malawi_Lil-20/1_1983  .............................................c.................................. [ 23920]
AY261360.1_Kenya_1950            ........c....................................c.......g.......................... [ 23920]
AM712240.1_OURT_88/3             -------------------------------------------------------------------------------- [ 23920]
AM712239.1_Benin_97/1            ......................a......................c.................................. [ 23920]
KM111295.1_Ken06.Bus             ............a........t.......................c....g...........................c. [ 23920]
```

```
KM111294.1_Ken05/Tk1          ...t..t...........t...................c......c.......g.......................... [ 23920]
KM262844.1_strain_L60         ......................a......................c.................................. [ 23920]
KM102979.1_26544/OG10_Italy   ......................a......................c.................................. [ 23920]
KM262845.1_strain_NHV         -------------------------------------------------------------------------------- [ 23920]
KX354450.1_47/Ss/2008         ......................a......................c.................................. [ 23920]
KP055815.1_strain_BA71        ----------------tc....c..cggg.cacca..c.t.c..tcac...c...------------------------- [ 23920]
MH025920.1_strain_R35         ............a........t.......................c....g...........................c. [ 23920]
MH025919.1_strain_N10         ............a........t.......................c....g...........................c. [ 23920]
MH025918.1_strain_R25         ............a........t.......................c....g...........................c. [ 23920]
MH025917.1_strain_R7          ............a........t.......................c....g...........................c. [ 23920]
MH025916.1_strain_R8          ............a........t.......................c....g...........................c. [ 23920]

FR682468.2_Georgia_2007/1     cactaggttctaatattttcttatgaagtattgaatgggggcttaaaagtcctttcttaaaaagaagtttcatcataaca [ 24000]
ASF_IND_20_CAD543             -------------------------------------------------------------------------------- [ 24000]
LR899193.1_Germany_2020/1     ................................................................................ [ 24000]
MN393477.1_Wuhan_2019         ................................................................................ [ 24000]
MW396979.1_Timor-Leste/2019/1 ................................................................................ [ 24000]
MK940252.1_CN/2019/InnerMongolia ................................................................................ [ 24000]
MK128995.1_China/2018/AnhuiXCGQ ................................................................................ [ 24000]
MH681419.1_ASFV/POL/2015      ................................................................................ [ 24000]
LS478113.1_Estonia_2014       ................................................................................ [ 24000]
KP843857.1_Odintsovo_02/14    ................................................................................ [ 24000]
FN557520.1_strain_E75         .........................................................................t...... [ 24000]
AY261366.1_Warthog            -------------------------------------------------------------------------------- [ 24000]
AY261365.1_Warmbaths          ...c...............t.....................................................t...... [ 24000]
AY261364.1_Tengani_62         -------------------------------------------------------------------------------- [ 24000]
AY261363.1_Pretoriuskop/96/4  ........................................................c................t...... [ 24000]
AY261362.1_Mkuzi_1979         ..t.........t............................................................t...... [ 24000]
AY261361.1_Malawi_Lil-20/1_1983 ...c...............t...................a.g.....................ag........t...... [ 24000]
AY261360.1_Kenya_1950         ................................................................t........t...t.. [ 24000]
AM712240.1_OURT_88/3          -------------------------------------------------------------------------------- [ 24000]
AM712239.1_Benin_97/1         .........................................................................t...... [ 24000]
KM111295.1_Ken06.Bus          ...................t...................................t........t........t...... [ 24000]
KM111294.1_Ken05/Tk1          ................................................................t........t...... [ 24000]
KM262844.1_strain_L60         .........................................................................t...... [ 24000]
KM102979.1_26544/OG10_Italy   .........................................................................t...... [ 24000]
KM262845.1_strain_NHV         -------------------------------------------------------------------------------- [ 24000]
KX354450.1_47/Ss/2008         .........................................................................t...... [ 24000]
KP055815.1_strain_BA71        -----------------.......a..tggcc..---------------------------------------------- [ 24000]
MH025920.1_strain_R35         ...................t...................................t........t........t...... [ 24000]
MH025919.1_strain_N10         ...................t...................................t........t........t...... [ 24000]
MH025918.1_strain_R25         ...................t...................................t........t........t...... [ 24000]
MH025917.1_strain_R7          ...................t...................................t........t........t...... [ 24000]
MH025916.1_strain_R8          ...................t...................................t........t........t...... [ 24000]
```

```
FR682468.2_Georgia_2007/1        ttcttttcttgtctaagaagagtttcttgta-ttttttttgtataaggattggcacccaaacttatacaaaaatgtacat [ 24080]
ASF_IND_20_CAD543                -------------------------------------------------------------------------------- [ 24080]
LR899193.1_Germany_2020/1        ...............................-................................................ [ 24080]
MN393477.1_Wuhan_2019            ...............................-................................................ [ 24080]
MW396979.1_Timor-Leste/2019/1    ...............................-................................................ [ 24080]
MK940252.1_CN/2019/InnerMongolia ...............................-................................................ [ 24080]
MK128995.1_China/2018/AnhuiXCGQ  ...............................-................................................ [ 24080]
MH681419.1_ASFV/POL/2015         ...............................-................................................ [ 24080]
LS478113.1_Estonia_2014          ...............................-................................................ [ 24080]
KP843857.1_Odintsovo_02/14       ...............................-................................................ [ 24080]
FN557520.1_strain_E75            ...............................-................................................ [ 24080]
AY261366.1_Warthog               -------------------------------------------------------------------------------- [ 24080]
AY261365.1_Warmbaths             ...............................-................................................ [ 24080]
AY261364.1_Tengani_62            -------------------------------------------------------------------------------- [ 24080]
AY261363.1_Pretoriuskop/96/4     ...............................-................................................ [ 24080]
AY261362.1_Mkuzi_1979            ...............................-................................................ [ 24080]
AY261361.1_Malawi_Lil-20/1_1983  ..........c....................-........a....................................... [ 24080]
AY261360.1_Kenya_1950            .a........c..........a.........-................................................ [ 24080]
AM712240.1_OURT_88/3             -------------------------------------------------------------------------------- [ 24080]
AM712239.1_Benin_97/1            ...............................-................................................ [ 24080]
KM111295.1_Ken06.Bus             ..........c....................-............c...g............................t.. [ 24080]
KM111294.1_Ken05/Tk1             ..........c....................-................................................ [ 24080]
KM262844.1_strain_L60            ...............................-................................................ [ 24080]
KM102979.1_26544/OG10_Italy      ...............................-................................................ [ 24080]
KM262845.1_strain_NHV            -------------------------------------------------------------------------------- [ 24080]
KX354450.1_47/Ss/2008            ...............................-................................................ [ 24080]
KP055815.1_strain_BA71           -----------------------cc.a.c..-....g...a..c...--------------------------------- [ 24080]
MH025920.1_strain_R35            ..........c....................t............c...g............................t.. [ 24080]
MH025919.1_strain_N10            ..........c....................t............c...g............................t.. [ 24080]
MH025918.1_strain_R25            ..........c....................t............c...g............................t.. [ 24080]
MH025917.1_strain_R7             ..........c....................t............c...g............................t.. [ 24080]
MH025916.1_strain_R8             ..........c....................t............c...g............................t.. [ 24080]

FR682468.2_Georgia_2007/1        tactccaaataccataatttgaaaagaaagttatttccctatttacttcatgattaatgaaacctatcaacgtctctaag [ 24160]
ASF_IND_20_CAD543                -------------------------------------------------------------------------------- [ 24160]
LR899193.1_Germany_2020/1        ................................................................................ [ 24160]
MN393477.1_Wuhan_2019            ................................................................................ [ 24160]
MW396979.1_Timor-Leste/2019/1    ................................................................................ [ 24160]
MK940252.1_CN/2019/InnerMongolia ................................................................................ [ 24160]
MK128995.1_China/2018/AnhuiXCGQ  ................................................................................ [ 24160]
MH681419.1_ASFV/POL/2015         ................................................................................ [ 24160]
LS478113.1_Estonia_2014          ................................................................................ [ 24160]
KP843857.1_Odintsovo_02/14       ................................................................................ [ 24160]
```

```
FN557520.1_strain_E75              ................................................................................ [ 24160]
AY261366.1_Warthog                 -------------------------------.....g.t.....gt.....------ca....g..g...ta.a.....a [ 24160]
AY261365.1_Warmbaths               ................................................................................ [ 24160]
AY261364.1_Tengani_62              -------------------------------------------------------------------------------- [ 24160]
AY261363.1_Pretoriuskop/96/4       .......................................c...................t...g......t......... [ 24160]
AY261362.1_Mkuzi_1979              ............................................c................................... [ 24160]
AY261361.1_Malawi_Lil-20/1_1983    ..........................................c....................g................ [ 24160]
AY261360.1_Kenya_1950              ..............................g.------.......g.................g........c....... [ 24160]
AM712240.1_OURT_88/3               -------------------------------------------------------------------------------- [ 24160]
AM712239.1_Benin_97/1              ................................................................................ [ 24160]
KM111295.1_Ken06.Bus               ...............................................................g................ [ 24160]
KM111294.1_Ken05/Tk1               ...............................................................g................ [ 24160]
KM262844.1_strain_L60              ................................................................................ [ 24160]
KM102979.1_26544/OG10_Italy        ................................................................................ [ 24160]
KM262845.1_strain_NHV              -------------------------------------------------------------------------------- [ 24160]
KX354450.1_47/Ss/2008              ................................................................................ [ 24160]
KP055815.1_strain_BA71             -------------------------------------------------------------------------------- [ 24160]
MH025920.1_strain_R35              ...............................................................g................ [ 24160]
MH025919.1_strain_N10              ...............................................................g................ [ 24160]
MH025918.1_strain_R25              ...............................................................g................ [ 24160]
MH025917.1_strain_R7               ...............................................................g................ [ 24160]
MH025916.1_strain_R8               ...............................................................g................ [ 24160]

FR682468.2_Georgia_2007/1          gccgtattgatatttgcgcctaaggcaaaacaatagtatatacccaatttattttgagggtacatacaagcaagcgacat [ 24240]
ASF_IND_20_CAD543                  -------------------------------------------------------------------------------- [ 24240]
LR899193.1_Germany_2020/1          ................................................................................ [ 24240]
MN393477.1_Wuhan_2019              ................................................................................ [ 24240]
MW396979.1_Timor-Leste/2019/1      ................................................................................ [ 24240]
MK940252.1_CN/2019/InnerMongolia   ................................................................................ [ 24240]
MK128995.1_China/2018/AnhuiXCGQ    ................................................................................ [ 24240]
MH681419.1_ASFV/POL/2015           ................................................................................ [ 24240]
LS478113.1_Estonia_2014            ................................................................................ [ 24240]
KP843857.1_Odintsovo_02/14         ................................................................................ [ 24240]
FN557520.1_strain_E75              ..a.........................................t........c.......................... [ 24240]
AY261366.1_Warthog                 ..at.g.ct....c.........a...........a........t........c..g..a.....g..g.....ta.... [ 24240]
AY261365.1_Warmbaths               ..a..............a..........................t........c......................g... [ 24240]
AY261364.1_Tengani_62              ----------------------------------------------------------------................ [ 24240]
AY261363.1_Pretoriuskop/96/4       ..a.........................................t........c......................g... [ 24240]
AY261362.1_Mkuzi_1979              ..a.........................................t........c.......................... [ 24240]
AY261361.1_Malawi_Lil-20/1_1983    ..a..............a..........................t................................... [ 24240]
AY261360.1_Kenya_1950              ..a..............c..........................t........c.....a.................... [ 24240]
AM712240.1_OURT_88/3               -------------------------------------------------------------------------------- [ 24240]
AM712239.1_Benin_97/1              ..a.........................................t........c.......................... [ 24240]
KM111295.1_Ken06.Bus               ..a.........................................t........c...................a...... [ 24240]
```

```
KM111294.1_Ken05/Tk1             ..a..............c..........................t..c................................ [ 24240]
KM262844.1_strain_L60            ..a.........................................t........c.......................... [ 24240]
KM102979.1_26544/OG10_Italy      ..a.........................................t........c.......................... [ 24240]
KM262845.1_strain_NHV            -------------------------------------------------------------------------------- [ 24240]
KX354450.1_47/Ss/2008            ..a.........................................t........c.......................... [ 24240]
KP055815.1_strain_BA71           -------------------------------------------------------------------------------- [ 24240]
MH025920.1_strain_R35            ..a.........................................t........c...................a...... [ 24240]
MH025919.1_strain_N10            ..a.........................................t........c...................a...... [ 24240]
MH025918.1_strain_R25            ..a.........................................t........c...................a...... [ 24240]
MH025917.1_strain_R7             ..a.........................................t........c...................a...... [ 24240]
MH025916.1_strain_R8             ..a.........................................t........c...................a...... [ 24240]

FR682468.2_Georgia_2007/1        catgtcatttggatctaaacgtatattttcctgaaaatatgcatgatggatttcatcaacattacctaagtatacagccg [ 24320]
ASF_IND_20_CAD543                -------------------------------------------------------------------------------- [ 24320]
LR899193.1_Germany_2020/1        ................................................................................ [ 24320]
MN393477.1_Wuhan_2019            ................................................................................ [ 24320]
MW396979.1_Timor-Leste/2019/1    ................................................................................ [ 24320]
MK940252.1_CN/2019/InnerMongolia ................................................................................ [ 24320]
MK128995.1_China/2018/AnhuiXCGQ  ................................................................................ [ 24320]
MH681419.1_ASFV/POL/2015         ................................................................................ [ 24320]
LS478113.1_Estonia_2014          ................................................................................ [ 24320]
KP843857.1_Odintsovo_02/14       ................................................................................ [ 24320]
FN557520.1_strain_E75            ................................................................................ [ 24320]
AY261366.1_Warthog               .......................................c...c....a.c........t..............t..... [ 24320]
AY261365.1_Warmbaths             ................................................................................ [ 24320]
AY261364.1_Tengani_62            .....................................................................a.......... [ 24320]
AY261363.1_Pretoriuskop/96/4     ................................................................................ [ 24320]
AY261362.1_Mkuzi_1979            ..............................t................................................. [ 24320]
AY261361.1_Malawi_Lil-20/1_1983  .....g.........................................................................a [ 24320]
AY261360.1_Kenya_1950            .....g.........................................................c................ [ 24320]
AM712240.1_OURT_88/3             -------------------------------------------------------------------------------- [ 24320]
AM712239.1_Benin_97/1            ................................................................................ [ 24320]
KM111295.1_Ken06.Bus             .....g........................t................................c................ [ 24320]
KM111294.1_Ken05/Tk1             .....t.......................................................................... [ 24320]
KM262844.1_strain_L60            ................................................................................ [ 24320]
KM102979.1_26544/OG10_Italy      t............................................................................... [ 24320]
KM262845.1_strain_NHV            -------------------------------------------------------------------------------- [ 24320]
KX354450.1_47/Ss/2008            t............................................................................... [ 24320]
KP055815.1_strain_BA71           -------------------------------------------------------------------------------- [ 24320]
MH025920.1_strain_R35            .....g........................t................................c................ [ 24320]
MH025919.1_strain_N10            .....g........................t................................c................ [ 24320]
MH025918.1_strain_R25            .....g........................t................................c................ [ 24320]
MH025917.1_strain_R7             .....g........................t................................c................ [ 24320]
MH025916.1_strain_R8             .....g........................t................................c................ [ 24320]
```

```
FR682468.2_Georgia_2007/1        tttttaaacgccaataatctaggtgaggaaatttcttactaagaaaacgaataggttttataagattaaactctatggcg [ 24400]
ASF_IND_20_CAD543                -------------------------------------------------------------------------------- [ 24400]
LR899193.1_Germany_2020/1        ................................................................................ [ 24400]
MN393477.1_Wuhan_2019            ................................................................................ [ 24400]
MW396979.1_Timor-Leste/2019/1    ................................................................................ [ 24400]
MK940252.1_CN/2019/InnerMongolia ................................................................................ [ 24400]
MK128995.1_China/2018/AnhuiXCGQ  ................................................................................ [ 24400]
MH681419.1_ASFV/POL/2015         ................................................................................ [ 24400]
LS478113.1_Estonia_2014          ................................................................................ [ 24400]
KP843857.1_Odintsovo_02/14       ................................................................................ [ 24400]
FN557520.1_strain_E75            ......................a......................................................... [ 24400]
AY261366.1_Warthog               c......g......at.....aa..g........t.........t...............c...t.....a......... [ 24400]
AY261365.1_Warmbaths             ........................a....................................................... [ 24400]
AY261364.1_Tengani_62            ...............c................................................................ [ 24400]
AY261363.1_Pretoriuskop/96/4     ...............c................................................................ [ 24400]
AY261362.1_Mkuzi_1979            ...............c........................................................g....... [ 24400]
AY261361.1_Malawi_Lil-20/1_1983  ...............c................................................................ [ 24400]
AY261360.1_Kenya_1950            ...............c.......................c....................c..................a [ 24400]
AM712240.1_OURT_88/3             -------------------------------------------------------------------------------- [ 24400]
AM712239.1_Benin_97/1            ......................a......................................................... [ 24400]
KM111295.1_Ken06.Bus             ...............c............................t....g..........c................... [ 24400]
KM111294.1_Ken05/Tk1             ...............g............................................................c... [ 24400]
KM262844.1_strain_L60            ......................a......................................................... [ 24400]
KM102979.1_26544/OG10_Italy      ......................a......................................................... [ 24400]
KM262845.1_strain_NHV            -------------------------------------------------------------------------------- [ 24400]
KX354450.1_47/Ss/2008            ......................a......................................................... [ 24400]
KP055815.1_strain_BA71           -------------------------------------------------------------------------------- [ 24400]
MH025920.1_strain_R35            ...............c............................t....g..........c................... [ 24400]
MH025919.1_strain_N10            ...............c............................t....g..........c................... [ 24400]
MH025918.1_strain_R25            ...............c............................t....g..........c................... [ 24400]
MH025917.1_strain_R7             ...............c............................t....g..........c................... [ 24400]
MH025916.1_strain_R8             ...............c............................t....g..........c................... [ 24400]

FR682468.2_Georgia_2007/1        atcttaaaccaaaattttaatacatatgtattttttatcattttt---tctttttcatctaaatttaagataaaacgatt [ 24480]
ASF_IND_20_CAD543                -------------------------------------------------------------------------------- [ 24480]
LR899193.1_Germany_2020/1        .............................................---................................ [ 24480]
MN393477.1_Wuhan_2019            .............................................---................................ [ 24480]
MW396979.1_Timor-Leste/2019/1    .............................................---................................ [ 24480]
MK940252.1_CN/2019/InnerMongolia .............................................---................................ [ 24480]
MK128995.1_China/2018/AnhuiXCGQ  .............................................---................................ [ 24480]
MH681419.1_ASFV/POL/2015         .............................................---................................ [ 24480]
LS478113.1_Estonia_2014          .............................................---................................ [ 24480]
KP843857.1_Odintsovo_02/14       .............................................---................................ [ 24480]
```

```
FN557520.1_strain_E75           .............................................---............................a... [ 24480]
AY261366.1_Warthog              .a....t.................ag..c....c.cg......c.tcg................a............... [ 24480]
AY261365.1_Warmbaths            .............................................---............................a... [ 24480]
AY261364.1_Tengani_62           .............................................---............................a... [ 24480]
AY261363.1_Pretoriuskop/96/4    .............................................---............................a... [ 24480]
AY261362.1_Mkuzi_1979           .a......................ag..........g......c.---..............g.a..g............ [ 24480]
AY261361.1_Malawi_Lil-20/1_1983 ........................ag.......c..g......c.---..............g.a..g....t....... [ 24480]
AY261360.1_Kenya_1950           .a....t.................ag..........g.t......---............................c... [ 24480]
AM712240.1_OURT_88/3            -------------------------------------------------------------------------------- [ 24480]
AM712239.1_Benin_97/1           .............................................---............................a... [ 24480]
KM111295.1_Ken06.Bus            .a......................a....g...c..g........---...................g............ [ 24480]
KM111294.1_Ken05/Tk1            .a....t.................ag.......c..g......c.---................................ [ 24480]
KM262844.1_strain_L60           .............................................---............................a... [ 24480]
KM102979.1_26544/OG10_Italy     .............................................---............................a... [ 24480]
KM262845.1_strain_NHV           -------------------------------------------------------------------------------- [ 24480]
KX354450.1_47/Ss/2008           .............................................---............................a... [ 24480]
KP055815.1_strain_BA71          -------------------------------------------------------------------------------- [ 24480]
MH025920.1_strain_R35           .a......................a....g...c..g........---...................g............ [ 24480]
MH025919.1_strain_N10           .a......................a....g...c..g........---...................g............ [ 24480]
MH025918.1_strain_R25           .a......................a....g...c..g........---...................g............ [ 24480]
MH025917.1_strain_R7            .a......................a....g...c..g........---...................g............ [ 24480]
MH025916.1_strain_R8            .a......................a....g...c..g........---...................g............ [ 24480]

FR682468.2_Georgia_2007/1       gtaaataaagtctatcaacacgtaaaaatcatggctatcaaaactgtcgagaatcgaaatattgtcataataaatatcta [ 24560]
ASF_IND_20_CAD543               -------------------------------------------------------------------------------- [ 24560]
LR899193.1_Germany_2020/1       ................................................................................ [ 24560]
MN393477.1_Wuhan_2019           ................................................................................ [ 24560]
MW396979.1_Timor-Leste/2019/1   ................................................................................ [ 24560]
MK940252.1_CN/2019/InnerMongolia ................................................................................ [ 24560]
MK128995.1_China/2018/AnhuiXCGQ ................................................................................ [ 24560]
MH681419.1_ASFV/POL/2015        ................................................................................ [ 24560]
LS478113.1_Estonia_2014         ................................................................................ [ 24560]
KP843857.1_Odintsovo_02/14      ................................................................................ [ 24560]
FN557520.1_strain_E75           ..................t.............................................a................ [ 24560]
AY261366.1_Warthog              .....c............t.g..g........a.........g..a.........a...................t..a. [ 24560]
AY261365.1_Warmbaths            ..........................................c....................a................ [ 24560]
AY261364.1_Tengani_62           ...c..............................a............................a................ [ 24560]
AY261363.1_Pretoriuskop/96/4    .................................a........c....................a................ [ 24560]
AY261362.1_Mkuzi_1979           ...c.......................................................c...a................ [ 24560]
AY261361.1_Malawi_Lil-20/1_1983 ...........t......tg..................................g.g...g..t................ [ 24560]
AY261360.1_Kenya_1950           ...............................................................a................ [ 24560]
AM712240.1_OURT_88/3            -------------------------------------------------------------------------------- [ 24560]
AM712239.1_Benin_97/1           ..................t.............................................a................ [ 24560]
KM111295.1_Ken06.Bus            ...........t.......g...g..............t...g............a.......a................ [ 24560]
```

KM111294.1_Ken05/Tk1 ...................gt.............tc.........a.................a................ [ 24560]
KM262844.1_strain_L60 ..................t............................................a................ [ 24560]
KM102979.1_26544/OG10_Italy ..................t............................................a................ [ 24560]
KM262845.1_strain_NHV -------------------------------------------------------------------------------- [ 24560]
KX354450.1_47/Ss/2008 ..................t............................................a................ [ 24560]
KP055815.1_strain_BA71 -------------------------------------------------------------------------------- [ 24560]
MH025920.1_strain_R35 ...........t.......g...g..............t...g............a.......a................ [ 24560]
MH025919.1_strain_N10 ...........t.......g...g..............t...g............a.......a................ [ 24560]
MH025918.1_strain_R25 ...........t.......g...g..............t...g............a.......a................ [ 24560]
MH025917.1_strain_R7 ...........t.......g...g..............t...g............a.......a................ [ 24560]
MH025916.1_strain_R8 ...........t.......g...g..............t...g............a.......a................ [ 24560]

FR682468.2_Georgia_2007/1 tagctaataagaccttttgttgtttaattagatcaacaaacatattatacaaccctacatctaaaaattttggatcagct [ 24640]
ASF_IND_20_CAD543 -------------------------------------------------------------------------------- [ 24640]
LR899193.1_Germany_2020/1 ................................................................................ [ 24640]
MN393477.1_Wuhan_2019 ................................................................................ [ 24640]
MW396979.1_Timor-Leste/2019/1 ................................................................................ [ 24640]
MK940252.1_CN/2019/InnerMongolia ................................................................................ [ 24640]
MK128995.1_China/2018/AnhuiXCGQ ................................................................................ [ 24640]
MH681419.1_ASFV/POL/2015 ................................................................................ [ 24640]
LS478113.1_Estonia_2014 ................................................................................ [ 24640]
KP843857.1_Odintsovo_02/14 ................................................................................ [ 24640]
FN557520.1_strain_E75 ....................................................t................c.....tg... [ 24640]
AY261366.1_Warthog .c..c.g...a.......t.c....g.cg....a...................................g......g... [ 24640]
AY261365.1_Warmbaths ................................................................................ [ 24640]
AY261364.1_Tengani_62 ................................................................................ [ 24640]
AY261363.1_Pretoriuskop/96/4 ..........................................a..................................... [ 24640]
AY261362.1_Mkuzi_1979 ...........................g.................................................... [ 24640]
AY261361.1_Malawi_Lil-20/1_1983 ...........................a................g.............c.........g........... [ 24640]
AY261360.1_Kenya_1950 ................................................................................ [ 24640]
AM712240.1_OURT_88/3 -------------------------------------------------------------------------------- [ 24640]
AM712239.1_Benin_97/1 ....................................................t................c.....tg... [ 24640]
KM111295.1_Ken06.Bus .g...............................t.................g.tt.tt...........c......g... [ 24640]
KM111294.1_Ken05/Tk1 .............t.............g.................................................... [ 24640]
KM262844.1_strain_L60 ....................................................t................c.....tg... [ 24640]
KM102979.1_26544/OG10_Italy ....................................................t................c.....tg... [ 24640]
KM262845.1_strain_NHV -------------------------------------------------------------------------------- [ 24640]
KX354450.1_47/Ss/2008 ....................................................t................c.....tg... [ 24640]
KP055815.1_strain_BA71 -------------------------------------------------------------------------------- [ 24640]
MH025920.1_strain_R35 .g...............................t.................g.tt.tt...........c......g... [ 24640]
MH025919.1_strain_N10 .g...............................t.................g.tt.tt...........c......g... [ 24640]
MH025918.1_strain_R25 .g...............................t.................g.tt.tt...........c......g... [ 24640]
MH025917.1_strain_R7 .g...............................t.................g.tt.tt...........c......g... [ 24640]
MH025916.1_strain_R8 .g...............................t.................g.tt.tt...........c......g... [ 24640]

```
FR682468.2_Georgia_2007/1        cctagttgaatacacagaactttcgtcctttccgtcttggcacatatgatgccataattaatgttggcaccccataaaac [ 24720]
ASF_IND_20_CAD543                -------------------------------------------------------------------------------- [ 24720]
LR899193.1_Germany_2020/1        ................................................................................ [ 24720]
MN393477.1_Wuhan_2019            ................................................................................ [ 24720]
MW396979.1_Timor-Leste/2019/1    ................................................................................ [ 24720]
MK940252.1_CN/2019/InnerMongolia ................................................................................ [ 24720]
MK128995.1_China/2018/AnhuiXCGQ  ................................................................................ [ 24720]
MH681419.1_ASFV/POL/2015         ................................................................................ [ 24720]
LS478113.1_Estonia_2014          ................................................................................ [ 24720]
KP843857.1_Odintsovo_02/14       ................................................................................ [ 24720]
FN557520.1_strain_E75            .......c.........gt............................................................. [ 24720]
AY261366.1_Warthog               ........g........gt....a...tgc.t...t........c.........a.......t.c............... [ 24720]
AY261365.1_Warmbaths             ..................g.....a.......t........g...................................... [ 24720]
AY261364.1_Tengani_62            ..............t............t.................................................... [ 24720]
AY261363.1_Pretoriuskop/96/4     ........g.....t................................................................. [ 24720]
AY261362.1_Mkuzi_1979            ...............................................................................t [ 24720]
AY261361.1_Malawi_Lil-20/1_1983  .................................a.........................g.................... [ 24720]
AY261360.1_Kenya_1950            ..............t.............................c.c.......a.......t.c...g........... [ 24720]
AM712240.1_OURT_88/3             -------------------------------------------------------------------------------- [ 24720]
AM712239.1_Benin_97/1            .......c.........gt............................................................. [ 24720]
KM111295.1_Ken06.Bus             .................gt..............a....a.....c..a................................ [ 24720]
KM111294.1_Ken05/Tk1             ..........g............t....................c..a.............c.................. [ 24720]
KM262844.1_strain_L60            .......c.........gt............................................................. [ 24720]
KM102979.1_26544/OG10_Italy      .......c.........gt............................................................. [ 24720]
KM262845.1_strain_NHV            -------------------------------------------------------------------------------- [ 24720]
KX354450.1_47/Ss/2008            .......c.........gt............................................................. [ 24720]
KP055815.1_strain_BA71           -----------------------------.g..a.......--------------------------------------- [ 24720]
MH025920.1_strain_R35            .................gt..............a....a.....c..a................................ [ 24720]
MH025919.1_strain_N10            .................gt..............a....a.....c..a................................ [ 24720]
MH025918.1_strain_R25            .................gt..............a....a.....c..a................................ [ 24720]
MH025917.1_strain_R7             .................gt..............a....a.....c..a................................ [ 24720]
MH025916.1_strain_R8             .................gt..............a....a.....c..a................................ [ 24720]

FR682468.2_Georgia_2007/1        aaataacttgattagatcagtctggtttttcttcacagccctcaccaaggctctgtcaagctcatagctgtcaacatcag [ 24800]
ASF_IND_20_CAD543                -------------------------------------------------------------------------------- [ 24800]
LR899193.1_Germany_2020/1        ................................................................................ [ 24800]
MN393477.1_Wuhan_2019            ................................................................................ [ 24800]
MW396979.1_Timor-Leste/2019/1    ................................................................................ [ 24800]
MK940252.1_CN/2019/InnerMongolia ................................................................................ [ 24800]
MK128995.1_China/2018/AnhuiXCGQ  ................................................................................ [ 24800]
MH681419.1_ASFV/POL/2015         ................................................................................ [ 24800]
LS478113.1_Estonia_2014          ................................................................................ [ 24800]
KP843857.1_Odintsovo_02/14       ................................................................................ [ 24800]
```

```
FN557520.1_strain_E75             ...c..............................................t................................g.... [ 24800]
AY261366.1_Warthog                .........a...................c.........tt.t..t......................a.....gt...... [ 24800]
AY261365.1_Warmbaths              ...........................................................................g.... [ 24800]
AY261364.1_Tengani_62             ..............a....................................................................g.... [ 24800]
AY261363.1_Pretoriuskop/96/4      ...........................................................................g.... [ 24800]
AY261362.1_Mkuzi_1979             ...........................................................................g.... [ 24800]
AY261361.1_Malawi_Lil-20/1_1983   ........................a........t......t..................................g.... [ 24800]
AY261360.1_Kenya_1950             .....................t.......c.........tt.t..t..........................g....... [ 24800]
AM712240.1_OURT_88/3              -------------------------------------------------------------------------------- [ 24800]
AM712239.1_Benin_97/1             ...c..............................................t................................g.... [ 24800]
KM111295.1_Ken06.Bus              ......ta.............t.......c.........tt.t..t.....c............................ [ 24800]
KM111294.1_Ken05/Tk1              ........................................g....................................... [ 24800]
KM262844.1_strain_L60             ...c..............................................t................................g.... [ 24800]
KM102979.1_26544/OG10_Italy       ...c..............................................t................................g.... [ 24800]
KM262845.1_strain_NHV             -------------------------------------------------------------------------------- [ 24800]
KX354450.1_47/Ss/2008             ...c..............................................t................................g.... [ 24800]
KP055815.1_strain_BA71            -----------------------------------------------..t.c.a.ca..at.g.c.a..t..c.tg.ag. [ 24800]
MH025920.1_strain_R35             ......ta.............t.......c.........tt.t..t.....c............................ [ 24800]
MH025919.1_strain_N10             ......ta.............t.......c.........tt.t..t.....c............................ [ 24800]
MH025918.1_strain_R25             ......ta.............t.......c.........tt.t..t.....c............................ [ 24800]
MH025917.1_strain_R7              ......ta.............t.......c.........tt.t..t.....c............................ [ 24800]
MH025916.1_strain_R8              ......ta.............t.......c.........tt.t..t.....c............................ [ 24800]

FR682468.2_Georgia_2007/1         aacatgacatagagccactggttaccattttacattgtttacaaaaacctatgggtccgttttcccaccataatccaagc [ 24880]
ASF_IND_20_CAD543                 ----------------------------------------------------------------................ [ 24880]
LR899193.1_Germany_2020/1         ................................................................................ [ 24880]
MN393477.1_Wuhan_2019             ................................................................................ [ 24880]
MW396979.1_Timor-Leste/2019/1     ................................................................................ [ 24880]
MK940252.1_CN/2019/InnerMongolia  ................................................................................ [ 24880]
MK128995.1_China/2018/AnhuiXCGQ   ................................................................................ [ 24880]
MH681419.1_ASFV/POL/2015          ................................................................................ [ 24880]
LS478113.1_Estonia_2014           ................................................................................ [ 24880]
KP843857.1_Odintsovo_02/14        ................................................................................ [ 24880]
FN557520.1_strain_E75             .g.....................................................g..a..a.g......c......... [ 24880]
AY261366.1_Warthog                .g.....................................................a..a..a.g................ [ 24880]
AY261365.1_Warmbaths              .g..........c......a...................................a................g....... [ 24880]
AY261364.1_Tengani_62             .g........g........ta..................................g..a..a.g........g....... [ 24880]
AY261363.1_Pretoriuskop/96/4      .g.....................................................g..a..a.g........g....... [ 24880]
AY261362.1_Mkuzi_1979             .g.....................................................g..a..a.g........g....... [ 24880]
AY261361.1_Malawi_Lil-20/1_1983   .g..................................................a.....................t.t... [ 24880]
AY261360.1_Kenya_1950             ..................c...................................................c.g....... [ 24880]
AM712240.1_OURT_88/3              -------------------------------------------------------------------------------- [ 24880]
AM712239.1_Benin_97/1             .g.....................................................g..a..a.g......c......... [ 24880]
KM111295.1_Ken06.Bus              .g........t......a.......t.............................g..............c.....t... [ 24880]
```

```
KM111294.1_Ken05/Tk1              .g.....................................................g..a.ca.g......c..c..t... [ 24880]
KM262844.1_strain_L60             .g.....................................................g..a..a.g......c......... [ 24880]
KM102979.1_26544/OG10_Italy       .g.....................................................g..a..a.g......c......... [ 24880]
KM262845.1_strain_NHV             -------------------------------------------------------------------------------- [ 24880]
KX354450.1_47/Ss/2008             .g.....................................................g..a..a.g......c......... [ 24880]
KP055815.1_strain_BA71            cg---------------------------------------------------------------.....gt..t.g..t [ 24880]
MH025920.1_strain_R35             .g........t......a.......t.............................g..............c.....t... [ 24880]
MH025919.1_strain_N10             .g........t......a.......t.............................g..............c.....t... [ 24880]
MH025918.1_strain_R25             .g........t......a.......t.............................g..............c.....t... [ 24880]
MH025917.1_strain_R7              .g........t......a.......t.............................g..............c.....t... [ 24880]
MH025916.1_strain_R8              .g........t......a.......t.............................g..............c.....t... [ 24880]

FR682468.2_Georgia_2007/1         tgctgtaaaataaaaatatcatcctcatgataatttgaaaaagccttgttttctatcaagactttttttgtaagaacctg [ 24960]
ASF_IND_20_CAD543                 ..y....r........................t.c.............k............................... [ 24960]
LR899193.1_Germany_2020/1         ................................................................................ [ 24960]
MN393477.1_Wuhan_2019             ................................................................................ [ 24960]
MW396979.1_Timor-Leste/2019/1     ................................................................................ [ 24960]
MK940252.1_CN/2019/InnerMongolia  ................................................................................ [ 24960]
MK128995.1_China/2018/AnhuiXCGQ   ................................................................................ [ 24960]
MH681419.1_ASFV/POL/2015          ................................................................................ [ 24960]
LS478113.1_Estonia_2014           ................................................................................ [ 24960]
KP843857.1_Odintsovo_02/14        ................................................................................ [ 24960]
FN557520.1_strain_E75             ................................t.c............................................. [ 24960]
AY261366.1_Warthog                ..t....................t........................g............................... [ 24960]
AY261365.1_Warmbaths              .......................t........t............................................... [ 24960]
AY261364.1_Tengani_62             ................................................................................ [ 24960]
AY261363.1_Pretoriuskop/96/4      .......................t........................................................ [ 24960]
AY261362.1_Mkuzi_1979             .......................t........t.c............................................. [ 24960]
AY261361.1_Malawi_Lil-20/1_1983   ................c......t........................................................ [ 24960]
AY261360.1_Kenya_1950             ...........g...c.......t........................................................ [ 24960]
AM712240.1_OURT_88/3              -------------------------------------------------------------------------------- [ 24960]
AM712239.1_Benin_97/1             ................................t.c............................................. [ 24960]
KM111295.1_Ken06.Bus              ...............c.......t.................g..............t....................... [ 24960]
KM111294.1_Ken05/Tk1              .........................................g......g............................... [ 24960]
KM262844.1_strain_L60             ................................t.c............................................. [ 24960]
KM102979.1_26544/OG10_Italy       ................................t.c............................................. [ 24960]
KM262845.1_strain_NHV             -------------------------------------------------------------------------------- [ 24960]
KX354450.1_47/Ss/2008             ................................t.c............................................. [ 24960]
KP055815.1_strain_BA71            .------------------------------------------------------------...cc.c..g.g..t...- [ 24960]
MH025920.1_strain_R35             ...............c.......t.................g..............t....................... [ 24960]
MH025919.1_strain_N10             ...............c.......t.................g..............t....................... [ 24960]
MH025918.1_strain_R25             ...............c.......t.................g..............t....................... [ 24960]
MH025917.1_strain_R7              ...............c.......t.................g..............t....................... [ 24960]
MH025916.1_strain_R8              ...............c.......t.................g..............t....................... [ 24960]
```

```
FR682468.2_Georgia_2007/1        taaagaattcatcgtattatcat-----gaatgaaagcagtaaatgtaatcaattataaaattgacttattgaagagaaa [ 25040]
ASF_IND_20_CAD543                ......r.............y..-----....r.m..s......y................................... [ 25040]
LR899193.1_Germany_2020/1        .......................-----.................................................... [ 25040]
MN393477.1_Wuhan_2019            .......................-----.................................................... [ 25040]
MW396979.1_Timor-Leste/2019/1    .......................-----.................................................... [ 25040]
MK940252.1_CN/2019/InnerMongolia .......................-----.................................................... [ 25040]
MK128995.1_China/2018/AnhuiXCGQ  .......................-----.................................................... [ 25040]
MH681419.1_ASFV/POL/2015         .......................-----.................................................... [ 25040]
LS478113.1_Estonia_2014          .......................-----.................................................... [ 25040]
KP843857.1_Odintsovo_02/14       .......................-----.................................................... [ 25040]
FN557520.1_strain_E75            .......................-----.g..a............................................... [ 25040]
AY261366.1_Warthog               .......................-----.g..a...a..a.....a.................................. [ 25040]
AY261365.1_Warmbaths             ....................t..-----.g..a...a........................................... [ 25040]
AY261364.1_Tengani_62            .......................-----.................................................... [ 25040]
AY261363.1_Pretoriuskop/96/4     .......................-----.g..a...a........................................... [ 25040]
AY261362.1_Mkuzi_1979            .......................ggcat.g..a............................................... [ 25040]
AY261361.1_Malawi_Lil-20/1_1983  .......................-----.g......a........................................... [ 25040]
AY261360.1_Kenya_1950            ......g..........c-.t..-----....a.c..a......c................g......t..tg...a.t. [ 25040]
AM712240.1_OURT_88/3             -------------------------------------------------------------------------------- [ 25040]
AM712239.1_Benin_97/1            .......................-----.g..a............................................... [ 25040]
KM111295.1_Ken06.Bus             ............t..........-----ag....tt......................................a...t. [ 25040]
KM111294.1_Ken05/Tk1             .............c...c.....-----.g..a...a........................c......c..a..a..... [ 25040]
KM262844.1_strain_L60            .......................-----.g..a............................................... [ 25040]
KM102979.1_26544/OG10_Italy      .......................-----.g..a............................................... [ 25040]
KM262845.1_strain_NHV            -------------------------------------------------------------------------------- [ 25040]
KX354450.1_47/Ss/2008            .......................-----.g..a............................................... [ 25040]
KP055815.1_strain_BA71           ------------------------------------------------------------------c.g...gtcga.gt [ 25040]
MH025920.1_strain_R35            ............t..........-----ag....tt......................................a...t. [ 25040]
MH025919.1_strain_N10            ............t..........-----ag....tt......................................a...t. [ 25040]
MH025918.1_strain_R25            ............t..........-----ag....tt......................................a...t. [ 25040]
MH025917.1_strain_R7             ............t..........-----ag....tt......................................a...t. [ 25040]
MH025916.1_strain_R8             ............t..........-----ag....tt......................................a...t. [ 25040]

FR682468.2_Georgia_2007/1        tgttaaatgagtgaaatcggtgtttatgatgatgtacatgatcatacgaagaaacacgttcactggtgtccatgatcaaa [ 25120]
ASF_IND_20_CAD543                ................................................................................ [ 25120]
LR899193.1_Germany_2020/1        ................................................................................ [ 25120]
MN393477.1_Wuhan_2019            ................................................................................ [ 25120]
MW396979.1_Timor-Leste/2019/1    ................................................................................ [ 25120]
MK940252.1_CN/2019/InnerMongolia ................................................................................ [ 25120]
MK128995.1_China/2018/AnhuiXCGQ  ................................................................................ [ 25120]
MH681419.1_ASFV/POL/2015         ................................................................................ [ 25120]
LS478113.1_Estonia_2014          ................................................................................ [ 25120]
KP843857.1_Odintsovo_02/14       ................................................................................ [ 25120]
```

```
FN557520.1_strain_E75          ..........................c........g....................g....................... [ 25120]
AY261366.1_Warthog             ...................t...............g........................t................... [ 25120]
AY261365.1_Warmbaths           ...................t...............g..........t...a......................a...... [ 25120]
AY261364.1_Tengani_62          ..........................c..................................................... [ 25120]
AY261363.1_Pretoriuskop/96/4   .........g.........a......c..................................................... [ 25120]
AY261362.1_Mkuzi_1979          ..........................c........g....................g....................... [ 25120]
AY261361.1_Malawi_Lil-20/1_1983 ...................a......c......a..........................................t... [ 25120]
AY261360.1_Kenya_1950          ct.g...------------------g....a....g..c-..t..............a.g..g..........a...... [ 25120]
AM712240.1_OURT_88/3           -------------------------------------------------------------------------------- [ 25120]
AM712239.1_Benin_97/1          ..........................c........g....................g....................... [ 25120]
KM111295.1_Ken06.Bus           .a...............t.a.....gc........g...........a.........a...................... [ 25120]
KM111294.1_Ken05/Tk1           ...................a......c........g...------.........t....ct................... [ 25120]
KM262844.1_strain_L60          ..........................c........g....................g....................... [ 25120]
KM102979.1_26544/OG10_Italy    ..........................c........g....................g....................... [ 25120]
KM262845.1_strain_NHV          -------------------------------------------------------------------------------- [ 25120]
KX354450.1_47/Ss/2008          ..........................c........g....................g....................... [ 25120]
KP055815.1_strain_BA71         ...cc..c....tg.c.t.aaacc.g------------------------------------------------------ [ 25120]
MH025920.1_strain_R35          .a...............t.a.....gc........g...........a.........a...................... [ 25120]
MH025919.1_strain_N10          .a...............t.a.....gc........g...........a.........a...................... [ 25120]
MH025918.1_strain_R25          .a...............t.a.....gc........g...........a.........a...................... [ 25120]
MH025917.1_strain_R7           .a...............t.a.....gc........g...........a.........a...................... [ 25120]
MH025916.1_strain_R8           .a...............t.a.....gc........g...........a.........a...................... [ 25120]

FR682468.2_Georgia_2007/1      atttaatgttttacgtaaaaagtacagatgttaactgtttagtttaaa-cataaatttaacctttagtttaaaccctagt [ 25200]
ASF_IND_20_CAD543              ............................y...................-...t.g....ca..aa..g....t.tt...a [ 25200]
LR899193.1_Germany_2020/1      ................................................-............................... [ 25200]
MN393477.1_Wuhan_2019          ................................................-............................... [ 25200]
MW396979.1_Timor-Leste/2019/1  ................................................-............................... [ 25200]
MK940252.1_CN/2019/InnerMongolia ................................................-............................... [ 25200]
MK128995.1_China/2018/AnhuiXCGQ ................................................-............................... [ 25200]
MH681419.1_ASFV/POL/2015       ................................................-............................... [ 25200]
LS478113.1_Estonia_2014        ................................................-............................... [ 25200]
KP843857.1_Odintsovo_02/14     ................................................-............................... [ 25200]
FN557520.1_strain_E75          .......a..c.................c...................-...t.g....ca..aa..g....t.tt...a [ 25200]
AY261366.1_Warthog             .......t..c..ta.............ca.......--.........-...t.g.....a..aa..g....t.tt...a [ 25200]
AY261365.1_Warmbaths           ..........c..t..............ca......a...........-tta.t......a..aa..a....t.tt...a [ 25200]
AY261364.1_Tengani_62          .......t.cc..t......g.......ca................g.-...t.g.c...a.............taa... [ 25200]
AY261363.1_Pretoriuskop/96/4   .......t.cc..t......g.......ca.......--.........-...t.g.....a..aa..g....t.tt...a [ 25200]
AY261362.1_Mkuzi_1979          .......a..c.................c...................-...t.g....ca..aa..g....t.tt...a [ 25200]
AY261361.1_Malawi_Lil-20/1_1983 .......t..c..ta.....-........a.......--.........-...t.g.....a.-------...t.tt..aa [ 25200]
AY261360.1_Kenya_1950          ..........c..t......g.......c....c......g.g...c.ttg.tt....-----.a.ac..t..ttaaccg [ 25200]
AM712240.1_OURT_88/3           -------------------------------------------------------------------------------- [ 25200]
AM712239.1_Benin_97/1          .......a..c.................c...................-...t.g....ca..aa..g....t.tt...a [ 25200]
KM111295.1_Ken06.Bus           .......t.cc..t.......a......c......ca....aa------...ttg.....a.a...........taa..g [ 25200]
```

```
KM111294.1_Ken05/Tk1          ..........c..t...............c........-------------------------............aa..a [ 25200]
KM262844.1_strain_L60         .......a..c..................c..................-...t.g....ca..aa..g....t.tt...a [ 25200]
KM102979.1_26544/OG10_Italy   .......a..c..................c..................-...t.g....ca..aa..g....t.tt...a [ 25200]
KM262845.1_strain_NHV         -------------------------------------------------------------------------------- [ 25200]
KX354450.1_47/Ss/2008         .......a..c..................c..................-...t.g....ca..aa..g....t.tt...a [ 25200]
KP055815.1_strain_BA71        -------------------------------------------------------------------------------- [ 25200]
MH025920.1_strain_R35         .......t.cc..t.......a.......c....tca....aa------...ttg.....a.a...........taa..g [ 25200]
MH025919.1_strain_N10         .......t.cc..t.......a.......c....tca....aa------...ttg.....a.a...........taa..g [ 25200]
MH025918.1_strain_R25         .......t.cc..t.......a.......c....tca....aa------...ttg.....a.a...........taa..g [ 25200]
MH025917.1_strain_R7          .......t.cc..t.......a.......c....tca....aa------...ttg.....a.a...........taa..g [ 25200]
MH025916.1_strain_R8          .......t.cc..t.......a.......c....tca....aa------...ttg.....a.a...........taa..g [ 25200]

FR682468.2_Georgia_2007/1     taatg--------------------atgtttaatatttcttctatactcattca----gggaagtg----taatgattct [ 25280]
ASF_IND_20_CAD543             .t.a.cct---agnnnnnnnnnnnnnnnnnnnnnnnnnnnnnnnnnnnnnnnnn----nnnnnnnn----nnnnnnnnnn [ 25280]
LR899193.1_Germany_2020/1     .....--------------------.............................----........----.......... [ 25280]
MN393477.1_Wuhan_2019         .....--------------------.............................----........----.......... [ 25280]
MW396979.1_Timor-Leste/2019/1 .....--------------------.............................----........----.......... [ 25280]
MK940252.1_CN/2019/InnerMongolia .....--------------------.............................----........----.......... [ 25280]
MK128995.1_China/2018/AnhuiXCGQ .....--------------------.............................----........----.......... [ 25280]
MH681419.1_ASFV/POL/2015      .....--------------------.............................----........----.......... [ 25280]
LS478113.1_Estonia_2014       .....--------------------.............................----........----.......... [ 25280]
KP843857.1_Odintsovo_02/14    .....--------------------.............................----........----.......... [ 25280]
FN557520.1_strain_E75         .t.a.cct---agtcaaca------......g.c........c..t..t..a.-----...gg.g.gtct....cg.... [ 25280]
AY261366.1_Warthog            .t.a.cct---agtcaaca------......g..........c..t..t..a.-----...gg.g.gttc....ag.... [ 25280]
AY261365.1_Warmbaths          .t.a.cct---agtcaaca------......g.c........c..t..t..c.c--cg...gg.g.gtcc....agc... [ 25280]
AY261364.1_Tengani_62         .t.actatttgagttaatg------.........c.......c..g..t..cag--gg...gg.g.gtcc....a...t. [ 25280]
AY261363.1_Pretoriuskop/96/4  .tga.cct---agtcaaca------......g.c........c..t..t..c.gaggg...gg.g.gtcc....agc... [ 25280]
AY261362.1_Mkuzi_1979         .t.a.cct---agtcaaca------......g.c........c..t..t..a.g----...gg.g.gtct....cg.... [ 25280]
AY261361.1_Malawi_Lil-20/1_1983 .t.a.cct---agtcaaca------......g.c........c..t..t..a.g----...gg.g.gtcc....ag.... [ 25280]
AY261360.1_Kenya_1950         .tta-------agttagcg------....c.g.ac.a.....c........c..--ag...gg.g.gtcc.c..agc... [ 25280]
AM712240.1_OURT_88/3          -------------------------------------------------------------------------------- [ 25280]
AM712239.1_Benin_97/1         .t.a.cct---agtcaaca------......g.c........c..t..t..a.-----...gg.g.gtct....cg.... [ 25280]
KM111295.1_Ken06.Bus          .t.aacct---agttaatg------.........gc......c...........--gg.aagt.------.....g.... [ 25280]
KM111294.1_Ken05/Tk1          .t.aacct---aatcaaca------......g.c.c......c..t..t..a.g--gg...ggt----cc....a...t. [ 25280]
KM262844.1_strain_L60         .t.a.cct---agtcaaca------......g.c........c..t..t..a.------..gg.g.gtct....cg.... [ 25280]
KM102979.1_26544/OG10_Italy   .t.a.cct---agtcaaca------......g.c........c..t..t..a.-----...gg.g.gtct....cg.... [ 25280]
KM262845.1_strain_NHV         -------------------------------------------------------------------------------- [ 25280]
KX354450.1_47/Ss/2008         .t.a.cct---agtcaaca------......g.c........c..t..t..a.-----...gg.g.gtct....cg.... [ 25280]
KP055815.1_strain_BA71        -------------------------------------------------------------------------------- [ 25280]
MH025920.1_strain_R35         .t.aacct---agttaatg------.........gc......c...........--gg.aagt.------.....g.... [ 25280]
MH025919.1_strain_N10         .t.aacct---agttaatg------.........gc......c...........--gg.aagt.------.....g.... [ 25280]
MH025918.1_strain_R25         .t.aacct---agttaatg------.........gc......c...........--gg.aagt.------.....g.... [ 25280]
MH025917.1_strain_R7          .t.aacct---agttaatg------.........gc......c...........--gg.aagt.------.....g.... [ 25280]
MH025916.1_strain_R8          .t.aacct---agttaatg------.........gc......c...........--gg.aagt.------.....g.... [ 25280]
```

```
FR682468.2_Georgia_2007/1        aatactgttgttatggattat------taatgaaaactttacagatgctg------gagggaataattttaatcat--ac [ 25360]
ASF_IND_20_CAD543                n.....a.....g......g.tgaaca.......g......t.......a------a.tca.tg...nnnnnnnnn--nn [ 25360]
LR899193.1_Germany_2020/1        .....................------.......................------....................--.. [ 25360]
MN393477.1_Wuhan_2019            .....................------.......................------....................--.. [ 25360]
MW396979.1_Timor-Leste/2019/1    .....................------.......................------....................--.. [ 25360]
MK940252.1_CN/2019/InnerMongolia .....................------.......................------....................--.. [ 25360]
MK128995.1_China/2018/AnhuiXCGQ  .....................------.......................------....................--.. [ 25360]
MH681419.1_ASFV/POL/2015         .....................------.......................------....................--.. [ 25360]
LS478113.1_Estonia_2014          .....................------.......................------....................--.. [ 25360]
KP843857.1_Odintsovo_02/14       .....................------.......................------....................--.. [ 25360]
FN557520.1_strain_E75            ......a.....g......g.tgaaca.......g......t.......a------a.tca.tg....a....a..ca.a [ 25360]
AY261366.1_Warthog               ......a.....g......g.tgaata.......g......t.......a------..ata.tg...ca..c.a.gg--- [ 25360]
AY261365.1_Warmbaths             ......a---.........g.tgaata..............t.......aaatcaca.tt.tt....ca..c-------- [ 25360]
AY261364.1_Tengani_62            ............g......g.------.g............t........------a.at.tt.....g..gag.aaata [ 25360]
AY261363.1_Pretoriuskop/96/4     ......a---.........g.tgaaca.......g......t.......a------t.att.tg...ac....a.ga--- [ 25360]
AY261362.1_Mkuzi_1979            ......a.....g......g.tgaaca.......g......t.......a------a.tca.tg....a....a..ca.a [ 25360]
AY261361.1_Malawi_Lil-20/1_1983  ......a---.........g.tgaaca.......g......t.......a------..tc..tg...a...c.agaa--- [ 25360]
AY261360.1_Kenya_1950            g.....a---.........g.tgaata............c.t......cc------.tatag.....aa....tg.a--- [ 25360]
AM712240.1_OURT_88/3             -------------------------------------------------------------------------------- [ 25360]
AM712239.1_Benin_97/1            ......a.....g......g.tgaaca.......g......t.......a------a.tca.tg....a....a..ca.a [ 25360]
KM111295.1_Ken06.Bus             ......a........a.....------.........t.............------.........g...a......a--- [ 25360]
KM111294.1_Ken05/Tk1             ............g......g.tgaata..............t.......a------a.tc.c.agc.c......g.a--- [ 25360]
KM262844.1_strain_L60            ......a.....g......g.tgaaca.......g......t.......a------a.tca.tg....a....a..ca.a [ 25360]
KM102979.1_26544/OG10_Italy      ......----------...g.tgaaca.......g......t.......a------a.tca.tg....a....a..ca.a [ 25360]
KM262845.1_strain_NHV            -------------------------------------------------------------------------------- [ 25360]
KX354450.1_47/Ss/2008            ......----------...g.tgaaca.......g......t.......a------a.tca.tg....a....a..ca.a [ 25360]
KP055815.1_strain_BA71           -------------------------------------------------------------------------------- [ 25360]
MH025920.1_strain_R35            ......a........a.....------.........t.............------.........g...a......a--- [ 25360]
MH025919.1_strain_N10            ......a........a.....------.........t.............------.........g...a......a--- [ 25360]
MH025918.1_strain_R25            ......a........a.....------.........t.............------.........g...a......a--- [ 25360]
MH025917.1_strain_R7             ......a........a.....------.........t.............------.........g...a......a--- [ 25360]
MH025916.1_strain_R8             ......a........a.....------.........t.............------.........g...a......a--- [ 25360]

FR682468.2_Georgia_2007/1        tgttttaatgtag------------------------------------------------------------------- [ 25440]
ASF_IND_20_CAD543                nnnnnnnnnnnnn-nnntataaatc------------------------------------------------------- [ 25440]
LR899193.1_Germany_2020/1        .............------------------------------------------------------------------- [ 25440]
MN393477.1_Wuhan_2019            .............------------------------------------------------------------------- [ 25440]
MW396979.1_Timor-Leste/2019/1    .............------------------------------------------------------------------- [ 25440]
MK940252.1_CN/2019/InnerMongolia .............------------------------------------------------------------------- [ 25440]
MK128995.1_China/2018/AnhuiXCGQ  .............------------------------------------------------------------------- [ 25440]
MH681419.1_ASFV/POL/2015         .............------------------------------------------------------------------- [ 25440]
LS478113.1_Estonia_2014          .............------------------------------------------------------------------- [ 25440]
KP843857.1_Odintsovo_02/14       .............------------------------------------------------------------------- [ 25440]
```

```
FN557520.1_strain_E75            ctg........ga---gctataaatc------------------------------------------------------ [ 25440]
AY261366.1_Warthog               ---------------agttataagtt------------------------------------------------------ [ 25440]
AY261365.1_Warmbaths             -------------------------------------------------------------------------------- [ 25440]
AY261364.1_Tengani_62            atac.a.t....ttacgttataagt------------------------------------------------------- [ 25440]
AY261363.1_Pretoriuskop/96/4     ---------------cgttataagtt------------------------------------------------------ [ 25440]
AY261362.1_Mkuzi_1979            ctg........ga---gctataaatc------------------------------------------------------ [ 25440]
AY261361.1_Malawi_Lil-20/1_1983  ---------------atatataagtt------------------------------------------------------ [ 25440]
AY261360.1_Kenya_1950            ctg........g---agatataagtttataaaaactttttgatcaaaatttaatttcttataaaaagagtacagacgtcg [ 25440]
AM712240.1_OURT_88/3             -------------------------------------------------------------------------------- [ 25440]
AM712239.1_Benin_97/1            ctg........ga---gctataaatc------------------------------------------------------ [ 25440]
KM111295.1_Ken06.Bus             ctg...t.---------atgtgagcttataaaagtattcatatcaaaatttaattttttttttataaaagtactgaaatta [ 25440]
KM111294.1_Ken05/Tk1             cag........--------------------------------------------------------------------- [ 25440]
KM262844.1_strain_L60            ctg........ga---gctataaatc------------------------------------------------------ [ 25440]
KM102979.1_26544/OG10_Italy      ctg........ga---gctataaatc------------------------------------------------------ [ 25440]
KM262845.1_strain_NHV            -------------------------------------------------------------------------------- [ 25440]
KX354450.1_47/Ss/2008            ctg........ga---gctataaatc------------------------------------------------------ [ 25440]
KP055815.1_strain_BA71           -------------------------------------------------------------------------------- [ 25440]
MH025920.1_strain_R35            ctg...t.---------atgtgagcttataaaagtattcatatcaaaatttaattttttttttataaaagtactgaaatta [ 25440]
MH025919.1_strain_N10            ctg...t.---------atgtgagcttataaaagtattcatatcaaaatttaattttttttttataaaagtactgaaatta [ 25440]
MH025918.1_strain_R25            ctg...t.---------atgtgagcttataaaagtattcatatcaaaatttaattttttttttataaaagtactgaaatta [ 25440]
MH025917.1_strain_R7             ctg...t.---------atgtgagcttataaaagtattcatatcaaaatttaattttttttttataaaagtactgaaatta [ 25440]
MH025916.1_strain_R8             ctg...t.---------atgtgagcttataaaagtattcatatcaaaatttaattttttttttataaaagtactgaaatta [ 25440]

FR682468.2_Georgia_2007/1        -------------------------------------------------------------------------------- [ 25520]
ASF_IND_20_CAD543                -------------------------------------------------------------------------------- [ 25520]
LR899193.1_Germany_2020/1        -------------------------------------------------------------------------------- [ 25520]
MN393477.1_Wuhan_2019            -------------------------------------------------------------------------------- [ 25520]
MW396979.1_Timor-Leste/2019/1    -------------------------------------------------------------------------------- [ 25520]
MK940252.1_CN/2019/InnerMongolia -------------------------------------------------------------------------------- [ 25520]
MK128995.1_China/2018/AnhuiXCGQ  -------------------------------------------------------------------------------- [ 25520]
MH681419.1_ASFV/POL/2015         -------------------------------------------------------------------------------- [ 25520]
LS478113.1_Estonia_2014          -------------------------------------------------------------------------------- [ 25520]
KP843857.1_Odintsovo_02/14       -------------------------------------------------------------------------------- [ 25520]
FN557520.1_strain_E75            -------------------------------------------------------------------------------- [ 25520]
AY261366.1_Warthog               -------------------------------------------------------------------------------- [ 25520]
AY261365.1_Warmbaths             -------------------------------------------------------------------------------- [ 25520]
AY261364.1_Tengani_62            -------------------------------------------------------------------------------- [ 25520]
AY261363.1_Pretoriuskop/96/4     -------------------------------------------------------------------------------- [ 25520]
AY261362.1_Mkuzi_1979            -------------------------------------------------------------------------------- [ 25520]
AY261361.1_Malawi_Lil-20/1_1983  -------------------------------------------------------------------------------- [ 25520]
AY261360.1_Kenya_1950            ---------cttgttta-------agcttcatcatgtttcattcattactttctacaattactggggggggggtccctca [ 25520]
AM712240.1_OURT_88/3             -------------------------------------------------------------------------------- [ 25520]
AM712239.1_Benin_97/1            -------------------------------------------------------------------------------- [ 25520]
KM111295.1_Ken06.Bus             aactaaagtttagttta--------------------------------------------------------------- [ 25520]
```

```
KM111294.1_Ken05/Tk1             -------------------------------------------------------------------------------- [ 25520]
KM262844.1_strain_L60            -------------------------------------------------------------------------------- [ 25520]
KM102979.1_26544/OG10_Italy      -------------------------------------------------------------------------------- [ 25520]
KM262845.1_strain_NHV            -------------------------------------------------------------------------------- [ 25520]
KX354450.1_47/Ss/2008            -------------------------------------------------------------------------------- [ 25520]
KP055815.1_strain_BA71           -------------------------------------------------------------------------------- [ 25520]
MH025920.1_strain_R35            aactaaagtttagttta--------------------------------------------------------------- [ 25520]
MH025919.1_strain_N10            aactaaagtttagtttagctatttgagttaatgatgtttaat---ctatcttccatgctcaatcagggggggggtcctaa [ 25520]
MH025918.1_strain_R25            aactaaagtttagttta--------------------------------------------------------------- [ 25520]
MH025917.1_strain_R7             aactaaagtttagttta--------------------------------------------------------------- [ 25520]
MH025916.1_strain_R8             aactaaagtttagttta--------------------------------------------------------------- [ 25520]

FR682468.2_Georgia_2007/1        -------------------------------------------------------------------------------- [ 25600]
ASF_IND_20_CAD543                -------------------------------------------------------------------------------- [ 25600]
LR899193.1_Germany_2020/1        -------------------------------------------------------------------------------- [ 25600]
MN393477.1_Wuhan_2019            -------------------------------------------------------------------------------- [ 25600]
MW396979.1_Timor-Leste/2019/1    -------------------------------------------------------------------------------- [ 25600]
MK940252.1_CN/2019/InnerMongolia -------------------------------------------------------------------------------- [ 25600]
MK128995.1_China/2018/AnhuiXCGQ  -------------------------------------------------------------------------------- [ 25600]
MH681419.1_ASFV/POL/2015         -------------------------------------------------------------------------------- [ 25600]
LS478113.1_Estonia_2014          -------------------------------------------------------------------------------- [ 25600]
KP843857.1_Odintsovo_02/14       -------------------------------------------------------------------------------- [ 25600]
FN557520.1_strain_E75            -------------------------------------------------------------------------------- [ 25600]
AY261366.1_Warthog               -------------------------------------------------------------------------------- [ 25600]
AY261365.1_Warmbaths             -------------------------------------------------------------------------------- [ 25600]
AY261364.1_Tengani_62            -------------------------------------------------------------------------------- [ 25600]
AY261363.1_Pretoriuskop/96/4     -------------------------------------------------------------------------------- [ 25600]
AY261362.1_Mkuzi_1979            -------------------------------------------------------------------------------- [ 25600]
AY261361.1_Malawi_Lil-20/1_1983  -------------------------------------------------------------------------------- [ 25600]
AY261360.1_Kenya_1950            tagctttagta---ttgctatggtttgctaattattatgtagaatttatagaagcatcaagaaataatattacttgtgct [ 25600]
AM712240.1_OURT_88/3             -------------------------------------------------------------------------------- [ 25600]
AM712239.1_Benin_97/1            -------------------------------------------------------------------------------- [ 25600]
KM111295.1_Ken06.Bus             -------------------------------------------------------------------------------- [ 25600]
KM111294.1_Ken05/Tk1             -------------------------------------------------------------------------------- [ 25600]
KM262844.1_strain_L60            -------------------------------------------------------------------------------- [ 25600]
KM102979.1_26544/OG10_Italy      -------------------------------------------------------------------------------- [ 25600]
KM262845.1_strain_NHV            -------------------------------------------------------------------------------- [ 25600]
KX354450.1_47/Ss/2008            -------------------------------------------------------------------------------- [ 25600]
KP055815.1_strain_BA71           -------------------------------------------------------------------------------- [ 25600]
MH025920.1_strain_R35            -------------------------------------------------------------------------------- [ 25600]
MH025919.1_strain_N10            tagttttaatactgttgctgtggattgttgaatataatgaaaactttatagatgc------------------------- [ 25600]
MH025918.1_strain_R25            -------------------------------------------------------------------------------- [ 25600]
MH025917.1_strain_R7             -------------------------------------------------------------------------------- [ 25600]
MH025916.1_strain_R8             -------------------------------------------------------------------------------- [ 25600]
```

```
FR682468.2_Georgia_2007/1         -------------------------------------------------------------------------------- [ 25680]
ASF_IND_20_CAD543                 -------------------------------------------------------------------------------- [ 25680]
LR899193.1_Germany_2020/1         -------------------------------------------------------------------------------- [ 25680]
MN393477.1_Wuhan_2019             -------------------------------------------------------------------------------- [ 25680]
MW396979.1_Timor-Leste/2019/1     -------------------------------------------------------------------------------- [ 25680]
MK940252.1_CN/2019/InnerMongolia  -------------------------------------------------------------------------------- [ 25680]
MK128995.1_China/2018/AnhuiXCGQ   -------------------------------------------------------------------------------- [ 25680]
MH681419.1_ASFV/POL/2015          -------------------------------------------------------------------------------- [ 25680]
LS478113.1_Estonia_2014           -------------------------------------------------------------------------------- [ 25680]
KP843857.1_Odintsovo_02/14        -------------------------------------------------------------------------------- [ 25680]
FN557520.1_strain_E75             -------------------------------------------------------------------------------- [ 25680]
AY261366.1_Warthog                -------------------------------------------------------------------------------- [ 25680]
AY261365.1_Warmbaths              -------------------------------------------------------------------------------- [ 25680]
AY261364.1_Tengani_62             -------------------------------------------------------------------------------- [ 25680]
AY261363.1_Pretoriuskop/96/4      -------------------------------------------------------------------------------- [ 25680]
AY261362.1_Mkuzi_1979             -------------------------------------------------------------------------------- [ 25680]
AY261361.1_Malawi_Lil-20/1_1983   -------------------------------------------------------------------------------- [ 25680]
AY261360.1_Kenya_1950             ctataagtttataaaaaatttcagtatattttttttatgaatagaacggaaatgatacaaaaataatttaatattgcaaa [ 25680]
AM712240.1_OURT_88/3              -------------------------------------------------------------------------------- [ 25680]
AM712239.1_Benin_97/1             -------------------------------------------------------------------------------- [ 25680]
KM111295.1_Ken06.Bus              -------------------------------------------------------------------------------- [ 25680]
KM111294.1_Ken05/Tk1              -------------------------------------------------------------------------------- [ 25680]
KM262844.1_strain_L60             -------------------------------------------------------------------------------- [ 25680]
KM102979.1_26544/OG10_Italy       -------------------------------------------------------------------------------- [ 25680]
KM262845.1_strain_NHV             -------------------------------------------------------------------------------- [ 25680]
KX354450.1_47/Ss/2008             -------------------------------------------------------------------------------- [ 25680]
KP055815.1_strain_BA71            -------------------------------------------------------------------------------- [ 25680]
MH025920.1_strain_R35             -------------------------------------------------------------------------------- [ 25680]
MH025919.1_strain_N10             -------------------------------------------------------------------------------- [ 25680]
MH025918.1_strain_R25             -------------------------------------------------------------------------------- [ 25680]
MH025917.1_strain_R7              -------------------------------------------------------------------------------- [ 25680]
MH025916.1_strain_R8              -------------------------------------------------------------------------------- [ 25680]

FR682468.2_Georgia_2007/1         -------------------------------------------------------------------------------- [ 25760]
ASF_IND_20_CAD543                 -------------------------------------------------------------------------------- [ 25760]
LR899193.1_Germany_2020/1         -------------------------------------------------------------------------------- [ 25760]
MN393477.1_Wuhan_2019             -------------------------------------------------------------------------------- [ 25760]
MW396979.1_Timor-Leste/2019/1     -------------------------------------------------------------------------------- [ 25760]
MK940252.1_CN/2019/InnerMongolia  -------------------------------------------------------------------------------- [ 25760]
MK128995.1_China/2018/AnhuiXCGQ   -------------------------------------------------------------------------------- [ 25760]
MH681419.1_ASFV/POL/2015          -------------------------------------------------------------------------------- [ 25760]
LS478113.1_Estonia_2014           -------------------------------------------------------------------------------- [ 25760]
KP843857.1_Odintsovo_02/14        -------------------------------------------------------------------------------- [ 25760]
```

```
FN557520.1_strain_E75            -------------------------------------------------------------------------------- [ 25760]
AY261366.1_Warthog               -------------------------------------------------------------------------------- [ 25760]
AY261365.1_Warmbaths             -------------------------------------------------------------------------------- [ 25760]
AY261364.1_Tengani_62            -------------------------------------------------------------------------------- [ 25760]
AY261363.1_Pretoriuskop/96/4     -------------------------------------------------------------------------------- [ 25760]
AY261362.1_Mkuzi_1979            -------------------------------------------------------------------------------- [ 25760]
AY261361.1_Malawi_Lil-20/1_1983  -------------------------------------------------------------------------------- [ 25760]
AY261360.1_Kenya_1950            aaaaattcataatgttgacatgaatatataataacgtgtattttataaactgactccttcaactaattattagatgaggt [ 25760]
AM712240.1_OURT_88/3             -------------------------------------------------------------------------------- [ 25760]
AM712239.1_Benin_97/1            -------------------------------------------------------------------------------- [ 25760]
KM111295.1_Ken06.Bus             -------------------------------------------------------------------------------- [ 25760]
KM111294.1_Ken05/Tk1             -------------------------------------------------------------------------------- [ 25760]
KM262844.1_strain_L60            -------------------------------------------------------------------------------- [ 25760]
KM102979.1_26544/OG10_Italy      -------------------------------------------------------------------------------- [ 25760]
KM262845.1_strain_NHV            -------------------------------------------------------------------------------- [ 25760]
KX354450.1_47/Ss/2008            -------------------------------------------------------------------------------- [ 25760]
KP055815.1_strain_BA71           -------------------------------------------------------------------------------- [ 25760]
MH025920.1_strain_R35            -------------------------------------------------------------------------------- [ 25760]
MH025919.1_strain_N10            --------------------------taaagcagcg-------------------------------------------- [ 25760]
MH025918.1_strain_R25            -------------------------------------------------------------------------------- [ 25760]
MH025917.1_strain_R7             -------------------------------------------------------------------------------- [ 25760]
MH025916.1_strain_R8             -------------------------------------------------------------------------------- [ 25760]

FR682468.2_Georgia_2007/1        -------------------------------------------------------------------------------- [ 25840]
ASF_IND_20_CAD543                -------------------------------------------------------------------------------- [ 25840]
LR899193.1_Germany_2020/1        -------------------------------------------------------------------------------- [ 25840]
MN393477.1_Wuhan_2019            -------------------------------------------------------------------------------- [ 25840]
MW396979.1_Timor-Leste/2019/1    -------------------------------------------------------------------------------- [ 25840]
MK940252.1_CN/2019/InnerMongolia -------------------------------------------------------------------------------- [ 25840]
MK128995.1_China/2018/AnhuiXCGQ  -------------------------------------------------------------------------------- [ 25840]
MH681419.1_ASFV/POL/2015         -------------------------------------------------------------------------------- [ 25840]
LS478113.1_Estonia_2014          -------------------------------------------------------------------------------- [ 25840]
KP843857.1_Odintsovo_02/14       -------------------------------------------------------------------------------- [ 25840]
FN557520.1_strain_E75            -------------------------------------------------------------------------------- [ 25840]
AY261366.1_Warthog               -------------------------------------------------------------------------------- [ 25840]
AY261365.1_Warmbaths             -------------------------------------------------------------------------------- [ 25840]
AY261364.1_Tengani_62            -------------------------------------------------------------------------------- [ 25840]
AY261363.1_Pretoriuskop/96/4     -------------------------------------------------------------------------------- [ 25840]
AY261362.1_Mkuzi_1979            -------------------------------------------------------------------------------- [ 25840]
AY261361.1_Malawi_Lil-20/1_1983  -------------------------------------------------------------------------------- [ 25840]
AY261360.1_Kenya_1950            accaacctatttatgatatgccgatgatagacattgtataccataaaacaaaattattttaaatgtattcatagatacat [ 25840]
AM712240.1_OURT_88/3             -------------------------------------------------------------------------------- [ 25840]
AM712239.1_Benin_97/1            -------------------------------------------------------------------------------- [ 25840]
KM111295.1_Ken06.Bus             -------------------------------------------------------------------------------- [ 25840]
```

```
KM111294.1_Ken05/Tk1                --------------------------------------------------------------------------------  [ 25840]
KM262844.1_strain_L60               --------------------------------------------------------------------------------  [ 25840]
KM102979.1_26544/OG10_Italy         --------------------------------------------------------------------------------  [ 25840]
KM262845.1_strain_NHV               --------------------------------------------------------------------------------  [ 25840]
KX354450.1_47/Ss/2008               --------------------------------------------------------------------------------  [ 25840]
KP055815.1_strain_BA71              --------------------------------------------------------------------------------  [ 25840]
MH025920.1_strain_R35               --------------------------------------------------------------------------------  [ 25840]
MH025919.1_strain_N10               ------------------------------------------------------------------------aaatatac  [ 25840]
MH025918.1_strain_R25               --------------------------------------------------------------------------------  [ 25840]
MH025917.1_strain_R7                --------------------------------------------------------------------------------  [ 25840]
MH025916.1_strain_R8                --------------------------------------------------------------------------------  [ 25840]

FR682468.2_Georgia_2007/1           --------------------------------------------------------------------------------  [ 25920]
ASF_IND_20_CAD543                   --------------------------------------------------------------------------------  [ 25920]
LR899193.1_Germany_2020/1           --------------------------------------------------------------------------------  [ 25920]
MN393477.1_Wuhan_2019               --------------------------------------------------------------------------------  [ 25920]
MW396979.1_Timor-Leste/2019/1       --------------------------------------------------------------------------------  [ 25920]
MK940252.1_CN/2019/InnerMongolia    --------------------------------------------------------------------------------  [ 25920]
MK128995.1_China/2018/AnhuiXCGQ     --------------------------------------------------------------------------------  [ 25920]
MH681419.1_ASFV/POL/2015            --------------------------------------------------------------------------------  [ 25920]
LS478113.1_Estonia_2014             --------------------------------------------------------------------------------  [ 25920]
KP843857.1_Odintsovo_02/14          --------------------------------------------------------------------------------  [ 25920]
FN557520.1_strain_E75               --------------------------------------------------------------------------------  [ 25920]
AY261366.1_Warthog                  --------------------------------------------------------------------------------  [ 25920]
AY261365.1_Warmbaths                --------------------------------------------------------------------------------  [ 25920]
AY261364.1_Tengani_62               --------------------------------------------------------------------------------  [ 25920]
AY261363.1_Pretoriuskop/96/4        --------------------------------------------------------------------------------  [ 25920]
AY261362.1_Mkuzi_1979               --------------------------------------------------------------------------------  [ 25920]
AY261361.1_Malawi_Lil-20/1_1983     --------------------------------------------------------------------------------  [ 25920]
AY261360.1_Kenya_1950               gtattcatgtatacatggacacatcataacatgtttaccgcaaattgtctctcagcgaagaaaatgaatgaaacgtttcc  [ 25920]
AM712240.1_OURT_88/3                --------------------------------------------------------------------------------  [ 25920]
AM712239.1_Benin_97/1               --------------------------------------------------------------------------------  [ 25920]
KM111295.1_Ken06.Bus                --------------------------------------------------------------------------------  [ 25920]
KM111294.1_Ken05/Tk1                --------------------------------------------------------------------------------  [ 25920]
KM262844.1_strain_L60               --------------------------------------------------------------------------------  [ 25920]
KM102979.1_26544/OG10_Italy         --------------------------------------------------------------------------------  [ 25920]
KM262845.1_strain_NHV               --------------------------------------------------------------------------------  [ 25920]
KX354450.1_47/Ss/2008               --------------------------------------------------------------------------------  [ 25920]
KP055815.1_strain_BA71              --------------------------------------------------------------------------------  [ 25920]
MH025920.1_strain_R35               --------------------------------------------------------------------------------  [ 25920]
MH025919.1_strain_N10               atgttc--------------------------------------------------------------------------  [ 25920]
MH025918.1_strain_R25               --------------------------------------------------------------------------------  [ 25920]
MH025917.1_strain_R7                --------------------------------------------------------------------------------  [ 25920]
MH025916.1_strain_R8                --------------------------------------------------------------------------------  [ 25920]
```

```
FR682468.2_Georgia_2007/1        -------------------------------------------------------------------------------- [ 26000]
ASF_IND_20_CAD543                -------------------------------------------------------------------------------- [ 26000]
LR899193.1_Germany_2020/1        -------------------------------------------------------------------------------- [ 26000]
MN393477.1_Wuhan_2019            -------------------------------------------------------------------------------- [ 26000]
MW396979.1_Timor-Leste/2019/1    -------------------------------------------------------------------------------- [ 26000]
MK940252.1_CN/2019/InnerMongolia -------------------------------------------------------------------------------- [ 26000]
MK128995.1_China/2018/AnhuiXCGQ  -------------------------------------------------------------------------------- [ 26000]
MH681419.1_ASFV/POL/2015         -------------------------------------------------------------------------------- [ 26000]
LS478113.1_Estonia_2014          -------------------------------------------------------------------------------- [ 26000]
KP843857.1_Odintsovo_02/14       -------------------------------------------------------------------------------- [ 26000]
FN557520.1_strain_E75            -------------------------------------------------------------------------------- [ 26000]
AY261366.1_Warthog               -------------------------------------------------------------------------------- [ 26000]
AY261365.1_Warmbaths             -------------------------------------------------------------------------------- [ 26000]
AY261364.1_Tengani_62            -------------------------------------------------------------------------------- [ 26000]
AY261363.1_Pretoriuskop/96/4     -------------------------------------------------------------------------------- [ 26000]
AY261362.1_Mkuzi_1979            -------------------------------------------------------------------------------- [ 26000]
AY261361.1_Malawi_Lil-20/1_1983  -------------------------------------------------------------------------------- [ 26000]
AY261360.1_Kenya_1950            gtatattcataggttgaaattattttacgcacctcactaggttctaatattttcttatgaagtattgaatgggggcctaa [ 26000]
AM712240.1_OURT_88/3             -------------------------------------------------------------------------------- [ 26000]
AM712239.1_Benin_97/1            -------------------------------------------------------------------------------- [ 26000]
KM111295.1_Ken06.Bus             -------------------------------------------------------------------------------- [ 26000]
KM111294.1_Ken05/Tk1             -------------------------------------------------------------------------------- [ 26000]
KM262844.1_strain_L60            -------------------------------------------------------------------------------- [ 26000]
KM102979.1_26544/OG10_Italy      -------------------------------------------------------------------------------- [ 26000]
KM262845.1_strain_NHV            -------------------------------------------------------------------------------- [ 26000]
KX354450.1_47/Ss/2008            -------------------------------------------------------------------------------- [ 26000]
KP055815.1_strain_BA71           -------------------------------------------------------------------------------- [ 26000]
MH025920.1_strain_R35            -------------------------------------------------------------------------------- [ 26000]
MH025919.1_strain_N10            --------------------------------------------------------------tgtttaatgtgagctata [ 26000]
MH025918.1_strain_R25            -------------------------------------------------------------------------------- [ 26000]
MH025917.1_strain_R7             -------------------------------------------------------------------------------- [ 26000]
MH025916.1_strain_R8             -------------------------------------------------------------------------------- [ 26000]

FR682468.2_Georgia_2007/1        -------------------------------------------------------------------------------- [ 26080]
ASF_IND_20_CAD543                -------------------------------------------------------------------------------- [ 26080]
LR899193.1_Germany_2020/1        -------------------------------------------------------------------------------- [ 26080]
MN393477.1_Wuhan_2019            -------------------------------------------------------------------------------- [ 26080]
MW396979.1_Timor-Leste/2019/1    -------------------------------------------------------------------------------- [ 26080]
MK940252.1_CN/2019/InnerMongolia -------------------------------------------------------------------------------- [ 26080]
MK128995.1_China/2018/AnhuiXCGQ  -------------------------------------------------------------------------------- [ 26080]
MH681419.1_ASFV/POL/2015         -------------------------------------------------------------------------------- [ 26080]
LS478113.1_Estonia_2014          -------------------------------------------------------------------------------- [ 26080]
KP843857.1_Odintsovo_02/14       -------------------------------------------------------------------------------- [ 26080]
```

```
FN557520.1_strain_E75          -------------------------------------------------------------------------------- [ 26080]
AY261366.1_Warthog             -------------------------------------------------------------------------------- [ 26080]
AY261365.1_Warmbaths           -------------------------------------------------------------------------------- [ 26080]
AY261364.1_Tengani_62          -------------------------------------------------------------------------------- [ 26080]
AY261363.1_Pretoriuskop/96/4   -------------------------------------------------------------------------------- [ 26080]
AY261362.1_Mkuzi_1979          -------------------------------------------------------------------------------- [ 26080]
AY261361.1_Malawi_Lil-20/1_1983 -------------------------------------------------------------------------------- [ 26080]
AY261360.1_Kenya_1950          aaatcctttcttaaaaagtagtttcattataacattcttttcttctctaagaagagtttcttgtattttttttgtatcag [ 26080]
AM712240.1_OURT_88/3           -------------------------------------------------------------------------------- [ 26080]
AM712239.1_Benin_97/1          -------------------------------------------------------------------------------- [ 26080]
KM111295.1_Ken06.Bus           -------------------------------------------------------------------------------- [ 26080]
KM111294.1_Ken05/Tk1           -------------------------------------------------------------------------------- [ 26080]
KM262844.1_strain_L60          -------------------------------------------------------------------------------- [ 26080]
KM102979.1_26544/OG10_Italy    -------------------------------------------------------------------------------- [ 26080]
KM262845.1_strain_NHV          -------------------------------------------------------------------------------- [ 26080]
KX354450.1_47/Ss/2008          -------------------------------------------------------------------------------- [ 26080]
KP055815.1_strain_BA71         -------------------------------------------------------------------------------- [ 26080]
MH025920.1_strain_R35          -------------------------------------------------------------------------------- [ 26080]
MH025919.1_strain_N10          aaatcttttcatcaaaa--------------------------------------------tttaatttttttttat--- [ 26080]
MH025918.1_strain_R25          -------------------------------------------------------------------------------- [ 26080]
MH025917.1_strain_R7           -------------------------------------------------------------------------------- [ 26080]
MH025916.1_strain_R8           -------------------------------------------------------------------------------- [ 26080]

FR682468.2_Georgia_2007/1      -------------------------------------------------------------------------------- [ 26160]
ASF_IND_20_CAD543              -------------------------------------------------------------------------------- [ 26160]
LR899193.1_Germany_2020/1      -------------------------------------------------------------------------------- [ 26160]
MN393477.1_Wuhan_2019          -------------------------------------------------------------------------------- [ 26160]
MW396979.1_Timor-Leste/2019/1  -------------------------------------------------------------------------------- [ 26160]
MK940252.1_CN/2019/InnerMongolia -------------------------------------------------------------------------------- [ 26160]
MK128995.1_China/2018/AnhuiXCGQ -------------------------------------------------------------------------------- [ 26160]
MH681419.1_ASFV/POL/2015       -------------------------------------------------------------------------------- [ 26160]
LS478113.1_Estonia_2014        -------------------------------------------------------------------------------- [ 26160]
KP843857.1_Odintsovo_02/14     -------------------------------------------------------------------------------- [ 26160]
FN557520.1_strain_E75          -------------------------------------------------------------------------------- [ 26160]
AY261366.1_Warthog             -------------------------------------------------------------------------------- [ 26160]
AY261365.1_Warmbaths           -------------------------------------------------------------------------------- [ 26160]
AY261364.1_Tengani_62          -------------------------------------------------------------------------------- [ 26160]
AY261363.1_Pretoriuskop/96/4   -------------------------------------------------------------------------------- [ 26160]
AY261362.1_Mkuzi_1979          -------------------------------------------------------------------------------- [ 26160]
AY261361.1_Malawi_Lil-20/1_1983 -------------------------------------------------------------------------------- [ 26160]
AY261360.1_Kenya_1950          ggttggctcccaaacttatacaaaaatgtatattactccaaataccataatttgaaaagaaagttatttccctatatact [ 26160]
AM712240.1_OURT_88/3           -------------------------------------------------------------------------------- [ 26160]
AM712239.1_Benin_97/1          -------------------------------------------------------------------------------- [ 26160]
KM111295.1_Ken06.Bus           -------------------------------------------------------------------------------- [ 26160]
```

```
KM111294.1_Ken05/Tk1          -------------------------------------------------------------------------------- [ 26160]
KM262844.1_strain_L60         -------------------------------------------------------------------------------- [ 26160]
KM102979.1_26544/OG10_Italy   -------------------------------------------------------------------------------- [ 26160]
KM262845.1_strain_NHV         -------------------------------------------------------------------------------- [ 26160]
KX354450.1_47/Ss/2008         -------------------------------------------------------------------------------- [ 26160]
KP055815.1_strain_BA71        -------------------------------------------------------------------------------- [ 26160]
MH025920.1_strain_R35         -------------------------------------------------------------------------------- [ 26160]
MH025919.1_strain_N10         -----------------------aaatgtaca------------------------------------------------ [ 26160]
MH025918.1_strain_R25         -------------------------------------------------------------------------------- [ 26160]
MH025917.1_strain_R7          -------------------------------------------------------------------------------- [ 26160]
MH025916.1_strain_R8          -------------------------------------------------------------------------------- [ 26160]

FR682468.2_Georgia_2007/1     -------------------------------------------------------------------------------- [ 26240]
ASF_IND_20_CAD543             -------------------------------------------------------------------------------- [ 26240]
LR899193.1_Germany_2020/1     -------------------------------------------------------------------------------- [ 26240]
MN393477.1_Wuhan_2019         -------------------------------------------------------------------------------- [ 26240]
MW396979.1_Timor-Leste/2019/1 -------------------------------------------------------------------------------- [ 26240]
MK940252.1_CN/2019/InnerMongolia -------------------------------------------------------------------------------- [ 26240]
MK128995.1_China/2018/AnhuiXCGQ -------------------------------------------------------------------------------- [ 26240]
MH681419.1_ASFV/POL/2015      -------------------------------------------------------------------------------- [ 26240]
LS478113.1_Estonia_2014       -------------------------------------------------------------------------------- [ 26240]
KP843857.1_Odintsovo_02/14    -------------------------------------------------------------------------------- [ 26240]
FN557520.1_strain_E75         -------------------------------------------------------------------------------- [ 26240]
AY261366.1_Warthog            -------------------------------------------------------------------------------- [ 26240]
AY261365.1_Warmbaths          -------------------------------------------------------------------------------- [ 26240]
AY261364.1_Tengani_62         -------------------------------------------------------------------------------- [ 26240]
AY261363.1_Pretoriuskop/96/4  -------------------------------------------------------------------------------- [ 26240]
AY261362.1_Mkuzi_1979         -------------------------------------------------------------------------------- [ 26240]
AY261361.1_Malawi_Lil-20/1_1983 -------------------------------------------------------------------------------- [ 26240]
AY261360.1_Kenya_1950         tcatgattaatgaaacgtatcaacgtctctaaggcagtattgatatttgcccctaaggcgaaacaatagtatatacctaa [ 26240]
AM712240.1_OURT_88/3          -------------------------------------------------------------------------------- [ 26240]
AM712239.1_Benin_97/1         -------------------------------------------------------------------------------- [ 26240]
KM111295.1_Ken06.Bus          -------------------------------------------------------------------------------- [ 26240]
KM111294.1_Ken05/Tk1          -------------------------------------------------------------------------------- [ 26240]
KM262844.1_strain_L60         -------------------------------------------------------------------------------- [ 26240]
KM102979.1_26544/OG10_Italy   -------------------------------------------------------------------------------- [ 26240]
KM262845.1_strain_NHV         -------------------------------------------------------------------------------- [ 26240]
KX354450.1_47/Ss/2008         -------------------------------------------------------------------------------- [ 26240]
KP055815.1_strain_BA71        -------------------------------------------------------------------------------- [ 26240]
MH025920.1_strain_R35         -------------------------------------------------------------------------------- [ 26240]
MH025919.1_strain_N10         -------------------------------------------------------------------------------- [ 26240]
MH025918.1_strain_R25         -------------------------------------------------------------------------------- [ 26240]
MH025917.1_strain_R7          -------------------------------------------------------------------------------- [ 26240]
MH025916.1_strain_R8          -------------------------------------------------------------------------------- [ 26240]
```

```
FR682468.2_Georgia_2007/1        -------------------------------------------------------------------------------- [ 26320]
ASF_IND_20_CAD543                -------------------------------------------------------------------------------- [ 26320]
LR899193.1_Germany_2020/1        -------------------------------------------------------------------------------- [ 26320]
MN393477.1_Wuhan_2019            -------------------------------------------------------------------------------- [ 26320]
MW396979.1_Timor-Leste/2019/1    -------------------------------------------------------------------------------- [ 26320]
MK940252.1_CN/2019/InnerMongolia -------------------------------------------------------------------------------- [ 26320]
MK128995.1_China/2018/AnhuiXCGQ  -------------------------------------------------------------------------------- [ 26320]
MH681419.1_ASFV/POL/2015         -------------------------------------------------------------------------------- [ 26320]
LS478113.1_Estonia_2014          -------------------------------------------------------------------------------- [ 26320]
KP843857.1_Odintsovo_02/14       -------------------------------------------------------------------------------- [ 26320]
FN557520.1_strain_E75            -------------------------------------------------------------------------------- [ 26320]
AY261366.1_Warthog               -------------------------------------------------------------------------------- [ 26320]
AY261365.1_Warmbaths             -------------------------------------------------------------------------------- [ 26320]
AY261364.1_Tengani_62            -------------------------------------------------------------------------------- [ 26320]
AY261363.1_Pretoriuskop/96/4     -------------------------------------------------------------------------------- [ 26320]
AY261362.1_Mkuzi_1979            -------------------------------------------------------------------------------- [ 26320]
AY261361.1_Malawi_Lil-20/1_1983  -------------------------------------------------------------------------------- [ 26320]
AY261360.1_Kenya_1950            tttattctgagggtacatacaagcaagcgacatcatgtgatttggttctaaacgtatattttcctgaaaatatgcatggt [ 26320]
AM712240.1_OURT_88/3             -------------------------------------------------------------------------------- [ 26320]
AM712239.1_Benin_97/1            -------------------------------------------------------------------------------- [ 26320]
KM111295.1_Ken06.Bus             -------------------------------------------------------------------------------- [ 26320]
KM111294.1_Ken05/Tk1             -------------------------------------------------------------------------------- [ 26320]
KM262844.1_strain_L60            -------------------------------------------------------------------------------- [ 26320]
KM102979.1_26544/OG10_Italy      -------------------------------------------------------------------------------- [ 26320]
KM262845.1_strain_NHV            -------------------------------------------------------------------------------- [ 26320]
KX354450.1_47/Ss/2008            -------------------------------------------------------------------------------- [ 26320]
KP055815.1_strain_BA71           -------------------------------------------------------------------------------- [ 26320]
MH025920.1_strain_R35            -------------------------------------------------------------------------------- [ 26320]
MH025919.1_strain_N10            -------------------------------------------------------------------------------- [ 26320]
MH025918.1_strain_R25            -------------------------------------------------------------------------------- [ 26320]
MH025917.1_strain_R7             -------------------------------------------------------------------------------- [ 26320]
MH025916.1_strain_R8             -------------------------------------------------------------------------------- [ 26320]

FR682468.2_Georgia_2007/1        -------------------------------------------------------------------------------- [ 26400]
ASF_IND_20_CAD543                -------------------------------------------------------------------------------- [ 26400]
LR899193.1_Germany_2020/1        -------------------------------------------------------------------------------- [ 26400]
MN393477.1_Wuhan_2019            -------------------------------------------------------------------------------- [ 26400]
MW396979.1_Timor-Leste/2019/1    -------------------------------------------------------------------------------- [ 26400]
MK940252.1_CN/2019/InnerMongolia -------------------------------------------------------------------------------- [ 26400]
MK128995.1_China/2018/AnhuiXCGQ  -------------------------------------------------------------------------------- [ 26400]
MH681419.1_ASFV/POL/2015         -------------------------------------------------------------------------------- [ 26400]
LS478113.1_Estonia_2014          -------------------------------------------------------------------------------- [ 26400]
KP843857.1_Odintsovo_02/14       -------------------------------------------------------------------------------- [ 26400]
```

```
FN557520.1_strain_E75          -------------------------------------------------------------------------------- [ 26400]
AY261366.1_Warthog             -------------------------------------------------------------------------------- [ 26400]
AY261365.1_Warmbaths           -------------------------------------------------------------------------------- [ 26400]
AY261364.1_Tengani_62          -------------------------------------------------------------------------------- [ 26400]
AY261363.1_Pretoriuskop/96/4   -------------------------------------------------------------------------------- [ 26400]
AY261362.1_Mkuzi_1979          -------------------------------------------------------------------------------- [ 26400]
AY261361.1_Malawi_Lil-20/1_1983 -------------------------------------------------------------------------------- [ 26400]
AY261360.1_Kenya_1950          ggatttcatcaacattccctaagtatacagccgtttttaaacgccaatcatctaggtgaggaaatttcttactaagaaaa [ 26400]
AM712240.1_OURT_88/3           -------------------------------------------------------------------------------- [ 26400]
AM712239.1_Benin_97/1          -------------------------------------------------------------------------------- [ 26400]
KM111295.1_Ken06.Bus           -------------------------------------------------------------------------------- [ 26400]
KM111294.1_Ken05/Tk1           -------------------------------------------------------------------------------- [ 26400]
KM262844.1_strain_L60          -------------------------------------------------------------------------------- [ 26400]
KM102979.1_26544/OG10_Italy    -------------------------------------------------------------------------------- [ 26400]
KM262845.1_strain_NHV          -------------------------------------------------------------------------------- [ 26400]
KX354450.1_47/Ss/2008          -------------------------------------------------------------------------------- [ 26400]
KP055815.1_strain_BA71         -------------------------------------------------------------------------------- [ 26400]
MH025920.1_strain_R35          -------------------------------------------------------------------------------- [ 26400]
MH025919.1_strain_N10          -------------------------------------------------------------------------------- [ 26400]
MH025918.1_strain_R25          -------------------------------------------------------------------------------- [ 26400]
MH025917.1_strain_R7           -------------------------------------------------------------------------------- [ 26400]
MH025916.1_strain_R8           -------------------------------------------------------------------------------- [ 26400]

FR682468.2_Georgia_2007/1      -------------------------------------------------------------------------------- [ 26480]
ASF_IND_20_CAD543              -------------------------------------------------------------------------------- [ 26480]
LR899193.1_Germany_2020/1      -------------------------------------------------------------------------------- [ 26480]
MN393477.1_Wuhan_2019          -------------------------------------------------------------------------------- [ 26480]
MW396979.1_Timor-Leste/2019/1  -------------------------------------------------------------------------------- [ 26480]
MK940252.1_CN/2019/InnerMongolia -------------------------------------------------------------------------------- [ 26480]
MK128995.1_China/2018/AnhuiXCGQ -------------------------------------------------------------------------------- [ 26480]
MH681419.1_ASFV/POL/2015       -------------------------------------------------------------------------------- [ 26480]
LS478113.1_Estonia_2014        -------------------------------------------------------------------------------- [ 26480]
KP843857.1_Odintsovo_02/14     -------------------------------------------------------------------------------- [ 26480]
FN557520.1_strain_E75          -------------------------------------------------------------------------------- [ 26480]
AY261366.1_Warthog             -------------------------------------------------------------------------------- [ 26480]
AY261365.1_Warmbaths           -------------------------------------------------------------------------------- [ 26480]
AY261364.1_Tengani_62          -------------------------------------------------------------------------------- [ 26480]
AY261363.1_Pretoriuskop/96/4   -------------------------------------------------------------------------------- [ 26480]
AY261362.1_Mkuzi_1979          -------------------------------------------------------------------------------- [ 26480]
AY261361.1_Malawi_Lil-20/1_1983 -------------------------------------------------------------------------------- [ 26480]
AY261360.1_Kenya_1950          cgaataggttttataagattaaactctatggccaacttataccaaaattttaatacaagtgtatttcttgtcattttttc [ 26480]
AM712240.1_OURT_88/3           -------------------------------------------------------------------------------- [ 26480]
AM712239.1_Benin_97/1          -------------------------------------------------------------------------------- [ 26480]
KM111295.1_Ken06.Bus           -------------------------------------------------------------------------------- [ 26480]
```

```
KM111294.1_Ken05/Tk1              -------------------------------------------------------------------------------- [ 26480]
KM262844.1_strain_L60             -------------------------------------------------------------------------------- [ 26480]
KM102979.1_26544/OG10_Italy       -------------------------------------------------------------------------------- [ 26480]
KM262845.1_strain_NHV             -------------------------------------------------------------------------------- [ 26480]
KX354450.1_47/Ss/2008             -------------------------------------------------------------------------------- [ 26480]
KP055815.1_strain_BA71            -------------------------------------------------------------------------------- [ 26480]
MH025920.1_strain_R35             -------------------------------------------------------------------------------- [ 26480]
MH025919.1_strain_N10             --------------aaaattaaact------------------------------------------------------- [ 26480]
MH025918.1_strain_R25             -------------------------------------------------------------------------------- [ 26480]
MH025917.1_strain_R7              -------------------------------------------------------------------------------- [ 26480]
MH025916.1_strain_R8              -------------------------------------------------------------------------------- [ 26480]

FR682468.2_Georgia_2007/1         -------------------------------------------------------------------------------- [ 26560]
ASF_IND_20_CAD543                 -------------------------------------------------------------------------------- [ 26560]
LR899193.1_Germany_2020/1         -------------------------------------------------------------------------------- [ 26560]
MN393477.1_Wuhan_2019             -------------------------------------------------------------------------------- [ 26560]
MW396979.1_Timor-Leste/2019/1     -------------------------------------------------------------------------------- [ 26560]
MK940252.1_CN/2019/InnerMongolia  -------------------------------------------------------------------------------- [ 26560]
MK128995.1_China/2018/AnhuiXCGQ   -------------------------------------------------------------------------------- [ 26560]
MH681419.1_ASFV/POL/2015          -------------------------------------------------------------------------------- [ 26560]
LS478113.1_Estonia_2014           -------------------------------------------------------------------------------- [ 26560]
KP843857.1_Odintsovo_02/14        -------------------------------------------------------------------------------- [ 26560]
FN557520.1_strain_E75             -------------------------------------------------------------------------------- [ 26560]
AY261366.1_Warthog                -------------------------------------------------------------------------------- [ 26560]
AY261365.1_Warmbaths              -------------------------------------------------------------------------------- [ 26560]
AY261364.1_Tengani_62             -------------------------------------------------------------------------------- [ 26560]
AY261363.1_Pretoriuskop/96/4      -------------------------------------------------------------------------------- [ 26560]
AY261362.1_Mkuzi_1979             -------------------------------------------------------------------------------- [ 26560]
AY261361.1_Malawi_Lil-20/1_1983   -------------------------------------------------------------------------------- [ 26560]
AY261360.1_Kenya_1950             tttttcatctaaatttaagatataaccattgtaaataaagtttatcaacgcgtgaaaatcatggctaccaaaactgtcga [ 26560]
AM712240.1_OURT_88/3              -------------------------------------------------------------------------------- [ 26560]
AM712239.1_Benin_97/1             -------------------------------------------------------------------------------- [ 26560]
KM111295.1_Ken06.Bus              -------------------------------------------------------------------------------- [ 26560]
KM111294.1_Ken05/Tk1              -------------------------------------------------------------------------------- [ 26560]
KM262844.1_strain_L60             -------------------------------------------------------------------------------- [ 26560]
KM102979.1_26544/OG10_Italy       -------------------------------------------------------------------------------- [ 26560]
KM262845.1_strain_NHV             -------------------------------------------------------------------------------- [ 26560]
KX354450.1_47/Ss/2008             -------------------------------------------------------------------------------- [ 26560]
KP055815.1_strain_BA71            -------------------------------------------------------------------------------- [ 26560]
MH025920.1_strain_R35             -------------------------------------------------------------------------------- [ 26560]
MH025919.1_strain_N10             -------------------------------------------------------------------------------- [ 26560]
MH025918.1_strain_R25             -------------------------------------------------------------------------------- [ 26560]
MH025917.1_strain_R7              -------------------------------------------------------------------------------- [ 26560]
MH025916.1_strain_R8              -------------------------------------------------------------------------------- [ 26560]
```

```
FR682468.2_Georgia_2007/1        -------------------------------------------------------------------------------- [ 26640]
ASF_IND_20_CAD543                -------------------------------------------------------------------------------- [ 26640]
LR899193.1_Germany_2020/1        -------------------------------------------------------------------------------- [ 26640]
MN393477.1_Wuhan_2019            -------------------------------------------------------------------------------- [ 26640]
MW396979.1_Timor-Leste/2019/1    -------------------------------------------------------------------------------- [ 26640]
MK940252.1_CN/2019/InnerMongolia -------------------------------------------------------------------------------- [ 26640]
MK128995.1_China/2018/AnhuiXCGQ  -------------------------------------------------------------------------------- [ 26640]
MH681419.1_ASFV/POL/2015         -------------------------------------------------------------------------------- [ 26640]
LS478113.1_Estonia_2014          -------------------------------------------------------------------------------- [ 26640]
KP843857.1_Odintsovo_02/14       -------------------------------------------------------------------------------- [ 26640]
FN557520.1_strain_E75            -------------------------------------------------------------------------------- [ 26640]
AY261366.1_Warthog               -------------------------------------------------------------------------------- [ 26640]
AY261365.1_Warmbaths             -------------------------------------------------------------------------------- [ 26640]
AY261364.1_Tengani_62            -------------------------------------------------------------------------------- [ 26640]
AY261363.1_Pretoriuskop/96/4     -------------------------------------------------------------------------------- [ 26640]
AY261362.1_Mkuzi_1979            -------------------------------------------------------------------------------- [ 26640]
AY261361.1_Malawi_Lil-20/1_1983  -------------------------------------------------------------------------------- [ 26640]
AY261360.1_Kenya_1950            gaatcaaaatattatcataataaatatctatagctaataagaccttttgttgcctaactagattaacaaacatattatat [ 26640]
AM712240.1_OURT_88/3             -------------------------------------------------------------------------------- [ 26640]
AM712239.1_Benin_97/1            -------------------------------------------------------------------------------- [ 26640]
KM111295.1_Ken06.Bus             -------------------------------------------------------------------------------- [ 26640]
KM111294.1_Ken05/Tk1             -------------------------------------------------------------------------------- [ 26640]
KM262844.1_strain_L60            -------------------------------------------------------------------------------- [ 26640]
KM102979.1_26544/OG10_Italy      -------------------------------------------------------------------------------- [ 26640]
KM262845.1_strain_NHV            -------------------------------------------------------------------------------- [ 26640]
KX354450.1_47/Ss/2008            -------------------------------------------------------------------------------- [ 26640]
KP055815.1_strain_BA71           -------------------------------------------------------------------------------- [ 26640]
MH025920.1_strain_R35            -------------------------------------------------------------------------------- [ 26640]
MH025919.1_strain_N10            -------------------------------------------------------------------------------- [ 26640]
MH025918.1_strain_R25            -------------------------------------------------------------------------------- [ 26640]
MH025917.1_strain_R7             -------------------------------------------------------------------------------- [ 26640]
MH025916.1_strain_R8             -------------------------------------------------------------------------------- [ 26640]

FR682468.2_Georgia_2007/1        -------------------------------------------------------------------------------- [ 26720]
ASF_IND_20_CAD543                -------------------------------------------------------------------------------- [ 26720]
LR899193.1_Germany_2020/1        -------------------------------------------------------------------------------- [ 26720]
MN393477.1_Wuhan_2019            -------------------------------------------------------------------------------- [ 26720]
MW396979.1_Timor-Leste/2019/1    -------------------------------------------------------------------------------- [ 26720]
MK940252.1_CN/2019/InnerMongolia -------------------------------------------------------------------------------- [ 26720]
MK128995.1_China/2018/AnhuiXCGQ  -------------------------------------------------------------------------------- [ 26720]
MH681419.1_ASFV/POL/2015         -------------------------------------------------------------------------------- [ 26720]
LS478113.1_Estonia_2014          -------------------------------------------------------------------------------- [ 26720]
KP843857.1_Odintsovo_02/14       -------------------------------------------------------------------------------- [ 26720]
```

```
FN557520.1_strain_E75            -------------------------------------------------------------------------------- [ 26720]
AY261366.1_Warthog               -------------------------------------------------------------------------------- [ 26720]
AY261365.1_Warmbaths             -------------------------------------------------------------------------------- [ 26720]
AY261364.1_Tengani_62            -------------------------------------------------------------------------------- [ 26720]
AY261363.1_Pretoriuskop/96/4     -------------------------------------------------------------------------------- [ 26720]
AY261362.1_Mkuzi_1979            -------------------------------------------------------------------------------- [ 26720]
AY261361.1_Malawi_Lil-20/1_1983  -------------------------------------------------------------------------------- [ 26720]
AY261360.1_Kenya_1950            agctttttattatctaaaaattctggattggctcctagttgaatacacagaactttcgtcctttccgtcttggcacacat [ 26720]
AM712240.1_OURT_88/3             -------------------------------------------------------------------------------- [ 26720]
AM712239.1_Benin_97/1            -------------------------------------------------------------------------------- [ 26720]
KM111295.1_Ken06.Bus             ------------------------------gctatttg------------------------------------------ [ 26720]
KM111294.1_Ken05/Tk1             -------------------------------------------------------------------------------- [ 26720]
KM262844.1_strain_L60            -------------------------------------------------------------------------------- [ 26720]
KM102979.1_26544/OG10_Italy      -------------------------------------------------------------------------------- [ 26720]
KM262845.1_strain_NHV            -------------------------------------------------------------------------------- [ 26720]
KX354450.1_47/Ss/2008            -------------------------------------------------------------------------------- [ 26720]
KP055815.1_strain_BA71           -------------------------------------------------------------------------------- [ 26720]
MH025920.1_strain_R35            ------------------------------gctatttg------------------------------------------ [ 26720]
MH025919.1_strain_N10            -----------------aaagtttaggttggctatttg------------------------------------------ [ 26720]
MH025918.1_strain_R25            ------------------------------gctatttg------------------------------------------ [ 26720]
MH025917.1_strain_R7             ------------------------------gctatttg------------------------------------------ [ 26720]
MH025916.1_strain_R8             ------------------------------gctatttg------------------------------------------ [ 26720]

FR682468.2_Georgia_2007/1        -------------------------------------------------------------------------------- [ 26800]
ASF_IND_20_CAD543                -------------------------------------------------------------------------------- [ 26800]
LR899193.1_Germany_2020/1        -------------------------------------------------------------------------------- [ 26800]
MN393477.1_Wuhan_2019            -------------------------------------------------------------------------------- [ 26800]
MW396979.1_Timor-Leste/2019/1    -------------------------------------------------------------------------------- [ 26800]
MK940252.1_CN/2019/InnerMongolia -------------------------------------------------------------------------------- [ 26800]
MK128995.1_China/2018/AnhuiXCGQ  -------------------------------------------------------------------------------- [ 26800]
MH681419.1_ASFV/POL/2015         -------------------------------------------------------------------------------- [ 26800]
LS478113.1_Estonia_2014          -------------------------------------------------------------------------------- [ 26800]
KP843857.1_Odintsovo_02/14       -------------------------------------------------------------------------------- [ 26800]
FN557520.1_strain_E75            -------------------------------------------------------------------------------- [ 26800]
AY261366.1_Warthog               -------------------------------------------------------------------------------- [ 26800]
AY261365.1_Warmbaths             -------------------------------------------------------------------------------- [ 26800]
AY261364.1_Tengani_62            -------------------------------------------------------------------------------- [ 26800]
AY261363.1_Pretoriuskop/96/4     -------------------------------------------------------------------------------- [ 26800]
AY261362.1_Mkuzi_1979            -------------------------------------------------------------------------------- [ 26800]
AY261361.1_Malawi_Lil-20/1_1983  -------------------------------------------------------------------------------- [ 26800]
AY261360.1_Kenya_1950            aatgccataattaatatcggcgttccataaaacaaataacttgattagatcagtctggttttccttcacagcttttacta [ 26800]
AM712240.1_OURT_88/3             -------------------------------------------------------------------------------- [ 26800]
AM712239.1_Benin_97/1            -------------------------------------------------------------------------------- [ 26800]
KM111295.1_Ken06.Bus             -------------------------------------------------------------------------------- [ 26800]
```

```
KM111294.1_Ken05/Tk1             -------------------------------------------------------------------------------- [ 26800]
KM262844.1_strain_L60            -------------------------------------------------------------------------------- [ 26800]
KM102979.1_26544/OG10_Italy      -------------------------------------------------------------------------------- [ 26800]
KM262845.1_strain_NHV            -------------------------------------------------------------------------------- [ 26800]
KX354450.1_47/Ss/2008            -------------------------------------------------------------------------------- [ 26800]
KP055815.1_strain_BA71           -------------------------------------------------------------------------------- [ 26800]
MH025920.1_strain_R35            -------------------------------------------------------------------------------- [ 26800]
MH025919.1_strain_N10            -------------------------------------------------------------------------------- [ 26800]
MH025918.1_strain_R25            -------------------------------------------------------------------------------- [ 26800]
MH025917.1_strain_R7             -------------------------------------------------------------------------------- [ 26800]
MH025916.1_strain_R8             -------------------------------------------------------------------------------- [ 26800]

FR682468.2_Georgia_2007/1        -------------------------------------------------------------------------------- [ 26880]
ASF_IND_20_CAD543                -------------------------------------------------------------------------------- [ 26880]
LR899193.1_Germany_2020/1        -------------------------------------------------------------------------------- [ 26880]
MN393477.1_Wuhan_2019            -------------------------------------------------------------------------------- [ 26880]
MW396979.1_Timor-Leste/2019/1    -------------------------------------------------------------------------------- [ 26880]
MK940252.1_CN/2019/InnerMongolia -------------------------------------------------------------------------------- [ 26880]
MK128995.1_China/2018/AnhuiXCGQ  -------------------------------------------------------------------------------- [ 26880]
MH681419.1_ASFV/POL/2015         -------------------------------------------------------------------------------- [ 26880]
LS478113.1_Estonia_2014          -------------------------------------------------------------------------------- [ 26880]
KP843857.1_Odintsovo_02/14       -------------------------------------------------------------------------------- [ 26880]
FN557520.1_strain_E75            -------------------------------------------------------------------------------- [ 26880]
AY261366.1_Warthog               -------------------------------------------------------------------------------- [ 26880]
AY261365.1_Warmbaths             -------------------------------------------------------------------------------- [ 26880]
AY261364.1_Tengani_62            -------------------------------------------------------------------------------- [ 26880]
AY261363.1_Pretoriuskop/96/4     -------------------------------------------------------------------------------- [ 26880]
AY261362.1_Mkuzi_1979            -------------------------------------------------------------------------------- [ 26880]
AY261361.1_Malawi_Lil-20/1_1983  -------------------------------------------------------------------------------- [ 26880]
AY261360.1_Kenya_1950            aggctctgtcaagctcatagctgtcgacatcagaacatgacatagatccgctgattaccattttacattgtttacaaaaa [ 26880]
AM712240.1_OURT_88/3             -------------------------------------------------------------------------------- [ 26880]
AM712239.1_Benin_97/1            -------------------------------------------------------------------------------- [ 26880]
KM111295.1_Ken06.Bus             -------------------------------------------------------------------------------- [ 26880]
KM111294.1_Ken05/Tk1             -------------------------------------------------------------------------------- [ 26880]
KM262844.1_strain_L60            -------------------------------------------------------------------------------- [ 26880]
KM102979.1_26544/OG10_Italy      -------------------------------------------------------------------------------- [ 26880]
KM262845.1_strain_NHV            -------------------------------------------------------------------------------- [ 26880]
KX354450.1_47/Ss/2008            -------------------------------------------------------------------------------- [ 26880]
KP055815.1_strain_BA71           -------------------------------------------------------------------------------- [ 26880]
MH025920.1_strain_R35            -------------------------------------------------------------------------------- [ 26880]
MH025919.1_strain_N10            -------------------------------------------------------------------------------- [ 26880]
MH025918.1_strain_R25            -------------------------------------------------------------------------------- [ 26880]
MH025917.1_strain_R7             -------------------------------------------------------------------------------- [ 26880]
MH025916.1_strain_R8             -------------------------------------------------------------------------------- [ 26880]
```

```
FR682468.2_Georgia_2007/1        -------------------------------------------------------------------------------- [ 26960]
ASF_IND_20_CAD543                -------------------------------------------------------------------------------- [ 26960]
LR899193.1_Germany_2020/1        -------------------------------------------------------------------------------- [ 26960]
MN393477.1_Wuhan_2019            -------------------------------------------------------------------------------- [ 26960]
MW396979.1_Timor-Leste/2019/1    -------------------------------------------------------------------------------- [ 26960]
MK940252.1_CN/2019/InnerMongolia -------------------------------------------------------------------------------- [ 26960]
MK128995.1_China/2018/AnhuiXCGQ  -------------------------------------------------------------------------------- [ 26960]
MH681419.1_ASFV/POL/2015         -------------------------------------------------------------------------------- [ 26960]
LS478113.1_Estonia_2014          -------------------------------------------------------------------------------- [ 26960]
KP843857.1_Odintsovo_02/14       -------------------------------------------------------------------------------- [ 26960]
FN557520.1_strain_E75            -------------------------------------------------------------------------------- [ 26960]
AY261366.1_Warthog               -------------------------------------------------------------------------------- [ 26960]
AY261365.1_Warmbaths             -------------------------------------------------------------------------------- [ 26960]
AY261364.1_Tengani_62            -------------------------------------------------------------------------------- [ 26960]
AY261363.1_Pretoriuskop/96/4     -------------------------------------------------------------------------------- [ 26960]
AY261362.1_Mkuzi_1979            -------------------------------------------------------------------------------- [ 26960]
AY261361.1_Malawi_Lil-20/1_1983  -------------------------------------------------------------------------------- [ 26960]
AY261360.1_Kenya_1950            cctatgggtccgttttcccaccacagtccaagctgctgtaaaataaaactatcatcttcatgataatttgaaaaagcctt [ 26960]
AM712240.1_OURT_88/3             -------------------------------------------------------------------------------- [ 26960]
AM712239.1_Benin_97/1            -------------------------------------------------------------------------------- [ 26960]
KM111295.1_Ken06.Bus             -------------------------------------------------------------------------------- [ 26960]
KM111294.1_Ken05/Tk1             -------------------------------------------------------------------------------- [ 26960]
KM262844.1_strain_L60            -------------------------------------------------------------------------------- [ 26960]
KM102979.1_26544/OG10_Italy      -------------------------------------------------------------------------------- [ 26960]
KM262845.1_strain_NHV            -------------------------------------------------------------------------------- [ 26960]
KX354450.1_47/Ss/2008            -------------------------------------------------------------------------------- [ 26960]
KP055815.1_strain_BA71           -------------------------------------------------------------------------------- [ 26960]
MH025920.1_strain_R35            -------------------------------------------------------------------------------- [ 26960]
MH025919.1_strain_N10            -------------------------------------------------------------------------------- [ 26960]
MH025918.1_strain_R25            -------------------------------------------------------------------------------- [ 26960]
MH025917.1_strain_R7             -------------------------------------------------------------------------------- [ 26960]
MH025916.1_strain_R8             -------------------------------------------------------------------------------- [ 26960]

FR682468.2_Georgia_2007/1        -------------------------------------------------------------------------------- [ 27040]
ASF_IND_20_CAD543                -------------------------------------------------------------------------------- [ 27040]
LR899193.1_Germany_2020/1        -------------------------------------------------------------------------------- [ 27040]
MN393477.1_Wuhan_2019            -------------------------------------------------------------------------------- [ 27040]
MW396979.1_Timor-Leste/2019/1    -------------------------------------------------------------------------------- [ 27040]
MK940252.1_CN/2019/InnerMongolia -------------------------------------------------------------------------------- [ 27040]
MK128995.1_China/2018/AnhuiXCGQ  -------------------------------------------------------------------------------- [ 27040]
MH681419.1_ASFV/POL/2015         -------------------------------------------------------------------------------- [ 27040]
LS478113.1_Estonia_2014          -------------------------------------------------------------------------------- [ 27040]
KP843857.1_Odintsovo_02/14       -------------------------------------------------------------------------------- [ 27040]
```

```
FN557520.1_strain_E75            -------------------------------------------------------------------------------- [ 27040]
AY261366.1_Warthog               -------------------------------------------------------------------------------- [ 27040]
AY261365.1_Warmbaths             -------------------------------------------------------------------------------- [ 27040]
AY261364.1_Tengani_62            -------------------------------------------------------------------------------- [ 27040]
AY261363.1_Pretoriuskop/96/4     -------------------------------------------------------------------------------- [ 27040]
AY261362.1_Mkuzi_1979            -------------------------------------------------------------------------------- [ 27040]
AY261361.1_Malawi_Lil-20/1_1983  -------------------------------------------------------------------------------- [ 27040]
AY261360.1_Kenya_1950            gttttctatcaagactttttttgtaagaacctgtaaagaattcatcctagtatcgtggatgaaagcagtaaatctaatca [ 27040]
AM712240.1_OURT_88/3             -------------------------------------------------------------------------------- [ 27040]
AM712239.1_Benin_97/1            -------------------------------------------------------------------------------- [ 27040]
KM111295.1_Ken06.Bus             -------------------------------------------------------------------------------- [ 27040]
KM111294.1_Ken05/Tk1             -------------------------------------------------------------------------------- [ 27040]
KM262844.1_strain_L60            -------------------------------------------------------------------------------- [ 27040]
KM102979.1_26544/OG10_Italy      -------------------------------------------------------------------------------- [ 27040]
KM262845.1_strain_NHV            -------------------------------------------------------------------------------- [ 27040]
KX354450.1_47/Ss/2008            -------------------------------------------------------------------------------- [ 27040]
KP055815.1_strain_BA71           -------------------------------------------------------------------------------- [ 27040]
MH025920.1_strain_R35            -------------------------------------------------------------------------------- [ 27040]
MH025919.1_strain_N10            -------------------------------------------------------------------------------- [ 27040]
MH025918.1_strain_R25            -------------------------------------------------------------------------------- [ 27040]
MH025917.1_strain_R7             -------------------------------------------------------------------------------- [ 27040]
MH025916.1_strain_R8             -------------------------------------------------------------------------------- [ 27040]

FR682468.2_Georgia_2007/1        -------------------------------------------------------------------------------- [ 27120]
ASF_IND_20_CAD543                -------------------------------------------------------------------------------- [ 27120]
LR899193.1_Germany_2020/1        -------------------------------------------------------------------------------- [ 27120]
MN393477.1_Wuhan_2019            -------------------------------------------------------------------------------- [ 27120]
MW396979.1_Timor-Leste/2019/1    -------------------------------------------------------------------------------- [ 27120]
MK940252.1_CN/2019/InnerMongolia -------------------------------------------------------------------------------- [ 27120]
MK128995.1_China/2018/AnhuiXCGQ  -------------------------------------------------------------------------------- [ 27120]
MH681419.1_ASFV/POL/2015         -------------------------------------------------------------------------------- [ 27120]
LS478113.1_Estonia_2014          -------------------------------------------------------------------------------- [ 27120]
KP843857.1_Odintsovo_02/14       -------------------------------------------------------------------------------- [ 27120]
FN557520.1_strain_E75            -------------------------------------------------------------------------------- [ 27120]
AY261366.1_Warthog               -------------------------------------------------------------------------------- [ 27120]
AY261365.1_Warmbaths             -------------------------------------------------------------------------------- [ 27120]
AY261364.1_Tengani_62            -------------------------------------------------------------------------------- [ 27120]
AY261363.1_Pretoriuskop/96/4     -------------------------------------------------------------------------------- [ 27120]
AY261362.1_Mkuzi_1979            -------------------------------------------------------------------------------- [ 27120]
AY261361.1_Malawi_Lil-20/1_1983  -------------------------------------------------------------------------------- [ 27120]
AY261360.1_Kenya_1950            attataaaattgacttattgaaaagatatattaaatgagtgaaattgatgtttgcgatgatgtgcatgatcatacgaaga [ 27120]
AM712240.1_OURT_88/3             -------------------------------------------------------------------------------- [ 27120]
AM712239.1_Benin_97/1            -------------------------------------------------------------------------------- [ 27120]
KM111295.1_Ken06.Bus             -------------------------------------------------------------------------------- [ 27120]
```

```
KM111294.1_Ken05/Tk1             -------------------------------------------------------------------------------- [ 27120]
KM262844.1_strain_L60            -------------------------------------------------------------------------------- [ 27120]
KM102979.1_26544/OG10_Italy      -------------------------------------------------------------------------------- [ 27120]
KM262845.1_strain_NHV            -------------------------------------------------------------------------------- [ 27120]
KX354450.1_47/Ss/2008            -------------------------------------------------------------------------------- [ 27120]
KP055815.1_strain_BA71           -------------------------------------------------------------------------------- [ 27120]
MH025920.1_strain_R35            -------------------------------------------------------------------------------- [ 27120]
MH025919.1_strain_N10            -------------------------------------------------------------------------------- [ 27120]
MH025918.1_strain_R25            -------------------------------------------------------------------------------- [ 27120]
MH025917.1_strain_R7             -------------------------------------------------------------------------------- [ 27120]
MH025916.1_strain_R8             -------------------------------------------------------------------------------- [ 27120]

FR682468.2_Georgia_2007/1        -------------------------------------------------------------------------------- [ 27200]
ASF_IND_20_CAD543                -------------------------------------------------------------------------------- [ 27200]
LR899193.1_Germany_2020/1        -------------------------------------------------------------------------------- [ 27200]
MN393477.1_Wuhan_2019            -------------------------------------------------------------------------------- [ 27200]
MW396979.1_Timor-Leste/2019/1    -------------------------------------------------------------------------------- [ 27200]
MK940252.1_CN/2019/InnerMongolia -------------------------------------------------------------------------------- [ 27200]
MK128995.1_China/2018/AnhuiXCGQ  -------------------------------------------------------------------------------- [ 27200]
MH681419.1_ASFV/POL/2015         -------------------------------------------------------------------------------- [ 27200]
LS478113.1_Estonia_2014          -------------------------------------------------------------------------------- [ 27200]
KP843857.1_Odintsovo_02/14       -------------------------------------------------------------------------------- [ 27200]
FN557520.1_strain_E75            -------------------------------------------------------------------------------- [ 27200]
AY261366.1_Warthog               -------------------------------------------------------------------------------- [ 27200]
AY261365.1_Warmbaths             -------------------------------------------------------------------------------- [ 27200]
AY261364.1_Tengani_62            -------------------------------------------------------------------------------- [ 27200]
AY261363.1_Pretoriuskop/96/4     -------------------------------------------------------------------------------- [ 27200]
AY261362.1_Mkuzi_1979            -------------------------------------------------------------------------------- [ 27200]
AY261361.1_Malawi_Lil-20/1_1983  -------------------------------------------------------------------------------- [ 27200]
AY261360.1_Kenya_1950            aacacgttcactggtgtccatgatcaaaatttaatgttctatgtaaaaagtacagacgttaaccctttaaacatttgttt [ 27200]
AM712240.1_OURT_88/3             -------------------------------------------------------------------------------- [ 27200]
AM712239.1_Benin_97/1            -------------------------------------------------------------------------------- [ 27200]
KM111295.1_Ken06.Bus             -------------------------------------------------------------------------------- [ 27200]
KM111294.1_Ken05/Tk1             -------------------------------------------------------------------------------- [ 27200]
KM262844.1_strain_L60            -------------------------------------------------------------------------------- [ 27200]
KM102979.1_26544/OG10_Italy      -------------------------------------------------------------------------------- [ 27200]
KM262845.1_strain_NHV            -------------------------------------------------------------------------------- [ 27200]
KX354450.1_47/Ss/2008            -------------------------------------------------------------------------------- [ 27200]
KP055815.1_strain_BA71           -------------------------------------------------------------------------------- [ 27200]
MH025920.1_strain_R35            -------------------------------------------------------------------------------- [ 27200]
MH025919.1_strain_N10            -------------------------------------------------------------------------------- [ 27200]
MH025918.1_strain_R25            -------------------------------------------------------------------------------- [ 27200]
MH025917.1_strain_R7             -------------------------------------------------------------------------------- [ 27200]
MH025916.1_strain_R8             -------------------------------------------------------------------------------- [ 27200]
```

```
FR682468.2_Georgia_2007/1        -------------------------------------------------------------------------------- [ 27280]
ASF_IND_20_CAD543                -------------------------------------------------------------------------------- [ 27280]
LR899193.1_Germany_2020/1        -------------------------------------------------------------------------------- [ 27280]
MN393477.1_Wuhan_2019            -------------------------------------------------------------------------------- [ 27280]
MW396979.1_Timor-Leste/2019/1    -------------------------------------------------------------------------------- [ 27280]
MK940252.1_CN/2019/InnerMongolia -------------------------------------------------------------------------------- [ 27280]
MK128995.1_China/2018/AnhuiXCGQ  -------------------------------------------------------------------------------- [ 27280]
MH681419.1_ASFV/POL/2015         -------------------------------------------------------------------------------- [ 27280]
LS478113.1_Estonia_2014          -------------------------------------------------------------------------------- [ 27280]
KP843857.1_Odintsovo_02/14       -------------------------------------------------------------------------------- [ 27280]
FN557520.1_strain_E75            -------------------------------------------------------------------------------- [ 27280]
AY261366.1_Warthog               -------------------------------------------------------------------------------- [ 27280]
AY261365.1_Warmbaths             -------------------------------------------------------------------------------- [ 27280]
AY261364.1_Tengani_62            -------------------------------------------------------------------------------- [ 27280]
AY261363.1_Pretoriuskop/96/4     -------------------------------------------------------------------------------- [ 27280]
AY261362.1_Mkuzi_1979            -------------------------------------------------------------------------------- [ 27280]
AY261361.1_Malawi_Lil-20/1_1983  -------------------------------------------------------------------------------- [ 27280]
AY261360.1_Kenya_1950            aaacattagtttaaactaaaggttaaacctagttaacaatgtttgacatttcttccattctt-atacggggggggggtcc [ 27280]
AM712240.1_OURT_88/3             -------------------------------------------------------------------------------- [ 27280]
AM712239.1_Benin_97/1            -------------------------------------------------------------------------------- [ 27280]
KM111295.1_Ken06.Bus             ------------------------------agttaatgatgtttaatctatcttccatgctcaatcagggggggggtcc [ 27280]
KM111294.1_Ken05/Tk1             -------------------------------------------------------------------------------- [ 27280]
KM262844.1_strain_L60            -------------------------------------------------------------------------------- [ 27280]
KM102979.1_26544/OG10_Italy      -------------------------------------------------------------------------------- [ 27280]
KM262845.1_strain_NHV            -------------------------------------------------------------------------------- [ 27280]
KX354450.1_47/Ss/2008            -------------------------------------------------------------------------------- [ 27280]
KP055815.1_strain_BA71           -------------------------------------------------------------------------------- [ 27280]
MH025920.1_strain_R35            ------------------------------agttaatgatgtttaatctatcttccatgctcaatcagggggggggtcc [ 27280]
MH025919.1_strain_N10            ------------------------------agttaatgatgtttaatcta------------------------------ [ 27280]
MH025918.1_strain_R25            ------------------------------agttaatgatgtttaatctatcttccatgctcaatcagggggggggtcc [ 27280]
MH025917.1_strain_R7             ------------------------------agttaatgatgtttaatctatcttccatgctcaatcagggggggggtcc [ 27280]
MH025916.1_strain_R8             ------------------------------agttaatgatgtttaatctatcttccatgctcaatcagggggggggtcc [ 27280]

FR682468.2_Georgia_2007/1        -------------------------------------------------------------------------------- [ 27360]
ASF_IND_20_CAD543                -------------------------------------------------------------------------------- [ 27360]
LR899193.1_Germany_2020/1        -------------------------------------------------------------------------------- [ 27360]
MN393477.1_Wuhan_2019            -------------------------------------------------------------------------------- [ 27360]
MW396979.1_Timor-Leste/2019/1    -------------------------------------------------------------------------------- [ 27360]
MK940252.1_CN/2019/InnerMongolia -------------------------------------------------------------------------------- [ 27360]
MK128995.1_China/2018/AnhuiXCGQ  -------------------------------------------------------------------------------- [ 27360]
MH681419.1_ASFV/POL/2015         -------------------------------------------------------------------------------- [ 27360]
LS478113.1_Estonia_2014          -------------------------------------------------------------------------------- [ 27360]
KP843857.1_Odintsovo_02/14       -------------------------------------------------------------------------------- [ 27360]
```

```
FN557520.1_strain_E75             -------------------------------------------------------------------------------- [ 27360]
AY261366.1_Warthog                -------------------------------------------------------------------------------- [ 27360]
AY261365.1_Warmbaths              -------------------------------------------------------------------------------- [ 27360]
AY261364.1_Tengani_62             -------------------------------------------------------------------------------- [ 27360]
AY261363.1_Pretoriuskop/96/4      -------------------------------------------------------------------------------- [ 27360]
AY261362.1_Mkuzi_1979             -------------------------------------------------------------------------------- [ 27360]
AY261361.1_Malawi_Lil-20/1_1983   -------------------------------------------------------------------------------- [ 27360]
AY261360.1_Kenya_1950             taataattttaatactgttgttgtggattgttg------atgaaaactttatagatgctaaagcaatgagttataataat [ 27360]
AM712240.1_OURT_88/3              -------------------------------------------------------------------------------- [ 27360]
AM712239.1_Benin_97/1             -------------------------------------------------------------------------------- [ 27360]
KM111295.1_Ken06.Bus              taatagttttaatactgttgctgtggattgttgaatataatgaaaactttatagatgctaaagcagcgaaatata---ca [ 27360]
KM111294.1_Ken05/Tk1              -------------------------------------------------------------------------------- [ 27360]
KM262844.1_strain_L60             -------------------------------------------------------------------------------- [ 27360]
KM102979.1_26544/OG10_Italy       -------------------------------------------------------------------------------- [ 27360]
KM262845.1_strain_NHV             -------------------------------------------------------------------------------- [ 27360]
KX354450.1_47/Ss/2008             -------------------------------------------------------------------------------- [ 27360]
KP055815.1_strain_BA71            -------------------------------------------------------------------------------- [ 27360]
MH025920.1_strain_R35             taatagttttaatactgttgctgtggattgttgaatataatgaaaactttatagatgctaaagcagcgaaatata---ca [ 27360]
MH025919.1_strain_N10             -------------------------------------------------------------------------------- [ 27360]
MH025918.1_strain_R25             taatagttttaatactgttgctgtggattgttgaatataatgaaaactttatagatgctaaagcagcgaaatata---ca [ 27360]
MH025917.1_strain_R7              taatagttttaatactgttgctgtggattgttgaatataatgaaaactttatagatgctaaagcagcgaaatata---ca [ 27360]
MH025916.1_strain_R8              taatagttttaatactgttgctgtggattgttgaatataatgaaaactttatagatgctaaagcagcgaaatata---ca [ 27360]

FR682468.2_Georgia_2007/1         ---------------------------ctatataagctttcatcaaaattt-aatttttttttataaaaatacacgaatt [ 27440]
ASF_IND_20_CAD543                 ---------------------------t....a...ta..t..........-.t........ata....g....g..... [ 27440]
LR899193.1_Germany_2020/1         ---------------------------........................-............................ [ 27440]
MN393477.1_Wuhan_2019             ---------------------------........................-............................ [ 27440]
MW396979.1_Timor-Leste/2019/1     ---------------------------........................-............................ [ 27440]
MK940252.1_CN/2019/InnerMongolia  ---------------------------........................-............................ [ 27440]
MK128995.1_China/2018/AnhuiXCGQ   ---------------------------........................-............................ [ 27440]
MH681419.1_ASFV/POL/2015          ---------------------------........................-............................ [ 27440]
LS478113.1_Estonia_2014           ---------------------------........................-............................ [ 27440]
KP843857.1_Odintsovo_02/14        ---------------------------........................-..-......................... [ 27440]
FN557520.1_strain_E75             ---------------------------t....a...ta..t..........-.t........ata....g....g..... [ 27440]
AY261366.1_Warthog                ---------------------------tata.a.c.t..............a.t..c.ca.aa.a....g.....atcaa [ 27440]
AY261365.1_Warmbaths              ---------------------------tgt.ta.t.t..............-.t........ata....g....g..... [ 27440]
AY261364.1_Tengani_62             --------------------------------------..g..........a.t..c.ca.--.a....g.....atcaa [ 27440]
AY261363.1_Pretoriuskop/96/4      ---------------------------tata.a.t.t..............a.t........ata.........a..... [ 27440]
AY261362.1_Mkuzi_1979             ---------------------------t....a...ta..t..........-.t........ata....g....g..... [ 27440]
AY261361.1_Malawi_Lil-20/1_1983   ---------------------------tata.a.t.t..............-.....c..a.a.a....g.....atcaa [ 27440]
AY261360.1_Kenya_1950             cgtactgtttaatgtgaactataagtttata.a.c.t..............-..........ata....g....a..... [ 27440]
AM712240.1_OURT_88/3              -------------------------------------------------------------------------------- [ 27440]
AM712239.1_Benin_97/1             ---------------------------t....a...ta..t..........-.t........ata....g....g..... [ 27440]
KM111295.1_Ken06.Bus              tgttctgtttaatgtgagc--------tata.a.tct..............-................tg....aa...- [ 27440]
```

KM111294.1_Ken05/Tk1 ---------------------------tata.a.t.t..............-..........ata....g....ga..a. [ 27440]
KM262844.1_strain_L60 ---------------------------t....a...ta..t..........-.t........ata....g....g..... [ 27440]
KM102979.1_26544/OG10_Italy ---------------------------t....a...ta..t..........-.t........ata....g....g..... [ 27440]
KM262845.1_strain_NHV -------------------------------------------------------------------------------- [ 27440]
KX354450.1_47/Ss/2008 ---------------------------t....a...ta..t..........-.t........ata....g....g..... [ 27440]
KP055815.1_strain_BA71 -------------------------------------------------------------------------------- [ 27440]
MH025920.1_strain_R35 tgttctgtttaatgtgagc--------tata.a.tct..............-................tg....aa...- [ 27440]
MH025919.1_strain_N10 -------------------------------------------------------------------------------- [ 27440]
MH025918.1_strain_R25 tgttctgtttaatgtgagc--------tata.a.tct..............-................tg....aa...- [ 27440]
MH025917.1_strain_R7 tgttctgtttaatgtgagc--------tata.a.tct..............-................tg....aa...- [ 27440]
MH025916.1_strain_R8 tgttctgtttaatgtgagc--------tata.a.tct..............-................tg....aa...- [ 27440]

FR682468.2_Georgia_2007/1 aaac-taaagtctaaact------------ttagtttgactatttgagttaatgatgcttaac---ttatcttccatgct [ 27520]
ASF_IND_20_CAD543 ..t.a.tt...t......------------........g.............nnnnnnnnnnn---nnnnnnnnnnnnnn [ 27520]
LR899193.1_Germany_2020/1 ....-.............------------.................................---.............. [ 27520]
MN393477.1_Wuhan_2019 ....-.............------------.................................---.............. [ 27520]
MW396979.1_Timor-Leste/2019/1 ....-.............------------.................................---.............. [ 27520]
MK940252.1_CN/2019/InnerMongolia ....-.............------------.................................---.............. [ 27520]
MK128995.1_China/2018/AnhuiXCGQ ....-.............------------.................................---.............. [ 27520]
MH681419.1_ASFV/POL/2015 ....-.............------------.................................---.............. [ 27520]
LS478113.1_Estonia_2014 ....-.............------------.................................---.............. [ 27520]
KP843857.1_Odintsovo_02/14 ....-.............------------.................................---.............. [ 27520]
FN557520.1_strain_E75 ..t.a.tt...t......------------........g.............c....t....t---a.t........... [ 27520]
AY261366.1_Warthog cgt.gcttgt.ta..--------------------------------t..c......t..g.ttca...ct.c.t.caa. [ 27520]
AY261365.1_Warmbaths ..t.a.tt...t.....--------------.......g.c.....gt..t......t....t---c............. [ 27520]
AY261364.1_Tengani_62 cgt.gctt.t.ta.g--------------------------------t..c......t..g.ttca...ct.c.t.caa. [ 27520]
AY261363.1_Pretoriuskop/96/4 ..t.a.tt...t......aaagtataaacg........g........a.........t....t---c...tc........ [ 27520]
AY261362.1_Mkuzi_1979 ..t.a.tt...t......------------........g.............c....t....t---a.t........... [ 27520]
AY261361.1_Malawi_Lil-20/1_1983 cgt.gcttgt.ta.g--------------------------------t..c......t...gttca...ct.c.t.caa. [ 27520]
AY261360.1_Kenya_1950 ..t.a.tt...t......aaagtttacact....a...gt.................t....ttca...ct...t.caa. [ 27520]
AM712240.1_OURT_88/3 ---------------------------------------------.......c....t....t---a.t........... [ 27520]
AM712239.1_Benin_97/1 ..t.a.tt...t......------------........g.............c....t....t---a.t........... [ 27520]
KM111295.1_Ken06.Bus ------------......aaag-------t....g...g..................t....t---c..........t.. [ 27520]
KM111294.1_Ken05/Tk1 ..t.a.tt...t......aaaagtt-tact........g..................t....t---.............. [ 27520]
KM262844.1_strain_L60 ..t.a.tt...t......------------........g.............c....t....t---a.t........... [ 27520]
KM102979.1_26544/OG10_Italy ..t.a.tt...t......------------........g.............c....t....t---a.t........... [ 27520]
KM262845.1_strain_NHV ---------------------------------------------.......c....t....t---a.t........... [ 27520]
KX354450.1_47/Ss/2008 ..t.a.tt...t......------------........g.............c....t....t---a.t........... [ 27520]
KP055815.1_strain_BA71 -----------------------------------------------------------------------------... [ 27520]
MH025920.1_strain_R35 ------------......aaag-------t....g...g..................t....t---c..........t.. [ 27520]
MH025919.1_strain_N10 ---------------------------------------------------------------------........t.. [ 27520]
MH025918.1_strain_R25 ------------......aaag-------t....g...g..................t....t---c..........t.. [ 27520]
MH025917.1_strain_R7 ------------......aaag-------t....g...g..................t....t---c..........t.. [ 27520]
MH025916.1_strain_R8 ------------......aaag-------t....g...g..................t....t---c..........t.. [ 27520]

```
FR682468.2_Georgia_2007/1        tatcaa--------ggggggggtcctaatagttttgatactattgttgtggattgttgaatataataaatactttataga [ 27600]
ASF_IND_20_CAD543                nnnnnnnnnnnnnnn.........t.......c.a.......................c.....g..a.......... [ 27600]
LR899193.1_Germany_2020/1        .....---------.................................................................. [ 27600]
MN393477.1_Wuhan_2019            .....---------.................................................................. [ 27600]
MW396979.1_Timor-Leste/2019/1    .....---------.................................................................. [ 27600]
MK940252.1_CN/2019/InnerMongolia .....---------.................................................................. [ 27600]
MK128995.1_China/2018/AnhuiXCGQ  .....---------.................................................................. [ 27600]
MH681419.1_ASFV/POL/2015         .....g-------................................................................... [ 27600]
LS478113.1_Estonia_2014          .....---------.................................................................. [ 27600]
KP843857.1_Odintsovo_02/14       .....---------.................................................................. [ 27600]
FN557520.1_strain_E75            .....ag----gg.........t....c...c.a.......................c.....g..a.......... [ 27600]
AY261366.1_Warthog               ..ct-gg-----gg.....atctt.....c...ag..---.....a...t...c...c...t..gt.g.a..c..... [ 27600]
AY261365.1_Warmbaths             ....gg-----gg...a.......a.....a....a.....g..a..a.......a..........g..a.......... [ 27600]
AY261364.1_Tengani_62            ..ctggg-----gg.......tctt.....aca..a...---..a......t...c...c...t..gt.g.aa...... [ 27600]
AY261363.1_Pretoriuskop/96/4     .....g-----gg..........a.....a....a.....g..a..a.......a..........g..a.......... [ 27600]
AY261362.1_Mkuzi_1979            .....a-----gg.........t....c...c.a.......................c.....g..a.......... [ 27600]
AY261361.1_Malawi_Lil-20/1_1983  ..c---------tg.......tctt.....c...ag..---.....a...c...c.a.c...t..gt.g.a..c..... [ 27600]
AY261360.1_Kenya_1950            ..ctggg-----gg.......tctt.....ac...ag..---..a..a...c...c.a.t...t..gt.g.a..c..... [ 27600]
AM712240.1_OURT_88/3             .....a------g.........t....c...c.a.......................c.....g..a.......... [ 27600]
AM712239.1_Benin_97/1            .....a------g.........t....c...c.a.......................c.....g..a.......... [ 27600]
KM111295.1_Ken06.Bus             cg..c.a-----gg.........t..........a...t.g..a.....................g..a.......... [ 27600]
KM111294.1_Ken05/Tk1             ...t..g-----gg.......a...............a........a.....................g..a........a. [ 27600]
KM262844.1_strain_L60            .....a-----gg.........t....c...c.a.......................c.....g..a.......... [ 27600]
KM102979.1_26544/OG10_Italy      .....ag----gg.........t....c...c.a.......................c.....g..a.......... [ 27600]
KM262845.1_strain_NHV            .....a------g.........t....c...c.a.......................c.....g..a.......... [ 27600]
KX354450.1_47/Ss/2008            .....a------g.........t....c...c.a.......................c.....g..a.......... [ 27600]
KP055815.1_strain_BA71           ggc..gaa----ggccaa.aat...c..a.aga.cacc.ac....c.ac..c.g.c....c----------------..c [ 27600]
MH025920.1_strain_R35            cg..c.ag----gg.........t..........a...t.g..a.....................g..a.......... [ 27600]
MH025919.1_strain_N10            cg..c.a------g.........t..........a...t.g..a.....................g..a.......... [ 27600]
MH025918.1_strain_R25            cg..c.ag----gg.........t..........a...t.g..a.....................g..a.......... [ 27600]
MH025917.1_strain_R7             cg..c.ag----gg.........t..........a...t.g..a.....................g..a.......... [ 27600]
MH025916.1_strain_R8             cg..c.ag----gg.........t..........a...t.g..a.....................g..a.......... [ 27600]

FR682468.2_Georgia_2007/1        tg-ctgaaatgtttgaaaataatagtacatcaat-------------------------gttgtaag----tttgatc-- [ 27680]
ASF_IND_20_CAD543                ..c..a....aa..nnnnnnnnnnnnnnnnnt..-------------------------ac.----------..a...-- [ 27680]
LR899193.1_Germany_2020/1        ..-...............................-------------------------........----.......-- [ 27680]
MN393477.1_Wuhan_2019            ..-...............................-------------------------........----.......-- [ 27680]
MW396979.1_Timor-Leste/2019/1    ..-...............................-------------------------........----.......-- [ 27680]
MK940252.1_CN/2019/InnerMongolia ..-...............................-------------------------........----.......-- [ 27680]
MK128995.1_China/2018/AnhuiXCGQ  ..-...............................-------------------------........----.......-- [ 27680]
MH681419.1_ASFV/POL/2015         ..-...............................-------------------------........----.......-- [ 27680]
LS478113.1_Estonia_2014          ..-...............................-------------------------........----.......-- [ 27680]
KP843857.1_Odintsovo_02/14       ..-...............................-------------------------........----.......-- [ 27680]
```

```
FN557520.1_strain_E75            ..-..a....aa.....tc...a.a..gtagt..-------------------------ac.----------..a...-- [ 27680]
AY261366.1_Warthog               a.-.a...t.aa.ga.t.t..c..ac.acaatgg----------------------tac...a....----.......-- [ 27680]
AY261365.1_Warmbaths             ..-..ag....ag..gtgc..t..t.cata.tg.ttaat----------------ttga.c.a...aatgt...c...-- [ 27680]
AY261364.1_Tengani_62            a.-.a.t...aga.a.ttg..c..a-------------------------------ggt...a....----.......-- [ 27680]
AY261363.1_Pretoriuskop/96/4     ..-..ag....ag..gtgc..t..t.cata.tg.ttaat----------------ttga.c.a...aatgt...c...-- [ 27680]
AY261362.1_Mkuzi_1979            ..-..a....aa.....tc...a.a..gtagt..-------------------------ac.----------..a...-- [ 27680]
AY261361.1_Malawi_Lil-20/1_1983  a.-.a.....aa.g..t.t..c..a..gcagtgc-------------------------a..a....----.......-- [ 27680]
AY261360.1_Kenya_1950            a.-.aa....aga.agt.cacc.gt.g.g-------------------------------g.a...c----.......-- [ 27680]
AM712240.1_OURT_88/3             ..-..a....aa.....tc...a.a..gtagt..-------------------------ac.----------..a...-- [ 27680]
AM712239.1_Benin_97/1            ..-..a....aa.....tc...a.a..gtagt..-------------------------ac.----------..a...-- [ 27680]
KM111295.1_Ken06.Bus             ..-..ac.cat...a.t.c..g.t..t.t-------------------------------gct...t----g.ga..tac [ 27680]
KM111294.1_Ken05/Tk1             ..-........ga...ttg....t....tcat.ctcaatgttgtcagctttaatgatttaaaaa...----.......-- [ 27680]
KM262844.1_strain_L60            ..-..-....aa.....tc...a.a..gtagt..-------------------------ac.----------..a...-- [ 27680]
KM102979.1_26544/OG10_Italy      ..-..a....aa.....tc...a.a..gtagt..-------------------------ac.----------..a...-- [ 27680]
KM262845.1_strain_NHV            ..-..a....aa.....tc...a.a..gtagt..-------------------------ac.----------..a...-- [ 27680]
KX354450.1_47/Ss/2008            ..-..a....aa.....tc...a.a..gtagt..-------------------------ac.----------..a...-- [ 27680]
KP055815.1_strain_BA71           ..-actg...agc.agcc----a.t..gcaatcc-------------------------ac.----------g.------ [ 27680]
MH025920.1_strain_R35            ..-..ac.cat...a.t.c..g.t..t.t-------------------------------gct...t----g.ga..tac [ 27680]
MH025919.1_strain_N10            ..-..ac.cat...a.t.c..g.t..t.t-------------------------------gct...t----g.ga..tac [ 27680]
MH025918.1_strain_R25            ..-..ac.cat...a.t.c..g.t..t.t-------------------------------gct...t----g.ga..tac [ 27680]
MH025917.1_strain_R7             ..-..ac.cat...a.t.c..g.t..t.t-------------------------------gct...t----g.ga..tac [ 27680]
MH025916.1_strain_R8             ..-..ac.cat...a.t.c..g.t..t.t-------------------------------gct...t----g.ga..tac [ 27680]

FR682468.2_Georgia_2007/1        -aaaatttaatttctcataaaaaaggta-cacatcaacattgctcatttaagtttcatgatgtttgattcattacttcct [ 27760]
ASF_IND_20_CAD543                -.......nnnnnnnnnnnnnnnnnnnn-nnnnnn.------t......................ag..........-.. [ 27760]
LR899193.1_Germany_2020/1        -...........................-................................................... [ 27760]
MN393477.1_Wuhan_2019            -...........................-................................................... [ 27760]
MW396979.1_Timor-Leste/2019/1    -...........................-................................................... [ 27760]
MK940252.1_CN/2019/InnerMongolia -...........................-................................................... [ 27760]
MK128995.1_China/2018/AnhuiXCGQ  -...........................-................................................... [ 27760]
MH681419.1_ASFV/POL/2015         -...........................-................................................... [ 27760]
LS478113.1_Estonia_2014          -...........................-................................................... [ 27760]
KP843857.1_Odintsovo_02/14       -...........................-................................................... [ 27760]
FN557520.1_strain_E75            -.......g...................-.......------t......................ag............. [ 27760]
AY261366.1_Warthog               -..........a...........g.t..-...........cat.......g..............a.............. [ 27760]
AY261365.1_Warmbaths             -............a..........a...-.........g.c...t------..........................t.. [ 27760]
AY261364.1_Tengani_62            -...........................-...........cat........a.............ag............. [ 27760]
AY261363.1_Pretoriuskop/96/4     -...........................-.........gcc...tg...........................g...... [ 27760]
AY261362.1_Mkuzi_1979            -.......g...................-.......------t......................ag............. [ 27760]
AY261361.1_Malawi_Lil-20/1_1983  -.......................a...-.......g.g.c...tg.................................. [ 27760]
AY261360.1_Kenya_1950            -.............at........atac--------....ca...gc..................a.............. [ 27760]
AM712240.1_OURT_88/3             -.......g...................-.......------t......................ag............. [ 27760]
AM712239.1_Benin_97/1            -.......g...................-.......------t......................ag............. [ 27760]
KM111295.1_Ken06.Bus             a....a..................aagta...........c.......c................c...........t.. [ 27760]
```

```
KM111294.1_Ken05/Tk1             -........................tac--.........----.......g.......c......a.............. [ 27760]
KM262844.1_strain_L60            -.......g...................-.......------t......................ag............. [ 27760]
KM102979.1_26544/OG10_Italy      -.......g...................-.......------t......................ag............. [ 27760]
KM262845.1_strain_NHV            -.......g...................-.......------t......................ag............. [ 27760]
KX354450.1_47/Ss/2008            -.......g...................-.......------t......................ag............. [ 27760]
KP055815.1_strain_BA71           ---------.c..t......g.---------------------.........a...g----------------------- [ 27760]
MH025920.1_strain_R35            a....a..................aagta...........c.......c................c...........t.. [ 27760]
MH025919.1_strain_N10            a....a..................aagta...........c.......c................c...........t.. [ 27760]
MH025918.1_strain_R25            a....a..................aagta...........c.......c................c...........t.. [ 27760]
MH025917.1_strain_R7             a....a..................aagta...........c.......c................c...........t.. [ 27760]
MH025916.1_strain_R8             a....a..................aagta...........c.......c................c...........t.. [ 27760]

FR682468.2_Georgia_2007/1        acaattactg---------------------------------------------------------------------- [ 27840]
ASF_IND_20_CAD543                ...nnnnnn----------------------------------------------------------------------- [ 27840]
LR899193.1_Germany_2020/1        .........----------------------------------------------------------------------- [ 27840]
MN393477.1_Wuhan_2019            .........----------------------------------------------------------------------- [ 27840]
MW396979.1_Timor-Leste/2019/1    .........----------------------------------------------------------------------- [ 27840]
MK940252.1_CN/2019/InnerMongolia .........----------------------------------------------------------------------- [ 27840]
MK128995.1_China/2018/AnhuiXCGQ  .........----------------------------------------------------------------------- [ 27840]
MH681419.1_ASFV/POL/2015         .........----------------------------------------------------------------------- [ 27840]
LS478113.1_Estonia_2014          .........----------------------------------------------------------------------- [ 27840]
KP843857.1_Odintsovo_02/14       .........----------------------------------------------------------------------- [ 27840]
FN557520.1_strain_E75            .........----------------------------------------------------------------------- [ 27840]
AY261366.1_Warthog               .........----------------------------------------------------------------------- [ 27840]
AY261365.1_Warmbaths             .........----------------------------------------------------------------------- [ 27840]
AY261364.1_Tengani_62            ......g..----------------------------------------------------------------------- [ 27840]
AY261363.1_Pretoriuskop/96/4     ..........ggggggggtctttaatcgctttagtattattatggcttgctgactattatgtagaattcatagaagcaat [ 27840]
AY261362.1_Mkuzi_1979            .........----------------------------------------------------------------------- [ 27840]
AY261361.1_Malawi_Lil-20/1_1983  .........----------------------------------------------------------------------- [ 27840]
AY261360.1_Kenya_1950            .g.......----------------------------------------------------------------------- [ 27840]
AM712240.1_OURT_88/3             .........----------------------------------------------------------------------- [ 27840]
AM712239.1_Benin_97/1            .........----------------------------------------------------------------------- [ 27840]
KM111295.1_Ken06.Bus             .........----------------------------------------------------------------------- [ 27840]
KM111294.1_Ken05/Tk1             .........----------------------------------------------------------------------- [ 27840]
KM262844.1_strain_L60            .........----------------------------------------------------------------------- [ 27840]
KM102979.1_26544/OG10_Italy      .........----------------------------------------------------------------------- [ 27840]
KM262845.1_strain_NHV            .........----------------------------------------------------------------------- [ 27840]
KX354450.1_47/Ss/2008            .........----------------------------------------------------------------------- [ 27840]
KP055815.1_strain_BA71           -------------------------------------------------------------------------------- [ 27840]
MH025920.1_strain_R35            .........----------------------------------------------------------------------- [ 27840]
MH025919.1_strain_N10            .........----------------------------------------------------------------------- [ 27840]
MH025918.1_strain_R25            .........----------------------------------------------------------------------- [ 27840]
MH025917.1_strain_R7             .........----------------------------------------------------------------------- [ 27840]
MH025916.1_strain_R8             .........----------------------------------------------------------------------- [ 27840]
```

```
FR682468.2_Georgia_2007/1        -------------------------------------------------------------------------------- [ 27920]
ASF_IND_20_CAD543                -------------------------------------------------------------------------------- [ 27920]
LR899193.1_Germany_2020/1        -------------------------------------------------------------------------------- [ 27920]
MN393477.1_Wuhan_2019            -------------------------------------------------------------------------------- [ 27920]
MW396979.1_Timor-Leste/2019/1    -------------------------------------------------------------------------------- [ 27920]
MK940252.1_CN/2019/InnerMongolia -------------------------------------------------------------------------------- [ 27920]
MK128995.1_China/2018/AnhuiXCGQ  -------------------------------------------------------------------------------- [ 27920]
MH681419.1_ASFV/POL/2015         -------------------------------------------------------------------------------- [ 27920]
LS478113.1_Estonia_2014          -------------------------------------------------------------------------------- [ 27920]
KP843857.1_Odintsovo_02/14       -------------------------------------------------------------------------------- [ 27920]
FN557520.1_strain_E75            -------------------------------------------------------------------------------- [ 27920]
AY261366.1_Warthog               -------------------------------------------------------------------------------- [ 27920]
AY261365.1_Warmbaths             -------------------------------------------------------------------------------- [ 27920]
AY261364.1_Tengani_62            -------------------------------------------------------------------------------- [ 27920]
AY261363.1_Pretoriuskop/96/4     aatagataatcatactaaggagttataagtttgatcaaaatttaatttcccataaaaaaggtacacatcaacatcattca [ 27920]
AY261362.1_Mkuzi_1979            -------------------------------------------------------------------------------- [ 27920]
AY261361.1_Malawi_Lil-20/1_1983  -------------------------------------------------------------------------------- [ 27920]
AY261360.1_Kenya_1950            -------------------------------------------------------------------------------- [ 27920]
AM712240.1_OURT_88/3             -------------------------------------------------------------------------------- [ 27920]
AM712239.1_Benin_97/1            -------------------------------------------------------------------------------- [ 27920]
KM111295.1_Ken06.Bus             -------------------------------------------------------------------------------- [ 27920]
KM111294.1_Ken05/Tk1             -------------------------------------------------------------------------------- [ 27920]
KM262844.1_strain_L60            -------------------------------------------------------------------------------- [ 27920]
KM102979.1_26544/OG10_Italy      -------------------------------------------------------------------------------- [ 27920]
KM262845.1_strain_NHV            -------------------------------------------------------------------------------- [ 27920]
KX354450.1_47/Ss/2008            -------------------------------------------------------------------------------- [ 27920]
KP055815.1_strain_BA71           -------------------------------------------------------------------------------- [ 27920]
MH025920.1_strain_R35            -------------------------------------------------------------------------------- [ 27920]
MH025919.1_strain_N10            -------------------------------------------------------------------------------- [ 27920]
MH025918.1_strain_R25            -------------------------------------------------------------------------------- [ 27920]
MH025917.1_strain_R7             -------------------------------------------------------------------------------- [ 27920]
MH025916.1_strain_R8             -------------------------------------------------------------------------------- [ 27920]

FR682468.2_Georgia_2007/1        -------------------------------------gggggggggggggggtctttaatagctttagcattgttatggt [ 28000]
ASF_IND_20_CAD543                --------------------------------------nnnnnnnnnnnnnnnnnnnnnnnnnnnnnnnnnnnnnnnnnn [ 28000]
LR899193.1_Germany_2020/1        ------------------------------------------...................................... [ 28000]
MN393477.1_Wuhan_2019            ---------------------------------------.........-............................... [ 28000]
MW396979.1_Timor-Leste/2019/1    ---------------------------------------.........-............................... [ 28000]
MK940252.1_CN/2019/InnerMongolia ---------------------------------------......................................... [ 28000]
MK128995.1_China/2018/AnhuiXCGQ  ---------------------------------------......................................... [ 28000]
MH681419.1_ASFV/POL/2015         -------------------------------------------..................................... [ 28000]
LS478113.1_Estonia_2014          -----------------------------------------....................................... [ 28000]
KP843857.1_Odintsovo_02/14       -------------------------------------........................................... [ 28000]
```

```
FN557520.1_strain_E75            ----------------------------------------..............cc.c..........t...a..g...c [ 28000]
AY261366.1_Warthog               --------------------------------------------..................ta...at...a..g.... [ 28000]
AY261365.1_Warmbaths             --------------------------------------------.......a................t........... [ 28000]
AY261364.1_Tengani_62            ------------------------------------g....................c..........t...a......c [ 28000]
AY261363.1_Pretoriuskop/96/4     tttaaatttcatgatgtttagttcattacttcctacaattact...........c.......ta...at...a..g.... [ 28000]
AY261362.1_Mkuzi_1979            --------------------------------------------........................t...a..g...c [ 28000]
AY261361.1_Malawi_Lil-20/1_1983  -----------------------------------------......................a...at...a..g.... [ 28000]
AY261360.1_Kenya_1950            --------------------------------------------.................a.....at.c......... [ 28000]
AM712240.1_OURT_88/3             --------------------------------------------..........cc.c..........t...a..g...c [ 28000]
AM712239.1_Benin_97/1            ---------------------------------------------.........cc.c..........t...a..g...c [ 28000]
KM111295.1_Ken06.Bus             -------------------------------------------..............c.........at.c......... [ 28000]
KM111294.1_Ken05/Tk1             -------------------------------------------...........cc.c..........t....c...... [ 28000]
KM262844.1_strain_L60            ---------------------------------------...............cc.c..........t...a..g...c [ 28000]
KM102979.1_26544/OG10_Italy      ----------------------------------------------........cc.c..........t...a..g...c [ 28000]
KM262845.1_strain_NHV            ------------------------------------------............cc.c..........t...a..g...c [ 28000]
KX354450.1_47/Ss/2008            ---------------------------------------------.........cc.c..........t...a..g...c [ 28000]
KP055815.1_strain_BA71           -------------------------------------------------------------------------------- [ 28000]
MH025920.1_strain_R35            ------------------------------------------...............c.........at.c......... [ 28000]
MH025919.1_strain_N10            -------------------------------------------..............c.........at.c......... [ 28000]
MH025918.1_strain_R25            ------------------------------------------...............c.........at.c......... [ 28000]
MH025917.1_strain_R7             ------------------------------------------...............c.........at.c......... [ 28000]
MH025916.1_strain_R8             -------------------------------------------..............c.........at.c......... [ 28000]

FR682468.2_Georgia_2007/1        ttgctgactattatgtagaattcatagaagcacgtttagatagtaata---tcactgcagtgtagattatgaaatacata [ 28080]
ASF_IND_20_CAD543                nnnnnnnnnnnnnnnnnnnnnnnnnnnnnnnnnnnnnnnnnnnnnc..---atg..a..t....a.....r......... [ 28080]
LR899193.1_Germany_2020/1        ................................................---............................. [ 28080]
MN393477.1_Wuhan_2019            ................................................---............................. [ 28080]
MW396979.1_Timor-Leste/2019/1    ................................................---............................. [ 28080]
MK940252.1_CN/2019/InnerMongolia ................................................---............................. [ 28080]
MK128995.1_China/2018/AnhuiXCGQ  ................................................---............................. [ 28080]
MH681419.1_ASFV/POL/2015         ................................................---............................. [ 28080]
LS478113.1_Estonia_2014          ................................................---............................. [ 28080]
KP843857.1_Odintsovo_02/14       ................................................---............................. [ 28080]
FN557520.1_strain_E75            ..a.............................acaa..t..gt..c..---atg..a..t....a.....a......... [ 28080]
AY261366.1_Warthog               ................................................---.t.......................t... [ 28080]
AY261365.1_Warmbaths             ................................gtaa..a.a.a..g..---cttg.c..a.................... [ 28080]
AY261364.1_Tengani_62            ................................aaa...t..ta..c..---atg..a..a....a.....a..----... [ 28080]
AY261363.1_Pretoriuskop/96/4     ................................aca..tt..ca..t..---atg..a..t.................... [ 28080]
AY261362.1_Mkuzi_1979            ..a.............................acaa..t..gt..c..---atg..a..t....a.....a......... [ 28080]
AY261361.1_Malawi_Lil-20/1_1983  ..a.........g...................gtaa.gt..ccatt..---at.gaa..a.................t.. [ 28080]
AY261360.1_Kenya_1950            .....a................t.........ga.a..tt..aat...ctaatg.ca..t.......c.........t.. [ 28080]
AM712240.1_OURT_88/3             ..a.............................acaa..t..gt..c..---atg..a..t....a.....a......... [ 28080]
AM712239.1_Benin_97/1            ..a.............................acaa..t..gt..c..---atg..a..t....a.....a......... [ 28080]
KM111295.1_Ken06.Bus             .....a................t.........gaaa.ga...a..g..----------gt....tg..g.a...------ [ 28080]
```

```
KM111294.1_Ken05/Tk1            .....a.t........................gaa..gt...a.tt..---at...a..t..............----.. [ 28080]
KM262844.1_strain_L60           ..a.............................acaa..t..gt..c..---atg..a..t....a.....a......... [ 28080]
KM102979.1_26544/OG10_Italy     ..a.............................acaa..t..gt..c..---atg..a..t....a.....a......... [ 28080]
KM262845.1_strain_NHV           ..a.............................acaa..t..gt..c..---atg..a..t....a.....a......... [ 28080]
KX354450.1_47/Ss/2008           ..a.............................acaa..t..gt..c..---atg..a..t....a.....a......... [ 28080]
KP055815.1_strain_BA71          -------------------------------------------------------------------------------- [ 28080]
MH025920.1_strain_R35           .....a................t.........gaaa.ga...a..g..----------gt....tg..g.a...------ [ 28080]
MH025919.1_strain_N10           .....a................t.........gaaa.ga...a..g..----------gt....tg..g.a...------ [ 28080]
MH025918.1_strain_R25           .....a................t.........gaaa.ga...a..g..----------gt....tg..g.a...------ [ 28080]
MH025917.1_strain_R7            .....a................t.........gaaa.ga...a..g..----------gt....tg..g.a...------ [ 28080]
MH025916.1_strain_R8            .....a................t.........gaaa.ga...a..g..----------gt....tg..g.a...------ [ 28080]

FR682468.2_Georgia_2007/1       ctaaactaatttcagtata--ttttttttgttcatataagttaaggtacaaaaatgat-taaacattgcaaaaaaagaa- [ 28160]
ASF_IND_20_CAD543               ..................---.....................................-......y.............- [ 28160]
LR899193.1_Germany_2020/1       ...................--.....................................-....................- [ 28160]
MN393477.1_Wuhan_2019           ...................--.....................................-....................- [ 28160]
MW396979.1_Timor-Leste/2019/1   ...................--.....................................-....................- [ 28160]
MK940252.1_CN/2019/InnerMongolia ...................--.....................................-....................- [ 28160]
MK128995.1_China/2018/AnhuiXCGQ ...................--.....................................-....................- [ 28160]
MH681419.1_ASFV/POL/2015        ...................--.....................................-....................- [ 28160]
LS478113.1_Estonia_2014         ...................--.....................................-....................- [ 28160]
KP843857.1_Odintsovo_02/14      ...................--.....................................-....................- [ 28160]
FN557520.1_strain_E75           ..................---.....................................-......c.............- [ 28160]
AY261366.1_Warthog              .-----.............--.................ga..................-....................a [ 28160]
AY261365.1_Warmbaths            .....t.............--.....................................-....................a [ 28160]
AY261364.1_Tengani_62           ...................-t.....................................-.................a.g- [ 28160]
AY261363.1_Pretoriuskop/96/4    ...................-t........................a............-....................a [ 28160]
AY261362.1_Mkuzi_1979           ...................--.....................................-......c.............- [ 28160]
AY261361.1_Malawi_Lil-20/1_1983 tg.................tt.....................................-.................a.ga [ 28160]
AY261360.1_Kenya_1950           ...................-t...........ac....c..c..at............t....a...............a [ 28160]
AM712240.1_OURT_88/3            ..................---.....................................-......c.............- [ 28160]
AM712239.1_Benin_97/1           ..................---.....................................-......c.............- [ 28160]
KM111295.1_Ken06.Bus            -.c..a.............-t...........a........----.............t....a.........----..a [ 28160]
KM111294.1_Ken05/Tk1            ...................-t...........a......c....a.............t....a....t.....---..a [ 28160]
KM262844.1_strain_L60           ..................---.....................................-......c.............- [ 28160]
KM102979.1_26544/OG10_Italy     ..................---.....................................-......c.............- [ 28160]
KM262845.1_strain_NHV           ..................---.....................................-......c.............- [ 28160]
KX354450.1_47/Ss/2008           ..................---.....................................-......c.............- [ 28160]
KP055815.1_strain_BA71          ----------g..gac..---.....c----------------------------------..t.g.ttgct.gg.a..- [ 28160]
MH025920.1_strain_R35           -.c..a.............--...........a........----.............t....a.........----..a [ 28160]
MH025919.1_strain_N10           -.c..a.............--...........a........----.............t....a.........----..a [ 28160]
MH025918.1_strain_R25           -.c..a.............--...........a........----.............t....a.........----..a [ 28160]
MH025917.1_strain_R7            -.c..a.............--...........a........----.............t....a.........----..a [ 28160]
MH025916.1_strain_R8            -.c..a.............--...........a........----.............t....a.........----..a [ 28160]
```

```
FR682468.2_Georgia_2007/1         ---aatcacaatgctatta---tacatagtgatcatagtggcttgtatcatttctaaactagttccaa---atgaatatt [ 28240]
ASF_IND_20_CAD543                 ---.....y..........---..............................................---......... [ 28240]
LR899193.1_Germany_2020/1         ---................---..............................................---......... [ 28240]
MN393477.1_Wuhan_2019             ---................---..............................................---......... [ 28240]
MW396979.1_Timor-Leste/2019/1     ---................---..............................................---......... [ 28240]
MK940252.1_CN/2019/InnerMongolia  ---................---..............................................---......... [ 28240]
MK128995.1_China/2018/AnhuiXCGQ   ---................---..............................................---......... [ 28240]
MH681419.1_ASFV/POL/2015          ---................---..............................................---......... [ 28240]
LS478113.1_Estonia_2014           ---................---..............................................---......... [ 28240]
KP843857.1_Odintsovo_02/14        ---................---..............................................---......... [ 28240]
FN557520.1_strain_E75             ---.....t..........---..............................................---......... [ 28240]
AY261366.1_Warthog                --g.....t..........---..............................................---......... [ 28240]
AY261365.1_Warmbaths              --g.....t..........---..............................................---......... [ 28240]
AY261364.1_Tengani_62             ---.....t..........---..............................................---......... [ 28240]
AY261363.1_Pretoriuskop/96/4      --g.....t..........---....................a..........t..t.t.........---......... [ 28240]
AY261362.1_Mkuzi_1979             ---.....t..........---..............................................---......... [ 28240]
AY261361.1_Malawi_Lil-20/1_1983   aag.....t..........---..............................................---......... [ 28240]
AY261360.1_Kenya_1950             --g................tta....ct.....g...t..a.a..........a......g......gata..a...... [ 28240]
AM712240.1_OURT_88/3              ---.....t..........---..............................................---......... [ 28240]
AM712239.1_Benin_97/1             ---.....t..........---..............................................---......... [ 28240]
KM111295.1_Ken06.Bus              --g................tta....ct..a..g...t..a.a..........a......g......gata..a...... [ 28240]
KM111294.1_Ken05/Tk1              --g.....t..........aca..t.ct.........t.aa.a......t...a........c....g---..a...... [ 28240]
KM262844.1_strain_L60             ---.....t..........---..............................................---......... [ 28240]
KM102979.1_26544/OG10_Italy       ---.....t..........---..............................................---......... [ 28240]
KM262845.1_strain_NHV             ---.....t..........---..............................................---......... [ 28240]
KX354450.1_47/Ss/2008             ---.....t..........---..............................................---......... [ 28240]
KP055815.1_strain_BA71            ---...ttt--------------------.a...t..ta.at.cac.c..c...-------------------------- [ 28240]
MH025920.1_strain_R35             --g................tta....ct..a..g...t..a.a..........a......g......gata..a...... [ 28240]
MH025919.1_strain_N10             --g................tta....ct..a..g...t..a.a..........a......g......gata..a.....g [ 28240]
MH025918.1_strain_R25             --g................tta....ct..a..g...t..a.a..........a......g......gata..a...... [ 28240]
MH025917.1_strain_R7              --g................tta....ct..a..g...t..a.a..........a......g......gata..a...... [ 28240]
MH025916.1_strain_R8              --g................tta....ct..a..g...t..a.a..........a......g......gata..a...... [ 28240]

FR682468.2_Georgia_2007/1         gggcaatacatctattttttatcattatgatttttatggtatatatgtatgaaaagttagatatacatcaaaaatctcag [ 28320]
ASF_IND_20_CAD543                 ...................y............................................................ [ 28320]
LR899193.1_Germany_2020/1         ................................................................................ [ 28320]
MN393477.1_Wuhan_2019             ................................................................................ [ 28320]
MW396979.1_Timor-Leste/2019/1     ................................................................................ [ 28320]
MK940252.1_CN/2019/InnerMongolia  ................................................................................ [ 28320]
MK128995.1_China/2018/AnhuiXCGQ   ................................................................................ [ 28320]
MH681419.1_ASFV/POL/2015          ................................................................................ [ 28320]
LS478113.1_Estonia_2014           ................................................................................ [ 28320]
KP843857.1_Odintsovo_02/14        ................................................................................ [ 28320]
```

```
FN557520.1_strain_E75            ...................c.......................................................a.... [ 28320]
AY261366.1_Warthog               ...................c.......................................................a.... [ 28320]
AY261365.1_Warmbaths             ...................c.............c.........c...........a......g.g..........a.... [ 28320]
AY261364.1_Tengani_62            ...................c......................................................aa.... [ 28320]
AY261363.1_Pretoriuskop/96/4     ...................c...................................a......g.g..........a.... [ 28320]
AY261362.1_Mkuzi_1979            ...................c.......................................................a.... [ 28320]
AY261361.1_Malawi_Lil-20/1_1983  ...................c..............................a........................a.... [ 28320]
AY261360.1_Kenya_1950            ..c........a.g.....ct.t..........ac..a..............................g..........a [ 28320]
AM712240.1_OURT_88/3             ...................c.......................................................a.... [ 28320]
AM712239.1_Benin_97/1            ...................c.......................................................a.... [ 28320]
KM111295.1_Ken06.Bus             ..c........a.g.....ct.t.....ag...ac..a..............................g..........a [ 28320]
KM111294.1_Ken05/Tk1             ..c.....t..a.g.....c..t..........ac..a...c..........................g..........a [ 28320]
KM262844.1_strain_L60            ...................c.......................................................a.... [ 28320]
KM102979.1_26544/OG10_Italy      ...................c.......................................................a.... [ 28320]
KM262845.1_strain_NHV            ...................c.......................................................a.... [ 28320]
KX354450.1_47/Ss/2008            ...................c.......................................................a.... [ 28320]
KP055815.1_strain_BA71           -----------------------------...c.c...c.-------..g.....aa..a.------------------- [ 28320]
MH025920.1_strain_R35            ..c........a.g.....ct.t.....ag...ac..a..............................g..........a [ 28320]
MH025919.1_strain_N10            ..c........a.g.....ct.t.....ag...ac..a..............................g..........a [ 28320]
MH025918.1_strain_R25            ..c........a.g.....ct.t.....ag...ac..a..............................g..........a [ 28320]
MH025917.1_strain_R7             ..c........a.g.....ct.t.....ag...ac..a..............................g..........a [ 28320]
MH025916.1_strain_R8             ..c........a.g.....ct.t.....ag...ac..a..............................g..........a [ 28320]

FR682468.2_Georgia_2007/1        ttctggaattataccatgtcaggcttatctggacataacgtacagg---taacatgtaagtgttact-----------aa [ 28400]
ASF_IND_20_CAD543                ..............................................---..................-----------.. [ 28400]
LR899193.1_Germany_2020/1        ..............................................---..................-----------.. [ 28400]
MN393477.1_Wuhan_2019            ..............................................---..................-----------.. [ 28400]
MW396979.1_Timor-Leste/2019/1    ..............................................---..................-----------.. [ 28400]
MK940252.1_CN/2019/InnerMongolia ..............................................---..................-----------.. [ 28400]
MK128995.1_China/2018/AnhuiXCGQ  ..............................................---..................-----------.. [ 28400]
MH681419.1_ASFV/POL/2015         ..............................................---..................-----------.. [ 28400]
LS478113.1_Estonia_2014          ..............................................---..................-----------.. [ 28400]
KP843857.1_Odintsovo_02/14       ..............................................---..................-----------.. [ 28400]
FN557520.1_strain_E75            ..............g...............................---...t..............-----------.. [ 28400]
AY261366.1_Warthog               ..............g..............................a---..................-----------.. [ 28400]
AY261365.1_Warmbaths             ..............t..............................a---..................-----------.. [ 28400]
AY261364.1_Tengani_62            ..............t..............................a---..................-----------.. [ 28400]
AY261363.1_Pretoriuskop/96/4     ..............t..........................t....---..tt..............----------a.. [ 28400]
AY261362.1_Mkuzi_1979            ..............g..............................a---..................-----------.. [ 28400]
AY261361.1_Malawi_Lil-20/1_1983  ..............t...............................---..................----------a.. [ 28400]
AY261360.1_Kenya_1950            ..............a...g..c.............ccc.....c.acaa...t......t.....a.ttttacaaaaa.. [ 28400]
AM712240.1_OURT_88/3             ..............g...............................---...t..............-----------.. [ 28400]
AM712239.1_Benin_97/1            ..............g...............................---...t..............-----------.. [ 28400]
KM111295.1_Ken06.Bus             .........c....a...g..c...........tgccc.....c.acaa...t......t.....a.tttttacaaaa.. [ 28400]
```

```
KM111294.1_Ken05/Tk1             .........g....g...g..c............gccc.....c..cat...tg..........tt.act------aa.. [ 28400]
KM262844.1_strain_L60            ..............g...............................---...t..............-----------.. [ 28400]
KM102979.1_26544/OG10_Italy      ..............g...............................---...t..............-----------.. [ 28400]
KM262845.1_strain_NHV            ..............g...............................---...t..............-----------.. [ 28400]
KX354450.1_47/Ss/2008            ..............g...............................---...t..............-----------.. [ 28400]
KP055815.1_strain_BA71           ---cca...ct...a....-------.......tc...t.aga.ac---..ga..tc.tc.t..ttc-----------.. [ 28400]
MH025920.1_strain_R35            .........c....a...g..c...........tgccc.....c.acaa...t......t.....a.tttttacaaaa.. [ 28400]
MH025919.1_strain_N10            .........c....a...g..c...........tgccc.....c.acaa...t......t.....a.tttttacaaaa.. [ 28400]
MH025918.1_strain_R25            .........c....a...g..c...........tgccc.....c.acaa...t......t.....a.tttttacaaaa.. [ 28400]
MH025917.1_strain_R7             .........c....a...g..c...........tgccc.....c.acaa...t......t.....a.tttttacaaaa.. [ 28400]
MH025916.1_strain_R8             .........c....a...g..c...........tgccc.....c.acaa...t......t.....a.tttttacaaaa.. [ 28400]

FR682468.2_Georgia_2007/1        atac--tatgaagtatctatttttttttgttgtaaaaaaaagaacttgatagtattttttaaaaa-----ataaaataa- [ 28480]
ASF_IND_20_CAD543                ....--.....................-----------------------------------.c.-----.c..------ [ 28480]
LR899193.1_Germany_2020/1        ....--...........................................................-----.........- [ 28480]
MN393477.1_Wuhan_2019            ....--...........................................................-----.........- [ 28480]
MW396979.1_Timor-Leste/2019/1    ....--...........................................................-----.........- [ 28480]
MK940252.1_CN/2019/InnerMongolia ....--...........................................................-----.........- [ 28480]
MK128995.1_China/2018/AnhuiXCGQ  ....--...........................................................-----.........- [ 28480]
MH681419.1_ASFV/POL/2015         ....--...........................................................-----.........- [ 28480]
LS478113.1_Estonia_2014          ....--...........................................................-----.........- [ 28480]
KP843857.1_Odintsovo_02/14       ....--.............-.............................................-----.........- [ 28480]
FN557520.1_strain_E75            ....--.....................acaac------------------------------------------------ [ 28480]
AY261366.1_Warthog               ....--.....................gt.gt-..................-.............-----.........- [ 28480]
AY261365.1_Warmbaths             ....--.....................gt.gta..................-.............ataac....g.c..- [ 28480]
AY261364.1_Tengani_62            ....--.....g-..............gt.gt--.................-.......c.....-----.........- [ 28480]
AY261363.1_Pretoriuskop/96/4     .c..--......----------.....at.gt--.............---.-........t....-----....g.c..- [ 28480]
AY261362.1_Mkuzi_1979            ....--.....................gt.gta..................-.............-----......c..g [ 28480]
AY261361.1_Malawi_Lil-20/1_1983  ....--....t.....................-t.......ac......................-----.........- [ 28480]
AY261360.1_Kenya_1950            .c..aa........g...-........ga.gt-----....t....c.g..-c..........-------....gg.--- [ 28480]
AM712240.1_OURT_88/3             ....--.....................acaac------------------------------------------------ [ 28480]
AM712239.1_Benin_97/1            ....--.....................acaac------------------------------------------------ [ 28480]
KM111295.1_Ken06.Bus             .c..aa........c.-.-........ga.gt-----..g--....c....-c.........----------.c.g.--- [ 28480]
KM111294.1_Ken05/Tk1             ....aa..........----.......ga.gt-----....-....ca...-cc.....a..---------....g.--- [ 28480]
KM262844.1_strain_L60            ....--.....................acaac------------------------------------------------ [ 28480]
KM102979.1_26544/OG10_Italy      ....--.....................acaac------------------------------------------------ [ 28480]
KM262845.1_strain_NHV            ....--.....................acaac------------------------------------------------ [ 28480]
KX354450.1_47/Ss/2008            ....--.....................acaac------------------------------------------------ [ 28480]
KP055815.1_strain_BA71           c.c.tt.c..g.a.g.t...cc.....cca-------------------------------------------------- [ 28480]
MH025920.1_strain_R35            .c..aa........c--.-........ga.gt-----..g--....c....-c.........----------.c.g.--- [ 28480]
MH025919.1_strain_N10            .c..aa........c.-.-........ga.gt-----..g--....c....-c.........----------.c.g.--- [ 28480]
MH025918.1_strain_R25            .c..aa........c--.-........ga.gt-----..g--....c....-c.........----------.c.g.--- [ 28480]
MH025917.1_strain_R7             .c..aa........c--.-........ga.gt-----..g--....c....-c.........----------.c.g.--- [ 28480]
MH025916.1_strain_R8             .c..aa........c--.-........ga.gt-----..g--....c....-c.........----------.c.g.--- [ 28480]
```

```
FR682468.2_Georgia_2007/1        ----ttaattgtacgtcaacttccttattttattcttt--aaaaa--taactcgtaagtattatttatctatttt-ttga [ 28560]
ASF_IND_20_CAD543                -------------------------------------------------------------------------------- [ 28560]
LR899193.1_Germany_2020/1        ----..................................--.....--............................-.... [ 28560]
MN393477.1_Wuhan_2019            ----..................................--.....--............................-.... [ 28560]
MW396979.1_Timor-Leste/2019/1    ----..................................--.....--............................-.... [ 28560]
MK940252.1_CN/2019/InnerMongolia ----..................................--.....--............................-.... [ 28560]
MK128995.1_China/2018/AnhuiXCGQ  ----..................................--.....--............................-.... [ 28560]
MH681419.1_ASFV/POL/2015         ----..................................--.....--............................-.... [ 28560]
LS478113.1_Estonia_2014          ----..................................--.....--............................-.... [ 28560]
KP843857.1_Odintsovo_02/14       ----..................................--.....--............................-.... [ 28560]
FN557520.1_strain_E75            -----------------------------------------------------------------------------aa. [ 28560]
AY261366.1_Warthog               ----..g.............a.................--.....tg...t...c.........c...t.-..cc-aaa. [ 28560]
AY261365.1_Warmbaths             ----..g.............a.......c.....t...-a.....tg.......c.........c...t.-..cc---a. [ 28560]
AY261364.1_Tengani_62            ------g...............................--.....tg.......c.........c...t.-..cc--aa. [ 28560]
AY261363.1_Pretoriuskop/96/4     ----..g.c...........a.................--.....tg..g....c.........c...t.-..cc-aaa. [ 28560]
AY261362.1_Mkuzi_1979            ataa..g...............................--.....tg...t...c.........c...t.-..cc-aaa. [ 28560]
AY261361.1_Malawi_Lil-20/1_1983  ----..g...............................--.....tg....a..c.........c...t.-..----..g [ 28560]
AY261360.1_Kenya_1950            ---t....cgtacaac...t.gt...........a...aa.....tg......ac.......c.c..ct.-..cc-aaa. [ 28560]
AM712240.1_OURT_88/3             -----------------------------------------------------------------------------aa. [ 28560]
AM712239.1_Benin_97/1            -----------------------------------------------------------------------------aa. [ 28560]
KM111295.1_Ken06.Bus             ---t..g..gtacaact..t.gt...........a...aa.....tg......ac.......c.c..ct.-...c-aaa. [ 28560]
KM111294.1_Ken05/Tk1             ---t..g.c------------gt...........a...aa.....tg......ac.......c.c..ct.-..cc-aaa. [ 28560]
KM262844.1_strain_L60            -----------------------------------------------------------------------------aa. [ 28560]
KM102979.1_26544/OG10_Italy      -----------------------------------------------------------------------------aa. [ 28560]
KM262845.1_strain_NHV            -----------------------------------------------------------------------------aa. [ 28560]
KX354450.1_47/Ss/2008            -----------------------------------------------------------------------------aa. [ 28560]
KP055815.1_strain_BA71           -------------------------------------------------------------------------------- [ 28560]
MH025920.1_strain_R35            ---t..g..gtacaact..t.gt...........a...aa.....tg......ac.......c.c..ct.-...caaaa. [ 28560]
MH025919.1_strain_N10            ---t..g..gtacaact..t.gt...........a...aa.....tg......ac.......c.c..ct.-...caaaa. [ 28560]
MH025918.1_strain_R25            ---t..g..gtacaact..t.gt...........a...aa.....tg......ac.......c.c..ct.-...caaaa. [ 28560]
MH025917.1_strain_R7             ---t..g..gtacaact..t.gt...........a...aa.....tg......ac.......c.c..ct.-...caaaa. [ 28560]
MH025916.1_strain_R8             ---t..g..gtacaact..t.gt...........a...aa.....tg......ac.......c.c..ct.-...caaaa. [ 28560]

FR682468.2_Georgia_2007/1        aaaaatagatgtaatcggtttcatcatttagg-tgtgtatttctttt--tagcatctatc-aagaattcattgtttagtg [ 28640]
ASF_IND_20_CAD543                ................................-..........--..--...........t................... [ 28640]
LR899193.1_Germany_2020/1        ................................-..............--...........-................... [ 28640]
MN393477.1_Wuhan_2019            ................................-..............--...........-................... [ 28640]
MW396979.1_Timor-Leste/2019/1    ................................-..............--...........-................... [ 28640]
MK940252.1_CN/2019/InnerMongolia ................................-..............--...........-................... [ 28640]
MK128995.1_China/2018/AnhuiXCGQ  ................................-..............--...........-................... [ 28640]
MH681419.1_ASFV/POL/2015         ................................-..............--...........-................... [ 28640]
LS478113.1_Estonia_2014          ................................-..............--...........-................... [ 28640]
KP843857.1_Odintsovo_02/14       ................................-..............--...........-................... [ 28640]
```

```
FN557520.1_strain_E75            .....................t..........-..........--..--........t..-.................c. [ 28640]
AY261366.1_Warthog               ................................-..............--a..t.......-................... [ 28640]
AY261365.1_Warmbaths             .....................t..........-.............----..........-................... [ 28640]
AY261364.1_Tengani_62            ..........a.....a...............-..............--a..t.......-................... [ 28640]
AY261363.1_Pretoriuskop/96/4     ................aa..............-..c...........-t...........-.........--........ [ 28640]
AY261362.1_Mkuzi_1979            ........................t.......-.........t--..--...........-................... [ 28640]
AY261361.1_Malawi_Lil-20/1_1983  ..............................------------------------------------------.....t.. [ 28640]
AY261360.1_Kenya_1950            .......a........a..c.t..........ac.ca..........tt..........g---att..gg.....a.a.a [ 28640]
AM712240.1_OURT_88/3             .....................t..........-..........--..--........t..-.................c. [ 28640]
AM712239.1_Benin_97/1            .....................t..........-..........--..--........t..-.................c. [ 28640]
KM111295.1_Ken06.Bus             ................a..c.tg.........ac.ca..........tt..........a---att..ag..c..a.a.a [ 28640]
KM111294.1_Ken05/Tk1             .......a........a..c.t..........acaca..........tt..........g---att..ag.....a.a.a [ 28640]
KM262844.1_strain_L60            .....................t..........-..........--..--........t..-.................c. [ 28640]
KM102979.1_26544/OG10_Italy      .....................t..........-..........--..--........t..-.................c. [ 28640]
KM262845.1_strain_NHV            .....................t..........-..........--..--........t..-.................c. [ 28640]
KX354450.1_47/Ss/2008            .....................t..........-..........--..--........t..-.................c. [ 28640]
KP055815.1_strain_BA71           ------------------------------------c.cc...--c.--.gca..t.tat-...g....cag.gc.--.t [ 28640]
MH025920.1_strain_R35            ................a..c.tg.........ac.ca..........tt..........a---att..ag..c..a.a.a [ 28640]
MH025919.1_strain_N10            ................a..c.tg.........ac.ca..........tt..........a---att..ag..c..a.a.a [ 28640]
MH025918.1_strain_R25            ................a..c.tg.........ac.ca..........tt..........a---att..ag..c..a.a.a [ 28640]
MH025917.1_strain_R7             ................a..c.tg.........ac.ca..........tt..........a---att..ag..c..a.a.a [ 28640]
MH025916.1_strain_R8             ................a..c.tg.........ac.ca..........tt..........a---att..ag..c..a.a.a [ 28640]

FR682468.2_Georgia_2007/1        atatgaaa-------acaatgaatgat-------cattatcttctattt---aacaaccacctaaataaatgaacgtctt [ 28720]
ASF_IND_20_CAD543                ........-------............-------...............---.....y.w...............r.... [ 28720]
LR899193.1_Germany_2020/1        ........-------............-------...............---............................ [ 28720]
MN393477.1_Wuhan_2019            ........-------............-------...............---............................ [ 28720]
MW396979.1_Timor-Leste/2019/1    ........-------............-------...............---............................ [ 28720]
MK940252.1_CN/2019/InnerMongolia ........-------............-------...............---............................ [ 28720]
MK128995.1_China/2018/AnhuiXCGQ  ........-------............-------...............---............................ [ 28720]
MH681419.1_ASFV/POL/2015         ........-------............-------...............---............................ [ 28720]
LS478113.1_Estonia_2014          ........-------............-------...............---............................ [ 28720]
KP843857.1_Odintsovo_02/14       ........-------............-------...............---............................ [ 28720]
FN557520.1_strain_E75            g...a...-------............-------...c..a........---.....t.t...............a.... [ 28720]
AY261366.1_Warthog               ....a...-------..........t.-------...c..a..t.....---.......................a.... [ 28720]
AY261365.1_Warmbaths             g...a...-------..........t.-------...c..a........---.....t.................a.... [ 28720]
AY261364.1_Tengani_62            ....a...-------..........t.-------...c..a..t.....---....t..................a--.. [ 28720]
AY261363.1_Pretoriuskop/96/4     g.......-------............-------...c..a........---.......................a.... [ 28720]
AY261362.1_Mkuzi_1979            g...a...-------............-------...--.a........---.....t.t...............a.... [ 28720]
AY261361.1_Malawi_Lil-20/1_1983  ....a...-------.....a......-------...c..a.....c..---.....t............---------- [ 28720]
AY261360.1_Kenya_1950            ga..atg.tg-----c..c.a..ca..gaatgat...c..a........---.....t.................a.... [ 28720]
AM712240.1_OURT_88/3             g...a...-------............-------...c..a........---.....t.t...............a.... [ 28720]
AM712239.1_Benin_97/1            g...a...-------............-------...c..a........---.....t.t...............a.... [ 28720]
KM111295.1_Ken06.Bus             ga..atg.tg-----c..c.a..ca..g-------...c.ac.......aac.......................----- [ 28720]
```

```
KM111294.1_Ken05/Tk1             ga..atg.tgatcatt..t.att.ag.g-----g...c..a........---.....t.................a.... [ 28720]
KM262844.1_strain_L60            g...a...-------............-------...c..a........---.....t.t...............a.... [ 28720]
KM102979.1_26544/OG10_Italy      g...a...-------............-------...c..a........---.....t.t...............a.... [ 28720]
KM262845.1_strain_NHV            g...a...-------............-------...c..a........---.....t.t...............a.... [ 28720]
KX354450.1_47/Ss/2008            g...a...-------............-------...c..a........---.....t.t...............a.... [ 28720]
KP055815.1_strain_BA71           ggg.c.g.-------...g.tc...c.a----tg...c..a........---.....t.t...............a.... [ 28720]
MH025920.1_strain_R35            ga..atg.tg-----c..c.a..ca..g-------...c.ac.......aac.......................----- [ 28720]
MH025919.1_strain_N10            ga..atg.tg-----c..c.a..ca..g-------...c.ac.......aac.......................----- [ 28720]
MH025918.1_strain_R25            ga..atg.tg-----c..c.a..ca..g-------...c.ac.......aac.......................----- [ 28720]
MH025917.1_strain_R7             ga..atg.tg-----c..c.a..ca..g-------...c.ac.......aac.......................----- [ 28720]
MH025916.1_strain_R8             ga..atg.tg-----c..c.a..ca..g-------...c.ac.......aac.......................----- [ 28720]

FR682468.2_Georgia_2007/1        tttcatcttaac--tgattaccaaaagttattttgcgaaaaggcatacatatgatcaatatcagacctacaatgaatatt [ 28800]
ASF_IND_20_CAD543                ......y.....-a....y..........................................................r.. [ 28800]
LR899193.1_Germany_2020/1        ............--.................................................................. [ 28800]
MN393477.1_Wuhan_2019            ............--.................................................................. [ 28800]
MW396979.1_Timor-Leste/2019/1    ............--.................................................................. [ 28800]
MK940252.1_CN/2019/InnerMongolia ............--.................................................................. [ 28800]
MK128995.1_China/2018/AnhuiXCGQ  ............--.................................................................. [ 28800]
MH681419.1_ASFV/POL/2015         ............--.................................................................. [ 28800]
LS478113.1_Estonia_2014          ............--.................................................................. [ 28800]
KP843857.1_Odintsovo_02/14       ............--.................................................................. [ 28800]
FN557520.1_strain_E75            ......t.....-a....c.....t...c................................................g.. [ 28800]
AY261366.1_Warthog               ............ta.a..c.....t...c...............................................ag.. [ 28800]
AY261365.1_Warmbaths             ............--....c.....t...c...............................................ag.. [ 28800]
AY261364.1_Tengani_62            ............--....c.....................................g....................g.. [ 28800]
AY261363.1_Pretoriuskop/96/4     ............--....c.........................................................ag.. [ 28800]
AY261362.1_Mkuzi_1979            ...t........-a....c.....t...c................................................g.. [ 28800]
AY261361.1_Malawi_Lil-20/1_1983  ---.........-a....c................t................................g........g.. [ 28800]
AY261360.1_Kenya_1950            ...t........--....c...................................g......................g.. [ 28800]
AM712240.1_OURT_88/3             ......t.....-a....c.....t...c................................................g.. [ 28800]
AM712239.1_Benin_97/1            ......t.....-a....c.....t...c................................................g.. [ 28800]
KM111295.1_Ken06.Bus             ----........--....c..........................................................g.. [ 28800]
KM111294.1_Ken05/Tk1             ...t........--....c..............................................t...........g.. [ 28800]
KM262844.1_strain_L60            ......t.....-a....c.....t...c................................................g.. [ 28800]
KM102979.1_26544/OG10_Italy      ......t.....-a....c.....t...c................................................g.. [ 28800]
KM262845.1_strain_NHV            ......t.....-a....c.....t...c................................................g.. [ 28800]
KX354450.1_47/Ss/2008            ......t.....-a....c.....t...c................................................g.. [ 28800]
KP055815.1_strain_BA71           ......t.....-a....c.....t...c................................................g.. [ 28800]
MH025920.1_strain_R35            ----........--....c..........................................................g.. [ 28800]
MH025919.1_strain_N10            ----........--....c..........................................................g.. [ 28800]
MH025918.1_strain_R25            ----........--....c..........................................................g.. [ 28800]
MH025917.1_strain_R7             ----........--....c..........................................................g.. [ 28800]
MH025916.1_strain_R8             ----........--....c..........................................................g.. [ 28800]
```

```
FR682468.2_Georgia_2007/1        tccataatatccctttattgtaataattctatttttgcattccgatatctcatcatctgtgctattatatgtttccataa [ 28880]
ASF_IND_20_CAD543                ................................................................................ [ 28880]
LR899193.1_Germany_2020/1        ................................................................................ [ 28880]
MN393477.1_Wuhan_2019            ................................................................................ [ 28880]
MW396979.1_Timor-Leste/2019/1    ................................................................................ [ 28880]
MK940252.1_CN/2019/InnerMongolia ................................................................................ [ 28880]
MK128995.1_China/2018/AnhuiXCGQ  ................................................................................ [ 28880]
MH681419.1_ASFV/POL/2015         ................................................................................ [ 28880]
LS478113.1_Estonia_2014          ................................................................................ [ 28880]
KP843857.1_Odintsovo_02/14       ................................................................................ [ 28880]
FN557520.1_strain_E75            ..................c.c........................................................... [ 28880]
AY261366.1_Warthog               .......c..........c.c........................................................... [ 28880]
AY261365.1_Warmbaths             .......c..........c.c.................................................a......... [ 28880]
AY261364.1_Tengani_62            ..................c.c........................................................... [ 28880]
AY261363.1_Pretoriuskop/96/4     .......c..........c.c........................................................... [ 28880]
AY261362.1_Mkuzi_1979            ..................c.c........................................................... [ 28880]
AY261361.1_Malawi_Lil-20/1_1983  .......c..........cac.......t..............a....t...........a...c.........g.g... [ 28880]
AY261360.1_Kenya_1950            ..................c.c.......t.............ta..........g.........c...........g... [ 28880]
AM712240.1_OURT_88/3             ..................c.c........................................................... [ 28880]
AM712239.1_Benin_97/1            ..................c.c........................................................... [ 28880]
KM111295.1_Ken06.Bus             .......c..........c.c.......t..............a....................c............... [ 28880]
KM111294.1_Ken05/Tk1             ..................c.c.......t..............a..gc......g.........c...........g... [ 28880]
KM262844.1_strain_L60            ..................c.c........................................................... [ 28880]
KM102979.1_26544/OG10_Italy      ..................c.c........................................................... [ 28880]
KM262845.1_strain_NHV            ..................c.c........................................................... [ 28880]
KX354450.1_47/Ss/2008            ..................c.c........................................................... [ 28880]
KP055815.1_strain_BA71           ..................c.c........................................................... [ 28880]
MH025920.1_strain_R35            .......c..........c.c.......t..............a....................c..........g.... [ 28880]
MH025919.1_strain_N10            .......c..........c.c.......t..............a....................c..........g.... [ 28880]
MH025918.1_strain_R25            .......c..........c.c.......t..............a....................c..........g.... [ 28880]
MH025917.1_strain_R7             .......c..........c.c.......t..............a....................c..........g.... [ 28880]
MH025916.1_strain_R8             .......c..........c.c.......t..............a....................c..........g.... [ 28880]

FR682468.2_Georgia_2007/1        ctgtttcatcatcaaacataaatcctgttaaataggcaaaagactttaatcccggatagatttttaccattttcctgaga [ 28960]
ASF_IND_20_CAD543                ........................................gcg.....................y........y...... [ 28960]
LR899193.1_Germany_2020/1        ................................................................................ [ 28960]
MN393477.1_Wuhan_2019            ................................................................................ [ 28960]
MW396979.1_Timor-Leste/2019/1    ................................................................................ [ 28960]
MK940252.1_CN/2019/InnerMongolia ................................................................................ [ 28960]
MK128995.1_China/2018/AnhuiXCGQ  ................................................................................ [ 28960]
MH681419.1_ASFV/POL/2015         ................................................................................ [ 28960]
LS478113.1_Estonia_2014          ................................................................................ [ 28960]
KP843857.1_Odintsovo_02/14       ................................................................................ [ 28960]
```

```
FN557520.1_strain_E75             ..........................a.......a.....gcg.....................c........t..a... [ 28960]
AY261366.1_Warthog                ................t..................a......g...............t..................... [ 28960]
AY261365.1_Warmbaths              ................t.........................gt..............t..................... [ 28960]
AY261364.1_Tengani_62             ................t.........................g...............t..............t..c... [ 28960]
AY261363.1_Pretoriuskop/96/4      ..c.......................................gt........t.....t..................... [ 28960]
AY261362.1_Mkuzi_1979             ................t.................a.....gcg...............t..................... [ 28960]
AY261361.1_Malawi_Lil-20/1_1983   ..........g...............a.......a.......g..................................... [ 28960]
AY261360.1_Kenya_1950             ..........................a...............gt....t.........t..................... [ 28960]
AM712240.1_OURT_88/3              ..........................a.......a.....gcg.....................c........t..a... [ 28960]
AM712239.1_Benin_97/1             ..........................a.......a.....gcg.....................c........t..a... [ 28960]
KM111295.1_Ken06.Bus              tg........................a.......a.......................t..............t...... [ 28960]
KM111294.1_Ken05/Tk1              ..........................a...............g.....t.........t..................... [ 28960]
KM262844.1_strain_L60             ..........................a.......a.....gcg.....................c........t..a... [ 28960]
KM102979.1_26544/OG10_Italy       ..........................a.......a.....gcg.....................c........t..a... [ 28960]
KM262845.1_strain_NHV             ..........................a.......a.....gcg.....................c........t..a... [ 28960]
KX354450.1_47/Ss/2008             ..........................a.......a.....gcg.....................c........t..a... [ 28960]
KP055815.1_strain_BA71            ..........................a.......a.....gcg.....................c........t..a... [ 28960]
MH025920.1_strain_R35             ta........................a.......a.......................t..............t...... [ 28960]
MH025919.1_strain_N10             ta........................a.......a.......................t..............t...... [ 28960]
MH025918.1_strain_R25             ta........................a.......a.......................t..............t...... [ 28960]
MH025917.1_strain_R7              ta........................a.......a.......................t..............t...... [ 28960]
MH025916.1_strain_R8              ta........................a.......a.......................t..............t...... [ 28960]

FR682468.2_Georgia_2007/1         gccgtgtatagcttgtaataaatggccaaaaatatgcaataaagcgtagaaagagagtaatttttggcataaaagatttt [ 29040]
ASF_IND_20_CAD543                 ............................r..............................................r.... [ 29040]
LR899193.1_Germany_2020/1         ................................................................................ [ 29040]
MN393477.1_Wuhan_2019             ................................................................................ [ 29040]
MW396979.1_Timor-Leste/2019/1     ................................................................................ [ 29040]
MK940252.1_CN/2019/InnerMongolia  ................................................................................ [ 29040]
MK128995.1_China/2018/AnhuiXCGQ   ................................................................................ [ 29040]
MH681419.1_ASFV/POL/2015          ................................................................................ [ 29040]
LS478113.1_Estonia_2014           ................................................................................ [ 29040]
KP843857.1_Odintsovo_02/14        ................................................................................ [ 29040]
FN557520.1_strain_E75             ...........t................g..............................................g.... [ 29040]
AY261366.1_Warthog                ............................g......a.......................................g.... [ 29040]
AY261365.1_Warmbaths              ...........................................................................g.... [ 29040]
AY261364.1_Tengani_62             ............................g......a............................................ [ 29040]
AY261363.1_Pretoriuskop/96/4      .........c..................g................................................... [ 29040]
AY261362.1_Mkuzi_1979             ...........................................................................g.... [ 29040]
AY261361.1_Malawi_Lil-20/1_1983   ...a.......t..a.............g......a.......................................g.... [ 29040]
AY261360.1_Kenya_1950             ........a...................g...............t..g................................ [ 29040]
AM712240.1_OURT_88/3              ............................g..............................................g.... [ 29040]
AM712239.1_Benin_97/1             ...........t................g..............................................g.... [ 29040]
KM111295.1_Ken06.Bus              ........a...................g..................g...........................g.... [ 29040]
```

```
KM111294.1_Ken05/Tk1           ..........................t.g...............t..............................g.... [ 29040]
KM262844.1_strain_L60          ............................g..............................................g.... [ 29040]
KM102979.1_26544/OG10_Italy    ...........t................g..............................................g.... [ 29040]
KM262845.1_strain_NHV          ............................g..............................................g.... [ 29040]
KX354450.1_47/Ss/2008          ...........t................g..............................................g.... [ 29040]
KP055815.1_strain_BA71         ............................g..............................................g.... [ 29040]
MH025920.1_strain_R35          ........a...................g..................g...........................g.... [ 29040]
MH025919.1_strain_N10          ........a...................g..................g...........................g.... [ 29040]
MH025918.1_strain_R25          ........a...................g..................g...........................g.... [ 29040]
MH025917.1_strain_R7           ........a...................g..................g...........................g.... [ 29040]
MH025916.1_strain_R8           ........a...................g..................g...........................g.... [ 29040]

FR682468.2_Georgia_2007/1      gaaggtttgatgaatggctaaatcgcatataatataagatacgattttaaagcgcacctgttcacgcagatttgttgaaa [ 29120]
ASF_IND_20_CAD543              ................................................................................ [ 29120]
LR899193.1_Germany_2020/1      ................................................................................ [ 29120]
MN393477.1_Wuhan_2019          ................................................................................ [ 29120]
MW396979.1_Timor-Leste/2019/1  ................................................................................ [ 29120]
MK940252.1_CN/2019/InnerMongolia ................................................................................ [ 29120]
MK128995.1_China/2018/AnhuiXCGQ ................................................................................ [ 29120]
MH681419.1_ASFV/POL/2015       ................................................................................ [ 29120]
LS478113.1_Estonia_2014        ................................................................................ [ 29120]
KP843857.1_Odintsovo_02/14     ................................................................................ [ 29120]
FN557520.1_strain_E75          ...........................g.................a.................................. [ 29120]
AY261366.1_Warthog             ...a..............c..........................a..........................g....... [ 29120]
AY261365.1_Warmbaths           ..................c.....a..g........t........a..................t.......a....... [ 29120]
AY261364.1_Tengani_62          ...............a...........g........t........a..........................g....... [ 29120]
AY261363.1_Pretoriuskop/96/4   ...........................g.................a..........................g....... [ 29120]
AY261362.1_Mkuzi_1979          ...........................g........t........a..........................a....... [ 29120]
AY261361.1_Malawi_Lil-20/1_1983 ...........................g.................a..............c...t............... [ 29120]
AY261360.1_Kenya_1950          ........................a..g.................a..............c.............c..... [ 29120]
AM712240.1_OURT_88/3           ...........................g.................a.................................. [ 29120]
AM712239.1_Benin_97/1          ...........................g.................a.................................. [ 29120]
KM111295.1_Ken06.Bus           ........................a..g........g........a..............c.............c..... [ 29120]
KM111294.1_Ken05/Tk1           ........................a..g..............a..a..............c...t............... [ 29120]
KM262844.1_strain_L60          ...........................g.................a.................................. [ 29120]
KM102979.1_26544/OG10_Italy    ...........................g.................a.................................. [ 29120]
KM262845.1_strain_NHV          ...........................g.................a.................................. [ 29120]
KX354450.1_47/Ss/2008          ...........................g.................a.................................. [ 29120]
KP055815.1_strain_BA71         ...........................g.................a.................................. [ 29120]
MH025920.1_strain_R35          ........................a..g........g........a..............c.............c..... [ 29120]
MH025919.1_strain_N10          ........................a..g........g........a..............c.............c..... [ 29120]
MH025918.1_strain_R25          ........................a..g........g........a..............c.............c..... [ 29120]
MH025917.1_strain_R7           ........................a..g........g........a..............c.............c..... [ 29120]
MH025916.1_strain_R8           ........................a..g........g........a..............c.............c..... [ 29120]
```

```
FR682468.2_Georgia_2007/1         aattcgtggaaagatttaacaaataaaaggttattaatagttgctcatcattccccttatacgacatcgtcagacgctct [ 29200]
ASF_IND_20_CAD543                 .......r........y............................................................... [ 29200]
LR899193.1_Germany_2020/1         ................................................................................ [ 29200]
MN393477.1_Wuhan_2019             ................................................................................ [ 29200]
MW396979.1_Timor-Leste/2019/1     ................................................................................ [ 29200]
MK940252.1_CN/2019/InnerMongolia  ................................................................................ [ 29200]
MK128995.1_China/2018/AnhuiXCGQ   ................................................................................ [ 29200]
MH681419.1_ASFV/POL/2015          ................................................................................ [ 29200]
LS478113.1_Estonia_2014           ................................................................................ [ 29200]
KP843857.1_Odintsovo_02/14        ................................................................................ [ 29200]
FN557520.1_strain_E75             .c.....a........c....................................t.......................... [ 29200]
AY261366.1_Warthog                ................c...................................tt................a......... [ 29200]
AY261365.1_Warmbaths              ...................t.t...............................t.......t.................. [ 29200]
AY261364.1_Tengani_62             ...................t.t...............................t.......................... [ 29200]
AY261363.1_Pretoriuskop/96/4      ................c....................................t.......................... [ 29200]
AY261362.1_Mkuzi_1979             ....t..............t.t.........c.....................t.......................... [ 29200]
AY261361.1_Malawi_Lil-20/1_1983   .c.....a........c...................................tt................a......... [ 29200]
AY261360.1_Kenya_1950             c......a........c....g...............................t.t..............a......... [ 29200]
AM712240.1_OURT_88/3              .c.....a........c....................................t.......................... [ 29200]
AM712239.1_Benin_97/1             .c.....a........c....................................t.......................... [ 29200]
KM111295.1_Ken06.Bus              c......a........c....g...............................t.t..............a......... [ 29200]
KM111294.1_Ken05/Tk1              c......a........c....g...............................t.t.....t........a......... [ 29200]
KM262844.1_strain_L60             .c.....a........c....................................t.......................... [ 29200]
KM102979.1_26544/OG10_Italy       .c.....a........c....................................t.......................... [ 29200]
KM262845.1_strain_NHV             .c.....a........c....................................t.......................... [ 29200]
KX354450.1_47/Ss/2008             .c.....a........c....................................t.......................... [ 29200]
KP055815.1_strain_BA71            .c.....a........c....................................t.......................... [ 29200]
MH025920.1_strain_R35             c......a........c....g...............................t.t..............a......... [ 29200]
MH025919.1_strain_N10             c......a........c....g...............................t.t..............a......... [ 29200]
MH025918.1_strain_R25             c......a........c....g...............................t.t..............a......... [ 29200]
MH025917.1_strain_R7              c......a........c....g...............................t.t..............a......... [ 29200]
MH025916.1_strain_R8              c......a........c....g...............................t.t..............a......... [ 29200]

FR682468.2_Georgia_2007/1         aatattttactactaggcacatctgccacatgttgaacatttaaagcctgttcttcttctgtgttacggcaaaagagccg [ 29280]
ASF_IND_20_CAD543                 ..............g.a............................................................... [ 29280]
LR899193.1_Germany_2020/1         ................................................................................ [ 29280]
MN393477.1_Wuhan_2019             ................................................................................ [ 29280]
MW396979.1_Timor-Leste/2019/1     ................................................................................ [ 29280]
MK940252.1_CN/2019/InnerMongolia  ................................................................................ [ 29280]
MK128995.1_China/2018/AnhuiXCGQ   ................................................................................ [ 29280]
MH681419.1_ASFV/POL/2015          ................................................................................ [ 29280]
LS478113.1_Estonia_2014           ................................................................................ [ 29280]
KP843857.1_Odintsovo_02/14        ................................................................................ [ 29280]
```

```
FN557520.1_strain_E75             ..g...........g.a...................................................a........... [ 29280]
AY261366.1_Warthog                ..g...........g.a..t................................................a........... [ 29280]
AY261365.1_Warmbaths              ..g...........g.a...................................................a........... [ 29280]
AY261364.1_Tengani_62             ..g.........tcg.............................................a.......a........... [ 29280]
AY261363.1_Pretoriuskop/96/4      ..g...........g.a............c......................................a........... [ 29280]
AY261362.1_Mkuzi_1979             ..g.................................................................a........... [ 29280]
AY261361.1_Malawi_Lil-20/1_1983   ..a...........g..t.t.................g.......................c...t..a........... [ 29280]
AY261360.1_Kenya_1950             ..g......tg.........c................g.....................c.................... [ 29280]
AM712240.1_OURT_88/3              ..g...........g.a...................................................a........... [ 29280]
AM712239.1_Benin_97/1             ..g...........g.a...................................................a........... [ 29280]
KM111295.1_Ken06.Bus              ..a......ta.........c...a...t........g..................a....................... [ 29280]
KM111294.1_Ken05/Tk1              ..g......t..t.g....t.........g.......g.......................................t.. [ 29280]
KM262844.1_strain_L60             ..g...........g.a...................................................a........... [ 29280]
KM102979.1_26544/OG10_Italy       ..g...........g.a...................................................a........... [ 29280]
KM262845.1_strain_NHV             ..g...........g.a...................................................a........... [ 29280]
KX354450.1_47/Ss/2008             ..g...........g.a...................................................a........... [ 29280]
KP055815.1_strain_BA71            ..g...........g.a...................................................a........... [ 29280]
MH025920.1_strain_R35             ..a......ta.........c...a...t........g..................a....................... [ 29280]
MH025919.1_strain_N10             ..a......ta.........c...a...t........g..................a....................... [ 29280]
MH025918.1_strain_R25             ..a......ta.........c...a...t........g..................a....................... [ 29280]
MH025917.1_strain_R7              ..a......ta.........c...a...t........g..................a....................... [ 29280]
MH025916.1_strain_R8              ..a......ta.........c...a...t........g..................a....................... [ 29280]

FR682468.2_Georgia_2007/1         tgcgtattcaggtgaagctccccaggataacaacgtccttgctacggctaaattttttttgacgatgacttttatcagaa [ 29360]
ASF_IND_20_CAD543                 .........................................tc..................................... [ 29360]
LR899193.1_Germany_2020/1         ................................................................................ [ 29360]
MN393477.1_Wuhan_2019             ................................................................................ [ 29360]
MW396979.1_Timor-Leste/2019/1     ................................................................................ [ 29360]
MK940252.1_CN/2019/InnerMongolia  ................................................................................ [ 29360]
MK128995.1_China/2018/AnhuiXCGQ   ................................................................................ [ 29360]
MH681419.1_ASFV/POL/2015          ................................................................................ [ 29360]
LS478113.1_Estonia_2014           ................................................................................ [ 29360]
KP843857.1_Odintsovo_02/14        ................................................................................ [ 29360]
FN557520.1_strain_E75             c..a..................................g..tc.............................c....... [ 29360]
AY261366.1_Warthog                c..a....................................................................c....... [ 29360]
AY261365.1_Warmbaths              c................................t....g...c.............................c....... [ 29360]
AY261364.1_Tengani_62             c.............................t.......g...c...a............................g.... [ 29360]
AY261363.1_Pretoriuskop/96/4      c.....................................g...........................a.....c....... [ 29360]
AY261362.1_Mkuzi_1979             c..a..................................g...c.............................c....... [ 29360]
AY261361.1_Malawi_Lil-20/1_1983   c........g..........................t................................c..c..g.... [ 29360]
AY261360.1_Kenya_1950             c.............................t.........................................c..g.... [ 29360]
AM712240.1_OURT_88/3              c..a..................................g..tc.............................c....... [ 29360]
AM712239.1_Benin_97/1             c..a..................................g..tc.............................c....... [ 29360]
KM111295.1_Ken06.Bus              c........g....................t..t......................................c..g.... [ 29360]
```

```
KM111294.1_Ken05/Tk1            c........g....................t..t......................................c..g.... [ 29360]
KM262844.1_strain_L60           c..a..................................g..tc.............................c....... [ 29360]
KM102979.1_26544/OG10_Italy     c..a..................................g..tc.............................c....... [ 29360]
KM262845.1_strain_NHV           c..a..................................g..tc.............................c....... [ 29360]
KX354450.1_47/Ss/2008           c..a..................................g..tc.............................c....... [ 29360]
KP055815.1_strain_BA71          c..a..................................g..tc.............................c....... [ 29360]
MH025920.1_strain_R35           c........g....................t..t......................................c..g.... [ 29360]
MH025919.1_strain_N10           c........g....................t..t......................................c..g.... [ 29360]
MH025918.1_strain_R25           c........g....................t..t......................................c..g.... [ 29360]
MH025917.1_strain_R7            c........g....................t..t......................................c..g.... [ 29360]
MH025916.1_strain_R8            c........g....................t..t......................................c..g.... [ 29360]

FR682468.2_Georgia_2007/1       ataagtctttatttttgcattgatcactatgcgaatttgtatagttgacgccgttgcattgagtacattgatataatgtt [ 29440]
ASF_IND_20_CAD543               ................................................................................ [ 29440]
LR899193.1_Germany_2020/1       ................................................................................ [ 29440]
MN393477.1_Wuhan_2019           ................................................................................ [ 29440]
MW396979.1_Timor-Leste/2019/1   ................................................................................ [ 29440]
MK940252.1_CN/2019/InnerMongolia ................................................................................ [ 29440]
MK128995.1_China/2018/AnhuiXCGQ ................................................................................ [ 29440]
MH681419.1_ASFV/POL/2015        ................................................................................ [ 29440]
LS478113.1_Estonia_2014         ................................................................................ [ 29440]
KP843857.1_Odintsovo_02/14      ................................................................................ [ 29440]
FN557520.1_strain_E75           ................................................................................ [ 29440]
AY261366.1_Warthog              ................................................t............................... [ 29440]
AY261365.1_Warmbaths            ................a..................................................c............ [ 29440]
AY261364.1_Tengani_62           ...g...c...............................................a........................ [ 29440]
AY261363.1_Pretoriuskop/96/4    ................................................................................ [ 29440]
AY261362.1_Mkuzi_1979           ................................................................................ [ 29440]
AY261361.1_Malawi_Lil-20/1_1983 .......c..............................a.......................a................. [ 29440]
AY261360.1_Kenya_1950           ...g...c........................a............................g.................. [ 29440]
AM712240.1_OURT_88/3            ................................................................................ [ 29440]
AM712239.1_Benin_97/1           ................................................................................ [ 29440]
KM111295.1_Ken06.Bus            ...g...c........................a............................................... [ 29440]
KM111294.1_Ken05/Tk1            ...g...c........................a............................................... [ 29440]
KM262844.1_strain_L60           ................................................................................ [ 29440]
KM102979.1_26544/OG10_Italy     ................................................................................ [ 29440]
KM262845.1_strain_NHV           ................................................................................ [ 29440]
KX354450.1_47/Ss/2008           ................................................................................ [ 29440]
KP055815.1_strain_BA71          ................................................................................ [ 29440]
MH025920.1_strain_R35           ...g...c..................t.....a............................................... [ 29440]
MH025919.1_strain_N10           ...g...c..................t.....a............................................... [ 29440]
MH025918.1_strain_R25           ...g...c..................t.....a............................................... [ 29440]
MH025917.1_strain_R7            ...g...c..................t.....a............................................... [ 29440]
MH025916.1_strain_R8            ...g...c..................t.....a............................................... [ 29440]
```

```
FR682468.2_Georgia_2007/1        ttacaattccagcgtagccctaaatggtataaaagaactgtattttcgacataagcatgctgattaacgatgtttttgag [ 29520]
ASF_IND_20_CAD543                ..................................................g............................. [ 29520]
LR899193.1_Germany_2020/1        ................................................................................ [ 29520]
MN393477.1_Wuhan_2019            ................................................................................ [ 29520]
MW396979.1_Timor-Leste/2019/1    ................................................................................ [ 29520]
MK940252.1_CN/2019/InnerMongolia ................................................................................ [ 29520]
MK128995.1_China/2018/AnhuiXCGQ  ................................................................................ [ 29520]
MH681419.1_ASFV/POL/2015         ................................................................................ [ 29520]
LS478113.1_Estonia_2014          ................................................................................ [ 29520]
KP843857.1_Odintsovo_02/14       ................................................................................ [ 29520]
FN557520.1_strain_E75            ..g................................g..............gc............................ [ 29520]
AY261366.1_Warthog               ..g...................................c........c....g........................... [ 29520]
AY261365.1_Warmbaths             ..g...................................c........c....g........................... [ 29520]
AY261364.1_Tengani_62            ..g..............t.............................................................. [ 29520]
AY261363.1_Pretoriuskop/96/4     ..g................................g...........c....g........................... [ 29520]
AY261362.1_Mkuzi_1979            ..g...................................c........c....g........................... [ 29520]
AY261361.1_Malawi_Lil-20/1_1983  ..g...................................g........c....g........................... [ 29520]
AY261360.1_Kenya_1950            ..g............................................t.g..g........................... [ 29520]
AM712240.1_OURT_88/3             ..g................................g..............gc............................ [ 29520]
AM712239.1_Benin_97/1            ..g................................g..............gc............................ [ 29520]
KM111295.1_Ken06.Bus             ..g............................................t.g..g........................... [ 29520]
KM111294.1_Ken05/Tk1             ..g...................................c........t.g..g........................a.. [ 29520]
KM262844.1_strain_L60            ..g................................g..............gc............................ [ 29520]
KM102979.1_26544/OG10_Italy      ..g................................g..............gc............................ [ 29520]
KM262845.1_strain_NHV            ..g................................g..............gc............................ [ 29520]
KX354450.1_47/Ss/2008            ..g................................g..............gc............................ [ 29520]
KP055815.1_strain_BA71           ..g................................g..............gc............................ [ 29520]
MH025920.1_strain_R35            ..g............................................t.g..g........................... [ 29520]
MH025919.1_strain_N10            ..g............................................t.g..g........................... [ 29520]
MH025918.1_strain_R25            ..g............................................t.g..g........................... [ 29520]
MH025917.1_strain_R7             ..g............................................t.g..g........................... [ 29520]
MH025916.1_strain_R8             ..g............................................t.g..g........................... [ 29520]

FR682468.2_Georgia_2007/1        acaacacgtcgttaaggacaccatattgt-ctccaatttgttagataaaagtctttactaaaaaaatagatttttagttt [ 29600]
ASF_IND_20_CAD543                .............................-.......................................r.......... [ 29600]
LR899193.1_Germany_2020/1        .............................-.................................................. [ 29600]
MN393477.1_Wuhan_2019            .............................-.................................................. [ 29600]
MW396979.1_Timor-Leste/2019/1    .............................-.................................................. [ 29600]
MK940252.1_CN/2019/InnerMongolia .............................-.................................................. [ 29600]
MK128995.1_China/2018/AnhuiXCGQ  .............................-.................................................. [ 29600]
MH681419.1_ASFV/POL/2015         .............................-.................................................. [ 29600]
LS478113.1_Estonia_2014          .............................-.................................................. [ 29600]
KP843857.1_Odintsovo_02/14       .............................-.................................................. [ 29600]
```

```
FN557520.1_strain_E75         .........t...................-............g..........................g.......... [ 29600]
AY261366.1_Warthog            .............................-.......................................g.......... [ 29600]
AY261365.1_Warmbaths          .............................-.......................................g.......... [ 29600]
AY261364.1_Tengani_62         ..................g...--.....-.......................................g.......... [ 29600]
AY261363.1_Pretoriuskop/96/4  .............................-......................................ag.......... [ 29600]
AY261362.1_Mkuzi_1979         .............................-.......................................g.......... [ 29600]
AY261361.1_Malawi_Lil-20/1_1983 ....t.t..a...................-...................................tat.t.......... [ 29600]
AY261360.1_Kenya_1950         ......t..a.c.......g.....g...c.c................g................tat.t.......... [ 29600]
AM712240.1_OURT_88/3          .........t...................-............g..........................g.......... [ 29600]
AM712239.1_Benin_97/1         .........t...................-............g..........................g.......... [ 29600]
KM111295.1_Ken06.Bus          ......t..a.........g.....g...-.c................g................tat.t.......... [ 29600]
KM111294.1_Ken05/Tk1          ......t..a.........g.....g...c.c................g................tat.t.......... [ 29600]
KM262844.1_strain_L60         .........t...................-............g..........................g.......... [ 29600]
KM102979.1_26544/OG10_Italy   .........t...................-............g..........................g.......... [ 29600]
KM262845.1_strain_NHV         .........t...................-............g..........................g.......... [ 29600]
KX354450.1_47/Ss/2008         .........t...................-............g..........................g.......... [ 29600]
KP055815.1_strain_BA71        .........t...................-............g..........................g.......... [ 29600]
MH025920.1_strain_R35         ......t..a.........g.....g...-.c................g................tat.t.......... [ 29600]
MH025919.1_strain_N10         ......t..a.........g.....g...-.c................g................tat.t.......... [ 29600]
MH025918.1_strain_R25         ......t..a.........g.....g...-.c................g................tat.t.......... [ 29600]
MH025917.1_strain_R7          ......t..a.........g.....g...-.c................g................tat.t.......... [ 29600]
MH025916.1_strain_R8          ......t..a.........g.....g...-.c................g................tat.t.......... [ 29600]

FR682468.2_Georgia_2007/1     taac-aatcgagattttattatttgg--------------------------------at----gcatcatcaa------ [ 29680]
ASF_IND_20_CAD543             ....-.....................--------------------------------..-ata..........------ [ 29680]
LR899193.1_Germany_2020/1     ....-.....................--------------------------------..----..........------ [ 29680]
MN393477.1_Wuhan_2019         ....-.....................--------------------------------..----..........------ [ 29680]
MW396979.1_Timor-Leste/2019/1 ....-.....................--------------------------------..----..........------ [ 29680]
MK940252.1_CN/2019/InnerMongolia ....-.....................--------------------------------..----..........------ [ 29680]
MK128995.1_China/2018/AnhuiXCGQ ....-.....................--------------------------------..----..........------ [ 29680]
MH681419.1_ASFV/POL/2015      ....-.....................--------------------------------..----..........------ [ 29680]
LS478113.1_Estonia_2014       ....-.....................--------------------------------..----..........------ [ 29680]
KP843857.1_Odintsovo_02/14    ....-.....................--------------------------------..----..........------ [ 29680]
FN557520.1_strain_E75         ....-.....................--------------------------------..-ata..........------ [ 29680]
AY261366.1_Warthog            ....a..ca..t..a...........--------------------------------..---a..........------ [ 29680]
AY261365.1_Warmbaths          ....-...t.................--------------------------------..----..........------ [ 29680]
AY261364.1_Tengani_62         ....-..a..................--------------------------------..-ataa.........------ [ 29680]
AY261363.1_Pretoriuskop/96/4  ....-...t.................--------------------------------..-ata..........------ [ 29680]
AY261362.1_Mkuzi_1979         ....-.......--.......c....aatgt-----------------------attc..-acat...t.gt..actcgt [ 29680]
AY261361.1_Malawi_Lil-20/1_1983 ....-.t........g.........aaatgtttccatacatcaacagtaaactcattt..-ataa.........------ [ 29680]
AY261360.1_Kenya_1950         ....-.t........g..........aatgtttccatacatcaatagtaaacttgttt.g-ataa.........------ [ 29680]
AM712240.1_OURT_88/3          ....-.....................--------------------------------..-ata..........------ [ 29680]
AM712239.1_Benin_97/1         ....-.....................--------------------------------..-ata..........------ [ 29680]
KM111295.1_Ken06.Bus          ....-.t........g..........aatgttttcatacatcaatagtaaactcgtttggaata.....c....------ [ 29680]
```

```
KM111294.1_Ken05/Tk1               ....-.t........g.........aaatgtttccatacatcaatag-aaactcatttga-ata.....c....------ [ 29680]
KM262844.1_strain_L60              ....-.....................--------------------------------..-ata..........------ [ 29680]
KM102979.1_26544/OG10_Italy        ....-.....................--------------------------------..-ata..........------ [ 29680]
KM262845.1_strain_NHV              ....-.....................--------------------------------..-ata..........------ [ 29680]
KX354450.1_47/Ss/2008              ....-.....................--------------------------------..-ata..........------ [ 29680]
KP055815.1_strain_BA71             ....-.....................--------------------------------..-ata..........------ [ 29680]
MH025920.1_strain_R35              ....-.t........g..........aatgttttcatacatcaatagtaaactcgtttggaata.....c....------ [ 29680]
MH025919.1_strain_N10              ....-.t........g..........aatgttttcatacatcaatagtaaactcgtttggaata.....c....------ [ 29680]
MH025918.1_strain_R25              ....-.t........g..........aatgttttcatacatcaatagtaaactcgtttggaata.....c....------ [ 29680]
MH025917.1_strain_R7               ....-.t........g..........aatgttttcatacatcaatagtaaactcgtttggaata.....c....------ [ 29680]
MH025916.1_strain_R8               ....-.t........g..........aatgttttcatacatcaatagtaaactcgtttggaata.....c....------ [ 29680]

FR682468.2_Georgia_2007/1          ----------aaagatttataagtat-------aagaggttgtataagaaaaaaaatgatgttatactatttatgttaaa [ 29760]
ASF_IND_20_CAD543                  ----------...-------------------------------------------------------............ [ 29760]
LR899193.1_Germany_2020/1          ----------................-------............................................... [ 29760]
MN393477.1_Wuhan_2019              ----------................-------............................................... [ 29760]
MW396979.1_Timor-Leste/2019/1      ----------................-------............................................... [ 29760]
MK940252.1_CN/2019/InnerMongolia   ----------................-------............................................... [ 29760]
MK128995.1_China/2018/AnhuiXCGQ    ----------................-------............................................... [ 29760]
MH681419.1_ASFV/POL/2015           ----------................-------............................................... [ 29760]
LS478113.1_Estonia_2014            ----------................-------............................................... [ 29760]
KP843857.1_Odintsovo_02/14         ----------................-------............................................... [ 29760]
FN557520.1_strain_E75              ----------........-------------------------------------------------------....... [ 29760]
AY261366.1_Warthog                 ----------.g..t...........aagacaa............................................... [ 29760]
AY261365.1_Warmbaths               ----------..............g.aggacaa.....................t.......cg................ [ 29760]
AY261364.1_Tengani_62              ----------........-------------------------------------------------------....... [ 29760]
AY261363.1_Pretoriuskop/96/4       ----------..............g.aggacaa.....................t........g................ [ 29760]
AY261362.1_Mkuzi_1979              ttggatcatc.g..........aa..aagactg........a............t.......cg.....g..g....... [ 29760]
AY261361.1_Malawi_Lil-20/1_1983    ----------.g....c.....aa.caagactg........a............t......---..g....cg....... [ 29760]
AY261360.1_Kenya_1950              ----------.g.a..c.....aa.caagactg........a............t......cag..g....c........ [ 29760]
AM712240.1_OURT_88/3               ----------........-------------------------------------------------------....... [ 29760]
AM712239.1_Benin_97/1              ----------........-------------------------------------------------------....... [ 29760]
KM111295.1_Ken06.Bus               ------------------------------------------------......t.....a.cg.......c........ [ 29760]
KM111294.1_Ken05/Tk1               ------------------------------------------------......t.......c-.......c........ [ 29760]
KM262844.1_strain_L60              ----------........-------------------------------------------------------....... [ 29760]
KM102979.1_26544/OG10_Italy        ----------........-------------------------------------------------------....... [ 29760]
KM262845.1_strain_NHV              ----------........-------------------------------------------------------....... [ 29760]
KX354450.1_47/Ss/2008              ----------........-------------------------------------------------------....... [ 29760]
KP055815.1_strain_BA71             ----------........-------------------------------------------------------....... [ 29760]
MH025920.1_strain_R35              ------------------------------------------------......t.....a.cg.......c........ [ 29760]
MH025919.1_strain_N10              ------------------------------------------------......t.....a.cg.......c........ [ 29760]
MH025918.1_strain_R25              ------------------------------------------------......t.....a.cg.......c........ [ 29760]
MH025917.1_strain_R7               ------------------------------------------------......t.....a.cg.......c........ [ 29760]
MH025916.1_strain_R8               ------------------------------------------------......t.....a.cg.......c........ [ 29760]
```

```
FR682468.2_Georgia_2007/1        atttaatttatcatataaaaagtacagatttaatcagttg-------------------------gttaaactatttagt [ 29840]
ASF_IND_20_CAD543                .........r..w..y..................-.a...-------------------------...........---- [ 29840]
LR899193.1_Germany_2020/1        ........................................-------------------------............... [ 29840]
MN393477.1_Wuhan_2019            ........................................-------------------------............... [ 29840]
MW396979.1_Timor-Leste/2019/1    ........................................-------------------------............... [ 29840]
MK940252.1_CN/2019/InnerMongolia ........................................-------------------------............... [ 29840]
MK128995.1_China/2018/AnhuiXCGQ  ........................................-------------------------............... [ 29840]
MH681419.1_ASFV/POL/2015         ........................................-------------------------............... [ 29840]
LS478113.1_Estonia_2014          ........................................-------------------------............... [ 29840]
KP843857.1_Odintsovo_02/14       ........................................-------------------------............... [ 29840]
FN557520.1_strain_E75            .........g..t..c..............c...-.a...-------------------------...t.......---- [ 29840]
AY261366.1_Warthog               ........................................-------------------------...------------ [ 29840]
AY261365.1_Warmbaths             ............t....................a.t..------------------------------------------ [ 29840]
AY261364.1_Tengani_62            .........g..t....................c---------------------------------------------- [ 29840]
AY261363.1_Pretoriuskop/96/4     ............t...........................-------------------------...t.....aa.... [ 29840]
AY261362.1_Mkuzi_1979            ............t...........................-------------------------...t.....aa---- [ 29840]
AY261361.1_Malawi_Lil-20/1_1983  .........t..t........a...........c..a...-------------------------......a..------ [ 29840]
AY261360.1_Kenya_1950            t...........t......g....................gtttaactaaatagttaactaaata...........---- [ 29840]
AM712240.1_OURT_88/3             .........g..t..c..............c...-.a...-------------------------...t.......---- [ 29840]
AM712239.1_Benin_97/1            .........g..t..c..............c...-.a...-------------------------...t.......---- [ 29840]
KM111295.1_Ken06.Bus             .........t.t.......g............t..ca...-------------------------a........c.---- [ 29840]
KM111294.1_Ken05/Tk1             .........t.........g............t..ca...-------------------------a........c.---- [ 29840]
KM262844.1_strain_L60            .........g..t..c..............c...-.a...-------------------------...t.......---- [ 29840]
KM102979.1_26544/OG10_Italy      .........g..t..c..............c...-.a...-------------------------...t.......---- [ 29840]
KM262845.1_strain_NHV            .........g..t..c..............c...-.a...-------------------------...t.......---- [ 29840]
KX354450.1_47/Ss/2008            .........g..t..c..............c...-.a...-------------------------...t.......---- [ 29840]
KP055815.1_strain_BA71           .........g..t..c..............c...-.a...-------------------------...t.......---- [ 29840]
MH025920.1_strain_R35            .........t.t.......g............t..ca...-------------------------a........c.---- [ 29840]
MH025919.1_strain_N10            .........t.t.......g............t..ca...-------------------------a........c.---- [ 29840]
MH025918.1_strain_R25            .........t.t.......g............t..ca...-------------------------a........c.---- [ 29840]
MH025917.1_strain_R7             .........t.t.......g............t..ca...-------------------------a........c.---- [ 29840]
MH025916.1_strain_R8             .........t.t.......g............t..ca...-------------------------a........c.---- [ 29840]

FR682468.2_Georgia_2007/1        ---------------------taattaaactaaatagtttaaccatttagtcagactac--------ttggttagcaatg [ 29920]
ASF_IND_20_CAD543                ----------------------------------.........t.......t.a.t...--------............. [ 29920]
LR899193.1_Germany_2020/1        ---------------------......................................--------............. [ 29920]
MN393477.1_Wuhan_2019            ---------------------......................................--------............. [ 29920]
MW396979.1_Timor-Leste/2019/1    ---------------------......................................--------............. [ 29920]
MK940252.1_CN/2019/InnerMongolia ---------------------......................................--------............. [ 29920]
MK128995.1_China/2018/AnhuiXCGQ  ---------------------......................................--------............. [ 29920]
MH681419.1_ASFV/POL/2015         ---------------------......................................--------............. [ 29920]
LS478113.1_Estonia_2014          ---------------------......................................--------............. [ 29920]
KP843857.1_Odintsovo_02/14       ---------------------......................................--------............. [ 29920]
```

```
FN557520.1_strain_E75           ----------------------------------.........t.......t.a.t...--------............. [ 29920]
AY261366.1_Warthog              --------------aactaaa..g...................................--------............. [ 29920]
AY261365.1_Warmbaths            ----------------------------------.........................-------t............. [ 29920]
AY261364.1_Tengani_62           -------------------------------------------.g......t.......--------............. [ 29920]
AY261363.1_Pretoriuskop/96/4    taactaaatagttaaactaaa..g...................................--------............. [ 29920]
AY261362.1_Mkuzi_1979           ----------------------------------.........t.ac............--------............. [ 29920]
AY261361.1_Malawi_Lil-20/1_1983 ----------------------------------a.....g..........tt......taattact.g.t...a..... [ 29920]
AY261360.1_Kenya_1950           ----------------------------------........tt.........ag....---tt-----........... [ 29920]
AM712240.1_OURT_88/3            ----------------------------------.........t.......t.a.t...--------............. [ 29920]
AM712239.1_Benin_97/1           ----------------------------------.........t.......t.a.t...--------............. [ 29920]
KM111295.1_Ken06.Bus            ----------------------------------......g..t.......tta.....---ttact.a.t......... [ 29920]
KM111294.1_Ken05/Tk1            ----------------------------------......g..t.......tta.t...---ttact.a.t......... [ 29920]
KM262844.1_strain_L60           ----------------------------------.........t.......t.a.t...--------............. [ 29920]
KM102979.1_26544/OG10_Italy     ----------------------------------.........t.......t.a.t...--------............. [ 29920]
KM262845.1_strain_NHV           ----------------------------------.........t.......t.a.t...--------............. [ 29920]
KX354450.1_47/Ss/2008           ----------------------------------.........t.......t.a.t...--------............. [ 29920]
KP055815.1_strain_BA71          ----------------------------------.........t.......t.a.t...--------............. [ 29920]
MH025920.1_strain_R35           ----------------------------------......g..t.......tta.....---ttact.a.t......... [ 29920]
MH025919.1_strain_N10           ----------------------------------......g..t.......tta.....---ttact.a.t......... [ 29920]
MH025918.1_strain_R25           ----------------------------------......g..t.......tta.....---ttact.a.t......... [ 29920]
MH025917.1_strain_R7            ----------------------------------......g..t.......tta.....---ttact.a.t......... [ 29920]
MH025916.1_strain_R8            ----------------------------------......g..t.......tta.....---ttact.a.t......... [ 29920]

FR682468.2_Georgia_2007/1       tttgagctttcttccattcttatcc-----------gggggggg---gtcctaatcgttctaatac---tattgtggata [ 30000]
ASF_IND_20_CAD543               .......................t.ga-----gggg........---........a...t.g....---......t.tgg [ 30000]
LR899193.1_Germany_2020/1       .........................------------.......---...................---........... [ 30000]
MN393477.1_Wuhan_2019           .........................g----------........---...................---........... [ 30000]
MW396979.1_Timor-Leste/2019/1   .........................g----------........---...................---........... [ 30000]
MK940252.1_CN/2019/InnerMongolia .........................-----------........---...................---........... [ 30000]
MK128995.1_China/2018/AnhuiXCGQ .........................g----------........---...................---........... [ 30000]
MH681419.1_ASFV/POL/2015        .........................g----------........---...................---........... [ 30000]
LS478113.1_Estonia_2014         .........................-----------........---...................---........... [ 30000]
KP843857.1_Odintsovo_02/14      .........................gg--------g........---...................---........... [ 30000]
FN557520.1_strain_E75           .......................t.ga------ggg........---........a...t.g....ttt.g........t [ 30000]
AY261366.1_Warthog              .........................gg-----gggg.-----------.......a.c........tgt.g........t [ 30000]
AY261365.1_Warmbaths            .........................ga-----gggg........---tc......a.c........tgt.g........t [ 30000]
AY261364.1_Tengani_62           .......................t.gg--ggggggg........tcc........a.c........tgt.g......... [ 30000]
AY261363.1_Pretoriuskop/96/4    .........................gg-----gggg......------.......a.........ttat.g........t [ 30000]
AY261362.1_Mkuzi_1979           ...............t.at.c....gg-----------......---............t......---........... [ 30000]
AY261361.1_Malawi_Lil-20/1_1983 ...a.t..a........a..c...aga-------gg........---...t....aac........tat.g........t [ 30000]
AY261360.1_Kenya_1950           .................a.......-----------........---...t.c..atc........---..........t [ 30000]
AM712240.1_OURT_88/3            .......................t.ga-------gg........---........a...t.g....ttt.g........t [ 30000]
AM712239.1_Benin_97/1           .......................t.ga----------.......---........a...t.g....ttt.g........t [ 30000]
KM111295.1_Ken06.Bus            .................a..c....----------g........---.....c..a.c...g....---..........t [ 30000]
```

```
KM111294.1_Ken05/Tk1             .................a..c....---------gg........---.....c..a.c...g....---..........t [ 30000]
KM262844.1_strain_L60            .......................t.ga------ggg........---........a...t.g....ttt.g........t [ 30000]
KM102979.1_26544/OG10_Italy      .......................t.gag-----ggg........---........a...t.g....ttt.g........t [ 30000]
KM262845.1_strain_NHV            .......................t.ga-------gg........---........a...t.g....ttt.g........t [ 30000]
KX354450.1_47/Ss/2008            .......................t.ga--------g........---........a...t.g....ttt.g........t [ 30000]
KP055815.1_strain_BA71           .......................t.gaggggggggg........---........a...t.g....ttt.g........t [ 30000]
MH025920.1_strain_R35            .................a..c....----------g........---.....c..a.c...g....---..........t [ 30000]
MH025919.1_strain_N10            .................a..c....----------g........---.....c..a.c...g....---..........t [ 30000]
MH025918.1_strain_R25            .................a..c....---------gg........---.....c..a.c...g....---..........t [ 30000]
MH025917.1_strain_R7             .................a..c....----------g........---.....c..a.c...g....---..........t [ 30000]
MH025916.1_strain_R8             .................a..c....g--------gg........---.....c..a.c...g....---..........t [ 30000]

FR682468.2_Georgia_2007/1        gttgaatataatgaagactttatagatgctataatgatg----------------------------------------- [ 30080]
ASF_IND_20_CAD543                a...---t.g.....a...g..c......nnnnnnn...gcatataatattaattnnnnnnnnnnnnnnnnnnnnnnnnn [ 30080]
LR899193.1_Germany_2020/1        .......................................----------------------------------------- [ 30080]
MN393477.1_Wuhan_2019            .......................................----------------------------------------- [ 30080]
MW396979.1_Timor-Leste/2019/1    .......................................----------------------------------------- [ 30080]
MK940252.1_CN/2019/InnerMongolia .......................................----------------------------------------- [ 30080]
MK128995.1_China/2018/AnhuiXCGQ  .......................................----------------------------------------- [ 30080]
MH681419.1_ASFV/POL/2015         .......................................----------------------------------------- [ 30080]
LS478113.1_Estonia_2014          .......................................----------------------------------------- [ 30080]
KP843857.1_Odintsovo_02/14       .......................................----------------------------------------- [ 30080]
FN557520.1_strain_E75            ....------.....a...g..c........a.gca...gcatataatattaattgtactgtttaatgtgagtttataag [ 30080]
AY261366.1_Warthog               a.....c........a...............a.gca...aat-------------------------------------- [ 30080]
AY261365.1_Warmbaths             a..............a...............g.g.a...aatccta---------------------------------- [ 30080]
AY261364.1_Tengani_62            ...............................g.g.a...gattgtaa---gaatattaccgtttaatatgag-------- [ 30080]
AY261363.1_Pretoriuskop/96/4     ....------.....a...............g...at.tgattgtag---taattgtactaactgttgtgag-------- [ 30080]
AY261362.1_Mkuzi_1979            ...............................a.......----------------------------------------- [ 30080]
AY261361.1_Malawi_Lil-20/1_1983  ...............a...............g.....gtggtgctat---tattcatactgtttaatttgag-------- [ 30080]
AY261360.1_Kenya_1950            ......c......t..........a.....tat.g.tatgattgtactaacaatggtactactaattgtact-------- [ 30080]
AM712240.1_OURT_88/3             ....------.....a...g..c........a.gca...gcatataatattaattgtactgtttaatgtgagtttataag [ 30080]
AM712239.1_Benin_97/1            ....------.....a...g..c........a.gca...gcatataatattaattgtactgtttaatgtgagtttataag [ 30080]
KM111295.1_Ken06.Bus             ...............................a...at.tgattgt------aattgtactcaatattgtgtg-------- [ 30080]
KM111294.1_Ken05/Tk1             ...............................a....t.tgattgt------aattgtactcaatgttgtgtg-------- [ 30080]
KM262844.1_strain_L60            ....------.....a...g..c........a.gca...gcatataatattaattgtactgtttaatgtgagtttataag [ 30080]
KM102979.1_26544/OG10_Italy      ....------.....a...g..c........a.gca...gcatataatattaattgtactgtttaatgtgagtttataag [ 30080]
KM262845.1_strain_NHV            ....------.....a...g..c........a.gca...gcatataatattaattgtactgtttaatgtgagtttataag [ 30080]
KX354450.1_47/Ss/2008            ....------.....a...g..c........a.gca...gcatataatattaattgtactgtttaatgtgagtttataag [ 30080]
KP055815.1_strain_BA71           ....------.....a...g..c........a.gca...gcatataatattaattgtactgtttaatgtgagtttataag [ 30080]
MH025920.1_strain_R35            ...............................a...at.tgattgt------aattgtactcaatattgtgtg-------- [ 30080]
MH025919.1_strain_N10            ...............................a...at.tgattgt------aattgtactcaatattgtgtg-------- [ 30080]
MH025918.1_strain_R25            ...............................a...at.tgattgt------aattgtactcaatattgtgtg-------- [ 30080]
MH025917.1_strain_R7             ...............................a...at.tgattgt------aattgtactcaatattgtgtg-------- [ 30080]
MH025916.1_strain_R8             ...............................a...at.tgattgt------aattgtactcaatattgtgtg-------- [ 30080]
```

```
FR682468.2_Georgia_2007/1        -------------------------------------------------------------------------------- [ 30160]
ASF_IND_20_CAD543                tatgtatcaaaatttatttttttttataaaagtannnnnnnnnnnnnnnnnnnnnnnnnnnnnnnnnnnnnnnnnnnnnn [ 30160]
LR899193.1_Germany_2020/1        -------------------------------------------------------------------------------- [ 30160]
MN393477.1_Wuhan_2019            -------------------------------------------------------------------------------- [ 30160]
MW396979.1_Timor-Leste/2019/1    -------------------------------------------------------------------------------- [ 30160]
MK940252.1_CN/2019/InnerMongolia -------------------------------------------------------------------------------- [ 30160]
MK128995.1_China/2018/AnhuiXCGQ  -------------------------------------------------------------------------------- [ 30160]
MH681419.1_ASFV/POL/2015         -------------------------------------------------------------------------------- [ 30160]
LS478113.1_Estonia_2014          -------------------------------------------------------------------------------- [ 30160]
KP843857.1_Odintsovo_02/14       -------------------------------------------------------------------------------- [ 30160]
FN557520.1_strain_E75            tatgtatcaaaatttatttttttttataaaagtacagaaattaatgttagcttaaactttagtttagctatttgagttaa [ 30160]
AY261366.1_Warthog               -------------------------------------------------------------------------------- [ 30160]
AY261365.1_Warmbaths             -------------------------------------------------------------------------------- [ 30160]
AY261364.1_Tengani_62            -------------------------------------------------------------------------------- [ 30160]
AY261363.1_Pretoriuskop/96/4     ---gtataaaga-------------------------------------------------------------------- [ 30160]
AY261362.1_Mkuzi_1979            ------------------------------------------------------aactgtagtgcagctgtataaaataa [ 30160]
AY261361.1_Malawi_Lil-20/1_1983  -------------------------------------------------------------------------------- [ 30160]
AY261360.1_Kenya_1950            -------------------------------------------------------------------------------- [ 30160]
AM712240.1_OURT_88/3             tatgtatcaaaatttatttttttttataaaagtacagaaattaatgttagcttaaactttagtttagctatttgagttaa [ 30160]
AM712239.1_Benin_97/1            tatgtatcaaaatttatttttttttataaaagtacagaaattaatgttagcttaaactttagtttagctatttgagttaa [ 30160]
KM111295.1_Ken06.Bus             -------------------------------------------------------------------------------- [ 30160]
KM111294.1_Ken05/Tk1             -------------------------------------------------------------------------------- [ 30160]
KM262844.1_strain_L60            tatgtatcaaaatttatttttttttataaaagtacagaaattaatgttagcttaaactttagtttagctatttgagttaa [ 30160]
KM102979.1_26544/OG10_Italy      tatgtatcaaaatttatttttttttataaaagtacagaaattaatgttagcttaaactttagtttagctatttgagttaa [ 30160]
KM262845.1_strain_NHV            tatgtatcaaaatttatttttttttataaaagtacagaaattaatgttagcttaaactttagtttagctatttgagttaa [ 30160]
KX354450.1_47/Ss/2008            tatgtatcaaaatttatttttttttataaaagtacagaaattaatgttagcttaaactttagtttagctatttgagttaa [ 30160]
KP055815.1_strain_BA71           tatgtatcaaaatttatttttttttataaaagtacagaaattaatgttagcttaaactttagtttagctatttgagttaa [ 30160]
MH025920.1_strain_R35            -------------------------------------------------------------------------------- [ 30160]
MH025919.1_strain_N10            -------------------------------------------------------------------------------- [ 30160]
MH025918.1_strain_R25            -------------------------------------------------------------------------------- [ 30160]
MH025917.1_strain_R7             -------------------------------------------------------------------------------- [ 30160]
MH025916.1_strain_R8             -------------------------------------------------------------------------------- [ 30160]

FR682468.2_Georgia_2007/1        -------------------------------------------------------------------------------- [ 30240]
ASF_IND_20_CAD543                nnnnnnnnnnntatcttccatgcttatcagnnnnnnnnnnnnnnnnnnnnnnnnnnnnnnataaatttgatcaaaattta [ 30240]
LR899193.1_Germany_2020/1        -------------------------------------------------------------------------------- [ 30240]
MN393477.1_Wuhan_2019            -------------------------------------------------------------------------------- [ 30240]
MW396979.1_Timor-Leste/2019/1    -------------------------------------------------------------------------------- [ 30240]
MK940252.1_CN/2019/InnerMongolia -------------------------------------------------------------------------------- [ 30240]
MK128995.1_China/2018/AnhuiXCGQ  -------------------------------------------------------------------------------- [ 30240]
MH681419.1_ASFV/POL/2015         -------------------------------------------------------------------------------- [ 30240]
LS478113.1_Estonia_2014          -------------------------------------------------------------------------------- [ 30240]
KP843857.1_Odintsovo_02/14       -------------------------------------------------------------------------------- [ 30240]
```

```
FN557520.1_strain_E75            tgatgtttaacctatcttccatgcttatcagggggggggggtcctaattgttttacgttataaatttgatcaaaattta [ 30240]
AY261366.1_Warthog               -------------------------------------------------------------------------------- [ 30240]
AY261365.1_Warmbaths             -------------------------------------------------------------------------------- [ 30240]
AY261364.1_Tengani_62            -------------------------------------------------------------------------------- [ 30240]
AY261363.1_Pretoriuskop/96/4     --------------------------------------------------------------------aatcaaaattta [ 30240]
AY261362.1_Mkuzi_1979            cctt-------------------------------------------------------------tttgatcaaaattta [ 30240]
AY261361.1_Malawi_Lil-20/1_1983  -------------------------------------------------------------------------------- [ 30240]
AY261360.1_Kenya_1950            -------------------------------------------------------------------------------- [ 30240]
AM712240.1_OURT_88/3             tgatgtttaacctatcttccatgcttatca---gggggggggtcctaattgttttacgttataaatttgatcaaaattta [ 30240]
AM712239.1_Benin_97/1            tgatgtttaacctatcttccatgcttatcagggggggggggtcctaattgttttacgttataaatttgatcaaaattta [ 30240]
KM111295.1_Ken06.Bus             -------------------------------------------------------------------------------- [ 30240]
KM111294.1_Ken05/Tk1             -------------------------------------------------------------------------------- [ 30240]
KM262844.1_strain_L60            tgatgtttaacctatcttccatgcttatcag--gggggggggtcctaattgttttacgttataaatttgatcaaaattta [ 30240]
KM102979.1_26544/OG10_Italy      tgatgtttaacctatcttccatgcttatca-----gggggggtcctaattgttttacgttataaatttgatcaaaattta [ 30240]
KM262845.1_strain_NHV            tgatgtttaacctatcttccatgcttatcag-ggggggggggtcctaattgttttacgttataaatttgatcaaaattta [ 30240]
KX354450.1_47/Ss/2008            tgatgtttaacctatcttccatgcttatca----ggggggggtcctaattgttttacgttataaatttgatcaaaattta [ 30240]
KP055815.1_strain_BA71           tgatgtttaacctatcttccatgcttatcag-ggggggggggtcctaattgttttacgttataaatttgatcaaaattta [ 30240]
MH025920.1_strain_R35            -------------------------------------------------------------------------------- [ 30240]
MH025919.1_strain_N10            -------------------------------------------------------------------------------- [ 30240]
MH025918.1_strain_R25            -------------------------------------------------------------------------------- [ 30240]
MH025917.1_strain_R7             -------------------------------------------------------------------------------- [ 30240]
MH025916.1_strain_R8             -------------------------------------------------------------------------------- [ 30240]

FR682468.2_Georgia_2007/1        -------------------------------------------------------------------------------- [ 30320]
ASF_IND_20_CAD543                atttctyataaaaaaggtacacatcaacattgctcatttaagtttcatratgtttgattcatnnnnnnnnnnnnnnnnnn [ 30320]
LR899193.1_Germany_2020/1        -------------------------------------------------------------------------------- [ 30320]
MN393477.1_Wuhan_2019            -------------------------------------------------------------------------------- [ 30320]
MW396979.1_Timor-Leste/2019/1    -------------------------------------------------------------------------------- [ 30320]
MK940252.1_CN/2019/InnerMongolia -------------------------------------------------------------------------------- [ 30320]
MK128995.1_China/2018/AnhuiXCGQ  -------------------------------------------------------------------------------- [ 30320]
MH681419.1_ASFV/POL/2015         -------------------------------------------------------------------------------- [ 30320]
LS478113.1_Estonia_2014          -------------------------------------------------------------------------------- [ 30320]
KP843857.1_Odintsovo_02/14       -------------------------------------------------------------------------------- [ 30320]
FN557520.1_strain_E75            atttcttataaaaaaggtacacatcaacattgctcatttaagtttcataatgtttgattcattacttcctacaattact- [ 30320]
AY261366.1_Warthog               -------------------------------------------------------------------------------- [ 30320]
AY261365.1_Warmbaths             -------------------------------------------------------------------------------- [ 30320]
AY261364.1_Tengani_62            -------------------------------------------------------------------------------- [ 30320]
AY261363.1_Pretoriuskop/96/4     atattttataaaaaaggtacagagtga-----------taagtt------------------------------------ [ 30320]
AY261362.1_Mkuzi_1979            a------------------------------------------------------------------------------- [ 30320]
AY261361.1_Malawi_Lil-20/1_1983  -------------------------------------------------------------------------------- [ 30320]
AY261360.1_Kenya_1950            -------------------------------------------------------------------------------- [ 30320]
AM712240.1_OURT_88/3             atttcttataaaaaaggtacacatcaacattgctcatttaagtttcataatgtttgattcattacttcctacaattact- [ 30320]
AM712239.1_Benin_97/1            atttcttataaaaaaggtacacatcaacattgctcatttaagtttcataatgtttgattcattacttcctacaattact- [ 30320]
KM111295.1_Ken06.Bus             -------------------------------------------------------------------------------- [ 30320]
```

```
KM111294.1_Ken05/Tk1             -------------------------------------------------------------------------------- [ 30320]
KM262844.1_strain_L60            atttcttataaaaaaggtacacatcaacattgctcatttaagtttcataatgtttgattcattacttcctacaattactg [ 30320]
KM102979.1_26544/OG10_Italy      atttcttataaaaaaggtacacatcaacattgctcatttaagtttcataatgtttgattcattacttcctacaattact- [ 30320]
KM262845.1_strain_NHV            atttcttataaaaaaggtacacatcaacattgctcatttaagtttcataatgtttgattcattacttcctacaattactg [ 30320]
KX354450.1_47/Ss/2008            atttcttataaaaaaggtacacatcaacattgctcatttaagtttcataatgtttgattcattacttcctacaattact- [ 30320]
KP055815.1_strain_BA71           atttcttataaaaaaggtacacatcaacattgctcatttaagtttcataatgtttgattcattacttcctacaattactg [ 30320]
MH025920.1_strain_R35            -------------------------------------------------------------------------------- [ 30320]
MH025919.1_strain_N10            -------------------------------------------------------------------------------- [ 30320]
MH025918.1_strain_R25            -------------------------------------------------------------------------------- [ 30320]
MH025917.1_strain_R7             -------------------------------------------------------------------------------- [ 30320]
MH025916.1_strain_R8             -------------------------------------------------------------------------------- [ 30320]

FR682468.2_Georgia_2007/1        -------------------------------------------------------------------------------- [ 30400]
ASF_IND_20_CAD543                -nnnnnnnnnntctttaatagcattaatattattatggtttnnnnnnnnnnnnnnnnnnnnnnnagcaaaaatagatact [ 30400]
LR899193.1_Germany_2020/1        -------------------------------------------------------------------------------- [ 30400]
MN393477.1_Wuhan_2019            -------------------------------------------------------------------------------- [ 30400]
MW396979.1_Timor-Leste/2019/1    -------------------------------------------------------------------------------- [ 30400]
MK940252.1_CN/2019/InnerMongolia -------------------------------------------------------------------------------- [ 30400]
MK128995.1_China/2018/AnhuiXCGQ  -------------------------------------------------------------------------------- [ 30400]
MH681419.1_ASFV/POL/2015         -------------------------------------------------------------------------------- [ 30400]
LS478113.1_Estonia_2014          -------------------------------------------------------------------------------- [ 30400]
KP843857.1_Odintsovo_02/14       -------------------------------------------------------------------------------- [ 30400]
FN557520.1_strain_E75            -ggggggggggtctttaatagcattaatattattatggtttgctgactatttggaatttatagaagcaaaaatagatact [ 30400]
AY261366.1_Warthog               -------------------------------------------------------------------------------- [ 30400]
AY261365.1_Warmbaths             -------------------------------------------------------------------------------- [ 30400]
AY261364.1_Tengani_62            -------------------------------------------------------------------------------- [ 30400]
AY261363.1_Pretoriuskop/96/4     -------------------------------------------------------------------------------- [ 30400]
AY261362.1_Mkuzi_1979            -------------------------------------------------------------------------------- [ 30400]
AY261361.1_Malawi_Lil-20/1_1983  -------------------------------------------------------------------------------- [ 30400]
AY261360.1_Kenya_1950            -------------------------------------------------------------------------------- [ 30400]
AM712240.1_OURT_88/3             -ggggggggggtctttaatagcattaatattattatggtttgctgactatttggaatttatagaagcaaaaatagatact [ 30400]
AM712239.1_Benin_97/1            --gggggggggtctttaatagcattaatattattatggtttgctgactatttggaatttatagaagcaaaaatagatact [ 30400]
KM111295.1_Ken06.Bus             -------------------------------------------------------------------------------- [ 30400]
KM111294.1_Ken05/Tk1             -------------------------------------------------------------------------------- [ 30400]
KM262844.1_strain_L60            -ggggggggggtctttaatagcattaatattattatggtttgctgactatttggaatttatagaagcaaaaatagatact [ 30400]
KM102979.1_26544/OG10_Italy      ---ggggggggtctttaatagcattaatattattatggtttgctgactatttggaatttatagaagcaaaaatagatact [ 30400]
KM262845.1_strain_NHV            -ggggggggggtctttaatagcattaatattattatggtttgctgactatttggaatttatagaagcaaaaatagatact [ 30400]
KX354450.1_47/Ss/2008            ---ggggggggtctttaatagcattaatattattatggtttgctgactatttggaatttatagaagcaaaaatagatact [ 30400]
KP055815.1_strain_BA71           gggggggggggtctttaatagcattaatattattatggtttgctgactatttggaatttatagaagcaaaaatagatact [ 30400]
MH025920.1_strain_R35            -------------------------------------------------------------------------------- [ 30400]
MH025919.1_strain_N10            -------------------------------------------------------------------------------- [ 30400]
MH025918.1_strain_R25            -------------------------------------------------------------------------------- [ 30400]
MH025917.1_strain_R7             -------------------------------------------------------------------------------- [ 30400]
MH025916.1_strain_R8             -------------------------------------------------------------------------------- [ 30400]
```

```
FR682468.2_Georgia_2007/1        -------------------------------------------------------------------------------- [ 30480]
ASF_IND_20_CAD543                tttnnnnatagcagtgcattataagtgatcaaaatttaannnnnnnnaaaaaaggtacacatcaacatcattcatttgtt [ 30480]
LR899193.1_Germany_2020/1        -------------------------------------------------------------------------------- [ 30480]
MN393477.1_Wuhan_2019            -------------------------------------------------------------------------------- [ 30480]
MW396979.1_Timor-Leste/2019/1    -------------------------------------------------------------------------------- [ 30480]
MK940252.1_CN/2019/InnerMongolia -------------------------------------------------------------------------------- [ 30480]
MK128995.1_China/2018/AnhuiXCGQ  -------------------------------------------------------------------------------- [ 30480]
MH681419.1_ASFV/POL/2015         -------------------------------------------------------------------------------- [ 30480]
LS478113.1_Estonia_2014          -------------------------------------------------------------------------------- [ 30480]
KP843857.1_Odintsovo_02/14       -------------------------------------------------------------------------------- [ 30480]
FN557520.1_strain_E75            tttactaatagcagtgcattataagtgatcaaaatttaatatctcataaaaaaggtacacatcaacatcattcatttgtt [ 30480]
AY261366.1_Warthog               -------------------------------------------------------------------------------- [ 30480]
AY261365.1_Warmbaths             -------------------------------------------------------------------------------- [ 30480]
AY261364.1_Tengani_62            -------------------------------------------------------------------------------- [ 30480]
AY261363.1_Pretoriuskop/96/4     ----------------------------------------------------------acatcaacg----taacttgtt [ 30480]
AY261362.1_Mkuzi_1979            -------------------------------------------------------------------------------- [ 30480]
AY261361.1_Malawi_Lil-20/1_1983  -------------------------------------------------------------------------------- [ 30480]
AY261360.1_Kenya_1950            -------------------------------------------------------------------------------- [ 30480]
AM712240.1_OURT_88/3             tttactaatagcagtgcattataagtgatcaaaatttaatatctcataaaaaaggtacacatcaacatcattcatttgtt [ 30480]
AM712239.1_Benin_97/1            tttactaatagcagtgcattataagtgatcaaaatttaatatctcataaaaaaggtacacatcaacatcattcatttgtt [ 30480]
KM111295.1_Ken06.Bus             -------------------------------------------------------------------------------- [ 30480]
KM111294.1_Ken05/Tk1             -------------------------------------------------------------------------------- [ 30480]
KM262844.1_strain_L60            tttactaatagcagtgcattataagtgatcaaaatttaatatctcataaaaaaggtacacatcaacatcattcatttgtt [ 30480]
KM102979.1_26544/OG10_Italy      tttactaatagcagtgcattataagtgatcaaaatttaatatctcataaaaaaggtacacatcaacatcattcatttgtt [ 30480]
KM262845.1_strain_NHV            tttactaatagcagtgcattataagtgatcaaaatttaatatctcataaaaaaggtacacatcaacatcattcatttgtt [ 30480]
KX354450.1_47/Ss/2008            tttactaatagcagtgcattataagtgatcaaaatttaatatctcataaaaaaggtacacatcaacatcattcatttgtt [ 30480]
KP055815.1_strain_BA71           tttactaatagcagtgcattataagtgatcaaaatttaatatctcataaaaaaggtacacatcaacatcattcatttgtt [ 30480]
MH025920.1_strain_R35            -------------------------------------------------------------------------------- [ 30480]
MH025919.1_strain_N10            -------------------------------------------------------------------------------- [ 30480]
MH025918.1_strain_R25            -------------------------------------------------------------------------------- [ 30480]
MH025917.1_strain_R7             -------------------------------------------------------------------------------- [ 30480]
MH025916.1_strain_R8             -------------------------------------------------------------------------------- [ 30480]

FR682468.2_Georgia_2007/1        -------------------------------------------------------------------------------- [ 30560]
ASF_IND_20_CAD543                tcatgatgtttgattcannnnnnnnnnnnnnnnnn----nnnnnnnnnnnnnnnnnnnnnnnnnnntattgctatggttt [ 30560]
LR899193.1_Germany_2020/1        -------------------------------------------------------------------------------- [ 30560]
MN393477.1_Wuhan_2019            -------------------------------------------------------------------------------- [ 30560]
MW396979.1_Timor-Leste/2019/1    -------------------------------------------------------------------------------- [ 30560]
MK940252.1_CN/2019/InnerMongolia -------------------------------------------------------------------------------- [ 30560]
MK128995.1_China/2018/AnhuiXCGQ  -------------------------------------------------------------------------------- [ 30560]
MH681419.1_ASFV/POL/2015         -------------------------------------------------------------------------------- [ 30560]
LS478113.1_Estonia_2014          -------------------------------------------------------------------------------- [ 30560]
KP843857.1_Odintsovo_02/14       -------------------------------------------------------------------------------- [ 30560]
```

```
FN557520.1_strain_E75           tcatgatgtttgattcattactttctacaatcact-------ggggggggtctctcatagctttagtattgctatggttt [ 30560]
AY261366.1_Warthog              -------------------------------------------------------------------------------- [ 30560]
AY261365.1_Warmbaths            -------------------------------------------------------------------------------- [ 30560]
AY261364.1_Tengani_62           -------------------------------------------------------------------------------- [ 30560]
AY261363.1_Pretoriuskop/96/4    ttatgatgtttgattcattactttctacaattact--------ggggggggtcctaatagctttagtattgttatggctt [ 30560]
AY261362.1_Mkuzi_1979           -------------------------------------------------------------------------------- [ 30560]
AY261361.1_Malawi_Lil-20/1_1983 -------------------------------------------------------------------------------- [ 30560]
AY261360.1_Kenya_1950           -------------------------------------------------------------------------------- [ 30560]
AM712240.1_OURT_88/3            tcatgatgtttgattcattactttctacaatcact----gggggggggggtctctcatagctttagtattgctatggttt [ 30560]
AM712239.1_Benin_97/1           tcatgatgtttgattcattactttctacaatcact-------ggggggggtctctcatagctttagtattgctatggttt [ 30560]
KM111295.1_Ken06.Bus            -------------------------------------------------------------------------------- [ 30560]
KM111294.1_Ken05/Tk1            -------------------------------------------------------------------------------- [ 30560]
KM262844.1_strain_L60           tcatgatgtttgattcattactttctacaatcactg-----gggggggggtctctcatagctttagtattgctatggttt [ 30560]
KM102979.1_26544/OG10_Italy     tcatgatgtttgattcattactttctacaatcact-------ggggggggtctctcatagctttagtattgctatggttt [ 30560]
KM262845.1_strain_NHV           tcatgatgtttgattcattactttctacaatcact-----ggggggggggtctctcatagctttagtattgctatggttt [ 30560]
KX354450.1_47/Ss/2008           tcatgatgtttgattcattactttctacaatcact------gggggggggtctctcatagctttagtattgctatggttt [ 30560]
KP055815.1_strain_BA71          tcatgatgtttgattcattactttctacaatcactgggggggggggggggtctctcatagctttagtattgctatggttt [ 30560]
MH025920.1_strain_R35           -------------------------------------------------------------------------------- [ 30560]
MH025919.1_strain_N10           -------------------------------------------------------------------------------- [ 30560]
MH025918.1_strain_R25           -------------------------------------------------------------------------------- [ 30560]
MH025917.1_strain_R7            -------------------------------------------------------------------------------- [ 30560]
MH025916.1_strain_R8            -------------------------------------------------------------------------------- [ 30560]

FR682468.2_Georgia_2007/1       -------------------------------------------------------------------------------- [ 30640]
ASF_IND_20_CAD543               actaattattatgtagaatttatagnnnnnn---nnnnnnnnnnnnnnnnnnnnnnnnnnnnnnnnnntaaa-------- [ 30640]
LR899193.1_Germany_2020/1       -------------------------------------------------------------------------------- [ 30640]
MN393477.1_Wuhan_2019           -------------------------------------------------------------------------------- [ 30640]
MW396979.1_Timor-Leste/2019/1   -------------------------------------------------------------------------------- [ 30640]
MK940252.1_CN/2019/InnerMongolia -------------------------------------------------------------------------------- [ 30640]
MK128995.1_China/2018/AnhuiXCGQ -------------------------------------------------------------------------------- [ 30640]
MH681419.1_ASFV/POL/2015        -------------------------------------------------------------------------------- [ 30640]
LS478113.1_Estonia_2014         -------------------------------------------------------------------------------- [ 30640]
KP843857.1_Odintsovo_02/14      -------------------------------------------------------------------------------- [ 30640]
FN557520.1_strain_E75           actaattattatgtagaatttataggagcag---atatatatggtggtaattgtactctataagtttataaa-------- [ 30640]
AY261366.1_Warthog              -------------------------------------------------------------------------------- [ 30640]
AY261365.1_Warmbaths            -------------------------------------------------------------------------------- [ 30640]
AY261364.1_Tengani_62           ----------------------------------------------------------tataagtttataaaaacta--- [ 30640]
AY261363.1_Pretoriuskop/96/4    gctgacaattctataacaattatagaagcagctattatgaaacctaataatacaacagtataagattttaaatgttaagc [ 30640]
AY261362.1_Mkuzi_1979           -------------------------------------------------------------------------------- [ 30640]
AY261361.1_Malawi_Lil-20/1_1983 -----------------------------------------------------------------ctataaaa------- [ 30640]
AY261360.1_Kenya_1950           ---------------------------------------------------------------gagttataa-------- [ 30640]
AM712240.1_OURT_88/3            actaattattatgtagaatttataggagcag---atatatatggtggtaattgtactctataagtttataaa-------- [ 30640]
AM712239.1_Benin_97/1           actaattattatgtagaatttataggagcag---atatatatggtggtaattgtactctataagtttataaa-------- [ 30640]
KM111295.1_Ken06.Bus            ---------------------------------------------------------------gtttaaaaaa------- [ 30640]
```

```
KM111294.1_Ken05/Tk1           ---------------------------------------------------------------gtttaaaaaa------- [ 30640]
KM262844.1_strain_L60          actaattattatgtagaatttataggagcag---atatatatggtggtaattgtactctataagtttataaa-------- [ 30640]
KM102979.1_26544/OG10_Italy    actaattattatgtagaatttataggagcag---atatatatggtggtaattgtactctataagtttataaa-------- [ 30640]
KM262845.1_strain_NHV          actaattattatgtagaatttataggagcag---atatatatggtggtaattgtactctataagtttataaa-------- [ 30640]
KX354450.1_47/Ss/2008          actaattattatgtagaatttataggagcag---atatatatggtggtaattgtactctataagtttataaa-------- [ 30640]
KP055815.1_strain_BA71         actaattattatgtagaatttataggagcag---atatatatggtggtaattgtactctataagtttataaa-------- [ 30640]
MH025920.1_strain_R35          ---------------------------------------------------------------gtttaaaaaa------- [ 30640]
MH025919.1_strain_N10          ---------------------------------------------------------------gtttaaaaaa------- [ 30640]
MH025918.1_strain_R25          ---------------------------------------------------------------gtttaaaaaa------- [ 30640]
MH025917.1_strain_R7           ---------------------------------------------------------------gtttaaaaaa------- [ 30640]
MH025916.1_strain_R8           ---------------------------------------------------------------gtttaaaaaa------- [ 30640]

FR682468.2_Georgia_2007/1      -aattctagtatgcct---------------------------------------------------------------- [ 30720]
ASF_IND_20_CAD543              -....tc.....att.tttttgttaatataannnnnnnnntttaaattacaaaaaaaagagaatcatcatgctattatac [ 30720]
LR899193.1_Germany_2020/1      -..............----------------------------------------------------------------- [ 30720]
MN393477.1_Wuhan_2019          -..............----------------------------------------------------------------- [ 30720]
MW396979.1_Timor-Leste/2019/1  -..............----------------------------------------------------------------- [ 30720]
MK940252.1_CN/2019/InnerMongolia -..............----------------------------------------------------------------- [ 30720]
MK128995.1_China/2018/AnhuiXCGQ  -..............----------------------------------------------------------------- [ 30720]
MH681419.1_ASFV/POL/2015       -..............----------------------------------------------------------------- [ 30720]
LS478113.1_Estonia_2014        -..............----------------------------------------------------------------- [ 30720]
KP843857.1_Odintsovo_02/14     -..............----------------------------------------------------------------- [ 30720]
FN557520.1_strain_E75          -....tc.....att.tttttgttaatataacagaaatgatttaaattacaaaaaaaagagaatcatcatgctattatac [ 30720]
AY261366.1_Warthog             -------------------------------------------------------------------------------- [ 30720]
AY261365.1_Warmbaths           -------------------------------------------------------------------------------- [ 30720]
AY261364.1_Tengani_62          -------------------------------------------------------------------------------- [ 30720]
AY261363.1_Pretoriuskop/96/4   t....tc.....att.tttttgttaatataacagaaatgatttaaattacaaaaaaaagagaatcatcatgctattatac [ 30720]
AY261362.1_Mkuzi_1979          -------------------------------------------------------------------------------- [ 30720]
AY261361.1_Malawi_Lil-20/1_1983 -------------------------------------------------------------------------------- [ 30720]
AY261360.1_Kenya_1950          -------------------------------------------------------------------------------- [ 30720]
AM712240.1_OURT_88/3           -....tc.....att.tttttgttaatataacagaaatgatttaaattacaaaaaaaagagaatcatcatgctattatac [ 30720]
AM712239.1_Benin_97/1          -....tc.....att.tttttgttaatataacagaaatgatttaaattacaaaaaaaagagaatcatcatgctattatac [ 30720]
KM111295.1_Ken06.Bus           -------------------------------------------------------------------------------- [ 30720]
KM111294.1_Ken05/Tk1           -------------------------------------------------------------------------------- [ 30720]
KM262844.1_strain_L60          -....tc.....att.tttttgttaatataacagaaatgatttaaattacaaaaaaaagagaatcatcatgctattatac [ 30720]
KM102979.1_26544/OG10_Italy    -....tc.....att.tttttgttaatataacagaaatgatttaaattacaaaaaaaagagaatcatcatgctattatac [ 30720]
KM262845.1_strain_NHV          -....tc.....att.tttttgttaatataacagaaatgatttaaattacaaaaaaaagagaatcatcatgctattatac [ 30720]
KX354450.1_47/Ss/2008          -....tc.....att.tttttgttaatataacagaaatgatttaaattacaaaaaaaagagaatcatcatgctattatac [ 30720]
KP055815.1_strain_BA71         -....tc.....att.tttttgttaatataacagaaatgatttaaattacaaaaaaaagagaatcatcatgctattatac [ 30720]
MH025920.1_strain_R35          -------------------------------------------------------------------------------- [ 30720]
MH025919.1_strain_N10          -------------------------------------------------------------------------------- [ 30720]
MH025918.1_strain_R25          -------------------------------------------------------------------------------- [ 30720]
MH025917.1_strain_R7           -------------------------------------------------------------------------------- [ 30720]
MH025916.1_strain_R8           -------------------------------------------------------------------------------- [ 30720]
```

```
FR682468.2_Georgia_2007/1       -------------------------------------------------------------------------------- [ 30800]
ASF_IND_20_CAD543               attgttattatagtagcatatgtaagttataaatnnnnnnnnnnnnnnnnnnnnnnnnnnnnnnnnnnnnnnnnnnnnnn [ 30800]
LR899193.1_Germany_2020/1       -------------------------------------------------------------------------------- [ 30800]
MN393477.1_Wuhan_2019           -------------------------------------------------------------------------------- [ 30800]
MW396979.1_Timor-Leste/2019/1   -------------------------------------------------------------------------------- [ 30800]
MK940252.1_CN/2019/InnerMongolia -------------------------------------------------------------------------------- [ 30800]
MK128995.1_China/2018/AnhuiXCGQ -------------------------------------------------------------------------------- [ 30800]
MH681419.1_ASFV/POL/2015        -------------------------------------------------------------------------------- [ 30800]
LS478113.1_Estonia_2014         -------------------------------------------------------------------------------- [ 30800]
KP843857.1_Odintsovo_02/14      -------------------------------------------------------------------------------- [ 30800]
FN557520.1_strain_E75           attgttattatagtagcatatgtaagttataaattggttccaaaacaatattggccaatactcatgtttatggcttacat [ 30800]
AY261366.1_Warthog              -------------------------------------------------------------------------------- [ 30800]
AY261365.1_Warmbaths            -------------------------------------------------------------------------------- [ 30800]
AY261364.1_Tengani_62           -------------------------------------------------------------------------------- [ 30800]
AY261363.1_Pretoriuskop/96/4    attgttattatagtagcatatgtaagttataaattggttccaaaacaatattggccaatactcatgtttatggcttacat [ 30800]
AY261362.1_Mkuzi_1979           -------------------------------------------------------------------------------- [ 30800]
AY261361.1_Malawi_Lil-20/1_1983 -------------------------------------------------------------------------------- [ 30800]
AY261360.1_Kenya_1950           -------------------------------------------------------------------------------- [ 30800]
AM712240.1_OURT_88/3            attgttattatagtagcatatgtaagttataaattggttccaaaacaatattggccaatactcatgtttatggcttacat [ 30800]
AM712239.1_Benin_97/1           attgttattatagtagcatatgtaagttataaattggttccaaaacaatattggccaatactcatgtttatggcttacat [ 30800]
KM111295.1_Ken06.Bus            -------------------------------------------------------------------------------- [ 30800]
KM111294.1_Ken05/Tk1            -------------------------------------------------------------------------------- [ 30800]
KM262844.1_strain_L60           attgttattatagtagcatatgtaagttataaattggttccaaaacaatattggccaatactcatgtttatggcttacat [ 30800]
KM102979.1_26544/OG10_Italy     attgttattatagtagcatatgtaagttataaattggttccaaaacaatattggccaatactcatgtttatggcttacat [ 30800]
KM262845.1_strain_NHV           attgttattatagtagcatatgtaagttataaattggttccaaaacaatattggccaatactcatgtttatggcttacat [ 30800]
KX354450.1_47/Ss/2008           attgttattatagtagcatatgtaagttataaattggttccaaaacaatattggccaatactcatgtttatggcttacat [ 30800]
KP055815.1_strain_BA71          attgttattatagtagcatatgtaagttataaattggttccaaaacaatattggccaatactcatgtttatggcttacat [ 30800]
MH025920.1_strain_R35           -------------------------------------------------------------------------------- [ 30800]
MH025919.1_strain_N10           -------------------------------------------------------------------------------- [ 30800]
MH025918.1_strain_R25           -------------------------------------------------------------------------------- [ 30800]
MH025917.1_strain_R7            -------------------------------------------------------------------------------- [ 30800]
MH025916.1_strain_R8            -------------------------------------------------------------------------------- [ 30800]

FR682468.2_Georgia_2007/1       -----------------------gtataa--------------------------------------------------- [ 30880]
ASF_IND_20_CAD543               nnnnnnnnnncatgaaaaattaga.....nnnnnnnnnnnnnnnnnnnnnnnnnnnnnnnnnnnnnnnnnnnnnnnnnnn [ 30880]
LR899193.1_Germany_2020/1       -----------------------......--------------------------------------------------- [ 30880]
MN393477.1_Wuhan_2019           -----------------------......--------------------------------------------------- [ 30880]
MW396979.1_Timor-Leste/2019/1   -----------------------......--------------------------------------------------- [ 30880]
MK940252.1_CN/2019/InnerMongolia -----------------------......--------------------------------------------------- [ 30880]
MK128995.1_China/2018/AnhuiXCGQ -----------------------......--------------------------------------------------- [ 30880]
MH681419.1_ASFV/POL/2015        -----------------------......--------------------------------------------------- [ 30880]
LS478113.1_Estonia_2014         -----------------------......--------------------------------------------------- [ 30880]
KP843857.1_Odintsovo_02/14      -----------------------......--------------------------------------------------- [ 30880]
```

```
FN557520.1_strain_E75           ggtatatacccatgaaaaattaga.....atgaaagatccggattttggaaatatattatagcaaaactattcagatgcc [ 30880]
AY261366.1_Warthog              ----tatacttatagcgtttaata.------------------------------------------------------- [ 30880]
AY261365.1_Warmbaths            ---gtatgcctgtataaaataat--------------------------------------------------------- [ 30880]
AY261364.1_Tengani_62           -------------------------------------------------------------------------------- [ 30880]
AY261363.1_Pretoriuskop/96/4    ggtatatacctatgaaaaattaga.....atgaaagatctggattttggaaatatattatagcaaaattattcagatgcc [ 30880]
AY261362.1_Mkuzi_1979           -------------------------------------------------------------------------------- [ 30880]
AY261361.1_Malawi_Lil-20/1_1983 -------------------------------------------------------------------------------- [ 30880]
AY261360.1_Kenya_1950           -------------------------------------------------------------------------------- [ 30880]
AM712240.1_OURT_88/3            ggtatatacccatgaaaaattaga.....atgaaagatccggattttggaaatatattatagcaaaactattcagatgcc [ 30880]
AM712239.1_Benin_97/1           ggtatatacccatgaaaaattaga.....atgaaagatccggattttggaaatatattatagcaaaactattcagatgcc [ 30880]
KM111295.1_Ken06.Bus            -------------------------------------------------------------------------------- [ 30880]
KM111294.1_Ken05/Tk1            -------------------------------------------------------------------------------- [ 30880]
KM262844.1_strain_L60           ggtatatacccatgaaaaattaga.....atgaaagatccggattttggaaatatattatagcaaaactattcagatgcc [ 30880]
KM102979.1_26544/OG10_Italy     ggtatatacccatgaaaaattaga.....atgaaagatccggattttggaaatatattatagcaaaactattcagatgcc [ 30880]
KM262845.1_strain_NHV           ggtatatacccatgaaaaattaga.....atgaaagatccggattttggaaatatattatagcaaaactattcagatgcc [ 30880]
KX354450.1_47/Ss/2008           ggtatatacccatgaaaaattaga.....atgaaagatccggattttggaaatatattatagcaaaactattcagatgcc [ 30880]
KP055815.1_strain_BA71          ggtatatacccatgaaaaattaga.....atgaaagatccggattttggaaatatattatagcaaaactattcagatgcc [ 30880]
MH025920.1_strain_R35           -------------------------------------------------------------------------------- [ 30880]
MH025919.1_strain_N10           -------------------------------------------------------------------------------- [ 30880]
MH025918.1_strain_R25           -------------------------------------------------------------------------------- [ 30880]
MH025917.1_strain_R7            -------------------------------------------------------------------------------- [ 30880]
MH025916.1_strain_R8            -------------------------------------------------------------------------------- [ 30880]

FR682468.2_Georgia_2007/1       -----------------------------------------------aataattaaccttttt----------------- [ 30960]
ASF_IND_20_CAD543               nnnnnnnnnnnnnnnnnnnnnnnnnnnnnnnnnnnnnnnnnnnnnnt.......g..ta....ttgnnnnnnnnnnnnnn [ 30960]
LR899193.1_Germany_2020/1       -----------------------------------------------................----------------- [ 30960]
MN393477.1_Wuhan_2019           -----------------------------------------------................----------------- [ 30960]
MW396979.1_Timor-Leste/2019/1   -----------------------------------------------................----------------- [ 30960]
MK940252.1_CN/2019/InnerMongolia -----------------------------------------------................----------------- [ 30960]
MK128995.1_China/2018/AnhuiXCGQ -----------------------------------------------................----------------- [ 30960]
MH681419.1_ASFV/POL/2015        -----------------------------------------------................----------------- [ 30960]
LS478113.1_Estonia_2014         -----------------------------------------------................----------------- [ 30960]
KP843857.1_Odintsovo_02/14      -----------------------------------------------................----------------- [ 30960]
FN557520.1_strain_E75           acggatgtgaaatatgtaagtgatattaaaaacataatacgaagtgt.......g..ta....ttgttgaatggaaactg [ 30960]
AY261366.1_Warthog              -------------------------------------------------------------------------------- [ 30960]
AY261365.1_Warmbaths            -------------------------------------------------------------------------------- [ 30960]
AY261364.1_Tengani_62           -------------------------------------------------------------------------------- [ 30960]
AY261363.1_Pretoriuskop/96/4    acggatgtggaatatgtaagtgatattaaaaacataatacgaagtgt.......g..ta....ttgttgaatggaaactg [ 30960]
AY261362.1_Mkuzi_1979           -------------------------------------------------------------------------------- [ 30960]
AY261361.1_Malawi_Lil-20/1_1983 -------------------------------------------------------------------------------- [ 30960]
AY261360.1_Kenya_1950           -------------------------------------------------------------------------------- [ 30960]
AM712240.1_OURT_88/3            acggatgtgaaatatgtaagtgatattaaaaacataatacgaagtgt.......g..ta....ttgttgaatggaaactg [ 30960]
AM712239.1_Benin_97/1           acggatgtgaaatatgtaagtgatattaaaaacataatacgaagtgt.......g..ta....ttgttgaatggaaactg [ 30960]
KM111295.1_Ken06.Bus            -------------------------------------------------------------------------------- [ 30960]
```

```
KM111294.1_Ken05/Tk1            -------------------------------------------------------------------------------- [ 30960]
KM262844.1_strain_L60           acggatgtgaaatatgtaagtgatattaaaaacataatacgaagtgt.......g..ta....ttgttgaatggaaactg [ 30960]
KM102979.1_26544/OG10_Italy     acggatgtgaaatatgtaagtgatattaaaaacataatacgaagtgt.......g..ta....ttgttgaatggaaactg [ 30960]
KM262845.1_strain_NHV           acggatgtgaaatatgtaagtgatattaaaaacataatacgaagtgt.......g..ta....ttgttgaatggaaactg [ 30960]
KX354450.1_47/Ss/2008           acggatgtgaaatatgtaagtgatattaaaaacataatacgaagtgt.......g..ta....ttgttgaatggaaactg [ 30960]
KP055815.1_strain_BA71          acggatgtgaaatatgtaagtgatattaaaaacataatacgaagtgt.......g..ta....ttgttgaatggaaactg [ 30960]
MH025920.1_strain_R35           -------------------------------------------------------------------------------- [ 30960]
MH025919.1_strain_N10           -------------------------------------------------------------------------------- [ 30960]
MH025918.1_strain_R25           -------------------------------------------------------------------------------- [ 30960]
MH025917.1_strain_R7            -------------------------------------------------------------------------------- [ 30960]
MH025916.1_strain_R8            -------------------------------------------------------------------------------- [ 30960]

FR682468.2_Georgia_2007/1       -------------------------------------------------------------------------------- [ 31040]
ASF_IND_20_CAD543               nnnnnnagtaatatgttgaacataannnnnnnnnnnnnnnnnnnnnnnnnnnnnttgtagtaattgtactaaatgnnnnnn [ 31040]
LR899193.1_Germany_2020/1       -------------------------------------------------------------------------------- [ 31040]
MN393477.1_Wuhan_2019           -------------------------------------------------------------------------------- [ 31040]
MW396979.1_Timor-Leste/2019/1   -------------------------------------------------------------------------------- [ 31040]
MK940252.1_CN/2019/InnerMongolia -------------------------------------------------------------------------------- [ 31040]
MK128995.1_China/2018/AnhuiXCGQ -------------------------------------------------------------------------------- [ 31040]
MH681419.1_ASFV/POL/2015        -------------------------------------------------------------------------------- [ 31040]
LS478113.1_Estonia_2014         -------------------------------------------------------------------------------- [ 31040]
KP843857.1_Odintsovo_02/14      -------------------------------------------------------------------------------- [ 31040]
FN557520.1_strain_E75           tttttgagtaatatgttgaacataatggtttttatagatgttaaatcatttgattgtagtaattgtactaaatgatgtga [ 31040]
AY261366.1_Warthog              -------------------------------------------------------------------------------- [ 31040]
AY261365.1_Warmbaths            -------------------------------------------------------------------------------- [ 31040]
AY261364.1_Tengani_62           -------------------------------------------------------------------------------- [ 31040]
AY261363.1_Pretoriuskop/96/4    tttttgagtaacatgttgaacataatagtttttatagatgttaaatcatttgattgtagtaattatactaaatgatgtga [ 31040]
AY261362.1_Mkuzi_1979           -------------------------------------------------------------------------------- [ 31040]
AY261361.1_Malawi_Lil-20/1_1983 -------------------------------------------------------------------------------- [ 31040]
AY261360.1_Kenya_1950           -------------------------------------------------------------------------------- [ 31040]
AM712240.1_OURT_88/3            tttttgagtaatatgttgaacataatggtttttatagatgttaaatcatttgattgtagtaattgtactaaatgatgtga [ 31040]
AM712239.1_Benin_97/1           tttttgagtaatatgttgaacataatggtttttatagatgttaaatcatttgattgtagtaattgtactaaatgatgtga [ 31040]
KM111295.1_Ken06.Bus            -------------------------------------------------------------------------------- [ 31040]
KM111294.1_Ken05/Tk1            -------------------------------------------------------------------------------- [ 31040]
KM262844.1_strain_L60           tttttgagtaatatgttgaacataatggtttttatagatgttaaatcatttgattgtagtaattgtactaaatgatgtga [ 31040]
KM102979.1_26544/OG10_Italy     tttttgagtaatatgttgaacataatggtttttatagatgttaaatcatttgattgtagtaattgtactaaatgatgtga [ 31040]
KM262845.1_strain_NHV           tttttgagtaatatgttgaacataatggtttttatagatgttaaatcatttgattgtagtaattgtactaaatgatgtga [ 31040]
KX354450.1_47/Ss/2008           tttttgagtaatatgttgaacataatggtttttatagatgttaaatcatttgattgtagtaattgtactaaatgatgtga [ 31040]
KP055815.1_strain_BA71          tttttgagtaatatgttgaacataatggtttttatagatgttaaatcatttgattgtagtaattgtactaaatgatgtga [ 31040]
MH025920.1_strain_R35           -------------------------------------------------------------------------------- [ 31040]
MH025919.1_strain_N10           -------------------------------------------------------------------------------- [ 31040]
MH025918.1_strain_R25           -------------------------------------------------------------------------------- [ 31040]
MH025917.1_strain_R7            -------------------------------------------------------------------------------- [ 31040]
MH025916.1_strain_R8            -------------------------------------------------------------------------------- [ 31040]
```

```
FR682468.2_Georgia_2007/1         ----------------gatcaaaatttaa-tttttttataaaaagctacagagtagtgttttattaaacgtggcttattt [ 31120]
ASF_IND_20_CAD543                 nnnnnnnnnnnnnnnnn............-.....................---.......................... [ 31120]
LR899193.1_Germany_2020/1         ----------------.............-.................................................. [ 31120]
MN393477.1_Wuhan_2019             ----------------.............-.................................................. [ 31120]
MW396979.1_Timor-Leste/2019/1     ----------------.............-.................................................. [ 31120]
MK940252.1_CN/2019/InnerMongolia  ----------------.............-.................................................. [ 31120]
MK128995.1_China/2018/AnhuiXCGQ   ----------------.............-.................................................. [ 31120]
MH681419.1_ASFV/POL/2015          ----------------.............-.................................................. [ 31120]
LS478113.1_Estonia_2014           ----------------.............-.................................................. [ 31120]
KP843857.1_Odintsovo_02/14        ----------------.............-.................................................. [ 31120]
FN557520.1_strain_E75             gatataaacctagttca............-.....................---.................a........ [ 31120]
AY261366.1_Warthog                ggactataaacattttc............-.......g.......-.......-t.g..............a.t...c.. [ 31120]
AY261365.1_Warmbaths              ------taaccttttt...t.........-.......................-t.a..............a........ [ 31120]
AY261364.1_Tengani_62             --------------ttc............t..............t........-t.g..............a........ [ 31120]
AY261363.1_Pretoriuskop/96/4      gatataaaactagttt.......t.....-.......g.......-.......-t.g......-.......a........ [ 31120]
AY261362.1_Mkuzi_1979             --------------------------------............aa...----------------......a....t... [ 31120]
AY261361.1_Malawi_Lil-20/1_1983   ----------tgttttc............-.g.gc.ata.....ag...g....ga.aga...catc.a..a...----- [ 31120]
AY261360.1_Kenya_1950             --------------ct.............---...........-.g.......-t.g..............a.......- [ 31120]
AM712240.1_OURT_88/3              gatataaacctagttca............-.....................---.................a........ [ 31120]
AM712239.1_Benin_97/1             gatataaacctagttca............-.....................---.................a........ [ 31120]
KM111295.1_Ken06.Bus              ----------tagttta............---...........cta.......-t.g..............a.......- [ 31120]
KM111294.1_Ken05/Tk1              ----------tagttt.............---...........-.g.......-t.g..............a.......- [ 31120]
KM262844.1_strain_L60             gatataaacctagttca............-.....................---.................a........ [ 31120]
KM102979.1_26544/OG10_Italy       gatataaacctagttca............-.....................---.................a........ [ 31120]
KM262845.1_strain_NHV             gatataaacctagttca............-.....................---.................a........ [ 31120]
KX354450.1_47/Ss/2008             gatataaacctagttca............-.....................---.................a........ [ 31120]
KP055815.1_strain_BA71            gatataaacctagttca............-.....................---.................a........ [ 31120]
MH025920.1_strain_R35             ----------tagttta............---...........cta.......-t.g..............a.......- [ 31120]
MH025919.1_strain_N10             ----------tagttta............---...........cta.......-t.g..............a.......- [ 31120]
MH025918.1_strain_R25             ----------tagttta............---...........cta.......-t.g..............a.......- [ 31120]
MH025917.1_strain_R7              ----------tagttta............---...........cta.......-t.g..............a.......- [ 31120]
MH025916.1_strain_R8              ----------tagttta............---...........cta.......-t.g..............a.......- [ 31120]

FR682468.2_Georgia_2007/1         -aaaagttacacaatgttaaaatctctacttactttaattctttgtggggttttattaactttatccatattatggctta [ 31200]
ASF_IND_20_CAD543                 a............................................................................... [ 31200]
LR899193.1_Germany_2020/1         -............................................................................... [ 31200]
MN393477.1_Wuhan_2019             -............................................................................... [ 31200]
MW396979.1_Timor-Leste/2019/1     -............................................................................... [ 31200]
MK940252.1_CN/2019/InnerMongolia  -............................................................................... [ 31200]
MK128995.1_China/2018/AnhuiXCGQ   -............................................................................... [ 31200]
MH681419.1_ASFV/POL/2015          -............................................................................... [ 31200]
LS478113.1_Estonia_2014           -............................................................................... [ 31200]
KP843857.1_Odintsovo_02/14        -............................................................................... [ 31200]
```

```
FN557520.1_strain_E75            a............................................................................... [ 31200]
AY261366.1_Warthog               ---.....t..tc............t.....t........a......t...........t.................... [ 31200]
AY261365.1_Warmbaths             a-.............................................................................. [ 31200]
AY261364.1_Tengani_62            -................................................a..........g................... [ 31200]
AY261363.1_Pretoriuskop/96/4     a..........................................................t.................... [ 31200]
AY261362.1_Mkuzi_1979            a..---.ct..tc...c........t.......................a................t............. [ 31200]
AY261361.1_Malawi_Lil-20/1_1983  -----a..t..tg............t.....t...............t............c..t................ [ 31200]
AY261360.1_Kenya_1950            ---....ct..tc............t.....t........a......t............c..t................ [ 31200]
AM712240.1_OURT_88/3             a............................................................................... [ 31200]
AM712239.1_Benin_97/1            a............................................................................... [ 31200]
KM111295.1_Ken06.Bus             ---....ct..tc.......g....t.....t.....t......................c..t................ [ 31200]
KM111294.1_Ken05/Tk1             ---....ct..tc............t.....t........a......t............c..t................ [ 31200]
KM262844.1_strain_L60            a............................................................................... [ 31200]
KM102979.1_26544/OG10_Italy      a............................................................................... [ 31200]
KM262845.1_strain_NHV            a............................................................................... [ 31200]
KX354450.1_47/Ss/2008            a............................................................................... [ 31200]
KP055815.1_strain_BA71           a............................................................................... [ 31200]
MH025920.1_strain_R35            ---....ct..tc.......g....t.....t.....t......................c..t................ [ 31200]
MH025919.1_strain_N10            ---....ct..tc.......g....t.....t.....t......................c..t................ [ 31200]
MH025918.1_strain_R25            ---....ct..tc.......g....t.....t.....t......................c..t................ [ 31200]
MH025917.1_strain_R7             ---....ct..tc.......g....t.....t.....t......................c..t................ [ 31200]
MH025916.1_strain_R8             ---....ct..tc.......g....t.....t.....t......................c..t................ [ 31200]

FR682468.2_Georgia_2007/1        ctacttaccatgtagaacttatagaggcaatagatgatttctacgactgaaatatagaat-----------agtccattt [ 31280]
ASF_IND_20_CAD543                y...........................................................-----------......... [ 31280]
LR899193.1_Germany_2020/1        ............................................................-----------......... [ 31280]
MN393477.1_Wuhan_2019            ............................................................-----------......... [ 31280]
MW396979.1_Timor-Leste/2019/1    ............................................................-----------......... [ 31280]
MK940252.1_CN/2019/InnerMongolia ............................................................-----------......... [ 31280]
MK128995.1_China/2018/AnhuiXCGQ  ............................................................-----------......... [ 31280]
MH681419.1_ASFV/POL/2015         ............................................................-----------......... [ 31280]
LS478113.1_Estonia_2014          ............................................................-----------......... [ 31280]
KP843857.1_Odintsovo_02/14       ............................................................-----------......... [ 31280]
FN557520.1_strain_E75            t........................a..................................-----------...t..... [ 31280]
AY261366.1_Warthog               ...tc..t....................................................-----------......... [ 31280]
AY261365.1_Warmbaths             ...tc..t....................................................-----------......... [ 31280]
AY261364.1_Tengani_62            ............................................................-----------......... [ 31280]
AY261363.1_Pretoriuskop/96/4     ...tc..t.....................c...gg.............a...........-----------.....---- [ 31280]
AY261362.1_Mkuzi_1979            ...tc..t...................................t................-----------......... [ 31280]
AY261361.1_Malawi_Lil-20/1_1983  .c..a..t.........t....g..a...c...gg.....................tc..aaattaatttc......... [ 31280]
AY261360.1_Kenya_1950            .c..g..t.........t...........c.g..g.....t.......a.......att.----------c.......g. [ 31280]
AM712240.1_OURT_88/3             t........................a..................................-----------...t..... [ 31280]
AM712239.1_Benin_97/1            t........................a..................................-----------...t..... [ 31280]
KM111295.1_Ken06.Bus             .c..g..t.........t...........c.g..g................--..tat..----------c....t..a. [ 31280]
```

```
KM111294.1_Ken05/Tk1              .c..g..t.........t...........t.g..g.....t.......a.......att.----------c.......g. [ 31280]
KM262844.1_strain_L60             t........................a..................................-----------...t..... [ 31280]
KM102979.1_26544/OG10_Italy       t........................a..................................-----------...t..... [ 31280]
KM262845.1_strain_NHV             t........................a..................................-----------...t..... [ 31280]
KX354450.1_47/Ss/2008             t........................a..................................-----------...t..... [ 31280]
KP055815.1_strain_BA71            t........................a..................................-----------...t..... [ 31280]
MH025920.1_strain_R35             .c..g..t.........t...........c.g..g................--..tat..----------c....t..a. [ 31280]
MH025919.1_strain_N10             .c..g..t.........t...........c.g..g................--..tat..----------c....t..a. [ 31280]
MH025918.1_strain_R25             .c..g..t.........t...........c.g..g................--..tat..----------c....t..a. [ 31280]
MH025917.1_strain_R7              .c..g..t.........t...........c.g..g................--..tat..----------c....t..a. [ 31280]
MH025916.1_strain_R8              .c..g..t.........t...........c.g..g................--..tat..----------c....t..a. [ 31280]

FR682468.2_Georgia_2007/1         tctatttgtaaaataatgatttatattctttcctaaaaat------gatactttatatggtt--tgaaaacaaatattaa [ 31360]
ASF_IND_20_CAD543                 ........................................acgatc...c............--................ [ 31360]
LR899193.1_Germany_2020/1         ........................................------................--................ [ 31360]
MN393477.1_Wuhan_2019             ........................................------................--................ [ 31360]
MW396979.1_Timor-Leste/2019/1     ........................................------................--................ [ 31360]
MK940252.1_CN/2019/InnerMongolia  ........................................------................--................ [ 31360]
MK128995.1_China/2018/AnhuiXCGQ   ........................................------................--................ [ 31360]
MH681419.1_ASFV/POL/2015          ........................................------................--................ [ 31360]
LS478113.1_Estonia_2014           ........................................------................--................ [ 31360]
KP843857.1_Odintsovo_02/14        ........................................------................--................ [ 31360]
FN557520.1_strain_E75             .............................c..........acgatc...c.........c..--.......g........ [ 31360]
AY261366.1_Warthog                ...........................t............a----c............aa..--.a.............. [ 31360]
AY261365.1_Warmbaths              ........................................a----ca...........aa..--.a.............. [ 31360]
AY261364.1_Tengani_62             .......a................................a----t............aa..--.a.....g........ [ 31360]
AY261363.1_Pretoriuskop/96/4      ---.....................................a----t....t.......aa..--.a.....g........ [ 31360]
AY261362.1_Mkuzi_1979             ........................................a----t............aa..--.a.............. [ 31360]
AY261361.1_Malawi_Lil-20/1_1983   ........................................ata--ac.c.........aa..--....g--t....c... [ 31360]
AY261360.1_Kenya_1950             ............----------.........t......taa---gcc...t....a..ca..-aaa....atg...c... [ 31360]
AM712240.1_OURT_88/3              .............................c..........acgatc...c.........c..--.......g........ [ 31360]
AM712239.1_Benin_97/1             .............................c..........acgatc...c.........c..--.......g........ [ 31360]
KM111295.1_Ken06.Bus              ----------.c----------...a............taa---gac.........g.ca..-tga....atg...c... [ 31360]
KM111294.1_Ken05/Tk1              ............----------.........t......taa---gac...t....g..ca..aaaa....atg...c... [ 31360]
KM262844.1_strain_L60             .............................c..........acgatc...c.........c..--.......g........ [ 31360]
KM102979.1_26544/OG10_Italy       .............................c..........acgatc...c.........c..--.......g........ [ 31360]
KM262845.1_strain_NHV             .............................c..........acgatc...c.........c..--.......g........ [ 31360]
KX354450.1_47/Ss/2008             .............................c..........acgatc...c.........c..--.......g........ [ 31360]
KP055815.1_strain_BA71            .............................c..........acgatc...c.........c..--.......g........ [ 31360]
MH025920.1_strain_R35             ----------.c----------...a............taa---gac.........g.ca..-tga....atg...c... [ 31360]
MH025919.1_strain_N10             ----------.c----------...a............taa---gac.........g.ca..-tga....atg...c... [ 31360]
MH025918.1_strain_R25             ----------.c----------...a............taa---gac.........g.ca..-tga....atg...c... [ 31360]
MH025917.1_strain_R7              ----------.c----------...a............taa---gac.........g.ca..-tga....atg...c... [ 31360]
MH025916.1_strain_R8              ----------.c----------...a............taa---gac.........g.ca..-tga....atg...c... [ 31360]
```

```
FR682468.2_Georgia_2007/1        caacttga---------tttttttttc----------------------tataaataaactataaatgaaaatagta--- [ 31440]
ASF_IND_20_CAD543                ........---------..........----------------------............................--- [ 31440]
LR899193.1_Germany_2020/1        ........---------..........----------------------............................--- [ 31440]
MN393477.1_Wuhan_2019            ........---------..........----------------------............................--- [ 31440]
MW396979.1_Timor-Leste/2019/1    ........---------..........----------------------............................--- [ 31440]
MK940252.1_CN/2019/InnerMongolia ........---------..........----------------------............................--- [ 31440]
MK128995.1_China/2018/AnhuiXCGQ  ........---------..........----------------------............................--- [ 31440]
MH681419.1_ASFV/POL/2015         ........---------..........----------------------............................--- [ 31440]
LS478113.1_Estonia_2014          ........---------..........----------------------............................--- [ 31440]
KP843857.1_Odintsovo_02/14       ........---------..........----------------------............................--- [ 31440]
FN557520.1_strain_E75            ........---------..........----------------------............................--- [ 31440]
AY261366.1_Warthog               ........---------..........----------------------............................--- [ 31440]
AY261365.1_Warmbaths             ........---------..........----------------------............................--- [ 31440]
AY261364.1_Tengani_62            t.......---------..........----------------------............................--- [ 31440]
AY261363.1_Pretoriuskop/96/4     ........---------..........----------------------............................--- [ 31440]
AY261362.1_Mkuzi_1979            ........----------.........----------------------............................--- [ 31440]
AY261361.1_Malawi_Lil-20/1_1983  ..c...a.taatgttta..........tttatagtttctataaagaaat.....g...t................a.atc [ 31440]
AY261360.1_Kenya_1950            .g..g.a.taatgtttc-.........-----------tataaagaaa-a..-..a..ct..............ta.ata [ 31440]
AM712240.1_OURT_88/3             ........---------..........----------------------............................--- [ 31440]
AM712239.1_Benin_97/1            ........---------..........----------------------............................--- [ 31440]
KM111295.1_Ken06.Bus             .g..g.a.taatgtttc..........-----------tataaagaaa-a.....a..ct..............ta.ata [ 31440]
KM111294.1_Ken05/Tk1             .g..g.a.taatgtttc-.........-----------tataaagaaa-a..-..a..ct...............a.ata [ 31440]
KM262844.1_strain_L60            ........---------..........----------------------............................--- [ 31440]
KM102979.1_26544/OG10_Italy      ........---------..........----------------------............................--- [ 31440]
KM262845.1_strain_NHV            ........---------..........----------------------............................--- [ 31440]
KX354450.1_47/Ss/2008            ........---------..........----------------------............................--- [ 31440]
KP055815.1_strain_BA71           ........---------..........----------------------............................--- [ 31440]
MH025920.1_strain_R35            .g..g.a.taatgtttc..........-----------tataaagaaa-a.....a..ct..............ta.ata [ 31440]
MH025919.1_strain_N10            .g..g.a.taatgtttc..........-----------tataaagaaa-a.....a..ct..............ta.ata [ 31440]
MH025918.1_strain_R25            .g..g.a.taatgtttc..........-----------tataaagaaa-a.....a..ct..............ta.ata [ 31440]
MH025917.1_strain_R7             .g..g.a.taatgtttc..........-----------tataaagaaa-a.....a..ct..............ta.ata [ 31440]
MH025916.1_strain_R8             .g..g.a.taatgtttc..........-----------tataaagaaa-a.....a..ct..............ta.ata [ 31440]

FR682468.2_Georgia_2007/1        ---aaactcata--gagtcttataagtgaacatcttcataatgttactcaaacgttggactattaaaaaatattccgtgt [ 31520]
ASF_IND_20_CAD543                ---.........--....................--...m........................................ [ 31520]
LR899193.1_Germany_2020/1        ---.........--.................................................................. [ 31520]
MN393477.1_Wuhan_2019            ---.........--.................................................................. [ 31520]
MW396979.1_Timor-Leste/2019/1    ---.........--.................................................................. [ 31520]
MK940252.1_CN/2019/InnerMongolia ---.........--.................................................................. [ 31520]
MK128995.1_China/2018/AnhuiXCGQ  ---.........--.................................................................. [ 31520]
MH681419.1_ASFV/POL/2015         ---.........--.................................................................. [ 31520]
LS478113.1_Estonia_2014          ---.........--.................................................................. [ 31520]
KP843857.1_Odintsovo_02/14       ---.........--.................................................................. [ 31520]
```

```
FN557520.1_strain_E75             ---.........--....................--...c............................g...c....... [ 31520]
AY261366.1_Warthog                ---......------........g........................................................ [ 31520]
AY261365.1_Warmbaths              ---......------........g............................................g........... [ 31520]
AY261364.1_Tengani_62             ---........----........g......t..---...c...............c............g........... [ 31520]
AY261363.1_Pretoriuskop/96/4      ---.......----t........g............................................g.......a... [ 31520]
AY261362.1_Mkuzi_1979             ---......------........g............................................g...c....... [ 31520]
AY261361.1_Malawi_Lil-20/1_1983   g-t......t..ta.........g..........a.................ga....t...c.....g.....t..... [ 31520]
AY261360.1_Kenya_1950             gta......t..ta.........g........a.a................cgaa...t...c.....g.....t..... [ 31520]
AM712240.1_OURT_88/3              ---.........--....................--...c............................g...c....... [ 31520]
AM712239.1_Benin_97/1             ---.........--....................--...c............................g...c....... [ 31520]
KM111295.1_Ken06.Bus              gta......t..ta.........g...a......a.................ga....t...c.....g.....t..... [ 31520]
KM111294.1_Ken05/Tk1              gta......t..ta.........g..........a................c.a....g.........g.........a. [ 31520]
KM262844.1_strain_L60             ---.........--....................--...c............................g...c....... [ 31520]
KM102979.1_26544/OG10_Italy       ---.........--....................--...c............................g...c....... [ 31520]
KM262845.1_strain_NHV             ---.........--....................--...c............................g...c....... [ 31520]
KX354450.1_47/Ss/2008             ---.........--....................--...c............................g...c....... [ 31520]
KP055815.1_strain_BA71            ---.........--....................--...c............................g...c....... [ 31520]
MH025920.1_strain_R35             gta......t..ta.........g...a......a.................ga....t...c.....g.....t..... [ 31520]
MH025919.1_strain_N10             gta......t..ta.........g...a......a.................ga....t...c.....g.....t..... [ 31520]
MH025918.1_strain_R25             gta......t..ta.........g...a......a.................ga....t...c.....g.....t..... [ 31520]
MH025917.1_strain_R7              gta......t..ta.........g...a......a.................ga....t...c.....g.....t..... [ 31520]
MH025916.1_strain_R8              gta......t..ta.........g...a......a.................ga....t...c.....g.....t..... [ 31520]

FR682468.2_Georgia_2007/1         -gcattattgcttttaatcagtatgattactttatacgaagccgctattaaaacgcttatcacacaccgaaaacaaattt [ 31600]
ASF_IND_20_CAD543                 -............................................................................... [ 31600]
LR899193.1_Germany_2020/1         -............................................................................... [ 31600]
MN393477.1_Wuhan_2019             -............................................................................... [ 31600]
MW396979.1_Timor-Leste/2019/1     -............................................................................... [ 31600]
MK940252.1_CN/2019/InnerMongolia  -............................................................................... [ 31600]
MK128995.1_China/2018/AnhuiXCGQ   -............................................................................... [ 31600]
MH681419.1_ASFV/POL/2015          -............................................................................... [ 31600]
LS478113.1_Estonia_2014           -............................................................................... [ 31600]
KP843857.1_Odintsovo_02/14        -............................................................................... [ 31600]
FN557520.1_strain_E75             -..........................................................................g.... [ 31600]
AY261366.1_Warthog                -..........................................................................g.... [ 31600]
AY261365.1_Warmbaths              -..........................................................................g.... [ 31600]
AY261364.1_Tengani_62             -..........................................t...............................g.... [ 31600]
AY261363.1_Pretoriuskop/96/4      -..........................................................................g.... [ 31600]
AY261362.1_Mkuzi_1979             -.......................................a........................................ [ 31600]
AY261361.1_Malawi_Lil-20/1_1983   g.........-..................................................................... [ 31600]
AY261360.1_Kenya_1950             -.........-..................................................................... [ 31600]
AM712240.1_OURT_88/3              -..........................................................................g.... [ 31600]
AM712239.1_Benin_97/1             -..........................................................................g.... [ 31600]
KM111295.1_Ken06.Bus              -.........-.......t..........................c.................................. [ 31600]
```

```
KM111294.1_Ken05/Tk1             -.........-................c.................................................... [ 31600]
KM262844.1_strain_L60            -..........................................................................g.... [ 31600]
KM102979.1_26544/OG10_Italy      -..........................................................................g.... [ 31600]
KM262845.1_strain_NHV            -..........................................................................g.... [ 31600]
KX354450.1_47/Ss/2008            -..........................................................................g.... [ 31600]
KP055815.1_strain_BA71           -..........................................................................g.... [ 31600]
MH025920.1_strain_R35            -.........-.......t..........................c.................................. [ 31600]
MH025919.1_strain_N10            -.........-.......t..........................c.................................. [ 31600]
MH025918.1_strain_R25            -.........-.......t..........................c.................................. [ 31600]
MH025917.1_strain_R7             -.........-.......t..........................c.................................. [ 31600]
MH025916.1_strain_R8             -.........-.......t..........................c.................................. [ 31600]

FR682468.2_Georgia_2007/1        taaaacaccccgatagccgtgaaattttactagctttggggttgtactgggataaaactcatattcttgttaaatgtcgt [ 31680]
ASF_IND_20_CAD543                ................................................................................ [ 31680]
LR899193.1_Germany_2020/1        ................................................................................ [ 31680]
MN393477.1_Wuhan_2019            ................................................................................ [ 31680]
MW396979.1_Timor-Leste/2019/1    ................................................................................ [ 31680]
MK940252.1_CN/2019/InnerMongolia ................................................................................ [ 31680]
MK128995.1_China/2018/AnhuiXCGQ  ................................................................................ [ 31680]
MH681419.1_ASFV/POL/2015         ................................................................................ [ 31680]
LS478113.1_Estonia_2014          ................................................................................ [ 31680]
KP843857.1_Odintsovo_02/14       ................................................................................ [ 31680]
FN557520.1_strain_E75            ................................................................................ [ 31680]
AY261366.1_Warthog               ....................................................................c....g..c.ac [ 31680]
AY261365.1_Warmbaths             ........................................a..a........................c....g..c.a. [ 31680]
AY261364.1_Tengani_62            ........................................a..a........................c....g..c.a. [ 31680]
AY261363.1_Pretoriuskop/96/4     ................................................................................ [ 31680]
AY261362.1_Mkuzi_1979            ...........................................a.................................... [ 31680]
AY261361.1_Malawi_Lil-20/1_1983  ..........t..........................a..a..a..............g.........c....g..c.ag [ 31680]
AY261360.1_Kenya_1950            ..........t...............c.............a.........a.................c....g..c.ac [ 31680]
AM712240.1_OURT_88/3             ................................................................................ [ 31680]
AM712239.1_Benin_97/1            ................................................................................ [ 31680]
KM111295.1_Ken06.Bus             ..........t...............c.............a..a......a.................c....g....ac [ 31680]
KM111294.1_Ken05/Tk1             ........................................a..a......a.................c....g..c.ac [ 31680]
KM262844.1_strain_L60            ................................................................................ [ 31680]
KM102979.1_26544/OG10_Italy      ................................................................................ [ 31680]
KM262845.1_strain_NHV            ................................................................................ [ 31680]
KX354450.1_47/Ss/2008            ................................................................................ [ 31680]
KP055815.1_strain_BA71           ................................................................................ [ 31680]
MH025920.1_strain_R35            ..........t...............c.............a..a......a.................c....g....ac [ 31680]
MH025919.1_strain_N10            ..........t...............c.............a..a......a.................c....g....ac [ 31680]
MH025918.1_strain_R25            ..........t...............c.............a..a......a.................c....g....ac [ 31680]
MH025917.1_strain_R7             ..........t...............c.............a..a......a.................c....g....ac [ 31680]
MH025916.1_strain_R8             ..........t...............c.............a..a......a.................c....g....ac [ 31680]
```

```
FR682468.2_Georgia_2007/1        gaatgtgggaatatgagtcttaccggaaaacacagtacaaaatgtattaacattaattgtctacttattcttgccata-a [ 31760]
ASF_IND_20_CAD543                ..............................................................................-. [ 31760]
LR899193.1_Germany_2020/1        ..............................................................................-. [ 31760]
MN393477.1_Wuhan_2019            ..............................................................................-. [ 31760]
MW396979.1_Timor-Leste/2019/1    ..............................................................................-. [ 31760]
MK940252.1_CN/2019/InnerMongolia ..............................................................................-. [ 31760]
MK128995.1_China/2018/AnhuiXCGQ  ..............................................................................-. [ 31760]
MH681419.1_ASFV/POL/2015         ..............................................................................-. [ 31760]
LS478113.1_Estonia_2014          ..............................................................................-. [ 31760]
KP843857.1_Odintsovo_02/14       .............................................................................--. [ 31760]
FN557520.1_strain_E75            ...........a..................................................................-. [ 31760]
AY261366.1_Warthog               ........a..a..............g.......................t...........................-. [ 31760]
AY261365.1_Warmbaths             ........a..a..............g.......................t...........................-. [ 31760]
AY261364.1_Tengani_62            ........a..a..............c.......................t...........................-. [ 31760]
AY261363.1_Pretoriuskop/96/4     ...........a...........t......................................................-. [ 31760]
AY261362.1_Mkuzi_1979            ...........a...........t......................................................-. [ 31760]
AY261361.1_Malawi_Lil-20/1_1983  .....c..a..a..a...................................t...........................a. [ 31760]
AY261360.1_Kenya_1950            ........a..a..a...................................t...........g...............-. [ 31760]
AM712240.1_OURT_88/3             ...........a..................................................................-. [ 31760]
AM712239.1_Benin_97/1            ...........a..................................................................-. [ 31760]
KM111295.1_Ken06.Bus             ........a..a..a...................................t...........g...........t...-. [ 31760]
KM111294.1_Ken05/Tk1             ........a..a..a...................................t...........g...............-. [ 31760]
KM262844.1_strain_L60            ...........a..................................................................-. [ 31760]
KM102979.1_26544/OG10_Italy      ...........a..................................................................-. [ 31760]
KM262845.1_strain_NHV            ...........a..................................................................-. [ 31760]
KX354450.1_47/Ss/2008            ...........a..................................................................-. [ 31760]
KP055815.1_strain_BA71           ...........a..................................................................-. [ 31760]
MH025920.1_strain_R35            ........a..a..a...................................t...........g...........t...-. [ 31760]
MH025919.1_strain_N10            ........a..a..a...................................t...........g...........t...-. [ 31760]
MH025918.1_strain_R25            ........a..a..a...................................t...........g...........t...-. [ 31760]
MH025917.1_strain_R7             ........a..a..a...................................t...........g...........t...-. [ 31760]
MH025916.1_strain_R8             ........a..a..a...................................t...........g...........t...-. [ 31760]

FR682468.2_Georgia_2007/1        aaaaaaagaataagcgtattgttgataccttgataggaatgggcgcggatgtaacatatatacatcttttaaagaataag [ 31840]
ASF_IND_20_CAD543                ................................................................................ [ 31840]
LR899193.1_Germany_2020/1        ................................................................................ [ 31840]
MN393477.1_Wuhan_2019            ................................................................................ [ 31840]
MW396979.1_Timor-Leste/2019/1    ................................................................................ [ 31840]
MK940252.1_CN/2019/InnerMongolia ................................................................................ [ 31840]
MK128995.1_China/2018/AnhuiXCGQ  ................................................................................ [ 31840]
MH681419.1_ASFV/POL/2015         ................................................................................ [ 31840]
LS478113.1_Estonia_2014          ................................................................................ [ 31840]
KP843857.1_Odintsovo_02/14       ................................................................................ [ 31840]
```

```
FN557520.1_strain_E75           ...................g...............a.......t.................................... [ 31840]
AY261366.1_Warthog              ...................g...............a.......t.................................... [ 31840]
AY261365.1_Warmbaths            ...................g...............a.......t..a..................t.............. [ 31840]
AY261364.1_Tengani_62           ...................g.......................t.............................a...... [ 31840]
AY261363.1_Pretoriuskop/96/4    ...................g...............a.......t.................................... [ 31840]
AY261362.1_Mkuzi_1979           ...................g...............a.......t.................................... [ 31840]
AY261361.1_Malawi_Lil-20/1_1983 ...........................................t..........a..........t.......t...... [ 31840]
AY261360.1_Kenya_1950           ...................g.................c.....t.....................t.............. [ 31840]
AM712240.1_OURT_88/3            ...................g...............a.......t.................................... [ 31840]
AM712239.1_Benin_97/1           ...................g...............a.......t.................................... [ 31840]
KM111295.1_Ken06.Bus            ...................g...........a.....c.....t.....................t.............. [ 31840]
KM111294.1_Ken05/Tk1            .................g.g.................c.....t.....................t.............. [ 31840]
KM262844.1_strain_L60           ...................g...............a.......t.................................... [ 31840]
KM102979.1_26544/OG10_Italy     ...................g...............a.......t.................................... [ 31840]
KM262845.1_strain_NHV           ...................g...............a.......t.................................... [ 31840]
KX354450.1_47/Ss/2008           ...................g...............a.......t.................................... [ 31840]
KP055815.1_strain_BA71          ...................g...............a.......t.................................... [ 31840]
MH025920.1_strain_R35           ...................g...........a.....c.....t.....................t.............. [ 31840]
MH025919.1_strain_N10           ...................g...........a.....c.....t.....................t.............. [ 31840]
MH025918.1_strain_R25           ...................g...........a.....c.....t.....................t.............. [ 31840]
MH025917.1_strain_R7            ...................g...........a.....c.....t.....................t.............. [ 31840]
MH025916.1_strain_R8            ...................g...........a.....c.....t.....................t.............. [ 31840]

FR682468.2_Georgia_2007/1       ataaaactgtcatacaaccagctgtctatgcttaaaagcaactcgcagatttcattgaaggagcttcatgctatatgcta [ 31920]
ASF_IND_20_CAD543               ................................................................................ [ 31920]
LR899193.1_Germany_2020/1       ................................................................................ [ 31920]
MN393477.1_Wuhan_2019           ................................................................................ [ 31920]
MW396979.1_Timor-Leste/2019/1   ................................................................................ [ 31920]
MK940252.1_CN/2019/InnerMongolia ................................................................................ [ 31920]
MK128995.1_China/2018/AnhuiXCGQ ................................................................................ [ 31920]
MH681419.1_ASFV/POL/2015        ................................................................................ [ 31920]
LS478113.1_Estonia_2014         ................................................................................ [ 31920]
KP843857.1_Odintsovo_02/14      ................................................................................ [ 31920]
FN557520.1_strain_E75           .....................................................g.......................... [ 31920]
AY261366.1_Warthog              ............................................a......g.g.......................t.. [ 31920]
AY261365.1_Warmbaths            ..................g.........................a........g.......................... [ 31920]
AY261364.1_Tengani_62           .....................t.............................g.g..a......t.......c.....t.. [ 31920]
AY261363.1_Pretoriuskop/96/4    .....................................................g.......................... [ 31920]
AY261362.1_Mkuzi_1979           .....................................................g.......................... [ 31920]
AY261361.1_Malawi_Lil-20/1_1983 .c......a.......................c.........c.c........g.........t.............t.. [ 31920]
AY261360.1_Kenya_1950           .c.........................gct.......................g.........t...g............ [ 31920]
AM712240.1_OURT_88/3            .....................................................g.......................... [ 31920]
AM712239.1_Benin_97/1           .....................................................g.......................... [ 31920]
KM111295.1_Ken06.Bus            .c......a..................gct.......................g.........t.............t.. [ 31920]
```

```
KM111294.1_Ken05/Tk1            .c.........................gct.......................g.........t...g............ [ 31920]
KM262844.1_strain_L60           .....................................................g.......................... [ 31920]
KM102979.1_26544/OG10_Italy     .....................................................g.......................... [ 31920]
KM262845.1_strain_NHV           .....................................................g.......................... [ 31920]
KX354450.1_47/Ss/2008           .....................................................g.......................... [ 31920]
KP055815.1_strain_BA71          .....................................................g.......................... [ 31920]
MH025920.1_strain_R35           .c......a..................gct.......................g.........t.............t.. [ 31920]
MH025919.1_strain_N10           .c......a..................gct.......................g.........t.............t.. [ 31920]
MH025918.1_strain_R25           .c......a..................gct.......................g.........t.............t.. [ 31920]
MH025917.1_strain_R7            .c......a..................gct.......................g.........t.............t.. [ 31920]
MH025916.1_strain_R8            .c......a..................gct.......................g.........t.............t.. [ 31920]

FR682468.2_Georgia_2007/1       tcttttatatggtcggcttcccaaaaaaattaaacaagggatgcgactgtgtaaaacaatggcgggactatgtggtgaac [ 32000]
ASF_IND_20_CAD543               ................................................................................ [ 32000]
LR899193.1_Germany_2020/1       ................................................................................ [ 32000]
MN393477.1_Wuhan_2019           ................................................................................ [ 32000]
MW396979.1_Timor-Leste/2019/1   ................................................................................ [ 32000]
MK940252.1_CN/2019/InnerMongolia ................................................................................ [ 32000]
MK128995.1_China/2018/AnhuiXCGQ ................................................................................ [ 32000]
MH681419.1_ASFV/POL/2015        ................................................................................ [ 32000]
LS478113.1_Estonia_2014         ................................................................................ [ 32000]
KP843857.1_Odintsovo_02/14      ................................................................................ [ 32000]
FN557520.1_strain_E75           c...........................................a................................... [ 32000]
AY261366.1_Warthog              c...........c...............................a................................... [ 32000]
AY261365.1_Warmbaths            c...........................................a................................... [ 32000]
AY261364.1_Tengani_62           c...........c...............................a.t.........................c....... [ 32000]
AY261363.1_Pretoriuskop/96/4    c...........................................a................................... [ 32000]
AY261362.1_Mkuzi_1979           c...........................................a................................... [ 32000]
AY261361.1_Malawi_Lil-20/1_1983 ca..........c..c..........................................c.............c....... [ 32000]
AY261360.1_Kenya_1950           ca........a.c..t..........................a.............................c..g.... [ 32000]
AM712240.1_OURT_88/3            c...........................................a................................... [ 32000]
AM712239.1_Benin_97/1           c...........................................a................................... [ 32000]
KM111295.1_Ken06.Bus            ca........a....t..........................a.....a.......................c..g.... [ 32000]
KM111294.1_Ken05/Tk1            c.........a.c..t.......................a....a...........................c....... [ 32000]
KM262844.1_strain_L60           c...........................................a................................... [ 32000]
KM102979.1_26544/OG10_Italy     c...........................................a................................... [ 32000]
KM262845.1_strain_NHV           c...........................................a................................... [ 32000]
KX354450.1_47/Ss/2008           c...........................................a................................... [ 32000]
KP055815.1_strain_BA71          c...........................................a................................... [ 32000]
MH025920.1_strain_R35           ca........a....t..........................a.....a.......................c..g.... [ 32000]
MH025919.1_strain_N10           ca........a....t..........................a.....a.......................c..g.... [ 32000]
MH025918.1_strain_R25           ca........a....t..........................a.....a.......................c..g.... [ 32000]
MH025917.1_strain_R7            ca........a....t..........................a.....a.......................c..g.... [ 32000]
MH025916.1_strain_R8            ca........a....t..........................a.....a.......................c..g.... [ 32000]
```

```
FR682468.2_Georgia_2007/1        ttttatgtgcatttttagctccgtaaatg-ataatatgtattt---------------------------aaaacaaaca [ 32080]
ASF_IND_20_CAD543                .............................t.at........a.---------------------------........y. [ 32080]
LR899193.1_Germany_2020/1        .............................-.............---------------------------.......... [ 32080]
MN393477.1_Wuhan_2019            .............................-.............---------------------------.......... [ 32080]
MW396979.1_Timor-Leste/2019/1    .............................-.............---------------------------.......... [ 32080]
MK940252.1_CN/2019/InnerMongolia .............................-.............---------------------------.......... [ 32080]
MK128995.1_China/2018/AnhuiXCGQ  .............................-.............---------------------------.......... [ 32080]
MH681419.1_ASFV/POL/2015         .............................-.............---------------------------.......... [ 32080]
LS478113.1_Estonia_2014          .............................-.............---------------------------.......... [ 32080]
KP843857.1_Odintsovo_02/14       .............................-.............---------------------------.......... [ 32080]
FN557520.1_strain_E75            .............................t.at........a.---------------------------........t. [ 32080]
AY261366.1_Warthog               .............................-...........a.---------------------------.......... [ 32080]
AY261365.1_Warmbaths             ..........................t..-.............---------------------------.......... [ 32080]
AY261364.1_Tengani_62            .................a...........-...........a.---------------------------.......... [ 32080]
AY261363.1_Pretoriuskop/96/4     .............................-.............---------------------------.......... [ 32080]
AY261362.1_Mkuzi_1979            .............................t.at........a.---------------------------........t. [ 32080]
AY261361.1_Malawi_Lil-20/1_1983  ................t............-......c....a.aaagtatataaagtatataaagtatat.......... [ 32080]
AY261360.1_Kenya_1950            ................t.........g..-......c....a.---------------------------.......... [ 32080]
AM712240.1_OURT_88/3             .............................t.at........a.---------------------------........t. [ 32080]
AM712239.1_Benin_97/1            .............................t.at........a.---------------------------........t. [ 32080]
KM111295.1_Ken06.Bus             ................t.........g..-......c....a.---------------------------.......... [ 32080]
KM111294.1_Ken05/Tk1             ................t.........g..-...........a.---------------------------.......... [ 32080]
KM262844.1_strain_L60            .............................t.at........a.---------------------------........t. [ 32080]
KM102979.1_26544/OG10_Italy      .............................t.at........a.---------------------------........t. [ 32080]
KM262845.1_strain_NHV            .............................t.at........a.---------------------------........t. [ 32080]
KX354450.1_47/Ss/2008            .............................t.at........a.---------------------------........t. [ 32080]
KP055815.1_strain_BA71           .............................t.at........a.---------------------------........t. [ 32080]
MH025920.1_strain_R35            ................t.........g..-......c....a.---------------------------.......... [ 32080]
MH025919.1_strain_N10            ................t.........g..-......c....a.---------------------------.......... [ 32080]
MH025918.1_strain_R25            ................t.........g..-......c....a.---------------------------.......... [ 32080]
MH025917.1_strain_R7             ................t.........g..-......c....a.---------------------------.......... [ 32080]
MH025916.1_strain_R8             ................t.........g..-......c....a.---------------------------.......... [ 32080]

FR682468.2_Georgia_2007/1        gatat--tacc-aaaatatattctatgtacataatatctgggaaattatttttttttctcatacccttaaatataaaaat [ 32160]
ASF_IND_20_CAD543                ....w--....-.................................................................... [ 32160]
LR899193.1_Germany_2020/1        .....--....-....................................-............................... [ 32160]
MN393477.1_Wuhan_2019            .....--....-.................................................................... [ 32160]
MW396979.1_Timor-Leste/2019/1    .....--....-.................................................................... [ 32160]
MK940252.1_CN/2019/InnerMongolia .....--....-.................................................................... [ 32160]
MK128995.1_China/2018/AnhuiXCGQ  .....--....-.................................................................... [ 32160]
MH681419.1_ASFV/POL/2015         .....--....-.................................................................... [ 32160]
LS478113.1_Estonia_2014          .....--....-.................................................................... [ 32160]
KP843857.1_Odintsovo_02/14       .....--....-....................................-............................... [ 32160]
```

```
FN557520.1_strain_E75            ....a--....-......................c...c..a...................................... [ 32160]
AY261366.1_Warthog               .....--....-.......................g....................-....................... [ 32160]
AY261365.1_Warmbaths             .g...--....a........a.............cg..c...g............---...................... [ 32160]
AY261364.1_Tengani_62            .....--....-..............t....c..c...c.....t.at........--c..................... [ 32160]
AY261363.1_Pretoriuskop/96/4     .g...--....-.......................g............................................ [ 32160]
AY261362.1_Mkuzi_1979            ....a--....-........a.....t........g....a......................t................ [ 32160]
AY261361.1_Malawi_Lil-20/1_1983  .....accg..-.......ca.............cg.tc....c...........---...................... [ 32160]
AY261360.1_Kenya_1950            .....ac..t.-.......ca.............c...c...gc....-.......--.t.................... [ 32160]
AM712240.1_OURT_88/3             ....a--....-......................c...c..a...................................... [ 32160]
AM712239.1_Benin_97/1            ....a--....-......................c...c..a...................................... [ 32160]
KM111295.1_Ken06.Bus             .....ac.g..-.......ca.............c...c...gc....-.......--...................... [ 32160]
KM111294.1_Ken05/Tk1             .....ac.g..-.......ca.............c...c...gc............--...................... [ 32160]
KM262844.1_strain_L60            ....a--....-......................c...c..a...................................... [ 32160]
KM102979.1_26544/OG10_Italy      ....a--....-......................c...c..a...................................... [ 32160]
KM262845.1_strain_NHV            ....a--....-......................c...c..a...................................... [ 32160]
KX354450.1_47/Ss/2008            ....a--....-......................c...c..a...................................... [ 32160]
KP055815.1_strain_BA71           ....a--....-......................c...c..a...................................... [ 32160]
MH025920.1_strain_R35            .....ac.g..-.......ca.............c...c...gc....-.......--...................... [ 32160]
MH025919.1_strain_N10            .....ac.g..-.......ca.............c...c...gc....-.......--...................... [ 32160]
MH025918.1_strain_R25            .....ac.g..-.......ca.............c...c...gc....-.......--...................... [ 32160]
MH025917.1_strain_R7             .....ac.g..-.......ca.............c...c...gc....-.......--...................... [ 32160]
MH025916.1_strain_R8             .....ac.g..-.......ca.............c...c...gc....-.......--...................... [ 32160]

FR682468.2_Georgia_2007/1        attgggtttcttcactaaactttagaggtaaaaatttttctttgttttgcaccatcatgtatgggtttaggctgtcccag [ 32240]
ASF_IND_20_CAD543                ................................................................................ [ 32240]
LR899193.1_Germany_2020/1        ................................................................................ [ 32240]
MN393477.1_Wuhan_2019            ................................................................................ [ 32240]
MW396979.1_Timor-Leste/2019/1    ................................................................................ [ 32240]
MK940252.1_CN/2019/InnerMongolia ................................................................................ [ 32240]
MK128995.1_China/2018/AnhuiXCGQ  ................................................................................ [ 32240]
MH681419.1_ASFV/POL/2015         ................................................................................ [ 32240]
LS478113.1_Estonia_2014          ................................................................................ [ 32240]
KP843857.1_Odintsovo_02/14       ................................................................................ [ 32240]
FN557520.1_strain_E75            ...................................................t...................t.a...... [ 32240]
AY261366.1_Warthog               ...................................................t...................t.a...... [ 32240]
AY261365.1_Warmbaths             ........................a....................................................... [ 32240]
AY261364.1_Tengani_62            ....a........g..........a...............c..............................tca...... [ 32240]
AY261363.1_Pretoriuskop/96/4     ...................................................t...................t.a...... [ 32240]
AY261362.1_Mkuzi_1979            ........................a................................................c...... [ 32240]
AY261361.1_Malawi_Lil-20/1_1983  .c......................a...............................g................t...... [ 32240]
AY261360.1_Kenya_1950            ........................a.......................c.....................a.cag..... [ 32240]
AM712240.1_OURT_88/3             ...................................................t...................t.a...... [ 32240]
AM712239.1_Benin_97/1            ...................................................t...................t.a...... [ 32240]
KM111295.1_Ken06.Bus             ........................a..........................t..................a.cag..... [ 32240]
```

```
KM111294.1_Ken05/Tk1              ........................a.......................c.....................a.cag..... [ 32240]
KM262844.1_strain_L60             ...................................................t...................t.a...... [ 32240]
KM102979.1_26544/OG10_Italy       ...................................................t...................t.a...... [ 32240]
KM262845.1_strain_NHV             ...................................................t...................t.a...... [ 32240]
KX354450.1_47/Ss/2008             ...................................................t...................t.a...... [ 32240]
KP055815.1_strain_BA71            ...................................................t...................t.a...... [ 32240]
MH025920.1_strain_R35             ........................a..........................t..................a.cag..... [ 32240]
MH025919.1_strain_N10             ........................a..........................t..................a.cag..... [ 32240]
MH025918.1_strain_R25             ........................a..........................t..................a.cag..... [ 32240]
MH025917.1_strain_R7              ........................a..........................t..................a.cag..... [ 32240]
MH025916.1_strain_R8              ........................a..........................t..................a.cag..... [ 32240]

FR682468.2_Georgia_2007/1         ggattgtttatttgaatatttcctaaataggaacacaacgccatgatcatatatctttcattctggtaagctttttgata [ 32320]
ASF_IND_20_CAD543                 ................................................................................ [ 32320]
LR899193.1_Germany_2020/1         ................................................................................ [ 32320]
MN393477.1_Wuhan_2019             ................................................................................ [ 32320]
MW396979.1_Timor-Leste/2019/1     ................................................................................ [ 32320]
MK940252.1_CN/2019/InnerMongolia  ................................................................................ [ 32320]
MK128995.1_China/2018/AnhuiXCGQ   ................................................................................ [ 32320]
MH681419.1_ASFV/POL/2015          ................................................................................ [ 32320]
LS478113.1_Estonia_2014           ................................................................................ [ 32320]
KP843857.1_Odintsovo_02/14        ................................................................................ [ 32320]
FN557520.1_strain_E75             ................................................................................ [ 32320]
AY261366.1_Warthog                ................................................................................ [ 32320]
AY261365.1_Warmbaths              ......................................t......................................... [ 32320]
AY261364.1_Tengani_62             ..................................................g............................. [ 32320]
AY261363.1_Pretoriuskop/96/4      ..............................................................t................. [ 32320]
AY261362.1_Mkuzi_1979             ......................................t...........ga............................ [ 32320]
AY261361.1_Malawi_Lil-20/1_1983   ......................................t...........ga..t..........a.............. [ 32320]
AY261360.1_Kenya_1950             .....................g..........g.....t...........ga.........aa.a.......c...t... [ 32320]
AM712240.1_OURT_88/3              ................................................................................ [ 32320]
AM712239.1_Benin_97/1             ................................................................................ [ 32320]
KM111295.1_Ken06.Bus              .....................g..........g.....t...........ga.........aa.a.......c...t... [ 32320]
KM111294.1_Ken05/Tk1              .....................g..........g.....t...........ga............................ [ 32320]
KM262844.1_strain_L60             ................................................................................ [ 32320]
KM102979.1_26544/OG10_Italy       ................................................................................ [ 32320]
KM262845.1_strain_NHV             ................................................................................ [ 32320]
KX354450.1_47/Ss/2008             ................................................................................ [ 32320]
KP055815.1_strain_BA71            ................................................................................ [ 32320]
MH025920.1_strain_R35             .....................g..........g.....t...........ga.........aa.a.......c...t... [ 32320]
MH025919.1_strain_N10             .....................g..........g.....t...........ga.........aa.a.......c...t... [ 32320]
MH025918.1_strain_R25             .....................g..........g.....t...........ga.........aa.a.......c...t... [ 32320]
MH025917.1_strain_R7              .....................g..........g.....t...........ga.........aa.a.......c...t... [ 32320]
MH025916.1_strain_R8              .....................g..........g.....t...........ga.........aa.a.......c...t... [ 32320]
```

```
FR682468.2_Georgia_2007/1        catcttcaaagatgccgtacctccgagtgtgtaacagcaaacaaacgtccgtacttttccatgggtcgcagcccattcca [ 32400]
ASF_IND_20_CAD543                ................................................................................ [ 32400]
LR899193.1_Germany_2020/1        ................................................................................ [ 32400]
MN393477.1_Wuhan_2019            ................................................................................ [ 32400]
MW396979.1_Timor-Leste/2019/1    ................................................................................ [ 32400]
MK940252.1_CN/2019/InnerMongolia ................................................................................ [ 32400]
MK128995.1_China/2018/AnhuiXCGQ  ................................................................................ [ 32400]
MH681419.1_ASFV/POL/2015         ................................................................................ [ 32400]
LS478113.1_Estonia_2014          ................................................................................ [ 32400]
KP843857.1_Odintsovo_02/14       ................................................................................ [ 32400]
FN557520.1_strain_E75            ................................................................................ [ 32400]
AY261366.1_Warthog               ................................................................a............... [ 32400]
AY261365.1_Warmbaths             .........g...................................................................... [ 32400]
AY261364.1_Tengani_62            .............................................t....................t............. [ 32400]
AY261363.1_Pretoriuskop/96/4     .............................................t................a................. [ 32400]
AY261362.1_Mkuzi_1979            ................a...................a....t...............c...................... [ 32400]
AY261361.1_Malawi_Lil-20/1_1983  ..a........................g........a....t...............c...................... [ 32400]
AY261360.1_Kenya_1950            ..g..c.....................g.....g.......t.........c.....c...........g.....c.... [ 32400]
AM712240.1_OURT_88/3             ................................................................................ [ 32400]
AM712239.1_Benin_97/1            ................................................................................ [ 32400]
KM111295.1_Ken06.Bus             ..g..c.....................g.....g.......t.........c.....c..g........g.......... [ 32400]
KM111294.1_Ken05/Tk1             .....c.....................g.....g.......t........ac.....c..g........g.....c.... [ 32400]
KM262844.1_strain_L60            ................................................................................ [ 32400]
KM102979.1_26544/OG10_Italy      ..............................................a................................. [ 32400]
KM262845.1_strain_NHV            ................................................................................ [ 32400]
KX354450.1_47/Ss/2008            ..............................................a................................. [ 32400]
KP055815.1_strain_BA71           ................................................................................ [ 32400]
MH025920.1_strain_R35            ..g..c.....................g.....g.......t.........c.....c..g........g.......... [ 32400]
MH025919.1_strain_N10            ..g..c.....................g.....g.......t.........c.....c..g........g.......... [ 32400]
MH025918.1_strain_R25            ..g..c.....................g.....g.......t.........c.....c..g........g.......... [ 32400]
MH025917.1_strain_R7             ..g..c.....................g.....g.......t.........c.....c..g........g.......... [ 32400]
MH025916.1_strain_R8             ..g..c.....................g.....g.......t.........c.....c..g........g.......... [ 32400]

FR682468.2_Georgia_2007/1        ttccgtagctcagcatcttttgctgtatttttttattcgctttataaaaaaagtttttcatccattccacgttctcataa [ 32480]
ASF_IND_20_CAD543                ................................................................................ [ 32480]
LR899193.1_Germany_2020/1        ................................................................................ [ 32480]
MN393477.1_Wuhan_2019            ................................................................................ [ 32480]
MW396979.1_Timor-Leste/2019/1    ................................................................................ [ 32480]
MK940252.1_CN/2019/InnerMongolia ................................................................................ [ 32480]
MK128995.1_China/2018/AnhuiXCGQ  ................................................................................ [ 32480]
MH681419.1_ASFV/POL/2015         ................................................................................ [ 32480]
LS478113.1_Estonia_2014          ................................................................................ [ 32480]
KP843857.1_Odintsovo_02/14       ................................................................................ [ 32480]
```

```
FN557520.1_strain_E75          .................c....................................a......................... [ 32480]
AY261366.1_Warthog             .......c.........c...................................c.......................... [ 32480]
AY261365.1_Warmbaths           .......c.........c...................................c.......................... [ 32480]
AY261364.1_Tengani_62          ...........................a..................g...t............................. [ 32480]
AY261363.1_Pretoriuskop/96/4   .......c.........c...................................c.......................... [ 32480]
AY261362.1_Mkuzi_1979          .................c............................g................................. [ 32480]
AY261361.1_Malawi_Lil-20/1_1983 ........t.......tc....t...............c......................................... [ 32480]
AY261360.1_Kenya_1950          ........t........c...t................t..............c.......................... [ 32480]
AM712240.1_OURT_88/3           .................c....................................a......................... [ 32480]
AM712239.1_Benin_97/1          .................c....................................a......................... [ 32480]
KM111295.1_Ken06.Bus           ........t........c...t................t..............c.......................... [ 32480]
KM111294.1_Ken05/Tk1           ........t........c...t................t..............c.......................... [ 32480]
KM262844.1_strain_L60          .................c....................................a......................... [ 32480]
KM102979.1_26544/OG10_Italy    .................c....................................a......................... [ 32480]
KM262845.1_strain_NHV          .................c....................................a......................... [ 32480]
KX354450.1_47/Ss/2008          .................c....................................a......................... [ 32480]
KP055815.1_strain_BA71         .................c....................................a......................... [ 32480]
MH025920.1_strain_R35          ........t........c...t................t..............c.......................... [ 32480]
MH025919.1_strain_N10          ........t........c...t................t..............c.......................... [ 32480]
MH025918.1_strain_R25          ........t........c...t................t..............c.......................... [ 32480]
MH025917.1_strain_R7           ........t........c...t................t..............c.......................... [ 32480]
MH025916.1_strain_R8           ........t........c...t................t..............c.......................... [ 32480]

FR682468.2_Georgia_2007/1      aaacaggcacttaaaaagagcactaggggtagtgtagtcttattatagaatgtaggaatgtatgttttagttattttttt [ 32560]
ASF_IND_20_CAD543              ................................................................................ [ 32560]
LR899193.1_Germany_2020/1      ................................................................................ [ 32560]
MN393477.1_Wuhan_2019          ................................................................................ [ 32560]
MW396979.1_Timor-Leste/2019/1  ................................................................................ [ 32560]
MK940252.1_CN/2019/InnerMongolia ................................................................................ [ 32560]
MK128995.1_China/2018/AnhuiXCGQ ................................................................................ [ 32560]
MH681419.1_ASFV/POL/2015       ................................................................................ [ 32560]
LS478113.1_Estonia_2014        ................................................................................ [ 32560]
KP843857.1_Odintsovo_02/14     ................................................................................ [ 32560]
FN557520.1_strain_E75          .......................c...tt................................................... [ 32560]
AY261366.1_Warthog             ............................t................................................... [ 32560]
AY261365.1_Warmbaths           ............................t................................................... [ 32560]
AY261364.1_Tengani_62          ............................t................................................... [ 32560]
AY261363.1_Pretoriuskop/96/4   ............................t................................................... [ 32560]
AY261362.1_Mkuzi_1979          .................a.........tt................................................... [ 32560]
AY261361.1_Malawi_Lil-20/1_1983 ...........................tt...c........................g..a................... [ 32560]
AY261360.1_Kenya_1950          .....t..g..................tt......c.......g...a...c......caa........a.......g.. [ 32560]
AM712240.1_OURT_88/3           .......................c...tt................................................... [ 32560]
AM712239.1_Benin_97/1          .......................c...tt................................................... [ 32560]
KM111295.1_Ken06.Bus           .....t..g..................tt......c.......g...a...c......c.g........a.......g.. [ 32560]
```

```
KM111294.1_Ken05/Tk1           .....t..g..................tt......c.......g...a..........c.a........a.......g.. [ 32560]
KM262844.1_strain_L60          .......................c...tt................................................... [ 32560]
KM102979.1_26544/OG10_Italy    .......................c...tt................................................... [ 32560]
KM262845.1_strain_NHV          .......................c...tt................................................... [ 32560]
KX354450.1_47/Ss/2008          .......................c...tt................................................... [ 32560]
KP055815.1_strain_BA71         .......................c...tt................................................... [ 32560]
MH025920.1_strain_R35          .....t..g..................tt......c.......g...a...c......c.g........a.......g.. [ 32560]
MH025919.1_strain_N10          .....t..g..................tt......c.......g...a...c......c.g........a.......g.. [ 32560]
MH025918.1_strain_R25          .....t..g..................tt......c.......g...a...c......c.g........a.......g.. [ 32560]
MH025917.1_strain_R7           .....t..g..................tt......c.......g...a...c......c.g........a.......g.. [ 32560]
MH025916.1_strain_R8           .....t..g..................tt......c.......g...a...c......c.g........a.......g.. [ 32560]

FR682468.2_Georgia_2007/1      caacgcgtgttccatactatgttttaccgccataaaaatacaaaaccaataccaactttttctataaaaggttttgctgt [ 32640]
ASF_IND_20_CAD543              ................................................................................ [ 32640]
LR899193.1_Germany_2020/1      ................................................................................ [ 32640]
MN393477.1_Wuhan_2019          ................................................................................ [ 32640]
MW396979.1_Timor-Leste/2019/1  ................................................................................ [ 32640]
MK940252.1_CN/2019/InnerMongolia ................................................................................ [ 32640]
MK128995.1_China/2018/AnhuiXCGQ ................................................................................ [ 32640]
MH681419.1_ASFV/POL/2015       ................................................................................ [ 32640]
LS478113.1_Estonia_2014        ................................................................................ [ 32640]
KP843857.1_Odintsovo_02/14     ................................................................................ [ 32640]
FN557520.1_strain_E75          ......c....t.....c.............................................................. [ 32640]
AY261366.1_Warthog             .................c.............................................................. [ 32640]
AY261365.1_Warmbaths           .................c.............................................................. [ 32640]
AY261364.1_Tengani_62          ...t.......................t....................g............t.................. [ 32640]
AY261363.1_Pretoriuskop/96/4   .................c.............................................................. [ 32640]
AY261362.1_Mkuzi_1979          ......c..........c.............................................................. [ 32640]
AY261361.1_Malawi_Lil-20/1_1983 .........c....................t.........................c.....c......c.......... [ 32640]
AY261360.1_Kenya_1950          ......c..a.g..............................................c..t.....g.c.....aa... [ 32640]
AM712240.1_OURT_88/3           ......c....t.....c.............................................................. [ 32640]
AM712239.1_Benin_97/1          ......c....t.....c.............................................................. [ 32640]
KM111295.1_Ken06.Bus           ......c..a.t..................t...........................c..t.....g.c......a... [ 32640]
KM111294.1_Ken05/Tk1           ......c..a.t.................................................t.......c.......... [ 32640]
KM262844.1_strain_L60          ......c....t.....c.............................................................. [ 32640]
KM102979.1_26544/OG10_Italy    ......c....t.....c.............................................................. [ 32640]
KM262845.1_strain_NHV          ......c....t.....c.............................................................. [ 32640]
KX354450.1_47/Ss/2008          ......c....t.....c.............................................................. [ 32640]
KP055815.1_strain_BA71         ......c....t.....c.............................................................. [ 32640]
MH025920.1_strain_R35          ......c..a.t..................t...........................c..t.....g.c......a... [ 32640]
MH025919.1_strain_N10          ......c..a.t..................t...........................c..t.....g.c......a... [ 32640]
MH025918.1_strain_R25          ......c..a.t..................t...........................c..t.....g.c......a... [ 32640]
MH025917.1_strain_R7           ......c..a.t..................t...........................c..t.....g.c......a... [ 32640]
MH025916.1_strain_R8           ......c..a.t..................t...........................c..t.....g.c......a... [ 32640]
```

```
FR682468.2_Georgia_2007/1        acacatataaacgagcaaaatatatttcaaactctatattctttttataaaaaaactcgagacagtcgtttatgttacga [ 32720]
ASF_IND_20_CAD543                ....................................w........................................... [ 32720]
LR899193.1_Germany_2020/1        ................................................................................ [ 32720]
MN393477.1_Wuhan_2019            ................................................................................ [ 32720]
MW396979.1_Timor-Leste/2019/1    ................................................................................ [ 32720]
MK940252.1_CN/2019/InnerMongolia ................................................................................ [ 32720]
MK128995.1_China/2018/AnhuiXCGQ  ................................................................................ [ 32720]
MH681419.1_ASFV/POL/2015         ................................................................................ [ 32720]
LS478113.1_Estonia_2014          ................................................................................ [ 32720]
KP843857.1_Odintsovo_02/14       ................................................................................ [ 32720]
FN557520.1_strain_E75            ......g.............................a..............................a............ [ 32720]
AY261366.1_Warthog               ....................................c......................................g.... [ 32720]
AY261365.1_Warmbaths             ....................................a......a.......................a............ [ 32720]
AY261364.1_Tengani_62            ..............a....................ga......a.......................a............ [ 32720]
AY261363.1_Pretoriuskop/96/4     ....................................c......................................g.... [ 32720]
AY261362.1_Mkuzi_1979            ......g.............................a......a.......................a............ [ 32720]
AY261361.1_Malawi_Lil-20/1_1983  ......g......g....................cg.g.................g.t.........a.......g.... [ 32720]
AY261360.1_Kenya_1950            ......g................t..........a......................t.........a............ [ 32720]
AM712240.1_OURT_88/3             ......g.............................a..............................a............ [ 32720]
AM712239.1_Benin_97/1            ......g.............................a..............................a............ [ 32720]
KM111295.1_Ken06.Bus             ......g...........................a......................t.........a............ [ 32720]
KM111294.1_Ken05/Tk1             ......g...........................a................................a............ [ 32720]
KM262844.1_strain_L60            ......g.............................a..............................a............ [ 32720]
KM102979.1_26544/OG10_Italy      ......g.............................a..............................a............ [ 32720]
KM262845.1_strain_NHV            ......g.............................a..............................a............ [ 32720]
KX354450.1_47/Ss/2008            ......g.............................a..............................a............ [ 32720]
KP055815.1_strain_BA71           ......g.............................a..............................a............ [ 32720]
MH025920.1_strain_R35            ......g...........................a......................t.........a............ [ 32720]
MH025919.1_strain_N10            ......g...........................a......................t.........a............ [ 32720]
MH025918.1_strain_R25            ......g...........................a......................t.........a............ [ 32720]
MH025917.1_strain_R7             ......g...........................a......................t.........a............ [ 32720]
MH025916.1_strain_R8             ......g...........................a......................t.........a............ [ 32720]

FR682468.2_Georgia_2007/1        ctttttctaaatacctcaaaaacagtaattaattcactgtcgctgtggaaatgttcgtaagctaactgtttaatgtcttt [ 32800]
ASF_IND_20_CAD543                .w.y.......w.................................................................... [ 32800]
LR899193.1_Germany_2020/1        ................................................................................ [ 32800]
MN393477.1_Wuhan_2019            ................................................................................ [ 32800]
MW396979.1_Timor-Leste/2019/1    ................................................................................ [ 32800]
MK940252.1_CN/2019/InnerMongolia ................................................................................ [ 32800]
MK128995.1_China/2018/AnhuiXCGQ  ................................................................................ [ 32800]
MH681419.1_ASFV/POL/2015         ................................................................................ [ 32800]
LS478113.1_Estonia_2014          ................................................................................ [ 32800]
KP843857.1_Odintsovo_02/14       ................................................................................ [ 32800]
```

```
FN557520.1_strain_E75            .a.c.......a..................................a.........a....................... [ 32800]
AY261366.1_Warthog               ...........a.................................................................... [ 32800]
AY261365.1_Warmbaths             .a.........a............................................a.................a..... [ 32800]
AY261364.1_Tengani_62            ...c.......a.................................................................... [ 32800]
AY261363.1_Pretoriuskop/96/4     .a.........a.................................................................... [ 32800]
AY261362.1_Mkuzi_1979            .a.........a............................................a....................... [ 32800]
AY261361.1_Malawi_Lil-20/1_1983  .a.........a..........t...................t.a........a....................c..... [ 32800]
AY261360.1_Kenya_1950            .a.c.......a..........t......................................................... [ 32800]
AM712240.1_OURT_88/3             .a.c.......a..................................a.........a....................... [ 32800]
AM712239.1_Benin_97/1            .a.c.......a..................................a.........a....................... [ 32800]
KM111295.1_Ken06.Bus             .a.c.......a..........tg........................................................ [ 32800]
KM111294.1_Ken05/Tk1             .a.c.......a..........t......................................................... [ 32800]
KM262844.1_strain_L60            .a.c.......a..................................a.........a....................... [ 32800]
KM102979.1_26544/OG10_Italy      .a.c.......a..................................a.........a....................... [ 32800]
KM262845.1_strain_NHV            .a.c.......a..................................a.........a....................... [ 32800]
KX354450.1_47/Ss/2008            .a.c.......a..................................a.........a....................... [ 32800]
KP055815.1_strain_BA71           .a.c.......a..................................a.........a....................... [ 32800]
MH025920.1_strain_R35            .a.c.......a..........tg........................................................ [ 32800]
MH025919.1_strain_N10            .a.c.......a..........tg........................................................ [ 32800]
MH025918.1_strain_R25            .a.c.......a..........tg........................................................ [ 32800]
MH025917.1_strain_R7             .a.c.......a..........tg........................................................ [ 32800]
MH025916.1_strain_R8             .a.c.......a..........tg........................................................ [ 32800]

FR682468.2_Georgia_2007/1        aggggtcaattctttttttgggagcagtggtttgagattcggcaaaggtcgtctaaagtagtgagcgaacttttcattcg [ 32880]
ASF_IND_20_CAD543                ................................................................................ [ 32880]
LR899193.1_Germany_2020/1        ................................................................................ [ 32880]
MN393477.1_Wuhan_2019            ................................................................................ [ 32880]
MW396979.1_Timor-Leste/2019/1    ................................................................................ [ 32880]
MK940252.1_CN/2019/InnerMongolia ................................................................................ [ 32880]
MK128995.1_China/2018/AnhuiXCGQ  ................................................................................ [ 32880]
MH681419.1_ASFV/POL/2015         ................................................................................ [ 32880]
LS478113.1_Estonia_2014          ................................................................................ [ 32880]
KP843857.1_Odintsovo_02/14       ................................................................................ [ 32880]
FN557520.1_strain_E75            ................................................................................ [ 32880]
AY261366.1_Warthog               ......................................c......c.................................. [ 32880]
AY261365.1_Warmbaths             ..................c....................t........................................ [ 32880]
AY261364.1_Tengani_62            ....c.........................c........t......................a................. [ 32880]
AY261363.1_Pretoriuskop/96/4     ......................c................t.....c..............a................... [ 32880]
AY261362.1_Mkuzi_1979            .....................ac................t........................................ [ 32880]
AY261361.1_Malawi_Lil-20/1_1983  .........................g.......a..g.....................c...t................. [ 32880]
AY261360.1_Kenya_1950            ......t..c.............t.........c....g......g................t...........t..... [ 32880]
AM712240.1_OURT_88/3             ................................................................................ [ 32880]
AM712239.1_Benin_97/1            ................................................................................ [ 32880]
KM111295.1_Ken06.Bus             .t....t..c.............t.........c....g.......................t................. [ 32880]
```

```
KM111294.1_Ken05/Tk1            ......t..c.............t.........c....g....................ga.tg.....gg......... [ 32880]
KM262844.1_strain_L60           ................................................................................ [ 32880]
KM102979.1_26544/OG10_Italy     ................................................................................ [ 32880]
KM262845.1_strain_NHV           ................................................................................ [ 32880]
KX354450.1_47/Ss/2008           ................................................................................ [ 32880]
KP055815.1_strain_BA71          ................................................................................ [ 32880]
MH025920.1_strain_R35           ......t..c.............t.........c....g.......................t................. [ 32880]
MH025919.1_strain_N10           ......t..c.............t.........c....g.......................t................. [ 32880]
MH025918.1_strain_R25           ......t..c.............t.........c....g.......................t................. [ 32880]
MH025917.1_strain_R7            ......t..c.............t.........c....g.......................t................. [ 32880]
MH025916.1_strain_R8            ......t..c.............t.........c....g.......................t................. [ 32880]

FR682468.2_Georgia_2007/1       ctccccaacacaaaagccgataagccagcatgtagttatcacgttttaccgcgtaaataagcaaatagtttatattgata [ 32960]
ASF_IND_20_CAD543               ................................................................................ [ 32960]
LR899193.1_Germany_2020/1       ................................................................................ [ 32960]
MN393477.1_Wuhan_2019           ................................................................................ [ 32960]
MW396979.1_Timor-Leste/2019/1   ................................................................................ [ 32960]
MK940252.1_CN/2019/InnerMongolia ................................................................................ [ 32960]
MK128995.1_China/2018/AnhuiXCGQ ................................................................................ [ 32960]
MH681419.1_ASFV/POL/2015        ................................................................................ [ 32960]
LS478113.1_Estonia_2014         ................................................................................ [ 32960]
KP843857.1_Odintsovo_02/14      ................................................................................ [ 32960]
FN557520.1_strain_E75           .........................t................a...............g..t.................. [ 32960]
AY261366.1_Warthog              ................................................................................ [ 32960]
AY261365.1_Warmbaths            ...........................a........................a........................... [ 32960]
AY261364.1_Tengani_62           ..................................a.......................g..t.................. [ 32960]
AY261363.1_Pretoriuskop/96/4    ..........................................................g..t.................. [ 32960]
AY261362.1_Mkuzi_1979           ...........................a.................................................... [ 32960]
AY261361.1_Malawi_Lil-20/1_1983 ........g.................................................g..t.................. [ 32960]
AY261360.1_Kenya_1950           .c......g.................................................g..t.................. [ 32960]
AM712240.1_OURT_88/3            .........................t................a...............g..t.................. [ 32960]
AM712239.1_Benin_97/1           .........................t................a...............g..t.................. [ 32960]
KM111295.1_Ken06.Bus            ........g......................a....................a.....g..t.................. [ 32960]
KM111294.1_Ken05/Tk1            ........g.................................................g..t.................. [ 32960]
KM262844.1_strain_L60           .........................t................a...............g..t.................. [ 32960]
KM102979.1_26544/OG10_Italy     .........................t................a...............g..t.................. [ 32960]
KM262845.1_strain_NHV           .........................t................a...............g..t.................. [ 32960]
KX354450.1_47/Ss/2008           .........................t................a...............g..t.................. [ 32960]
KP055815.1_strain_BA71          .........................t................a...............g..t.................. [ 32960]
MH025920.1_strain_R35           ........g......................a....................a.....g..t.................. [ 32960]
MH025919.1_strain_N10           ........g......................a....................a.....g..t.................. [ 32960]
MH025918.1_strain_R25           ........g......................a....................a.....g..t.................. [ 32960]
MH025917.1_strain_R7            ........g......................a....................a.....g..t.................. [ 32960]
MH025916.1_strain_R8            ........g......................a....................a.....g..t.................. [ 32960]
```

```
FR682468.2_Georgia_2007/1        catgt---accatgttgctgcccgtttggacatatgttgccgcattctgaacacttatgaatgagatcatagttcttaca [ 33040]
ASF_IND_20_CAD543                .....---........................................................................ [ 33040]
LR899193.1_Germany_2020/1        .....---........................................................................ [ 33040]
MN393477.1_Wuhan_2019            .....---........................................................................ [ 33040]
MW396979.1_Timor-Leste/2019/1    .....---........................................................................ [ 33040]
MK940252.1_CN/2019/InnerMongolia .....---........................................................................ [ 33040]
MK128995.1_China/2018/AnhuiXCGQ  .....---........................................................................ [ 33040]
MH681419.1_ASFV/POL/2015         .....---........................................................................ [ 33040]
LS478113.1_Estonia_2014          .....---........................................................................ [ 33040]
KP843857.1_Odintsovo_02/14       .....---........................................................................ [ 33040]
FN557520.1_strain_E75            .....---............t.t......g.................................................. [ 33040]
AY261366.1_Warthog               ....c---............t........................................................... [ 33040]
AY261365.1_Warmbaths             ....c---............t........................................................... [ 33040]
AY261364.1_Tengani_62            .....---.........t..t.......a.................................a.t.........t..... [ 33040]
AY261363.1_Pretoriuskop/96/4     .....---............t........................................................... [ 33040]
AY261362.1_Mkuzi_1979            ....c---............t........................................................... [ 33040]
AY261361.1_Malawi_Lil-20/1_1983  .....atc............t.....................................c.....t.a......c...... [ 33040]
AY261360.1_Kenya_1950            ....---c............t..............a.................t....c.....t..t............ [ 33040]
AM712240.1_OURT_88/3             .....---............t.t......g.................................................. [ 33040]
AM712239.1_Benin_97/1            .....---............t.t......g.................................................. [ 33040]
KM111295.1_Ken06.Bus             ....---c............t..............a......................c.....t..t............ [ 33040]
KM111294.1_Ken05/Tk1             .....atc............t.......a.............................c.....t.at............ [ 33040]
KM262844.1_strain_L60            .....---............t.t......g.................................................. [ 33040]
KM102979.1_26544/OG10_Italy      .....---............t.t......g.................................................. [ 33040]
KM262845.1_strain_NHV            .....---............t.t......g.................................................. [ 33040]
KX354450.1_47/Ss/2008            .....---............t.t......g.................................................. [ 33040]
KP055815.1_strain_BA71           .....---............t.t......g.................................................. [ 33040]
MH025920.1_strain_R35            ....---c............t..............a......................c.....t..t............ [ 33040]
MH025919.1_strain_N10            ....---c............t..............a......................c.....t..t............ [ 33040]
MH025918.1_strain_R25            ....---c............t..............a......................c.....t..t............ [ 33040]
MH025917.1_strain_R7             ....---c............t..............a......................c.....t..t............ [ 33040]
MH025916.1_strain_R8             ....---c............t..............a......................c.....t..t............ [ 33040]

FR682468.2_Georgia_2007/1        acataaccccaaacgggttagtacttctttgtcacgttttaaaaactcgacatgattctttaatgttaatgctttgagcg [ 33120]
ASF_IND_20_CAD543                .......................tg.........y..............................------......... [ 33120]
LR899193.1_Germany_2020/1        ................................................................................ [ 33120]
MN393477.1_Wuhan_2019            ................................................................................ [ 33120]
MW396979.1_Timor-Leste/2019/1    ................................................................................ [ 33120]
MK940252.1_CN/2019/InnerMongolia ................................................................................ [ 33120]
MK128995.1_China/2018/AnhuiXCGQ  ................................................................................ [ 33120]
MH681419.1_ASFV/POL/2015         ................................................................................ [ 33120]
LS478113.1_Estonia_2014          ................................................................................ [ 33120]
KP843857.1_Odintsovo_02/14       ................................................................................ [ 33120]
```

```
FN557520.1_strain_E75           .............t......a..tg.........t...............t...............------........ [ 33120]
AY261366.1_Warthog              ................................................................................ [ 33120]
AY261365.1_Warmbaths            ................................................................................ [ 33120]
AY261364.1_Tengani_62           ...........g.t.......................................................c.....t.... [ 33120]
AY261363.1_Pretoriuskop/96/4    .............t....................t.............t....................c.......... [ 33120]
AY261362.1_Mkuzi_1979           .............t.........t..t..................................................... [ 33120]
AY261361.1_Malawi_Lil-20/1_1983 .............t.........t.............................t..gt..c.t.a..........t.... [ 33120]
AY261360.1_Kenya_1950           ......t................t.....................t...t..ctc....................t.... [ 33120]
AM712240.1_OURT_88/3            .............t......a..tg.........t...............t...............------........ [ 33120]
AM712239.1_Benin_97/1           .............t......a..tg.........t...............t...............------........ [ 33120]
KM111295.1_Ken06.Bus            ......t................t.....................t...t..ctc....................t.... [ 33120]
KM111294.1_Ken05/Tk1            ......t................t.....................t...t..ctc....................t.... [ 33120]
KM262844.1_strain_L60           .............t......a..tg.........t...............t...............------........ [ 33120]
KM102979.1_26544/OG10_Italy     .............t......a..tg.........t...............t...............------........ [ 33120]
KM262845.1_strain_NHV           .............t......a..tg.........t...............t...............------........ [ 33120]
KX354450.1_47/Ss/2008           .............t......a..tg.........t...............t...............------........ [ 33120]
KP055815.1_strain_BA71          .............t......a..tg.........t...............t...............------........ [ 33120]
MH025920.1_strain_R35           ......t................t.....................t...t..ctc....................t.... [ 33120]
MH025919.1_strain_N10           ......t................t.....................t...t..ctc....................t.... [ 33120]
MH025918.1_strain_R25           ......t................t.....................t...t..ctc....................t.... [ 33120]
MH025917.1_strain_R7            ......t................t.....................t...t..ctc....................t.... [ 33120]
MH025916.1_strain_R8            ......t................t.....................t...t..ctc....................t.... [ 33120]

FR682468.2_Georgia_2007/1       caatgttaaataaactctgcattttattaaa---atgagg-----------------ttagtatcatg------ttttag [ 33200]
ASF_IND_20_CAD543               ..........................w....---......-------------tgtt.......y...-------..... [ 33200]
LR899193.1_Germany_2020/1       ...............................---......-----------------...........------...... [ 33200]
MN393477.1_Wuhan_2019           ...............................---......-----------------...........------...... [ 33200]
MW396979.1_Timor-Leste/2019/1   ...............................---......-----------------...........------...... [ 33200]
MK940252.1_CN/2019/InnerMongolia ...............................---......-----------------...........------...... [ 33200]
MK128995.1_China/2018/AnhuiXCGQ ...............................---......-----------------...........------...... [ 33200]
MH681419.1_ASFV/POL/2015        ...............................---......-----------------...........------...... [ 33200]
LS478113.1_Estonia_2014         ...............................---......-----------------...........------...... [ 33200]
KP843857.1_Odintsovo_02/14      ...............................---......-----------------...........------...... [ 33200]
FN557520.1_strain_E75           .................a........a....---......-------------tgtt.......t...-------..... [ 33200]
AY261366.1_Warthog              ...............................---......-----------------...........------...... [ 33200]
AY261365.1_Warmbaths            ...............................atg......-----------------.....-.....------...... [ 33200]
AY261364.1_Tengani_62           .........................t.....a--.c...a-------------cg--------------------..... [ 33200]
AY261363.1_Pretoriuskop/96/4    .................a.............---.....a-------------tgtt..ca.......------....g. [ 33200]
AY261362.1_Mkuzi_1979           ...............................---....t.-------------aggt.agt.catg.--------..... [ 33200]
AY261361.1_Malawi_Lil-20/1_1983 .............g...a.........gt..---......-------------tgtt...a...gt..ttagtg...... [ 33200]
AY261360.1_Kenya_1950           .............g...a.........g...---......tgttttagtataatgtt.......a...ttagtg.....a [ 33200]
AM712240.1_OURT_88/3            .................a........a....---......-------------tgtt.......t...-------..... [ 33200]
AM712239.1_Benin_97/1           .................a........a....---......-------------tgtt.......t...-------..... [ 33200]
KM111295.1_Ken06.Bus            .............g...a.........g...---......-------------tgtt.......at..ttagtg...... [ 33200]
```

```
KM111294.1_Ken05/Tk1             .............g...a.........g...---....a.tgttttaacaagatgtt....c.aa...ttaatg.....a [ 33200]
KM262844.1_strain_L60            .................a........a....---......-------------tgtt.......t...-------..... [ 33200]
KM102979.1_26544/OG10_Italy      .................a........a....---......-------------tgtt.......t...-------..... [ 33200]
KM262845.1_strain_NHV            .................a........a....---......-------------tgtt.......t...-------..... [ 33200]
KX354450.1_47/Ss/2008            .................a........a....---......-------------tgtt.......t...-------..... [ 33200]
KP055815.1_strain_BA71           .................a........a....---......-------------tgtt.......t...-------..... [ 33200]
MH025920.1_strain_R35            .............g...a.........g...---......-------------tgtt.......at..ttagtg...... [ 33200]
MH025919.1_strain_N10            .............g...a.........g...---......-------------tgtt.......at..ttagtg...... [ 33200]
MH025918.1_strain_R25            .............g...a.........g...---......-------------tgtt.......at..ttagtg...... [ 33200]
MH025917.1_strain_R7             .............g...a.........g...---......-------------tgtt.......at..ttagtg...... [ 33200]
MH025916.1_strain_R8             .............g...a.........g...---......-------------tgtt.......at..ttagtg...... [ 33200]

FR682468.2_Georgia_2007/1        tataaaattta---------------------------------------------------------gcggc-tgttt- [ 33280]
ASF_IND_20_CAD543                ...........---------------------------------------------------------....ta.....- [ 33280]
LR899193.1_Germany_2020/1        ...........---------------------------------------------------------.....-.....- [ 33280]
MN393477.1_Wuhan_2019            ...........---------------------------------------------------------.....-.....- [ 33280]
MW396979.1_Timor-Leste/2019/1    ...........---------------------------------------------------------.....-.....- [ 33280]
MK940252.1_CN/2019/InnerMongolia ...........---------------------------------------------------------.....-.....- [ 33280]
MK128995.1_China/2018/AnhuiXCGQ  ...........---------------------------------------------------------.....-.....- [ 33280]
MH681419.1_ASFV/POL/2015         ...........---------------------------------------------------------.....-.....- [ 33280]
LS478113.1_Estonia_2014          ...........---------------------------------------------------------.....-.....- [ 33280]
KP843857.1_Odintsovo_02/14       ...........---------------------------------------------------------.....-.....- [ 33280]
FN557520.1_strain_E75            ...........---------------------------------------------------------....ta.....- [ 33280]
AY261366.1_Warthog               ...........---------------------------------------------------------.....-.....- [ 33280]
AY261365.1_Warmbaths             ...........---------------------------------------------------------.....-.....- [ 33280]
AY261364.1_Tengani_62            ...........---------------------------------------------------------a.a.a-...c.- [ 33280]
AY261363.1_Pretoriuskop/96/4     .........c.---------------------------------------------------------.....ac..c.- [ 33280]
AY261362.1_Mkuzi_1979            ...........---------------------------------------------------------.....-.....- [ 33280]
AY261361.1_Malawi_Lil-20/1_1983  ...c.tg..a.tacatgatgttaatacatgatgttaatata--taatgttaatatatgatgttaatatat.atg.taa.- [ 33280]
AY261360.1_Kenya_1950            .....tg...t------------------------agtataatgtatgttagtgtttta---------.taaaa.....- [ 33280]
AM712240.1_OURT_88/3             ...........---------------------------------------------------------....ta.....- [ 33280]
AM712239.1_Benin_97/1            ...........---------------------------------------------------------....ta.....- [ 33280]
KM111295.1_Ken06.Bus             ....gtg...t------------------------agtata----atgttagtgtttta---------.taaaa.....a [ 33280]
KM111294.1_Ken05/Tk1             .....tg..----------------------------tgta--------------ttta---------.....a...c.- [ 33280]
KM262844.1_strain_L60            ...........---------------------------------------------------------....ta.....- [ 33280]
KM102979.1_26544/OG10_Italy      ...........---------------------------------------------------------....ta.....- [ 33280]
KM262845.1_strain_NHV            ...........---------------------------------------------------------....ta.....- [ 33280]
KX354450.1_47/Ss/2008            ...........---------------------------------------------------------....ta.....- [ 33280]
KP055815.1_strain_BA71           ...........---------------------------------------------------------....ta.....- [ 33280]
MH025920.1_strain_R35            ....gtg...t------------------------agtata----atgttagtgtttta---------.taaaa.....a [ 33280]
MH025919.1_strain_N10            ....gtg...t------------------------agtata----atgttagtgtttta---------.taaaa.....a [ 33280]
MH025918.1_strain_R25            ....gtg...t------------------------agtata----atgttagtgtttta---------.taaaa.....a [ 33280]
MH025917.1_strain_R7             ....gtg...t------------------------agtata----atgttagtgtttta---------.taaaa.....a [ 33280]
MH025916.1_strain_R8             ....gtg...t------------------------agtata----atgttagtgtttta---------.taaaa.....a [ 33280]
```

```
FR682468.2_Georgia_2007/1        ------------acataatgctaaataaacttaacgttcctactaaacc-aaaaaaaaatcaaattgactaagtcataga [ 33360]
ASF_IND_20_CAD543                ------------.....................................-.............................. [ 33360]
LR899193.1_Germany_2020/1        ------------.....................................-.............................. [ 33360]
MN393477.1_Wuhan_2019            ------------.....................................-.............................. [ 33360]
MW396979.1_Timor-Leste/2019/1    ------------.....................................-.............................. [ 33360]
MK940252.1_CN/2019/InnerMongolia ------------.....................................-.............................. [ 33360]
MK128995.1_China/2018/AnhuiXCGQ  ------------.....................................-.............................. [ 33360]
MH681419.1_ASFV/POL/2015         ------------.....................................-.............................. [ 33360]
LS478113.1_Estonia_2014          ------------.....................................-.............................. [ 33360]
KP843857.1_Odintsovo_02/14       ------------.....................................--............................. [ 33360]
FN557520.1_strain_E75            ------------g...g................................-....................g.c....... [ 33360]
AY261366.1_Warthog               ------------.....................................-...............----.g.c....... [ 33360]
AY261365.1_Warmbaths             ------------.....................................-....................g.c....... [ 33360]
AY261364.1_Tengani_62            ------------....g..................a.............-...............----.g.c.....a. [ 33360]
AY261363.1_Pretoriuskop/96/4     ------------....g.........................t......-...............----.g.c....... [ 33360]
AY261362.1_Mkuzi_1979            ------------.....................................-....................g.c....... [ 33360]
AY261361.1_Malawi_Lil-20/1_1983  ------------.t..g...t.................g.......g..-...............----...c....c.. [ 33360]
AY261360.1_Kenya_1950            --attgcagccgggc.gcc...........g...............g..-...............----.g.c....c.. [ 33360]
AM712240.1_OURT_88/3             ------------g...g................................-....................g.c....... [ 33360]
AM712239.1_Benin_97/1            ------------g...g................................-....................g.c....... [ 33360]
KM111295.1_Ken06.Bus             gaagcatgcgcgggc.gcc...........g...............g..-...............----.g.c....c.. [ 33360]
KM111294.1_Ken05/Tk1             ----------------------.c..g.tg...................a...............----.g.c....-.. [ 33360]
KM262844.1_strain_L60            ------------g...g................................-....................g.c....... [ 33360]
KM102979.1_26544/OG10_Italy      ------------g...g................................-....................g.c....... [ 33360]
KM262845.1_strain_NHV            ------------g...g................................-....................g.c....... [ 33360]
KX354450.1_47/Ss/2008            ------------g...g................................-....................g.c....... [ 33360]
KP055815.1_strain_BA71           ------------g...g................................-....................g.c....... [ 33360]
MH025920.1_strain_R35            gaagcatgcgcgggc.gcc...........g...............g..-...............----.g.c....c.. [ 33360]
MH025919.1_strain_N10            gaagcatgcgcgggc.gcc...........g...............g..-...............----.g.c....c.. [ 33360]
MH025918.1_strain_R25            gaagcatgcgcgggc.gcc...........g...............g..-...............----.g.c....c.. [ 33360]
MH025917.1_strain_R7             gaagcatgcgcgggc.gcc...........g...............g..-...............----.g.c....c.. [ 33360]
MH025916.1_strain_R8             gaagcatgcgcgggc.gcc...........g...............g..-...............----.g.c....c.. [ 33360]

FR682468.2_Georgia_2007/1        gaatt------tgacgatgttggtaggtaatttttaacatggtatatattttttttagggtcggttatattaggtaataa [ 33440]
ASF_IND_20_CAD543                .....------..................................................................... [ 33440]
LR899193.1_Germany_2020/1        .....------..................................................................... [ 33440]
MN393477.1_Wuhan_2019            .....------..................................................................... [ 33440]
MW396979.1_Timor-Leste/2019/1    .....------..................................................................... [ 33440]
MK940252.1_CN/2019/InnerMongolia .....------..................................................................... [ 33440]
MK128995.1_China/2018/AnhuiXCGQ  .....------..................................................................... [ 33440]
MH681419.1_ASFV/POL/2015         .....------..................................................................... [ 33440]
LS478113.1_Estonia_2014          .....------..................................................................... [ 33440]
KP843857.1_Odintsovo_02/14       .....------..................................................................... [ 33440]
```

```
FN557520.1_strain_E75              .....------................................................a..c................. [ 33440]
AY261366.1_Warthog                 .....------...................................................a................. [ 33440]
AY261365.1_Warmbaths               .....------...................................................a................. [ 33440]
AY261364.1_Tengani_62              .....------................................................a..a................. [ 33440]
AY261363.1_Pretoriuskop/96/4       .....------...................................................a................. [ 33440]
AY261362.1_Mkuzi_1979              .....------................................................a..a................. [ 33440]
AY261361.1_Malawi_Lil-20/1_1983    .....------............c...................................a..a................. [ 33440]
AY261360.1_Kenya_1950              .....------.......a........................................a................g... [ 33440]
AM712240.1_OURT_88/3               .....------................................................a..c................. [ 33440]
AM712239.1_Benin_97/1              .....------................................................a..c................. [ 33440]
KM111295.1_Ken06.Bus               .....------.......a........................................a................g... [ 33440]
KM111294.1_Ken05/Tk1               .....ataaac.......a.....t..................................a.................... [ 33440]
KM262844.1_strain_L60              .....------................................................a..c................. [ 33440]
KM102979.1_26544/OG10_Italy        .....------................................................a..c................. [ 33440]
KM262845.1_strain_NHV              .....------................................................a..c................. [ 33440]
KX354450.1_47/Ss/2008              .....------................................................a..c................. [ 33440]
KP055815.1_strain_BA71             .....------................................................a..c................. [ 33440]
MH025920.1_strain_R35              .....------.......a........................................a................g... [ 33440]
MH025919.1_strain_N10              .....------.......a........................................a................g... [ 33440]
MH025918.1_strain_R25              .....------.......a........................................a................g... [ 33440]
MH025917.1_strain_R7               .....------.......a........................................a................g... [ 33440]
MH025916.1_strain_R8               .....------.......a........................................a................g... [ 33440]

FR682468.2_Georgia_2007/1          aagaggacgtgccgttaaagtattttg---cttaagatcctttagatccttacaaaaatatagattgttcgtctgatgat [ 33520]
ASF_IND_20_CAD543                  ...........................---..............r................................... [ 33520]
LR899193.1_Germany_2020/1          ...........................---.................................................. [ 33520]
MN393477.1_Wuhan_2019              ...........................---.................................................. [ 33520]
MW396979.1_Timor-Leste/2019/1      ...........................---.................................................. [ 33520]
MK940252.1_CN/2019/InnerMongolia   ...........................---.................................................. [ 33520]
MK128995.1_China/2018/AnhuiXCGQ    ...........................---.................................................. [ 33520]
MH681419.1_ASFV/POL/2015           ...........................---.................................................. [ 33520]
LS478113.1_Estonia_2014            ...........................---.................................................. [ 33520]
KP843857.1_Odintsovo_02/14         ...........................---.................................................. [ 33520]
FN557520.1_strain_E75              ...........................---..............a...........................a....a.. [ 33520]
AY261366.1_Warthog                 .........a.........a.......---.................................................. [ 33520]
AY261365.1_Warmbaths               .........a.........a.......---.................................................. [ 33520]
AY261364.1_Tengani_62              .........a.........a.......---t.......at........................................ [ 33520]
AY261363.1_Pretoriuskop/96/4       .........a.................---.................................................. [ 33520]
AY261362.1_Mkuzi_1979              ...........................---.................................................. [ 33520]
AY261361.1_Malawi_Lil-20/1_1983    .........a.........ac......---.........t..................c...t................. [ 33520]
AY261360.1_Kenya_1950              .........c.........cc......ttct..t.a...t..................c..................... [ 33520]
AM712240.1_OURT_88/3               ...........................---..............a...........................a....a.. [ 33520]
AM712239.1_Benin_97/1              ...........................---..............a...........................a....a.. [ 33520]
KM111295.1_Ken06.Bus               ......g..c.........cc......ctc...t.a.c.t....................c................... [ 33520]
```

```
KM111294.1_Ken05/Tk1           .........a.........a.......---t...................................a..t........c. [ 33520]
KM262844.1_strain_L60          ...........................---..............a...........................a....a.. [ 33520]
KM102979.1_26544/OG10_Italy    ...........................---..............a...........................a....a.. [ 33520]
KM262845.1_strain_NHV          ...........................---..............a...........................a....a.. [ 33520]
KX354450.1_47/Ss/2008          ...........................---..............a...........................a....a.. [ 33520]
KP055815.1_strain_BA71         ...........................---..............a...........................a....a.. [ 33520]
MH025920.1_strain_R35          ......g..c.........cc......ctc...t.a.c.t....................c................... [ 33520]
MH025919.1_strain_N10          ......g..c.........cc......ctc...t.a.c......................c................... [ 33520]
MH025918.1_strain_R25          ......g..c.........cc......ctc...t.a.c.t....................c................... [ 33520]
MH025917.1_strain_R7           ......g..c.........cc......ctc...t.a.c.t....................c................... [ 33520]
MH025916.1_strain_R8           ......g..c.........cc......ctc...t.a.c.t....................c................... [ 33520]

FR682468.2_Georgia_2007/1      gccactgtgttgcagtgatggcttgatcaatatcac-ctcccaagacaaaacagtagtatatcgttaaaaagttgtaatc [ 33600]
ASF_IND_20_CAD543              ....................................-........................................... [ 33600]
LR899193.1_Germany_2020/1      ....................................-........................................... [ 33600]
MN393477.1_Wuhan_2019          ....................................-........................................... [ 33600]
MW396979.1_Timor-Leste/2019/1  ....................................-........................................... [ 33600]
MK940252.1_CN/2019/InnerMongolia ....................................-........................................... [ 33600]
MK128995.1_China/2018/AnhuiXCGQ ....................................-........................................... [ 33600]
MH681419.1_ASFV/POL/2015       ....................................-........................................... [ 33600]
LS478113.1_Estonia_2014        ....................................-........................................... [ 33600]
KP843857.1_Odintsovo_02/14     ....................................-........................................... [ 33600]
FN557520.1_strain_E75          .....c.....a.......................g-........................................... [ 33600]
AY261366.1_Warthog             ...........a............t...........-........................................... [ 33600]
AY261365.1_Warmbaths           ...........a............t...........-........................................... [ 33600]
AY261364.1_Tengani_62          ...........a.......................g-.....................................c.g... [ 33600]
AY261363.1_Pretoriuskop/96/4   ...........a........................-........................................... [ 33600]
AY261362.1_Mkuzi_1979          ...........a.......................g-........................................... [ 33600]
AY261361.1_Malawi_Lil-20/1_1983 .........a.a..t.t...................-.............................g............. [ 33600]
AY261360.1_Kenya_1950          .....aa....a..t.t..a...............g-.c....g......g.........................c... [ 33600]
AM712240.1_OURT_88/3           ...........a.......................g-........................................... [ 33600]
AM712239.1_Benin_97/1          .....c.....a.......................g-........................................... [ 33600]
KM111295.1_Ken06.Bus           .....aa....a..t.t......c...........gc.c...........g...........t.............c... [ 33600]
KM111294.1_Ken05/Tk1           .....ga..a.a.gc.t...................-.c..tgca.................t........a........ [ 33600]
KM262844.1_strain_L60          ...........a.......................g-........................................... [ 33600]
KM102979.1_26544/OG10_Italy    .....c.....a.......................g-........................................... [ 33600]
KM262845.1_strain_NHV          ...........a.......................g-........................................... [ 33600]
KX354450.1_47/Ss/2008          .....c.....a.......................g-........................................... [ 33600]
KP055815.1_strain_BA71         ...........a.......................g-........................................... [ 33600]
MH025920.1_strain_R35          .....aa....a..t.t......c...........g-.c...........g...........t.............c... [ 33600]
MH025919.1_strain_N10          .....aa....a..t.t......c...........g-.c...........g...........t.............c... [ 33600]
MH025918.1_strain_R25          .....aa....a..t.t......c...........g-.c...........g...........t.............c... [ 33600]
MH025917.1_strain_R7           .....aa....a..t.t......c...........g-.c...........g...........t.............c... [ 33600]
MH025916.1_strain_R8           .....aa....a..t.t......c...........g-.c...........g...........t.............c... [ 33600]
```

```
FR682468.2_Georgia_2007/1        tttcatacaagccaactgcatcattttatcgatgtccatatgaacgatcttttgctcgtatatttcatgaaggtcaaata [ 33680]
ASF_IND_20_CAD543                ................................................................................ [ 33680]
LR899193.1_Germany_2020/1        ................................................................................ [ 33680]
MN393477.1_Wuhan_2019            ................................................................................ [ 33680]
MW396979.1_Timor-Leste/2019/1    ................................................................................ [ 33680]
MK940252.1_CN/2019/InnerMongolia ................................................................................ [ 33680]
MK128995.1_China/2018/AnhuiXCGQ  ................................................................................ [ 33680]
MH681419.1_ASFV/POL/2015         ................................................................................ [ 33680]
LS478113.1_Estonia_2014          ................................................................................ [ 33680]
KP843857.1_Odintsovo_02/14       ................................................................................ [ 33680]
FN557520.1_strain_E75            ..g............................................................................. [ 33680]
AY261366.1_Warthog               ..g..............t.............................................................. [ 33680]
AY261365.1_Warmbaths             ..g..............t.............................................................. [ 33680]
AY261364.1_Tengani_62            ..g......................................................a....g................. [ 33680]
AY261363.1_Pretoriuskop/96/4     ..g..............t.............................................................. [ 33680]
AY261362.1_Mkuzi_1979            ..g............................................................................. [ 33680]
AY261361.1_Malawi_Lil-20/1_1983  .cg...g........t..........c......a.........g.............a...................... [ 33680]
AY261360.1_Kenya_1950            ..c............a......................................t..a...................... [ 33680]
AM712240.1_OURT_88/3             ..g............................................................................. [ 33680]
AM712239.1_Benin_97/1            ..g............................................................................. [ 33680]
KM111295.1_Ken06.Bus             ..g............a......................................t..a...................... [ 33680]
KM111294.1_Ken05/Tk1             ..g..a.........a..........c.............................t...c.gc................ [ 33680]
KM262844.1_strain_L60            ..g............................................................................. [ 33680]
KM102979.1_26544/OG10_Italy      ..g............................................................................. [ 33680]
KM262845.1_strain_NHV            ..g............................................................................. [ 33680]
KX354450.1_47/Ss/2008            ..g............................................................................. [ 33680]
KP055815.1_strain_BA71           ..g............................................................................. [ 33680]
MH025920.1_strain_R35            ..g............a......................................t..a...................... [ 33680]
MH025919.1_strain_N10            ..g............a......................................t..a...................... [ 33680]
MH025918.1_strain_R25            ..g............a......................................t..a...................... [ 33680]
MH025917.1_strain_R7             ..g............a......................................t..a...................... [ 33680]
MH025916.1_strain_R8             ..g............a......................................t..a...................... [ 33680]

FR682468.2_Georgia_2007/1        cattgttgaagtaaatggcgcacatgagtcgccacatactaaggtgcccatatgtttgatagaaaaaggagatagctctt [ 33760]
ASF_IND_20_CAD543                ................................................................................ [ 33760]
LR899193.1_Germany_2020/1        ................................................................................ [ 33760]
MN393477.1_Wuhan_2019            ................................................................................ [ 33760]
MW396979.1_Timor-Leste/2019/1    ................................................................................ [ 33760]
MK940252.1_CN/2019/InnerMongolia ................................................................................ [ 33760]
MK128995.1_China/2018/AnhuiXCGQ  ................................................................................ [ 33760]
MH681419.1_ASFV/POL/2015         ................................................................................ [ 33760]
LS478113.1_Estonia_2014          ................................................................................ [ 33760]
KP843857.1_Odintsovo_02/14       ................................................................................ [ 33760]
```

```
FN557520.1_strain_E75             ....................................................................a........... [ 33760]
AY261366.1_Warthog                ................................................................................ [ 33760]
AY261365.1_Warmbaths              ................................................................................ [ 33760]
AY261364.1_Tengani_62             .............................................a.................................. [ 33760]
AY261363.1_Pretoriuskop/96/4      ...........a.................................................................... [ 33760]
AY261362.1_Mkuzi_1979             ................................................................................ [ 33760]
AY261361.1_Malawi_Lil-20/1_1983  t...a.................................t........................................c [ 33760]
AY261360.1_Kenya_1950             ...........g..........................t.......gt.g.............................c [ 33760]
AM712240.1_OURT_88/3              ....................................................................a........... [ 33760]
AM712239.1_Benin_97/1             ....................................................................a........... [ 33760]
KM111295.1_Ken06.Bus              ...........g..........................t.......gt.g.........a...................c [ 33760]
KM111294.1_Ken05/Tk1              ......................................t........t...............................c [ 33760]
KM262844.1_strain_L60             ....................................................................a........... [ 33760]
KM102979.1_26544/OG10_Italy       ....................................................................a........... [ 33760]
KM262845.1_strain_NHV             ....................................................................a........... [ 33760]
KX354450.1_47/Ss/2008             ....................................................................a........... [ 33760]
KP055815.1_strain_BA71            ....................................................................a........... [ 33760]
MH025920.1_strain_R35             ...........g..........................t.......gt.g.........a...................c [ 33760]
MH025919.1_strain_N10             ...........g..........................t.......gt.g.........a...................c [ 33760]
MH025918.1_strain_R25             ...........g..........................t.......gt.g.........a...................c [ 33760]
MH025917.1_strain_R7              ...........g..........................t.......gt.g.........a...................c [ 33760]
MH025916.1_strain_R8              ...........g..........................t.......gt.g.........a...................c [ 33760]

FR682468.2_Georgia_2007/1         ttaagcttatattttactgctatggcatagcagtatttaacgaatacgttcatgggtacattatctaagatataaaatat [ 33840]
ASF_IND_20_CAD543                 ................................................................................ [ 33840]
LR899193.1_Germany_2020/1         ................................................................................ [ 33840]
MN393477.1_Wuhan_2019             ................................................................................ [ 33840]
MW396979.1_Timor-Leste/2019/1     ................................................................................ [ 33840]
MK940252.1_CN/2019/InnerMongolia  ................................................................................ [ 33840]
MK128995.1_China/2018/AnhuiXCGQ   ................................................................................ [ 33840]
MH681419.1_ASFV/POL/2015          ................................................................................ [ 33840]
LS478113.1_Estonia_2014           ................................................................................ [ 33840]
KP843857.1_Odintsovo_02/14        ................................................................................ [ 33840]
FN557520.1_strain_E75             .............................c................t..................c.............. [ 33840]
AY261366.1_Warthog                .............................c..........tt.......................c.............. [ 33840]
AY261365.1_Warmbaths              .............................c..........tt.......................c.............. [ 33840]
AY261364.1_Tengani_62             .............................c...................................c.............c [ 33840]
AY261363.1_Pretoriuskop/96/4      ....................c........c...........t.......................c.............. [ 33840]
AY261362.1_Mkuzi_1979             .................................................................c.............. [ 33840]
AY261361.1_Malawi_Lil-20/1_1983   ................tc...........c...........t........a..............c.............. [ 33840]
AY261360.1_Kenya_1950             .................c...........c........ttt....gt...g..............c.............. [ 33840]
AM712240.1_OURT_88/3              .............................c................t..................c.............. [ 33840]
AM712239.1_Benin_97/1             .............................c................t..................c.............. [ 33840]
KM111295.1_Ken06.Bus              .................c...........c...........t...gt...g.....c........c.............. [ 33840]
```

```
KM111294.1_Ken05/Tk1             .................c........g..c...a.......t...gt...a..........g.................. [ 33840]
KM262844.1_strain_L60            .............................c................t..................c.............. [ 33840]
KM102979.1_26544/OG10_Italy      .............................c................t..................c.............. [ 33840]
KM262845.1_strain_NHV            .............................c................t..................c.............. [ 33840]
KX354450.1_47/Ss/2008            .............................c................t..................c.............. [ 33840]
KP055815.1_strain_BA71           .............................c................t..................c.............. [ 33840]
MH025920.1_strain_R35            .................c...........c...........t...gt...g.....c........c.............. [ 33840]
MH025919.1_strain_N10            .................c...........c...........t...gt...g.....c........c.............. [ 33840]
MH025918.1_strain_R25            .................c...........c...........t...gt...g.....c........c.............. [ 33840]
MH025917.1_strain_R7             .................c...........c...........t...gt...g.....c........c.............. [ 33840]
MH025916.1_strain_R8             .................c...........c...........t...gt...g.....c........c.............. [ 33840]

FR682468.2_Georgia_2007/1        gaaaaactttaactctcgatgaatctcttcccccatttcctgtacatttagagcttccaacataggatttttatcaaata [ 33920]
ASF_IND_20_CAD543                .....r.......................................................................... [ 33920]
LR899193.1_Germany_2020/1        ................................................................................ [ 33920]
MN393477.1_Wuhan_2019            ................................................................................ [ 33920]
MW396979.1_Timor-Leste/2019/1    ................................................................................ [ 33920]
MK940252.1_CN/2019/InnerMongolia ................................................................................ [ 33920]
MK128995.1_China/2018/AnhuiXCGQ  ................................................................................ [ 33920]
MH681419.1_ASFV/POL/2015         ................................................................................ [ 33920]
LS478113.1_Estonia_2014          ................................................................................ [ 33920]
KP843857.1_Odintsovo_02/14       ................................................................................ [ 33920]
FN557520.1_strain_E75            .....g.....................c..t...........c..................................... [ 33920]
AY261366.1_Warthog               .....g.....................c......................a..t.......................... [ 33920]
AY261365.1_Warmbaths             .....g.....................c.........................t.......................... [ 33920]
AY261364.1_Tengani_62            .....g.....................c.................................................... [ 33920]
AY261363.1_Pretoriuskop/96/4     a....g.....................c.................t.......t........................a. [ 33920]
AY261362.1_Mkuzi_1979            .....g.....................c.................................................... [ 33920]
AY261361.1_Malawi_Lil-20/1_1983  ...............c.....g.....c.....a.aa........tg......t......t........a....t..... [ 33920]
AY261360.1_Kenya_1950            ..g.tg.........c...........c.....a.ag.....c..t......atc..t................t..... [ 33920]
AM712240.1_OURT_88/3             .....g.....................c..t...........c..................................... [ 33920]
AM712239.1_Benin_97/1            .....g.....................c..t...........c..................................... [ 33920]
KM111295.1_Ken06.Bus             ..g.tg.........c...........cc..g...a......c..t......atgc.t................t..... [ 33920]
KM111294.1_Ken05/Tk1             t...tg.........ct....g.....cc..a.t...........tg......t...t......g.........t..... [ 33920]
KM262844.1_strain_L60            .....g.....................c..t...........c..................................... [ 33920]
KM102979.1_26544/OG10_Italy      .....g.....................c..t...........c..................................... [ 33920]
KM262845.1_strain_NHV            .....g.....................c..t...........c..................................... [ 33920]
KX354450.1_47/Ss/2008            .....g.....................c..t...........c..................................... [ 33920]
KP055815.1_strain_BA71           .....g.....................c..t...........c..................................... [ 33920]
MH025920.1_strain_R35            ..g.tg.........c...........cc..g...a......c..t......atgc.t................t..... [ 33920]
MH025919.1_strain_N10            ..g.tg.........c...........cc..g...a......c..t......atgc.t................t..... [ 33920]
MH025918.1_strain_R25            ..g.tg.........c...........cc..g...a......c..t......atgc.t................t..... [ 33920]
MH025917.1_strain_R7             ..g.tg.........c...........cc..g...a......c..t......atgc.t................t..... [ 33920]
MH025916.1_strain_R8             ..g.tg.........c...........cc..g...a......c..t......atgc.t................t..... [ 33920]
```

```
FR682468.2_Georgia_2007/1        tttcatgacataaaataatgttattgctcgttttatgacgcattaaaccggtgaaaatttccttattatttaaactatct [ 34000]
ASF_IND_20_CAD543                ...........................................................................y.... [ 34000]
LR899193.1_Germany_2020/1        ................................................................................ [ 34000]
MN393477.1_Wuhan_2019            ................................................................................ [ 34000]
MW396979.1_Timor-Leste/2019/1    ................................................................................ [ 34000]
MK940252.1_CN/2019/InnerMongolia ................................................................................ [ 34000]
MK128995.1_China/2018/AnhuiXCGQ  ................................................................................ [ 34000]
MH681419.1_ASFV/POL/2015         ................................................................................ [ 34000]
LS478113.1_Estonia_2014          ................................................................................ [ 34000]
KP843857.1_Odintsovo_02/14       ................................................................................ [ 34000]
FN557520.1_strain_E75            .........................a.........................c.......................c.... [ 34000]
AY261366.1_Warthog               ......................................................................c......... [ 34000]
AY261365.1_Warmbaths             ...................a..............................a...................c......... [ 34000]
AY261364.1_Tengani_62            .........................a........................a........................gt... [ 34000]
AY261363.1_Pretoriuskop/96/4     ......................................................................c......... [ 34000]
AY261362.1_Mkuzi_1979            .........................a...................................................... [ 34000]
AY261361.1_Malawi_Lil-20/1_1983  .............t............................g...c..aa.a....................t.c.... [ 34000]
AY261360.1_Kenya_1950            .c.....g..............................t...a.....t.a.......g.....t....c.....c.... [ 34000]
AM712240.1_OURT_88/3             .........................a.........................c.......................c.... [ 34000]
AM712239.1_Benin_97/1            .........................a.........................c.......................c.... [ 34000]
KM111295.1_Ken06.Bus             .c.....g..................................a.......a.......g.....t....c.....c.... [ 34000]
KM111294.1_Ken05/Tk1             ..................ca.....a..........tg....g...c..aa...........c............ct... [ 34000]
KM262844.1_strain_L60            .........................a.........................c.......................c.... [ 34000]
KM102979.1_26544/OG10_Italy      .........................a.........................c.......................c.... [ 34000]
KM262845.1_strain_NHV            .........................a.........................c.......................c.... [ 34000]
KX354450.1_47/Ss/2008            .........................a.........................c.......................c.... [ 34000]
KP055815.1_strain_BA71           .........................a.........................c.......................c.... [ 34000]
MH025920.1_strain_R35            .c.....g..................................a.......a.......g.....t....c.....c.... [ 34000]
MH025919.1_strain_N10            .c.....g..................................a.......a.......g.....t....c.....c.... [ 34000]
MH025918.1_strain_R25            .c.....g..................................a.......a.......g.....t....c.....c.... [ 34000]
MH025917.1_strain_R7             .c.....g..................................a.......a.......g.....t....c.....c.... [ 34000]
MH025916.1_strain_R8             .c.....g..................................a.......a.......g.....t....c.....c.... [ 34000]

FR682468.2_Georgia_2007/1        ttagctcctaactttcgacacagctcctgagtttgttccgtcctagcacaggtcagcccataataaatgtttgctcccca [ 34080]
ASF_IND_20_CAD543                ................................................................................ [ 34080]
LR899193.1_Germany_2020/1        ................................................................................ [ 34080]
MN393477.1_Wuhan_2019            ................................................................................ [ 34080]
MW396979.1_Timor-Leste/2019/1    ................................................................................ [ 34080]
MK940252.1_CN/2019/InnerMongolia ................................................................................ [ 34080]
MK128995.1_China/2018/AnhuiXCGQ  ................................................................................ [ 34080]
MH681419.1_ASFV/POL/2015         ................................................................................ [ 34080]
LS478113.1_Estonia_2014          ................................................................................ [ 34080]
KP843857.1_Odintsovo_02/14       ................................................................................ [ 34080]
```

```
FN557520.1_strain_E75            ................................................................................ [ 34080]
AY261366.1_Warthog               .............g.................................................................. [ 34080]
AY261365.1_Warmbaths             ................................................................................ [ 34080]
AY261364.1_Tengani_62            .....................................................t.......................... [ 34080]
AY261363.1_Pretoriuskop/96/4     .........................t................................................c..... [ 34080]
AY261362.1_Mkuzi_1979            ................................................................................ [ 34080]
AY261361.1_Malawi_Lil-20/1_1983  .............c...g....a......................................................... [ 34080]
AY261360.1_Kenya_1950            ......................a.........a.a...............ac....a..g.................... [ 34080]
AM712240.1_OURT_88/3             ................................................................................ [ 34080]
AM712239.1_Benin_97/1            ................................................................................ [ 34080]
KM111295.1_Ken06.Bus             ................................a.a...t...........ac....a..g.................... [ 34080]
KM111294.1_Ken05/Tk1             ............g................................................................... [ 34080]
KM262844.1_strain_L60            ................................................................................ [ 34080]
KM102979.1_26544/OG10_Italy      ................................................................................ [ 34080]
KM262845.1_strain_NHV            ................................................................................ [ 34080]
KX354450.1_47/Ss/2008            ................................................................................ [ 34080]
KP055815.1_strain_BA71           ................................................................................ [ 34080]
MH025920.1_strain_R35            ................................a.a...t...........ac....a..g.................... [ 34080]
MH025919.1_strain_N10            ................................a.a...t...........ac....a..g.................... [ 34080]
MH025918.1_strain_R25            ................................a.a...t...........ac....a..g.................... [ 34080]
MH025917.1_strain_R7             ................................a.a...t...........ac....a..g.................... [ 34080]
MH025916.1_strain_R8             ................................a.a...t...........ac....a..g.................... [ 34080]

FR682468.2_Georgia_2007/1        ctcggtgaacagccttattacgtcatagttattttcttttatggccatgattaatgccacatcaagatgaagaagttccc [ 34160]
ASF_IND_20_CAD543                .....................r.......................................................... [ 34160]
LR899193.1_Germany_2020/1        ................................................................................ [ 34160]
MN393477.1_Wuhan_2019            ................................................................................ [ 34160]
MW396979.1_Timor-Leste/2019/1    ................................................................................ [ 34160]
MK940252.1_CN/2019/InnerMongolia ................................................................................ [ 34160]
MK128995.1_China/2018/AnhuiXCGQ  ................................................................................ [ 34160]
MH681419.1_ASFV/POL/2015         ................................................................................ [ 34160]
LS478113.1_Estonia_2014          ................................................................................ [ 34160]
KP843857.1_Odintsovo_02/14       ................................................................................ [ 34160]
FN557520.1_strain_E75            t....................a................................c......................... [ 34160]
AY261366.1_Warthog               .....................a..g.................a..................................... [ 34160]
AY261365.1_Warmbaths             .....................a..g..............g..a..................................... [ 34160]
AY261364.1_Tengani_62            .....................a..g....................................................... [ 34160]
AY261363.1_Pretoriuskop/96/4     .....................a..g.................a...........c......................... [ 34160]
AY261362.1_Mkuzi_1979            ......a..............a................................c......................... [ 34160]
AY261361.1_Malawi_Lil-20/1_1983  .....................a..g..............g........................................ [ 34160]
AY261360.1_Kenya_1950            ............t.....c.aa..g.g.....................a...................---......... [ 34160]
AM712240.1_OURT_88/3             t....................a................................c......................... [ 34160]
AM712239.1_Benin_97/1            t....................a................................c......................... [ 34160]
KM111295.1_Ken06.Bus             ............t.....c.aa..g.g.........................................---......... [ 34160]
```

```
KM111294.1_Ken05/Tk1              ....................aa..g..............................a.t...................... [ 34160]
KM262844.1_strain_L60             t....................a................................c......................... [ 34160]
KM102979.1_26544/OG10_Italy       t....................a................................c......................... [ 34160]
KM262845.1_strain_NHV             t....................a................................c......................... [ 34160]
KX354450.1_47/Ss/2008             t....................a................................c......................... [ 34160]
KP055815.1_strain_BA71            t....................a................................c......................... [ 34160]
MH025920.1_strain_R35             ............t.....c.aa..g.g.........................................---......... [ 34160]
MH025919.1_strain_N10             ............t.....c.aa..g.g.........................................---......... [ 34160]
MH025918.1_strain_R25             ............t.....c.aa..g.g.........................................---......... [ 34160]
MH025917.1_strain_R7              ............t.....c.aa..g.g.........................................---......... [ 34160]
MH025916.1_strain_R8              ............t.....c.aa..g.g.........................................---......... [ 34160]

FR682468.2_Georgia_2007/1         ccttaaagggggttgagcttaaaataacgtaattacagtagtgacataagctaatgggcttgttttgccaccataagcca [ 34240]
ASF_IND_20_CAD543                 ................................................................................ [ 34240]
LR899193.1_Germany_2020/1         ................................................................................ [ 34240]
MN393477.1_Wuhan_2019             ................................................................................ [ 34240]
MW396979.1_Timor-Leste/2019/1     ................................................................................ [ 34240]
MK940252.1_CN/2019/InnerMongolia  ................................................................................ [ 34240]
MK128995.1_China/2018/AnhuiXCGQ   ................................................................................ [ 34240]
MH681419.1_ASFV/POL/2015          ................................................................................ [ 34240]
LS478113.1_Estonia_2014           ................................................................................ [ 34240]
KP843857.1_Odintsovo_02/14        ................................................................................ [ 34240]
FN557520.1_strain_E75             ................................................................................ [ 34240]
AY261366.1_Warthog                ........................................a....................a.................. [ 34240]
AY261365.1_Warmbaths              ................................................................................ [ 34240]
AY261364.1_Tengani_62             ................................................................................ [ 34240]
AY261363.1_Pretoriuskop/96/4      ...............................................................................g [ 34240]
AY261362.1_Mkuzi_1979             ................................................................................ [ 34240]
AY261361.1_Malawi_Lil-20/1_1983   t.......a........t.........t...............g..c..a..........................a... [ 34240]
AY261360.1_Kenya_1950             .................t...c....................c......atg...a........................ [ 34240]
AM712240.1_OURT_88/3              ................................................................................ [ 34240]
AM712239.1_Benin_97/1             ................................................................................ [ 34240]
KM111295.1_Ken06.Bus              .................t...c....................c...c..atg...a........................ [ 34240]
KM111294.1_Ken05/Tk1              ...........................t.............c....c..a.............c................ [ 34240]
KM262844.1_strain_L60             ................................................................................ [ 34240]
KM102979.1_26544/OG10_Italy       ................................................................................ [ 34240]
KM262845.1_strain_NHV             ................................................................................ [ 34240]
KX354450.1_47/Ss/2008             ................................................................................ [ 34240]
KP055815.1_strain_BA71            ................................................................................ [ 34240]
MH025920.1_strain_R35             .................t...c....................c...c..atg...a........................ [ 34240]
MH025919.1_strain_N10             .................t...c....................c...c..atg...a........................ [ 34240]
MH025918.1_strain_R25             .................t...c....................c...c..atg...a........................ [ 34240]
MH025917.1_strain_R7              .................t...c....................c...c..atg...a........................ [ 34240]
MH025916.1_strain_R8              .................t...c....................c...c..atg...a........................ [ 34240]
```

```
FR682468.2_Georgia_2007/1        caatattttaaaatataatgatactcctcaggcacgctctgtttggccacagcctttttggccagggtttgcaaggagag [ 34320]
ASF_IND_20_CAD543                ................................................................................ [ 34320]
LR899193.1_Germany_2020/1        ................................................................................ [ 34320]
MN393477.1_Wuhan_2019            ................................................................................ [ 34320]
MW396979.1_Timor-Leste/2019/1    ................................................................................ [ 34320]
MK940252.1_CN/2019/InnerMongolia ................................................................................ [ 34320]
MK128995.1_China/2018/AnhuiXCGQ  ................................................................................ [ 34320]
MH681419.1_ASFV/POL/2015         ................................................................................ [ 34320]
LS478113.1_Estonia_2014          ................................................................................ [ 34320]
KP843857.1_Odintsovo_02/14       ................................................................................ [ 34320]
FN557520.1_strain_E75            ................................................................................ [ 34320]
AY261366.1_Warthog               ................................................................................ [ 34320]
AY261365.1_Warmbaths             ...........................................................a.................... [ 34320]
AY261364.1_Tengani_62            ..................................................c............................. [ 34320]
AY261363.1_Pretoriuskop/96/4     ..........................t....................g......c......................... [ 34320]
AY261362.1_Mkuzi_1979            ................................................................................ [ 34320]
AY261361.1_Malawi_Lil-20/1_1983  ..........................t.....................................a.........t..... [ 34320]
AY261360.1_Kenya_1950            ..g..............g........t......................g............t...........a..... [ 34320]
AM712240.1_OURT_88/3             ................................................................................ [ 34320]
AM712239.1_Benin_97/1            ................................................................................ [ 34320]
KM111295.1_Ken06.Bus             ..g..............g........t......................g........................a..... [ 34320]
KM111294.1_Ken05/Tk1             ................................................................a.........a..... [ 34320]
KM262844.1_strain_L60            ................................................................................ [ 34320]
KM102979.1_26544/OG10_Italy      ................................................................................ [ 34320]
KM262845.1_strain_NHV            ................................................................................ [ 34320]
KX354450.1_47/Ss/2008            ................................................................................ [ 34320]
KP055815.1_strain_BA71           ................................................................................ [ 34320]
MH025920.1_strain_R35            ..g..............g........t......................g........................a..... [ 34320]
MH025919.1_strain_N10            ..g..............g........t......................g........................a..... [ 34320]
MH025918.1_strain_R25            ..g..............g........t......................g........................a..... [ 34320]
MH025917.1_strain_R7             ..g..............g........t......................g........................a..... [ 34320]
MH025916.1_strain_R8             ..g..............g........t......................g........................a..... [ 34320]

FR682468.2_Georgia_2007/1        catgataacttct-tg----aaaaaaaaactcaaattaagttcctacttttttaaaatatta------gtat-------- [ 34400]
ASF_IND_20_CAD543                .............-..----..........................................------....-------- [ 34400]
LR899193.1_Germany_2020/1        .............-..----..........................................------....-------- [ 34400]
MN393477.1_Wuhan_2019            .............-..----..........................................------....-------- [ 34400]
MW396979.1_Timor-Leste/2019/1    .............-..----..........................................------....-------- [ 34400]
MK940252.1_CN/2019/InnerMongolia .............-..----..........................................------....-------- [ 34400]
MK128995.1_China/2018/AnhuiXCGQ  .............-..----..........................................------....-------- [ 34400]
MH681419.1_ASFV/POL/2015         .............-..----..........................................------....-------- [ 34400]
LS478113.1_Estonia_2014          .............-..----..........................................------....-------- [ 34400]
KP843857.1_Odintsovo_02/14       .............-..----..........................................------....-------- [ 34400]
```

```
FN557520.1_strain_E75            .............-..----..........................................------....-------- [ 34400]
AY261366.1_Warthog               ....t......t.t.t----g.....................t...................-----t....-------- [ 34400]
AY261365.1_Warmbaths             ....t......t.t.t----g.....................t..........-...c....-----t....-------- [ 34400]
AY261364.1_Tengani_62            ...a.........-..----..........................................t--tat....-------- [ 34400]
AY261363.1_Pretoriuskop/96/4     ....t......t.t.t----g.....................t...................-----t....-------- [ 34400]
AY261362.1_Mkuzi_1979            .............-..----..........................................------....-------- [ 34400]
AY261361.1_Malawi_Lil-20/1_1983  ...........t.-.t----g........a.........a.....g.......-...c....tgttat....-------- [ 34400]
AY261360.1_Kenya_1950            ...........t.-.t----cg....................t..gt......t...c..g.-----t....-------- [ 34400]
AM712240.1_OURT_88/3             .............-..----..........................................------....-------- [ 34400]
AM712239.1_Benin_97/1            .............-..----..........................................------....-------- [ 34400]
KM111295.1_Ken06.Bus             ...........t.-.ttcg-......................t..............cg...-----t....-------- [ 34400]
KM111294.1_Ken05/Tk1             ...........t.-.ttg--.........g.............t.gt..........c....-----t..t.tgtatata [ 34400]
KM262844.1_strain_L60            .............-..----..........................................------....-------- [ 34400]
KM102979.1_26544/OG10_Italy      .............-..----..........................................------....-------- [ 34400]
KM262845.1_strain_NHV            .............-..----..........................................------....-------- [ 34400]
KX354450.1_47/Ss/2008            .............-..----..........................................------....-------- [ 34400]
KP055815.1_strain_BA71           .............-..----..........................................------....-------- [ 34400]
MH025920.1_strain_R35            ...........t.-.ttcga......................t..............cg...-----t....-------- [ 34400]
MH025919.1_strain_N10            ...........t.-.ttcga......................t..............cg...-----t....-------- [ 34400]
MH025918.1_strain_R25            ...........t.-.ttcga......................t..............cg...-----t....-------- [ 34400]
MH025917.1_strain_R7             ...........t.-.ttcga......................t..............cg...-----t....-------- [ 34400]
MH025916.1_strain_R8             ...........t.-.ttcga......................t..............cg...-----t....-------- [ 34400]

FR682468.2_Georgia_2007/1        ggacagatc-------------------------taccatc--atatgaaggaa---tt--------------------- [ 34480]
ASF_IND_20_CAD543                .........-------------------------.......--...........---..--------------------- [ 34480]
LR899193.1_Germany_2020/1        .........-------------------------.......--...........---..--------------------- [ 34480]
MN393477.1_Wuhan_2019            .........-------------------------.......--...........---..--------------------- [ 34480]
MW396979.1_Timor-Leste/2019/1    .........-------------------------.......--...........---..--------------------- [ 34480]
MK940252.1_CN/2019/InnerMongolia .........-------------------------.......--...........---..--------------------- [ 34480]
MK128995.1_China/2018/AnhuiXCGQ  .........-------------------------.......--...........---..--------------------- [ 34480]
MH681419.1_ASFV/POL/2015         .........-------------------------.......--...........---..--------------------- [ 34480]
LS478113.1_Estonia_2014          .........-------------------------.......--...........---..--------------------- [ 34480]
KP843857.1_Odintsovo_02/14       .........-------------------------.......--...........---..--------------------- [ 34480]
FN557520.1_strain_E75            .........-------------------------.......--...........---..--------------------- [ 34480]
AY261366.1_Warthog               a.g......cgccg-------------------a.......--.....g.....---..--------------------- [ 34480]
AY261365.1_Warmbaths             ..g......cgccg-------------------a.......--........a..---..--------------------- [ 34480]
AY261364.1_Tengani_62            a.g......tgccg-------------------c.......at...........---..--------------------- [ 34480]
AY261363.1_Pretoriuskop/96/4     a.g......cgccg-------------------a.......--...........---..--------------------- [ 34480]
AY261362.1_Mkuzi_1979            ..g......-------------------------.......--...........---..--------------------- [ 34480]
AY261361.1_Malawi_Lil-20/1_1983  a...g...tcgccgattat-------------ct.......--c..........---..cttctttcatcgttaaatact [ 34480]
AY261360.1_Kenya_1950            ..g.c....cgccggagatctctttcagaagaccc..t..t--c......a...---..--------------------- [ 34480]
AM712240.1_OURT_88/3             .........-------------------------.......--...........---..--------------------- [ 34480]
AM712239.1_Benin_97/1            .........-------------------------.......--...........---..--------------------- [ 34480]
KM111295.1_Ken06.Bus             .ag.c....cgccggagatccctttcagaagacc....g.t--c......a...---..--------------------- [ 34480]
```

```
KM111294.1_Ken05/Tk1             tagat....caccgatgat-------------cc..t....--c..........ttc..--------------------- [ 34480]
KM262844.1_strain_L60            .........-------------------------.......--...........---..--------------------- [ 34480]
KM102979.1_26544/OG10_Italy      .........-------------------------.......--...........---..--------------------- [ 34480]
KM262845.1_strain_NHV            .........-------------------------.......--...........---..--------------------- [ 34480]
KX354450.1_47/Ss/2008            .........-------------------------.......--...........---..--------------------- [ 34480]
KP055815.1_strain_BA71           .........-------------------------.......--...........---..--------------------- [ 34480]
MH025920.1_strain_R35            .ag.c....cgccggagatccctttcagaagacc....g.t--c......a...---..--------------------- [ 34480]
MH025919.1_strain_N10            .ag.c....cgccggagatccctttcagaagacc....g.t--c......a...---..--------------------- [ 34480]
MH025918.1_strain_R25            .ag.c....cgccggagatccctttcagaagacc....g.t--c......a...---..--------------------- [ 34480]
MH025917.1_strain_R7             .ag.c....cgccggagatccctttcagaagacc....g.t--c......a...---..--------------------- [ 34480]
MH025916.1_strain_R8             .ag.c....cgccggagatccctttcagaagacc....g.t--c......a...---..--------------------- [ 34480]

FR682468.2_Georgia_2007/1        -----------------------------------------------------------------------ctttcatcg [ 34560]
ASF_IND_20_CAD543                -----------------------------------------------------------------------......... [ 34560]
LR899193.1_Germany_2020/1        -----------------------------------------------------------------------......... [ 34560]
MN393477.1_Wuhan_2019            -----------------------------------------------------------------------......... [ 34560]
MW396979.1_Timor-Leste/2019/1    -----------------------------------------------------------------------......... [ 34560]
MK940252.1_CN/2019/InnerMongolia -----------------------------------------------------------------------......... [ 34560]
MK128995.1_China/2018/AnhuiXCGQ  -----------------------------------------------------------------------......... [ 34560]
MH681419.1_ASFV/POL/2015         -----------------------------------------------------------------------......... [ 34560]
LS478113.1_Estonia_2014          -----------------------------------------------------------------------......... [ 34560]
KP843857.1_Odintsovo_02/14       -----------------------------------------------------------------------......... [ 34560]
FN557520.1_strain_E75            -----------------------------------------------------------------------......... [ 34560]
AY261366.1_Warthog               -----------------------------------------------------------------------....g.... [ 34560]
AY261365.1_Warmbaths             -----------------------------------------------------------------------....g.... [ 34560]
AY261364.1_Tengani_62            -----------------------------------------------------------------------......... [ 34560]
AY261363.1_Pretoriuskop/96/4     -----------------------------------------------------------------------....g.... [ 34560]
AY261362.1_Mkuzi_1979            -----------------------------------------------------------------------......... [ 34560]
AY261361.1_Malawi_Lil-20/1_1983  gaagagataatactttcatcgttaaatactgaagagataatactttcatcattaaatactgaagagtaata......... [ 34560]
AY261360.1_Kenya_1950            -----------------------------------------------------------------------t........ [ 34560]
AM712240.1_OURT_88/3             -----------------------------------------------------------------------......... [ 34560]
AM712239.1_Benin_97/1            -----------------------------------------------------------------------......... [ 34560]
KM111295.1_Ken06.Bus             -----------------------------------------------------------------------t........ [ 34560]
KM111294.1_Ken05/Tk1             -----------------------------------------------------------------------.......t. [ 34560]
KM262844.1_strain_L60            -----------------------------------------------------------------------......... [ 34560]
KM102979.1_26544/OG10_Italy      -----------------------------------------------------------------------......... [ 34560]
KM262845.1_strain_NHV            -----------------------------------------------------------------------......... [ 34560]
KX354450.1_47/Ss/2008            -----------------------------------------------------------------------......... [ 34560]
KP055815.1_strain_BA71           -----------------------------------------------------------------------......... [ 34560]
MH025920.1_strain_R35            -----------------------------------------------------------------------t........ [ 34560]
MH025919.1_strain_N10            -----------------------------------------------------------------------t........ [ 34560]
MH025918.1_strain_R25            -----------------------------------------------------------------------t........ [ 34560]
MH025917.1_strain_R7             -----------------------------------------------------------------------t........ [ 34560]
MH025916.1_strain_R8             -----------------------------------------------------------------------t........ [ 34560]
```

```
FR682468.2_Georgia_2007/1        ttaaacactgaagag--ataatactttcatcgtatagagaatatcatgtcaatccatatattgaatgttatatatcatta [ 34640]
ASF_IND_20_CAD543                ...............--............................................................... [ 34640]
LR899193.1_Germany_2020/1        ...............--............................................................... [ 34640]
MN393477.1_Wuhan_2019            ...............--............................................................... [ 34640]
MW396979.1_Timor-Leste/2019/1    ...............--............................................................... [ 34640]
MK940252.1_CN/2019/InnerMongolia ...............--............................................................... [ 34640]
MK128995.1_China/2018/AnhuiXCGQ  ...............--............................................................... [ 34640]
MH681419.1_ASFV/POL/2015         ...............--............................................................... [ 34640]
LS478113.1_Estonia_2014          ...............--............................................................... [ 34640]
KP843857.1_Odintsovo_02/14       ...............--............................................................... [ 34640]
FN557520.1_strain_E75            ...............--............................................................... [ 34640]
AY261366.1_Warthog               ...............at..............................a................................ [ 34640]
AY261365.1_Warmbaths             ...............at............................--a................................ [ 34640]
AY261364.1_Tengani_62            ............a..--......t........................................................ [ 34640]
AY261363.1_Pretoriuskop/96/4     ...............at.............a................a................................ [ 34640]
AY261362.1_Mkuzi_1979            ...............--............................................................... [ 34640]
AY261361.1_Malawi_Lil-20/1_1983  .....t.........--..............................a......----------------.....t..c. [ 34640]
AY261360.1_Kenya_1950            ..g.....a......--..........t................t.........-------------a.cc....t.... [ 34640]
AM712240.1_OURT_88/3             ...............--............................................................... [ 34640]
AM712239.1_Benin_97/1            ...............--............................................................... [ 34640]
KM111295.1_Ken06.Bus             ..g.....g......--.....................................-------------a.cc....t.... [ 34640]
KM111294.1_Ken05/Tk1             .....t.........--.....................................-------------a.cc....g.... [ 34640]
KM262844.1_strain_L60            ...............--............................................................... [ 34640]
KM102979.1_26544/OG10_Italy      ...............--............................................................... [ 34640]
KM262845.1_strain_NHV            ...............--............................................................... [ 34640]
KX354450.1_47/Ss/2008            ...............--............................................................... [ 34640]
KP055815.1_strain_BA71           ...............--............................................................... [ 34640]
MH025920.1_strain_R35            ..g.....g......--.....................................-------------a.cc....t.... [ 34640]
MH025919.1_strain_N10            ..g.....g......--.....................................-------------a.cc....t.... [ 34640]
MH025918.1_strain_R25            ..g.....g......--.....................................-------------a.cc....t.... [ 34640]
MH025917.1_strain_R7             ..g.....g......--.....................................-------------a.cc....t.... [ 34640]
MH025916.1_strain_R8             ..g.....g......--.....................................-------------a.cc....t.... [ 34640]

FR682468.2_Georgia_2007/1        aacccatcattaatatagtgtttatgtgctatggacaggttttttgaatgataatcttttaacatacgttttataacttc [ 34720]
ASF_IND_20_CAD543                ................................................................................ [ 34720]
LR899193.1_Germany_2020/1        ................................................................................ [ 34720]
MN393477.1_Wuhan_2019            ................................................................................ [ 34720]
MW396979.1_Timor-Leste/2019/1    ................................................................................ [ 34720]
MK940252.1_CN/2019/InnerMongolia ................................................................................ [ 34720]
MK128995.1_China/2018/AnhuiXCGQ  ................................................................................ [ 34720]
MH681419.1_ASFV/POL/2015         ................................................................................ [ 34720]
LS478113.1_Estonia_2014          ................................................................................ [ 34720]
KP843857.1_Odintsovo_02/14       ................................................................................ [ 34720]
```

```
FN557520.1_strain_E75          ................................................................................ [ 34720]
AY261366.1_Warthog             .......gt..........a.........g.......t.......................................... [ 34720]
AY261365.1_Warmbaths           .......gt..........a.................t.......................................... [ 34720]
AY261364.1_Tengani_62          ................................................................................ [ 34720]
AY261363.1_Pretoriuskop/96/4   .......gt..........a.................t.......................................... [ 34720]
AY261362.1_Mkuzi_1979          .................................a...a.......................................... [ 34720]
AY261361.1_Malawi_Lil-20/1_1983 .......ac..........a.........c.......ta......................................... [ 34720]
AY261360.1_Kenya_1950          .......ac..........a.........c.......t.................t......................at [ 34720]
AM712240.1_OURT_88/3           ................................................................................ [ 34720]
AM712239.1_Benin_97/1          ................................................................................ [ 34720]
KM111295.1_Ken06.Bus           .......ac..........a.........c.......ta................t......................at [ 34720]
KM111294.1_Ken05/Tk1           .......ac..........a.........c.......ta.......................................at [ 34720]
KM262844.1_strain_L60          ................................................................................ [ 34720]
KM102979.1_26544/OG10_Italy    ................................................................................ [ 34720]
KM262845.1_strain_NHV          ................................................................................ [ 34720]
KX354450.1_47/Ss/2008          ................................................................................ [ 34720]
KP055815.1_strain_BA71         ................................................................................ [ 34720]
MH025920.1_strain_R35          .......ac..........a.........c.......ta................t......................at [ 34720]
MH025919.1_strain_N10          .......ac..........a.........c.......ta................t......................at [ 34720]
MH025918.1_strain_R25          .......ac..........a.........c.......ta................t......................at [ 34720]
MH025917.1_strain_R7           .......ac..........a.........c.......ta................t......................at [ 34720]
MH025916.1_strain_R8           .......ac..........a.........c.......ta................t......................at [ 34720]

FR682468.2_Georgia_2007/1      gggatcagtttcttttaaagataaagaatcattcatgttataacaatttaatgataacatgctggcaatgaacgagttgt [ 34800]
ASF_IND_20_CAD543              ................................................................................ [ 34800]
LR899193.1_Germany_2020/1      ................................................................................ [ 34800]
MN393477.1_Wuhan_2019          ................................................................................ [ 34800]
MW396979.1_Timor-Leste/2019/1  ................................................................................ [ 34800]
MK940252.1_CN/2019/InnerMongolia ................................................................................ [ 34800]
MK128995.1_China/2018/AnhuiXCGQ ................................................................................ [ 34800]
MH681419.1_ASFV/POL/2015       ................................................................................ [ 34800]
LS478113.1_Estonia_2014        ................................................................................ [ 34800]
KP843857.1_Odintsovo_02/14     ................................................................................ [ 34800]
FN557520.1_strain_E75          .....................................c.......................................... [ 34800]
AY261366.1_Warthog             ......g.......................g......c............t.....ga.g...............t.... [ 34800]
AY261365.1_Warmbaths           ......g..............................c..................ga.................t.... [ 34800]
AY261364.1_Tengani_62          ...............a.................g...c..................g....................... [ 34800]
AY261363.1_Pretoriuskop/96/4   ......g.......................g......c............t.....ga.g...............t.... [ 34800]
AY261362.1_Mkuzi_1979          .....................................c.......................................... [ 34800]
AY261361.1_Malawi_Lil-20/1_1983 ......g.......................g...t.ac..................ga.................t.... [ 34800]
AY261360.1_Kenya_1950          ...g..g.....a........................c..................ga.................t.... [ 34800]
AM712240.1_OURT_88/3           .....................................c.......................................... [ 34800]
AM712239.1_Benin_97/1          .....................................c.......................................... [ 34800]
KM111295.1_Ken06.Bus           ...g..ga....a........................c..................ga.................t.... [ 34800]
```

```
KM111294.1_Ken05/Tk1             ...g..g.....a........................c...................a.................t.... [ 34800]
KM262844.1_strain_L60            .....................................c.......................................... [ 34800]
KM102979.1_26544/OG10_Italy      .....................................c.......................................... [ 34800]
KM262845.1_strain_NHV            .....................................c.......................................... [ 34800]
KX354450.1_47/Ss/2008            .....................................c.......................................... [ 34800]
KP055815.1_strain_BA71           .....................................c.......................................... [ 34800]
MH025920.1_strain_R35            ...g..ga....a........................c..................ga.................t.... [ 34800]
MH025919.1_strain_N10            ...g..ga....a........................c..................ga.................t.... [ 34800]
MH025918.1_strain_R25            ...g..ga....a........................c..................ga.................t.... [ 34800]
MH025917.1_strain_R7             ...g..ga....a........................c..................ga.................t.... [ 34800]
MH025916.1_strain_R8             ...g..ga....a........................c..................ga.................t.... [ 34800]

FR682468.2_Georgia_2007/1        ctttttgatgcgctagagtctttccctcctcaaaggcattggcgcctaagtctatacaaaagaatatgtttccgatatta [ 34880]
ASF_IND_20_CAD543                ................................................................................ [ 34880]
LR899193.1_Germany_2020/1        ................................................................................ [ 34880]
MN393477.1_Wuhan_2019            ................................................................................ [ 34880]
MW396979.1_Timor-Leste/2019/1    ................................................................................ [ 34880]
MK940252.1_CN/2019/InnerMongolia ................................................................................ [ 34880]
MK128995.1_China/2018/AnhuiXCGQ  ................................................................................ [ 34880]
MH681419.1_ASFV/POL/2015         ................................................................................ [ 34880]
LS478113.1_Estonia_2014          ................................................................................ [ 34880]
KP843857.1_Odintsovo_02/14       ................................................................................ [ 34880]
FN557520.1_strain_E75            .......g........................................................................ [ 34880]
AY261366.1_Warthog               ........a................t.......t..............................c............... [ 34880]
AY261365.1_Warmbaths             ........a................t.......t..............................c............... [ 34880]
AY261364.1_Tengani_62            ................................................................................ [ 34880]
AY261363.1_Pretoriuskop/96/4     ........a................t.......t..............................c............... [ 34880]
AY261362.1_Mkuzi_1979            ................................................................................ [ 34880]
AY261361.1_Malawi_Lil-20/1_1983  .g.......a.....c.........t.............c...................................g.... [ 34880]
AY261360.1_Kenya_1950            .a.......a.....a.......................c........................c..........g.... [ 34880]
AM712240.1_OURT_88/3             .......g........................................................................ [ 34880]
AM712239.1_Benin_97/1            .......g........................................................................ [ 34880]
KM111295.1_Ken06.Bus             .a.......a.............................c........................c..........g.... [ 34880]
KM111294.1_Ken05/Tk1             .a.......a.....a.......................c........................c..........g.... [ 34880]
KM262844.1_strain_L60            .......g........................................................................ [ 34880]
KM102979.1_26544/OG10_Italy      .......g........................................................................ [ 34880]
KM262845.1_strain_NHV            .......g........................................................................ [ 34880]
KX354450.1_47/Ss/2008            .......g........................................................................ [ 34880]
KP055815.1_strain_BA71           .......g........................................................................ [ 34880]
MH025920.1_strain_R35            .a.......a.............................c........................c..........g.... [ 34880]
MH025919.1_strain_N10            .a.......a.............................c........................c..........g.... [ 34880]
MH025918.1_strain_R25            .a.......a.............................c........................c..........g.... [ 34880]
MH025917.1_strain_R7             .a.......a.............................c........................c..........g.... [ 34880]
MH025916.1_strain_R8             .a.......a.............................c........................c..........g.... [ 34880]
```

```
FR682468.2_Georgia_2007/1        tagaactgaatagaatgaaacatggcctgattgatatcagcccctaagacgacgcaacagtaataaatcgttaaatagtt [ 34960]
ASF_IND_20_CAD543                ................................................................................ [ 34960]
LR899193.1_Germany_2020/1        ................................................................................ [ 34960]
MN393477.1_Wuhan_2019            ................................................................................ [ 34960]
MW396979.1_Timor-Leste/2019/1    ................................................................................ [ 34960]
MK940252.1_CN/2019/InnerMongolia ................................................................................ [ 34960]
MK128995.1_China/2018/AnhuiXCGQ  ................................................................................ [ 34960]
MH681419.1_ASFV/POL/2015         ................................................................................ [ 34960]
LS478113.1_Estonia_2014          ................................................................................ [ 34960]
KP843857.1_Odintsovo_02/14       ................................................................................ [ 34960]
FN557520.1_strain_E75            ................................................g............................... [ 34960]
AY261366.1_Warthog               ....................................................t...g....................... [ 34960]
AY261365.1_Warmbaths             ..................................................a.t...g....................... [ 34960]
AY261364.1_Tengani_62            ................................................................................ [ 34960]
AY261363.1_Pretoriuskop/96/4     ....................................................t...g....................... [ 34960]
AY261362.1_Mkuzi_1979            ................................................................................ [ 34960]
AY261361.1_Malawi_Lil-20/1_1983  ............................t.....gg..............a.t...g....................... [ 34960]
AY261360.1_Kenya_1950            ......a.....................t.....g.................t...g....................... [ 34960]
AM712240.1_OURT_88/3             ................................................g............................... [ 34960]
AM712239.1_Benin_97/1            ................................................g............................... [ 34960]
KM111295.1_Ken06.Bus             ......a.....................t.....gg................t...g....................... [ 34960]
KM111294.1_Ken05/Tk1             ......a.....................t.....g...............a.t...g....................... [ 34960]
KM262844.1_strain_L60            ................................................g............................... [ 34960]
KM102979.1_26544/OG10_Italy      ................................................g............................... [ 34960]
KM262845.1_strain_NHV            ................................................g............................... [ 34960]
KX354450.1_47/Ss/2008            ................................................g............................... [ 34960]
KP055815.1_strain_BA71           ................................................g............................... [ 34960]
MH025920.1_strain_R35            ......a.....................t.....gg................t...g....................... [ 34960]
MH025919.1_strain_N10            ......a.....................t.....gg................t...g....................... [ 34960]
MH025918.1_strain_R25            ......a.....................t.....gg................t...g....................... [ 34960]
MH025917.1_strain_R7             ......a.....................t.....gg................t...g....................... [ 34960]
MH025916.1_strain_R8             ......a.....................t.....gg................t...g....................... [ 34960]

FR682468.2_Georgia_2007/1        atagttcttgcgacaggcccactttagcatttcattcatgtctatgcgaatcctctccttttcgtacacttcgtgaagtt [ 35040]
ASF_IND_20_CAD543                ..................................................c.t...............gc........k. [ 35040]
LR899193.1_Germany_2020/1        ................................................................................ [ 35040]
MN393477.1_Wuhan_2019            ................................................................................ [ 35040]
MW396979.1_Timor-Leste/2019/1    ................................................................................ [ 35040]
MK940252.1_CN/2019/InnerMongolia ................................................................................ [ 35040]
MK128995.1_China/2018/AnhuiXCGQ  ................................................................................ [ 35040]
MH681419.1_ASFV/POL/2015         ................................................................................ [ 35040]
LS478113.1_Estonia_2014          ................................................................................ [ 35040]
KP843857.1_Odintsovo_02/14       ................................................................................ [ 35040]
```

```
FN557520.1_strain_E75            ...a..................c............c......c..a....c.t........g......gc........g. [ 35040]
AY261366.1_Warthog               ...................................c......c..a......t........g......gc........g. [ 35040]
AY261365.1_Warmbaths             ...................................c......c..a......t........g......gc........g. [ 35040]
AY261364.1_Tengani_62            ...................................c......c..a....c.t..........a..............g. [ 35040]
AY261363.1_Pretoriuskop/96/4     ...................................c......c..a......t........g......gc........g. [ 35040]
AY261362.1_Mkuzi_1979            ......................c............c......c..a....c.t........g......gc........g. [ 35040]
AY261361.1_Malawi_Lil-20/1_1983  ........................c..........c......c..a....c.t.......cgt.....gc........g. [ 35040]
AY261360.1_Kenya_1950            ........................c.......t..c......c..at...ctt........g..c...gc........g. [ 35040]
AM712240.1_OURT_88/3             ...a..................c............c......c..a....c.t........g......gc........g. [ 35040]
AM712239.1_Benin_97/1            ...a..................c............c......c..a....c.t........g......gc........g. [ 35040]
KM111295.1_Ken06.Bus             ........................c.......t..c......c..at...ctt........g..c...gc........g. [ 35040]
KM111294.1_Ken05/Tk1             ........................c.......t..c......c..at...ctt........g..c...gc........g. [ 35040]
KM262844.1_strain_L60            ...a..................c............c......c..a....c.t........g......gc........g. [ 35040]
KM102979.1_26544/OG10_Italy      ...a..................c............c......c..a....c.t........g......gc........g. [ 35040]
KM262845.1_strain_NHV            ...a..................c............c......c..a....c.t........g......gc........g. [ 35040]
KX354450.1_47/Ss/2008            ...a..................c............c......c..a....c.t........g......gc........g. [ 35040]
KP055815.1_strain_BA71           ...a..................c............c......c..a....c.t........g......gc........g. [ 35040]
MH025920.1_strain_R35            ........................c.......t..c......c..at...ctt........g..c...gc........g. [ 35040]
MH025919.1_strain_N10            ........................c.......t..c......c..at...ctt........g..c...gc........g. [ 35040]
MH025918.1_strain_R25            ........................c.......t..c......c..at...ctt........g..c...gc........g. [ 35040]
MH025917.1_strain_R7             ........................c.......t..c......c..at...ctt........g..c...gc........g. [ 35040]
MH025916.1_strain_R8             ........................c.......t..c......c..at...ctt........g..c...gc........g. [ 35040]

FR682468.2_Georgia_2007/1        caaacacattattgtaaaaaagggcgcacataagccgccaccgatgtagatgagcatatctctgataaaaatagcaaatc [ 35120]
ASF_IND_20_CAD543                .r.....r..r...............------------------------------------------------------ [ 35120]
LR899193.1_Germany_2020/1        ................................................................................ [ 35120]
MN393477.1_Wuhan_2019            ................................................................................ [ 35120]
MW396979.1_Timor-Leste/2019/1    ................................................................................ [ 35120]
MK940252.1_CN/2019/InnerMongolia ................................................................................ [ 35120]
MK128995.1_China/2018/AnhuiXCGQ  ................................................................................ [ 35120]
MH681419.1_ASFV/POL/2015         ................................................................................ [ 35120]
LS478113.1_Estonia_2014          ................................................................................ [ 35120]
KP843857.1_Odintsovo_02/14       ................................................................................ [ 35120]
FN557520.1_strain_E75            .g.............................................................................. [ 35120]
AY261366.1_Warthog               .gc....g..g...............t....g.........t.g.........a....g.g...g.....g...tgg..g [ 35120]
AY261365.1_Warmbaths             .gc....g..g...............t....g.........t.g.........a....g.g...g.....g...tgg..g [ 35120]
AY261364.1_Tengani_62            .g.............................................................................. [ 35120]
AY261363.1_Pretoriuskop/96/4     .gc....g..g...............t....g.........t.g.........a....g.g...g.....g...tgg..g [ 35120]
AY261362.1_Mkuzi_1979            .g.............................................................................. [ 35120]
AY261361.1_Malawi_Lil-20/1_1983  .g.....g..g....................g.........t.g..............g.g...g.........tgg..g [ 35120]
AY261360.1_Kenya_1950            .......g..g..a.................g.........tcg..............g.g...g.....g...tgg..g [ 35120]
AM712240.1_OURT_88/3             .g.............................................................................. [ 35120]
AM712239.1_Benin_97/1            .g.............................................................................. [ 35120]
KM111295.1_Ken06.Bus             .......g..g....................g.........tcg.........t....g.g...g.....g...tgg..g [ 35120]
```

```
KM111294.1_Ken05/Tk1             .......g..g..a.................g.........tcg..............g.g...g.....g...tgg..g [ 35120]
KM262844.1_strain_L60            .g.............................................................................. [ 35120]
KM102979.1_26544/OG10_Italy      .g.............................................................................. [ 35120]
KM262845.1_strain_NHV            .g.............................................................................. [ 35120]
KX354450.1_47/Ss/2008            .g.............................................................................. [ 35120]
KP055815.1_strain_BA71           .g.............................................................................. [ 35120]
MH025920.1_strain_R35            .......g..g....................g.........tcg.........t....g.g...g.....g...tgg..g [ 35120]
MH025919.1_strain_N10            .......g..g....................g.........tcg.........t....g.g...g.....g...tgg..g [ 35120]
MH025918.1_strain_R25            .......g..g....................g.........tcg.........t....g.g...g.....g...tgg..g [ 35120]
MH025917.1_strain_R7             .......g..g....................g.........tcg.........t....g.g...g.....g...tgg..g [ 35120]
MH025916.1_strain_R8             .......g..g....................g.........tcg.........t....g.g...g.....g...tgg..g [ 35120]

FR682468.2_Georgia_2007/1        gcctccttaaggttacattctattgccatcgcgtaccaatatttagtaaacatctcgcttaatatatcggtttctaccat [ 35200]
ASF_IND_20_CAD543                -------------------------------------------------------------------------------- [ 35200]
LR899193.1_Germany_2020/1        ................................................................................ [ 35200]
MN393477.1_Wuhan_2019            ................................................................................ [ 35200]
MW396979.1_Timor-Leste/2019/1    ................................................................................ [ 35200]
MK940252.1_CN/2019/InnerMongolia ................................................................................ [ 35200]
MK128995.1_China/2018/AnhuiXCGQ  ................................................................................ [ 35200]
MH681419.1_ASFV/POL/2015         ................................................................................ [ 35200]
LS478113.1_Estonia_2014          ................................................................................ [ 35200]
KP843857.1_Odintsovo_02/14       ................................................................................ [ 35200]
FN557520.1_strain_E75            ...............a................................................................ [ 35200]
AY261366.1_Warthog               ....ttg........a.c.....g..t..ac..........c..t..t.g..a......c........c..a.t.t.... [ 35200]
AY261365.1_Warmbaths             ....ttg........a.c.....g..t..ac..........c..t..t.g..a......c........c..a.t.t.... [ 35200]
AY261364.1_Tengani_62            ................................................................................ [ 35200]
AY261363.1_Pretoriuskop/96/4     ....ttg........a.c.....g..t..ac..........c..t..t.g..a......c........c..a.t.t.... [ 35200]
AY261362.1_Mkuzi_1979            ...............a................................................................ [ 35200]
AY261361.1_Malawi_Lil-20/1_1983  ....ttg........a.c....cg.....ac..........c..t..t.g..gt........g.....c..a.t.t.... [ 35200]
AY261360.1_Kenya_1950            ....ttg.c.....ga.c....cg.....ac.............t..t.g..a...............c..a.t.t.... [ 35200]
AM712240.1_OURT_88/3             ...............a................................................................ [ 35200]
AM712239.1_Benin_97/1            ...............a................................................................ [ 35200]
KM111295.1_Ken06.Bus             ....ttg........a.c....cg.....ac.............t..t.g..a...........c...c..a.t.t.... [ 35200]
KM111294.1_Ken05/Tk1             ....ttg.c.....ga.c....cg.....ac.............t..t.g..a...............c..a.t.t.... [ 35200]
KM262844.1_strain_L60            ...............a................................................................ [ 35200]
KM102979.1_26544/OG10_Italy      ...............a................................................................ [ 35200]
KM262845.1_strain_NHV            ...............a................................................................ [ 35200]
KX354450.1_47/Ss/2008            ...............a................................................................ [ 35200]
KP055815.1_strain_BA71           ...............a................................................................ [ 35200]
MH025920.1_strain_R35            ....ttg........a.c....cg.....ac.............t..t.g..a...........c...c..a.t.t.... [ 35200]
MH025919.1_strain_N10            ....ttg........a.c....cg.....ac.............t..t.g..a...........c...c..a.t.t.... [ 35200]
MH025918.1_strain_R25            ....ttg........a.c....cg.....ac.............t..t.g..a...........c...c..a.t.t.... [ 35200]
MH025917.1_strain_R7             ....ttg........a.c....cg.....ac.............t..t.g..a...........c...c..a.t.t.... [ 35200]
MH025916.1_strain_R8             ....ttg........a.c....cg.....ac.............t..t.g..a...........c...c..a.t.t.... [ 35200]
```

```
FR682468.2_Georgia_2007/1        taatccctccagttgttcataaatcattccctttacttcaaaacgatttatggtatctaaaatgggattattagaaaata [ 35280]
ASF_IND_20_CAD543                -------------------------------------------------------------------------------- [ 35280]
LR899193.1_Germany_2020/1        ................................................................................ [ 35280]
MN393477.1_Wuhan_2019            ................................................................................ [ 35280]
MW396979.1_Timor-Leste/2019/1    ................................................................................ [ 35280]
MK940252.1_CN/2019/InnerMongolia ................................................................................ [ 35280]
MK128995.1_China/2018/AnhuiXCGQ  ................................................................................ [ 35280]
MH681419.1_ASFV/POL/2015         ................................................................................ [ 35280]
LS478113.1_Estonia_2014          ................................................................................ [ 35280]
KP843857.1_Odintsovo_02/14       ................................................................................ [ 35280]
FN557520.1_strain_E75            ................................................................................ [ 35280]
AY261366.1_Warthog               ....g.t.....c...........g......c...ac.tggt.........t..t.....g..........g......c. [ 35280]
AY261365.1_Warmbaths             ....g.t.....c...........g......c...ac.tggt.........t..t.....g..........g......c. [ 35280]
AY261364.1_Tengani_62            ................................................................................ [ 35280]
AY261363.1_Pretoriuskop/96/4     ....g.t....ac...........g......c...ac.tggt.........t..t.....g..........g......c. [ 35280]
AY261362.1_Mkuzi_1979            ...................................................a............................ [ 35280]
AY261361.1_Malawi_Lil-20/1_1983  ....g.t.....c.....g.....g......c...a..tggt.........c..t.....g..........g......c. [ 35280]
AY261360.1_Kenya_1950            ....g.t.....c...........g......c...a..tggt.........c..t.....g..a.a.....g......c. [ 35280]
AM712240.1_OURT_88/3             ................................................................................ [ 35280]
AM712239.1_Benin_97/1            ................................................................................ [ 35280]
KM111295.1_Ken06.Bus             ....g.t.....c.....g.....g......c...a..tggt........cc..t.....g....a.....g......c. [ 35280]
KM111294.1_Ken05/Tk1             ....g.t.....c...........g......c...a..tggt.........c..t.....g..a.a.....g......c. [ 35280]
KM262844.1_strain_L60            ................................................................................ [ 35280]
KM102979.1_26544/OG10_Italy      ................................................................................ [ 35280]
KM262845.1_strain_NHV            ................................................................................ [ 35280]
KX354450.1_47/Ss/2008            ................................................................................ [ 35280]
KP055815.1_strain_BA71           ................................................................................ [ 35280]
MH025920.1_strain_R35            ....g.t.....c.....g.....g......c...a..tggt........cc..t.....g....a.....g......c. [ 35280]
MH025919.1_strain_N10            ....g.t.....c.....g.....g......c...a..tggt........cc..t.....g....a.....g......c. [ 35280]
MH025918.1_strain_R25            ....g.t.....c.....g.....g......c...a..tggt........cc..t.....g....a.....g......c. [ 35280]
MH025917.1_strain_R7             ....g.t.....c.....g.....g......c...a..tggt........cc..t.....g....a.....g......c. [ 35280]
MH025916.1_strain_R8             ....g.t.....c.....g.....g......c...a..tggt........cc..t.....g....a.....g......c. [ 35280]

FR682468.2_Georgia_2007/1        cctcatggcagaaaatgatgttactgctagttagatcacgtttcaatgtgtaaaaaaatcgtaaaatttcctggtcattt [ 35360]
ASF_IND_20_CAD543                -------------------------------------------------------------------------------- [ 35360]
LR899193.1_Germany_2020/1        ................................................................................ [ 35360]
MN393477.1_Wuhan_2019            ................................................................................ [ 35360]
MW396979.1_Timor-Leste/2019/1    ................................................................................ [ 35360]
MK940252.1_CN/2019/InnerMongolia ................................................................................ [ 35360]
MK128995.1_China/2018/AnhuiXCGQ  ................................................................................ [ 35360]
MH681419.1_ASFV/POL/2015         ................................................................................ [ 35360]
LS478113.1_Estonia_2014          ................................................................................ [ 35360]
KP843857.1_Odintsovo_02/14       ................................................................................ [ 35360]
```

```
FN557520.1_strain_E75            ................................................................................ [ 35360]
AY261366.1_Warthog               ....g.....c.....a...........g.......t......t.t.a...t....t.ata.....c.....ct.gc... [ 35360]
AY261365.1_Warmbaths             ....g.....c.....a...........g.......t......t.t.a...t....t.ata.....c.....ct.gc... [ 35360]
AY261364.1_Tengani_62            ................................................................................ [ 35360]
AY261363.1_Pretoriuskop/96/4     ....g.....c.....a...........g.......t......t.t.....t....t.ata.....c.....ct.gc... [ 35360]
AY261362.1_Mkuzi_1979            ................................................................................ [ 35360]
AY261361.1_Malawi_Lil-20/1_1983  ....g.....c.....a..........cg..............t.cca...t......at.c..c.c......t.gc... [ 35360]
AY261360.1_Kenya_1950            ....g.....c.....a...........g..............t.cca...t......at.c...........t.g.... [ 35360]
AM712240.1_OURT_88/3             ................................................................................ [ 35360]
AM712239.1_Benin_97/1            ................................................................................ [ 35360]
KM111295.1_Ken06.Bus             ....g.....c.....a..........cg..............t.cca...t......at.c...........t.g.... [ 35360]
KM111294.1_Ken05/Tk1             ....g.....c.....a...........g..............t.cca...t......at.c...........t.g.... [ 35360]
KM262844.1_strain_L60            ................................................................................ [ 35360]
KM102979.1_26544/OG10_Italy      ................................................................................ [ 35360]
KM262845.1_strain_NHV            ................................................................................ [ 35360]
KX354450.1_47/Ss/2008            ................................................................................ [ 35360]
KP055815.1_strain_BA71           ................................................................................ [ 35360]
MH025920.1_strain_R35            ....g.....c.....a..........cg..............t.cca...t......at.c...........t.g.... [ 35360]
MH025919.1_strain_N10            ....g.....c.....a..........cg..............t.cca...t......at.c...........t.g.... [ 35360]
MH025918.1_strain_R25            ....g.....c.....a..........cg..............t.cca...t......at.c...........t.g.... [ 35360]
MH025917.1_strain_R7             ....g.....c.....a..........cg..............t.cca...t......at.c...........t.g.... [ 35360]
MH025916.1_strain_R8             ....g.....c.....a..........cg..............t.cca...t......at.c...........t.g.... [ 35360]

FR682468.2_Georgia_2007/1        aactgttctttggcacctagctgcctgcacaggtctcgggtgtgctccgtgttgacagaaagcaaaccgtagttgatgtt [ 35440]
ASF_IND_20_CAD543                -------------------------------------------------------------------------------- [ 35440]
LR899193.1_Germany_2020/1        ................................................................................ [ 35440]
MN393477.1_Wuhan_2019            ................................................................................ [ 35440]
MW396979.1_Timor-Leste/2019/1    ................................................................................ [ 35440]
MK940252.1_CN/2019/InnerMongolia ................................................................................ [ 35440]
MK128995.1_China/2018/AnhuiXCGQ  ................................................................................ [ 35440]
MH681419.1_ASFV/POL/2015         ................................................................................ [ 35440]
LS478113.1_Estonia_2014          ................................................................................ [ 35440]
KP843857.1_Odintsovo_02/14       ................................................................................ [ 35440]
FN557520.1_strain_E75            ..........................a..................................................... [ 35440]
AY261366.1_Warthog               ...........a..g.......ct........a.....a...c...................t....ta....c...... [ 35440]
AY261365.1_Warmbaths             ...........a..g.......ct........a.....a...c...................t....ta....c...... [ 35440]
AY261364.1_Tengani_62            ................................................................................ [ 35440]
AY261363.1_Pretoriuskop/96/4     ...........a..g.......ct........a.....a...c...................t....ta....c...... [ 35440]
AY261362.1_Mkuzi_1979            ..........................a..................................................... [ 35440]
AY261361.1_Malawi_Lil-20/1_1983  ..............g.......ct.......aa.........ca...................t....t.....c...... [ 35440]
AY261360.1_Kenya_1950            ..............g....a..ct........a..........a..............c...t....t.....c...... [ 35440]
AM712240.1_OURT_88/3             ..........................a..................................................... [ 35440]
AM712239.1_Benin_97/1            ..........................a..................................................... [ 35440]
KM111295.1_Ken06.Bus             ..............g....a..ct........a.....a....a..............c...t....t.....c...... [ 35440]
```

```
KM111294.1_Ken05/Tk1            ..............g....a..ct........a..........a..............c...t....t.....c...... [ 35440]
KM262844.1_strain_L60           ..........................a..................................................... [ 35440]
KM102979.1_26544/OG10_Italy     ..........................a..................................................... [ 35440]
KM262845.1_strain_NHV           ..........................a..................................................... [ 35440]
KX354450.1_47/Ss/2008           ..........................a..................................................... [ 35440]
KP055815.1_strain_BA71          ..........................a..................................................... [ 35440]
MH025920.1_strain_R35           ..............g....a..ct........a.....a....a..............c...t....t.....c...... [ 35440]
MH025919.1_strain_N10           ..............g....a..ct........a.....a....a..............c...t....t.....c...... [ 35440]
MH025918.1_strain_R25           ..............g....a..ct........a.....a....a..............c...t....t.....c...... [ 35440]
MH025917.1_strain_R7            ..............g....a..ct........a.....a....a..............c...t....t.....c...... [ 35440]
MH025916.1_strain_R8            ..............g....a..ct........a.....a....a..............c...t....t.....c...... [ 35440]

FR682468.2_Georgia_2007/1       tgcaccccactcggtgaacaattctattagatcgtgattgttttcctccacagctttcaccaaggccgcgttaagatttg [ 35520]
ASF_IND_20_CAD543               -------------------------------------------------------------------------------- [ 35520]
LR899193.1_Germany_2020/1       ................................................................................ [ 35520]
MN393477.1_Wuhan_2019           ................................................................................ [ 35520]
MW396979.1_Timor-Leste/2019/1   ................................................................................ [ 35520]
MK940252.1_CN/2019/InnerMongolia ................................................................................ [ 35520]
MK128995.1_China/2018/AnhuiXCGQ ................................................................................ [ 35520]
MH681419.1_ASFV/POL/2015        ................................................................................ [ 35520]
LS478113.1_Estonia_2014         ................................................................................ [ 35520]
KP843857.1_Odintsovo_02/14      ................................................................................ [ 35520]
FN557520.1_strain_E75           ...g............................................................................ [ 35520]
AY261366.1_Warthog              c..............a.......t......g..a.a...a.....ag....g...........c.....a....t..gca [ 35520]
AY261365.1_Warmbaths            c..............a.......t......g..a.a...a.....ag....g...........c.....a....t..gca [ 35520]
AY261364.1_Tengani_62           ...g.............................a.............................................. [ 35520]
AY261363.1_Pretoriuskop/96/4    c..............a.......t......g..a.a...a.....ag.tg.g...........c.....a....t..gca [ 35520]
AY261362.1_Mkuzi_1979           ...g............................................................................ [ 35520]
AY261361.1_Malawi_Lil-20/1_1983 c..g...........a......a....c.a...a.a...........t...g...........c.....a....t...ca [ 35520]
AY261360.1_Kenya_1950           c..g...........a.......t......g..a.a...a.....t.t...g...........c.....a.....cg.ca [ 35520]
AM712240.1_OURT_88/3            ...g............................................................................ [ 35520]
AM712239.1_Benin_97/1           ...g.....t...................................................................... [ 35520]
KM111295.1_Ken06.Bus            c..g...........a.......t......g..a.a...a.....t.t...g...........c.....a.....cg.ca [ 35520]
KM111294.1_Ken05/Tk1            c..g...........a.......t......g..a.a...a.....t.t...g...........c.....a.....cg.ca [ 35520]
KM262844.1_strain_L60           ...g............................................................................ [ 35520]
KM102979.1_26544/OG10_Italy     ...g............................................................................ [ 35520]
KM262845.1_strain_NHV           ...g............................................................................ [ 35520]
KX354450.1_47/Ss/2008           ...g............................................................................ [ 35520]
KP055815.1_strain_BA71          ...g............................................................................ [ 35520]
MH025920.1_strain_R35           c..g...........a.......t......g..a.a...a.....t.t...g...........c.....a.....cg.ca [ 35520]
MH025919.1_strain_N10           c..g...........a.......t......g..a.a...a.....t.t...g...........c.....a.....cg.ca [ 35520]
MH025918.1_strain_R25           c..g...........a.......t......g..a.a...a.....t.t...g...........c.....a.....cg.ca [ 35520]
MH025917.1_strain_R7            c..g...........a.......t......g..a.a...a.....t.t...g...........c.....a.....cg.ca [ 35520]
MH025916.1_strain_R8            c..g...........a.......t......g..a.a...a.....t.t...g...........c.....a.....cg.ca [ 35520]
```

```
FR682468.2_Georgia_2007/1        tgccgttcttaaaatacggcgtccatattttcttt------tgatgatacatgatagggccattatgccaccatagaccg [ 35600]
ASF_IND_20_CAD543                -------------------------------------------------------------------------------- [ 35600]
LR899193.1_Germany_2020/1        ...................................------....................................... [ 35600]
MN393477.1_Wuhan_2019            ...................................------....................................... [ 35600]
MW396979.1_Timor-Leste/2019/1    ...................................------....................................... [ 35600]
MK940252.1_CN/2019/InnerMongolia ...................................------....................................... [ 35600]
MK128995.1_China/2018/AnhuiXCGQ  ...................................------....................................... [ 35600]
MH681419.1_ASFV/POL/2015         ...................................------....................................... [ 35600]
LS478113.1_Estonia_2014          ...................................------....................................... [ 35600]
KP843857.1_Odintsovo_02/14       ...................................------....................................... [ 35600]
FN557520.1_strain_E75            ...................................------....................................... [ 35600]
AY261366.1_Warthog               c...a.ctg.....at.....a.........t...tggttt....c.a.t..c..g..a.....g...........g..a [ 35600]
AY261365.1_Warmbaths             c...a.ctg.....at.....a.........t...tggttt....c.a.t..c..g..a.....g...........g..a [ 35600]
AY261364.1_Tengani_62            ...................................------....................................... [ 35600]
AY261363.1_Pretoriuskop/96/4     c...a.ctg.....at.....a.........t...tggttc....c.a.t..c..g..a.....g...........g..a [ 35600]
AY261362.1_Mkuzi_1979            ...................................------....................................... [ 35600]
AY261361.1_Malawi_Lil-20/1_1983  c...a.ctg.....a...at.a.............tggttt..........c......a....c................ [ 35600]
AY261360.1_Kenya_1950            c...a.c.g.....at.....a.............tggttt..........c...g..a..g.c................ [ 35600]
AM712240.1_OURT_88/3             ...................................------....................................... [ 35600]
AM712239.1_Benin_97/1            ...................................------....................................... [ 35600]
KM111295.1_Ken06.Bus             c...a.ctg.....at.....a.............tggttt..........c...g..a..g.c................ [ 35600]
KM111294.1_Ken05/Tk1             c...a.c.g.....at.....a.............tggttt..........c...g..a..g.c................ [ 35600]
KM262844.1_strain_L60            ...................................------....................................... [ 35600]
KM102979.1_26544/OG10_Italy      ...................................------....................................... [ 35600]
KM262845.1_strain_NHV            ...................................------....................................... [ 35600]
KX354450.1_47/Ss/2008            ...................................------....................................... [ 35600]
KP055815.1_strain_BA71           ...................................------....................................... [ 35600]
MH025920.1_strain_R35            c...a.ctg.....at.....a.............tggttt..........c...g..a..g.c................ [ 35600]
MH025919.1_strain_N10            c...a.ctg.....at.....a.............tggttt..........c...g..a..g.c................ [ 35600]
MH025918.1_strain_R25            c...a.ctg.....at.....a.............tggttt..........c...g..a..g.c................ [ 35600]
MH025917.1_strain_R7             c...a.ctg.....at.....a.............tggttt..........c...g..a..g.c................ [ 35600]
MH025916.1_strain_R8             c...a.ctg.....at.....a.............tggttt..........c...g..a..g.c................ [ 35600]

FR682468.2_Georgia_2007/1        cagcacttcaaaaaatgaggatggcatttggccggatactggctggccagcacctttttggtgagagtctgcagagagag [ 35680]
ASF_IND_20_CAD543                -------------------------------------------------------------------------------- [ 35680]
LR899193.1_Germany_2020/1        ................................................................................ [ 35680]
MN393477.1_Wuhan_2019            ................................................................................ [ 35680]
MW396979.1_Timor-Leste/2019/1    ................................................................................ [ 35680]
MK940252.1_CN/2019/InnerMongolia ................................................................................ [ 35680]
MK128995.1_China/2018/AnhuiXCGQ  ................................................................................ [ 35680]
MH681419.1_ASFV/POL/2015         ................................................................................ [ 35680]
LS478113.1_Estonia_2014          ................................................................................ [ 35680]
KP843857.1_Odintsovo_02/14       ................................................................................ [ 35680]
```

```
FN557520.1_strain_E75          ................................................................................ [ 35680]
AY261366.1_Warthog             t.a..t..t...........g..........t...gcgt..t.c...t.a...t.......c...g.............. [ 35680]
AY261365.1_Warmbaths           t.a..t..t...........g..........t...gcgt..t.c...t.a...t.......c...g.............. [ 35680]
AY261364.1_Tengani_62          .....................................g.......................................... [ 35680]
AY261363.1_Pretoriuskop/96/4   t.a..t..t...........g..........t...gcgt..t.c...t.a...t.......c...g.............. [ 35680]
AY261362.1_Mkuzi_1979          ................................................................................ [ 35680]
AY261361.1_Malawi_Lil-20/1_1983 ........t.........t.........g............t...................................... [ 35680]
AY261360.1_Kenya_1950          ........t.........t.........c............t........t..............c.............. [ 35680]
AM712240.1_OURT_88/3           ................................................-------------------------------- [ 35680]
AM712239.1_Benin_97/1          ................................................................................ [ 35680]
KM111295.1_Ken06.Bus           ........t.........t.........c............t.......a...............c.............. [ 35680]
KM111294.1_Ken05/Tk1           ........t.........t....a....c............t.......................c.............. [ 35680]
KM262844.1_strain_L60          ................................................................................ [ 35680]
KM102979.1_26544/OG10_Italy    ................................................................................ [ 35680]
KM262845.1_strain_NHV          ................................................-------------------------------- [ 35680]
KX354450.1_47/Ss/2008          ................................................................................ [ 35680]
KP055815.1_strain_BA71         ................................................................................ [ 35680]
MH025920.1_strain_R35          ........t.........t.........c............t.......a...............c.............. [ 35680]
MH025919.1_strain_N10          ........t.........t.........c............t.......a...............c.............. [ 35680]
MH025918.1_strain_R25          ........t.........t.........c............t.......a...............c.............. [ 35680]
MH025917.1_strain_R7           ........t.........t.........c............t.......a...............c.............. [ 35680]
MH025916.1_strain_R8           ........t.........t.........c............t.......a...............c.............. [ 35680]

FR682468.2_Georgia_2007/1      gaccatatttct--tttttttgaaaaaatcaaatt-aaaaa----------------aatc-atgcttgtttagcataca [ 35760]
ASF_IND_20_CAD543              -------------------------------------------------------------------------------- [ 35760]
LR899193.1_Germany_2020/1      ............--.....................-.....----------------....-.................. [ 35760]
MN393477.1_Wuhan_2019          ............--.....................-.....----------------....-.................. [ 35760]
MW396979.1_Timor-Leste/2019/1  ............--.....................-.....----------------....-.................. [ 35760]
MK940252.1_CN/2019/InnerMongolia ............--.....................-.....----------------....-.................. [ 35760]
MK128995.1_China/2018/AnhuiXCGQ ............--.....................-.....----------------....-.................. [ 35760]
MH681419.1_ASFV/POL/2015       ............--.....................-.....----------------....-.................. [ 35760]
LS478113.1_Estonia_2014        ............--.....................-.....----------------....-.................. [ 35760]
KP843857.1_Odintsovo_02/14     ............--.....................-.....----------------....-.................. [ 35760]
FN557520.1_strain_E75          ...........---.....................a.....----------------..a.-..........g....... [ 35760]
AY261366.1_Warthog             ......g....c--.....................a.....---------------------..aa..t......----- [ 35760]
AY261365.1_Warmbaths           ......g....c--.....................a.....a--------------t....a..aa..t......----- [ 35760]
AY261364.1_Tengani_62          ............--.....................a.....----------------....-.................. [ 35760]
AY261363.1_Pretoriuskop/96/4   ......g....c-t.....................a.....----------------....-.................. [ 35760]
AY261362.1_Mkuzi_1979          ...........---.....................-.....----------------..a.-..........g.....t. [ 35760]
AY261361.1_Malawi_Lil-20/1_1983 ......g...tc-t......atg............a.....---------------------.ata..t...g..----- [ 35760]
AY261360.1_Kenya_1950          a.....g.c..ctt.....................a.....ac----ataaaaaaca..at-.aatc.a........... [ 35760]
AM712240.1_OURT_88/3           -------------------------------------------------------------------------------- [ 35760]
AM712239.1_Benin_97/1          ...........---.....................a.....----------------..a.-..........g....... [ 35760]
KM111295.1_Ken06.Bus           ......g.c..ctt.....................a.....actttaaaaaaaaaca..at-.aatc.a........... [ 35760]
```

```
KM111294.1_Ken05/Tk1             ......g.c..ctt.......tc.....atc....t.....actttaaaaaaaaaca..at-.aatc.a........... [ 35760]
KM262844.1_strain_L60            ...........---.....................a.....----------------..a.-..........g....... [ 35760]
KM102979.1_26544/OG10_Italy      ...........---.....................-.....----------------..a.-..........g....... [ 35760]
KM262845.1_strain_NHV            -------------------------------------------------------------------------------- [ 35760]
KX354450.1_47/Ss/2008            ...........---.....................-.....----------------..a.-..........g....... [ 35760]
KP055815.1_strain_BA71           ...........---.....................a.....----------------..a.-..........g....... [ 35760]
MH025920.1_strain_R35            ......g.c..ctt.....................a.....actttaaaaaaaaaca..at-.aatc.a........... [ 35760]
MH025919.1_strain_N10            ......g.c..ctt.....................a.....actttaaaaaaaaaca..at-.aatc.a........... [ 35760]
MH025918.1_strain_R25            ......g.c..ctt.....................a.....actttaaaaaaaaaca..at-.aatc.a........... [ 35760]
MH025917.1_strain_R7             ......g.c..ctt.....................a.....actttaaaaaaaaaca..at-.aatc.a........... [ 35760]
MH025916.1_strain_R8             ......g.c..ctt.....................a.....actttaaaaaaaaaca..at-.aatc.a........... [ 35760]

FR682468.2_Georgia_2007/1        tgtaatattgttataattacgttataattacgttataattacgttataactatattataacaatggtataacaatggtat [ 35840]
ASF_IND_20_CAD543                -------------------------------------------------------------------------------- [ 35840]
LR899193.1_Germany_2020/1        ................................................................................ [ 35840]
MN393477.1_Wuhan_2019            ................................................................................ [ 35840]
MW396979.1_Timor-Leste/2019/1    ................................................................................ [ 35840]
MK940252.1_CN/2019/InnerMongolia ................................................................................ [ 35840]
MK128995.1_China/2018/AnhuiXCGQ  ................................................................................ [ 35840]
MH681419.1_ASFV/POL/2015         ................................................................................ [ 35840]
LS478113.1_Estonia_2014          ................................................................................ [ 35840]
KP843857.1_Odintsovo_02/14       ................................................................................ [ 35840]
FN557520.1_strain_E75            .....ct--------------------------------------------..g.......c...t.......t..t... [ 35840]
AY261366.1_Warthog               -------------------------------------------------------------------------------- [ 35840]
AY261365.1_Warmbaths             -------------------------------------------------------------------------------- [ 35840]
AY261364.1_Tengani_62            ..................----------------------.........t..cg.......t..at......g....... [ 35840]
AY261363.1_Pretoriuskop/96/4     ....gct-------------------------------------------------------...t........c.t... [ 35840]
AY261362.1_Mkuzi_1979            .....cta.........a----------------------.t........at.g......t.t..t.....t.t..t... [ 35840]
AY261361.1_Malawi_Lil-20/1_1983  -------------------------------------------------------------------------------- [ 35840]
AY261360.1_Kenya_1950            c.------------------------------------------------------------------------------ [ 35840]
AM712240.1_OURT_88/3             -------------------------------------------------------------------------------- [ 35840]
AM712239.1_Benin_97/1            .....cta---------------------------------t........c..g.......c...t.......t..t... [ 35840]
KM111295.1_Ken06.Bus             c.------------------------------------------------------------------------------ [ 35840]
KM111294.1_Ken05/Tk1             -------------------------------------------------------------------------------- [ 35840]
KM262844.1_strain_L60            .....ct--------------------------------------------..g.......c...t.......t..t... [ 35840]
KM102979.1_26544/OG10_Italy      .....ct--------------------------------------------..g.......c...t.......t..t... [ 35840]
KM262845.1_strain_NHV            -------------------------------------------------------------------------------- [ 35840]
KX354450.1_47/Ss/2008            .....ct--------------------------------------------..g.......c...t.......t..t... [ 35840]
KP055815.1_strain_BA71           .....ct--------------------------------------------..g.......c...t.......t..t... [ 35840]
MH025920.1_strain_R35            c.------------------------------------------------------------------------------ [ 35840]
MH025919.1_strain_N10            c.------------------------------------------------------------------------------ [ 35840]
MH025918.1_strain_R25            c.------------------------------------------------------------------------------ [ 35840]
MH025917.1_strain_R7             c.------------------------------------------------------------------------------ [ 35840]
MH025916.1_strain_R8             c.------------------------------------------------------------------------------ [ 35840]
```

```
FR682468.2_Georgia_2007/1        aacaatgttataacaatgttataacgatgtatcattgatgtcatcattcaactaggccaacatactttttaatttatagt [ 35920]
ASF_IND_20_CAD543                -------------------------------------------------------------------------------- [ 35920]
LR899193.1_Germany_2020/1        ................................................................................ [ 35920]
MN393477.1_Wuhan_2019            ................................................................................ [ 35920]
MW396979.1_Timor-Leste/2019/1    ................................................................................ [ 35920]
MK940252.1_CN/2019/InnerMongolia ................................................................................ [ 35920]
MK128995.1_China/2018/AnhuiXCGQ  ................................................................................ [ 35920]
MH681419.1_ASFV/POL/2015         ................................................................................ [ 35920]
LS478113.1_Estonia_2014          ................................................................................ [ 35920]
KP843857.1_Odintsovo_02/14       ................................................................................ [ 35920]
FN557520.1_strain_E75            ..tg...g.......ta...........................a......a..................g......... [ 35920]
AY261366.1_Warthog               -------------------c..c.tat.....------ca....gg..a..a..t...gt....t.....g......... [ 35920]
AY261365.1_Warmbaths             -------------------c..c.tat.....------ca....gg..a..a..t...gt....t.....g......... [ 35920]
AY261364.1_Tengani_62            ...............t.........a.............a...........a............t.....g......... [ 35920]
AY261363.1_Pretoriuskop/96/4     ....tc..........c.........t.....t.....c.....a......a........t.........g......... [ 35920]
AY261362.1_Mkuzi_1979            ..tg...g.......ta...........................a......a..................g......... [ 35920]
AY261361.1_Malawi_Lil-20/1_1983  -------------------c..t..ac...g....gccca....ag.ca..a..ta...t....t.....g......... [ 35920]
AY261360.1_Kenya_1950            ----------....g......cg...t.........a....g..a..c...a......g...........g......... [ 35920]
AM712240.1_OURT_88/3             -------------------------------------------------------------------------------- [ 35920]
AM712239.1_Benin_97/1            ..tg...g.......ta...........................a......a..................g......... [ 35920]
KM111295.1_Ken06.Bus             ----------....t......cg...t..............g..a..c...a..................g......... [ 35920]
KM111294.1_Ken05/Tk1             -----------gc.t......cg...t.........at...g..a..c...a......g...........g.......a. [ 35920]
KM262844.1_strain_L60            ..tg...g.......ta...........................a......a..................g......... [ 35920]
KM102979.1_26544/OG10_Italy      ..tg...g.......ta...........................a......a..................g......... [ 35920]
KM262845.1_strain_NHV            -------------------------------------------------------------------------------- [ 35920]
KX354450.1_47/Ss/2008            ..tg...g.......ta...........................a......a..................g......... [ 35920]
KP055815.1_strain_BA71           ..tg...g.......ta...........................a......a..................g......... [ 35920]
MH025920.1_strain_R35            ----------....t......cg...t..............g..a..c...a..................g......... [ 35920]
MH025919.1_strain_N10            ----------....t......cg...t..............g..a..c...a..................g......... [ 35920]
MH025918.1_strain_R25            ----------....t......cg...t..............g..a..c...a..................g......... [ 35920]
MH025917.1_strain_R7             ----------....t......cg...t..............g..a..c...a..................g......... [ 35920]
MH025916.1_strain_R8             ----------....t......cg...t..............g..a..c...a..................g......... [ 35920]

FR682468.2_Georgia_2007/1        tttttaatagatgatatattttgttaggatctgcttcttttaacgttaatagcgaggagtctgcactataaatgtctaat [ 36000]
ASF_IND_20_CAD543                -------------------------------------------------------------------------------- [ 36000]
LR899193.1_Germany_2020/1        ................................................................................ [ 36000]
MN393477.1_Wuhan_2019            .......................c........................................................ [ 36000]
MW396979.1_Timor-Leste/2019/1    .......................c........................................................ [ 36000]
MK940252.1_CN/2019/InnerMongolia .......................c........................................................ [ 36000]
MK128995.1_China/2018/AnhuiXCGQ  .......................c........................................................ [ 36000]
MH681419.1_ASFV/POL/2015         ................................................................................ [ 36000]
LS478113.1_Estonia_2014          ................................................................................ [ 36000]
KP843857.1_Odintsovo_02/14       ................................................................................ [ 36000]
```

FN557520.1_strain_E75 ...........c...................................................g................ [ 36000]
AY261366.1_Warthog ............................g..c...............................g................ [ 36000]
AY261365.1_Warmbaths ............................g..c...............................g................ [ 36000]
AY261364.1_Tengani_62 ...............t.................t...................a.........gct.............. [ 36000]
AY261363.1_Pretoriuskop/96/4 ...............................c...................a......a....g................ [ 36000]
AY261362.1_Mkuzi_1979 ...........c...................................................g................ [ 36000]
AY261361.1_Malawi_Lil-20/1_1983 ......................t........................................g.........a....t. [ 36000]
AY261360.1_Kenya_1950 .g...........g........t.....g.tc..........g...g.....a....ga....g.tg...g..a...... [ 36000]
AM712240.1_OURT_88/3 -------------------------------------------------------------------------------- [ 36000]
AM712239.1_Benin_97/1 ...........c...................................................g................ [ 36000]
KM111295.1_Ken06.Bus .g...........g........t.....g.tc..........g...g.....a....ga....g.tg...g..a...... [ 36000]
KM111294.1_Ken05/Tk1 .g...........g........t.....g.tc..........g...g.....a....ga....g.tg...g..a...... [ 36000]
KM262844.1_strain_L60 ...........c...................................................g................ [ 36000]
KM102979.1_26544/OG10_Italy ...........c...................................................g................ [ 36000]
KM262845.1_strain_NHV -------------------------------------------------------------------------------- [ 36000]
KX354450.1_47/Ss/2008 ...........c...................................................g................ [ 36000]
KP055815.1_strain_BA71 ...........c...................................................g................ [ 36000]
MH025920.1_strain_R35 .g...........g........t.....g.tc..........g...g.....a....ga....g.tg...g..a...... [ 36000]
MH025919.1_strain_N10 .g...........g........t.....g.tc..........g...g.....a....ga....g.tg...g..a...... [ 36000]
MH025918.1_strain_R25 .g...........g........t.....g.tc..........g...g.....a....ga....g.tg...g..a...... [ 36000]
MH025917.1_strain_R7 .g...........g........t.....g.tc..........g...g.....a....ga....g.tg...g..a...... [ 36000]
MH025916.1_strain_R8 .g...........g........t.....g.tc..........g...g.....a....ga....g.tg...g..a...... [ 36000]

FR682468.2_Georgia_2007/1 gataaatgatgagatatcaaatagtaattccgttgctctgctagggcctttgcctcttcaaaggcgtcggctcccagatc [ 36080]
ASF_IND_20_CAD543 -------------------------------------------------------------------------------- [ 36080]
LR899193.1_Germany_2020/1 ................................................................................ [ 36080]
MN393477.1_Wuhan_2019 ................................................................................ [ 36080]
MW396979.1_Timor-Leste/2019/1 ................................................................................ [ 36080]
MK940252.1_CN/2019/InnerMongolia ................................................................................ [ 36080]
MK128995.1_China/2018/AnhuiXCGQ ................................................................................ [ 36080]
MH681419.1_ASFV/POL/2015 ................................................................................ [ 36080]
LS478113.1_Estonia_2014 ................................................................................ [ 36080]
KP843857.1_Odintsovo_02/14 ................................................................................ [ 36080]
FN557520.1_strain_E75 ......c.......................t..........c.a...t.................a.....c........ [ 36080]
AY261366.1_Warthog ......c..............a...................c.....t.................a.....c....ag.. [ 36080]
AY261365.1_Warmbaths ......c..............a...................c..a..........................c.....g.. [ 36080]
AY261364.1_Tengani_62 ......c..........t.......................c.....t................................ [ 36080]
AY261363.1_Pretoriuskop/96/4 ......ca.............a...................c.....t........c........a.....c.....g.. [ 36080]
AY261362.1_Mkuzi_1979 ......c.......................t..........c.a.....................a.....c........ [ 36080]
AY261361.1_Malawi_Lil-20/1_1983 ......c.....t.c......a..........c..g.....c.....t.................a...........g.. [ 36080]
AY261360.1_Kenya_1950 ..c...c....t.....t.....a..g.c...c..............t...........g.....a.t...c....ag.. [ 36080]
AM712240.1_OURT_88/3 -------------------------------------------------------------------------------- [ 36080]
AM712239.1_Benin_97/1 ......c.......................t..........c.a...t.................a.....c........ [ 36080]
KM111295.1_Ken06.Bus ..c...c.t..t.....t.....a..g.c...c..............t...........g.....a.t........ag.. [ 36080]

```
KM111294.1_Ken05/Tk1           ..c...c....t.....t.....a..g.c...c..............t...........g.....a.t...c....ag.. [ 36080]
KM262844.1_strain_L60          ......c.......................t..........c.a...t.................a.....c........ [ 36080]
KM102979.1_26544/OG10_Italy    ......c.......................t..........c.a...t.................a.....c........ [ 36080]
KM262845.1_strain_NHV          -------------------------------------------------------------------------------- [ 36080]
KX354450.1_47/Ss/2008          ......c.......................t..........c.a...t.................a.....c........ [ 36080]
KP055815.1_strain_BA71         ......c.......................t..........c.a...t.................a.....c........ [ 36080]
MH025920.1_strain_R35          ..c...c.t..t.....t.....a..g.c...c..............t...........g.....a.t........ag.. [ 36080]
MH025919.1_strain_N10          ..c...c.t..t.....t.....a..g.c...c..............t...........g.....a.t........ag.. [ 36080]
MH025918.1_strain_R25          ..c...c.t..t.....t.....a..g.c...c..............t...........g.....a.t........ag.. [ 36080]
MH025917.1_strain_R7           ..c...c.t..t.....t.....a..g.c...c..............t...........g.....a.t........ag.. [ 36080]
MH025916.1_strain_R8           ..c...c.t..t.....t.....a..g.c...c..............t...........g.....a.t........ag.. [ 36080]

FR682468.2_Georgia_2007/1      tatacaaaagaacaagttatccatattataaaatcgtacgcaggcaagcatagctgaattaatattagctcctaagagaa [ 36160]
ASF_IND_20_CAD543              -------------------------------------------------------------------------------- [ 36160]
LR899193.1_Germany_2020/1      ................................................................................ [ 36160]
MN393477.1_Wuhan_2019          ................................................................................ [ 36160]
MW396979.1_Timor-Leste/2019/1  ................................................................................ [ 36160]
MK940252.1_CN/2019/InnerMongolia ................................................................................ [ 36160]
MK128995.1_China/2018/AnhuiXCGQ ................................................................................ [ 36160]
MH681419.1_ASFV/POL/2015       ................................................................................ [ 36160]
LS478113.1_Estonia_2014        ................................................................................ [ 36160]
KP843857.1_Odintsovo_02/14     ................................................................................ [ 36160]
FN557520.1_strain_E75          ..............g...............g.......t.g......t...g..ca.....................t.. [ 36160]
AY261366.1_Warthog             ..............g.......................t.g......t...g..ca.....................t.. [ 36160]
AY261365.1_Warmbaths           ..............g...............g.......t.g......t...g..ca.....................t.. [ 36160]
AY261364.1_Tengani_62          ................................................................................ [ 36160]
AY261363.1_Pretoriuskop/96/4   ............t........t..g.....g...t...t.g......t.......a.....................t.. [ 36160]
AY261362.1_Mkuzi_1979          ..............g...............g.......t.g......t...g..ca.....................t.. [ 36160]
AY261361.1_Malawi_Lil-20/1_1983 ..............................g...t...................ctg....................... [ 36160]
AY261360.1_Kenya_1950          ..............ga............a.g..ct...t.gca........g..cat...g....cg..c.....a.t.. [ 36160]
AM712240.1_OURT_88/3           -------------------------------------------------------------------------------- [ 36160]
AM712239.1_Benin_97/1          ..............g...............g.......t.g......t...g..ca.....................t.. [ 36160]
KM111295.1_Ken06.Bus           ..............ga.....t......a.g..ct...t.gca........g..cat...g....cg..c.....a.t.. [ 36160]
KM111294.1_Ken05/Tk1           ..............ga.....t......a....ct...t.gca........c..cat...g....cg..c.....a.t.. [ 36160]
KM262844.1_strain_L60          ..............g...............g.......t.g......t...g..ca.....................t.. [ 36160]
KM102979.1_26544/OG10_Italy    ..............g...............g.......t.g......t...g..ca.....................t.. [ 36160]
KM262845.1_strain_NHV          -------------------------------------------------------------------------------- [ 36160]
KX354450.1_47/Ss/2008          ..............g...............g.......t.g......t...g..ca.....................t.. [ 36160]
KP055815.1_strain_BA71         ..............g...............g.......t.g......t...g..ca.....................t.. [ 36160]
MH025920.1_strain_R35          ..............ga.....t......a.g..ct...t.gca........g..cat...g....cg..c.....a.t.. [ 36160]
MH025919.1_strain_N10          ..............ga.....t......a.g..ct...t.gca........g..cat...g....cg..c.....a.t.. [ 36160]
MH025918.1_strain_R25          ..............ga.....t......a.g..ct...t.gca........g..cat...g....cg..c.....a.t.. [ 36160]
MH025917.1_strain_R7           ..............ga.....t......a.g..ct...t.gca........g..cat...g....cg..c.....a.t.. [ 36160]
MH025916.1_strain_R8           ..............ga.....t......a.g..ct...t.gca........g..cat...g....cg..c.....a.t.. [ 36160]
```

```
FR682468.2_Georgia_2007/1        aacaataatatatggttaaaaaattgttatcttttgtgcaggccatccgcatcatttcatccacgtccatgcggatcttt [ 36240]
ASF_IND_20_CAD543                -------------------------------------------------------------------------------- [ 36240]
LR899193.1_Germany_2020/1        ................................................................................ [ 36240]
MN393477.1_Wuhan_2019            ................................................................................ [ 36240]
MW396979.1_Timor-Leste/2019/1    ................................................................................ [ 36240]
MK940252.1_CN/2019/InnerMongolia ................................................................................ [ 36240]
MK128995.1_China/2018/AnhuiXCGQ  ................................................................................ [ 36240]
MH681419.1_ASFV/POL/2015         ................................................................................ [ 36240]
LS478113.1_Estonia_2014          ................................................................................ [ 36240]
KP843857.1_Odintsovo_02/14       ................................................................................ [ 36240]
FN557520.1_strain_E75            .............a.............................t...................t......a.....t... [ 36240]
AY261366.1_Warthog               .............a.............................t...................t............t... [ 36240]
AY261365.1_Warmbaths             .............a.............................t...................t............t... [ 36240]
AY261364.1_Tengani_62            .............a.................................................................. [ 36240]
AY261363.1_Pretoriuskop/96/4     ...........................................t...................t................ [ 36240]
AY261362.1_Mkuzi_1979            .............a.............................t...................t......a.....t... [ 36240]
AY261361.1_Malawi_Lil-20/1_1983  ....g.................g...........a........t..g.a..............t................ [ 36240]
AY261360.1_Kenya_1950            ..........c..c........g...........c.....a..t..g.............ta.t...t.....a..t... [ 36240]
AM712240.1_OURT_88/3             -------------------------------------------------------------------------------- [ 36240]
AM712239.1_Benin_97/1            .............a.............................t...................t......a.....t... [ 36240]
KM111295.1_Ken06.Bus             ..........c..c........g..a........c.....a..t..g.............ta.t...t.....a...... [ 36240]
KM111294.1_Ken05/Tk1             ..........c..c........g...........c.....a..t..g.............ta.t...t.....a..t... [ 36240]
KM262844.1_strain_L60            .............a.............................t...................t......a.....t... [ 36240]
KM102979.1_26544/OG10_Italy      .............a.............................t...................t......a.....t... [ 36240]
KM262845.1_strain_NHV            -------------------------------------------------------------------------------- [ 36240]
KX354450.1_47/Ss/2008            .............a.............................t...................t......a.....t... [ 36240]
KP055815.1_strain_BA71           .............a.............................t...................t......a.....t... [ 36240]
MH025920.1_strain_R35            ..........c..c........g...........c.....a..t..g.............ta.t...t.....a...... [ 36240]
MH025919.1_strain_N10            ..........c..c........g...........c.....a..t..g.............ta.t...t.....a...... [ 36240]
MH025918.1_strain_R25            ..........c..c........g...........c.....a..t..g.............ta.t...t.....a...... [ 36240]
MH025917.1_strain_R7             ..........c..c........g...........c.....a..t..g.............ta.t...t.....a...... [ 36240]
MH025916.1_strain_R8             ..........c..c........g...........c.....a..t..g.............ta.t...t.....a...... [ 36240]

FR682468.2_Georgia_2007/1        tccttttcatacaaattatgtaggtcaaacagcttattaaaacaaagagcacagattaaccaccacgtatttagatactt [ 36320]
ASF_IND_20_CAD543                -------------------------------------------------------------------------------- [ 36320]
LR899193.1_Germany_2020/1        ................................................................................ [ 36320]
MN393477.1_Wuhan_2019            ................................................................................ [ 36320]
MW396979.1_Timor-Leste/2019/1    ................................................................................ [ 36320]
MK940252.1_CN/2019/InnerMongolia ................................................................................ [ 36320]
MK128995.1_China/2018/AnhuiXCGQ  ................................................................................ [ 36320]
MH681419.1_ASFV/POL/2015         ................................................................................ [ 36320]
LS478113.1_Estonia_2014          ................................................................................ [ 36320]
KP843857.1_Odintsovo_02/14       ................................................................................ [ 36320]
```

```
FN557520.1_strain_E75          ..t.....g..............................t.g...........tg................c........ [ 36320]
AY261366.1_Warthog             ........g.................................................g............c.....t.. [ 36320]
AY261365.1_Warmbaths           ........g.................................................g.......a....c........ [ 36320]
AY261364.1_Tengani_62          ................................................................................ [ 36320]
AY261363.1_Pretoriuskop/96/4   ................g..............c.......t.g......a....tg................c........ [ 36320]
AY261362.1_Mkuzi_1979          ..t.....g..............................t.g...........tg................c.a...... [ 36320]
AY261361.1_Malawi_Lil-20/1_1983 ................c......................................................c....g... [ 36320]
AY261360.1_Kenya_1950          ....gc.....g.g..a...g...........t...........c..........................c..g.g... [ 36320]
AM712240.1_OURT_88/3           -------------------------------------------------------------------------------- [ 36320]
AM712239.1_Benin_97/1          ..t.....g..............................t.g...........tg................c........ [ 36320]
KM111295.1_Ken06.Bus           ....gc.....g.g..a...g......g...t.........t.c..........................c.tg.g... [ 36320]
KM111294.1_Ken05/Tk1           ....gc.....g.g..a...g...........t...........c..........................c..g.g... [ 36320]
KM262844.1_strain_L60          ..t.....g..............................t.g...........tg................c........ [ 36320]
KM102979.1_26544/OG10_Italy    ..t.....g..............................t.g...........tg................c........ [ 36320]
KM262845.1_strain_NHV          -------------------------------------------------------------------------------- [ 36320]
KX354450.1_47/Ss/2008          ..t.....g..............................t.g...........tg................c........ [ 36320]
KP055815.1_strain_BA71         ..t.....g..............................t.g...........tg................c........ [ 36320]
MH025920.1_strain_R35          ....gc.....g.g..a...g......g...t...........c..........................c.tg.g... [ 36320]
MH025919.1_strain_N10          ....gc.....g.g..a...g......g...t...........c..........................c.tg.g... [ 36320]
MH025918.1_strain_R25          ....gc.....g.g..a...g......g...t...........c..........................c.tg.g... [ 36320]
MH025917.1_strain_R7           ....gc.....g.g..a...g......g...t...........c..........................c.tg.g... [ 36320]
MH025916.1_strain_R8           ....gc.....g.g..a...g......g...t...........c..........................c.tg.g... [ 36320]

FR682468.2_Georgia_2007/1      aaaatgttggtaaacataagaaatggcctccctaagattatcctgcaatgccactataaaacagtatatcgttaacatat [ 36400]
ASF_IND_20_CAD543              -------------------------------------------------------------------------------- [ 36400]
LR899193.1_Germany_2020/1      ................................................................................ [ 36400]
MN393477.1_Wuhan_2019          ................................................................................ [ 36400]
MW396979.1_Timor-Leste/2019/1  ................................................................................ [ 36400]
MK940252.1_CN/2019/InnerMongolia ................................................................................ [ 36400]
MK128995.1_China/2018/AnhuiXCGQ ................................................................................ [ 36400]
MH681419.1_ASFV/POL/2015       ................................................................................ [ 36400]
LS478113.1_Estonia_2014        ................................................................................ [ 36400]
KP843857.1_Odintsovo_02/14     ................................................................................ [ 36400]
FN557520.1_strain_E75          ......................................................g..............a.......... [ 36400]
AY261366.1_Warthog             ......................................................c......................... [ 36400]
AY261365.1_Warmbaths           ......................................................c......................... [ 36400]
AY261364.1_Tengani_62          ......................................................................a......... [ 36400]
AY261363.1_Pretoriuskop/96/4   ......................................................g..............a.......... [ 36400]
AY261362.1_Mkuzi_1979          ......................................................g..............a.......... [ 36400]
AY261361.1_Malawi_Lil-20/1_1983 ..............a......................................gg..............a..c....... [ 36400]
AY261360.1_Kenya_1950          ..............a...c..g.c..............c..t......g.....c........a....aa....g..... [ 36400]
AM712240.1_OURT_88/3           -------------------------------------------------------------------------------- [ 36400]
AM712239.1_Benin_97/1          ......................................................g..............a.......... [ 36400]
KM111295.1_Ken06.Bus           ........c.....a...c..g.c..............c..t......g.....c........a.....a....g..... [ 36400]
```

```
KM111294.1_Ken05/Tk1              ..............a...c..g.c.............cc..t......g.....c........a....aa....g..... [ 36400]
KM262844.1_strain_L60             ......................................................g..............a.......... [ 36400]
KM102979.1_26544/OG10_Italy       ......................................................g..............a.......... [ 36400]
KM262845.1_strain_NHV             -------------------------------------------------------------------------------- [ 36400]
KX354450.1_47/Ss/2008             ......................................................g..............a.......... [ 36400]
KP055815.1_strain_BA71            ......................................................g..............a.......... [ 36400]
MH025920.1_strain_R35             ........c.....a...c..g.c..............c..t......g.....c........a.....a....g..... [ 36400]
MH025919.1_strain_N10             ........c.....a...c..g.c..............c..t......g.....c........a.....a....g..... [ 36400]
MH025918.1_strain_R25             ........c.....a...c..g.c..............c..t......g.....c........a.....a....g..... [ 36400]
MH025917.1_strain_R7              ........c.....a...c..g.c..............c..t......g.....c........a.....a....g..... [ 36400]
MH025916.1_strain_R8              ........c.....a...c..g.c..............c..t......g.....c........a.....a....g..... [ 36400]

FR682468.2_Georgia_2007/1         caccatccgacatattacttaatatgtcggtgtcttctactaaccttttcaacttccaatatatggatgaccttatttcc [ 36480]
ASF_IND_20_CAD543                 -------------------------------------------------------------------------------- [ 36480]
LR899193.1_Germany_2020/1         ................................................................................ [ 36480]
MN393477.1_Wuhan_2019             ................................................................................ [ 36480]
MW396979.1_Timor-Leste/2019/1     ................................................................................ [ 36480]
MK940252.1_CN/2019/InnerMongolia  ................................................................................ [ 36480]
MK128995.1_China/2018/AnhuiXCGQ   ................................................................................ [ 36480]
MH681419.1_ASFV/POL/2015          ................................................................................ [ 36480]
LS478113.1_Estonia_2014           ................................................................................ [ 36480]
KP843857.1_Odintsovo_02/14        ................................................................................ [ 36480]
FN557520.1_strain_E75             ..........a.............c..t...a......g....at....t.g........g...ac.c...t........ [ 36480]
AY261366.1_Warthog                ..........a.............c...................t......g...................t........ [ 36480]
AY261365.1_Warmbaths              .......t..a....c........c...................t..c...g...................t........ [ 36480]
AY261364.1_Tengani_62             ...................................................g................at.......... [ 36480]
AY261363.1_Pretoriuskop/96/4      .t........a.............c..t...a......g.....t....t.g........g...ac.c...t........ [ 36480]
AY261362.1_Mkuzi_1979             ..........a.............c..t...a......g....at....t.g........g...ac.c...t........ [ 36480]
AY261361.1_Malawi_Lil-20/1_1983   ..t.........................a..................c...g............a..c...t........ [ 36480]
AY261360.1_Kenya_1950             .gt.g.....a.c..cg.g..g..g...ttc............t..cc...g.c.......a.ga..c...t........ [ 36480]
AM712240.1_OURT_88/3              -------------------------------------------------------------------------------- [ 36480]
AM712239.1_Benin_97/1             ..........a.............c..t...a......g....at....t.g........g...ac.c...t........ [ 36480]
KM111295.1_Ken06.Bus              .g..g.....a.c..c..g..g..g...ttca..............cc...g.c.......a.ga..c...t........ [ 36480]
KM111294.1_Ken05/Tk1              .g..g.....a.c..cg.g..g..g...ttc...............cc...g.c.......a.ga..c...t........ [ 36480]
KM262844.1_strain_L60             ..........a.............c..t...a......g....at....t.g........g...ac.c...t........ [ 36480]
KM102979.1_26544/OG10_Italy       ..........a.............c..t...a......g....at....t.g........g...ac.c...t........ [ 36480]
KM262845.1_strain_NHV             -------------------------------------------------------------------------------- [ 36480]
KX354450.1_47/Ss/2008             ..........a.............c..t...a......g....at....t.g........g...ac.c...t........ [ 36480]
KP055815.1_strain_BA71            ..........a.............c..t...a......g....at....t.g........g...ac.c...t........ [ 36480]
MH025920.1_strain_R35             .g..g.....a.c..c..g..g..g...ttca..............cc...g.c.......a.ga..c...t........ [ 36480]
MH025919.1_strain_N10             .g..g.....a.c..c..g..g..g...ttca..............cc...g.c.......a.ga..c...t........ [ 36480]
MH025918.1_strain_R25             .g..g.....a.c..c..g..g..g...ttca..............cc...g.c.......a.ga..c...t........ [ 36480]
MH025917.1_strain_R7              .g..g.....a.c..c..g..g..g...ttca..............cc...g.c.......a.ga..c...t........ [ 36480]
MH025916.1_strain_R8              .g..g.....a.c..c..g..g..g...ttca..............cc...g.c.......a.ga..c...t........ [ 36480]
```

```
FR682468.2_Georgia_2007/1        cttataatgacataggctggaaagggattatcattaaaaagtttaagacataagataatattactgctagtagtgccagg [ 36560]
ASF_IND_20_CAD543                -------------------------------------------------------------------------------- [ 36560]
LR899193.1_Germany_2020/1        ................................................................................ [ 36560]
MN393477.1_Wuhan_2019            ................................................................................ [ 36560]
MW396979.1_Timor-Leste/2019/1    ................................................................................ [ 36560]
MK940252.1_CN/2019/InnerMongolia ................................................................................ [ 36560]
MK128995.1_China/2018/AnhuiXCGQ  ................................................................................ [ 36560]
MH681419.1_ASFV/POL/2015         ...............................................g................................ [ 36560]
LS478113.1_Estonia_2014          ................................................................................ [ 36560]
KP843857.1_Odintsovo_02/14       ................................................................................ [ 36560]
FN557520.1_strain_E75            ...................a.........g..........a......................t.a.........tt... [ 36560]
AY261366.1_Warthog               ...................a......g..................................................... [ 36560]
AY261365.1_Warmbaths             ...................a......g..................................................... [ 36560]
AY261364.1_Tengani_62            ................................................................................ [ 36560]
AY261363.1_Pretoriuskop/96/4     ...................a.........g...................................a..........t... [ 36560]
AY261362.1_Mkuzi_1979            ...................a.........g..........a........................a..........t... [ 36560]
AY261361.1_Malawi_Lil-20/1_1983  ................t......a.......t.................................a............t. [ 36560]
AY261360.1_Kenya_1950            ........c.......ta.....a.....c...........c...t...............actga..c..tt..tt--- [ 36560]
AM712240.1_OURT_88/3             -------------------------------------------------------------------------------- [ 36560]
AM712239.1_Benin_97/1            ...................a.........g..........a......................t.a.........tt... [ 36560]
KM111295.1_Ken06.Bus             ........c.......ta.....a.....c...........c...t...............attga..t..tt..tt--- [ 36560]
KM111294.1_Ken05/Tk1             ........c.......ta.....a.....c...........c...t...............attga..c..tt..tt--- [ 36560]
KM262844.1_strain_L60            ...................a.........g..........a......................t.a.........tt... [ 36560]
KM102979.1_26544/OG10_Italy      ...................a.........g..........a......................t.a.........tt... [ 36560]
KM262845.1_strain_NHV            -------------------------------------------------------------------------------- [ 36560]
KX354450.1_47/Ss/2008            ...................a.........g..........a......................t.a.........tt... [ 36560]
KP055815.1_strain_BA71           ...................a.........g..........a......................t.a.........tt... [ 36560]
MH025920.1_strain_R35            ........c.......ta.....a.....c...........c...t...............attga..t..tt..tt--- [ 36560]
MH025919.1_strain_N10            ........c.......ta.....a.....c...........c...t...............attga..t..tt..tt--- [ 36560]
MH025918.1_strain_R25            ........c.......ta.....a.....c...........c...t...............attga..t..tt..tt--- [ 36560]
MH025917.1_strain_R7             ........c.......ta.....a.....c...........c...t...............attga..t..tt..tt--- [ 36560]
MH025916.1_strain_R8             ........c.......ta.....a.....c...........c...t...............attga..t..tt..tt--- [ 36560]

FR682468.2_Georgia_2007/1        gtgtattaatttaaagaacatgtgcataatcttctttttatccacgcggtacttggctcctaattcccagcaaaattctc [ 36640]
ASF_IND_20_CAD543                -------------------------------------------------------------------------------- [ 36640]
LR899193.1_Germany_2020/1        ................................................................................ [ 36640]
MN393477.1_Wuhan_2019            ................................................................................ [ 36640]
MW396979.1_Timor-Leste/2019/1    ................................................................................ [ 36640]
MK940252.1_CN/2019/InnerMongolia ................................................................................ [ 36640]
MK128995.1_China/2018/AnhuiXCGQ  ................................................................................ [ 36640]
MH681419.1_ASFV/POL/2015         ................................................................................ [ 36640]
LS478113.1_Estonia_2014          ................................................................................ [ 36640]
KP843857.1_Odintsovo_02/14       ................................................................................ [ 36640]
```

```
FN557520.1_strain_E75            a.................t..........................tt................................. [ 36640]
AY261366.1_Warthog               a.............................................................g.........c....... [ 36640]
AY261365.1_Warmbaths             a.............................................................g.........c....... [ 36640]
AY261364.1_Tengani_62            a.......................t.........................g............................. [ 36640]
AY261363.1_Pretoriuskop/96/4     a.................t..........................tt................................. [ 36640]
AY261362.1_Mkuzi_1979            a.................t..........................tt...g............................. [ 36640]
AY261361.1_Malawi_Lil-20/1_1983  a..............a.....a.......c...a...........tt................................. [ 36640]
AY261360.1_Kenya_1950            c.c...a........a.....a.....c.c...t........t.a...a..g..t..............a.....c.... [ 36640]
AM712240.1_OURT_88/3             -------------------------------------------------------------------------------- [ 36640]
AM712239.1_Benin_97/1            a.................t..........................tt................................. [ 36640]
KM111295.1_Ken06.Bus             c.c...a.................t..c.c...t..........t...a..g..c..............a.....c.... [ 36640]
KM111294.1_Ken05/Tk1             c.c...a........a.....a.....c.c...t........t.a...a..g..c..............a.....c.... [ 36640]
KM262844.1_strain_L60            a.................t..........................tt................................. [ 36640]
KM102979.1_26544/OG10_Italy      a.................t..............t...........tt................................. [ 36640]
KM262845.1_strain_NHV            -------------------------------------------------------------------------------- [ 36640]
KX354450.1_47/Ss/2008            a.................t..............t...........tt................................. [ 36640]
KP055815.1_strain_BA71           a.................t..........................tt................................. [ 36640]
MH025920.1_strain_R35            c.c...a.................t..c.c...t..........t...a..g..c..............a.....c.... [ 36640]
MH025919.1_strain_N10            c.c...a.................t..c.c...t..........t...a..g..c..............a.....c.... [ 36640]
MH025918.1_strain_R25            c.c...a.................t..c.c...t..........t...a..g..c..............a.....c.... [ 36640]
MH025917.1_strain_R7             c.c...a.................t..c.c...t..........t...a..g..c..............a.....c.... [ 36640]
MH025916.1_strain_R8             c.c...a.................t..c.c...t..........t...a..g..c..............a.....c.... [ 36640]

FR682468.2_Georgia_2007/1        gaacaggcggcgtattggcgcaaattaacccatagttgatgtctgcgccccattctgtaaacagttttattaactgatag [ 36720]
ASF_IND_20_CAD543                -------------------------------------------------------------------------------- [ 36720]
LR899193.1_Germany_2020/1        ................................................................................ [ 36720]
MN393477.1_Wuhan_2019            ................................................................................ [ 36720]
MW396979.1_Timor-Leste/2019/1    ................................................................................ [ 36720]
MK940252.1_CN/2019/InnerMongolia ................................................................................ [ 36720]
MK128995.1_China/2018/AnhuiXCGQ  ................................................................................ [ 36720]
MH681419.1_ASFV/POL/2015         ................................................................................ [ 36720]
LS478113.1_Estonia_2014          ................................................................................ [ 36720]
KP843857.1_Odintsovo_02/14       ................................................................................ [ 36720]
FN557520.1_strain_E75            ...t...t........c...................................a..c.......t.........t...... [ 36720]
AY261366.1_Warthog               .......................................................c........................ [ 36720]
AY261365.1_Warmbaths             ..........t............................................c........................ [ 36720]
AY261364.1_Tengani_62            ......a................................................c........................ [ 36720]
AY261363.1_Pretoriuskop/96/4     .......t........c...................................a..c.......t.........t...... [ 36720]
AY261362.1_Mkuzi_1979            ...t...t........c...................................a..c.......t.........t...... [ 36720]
AY261361.1_Malawi_Lil-20/1_1983  ................a...........t..............c...........c........................ [ 36720]
AY261360.1_Kenya_1950            ..gt....t.t........a.....gt.t.............t...a.....c..c..g..............t...... [ 36720]
AM712240.1_OURT_88/3             -------------------------------------------------------------------------------- [ 36720]
AM712239.1_Benin_97/1            ...t...t........c...................................a..c.......t.........t...... [ 36720]
KM111295.1_Ken06.Bus             .ggt....t.t........a.....gt.t.............t...a.....c..c..g..t...........t...... [ 36720]
```

```
KM111294.1_Ken05/Tk1            ..gt....t.t........a.....gt.t.............t...a.....c..c..g..............t...... [ 36720]
KM262844.1_strain_L60           ...t...t........c...................................a..c.......t.........t...... [ 36720]
KM102979.1_26544/OG10_Italy     ...t...t........c...................................a..c.......t.........t...... [ 36720]
KM262845.1_strain_NHV           -------------------------------------------------------------------------------- [ 36720]
KX354450.1_47/Ss/2008           ...t...t........c...................................a..c.......t.........t...... [ 36720]
KP055815.1_strain_BA71          ...t...t........c...................................a..c.......t.........t...... [ 36720]
MH025920.1_strain_R35           .ggt....t.t........a.....gt.t.t...........t...a.....c..c..g..............t...... [ 36720]
MH025919.1_strain_N10           .ggt....t.t........a.....gt.t.t...........t...a.....c..c..g..............t...... [ 36720]
MH025918.1_strain_R25           .ggt....t.t........a.....gt.t.t...........t...a.....c..c..g..............t...... [ 36720]
MH025917.1_strain_R7            .ggt....t.t........a.....gt.t.t...........t...a.....c..c..g..............t...... [ 36720]
MH025916.1_strain_R8            .ggt....t.t........a.....gt.t.t...........t...a.....c..c..g..............t...... [ 36720]

FR682468.2_Georgia_2007/1       ttgttttcctttgtagccaacattagtgccgtattaaggtccaagccgtctgcaaagcttggcagctttatcagcatatg [ 36800]
ASF_IND_20_CAD543               -------------------------------------------------------------------------------- [ 36800]
LR899193.1_Germany_2020/1       ................................................................................ [ 36800]
MN393477.1_Wuhan_2019           ................................................................................ [ 36800]
MW396979.1_Timor-Leste/2019/1   ................................................................................ [ 36800]
MK940252.1_CN/2019/InnerMongolia ................................................................................ [ 36800]
MK128995.1_China/2018/AnhuiXCGQ ................................................................................ [ 36800]
MH681419.1_ASFV/POL/2015        ................................................................................ [ 36800]
LS478113.1_Estonia_2014         ................................................................................ [ 36800]
KP843857.1_Odintsovo_02/14      ................................................................................ [ 36800]
FN557520.1_strain_E75           ...............................................a.........t...................... [ 36800]
AY261366.1_Warthog              ...............................................a................................ [ 36800]
AY261365.1_Warmbaths            ...............................................a................................ [ 36800]
AY261364.1_Tengani_62           ..a............................................................................. [ 36800]
AY261363.1_Pretoriuskop/96/4    ...............................................a.........t...................... [ 36800]
AY261362.1_Mkuzi_1979           ...............................................a.........t...................... [ 36800]
AY261361.1_Malawi_Lil-20/1_1983 ...............................................a.......................t.a...... [ 36800]
AY261360.1_Kenya_1950           .............c...t.t...c.a.........g.......g...a..c......ta.....a.c....t.a...g.. [ 36800]
AM712240.1_OURT_88/3            -------------------------------------------------------------------------------- [ 36800]
AM712239.1_Benin_97/1           ...............................................a.........t...................... [ 36800]
KM111295.1_Ken06.Bus            .............c...t.t...c.a.........g..a....g...a..c......ta.....a.c....t.a...g.. [ 36800]
KM111294.1_Ken05/Tk1            .............c...t.t...c.a.........g.......g...a..c......ta.....a.c....t.a...g.. [ 36800]
KM262844.1_strain_L60           ...............................................a.........t...................... [ 36800]
KM102979.1_26544/OG10_Italy     ...............................................a.........t...................... [ 36800]
KM262845.1_strain_NHV           -------------------------------------------------------------------------------- [ 36800]
KX354450.1_47/Ss/2008           ...............................................a.........t...................... [ 36800]
KP055815.1_strain_BA71          ...............................................a.........t...................... [ 36800]
MH025920.1_strain_R35           .............c...t.t...c.a.........g..a....g...a..c......ta.....a.c....t.a...g.. [ 36800]
MH025919.1_strain_N10           .............c...t.t...c.a.........g..a....g...a..c......ta.....a.c....t.a...g.. [ 36800]
MH025918.1_strain_R25           .............c...t.t...c.a.........g..a....g...a..c......ta.....a.c....t.a...g.. [ 36800]
MH025917.1_strain_R7            .............c...t.t...c.a.........g..a....g...a..c......ta.....a.c....t.a...g.. [ 36800]
MH025916.1_strain_R8            .............c...t.t...c.a.........g..a....g...a..c......ta.....a.c....t.a...g.. [ 36800]
```

```
FR682468.2_Georgia_2007/1        tttgcaatcaagggaaattggggccttataccaccatagtccgcagcgttctaagataacatggtactcaatag---ata [ 36880]
ASF_IND_20_CAD543                -------------------------------------------------------------------------------- [ 36880]
LR899193.1_Germany_2020/1        ..........................................................................---... [ 36880]
MN393477.1_Wuhan_2019            ..........................................................................---... [ 36880]
MW396979.1_Timor-Leste/2019/1    ..........................................................................---... [ 36880]
MK940252.1_CN/2019/InnerMongolia ..........................................................................---... [ 36880]
MK128995.1_China/2018/AnhuiXCGQ  ..........................................................................---... [ 36880]
MH681419.1_ASFV/POL/2015         ..........................................................................---... [ 36880]
LS478113.1_Estonia_2014          ..........................................................................---... [ 36880]
KP843857.1_Odintsovo_02/14       ..........................................................................---... [ 36880]
FN557520.1_strain_E75            ............t........a.................c..........................a.......---... [ 36880]
AY261366.1_Warthog               .....................a............................................a.......---... [ 36880]
AY261365.1_Warmbaths             ............t........a............................................a.......---... [ 36880]
AY261364.1_Tengani_62            ......g.....t........a.......................a............................---... [ 36880]
AY261363.1_Pretoriuskop/96/4     ............t........a.................c...................t.....ga.....g.---... [ 36880]
AY261362.1_Mkuzi_1979            ............t........a.................c.........................ga.......---... [ 36880]
AY261361.1_Malawi_Lil-20/1_1983  ............t........a.......................at...t.......ta...a..a.....g.---... [ 36880]
AY261360.1_Kenya_1950            ...a........t........a................ac........c.tc..a...tg..a...g..ttc..gcagc. [ 36880]
AM712240.1_OURT_88/3             -------------------------------------------------------------------------------- [ 36880]
AM712239.1_Benin_97/1            ............t........a.................c..........................a.......---... [ 36880]
KM111295.1_Ken06.Bus             ............t........a...g............ac......ta..tc..a...tg..a...g..ttc..gcagc. [ 36880]
KM111294.1_Ken05/Tk1             ...a........t........a................ac........c.tc..a...tg..a...g..ttc..gcagc. [ 36880]
KM262844.1_strain_L60            ............t........a.................c..........................a.......---... [ 36880]
KM102979.1_26544/OG10_Italy      ............t........a.................c..........................a.......---... [ 36880]
KM262845.1_strain_NHV            -------------------------------------------------------------------------------- [ 36880]
KX354450.1_47/Ss/2008            ............t........a.................c..........................a.......---... [ 36880]
KP055815.1_strain_BA71           ............t........a.................c..........................a.......---... [ 36880]
MH025920.1_strain_R35            ............t........a...g............ac......ta..tc..a...tg..a...g..ttc..gcagc. [ 36880]
MH025919.1_strain_N10            ............t........a...g............ac......ta..tc..a...tg..a...g..ttc..gcagc. [ 36880]
MH025918.1_strain_R25            ............t........a...g............ac......ta..tc..a...tg..a...g..ttc..gcagc. [ 36880]
MH025917.1_strain_R7             ............t........a...g............ac......ta..tc..a...tg..a...g..ttc..gcagc. [ 36880]
MH025916.1_strain_R8             ............t........a...g............ac......ta..tc..a...tg..a...g..ttc..gcagc. [ 36880]

FR682468.2_Georgia_2007/1        cttgctgtctggctagtacctttttggcgaaggattgtaaggaaggaaacatcctgtttct-tttttttttaaaaatcaa [ 36960]
ASF_IND_20_CAD543                -------------------------------------------------------------------------------- [ 36960]
LR899193.1_Germany_2020/1        .............................................................-.................. [ 36960]
MN393477.1_Wuhan_2019            .............................................................-.................. [ 36960]
MW396979.1_Timor-Leste/2019/1    .............................................................-.................. [ 36960]
MK940252.1_CN/2019/InnerMongolia .............................................................-.................. [ 36960]
MK128995.1_China/2018/AnhuiXCGQ  .............................................................-.................. [ 36960]
MH681419.1_ASFV/POL/2015         .............................................................-.................. [ 36960]
LS478113.1_Estonia_2014          .............................................................-.................. [ 36960]
KP843857.1_Odintsovo_02/14       .............................................................--................. [ 36960]
```

```
FN557520.1_strain_E75          .a.....a.......a.....................c..........c...........---................. [ 36960]
AY261366.1_Warthog             .a..........................................................--........a......... [ 36960]
AY261365.1_Warmbaths           .a..........................................................--........a......... [ 36960]
AY261364.1_Tengani_62          ............................................................--.................. [ 36960]
AY261363.1_Pretoriuskop/96/4   .a...........................................................-........a..t...... [ 36960]
AY261362.1_Mkuzi_1979          .a.....a.......a.....................c......................---........t...-.... [ 36960]
AY261361.1_Malawi_Lil-20/1_1983 .a..a.....a....a........................a....................t.................. [ 36960]
AY261360.1_Kenya_1950          .a.....g.......a..t............a.....c.g.................c..--........a....t.... [ 36960]
AM712240.1_OURT_88/3           -------------------------------------------------------------------------------- [ 36960]
AM712239.1_Benin_97/1          .a.....a.......a.....................c..........c...........---................. [ 36960]
KM111295.1_Ken06.Bus           .a.....a.......a..t............a..c.....a................c..--........a....t.... [ 36960]
KM111294.1_Ken05/Tk1           .a.....g.......a..t............a.....c.g.................c..--.............t.... [ 36960]
KM262844.1_strain_L60          .a.....a.......a.....................c..........c...........---................. [ 36960]
KM102979.1_26544/OG10_Italy    .a.....a.......a.....................c..........c...........---................. [ 36960]
KM262845.1_strain_NHV          -------------------------------------------------------------------------------- [ 36960]
KX354450.1_47/Ss/2008          .a.....a.......a.....................c..........c...........---................. [ 36960]
KP055815.1_strain_BA71         .a.....a.......a.....................c..........c...........---................. [ 36960]
MH025920.1_strain_R35          .a.....a.......a..t............a..c.....a................c..--........a....t.... [ 36960]
MH025919.1_strain_N10          .a.....a.......a..t............a..c.....a................c..--........a....t.... [ 36960]
MH025918.1_strain_R25          .a.....a.......a..t............a..c.....a................c..--........a....t.... [ 36960]
MH025917.1_strain_R7           .a.....a.......a..t............a..c.....a................c..--........a....t.... [ 36960]
MH025916.1_strain_R8           .a.....a.......a..t............a..c.....a................c..--........a....t.... [ 36960]

FR682468.2_Georgia_2007/1      ttatctttgttcataatcaagaaaaatccccatatttattgagtgataattttttaacatgcaatttattttttcagggt [ 37040]
ASF_IND_20_CAD543              -------------------------------------------------------------------------------- [ 37040]
LR899193.1_Germany_2020/1      ................................................................................ [ 37040]
MN393477.1_Wuhan_2019          ................................................................................ [ 37040]
MW396979.1_Timor-Leste/2019/1  ................................................................................ [ 37040]
MK940252.1_CN/2019/InnerMongolia ................................................................................ [ 37040]
MK128995.1_China/2018/AnhuiXCGQ ................................................................................ [ 37040]
MH681419.1_ASFV/POL/2015       ................................................................................ [ 37040]
LS478113.1_Estonia_2014        ................................................................................ [ 37040]
KP843857.1_Odintsovo_02/14     ................................................................................ [ 37040]
FN557520.1_strain_E75          .............................................................t.................. [ 37040]
AY261366.1_Warthog             ..............................................................g................. [ 37040]
AY261365.1_Warmbaths           ..............................................................g................. [ 37040]
AY261364.1_Tengani_62          ..........................................................................t..... [ 37040]
AY261363.1_Pretoriuskop/96/4   ..............................................................g................. [ 37040]
AY261362.1_Mkuzi_1979          .............................................................t.................. [ 37040]
AY261361.1_Malawi_Lil-20/1_1983 .......c..................c..t.................................................. [ 37040]
AY261360.1_Kenya_1950          ......gc..........g.......c.a.......ct......................ag.......c.....t.... [ 37040]
AM712240.1_OURT_88/3           -------------------------------------------------------------------------------- [ 37040]
AM712239.1_Benin_97/1          .............................................................t.................. [ 37040]
KM111295.1_Ken06.Bus           ......gc..........g.......c.a.......ct......................ag.......c.....c.... [ 37040]
```

```
KM111294.1_Ken05/Tk1            ......gc..........g.......c.a.......ct......................ag.......c.....t.... [ 37040]
KM262844.1_strain_L60           .............................................................t.................. [ 37040]
KM102979.1_26544/OG10_Italy     .............................................................t.................. [ 37040]
KM262845.1_strain_NHV           -------------------------------------------------------------------------------- [ 37040]
KX354450.1_47/Ss/2008           .............................................................t.................. [ 37040]
KP055815.1_strain_BA71          .............................................................t.................. [ 37040]
MH025920.1_strain_R35           ......gc..........g.......c.a.......ct......................ag.......c.....c.... [ 37040]
MH025919.1_strain_N10           ......gc..........g.......c.a.......ct......................ag.......c.....c.... [ 37040]
MH025918.1_strain_R25           ......gc..........g.......c.a.......ct......................ag.......c.....c.... [ 37040]
MH025917.1_strain_R7            ......gc..........g.......c.a.......ct......................ag.......c.....c.... [ 37040]
MH025916.1_strain_R8            ......gc..........g.......c.a.......ct......................ag.......c.....c.... [ 37040]

FR682468.2_Georgia_2007/1       ccgtaacgatcgacaacagagaaataaccggattgtaatgctttaatgataaggcatgggctatcagataattttccttt [ 37120]
ASF_IND_20_CAD543               -------------------------------------------------------------------------------- [ 37120]
LR899193.1_Germany_2020/1       ................................................................................ [ 37120]
MN393477.1_Wuhan_2019           ................................................................................ [ 37120]
MW396979.1_Timor-Leste/2019/1   ................................................................................ [ 37120]
MK940252.1_CN/2019/InnerMongolia ................................................................................ [ 37120]
MK128995.1_China/2018/AnhuiXCGQ ................................................................................ [ 37120]
MH681419.1_ASFV/POL/2015        ................................................................................ [ 37120]
LS478113.1_Estonia_2014         ................................................................................ [ 37120]
KP843857.1_Odintsovo_02/14      ................................................................................ [ 37120]
FN557520.1_strain_E75           ..a.....t................g..t................................................... [ 37120]
AY261366.1_Warthog              .......c.c...............g...............................t................c..... [ 37120]
AY261365.1_Warmbaths            .......c.c...............g...............................t................c..... [ 37120]
AY261364.1_Tengani_62           .t.......c...............g...............................c................c..... [ 37120]
AY261363.1_Pretoriuskop/96/4    .......c.c...............g...............................t................c..... [ 37120]
AY261362.1_Mkuzi_1979           ..a......................g................................................c..... [ 37120]
AY261361.1_Malawi_Lil-20/1_1983 ..a...t..c...............g...............................c...................... [ 37120]
AY261360.1_Kenya_1950           .g..c...gc.........g.....c.....g..a.....t................c........a............. [ 37120]
AM712240.1_OURT_88/3            -------------------------------------------------------------------------------- [ 37120]
AM712239.1_Benin_97/1           ..a.....t................g..t................................................... [ 37120]
KM111295.1_Ken06.Bus            .g..c...gc.........g.....c.....g..a.....t................c........a............. [ 37120]
KM111294.1_Ken05/Tk1            .g..c...gc.........g.....c.....g..a.....t................c........a............. [ 37120]
KM262844.1_strain_L60           ..a.....t................g..t................................................... [ 37120]
KM102979.1_26544/OG10_Italy     ..a.....t................g..t................................................... [ 37120]
KM262845.1_strain_NHV           -------------------------------------------------------------------------------- [ 37120]
KX354450.1_47/Ss/2008           ..a.....t................g..t................................................... [ 37120]
KP055815.1_strain_BA71          ..a.....t................g..t................................................... [ 37120]
MH025920.1_strain_R35           .g..c...gc.........g.....c.....g..a.....t................c........a............. [ 37120]
MH025919.1_strain_N10           .g..c...gc.........g.....c.....g..a.....t................c........a............. [ 37120]
MH025918.1_strain_R25           .g..c...gc.........g.....c.....g..a.....t................c........a............. [ 37120]
MH025917.1_strain_R7            .g..c...gc.........g.....c.....g..a.....t................c........a............. [ 37120]
MH025916.1_strain_R8            .g..c...gc.........g.....c.....g..a.....t................c........a............. [ 37120]
```

```
FR682468.2_Georgia_2007/1        tgttctgccaaagctttgccctcctcaaaggcatcggcacccaggtctatacaaaagaacaggtttccaagattatagtt [ 37200]
ASF_IND_20_CAD543                -------------------------------------------------------------------------------- [ 37200]
LR899193.1_Germany_2020/1        ................................................................................ [ 37200]
MN393477.1_Wuhan_2019            ................................................................................ [ 37200]
MW396979.1_Timor-Leste/2019/1    ................................................................................ [ 37200]
MK940252.1_CN/2019/InnerMongolia ................................................................................ [ 37200]
MK128995.1_China/2018/AnhuiXCGQ  ................................................................................ [ 37200]
MH681419.1_ASFV/POL/2015         ................................................................................ [ 37200]
LS478113.1_Estonia_2014          ................................................................................ [ 37200]
KP843857.1_Odintsovo_02/14       ................................................................................ [ 37200]
FN557520.1_strain_E75            ................................g.t...g......................................... [ 37200]
AY261366.1_Warthog               ................................g.t...c..........................c.............. [ 37200]
AY261365.1_Warmbaths             ................................g.t..............................c.............. [ 37200]
AY261364.1_Tengani_62            ....................t...........g.t...t..........................c.............. [ 37200]
AY261363.1_Pretoriuskop/96/4     .......................t........g.t..............................c.............. [ 37200]
AY261362.1_Mkuzi_1979            ................................g.t...g......................................... [ 37200]
AY261361.1_Malawi_Lil-20/1_1983  .................c..............g.t...c.....a................t...c.............. [ 37200]
AY261360.1_Kenya_1950            ...at...t.gg.t......t..t........g.t..........................t...c..g........... [ 37200]
AM712240.1_OURT_88/3             -------------------------------------------------------------------------------- [ 37200]
AM712239.1_Benin_97/1            ................................g.t...g......................................... [ 37200]
KM111295.1_Ken06.Bus             ...at...t.gg........t..t........g.t..........................t...c..g........... [ 37200]
KM111294.1_Ken05/Tk1             ...at...t.gg........t..t........g.t..........................t...c..g........... [ 37200]
KM262844.1_strain_L60            ................................g.t...g......................................... [ 37200]
KM102979.1_26544/OG10_Italy      ................................g.t...g......................................... [ 37200]
KM262845.1_strain_NHV            -------------------------------------------------------------------------------- [ 37200]
KX354450.1_47/Ss/2008            ................................g.t...g......................................... [ 37200]
KP055815.1_strain_BA71           ................................g.t...g......................................... [ 37200]
MH025920.1_strain_R35            ...at...t.gg........t..t........g.t..........................t...c..g........... [ 37200]
MH025919.1_strain_N10            ...at...t.gg........t..t........g.t..........................t...c..g........... [ 37200]
MH025918.1_strain_R25            ...at...t.gg........t..t........g.t..........................t...c..g........... [ 37200]
MH025917.1_strain_R7             ...at...t.gg........t..t........g.t..........................t...c..g........... [ 37200]
MH025916.1_strain_R8             ...at...t.gg........t..t........g.t..........................t...c..g........... [ 37200]

FR682468.2_Georgia_2007/1        ttgtatggaaacaagcatggcttgattgatgttggctcccatgataaaacagtagtaaatggccgaatagctataatctt [ 37280]
ASF_IND_20_CAD543                -------------------------------------------------------------------------------- [ 37280]
LR899193.1_Germany_2020/1        ................................................................................ [ 37280]
MN393477.1_Wuhan_2019            ................................................................................ [ 37280]
MW396979.1_Timor-Leste/2019/1    ................................................................................ [ 37280]
MK940252.1_CN/2019/InnerMongolia ................................................................................ [ 37280]
MK128995.1_China/2018/AnhuiXCGQ  ................................................................................ [ 37280]
MH681419.1_ASFV/POL/2015         ................................................................................ [ 37280]
LS478113.1_Estonia_2014          ................................................................................ [ 37280]
KP843857.1_Odintsovo_02/14       ................................................................................ [ 37280]
```

```
FN557520.1_strain_E75          ............c................................................................... [ 37280]
AY261366.1_Warthog             ............c..................................................t................ [ 37280]
AY261365.1_Warmbaths           ............c............................................g.....t................ [ 37280]
AY261364.1_Tengani_62          ............c................................................................... [ 37280]
AY261363.1_Pretoriuskop/96/4   ............c..................................................t................ [ 37280]
AY261362.1_Mkuzi_1979          ............c.....a..........................................a.................. [ 37280]
AY261361.1_Malawi_Lil-20/1_1983 ......a.............................c.................a.....a....c.............. [ 37280]
AY261360.1_Kenya_1950          ..............a.....................c....................g..a..t................ [ 37280]
AM712240.1_OURT_88/3           -------------------------------------------------------------------------------- [ 37280]
AM712239.1_Benin_97/1          ............c................................................................... [ 37280]
KM111295.1_Ken06.Bus           ..............a.....................a.......................a.tt................ [ 37280]
KM111294.1_Ken05/Tk1           ..............a.....................c.......................a..t................ [ 37280]
KM262844.1_strain_L60          ............c................................................................... [ 37280]
KM102979.1_26544/OG10_Italy    ............c................................................................... [ 37280]
KM262845.1_strain_NHV          -------------------------------------------------------------------------------- [ 37280]
KX354450.1_47/Ss/2008          ............c................................................................... [ 37280]
KP055815.1_strain_BA71         ............c................................................................... [ 37280]
MH025920.1_strain_R35          ..............a.....................a.......................a.tt................ [ 37280]
MH025919.1_strain_N10          ..............a.....................a.......................a.tt................ [ 37280]
MH025918.1_strain_R25          ..............a.....................a.......................a.tt................ [ 37280]
MH025917.1_strain_R7           ..............a.....................a.......................a.tt................ [ 37280]
MH025916.1_strain_R8           ..............a.....................a.......................a.tt................ [ 37280]

FR682468.2_Georgia_2007/1      ggatgcaggctatgtgcatcatttcatcaatatccatgcggaccctttctatttcgtacagctcgtgaaggtcgaacacg [ 37360]
ASF_IND_20_CAD543              -------------------------------------------------------------------------------- [ 37360]
LR899193.1_Germany_2020/1      ................................................................................ [ 37360]
MN393477.1_Wuhan_2019          ................................................................................ [ 37360]
MW396979.1_Timor-Leste/2019/1  ................................................................................ [ 37360]
MK940252.1_CN/2019/InnerMongolia ................................................................................ [ 37360]
MK128995.1_China/2018/AnhuiXCGQ ................................................................................ [ 37360]
MH681419.1_ASFV/POL/2015       ................................................................................ [ 37360]
LS478113.1_Estonia_2014        ................................................................................ [ 37360]
KP843857.1_Odintsovo_02/14     ................................................................................ [ 37360]
FN557520.1_strain_E75          ............................g..........................a........................ [ 37360]
AY261366.1_Warthog             ...c...........................g.......................a........................ [ 37360]
AY261365.1_Warmbaths           ...c...........................g.......................a........................ [ 37360]
AY261364.1_Tengani_62          ...c...........................g................................................ [ 37360]
AY261363.1_Pretoriuskop/96/4   ...c...........................g.......................a........................ [ 37360]
AY261362.1_Mkuzi_1979          .......................................................a........................ [ 37360]
AY261361.1_Malawi_Lil-20/1_1983 ...c.................................................a.a........................ [ 37360]
AY261360.1_Kenya_1950          ......................c.....g..........................a.......................a [ 37360]
AM712240.1_OURT_88/3           -------------------------------------------------------------------------------- [ 37360]
AM712239.1_Benin_97/1          ............................g..........................a........................ [ 37360]
KM111295.1_Ken06.Bus           ......................c.....g.............t............a.......................a [ 37360]
```

```
KM111294.1_Ken05/Tk1             ......................c.....g..........................a.......................a [ 37360]
KM262844.1_strain_L60            ............................g..........................a........................ [ 37360]
KM102979.1_26544/OG10_Italy      ............................g..........................a........................ [ 37360]
KM262845.1_strain_NHV            -------------------------------------------------------------------------------- [ 37360]
KX354450.1_47/Ss/2008            ............................g..........................a........................ [ 37360]
KP055815.1_strain_BA71           ............................g..........................a........................ [ 37360]
MH025920.1_strain_R35            ......................c.....g.............t............a.......................a [ 37360]
MH025919.1_strain_N10            ......................c.....g.............t............a.......................a [ 37360]
MH025918.1_strain_R25            ......................c.....g.............t............a.......................a [ 37360]
MH025917.1_strain_R7             ......................c.....g.............t............a.......................a [ 37360]
MH025916.1_strain_R8             ......................c.....g.............t............a.......................a [ 37360]

FR682468.2_Georgia_2007/1        ttgttgtaaaaaagggcgcacatgagccgccacctatgtagacgcgggtatttctggtaaaagtagcggatagcatcttt [ 37440]
ASF_IND_20_CAD543                -------------------------------------------------------------------------------- [ 37440]
LR899193.1_Germany_2020/1        ................................................................................ [ 37440]
MN393477.1_Wuhan_2019            ................................................................................ [ 37440]
MW396979.1_Timor-Leste/2019/1    ................................................................................ [ 37440]
MK940252.1_CN/2019/InnerMongolia ................................................................................ [ 37440]
MK128995.1_China/2018/AnhuiXCGQ  ................................................................................ [ 37440]
MH681419.1_ASFV/POL/2015         ................................................................................ [ 37440]
LS478113.1_Estonia_2014          ................................................................................ [ 37440]
KP843857.1_Odintsovo_02/14       ................................................................................ [ 37440]
FN557520.1_strain_E75            ..................................g......g..........................a........... [ 37440]
AY261366.1_Warthog               ..............a..........................c...................................... [ 37440]
AY261365.1_Warmbaths             .........................................c...................................... [ 37440]
AY261364.1_Tengani_62            .......................................................................c........ [ 37440]
AY261363.1_Pretoriuskop/96/4     .........................................c...................................... [ 37440]
AY261362.1_Mkuzi_1979            .........................................g..........................a........... [ 37440]
AY261361.1_Malawi_Lil-20/1_1983  .......................a........gtgg......t.g.ta..cc....a.....a..at.......c.t... [ 37440]
AY261360.1_Kenya_1950            .....a.................a..........g.......t.g.ta..c.....a.....a...t....g..c.t... [ 37440]
AM712240.1_OURT_88/3             -------------------------------------------------------------------------------- [ 37440]
AM712239.1_Benin_97/1            ..................................g......g..........................a........... [ 37440]
KM111295.1_Ken06.Bus             .....a............................g.......t.g.ta..c.....a.....a...t....g..c.t... [ 37440]
KM111294.1_Ken05/Tk1             .....a.................a..........g.......t.g.ta..c.....a.....a...t....g..c.t... [ 37440]
KM262844.1_strain_L60            ..................................g......g..........................a........... [ 37440]
KM102979.1_26544/OG10_Italy      ..................................g......g..........................a........... [ 37440]
KM262845.1_strain_NHV            -------------------------------------------------------------------------------- [ 37440]
KX354450.1_47/Ss/2008            ..................................g......g..........................a........... [ 37440]
KP055815.1_strain_BA71           ..................................g......g..........................a........... [ 37440]
MH025920.1_strain_R35            .....a............................g.......t.g.ta..c.....a.....a...t....g..c.t... [ 37440]
MH025919.1_strain_N10            .....a............................g.......t.g.ta..c.....a.....a...t....g..c.t... [ 37440]
MH025918.1_strain_R25            .....a............................g.......t.g.ta..c.....a.....a...t....g..c.t... [ 37440]
MH025917.1_strain_R7             .....a............................g.......t.g.ta..c.....a.....a...t....g..c.t... [ 37440]
MH025916.1_strain_R8             .....a............................g.......t.g.ta..c.....a.....a...t....g..c.t... [ 37440]
```

```
FR682468.2_Georgia_2007/1        gaggtcatagtccaccgctatcgcgtaccagtatttggttaaaacagtgctaaagctatcatcatggtccagcatgaagg [ 37520]
ASF_IND_20_CAD543                -------------------------------------------------------------------------------- [ 37520]
LR899193.1_Germany_2020/1        ................................................................................ [ 37520]
MN393477.1_Wuhan_2019            ................................................................................ [ 37520]
MW396979.1_Timor-Leste/2019/1    ................................................................................ [ 37520]
MK940252.1_CN/2019/InnerMongolia ................................................................................ [ 37520]
MK128995.1_China/2018/AnhuiXCGQ  ................................................................................ [ 37520]
MH681419.1_ASFV/POL/2015         ................................................................................ [ 37520]
LS478113.1_Estonia_2014          ................................................................................ [ 37520]
KP843857.1_Odintsovo_02/14       ................................................................................ [ 37520]
FN557520.1_strain_E75            .....................a..a............................................t.......... [ 37520]
AY261366.1_Warthog               ................................................a............................... [ 37520]
AY261365.1_Warmbaths             ................................................................................ [ 37520]
AY261364.1_Tengani_62            ................................................................................ [ 37520]
AY261363.1_Pretoriuskop/96/4     ................................................................................ [ 37520]
AY261362.1_Mkuzi_1979            .....................a..a............................................t.......... [ 37520]
AY261361.1_Malawi_Lil-20/1_1983  ....c.......t........g....................c.t.t........gag.tg..........a....t... [ 37520]
AY261360.1_Kenya_1950            ....c.g.....t........g....................c.t.t....g...gag.tg..........a....t... [ 37520]
AM712240.1_OURT_88/3             -------------------------------------------------------------------------------- [ 37520]
AM712239.1_Benin_97/1            .....................a..a............................................t.......... [ 37520]
KM111295.1_Ken06.Bus             ....c.g.....t........g....................c.tgt....g...gag.tg..........a....t... [ 37520]
KM111294.1_Ken05/Tk1             ....c.g.....t........g....................c.t.t....g...gag.tg..........a....t... [ 37520]
KM262844.1_strain_L60            .....................a..a............................................t.......... [ 37520]
KM102979.1_26544/OG10_Italy      .....................a..a............................................t.......... [ 37520]
KM262845.1_strain_NHV            -------------------------------------------------------------------------------- [ 37520]
KX354450.1_47/Ss/2008            .....................a..a............................................t.......... [ 37520]
KP055815.1_strain_BA71           .....................a..a............................................t.......... [ 37520]
MH025920.1_strain_R35            ....c.g.....t........g....................c.tgt....g...gag.tg..........a....t... [ 37520]
MH025919.1_strain_N10            ....c.g.....t........g....................c.tgt....g...gag.tg..........a....t... [ 37520]
MH025918.1_strain_R25            ....c.g.....t........g....................c.tgt....g...gag.tg..........a....t... [ 37520]
MH025917.1_strain_R7             ....c.g.....t........g....................c.tgt....g...gag.tg..........a....t... [ 37520]
MH025916.1_strain_R8             ....c.g.....t........g....................c.tgt....g...gag.tg..........a....t... [ 37520]

FR682468.2_Georgia_2007/1        ttatctccatgagccctcttaactcccacatgatttcccccctcagatccagattatctataatccttaaattggggtta [ 37600]
ASF_IND_20_CAD543                -------------------------------------------------------------------------------- [ 37600]
LR899193.1_Germany_2020/1        ................................................................................ [ 37600]
MN393477.1_Wuhan_2019            ................................................................................ [ 37600]
MW396979.1_Timor-Leste/2019/1    ................................................................................ [ 37600]
MK940252.1_CN/2019/InnerMongolia ................................................................................ [ 37600]
MK128995.1_China/2018/AnhuiXCGQ  ................................................................................ [ 37600]
MH681419.1_ASFV/POL/2015         ................................................................................ [ 37600]
LS478113.1_Estonia_2014          ................................................................................ [ 37600]
KP843857.1_Odintsovo_02/14       ................................................................................ [ 37600]
```

```
FN557520.1_strain_E75              ................................................................................ [ 37600]
AY261366.1_Warthog                 .....................................t.....t.................................... [ 37600]
AY261365.1_Warmbaths               ................................................................................ [ 37600]
AY261364.1_Tengani_62              ................................................................................ [ 37600]
AY261363.1_Pretoriuskop/96/4       ................................................................................ [ 37600]
AY261362.1_Mkuzi_1979              .....................................t.......................................... [ 37600]
AY261361.1_Malawi_Lil-20/1_1983    .............t..c....g................t............a......c.c.t.tt.c.gg......... [ 37600]
AY261360.1_Kenya_1950              .............t..c....g................t............a..c...c.c.tgtt.c.g.......... [ 37600]
AM712240.1_OURT_88/3               -------------------------------------------------------------------------------- [ 37600]
AM712239.1_Benin_97/1              ................................................................................ [ 37600]
KM111295.1_Ken06.Bus               .............t..c....g................t.......g....a..c...c.c.tgtt.c.g.......... [ 37600]
KM111294.1_Ken05/Tk1               ....t........t..c....g................t............a..c...c.c.tgtt.c.g.....c.... [ 37600]
KM262844.1_strain_L60              ................................................................................ [ 37600]
KM102979.1_26544/OG10_Italy        ................................................................................ [ 37600]
KM262845.1_strain_NHV              -------------------------------------------------------------------------------- [ 37600]
KX354450.1_47/Ss/2008              ................................................................................ [ 37600]
KP055815.1_strain_BA71             ................................................................................ [ 37600]
MH025920.1_strain_R35              .............t..c....g................t.......g....a..c...c.c.tgtt.c.g.......... [ 37600]
MH025919.1_strain_N10              .............t..c....g................t.......g....a..c...c.c.tgtt.c.g.......... [ 37600]
MH025918.1_strain_R25              .............t..c....g................t.......g....a..c...c.c.tgtt.c.g.......... [ 37600]
MH025917.1_strain_R7               .............t..c....g................t.......g....a..c...c.c.tgtt.c.g.......... [ 37600]
MH025916.1_strain_R8               .............t..c....g................t.......g....a..c...c.c.tgtt.c.g.......... [ 37600]

FR682468.2_Georgia_2007/1          ttggaaaacacctcgtggcaaaagataatattgctactggttttatcgcgcgttgtatcaaagaaaatttttaaaatata [ 37680]
ASF_IND_20_CAD543                  -------------------------------------------------------------------------------- [ 37680]
LR899193.1_Germany_2020/1          ................................................................................ [ 37680]
MN393477.1_Wuhan_2019              ................................................................................ [ 37680]
MW396979.1_Timor-Leste/2019/1      ................................................................................ [ 37680]
MK940252.1_CN/2019/InnerMongolia   ................................................................................ [ 37680]
MK128995.1_China/2018/AnhuiXCGQ    ................................................................................ [ 37680]
MH681419.1_ASFV/POL/2015           ................................................................................ [ 37680]
LS478113.1_Estonia_2014            ................................................................................ [ 37680]
KP843857.1_Odintsovo_02/14         ................................................................................ [ 37680]
FN557520.1_strain_E75              .....................................................c.......................... [ 37680]
AY261366.1_Warthog                 .................................t.............................................. [ 37680]
AY261365.1_Warmbaths               .................................t.............................................. [ 37680]
AY261364.1_Tengani_62              .................................t...................c.......................... [ 37680]
AY261363.1_Pretoriuskop/96/4       .................................t.............................................. [ 37680]
AY261362.1_Mkuzi_1979              .....................................................c.......................... [ 37680]
AY261361.1_Malawi_Lil-20/1_1983    ..........tt........t...........a..gtc......g.t......c.......................... [ 37680]
AY261360.1_Kenya_1950              ..........tt........t...........a..gtc......g.t......c.......................... [ 37680]
AM712240.1_OURT_88/3               -------------------------------------------------------------------------------- [ 37680]
AM712239.1_Benin_97/1              .....................................................c.......................... [ 37680]
KM111295.1_Ken06.Bus               ..........tt........t...........a..gtc......g.t......c.......................... [ 37680]
```

```
KM111294.1_Ken05/Tk1             ..........tt........t...........a..gtc......g.t......c.......................... [ 37680]
KM262844.1_strain_L60            .....................................................c.......................... [ 37680]
KM102979.1_26544/OG10_Italy      .....................................................c.......................... [ 37680]
KM262845.1_strain_NHV            -------------------------------------------------------------------------------- [ 37680]
KX354450.1_47/Ss/2008            .....................................................c.......................... [ 37680]
KP055815.1_strain_BA71           .....................................................c.......................... [ 37680]
MH025920.1_strain_R35            ..........tt........t...........a..gtc......g.t......c.......................... [ 37680]
MH025919.1_strain_N10            ..........tt........t...........a..gtc......g.t......c.......................... [ 37680]
MH025918.1_strain_R25            ..........tt........t...........a..gtc......g.t......c.......................... [ 37680]
MH025917.1_strain_R7             ..........tt........t...........a..gtc......g.t......c.......................... [ 37680]
MH025916.1_strain_R8             ..........tt........t...........a..gtc......g.t......c.......................... [ 37680]

FR682468.2_Georgia_2007/1        ctctctttctaaatattctttggctcccagctctttgcacagatcacgggtattttccgtgagagcacaaatcattccat [ 37760]
ASF_IND_20_CAD543                -------------------------------------------------------------------------------- [ 37760]
LR899193.1_Germany_2020/1        ................................................................................ [ 37760]
MN393477.1_Wuhan_2019            ................................................................................ [ 37760]
MW396979.1_Timor-Leste/2019/1    ................................................................................ [ 37760]
MK940252.1_CN/2019/InnerMongolia ................................................................................ [ 37760]
MK128995.1_China/2018/AnhuiXCGQ  ................................................................................ [ 37760]
MH681419.1_ASFV/POL/2015         ................................................................................ [ 37760]
LS478113.1_Estonia_2014          ................................................................................ [ 37760]
KP843857.1_Odintsovo_02/14       ................................................................................ [ 37760]
FN557520.1_strain_E75            ................................................................................ [ 37760]
AY261366.1_Warthog               ................................................................................ [ 37760]
AY261365.1_Warmbaths             ................................................................................ [ 37760]
AY261364.1_Tengani_62            ........................a....................................................... [ 37760]
AY261363.1_Pretoriuskop/96/4     ................................................................................ [ 37760]
AY261362.1_Mkuzi_1979            ................................................................................ [ 37760]
AY261361.1_Malawi_Lil-20/1_1983  a.....a......cga...........t..t...c....t....t......g.gc.....at.g..........g..... [ 37760]
AY261360.1_Kenya_1950            ......a......cgg...........a..t...c....t....tg.....g.gc....aat.........c..g..... [ 37760]
AM712240.1_OURT_88/3             -------------------------------------------------------------------------------- [ 37760]
AM712239.1_Benin_97/1            ................................................................................ [ 37760]
KM111295.1_Ken06.Bus             ....t.a......cgg........c..t..t...c....t.....g.....g.gc....aat.........c..g..... [ 37760]
KM111294.1_Ken05/Tk1             ......a......cgg...........a..t...c.a..t....tg.....g.gc....aat.........c..g..... [ 37760]
KM262844.1_strain_L60            ................................................................................ [ 37760]
KM102979.1_26544/OG10_Italy      ................................................................................ [ 37760]
KM262845.1_strain_NHV            -------------------------------------------------------------------------------- [ 37760]
KX354450.1_47/Ss/2008            ................................................................................ [ 37760]
KP055815.1_strain_BA71           ................................................................................ [ 37760]
MH025920.1_strain_R35            ....t.a......cgg........c..t..t...c....t.....g.....g.gc....aat.........c..g..... [ 37760]
MH025919.1_strain_N10            ....t.a......cgg........c..t..t...c....t.....g.....g.gc....aat.........c..g..... [ 37760]
MH025918.1_strain_R25            ....t.a......cgg........c..t..t...c....t.....g.....g.gc....aat.........c..g..... [ 37760]
MH025917.1_strain_R7             ....t.a......cgg........c..t..t...c....t.....g.....g.gc....aat.........c..g..... [ 37760]
MH025916.1_strain_R8             ....t.a......cgg........c..t..t...c....t.....g.....g.gc....aat.........c..g..... [ 37760]
```

```
FR682468.2_Georgia_2007/1        agttaatatctgcaccccattcagtaaacagctttatcaagtcatgattattctccttcacggctttcatcagtcctatg [ 37840]
ASF_IND_20_CAD543                -------------------------------------------------------------------------------- [ 37840]
LR899193.1_Germany_2020/1        ................................................................................ [ 37840]
MN393477.1_Wuhan_2019            ................................................................................ [ 37840]
MW396979.1_Timor-Leste/2019/1    ................................................................................ [ 37840]
MK940252.1_CN/2019/InnerMongolia ................................................................................ [ 37840]
MK128995.1_China/2018/AnhuiXCGQ  ................................................................................ [ 37840]
MH681419.1_ASFV/POL/2015         ................................................................................ [ 37840]
LS478113.1_Estonia_2014          ................................................................................ [ 37840]
KP843857.1_Odintsovo_02/14       ................................................................................ [ 37840]
FN557520.1_strain_E75            ................................................................................ [ 37840]
AY261366.1_Warthog               ................................................................................ [ 37840]
AY261365.1_Warmbaths             ................................................................................ [ 37840]
AY261364.1_Tengani_62            ................................................................................ [ 37840]
AY261363.1_Pretoriuskop/96/4     ................................................................................ [ 37840]
AY261362.1_Mkuzi_1979            ................................................................................ [ 37840]
AY261361.1_Malawi_Lil-20/1_1983  .a........g........c..g.c.......c.............g..g..t......g..............a.gg.. [ 37840]
AY261360.1_Kenya_1950            .a........g........c...ac.......c................g..t......g..............a.gg.. [ 37840]
AM712240.1_OURT_88/3             -------------------------------------------------------------------------------- [ 37840]
AM712239.1_Benin_97/1            ................................................................................ [ 37840]
KM111295.1_Ken06.Bus             .a........g........c........t.at.....t.g.........g..t......g..............a.gg.. [ 37840]
KM111294.1_Ken05/Tk1             .a........g........c...ac.......c................g..t......g..............a.gg.. [ 37840]
KM262844.1_strain_L60            ................................................................................ [ 37840]
KM102979.1_26544/OG10_Italy      ................................................................................ [ 37840]
KM262845.1_strain_NHV            -------------------------------------------------------------------------------- [ 37840]
KX354450.1_47/Ss/2008            ................................................................................ [ 37840]
KP055815.1_strain_BA71           ................................................................................ [ 37840]
MH025920.1_strain_R35            .a........g........c........t.at.....t.g.........g..t......g..............a.gg.. [ 37840]
MH025919.1_strain_N10            .a........g........c........t.at.....t.g.........g..t......g..............a.gg.. [ 37840]
MH025918.1_strain_R25            .a........g........c........t.at.....t.g.........g..t......g..............a.gg.. [ 37840]
MH025917.1_strain_R7             .a........g........c........t.at.....t.g.........g..t......g..............a.gg.. [ 37840]
MH025916.1_strain_R8             .a........g........c........t.at.....t.g.........g..t......g..............a.gg.. [ 37840]

FR682468.2_Georgia_2007/1        tttaactcgataccttgactaaaacaggttgaccttataaataatttattgcgtcgaatatgaagcataatggggccatt [ 37920]
ASF_IND_20_CAD543                -------------------------------------------------------------------------------- [ 37920]
LR899193.1_Germany_2020/1        ................................................................................ [ 37920]
MN393477.1_Wuhan_2019            ................................................................................ [ 37920]
MW396979.1_Timor-Leste/2019/1    ................................................................................ [ 37920]
MK940252.1_CN/2019/InnerMongolia ................................................................................ [ 37920]
MK128995.1_China/2018/AnhuiXCGQ  ................................................................................ [ 37920]
MH681419.1_ASFV/POL/2015         ................................................................................ [ 37920]
LS478113.1_Estonia_2014          ................................................................................ [ 37920]
KP843857.1_Odintsovo_02/14       ................................................................................ [ 37920]
```

```
FN557520.1_strain_E75            .......................g...................g......a............................. [ 37920]
AY261366.1_Warthog               .......................g...................t.c....a.................t........... [ 37920]
AY261365.1_Warmbaths             .......................g...................t.c....a.................t........... [ 37920]
AY261364.1_Tengani_62            .......................g...................g......a........c..............t..... [ 37920]
AY261363.1_Pretoriuskop/96/4     .......................g...................t.c....a.................t........... [ 37920]
AY261362.1_Mkuzi_1979            .......................g...................g......a............................. [ 37920]
AY261361.1_Malawi_Lil-20/1_1983  ..c..t..a.....c.cg.......................c.t...g.ga..ata..cg.a..atg....a.......c [ 37920]
AY261360.1_Kenya_1950            ..c..t..a.....c.cg...............t.......c.t...g..a..atacg...a..atg....a.......c [ 37920]
AM712240.1_OURT_88/3             -------------------------------------------------------------------------------- [ 37920]
AM712239.1_Benin_97/1            .......................g...................g......a............................. [ 37920]
KM111295.1_Ken06.Bus             ..c..t..a.....c.cg...............t.......c.t...g..a..atacg...a..atg....a.......c [ 37920]
KM111294.1_Ken05/Tk1             ..c..t..a.....c.cg...............t.......c.t...g..a..atacg...a..atg....a.......c [ 37920]
KM262844.1_strain_L60            .......................g...................g......a............................. [ 37920]
KM102979.1_26544/OG10_Italy      .......................g...................g......a............................. [ 37920]
KM262845.1_strain_NHV            -------------------------------------------------------------------------------- [ 37920]
KX354450.1_47/Ss/2008            .......................g...................g......a............................. [ 37920]
KP055815.1_strain_BA71           .......................g...................g......a............................. [ 37920]
MH025920.1_strain_R35            ..c..t..a.....c.cg...............t.......c.t...g..a..atacg...a..atg....a.......c [ 37920]
MH025919.1_strain_N10            ..c..t..a.....c.cg...............t.......c.t...g..a..atacg...a..atg....a.......c [ 37920]
MH025918.1_strain_R25            ..c..t..a.....c.cg...............t.......c.t...g..a..atacg...a..atg....a.......c [ 37920]
MH025917.1_strain_R7             ..c..t..a.....c.cg...............t.......c.t...g..a..atacg...a..atg....a.......c [ 37920]
MH025916.1_strain_R8             ..c..t..a.....c.cg...............t.......c.t...g..a..atacg...a..atg....a.......c [ 37920]

FR682468.2_Georgia_2007/1        atgccaccacaggccacaacacttcaggacatgatattgatctaccggtatacactgcccggccagtactttcttcgtga [ 38000]
ASF_IND_20_CAD543                -------------------------------------------------------------------------------- [ 38000]
LR899193.1_Germany_2020/1        ................................................................................ [ 38000]
MN393477.1_Wuhan_2019            ................................................................................ [ 38000]
MW396979.1_Timor-Leste/2019/1    ................................................................................ [ 38000]
MK940252.1_CN/2019/InnerMongolia ................................................................................ [ 38000]
MK128995.1_China/2018/AnhuiXCGQ  ................................................................................ [ 38000]
MH681419.1_ASFV/POL/2015         ................................................................................ [ 38000]
LS478113.1_Estonia_2014          ................................................................................ [ 38000]
KP843857.1_Odintsovo_02/14       ................................................................................ [ 38000]
FN557520.1_strain_E75            .............................t......a.....c....ac.cg............................ [ 38000]
AY261366.1_Warthog               .............................t......a.....c....a..cg............................ [ 38000]
AY261365.1_Warmbaths             .............................t......a.....c....ac.cg............................ [ 38000]
AY261364.1_Tengani_62            .............................t......a.....c....ac.cg............................ [ 38000]
AY261363.1_Pretoriuskop/96/4     .............................t......a..........ac.cg........a................... [ 38000]
AY261362.1_Mkuzi_1979            .............................t......a.....c....ac.cg............................ [ 38000]
AY261361.1_Malawi_Lil-20/1_1983  ..a........ac..g..g.....t.a.......................c......ttt......c........a..t. [ 38000]
AY261360.1_Kenya_1950            ..a........ac..g..g..t.......t...........................ttt......c........g.at. [ 38000]
AM712240.1_OURT_88/3             -------------------------------------------------------------------------------- [ 38000]
AM712239.1_Benin_97/1            .............................t......a.....c....ac.cg............................ [ 38000]
KM111295.1_Ken06.Bus             ..a........ac..g....tt.......t...........................ttt......c.....t..g.at. [ 38000]
```

```
KM111294.1_Ken05/Tk1           ..a........ac..g..g..t.......t...........................ttt......c........g.at. [ 38000]
KM262844.1_strain_L60          .............................t......a.....c....ac.cg............................ [ 38000]
KM102979.1_26544/OG10_Italy    .............................t......a.....c....ac.cg............................ [ 38000]
KM262845.1_strain_NHV          -------------------------------------------------------------------------------- [ 38000]
KX354450.1_47/Ss/2008          .............................t......a.....c....ac.cg............................ [ 38000]
KP055815.1_strain_BA71         .............................t......a.....c....ac.cg............................ [ 38000]
MH025920.1_strain_R35          ..a........ac..g....tt.......t...........................ttt......c.....t..g.at. [ 38000]
MH025919.1_strain_N10          ..a........ac..g....tt.......t...........................ttt......c.....t..g.at. [ 38000]
MH025918.1_strain_R25          ..a........ac..g....tt.......t...........................ttt......c.....t..g.at. [ 38000]
MH025917.1_strain_R7           ..a........ac..g....tt.......t...........................ttt......c.....t..g.at. [ 38000]
MH025916.1_strain_R8           ..a........ac..g....tt.......t...........................ttt......c.....t..g.at. [ 38000]

FR682468.2_Georgia_2007/1      gggattgcagggaaggcaacatgcctt-tccatcctttgacggaaatcaaattatctactaataactatcagtgtttata [ 38080]
ASF_IND_20_CAD543              -------------------------------------------------------------------------------- [ 38080]
LR899193.1_Germany_2020/1      ...........................-.................................................... [ 38080]
MN393477.1_Wuhan_2019          ...........................-.................................................... [ 38080]
MW396979.1_Timor-Leste/2019/1  ...........................-.................................................... [ 38080]
MK940252.1_CN/2019/InnerMongolia ...........................-.................................................... [ 38080]
MK128995.1_China/2018/AnhuiXCGQ ...........................-.................................................... [ 38080]
MH681419.1_ASFV/POL/2015       ...........................-.................................................... [ 38080]
LS478113.1_Estonia_2014        ...........................-.................................................... [ 38080]
KP843857.1_Odintsovo_02/14     ...........................-.................................................... [ 38080]
FN557520.1_strain_E75          ...........................-............t.................t.......c............. [ 38080]
AY261366.1_Warthog             .......................t...-............t..g..............---------------------. [ 38080]
AY261365.1_Warmbaths           .......................g...-............t.................t.......c..t.......... [ 38080]
AY261364.1_Tengani_62          ......................at...-............t.................t.....t.c..t.......... [ 38080]
AY261363.1_Pretoriuskop/96/4   .......................t...-............t.................---------------------. [ 38080]
AY261362.1_Mkuzi_1979          ...........................-............t.................t.......c............. [ 38080]
AY261361.1_Malawi_Lil-20/1_1983 aa.....t.a.................t..............ag..............t.-----t..aa.t......-- [ 38080]
AY261360.1_Kenya_1950          .........a.............t.c.t.t..........t.................t.------..aa.a.....ta. [ 38080]
AM712240.1_OURT_88/3           -------------------------------------------------------------------------------- [ 38080]
AM712239.1_Benin_97/1          ...........................-............t.................t.......c............. [ 38080]
KM111295.1_Ken06.Bus           .........a.............t.c.t.t..........t.................t.------..aa.a.....ta. [ 38080]
KM111294.1_Ken05/Tk1           .........a.............t.c.t.t...t......t.................t.------..aa.a.....ta. [ 38080]
KM262844.1_strain_L60          ...........................-............t.................t.......c............. [ 38080]
KM102979.1_26544/OG10_Italy    ...........................-............t.................t.......c............. [ 38080]
KM262845.1_strain_NHV          -------------------------------------------------------------------------------- [ 38080]
KX354450.1_47/Ss/2008          ...........................-............t.................t.......c............. [ 38080]
KP055815.1_strain_BA71         ...........................-............t.................t.......c............. [ 38080]
MH025920.1_strain_R35          .......t.a.............t.c.t.t..........t.................t.------..aa.a.....ta. [ 38080]
MH025919.1_strain_N10          .........a.............t.c.t.t..........t.................t.------..aa.a.....ta. [ 38080]
MH025918.1_strain_R25          .......t.a.............t.c.t.t..........t.................t.------..aa.a.....ta. [ 38080]
MH025917.1_strain_R7           .......t.a.............t.c.t.t..........t.................t.------..aa.a.....ta. [ 38080]
MH025916.1_strain_R8           .......t.a.............t.c.t.t..........t.................t.------..aa.a.....ta. [ 38080]
```

```
FR682468.2_Georgia_2007/1        ttaagtatttagatattatcccgggctggatacgtagtatcgctattcacatgtacttccaactctagccggagcctgca [ 38160]
ASF_IND_20_CAD543                -------------------------------------------------------------------------------- [ 38160]
LR899193.1_Germany_2020/1        ................................................................................ [ 38160]
MN393477.1_Wuhan_2019            ................................................................................ [ 38160]
MW396979.1_Timor-Leste/2019/1    ................................................................................ [ 38160]
MK940252.1_CN/2019/InnerMongolia ................................................................................ [ 38160]
MK128995.1_China/2018/AnhuiXCGQ  ................................................................................ [ 38160]
MH681419.1_ASFV/POL/2015         ................................................................................ [ 38160]
LS478113.1_Estonia_2014          ................................................................................ [ 38160]
KP843857.1_Odintsovo_02/14       ................................................................................ [ 38160]
FN557520.1_strain_E75            ...g......g...........a.................................................c....... [ 38160]
AY261366.1_Warthog               ..t.aa.a..gttg.a..c.taa..t.atg..aaa.a....------------------------------------... [ 38160]
AY261365.1_Warmbaths             ...g......g........................g..................g......gt..------.c....... [ 38160]
AY261364.1_Tengani_62            ...g...........................gg..g..............t...gt.....gt..------.c....... [ 38160]
AY261363.1_Pretoriuskop/96/4     ..t.aa.a..gttg.a..c.taa..t.atg..aaa.a....------------------------------------... [ 38160]
AY261362.1_Mkuzi_1979            ...--------.............a......................................................g [ 38160]
AY261361.1_Malawi_Lil-20/1_1983  -----------------------------------------------------------------------------.t. [ 38160]
AY261360.1_Kenya_1950            .agga..---------------a..t.att.-aaa.a......a---------------------------------.at [ 38160]
AM712240.1_OURT_88/3             -------------------------------------------------------------------------------- [ 38160]
AM712239.1_Benin_97/1            ...g......g...........a.................................................c....... [ 38160]
KM111295.1_Ken06.Bus             .ggg...---------------a..t..tt.-aaa.a.g...ta---------------------------------.ac [ 38160]
KM111294.1_Ken05/Tk1             .agga..---------------a..t.att..aaa.a......a---------------------------------.at [ 38160]
KM262844.1_strain_L60            ...g......g...........a.................................................c....... [ 38160]
KM102979.1_26544/OG10_Italy      ...g......g...........a.................................................c....... [ 38160]
KM262845.1_strain_NHV            -------------------------------------------------------------------------------- [ 38160]
KX354450.1_47/Ss/2008            ...g......g...........a.................................................c....... [ 38160]
KP055815.1_strain_BA71           ...g......g...........a.................................................c....... [ 38160]
MH025920.1_strain_R35            .ggg...---------------a..t..tt.-aaa.a.g...ta---------------------------------.ac [ 38160]
MH025919.1_strain_N10            .ggg...---------------a..t..tt.-aaa.a.g...ta---------------------------------.ac [ 38160]
MH025918.1_strain_R25            .ggg...---------------a..t..tt.-aaa.a.g...ta---------------------------------.ac [ 38160]
MH025917.1_strain_R7             .ggg...---------------a..t..tt.-aaa.a.g...ta---------------------------------.ac [ 38160]
MH025916.1_strain_R8             .ggg...---------------a..t..tt.-aaa.a.g...ta---------------------------------.ac [ 38160]

FR682468.2_Georgia_2007/1        gggtcatttatttttaat-a--------------------------------ttgattc-tttttgtatttaatcattta [ 38240]
ASF_IND_20_CAD543                -------------------------------------------------------------------------------- [ 38240]
LR899193.1_Germany_2020/1        ..................-.--------------------------------.......-.................... [ 38240]
MN393477.1_Wuhan_2019            ..................-.--------------------------------.......-.................... [ 38240]
MW396979.1_Timor-Leste/2019/1    ..................-.--------------------------------.......-.................... [ 38240]
MK940252.1_CN/2019/InnerMongolia ..................-.--------------------------------.......-.................... [ 38240]
MK128995.1_China/2018/AnhuiXCGQ  ..................-.--------------------------------.......-.................... [ 38240]
MH681419.1_ASFV/POL/2015         ..................-.--------------------------------.......-.................... [ 38240]
LS478113.1_Estonia_2014          ..................-.--------------------------------.......-.................... [ 38240]
KP843857.1_Odintsovo_02/14       ..................-.--------------------------------.......-.................... [ 38240]
```

```
FN557520.1_strain_E75          ..-..............-.-------------------------------......-..................... [ 38240]
AY261366.1_Warthog             .atatcc...a......c-.a------------taaaac----taataaat.....c.-..............t..... [ 38240]
AY261365.1_Warmbaths           ...c.............-.-------------------------------......-..................... [ 38240]
AY261364.1_Tengani_62          ...c.............-.-------------------------------.......t.................... [ 38240]
AY261363.1_Pretoriuskop/96/4   .atatcc...a......c-ga------------taaaacagtgtaataaat.......t.............t..... [ 38240]
AY261362.1_Mkuzi_1979          ..........a......-.-------------------------------......-..................... [ 38240]
AY261361.1_Malawi_Lil-20/1_1983 cacaacc...a.......-ga------------taaaacagtgcgataaaa.....c.t......gg.....t..... [ 38240]
AY261360.1_Kenya_1950          ata..cc..ga...g..c-gataaaaactcccataaaactgtgccataaaa.....c.-......gg......t.c..g [ 38240]
AM712240.1_OURT_88/3           ------------------------------------------------------------------------------ [ 38240]
AM712239.1_Benin_97/1          ..-..............-.-------------------------------......-..................... [ 38240]
KM111295.1_Ken06.Bus           ata..cc..ga...g..cggataaatactgccataaaactgtgccataaaa.....c.-......gg......t.c..g [ 38240]
KM111294.1_Ken05/Tk1           ata..cc..ga...g..c-gataaaaactcccataaaactgtgccataaaa.....c.-......gg......t.c..g [ 38240]
KM262844.1_strain_L60          ..-..............-.-------------------------------......-..................... [ 38240]
KM102979.1_26544/OG10_Italy    ..-..............-.-------------------------------......-..................... [ 38240]
KM262845.1_strain_NHV          ------------------------------------------------------------------------------ [ 38240]
KX354450.1_47/Ss/2008          ..-..............-.-------------------------------......-..................... [ 38240]
KP055815.1_strain_BA71         ..-..............-.-------------------------------......-..................... [ 38240]
MH025920.1_strain_R35          ata..cc..ga...g..cggataaatactgccataaaactgtgccataaaa.....c.-......gg......t.c..g [ 38240]
MH025919.1_strain_N10          ata..cc..ga...g..cggataaatactgccataaaactgtgccataaaa.....c.-......gg......t.c..g [ 38240]
MH025918.1_strain_R25          ata..cc..ga...g..cggataaatactgccataaaactgtgccataaaa.....c.-......gg......t.c..g [ 38240]
MH025917.1_strain_R7           ata..cc..ga...g..cggataaatactgccataaaactgtgccataaaa.....c.-......gg......t.c..g [ 38240]
MH025916.1_strain_R8           ata..cc..ga...g..cggataaatactgccataaaactgtgccataaaa.....c.-......gg......t.c..g [ 38240]

FR682468.2_Georgia_2007/1      g----agaaggtcatcataggagccagatgttctctctccagaacttatgtcgaaaaacattacctaaccgtaaacttcc [ 38320]
ASF_IND_20_CAD543              -------------------------------------------------------------------------------- [ 38320]
LR899193.1_Germany_2020/1      .----........................................................................... [ 38320]
MN393477.1_Wuhan_2019          .----........................................................................... [ 38320]
MW396979.1_Timor-Leste/2019/1  .----........................................................................... [ 38320]
MK940252.1_CN/2019/InnerMongolia .----........................................................................... [ 38320]
MK128995.1_China/2018/AnhuiXCGQ .----........................................................................... [ 38320]
MH681419.1_ASFV/POL/2015       .----........................................................................... [ 38320]
LS478113.1_Estonia_2014        .----........................................................................... [ 38320]
KP843857.1_Odintsovo_02/14     .----........................................................................... [ 38320]
FN557520.1_strain_E75          .----...........................................................g..t........... [ 38320]
AY261366.1_Warthog             aaa-t..t..a...ctgc.............t.............g.................g..t........... [ 38320]
AY261365.1_Warmbaths           .----...................................c...........t.....g..t........... [ 38320]
AY261364.1_Tengani_62          .----...........................t......c.......................g..t........... [ 38320]
AY261363.1_Pretoriuskop/96/4   aaa-t..t..a...cggc..............................................g..t........... [ 38320]
AY261362.1_Mkuzi_1979          .----...........................................................g.tt........... [ 38320]
AY261361.1_Malawi_Lil-20/1_1983 aaatagtg..aa..ctgc..............................................g..t........... [ 38320]
AY261360.1_Kenya_1950          aaa-t..tg.....ctgc......t.......t......................c......g.............. [ 38320]
AM712240.1_OURT_88/3           -------------------------------------------------------------------------------- [ 38320]
AM712239.1_Benin_97/1          .----...........................................................g..t........... [ 38320]
KM111295.1_Ken06.Bus           aaa-t..tg.....ctgc.a...................................c......g..t........... [ 38320]
```

```
KM111294.1_Ken05/Tk1             aaa-t..tg.....ctgc......t................a.................c......g..t.......... [ 38320]
KM262844.1_strain_L60            .----.............................................................g..t.......... [ 38320]
KM102979.1_26544/OG10_Italy      .----.............................................................g..t.......... [ 38320]
KM262845.1_strain_NHV            -------------------------------------------------------------------------------- [ 38320]
KX354450.1_47/Ss/2008            .----.............................................................g..t.......... [ 38320]
KP055815.1_strain_BA71           .----.............................................................g..t.......... [ 38320]
MH025920.1_strain_R35            aaa-t..tg.....ctgc.a.......................................c......g..t.......... [ 38320]
MH025919.1_strain_N10            aaa-t..tg.....ctgc.a.......................................c......g..t.......... [ 38320]
MH025918.1_strain_R25            aaa-t..tg.....ctgc.a.......................................c......g..t.......... [ 38320]
MH025917.1_strain_R7             aaa-t..tg.....ctgc.a.......................................c......g..t.......... [ 38320]
MH025916.1_strain_R8             aaa-t..tg.....ctgc.a.......................................c......g..t.......... [ 38320]

FR682468.2_Georgia_2007/1        tgaattttttgacgaatatatattacaactgctgggattatactgggaaaaccatggaactattcaacgagcaggaaaca [ 38400]
ASF_IND_20_CAD543                -------------------------------------------------------------------------------- [ 38400]
LR899193.1_Germany_2020/1        ................................................................................ [ 38400]
MN393477.1_Wuhan_2019            ................................................................................ [ 38400]
MW396979.1_Timor-Leste/2019/1    ................................................................................ [ 38400]
MK940252.1_CN/2019/InnerMongolia ................................................................................ [ 38400]
MK128995.1_China/2018/AnhuiXCGQ  ................................................................................ [ 38400]
MH681419.1_ASFV/POL/2015         ................................................................................ [ 38400]
LS478113.1_Estonia_2014          ................................................................................ [ 38400]
KP843857.1_Odintsovo_02/14       ................................................................................ [ 38400]
FN557520.1_strain_E75            ...............t.....................................................g.......... [ 38400]
AY261366.1_Warthog               ................................................................................ [ 38400]
AY261365.1_Warmbaths             ................................................................................ [ 38400]
AY261364.1_Tengani_62            ................................................................................ [ 38400]
AY261363.1_Pretoriuskop/96/4     ................................................................................ [ 38400]
AY261362.1_Mkuzi_1979            ...............t.....................................................g.......... [ 38400]
AY261361.1_Malawi_Lil-20/1_1983  ................................................................................ [ 38400]
AY261360.1_Kenya_1950            ...............t...g........................................g................... [ 38400]
AM712240.1_OURT_88/3             -------------------------------------------------------------------------------- [ 38400]
AM712239.1_Benin_97/1            ...............t.....................................................g.......... [ 38400]
KM111295.1_Ken06.Bus             ...g...........t...g................c..............t.....t..g................... [ 38400]
KM111294.1_Ken05/Tk1             ...............t...g........................................g................... [ 38400]
KM262844.1_strain_L60            ...............t.....................................................g.......... [ 38400]
KM102979.1_26544/OG10_Italy      ...............t.....................................................g.......... [ 38400]
KM262845.1_strain_NHV            -------------------------------------------------------------------------------- [ 38400]
KX354450.1_47/Ss/2008            ...............t.....................................................g.......... [ 38400]
KP055815.1_strain_BA71           ...............t.....................................................g.......... [ 38400]
MH025920.1_strain_R35            ...g...........t...g................c..............t.....t..g................... [ 38400]
MH025919.1_strain_N10            ...g...........t...g................c..............t.....t..g................... [ 38400]
MH025918.1_strain_R25            ...g...........t...g................c..............t.....t..g................... [ 38400]
MH025917.1_strain_R7             ...g...........t...g................c..............t.....t..g................... [ 38400]
MH025916.1_strain_R8             ...g...........t...g................c..............t.....t..g................... [ 38400]
```

```
FR682468.2_Georgia_2007/1        actgtgtgcttatacagcaacataccctcattcccgtaaatgaagccctgagaacagcagcatctgaagaaaattatgag [ 38480]
ASF_IND_20_CAD543                -------------------------------------------------------------------------------- [ 38480]
LR899193.1_Germany_2020/1        ................................................................................ [ 38480]
MN393477.1_Wuhan_2019            ................................................................................ [ 38480]
MW396979.1_Timor-Leste/2019/1    ................................................................................ [ 38480]
MK940252.1_CN/2019/InnerMongolia ................................................................................ [ 38480]
MK128995.1_China/2018/AnhuiXCGQ  ................................................................................ [ 38480]
MH681419.1_ASFV/POL/2015         ................................................................................ [ 38480]
LS478113.1_Estonia_2014          ................................................................................ [ 38480]
KP843857.1_Odintsovo_02/14       ................................................................................ [ 38480]
FN557520.1_strain_E75            ................a................................a....t......................... [ 38480]
AY261366.1_Warthog               ........................................c.............t......................... [ 38480]
AY261365.1_Warmbaths             .....................g................................t......................... [ 38480]
AY261364.1_Tengani_62            ........................a........................a....t......................... [ 38480]
AY261363.1_Pretoriuskop/96/4     ...........g....a.......................c.............t......g.................. [ 38480]
AY261362.1_Mkuzi_1979            ............................t......a............................................ [ 38480]
AY261361.1_Malawi_Lil-20/1_1983  .............c..........a...t....................a....t......................... [ 38480]
AY261360.1_Kenya_1950            .......a.....c..a.......a........................a....t......................... [ 38480]
AM712240.1_OURT_88/3             -------------------------------------------------------------------------------- [ 38480]
AM712239.1_Benin_97/1            ................a................................a....t......................... [ 38480]
KM111295.1_Ken06.Bus             .......a.....c..a.......a........................a....t............g............ [ 38480]
KM111294.1_Ken05/Tk1             .......a.....c..a.......a........................a....t......................... [ 38480]
KM262844.1_strain_L60            ................a................................a....t......................... [ 38480]
KM102979.1_26544/OG10_Italy      ................a................................a....t......................... [ 38480]
KM262845.1_strain_NHV            -------------------------------------------------------------------------------- [ 38480]
KX354450.1_47/Ss/2008            ................a................................a....t......................... [ 38480]
KP055815.1_strain_BA71           ................a................................a....t......................... [ 38480]
MH025920.1_strain_R35            .......a.....c..a.......a........................a....t............g............ [ 38480]
MH025919.1_strain_N10            .......a.....c..a.......a........................a....t............g............ [ 38480]
MH025918.1_strain_R25            .......a.....c..a.......a........................a....t............g............ [ 38480]
MH025917.1_strain_R7             .......a.....c..a.......a........................a....t............g............ [ 38480]
MH025916.1_strain_R8             .......a.....c..a.......a........................a....t............g............ [ 38480]

FR682468.2_Georgia_2007/1        atcgtgagccttttattagcgtgggaggggaacctttactatgctattataggggctctagagggcaaccgccacgactt [ 38560]
ASF_IND_20_CAD543                -------------------------------------------------------------------------------- [ 38560]
LR899193.1_Germany_2020/1        ................................................................................ [ 38560]
MN393477.1_Wuhan_2019            ................................................................................ [ 38560]
MW396979.1_Timor-Leste/2019/1    ................................................................................ [ 38560]
MK940252.1_CN/2019/InnerMongolia ................................................................................ [ 38560]
MK128995.1_China/2018/AnhuiXCGQ  ................................................................................ [ 38560]
MH681419.1_ASFV/POL/2015         ................................................................................ [ 38560]
LS478113.1_Estonia_2014          ................................................................................ [ 38560]
KP843857.1_Odintsovo_02/14       ................................................................................ [ 38560]
```

```
FN557520.1_strain_E75          ......g..........................................................t......t..a.... [ 38560]
AY261366.1_Warthog             .............................a.......................a.......................... [ 38560]
AY261365.1_Warmbaths           ......g..............................................a.......................... [ 38560]
AY261364.1_Tengani_62          ........t...........a........................................................... [ 38560]
AY261363.1_Pretoriuskop/96/4   .............................a.......................a.......................... [ 38560]
AY261362.1_Mkuzi_1979          ....................a...........t............................................... [ 38560]
AY261361.1_Malawi_Lil-20/1_1983 .............................a...........................................c...... [ 38560]
AY261360.1_Kenya_1950          ....................t............................................t..t......a.... [ 38560]
AM712240.1_OURT_88/3           -------------------------------------------------------------------------------- [ 38560]
AM712239.1_Benin_97/1          ......g..........................................................t......t..a.... [ 38560]
KM111295.1_Ken06.Bus           .................................................................g....a....a.... [ 38560]
KM111294.1_Ken05/Tk1           .................................................................t.........a.... [ 38560]
KM262844.1_strain_L60          ......g..........................................................t......t..a.... [ 38560]
KM102979.1_26544/OG10_Italy    ......g..........................................................t......t..a.... [ 38560]
KM262845.1_strain_NHV          -------------------------------------------------------------------------------- [ 38560]
KX354450.1_47/Ss/2008          ......g..........................................................t......t..a.... [ 38560]
KP055815.1_strain_BA71         ......g..........................................................t......t..a.... [ 38560]
MH025920.1_strain_R35          ..........................a......................................g....a....a.... [ 38560]
MH025919.1_strain_N10          ..........................a......................................g....a....a.... [ 38560]
MH025918.1_strain_R25          ..........................a......................................g....a....a.... [ 38560]
MH025917.1_strain_R7           ..........................a......................................g....a....a.... [ 38560]
MH025916.1_strain_R8           ..........................a......................................g....a....a.... [ 38560]

FR682468.2_Georgia_2007/1      aattcgtaaatatgatgaccaaatcaaggaccatcatgaaattctgccattcattgacgatccagtcatatttcacaaat [ 38640]
ASF_IND_20_CAD543              -------------------------------------------------------------------------------- [ 38640]
LR899193.1_Germany_2020/1      ................................................................................ [ 38640]
MN393477.1_Wuhan_2019          ................................................................................ [ 38640]
MW396979.1_Timor-Leste/2019/1  ................................................................................ [ 38640]
MK940252.1_CN/2019/InnerMongolia ................................................................................ [ 38640]
MK128995.1_China/2018/AnhuiXCGQ ................................................................................ [ 38640]
MH681419.1_ASFV/POL/2015       ................................................................................ [ 38640]
LS478113.1_Estonia_2014        ................................................................................ [ 38640]
KP843857.1_Odintsovo_02/14     ................................................................................ [ 38640]
FN557520.1_strain_E75          ..................t....................c.................t......a............... [ 38640]
AY261366.1_Warthog             ................................................................................ [ 38640]
AY261365.1_Warmbaths           ................................................................................ [ 38640]
AY261364.1_Tengani_62          .........g........................................................a............. [ 38640]
AY261363.1_Pretoriuskop/96/4   ..........................................................a..................... [ 38640]
AY261362.1_Mkuzi_1979          ..................................................................a............. [ 38640]
AY261361.1_Malawi_Lil-20/1_1983 ...........................................t..........c.........a............... [ 38640]
AY261360.1_Kenya_1950          ...................................................t............................ [ 38640]
AM712240.1_OURT_88/3           -------------------------------------------------------------------------------- [ 38640]
AM712239.1_Benin_97/1          ..................t....................c.................t......a............... [ 38640]
KM111295.1_Ken06.Bus           ............c......................................t............................ [ 38640]
```

```
KM111294.1_Ken05/Tk1             ...................................................t..............t............. [ 38640]
KM262844.1_strain_L60            ..................t....................c.................t......a............... [ 38640]
KM102979.1_26544/OG10_Italy      ..................t....................c.................t......a............... [ 38640]
KM262845.1_strain_NHV            -------------------------------------------------------------------------------- [ 38640]
KX354450.1_47/Ss/2008            ..................t....................c.................t......a............... [ 38640]
KP055815.1_strain_BA71           ..................t....................c.................t......a............... [ 38640]
MH025920.1_strain_R35            ............c......................................t............................ [ 38640]
MH025919.1_strain_N10            ............c......................................t............................ [ 38640]
MH025918.1_strain_R25            ............c......................................t............................ [ 38640]
MH025917.1_strain_R7             ............c......................................t............................ [ 38640]
MH025916.1_strain_R8             ............c......................................t............................ [ 38640]

FR682468.2_Georgia_2007/1        gccatatcatgcggcaatgcttttttgattgtattttatatcaagctgtaaaatatagtaagtttcgcgttcttctttac [ 38720]
ASF_IND_20_CAD543                -------------------------------------------------------------------------------- [ 38720]
LR899193.1_Germany_2020/1        ................................................................................ [ 38720]
MN393477.1_Wuhan_2019            ................................................................................ [ 38720]
MW396979.1_Timor-Leste/2019/1    ................................................................................ [ 38720]
MK940252.1_CN/2019/InnerMongolia ................................................................................ [ 38720]
MK128995.1_China/2018/AnhuiXCGQ  ................................................................................ [ 38720]
MH681419.1_ASFV/POL/2015         ................................................................................ [ 38720]
LS478113.1_Estonia_2014          ................................................................................ [ 38720]
KP843857.1_Odintsovo_02/14       ................................................................................ [ 38720]
FN557520.1_strain_E75            ...............g...................................................t...........t [ 38720]
AY261366.1_Warthog               ...............g...............................................................t [ 38720]
AY261365.1_Warmbaths             .......a..c..a.g..........a..................................................... [ 38720]
AY261364.1_Tengani_62            ..............................................................................ct [ 38720]
AY261363.1_Pretoriuskop/96/4     .......a.......g...............................................................t [ 38720]
AY261362.1_Mkuzi_1979            ...............g...............................................................t [ 38720]
AY261361.1_Malawi_Lil-20/1_1983  .t..c..a.......g..........a........................................t...........t [ 38720]
AY261360.1_Kenya_1950            .t.....g.......g..........a..........g...........................a..........a..t [ 38720]
AM712240.1_OURT_88/3             -------------------------------------------------------------------------------- [ 38720]
AM712239.1_Benin_97/1            ...............g...................................................t...........t [ 38720]
KM111295.1_Ken06.Bus             .t....ca.......g..........a.....................................ca..........a... [ 38720]
KM111294.1_Ken05/Tk1             .t.....a.......g..........a.......g..g...........................a..........a..t [ 38720]
KM262844.1_strain_L60            ...............g...................................................t...........t [ 38720]
KM102979.1_26544/OG10_Italy      ...............g...................................................t...........t [ 38720]
KM262845.1_strain_NHV            -------------------------------------------------------------------------------- [ 38720]
KX354450.1_47/Ss/2008            ...............g...................................................t...........t [ 38720]
KP055815.1_strain_BA71           ...............g...................................................t...........t [ 38720]
MH025920.1_strain_R35            .t....ca.......g..........a.....................................ca..........a... [ 38720]
MH025919.1_strain_N10            .t....ca.......g..........a.....................................ca..........a... [ 38720]
MH025918.1_strain_R25            .t....ca.......g..........a.....................................ca..........a... [ 38720]
MH025917.1_strain_R7             .t....ca.......g..........a.....................................ca..........a... [ 38720]
MH025916.1_strain_R8             .t....ca.......g..........a.....................................ca..........a... [ 38720]
```

```
FR682468.2_Georgia_2007/1        tttaaacatagattagaggatgatttgcccttcactcatttacttattgaaaaggcatgtaaagatcataattatgaagt [ 38800]
ASF_IND_20_CAD543                -------------------------------------------------------------------------------- [ 38800]
LR899193.1_Germany_2020/1        ................................................................................ [ 38800]
MN393477.1_Wuhan_2019            ................................................................................ [ 38800]
MW396979.1_Timor-Leste/2019/1    ................................................................................ [ 38800]
MK940252.1_CN/2019/InnerMongolia ................................................................................ [ 38800]
MK128995.1_China/2018/AnhuiXCGQ  ................................................................................ [ 38800]
MH681419.1_ASFV/POL/2015         ................................................................................ [ 38800]
LS478113.1_Estonia_2014          ................................................................................ [ 38800]
KP843857.1_Odintsovo_02/14       ................................................................................ [ 38800]
FN557520.1_strain_E75            ......t...c...................c..gt.........................g................... [ 38800]
AY261366.1_Warthog               ......t....g......a...........c..g..............a...........g................... [ 38800]
AY261365.1_Warmbaths             ..............................c.............................g................... [ 38800]
AY261364.1_Tengani_62            ...............ag............................................................... [ 38800]
AY261363.1_Pretoriuskop/96/4     ......t....g......a...........c..g...........g..............g................... [ 38800]
AY261362.1_Mkuzi_1979            ......t....g..............a...c.............................g..a................ [ 38800]
AY261361.1_Malawi_Lil-20/1_1983  ...........g....g.............c.............................g................... [ 38800]
AY261360.1_Kenya_1950            ......t...t....a....ga........c..gt.....c...............g...g................... [ 38800]
AM712240.1_OURT_88/3             -------------------------------------------------------------------------------- [ 38800]
AM712239.1_Benin_97/1            ......t...c...................c..gt.........................g................... [ 38800]
KM111295.1_Ken06.Bus             ......t...t....a....ga........c..gtc..c.c...............g.a....t.............g.. [ 38800]
KM111294.1_Ken05/Tk1             ......t...t....a....aa........c..gtc..c.c...c........a..g.a....t.............g.. [ 38800]
KM262844.1_strain_L60            ......t...c...................c..gt.........................g................... [ 38800]
KM102979.1_26544/OG10_Italy      ......t...c...................c..gt.........................g................... [ 38800]
KM262845.1_strain_NHV            -------------------------------------------------------------------------------- [ 38800]
KX354450.1_47/Ss/2008            ......t...c...................c..gt.........................g................... [ 38800]
KP055815.1_strain_BA71           ......t...c...................c..gt.........................g................... [ 38800]
MH025920.1_strain_R35            ......t...t....a....ga........c..gtc..c.c...............g.a....t.............g.. [ 38800]
MH025919.1_strain_N10            ......t...t....a....ga........c..gtc..c.c...............g.a....t.............g.. [ 38800]
MH025918.1_strain_R25            ......t...t....a....ga........c..gtc..c.c...............g.a....t.............g.. [ 38800]
MH025917.1_strain_R7             ......t...t....a....ga........c..gtc..c.c...............g.a....t.............g.. [ 38800]
MH025916.1_strain_R8             ......t...t....a....ga........c..gtc..c.c...............g.a....t.............g.. [ 38800]

FR682468.2_Georgia_2007/1        tattaaatggatatatgaaaacctacatatctacaatatgatagatacctttgaatgtgctattgcccataaggatctac [ 38880]
ASF_IND_20_CAD543                -------------------------------------------------------------------------------- [ 38880]
LR899193.1_Germany_2020/1        ................................................................................ [ 38880]
MN393477.1_Wuhan_2019            ................................................................................ [ 38880]
MW396979.1_Timor-Leste/2019/1    ................................................................................ [ 38880]
MK940252.1_CN/2019/InnerMongolia ................................................................................ [ 38880]
MK128995.1_China/2018/AnhuiXCGQ  ................................................................................ [ 38880]
MH681419.1_ASFV/POL/2015         ................................................................................ [ 38880]
LS478113.1_Estonia_2014          ................................................................................ [ 38880]
KP843857.1_Odintsovo_02/14       ................................................................................ [ 38880]
```

```
FN557520.1_strain_E75           ............................g...g.c....a..............c.................a....... [ 38880]
AY261366.1_Warthog              .......................................a..g..........g.......................... [ 38880]
AY261365.1_Warmbaths            .............................c....g....a..g..................................... [ 38880]
AY261364.1_Tengani_62           a............................................................................... [ 38880]
AY261363.1_Pretoriuskop/96/4    ..........t............................a..g..g.................................. [ 38880]
AY261362.1_Mkuzi_1979           .....g.......................................................................... [ 38880]
AY261361.1_Malawi_Lil-20/1_1983 ............c................g.........a..g..................................... [ 38880]
AY261360.1_Kenya_1950           ............c....................tg.a..a..g.........a...................a....... [ 38880]
AM712240.1_OURT_88/3            -------------------------------------------------------------------------------- [ 38880]
AM712239.1_Benin_97/1           ............................g...g.c....a..............c.................a....... [ 38880]
KM111295.1_Ken06.Bus            ............c....................tg....c...a....t...a...................a....... [ 38880]
KM111294.1_Ken05/Tk1            ............c........t...........tg....c...a....t...a...a...............a....... [ 38880]
KM262844.1_strain_L60           ............................g...g.c....a..............c.................a....... [ 38880]
KM102979.1_26544/OG10_Italy     ............................g...g.c....a..............c.................a....... [ 38880]
KM262845.1_strain_NHV           -------------------------------------------------------------------------------- [ 38880]
KX354450.1_47/Ss/2008           ............................g...g.c....a..............c.................a....... [ 38880]
KP055815.1_strain_BA71          ............................g...g.c....a..............c.................a....... [ 38880]
MH025920.1_strain_R35           ............c....................tg....c...a....t...a...................a....... [ 38880]
MH025919.1_strain_N10           ............c....................tg....c...a....t...a...................a....... [ 38880]
MH025918.1_strain_R25           ............c....................tg....c...a....t...a...................a....... [ 38880]
MH025917.1_strain_R7            ............c....................tg....c...a....t...a...................a....... [ 38880]
MH025916.1_strain_R8            ............c....................tg....c...a....t...a...................a....... [ 38880]

FR682468.2_Georgia_2007/1       atctatattgtttggggtatagatttatatataacagaatcgtacccgataagtatcatcatttagatattcgcatgctt [ 38960]
ASF_IND_20_CAD543               -------------------------------------------------------------------------------- [ 38960]
LR899193.1_Germany_2020/1       ................................................................................ [ 38960]
MN393477.1_Wuhan_2019           ................................................................................ [ 38960]
MW396979.1_Timor-Leste/2019/1   ................................................................................ [ 38960]
MK940252.1_CN/2019/InnerMongolia ................................................................................ [ 38960]
MK128995.1_China/2018/AnhuiXCGQ ................................................................................ [ 38960]
MH681419.1_ASFV/POL/2015        ................................................................................ [ 38960]
LS478113.1_Estonia_2014         ................................................................................ [ 38960]
KP843857.1_Odintsovo_02/14      ................................................................................ [ 38960]
FN557520.1_strain_E75           g.t..................c.........c.....g..t......t........................t...a... [ 38960]
AY261366.1_Warthog              g.......c............c...............g.........t...........t........g...t...t... [ 38960]
AY261365.1_Warmbaths            g....................c...............g.........t..g.....................t...t... [ 38960]
AY261364.1_Tengani_62           g....................................g.......................................... [ 38960]
AY261363.1_Pretoriuskop/96/4    g.......c............c...............g.........t...........t........g...t...t... [ 38960]
AY261362.1_Mkuzi_1979           g....................c...............g.........a..............a.........t...t... [ 38960]
AY261361.1_Malawi_Lil-20/1_1983 g....................c............t..g.........t.......................t....a... [ 38960]
AY261360.1_Kenya_1950           ..t.....a....a..a....c............t..g.........t............................a... [ 38960]
AM712240.1_OURT_88/3            -------------------------------------------------------------------------------- [ 38960]
AM712239.1_Benin_97/1           g.t..................c.........c.....g..t......t........................t...a... [ 38960]
KM111295.1_Ken06.Bus            g.t..........a.......c....g.......t............t........t...................c... [ 38960]
```

```
KM111294.1_Ken05/Tk1             g.t..........a.......c....g.......t..g.........t........t...................c... [ 38960]
KM262844.1_strain_L60            g.t..................c.........c.....g..t......t........................t...a... [ 38960]
KM102979.1_26544/OG10_Italy      g.t..................c.........c.....g..t......t........................t...a... [ 38960]
KM262845.1_strain_NHV            -------------------------------------------------------------------------------- [ 38960]
KX354450.1_47/Ss/2008            g.t..................c.........c.....g..t......t........................t...a... [ 38960]
KP055815.1_strain_BA71           g.t..................c.........c.....g..t......t........................t...a... [ 38960]
MH025920.1_strain_R35            g.t..........a.......c....g.......t............t........t...................c... [ 38960]
MH025919.1_strain_N10            g.t..........a.......c....g.......t............t........t...................c... [ 38960]
MH025918.1_strain_R25            g.t..........a.......c....g.......t............t........t...................c... [ 38960]
MH025917.1_strain_R7             g.t..........a.......c....g.......t............t........t...................c... [ 38960]
MH025916.1_strain_R8             g.t..........a.......c....g.......t............t........t...................c... [ 38960]

FR682468.2_Georgia_2007/1        tcaagcctacaactcctacataaggtggcagccaaaggatacttagattttatcctagaaaccttaaagtatgatcataa [ 39040]
ASF_IND_20_CAD543                -------------------------------------------------------------------------------- [ 39040]
LR899193.1_Germany_2020/1        ................................................................................ [ 39040]
MN393477.1_Wuhan_2019            ................................................................................ [ 39040]
MW396979.1_Timor-Leste/2019/1    ................................................................................ [ 39040]
MK940252.1_CN/2019/InnerMongolia ................................................................................ [ 39040]
MK128995.1_China/2018/AnhuiXCGQ  ................................................................................ [ 39040]
MH681419.1_ASFV/POL/2015         ................................................................................ [ 39040]
LS478113.1_Estonia_2014          ................................................................................ [ 39040]
KP843857.1_Odintsovo_02/14       ................................................................................ [ 39040]
FN557520.1_strain_E75            .............................g.................................................. [ 39040]
AY261366.1_Warthog               ...g.......t.....gt............................................................. [ 39040]
AY261365.1_Warmbaths             .................g.............................................................. [ 39040]
AY261364.1_Tengani_62            .................g.............................................................. [ 39040]
AY261363.1_Pretoriuskop/96/4     ...g.......t.....gt............................................................. [ 39040]
AY261362.1_Mkuzi_1979            .................g.............................c................................ [ 39040]
AY261361.1_Malawi_Lil-20/1_1983  .................g..........................g................................... [ 39040]
AY261360.1_Kenya_1950            .....g........t............a................g..c................................ [ 39040]
AM712240.1_OURT_88/3             -------------------------------------------------------------------------------- [ 39040]
AM712239.1_Benin_97/1            .............................g.................................................. [ 39040]
KM111295.1_Ken06.Bus             .t...g........t............a................g................................... [ 39040]
KM111294.1_Ken05/Tk1             .t...g........t............a................g................................... [ 39040]
KM262844.1_strain_L60            .............................g.................................................. [ 39040]
KM102979.1_26544/OG10_Italy      .............................g.................................................. [ 39040]
KM262845.1_strain_NHV            -------------------------------------------------------------------------------- [ 39040]
KX354450.1_47/Ss/2008            .............................g.................................................. [ 39040]
KP055815.1_strain_BA71           .............................g.................................................. [ 39040]
MH025920.1_strain_R35            .t...g........t............a................g................................... [ 39040]
MH025919.1_strain_N10            .t...g........t............a................g................................... [ 39040]
MH025918.1_strain_R25            .t...g........t............a................g................................... [ 39040]
MH025917.1_strain_R7             .t...g........t............a................g................................... [ 39040]
MH025916.1_strain_R8             .t...g........t............a................g................................... [ 39040]
```

```
FR682468.2_Georgia_2007/1        taaagataatataaatattattctaacacaagctgcaacctataaccatagaaaaattttaatctatttcattcctcaat [ 39120]
ASF_IND_20_CAD543                -------------------------------------------------------------------------------- [ 39120]
LR899193.1_Germany_2020/1        ................................................................................ [ 39120]
MN393477.1_Wuhan_2019            ................................................................................ [ 39120]
MW396979.1_Timor-Leste/2019/1    ................................................................................ [ 39120]
MK940252.1_CN/2019/InnerMongolia ................................................................................ [ 39120]
MK128995.1_China/2018/AnhuiXCGQ  ................................................................................ [ 39120]
MH681419.1_ASFV/POL/2015         ................................................................................ [ 39120]
LS478113.1_Estonia_2014          ................................................................................ [ 39120]
KP843857.1_Odintsovo_02/14       ................................................................................ [ 39120]
FN557520.1_strain_E75            ..t.......t..g..g......................a......................c......t.......... [ 39120]
AY261366.1_Warthog               c..t......t..g.........................a......................c.....a........... [ 39120]
AY261365.1_Warmbaths             c..t......t..g.........................a......................c................. [ 39120]
AY261364.1_Tengani_62            ................................................................................ [ 39120]
AY261363.1_Pretoriuskop/96/4     c..t......t..g.....................t...a......................c.....a........... [ 39120]
AY261362.1_Mkuzi_1979            c..t......t..g.........................a......................c......t.......... [ 39120]
AY261361.1_Malawi_Lil-20/1_1983  c.t.a........g................................................c................. [ 39120]
AY261360.1_Kenya_1950            ........................g.............................c.......c................. [ 39120]
AM712240.1_OURT_88/3             -------------------------------------------------------------------------------- [ 39120]
AM712239.1_Benin_97/1            ..t.......t..g..g......................a......................c......t.......... [ 39120]
KM111295.1_Ken06.Bus             ..c.a.c......g.......c........g.......................c.......c................. [ 39120]
KM111294.1_Ken05/Tk1             ..c.a.c......g.......c........g.......................c.......c................. [ 39120]
KM262844.1_strain_L60            ..t.......t..g..g......................a......................c......t.......... [ 39120]
KM102979.1_26544/OG10_Italy      ..t.......t..g..g.................a....a......................c......t.......... [ 39120]
KM262845.1_strain_NHV            -------------------------------------------------------------------------------- [ 39120]
KX354450.1_47/Ss/2008            ..t.......t..g..g.................a....a......................c......t.......... [ 39120]
KP055815.1_strain_BA71           ..t.......t..g..g......................a......................c......t.......... [ 39120]
MH025920.1_strain_R35            ..c.a.c......g.......c........g.......................c.......c................. [ 39120]
MH025919.1_strain_N10            ..c.a.c......g.......c........g.......................c.......c................. [ 39120]
MH025918.1_strain_R25            ..c.a.c......g.......c........g.......................c.......c................. [ 39120]
MH025917.1_strain_R7             ..c.a.c......g.......c........g.......................c.......c................. [ 39120]
MH025916.1_strain_R8             ..c.a.c......g.......c........g.......................c.......c................. [ 39120]

FR682468.2_Georgia_2007/1        caacccacgcacagatagaacaatgtttactagtggcgataaaagcaaaatcttccaggaaaaccttgaacttactactg [ 39200]
ASF_IND_20_CAD543                -------------------------------------------------------------------------------- [ 39200]
LR899193.1_Germany_2020/1        ................................................................................ [ 39200]
MN393477.1_Wuhan_2019            ................................................................................ [ 39200]
MW396979.1_Timor-Leste/2019/1    ................................................................................ [ 39200]
MK940252.1_CN/2019/InnerMongolia ................................................................................ [ 39200]
MK128995.1_China/2018/AnhuiXCGQ  ................................................................................ [ 39200]
MH681419.1_ASFV/POL/2015         ................................................................................ [ 39200]
LS478113.1_Estonia_2014          ................................................................................ [ 39200]
KP843857.1_Odintsovo_02/14       ................................................................................ [ 39200]
```

```
FN557520.1_strain_E75          .....t.......a..............gt.c............a............a...................... [ 39200]
AY261366.1_Warthog             t....t.t.....a..............gt.ca.......t...aa...........a...................... [ 39200]
AY261365.1_Warmbaths           t....t.......a..............gt.ca...........ag...........a...................... [ 39200]
AY261364.1_Tengani_62          ............................................a............a...................... [ 39200]
AY261363.1_Pretoriuskop/96/4   t....t.t.....a..............gt.ca.......t...aa...........a...................... [ 39200]
AY261362.1_Mkuzi_1979          .....t.......a..............ga.g............a............a............t......t.. [ 39200]
AY261361.1_Malawi_Lil-20/1_1983 t....t......................................a.....g......a...................... [ 39200]
AY261360.1_Kenya_1950          .....t....g..............c...t.c.....a......a....cg......a............t......t.a [ 39200]
AM712240.1_OURT_88/3           -------------------------------------------------------------------------------- [ 39200]
AM712239.1_Benin_97/1          .....t.......a..............gt.c............a............a...................... [ 39200]
KM111295.1_Ken06.Bus           .....t.......a...............t.c............a....tg......a............t......t.a [ 39200]
KM111294.1_Ken05/Tk1           .....t.......a...........c...t.c.....a......a....cg......aa...........t......t.a [ 39200]
KM262844.1_strain_L60          .....t.......a..............gt.c............a............a...................... [ 39200]
KM102979.1_26544/OG10_Italy    .....t.......a..............gt.c............a............a...................... [ 39200]
KM262845.1_strain_NHV          -------------------------------------------------------------------------------- [ 39200]
KX354450.1_47/Ss/2008          .....t.......a..............gt.c............a............a...................... [ 39200]
KP055815.1_strain_BA71         .....t.......a..............gt.c............a............a...................... [ 39200]
MH025920.1_strain_R35          .....t.......a...............t.c............a....tg......a............t......t.a [ 39200]
MH025919.1_strain_N10          .....t.......a...............t.c............a....tg......a............t......t.a [ 39200]
MH025918.1_strain_R25          .....t.......a...............t.c............a....tg......a............t......t.a [ 39200]
MH025917.1_strain_R7           .....t.......a...............t.c............a....tg......a............t......t.a [ 39200]
MH025916.1_strain_R8           .....t.......a...............t.c............a....tg......a............t......t.a [ 39200]

FR682468.2_Georgia_2007/1      tctcacctaaacctttccatcaacctcatcaaaaaaataagccattatgttgccacttacaattcaacaaatataatagg [ 39280]
ASF_IND_20_CAD543              -------------------------------------------------------------------------------- [ 39280]
LR899193.1_Germany_2020/1      ................................................................................ [ 39280]
MN393477.1_Wuhan_2019          ................................................................................ [ 39280]
MW396979.1_Timor-Leste/2019/1  ................................................................................ [ 39280]
MK940252.1_CN/2019/InnerMongolia ................................................................................ [ 39280]
MK128995.1_China/2018/AnhuiXCGQ ................................................................................ [ 39280]
MH681419.1_ASFV/POL/2015       ................................................................................ [ 39280]
LS478113.1_Estonia_2014        ................................................................................ [ 39280]
KP843857.1_Odintsovo_02/14     ................................................................................ [ 39280]
FN557520.1_strain_E75          .......................a......c.......c.....a.............t..................... [ 39280]
AY261366.1_Warthog             ...........g...........a....................a................................... [ 39280]
AY261365.1_Warmbaths           ..........c..........g.a....................a.......t........................... [ 39280]
AY261364.1_Tengani_62          .......................t........................................................ [ 39280]
AY261363.1_Pretoriuskop/96/4   ...........g...........a....................a................................... [ 39280]
AY261362.1_Mkuzi_1979          ..c....................a....................a.......t........................... [ 39280]
AY261361.1_Malawi_Lil-20/1_1983 .......................a....................a.......t.....................c...a. [ 39280]
AY261360.1_Kenya_1950          .................t.....a...g........t.......a.......ttg.......g...........c...a. [ 39280]
AM712240.1_OURT_88/3           -------------------------------------------------------------------------------- [ 39280]
AM712239.1_Benin_97/1          .......................a......c.......c.....a.............t..................... [ 39280]
KM111295.1_Ken06.Bus           ...........t.....t.....a...g........t.......a.......t.g.......g...........c...a. [ 39280]
```

```
KM111294.1_Ken05/Tk1             .................t.....a...g........t.......a.......t.g.......g...........c...a. [ 39280]
KM262844.1_strain_L60            .......................a......c.......c.....a.............t..................... [ 39280]
KM102979.1_26544/OG10_Italy      ...........t...........a......c.......c.....a.............t..................... [ 39280]
KM262845.1_strain_NHV            -------------------------------------------------------------------------------- [ 39280]
KX354450.1_47/Ss/2008            ...........t...........a......c.......c.....a.............t..................... [ 39280]
KP055815.1_strain_BA71           .......................a......c.......c.....a.............t..................... [ 39280]
MH025920.1_strain_R35            ...........t.....t.....a...g........t.......a.......t.g.......g...........c...a. [ 39280]
MH025919.1_strain_N10            ...........t.....t.....a...g........t.......a.......t.g.......g...........c...a. [ 39280]
MH025918.1_strain_R25            ...........t.....t.....a...g........t.......a.......t.g.......g...........c...a. [ 39280]
MH025917.1_strain_R7             ...........t.....t.....a...g........t.......a.......t.g.......g...........c...a. [ 39280]
MH025916.1_strain_R8             ...........t.....t.....a...g........t.......a.......t.g.......g...........c...a. [ 39280]

FR682468.2_Georgia_2007/1        cattctgagtatgcggcggaaaaagaagatatatttagatatcatattgacaaaatttgtaaaaaaagctatttttaata [ 39360]
ASF_IND_20_CAD543                -------------------------------------------------------------------------------- [ 39360]
LR899193.1_Germany_2020/1        ................................................................................ [ 39360]
MN393477.1_Wuhan_2019            ................................................................................ [ 39360]
MW396979.1_Timor-Leste/2019/1    ................................................................................ [ 39360]
MK940252.1_CN/2019/InnerMongolia ................................................................................ [ 39360]
MK128995.1_China/2018/AnhuiXCGQ  ................................................................................ [ 39360]
MH681419.1_ASFV/POL/2015         ................................................................................ [ 39360]
LS478113.1_Estonia_2014          ................................................................................ [ 39360]
KP843857.1_Odintsovo_02/14       ................................................................................ [ 39360]
FN557520.1_strain_E75            .............aa.....................g.............................t............. [ 39360]
AY261366.1_Warthog               ........a....................................................................... [ 39360]
AY261365.1_Warmbaths             .............aa..............................g..........a.......t.c............. [ 39360]
AY261364.1_Tengani_62            ................................................................................ [ 39360]
AY261363.1_Pretoriuskop/96/4     ........a....aa.................................................t.c............. [ 39360]
AY261362.1_Mkuzi_1979            .............aa...a........................t.g....................t............. [ 39360]
AY261361.1_Malawi_Lil-20/1_1983  .............................................t......g.g...........c............. [ 39360]
AY261360.1_Kenya_1950            ..............a..aa..........................t........gg..........c..........t.. [ 39360]
AM712240.1_OURT_88/3             -------------------------------------------------------------------------------- [ 39360]
AM712239.1_Benin_97/1            .............aa.....................g.............................t............. [ 39360]
KM111295.1_Ken06.Bus             ...........g..a..aa..........................t........gg..........t...g......... [ 39360]
KM111294.1_Ken05/Tk1             ..............a..aa..........................t........gg..........c...g......t.. [ 39360]
KM262844.1_strain_L60            .............aa.....................g.............................t............. [ 39360]
KM102979.1_26544/OG10_Italy      .............aa.....................g.............................t............. [ 39360]
KM262845.1_strain_NHV            -------------------------------------------------------------------------------- [ 39360]
KX354450.1_47/Ss/2008            .............aa.....................g.............................t............. [ 39360]
KP055815.1_strain_BA71           .............aa.....................g.............................t............. [ 39360]
MH025920.1_strain_R35            ...........g..a..aa..........................t........gg..........t...g......... [ 39360]
MH025919.1_strain_N10            ...........g..a..aa..........................t........gg..........t...g......... [ 39360]
MH025918.1_strain_R25            ...........g..a..aa..........................t........gg..........t...g......... [ 39360]
MH025917.1_strain_R7             ...........g..a..aa..........................t........gg..........t...g......... [ 39360]
MH025916.1_strain_R8             ...........g..a..aa..........................t........gg..........t...g......... [ 39360]
```

```
FR682468.2_Georgia_2007/1        agtttgtcgttcgatgtatggatacattttctataaacccggaaagaatccttaaaatagccgcgcgaataaataggatg [ 39440]
ASF_IND_20_CAD543                -------------------------------------------------------------------------------- [ 39440]
LR899193.1_Germany_2020/1        ................................................................................ [ 39440]
MN393477.1_Wuhan_2019            ................................................................................ [ 39440]
MW396979.1_Timor-Leste/2019/1    ................................................................................ [ 39440]
MK940252.1_CN/2019/InnerMongolia ................................................................................ [ 39440]
MK128995.1_China/2018/AnhuiXCGQ  ................................................................................ [ 39440]
MH681419.1_ASFV/POL/2015         ................................................................................ [ 39440]
LS478113.1_Estonia_2014          ................................................................................ [ 39440]
KP843857.1_Odintsovo_02/14       ................................................................................ [ 39440]
FN557520.1_strain_E75            .a....................g.g.........................g.c.....g..t.....t.......a.... [ 39440]
AY261366.1_Warthog               .a.............................c..................a.c.....g................a.... [ 39440]
AY261365.1_Warmbaths             .a.a....................g......c..................a.c.....g..................... [ 39440]
AY261364.1_Tengani_62            .a............................................................................c. [ 39440]
AY261363.1_Pretoriuskop/96/4     .a.a.........t.................c..................a.c.....g................a.... [ 39440]
AY261362.1_Mkuzi_1979            .a.............a.........g...................a....g.c.....g................a.... [ 39440]
AY261361.1_Malawi_Lil-20/1_1983  .a.............................c.................tg.c.....g.....a............... [ 39440]
AY261360.1_Kenya_1950            .a......a..g.........t.........c........a........tg.c.....g..............a.a.... [ 39440]
AM712240.1_OURT_88/3             -------------------------------------------------------------------------------- [ 39440]
AM712239.1_Benin_97/1            .a....................g.g.........................g.c.....g..t.....t.......a.... [ 39440]
KM111295.1_Ken06.Bus             .a......a..g.........t.........c........a........tg.c.....g..............a.a.... [ 39440]
KM111294.1_Ken05/Tk1             .a......a..g.........t.........c........a........tg.c.....g..............a.a.... [ 39440]
KM262844.1_strain_L60            .a....................g.g.........................g.c.....g..t.....t.......a.... [ 39440]
KM102979.1_26544/OG10_Italy      .a....................g.g.........................g.c.....g..t.....t.......a.... [ 39440]
KM262845.1_strain_NHV            -------------------------------------------------------------------------------- [ 39440]
KX354450.1_47/Ss/2008            .a....................g.g.........................g.c.....g..t.....t.......a.... [ 39440]
KP055815.1_strain_BA71           .a....................g.g.........................g.c.....g..t.....t.......a.... [ 39440]
MH025920.1_strain_R35            .a......a..g.........t.........c........a........tg.c.....g..............a.a.... [ 39440]
MH025919.1_strain_N10            .a......a..g.........t.........c........a........tg.c.....g..............a.a.... [ 39440]
MH025918.1_strain_R25            .a......a..g.........t.........c........a........tg.c.....g..............a.a.... [ 39440]
MH025917.1_strain_R7             .a......a..g.........t.........c........a........tg.c.....g..............a.a.... [ 39440]
MH025916.1_strain_R8             .a......a..g.........t.........c........a........tg.c.....g..............a.a.... [ 39440]

FR682468.2_Georgia_2007/1        atgttagtgaaaaaaatatctgaacatgtttggaaaaatcatgcggttagacttaaataccttaaacatgcggtacacac [ 39520]
ASF_IND_20_CAD543                -------------------------------------------------------------------------------- [ 39520]
LR899193.1_Germany_2020/1        ................................................................................ [ 39520]
MN393477.1_Wuhan_2019            ................................................................................ [ 39520]
MW396979.1_Timor-Leste/2019/1    ................................................................................ [ 39520]
MK940252.1_CN/2019/InnerMongolia ................................................................................ [ 39520]
MK128995.1_China/2018/AnhuiXCGQ  ................................................................................ [ 39520]
MH681419.1_ASFV/POL/2015         ................................................................................ [ 39520]
LS478113.1_Estonia_2014          ................................................................................ [ 39520]
KP843857.1_Odintsovo_02/14       ................................................................................ [ 39520]
```

```
FN557520.1_strain_E75          ..............................................c..........c...................... [ 39520]
AY261366.1_Warthog             t...........................c.................c..........c.......g.........t.... [ 39520]
AY261365.1_Warmbaths           ..............................................c..........c.......g.............. [ 39520]
AY261364.1_Tengani_62          ................................................................................ [ 39520]
AY261363.1_Pretoriuskop/96/4   t....................c......c.................c..........c.......g.........t.... [ 39520]
AY261362.1_Mkuzi_1979          ...........................a..................c.......g..c...................... [ 39520]
AY261361.1_Malawi_Lil-20/1_1983 ..............t..........g.......................a.......c.......g...........t.. [ 39520]
AY261360.1_Kenya_1950          .a............t...............................c..a.......c.......g.............. [ 39520]
AM712240.1_OURT_88/3           -------------------------------------------------------------------------------- [ 39520]
AM712239.1_Benin_97/1          ..............................................c..........c...................... [ 39520]
KM111295.1_Ken06.Bus           .a......a.....t..................................a.........t.....g.............. [ 39520]
KM111294.1_Ken05/Tk1           .a......a.....t...............................c..a.......c.t.....g.............. [ 39520]
KM262844.1_strain_L60          ..............................................c..........c...................... [ 39520]
KM102979.1_26544/OG10_Italy    ..............................................c..........c...................... [ 39520]
KM262845.1_strain_NHV          -------------------------------------------------------------------------------- [ 39520]
KX354450.1_47/Ss/2008          ..............................................c..........c...................... [ 39520]
KP055815.1_strain_BA71         ..............................................c..........c...................... [ 39520]
MH025920.1_strain_R35          .a......a.....t..................................a.......c.......g.............. [ 39520]
MH025919.1_strain_N10          .a......a.....t..................................a.......c.......g.............. [ 39520]
MH025918.1_strain_R25          .a......a.....t..................................a.......c.......g.............. [ 39520]
MH025917.1_strain_R7           .a......a.....t..................................a.......c.......g.............. [ 39520]
MH025916.1_strain_R8           .a......a.....t..................................a.......c.......g.............. [ 39520]

FR682468.2_Georgia_2007/1      gatgaagcataaagatgggaaaaatagactcatgaactttatctatgatcgctgttattaccatatgcaaggggaagaaa [ 39600]
ASF_IND_20_CAD543              -------------------------------------------------------------------------------- [ 39600]
LR899193.1_Germany_2020/1      ................................................................................ [ 39600]
MN393477.1_Wuhan_2019          ................................................................................ [ 39600]
MW396979.1_Timor-Leste/2019/1  ................................................................................ [ 39600]
MK940252.1_CN/2019/InnerMongolia ................................................................................ [ 39600]
MK128995.1_China/2018/AnhuiXCGQ ................................................................................ [ 39600]
MH681419.1_ASFV/POL/2015       ................................................................................ [ 39600]
LS478113.1_Estonia_2014        ................................................................................ [ 39600]
KP843857.1_Odintsovo_02/14     ................................................................................ [ 39600]
FN557520.1_strain_E75          ................................................a.a...c......................... [ 39600]
AY261366.1_Warthog             ......a................................g..........a..ac......................... [ 39600]
AY261365.1_Warmbaths           ......................................................c......................... [ 39600]
AY261364.1_Tengani_62          ................................................................................ [ 39600]
AY261363.1_Pretoriuskop/96/4   ......a................................g..........a..ac......................... [ 39600]
AY261362.1_Mkuzi_1979          ................................................gta...c......................... [ 39600]
AY261361.1_Malawi_Lil-20/1_1983 ..........c....a..a...............................a...c........c..............g. [ 39600]
AY261360.1_Kenya_1950          ..........c....a..a.........t.a...................a............c........a.....g. [ 39600]
AM712240.1_OURT_88/3           -------------------------------------------------------------------------------- [ 39600]
AM712239.1_Benin_97/1          ................................................a.a...c......................... [ 39600]
KM111295.1_Ken06.Bus           ..........c....a..a.........t.a...................a............c.......aa.....g. [ 39600]
```

```
KM111294.1_Ken05/Tk1           ..........c....a..a...............................a...c........c........a.....g. [ 39600]
KM262844.1_strain_L60          ................................................a.a...c......................... [ 39600]
KM102979.1_26544/OG10_Italy    ................................................a.a...c......................... [ 39600]
KM262845.1_strain_NHV          -------------------------------------------------------------------------------- [ 39600]
KX354450.1_47/Ss/2008          ................................................a.a...c......................... [ 39600]
KP055815.1_strain_BA71         ................................................a.a...c......................... [ 39600]
MH025920.1_strain_R35          ..........c....a..a.........t.a...................a...c........c.......aa.....g. [ 39600]
MH025919.1_strain_N10          ..........c....a..a.........t.a...................a...c........c.......aa.....g. [ 39600]
MH025918.1_strain_R25          ..........c....a..a.........t.a...................a...c........c.......aa.....g. [ 39600]
MH025917.1_strain_R7           ..........c....a..a.........t.a...................a...c........c.......aa.....g. [ 39600]
MH025916.1_strain_R8           ..........c....a..a.........t.a...................a...c........c.......aa.....g. [ 39600]

FR682468.2_Georgia_2007/1      tctttagcctcgcaagattttatgcaatccatcatgcaccaaagttgtttgacgttttttatgattgttgtatcctagat [ 39680]
ASF_IND_20_CAD543              -------------------------------------------------------------------------------- [ 39680]
LR899193.1_Germany_2020/1      ................................................................................ [ 39680]
MN393477.1_Wuhan_2019          ................................................................................ [ 39680]
MW396979.1_Timor-Leste/2019/1  ................................................................................ [ 39680]
MK940252.1_CN/2019/InnerMongolia ................................................................................ [ 39680]
MK128995.1_China/2018/AnhuiXCGQ ................................................................................ [ 39680]
MH681419.1_ASFV/POL/2015       ................................................................................ [ 39680]
LS478113.1_Estonia_2014        ................................................................................ [ 39680]
KP843857.1_Odintsovo_02/14     ................................................................................ [ 39680]
FN557520.1_strain_E75          .t...............................................c............a................. [ 39680]
AY261366.1_Warthog             .......................................................................c........ [ 39680]
AY261365.1_Warmbaths           ................................................................................ [ 39680]
AY261364.1_Tengani_62          ................................................................................ [ 39680]
AY261363.1_Pretoriuskop/96/4   .......................................................................c........ [ 39680]
AY261362.1_Mkuzi_1979          .t...............................................c............a................. [ 39680]
AY261361.1_Malawi_Lil-20/1_1983 .....g..................................c..........................c..c..g...... [ 39680]
AY261360.1_Kenya_1950          .......t..t..........................g....a.....c.............................. [ 39680]
AM712240.1_OURT_88/3           -------------------------------------------------------------------------------- [ 39680]
AM712239.1_Benin_97/1          .t...............................................c............a................. [ 39680]
KM111295.1_Ken06.Bus           .....................................g....a.....c.............................. [ 39680]
KM111294.1_Ken05/Tk1           .......t..t..........................g....a.....c.............................. [ 39680]
KM262844.1_strain_L60          .t...............................................c............a................. [ 39680]
KM102979.1_26544/OG10_Italy    .t...............................................c............a................. [ 39680]
KM262845.1_strain_NHV          -------------------------------------------------------------------------------- [ 39680]
KX354450.1_47/Ss/2008          .t...............................................c............a................. [ 39680]
KP055815.1_strain_BA71         .t...............................................c............a................. [ 39680]
MH025920.1_strain_R35          .....................................g....a.....c.............................. [ 39680]
MH025919.1_strain_N10          .....................................g....a.....c.............................. [ 39680]
MH025918.1_strain_R25          .....................................g....a.....c.............................. [ 39680]
MH025917.1_strain_R7           .....................................g....a.....c.............................. [ 39680]
MH025916.1_strain_R8           .....................................g....a.....c.............................. [ 39680]
```

```
FR682468.2_Georgia_2007/1        acgatacgattcaaaagccttcttttagattgttcacatatcataggtaaaaacgctcatgatgctaccaatatcaacat [ 39760]
ASF_IND_20_CAD543                -------------------------------------------------------------------------------- [ 39760]
LR899193.1_Germany_2020/1        ................................................................................ [ 39760]
MN393477.1_Wuhan_2019            ................................................................................ [ 39760]
MW396979.1_Timor-Leste/2019/1    ................................................................................ [ 39760]
MK940252.1_CN/2019/InnerMongolia ................................................................................ [ 39760]
MK128995.1_China/2018/AnhuiXCGQ  ................................................................................ [ 39760]
MH681419.1_ASFV/POL/2015         ................................................................................ [ 39760]
LS478113.1_Estonia_2014          ................................................................................ [ 39760]
KP843857.1_Odintsovo_02/14       ................................................................................ [ 39760]
FN557520.1_strain_E75            ....................................................................t........... [ 39760]
AY261366.1_Warthog               ..t........t....a.....................c.............................t.......ct.. [ 39760]
AY261365.1_Warmbaths             ..t....a........................................................................ [ 39760]
AY261364.1_Tengani_62            ................................................................................ [ 39760]
AY261363.1_Pretoriuskop/96/4     ..t........t..........................c.............................t.......ct.. [ 39760]
AY261362.1_Mkuzi_1979            ....................................................................t........... [ 39760]
AY261361.1_Malawi_Lil-20/1_1983  g.t.c......t........g............c.......................t........---gg.......c. [ 39760]
AY261360.1_Kenya_1950            ...........t........g...............t.........c...................---.g...t..... [ 39760]
AM712240.1_OURT_88/3             -------------------------------------------------------------------------------- [ 39760]
AM712239.1_Benin_97/1            ....................................................................t........... [ 39760]
KM111295.1_Ken06.Bus             ...........t........g.........................t...................---.g...t..... [ 39760]
KM111294.1_Ken05/Tk1             ...........t........g....................t....c...................---.g...t..... [ 39760]
KM262844.1_strain_L60            ....................................................................t........... [ 39760]
KM102979.1_26544/OG10_Italy      ....................................................................t........... [ 39760]
KM262845.1_strain_NHV            -------------------------------------------------------------------------------- [ 39760]
KX354450.1_47/Ss/2008            ....................................................................t........... [ 39760]
KP055815.1_strain_BA71           ....................................................................t........... [ 39760]
MH025920.1_strain_R35            ...........t........g.........................t...................---.g...t..... [ 39760]
MH025919.1_strain_N10            ...........t........g.........................t...................---.g...t..... [ 39760]
MH025918.1_strain_R25            ...........t........g.........................t...................---.g...t..... [ 39760]
MH025917.1_strain_R7             ...........t........g.........................t...................---.g...t..... [ 39760]
MH025916.1_strain_R8             ...........t........g.........................t...................---.g...t..... [ 39760]

FR682468.2_Georgia_2007/1        cgtgaacaagtatatcggcaacctgtttgttatgggagttcttagcaaaaaagaaatcttacaggactatccatccattt [ 39840]
ASF_IND_20_CAD543                -------------------------------------------------------------------------------- [ 39840]
LR899193.1_Germany_2020/1        ................................................................................ [ 39840]
MN393477.1_Wuhan_2019            ................................................................................ [ 39840]
MW396979.1_Timor-Leste/2019/1    ................................................................................ [ 39840]
MK940252.1_CN/2019/InnerMongolia ................................................................................ [ 39840]
MK128995.1_China/2018/AnhuiXCGQ  ................................................................................ [ 39840]
MH681419.1_ASFV/POL/2015         ................................................................................ [ 39840]
LS478113.1_Estonia_2014          ................................................................................ [ 39840]
KP843857.1_Odintsovo_02/14       ................................................................................ [ 39840]
```

```
FN557520.1_strain_E75          ...............t.............c................................................c. [ 39840]
AY261366.1_Warthog             ...............t.............c................................................c. [ 39840]
AY261365.1_Warmbaths           ...............t...g.........c................................................c. [ 39840]
AY261364.1_Tengani_62          ..............................................................................c. [ 39840]
AY261363.1_Pretoriuskop/96/4   ...............t.............c................................................c. [ 39840]
AY261362.1_Mkuzi_1979          ...............t.a...........c.................................................. [ 39840]
AY261361.1_Malawi_Lil-20/1_1983 .........a.....t........a....c......g......................................t..c. [ 39840]
AY261360.1_Kenya_1950          ......t........t.....t..a....c......g..........................a..............c. [ 39840]
AM712240.1_OURT_88/3           -------------------------------------------------------------------------------- [ 39840]
AM712239.1_Benin_97/1          ...............t.............c................................................c. [ 39840]
KM111295.1_Ken06.Bus           ...............t.....t..a....c.................................a...........t..c. [ 39840]
KM111294.1_Ken05/Tk1           ......t........t.....t..a....c.................................a..............c. [ 39840]
KM262844.1_strain_L60          ...............t.............c................................................c. [ 39840]
KM102979.1_26544/OG10_Italy    ...............t.............c................................................c. [ 39840]
KM262845.1_strain_NHV          -------------------------------------------------------------------------------- [ 39840]
KX354450.1_47/Ss/2008          ...............t.............c................................................c. [ 39840]
KP055815.1_strain_BA71         ...............t.............c................................................c. [ 39840]
MH025920.1_strain_R35          ...............t.....t..a....c.................................a...........t..c. [ 39840]
MH025919.1_strain_N10          ...............t.....t..a....c.................................a...........t..c. [ 39840]
MH025918.1_strain_R25          ...............t.....t..a....c.................................a...........t..c. [ 39840]
MH025917.1_strain_R7           ...............t.....t..a....c.................................a...........t..c. [ 39840]
MH025916.1_strain_R8           ...............t.....t..a....c.................................a...........t..c. [ 39840]

FR682468.2_Georgia_2007/1      attctaaacaatacatgccttagttta-ttttt-tttg-cggccgaaacattattcttaccct-agaaaacgcttatagt [ 39920]
ASF_IND_20_CAD543              -------------------------------------------------------------------------------- [ 39920]
LR899193.1_Germany_2020/1      ...........................-.....-....-........................-................ [ 39920]
MN393477.1_Wuhan_2019          ...........................-.....-....-........................-................ [ 39920]
MW396979.1_Timor-Leste/2019/1  ...........................-.....-....-........................-................ [ 39920]
MK940252.1_CN/2019/InnerMongolia ...........................-.....-....-........................-................ [ 39920]
MK128995.1_China/2018/AnhuiXCGQ ...........................-.....-....-........................-................ [ 39920]
MH681419.1_ASFV/POL/2015       ...........................-.....-....-........................-................ [ 39920]
LS478113.1_Estonia_2014        ...........................-.....-....-........................-................ [ 39920]
KP843857.1_Odintsovo_02/14     ...........................-.....-....-........................-................ [ 39920]
FN557520.1_strain_E75          ..........t..t.............-.....---..-....t.......c.......-...-......t......... [ 39920]
AY261366.1_Warthog             ..........t................-.....t....-.a..t....t..............-................ [ 39920]
AY261365.1_Warmbaths           ..........t.......t........---...t....-....t----...............-.a.............. [ 39920]
AY261364.1_Tengani_62          ..........t................-.....t....-..........c.............-................ [ 39920]
AY261363.1_Pretoriuskop/96/4   ..........t................-.....t....-.a..t....t..............-................ [ 39920]
AY261362.1_Mkuzi_1979          ..........t..t.............-.....t....-....t.......c.......-...-......t..a...... [ 39920]
AY261361.1_Malawi_Lil-20/1_1983 ..........tg.t...tt........-.....t....-t...t...............-...-.......a..----.. [ 39920]
AY261360.1_Kenya_1950          .......gg.t.......t........-.....t...aga.c.t...c..........t..t.c.......a..---... [ 39920]
AM712240.1_OURT_88/3           -------------------------------------------------------------------------------- [ 39920]
AM712239.1_Benin_97/1          ..........t..t.............-.....---..-....t.......c.......-...-......t......... [ 39920]
KM111295.1_Ken06.Bus           .......gg.t.......t........-.....t...a-a.c.t...c..a.......t..t.c.a.....a..---... [ 39920]
```

```
KM111294.1_Ken05/Tk1          ........t.tg....a.t........t.....t...aaa.c.t...cg.a.......t..t.c.......a..---... [ 39920]
KM262844.1_strain_L60         ..........t..t.............-.....---..-....t.......c.......-...-......t......... [ 39920]
KM102979.1_26544/OG10_Italy   ..........t..t.............-.....---..-....t.......c.......-...-......t......... [ 39920]
KM262845.1_strain_NHV         -------------------------------------------------------------------------------- [ 39920]
KX354450.1_47/Ss/2008         ..........t..t.............-.....---..-....t.......c.......-...-......t......... [ 39920]
KP055815.1_strain_BA71        ..........t..t.............-.....---..-....t.......c.......-...-......t......... [ 39920]
MH025920.1_strain_R35         .......gg.t.......t........-.....t...a-aac.t...c..a.......t..t.c.......a..---... [ 39920]
MH025919.1_strain_N10         .......gg.t.......t........-.....t...a-aac.t...c..a.......t..t.c.......a..---... [ 39920]
MH025918.1_strain_R25         .......gg.t.......t........-.....t...a-aac.t...c..a.......t..t.c.......a..---... [ 39920]
MH025917.1_strain_R7          .......gg.t.......t........-.....t...a-aac.t...c..a.......t..t.c.......a..---... [ 39920]
MH025916.1_strain_R8          .......gg.t.......t........-.....t...a-aac.t...c..a.......t..t.c.......a..---... [ 39920]

FR682468.2_Georgia_2007/1     catcttaaatcataggtaaggaagatcatcatattttttgaaacgtaattttttaacgcatgatctatgatttcagggtc [ 40000]
ASF_IND_20_CAD543             -------------------------------------------------------------------------------- [ 40000]
LR899193.1_Germany_2020/1     ................................................................................ [ 40000]
MN393477.1_Wuhan_2019         ................................................................................ [ 40000]
MW396979.1_Timor-Leste/2019/1 ................................................................................ [ 40000]
MK940252.1_CN/2019/InnerMongolia ................................................................................ [ 40000]
MK128995.1_China/2018/AnhuiXCGQ ................................................................................ [ 40000]
MH681419.1_ASFV/POL/2015      ................................................................................ [ 40000]
LS478113.1_Estonia_2014       ................................................................................ [ 40000]
KP843857.1_Odintsovo_02/14    ................................................................................ [ 40000]
FN557520.1_strain_E75         ................a..........................a.................................... [ 40000]
AY261366.1_Warthog            ...........g........................................................a........... [ 40000]
AY261365.1_Warmbaths          ...........g...........................................c........................ [ 40000]
AY261364.1_Tengani_62         ................................................................................ [ 40000]
AY261363.1_Pretoriuskop/96/4  ...........g........................................................a........... [ 40000]
AY261362.1_Mkuzi_1979         ................a..........................a.................................... [ 40000]
AY261361.1_Malawi_Lil-20/1_1983 ...........g....a.......g................ttta............a..........t........... [ 40000]
AY261360.1_Kenya_1950         .......g......a..........c.................a.............t..........a........... [ 40000]
AM712240.1_OURT_88/3          -------------------------------------------------------------------------------- [ 40000]
AM712239.1_Benin_97/1         ................a..........................a.................................... [ 40000]
KM111295.1_Ken06.Bus          .......g......a..........c.................a........................a........... [ 40000]
KM111294.1_Ken05/Tk1          .......g......a..........c.................a.............t..........a........... [ 40000]
KM262844.1_strain_L60         ................a..........................a.................................... [ 40000]
KM102979.1_26544/OG10_Italy   ................a..........................a.................................... [ 40000]
KM262845.1_strain_NHV         -------------------------------------------------------------------------------- [ 40000]
KX354450.1_47/Ss/2008         ................a..........................a.................................... [ 40000]
KP055815.1_strain_BA71        ................a..........................a.................................... [ 40000]
MH025920.1_strain_R35         .......g......a..........c.................a........................a........... [ 40000]
MH025919.1_strain_N10         .......g......a..........c.................a...............g........a........... [ 40000]
MH025918.1_strain_R25         .......g......a..........c.................a........................a........... [ 40000]
MH025917.1_strain_R7          .......g......a..........c.................a........................a........... [ 40000]
MH025916.1_strain_R8          .......g......a..........c.................a........................a........... [ 40000]
```

```
FR682468.2_Georgia_2007/1        cgtgcttttaggcaacggggtggtggccggactataaatctttagggataaaatgttctttataagctcatacccttccc [ 40080]
ASF_IND_20_CAD543                -------------------------------------------------------------------------------- [ 40080]
LR899193.1_Germany_2020/1        ................................................................................ [ 40080]
MN393477.1_Wuhan_2019            ................................................................................ [ 40080]
MW396979.1_Timor-Leste/2019/1    ................................................................................ [ 40080]
MK940252.1_CN/2019/InnerMongolia ................................................................................ [ 40080]
MK128995.1_China/2018/AnhuiXCGQ  ................................................................................ [ 40080]
MH681419.1_ASFV/POL/2015         ................................................................................ [ 40080]
LS478113.1_Estonia_2014          ................................................................................ [ 40080]
KP843857.1_Odintsovo_02/14       ................................................................................ [ 40080]
FN557520.1_strain_E75            ...................a......t..................a...........t...................... [ 40080]
AY261366.1_Warthog               ....................................................................t........... [ 40080]
AY261365.1_Warmbaths             ....................................................................t........... [ 40080]
AY261364.1_Tengani_62            .a.............................................................................. [ 40080]
AY261363.1_Pretoriuskop/96/4     ....................................................................t........... [ 40080]
AY261362.1_Mkuzi_1979            ...................a......t..................a...........t...................... [ 40080]
AY261361.1_Malawi_Lil-20/1_1983  ....t.......t.......ca....a.........g.....................c..................... [ 40080]
AY261360.1_Kenya_1950            ..............................g.......................a...c.....g..............t [ 40080]
AM712240.1_OURT_88/3             -------------------------------------------------------------------------------- [ 40080]
AM712239.1_Benin_97/1            ...................a......t..................a...........t...................... [ 40080]
KM111295.1_Ken06.Bus             ...............t.....a........g.......................a...c.....g..............t [ 40080]
KM111294.1_Ken05/Tk1             ..............................g.......................a...c.....g.......t......t [ 40080]
KM262844.1_strain_L60            ...................a......t..................a...........t...................... [ 40080]
KM102979.1_26544/OG10_Italy      ...................a......t..................a...........t...................... [ 40080]
KM262845.1_strain_NHV            -------------------------------------------------------------------------------- [ 40080]
KX354450.1_47/Ss/2008            ...................a......t..................a...........t...................... [ 40080]
KP055815.1_strain_BA71           ...................a......t..................a...........t...................... [ 40080]
MH025920.1_strain_R35            ...............t.....a........g.......................a...c.....g..............t [ 40080]
MH025919.1_strain_N10            ...............t.....a........g.......................a...c....................t [ 40080]
MH025918.1_strain_R25            ...............t.....a........g.......................a...c.....g..............t [ 40080]
MH025917.1_strain_R7             ...............t.....a........g.......................a...c.....g..............t [ 40080]
MH025916.1_strain_R8             ...............t.....a........g.......................a...c.....g..............t [ 40080]

FR682468.2_Georgia_2007/1        ------ctaaagctgtagtaccctcttcgaaaacatcagcccccagatctatacaaaagaacatgttttctatattatag [ 40160]
ASF_IND_20_CAD543                -------------------------------------------------------------------------------- [ 40160]
LR899193.1_Germany_2020/1        ------.......................................................................... [ 40160]
MN393477.1_Wuhan_2019            ------.......................................................................... [ 40160]
MW396979.1_Timor-Leste/2019/1    ------.......................................................................... [ 40160]
MK940252.1_CN/2019/InnerMongolia ------.......................................................................... [ 40160]
MK128995.1_China/2018/AnhuiXCGQ  ------.......................................................................... [ 40160]
MH681419.1_ASFV/POL/2015         ------.......................................................................... [ 40160]
LS478113.1_Estonia_2014          ------.......................................................................... [ 40160]
KP843857.1_Odintsovo_02/14       ------.......................................................................... [ 40160]
```

```
FN557520.1_strain_E75              ------...........................................c.............................. [ 40160]
AY261366.1_Warthog                 ------........a....g............g................c.............................. [ 40160]
AY261365.1_Warmbaths               ------........a.................g................c......................c....... [ 40160]
AY261364.1_Tengani_62              ------.......................................................................... [ 40160]
AY261363.1_Pretoriuskop/96/4       ------........a....g............g................c.............................. [ 40160]
AY261362.1_Mkuzi_1979              ------...........................................c.............................. [ 40160]
AY261361.1_Malawi_Lil-20/1_1983    ------...c..gca..---tt...........................c.............................. [ 40160]
AY261360.1_Kenya_1950              gtccca...g...ca..---........a...g....g...........c...........t.................. [ 40160]
AM712240.1_OURT_88/3               -------------------------------------------------------------------------------- [ 40160]
AM712239.1_Benin_97/1              ------...........................................c.............................. [ 40160]
KM111295.1_Ken06.Bus               gtccca...g...ca..---........a...g....g...........c...........t.................. [ 40160]
KM111294.1_Ken05/Tk1               gtccca...g...ca..---........a...g....g...........c...........t.................. [ 40160]
KM262844.1_strain_L60              ------...........................................c.............................. [ 40160]
KM102979.1_26544/OG10_Italy        ------...........................................c.............................. [ 40160]
KM262845.1_strain_NHV              -------------------------------------------------------------------------------- [ 40160]
KX354450.1_47/Ss/2008              ------...........................................c.............................. [ 40160]
KP055815.1_strain_BA71             ------...........................................c.............................. [ 40160]
MH025920.1_strain_R35              gtccca...g...ca..---........a...g....g...........c...........t.................. [ 40160]
MH025919.1_strain_N10              gtccca...g...ca..---........a...g....g...........c...........t.................. [ 40160]
MH025918.1_strain_R25              gtccca...g...ca..---........a...g....g...........c...........t.................. [ 40160]
MH025917.1_strain_R7               gtccca...g...ca..---........a...g....g...........c...........t.................. [ 40160]
MH025916.1_strain_R8               gtccca...g...ca..---........a...g....g...........c...........t.................. [ 40160]

FR682468.2_Georgia_2007/1          tactgtattgagctaagcatggcttgattgatgttggcgcccaggacatagcagtagtacatggttgaaaggttgtggtc [ 40240]
ASF_IND_20_CAD543                  -------------------------------------------------------------------------------- [ 40240]
LR899193.1_Germany_2020/1          ................................................................................ [ 40240]
MN393477.1_Wuhan_2019              ................................................................................ [ 40240]
MW396979.1_Timor-Leste/2019/1      ................................................................................ [ 40240]
MK940252.1_CN/2019/InnerMongolia   ................................................................................ [ 40240]
MK128995.1_China/2018/AnhuiXCGQ    ................................................................................ [ 40240]
MH681419.1_ASFV/POL/2015           ................................................................................ [ 40240]
LS478113.1_Estonia_2014            ................................................................................ [ 40240]
KP843857.1_Odintsovo_02/14         ................................................................................ [ 40240]
FN557520.1_strain_E75              ............g..........c...................ac.t................................. [ 40240]
AY261366.1_Warthog                 ............g..........c........................................................ [ 40240]
AY261365.1_Warmbaths               ............g..........c....................a................................... [ 40240]
AY261364.1_Tengani_62              .......................c........................................................ [ 40240]
AY261363.1_Pretoriuskop/96/4       ............g..........c............c.c......................................... [ 40240]
AY261362.1_Mkuzi_1979              ............g..........c...................ac.t................................. [ 40240]
AY261361.1_Malawi_Lil-20/1_1983    ............gc.........c.....a.............................t..c................. [ 40240]
AY261360.1_Kenya_1950              ............g..........c....................c.t............t..c.........c....... [ 40240]
AM712240.1_OURT_88/3               -------------------------------------------------------------------------------- [ 40240]
AM712239.1_Benin_97/1              ............g..........c...................ac.t................................. [ 40240]
KM111295.1_Ken06.Bus               ............g..........c....................c.t............t..t................. [ 40240]
```

```
KM111294.1_Ken05/Tk1             ............g..........c...................ac.t............t..c.........c....... [ 40240]
KM262844.1_strain_L60            ............g..........c...................ac.t................................. [ 40240]
KM102979.1_26544/OG10_Italy      ............g..........c...................ac.t................................. [ 40240]
KM262845.1_strain_NHV            -------------------------------------------------------------------------------- [ 40240]
KX354450.1_47/Ss/2008            ............g..........c...................ac.t................................. [ 40240]
KP055815.1_strain_BA71           ............g..........c...................ac.t................................. [ 40240]
MH025920.1_strain_R35            ............g..........c....................c.t............t..t................. [ 40240]
MH025919.1_strain_N10            ............g..........c....................c.t............t..t................. [ 40240]
MH025918.1_strain_R25            ............g..........c....................c.t............t..t................. [ 40240]
MH025917.1_strain_R7             ............g..........c....................c.t............t..t................. [ 40240]
MH025916.1_strain_R8             ............g..........c....................c.t............t..t................. [ 40240]

FR682468.2_Georgia_2007/1        tttgatgcaggcgatccgcatcatctcttctatgtccatatggatcttgtccttttcatacgcctcatgaaggtcaaaca [ 40320]
ASF_IND_20_CAD543                -------------------------------------------------------------------------------- [ 40320]
LR899193.1_Germany_2020/1        ................................................................................ [ 40320]
MN393477.1_Wuhan_2019            ................................................................................ [ 40320]
MW396979.1_Timor-Leste/2019/1    ................................................................................ [ 40320]
MK940252.1_CN/2019/InnerMongolia ................................................................................ [ 40320]
MK128995.1_China/2018/AnhuiXCGQ  ................................................................................ [ 40320]
MH681419.1_ASFV/POL/2015         ................................................................................ [ 40320]
LS478113.1_Estonia_2014          ................................................................................ [ 40320]
KP843857.1_Odintsovo_02/14       ................................................................................ [ 40320]
FN557520.1_strain_E75            ............a........................................................g.......... [ 40320]
AY261366.1_Warthog               ................................................................................ [ 40320]
AY261365.1_Warmbaths             .........................................a...................................... [ 40320]
AY261364.1_Tengani_62            ................................................................................ [ 40320]
AY261363.1_Pretoriuskop/96/4     ..........................................a..................................... [ 40320]
AY261362.1_Mkuzi_1979            ...a.....................................a...........................g.......... [ 40320]
AY261361.1_Malawi_Lil-20/1_1983  ............t.ct................a........a...................................... [ 40320]
AY261360.1_Kenya_1950            ............t...................aa......ca...................................... [ 40320]
AM712240.1_OURT_88/3             -------------------------------------------------------------------------------- [ 40320]
AM712239.1_Benin_97/1            ............a........................................................g.......... [ 40320]
KM111295.1_Ken06.Bus             ............t...................aa......ca...................................... [ 40320]
KM111294.1_Ken05/Tk1             ............t...................aa......c....................................... [ 40320]
KM262844.1_strain_L60            ............a........................................................g.......... [ 40320]
KM102979.1_26544/OG10_Italy      ............a........................................................g.......... [ 40320]
KM262845.1_strain_NHV            -------------------------------------------------------------------------------- [ 40320]
KX354450.1_47/Ss/2008            ............a........................................................g.......... [ 40320]
KP055815.1_strain_BA71           ............a........................................................g.......... [ 40320]
MH025920.1_strain_R35            ............t...................aa......ca...................................... [ 40320]
MH025919.1_strain_N10            ............t...................aa......ca...................................... [ 40320]
MH025918.1_strain_R25            ............t...................aa......ca...................................... [ 40320]
MH025917.1_strain_R7             ............t...................aa......ca...................................... [ 40320]
MH025916.1_strain_R8             ............t...................aa......ca...................................... [ 40320]
```

```
FR682468.2_Georgia_2007/1        cattattaaaacaaagagcacatgttaaccgccacgtattcaggtgtgtatatttttggtaaaaatactgtatggcctct [ 40400]
ASF_IND_20_CAD543                -------------------------------------------------------------------------------- [ 40400]
LR899193.1_Germany_2020/1        ................................................................................ [ 40400]
MN393477.1_Wuhan_2019            ................................................................................ [ 40400]
MW396979.1_Timor-Leste/2019/1    ................................................................................ [ 40400]
MK940252.1_CN/2019/InnerMongolia ................................................................................ [ 40400]
MK128995.1_China/2018/AnhuiXCGQ  ................................................................................ [ 40400]
MH681419.1_ASFV/POL/2015         ................................................................................ [ 40400]
LS478113.1_Estonia_2014          ................................................................................ [ 40400]
KP843857.1_Odintsovo_02/14       ................................................................................ [ 40400]
FN557520.1_strain_E75            ................................................................................ [ 40400]
AY261366.1_Warthog               ................................................................................ [ 40400]
AY261365.1_Warmbaths             ..................................t............................................. [ 40400]
AY261364.1_Tengani_62            ................................................................................ [ 40400]
AY261363.1_Pretoriuskop/96/4     ................................................................................ [ 40400]
AY261362.1_Mkuzi_1979            ............................................................................t... [ 40400]
AY261361.1_Malawi_Lil-20/1_1983  ..........................................................a..................... [ 40400]
AY261360.1_Kenya_1950            .g..........................t.................c................................. [ 40400]
AM712240.1_OURT_88/3             -------------------------------------------------------------------------------- [ 40400]
AM712239.1_Benin_97/1            ..........................................t..................................... [ 40400]
KM111295.1_Ken06.Bus             .g.............................................................................. [ 40400]
KM111294.1_Ken05/Tk1             .g.............................................................................. [ 40400]
KM262844.1_strain_L60            ................................................................................ [ 40400]
KM102979.1_26544/OG10_Italy      ................................................................................ [ 40400]
KM262845.1_strain_NHV            -------------------------------------------------------------------------------- [ 40400]
KX354450.1_47/Ss/2008            ................................................................................ [ 40400]
KP055815.1_strain_BA71           ................................................................................ [ 40400]
MH025920.1_strain_R35            .g.............................................................................. [ 40400]
MH025919.1_strain_N10            .g.............................................................................. [ 40400]
MH025918.1_strain_R25            .g.............................................................................. [ 40400]
MH025917.1_strain_R7             .g.............................................................................. [ 40400]
MH025916.1_strain_R8             .g.............................................................................. [ 40400]

FR682468.2_Georgia_2007/1        ttcaggttatagcgtatggctatagcgtaccagtatttgagtagtaatgtactgagcgaaaactcattatttagcagatc [ 40480]
ASF_IND_20_CAD543                -------------------------------------------------------------------------------- [ 40480]
LR899193.1_Germany_2020/1        ................................................................................ [ 40480]
MN393477.1_Wuhan_2019            ................................................................................ [ 40480]
MW396979.1_Timor-Leste/2019/1    ................................................................................ [ 40480]
MK940252.1_CN/2019/InnerMongolia ................................................................................ [ 40480]
MK128995.1_China/2018/AnhuiXCGQ  ................................................................................ [ 40480]
MH681419.1_ASFV/POL/2015         ................................................................................ [ 40480]
LS478113.1_Estonia_2014          ................................................................................ [ 40480]
KP843857.1_Odintsovo_02/14       ................................................................................ [ 40480]
```

```
FN557520.1_strain_E75          ......c.........c........................................a...................... [ 40480]
AY261366.1_Warthog             .................................................c.............................. [ 40480]
AY261365.1_Warmbaths           ................................................................................ [ 40480]
AY261364.1_Tengani_62          ................................................................................ [ 40480]
AY261363.1_Pretoriuskop/96/4   ......................................a......................................... [ 40480]
AY261362.1_Mkuzi_1979          ............................................c................................... [ 40480]
AY261361.1_Malawi_Lil-20/1_1983 ................c..........................a.......t....t................a...... [ 40480]
AY261360.1_Kenya_1950          ....a...g.......c..........................a..g....t............................ [ 40480]
AM712240.1_OURT_88/3           -------------------------------------------------------------------------------- [ 40480]
AM712239.1_Benin_97/1          ......c.........c........................................a...................... [ 40480]
KM111295.1_Ken06.Bus           ....a...g.......c..........................a..g....t............................ [ 40480]
KM111294.1_Ken05/Tk1           ................c..........................a..g....t.a.......................... [ 40480]
KM262844.1_strain_L60          ......c.........c........................................a...................... [ 40480]
KM102979.1_26544/OG10_Italy    ......c...t.....c........................................a...................... [ 40480]
KM262845.1_strain_NHV          -------------------------------------------------------------------------------- [ 40480]
KX354450.1_47/Ss/2008          ......c...t.....c........................................a...................... [ 40480]
KP055815.1_strain_BA71         ......c.........c........................................a...................... [ 40480]
MH025920.1_strain_R35          ....a...g.......c..........................a..g....t............................ [ 40480]
MH025919.1_strain_N10          ....a...g.......c..........................a..g....t............................ [ 40480]
MH025918.1_strain_R25          ....a...g.......c..........................a..g....t............................ [ 40480]
MH025917.1_strain_R7           ....a...g.......c..........................a..g....t............................ [ 40480]
MH025916.1_strain_R8           ....a...g.......c..........................a..g....t............................ [ 40480]

FR682468.2_Georgia_2007/1      ggtttttactattaactcccttaactcccagaaaatttctatcctcatttttatattatttactttttgtaatatcggat [ 40560]
ASF_IND_20_CAD543              -------------------------------------------------------------------------------- [ 40560]
LR899193.1_Germany_2020/1      ................................................................................ [ 40560]
MN393477.1_Wuhan_2019          ................................................................................ [ 40560]
MW396979.1_Timor-Leste/2019/1  ................................................................................ [ 40560]
MK940252.1_CN/2019/InnerMongolia ................................................................................ [ 40560]
MK128995.1_China/2018/AnhuiXCGQ ................................................................................ [ 40560]
MH681419.1_ASFV/POL/2015       ................................................................................ [ 40560]
LS478113.1_Estonia_2014        ................................................................................ [ 40560]
KP843857.1_Odintsovo_02/14     ................................................................................ [ 40560]
FN557520.1_strain_E75          .......t.......................................a...c.g.......................... [ 40560]
AY261366.1_Warthog             .......t........................................................................ [ 40560]
AY261365.1_Warmbaths           .......t........................................................................ [ 40560]
AY261364.1_Tengani_62          .......t........................................................................ [ 40560]
AY261363.1_Pretoriuskop/96/4   .......t........................................................................ [ 40560]
AY261362.1_Mkuzi_1979          .......t.....................a................................................. [ 40560]
AY261361.1_Malawi_Lil-20/1_1983 .......tt..........................................c.g....................a..... [ 40560]
AY261360.1_Kenya_1950          .......t......t...............a....................c.gg........c....t...a.gta... [ 40560]
AM712240.1_OURT_88/3           -------------------------------------------------------------------------------- [ 40560]
AM712239.1_Benin_97/1          .......t.......................................a...c.g.......................... [ 40560]
KM111295.1_Ken06.Bus           .......t......t...............a....................c.gg........cc...t...a.gtaa.. [ 40560]
```

```
KM111294.1_Ken05/Tk1          .......t......t...............a......................gg........c....t...a.gta... [ 40560]
KM262844.1_strain_L60         .......t.......................................a...c.g.......................... [ 40560]
KM102979.1_26544/OG10_Italy   .......t.......................................a...c.g.......................... [ 40560]
KM262845.1_strain_NHV         -------------------------------------------------------------------------------- [ 40560]
KX354450.1_47/Ss/2008         .......t.......................................a...c.g.......................... [ 40560]
KP055815.1_strain_BA71        .......t.......................................a...c.g.......................... [ 40560]
MH025920.1_strain_R35         .......t......t...............a....................c.gg........cc...t...a.gtaa.. [ 40560]
MH025919.1_strain_N10         .......tt.....t...............a....................c.gg........cc...t...a.gtaa.. [ 40560]
MH025918.1_strain_R25         .......t......t...............a....................c.gg........cc...t...a.gtaa.. [ 40560]
MH025917.1_strain_R7          .......t......t...............a....................c.gg........cc...t...a.gtaa.. [ 40560]
MH025916.1_strain_R8          .......t......t...............a....................c.gg........cc...t...a.gtaa.. [ 40560]

FR682468.2_Georgia_2007/1     tgttggaaaacacctcatggcataaaataatgttactactagttttatgaaactttagatctataaaaatttgtaaaatt [ 40640]
ASF_IND_20_CAD543             -------------------------------------------------------------------------------- [ 40640]
LR899193.1_Germany_2020/1     ................................................................................ [ 40640]
MN393477.1_Wuhan_2019         ................................................................................ [ 40640]
MW396979.1_Timor-Leste/2019/1 ................................................................................ [ 40640]
MK940252.1_CN/2019/InnerMongolia ................................................................................ [ 40640]
MK128995.1_China/2018/AnhuiXCGQ ................................................................................ [ 40640]
MH681419.1_ASFV/POL/2015      ................................................................................ [ 40640]
LS478113.1_Estonia_2014       ................................................................................ [ 40640]
KP843857.1_Odintsovo_02/14    ................................................................................ [ 40640]
FN557520.1_strain_E75         ................................................................g............... [ 40640]
AY261366.1_Warthog            ................................................................................ [ 40640]
AY261365.1_Warmbaths          ................................................................................ [ 40640]
AY261364.1_Tengani_62         ................................................................................ [ 40640]
AY261363.1_Pretoriuskop/96/4  ................................................................................ [ 40640]
AY261362.1_Mkuzi_1979         ................................................................................ [ 40640]
AY261361.1_Malawi_Lil-20/1_1983 ................................................a............g..g......c........ [ 40640]
AY261360.1_Kenya_1950         .a.c...............a.....g..............g................a......g..............a [ 40640]
AM712240.1_OURT_88/3          -------------------------------------------------------------------------------- [ 40640]
AM712239.1_Benin_97/1         ................................................................g............... [ 40640]
KM111295.1_Ken06.Bus          .a.c...............a.....g..............g.......a...t....a......g..............a [ 40640]
KM111294.1_Ken05/Tk1          .a.c...............a.....g.....a........g................a......g..............a [ 40640]
KM262844.1_strain_L60         ................................................................g............... [ 40640]
KM102979.1_26544/OG10_Italy   ................................................................g............... [ 40640]
KM262845.1_strain_NHV         -------------------------------------------------------------------------------- [ 40640]
KX354450.1_47/Ss/2008         ................................................................g............... [ 40640]
KP055815.1_strain_BA71        ................................................................g............... [ 40640]
MH025920.1_strain_R35         .a.c...............a.....g..............g.......a...t....a......g..............a [ 40640]
MH025919.1_strain_N10         .a.c...............a.....g..............g.......a...t....a......g..............a [ 40640]
MH025918.1_strain_R25         .a.c...............a.....g..............g.......a...t....a......g..............a [ 40640]
MH025917.1_strain_R7          .a.c...............a.....g..............g.......a...t....a......g..............a [ 40640]
MH025916.1_strain_R8          .a.c...............a.....g..............g.......a...t....a......g..............a [ 40640]
```

```
FR682468.2_Georgia_2007/1        tcttcttcattcaaggtttccttggcacctagctctcgacagaggtcccaggtgtgctccgtgttgacagataccagccc [ 40720]
ASF_IND_20_CAD543                -------------------------------------------------------------------------------- [ 40720]
LR899193.1_Germany_2020/1        ................................................................................ [ 40720]
MN393477.1_Wuhan_2019            ................................................................................ [ 40720]
MW396979.1_Timor-Leste/2019/1    ................................................................................ [ 40720]
MK940252.1_CN/2019/InnerMongolia ................................................................................ [ 40720]
MK128995.1_China/2018/AnhuiXCGQ  ................................................................................ [ 40720]
MH681419.1_ASFV/POL/2015         ................................................................................ [ 40720]
LS478113.1_Estonia_2014          ................................................................................ [ 40720]
KP843857.1_Odintsovo_02/14       ................................................................................ [ 40720]
FN557520.1_strain_E75            ..c............a.c.............a....t............g...a.......................... [ 40720]
AY261366.1_Warthog               .................c...............................g.............................. [ 40720]
AY261365.1_Warmbaths             .................c...............................g.............................. [ 40720]
AY261364.1_Tengani_62            .................c.............................................................. [ 40720]
AY261363.1_Pretoriuskop/96/4     .................c.............................................................. [ 40720]
AY261362.1_Mkuzi_1979            ....t............c...........................................a.c................ [ 40720]
AY261361.1_Malawi_Lil-20/1_1983  ..c........t.....c........g..........tg..........g.............................. [ 40720]
AY261360.1_Kenya_1950            ..cc.......t.....c.t......g..............................................t...... [ 40720]
AM712240.1_OURT_88/3             -------------------------------------------------------------------------------- [ 40720]
AM712239.1_Benin_97/1            ..c............a.c.............a....t............g...a.......................... [ 40720]
KM111295.1_Ken06.Bus             ..cc.......t.....c.t......g..............................................t...... [ 40720]
KM111294.1_Ken05/Tk1             ..cc.......t.....c.t......g.................a............................t...... [ 40720]
KM262844.1_strain_L60            ..c............a.c.............a....t............g...a.......................... [ 40720]
KM102979.1_26544/OG10_Italy      ..c............a.c.............a....t............g...a.......................... [ 40720]
KM262845.1_strain_NHV            -------------------------------------------------------------------------------- [ 40720]
KX354450.1_47/Ss/2008            ..c............a.c.............a....t............g...a.......................... [ 40720]
KP055815.1_strain_BA71           ..c............a.c.............a....t............g...a.......................... [ 40720]
MH025920.1_strain_R35            ..cc.......t.....c.t......g..............................................t...... [ 40720]
MH025919.1_strain_N10            ..cc.......t.....c.t......g..............................................t...... [ 40720]
MH025918.1_strain_R25            ..cc.......t.....c.t......g..............................................t...... [ 40720]
MH025917.1_strain_R7             ..cc.......t.....c.t......g..............................................t...... [ 40720]
MH025916.1_strain_R8             ..cc.......t.....c.t......g..............................................t...... [ 40720]

FR682468.2_Georgia_2007/1        gtagttgatgtccgccccccactctgcaaacagttttataaggttgtagttgttttcccttacagccttcactaacgccg [ 40800]
ASF_IND_20_CAD543                -------------------------------------------------------------------------------- [ 40800]
LR899193.1_Germany_2020/1        ................................................................................ [ 40800]
MN393477.1_Wuhan_2019            ................................................................................ [ 40800]
MW396979.1_Timor-Leste/2019/1    ................................................................................ [ 40800]
MK940252.1_CN/2019/InnerMongolia ................................................................................ [ 40800]
MK128995.1_China/2018/AnhuiXCGQ  ................................................................................ [ 40800]
MH681419.1_ASFV/POL/2015         ................................................................................ [ 40800]
LS478113.1_Estonia_2014          ................................................................................ [ 40800]
KP843857.1_Odintsovo_02/14       ................................................................................ [ 40800]
```

```
FN557520.1_strain_E75           a.............-...........t...t............c.........................t.......... [ 40800]
AY261366.1_Warthog              a..........t....................................a............................... [ 40800]
AY261365.1_Warmbaths            a............................................................................... [ 40800]
AY261364.1_Tengani_62           .....................................................................t.......... [ 40800]
AY261363.1_Pretoriuskop/96/4    a..........t.................................................................... [ 40800]
AY261362.1_Mkuzi_1979           a...............................a.........................t.............c....... [ 40800]
AY261361.1_Malawi_Lil-20/1_1983 ..........................t...a..c...................................t.......... [ 40800]
AY261360.1_Kenya_1950           ...........t...t........c.t...t...........ac..............t................t...a [ 40800]
AM712240.1_OURT_88/3            -------------------------------------------------------------------------------- [ 40800]
AM712239.1_Benin_97/1           a.........................t...t............c.........................t.......... [ 40800]
KM111295.1_Ken06.Bus            ......a....t............c.t...t...........ac..............t................t...a [ 40800]
KM111294.1_Ken05/Tk1            ...........t...t........c.t...t...........ac..............t................t...a [ 40800]
KM262844.1_strain_L60           a.........................t...t............c.........................t.......... [ 40800]
KM102979.1_26544/OG10_Italy     a.........................t...t............c.........................t.......... [ 40800]
KM262845.1_strain_NHV           -------------------------------------------------------------------------------- [ 40800]
KX354450.1_47/Ss/2008           a.........................t...t............c.........................t.......... [ 40800]
KP055815.1_strain_BA71          a.........................t...t............c.........................t.......... [ 40800]
MH025920.1_strain_R35           ......a....t............c.t...t...........ac..............t................t...a [ 40800]
MH025919.1_strain_N10           ......a....t............c.t...t...........ac..............t................t...a [ 40800]
MH025918.1_strain_R25           ......a....t............c.t...t...........ac..............t................t...a [ 40800]
MH025917.1_strain_R7            ......a....t............c.t...t...........ac..............t................t...a [ 40800]
MH025916.1_strain_R8            ......a....t............c.t...t...........ac..............t................t...a [ 40800]

FR682468.2_Georgia_2007/1       tatttaggtttaagccctctttaatacctgctgattttatgagccttaggttatgatcaaacgtgatcggagcatcatgc [ 40880]
ASF_IND_20_CAD543               -------------------------------------------------------------------------------- [ 40880]
LR899193.1_Germany_2020/1       ................................................................................ [ 40880]
MN393477.1_Wuhan_2019           ................................................................................ [ 40880]
MW396979.1_Timor-Leste/2019/1   ................................................................................ [ 40880]
MK940252.1_CN/2019/InnerMongolia ................................................................................ [ 40880]
MK128995.1_China/2018/AnhuiXCGQ ................................................................................ [ 40880]
MH681419.1_ASFV/POL/2015        ................................................................................ [ 40880]
LS478113.1_Estonia_2014         ................................................................................ [ 40880]
KP843857.1_Odintsovo_02/14      ................................................................................ [ 40880]
FN557520.1_strain_E75           .........c...............g....a............................................t..a. [ 40880]
AY261366.1_Warthog              .........c...............g..................t..........................c........ [ 40880]
AY261365.1_Warmbaths            .........c...............g..................t..........................c........ [ 40880]
AY261364.1_Tengani_62           ................................................................................ [ 40880]
AY261363.1_Pretoriuskop/96/4    .........c...............g..................t..........................c........ [ 40880]
AY261362.1_Mkuzi_1979           .........c...............g...................................................... [ 40880]
AY261361.1_Malawi_Lil-20/1_1983 .........c......a........g..............c....c..........................a.....a. [ 40880]
AY260360.1_Kenya_1950           .g.......c......t....g..gg..............c..................................t..t. [ 40880]
AM712240.1_OURT_88/3            -------------------------------------------------------------------------------- [ 40880]
AM712239.1_Benin_97/1           .........c...............g....a............................................t..a. [ 40880]
KM111295.1_Ken06.Bus            .g.......c......t....g...g......a.......c..................................t..t. [ 40880]
```

KM111294.1_Ken05/Tk1 .g.......c......t....g..gg......a.......c..................................t..t. [ 40880]
KM262844.1_strain_L60 .........c...............g....a............................................t..a. [ 40880]
KM102979.1_26544/OG10_Italy .........c...............g....a............................................t..a. [ 40880]
KM262845.1_strain_NHV -------------------------------------------------------------------------------- [ 40880]
KX354450.1_47/Ss/2008 .........c...............g....a............................................t..a. [ 40880]
KP055815.1_strain_BA71 .........c...............g....a............................................t..a. [ 40880]
MH025920.1_strain_R35 .g.......c......t....g...g......a.......c..................................t..t. [ 40880]
MH025919.1_strain_N10 .g.......c......t....g...g......a.......c..................................t..t. [ 40880]
MH025918.1_strain_R25 .g.......c......t....g...g......a.......c..................................t..t. [ 40880]
MH025917.1_strain_R7 .g.......c......t....g...g......a.......c..................................t..t. [ 40880]
MH025916.1_strain_R8 .g.......c......t....g...g......a.......c..................................t..t. [ 40880]

FR682468.2_Georgia_2007/1 caccataggtcataacactttaaaagataatgttggttcgtgggcacgcattgtccagccaacaccttttttggtcagaga [ 40960]
ASF_IND_20_CAD543 -------------------------------------------------------------------------------- [ 40960]
LR899193.1_Germany_2020/1 ................................................................................ [ 40960]
MN393477.1_Wuhan_2019 ................................................................................ [ 40960]
MW396979.1_Timor-Leste/2019/1 ................................................................................ [ 40960]
MK940252.1_CN/2019/InnerMongolia ................................................................................ [ 40960]
MK128995.1_China/2018/AnhuiXCGQ ................................................................................ [ 40960]
MH681419.1_ASFV/POL/2015 ................................................................................ [ 40960]
LS478113.1_Estonia_2014 ................................................................................ [ 40960]
KP843857.1_Odintsovo_02/14 ................................................................................ [ 40960]
FN557520.1_strain_E75 ........a......t.t..........................................................a... [ 40960]
AY261366.1_Warthog ................................................................................ [ 40960]
AY261365.1_Warmbaths ................................................................................ [ 40960]
AY261364.1_Tengani_62 ................................................................................ [ 40960]
AY261363.1_Pretoriuskop/96/4 ...........g..........................t......................................... [ 40960]
AY261362.1_Mkuzi_1979 ...............t.t......................................g...................a... [ 40960]
AY261361.1_Malawi_Lil-20/1_1983 ...........g...................a.....c...c....g..............gt................. [ 40960]
AY261360.1_Kenya_1950 ........a..g.........................c.tc.....ga.............g.................. [ 40960]
AM712240.1_OURT_88/3 -------------------------------------------------------------------------------- [ 40960]
AM712239.1_Benin_97/1 ........a......t.t..........................................................a... [ 40960]
KM111295.1_Ken06.Bus ........a..g.........................c.tc.....ga.......t.....g..............a... [ 40960]
KM111294.1_Ken05/Tk1 ........a..g.........................c.tc.....ga.............g.................. [ 40960]
KM262844.1_strain_L60 ........a......t.t..........................................................a... [ 40960]
KM102979.1_26544/OG10_Italy ........a......t.t..........................................................a... [ 40960]
KM262845.1_strain_NHV -------------------------------------------------------------------------------- [ 40960]
KX354450.1_47/Ss/2008 ........a......t.t..........................................................a... [ 40960]
KP055815.1_strain_BA71 ........a......t.t..........................................................a... [ 40960]
MH025920.1_strain_R35 ........a..g.........................c.tc.....ga.......t.....g..............a... [ 40960]
MH025919.1_strain_N10 ........a..g.........................c.tc.....ga.......t.....g..............a... [ 40960]
MH025918.1_strain_R25 ........a..g.........................c.tc.....ga.......t.....g..............a... [ 40960]
MH025917.1_strain_R7 ........a..g.........................c.tc.....ga.......t.....g..............a... [ 40960]
MH025916.1_strain_R8 ........a..g.........................c.tc.....ga.......t.....g..............a... [ 40960]

```
FR682468.2_Georgia_2007/1        ttgcagggaaggcaacatgtctcttcatcttttaaaaaaaaatcaaattaattag-ccgaataaatttttctttcgaggg [ 41040]
ASF_IND_20_CAD543                -------------------------------------------------------------------------------- [ 41040]
LR899193.1_Germany_2020/1        .......................................................-........................ [ 41040]
MN393477.1_Wuhan_2019            .......................................................-........................ [ 41040]
MW396979.1_Timor-Leste/2019/1    .......................................................-........................ [ 41040]
MK940252.1_CN/2019/InnerMongolia .......................................................-........................ [ 41040]
MK128995.1_China/2018/AnhuiXCGQ  .......................................................-........................ [ 41040]
MH681419.1_ASFV/POL/2015         .......................................................-........................ [ 41040]
LS478113.1_Estonia_2014          .......................................................-........................ [ 41040]
KP843857.1_Odintsovo_02/14       .......................................................-........................ [ 41040]
FN557520.1_strain_E75            ...................................................c...-.........a.............. [ 41040]
AY261366.1_Warthog               ...................................................c...-.....................aa. [ 41040]
AY261365.1_Warmbaths             ...................................................c...-.....................a.. [ 41040]
AY261364.1_Tengani_62            .......................................................-........................ [ 41040]
AY261363.1_Pretoriuskop/96/4     ...................................................c...-.....................a.. [ 41040]
AY261362.1_Mkuzi_1979            ...................................................c...-.........a.............. [ 41040]
AY261361.1_Malawi_Lil-20/1_1983  ................................a.....................c-....................ta.a [ 41040]
AY261360.1_Kenya_1950            ...................................................c.tct...g...............ac..a [ 41040]
AM712240.1_OURT_88/3             -------------------------------------------------------------------------------- [ 41040]
AM712239.1_Benin_97/1            ...................................................c...-.........a.............. [ 41040]
KM111295.1_Ken06.Bus             ................................a..................c.tct...g...............ac..a [ 41040]
KM111294.1_Ken05/Tk1             ...................................................c.tct...g...............ac..a [ 41040]
KM262844.1_strain_L60            ...................................................c...-.........a.............. [ 41040]
KM102979.1_26544/OG10_Italy      ...................................................c...-.........a.............. [ 41040]
KM262845.1_strain_NHV            -------------------------------------------------------------------------------- [ 41040]
KX354450.1_47/Ss/2008            ...................................................c...-.........a.............. [ 41040]
KP055815.1_strain_BA71           ...................................................c...-.........a.............. [ 41040]
MH025920.1_strain_R35            ...................................................c.tct...g...............ac..a [ 41040]
MH025919.1_strain_N10            ...................................................c.tct...g...............ac..a [ 41040]
MH025918.1_strain_R25            ...................................................c.tct...g...............ac..a [ 41040]
MH025917.1_strain_R7             ...................................................c.tct...g...............ac..a [ 41040]
MH025916.1_strain_R8             ...................................................c.tct...g...............ac..a [ 41040]

FR682468.2_Georgia_2007/1        ctttt----------------------------t--aaaagagctcttt----aag----------agctctttaagagc [ 41120]
ASF_IND_20_CAD543                -------------------------------------------------------------------------------- [ 41120]
LR899193.1_Germany_2020/1        .....----------------------------.--.............----...----------.............. [ 41120]
MN393477.1_Wuhan_2019            .....----------------------------.--.............----...----------.............. [ 41120]
MW396979.1_Timor-Leste/2019/1    .....----------------------------.--.............----...----------.............. [ 41120]
MK940252.1_CN/2019/InnerMongolia .....----------------------------.--.............----...----------.............. [ 41120]
MK128995.1_China/2018/AnhuiXCGQ  .....----------------------------.--.............----...----------.............. [ 41120]
MH681419.1_ASFV/POL/2015         .....----------------------------.--.............----...----------.............. [ 41120]
LS478113.1_Estonia_2014          .....----------------------------.--.............----...----------.............. [ 41120]
KP843857.1_Odintsovo_02/14       .....----------------------------.--.............----...----------.............. [ 41120]
```

```
FN557520.1_strain_E75           .....----------------------------.ta......a......----...agcttttaaa....t..a..a... [ 41120]
AY261366.1_Warthog              .....taaaagagagctctttaagagatatatc.tt.......a.....----..aag--------.ta..........a [ 41120]
AY261365.1_Warmbaths            ...------------------------------.tt...g.........----...ag--------.ta..........a [ 41120]
AY261364.1_Tengani_62           ...------------------------------.tt.............----...----------.............. [ 41120]
AY261363.1_Pretoriuskop/96/4    ...------------------------------.tt.......a.a.c.ttaa...at--------.ta..........a [ 41120]
AY261362.1_Mkuzi_1979           .....----------------------------.ta......a......----...----------....t.....a... [ 41120]
AY261361.1_Malawi_Lil-20/1_1983 .....----------------------------.ca...---------------------------....t.....a... [ 41120]
AY261360.1_Kenya_1950           ..g..----------------------------.ta.........t..c----..aa---------....t--------- [ 41120]
AM712240.1_OURT_88/3            -------------------------------------------------------------------------------- [ 41120]
AM712239.1_Benin_97/1           .....----------------------------.ta......a......----...agcttttaaa....t..a..a... [ 41120]
KM111295.1_Ken06.Bus            ..g..----------------------------.ta.........t..c----..aa---------....t..c..a.ag [ 41120]
KM111294.1_Ken05/Tk1            ..g..----------------------------.ta.........t..c----..aa---------....t--------- [ 41120]
KM262844.1_strain_L60           .....----------------------------.ta......a......----...agcttttaaa....t..a..a... [ 41120]
KM102979.1_26544/OG10_Italy     .....----------------------------.ta......a......----...agcttttaaa....t..a..a... [ 41120]
KM262845.1_strain_NHV           -------------------------------------------------------------------------------- [ 41120]
KX354450.1_47/Ss/2008           .....----------------------------.ta......a......----...agcttttaaa....t..a..a... [ 41120]
KP055815.1_strain_BA71          .....----------------------------.ta......a......----...agcttttaaa....t..a..a... [ 41120]
MH025920.1_strain_R35           ..g..----------------------------.ta.........t..c----..aa---------....t..c..a.ag [ 41120]
MH025919.1_strain_N10           ..g..----------------------------.ta.........t..c----..aa---------....t..c..a.ag [ 41120]
MH025918.1_strain_R25           ..g..----------------------------.ta.........t..c----..aa---------....t..c..a.ag [ 41120]
MH025917.1_strain_R7            ..g..----------------------------.ta.........t..c----..aa---------....t..c..a.ag [ 41120]
MH025916.1_strain_R8            ..g..----------------------------.ta.........t..c----..aa---------....t..c..a.ag [ 41120]

FR682468.2_Georgia_2007/1       tt--tttaagagattaaaaa--attattcttgctggcattctgccaagtatgcgg------------------------- [ 41200]
ASF_IND_20_CAD543               -------------------------------------------------------------------------------- [ 41200]
LR899193.1_Germany_2020/1       ..--................--.................................------------------------- [ 41200]
MN393477.1_Wuhan_2019           ..--................--.................................------------------------- [ 41200]
MW396979.1_Timor-Leste/2019/1   ..--................--.................................------------------------- [ 41200]
MK940252.1_CN/2019/InnerMongolia ..--................--.................................------------------------- [ 41200]
MK128995.1_China/2018/AnhuiXCGQ ..--................--.................................------------------------- [ 41200]
MH681419.1_ASFV/POL/2015        ..--................--.................................------------------------- [ 41200]
LS478113.1_Estonia_2014         ..--................--.................................------------------------- [ 41200]
KP843857.1_Odintsovo_02/14      ..--................--.................................------------------------- [ 41200]
FN557520.1_strain_E75           ..--...............---.....c............t.......c......------------------------- [ 41200]
AY261366.1_Warthog              .c--...............----....c............t........g...a-------------------------- [ 41200]
AY261365.1_Warmbaths            ..-----------aa....----...cc............t..........a...------------------------- [ 41200]
AY261364.1_Tengani_62           ..--................--..g...............t..............------------------------- [ 41200]
AY261363.1_Pretoriuskop/96/4    ..-------------....----....c............t...........t..ca----------------------- [ 41200]
AY261362.1_Mkuzi_1979           ..--...............----....c............t.......c......------------------------- [ 41200]
AY261361.1_Malawi_Lil-20/1_1983 ..--..--.a.ag...g...--.a.cac...c........t........-.....caaacatatgtttgcaaacatatag [ 41200]
AY261360.1_Kenya_1950           ----...tta..t...g..---..c..c...t........t....gca.------------------------------- [ 41200]
AM712240.1_OURT_88/3            -------------------------------------------------------------------------------- [ 41200]
AM712239.1_Benin_97/1           ..--...............---.....c............t.......c......------------------------- [ 41200]
KM111295.1_Ken06.Bus            c.gc...tta..t...g..---..cg.c...c........t....gca.------------------------------- [ 41200]
```

```
KM111294.1_Ken05/Tk1                ----...tta..t...g...tc..c..c...c........t....gca.------------------------------- [ 41200]
KM262844.1_strain_L60               ..--...............---.....c............t.......c......------------------------- [ 41200]
KM102979.1_26544/OG10_Italy         ..--...............---.....c............t.......c......------------------------- [ 41200]
KM262845.1_strain_NHV               -------------------------------------------------------------------------------- [ 41200]
KX354450.1_47/Ss/2008               ..--...............---.....c............t.......c......------------------------- [ 41200]
KP055815.1_strain_BA71              ..--...............---.....c............t.......c......------------------------- [ 41200]
MH025920.1_strain_R35               c.gc...tta..t...g..---..cg.c...c........t....gca.------------------------------- [ 41200]
MH025919.1_strain_N10               c.gc...tta..t...g..---..cg.c...c........t....gca.------------------------------- [ 41200]
MH025918.1_strain_R25               c.gc...tta..t...g..---..cg.c...c........t....gca.------------------------------- [ 41200]
MH025917.1_strain_R7                c.gc...tta..t...g..---..cg.c...c........t....gca.------------------------------- [ 41200]
MH025916.1_strain_R8                c.gc...tta..t...g..---..cg.c...c........t....gca.------------------------------- [ 41200]

FR682468.2_Georgia_2007/1           ---------------cattc-ctatcatctatagtatattatgagaatattcccaaatgatggataagttttttgattta [ 41280]
ASF_IND_20_CAD543                   -------------------------------------------------------------------------------- [ 41280]
LR899193.1_Germany_2020/1           ---------------.....-........................................................... [ 41280]
MN393477.1_Wuhan_2019               ---------------.....-........................................................... [ 41280]
MW396979.1_Timor-Leste/2019/1       ---------------.....-........................................................... [ 41280]
MK940252.1_CN/2019/InnerMongolia    ---------------.....-........................................................... [ 41280]
MK128995.1_China/2018/AnhuiXCGQ     ---------------.....-........................................................... [ 41280]
MH681419.1_ASFV/POL/2015            ---------------.....-........................................................... [ 41280]
LS478113.1_Estonia_2014             ---------------.....-........................................................... [ 41280]
KP843857.1_Odintsovo_02/14          ---------------.....-........................................................... [ 41280]
FN557520.1_strain_E75               ---------------.....-............a.............................................. [ 41280]
AY261366.1_Warthog                  ----------------ca..-........................................................... [ 41280]
AY261365.1_Warmbaths                ---------------.....-........................................................... [ 41280]
AY261364.1_Tengani_62               ---------------.....-........................................................... [ 41280]
AY261363.1_Pretoriuskop/96/4        --------------g....g-t....................................t..................... [ 41280]
AY261362.1_Mkuzi_1979               ---------------.....-............a.............................................. [ 41280]
AY261361.1_Malawi_Lil-20/1_1983     ctatatgtagagcta...a.a....g..t..g..............g..a...a.............a............ [ 41280]
AY261360.1_Kenya_1950               ---------------tt.g.-.acg.gct..g.....t........g..a...ag............a............ [ 41280]
AM712240.1_OURT_88/3                -------------------------------------------------------------------------------- [ 41280]
AM712239.1_Benin_97/1               ---------------.....-............a.............................................. [ 41280]
KM111295.1_Ken06.Bus                ---------------tt.g.-.acg.gct..g.....t........g..a...ag............a............ [ 41280]
KM111294.1_Ken05/Tk1                ---------------tt.g.-.acg.gct..g.....t........g..a...ag............a............ [ 41280]
KM262844.1_strain_L60               ---------------.....-............a.............................................. [ 41280]
KM102979.1_26544/OG10_Italy         ---------------.....-............a.............................................. [ 41280]
KM262845.1_strain_NHV               -------------------------------------------------------------------------------- [ 41280]
KX354450.1_47/Ss/2008               ---------------.....-............a.............................................. [ 41280]
KP055815.1_strain_BA71              ---------------.....-............a.............................................. [ 41280]
MH025920.1_strain_R35               ---------------tt.g.-.acg.gct..g.....t........g..a...ag............a............ [ 41280]
MH025919.1_strain_N10               ---------------tt.g.-.acg.gct..g.....t........g..a...ag............a............ [ 41280]
MH025918.1_strain_R25               ---------------tt.g.-.acg.gct..g.....t........g..a...ag............a............ [ 41280]
MH025917.1_strain_R7                ---------------tt.g.-.acg.gct..g.....t........g..a...ag............a............ [ 41280]
MH025916.1_strain_R8                ---------------tt.g.-.acg.gct..g.....t........g..a...ag............a............ [ 41280]
```

```
FR682468.2_Georgia_2007/1        taatcttttaataaactgcttatttcttcggggtcctttaagtttagtggcaaggaagcatctgagctgtaaatatccaa [ 41360]
ASF_IND_20_CAD543                -------------------------------------------------------------------------------- [ 41360]
LR899193.1_Germany_2020/1        ................................................................................ [ 41360]
MN393477.1_Wuhan_2019            ................................................................................ [ 41360]
MW396979.1_Timor-Leste/2019/1    ................................................................................ [ 41360]
MK940252.1_CN/2019/InnerMongolia ................................................................................ [ 41360]
MK128995.1_China/2018/AnhuiXCGQ  ................................................................................ [ 41360]
MH681419.1_ASFV/POL/2015         ................................................................................ [ 41360]
LS478113.1_Estonia_2014          ................................................................................ [ 41360]
KP843857.1_Odintsovo_02/14       ................................................................................ [ 41360]
FN557520.1_strain_E75            .........................t...................a..................a.............. [ 41360]
AY261366.1_Warthog               .....c.......................................................................... [ 41360]
AY261365.1_Warmbaths             ...c.c.......................................................................... [ 41360]
AY261364.1_Tengani_62            ................................................................................ [ 41360]
AY261363.1_Pretoriuskop/96/4     .....c.....a.........................................................a.......... [ 41360]
AY261362.1_Mkuzi_1979            ..............................................a..................a.............. [ 41360]
AY261361.1_Malawi_Lil-20/1_1983  ....t......c..................................a...............c.....a.......t... [ 41360]
AY261360.1_Kenya_1950            ....t...........................aa............a.....................a.......t... [ 41360]
AM712240.1_OURT_88/3             -------------------------------------------------------------------------------- [ 41360]
AM712239.1_Benin_97/1            .........................t...................a..................a.............. [ 41360]
KM111295.1_Ken06.Bus             ....t...........................a.............a.....................a.......t... [ 41360]
KM111294.1_Ken05/Tk1             ....t...........................aa............a.....................a.......t... [ 41360]
KM262844.1_strain_L60            .........................t...................a..................a.............. [ 41360]
KM102979.1_26544/OG10_Italy      .........................t...................a..................a.............. [ 41360]
KM262845.1_strain_NHV            -------------------------------------------------------------------------------- [ 41360]
KX354450.1_47/Ss/2008            .........................t...................a..................a.............. [ 41360]
KP055815.1_strain_BA71           .........................t...................a..................a.............. [ 41360]
MH025920.1_strain_R35            ....t...........................a.............a.....................a.......t... [ 41360]
MH025919.1_strain_N10            ....t...........................a.............a.....................a.......t... [ 41360]
MH025918.1_strain_R25            ....t...........................a.............a.....................a.......t... [ 41360]
MH025917.1_strain_R7             ....t...........................a.............a.....................a.......t... [ 41360]
MH025916.1_strain_R8             ....t...........................a.............a.....................a.......t... [ 41360]

FR682468.2_Georgia_2007/1        agccaaactatggctcagaaaattataacctttttgttccgctatggcacgaccctcttcaaaggcattaccacccaaat [ 41440]
ASF_IND_20_CAD543                -------------------------------------------------------------------------------- [ 41440]
LR899193.1_Germany_2020/1        ................................................................................ [ 41440]
MN393477.1_Wuhan_2019            ................................................................................ [ 41440]
MW396979.1_Timor-Leste/2019/1    ................................................................................ [ 41440]
MK940252.1_CN/2019/InnerMongolia ................................................................................ [ 41440]
MK128995.1_China/2018/AnhuiXCGQ  ................................................................................ [ 41440]
MH681419.1_ASFV/POL/2015         ................................................................................ [ 41440]
LS478113.1_Estonia_2014          ................................................................................ [ 41440]
KP843857.1_Odintsovo_02/14       ................................................................................ [ 41440]
```

```
FN557520.1_strain_E75           ..t...g........t.a.............................................................. [ 41440]
AY261366.1_Warthog              ..t............t.......g.....................a.................................. [ 41440]
AY261365.1_Warmbaths            ...............t.......g........................................................ [ 41440]
AY261364.1_Tengani_62           ................................................................................ [ 41440]
AY261363.1_Pretoriuskop/96/4    ...............t................................................................ [ 41440]
AY261362.1_Mkuzi_1979           ..t...g........t.a......................................................c....... [ 41440]
AY261361.1_Malawi_Lil-20/1_1983 .a.............t.....................ct...........t............................. [ 41440]
AY261360.1_Kenya_1950           .att..........at.a............a......ct...........t............................. [ 41440]
AM712240.1_OURT_88/3            -------------------------------------------------------------------------------- [ 41440]
AM712239.1_Benin_97/1           ..t...g........t.a.............................................................. [ 41440]
KM111295.1_Ken06.Bus            .a.............t.a............a......ct...........t............................. [ 41440]
KM111294.1_Ken05/Tk1            .at...........at.a............a......ct...........t............................. [ 41440]
KM262844.1_strain_L60           ..t...g........t.a.............................................................. [ 41440]
KM102979.1_26544/OG10_Italy     ..t...g........t.a.............................................................. [ 41440]
KM262845.1_strain_NHV           -------------------------------------------------------------------------------- [ 41440]
KX354450.1_47/Ss/2008           ..t...g........t.a.............................................................. [ 41440]
KP055815.1_strain_BA71          ..t...g........t.a.............................................................. [ 41440]
MH025920.1_strain_R35           .a.............t.a............a......ct...........t............................. [ 41440]
MH025919.1_strain_N10           .a.............t.a............a......ct...........t............................. [ 41440]
MH025918.1_strain_R25           .a.............t.a............a......ct...........t............................. [ 41440]
MH025917.1_strain_R7            .a.............t.a............a......ct...........t............................. [ 41440]
MH025916.1_strain_R8            .a.............t.a............a......ct...........t............................. [ 41440]

FR682468.2_Georgia_2007/1       ctatacagaaaaatatattaccgatgttataatattgtactgaagtaagcatagcttggttgatgttgccccccagcgcg [ 41520]
ASF_IND_20_CAD543               -------------------------------------------------------------------------------- [ 41520]
LR899193.1_Germany_2020/1       ................................................................................ [ 41520]
MN393477.1_Wuhan_2019           ................................................................................ [ 41520]
MW396979.1_Timor-Leste/2019/1   ................................................................................ [ 41520]
MK940252.1_CN/2019/InnerMongolia ................................................................................ [ 41520]
MK128995.1_China/2018/AnhuiXCGQ ................................................................................ [ 41520]
MH681419.1_ASFV/POL/2015        ................................................................................ [ 41520]
LS478113.1_Estonia_2014         ................................................................................ [ 41520]
KP843857.1_Odintsovo_02/14      ................................................................................ [ 41520]
FN557520.1_strain_E75           ........................c.....................g.............................g... [ 41520]
AY261366.1_Warthog              ............................................................................g... [ 41520]
AY261365.1_Warmbaths            ............................................................................g... [ 41520]
AY261364.1_Tengani_62           ................................................................................ [ 41520]
AY261363.1_Pretoriuskop/96/4    .......................................t....................................g... [ 41520]
AY261362.1_Mkuzi_1979           ..............................................g......................t......g... [ 41520]
AY261361.1_Malawi_Lil-20/1_1983 ...........t.........................c......................................g..a [ 41520]
AY261360.1_Kenya_1950           ...........t................................................................g..a [ 41520]
AM712240.1_OURT_88/3            -------------------------------------------------------------------------------- [ 41520]
AM712239.1_Benin_97/1           ........................c.....................g.............................g... [ 41520]
KM111295.1_Ken06.Bus            ...........t..............................................a.................g..a [ 41520]
```

```
KM111294.1_Ken05/Tk1             ...........t................................................................ga.a [ 41520]
KM262844.1_strain_L60            ........................c.....................g.............................g... [ 41520]
KM102979.1_26544/OG10_Italy      ........................c.....................g.............................g... [ 41520]
KM262845.1_strain_NHV            -------------------------------------------------------------------------------- [ 41520]
KX354450.1_47/Ss/2008            ........................c.....................g.............................g... [ 41520]
KP055815.1_strain_BA71           ........................c.....................g.............................g... [ 41520]
MH025920.1_strain_R35            ...........t..............................................a.................g..a [ 41520]
MH025919.1_strain_N10            ...........t..............................................a.................g..a [ 41520]
MH025918.1_strain_R25            ...........t..............................................a.................g..a [ 41520]
MH025917.1_strain_R7             ...........t..............................................a.................g..a [ 41520]
MH025916.1_strain_R8             ...........t..............................................a.................g..a [ 41520]

FR682468.2_Georgia_2007/1        taacagtaatatattgttaatggattgttatccttggtagaagccagacatatcatgtcatggacgtctatttggatgtt [ 41600]
ASF_IND_20_CAD543                -------------------------------------------------------------------------------- [ 41600]
LR899193.1_Germany_2020/1        ................................................................................ [ 41600]
MN393477.1_Wuhan_2019            ................................................................................ [ 41600]
MW396979.1_Timor-Leste/2019/1    ................................................................................ [ 41600]
MK940252.1_CN/2019/InnerMongolia ................................................................................ [ 41600]
MK128995.1_China/2018/AnhuiXCGQ  ................................................................................ [ 41600]
MH681419.1_ASFV/POL/2015         ................................................................................ [ 41600]
LS478113.1_Estonia_2014          ................................................................................ [ 41600]
KP843857.1_Odintsovo_02/14       ................................................................................ [ 41600]
FN557520.1_strain_E75            .....................a...............c.c....................................ct.. [ 41600]
AY261366.1_Warthog               .....a......................................................................ct.. [ 41600]
AY261365.1_Warmbaths             ............................................................................ct.. [ 41600]
AY261364.1_Tengani_62            .......................g........................................................ [ 41600]
AY261363.1_Pretoriuskop/96/4     ............................................................................ct.. [ 41600]
AY261362.1_Mkuzi_1979            .....................a...............c.c....................................ct.. [ 41600]
AY261361.1_Malawi_Lil-20/1_1983  .................................c..a..c.......gg...a...t....ta..............t.. [ 41600]
AY261360.1_Kenya_1950            .....a...............a...........c..a..c........g...a...t....t...............t.. [ 41600]
AM712240.1_OURT_88/3             -------------------------------------------------------------------------------- [ 41600]
AM712239.1_Benin_97/1            .....................a...............c.c....................................ct.. [ 41600]
KM111295.1_Ken06.Bus             .....a...............a...........c..a..c........g...a...t....t...............t.. [ 41600]
KM111294.1_Ken05/Tk1             .....a...............a...........c..a..c........g...a...t....t...............t.. [ 41600]
KM262844.1_strain_L60            .....................a...............c.c....................................ct.. [ 41600]
KM102979.1_26544/OG10_Italy      .....................a...............c.c....................................ct.. [ 41600]
KM262845.1_strain_NHV            -------------------------------------------------------------------------------- [ 41600]
KX354450.1_47/Ss/2008            .....................a...............c.c....................................ct.. [ 41600]
KP055815.1_strain_BA71           .....................a...............c.c....................................ct.. [ 41600]
MH025920.1_strain_R35            .....a...............a...........c..a..c........g...a...t....t...............t.. [ 41600]
MH025919.1_strain_N10            .....a...............a...........c..a..c........g...a...t....t...............t.. [ 41600]
MH025918.1_strain_R25            .....a...............a...........c..a..c........g...a...t....t...............t.. [ 41600]
MH025917.1_strain_R7             .....a...............a...........c..a..c........g...a...t....t...............t.. [ 41600]
MH025916.1_strain_R8             .....a...............a...........c..a..c........g...a...t....t...............t.. [ 41600]
```

```
FR682468.2_Georgia_2007/1        ttccttgtggtacatctcatgaagctcatatattttgttataatacaggagacattttaatcgccattcattaagatccg [ 41680]
ASF_IND_20_CAD543                -------------------------------------------------------------------------------- [ 41680]
LR899193.1_Germany_2020/1        ................................................................................ [ 41680]
MN393477.1_Wuhan_2019            ................................................................................ [ 41680]
MW396979.1_Timor-Leste/2019/1    ................................................................................ [ 41680]
MK940252.1_CN/2019/InnerMongolia ................................................................................ [ 41680]
MK128995.1_China/2018/AnhuiXCGQ  ................................................................................ [ 41680]
MH681419.1_ASFV/POL/2015         ................................................................................ [ 41680]
LS478113.1_Estonia_2014          ................................................................................ [ 41680]
KP843857.1_Odintsovo_02/14       ................................................................................ [ 41680]
FN557520.1_strain_E75            ..t...............................................................c..g.......... [ 41680]
AY261366.1_Warthog               ................................................................................ [ 41680]
AY261365.1_Warmbaths             a...........................................................................c... [ 41680]
AY261364.1_Tengani_62            ................................................................................ [ 41680]
AY261363.1_Pretoriuskop/96/4     ........................................a....................................... [ 41680]
AY261362.1_Mkuzi_1979            ..t..................................................................g.......... [ 41680]
AY261361.1_Malawi_Lil-20/1_1983  ............t..a.............................t...ta............................. [ 41680]
AY261360.1_Kenya_1950            ..t.........t..a...........................c.t...ta....................c........ [ 41680]
AM712240.1_OURT_88/3             -------------------------------------------------------------------------------- [ 41680]
AM712239.1_Benin_97/1            ..t...............................................................c..g.......... [ 41680]
KM111295.1_Ken06.Bus             ............t..a...........................c.t...ta...........a........c........ [ 41680]
KM111294.1_Ken05/Tk1             ............t..a...........................c.t...ta....................c........ [ 41680]
KM262844.1_strain_L60            ..t...............................................................c..g.......... [ 41680]
KM102979.1_26544/OG10_Italy      ..t...............................................................c..g.......... [ 41680]
KM262845.1_strain_NHV            -------------------------------------------------------------------------------- [ 41680]
KX354450.1_47/Ss/2008            ..t...............................................................c..g.......... [ 41680]
KP055815.1_strain_BA71           ..t...............................................................c..g.......... [ 41680]
MH025920.1_strain_R35            .c..........t..a...........................c.t...ta....................c........ [ 41680]
MH025919.1_strain_N10            .c..........t..a...........................c.t...ta....................c........ [ 41680]
MH025918.1_strain_R25            .c..........t..a...........................c.t...ta....................c........ [ 41680]
MH025917.1_strain_R7             .c..........t..a...........................c.t...ta....................c........ [ 41680]
MH025916.1_strain_R8             .c..........t..a...........................c.t...ta....................c........ [ 41680]

FR682468.2_Georgia_2007/1        tatatttctcatctagaaaacaaatggcgtccttacaatcgtattgtactgctttggcgtaccaatacttcactagtaaa [ 41760]
ASF_IND_20_CAD543                -------------------------------------------------------------------------------- [ 41760]
LR899193.1_Germany_2020/1        ................................................................................ [ 41760]
MN393477.1_Wuhan_2019            ................................................................................ [ 41760]
MW396979.1_Timor-Leste/2019/1    ................................................................................ [ 41760]
MK940252.1_CN/2019/InnerMongolia ................................................................................ [ 41760]
MK128995.1_China/2018/AnhuiXCGQ  ................................................................................ [ 41760]
MH681419.1_ASFV/POL/2015         ................................................................................ [ 41760]
LS478113.1_Estonia_2014          ................................................................................ [ 41760]
KP843857.1_Odintsovo_02/14       ................................................................................ [ 41760]
```

```
FN557520.1_strain_E75            ...................g..........t.......a...................a..................... [ 41760]
AY261366.1_Warthog               ...................g.................c.......................................... [ 41760]
AY261365.1_Warmbaths             ...................g.................c.......................................... [ 41760]
AY261364.1_Tengani_62            ...................g.................c.......................................... [ 41760]
AY261363.1_Pretoriuskop/96/4     ..........t........g.................c.......................................... [ 41760]
AY261362.1_Mkuzi_1979            ...................g..........t.......a...................a..................... [ 41760]
AY261361.1_Malawi_Lil-20/1_1983  .............c.....g........a..t...t...............................t...........t [ 41760]
AY261360.1_Kenya_1950            .......t.....c.....g........a..t...tg...............a..............t...........t [ 41760]
AM712240.1_OURT_88/3             -------------------------------------------------------------------------------- [ 41760]
AM712239.1_Benin_97/1            ...................g..........t.......a...................a..................... [ 41760]
KM111295.1_Ken06.Bus             .......t.....c.....g........a..t...tgg..............a..............t........c..t [ 41760]
KM111294.1_Ken05/Tk1             .......t.....c.....g........a..t...tg...............a..............t...........t [ 41760]
KM262844.1_strain_L60            ...................g..........t.......a...................a..................... [ 41760]
KM102979.1_26544/OG10_Italy      ...................g..........t.......a...................a..................... [ 41760]
KM262845.1_strain_NHV            -------------------------------------------------------------------------------- [ 41760]
KX354450.1_47/Ss/2008            ...................g..........t.......a...................a..................... [ 41760]
KP055815.1_strain_BA71           ...................g..........t.......a...................a..................... [ 41760]
MH025920.1_strain_R35            .......t.....c.....g........a..t...tgg..............a..............t........c..t [ 41760]
MH025919.1_strain_N10            .......t.....c.....g........a..t...tgg..............a..............t........c..t [ 41760]
MH025918.1_strain_R25            .......t.....c.....g........a..t...tgg..............a..............t........c..t [ 41760]
MH025917.1_strain_R7             .......t.....c.....g........a..t...tgg..............a..............t........c..t [ 41760]
MH025916.1_strain_R8             .......t.....c.....g........a..t...tgg..............a..............t........c..t [ 41760]

FR682468.2_Georgia_2007/1        ccatttaactcgtccgtttcttttatttctatgagcccccatagtcttttataaattaagccccttaattgtataacaaa [ 41840]
ASF_IND_20_CAD543                -------------------------------------------------------------------------------- [ 41840]
LR899193.1_Germany_2020/1        ................................................................................ [ 41840]
MN393477.1_Wuhan_2019            ................................................................................ [ 41840]
MW396979.1_Timor-Leste/2019/1    ................................................................................ [ 41840]
MK940252.1_CN/2019/InnerMongolia ................................................................................ [ 41840]
MK128995.1_China/2018/AnhuiXCGQ  ................................................................................ [ 41840]
MH681419.1_ASFV/POL/2015         ................................................................................ [ 41840]
LS478113.1_Estonia_2014          ................................................................................ [ 41840]
KP843857.1_Odintsovo_02/14       ................................................................................ [ 41840]
FN557520.1_strain_E75            t............................................................................... [ 41840]
AY261366.1_Warthog               t.g........a.......................................c.......................gt... [ 41840]
AY261365.1_Warmbaths             t.g........a.........................t.............c.......................gt... [ 41840]
AY261364.1_Tengani_62            ..g........a...............................................................gt... [ 41840]
AY261363.1_Pretoriuskop/96/4     t.g........a.......................................c.......................gt... [ 41840]
AY261362.1_Mkuzi_1979            t.........t..................................................................... [ 41840]
AY261361.1_Malawi_Lil-20/1_1983  t..c............................a.....g......................g..........gc.g.... [ 41840]
AY261360.1_Kenya_1950            t.gc............................a.....g....a.................g..........gc.g.... [ 41840]
AM712240.1_OURT_88/3             -------------------------------------------------------------------------------- [ 41840]
AM712239.1_Benin_97/1            t............................................................................... [ 41840]
KM111295.1_Ken06.Bus             t.gc............................a.....g......................g..........gc.g.... [ 41840]
```

```
KM111294.1_Ken05/Tk1              t.gc............................a.....g.........c............g..........gc.g.... [ 41840]
KM262844.1_strain_L60             t............................................................................... [ 41840]
KM102979.1_26544/OG10_Italy       t............................................................................... [ 41840]
KM262845.1_strain_NHV             -------------------------------------------------------------------------------- [ 41840]
KX354450.1_47/Ss/2008             t............................................................................... [ 41840]
KP055815.1_strain_BA71            t............................................................................... [ 41840]
MH025920.1_strain_R35             t.gc............................a.....g......................g..........gc.g.... [ 41840]
MH025919.1_strain_N10             t.gc............................a.....g......................g..........gc.g.... [ 41840]
MH025918.1_strain_R25             t.gc............................a.....g......................g..........gc.g.... [ 41840]
MH025917.1_strain_R7              t.gc............................a.....g......................g..........gc.g.... [ 41840]
MH025916.1_strain_R8              t.gc............................a.....g......................g..........gc.g.... [ 41840]

FR682468.2_Georgia_2007/1         tttgttttctaaaataggattattcataaaaatttcatggcacaaaataatactgccgctggttttattgtgcattatcc [ 41920]
ASF_IND_20_CAD543                 -------------------------------------------------------------------------------- [ 41920]
LR899193.1_Germany_2020/1         ................................................................................ [ 41920]
MN393477.1_Wuhan_2019             ................................................................................ [ 41920]
MW396979.1_Timor-Leste/2019/1     ................................................................................ [ 41920]
MK940252.1_CN/2019/InnerMongolia  ................................................................................ [ 41920]
MK128995.1_China/2018/AnhuiXCGQ   ................................................................................ [ 41920]
MH681419.1_ASFV/POL/2015          ................................................................................ [ 41920]
LS478113.1_Estonia_2014           ................................................................................ [ 41920]
KP843857.1_Odintsovo_02/14        ................................................................................ [ 41920]
FN557520.1_strain_E75             ............c................................................................... [ 41920]
AY261366.1_Warthog                ...........t.t................t..c...................g.........................t [ 41920]
AY261365.1_Warmbaths              ...........t.t................t..c...................g.........................t [ 41920]
AY261364.1_Tengani_62             ...a.c.....t.t................t..a.............................................t [ 41920]
AY261363.1_Pretoriuskop/96/4      ...........t.t................t..a...................g.........................t [ 41920]
AY261362.1_Mkuzi_1979             ................................................................................ [ 41920]
AY261361.1_Malawi_Lil-20/1_1983   .............t..........t.....t..c................c..................a.........t [ 41920]
AY261360.1_Kenya_1950             g...........tt..........t.....t..c..........................a........a........tt [ 41920]
AM712240.1_OURT_88/3              -------------------------------------------------------------------------------- [ 41920]
AM712239.1_Benin_97/1             ............c................................................................... [ 41920]
KM111295.1_Ken06.Bus              g............t..........t.....t..c..........................a........a.........t [ 41920]
KM111294.1_Ken05/Tk1              .............t..........t.....t..c..........................ac.......a........tt [ 41920]
KM262844.1_strain_L60             ............c................................................................... [ 41920]
KM102979.1_26544/OG10_Italy       ............c................................................................... [ 41920]
KM262845.1_strain_NHV             -------------------------------------------------------------------------------- [ 41920]
KX354450.1_47/Ss/2008             ............c................................................................... [ 41920]
KP055815.1_strain_BA71            ............c................................................................... [ 41920]
MH025920.1_strain_R35             g............t..........t.....t..c..........................a........a.........t [ 41920]
MH025919.1_strain_N10             g............t..........t.....t..c..........................a........a.........t [ 41920]
MH025918.1_strain_R25             g............t..........t.....t..c..........................a........a.........t [ 41920]
MH025917.1_strain_R7              g............t..........t.....t..c..........................a........a.........t [ 41920]
MH025916.1_strain_R8              g............t..........t.....t..c..........................a........a.........t [ 41920]
```

```
FR682468.2_Georgia_2007/1        tggtaaaaatacggaaaatatcgttgtcctctagagtttctttggcgcctagctgtctacacaactctcggatgtgcttc [ 42000]
ASF_IND_20_CAD543                -------------------------------------------------------------------------------- [ 42000]
LR899193.1_Germany_2020/1        ................................................................................ [ 42000]
MN393477.1_Wuhan_2019            ................................................................................ [ 42000]
MW396979.1_Timor-Leste/2019/1    ................................................................................ [ 42000]
MK940252.1_CN/2019/InnerMongolia ................................................................................ [ 42000]
MK128995.1_China/2018/AnhuiXCGQ  ................................................................................ [ 42000]
MH681419.1_ASFV/POL/2015         ................................................................................ [ 42000]
LS478113.1_Estonia_2014          ................................................................................ [ 42000]
KP843857.1_Odintsovo_02/14       ................................................................................ [ 42000]
FN557520.1_strain_E75            ....g........................................................................... [ 42000]
AY261366.1_Warthog               ....g.........................................a...............................c. [ 42000]
AY261365.1_Warmbaths             ....g..................................t......a.............................t.c. [ 42000]
AY261364.1_Tengani_62            ....g...................c...t................................................... [ 42000]
AY261363.1_Pretoriuskop/96/4     ....g.............c.....c...t.................................................c. [ 42000]
AY261362.1_Mkuzi_1979            ....g........................................................................... [ 42000]
AY261361.1_Malawi_Lil-20/1_1983  ..t.g.......a...........a......c................................a.......c....... [ 42000]
AY261360.1_Kenya_1950            .at.g.......a...........a...g...................................a......gc....... [ 42000]
AM712240.1_OURT_88/3             -------------------------------------------------------------------------------- [ 42000]
AM712239.1_Benin_97/1            ....g........................................................................... [ 42000]
KM111295.1_Ken06.Bus             ..t.g.......a...........a...g...................................a......gc....... [ 42000]
KM111294.1_Ken05/Tk1             .at.g.......a...........a...g...................................a......gc....... [ 42000]
KM262844.1_strain_L60            ....g........................................................................... [ 42000]
KM102979.1_26544/OG10_Italy      ....g........................................................................... [ 42000]
KM262845.1_strain_NHV            -------------------------------------------------------------------------------- [ 42000]
KX354450.1_47/Ss/2008            ....g........................................................................... [ 42000]
KP055815.1_strain_BA71           ....g........................................................................... [ 42000]
MH025920.1_strain_R35            ..t.g.......a...........a...g...................................a......gc....... [ 42000]
MH025919.1_strain_N10            ..t.g.......a...........a...g...................................a......gc....... [ 42000]
MH025918.1_strain_R25            ..t.g.......a...........a...g...................................a......gc....... [ 42000]
MH025917.1_strain_R7             ..t.g.......a...........a...g...................................a......gc....... [ 42000]
MH025916.1_strain_R8             ..t.g.......a...........a...g...................................a......gc....... [ 42000]

FR682468.2_Georgia_2007/1        gtattgatagaaagcaaaccatagttgatatttgcgccccactctgtaaagagctttatcagactatagttgttttcctt [ 42080]
ASF_IND_20_CAD543                -------------------------------------------------------------------------------- [ 42080]
LR899193.1_Germany_2020/1        ................................................................................ [ 42080]
MN393477.1_Wuhan_2019            ................................................................................ [ 42080]
MW396979.1_Timor-Leste/2019/1    ................................................................................ [ 42080]
MK940252.1_CN/2019/InnerMongolia ................................................................................ [ 42080]
MK128995.1_China/2018/AnhuiXCGQ  ................................................................................ [ 42080]
MH681419.1_ASFV/POL/2015         ................................................................................ [ 42080]
LS478113.1_Estonia_2014          ................................................................................ [ 42080]
KP843857.1_Odintsovo_02/14       ................................................................................ [ 42080]
```

```
FN557520.1_strain_E75            ...................t............................................................ [ 42080]
AY261366.1_Warthog               ...................t...........g.........t...................................... [ 42080]
AY261365.1_Warmbaths             ...................t...........g.........t...................................... [ 42080]
AY261364.1_Tengani_62            ...................t............................................................ [ 42080]
AY261363.1_Pretoriuskop/96/4     .....a.............t...........g.........t...................................... [ 42080]
AY261362.1_Mkuzi_1979            ...................t............................................................ [ 42080]
AY261361.1_Malawi_Lil-20/1_1983  ................g..t...................a..........a........t...t................ [ 42080]
AY261360.1_Kenya_1950            ................g..t...........a..................t.a..................a........ [ 42080]
AM712240.1_OURT_88/3             -------------------------------------------------------------------------------- [ 42080]
AM712239.1_Benin_97/1            ...................t............................................................ [ 42080]
KM111295.1_Ken06.Bus             ................g..t...........a..................t.a........t.................. [ 42080]
KM111294.1_Ken05/Tk1             ..............t.g..t...........a..................t.a..................a........ [ 42080]
KM262844.1_strain_L60            ...................t............................................................ [ 42080]
KM102979.1_26544/OG10_Italy      ...................t............................................................ [ 42080]
KM262845.1_strain_NHV            -------------------------------------------------------------------------------- [ 42080]
KX354450.1_47/Ss/2008            ...................t............................................................ [ 42080]
KP055815.1_strain_BA71           ...................t............................................................ [ 42080]
MH025920.1_strain_R35            ................g..t...........a..................t.a........t.................. [ 42080]
MH025919.1_strain_N10            ................g..t...........a..................t.a........t.................. [ 42080]
MH025918.1_strain_R25            ................g..t...........a..................t.a........t.................. [ 42080]
MH025917.1_strain_R7             ................g..t...........a..................t.a........t.................. [ 42080]
MH025916.1_strain_R8             ................g..t...........a..................t.a........t.................. [ 42080]

FR682468.2_Georgia_2007/1        aacagctattattaatgccacacgaaggtctatatcttctcctaaaaatcctgattttatttgtattcggccacgatcca [ 42160]
ASF_IND_20_CAD543                -------------------------------------------------------------------------------- [ 42160]
LR899193.1_Germany_2020/1        ................................................................................ [ 42160]
MN393477.1_Wuhan_2019            ................................................................................ [ 42160]
MW396979.1_Timor-Leste/2019/1    ................................................................................ [ 42160]
MK940252.1_CN/2019/InnerMongolia ................................................................................ [ 42160]
MK128995.1_China/2018/AnhuiXCGQ  ................................................................................ [ 42160]
MH681419.1_ASFV/POL/2015         ................................................................................ [ 42160]
LS478113.1_Estonia_2014          ................................................................................ [ 42160]
KP843857.1_Odintsovo_02/14       ................................................................................ [ 42160]
FN557520.1_strain_E75            ................................................................................ [ 42160]
AY261366.1_Warthog               ..................t.t.t.g..........................................t............ [ 42160]
AY261365.1_Warmbaths             ..................t...t.g..a.......................................t............ [ 42160]
AY261364.1_Tengani_62            ..................t.t.t.g..........................................t............ [ 42160]
AY261363.1_Pretoriuskop/96/4     ..................t.t.t.g...........c..............................t............ [ 42160]
AY261362.1_Mkuzi_1979            ................................................................................ [ 42160]
AY261361.1_Malawi_Lil-20/1_1983  .......................a...........................c............................ [ 42160]
AY261360.1_Kenya_1950            .......................a....................gg.....c........g.a......t........t. [ 42160]
AM712240.1_OURT_88/3             -------------------------------------------------------------------------------- [ 42160]
AM712239.1_Benin_97/1            ................................................................................ [ 42160]
KM111295.1_Ken06.Bus             .......................a....................gg.....c........g.a......c........t. [ 42160]
```

```
KM111294.1_Ken05/Tk1            .......................a....................gg.....c........g.a......c........t. [ 42160]
KM262844.1_strain_L60           ................................................................................ [ 42160]
KM102979.1_26544/OG10_Italy     ................................................................................ [ 42160]
KM262845.1_strain_NHV           -------------------------------------------------------------------------------- [ 42160]
KX354450.1_47/Ss/2008           ................................................................................ [ 42160]
KP055815.1_strain_BA71          ................................................................................ [ 42160]
MH025920.1_strain_R35           .......................a....................gg.....c........g.a......c........t. [ 42160]
MH025919.1_strain_N10           .......................a....................gg.....c........g.a......c........t. [ 42160]
MH025918.1_strain_R25           .......................a....................gg.....c........g.a......c........t. [ 42160]
MH025917.1_strain_R7            .......................a....................gg.....c........g.a......c........t. [ 42160]
MH025916.1_strain_R8            .......................a....................gg.....c........g.a......c........t. [ 42160]

FR682468.2_Georgia_2007/1       tacaaagcttgagaggagcatcatgccaccataggccacaatatttcaaaatgcagtgttcatctattgacaaacactg- [ 42240]
ASF_IND_20_CAD543               -------------------------------------------------------------------------------- [ 42240]
LR899193.1_Germany_2020/1       ...............................................................................- [ 42240]
MN393477.1_Wuhan_2019           ...............................................................................- [ 42240]
MW396979.1_Timor-Leste/2019/1   ...............................................................................- [ 42240]
MK940252.1_CN/2019/InnerMongolia ...............................................................................- [ 42240]
MK128995.1_China/2018/AnhuiXCGQ ...............................................................................- [ 42240]
MH681419.1_ASFV/POL/2015        ...............................................................................- [ 42240]
LS478113.1_Estonia_2014         ...............................................................................- [ 42240]
KP843857.1_Odintsovo_02/14      ...............................................................................- [ 42240]
FN557520.1_strain_E75           ........................................g......................................- [ 42240]
AY261366.1_Warthog              .............g..ga......................g.g.....g................g.c...........- [ 42240]
AY261365.1_Warmbaths            .............g..ga......................g.g.....g................g.c...........- [ 42240]
AY261364.1_Tengani_62           .............g..ga......................g.g.....g..................c......g....- [ 42240]
AY261363.1_Pretoriuskop/96/4    .............g..ga......................g.g.....g..................c...........- [ 42240]
AY261362.1_Mkuzi_1979           ........................................g......................................- [ 42240]
AY261361.1_Malawi_Lil-20/1_1983 ..t.............g................a.t....g...........---............c........tgat [ 42240]
AY261360.1_Kenya_1950           .gt..........g...................a......g...........---............c........tgat [ 42240]
AM712240.1_OURT_88/3            -------------------------------------------------------------------------------- [ 42240]
AM712239.1_Benin_97/1           ...............................................................................- [ 42240]
KM111295.1_Ken06.Bus            .gt..........g...................a......g...........---............c........tgat [ 42240]
KM111294.1_Ken05/Tk1            .gt..........g...................a......g...........---............c........tgat [ 42240]
KM262844.1_strain_L60           ........................................g......................................- [ 42240]
KM102979.1_26544/OG10_Italy     ........................................g......................................- [ 42240]
KM262845.1_strain_NHV           -------------------------------------------------------------------------------- [ 42240]
KX354450.1_47/Ss/2008           ........................................g......................................- [ 42240]
KP055815.1_strain_BA71          ........................................g......................................- [ 42240]
MH025920.1_strain_R35           .gt..........g...................a......g...........---............c........tgat [ 42240]
MH025919.1_strain_N10           .gt..........g..............t....a......g...........---............c........tgat [ 42240]
MH025918.1_strain_R25           .gt..........g...................a......g...........---............c........tgat [ 42240]
MH025917.1_strain_R7            .gt..........g...................a......g...........---............c........tgat [ 42240]
MH025916.1_strain_R8            .gt..........g...................a......g...........---............c........tgat [ 42240]
```

```
FR682468.2_Georgia_2007/1        --gctggctatcgtctttttgacgagggtctgcagagagagcggcaacgacatgtttctttttcac-----------caa [ 42320]
ASF_IND_20_CAD543                -------------------------------------------------------------------------------- [ 42320]
LR899193.1_Germany_2020/1        --................................................................-----------... [ 42320]
MN393477.1_Wuhan_2019            --................................................................-----------... [ 42320]
MW396979.1_Timor-Leste/2019/1    --................................................................-----------... [ 42320]
MK940252.1_CN/2019/InnerMongolia --................................................................-----------... [ 42320]
MK128995.1_China/2018/AnhuiXCGQ  --................................................................-----------... [ 42320]
MH681419.1_ASFV/POL/2015         --................................................................-----------... [ 42320]
LS478113.1_Estonia_2014          --................................................................-----------... [ 42320]
KP843857.1_Odintsovo_02/14       --................................................................-----------.-. [ 42320]
FN557520.1_strain_E75            --........c.......................................................-----------... [ 42320]
AY261366.1_Warthog               --........c....................................................tca-----------.c. [ 42320]
AY261365.1_Warmbaths             --........c...t................................................tca-----------.c. [ 42320]
AY261364.1_Tengani_62            --.t......c.................................t..t...............tca-----------.c. [ 42320]
AY261363.1_Pretoriuskop/96/4     --........c....................................................tca-----------.c. [ 42320]
AY261362.1_Mkuzi_1979            --........c.......................................................-----------... [ 42320]
AY261361.1_Malawi_Lil-20/1_1983  gta.......c..................t..t............gc.............c..tgaaataattattt.c. [ 42320]
AY261360.1_Kenya_1950            gta.......c..................................g..............c..tgaaataattattt.c. [ 42320]
AM712240.1_OURT_88/3             -------------------------------------------------------------------------------- [ 42320]
AM712239.1_Benin_97/1            --........c.......................................................-----------... [ 42320]
KM111295.1_Ken06.Bus             gta.c.....c..................................g..............c..tgaaataattattt.c. [ 42320]
KM111294.1_Ken05/Tk1             gta.......c..................................g..............c..tgaaataattattt.c. [ 42320]
KM262844.1_strain_L60            --........c.......................................................-----------... [ 42320]
KM102979.1_26544/OG10_Italy      --........c.......................................................-----------... [ 42320]
KM262845.1_strain_NHV            -------------------------------------------------------------------------------- [ 42320]
KX354450.1_47/Ss/2008            --........c.......................................................-----------... [ 42320]
KP055815.1_strain_BA71           --........c.......................................................-----------... [ 42320]
MH025920.1_strain_R35            gta.c.....c..................................g..............c..tgaaataattattt.c. [ 42320]
MH025919.1_strain_N10            gta.c.....c..................................g..............c..tgaaataattattt.c. [ 42320]
MH025918.1_strain_R25            gta.c.....c..................................g..............c..tgaaataattattt.c. [ 42320]
MH025917.1_strain_R7             gta.c.....c..................................g..............c..tgaaataattattt.c. [ 42320]
MH025916.1_strain_R8             gta.c.....c..................................g..............c..tgaaataattattt.c. [ 42320]

FR682468.2_Georgia_2007/1        aaaaaaatcaaatgttc--tcgtcttt--aaaggt------------taattcatgttcttaaaatgttcattt-cat-- [ 42400]
ASF_IND_20_CAD543                -------------------------------------------------------------------------------- [ 42400]
LR899193.1_Germany_2020/1        .................--........--......------------...........................-...-- [ 42400]
MN393477.1_Wuhan_2019            .................--........--......------------...........................-...-- [ 42400]
MW396979.1_Timor-Leste/2019/1    .................--........--......------------...........................-...-- [ 42400]
MK940252.1_CN/2019/InnerMongolia .................--........--......------------...........................-...-- [ 42400]
MK128995.1_China/2018/AnhuiXCGQ  .................--........--......------------...........................-...-- [ 42400]
MH681419.1_ASFV/POL/2015         .................--........--......------------...........................-...-- [ 42400]
LS478113.1_Estonia_2014          .................--........--......------------...........................-...-- [ 42400]
KP843857.1_Odintsovo_02/14       .................--........--......------------...........................-...-- [ 42400]
```

```
FN557520.1_strain_E75           .................--........--......------------...........................t...-- [ 42400]
AY261366.1_Warthog              .................--.ac.a...--.....cgtaattcatgtac.tgaa..a.a.a..............-...-- [ 42400]
AY261365.1_Warmbaths            .................--.ac.a...--.....cgtaattcatgtgc.t......ca.a.......a.g....-...-- [ 42400]
AY261364.1_Tengani_62           .................--.gc.a...--.....cgtaattc----------------------...ca.....-...-- [ 42400]
AY261363.1_Pretoriuskop/96/4    .................--.ac.a...--.....cgtaattcatgtac.t.......a.a..............-...-- [ 42400]
AY261362.1_Mkuzi_1979           .................--........--......------------...........................a...-- [ 42400]
AY261361.1_Malawi_Lil-20/1_1983 .................--.a.....g--.....c-----------t........ca..a.......a......-t..at [ 42400]
AY261360.1_Kenya_1950           .................--.a......aa.....c-----------t........---.a.......a....a.tt.ata [ 42400]
AM712240.1_OURT_88/3            -------------------------------------------------------------------------------- [ 42400]
AM712239.1_Benin_97/1           .................--........--......------------...........................t...-- [ 42400]
KM111295.1_Ken06.Bus            .................--.a..t...--.....c-----------t........---.a...c...a....a.tt.ata [ 42400]
KM111294.1_Ken05/Tk1            .................ta.a......aa.....c-----------t........---.a.......a....a.tt.ata [ 42400]
KM262844.1_strain_L60           .................--........--......------------...........................t...-- [ 42400]
KM102979.1_26544/OG10_Italy     .................--........--......------------...........................t...-- [ 42400]
KM262845.1_strain_NHV           -------------------------------------------------------------------------------- [ 42400]
KX354450.1_47/Ss/2008           .................--........--......------------...........................t...-- [ 42400]
KP055815.1_strain_BA71          .................--........--......------------...........................t...-- [ 42400]
MH025920.1_strain_R35           .................--.a..t...--.....c-----------t........---.a...c...a....a.tt.ata [ 42400]
MH025919.1_strain_N10           .................--.a..t...--.....c-----------t........---.a...c...a....a.tt.ata [ 42400]
MH025918.1_strain_R25           .................--.a..t...--.....c-----------t........---.a...c...a....a.tt.ata [ 42400]
MH025917.1_strain_R7            .................--.a..t...--.....c-----------t........---.a...c...a....a.tt.ata [ 42400]
MH025916.1_strain_R8            .................--.a..t...--.....c-----------t........---.a...c...a....a.tt.ata [ 42400]

FR682468.2_Georgia_2007/1       -------------------------------------------------------------------------------- [ 42480]
ASF_IND_20_CAD543               -------------------------------------------------------------------------------- [ 42480]
LR899193.1_Germany_2020/1       -------------------------------------------------------------------------------- [ 42480]
MN393477.1_Wuhan_2019           -------------------------------------------------------------------------------- [ 42480]
MW396979.1_Timor-Leste/2019/1   -------------------------------------------------------------------------------- [ 42480]
MK940252.1_CN/2019/InnerMongolia -------------------------------------------------------------------------------- [ 42480]
MK128995.1_China/2018/AnhuiXCGQ -------------------------------------------------------------------------------- [ 42480]
MH681419.1_ASFV/POL/2015        -------------------------------------------------------------------------------- [ 42480]
LS478113.1_Estonia_2014         -------------------------------------------------------------------------------- [ 42480]
KP843857.1_Odintsovo_02/14      -------------------------------------------------------------------------------- [ 42480]
FN557520.1_strain_E75           -------------------------------------------------------------------------------- [ 42480]
AY261366.1_Warthog              -------------------------------------------------------------------------------- [ 42480]
AY261365.1_Warmbaths            -------------------------------------------------------------------------------- [ 42480]
AY261364.1_Tengani_62           -------------------------------------------------------------------------------- [ 42480]
AY261363.1_Pretoriuskop/96/4    -------------------------------------------------------------------------------- [ 42480]
AY261362.1_Mkuzi_1979           -------------------------------------------------------------------------------- [ 42480]
AY261361.1_Malawi_Lil-20/1_1983 att----------------------------------------------------------------------------- [ 42480]
AY261360.1_Kenya_1950           acgcaaacacagtttaataacacaaacacagtttaataacacaaacacagtttaataacgcaaacacagtttaataacac [ 42480]
AM712240.1_OURT_88/3            -------------------------------------------------------------------------------- [ 42480]
AM712239.1_Benin_97/1           -------------------------------------------------------------------------------- [ 42480]
KM111295.1_Ken06.Bus            acg----------------------------------------------------------------------------- [ 42480]
```

```
KM111294.1_Ken05/Tk1           atg----------------------------------------------------------------------------c [ 42480]
KM262844.1_strain_L60          -------------------------------------------------------------------------------- [ 42480]
KM102979.1_26544/OG10_Italy    -------------------------------------------------------------------------------- [ 42480]
KM262845.1_strain_NHV          -------------------------------------------------------------------------------- [ 42480]
KX354450.1_47/Ss/2008          -------------------------------------------------------------------------------- [ 42480]
KP055815.1_strain_BA71         -------------------------------------------------------------------------------- [ 42480]
MH025920.1_strain_R35          acg----------------------------------------------------------------------------- [ 42480]
MH025919.1_strain_N10          acg----------------------------------------------------------------------------- [ 42480]
MH025918.1_strain_R25          acg----------------------------------------------------------------------------- [ 42480]
MH025917.1_strain_R7           acg----------------------------------------------------------------------------- [ 42480]
MH025916.1_strain_R8           acg----------------------------------------------------------------------------- [ 42480]

FR682468.2_Georgia_2007/1      --------------------gatagtgattaataatatggtt-------------taataacgctagaaggcttgttta- [ 42560]
ASF_IND_20_CAD543              -------------------------------------------------------------------------------- [ 42560]
LR899193.1_Germany_2020/1      --------------------......................-------------........................- [ 42560]
MN393477.1_Wuhan_2019          --------------------......................-------------........................- [ 42560]
MW396979.1_Timor-Leste/2019/1  --------------------......................-------------........................- [ 42560]
MK940252.1_CN/2019/InnerMongolia --------------------......................-------------........................- [ 42560]
MK128995.1_China/2018/AnhuiXCGQ  --------------------......................-------------........................- [ 42560]
MH681419.1_ASFV/POL/2015       --------------------......................-------------........................- [ 42560]
LS478113.1_Estonia_2014        --------------------......................-------------........................- [ 42560]
KP843857.1_Odintsovo_02/14     --------------------......................-------------........................- [ 42560]
FN557520.1_strain_E75          --------------------............-.........-------------........................- [ 42560]
AY261366.1_Warthog             --------------------a...c..t..c..g...c....-------------......t.........t.------- [ 42560]
AY261365.1_Warmbaths           --------------------.......t.......c.c....-------------..............a......c..t [ 42560]
AY261364.1_Tengani_62          --------------------....c..t..c...g..c....-------------............-........a..- [ 42560]
AY261363.1_Pretoriuskop/96/4   --------------------....c..t..c..g...c....-------------................t.------- [ 42560]
AY261362.1_Mkuzi_1979          --------------------............-.........taataatacggtt........................- [ 42560]
AY261361.1_Malawi_Lil-20/1_1983 ----------taataagacaa.c.cc.t.......c.caaac-------accgtt......t..---..act..taa..t [ 42560]
AY261360.1_Kenya_1950          aaacacagtttaataacacaa.c.ca.t........gcaaac-------acagtt.........a.ac.t.a..taa..a [ 42560]
AM712240.1_OURT_88/3           -------------------------------------------------------------------------------- [ 42560]
AM712239.1_Benin_97/1          --------------------............-.........-------------........................- [ 42560]
KM111295.1_Ken06.Bus           ------------------taa.c.ca.t........g.aaac-------acagtt....c.t..a.ac.t.g..taa..a [ 42560]
KM111294.1_Ken05/Tk1           aaacacagtttaataatgcaa.c.caat........gcaaac-------acagtt.........a.ac.t.a..caa..a [ 42560]
KM262844.1_strain_L60          --------------------............-.........-------------........................- [ 42560]
KM102979.1_26544/OG10_Italy    --------------------............-.........-------------........................- [ 42560]
KM262845.1_strain_NHV          -------------------------------------------------------------------------------- [ 42560]
KX354450.1_47/Ss/2008          --------------------............-.........-------------........................- [ 42560]
KP055815.1_strain_BA71         --------------------............-.........-------------........................- [ 42560]
MH025920.1_strain_R35          ------------------taa.c.ca.t........g.aaac-------acagtt....c.t..a.ac.t.g..taa..a [ 42560]
MH025919.1_strain_N10          ------------------taa.c.ca.t........g.aaac-------acagtt....c.t..a.ac.t.g..taa..a [ 42560]
MH025918.1_strain_R25          ------------------taa.c.ca.t........g.aaac-------acagtt....c.t..a.ac.t.g..taa..a [ 42560]
MH025917.1_strain_R7           ------------------taa.c.ca.t........g.aaac-------acagtt....c.t..a.ac.t.g..taa..a [ 42560]
MH025916.1_strain_R8           ------------------taa.c.ca.t........g.aaac-------acagtt....c.t..a.ac.t.g..taa..a [ 42560]
```

```
FR682468.2_Georgia_2007/1        --------------------taagacagtc-ataag-c-agtctataagac--agtctataa-gc-agtctataagacag [ 42640]
ASF_IND_20_CAD543                -------------------------------------------------------------------------------- [ 42640]
LR899193.1_Germany_2020/1        --------------------..........-.....-.-............--.........-..-.............. [ 42640]
MN393477.1_Wuhan_2019            --------------------..........-.....-.-............--.........-..-.............. [ 42640]
MW396979.1_Timor-Leste/2019/1    --------------------..........-.....-.-............--.........-..-.............. [ 42640]
MK940252.1_CN/2019/InnerMongolia --------------------..........-.....-.-............--.........-..-.............. [ 42640]
MK128995.1_China/2018/AnhuiXCGQ  --------------------..........-.....-.-............--.........-..-.............. [ 42640]
MH681419.1_ASFV/POL/2015         --------------------..........-.....-.-............--.........-..-.............. [ 42640]
LS478113.1_Estonia_2014          --------------------..........-.....-.-............--.........-..-.............. [ 42640]
KP843857.1_Odintsovo_02/14       --------------------..........-.....-.-............--.........-..-.............. [ 42640]
FN557520.1_strain_E75            --------------------.....-....t.....-.-............--.......g.---------------... [ 42640]
AY261366.1_Warthog               ----------------------------------------..t.....a..--.....---------........----- [ 42640]
AY261365.1_Warmbaths             aa----------acagcctg....g.....tg....---.........a..--....------------......----- [ 42640]
AY261364.1_Tengani_62            --------------------..........t.....-.-...t....g...tt.a.......ga.t.a...tat.----- [ 42640]
AY261363.1_Pretoriuskop/96/4     ----------------------------------------..t.....a..--.....---------........----- [ 42640]
AY261362.1_Mkuzi_1979            --------------------..........t.....-.-............--.........ga.-.......g.ctt.. [ 42640]
AY261361.1_Malawi_Lil-20/1_1983  g----------------ctaa...g.t...t.....a.-c...........--c........ga.-g..t.....----- [ 42640]
AY261360.1_Kenya_1950            cataaatttttaatatgttaa.....t...ttat..a.-t..........ttt.........------------------ [ 42640]
AM712240.1_OURT_88/3             -------------------------------------------------------------------------------- [ 42640]
AM712239.1_Benin_97/1            --------------------.....-....t.....-.-............--.......g.---------------... [ 42640]
KM111295.1_Ken06.Bus             cataaatttttaatacgctaa...g.t...t.....a.ttagt.......ttt.a.......------------------ [ 42640]
KM111294.1_Ken05/Tk1             cataaatttttaatacgctaa.....t...t.....a.-------------tt.........------------------ [ 42640]
KM262844.1_strain_L60            --------------------.....-....t.....-.-............--.......g.---------------... [ 42640]
KM102979.1_26544/OG10_Italy      --------------------.....-....t.....-.-............--.......g.---------------... [ 42640]
KM262845.1_strain_NHV            -------------------------------------------------------------------------------- [ 42640]
KX354450.1_47/Ss/2008            --------------------.....-....t.....-.-............--.......g.---------------... [ 42640]
KP055815.1_strain_BA71           --------------------.....-....t.....-.-............--.......g.---------------... [ 42640]
MH025920.1_strain_R35            cataaatttttaatacgctaa...g.t...t.....a.ttagt.......ttt.a.......------------------ [ 42640]
MH025919.1_strain_N10            cataaatttttaatacgctaa...g.t...t.....a.ttagt.......ttt.a.......------------------ [ 42640]
MH025918.1_strain_R25            cataaatttttaatacgctaa...g.t...t.....a.ttagt.......ttt.a.......------------------ [ 42640]
MH025917.1_strain_R7             cataaatttttaatacgctaa...g.t...t.....a.ttagt.......ttt.a.......------------------ [ 42640]
MH025916.1_strain_R8             cataaatttttaatacgctaa...g.t...t.....a.ttagt.......ttt.a.......------------------ [ 42640]

FR682468.2_Georgia_2007/1        tctatgacttagtctataactataatttctggatgggctgtaagatactcttcggctcgtttcagattttttgaagtata [ 42720]
ASF_IND_20_CAD543                -------------------------------------------------------------------------------- [ 42720]
LR899193.1_Germany_2020/1        ................................................................................ [ 42720]
MN393477.1_Wuhan_2019            ................................................................................ [ 42720]
MW396979.1_Timor-Leste/2019/1    ................................................................................ [ 42720]
MK940252.1_CN/2019/InnerMongolia ................................................................................ [ 42720]
MK128995.1_China/2018/AnhuiXCGQ  ................................................................................ [ 42720]
MH681419.1_ASFV/POL/2015         ................................................................................ [ 42720]
LS478113.1_Estonia_2014          ................................................................................ [ 42720]
KP843857.1_Odintsovo_02/14       ................................................................................ [ 42720]
```

```
FN557520.1_strain_E75          ................................................................t............... [ 42720]
AY261366.1_Warthog             -----........................................................................... [ 42720]
AY261365.1_Warmbaths           -----...........................................................t............... [ 42720]
AY261364.1_Tengani_62          -----................................................a.......................... [ 42720]
AY261363.1_Pretoriuskop/96/4   -----........................................................................... [ 42720]
AY261362.1_Mkuzi_1979          ........................................................c.......t............... [ 42720]
AY261361.1_Malawi_Lil-20/1_1983 -----.................c....................a.............t....t...........c..... [ 42720]
AY261360.1_Kenya_1950          -----..t..............c....................a..................t...........c..... [ 42720]
AM712240.1_OURT_88/3           -------------------------------------------------------------------------------- [ 42720]
AM712239.1_Benin_97/1          ................................................................t............... [ 42720]
KM111295.1_Ken06.Bus           -----..t............g.c....................a..................t...........c..... [ 42720]
KM111294.1_Ken05/Tk1           -----.................c....................a..................t...........c..... [ 42720]
KM262844.1_strain_L60          ................................................................t............... [ 42720]
KM102979.1_26544/OG10_Italy    ................................................................t............... [ 42720]
KM262845.1_strain_NHV          -------------------------------------------------------------------------------- [ 42720]
KX354450.1_47/Ss/2008          ................................................................t............... [ 42720]
KP055815.1_strain_BA71         ................................................................t............... [ 42720]
MH025920.1_strain_R35          -----..t............g.c....................a..................t...........c..... [ 42720]
MH025919.1_strain_N10          -----..t............g.c....................a..................t...........c..... [ 42720]
MH025918.1_strain_R25          -----..t............g.c....................a..................t...........c..... [ 42720]
MH025917.1_strain_R7           -----..t............g.c....................a..................t...........c..... [ 42720]
MH025916.1_strain_R8           -----..t............g.c....................a..................t...........c..... [ 42720]

FR682468.2_Georgia_2007/1      tgtctttagcatatcatatatttcctggggttcggttacatctaataccaaggtcacatcacggctgaaaagctgcttta [ 42800]
ASF_IND_20_CAD543              -------------------------------------------------------------------------------- [ 42800]
LR899193.1_Germany_2020/1      ................................................................................ [ 42800]
MN393477.1_Wuhan_2019          ................................................................................ [ 42800]
MW396979.1_Timor-Leste/2019/1  ................................................................................ [ 42800]
MK940252.1_CN/2019/InnerMongolia ................................................................................ [ 42800]
MK128995.1_China/2018/AnhuiXCGQ ................................................................................ [ 42800]
MH681419.1_ASFV/POL/2015       ................................................................................ [ 42800]
LS478113.1_Estonia_2014        ................................................................................ [ 42800]
KP843857.1_Odintsovo_02/14     ................................................................................ [ 42800]
FN557520.1_strain_E75          ................................................................................ [ 42800]
AY261366.1_Warthog             ..a...........................................g................................. [ 42800]
AY261365.1_Warmbaths           ..c.......c...................................g................................. [ 42800]
AY261364.1_Tengani_62          ..a...........................................g................................. [ 42800]
AY261363.1_Pretoriuskop/96/4   ..a.......c...................................g................................. [ 42800]
AY261362.1_Mkuzi_1979          ................................................................................ [ 42800]
AY261361.1_Malawi_Lil-20/1_1983 ...a..................g.......c..a....g........t....c......................t.... [ 42800]
AY261360.1_Kenya_1950          ......................g...............t..t..........c..g...................t.... [ 42800]
AM712240.1_OURT_88/3           -------------------------------------------------------------------------------- [ 42800]
AM712239.1_Benin_97/1          ................................................................................ [ 42800]
KM111295.1_Ken06.Bus           ......................g...............t..t..........c..g...................t.... [ 42800]
```

```
KM111294.1_Ken05/Tk1              ......................g...............t..t..........c..g...................t.... [ 42800]
KM262844.1_strain_L60             ................................................................................ [ 42800]
KM102979.1_26544/OG10_Italy       ................................................................................ [ 42800]
KM262845.1_strain_NHV             -------------------------------------------------------------------------------- [ 42800]
KX354450.1_47/Ss/2008             ................................................................................ [ 42800]
KP055815.1_strain_BA71            ................................................................................ [ 42800]
MH025920.1_strain_R35             ......................g...............t..t..........c..g...................t.... [ 42800]
MH025919.1_strain_N10             ......................g...............t..t..........c..g...................t.... [ 42800]
MH025918.1_strain_R25             ......................g...............t..t..........c..g...................t.... [ 42800]
MH025917.1_strain_R7              ......................g...............t..t..........c..g...................t.... [ 42800]
MH025916.1_strain_R8              ......................g...............t..t..........c..g...................t.... [ 42800]

FR682468.2_Georgia_2007/1         ctaagaaaatgttgctcaagttatacatataagctttgtgcgcaatgagttgtgccctatcaaaatcggcag-cccccaa [ 42880]
ASF_IND_20_CAD543                 -------------------------------------------------------------------------------- [ 42880]
LR899193.1_Germany_2020/1         ........................................................................-....... [ 42880]
MN393477.1_Wuhan_2019             ........................................................................c....... [ 42880]
MW396979.1_Timor-Leste/2019/1     ........................................................................c....... [ 42880]
MK940252.1_CN/2019/InnerMongolia  ........................................................................-....... [ 42880]
MK128995.1_China/2018/AnhuiXCGQ   ........................................................................-....... [ 42880]
MH681419.1_ASFV/POL/2015          ........................................................................-....... [ 42880]
LS478113.1_Estonia_2014           ........................................................................-....... [ 42880]
KP843857.1_Odintsovo_02/14        ........................................................................-....... [ 42880]
FN557520.1_strain_E75             ......................................................................g.-....... [ 42880]
AY261366.1_Warthog                ....a............................t.............c...................a.t..-....... [ 42880]
AY261365.1_Warmbaths              ....a.g........................................c...................a.t..-....... [ 42880]
AY261364.1_Tengani_62             ...............................................c...................a.t..-....... [ 42880]
AY261363.1_Pretoriuskop/96/4      ...............................................t...................a.t..-....... [ 42880]
AY261362.1_Mkuzi_1979             ........................................................................-....... [ 42880]
AY261361.1_Malawi_Lil-20/1_1983   ....a....g......................a..c.ac...........c.c...........g.ta.t..-....... [ 42880]
AY261360.1_Kenya_1950             ....a...........................a..c.a............c.c..t........g.ta.t..-....... [ 42880]
AM712240.1_OURT_88/3              -------------------------------------------------------------------------------- [ 42880]
AM712239.1_Benin_97/1             ......................................................................g.-....... [ 42880]
KM111295.1_Ken06.Bus              ....a...........................a..c.a............c.c..t........g.ta.t..-....... [ 42880]
KM111294.1_Ken05/Tk1              ....a...........................a..c.a............c.c..t........g.ta.t..-....... [ 42880]
KM262844.1_strain_L60             ......................................................................g.-....... [ 42880]
KM102979.1_26544/OG10_Italy       ......................................................................g.-....... [ 42880]
KM262845.1_strain_NHV             -------------------------------------------------------------------------------- [ 42880]
KX354450.1_47/Ss/2008             ......................................................................g.-....... [ 42880]
KP055815.1_strain_BA71            ......................................................................g.-....... [ 42880]
MH025920.1_strain_R35             ....a...........................a..c.a............c.c..t........g.ta.t..-....... [ 42880]
MH025919.1_strain_N10             ....a...........................a..c.a............c.c..t........g.ta.t..-....... [ 42880]
MH025918.1_strain_R25             ....a...........................a..c.a............c.c..t........g.ta.t..-....... [ 42880]
MH025917.1_strain_R7              ....a...........................a..c.a............c.c..t........g.ta.t..-....... [ 42880]
MH025916.1_strain_R8              ....a...........................a..c.a............c.c..t........g.ta.t..-....... [ 42880]
```

```
FR682468.2_Georgia_2007/1        atcaatacagaaaaacatgtttaaagtattattgttatagatagaaagattcatgccataatcgagactagcccccaacc [ 42960]
ASF_IND_20_CAD543                -------------------------------------------------------------------------------- [ 42960]
LR899193.1_Germany_2020/1        ................................................................................ [ 42960]
MN393477.1_Wuhan_2019            ................................................................................ [ 42960]
MW396979.1_Timor-Leste/2019/1    ................................................................................ [ 42960]
MK940252.1_CN/2019/InnerMongolia ................................................................................ [ 42960]
MK128995.1_China/2018/AnhuiXCGQ  ................................................................................ [ 42960]
MH681419.1_ASFV/POL/2015         ................................................................................ [ 42960]
LS478113.1_Estonia_2014          ................................................................................ [ 42960]
KP843857.1_Odintsovo_02/14       ................................................................................ [ 42960]
FN557520.1_strain_E75            ................................................................................ [ 42960]
AY261366.1_Warthog               ...g..g....................................................................t.... [ 42960]
AY261365.1_Warmbaths             ...g..g..........g........a.............................................t....... [ 42960]
AY261364.1_Tengani_62            ...g..g....................................................................t.... [ 42960]
AY261363.1_Pretoriuskop/96/4     ...g..g....................................................................t.... [ 42960]
AY261362.1_Mkuzi_1979            ................................................................................ [ 42960]
AY261361.1_Malawi_Lil-20/1_1983  ......g........t.g...........................t.................a...........t..a [ 42960]
AY261360.1_Kenya_1950            ...............t.g...........................t..................g..............a [ 42960]
AM712240.1_OURT_88/3             -------------------------------------------------------------------------------- [ 42960]
AM712239.1_Benin_97/1            ................................................................................ [ 42960]
KM111295.1_Ken06.Bus             ......g........t.g...........................t..................g..............a [ 42960]
KM111294.1_Ken05/Tk1             ......g........t.g.............g..........g..t.................a...............a [ 42960]
KM262844.1_strain_L60            ................................................................................ [ 42960]
KM102979.1_26544/OG10_Italy      ................................................................................ [ 42960]
KM262845.1_strain_NHV            -------------------------------------------------------------------------------- [ 42960]
KX354450.1_47/Ss/2008            ................................................................................ [ 42960]
KP055815.1_strain_BA71           ................................................................................ [ 42960]
MH025920.1_strain_R35            ......g........t.g...........................t..................g..............a [ 42960]
MH025919.1_strain_N10            ......g........t.g...........................t..................g..............a [ 42960]
MH025918.1_strain_R25            ......g........t.g...........................t..................g..............a [ 42960]
MH025917.1_strain_R7             ......g........t.g...........................t..................g..............a [ 42960]
MH025916.1_strain_R8             ......g........t.g...........................t..................g..............a [ 42960]

FR682468.2_Georgia_2007/1        tatgacagtaataaatggccgcgtaatttttttcccgcaagcaagcaaatttcatcatcagattagggctgatgcaaatc [ 43040]
ASF_IND_20_CAD543                -------------------------------------------------------------------------------- [ 43040]
LR899193.1_Germany_2020/1        ................................................................................ [ 43040]
MN393477.1_Wuhan_2019            ................................................................................ [ 43040]
MW396979.1_Timor-Leste/2019/1    ................................................................................ [ 43040]
MK940252.1_CN/2019/InnerMongolia ................................................................................ [ 43040]
MK128995.1_China/2018/AnhuiXCGQ  ................................................................................ [ 43040]
MH681419.1_ASFV/POL/2015         ................................................................................ [ 43040]
LS478113.1_Estonia_2014          ................................................................................ [ 43040]
KP843857.1_Odintsovo_02/14       ................................................................................ [ 43040]
```

```
FN557520.1_strain_E75            ......................................................................a......... [ 43040]
AY261366.1_Warthog               ...a..................a..........t.......................................a...... [ 43040]
AY261365.1_Warmbaths             ...a..................................................................a......... [ 43040]
AY261364.1_Tengani_62            ...a..................a..........t.t............................................ [ 43040]
AY261363.1_Pretoriuskop/96/4     ...a..................a..........t.t............................................ [ 43040]
AY261362.1_Mkuzi_1979            ......................................................................a......... [ 43040]
AY261361.1_Malawi_Lil-20/1_1983  ...................t..a.......c.......................................a......... [ 43040]
AY261360.1_Kenya_1950            ...................t..a.......c........g..................t...........a......... [ 43040]
AM712240.1_OURT_88/3             -------------------------------------------------------------------------------- [ 43040]
AM712239.1_Benin_97/1            ......................................................................a......... [ 43040]
KM111295.1_Ken06.Bus             ...................t..a.......c........g..................t...........a......... [ 43040]
KM111294.1_Ken05/Tk1             ...................t..a.......c........ga.................t...........a......... [ 43040]
KM262844.1_strain_L60            ......................................................................a......... [ 43040]
KM102979.1_26544/OG10_Italy      ......................................................................a......... [ 43040]
KM262845.1_strain_NHV            -------------------------------------------------------------------------------- [ 43040]
KX354450.1_47/Ss/2008            ...................................................a..................a......... [ 43040]
KP055815.1_strain_BA71           ......................................................................a......... [ 43040]
MH025920.1_strain_R35            ...................t..a.......c........g..................t...........a......... [ 43040]
MH025919.1_strain_N10            ...................t..a.......c........g..................t...........a......... [ 43040]
MH025918.1_strain_R25            ...................t..a.......c........g..................t...........a......... [ 43040]
MH025917.1_strain_R7             ...................t..a.......c........g..................t...........a......... [ 43040]
MH025916.1_strain_R8             ...................t..a.......c........g..................t...........a......... [ 43040]

FR682468.2_Georgia_2007/1        tctttttcacgacacaactcgtgtatgtcaaaaatgttattaaaataaaggctacaagctacccgccaatagaggtgatt [ 43120]
ASF_IND_20_CAD543                -------------------------------------------------------------------------------- [ 43120]
LR899193.1_Germany_2020/1        ................................................................................ [ 43120]
MN393477.1_Wuhan_2019            ................................................................................ [ 43120]
MW396979.1_Timor-Leste/2019/1    ................................................................................ [ 43120]
MK940252.1_CN/2019/InnerMongolia ................................................................................ [ 43120]
MK128995.1_China/2018/AnhuiXCGQ  ................................................................................ [ 43120]
MH681419.1_ASFV/POL/2015         ................................................................................ [ 43120]
LS478113.1_Estonia_2014          ................................................................................ [ 43120]
KP843857.1_Odintsovo_02/14       ................................................................................ [ 43120]
FN557520.1_strain_E75            ................g.........................................................c..... [ 43120]
AY261366.1_Warthog               ................g............................................................... [ 43120]
AY261365.1_Warmbaths             ................g............................................................... [ 43120]
AY261364.1_Tengani_62            ................................................................................ [ 43120]
AY261363.1_Pretoriuskop/96/4     ................g............................................................... [ 43120]
AY261362.1_Mkuzi_1979            ................g.........................................................c..... [ 43120]
AY261361.1_Malawi_Lil-20/1_1983  .......t..a.....g...a..c.a............g...................tc.gt...........c..... [ 43120]
AY261360.1_Kenya_1950            .......t.ga.....g...a..c.a.........a.......................c.g............c..... [ 43120]
AM712240.1_OURT_88/3             -------------------------------------------------------------------------------- [ 43120]
AM712239.1_Benin_97/1            ................g.........................................................c..... [ 43120]
KM111295.1_Ken06.Bus             .......t.ga.....g...a..c.a................................t..g.........c..c..... [ 43120]
```

```
KM111294.1_Ken05/Tk1            .......t.ga.....g...a..c.a.........a......................t..g.........c..c..... [ 43120]
KM262844.1_strain_L60           ................g.........................................................c..... [ 43120]
KM102979.1_26544/OG10_Italy     ................g.........................................................c..... [ 43120]
KM262845.1_strain_NHV           -------------------------------------------------------------------------------- [ 43120]
KX354450.1_47/Ss/2008           ................g.........................................................c..... [ 43120]
KP055815.1_strain_BA71          ................g.........................................................c..... [ 43120]
MH025920.1_strain_R35           .......t.ga.....g...a..c.a................................t..g.........c..c..... [ 43120]
MH025919.1_strain_N10           .......t.ga.....g...a..c.a................................t..g.........c..c..... [ 43120]
MH025918.1_strain_R25           .......t.ga.....g...a..c.a................................t..g.........c..c..... [ 43120]
MH025917.1_strain_R7            .......t.ga.....g...a..c.a................................t..g.........c..c..... [ 43120]
MH025916.1_strain_R8            .......t.ga.....g...a..c.a................................t..g.........c..c..... [ 43120]

FR682468.2_Georgia_2007/1       tttatgccttttatagaaatagtgaatagcctttgtaaaattatgtcgtaatgccagggcaaaccaaaactttgttaata [ 43200]
ASF_IND_20_CAD543               -------------------------------------------------------------------------------- [ 43200]
LR899193.1_Germany_2020/1       ................................................................................ [ 43200]
MN393477.1_Wuhan_2019           ................................................................................ [ 43200]
MW396979.1_Timor-Leste/2019/1   ................................................................................ [ 43200]
MK940252.1_CN/2019/InnerMongolia ................................................................................ [ 43200]
MK128995.1_China/2018/AnhuiXCGQ ................................................................................ [ 43200]
MH681419.1_ASFV/POL/2015        ................................................................................ [ 43200]
LS478113.1_Estonia_2014         ................................................................................ [ 43200]
KP843857.1_Odintsovo_02/14      ................................................................................ [ 43200]
FN557520.1_strain_E75           ...............a........g....................................................... [ 43200]
AY261366.1_Warthog              ...............a................................................................ [ 43200]
AY261365.1_Warmbaths            .........................................................c...................... [ 43200]
AY261364.1_Tengani_62           ................................................................................ [ 43200]
AY261363.1_Pretoriuskop/96/4    ................................................................................ [ 43200]
AY261362.1_Mkuzi_1979           ...............a........g....................................................... [ 43200]
AY261361.1_Malawi_Lil-20/1_1983 .......t...cg.................t...........................................a..... [ 43200]
AY261360.1_Kenya_1950           .......t...c...............c..t.........................................c....... [ 43200]
AM712240.1_OURT_88/3            -------------------------------------------------------------------------------- [ 43200]
AM712239.1_Benin_97/1           ...............a........g....................................................... [ 43200]
KM111295.1_Ken06.Bus            ......tt...cg..............c..t..............c..........................c....... [ 43200]
KM111294.1_Ken05/Tk1            .......t...c..................t................................................. [ 43200]
KM262844.1_strain_L60           ...............a........g....................................................... [ 43200]
KM102979.1_26544/OG10_Italy     ...............a........g....................................................... [ 43200]
KM262845.1_strain_NHV           -------------------------------------------------------------------------------- [ 43200]
KX354450.1_47/Ss/2008           ...............a........g....................................................... [ 43200]
KP055815.1_strain_BA71          ...............a........g....................................................... [ 43200]
MH025920.1_strain_R35           ......tt...cg..............c..t..............c..........................c....... [ 43200]
MH025919.1_strain_N10           ......tt...cg..............c..t..............c..........................c....... [ 43200]
MH025918.1_strain_R25           ......tt...cg..............c..t..............c..........................c....... [ 43200]
MH025917.1_strain_R7            ......tt...cg..............c..t..............c..........................c....... [ 43200]
MH025916.1_strain_R8            ......tt...cg..............c..t..............c..........................c....... [ 43200]
```

```
FR682468.2_Georgia_2007/1        ggtggtgcgccgtatcccccgtcaacggaatgtttgaacaggtgtacgtaactgtgtctaaagtggttctagttacggtt [ 43280]
ASF_IND_20_CAD543                -------------------------------------------------------------------------------- [ 43280]
LR899193.1_Germany_2020/1        ................................................................................ [ 43280]
MN393477.1_Wuhan_2019            ................................................................................ [ 43280]
MW396979.1_Timor-Leste/2019/1    ................................................................................ [ 43280]
MK940252.1_CN/2019/InnerMongolia ................................................................................ [ 43280]
MK128995.1_China/2018/AnhuiXCGQ  ................................................................................ [ 43280]
MH681419.1_ASFV/POL/2015         ................................................................................ [ 43280]
LS478113.1_Estonia_2014          ................................................................................ [ 43280]
KP843857.1_Odintsovo_02/14       ................................................................................ [ 43280]
FN557520.1_strain_E75            ..a............................................................................. [ 43280]
AY261366.1_Warthog               ................................................................................ [ 43280]
AY261365.1_Warmbaths             ....................a..........t.........c.....a...........................t.... [ 43280]
AY261364.1_Tengani_62            ................................................................................ [ 43280]
AY261363.1_Pretoriuskop/96/4     ..a............................................................................. [ 43280]
AY261362.1_Mkuzi_1979            ..a............................................................................. [ 43280]
AY261361.1_Malawi_Lil-20/1_1983  ..a............................t.......g.......a.............g...........c...... [ 43280]
AY261360.1_Kenya_1950            ..a............................t.......g.......a....g....t...............c...... [ 43280]
AM712240.1_OURT_88/3             -------------------------------------------------------------------------------- [ 43280]
AM712239.1_Benin_97/1            ..a............................................................................. [ 43280]
KM111295.1_Ken06.Bus             ..a............................t.......g...a...a.........t...............c...... [ 43280]
KM111294.1_Ken05/Tk1             ..a............................t.......g.......a.........................c...... [ 43280]
KM262844.1_strain_L60            ..a............................................................................. [ 43280]
KM102979.1_26544/OG10_Italy      ..a............................................................................. [ 43280]
KM262845.1_strain_NHV            -------------------------------------------------------------------------------- [ 43280]
KX354450.1_47/Ss/2008            ..a............................................................................. [ 43280]
KP055815.1_strain_BA71           ..a............................................................................. [ 43280]
MH025920.1_strain_R35            ..a............................t.......g...a...a.........t...............c...... [ 43280]
MH025919.1_strain_N10            ..a............................t.......g...a...a.........t...............c...... [ 43280]
MH025918.1_strain_R25            ..a............................t.......g...a...a.........t...............c...... [ 43280]
MH025917.1_strain_R7             ..a............................t.......g...a...a.........t...............c...... [ 43280]
MH025916.1_strain_R8             ..a............................t.......g...a...a.........t...............c...... [ 43280]

FR682468.2_Georgia_2007/1        tccaagagtggattatgacaaaacatgtcataacccagcagaactcctgcacaggattttagcctggccacttcttttaa [ 43360]
ASF_IND_20_CAD543                -------------------------------------------------------------------------------- [ 43360]
LR899193.1_Germany_2020/1        ................................................................................ [ 43360]
MN393477.1_Wuhan_2019            ................................................................................ [ 43360]
MW396979.1_Timor-Leste/2019/1    ................................................................................ [ 43360]
MK940252.1_CN/2019/InnerMongolia ................................................................................ [ 43360]
MK128995.1_China/2018/AnhuiXCGQ  ................................................................................ [ 43360]
MH681419.1_ASFV/POL/2015         ................................................................................ [ 43360]
LS478113.1_Estonia_2014          ................................................................................ [ 43360]
KP843857.1_Odintsovo_02/14       ................................................................................ [ 43360]
```

```
FN557520.1_strain_E75            ......................................t.......................gt................ [ 43360]
AY261366.1_Warthog               ......................................t........................t................ [ 43360]
AY261365.1_Warmbaths             ...........g....a.....................t.......................gt................ [ 43360]
AY261364.1_Tengani_62            ................................................................................ [ 43360]
AY261363.1_Pretoriuskop/96/4     ......................................t........................t................ [ 43360]
AY261362.1_Mkuzi_1979            ......................................t.......................gt................ [ 43360]
AY261361.1_Malawi_Lil-20/1_1983  ......................................t....................c..atc............... [ 43360]
AY261360.1_Kenya_1950            ......................................t....................c..atc............... [ 43360]
AM712240.1_OURT_88/3             -------------------------------------------------------------------------------- [ 43360]
AM712239.1_Benin_97/1            ......................................t.......................gt................ [ 43360]
KM111295.1_Ken06.Bus             ....ta................................t....................c..atc............... [ 43360]
KM111294.1_Ken05/Tk1             ......................................t....................c..atc............... [ 43360]
KM262844.1_strain_L60            ......................................t.......................gt................ [ 43360]
KM102979.1_26544/OG10_Italy      ......................................t.......................gt................ [ 43360]
KM262845.1_strain_NHV            -------------------------------------------------------------------------------- [ 43360]
KX354450.1_47/Ss/2008            ......................................t.......................gt................ [ 43360]
KP055815.1_strain_BA71           ......................................t.......................gt................ [ 43360]
MH025920.1_strain_R35            ....ta................................t....................c..atc............... [ 43360]
MH025919.1_strain_N10            ....ta................................t....................c..atc............... [ 43360]
MH025918.1_strain_R25            ....ta................................t....................c..atc............... [ 43360]
MH025917.1_strain_R7             ....ta................................t....................c..atc............... [ 43360]
MH025916.1_strain_R8             ....ta................................t....................c..atc............... [ 43360]

FR682468.2_Georgia_2007/1        aatttccagaagacggggttcggatacaggcgttaagcctcccagttccgcacacagccgctttagatacacggcaggaa [ 43440]
ASF_IND_20_CAD543                -------------------------------------------------------------------------------- [ 43440]
LR899193.1_Germany_2020/1        ................................................................................ [ 43440]
MN393477.1_Wuhan_2019            ................................................................................ [ 43440]
MW396979.1_Timor-Leste/2019/1    ................................................................................ [ 43440]
MK940252.1_CN/2019/InnerMongolia ................................................................................ [ 43440]
MK128995.1_China/2018/AnhuiXCGQ  ................................................................................ [ 43440]
MH681419.1_ASFV/POL/2015         ................................................................................ [ 43440]
LS478113.1_Estonia_2014          ................................................................................ [ 43440]
KP843857.1_Odintsovo_02/14       ................................................................................ [ 43440]
FN557520.1_strain_E75            .................................c.............................................. [ 43440]
AY261366.1_Warthog               ..................................................tg................g........... [ 43440]
AY261365.1_Warmbaths             ..............................t.................t.tg................g.....t..... [ 43440]
AY261364.1_Tengani_62            .................a.....................a..........tg................g........... [ 43440]
AY261363.1_Pretoriuskop/96/4     .......................t..........................tg................g........... [ 43440]
AY261362.1_Mkuzi_1979            ................................................................................ [ 43440]
AY261361.1_Malawi_Lil-20/1_1983  ...c...........a.a............t...........t......atg................g........... [ 43440]
AY261360.1_Kenya_1950            ...............a.a......g.....t...........t.....t.tg........t.......c........... [ 43440]
AM712240.1_OURT_88/3             -------------------------------------------------------------------------------- [ 43440]
AM712239.1_Benin_97/1            .................................c.............................................. [ 43440]
KM111295.1_Ken06.Bus             ...............a.a......g.....t...........t.......tg........t.......g........... [ 43440]
```

```
KM111294.1_Ken05/Tk1            ...............a.a...a..g.....t...........t.......tg................g.t....g.... [ 43440]
KM262844.1_strain_L60           .................................c.............................................. [ 43440]
KM102979.1_26544/OG10_Italy     .................................c.............................................. [ 43440]
KM262845.1_strain_NHV           -------------------------------------------------------------------------------- [ 43440]
KX354450.1_47/Ss/2008           .................................c.............................................. [ 43440]
KP055815.1_strain_BA71          .................................c.............................................. [ 43440]
MH025920.1_strain_R35           ...............a.a......g.....t...........t.......tg........t.......g........... [ 43440]
MH025919.1_strain_N10           ...............a.a......g.....t...........t.......tg........t.......g........... [ 43440]
MH025918.1_strain_R25           ...............a.a......g.....t...........t.......tg........t.......g........... [ 43440]
MH025917.1_strain_R7            ...............a.a......g.....t...........t.......tg........t.......g........... [ 43440]
MH025916.1_strain_R8            ...............a.a......g.....t...........t.......tg........t.......g........... [ 43440]

FR682468.2_Georgia_2007/1       cacgtataagcccatattcaggatttgcgccccaatccacaaataaacgtataagttcaagattatcgctcttcacggcc [ 43520]
ASF_IND_20_CAD543               -------------------------------------------------------------------------------- [ 43520]
LR899193.1_Germany_2020/1       ................................................................................ [ 43520]
MN393477.1_Wuhan_2019           ................................................................................ [ 43520]
MW396979.1_Timor-Leste/2019/1   ................................................................................ [ 43520]
MK940252.1_CN/2019/InnerMongolia ................................................................................ [ 43520]
MK128995.1_China/2018/AnhuiXCGQ ................................................................................ [ 43520]
MH681419.1_ASFV/POL/2015        ................................................................................ [ 43520]
LS478113.1_Estonia_2014         ................................................................................ [ 43520]
KP843857.1_Odintsovo_02/14      ................................................................................ [ 43520]
FN557520.1_strain_E75           ................................................................................ [ 43520]
AY261366.1_Warthog              ..........t............................................c........................ [ 43520]
AY261365.1_Warmbaths            ...a......t.................a..........................c...........c............ [ 43520]
AY261364.1_Tengani_62           ..........t............................................c........................ [ 43520]
AY261363.1_Pretoriuskop/96/4    ..........t............................................c........................ [ 43520]
AY261362.1_Mkuzi_1979           ................................................................................ [ 43520]
AY261361.1_Malawi_Lil-20/1_1983 ..........t................................c..t........c........................ [ 43520]
AY261360.1_Kenya_1950           ..........t................................c..t........c........................ [ 43520]
AM712240.1_OURT_88/3            -------------------------------------------------------------------------------- [ 43520]
AM712239.1_Benin_97/1           ................................................................................ [ 43520]
KM111295.1_Ken06.Bus            ..........t................................c..t........c........................ [ 43520]
KM111294.1_Ken05/Tk1            .g.a......tg.......g.......................c..t........c........................ [ 43520]
KM262844.1_strain_L60           ................................................................................ [ 43520]
KM102979.1_26544/OG10_Italy     ................................................................................ [ 43520]
KM262845.1_strain_NHV           -------------------------------------------------------------------------------- [ 43520]
KX354450.1_47/Ss/2008           ................................................................................ [ 43520]
KP055815.1_strain_BA71          ................................................................................ [ 43520]
MH025920.1_strain_R35           ..........t................................c..t........c........................ [ 43520]
MH025919.1_strain_N10           ..........t................................c..t........c........................ [ 43520]
MH025918.1_strain_R25           ..........t................................c..t........c........................ [ 43520]
MH025917.1_strain_R7            ..........t................................c..t........c........................ [ 43520]
MH025916.1_strain_R8            ..........t................................c..t........c........................ [ 43520]
```

```
FR682468.2_Georgia_2007/1        tttactagcgccgcttcgagacaaagatcatcctcagaaaaacactgtaaatgtttatacgaaaaaacttgcttacaatt [ 43600]
ASF_IND_20_CAD543                -------------------------------------------------------------------------------- [ 43600]
LR899193.1_Germany_2020/1        ................................................................................ [ 43600]
MN393477.1_Wuhan_2019            ................................................................................ [ 43600]
MW396979.1_Timor-Leste/2019/1    ................................................................................ [ 43600]
MK940252.1_CN/2019/InnerMongolia ................................................................................ [ 43600]
MK128995.1_China/2018/AnhuiXCGQ  ................................................................................ [ 43600]
MH681419.1_ASFV/POL/2015         ................................................................................ [ 43600]
LS478113.1_Estonia_2014          ................................................................................ [ 43600]
KP843857.1_Odintsovo_02/14       ................................................................................ [ 43600]
FN557520.1_strain_E75            .................a.............................................................. [ 43600]
AY261366.1_Warthog               .................a.................................................t............ [ 43600]
AY261365.1_Warmbaths             .................a.....g...........................................t............ [ 43600]
AY261364.1_Tengani_62            .................a...............................c.................t............ [ 43600]
AY261363.1_Pretoriuskop/96/4     .................a.................................................t............ [ 43600]
AY261362.1_Mkuzi_1979            .................a.............................................................. [ 43600]
AY261361.1_Malawi_Lil-20/1_1983  .................a.........................................t.......t.........tg. [ 43600]
AY261360.1_Kenya_1950            ..............g..a.............................c.......g...........t.........tg. [ 43600]
AM712240.1_OURT_88/3             -------------------------------------------------------------------------------- [ 43600]
AM712239.1_Benin_97/1            .................a.............................................................. [ 43600]
KM111295.1_Ken06.Bus             ..............g..a...............a.............c.......g...........t.........tg. [ 43600]
KM111294.1_Ken05/Tk1             ..............g..a.............................c.......g...........t.........tg. [ 43600]
KM262844.1_strain_L60            .................a.............................................................. [ 43600]
KM102979.1_26544/OG10_Italy      .................a.............................................................. [ 43600]
KM262845.1_strain_NHV            -------------------------------------------------------------------------------- [ 43600]
KX354450.1_47/Ss/2008            .................a.............................................................. [ 43600]
KP055815.1_strain_BA71           .................a.............................................................. [ 43600]
MH025920.1_strain_R35            ..............g..a...............a.............c.......g...........t.........tg. [ 43600]
MH025919.1_strain_N10            ..............g..a...............a.............c.......g...........t.........tg. [ 43600]
MH025918.1_strain_R25            ..............g..a...............a.............c.......g...........t.........tg. [ 43600]
MH025917.1_strain_R7             ..............g..a...............a.............c.......g...........t.........tg. [ 43600]
MH025916.1_strain_R8             ..............g..a...............a.............c.......g...........t.........tg. [ 43600]

FR682468.2_Georgia_2007/1        gttacataggtgaataggacctaaatcccaccacaaaccaaaacgctgcaacgtataatcatagtcacttgaaagataat [ 43680]
ASF_IND_20_CAD543                -------------------------------------------------------------------------------- [ 43680]
LR899193.1_Germany_2020/1        ................................................................................ [ 43680]
MN393477.1_Wuhan_2019            ................................................................................ [ 43680]
MW396979.1_Timor-Leste/2019/1    ................................................................................ [ 43680]
MK940252.1_CN/2019/InnerMongolia ................................................................................ [ 43680]
MK128995.1_China/2018/AnhuiXCGQ  ................................................................................ [ 43680]
MH681419.1_ASFV/POL/2015         ................................................................................ [ 43680]
LS478113.1_Estonia_2014          ................................................................................ [ 43680]
KP843857.1_Odintsovo_02/14       ................................................................................ [ 43680]
```

```
FN557520.1_strain_E75            ................................................................................ [ 43680]
AY261366.1_Warthog               ....................................................a........................... [ 43680]
AY261365.1_Warmbaths             ......c.............................................a........................... [ 43680]
AY261364.1_Tengani_62            a............................................................................... [ 43680]
AY261363.1_Pretoriuskop/96/4     ......c......................................................................... [ 43680]
AY261362.1_Mkuzi_1979            ................................................................................ [ 43680]
AY261361.1_Malawi_Lil-20/1_1983  a.......a.........................................g............a................ [ 43680]
AY261360.1_Kenya_1950            a.......a.........................................g............................. [ 43680]
AM712240.1_OURT_88/3             -------------------------------------------------------------------------------- [ 43680]
AM712239.1_Benin_97/1            ................................................................................ [ 43680]
KM111295.1_Ken06.Bus             a.......a.........................................g............................. [ 43680]
KM111294.1_Ken05/Tk1             a.......a.........................................g............................. [ 43680]
KM262844.1_strain_L60            ................................................................................ [ 43680]
KM102979.1_26544/OG10_Italy      ................................................................................ [ 43680]
KM262845.1_strain_NHV            -------------------------------------------------------------------------------- [ 43680]
KX354450.1_47/Ss/2008            ................................................................................ [ 43680]
KP055815.1_strain_BA71           ................................................................................ [ 43680]
MH025920.1_strain_R35            a.......a.........................................g............................. [ 43680]
MH025919.1_strain_N10            a.......a.........................................g............................. [ 43680]
MH025918.1_strain_R25            a.......a.........................................g............................. [ 43680]
MH025917.1_strain_R7             a.......a.........................................g............................. [ 43680]
MH025916.1_strain_R8             a.......a.........................................g............................. [ 43680]

FR682468.2_Georgia_2007/1        tgcatgccacaacttttttggccaacgtttgtaaagacaacatactaagtttaaaacatcttaaatctaagctagc---t [ 43760]
ASF_IND_20_CAD543                -------------------.........................................................---. [ 43760]
LR899193.1_Germany_2020/1        ............................................................................---. [ 43760]
MN393477.1_Wuhan_2019            ............................................................................---. [ 43760]
MW396979.1_Timor-Leste/2019/1    ............................................................................---. [ 43760]
MK940252.1_CN/2019/InnerMongolia ............................................................................---. [ 43760]
MK128995.1_China/2018/AnhuiXCGQ  ............................................................................---. [ 43760]
MH681419.1_ASFV/POL/2015         ............................................................................---. [ 43760]
LS478113.1_Estonia_2014          ............................................................................---. [ 43760]
KP843857.1_Odintsovo_02/14       ............................................................................---. [ 43760]
FN557520.1_strain_E75            ...........................................g................................---. [ 43760]
AY261366.1_Warthog               .............c...........................................ga.................---. [ 43760]
AY261365.1_Warmbaths             .............c...........................................ga.................---. [ 43760]
AY261364.1_Tengani_62            .............c...........t..................................................---- [ 43760]
AY261363.1_Pretoriuskop/96/4     .............c................................t.............................---. [ 43760]
AY261362.1_Mkuzi_1979            ............................................................................---. [ 43760]
AY261361.1_Malawi_Lil-20/1_1983  ........................g...............................t.g......c..........---. [ 43760]
AY261360.1_Kenya_1950            .............c.....................................----------....c....at....tac. [ 43760]
AM712240.1_OURT_88/3             -------------------------------------------------------------------------------- [ 43760]
AM712239.1_Benin_97/1            ...........................................g................................---. [ 43760]
KM111295.1_Ken06.Bus             ....c.....................................................g-c....c....at....tac. [ 43760]
```

```
KM111294.1_Ken05/Tk1             .............c................................t...........g......c....at....tac. [ 43760]
KM262844.1_strain_L60            ...........................................g................................---. [ 43760]
KM102979.1_26544/OG10_Italy      ...........................................g................................---. [ 43760]
KM262845.1_strain_NHV            -------------------------------------------------------------------------------- [ 43760]
KX354450.1_47/Ss/2008            ...........................................g................................---. [ 43760]
KP055815.1_strain_BA71           ...........................................g................................---. [ 43760]
MH025920.1_strain_R35            ....c.....................................................g-c....c....at....tac. [ 43760]
MH025919.1_strain_N10            ....c.....................................................g-c....c....at....tac. [ 43760]
MH025918.1_strain_R25            ....c.....................................................g-c....c....at....tac. [ 43760]
MH025917.1_strain_R7             ....c.....................................................g-c....c....at....tac. [ 43760]
MH025916.1_strain_R8             ....c.....................................................g-c....c....at....tac. [ 43760]

FR682468.2_Georgia_2007/1        aactttcaagaaaaccctctatccctaagaatatatcttataactagacttatagcagtaaaaatcaactttggttattc [ 43840]
ASF_IND_20_CAD543                ................................................................................ [ 43840]
LR899193.1_Germany_2020/1        ................................................................................ [ 43840]
MN393477.1_Wuhan_2019            ................................................................................ [ 43840]
MW396979.1_Timor-Leste/2019/1    ................................................................................ [ 43840]
MK940252.1_CN/2019/InnerMongolia ................................................................................ [ 43840]
MK128995.1_China/2018/AnhuiXCGQ  ................................................................................ [ 43840]
MH681419.1_ASFV/POL/2015         ................................................................................ [ 43840]
LS478113.1_Estonia_2014          ................................................................................ [ 43840]
KP843857.1_Odintsovo_02/14       ................................................................................ [ 43840]
FN557520.1_strain_E75            ..............t................................................................. [ 43840]
AY261366.1_Warthog               ...c..........t................................................................. [ 43840]
AY261365.1_Warmbaths             ...c..........t...............................a................................. [ 43840]
AY261364.1_Tengani_62            g.........-...t................................................................. [ 43840]
AY261363.1_Pretoriuskop/96/4     ..........-...t................................................................. [ 43840]
AY261362.1_Mkuzi_1979            ..............t................................................................. [ 43840]
AY261361.1_Malawi_Lil-20/1_1983  ..t..ct.......t...-----.....ag.....c.............a...g...a....................ag [ 43840]
AY261360.1_Kenya_1950            .....c........t.....t........g-..................a..ag..ga....................a. [ 43840]
AM712240.1_OURT_88/3             -------------------------------------------------------------------------------- [ 43840]
AM712239.1_Benin_97/1            ..............t................................................................. [ 43840]
KM111295.1_Ken06.Bus             .....c........t.t...t........g...................a..ag...a....................a. [ 43840]
KM111294.1_Ken05/Tk1             .....c........t.....t........g-..................a..ag..ga....................a. [ 43840]
KM262844.1_strain_L60            ..............t................................................................. [ 43840]
KM102979.1_26544/OG10_Italy      ..............t................................................................. [ 43840]
KM262845.1_strain_NHV            -------------------------------------------------------------------------------- [ 43840]
KX354450.1_47/Ss/2008            ..............t................................................................. [ 43840]
KP055815.1_strain_BA71           ..............t................................................................. [ 43840]
MH025920.1_strain_R35            .....c........t.t...t........g...................a..ag...a....................a. [ 43840]
MH025919.1_strain_N10            .....c........t.t...t........g...................a..ag...a....................a. [ 43840]
MH025918.1_strain_R25            .....c........t.t...t........g...................a..ag...a....................a. [ 43840]
MH025917.1_strain_R7             .....c........t.t...t........g...................a..ag...a....................a. [ 43840]
MH025916.1_strain_R8             .....c........t.t...t........g...................a..ag...a....................a. [ 43840]
```

```
FR682468.2_Georgia_2007/1        tttttaatataaaacgtctaattacttgcaaaggactata---------------------------------------- [ 43920]
ASF_IND_20_CAD543                .................................k.y...m---------------------------------------- [ 43920]
LR899193.1_Germany_2020/1        ........................................---------------------------------------- [ 43920]
MN393477.1_Wuhan_2019            ........................................---------------------------------------- [ 43920]
MW396979.1_Timor-Leste/2019/1    ........................................---------------------------------------- [ 43920]
MK940252.1_CN/2019/InnerMongolia ........................................---------------------------------------- [ 43920]
MK128995.1_China/2018/AnhuiXCGQ  ........................................---------------------------------------- [ 43920]
MH681419.1_ASFV/POL/2015         ........................................---------------------------------------- [ 43920]
LS478113.1_Estonia_2014          ........................................---------------------------------------- [ 43920]
KP843857.1_Odintsovo_02/14       ........................................---------------------------------------- [ 43920]
FN557520.1_strain_E75            .................................t.t...c---------------------------------------- [ 43920]
AY261366.1_Warthog               ...............................---.g..cta--------------------------------------- [ 43920]
AY261365.1_Warmbaths             ................c................t.g..cta--------------------------------------- [ 43920]
AY261364.1_Tengani_62            .........................c..............---------------------------------------- [ 43920]
AY261363.1_Pretoriuskop/96/4     .............................t.ta----------------------------------------------- [ 43920]
AY261362.1_Mkuzi_1979            .................................t.t....---------------------------------------- [ 43920]
AY261361.1_Malawi_Lil-20/1_1983  ..........g.....................atc..tgcaaaagtccttgcaaagttcttgcaaagttcttgcaaagac [ 43920]
AY261360.1_Kenya_1950            ..........g..........g...........tc..tgcaa-------------------------------------- [ 43920]
AM712240.1_OURT_88/3             -------------------------------------------------------------------------------- [ 43920]
AM712239.1_Benin_97/1            .................................t.t...c---------------------------------------- [ 43920]
KM111295.1_Ken06.Bus             ..........g..........g...........t...tgcaa-------------------------------------- [ 43920]
KM111294.1_Ken05/Tk1             ..........g..........g...........t...tgcaa-------------------------------------- [ 43920]
KM262844.1_strain_L60            .................................t.t...c---------------------------------------- [ 43920]
KM102979.1_26544/OG10_Italy      .................................t.t...c---------------------------------------- [ 43920]
KM262845.1_strain_NHV            -------------------------------------------------------------------------------- [ 43920]
KX354450.1_47/Ss/2008            .................................t.t...c---------------------------------------- [ 43920]
KP055815.1_strain_BA71           .................................t.t...c---------------------------------------- [ 43920]
MH025920.1_strain_R35            ..........g..........g...........t...tgcaa-------------------------------------- [ 43920]
MH025919.1_strain_N10            ..........g..........g...........t...tgcaa-------------------------------------- [ 43920]
MH025918.1_strain_R25            ..........g..........g...........t...tgcaa-------------------------------------- [ 43920]
MH025917.1_strain_R7             ..........g..........g...........t...tgcaa-------------------------------------- [ 43920]
MH025916.1_strain_R8             ..........g..........g...........t...tgcaa-------------------------------------- [ 43920]

FR682468.2_Georgia_2007/1        -----------------aagcccattttcctcagctagaattttta-----------------ttttttaatgaagta-g [ 44000]
ASF_IND_20_CAD543                -----------------.............................-----------------...............-. [ 44000]
LR899193.1_Germany_2020/1        -----------------.............................-----------------...............-. [ 44000]
MN393477.1_Wuhan_2019            -----------------.............................-----------------...............-. [ 44000]
MW396979.1_Timor-Leste/2019/1    -----------------.............................-----------------...............-. [ 44000]
MK940252.1_CN/2019/InnerMongolia -----------------.............................-----------------...............-. [ 44000]
MK128995.1_China/2018/AnhuiXCGQ  -----------------.............................-----------------...............-. [ 44000]
MH681419.1_ASFV/POL/2015         -----------------.............................-----------------...............-. [ 44000]
LS478113.1_Estonia_2014          -----------------.............................-----------------...............-. [ 44000]
KP843857.1_Odintsovo_02/14       -----------------.............................-----------------...............-. [ 44000]
```

```
FN557520.1_strain_E75           -----------------.............................-----------------...............-. [ 44000]
AY261366.1_Warthog              ----------------t.................a...........-----------------...........g...-. [ 44000]
AY261365.1_Warmbaths            ----------------t.................a...........-----------------...........g...g. [ 44000]
AY261364.1_Tengani_62           -----------------.....t...........ac..........-----------------...........g...g. [ 44000]
AY261363.1_Pretoriuskop/96/4    -----------------g....t...........ac..........-----------------...........g...-- [ 44000]
AY261362.1_Mkuzi_1979           -----------------g............................-----------------...............-. [ 44000]
AY261361.1_Malawi_Lil-20/1_1983 cttgcatagtctttgca.....gtca.......ta.........c.atgccatgaatttttat...........g...-. [ 44000]
AY261360.1_Kenya_1950           -------agactttgca.....ttca.......ta..t......c.atgacatgaattttta-...........g...-. [ 44000]
AM712240.1_OURT_88/3            -------------------------------------------------------------------------------- [ 44000]
AM712239.1_Benin_97/1           -----------------.............................-----------------...............-. [ 44000]
KM111295.1_Ken06.Bus            -------agtccttgca.....ttca.......ta..t......c.atgacatgaattttta-...........g...-. [ 44000]
KM111294.1_Ken05/Tk1            -------agtccttgca....-------.....ta.........c.atgacatgaattttta-...........g...-. [ 44000]
KM262844.1_strain_L60           -----------------.............................-----------------...............-. [ 44000]
KM102979.1_26544/OG10_Italy     -----------------.............................-----------------...............-. [ 44000]
KM262845.1_strain_NHV           -------------------------------------------------------------------------------- [ 44000]
KX354450.1_47/Ss/2008           -----------------.............................-----------------...............-. [ 44000]
KP055815.1_strain_BA71          -----------------.............................-----------------...............-. [ 44000]
MH025920.1_strain_R35           -------agtccttgca.....ttca.......ta..t......c.atgacatgaattttta-...........g...-. [ 44000]
MH025919.1_strain_N10           -------agtccttgca.....ttca.......ta..t......c.atgacatgaattttta-...........g...-. [ 44000]
MH025918.1_strain_R25           -------agtccttgca.....ttca.......ta..t......c.atgacatgaattttta-...........g...-. [ 44000]
MH025917.1_strain_R7            -------agtccttgca.....ttca.......ta..t......c.atgacatgaattttta-...........g...-. [ 44000]
MH025916.1_strain_R8            -------agtccttgca.....ttca.......ta..t......c.atgacatgaattttta-...........g...-. [ 44000]

FR682468.2_Georgia_2007/1       ggggatatgttttcccttcaagacctttgccgaaagcatctttttattcttcccgatgtttttggcgagcatgtactaca [ 44080]
ASF_IND_20_CAD543               ................................................................................ [ 44080]
LR899193.1_Germany_2020/1       ................................................................................ [ 44080]
MN393477.1_Wuhan_2019           ................................................................................ [ 44080]
MW396979.1_Timor-Leste/2019/1   ................................................................................ [ 44080]
MK940252.1_CN/2019/InnerMongolia ................................................................................ [ 44080]
MK128995.1_China/2018/AnhuiXCGQ ................................................................................ [ 44080]
MH681419.1_ASFV/POL/2015        ................................................................................ [ 44080]
LS478113.1_Estonia_2014         ................................................................................ [ 44080]
KP843857.1_Odintsovo_02/14      ................................................................................ [ 44080]
FN557520.1_strain_E75           ................................................................................ [ 44080]
AY261366.1_Warthog              .....c.....................................................................t.... [ 44080]
AY261365.1_Warmbaths            ...........................................................................t.... [ 44080]
AY261364.1_Tengani_62           ...a.c.......................................................................... [ 44080]
AY261363.1_Pretoriuskop/96/4    ...a............................................................................ [ 44080]
AY261362.1_Mkuzi_1979           ................................................................................ [ 44080]
AY261361.1_Malawi_Lil-20/1_1983 ...a.a.................t........................................................ [ 44080]
AY261360.1_Kenya_1950           ...-ga.................................................................c........ [ 44080]
AM712240.1_OURT_88/3            -------------------------------------------------------------------------------- [ 44080]
AM712239.1_Benin_97/1           ................................................................................ [ 44080]
KM111295.1_Ken06.Bus            ...-ga.......................................................................... [ 44080]
```

```
KM111294.1_Ken05/Tk1             ...a.a...................a...................................................... [ 44080]
KM262844.1_strain_L60            ................................................................................ [ 44080]
KM102979.1_26544/OG10_Italy      ................................................................................ [ 44080]
KM262845.1_strain_NHV            -------------------------------------------------------------------------------- [ 44080]
KX354450.1_47/Ss/2008            ................................................................................ [ 44080]
KP055815.1_strain_BA71           ................................................................................ [ 44080]
MH025920.1_strain_R35            ...-ga.......................................................................... [ 44080]
MH025919.1_strain_N10            ...-ga.......................................................................... [ 44080]
MH025918.1_strain_R25            ...-ga.......................................................................... [ 44080]
MH025917.1_strain_R7             ...-ga.......................................................................... [ 44080]
MH025916.1_strain_R8             ...-ga.......................................................................... [ 44080]

FR682468.2_Georgia_2007/1        acgattaggactgtattggagatgtcacggctcccttcaacgcataggagacgaccacatactcatacgacgggatctca [ 44160]
ASF_IND_20_CAD543                ................................................................................ [ 44160]
LR899193.1_Germany_2020/1        ................................................................................ [ 44160]
MN393477.1_Wuhan_2019            ................................................................................ [ 44160]
MW396979.1_Timor-Leste/2019/1    ................................................................................ [ 44160]
MK940252.1_CN/2019/InnerMongolia ................................................................................ [ 44160]
MK128995.1_China/2018/AnhuiXCGQ  ................................................................................ [ 44160]
MH681419.1_ASFV/POL/2015         ................................................................................ [ 44160]
LS478113.1_Estonia_2014          ................................................................................ [ 44160]
KP843857.1_Odintsovo_02/14       ................................................................................ [ 44160]
FN557520.1_strain_E75            ................................................................................ [ 44160]
AY261366.1_Warthog               ....................a................................................ga......... [ 44160]
AY261365.1_Warmbaths             ...........................................g.........................g.......... [ 44160]
AY261364.1_Tengani_62            ...........................................g.........................g.......... [ 44160]
AY261363.1_Pretoriuskop/96/4     ...........................................g................c........g.......... [ 44160]
AY261362.1_Mkuzi_1979            ................................................................................ [ 44160]
AY261361.1_Malawi_Lil-20/1_1983  ..ag...........c....a.c..........t...................................g.....c.... [ 44160]
AY261360.1_Kenya_1950            ..a...g..g.....c...ga.aaa........t........a.....ga....t..............g.......... [ 44160]
AM712240.1_OURT_88/3             -------------------------------------------------------------------------------- [ 44160]
AM712239.1_Benin_97/1            ................................................................................ [ 44160]
KM111295.1_Ken06.Bus             ..a...g..g.....c...ga.aaa........t........a.....ga....t..............g.......... [ 44160]
KM111294.1_Ken05/Tk1             ...g...........c...g..c..........t........a..c..g.....t..............g.....c.... [ 44160]
KM262844.1_strain_L60            ................................................................................ [ 44160]
KM102979.1_26544/OG10_Italy      ................................................................................ [ 44160]
KM262845.1_strain_NHV            -------------------------------------------------------------------------------- [ 44160]
KX354450.1_47/Ss/2008            ................................................................................ [ 44160]
KP055815.1_strain_BA71           ................................................................................ [ 44160]
MH025920.1_strain_R35            ..a...g..g.....c...ga.aaa........t........a.....ga....t..............g.......... [ 44160]
MH025919.1_strain_N10            ..a...g..g.....c...ga.aaa........t........a.....ga....t..............g.......... [ 44160]
MH025918.1_strain_R25            ..a...g..g.....c...ga.aaa........t........a.....ga....t..............g.......... [ 44160]
MH025917.1_strain_R7             ..a...g..g.....c...ga.aaa........t........a.....ga....t..............g.......... [ 44160]
MH025916.1_strain_R8             ..a...g..g.....c...ga.aaa........t........a.....ga....t..............g.......... [ 44160]
```

```
FR682468.2_Georgia_2007/1        tcctttccaccaacgaggccttaagaatggcgggagaggaaggaaacaatgaagtagtaaagctcttgttactgtggaag [ 44240]
ASF_IND_20_CAD543                ................................................................................ [ 44240]
LR899193.1_Germany_2020/1        ................................................................................ [ 44240]
MN393477.1_Wuhan_2019            ................................................................................ [ 44240]
MW396979.1_Timor-Leste/2019/1    ................................................................................ [ 44240]
MK940252.1_CN/2019/InnerMongolia ................................................................................ [ 44240]
MK128995.1_China/2018/AnhuiXCGQ  ................................................................................ [ 44240]
MH681419.1_ASFV/POL/2015         ................................................................................ [ 44240]
LS478113.1_Estonia_2014          ................................................................................ [ 44240]
KP843857.1_Odintsovo_02/14       ................................................................................ [ 44240]
FN557520.1_strain_E75            ................................................................................ [ 44240]
AY261366.1_Warthog               .......t.............................a.......................................... [ 44240]
AY261365.1_Warmbaths             .....................................a.......................................... [ 44240]
AY261364.1_Tengani_62            ................................................................................ [ 44240]
AY261363.1_Pretoriuskop/96/4     .....................................a.......................................... [ 44240]
AY261362.1_Mkuzi_1979            ................................................................................ [ 44240]
AY261361.1_Malawi_Lil-20/1_1983  ........................a....................................................g.. [ 44240]
AY261360.1_Kenya_1950            .......t.t..............a...a....c..............................t........a...... [ 44240]
AM712240.1_OURT_88/3             -------------------------------------------------------------------------------- [ 44240]
AM712239.1_Benin_97/1            ................................................................................ [ 44240]
KM111295.1_Ken06.Bus             .......t.t..............a...a....c.............................................. [ 44240]
KM111294.1_Ken05/Tk1             ........................a............a.......................................g.. [ 44240]
KM262844.1_strain_L60            ................................................................................ [ 44240]
KM102979.1_26544/OG10_Italy      ................................................................................ [ 44240]
KM262845.1_strain_NHV            -------------------------------------------------------------------------------- [ 44240]
KX354450.1_47/Ss/2008            ................................................................................ [ 44240]
KP055815.1_strain_BA71           ................................................................................ [ 44240]
MH025920.1_strain_R35            .......t.t..............a...a....c.............................................. [ 44240]
MH025919.1_strain_N10            .......t.t..............a...a....c.............................................. [ 44240]
MH025918.1_strain_R25            .......t.t..............a...a....c.............................................. [ 44240]
MH025917.1_strain_R7             .......t.t..............a...a....c.............................................. [ 44240]
MH025916.1_strain_R8             .......t.t..............a...a....c.............................................. [ 44240]

FR682468.2_Georgia_2007/1        ggaaatcttcattacgccgtcataggagccttgcagggtgatcaatatgacctgatccataagtatgaaaaccaaatcgg [ 44320]
ASF_IND_20_CAD543                ................................................................................ [ 44320]
LR899193.1_Germany_2020/1        ................................................................................ [ 44320]
MN393477.1_Wuhan_2019            ................................................................................ [ 44320]
MW396979.1_Timor-Leste/2019/1    ................................................................................ [ 44320]
MK940252.1_CN/2019/InnerMongolia ................................................................................ [ 44320]
MK128995.1_China/2018/AnhuiXCGQ  ................................................................................ [ 44320]
MH681419.1_ASFV/POL/2015         ................................................................................ [ 44320]
LS478113.1_Estonia_2014          ................................................................................ [ 44320]
KP843857.1_Odintsovo_02/14       ................................................................................ [ 44320]
```

```
FN557520.1_strain_E75            ..............................c................................................. [ 44320]
AY261366.1_Warthog               ..................a..................................a.......................... [ 44320]
AY261365.1_Warmbaths             ..................a..................................a.......................... [ 44320]
AY261364.1_Tengani_62            ................................................................................ [ 44320]
AY261363.1_Pretoriuskop/96/4     ..................a............................................................. [ 44320]
AY261362.1_Mkuzi_1979            ................................................................................ [ 44320]
AY261361.1_Malawi_Lil-20/1_1983  .....c........t...a.......g.....a.................t............................a [ 44320]
AY261360.1_Kenya_1950            ..............t...a..........t...g.......c...............t...c...............t.a [ 44320]
AM712240.1_OURT_88/3             -------------------------------------------------------------------------------- [ 44320]
AM712239.1_Benin_97/1            ..............................c................................................. [ 44320]
KM111295.1_Ken06.Bus             ..............t...a......................c...........c.......c...c.............a [ 44320]
KM111294.1_Ken05/Tk1             ..............t..ta.......g..t..a....................a.........................a [ 44320]
KM262844.1_strain_L60            ..............................c................................................. [ 44320]
KM102979.1_26544/OG10_Italy      ..............................c................................................. [ 44320]
KM262845.1_strain_NHV            -------------------------------------------------------------------------------- [ 44320]
KX354450.1_47/Ss/2008            ..............................c................................................. [ 44320]
KP055815.1_strain_BA71           ..............................c................................................. [ 44320]
MH025920.1_strain_R35            ..............t...a......................c...........c.......c...c.............a [ 44320]
MH025919.1_strain_N10            ..............t...a......................c...........c.......c...c.............a [ 44320]
MH025918.1_strain_R25            ..............t...a......................c...........c.......c...c.............a [ 44320]
MH025917.1_strain_R7             ..............t...a......................c...........c.......c...c.............a [ 44320]
MH025916.1_strain_R8             ..............t...a......................c...........c.......c...c.............a [ 44320]

FR682468.2_Georgia_2007/1        cgactttcattttatcttaccattgattcaagacgcgaatacgtttgaaaaatgccacgctttagaacgtttttgtggtg [ 44400]
ASF_IND_20_CAD543                ................................................................................ [ 44400]
LR899193.1_Germany_2020/1        ................................................................................ [ 44400]
MN393477.1_Wuhan_2019            ................................................................................ [ 44400]
MW396979.1_Timor-Leste/2019/1    ................................................................................ [ 44400]
MK940252.1_CN/2019/InnerMongolia ................................................................................ [ 44400]
MK128995.1_China/2018/AnhuiXCGQ  ................................................................................ [ 44400]
MH681419.1_ASFV/POL/2015         ................................................................................ [ 44400]
LS478113.1_Estonia_2014          ................................................................................ [ 44400]
KP843857.1_Odintsovo_02/14       ................................................................................ [ 44400]
FN557520.1_strain_E75            ................................................................................ [ 44400]
AY261366.1_Warthog               ..........c......................t.....a........................................ [ 44400]
AY261365.1_Warmbaths             ..........c......................t.....a........................................ [ 44400]
AY261364.1_Tengani_62            ..........c............................a.................t...................... [ 44400]
AY261363.1_Pretoriuskop/96/4     ..........c......................t.....a........................................ [ 44400]
AY261362.1_Mkuzi_1979            ................................................................................ [ 44400]
AY261361.1_Malawi_Lil-20/1_1983  a....a....ca......g..............t.....a....................c................a.. [ 44400]
AY261360.1_Kenya_1950            a....a....ca......g..............t.....a....................c..g.............a.. [ 44400]
AM712240.1_OURT_88/3             -------------------------------------------------------------------------------- [ 44400]
AM712239.1_Benin_97/1            ................................................................................ [ 44400]
KM111295.1_Ken06.Bus             a....a....ca......g..............t.....a....................c..g.............a.. [ 44400]
```

```
KM111294.1_Ken05/Tk1           a....a....ca.....................t...g.a....................c................a.. [ 44400]
KM262844.1_strain_L60          ................................................................................ [ 44400]
KM102979.1_26544/OG10_Italy    ................................................................................ [ 44400]
KM262845.1_strain_NHV          -------------------------------------------------------------------------------- [ 44400]
KX354450.1_47/Ss/2008          ................................................................................ [ 44400]
KP055815.1_strain_BA71         ................................................................................ [ 44400]
MH025920.1_strain_R35          a....a....ca......g..............t.....a....................c..g.............a.. [ 44400]
MH025919.1_strain_N10          a....a....ca......g..............t.....a....................c..g.............a.. [ 44400]
MH025918.1_strain_R25          a....a....ca......g..............t.....a....................c..g.............a.. [ 44400]
MH025917.1_strain_R7           a....a....ca......g..............t.....a....................c..g.............a.. [ 44400]
MH025916.1_strain_R8           a....a....ca......g..............t.....a....................c..g.............a.. [ 44400]

FR682468.2_Georgia_2007/1      tttcatgtctgctaaaacatgctacaaaatacaacatgctccctattctccaaaaataccaagaagagctgtctatgaga [ 44480]
ASF_IND_20_CAD543              ................................................................................ [ 44480]
LR899193.1_Germany_2020/1      ................................................................................ [ 44480]
MN393477.1_Wuhan_2019          ................................................................................ [ 44480]
MW396979.1_Timor-Leste/2019/1  ................................................................................ [ 44480]
MK940252.1_CN/2019/InnerMongolia ................................................................................ [ 44480]
MK128995.1_China/2018/AnhuiXCGQ ................................................................................ [ 44480]
MH681419.1_ASFV/POL/2015       ................................................................................ [ 44480]
LS478113.1_Estonia_2014        ................................................................................ [ 44480]
KP843857.1_Odintsovo_02/14     ................................................................................ [ 44480]
FN557520.1_strain_E75          ................................................................................ [ 44480]
AY261366.1_Warthog             ........................................t.............c......................... [ 44480]
AY261365.1_Warmbaths           ......................................................c......................... [ 44480]
AY261364.1_Tengani_62          ............................................g............................c...... [ 44480]
AY261363.1_Pretoriuskop/96/4   ..............g.......................................c......................... [ 44480]
AY261362.1_Mkuzi_1979          ................................................................................ [ 44480]
AY261361.1_Malawi_Lil-20/1_1983 ..c...........g..............c..............................................a... [ 44480]
AY261360.1_Kenya_1950          ..c....c......g.g.......t....c............................t........a....t...a... [ 44480]
AM712240.1_OURT_88/3           -------------------------------------------------------------------------------- [ 44480]
AM712239.1_Benin_97/1          ................................................................................ [ 44480]
KM111295.1_Ken06.Bus           ..c....c..a...g..............c..........................................t...a... [ 44480]
KM111294.1_Ken05/Tk1           ..c...........g..............c............................t.........t.......a... [ 44480]
KM262844.1_strain_L60          ................................................................................ [ 44480]
KM102979.1_26544/OG10_Italy    ................................................................................ [ 44480]
KM262845.1_strain_NHV          -------------------------------------------------------------------------------- [ 44480]
KX354450.1_47/Ss/2008          ................................................................................ [ 44480]
KP055815.1_strain_BA71         ................................................................................ [ 44480]
MH025920.1_strain_R35          ..c....c..a...g..............c..........................................t...a... [ 44480]
MH025919.1_strain_N10          ..c....c..a...g..............c..........................................t...a... [ 44480]
MH025918.1_strain_R25          ..c....c..a...g..............c..........................................t...a... [ 44480]
MH025917.1_strain_R7           ..c....c..a...g..............c..........................................t...a... [ 44480]
MH025916.1_strain_R8           ..c....c..a...g..............c..........................................t...a... [ 44480]
```

```
FR682468.2_Georgia_2007/1        gcgtatcttcacgaaaccctatttgaactagcatgcctatggcagaggtatgatgtccttaaatggatagagcaaaccat [ 44560]
ASF_IND_20_CAD543                ................................................................................ [ 44560]
LR899193.1_Germany_2020/1        ................................................................................ [ 44560]
MN393477.1_Wuhan_2019            ................................................................................ [ 44560]
MW396979.1_Timor-Leste/2019/1    ................................................................................ [ 44560]
MK940252.1_CN/2019/InnerMongolia ................................................................................ [ 44560]
MK128995.1_China/2018/AnhuiXCGQ  ................................................................................ [ 44560]
MH681419.1_ASFV/POL/2015         ................................................................................ [ 44560]
LS478113.1_Estonia_2014          ................................................................................ [ 44560]
KP843857.1_Odintsovo_02/14       ................................................................................ [ 44560]
FN557520.1_strain_E75            ..........................................................................g..... [ 44560]
AY261366.1_Warthog               .ta.......g..................................................................... [ 44560]
AY261365.1_Warmbaths             .ta.......g..................................................................... [ 44560]
AY261364.1_Tengani_62            ..a......tg..................................................................... [ 44560]
AY261363.1_Pretoriuskop/96/4     ..a.......g.....................g.....g......................................... [ 44560]
AY261362.1_Mkuzi_1979            ..........................................................................g..... [ 44560]
AY261361.1_Malawi_Lil-20/1_1983  .t........g............c................................t....................... [ 44560]
AY261360.1_Kenya_1950            .t......c.g...........................................a.t....................... [ 44560]
AM712240.1_OURT_88/3             -------------------------------------------------------------------------------- [ 44560]
AM712239.1_Benin_97/1            ..........................................................................g..... [ 44560]
KM111295.1_Ken06.Bus             .t......c.g...........................................a......................... [ 44560]
KM111294.1_Ken05/Tk1             .t......a.g............c..............................a.t....................... [ 44560]
KM262844.1_strain_L60            ..........................................................................g..... [ 44560]
KM102979.1_26544/OG10_Italy      ..........................................................................g..... [ 44560]
KM262845.1_strain_NHV            -------------------------------------------------------------------------------- [ 44560]
KX354450.1_47/Ss/2008            ..........................................................................g..... [ 44560]
KP055815.1_strain_BA71           ..........................................................................g..... [ 44560]
MH025920.1_strain_R35            .t......c.g...........................................a......................... [ 44560]
MH025919.1_strain_N10            .t......c.g...........................................a......................... [ 44560]
MH025918.1_strain_R25            .t......c.g...........................................a......................... [ 44560]
MH025917.1_strain_R7             .t......c.g...........................................a......................... [ 44560]
MH025916.1_strain_R8             .t......c.g...........................................a......................... [ 44560]

FR682468.2_Georgia_2007/1        acatgtttacgacctaaagattatgtttaatattgccatctccaagagggatctgactatgtactccttaggatatattt [ 44640]
ASF_IND_20_CAD543                ................................................................................ [ 44640]
LR899193.1_Germany_2020/1        ................................................................................ [ 44640]
MN393477.1_Wuhan_2019            ................................................................................ [ 44640]
MW396979.1_Timor-Leste/2019/1    ................................................................................ [ 44640]
MK940252.1_CN/2019/InnerMongolia ................................................................................ [ 44640]
MK128995.1_China/2018/AnhuiXCGQ  ................................................................................ [ 44640]
MH681419.1_ASFV/POL/2015         ................................................................................ [ 44640]
LS478113.1_Estonia_2014          ................................................................................ [ 44640]
KP843857.1_Odintsovo_02/14       ................................................................................ [ 44640]
```

```
FN557520.1_strain_E75              g............................................................................... [ 44640]
AY261366.1_Warthog                 g..................g...........................................t...............c [ 44640]
AY261365.1_Warmbaths               g..................g............................a..............t...............c [ 44640]
AY261364.1_Tengani_62              g............................................................................... [ 44640]
AY261363.1_Pretoriuskop/96/4       g........................................................g.....t...............c [ 44640]
AY261362.1_Mkuzi_1979              g............................................................................... [ 44640]
AY261361.1_Malawi_Lil-20/1_1983    g.....................................................a.gc..................g..c [ 44640]
AY261360.1_Kenya_1950              g...........t.....a.....a.............................a.gc..................g..c [ 44640]
AM712240.1_OURT_88/3               -------------------------------------------------------------------------------- [ 44640]
AM712239.1_Benin_97/1              g............................................................................... [ 44640]
KM111295.1_Ken06.Bus               g.....................................................a.gc........t....a.......c [ 44640]
KM111294.1_Ken05/Tk1               g...........t.....a.....a.............................a.gc..................g..c [ 44640]
KM262844.1_strain_L60              g............................................................................... [ 44640]
KM102979.1_26544/OG10_Italy        g............................................................................... [ 44640]
KM262845.1_strain_NHV              -------------------------------------------------------------------------------- [ 44640]
KX354450.1_47/Ss/2008              g............................................................................... [ 44640]
KP055815.1_strain_BA71             g............................................................................... [ 44640]
MH025920.1_strain_R35              g.....................................................a.gc........t............c [ 44640]
MH025919.1_strain_N10              g.....................................................a.gc........t............c [ 44640]
MH025918.1_strain_R25              g.....................................................a.gc........t............c [ 44640]
MH025917.1_strain_R7               g.....................................................a.gc........t............c [ 44640]
MH025916.1_strain_R8               g.....................................................a.gc........t............c [ 44640]

FR682468.2_Georgia_2007/1          tcctttttgatagagggaacaccga---------agctacgttgctaacgcaacatctcaagaagacagcggccaaaggg [ 44720]
ASF_IND_20_CAD543                  .........................---------.............................................. [ 44720]
LR899193.1_Germany_2020/1          .........................---------.............................................. [ 44720]
MN393477.1_Wuhan_2019              .........................---------.............................................. [ 44720]
MW396979.1_Timor-Leste/2019/1      .........................---------.............................................. [ 44720]
MK940252.1_CN/2019/InnerMongolia   .........................---------.............................................. [ 44720]
MK128995.1_China/2018/AnhuiXCGQ    .........................---------.............................................. [ 44720]
MH681419.1_ASFV/POL/2015           .........................---------.............................................. [ 44720]
LS478113.1_Estonia_2014            .........................---------.............................................. [ 44720]
KP843857.1_Odintsovo_02/14         .........................---------.............................................. [ 44720]
FN557520.1_strain_E75              .........................---------.........................g.................... [ 44720]
AY261366.1_Warthog                 .t.............a.........---------.........................g....a...........g... [ 44720]
AY261365.1_Warmbaths               .t.............a.........---------.........................g....a...........g... [ 44720]
AY261364.1_Tengani_62              .........................---------...................t.....g.................... [ 44720]
AY261363.1_Pretoriuskop/96/4       .t.............a.........---------..........t....a.........g.................... [ 44720]
AY261362.1_Mkuzi_1979              .........................---------.........................g.................... [ 44720]
AY261361.1_Malawi_Lil-20/1_1983    ................a....t...---------.........c...............g.................... [ 44720]
AY261360.1_Kenya_1950              .........................---------......c...t..........c...g................g... [ 44720]
AM712240.1_OURT_88/3               -------------------------------------------------------------------------------- [ 44720]
AM712239.1_Benin_97/1              .........................---------.........................g.................... [ 44720]
KM111295.1_Ken06.Bus               ..t.....a................tgttgcgtt...a.....................g................g... [ 44720]
```

```
KM111294.1_Ken05/Tk1            .........................---------......c...t..........c...g................g... [ 44720]
KM262844.1_strain_L60           .........................---------.........................g.................... [ 44720]
KM102979.1_26544/OG10_Italy     .........................---------.........................g.................... [ 44720]
KM262845.1_strain_NHV           -------------------------------------------------------------------------------- [ 44720]
KX354450.1_47/Ss/2008           .........................---------.........................g.................... [ 44720]
KP055815.1_strain_BA71          .........................---------.........................g.................... [ 44720]
MH025920.1_strain_R35           ........a................tgttgcgtt...a.....................g................g... [ 44720]
MH025919.1_strain_N10           ........a................tgttgcgtt...a.....................g................g... [ 44720]
MH025918.1_strain_R25           ........a................tgttgcgtt...a.....................g................g... [ 44720]
MH025917.1_strain_R7            ........a................tgttgcgtt...a.....................g................g... [ 44720]
MH025916.1_strain_R8            ........a................tgttgcgtt...a.....................g................g... [ 44720]

FR682468.2_Georgia_2007/1       ctcctccactttgtgctagaaacgttaaaatacggcggcaacatagataccgtcctgacccaagccgtaaagtacaatca [ 44800]
ASF_IND_20_CAD543               .............................................................................y.. [ 44800]
LR899193.1_Germany_2020/1       ................................................................................ [ 44800]
MN393477.1_Wuhan_2019           ................................................................................ [ 44800]
MW396979.1_Timor-Leste/2019/1   ................................................................................ [ 44800]
MK940252.1_CN/2019/InnerMongolia ................................................................................ [ 44800]
MK128995.1_China/2018/AnhuiXCGQ ................................................................................ [ 44800]
MH681419.1_ASFV/POL/2015        ................................................................................ [ 44800]
LS478113.1_Estonia_2014         ................................................................................ [ 44800]
KP843857.1_Odintsovo_02/14      ................................................................................ [ 44800]
FN557520.1_strain_E75           .............................................................................c.. [ 44800]
AY261366.1_Warthog              ......................................t..........t.......t.t.................... [ 44800]
AY261365.1_Warmbaths            ......................................t..........t.......t...................... [ 44800]
AY261364.1_Tengani_62           ......................................t..........tt............................. [ 44800]
AY261363.1_Pretoriuskop/96/4    ......t...............................t..........t.........t.................... [ 44800]
AY261362.1_Mkuzi_1979           ................................................................................ [ 44800]
AY261361.1_Malawi_Lil-20/1_1983 ......................................t...t..a...t.......t.............a........ [ 44800]
AY261360.1_Kenya_1950           ......................................t......a.c.t......at.............a........ [ 44800]
AM712240.1_OURT_88/3            -------------------------------------------------------------------------------- [ 44800]
AM712239.1_Benin_97/1           .............................................................................c.. [ 44800]
KM111295.1_Ken06.Bus            ......................................t......a.c.t......ct.............a........ [ 44800]
KM111294.1_Ken05/Tk1            ......................................t...t..a...t.......tt............a........ [ 44800]
KM262844.1_strain_L60           .............................................................................c.. [ 44800]
KM102979.1_26544/OG10_Italy     .............................................................................c.. [ 44800]
KM262845.1_strain_NHV           -------------------------------------------------------------------------------- [ 44800]
KX354450.1_47/Ss/2008           .............................................................................c.. [ 44800]
KP055815.1_strain_BA71          .............................................................................c.. [ 44800]
MH025920.1_strain_R35           ......................................t......a.c.t......ct.............a........ [ 44800]
MH025919.1_strain_N10           ......................................t......a.c.t......ct.............a........ [ 44800]
MH025918.1_strain_R25           ......................................t......a.c.t......ct.............a........ [ 44800]
MH025917.1_strain_R7            ......................................t......a.c.t......ct.............a........ [ 44800]
MH025916.1_strain_R8            ......................................t......a.c.t......ct.............a........ [ 44800]
```

```
FR682468.2_Georgia_2007/1        tagaaaacttttagattattttctgcgtcaactacctcgtaaacatattgaaaaacttttgttgctggccgtgcaggaaa [ 44880]
ASF_IND_20_CAD543                ................................................................................ [ 44880]
LR899193.1_Germany_2020/1        ................................................................................ [ 44880]
MN393477.1_Wuhan_2019            ................................................................................ [ 44880]
MW396979.1_Timor-Leste/2019/1    ................................................................................ [ 44880]
MK940252.1_CN/2019/InnerMongolia ................................................................................ [ 44880]
MK128995.1_China/2018/AnhuiXCGQ  ................................................................................ [ 44880]
MH681419.1_ASFV/POL/2015         ................................................................................ [ 44880]
LS478113.1_Estonia_2014          ................................................................................ [ 44880]
KP843857.1_Odintsovo_02/14       ................................................................................ [ 44880]
FN557520.1_strain_E75            ......................................................g..............t..t....... [ 44880]
AY261366.1_Warthog               ......................................................gt........................ [ 44880]
AY261365.1_Warmbaths             ...........................................a.........................t..t....... [ 44880]
AY261364.1_Tengani_62            ......................................................g......................... [ 44880]
AY261363.1_Pretoriuskop/96/4     ......................................................g......................... [ 44880]
AY261362.1_Mkuzi_1979            .....................................................................t..t....... [ 44880]
AY261361.1_Malawi_Lil-20/1_1983  ............g..............................a.................................... [ 44880]
AY261360.1_Kenya_1950            ...g.......................................a....................t.a....a....... [ 44880]
AM712240.1_OURT_88/3             -------------------------------------------------------------------------------- [ 44880]
AM712239.1_Benin_97/1            ......................................................g..............t..t....... [ 44880]
KM111295.1_Ken06.Bus             ...........................................a............................a....... [ 44880]
KM111294.1_Ken05/Tk1             ....................................c......a.........................t.......... [ 44880]
KM262844.1_strain_L60            ......................................................g..............t..t....... [ 44880]
KM102979.1_26544/OG10_Italy      ......................................................g..............t..t....... [ 44880]
KM262845.1_strain_NHV            -------------------------------------------------------------------------------- [ 44880]
KX354450.1_47/Ss/2008            ......................................................g..............t..t....... [ 44880]
KP055815.1_strain_BA71           ......................................................g..............t..t....... [ 44880]
MH025920.1_strain_R35            ...........................................a............................a....... [ 44880]
MH025919.1_strain_N10            ...........................................a............................a....... [ 44880]
MH025918.1_strain_R25            ...........................................a............................a....... [ 44880]
MH025917.1_strain_R7             ...........................................a............................a....... [ 44880]
MH025916.1_strain_R8             ...........................................a............................a....... [ 44880]

FR682468.2_Georgia_2007/1        aggcttctaaaaaaacattgaacttactgttgtcacatttaaactactccgtgaaacgcatcaaaaaactaccgcgctat [ 44960]
ASF_IND_20_CAD543                ........................................................................y....... [ 44960]
LR899193.1_Germany_2020/1        ................................................................................ [ 44960]
MN393477.1_Wuhan_2019            ................................................................................ [ 44960]
MW396979.1_Timor-Leste/2019/1    ................................................................................ [ 44960]
MK940252.1_CN/2019/InnerMongolia ................................................................................ [ 44960]
MK128995.1_China/2018/AnhuiXCGQ  ................................................................................ [ 44960]
MH681419.1_ASFV/POL/2015         ................................................................................ [ 44960]
LS478113.1_Estonia_2014          ................................................................................ [ 44960]
KP843857.1_Odintsovo_02/14       ................................................................................ [ 44960]
```

```
FN557520.1_strain_E75            ......................................................................g.t....... [ 44960]
AY261366.1_Warthog               ......................................................................g.t....... [ 44960]
AY261365.1_Warmbaths             ..................................c...................................g.t....... [ 44960]
AY261364.1_Tengani_62            ......................................................................g.t....... [ 44960]
AY261363.1_Pretoriuskop/96/4     ..................................c...c...................t...........g.t....... [ 44960]
AY261362.1_Mkuzi_1979            ......................................................................g.t....... [ 44960]
AY261361.1_Malawi_Lil-20/1_1983  .......................c.............c...................a............g.t....... [ 44960]
AY261360.1_Kenya_1950            ..........g.......................tt..................................g.t....... [ 44960]
AM712240.1_OURT_88/3             -------------------------------------------------------------------------------- [ 44960]
AM712239.1_Benin_97/1            ......................................................................g.t....... [ 44960]
KM111295.1_Ken06.Bus             g.........g.......................tt..................................g.t....... [ 44960]
KM111294.1_Ken05/Tk1             .a........g...........................................................g.t....... [ 44960]
KM262844.1_strain_L60            ......................................................................g.t....... [ 44960]
KM102979.1_26544/OG10_Italy      ......................................................................g.t....... [ 44960]
KM262845.1_strain_NHV            -------------------------------------------------------------------------------- [ 44960]
KX354450.1_47/Ss/2008            ......................................................................g.t....... [ 44960]
KP055815.1_strain_BA71           ......................................................................g.t....... [ 44960]
MH025920.1_strain_R35            g.........g.......................tt..................................g.t....... [ 44960]
MH025919.1_strain_N10            g.........g.......................tt..................................g.t....... [ 44960]
MH025918.1_strain_R25            g.........g.......................tt..................................g.t....... [ 44960]
MH025917.1_strain_R7             g.........g.......................tt..................................g.t....... [ 44960]
MH025916.1_strain_R8             g.........g.......................tt..................................g.t....... [ 44960]

FR682468.2_Georgia_2007/1        gtgatagagtacgagtccaccttggtgataaagattttatt-aaaaaaaagagtgaacctgatagatgccatgttggaaa [ 45040]
ASF_IND_20_CAD543                .........................................-...................................... [ 45040]
LR899193.1_Germany_2020/1        .........................................-...................................... [ 45040]
MN393477.1_Wuhan_2019            .........................................-...................................... [ 45040]
MW396979.1_Timor-Leste/2019/1    .........................................-...................................... [ 45040]
MK940252.1_CN/2019/InnerMongolia .........................................-...................................... [ 45040]
MK128995.1_China/2018/AnhuiXCGQ  .........................................-...................................... [ 45040]
MH681419.1_ASFV/POL/2015         .........................................-...................................... [ 45040]
LS478113.1_Estonia_2014          .........................................-...................................... [ 45040]
KP843857.1_Odintsovo_02/14       .........................................--..................................... [ 45040]
FN557520.1_strain_E75            ...........t..........................g..a............a......................... [ 45040]
AY261366.1_Warthog               ...........t.............................-............a......................... [ 45040]
AY261365.1_Warmbaths             .........................................-............a......................... [ 45040]
AY261364.1_Tengani_62            .........................................-............a......................... [ 45040]
AY261363.1_Pretoriuskop/96/4     .........................................-............a......................... [ 45040]
AY261362.1_Mkuzi_1979            ...........t..........................g..-............a......................... [ 45040]
AY261361.1_Malawi_Lil-20/1_1983  .................t..............ac.......-g...........a...........c...g......... [ 45040]
AY261360.1_Kenya_1950            ....................ac.........g.........-..........a.a.....a.....c...g......... [ 45040]
AM712240.1_OURT_88/3             -------------------------------------------------------------------------------- [ 45040]
AM712239.1_Benin_97/1            ...........t..........................g..-............a......................... [ 45040]
KM111295.1_Ken06.Bus             .........c.t........ac.........g.........-..........a.a.....a.....c...g......... [ 45040]
```

```
KM111294.1_Ken05/Tk1             .................t.......................-............a.....a.....c...g......... [ 45040]
KM262844.1_strain_L60            ...........t..........................g..-............a......................... [ 45040]
KM102979.1_26544/OG10_Italy      ...........t..........................g..-............a......................... [ 45040]
KM262845.1_strain_NHV            -------------------------------------------------------------------------------- [ 45040]
KX354450.1_47/Ss/2008            ...........t..........................g..-............a......................... [ 45040]
KP055815.1_strain_BA71           ...........t..........................g..-............a......................... [ 45040]
MH025920.1_strain_R35            .........c.t........ac.........g.........-..........a.a.....a.....c...g......... [ 45040]
MH025919.1_strain_N10            .........c.t........ac.........g.........-..........a.a.....a.....c...g......... [ 45040]
MH025918.1_strain_R25            .........c.t........ac.........g.........-..........a.a.....a.....c...g......... [ 45040]
MH025917.1_strain_R7             .........c.t........ac.........g.........-..........a.a.....a.....c...g......... [ 45040]
MH025916.1_strain_R8             .........c.t........ac.........g.........-..........a.a.....a.....c...g......... [ 45040]

FR682468.2_Georgia_2007/1        agatggtaagatatttttctgcgacgaaagtgaggacgatcatggatgagctttcgattagtccggaaagagtcattaag [ 45120]
ASF_IND_20_CAD543                ................................................................................ [ 45120]
LR899193.1_Germany_2020/1        ................................................................................ [ 45120]
MN393477.1_Wuhan_2019            ................................................................................ [ 45120]
MW396979.1_Timor-Leste/2019/1    ................................................................................ [ 45120]
MK940252.1_CN/2019/InnerMongolia ................................................................................ [ 45120]
MK128995.1_China/2018/AnhuiXCGQ  ................................................................................ [ 45120]
MH681419.1_ASFV/POL/2015         ................................................................................ [ 45120]
LS478113.1_Estonia_2014          ................................................................................ [ 45120]
KP843857.1_Odintsovo_02/14       ................................................................................ [ 45120]
FN557520.1_strain_E75            ................................................................................ [ 45120]
AY261366.1_Warthog               .................................a.....................a........................ [ 45120]
AY261365.1_Warmbaths             ................................................................................ [ 45120]
AY261364.1_Tengani_62            ......................................................t......................... [ 45120]
AY261363.1_Pretoriuskop/96/4     ......................................................t......................... [ 45120]
AY261362.1_Mkuzi_1979            .........................a...................................................... [ 45120]
AY261361.1_Malawi_Lil-20/1_1983  ...at...................tt...........t..t....................................... [ 45120]
AY261360.1_Kenya_1950            ...ct................a.........a.a...t..t...................a........a...t...... [ 45120]
AM712240.1_OURT_88/3             -------------------------------------------------------------------------------- [ 45120]
AM712239.1_Benin_97/1            ................................................................................ [ 45120]
KM111295.1_Ken06.Bus             ...ct...............at...t..................................a........a.........a [ 45120]
KM111294.1_Ken05/Tk1             ...ct................a.........a.a...t..t...................a........a.......... [ 45120]
KM262844.1_strain_L60            ................................................................................ [ 45120]
KM102979.1_26544/OG10_Italy      ................................................................................ [ 45120]
KM262845.1_strain_NHV            -------------------------------------------------------------------------------- [ 45120]
KX354450.1_47/Ss/2008            ................................................................................ [ 45120]
KP055815.1_strain_BA71           ................................................................................ [ 45120]
MH025920.1_strain_R35            ...ct...............at...t..................................a........a.........a [ 45120]
MH025919.1_strain_N10            ...ct...............at...t..................................a........a.........a [ 45120]
MH025918.1_strain_R25            ...ct...............at...t..................................a........a.........a [ 45120]
MH025917.1_strain_R7             ...ct...............at...t..................................a........a.........a [ 45120]
MH025916.1_strain_R8             ...ct...............at...t..................................a........a.........a [ 45120]
```

```
FR682468.2_Georgia_2007/1        atggctatacagaaaatgagaacggatatcgtaatccatacttcttatgtttgggaggatgatctagaacgtcttactcg [ 45200]
ASF_IND_20_CAD543                ................................................................................ [ 45200]
LR899193.1_Germany_2020/1        ................................................................................ [ 45200]
MN393477.1_Wuhan_2019            ................................................................................ [ 45200]
MW396979.1_Timor-Leste/2019/1    ................................................................................ [ 45200]
MK940252.1_CN/2019/InnerMongolia ................................................................................ [ 45200]
MK128995.1_China/2018/AnhuiXCGQ  ................................................................................ [ 45200]
MH681419.1_ASFV/POL/2015         ................................................................................ [ 45200]
LS478113.1_Estonia_2014          ................................................................................ [ 45200]
KP843857.1_Odintsovo_02/14       ................................................................................ [ 45200]
FN557520.1_strain_E75            ................................................................................ [ 45200]
AY261366.1_Warthog               ................................................a......................a........ [ 45200]
AY261365.1_Warmbaths             .......................a...............................................a........ [ 45200]
AY261364.1_Tengani_62            ..................................................a....................a........ [ 45200]
AY261363.1_Pretoriuskop/96/4     .......................................................................a........ [ 45200]
AY261362.1_Mkuzi_1979            ........................................................................t....... [ 45200]
AY261361.1_Malawi_Lil-20/1_1983  .....c.......................t..g..t..g.........a....................a.a....t... [ 45200]
AY261360.1_Kenya_1950            .....c.......................t..g.....a.........a....................a.a....t... [ 45200]
AM712240.1_OURT_88/3             -------------------------------------------------------------------------------- [ 45200]
AM712239.1_Benin_97/1            ................................................................................ [ 45200]
KM111295.1_Ken06.Bus             .....cg.g.......................g....................................a.a....t... [ 45200]
KM111294.1_Ken05/Tk1             .....c.......................t..g.....a.........a....................a.a....t... [ 45200]
KM262844.1_strain_L60            ................................................................................ [ 45200]
KM102979.1_26544/OG10_Italy      ................................................................................ [ 45200]
KM262845.1_strain_NHV            -------------------------------------------------------------------------------- [ 45200]
KX354450.1_47/Ss/2008            ................................................................................ [ 45200]
KP055815.1_strain_BA71           ................................................................................ [ 45200]
MH025920.1_strain_R35            .....cg.g.......................g....................................a.a....t... [ 45200]
MH025919.1_strain_N10            .....cg.g.......................g....................................a.a....t... [ 45200]
MH025918.1_strain_R25            .....cg.g.......................g....................................a.a....t... [ 45200]
MH025917.1_strain_R7             .....cg.g.......................g....................................a.a....t... [ 45200]
MH025916.1_strain_R8             .....cg.g.......................g....................................a.a....t... [ 45200]

FR682468.2_Georgia_2007/1        tcttaaaaatatggtatacaccataaagtacgaacatgggaaaaaaatgttaattaaagtcatgcacggcatatacaaaa [ 45280]
ASF_IND_20_CAD543                ................................................................................ [ 45280]
LR899193.1_Germany_2020/1        ................................................................................ [ 45280]
MN393477.1_Wuhan_2019            ................................................................................ [ 45280]
MW396979.1_Timor-Leste/2019/1    ................................................................................ [ 45280]
MK940252.1_CN/2019/InnerMongolia ................................................................................ [ 45280]
MK128995.1_China/2018/AnhuiXCGQ  ................................................................................ [ 45280]
MH681419.1_ASFV/POL/2015         ................................................................................ [ 45280]
LS478113.1_Estonia_2014          ................................................................................ [ 45280]
KP843857.1_Odintsovo_02/14       ................................................................................ [ 45280]
```

```
FN557520.1_strain_E75            ................................................................................ [ 45280]
AY261366.1_Warthog               .......g......................t.....c........................................... [ 45280]
AY261365.1_Warmbaths             ...................g..........t.......................g......................... [ 45280]
AY261364.1_Tengani_62            .......g..............g........................................a................ [ 45280]
AY261363.1_Pretoriuskop/96/4     ................................................................................ [ 45280]
AY261362.1_Mkuzi_1979            ................................................................................ [ 45280]
AY261361.1_Malawi_Lil-20/1_1983  ..............................t.............................t..t................ [ 45280]
AY261360.1_Kenya_1950            ..............................t.......................g.....t..t................ [ 45280]
AM712240.1_OURT_88/3             -------------------------------------------------------------------------------- [ 45280]
AM712239.1_Benin_97/1            ................................................................................ [ 45280]
KM111295.1_Ken06.Bus             .............c.......t........t..g.....a.........c.............t................ [ 45280]
KM111294.1_Ken05/Tk1             ..............................t.......................g.....t..t................ [ 45280]
KM262844.1_strain_L60            ................................................................................ [ 45280]
KM102979.1_26544/OG10_Italy      ................................................................................ [ 45280]
KM262845.1_strain_NHV            -------------------------------------------------------------------------------- [ 45280]
KX354450.1_47/Ss/2008            ................................................................................ [ 45280]
KP055815.1_strain_BA71           ................................................................................ [ 45280]
MH025920.1_strain_R35            .............c.......t........t..g....aa.........c.............t................ [ 45280]
MH025919.1_strain_N10            .............c.......t........t..g....aa.........c.............t................ [ 45280]
MH025918.1_strain_R25            .............c.......t........t..g....aa.........c.............t................ [ 45280]
MH025917.1_strain_R7             .............c.......t........t..g....aa.........c.............t................ [ 45280]
MH025916.1_strain_R8             .............c.......t........t..g....aa.........c.............t................ [ 45280]

FR682468.2_Georgia_2007/1        acttattatacggcgaaagggaaaaagtcatgttttatttagccaagctctatgttgctcaaaacgcggccacccaattc [ 45360]
ASF_IND_20_CAD543                ................................................................................ [ 45360]
LR899193.1_Germany_2020/1        ................................................................................ [ 45360]
MN393477.1_Wuhan_2019            ................................................................................ [ 45360]
MW396979.1_Timor-Leste/2019/1    ................................................................................ [ 45360]
MK940252.1_CN/2019/InnerMongolia ................................................................................ [ 45360]
MK128995.1_China/2018/AnhuiXCGQ  ................................................................................ [ 45360]
MH681419.1_ASFV/POL/2015         ................................................................................ [ 45360]
LS478113.1_Estonia_2014          ................................................................................ [ 45360]
KP843857.1_Odintsovo_02/14       ................................................................................ [ 45360]
FN557520.1_strain_E75            ...................................c............................t............... [ 45360]
AY261366.1_Warthog               ............a......a.....g.........c............................................ [ 45360]
AY261365.1_Warmbaths             ...................a.....g.........c............................................ [ 45360]
AY261364.1_Tengani_62            ..................aa.....g.........c............................................ [ 45360]
AY261363.1_Pretoriuskop/96/4     ............a......a.....g.........c............................................ [ 45360]
AY261362.1_Mkuzi_1979            ...................................c............................t............... [ 45360]
AY261361.1_Malawi_Lil-20/1_1983  ..................aa...............c..........a................................. [ 45360]
AY261360.1_Kenya_1950            ........c...at....aa...............c...........t.....a.......................... [ 45360]
AM712240.1_OURT_88/3             -------------------------------------------------------------------------------- [ 45360]
AM712239.1_Benin_97/1            ...................................c............................t............... [ 45360]
KM111295.1_Ken06.Bus             ......---.........aa...............a....g......t......c......................... [ 45360]
```

```
KM111294.1_Ken05/Tk1             ........c...at....aa...............c...........t.....a.......................... [ 45360]
KM262844.1_strain_L60            ...................................c............................t............... [ 45360]
KM102979.1_26544/OG10_Italy      ...................................c............................t............... [ 45360]
KM262845.1_strain_NHV            -------------------------------------------------------------------------------- [ 45360]
KX354450.1_47/Ss/2008            ...................................c............................t............... [ 45360]
KP055815.1_strain_BA71           ...................................c............................t............... [ 45360]
MH025920.1_strain_R35            ......---.........aa...............a....g......t......c......................... [ 45360]
MH025919.1_strain_N10            ......---.........aa...............a....g......t......c......................... [ 45360]
MH025918.1_strain_R25            ......---.........aa...............a....g......t......c......................... [ 45360]
MH025917.1_strain_R7             ......---.........aa...............a....g......t......c......................... [ 45360]
MH025916.1_strain_R8             ......---.........aa...............a....g......t......c......................... [ 45360]

FR682468.2_Georgia_2007/1        agagacatttgtaaggactgttacaaactggatgtggcacggtttaaaccgcggtttaagcaactaatattagactgttt [ 45440]
ASF_IND_20_CAD543                ................................................................................ [ 45440]
LR899193.1_Germany_2020/1        ................................................................................ [ 45440]
MN393477.1_Wuhan_2019            ................................................................................ [ 45440]
MW396979.1_Timor-Leste/2019/1    ................................................................................ [ 45440]
MK940252.1_CN/2019/InnerMongolia ................................................................................ [ 45440]
MK128995.1_China/2018/AnhuiXCGQ  ................................................................................ [ 45440]
MH681419.1_ASFV/POL/2015         ................................................................................ [ 45440]
LS478113.1_Estonia_2014          ................................................................................ [ 45440]
KP843857.1_Odintsovo_02/14       ................................................................................ [ 45440]
FN557520.1_strain_E75            ................................................................................ [ 45440]
AY261366.1_Warthog               .........................g.......t.........................a........g........... [ 45440]
AY261365.1_Warmbaths             .......................................------------................cg...a....... [ 45440]
AY261364.1_Tengani_62            ....................................................................g........... [ 45440]
AY261363.1_Pretoriuskop/96/4     ....................................................................g........... [ 45440]
AY261362.1_Mkuzi_1979            ................................................................................ [ 45440]
AY261361.1_Malawi_Lil-20/1_1983  ..g...................g...............g....................a........t........... [ 45440]
AY261360.1_Kenya_1950            ..............a.......g...............g....................a........t........... [ 45440]
AM712240.1_OURT_88/3             -------------------------------------------------------------------------------- [ 45440]
AM712239.1_Benin_97/1            ................................................................................ [ 45440]
KM111295.1_Ken06.Bus             .a.....c......a.......g..............------------..........a........t........... [ 45440]
KM111294.1_Ken05/Tk1             ..............a.......g...............g....................a........t........... [ 45440]
KM262844.1_strain_L60            ................................................................................ [ 45440]
KM102979.1_26544/OG10_Italy      ................................................................................ [ 45440]
KM262845.1_strain_NHV            -------------------------------------------------------------------------------- [ 45440]
KX354450.1_47/Ss/2008            ................................................................................ [ 45440]
KP055815.1_strain_BA71           ................................................................................ [ 45440]
MH025920.1_strain_R35            .a.....c......a.......g..............------------..........a........t........... [ 45440]
MH025919.1_strain_N10            .a.....c......a.......g..............------------..........a........t........... [ 45440]
MH025918.1_strain_R25            .a.....c......a.......g..............------------..........a........t........... [ 45440]
MH025917.1_strain_R7             .a.....c......a.......g..............------------..........a........t........... [ 45440]
MH025916.1_strain_R8             .a.....c......a.......g..............------------..........a........t........... [ 45440]
```

```
FR682468.2_Georgia_2007/1        agaaattattactaaaaaatcttgctatagtatcctggaaatcttagaaaaacatattatttccctgtttactatgaaag [ 45520]
ASF_IND_20_CAD543                .............-....................................................r............. [ 45520]
LR899193.1_Germany_2020/1        ................................................................................ [ 45520]
MN393477.1_Wuhan_2019            ................................................................................ [ 45520]
MW396979.1_Timor-Leste/2019/1    ................................................................................ [ 45520]
MK940252.1_CN/2019/InnerMongolia ................................................................................ [ 45520]
MK128995.1_China/2018/AnhuiXCGQ  ................................................................................ [ 45520]
MH681419.1_ASFV/POL/2015         ................................................................................ [ 45520]
LS478113.1_Estonia_2014          ................................................................................ [ 45520]
KP843857.1_Odintsovo_02/14       ................................................................................ [ 45520]
FN557520.1_strain_E75            .......g..........................................................a............. [ 45520]
AY261366.1_Warthog               .......g..........................................................a............. [ 45520]
AY261365.1_Warmbaths             ...........................................c......................a............. [ 45520]
AY261364.1_Tengani_62            .......g.........................t.......................g........a............. [ 45520]
AY261363.1_Pretoriuskop/96/4     .......g..........................................................a............. [ 45520]
AY261362.1_Mkuzi_1979            .......g..........................................................a............. [ 45520]
AY261361.1_Malawi_Lil-20/1_1983  ......gg..................t......ta.a..............ct.............a...gtg....... [ 45520]
AY261360.1_Kenya_1950            ...t...............a.g...ctc.a...ta................t...........g..a...g........a [ 45520]
AM712240.1_OURT_88/3             -------------------------------------------------------------------------------- [ 45520]
AM712239.1_Benin_97/1            .......g..........................................................a............. [ 45520]
KM111295.1_Ken06.Bus             ...c...............a.g...ctc......a................t...........g..a...g........a [ 45520]
KM111294.1_Ken05/Tk1             .......g..................t......ta.a.....t........c..............a....tg....... [ 45520]
KM262844.1_strain_L60            .......g..........................................................a............. [ 45520]
KM102979.1_26544/OG10_Italy      .......g..........................................................a............. [ 45520]
KM262845.1_strain_NHV            -------------------------------------------------------------------------------- [ 45520]
KX354450.1_47/Ss/2008            .......g..........................................................a............. [ 45520]
KP055815.1_strain_BA71           .......g..........................................................a............. [ 45520]
MH025920.1_strain_R35            ...c...............a.g...ctc......a................t...........g..a...g........a [ 45520]
MH025919.1_strain_N10            ...c...............a.g...ctc......a................t...........g..a...g........a [ 45520]
MH025918.1_strain_R25            ...c...............a.g...ctc......a................t...........g..a...g........a [ 45520]
MH025917.1_strain_R7             ...c...............a.g...ctc......a................t...........g..a...g........a [ 45520]
MH025916.1_strain_R8             ...c...............a.g...ctc......a................t...........g..a...g........a [ 45520]

FR682468.2_Georgia_2007/1        ttatgactgaagaagaaaaaaacctatgtttagaaatattatataaagtaattcattataaaacaatacaatgttaaaat [ 45600]
ASF_IND_20_CAD543                ................................................................................ [ 45600]
LR899193.1_Germany_2020/1        ................................................................................ [ 45600]
MN393477.1_Wuhan_2019            ................................................................................ [ 45600]
MW396979.1_Timor-Leste/2019/1    ................................................................................ [ 45600]
MK940252.1_CN/2019/InnerMongolia ................................................................................ [ 45600]
MK128995.1_China/2018/AnhuiXCGQ  ................................................................................ [ 45600]
MH681419.1_ASFV/POL/2015         ................................................................................ [ 45600]
LS478113.1_Estonia_2014          ................................................................................ [ 45600]
KP843857.1_Odintsovo_02/14       ................................................................................ [ 45600]
```

```
FN557520.1_strain_E75            ................................................................................ [ 45600]
AY261366.1_Warthog               ............................................-................t.t................ [ 45600]
AY261365.1_Warmbaths             .............................................................t.................. [ 45600]
AY261364.1_Tengani_62            ............................................-..............-----................ [ 45600]
AY261363.1_Pretoriuskop/96/4     .............g..............................-................t.................. [ 45600]
AY261362.1_Mkuzi_1979            ................................................................................ [ 45600]
AY261361.1_Malawi_Lil-20/1_1983  .c..c..............................c.................ag...c.....g...t........... [ 45600]
AY261360.1_Kenya_1950            .g...........t............g........t.................ag...c....tg...tc..a....... [ 45600]
AM712240.1_OURT_88/3             -------------------------------------------------------------------------------- [ 45600]
AM712239.1_Benin_97/1            ................................................................................ [ 45600]
KM111295.1_Ken06.Bus             .g.....g..g..t.........t..g..........................ag...c....tg...tc..a....... [ 45600]
KM111294.1_Ken05/Tk1             ....c........t............g........t.................ag...c....tg...tc..a....... [ 45600]
KM262844.1_strain_L60            ................................................................................ [ 45600]
KM102979.1_26544/OG10_Italy      ................................................................................ [ 45600]
KM262845.1_strain_NHV            -------------------------------------------------------------------------------- [ 45600]
KX354450.1_47/Ss/2008            ................................................................................ [ 45600]
KP055815.1_strain_BA71           ................................................................................ [ 45600]
MH025920.1_strain_R35            .g.....g..g..t.........t..g..........................ag...c....tg...tc..a....... [ 45600]
MH025919.1_strain_N10            .g.....g..g..t.........t..g..........................ag...c....tg...tc..a....... [ 45600]
MH025918.1_strain_R25            .g.....g..g..t.........t..g..........................ag...c....tg...tc..a....... [ 45600]
MH025917.1_strain_R7             .g.....g..g..t.........t..g..........................ag...c....tg...tc..a....... [ 45600]
MH025916.1_strain_R8             .g.....g..g..t.........t..g..........................ag...c....tg...tc..a....... [ 45600]

FR682468.2_Georgia_2007/1        tcaatagatatccatcattaatattgattatattttcgaatattatcttctatggtgcaagataatcatctagcgcgtga [ 45680]
ASF_IND_20_CAD543                ...........m.................................................................... [ 45680]
LR899193.1_Germany_2020/1        ................................................................................ [ 45680]
MN393477.1_Wuhan_2019            ................................................................................ [ 45680]
MW396979.1_Timor-Leste/2019/1    ................................................................................ [ 45680]
MK940252.1_CN/2019/InnerMongolia ................................................................................ [ 45680]
MK128995.1_China/2018/AnhuiXCGQ  ................................................................................ [ 45680]
MH681419.1_ASFV/POL/2015         ................................................................................ [ 45680]
LS478113.1_Estonia_2014          ................................................................................ [ 45680]
KP843857.1_Odintsovo_02/14       ................................................................................ [ 45680]
FN557520.1_strain_E75            ...........a.................................................................... [ 45680]
AY261366.1_Warthog               ..........cgt........c.....................................t...c................ [ 45680]
AY261365.1_Warmbaths             ...........gt..............................................t...c................ [ 45680]
AY261364.1_Tengani_62            ...........at..............................................t...c..........a..... [ 45680]
AY261363.1_Pretoriuskop/96/4     ...........gt..............................................t...c................ [ 45680]
AY261362.1_Mkuzi_1979            ..........aa.................................................................... [ 45680]
AY261361.1_Malawi_Lil-20/1_1983  ....c......gt..t...........................................t..........a...a..... [ 45680]
AY261360.1_Kenya_1950            ....c...a..gt..a.c.....................t...................t..............a..... [ 45680]
AM712240.1_OURT_88/3             -------------------------------------------------------------------------------- [ 45680]
AM712239.1_Benin_97/1            ...........a.................................................................... [ 45680]
KM111295.1_Ken06.Bus             ....c......gt..a.c.....................t...................t..............a..... [ 45680]
```

```
KM111294.1_Ken05/Tk1             ....c......gt..a.c.....................t...................t..............a..... [ 45680]
KM262844.1_strain_L60            ...........a.................................................................... [ 45680]
KM102979.1_26544/OG10_Italy      ...........a.................................................................... [ 45680]
KM262845.1_strain_NHV            -------------------------------------------------------------------------------- [ 45680]
KX354450.1_47/Ss/2008            ...........a.................................................................... [ 45680]
KP055815.1_strain_BA71           ...........a.................................................................... [ 45680]
MH025920.1_strain_R35            ....c......gt..a.c.....................t...................t..............a..... [ 45680]
MH025919.1_strain_N10            ....c......gt..a.c.....................t...................t..............a..... [ 45680]
MH025918.1_strain_R25            ....c......gt..a.c.....................t...................t..............a..... [ 45680]
MH025917.1_strain_R7             ....c......gt..a.c.....................t...................t..............a..... [ 45680]
MH025916.1_strain_R8             ....c......gt..a.c.....................t...................t..............a..... [ 45680]

FR682468.2_Georgia_2007/1        aacatgtcctct---tctcttcaggaactttgtcgaaaaaagctgcctgactgcatacttccagagttttttgacgacta [ 45760]
ASF_IND_20_CAD543                ............---..y.............................................................. [ 45760]
LR899193.1_Germany_2020/1        ............---................................................................. [ 45760]
MN393477.1_Wuhan_2019            ............---................................................................. [ 45760]
MW396979.1_Timor-Leste/2019/1    ............---................................................................. [ 45760]
MK940252.1_CN/2019/InnerMongolia ............---................................................................. [ 45760]
MK128995.1_China/2018/AnhuiXCGQ  ............---................................................................. [ 45760]
MH681419.1_ASFV/POL/2015         ............---................................................................. [ 45760]
LS478113.1_Estonia_2014          ............---................................................................. [ 45760]
KP843857.1_Odintsovo_02/14       ............---................................................................. [ 45760]
FN557520.1_strain_E75            ............---..c.............................................................. [ 45760]
AY261366.1_Warthog               ............---..c..............c......................................c........ [ 45760]
AY261365.1_Warmbaths             ............---..c..............c......................................c........ [ 45760]
AY261364.1_Tengani_62            ............tcc..c..............c.......................c..............c........ [ 45760]
AY261363.1_Pretoriuskop/96/4     ........t...---..c.............................................................. [ 45760]
AY261362.1_Mkuzi_1979            ............---..c.............................................................. [ 45760]
AY261361.1_Malawi_Lil-20/1_1983  g.....---------..c..............c........c.....c......ga............c........... [ 45760]
AY261360.1_Kenya_1950            ............---..c..............c........ct.a..c...ga...............c........... [ 45760]
AM712240.1_OURT_88/3             -------------------------------------------------------------------------------- [ 45760]
AM712239.1_Benin_97/1            ............---..c.............................................................. [ 45760]
KM111295.1_Ken06.Bus             ............---..c..............c........ct.a..c...ca...............c........... [ 45760]
KM111294.1_Ken05/Tk1             ............---..c..............c........ct.a..c....a...............c........... [ 45760]
KM262844.1_strain_L60            ............---..c.............................................................. [ 45760]
KM102979.1_26544/OG10_Italy      ............---..c.............................................................. [ 45760]
KM262845.1_strain_NHV            -------------------------------------------------------------------------------- [ 45760]
KX354450.1_47/Ss/2008            ............---..c.............................................................. [ 45760]
KP055815.1_strain_BA71           ............---..c.............................................................. [ 45760]
MH025920.1_strain_R35            ............---..c..............c........ct.a..c...ca...............c........... [ 45760]
MH025919.1_strain_N10            ............---..c..............c........ct.a..c...ca...............c........... [ 45760]
MH025918.1_strain_R25            ............---..c..............c........ct.a..c...ca...............c........... [ 45760]
MH025917.1_strain_R7             ............---..c..............c........ct.a..c...ca...............c........... [ 45760]
MH025916.1_strain_R8             ............---..c..............c........ct.a..c...ca...............c........... [ 45760]
```

```
FR682468.2_Georgia_2007/1        tgtattgcaactgttaggactgcactggcaagatcatggttcccttcagcgtatcgagaagaaccagatacttgttcaac [ 45840]
ASF_IND_20_CAD543                ................................................................................ [ 45840]
LR899193.1_Germany_2020/1        ................................................................................ [ 45840]
MN393477.1_Wuhan_2019            ................................................................................ [ 45840]
MW396979.1_Timor-Leste/2019/1    ................................................................................ [ 45840]
MK940252.1_CN/2019/InnerMongolia ................................................................................ [ 45840]
MK128995.1_China/2018/AnhuiXCGQ  ................................................................................ [ 45840]
MH681419.1_ASFV/POL/2015         ................................................................................ [ 45840]
LS478113.1_Estonia_2014          ................................................................................ [ 45840]
KP843857.1_Odintsovo_02/14       ................................................................................ [ 45840]
FN557520.1_strain_E75            ................................................................................ [ 45840]
AY261366.1_Warthog               ...................t....t.................t..................................... [ 45840]
AY261365.1_Warmbaths             ..........................................t..................................... [ 45840]
AY261364.1_Tengani_62            ..........................................t..................................... [ 45840]
AY261363.1_Pretoriuskop/96/4     ..........................................t..................................... [ 45840]
AY261362.1_Mkuzi_1979            ................................................................................ [ 45840]
AY261361.1_Malawi_Lil-20/1_1983  ...................t......................t..........c..g......t...g............ [ 45840]
AY261360.1_Kenya_1950            ...g........a........a....................t.....a....c..ga.....t...g............ [ 45840]
AM712240.1_OURT_88/3             -------------------------------------------------------------------------------- [ 45840]
AM712239.1_Benin_97/1            ................................................................................ [ 45840]
KM111295.1_Ken06.Bus             .a..........a.............................t..........c.........t...g............ [ 45840]
KM111294.1_Ken05/Tk1             ...g........a........a....................t.....a....c..ga.....t...g............ [ 45840]
KM262844.1_strain_L60            ................................................................................ [ 45840]
KM102979.1_26544/OG10_Italy      ................................................................................ [ 45840]
KM262845.1_strain_NHV            -------------------------------------------------------------------------------- [ 45840]
KX354450.1_47/Ss/2008            ................................................................................ [ 45840]
KP055815.1_strain_BA71           ................................................................................ [ 45840]
MH025920.1_strain_R35            .a..........a.............................t..........c.........t...g............ [ 45840]
MH025919.1_strain_N10            .a..........a.............................t..........c.........t...g............ [ 45840]
MH025918.1_strain_R25            .a..........a.............................t..........c.........t...g............ [ 45840]
MH025917.1_strain_R7             .a..........a.............................t..........c.........t...g............ [ 45840]
MH025916.1_strain_R8             .a..........a.............................t..........c.........t...g............ [ 45840]

FR682468.2_Georgia_2007/1        aggaacccatccatatcaatgaagcactcaaagtagcagcatcggaagggaactatgaaatcgtagagctgttgttgtca [ 45920]
ASF_IND_20_CAD543                ................................................................................ [ 45920]
LR899193.1_Germany_2020/1        ................................................................................ [ 45920]
MN393477.1_Wuhan_2019            ................................................................................ [ 45920]
MW396979.1_Timor-Leste/2019/1    ................................................................................ [ 45920]
MK940252.1_CN/2019/InnerMongolia ................................................................................ [ 45920]
MK128995.1_China/2018/AnhuiXCGQ  ................................................................................ [ 45920]
MH681419.1_ASFV/POL/2015         ................................................................................ [ 45920]
LS478113.1_Estonia_2014          ................................................................................ [ 45920]
KP843857.1_Odintsovo_02/14       ................................................................................ [ 45920]
```

```
FN557520.1_strain_E75             ................................................................................ [ 45920]
AY261366.1_Warthog                ................................................................................ [ 45920]
AY261365.1_Warmbaths              ................................................................................ [ 45920]
AY261364.1_Tengani_62             ................................................................................ [ 45920]
AY261363.1_Pretoriuskop/96/4      ................................................................................ [ 45920]
AY261362.1_Mkuzi_1979             ................................................................................ [ 45920]
AY261361.1_Malawi_Lil-20/1_1983   ............................a...agt..g.....a...........................c........ [ 45920]
AY261360.1_Kenya_1950             ..........t.....t...........a.....g........a..g..a....t............a............ [ 45920]
AM712240.1_OURT_88/3              -------------------------------------------------------------------------------- [ 45920]
AM712239.1_Benin_97/1             ................................................................................ [ 45920]
KM111295.1_Ken06.Bus              ............................a.....g........a.....a.....c.....a.................. [ 45920]
KM111294.1_Ken05/Tk1              ..........t.................a.....g........a..g..a....t............a............ [ 45920]
KM262844.1_strain_L60             ................................................................................ [ 45920]
KM102979.1_26544/OG10_Italy       ................................................................................ [ 45920]
KM262845.1_strain_NHV             -------------------------------------------------------------------------------- [ 45920]
KX354450.1_47/Ss/2008             ................................................................................ [ 45920]
KP055815.1_strain_BA71            ................................................................................ [ 45920]
MH025920.1_strain_R35             ............................a.....g........a.....a.....c.....a.................. [ 45920]
MH025919.1_strain_N10             ............................a.....g........a.....a.....c.....a.................. [ 45920]
MH025918.1_strain_R25             ............................a.....g........a.....a.....c.....a.................. [ 45920]
MH025917.1_strain_R7              ............................a.....g........a.....a.....c.....a.................. [ 45920]
MH025916.1_strain_R8              ............................a.....g........a.....a.....c.....a.................. [ 45920]

FR682468.2_Georgia_2007/1         tgggaggcagatccccgctacgccgtcgtaggagccctagaaagcaaatactatgacctggtttacaaatactatgacca [ 46000]
ASF_IND_20_CAD543                 ................................................................................ [ 46000]
LR899193.1_Germany_2020/1         ................................................................................ [ 46000]
MN393477.1_Wuhan_2019             ................................................................................ [ 46000]
MW396979.1_Timor-Leste/2019/1     ................................................................................ [ 46000]
MK940252.1_CN/2019/InnerMongolia  ................................................................................ [ 46000]
MK128995.1_China/2018/AnhuiXCGQ   ................................................................................ [ 46000]
MH681419.1_ASFV/POL/2015          ................................................................................ [ 46000]
LS478113.1_Estonia_2014           ................................................................................ [ 46000]
KP843857.1_Odintsovo_02/14        ................................................................................ [ 46000]
FN557520.1_strain_E75             ...................................t...........................................t [ 46000]
AY261366.1_Warthog                ...................................t..c........................c............g... [ 46000]
AY261365.1_Warmbaths              ...................................t..c........................c............g... [ 46000]
AY261364.1_Tengani_62             ...................................t..c........................c................ [ 46000]
AY261363.1_Pretoriuskop/96/4      ...................................t..c........................c............g... [ 46000]
AY261362.1_Mkuzi_1979             ...................................t............................................ [ 46000]
AY261361.1_Malawi_Lil-20/1_1983   .......................t.......................t...............c.......t........ [ 46000]
AY261360.1_Kenya_1950             ...a...................t...........g.......................c........g..t..ca...t [ 46000]
AM712240.1_OURT_88/3              -------------------------------------------------------------------------------- [ 46000]
AM712239.1_Benin_97/1             ...................................t...........................................t [ 46000]
KM111295.1_Ken06.Bus              ...a.......c.....a.....t...........g.......................c...........t..c..... [ 46000]
```

```
KM111294.1_Ken05/Tk1             ...a.......c...........t...........g.......................c........g..t..ca...t [ 46000]
KM262844.1_strain_L60            ...................................t...........................................t [ 46000]
KM102979.1_26544/OG10_Italy      ...................................t...............c...........................t [ 46000]
KM262845.1_strain_NHV            -------------------------------------------------------------------------------- [ 46000]
KX354450.1_47/Ss/2008            ...................................t...............c.......a...................t [ 46000]
KP055815.1_strain_BA71           ...................................t...........................................t [ 46000]
MH025920.1_strain_R35            ...a.......c.....a.....t...........g.......................c...........t..c..... [ 46000]
MH025919.1_strain_N10            ...a.......c.....a.....t...........g.......................c...........t..c..... [ 46000]
MH025918.1_strain_R25            ...a.......c.....a.....t...........g.......................c...........t..c..... [ 46000]
MH025917.1_strain_R7             ...a.......c.....a.....t...........g.......................c...........t..c..... [ 46000]
MH025916.1_strain_R8             ...a.......c.....a.....t...........g.......................c...........t..c..... [ 46000]

FR682468.2_Georgia_2007/1        agttaaagactgccatgatatcttgccgctgattcaaaatccggaaacattcgaaagatgtcatgagttaaacagcacct [ 46080]
ASF_IND_20_CAD543                ................................................................................ [ 46080]
LR899193.1_Germany_2020/1        ................................................................................ [ 46080]
MN393477.1_Wuhan_2019            ................................................................................ [ 46080]
MW396979.1_Timor-Leste/2019/1    ................................................................................ [ 46080]
MK940252.1_CN/2019/InnerMongolia ................................................................................ [ 46080]
MK128995.1_China/2018/AnhuiXCGQ  ................................................................................ [ 46080]
MH681419.1_ASFV/POL/2015         ................................................................................ [ 46080]
LS478113.1_Estonia_2014          ................................................................................ [ 46080]
KP843857.1_Odintsovo_02/14       ................................................................................ [ 46080]
FN557520.1_strain_E75            g..................................................t....a.................a.c... [ 46080]
AY261366.1_Warthog               ...................................................t....ag...................... [ 46080]
AY261365.1_Warmbaths             ........................................................a....................... [ 46080]
AY261364.1_Tengani_62            ........................................................ag...................... [ 46080]
AY261363.1_Pretoriuskop/96/4     ........................................................a....................... [ 46080]
AY261362.1_Mkuzi_1979            ...................................................t....a.................a.c... [ 46080]
AY261361.1_Malawi_Lil-20/1_1983  .....................gc.a..a.....c.............tg.......a.................a..... [ 46080]
AY261360.1_Kenya_1950            .a..g.....c..........g...........c......t.a.....g.........................a---.. [ 46080]
AM712240.1_OURT_88/3             -------------------------------------------------------------------------------- [ 46080]
AM712239.1_Benin_97/1            g..................................................t....a.................a.c... [ 46080]
KM111295.1_Ken06.Bus             .................................c......t............................g....a---t. [ 46080]
KM111294.1_Ken05/Tk1             ....g.....c..........g...........c......t.a.....g.........................a---.. [ 46080]
KM262844.1_strain_L60            g..................................................t....a.................a.c... [ 46080]
KM102979.1_26544/OG10_Italy      g..................................................t....a.................a.c... [ 46080]
KM262845.1_strain_NHV            -------------------------------------------------------------------------------- [ 46080]
KX354450.1_47/Ss/2008            g..................................................t....a.................a.c... [ 46080]
KP055815.1_strain_BA71           g..................................................t....a.................a.c... [ 46080]
MH025920.1_strain_R35            .................................c......t............................g....a---t. [ 46080]
MH025919.1_strain_N10            .................................c......t............................g....a---t. [ 46080]
MH025918.1_strain_R25            .................................c......t............................g....a---t. [ 46080]
MH025917.1_strain_R7             .................................c......t............................g....a---t. [ 46080]
MH025916.1_strain_R8             .................................c......t............................g....a---t. [ 46080]
```

```
FR682468.2_Georgia_2007/1        gttcactgaaatgcttattcaagcatgctgtgataaatgacatgctgccgattcttcaaaaatatacagactatctggat [ 46160]
ASF_IND_20_CAD543                ....................................................................k...w....... [ 46160]
LR899193.1_Germany_2020/1        ................................................................................ [ 46160]
MN393477.1_Wuhan_2019            ................................................................................ [ 46160]
MW396979.1_Timor-Leste/2019/1    ................................................................................ [ 46160]
MK940252.1_CN/2019/InnerMongolia ................................................................................ [ 46160]
MK128995.1_China/2018/AnhuiXCGQ  ................................................................................ [ 46160]
MH681419.1_ASFV/POL/2015         ................................................................................ [ 46160]
LS478113.1_Estonia_2014          ................................................................................ [ 46160]
KP843857.1_Odintsovo_02/14       ................................................................................ [ 46160]
FN557520.1_strain_E75            ....t..t...........................c................................t...t....... [ 46160]
AY261366.1_Warthog               ..................................g.....................g....................... [ 46160]
AY261365.1_Warmbaths             ..................................g.....................g....................... [ 46160]
AY261364.1_Tengani_62            ................................................................................ [ 46160]
AY261363.1_Pretoriuskop/96/4     ..................................g.....................g....................... [ 46160]
AY261362.1_Mkuzi_1979            ....t..t...........................c................................t...t....... [ 46160]
AY261361.1_Malawi_Lil-20/1_1983  ....t..t..............a............c.............................t........t..... [ 46160]
AY261360.1_Kenya_1950            ....t..t...........................t......a............a.........g........t....c [ 46160]
AM712240.1_OURT_88/3             -------------------------------------------------------------------------------- [ 46160]
AM712239.1_Benin_97/1            ....t..t...........................c................................t...t....... [ 46160]
KM111295.1_Ken06.Bus             ....t..t...........................t......a......................g..ag.......... [ 46160]
KM111294.1_Ken05/Tk1             ....t..t.......................a...t......a............a.........g........t....c [ 46160]
KM262844.1_strain_L60            ....t..t...........................c................................t...t....... [ 46160]
KM102979.1_26544/OG10_Italy      ....t..t...........................c................................t...t....... [ 46160]
KM262845.1_strain_NHV            -------------------------------------------------------------------------------- [ 46160]
KX354450.1_47/Ss/2008            ....t..t...........................c................................t...t....... [ 46160]
KP055815.1_strain_BA71           ....t..t...........................c................................t...t....... [ 46160]
MH025920.1_strain_R35            ....t..t...........................t......a......................g..ag.......... [ 46160]
MH025919.1_strain_N10            ....t..t...........................t......a......................g..ag.......... [ 46160]
MH025918.1_strain_R25            ....t..t...........................t......a......................g..ag.......... [ 46160]
MH025917.1_strain_R7             ....t..t...........................t......a......................g..ag.......... [ 46160]
MH025916.1_strain_R8             ....t..t...........................t......a......................g..ag.......... [ 46160]

FR682468.2_Georgia_2007/1        aggtgggagtattgcagccagatgctgttcgaactggcatgtagtaaaaaaaaatatgagatggttgtgtggatagaggg [ 46240]
ASF_IND_20_CAD543                ................................................................................ [ 46240]
LR899193.1_Germany_2020/1        ................................................................................ [ 46240]
MN393477.1_Wuhan_2019            ................................................................................ [ 46240]
MW396979.1_Timor-Leste/2019/1    ................................................................................ [ 46240]
MK940252.1_CN/2019/InnerMongolia ................................................................................ [ 46240]
MK128995.1_China/2018/AnhuiXCGQ  ................................................................................ [ 46240]
MH681419.1_ASFV/POL/2015         ................................................................................ [ 46240]
LS478113.1_Estonia_2014          ................................................................................ [ 46240]
KP843857.1_Odintsovo_02/14       ................................................................................ [ 46240]
```

```
FN557520.1_strain_E75            g...............a............................................................... [ 46240]
AY261366.1_Warthog               .................................t..........a................................... [ 46240]
AY261365.1_Warmbaths             .................................t..........a................................... [ 46240]
AY261364.1_Tengani_62            ..............t..................t..........a........g.......................... [ 46240]
AY261363.1_Pretoriuskop/96/4     .................................t.......c..a................................... [ 46240]
AY261362.1_Mkuzi_1979            g................................t......................c....................... [ 46240]
AY261361.1_Malawi_Lil-20/1_1983  g.....c.........at.....a.........t.........aa.g.c........a.t.................... [ 46240]
AY261360.1_Kenya_1950            g.....cc........a...............gt.........aa...c........a.c.................... [ 46240]
AM712240.1_OURT_88/3             -----------------------------------............................................. [ 46240]
AM712239.1_Benin_97/1            g...............a............................................................... [ 46240]
KM111295.1_Ken06.Bus             g.....cc........a...............gt.........aa...c........a.c....c...a........... [ 46240]
KM111294.1_Ken05/Tk1             g.....cc........a...............gt.........aa...c........a.c.................... [ 46240]
KM262844.1_strain_L60            g...............a............................................................... [ 46240]
KM102979.1_26544/OG10_Italy      g...............a............................................................... [ 46240]
KM262845.1_strain_NHV            -----------------------------------............................................. [ 46240]
KX354450.1_47/Ss/2008            g...............a............................................................... [ 46240]
KP055815.1_strain_BA71           g...............a............................................................... [ 46240]
MH025920.1_strain_R35            g.....cc........a...............gt.........aa...c........a.c....c...a........... [ 46240]
MH025919.1_strain_N10            g.....cc........a...............gt.........aa...c........a.c....c...a........... [ 46240]
MH025918.1_strain_R25            g.....cc........a...............gt.........aa...c........a.c....c...a........... [ 46240]
MH025917.1_strain_R7             g.....cc........a...............gt.........aa...c........a.c....c...a........... [ 46240]
MH025916.1_strain_R8             g.....cc........a...............gt.........aa...c........a.c....c...a........... [ 46240]

FR682468.2_Georgia_2007/1        agttctaggcgtcggcaaagttacatctcttttcaccattgcgattagcaacagagacctacagctgtattctctgggct [ 46320]
ASF_IND_20_CAD543                ................................................................................ [ 46320]
LR899193.1_Germany_2020/1        ................................................................................ [ 46320]
MN393477.1_Wuhan_2019            ................................................................................ [ 46320]
MW396979.1_Timor-Leste/2019/1    ................................................................................ [ 46320]
MK940252.1_CN/2019/InnerMongolia ................................................................................ [ 46320]
MK128995.1_China/2018/AnhuiXCGQ  ................................................................................ [ 46320]
MH681419.1_ASFV/POL/2015         ................................................................................ [ 46320]
LS478113.1_Estonia_2014          ................................................................................ [ 46320]
KP843857.1_Odintsovo_02/14       ................................................................................ [ 46320]
FN557520.1_strain_E75            ..........a.........................t.......................g..c........c......c [ 46320]
AY261366.1_Warthog               ........................g....................................................... [ 46320]
AY261365.1_Warmbaths             ..........a.............gc...................................................... [ 46320]
AY261364.1_Tengani_62            ..c...g.....................................................g................... [ 46320]
AY261363.1_Pretoriuskop/96/4     .......t................g....................................................... [ 46320]
AY261362.1_Mkuzi_1979            ..........a.........................t.......................g..c........c......c [ 46320]
AY261361.1_Malawi_Lil-20/1_1983  ..................ct...a.at...............c....a.........t....................g. [ 46320]
AY261360.1_Kenya_1950            ...c..g...........ct.c...at.........g..........t..a..g.................c....... [ 46320]
AM712240.1_OURT_88/3             ..........a.........................t.......................g..c........c......c [ 46320]
AM712239.1_Benin_97/1            ..........a.........................t.......................g..c........c......c [ 46320]
KM111295.1_Ken06.Bus             ...c..g...........ct.c...at.........g..........t..a....................c....... [ 46320]
```

```
KM111294.1_Ken05/Tk1           ...c..g...........ct.c...at.........a..........t...a..g.................c....... [ 46320]
KM262844.1_strain_L60          ..........a.........................t.......................g..c........c......c [ 46320]
KM102979.1_26544/OG10_Italy    ..........a.........................t.......................g..c........c......c [ 46320]
KM262845.1_strain_NHV          ..........a.........................t.......................g..c........c......c [ 46320]
KX354450.1_47/Ss/2008          ..........a.........................t.......................g..c........c......c [ 46320]
KP055815.1_strain_BA71         ..........a.........................t.......................g..c........c......c [ 46320]
MH025920.1_strain_R35          ...c..g...........ct.c...at.........g..........t...a.....................c....... [ 46320]
MH025919.1_strain_N10          ...c..g...........ct.c...at.........g..........t...a.....................c....... [ 46320]
MH025918.1_strain_R25          ...c..g...........ct.c...at.........g..........t...a.....................c....... [ 46320]
MH025917.1_strain_R7           ...c..g...........ct.c...at.........g..........t...a.....................c....... [ 46320]
MH025916.1_strain_R8           ...c..g...........ct.c...at.........g..........t...a.....................c....... [ 46320]

FR682468.2_Georgia_2007/1      actcaattatccttgagaatttgtactcctgtggacaggaccccaagtttttactaaatcatttcctgcgagacgtttca [ 46400]
ASF_IND_20_CAD543              ..................gaa..c.g...................................................... [ 46400]
LR899193.1_Germany_2020/1      ................................................................................ [ 46400]
MN393477.1_Wuhan_2019          ................................................................................ [ 46400]
MW396979.1_Timor-Leste/2019/1  ................................................................................ [ 46400]
MK940252.1_CN/2019/InnerMongolia ................................................................................ [ 46400]
MK128995.1_China/2018/AnhuiXCGQ ................................................................................ [ 46400]
MH681419.1_ASFV/POL/2015       ................................................................................ [ 46400]
LS478113.1_Estonia_2014        ................................................................................ [ 46400]
KP843857.1_Odintsovo_02/14     ................................................................................ [ 46400]
FN557520.1_strain_E75          ...t...c..t.......gaa..c.g...........a.......c.................................. [ 46400]
AY261366.1_Warthog             t.........t........a.........................c.................................g [ 46400]
AY261365.1_Warmbaths           ..........t.......gaa....t.......a......t....c.................................g [ 46400]
AY261364.1_Tengani_62          .......g..t........a.........................c.................................g [ 46400]
AY261363.1_Pretoriuskop/96/4   ..........t........aa...........c............c.................................g [ 46400]
AY261362.1_Mkuzi_1979          ...t...c..t.......gaa..c.g...........a...........................t.............. [ 46400]
AY261361.1_Malawi_Lil-20/1_1983 ...t...c..t.....a.ga.................a.......c............c........a............ [ 46400]
AY261360.1_Kenya_1950          ....c..c..........gaa..............t.c.......c...............c.at........tg...... [ 46400]
AM712240.1_OURT_88/3           ...t...c..t.......gaa..c.g...........a.......c.................................. [ 46400]
AM712239.1_Benin_97/1          ...t...c..t.......gaa..c.g...........a.......c.................................. [ 46400]
KM111295.1_Ken06.Bus           .......g..t........aa..............t.c.....t.c.................at......at....... [ 46400]
KM111294.1_Ken05/Tk1           ....c..c..........gaa....t.........t.c.......c...............c.at........tg...... [ 46400]
KM262844.1_strain_L60          ...t...c..t.......gaa..c.g...........a.......c.................................. [ 46400]
KM102979.1_26544/OG10_Italy    ...t...c..t.......gaa..c.g...........a.......c.................................. [ 46400]
KM262845.1_strain_NHV          ...t...c..t.......gaa..c.g...........a.......c.................................. [ 46400]
KX354450.1_47/Ss/2008          ...t...c..t.......gaa..c.g...........a.......c.................................. [ 46400]
KP055815.1_strain_BA71         ...t...c..t.......gaa..c.g...........a.......c.................................. [ 46400]
MH025920.1_strain_R35          .......g..t........aa..............t.c.....t.c.................at......at....... [ 46400]
MH025919.1_strain_N10          .......g..t........aa..............t.c.....t.c.................at......at....... [ 46400]
MH025918.1_strain_R25          .......g..t........aa..............t.c.....t.c.................at......at....... [ 46400]
MH025917.1_strain_R7           .......g..t........aa..............t.c.....t.c.................at......at....... [ 46400]
MH025916.1_strain_R8           .......g..t........aa..............t.c.....t.c.................at......at....... [ 46400]
```

```
FR682468.2_Georgia_2007/1        ataaaagggcttctaccctttgtaatcaaaaccatagaatatggtggaagcaaggagatagccataactctggctaaaaa [ 46480]
ASF_IND_20_CAD543                ..............c..t......c.g..................................................... [ 46480]
LR899193.1_Germany_2020/1        ................................................................................ [ 46480]
MN393477.1_Wuhan_2019            ................................................................................ [ 46480]
MW396979.1_Timor-Leste/2019/1    ................................................................................ [ 46480]
MK940252.1_CN/2019/InnerMongolia ................................................................................ [ 46480]
MK128995.1_China/2018/AnhuiXCGQ  ................................................................................ [ 46480]
MH681419.1_ASFV/POL/2015         ................................................................................ [ 46480]
LS478113.1_Estonia_2014          ................................................................................ [ 46480]
KP843857.1_Odintsovo_02/14       ................................................................................ [ 46480]
FN557520.1_strain_E75            ..............c..t......c.g..................................................... [ 46480]
AY261366.1_Warthog               .....g........g.........c....................................................... [ 46480]
AY261365.1_Warmbaths             .....g........g.........c.g..........................a.......................... [ 46480]
AY261364.1_Tengani_62            .....g........g.........c....................................................... [ 46480]
AY261363.1_Pretoriuskop/96/4     .....g........g.........c.g..................................................... [ 46480]
AY261362.1_Mkuzi_1979            ..............c.........c.g..................................................... [ 46480]
AY261361.1_Malawi_Lil-20/1_1983  ..g..g........g.........c.g.............t.........t............................. [ 46480]
AY261360.1_Kenya_1950            .c............g..t......c.g..........................a...........c...t.......... [ 46480]
AM712240.1_OURT_88/3             ..............c..t......c.g..................................................... [ 46480]
AM712239.1_Benin_97/1            ..............c..t......c.g..................................................... [ 46480]
KM111295.1_Ken06.Bus             .c...g........g........gc.g..........................a...........c...t.......... [ 46480]
KM111294.1_Ken05/Tk1             .c............g..t......c.g..........................a.........t.....t.......... [ 46480]
KM262844.1_strain_L60            ..............c..t......c.g..................................................... [ 46480]
KM102979.1_26544/OG10_Italy      ..............c..t......c.g..................................................... [ 46480]
KM262845.1_strain_NHV            ..............c..t......c.g..................................................... [ 46480]
KX354450.1_47/Ss/2008            ..............c..t......c.g..................................................... [ 46480]
KP055815.1_strain_BA71           ..............c..t......c.g..................................................... [ 46480]
MH025920.1_strain_R35            .c...g........g........gc.g..........................a...........c...t.......... [ 46480]
MH025919.1_strain_N10            .c...g........g........gc.g..........................a...........c...t.......... [ 46480]
MH025918.1_strain_R25            .c...g........g........gc.g..........................a...........c...t.......... [ 46480]
MH025917.1_strain_R7             .c...g........g........gc.g..........................a...........c...t.......... [ 46480]
MH025916.1_strain_R8             .c...g........g........gc.g..........................a...........c...t.......... [ 46480]

FR682468.2_Georgia_2007/1        atatcagcataaacatattttgaaatacttcgaaacctgggaaagctagg---------ttcagtatggtgtactca-ct [ 46560]
ASF_IND_20_CAD543                ....................................nnnnnnnnnnnnnn---------nnnnnnnnnnnnnnnnnn-nn [ 46560]
LR899193.1_Germany_2020/1        ..................................................---------..................-.. [ 46560]
MN393477.1_Wuhan_2019            ..................................................---------..................-.. [ 46560]
MW396979.1_Timor-Leste/2019/1    ..................................................---------..................-.. [ 46560]
MK940252.1_CN/2019/InnerMongolia ..................................................---------..................-.. [ 46560]
MK128995.1_China/2018/AnhuiXCGQ  ..................................................---------..................-.. [ 46560]
MH681419.1_ASFV/POL/2015         ..................................................---------..................-.. [ 46560]
LS478113.1_Estonia_2014          ..................................................---------..................-.. [ 46560]
KP843857.1_Odintsovo_02/14       ..................................................---------..................-.. [ 46560]
```

```
FN557520.1_strain_E75          .....................................g..a..t....aa---------.g.g.....a.....c..--- [ 46560]
AY261366.1_Warthog             ..................................................---------...............c..-.. [ 46560]
AY261365.1_Warmbaths           ....................................a...........a.---------..................c.. [ 46560]
AY261364.1_Tengani_62          ....................................t.............---------..........c.......-.g [ 46560]
AY261363.1_Pretoriuskop/96/4   ..................................................---------...............c..-.. [ 46560]
AY261362.1_Mkuzi_1979          .....................................g..a..t....aa---------.g.g.....a.....c..--- [ 46560]
AY261361.1_Malawi_Lil-20/1_1983 ........................g..t.........gaa...t....a.gttgttaaa.a....g........c.ca.. [ 46560]
AY261360.1_Kenya_1950          g.........g..ac........g.............a..a..tc.c...-------ag.g.taa..aca.c.ac.--.. [ 46560]
AM712240.1_OURT_88/3           .....................................g..a..t....aa---------.g.g.....a.....c..--- [ 46560]
AM712239.1_Benin_97/1          .....................................g..a..t....aa---------.g.g.....a.....c..--- [ 46560]
KM111295.1_Ken06.Bus           g.........g..a.........g.............a..a...c.c.a.-------ag.g.taa..aca.c.ac.--.. [ 46560]
KM111294.1_Ken05/Tk1           ..........g..ac........g.............a..a..tc.c...-------ag.g.taa..aca.c.ac.--.. [ 46560]
KM262844.1_strain_L60          .....................................g..a..t....aa---------.g.g.....a.....c..--- [ 46560]
KM102979.1_26544/OG10_Italy    .....................................g..a..t....aa---------.g.g.....a.....c..--- [ 46560]
KM262845.1_strain_NHV          .....................................g..a..t....aa---------.g.g.....a.....c..--- [ 46560]
KX354450.1_47/Ss/2008          .....................................g..a..t....aa---------.g.g.....a.....c..--- [ 46560]
KP055815.1_strain_BA71         .....................................g..a..t....aa---------.g.g.....a.....c..--- [ 46560]
MH025920.1_strain_R35          g.........g..a.........g.............a..a...c.c.a.-------ag.g.taa..aca.c.ac.--.. [ 46560]
MH025919.1_strain_N10          g.........g..a.........g.............a..a...c.c.a.-------ag.g.taa..aca.c.ac.--.. [ 46560]
MH025918.1_strain_R25          g.........g..a.........g.............a..a...c.c.a.-------ag.g.taa..aca.c.ac.--.. [ 46560]
MH025917.1_strain_R7           g.........g..a.........g.............a..a...c.c.a.-------ag.g.taa..aca.c.ac.--.. [ 46560]
MH025916.1_strain_R8           g.........g..a.........g.............a..a...c.c.a.-------ag.g.taa..aca.c.ac.--.. [ 46560]

FR682468.2_Georgia_2007/1      attgtagtgaatcgta---------------------------------------------------------------- [ 46640]
ASF_IND_20_CAD543              nnnnnnnnnnnnnn.----------------------------------------------------------------- [ 46640]
LR899193.1_Germany_2020/1      ................---------------------------------------------------------------- [ 46640]
MN393477.1_Wuhan_2019          ................---------------------------------------------------------------- [ 46640]
MW396979.1_Timor-Leste/2019/1  ................---------------------------------------------------------------- [ 46640]
MK940252.1_CN/2019/InnerMongolia ................---------------------------------------------------------------- [ 46640]
MK128995.1_China/2018/AnhuiXCGQ ................---------------------------------------------------------------- [ 46640]
MH681419.1_ASFV/POL/2015       ................---------------------------------------------------------------- [ 46640]
LS478113.1_Estonia_2014        ................---------------------------------------------------------------- [ 46640]
KP843857.1_Odintsovo_02/14     ................---------------------------------------------------------------- [ 46640]
FN557520.1_strain_E75          ......a.........---------------------------------------------------------------- [ 46640]
AY261366.1_Warthog             ......a........----------------------------------------------------------------- [ 46640]
AY261365.1_Warmbaths           ................cct------------------------------------------------------------- [ 46640]
AY261364.1_Tengani_62          ......a..g...a..---------------------------------------------------------------- [ 46640]
AY261363.1_Pretoriuskop/96/4   ......a........----------------------------------------------------------------- [ 46640]
AY261362.1_Mkuzi_1979          ......a.........---------------------------------------------------------------- [ 46640]
AY261361.1_Malawi_Lil-20/1_1983 ......a.....t...tattttatgtgtacctactattgtattctgtatattttatgtgtaacctactct---------- [ 46640]
AY261360.1_Kenya_1950          .....ga.....a...-------------------actatattgtatacgttgtatacgttgtatacgttgtatacgttg [ 46640]
AM712240.1_OURT_88/3           ......a.........---------------------------------------------------------------- [ 46640]
AM712239.1_Benin_97/1          ......a.........---------------------------------------------------------------- [ 46640]
KM111295.1_Ken06.Bus           .....ga..t..a...-------------------actatattgtatacgttgtatacgttgtatatgttg--------- [ 46640]
```

```
KM111294.1_Ken05/Tk1             .....ga.....a...-------------------actatattgtatacgttgtaaacgttg------------------ [ 46640]
KM262844.1_strain_L60            ......a.........---------------------------------------------------------------- [ 46640]
KM102979.1_26544/OG10_Italy      ......a.........---------------------------------------------------------------- [ 46640]
KM262845.1_strain_NHV            ......a.........---------------------------------------------------------------- [ 46640]
KX354450.1_47/Ss/2008            ......a.........---------------------------------------------------------------- [ 46640]
KP055815.1_strain_BA71           ......a.........---------------------------------------------------------------- [ 46640]
MH025920.1_strain_R35            .....ga..t..a...-------------------actatattgtatacgttgtatacgttgtatatgttg--------- [ 46640]
MH025919.1_strain_N10            .....ga..t..a...-------------------actatattgtatacgttgtatacgttgtatatgttg--------- [ 46640]
MH025918.1_strain_R25            .....ga..t..a...-------------------actatattgtatacgttgtatacgttgtatatgttg--------- [ 46640]
MH025917.1_strain_R7             .....ga..t..a...-------------------actatattgtatacgttgtatacgttgtatatgttg--------- [ 46640]
MH025916.1_strain_R8             .....ga..t..a...-------------------actatattgtatacgttgtatacgttgtatatgttg--------- [ 46640]

FR682468.2_Georgia_2007/1        -------------------------------------------------------------------------------- [ 46720]
ASF_IND_20_CAD543                -------------------------------------------------------------------------------- [ 46720]
LR899193.1_Germany_2020/1        -------------------------------------------------------------------------------- [ 46720]
MN393477.1_Wuhan_2019            -------------------------------------------------------------------------------- [ 46720]
MW396979.1_Timor-Leste/2019/1    -------------------------------------------------------------------------------- [ 46720]
MK940252.1_CN/2019/InnerMongolia -------------------------------------------------------------------------------- [ 46720]
MK128995.1_China/2018/AnhuiXCGQ  -------------------------------------------------------------------------------- [ 46720]
MH681419.1_ASFV/POL/2015         -------------------------------------------------------------------------------- [ 46720]
LS478113.1_Estonia_2014          -------------------------------------------------------------------------------- [ 46720]
KP843857.1_Odintsovo_02/14       -------------------------------------------------------------------------------- [ 46720]
FN557520.1_strain_E75            -------------------------------------------------------------------------------- [ 46720]
AY261366.1_Warthog               -------------------------------------------------------------------------------- [ 46720]
AY261365.1_Warmbaths             -------------------------------------------------------------------------------- [ 46720]
AY261364.1_Tengani_62            -------------------------------------------------------------------------------- [ 46720]
AY261363.1_Pretoriuskop/96/4     -------------------------------------------------------------------------------- [ 46720]
AY261362.1_Mkuzi_1979            -------------------------------------------------------------------------------- [ 46720]
AY261361.1_Malawi_Lil-20/1_1983  -----------------------------------------------------------------tgtattctgtatatt [ 46720]
AY261360.1_Kenya_1950            tatatgttgtaaacgttgtatacgttgtatacgctgtaaacgttgtatacgctgtatacgttgtatatacgttgtacctt [ 46720]
AM712240.1_OURT_88/3             -------------------------------------------------------------------------------- [ 46720]
AM712239.1_Benin_97/1            -------------------------------------------------------------------------------- [ 46720]
KM111295.1_Ken06.Bus             -----------------------------------------------------------------tatacgttgtacctt [ 46720]
KM111294.1_Ken05/Tk1             -----------------------------------------------------------------tatacattgtacctt [ 46720]
KM262844.1_strain_L60            -------------------------------------------------------------------------------- [ 46720]
KM102979.1_26544/OG10_Italy      -------------------------------------------------------------------------------- [ 46720]
KM262845.1_strain_NHV            -------------------------------------------------------------------------------- [ 46720]
KX354450.1_47/Ss/2008            -------------------------------------------------------------------------------- [ 46720]
KP055815.1_strain_BA71           -------------------------------------------------------------------------------- [ 46720]
MH025920.1_strain_R35            -----------------------------------------------------------------tatacgttgtacctt [ 46720]
MH025919.1_strain_N10            -----------------------------------------------------------------tatacgttgtacctt [ 46720]
MH025918.1_strain_R25            -----------------------------------------------------------------tatacgttgtacctt [ 46720]
MH025917.1_strain_R7             -----------------------------------------------------------------tatacgttgtacctt [ 46720]
MH025916.1_strain_R8             -----------------------------------------------------------------tatacgttgtacctt [ 46720]
```

```
FR682468.2_Georgia_2007/1        -------------------------------tcctgtaaattttgtaaa---------------------------aaag [ 46800]
ASF_IND_20_CAD543                -------------------------------ctt.a.....g......-----------------------------.nn [ 46800]
LR899193.1_Germany_2020/1        -------------------------------..................---------------------------.... [ 46800]
MN393477.1_Wuhan_2019            -------------------------------..................---------------------------.... [ 46800]
MW396979.1_Timor-Leste/2019/1    -------------------------------..................---------------------------.... [ 46800]
MK940252.1_CN/2019/InnerMongolia -------------------------------..................---------------------------.... [ 46800]
MK128995.1_China/2018/AnhuiXCGQ  -------------------------------..................---------------------------.... [ 46800]
MH681419.1_ASFV/POL/2015         -------------------------------..................---------------------------.... [ 46800]
LS478113.1_Estonia_2014          -------------------------------..................---------------------------.... [ 46800]
KP843857.1_Odintsovo_02/14       -------------------------------..................---------------------------.... [ 46800]
FN557520.1_strain_E75            ------------------------------tctt.a.....g......-----------------------------..a [ 46800]
AY261366.1_Warthog               ------------------------------a..................---------------------------..g. [ 46800]
AY261365.1_Warmbaths             -----------------gtaaattttgtaaa.tt..........c....---------------------------.... [ 46800]
AY261364.1_Tengani_62            -------------------------------..........-......-------------------------------- [ 46800]
AY261363.1_Pretoriuskop/96/4     ---------------------atcctgtaaa.tt..........a....---------------------------.gg. [ 46800]
AY261362.1_Mkuzi_1979            ------------------------------tctt.a.....g......-----------------------------..a [ 46800]
AY261361.1_Malawi_Lil-20/1_1983  ttatgtataacctactattgtattctgtata.tt...gtt.........---------------------------.... [ 46800]
AY261360.1_Kenya_1950            aaacattttacccgc------atcatata-a.tt.ag....c..t...gccactgaacaatactattatgatgtat.g.. [ 46800]
AM712240.1_OURT_88/3             ------------------------------tctt.a.....g......-----------------------------..a [ 46800]
AM712239.1_Benin_97/1            ------------------------------tctt.a.....g......-----------------------------..a [ 46800]
KM111295.1_Ken06.Bus             aaacattttacccac------atcatatg-a.tt.ag....c..t...gccactgaacaatactattatgatgtat.g.. [ 46800]
KM111294.1_Ken05/Tk1             aaacattttacccac------atcatatg-a.tt.ag....c..t...gccactaaacaatactattatgatgtat.g.. [ 46800]
KM262844.1_strain_L60            ------------------------------tctt.a.....g......-----------------------------..a [ 46800]
KM102979.1_26544/OG10_Italy      ------------------------------tctt.a.....g......-----------------------------..a [ 46800]
KM262845.1_strain_NHV            ------------------------------tctt.a.....g......-----------------------------..a [ 46800]
KX354450.1_47/Ss/2008            ------------------------------tctt.a.....g......-----------------------------..a [ 46800]
KP055815.1_strain_BA71           ------------------------------tctt.a.....g......-----------------------------..a [ 46800]
MH025920.1_strain_R35            aaacattttacccac------atcatatg-a.tt.ag....c..t...gccactgaacaatactattatgatgtat.g.. [ 46800]
MH025919.1_strain_N10            aaacattttacccac------atcatatg-a.tt.ag....c..t...gccactgaacaatactattatgatgtat.g.. [ 46800]
MH025918.1_strain_R25            aaacattttacccac------atcatatg-a.tt.ag....c..t...gccactgaacaatactattatgatgtat.g.. [ 46800]
MH025917.1_strain_R7             aaacattttacccac------atcatatg-a.tt.ag....c..t...gccactgaacaatactattatgatgtat.g.. [ 46800]
MH025916.1_strain_R8             aaacattttacccac------atcatatg-a.tt.ag....c..t...gccactgaacaatactattatgatgtat.g.. [ 46800]

FR682468.2_Georgia_2007/1        --------------------------cttaaact-tttgaccacatcatattgttttagaaatctcaaaccagtgaacaa [ 46880]
ASF_IND_20_CAD543                --------------------------nnnnnnnn-................ga..........y....g.tgc....... [ 46880]
LR899193.1_Germany_2020/1        --------------------------........-............................................. [ 46880]
MN393477.1_Wuhan_2019            --------------------------........-............................................. [ 46880]
MW396979.1_Timor-Leste/2019/1    --------------------------........-............................................. [ 46880]
MK940252.1_CN/2019/InnerMongolia --------------------------........-............................................. [ 46880]
MK128995.1_China/2018/AnhuiXCGQ  --------------------------........-............................................. [ 46880]
MH681419.1_ASFV/POL/2015         --------------------------........-............................................. [ 46880]
LS478113.1_Estonia_2014          --------------------------........-............................................. [ 46880]
KP843857.1_Odintsovo_02/14       --------------------------........-............................................. [ 46880]
```

```
FN557520.1_strain_E75             --------------------------.....cta-................ga..........t....g.tgc....... [ 46880]
AY261366.1_Warthog                --------------------------.......a-.................aa....a.........g..g........ [ 46880]
AY261365.1_Warmbaths              --------------------------.......a-.................ag.................g........ [ 46880]
AY261364.1_Tengani_62             -----------------------------...ta-.................a.....a.........g........... [ 46880]
AY261363.1_Pretoriuskop/96/4      --------------------------.......a-.................aa..............g..g........ [ 46880]
AY261362.1_Mkuzi_1979             --------------------------.....cta-................ga...............g.tgc....... [ 46880]
AY261361.1_Malawi_Lil-20/1_1983   --------------------------.......a-...at............a...............g..g........ [ 46880]
AY261360.1_Kenya_1950             --taacccctattgtatatgttgtac......tat...ac.t.t.....g.ga............t..g..g........ [ 46880]
AM712240.1_OURT_88/3              --------------------------.....cta-................ga..........t....g.tgc....... [ 46880]
AM712239.1_Benin_97/1             --------------------------.....cta-................ga..........t....g.tgc....... [ 46880]
KM111295.1_Ken06.Bus              --taacccctattgtatatgttgtac......ta-...ac...........ga........t...t..g...a....... [ 46880]
KM111294.1_Ken05/Tk1              tataacccctattgtatatgttgtac......tat...ac.t.t.....g.ga............t..g..g........ [ 46880]
KM262844.1_strain_L60             --------------------------.....cta-................ga..........t....g.tgc....... [ 46880]
KM102979.1_26544/OG10_Italy       --------------------------.....cta-................ga..........t....g.tgc....... [ 46880]
KM262845.1_strain_NHV             --------------------------.....cta-................ga..........t....g.tgc....... [ 46880]
KX354450.1_47/Ss/2008             --------------------------.....cta-................ga..........t....g.tgc....... [ 46880]
KP055815.1_strain_BA71            --------------------------.....cta-................ga..........t....g.tgc....... [ 46880]
MH025920.1_strain_R35             --taacccctattgtatatgttgtac......ta-...ac...........ga........t...t..g...a....... [ 46880]
MH025919.1_strain_N10             --taacccctattgtatatgttgtac......ta-...ac...........ga........t...t..g...a....... [ 46880]
MH025918.1_strain_R25             --taacccctattgtatatgttgtac......ta-...ac...........ga........t...t..g...a....... [ 46880]
MH025917.1_strain_R7              --taacccctattgtatatgttgtac......ta-...ac...........ga........t...t..g...a....... [ 46880]
MH025916.1_strain_R8              --taacccctattgtatatgttgtac......ta-...ac...........ga........t...t..g...a....... [ 46880]

FR682468.2_Georgia_2007/1         cagtcttatcatacattaaaattccagtaaaatttatattttttttggtaaacaaatgttttctcttcaagacatctgtc [ 46960]
ASF_IND_20_CAD543                 ....y........................................................................... [ 46960]
LR899193.1_Germany_2020/1         ................................................................................ [ 46960]
MN393477.1_Wuhan_2019             ................................................................................ [ 46960]
MW396979.1_Timor-Leste/2019/1     ................................................................................ [ 46960]
MK940252.1_CN/2019/InnerMongolia  ................................................................................ [ 46960]
MK128995.1_China/2018/AnhuiXCGQ   ................................................................................ [ 46960]
MH681419.1_ASFV/POL/2015          ................................................................................ [ 46960]
LS478113.1_Estonia_2014           ................................................................................ [ 46960]
KP843857.1_Odintsovo_02/14        ...........................................-.................................... [ 46960]
FN557520.1_strain_E75             ....t.............................................................c............. [ 46960]
AY261366.1_Warthog                .g......................t......................c................................ [ 46960]
AY261365.1_Warmbaths              ........................t......................c................................ [ 46960]
AY261364.1_Tengani_62             ............g................................................................... [ 46960]
AY261363.1_Pretoriuskop/96/4      ........................t......................t................................ [ 46960]
AY261362.1_Mkuzi_1979             ..................................................................c............. [ 46960]
AY261361.1_Malawi_Lil-20/1_1983   tc..a........t..g......t.........................g..t........................... [ 46960]
AY261360.1_Kenya_1950             ....g...................t....................................................... [ 46960]
AM712240.1_OURT_88/3              ....t.............................................................c............. [ 46960]
AM712239.1_Benin_97/1             ....t.............................................................c............. [ 46960]
KM111295.1_Ken06.Bus              g...g.....c.............t........................g..t........................... [ 46960]
```

```
KM111294.1_Ken05/Tk1              ....g...................t.......................cg..t........................... [ 46960]
KM262844.1_strain_L60             ....t.............................................................c............. [ 46960]
KM102979.1_26544/OG10_Italy       ....t.............................................................c............. [ 46960]
KM262845.1_strain_NHV             ....t.............................................................c............. [ 46960]
KX354450.1_47/Ss/2008             ....t.............................................................c............. [ 46960]
KP055815.1_strain_BA71            ....t.............................................................c............. [ 46960]
MH025920.1_strain_R35             g...g.....c.............t........................g..t........................... [ 46960]
MH025919.1_strain_N10             g...g.....c.............t........................g..t........................... [ 46960]
MH025918.1_strain_R25             g...g.....c.............t........................g..t........................... [ 46960]
MH025917.1_strain_R7              g...g.....c.............t........................g..t........................... [ 46960]
MH025916.1_strain_R8              g...g.....c.............t........................g..t........................... [ 46960]

FR682468.2_Georgia_2007/1         ggaaacatctttttcaacttcctgacgcttttgatgaatatatattacaagcgctaggactatactgggaaaaacacgga [ 47040]
ASF_IND_20_CAD543                 ..........................................cg.................................... [ 47040]
LR899193.1_Germany_2020/1         ................................................................................ [ 47040]
MN393477.1_Wuhan_2019             ................................................................................ [ 47040]
MW396979.1_Timor-Leste/2019/1     ................................................................................ [ 47040]
MK940252.1_CN/2019/InnerMongolia  ................................................................................ [ 47040]
MK128995.1_China/2018/AnhuiXCGQ   ................................................................................ [ 47040]
MH681419.1_ASFV/POL/2015          ................................................................................ [ 47040]
LS478113.1_Estonia_2014           ................................................................................ [ 47040]
KP843857.1_Odintsovo_02/14        ................................................................................ [ 47040]
FN557520.1_strain_E75             .....t....................t...............cg.......t......g..................... [ 47040]
AY261366.1_Warthog                ................................................................t............... [ 47040]
AY261365.1_Warmbaths              .............c.............................g...........................g........ [ 47040]
AY261364.1_Tengani_62             .......................................t........................t............... [ 47040]
AY261363.1_Pretoriuskop/96/4      .................g.............................................................. [ 47040]
AY261362.1_Mkuzi_1979             ..........................................cg.......t......g.................t... [ 47040]
AY261361.1_Malawi_Lil-20/1_1983   ...............tg......agtt.........................a........................... [ 47040]
AY261360.1_Kenya_1950             ...............tg......agtt.....a..................t.........c.................. [ 47040]
AM712240.1_OURT_88/3              .....t....................t...............cg.......t......g..................... [ 47040]
AM712239.1_Benin_97/1             .....t....................t...............cg.......t......g..................... [ 47040]
KM111295.1_Ken06.Bus              ...............tg......agtt........................t.........c.................. [ 47040]
KM111294.1_Ken05/Tk1              ..............tt.......agtt..................................................... [ 47040]
KM262844.1_strain_L60             .....t....................t...............cg.......t......g..................... [ 47040]
KM102979.1_26544/OG10_Italy       .....t....................t...............cg.......t......g..................... [ 47040]
KM262845.1_strain_NHV             .....t....................t...............cg.......t......g..................... [ 47040]
KX354450.1_47/Ss/2008             .....t....................t...............cg.......t......g..................... [ 47040]
KP055815.1_strain_BA71            .....t....................t...............cg.......t......g..................... [ 47040]
MH025920.1_strain_R35             ...............tg......agtt........................t.........c.................. [ 47040]
MH025919.1_strain_N10             ...............tg......agtt........................t.........c.................. [ 47040]
MH025918.1_strain_R25             ...............tg......agtt........................t.........c.................. [ 47040]
MH025917.1_strain_R7              ...............tg......agtt........................t.........c.................. [ 47040]
MH025916.1_strain_R8              ...............tg......agtt........................t.........c.................. [ 47040]
```

```
FR682468.2_Georgia_2007/1        tctcttcaacgaataagaaaggacgctgtgtttgtacagcgaaa-catcgtcctttctaccaatgaggccctgagaatcg [ 47120]
ASF_IND_20_CAD543                ............................................-tc..a..a................tt.a....... [ 47120]
LR899193.1_Germany_2020/1        ............................................-................................... [ 47120]
MN393477.1_Wuhan_2019            ............................................-................................... [ 47120]
MW396979.1_Timor-Leste/2019/1    ............................................-................................... [ 47120]
MK940252.1_CN/2019/InnerMongolia ............................................-................................... [ 47120]
MK128995.1_China/2018/AnhuiXCGQ  ............................................-................................... [ 47120]
MH681419.1_ASFV/POL/2015         ............................................-................................... [ 47120]
LS478113.1_Estonia_2014          ............................................-................................... [ 47120]
KP843857.1_Odintsovo_02/14       ............................................-................................... [ 47120]
FN557520.1_strain_E75            ............................................-tc..a..a......t.........tt.a....... [ 47120]
AY261366.1_Warthog               ............................................-tc..a..a........................... [ 47120]
AY261365.1_Warmbaths             .....................................g......-tc..a..a........................... [ 47120]
AY261364.1_Tengani_62            ............................................-............c...................... [ 47120]
AY261363.1_Pretoriuskop/96/4     ............................................-tc..ac.a........................... [ 47120]
AY261362.1_Mkuzi_1979            ............................................-tc..a..a......t.........tt.a....... [ 47120]
AY261361.1_Malawi_Lil-20/1_1983  ......................................a.....-......t.....c..............a....... [ 47120]
AY261360.1_Kenya_1950            .......................t....................-t...ac......c..g.........t......... [ 47120]
AM712240.1_OURT_88/3             ............................................-tc..a..a......t.........tt.a....... [ 47120]
AM712239.1_Benin_97/1            ............................................-tc..a..a......t.........tt.a....... [ 47120]
KM111295.1_Ken06.Bus             .......................t......a.............-t..tac......c......c.....t......... [ 47120]
KM111294.1_Ken05/Tk1             ...........................................g-............c...................... [ 47120]
KM262844.1_strain_L60            ............................................-tc..a..a......t.........tt.a....... [ 47120]
KM102979.1_26544/OG10_Italy      ............................................-tc..a..a......t.........tt.a....... [ 47120]
KM262845.1_strain_NHV            ............................................-tc..a..a......t.........tt.a....... [ 47120]
KX354450.1_47/Ss/2008            ............................................-tc..a..a......t.........tt.a....... [ 47120]
KP055815.1_strain_BA71           ............................................-tc..a..a......t.........tt.a....... [ 47120]
MH025920.1_strain_R35            .......................t......a.............-t..tac......c......c.....t......... [ 47120]
MH025919.1_strain_N10            .......................t......a.............tt..tac......c......c.....t......... [ 47120]
MH025918.1_strain_R25            .......................t......a.............-t..tac......c......c.....t......... [ 47120]
MH025917.1_strain_R7             .......................t......a.............-t..tac......c......c.....t......... [ 47120]
MH025916.1_strain_R8             .......................t......a.............-t..tac......c......c.....t......... [ 47120]

FR682468.2_Georgia_2007/1        cagcctcagagggaaacgaaagggtaataaaacttctgttatcatgggagggaaattttcattatgtgatcataggagct [ 47200]
ASF_IND_20_CAD543                ..................gt............................................................ [ 47200]
LR899193.1_Germany_2020/1        ................................................................................ [ 47200]
MN393477.1_Wuhan_2019            ................................................................................ [ 47200]
MW396979.1_Timor-Leste/2019/1    ................................................................................ [ 47200]
MK940252.1_CN/2019/InnerMongolia ................................................................................ [ 47200]
MK128995.1_China/2018/AnhuiXCGQ  ................................................................................ [ 47200]
MH681419.1_ASFV/POL/2015         ................................................................................ [ 47200]
LS478113.1_Estonia_2014          ................................................................................ [ 47200]
KP843857.1_Odintsovo_02/14       ................................................................................ [ 47200]
```

```
FN557520.1_strain_E75            ....t........c....gt......g......................a.............................. [ 47200]
AY261366.1_Warthog               ...............................t.................a.............................. [ 47200]
AY261365.1_Warmbaths             ................................................................................ [ 47200]
AY261364.1_Tengani_62            ...............................t.................a.............................. [ 47200]
AY261363.1_Pretoriuskop/96/4     ...............................t.....c...........a.............................. [ 47200]
AY261362.1_Mkuzi_1979            ....t........c.....t.a....g......................a.............................. [ 47200]
AY261361.1_Malawi_Lil-20/1_1983  ...............................g.....................g.......................... [ 47200]
AY261360.1_Kenya_1950            ..........................g.................................................g... [ 47200]
AM712240.1_OURT_88/3             ....t........c....gt......g......................a.............................. [ 47200]
AM712239.1_Benin_97/1            ....t........c....gt......g......................a.............................. [ 47200]
KM111295.1_Ken06.Bus             ................t.........g.................................................g... [ 47200]
KM111294.1_Ken05/Tk1             ............................................................................g... [ 47200]
KM262844.1_strain_L60            ....t........c....gt......g......................a.............................. [ 47200]
KM102979.1_26544/OG10_Italy      ....t........c....gt......g......................a.............................. [ 47200]
KM262845.1_strain_NHV            ....t........c....gt......g......................a.............................. [ 47200]
KX354450.1_47/Ss/2008            ....t........c....gt......g......................a.............................. [ 47200]
KP055815.1_strain_BA71           ....t........c....gt......g......................a.............................. [ 47200]
MH025920.1_strain_R35            ................t.........g.................................................g... [ 47200]
MH025919.1_strain_N10            ................t.........g.................................................g... [ 47200]
MH025918.1_strain_R25            ................t.........g.................................................g... [ 47200]
MH025917.1_strain_R7             ................t.........g.................................................g... [ 47200]
MH025916.1_strain_R8             ................t.........g.................................................g... [ 47200]

FR682468.2_Georgia_2007/1        ctagagggtgaccaatatgacctaattcataagtatgatagtcaaattaaagactaccacatgattttatcattgatcca [ 47280]
ASF_IND_20_CAD543                .........................................................................y...... [ 47280]
LR899193.1_Germany_2020/1        ................................................................................ [ 47280]
MN393477.1_Wuhan_2019            ................................................................................ [ 47280]
MW396979.1_Timor-Leste/2019/1    ................................................................................ [ 47280]
MK940252.1_CN/2019/InnerMongolia ................................................................................ [ 47280]
MK128995.1_China/2018/AnhuiXCGQ  ................................................................................ [ 47280]
MH681419.1_ASFV/POL/2015         ................................................................................ [ 47280]
LS478113.1_Estonia_2014          ................................................................................ [ 47280]
KP843857.1_Odintsovo_02/14       ................................................................................ [ 47280]
FN557520.1_strain_E75            ..............t..c.....g.............gc........cg........................cc..... [ 47280]
AY261366.1_Warthog               .........a...gt..c.....g.............g...................................cc..... [ 47280]
AY261365.1_Warmbaths             .....................................g........................a................. [ 47280]
AY261364.1_Tengani_62            .........a...gt..c.....g.............g...................................cc..... [ 47280]
AY261363.1_Pretoriuskop/96/4     .........a...gt..c.....g.............g...................................cc..... [ 47280]
AY261362.1_Mkuzi_1979            ..............t..c.....g.............gc........cg........................cc..... [ 47280]
AY261361.1_Malawi_Lil-20/1_1983  t............g.......................t.....................t.at................. [ 47280]
AY261360.1_Kenya_1950            .......................g............a...................t...g............ct..... [ 47280]
AM712240.1_OURT_88/3             ..............t..c.....g.............gc........cg........................cc..... [ 47280]
AM712239.1_Benin_97/1            ..............t..c.....g.............gc........cg........................cc..... [ 47280]
KM111295.1_Ken06.Bus             .......................g............a...................t...g.t..........ct..... [ 47280]
```

```
KM111294.1_Ken05/Tk1             .......................g............a...................t...g................... [ 47280]
KM262844.1_strain_L60            ..............t..c.....g.............gc........cg........................cc..... [ 47280]
KM102979.1_26544/OG10_Italy      ..............t..c.....g.............gc........cg........................cc..... [ 47280]
KM262845.1_strain_NHV            ..............t..c.....g.............gc........cg........................cc..... [ 47280]
KX354450.1_47/Ss/2008            ..............t..c.....g.............gc........cg........................cc..... [ 47280]
KP055815.1_strain_BA71           ..............t..c.....g.............gc........cg........................cc..... [ 47280]
MH025920.1_strain_R35            .......................g............a...................t...g.t..........ct..... [ 47280]
MH025919.1_strain_N10            .......................g............a...................t...g.t..........ct..... [ 47280]
MH025918.1_strain_R25            .......................g............a...................t...g.t..........ct..... [ 47280]
MH025917.1_strain_R7             .......................g............a...................t...g.t..........ct..... [ 47280]
MH025916.1_strain_R8             .......................g............a...................t...g.t..........ct..... [ 47280]

FR682468.2_Georgia_2007/1        aaatgcaaatacctttgaaaagtgtcatcagttatccaatagtaatatgtggtgtcttatacagaatgctataaaatata [ 47360]
ASF_IND_20_CAD543                .........y...........r......s................................................... [ 47360]
LR899193.1_Germany_2020/1        ................................................................................ [ 47360]
MN393477.1_Wuhan_2019            ................................................................................ [ 47360]
MW396979.1_Timor-Leste/2019/1    ................................................................................ [ 47360]
MK940252.1_CN/2019/InnerMongolia ................................................................................ [ 47360]
MK128995.1_China/2018/AnhuiXCGQ  ................................................................................ [ 47360]
MH681419.1_ASFV/POL/2015         ................................................................................ [ 47360]
LS478113.1_Estonia_2014          ................................................................................ [ 47360]
KP843857.1_Odintsovo_02/14       ................................................................................ [ 47360]
FN557520.1_strain_E75            t........c...........a......g...........t..g.................................... [ 47360]
AY261366.1_Warthog               ...c.................a......g...........t..g.................g.c................ [ 47360]
AY261365.1_Warmbaths             .....................a..............................c........................... [ 47360]
AY261364.1_Tengani_62            ...c.................a......g...........t..g.................g.c................ [ 47360]
AY261363.1_Pretoriuskop/96/4     ...c.................a......g...........t..g.................g.c................ [ 47360]
AY261362.1_Mkuzi_1979            t....................a......g......a....t..g.................................... [ 47360]
AY261361.1_Malawi_Lil-20/1_1983  ..............................c................................................. [ 47360]
AY261360.1_Kenya_1950            .....................a.............a....t...c.......c.....g..................... [ 47360]
AM712240.1_OURT_88/3             t........c...........a......g...........t..g.................................... [ 47360]
AM712239.1_Benin_97/1            t........c...........a......g...........t..g.................................... [ 47360]
KM111295.1_Ken06.Bus             .....................a.............a....t...c.......c.....g..................... [ 47360]
KM111294.1_Ken05/Tk1             .....................a.............a....t...c.......c.....g..................... [ 47360]
KM262844.1_strain_L60            t........c...........a......g...........t..g.................................... [ 47360]
KM102979.1_26544/OG10_Italy      t........c...........a......g...........t..g.................................... [ 47360]
KM262845.1_strain_NHV            t........c...........a......g...........t..g.................................... [ 47360]
KX354450.1_47/Ss/2008            t........c...........a......g...........t..g.................................... [ 47360]
KP055815.1_strain_BA71           t........c...........a......g...........t..g.................................... [ 47360]
MH025920.1_strain_R35            .....................a.............a....t...c.......c.....g..................... [ 47360]
MH025919.1_strain_N10            .....................a.............a....t...c.......c.....g..................... [ 47360]
MH025918.1_strain_R25            .....................a.............a....t...c.......c.....g..................... [ 47360]
MH025917.1_strain_R7             .....................a.............a....t...c.......c.....g..................... [ 47360]
MH025916.1_strain_R8             .....................a.............a....t...c.......c.....g..................... [ 47360]
```

```
FR682468.2_Georgia_2007/1        atatgctccctattctccaaaaacacagaaatattctgacacatgagggagagaatcaggaattgtttgagatggcatgt [ 47440]
ASF_IND_20_CAD543                ................................................................................ [ 47440]
LR899193.1_Germany_2020/1        ................................................................................ [ 47440]
MN393477.1_Wuhan_2019            ................................................................................ [ 47440]
MW396979.1_Timor-Leste/2019/1    ................................................................................ [ 47440]
MK940252.1_CN/2019/InnerMongolia ................................................................................ [ 47440]
MK128995.1_China/2018/AnhuiXCGQ  ................................................................................ [ 47440]
MH681419.1_ASFV/POL/2015         ................................................................................ [ 47440]
LS478113.1_Estonia_2014          ................................................................................ [ 47440]
KP843857.1_Odintsovo_02/14       ................................................................................ [ 47440]
FN557520.1_strain_E75            ................................................................................ [ 47440]
AY261366.1_Warthog               .......tt.......................t........g.a...ttg..............t.......c....... [ 47440]
AY261365.1_Warmbaths             ................................................................................ [ 47440]
AY261364.1_Tengani_62            .......tt.......................t........g.a...ttg..............t.......c....... [ 47440]
AY261363.1_Pretoriuskop/96/4     .......tt.......................t........g.a...ttg..............t.......c....... [ 47440]
AY261362.1_Mkuzi_1979            ................................................................................ [ 47440]
AY261361.1_Malawi_Lil-20/1_1983  .......tt..................a....tg.............................................. [ 47440]
AY261360.1_Kenya_1950            .......tt.......................c........g..........c...................a....... [ 47440]
AM712240.1_OURT_88/3             ................................................................................ [ 47440]
AM712239.1_Benin_97/1            ................................................................................ [ 47440]
KM111295.1_Ken06.Bus             .......tt.......................c........g..........c...........a.......a....... [ 47440]
KM111294.1_Ken05/Tk1             .......tt.......................c........g..........c...................a....... [ 47440]
KM262844.1_strain_L60            ................................................................................ [ 47440]
KM102979.1_26544/OG10_Italy      ................................................................................ [ 47440]
KM262845.1_strain_NHV            ................................................................................ [ 47440]
KX354450.1_47/Ss/2008            ................................................................................ [ 47440]
KP055815.1_strain_BA71           ................................................................................ [ 47440]
MH025920.1_strain_R35            .......tt.......................c........g..........c...........a.......a....... [ 47440]
MH025919.1_strain_N10            .......tt.......................c........g..........c...........a.......a....... [ 47440]
MH025918.1_strain_R25            .......tt.......................c........g..........c...........a.......a....... [ 47440]
MH025917.1_strain_R7             .......tt.......................c........g..........c...........a.......a....... [ 47440]
MH025916.1_strain_R8             .......tt.......................c........g..........c...........a.......a....... [ 47440]

FR682468.2_Georgia_2007/1        gaggaacagaaatatgacatagttttatggataggacaaaccctaatgttaaatgagccggagtttatttttgatatcgc [ 47520]
ASF_IND_20_CAD543                ................................................................................ [ 47520]
LR899193.1_Germany_2020/1        ................................................................................ [ 47520]
MN393477.1_Wuhan_2019            ................................................................................ [ 47520]
MW396979.1_Timor-Leste/2019/1    ................................................................................ [ 47520]
MK940252.1_CN/2019/InnerMongolia ................................................................................ [ 47520]
MK128995.1_China/2018/AnhuiXCGQ  ................................................................................ [ 47520]
MH681419.1_ASFV/POL/2015         ................................................................................ [ 47520]
LS478113.1_Estonia_2014          ................................................................................ [ 47520]
KP843857.1_Odintsovo_02/14       ................................................................................ [ 47520]
```

```
FN557520.1_strain_E75             ................................................................................ [ 47520]
AY261366.1_Warthog                .................t.......................t...t.a...g........a.a.............ct.. [ 47520]
AY261365.1_Warmbaths              .................t.................................g............................ [ 47520]
AY261364.1_Tengani_62             .................t.......................t.........g........a.a.............ct.. [ 47520]
AY261363.1_Pretoriuskop/96/4      .................t..............g........t...t.a...g........a.a.............ct.. [ 47520]
AY261362.1_Mkuzi_1979             ................................................................................ [ 47520]
AY261361.1_Malawi_Lil-20/1_1983   .................t..............................g...............c...........c... [ 47520]
AY261360.1_Kenya_1950             .................t..c..............................g........a...c............... [ 47520]
AM712240.1_OURT_88/3              ................................................................................ [ 47520]
AM712239.1_Benin_97/1             ................................................................................ [ 47520]
KM111295.1_Ken06.Bus              .................t..ca.............................g............g...........c... [ 47520]
KM111294.1_Ken05/Tk1              .................t..c..............................g............c............... [ 47520]
KM262844.1_strain_L60             ................................................................................ [ 47520]
KM102979.1_26544/OG10_Italy       ................................................................................ [ 47520]
KM262845.1_strain_NHV             ................................................................................ [ 47520]
KX354450.1_47/Ss/2008             ................................................................................ [ 47520]
KP055815.1_strain_BA71            ................................................................................ [ 47520]
MH025920.1_strain_R35             .................t..ca.............................g............g...........c... [ 47520]
MH025919.1_strain_N10             .................t..ca.............................g............g...........c... [ 47520]
MH025918.1_strain_R25             .................t..ca.............................g............g...........c... [ 47520]
MH025917.1_strain_R7              .................t..ca.............................g............g...........c... [ 47520]
MH025916.1_strain_R8              .................t..ca.............................g............g...........c... [ 47520]

FR682468.2_Georgia_2007/1         cttcgaacggatagatttttctttattaacaatgggttatagccttctttttgataacaagatgagtagtatagacattc [ 47600]
ASF_IND_20_CAD543                 ....................................................r........................... [ 47600]
LR899193.1_Germany_2020/1         ................................................................................ [ 47600]
MN393477.1_Wuhan_2019             ................................................................................ [ 47600]
MW396979.1_Timor-Leste/2019/1     ................................................................................ [ 47600]
MK940252.1_CN/2019/InnerMongolia  ................................................................................ [ 47600]
MK128995.1_China/2018/AnhuiXCGQ   ................................................................................ [ 47600]
MH681419.1_ASFV/POL/2015          ................................................................................ [ 47600]
LS478113.1_Estonia_2014           ................................................................................ [ 47600]
KP843857.1_Odintsovo_02/14        ................................................................................ [ 47600]
FN557520.1_strain_E75             ....................................................a........................... [ 47600]
AY261366.1_Warthog                ....a..at.g.............g.....t..a..c....................t......---------....... [ 47600]
AY261365.1_Warmbaths              .c..a........................................................................... [ 47600]
AY261364.1_Tengani_62             .......at.g.............g.....t.....c....................t......---ga........... [ 47600]
AY261363.1_Pretoriuskop/96/4      ....a..at.g.............g.....t..a..c....................t......---------....... [ 47600]
AY261362.1_Mkuzi_1979             ................................................................................ [ 47600]
AY261361.1_Malawi_Lil-20/1_1983   .c...c...........................c........g.........................t........... [ 47600]
AY261360.1_Kenya_1950             .c.t.........................t.......................c.......................... [ 47600]
AM712240.1_OURT_88/3              ....................................................a........................... [ 47600]
AM712239.1_Benin_97/1             ....................................................a........................... [ 47600]
KM111295.1_Ken06.Bus              .c.t...........................g................................................ [ 47600]
```

```
KM111294.1_Ken05/Tk1            .c.t.........................t.......................c.......................... [ 47600]
KM262844.1_strain_L60           ....................................................a........................... [ 47600]
KM102979.1_26544/OG10_Italy     ....................................................a........................... [ 47600]
KM262845.1_strain_NHV           ....................................................a........................... [ 47600]
KX354450.1_47/Ss/2008           ....................................................a........................... [ 47600]
KP055815.1_strain_BA71          ....................................................a........................... [ 47600]
MH025920.1_strain_R35           .c.t...........................g................................................ [ 47600]
MH025919.1_strain_N10           .c.t...........................g................................................ [ 47600]
MH025918.1_strain_R25           .c.t...........................g................................................ [ 47600]
MH025917.1_strain_R7            .c.t...........................g................................................ [ 47600]
MH025916.1_strain_R8            .c.t...........................g................................................ [ 47600]

FR682468.2_Georgia_2007/1       atgatgaagaagatcttacttcattaccaacagaacacctcgaaaaagcagccactaagggatgtttcttctttatgcta [ 47680]
ASF_IND_20_CAD543               ................................................................................ [ 47680]
LR899193.1_Germany_2020/1       ................................................................................ [ 47680]
MN393477.1_Wuhan_2019           ................................................................................ [ 47680]
MW396979.1_Timor-Leste/2019/1   ................................................................................ [ 47680]
MK940252.1_CN/2019/InnerMongolia ...................................................................t............ [ 47680]
MK128995.1_China/2018/AnhuiXCGQ ................................................................................ [ 47680]
MH681419.1_ASFV/POL/2015        ................................................................................ [ 47680]
LS478113.1_Estonia_2014         ................................................................................ [ 47680]
KP843857.1_Odintsovo_02/14      ................................................................................ [ 47680]
FN557520.1_strain_E75           ..................t........t.................................................... [ 47680]
AY261366.1_Warthog              ------....t................t.....................g..tg........cag..g.a.......t.. [ 47680]
AY261365.1_Warmbaths            ...........................t..................................cag..g.........t.. [ 47680]
AY261364.1_Tengani_62           .a.....g....c........t.....tc.tgc....t.......t...g..tg........cag..g.a.......t.. [ 47680]
AY261363.1_Pretoriuskop/96/4    ------....t................t.................t...g..tg........cag..g.a.......t.. [ 47680]
AY261362.1_Mkuzi_1979           ..................t........t.................................................... [ 47680]
AY261361.1_Malawi_Lil-20/1_1983 ........................c..t..........................t..................c...... [ 47680]
AY261360.1_Kenya_1950           ...........................t............t...........t............c............a. [ 47680]
AM712240.1_OURT_88/3            ..................t........t.................................................... [ 47680]
AM712239.1_Benin_97/1           ..................t........t.................................................... [ 47680]
KM111295.1_Ken06.Bus            ...........................t............ta..........t.....a......c............a. [ 47680]
KM111294.1_Ken05/Tk1            ...........................t............t...........t............c............a. [ 47680]
KM262844.1_strain_L60           ..................t........t.................................................... [ 47680]
KM102979.1_26544/OG10_Italy     ..................t........t.................................................... [ 47680]
KM262845.1_strain_NHV           ..................t........t.................................................... [ 47680]
KX354450.1_47/Ss/2008           ..................t........t.................................................... [ 47680]
KP055815.1_strain_BA71          ..................t........t.................................................... [ 47680]
MH025920.1_strain_R35           ...........................t............ta..........t.....a......c............a. [ 47680]
MH025919.1_strain_N10           ...........................t............ta..........t.....a......c............a. [ 47680]
MH025918.1_strain_R25           ...........................t............ta..........t.....a......c............a. [ 47680]
MH025917.1_strain_R7            ...........................t............ta..........t.....a......c............a. [ 47680]
MH025916.1_strain_R8            ...........................t............ta..........t.....a......c............a. [ 47680]
```

```
FR682468.2_Georgia_2007/1        gaaactttaaaacatggtggaaatgtaaatatggcagtcttatctaaagctgttgagtataatcatagaaaaattttaga [ 47760]
ASF_IND_20_CAD543                ...........................................................................y.... [ 47760]
LR899193.1_Germany_2020/1        ................................................................................ [ 47760]
MN393477.1_Wuhan_2019            ................................................................................ [ 47760]
MW396979.1_Timor-Leste/2019/1    ................................................................................ [ 47760]
MK940252.1_CN/2019/InnerMongolia ................................................................................ [ 47760]
MK128995.1_China/2018/AnhuiXCGQ  ................................................................................ [ 47760]
MH681419.1_ASFV/POL/2015         ................................................................................ [ 47760]
LS478113.1_Estonia_2014          ................................................................................ [ 47760]
KP843857.1_Odintsovo_02/14       ................................................................................ [ 47760]
FN557520.1_strain_E75            ...........................................................................c.... [ 47760]
AY261366.1_Warthog               .....................g........t.c.ag..t..............c...a..g..............c.... [ 47760]
AY261365.1_Warmbaths             ..............................t....g..t..............c.....................c.... [ 47760]
AY261364.1_Tengani_62            ..............................t.c.ag..t..............c...a..g..............c.... [ 47760]
AY261363.1_Pretoriuskop/96/4     .....................g........t.c.ag..t..............c...a..g..............c.... [ 47760]
AY261362.1_Mkuzi_1979            ...........................................................................c.... [ 47760]
AY261361.1_Malawi_Lil-20/1_1983  ..................................................c............................. [ 47760]
AY261360.1_Kenya_1950            .................c..............a..g............................................ [ 47760]
AM712240.1_OURT_88/3             ...........................................................................c.... [ 47760]
AM712239.1_Benin_97/1            ...........................................................................c.... [ 47760]
KM111295.1_Ken06.Bus             ....t.......t.......g...........a..g.................ca......................... [ 47760]
KM111294.1_Ken05/Tk1             .................c..............a..g............................................ [ 47760]
KM262844.1_strain_L60            ...........................................................................c.... [ 47760]
KM102979.1_26544/OG10_Italy      ...........................................................................c.... [ 47760]
KM262845.1_strain_NHV            ...........................................................................c.... [ 47760]
KX354450.1_47/Ss/2008            ...........................................................................c.... [ 47760]
KP055815.1_strain_BA71           ...........................................................................c.... [ 47760]
MH025920.1_strain_R35            ....t.......t.......g...........a..g.................ca......................... [ 47760]
MH025919.1_strain_N10            ....t.......t.......g...........a..g.................ca......................... [ 47760]
MH025918.1_strain_R25            ....t.......t.......g...........a..g.................ca......................... [ 47760]
MH025917.1_strain_R7             ....t.......t.......g...........a..g.................ca......................... [ 47760]
MH025916.1_strain_R8             ....t.......t.......g...........a..g.................ca......................... [ 47760]

FR682468.2_Georgia_2007/1        ccattttattcggcggcaaaaatgtttatcacgtgaagagattgaaaacctattattaaccgccataaccaattgtgcat [ 47840]
ASF_IND_20_CAD543                .y.......---a....................yr....k..................gta................... [ 47840]
LR899193.1_Germany_2020/1        ................................................................................ [ 47840]
MN393477.1_Wuhan_2019            ................................................................................ [ 47840]
MW396979.1_Timor-Leste/2019/1    ................................................................................ [ 47840]
MK940252.1_CN/2019/InnerMongolia ................................................................................ [ 47840]
MK128995.1_China/2018/AnhuiXCGQ  ................................................................................ [ 47840]
MH681419.1_ASFV/POL/2015         ................................................................................ [ 47840]
LS478113.1_Estonia_2014          ................................................................................ [ 47840]
KP843857.1_Odintsovo_02/14       ................................................................................ [ 47840]
```

```
FN557520.1_strain_E75            .t.......---a....................ca....t........a.........gta......t.....a...... [ 47840]
AY261366.1_Warthog               .t.................c...................t..................t..................... [ 47840]
AY261365.1_Warmbaths             .t.................c...................t..c...............t..................... [ 47840]
AY261364.1_Tengani_62            .t.................c...................t................................a....... [ 47840]
AY261363.1_Pretoriuskop/96/4     .t.................c...a...............t..................t..................... [ 47840]
AY261362.1_Mkuzi_1979            .t.......---a....................ca....t........a.........gta......t.....a...... [ 47840]
AY261361.1_Malawi_Lil-20/1_1983  .t............................g.................a.........t.a.g....t.t...g.c.... [ 47840]
AY261360.1_Kenya_1950            .t...............................ca....t........a..........ta..t..t.aag..g.c.g.. [ 47840]
AM712240.1_OURT_88/3             .t.......---a....................ca....t........a.........gta......t.....a...... [ 47840]
AM712239.1_Benin_97/1            .t.......---a....................ca....t........a.........gta......t.....a...... [ 47840]
KM111295.1_Ken06.Bus             .t...............................c....gt........a...ac.....ta.....t.aatg.g.c.g.. [ 47840]
KM111294.1_Ken05/Tk1             .t...............................ca....t........a..........ta..t..t.aac..g.c.g.. [ 47840]
KM262844.1_strain_L60            .t.......---a....................ca....t........a.........gta......t.....a...... [ 47840]
KM102979.1_26544/OG10_Italy      .t.......---a....................ca....t........a.........gta......t.....a...... [ 47840]
KM262845.1_strain_NHV            .t.......---a....................ca....t........a.........gta......t.....a...... [ 47840]
KX354450.1_47/Ss/2008            .t.......---a....................ca....t........a.........gta......t.....a...... [ 47840]
KP055815.1_strain_BA71           .t.......---a....................ca....t........a.........gta......t.....a...... [ 47840]
MH025920.1_strain_R35            .t...............................c....gt........a...ac.....ta.....t.aatg.g.c.g.. [ 47840]
MH025919.1_strain_N10            .t...............................c....gt........a...ac.....ta.....t.aatg.g.c.g.. [ 47840]
MH025918.1_strain_R25            .t...............................c....gt........a...ac.....ta.....t.aatg.g.c.g.. [ 47840]
MH025917.1_strain_R7             .t...............................c....gt........a...ac.....ta.....t.aatg.g.c.g.. [ 47840]
MH025916.1_strain_R8             .t...............................c....gt........a...ac.....ta.....t.aatg.g.c.g.. [ 47840]

FR682468.2_Georgia_2007/1        ccataaaaacgttaaacttactcttgtcttacctaaactattccgtaaaaaatatcattggaaaaatagtacaacatgtc [ 47920]
ASF_IND_20_CAD543                ..........................................................................tc...t [ 47920]
LR899193.1_Germany_2020/1        ................................................................................ [ 47920]
MN393477.1_Wuhan_2019            ................................................................................ [ 47920]
MW396979.1_Timor-Leste/2019/1    ................................................................................ [ 47920]
MK940252.1_CN/2019/InnerMongolia ................................................................................ [ 47920]
MK128995.1_China/2018/AnhuiXCGQ  ................................................................................ [ 47920]
MH681419.1_ASFV/POL/2015         ................................................................................ [ 47920]
LS478113.1_Estonia_2014          ................................................................................ [ 47920]
KP843857.1_Odintsovo_02/14       ................................................................................ [ 47920]
FN557520.1_strain_E75            ...a......t...........................c...................................tc...t [ 47920]
AY261366.1_Warthog               ...a......t....................t............a............c..a................... [ 47920]
AY261365.1_Warmbaths             ...a...........................t......c.....a................................... [ 47920]
AY261364.1_Tengani_62            ...a.......................t...tt...........a.............................a..a.. [ 47920]
AY261363.1_Pretoriuskop/96/4     ...a......t....................t............a............c..a................... [ 47920]
AY261362.1_Mkuzi_1979            ...a......t...........................c...................................tc.a.t [ 47920]
AY261361.1_Malawi_Lil-20/1_1983  .........................a.....t..................c....t...a..............t..... [ 47920]
AY261360.1_Kenya_1950            ...a.............c.......a.....t...............c.......g...............g..gc.... [ 47920]
AM712240.1_OURT_88/3             ...a......t...........................c...................................tc...t [ 47920]
AM712239.1_Benin_97/1            ...a......t...........................c...................................tc...t [ 47920]
KM111295.1_Ken06.Bus             .t.a.............c.............t...............c.......g...............g..gc.... [ 47920]
```

```
KM111294.1_Ken05/Tk1           ...a.............c.......a.....t...............c.......g...............g..gc.... [ 47920]
KM262844.1_strain_L60          ...a......t...........................c...................................tc...t [ 47920]
KM102979.1_26544/OG10_Italy    ...a......t...........................c...................................tc...t [ 47920]
KM262845.1_strain_NHV          ...a......t...........................c...................................tc...t [ 47920]
KX354450.1_47/Ss/2008          ...a......t...........................c...................................tc...t [ 47920]
KP055815.1_strain_BA71         ...a......t...........................c...................................tc...t [ 47920]
MH025920.1_strain_R35          .t.a.............c.............t...............c.......g...............g..gc.... [ 47920]
MH025919.1_strain_N10          .t.a.............c.............t...............c.......g...............g..gc.... [ 47920]
MH025918.1_strain_R25          .t.a.............c.............t...............c.......g...............g..gc.... [ 47920]
MH025917.1_strain_R7           .t.a.............c.............t...............c.......g...............g..gc.... [ 47920]
MH025916.1_strain_R8           .t.a.............c.............t...............c.......g...............g..gc.... [ 47920]

FR682468.2_Georgia_2007/1      ataaaagatggtgattataccatcatattacttttaaaaaaaaagaaaataaacctagtggaacctgttttaacaggttt [ 48000]
ASF_IND_20_CAD543              t.....r.........w..........w..t................................................. [ 48000]
LR899193.1_Germany_2020/1      ................................................................................ [ 48000]
MN393477.1_Wuhan_2019          ................................................................................ [ 48000]
MW396979.1_Timor-Leste/2019/1  ................................................................................ [ 48000]
MK940252.1_CN/2019/InnerMongolia ................................................................................ [ 48000]
MK128995.1_China/2018/AnhuiXCGQ ................................................................................ [ 48000]
MH681419.1_ASFV/POL/2015       ................................................................................ [ 48000]
LS478113.1_Estonia_2014        ................................................................................ [ 48000]
KP843857.1_Odintsovo_02/14     ................................................................................ [ 48000]
FN557520.1_strain_E75          t.....a.........t..........a..t....................................c...g.t...... [ 48000]
AY261366.1_Warthog             .......g...................a...........................................g........ [ 48000]
AY261365.1_Warmbaths           ................t..........a...........................................g........ [ 48000]
AY261364.1_Tengani_62          .......g...................a...........................................g........ [ 48000]
AY261363.1_Pretoriuskop/96/4   .......g...................a...........................................g........ [ 48000]
AY261362.1_Mkuzi_1979          t.....a.........t.t........a..t....................................c...g.t...... [ 48000]
AY261361.1_Malawi_Lil-20/1_1983 ........a..c....t.........cg..g.........g...a.................g........gt....... [ 48000]
AY261360.1_Kenya_1950          .....t..........t..........a.ca.......................t.......ct.c.....gg.g..... [ 48000]
AM712240.1_OURT_88/3           t.....a.........t..........a..t....................................c...g.t...... [ 48000]
AM712239.1_Benin_97/1          t.....a.........t..........a..t....................................c...g.t...... [ 48000]
KM111295.1_Ken06.Bus           .....t..........t..........a.ca.......................t.......c..c.....ggtg..... [ 48000]
KM111294.1_Ken05/Tk1           .....t..........t..........a.ca.......................t.......ct.c.....gg.g..... [ 48000]
KM262844.1_strain_L60          t.....a.........t..........a..t....................................c...g.t...... [ 48000]
KM102979.1_26544/OG10_Italy    t.....a.........t..........a..t....................................c...g.t...... [ 48000]
KM262845.1_strain_NHV          t.....a.........t..........a..t....................................c...g.t...... [ 48000]
KX354450.1_47/Ss/2008          t.....a.........t..........a..t....................................c...g.t...... [ 48000]
KP055815.1_strain_BA71         t.....a.........t..........a..t....................................c...g.t...... [ 48000]
MH025920.1_strain_R35          .....t..........t..........a.ca.......................t.......c..c.....gg.g..... [ 48000]
MH025919.1_strain_N10          .....t..........t..........a.ca.......................t.......c..c.....gg.g..... [ 48000]
MH025918.1_strain_R25          .....t..........t..........a.ca.......................t.......c..c.....gg.g..... [ 48000]
MH025917.1_strain_R7           .....t..........t..........a.ca.......................t.......c..c.....gg.g..... [ 48000]
MH025916.1_strain_R8           .....t..........t..........a.ca.......................t.......c..c.....gg.g..... [ 48000]
```

```
FR682468.2_Georgia_2007/1        tatagattattactatagctattgttttataaaacattttatccaagagtttgctattcgtccggaaaaactgattaaaa [ 48080]
ASF_IND_20_CAD543                ............................w..r....w........w.......ac...............m......... [ 48080]
LR899193.1_Germany_2020/1        ................................................................................ [ 48080]
MN393477.1_Wuhan_2019            ................................................................................ [ 48080]
MW396979.1_Timor-Leste/2019/1    ................................................................................ [ 48080]
MK940252.1_CN/2019/InnerMongolia ................................................................................ [ 48080]
MK128995.1_China/2018/AnhuiXCGQ  ................................................................................ [ 48080]
MH681419.1_ASFV/POL/2015         ................................................................................ [ 48080]
LS478113.1_Estonia_2014          ................................................................................ [ 48080]
KP843857.1_Odintsovo_02/14       ................................................................................ [ 48080]
FN557520.1_strain_E75            ....a.......................t..g....a........t.......ac...............a......... [ 48080]
AY261366.1_Warthog               .......c.g......................g..a.........gt.........c.............g........... [ 48080]
AY261365.1_Warmbaths             .......c...........a...........g..a.g........t........c.......................... [ 48080]
AY261364.1_Tengani_62            .......c...........ac..........g..a..........t........c...............g....c.... [ 48080]
AY261363.1_Pretoriuskop/96/4     .......c........g..a...........g..a.........gt..a.....c...............g......... [ 48080]
AY261362.1_Mkuzi_1979            ....a.......................t..g....a........t.......ac...............a......... [ 48080]
AY261361.1_Malawi_Lil-20/1_1983  ...g...g.............................g......g.t.......ac...............a.a....... [ 48080]
AY261360.1_Kenya_1950            .t.....c...ttc...c..................gt.........t........................a.a....... [ 48080]
AM712240.1_OURT_88/3             ....a.......................t..g....a........t.......ac...............a......... [ 48080]
AM712239.1_Benin_97/1            ....a.......................t..g....a........t.......ac...............a......... [ 48080]
KM111295.1_Ken06.Bus             .t.g...ct..t.c......................gt.........t........................a.a....... [ 48080]
KM111294.1_Ken05/Tk1             .t.....c...t.c...c..................gt.........t........................a.a....... [ 48080]
KM262844.1_strain_L60            ....a.......................t..g....a........t.......ac...............a......... [ 48080]
KM102979.1_26544/OG10_Italy      ....a.......................t..g....a........t.......ac...............a......... [ 48080]
KM262845.1_strain_NHV            ....a.......................t..g....a........t.......ac...............a......... [ 48080]
KX354450.1_47/Ss/2008            ....a.......................t..g....a........t.......ac...............a......... [ 48080]
KP055815.1_strain_BA71           ....a.......................t..g....a........t.......ac...............a......... [ 48080]
MH025920.1_strain_R35            .t.g...ct..t.c......................gt.........t........................a.a....... [ 48080]
MH025919.1_strain_N10            .t.g...ct..t.c......................gt.........t........................a.a....... [ 48080]
MH025918.1_strain_R25            .t.g...ct..t.c......................gt.........t........................a.a....... [ 48080]
MH025917.1_strain_R7             .t.g...ct..t.c......................gt.........t........................a.a....... [ 48080]
MH025916.1_strain_R8             .t.g...ct..t.c......................gt.........t........................a.a....... [ 48080]

FR682468.2_Georgia_2007/1        tggccgcgcgaaaaggtaaactaaatatgattatcgaattccttaacgaaaaatatgttcataaagatgatcttggaact [ 48160]
ASF_IND_20_CAD543                ...............................y........tt.a..t..............................r.y [ 48160]
LR899193.1_Germany_2020/1        ................................................................................ [ 48160]
MN393477.1_Wuhan_2019            ................................................................................ [ 48160]
MW396979.1_Timor-Leste/2019/1    ................................................................................ [ 48160]
MK940252.1_CN/2019/InnerMongolia ................................................................................ [ 48160]
MK128995.1_China/2018/AnhuiXCGQ  ................................................................................ [ 48160]
MH681419.1_ASFV/POL/2015         ................................................................................ [ 48160]
LS478113.1_Estonia_2014          ................................................................................ [ 48160]
KP843857.1_Odintsovo_02/14       ................................................................................ [ 48160]
```

```
FN557520.1_strain_E75             ...........................g...........c........tt.a..t..............................g.c [ 48160]
AY261366.1_Warthog                ....t..a......................................t................................. [ 48160]
AY261365.1_Warmbaths              ....t........................................................................... [ 48160]
AY261364.1_Tengani_62             ....t........................................................c.................. [ 48160]
AY261363.1_Pretoriuskop/96/4      ....t........................................................................... [ 48160]
AY261362.1_Mkuzi_1979             ...................g...........c........tt.a..t..............................g.c [ 48160]
AY261361.1_Malawi_Lil-20/1_1983   ....t...............t...................tt....t....t....cc................aa.... [ 48160]
AY261360.1_Kenya_1950             ...............a.......g....a.....t.....tt....t..t.t....cc...................... [ 48160]
AM712240.1_OURT_88/3              ...................g...........c........tt.a..t..............................g.c [ 48160]
AM712239.1_Benin_97/1             ...................g...........c........tt.a..t..............................g.c [ 48160]
KM111295.1_Ken06.Bus              ...............a.......g....a.....t.....tt....t..t.t....cc.................a.... [ 48160]
KM111294.1_Ken05/Tk1              ...............a.......g....a.....t.....tt....t..t.t....cc...................... [ 48160]
KM262844.1_strain_L60             ...................g...........c........tt.a..t..............................g.c [ 48160]
KM102979.1_26544/OG10_Italy       ...................g...........c........tt.a..t..............................g.c [ 48160]
KM262845.1_strain_NHV             ...................g...........c........tt.a..t..............................g.c [ 48160]
KX354450.1_47/Ss/2008             ...................g...........c........tt.a..t..............................g.c [ 48160]
KP055815.1_strain_BA71            ...................g...........c........tt.a..t..............................g.c [ 48160]
MH025920.1_strain_R35             ...............a.......g....a.....t.....tt....t..t.t....cc.................a.... [ 48160]
MH025919.1_strain_N10             ...............a.......g....a.....t.....tt....t..t.t....cc.................a.... [ 48160]
MH025918.1_strain_R25             ...............a.......g....a.....t.....tt....t..t.t....cc.................a.... [ 48160]
MH025917.1_strain_R7              ...............a.......g....a.....t.....tt....t..t.t....cc.................a.... [ 48160]
MH025916.1_strain_R8              ...............a.......g....a.....t.....tt....t..t.t....cc.................a.... [ 48160]

FR682468.2_Georgia_2007/1         atatttaaatatctcaaaaccctagtatgtaccatgaaacataaaaaaggaaaagagacattaattgttcttattcataa [ 48240]
ASF_IND_20_CAD543                 ..........w..................................................................... [ 48240]
LR899193.1_Germany_2020/1         ................................................................................ [ 48240]
MN393477.1_Wuhan_2019             ................................................................................ [ 48240]
MW396979.1_Timor-Leste/2019/1     ................................................................................ [ 48240]
MK940252.1_CN/2019/InnerMongolia  ................................................................................ [ 48240]
MK128995.1_China/2018/AnhuiXCGQ   ................................................................................ [ 48240]
MH681419.1_ASFV/POL/2015          ................................................................................ [ 48240]
LS478113.1_Estonia_2014           ................................................................................ [ 48240]
KP843857.1_Odintsovo_02/14        ................................................................................ [ 48240]
FN557520.1_strain_E75             ..........t........a............................................................ [ 48240]
AY261366.1_Warthog                .........ct........a..............................................a............. [ 48240]
AY261365.1_Warmbaths              .........ct........at.....................................g......a............. [ 48240]
AY261364.1_Tengani_62             .........ct........a..............................................a...........c. [ 48240]
AY261363.1_Pretoriuskop/96/4      .........ct........a...........................................c..a...........g. [ 48240]
AY261362.1_Mkuzi_1979             ..........t........a............................................................ [ 48240]
AY261361.1_Malawi_Lil-20/1_1983   ..g.......t........a........a............................gtg.....c.g............ [ 48240]
AY261360.1_Kenya_1950             .....a..tat........a.a.....aa................g...........gt........g............ [ 48240]
AM712240.1_OURT_88/3              ..........t........a............................................................ [ 48240]
AM712239.1_Benin_97/1             ..........t........a............................................................ [ 48240]
KM111295.1_Ken06.Bus              g.......tat........a.a.....aa................g............t........g............ [ 48240]
```

| | | |
|---|---|---|
| KM111294.1_Ken05/Tk1 | .....a..tat........ata.....aa................g...........gt........g............ | [ 48240] |
| KM262844.1_strain_L60 | ..........t........a............................................................ | [ 48240] |
| KM102979.1_26544/OG10_Italy | ..........t........a............................................................ | [ 48240] |
| KM262845.1_strain_NHV | ..........t........a............................................................ | [ 48240] |
| KX354450.1_47/Ss/2008 | ..........t........a............................................................ | [ 48240] |
| KP055815.1_strain_BA71 | ..........t........a............................................................ | [ 48240] |
| MH025920.1_strain_R35 | g.......tat........a.a.....aa................g............t........g............ | [ 48240] |
| MH025919.1_strain_N10 | g.......tat........a.a.....aa................g............t........g............ | [ 48240] |
| MH025918.1_strain_R25 | g.......tat........a.a.....aa................g............t........g............ | [ 48240] |
| MH025917.1_strain_R7 | g.......tat........a.a.....aa................g............t........g............ | [ 48240] |
| MH025916.1_strain_R8 | g.......tat........a.a.....aa................g............t........g............ | [ 48240] |

| | | |
|---|---|---|
| FR682468.2_Georgia_2007/1 | aatatatcaagatattcatctggagactaaagaaaaatttaaattattaagattttatgtcatgcatgatgcaactatcc | [ 48320] |
| ASF_IND_20_CAD543 | ................................................................................ | [ 48320] |
| LR899193.1_Germany_2020/1 | ................................................................................ | [ 48320] |
| MN393477.1_Wuhan_2019 | ................................................................................ | [ 48320] |
| MW396979.1_Timor-Leste/2019/1 | ................................................................................ | [ 48320] |
| MK940252.1_CN/2019/InnerMongolia | ................................................................................ | [ 48320] |
| MK128995.1_China/2018/AnhuiXCGQ | ................................................................................ | [ 48320] |
| MH681419.1_ASFV/POL/2015 | ................................................................................ | [ 48320] |
| LS478113.1_Estonia_2014 | ................................................................................ | [ 48320] |
| KP843857.1_Odintsovo_02/14 | ................................................................................ | [ 48320] |
| FN557520.1_strain_E75 | ...........t......a......................................c...................... | [ 48320] |
| AY261366.1_Warthog | ..................................................a......c...................... | [ 48320] |
| AY261365.1_Warmbaths | ...........c............t................................c...................... | [ 48320] |
| AY261364.1_Tengani_62 | .........................................................c...................... | [ 48320] |
| AY261363.1_Pretoriuskop/96/4 | ........g..c............t.....................................................t. | [ 48320] |
| AY261362.1_Mkuzi_1979 | ...........t......a......................................c...................... | [ 48320] |
| AY261361.1_Malawi_Lil-20/1_1983 | ..........tc.........a....a........tg.....t.......a......c......t..a......a..... | [ 48320] |
| AY261360.1_Kenya_1950 | ........g.at..............a........tg.....t..gg..c.......ca.t......a......a..... | [ 48320] |
| AM712240.1_OURT_88/3 | ...........t......a......................................c...................... | [ 48320] |
| AM712239.1_Benin_97/1 | ...........t......a......................................c...................... | [ 48320] |
| KM111295.1_Ken06.Bus | ........g.at..............a........tg.....g..gg..c.......c.........a......a..... | [ 48320] |
| KM111294.1_Ken05/Tk1 | ........g.at..............a........tg.....t..gg..c.......ca........a......a..... | [ 48320] |
| KM262844.1_strain_L60 | ...........t......a......................................c...................... | [ 48320] |
| KM102979.1_26544/OG10_Italy | ...........t......a......................................c...................... | [ 48320] |
| KM262845.1_strain_NHV | ...........t......a......................................c...................... | [ 48320] |
| KX354450.1_47/Ss/2008 | ...........t......a......................................c...................... | [ 48320] |
| KP055815.1_strain_BA71 | ...........t......a......................................c...................... | [ 48320] |
| MH025920.1_strain_R35 | ........g.at..............a........tg.....g..gg..c.......c.........a......a..... | [ 48320] |
| MH025919.1_strain_N10 | ........g.at..............a........tg.....g..gg..c.......c.........a......a..... | [ 48320] |
| MH025918.1_strain_R25 | ........g.at..............a........tg.....g..gg..c.......c.........a......a..... | [ 48320] |
| MH025917.1_strain_R7 | ........g.at..............a........tg.....g..gg..c.......c.........a......a..... | [ 48320] |
| MH025916.1_strain_R8 | ........g.at..............a........tg.....g..gg..c.......c.........a......a..... | [ 48320] |

```
FR682468.2_Georgia_2007/1        aatttctatctatgtgcaaagactgttttaatttagccggttttaaaccatttgttttagaatgtttggatattgctatt [ 48400]
ASF_IND_20_CAD543                ...............r.....................................s.......................... [ 48400]
LR899193.1_Germany_2020/1        ................................................................................ [ 48400]
MN393477.1_Wuhan_2019            ................................................................................ [ 48400]
MW396979.1_Timor-Leste/2019/1    ................................................................................ [ 48400]
MK940252.1_CN/2019/InnerMongolia ................................................................................ [ 48400]
MK128995.1_China/2018/AnhuiXCGQ  ................................................................................ [ 48400]
MH681419.1_ASFV/POL/2015         ................................................................................ [ 48400]
LS478113.1_Estonia_2014          ................................................................................ [ 48400]
KP843857.1_Odintsovo_02/14       ................................................................................ [ 48400]
FN557520.1_strain_E75            .....a.........a.....................................c.......................... [ 48400]
AY261366.1_Warthog               ..................................g............................................. [ 48400]
AY261365.1_Warmbaths             ................................................................................ [ 48400]
AY261364.1_Tengani_62            ..................................gc.................a.......................... [ 48400]
AY261363.1_Pretoriuskop/96/4     ................................................................................ [ 48400]
AY261362.1_Mkuzi_1979            .....a...............................................c.......................... [ 48400]
AY261361.1_Malawi_Lil-20/1_1983  .....a..g..c.t.................a.................g...c.......................... [ 48400]
AY261360.1_Kenya_1950            .....a....g..t.......................t...............c...c...................... [ 48400]
AM712240.1_OURT_88/3             .....a.........a.....................................c.......................... [ 48400]
AM712239.1_Benin_97/1            .....a.........a.....................................c.......................... [ 48400]
KM111295.1_Ken06.Bus             .....a.......t.......................................c...c...................... [ 48400]
KM111294.1_Ken05/Tk1             .....a....g..t.......................................c...c...................... [ 48400]
KM262844.1_strain_L60            .....a.........a.....................................c.......................... [ 48400]
KM102979.1_26544/OG10_Italy      .....a.........a.....................................c.......................... [ 48400]
KM262845.1_strain_NHV            .....a.........a.....................................c.......................... [ 48400]
KX354450.1_47/Ss/2008            .....a.........a.....................................c.......................... [ 48400]
KP055815.1_strain_BA71           .....a.........a.....................................c.......................... [ 48400]
MH025920.1_strain_R35            .....a.......t.......................................c...c...................... [ 48400]
MH025919.1_strain_N10            .....a.......t.......................................c...c...................... [ 48400]
MH025918.1_strain_R25            .....a.......t.......................................c...c...................... [ 48400]
MH025917.1_strain_R7             .....a.......t.......................................c...c...................... [ 48400]
MH025916.1_strain_R8             .....a.......t.......................................c...c...................... [ 48400]

FR682468.2_Georgia_2007/1        aaaaaaaattaccctgatatgatacaatatatagaaattctatcgaaatctgagtaaaatttatttttttgatcagagta [ 48480]
ASF_IND_20_CAD543                .........................g.a.........y...r.y.....s.............................. [ 48480]
LR899193.1_Germany_2020/1        ................................................................................ [ 48480]
MN393477.1_Wuhan_2019            ................................................................................ [ 48480]
MW396979.1_Timor-Leste/2019/1    ................................................................................ [ 48480]
MK940252.1_CN/2019/InnerMongolia ................................................................................ [ 48480]
MK128995.1_China/2018/AnhuiXCGQ  ................................................................................ [ 48480]
MH681419.1_ASFV/POL/2015         ................................................................................ [ 48480]
LS478113.1_Estonia_2014          ................................................................................ [ 48480]
KP843857.1_Odintsovo_02/14       -............................................................................... [ 48480]
```

```
FN557520.1_strain_E75            .........................g.a.........c...g.t.....g.............................. [ 48480]
AY261366.1_Warthog               .........c.....a...........a.............g.......a.............................. [ 48480]
AY261365.1_Warmbaths             .........c.....a...........a.............g.......a.............................. [ 48480]
AY261364.1_Tengani_62            .........c.....a...........a.................................................... [ 48480]
AY261363.1_Pretoriuskop/96/4     .........c.....a........g..a.............g.......a.............................. [ 48480]
AY261362.1_Mkuzi_1979            .........................g.a.............g.t.....g.............................. [ 48480]
AY261361.1_Malawi_Lil-20/1_1983  ....g...............t......a.....a..c......ta....a.............................. [ 48480]
AY261360.1_Kenya_1950            ..............c.....tg...g.a.....a......a..t.....a........................t..... [ 48480]
AM712240.1_OURT_88/3             .........................g.a.........c...g.t.....g.............................. [ 48480]
AM712239.1_Benin_97/1            .........................g.a.........c...g.t.....g.............................. [ 48480]
KM111295.1_Ken06.Bus             ..............c.....tg...g.a.....a......a..t.....a........................t..... [ 48480]
KM111294.1_Ken05/Tk1             ..............c.....tg...g.a.....a......a..t.....a........................t..... [ 48480]
KM262844.1_strain_L60            .........................g.a.........c...g.t.....g.............................. [ 48480]
KM102979.1_26544/OG10_Italy      .........................g.a.........c...g.t.....g.............................. [ 48480]
KM262845.1_strain_NHV            .........................g.a.........c...g.t.....g.............................. [ 48480]
KX354450.1_47/Ss/2008            .........................g.a.........c...g.t.....g.............................. [ 48480]
KP055815.1_strain_BA71           .........................g.a.........c...g.t.....g.............................. [ 48480]
MH025920.1_strain_R35            ..............c.....tg...g.a.....a......a..t.....a........................t..... [ 48480]
MH025919.1_strain_N10            ..............c.....tg...g.a.....a......a..t.....a........................t..... [ 48480]
MH025918.1_strain_R25            ..............c.....tg...g.a.....a......a..t.....a........................t..... [ 48480]
MH025917.1_strain_R7             ..............c.....tg...g.a.....a......a..t.....a........................t..... [ 48480]
MH025916.1_strain_R8             ..............c.....tg...g.a.....a......a..t.....a........................t..... [ 48480]

FR682468.2_Georgia_2007/1        agaaaatgttctccctccaggagatctgtcgaaagaacatctactttctacctgactggctcggtgagcatgtgattcag [ 48560]
ASF_IND_20_CAD543                ................................................................................ [ 48560]
LR899193.1_Germany_2020/1        ................................................................................ [ 48560]
MN393477.1_Wuhan_2019            ................................................................................ [ 48560]
MW396979.1_Timor-Leste/2019/1    ................................................................................ [ 48560]
MK940252.1_CN/2019/InnerMongolia ................................................................................ [ 48560]
MK128995.1_China/2018/AnhuiXCGQ  ................................................................................ [ 48560]
MH681419.1_ASFV/POL/2015         ................................................................................ [ 48560]
LS478113.1_Estonia_2014          ................................................................................ [ 48560]
KP843857.1_Odintsovo_02/14       ................................................................................ [ 48560]
FN557520.1_strain_E75            ..............................................................a................. [ 48560]
AY261366.1_Warthog               ..............................................................a................. [ 48560]
AY261365.1_Warmbaths             ..............................................................a................. [ 48560]
AY261364.1_Tengani_62            ................................................................................ [ 48560]
AY261363.1_Pretoriuskop/96/4     ..............................................................a................. [ 48560]
AY261362.1_Mkuzi_1979            .........................t....................................a................. [ 48560]
AY261361.1_Malawi_Lil-20/1_1983  ...........a.............t........a..............t...........taa..........gc...a [ 48560]
AY261360.1_Kenya_1950            c...............a.................a........tc....t..a........t.....a..c...g....a [ 48560]
AM712240.1_OURT_88/3             ..............................................................a................. [ 48560]
AM712239.1_Benin_97/1            ..............................................................a................. [ 48560]
KM111295.1_Ken06.Bus             c...............a.................a........tc....t..a........t.....a......g....a [ 48560]
```

```
KM111294.1_Ken05/Tk1             c...............a.................a...c..........t...........taa.......a..gc...a [ 48560]
KM262844.1_strain_L60            ..............................................................a................. [ 48560]
KM102979.1_26544/OG10_Italy      ..............................................................a................. [ 48560]
KM262845.1_strain_NHV            ..............................................................a................. [ 48560]
KX354450.1_47/Ss/2008            ..............................................................a................. [ 48560]
KP055815.1_strain_BA71           ..............................................................a................. [ 48560]
MH025920.1_strain_R35            c...............a.................a........tc....t..a........t.....a......g....a [ 48560]
MH025919.1_strain_N10            c...............a.................a........tc....t..a........t.....a......g....a [ 48560]
MH025918.1_strain_R25            c...............a.................a........tc....t..a........t.....a......g....a [ 48560]
MH025917.1_strain_R7             c...............a.................a........tc....t..a........t.....a......g....a [ 48560]
MH025916.1_strain_R8             c...............a.................a........tc....t..a........t.....a......g....a [ 48560]

FR682468.2_Georgia_2007/1        cgactaggtctgtactgggaaaaacatggttctcttcagcgaatcggagacaactatgtacttatacaacaggacctcat [ 48640]
ASF_IND_20_CAD543                ..........................................................................y..... [ 48640]
LR899193.1_Germany_2020/1        ................................................................................ [ 48640]
MN393477.1_Wuhan_2019            ................................................................................ [ 48640]
MW396979.1_Timor-Leste/2019/1    ................................................................................ [ 48640]
MK940252.1_CN/2019/InnerMongolia ................................................................................ [ 48640]
MK128995.1_China/2018/AnhuiXCGQ  ................................................................................ [ 48640]
MH681419.1_ASFV/POL/2015         ................................................................................ [ 48640]
LS478113.1_Estonia_2014          ................................................................................ [ 48640]
KP843857.1_Odintsovo_02/14       ................................................................................ [ 48640]
FN557520.1_strain_E75            ...................................................g......................t..... [ 48640]
AY261366.1_Warthog               ...................................................g............................ [ 48640]
AY261365.1_Warmbaths             ...................................................g......................t..... [ 48640]
AY261364.1_Tengani_62            ................................................................................ [ 48640]
AY261363.1_Pretoriuskop/96/4     ...................................................g............................ [ 48640]
AY261362.1_Mkuzi_1979            ...................................................g......................t..... [ 48640]
AY261361.1_Malawi_Lil-20/1_1983  ........a.............................a.....a......g......................t..... [ 48640]
AY261360.1_Kenya_1950            ........gt....t...a.......c...a.......a..g..a......g............................ [ 48640]
AM712240.1_OURT_88/3             ...................................................g......................t..... [ 48640]
AM712239.1_Benin_97/1            ...................................................g......................t..... [ 48640]
KM111295.1_Ken06.Bus             ........gt....t...a.......c...a.......a..g..a.....tg............................ [ 48640]
KM111294.1_Ken05/Tk1             ........a.............................a...g.aac...tg.......g...g.c........t..... [ 48640]
KM262844.1_strain_L60            ...................................................g......................t..... [ 48640]
KM102979.1_26544/OG10_Italy      ...................................................g......................t..... [ 48640]
KM262845.1_strain_NHV            ...................................................g......................t..... [ 48640]
KX354450.1_47/Ss/2008            ...................................................g......................t..... [ 48640]
KP055815.1_strain_BA71           ...................................................g......................t..... [ 48640]
MH025920.1_strain_R35            ........gt....t...a.......c...a.......a..g..a.....tg............................ [ 48640]
MH025919.1_strain_N10            ........gt....t...a.......c...a.......a..g..a.....tg............................ [ 48640]
MH025918.1_strain_R25            ........gt....t...a.......c...a.......a..g..a.....tg............................ [ 48640]
MH025917.1_strain_R7             ........gt....t...a.......c...a.......a..g..a.....tg............................ [ 48640]
MH025916.1_strain_R8             ........gt....t...a.......c...a.......a..g..a.....tg............................ [ 48640]
```

```
FR682468.2_Georgia_2007/1        catccccatcaatgaagccctaagaatggcaggggaggaggggaatgatgaggtggtacaactcctattactatgggagg [ 48720]
ASF_IND_20_CAD543                ................................................................................ [ 48720]
LR899193.1_Germany_2020/1        ................................................................................ [ 48720]
MN393477.1_Wuhan_2019            ................................................................................ [ 48720]
MW396979.1_Timor-Leste/2019/1    ................................................................................ [ 48720]
MK940252.1_CN/2019/InnerMongolia ................................................................................ [ 48720]
MK128995.1_China/2018/AnhuiXCGQ  ................................................................................ [ 48720]
MH681419.1_ASFV/POL/2015         ................................................................................ [ 48720]
LS478113.1_Estonia_2014          ................................................................................ [ 48720]
KP843857.1_Odintsovo_02/14       ................................................................................ [ 48720]
FN557520.1_strain_E75            ..................t............................................................. [ 48720]
AY261366.1_Warthog               ..................t..............a..........g...................t............... [ 48720]
AY261365.1_Warmbaths             ..................t..............a..........g...................t............... [ 48720]
AY261364.1_Tengani_62            ..................t..............a..........g...................t............... [ 48720]
AY261363.1_Pretoriuskop/96/4     ......t...........t..............a.............................tt............... [ 48720]
AY261362.1_Mkuzi_1979            ..................t.............c...............................t............... [ 48720]
AY261361.1_Malawi_Lil-20/1_1983  t..tt.............t..............a..............................t..c...a........ [ 48720]
AY261360.1_Kenya_1950            t..t..t...........t..g...........a.....................a..g.g...t.gc............ [ 48720]
AM712240.1_OURT_88/3             ..................t............................................................. [ 48720]
AM712239.1_Benin_97/1            ..................t............................................................. [ 48720]
KM111295.1_Ken06.Bus             t..t..............t..g...........a.....................a..g.g...t....g......a... [ 48720]
KM111294.1_Ken05/Tk1             t..tt.............t..............a.....................a..g.g...t..c............ [ 48720]
KM262844.1_strain_L60            ..................t............................................................. [ 48720]
KM102979.1_26544/OG10_Italy      ..................t............................................................. [ 48720]
KM262845.1_strain_NHV            ..................t............................................................. [ 48720]
KX354450.1_47/Ss/2008            ..................t............................................................. [ 48720]
KP055815.1_strain_BA71           ..................t............................................................. [ 48720]
MH025920.1_strain_R35            t..t..............t..g...........a.....................a..g.g...t....g......a... [ 48720]
MH025919.1_strain_N10            t..t..............t..g...........a.....................a..g.g...t....g......a... [ 48720]
MH025918.1_strain_R25            t..t..............t..g...........a.....................a..g.g...t....g......a... [ 48720]
MH025917.1_strain_R7             t..t..............t..g...........a.....................a..g.g...t....g......a... [ 48720]
MH025916.1_strain_R8             t..t..............t..g...........a.....................a..g.g...t....g......a... [ 48720]

FR682468.2_Georgia_2007/1        gaaacattcattatgccatcataggagctttggagagtgaccattatagcctaatacgtaagctctatgaccaaatcgaa [ 48800]
ASF_IND_20_CAD543                ................................................................................ [ 48800]
LR899193.1_Germany_2020/1        ................................................................................ [ 48800]
MN393477.1_Wuhan_2019            ................................................................................ [ 48800]
MW396979.1_Timor-Leste/2019/1    ................................................................................ [ 48800]
MK940252.1_CN/2019/InnerMongolia ................................................................................ [ 48800]
MK128995.1_China/2018/AnhuiXCGQ  ................................................................................ [ 48800]
MH681419.1_ASFV/POL/2015         ................................................................................ [ 48800]
LS478113.1_Estonia_2014          ................................................................................ [ 48800]
KP843857.1_Odintsovo_02/14       ................................................................................ [ 48800]
```

```
FN557520.1_strain_E75            ................................................................................ [ 48800]
AY261366.1_Warthog               ...................................g............................................ [ 48800]
AY261365.1_Warmbaths             ....................g..............g...................c........................ [ 48800]
AY261364.1_Tengani_62            ............................c......g.c.......................................... [ 48800]
AY261363.1_Pretoriuskop/96/4     ...................................g.........................................a.. [ 48800]
AY261362.1_Mkuzi_1979            ................................................................................ [ 48800]
AY261361.1_Malawi_Lil-20/1_1983  ........tt.....................a...g....t...g.......ggc.ta..........ag.......ag. [ 48800]
AY261360.1_Kenya_1950            ........t................g.....a...g........g........gc.ta..........agt......ag. [ 48800]
AM712240.1_OURT_88/3             ..............................................................................g. [ 48800]
AM712239.1_Benin_97/1            ................................................................................ [ 48800]
KM111295.1_Ken06.Bus             .....g.............................g....t...g...at..g........................... [ 48800]
KM111294.1_Ken05/Tk1             ........t......................a...g....t...g........gc.ta..........ag.......a.. [ 48800]
KM262844.1_strain_L60            ................................................................................ [ 48800]
KM102979.1_26544/OG10_Italy      ................................................................................ [ 48800]
KM262845.1_strain_NHV            ..............................................................................g. [ 48800]
KX354450.1_47/Ss/2008            ................................................................................ [ 48800]
KP055815.1_strain_BA71           ................................................................................ [ 48800]
MH025920.1_strain_R35            .....g.............................g....t...g...at..g........................... [ 48800]
MH025919.1_strain_N10            .....g.............................g....t...g...at..g........................... [ 48800]
MH025918.1_strain_R25            .....g.............................g....t...g...at..g........................... [ 48800]
MH025917.1_strain_R7             .....g.............................g....t...g...at..g........................... [ 48800]
MH025916.1_strain_R8             .....g.............................g....t...g...at..g........................... [ 48800]

FR682468.2_Georgia_2007/1        gactgtcacgacatccttcccttgattcaagacccaaaactctttgaaaaatgccatgaattagataaatcttgtaacat [ 48880]
ASF_IND_20_CAD543                .......................................m..............................y......... [ 48880]
LR899193.1_Germany_2020/1        ................................................................................ [ 48880]
MN393477.1_Wuhan_2019            ................................................................................ [ 48880]
MW396979.1_Timor-Leste/2019/1    ................................................................................ [ 48880]
MK940252.1_CN/2019/InnerMongolia ................................................................................ [ 48880]
MK128995.1_China/2018/AnhuiXCGQ  ................................................................................ [ 48880]
MH681419.1_ASFV/POL/2015         ................................................................................ [ 48880]
LS478113.1_Estonia_2014          ................................................................................ [ 48880]
KP843857.1_Odintsovo_02/14       ................................................................................ [ 48880]
FN557520.1_strain_E75            .......................a...............a..............................t......... [ 48880]
AY261366.1_Warthog               .......................a...............a........................................ [ 48880]
AY261365.1_Warmbaths             .......................a...............a.............t.................c.....t.. [ 48880]
AY261364.1_Tengani_62            .......................a...............a.............t........g.............c... [ 48880]
AY261363.1_Pretoriuskop/96/4     ..t......a.............a...............ac.........g.....c.........g....c.....t.. [ 48880]
AY261362.1_Mkuzi_1979            .......................a...............a..............................t......... [ 48880]
AY261361.1_Malawi_Lil-20/1_1983  ..t.....tac...t........a...............a..........g.....c..c......g....c........ [ 48880]
AY261360.1_Kenya_1950            ..t.....ta....t........a...............a..........g.....c..c......g....c........ [ 48880]
AM712240.1_OURT_88/3             .......................a...............a..............................t......... [ 48880]
AM712239.1_Benin_97/1            .......................a...............a..............................t......... [ 48880]
KM111295.1_Ken06.Bus             ..t......ag............a...............act........g..t..c.....g...g.g..c.....t.. [ 48880]
```

```
KM111294.1_Ken05/Tk1           ..t.....ta....t........a...............a..........g.....c..c......g....c........ [ 48880]
KM262844.1_strain_L60          .......................a...............a..............................t......... [ 48880]
KM102979.1_26544/OG10_Italy    .......................a...............a..............................t......... [ 48880]
KM262845.1_strain_NHV          .......................a...............a..............................t......... [ 48880]
KX354450.1_47/Ss/2008          .......................a...............a..............................t......... [ 48880]
KP055815.1_strain_BA71         .......................a...............a..............................t......... [ 48880]
MH025920.1_strain_R35          ..t......ag............a...............act........g..t..c.....g...g.g..c.....t.. [ 48880]
MH025919.1_strain_N10          ..t......ag............a...............act........g..t..c.....g...g.g..c.....t.. [ 48880]
MH025918.1_strain_R25          ..t......ag............a...............act........g..t..c.....g...g.g..c.....t.. [ 48880]
MH025917.1_strain_R7           ..t......ag............a...............act........g..t..c.....g...g.g..c.....t.. [ 48880]
MH025916.1_strain_R8           ..t......ag............a...............act........g..t..c.....g...g.g..c.....t.. [ 48880]

FR682468.2_Georgia_2007/1      tttatgtctcgtattacacgccgtaaaaaacgatatgctttgcattcttcaagagtataaaatgcatctaagtggagagg [ 48960]
ASF_IND_20_CAD543              ................................................................................ [ 48960]
LR899193.1_Germany_2020/1      ................................................................................ [ 48960]
MN393477.1_Wuhan_2019          ................................................................................ [ 48960]
MW396979.1_Timor-Leste/2019/1  ................................................................................ [ 48960]
MK940252.1_CN/2019/InnerMongolia ................................................................................ [ 48960]
MK128995.1_China/2018/AnhuiXCGQ ................................................................................ [ 48960]
MH681419.1_ASFV/POL/2015       ................................................................................ [ 48960]
LS478113.1_Estonia_2014        ................................................................................ [ 48960]
KP843857.1_Odintsovo_02/14     ................................................................................ [ 48960]
FN557520.1_strain_E75          ................................................................................ [ 48960]
AY261366.1_Warthog             ......................................c...............t.......c................. [ 48960]
AY261365.1_Warmbaths           ..c...................................................a..........g...........g.. [ 48960]
AY261364.1_Tengani_62          ...t..................................................t.......c................. [ 48960]
AY261363.1_Pretoriuskop/96/4   ..c...................................................a..........g...........g.. [ 48960]
AY261362.1_Mkuzi_1979          ................................................................................ [ 48960]
AY261361.1_Malawi_Lil-20/1_1983 ..c.............a.......g...c.t.......................a..........t.........g.g.. [ 48960]
AY261360.1_Kenya_1950          ..c.............a...........c.t.......................a..........t...........g.. [ 48960]
AM712240.1_OURT_88/3           ................................................................................ [ 48960]
AM712239.1_Benin_97/1          ................................................................................ [ 48960]
KM111295.1_Ken06.Bus           ..c.............t....................t....t...........c....t.....g.........g.g.. [ 48960]
KM111294.1_Ken05/Tk1           ..c.............a...........c.t.......................a..........t...........g.. [ 48960]
KM262844.1_strain_L60          ................................................................................ [ 48960]
KM102979.1_26544/OG10_Italy    ................................................................................ [ 48960]
KM262845.1_strain_NHV          ................................................................................ [ 48960]
KX354450.1_47/Ss/2008          ................................................................................ [ 48960]
KP055815.1_strain_BA71         ................................................................................ [ 48960]
MH025920.1_strain_R35          ..c.............t....................t....t...........c....t.....g.........g.g.. [ 48960]
MH025919.1_strain_N10          ..c.............t....................t....t...........c....t.....g.........g.g.. [ 48960]
MH025918.1_strain_R25          ..c.............t....................t....t...........c....t.....g.........g.g.. [ 48960]
MH025917.1_strain_R7           ..c.............t....................t....t...........c....t.....g.........g.g.. [ 48960]
MH025916.1_strain_R8           ..c.............t....................t....t...........c....t.....g.........g.g.. [ 48960]
```

```
FR682468.2_Georgia_2007/1        atattcaagtggtgtttgaaacagcatgccgttcacaaaaaaacgatattgtgtcatggatgggacaaaatattgcaata [ 49040]
ASF_IND_20_CAD543                ................................................................................ [ 49040]
LR899193.1_Germany_2020/1        ................................................................................ [ 49040]
MN393477.1_Wuhan_2019            ................................................................................ [ 49040]
MW396979.1_Timor-Leste/2019/1    ................................................................................ [ 49040]
MK940252.1_CN/2019/InnerMongolia ................................................................................ [ 49040]
MK128995.1_China/2018/AnhuiXCGQ  ................................................................................ [ 49040]
MH681419.1_ASFV/POL/2015         ................................................................................ [ 49040]
LS478113.1_Estonia_2014          ................................................................................ [ 49040]
KP843857.1_Odintsovo_02/14       ................................................................................ [ 49040]
FN557520.1_strain_E75            ................................................................................ [ 49040]
AY261366.1_Warthog               ....c.............................g......t...........ag......................... [ 49040]
AY261365.1_Warmbaths             ....c....aa.......................g......t...........a...........g.......c...... [ 49040]
AY261364.1_Tengani_62            ....c....................................t...........ag.........g............... [ 49040]
AY261363.1_Pretoriuskop/96/4     ....c....aa.......................g......t...........a..........gg.......c...... [ 49040]
AY261362.1_Mkuzi_1979            .........c...................................................................... [ 49040]
AY261361.1_Malawi_Lil-20/1_1983  ....c....aa..........t..............t....t.......c..ta...................c...... [ 49040]
AY261360.1_Kenya_1950            ....c....aa..............................t..........ta...................c...... [ 49040]
AM712240.1_OURT_88/3             ................................................................................ [ 49040]
AM712239.1_Benin_97/1            ................................................................................ [ 49040]
KM111295.1_Ken06.Bus             ....c..g.aa.................t....t.......t.......c..ta...................c...... [ 49040]
KM111294.1_Ken05/Tk1             ....c....aa..............................t..........ta...................c...... [ 49040]
KM262844.1_strain_L60            ................................................................................ [ 49040]
KM102979.1_26544/OG10_Italy      ................................................................................ [ 49040]
KM262845.1_strain_NHV            ................................................................................ [ 49040]
KX354450.1_47/Ss/2008            ................................................................................ [ 49040]
KP055815.1_strain_BA71           ................................................................................ [ 49040]
MH025920.1_strain_R35            ....c..g.aa.................t....t.......t.......c..ta...................c...... [ 49040]
MH025919.1_strain_N10            ....c..g.aa.................t....t.......t.......c..ta...................c...... [ 49040]
MH025918.1_strain_R25            ....c..g.aa.................t....t.......t.......c..ta...................c...... [ 49040]
MH025917.1_strain_R7             ....c..g.aa.................t....t.......t.......c..ta...................c...... [ 49040]
MH025916.1_strain_R8             ....c..g.aa.................t....t.......t.......c..ta...................c...... [ 49040]

FR682468.2_Georgia_2007/1        tacaaccccgaagttatttttgatattgcctttgataagatgaatgtgtccttattatctatagggtatacgcttctttt [ 49120]
ASF_IND_20_CAD543                ..........r..................................................................... [ 49120]
LR899193.1_Germany_2020/1        ................................................................................ [ 49120]
MN393477.1_Wuhan_2019            ................................................................................ [ 49120]
MW396979.1_Timor-Leste/2019/1    ................................................................................ [ 49120]
MK940252.1_CN/2019/InnerMongolia ................................................................................ [ 49120]
MK128995.1_China/2018/AnhuiXCGQ  ................................................................................ [ 49120]
MH681419.1_ASFV/POL/2015         ................................................................................ [ 49120]
LS478113.1_Estonia_2014          ................................................................................ [ 49120]
KP843857.1_Odintsovo_02/14       ................................................................................ [ 49120]
```

```
FN557520.1_strain_E75            ......t...g.........................................................c........... [ 49120]
AY261366.1_Warthog               .....................a...................a..cac.a......................a........ [ 49120]
AY261365.1_Warmbaths             ........g.g...cg....................ct...a.....a..........................a..... [ 49120]
AY261364.1_Tengani_62            .....................a...................a..ca..a......................a........ [ 49120]
AY261363.1_Pretoriuskop/96/4     ........g.g...cg....................ct.......................................... [ 49120]
AY261362.1_Mkuzi_1979            ................................................................................ [ 49120]
AY261361.1_Malawi_Lil-20/1_1983  ..........g..................................................................... [ 49120]
AY261360.1_Kenya_1950            ..........gg..c.......................................c..........a.............. [ 49120]
AM712240.1_OURT_88/3             ......t...g.........................................................c........... [ 49120]
AM712239.1_Benin_97/1            ......t...g.........................................................c........... [ 49120]
KM111295.1_Ken06.Bus             ...t......gg..c................................a......c..........t.............. [ 49120]
KM111294.1_Ken05/Tk1             ..........gg..c................................a........g....................... [ 49120]
KM262844.1_strain_L60            ......t...g.........................................................c........... [ 49120]
KM102979.1_26544/OG10_Italy      ......t...g.........................................................c........... [ 49120]
KM262845.1_strain_NHV            ......t...g.........................................................c........... [ 49120]
KX354450.1_47/Ss/2008            ......t...g.........................................................c........... [ 49120]
KP055815.1_strain_BA71           ......t...g.........................................................c........... [ 49120]
MH025920.1_strain_R35            ...t......gg..c................................a......c..........t.............. [ 49120]
MH025919.1_strain_N10            ...t......gg..c................................a......c..........t.............. [ 49120]
MH025918.1_strain_R25            ...t......gg..c................................a......c..........t.............. [ 49120]
MH025917.1_strain_R7             ...t......gg..c................................a......c..........t.............. [ 49120]
MH025916.1_strain_R8             ...t......gg..c................................a......c..........t.............. [ 49120]

FR682468.2_Georgia_2007/1        caatcatcatataaataatacgaacga---------aaatattaattctttattgacacaacatcttgaatgggctgccg [ 49200]
ASF_IND_20_CAD543                ...........................---------............................................ [ 49200]
LR899193.1_Germany_2020/1        ...........................---------............................................ [ 49200]
MN393477.1_Wuhan_2019            ...........................---------............................................ [ 49200]
MW396979.1_Timor-Leste/2019/1    ...........................---------............................................ [ 49200]
MK940252.1_CN/2019/InnerMongolia ...........................---------............................................ [ 49200]
MK128995.1_China/2018/AnhuiXCGQ  ...........................---------............................................ [ 49200]
MH681419.1_ASFV/POL/2015         ...........................---------............................................ [ 49200]
LS478113.1_Estonia_2014          ...........................---------............................................ [ 49200]
KP843857.1_Odintsovo_02/14       ...........................---------............................................ [ 49200]
FN557520.1_strain_E75            ...........................---------............................................ [ 49200]
AY261366.1_Warthog               ........g...........t...t..---------.......g.................................... [ 49200]
AY261365.1_Warmbaths             .g..................t...t..---------.......g...........g........................ [ 49200]
AY261364.1_Tengani_62            ....................t...t..---------.......g.................................... [ 49200]
AY261363.1_Pretoriuskop/96/4     ....................t...t..---------.......g...........g........................ [ 49200]
AY261362.1_Mkuzi_1979            ........g..................---------............................................ [ 49200]
AY261361.1_Malawi_Lil-20/1_1983  ................c...ta..tc.tgaaaatat.g..g.....................................t. [ 49200]
AY261360.1_Kenya_1950            .g.a........c.......t..tt..aaacaatatgg...ca.............tg....................ta [ 49200]
AM712240.1_OURT_88/3             ...........................---------............................................ [ 49200]
AM712239.1_Benin_97/1            ...........................---------............................................ [ 49200]
KM111295.1_Ken06.Bus             .g.a........c.......t..tt..aaataatatgg...ca.............tg....................ta [ 49200]
```

```
KM111294.1_Ken05/Tk1              .g.a........c.......t..tt..aaacaatatgg...ca.............tg....................ta [ 49200]
KM262844.1_strain_L60             ...........................---------............................................ [ 49200]
KM102979.1_26544/OG10_Italy       ...........................---------............................................ [ 49200]
KM262845.1_strain_NHV             ...........................---------............................................ [ 49200]
KX354450.1_47/Ss/2008             ...........................---------............................................ [ 49200]
KP055815.1_strain_BA71            ...........................---------............................................ [ 49200]
MH025920.1_strain_R35             .g.a........c.......t..tt..aaataatatgg...ca.............tg....................ta [ 49200]
MH025919.1_strain_N10             .g.a........c.......t..tt..aaataatatgg...ca.............tg....................ta [ 49200]
MH025918.1_strain_R25             .g.a........c.......t..tt..aaataatatgg...ca.............tg....................ta [ 49200]
MH025917.1_strain_R7              .g.a........c.......t..tt..aaataatatgg...ca.............tg....................ta [ 49200]
MH025916.1_strain_R8              .g.a........c.......t..tt..aaataatatgg...ca.............tg....................ta [ 49200]

FR682468.2_Georgia_2007/1         gcatgggccttcttcattttatgctggaaactttaaagtatggcggggatgtaacgataatagtcttgtctgaggccgta [ 49280]
ASF_IND_20_CAD543                 ................................................................................ [ 49280]
LR899193.1_Germany_2020/1         ................................................................................ [ 49280]
MN393477.1_Wuhan_2019             ................................................................................ [ 49280]
MW396979.1_Timor-Leste/2019/1     ................................................................................ [ 49280]
MK940252.1_CN/2019/InnerMongolia  ................................................................................ [ 49280]
MK128995.1_China/2018/AnhuiXCGQ   ................................................................................ [ 49280]
MH681419.1_ASFV/POL/2015          ................................................................................ [ 49280]
LS478113.1_Estonia_2014           ................................................................................ [ 49280]
KP843857.1_Odintsovo_02/14        ................................................................................ [ 49280]
FN557520.1_strain_E75             ................................................................t............... [ 49280]
AY261366.1_Warthog                .............c.........t...................................gc................... [ 49280]
AY261365.1_Warmbaths              .............c.........t....................................c................... [ 49280]
AY261364.1_Tengani_62             .............c.........t...................t............c....................... [ 49280]
AY261363.1_Pretoriuskop/96/4      .............c.........t....................................c................... [ 49280]
AY261362.1_Mkuzi_1979             ................................................................t............... [ 49280]
AY261361.1_Malawi_Lil-20/1_1983   .......................t...........................................a....c...g... [ 49280]
AY261360.1_Kenya_1950             ...ca...t..............t.......g........c..............a.......agc.a..g.c...g... [ 49280]
AM712240.1_OURT_88/3              ................................................................t............... [ 49280]
AM712239.1_Benin_97/1             ................................................................t............... [ 49280]
KM111295.1_Ken06.Bus              ...ca...t..............t.......g.......................a.......ag..a..g.c...g... [ 49280]
KM111294.1_Ken05/Tk1              ...ca...t..............t.......g........c..............a.......agc.a..g.c...g... [ 49280]
KM262844.1_strain_L60             ................................................................t............... [ 49280]
KM102979.1_26544/OG10_Italy       ................................................................t............... [ 49280]
KM262845.1_strain_NHV             ................................................................t............... [ 49280]
KX354450.1_47/Ss/2008             ................................................................t............... [ 49280]
KP055815.1_strain_BA71            ................................................................t............... [ 49280]
MH025920.1_strain_R35             ...ca...t..............t.......g.......................a.......ag..a..g.c...g... [ 49280]
MH025919.1_strain_N10             ...ca...t..............t.......g.......................a.......ag..a..g.c...g... [ 49280]
MH025918.1_strain_R25             ...ca...t..............t.......g.......................a.......ag..a..g.c...g... [ 49280]
MH025917.1_strain_R7              ...ca...t..............t.......g.......................a.......ag..a..g.c...g... [ 49280]
MH025916.1_strain_R8              ...ca...t..............t.......g.......................a.......ag..a..g.c...g... [ 49280]
```

```
FR682468.2_Georgia_2007/1         aaatatgaccacagaaagattttagattattttctccgtcgaaaaaacttgtaccaagaagatcttgaagaactattatt [ 49360]
ASF_IND_20_CAD543                 ................................................................................ [ 49360]
LR899193.1_Germany_2020/1         ................................................................................ [ 49360]
MN393477.1_Wuhan_2019             ................................................................................ [ 49360]
MW396979.1_Timor-Leste/2019/1     ................................................................................ [ 49360]
MK940252.1_CN/2019/InnerMongolia  ................................................................................ [ 49360]
MK128995.1_China/2018/AnhuiXCGQ   ................................................................................ [ 49360]
MH681419.1_ASFV/POL/2015          ................................................................................ [ 49360]
LS478113.1_Estonia_2014           ................................................................................ [ 49360]
KP843857.1_Odintsovo_02/14        ................................................................................ [ 49360]
FN557520.1_strain_E75             ................................................................................ [ 49360]
AY261366.1_Warthog                ..................g...........................................c................. [ 49360]
AY261365.1_Warmbaths              ..................g...........................................c................. [ 49360]
AY261364.1_Tengani_62             ..................g...........................................c................. [ 49360]
AY261363.1_Pretoriuskop/96/4      ..................g...........................................c................. [ 49360]
AY261362.1_Mkuzi_1979             ..g...............g..................................t.......................... [ 49360]
AY261361.1_Malawi_Lil-20/1_1983   ........t..t......g............................t.......g.......a................ [ 49360]
AY261360.1_Kenya_1950             ...........t......g............................g...cc..g....acc.....ga.......... [ 49360]
AM712240.1_OURT_88/3              ................................................................................ [ 49360]
AM712239.1_Benin_97/1             ................................................................................ [ 49360]
KM111295.1_Ken06.Bus              ........t..t......g............................g...cc..g....acc.....ga.......... [ 49360]
KM111294.1_Ken05/Tk1              ...........t......g............................g...cc..g....acc.....ga.......... [ 49360]
KM262844.1_strain_L60             ................................................................................ [ 49360]
KM102979.1_26544/OG10_Italy       ................................................................................ [ 49360]
KM262845.1_strain_NHV             ................................................................................ [ 49360]
KX354450.1_47/Ss/2008             ................................................................................ [ 49360]
KP055815.1_strain_BA71            ................................................................................ [ 49360]
MH025920.1_strain_R35             ........t..t......g............................g...cc..g....acc.....ga.......... [ 49360]
MH025919.1_strain_N10             ........t..t......g............................g...cc..g....acc.....ga.......... [ 49360]
MH025918.1_strain_R25             ........t..t......g............................g...cc..g....acc.....ga.......... [ 49360]
MH025917.1_strain_R7              ........t..t......g............................g...cc..g....acc.....ga.......... [ 49360]
MH025916.1_strain_R8              ........t..t......g............................g...cc..g....acc.....ga.......... [ 49360]

FR682468.2_Georgia_2007/1         gttggcgatacgtgcagattgttctaaaaagaccttaaacttgttattatcttacttaaactattccataaacaatatcc [ 49440]
ASF_IND_20_CAD543                 ................................................................................ [ 49440]
LR899193.1_Germany_2020/1         ................................................................................ [ 49440]
MN393477.1_Wuhan_2019             ................................................................................ [ 49440]
MW396979.1_Timor-Leste/2019/1     ................................................................................ [ 49440]
MK940252.1_CN/2019/InnerMongolia  ................................................................................ [ 49440]
MK128995.1_China/2018/AnhuiXCGQ   ................................................................................ [ 49440]
MH681419.1_ASFV/POL/2015          ................................................................................ [ 49440]
LS478113.1_Estonia_2014           ................................................................................ [ 49440]
KP843857.1_Odintsovo_02/14        ................................................................................ [ 49440]
```

```
FN557520.1_strain_E75          ................................................................................ [ 49440]
AY261366.1_Warthog             ......................................................t........c................ [ 49440]
AY261365.1_Warmbaths           ................................................................................ [ 49440]
AY261364.1_Tengani_62          a.........................g..................................................... [ 49440]
AY261363.1_Pretoriuskop/96/4   a............................................................................... [ 49440]
AY261362.1_Mkuzi_1979          ................................................................................ [ 49440]
AY261361.1_Malawi_Lil-20/1_1983 ..........a.......................................................t...g......... [ 49440]
AY261360.1_Kenya_1950          .c.a..t...t...a..c.................................c........t......g.gg......... [ 49440]
AM712240.1_OURT_88/3           ................................................................................ [ 49440]
AM712239.1_Benin_97/1          ................................................................................ [ 49440]
KM111295.1_Ken06.Bus           .g...t...t...a..c.................................c........t......g.g.......... [ 49440]
KM111294.1_Ken05/Tk1           .c.a..t...t...a..c.................................c........t......g.g.......... [ 49440]
KM262844.1_strain_L60          ................................................................................ [ 49440]
KM102979.1_26544/OG10_Italy    ................................................................................ [ 49440]
KM262845.1_strain_NHV          ................................................................................ [ 49440]
KX354450.1_47/Ss/2008          ................................................................................ [ 49440]
KP055815.1_strain_BA71         ................................................................................ [ 49440]
MH025920.1_strain_R35          .g...t...t...a..c.................................c........t......g.g.......... [ 49440]
MH025919.1_strain_N10          .g...t...t...a..c.................................c........t......g.g.......... [ 49440]
MH025918.1_strain_R25          .g...t...t...a..c.................................c........t......g.g.......... [ 49440]
MH025917.1_strain_R7           .g...t...t...a..c.................................c........t......g.g.......... [ 49440]
MH025916.1_strain_R8           .g...t...t...a..c.................................c........t......g.g.......... [ 49440]

FR682468.2_Georgia_2007/1      gtaaaaaaatattacaatgtgtaaaagaatatgaaacgaccgttattataaaaattttacggaaaagaaag---ataaat [ 49520]
ASF_IND_20_CAD543              ........................................................y..y...........---...... [ 49520]
LR899193.1_Germany_2020/1      .........................................a.............................---...... [ 49520]
MN393477.1_Wuhan_2019          .......................................................................---...... [ 49520]
MW396979.1_Timor-Leste/2019/1  .......................................................................---...... [ 49520]
MK940252.1_CN/2019/InnerMongolia .......................................................................---...... [ 49520]
MK128995.1_China/2018/AnhuiXCGQ .......................................................................---...... [ 49520]
MH681419.1_ASFV/POL/2015       .......................................................................---...... [ 49520]
LS478113.1_Estonia_2014        .......................................................................---...... [ 49520]
KP843857.1_Odintsovo_02/14     .......................................................................---...... [ 49520]
FN557520.1_strain_E75          ........................................................c..t...........---...... [ 49520]
AY261366.1_Warthog             .........................................a.................t...........---...... [ 49520]
AY261365.1_Warmbaths           .........................................a.................t...........---...... [ 49520]
AY261364.1_Tengani_62          .........................................a.................t...........---...... [ 49520]
AY261363.1_Pretoriuskop/96/4   .........................................a.................t...........---...... [ 49520]
AY261362.1_Mkuzi_1979          ...........................................................t...........---...... [ 49520]
AY261361.1_Malawi_Lil-20/1_1983 ..................a...c.................tc.................t......a....aaa...... [ 49520]
AY261360.1_Kenya_1950          ..........c.......a.......c..........a..tc.................t......a..gaaag...... [ 49520]
AM712240.1_OURT_88/3           ........................................................c..t...........---...... [ 49520]
AM712239.1_Benin_97/1          ........................................................c..t...........---...... [ 49520]
KM111295.1_Ken06.Bus           ..........ca......a.......c..........a..tc.................t......a..gaaag...... [ 49520]
```

KM111294.1_Ken05/Tk1 ..........c.......a.......c..........a..tc.................t......a..gaaag...... [ 49520]
KM262844.1_strain_L60 ........................................................c..t...........---...... [ 49520]
KM102979.1_26544/OG10_Italy ........................................................c..t...........---...... [ 49520]
KM262845.1_strain_NHV ........................................................c..t...........---...... [ 49520]
KX354450.1_47/Ss/2008 ........................................................c..t...........---...... [ 49520]
KP055815.1_strain_BA71 ........................................................c..t...........---...... [ 49520]
MH025920.1_strain_R35 ..........ca......a.......c..........a..tc.................t......a..gaaag...... [ 49520]
MH025919.1_strain_N10 ..........ca......a.......c..........a..tc.................t......a..gaaag...... [ 49520]
MH025918.1_strain_R25 ..........ca......a.......c..........a..tc.................t......a..gaaag...... [ 49520]
MH025917.1_strain_R7 ..........ca......a.......c..........a..tc.................t......a..gaaag...... [ 49520]
MH025916.1_strain_R8 ..........ca......a.......c..........a..tc.................t......a..gaaag...... [ 49520]

FR682468.2_Georgia_2007/1 ctgatagagcccattttggcagactttataggatatcatagctatacctatatggtagattttatgcgtgagttttccat [ 49600]
ASF_IND_20_CAD543 ................................................................................ [ 49600]
LR899193.1_Germany_2020/1 ................................................................................ [ 49600]
MN393477.1_Wuhan_2019 ................................................................................ [ 49600]
MW396979.1_Timor-Leste/2019/1 ................................................................................ [ 49600]
MK940252.1_CN/2019/InnerMongolia ................................................................................ [ 49600]
MK128995.1_China/2018/AnhuiXCGQ ................................................................................ [ 49600]
MH681419.1_ASFV/POL/2015 ................................................................................ [ 49600]
LS478113.1_Estonia_2014 ................................................................................ [ 49600]
KP843857.1_Odintsovo_02/14 ................................................................................ [ 49600]
FN557520.1_strain_E75 ....................................................................c........... [ 49600]
AY261366.1_Warthog .......ga....................................................................... [ 49600]
AY261365.1_Warmbaths .......ga....................................................................... [ 49600]
AY261364.1_Tengani_62 .......ga....................................................................... [ 49600]
AY261363.1_Pretoriuskop/96/4 .......ga....................................................................... [ 49600]
AY261362.1_Mkuzi_1979 ....................................................................c........... [ 49600]
AY261361.1_Malawi_Lil-20/1_1983 ...........t..c.......................................aa..g............t..c..... [ 49600]
AY261360.1_Kenya_1950 ........t..t...............g.......................t..a..a...................... [ 49600]
AM712240.1_OURT_88/3 ....................................................................c........... [ 49600]
AM712239.1_Benin_97/1 ....................................................................c........... [ 49600]
KM111295.1_Ken06.Bus ........t..t...............g............a..........t..a..a...................... [ 49600]
KM111294.1_Ken05/Tk1 ........t..t...............g.............t.........t..a..a...................... [ 49600]
KM262844.1_strain_L60 ....................................................................c........... [ 49600]
KM102979.1_26544/OG10_Italy ....................................................................c........... [ 49600]
KM262845.1_strain_NHV ....................................................................c........... [ 49600]
KX354450.1_47/Ss/2008 ....................................................................c........... [ 49600]
KP055815.1_strain_BA71 ....................................................................c........... [ 49600]
MH025920.1_strain_R35 ........t..t...............g............a..........t..a..a...................... [ 49600]
MH025919.1_strain_N10 ........t..t...............g............a..........t..a..a...................... [ 49600]
MH025918.1_strain_R25 ........t..t...............g............a..........t..a..a...................... [ 49600]
MH025917.1_strain_R7 ........t..t...............g............a..........t..a..a...................... [ 49600]
MH025916.1_strain_R8 ........t..t...............g............a..........t..a..a...................... [ 49600]

```
FR682468.2_Georgia_2007/1        ccatccggaaaaaatgatcaaaatggctgcacgagaatcgagggaggacttgatcataaaattttccaaaaaagtttgca [ 49680]
ASF_IND_20_CAD543                ...................................................r............................ [ 49680]
LR899193.1_Germany_2020/1        ................................................................................ [ 49680]
MN393477.1_Wuhan_2019            ................................................................................ [ 49680]
MW396979.1_Timor-Leste/2019/1    ................................................................................ [ 49680]
MK940252.1_CN/2019/InnerMongolia ................................................................................ [ 49680]
MK128995.1_China/2018/AnhuiXCGQ  ................................................................................ [ 49680]
MH681419.1_ASFV/POL/2015         ................................................................................ [ 49680]
LS478113.1_Estonia_2014          ................................................................................ [ 49680]
KP843857.1_Odintsovo_02/14       ................................................................................ [ 49680]
FN557520.1_strain_E75            ..............................g....................a............................ [ 49680]
AY261366.1_Warthog               ........................................................................t....... [ 49680]
AY261365.1_Warmbaths             ........................................................................t....... [ 49680]
AY261364.1_Tengani_62            ........................................................................t....... [ 49680]
AY261363.1_Pretoriuskop/96/4     ........................................................................t....a.. [ 49680]
AY261362.1_Mkuzi_1979            ..............................g....................a............................ [ 49680]
AY261361.1_Malawi_Lil-20/1_1983  ...........g...a.............................a.....a....................t....... [ 49680]
AY261360.1_Kenya_1950            .t.........g...a..t................t.g.......a.....ag.t......................... [ 49680]
AM712240.1_OURT_88/3             ..............................g....................a............................ [ 49680]
AM712239.1_Benin_97/1            ..............................g....................a............................ [ 49680]
KM111295.1_Ken06.Bus             .t.........g...a..t................t.g.......a.....ag.t......................... [ 49680]
KM111294.1_Ken05/Tk1             .t.........g...a..t................t.g.......a.....ag.t......................... [ 49680]
KM262844.1_strain_L60            ..............................g....................a............................ [ 49680]
KM102979.1_26544/OG10_Italy      ..............................g....................a............................ [ 49680]
KM262845.1_strain_NHV            ..............................g....................a............................ [ 49680]
KX354450.1_47/Ss/2008            ..............................g....................a............................ [ 49680]
KP055815.1_strain_BA71           ..............................g....................a............................ [ 49680]
MH025920.1_strain_R35            .t.........g...a..t................t.g.......a.....ag.t......................... [ 49680]
MH025919.1_strain_N10            .t.........g...a..t................t.g.......a.....ag.t......................... [ 49680]
MH025918.1_strain_R25            .t.........g...a..t................t.g.......a.....ag.t......................... [ 49680]
MH025917.1_strain_R7             .t.........g...a..t................t.g.......a.....ag.t......................... [ 49680]
MH025916.1_strain_R8             .t.........g...a..t................t.g.......a.....ag.t......................... [ 49680]

FR682468.2_Georgia_2007/1        aagagcctaaagatagacttcactatctcaaaagcttagtgtatactatgcgacataaagaaggcaaacaactgttaatt [ 49760]
ASF_IND_20_CAD543                ................................................................................ [ 49760]
LR899193.1_Germany_2020/1        ................................................................................ [ 49760]
MN393477.1_Wuhan_2019            ................................................................................ [ 49760]
MW396979.1_Timor-Leste/2019/1    ................................................................................ [ 49760]
MK940252.1_CN/2019/InnerMongolia ................................................................................ [ 49760]
MK128995.1_China/2018/AnhuiXCGQ  ................................................................................ [ 49760]
MH681419.1_ASFV/POL/2015         ................................................................................ [ 49760]
LS478113.1_Estonia_2014          ................................................................................ [ 49760]
KP843857.1_Odintsovo_02/14       ................................................................................ [ 49760]
```

FN557520.1_strain_E75 ................................................................................ [ 49760]
AY261366.1_Warthog ................................................................................ [ 49760]
AY261365.1_Warmbaths ................................................................................ [ 49760]
AY261364.1_Tengani_62 ................................................................................ [ 49760]
AY261363.1_Pretoriuskop/96/4 ................................................................................ [ 49760]
AY261362.1_Mkuzi_1979 ................................................................................ [ 49760]
AY261361.1_Malawi_Lil-20/1_1983 .................................a................aa........g................... [ 49760]
AY261360.1_Kenya_1950 ....c....t.........ga.......g....c................a.........c........g.g........ [ 49760]
AM712240.1_OURT_88/3 ................................................................................ [ 49760]
AM712239.1_Benin_97/1 ................................................................................ [ 49760]
KM111295.1_Ken06.Bus ....c..c.t.........ga.......g....c................aa........c........g.a........ [ 49760]
KM111294.1_Ken05/Tk1 ....c....t.........ga.......g....c................a.........c........g.g........ [ 49760]
KM262844.1_strain_L60 ................................................................................ [ 49760]
KM102979.1_26544/OG10_Italy ................................................................................ [ 49760]
KM262845.1_strain_NHV ................................................................................ [ 49760]
KX354450.1_47/Ss/2008 ................................................................................ [ 49760]
KP055815.1_strain_BA71 ................................................................................ [ 49760]
MH025920.1_strain_R35 ....c..c.t.........ga.......g....c................aa........c........g.a........ [ 49760]
MH025919.1_strain_N10 ....c..c.t.........ga.......g....c................aa........c........g.a........ [ 49760]
MH025918.1_strain_R25 ....c..c.t.........ga.......g....c................aa........c........g.a........ [ 49760]
MH025917.1_strain_R7 ....c..c.t.........ga.......g....c................aa........c........g.a........ [ 49760]
MH025916.1_strain_R8 ....c..c.t.........ga.......g....c................aa........c........g.a........ [ 49760]

FR682468.2_Georgia_2007/1 tatacaatccataacttatacaaagcttgtcatctagagagtaaagaaatgtttaatttggcacgattttatgcacggca [ 49840]
ASF_IND_20_CAD543 ................................................................................ [ 49840]
LR899193.1_Germany_2020/1 ................................................................................ [ 49840]
MN393477.1_Wuhan_2019 ................................................................................ [ 49840]
MW396979.1_Timor-Leste/2019/1 ................................................................................ [ 49840]
MK940252.1_CN/2019/InnerMongolia ................................................................................ [ 49840]
MK128995.1_China/2018/AnhuiXCGQ ................................................................................ [ 49840]
MH681419.1_ASFV/POL/2015 ................................................................................ [ 49840]
LS478113.1_Estonia_2014 ................................................................................ [ 49840]
KP843857.1_Odintsovo_02/14 ................................................................................ [ 49840]
FN557520.1_strain_E75 ................................................................................ [ 49840]
AY261366.1_Warthog ........................................................c....................a.. [ 49840]
AY261365.1_Warmbaths ........................................................c....................a.. [ 49840]
AY261364.1_Tengani_62 .............................................................................a.. [ 49840]
AY261363.1_Pretoriuskop/96/4 .............................................................................a.. [ 49840]
AY261362.1_Mkuzi_1979 .............................................................................a.. [ 49840]
AY261361.1_Malawi_Lil-20/1_1983 ...........................a..tg........................ac................c..... [ 49840]
AY261360.1_Kenya_1950 ........t...........t.....ca..t....g....................g....................... [ 49840]
AM712240.1_OURT_88/3 ................................................................................ [ 49840]
AM712239.1_Benin_97/1 ................................................................................ [ 49840]
KM111295.1_Ken06.Bus .....g..t...........t.....ca..t....g....................g....................... [ 49840]

```
KM111294.1_Ken05/Tk1             ........t...........t...a.ca..t....g....................g....................... [ 49840]
KM262844.1_strain_L60            ................................................................................ [ 49840]
KM102979.1_26544/OG10_Italy      ................................................................................ [ 49840]
KM262845.1_strain_NHV            ................................................................................ [ 49840]
KX354450.1_47/Ss/2008            ................................................................................ [ 49840]
KP055815.1_strain_BA71           ................................................................................ [ 49840]
MH025920.1_strain_R35            .....g..t...........t.....ca..t....g....................g....................... [ 49840]
MH025919.1_strain_N10            .....g..t...........t.....ca..t....g....................g....................... [ 49840]
MH025918.1_strain_R25            .....g..t...........t.....ca..t....g....................g....................... [ 49840]
MH025917.1_strain_R7             .....g..t...........t.....ca..t....g....................g....................... [ 49840]
MH025916.1_strain_R8             .....g..t...........t.....ca..t....g....................g....................... [ 49840]

FR682468.2_Georgia_2007/1        taatgcagtgatccagttcaaatcgatttgccacgatctctccaagctcaatattaatatcaaaaacttgttgttagaat [ 49920]
ASF_IND_20_CAD543                ................................................................................ [ 49920]
LR899193.1_Germany_2020/1        ................................................................................ [ 49920]
MN393477.1_Wuhan_2019            ................................................................................ [ 49920]
MW396979.1_Timor-Leste/2019/1    ................................................................................ [ 49920]
MK940252.1_CN/2019/InnerMongolia ................................................................................ [ 49920]
MK128995.1_China/2018/AnhuiXCGQ  ................................................................................ [ 49920]
MH681419.1_ASFV/POL/2015         ................................................................................ [ 49920]
LS478113.1_Estonia_2014          ................................................................................ [ 49920]
KP843857.1_Odintsovo_02/14       ................................................................................ [ 49920]
FN557520.1_strain_E75            ..............................t.................g............................... [ 49920]
AY261366.1_Warthog               .........a........t...........t..g..............g..c..................c.t....... [ 49920]
AY261365.1_Warmbaths             .........a........t...........t.................g.....................c.t....... [ 49920]
AY261364.1_Tengani_62            ..................t...........tg..........t.....g.....................c.t.....t. [ 49920]
AY261363.1_Pretoriuskop/96/4     ..................t...........t.................g.....................c.t....... [ 49920]
AY261362.1_Mkuzi_1979            ..............................t.................g............................... [ 49920]
AY261361.1_Malawi_Lil-20/1_1983  .g.....ag.........t.....t.....t................ag...........t................... [ 49920]
AY261360.1_Kenya_1950            .g.....actt.......t.t...t.....t..t..............a......g....t.........c.t.c..... [ 49920]
AM712240.1_OURT_88/3             ..............................t.................g............................... [ 49920]
AM712239.1_Benin_97/1            ..............................t.................g............................... [ 49920]
KM111295.1_Ken06.Bus             .g.....actt.......t.t...t.....t...........t.....a......g....t.........c.t.c..... [ 49920]
KM111294.1_Ken05/Tk1             .g.....actt.......t.t...t.....t..t..............a......g....t.........c.t.c..... [ 49920]
KM262844.1_strain_L60            ..............................t.................g............................... [ 49920]
KM102979.1_26544/OG10_Italy      ..............................t.................g............................... [ 49920]
KM262845.1_strain_NHV            ..............................t.................g............................... [ 49920]
KX354450.1_47/Ss/2008            ..............................t.................g............................... [ 49920]
KP055815.1_strain_BA71           ..............................t.................g............................... [ 49920]
MH025920.1_strain_R35            .g.....actt.......t.t...t.....t...........t.....a......g....t.........c.t.c..... [ 49920]
MH025919.1_strain_N10            .g.....actt.......t.t...t.....t...........t.....a......g....t.........c.t.c..... [ 49920]
MH025918.1_strain_R25            .g.....actt.......t.t...t.....t...........t.....a......g....t.........c.t.c..... [ 49920]
MH025917.1_strain_R7             .g.....actt.......t.t...t.....t...........t.....a......g....t.........c.t.c..... [ 49920]
MH025916.1_strain_R8             .g.....actt.......t.t...t.....t...........t.....a......g....t.........c.t.c..... [ 49920]
```

```
FR682468.2_Georgia_2007/1        gtttaggtattgctattaaaaaaaattactttcaacttatcaaaacaatagaaacggatatgcgttatgagtaacatttt [ 50000]
ASF_IND_20_CAD543                ..........................................................................m..... [ 50000]
LR899193.1_Germany_2020/1        ................................................................................ [ 50000]
MN393477.1_Wuhan_2019            ................................................................................ [ 50000]
MW396979.1_Timor-Leste/2019/1    ................................................................................ [ 50000]
MK940252.1_CN/2019/InnerMongolia ................................................................................ [ 50000]
MK128995.1_China/2018/AnhuiXCGQ  ................................................................................ [ 50000]
MH681419.1_ASFV/POL/2015         ................................................................................ [ 50000]
LS478113.1_Estonia_2014          ................................................................................ [ 50000]
KP843857.1_Odintsovo_02/14       ................................................................................ [ 50000]
FN557520.1_strain_E75            ..........................................................................a..... [ 50000]
AY261366.1_Warthog               .....a.g.....................cc...........g.......a...........a.............-... [ 50000]
AY261365.1_Warmbaths             .....a.g.....................cc...................a...........a.............-... [ 50000]
AY261364.1_Tengani_62            .....a.g......................c...........g.......a...........a...........ac.... [ 50000]
AY261363.1_Pretoriuskop/96/4     .....a.g.....................cc...........g.......a...........a...........a..... [ 50000]
AY261362.1_Mkuzi_1979            ....................................................g..........a............-... [ 50000]
AY261361.1_Malawi_Lil-20/1_1983  ..c......................................g....c.......t.ac....aa...c......a..... [ 50000]
AY261360.1_Kenya_1950            .c....aa..............t.....tcc.............g.t...a...t.......aa..........a..g.. [ 50000]
AM712240.1_OURT_88/3             ..........................................................................a..... [ 50000]
AM712239.1_Benin_97/1            ..........................................................................a..... [ 50000]
KM111295.1_Ken06.Bus             .c....aa..............t.....tcc..g.a........g.t...a...t.......aa..........a..g.. [ 50000]
KM111294.1_Ken05/Tk1             .c....aa..............t.....tcc..g..........g.t...a...t.......aa..........a..g.. [ 50000]
KM262844.1_strain_L60            ..........................................................................a..... [ 50000]
KM102979.1_26544/OG10_Italy      ..........................................................................a..... [ 50000]
KM262845.1_strain_NHV            ..........................................................................a..... [ 50000]
KX354450.1_47/Ss/2008            ..........................................................................a..... [ 50000]
KP055815.1_strain_BA71           ..........................................................................a..... [ 50000]
MH025920.1_strain_R35            .c....aa..............t.....tcc..g.a........g.t...a...t.......aa..........a..g.. [ 50000]
MH025919.1_strain_N10            .c....aa..............t.....tcc..g.a........g.t...a...t.......aa..........a..g.. [ 50000]
MH025918.1_strain_R25            .c....aa..............t.....tcc..g.a........g.t...a...t.......aa..........a..g.. [ 50000]
MH025917.1_strain_R7             .c....aa..............t.....tcc..g.a........g.t...a...t.......aa..........a..g.. [ 50000]
MH025916.1_strain_R8             .c....aa..............t.....tcc..g.a........g.t...a...t.......aa..........a..g.. [ 50000]

FR682468.2_Georgia_2007/1        taga----------tgagggaagattctaccaaactaactaagacctttcgctagaatgtatcttattgttaatatagat [ 50080]
ASF_IND_20_CAD543                .r..----------.................................................................. [ 50080]
LR899193.1_Germany_2020/1        ....----------.................................................................. [ 50080]
MN393477.1_Wuhan_2019            ....----------.................................................................. [ 50080]
MW396979.1_Timor-Leste/2019/1    ....----------.................................................................. [ 50080]
MK940252.1_CN/2019/InnerMongolia ....----------.................................................................. [ 50080]
MK128995.1_China/2018/AnhuiXCGQ  ....----------.................................................................. [ 50080]
MH681419.1_ASFV/POL/2015         ....----------.................................................................. [ 50080]
LS478113.1_Estonia_2014          ....----------.................................................................. [ 50080]
KP843857.1_Odintsovo_02/14       ....----------.................................................................. [ 50080]
```

```
FN557520.1_strain_E75            .g..----------.a................................................................ [ 50080]
AY261366.1_Warthog               .g..----------.a..a......................................c.c......a...---------- [ 50080]
AY261365.1_Warmbaths             .g..----------.a..................................................---.---------- [ 50080]
AY261364.1_Tengani_62            .g..----------.a........-.............-----ga.c..t.............................. [ 50080]
AY261363.1_Pretoriuskop/96/4     ....----------.a...........................................c......a...---------- [ 50080]
AY261362.1_Mkuzi_1979            .g..----------.a................................................................ [ 50080]
AY261361.1_Malawi_Lil-20/1_1983  .g..----------.a.........c............----........a.................a........... [ 50080]
AY261360.1_Kenya_1950            .t.gatatcaacat.ag.cag.a..c.........at.---...t.............ac.at........--------- [ 50080]
AM712240.1_OURT_88/3             .g..----------.a................................................................ [ 50080]
AM712239.1_Benin_97/1            .g..----------.a................................................................ [ 50080]
KM111295.1_Ken06.Bus             .t.gatatcaacat.a..cag.a..c.........at.---...tt............ac..t........--------- [ 50080]
KM111294.1_Ken05/Tk1             .t.gatatcaacat.ag.cag.a..c........tat.---...t.............ac.at........--------- [ 50080]
KM262844.1_strain_L60            .g..----------.a................................................................ [ 50080]
KM102979.1_26544/OG10_Italy      .g..----------.a................................................................ [ 50080]
KM262845.1_strain_NHV            .g..----------.a................................................................ [ 50080]
KX354450.1_47/Ss/2008            .g..----------.a................................................................ [ 50080]
KP055815.1_strain_BA71           .g..----------.a................................................................ [ 50080]
MH025920.1_strain_R35            .t.gatatcaacat.a..cag.a..c.........at.---...t.............ac..t........--------- [ 50080]
MH025919.1_strain_N10            .t.gatatcaacat.a..cag.a..c.........at.---...t.............ac..t........--------- [ 50080]
MH025918.1_strain_R25            .t.gatatcaacat.a..cag.a..c.........at.---...t.............ac..t........--------- [ 50080]
MH025917.1_strain_R7             .t.gatatcaacat.a..cag.a..c.........at.---...t.............ac..t........--------- [ 50080]
MH025916.1_strain_R8             .t.gatatcaacat.a..cag.a..c.........at.---...t.............ac..t........--------- [ 50080]

FR682468.2_Georgia_2007/1        gagatatgtcattgtgaaaaaatagattaggtaggttgtgaaaaacagattaaacttaaaattatgtgtattatgtaaaa [ 50160]
ASF_IND_20_CAD543                ..........---................................................................... [ 50160]
LR899193.1_Germany_2020/1        ................................................................................ [ 50160]
MN393477.1_Wuhan_2019            ................................................................................ [ 50160]
MW396979.1_Timor-Leste/2019/1    ................................................................................ [ 50160]
MK940252.1_CN/2019/InnerMongolia ................................................................................ [ 50160]
MK128995.1_China/2018/AnhuiXCGQ  ................................................................................ [ 50160]
MH681419.1_ASFV/POL/2015         ................................................................................ [ 50160]
LS478113.1_Estonia_2014          ................................................................................ [ 50160]
KP843857.1_Odintsovo_02/14       ................................................................................ [ 50160]
FN557520.1_strain_E75            t..g....a.---................................................................... [ 50160]
AY261366.1_Warthog               ------------------.......t...................................................... [ 50160]
AY261365.1_Warmbaths             ------------------g......t...................................................... [ 50160]
AY261364.1_Tengani_62            ........a........................a.............................................. [ 50160]
AY261363.1_Pretoriuskop/96/4     ------------------.......t.....................................................t [ 50160]
AY261362.1_Mkuzi_1979            t..g....a....................................................................... [ 50160]
AY261361.1_Malawi_Lil-20/1_1983  t..g....a........................ta..........a..............---------........... [ 50160]
AY261360.1_Kenya_1950            ------------------........g......ca....a.....a................a................t [ 50160]
AM712240.1_OURT_88/3             t..g....a.---................................................................... [ 50160]
AM712239.1_Benin_97/1            t..g....a.---................................................................... [ 50160]
KM111295.1_Ken06.Bus             ------------------........a......ca....a.....a........t.......a................t [ 50160]
```

```
KM111294.1_Ken05/Tk1             ------------------........g......ca....a.....a................a................t [ 50160]
KM262844.1_strain_L60            t..g....a.---................................................................... [ 50160]
KM102979.1_26544/OG10_Italy      t..g....a.---................................................................... [ 50160]
KM262845.1_strain_NHV            t..g....a.---................................................................... [ 50160]
KX354450.1_47/Ss/2008            t..g....a.---................................................................... [ 50160]
KP055815.1_strain_BA71           t..g....a.---................................................................... [ 50160]
MH025920.1_strain_R35            ------------------........a......ca....a.....a................a................t [ 50160]
MH025919.1_strain_N10            ------------------........a......ca....a.....a................a................t [ 50160]
MH025918.1_strain_R25            ------------------........a......ca....a.....a................a................t [ 50160]
MH025917.1_strain_R7             ------------------........a......ca....a.....a................a................t [ 50160]
MH025916.1_strain_R8             ------------------........a......ca....a.....a................a................t [ 50160]

FR682468.2_Georgia_2007/1        ttttagaaataaaaattta-tttttttttattga-gggtacggaaaatgttctccctacaggacctctgtcggaagaaca [ 50240]
ASF_IND_20_CAD543                ...................-..............-.....s................w...................... [ 50240]
LR899193.1_Germany_2020/1        ...................-..............-............................................. [ 50240]
MN393477.1_Wuhan_2019            ...................-..............-............................................. [ 50240]
MW396979.1_Timor-Leste/2019/1    ...................-..............-............................................. [ 50240]
MK940252.1_CN/2019/InnerMongolia ...................-..............-............................................. [ 50240]
MK128995.1_China/2018/AnhuiXCGQ  ...................-..............-............................................. [ 50240]
MH681419.1_ASFV/POL/2015         ...................t..............-............................................. [ 50240]
LS478113.1_Estonia_2014          ...................-..............-............................................. [ 50240]
KP843857.1_Odintsovo_02/14       ...................--.............-............................................. [ 50240]
FN557520.1_strain_E75            ...................-..............-............................................. [ 50240]
AY261366.1_Warthog               ...................-............a.-......................t...................... [ 50240]
AY261365.1_Warmbaths             ...................-............a.-............................................. [ 50240]
AY261364.1_Tengani_62            ...................-..............-......................t...................... [ 50240]
AY261363.1_Pretoriuskop/96/4     ...................-............a.-......................t...................... [ 50240]
AY261362.1_Mkuzi_1979            ...................-..............-............................................. [ 50240]
AY261361.1_Malawi_Lil-20/1_1983  ...................-..............-............................................. [ 50240]
AY261360.1_Kenya_1950            ...................-..............g...........................................t. [ 50240]
AM712240.1_OURT_88/3             ...................--.............-............................................. [ 50240]
AM712239.1_Benin_97/1            ...................-..............-............................................. [ 50240]
KM111295.1_Ken06.Bus             ...................--.............g............................................. [ 50240]
KM111294.1_Ken05/Tk1             ...................-..............g...........................................t. [ 50240]
KM262844.1_strain_L60            ...................-..............-............................................. [ 50240]
KM102979.1_26544/OG10_Italy      ...................-..............-............................................. [ 50240]
KM262845.1_strain_NHV            ...................--.............-............................................. [ 50240]
KX354450.1_47/Ss/2008            ...................-..............-............................................. [ 50240]
KP055815.1_strain_BA71           ...................-..............-............................................. [ 50240]
MH025920.1_strain_R35            ...................--.............g............................................. [ 50240]
MH025919.1_strain_N10            ...................--.............g............................................. [ 50240]
MH025918.1_strain_R25            ...................--.............g............................................. [ 50240]
MH025917.1_strain_R7             ...................--.............g............................................. [ 50240]
MH025916.1_strain_R8             ...................--.............g............................................. [ 50240]
```

```
FR682468.2_Georgia_2007/1        ttttcttccttccaaatgattttagcaagcataccctacaatggctgggattatattggaaagagcatggatccgtccat [ 50320]
ASF_IND_20_CAD543                yy.............r................................................................ [ 50320]
LR899193.1_Germany_2020/1        ................................................................................ [ 50320]
MN393477.1_Wuhan_2019            ................................................................................ [ 50320]
MW396979.1_Timor-Leste/2019/1    ................................................................................ [ 50320]
MK940252.1_CN/2019/InnerMongolia ................................................................................ [ 50320]
MK128995.1_China/2018/AnhuiXCGQ  ................................................................................ [ 50320]
MH681419.1_ASFV/POL/2015         ................................................................................ [ 50320]
LS478113.1_Estonia_2014          ................................................................................ [ 50320]
KP843857.1_Odintsovo_02/14       ................................................................................ [ 50320]
FN557520.1_strain_E75            c.................................t......c.......g..g........................... [ 50320]
AY261366.1_Warthog               .................................................g.............................. [ 50320]
AY261365.1_Warmbaths             ................c................................g.............................. [ 50320]
AY261364.1_Tengani_62            c........................................c.......g.............................. [ 50320]
AY261363.1_Pretoriuskop/96/4     ................c................................g.............................. [ 50320]
AY261362.1_Mkuzi_1979            c...........................t............c.......g.............................. [ 50320]
AY261361.1_Malawi_Lil-20/1_1983  c.................g.....a................ct......g.............................. [ 50320]
AY261360.1_Kenya_1950            c.......................ag..a........g..gct..............................t...... [ 50320]
AM712240.1_OURT_88/3             c.................................t......c.......g..g........................... [ 50320]
AM712239.1_Benin_97/1            c.................................t......c.......g..g........................... [ 50320]
KM111295.1_Ken06.Bus             c...........................a..c.....g..gct......g.............................. [ 50320]
KM111294.1_Ken05/Tk1             c.......................ag..a........g..gct..............................t...... [ 50320]
KM262844.1_strain_L60            c.................................t......c.......g..g........................... [ 50320]
KM102979.1_26544/OG10_Italy      c.................................t......c.......g..g........................... [ 50320]
KM262845.1_strain_NHV            c.................................t......c.......g..g........................... [ 50320]
KX354450.1_47/Ss/2008            c.................................t......c.......g..g........................... [ 50320]
KP055815.1_strain_BA71           c.................................t......c.......g..g........................... [ 50320]
MH025920.1_strain_R35            c...........................a..c.....g..gct......g.............................. [ 50320]
MH025919.1_strain_N10            c...........................a..c.....g..gct......g.............................. [ 50320]
MH025918.1_strain_R25            c...........................a..c.....g..gct......g.............................. [ 50320]
MH025917.1_strain_R7             c...........................a..c.....g..gct......g.............................. [ 50320]
MH025916.1_strain_R8             c...........................a..c.....g..gct......g.............................. [ 50320]

FR682468.2_Georgia_2007/1        cgagcagaaaaagacagcataatgatacagaatgaattggttctttctatcaatgatgctttacagcttgcaggagagga [ 50400]
ASF_IND_20_CAD543                ...at........................................................................... [ 50400]
LR899193.1_Germany_2020/1        ................................................................................ [ 50400]
MN393477.1_Wuhan_2019            ................................................................................ [ 50400]
MW396979.1_Timor-Leste/2019/1    ................................................................................ [ 50400]
MK940252.1_CN/2019/InnerMongolia ................................................................................ [ 50400]
MK128995.1_China/2018/AnhuiXCGQ  ................................................................................ [ 50400]
MH681419.1_ASFV/POL/2015         ................................................................................ [ 50400]
LS478113.1_Estonia_2014          ................................................................................ [ 50400]
KP843857.1_Odintsovo_02/14       ................................................................................ [ 50400]
```

```
FN557520.1_strain_E75            ...at......g....................................................................  [ 50400]
AY261366.1_Warthog               ............a........g..................................................g..a..  [ 50400]
AY261365.1_Warmbaths             ................................................................t...............  [ 50400]
AY261364.1_Tengani_62            ................................................................................  [ 50400]
AY261363.1_Pretoriuskop/96/4     ................................................................t...............  [ 50400]
AY261362.1_Mkuzi_1979            ...at......g.....................................................t..............  [ 50400]
AY261361.1_Malawi_Lil-20/1_1983  ....t......g....a...............................................t...............  [ 50400]
AY261360.1_Kenya_1950            ................a............a......c..a.....a..g...............................  [ 50400]
AM712240.1_OURT_88/3             ...at......g....................................................................  [ 50400]
AM712239.1_Benin_97/1            ...at......g....................................................................  [ 50400]
KM111295.1_Ken06.Bus             ...a............a...................c..a.....a..g...............................  [ 50400]
KM111294.1_Ken05/Tk1             ................a............a......c..a.....a..g...............................  [ 50400]
KM262844.1_strain_L60            ...at......g....................................................................  [ 50400]
KM102979.1_26544/OG10_Italy      ...at......g....................................................................  [ 50400]
KM262845.1_strain_NHV            ...at......g....................................................................  [ 50400]
KX354450.1_47/Ss/2008            ...at......g....................................................................  [ 50400]
KP055815.1_strain_BA71           ...at......g....................................................................  [ 50400]
MH025920.1_strain_R35            ...a............a...................c..a.....a..g...............................  [ 50400]
MH025919.1_strain_N10            ...a............a...................c..a.....a..g...............................  [ 50400]
MH025918.1_strain_R25            ...a............a...................c..a.....a..g...............................  [ 50400]
MH025917.1_strain_R7             ...a............a...................c..a.....a..g...............................  [ 50400]
MH025916.1_strain_R8             ...a............a...................c..a.....a..g...............................  [ 50400]

FR682468.2_Georgia_2007/1        gggggacacagatgtagtacagctcttgttattatgggagggaaatctgcattatgccatcataggagccttgaagactg  [ 50480]
ASF_IND_20_CAD543                ...............................y.....................................y....n.....  [ 50480]
LR899193.1_Germany_2020/1        ................................................................................  [ 50480]
MN393477.1_Wuhan_2019            ................................................................................  [ 50480]
MW396979.1_Timor-Leste/2019/1    ................................................................................  [ 50480]
MK940252.1_CN/2019/InnerMongolia ................................................................................  [ 50480]
MK128995.1_China/2018/AnhuiXCGQ  ................................................................................  [ 50480]
MH681419.1_ASFV/POL/2015         ................................................................................  [ 50480]
LS478113.1_Estonia_2014          ................................................................................  [ 50480]
KP843857.1_Odintsovo_02/14       ................................................................................  [ 50480]
FN557520.1_strain_E75            ...............g...............c.....................................t..........  [ 50480]
AY261366.1_Warthog               ................................................................................  [ 50480]
AY261365.1_Warmbaths             ........t.................................t.....................................  [ 50480]
AY261364.1_Tengani_62            ...........................a...........a........t...............................  [ 50480]
AY261363.1_Pretoriuskop/96/4     ..............................gc.......a........................................  [ 50480]
AY261362.1_Mkuzi_1979            ....a...........................................................................  [ 50480]
AY261361.1_Malawi_Lil-20/1_1983  ......t........g.....a.....a...c............................t...........a.......  [ 50480]
AY261360.1_Kenya_1950            ......t........g.........c.....c..............t.............................g...  [ 50480]
AM712240.1_OURT_88/3             ...............g...............c.....................................t..........  [ 50480]
AM712239.1_Benin_97/1            ...............g...............c.....................................t..........  [ 50480]
KM111295.1_Ken06.Bus             .............a.g...g.a.........c................a............................a..  [ 50480]
```

```
KM111294.1_Ken05/Tk1            ......t........g.........c.....c..............t.............................g... [ 50480]
KM262844.1_strain_L60           ...............g...............c.....................................t.......... [ 50480]
KM102979.1_26544/OG10_Italy     ...............g...............c.....................................t.......... [ 50480]
KM262845.1_strain_NHV           ...............g...............c.....................................t.......... [ 50480]
KX354450.1_47/Ss/2008           ...............g...............c.....................................t.......... [ 50480]
KP055815.1_strain_BA71          ...............g...............c.....................................t.......... [ 50480]
MH025920.1_strain_R35           .............a.g...g.a.........c................a............................a.. [ 50480]
MH025919.1_strain_N10           .............a.g...g.a.........c................a............................a.. [ 50480]
MH025918.1_strain_R25           .............a.g...g.a.........c................a............................a.. [ 50480]
MH025917.1_strain_R7            .............a.g...g.a.........c................a............................a.. [ 50480]
MH025916.1_strain_R8            .............a.g...g.a.........c................a............................a.. [ 50480]

FR682468.2_Georgia_2007/1       aaaaatataacctaatatgtgagtatcatagccaaattcaggactggcatattctcctacccatgattcaagatccagaa [ 50560]
ASF_IND_20_CAD543               .r............r..........y.....y..................r...........w................. [ 50560]
LR899193.1_Germany_2020/1       ................................................................................ [ 50560]
MN393477.1_Wuhan_2019           ................................................................................ [ 50560]
MW396979.1_Timor-Leste/2019/1   ................................................................................ [ 50560]
MK940252.1_CN/2019/InnerMongolia ................................................................................ [ 50560]
MK128995.1_China/2018/AnhuiXCGQ ................................................................................ [ 50560]
MH681419.1_ASFV/POL/2015        ................................................................................ [ 50560]
LS478113.1_Estonia_2014         ................................................................................ [ 50560]
KP843857.1_Odintsovo_02/14      ................................................................................ [ 50560]
FN557520.1_strain_E75           .g..t.....t...g..........c....................................t................. [ 50560]
AY261366.1_Warthog              .g........t..............c........................g.......c...t................. [ 50560]
AY261365.1_Warmbaths            .g.......................c...................................................... [ 50560]
AY261364.1_Tengani_62           .g................a......c........................g.....t....................... [ 50560]
AY261363.1_Pretoriuskop/96/4    .g.......g....g..........c........................g.......c...t................. [ 50560]
AY261362.1_Mkuzi_1979           .g.......................c...............a........g.......c...t................. [ 50560]
AY261361.1_Malawi_Lil-20/1_1983 ...........t.............c............g......................................... [ 50560]
AY261360.1_Kenya_1950           .g..t........c...........c............g...........gcc.........t..........c...... [ 50560]
AM712240.1_OURT_88/3            .g..t.....t...g..........c....................................t................. [ 50560]
AM712239.1_Benin_97/1           .g..t.....t...g..........c....................................t................. [ 50560]
KM111295.1_Ken06.Bus            .gc.t........c...........c............g...........gcc.........t..........c...... [ 50560]
KM111294.1_Ken05/Tk1            ....t........c...........c............g...........gcc.........t..........c...... [ 50560]
KM262844.1_strain_L60           .g..t.....t...g..........c....................................t................. [ 50560]
KM102979.1_26544/OG10_Italy     .g..t.....t...g..........c....................................t................. [ 50560]
KM262845.1_strain_NHV           .g..t.....t...g..........c....................................t................. [ 50560]
KX354450.1_47/Ss/2008           .g..t.....t...g..........c....................................t................. [ 50560]
KP055815.1_strain_BA71          .g..t.....t...g..........c....................................t................. [ 50560]
MH025920.1_strain_R35           .gc.t........c...........c............g...........gcc.........t..........c...... [ 50560]
MH025919.1_strain_N10           .gc.t........c...........c............g...........gcc.........t..........c...... [ 50560]
MH025918.1_strain_R25           .gc.t........c...........c............g...........gcc.........t..........c...... [ 50560]
MH025917.1_strain_R7            .gc.t........c...........c............g...........gcc.........t..........c...... [ 50560]
MH025916.1_strain_R8            .gc.t........c...........c............g...........gcc.........t..........c...... [ 50560]
```

```
FR682468.2_Georgia_2007/1        acattcgaaaaatgtcatgatttaagccttggatgtgactttatttgccttctccaacatgctgtaaaatacaacatgct [ 50640]
ASF_IND_20_CAD543                ...............................r......yy....w.........................gtg....... [ 50640]
LR899193.1_Germany_2020/1        ................................................................................ [ 50640]
MN393477.1_Wuhan_2019            ................................................................................ [ 50640]
MW396979.1_Timor-Leste/2019/1    ................................................................................ [ 50640]
MK940252.1_CN/2019/InnerMongolia ................................................................................ [ 50640]
MK128995.1_China/2018/AnhuiXCGQ  ................................................................................ [ 50640]
MH681419.1_ASFV/POL/2015         ................................................................................ [ 50640]
LS478113.1_Estonia_2014          ................................................................................ [ 50640]
KP843857.1_Odintsovo_02/14       ................................................................................ [ 50640]
FN557520.1_strain_E75            .......................................c....a.........................gtg.t..... [ 50640]
AY261366.1_Warthog               ...............................c............a........a.................t........ [ 50640]
AY261365.1_Warmbaths             ...............................c............a..........................t........ [ 50640]
AY261364.1_Tengani_62            ...............................c............a..................a.......t........ [ 50640]
AY261363.1_Pretoriuskop/96/4     ...............................a............a........a................gtg....... [ 50640]
AY261362.1_Mkuzi_1979            ......................................................................gtg....... [ 50640]
AY261361.1_Malawi_Lil-20/1_1983  .....t........................ca............a........a.........a.......t....a... [ 50640]
AY261360.1_Kenya_1950            ..c..t........c.....a.....t............c..t.a..t.....t.........a......gt........ [ 50640]
AM712240.1_OURT_88/3             .......................................c....a.........................gtg.t..... [ 50640]
AM712239.1_Benin_97/1            .......................................c....a.........................gtg.t..... [ 50640]
KM111295.1_Ken06.Bus             ..c..t........c.....a.....t...........tc..tca........a...............c.t........ [ 50640]
KM111294.1_Ken05/Tk1             ..c..t........c.....a.....t............c..t.a..t.....t................gt........ [ 50640]
KM262844.1_strain_L60            .......................................c....a.........................gtg.t..... [ 50640]
KM102979.1_26544/OG10_Italy      .......................................c....a.........................gtg.t..... [ 50640]
KM262845.1_strain_NHV            .......................................c....a.........................gtg.t..... [ 50640]
KX354450.1_47/Ss/2008            .......................................c....a.........................gtg.t..... [ 50640]
KP055815.1_strain_BA71           .......................................c....a.........................gtg.t..... [ 50640]
MH025920.1_strain_R35            ..c..t........c.....a.....t...........tc..tca........a...............c.t........ [ 50640]
MH025919.1_strain_N10            ..c..t........c.....a.....t...........tc..tca........a...............c.t........ [ 50640]
MH025918.1_strain_R25            ..c..t........c.....a.....t...........tc..tca........a...............c.t........ [ 50640]
MH025917.1_strain_R7             ..c..t........c.....a.....t...........tc..tca........a...............c.t........ [ 50640]
MH025916.1_strain_R8             ..c..t........c.....a.....t...........tc..tca........a...............c.t........ [ 50640]

FR682468.2_Georgia_2007/1        ttctattcttgtcaaatataaggaggatctactaaatgcaaggattaggcatcgtatccaatccctgtttgttttggcat [ 50720]
ASF_IND_20_CAD543                ...k........y........r......y.........y......................................... [ 50720]
LR899193.1_Germany_2020/1        ................................................................................ [ 50720]
MN393477.1_Wuhan_2019            ................................................................................ [ 50720]
MW396979.1_Timor-Leste/2019/1    ................................................................................ [ 50720]
MK940252.1_CN/2019/InnerMongolia ................................................................................ [ 50720]
MK128995.1_China/2018/AnhuiXCGQ  ................................................................................ [ 50720]
MH681419.1_ASFV/POL/2015         ................................................................................ [ 50720]
LS478113.1_Estonia_2014          ................................................................................ [ 50720]
KP843857.1_Odintsovo_02/14       ................................................................................ [ 50720]
```

```
FN557520.1_strain_E75          ............t...............t.........t.................c....................... [ 50720]
AY261366.1_Warthog             ......c.................a..ct.........t..........t.............................. [ 50720]
AY261365.1_Warmbaths           ......c.................a..ct.........t..........t.............................. [ 50720]
AY261364.1_Tengani_62          ......c.................a..ct.........t..........t.............................. [ 50720]
AY261363.1_Pretoriuskop/96/4   ............t........a................t......................................... [ 50720]
AY261362.1_Mkuzi_1979          ............t................................................................... [ 50720]
AY261361.1_Malawi_Lil-20/1_1983 ...c..c.....t........a.....ct........at................g.t........c............. [ 50720]
AY261360.1_Kenya_1950          ......c.....g..g........a...t..t.........................t.....t................ [ 50720]
AM712240.1_OURT_88/3           ............t...............t.........t.................c....................... [ 50720]
AM712239.1_Benin_97/1          ............t...............t.........t.................c....................... [ 50720]
KM111295.1_Ken06.Bus           ..........a.g..g........a...t..t.....at.ttat.............t...................... [ 50720]
KM111294.1_Ken05/Tk1           ......c.....g..g........a...t..t.........................t.....t................ [ 50720]
KM262844.1_strain_L60          ............t...............t.........t.................c....................... [ 50720]
KM102979.1_26544/OG10_Italy    ............t...............t.........t.................c....................... [ 50720]
KM262845.1_strain_NHV          ............t...............t.........t.................c....................... [ 50720]
KX354450.1_47/Ss/2008          ............t...............t.........t.................c....................... [ 50720]
KP055815.1_strain_BA71         ............t...............t.........t.................c....................... [ 50720]
MH025920.1_strain_R35          ..........a.g..g........a...t..t.....at.ttat.............t...................... [ 50720]
MH025919.1_strain_N10          ..........a.g..g........a...t..t.....at.ttat.............t...................... [ 50720]
MH025918.1_strain_R25          ..........a.g..g........a...t..t.....at.ttat.............t...................... [ 50720]
MH025917.1_strain_R7           ..........a.g..g........a...t..t.....at.ttat.............t...................... [ 50720]
MH025916.1_strain_R8           ..........a.g..g........a...t..t.....at.ttat.............t...................... [ 50720]

FR682468.2_Georgia_2007/1      gcgaaaatcggagaattgaaattattgattggataggccaaaatctgccaattcctgaacctgatgccatttttagcatt [ 50800]
ASF_IND_20_CAD543              ................................................................................ [ 50800]
LR899193.1_Germany_2020/1      ................................................................................ [ 50800]
MN393477.1_Wuhan_2019          ................................................................................ [ 50800]
MW396979.1_Timor-Leste/2019/1  ................................................................................ [ 50800]
MK940252.1_CN/2019/InnerMongolia ................................................................................ [ 50800]
MK128995.1_China/2018/AnhuiXCGQ ................................................................................ [ 50800]
MH681419.1_ASFV/POL/2015       ................................................................................ [ 50800]
LS478113.1_Estonia_2014        ................................................................................ [ 50800]
KP843857.1_Odintsovo_02/14     ................................................................................ [ 50800]
FN557520.1_strain_E75          ..............t....g........a........t.........t................g............... [ 50800]
AY261366.1_Warthog             .t.....c...........g............................................................ [ 50800]
AY261365.1_Warmbaths           .......c...........g......c..................................................... [ 50800]
AY261364.1_Tengani_62          .t.....c...........g........a........t.......................................... [ 50800]
AY261363.1_Pretoriuskop/96/4   ..............t....g........a........t..........................g............... [ 50800]
AY261362.1_Mkuzi_1979          ..............t....g........a........t..........................g............... [ 50800]
AY261361.1_Malawi_Lil-20/1_1983 .t.....c..a....c...g........a..................t................................ [ 50800]
AY261360.1_Kenya_1950          ......cc...........g.....ccta........t..........................ga.t............ [ 50800]
AM712240.1_OURT_88/3           ..............t....g........a........t.........t................g............... [ 50800]
AM712239.1_Benin_97/1          ..............t....g........a........t.........t................g............... [ 50800]
KM111295.1_Ken06.Bus           .......c...........g.....catg...................................g..t............ [ 50800]
```

```
KM111294.1_Ken05/Tk1            .......c...........g.....ccta........t..........................ga.t............ [ 50800]
KM262844.1_strain_L60           ..............t....g........a........t.........t................g............... [ 50800]
KM102979.1_26544/OG10_Italy     ..............t....g........a........t.........t................g............... [ 50800]
KM262845.1_strain_NHV           ..............t....g........a........t.........t................g............... [ 50800]
KX354450.1_47/Ss/2008           ..............t....g........a........t.........t................g............... [ 50800]
KP055815.1_strain_BA71          ..............t....g........a........t.........t................g............... [ 50800]
MH025920.1_strain_R35           .......c...........g.....catg...................................g..t............ [ 50800]
MH025919.1_strain_N10           .......c...........g.....catg...................................g..t............ [ 50800]
MH025918.1_strain_R25           .......c...........g.....catg...................................g..t............ [ 50800]
MH025917.1_strain_R7            .......c...........g.....catg...................................g..t............ [ 50800]
MH025916.1_strain_R8            .......c...........g.....catg...................................g..t............ [ 50800]

FR682468.2_Georgia_2007/1       gctgttgctacaagagatttagaactgttttccttagggtacaagattatttttgattacatgcaaagacagggaatcat [ 50880]
ASF_IND_20_CAD543               .........................................y..r................................tt. [ 50880]
LR899193.1_Germany_2020/1       ................................................................................ [ 50880]
MN393477.1_Wuhan_2019           ................................................................................ [ 50880]
MW396979.1_Timor-Leste/2019/1   ................................................................................ [ 50880]
MK940252.1_CN/2019/InnerMongolia ................................................................................ [ 50880]
MK128995.1_China/2018/AnhuiXCGQ ................................................................................ [ 50880]
MH681419.1_ASFV/POL/2015        ................................................................................ [ 50880]
LS478113.1_Estonia_2014         ................................................................................ [ 50880]
KP843857.1_Odintsovo_02/14      ................................................................................ [ 50880]
FN557520.1_strain_E75           ...a...t.....a....g...................a..t..a..........................a.....tt. [ 50880]
AY261366.1_Warthog              ..............................................................................t. [ 50880]
AY261365.1_Warmbaths            ..............................................................................t. [ 50880]
AY261364.1_Tengani_62           ..............................................................................t. [ 50880]
AY261363.1_Pretoriuskop/96/4    ...a...t.....a....a...................a......c.........................a....ctt. [ 50880]
AY261362.1_Mkuzi_1979           ...a...t.....a....a..................c.......c.........................a....ctt. [ 50880]
AY261361.1_Malawi_Lil-20/1_1983 ...a.........a........................a.......................................c. [ 50880]
AY261360.1_Kenya_1950           ......at.....a........................a........a.......................a.....tt. [ 50880]
AM712240.1_OURT_88/3            ...a...t.....a....g...................a..t..a..........................a.....tt. [ 50880]
AM712239.1_Benin_97/1           ...a...t.....a....g...................a..t..a..........................a.....tt. [ 50880]
KM111295.1_Ken06.Bus            .......t.....a........................a.....a..........................a.....tt. [ 50880]
KM111294.1_Ken05/Tk1            ......at.....a........................a........a.......................a.....tt. [ 50880]
KM262844.1_strain_L60           ...a...t.....a....g...................a..t..a..........................a.....tt. [ 50880]
KM102979.1_26544/OG10_Italy     ...a...t.....a....g...................a..t..a..........................a.....tt. [ 50880]
KM262845.1_strain_NHV           ...a...t.....a....g...................a..t..a..........................a.....tt. [ 50880]
KX354450.1_47/Ss/2008           ...a...t.....a....g...................a..t..a..........................a.....tt. [ 50880]
KP055815.1_strain_BA71          ...a...t.....a....g...................a..t..a..........................a.....tt. [ 50880]
MH025920.1_strain_R35           .......t.....a........................a.....a..........................a.....tt. [ 50880]
MH025919.1_strain_N10           .......t.....a........................a.....a..........................a.....tt. [ 50880]
MH025918.1_strain_R25           .......t.....a........................a.....a..........................a.....tt. [ 50880]
MH025917.1_strain_R7            .......t.....a........................a.....a..........................a.....tt. [ 50880]
MH025916.1_strain_R8            .......t.....a........................a.....a..........................a.....tt. [ 50880]
```

```
FR682468.2_Georgia_2007/1        tcaattaaccaatggagttcgcatggttgtgctaaatcgtcacattagcatggcaatagataatggtcttttaccttttg [ 50960]
ASF_IND_20_CAD543                ................................................y.................w............r [ 50960]
LR899193.1_Germany_2020/1        ................................................................................ [ 50960]
MN393477.1_Wuhan_2019            ................................................................................ [ 50960]
MW396979.1_Timor-Leste/2019/1    ................................................................................ [ 50960]
MK940252.1_CN/2019/InnerMongolia ................................................................................ [ 50960]
MK128995.1_China/2018/AnhuiXCGQ  ................................................................................ [ 50960]
MH681419.1_ASFV/POL/2015         ................................................................................ [ 50960]
LS478113.1_Estonia_2014          ................................................................................ [ 50960]
KP843857.1_Odintsovo_02/14       ................................................................................ [ 50960]
FN557520.1_strain_E75            ..............t..........c..c.............t...g.t...........a..a..a............a [ 50960]
AY261366.1_Warthog               .......t..........................................................a........c.... [ 50960]
AY261365.1_Warmbaths             .......t..........................................................a........c.... [ 50960]
AY261364.1_Tengani_62            ...........................................................................c.... [ 50960]
AY261363.1_Pretoriuskop/96/4     .............atg.........c..c...........t.....g.t...........a..a..c........c...a [ 50960]
AY261362.1_Mkuzi_1979            .............atg..c......c..c..........gt.....g.t...........a..a..a........c...a [ 50960]
AY261361.1_Malawi_Lil-20/1_1983  .......t........c.................................................g........c.... [ 50960]
AY261360.1_Kenya_1950            .......t..g.g.t........a.a..t.................gat......g..a....a..a........c.... [ 50960]
AM712240.1_OURT_88/3             ..............t..........c..c.............t...g.t...........a..a..a............a [ 50960]
AM712239.1_Benin_97/1            ..............t..........c..c.............t...g.t...........a..a..a............a [ 50960]
KM111295.1_Ken06.Bus             c..gc..t.....cc.............ac................g.t....t.........a..a........c.... [ 50960]
KM111294.1_Ken05/Tk1             .......t..g.a.t........a.a..t.................gat......g..a....a..a........c.... [ 50960]
KM262844.1_strain_L60            ..............t..........c..c.............t...g.t...........a..a..a............a [ 50960]
KM102979.1_26544/OG10_Italy      ..............t..........c..c.............t...g.t...........a..a..a............a [ 50960]
KM262845.1_strain_NHV            ..............t..........c..c.............t...g.t...........a..a..a............a [ 50960]
KX354450.1_47/Ss/2008            ..............t..........c..c.............t...g.t...........a..a..a............a [ 50960]
KP055815.1_strain_BA71           ..............t..........c..c.............t...g.t...........a..a..a............a [ 50960]
MH025920.1_strain_R35            c..gc..t.....cc.............ac................g.t..............a..a........c.... [ 50960]
MH025919.1_strain_N10            c..gc..t.....cc.............ac................g.t..............a..a........c.... [ 50960]
MH025918.1_strain_R25            c..gc..t.....cc.............ac................g.t..............a..a........c.... [ 50960]
MH025917.1_strain_R7             c..gc..t.....cc.............ac................g.t..............a..a........c.... [ 50960]
MH025916.1_strain_R8             c..gc..t.....cc.............ac................g.t..............a..a........c.... [ 50960]

FR682468.2_Georgia_2007/1        ttctggaaactttaaaacatggtgggaatatacatagagccttatcttatgcagtaacacacaatagaagaaaaattctg [ 51040]
ASF_IND_20_CAD543                .................y.......t.g.g...........................t.g.t....a..........a.a [ 51040]
LR899193.1_Germany_2020/1        ................................................................................ [ 51040]
MN393477.1_Wuhan_2019            ................................................................................ [ 51040]
MW396979.1_Timor-Leste/2019/1    ................................................................................ [ 51040]
MK940252.1_CN/2019/InnerMongolia ................................................................................ [ 51040]
MK128995.1_China/2018/AnhuiXCGQ  ................................................................................ [ 51040]
MH681419.1_ASFV/POL/2015         ................................................................................ [ 51040]
LS478113.1_Estonia_2014          ................................................................................ [ 51040]
KP843857.1_Odintsovo_02/14       ................................................................................ [ 51040]
```

```
FN557520.1_strain_E75          .................t.......t.g.g..a.a.....t................t.g.t....a..........a.a [ 51040]
AY261366.1_Warthog             ................................................................................ [ 51040]
AY261365.1_Warmbaths           ................................................................................ [ 51040]
AY261364.1_Tengani_62          ..................................................................a............. [ 51040]
AY261363.1_Pretoriuskop/96/4   .cg..............t.......t.g.g..a.......t..............c.t.g.t....a..........a.a [ 51040]
AY261362.1_Mkuzi_1979          .................t.......t.g.g..a.......t................t.g.t....a..........a.a [ 51040]
AY261361.1_Malawi_Lil-20/1_1983 .........................t..............t...............t....t...............t.. [ 51040]
AY261360.1_Kenya_1950          .g...........t...t.......t.g.g..a.a.............g........tgg......a.......g..a.a [ 51040]
AM712240.1_OURT_88/3           .................t.......t.g.g..a.a.....t................t.g.t....a..........a.a [ 51040]
AM712239.1_Benin_97/1          .................t.......t.g.g..a.a.....t................t.g.t....a..........a.a [ 51040]
KM111295.1_Ken06.Bus           .g...........t...........t.g.g..a.a......................t.a......a..........a.a [ 51040]
KM111294.1_Ken05/Tk1           .g...........t...t.......t.g....a.a.............g........tgg......a.......g..a.a [ 51040]
KM262844.1_strain_L60          .................t.......t.g.g..a.a.....t................t.g.t....a..........a.a [ 51040]
KM102979.1_26544/OG10_Italy    .................t.......t.g.g..a.a.....t................t.g.t....a..........a.a [ 51040]
KM262845.1_strain_NHV          .................t.......t.g.g..a.a.....t................t.g.t....a..........a.a [ 51040]
KX354450.1_47/Ss/2008          .................t.......t.g.g..a.a.....t................t.g.t....a..........a.a [ 51040]
KP055815.1_strain_BA71         .................t.......t.g.g..a.a.....t................t.g.t....a..........a.a [ 51040]
MH025920.1_strain_R35          .g...........t...........t.g.g..a.a......................t.a......a..........a.a [ 51040]
MH025919.1_strain_N10          .g...........t...........t.g.g..a.a......................t.a......a..........a.a [ 51040]
MH025918.1_strain_R25          .g...........t...........t.g.g..a.a......................t.a......a..........a.a [ 51040]
MH025917.1_strain_R7           .g...........t...........t.g.g..a.a......................t.a......a..........a.a [ 51040]
MH025916.1_strain_R8           .g...........t...........t.g.g..a.a......................t.a......a..........a.a [ 51040]

FR682468.2_Georgia_2007/1      gattatcttattcgccagaaaaatatagcccctaatacaattgaaagacttttatatctggccgtgaaaaatcaatcttc [ 51120]
ASF_IND_20_CAD543              .nnnnnnnnnnnnnnn.tg........c...gagga..t..............gc........................- [ 51120]
LR899193.1_Germany_2020/1      ................................................................................ [ 51120]
MN393477.1_Wuhan_2019          ................................................................................ [ 51120]
MW396979.1_Timor-Leste/2019/1  ................................................................................ [ 51120]
MK940252.1_CN/2019/InnerMongolia ................................................................................ [ 51120]
MK128995.1_China/2018/AnhuiXCGQ ................................................................................ [ 51120]
MH681419.1_ASFV/POL/2015       ................................................................................ [ 51120]
LS478113.1_Estonia_2014        ................................................................................ [ 51120]
KP843857.1_Odintsovo_02/14     ................................................................................ [ 51120]
FN557520.1_strain_E75          ..c......g.a.....tg........c...gagga..t..............gc....a..t........a........ [ 51120]
AY261366.1_Warthog             ................................................................................ [ 51120]
AY261365.1_Warmbaths           ................................................................................ [ 51120]
AY261364.1_Tengani_62          .............................................................................a.. [ 51120]
AY261363.1_Pretoriuskop/96/4   ..c......g.a..t..tg........c...g.gga..t..............gc....a..t........a........ [ 51120]
AY261362.1_Mkuzi_1979          ..c......g.a.....tg........c...g.gga..t..............gc....a..t........a........ [ 51120]
AY261361.1_Malawi_Lil-20/1_1983 .....c.....................c....c............................................... [ 51120]
AY261360.1_Kenya_1950          ..c......g.......tg........t...a.c.a..c....................a..t........a..t..... [ 51120]
AM712240.1_OURT_88/3           ..c......g.a.....tg........c...gagga..t..............gc....a..t........a........ [ 51120]
AM712239.1_Benin_97/1          ..c......g.a.....tg........c...gagga..t..............gc....a..t........a........ [ 51120]
KM111295.1_Ken06.Bus           ..c......g....t..tg.....g..t...a.c.a..c................g...a..t........c..tg.... [ 51120]
```

```
KM111294.1_Ken05/Tk1            ..c......g.......tg........t...a.c.a..c....................a..t........a..t..... [ 51120]
KM262844.1_strain_L60           ..c......g.a.....tg........c...gagga..t..............gc....a..t........a........ [ 51120]
KM102979.1_26544/OG10_Italy     ..c......g.a.....tg........c...gagga..t..............gc....a..t........a........ [ 51120]
KM262845.1_strain_NHV           ..c......g.a.....tg........c...gagga..t..............gc....a..t........a........ [ 51120]
KX354450.1_47/Ss/2008           ..c......g.a.....tg........c...gagga..t..............gc....a..t........a........ [ 51120]
KP055815.1_strain_BA71          ..c......g.a.....tg........c...gagga..t..............gc....a..t........a........ [ 51120]
MH025920.1_strain_R35           ..c......g....t..tg.....g..t...a.c.a..c................g...a..t........c..tg.... [ 51120]
MH025919.1_strain_N10           ..c......g....t..tg.....g..t...a.c.a..c................g...a..t........c..tg.... [ 51120]
MH025918.1_strain_R25           ..c......g....t..tg.....g..t...a.c.a..c................g...a..t........c..tg.... [ 51120]
MH025917.1_strain_R7            ..c......g....t..tg.....g..t...a.c.a..c................g...a..t........c..tg.... [ 51120]
MH025916.1_strain_R8            ..c......g....t..tg.....g..t...a.c.a..c................g...a..t........c..tg.... [ 51120]

FR682468.2_Georgia_2007/1       caggaaaactttgaacttgttgctatcttacataaattacaaggtgaaaaatgttaaaaagctggtagagcatgtagtaa [ 51200]
ASF_IND_20_CAD543               -------------------------------------------------------------------------------- [ 51200]
LR899193.1_Germany_2020/1       ................................................................................ [ 51200]
MN393477.1_Wuhan_2019           ................................................................................ [ 51200]
MW396979.1_Timor-Leste/2019/1   ................................................................................ [ 51200]
MK940252.1_CN/2019/InnerMongolia ................................................................................ [ 51200]
MK128995.1_China/2018/AnhuiXCGQ ................................................................................ [ 51200]
MH681419.1_ASFV/POL/2015        ................................................................................ [ 51200]
LS478113.1_Estonia_2014         ................................................................................ [ 51200]
KP843857.1_Odintsovo_02/14      ................................................................................ [ 51200]
FN557520.1_strain_E75           ...............................................................................g [ 51200]
AY261366.1_Warthog              ...............................................................a................ [ 51200]
AY261365.1_Warmbaths            ................................................................................ [ 51200]
AY261364.1_Tengani_62           ...a.................a.......................................................... [ 51200]
AY261363.1_Pretoriuskop/96/4    ...a................................c..........................................g [ 51200]
AY261362.1_Mkuzi_1979           ...a...........................................................................g [ 51200]
AY261361.1_Malawi_Lil-20/1_1983 ......g........................................................a...............g [ 51200]
AY261360.1_Kenya_1950           ....................................................a...........t....t.....tc..g [ 51200]
AM712240.1_OURT_88/3            ...............................................................................g [ 51200]
AM712239.1_Benin_97/1           ...............................................................................g [ 51200]
KM111295.1_Ken06.Bus            ....................................................a...........t....tt.....c..g [ 51200]
KM111294.1_Ken05/Tk1            ..a.................................................a...........t....t.....tc..g [ 51200]
KM262844.1_strain_L60           ...............................................................................g [ 51200]
KM102979.1_26544/OG10_Italy     ...............................................................................g [ 51200]
KM262845.1_strain_NHV           ...............................................................................g [ 51200]
KX354450.1_47/Ss/2008           ...............................................................................g [ 51200]
KP055815.1_strain_BA71          ...............................................................................g [ 51200]
MH025920.1_strain_R35           ....................................................a...........t....tt.....c..g [ 51200]
MH025919.1_strain_N10           ....................................................a...........t....tt.....c..g [ 51200]
MH025918.1_strain_R25           ....................................................a...........t....tt.....c..g [ 51200]
MH025917.1_strain_R7            ....................................................a...........t....tt.....c..g [ 51200]
MH025916.1_strain_R8            ....................................................a...........t....tt.....c..g [ 51200]
```

```
FR682468.2_Georgia_2007/1        atgagaaatccactcttgtgttaaaaattttattagaaaaaaaggaaaatctagtggatgctgttttaacaagacttgta [ 51280]
ASF_IND_20_CAD543                -------------------------------------------------------------------------------- [ 51280]
LR899193.1_Germany_2020/1        ................................................................................ [ 51280]
MN393477.1_Wuhan_2019            ................................................................................ [ 51280]
MW396979.1_Timor-Leste/2019/1    ................................................................................ [ 51280]
MK940252.1_CN/2019/InnerMongolia ................................................................................ [ 51280]
MK128995.1_China/2018/AnhuiXCGQ  ................................................................................ [ 51280]
MH681419.1_ASFV/POL/2015         ................................................................................ [ 51280]
LS478113.1_Estonia_2014          ................................................................................ [ 51280]
KP843857.1_Odintsovo_02/14       ................................................................................ [ 51280]
FN557520.1_strain_E75            ..c.c.............................g............................................. [ 51280]
AY261366.1_Warthog               ..................................g............................................. [ 51280]
AY261365.1_Warmbaths             ..................................g............................................. [ 51280]
AY261364.1_Tengani_62            ................................................................................ [ 51280]
AY261363.1_Pretoriuskop/96/4     ..c.c..c..t..c....................g...............t............................. [ 51280]
AY261362.1_Mkuzi_1979            ..c.c..c..t.......................g............................................. [ 51280]
AY261361.1_Malawi_Lil-20/1_1983  ..c.c..c............g...c.........g.........at.......a.a........................ [ 51280]
AY261360.1_Kenya_1950            g..gc..c..........................g.........c....c..g..........c................ [ 51280]
AM712240.1_OURT_88/3             ..c.c.............................g............................................. [ 51280]
AM712239.1_Benin_97/1            ..c.c.............................g............................................. [ 51280]
KM111295.1_Ken06.Bus             g...c..c..........................g.........c....c.............c................ [ 51280]
KM111294.1_Ken05/Tk1             g..gc..c..........................g.........c....c..g..........c................ [ 51280]
KM262844.1_strain_L60            ..c.c.............................g............................................. [ 51280]
KM102979.1_26544/OG10_Italy      ..c.c.............................g............................................. [ 51280]
KM262845.1_strain_NHV            ..c.c.............................g............................................. [ 51280]
KX354450.1_47/Ss/2008            ..c.c.............................g............................................. [ 51280]
KP055815.1_strain_BA71           ..c.c.............................g............................................. [ 51280]
MH025920.1_strain_R35            g...c..c..........................g.........c....c.............c................ [ 51280]
MH025919.1_strain_N10            g...c..c..........................g.........c....c.............c................ [ 51280]
MH025918.1_strain_R25            g...c..c..........................g.........c....c.............c................ [ 51280]
MH025917.1_strain_R7             g...c..c..........................g.........c....c.............c................ [ 51280]
MH025916.1_strain_R8             g...c..c..........................g.........c....c.............c................ [ 51280]

FR682468.2_Georgia_2007/1        aaacattctacatatttccaggtgagagaatttatccaggagttttccatcagcccagaaaaattcattaaaatagctgt [ 51360]
ASF_IND_20_CAD543                -------------------------------------------------------------------------------- [ 51360]
LR899193.1_Germany_2020/1        ................................................................................ [ 51360]
MN393477.1_Wuhan_2019            ................................................................................ [ 51360]
MW396979.1_Timor-Leste/2019/1    ................................................................................ [ 51360]
MK940252.1_CN/2019/InnerMongolia ................................................................................ [ 51360]
MK128995.1_China/2018/AnhuiXCGQ  ................................................................................ [ 51360]
MH681419.1_ASFV/POL/2015         ................................................................................ [ 51360]
LS478113.1_Estonia_2014          ................................................................................ [ 51360]
KP843857.1_Odintsovo_02/14       ................................................................................ [ 51360]
```

```
FN557520.1_strain_E75            ................................................................................ [ 51360]
AY261366.1_Warthog               ................................................................................ [ 51360]
AY261365.1_Warmbaths             ................................................................................ [ 51360]
AY261364.1_Tengani_62            ................................................................................ [ 51360]
AY261363.1_Pretoriuskop/96/4     ...................g............................................................ [ 51360]
AY261362.1_Mkuzi_1979            ...................g...a........................................................ [ 51360]
AY261361.1_Malawi_Lil-20/1_1983  ...................g.....................................----------------------- [ 51360]
AY261360.1_Kenya_1950            ...................g.....a..........g....c...............----------------------- [ 51360]
AM712240.1_OURT_88/3             ................................................................................ [ 51360]
AM712239.1_Benin_97/1            ................................................................................ [ 51360]
KM111295.1_Ken06.Bus             ...................g................g....c...............----------------------- [ 51360]
KM111294.1_Ken05/Tk1             ...................g.....a..........g....c...............----------------------- [ 51360]
KM262844.1_strain_L60            ................................................................................ [ 51360]
KM102979.1_26544/OG10_Italy      ................................................................................ [ 51360]
KM262845.1_strain_NHV            ................................................................................ [ 51360]
KX354450.1_47/Ss/2008            ................................................................................ [ 51360]
KP055815.1_strain_BA71           ................................................................................ [ 51360]
MH025920.1_strain_R35            ...................g................g....c...............----------------------- [ 51360]
MH025919.1_strain_N10            ...................g................g....c...............----------------------- [ 51360]
MH025918.1_strain_R25            ...................g................g....c...............----------------------- [ 51360]
MH025917.1_strain_R7             ...................g................g....c...............----------------------- [ 51360]
MH025916.1_strain_R8             ...................g................g....c...............----------------------- [ 51360]

FR682468.2_Georgia_2007/1        gcgggaaaagaaaaatgtgttaatcgaggctatttctgaagatatttgggaaaatcccacagaaagaattacttatctca [ 51440]
ASF_IND_20_CAD543                -------------------------------------------------------------------------------- [ 51440]
LR899193.1_Germany_2020/1        ................................................................................ [ 51440]
MN393477.1_Wuhan_2019            ................................................................................ [ 51440]
MW396979.1_Timor-Leste/2019/1    ................................................................................ [ 51440]
MK940252.1_CN/2019/InnerMongolia ................................................................................ [ 51440]
MK128995.1_China/2018/AnhuiXCGQ  ................................................................................ [ 51440]
MH681419.1_ASFV/POL/2015         ................................................................................ [ 51440]
LS478113.1_Estonia_2014          ................................................................................ [ 51440]
KP843857.1_Odintsovo_02/14       ................................................................................ [ 51440]
FN557520.1_strain_E75            ................................................................................ [ 51440]
AY261366.1_Warthog               ................................................................................ [ 51440]
AY261365.1_Warmbaths             ................................................................................ [ 51440]
AY261364.1_Tengani_62            .....................g.......................................................... [ 51440]
AY261363.1_Pretoriuskop/96/4     .......c........................................................................ [ 51440]
AY261362.1_Mkuzi_1979            ................................................................................ [ 51440]
AY261361.1_Malawi_Lil-20/1_1983  -------------------------------------------------------------------------------- [ 51440]
AY261360.1_Kenya_1950            -------------------------------------------------------------------------------- [ 51440]
AM712240.1_OURT_88/3             ................................................................................ [ 51440]
AM712239.1_Benin_97/1            ................................................................................ [ 51440]
KM111295.1_Ken06.Bus             -------------------------------------------------------------------------------- [ 51440]
```

```
KM111294.1_Ken05/Tk1              -------------------------------------------------------------------------------- [ 51440]
KM262844.1_strain_L60             ................................................................................ [ 51440]
KM102979.1_26544/OG10_Italy       ................................................................................ [ 51440]
KM262845.1_strain_NHV             ................................................................................ [ 51440]
KX354450.1_47/Ss/2008             ................................................................................ [ 51440]
KP055815.1_strain_BA71            ................................................................................ [ 51440]
MH025920.1_strain_R35             -------------------------------------------------------------------------------- [ 51440]
MH025919.1_strain_N10             -------------------------------------------------------------------------------- [ 51440]
MH025918.1_strain_R25             -------------------------------------------------------------------------------- [ 51440]
MH025917.1_strain_R7              -------------------------------------------------------------------------------- [ 51440]
MH025916.1_strain_R8              -------------------------------------------------------------------------------- [ 51440]

FR682468.2_Georgia_2007/1         aacagatagtgcacaccataaaatatgaaagtggaaggcgatttttggtagacatcattcacagcatttaccaaagttac [ 51520]
ASF_IND_20_CAD543                 -------------------------------------------------------------------------------- [ 51520]
LR899193.1_Germany_2020/1         ................................................................................ [ 51520]
MN393477.1_Wuhan_2019             ................................................................................ [ 51520]
MW396979.1_Timor-Leste/2019/1     ................................................................................ [ 51520]
MK940252.1_CN/2019/InnerMongolia  ................................................................................ [ 51520]
MK128995.1_China/2018/AnhuiXCGQ   ................................................................................ [ 51520]
MH681419.1_ASFV/POL/2015          ................................................................................ [ 51520]
LS478113.1_Estonia_2014           ................................................................................ [ 51520]
KP843857.1_Odintsovo_02/14        ................................................................................ [ 51520]
FN557520.1_strain_E75             ........................................g......a................................ [ 51520]
AY261366.1_Warthog                .....................................a.........a................................ [ 51520]
AY261365.1_Warmbaths              .....................................a.........a....g........................... [ 51520]
AY261364.1_Tengani_62             ................................................................................ [ 51520]
AY261363.1_Pretoriuskop/96/4      ........................................g......a................................ [ 51520]
AY261362.1_Mkuzi_1979             ........................................g......a...............................t [ 51520]
AY261361.1_Malawi_Lil-20/1_1983   -------------------------------------------------------------------------------- [ 51520]
AY261360.1_Kenya_1950             -------------------------------------------------------------------------------- [ 51520]
AM712240.1_OURT_88/3              ........................................g......a................................ [ 51520]
AM712239.1_Benin_97/1             ........................................g......a................................ [ 51520]
KM111295.1_Ken06.Bus              -------------------------------------------------------------------------------- [ 51520]
KM111294.1_Ken05/Tk1              -------------------------------------------------------------------------------- [ 51520]
KM262844.1_strain_L60             ........................................g......a................................ [ 51520]
KM102979.1_26544/OG10_Italy       ........................................g......a................................ [ 51520]
KM262845.1_strain_NHV             ........................................g......a................................ [ 51520]
KX354450.1_47/Ss/2008             ........................................g......a................................ [ 51520]
KP055815.1_strain_BA71            ........................................g......a................................ [ 51520]
MH025920.1_strain_R35             -------------------------------------------------------------------------------- [ 51520]
MH025919.1_strain_N10             -------------------------------------------------------------------------------- [ 51520]
MH025918.1_strain_R25             -------------------------------------------------------------------------------- [ 51520]
MH025917.1_strain_R7              -------------------------------------------------------------------------------- [ 51520]
MH025916.1_strain_R8              -------------------------------------------------------------------------------- [ 51520]
```

```
FR682468.2_Georgia_2007/1        tcactaaaacacgaagatattcttaaactggcaacattttatgtcaaacacaatgcaatcacccattttaaagacctctg [ 51600]
ASF_IND_20_CAD543                -------------------------------------------------------------------------------- [ 51600]
LR899193.1_Germany_2020/1        ................................................................................ [ 51600]
MN393477.1_Wuhan_2019            ................................................................................ [ 51600]
MW396979.1_Timor-Leste/2019/1    ................................................................................ [ 51600]
MK940252.1_CN/2019/InnerMongolia ................................................................................ [ 51600]
MK128995.1_China/2018/AnhuiXCGQ  ................................................................................ [ 51600]
MH681419.1_ASFV/POL/2015         ................................................................................ [ 51600]
LS478113.1_Estonia_2014          ................................................................................ [ 51600]
KP843857.1_Odintsovo_02/14       ................................................................................ [ 51600]
FN557520.1_strain_E75            ..........................................................................t..... [ 51600]
AY261366.1_Warthog               ................................................t.........................t..... [ 51600]
AY261365.1_Warmbaths             ..........................................................c...............t..... [ 51600]
AY261364.1_Tengani_62            ..........................................................c...............t..... [ 51600]
AY261363.1_Pretoriuskop/96/4     ..................................t.............t.........................t..... [ 51600]
AY261362.1_Mkuzi_1979            ...........t.........t....................................................t..... [ 51600]
AY261361.1_Malawi_Lil-20/1_1983  -------------------------------------------------------------------------------- [ 51600]
AY261360.1_Kenya_1950            -------------------------------------------------------------------------------- [ 51600]
AM712240.1_OURT_88/3             ..........................................................................t..... [ 51600]
AM712239.1_Benin_97/1            ..........................................................................t..... [ 51600]
KM111295.1_Ken06.Bus             -------------------------------------------------------------------------------- [ 51600]
KM111294.1_Ken05/Tk1             -------------------------------------------------------------------------------- [ 51600]
KM262844.1_strain_L60            ..........................................................................t..... [ 51600]
KM102979.1_26544/OG10_Italy      ..........................................................................t..... [ 51600]
KM262845.1_strain_NHV            ..........................................................................t..... [ 51600]
KX354450.1_47/Ss/2008            ..........................................................................t..... [ 51600]
KP055815.1_strain_BA71           ..........................................................................t..... [ 51600]
MH025920.1_strain_R35            -------------------------------------------------------------------------------- [ 51600]
MH025919.1_strain_N10            -------------------------------------------------------------------------------- [ 51600]
MH025918.1_strain_R25            -------------------------------------------------------------------------------- [ 51600]
MH025917.1_strain_R7             -------------------------------------------------------------------------------- [ 51600]
MH025916.1_strain_R8             -------------------------------------------------------------------------------- [ 51600]

FR682468.2_Georgia_2007/1        caaatatctttggctgaacagaggaacagaaagtaagaaactgtttttagagtgtttagaaattgctgatgagaaggagt [ 51680]
ASF_IND_20_CAD543                -------------------------------------------------------------------------------- [ 51680]
LR899193.1_Germany_2020/1        ................................................................................ [ 51680]
MN393477.1_Wuhan_2019            ................................................................................ [ 51680]
MW396979.1_Timor-Leste/2019/1    ................................................................................ [ 51680]
MK940252.1_CN/2019/InnerMongolia ................................................................................ [ 51680]
MK128995.1_China/2018/AnhuiXCGQ  ................................................................................ [ 51680]
MH681419.1_ASFV/POL/2015         ................................................................................ [ 51680]
LS478113.1_Estonia_2014          ................................................................................ [ 51680]
KP843857.1_Odintsovo_02/14       ................................................................................ [ 51680]
```

```
FN557520.1_strain_E75              ......................................................c..g...................... [ 51680]
AY261366.1_Warthog                 ................................................................................ [ 51680]
AY261365.1_Warmbaths               ................................................................................ [ 51680]
AY261364.1_Tengani_62              ....................................a.....c..................................... [ 51680]
AY261363.1_Pretoriuskop/96/4       ......................................................c..g...................... [ 51680]
AY261362.1_Mkuzi_1979              ......................................................c..g...................... [ 51680]
AY261361.1_Malawi_Lil-20/1_1983    -------------------------------------------------------------------------------- [ 51680]
AY261360.1_Kenya_1950              -------------------------------------------------------------------------------- [ 51680]
AM712240.1_OURT_88/3               ......................................................c..g...................... [ 51680]
AM712239.1_Benin_97/1              ......................................................c..g...................... [ 51680]
KM111295.1_Ken06.Bus               -------------------------------------------------------------------------------- [ 51680]
KM111294.1_Ken05/Tk1               -------------------------------------------------------------------------------- [ 51680]
KM262844.1_strain_L60              ......................................................c..g...................... [ 51680]
KM102979.1_26544/OG10_Italy        ......................................................c..g...................... [ 51680]
KM262845.1_strain_NHV              ......................................................c..g...................... [ 51680]
KX354450.1_47/Ss/2008              ......................................................c..g...................... [ 51680]
KP055815.1_strain_BA71             ......................................................c..g...................... [ 51680]
MH025920.1_strain_R35              -------------------------------------------------------------------------------- [ 51680]
MH025919.1_strain_N10              -------------------------------------------------------------------------------- [ 51680]
MH025918.1_strain_R25              -------------------------------------------------------------------------------- [ 51680]
MH025917.1_strain_R7               -------------------------------------------------------------------------------- [ 51680]
MH025916.1_strain_R8               -------------------------------------------------------------------------------- [ 51680]

FR682468.2_Georgia_2007/1          ttcctgatattaaaagtattgtgagtgaatatattaactacttgtttactgcaggagctattaccaaggaagaaatcatg [ 51760]
ASF_IND_20_CAD543                  -------------------------------------------------------------------------------- [ 51760]
LR899193.1_Germany_2020/1          ................................................................................ [ 51760]
MN393477.1_Wuhan_2019              ................................................................................ [ 51760]
MW396979.1_Timor-Leste/2019/1      ................................................................................ [ 51760]
MK940252.1_CN/2019/InnerMongolia   ................................................................................ [ 51760]
MK128995.1_China/2018/AnhuiXCGQ    ................................................................................ [ 51760]
MH681419.1_ASFV/POL/2015           ................................................................................ [ 51760]
LS478113.1_Estonia_2014            ................................................................................ [ 51760]
KP843857.1_Odintsovo_02/14         ................................................................................ [ 51760]
FN557520.1_strain_E75              ................................................................................ [ 51760]
AY261366.1_Warthog                 ................c............................................................... [ 51760]
AY261365.1_Warmbaths               ........................................................a....................... [ 51760]
AY261364.1_Tengani_62              ................c.......................................................-------- [ 51760]
AY261363.1_Pretoriuskop/96/4       ................c...............................................t..............a [ 51760]
AY261362.1_Mkuzi_1979              ................c............................................................... [ 51760]
AY261361.1_Malawi_Lil-20/1_1983    -------------------------------------------------------------------------------- [ 51760]
AY261360.1_Kenya_1950              -------------------------------------------------------------------------------- [ 51760]
AM712240.1_OURT_88/3               ................................................................................ [ 51760]
AM712239.1_Benin_97/1              ...............a................................................................ [ 51760]
KM111295.1_Ken06.Bus               -------------------------------------------------------------------------------- [ 51760]
```

```
KM111294.1_Ken05/Tk1             -------------------------------------------------------------------------------- [ 51760]
KM262844.1_strain_L60            ................................................................................ [ 51760]
KM102979.1_26544/OG10_Italy      ................................................................................ [ 51760]
KM262845.1_strain_NHV            ................................................................................ [ 51760]
KX354450.1_47/Ss/2008            ................................................................................ [ 51760]
KP055815.1_strain_BA71           ................................................................................ [ 51760]
MH025920.1_strain_R35            -------------------------------------------------------------------------------- [ 51760]
MH025919.1_strain_N10            -------------------------------------------------------------------------------- [ 51760]
MH025918.1_strain_R25            -------------------------------------------------------------------------------- [ 51760]
MH025917.1_strain_R7             -------------------------------------------------------------------------------- [ 51760]
MH025916.1_strain_R8             -------------------------------------------------------------------------------- [ 51760]

FR682468.2_Georgia_2007/1        caagcctat-gatgctttagagta-gccatgtattaacattctgaaagtagaataaaatatactatatactaaaaa--cc [ 51840]
ASF_IND_20_CAD543                -------------------------------------------------------------------------------- [ 51840]
LR899193.1_Germany_2020/1        .........-..............-...................................................--.. [ 51840]
MN393477.1_Wuhan_2019            .........-..............-...................................................--.. [ 51840]
MW396979.1_Timor-Leste/2019/1    .........-..............-...................................................--.. [ 51840]
MK940252.1_CN/2019/InnerMongolia .........-..............-...................................................--.. [ 51840]
MK128995.1_China/2018/AnhuiXCGQ  .........-..............-...................................................--.. [ 51840]
MH681419.1_ASFV/POL/2015         .........-..............-...................................................--.. [ 51840]
LS478113.1_Estonia_2014          .........-..............-...................................................--.. [ 51840]
KP843857.1_Odintsovo_02/14       .........-..............-...................................................--.. [ 51840]
FN557520.1_strain_E75            .....t.----.............t.a..........at.......g.....c................t......at.. [ 51840]
AY261366.1_Warthog               .........-..............-...........................c..........a..g.........--.. [ 51840]
AY261365.1_Warmbaths             .........-..............-...........................c.............g..t......--.. [ 51840]
AY261364.1_Tengani_62            -........g..............-...........................c.............g.........--.. [ 51840]
AY261363.1_Pretoriuskop/96/4     t........-...........a..-...........................c.............g.........--.. [ 51840]
AY261362.1_Mkuzi_1979            .........-a.............-.....a..................g..c.............g.........--.. [ 51840]
AY261361.1_Malawi_Lil-20/1_1983  -------------------------------------------------------------------------------- [ 51840]
AY261360.1_Kenya_1950            -------------------------------------------------------------------------------- [ 51840]
AM712240.1_OURT_88/3             .....t.----.............t.a..........at.......g.....c................t......at.. [ 51840]
AM712239.1_Benin_97/1            .....t.----.............t.a..........at.......g.....c................t......at.. [ 51840]
KM111295.1_Ken06.Bus             -------------------------------------------------------------------------------- [ 51840]
KM111294.1_Ken05/Tk1             -------------------------------------------------------------------------------- [ 51840]
KM262844.1_strain_L60            .....t.----.............t.a..........at.......g.....c................t......at.. [ 51840]
KM102979.1_26544/OG10_Italy      .....t.----.............t.a..........at.......g.....c................t......at.. [ 51840]
KM262845.1_strain_NHV            .....t.----.............t.a..........at.......g.....c................t......at.. [ 51840]
KX354450.1_47/Ss/2008            .....t.----.............t.a..........at.......g.....c................t......at.. [ 51840]
KP055815.1_strain_BA71           .....t.----.............t.a..........at.......g.....c................t......at.. [ 51840]
MH025920.1_strain_R35            -------------------------------------------------------------------------------- [ 51840]
MH025919.1_strain_N10            -------------------------------------------------------------------------------- [ 51840]
MH025918.1_strain_R25            -------------------------------------------------------------------------------- [ 51840]
MH025917.1_strain_R7             -------------------------------------------------------------------------------- [ 51840]
MH025916.1_strain_R8             -------------------------------------------------------------------------------- [ 51840]
```

```
FR682468.2_Georgia_2007/1        aaattagcca-tttttaactatcttcttcttaaaaactctggataaaaatttatttttttttaatttgggtagggaaaat [ 51920]
ASF_IND_20_CAD543                -------------------------------------------------------------------------------- [ 51920]
LR899193.1_Germany_2020/1        ..........-..................................................................... [ 51920]
MN393477.1_Wuhan_2019            ..........-..................................................................... [ 51920]
MW396979.1_Timor-Leste/2019/1    ..........-..................................................................... [ 51920]
MK940252.1_CN/2019/InnerMongolia ..........-..................................................................... [ 51920]
MK128995.1_China/2018/AnhuiXCGQ  ..........-..................................................................... [ 51920]
MH681419.1_ASFV/POL/2015         ..........-..................................................................... [ 51920]
LS478113.1_Estonia_2014          ..........-..................................................................... [ 51920]
KP843857.1_Odintsovo_02/14       ..........-..........................................-.......................... [ 51920]
FN557520.1_strain_E75            ...ac.....-..........a..............................................a........... [ 51920]
AY261366.1_Warthog               ..........t................c.................................................... [ 51920]
AY261365.1_Warmbaths             ..........t................c.................................................... [ 51920]
AY261364.1_Tengani_62            ..........-..................................................................... [ 51920]
AY261363.1_Pretoriuskop/96/4     ..........t..................................................................... [ 51920]
AY261362.1_Mkuzi_1979            ..........-..........a.......................................................... [ 51920]
AY261361.1_Malawi_Lil-20/1_1983  -------------------------------------------------------------------------------- [ 51920]
AY261360.1_Kenya_1950            -------------------------------------------------------------------------------- [ 51920]
AM712240.1_OURT_88/3             ...ac.....-..........a...............................-..............a........... [ 51920]
AM712239.1_Benin_97/1            ...ac.....-..........a..............................................a........... [ 51920]
KM111295.1_Ken06.Bus             -------------------------------------------------------------------------------- [ 51920]
KM111294.1_Ken05/Tk1             -------------------------------------------------------------------------------- [ 51920]
KM262844.1_strain_L60            ...ac.....-..........a..............................................a........... [ 51920]
KM102979.1_26544/OG10_Italy      ...ac.....-..........a..............................................a........... [ 51920]
KM262845.1_strain_NHV            ...ac.....-..........a...............................-..............a........... [ 51920]
KX354450.1_47/Ss/2008            ...ac.....-..........a..............................................a........... [ 51920]
KP055815.1_strain_BA71           ...ac.....-..........a..............................................a........... [ 51920]
MH025920.1_strain_R35            -------------------------------------------------------------------------------- [ 51920]
MH025919.1_strain_N10            -------------------------------------------------------------------------------- [ 51920]
MH025918.1_strain_R25            -------------------------------------------------------------------------------- [ 51920]
MH025917.1_strain_R7             -------------------------------------------------------------------------------- [ 51920]
MH025916.1_strain_R8             -------------------------------------------------------------------------------- [ 51920]

FR682468.2_Georgia_2007/1        gttctcccttcaggacctctgtcggaagaacaccttcttccttccaagtgattttagcaagcataccctgcatttgctgg [ 52000]
ASF_IND_20_CAD543                -------------------------------------------------------------------------------- [ 52000]
LR899193.1_Germany_2020/1        ................................................................................ [ 52000]
MN393477.1_Wuhan_2019            ................................................................................ [ 52000]
MW396979.1_Timor-Leste/2019/1    ................................................................................ [ 52000]
MK940252.1_CN/2019/InnerMongolia ................................................................................ [ 52000]
MK128995.1_China/2018/AnhuiXCGQ  ................................................................................ [ 52000]
MH681419.1_ASFV/POL/2015         ................................................................................ [ 52000]
LS478113.1_Estonia_2014          ................................................................................ [ 52000]
KP843857.1_Odintsovo_02/14       ................................................................................ [ 52000]
```

```
FN557520.1_strain_E75            ..............................................ga.a....................t......... [ 52000]
AY261366.1_Warthog               ..................................................................t............. [ 52000]
AY261365.1_Warmbaths             ................................................................................ [ 52000]
AY261364.1_Tengani_62            ................................................................................ [ 52000]
AY261363.1_Pretoriuskop/96/4     .................................................a.............................. [ 52000]
AY261362.1_Mkuzi_1979            ......t.......................................ga.a.............................. [ 52000]
AY261361.1_Malawi_Lil-20/1_1983  -------------------------------------------------------------------------------- [ 52000]
AY261360.1_Kenya_1950            -------------------------------------------------------------------------------- [ 52000]
AM712240.1_OURT_88/3             ..............................................ga.a....................t......... [ 52000]
AM712239.1_Benin_97/1            ..............................................ga.a....................t......... [ 52000]
KM111295.1_Ken06.Bus             -------------------------------------------------------------------------------- [ 52000]
KM111294.1_Ken05/Tk1             -------------------------------------------------------------------------------- [ 52000]
KM262844.1_strain_L60            ..............................................ga.a....................t......... [ 52000]
KM102979.1_26544/OG10_Italy      ..............................................ga.a....................t......... [ 52000]
KM262845.1_strain_NHV            ..............................................ga.a....................t......... [ 52000]
KX354450.1_47/Ss/2008            ..............................................ga.a....................t......... [ 52000]
KP055815.1_strain_BA71           ..............................................ga.a....................t......... [ 52000]
MH025920.1_strain_R35            -------------------------------------------------------------------------------- [ 52000]
MH025919.1_strain_N10            -------------------------------------------------------------------------------- [ 52000]
MH025918.1_strain_R25            -------------------------------------------------------------------------------- [ 52000]
MH025917.1_strain_R7             -------------------------------------------------------------------------------- [ 52000]
MH025916.1_strain_R8             -------------------------------------------------------------------------------- [ 52000]

FR682468.2_Georgia_2007/1        ggttatactggaaggggcatggatctatccaaaggataaagaatgatggtgtgcttatagagcatgatcttactctttcc [ 52080]
ASF_IND_20_CAD543                -------------------------------------------------------------------------------- [ 52080]
LR899193.1_Germany_2020/1        ................................................................................ [ 52080]
MN393477.1_Wuhan_2019            ................................................................................ [ 52080]
MW396979.1_Timor-Leste/2019/1    ................................................................................ [ 52080]
MK940252.1_CN/2019/InnerMongolia ................................................................................ [ 52080]
MK128995.1_China/2018/AnhuiXCGQ  ................................................................................ [ 52080]
MH681419.1_ASFV/POL/2015         ................................................................................ [ 52080]
LS478113.1_Estonia_2014          ................................................................................ [ 52080]
KP843857.1_Odintsovo_02/14       ................................................................................ [ 52080]
FN557520.1_strain_E75            ................a.................a.c..t............a..g.........a......a....... [ 52080]
AY261366.1_Warthog               .......................................................................ct....... [ 52080]
AY261365.1_Warmbaths             .......................................................................ct....... [ 52080]
AY261364.1_Tengani_62            ................................................................................ [ 52080]
AY261363.1_Pretoriuskop/96/4     ................................................................................ [ 52080]
AY261362.1_Mkuzi_1979            ................................................................................ [ 52080]
AY261361.1_Malawi_Lil-20/1_1983  -------------------------------------------------------------------------------- [ 52080]
AY261360.1_Kenya_1950            -------------------------------------------------------------------------------- [ 52080]
AM712240.1_OURT_88/3             ................a.................a.c..t.....t......a..g.........a......a....... [ 52080]
AM712239.1_Benin_97/1            ................a.................a.c..t............a..g.........a......a....... [ 52080]
KM111295.1_Ken06.Bus             -------------------------------------------------------------------------------- [ 52080]
```

```
KM111294.1_Ken05/Tk1            -------------------------------------------------------------------------------- [ 52080]
KM262844.1_strain_L60           ................a.................a.c..t............a.g..........a......a....... [ 52080]
KM102979.1_26544/OG10_Italy     ................a.................a.c..t............a.g..........a......a....... [ 52080]
KM262845.1_strain_NHV           ................a.................a.c..t.....t......a.g..........a......a....... [ 52080]
KX354450.1_47/Ss/2008           ................a.................a.c..t............a.g..........a......a....... [ 52080]
KP055815.1_strain_BA71          ................a.................a.c..t............a.g..........a......a....... [ 52080]
MH025920.1_strain_R35           -------------------------------------------------------------------------------- [ 52080]
MH025919.1_strain_N10           -------------------------------------------------------------------------------- [ 52080]
MH025918.1_strain_R25           -------------------------------------------------------------------------------- [ 52080]
MH025917.1_strain_R7            -------------------------------------------------------------------------------- [ 52080]
MH025916.1_strain_R8            -------------------------------------------------------------------------------- [ 52080]

FR682468.2_Georgia_2007/1       atcaatgaagccttaattcttgcaggagaagagggaaacaatgaagtagtaaagctcttgttactatgggaaggaaatct [ 52160]
ASF_IND_20_CAD543               -------------------------------------------------------------------------------- [ 52160]
LR899193.1_Germany_2020/1       ................................................................................ [ 52160]
MN393477.1_Wuhan_2019           ................................................................................ [ 52160]
MW396979.1_Timor-Leste/2019/1   ................................................................................ [ 52160]
MK940252.1_CN/2019/InnerMongolia ................................................................................ [ 52160]
MK128995.1_China/2018/AnhuiXCGQ ................................................................................ [ 52160]
MH681419.1_ASFV/POL/2015        ................................................................................ [ 52160]
LS478113.1_Estonia_2014         ................................................................................ [ 52160]
KP843857.1_Odintsovo_02/14      ................................................................................ [ 52160]
FN557520.1_strain_E75           .................c..........................t......c.a.....a..g................. [ 52160]
AY261366.1_Warthog              .................c..........................t......c..........g................. [ 52160]
AY261365.1_Warmbaths            .................c..........................ta.....c..........g...........t..... [ 52160]
AY261364.1_Tengani_62           ................................................................................ [ 52160]
AY261363.1_Pretoriuskop/96/4    .................c..........................ta.....c..........g................. [ 52160]
AY261362.1_Mkuzi_1979           .................c..........................ta.g...c....t.....g................. [ 52160]
AY261361.1_Malawi_Lil-20/1_1983 -------------------------------------------------------------------------------- [ 52160]
AY261360.1_Kenya_1950           -------------------------------------------------------------------------------- [ 52160]
AM712240.1_OURT_88/3            .................c..........................t......c.a.....a..g................. [ 52160]
AM712239.1_Benin_97/1           .................c..........................t......c.a.....a..g................. [ 52160]
KM111295.1_Ken06.Bus            -------------------------------------------------------------------------------- [ 52160]
KM111294.1_Ken05/Tk1            -------------------------------------------------------------------------------- [ 52160]
KM262844.1_strain_L60           .................c..........................t......c.a.....a..g................. [ 52160]
KM102979.1_26544/OG10_Italy     .................c..........................t......c.a.....a..g................. [ 52160]
KM262845.1_strain_NHV           .................c..........................t......c.a.....a..g................. [ 52160]
KX354450.1_47/Ss/2008           .................c..........................t......c.a.....a..g................. [ 52160]
KP055815.1_strain_BA71          .................c..........................t......c.a.....a..g................. [ 52160]
MH025920.1_strain_R35           -------------------------------------------------------------------------------- [ 52160]
MH025919.1_strain_N10           -------------------------------------------------------------------------------- [ 52160]
MH025918.1_strain_R25           -------------------------------------------------------------------------------- [ 52160]
MH025917.1_strain_R7            -------------------------------------------------------------------------------- [ 52160]
MH025916.1_strain_R8            -------------------------------------------------------------------------------- [ 52160]
```

```
FR682468.2_Georgia_2007/1        tcattatgccatcataggagctttgaggactgagaactataacctagtatgtgagtaccatagtcaaattcaggactggc [ 52240]
ASF_IND_20_CAD543                -------------------------------------------------------------------------------- [ 52240]
LR899193.1_Germany_2020/1        ................................................................................ [ 52240]
MN393477.1_Wuhan_2019            ................................................................................ [ 52240]
MW396979.1_Timor-Leste/2019/1    ................................................................................ [ 52240]
MK940252.1_CN/2019/InnerMongolia ................................................................................ [ 52240]
MK128995.1_China/2018/AnhuiXCGQ  ................................................................................ [ 52240]
MH681419.1_ASFV/POL/2015         ................................................................................ [ 52240]
LS478113.1_Estonia_2014          ................................................................................ [ 52240]
KP843857.1_Odintsovo_02/14       ................................................................................ [ 52240]
FN557520.1_strain_E75            ..........................a.........a...gg.t..a................c................ [ 52240]
AY261366.1_Warthog               g...................................a...gg.t..a................c................ [ 52240]
AY261365.1_Warmbaths             g...................................a...gg.t..a..........t.....c................ [ 52240]
AY261364.1_Tengani_62            ................................................................................ [ 52240]
AY261363.1_Pretoriuskop/96/4     g....................c....a.........a....g.....................c................ [ 52240]
AY261362.1_Mkuzi_1979            ..........................a.........a...gg.t..a................c................ [ 52240]
AY261361.1_Malawi_Lil-20/1_1983  -------------------------------------------------------------------------------- [ 52240]
AY261360.1_Kenya_1950            -------------------------------------------------------------------------------- [ 52240]
AM712240.1_OURT_88/3             ..........................a.........a...gg.t..a................c................ [ 52240]
AM712239.1_Benin_97/1            ..........................a.........a...gg.t..a................c................ [ 52240]
KM111295.1_Ken06.Bus             -------------------------------------------------------------------------------- [ 52240]
KM111294.1_Ken05/Tk1             -------------------------------------------------------------------------------- [ 52240]
KM262844.1_strain_L60            ..........................a.........a...gg.t..a................c................ [ 52240]
KM102979.1_26544/OG10_Italy      ..........................a.........a...gg.t..a................c................ [ 52240]
KM262845.1_strain_NHV            ..........................a.........a...gg.t..a................c................ [ 52240]
KX354450.1_47/Ss/2008            ..........................a.........a...gg.t..a................c................ [ 52240]
KP055815.1_strain_BA71           ..........................a.........a...gg.t..a................c................ [ 52240]
MH025920.1_strain_R35            -------------------------------------------------------------------------------- [ 52240]
MH025919.1_strain_N10            -------------------------------------------------------------------------------- [ 52240]
MH025918.1_strain_R25            -------------------------------------------------------------------------------- [ 52240]
MH025917.1_strain_R7             -------------------------------------------------------------------------------- [ 52240]
MH025916.1_strain_R8             -------------------------------------------------------------------------------- [ 52240]

FR682468.2_Georgia_2007/1        atgttctcctccctttgattcaagatccagaaacattcgaaaaatgtcatgatttaagccttgaatgtgatctttcatgc [ 52320]
ASF_IND_20_CAD543                -------------------------------------------------------------------------------- [ 52320]
LR899193.1_Germany_2020/1        ................................................................................ [ 52320]
MN393477.1_Wuhan_2019            ................................................................................ [ 52320]
MW396979.1_Timor-Leste/2019/1    ................................................................................ [ 52320]
MK940252.1_CN/2019/InnerMongolia ................................................................................ [ 52320]
MK128995.1_China/2018/AnhuiXCGQ  ................................................................................ [ 52320]
MH681419.1_ASFV/POL/2015         ................................................................................ [ 52320]
LS478113.1_Estonia_2014          ................................................................................ [ 52320]
KP843857.1_Odintsovo_02/14       ................................................................................ [ 52320]
```

```
FN557520.1_strain_E75           .............c.................................................................. [ 52320]
AY261366.1_Warthog              .............c.................................................................. [ 52320]
AY261365.1_Warmbaths            .............c.................................................................. [ 52320]
AY261364.1_Tengani_62           ................................................................................ [ 52320]
AY261363.1_Pretoriuskop/96/4    .............c.................................................................. [ 52320]
AY261362.1_Mkuzi_1979           .............c.................................................................. [ 52320]
AY261361.1_Malawi_Lil-20/1_1983 -------------------------------------------------------------------------------- [ 52320]
AY261360.1_Kenya_1950           -------------------------------------------------------------------------------- [ 52320]
AM712240.1_OURT_88/3            .............c.................................................................. [ 52320]
AM712239.1_Benin_97/1           .............c.................................................................. [ 52320]
KM111295.1_Ken06.Bus            -------------------------------------------------------------------------------- [ 52320]
KM111294.1_Ken05/Tk1            -------------------------------------------------------------------------------- [ 52320]
KM262844.1_strain_L60           .............c.................................................................. [ 52320]
KM102979.1_26544/OG10_Italy     .............c.................................................................. [ 52320]
KM262845.1_strain_NHV           .............c.................................................................. [ 52320]
KX354450.1_47/Ss/2008           .............c.................................................................. [ 52320]
KP055815.1_strain_BA71          .............c.................................................................. [ 52320]
MH025920.1_strain_R35           -------------------------------------------------------------------------------- [ 52320]
MH025919.1_strain_N10           -------------------------------------------------------------------------------- [ 52320]
MH025918.1_strain_R25           -------------------------------------------------------------------------------- [ 52320]
MH025917.1_strain_R7            -------------------------------------------------------------------------------- [ 52320]
MH025916.1_strain_R8            -------------------------------------------------------------------------------- [ 52320]

FR682468.2_Georgia_2007/1       cttctccaacatgctgtaaaatataacatgctttcgattcttgttaaatataaagaggatctactaaatgtactatttag [ 52400]
ASF_IND_20_CAD543               -------------------------------------------------------------------------------- [ 52400]
LR899193.1_Germany_2020/1       ................................................................................ [ 52400]
MN393477.1_Wuhan_2019           ................................................................................ [ 52400]
MW396979.1_Timor-Leste/2019/1   ................................................................................ [ 52400]
MK940252.1_CN/2019/InnerMongolia ................................................................................ [ 52400]
MK128995.1_China/2018/AnhuiXCGQ ................................................................................ [ 52400]
MH681419.1_ASFV/POL/2015        ................................................................................ [ 52400]
LS478113.1_Estonia_2014         ................................................................................ [ 52400]
KP843857.1_Odintsovo_02/14      ................................................................................ [ 52400]
FN557520.1_strain_E75           ...................................t.....................a......t................ [ 52400]
AY261366.1_Warthog              ...................................t...................................g........ [ 52400]
AY261365.1_Warmbaths            ...................................t...................................g........ [ 52400]
AY261364.1_Tengani_62           ...................................t............................................ [ 52400]
AY261363.1_Pretoriuskop/96/4    ...................................t...................................g........ [ 52400]
AY261362.1_Mkuzi_1979           ...................................t...........................t................ [ 52400]
AY261361.1_Malawi_Lil-20/1_1983 -------------------------------------------------------------------------------- [ 52400]
AY261360.1_Kenya_1950           -------------------------------------------------------------------------------- [ 52400]
AM712240.1_OURT_88/3            ...................................t...........................t................ [ 52400]
AM712239.1_Benin_97/1           ...................................t...........................t................ [ 52400]
KM111295.1_Ken06.Bus            -------------------------------------------------------------------------------- [ 52400]
```

```
KM111294.1_Ken05/Tk1             -------------------------------------------------------------------------------- [ 52400]
KM262844.1_strain_L60            ...................................t...........................t................ [ 52400]
KM102979.1_26544/OG10_Italy      ...................................t...........................t................ [ 52400]
KM262845.1_strain_NHV            ...................................t...........................t................ [ 52400]
KX354450.1_47/Ss/2008            ...................................t...........................t................ [ 52400]
KP055815.1_strain_BA71           ...................................t...........................t................ [ 52400]
MH025920.1_strain_R35            -------------------------------------------------------------------------------- [ 52400]
MH025919.1_strain_N10            -------------------------------------------------------------------------------- [ 52400]
MH025918.1_strain_R25            -------------------------------------------------------------------------------- [ 52400]
MH025917.1_strain_R7             -------------------------------------------------------------------------------- [ 52400]
MH025916.1_strain_R8             -------------------------------------------------------------------------------- [ 52400]

FR682468.2_Georgia_2007/1        gcaacaaattcaaggactatttattttagcatgtgaaaatcggaagcttgagattcttacgtggatgggtcaaaatctgc [ 52480]
ASF_IND_20_CAD543                -------------------------------------------------------------------------------- [ 52480]
LR899193.1_Germany_2020/1        ................................................................................ [ 52480]
MN393477.1_Wuhan_2019            ................................................................................ [ 52480]
MW396979.1_Timor-Leste/2019/1    ................................................................................ [ 52480]
MK940252.1_CN/2019/InnerMongolia ................................................................................ [ 52480]
MK128995.1_China/2018/AnhuiXCGQ  ................................................................................ [ 52480]
MH681419.1_ASFV/POL/2015         ................................................................................ [ 52480]
LS478113.1_Estonia_2014          ................................................................................ [ 52480]
KP843857.1_Odintsovo_02/14       ................................................................................ [ 52480]
FN557520.1_strain_E75            .....................................t......g.a................................. [ 52480]
AY261366.1_Warthog               ..............................................t.a............................... [ 52480]
AY261365.1_Warmbaths             ..............................................t................................. [ 52480]
AY261364.1_Tengani_62            ................................................................................ [ 52480]
AY261363.1_Pretoriuskop/96/4     ............................................t.t................................. [ 52480]
AY261362.1_Mkuzi_1979            .....................................c......t.t................................. [ 52480]
AY261361.1_Malawi_Lil-20/1_1983  -------------------------------------------------------------------------------- [ 52480]
AY261360.1_Kenya_1950            -------------------------------------------------------------------------------- [ 52480]
AM712240.1_OURT_88/3             .....................................c......g.a................................. [ 52480]
AM712239.1_Benin_97/1            .....................................c......g.a................................. [ 52480]
KM111295.1_Ken06.Bus             -------------------------------------------------------------------------------- [ 52480]
KM111294.1_Ken05/Tk1             -------------------------------------------------------------------------------- [ 52480]
KM262844.1_strain_L60            .....................................c......g.a................................. [ 52480]
KM102979.1_26544/OG10_Italy      .....................................c......g.a................................. [ 52480]
KM262845.1_strain_NHV            .....................................c......g.a................................. [ 52480]
KX354450.1_47/Ss/2008            .....................................c......g.a................................. [ 52480]
KP055815.1_strain_BA71           .....................................c......g.a................................. [ 52480]
MH025920.1_strain_R35            -------------------------------------------------------------------------------- [ 52480]
MH025919.1_strain_N10            -------------------------------------------------------------------------------- [ 52480]
MH025918.1_strain_R25            -------------------------------------------------------------------------------- [ 52480]
MH025917.1_strain_R7             -------------------------------------------------------------------------------- [ 52480]
MH025916.1_strain_R8             -------------------------------------------------------------------------------- [ 52480]
```

```
FR682468.2_Georgia_2007/1        caattcctgatcctgagcctatttttagcattgctgttgtcacaaaagatttagaaatgttttccttagggtacaagatt [ 52560]
ASF_IND_20_CAD543                -------------------------------------------------------------------------------- [ 52560]
LR899193.1_Germany_2020/1        ................................................................................ [ 52560]
MN393477.1_Wuhan_2019            ................................................................................ [ 52560]
MW396979.1_Timor-Leste/2019/1    ................................................................................ [ 52560]
MK940252.1_CN/2019/InnerMongolia ................................................................................ [ 52560]
MK128995.1_China/2018/AnhuiXCGQ  ................................................................................ [ 52560]
MH681419.1_ASFV/POL/2015         ................................................................................ [ 52560]
LS478113.1_Estonia_2014          ................................................................................ [ 52560]
KP843857.1_Odintsovo_02/14       ................................................................................ [ 52560]
FN557520.1_strain_E75            ................................................................................ [ 52560]
AY261366.1_Warthog               ...........................a.................................................... [ 52560]
AY261365.1_Warmbaths             ................................................................................ [ 52560]
AY261364.1_Tengani_62            ................................................................................ [ 52560]
AY261363.1_Pretoriuskop/96/4     ................................................................................ [ 52560]
AY261362.1_Mkuzi_1979            ................................................................................ [ 52560]
AY261361.1_Malawi_Lil-20/1_1983  -------------------------------------------------------------------------------- [ 52560]
AY261360.1_Kenya_1950            -------------------------------------------------------------------------------- [ 52560]
AM712240.1_OURT_88/3             ................................................................................ [ 52560]
AM712239.1_Benin_97/1            ................................................................................ [ 52560]
KM111295.1_Ken06.Bus             -------------------------------------------------------------------------------- [ 52560]
KM111294.1_Ken05/Tk1             -------------------------------------------------------------------------------- [ 52560]
KM262844.1_strain_L60            ................................................................................ [ 52560]
KM102979.1_26544/OG10_Italy      ................................................................................ [ 52560]
KM262845.1_strain_NHV            ................................................................................ [ 52560]
KX354450.1_47/Ss/2008            ................................................................................ [ 52560]
KP055815.1_strain_BA71           ................................................................................ [ 52560]
MH025920.1_strain_R35            -------------------------------------------------------------------------------- [ 52560]
MH025919.1_strain_N10            -------------------------------------------------------------------------------- [ 52560]
MH025918.1_strain_R25            -------------------------------------------------------------------------------- [ 52560]
MH025917.1_strain_R7             -------------------------------------------------------------------------------- [ 52560]
MH025916.1_strain_R8             -------------------------------------------------------------------------------- [ 52560]

FR682468.2_Georgia_2007/1        gtttttgaatacatggaaaaccaaggac---ttcatttaacccaggtagttcgtatggttatgctaaatcatcactttgg [ 52640]
ASF_IND_20_CAD543                -------------------------------------------------------------------------------- [ 52640]
LR899193.1_Germany_2020/1        ............................---................................................. [ 52640]
MN393477.1_Wuhan_2019            ............................---................................................. [ 52640]
MW396979.1_Timor-Leste/2019/1    ............................---................................................. [ 52640]
MK940252.1_CN/2019/InnerMongolia ............................---................................................. [ 52640]
MK128995.1_China/2018/AnhuiXCGQ  ............................---................................................. [ 52640]
MH681419.1_ASFV/POL/2015         ............................---................................................. [ 52640]
LS478113.1_Estonia_2014          ............................---................................................. [ 52640]
KP843857.1_Odintsovo_02/14       ............................---................................................. [ 52640]
```

```
FN557520.1_strain_E75            ....................t.......tat............................................c.... [ 52640]
AY261366.1_Warthog               ............................tgt................................a......g.......a. [ 52640]
AY261365.1_Warmbaths             ............................tgt................................a......g.......a. [ 52640]
AY261364.1_Tengani_62            ............................---................................................. [ 52640]
AY261363.1_Pretoriuskop/96/4     ....................t.......tgt.......................................g.......a. [ 52640]
AY261362.1_Mkuzi_1979            ....................t.......tgt................................................a [ 52640]
AY261361.1_Malawi_Lil-20/1_1983  -------------------------------------------------------------------------------- [ 52640]
AY261360.1_Kenya_1950            -------------------------------------------------------------------------------- [ 52640]
AM712240.1_OURT_88/3             ....................t.......tat............................................c.... [ 52640]
AM712239.1_Benin_97/1            ....................t.......tat............................................c.... [ 52640]
KM111295.1_Ken06.Bus             -------------------------------------------------------------------------------- [ 52640]
KM111294.1_Ken05/Tk1             -------------------------------------------------------------------------------- [ 52640]
KM262844.1_strain_L60            ....................t.......tat............................................c.... [ 52640]
KM102979.1_26544/OG10_Italy      ....................t.......tat............................................c.... [ 52640]
KM262845.1_strain_NHV            ....................t.......tat............................................c.... [ 52640]
KX354450.1_47/Ss/2008            ....................t.......tat............................................c.... [ 52640]
KP055815.1_strain_BA71           ....................t.......tat............................................c.... [ 52640]
MH025920.1_strain_R35            -------------------------------------------------------------------------------- [ 52640]
MH025919.1_strain_N10            -------------------------------------------------------------------------------- [ 52640]
MH025918.1_strain_R25            -------------------------------------------------------------------------------- [ 52640]
MH025917.1_strain_R7             -------------------------------------------------------------------------------- [ 52640]
MH025916.1_strain_R8             -------------------------------------------------------------------------------- [ 52640]

FR682468.2_Georgia_2007/1        catggtaataaataaaggacttttaccctttgtgctggaaattttaaattatggtgggaatgtaaatagagccttatctt [ 52720]
ASF_IND_20_CAD543                -------------------------------------------------------------------------------- [ 52720]
LR899193.1_Germany_2020/1        ................................................................................ [ 52720]
MN393477.1_Wuhan_2019            ................................................................................ [ 52720]
MW396979.1_Timor-Leste/2019/1    ................................................................................ [ 52720]
MK940252.1_CN/2019/InnerMongolia ................................................................................ [ 52720]
MK128995.1_China/2018/AnhuiXCGQ  ................................................................................ [ 52720]
MH681419.1_ASFV/POL/2015         ................................................................................ [ 52720]
LS478113.1_Estonia_2014          ................................................................................ [ 52720]
KP843857.1_Odintsovo_02/14       ................................................................................ [ 52720]
FN557520.1_strain_E75            .........................................c......ac.............................. [ 52720]
AY261366.1_Warthog               ................................................ac........c..................... [ 52720]
AY261365.1_Warmbaths             ................................................ac........c..................... [ 52720]
AY261364.1_Tengani_62            ................................................a............................... [ 52720]
AY261363.1_Pretoriuskop/96/4     ................................................ac.............................. [ 52720]
AY261362.1_Mkuzi_1979            .........................................c......ac.............................. [ 52720]
AY261361.1_Malawi_Lil-20/1_1983  -------------------------------------------------------------------------------- [ 52720]
AY261360.1_Kenya_1950            -------------------------------------------------------------------------------- [ 52720]
AM712240.1_OURT_88/3             .........................................c......ac.............................. [ 52720]
AM712239.1_Benin_97/1            .........................................c......ac.............................. [ 52720]
KM111295.1_Ken06.Bus             -------------------------------------------------------------------------------- [ 52720]
```

```
KM111294.1_Ken05/Tk1              -------------------------------------------------------------------------------- [ 52720]
KM262844.1_strain_L60             .........................................c......ac.............................. [ 52720]
KM102979.1_26544/OG10_Italy       .........................................c......ac.............................. [ 52720]
KM262845.1_strain_NHV             .........................................c......ac.............................. [ 52720]
KX354450.1_47/Ss/2008             .........................................c......ac.............................. [ 52720]
KP055815.1_strain_BA71            .........................................c......ac.............................. [ 52720]
MH025920.1_strain_R35             -------------------------------------------------------------------------------- [ 52720]
MH025919.1_strain_N10             -------------------------------------------------------------------------------- [ 52720]
MH025918.1_strain_R25             -------------------------------------------------------------------------------- [ 52720]
MH025917.1_strain_R7              -------------------------------------------------------------------------------- [ 52720]
MH025916.1_strain_R8              -------------------------------------------------------------------------------- [ 52720]

FR682468.2_Georgia_2007/1         atgctgtcacacaaaataaaagaaagattttagaccatgttgttcgccaaaagaatataccccataaaaccattgaaaga [ 52800]
ASF_IND_20_CAD543                 -------------------------------------------------------------------------------- [ 52800]
LR899193.1_Germany_2020/1         ................................................................................ [ 52800]
MN393477.1_Wuhan_2019             ................................................................................ [ 52800]
MW396979.1_Timor-Leste/2019/1     ................................................................................ [ 52800]
MK940252.1_CN/2019/InnerMongolia  ................................................................................ [ 52800]
MK128995.1_China/2018/AnhuiXCGQ   ................................................................................ [ 52800]
MH681419.1_ASFV/POL/2015          ................................................................................ [ 52800]
LS478113.1_Estonia_2014           ................................................................................ [ 52800]
KP843857.1_Odintsovo_02/14        ................................................................................ [ 52800]
FN557520.1_strain_E75             ................c............................................................... [ 52800]
AY261366.1_Warthog                ........................................................g....................... [ 52800]
AY261365.1_Warmbaths              ..........................g.............................g....................... [ 52800]
AY261364.1_Tengani_62             ................................................................................ [ 52800]
AY261363.1_Pretoriuskop/96/4      ........................................................g....................... [ 52800]
AY261362.1_Mkuzi_1979             ................c............................................................... [ 52800]
AY261361.1_Malawi_Lil-20/1_1983   -------------------------------------------------------------------------------- [ 52800]
AY261360.1_Kenya_1950             -------------------------------------------------------------------------------- [ 52800]
AM712240.1_OURT_88/3              ................c............................................................... [ 52800]
AM712239.1_Benin_97/1             ................c............................................................... [ 52800]
KM111295.1_Ken06.Bus              -------------------------------------------------------------------------------- [ 52800]
KM111294.1_Ken05/Tk1              -------------------------------------------------------------------------------- [ 52800]
KM262844.1_strain_L60             ................c............................................................... [ 52800]
KM102979.1_26544/OG10_Italy       ................c............................................................... [ 52800]
KM262845.1_strain_NHV             ................c............................................................... [ 52800]
KX354450.1_47/Ss/2008             ................c............................................................... [ 52800]
KP055815.1_strain_BA71            ................c............................................................... [ 52800]
MH025920.1_strain_R35             -------------------------------------------------------------------------------- [ 52800]
MH025919.1_strain_N10             -------------------------------------------------------------------------------- [ 52800]
MH025918.1_strain_R25             -------------------------------------------------------------------------------- [ 52800]
MH025917.1_strain_R7              -------------------------------------------------------------------------------- [ 52800]
MH025916.1_strain_R8              -------------------------------------------------------------------------------- [ 52800]
```

```
FR682468.2_Georgia_2007/1        atgttgcatctggctgtaaaaaagcatgctcccaggaaaactctgaacttgttactatcttacataaattacaaggtgaa [ 52880]
ASF_IND_20_CAD543                -------------------------------...........y..........r.......................... [ 52880]
LR899193.1_Germany_2020/1        ................................................................................ [ 52880]
MN393477.1_Wuhan_2019            ................................................................................ [ 52880]
MW396979.1_Timor-Leste/2019/1    ................................................................................ [ 52880]
MK940252.1_CN/2019/InnerMongolia ................................................................................ [ 52880]
MK128995.1_China/2018/AnhuiXCGQ  ................................................................................ [ 52880]
MH681419.1_ASFV/POL/2015         ................................................................................ [ 52880]
LS478113.1_Estonia_2014          ................................................................................ [ 52880]
KP843857.1_Odintsovo_02/14       ................................................................................ [ 52880]
FN557520.1_strain_E75            ................................................................................ [ 52880]
AY261366.1_Warthog               ................................................................................ [ 52880]
AY261365.1_Warmbaths             ................................t............................................... [ 52880]
AY261364.1_Tengani_62            ................................................................................ [ 52880]
AY261363.1_Pretoriuskop/96/4     ................................................................................ [ 52880]
AY261362.1_Mkuzi_1979            ................................................................................ [ 52880]
AY261361.1_Malawi_Lil-20/1_1983  -------------------------------------------------------------------------------- [ 52880]
AY261360.1_Kenya_1950            -------------------------------------------------------------------------------- [ 52880]
AM712240.1_OURT_88/3             ................................................................................ [ 52880]
AM712239.1_Benin_97/1            ................................................................................ [ 52880]
KM111295.1_Ken06.Bus             -------------------------------------------------------------------------------- [ 52880]
KM111294.1_Ken05/Tk1             -------------------------------------------------------------------------------- [ 52880]
KM262844.1_strain_L60            ................................................................................ [ 52880]
KM102979.1_26544/OG10_Italy      ................................................................................ [ 52880]
KM262845.1_strain_NHV            ................................................................................ [ 52880]
KX354450.1_47/Ss/2008            ................................................................................ [ 52880]
KP055815.1_strain_BA71           ................................................................................ [ 52880]
MH025920.1_strain_R35            -------------------------------------------------------------------------------- [ 52880]
MH025919.1_strain_N10            -------------------------------------------------------------------------------- [ 52880]
MH025918.1_strain_R25            -------------------------------------------------------------------------------- [ 52880]
MH025917.1_strain_R7             -------------------------------------------------------------------------------- [ 52880]
MH025916.1_strain_R8             -------------------------------------------------------------------------------- [ 52880]

FR682468.2_Georgia_2007/1        aaatgttaaaaagttgttagaacatgtagtgaaatacaactctactcttgtgataagactcttgttagaaaaaaagaaaa [ 52960]
ASF_IND_20_CAD543                .............y..k....r.......................................................... [ 52960]
LR899193.1_Germany_2020/1        ................................................................................ [ 52960]
MN393477.1_Wuhan_2019            ................................................................................ [ 52960]
MW396979.1_Timor-Leste/2019/1    ................................................................................ [ 52960]
MK940252.1_CN/2019/InnerMongolia ................................................................................ [ 52960]
MK128995.1_China/2018/AnhuiXCGQ  ................................................................................ [ 52960]
MH681419.1_ASFV/POL/2015         ................................................................................ [ 52960]
LS478113.1_Estonia_2014          ................................................................................ [ 52960]
KP843857.1_Odintsovo_02/14       ................................................................................ [ 52960]
```

```
FN557520.1_strain_E75              ..........................................................a..................... [ 52960]
AY261366.1_Warthog                 ..........................................................a..................... [ 52960]
AY261365.1_Warmbaths               ...............a..........................................a..................... [ 52960]
AY261364.1_Tengani_62              ................................................................................ [ 52960]
AY261363.1_Pretoriuskop/96/4       ..........................................................a..................... [ 52960]
AY261362.1_Mkuzi_1979              ..........................................................a.t................... [ 52960]
AY261361.1_Malawi_Lil-20/1_1983    -------------------------------------------------------------------------------- [ 52960]
AY261360.1_Kenya_1950              -------------------------------------------------------------------------------- [ 52960]
AM712240.1_OURT_88/3               ..........................................................a..................... [ 52960]
AM712239.1_Benin_97/1              ..........................................................a..................... [ 52960]
KM111295.1_Ken06.Bus               -------------------------------------------------------------------------------- [ 52960]
KM111294.1_Ken05/Tk1               -------------------------------------------------------------------------------- [ 52960]
KM262844.1_strain_L60              ..........................................................a..................... [ 52960]
KM102979.1_26544/OG10_Italy        ..........................................................a..................... [ 52960]
KM262845.1_strain_NHV              ..........................................................a..................... [ 52960]
KX354450.1_47/Ss/2008              ..........................................................a..................... [ 52960]
KP055815.1_strain_BA71             ..........................................................a..................... [ 52960]
MH025920.1_strain_R35              -------------------------------------------------------------------------------- [ 52960]
MH025919.1_strain_N10              -------------------------------------------------------------------------------- [ 52960]
MH025918.1_strain_R25              -------------------------------------------------------------------------------- [ 52960]
MH025917.1_strain_R7               -------------------------------------------------------------------------------- [ 52960]
MH025916.1_strain_R8               -------------------------------------------------------------------------------- [ 52960]

FR682468.2_Georgia_2007/1          acctgctggatgctactttgacaagatatgtcaaagattctacatactttcaggtgaaagaatttatgcaagacttctcc [ 53040]
ASF_IND_20_CAD543                  .........................................................r.........s........y... [ 53040]
LR899193.1_Germany_2020/1          ................................................................................ [ 53040]
MN393477.1_Wuhan_2019              ................................................................................ [ 53040]
MW396979.1_Timor-Leste/2019/1      ................................................................................ [ 53040]
MK940252.1_CN/2019/InnerMongolia   ................................................................................ [ 53040]
MK128995.1_China/2018/AnhuiXCGQ    ................................................................................ [ 53040]
MH681419.1_ASFV/POL/2015           ................................................................................ [ 53040]
LS478113.1_Estonia_2014            ................................................................................ [ 53040]
KP843857.1_Odintsovo_02/14         ................................................................................ [ 53040]
FN557520.1_strain_E75              ................................................................................ [ 53040]
AY261366.1_Warthog                 ...................................................g.....................t...... [ 53040]
AY261365.1_Warmbaths               ...................................................g............................ [ 53040]
AY261364.1_Tengani_62              ................................................................................ [ 53040]
AY261363.1_Pretoriuskop/96/4       ...................................................g...a.g.........c..g..g..t... [ 53040]
AY261362.1_Mkuzi_1979              .......a...........................................g............................ [ 53040]
AY261361.1_Malawi_Lil-20/1_1983    -------------------------------------------------------------------------------- [ 53040]
AY261360.1_Kenya_1950              -------------------------------------------------------------------------------- [ 53040]
AM712240.1_OURT_88/3               ................................................................................ [ 53040]
AM712239.1_Benin_97/1              ................................................................................ [ 53040]
KM111295.1_Ken06.Bus               -------------------------------------------------------------------------------- [ 53040]
```

```
KM111294.1_Ken05/Tk1            -------------------------------------------------------------------------------- [ 53040]
KM262844.1_strain_L60           ................................................................................ [ 53040]
KM102979.1_26544/OG10_Italy     ................................................................................ [ 53040]
KM262845.1_strain_NHV           ................................................................................ [ 53040]
KX354450.1_47/Ss/2008           ................................................................................ [ 53040]
KP055815.1_strain_BA71          ................................................................................ [ 53040]
MH025920.1_strain_R35           -------------------------------------------------------------------------------- [ 53040]
MH025919.1_strain_N10           -------------------------------------------------------------------------------- [ 53040]
MH025918.1_strain_R25           -------------------------------------------------------------------------------- [ 53040]
MH025917.1_strain_R7            -------------------------------------------------------------------------------- [ 53040]
MH025916.1_strain_R8            -------------------------------------------------------------------------------- [ 53040]

FR682468.2_Georgia_2007/1       atcagcccagaaaaattcattaaaatagctgtgcgggaaaagagaaatgtgttgatcaagggtatttctgaagatatttg [ 53120]
ASF_IND_20_CAD543               ............................................r........r...r...s.................. [ 53120]
LR899193.1_Germany_2020/1       ................................................................................ [ 53120]
MN393477.1_Wuhan_2019           ................................................................................ [ 53120]
MW396979.1_Timor-Leste/2019/1   ................................................................................ [ 53120]
MK940252.1_CN/2019/InnerMongolia ................................................................................ [ 53120]
MK128995.1_China/2018/AnhuiXCGQ ................................................................................ [ 53120]
MH681419.1_ASFV/POL/2015        ................................................................................ [ 53120]
LS478113.1_Estonia_2014         ................................................................................ [ 53120]
KP843857.1_Odintsovo_02/14      ................................................................................ [ 53120]
FN557520.1_strain_E75           ............................................g................................... [ 53120]
AY261366.1_Warthog              .......................................c...a.................c.....g............ [ 53120]
AY261365.1_Warmbaths            .......................................c...a.................c.....g............ [ 53120]
AY261364.1_Tengani_62           ................................................................................ [ 53120]
AY261363.1_Pretoriuskop/96/4    .......................................c...a.................c.....g............ [ 53120]
AY261362.1_Mkuzi_1979           ................................................................................ [ 53120]
AY261361.1_Malawi_Lil-20/1_1983 ---------........t.................................a.a.......................... [ 53120]
AY261360.1_Kenya_1950           ---------..............................c...a.......a.c.......t.....g............ [ 53120]
AM712240.1_OURT_88/3            ............................................g................................... [ 53120]
AM712239.1_Benin_97/1           ............................................g................................... [ 53120]
KM111295.1_Ken06.Bus            ---------..............................c...a.......a.c.......t.....g............ [ 53120]
KM111294.1_Ken05/Tk1            ---------..............................c...a.......a.c.......t.....g............ [ 53120]
KM262844.1_strain_L60           ............................................g................................... [ 53120]
KM102979.1_26544/OG10_Italy     ............................................g................................... [ 53120]
KM262845.1_strain_NHV           ............................................g................................... [ 53120]
KX354450.1_47/Ss/2008           ............................................g................................... [ 53120]
KP055815.1_strain_BA71          ............................................g................................... [ 53120]
MH025920.1_strain_R35           ---------..............................c...a.......a.c.......t.....g............ [ 53120]
MH025919.1_strain_N10           ---------..............................c...a.......a.c.......t.....g............ [ 53120]
MH025918.1_strain_R25           ---------..............................c...a.......a.c.......t.....g............ [ 53120]
MH025917.1_strain_R7            ---------..............................c...a.......a.c.......t.....g............ [ 53120]
MH025916.1_strain_R8            ---------..............................c...a.......a.c.......t.....g............ [ 53120]
```

```
FR682468.2_Georgia_2007/1         ggaaaatcccgcggaaagaatcaggaatcttaagcagatagtgtgtaccataaaatatgaaagtggaagacaattcctga [ 53200]
ASF_IND_20_CAD543                 ..........r.r.................y..r...................................r.r.......r [ 53200]
LR899193.1_Germany_2020/1         ................................................................................ [ 53200]
MN393477.1_Wuhan_2019             ................................................................................ [ 53200]
MW396979.1_Timor-Leste/2019/1     ................................................................................ [ 53200]
MK940252.1_CN/2019/InnerMongolia  ................................................................................ [ 53200]
MK128995.1_China/2018/AnhuiXCGQ   ................................................................................ [ 53200]
MH681419.1_ASFV/POL/2015          ................................................................................ [ 53200]
LS478113.1_Estonia_2014           ................................................................................ [ 53200]
KP843857.1_Odintsovo_02/14        ................................................................................ [ 53200]
FN557520.1_strain_E75             ................................................................................ [ 53200]
AY261366.1_Warthog                ............a................................................................... [ 53200]
AY261365.1_Warmbaths              ............a................................................................... [ 53200]
AY261364.1_Tengani_62             ................................................................................ [ 53200]
AY261363.1_Pretoriuskop/96/4      ............a................................................................... [ 53200]
AY261362.1_Mkuzi_1979             ................................................................................ [ 53200]
AY261361.1_Malawi_Lil-20/1_1983   ......c.....a........t.ctt....c..a.........cac........................g......... [ 53200]
AY261360.1_Kenya_1950             ......c.....a............t....g...........aa.c.g.....................g.....t.... [ 53200]
AM712240.1_OURT_88/3              ................................................................................ [ 53200]
AM712239.1_Benin_97/1             ................................................................................ [ 53200]
KM111295.1_Ken06.Bus              ......c.....a............t....g...........aa.c.g.....................g.....t.... [ 53200]
KM111294.1_Ken05/Tk1              ......c.....a............t....g...........aa.c.g...........................t.... [ 53200]
KM262844.1_strain_L60             ................................................................................ [ 53200]
KM102979.1_26544/OG10_Italy       ................................................................................ [ 53200]
KM262845.1_strain_NHV             ................................................................................ [ 53200]
KX354450.1_47/Ss/2008             ................................................................................ [ 53200]
KP055815.1_strain_BA71            ................................................................................ [ 53200]
MH025920.1_strain_R35             ......c.....a............t....g...........aa.c.g.....................g.....t.... [ 53200]
MH025919.1_strain_N10             ......c.....a............t....g...........aa.c.g.....................g.....t.... [ 53200]
MH025918.1_strain_R25             ......c.....a............t....g...........aa.c.g.....................g.....t.... [ 53200]
MH025917.1_strain_R7              ......c.....a............t....g...........aa.c.g.....................g.....t.... [ 53200]
MH025916.1_strain_R8              ......c.....a............t....g...........aa.c.g.....................g.....t.... [ 53200]

FR682468.2_Georgia_2007/1         taaatatcattcacaccatttaccagagttattctttgaaacctgaagaaattcttaaattggcaacattttatgtcaaa [ 53280]
ASF_IND_20_CAD543                 ..r.y..........s.........r.......................w.........y.................... [ 53280]
LR899193.1_Germany_2020/1         ................................................................................ [ 53280]
MN393477.1_Wuhan_2019             ................................................................................ [ 53280]
MW396979.1_Timor-Leste/2019/1     ................................................................................ [ 53280]
MK940252.1_CN/2019/InnerMongolia  ................................................................................ [ 53280]
MK128995.1_China/2018/AnhuiXCGQ   ................................................................................ [ 53280]
MH681419.1_ASFV/POL/2015          ................................................................................ [ 53280]
LS478113.1_Estonia_2014           ................................................................................ [ 53280]
KP843857.1_Odintsovo_02/14        ................................................................................ [ 53280]
```

```
FN557520.1_strain_E75           ...........................................................c.................... [ 53280]
AY261366.1_Warthog              ................................................................................ [ 53280]
AY261365.1_Warmbaths            ................................................................................ [ 53280]
AY261364.1_Tengani_62           ................................................................................ [ 53280]
AY261363.1_Pretoriuskop/96/4    ..................................................g............................. [ 53280]
AY261362.1_Mkuzi_1979           ......................................................................c......... [ 53280]
AY261361.1_Malawi_Lil-20/1_1983 ..................................c...............................g.........g... [ 53280]
AY261360.1_Kenya_1950           ....c...........t............c....cc.................t.......................... [ 53280]
AM712240.1_OURT_88/3            ...........................................................c.................... [ 53280]
AM712239.1_Benin_97/1           ...........................................................c.................... [ 53280]
KM111295.1_Ken06.Bus            ....c...........t............c....cc.................t.......................... [ 53280]
KM111294.1_Ken05/Tk1            ....c...........t............c....cc.................t.......................... [ 53280]
KM262844.1_strain_L60           ...........................................................c.................... [ 53280]
KM102979.1_26544/OG10_Italy     ...........................................................c.................... [ 53280]
KM262845.1_strain_NHV           ...........................................................c.................... [ 53280]
KX354450.1_47/Ss/2008           ...........................................................c.................... [ 53280]
KP055815.1_strain_BA71          ...........................................................c........-........... [ 53280]
MH025920.1_strain_R35           ....c...........t............c....cc.................t.......................... [ 53280]
MH025919.1_strain_N10           ....c...........t............c....cc.................t.......................... [ 53280]
MH025918.1_strain_R25           ....c...........t............c....cc.................t.......................... [ 53280]
MH025917.1_strain_R7            ....c...........t............c....cc.................t.......................... [ 53280]
MH025916.1_strain_R8            ....c...........t............c....cc.................t.......................... [ 53280]

FR682468.2_Georgia_2007/1       cacaatgcaaccacccattttaaagatctctgcaaatatctttggctgaacagaagaacagaaagtaagaaactgttttt [ 53360]
ASF_IND_20_CAD543               ..........y...............y...........................r......................... [ 53360]
LR899193.1_Germany_2020/1       ................................................................................ [ 53360]
MN393477.1_Wuhan_2019           ................................................................................ [ 53360]
MW396979.1_Timor-Leste/2019/1   ................................................................................ [ 53360]
MK940252.1_CN/2019/InnerMongolia ................................................................................ [ 53360]
MK128995.1_China/2018/AnhuiXCGQ ................................................................................ [ 53360]
MH681419.1_ASFV/POL/2015        ................................................................................ [ 53360]
LS478113.1_Estonia_2014         ................................................................................ [ 53360]
KP843857.1_Odintsovo_02/14      ................................................................................ [ 53360]
FN557520.1_strain_E75           ......................................................g......................... [ 53360]
AY261366.1_Warthog              ......................................................g......................... [ 53360]
AY261365.1_Warmbaths            ......................................................g......................... [ 53360]
AY261364.1_Tengani_62           ................................................................................ [ 53360]
AY261363.1_Pretoriuskop/96/4    ..t...................................................g......................... [ 53360]
AY261362.1_Mkuzi_1979           ......................................................g......................... [ 53360]
AY261361.1_Malawi_Lil-20/1_1983 ...........a..t..............t...............t........g......................... [ 53360]
AY261360.1_Kenya_1950           ..t......gt.........c........t.......t................g.g................ac..c.. [ 53360]
AM712240.1_OURT_88/3            ......................................................g......................... [ 53360]
AM712239.1_Benin_97/1           ......................................................g......................... [ 53360]
KM111295.1_Ken06.Bus            ..t......gt.........c........t.......t................g.g................ac..c.. [ 53360]
```

```
KM111294.1_Ken05/Tk1            ..t......gt.........c........t.......t............t...g.g.t..............ac..c.. [ 53360]
KM262844.1_strain_L60           ......................................................g......................... [ 53360]
KM102979.1_26544/OG10_Italy     ......................................................g......................... [ 53360]
KM262845.1_strain_NHV           ......................................................g......................... [ 53360]
KX354450.1_47/Ss/2008           ............................................a.........g......................... [ 53360]
KP055815.1_strain_BA71          ......................................................g......................... [ 53360]
MH025920.1_strain_R35           ..t......gt.........c........t.......t................g.g................ac..c.. [ 53360]
MH025919.1_strain_N10           ..t......gt.........c........t.......t................g.g................ac..c.. [ 53360]
MH025918.1_strain_R25           ..t......gt.........c........t.......t................g.g................ac..c.. [ 53360]
MH025917.1_strain_R7            ..t......gt.........c........t.......t................g.g................ac..c.. [ 53360]
MH025916.1_strain_R8            ..t......gt.........c........t.......t................g.g................ac..c.. [ 53360]

FR682468.2_Georgia_2007/1       agagtgcttggaaattgctgataagaaggagtttcctgatattaaaagtattgtgagtgaatacattaactatttgttta [ 53440]
ASF_IND_20_CAD543               ......y..r............r........................................y........c....... [ 53440]
LR899193.1_Germany_2020/1       ................................................................................ [ 53440]
MN393477.1_Wuhan_2019           ................................................................................ [ 53440]
MW396979.1_Timor-Leste/2019/1   ................................................................................ [ 53440]
MK940252.1_CN/2019/InnerMongolia ................................................................................ [ 53440]
MK128995.1_China/2018/AnhuiXCGQ ................................................................................ [ 53440]
MH681419.1_ASFV/POL/2015        ................................................................................ [ 53440]
LS478113.1_Estonia_2014         ................................................................................ [ 53440]
KP843857.1_Odintsovo_02/14      ................................................................................ [ 53440]
FN557520.1_strain_E75           ......................g........................................t........c....... [ 53440]
AY261366.1_Warthog              ................................................................................ [ 53440]
AY261365.1_Warmbaths            ................................................................................ [ 53440]
AY261364.1_Tengani_62           ................................................................................ [ 53440]
AY261363.1_Pretoriuskop/96/4    ................................................................................ [ 53440]
AY261362.1_Mkuzi_1979           ......................g.........................c..............t........c....... [ 53440]
AY261361.1_Malawi_Lil-20/1_1983 .........a....................t.......c........a................................ [ 53440]
AY261360.1_Kenya_1950           .a....t...............g.......t.....c.cc.................................a...... [ 53440]
AM712240.1_OURT_88/3            ......................g........................................t........c....... [ 53440]
AM712239.1_Benin_97/1           ......................g........................................t........c....... [ 53440]
KM111295.1_Ken06.Bus            .ag...t...............g.....a.t.....c.cc.................................a...... [ 53440]
KM111294.1_Ken05/Tk1            .ag...t...............g.......t.....c.cc.................................a...... [ 53440]
KM262844.1_strain_L60           ......................g........................................t........c....... [ 53440]
KM102979.1_26544/OG10_Italy     ......................g........................................t........c....... [ 53440]
KM262845.1_strain_NHV           ......................g........................................t........c....... [ 53440]
KX354450.1_47/Ss/2008           ......................g........................................t........c....... [ 53440]
KP055815.1_strain_BA71          ......................g........................................t........c....... [ 53440]
MH025920.1_strain_R35           .ag...t...............g.....a.t.....c.cc.................................a...... [ 53440]
MH025919.1_strain_N10           .ag...t...............g.....a.t.....c.cc.................................a...... [ 53440]
MH025918.1_strain_R25           .ag...t...............g.....a.t.....c.cc.................................a...... [ 53440]
MH025917.1_strain_R7            .ag...t...............g.....a.t.....c.cc.................................a...... [ 53440]
MH025916.1_strain_R8            .ag...t...............g.....a.t.....c.cc.................................a...... [ 53440]
```

```
FR682468.2_Georgia_2007/1         ctgcaggagctattaccaaggaagaaatcatgcaagcctatgctttggagtatgccatgtattaa----atttctgaatc [ 53520]
ASF_IND_20_CAD543                 ....................................y.........r..................----........... [ 53520]
LR899193.1_Germany_2020/1         .................................................................----........... [ 53520]
MN393477.1_Wuhan_2019             .................................................................----........... [ 53520]
MW396979.1_Timor-Leste/2019/1     .................................................................----........... [ 53520]
MK940252.1_CN/2019/InnerMongolia  .................................................................----........... [ 53520]
MK128995.1_China/2018/AnhuiXCGQ   .................................................................----........... [ 53520]
MH681419.1_ASFV/POL/2015          .................................................................----........... [ 53520]
LS478113.1_Estonia_2014           .................................................................----........... [ 53520]
KP843857.1_Odintsovo_02/14        .................................................................----........... [ 53520]
FN557520.1_strain_E75             ....................................t............................----..........a [ 53520]
AY261366.1_Warthog                .................................g..t............................----.........c. [ 53520]
AY261365.1_Warmbaths              ....................................t..................t.........----........... [ 53520]
AY261364.1_Tengani_62             .................................................................----........... [ 53520]
AY261363.1_Pretoriuskop/96/4      ....................................t..................t.........----........... [ 53520]
AY261362.1_Mkuzi_1979             .................................................................----..........a [ 53520]
AY261361.1_Malawi_Lil-20/1_1983   ....................................t....a.......................----..........a [ 53520]
AY261360.1_Kenya_1950             ........a.c..g........t......................c.......a......g....gtttg.........a [ 53520]
AM712240.1_OURT_88/3              ....................................t............................----..........a [ 53520]
AM712239.1_Benin_97/1             ....................................t............................----..........a [ 53520]
KM111295.1_Ken06.Bus              ........a.c..g........t......................ct..................----g.........a [ 53520]
KM111294.1_Ken05/Tk1              ........a.c..g........t......................c..............g....----g.........a [ 53520]
KM262844.1_strain_L60             ....................................t............................----..........a [ 53520]
KM102979.1_26544/OG10_Italy       ....................................t............................----..........a [ 53520]
KM262845.1_strain_NHV             ....................................t............................----..........a [ 53520]
KX354450.1_47/Ss/2008             ....................................t............................----..........a [ 53520]
KP055815.1_strain_BA71            ....................................t............................----..........a [ 53520]
MH025920.1_strain_R35             ........a.c..g........t......................ct..................----g.........a [ 53520]
MH025919.1_strain_N10             ........a.c..g........t......................ct..................----g.........a [ 53520]
MH025918.1_strain_R25             ........a.c..g........t......................ct..................----g.........a [ 53520]
MH025917.1_strain_R7              ........a.c..g........t......................ct..................----g.........a [ 53520]
MH025916.1_strain_R8              ........a.c..g........t......................ct..................----g.........a [ 53520]

FR682468.2_Georgia_2007/1         agtaagcaata--------------------------------------------------------------------- [ 53600]
ASF_IND_20_CAD543                 .....ky....--------------------------------------------------------------------- [ 53600]
LR899193.1_Germany_2020/1         ...........--------------------------------------------------------------------- [ 53600]
MN393477.1_Wuhan_2019             ...........--------------------------------------------------------------------- [ 53600]
MW396979.1_Timor-Leste/2019/1     ...........--------------------------------------------------------------------- [ 53600]
MK940252.1_CN/2019/InnerMongolia  ...........--------------------------------------------------------------------- [ 53600]
MK128995.1_China/2018/AnhuiXCGQ   ...........--------------------------------------------------------------------- [ 53600]
MH681419.1_ASFV/POL/2015          ...........--------------------------------------------------------------------- [ 53600]
LS478113.1_Estonia_2014           ...........--------------------------------------------------------------------- [ 53600]
KP843857.1_Odintsovo_02/14        ...........--------------------------------------------------------------------- [ 53600]
```

```
FN557520.1_strain_E75             ...........--------------------------------------------------------------------- [ 53600]
AY261366.1_Warthog                t..........--------------------------------------------------------------------- [ 53600]
AY261365.1_Warmbaths              .....t.....--------------------------------------------------------------------- [ 53600]
AY261364.1_Tengani_62             ......t...---------------------------------------------------------------------- [ 53600]
AY261363.1_Pretoriuskop/96/4      ...........--------------------------------------------------------------------- [ 53600]
AY261362.1_Mkuzi_1979             ...........--------------------------------------------------------------------- [ 53600]
AY261361.1_Malawi_Lil-20/1_1983   ...........gattttagaatatgctgtattaagttagtttctgaataagtaagcaatagatagattttagaatatgct [ 53600]
AY261360.1_Kenya_1950             ...c.ttt...-------------------------------------------------agtagattttagaataggtt [ 53600]
AM712240.1_OURT_88/3              ...........--------------------------------------------------------------------- [ 53600]
AM712239.1_Benin_97/1             ...........--------------------------------------------------------------------- [ 53600]
KM111295.1_Ken06.Bus              ...c.ttc...-------------------------------------------------ggtagattttagaatatgct [ 53600]
KM111294.1_Ken05/Tk1              ...c.ttc...-------------------------------------------------ggtagattttagaataggtt [ 53600]
KM262844.1_strain_L60             ...........--------------------------------------------------------------------- [ 53600]
KM102979.1_26544/OG10_Italy       ...........--------------------------------------------------------------------- [ 53600]
KM262845.1_strain_NHV             ...........--------------------------------------------------------------------- [ 53600]
KX354450.1_47/Ss/2008             ...........--------------------------------------------------------------------- [ 53600]
KP055815.1_strain_BA71            ...........--------------------------------------------------------------------- [ 53600]
MH025920.1_strain_R35             ...c.ttc...-------------------------------------------------ggtagattttagaatatgct [ 53600]
MH025919.1_strain_N10             ...c.ttc...-------------------------------------------------ggtagattttagaatatgct [ 53600]
MH025918.1_strain_R25             ...c.ttc...-------------------------------------------------ggtagattttagaatatgct [ 53600]
MH025917.1_strain_R7              ...c.ttc...-------------------------------------------------ggtagattttagaatatgct [ 53600]
MH025916.1_strain_R8              ...c.ttc...-------------------------------------------------ggtagattttagaatatgct [ 53600]

FR682468.2_Georgia_2007/1         ---------------------------------gatagattttagaatatgctgtattaagttagtttctgaataagtaa [ 53680]
ASF_IND_20_CAD543                 ---------------------------------............................................... [ 53680]
LR899193.1_Germany_2020/1         ---------------------------------............................................... [ 53680]
MN393477.1_Wuhan_2019             ---------------------------------............................................... [ 53680]
MW396979.1_Timor-Leste/2019/1     ---------------------------------............................................... [ 53680]
MK940252.1_CN/2019/InnerMongolia  ---------------------------------............................................... [ 53680]
MK128995.1_China/2018/AnhuiXCGQ   ---------------------------------............................................... [ 53680]
MH681419.1_ASFV/POL/2015          ---------------------------------............................................... [ 53680]
LS478113.1_Estonia_2014           ---------------------------------............................................... [ 53680]
KP843857.1_Odintsovo_02/14        ---------------------------------............................................... [ 53680]
FN557520.1_strain_E75             ---------------------------------.....g......................................... [ 53680]
AY261366.1_Warthog                ---------------------------------............................................... [ 53680]
AY261365.1_Warmbaths              ---------------------------------............................................... [ 53680]
AY261364.1_Tengani_62             ----------------------------------.............................................. [ 53680]
AY261363.1_Pretoriuskop/96/4      ---------------------------------............................................... [ 53680]
AY261362.1_Mkuzi_1979             ---------------------------------.....g......................................... [ 53680]
AY261361.1_Malawi_Lil-20/1_1983   gtattaagttagtttctgaataagtaagcaata.............................................t. [ 53680]
AY261360.1_Kenya_1950             gtattg---------------aagttattgata.g....g...................----..g..a.........c. [ 53680]
AM712240.1_OURT_88/3              ---------------------------------.....g......................................... [ 53680]
AM712239.1_Benin_97/1             ---------------------------------.....g......................................... [ 53680]
KM111295.1_Ken06.Bus              gtattg---------------aagttattgata.g....g....a..g...........----..c..........a.c. [ 53680]
```

```
KM111294.1_Ken05/Tk1            gcattg---------------aagttattgata.g....g...................----..g..a.........c. [ 53680]
KM262844.1_strain_L60           ---------------------------------.....g......................................... [ 53680]
KM102979.1_26544/OG10_Italy     ---------------------------------.....g......................................... [ 53680]
KM262845.1_strain_NHV           ---------------------------------.....g......................................... [ 53680]
KX354450.1_47/Ss/2008           ---------------------------------.....g......................................... [ 53680]
KP055815.1_strain_BA71          ---------------------------------.....g......................................... [ 53680]
MH025920.1_strain_R35           gtattg---------------aagttattgata.g....g....a..g...........----..c..........a.c. [ 53680]
MH025919.1_strain_N10           gtattg---------------aagttattgata.g....g....a..g...........----..c..........a.c. [ 53680]
MH025918.1_strain_R25           gtattg---------------aagttattgata.g....g....a..g...........----..c..........a.c. [ 53680]
MH025917.1_strain_R7            gtattg---------------aagttattgata.g....g....a..g...........----..c..........a.c. [ 53680]
MH025916.1_strain_R8            gtattg---------------aagttattgata.g....g....a..g...........----..c..........a.c. [ 53680]

FR682468.2_Georgia_2007/1       ttaatagatagattttagtttatgtaaaaatgttaacatttgt------------------------------------- [ 53760]
ASF_IND_20_CAD543               ...........................................------------------------------------- [ 53760]
LR899193.1_Germany_2020/1       ...........................................------------------------------------- [ 53760]
MN393477.1_Wuhan_2019           ...........................................------------------------------------- [ 53760]
MW396979.1_Timor-Leste/2019/1   ...........................................------------------------------------- [ 53760]
MK940252.1_CN/2019/InnerMongolia ...........................................------------------------------------- [ 53760]
MK128995.1_China/2018/AnhuiXCGQ ...........................................------------------------------------- [ 53760]
MH681419.1_ASFV/POL/2015        ...........................................------------------------------------- [ 53760]
LS478113.1_Estonia_2014         ...........................................------------------------------------- [ 53760]
KP843857.1_Odintsovo_02/14      ...........................................------------------------------------- [ 53760]
FN557520.1_strain_E75           ...........................................------------------------------------- [ 53760]
AY261366.1_Warthog              .c.........................................ta----------------------------------- [ 53760]
AY261365.1_Warmbaths            .c.........................................ta----------------------------------- [ 53760]
AY261364.1_Tengani_62           .c.........................................------------------------------------- [ 53760]
AY261363.1_Pretoriuskop/96/4    .c.........t...............................------------------------------------- [ 53760]
AY261362.1_Mkuzi_1979           ...........................................------------------------------------- [ 53760]
AY261361.1_Malawi_Lil-20/1_1983 gc.................g.....t.................------------------------------------- [ 53760]
AY261360.1_Kenya_1950           .a....t...--c............-.........g.......taatatgttttaaatagttaccattttagataagtaa [ 53760]
AM712240.1_OURT_88/3            ...........................................------------------------------------- [ 53760]
AM712239.1_Benin_97/1           ...........................................------------------------------------- [ 53760]
KM111295.1_Ken06.Bus            .ag...t...--c.......-------........g.......taatatgttttagataattaccattttagataattac [ 53760]
KM111294.1_Ken05/Tk1            ..c....g.------------------.g..t...g-------------------------------------------- [ 53760]
KM262844.1_strain_L60           ...........................................------------------------------------- [ 53760]
KM102979.1_26544/OG10_Italy     ...........................................------------------------------------- [ 53760]
KM262845.1_strain_NHV           ...........................................------------------------------------- [ 53760]
KX354450.1_47/Ss/2008           ...........................................------------------------------------- [ 53760]
KP055815.1_strain_BA71          ...........................................------------------------------------- [ 53760]
MH025920.1_strain_R35           .ag...t...--c.......-------........g.......taatatgttttagataattaccattttagataattac [ 53760]
MH025919.1_strain_N10           .ag...t...--c.......-------........g.......taatatgttttagataattacctttttagataattac [ 53760]
MH025918.1_strain_R25           .ag...t...--c.......-------........g.......taatatgttttagataattaccattttagataattac [ 53760]
MH025917.1_strain_R7            .ag...t...--c.......-------........g.......taatatgttttagataattaccattttagataattac [ 53760]
MH025916.1_strain_R8            .ag...t...--c.......-------........g.......taatatgttttagataattaccattttagataattac [ 53760]
```

```
FR682468.2_Georgia_2007/1        --------------------------------------tcataag-----ttttagata----ccattttag-------- [ 53840]
ASF_IND_20_CAD543                --------------------------------------.m.....-----.........atta.........-------- [ 53840]
LR899193.1_Germany_2020/1        --------------------------------------.......-----.........----.........-------- [ 53840]
MN393477.1_Wuhan_2019            --------------------------------------.......-----.........----.........-------- [ 53840]
MW396979.1_Timor-Leste/2019/1    --------------------------------------.......-----.........----.........-------- [ 53840]
MK940252.1_CN/2019/InnerMongolia --------------------------------------.......-----.........----.........-------- [ 53840]
MK128995.1_China/2018/AnhuiXCGQ  --------------------------------------.......-----.........----.........-------- [ 53840]
MH681419.1_ASFV/POL/2015         --------------------------------------.......-----.........----.........-------- [ 53840]
LS478113.1_Estonia_2014          --------------------------------------.......-----.........----.........-------- [ 53840]
KP843857.1_Odintsovo_02/14       --------------------------------------.......-----.........----.........-------- [ 53840]
FN557520.1_strain_E75            --------------------------------------.a.....-----.........atta.t.......-------- [ 53840]
AY261366.1_Warthog               -------------------------------------a.a.....-----.........----.........-------- [ 53840]
AY261365.1_Warmbaths             -------------------------------------a.a.....-----.........----.........-------- [ 53840]
AY261364.1_Tengani_62            --------------------------------------.......-----.........----.........-------- [ 53840]
AY261363.1_Pretoriuskop/96/4     --------------------------------------.a.....-----.........----.........-------- [ 53840]
AY261362.1_Mkuzi_1979            --------------------------------------.a.....-----.......----ta.........-------- [ 53840]
AY261361.1_Malawi_Lil-20/1_1983  --------------------------------------.a.....-----.........gtta.......---------- [ 53840]
AY261360.1_Kenya_1950            tattttagataggtaccattttagataagtaatgttttaa.....taatg.....a...gtta.........ataagtaa [ 53840]
AM712240.1_OURT_88/3             --------------------------------------.a.....-----.........atta.t.......-------- [ 53840]
AM712239.1_Benin_97/1            --------------------------------------.a.....-----.........atta.t.......-------- [ 53840]
KM111295.1_Ken06.Bus             tattttagataagtattattttagataggtactattttag...g.tacta.........atta.tc-------------- [ 53840]
KM111294.1_Ken05/Tk1             -------aataggttgcattgaag----------ttat.g...g.tacta.........atta.t.......ataagtat [ 53840]
KM262844.1_strain_L60            --------------------------------------.a.....-----.........atta.t.......-------- [ 53840]
KM102979.1_26544/OG10_Italy      --------------------------------------.a.....-----.........atta.t.......-------- [ 53840]
KM262845.1_strain_NHV            --------------------------------------.a.....-----.........atta.t.......-------- [ 53840]
KX354450.1_47/Ss/2008            --------------------------------------.a.....-----.........atta.t.......-------- [ 53840]
KP055815.1_strain_BA71           --------------------------------------.a.....-----.........atta.t.......-------- [ 53840]
MH025920.1_strain_R35            tattttagataagtattattttagataggtactattttag...g.tacta.........atta.tcc.....ataagtat [ 53840]
MH025919.1_strain_N10            tattttagataagtattattttagataggtactattttag...g.tacta.........atta.tcc.....ataagtat [ 53840]
MH025918.1_strain_R25            tattttagataagtattattttagataggtactattttag...g.tacta.........atta.tcc.....ataagtat [ 53840]
MH025917.1_strain_R7             tattttagataagtattattttagataggtactattttag...g.tacta.........atta.tcc.....ataagtat [ 53840]
MH025916.1_strain_R8             tattttagataagtattattttagataggtactattttag...g.tacta.........atta.tcc.....ataagtat [ 53840]

FR682468.2_Georgia_2007/1        --------------------------------------------------------------------------agttac [ 53920]
ASF_IND_20_CAD543                --------------------------------------------------------------------------...... [ 53920]
LR899193.1_Germany_2020/1        --------------------------------------------------------------------------...... [ 53920]
MN393477.1_Wuhan_2019            --------------------------------------------------------------------------...... [ 53920]
MW396979.1_Timor-Leste/2019/1    --------------------------------------------------------------------------...... [ 53920]
MK940252.1_CN/2019/InnerMongolia --------------------------------------------------------------------------...... [ 53920]
MK128995.1_China/2018/AnhuiXCGQ  --------------------------------------------------------------------------...... [ 53920]
MH681419.1_ASFV/POL/2015         --------------------------------------------------------------------------...... [ 53920]
LS478113.1_Estonia_2014          --------------------------------------------------------------------------...... [ 53920]
KP843857.1_Odintsovo_02/14       --------------------------------------------------------------------------...... [ 53920]
```

```
FN557520.1_strain_E75          --------------------------------------------------------------------------...... [ 53920]
AY261366.1_Warthog             --------------------------------------------------------------------------...... [ 53920]
AY261365.1_Warmbaths           --------------------------------------------------------------------------...c.. [ 53920]
AY261364.1_Tengani_62          --------------------------------------------------------------------------...... [ 53920]
AY261363.1_Pretoriuskop/96/4   --------------------------------------------------------------------------...... [ 53920]
AY261362.1_Mkuzi_1979          --------------------------------------------------------------------------...... [ 53920]
AY261361.1_Malawi_Lil-20/1_1983 ---------------------------------------------------------------------------..... [ 53920]
AY261360.1_Kenya_1950          tatttta------------------------------------------------gatagatactattttagat..g... [ 53920]
AM712240.1_OURT_88/3           --------------------------------------------------------------------------...... [ 53920]
AM712239.1_Benin_97/1          --------------------------------------------------------------------------...... [ 53920]
KM111295.1_Ken06.Bus           ------------------cttta--------------------------------gataagtattattttagat.a.... [ 53920]
KM111294.1_Ken05/Tk1           tatttta------------------------------------------------gataagtattattttagat..a... [ 53920]
KM262844.1_strain_L60          --------------------------------------------------------------------------...... [ 53920]
KM102979.1_26544/OG10_Italy    --------------------------------------------------------------------------...... [ 53920]
KM262845.1_strain_NHV          --------------------------------------------------------------------------...... [ 53920]
KX354450.1_47/Ss/2008          --------------------------------------------------------------------------...... [ 53920]
KP055815.1_strain_BA71         --------------------------------------------------------------------------...... [ 53920]
MH025920.1_strain_R35          tattttagataattacttctttaaataagtattattttagataattactcctttaaataagtattattttagat.a.... [ 53920]
MH025919.1_strain_N10          tattttagataattacttctttaaataagtattattttagataattactcctttaaataagtattattttagat.a.... [ 53920]
MH025918.1_strain_R25          tattttagataattacttctttaaataagtattattttagataattactcctttaaataagtattattttagat.a.... [ 53920]
MH025917.1_strain_R7           tattttagataattacttctttaaataagtattattttagataattactcctttaaataagtattattttagat.a.... [ 53920]
MH025916.1_strain_R8           tattttagataattacttctttaaataagtattattttagataattactcctttaaataagtattattttagat.a.... [ 53920]

FR682468.2_Georgia_2007/1      ttttttagata-ttactattttagcca-ttattatcttaaataatcactattttagataggtccccgtattaaaaaccaa [ 54000]
ASF_IND_20_CAD543              ...........-....y........y.-.................................................... [ 54000]
LR899193.1_Germany_2020/1      ...........-...............-.................................................... [ 54000]
MN393477.1_Wuhan_2019          ...........-...............-.................................................... [ 54000]
MW396979.1_Timor-Leste/2019/1  ...........-...............-.................................................... [ 54000]
MK940252.1_CN/2019/InnerMongolia ...........-...............-.................................................... [ 54000]
MK128995.1_China/2018/AnhuiXCGQ ...........-...............-.................................................... [ 54000]
MH681419.1_ASFV/POL/2015       ...........-...............-.................................................... [ 54000]
LS478113.1_Estonia_2014        ...........-...............-.................................................... [ 54000]
KP843857.1_Odintsovo_02/14     ...........-...............-.................................................... [ 54000]
FN557520.1_strain_E75          .a........t-....c........t.-.................................................... [ 54000]
AY261366.1_Warthog             ...........-...............-...................................t..............g. [ 54000]
AY261365.1_Warmbaths           .a.........-...............-........................................------------ [ 54000]
AY261364.1_Tengani_62          ...........-...............-.................................................... [ 54000]
AY261363.1_Pretoriuskop/96/4   ...........-...............-..................................................g. [ 54000]
AY261362.1_Mkuzi_1979          .a.........-....c........t.-.................................................... [ 54000]
AY261361.1_Malawi_Lil-20/1_1983 .a.........-.............t.-........c........t....................t.........a... [ 54000]
AY261360.1_Kenya_1950          ca.........ga...........at.gg..cc..t...g....gt.t...........at.a..atgta.t........ [ 54000]
AM712240.1_OURT_88/3           .a........t-....c........t.-.................................................... [ 54000]
AM712239.1_Benin_97/1          .a........t-....c........t.-.................................................... [ 54000]
KM111295.1_Ken06.Bus           ..c....a...ag..t........at.a...c.cct........gt.t...........at.a..atgta.t........ [ 54000]
```

```
KM111294.1_Ken05/Tk1           .a.........a............at.ag......t...g...gat.............at.a..atgta.t........ [ 54000]
KM262844.1_strain_L60          .a........t-....c........t.-.................................................... [ 54000]
KM102979.1_26544/OG10_Italy    .a........t-....c........t.-.................................................... [ 54000]
KM262845.1_strain_NHV          .a........t-....c........t.-.................................................... [ 54000]
KX354450.1_47/Ss/2008          .a........t-....c........t.-.................................................... [ 54000]
KP055815.1_strain_BA71         .a........t-....c........t.-.................................................... [ 54000]
MH025920.1_strain_R35          ..c....a...ag..t........at.a...c.cct........gt.t...........at.a..atgta.t........ [ 54000]
MH025919.1_strain_N10          ..c....a...ag..t........at.a...c.cct........gt.t...........at.a..atgta.t........ [ 54000]
MH025918.1_strain_R25          ..c....a...ag..t........at.a...c.cct........gt.t...........at.a..atgta.t........ [ 54000]
MH025917.1_strain_R7           ..c....a...ag..t........at.a...c.cct........gt.t...........at.a..atgta.t........ [ 54000]
MH025916.1_strain_R8           ..c....a...ag..t........at.a...c.cct........gt.t...........at.a..atgta.t........ [ 54000]

FR682468.2_Georgia_2007/1      attaaccattatctatgtttttaataatactttttaaaaaccctccataaaaatttattttttttttcataaaagtagaga [ 54080]
ASF_IND_20_CAD543              ................................................................................ [ 54080]
LR899193.1_Germany_2020/1      ................................................................................ [ 54080]
MN393477.1_Wuhan_2019          ................................................................................ [ 54080]
MW396979.1_Timor-Leste/2019/1  ................................................................................ [ 54080]
MK940252.1_CN/2019/InnerMongolia ................................................................................ [ 54080]
MK128995.1_China/2018/AnhuiXCGQ ................................................................................ [ 54080]
MH681419.1_ASFV/POL/2015       ................................................................................ [ 54080]
LS478113.1_Estonia_2014        ................................................................................ [ 54080]
KP843857.1_Odintsovo_02/14     .........................................................-...................... [ 54080]
FN557520.1_strain_E75          ................................................................................ [ 54080]
AY261366.1_Warthog             ........c....................................................................... [ 54080]
AY261365.1_Warmbaths           ................................................................................ [ 54080]
AY261364.1_Tengani_62          ...................a............................................................ [ 54080]
AY261363.1_Pretoriuskop/96/4   ........c....................................................................... [ 54080]
AY261362.1_Mkuzi_1979          ................................................................................ [ 54080]
AY261361.1_Malawi_Lil-20/1_1983 .........ct.t................................................................... [ 54080]
AY261360.1_Kenya_1950          ....g....c..........a...........c.........t.ag.................................. [ 54080]
AM712240.1_OURT_88/3           ................................................................................ [ 54080]
AM712239.1_Benin_97/1          ................................................................................ [ 54080]
KM111295.1_Ken06.Bus           ....g.t..c..........a...........c.........t.ag.................................. [ 54080]
KM111294.1_Ken05/Tk1           ....g....c..........a...........c.........t.ag.................................. [ 54080]
KM262844.1_strain_L60          ................................................................................ [ 54080]
KM102979.1_26544/OG10_Italy    ................................................................................ [ 54080]
KM262845.1_strain_NHV          .........................................................-...................... [ 54080]
KX354450.1_47/Ss/2008          ................................................................................ [ 54080]
KP055815.1_strain_BA71         ................................................................................ [ 54080]
MH025920.1_strain_R35          ....g.t..c..........a...........c.........t.ag.................................. [ 54080]
MH025919.1_strain_N10          ....g.t..c..........a...........c.........t.ag.................................. [ 54080]
MH025918.1_strain_R25          ....g.t..c..........a...........c.........t.ag.................................. [ 54080]
MH025917.1_strain_R7           ....g.t..c..........a...........c.........t.ag.................................. [ 54080]
MH025916.1_strain_R8           ....g.t..c..........a...........c.........t.ag.................................. [ 54080]
```

```
FR682468.2_Georgia_2007/1        aaatgttctccctacaggatctctgtcggaagaacctttttcttccacttgagcccttaggcaagcatgtggttcaacgg [ 54160]
ASF_IND_20_CAD543                ................................................................................ [ 54160]
LR899193.1_Germany_2020/1        ................................................................................ [ 54160]
MN393477.1_Wuhan_2019            ................................................................................ [ 54160]
MW396979.1_Timor-Leste/2019/1    ................................................................................ [ 54160]
MK940252.1_CN/2019/InnerMongolia ................................................................................ [ 54160]
MK128995.1_China/2018/AnhuiXCGQ  ................................................................................ [ 54160]
MH681419.1_ASFV/POL/2015         ................................................................................ [ 54160]
LS478113.1_Estonia_2014          ................................................................................ [ 54160]
KP843857.1_Odintsovo_02/14       ................................................................................ [ 54160]
FN557520.1_strain_E75            ................................................................................ [ 54160]
AY261366.1_Warthog               ...................c............................................................ [ 54160]
AY261365.1_Warmbaths             ...............................................g................................ [ 54160]
AY261364.1_Tengani_62            ................................................................................ [ 54160]
AY261363.1_Pretoriuskop/96/4     ................................................................................ [ 54160]
AY261362.1_Mkuzi_1979            ................................................................................ [ 54160]
AY261361.1_Malawi_Lil-20/1_1983  ................................................................................ [ 54160]
AY261360.1_Kenya_1950            .............t.....c.....................ac...c.....a...........a............... [ 54160]
AM712240.1_OURT_88/3             ................................................................................ [ 54160]
AM712239.1_Benin_97/1            ................................................................................ [ 54160]
KM111295.1_Ken06.Bus             .............t.....c.....................ac...c.....a...........a............... [ 54160]
KM111294.1_Ken05/Tk1             .............t.....c.....................ac...c.....a...........a............... [ 54160]
KM262844.1_strain_L60            ................................................................................ [ 54160]
KM102979.1_26544/OG10_Italy      ................................................................................ [ 54160]
KM262845.1_strain_NHV            ................................................................................ [ 54160]
KX354450.1_47/Ss/2008            ................................................................................ [ 54160]
KP055815.1_strain_BA71           ................................................................................ [ 54160]
MH025920.1_strain_R35            .............t.....c.....................ac...c.....a...........a............... [ 54160]
MH025919.1_strain_N10            .............t.....c.....................ac...c.....a...........a............... [ 54160]
MH025918.1_strain_R25            .............t.....c.....................ac...c.....a...........a............... [ 54160]
MH025917.1_strain_R7             .............t.....c.....................ac...c.....a...........a............... [ 54160]
MH025916.1_strain_R8             .............t.....c.....................ac...c.....a...........a............... [ 54160]

FR682468.2_Georgia_2007/1        ctgggattatactgggaaggccatggttcagttaaacgagtgggtgattgctttatatgtgtagaccagatttggatgct [ 54240]
ASF_IND_20_CAD543                ................................................................................ [ 54240]
LR899193.1_Germany_2020/1        ................................................................................ [ 54240]
MN393477.1_Wuhan_2019            ................................................................................ [ 54240]
MW396979.1_Timor-Leste/2019/1    ................................................................................ [ 54240]
MK940252.1_CN/2019/InnerMongolia ................................................................................ [ 54240]
MK128995.1_China/2018/AnhuiXCGQ  ................................................................................ [ 54240]
MH681419.1_ASFV/POL/2015         ................................................................................ [ 54240]
LS478113.1_Estonia_2014          ................................................................................ [ 54240]
KP843857.1_Odintsovo_02/14       ................................................................................ [ 54240]
```

```
FN557520.1_strain_E75          ..............................c...................................a..........c.. [ 54240]
AY261366.1_Warthog             ......................................................g....................g.c.. [ 54240]
AY261365.1_Warmbaths           ......................................................g....................g.c.. [ 54240]
AY261364.1_Tengani_62          ......................................................g....................g.c.. [ 54240]
AY261363.1_Pretoriuskop/96/4   ....................t........................................................c.. [ 54240]
AY261362.1_Mkuzi_1979          ..............................c....................................g.........c.. [ 54240]
AY261361.1_Malawi_Lil-20/1_1983 ...............a..............c......a.a.....c.a......g.g...a.........c......t.. [ 54240]
AY261360.1_Kenya_1950          ..a.....g....................gc.......ga....gc........g.g.....g.........cat..t.. [ 54240]
AM712240.1_OURT_88/3           ..............................c...................................a..........c.. [ 54240]
AM712239.1_Benin_97/1          ..............................c...................................a..........c.. [ 54240]
KM111295.1_Ken06.Bus           ..a.....g....................gc.......ga....gc........g.g.....g.........cat..t.. [ 54240]
KM111294.1_Ken05/Tk1           ..a.....g....................gc.......ga....gc........g.g.....g.........cat..t.. [ 54240]
KM262844.1_strain_L60          ..............................c...................................a..........c.. [ 54240]
KM102979.1_26544/OG10_Italy    ..............................c...................................a..........c.. [ 54240]
KM262845.1_strain_NHV          ..............................c...................................a..........c.. [ 54240]
KX354450.1_47/Ss/2008          ..............................c...................................a..........c.. [ 54240]
KP055815.1_strain_BA71         ..............................c...................................a..........c.. [ 54240]
MH025920.1_strain_R35          ..a.....g....................gc.......ga....gc........g.g.....g.........cat..t.. [ 54240]
MH025919.1_strain_N10          ..a.....g....................gc.......ga....gc........g.g.....g.........cat..t.. [ 54240]
MH025918.1_strain_R25          ..a.....g....................gc.......ga....gc........g.g.....g.........cat..t.. [ 54240]
MH025917.1_strain_R7           ..a.....g....................gc.......ga....gc........g.g.....g.........cat..t.. [ 54240]
MH025916.1_strain_R8           ..a.....g....................gc.......ga....gc........g.g.....g.........cat..t.. [ 54240]

FR682468.2_Georgia_2007/1      atcaatccataaggctatacaaattgcagcctcggaaggaaatgagaacattgtcaagcttttcttactatggaagggga [ 54320]
ASF_IND_20_CAD543              ................................................................................ [ 54320]
LR899193.1_Germany_2020/1      ................................................................................ [ 54320]
MN393477.1_Wuhan_2019          ................................................................................ [ 54320]
MW396979.1_Timor-Leste/2019/1  ................................................................................ [ 54320]
MK940252.1_CN/2019/InnerMongolia ................................................................................ [ 54320]
MK128995.1_China/2018/AnhuiXCGQ ................................................................................ [ 54320]
MH681419.1_ASFV/POL/2015       ................................................................................ [ 54320]
LS478113.1_Estonia_2014        ................................................................................ [ 54320]
KP843857.1_Odintsovo_02/14     ................................................................................ [ 54320]
FN557520.1_strain_E75          ...c.................................................................g.......... [ 54320]
AY261366.1_Warthog             .c...c..........g.............................g......................g.......... [ 54320]
AY261365.1_Warmbaths           .c..............g.............................g.g....................g.......... [ 54320]
AY261364.1_Tengani_62          .c...c..........g.............................g.g....................g.......... [ 54320]
AY261363.1_Pretoriuskop/96/4   ................g.............................g......................g.....a.... [ 54320]
AY261362.1_Mkuzi_1979          ...c.................................................................g.......... [ 54320]
AY261361.1_Malawi_Lil-20/1_1983 ......t.........t......................g......g.t........a.....ga..........t..a. [ 54320]
AY261360.1_Kenya_1950          .......a.ctt...c...a...........g.......g......g.g.....t........a.....g...gg...a. [ 54320]
AM712240.1_OURT_88/3           ...c.................................................................g.......... [ 54320]
AM712239.1_Benin_97/1          ...c.................................................................g.......... [ 54320]
KM111295.1_Ken06.Bus           .......a.ctt...c...a...........g.......g......g.g.....t........a.....g...gg...a. [ 54320]
```

```
KM111294.1_Ken05/Tk1             .......a.ctt...c...a...........g.......g......g.g.....t........a.....g...gg...a. [ 54320]
KM262844.1_strain_L60            ...c.................................................................g.......... [ 54320]
KM102979.1_26544/OG10_Italy      ...c.................................................................g.......... [ 54320]
KM262845.1_strain_NHV            ...c.................................................................g.......... [ 54320]
KX354450.1_47/Ss/2008            ...c.................................................................g.......... [ 54320]
KP055815.1_strain_BA71           ...c.................................................................g.......... [ 54320]
MH025920.1_strain_R35            .......a.ctt...c...a...........g.......g......g.g.....t........a.....g...gg...a. [ 54320]
MH025919.1_strain_N10            .......a.ctt...c...a...........g.......g......g.g.....t........a.....g...gg...a. [ 54320]
MH025918.1_strain_R25            .......a.ctt...c...a...........g.......g......g.g.....t........a.....g...gg...a. [ 54320]
MH025917.1_strain_R7             .......a.ctt...c...a...........g.......g......g.g.....t........a.....g...gg...a. [ 54320]
MH025916.1_strain_R8             .......a.ctt...c...a...........g.......g......g.g.....t........a.....g...gg...a. [ 54320]

FR682468.2_Georgia_2007/1        gtctacaatatgccatcataggagccttagagggcaggcaatatgatctgattcaaaaatattacaaccaaattggggac [ 54400]
ASF_IND_20_CAD543                ................................................................................ [ 54400]
LR899193.1_Germany_2020/1        ................................................................................ [ 54400]
MN393477.1_Wuhan_2019            ................................................................................ [ 54400]
MW396979.1_Timor-Leste/2019/1    ................................................................................ [ 54400]
MK940252.1_CN/2019/InnerMongolia ................................................................................ [ 54400]
MK128995.1_China/2018/AnhuiXCGQ  ................................................................................ [ 54400]
MH681419.1_ASFV/POL/2015         ................................................................................ [ 54400]
LS478113.1_Estonia_2014          ................................................................................ [ 54400]
KP843857.1_Odintsovo_02/14       ................................................................................ [ 54400]
FN557520.1_strain_E75            ................................................................................ [ 54400]
AY261366.1_Warthog               .....................................a....................g..............c...... [ 54400]
AY261365.1_Warmbaths             .....................................a....................g..................... [ 54400]
AY261364.1_Tengani_62            .....................................a.......................................... [ 54400]
AY261363.1_Pretoriuskop/96/4     .....................................a....................g..................... [ 54400]
AY261362.1_Mkuzi_1979            ..............g................................................................. [ 54400]
AY261361.1_Malawi_Lil-20/1_1983  a....a...................t..g..a.....a..t......................................t [ 54400]
AY261360.1_Kenya_1950            a...g..t........................a....a........g.....c.tg.t....g.g..t............ [ 54400]
AM712240.1_OURT_88/3             ................................................................................ [ 54400]
AM712239.1_Benin_97/1            ................................................................................ [ 54400]
KM111295.1_Ken06.Bus             ac..g..t........................a....a........g.....c.tt.tg...g.a..t............ [ 54400]
KM111294.1_Ken05/Tk1             a...g..t........................a....a........g.....c.tg.t....g.g..t............ [ 54400]
KM262844.1_strain_L60            ................................................................................ [ 54400]
KM102979.1_26544/OG10_Italy      ................................................................................ [ 54400]
KM262845.1_strain_NHV            ................................................................................ [ 54400]
KX354450.1_47/Ss/2008            ................................................................................ [ 54400]
KP055815.1_strain_BA71           ................................................................................ [ 54400]
MH025920.1_strain_R35            ac..g..t........................a....a........g.....c.tt.tg...g.a..t............ [ 54400]
MH025919.1_strain_N10            ac..g..t........................a....a........g.....c.tt.tg...g.a..t............ [ 54400]
MH025918.1_strain_R25            ac..g..t........................a....a........g.....c.tt.tg...g.a..t............ [ 54400]
MH025917.1_strain_R7             ac..g..t........................a....a........g.....c.tt.tg...g.a..t............ [ 54400]
MH025916.1_strain_R8             ac..g..t........................a....a........g.....c.tt.tg...g.a..t............ [ 54400]
```

```
FR682468.2_Georgia_2007/1        tgccatcagattctaccactgattcaagatccagaaatttacgaaagatgtcatgaattaaatgttacatgtacctttca [ 54480]
ASF_IND_20_CAD543                ................................................................................ [ 54480]
LR899193.1_Germany_2020/1        ................................................................................ [ 54480]
MN393477.1_Wuhan_2019            ................................................................................ [ 54480]
MW396979.1_Timor-Leste/2019/1    ................................................................................ [ 54480]
MK940252.1_CN/2019/InnerMongolia ................................................................................ [ 54480]
MK128995.1_China/2018/AnhuiXCGQ  ................................................................................ [ 54480]
MH681419.1_ASFV/POL/2015         ................................................................................ [ 54480]
LS478113.1_Estonia_2014          ................................................................................ [ 54480]
KP843857.1_Odintsovo_02/14       ................................................................................ [ 54480]
FN557520.1_strain_E75            ......g...........................................c............................. [ 54480]
AY261366.1_Warthog               ................................................................................ [ 54480]
AY261365.1_Warmbaths             ................................................................................ [ 54480]
AY261364.1_Tengani_62            ................................................................................ [ 54480]
AY261363.1_Pretoriuskop/96/4     ................................................................................ [ 54480]
AY261362.1_Mkuzi_1979            ......g...........................................c............................. [ 54480]
AY261361.1_Malawi_Lil-20/1_1983  ......g..........c...............a....c......................................... [ 54480]
AY261360.1_Kenya_1950            ..g...g.c..c...t.c..c....g........tg..c..t........c............................. [ 54480]
AM712240.1_OURT_88/3             ......g...........................................c............................. [ 54480]
AM712239.1_Benin_97/1            ......g...........................................c............................. [ 54480]
KM111295.1_Ken06.Bus             .at...g.c..c...t.c..c....g........tg..c..t........c............................. [ 54480]
KM111294.1_Ken05/Tk1             .at...g.c..c...t.c..c....g........tg..c..t........c............................. [ 54480]
KM262844.1_strain_L60            ......g...........................................c............................. [ 54480]
KM102979.1_26544/OG10_Italy      ......g.....t.....................................c............................. [ 54480]
KM262845.1_strain_NHV            ......g...........................................c............................. [ 54480]
KX354450.1_47/Ss/2008            ......g.....t.....................................c............................. [ 54480]
KP055815.1_strain_BA71           ......g...........................................c............................. [ 54480]
MH025920.1_strain_R35            .at...g.c..c...t.c..c....g........tg..c..t........c............................. [ 54480]
MH025919.1_strain_N10            .at...g.c..c...t.c..c....g...c....tg..c..t........c............................. [ 54480]
MH025918.1_strain_R25            .at...g.c..c...t.c..c....g........tg..c..t........c............................. [ 54480]
MH025917.1_strain_R7             .at...g.c..c...t.c..c....g........tg..c..t........c............................. [ 54480]
MH025916.1_strain_R8             .at...g.c..c...t.c..c....g........tg..c..t........c............................. [ 54480]

FR682468.2_Georgia_2007/1        atgcttatttcaacatgctataagagataacatgctgcccattttccaaaaatatggagaagatctgaatggaaacagga [ 54560]
ASF_IND_20_CAD543                ..............................................................................r. [ 54560]
LR899193.1_Germany_2020/1        ................................................................................ [ 54560]
MN393477.1_Wuhan_2019            ................................................................................ [ 54560]
MW396979.1_Timor-Leste/2019/1    ................................................................................ [ 54560]
MK940252.1_CN/2019/InnerMongolia ................................................................................ [ 54560]
MK128995.1_China/2018/AnhuiXCGQ  ................................................................................ [ 54560]
MH681419.1_ASFV/POL/2015         ................................................................................ [ 54560]
LS478113.1_Estonia_2014          ................................................................................ [ 54560]
KP843857.1_Odintsovo_02/14       ................................................................................ [ 54560]
```

```
FN557520.1_strain_E75           ..............................................................................a. [ 54560]
AY261366.1_Warthog              .............................................t................................ag [ 54560]
AY261365.1_Warmbaths            .............................................t................................a. [ 54560]
AY261364.1_Tengani_62           ..............................................................................a. [ 54560]
AY261363.1_Pretoriuskop/96/4    .............................................t................................ag [ 54560]
AY261362.1_Mkuzi_1979           ..............................................................................a. [ 54560]
AY261361.1_Malawi_Lil-20/1_1983 ...t.................t.a...................c.t............c...................a. [ 54560]
AY261360.1_Kenya_1950           ...t.................t...c....t...g.tt.....c.t.........a.......c..agc.aac.....a. [ 54560]
AM712240.1_OURT_88/3            ..............................................................................a. [ 54560]
AM712239.1_Benin_97/1           ..............................................................................a. [ 54560]
KM111295.1_Ken06.Bus            ...t.................t...c....t...g.tt.....c.t.........a.......c..agc.cat.....a. [ 54560]
KM111294.1_Ken05/Tk1            ...t.................t...c....t...g.tt.....c.t.........a.......c..agc.aat.....a. [ 54560]
KM262844.1_strain_L60           ..............................................................................a. [ 54560]
KM102979.1_26544/OG10_Italy     ..............................................................................a. [ 54560]
KM262845.1_strain_NHV           ..............................................................................a. [ 54560]
KX354450.1_47/Ss/2008           ..............................................................................a. [ 54560]
KP055815.1_strain_BA71          ..............................................................................a. [ 54560]
MH025920.1_strain_R35           ...t.................t...c....t...g.tt.....c.t.........a.......c..agc.cat.....a. [ 54560]
MH025919.1_strain_N10           ...t.................t...c....t...g.tt.....c.t.........a.......c..agc.cat.....a. [ 54560]
MH025918.1_strain_R25           ...t.................t...c....t...g.tt.....c.t.........a.......c..agc.cat.....a. [ 54560]
MH025917.1_strain_R7            ...t.................t...c....t...g.tt.....c.t.........a.......c..agc.cat.....a. [ 54560]
MH025916.1_strain_R8            ...t.................t...c....t...g.tt.....c.t.........a.......c..agc.cat.....a. [ 54560]

FR682468.2_Georgia_2007/1       gaatggttcaacttctgtatgagatggcatgccgattacaaaattatgatatcatcaaatggataggatctaacctgcat [ 54640]
ASF_IND_20_CAD543               ................................................................................ [ 54640]
LR899193.1_Germany_2020/1       ................................................................................ [ 54640]
MN393477.1_Wuhan_2019           ................................................................................ [ 54640]
MW396979.1_Timor-Leste/2019/1   ................................................................................ [ 54640]
MK940252.1_CN/2019/InnerMongolia ................................................................................ [ 54640]
MK128995.1_China/2018/AnhuiXCGQ ................................................................................ [ 54640]
MH681419.1_ASFV/POL/2015        ................................................................................ [ 54640]
LS478113.1_Estonia_2014         ................................................................................ [ 54640]
KP843857.1_Odintsovo_02/14      ................................................................................ [ 54640]
FN557520.1_strain_E75           ................a.....a..............................................t.......... [ 54640]
AY261366.1_Warthog              .....................................................................t.......... [ 54640]
AY261365.1_Warmbaths            .....................................................................t.......... [ 54640]
AY261364.1_Tengani_62           ................................................................................ [ 54640]
AY261363.1_Pretoriuskop/96/4    ....................................................t................t.......... [ 54640]
AY261362.1_Mkuzi_1979           ................a.....a..............................................t.......... [ 54640]
AY261361.1_Malawi_Lil-20/1_1983 .....................................................................t......a..a [ 54640]
AY261360.1_Kenya_1950           .....a.c..............a................................t.cc.......c.cg...ttg.... [ 54640]
AM712240.1_OURT_88/3            ................a.....a..............................................t.......... [ 54640]
AM712239.1_Benin_97/1           ................a.....a..............................................t.......... [ 54640]
KM111295.1_Ken06.Bus            .....a.c..............a................................t.cc.......cgcg....tg.... [ 54640]
```

```
KM111294.1_Ken05/Tk1             .....a.c..............a................................t.cc.......cgcg....tg.... [ 54640]
KM262844.1_strain_L60            ................a.....a..............................................t.......... [ 54640]
KM102979.1_26544/OG10_Italy      ................a.....a..............................................t.......... [ 54640]
KM262845.1_strain_NHV            ................a.....a..............................................t.......... [ 54640]
KX354450.1_47/Ss/2008            ................a.....a..............................................t.......... [ 54640]
KP055815.1_strain_BA71           ................a.....a..............................................t.......... [ 54640]
MH025920.1_strain_R35            .....a.c..............a................................t.cc.......cgcg....tg.... [ 54640]
MH025919.1_strain_N10            .....a.c..............a................................t.cc.......cgcg....tg.... [ 54640]
MH025918.1_strain_R25            .....a.c..............a................................t.cc.......cgcg....tg.... [ 54640]
MH025917.1_strain_R7             .....a.c..............a................................t.cc.......cgcg....tg.... [ 54640]
MH025916.1_strain_R8             .....a.c..............a................................t.cc.......cgcg....tg.... [ 54640]

FR682468.2_Georgia_2007/1        gtttataacttggaagccatttttagcattgcttttgttagaaaggatttaactttgtattctttaggctacatgcttct [ 54720]
ASF_IND_20_CAD543                ................................................................................ [ 54720]
LR899193.1_Germany_2020/1        ................................................................................ [ 54720]
MN393477.1_Wuhan_2019            ................................................................................ [ 54720]
MW396979.1_Timor-Leste/2019/1    ................................................................................ [ 54720]
MK940252.1_CN/2019/InnerMongolia ................................................................................ [ 54720]
MK128995.1_China/2018/AnhuiXCGQ  ................................................................................ [ 54720]
MH681419.1_ASFV/POL/2015         ................................................................................ [ 54720]
LS478113.1_Estonia_2014          ................................................................................ [ 54720]
KP843857.1_Odintsovo_02/14       ................................................................................ [ 54720]
FN557520.1_strain_E75            ................................................................................ [ 54720]
AY261366.1_Warthog               ............................................a................................... [ 54720]
AY261365.1_Warmbaths             .....................................................a.......................... [ 54720]
AY261364.1_Tengani_62            ................................................................................ [ 54720]
AY261363.1_Pretoriuskop/96/4     ............................................a................................... [ 54720]
AY261362.1_Mkuzi_1979            ................................................................................ [ 54720]
AY261361.1_Malawi_Lil-20/1_1983  .....................................................c.......................... [ 54720]
AY261360.1_Kenya_1950            ....t...ta.a..g........c............a.........................c.....g......t.... [ 54720]
AM712240.1_OURT_88/3             ................................................................................ [ 54720]
AM712239.1_Benin_97/1            ................................................................................ [ 54720]
KM111295.1_Ken06.Bus             ........ta.a..g.....................a.........................c.....g.....at.... [ 54720]
KM111294.1_Ken05/Tk1             ........ta.a..g........c............a.........................c.....g......t.... [ 54720]
KM262844.1_strain_L60            ................................................................................ [ 54720]
KM102979.1_26544/OG10_Italy      ................................................................................ [ 54720]
KM262845.1_strain_NHV            ................................................................................ [ 54720]
KX354450.1_47/Ss/2008            ................................................................................ [ 54720]
KP055815.1_strain_BA71           ................................................................................ [ 54720]
MH025920.1_strain_R35            ........ta.a..g.....................a.........................c.....g.....at.... [ 54720]
MH025919.1_strain_N10            ........ta.a..g.....................a.........................c.....g.....at.... [ 54720]
MH025918.1_strain_R25            ........ta.a..g.....................a.........................c.....g.....at.... [ 54720]
MH025917.1_strain_R7             ........ta.a..g.....................a.........................c.....g.....at.... [ 54720]
MH025916.1_strain_R8             ........ta.a..g.....................a.........................c.....g.....at.... [ 54720]
```

```
FR682468.2_Georgia_2007/1        tctgggtagaatgagtactgaagatagaaactttatctcaatcataacacgccatcttgaatacgcatcaaaaaagggac [ 54800]
ASF_IND_20_CAD543                ................................................................................ [ 54800]
LR899193.1_Germany_2020/1        ................................................................................ [ 54800]
MN393477.1_Wuhan_2019            ................................................................................ [ 54800]
MW396979.1_Timor-Leste/2019/1    ................................................................................ [ 54800]
MK940252.1_CN/2019/InnerMongolia ................................................................................ [ 54800]
MK128995.1_China/2018/AnhuiXCGQ  ................................................................................ [ 54800]
MH681419.1_ASFV/POL/2015         ................................................................................ [ 54800]
LS478113.1_Estonia_2014          ................................................................................ [ 54800]
KP843857.1_Odintsovo_02/14       ................................................................................ [ 54800]
FN557520.1_strain_E75            ....................................t........................................... [ 54800]
AY261366.1_Warthog               ....................t........................................................... [ 54800]
AY261365.1_Warmbaths             ................................................................................ [ 54800]
AY261364.1_Tengani_62            .....................................................................g.......... [ 54800]
AY261363.1_Pretoriuskop/96/4     .....a..............t........................................................... [ 54800]
AY261362.1_Mkuzi_1979            ................................................................................ [ 54800]
AY261361.1_Malawi_Lil-20/1_1983  ....aa.......................................................................... [ 54800]
AY261360.1_Kenya_1950            .....a.........c.t.................aa.................c........t..gg.c.......... [ 54800]
AM712240.1_OURT_88/3             ....................................t........................................... [ 54800]
AM712239.1_Benin_97/1            ....................................t........................................... [ 54800]
KM111295.1_Ken06.Bus             .....a.........c.t.................aa.................c........t..gg.c.......... [ 54800]
KM111294.1_Ken05/Tk1             .....a.........c.t.................aa.................c........t..gg.c.......... [ 54800]
KM262844.1_strain_L60            ....................................t........................................... [ 54800]
KM102979.1_26544/OG10_Italy      ....................................t........................................... [ 54800]
KM262845.1_strain_NHV            ....................................t........................................... [ 54800]
KX354450.1_47/Ss/2008            ....................................t........................................... [ 54800]
KP055815.1_strain_BA71           ....................................t........................................... [ 54800]
MH025920.1_strain_R35            .....a.........c.t.................aa.................c........t..gg.c.......... [ 54800]
MH025919.1_strain_N10            .....a.........c.t.................aa.................c........t..gg.c.......... [ 54800]
MH025918.1_strain_R25            .....a.........c.t.................aa.................c........t..gg.c.......... [ 54800]
MH025917.1_strain_R7             .....a.........c.t.................aa.................c........t..gg.c.......... [ 54800]
MH025916.1_strain_R8             .....a.........c.t.................aa.................c........t..gg.c.......... [ 54800]

FR682468.2_Georgia_2007/1        tttttgactttgtactagaatctttgaaatatggaggtcaagtggatacagtgttgtttcaggctgtaaaatacaaccat [ 54880]
ASF_IND_20_CAD543                ................................................................................ [ 54880]
LR899193.1_Germany_2020/1        ................................................................................ [ 54880]
MN393477.1_Wuhan_2019            ................................................................................ [ 54880]
MW396979.1_Timor-Leste/2019/1    ................................................................................ [ 54880]
MK940252.1_CN/2019/InnerMongolia ................................................................................ [ 54880]
MK128995.1_China/2018/AnhuiXCGQ  ................................................................................ [ 54880]
MH681419.1_ASFV/POL/2015         ................................................................................ [ 54880]
LS478113.1_Estonia_2014          ................................................................................ [ 54880]
KP843857.1_Odintsovo_02/14       ................................................................................ [ 54880]
```

```
FN557520.1_strain_E75            ...............................c................................................ [ 54880]
AY261366.1_Warthog               ................................................................................ [ 54880]
AY261365.1_Warmbaths             ...............................c................................................ [ 54880]
AY261364.1_Tengani_62            ...............................c................................................ [ 54880]
AY261363.1_Pretoriuskop/96/4     ................................................................................ [ 54880]
AY261362.1_Mkuzi_1979            ...............................c................................................ [ 54880]
AY261361.1_Malawi_Lil-20/1_1983  ....g..........................c....................a........................... [ 54880]
AY261360.1_Kenya_1950            .............c.........c.............c.....a.................................... [ 54880]
AM712240.1_OURT_88/3             ...............................c................................................ [ 54880]
AM712239.1_Benin_97/1            ...............................c................................................ [ 54880]
KM111295.1_Ken06.Bus             .............c.........c.............c.....a................................t... [ 54880]
KM111294.1_Ken05/Tk1             .............c.........c.............c.....a.................................... [ 54880]
KM262844.1_strain_L60            ...............................c................................................ [ 54880]
KM102979.1_26544/OG10_Italy      ...............................c................................................ [ 54880]
KM262845.1_strain_NHV            ...............................c................................................ [ 54880]
KX354450.1_47/Ss/2008            ...............................c................................................ [ 54880]
KP055815.1_strain_BA71           ...............................c................................................ [ 54880]
MH025920.1_strain_R35            .............c.........c.............c.....a................................t... [ 54880]
MH025919.1_strain_N10            .............c.........c.............c.....a................................t... [ 54880]
MH025918.1_strain_R25            .............c.........c.............c.....a................................t... [ 54880]
MH025917.1_strain_R7             .............c.........c.............c.....a................................t... [ 54880]
MH025916.1_strain_R8             .............c.........c.............c.....a................................t... [ 54880]

FR682468.2_Georgia_2007/1        aggaaaattttggcccattttattcatgaaattccccgtgaaacggttgaaaagctgatactccatgctgtggagtcacg [ 54960]
ASF_IND_20_CAD543                ..r............................................................................. [ 54960]
LR899193.1_Germany_2020/1        ................................................................................ [ 54960]
MN393477.1_Wuhan_2019            ................................................................................ [ 54960]
MW396979.1_Timor-Leste/2019/1    ................................................................................ [ 54960]
MK940252.1_CN/2019/InnerMongolia ................................................................................ [ 54960]
MK128995.1_China/2018/AnhuiXCGQ  ................................................................................ [ 54960]
MH681419.1_ASFV/POL/2015         ................................................................................ [ 54960]
LS478113.1_Estonia_2014          ................................................................................ [ 54960]
KP843857.1_Odintsovo_02/14       ................................................................................ [ 54960]
FN557520.1_strain_E75            ..a................................g........a..........................a..a..g.. [ 54960]
AY261366.1_Warthog               ................................................................................ [ 54960]
AY261365.1_Warmbaths             .............................................................................g.. [ 54960]
AY261364.1_Tengani_62            ................................................................................ [ 54960]
AY261363.1_Pretoriuskop/96/4     .............................................................................g.. [ 54960]
AY261362.1_Mkuzi_1979            ..a................................g........a................................g.. [ 54960]
AY261361.1_Malawi_Lil-20/1_1983  ...........................c.tg............tc................................g.. [ 54960]
AY261360.1_Kenya_1950            ..a......c.....t.............g.cc......a.................t....t.....c........g.. [ 54960]
AM712240.1_OURT_88/3             ..a................................g........a..........................a..a..g.. [ 54960]
AM712239.1_Benin_97/1            ..a................................g........a..........................a..a..g.. [ 54960]
KM111295.1_Ken06.Bus             ..a......c.....t.............g.cc......a.................t....t.....c........g.. [ 54960]
```

```
KM111294.1_Ken05/Tk1            ..a......c.....t.............g.cc......a.................t....t.....c........g.. [ 54960]
KM262844.1_strain_L60           ..a................................g........a..........................a..a..g.. [ 54960]
KM102979.1_26544/OG10_Italy     ..a................................g........a..........................a..a..g.. [ 54960]
KM262845.1_strain_NHV           ..a................................g........a..........................a..a..g.. [ 54960]
KX354450.1_47/Ss/2008           ..a................................g........a..........................a..a..g.. [ 54960]
KP055815.1_strain_BA71          ..a................................g........a..........................a..a..g.. [ 54960]
MH025920.1_strain_R35           ..a......c.....t.............g.cc......a.................t....t.....c........g.. [ 54960]
MH025919.1_strain_N10           ..a......c.....t.............g.cc......a.................t....t.....c........g.. [ 54960]
MH025918.1_strain_R25           ..a......c.....t.............g.cc......a.................t....t.....c........g.. [ 54960]
MH025917.1_strain_R7            ..a......c.....t.............g.cc......a.................t....t.....c........g.. [ 54960]
MH025916.1_strain_R8            ..a......c.....t.............g.cc......a.................t....t.....c........g.. [ 54960]

FR682468.2_Georgia_2007/1       ggcctccagaaaaacattcaacctgcttttatcttccataaactactgtgtgaacccttttgtcaaaaaactactgcacg [ 55040]
ASF_IND_20_CAD543               ................................................................................ [ 55040]
LR899193.1_Germany_2020/1       ................................................................................ [ 55040]
MN393477.1_Wuhan_2019           ................................................................................ [ 55040]
MW396979.1_Timor-Leste/2019/1   ................................................................................ [ 55040]
MK940252.1_CN/2019/InnerMongolia ................................................................................ [ 55040]
MK128995.1_China/2018/AnhuiXCGQ ................................................................................ [ 55040]
MH681419.1_ASFV/POL/2015        ................................................................................ [ 55040]
LS478113.1_Estonia_2014         ................................................................................ [ 55040]
KP843857.1_Odintsovo_02/14      ................................................................................ [ 55040]
FN557520.1_strain_E75           .........g.........................................a...........................a [ 55040]
AY261366.1_Warthog              .........g............................................t......................... [ 55040]
AY261365.1_Warmbaths            ........ag...................................................................... [ 55040]
AY261364.1_Tengani_62           .........g...................................................................... [ 55040]
AY261363.1_Pretoriuskop/96/4    .........g...................................................................... [ 55040]
AY261362.1_Mkuzi_1979           .........g...................................................................... [ 55040]
AY261361.1_Malawi_Lil-20/1_1983 .........g.............................t.........a.......................t....t. [ 55040]
AY261360.1_Kenya_1950           ...t..t.ag.....ca.g...........g.....tc.g.......c.a.ac.tt.ca.ca.t.....g..c...t... [ 55040]
AM712240.1_OURT_88/3            .........g.........................................a...........................a [ 55040]
AM712239.1_Benin_97/1           .........g.........................................a...........................a [ 55040]
KM111295.1_Ken06.Bus            ...t..t.ag.....ca.g...........g......c.........c.a.ac.tt.ca.ca.t.....g..c...t... [ 55040]
KM111294.1_Ken05/Tk1            a..t..t.ag.....ca.g...........g.....tc.g.......c.a.ac.tt.ca.ca.t.....g..c...t... [ 55040]
KM262844.1_strain_L60           .........g.........................................a...........................a [ 55040]
KM102979.1_26544/OG10_Italy     .........g.........................................a...........................a [ 55040]
KM262845.1_strain_NHV           .........g.........................................a...........................a [ 55040]
KX354450.1_47/Ss/2008           .........g.........................................a...........................a [ 55040]
KP055815.1_strain_BA71          .........g.........................................a...........................a [ 55040]
MH025920.1_strain_R35           ...t..t.ag.....ca.g...........g......c.........c.a.ac.tt.ca.ca.t.....g..c...t... [ 55040]
MH025919.1_strain_N10           ...t..t.ag.....ca.g...........g......c.........c.a.ac.tt.ca.ca.t.....g..c...t... [ 55040]
MH025918.1_strain_R25           ...t..t.ag.....ca.g...........g......c.........c.a.ac.tt.ca.ca.t.....g..c...t... [ 55040]
MH025917.1_strain_R7            ...t..t.ag.....ca.g...........g......c.........c.a.ac.tt.ca.ca.t.....g..c...t... [ 55040]
MH025916.1_strain_R8            ...t..t.ag.....ca.g...........g......c.........c.a.ac.tt.ca.ca.t.....g..c...t... [ 55040]
```

```
FR682468.2_Georgia_2007/1        ctgtggtgaaacacaagtacatgcttatcataaagcttttgctcgagcggcccaaaaagaagataaacctggtagatgct [ 55120]
ASF_IND_20_CAD543                ........r....................................................................... [ 55120]
LR899193.1_Germany_2020/1        ........g....................................................................... [ 55120]
MN393477.1_Wuhan_2019            ........g....................................................................... [ 55120]
MW396979.1_Timor-Leste/2019/1    ........g....................................................................... [ 55120]
MK940252.1_CN/2019/InnerMongolia ........g....................................................................... [ 55120]
MK128995.1_China/2018/AnhuiXCGQ  ........g....................................................................... [ 55120]
MH681419.1_ASFV/POL/2015         ................................................................................ [ 55120]
LS478113.1_Estonia_2014          ........g....................................................................... [ 55120]
KP843857.1_Odintsovo_02/14       ........g....................................................................... [ 55120]
FN557520.1_strain_E75            .c.............................................................................. [ 55120]
AY261366.1_Warthog               ................................................................................ [ 55120]
AY261365.1_Warmbaths             ................................................a............................... [ 55120]
AY261364.1_Tengani_62            ................................................................................ [ 55120]
AY261363.1_Pretoriuskop/96/4     ................................................................................ [ 55120]
AY261362.1_Mkuzi_1979            ................................................a............................... [ 55120]
AY261361.1_Malawi_Lil-20/1_1983  .......a........a.t.................................................t.a...a..... [ 55120]
AY261360.1_Kenya_1950            ...................t......g....................a......................a......... [ 55120]
AM712240.1_OURT_88/3             .c.............................................................................. [ 55120]
AM712239.1_Benin_97/1            .c.............................................................................. [ 55120]
KM111295.1_Ken06.Bus             ...................t......g.................ac......a.........t.......a......... [ 55120]
KM111294.1_Ken05/Tk1             ...................t......g..................c......a.........t.......a......... [ 55120]
KM262844.1_strain_L60            .c.............................................................................. [ 55120]
KM102979.1_26544/OG10_Italy      .c.............................................................................. [ 55120]
KM262845.1_strain_NHV            .c.............................................................................. [ 55120]
KX354450.1_47/Ss/2008            .c.............................................................................. [ 55120]
KP055815.1_strain_BA71           .c.............................................................................. [ 55120]
MH025920.1_strain_R35            ...................t......g.................ac......a.........t.......a......... [ 55120]
MH025919.1_strain_N10            ...................t......g.................ac......a.........t.......a......... [ 55120]
MH025918.1_strain_R25            ...................t......g.................ac......a.........t.......a......... [ 55120]
MH025917.1_strain_R7             ...................t......g.................ac......a.........t.......a......... [ 55120]
MH025916.1_strain_R8             ...................t......g.................ac......a.........t.......a......... [ 55120]

FR682468.2_Georgia_2007/1        gctctattcaaacttgtaaaatactctacttatacagaaatagtaaaatacatgggtgagttttctgtggacccaaaaag [ 55200]
ASF_IND_20_CAD543                .................................................w....aaa....................... [ 55200]
LR899193.1_Germany_2020/1        ................................................................................ [ 55200]
MN393477.1_Wuhan_2019            ................................................................................ [ 55200]
MW396979.1_Timor-Leste/2019/1    ................................................................................ [ 55200]
MK940252.1_CN/2019/InnerMongolia ................................................................................ [ 55200]
MK128995.1_China/2018/AnhuiXCGQ  ................................................................................ [ 55200]
MH681419.1_ASFV/POL/2015         ................................................................................ [ 55200]
LS478113.1_Estonia_2014          ................................................................................ [ 55200]
KP843857.1_Odintsovo_02/14       ................................................................................ [ 55200]
```

```
FN557520.1_strain_E75           .................................g...............t....aaa..................g.... [ 55200]
AY261366.1_Warthog              ..................................a....................a........................ [ 55200]
AY261365.1_Warmbaths            ..................................a....................a.............a....gg.... [ 55200]
AY261364.1_Tengani_62           ................................................................................ [ 55200]
AY261363.1_Pretoriuskop/96/4    ..................................a....................a........................ [ 55200]
AY261362.1_Mkuzi_1979           ..................................a....................a.............a....gg.... [ 55200]
AY261361.1_Malawi_Lil-20/1_1983 ..........................c......c....................aag............a....gg...t [ 55200]
AY261360.1_Kenya_1950           .t.....at.g..........c....c..aa..g.............g.tt...aa.............ag...gg.... [ 55200]
AM712240.1_OURT_88/3            .................................g...............t....aaa..................g.... [ 55200]
AM712239.1_Benin_97/1           .................................g...............t....aaa..................g.... [ 55200]
KM111295.1_Ken06.Bus            .t.....at.g..........c....c..ga..g.............g.tt...aa.............ag...gg.... [ 55200]
KM111294.1_Ken05/Tk1            .t.....at.g..........c....c..ga..g.............g.tt...aa.............ag...gg.... [ 55200]
KM262844.1_strain_L60           .................................g...............t....aaa..................g.... [ 55200]
KM102979.1_26544/OG10_Italy     .................................g...............t....aaa..................g.... [ 55200]
KM262845.1_strain_NHV           .................................g...............t....aaa..................g.... [ 55200]
KX354450.1_47/Ss/2008           .................................g...............t....aaa..................g.... [ 55200]
KP055815.1_strain_BA71          .................................g...............t....aaa..................g.... [ 55200]
MH025920.1_strain_R35           .t.....at.g..........c....c..ga..g.............g.tt...aa.............ag...gg.... [ 55200]
MH025919.1_strain_N10           .t.....at.g..........c....c..ga..g.............g.tt...aa.............ag...gg.... [ 55200]
MH025918.1_strain_R25           .t.....at.g..........c....c..ga..g.............g.tt...aa.............ag...gg.... [ 55200]
MH025917.1_strain_R7            .t.....at.g..........c....c..ga..g.............g.tt...aa.............ag...gg.... [ 55200]
MH025916.1_strain_R8            .t.....at.g..........c....c..ga..g.............g.tt...aa.............ag...gg.... [ 55200]

FR682468.2_Georgia_2007/1       ggtggtcaaaatggcagcacgactcatgagagtggacctgattaaaaagatttctaatgatgcatgggaagataaactag [ 55280]
ASF_IND_20_CAD543               ......................................................................r......... [ 55280]
LR899193.1_Germany_2020/1       ................................................................................ [ 55280]
MN393477.1_Wuhan_2019           ................................................................................ [ 55280]
MW396979.1_Timor-Leste/2019/1   ................................................................................ [ 55280]
MK940252.1_CN/2019/InnerMongolia ................................................................................ [ 55280]
MK128995.1_China/2018/AnhuiXCGQ ................................................................................ [ 55280]
MH681419.1_ASFV/POL/2015        ................................................................................ [ 55280]
LS478113.1_Estonia_2014         ................................................................................ [ 55280]
KP843857.1_Odintsovo_02/14      ................................................................................ [ 55280]
FN557520.1_strain_E75           .........................................................c............a......... [ 55280]
AY261366.1_Warthog              ................................................................................ [ 55280]
AY261365.1_Warmbaths            ................................................................................ [ 55280]
AY261364.1_Tengani_62           ................................................................................ [ 55280]
AY261363.1_Pretoriuskop/96/4    ................................................................................ [ 55280]
AY261362.1_Mkuzi_1979           ................................................a............................... [ 55280]
AY261361.1_Malawi_Lil-20/1_1983 ....a....................................................a............a......... [ 55280]
AY261360.1_Kenya_1950           ....a.....g.a.........t.a.......................c........a....t.......a......... [ 55280]
AM712240.1_OURT_88/3            .........................................................c............a......... [ 55280]
AM712239.1_Benin_97/1           .........................................................c............a......... [ 55280]
KM111295.1_Ken06.Bus            ....a.................t.a................................a...a........a......... [ 55280]
```

```
KM111294.1_Ken05/Tk1            ....a.....g.a.........t.a................................a....t.......a......... [ 55280]
KM262844.1_strain_L60           .........................................................c............a......... [ 55280]
KM102979.1_26544/OG10_Italy     .........................................................c............a......... [ 55280]
KM262845.1_strain_NHV           .........................................................c............a......... [ 55280]
KX354450.1_47/Ss/2008           .........................................................c............a......... [ 55280]
KP055815.1_strain_BA71          .........................................................c............a......... [ 55280]
MH025920.1_strain_R35           ....a.................t.a................................a...a........a......... [ 55280]
MH025919.1_strain_N10           ....a.................t.a................................a...a........a......... [ 55280]
MH025918.1_strain_R25           ....a.................t.a................................a...a........a......... [ 55280]
MH025917.1_strain_R7            ....a.................t.a................................a...a........a......... [ 55280]
MH025916.1_strain_R8            ....a.................t.a................................a...a........a......... [ 55280]

FR682468.2_Georgia_2007/1       agagaatcaagcaccttaaacagatggtaaataccatgaaccacagaaatggaaaaaatctattgatgtacaatattcac [ 55360]
ASF_IND_20_CAD543               ................................................................................ [ 55360]
LR899193.1_Germany_2020/1       ................................................................................ [ 55360]
MN393477.1_Wuhan_2019           ................................................................................ [ 55360]
MW396979.1_Timor-Leste/2019/1   ................................................................................ [ 55360]
MK940252.1_CN/2019/InnerMongolia ................................................................................ [ 55360]
MK128995.1_China/2018/AnhuiXCGQ ................................................................................ [ 55360]
MH681419.1_ASFV/POL/2015        ................................................................................ [ 55360]
LS478113.1_Estonia_2014         ................................................................................ [ 55360]
KP843857.1_Odintsovo_02/14      ................................................................................ [ 55360]
FN557520.1_strain_E75           ................................................................................ [ 55360]
AY261366.1_Warthog              ................................................................................ [ 55360]
AY261365.1_Warmbaths            ................................................................................ [ 55360]
AY261364.1_Tengani_62           ................................................................................ [ 55360]
AY261363.1_Pretoriuskop/96/4    ................................................................................ [ 55360]
AY261362.1_Mkuzi_1979           ................................................................................ [ 55360]
AY261361.1_Malawi_Lil-20/1_1983 .............t..............tc..........t.....................a.a............t.t [ 55360]
AY261360.1_Kenya_1950           ..........a...........a..a...t..........g.....................c..............t.. [ 55360]
AM712240.1_OURT_88/3            ................................................................................ [ 55360]
AM712239.1_Benin_97/1           ................................................................................ [ 55360]
KM111295.1_Ken06.Bus            ..........a...........a......t..........g........c............c..............t.. [ 55360]
KM111294.1_Ken05/Tk1            ..........a...........a..a...t..........g.....................c..............t.. [ 55360]
KM262844.1_strain_L60           ................................................................................ [ 55360]
KM102979.1_26544/OG10_Italy     ................................................................................ [ 55360]
KM262845.1_strain_NHV           ................................................................................ [ 55360]
KX354450.1_47/Ss/2008           ................................................................................ [ 55360]
KP055815.1_strain_BA71          ................................................................................ [ 55360]
MH025920.1_strain_R35           ..........a...........a......t..........g.....................c..............t.. [ 55360]
MH025919.1_strain_N10           ..........a...........a......t..........g.....................c..............t.. [ 55360]
MH025918.1_strain_R25           ..........a...........a......t..........g.....................c..............t.. [ 55360]
MH025917.1_strain_R7            ..........a...........a......t..........g.....................c..............t.. [ 55360]
MH025916.1_strain_R8            ..........a...........a......t..........g.....................c..............t.. [ 55360]
```

```
FR682468.2_Georgia_2007/1        aatattactggatatacctatctgaacaccaaagaagcatttaacttaacaagattttatgctgtccacaatgcaacatg [ 55440]
ASF_IND_20_CAD543                ...................gct.......................................................... [ 55440]
LR899193.1_Germany_2020/1        ................................................................................ [ 55440]
MN393477.1_Wuhan_2019            ................................................................................ [ 55440]
MW396979.1_Timor-Leste/2019/1    ................................................................................ [ 55440]
MK940252.1_CN/2019/InnerMongolia ................................................................................ [ 55440]
MK128995.1_China/2018/AnhuiXCGQ  ................................................................................ [ 55440]
MH681419.1_ASFV/POL/2015         ................................................................................ [ 55440]
LS478113.1_Estonia_2014          ................................................................................ [ 55440]
KP843857.1_Odintsovo_02/14       ................................................................................ [ 55440]
FN557520.1_strain_E75            ...................gct.......................................................... [ 55440]
AY261366.1_Warthog               ..................c.c...............................a........................... [ 55440]
AY261365.1_Warmbaths             ....................c...............................a........................... [ 55440]
AY261364.1_Tengani_62            ....................c......t...................................a................ [ 55440]
AY261363.1_Pretoriuskop/96/4     ..................c.c...............................a........................... [ 55440]
AY261362.1_Mkuzi_1979            ....................c........................................................... [ 55440]
AY261361.1_Malawi_Lil-20/1_1983  g...................c.....tg................t................................... [ 55440]
AY261360.1_Kenya_1950            ....................ca......t.......................a........................... [ 55440]
AM712240.1_OURT_88/3             ...................gct.......................................................... [ 55440]
AM712239.1_Benin_97/1            ...................gct.......................................................... [ 55440]
KM111295.1_Ken06.Bus             ................t...c.....tg....................g............................... [ 55440]
KM111294.1_Ken05/Tk1             ...g................ca......t.......................a........................... [ 55440]
KM262844.1_strain_L60            ...................gct.......................................................... [ 55440]
KM102979.1_26544/OG10_Italy      ...................gct.......................................................... [ 55440]
KM262845.1_strain_NHV            ...................gct.......................................................... [ 55440]
KX354450.1_47/Ss/2008            ...................gct.......................................................... [ 55440]
KP055815.1_strain_BA71           ...................gct.......................................................... [ 55440]
MH025920.1_strain_R35            ................t...c.....tg....................g............................... [ 55440]
MH025919.1_strain_N10            ................t...c.....tg....................g............................... [ 55440]
MH025918.1_strain_R25            ................t...c.....tg....................g............................... [ 55440]
MH025917.1_strain_R7             ................t...c.....tg....................g............................... [ 55440]
MH025916.1_strain_R8             ................t...c.....tg....................g............................... [ 55440]

FR682468.2_Georgia_2007/1        tttgtttaaagaaatgtgtaaaagctgttttgtac---atgataaaatacagctcagagaattgcttgaagattgtttac [ 55520]
ASF_IND_20_CAD543                ...................................---.......................................... [ 55520]
LR899193.1_Germany_2020/1        ...................................---.......................................... [ 55520]
MN393477.1_Wuhan_2019            ...................................---.......................................... [ 55520]
MW396979.1_Timor-Leste/2019/1    ...................................---.......................................... [ 55520]
MK940252.1_CN/2019/InnerMongolia ...................................---.......................................... [ 55520]
MK128995.1_China/2018/AnhuiXCGQ  ...................................---.......................................... [ 55520]
MH681419.1_ASFV/POL/2015         ...................................---.......................................... [ 55520]
LS478113.1_Estonia_2014          ...................................---.......................................... [ 55520]
KP843857.1_Odintsovo_02/14       ...................................---.......................................... [ 55520]
```

```
FN557520.1_strain_E75          ...................................---..............t........................... [ 55520]
AY261366.1_Warthog             ................................a..---..............t........................... [ 55520]
AY261365.1_Warmbaths           ...................................---......................................c... [ 55520]
AY261364.1_Tengani_62          ...................................---......................................c... [ 55520]
AY261363.1_Pretoriuskop/96/4   ................................a..---......................................... [ 55520]
AY261362.1_Mkuzi_1979          ...................................---......................................c... [ 55520]
AY261361.1_Malawi_Lil-20/1_1983 c......................a...........---.....tt.........t......................... [ 55520]
AY261360.1_Kenya_1950          c......................a...........---.....tt.........t......................... [ 55520]
AM712240.1_OURT_88/3           ...................................---..............t........................... [ 55520]
AM712239.1_Benin_97/1          ...................................---..............t........................... [ 55520]
KM111295.1_Ken06.Bus           c..............a...................atg..............t.t......a.a...a............ [ 55520]
KM111294.1_Ken05/Tk1           c......................a...........---.....tt.........t......................... [ 55520]
KM262844.1_strain_L60          ...................................---..............t........................... [ 55520]
KM102979.1_26544/OG10_Italy    ...................................---..............t........................... [ 55520]
KM262845.1_strain_NHV          ...................................---..............t........................... [ 55520]
KX354450.1_47/Ss/2008          ...................................---..............t........................... [ 55520]
KP055815.1_strain_BA71         ...................................---..............t........................... [ 55520]
MH025920.1_strain_R35          c..............a...................atg..............t.t......a.a...a............ [ 55520]
MH025919.1_strain_N10          c..............a...................atg..............t.t......a.a...a............ [ 55520]
MH025918.1_strain_R25          c..............a...................atg..............t.t......a.a...a............ [ 55520]
MH025917.1_strain_R7           c..............a...................atg..............t.t......a.a...a............ [ 55520]
MH025916.1_strain_R8           c..............a...................atg..............t.t......a.a...a............ [ 55520]

FR682468.2_Georgia_2007/1      atattgctaataggcatgattatatccagattgcagaaaccgcagatgaatgtatcaaatatatagat------cttatt [ 55600]
ASF_IND_20_CAD543              ....................................................................------...... [ 55600]
LR899193.1_Germany_2020/1      ....................................................................------...... [ 55600]
MN393477.1_Wuhan_2019          ....................................................................------...... [ 55600]
MW396979.1_Timor-Leste/2019/1  ....................................................................------...... [ 55600]
MK940252.1_CN/2019/InnerMongolia ....................................................................------...... [ 55600]
MK128995.1_China/2018/AnhuiXCGQ ....................................................................------...... [ 55600]
MH681419.1_ASFV/POL/2015       ....................................................................------...... [ 55600]
LS478113.1_Estonia_2014        ....................................................................------...... [ 55600]
KP843857.1_Odintsovo_02/14     ....................................................................------...... [ 55600]
FN557520.1_strain_E75          ....................................................................------...... [ 55600]
AY261366.1_Warthog             ..................c.........a.......................................------...... [ 55600]
AY261365.1_Warmbaths           ............................a.......................................------.....c [ 55600]
AY261364.1_Tengani_62          .............c..............a...........t..........a................------...... [ 55600]
AY261363.1_Pretoriuskop/96/4   ..................c.........a.......................................------...... [ 55600]
AY261362.1_Mkuzi_1979          ............................a.......................................------...c.. [ 55600]
AY261361.1_Malawi_Lil-20/1_1983 .............a....c.........a...........t...a......a...t............------ga...a [ 55600]
AY261360.1_Kenya_1950          ............a.....c......t............g.....a......a................gatatata...a [ 55600]
AM712240.1_OURT_88/3           ....................................................................------...... [ 55600]
AM712239.1_Benin_97/1          ....................................................................------...... [ 55600]
KM111295.1_Ken06.Bus           ..................c.........a..............g.......a........t...g...------.....g [ 55600]
```

```
KM111294.1_Ken05/Tk1           ............a.....c......t............g.....a......a................gatatata...a [ 55600]
KM262844.1_strain_L60          ....................................................................------...... [ 55600]
KM102979.1_26544/OG10_Italy    ....................................................................------...... [ 55600]
KM262845.1_strain_NHV          ....................................................................------...... [ 55600]
KX354450.1_47/Ss/2008          ....................................................................------...... [ 55600]
KP055815.1_strain_BA71         ....................................................................------...... [ 55600]
MH025920.1_strain_R35          ..................c.........a..............g.......a........t...g...------.....g [ 55600]
MH025919.1_strain_N10          ..................c.........a..............g.......a........t...g...------.....g [ 55600]
MH025918.1_strain_R25          ..................c.........a..............g.......a........t...g...------.....g [ 55600]
MH025917.1_strain_R7           ..................c.........a..............g.......a........t...g...------.....g [ 55600]
MH025916.1_strain_R8           ..................c.........a..............g.......a........t...g...------.....g [ 55600]

FR682468.2_Georgia_2007/1      acatttaagtaaaccatg--------------------tatatatcaagtaaatccagattaaatcaggctaattgtaaa [ 55680]
ASF_IND_20_CAD543              ..................aa------------------a......................................... [ 55680]
LR899193.1_Germany_2020/1      ..................--------------------.......................................... [ 55680]
MN393477.1_Wuhan_2019          ..................--------------------.......................................... [ 55680]
MW396979.1_Timor-Leste/2019/1  ..................--------------------.......................................... [ 55680]
MK940252.1_CN/2019/InnerMongolia ..................--------------------.......................................... [ 55680]
MK128995.1_China/2018/AnhuiXCGQ ..................--------------------.......................................... [ 55680]
MH681419.1_ASFV/POL/2015       ..................--------------------.......................................... [ 55680]
LS478113.1_Estonia_2014        ..................--------------------.......................................... [ 55680]
KP843857.1_Odintsovo_02/14     ..................--------------------.......................................... [ 55680]
FN557520.1_strain_E75          ...cc........t....aa------------------a......................................... [ 55680]
AY261366.1_Warthog             ...cc........t....aa------------------a....................a.................... [ 55680]
AY261365.1_Warmbaths           ..................tatatatcaagtaaaccatg.......................................... [ 55680]
AY261364.1_Tengani_62          ...g..............--------------------..........................a............... [ 55680]
AY261363.1_Pretoriuskop/96/4   ...cc........t....aa------------------a....................a.................... [ 55680]
AY261362.1_Mkuzi_1979          .t.cc........tt...aa------------------a......................................... [ 55680]
AY261361.1_Malawi_Lil-20/1_1983 ...cc........t..a.--------------------.........................g................ [ 55680]
AY261360.1_Kenya_1950          t..c.........t....--------------------......................g..................- [ 55680]
AM712240.1_OURT_88/3           ...cc........t....aa------------------a......................................... [ 55680]
AM712239.1_Benin_97/1          ...cc........t....aa------------------a......................................... [ 55680]
KM111295.1_Ken06.Bus           ttgcc........t....--------------------..............c.tt....g................... [ 55680]
KM111294.1_Ken05/Tk1           t..c.........t....--------------------......................g..................- [ 55680]
KM262844.1_strain_L60          ...cc........t....aa------------------a......................................... [ 55680]
KM102979.1_26544/OG10_Italy    ...cc........t....aa------------------a......................................... [ 55680]
KM262845.1_strain_NHV          ...cc........t....aa------------------a......................................... [ 55680]
KX354450.1_47/Ss/2008          ...cc........t....aa------------------a......................................... [ 55680]
KP055815.1_strain_BA71         ...cc........t....aa------------------a......................................... [ 55680]
MH025920.1_strain_R35          ttgcc........t....--------------------..............c.tt....g................... [ 55680]
MH025919.1_strain_N10          ttgcc........t....--------------------..............c.tt....g................... [ 55680]
MH025918.1_strain_R25          ttgcc........t....--------------------..............c.tt....g................... [ 55680]
MH025917.1_strain_R7           ttgcc........t....--------------------..............c.tt....g................... [ 55680]
MH025916.1_strain_R8           ttgcc........t....--------------------..............c.tt....g................... [ 55680]
```

```
FR682468.2_Georgia_2007/1        tagttgtagataccatataatgaatgttttattaggatagtagttcagttaaga-----------------tagtagttt [ 55760]
ASF_IND_20_CAD543                ............................................---------------------------..a...... [ 55760]
LR899193.1_Germany_2020/1        ......................................................-----------------......... [ 55760]
MN393477.1_Wuhan_2019            ......................................................-----------------......... [ 55760]
MW396979.1_Timor-Leste/2019/1    ......................................................-----------------......... [ 55760]
MK940252.1_CN/2019/InnerMongolia ......................................................-----------------......... [ 55760]
MK128995.1_China/2018/AnhuiXCGQ  ......................................................-----------------......... [ 55760]
MH681419.1_ASFV/POL/2015         ......................................................-----------------......... [ 55760]
LS478113.1_Estonia_2014          ......................................................-----------------......... [ 55760]
KP843857.1_Odintsovo_02/14       ......................................................-----------------......... [ 55760]
FN557520.1_strain_E75            ............................................---------------------------..a...... [ 55760]
AY261366.1_Warthog               .............................................g........cagtagtttagttaagac........ [ 55760]
AY261365.1_Warmbaths             ......................................................cagtagttca---------------. [ 55760]
AY261364.1_Tengani_62            .............................................g.....g..tagtagttgagttagga........c [ 55760]
AY261363.1_Pretoriuskop/96/4     ......................................................tagtagttcagttaagac.......c [ 55760]
AY261362.1_Mkuzi_1979            .............................................t.....t..tagtggtttagttatga....g.... [ 55760]
AY261361.1_Malawi_Lil-20/1_1983  ..........................a.......a....a.............tcaatagttcagttaagtc.a.....c [ 55760]
AY261360.1_Kenya_1950            -----a....................a.......a..cg......tca......cagtagttctattaaga..a.....c [ 55760]
AM712240.1_OURT_88/3             ............................................---------------------------..a...... [ 55760]
AM712239.1_Benin_97/1            ............................................---------------------------..a...... [ 55760]
KM111295.1_Ken06.Bus             .....a.......t............a....------------------------------------------------- [ 55760]
KM111294.1_Ken05/Tk1             -----a....................a.......a..cg......tca......cagtagttctattaaga..a.....c [ 55760]
KM262844.1_strain_L60            ............................................---------------------------..a...... [ 55760]
KM102979.1_26544/OG10_Italy      ............................................---------------------------..a...... [ 55760]
KM262845.1_strain_NHV            ............................................---------------------------..a...... [ 55760]
KX354450.1_47/Ss/2008            ............................................---------------------------..a...... [ 55760]
KP055815.1_strain_BA71           ............................................---------------------------..a...... [ 55760]
MH025920.1_strain_R35            .....a.......t............a....------------------------------------------------- [ 55760]
MH025919.1_strain_N10            .....a.......t............a....------------------------------------------------- [ 55760]
MH025918.1_strain_R25            .....a.......t............a....------------------------------------------------- [ 55760]
MH025917.1_strain_R7             .....a.......t............a....------------------------------------------------- [ 55760]
MH025916.1_strain_R8             .....a.......t............a....------------------------------------------------- [ 55760]

FR682468.2_Georgia_2007/1        agttaagatagtagtttagttaagatagtag------------------------------------------------- [ 55840]
ASF_IND_20_CAD543                ................c-------------t------------------------------------------------- [ 55840]
LR899193.1_Germany_2020/1        ...............................------------------------------------------------- [ 55840]
MN393477.1_Wuhan_2019            ...............................------------------------------------------------- [ 55840]
MW396979.1_Timor-Leste/2019/1    ...............................------------------------------------------------- [ 55840]
MK940252.1_CN/2019/InnerMongolia ...............................------------------------------------------------- [ 55840]
MK128995.1_China/2018/AnhuiXCGQ  ...............................------------------------------------------------- [ 55840]
MH681419.1_ASFV/POL/2015         ...............................------------------------------------------------- [ 55840]
LS478113.1_Estonia_2014          ...............................------------------------------------------------- [ 55840]
KP843857.1_Odintsovo_02/14       ...............................------------------------------------------------- [ 55840]
```

```
FN557520.1_strain_E75            ........c.......ct-------------------------------------------------------------- [ 55840]
AY261366.1_Warthog               ........c.......ct............a------------------------------------------------- [ 55840]
AY261365.1_Warmbaths             ................ct-------------------------------------------------------------- [ 55840]
AY261364.1_Tengani_62            ........c.a..............c.....------------------------------------------------- [ 55840]
AY261363.1_Pretoriuskop/96/4     t...............c........c.....------------------------------------------------- [ 55840]
AY261362.1_Mkuzi_1979            ........c.......ct.............t------------------------------------------------ [ 55840]
AY261361.1_Malawi_Lil-20/1_1983  ..........a.....c.......tc.a...ttcagttaagtcaatagttcagttaagtcaatagttcagttaagacagt [ 55840]
AY261360.1_Kenya_1950            t.......c........t.......c.....------------------------------------------------- [ 55840]
AM712240.1_OURT_88/3             ........c.......ct-------------------------------------------------------------- [ 55840]
AM712239.1_Benin_97/1            ........c.......ct-------------------------------------------------------------- [ 55840]
KM111295.1_Ken06.Bus             -------------------------------------------------------------------------------- [ 55840]
KM111294.1_Ken05/Tk1             t................t......-------------------------------------------------------- [ 55840]
KM262844.1_strain_L60            ........c.......ct-------------------------------------------------------------- [ 55840]
KM102979.1_26544/OG10_Italy      ........c.......ct-------------------------------------------------------------- [ 55840]
KM262845.1_strain_NHV            ........c.......ct-------------------------------------------------------------- [ 55840]
KX354450.1_47/Ss/2008            ........c.......ct-------------------------------------------------------------- [ 55840]
KP055815.1_strain_BA71           ........c.......ct-------------------------------------------------------------- [ 55840]
MH025920.1_strain_R35            -------------------------------------------------------------------------------- [ 55840]
MH025919.1_strain_N10            -------------------------------------------------------------------------------- [ 55840]
MH025918.1_strain_R25            -------------------------------------------------------------------------------- [ 55840]
MH025917.1_strain_R7             -------------------------------------------------------------------------------- [ 55840]
MH025916.1_strain_R8             -------------------------------------------------------------------------------- [ 55840]

FR682468.2_Georgia_2007/1        -------------------------------------------------------------------------------- [ 55920]
ASF_IND_20_CAD543                -------------------------------------------------------------------------------- [ 55920]
LR899193.1_Germany_2020/1        -------------------------------------------------------------------------------- [ 55920]
MN393477.1_Wuhan_2019            -------------------------------------------------------------------------------- [ 55920]
MW396979.1_Timor-Leste/2019/1    -------------------------------------------------------------------------------- [ 55920]
MK940252.1_CN/2019/InnerMongolia -------------------------------------------------------------------------------- [ 55920]
MK128995.1_China/2018/AnhuiXCGQ  -------------------------------------------------------------------------------- [ 55920]
MH681419.1_ASFV/POL/2015         -------------------------------------------------------------------------------- [ 55920]
LS478113.1_Estonia_2014          -------------------------------------------------------------------------------- [ 55920]
KP843857.1_Odintsovo_02/14       -------------------------------------------------------------------------------- [ 55920]
FN557520.1_strain_E75            -------------------------------------------------------------------------------- [ 55920]
AY261366.1_Warthog               -------------------------------------------------------------------------------- [ 55920]
AY261365.1_Warmbaths             -------------------------------------------------------------------------------- [ 55920]
AY261364.1_Tengani_62            -------------------------------------------------------------------------------- [ 55920]
AY261363.1_Pretoriuskop/96/4     -------------------------------------------------------------------------------- [ 55920]
AY261362.1_Mkuzi_1979            -----------------------------------------------------------------------ttagttatg [ 55920]
AY261361.1_Malawi_Lil-20/1_1983  agttcaattaagacagtagttcagttaagtcaatagttcagttaagtcaatagttcagttaagacagtagttcagttaag [ 55920]
AY261360.1_Kenya_1950            ----------------------------------------------------------------------ttctattaag [ 55920]
AM712240.1_OURT_88/3             -------------------------------------------------------------------------------- [ 55920]
AM712239.1_Benin_97/1            -------------------------------------------------------------------------------- [ 55920]
KM111295.1_Ken06.Bus             -------------------------------------------------------------------------------- [ 55920]
```

```
KM111294.1_Ken05/Tk1             -------------------------------------------------------------------------------- [ 55920]
KM262844.1_strain_L60            -------------------------------------------------------------------------------- [ 55920]
KM102979.1_26544/OG10_Italy      -------------------------------------------------------------------------------- [ 55920]
KM262845.1_strain_NHV            -------------------------------------------------------------------------------- [ 55920]
KX354450.1_47/Ss/2008            -------------------------------------------------------------------------------- [ 55920]
KP055815.1_strain_BA71           -------------------------------------------------------------------------------- [ 55920]
MH025920.1_strain_R35            -------------------------------------------------------------------------------- [ 55920]
MH025919.1_strain_N10            -------------------------------------------------------------------------------- [ 55920]
MH025918.1_strain_R25            -------------------------------------------------------------------------------- [ 55920]
MH025917.1_strain_R7             -------------------------------------------------------------------------------- [ 55920]
MH025916.1_strain_R8             -------------------------------------------------------------------------------- [ 55920]

FR682468.2_Georgia_2007/1        ------------------------ttatgttaagatagtagttctgttaagataatagtttagttaaaac--tagttcat [ 56000]
ASF_IND_20_CAD543                ------------------------..c.................w.........r............g.----------- [ 56000]
LR899193.1_Germany_2020/1        ------------------------..............................................--........ [ 56000]
MN393477.1_Wuhan_2019            ------------------------..............................................--........ [ 56000]
MW396979.1_Timor-Leste/2019/1    ------------------------..............................................--........ [ 56000]
MK940252.1_CN/2019/InnerMongolia ------------------------..............................................--........ [ 56000]
MK128995.1_China/2018/AnhuiXCGQ  ------------------------..............................................--........ [ 56000]
MH681419.1_ASFV/POL/2015         ------------------------..............................................--........ [ 56000]
LS478113.1_Estonia_2014          ------------------------..............................................--........ [ 56000]
KP843857.1_Odintsovo_02/14       ------------------------..............................................--........ [ 56000]
FN557520.1_strain_E75            ------------------------..c...........................g...........tg.----------- [ 56000]
AY261366.1_Warthog               ------------------------..c.............a..........................g.----------- [ 56000]
AY261365.1_Warmbaths             --------------------------------------------------------------.....g.----------- [ 56000]
AY261364.1_Tengani_62            ------------------------..ta.......c.......ta...............c.........--........ [ 56000]
AY261363.1_Pretoriuskop/96/4     ------------------------..ca.......................................g.----------- [ 56000]
AY261362.1_Mkuzi_1979            atagtggtttagttaagacaatag..t........c..................g............gt----------- [ 56000]
AY261361.1_Malawi_Lil-20/1_1983  acagtagttcagttaagtcaatag..ca.......c........a.........g.....c......gt.aa.......- [ 56000]
AY261360.1_Kenya_1950            ataatagttctgttaagacagtag..t........c.......t........c.g......t.....g.----------- [ 56000]
AM712240.1_OURT_88/3             ------------------------..c...........................g...........tg.----------- [ 56000]
AM712239.1_Benin_97/1            ------------------------..c...........................g...........tg.----------- [ 56000]
KM111295.1_Ken06.Bus             ---------------------------------------------------------------....g.----------- [ 56000]
KM111294.1_Ken05/Tk1             ----------------------------------.........t..........g......ca....g.----------- [ 56000]
KM262844.1_strain_L60            ------------------------..c...........................g...........tg.----------- [ 56000]
KM102979.1_26544/OG10_Italy      ------------------------..c...........................g...........tg.----------- [ 56000]
KM262845.1_strain_NHV            ------------------------..c...........................g...........tg.----------- [ 56000]
KX354450.1_47/Ss/2008            ------------------------..c...........................g...........tg.----------- [ 56000]
KP055815.1_strain_BA71           ------------------------..c...........................g...........tg.----------- [ 56000]
MH025920.1_strain_R35            ---------------------------------------------------------------....g.----------- [ 56000]
MH025919.1_strain_N10            ---------------------------------------------------------------....g.----------- [ 56000]
MH025918.1_strain_R25            ---------------------------------------------------------------....g.----------- [ 56000]
MH025917.1_strain_R7             ---------------------------------------------------------------....g.----------- [ 56000]
MH025916.1_strain_R8             ---------------------------------------------------------------....g.----------- [ 56000]
```

```
FR682468.2_Georgia_2007/1        gttaagttaatagttttgttaagacaatagttcatttaagtcaatagttcagttaagtcaatagttttgttaagtcaa-- [ 56080]
ASF_IND_20_CAD543                ----------------------------------------------------------t.g......a......wy..-- [ 56080]
LR899193.1_Germany_2020/1        ..............................................................................-- [ 56080]
MN393477.1_Wuhan_2019            ..............................................................................-- [ 56080]
MW396979.1_Timor-Leste/2019/1    ..............................................................................-- [ 56080]
MK940252.1_CN/2019/InnerMongolia ..............................................................................-- [ 56080]
MK128995.1_China/2018/AnhuiXCGQ  ..............................................................................-- [ 56080]
MH681419.1_ASFV/POL/2015         ..............................................................................-- [ 56080]
LS478113.1_Estonia_2014          ..............................................................................-- [ 56080]
KP843857.1_Odintsovo_02/14       ..............................................................................-- [ 56080]
FN557520.1_strain_E75            ----------------------------------------------------------t.g.g....a......a...-- [ 56080]
AY261366.1_Warthog               ----------------------------------------------------------t.......ca.....aa.t.g- [ 56080]
AY261365.1_Warmbaths             ----------------------------------------------------------t........a.....aa.t.g- [ 56080]
AY261364.1_Tengani_62            ......a---------------------------------------------------t.t.............at.t-- [ 56080]
AY261363.1_Pretoriuskop/96/4     ----------------------------------------------------------t.......ca.....aa.t.g- [ 56080]
AY261362.1_Mkuzi_1979            ----------------------------------------------------------..g.....ca..........-- [ 56080]
AY261361.1_Malawi_Lil-20/1_1983  ......ac.g.....ca.........g.......a.....a..g......................ca......a..gta [ 56080]
AY261360.1_Kenya_1950            ----------------------------------------------------------t.g.....cca.....at.g-- [ 56080]
AM712240.1_OURT_88/3             ----------------------------------------------------------t.g.g....a......a...-- [ 56080]
AM712239.1_Benin_97/1            ----------------------------------------------------------t.g.g....a......a...-- [ 56080]
KM111295.1_Ken06.Bus             ----------------------------------------------------------t.......c.......a..g-- [ 56080]
KM111294.1_Ken05/Tk1             ----------------------------------------------------------t.g......ca.....at.g-- [ 56080]
KM262844.1_strain_L60            ----------------------------------------------------------t.g.g....a......a...-- [ 56080]
KM102979.1_26544/OG10_Italy      ----------------------------------------------------------t.g.g....a......a...-- [ 56080]
KM262845.1_strain_NHV            ----------------------------------------------------------t.g.g....a......a...-- [ 56080]
KX354450.1_47/Ss/2008            ----------------------------------------------------------t.g.g....a......a...-- [ 56080]
KP055815.1_strain_BA71           ----------------------------------------------------------t.g.g....a......a...-- [ 56080]
MH025920.1_strain_R35            ----------------------------------------------------------t.......c.......a..g-- [ 56080]
MH025919.1_strain_N10            ----------------------------------------------------------t.......c.......a..g-- [ 56080]
MH025918.1_strain_R25            ----------------------------------------------------------t.......c.......a..g-- [ 56080]
MH025917.1_strain_R7             ----------------------------------------------------------t.......c.......a..g-- [ 56080]
MH025916.1_strain_R8             ----------------------------------------------------------t.......c.......a..g-- [ 56080]

FR682468.2_Georgia_2007/1        ---------------------------------tagtttagttaagtcaatagtttagttaagtcaatagtttagttaag [ 56160]
ASF_IND_20_CAD543                ---------------------------------......w......wy.r.....ct...............y....... [ 56160]
LR899193.1_Germany_2020/1        ---------------------------------............................................... [ 56160]
MN393477.1_Wuhan_2019            ---------------------------------............................................... [ 56160]
MW396979.1_Timor-Leste/2019/1    ---------------------------------............................................... [ 56160]
MK940252.1_CN/2019/InnerMongolia ---------------------------------............................................... [ 56160]
MK128995.1_China/2018/AnhuiXCGQ  ---------------------------------............................................... [ 56160]
MH681419.1_ASFV/POL/2015         ---------------------------------............................................... [ 56160]
LS478113.1_Estonia_2014          ---------------------------------............................................... [ 56160]
KP843857.1_Odintsovo_02/14       ---------------------------------............................................... [ 56160]
```

```
FN557520.1_strain_E75             ---------------------------------......t.....a..g....ct..............c....... [ 56160]
AY261366.1_Warthog                ------------------ttcatgttaaaatat......t....a.........t.....a.......c....... [ 56160]
AY261365.1_Warmbaths              ------------------ttca----------------t......t.......t.....a.......c.t..... [ 56160]
AY261364.1_Tengani_62             ---------------------------------......t.....a........t.....a.......c....... [ 56160]
AY261363.1_Pretoriuskop/96/4      ------------------ttca----------------t.....at.t......t..............c....... [ 56160]
AY261362.1_Mkuzi_1979             ---------------------------------......t..............t..............t...... [ 56160]
AY261361.1_Malawi_Lil-20/1_1983  gttcagttaagacagtagttca-gttaagtcaa.....c.......a..g....c......at.g....c....... [ 56160]
AY261360.1_Kenya_1950             ---------------------------------.....cca....at.g....cca....at.g.....ca..... [ 56160]
AM712240.1_OURT_88/3              ---------------------------------......t.....a..g....ct..............c....... [ 56160]
AM712239.1_Benin_97/1             ---------------------------------......t.....a..g....ct..............c....... [ 56160]
KM111295.1_Ken06.Bus              ---------------------------------.....cta....at.......ca....at.g.....ca..... [ 56160]
KM111294.1_Ken05/Tk1              ---------------------------------......t.....a..g....cca....at.g.....ca..... [ 56160]
KM262844.1_strain_L60             ---------------------------------......t.....a..g....ct..............c....... [ 56160]
KM102979.1_26544/OG10_Italy       ---------------------------------......t.....a..g....ct..............c....... [ 56160]
KM262845.1_strain_NHV             ---------------------------------......t.....a..g....ct..............c....... [ 56160]
KX354450.1_47/Ss/2008             ---------------------------------......t.....a..g....ct..............c....... [ 56160]
KP055815.1_strain_BA71            ---------------------------------......t.....a..g....ct..............c....... [ 56160]
MH025920.1_strain_R35             ---------------------------------.....cta....at.......ca....at.g.....ca..... [ 56160]
MH025919.1_strain_N10             ---------------------------------.....cta....at.......ca....at.g.....ca..... [ 56160]
MH025918.1_strain_R25             ---------------------------------.....cta....at.......ca....at.g.....ca..... [ 56160]
MH025917.1_strain_R7              ---------------------------------.....cta....at.......ca....at.g.....ca..... [ 56160]
MH025916.1_strain_R8              ---------------------------------.....cta....at.......ca....at.g.....ca..... [ 56160]

FR682468.2_Georgia_2007/1         tcaatagttatattaagacattagttctgctaatacattagttt------------------------------------ [ 56240]
ASF_IND_20_CAD543                 .........w.r.....w..w...................y------------------------------------ [ 56240]
LR899193.1_Germany_2020/1         ............................................------------------------------------ [ 56240]
MN393477.1_Wuhan_2019             ............................................------------------------------------ [ 56240]
MW396979.1_Timor-Leste/2019/1     ............................................------------------------------------ [ 56240]
MK940252.1_CN/2019/InnerMongolia ............................................------------------------------------ [ 56240]
MK128995.1_China/2018/AnhuiXCGQ   ............................................------------------------------------ [ 56240]
MH681419.1_ASFV/POL/2015          ............................................------------------------------------ [ 56240]
LS478113.1_Estonia_2014           ............................................------------------------------------ [ 56240]
KP843857.1_Odintsovo_02/14        ............................................------------------------------------ [ 56240]
FN557520.1_strain_E75             .........t.g.....t..a...................c------------------------------------ [ 56240]
AY261366.1_Warthog                ..............................t...........gc------------------------------------ [ 56240]
AY261365.1_Warmbaths              ..............................t...........gc------------------------------------ [ 56240]
AY261364.1_Tengani_62             ..............................t...........gc------------------------------------ [ 56240]
AY261363.1_Pretoriuskop/96/4      .............................at...........gc------------------------------------ [ 56240]
AY261362.1_Mkuzi_1979             ..............................t............c------------------------------------ [ 56240]
AY261361.1_Malawi_Lil-20/1_1983  .........cag.......g.....a.t...g...g.....cagttaagacagtagttcagttaagtcaatagttcag [ 56240]
AY261360.1_Kenya_1950             at.......c.g.....g..g..a...a.t...g.t.a.....c------------------------------------ [ 56240]
AM712240.1_OURT_88/3              .........t.g.....t..a...................c------------------------------------ [ 56240]
AM712239.1_Benin_97/1             .........t.g.....t..a...................c------------------------------------ [ 56240]
KM111295.1_Ken06.Bus              at.......c.g.....c..a..a...a.t...a.t.a.....c------------------------------------ [ 56240]
```

```
KM111294.1_Ken05/Tk1             at.......c.g.....g..g..a...a.t...g.t.a.....c------------------------------------ [ 56240]
KM262844.1_strain_L60            .........t.g.....t..a......................c------------------------------------ [ 56240]
KM102979.1_26544/OG10_Italy      .........t.g.....t..a......................c------------------------------------ [ 56240]
KM262845.1_strain_NHV            .........t.g.....t..a......................c------------------------------------ [ 56240]
KX354450.1_47/Ss/2008            .........t.g.....t..a......................c------------------------------------ [ 56240]
KP055815.1_strain_BA71           .........t.g.....t..a......................c------------------------------------ [ 56240]
MH025920.1_strain_R35            at.......c.g.....c..a..a...a.t...a.t.a.....c------------------------------------ [ 56240]
MH025919.1_strain_N10            at.......c.g.....c..a..a...a.t...a.t.a.....c------------------------------------ [ 56240]
MH025918.1_strain_R25            at.......c.g.....c..a..a...a.t...a.t.a.....c------------------------------------ [ 56240]
MH025917.1_strain_R7             at.......c.g.....c..a..a...a.t...a.t.a.....c------------------------------------ [ 56240]
MH025916.1_strain_R8             at.......c.g.....c..a..a...a.t...a.t.a.....c------------------------------------ [ 56240]

FR682468.2_Georgia_2007/1        ------------tgttaagataataaaaatttatttttttttcatcagggtagagaaaatgttctccctacaggagctct [ 56320]
ASF_IND_20_CAD543                ------------...................................r................................ [ 56320]
LR899193.1_Germany_2020/1        ------------.................................................................... [ 56320]
MN393477.1_Wuhan_2019            ------------.................................................................... [ 56320]
MW396979.1_Timor-Leste/2019/1    ------------.................................................................... [ 56320]
MK940252.1_CN/2019/InnerMongolia ------------.................................................................... [ 56320]
MK128995.1_China/2018/AnhuiXCGQ  ------------.................................................................... [ 56320]
MH681419.1_ASFV/POL/2015         ------------.................................................................... [ 56320]
LS478113.1_Estonia_2014          ------------.................................................................... [ 56320]
KP843857.1_Odintsovo_02/14       ------------.............................-...................................... [ 56320]
FN557520.1_strain_E75            ------------...................................a.....................t.......... [ 56320]
AY261366.1_Warthog               ------------.....................-.............a................................ [ 56320]
AY261365.1_Warmbaths             ------------.....................-.............a................................ [ 56320]
AY261364.1_Tengani_62            ------------.....................-.............a.....................t.......... [ 56320]
AY261363.1_Pretoriuskop/96/4     ------------.....................-.............a................................ [ 56320]
AY261362.1_Mkuzi_1979            ------------...................................a.....................t.......... [ 56320]
AY261361.1_Malawi_Lil-20/1_1983  ttaagtcaataga....................-.............a..........-..........t..a....... [ 56320]
AY261360.1_Kenya_1950            ------------.................................a.aa....................t.......... [ 56320]
AM712240.1_OURT_88/3             ------------.....................-.............a.....................t.......... [ 56320]
AM712239.1_Benin_97/1            ------------...................................a.....................t.......... [ 56320]
KM111295.1_Ken06.Bus             ------------.................................a.aa....................t.......... [ 56320]
KM111294.1_Ken05/Tk1             ------------.................................a.aa....................t.......... [ 56320]
KM262844.1_strain_L60            ------------...................................a.....................t.......... [ 56320]
KM102979.1_26544/OG10_Italy      ------------...................................a.....................t.......... [ 56320]
KM262845.1_strain_NHV            ------------.....................-.............a.....................t.......... [ 56320]
KX354450.1_47/Ss/2008            ------------...................................a.....................t.......... [ 56320]
KP055815.1_strain_BA71           ------------...................................a.....................t.......... [ 56320]
MH025920.1_strain_R35            ------------.....................-...........a.aa....................t.......... [ 56320]
MH025919.1_strain_N10            ------------.....................-...........a.aa....................t.......... [ 56320]
MH025918.1_strain_R25            ------------.....................-...........a.aa....................t.......... [ 56320]
MH025917.1_strain_R7             ------------.....................-...........a.aa....................t.......... [ 56320]
MH025916.1_strain_R8             ------------.....................-...........a.aa....................t.......... [ 56320]
```

```
FR682468.2_Georgia_2007/1        gccggaagaacatttacattcttccttaccccttggctaagcatgtacttcaacaactagggctgtactggaagggacat [ 56400]
ASF_IND_20_CAD543                ................................................................................ [ 56400]
LR899193.1_Germany_2020/1        ................................................................................ [ 56400]
MN393477.1_Wuhan_2019            ................................................................................ [ 56400]
MW396979.1_Timor-Leste/2019/1    ................................................................................ [ 56400]
MK940252.1_CN/2019/InnerMongolia ................................................................................ [ 56400]
MK128995.1_China/2018/AnhuiXCGQ  ................................................................................ [ 56400]
MH681419.1_ASFV/POL/2015         ................................................................................ [ 56400]
LS478113.1_Estonia_2014          ................................................................................ [ 56400]
KP843857.1_Odintsovo_02/14       ................................................................................ [ 56400]
FN557520.1_strain_E75            ....................................g........................................... [ 56400]
AY261366.1_Warthog               ....................................g........................................... [ 56400]
AY261365.1_Warmbaths             ....................................g........................................... [ 56400]
AY261364.1_Tengani_62            ....................................g........................................... [ 56400]
AY261363.1_Pretoriuskop/96/4     ....................................g........................................... [ 56400]
AY261362.1_Mkuzi_1979            ................................................................................ [ 56400]
AY261361.1_Malawi_Lil-20/1_1983  ...............................tc...g.g......................................t.. [ 56400]
AY261360.1_Kenya_1950            .............................t.t....gc.................t........a......g..aa...c [ 56400]
AM712240.1_OURT_88/3             ....................................g........................................... [ 56400]
AM712239.1_Benin_97/1            ....................................g........................................... [ 56400]
KM111295.1_Ken06.Bus             .............................t.t....gc.................t........a......g..aa...c [ 56400]
KM111294.1_Ken05/Tk1             .............................t.t....gc.................t........a......g.aaa.... [ 56400]
KM262844.1_strain_L60            ....................................g........................................... [ 56400]
KM102979.1_26544/OG10_Italy      ....................................g........................................... [ 56400]
KM262845.1_strain_NHV            ....................................g........................................... [ 56400]
KX354450.1_47/Ss/2008            ....................................g........................................... [ 56400]
KP055815.1_strain_BA71           ....................................g........................................... [ 56400]
MH025920.1_strain_R35            .............................t.t....gc.................t........a......g..aa...c [ 56400]
MH025919.1_strain_N10            .............................t.t....gc.................t........a......g..aa...c [ 56400]
MH025918.1_strain_R25            .............................t.t....gc.................t........a......g..aa...c [ 56400]
MH025917.1_strain_R7             .............................t.t....gc.................t........a......g..aa...c [ 56400]
MH025916.1_strain_R8             .............................t.t....gc.................t........a......g..aa...c [ 56400]

FR682468.2_Georgia_2007/1        ggatctcttcaacgaatcggagatgaccatgtactcttacagcaggacctgatcttttccatcaacgaggccttaagaat [ 56480]
ASF_IND_20_CAD543                ................................................................................ [ 56480]
LR899193.1_Germany_2020/1        ................................................................................ [ 56480]
MN393477.1_Wuhan_2019            ................................................................................ [ 56480]
MW396979.1_Timor-Leste/2019/1    ................................................................................ [ 56480]
MK940252.1_CN/2019/InnerMongolia ................................................................................ [ 56480]
MK128995.1_China/2018/AnhuiXCGQ  ................................................................................ [ 56480]
MH681419.1_ASFV/POL/2015         ................................................................................ [ 56480]
LS478113.1_Estonia_2014          ................................................................................ [ 56480]
KP843857.1_Odintsovo_02/14       ................................................................................ [ 56480]
```

```
FN557520.1_strain_E75            .........................................a...................................... [ 56480]
AY261366.1_Warthog               ................................................................................ [ 56480]
AY261365.1_Warmbaths             ............................................a................................... [ 56480]
AY261364.1_Tengani_62            ................................................................................ [ 56480]
AY261363.1_Pretoriuskop/96/4     ................................................................................ [ 56480]
AY261362.1_Mkuzi_1979            ................................................................................ [ 56480]
AY261361.1_Malawi_Lil-20/1_1983  .........................................a........c..............t.............. [ 56480]
AY261360.1_Kenya_1950            ..t.......................tt.............a..a..t..c.........g....t.............. [ 56480]
AM712240.1_OURT_88/3             .........................................a.......................t.............. [ 56480]
AM712239.1_Benin_97/1            .........................................a...................................... [ 56480]
KM111295.1_Ken06.Bus             ..c..............a........tt.............a..a..t..c.........g....t...........g.. [ 56480]
KM111294.1_Ken05/Tk1             ..t.......................tt.............a..a..t..c.........g....t...........g.. [ 56480]
KM262844.1_strain_L60            .........................................a...................................... [ 56480]
KM102979.1_26544/OG10_Italy      .........................................a...................................... [ 56480]
KM262845.1_strain_NHV            .........................................a.......................t.............. [ 56480]
KX354450.1_47/Ss/2008            .........................................a...................................... [ 56480]
KP055815.1_strain_BA71           .........................................a...................................... [ 56480]
MH025920.1_strain_R35            ..c..............a........tt.............a..a..t..c.........g....t...........g.. [ 56480]
MH025919.1_strain_N10            ..c..............a........tt.............a..a..t..c.........g....t...........g.. [ 56480]
MH025918.1_strain_R25            ..c..............a........tt.............a..a..t..c.........g....t...........g.. [ 56480]
MH025917.1_strain_R7             ..c..............a........tt.............a..a..t..c.........g....t...........g.. [ 56480]
MH025916.1_strain_R8             ..c..............a........tt.............a..a..t..c.........g....t...........g.. [ 56480]

FR682468.2_Georgia_2007/1        ggcaggagaggaaggaaacaatgaagtagtaaagctcttgttactatgggagggaaaccttcattatgccatcataggag [ 56560]
ASF_IND_20_CAD543                ...g.c...r..r......................................r.....y...................... [ 56560]
LR899193.1_Germany_2020/1        ................................................................................ [ 56560]
MN393477.1_Wuhan_2019            ................................................................................ [ 56560]
MW396979.1_Timor-Leste/2019/1    ................................................................................ [ 56560]
MK940252.1_CN/2019/InnerMongolia ................................................................................ [ 56560]
MK128995.1_China/2018/AnhuiXCGQ  ................................................................................ [ 56560]
MH681419.1_ASFV/POL/2015         ................................................................................ [ 56560]
LS478113.1_Estonia_2014          ................................................................................ [ 56560]
KP843857.1_Odintsovo_02/14       ................................................................................ [ 56560]
FN557520.1_strain_E75            ...g.c...a...................................g.................................. [ 56560]
AY261366.1_Warthog               ................................................................................ [ 56560]
AY261365.1_Warmbaths             ................................................................................ [ 56560]
AY261364.1_Tengani_62            ................................................................................ [ 56560]
AY261363.1_Pretoriuskop/96/4     ................................................................................ [ 56560]
AY261362.1_Mkuzi_1979            .............................................g.................................. [ 56560]
AY261361.1_Malawi_Lil-20/1_1983  ...g.............................a...........g.................................. [ 56560]
AY261360.1_Kenya_1950            ...g.c.......................................g.......................t.......... [ 56560]
AM712240.1_OURT_88/3             ...g.c...a...................................g.................................. [ 56560]
AM712239.1_Benin_97/1            ...g.c...a...................................g..................c............... [ 56560]
KM111295.1_Ken06.Bus             ...g.c.......................................g.................................. [ 56560]
```

```
KM111294.1_Ken05/Tk1           ...g.c.......................................g.................................. [ 56560]
KM262844.1_strain_L60          ...g.c...a...................................g.................................. [ 56560]
KM102979.1_26544/OG10_Italy    ...g.c...a...................................g.................................. [ 56560]
KM262845.1_strain_NHV          ...g.c...a...................................g.................................. [ 56560]
KX354450.1_47/Ss/2008          ...g.c...a...................................g.................................. [ 56560]
KP055815.1_strain_BA71         ...g.c...a...................................g.................................. [ 56560]
MH025920.1_strain_R35          ...g.c.......................................g.................................. [ 56560]
MH025919.1_strain_N10          ...g.c.......................................g.................................. [ 56560]
MH025918.1_strain_R25          ...g.c.......................................g.................................. [ 56560]
MH025917.1_strain_R7           ...g.c.......................................g.................................. [ 56560]
MH025916.1_strain_R8           ...g.c.......................................g.................................. [ 56560]

FR682468.2_Georgia_2007/1      ctttagagggcgaccgatatgaccttatccataaatattatgatcaaattggggactgccacaagattcttcctttaatc [ 56640]
ASF_IND_20_CAD543              ................................................................................ [ 56640]
LR899193.1_Germany_2020/1      ................................................................................ [ 56640]
MN393477.1_Wuhan_2019          ................................................................................ [ 56640]
MW396979.1_Timor-Leste/2019/1  ................................................................................ [ 56640]
MK940252.1_CN/2019/InnerMongolia ................................................................................ [ 56640]
MK128995.1_China/2018/AnhuiXCGQ ................................................................................ [ 56640]
MH681419.1_ASFV/POL/2015       ................................................................................ [ 56640]
LS478113.1_Estonia_2014        ................................................................................ [ 56640]
KP843857.1_Odintsovo_02/14     ................................................................................ [ 56640]
FN557520.1_strain_E75          ...................c.......................a.................................... [ 56640]
AY261366.1_Warthog             ...................c............................................................ [ 56640]
AY261365.1_Warmbaths           ...................c............................................................ [ 56640]
AY261364.1_Tengani_62          ...................c............................................................ [ 56640]
AY261363.1_Pretoriuskop/96/4   ...................c............................................................ [ 56640]
AY261362.1_Mkuzi_1979          ...................c............................................................ [ 56640]
AY261361.1_Malawi_Lil-20/1_1983 ...................c.....c................gc.....c..a...........a........c.....t [ 56640]
AY261360.1_Kenya_1950          ....g......a.......c.....c.................c.............................c...... [ 56640]
AM712240.1_OURT_88/3           ...................c.......................a.................................... [ 56640]
AM712239.1_Benin_97/1          ...................c.......................a.................................... [ 56640]
KM111295.1_Ken06.Bus           ....g......a.......c.....c.................c.....c.......................c...... [ 56640]
KM111294.1_Ken05/Tk1           ....g......a.......c.....c.................c.............................c...... [ 56640]
KM262844.1_strain_L60          ...................c.......................a.................................... [ 56640]
KM102979.1_26544/OG10_Italy    ...................c.......................a.................................... [ 56640]
KM262845.1_strain_NHV          ...................c.......................a.................................... [ 56640]
KX354450.1_47/Ss/2008          ...................c.......................a.................................... [ 56640]
KP055815.1_strain_BA71         ...................c.......................a.................................... [ 56640]
MH025920.1_strain_R35          ....g......a.......c.....c.................c.....c.......................c...... [ 56640]
MH025919.1_strain_N10          ....g......a.......c.....c.................c.....c.......................c...... [ 56640]
MH025918.1_strain_R25          ....g......a.......c.....c.................c.....c.......................c...... [ 56640]
MH025917.1_strain_R7           ....g......a.......c.....c.................c.....c.......................c...... [ 56640]
MH025916.1_strain_R8           ....g......a.......c.....c.................c.....c.......................c...... [ 56640]
```

```
FR682468.2_Georgia_2007/1         caagacccgcaaatctttgaaaaatgccatgaattgagtaactcctgtaatattcgatgccttttagaacatgcagtaaa [ 56720]
ASF_IND_20_CAD543                 ................................................nnn............................. [ 56720]
LR899193.1_Germany_2020/1         ................................................................................ [ 56720]
MN393477.1_Wuhan_2019             ................................................................................ [ 56720]
MW396979.1_Timor-Leste/2019/1     ................................................................................ [ 56720]
MK940252.1_CN/2019/InnerMongolia  ................................................................................ [ 56720]
MK128995.1_China/2018/AnhuiXCGQ   ................................................................................ [ 56720]
MH681419.1_ASFV/POL/2015          ................................................................................ [ 56720]
LS478113.1_Estonia_2014           ................................................................................ [ 56720]
KP843857.1_Odintsovo_02/14        ................................................................................ [ 56720]
FN557520.1_strain_E75             ..................................................c............................. [ 56720]
AY261366.1_Warthog                ..................................................c............................. [ 56720]
AY261365.1_Warmbaths              ..................................................c............................. [ 56720]
AY261364.1_Tengani_62             ........a............................a............c............................. [ 56720]
AY261363.1_Pretoriuskop/96/4      ..................................................c............................. [ 56720]
AY261362.1_Mkuzi_1979             ..................................................c............................. [ 56720]
AY261361.1_Malawi_Lil-20/1_1983   .....t..................................c.........c..........................g.. [ 56720]
AY261360.1_Kenya_1950             .....t..................................ct........c.......................g..g.. [ 56720]
AM712240.1_OURT_88/3              ..................................................c............................. [ 56720]
AM712239.1_Benin_97/1             ..................................................c............................. [ 56720]
KM111295.1_Ken06.Bus              .....t..................................ct........c..........................g.. [ 56720]
KM111294.1_Ken05/Tk1              .....t..................................ct........c.......................g..g.. [ 56720]
KM262844.1_strain_L60             ..................................................c............................. [ 56720]
KM102979.1_26544/OG10_Italy       ..................................................c............................. [ 56720]
KM262845.1_strain_NHV             ..................................................c............................. [ 56720]
KX354450.1_47/Ss/2008             ..................................................c............................. [ 56720]
KP055815.1_strain_BA71            ..................................................c............................. [ 56720]
MH025920.1_strain_R35             .....t..................................ct........c..........................g.. [ 56720]
MH025919.1_strain_N10             .....t..................................ct........c..........................g.. [ 56720]
MH025918.1_strain_R25             .....t..................................ct........c..........................g.. [ 56720]
MH025917.1_strain_R7              .....t..................................ct........c..........................g.. [ 56720]
MH025916.1_strain_R8              .....t..................................ct........c..........................g.. [ 56720]

FR682468.2_Georgia_2007/1         acacgacatgctttctattcttcaaaaacacaaggagcaaataagattacacatggcattaacccaaatactatttgaat [ 56800]
ASF_IND_20_CAD543                 ....r.........................y.....k........................................... [ 56800]
LR899193.1_Germany_2020/1         ................................................................................ [ 56800]
MN393477.1_Wuhan_2019             ................................................................................ [ 56800]
MW396979.1_Timor-Leste/2019/1     ................................................................................ [ 56800]
MK940252.1_CN/2019/InnerMongolia  ................................................................................ [ 56800]
MK128995.1_China/2018/AnhuiXCGQ   ................................................................................ [ 56800]
MH681419.1_ASFV/POL/2015          ................................................................................ [ 56800]
LS478113.1_Estonia_2014           ................................................................................ [ 56800]
KP843857.1_Odintsovo_02/14        ................................................................................ [ 56800]
```

```
FN557520.1_strain_E75              ....a.....................t.....t........................................... [ 56800]
AY261366.1_Warthog                 ................................g...................t........................... [ 56800]
AY261365.1_Warmbaths               ................................................................................ [ 56800]
AY261364.1_Tengani_62              ....a.t..........................a..................t........................... [ 56800]
AY261363.1_Pretoriuskop/96/4       ....................................................t........................... [ 56800]
AY261362.1_Mkuzi_1979              ................................................................................ [ 56800]
AY261361.1_Malawi_Lil-20/1_1983    ....a.....................t.....c........g..t...t.........................g. [ 56800]
AY261360.1_Kenya_1950              ....a.t...................t.....c........g......t.........t....g..........g. [ 56800]
AM712240.1_OURT_88/3               ....a.....................t.....t........................................... [ 56800]
AM712239.1_Benin_97/1              ....a.....................t.....t........................................... [ 56800]
KM111295.1_Ken06.Bus               ....a.t...................t.....t..t.....g......t.........t....g..........g. [ 56800]
KM111294.1_Ken05/Tk1               ....a.t...................t.....c........g......t.........t....g..........g. [ 56800]
KM262844.1_strain_L60              ....a.....................t.....t........................................... [ 56800]
KM102979.1_26544/OG10_Italy        ....a.....................t.....t........................................... [ 56800]
KM262845.1_strain_NHV              ....a.....................t.....t........................................... [ 56800]
KX354450.1_47/Ss/2008              ....a.....................t.....t........................................... [ 56800]
KP055815.1_strain_BA71             ....a.....................t.....t........................................... [ 56800]
MH025920.1_strain_R35              ....a.t...................t.....t..t.....g......t.........t....g..........g. [ 56800]
MH025919.1_strain_N10              ....a.t...................t.....t..t.....g......t.........t....g..........g. [ 56800]
MH025918.1_strain_R25              ....a.t...................t.....t..t.....g......t.........t....g..........g. [ 56800]
MH025917.1_strain_R7               ....a.t...................t.....t..t.....g......t.........t....g..........g. [ 56800]
MH025916.1_strain_R8               ....a.t...................t.....t..t.....g......t.........t....g..........g. [ 56800]

FR682468.2_Georgia_2007/1          tggcgtgtcatgaacgtaaaaatgacatcattagatggatcggttattccctgcacatacaccatctagagactattttt [ 56880]
ASF_IND_20_CAD543                  ...........................................................y.................... [ 56880]
LR899193.1_Germany_2020/1          ................................................................................ [ 56880]
MN393477.1_Wuhan_2019              ................................................................................ [ 56880]
MW396979.1_Timor-Leste/2019/1      ................................................................................ [ 56880]
MK940252.1_CN/2019/InnerMongolia   ................................................................................ [ 56880]
MK128995.1_China/2018/AnhuiXCGQ    ................................................................................ [ 56880]
MH681419.1_ASFV/POL/2015           ................................................................................ [ 56880]
LS478113.1_Estonia_2014            ................................................................................ [ 56880]
KP843857.1_Odintsovo_02/14         ................................................................................ [ 56880]
FN557520.1_strain_E75              .......c...........g.......................................t.................... [ 56880]
AY261366.1_Warthog                 ...................g............................................................ [ 56880]
AY261365.1_Warmbaths               ...................g............................................................ [ 56880]
AY261364.1_Tengani_62              ...................g............................................................ [ 56880]
AY261363.1_Pretoriuskop/96/4       ...................g............................................................ [ 56880]
AY261362.1_Mkuzi_1979              ...................g............................................................ [ 56880]
AY261361.1_Malawi_Lil-20/1_1983    ...................g.......................................t....a..g............ [ 56880]
AY261360.1_Kenya_1950              .........g.....a.............g..c.g........c.........t.......t.................. [ 56880]
AM712240.1_OURT_88/3               .......c...........g.......................................t.................... [ 56880]
AM712239.1_Benin_97/1              .......c...........g.......................................t.................... [ 56880]
KM111295.1_Ken06.Bus               ...............a.............g..c..........c........a........t.................. [ 56880]
```

```
KM111294.1_Ken05/Tk1             .........g.....a.............g..c.g........c.........t.......t.................. [ 56880]
KM262844.1_strain_L60            .......c...........g.......................................t.................... [ 56880]
KM102979.1_26544/OG10_Italy      .......c...........g.......................................t.................... [ 56880]
KM262845.1_strain_NHV            .......c...........g.......................................t.................... [ 56880]
KX354450.1_47/Ss/2008            .......c...........g.......................................t.................... [ 56880]
KP055815.1_strain_BA71           .......c...........g.......................................t.................... [ 56880]
MH025920.1_strain_R35            ...............a.............g..c..........c........a........t.................. [ 56880]
MH025919.1_strain_N10            ...............a.............g..c..........c........a........t.................. [ 56880]
MH025918.1_strain_R25            ...............a.............g..c..........c........a........t.................. [ 56880]
MH025917.1_strain_R7             ...............a.............g..c..........c........a........t.................. [ 56880]
MH025916.1_strain_R8             ...............a.............g..c..........c........a........t.................. [ 56880]

FR682468.2_Georgia_2007/1        gatgttgcattcgcccataaaaatttatccttatacgttttagggtatgaacttctcatgcacaaagtaaatacagaggc [ 56960]
ASF_IND_20_CAD543                ................................................................................ [ 56960]
LR899193.1_Germany_2020/1        ................................................................................ [ 56960]
MN393477.1_Wuhan_2019            ................................................................................ [ 56960]
MW396979.1_Timor-Leste/2019/1    ................................................................................ [ 56960]
MK940252.1_CN/2019/InnerMongolia ................................................................................ [ 56960]
MK128995.1_China/2018/AnhuiXCGQ  ................................................................................ [ 56960]
MH681419.1_ASFV/POL/2015         ................................................................................ [ 56960]
LS478113.1_Estonia_2014          ................................................................................ [ 56960]
KP843857.1_Odintsovo_02/14       ................................................................................ [ 56960]
FN557520.1_strain_E75            ................................................................................ [ 56960]
AY261366.1_Warthog               .......................................................................c........ [ 56960]
AY261365.1_Warmbaths             ................................................................................ [ 56960]
AY261364.1_Tengani_62            ................................................................................ [ 56960]
AY261363.1_Pretoriuskop/96/4     ................................................................................ [ 56960]
AY261362.1_Mkuzi_1979            ................................................................................ [ 56960]
AY261361.1_Malawi_Lil-20/1_1983  ....................g..............t....................t....................... [ 56960]
AY261360.1_Kenya_1950            ........cc.gct.....................t....................t...........g........... [ 56960]
AM712240.1_OURT_88/3             ................................................................................ [ 56960]
AM712239.1_Benin_97/1            ................................................................................ [ 56960]
KM111295.1_Ken06.Bus             ...acc..cc.gct.....................t....................t........g..g........... [ 56960]
KM111294.1_Ken05/Tk1             ........cc.gct.....................t....................t...........g........... [ 56960]
KM262844.1_strain_L60            ................................................................................ [ 56960]
KM102979.1_26544/OG10_Italy      ................................................................................ [ 56960]
KM262845.1_strain_NHV            ................................................................................ [ 56960]
KX354450.1_47/Ss/2008            ................................................................................ [ 56960]
KP055815.1_strain_BA71           ................................................................................ [ 56960]
MH025920.1_strain_R35            ...acc..cc.gct.....................t....................t........g..g........... [ 56960]
MH025919.1_strain_N10            ...acc..cc.gct.....................t....................t........g..g........... [ 56960]
MH025918.1_strain_R25            ...acc..cc.gct.....................t....................t........g..g........... [ 56960]
MH025917.1_strain_R7             ...acc..cc.gct.....................t....................t........g..g........... [ 56960]
MH025916.1_strain_R8             ...acc..cc.gct.....................t....................t........g..g........... [ 56960]
```

```
FR682468.2_Georgia_2007/1        tgcatatatagaattacccaatttgctatcatatcaccttcgaactgcggcggcaggaggtcttcttaactttatgttag [ 57040]
ASF_IND_20_CAD543                ................................................................................ [ 57040]
LR899193.1_Germany_2020/1        ................................................................................ [ 57040]
MN393477.1_Wuhan_2019            ................................................................................ [ 57040]
MW396979.1_Timor-Leste/2019/1    ................................................................................ [ 57040]
MK940252.1_CN/2019/InnerMongolia ................................................................................ [ 57040]
MK128995.1_China/2018/AnhuiXCGQ  ................................................................................ [ 57040]
MH681419.1_ASFV/POL/2015         ................................................................................ [ 57040]
LS478113.1_Estonia_2014          ................................................................................ [ 57040]
KP843857.1_Odintsovo_02/14       ................................................................................ [ 57040]
FN557520.1_strain_E75            ....a.......t........................................................t.......... [ 57040]
AY261366.1_Warthog               ....................................t........................................... [ 57040]
AY261365.1_Warmbaths             ................................................................................ [ 57040]
AY261364.1_Tengani_62            .....................................................................t.......... [ 57040]
AY261363.1_Pretoriuskop/96/4     ................................................................................ [ 57040]
AY261362.1_Mkuzi_1979            ................................................................................ [ 57040]
AY261361.1_Malawi_Lil-20/1_1983  ....a.......t....................c.................c...........c...c.t........t. [ 57040]
AY261360.1_Kenya_1950            ....a.......tg..t..g........a..c.g..........gc.....a...........c...c.t.......... [ 57040]
AM712240.1_OURT_88/3             ....a.......t........................................................t.......... [ 57040]
AM712239.1_Benin_97/1            ....a.......t........................................................t.......... [ 57040]
KM111295.1_Ken06.Bus             ....a.......tg.....g........a..c.g..........gc.....a...........c...c.t.......... [ 57040]
KM111294.1_Ken05/Tk1             ....a.......tg..t..g........a..c.g..........gc.....a...........c...c.t.......... [ 57040]
KM262844.1_strain_L60            ....a.......t........................................................t.......... [ 57040]
KM102979.1_26544/OG10_Italy      ....a.......t........................................................t.......... [ 57040]
KM262845.1_strain_NHV            ....a.......t........................................................t.......... [ 57040]
KX354450.1_47/Ss/2008            ....a.......t........................................................t.......... [ 57040]
KP055815.1_strain_BA71           ....a.......t........................................................t.......... [ 57040]
MH025920.1_strain_R35            ....a.......tg.....g........a..c.g..........gc.....a...........c...c.t.......... [ 57040]
MH025919.1_strain_N10            ....a.......tg.....g........a..c.g..........gc.....a...........c...c.t.......... [ 57040]
MH025918.1_strain_R25            ....a.......tg.....g........a..c.g..........gc.....a...........c...c.t.......... [ 57040]
MH025917.1_strain_R7             ....a.......tg.....g........a..c.g..........gc.....a...........c...c.t.......... [ 57040]
MH025916.1_strain_R8             ....a.......tg.....g........a..c.g..........gc.....a...........c...c.t.......... [ 57040]

FR682468.2_Georgia_2007/1        aaacaataaagcatggtggatatctggataaaacggttttatccgcggctatcaggtacaagcataggaaaattgtggct [ 57120]
ASF_IND_20_CAD543                ...................r.r.s........................................................ [ 57120]
LR899193.1_Germany_2020/1        ................................................................................ [ 57120]
MN393477.1_Wuhan_2019            ................................................................................ [ 57120]
MW396979.1_Timor-Leste/2019/1    ................................................................................ [ 57120]
MK940252.1_CN/2019/InnerMongolia ................................................................................ [ 57120]
MK128995.1_China/2018/AnhuiXCGQ  ................................................................................ [ 57120]
MH681419.1_ASFV/POL/2015         ................................................................................ [ 57120]
LS478113.1_Estonia_2014          ................................................................................ [ 57120]
KP843857.1_Odintsovo_02/14       ................................................................................ [ 57120]
```

```
FN557520.1_strain_E75            ...................g.g.g..........c...c......................................... [ 57120]
AY261366.1_Warthog               .......................g..........c............................................. [ 57120]
AY261365.1_Warmbaths             ................................................................................ [ 57120]
AY261364.1_Tengani_62            ......................................c......................................... [ 57120]
AY261363.1_Pretoriuskop/96/4     .......................g..........c............................................. [ 57120]
AY261362.1_Mkuzi_1979            ................................................................................ [ 57120]
AY261361.1_Malawi_Lil-20/1_1983  .g...t.................g........g.c....................t.....a.........g........ [ 57120]
AY261360.1_Kenya_1950            .g...t.....t.......g.g.g..................t...a..g.....t.....a.................. [ 57120]
AM712240.1_OURT_88/3             ...................g.g.g..........c...c......................................... [ 57120]
AM712239.1_Benin_97/1            ...................g.g.g..........c...c......................................... [ 57120]
KM111295.1_Ken06.Bus             .g...t.....t.......g.g.g..................t...a..g.....t.....a.................. [ 57120]
KM111294.1_Ken05/Tk1             .g...t.....t.......g.g.g..................t...a..g.....t.....a.................. [ 57120]
KM262844.1_strain_L60            ...................g.g.g..........c...c......................................... [ 57120]
KM102979.1_26544/OG10_Italy      ...................g.g.g..........c...c......................................... [ 57120]
KM262845.1_strain_NHV            ...................g.g.g..........c...c......................................... [ 57120]
KX354450.1_47/Ss/2008            ...................g.g.g..........c...c......................................... [ 57120]
KP055815.1_strain_BA71           ...................g.g.g..........c...c......................................... [ 57120]
MH025920.1_strain_R35            .g...t.....t.......g.g.g..................t...a..g.....t.....a.................. [ 57120]
MH025919.1_strain_N10            .g...t.....t.......g.g.g..................t...a..g.....t.....a.................. [ 57120]
MH025918.1_strain_R25            .g...t.....t.......g.g.g..................t...a..g.....t.....a.................. [ 57120]
MH025917.1_strain_R7             .g...t.....t.......g.g.g..................t...a..g.....t.....a.................. [ 57120]
MH025916.1_strain_R8             .g...t.....t.......g.g.g..................t...a..g.....t.....a.................. [ 57120]

FR682468.2_Georgia_2007/1        cattttattcatcaggttccccgtaaaaccgttaaaaaactgttactctatgctgtgcaggctcgggcccccaaaaaaac [ 57200]
ASF_IND_20_CAD543                ................................................................................ [ 57200]
LR899193.1_Germany_2020/1        ................................................................................ [ 57200]
MN393477.1_Wuhan_2019            ................................................................................ [ 57200]
MW396979.1_Timor-Leste/2019/1    ................................................................................ [ 57200]
MK940252.1_CN/2019/InnerMongolia ................................................................................ [ 57200]
MK128995.1_China/2018/AnhuiXCGQ  ................................................................................ [ 57200]
MH681419.1_ASFV/POL/2015         ................................................................................ [ 57200]
LS478113.1_Estonia_2014          ................................................................................ [ 57200]
KP843857.1_Odintsovo_02/14       ................................................................................ [ 57200]
FN557520.1_strain_E75            ..........................................c.............................-....... [ 57200]
AY261366.1_Warthog               ..............a...........................c..................................... [ 57200]
AY261365.1_Warmbaths             ..........................................c..................t.................. [ 57200]
AY261364.1_Tengani_62            ..........................................c..................................... [ 57200]
AY261363.1_Pretoriuskop/96/4     ..........................................c..................................... [ 57200]
AY261362.1_Mkuzi_1979            ................................................................................ [ 57200]
AY261361.1_Malawi_Lil-20/1_1983  ..............a..............g...g........c.....c....c......a................... [ 57200]
AY261360.1_Kenya_1950            ..............a..............g...g..g....ac.....c....c......a................... [ 57200]
AM712240.1_OURT_88/3             ..........................................c..................................... [ 57200]
AM712239.1_Benin_97/1            ..........................................c..................................... [ 57200]
KM111295.1_Ken06.Bus             .........t....a..............a...g..g....ac.....c....c......a................... [ 57200]
```

```
KM111294.1_Ken05/Tk1             .............ga..............g...g..g....a......c....c......a.c................. [ 57200]
KM262844.1_strain_L60            ..........................................c..................................... [ 57200]
KM102979.1_26544/OG10_Italy      ..........................................c..................................... [ 57200]
KM262845.1_strain_NHV            ..........................................c..................................... [ 57200]
KX354450.1_47/Ss/2008            ..........................................c..................................... [ 57200]
KP055815.1_strain_BA71           ..........................................c..................................... [ 57200]
MH025920.1_strain_R35            ..............a..............a...g..g....ac.....c....c......a................... [ 57200]
MH025919.1_strain_N10            ..............a..............a...g..g....ac.....c....c......a................... [ 57200]
MH025918.1_strain_R25            ..............a..............a...g..g....ac.....c....c......a................... [ 57200]
MH025917.1_strain_R7             ..............a..............a...g..g....ac.....c....c......a................... [ 57200]
MH025916.1_strain_R8             ..............a..............a...g..g....ac.....c....c......a................... [ 57200]

FR682468.2_Georgia_2007/1        actgaacctacttttatcttccttaaactactccgtgcacaccatcaccaaacaactcgtacacaatgtcgtcatctaca [ 57280]
ASF_IND_20_CAD543                ...r..y.......................tg................................................ [ 57280]
LR899193.1_Germany_2020/1        ................................................................................ [ 57280]
MN393477.1_Wuhan_2019            ................................................................................ [ 57280]
MW396979.1_Timor-Leste/2019/1    ................................................................................ [ 57280]
MK940252.1_CN/2019/InnerMongolia ................................................................................ [ 57280]
MK128995.1_China/2018/AnhuiXCGQ  ................................................................................ [ 57280]
MH681419.1_ASFV/POL/2015         ................................................................................ [ 57280]
LS478113.1_Estonia_2014          ................................................................................ [ 57280]
KP843857.1_Odintsovo_02/14       ................................................................................ [ 57280]
FN557520.1_strain_E75            ...a..t.......................tg......................................a...a..... [ 57280]
AY261366.1_Warthog               ................................................................................ [ 57280]
AY261365.1_Warmbaths             ................................................................................ [ 57280]
AY261364.1_Tengani_62            ........................................t....................................... [ 57280]
AY261363.1_Pretoriuskop/96/4     ................................................................................ [ 57280]
AY261362.1_Mkuzi_1979            ................................................................................ [ 57280]
AY261361.1_Malawi_Lil-20/1_1983  ........................................c.....................g..........gat..t. [ 57280]
AY261360.1_Kenya_1950            t..............................................t..............g..g...t.c........ [ 57280]
AM712240.1_OURT_88/3             ...a..t.......................tg......................................a...a..... [ 57280]
AM712239.1_Benin_97/1            ...a..t.......................tg......................................a...a..... [ 57280]
KM111295.1_Ken06.Bus             t..............................................t..............g..g...t.c........ [ 57280]
KM111294.1_Ken05/Tk1             t..............................................t..............g..g...t.c........ [ 57280]
KM262844.1_strain_L60            ...a..t.......................tg......................................a...a..... [ 57280]
KM102979.1_26544/OG10_Italy      ...a..t.......................tg......................................a...a..... [ 57280]
KM262845.1_strain_NHV            ...a..t.......................tg......................................a...a..... [ 57280]
KX354450.1_47/Ss/2008            ...a..t.......................tg......................................a...a..... [ 57280]
KP055815.1_strain_BA71           ...a..t.......................tg......................................a...a..... [ 57280]
MH025920.1_strain_R35            t..............................................t..............g..g...t.c........ [ 57280]
MH025919.1_strain_N10            t..............................................t..............g..g...t.c........ [ 57280]
MH025918.1_strain_R25            t..............................................t..............g..g...t.c........ [ 57280]
MH025917.1_strain_R7             t..............................................t..............g..g...t.c........ [ 57280]
MH025916.1_strain_R8             t..............................................t..............g..g...t.c........ [ 57280]
```

```
FR682468.2_Georgia_2007/1        gttccacgcttatcgtaaagcttttactcatgcggcgaaaaaacaagttaaacctagtagatgccgttttagccagactt [ 57360]
ASF_IND_20_CAD543                ................................................................................ [ 57360]
LR899193.1_Germany_2020/1        ................................................................................ [ 57360]
MN393477.1_Wuhan_2019            ................................................................................ [ 57360]
MW396979.1_Timor-Leste/2019/1    ................................................................................ [ 57360]
MK940252.1_CN/2019/InnerMongolia ................................................................................ [ 57360]
MK128995.1_China/2018/AnhuiXCGQ  ................................................................................ [ 57360]
MH681419.1_ASFV/POL/2015         ................................................................................ [ 57360]
LS478113.1_Estonia_2014          ................................................................................ [ 57360]
KP843857.1_Odintsovo_02/14       ................................................................................ [ 57360]
FN557520.1_strain_E75            ...........g..............................gg.................................... [ 57360]
AY261366.1_Warthog               ...........g.................................................................... [ 57360]
AY261365.1_Warmbaths             ...........g.....................................................a.............. [ 57360]
AY261364.1_Tengani_62            ...........g...........................................g........................ [ 57360]
AY261363.1_Pretoriuskop/96/4     ................................................................................ [ 57360]
AY261362.1_Mkuzi_1979            ................................................................................ [ 57360]
AY261361.1_Malawi_Lil-20/1_1983  .g........................................ga....................t............... [ 57360]
AY261360.1_Kenya_1950            .g.........g.............g.................g............a.......t............... [ 57360]
AM712240.1_OURT_88/3             ...........g..............................gg.................................... [ 57360]
AM712239.1_Benin_97/1            ...........g..............................gg.................................... [ 57360]
KM111295.1_Ken06.Bus             .g.........g.............g.................g............a.......t............... [ 57360]
KM111294.1_Ken05/Tk1             .g.........g.............g.................g............a.......t............... [ 57360]
KM262844.1_strain_L60            ...........g..............................gg.................................... [ 57360]
KM102979.1_26544/OG10_Italy      ...........g..............................gg.................................... [ 57360]
KM262845.1_strain_NHV            ...........g..............................gg.................................... [ 57360]
KX354450.1_47/Ss/2008            ...........g..............................gg.................................... [ 57360]
KP055815.1_strain_BA71           ...........g..............................gg.................................... [ 57360]
MH025920.1_strain_R35            .g.........g.............g.................g............a.......t............... [ 57360]
MH025919.1_strain_N10            .g.........g.............g.................g............a.......t............... [ 57360]
MH025918.1_strain_R25            .g.........g.............g.................g............a.......t............... [ 57360]
MH025917.1_strain_R7             .g.........g.............g.................g............a.......t............... [ 57360]
MH025916.1_strain_R8             .g.........g.............g.................g............a.......t............... [ 57360]

FR682468.2_Georgia_2007/1        gtaaaatattccacctatacagacattgtacaattcatgggtgagttttctgtgagcccagaaagggtgatcaaaatggc [ 57440]
ASF_IND_20_CAD543                ................................................................................ [ 57440]
LR899193.1_Germany_2020/1        ................................................................................ [ 57440]
MN393477.1_Wuhan_2019            ................................................................................ [ 57440]
MW396979.1_Timor-Leste/2019/1    ................................................................................ [ 57440]
MK940252.1_CN/2019/InnerMongolia ................................................................................ [ 57440]
MK128995.1_China/2018/AnhuiXCGQ  ................................................................................ [ 57440]
MH681419.1_ASFV/POL/2015         ................................................................................ [ 57440]
LS478113.1_Estonia_2014          ................................................................................ [ 57440]
KP843857.1_Odintsovo_02/14       ................................................................................ [ 57440]
```

```
FN557520.1_strain_E75          ................................................................................ [ 57440]
AY261366.1_Warthog             .......................t...............a...................g.................... [ 57440]
AY261365.1_Warmbaths           .......................................a...................g....a............... [ 57440]
AY261364.1_Tengani_62          ...........................................................g.................... [ 57440]
AY261363.1_Pretoriuskop/96/4   ..............t........t...............a...................g.................... [ 57440]
AY261362.1_Mkuzi_1979          ................................................................................ [ 57440]
AY261361.1_Malawi_Lil-20/1_1983 ....g....................c....a............................g....a...t........... [ 57440]
AY261360.1_Kenya_1950          ........c.g...................a.......ac...................g.................... [ 57440]
AM712240.1_OURT_88/3           ................................................................................ [ 57440]
AM712239.1_Benin_97/1          ................................................................................ [ 57440]
KM111295.1_Ken06.Bus           ........c.g...................a.......ac...................g.................... [ 57440]
KM111294.1_Ken05/Tk1           ........c.g...................a.......ac...................g.................... [ 57440]
KM262844.1_strain_L60          ................................................................................ [ 57440]
KM102979.1_26544/OG10_Italy    ................................................................................ [ 57440]
KM262845.1_strain_NHV          ................................................................................ [ 57440]
KX354450.1_47/Ss/2008          ................................................................................ [ 57440]
KP055815.1_strain_BA71         ................................................................................ [ 57440]
MH025920.1_strain_R35          ........c.g...................a.......ac...................g.................... [ 57440]
MH025919.1_strain_N10          ........c.g...................a.......ac...................g.................... [ 57440]
MH025918.1_strain_R25          ........c.g...................a.......ac...................g.................... [ 57440]
MH025917.1_strain_R7           ........c.g...................a.......ac...................g.................... [ 57440]
MH025916.1_strain_R8           ........c.g...................a.......ac...................g.................... [ 57440]

FR682468.2_Georgia_2007/1      tgcacgggaatccaggacctttctgattgaaatgatctccaaagctgcttggggaaatcacccacagacgttgattcatc [ 57520]
ASF_IND_20_CAD543              ................................................................................ [ 57520]
LR899193.1_Germany_2020/1      ................................................................................ [ 57520]
MN393477.1_Wuhan_2019          ................................................................................ [ 57520]
MW396979.1_Timor-Leste/2019/1  ................................................................................ [ 57520]
MK940252.1_CN/2019/InnerMongolia ................................................................................ [ 57520]
MK128995.1_China/2018/AnhuiXCGQ ................................................................................ [ 57520]
MH681419.1_ASFV/POL/2015       ................................................................................ [ 57520]
LS478113.1_Estonia_2014        ................................................................................ [ 57520]
KP843857.1_Odintsovo_02/14     ................................................................................ [ 57520]
FN557520.1_strain_E75          ................................................................................ [ 57520]
AY261366.1_Warthog             ............t................................................................... [ 57520]
AY261365.1_Warmbaths           ............t................................................................... [ 57520]
AY261364.1_Tengani_62          ......................................................................a.......c. [ 57520]
AY261363.1_Pretoriuskop/96/4   ............t................................................................... [ 57520]
AY261362.1_Mkuzi_1979          ................................................................................ [ 57520]
AY261361.1_Malawi_Lil-20/1_1983 ...c........t.........a....c...................t....aa................a......... [ 57520]
AY261360.1_Kenya_1950          ...........................c........................a.....g...........a.......c. [ 57520]
AM712240.1_OURT_88/3           ................................................................................ [ 57520]
AM712239.1_Benin_97/1          ................................................................................ [ 57520]
KM111295.1_Ken06.Bus           ............t..............c........................a.................a.......c. [ 57520]
```

```
KM111294.1_Ken05/Tk1             ...........................c........................a.....g...........a.......c. [ 57520]
KM262844.1_strain_L60            ................................................................................ [ 57520]
KM102979.1_26544/OG10_Italy      ................................................................................ [ 57520]
KM262845.1_strain_NHV            ................................................................................ [ 57520]
KX354450.1_47/Ss/2008            ................................................................................ [ 57520]
KP055815.1_strain_BA71           ................................................................................ [ 57520]
MH025920.1_strain_R35            ............t..............c........................a.................a.......c. [ 57520]
MH025919.1_strain_N10            ............t..............c........................a.................a.......c. [ 57520]
MH025918.1_strain_R25            ............t..............c........................a.................a.......c. [ 57520]
MH025917.1_strain_R7             ............t..............c........................a.................a.......c. [ 57520]
MH025916.1_strain_R8             ............t..............c........................a.................a.......c. [ 57520]

FR682468.2_Georgia_2007/1        atctcaaacaactaaccaataccatgaagcctcaatctggaaaagaccacatcatatataccatccactatatttatcta [ 57600]
ASF_IND_20_CAD543                ................................................................................ [ 57600]
LR899193.1_Germany_2020/1        ................................................................................ [ 57600]
MN393477.1_Wuhan_2019            ................................................................................ [ 57600]
MW396979.1_Timor-Leste/2019/1    ................................................................................ [ 57600]
MK940252.1_CN/2019/InnerMongolia ................................................................................ [ 57600]
MK128995.1_China/2018/AnhuiXCGQ  ................................................................................ [ 57600]
MH681419.1_ASFV/POL/2015         ................................................................................ [ 57600]
LS478113.1_Estonia_2014          ................................................................................ [ 57600]
KP843857.1_Odintsovo_02/14       ................................................................................ [ 57600]
FN557520.1_strain_E75            ..........c.....................................t............................... [ 57600]
AY261366.1_Warthog               ..............g..c...........t..................t............................... [ 57600]
AY261365.1_Warmbaths             ..............g..c...........t..................t............................... [ 57600]
AY261364.1_Tengani_62            ..............g..c...........t..................t.............................a. [ 57600]
AY261363.1_Pretoriuskop/96/4     ..............g..c...........t..................t............................... [ 57600]
AY261362.1_Mkuzi_1979            ..........c.....................................t...............t............... [ 57600]
AY261361.1_Malawi_Lil-20/1_1983  ..............g..g..............................t.............................a. [ 57600]
AY261360.1_Kenya_1950            .................t..............g...............t.c.........tg................a. [ 57600]
AM712240.1_OURT_88/3             ..........c.....................................t............................... [ 57600]
AM712239.1_Benin_97/1            ..........c.....................................t............................... [ 57600]
KM111295.1_Ken06.Bus             .................t..............g...............ttc.........tg................a. [ 57600]
KM111294.1_Ken05/Tk1             .................t..............g...............t.c.........tg................a. [ 57600]
KM262844.1_strain_L60            ..........c.....................................t............................... [ 57600]
KM102979.1_26544/OG10_Italy      ..........c.....................................t............................... [ 57600]
KM262845.1_strain_NHV            ..........c.....................................t............................... [ 57600]
KX354450.1_47/Ss/2008            ..........c.....................................t............................... [ 57600]
KP055815.1_strain_BA71           ..........c.....................................t............................... [ 57600]
MH025920.1_strain_R35            .................t..............g...............ttc.........tg................a. [ 57600]
MH025919.1_strain_N10            .................t..............g...............ttc.........tg................a. [ 57600]
MH025918.1_strain_R25            .................t..............g...............ttc.........tg................a. [ 57600]
MH025917.1_strain_R7             .................t..............g...............ttc.........tg................a. [ 57600]
MH025916.1_strain_R8             .................t..............g...............ttc.........tg................a. [ 57600]
```

```
FR682468.2_Georgia_2007/1        aactctaatatgctggtagcggaggaggaaaaaaatattttttaaattagcaaaattttatgcgaatcataatgcggtaaa [ 57680]
ASF_IND_20_CAD543                ................................................................................ [ 57680]
LR899193.1_Germany_2020/1        ................................................................................ [ 57680]
MN393477.1_Wuhan_2019            ................................................................................ [ 57680]
MW396979.1_Timor-Leste/2019/1    ................................................................................ [ 57680]
MK940252.1_CN/2019/InnerMongolia ................................................................................ [ 57680]
MK128995.1_China/2018/AnhuiXCGQ  ................................................................................ [ 57680]
MH681419.1_ASFV/POL/2015         ................................................................................ [ 57680]
LS478113.1_Estonia_2014          ................................................................................ [ 57680]
KP843857.1_Odintsovo_02/14       ................................................................................ [ 57680]
FN557520.1_strain_E75            .......................................................................c........ [ 57680]
AY261366.1_Warthog               .......................................................................c..a..... [ 57680]
AY261365.1_Warmbaths             .......................................................................c..a..... [ 57680]
AY261364.1_Tengani_62            .g.......t.........t................................c..................c..a..... [ 57680]
AY261363.1_Pretoriuskop/96/4     .......................................................................c........ [ 57680]
AY261362.1_Mkuzi_1979            .......................a.......................................c................ [ 57680]
AY261361.1_Malawi_Lil-20/1_1983  .g.......t.....................................g.......................ct....... [ 57680]
AY261360.1_Kenya_1950            .c.......t.....................................g....ct...........g.....ct....... [ 57680]
AM712240.1_OURT_88/3             .......................................................................c........ [ 57680]
AM712239.1_Benin_97/1            .......................................................................c........ [ 57680]
KM111295.1_Ken06.Bus             .c.......t...................................c.g....ct...........g.....ct....... [ 57680]
KM111294.1_Ken05/Tk1             .c.......t.....................................g....ct...........g.....ct....... [ 57680]
KM262844.1_strain_L60            .......................................................................c........ [ 57680]
KM102979.1_26544/OG10_Italy      .......................................................................c........ [ 57680]
KM262845.1_strain_NHV            .......................................................................c........ [ 57680]
KX354450.1_47/Ss/2008            .......................................................................c........ [ 57680]
KP055815.1_strain_BA71           .......................................................................c........ [ 57680]
MH025920.1_strain_R35            .c.......t...................................c.g....ct...........g.....ct....... [ 57680]
MH025919.1_strain_N10            .c.......t...................................c.g....ct...........g.....ct....... [ 57680]
MH025918.1_strain_R25            .c.......t...................................c.g....ct...........g.....ct....... [ 57680]
MH025917.1_strain_R7             .c.......t...................................c.g....ct...........g.....ct....... [ 57680]
MH025916.1_strain_R8             .c.......t...................................c.g....ct...........g.....ct....... [ 57680]

FR682468.2_Georgia_2007/1        caggtttaaacaaatttgtgaagactattatatat------tagatgcacgatttaaaacacttattttagaatgttttg [ 57760]
ASF_IND_20_CAD543                ...................................------....................................... [ 57760]
LR899193.1_Germany_2020/1        ...................................------....................................... [ 57760]
MN393477.1_Wuhan_2019            ...................................------....................................... [ 57760]
MW396979.1_Timor-Leste/2019/1    ...................................------....................................... [ 57760]
MK940252.1_CN/2019/InnerMongolia ...................................------....................................... [ 57760]
MK128995.1_China/2018/AnhuiXCGQ  ...................................------....................................... [ 57760]
MH681419.1_ASFV/POL/2015         ...................................------....................................... [ 57760]
LS478113.1_Estonia_2014          ...................................------....................................... [ 57760]
KP843857.1_Odintsovo_02/14       ...................................------....................................... [ 57760]
```

```
FN557520.1_strain_E75              ...................................------....................................... [ 57760]
AY261366.1_Warthog                 .............g.................gc.ctagatg..........g............................ [ 57760]
AY261365.1_Warmbaths               .............g.................gc.ctagatg..........g............................ [ 57760]
AY261364.1_Tengani_62              .............g.................gc.ctagatg..........g...........c................ [ 57760]
AY261363.1_Pretoriuskop/96/4       .............g.................gc.ctagatg..........g............................ [ 57760]
AY261362.1_Mkuzi_1979              ...................................------....................................... [ 57760]
AY261361.1_Malawi_Lil-20/1_1983    ...............a........t.......c..tagacg.....a....g.......................c.... [ 57760]
AY261360.1_Kenya_1950              ...............a...............gc.ctagatg..........g.......................c.... [ 57760]
AM712240.1_OURT_88/3               .................................g.------....................................... [ 57760]
AM712239.1_Benin_97/1              ...................................------....................................... [ 57760]
KM111295.1_Ken06.Bus               ...............a...............gc.ctagacg..........g.......................c.... [ 57760]
KM111294.1_Ken05/Tk1               ...............a...............gc.ctagatg..........g.......................c.... [ 57760]
KM262844.1_strain_L60              ...................................------....................................... [ 57760]
KM102979.1_26544/OG10_Italy        ...................a..a............------....................................... [ 57760]
KM262845.1_strain_NHV              .................................g.------....................................... [ 57760]
KX354450.1_47/Ss/2008              ...................a..a............------....................................... [ 57760]
KP055815.1_strain_BA71             ...................................------....................................... [ 57760]
MH025920.1_strain_R35              ...............a...............gc.ctagacg..........g.......................c.... [ 57760]
MH025919.1_strain_N10              ...............a...............gc.ctagacg..........g.......................c.... [ 57760]
MH025918.1_strain_R25              ...............a...............gc.ctagacg..........g.......................c.... [ 57760]
MH025917.1_strain_R7               ...............a...............gc.ctagacg..........g.......................c.... [ 57760]
MH025916.1_strain_R8               ...............a...............gc.ctagacg..........g.......................c.... [ 57760]

FR682468.2_Georgia_2007/1          aaattgccgtccagaaaaactatcctagaattgcaaatattgtggatgactatattcgattccttttttacaggggaaat [ 57840]
ASF_IND_20_CAD543                  ................................................................................ [ 57840]
LR899193.1_Germany_2020/1          ................................................................................ [ 57840]
MN393477.1_Wuhan_2019              ................................................................................ [ 57840]
MW396979.1_Timor-Leste/2019/1      ................................................................................ [ 57840]
MK940252.1_CN/2019/InnerMongolia   ................................................................................ [ 57840]
MK128995.1_China/2018/AnhuiXCGQ    ................................................................................ [ 57840]
MH681419.1_ASFV/POL/2015           ................................................................................ [ 57840]
LS478113.1_Estonia_2014            ................................................................................ [ 57840]
KP843857.1_Odintsovo_02/14         ................................................................................ [ 57840]
FN557520.1_strain_E75              ................................................................................ [ 57840]
AY261366.1_Warthog                 .....................................................g.......................g.. [ 57840]
AY261365.1_Warmbaths               .....................................................g.......................g.. [ 57840]
AY261364.1_Tengani_62              .....................................................g.......................g.. [ 57840]
AY261363.1_Pretoriuskop/96/4       .....................................................g.......................g.. [ 57840]
AY261362.1_Mkuzi_1979              ................................................................................ [ 57840]
AY261361.1_Malawi_Lil-20/1_1983    ..........g..a......................c..............t..................t.aa...g.. [ 57840]
AY261360.1_Kenya_1950              .............a......................g..............t..................t.a....g.. [ 57840]
AM712240.1_OURT_88/3               ................................................................................ [ 57840]
AM712239.1_Benin_97/1              ................................................................................ [ 57840]
KM111295.1_Ken06.Bus               .............a.....................................t..................t.a....g.. [ 57840]
```

```
KM111294.1_Ken05/Tk1            .............a......................g..............t..................t.a....g.. [ 57840]
KM262844.1_strain_L60           ................................................................................ [ 57840]
KM102979.1_26544/OG10_Italy     ................................................................................ [ 57840]
KM262845.1_strain_NHV           ................................................................................ [ 57840]
KX354450.1_47/Ss/2008           ................................................................................ [ 57840]
KP055815.1_strain_BA71          ................................................................................ [ 57840]
MH025920.1_strain_R35           .............a.....................................t..................t.a....g.. [ 57840]
MH025919.1_strain_N10           .............a.....................................t..................t.a....g.. [ 57840]
MH025918.1_strain_R25           .............a.....................................t..................t.a....g.. [ 57840]
MH025917.1_strain_R7            .............a.....................................t..................t.a....g.. [ 57840]
MH025916.1_strain_R8            .............a.....................................t..................t.a....g.. [ 57840]

FR682468.2_Georgia_2007/1       ataaccgaggaagaaattcgtgaagcctattctttaaaagatgctgaggtttatgtagatttaaaatggttacaac---- [ 57920]
ASF_IND_20_CAD543               ............................................................................---- [ 57920]
LR899193.1_Germany_2020/1       ............................................................................---- [ 57920]
MN393477.1_Wuhan_2019           ............................................................................---- [ 57920]
MW396979.1_Timor-Leste/2019/1   ............................................................................---- [ 57920]
MK940252.1_CN/2019/InnerMongolia ............................................................................---- [ 57920]
MK128995.1_China/2018/AnhuiXCGQ ............................................................................---- [ 57920]
MH681419.1_ASFV/POL/2015        ............................................................................---- [ 57920]
LS478113.1_Estonia_2014         ............................................................................---- [ 57920]
KP843857.1_Odintsovo_02/14      ............................................................................---- [ 57920]
FN557520.1_strain_E75           ............................................................................---- [ 57920]
AY261366.1_Warthog              ...................a............................t...........................atga [ 57920]
AY261365.1_Warmbaths            ........a.......................................t...........................---- [ 57920]
AY261364.1_Tengani_62           ................................................t...........................atga [ 57920]
AY261363.1_Pretoriuskop/96/4    ........a..........a............................t...........................atga [ 57920]
AY261362.1_Mkuzi_1979           ............................................................................---- [ 57920]
AY261361.1_Malawi_Lil-20/1_1983 ..................a.c..............g..ga........c...........................aaga [ 57920]
AY261360.1_Kenya_1950           ......a...........a.c..............g..ga........t.a........c................aag- [ 57920]
AM712240.1_OURT_88/3            ............................................................................---- [ 57920]
AM712239.1_Benin_97/1           ............................................................................---- [ 57920]
KM111295.1_Ken06.Bus            ......a...........a.c..............g..g.........t.a........c................aag- [ 57920]
KM111294.1_Ken05/Tk1            ......a...........a.c..............g..g.........t.a........c................aag- [ 57920]
KM262844.1_strain_L60           ............................................................................---- [ 57920]
KM102979.1_26544/OG10_Italy     ............................................................................---- [ 57920]
KM262845.1_strain_NHV           ............................................................................---- [ 57920]
KX354450.1_47/Ss/2008           ............................................................................---- [ 57920]
KP055815.1_strain_BA71          ............................................................................---- [ 57920]
MH025920.1_strain_R35           ......a...........a.c..............g..g.........t.a........c................aag- [ 57920]
MH025919.1_strain_N10           ......a...........a.c..............g..g.........t.a........c................aag- [ 57920]
MH025918.1_strain_R25           ......a...........a.c..............g..g.........t.a........c................aag- [ 57920]
MH025917.1_strain_R7            ......a...........a.c..............g..g.........t.a........c................aag- [ 57920]
MH025916.1_strain_R8            ......a...........a.c..............g..g.........t.a........c................aag- [ 57920]
```

```
FR682468.2_Georgia_2007/1        ---------aaggagaaatggtttaaaccaaatc----------------cggtttaaactaaatcca-----atttaaa [ 58000]
ASF_IND_20_CAD543                ---------.........................----------------..................-----....... [ 58000]
LR899193.1_Germany_2020/1        ---------.........................----------------..................-----....... [ 58000]
MN393477.1_Wuhan_2019            ---------.........................----------------..................-----....... [ 58000]
MW396979.1_Timor-Leste/2019/1    ---------.........................----------------..................-----....... [ 58000]
MK940252.1_CN/2019/InnerMongolia ---------.........................----------------..................-----....... [ 58000]
MK128995.1_China/2018/AnhuiXCGQ  ---------.........................----------------..................-----....... [ 58000]
MH681419.1_ASFV/POL/2015         ---------.........................----------------..................-----....... [ 58000]
LS478113.1_Estonia_2014          ---------.........................----------------..................-----....... [ 58000]
KP843857.1_Odintsovo_02/14       ---------.........................----------------..................-----....... [ 58000]
FN557520.1_strain_E75            ---------...................t.....----------------...............t.g-----g...... [ 58000]
AY261366.1_Warthog               tttgtttta..ta...gt.a........t.....tgatttaaactacattt...........c..ttg-----g...... [ 58000]
AY261365.1_Warmbaths             ---------..........a........t.....----------------ta..............t.-----g...... [ 58000]
AY261364.1_Tengani_62            tttgtttta..ta...gt.a........t.....--------------------------------tg-----.c..... [ 58000]
AY261363.1_Pretoriuskop/96/4     tttgtttta..ta...gt.a........t.....----------------t.a.........c..ttg-----g...... [ 58000]
AY261362.1_Mkuzi_1979            ---------..........a........t.....----------------.................g-----g...... [ 58000]
AY261361.1_Malawi_Lil-20/1_1983  agagaatta..cc.c.tt..t.........c..t----------------t..c........c..ttg-----gc..... [ 58000]
AY261360.1_Kenya_1950            ---------..taga..tg.t.......a.gg.ttagtt----------------.....a.g----------g...... [ 58000]
AM712240.1_OURT_88/3             ---------...................t.....----------------...............t.g-----g...... [ 58000]
AM712239.1_Benin_97/1            ---------...................t.....----------------...............t.g-----g...... [ 58000]
KM111295.1_Ken06.Bus             ---------..taga..tg.t.......a.gg.ttagtttaaa-------.a..g.....a..----------g...... [ 58000]
KM111294.1_Ken05/Tk1             ---------..taga..tg.t.......a.gg.tta-----aa-------.aa.......a..g.tt.aacagg...... [ 58000]
KM262844.1_strain_L60            ---------...................t.....----------------...............t.g-----g...... [ 58000]
KM102979.1_26544/OG10_Italy      ---------...................t.....----------------...............t.g-----g...... [ 58000]
KM262845.1_strain_NHV            ---------...................t.....----------------...............t.g-----g...... [ 58000]
KX354450.1_47/Ss/2008            ---------...................t.....----------------...............t.g-----g...... [ 58000]
KP055815.1_strain_BA71           ---------...................t.....----------------...............t.g-----g...... [ 58000]
MH025920.1_strain_R35            ---------..taga..tg.t.......a.gg.ttagtttaaa-------.a..g.....a..----------g...... [ 58000]
MH025919.1_strain_N10            ---------..taga..tg.t.......a.gg.ttagtttaaa-------.a..g.....a..----------g...... [ 58000]
MH025918.1_strain_R25            ---------..taga..tg.t.......a.gg.ttagtttaaa-------.a..g.....a..----------g...... [ 58000]
MH025917.1_strain_R7             ---------..taga..tg.t.......a.gg.ttagtttaaa-------.a..g.....a..----------g...... [ 58000]
MH025916.1_strain_R8             ---------..taga..tg.t.......a.gg.ttagtttaaa-------.a..g.....a..----------g...... [ 58000]

FR682468.2_Georgia_2007/1        ctacatttggtttatcattagtcattgaaa-ccatcg-aaaaaaaagctatttgtttatccccataaactcatc-t-ttt [ 58080]
ASF_IND_20_CAD543                ..............................-......-.........-..........................-.-... [ 58080]
LR899193.1_Germany_2020/1        ..............................-......-....................................-.-... [ 58080]
MN393477.1_Wuhan_2019            ..............................-......-....................................-.-... [ 58080]
MW396979.1_Timor-Leste/2019/1    ..............................-......-....................................-.-... [ 58080]
MK940252.1_CN/2019/InnerMongolia ..............................-......-....................................-.-... [ 58080]
MK128995.1_China/2018/AnhuiXCGQ  ..............................-......-....................................-.-... [ 58080]
MH681419.1_ASFV/POL/2015         ..............................-......-....................................-.-... [ 58080]
LS478113.1_Estonia_2014          ..............................-......-....................................-.-... [ 58080]
KP843857.1_Odintsovo_02/14       ..............................-......--...................................-.-... [ 58080]
```

```
FN557520.1_strain_E75          ..............................-......-.........-..........................-.t... [ 58080]
AY261366.1_Warthog             ....................t.........-......a.......................t............-.-... [ 58080]
AY261365.1_Warmbaths           ....................t.........-......-.......................t...........--.-... [ 58080]
AY261364.1_Tengani_62          ....................t.........-......-..........----.........t...........--.-... [ 58080]
AY261363.1_Pretoriuskop/96/4   ....................t.........-......a..........----.........t..........---.-... [ 58080]
AY261362.1_Mkuzi_1979          ..............................-......-....................................-.-... [ 58080]
AY261361.1_Malawi_Lil-20/1_1983 .c................g.t.........-t.....-.......................t...........----... [ 58080]
AY261360.1_Kenya_1950          .a.g....a----.a.c...t.........t......-...........g...........-............t.t... [ 58080]
AM712240.1_OURT_88/3           ..............................-......--........-..........................-.-... [ 58080]
AM712239.1_Benin_97/1          ..............................-......-.........-..........................-.-... [ 58080]
KM111295.1_Ken06.Bus           .a.gg...a----.a.c...t.........t......-...........g...........-............t.t... [ 58080]
KM111294.1_Ken05/Tk1           .a.g....a----.a.c...t.........t......-...........g...........-............-.t... [ 58080]
KM262844.1_strain_L60          ..............................-......-.........-..........................-.-... [ 58080]
KM102979.1_26544/OG10_Italy    ..............................-......-.........-..........................-.-... [ 58080]
KM262845.1_strain_NHV          ..............................-......-.........-..........................-.-... [ 58080]
KX354450.1_47/Ss/2008          ..............................-......-.........-..........................-.-... [ 58080]
KP055815.1_strain_BA71         ..............................-......-.........-..........................-.-... [ 58080]
MH025920.1_strain_R35          .a.gg...a----.a.c...t.........t......-...........g...........-............-.t... [ 58080]
MH025919.1_strain_N10          .a.gg...a----.a.c...t.........t......-...........g...........-............-.t... [ 58080]
MH025918.1_strain_R25          .a.gg...a----.a.c...t.........t......-...........g...........-............-.t... [ 58080]
MH025917.1_strain_R7           .a.gg...a----.a.c...t.........t......-...........g...........-............-.t... [ 58080]
MH025916.1_strain_R8           .a.gg...a----.a.c...t.........t......-...........g...........-............-.t... [ 58080]

FR682468.2_Georgia_2007/1      ttttgtctcaaagtttgacactaaaattcagtgttttatagtgtttataattaagtgttttgcatgcattgcagaaattt [ 58160]
ASF_IND_20_CAD543              ................................................................................ [ 58160]
LR899193.1_Germany_2020/1      ................................................................................ [ 58160]
MN393477.1_Wuhan_2019          ................................................................................ [ 58160]
MW396979.1_Timor-Leste/2019/1  ................................................................................ [ 58160]
MK940252.1_CN/2019/InnerMongolia ................................................................................ [ 58160]
MK128995.1_China/2018/AnhuiXCGQ ................................................................................ [ 58160]
MH681419.1_ASFV/POL/2015       ................................................................................ [ 58160]
LS478113.1_Estonia_2014        ................................................................................ [ 58160]
KP843857.1_Odintsovo_02/14     ................................................................................ [ 58160]
FN557520.1_strain_E75          ................................................................................ [ 58160]
AY261366.1_Warthog             ................................................................................ [ 58160]
AY261365.1_Warmbaths           ................................................................................ [ 58160]
AY261364.1_Tengani_62          ................................................................................ [ 58160]
AY261363.1_Pretoriuskop/96/4   ................................................................................ [ 58160]
AY261362.1_Mkuzi_1979          ................................................................................ [ 58160]
AY261361.1_Malawi_Lil-20/1_1983 ................................a........................g...................... [ 58160]
AY261360.1_Kenya_1950          ...g..t.........t.t.........t..ca.......a.a.........................c..........c [ 58160]
AM712240.1_OURT_88/3           ................................................................................ [ 58160]
AM712239.1_Benin_97/1          ................................................................................ [ 58160]
KM111295.1_Ken06.Bus           ...g..t.........t.t.........t..ca.......a.a.........................c..........c [ 58160]
```

```
KM111294.1_Ken05/Tk1            ...g..t.........t.t.........t..ca.......a.a.........................c..........c [ 58160]
KM262844.1_strain_L60           ................................................................................ [ 58160]
KM102979.1_26544/OG10_Italy     ................................................................................ [ 58160]
KM262845.1_strain_NHV           ................................................................................ [ 58160]
KX354450.1_47/Ss/2008           ................................................................................ [ 58160]
KP055815.1_strain_BA71          ................................................................................ [ 58160]
MH025920.1_strain_R35           ...g..t.........t.t.........t..ca.......a.a.........................c..........c [ 58160]
MH025919.1_strain_N10           ...g..t.........t.t.........t..ca.......a.a.........................c..........c [ 58160]
MH025918.1_strain_R25           ...g..t.........t.t.........t..ca.......a.a.........................c..........c [ 58160]
MH025917.1_strain_R7            ...g..t.........t.t.........t..ca.......a.a.........................c..........c [ 58160]
MH025916.1_strain_R8            ...g..t.........t.t.........t..ca.......a.a.........................c..........c [ 58160]

FR682468.2_Georgia_2007/1       tcatcttttttaattggttcaataccacatgtcatacaatatgttgtttgattatcaagattaactttatgaaaggaaag [ 58240]
ASF_IND_20_CAD543               ................................................................................ [ 58240]
LR899193.1_Germany_2020/1       ................................................................................ [ 58240]
MN393477.1_Wuhan_2019           ................................................................................ [ 58240]
MW396979.1_Timor-Leste/2019/1   ................................................................................ [ 58240]
MK940252.1_CN/2019/InnerMongolia ................................................................................ [ 58240]
MK128995.1_China/2018/AnhuiXCGQ ................................................................................ [ 58240]
MH681419.1_ASFV/POL/2015        ................................................................................ [ 58240]
LS478113.1_Estonia_2014         ................................................................................ [ 58240]
KP843857.1_Odintsovo_02/14      ................................................................................ [ 58240]
FN557520.1_strain_E75           ................................................................................ [ 58240]
AY261366.1_Warthog              ................................................................................ [ 58240]
AY261365.1_Warmbaths            .......................g........................................................ [ 58240]
AY261364.1_Tengani_62           ................................................................................ [ 58240]
AY261363.1_Pretoriuskop/96/4    ................................................................................ [ 58240]
AY261362.1_Mkuzi_1979           ................................................................................ [ 58240]
AY261361.1_Malawi_Lil-20/1_1983 ........a......t.............................a....g............................. [ 58240]
AY261360.1_Kenya_1950           ........a......t..................................g............................. [ 58240]
AM712240.1_OURT_88/3            ................................................................................ [ 58240]
AM712239.1_Benin_97/1           ................................................................................ [ 58240]
KM111295.1_Ken06.Bus            ........a......t..................................g............................. [ 58240]
KM111294.1_Ken05/Tk1            ........a......t..................................g........t.................... [ 58240]
KM262844.1_strain_L60           ................................................................................ [ 58240]
KM102979.1_26544/OG10_Italy     ................................................................................ [ 58240]
KM262845.1_strain_NHV           ................................................................................ [ 58240]
KX354450.1_47/Ss/2008           ................................................................................ [ 58240]
KP055815.1_strain_BA71          ................................................................................ [ 58240]
MH025920.1_strain_R35           ........a......t..................................g............................. [ 58240]
MH025919.1_strain_N10           ........a......t..................................g............................. [ 58240]
MH025918.1_strain_R25           ........a......t..................................g............................. [ 58240]
MH025917.1_strain_R7            ........a......t..................................g............................. [ 58240]
MH025916.1_strain_R8            ........a......t..................................g............................. [ 58240]
```

```
FR682468.2_Georgia_2007/1        taagtgagccgcaaatttaaaagtaaaatatctttcatttaaaatgatcttatgaatgtattttcgataaggaggaatga [ 58320]
ASF_IND_20_CAD543                ..............................................................................r. [ 58320]
LR899193.1_Germany_2020/1        ................................................................................ [ 58320]
MN393477.1_Wuhan_2019            ................................................................................ [ 58320]
MW396979.1_Timor-Leste/2019/1    ................................................................................ [ 58320]
MK940252.1_CN/2019/InnerMongolia ................................................................................ [ 58320]
MK128995.1_China/2018/AnhuiXCGQ  ................................................................................ [ 58320]
MH681419.1_ASFV/POL/2015         ................................................................................ [ 58320]
LS478113.1_Estonia_2014          ................................................................................ [ 58320]
KP843857.1_Odintsovo_02/14       ................................................................................ [ 58320]
FN557520.1_strain_E75            ..............................................................................a. [ 58320]
AY261366.1_Warthog               .................................g.......................a...................... [ 58320]
AY261365.1_Warmbaths             ................................gg...........a...........a...................... [ 58320]
AY261364.1_Tengani_62            ..g..a...........................g.......................a...................... [ 58320]
AY261363.1_Pretoriuskop/96/4     .................................g.......................a...................... [ 58320]
AY261362.1_Mkuzi_1979            ..............................................................................a. [ 58320]
AY261361.1_Malawi_Lil-20/1_1983  c.g..a...ta..................................a...........a.................c.... [ 58320]
AY261360.1_Kenya_1950            c.g..a...a....................c..............a...........a.................c.... [ 58320]
AM712240.1_OURT_88/3             ..............................................................................a. [ 58320]
AM712239.1_Benin_97/1            ..............................................................................a. [ 58320]
KM111295.1_Ken06.Bus             c.g..a...a....................c..............a...........a.................c.... [ 58320]
KM111294.1_Ken05/Tk1             c.g..a...a....................c..............a...........a........g........c.... [ 58320]
KM262844.1_strain_L60            ..............................................................................a. [ 58320]
KM102979.1_26544/OG10_Italy      ..............................................................................a. [ 58320]
KM262845.1_strain_NHV            ..............................................................................a. [ 58320]
KX354450.1_47/Ss/2008            ..............................................................................a. [ 58320]
KP055815.1_strain_BA71           ..............................................................................a. [ 58320]
MH025920.1_strain_R35            c.g..a...a....................c..............a...........a.................c.... [ 58320]
MH025919.1_strain_N10            c.g..a...a....................c..............a...........a.................c.... [ 58320]
MH025918.1_strain_R25            c.g..a...a....................c..............a...........a.................c.... [ 58320]
MH025917.1_strain_R7             c.g..a...a....................c..............a...........a.................c.... [ 58320]
MH025916.1_strain_R8             c.g..a...a....................c..............a...........a.................c.... [ 58320]

FR682468.2_Georgia_2007/1        aagcatttgccaaaataaatcgcataaaaggcttggaaaaacccatatcttctaatcttttgtgggtataaaccctattt [ 58400]
ASF_IND_20_CAD543                ................................................................................ [ 58400]
LR899193.1_Germany_2020/1        ................................................................................ [ 58400]
MN393477.1_Wuhan_2019            ................................................................................ [ 58400]
MW396979.1_Timor-Leste/2019/1    ................................................................................ [ 58400]
MK940252.1_CN/2019/InnerMongolia ................................................................................ [ 58400]
MK128995.1_China/2018/AnhuiXCGQ  ................................................................................ [ 58400]
MH681419.1_ASFV/POL/2015         ................................................................................ [ 58400]
LS478113.1_Estonia_2014          ................................................................................ [ 58400]
KP843857.1_Odintsovo_02/14       ................................................................................ [ 58400]
```

```
FN557520.1_strain_E75            ............................g................................................... [ 58400]
AY261366.1_Warthog               ....................t.......g...........c...............t....................... [ 58400]
AY261365.1_Warmbaths             ....................t.......g...........c....................................... [ 58400]
AY261364.1_Tengani_62            ....................t.......g...........g....................................... [ 58400]
AY261363.1_Pretoriuskop/96/4     ....................t.......g...........c...............t....................... [ 58400]
AY261362.1_Mkuzi_1979            ............................g................................................... [ 58400]
AY261361.1_Malawi_Lil-20/1_1983  ..............................a........................c...........g.....t...... [ 58400]
AY261360.1_Kenya_1950            ............................gca........................c.................t...... [ 58400]
AM712240.1_OURT_88/3             ............................g................................................... [ 58400]
AM712239.1_Benin_97/1            ............................g................................................... [ 58400]
KM111295.1_Ken06.Bus             .........................g..gca........................c.................t...... [ 58400]
KM111294.1_Ken05/Tk1             ............................gca...a....................c.................t...... [ 58400]
KM262844.1_strain_L60            ............................g................................................... [ 58400]
KM102979.1_26544/OG10_Italy      ............................g................................................... [ 58400]
KM262845.1_strain_NHV            ............................g................................................... [ 58400]
KX354450.1_47/Ss/2008            ............................g................................................... [ 58400]
KP055815.1_strain_BA71           ............................g................................................... [ 58400]
MH025920.1_strain_R35            .........................g..gca........................c.................t...... [ 58400]
MH025919.1_strain_N10            .........................g..gca........................c.................t...... [ 58400]
MH025918.1_strain_R25            .........................g..gca........................c.................t...... [ 58400]
MH025917.1_strain_R7             .........................g..gca........................c.................t...... [ 58400]
MH025916.1_strain_R8             .........................g..gca........................c.................t...... [ 58400]

FR682468.2_Georgia_2007/1        tggtgttttacaaaaacttcattgttataatagtcgttatagctatcaatcatttttttaagtcctataatgcccaaggt [ 58480]
ASF_IND_20_CAD543                .k.............................................................................. [ 58480]
LR899193.1_Germany_2020/1        ................................................................................ [ 58480]
MN393477.1_Wuhan_2019            ................................................................................ [ 58480]
MW396979.1_Timor-Leste/2019/1    ................................................................................ [ 58480]
MK940252.1_CN/2019/InnerMongolia ................................................................................ [ 58480]
MK128995.1_China/2018/AnhuiXCGQ  ................................................................................ [ 58480]
MH681419.1_ASFV/POL/2015         ................................................................................ [ 58480]
LS478113.1_Estonia_2014          ................................................................................ [ 58480]
KP843857.1_Odintsovo_02/14       ................................................................................ [ 58480]
FN557520.1_strain_E75            .t.............................................................................. [ 58480]
AY261366.1_Warthog               .t.........................................................t.................... [ 58480]
AY261365.1_Warmbaths             .t.............................................................................. [ 58480]
AY261364.1_Tengani_62            .t.........................................................t.................... [ 58480]
AY261363.1_Pretoriuskop/96/4     .t.............................................................................. [ 58480]
AY261362.1_Mkuzi_1979            .t.............................................................................. [ 58480]
AY261361.1_Malawi_Lil-20/1_1983  .t..........g...........g..a.......a.....a...................................... [ 58480]
AY261360.1_Kenya_1950            .t..........g.........g.g..a..................................c................. [ 58480]
AM712240.1_OURT_88/3             .t.............................................................................. [ 58480]
AM712239.1_Benin_97/1            .t.............................................................................. [ 58480]
KM111295.1_Ken06.Bus             .t..........g.........g.g..a..................................c................. [ 58480]
```

```
KM111294.1_Ken05/Tk1             .t..........g.........g.g..a..................................c................. [ 58480]
KM262844.1_strain_L60            .t.............................................................................. [ 58480]
KM102979.1_26544/OG10_Italy      .t.............................................................................. [ 58480]
KM262845.1_strain_NHV            .t.............................................................................. [ 58480]
KX354450.1_47/Ss/2008            .t.............................................................................. [ 58480]
KP055815.1_strain_BA71           .t.............................................................................. [ 58480]
MH025920.1_strain_R35            .t..........g.........g.g..a..................................c................. [ 58480]
MH025919.1_strain_N10            .t..........g.........g.g..a..................................c................. [ 58480]
MH025918.1_strain_R25            .t..........g.........g.g..a..................................c................. [ 58480]
MH025917.1_strain_R7             .t..........g.........g.g..a..................................c................. [ 58480]
MH025916.1_strain_R8             .t..........g.........g.g..a..................................c................. [ 58480]

FR682468.2_Georgia_2007/1        tgcacgcataaagccacagtttctgctccaaaaagcatgcacctgtaaagggtgcttttcatataaccaattacaaaatt [ 58560]
ASF_IND_20_CAD543                .................................gc............................................. [ 58560]
LR899193.1_Germany_2020/1        ................................................................................ [ 58560]
MN393477.1_Wuhan_2019            ................................................................................ [ 58560]
MW396979.1_Timor-Leste/2019/1    ................................................................................ [ 58560]
MK940252.1_CN/2019/InnerMongolia ................................................................................ [ 58560]
MK128995.1_China/2018/AnhuiXCGQ  ................................................................................ [ 58560]
MH681419.1_ASFV/POL/2015         ................................................................................ [ 58560]
LS478113.1_Estonia_2014          ................................................................................ [ 58560]
KP843857.1_Odintsovo_02/14       ................................................................................ [ 58560]
FN557520.1_strain_E75            ...............g........a........gc............................................. [ 58560]
AY261366.1_Warthog               ...............g..................c............................................. [ 58560]
AY261365.1_Warmbaths             ...............g................................................................ [ 58560]
AY261364.1_Tengani_62            ...............g..................c..................a.......................... [ 58560]
AY261363.1_Pretoriuskop/96/4     ...............g..................c............................................. [ 58560]
AY261362.1_Mkuzi_1979            ...............g........a........gc............................................. [ 58560]
AY261361.1_Malawi_Lil-20/1_1983  ..............................g..gc..............................t.............. [ 58560]
AY261360.1_Kenya_1950            ..............................g..gc................a............................ [ 58560]
AM712240.1_OURT_88/3             ...............g........a........gc............................................. [ 58560]
AM712239.1_Benin_97/1            ...............g........a........gc............................................. [ 58560]
KM111295.1_Ken06.Bus             ..............................g..gc................a............................ [ 58560]
KM111294.1_Ken05/Tk1             ..............................g..gc................a............................ [ 58560]
KM262844.1_strain_L60            ...............g........a........gc............................................. [ 58560]
KM102979.1_26544/OG10_Italy      ...............g........a........gc............................................. [ 58560]
KM262845.1_strain_NHV            ...............g........a........gc............................................. [ 58560]
KX354450.1_47/Ss/2008            ...............g........a........gc............................................. [ 58560]
KP055815.1_strain_BA71           ...............g........a........gc............................................. [ 58560]
MH025920.1_strain_R35            ..............................g..gc................a..t......................... [ 58560]
MH025919.1_strain_N10            ..............................g..gc................a..t......................... [ 58560]
MH025918.1_strain_R25            ..............................g..gc................a..t......................... [ 58560]
MH025917.1_strain_R7             ..............................g..gc................a..t......................... [ 58560]
MH025916.1_strain_R8             ..............................g..gc................a..t......................... [ 58560]
```

```
FR682468.2_Georgia_2007/1        tcattccgcaacagtagcatgttatttcagtgggggatgtatagaataatccggcattcgaaaatttttcataatttttt [ 58640]
ASF_IND_20_CAD543                ................................................................................ [ 58640]
LR899193.1_Germany_2020/1        ................................................................................ [ 58640]
MN393477.1_Wuhan_2019            ................................................................................ [ 58640]
MW396979.1_Timor-Leste/2019/1    ................................................................................ [ 58640]
MK940252.1_CN/2019/InnerMongolia ................................................................................ [ 58640]
MK128995.1_China/2018/AnhuiXCGQ  ................................................................................ [ 58640]
MH681419.1_ASFV/POL/2015         ................................................................................ [ 58640]
LS478113.1_Estonia_2014          ................................................................................ [ 58640]
KP843857.1_Odintsovo_02/14       ................................................................................ [ 58640]
FN557520.1_strain_E75            ..............................................a................................. [ 58640]
AY261366.1_Warthog               ................................................................................ [ 58640]
AY261365.1_Warmbaths             ................................................................................ [ 58640]
AY261364.1_Tengani_62            ........................................g....................................... [ 58640]
AY261363.1_Pretoriuskop/96/4     ................................................................................ [ 58640]
AY261362.1_Mkuzi_1979            ................................................................................ [ 58640]
AY261361.1_Malawi_Lil-20/1_1983  ..........................................................a.....c............... [ 58640]
AY261360.1_Kenya_1950            ................a.................t.......................a.....c............... [ 58640]
AM712240.1_OURT_88/3             ..............................................a................................. [ 58640]
AM712239.1_Benin_97/1            ..............................................a................................. [ 58640]
KM111295.1_Ken06.Bus             ................a.................t.......................a.....c............... [ 58640]
KM111294.1_Ken05/Tk1             ................a.................t.......................a.....c............... [ 58640]
KM262844.1_strain_L60            ..............................................a................................. [ 58640]
KM102979.1_26544/OG10_Italy      ..............................................a................................. [ 58640]
KM262845.1_strain_NHV            ..............................................a................................. [ 58640]
KX354450.1_47/Ss/2008            ..............................................a................................. [ 58640]
KP055815.1_strain_BA71           ..............................................a................................. [ 58640]
MH025920.1_strain_R35            ................a.................t.......................a.....c............... [ 58640]
MH025919.1_strain_N10            ................a.................t.......................a.....c............... [ 58640]
MH025918.1_strain_R25            ................a.................t.......................a.....c............... [ 58640]
MH025917.1_strain_R7             ................a.................t.......................a.....c............... [ 58640]
MH025916.1_strain_R8             ................a.................t.......................a.....c............... [ 58640]

FR682468.2_Georgia_2007/1        atgtcatggattgcgaagctttgatttcgtgcatctatggagctatagcctacatatttaggttttacttcaaataatcg [ 58720]
ASF_IND_20_CAD543                ................................................................................ [ 58720]
LR899193.1_Germany_2020/1        ................................................................................ [ 58720]
MN393477.1_Wuhan_2019            ................................................................................ [ 58720]
MW396979.1_Timor-Leste/2019/1    ................................................................................ [ 58720]
MK940252.1_CN/2019/InnerMongolia ................................................................................ [ 58720]
MK128995.1_China/2018/AnhuiXCGQ  ................................................................................ [ 58720]
MH681419.1_ASFV/POL/2015         ................................................................................ [ 58720]
LS478113.1_Estonia_2014          ................................................................................ [ 58720]
KP843857.1_Odintsovo_02/14       ................................................................................ [ 58720]
```

```
FN557520.1_strain_E75              ...c............................................................................ [ 58720]
AY261366.1_Warthog                 ..a..................................c......g................................... [ 58720]
AY261365.1_Warmbaths               ..a..................................c......g................................... [ 58720]
AY261364.1_Tengani_62              ..a..................................c......g...t............................... [ 58720]
AY261363.1_Pretoriuskop/96/4       ..a..................................c......g................................... [ 58720]
AY261362.1_Mkuzi_1979              ................................................................................ [ 58720]
AY261361.1_Malawi_Lil-20/1_1983    .....g........a.......tg...........c.c...........................g.............. [ 58720]
AY261360.1_Kenya_1950              ......................tg...........c.c.............g.............g.............. [ 58720]
AM712240.1_OURT_88/3               ...c............................................................................ [ 58720]
AM712239.1_Benin_97/1              ...c............................................................................ [ 58720]
KM111295.1_Ken06.Bus               ......................tg...........c.c.............g.............g.............. [ 58720]
KM111294.1_Ken05/Tk1               ......................tg...........c.c.............g.............g.............. [ 58720]
KM262844.1_strain_L60              ...c............................................................................ [ 58720]
KM102979.1_26544/OG10_Italy        ...c............................................................................ [ 58720]
KM262845.1_strain_NHV              ...c............................................................................ [ 58720]
KX354450.1_47/Ss/2008              ...c............................................................................ [ 58720]
KP055815.1_strain_BA71             ...c............................................................................ [ 58720]
MH025920.1_strain_R35              ......................tg...........c.c.............g.............g.............. [ 58720]
MH025919.1_strain_N10              ......................tg...........c.c.............g.............g.............. [ 58720]
MH025918.1_strain_R25              ......................tg...........c.c.............g.............g.............. [ 58720]
MH025917.1_strain_R7               ......................tg...........c.c.............g.............g.............. [ 58720]
MH025916.1_strain_R8               ......................tg...........c.c.............g.............g.............. [ 58720]

FR682468.2_Georgia_2007/1          caaagagatgtatggatctatcgtatttattttaggaaacatttcataattttaaattcttatatataatat-aaaaaaa [ 58800]
ASF_IND_20_CAD543                  .....................y..................................................-....... [ 58800]
LR899193.1_Germany_2020/1          ........................................................................-....... [ 58800]
MN393477.1_Wuhan_2019              ........................................................................-....... [ 58800]
MW396979.1_Timor-Leste/2019/1      ........................................................................-....... [ 58800]
MK940252.1_CN/2019/InnerMongolia   ........................................................................-....... [ 58800]
MK128995.1_China/2018/AnhuiXCGQ    ........................................................................-....... [ 58800]
MH681419.1_ASFV/POL/2015           ........................................................................-....... [ 58800]
LS478113.1_Estonia_2014            ........................................................................-....... [ 58800]
KP843857.1_Odintsovo_02/14         ........................................................................-....... [ 58800]
FN557520.1_strain_E75              .....................t..................................................-....... [ 58800]
AY261366.1_Warthog                 t....................t..................................................-....... [ 58800]
AY261365.1_Warmbaths               t....................t..................................................-....... [ 58800]
AY261364.1_Tengani_62              t.......................................................................-....... [ 58800]
AY261363.1_Pretoriuskop/96/4       t....................t..................................................-....... [ 58800]
AY261362.1_Mkuzi_1979              ........................................................................-....... [ 58800]
AY261361.1_Malawi_Lil-20/1_1983    t........a....t......................t.........-------..................a....... [ 58800]
AY261360.1_Kenya_1950              t........a....t......................t.........-------..................-....... [ 58800]
AM712240.1_OURT_88/3               .....................t..................................................-....... [ 58800]
AM712239.1_Benin_97/1              .....................t..................................................-....... [ 58800]
KM111295.1_Ken06.Bus               t........a....t......................t.........-------..................-....... [ 58800]
```

```
KM111294.1_Ken05/Tk1            t........a....t......................t.........-------..................-....... [ 58800]
KM262844.1_strain_L60           .....................t..................................................-....... [ 58800]
KM102979.1_26544/OG10_Italy     .....................t..................................................-....... [ 58800]
KM262845.1_strain_NHV           .....................t..................................................-....... [ 58800]
KX354450.1_47/Ss/2008           .....................t..................................................-....... [ 58800]
KP055815.1_strain_BA71          .....................t..................................................-....... [ 58800]
MH025920.1_strain_R35           t........a....t......................t.........-------..................-....... [ 58800]
MH025919.1_strain_N10           t........a....t......................t.........-------..................-....... [ 58800]
MH025918.1_strain_R25           t........a....t......................t.........-------..................-....... [ 58800]
MH025917.1_strain_R7            t........a....t......................t.........-------..................-....... [ 58800]
MH025916.1_strain_R8            t........a....t......................t.........-------..................-....... [ 58800]

FR682468.2_Georgia_2007/1       attacaaacatttgtaatgatcatcctcaattgaaggctgagttgtaggctttatttttctaattatacgaagaaggtag [ 58880]
ASF_IND_20_CAD543               ................................................................................ [ 58880]
LR899193.1_Germany_2020/1       ................................................................................ [ 58880]
MN393477.1_Wuhan_2019           ................................................................................ [ 58880]
MW396979.1_Timor-Leste/2019/1   ................................................................................ [ 58880]
MK940252.1_CN/2019/InnerMongolia ................................................................................ [ 58880]
MK128995.1_China/2018/AnhuiXCGQ ................................................................................ [ 58880]
MH681419.1_ASFV/POL/2015        ................................................................................ [ 58880]
LS478113.1_Estonia_2014         ................................................................................ [ 58880]
KP843857.1_Odintsovo_02/14      ................................................................................ [ 58880]
FN557520.1_strain_E75           ...................................a......c..c.................................. [ 58880]
AY261366.1_Warthog              ........t.............t...........gc.......................................aa... [ 58880]
AY261365.1_Warmbaths            ......................t...........g.........................................a... [ 58880]
AY261364.1_Tengani_62           ........t.............t...........ga........................................a... [ 58880]
AY261363.1_Pretoriuskop/96/4    ........t.............t...........g........................................aa... [ 58880]
AY261362.1_Mkuzi_1979           ...................................a......c..c.................................. [ 58880]
AY261361.1_Malawi_Lil-20/1_1983 ...............................a..t.............................a.c.ta.....ca... [ 58880]
AY261360.1_Kenya_1950           ...................c...........a..ca......c.....................a.c.t....g.ca... [ 58880]
AM712240.1_OURT_88/3            ...................................a............a............................... [ 58880]
AM712239.1_Benin_97/1           ...................................a......c..c.................................. [ 58880]
KM111295.1_Ken06.Bus            ...................c...........a..ca..c.........................a.c.t......ca... [ 58880]
KM111294.1_Ken05/Tk1            ...................c...........a..ca............................a.c.t....g.ca... [ 58880]
KM262844.1_strain_L60           ...................................a......c..c.................................. [ 58880]
KM102979.1_26544/OG10_Italy     ...................................a......c..c.................................. [ 58880]
KM262845.1_strain_NHV           ...................................a............a............................... [ 58880]
KX354450.1_47/Ss/2008           ...................................ac.....c..c.................................. [ 58880]
KP055815.1_strain_BA71          ...................................a......c..c.................................. [ 58880]
MH025920.1_strain_R35           ...................c...........a..ca..c.........................a.c.t......ca... [ 58880]
MH025919.1_strain_N10           ...................c...........a..ca..c.........................a.c.t......ca... [ 58880]
MH025918.1_strain_R25           ...................c...........a..ca..c.........................a.c.t......ca... [ 58880]
MH025917.1_strain_R7            ...................c...........a..ca..c.........................a.c.t......ca... [ 58880]
MH025916.1_strain_R8            ...................c...........a..ca..c.........................a.c.t......ca... [ 58880]
```

```
FR682468.2_Georgia_2007/1        gttctcataaagccttcaagatgactattgatgtttccaatacattttctcaatgagttcataaacccagacattttgct [ 58960]
ASF_IND_20_CAD543                ................................................................................ [ 58960]
LR899193.1_Germany_2020/1        ................................................................................ [ 58960]
MN393477.1_Wuhan_2019            ................................................................................ [ 58960]
MW396979.1_Timor-Leste/2019/1    ................................................................................ [ 58960]
MK940252.1_CN/2019/InnerMongolia ................................................................................ [ 58960]
MK128995.1_China/2018/AnhuiXCGQ  ................................................................................ [ 58960]
MH681419.1_ASFV/POL/2015         ................................................................................ [ 58960]
LS478113.1_Estonia_2014          ................................................................................ [ 58960]
KP843857.1_Odintsovo_02/14       ................................................................................ [ 58960]
FN557520.1_strain_E75            ................................................................................ [ 58960]
AY261366.1_Warthog               ................................................................................ [ 58960]
AY261365.1_Warmbaths             ................................................................................ [ 58960]
AY261364.1_Tengani_62            ................................................................................ [ 58960]
AY261363.1_Pretoriuskop/96/4     ................................................................................ [ 58960]
AY261362.1_Mkuzi_1979            ................................................................................ [ 58960]
AY261361.1_Malawi_Lil-20/1_1983  ...................tt..g.........a............................g.....t.c..-...... [ 58960]
AY261360.1_Kenya_1950            t.........ga........tc.gt.......ca........................g...g..g..t.g......... [ 58960]
AM712240.1_OURT_88/3             ................................................................................ [ 58960]
AM712239.1_Benin_97/1            ................................................................................ [ 58960]
KM111295.1_Ken06.Bus             t.........ga..c.....tc.gt.......ca........................g...g..g..t.gt........ [ 58960]
KM111294.1_Ken05/Tk1             t.........ga........tc.gt.......ca........................g...g..g..t.g......... [ 58960]
KM262844.1_strain_L60            ................................................................................ [ 58960]
KM102979.1_26544/OG10_Italy      ................................................................................ [ 58960]
KM262845.1_strain_NHV            ................................................................................ [ 58960]
KX354450.1_47/Ss/2008            ................................................................................ [ 58960]
KP055815.1_strain_BA71           ................................................................................ [ 58960]
MH025920.1_strain_R35            t.........ga..c.....tcagt.......ca........................g...g..g..t.g......... [ 58960]
MH025919.1_strain_N10            t.........ga..c.....tc.gt.......ca........................g...g..g..t.g......... [ 58960]
MH025918.1_strain_R25            t.........ga..c.....tcagt.......ca........................g...g..g..t.g......... [ 58960]
MH025917.1_strain_R7             t.........ga..c.....tcagt.......ca........................g...g..g..t.g......... [ 58960]
MH025916.1_strain_R8             t.........ga..c.....tcagt.......ca........................g...g..g..t.g......... [ 58960]

FR682468.2_Georgia_2007/1        aatggcttggcaaagtgccaacaagttgtccac--aaagta-ctggtagattgcc-----------actagctatag--- [ 59040]
ASF_IND_20_CAD543                .................................--......-.............tactagctatag..........--- [ 59040]
LR899193.1_Germany_2020/1        .................................--......-.............-----------...........--- [ 59040]
MN393477.1_Wuhan_2019            .................................--......-.............-----------...........--- [ 59040]
MW396979.1_Timor-Leste/2019/1    .................................--......-.............-----------...........--- [ 59040]
MK940252.1_CN/2019/InnerMongolia .................................--......-.............-----------...........--- [ 59040]
MK128995.1_China/2018/AnhuiXCGQ  .................................--......-.............-----------...........--- [ 59040]
MH681419.1_ASFV/POL/2015         .................................--......-.............-----------...........--- [ 59040]
LS478113.1_Estonia_2014          .................................--......-.............-----------...........--- [ 59040]
KP843857.1_Odintsovo_02/14       .................................--......-.............-----------...........--- [ 59040]
```

```
FN557520.1_strain_E75          .................................--......-.............t----------...........--- [ 59040]
AY261366.1_Warthog             .........a.........gg..ga..c.....c-......-...a.........t----------...........--- [ 59040]
AY261365.1_Warmbaths           .........a.........gg..ga........t-......-.............t----------...........--- [ 59040]
AY261364.1_Tengani_62          .........a.........gg..ga..c.....c-......-..aa.........t----------....---------- [ 59040]
AY261363.1_Pretoriuskop/96/4   .........a.........gg..ga........c-......-...a.........t----------...........--- [ 59040]
AY261362.1_Mkuzi_1979          .................................--......-.............t----------...........c-- [ 59040]
AY261361.1_Malawi_Lil-20/1_1983 ...t.t.....gg..........ga...c....ta......-tcaa......a..ta---------ct......ataggt [ 59040]
AY261360.1_Kenya_1950          ...t........g..........ga...c....t-......t..aa......a..ta----------------------- [ 59040]
AM712240.1_OURT_88/3           .................................--......-.............t----------...........--- [ 59040]
AM712239.1_Benin_97/1          .................................--......-.............t----------...........--- [ 59040]
KM111295.1_Ken06.Bus           ...t........g..........ga...c....t-......t..aa......a..ta----------------------- [ 59040]
KM111294.1_Ken05/Tk1           ...t........g..........ga...c....t-......t..aa......a..ta----------------------- [ 59040]
KM262844.1_strain_L60          .................................--......-.............t----------...........--- [ 59040]
KM102979.1_26544/OG10_Italy    .................................--......-.............t----------...........--- [ 59040]
KM262845.1_strain_NHV          .................................--......-.............t----------...........--- [ 59040]
KX354450.1_47/Ss/2008          .................................--......-.............t----------...........--- [ 59040]
KP055815.1_strain_BA71         .................................--......-.............t----------...........--- [ 59040]
MH025920.1_strain_R35          ...t........g..........ga...c....t-......t..aa......a..ta----------------------- [ 59040]
MH025919.1_strain_N10          ...t........g..........ga...c....t-......t..aa......a..ta----------------------- [ 59040]
MH025918.1_strain_R25          ...t........g..........ga...c....t-......t..aa......a..ta----------------------- [ 59040]
MH025917.1_strain_R7           ...t........g..........ga...c....t-......t..aa......a..ta----------------------- [ 59040]
MH025916.1_strain_R8           ...t........g..........ga...c....t-......t..aa......a..ta----------------------- [ 59040]

FR682468.2_Georgia_2007/1      -ctagctatagtgagccaacctctctgtatgtattttatatatttca-ttttttaatagatttaatattttta-taaaaa [ 59120]
ASF_IND_20_CAD543              -.............................k................-.........................-...... [ 59120]
LR899193.1_Germany_2020/1      -..............................................-.........................-...... [ 59120]
MN393477.1_Wuhan_2019          -..............................................-.........................-...... [ 59120]
MW396979.1_Timor-Leste/2019/1  -..............................................-.........................-...... [ 59120]
MK940252.1_CN/2019/InnerMongolia -..............................................-.........................-...... [ 59120]
MK128995.1_China/2018/AnhuiXCGQ -..............................................-.........................-...... [ 59120]
MH681419.1_ASFV/POL/2015       -..............................................-.........................-...... [ 59120]
LS478113.1_Estonia_2014        -..............................................-.........................-...... [ 59120]
KP843857.1_Odintsovo_02/14     -..............................................-.........................-...... [ 59120]
FN557520.1_strain_E75          -.............................t................-.........................-...... [ 59120]
AY261366.1_Warthog             -.............................t...a............-.........................-...... [ 59120]
AY261365.1_Warmbaths           -.............................t...g............-.........................-...... [ 59120]
AY261364.1_Tengani_62          -.............................t...a............-.........................-...... [ 59120]
AY261363.1_Pretoriuskop/96/4   -.............................t...a............-.........................-...... [ 59120]
AY261362.1_Mkuzi_1979          t.............................t................-.........................-...... [ 59120]
AY261361.1_Malawi_Lil-20/1_1983 a..................t......a.............c......-................t..-.....-...... [ 59120]
AY261360.1_Kenya_1950          -.....g...................a...t.........ct.....t................t......at-a..... [ 59120]
AM712240.1_OURT_88/3           -.............................t................-.........................-...... [ 59120]
AM712239.1_Benin_97/1          -.............................t................-.........................-...... [ 59120]
KM111295.1_Ken06.Bus           -...............t.........a...t.........ct.....-.......................at-a..... [ 59120]
```

```
KM111294.1_Ken05/Tk1            -.........................a...t.........ct.....-.......................at-a..... [ 59120]
KM262844.1_strain_L60           -.............................t................-.........................-...... [ 59120]
KM102979.1_26544/OG10_Italy     -.............................t................-.........................-...... [ 59120]
KM262845.1_strain_NHV           -.............................t................-.........................-...... [ 59120]
KX354450.1_47/Ss/2008           -.............................t................-.........................-...... [ 59120]
KP055815.1_strain_BA71          -.............................t................-.........................-...... [ 59120]
MH025920.1_strain_R35           -...............t.........a...t.........ct.....-.......................ataa..... [ 59120]
MH025919.1_strain_N10           -...............t.........a...t.........ct.....-.......................ataa..... [ 59120]
MH025918.1_strain_R25           -...............t.........a...t.........ct.....-.......................ataa..... [ 59120]
MH025917.1_strain_R7            -...............t.........a...t.........ct.....-.......................ataa..... [ 59120]
MH025916.1_strain_R8            -...............t.........a...t.........ct.....-.......................ataa..... [ 59120]

FR682468.2_Georgia_2007/1       aatatttagttttttatacaagaatgtcgacaaaaaaaaagcccacaattaccaagcaagagctttactccttagtagcg [ 59200]
ASF_IND_20_CAD543               ................................................................................ [ 59200]
LR899193.1_Germany_2020/1       ................................................................................ [ 59200]
MN393477.1_Wuhan_2019           ................................................................................ [ 59200]
MW396979.1_Timor-Leste/2019/1   ................................................................................ [ 59200]
MK940252.1_CN/2019/InnerMongolia ................................................................................ [ 59200]
MK128995.1_China/2018/AnhuiXCGQ ................................................................................ [ 59200]
MH681419.1_ASFV/POL/2015        ................................................................................ [ 59200]
LS478113.1_Estonia_2014         ................................................................................ [ 59200]
KP843857.1_Odintsovo_02/14      ................................................................................ [ 59200]
FN557520.1_strain_E75           ................................................................................ [ 59200]
AY261366.1_Warthog              .............................g.................................................. [ 59200]
AY261365.1_Warmbaths            .............................g.................................................. [ 59200]
AY261364.1_Tengani_62           ................................................................................ [ 59200]
AY261363.1_Pretoriuskop/96/4    ................................................................................ [ 59200]
AY261362.1_Mkuzi_1979           ................................................................................ [ 59200]
AY261361.1_Malawi_Lil-20/1_1983 ................................................................................ [ 59200]
AY261360.1_Kenya_1950           ..............................................g........................c....g... [ 59200]
AM712240.1_OURT_88/3            ................................................................................ [ 59200]
AM712239.1_Benin_97/1           ................................................................................ [ 59200]
KM111295.1_Ken06.Bus            ..............................................g........................c........ [ 59200]
KM111294.1_Ken05/Tk1            ..............................................g........................c....g... [ 59200]
KM262844.1_strain_L60           ................................................................................ [ 59200]
KM102979.1_26544/OG10_Italy     ................................................................................ [ 59200]
KM262845.1_strain_NHV           ................................................................................ [ 59200]
KX354450.1_47/Ss/2008           ................................................................................ [ 59200]
KP055815.1_strain_BA71          ................................................................................ [ 59200]
MH025920.1_strain_R35           ..............................................g........................c........ [ 59200]
MH025919.1_strain_N10           ..............................................g........................c........ [ 59200]
MH025918.1_strain_R25           ..............................................g........................c........ [ 59200]
MH025917.1_strain_R7            ..............................................g........................c........ [ 59200]
MH025916.1_strain_R8            ..............................................g........................c........ [ 59200]
```

```
FR682468.2_Georgia_2007/1        gcagatacccagttaaataaagcattgattgaaagaatctttacaagtcagcaaaaaataatacaaaatgctttaaagca [ 59280]
ASF_IND_20_CAD543                ..............................................................m................. [ 59280]
LR899193.1_Germany_2020/1        ................................................................................ [ 59280]
MN393477.1_Wuhan_2019            ................................................................................ [ 59280]
MW396979.1_Timor-Leste/2019/1    ................................................................................ [ 59280]
MK940252.1_CN/2019/InnerMongolia ................................................................................ [ 59280]
MK128995.1_China/2018/AnhuiXCGQ  ................................................................................ [ 59280]
MH681419.1_ASFV/POL/2015         ................................................................................ [ 59280]
LS478113.1_Estonia_2014          ................................................................................ [ 59280]
KP843857.1_Odintsovo_02/14       ................................................................................ [ 59280]
FN557520.1_strain_E75            ...............................................c..............c................. [ 59280]
AY261366.1_Warthog               ..............................................................c................. [ 59280]
AY261365.1_Warmbaths             ..............................................................c................. [ 59280]
AY261364.1_Tengani_62            ..................................................a...........c................. [ 59280]
AY261363.1_Pretoriuskop/96/4     ...............................................c..............c................. [ 59280]
AY261362.1_Mkuzi_1979            ..............................................................c................. [ 59280]
AY261361.1_Malawi_Lil-20/1_1983  ..............................................................c........c........ [ 59280]
AY261360.1_Kenya_1950            .......................t..............t.......................c........c.....a.. [ 59280]
AM712240.1_OURT_88/3             ...............................................c..............c................. [ 59280]
AM712239.1_Benin_97/1            ...............................................c..............c................. [ 59280]
KM111295.1_Ken06.Bus             .......................t..........................a...........c........c.....a.. [ 59280]
KM111294.1_Ken05/Tk1             .......................t..............t.......................c........c.....a.. [ 59280]
KM262844.1_strain_L60            ...............................................c..............c................. [ 59280]
KM102979.1_26544/OG10_Italy      ...............................................c..............c................. [ 59280]
KM262845.1_strain_NHV            ...............................................c..............c................. [ 59280]
KX354450.1_47/Ss/2008            ...............................................c..............c................. [ 59280]
KP055815.1_strain_BA71           ...............................................c..............c................. [ 59280]
MH025920.1_strain_R35            .......................t..........................a...........c........c.....a.. [ 59280]
MH025919.1_strain_N10            .......................t..........................a...........c........c.....a.. [ 59280]
MH025918.1_strain_R25            .......................t..........................a...........c........c.....a.. [ 59280]
MH025917.1_strain_R7             .......................t..........................a...........c........c.....a.. [ 59280]
MH025916.1_strain_R8             .......................t..........................a...........c........c.....a.. [ 59280]

FR682468.2_Georgia_2007/1        caatcaagaagttattataccacccggaatcaagttcaccgtcgttacggtgaaagctaaacctgctcgccagggccata [ 59360]
ASF_IND_20_CAD543                ................................................................................ [ 59360]
LR899193.1_Germany_2020/1        ................................................................................ [ 59360]
MN393477.1_Wuhan_2019            ................................................................................ [ 59360]
MW396979.1_Timor-Leste/2019/1    ................................................................................ [ 59360]
MK940252.1_CN/2019/InnerMongolia ................................................................................ [ 59360]
MK128995.1_China/2018/AnhuiXCGQ  ................................................................................ [ 59360]
MH681419.1_ASFV/POL/2015         ................................................................................ [ 59360]
LS478113.1_Estonia_2014          ................................................................................ [ 59360]
KP843857.1_Odintsovo_02/14       ................................................................................ [ 59360]
```

| | | |
|---|---|---|
| FN557520.1_strain_E75 | ................................................a............................... | [ 59360] |
| AY261366.1_Warthog | ................................................a............................... | [ 59360] |
| AY261365.1_Warmbaths | t...............................................a............................... | [ 59360] |
| AY261364.1_Tengani_62 | ................................................a............................... | [ 59360] |
| AY261363.1_Pretoriuskop/96/4 | ................................................a.................a............. | [ 59360] |
| AY261362.1_Mkuzi_1979 | ................................................a............................... | [ 59360] |
| AY261361.1_Malawi_Lil-20/1_1983 | ............g.............................t..g.................................. | [ 59360] |
| AY261360.1_Kenya_1950 | ............g...................................a..a....................a....... | [ 59360] |
| AM712240.1_OURT_88/3 | ................................................a............................... | [ 59360] |
| AM712239.1_Benin_97/1 | ................................................a............................... | [ 59360] |
| KM111295.1_Ken06.Bus | ............g...................................a..a....................a....... | [ 59360] |
| KM111294.1_Ken05/Tk1 | ............g...................................a..a....................a....... | [ 59360] |
| KM262844.1_strain_L60 | ................................................a............................... | [ 59360] |
| KM102979.1_26544/OG10_Italy | ................................................a............................... | [ 59360] |
| KM262845.1_strain_NHV | ................................................a............................... | [ 59360] |
| KX354450.1_47/Ss/2008 | ................................................a............................... | [ 59360] |
| KP055815.1_strain_BA71 | ................................................a............................... | [ 59360] |
| MH025920.1_strain_R35 | ............g...................................a..a....................a....... | [ 59360] |
| MH025919.1_strain_N10 | ............g...................................a..a....................a....... | [ 59360] |
| MH025918.1_strain_R25 | ............g...................................a..a....................a....... | [ 59360] |
| MH025917.1_strain_R7 | ............g...................................a..a....................a....... | [ 59360] |
| MH025916.1_strain_R8 | ............g...................................a..a....................a....... | [ 59360] |

| | | |
|---|---|---|
| FR682468.2_Georgia_2007/1 | atcccgccacaggagagcctattcaaattaaagctaaacctgaacataaagccgtaaagatacgagcattgaaacctgtc | [ 59440] |
| ASF_IND_20_CAD543 | ................................................................................ | [ 59440] |
| LR899193.1_Germany_2020/1 | ................................................................................ | [ 59440] |
| MN393477.1_Wuhan_2019 | ................................................................................ | [ 59440] |
| MW396979.1_Timor-Leste/2019/1 | ................................................................................ | [ 59440] |
| MK940252.1_CN/2019/InnerMongolia | ................................................................................ | [ 59440] |
| MK128995.1_China/2018/AnhuiXCGQ | ................................................................................ | [ 59440] |
| MH681419.1_ASFV/POL/2015 | ................................................................................ | [ 59440] |
| LS478113.1_Estonia_2014 | ................................................................................ | [ 59440] |
| KP843857.1_Odintsovo_02/14 | ................................................................................ | [ 59440] |
| FN557520.1_strain_E75 | .c.....a..........................c..g.........................................t | [ 59440] |
| AY261366.1_Warthog | .c.....a..........................c..g.......................................... | [ 59440] |
| AY261365.1_Warmbaths | .c.....a..........................c..g.......................................... | [ 59440] |
| AY261364.1_Tengani_62 | .c.....a........a.................c.....a....................................... | [ 59440] |
| AY261363.1_Pretoriuskop/96/4 | .c.....a..........................c..g.......................................... | [ 59440] |
| AY261362.1_Mkuzi_1979 | .c.....a..........................c..g.......................................... | [ 59440] |
| AY261361.1_Malawi_Lil-20/1_1983 | .c.....a................................a................................g...... | [ 59440] |
| AY261360.1_Kenya_1950 | ....g..a................................c...........g.....a...........a........a | [ 59440] |
| AM712240.1_OURT_88/3 | .c.....a..........................c..g.........................................t | [ 59440] |
| AM712239.1_Benin_97/1 | .c.....a..........................c..g.........................................t | [ 59440] |
| KM111295.1_Ken06.Bus | ....g..a................................c...........g.....a....................a | [ 59440] |

```
KM111294.1_Ken05/Tk1            ....g..a................................c...........g.....a....................a [ 59440]
KM262844.1_strain_L60           .c.....a..........................c..g.........................................t [ 59440]
KM102979.1_26544/OG10_Italy     .c.....a..........................c..g.........................................t [ 59440]
KM262845.1_strain_NHV           .c.....a..........................c..g.........................................t [ 59440]
KX354450.1_47/Ss/2008           .c.....a..........................c..g.........................................t [ 59440]
KP055815.1_strain_BA71          .c.....a..........................c..g.........................................t [ 59440]
MH025920.1_strain_R35           ....g..a................................c...........g.....a....................a [ 59440]
MH025919.1_strain_N10           ....g..a................................c...........g.....a....................a [ 59440]
MH025918.1_strain_R25           ....g..a................................c...........g.....a....................a [ 59440]
MH025917.1_strain_R7            ....g..a................................c...........g.....a....................a [ 59440]
MH025916.1_strain_R8            ....g..a................................c...........g.....a....................a [ 59440]

FR682468.2_Georgia_2007/1       catgatatgttaaactaaactataaa------------------------------------------------------ [ 59520]
ASF_IND_20_CAD543               ..............y....ta.....gc--------------catcatcat----------------------------- [ 59520]
LR899193.1_Germany_2020/1       ..........................------------------------------------------------------ [ 59520]
MN393477.1_Wuhan_2019           ..........................------------------------------------------------------ [ 59520]
MW396979.1_Timor-Leste/2019/1   ..........................------------------------------------------------------ [ 59520]
MK940252.1_CN/2019/InnerMongolia ..........................------------------------------------------------------ [ 59520]
MK128995.1_China/2018/AnhuiXCGQ ..........................------------------------------------------------------ [ 59520]
MH681419.1_ASFV/POL/2015        ..........................------------------------------------------------------ [ 59520]
LS478113.1_Estonia_2014         ..........................------------------------------------------------------ [ 59520]
KP843857.1_Odintsovo_02/14      ..........................------------------------------------------------------ [ 59520]
FN557520.1_strain_E75           ..............t....ta.....gc--------------catcatcat----------------------------- [ 59520]
AY261366.1_Warthog              ..c.................g.....g-----------------tcatcat----------------------------- [ 59520]
AY261365.1_Warmbaths            ...................tg.....gc--------------catcatcat----------------------------- [ 59520]
AY261364.1_Tengani_62           ...................tg...tgttaaactaaatgataaagtcattat----------------------------- [ 59520]
AY261363.1_Pretoriuskop/96/4    ...................tg.....g-----------------tcatcat----------------------------- [ 59520]
AY261362.1_Mkuzi_1979           ...................tg.....gc--------------catcatcat----------------------------- [ 59520]
AY261361.1_Malawi_Lil-20/1_1983 ..........................g-----------------ttgttat----------------------------- [ 59520]
AY261360.1_Kenya_1950           .....c............tt.....ttcatattcctcatctataccatcattttcttcaatgcatttttgccaatcaaaa [ 59520]
AM712240.1_OURT_88/3            ..............t....ta.....gc--------------catcatcat----------------------------- [ 59520]
AM712239.1_Benin_97/1           ..............t....ta.....gc--------------catcatcat----------------------------- [ 59520]
KM111295.1_Ken06.Bus            .....c...----------------------------------------------------------------------- [ 59520]
KM111294.1_Ken05/Tk1            .....c............tt.....ttcatattcctcatctataccatcatcttcttcaatgcatttttgccaatcaaaa [ 59520]
KM262844.1_strain_L60           ..............t....ta.....gc--------------catcatcat----------------------------- [ 59520]
KM102979.1_26544/OG10_Italy     ..............t....ta.....gc--------------catcatcat----------------------------- [ 59520]
KM262845.1_strain_NHV           ..............t....ta.....gc--------------catcatcat----------------------------- [ 59520]
KX354450.1_47/Ss/2008           ..............t....ta.....gc--------------catcatcat----------------------------- [ 59520]
KP055815.1_strain_BA71          ..............t....ta.....gc--------------catcatcat----------------------------- [ 59520]
MH025920.1_strain_R35           .....c...----------------------------------------------------------------------- [ 59520]
MH025919.1_strain_N10           .....c...----------------------------------------------------------------------- [ 59520]
MH025918.1_strain_R25           .....c...----------------------------------------------------------------------- [ 59520]
MH025917.1_strain_R7            .....c...----------------------------------------------------------------------- [ 59520]
MH025916.1_strain_R8            .....c...----------------------------------------------------------------------- [ 59520]
```

```
FR682468.2_Georgia_2007/1        -------------------------------------------gtcatattcttctttatcgttattatcttcaatatat [ 59600]
ASF_IND_20_CAD543                ------------------------------------------c...g..............r...............y.. [ 59600]
LR899193.1_Germany_2020/1        -------------------------------------------..................................... [ 59600]
MN393477.1_Wuhan_2019            -------------------------------------------..................................... [ 59600]
MW396979.1_Timor-Leste/2019/1    -------------------------------------------..................................... [ 59600]
MK940252.1_CN/2019/InnerMongolia -------------------------------------------..................................... [ 59600]
MK128995.1_China/2018/AnhuiXCGQ  -------------------------------------------..................................... [ 59600]
MH681419.1_ASFV/POL/2015         -------------------------------------------..................................... [ 59600]
LS478113.1_Estonia_2014          -------------------------------------------..................................... [ 59600]
KP843857.1_Odintsovo_02/14       -------------------------------------------..................................... [ 59600]
FN557520.1_strain_E75            ------------------------------------------c...g..............a...............c.. [ 59600]
AY261366.1_Warthog               ------------------------------------------c..................a...............c.. [ 59600]
AY261365.1_Warmbaths             ------------------------------------------c..................a...............c.. [ 59600]
AY261364.1_Tengani_62            ------------------------------------------ca................ta.c.a......t....c.. [ 59600]
AY261363.1_Pretoriuskop/96/4     ------------------------------------------c..................a...............c.. [ 59600]
AY261362.1_Mkuzi_1979            ------------------------------------------c..................a...............c.. [ 59600]
AY261361.1_Malawi_Lil-20/1_1983  ------------------------------------------ca.................a...............c.. [ 59600]
AY261360.1_Kenya_1950            cggactacatccaggtcatgaacattaaactaatttataatcc..at.cc..a...a.gc.a.c..ct........gc.. [ 59600]
AM712240.1_OURT_88/3             ------------------------------------------c...g..............a...............c.. [ 59600]
AM712239.1_Benin_97/1            ------------------------------------------c...g..............a...............c.. [ 59600]
KM111295.1_Ken06.Bus             ------------------------ttaaattaatttataatcc..gt.cc..a...a.gc.a.c..ct.........c.. [ 59600]
KM111294.1_Ken05/Tk1             cggactacatctaggtcatgaacattaaactaatttataatcc..at.cc..a...a.gc.a.c..ct........gc.. [ 59600]
KM262844.1_strain_L60            ------------------------------------------c...g..............a...............c.. [ 59600]
KM102979.1_26544/OG10_Italy      ------------------------------------------c...g..............a...............c.. [ 59600]
KM262845.1_strain_NHV            ------------------------------------------c...g..............a...............c.. [ 59600]
KX354450.1_47/Ss/2008            ------------------------------------------c...g..............a...............c.. [ 59600]
KP055815.1_strain_BA71           ------------------------------------------c...g..............a...............c.. [ 59600]
MH025920.1_strain_R35            ------------------------ttaaattaatttataatcc..gt.cc..a...a.gc.a.c..ct.........c.. [ 59600]
MH025919.1_strain_N10            ------------------------ttaaattaatttataatcc..gt.cc..a...a.gc.a.c..ct.........c.. [ 59600]
MH025918.1_strain_R25            ------------------------ttaaattaatttataatcc..gt.cc..a...a.gc.a.c..ct.........c.. [ 59600]
MH025917.1_strain_R7             ------------------------ttaaattaatttataatcc..gt.cc..a...a.gc.a.c..ct.........c.. [ 59600]
MH025916.1_strain_R8             ------------------------ttaaattaatttataatcc..gt.cc..a...a.gc.a.c..ct.........c.. [ 59600]

FR682468.2_Georgia_2007/1        ttttgccaatcgaaatcgaataaattcagatcctggacatttaaatacttatcatcgtacattttaatataatttaaaca [ 59680]
ASF_IND_20_CAD543                ................................................................................ [ 59680]
LR899193.1_Germany_2020/1        ................................................................................ [ 59680]
MN393477.1_Wuhan_2019            ................................................................................ [ 59680]
MW396979.1_Timor-Leste/2019/1    ................................................................................ [ 59680]
MK940252.1_CN/2019/InnerMongolia ................................................................................ [ 59680]
MK128995.1_China/2018/AnhuiXCGQ  ................................................................................ [ 59680]
MH681419.1_ASFV/POL/2015         ................................................................................ [ 59680]
LS478113.1_Estonia_2014          ................................................................................ [ 59680]
KP843857.1_Odintsovo_02/14       ................................................................................ [ 59680]
```

```
FN557520.1_strain_E75          ................................................................................ [ 59680]
AY261366.1_Warthog             ........t....................t..t..a......................................c..... [ 59680]
AY261365.1_Warmbaths           .............................t..t..a......................................c..... [ 59680]
AY261364.1_Tengani_62          ............................a......a...........................c............t... [ 59680]
AY261363.1_Pretoriuskop/96/4   ........t..a.................t..t..a......................................c..... [ 59680]
AY261362.1_Mkuzi_1979          ................................................................................ [ 59680]
AY261361.1_Malawi_Lil-20/1_1983 ...................t.........g..t..a.....a.g...........................t..c..... [ 59680]
AY261360.1_Kenya_1950          ...........a....g..t..g..ct.ag..g..a.....a......a..c....a..............g.c....tt [ 59680]
AM712240.1_OURT_88/3           ................................................................................ [ 59680]
AM712239.1_Benin_97/1          ................................................................................ [ 59680]
KM111295.1_Ken06.Bus           ...........a...cg..t.....ct.ag..g..a.t...a......a..c....a..............g.c....tt [ 59680]
KM111294.1_Ken05/Tk1           ...........a...cg..t..g..ct.ag..g..a.t...a......a..c....a..............g.c....tt [ 59680]
KM262844.1_strain_L60          ................................................................................ [ 59680]
KM102979.1_26544/OG10_Italy    ................................................................................ [ 59680]
KM262845.1_strain_NHV          ................................................................................ [ 59680]
KX354450.1_47/Ss/2008          ................................................................................ [ 59680]
KP055815.1_strain_BA71         ..................................a............................................. [ 59680]
MH025920.1_strain_R35          ...........a...cg..t.....ct.ag..g..a.t...a......a..c....a..............g.c....tt [ 59680]
MH025919.1_strain_N10          ...........a...cg..t.....ct.ag..g..a.t...a......a..c....a..............g.c....tt [ 59680]
MH025918.1_strain_R25          ...........a...cg..t.....ct.ag..g..a.t...a......a..c....a..............g.c....tt [ 59680]
MH025917.1_strain_R7           ...........a...cg..t.....ct.ag..g..a.t...a......a..c....a..............g.c....tt [ 59680]
MH025916.1_strain_R8           ...........a...cg..t.....ct.ag..g..a.t...a......a..c....a..............g.c....tt [ 59680]

FR682468.2_Georgia_2007/1      tgagttgttgtcaaaaacttttagcgtttttgttaaaattatcatatgaataatttccttattaagagttgccggaataa [ 59760]
ASF_IND_20_CAD543              .......................................c..tg.................................... [ 59760]
LR899193.1_Germany_2020/1      ................................................................................ [ 59760]
MN393477.1_Wuhan_2019          ................................................................................ [ 59760]
MW396979.1_Timor-Leste/2019/1  ................................................................................ [ 59760]
MK940252.1_CN/2019/InnerMongolia ................................................................................ [ 59760]
MK128995.1_China/2018/AnhuiXCGQ ................................................................................ [ 59760]
MH681419.1_ASFV/POL/2015       ................................................................................ [ 59760]
LS478113.1_Estonia_2014        ................................................................................ [ 59760]
KP843857.1_Odintsovo_02/14     ................................................................................ [ 59760]
FN557520.1_strain_E75          .......................t...............c..tg.................................... [ 59760]
AY261366.1_Warthog             .......................t...............c..tg...................................g [ 59760]
AY261365.1_Warmbaths           .......................t...............c..tg.................................... [ 59760]
AY261364.1_Tengani_62          .........c.............t..................tg.......g............................ [ 59760]
AY261363.1_Pretoriuskop/96/4   .......................t...............c..tg.................................... [ 59760]
AY261362.1_Mkuzi_1979          .......................t...............c..tg.................................... [ 59760]
AY261361.1_Malawi_Lil-20/1_1983 .......................a..............cg..at...a...................tg.........g. [ 59760]
AY261360.1_Kenya_1950          .a....t..c.............t........ag.....g...t...c...t...............t....a......g [ 59760]
AM712240.1_OURT_88/3           .......................t...............c..tg.................................... [ 59760]
AM712239.1_Benin_97/1          .......................t...............c..tg.................................... [ 59760]
KM111295.1_Ken06.Bus           .a....t..c.............t........ag.....g...t...c...t...............t....a....... [ 59760]
```

```
KM111294.1_Ken05/Tk1             .a....t..c.............t........ag.....g...t...c...t...............t....a......g [ 59760]
KM262844.1_strain_L60            .......................t...............c..tg.................................... [ 59760]
KM102979.1_26544/OG10_Italy      .......................t...............c..tg.................................... [ 59760]
KM262845.1_strain_NHV            .......................t...............c..tg.................................... [ 59760]
KX354450.1_47/Ss/2008            .......................t...............c..tg.................................... [ 59760]
KP055815.1_strain_BA71           .......................t...............c..tg.................................... [ 59760]
MH025920.1_strain_R35            .a....t..c.............t........ag.....g...t...c...t...............t....a....... [ 59760]
MH025919.1_strain_N10            .a....t..c.............t........ag.....g...t...c...t...............t....a....... [ 59760]
MH025918.1_strain_R25            .a....t..c.............t........ag.....g...t...c...t...............t....a....... [ 59760]
MH025917.1_strain_R7             .a....t..c.............t........ag.....g...t...c...t...............t....a....... [ 59760]
MH025916.1_strain_R8             .a....t..c.............t........ag.....g...t...c...t...............t....a....... [ 59760]

FR682468.2_Georgia_2007/1        tacaaaacctatttttaggtacatcatccatgataatagtaaaattagtaaaaattgtttcttgtttttcttttgtttca [ 59840]
ASF_IND_20_CAD543                ................................................................................ [ 59840]
LR899193.1_Germany_2020/1        ................................................................................ [ 59840]
MN393477.1_Wuhan_2019            ................................................................................ [ 59840]
MW396979.1_Timor-Leste/2019/1    ................................................................................ [ 59840]
MK940252.1_CN/2019/InnerMongolia ................................................................................ [ 59840]
MK128995.1_China/2018/AnhuiXCGQ  ................................................................................ [ 59840]
MH681419.1_ASFV/POL/2015         ................................................................................ [ 59840]
LS478113.1_Estonia_2014          ................................................................................ [ 59840]
KP843857.1_Odintsovo_02/14       ................................................................................ [ 59840]
FN557520.1_strain_E75            ......g....................t.................................................... [ 59840]
AY261366.1_Warthog               ......g....................t.................................................... [ 59840]
AY261365.1_Warmbaths             ......g....................t.................................................... [ 59840]
AY261364.1_Tengani_62            ..t...g....................t.................................................... [ 59840]
AY261363.1_Pretoriuskop/96/4     ......g....................t..................c................................. [ 59840]
AY261362.1_Mkuzi_1979            ......g....................t.................................................... [ 59840]
AY261361.1_Malawi_Lil-20/1_1983  .......tt..................t.......g.ga........t........a..........a.....ctc.... [ 59840]
AY261360.1_Kenya_1950            c......tt.............t....tt......t..a....t.g..........a...............c....... [ 59840]
AM712240.1_OURT_88/3             ......g....................t.................................................... [ 59840]
AM712239.1_Benin_97/1            ......g....................t.................................................... [ 59840]
KM111295.1_Ken06.Bus             ........t.............t....tt......t..a....t.g..........a...............c....... [ 59840]
KM111294.1_Ken05/Tk1             c......tt...c.........t....tt......t..a....t.g..........a...............c....... [ 59840]
KM262844.1_strain_L60            ......g....................t.................................................... [ 59840]
KM102979.1_26544/OG10_Italy      ......g....................t.................................................... [ 59840]
KM262845.1_strain_NHV            ......g....................t.................................................... [ 59840]
KX354450.1_47/Ss/2008            ......g....................t.................................................... [ 59840]
KP055815.1_strain_BA71           ......g....................t.................................................... [ 59840]
MH025920.1_strain_R35            ........t.............t....tt......t..a....t.g..........a...............c....... [ 59840]
MH025919.1_strain_N10            ........t.............t....tt......t..a....t.g..........a...............c....... [ 59840]
MH025918.1_strain_R25            ........t.............t....tt......t..a....t.g..........a...............c....... [ 59840]
MH025917.1_strain_R7             ........t.............t....tt......t..a....t.g..........a...............c....... [ 59840]
MH025916.1_strain_R8             ........t.............t....tt......t..a....t.g..........a...............c....... [ 59840]
```

```
FR682468.2_Georgia_2007/1        aataaacgttgtaaggttaaaggtttctcgttcaatggtttctttgaagataaaaagaatgtataatctggtttaaaggt [ 59920]
ASF_IND_20_CAD543                ..........................y..r.................................................. [ 59920]
LR899193.1_Germany_2020/1        ................................................................................ [ 59920]
MN393477.1_Wuhan_2019            ................................................................................ [ 59920]
MW396979.1_Timor-Leste/2019/1    ................................................................................ [ 59920]
MK940252.1_CN/2019/InnerMongolia ................................................................................ [ 59920]
MK128995.1_China/2018/AnhuiXCGQ  ................................................................................ [ 59920]
MH681419.1_ASFV/POL/2015         ................................................................................ [ 59920]
LS478113.1_Estonia_2014          ................................................................................ [ 59920]
KP843857.1_Odintsovo_02/14       ................................................................................ [ 59920]
FN557520.1_strain_E75            ..........................t..a......................................c.........a. [ 59920]
AY261366.1_Warthog               ...........c..............t..a......................................c........... [ 59920]
AY261365.1_Warmbaths             ..........................t..a......................................c.........a. [ 59920]
AY261364.1_Tengani_62            ..........................t..................................................... [ 59920]
AY261363.1_Pretoriuskop/96/4     ..........................t..a......................................c.........a. [ 59920]
AY261362.1_Mkuzi_1979            ..........................t..a......................................c.........a. [ 59920]
AY261361.1_Malawi_Lil-20/1_1983  .....t.......t.t.a..g.....t..t...........g..........................c........a.. [ 59920]
AY261360.1_Kenya_1950            t.c..t........atg...g.....t..a.......a...g...g...............g.....c..g..g..... [ 59920]
AM712240.1_OURT_88/3             ..........................t..a......................................c.........a. [ 59920]
AM712239.1_Benin_97/1            ..........................t..a......................................c.........a. [ 59920]
KM111295.1_Ken06.Bus             t.c..t........atg...g.....t..a..t........g....................g.....c..a..g..a.. [ 59920]
KM111294.1_Ken05/Tk1             t....t........atg...g.....t..a...........g....................g........g..g..... [ 59920]
KM262844.1_strain_L60            ..........................t..a......................................c.........a. [ 59920]
KM102979.1_26544/OG10_Italy      ..........................t..a......................................c.........a. [ 59920]
KM262845.1_strain_NHV            ..........................t..a......................................c.........a. [ 59920]
KX354450.1_47/Ss/2008            ..........................t..a......................................c.........a. [ 59920]
KP055815.1_strain_BA71           ..........................t..a......................................c.........a. [ 59920]
MH025920.1_strain_R35            t.c..t........atg...g.....t..a..t........g....................g.....c..a..g..a.. [ 59920]
MH025919.1_strain_N10            t.c..t........atg...g.....t..a..t........g....................g.....c..a..g..a.. [ 59920]
MH025918.1_strain_R25            t.c..t........atg...g.....t..a..t........g....................g.....c..a..g..a.. [ 59920]
MH025917.1_strain_R7             t.c..t........atg...g.....t..a..t........g....................g.....c..a..g..a.. [ 59920]
MH025916.1_strain_R8             t.c..t........atg...g.....t..a..t........g....................g.....c..a..g..a.. [ 59920]

FR682468.2_Georgia_2007/1        atttttggtttcaatcgtgattccatctgcttgagcatatactaaaccagaccaaatataacggtccactattacaatat [ 60000]
ASF_IND_20_CAD543                ...............................................................................w [ 60000]
LR899193.1_Germany_2020/1        ................................................................................ [ 60000]
MN393477.1_Wuhan_2019            ................................................................................ [ 60000]
MW396979.1_Timor-Leste/2019/1    ................................................................................ [ 60000]
MK940252.1_CN/2019/InnerMongolia ................................................................................ [ 60000]
MK128995.1_China/2018/AnhuiXCGQ  ................................................................................ [ 60000]
MH681419.1_ASFV/POL/2015         ................................................................................ [ 60000]
LS478113.1_Estonia_2014          ................................................................................ [ 60000]
KP843857.1_Odintsovo_02/14       ................................................................................ [ 60000]
```

```
FN557520.1_strain_E75              ........................................g....................t.................a [ 60000]
AY261366.1_Warthog                 .......................................cg......................................a [ 60000]
AY261365.1_Warmbaths               .......c................................g......................................a [ 60000]
AY261364.1_Tengani_62              ........................................g....c.................................. [ 60000]
AY261363.1_Pretoriuskop/96/4       .......................................cg......................................a [ 60000]
AY261362.1_Mkuzi_1979              ........................................g......................................a [ 60000]
AY261361.1_Malawi_Lil-20/1_1983    ..c.....................................g.c..t..g....................a.........a [ 60000]
AY261360.1_Kenya_1950              .........a......a.t....tg...............g.c..........................a.......... [ 60000]
AM712240.1_OURT_88/3               ........................................g....................t.................a [ 60000]
AM712239.1_Benin_97/1              ........................................g....................t.................a [ 60000]
KM111295.1_Ken06.Bus               .........a......a.t....tg...............g.c..........................a.........a [ 60000]
KM111294.1_Ken05/Tk1               .........a......a.t....tg...............g.c..........................a.........a [ 60000]
KM262844.1_strain_L60              ........................................g....................t.................a [ 60000]
KM102979.1_26544/OG10_Italy        ........................................g....................t.................a [ 60000]
KM262845.1_strain_NHV              ........................................g....................t.................a [ 60000]
KX354450.1_47/Ss/2008              ........................................g....................t.................a [ 60000]
KP055815.1_strain_BA71             ........................................g....................t.................a [ 60000]
MH025920.1_strain_R35              .........a......a.t....tg...............g.c..........................a.........a [ 60000]
MH025919.1_strain_N10              .........a......a.t....tg...............g.c..........................a.........a [ 60000]
MH025918.1_strain_R25              .........a......a.t....tg...............g.c..........................a.........a [ 60000]
MH025917.1_strain_R7               .........a......a.t....tg...............g.c..........................a.........a [ 60000]
MH025916.1_strain_R8               .........a......a.t....tg...............g.c..........................a.........a [ 60000]

FR682468.2_Georgia_2007/1          aatttagcttaagtagcactgcaatttctgcgataaattcactacgatgttttgtaaataatttatgtaattgttccgat [ 60080]
ASF_IND_20_CAD543                  .......y........................................................................ [ 60080]
LR899193.1_Germany_2020/1          ................................................................................ [ 60080]
MN393477.1_Wuhan_2019              ................................................................................ [ 60080]
MW396979.1_Timor-Leste/2019/1      ................................................................................ [ 60080]
MK940252.1_CN/2019/InnerMongolia   ................................................................................ [ 60080]
MK128995.1_China/2018/AnhuiXCGQ    ................................................................................ [ 60080]
MH681419.1_ASFV/POL/2015           ................................................................................ [ 60080]
LS478113.1_Estonia_2014            ................................................................................ [ 60080]
KP843857.1_Odintsovo_02/14         ................................................................................ [ 60080]
FN557520.1_strain_E75              .......t........tg..............c...........t......................c............ [ 60080]
AY261366.1_Warthog                 .......t........tg...............c...........a.....................c............ [ 60080]
AY261365.1_Warmbaths               .......t........tg..............c..................................c............ [ 60080]
AY261364.1_Tengani_62              ...................................................................c............ [ 60080]
AY261363.1_Pretoriuskop/96/4       .......t........tg...............c...........ag....................c............ [ 60080]
AY261362.1_Mkuzi_1979              .......t........tg..............c..................................c............ [ 60080]
AY261361.1_Malawi_Lil-20/1_1983    ....c..t.......atg............t..a.................................c............ [ 60080]
AY261360.1_Kenya_1950              .......t........tg.............cg.........at...........g.......c...c........t... [ 60080]
AM712240.1_OURT_88/3               .......t........tg..............c...........t......................c............ [ 60080]
AM712239.1_Benin_97/1              .......t........tg..............c...........t......................c............ [ 60080]
KM111295.1_Ken06.Bus               .......t........tg.............cg.........at...........g......ccg..c....c...t... [ 60080]
```

```
KM111294.1_Ken05/Tk1          .......t........tg.............cg.........at...........g.......c...c........t... [ 60080]
KM262844.1_strain_L60         .......t........tg..............c...........t......................c............ [ 60080]
KM102979.1_26544/OG10_Italy   .......t........tg..............c...........t......................c............ [ 60080]
KM262845.1_strain_NHV         .......t........tg..............c...........t......................c............ [ 60080]
KX354450.1_47/Ss/2008         .......t........tg..............c...........t......................c............ [ 60080]
KP055815.1_strain_BA71        .......t........tg..............c...........t......................c............ [ 60080]
MH025920.1_strain_R35         .......t........tg.............cg.........at...........g......ccg..c....c...t... [ 60080]
MH025919.1_strain_N10         .......t........tg.............cg.........at...........g......ccg..c....c...t... [ 60080]
MH025918.1_strain_R25         .......t........tg.............cg.........at...........g......ccg..c....c...t... [ 60080]
MH025917.1_strain_R7          .......t........tg.............cg.........at...........g......ccg..c....c...t... [ 60080]
MH025916.1_strain_R8          .......t........tg.............cg.........at...........g......ccg..c....c...t... [ 60080]

FR682468.2_Georgia_2007/1     gacatttctatggttttatttaacacctgcaatataagatcaccggtggtcgtgtctggattaggaaaatg----tatac [ 60160]
ASF_IND_20_CAD543             .........g.ca..........................................................----..... [ 60160]
LR899193.1_Germany_2020/1     .......................................................................----..... [ 60160]
MN393477.1_Wuhan_2019         .......................................................................----..... [ 60160]
MW396979.1_Timor-Leste/2019/1 .......................................................................----..... [ 60160]
MK940252.1_CN/2019/InnerMongolia .......................................................................----..... [ 60160]
MK128995.1_China/2018/AnhuiXCGQ .......................................................................----..... [ 60160]
MH681419.1_ASFV/POL/2015      .......................................................................----..... [ 60160]
LS478113.1_Estonia_2014       .......................................................................----..... [ 60160]
KP843857.1_Odintsovo_02/14    .......................................................................----..... [ 60160]
FN557520.1_strain_E75         .........g.ca.........................gc.............................c.----..... [ 60160]
AY261366.1_Warthog            ...........ca.........................g..............................c.----..... [ 60160]
AY261365.1_Warmbaths          ...........ca.........................g..............................c.----..... [ 60160]
AY261364.1_Tengani_62         .......................................................................----..... [ 60160]
AY261363.1_Pretoriuskop/96/4  ...........ca.........................g..............................c.----..... [ 60160]
AY261362.1_Mkuzi_1979         ..................c...................g.t..............................----..... [ 60160]
AY261361.1_Malawi_Lil-20/1_1983 .....c....ca.......c...t.....g........g.t.......ac...a.................cata....t [ 60160]
AY261360.1_Kenya_1950         .g.....t..c.a..........t..a.c.........c.t......atc.....t....c.g........----c.a.t [ 60160]
AM712240.1_OURT_88/3          .........g.ca.........................gc.............................c.----..... [ 60160]
AM712239.1_Benin_97/1         .........g.ca.........................gc.............................c.----..... [ 60160]
KM111295.1_Ken06.Bus          .g.....t..c.a..........t..a.c.........c.t.......tc.....t....c.g..-.....----c...t [ 60160]
KM111294.1_Ken05/Tk1          .g.....t..c.a..........t..a.c.........c.t......atc.....t....c.g........----c...t [ 60160]
KM262844.1_strain_L60         .........g.ca.........................gc.............................c.----..... [ 60160]
KM102979.1_26544/OG10_Italy   .........g.ca.........................gc.............................c.----..... [ 60160]
KM262845.1_strain_NHV         .........g.ca.........................gc.............................c.----..... [ 60160]
KX354450.1_47/Ss/2008         .........g.ca.........................gc.............................c.----..... [ 60160]
KP055815.1_strain_BA71        .........g.ca.........................gc.............................c.----..... [ 60160]
MH025920.1_strain_R35         .g.....t..c.a..........t..a.c.........c.t.......tc.....t....c.g........----c...t [ 60160]
MH025919.1_strain_N10         .g.....t..c.a..........t..a.c.........c.t.......tc.....t....c.g........----c...t [ 60160]
MH025918.1_strain_R25         .g.....t..c.a..........t..a.c.........c.t.......tc.....t....c.g........----c...t [ 60160]
MH025917.1_strain_R7          .g.....t..c.a..........t..a.c.........c.t.......tc.....t....c.g........----c...t [ 60160]
MH025916.1_strain_R8          .g.....t..c.a..........t..a.c.........c.t.......tc.....t....c.g........----c...t [ 60160]
```

```
FR682468.2_Georgia_2007/1        atatagcattataatccatgcattccaatgtttcttttaatttcattgcctgtgtgctttttcccacaccattgattccc [ 60240]
ASF_IND_20_CAD543                ..............................cc.t.......y...................................... [ 60240]
LR899193.1_Germany_2020/1        ................................................................................ [ 60240]
MN393477.1_Wuhan_2019            ................................................................................ [ 60240]
MW396979.1_Timor-Leste/2019/1    ................................................................................ [ 60240]
MK940252.1_CN/2019/InnerMongolia ................................................................................ [ 60240]
MK128995.1_China/2018/AnhuiXCGQ  ................................................................................ [ 60240]
MH681419.1_ASFV/POL/2015         ................................................................................ [ 60240]
LS478113.1_Estonia_2014          ................................................................................ [ 60240]
KP843857.1_Odintsovo_02/14       ................................................................................ [ 60240]
FN557520.1_strain_E75            ...c.....................t....cc.t.......c...................................... [ 60240]
AY261366.1_Warthog               ...c.....................t....cc.t.......c...................................... [ 60240]
AY261365.1_Warmbaths             ...c.....................t....cc.t.......c....g.....c........................... [ 60240]
AY261364.1_Tengani_62            .........................................................................a...... [ 60240]
AY261363.1_Pretoriuskop/96/4     ...c.....................t....cc.t.......c...........................g.....c.... [ 60240]
AY261362.1_Mkuzi_1979            ............................c.cc.t.......c....g.....c........................... [ 60240]
AY261361.1_Malawi_Lil-20/1_1983  .............t................ac.t.......c....c................................. [ 60240]
AY261360.1_Kenya_1950            ....ca...c...tct.t.att........ca.t.....gcc.t.................ct.......g........t [ 60240]
AM712240.1_OURT_88/3             ...c.....................t....cc.t.......c...................................... [ 60240]
AM712239.1_Benin_97/1            ...c.....................t....cc.t.......c...................................... [ 60240]
KM111295.1_Ken06.Bus             ....ca...c...tct.t.att........ca.t.....gc..tc....t...........c.................t [ 60240]
KM111294.1_Ken05/Tk1             ....ca...c...tct.t.att........ca.t......cc.t.................c........g........t [ 60240]
KM262844.1_strain_L60            ...c.....................t....cc.t.......c...................................... [ 60240]
KM102979.1_26544/OG10_Italy      ...c.....................t....cc.t.......c...................................... [ 60240]
KM262845.1_strain_NHV            ...c.....................t....cc.t.......c...................................... [ 60240]
KX354450.1_47/Ss/2008            ...c.....................t....cc.t.......c...................................... [ 60240]
KP055815.1_strain_BA71           ...c.....................t....cc.t.......c...................................... [ 60240]
MH025920.1_strain_R35            ....ca...c...tct.t.att........ca.t.....gc..tc....t...........c.................t [ 60240]
MH025919.1_strain_N10            ....ca...c...tct.t.att........ca.t.....gc..tc....t...........c.................t [ 60240]
MH025918.1_strain_R25            ....ca...c...tct.t.att........ca.t.....gc..tc....t...........c.................t [ 60240]
MH025917.1_strain_R7             ....ca...c...tct.t.att........ca.t.....gc..tc....t...........c.................t [ 60240]
MH025916.1_strain_R8             ....ca...c...tct.t.att........ca.t.....gc..tc....t...........c.................t [ 60240]

FR682468.2_Georgia_2007/1        tcgatggcaatgagtattccacgcatgattaataaaggaaaaaaaagaattcagtttttaacatttcttacaaatctttt [ 60320]
ASF_IND_20_CAD543                ................................................................................ [ 60320]
LR899193.1_Germany_2020/1        ................................................................................ [ 60320]
MN393477.1_Wuhan_2019            ................................................................................ [ 60320]
MW396979.1_Timor-Leste/2019/1    ................................................................................ [ 60320]
MK940252.1_CN/2019/InnerMongolia ................................................................................ [ 60320]
MK128995.1_China/2018/AnhuiXCGQ  ................................................................................ [ 60320]
MH681419.1_ASFV/POL/2015         ................................................................................ [ 60320]
LS478113.1_Estonia_2014          ................................................................................ [ 60320]
KP843857.1_Odintsovo_02/14       ................................................................................ [ 60320]
```

```
FN557520.1_strain_E75            .......t...a........................t..gt.......g............................a.. [ 60320]
AY261366.1_Warthog               ...........a........................t..gtg.................--------..........a.. [ 60320]
AY261365.1_Warmbaths             ...........a........................t..gt..................--------.------------ [ 60320]
AY261364.1_Tengani_62            ..................................c.t..gcg...................................a.. [ 60320]
AY261363.1_Pretoriuskop/96/4     ...........a......................c.t..gc.................c--------..........a.. [ 60320]
AY261362.1_Mkuzi_1979            ...........a........................t..gt..................--------..........a.. [ 60320]
AY261361.1_Malawi_Lil-20/1_1983  ...........a...............g........t..-t............a..c..----------......----- [ 60320]
AY261360.1_Kenya_1950            ...........a........................ta.gt.....a......t................t....----- [ 60320]
AM712240.1_OURT_88/3             .......t...a........................t..gt.......g............................a.. [ 60320]
AM712239.1_Benin_97/1            .......t...a........................t..gt.......g............................a.. [ 60320]
KM111295.1_Ken06.Bus             ...........a....c...................c.agt............t................t....----- [ 60320]
KM111294.1_Ken05/Tk1             ...........a........................ta.gt.....-......t................t....----- [ 60320]
KM262844.1_strain_L60            .......t...a........................t..gt.......g............................a.. [ 60320]
KM102979.1_26544/OG10_Italy      .......t...a........................t..gt.......g............................a.. [ 60320]
KM262845.1_strain_NHV            .......t...a........................t..gt.......g............................a.. [ 60320]
KX354450.1_47/Ss/2008            .......t...a........................t..gt.......g............................a.. [ 60320]
KP055815.1_strain_BA71           .......t...a........................t..gt.......g............................a.. [ 60320]
MH025920.1_strain_R35            ...........a....c...................c.agt............t................t....----- [ 60320]
MH025919.1_strain_N10            ...........a....c...................c.agt............t................t....----- [ 60320]
MH025918.1_strain_R25            ...........a....c...................c.agt............t................t....----- [ 60320]
MH025917.1_strain_R7             ...........a....c...................c.agt............t................t....----- [ 60320]
MH025916.1_strain_R8             ...........a....c...................c.agt............t................t....----- [ 60320]

FR682468.2_Georgia_2007/1        tt--------------------tatacaacattgtacaacactgcattagcggtatatgatgttatagcttcatt----a [ 60400]
ASF_IND_20_CAD543                ..--------------------...........................a.c.c................n....----. [ 60400]
LR899193.1_Germany_2020/1        ..--------------------.....................................................----. [ 60400]
MN393477.1_Wuhan_2019            ..--------------------.....................................................----. [ 60400]
MW396979.1_Timor-Leste/2019/1    ..--------------------.....................................................----. [ 60400]
MK940252.1_CN/2019/InnerMongolia ..--------------------.....................................................----. [ 60400]
MK128995.1_China/2018/AnhuiXCGQ  ..--------------------.....................................................----. [ 60400]
MH681419.1_ASFV/POL/2015         ..--------------------.....................................................----. [ 60400]
LS478113.1_Estonia_2014          ..--------------------.....................................................----. [ 60400]
KP843857.1_Odintsovo_02/14       ..--------------------.....................................................----. [ 60400]
FN557520.1_strain_E75            ..--------------------.....................a.....a.c.c...c.................----. [ 60400]
AY261366.1_Warthog               ..--------------------.....................a.....a.c.c...c.................----. [ 60400]
AY261365.1_Warmbaths             ------------------------...................a.....a.c.c...c................c----. [ 60400]
AY261364.1_Tengani_62            ..tgtacaacagtgtacaacat.g...................a.....a.c.c...c.............t...----. [ 60400]
AY261363.1_Pretoriuskop/96/4     ..--------------------.....................a...g.a.c.c...c.................----. [ 60400]
AY261362.1_Mkuzi_1979            ..--------------------.....................a.....a.c.c...c................c----. [ 60400]
AY261361.1_Malawi_Lil-20/1_1983  ---------------------g.....g.-----...gt.g..........c.c.a.c...........c.....----. [ 60400]
AY261360.1_Kenya_1950            ---------------------------c.t.................c.a.c.c----------...........aaaa. [ 60400]
AM712240.1_OURT_88/3             ..--------------------.....................a.....a.c.c...c.................----. [ 60400]
AM712239.1_Benin_97/1            ..--------------------.....................a.....a.c.c...c.................----. [ 60400]
KM111295.1_Ken06.Bus             --------------------------ac.t.................c...c.c----------........g..-aaa. [ 60400]
```

```
KM111294.1_Ken05/Tk1            ---------------------------c.t.................c.a.c.c----------...........-aaa. [ 60400]
KM262844.1_strain_L60           ..--------------------.....................a.....a.c.c...c.................----. [ 60400]
KM102979.1_26544/OG10_Italy     ..--------------------.....................a.....a.c.c...c.................----. [ 60400]
KM262845.1_strain_NHV           ..--------------------.....................a.....a.c.c...c.................----. [ 60400]
KX354450.1_47/Ss/2008           ..--------------------.....................a.....a.c.c...c.................----. [ 60400]
KP055815.1_strain_BA71          ..--------------------.....................a.....a.c.c...c.................----. [ 60400]
MH025920.1_strain_R35           --------------------------ac.t.................c...c.c----------........g..-aaa. [ 60400]
MH025919.1_strain_N10           --------------------------ac.t.................c...c.c----------........g..-aaa. [ 60400]
MH025918.1_strain_R25           --------------------------ac.t.................c...c.c----------........g..-aaa. [ 60400]
MH025917.1_strain_R7            --------------------------ac.t.................c...c.c----------........g..-aaa. [ 60400]
MH025916.1_strain_R8            --------------------------ac.t.................c...c.c----------........g..-aaa. [ 60400]

FR682468.2_Georgia_2007/1       aatatt------tgc-ttttatataatctttaccaacctata------------tttggtag-----atcactgcagat- [ 60480]
ASF_IND_20_CAD543               ......------...-..........................------------........-----............- [ 60480]
LR899193.1_Germany_2020/1       ......------...-..........................------------........-----............- [ 60480]
MN393477.1_Wuhan_2019           ......------...-..........................------------........-----............- [ 60480]
MW396979.1_Timor-Leste/2019/1   ......------...-..........................------------........-----............- [ 60480]
MK940252.1_CN/2019/InnerMongolia ......------...-..........................------------........-----............- [ 60480]
MK128995.1_China/2018/AnhuiXCGQ ......------...-..........................------------........-----............- [ 60480]
MH681419.1_ASFV/POL/2015        ......------...-..........................------------........-----............- [ 60480]
LS478113.1_Estonia_2014         ......------...-..........................------------........-----............- [ 60480]
KP843857.1_Odintsovo_02/14      ......------...-..........................------------........-----............- [ 60480]
FN557520.1_strain_E75           ......------...-..................g.......------------.....c..-----............- [ 60480]
AY261366.1_Warthog              ......------...t...--............tg.......------------.....c..-----............- [ 60480]
AY261365.1_Warmbaths            ......------...t............c.....g.......------------....ac..a--tc............- [ 60480]
AY261364.1_Tengani_62           ......------...-..........................------------.....c..-----............- [ 60480]
AY261363.1_Pretoriuskop/96/4    ......------...t...-..............g.......------------.....c..-----............- [ 60480]
AY261362.1_Mkuzi_1979           ......------...t............c.....g.......------------....ac..a--tc............- [ 60480]
AY261361.1_Malawi_Lil-20/1_1983 .c....ttgcga...-..........c..a..tt..t..t..------------ccaac..tacatata.ttg......- [ 60480]
AY261360.1_Kenya_1950           ...g..gcg-ga...c....g.....c..a..ttc.t..t..ccaacatatatac...ac..-----.cag.cag.t..g [ 60480]
AM712240.1_OURT_88/3            ......------...-..................g.......------------.....c..-----............- [ 60480]
AM712239.1_Benin_97/1           ......------...-..................g.......------------.....c..-----............- [ 60480]
KM111295.1_Ken06.Bus            ...g.agcg-ga...t.......c..c..a..ttc.t..t..-caacatatatac....c.c-----.gag.cat.tc.- [ 60480]
KM111294.1_Ken05/Tk1            ...g..gcg-ga...c....g.....c..a..ttc.t..t..ccaacatatatac....c..-----.cag.cag.t..- [ 60480]
KM262844.1_strain_L60           ......------...-..................g.......------------.....c..-----............- [ 60480]
KM102979.1_26544/OG10_Italy     ......------...-..................g.......------------.....c..-----............- [ 60480]
KM262845.1_strain_NHV           ......------...-..................g.......------------.....c..-----............- [ 60480]
KX354450.1_47/Ss/2008           ......------...-..................g.......------------.....c..-----............- [ 60480]
KP055815.1_strain_BA71          ......------...-..................g.......------------.....c..-----............- [ 60480]
MH025920.1_strain_R35           ...g.agcg-ga...t.......c..c..a..ttc.t..t..-caacatatatac....c.c-----.gag.cat.tc.- [ 60480]
MH025919.1_strain_N10           ...g.agcg-ga...t.......c..c..a..ttc.t..t..-caacatatatac....c.c-----.gag.cat.tc.- [ 60480]
MH025918.1_strain_R25           ...g.agcg-ga...t.......c..c..a..ttc.t..t..-caacatatatac....c.c-----.gag.cat.tc.- [ 60480]
MH025917.1_strain_R7            ...g.agcg-ga...t.......c..c..a..ttc.t..t..-caacatatatac....c.c-----.gag.cat.tc.- [ 60480]
MH025916.1_strain_R8            ...g.agcg-ga...t.......c..c..a..ttc.t..t..-caacatatatac....c.c-----.gag.cat.tc.- [ 60480]
```

```
FR682468.2_Georgia_2007/1        -----------gg---tcataaataggccataactaagataaaaattatttcagacgctactacggtagtattattaaaa [ 60560]
ASF_IND_20_CAD543                -------------------------..............................y........................ [ 60560]
LR899193.1_Germany_2020/1        -----------..---................................................................ [ 60560]
MN393477.1_Wuhan_2019            -----------..---................................................................ [ 60560]
MW396979.1_Timor-Leste/2019/1    -----------..---................................................................ [ 60560]
MK940252.1_CN/2019/InnerMongolia -----------..---................................................................ [ 60560]
MK128995.1_China/2018/AnhuiXCGQ  -----------..---................................................................ [ 60560]
MH681419.1_ASFV/POL/2015         -----------..---................................................................ [ 60560]
LS478113.1_Estonia_2014          -----------..---................................................................ [ 60560]
KP843857.1_Odintsovo_02/14       -----------..---................................................................ [ 60560]
FN557520.1_strain_E75            -------------------------..............................t......g...c............. [ 60560]
AY261366.1_Warthog               -----------..---......c................................t......g...c............. [ 60560]
AY261365.1_Warmbaths             -------------------------..............................t......g...c............. [ 60560]
AY261364.1_Tengani_62            -----------..tca......c...............................c.......g...c............. [ 60560]
AY261363.1_Pretoriuskop/96/4     -----------..---..g...c.......................g........t......g...c............. [ 60560]
AY261362.1_Mkuzi_1979            -------------------------..............................t......g...c............. [ 60560]
AY261361.1_Malawi_Lil-20/1_1983  -----------..---c.....t.g..........................t....a.gg..g.a.cg.---..g..... [ 60560]
AY261360.1_Kenya_1950            ggcccatatct..gc-c....tc.g........t.................t.......g....a.cga---a..c..c. [ 60560]
AM712240.1_OURT_88/3             -------------------------..............................t......g...c............. [ 60560]
AM712239.1_Benin_97/1            -------------------------..............................t......g...c............. [ 60560]
KM111295.1_Ken06.Bus             ------------.gc-c....tc.g........t.................t.......g....a.cga---a..c..c. [ 60560]
KM111294.1_Ken05/Tk1             -----------..gc-c....tc.g........t.................t.......g....a.cga---a..c..c. [ 60560]
KM262844.1_strain_L60            -------------------------..............................t......g...c............. [ 60560]
KM102979.1_26544/OG10_Italy      -------------------------..............................t......g...c............. [ 60560]
KM262845.1_strain_NHV            -------------------------..............................t......g...c............. [ 60560]
KX354450.1_47/Ss/2008            -------------------------..............................t......g...c............. [ 60560]
KP055815.1_strain_BA71           -------------------------..............................t......g...c............. [ 60560]
MH025920.1_strain_R35            ------------.gc-c....tc.g........t.................t.......g....a.cga---a..c..c. [ 60560]
MH025919.1_strain_N10            ------------.gc-c....tc.g........t.................t.......g....a.cga---a..c..c. [ 60560]
MH025918.1_strain_R25            ------------.gc-c....tc.g........t.................t.......g....a.cga---a..c..c. [ 60560]
MH025917.1_strain_R7             ------------.gc-c....tc.g........t.................t.......g....a.cga---a..c..c. [ 60560]
MH025916.1_strain_R8             ------------.gc-c....tc.g........t.................t.......g....a.cga---a..c..c. [ 60560]

FR682468.2_Georgia_2007/1        tcatgtgtgg---------------------------------------------------------------------- [ 60640]
ASF_IND_20_CAD543                ..........---------------------------------------------------------------------- [ 60640]
LR899193.1_Germany_2020/1        ..........---------------------------------------------------------------------- [ 60640]
MN393477.1_Wuhan_2019            ..........---------------------------------------------------------------------- [ 60640]
MW396979.1_Timor-Leste/2019/1    ..........---------------------------------------------------------------------- [ 60640]
MK940252.1_CN/2019/InnerMongolia ..........---------------------------------------------------------------------- [ 60640]
MK128995.1_China/2018/AnhuiXCGQ  ..........---------------------------------------------------------------------- [ 60640]
MH681419.1_ASFV/POL/2015         ..........---------------------------------------------------------------------- [ 60640]
LS478113.1_Estonia_2014          ..........---------------------------------------------------------------------- [ 60640]
KP843857.1_Odintsovo_02/14       ..........---------------------------------------------------------------------- [ 60640]
```

```
FN557520.1_strain_E75            .......c..---------------------------------------------------------------------- [ 60640]
AY261366.1_Warthog               .......c..---------------------------------------------------------------------- [ 60640]
AY261365.1_Warmbaths             .......c..---------------------------------------------------------------------- [ 60640]
AY261364.1_Tengani_62            .......c..---------------------------------------------------------------------- [ 60640]
AY261363.1_Pretoriuskop/96/4     ..........---------------------------------------------------------------------- [ 60640]
AY261362.1_Mkuzi_1979            .......c..---------------------------------------------------------------------- [ 60640]
AY261361.1_Malawi_Lil-20/1_1983  ..........c-acatag--------------------------tgctgcacatagtgccgcacatagtgccgtattata [ 60640]
AY261360.1_Kenya_1950            .t....act.cggcatagcatgtggcatatcatgcggcatagcatgcggcatagcatgcggcatagcatgcc----cata [ 60640]
AM712240.1_OURT_88/3             .......c..---------------------------------------------------------------------- [ 60640]
AM712239.1_Benin_97/1            .......c..---------------------------------------------------------------------- [ 60640]
KM111295.1_Ken06.Bus             .t....ac------------------------------------------------tgcggcatattatgcc----cata [ 60640]
KM111294.1_Ken05/Tk1             .t....ac------------------------------------tgcggcatagcatgcggcatagcatgcc----cata [ 60640]
KM262844.1_strain_L60            .......c..---------------------------------------------------------------------- [ 60640]
KM102979.1_26544/OG10_Italy      .......c..---------------------------------------------------------------------- [ 60640]
KM262845.1_strain_NHV            .......c..---------------------------------------------------------------------- [ 60640]
KX354450.1_47/Ss/2008            .......c..---------------------------------------------------------------------- [ 60640]
KP055815.1_strain_BA71           .......c..---------------------------------------------------------------------- [ 60640]
MH025920.1_strain_R35            .t....ac------------------------------------------------tgcggcatattatgcc----cata [ 60640]
MH025919.1_strain_N10            .t....ac------------------------------------------------tgcggcatattatgcc----cata [ 60640]
MH025918.1_strain_R25            .t....ac------------------------------------------------tgcggcatattatgcc----cata [ 60640]
MH025917.1_strain_R7             .t....ac------------------------------------------------tgcggcatattatgcc----cata [ 60640]
MH025916.1_strain_R8             .t....ac------------------------------------------------tgcggcatattatgcc----cata [ 60640]

FR682468.2_Georgia_2007/1        ---------caatgtatgacgtcttaatagataaaacatttaaggaaaacaaatttgaataaaaaaaataattgttatga [ 60720]
ASF_IND_20_CAD543                ---------...........................m.......k...........................w....... [ 60720]
LR899193.1_Germany_2020/1        ---------....................................................................... [ 60720]
MN393477.1_Wuhan_2019            ---------....................................................................... [ 60720]
MW396979.1_Timor-Leste/2019/1    ---------....................................................................... [ 60720]
MK940252.1_CN/2019/InnerMongolia ---------....................................................................... [ 60720]
MK128995.1_China/2018/AnhuiXCGQ  ---------....................................................................... [ 60720]
MH681419.1_ASFV/POL/2015         ---------....................................................................... [ 60720]
LS478113.1_Estonia_2014          ---------....................................................................... [ 60720]
KP843857.1_Odintsovo_02/14       ---------...................................................-................... [ 60720]
FN557520.1_strain_E75            ---------...........................a.......t...........................a....... [ 60720]
AY261366.1_Warthog               ---------...........................a.......t...........................a....... [ 60720]
AY261365.1_Warmbaths             ---------...........................a......at...........................a....... [ 60720]
AY261364.1_Tengani_62            ---------....................................................................... [ 60720]
AY261363.1_Pretoriuskop/96/4     ---------...................................t................................... [ 60720]
AY261362.1_Mkuzi_1979            ---------...........................a......at...........................a....... [ 60720]
AY261361.1_Malawi_Lil-20/1_1983  ggcaaccta....a.......c...g..................t................................... [ 60720]
AY261360.1_Kenya_1950            agcaatcta....a...a...-......gc..............t.................................a. [ 60720]
AM712240.1_OURT_88/3             ---------...........................a.......t...........................a....... [ 60720]
AM712239.1_Benin_97/1            ---------...........................a.......t...........................a....... [ 60720]
KM111295.1_Ken06.Bus             agtaatctg...................gc..............t.................................a. [ 60720]
```

```
KM111294.1_Ken05/Tk1             agcaatcta........a...-......gc..............t.................................a. [ 60720]
KM262844.1_strain_L60            ---------...........................a.......t...........................a....... [ 60720]
KM102979.1_26544/OG10_Italy      ---------...........................a.......t...........................a....... [ 60720]
KM262845.1_strain_NHV            ---------...........................a.......t...........................a....... [ 60720]
KX354450.1_47/Ss/2008            ---------...........................a.......t...........................a....... [ 60720]
KP055815.1_strain_BA71           ---------...........................a.......t...........................a....... [ 60720]
MH025920.1_strain_R35            agtaatctg...................gc..............t.................................a. [ 60720]
MH025919.1_strain_N10            agtaatctg...................gc..............t.................................a. [ 60720]
MH025918.1_strain_R25            agtaatctg...................gc..............t.................................a. [ 60720]
MH025917.1_strain_R7             agtaatctg...................gc..............t.................................a. [ 60720]
MH025916.1_strain_R8             agtaatctg...................gc..............t.................................a. [ 60720]

FR682468.2_Georgia_2007/1        tggcgttgttacacaaagaaaagcttatagagtgcatctatcatgagctagaaaatggcgggacaatattgcttctaaca [ 60800]
ASF_IND_20_CAD543                ......................................g.aa.....g.gctt.gc..t..t...g.......c...gt. [ 60800]
LR899193.1_Germany_2020/1        ................................................................................ [ 60800]
MN393477.1_Wuhan_2019            ................................................................................ [ 60800]
MW396979.1_Timor-Leste/2019/1    ................................................................................ [ 60800]
MK940252.1_CN/2019/InnerMongolia ................................................................................ [ 60800]
MK128995.1_China/2018/AnhuiXCGQ  ................................................................................ [ 60800]
MH681419.1_ASFV/POL/2015         ................................................................................ [ 60800]
LS478113.1_Estonia_2014          ................................................................................ [ 60800]
KP843857.1_Odintsovo_02/14       ................................................................................ [ 60800]
FN557520.1_strain_E75            ......................................g.aa.....g.gctt.gc..t..t...g.......c...gt. [ 60800]
AY261366.1_Warthog               ......................................g.aa.....g.gctt.gc.....t...g...........gt. [ 60800]
AY261365.1_Warmbaths             ......................................g.aa.....g.gctt.gc..t..t...g...........gt. [ 60800]
AY261364.1_Tengani_62            ......................................g.aa.....g.gctt..c.....t..gg.....t.....gt. [ 60800]
AY261363.1_Pretoriuskop/96/4     ......................................g.aa.....g.gctt.gc..t..t...g...........gt. [ 60800]
AY261362.1_Mkuzi_1979            ......................................g.aa.....g.gctt.gc..t..t...g...........gt. [ 60800]
AY261361.1_Malawi_Lil-20/1_1983  .........c.t.t.......................tg..a.......gc....c..t..a...t.............. [ 60800]
AY261360.1_Kenya_1950            ...............................a.....tg..a.......gc....ca.t..a..gc.............. [ 60800]
AM712240.1_OURT_88/3             ......................................g.aa.....g.gctt.gc..t..t...g.......c...gt. [ 60800]
AM712239.1_Benin_97/1            .................a....................g.aa.....g.gctt.gc..t..t...g.......c...gt. [ 60800]
KM111295.1_Ken06.Bus             ...............................a.....tg..a.......gc....ca.t..a..gc....a......... [ 60800]
KM111294.1_Ken05/Tk1             ...............................a.....tg..a.......gc....ca.t..a..gc.............. [ 60800]
KM262844.1_strain_L60            ......................................g.aa.....g.gctt.gc..t..t...g.......c...gt. [ 60800]
KM102979.1_26544/OG10_Italy      ......................................g.aa.....g.gctt.gc..t..t...g.......c...gt. [ 60800]
KM262845.1_strain_NHV            ......................................g.aa.....g.gctt.gc..t..t...g.......c...gt. [ 60800]
KX354450.1_47/Ss/2008            ......................................g.aa.....g.gctt.gc..t..t...g.......c...gt. [ 60800]
KP055815.1_strain_BA71           ......................................g.aa.....g.gctt.gc..t..t...g.......c...gt. [ 60800]
MH025920.1_strain_R35            ...............................a.....tg..a.......gc....ca.t..a..gc....a......... [ 60800]
MH025919.1_strain_N10            ...............................a.....tg..a.......gc....ca.t..a..gc....a......... [ 60800]
MH025918.1_strain_R25            ...............................a.....tg..a.......gc....ca.t..a..gc....a......... [ 60800]
MH025917.1_strain_R7             ...............................a.....tg..a.......gc....ca.t..a..gc....a......... [ 60800]
MH025916.1_strain_R8             ...............................a.....tg..a.......gc....ca.t..a..gc....a......... [ 60800]
```

```
FR682468.2_Georgia_2007/1        aaaaatattgttgtgtcagaaatttcatacattggcaatacttataaatattttacctttaatgacaatcatgatctgat [ 60880]
ASF_IND_20_CAD543                ..g............................................................................w [ 60880]
LR899193.1_Germany_2020/1        ................................................................................ [ 60880]
MN393477.1_Wuhan_2019            ................................................................................ [ 60880]
MW396979.1_Timor-Leste/2019/1    ................................................................................ [ 60880]
MK940252.1_CN/2019/InnerMongolia ................................................................................ [ 60880]
MK128995.1_China/2018/AnhuiXCGQ  ................................................................................ [ 60880]
MH681419.1_ASFV/POL/2015         ................................................................................ [ 60880]
LS478113.1_Estonia_2014          ................................................................................ [ 60880]
KP843857.1_Odintsovo_02/14       ................................................................................ [ 60880]
FN557520.1_strain_E75            ..g...............................a.....g.......................c..............a [ 60880]
AY261366.1_Warthog               ................................g..tg...g.......................c..............a [ 60880]
AY261365.1_Warmbaths             ..g.............................g...g...g.......................c..............a [ 60880]
AY261364.1_Tengani_62            ........................................g.......................c..............a [ 60880]
AY261363.1_Pretoriuskop/96/4     ..g.............................g...g...g.......................c..............a [ 60880]
AY261362.1_Mkuzi_1979            ..g.............................g...g...g.......................c..............a [ 60880]
AY261361.1_Malawi_Lil-20/1_1983  ..t....a......................ta...a...gg...............t......ac.........ct.... [ 60880]
AY261360.1_Kenya_1950            ..............a................a.......ga........................g........ct.... [ 60880]
AM712240.1_OURT_88/3             ..g...............................a.....g.......................c..............a [ 60880]
AM712239.1_Benin_97/1            ..g...............................a.....g.......................c..............a [ 60880]
KM111295.1_Ken06.Bus             ..............a................a.......ga.................................c..... [ 60880]
KM111294.1_Ken05/Tk1             ..............a................a.......ga.................................ct.... [ 60880]
KM262844.1_strain_L60            ..g...............................a.....g.......................c..............a [ 60880]
KM102979.1_26544/OG10_Italy      ..g...............................a.....g.......................c..............a [ 60880]
KM262845.1_strain_NHV            ..g...............................a.....g.......................c..............a [ 60880]
KX354450.1_47/Ss/2008            ..g...............................a.....g.......................c..............a [ 60880]
KP055815.1_strain_BA71           ..g...............................a.....g.......................c..............a [ 60880]
MH025920.1_strain_R35            ..............a................a.......ga.................................c..... [ 60880]
MH025919.1_strain_N10            ..............a................a.......ga.................................c..... [ 60880]
MH025918.1_strain_R25            ..............a................a.......ga.................................c..... [ 60880]
MH025917.1_strain_R7             ..............a................a.......ga.................................c..... [ 60880]
MH025916.1_strain_R8             ..............a................a.......ga.................................c..... [ 60880]

FR682468.2_Georgia_2007/1        aagcaaagaagatcttaaaggagcaacatccaaaaacattgctaaaatgatttataattggattataaaaaatcctcaaa [ 60960]
ASF_IND_20_CAD543                ................................................................................ [ 60960]
LR899193.1_Germany_2020/1        ................................................................................ [ 60960]
MN393477.1_Wuhan_2019            ................................................................................ [ 60960]
MW396979.1_Timor-Leste/2019/1    ................................................................................ [ 60960]
MK940252.1_CN/2019/InnerMongolia ................................................................................ [ 60960]
MK128995.1_China/2018/AnhuiXCGQ  ................................................................................ [ 60960]
MH681419.1_ASFV/POL/2015         ................................................................................ [ 60960]
LS478113.1_Estonia_2014          ................................................................................ [ 60960]
KP843857.1_Odintsovo_02/14       ................................................................................ [ 60960]
```

```
FN557520.1_strain_E75          .................................c.............................................. [ 60960]
AY261366.1_Warthog             ...t.............................g.............................................. [ 60960]
AY261365.1_Warmbaths           .c...............................c.............................................. [ 60960]
AY261364.1_Tengani_62          ....c.......a....................g.............................................. [ 60960]
AY261363.1_Pretoriuskop/96/4   .................................g.............................................. [ 60960]
AY261362.1_Mkuzi_1979          .c...............................c.............................................. [ 60960]
AY261361.1_Malawi_Lil-20/1_1983 ....c.g.....a....................c..........g..................c..g............. [ 60960]
AY261360.1_Kenya_1950          .g..c.....a......................c...........g..g.............cgcg............. [ 60960]
AM712240.1_OURT_88/3           .................................c.............................................. [ 60960]
AM712239.1_Benin_97/1          .................................c.............................................. [ 60960]
KM111295.1_Ken06.Bus           .g..c.....a......................c...........g..g.............cgcg............. [ 60960]
KM111294.1_Ken05/Tk1           .g..c.....a......................c...........g..g.............cgcg............. [ 60960]
KM262844.1_strain_L60          .................................c.............................................. [ 60960]
KM102979.1_26544/OG10_Italy    .................................c.............................................. [ 60960]
KM262845.1_strain_NHV          .................................c.............................................. [ 60960]
KX354450.1_47/Ss/2008          .................................c.............................................. [ 60960]
KP055815.1_strain_BA71         .................................c.............................................. [ 60960]
MH025920.1_strain_R35          .g..c.....a......................c...........g..g.............cgcg............. [ 60960]
MH025919.1_strain_N10          .g..c.....a......................c...........g..g.............cgcg............. [ 60960]
MH025918.1_strain_R25          .g..c.....a......................c...........g..g.............cgcg............. [ 60960]
MH025917.1_strain_R7           .g..c.....a......................c...........g..g.............cgcg............. [ 60960]
MH025916.1_strain_R8           .g..c.....a......................c...........g..g.............cgcg............. [ 60960]

FR682468.2_Georgia_2007/1      ataataagatttggagtggtgagccgcgtactcaaatttattttgaaaatgatttatatcatacaaattacaatcataaa [ 61040]
ASF_IND_20_CAD543              ................................................................................ [ 61040]
LR899193.1_Germany_2020/1      ................................................................................ [ 61040]
MN393477.1_Wuhan_2019          ................................................................................ [ 61040]
MW396979.1_Timor-Leste/2019/1  ................................................................................ [ 61040]
MK940252.1_CN/2019/InnerMongolia ................................................................................ [ 61040]
MK128995.1_China/2018/AnhuiXCGQ ................................................................................ [ 61040]
MH681419.1_ASFV/POL/2015       ................................................................................ [ 61040]
LS478113.1_Estonia_2014        ................................................................................ [ 61040]
KP843857.1_Odintsovo_02/14     ................................................................................ [ 61040]
FN557520.1_strain_E75          ............................c...................................g.....t......g.. [ 61040]
AY261366.1_Warthog             ............................c...................................g............g.. [ 61040]
AY261365.1_Warmbaths           ............................c...................................g............g.. [ 61040]
AY261364.1_Tengani_62          ............................c.......................c........................g.. [ 61040]
AY261363.1_Pretoriuskop/96/4   ............................c...................................g.....t......g.. [ 61040]
AY261362.1_Mkuzi_1979          ............................c...................................g............g.. [ 61040]
AY261361.1_Malawi_Lil-20/1_1983 ...............a...........aa....g...c............a..g.......c..g........c...g.. [ 61040]
AY261360.1_Kenya_1950          c.......g.....gtc..c..a.....a.t..gc.........a.....a.............t...a.t..c...gt. [ 61040]
AM712240.1_OURT_88/3           ............................c...................................g.....t......g.. [ 61040]
AM712239.1_Benin_97/1          ............................c...................................g.....t......g.. [ 61040]
KM111295.1_Ken06.Bus           c.............gt......a.....a.t..gc.........a....ca.............t...a.t..c...gt. [ 61040]
```

| | | |
|---|---|---|
| KM111294.1_Ken05/Tk1 | c.......g.....gtc..c..a.....a.t..gc.........a.....a.............t...a.t..c...gt. | [ 61040] |
| KM262844.1_strain_L60 | ............................c...................................g.....t......g.. | [ 61040] |
| KM102979.1_26544/OG10_Italy | ............................c...................................g.....t......g.. | [ 61040] |
| KM262845.1_strain_NHV | ............................c...................................g.....t......g.. | [ 61040] |
| KX354450.1_47/Ss/2008 | ............................c...................................g.....t......g.. | [ 61040] |
| KP055815.1_strain_BA71 | ............................c...................................g.....t......g.. | [ 61040] |
| MH025920.1_strain_R35 | c.............gt......a.....a.t..gc.........a....ca.............t...a.t..c...gt. | [ 61040] |
| MH025919.1_strain_N10 | c.............gt......a.....a.t..gc.........a....ca.............t...a.t..c...gt. | [ 61040] |
| MH025918.1_strain_R25 | c.............gt......a.....a.t..gc.........a....ca.............t...a.t..c...gt. | [ 61040] |
| MH025917.1_strain_R7 | c.............gt......a.....a.t..gc.........a....ca.............t...a.t..c...gt. | [ 61040] |
| MH025916.1_strain_R8 | c.............gt......a.....a.t..gc.........a....ca.............t...a.t..c...gt. | [ 61040] |

| | | |
|---|---|---|
| FR682468.2_Georgia_2007/1 | tgtataaaagatttttggaatgtttcaacttcagtcggtcctcatatctttaatgatcgtagcatttggtgtactaaatg | [ 61120] |
| ASF_IND_20_CAD543 | ..........................................tg.................................... | [ 61120] |
| LR899193.1_Germany_2020/1 | ................................................................................ | [ 61120] |
| MN393477.1_Wuhan_2019 | ................................................................................ | [ 61120] |
| MW396979.1_Timor-Leste/2019/1 | ................................................................................ | [ 61120] |
| MK940252.1_CN/2019/InnerMongolia | ................................................................................ | [ 61120] |
| MK128995.1_China/2018/AnhuiXCGQ | ................................................................................ | [ 61120] |
| MH681419.1_ASFV/POL/2015 | ................................................................................ | [ 61120] |
| LS478113.1_Estonia_2014 | ................................................................................ | [ 61120] |
| KP843857.1_Odintsovo_02/14 | ................................................................................ | [ 61120] |
| FN557520.1_strain_E75 | ..................g..........c............tg.................................... | [ 61120] |
| AY261366.1_Warthog | .............................c............tg.................................... | [ 61120] |
| AY261365.1_Warmbaths | .............................c............tg.................................... | [ 61120] |
| AY261364.1_Tengani_62 | ...................................t......tg...................g................ | [ 61120] |
| AY261363.1_Pretoriuskop/96/4 | .............................c............tg.................................... | [ 61120] |
| AY261362.1_Mkuzi_1979 | .............................c............tg...t................................ | [ 61120] |
| AY261361.1_Malawi_Lil-20/1_1983 | ..................g.a.....g...aa..........tg.................................... | [ 61120] |
| AY261360.1_Kenya_1950 | ..............c.at..g.......tc.....ta.....a..................................... | [ 61120] |
| AM712240.1_OURT_88/3 | ..................g..........c............tg.................................... | [ 61120] |
| AM712239.1_Benin_97/1 | ..................g..........c............tg.................................... | [ 61120] |
| KM111295.1_Ken06.Bus | ..............c.at..g........c.....t......a..................................... | [ 61120] |
| KM111294.1_Ken05/Tk1 | ..............c.at..g........c.....t......a..................................... | [ 61120] |
| KM262844.1_strain_L60 | ..................g..........c............tg.................................... | [ 61120] |
| KM102979.1_26544/OG10_Italy | ..................g..........c............tg.................................... | [ 61120] |
| KM262845.1_strain_NHV | ..................g..........c............tg.................................... | [ 61120] |
| KX354450.1_47/Ss/2008 | ..................g..........c............tg.................................... | [ 61120] |
| KP055815.1_strain_BA71 | ..................g..........c............tg.................................... | [ 61120] |
| MH025920.1_strain_R35 | ..............c.at..g........c.....t......a..................................... | [ 61120] |
| MH025919.1_strain_N10 | ..............c.at..g........c.....t......a..................................... | [ 61120] |
| MH025918.1_strain_R25 | ..............c.at..g........c.....t......a..................................... | [ 61120] |
| MH025917.1_strain_R7 | ..............c.at..g........c.....t......a..................................... | [ 61120] |
| MH025916.1_strain_R8 | ..............c.at..g........c.....t......a..................................... | [ 61120] |

```
FR682468.2_Georgia_2007/1        cacatccttttacccatttaccaacattatgtcgcccaatatat-tccaataaattagatatctt----tgctattaaaa [ 61200]
ASF_IND_20_CAD543                ............................................-....................ttac.a......... [ 61200]
LR899193.1_Germany_2020/1        ............................................-....................----........... [ 61200]
MN393477.1_Wuhan_2019            ............................................-....................----........... [ 61200]
MW396979.1_Timor-Leste/2019/1    ............................................-....................----........... [ 61200]
MK940252.1_CN/2019/InnerMongolia ............................................-....................----........... [ 61200]
MK128995.1_China/2018/AnhuiXCGQ  ............................................-....................----........... [ 61200]
MH681419.1_ASFV/POL/2015         ............................................-....................----........... [ 61200]
LS478113.1_Estonia_2014          ............................................-....................----........... [ 61200]
KP843857.1_Odintsovo_02/14       ............................................-....................----........... [ 61200]
FN557520.1_strain_E75            ............................................-....................ttac.a......... [ 61200]
AY261366.1_Warthog               .................................a..........-....................ttat.a......... [ 61200]
AY261365.1_Warmbaths             ........................................g...-....................ttac.a......... [ 61200]
AY261364.1_Tengani_62            ..................c...........a.......g.....ac...................ttac.a......... [ 61200]
AY261363.1_Pretoriuskop/96/4     .................................a..........-....................ttat.a......... [ 61200]
AY261362.1_Mkuzi_1979            ............................................-....................ttac.a......... [ 61200]
AY261361.1_Malawi_Lil-20/1_1983  ....a.......t.......g.......t.............g.-.gg.................ttac.........g. [ 61200]
AY261360.1_Kenya_1950            ............t.....c.g..g....t.a..a......t...-................ca..ttac.a.g....... [ 61200]
AM712240.1_OURT_88/3             ..............................a.............-....................ttac.a......... [ 61200]
AM712239.1_Benin_97/1            ............................................-....................ttac.a......... [ 61200]
KM111295.1_Ken06.Bus             ............t.....c.g..g....t.a..a......t...-................ca..ttac.a.g.....g. [ 61200]
KM111294.1_Ken05/Tk1             ............t.....c.g..g....t.a..a......t...-................ca..ttac.a.g....... [ 61200]
KM262844.1_strain_L60            ............................................-....................ttac.a......... [ 61200]
KM102979.1_26544/OG10_Italy      ............................................-....................ttac.a......... [ 61200]
KM262845.1_strain_NHV            ..............................a.............-....................ttac.a......... [ 61200]
KX354450.1_47/Ss/2008            ............................................-....................ttac.a......... [ 61200]
KP055815.1_strain_BA71           ............................................-....................ttac.a......... [ 61200]
MH025920.1_strain_R35            ............t.....c.g..g....t.a..a......t...-................ca..ttac.a.g.....g. [ 61200]
MH025919.1_strain_N10            ............t.....c.g..g....t.a..a......t...-................ca..ttac.a.g.....g. [ 61200]
MH025918.1_strain_R25            ............t.....c.g..g....t.a..a......t...-................ca..ttac.a.g.....g. [ 61200]
MH025917.1_strain_R7             ............t.....c.g..g....t.a..a......t...-................ca..ttac.a.g.....g. [ 61200]
MH025916.1_strain_R8             ............t.....c.g..g....t.a..a......t...-................ca..ttac.a.g.....g. [ 61200]

FR682468.2_Georgia_2007/1        tagttaa--aaaccttataggataattaggtac----------------------------------tttattacgataa [ 61280]
ASF_IND_20_CAD543                .......--........................----------------------------------...........g. [ 61280]
LR899193.1_Germany_2020/1        .......--........................----------------------------------............. [ 61280]
MN393477.1_Wuhan_2019            .......--........................----------------------------------............. [ 61280]
MW396979.1_Timor-Leste/2019/1    .......--........................----------------------------------............. [ 61280]
MK940252.1_CN/2019/InnerMongolia .......--........................----------------------------------............. [ 61280]
MK128995.1_China/2018/AnhuiXCGQ  .......--........................----------------------------------............. [ 61280]
MH681419.1_ASFV/POL/2015         .......--........................----------------------------------............. [ 61280]
LS478113.1_Estonia_2014          .......--........................----------------------------------............. [ 61280]
KP843857.1_Odintsovo_02/14       .......--........................----------------------------------............. [ 61280]
```

```
FN557520.1_strain_E75           .......--t.......................----------------------------------...........g. [ 61280]
AY261366.1_Warthog              .......--t...a...............a..-------------------------------------.........g. [ 61280]
AY261365.1_Warmbaths            .......--t...a...............a..-------------------------------------.........g. [ 61280]
AY261364.1_Tengani_62           .......--t...a.............ta...t----------------------------------.......ta..tt [ 61280]
AY261363.1_Pretoriuskop/96/4    .......--t...a...............a..-------------------------------------.........g. [ 61280]
AY261362.1_Mkuzi_1979           .......aat...a...............a..-------------------------------------.........g. [ 61280]
AY261361.1_Malawi_Lil-20/1_1983 ..a..tgt------.at..........t.t..ttaagataactaattgcattcttatgattcgttac.......t...tc [ 61280]
AY261360.1_Kenya_1950           ..a.g..---------------------------------------------------------tgca......t..c.. [ 61280]
AM712240.1_OURT_88/3            .......--t.......................----------------------------------...........g. [ 61280]
AM712239.1_Benin_97/1           .......--t.......................----------------------------------...........g. [ 61280]
KM111295.1_Ken06.Bus            ..a.g..--------------------------------------------------------ttgca......t...-. [ 61280]
KM111294.1_Ken05/Tk1            ..a.g..---------------------------------------------------------tgca......t..c.. [ 61280]
KM262844.1_strain_L60           .......--t.......................----------------------------------...........g. [ 61280]
KM102979.1_26544/OG10_Italy     .......--t.......................----------------------------------...........g. [ 61280]
KM262845.1_strain_NHV           .......--t.......................----------------------------------...........g. [ 61280]
KX354450.1_47/Ss/2008           .......--t.......................----------------------------------...........g. [ 61280]
KP055815.1_strain_BA71          .......--t.......................----------------------------------...........g. [ 61280]
MH025920.1_strain_R35           ..a.g..--------------------------------------------------------ttgca......t...-. [ 61280]
MH025919.1_strain_N10           ..a.g..--------------------------------------------------------ttgca......t...-. [ 61280]
MH025918.1_strain_R25           ..a.g..--------------------------------------------------------ttgca......t...-. [ 61280]
MH025917.1_strain_R7            ..a.g..--------------------------------------------------------ttgca......t...-. [ 61280]
MH025916.1_strain_R8            ..a.g..--------------------------------------------------------ttgca......t...-. [ 61280]

FR682468.2_Georgia_2007/1       a--------------------------------------------------------------ttatgatattttataat [ 61360]
ASF_IND_20_CAD543               .ttagtttat--------gwtattttataatt-----ag-ktactttattacgatgaattagt................. [ 61360]
LR899193.1_Germany_2020/1       .--------------------------------------------------------------................. [ 61360]
MN393477.1_Wuhan_2019           .--------------------------------------------------------------................. [ 61360]
MW396979.1_Timor-Leste/2019/1   .--------------------------------------------------------------................. [ 61360]
MK940252.1_CN/2019/InnerMongolia .--------------------------------------------------------------................. [ 61360]
MK128995.1_China/2018/AnhuiXCGQ .--------------------------------------------------------------................. [ 61360]
MH681419.1_ASFV/POL/2015        .--------------------------------------------------------------................. [ 61360]
LS478113.1_Estonia_2014         .--------------------------------------------------------------................. [ 61360]
KP843857.1_Odintsovo_02/14      .--------------------------------------------------------------................. [ 61360]
FN557520.1_strain_E75           .ttagtttat--------gttattttataatt-----ag-gtactttattacgatgaattagt.....t........... [ 61360]
AY261366.1_Warthog              .ttagtttat--------gttattttataagataattag-ttactttattatggttaatctct...------..a.cg.. [ 61360]
AY261365.1_Warmbaths            .ttagtttat--------gttattttataaga---------tactttattatggttaatctct...------..a..g.. [ 61360]
AY261364.1_Tengani_62           .------------------gtattttataagataattag-ttactttattatgattaatctct...-------....g.. [ 61360]
AY261363.1_Pretoriuskop/96/4    .ttagtttat--------gttattttataaga---------tactttattatggttaatctct...------..a.cg.. [ 61360]
AY261362.1_Mkuzi_1979           .ttagtttat--------gttattttata--------ag-atactttattatggttaatctct...------..a..g.. [ 61360]
AY261361.1_Malawi_Lil-20/1_1983 .ttactttattataataaactattttataaaataattagcttactttattgcgattaactgca...-------...cg.. [ 61360]
AY261360.1_Kenya_1950           .------------------------------------aaaatgcattattacggctggttacta..-------..g.g.. [ 61360]
AM712240.1_OURT_88/3            .ttagtttat--------gttattttataatt-----ag-gtactttattacgatgaattagt.....t........... [ 61360]
AM712239.1_Benin_97/1           .ttagtttat--------gttattttataatt-----ag-gtactttattacgatgaattagt.....t........... [ 61360]
KM111295.1_Ken06.Bus            .------------------------------------aaattgcattattacggctgatt-------------------- [ 61360]
```

```
KM111294.1_Ken05/Tk1             .------------------------------------aaaatgcattattacggctggttacta..-------..gcg.. [ 61360]
KM262844.1_strain_L60            .ttagtttat--------gttattttataatt-----ag-gtactttattacgatgaattagt.....t........... [ 61360]
KM102979.1_26544/OG10_Italy      .ttagtttat--------gttattttataatt-----ag-gtactttattacgatgaattagt.....t........... [ 61360]
KM262845.1_strain_NHV            .ttagtttat--------gttattttataatt-----ag-gtactttattacgatgaattagt.....t........... [ 61360]
KX354450.1_47/Ss/2008            .ttagtttat--------gttattttataatt-----ag-gtactttattacgatgaattagt.....t........... [ 61360]
KP055815.1_strain_BA71           .ttagtttat--------gttattttataatt-----ag-gtactttattacgatgaattagt.....t........... [ 61360]
MH025920.1_strain_R35            .------------------------------------aaattgcattattacggctgatt-------------------- [ 61360]
MH025919.1_strain_N10            .------------------------------------aaattgcattattacggctgatt-------------------- [ 61360]
MH025918.1_strain_R25            .------------------------------------aaattgcattattacggctgatt-------------------- [ 61360]
MH025917.1_strain_R7             .------------------------------------aaattgcattattacggctgatt-------------------- [ 61360]
MH025916.1_strain_R8             .------------------------------------aaattgcattattacggctgatt-------------------- [ 61360]

FR682468.2_Georgia_2007/1        t---------------agttactttatta-taattaatct-------------ctttattaatgaatta----------- [ 61440]
ASF_IND_20_CAD543                .---------------.............-cg..g.....ttattaacgattct...............----------- [ 61440]
LR899193.1_Germany_2020/1        .---------------.............-..........-------------................----------- [ 61440]
MN393477.1_Wuhan_2019            .---------------.............-..........-------------................----------- [ 61440]
MW396979.1_Timor-Leste/2019/1    .---------------.............-..........-------------................----------- [ 61440]
MK940252.1_CN/2019/InnerMongolia .---------------.............-..........-------------................----------- [ 61440]
MK128995.1_China/2018/AnhuiXCGQ  .---------------.............-..........-------------................----------- [ 61440]
MH681419.1_ASFV/POL/2015         .---------------.............-..........-------------................----------- [ 61440]
LS478113.1_Estonia_2014          .---------------.............-..........-------------................----------- [ 61440]
KP843857.1_Odintsovo_02/14       .---------------.............-..........-------------................----------- [ 61440]
FN557520.1_strain_E75            .---------------.............-cg..g.....ttattaacgattct...............----------- [ 61440]
AY261366.1_Warthog               .---------------.a.ct....-------........-------------t...............----------- [ 61440]
AY261365.1_Warmbaths             .---------------.a.ctt.......a.g........-------------t...............----------- [ 61440]
AY261364.1_Tengani_62            .---------------.a.ct........acg......t.-------------................----------- [ 61440]
AY261363.1_Pretoriuskop/96/4     .---------------.a.ct....-------........-------------t...............----------- [ 61440]
AY261362.1_Mkuzi_1979            .---------------.a.ctt.......-------------acgattaatctt...............----------- [ 61440]
AY261361.1_Malawi_Lil-20/1_1983  .---------------g..a.........-....a..cta-------------t.....a..at.....gcttactttat [ 61440]
AY261360.1_Kenya_1950            .catcattttatgataca.c.t.....g.-..ca.t....--ataggataatt.a....ct..agga..acgagatat-- [ 61440]
AM712240.1_OURT_88/3             .---------------.............-cg..g.....ttattaacgattct...............----------- [ 61440]
AM712239.1_Benin_97/1            .---------------.............-cg..g.....ttattaacgattct...............----------- [ 61440]
KM111295.1_Ken06.Bus             ----------------ca.c.t.....g.-...a.t....--atag---------------------------gatat-- [ 61440]
KM111294.1_Ken05/Tk1             .---------------ca.c.t.....g.-..ca.t....--ataggataattta....ct..agga..tctcgttat-- [ 61440]
KM262844.1_strain_L60            .---------------.............-cg..g.....ttattaacgattct...............----------- [ 61440]
KM102979.1_26544/OG10_Italy      .---------------.............-cg..g.....ttattaacgattct...............----------- [ 61440]
KM262845.1_strain_NHV            .---------------.............-cg..g.....ttattaacgattct...............----------- [ 61440]
KX354450.1_47/Ss/2008            .---------------.............-cg..g.....ttattaacgattct...............----------- [ 61440]
KP055815.1_strain_BA71           .---------------.............-cg..g.....ttattaacgattct...............----------- [ 61440]
MH025920.1_strain_R35            ----------------ca.c.t.....g.-...a.t....--atag---------------------------gatat-- [ 61440]
MH025919.1_strain_N10            ----------------ca.c.t.....g.-...a.t....--atag---------------------------gatat-- [ 61440]
MH025918.1_strain_R25            ----------------ca.c.t.....g.-...a.t....--atag---------------------------gatat-- [ 61440]
MH025917.1_strain_R7             ----------------ca.c.t.....g.-...a.t....--atag---------------------------gatat-- [ 61440]
MH025916.1_strain_R8             ----------------ca.c.t.....g.-...a.t....--atag---------------------------gatat-- [ 61440]
```

```
FR682468.2_Georgia_2007/1        ----------------------------------------tcataagataactaattatttttttc--catatatcagat [ 61520]
ASF_IND_20_CAD543                ----------------------------------------...........m..............tc........gc.. [ 61520]
LR899193.1_Germany_2020/1        ----------------------------------------..........................--............ [ 61520]
MN393477.1_Wuhan_2019            ----------------------------------------..........................--............ [ 61520]
MW396979.1_Timor-Leste/2019/1    ----------------------------------------..........................--............ [ 61520]
MK940252.1_CN/2019/InnerMongolia ----------------------------------------..........................--............ [ 61520]
MK128995.1_China/2018/AnhuiXCGQ  ----------------------------------------..........................--............ [ 61520]
MH681419.1_ASFV/POL/2015         ----------------------------------------..........................--............ [ 61520]
LS478113.1_Estonia_2014          ----------------------------------------..........................--............ [ 61520]
KP843857.1_Odintsovo_02/14       ----------------------------------------..........................--............ [ 61520]
FN557520.1_strain_E75            ----------------------------------------...........a..............tc........gc.. [ 61520]
AY261366.1_Warthog               ----------------------------------------...........a.............t-ct........c.. [ 61520]
AY261365.1_Warmbaths             ----------------------------------------...........a.............t-ct.......gc.. [ 61520]
AY261364.1_Tengani_62            ----------------------------------------...........a..............tc...c....gc.. [ 61520]
AY261363.1_Pretoriuskop/96/4     ----------------------------------------...........a.............t-ct........c.. [ 61520]
AY261362.1_Mkuzi_1979            ----------------------------------------...........a..............tc........gc.. [ 61520]
AY261361.1_Malawi_Lil-20/1_1983  tgcgattaactgcattattacgattggtaactttattatc.t........................-tt........c.. [ 61520]
AY261360.1_Kenya_1950            ----------------------------------------c.tat......tg............atct........... [ 61520]
AM712240.1_OURT_88/3             ----------------------------------------...........a..............tc........gc.. [ 61520]
AM712239.1_Benin_97/1            ----------------------------------------...........a..............tc........gc.. [ 61520]
KM111295.1_Ken06.Bus             ----------------------------------------c.tat......tg............atct........c.. [ 61520]
KM111294.1_Ken05/Tk1             ----------------------------------------c.tat......tg............atct........c.. [ 61520]
KM262844.1_strain_L60            ----------------------------------------...........a..............tc........gc.. [ 61520]
KM102979.1_26544/OG10_Italy      ----------------------------------------...........a..............tc........gc.. [ 61520]
KM262845.1_strain_NHV            ----------------------------------------...........a..............tc........gc.. [ 61520]
KX354450.1_47/Ss/2008            ----------------------------------------...........a..............tc........gc.. [ 61520]
KP055815.1_strain_BA71           ----------------------------------------...........a..............tc........gc.. [ 61520]
MH025920.1_strain_R35            ----------------------------------------c.tat......tg............atct........c.. [ 61520]
MH025919.1_strain_N10            ----------------------------------------c.tat......tg............atct........c.. [ 61520]
MH025918.1_strain_R25            ----------------------------------------c.tat......tg............atct........c.. [ 61520]
MH025917.1_strain_R7             ----------------------------------------c.tat......tg............atct........c.. [ 61520]
MH025916.1_strain_R8             ----------------------------------------c.tat......tg............atct........c.. [ 61520]

FR682468.2_Georgia_2007/1        aataaatctgatatgggctaaaagtatgtttcaa---------------------------------------------- [ 61600]
ASF_IND_20_CAD543                ..................................---------------------------------------------- [ 61600]
LR899193.1_Germany_2020/1        ..................................---------------------------------------------- [ 61600]
MN393477.1_Wuhan_2019            ..................................---------------------------------------------- [ 61600]
MW396979.1_Timor-Leste/2019/1    ..................................---------------------------------------------- [ 61600]
MK940252.1_CN/2019/InnerMongolia ..................................---------------------------------------------- [ 61600]
MK128995.1_China/2018/AnhuiXCGQ  ..................................---------------------------------------------- [ 61600]
MH681419.1_ASFV/POL/2015         ..................................---------------------------------------------- [ 61600]
LS478113.1_Estonia_2014          ..................................---------------------------------------------- [ 61600]
KP843857.1_Odintsovo_02/14       ..................................---------------------------------------------- [ 61600]
```

```
FN557520.1_strain_E75          ....gg....g.......................---------------------------------------------- [ 61600]
AY261366.1_Warthog             ....g..................--------------------------------------------------------- [ 61600]
AY261365.1_Warmbaths           ....gg...........t................---------------------------------------------- [ 61600]
AY261364.1_Tengani_62          ....ggc....g......................---------------------------------------------- [ 61600]
AY261363.1_Pretoriuskop/96/4   ....g.............................---------------------------------------------- [ 61600]
AY261362.1_Mkuzi_1979          ....gg............................---------------------------------------------- [ 61600]
AY261361.1_Malawi_Lil-20/1_1983 .g..g...................c......a..ctctatttataatagaaattgttatatatcacatagtagatctgat [ 61600]
AY261360.1_Kenya_1950          ..-.g.......................c.ca..c--------------------------------------------- [ 61600]
AM712240.1_OURT_88/3           ....gg....g.......................---------------------------------------------- [ 61600]
AM712239.1_Benin_97/1          ....gg....g.......................---------------------------------------------- [ 61600]
KM111295.1_Ken06.Bus           ..-.g......................a..ca..c--------------------------------------------- [ 61600]
KM111294.1_Ken05/Tk1           ..-.g.........................ca..c--------------------------------------------- [ 61600]
KM262844.1_strain_L60          ....gg....g.......................---------------------------------------------- [ 61600]
KM102979.1_26544/OG10_Italy    ....gg....g.......................---------------------------------------------- [ 61600]
KM262845.1_strain_NHV          ....gg....g.......................---------------------------------------------- [ 61600]
KX354450.1_47/Ss/2008          ....gg....g.......................---------------------------------------------- [ 61600]
KP055815.1_strain_BA71         ....gg....g.......................---------------------------------------------- [ 61600]
MH025920.1_strain_R35          ..-.g......................a..ca..c--------------------------------------------- [ 61600]
MH025919.1_strain_N10          ..-.g......................a..ca..c--------------------------------------------- [ 61600]
MH025918.1_strain_R25          ..-.g......................a..ca..c--------------------------------------------- [ 61600]
MH025917.1_strain_R7           ..-.g......................a..ca..c--------------------------------------------- [ 61600]
MH025916.1_strain_R8           ..-.g......................a..ca..c--------------------------------------------- [ 61600]

FR682468.2_Georgia_2007/1      -----------------------actatttacaatagaatttctgttaag----------aaaacatacataatttgaat [ 61680]
ASF_IND_20_CAD543              -----------------------...........................----------.................... [ 61680]
LR899193.1_Germany_2020/1      -----------------------...........................----------.................... [ 61680]
MN393477.1_Wuhan_2019          -----------------------...........................----------.................... [ 61680]
MW396979.1_Timor-Leste/2019/1  -----------------------...........................----------.................... [ 61680]
MK940252.1_CN/2019/InnerMongolia -----------------------...........................----------.................... [ 61680]
MK128995.1_China/2018/AnhuiXCGQ -----------------------...........................----------.................... [ 61680]
MH681419.1_ASFV/POL/2015       -----------------------...........................----------.................... [ 61680]
LS478113.1_Estonia_2014        -----------------------...........................----------.................... [ 61680]
KP843857.1_Odintsovo_02/14     -----------------------...........................----------.................... [ 61680]
FN557520.1_strain_E75          -----------------------...........................----------.................... [ 61680]
AY261366.1_Warthog             -------------------------................-........----------.................... [ 61680]
AY261365.1_Warmbaths           -----------------------..................--.......----------.................... [ 61680]
AY261364.1_Tengani_62          -----------------------..................-........----------.................... [ 61680]
AY261363.1_Pretoriuskop/96/4   -----------------------..................-........----------.................... [ 61680]
AY261362.1_Mkuzi_1979          -----------------------........t..................----------.................... [ 61680]
AY261361.1_Malawi_Lil-20/1_1983 atgggctaaaagcatgtttaaact.......t.......g.--......acatcatacat.g.ttgg.t....a...... [ 61680]
AY261360.1_Kenya_1950          -----------------------tt......tc...t.gg.-.......a-aacatacat.ggtt...t....a...... [ 61680]
AM712240.1_OURT_88/3           -----------------------...........................----------.................... [ 61680]
AM712239.1_Benin_97/1          -----------------------...........................----------.................... [ 61680]
KM111295.1_Ken06.Bus           -------------------------......tc...t.gg.-.......a-aacatacat.ggtt..tt....a...... [ 61680]
```

```
KM111294.1_Ken05/Tk1           -----------------------tt......tc...t.gg.-.......a-aacatacat.ggtt...t....a...... [ 61680]
KM262844.1_strain_L60          -----------------------...........................----------.................... [ 61680]
KM102979.1_26544/OG10_Italy    -----------------------...........................----------.................... [ 61680]
KM262845.1_strain_NHV          -----------------------...........................----------.................... [ 61680]
KX354450.1_47/Ss/2008          -----------------------...........................----------.................... [ 61680]
KP055815.1_strain_BA71         -----------------------...........................----------.................... [ 61680]
MH025920.1_strain_R35          -------------------------......tc...t.gg.-.......a-aacatacat.ggtt..tt....a...... [ 61680]
MH025919.1_strain_N10          -------------------------......tc...t.gg.-.......a-aacatacat.ggtt..tt....a...... [ 61680]
MH025918.1_strain_R25          -------------------------......tc...t.gg.-.......a-aacatacat.ggtt..tt....a...... [ 61680]
MH025917.1_strain_R7           -------------------------......tc...t.gg.-.......a-aacatacat.ggtt..tt....a...... [ 61680]
MH025916.1_strain_R8           -------------------------......tc...t.gg.-.......a-aacatacat.ggtt..tt....a...... [ 61680]

FR682468.2_Georgia_2007/1      aaaatttttttaaatatcaccgaaacaatcaacatggtgttaatagagtttttaacaggtttcttctatttatatggaaa [ 61760]
ASF_IND_20_CAD543              ...........................tc................................................... [ 61760]
LR899193.1_Germany_2020/1      ................................................................................ [ 61760]
MN393477.1_Wuhan_2019          ................................................................................ [ 61760]
MW396979.1_Timor-Leste/2019/1  ................................................................................ [ 61760]
MK940252.1_CN/2019/InnerMongolia ................................................................................ [ 61760]
MK128995.1_China/2018/AnhuiXCGQ ................................................................................ [ 61760]
MH681419.1_ASFV/POL/2015       ................................................................................ [ 61760]
LS478113.1_Estonia_2014        ................................................................................ [ 61760]
KP843857.1_Odintsovo_02/14     ................................................................................ [ 61760]
FN557520.1_strain_E75          ...........................tc................................................... [ 61760]
AY261366.1_Warthog             ...t.......................t.................................................... [ 61760]
AY261365.1_Warmbaths           ...........................t.................................................... [ 61760]
AY261364.1_Tengani_62          ...........................t...................a................................ [ 61760]
AY261363.1_Pretoriuskop/96/4   ...........................t..............g..................................... [ 61760]
AY261362.1_Mkuzi_1979          c..........................tc................................................... [ 61760]
AY261361.1_Malawi_Lil-20/1_1983 ......................g....t..............g.....................c............t.. [ 61760]
AY261360.1_Kenya_1950          ...........................t.............g...c...............................t.. [ 61760]
AM712240.1_OURT_88/3           ...........................tc................................................... [ 61760]
AM712239.1_Benin_97/1          ...........................tc................................................... [ 61760]
KM111295.1_Ken06.Bus           ......................g....t.............g...c...............................t.. [ 61760]
KM111294.1_Ken05/Tk1           ...........................t.............g...c...............................t.. [ 61760]
KM262844.1_strain_L60          ...........................tc................................................... [ 61760]
KM102979.1_26544/OG10_Italy    ...........................tc................................................... [ 61760]
KM262845.1_strain_NHV          ...........................tc................................................... [ 61760]
KX354450.1_47/Ss/2008          ...........................tc................................................... [ 61760]
KP055815.1_strain_BA71         ...........................tc................................................... [ 61760]
MH025920.1_strain_R35          ......................g....t.............g...c...............................t.. [ 61760]
MH025919.1_strain_N10          ......................g....t.............g...c...............................t.. [ 61760]
MH025918.1_strain_R25          ......................g....t.............g...c...............................t.. [ 61760]
MH025917.1_strain_R7           ......................g....t.............g...c...............................t.. [ 61760]
MH025916.1_strain_R8           ......................g....t.............g...c...............................t.. [ 61760]
```

```
FR682468.2_Georgia_2007/1        gagactgttttccattagtaaagtcatggacatgatatgtctagactattataccattattcctgctcctctggcgatga [ 61840]
ASF_IND_20_CAD543                ................................................................................ [ 61840]
LR899193.1_Germany_2020/1        ................................................................................ [ 61840]
MN393477.1_Wuhan_2019            ................................................................................ [ 61840]
MW396979.1_Timor-Leste/2019/1    ................................................................................ [ 61840]
MK940252.1_CN/2019/InnerMongolia ................................................................................ [ 61840]
MK128995.1_China/2018/AnhuiXCGQ  ................................................................................ [ 61840]
MH681419.1_ASFV/POL/2015         ................................................................................ [ 61840]
LS478113.1_Estonia_2014          ................................................................................ [ 61840]
KP843857.1_Odintsovo_02/14       ................................................................................ [ 61840]
FN557520.1_strain_E75            ..................c............................................................. [ 61840]
AY261366.1_Warthog               ..............................t................................................. [ 61840]
AY261365.1_Warmbaths             ................................................................................ [ 61840]
AY261364.1_Tengani_62            ................................................................................ [ 61840]
AY261363.1_Pretoriuskop/96/4     ..............................t................................................. [ 61840]
AY261362.1_Mkuzi_1979            ..................c............................................................. [ 61840]
AY261361.1_Malawi_Lil-20/1_1983  ..a.........ag........a..............................................c.......... [ 61840]
AY261360.1_Kenya_1950            ............ag..........t.................................c..................... [ 61840]
AM712240.1_OURT_88/3             ..................c............................................................. [ 61840]
AM712239.1_Benin_97/1            ..................c............................................................. [ 61840]
KM111295.1_Ken06.Bus             .........c..a.....c.....t.................................c..................... [ 61840]
KM111294.1_Ken05/Tk1             ............ag..........t.................................c..................... [ 61840]
KM262844.1_strain_L60            ..................c............................................................. [ 61840]
KM102979.1_26544/OG10_Italy      ..................c............................................................. [ 61840]
KM262845.1_strain_NHV            ..................c............................................................. [ 61840]
KX354450.1_47/Ss/2008            ..................c............................................................. [ 61840]
KP055815.1_strain_BA71           ..................c............................................................. [ 61840]
MH025920.1_strain_R35            .........c..a.....c.....t.................................c..................... [ 61840]
MH025919.1_strain_N10            .........c..a.....c.....t.................................c..................... [ 61840]
MH025918.1_strain_R25            .........c..a.....c.....t.................................c..................... [ 61840]
MH025917.1_strain_R7             .........c..a.....c.....t.................................c..................... [ 61840]
MH025916.1_strain_R8             .........c..a.....c.....t.................................c..................... [ 61840]

FR682468.2_Georgia_2007/1        tgttagcggcaagactaaaaaactatgacctcatgaaacgactgcacgaatgggaaatctctattgactacgctctactt [ 61920]
ASF_IND_20_CAD543                ................................................................................ [ 61920]
LR899193.1_Germany_2020/1        ................................................................................ [ 61920]
MN393477.1_Wuhan_2019            ................................................................................ [ 61920]
MW396979.1_Timor-Leste/2019/1    ................................................................................ [ 61920]
MK940252.1_CN/2019/InnerMongolia ................................................................................ [ 61920]
MK128995.1_China/2018/AnhuiXCGQ  ................................................................................ [ 61920]
MH681419.1_ASFV/POL/2015         ................................................................................ [ 61920]
LS478113.1_Estonia_2014          ................................................................................ [ 61920]
KP843857.1_Odintsovo_02/14       ................................................................................ [ 61920]
```

```
FN557520.1_strain_E75             ..............a...............................................g................. [ 61920]
AY261366.1_Warthog                .......a......................................................g...........t..... [ 61920]
AY261365.1_Warmbaths              .......a......a...............................................g................. [ 61920]
AY261364.1_Tengani_62             .......a........................................................................ [ 61920]
AY261363.1_Pretoriuskop/96/4      .......a.....................................................cg...........t..... [ 61920]
AY261362.1_Mkuzi_1979             .......a......a...............................................g...........t..... [ 61920]
AY261361.1_Malawi_Lil-20/1_1983   .......a..t...g........................................c..................t..... [ 61920]
AY261360.1_Kenya_1950             ..............g.......................aa...................c..g...........t..... [ 61920]
AM712240.1_OURT_88/3              ..............a...............................................g................. [ 61920]
AM712239.1_Benin_97/1             ..............a...............................................g................. [ 61920]
KM111295.1_Ken06.Bus              ..........g...g...................a...aa...................c..g.......t...t..... [ 61920]
KM111294.1_Ken05/Tk1              ..............g................t......aa...................c..g...........t..... [ 61920]
KM262844.1_strain_L60             ..............a...............................................g................. [ 61920]
KM102979.1_26544/OG10_Italy       ..............a...............................................g................. [ 61920]
KM262845.1_strain_NHV             ..............a...............................................g................. [ 61920]
KX354450.1_47/Ss/2008             ..............a...............................................g................. [ 61920]
KP055815.1_strain_BA71            ..............a...............................................g................. [ 61920]
MH025920.1_strain_R35             ..........g...g...................a...aa...................c..g.......t...t..... [ 61920]
MH025919.1_strain_N10             ..........g...g...................a...aa...................c..g.......t...t..... [ 61920]
MH025918.1_strain_R25             ..........g...g...................a...aa...................c..g.......t...t..... [ 61920]
MH025917.1_strain_R7              ..........g...g...................a...aa...................c..g.......t...t..... [ 61920]
MH025916.1_strain_R8              ..........g...g...................a...aa...................c..g.......t...t..... [ 61920]

FR682468.2_Georgia_2007/1         gtagtagatgatgtgccgtctattgactattgcttaagtcttggcgctagatccccgactagagcacaaaaaagagaact [ 62000]
ASF_IND_20_CAD543                 ................................................................................ [ 62000]
LR899193.1_Germany_2020/1         ................................................................................ [ 62000]
MN393477.1_Wuhan_2019             ................................................................................ [ 62000]
MW396979.1_Timor-Leste/2019/1     ................................................................................ [ 62000]
MK940252.1_CN/2019/InnerMongolia  ................................................................................ [ 62000]
MK128995.1_China/2018/AnhuiXCGQ   ................................................................................ [ 62000]
MH681419.1_ASFV/POL/2015          ................................................................................ [ 62000]
LS478113.1_Estonia_2014           ................................................................................ [ 62000]
KP843857.1_Odintsovo_02/14        ................................................................................ [ 62000]
FN557520.1_strain_E75             ..................ag...c....t....................a.........................c.... [ 62000]
AY261366.1_Warthog                ..................ag...c....t....................a.............................. [ 62000]
AY261365.1_Warmbaths              ..................ag...c....t....................a.........................c.... [ 62000]
AY261364.1_Tengani_62             ................................................................................ [ 62000]
AY261363.1_Pretoriuskop/96/4      ..................ag...c....t....................a.............................. [ 62000]
AY261362.1_Mkuzi_1979             ..................a....c............g............a...t.....................c.... [ 62000]
AY261361.1_Malawi_Lil-20/1_1983   ..g..............aa....c.............................t..a..................c.... [ 62000]
AY261360.1_Kenya_1950             ..g...............a.c..c.....c........c..........ac..t.....................cg... [ 62000]
AM712240.1_OURT_88/3              ..................ag...c....t....................a.........................c.... [ 62000]
AM712239.1_Benin_97/1             ..................ag...c....t....................a.........................c.... [ 62000]
KM111295.1_Ken06.Bus              ..................a....c.....c........c........c.tc..t.....................cg... [ 62000]
```

```
KM111294.1_Ken05/Tk1             ..g...............a.c..c.....c........c..........ac..t.....................cg... [ 62000]
KM262844.1_strain_L60            ..................ag...c....t....................a.........................c.... [ 62000]
KM102979.1_26544/OG10_Italy      ..................ag...c....t....................a.........................c.... [ 62000]
KM262845.1_strain_NHV            ..................ag...c....t....................a.........................c.... [ 62000]
KX354450.1_47/Ss/2008            ..................ag...c....t....................a.........................c.... [ 62000]
KP055815.1_strain_BA71           ..................ag...c....t....................a.........................c.... [ 62000]
MH025920.1_strain_R35            ..................a....c.....c........c........c.tc..t.....................cg... [ 62000]
MH025919.1_strain_N10            ..................a....c.....c........c........c.tc..t.....................cg... [ 62000]
MH025918.1_strain_R25            ..................a....c.....c........c........c.tc..t.....................cg... [ 62000]
MH025917.1_strain_R7             ..................a....c.....c........c........c.tc..t.....................cg... [ 62000]
MH025916.1_strain_R8             ..................a....c.....c........c........c.tc..t.....................cg... [ 62000]

FR682468.2_Georgia_2007/1        gctgagggacaacacgtttaatcccgtgtataagtatcttatgaactgttccggcttcccaacaaagagagaaaaaaaca [ 62080]
ASF_IND_20_CAD543                .................................................................ga............. [ 62080]
LR899193.1_Germany_2020/1        ................................................................................ [ 62080]
MN393477.1_Wuhan_2019            ................................................................................ [ 62080]
MW396979.1_Timor-Leste/2019/1    ................................................................................ [ 62080]
MK940252.1_CN/2019/InnerMongolia ................................................................................ [ 62080]
MK128995.1_China/2018/AnhuiXCGQ  ................................................................................ [ 62080]
MH681419.1_ASFV/POL/2015         ................................................................................ [ 62080]
LS478113.1_Estonia_2014          ................................................................................ [ 62080]
KP843857.1_Odintsovo_02/14       ................................................................................ [ 62080]
FN557520.1_strain_E75            .t....................................................t..t.......ga............. [ 62080]
AY261366.1_Warthog               ......................................................t..t........a............. [ 62080]
AY261365.1_Warmbaths             ......................................................t..t.......ga............. [ 62080]
AY261364.1_Tengani_62            .........................................................t...................... [ 62080]
AY261363.1_Pretoriuskop/96/4     ......................................................t..t........a............. [ 62080]
AY261362.1_Mkuzi_1979            ......................................................t..t.......ga............. [ 62080]
AY261361.1_Malawi_Lil-20/1_1983  ...ag.a.....t.....c.gc.....t.............................t....................t. [ 62080]
AY261360.1_Kenya_1950            .t.a.......c...a..c..c.....t....................c........t...................... [ 62080]
AM712240.1_OURT_88/3             .t....................................................t..t.......ga............. [ 62080]
AM712239.1_Benin_97/1            .t....................................................t..t.......ga............. [ 62080]
KM111295.1_Ken06.Bus             ...a.......c...a..c..c...a.t....................c........t...................... [ 62080]
KM111294.1_Ken05/Tk1             ...a.......c...a..c..c.....t....................c........t...................... [ 62080]
KM262844.1_strain_L60            .t....................................................t..t.......ga............. [ 62080]
KM102979.1_26544/OG10_Italy      .t....................................................t..t.......ga............. [ 62080]
KM262845.1_strain_NHV            .t....................................................t..t.......ga............. [ 62080]
KX354450.1_47/Ss/2008            .t....................................................t..t.......ga............. [ 62080]
KP055815.1_strain_BA71           .t....................................................t..t.......ga............. [ 62080]
MH025920.1_strain_R35            ...a.......c...a..c..c...a.t....................c........t...................... [ 62080]
MH025919.1_strain_N10            ...a.......c...a..c..c...a.t....................c........t...................... [ 62080]
MH025918.1_strain_R25            ...a.......c...a..c..c...a.t....................c........t...................... [ 62080]
MH025917.1_strain_R7             ...a.......c...a..c..c...a.t....................c........t...................... [ 62080]
MH025916.1_strain_R8             ...a.......c...a..c..c...a.t....................c........t...................... [ 62080]
```

```
FR682468.2_Georgia_2007/1        ttccttgtgatgttcaatgcgaaagactgcaaaaaaacattataaaagaactggtatttaactgctctgtactgcttgaa [ 62160]
ASF_IND_20_CAD543                ...................................................................c..g......... [ 62160]
LR899193.1_Germany_2020/1        ................................................................................ [ 62160]
MN393477.1_Wuhan_2019            ................................................................................ [ 62160]
MW396979.1_Timor-Leste/2019/1    ................................................................................ [ 62160]
MK940252.1_CN/2019/InnerMongolia ................................................................................ [ 62160]
MK128995.1_China/2018/AnhuiXCGQ  ................................................................................ [ 62160]
MH681419.1_ASFV/POL/2015         ................................................................................ [ 62160]
LS478113.1_Estonia_2014          ................................................................................ [ 62160]
KP843857.1_Odintsovo_02/14       ................................................................................ [ 62160]
FN557520.1_strain_E75            ..........................................................c........c..g......... [ 62160]
AY261366.1_Warthog               ...................................................................c..g......... [ 62160]
AY261365.1_Warmbaths             ...................................................................c..g......... [ 62160]
AY261364.1_Tengani_62            ................................................................................ [ 62160]
AY261363.1_Pretoriuskop/96/4     ...................................................................c..g......... [ 62160]
AY261362.1_Mkuzi_1979            ...................................................................c..g......... [ 62160]
AY261361.1_Malawi_Lil-20/1_1983  ......c............t................ta..c............a.............c............ [ 62160]
AY261360.1_Kenya_1950            .............a......................c...c..........................c............ [ 62160]
AM712240.1_OURT_88/3             ..........................................................c........c..g......... [ 62160]
AM712239.1_Benin_97/1            ..........................................................c........c..g......... [ 62160]
KM111295.1_Ken06.Bus             .............g......................c...c..........................c............ [ 62160]
KM111294.1_Ken05/Tk1             .............a......................c...c..........................c............ [ 62160]
KM262844.1_strain_L60            ..........................................................c........c..g......... [ 62160]
KM102979.1_26544/OG10_Italy      ..........................................................c........c..g......... [ 62160]
KM262845.1_strain_NHV            ..........................................................c........c..g......... [ 62160]
KX354450.1_47/Ss/2008            ..........................................................c........c..g......... [ 62160]
KP055815.1_strain_BA71           ..........................................................c........c..g......... [ 62160]
MH025920.1_strain_R35            .............g......................c...c..........................c............ [ 62160]
MH025919.1_strain_N10            .............g......................c...c..........................c............ [ 62160]
MH025918.1_strain_R25            .............g......................c...c..........................c............ [ 62160]
MH025917.1_strain_R7             .............g......................c...c..........................c............ [ 62160]
MH025916.1_strain_R8             .............g......................c...c..........................c............ [ 62160]

FR682468.2_Georgia_2007/1        atggtactgcacacagaaagagaatatgcatacgccctacactgtgctgcaaaacataaccaattgcccatcctcatgta [ 62240]
ASF_IND_20_CAD543                ................................................................................ [ 62240]
LR899193.1_Germany_2020/1        ................................................................................ [ 62240]
MN393477.1_Wuhan_2019            ................................................................................ [ 62240]
MW396979.1_Timor-Leste/2019/1    ................................................................................ [ 62240]
MK940252.1_CN/2019/InnerMongolia ................................................................................ [ 62240]
MK128995.1_China/2018/AnhuiXCGQ  ................................................................................ [ 62240]
MH681419.1_ASFV/POL/2015         ................................................................................ [ 62240]
LS478113.1_Estonia_2014          ................................................................................ [ 62240]
KP843857.1_Odintsovo_02/14       ................................................................................ [ 62240]
```

```
FN557520.1_strain_E75          ................................................................................ [ 62240]
AY261366.1_Warthog             ..........................................................................t..... [ 62240]
AY261365.1_Warmbaths           ................................................................................ [ 62240]
AY261364.1_Tengani_62          ................................................................................ [ 62240]
AY261363.1_Pretoriuskop/96/4   ..........................................................................t..... [ 62240]
AY261362.1_Mkuzi_1979          ..................................................................t.t........... [ 62240]
AY261361.1_Malawi_Lil-20/1_1983 .........ttg...............................a...a......t........a....t........... [ 62240]
AY261360.1_Kenya_1950          ...a......t.t......a.......................a...a......t.............t........... [ 62240]
AM712240.1_OURT_88/3           ................................................................................ [ 62240]
AM712239.1_Benin_97/1          ................................................................................ [ 62240]
KM111295.1_Ken06.Bus           ...a......t.t......a.......................a...a......t.............t.....g..... [ 62240]
KM111294.1_Ken05/Tk1           ...a......t.t......a.......................a...a......t.............t........... [ 62240]
KM262844.1_strain_L60          ................................................................................ [ 62240]
KM102979.1_26544/OG10_Italy    ................................................................................ [ 62240]
KM262845.1_strain_NHV          ................................................................................ [ 62240]
KX354450.1_47/Ss/2008          ................................................................................ [ 62240]
KP055815.1_strain_BA71         ................................................................................ [ 62240]
MH025920.1_strain_R35          ...a......t.t......a.......a...............a...a......t.............t.....g..... [ 62240]
MH025919.1_strain_N10          ...a......t.t......a.......a...............a...a......t.............t.....g..... [ 62240]
MH025918.1_strain_R25          ...a......t.t......a.......a...............a...a......t.............t.....g..... [ 62240]
MH025917.1_strain_R7           ...a......t.t......a.......a...............a...a......t.............t.....g..... [ 62240]
MH025916.1_strain_R8           ...a......t.t......a.......a...............a...a......t.............t.....g..... [ 62240]

FR682468.2_Georgia_2007/1      ttgttggcaacaatccacagacgcggaatctattttgttgaaaacctgctgttctgataagaacatcaattgttttaact [ 62320]
ASF_IND_20_CAD543              ..............................................................................y. [ 62320]
LR899193.1_Germany_2020/1      ................................................................................ [ 62320]
MN393477.1_Wuhan_2019          ................................................................................ [ 62320]
MW396979.1_Timor-Leste/2019/1  ................................................................................ [ 62320]
MK940252.1_CN/2019/InnerMongolia ................................................................................ [ 62320]
MK128995.1_China/2018/AnhuiXCGQ ................................................................................ [ 62320]
MH681419.1_ASFV/POL/2015       ................................................................................ [ 62320]
LS478113.1_Estonia_2014        ................................................................................ [ 62320]
KP843857.1_Odintsovo_02/14     ................................................................................ [ 62320]
FN557520.1_strain_E75          ..............................c...............................................t. [ 62320]
AY261366.1_Warthog             ..............................c...............................................t. [ 62320]
AY261365.1_Warmbaths           ..............................c...............................................t. [ 62320]
AY261364.1_Tengani_62          ................................................................................ [ 62320]
AY261363.1_Pretoriuskop/96/4   ..............................c...............................................t. [ 62320]
AY261362.1_Mkuzi_1979          ..............................c...............................................t. [ 62320]
AY261361.1_Malawi_Lil-20/1_1983 ......................t.......cg.......a........t.............................tc [ 62320]
AY261360.1_Kenya_1950          ...c...........a...a.t........c........a........t..c..g.......................tc [ 62320]
AM712240.1_OURT_88/3           ..............................c...............................................t. [ 62320]
AM712239.1_Benin_97/1          ..............................c...............................................t. [ 62320]
KM111295.1_Ken06.Bus           ...c...........a...a...........g.......a...........c........a.................tc [ 62320]
```

```
KM111294.1_Ken05/Tk1             ...c...........a...a.t........c........a........t..c..g.......................tc [ 62320]
KM262844.1_strain_L60            ..............................c...............................................t. [ 62320]
KM102979.1_26544/OG10_Italy      ..............................c...............................................t. [ 62320]
KM262845.1_strain_NHV            ..............................c...............................................t. [ 62320]
KX354450.1_47/Ss/2008            ..............................c...............................................t. [ 62320]
KP055815.1_strain_BA71           ..............................c...............................................t. [ 62320]
MH025920.1_strain_R35            ...c...........a...a...........g.......a...........c........a.................tc [ 62320]
MH025919.1_strain_N10            ...c...........a...a...........g.......a...........c........a.................tc [ 62320]
MH025918.1_strain_R25            ...c...........a...a...........g.......a...........c........a.................tc [ 62320]
MH025917.1_strain_R7             ...c...........a...a...........g.......a...........c........a.................tc [ 62320]
MH025916.1_strain_R8             ...c...........a...a...........g.......a...........c........a.................tc [ 62320]

FR682468.2_Georgia_2007/1        attgtattctatatggcggcgcccaaaatttggatgctgcaatggtggaagcggcaaagcacgatgcccggatgctgata [ 62400]
ASF_IND_20_CAD543                ................................................................................ [ 62400]
LR899193.1_Germany_2020/1        ................................................................................ [ 62400]
MN393477.1_Wuhan_2019            ................................................................................ [ 62400]
MW396979.1_Timor-Leste/2019/1    ................................................................................ [ 62400]
MK940252.1_CN/2019/InnerMongolia ................................................................................ [ 62400]
MK128995.1_China/2018/AnhuiXCGQ  ................................................................................ [ 62400]
MH681419.1_ASFV/POL/2015         ................................................................................ [ 62400]
LS478113.1_Estonia_2014          ................................................................................ [ 62400]
KP843857.1_Odintsovo_02/14       ................................................................................ [ 62400]
FN557520.1_strain_E75            .........................g......a............................t.................. [ 62400]
AY261366.1_Warthog               .........................g...................................................... [ 62400]
AY261365.1_Warmbaths             .........................g......a............................................... [ 62400]
AY261364.1_Tengani_62            ................t........g......a............................................... [ 62400]
AY261363.1_Pretoriuskop/96/4     .........................g...................................................... [ 62400]
AY261362.1_Mkuzi_1979            .........................g......a............................................... [ 62400]
AY261361.1_Malawi_Lil-20/1_1983  .........................g.....c..........................a..t.........c........ [ 62400]
AY261360.1_Kenya_1950            .......c.....c........g..g...c..a...........a.a...........a...........a.....a..c [ 62400]
AM712240.1_OURT_88/3             .........................g......a............................t.................. [ 62400]
AM712239.1_Benin_97/1            .........................g......a............................t.................. [ 62400]
KM111295.1_Ken06.Bus             .......ca.............g..g...c..............a.a...........a...........a.....a..c [ 62400]
KM111294.1_Ken05/Tk1             .......c.....c........g..g...c..............a.a...........a...........a.....a..c [ 62400]
KM262844.1_strain_L60            .........................g......a............................t.................. [ 62400]
KM102979.1_26544/OG10_Italy      .........................g......a............................t.................. [ 62400]
KM262845.1_strain_NHV            .........................g......a............................t.................. [ 62400]
KX354450.1_47/Ss/2008            .........................g......a............................t.................. [ 62400]
KP055815.1_strain_BA71           .........................g......a............................t.................. [ 62400]
MH025920.1_strain_R35            .......ca.............g..g...c..............a.a...........a...........a.....a..c [ 62400]
MH025919.1_strain_N10            .......ca.............g..g...c..............a.a...........a...........a.....a..c [ 62400]
MH025918.1_strain_R25            .......ca.............g..g...c..............a.a...........a...........a.....a..c [ 62400]
MH025917.1_strain_R7             .......ca.............g..g...c..............a.a...........a...........a.....a..c [ 62400]
MH025916.1_strain_R8             .......ca.............g..g...c..............a.a...........a...........a.....a..c [ 62400]
```

```
FR682468.2_Georgia_2007/1        aactactgtgtcatgcttggtggaagatccttaaacgaagcaaaagaaacggctgccatgtttggacacattgaatgcgc [ 62480]
ASF_IND_20_CAD543                ................................................................................ [ 62480]
LR899193.1_Germany_2020/1        ................................................................................ [ 62480]
MN393477.1_Wuhan_2019            ................................................................................ [ 62480]
MW396979.1_Timor-Leste/2019/1    ................................................................................ [ 62480]
MK940252.1_CN/2019/InnerMongolia ................................................................................ [ 62480]
MK128995.1_China/2018/AnhuiXCGQ  ................................................................................ [ 62480]
MH681419.1_ASFV/POL/2015         ................................................................................ [ 62480]
LS478113.1_Estonia_2014          ................................................................................ [ 62480]
KP843857.1_Odintsovo_02/14       ................................................................................ [ 62480]
FN557520.1_strain_E75            ................................................................................ [ 62480]
AY261366.1_Warthog               ................................................................................ [ 62480]
AY261365.1_Warmbaths             ...................................t............................................ [ 62480]
AY261364.1_Tengani_62            ..........................t..................................................... [ 62480]
AY261363.1_Pretoriuskop/96/4     ....................................c........................................... [ 62480]
AY261362.1_Mkuzi_1979            .................................................t.............................. [ 62480]
AY261361.1_Malawi_Lil-20/1_1983  .................c..c..............t.......g............tgca.........c.......t.. [ 62480]
AY261360.1_Kenya_1950            .........................ag...c............g.........catt..a.................... [ 62480]
AM712240.1_OURT_88/3             ................................................................................ [ 62480]
AM712239.1_Benin_97/1            ................................................................................ [ 62480]
KM111295.1_Ken06.Bus             ..............................c............g.........catt..a.................... [ 62480]
KM111294.1_Ken05/Tk1             ..........................g...c............g.........catt..a.................... [ 62480]
KM262844.1_strain_L60            ................................................................................ [ 62480]
KM102979.1_26544/OG10_Italy      ................................................................................ [ 62480]
KM262845.1_strain_NHV            ................................................................................ [ 62480]
KX354450.1_47/Ss/2008            ................................................................................ [ 62480]
KP055815.1_strain_BA71           ................................................................................ [ 62480]
MH025920.1_strain_R35            ..............................c............g.........catt..a.................... [ 62480]
MH025919.1_strain_N10            ..............................c............g.........catt..a.................... [ 62480]
MH025918.1_strain_R25            ..............................c............g.........catt..a.................... [ 62480]
MH025917.1_strain_R7             ..............................c............g.........catt..a.................... [ 62480]
MH025916.1_strain_R8             ..............................c............g.........catt..a.................... [ 62480]

FR682468.2_Georgia_2007/1        acaacactgttttaaa---------------------------------------------------------------- [ 62560]
ASF_IND_20_CAD543                ................---------------------------------------------------------------- [ 62560]
LR899193.1_Germany_2020/1        ................---------------------------------------------------------------- [ 62560]
MN393477.1_Wuhan_2019            ................---------------------------------------------------------------- [ 62560]
MW396979.1_Timor-Leste/2019/1    ................---------------------------------------------------------------- [ 62560]
MK940252.1_CN/2019/InnerMongolia ................---------------------------------------------------------------- [ 62560]
MK128995.1_China/2018/AnhuiXCGQ  ................---------------------------------------------------------------- [ 62560]
MH681419.1_ASFV/POL/2015         ................---------------------------------------------------------------- [ 62560]
LS478113.1_Estonia_2014          ................---------------------------------------------------------------- [ 62560]
KP843857.1_Odintsovo_02/14       ................---------------------------------------------------------------- [ 62560]
```

```
FN557520.1_strain_E75            .............g..---------------------------------------------------------------- [ 62560]
AY261366.1_Warthog               ................---------------------------------------------------------------- [ 62560]
AY261365.1_Warmbaths             ................ctgcagtcttatgtcat----------------------------------------------- [ 62560]
AY261364.1_Tengani_62            ................ctgcagtcttacgtcgtggacacatcgaatgcggacgacactgattaaagctgcagatttacgt [ 62560]
AY261363.1_Pretoriuskop/96/4     ................---------------------------------------------------------------- [ 62560]
AY261362.1_Mkuzi_1979            ................---------------------------------------------------------------- [ 62560]
AY261361.1_Malawi_Lil-20/1_1983  ............a.ggctgcaatcttacattgt----------------------------------------------- [ 62560]
AY261360.1_Kenya_1950            ......t..c.cg.g.---------------------------------------------------------------- [ 62560]
AM712240.1_OURT_88/3             .............g..---------------------------------------------------------------- [ 62560]
AM712239.1_Benin_97/1            .............g..---------------------------------------------------------------- [ 62560]
KM111295.1_Ken06.Bus             ...t..t..c.cg.g.---------------------------------------------------------------- [ 62560]
KM111294.1_Ken05/Tk1             ......t..c.cg.g.---------------------------------------------------------------- [ 62560]
KM262844.1_strain_L60            .............g..---------------------------------------------------------------- [ 62560]
KM102979.1_26544/OG10_Italy      .............g..---------------------------------------------------------------- [ 62560]
KM262845.1_strain_NHV            .............g..---------------------------------------------------------------- [ 62560]
KX354450.1_47/Ss/2008            .............g..---------------------------------------------------------------- [ 62560]
KP055815.1_strain_BA71           .............g..---------------------------------------------------------------- [ 62560]
MH025920.1_strain_R35            ...t..t..c.cg.g.---------------------------------------------------------------- [ 62560]
MH025919.1_strain_N10            ...t..t..c.cg.g.---------------------------------------------------------------- [ 62560]
MH025918.1_strain_R25            ...t..t..c.cg.g.---------------------------------------------------------------- [ 62560]
MH025917.1_strain_R7             ...t..t..c.cg.g.---------------------------------------------------------------- [ 62560]
MH025916.1_strain_R8             ...t..t..c.cg.g.---------------------------------------------------------------- [ 62560]

FR682468.2_Georgia_2007/1        -------------------------------------------------------------------------------- [ 62640]
ASF_IND_20_CAD543                -------------------------------------------------------------------------------- [ 62640]
LR899193.1_Germany_2020/1        -------------------------------------------------------------------------------- [ 62640]
MN393477.1_Wuhan_2019            -------------------------------------------------------------------------------- [ 62640]
MW396979.1_Timor-Leste/2019/1    -------------------------------------------------------------------------------- [ 62640]
MK940252.1_CN/2019/InnerMongolia -------------------------------------------------------------------------------- [ 62640]
MK128995.1_China/2018/AnhuiXCGQ  -------------------------------------------------------------------------------- [ 62640]
MH681419.1_ASFV/POL/2015         -------------------------------------------------------------------------------- [ 62640]
LS478113.1_Estonia_2014          -------------------------------------------------------------------------------- [ 62640]
KP843857.1_Odintsovo_02/14       -------------------------------------------------------------------------------- [ 62640]
FN557520.1_strain_E75            -------------------------------------------------------------------------------- [ 62640]
AY261366.1_Warthog               -------------------------------------------------------------------------------- [ 62640]
AY261365.1_Warmbaths             -----------------------------------------------------ggacgcattgaatgcggacgacactaa [ 62640]
AY261364.1_Tengani_62            cacgaatgcattgaatgcggacgacactgattaaagctgcagacttacgtcacggacgcattgaatgcggacgacactga [ 62640]
AY261363.1_Pretoriuskop/96/4     -------------------------------------------------------------------------------- [ 62640]
AY261362.1_Mkuzi_1979            -------------------------------------------------------------------------------- [ 62640]
AY261361.1_Malawi_Lil-20/1_1983  ------------------ggacgatactgattaaagctgcaaacttacatcatggacatattgaatgtgaacaacactgt [ 62640]
AY261360.1_Kenya_1950            -------------------------------------------------------------------------------- [ 62640]
AM712240.1_OURT_88/3             -------------------------------------------------------------------------------- [ 62640]
AM712239.1_Benin_97/1            -------------------------------------------------------------------------------- [ 62640]
KM111295.1_Ken06.Bus             -------------------------------------------------------------------------------- [ 62640]
```

```
KM111294.1_Ken05/Tk1            -------------------------------------------------------------------------------- [ 62640]
KM262844.1_strain_L60           -------------------------------------------------------------------------------- [ 62640]
KM102979.1_26544/OG10_Italy     -------------------------------------------------------------------------------- [ 62640]
KM262845.1_strain_NHV           -------------------------------------------------------------------------------- [ 62640]
KX354450.1_47/Ss/2008           -------------------------------------------------------------------------------- [ 62640]
KP055815.1_strain_BA71          -------------------------------------------------------------------------------- [ 62640]
MH025920.1_strain_R35           -------------------------------------------------------------------------------- [ 62640]
MH025919.1_strain_N10           -------------------------------------------------------------------------------- [ 62640]
MH025918.1_strain_R25           -------------------------------------------------------------------------------- [ 62640]
MH025917.1_strain_R7            -------------------------------------------------------------------------------- [ 62640]
MH025916.1_strain_R8            -------------------------------------------------------------------------------- [ 62640]

FR682468.2_Georgia_2007/1       --------------------------------------------------------ctgcagtcttacgtcgtggacaca [ 62720]
ASF_IND_20_CAD543               --------------------------------------------------------........................ [ 62720]
LR899193.1_Germany_2020/1       --------------------------------------------------------........................ [ 62720]
MN393477.1_Wuhan_2019           --------------------------------------------------------........................ [ 62720]
MW396979.1_Timor-Leste/2019/1   --------------------------------------------------------........................ [ 62720]
MK940252.1_CN/2019/InnerMongolia --------------------------------------------------------........................ [ 62720]
MK128995.1_China/2018/AnhuiXCGQ --------------------------------------------------------........................ [ 62720]
MH681419.1_ASFV/POL/2015        --------------------------------------------------------........................ [ 62720]
LS478113.1_Estonia_2014         --------------------------------------------------------........................ [ 62720]
KP843857.1_Odintsovo_02/14      --------------------------------------------------------........................ [ 62720]
FN557520.1_strain_E75           --------------------------------------------------------...........t...a.....g.. [ 62720]
AY261366.1_Warthog              ------ctgcagtcttacgtcatggacacattgaatgtggacgagactgattaaag.....a.....t...ac....... [ 62720]
AY261365.1_Warmbaths            ttaaagctgcagtcttacgtcatggacacattgaatgcggacgacactaattaaag...........t...ac....... [ 62720]
AY261364.1_Tengani_62           ttaaagctgcaatcttacgtcacggtcgcattgaatgcagacgatactgattaaag.....a.........ac..t.g.. [ 62720]
AY261363.1_Pretoriuskop/96/4    ------ctgcagtcttacgtcatggacacattgaatgtggacgacactgattaaag.....a..g..t...aca...... [ 62720]
AY261362.1_Mkuzi_1979           --------------------------------------------------------...........t...a.....g.. [ 62720]
AY261361.1_Malawi_Lil-20/1_1983 ttaaagctgcaaacttacatcatggacatattgaatgtaaacgacactgattaaag.....aa.....a..a......t. [ 62720]
AY261360.1_Kenya_1950           -------------------------------------------------------------------------------- [ 62720]
AM712240.1_OURT_88/3            --------------------------------------------------------...........t...a.....g.. [ 62720]
AM712239.1_Benin_97/1           --------------------------------------------------------...........t...a.....g.. [ 62720]
KM111295.1_Ken06.Bus            -------------------------------------------------------------------------------- [ 62720]
KM111294.1_Ken05/Tk1            -------------------------------------------------------------------------------- [ 62720]
KM262844.1_strain_L60           --------------------------------------------------------...........t...a.....g.. [ 62720]
KM102979.1_26544/OG10_Italy     --------------------------------------------------------...........t...a.....g.. [ 62720]
KM262845.1_strain_NHV           --------------------------------------------------------...........t...a.....g.. [ 62720]
KX354450.1_47/Ss/2008           --------------------------------------------------------...........t...a.....g.. [ 62720]
KP055815.1_strain_BA71          --------------------------------------------------------...........t...a.....g.. [ 62720]
MH025920.1_strain_R35           -------------------------------------------------------------------------------- [ 62720]
MH025919.1_strain_N10           -------------------------------------------------------------------------------- [ 62720]
MH025918.1_strain_R25           -------------------------------------------------------------------------------- [ 62720]
MH025917.1_strain_R7            -------------------------------------------------------------------------------- [ 62720]
MH025916.1_strain_R8            -------------------------------------------------------------------------------- [ 62720]
```

```
FR682468.2_Georgia_2007/1        tcgaatacagacgacactgattaaagcgacaatcttacgtcatgaac---------------gact--gtcttttgag-- [ 62800]
ASF_IND_20_CAD543                ...............................................---------------....--..........-- [ 62800]
LR899193.1_Germany_2020/1        ...............................................---------------....--..........-- [ 62800]
MN393477.1_Wuhan_2019            ...............................................---------------....--..........-- [ 62800]
MW396979.1_Timor-Leste/2019/1    ...............................................---------------....--..........-- [ 62800]
MK940252.1_CN/2019/InnerMongolia ...............................................---------------....--..........-- [ 62800]
MK128995.1_China/2018/AnhuiXCGQ  ...............................................---------------....--..........-- [ 62800]
MH681419.1_ASFV/POL/2015         ...............................................---------------....--..........-- [ 62800]
LS478113.1_Estonia_2014          ...............................................---------------....--..........-- [ 62800]
KP843857.1_Odintsovo_02/14       ...............................................---------------....--..........-- [ 62800]
FN557520.1_strain_E75            .t....g.g......g..............................t---------------....--..........-- [ 62800]
AY261366.1_Warthog               .t....g.g...a..............tg...............g..---------------....aa..........ta [ 62800]
AY261365.1_Warmbaths             .t....g.g.........a........tg..............a...---------------....aa..........-- [ 62800]
AY261364.1_Tengani_62            .t....g.......t............tg...c..............---------------...gaa..........-- [ 62800]
AY261363.1_Pretoriuskop/96/4     .t....gtg..................tg...............g.t---------------....aa..t.......ta [ 62800]
AY261362.1_Mkuzi_1979            .t....g.g......g..............................t---------------....--..........-- [ 62800]
AY261361.1_Malawi_Lil-20/1_1983  .t....gt.a.t...............tg...a.....a.....g..atattgaatgtggaa....--.........a-- [ 62800]
AY261360.1_Kenya_1950            --------------------------.t................cg.---------------....aaa....ga.ga-- [ 62800]
AM712240.1_OURT_88/3             .t....g.g......g..............................t---------------....--..........-- [ 62800]
AM712239.1_Benin_97/1            .t....g.g......g..............................t---------------....--..........-- [ 62800]
KM111295.1_Ken06.Bus             --------------------------ttg..........c....cg.---------------....aaa....ga.ga-- [ 62800]
KM111294.1_Ken05/Tk1             --------------------------.tg...............cg.---------------....aaa....ga.ga-- [ 62800]
KM262844.1_strain_L60            .t....g.g......g..............................t---------------....--..........-- [ 62800]
KM102979.1_26544/OG10_Italy      .t....g.g......g..............................t---------------....--..........-- [ 62800]
KM262845.1_strain_NHV            .t....g.g......g..............................t---------------....--..........-- [ 62800]
KX354450.1_47/Ss/2008            .t....g.g......g..............................t---------------....--..........-- [ 62800]
KP055815.1_strain_BA71           .t....g.g......g..............................t---------------....--..........-- [ 62800]
MH025920.1_strain_R35            --------------------------ttg..........c....cg.---------------....aaa....ga.ga-- [ 62800]
MH025919.1_strain_N10            --------------------------ttg..........c....cg.---------------....aaa....ga.ga-- [ 62800]
MH025918.1_strain_R25            --------------------------ttg..........c....cg.---------------....aaa....ga.ga-- [ 62800]
MH025917.1_strain_R7             --------------------------ttg..........c....cg.---------------....aaa....ga.ga-- [ 62800]
MH025916.1_strain_R8             --------------------------ttg..........c....cg.---------------....aaa....ga.ga-- [ 62800]

FR682468.2_Georgia_2007/1        ------------tatctatacttacattatatttttttatg-aaaaaaatataaaggttgtat----------------- [ 62880]
ASF_IND_20_CAD543                ------------.............................-.....................----------------- [ 62880]
LR899193.1_Germany_2020/1        ------------.............................-.....................----------------- [ 62880]
MN393477.1_Wuhan_2019            ------------.............................-.....................----------------- [ 62880]
MW396979.1_Timor-Leste/2019/1    ------------.............................-.....................----------------- [ 62880]
MK940252.1_CN/2019/InnerMongolia ------------.............................-.....................----------------- [ 62880]
MK128995.1_China/2018/AnhuiXCGQ  ------------.............................-.....................----------------- [ 62880]
MH681419.1_ASFV/POL/2015         ------------.............................-.....................----------------- [ 62880]
LS478113.1_Estonia_2014          ------------.............................-.....................----------------- [ 62880]
KP843857.1_Odintsovo_02/14       ------------.............................-.....................----------------- [ 62880]
```

```
FN557520.1_strain_E75           ------------.............................--....................----------------- [ 62880]
AY261366.1_Warthog              ttgttttttgaa........a.a..c...........gg..-............g..----------------------- [ 62880]
AY261365.1_Warmbaths            ------------..........................cat-............g.a......----------------- [ 62880]
AY261364.1_Tengani_62           ------------........a....................--...........g........----------------- [ 62880]
AY261363.1_Pretoriuskop/96/4    ttgttttttgaa........a.a..c...........g...-............g........----------------- [ 62880]
AY261362.1_Mkuzi_1979           ------------.............................-.....................----------------- [ 62880]
AY261361.1_Malawi_Lil-20/1_1983 ------------........a.................g..-............g....a...atacaaaagcttgtata [ 62880]
AY261360.1_Kenya_1950           -----------g........t.................gaca............g.t......--acaaagatctgtata [ 62880]
AM712240.1_OURT_88/3            ------------.............................-.....................----------------- [ 62880]
AM712239.1_Benin_97/1           ------------.............................-.....................----------------- [ 62880]
KM111295.1_Ken06.Bus            -----------g..--....t...t.............gac-............g.t......--acaaagatctatata [ 62880]
KM111294.1_Ken05/Tk1            -----------g........t...................ca............g.t......--acaaagatctgtata [ 62880]
KM262844.1_strain_L60           ------------.............................-.....................----------------- [ 62880]
KM102979.1_26544/OG10_Italy     ------------.............................-.....................----------------- [ 62880]
KM262845.1_strain_NHV           ------------.............................-.....................----------------- [ 62880]
KX354450.1_47/Ss/2008           ------------.............................-.....................----------------- [ 62880]
KP055815.1_strain_BA71          ------------.............................-.....................----------------- [ 62880]
MH025920.1_strain_R35           -----------g..--....t...t.............gac-............g.t......--acaaagatctgtata [ 62880]
MH025919.1_strain_N10           -----------g..--....t...t.............gac-............g.t......--acaaagatctgtata [ 62880]
MH025918.1_strain_R25           -----------g..--....t...t.............gac-............g.t......--acaaagatctgtata [ 62880]
MH025917.1_strain_R7            -----------g..--....t...t.............gac-............g.t......--acaaagatctgtata [ 62880]
MH025916.1_strain_R8            -----------g..--....t...t.............gac-............g.t......--acaaagatctgtata [ 62880]

FR682468.2_Georgia_2007/1       ----------------------------acaaacctttgtatacaagaaatttggatcatt-aaacaataattaatttgg [ 62960]
ASF_IND_20_CAD543               ----------------------------.....----g.......................-.................. [ 62960]
LR899193.1_Germany_2020/1       ----------------------------.................................-.................. [ 62960]
MN393477.1_Wuhan_2019           ----------------------------.................................-.................. [ 62960]
MW396979.1_Timor-Leste/2019/1   ----------------------------.................................-.................. [ 62960]
MK940252.1_CN/2019/InnerMongolia ----------------------------.................................-.................. [ 62960]
MK128995.1_China/2018/AnhuiXCGQ ----------------------------.................................-.................. [ 62960]
MH681419.1_ASFV/POL/2015        ----------------------------.................................-.................. [ 62960]
LS478113.1_Estonia_2014         ----------------------------.................................-.................. [ 62960]
KP843857.1_Odintsovo_02/14      ----------------------------.................................-.................. [ 62960]
FN557520.1_strain_E75           ----------------------------.....g----.......................-................a. [ 62960]
AY261366.1_Warthog              ------------------------------------.................a...g...-.................. [ 62960]
AY261365.1_Warmbaths            ----------------------------..................a..........a...-................a. [ 62960]
AY261364.1_Tengani_62           ----------------------------.....gg...........a......a...a...------------....... [ 62960]
AY261363.1_Pretoriuskop/96/4    ----------------------------.......-.....................a...a...g.......g...... [ 62960]
AY261362.1_Mkuzi_1979           ----------------------------.....gg..........................-................a. [ 62960]
AY261361.1_Malawi_Lil-20/1_1983 caaaaaatttgtatacaaaagcttgtaa.....agc..................t..a...-...........g...... [ 62960]
AY261360.1_Kenya_1950           ca---------------aagatttgtat.....ga......c....a.gg.-----.a...-...........g-..... [ 62960]
AM712240.1_OURT_88/3            ----------------------------.....g----.......................-................a. [ 62960]
AM712239.1_Benin_97/1           ----------------------------.....g----.......................-................a. [ 62960]
KM111295.1_Ken06.Bus            t-----------------agatttgtat.....ga......c....a.gg.-----.a...-...........g-..... [ 62960]
```

```
KM111294.1_Ken05/Tk1            caa-aaatctgtatacaaagatttgtat.....ga......c....a.gg.-----.a...-...........g-..... [ 62960]
KM262844.1_strain_L60           ----------------------------.....g---........................-................a. [ 62960]
KM102979.1_26544/OG10_Italy     ----------------------------.....g---........................-................a. [ 62960]
KM262845.1_strain_NHV           ----------------------------.....g---........................-................a. [ 62960]
KX354450.1_47/Ss/2008           ----------------------------.....g---........................-................a. [ 62960]
KP055815.1_strain_BA71          ----------------------------.....g---........................-................a. [ 62960]
MH025920.1_strain_R35           t-----------------agatttgtat.....ga......c....a.gg.-----.a...-...........g-..... [ 62960]
MH025919.1_strain_N10           t-----------------agatttgtat.....ga......c....a.gg.-----.a...-...........g-..... [ 62960]
MH025918.1_strain_R25           t-----------------agatttgtat.....ga......c....a.gg.-----.a...-...........g-..... [ 62960]
MH025917.1_strain_R7            t-----------------agatttgtat.....ga......c....a.gg.-----.a...-...........g-..... [ 62960]
MH025916.1_strain_R8            t-----------------agatttgtat.....ga......c....a.gg.-----.a...-...........g-..... [ 62960]

FR682468.2_Georgia_2007/1       acacagg-aaacgatctagatcgatcaaaaagcta--ttttttttgcaca------------cagaacatttagataatt [ 63040]
ASF_IND_20_CAD543               .......a...........................---............------------.................. [ 63040]
LR899193.1_Germany_2020/1       .......-...........................--.............------------.................. [ 63040]
MN393477.1_Wuhan_2019           .......-...........................--.............------------.................. [ 63040]
MW396979.1_Timor-Leste/2019/1   .......-...........................--.............------------.................. [ 63040]
MK940252.1_CN/2019/InnerMongolia .......-...........................--.............------------.................. [ 63040]
MK128995.1_China/2018/AnhuiXCGQ .......-...........................--.............------------.................. [ 63040]
MH681419.1_ASFV/POL/2015        .......-...........................--.............------------.................. [ 63040]
LS478113.1_Estonia_2014         .......-...........................--.............------------.................. [ 63040]
KP843857.1_Odintsovo_02/14      .......-...........................--.............------------.................. [ 63040]
FN557520.1_strain_E75           .......a...........................---............------------.....t............ [ 63040]
AY261366.1_Warthog              g......a..............ag.a..t......--.............------------.................. [ 63040]
AY261365.1_Warmbaths            .......a...g...t......ag.a..t......--.............------------.................. [ 63040]
AY261364.1_Tengani_62           g......a.............t.g.a..t......t-.............------------t................. [ 63040]
AY261363.1_Pretoriuskop/96/4    .......-..............ag.a..t......--.............------------...............--- [ 63040]
AY261362.1_Mkuzi_1979           .......a...........................--.............------------.................. [ 63040]
AY261361.1_Malawi_Lil-20/1_1983 gtg...aa..t.a.....a..t.g.a..t.....tat...........t.------------.................c [ 63040]
AY261360.1_Kenya_1950           ......aa..t.a........-ag.a..t.....ta--..........t.atacagacaaatt...t....a....c..c [ 63040]
AM712240.1_OURT_88/3            .......a...........................---............------------.....t............ [ 63040]
AM712239.1_Benin_97/1           .......a...........................---............------------.....t............ [ 63040]
KM111295.1_Ken06.Bus            ......aa..t.a......cg-.g.a..t.....tat...........t.atacagagaattt...t....a....c..c [ 63040]
KM111294.1_Ken05/Tk1            ......aa..t.a........-ag.a..t.....ta--..........t.atacagacaaatt...t...ca....c..c [ 63040]
KM262844.1_strain_L60           .......a...........................---............------------.....t............ [ 63040]
KM102979.1_26544/OG10_Italy     .......a...........................---............------------.....t............ [ 63040]
KM262845.1_strain_NHV           .......a...........................---............------------.....t............ [ 63040]
KX354450.1_47/Ss/2008           .......a...........................---............------------.....t............ [ 63040]
KP055815.1_strain_BA71          .......a...........................---............------------.....t............ [ 63040]
MH025920.1_strain_R35           ......aa..t.a......cg-.g.a..t.....tat...........t.atacagagaattt...t....a....c..c [ 63040]
MH025919.1_strain_N10           ......aa..t.a......cg-.g.a..t.....tat...........t.atacagagaattt...t....a....c..c [ 63040]
MH025918.1_strain_R25           ......aa..t.a......cg-.g.a..t.....tat...........t.atacagagaattt...t....a....c..c [ 63040]
MH025917.1_strain_R7            ......aa..t.a......cg-.g.a..t.....tat...........t.atacagagaattt...t....a....c..c [ 63040]
MH025916.1_strain_R8            ......aa..t.a......cg-.g.a..t.....tat...........t.atacagagaattt...t....a....c..c [ 63040]
```

```
FR682468.2_Georgia_2007/1        g---------agagattactttccatacttgttaagcttttttacacacaggaactttggattctgttcaggaagttttt [ 63120]
ASF_IND_20_CAD543                .---------...................................................................... [ 63120]
LR899193.1_Germany_2020/1        .---------...................................................................... [ 63120]
MN393477.1_Wuhan_2019            .---------...................................................................... [ 63120]
MW396979.1_Timor-Leste/2019/1    .---------...................................................................... [ 63120]
MK940252.1_CN/2019/InnerMongolia .---------...................................................................... [ 63120]
MK128995.1_China/2018/AnhuiXCGQ  .---------...................................................................... [ 63120]
MH681419.1_ASFV/POL/2015         .---------...................................................................... [ 63120]
LS478113.1_Estonia_2014          .---------...................................................................... [ 63120]
KP843857.1_Odintsovo_02/14       .---------...................................................................... [ 63120]
FN557520.1_strain_E75            a---------.......................c..t.................g......................... [ 63120]
AY261366.1_Warthog               a---------.......................c..t...........t....g......................... [ 63120]
AY261365.1_Warmbaths             a---------.......................c..t...........t....g......................... [ 63120]
AY261364.1_Tengani_62            a---------..........................t.................g......................... [ 63120]
AY261363.1_Pretoriuskop/96/4     --------------...................c..t...........t....g......................... [ 63120]
AY261362.1_Mkuzi_1979            a---------.......................c..t...........t....g......................... [ 63120]
AY261361.1_Malawi_Lil-20/1_1983  a---------..........................t...........t..a..g....a.................... [ 63120]
AY261360.1_Kenya_1950            aagatcatca.....c.....a..............t...........t..a..g......................... [ 63120]
AM712240.1_OURT_88/3             a---------.......................c..t.................g......................... [ 63120]
AM712239.1_Benin_97/1            a---------.......................c..t.................g......................... [ 63120]
KM111295.1_Ken06.Bus             aag-------...t.......a..............t...........t..a..g......................... [ 63120]
KM111294.1_Ken05/Tk1             a---------...........a..............t...........t..a..g......................... [ 63120]
KM262844.1_strain_L60            a---------.......................c..t.................g......................... [ 63120]
KM102979.1_26544/OG10_Italy      a---------.......................c..t.................g......................... [ 63120]
KM262845.1_strain_NHV            a---------.......................c..t.................g......................... [ 63120]
KX354450.1_47/Ss/2008            a---------.......................c..t.................g......................... [ 63120]
KP055815.1_strain_BA71           a---------.......................c..t.................g......................... [ 63120]
MH025920.1_strain_R35            aag-------...t.......a..............t...........t..a..g......................... [ 63120]
MH025919.1_strain_N10            aag-------...t.......a..............t...........t..a..g......................... [ 63120]
MH025918.1_strain_R25            aag-------...t.......a..............t...........t..a..g......................... [ 63120]
MH025917.1_strain_R7             aag-------...t.......a..............t...........t..a..g......................... [ 63120]
MH025916.1_strain_R8             aag-------...t.......a..............t...........t..a..g......................... [ 63120]

FR682468.2_Georgia_2007/1        catagacattatgtttacagccagtaataataattttgggctttttcttaaaccaccggtggaaaacatccagcttgtaa [ 63200]
ASF_IND_20_CAD543                ............k................................................................... [ 63200]
LR899193.1_Germany_2020/1        ................................................................................ [ 63200]
MN393477.1_Wuhan_2019            ................................................................................ [ 63200]
MW396979.1_Timor-Leste/2019/1    ................................................................................ [ 63200]
MK940252.1_CN/2019/InnerMongolia ................................................................................ [ 63200]
MK128995.1_China/2018/AnhuiXCGQ  ................................................................................ [ 63200]
MH681419.1_ASFV/POL/2015         ................................................................................ [ 63200]
LS478113.1_Estonia_2014          ................................................................................ [ 63200]
KP843857.1_Odintsovo_02/14       ................................................................................ [ 63200]
```

```
FN557520.1_strain_E75           ............t................................................................... [ 63200]
AY261366.1_Warthog              ............t................................................................... [ 63200]
AY261365.1_Warmbaths            ............t................................................................... [ 63200]
AY261364.1_Tengani_62           ............c..c................................................................ [ 63200]
AY261363.1_Pretoriuskop/96/4    ............t................................................................... [ 63200]
AY261362.1_Mkuzi_1979           ............t................................................................... [ 63200]
AY261361.1_Malawi_Lil-20/1_1983 ..........g.c..c........................t........g....tg..a..................... [ 63200]
AY261360.1_Kenya_1950           ............c..c.....t...g...........a..t........gtg..tg..a..............t...... [ 63200]
AM712240.1_OURT_88/3            ............t................................................................... [ 63200]
AM712239.1_Benin_97/1           ............t................................................................... [ 63200]
KM111295.1_Ken06.Bus            ............c..c.....t...g...........a..t.........tg..tg..a..............t...... [ 63200]
KM111294.1_Ken05/Tk1            ............c..c.....t...g...........a..t........gtg..tg..a..............t...... [ 63200]
KM262844.1_strain_L60           ............t................................................................... [ 63200]
KM102979.1_26544/OG10_Italy     ............t................................................................... [ 63200]
KM262845.1_strain_NHV           ............t................................................................... [ 63200]
KX354450.1_47/Ss/2008           ............t................................................................... [ 63200]
KP055815.1_strain_BA71          ............t................................................................... [ 63200]
MH025920.1_strain_R35           ............c..c.....t...g...........a..t.........tg..tg..a..............t...... [ 63200]
MH025919.1_strain_N10           ............c..c.....t...g...........a..t.........tg..tg..a..............t...... [ 63200]
MH025918.1_strain_R25           ............c..c.....t...g...........a..t.........tg..tg..a..............t...... [ 63200]
MH025917.1_strain_R7            ............c..c.....t...g...........a..t.........tg..tg..a..............t...... [ 63200]
MH025916.1_strain_R8            ............c..c.....t...g...........a..t.........tg..tg..a..............t...... [ 63200]

FR682468.2_Georgia_2007/1       agagggaaatgcatgtagagaggttt-tggtagtcatggttaagagatttgactaactccatgtttcctgtaaagactgc [ 63280]
ASF_IND_20_CAD543               ..........................t..................................................... [ 63280]
LR899193.1_Germany_2020/1       ..........................-..................................................... [ 63280]
MN393477.1_Wuhan_2019           ..........................-..................................................... [ 63280]
MW396979.1_Timor-Leste/2019/1   ..........................-..................................................... [ 63280]
MK940252.1_CN/2019/InnerMongolia ..........................-..................................................... [ 63280]
MK128995.1_China/2018/AnhuiXCGQ ..........................-..................................................... [ 63280]
MH681419.1_ASFV/POL/2015        ..........................-..................................................... [ 63280]
LS478113.1_Estonia_2014         ..........................-..................................................... [ 63280]
KP843857.1_Odintsovo_02/14      ..........................-..................................................... [ 63280]
FN557520.1_strain_E75           ..........................t...................................a................. [ 63280]
AY261366.1_Warthog              ..........................t..................................................... [ 63280]
AY261365.1_Warmbaths            ..........................t..................................................... [ 63280]
AY261364.1_Tengani_62           ...............a..........t..................................................... [ 63280]
AY261363.1_Pretoriuskop/96/4    ..........................t..................................................... [ 63280]
AY261362.1_Mkuzi_1979           ..........................t...................................a................. [ 63280]
AY261361.1_Malawi_Lil-20/1_1983 .....a.........a....g.....t......c.c.t........t.........a..tgc..g............... [ 63280]
AY261360.1_Kenya_1950           ...............a....g.....t........c.t...c........a.....a..tgc..g............... [ 63280]
AM712240.1_OURT_88/3            ..........................t...................................a................. [ 63280]
AM712239.1_Benin_97/1           ..........................t...................................a................. [ 63280]
KM111295.1_Ken06.Bus            ...............a....g.....t........c.t...c..............a..tgca.g.........a..... [ 63280]
```

```
KM111294.1_Ken05/Tk1           ...............a....g.....t........c.t...c........a.....a..tgc..g............... [ 63280]
KM262844.1_strain_L60          ..........................t...................................a................. [ 63280]
KM102979.1_26544/OG10_Italy    ..........................t...................................a................. [ 63280]
KM262845.1_strain_NHV          ..........................t...................................a................. [ 63280]
KX354450.1_47/Ss/2008          ..........................t...................................a................. [ 63280]
KP055815.1_strain_BA71         ..........................t...................................a................. [ 63280]
MH025920.1_strain_R35          ...............a....g.....t........c.t...c..............a..tgca.g.........a..... [ 63280]
MH025919.1_strain_N10          ...............a....g.....t........c.t...c..............a..tgca.g.........a..... [ 63280]
MH025918.1_strain_R25          ...............a....g.....t........c.t...c..............a..tgca.g.........a..... [ 63280]
MH025917.1_strain_R7           ...............a....g.....t........c.t...c..............a..tgca.g.........a..... [ 63280]
MH025916.1_strain_R8           ...............a....g.....t........c.t...c..............a..tgca.g.........a..... [ 63280]

FR682468.2_Georgia_2007/1      ccagtcccaagcagtaaaacctctatgatagtctttttgagtcggatctgctccaaattttatgagagaaagcatattta [ 63360]
ASF_IND_20_CAD543              ..................r............................................................. [ 63360]
LR899193.1_Germany_2020/1      ................................................................................ [ 63360]
MN393477.1_Wuhan_2019          ................................................................................ [ 63360]
MW396979.1_Timor-Leste/2019/1  ................................................................................ [ 63360]
MK940252.1_CN/2019/InnerMongolia ................................................................................ [ 63360]
MK128995.1_China/2018/AnhuiXCGQ ................................................................................ [ 63360]
MH681419.1_ASFV/POL/2015       ................................................................................ [ 63360]
LS478113.1_Estonia_2014        ................................................................................ [ 63360]
KP843857.1_Odintsovo_02/14     ................................................................................ [ 63360]
FN557520.1_strain_E75          ..................g.......................................................c....g [ 63360]
AY261366.1_Warthog             ..................g........................................................g.... [ 63360]
AY261365.1_Warmbaths           ..................g.....................................................t.c..... [ 63360]
AY261364.1_Tengani_62          ...a..............g....................g..t..................................... [ 63360]
AY261363.1_Pretoriuskop/96/4   ...a..............g.......................t..................................... [ 63360]
AY261362.1_Mkuzi_1979          ..................g.......................................................c..... [ 63360]
AY261361.1_Malawi_Lil-20/1_1983 ...a.......t......c...t..a...........c.gt.t....t...c...t......a.........g.c.c... [ 63360]
AY261360.1_Kenya_1950          ...a.......t.t....t...t..a.............gt......t.......t................g.c.c... [ 63360]
AM712240.1_OURT_88/3           ..................g.......................................................c....g [ 63360]
AM712239.1_Benin_97/1          ..................g.......................................................c....g [ 63360]
KM111295.1_Ken06.Bus           ...a.......t.t....t...t..a...........c.gt......t.......t................g.c.c... [ 63360]
KM111294.1_Ken05/Tk1           ...a.......t.t....t...t..a.............gt......t.......t................g.c.c... [ 63360]
KM262844.1_strain_L60          ..................g.......................................................c....g [ 63360]
KM102979.1_26544/OG10_Italy    ..................g.......................................................c....g [ 63360]
KM262845.1_strain_NHV          ..................g.......................................................c....g [ 63360]
KX354450.1_47/Ss/2008          ..................g.......................................................c....g [ 63360]
KP055815.1_strain_BA71         ..................g.......................................................c....g [ 63360]
MH025920.1_strain_R35          ...a.......t.t....t...t..a...........c.gt......t.......t................g.c.c... [ 63360]
MH025919.1_strain_N10          ...a.......t.t....t...t..a...........c.gt......t.......t................g.c.c... [ 63360]
MH025918.1_strain_R25          ...a.......t.t....t...t..a...........c.gt......t.......t................g.c.c... [ 63360]
MH025917.1_strain_R7           ...a.......t.t....t...t..a...........c.gt......t.......t................g.c.c... [ 63360]
MH025916.1_strain_R8           ...a.......t.t....t...t..a...........c.gt......t.......t................g.c.c... [ 63360]
```

```
FR682468.2_Georgia_2007/1        aagaacggccccgtattgcggccttcatcacaggagtcatcccattaaaattcggtaaacaaattctggtcccatttttt [ 63440]
ASF_IND_20_CAD543                .................................................................y.............. [ 63440]
LR899193.1_Germany_2020/1        ................................................................................ [ 63440]
MN393477.1_Wuhan_2019            ................................................................................ [ 63440]
MW396979.1_Timor-Leste/2019/1    ................................................................................ [ 63440]
MK940252.1_CN/2019/InnerMongolia ................................................................................ [ 63440]
MK128995.1_China/2018/AnhuiXCGQ  ................................................................................ [ 63440]
MH681419.1_ASFV/POL/2015         ................................................................................ [ 63440]
LS478113.1_Estonia_2014          ................................................................................ [ 63440]
KP843857.1_Odintsovo_02/14       ................................................................................ [ 63440]
FN557520.1_strain_E75            ...t.............................................................t.............. [ 63440]
AY261366.1_Warthog               .............c.......................t.......................................... [ 63440]
AY261365.1_Warmbaths             ...tg........c..............t.............................g..................... [ 63440]
AY261364.1_Tengani_62            .............c..g..............................................................g [ 63440]
AY261363.1_Pretoriuskop/96/4     ...t.........c............................................g..................... [ 63440]
AY261362.1_Mkuzi_1979            ...tg.....................................................g..................... [ 63440]
AY261361.1_Malawi_Lil-20/1_1983  ...t...a.....c.....a........a.t......a.c...............a.......c....a.g......... [ 63440]
AY261360.1_Kenya_1950            g..t...a.....c.....a........a..g..c..a...............c.a.......c.t..a.g...a..... [ 63440]
AM712240.1_OURT_88/3             ...t.............................................................t.............. [ 63440]
AM712239.1_Benin_97/1            ...t.............................................................t.............. [ 63440]
KM111295.1_Ken06.Bus             g..t...a.....c.....a........a.tg..c..a...............c.a.......c.t..a.g...a..... [ 63440]
KM111294.1_Ken05/Tk1             g..t...a.....c.....a........a..g..c..a...............c.a.......c.t..a.g...a..... [ 63440]
KM262844.1_strain_L60            ...t.............................................................t.............. [ 63440]
KM102979.1_26544/OG10_Italy      ...t.............................................................t.............. [ 63440]
KM262845.1_strain_NHV            ...t.............................................................t.............. [ 63440]
KX354450.1_47/Ss/2008            ...t.............................................................t.............. [ 63440]
KP055815.1_strain_BA71           ...t.............................................................t.............. [ 63440]
MH025920.1_strain_R35            g..t...a.....c.....a........a.tg..c..a...............c.a.......c.t..a.g...a..... [ 63440]
MH025919.1_strain_N10            g..t...a.....c.....a........a.tg..c..a...............c.a.......c.t..a.g...a..... [ 63440]
MH025918.1_strain_R25            g..t...a.....c.....a........a.tg..c..a...............c.a.......c.t..a.g...a..... [ 63440]
MH025917.1_strain_R7             g..t...a.....c.....a........a.tg..c..a...............c.a.......c.t..a.g...a..... [ 63440]
MH025916.1_strain_R8             g..t...a.....c.....a........a.tg..c..a...............c.a.......c.t..a.g...a..... [ 63440]

FR682468.2_Georgia_2007/1        ccgaaatagcccaacaccccttccaggattaaatgattttttttctcagctaaataatgtaaagcagagtttccatcttt [ 63520]
ASF_IND_20_CAD543                ..................y............................................................. [ 63520]
LR899193.1_Germany_2020/1        ................................................................................ [ 63520]
MN393477.1_Wuhan_2019            ................................................................................ [ 63520]
MW396979.1_Timor-Leste/2019/1    ................................................................................ [ 63520]
MK940252.1_CN/2019/InnerMongolia ................................................................................ [ 63520]
MK128995.1_China/2018/AnhuiXCGQ  ................................................................................ [ 63520]
MH681419.1_ASFV/POL/2015         ................................................................................ [ 63520]
LS478113.1_Estonia_2014          ................................................................................ [ 63520]
KP843857.1_Odintsovo_02/14       ................................................................................ [ 63520]
```

```
FN557520.1_strain_E75            ..................t............................................................. [ 63520]
AY261366.1_Warthog               ..a...............t....t..a.......................c............................. [ 63520]
AY261365.1_Warmbaths             .....g...t........t......a........................c............................. [ 63520]
AY261364.1_Tengani_62            ..................t............................................................. [ 63520]
AY261363.1_Pretoriuskop/96/4     ..................t..........................................................g.. [ 63520]
AY261362.1_Mkuzi_1979            .....g...t........t......a........................c............................. [ 63520]
AY261361.1_Malawi_Lil-20/1_1983  ........at...g...tt......aa.gg.g......g.....t.....c........cc.......a........... [ 63520]
AY261360.1_Kenya_1950            ..a.....tt...g...tt......aa.gg.g......c.....t..c..c........cc..a....a........... [ 63520]
AM712240.1_OURT_88/3             ..................t............................................................. [ 63520]
AM712239.1_Benin_97/1            ..................t............................................................. [ 63520]
KM111295.1_Ken06.Bus             ..a.....tt...g...tt.c....aa.gg.g......c.....t..c..c........cc..a....a........... [ 63520]
KM111294.1_Ken05/Tk1             ........tt...g...tt......aa.gg.g......c.....t..c..c........cc..a....a........... [ 63520]
KM262844.1_strain_L60            ..................t............................................................. [ 63520]
KM102979.1_26544/OG10_Italy      ..................t............................................................. [ 63520]
KM262845.1_strain_NHV            ..................t............................................................. [ 63520]
KX354450.1_47/Ss/2008            ..................t............................................................. [ 63520]
KP055815.1_strain_BA71           ..................t............................................................. [ 63520]
MH025920.1_strain_R35            ..a.....tt...g...tt.c....aa.gg.g......c.....t..c..c........cc..a....a........... [ 63520]
MH025919.1_strain_N10            ..a.....tt...g...tt.c....aa.gg.g......c.....t..c..c........cc..a....a........... [ 63520]
MH025918.1_strain_R25            ..a.....tt...g...tt.c....aa.gg.g......c.....t..c..c........cc..a....a........... [ 63520]
MH025917.1_strain_R7             ..a.....tt...g...tt.c....aa.gg.g......c.....t..c..c........cc..a....a........... [ 63520]
MH025916.1_strain_R8             ..a.....tt...g...tt.c....aa.gg.g......c.....t..c..c........cc..a....a........... [ 63520]

FR682468.2_Georgia_2007/1        atccctcctatgagggttaattatttctccaggataagattcttgttcaaaaagaaattttaaaaagtctatacgtccgt [ 63600]
ASF_IND_20_CAD543                ................................................................................ [ 63600]
LR899193.1_Germany_2020/1        ................................................................................ [ 63600]
MN393477.1_Wuhan_2019            ................................................................................ [ 63600]
MW396979.1_Timor-Leste/2019/1    ................................................................................ [ 63600]
MK940252.1_CN/2019/InnerMongolia ................................................................................ [ 63600]
MK128995.1_China/2018/AnhuiXCGQ  ................................................................................ [ 63600]
MH681419.1_ASFV/POL/2015         ................................................................................ [ 63600]
LS478113.1_Estonia_2014          ................................................................................ [ 63600]
KP843857.1_Odintsovo_02/14       ................................................................................ [ 63600]
FN557520.1_strain_E75            ..............................g................................................. [ 63600]
AY261366.1_Warthog               ..............................g................t................................ [ 63600]
AY261365.1_Warmbaths             ..............................g................t................................ [ 63600]
AY261364.1_Tengani_62            ..............................g................t................................ [ 63600]
AY261363.1_Pretoriuskop/96/4     .............t................g................t................................ [ 63600]
AY261362.1_Mkuzi_1979            ..............................g................................................. [ 63600]
AY261361.1_Malawi_Lil-20/1_1983  ......t....a.t.a..c.c......c..g.....g.....c....t...g......c............a...c.... [ 63600]
AY261360.1_Kenya_1950            ......t....a.t..c.t........c..g.....g..........t..........c............a......a. [ 63600]
AM712240.1_OURT_88/3             ..............................g................................................. [ 63600]
AM712239.1_Benin_97/1            ..............................g................................................. [ 63600]
KM111295.1_Ken06.Bus             ......t....a.ta.c.t........c..g.....g..........t..........c............a......a. [ 63600]
```

```
KM111294.1_Ken05/Tk1            ......t....a.t..c.t........c..g.....g..........t..........c............a......a. [ 63600]
KM262844.1_strain_L60           ..............................g................................................. [ 63600]
KM102979.1_26544/OG10_Italy     ..............................g................................................. [ 63600]
KM262845.1_strain_NHV           ..............................g................................................. [ 63600]
KX354450.1_47/Ss/2008           ..............................g................................................. [ 63600]
KP055815.1_strain_BA71          ..............................g................................................. [ 63600]
MH025920.1_strain_R35           ......t....a.ta.c.t........c..g.....g..........t..........c............a......a. [ 63600]
MH025919.1_strain_N10           ......t....a.ta.c.t........c..g.....g..........t..........c............a......a. [ 63600]
MH025918.1_strain_R25           ......t....a.ta.c.t........c..g.....g..........t..........c............a......a. [ 63600]
MH025917.1_strain_R7            ......t....a.ta.c.t........c..g.....g..........t..........c............a......a. [ 63600]
MH025916.1_strain_R8            ......t....a.ta.c.t........c..g.....g..........t..........c............a......a. [ 63600]

FR682468.2_Georgia_2007/1       agatgcatatccacatgaataccgaggatccatttttatcgcatctattgacaatccacggatctgttttaaaaaattcc [ 63680]
ASF_IND_20_CAD543               ................................................................................ [ 63680]
LR899193.1_Germany_2020/1       ................................................................................ [ 63680]
MN393477.1_Wuhan_2019           ................................................................................ [ 63680]
MW396979.1_Timor-Leste/2019/1   ................................................................................ [ 63680]
MK940252.1_CN/2019/InnerMongolia ................................................................................ [ 63680]
MK128995.1_China/2018/AnhuiXCGQ ................................................................................ [ 63680]
MH681419.1_ASFV/POL/2015        ................................................................................ [ 63680]
LS478113.1_Estonia_2014         ................................................................................ [ 63680]
KP843857.1_Odintsovo_02/14      ................................................................................ [ 63680]
FN557520.1_strain_E75           ................................................................................ [ 63680]
AY261366.1_Warthog              ................................................................................ [ 63680]
AY261365.1_Warmbaths            ................................................................................ [ 63680]
AY261364.1_Tengani_62           ................................................................................ [ 63680]
AY261363.1_Pretoriuskop/96/4    ................................................................................ [ 63680]
AY261362.1_Mkuzi_1979           ................................................................................ [ 63680]
AY261361.1_Malawi_Lil-20/1_1983 ...........a....a.......taat.tt........t..................t.......ca.......g.gtt [ 63680]
AY261360.1_Kenya_1950           ................a.......t.at.tt.c..cg.....................t.......c..........ct. [ 63680]
AM712240.1_OURT_88/3            ................................................................................ [ 63680]
AM712239.1_Benin_97/1           ..........................................................t..................... [ 63680]
KM111295.1_Ken06.Bus            .a..............a.......t.at.tt.c...g.....................t.......c..........ct. [ 63680]
KM111294.1_Ken05/Tk1            ................a.......t.at.tt.c...g.....................t.......c..........ct. [ 63680]
KM262844.1_strain_L60           ................................................................................ [ 63680]
KM102979.1_26544/OG10_Italy     ................................................................................ [ 63680]
KM262845.1_strain_NHV           ................................................................................ [ 63680]
KX354450.1_47/Ss/2008           ................................................................................ [ 63680]
KP055815.1_strain_BA71          ................................................................................ [ 63680]
MH025920.1_strain_R35           .a..............a.......t.at.tt.c...g.....................t.......c..........ct. [ 63680]
MH025919.1_strain_N10           .a..............a.......t.at.tt.c...g.....................t.......c..........ct. [ 63680]
MH025918.1_strain_R25           .a..............a.......t.at.tt.c...g.....................t.......c..........ct. [ 63680]
MH025917.1_strain_R7            .a..............a.......t.at.tt.c...g.....................t.......c..........ct. [ 63680]
MH025916.1_strain_R8            .a..............a.......t.at.tt.c...g.....................t.......c..........ct. [ 63680]
```

```
FR682468.2_Georgia_2007/1        tcaaatagtgtaagattcccatttc---taatatgttttttaatccatttaacaaacaagttttctatctccctttctgg [ 63760]
ASF_IND_20_CAD543                .........................---.r.................................................. [ 63760]
LR899193.1_Germany_2020/1        .........................---.................................................... [ 63760]
MN393477.1_Wuhan_2019            .........................---.................................................... [ 63760]
MW396979.1_Timor-Leste/2019/1    .........................---.................................................... [ 63760]
MK940252.1_CN/2019/InnerMongolia .........................---.................................................... [ 63760]
MK128995.1_China/2018/AnhuiXCGQ  .........................---.................................................... [ 63760]
MH681419.1_ASFV/POL/2015         .........................---.................................................... [ 63760]
LS478113.1_Estonia_2014          .........................---.................................................... [ 63760]
KP843857.1_Odintsovo_02/14       .........................---.................................................... [ 63760]
FN557520.1_strain_E75            .........................---.g.................................................. [ 63760]
AY261366.1_Warthog               .........................---.g..................cc....................gat....... [ 63760]
AY261365.1_Warmbaths             .........................tga.g..................cc....................gat....c.. [ 63760]
AY261364.1_Tengani_62            .........................---.g..................cc....................g.t....... [ 63760]
AY261363.1_Pretoriuskop/96/4     ................ct.......---.g..................cc...............g....gat....... [ 63760]
AY261362.1_Mkuzi_1979            .........................tga.g.................................................. [ 63760]
AY261361.1_Malawi_Lil-20/1_1983  ........g.......c...t...t---.....ca........c...c.c...........g....gct..g---a...a [ 63760]
AY261360.1_Kenya_1950            ..............g.ct..t...t---.g.................c.c..t........g.....c...g---g...a [ 63760]
AM712240.1_OURT_88/3             .........................---.g.................................................. [ 63760]
AM712239.1_Benin_97/1            .........................---.g.................................................. [ 63760]
KM111295.1_Ken06.Bus             ..............g.ct..t...t---.g...a.a...........g....t........g.....c...g---g...a [ 63760]
KM111294.1_Ken05/Tk1             ..............g.ct..t...t---.g.................c.c..t........g.....ct..g---g...a [ 63760]
KM262844.1_strain_L60            .........................---.g.................................................. [ 63760]
KM102979.1_26544/OG10_Italy      .........................---.g.................................................. [ 63760]
KM262845.1_strain_NHV            .........................---.g.................................................. [ 63760]
KX354450.1_47/Ss/2008            .........................---.g.................................................. [ 63760]
KP055815.1_strain_BA71           .................t.......---.g.................................................. [ 63760]
MH025920.1_strain_R35            ..............g.ct..t...t---.g...a.a...........g....t........g.....c...g---g...a [ 63760]
MH025919.1_strain_N10            ..............g.ct..t...t---.g...a.a...........g....t........g.....c...g---g...a [ 63760]
MH025918.1_strain_R25            ..............g.ct..t...t---.g...a.a...........g....t........g.....c...g---g...a [ 63760]
MH025917.1_strain_R7             ..............g.ct..t...t---.g...a.a...........g....t........g.....c...g---g...a [ 63760]
MH025916.1_strain_R8             ..............g.ct..t...t---.g...a.a...........g....t........g.....c...g---g...a [ 63760]

FR682468.2_Georgia_2007/1        aaacatgtgt------tccattttgaatgtcgcccc--tactccactatatgattt-tactcctttaatttttaatgtcc [ 63840]
ASF_IND_20_CAD543                ..........------....................---.................-....................... [ 63840]
LR899193.1_Germany_2020/1        ..........------....................--..................-....................... [ 63840]
MN393477.1_Wuhan_2019            ..........------....................--..................-....................... [ 63840]
MW396979.1_Timor-Leste/2019/1    ..........------....................--..................-....................... [ 63840]
MK940252.1_CN/2019/InnerMongolia ..........------....................--..................-....................... [ 63840]
MK128995.1_China/2018/AnhuiXCGQ  ..........------....................--..................-....................... [ 63840]
MH681419.1_ASFV/POL/2015         ..........------....................--..................-....................... [ 63840]
LS478113.1_Estonia_2014          ..........------....................--..................-....................... [ 63840]
KP843857.1_Odintsovo_02/14       ..........------....................--..................-....................... [ 63840]
```

```
FN557520.1_strain_E75             ..........------....................---..c...t.....a....-....................... [ 63840]
AY261366.1_Warthog                ..........------...............a-...--...c...t.....a....-...c..............c.... [ 63840]
AY261365.1_Warmbaths              ..........------................-...--...c...t.....a....-...c..............c.... [ 63840]
AY261364.1_Tengani_62             ..........------...................t---..c...t.....a.c..-...c..............c.... [ 63840]
AY261363.1_Pretoriuskop/96/4      ..........------..............g.....--...c...t..........-...c................... [ 63840]
AY261362.1_Mkuzi_1979             ..........------...................t---..c...t.....a....-....................... [ 63840]
AY261361.1_Malawi_Lil-20/1_1983   .....g.cc.atcgta........t....gat..t.gtgctct....c...a.agag...ct.............c.... [ 63840]
AY261360.1_Kenya_1950             .....g.cc.accgta........t.ggtagt..t.gtactct....t...a.ggag..t.t.............c.... [ 63840]
AM712240.1_OURT_88/3              ..........------....................---..c...t.....a....-....................... [ 63840]
AM712239.1_Benin_97/1             ..........------....................---..c...t.....a....-....................... [ 63840]
KM111295.1_Ken06.Bus              .....g.cc.actgta........t.ggtagt..t.gtactct....t...a.ggag..t.t.............c.... [ 63840]
KM111294.1_Ken05/Tk1              .....g.cc.accgta........t.ggtagt..t.gtactct....t...a.ggag..t.t.............c.... [ 63840]
KM262844.1_strain_L60             ..........------....................---..c...t.....a....-....................... [ 63840]
KM102979.1_26544/OG10_Italy       ..........------....................---..c...t.....a....-....................... [ 63840]
KM262845.1_strain_NHV             ..........------....................---..c...t.....a....-....................... [ 63840]
KX354450.1_47/Ss/2008             ..........------....................---..c...t.....a....-....................... [ 63840]
KP055815.1_strain_BA71            ..........------....................---..c...t.....a....-....................... [ 63840]
MH025920.1_strain_R35             .....g.cc.actgta........t.ggtagt..t.gtactct....t...a.ggag..t.t.............c.... [ 63840]
MH025919.1_strain_N10             .....g.cc.actgta........t.ggtagt..t.gtactct....t...a.ggag..t.t.............c.... [ 63840]
MH025918.1_strain_R25             .....g.cc.actgta........t.ggtagt..t.gtactct....t...a.ggag..t.t.............c.... [ 63840]
MH025917.1_strain_R7              .....g.cc.actgta........t.ggtagt..t.gtactct....t...a.ggag..t.t.............c.... [ 63840]
MH025916.1_strain_R8              .....g.cc.actgta........t.ggtagt..t.gtactct....t...a.ggag..t.t.............c.... [ 63840]

FR682468.2_Georgia_2007/1         tttttttt-cggacttctttggataagctgtttattaccatctttaaatgccttatagcggggaggagccaggccctttt [ 63920]
ASF_IND_20_CAD543                 ........-..................................................................gc... [ 63920]
LR899193.1_Germany_2020/1         ........-....................................................................... [ 63920]
MN393477.1_Wuhan_2019             ........-....................................................................... [ 63920]
MW396979.1_Timor-Leste/2019/1     ........-....................................................................... [ 63920]
MK940252.1_CN/2019/InnerMongolia  ........-....................................................................... [ 63920]
MK128995.1_China/2018/AnhuiXCGQ   ........-....................................................................... [ 63920]
MH681419.1_ASFV/POL/2015          ........-....................................................................... [ 63920]
LS478113.1_Estonia_2014           ........-....................................................................... [ 63920]
KP843857.1_Odintsovo_02/14        ........-....................................................................... [ 63920]
FN557520.1_strain_E75             ........-..................................................................gc... [ 63920]
AY261366.1_Warthog                ........-................................................t...................... [ 63920]
AY261365.1_Warmbaths              ........-................................................t...................... [ 63920]
AY261364.1_Tengani_62             ........t....................................................................... [ 63920]
AY261363.1_Pretoriuskop/96/4      ........-....................................................................... [ 63920]
AY261362.1_Mkuzi_1979             ........-..................................................................t.... [ 63920]
AY261361.1_Malawi_Lil-20/1_1983   ........-..................................................................gc... [ 63920]
AY261360.1_Kenya_1950             ........-...................c...............................a..............ac... [ 63920]
AM712240.1_OURT_88/3              ........-..................................................................gc... [ 63920]
AM712239.1_Benin_97/1             ........-..................................................................gc... [ 63920]
KM111295.1_Ken06.Bus              ........-...................c............................a..a..............ac... [ 63920]
```

```
KM111294.1_Ken05/Tk1             ........-...................c...............................a..............gc... [ 63920]
KM262844.1_strain_L60            ........-..................................................................gc... [ 63920]
KM102979.1_26544/OG10_Italy      ........-..................................................................gc... [ 63920]
KM262845.1_strain_NHV            ........-..................................................................gc... [ 63920]
KX354450.1_47/Ss/2008            ........-..................................................................gc... [ 63920]
KP055815.1_strain_BA71           ........-..................................................................gc... [ 63920]
MH025920.1_strain_R35            ........-...................c............................a..a..............ac... [ 63920]
MH025919.1_strain_N10            ........-...................c............................a..a..............ac... [ 63920]
MH025918.1_strain_R25            ........-...................c............................a..a..............ac... [ 63920]
MH025917.1_strain_R7             ........-...................c............................a..a..............ac... [ 63920]
MH025916.1_strain_R8             ........-...................c............................a..a..............ac... [ 63920]

FR682468.2_Georgia_2007/1        cccatatgtgcggtaattcttggtgtttatgcttgcctttggcataaccaggccagtatttttcgatatattcagggttt [ 64000]
ASF_IND_20_CAD543                .........................................................................g..a... [ 64000]
LR899193.1_Germany_2020/1        ................................................................................ [ 64000]
MN393477.1_Wuhan_2019            ................................................................................ [ 64000]
MW396979.1_Timor-Leste/2019/1    ................................................................................ [ 64000]
MK940252.1_CN/2019/InnerMongolia ................................................................................ [ 64000]
MK128995.1_China/2018/AnhuiXCGQ  ................................................................................ [ 64000]
MH681419.1_ASFV/POL/2015         ................................................................................ [ 64000]
LS478113.1_Estonia_2014          ................................................................................ [ 64000]
KP843857.1_Odintsovo_02/14       ................................................................................ [ 64000]
FN557520.1_strain_E75            ..............................................t..........................g..a... [ 64000]
AY261366.1_Warthog               ......................................................................g......... [ 64000]
AY261365.1_Warmbaths             ................................................................................ [ 64000]
AY261364.1_Tengani_62            ..............................................c................................. [ 64000]
AY261363.1_Pretoriuskop/96/4     ................................................................................ [ 64000]
AY261362.1_Mkuzi_1979            ................................................................................ [ 64000]
AY261361.1_Malawi_Lil-20/1_1983  ..........t........................................................g............ [ 64000]
AY261360.1_Kenya_1950            ...................................................................g............ [ 64000]
AM712240.1_OURT_88/3             ..............................................t..........................g..a... [ 64000]
AM712239.1_Benin_97/1            ..............................................t..........................g..a... [ 64000]
KM111295.1_Ken06.Bus             ...................................................................g............ [ 64000]
KM111294.1_Ken05/Tk1             ...................................................................g............ [ 64000]
KM262844.1_strain_L60            ..............................................t..........................g..a... [ 64000]
KM102979.1_26544/OG10_Italy      ..............................................t..........................g..a... [ 64000]
KM262845.1_strain_NHV            ..............................................t..........................g..a... [ 64000]
KX354450.1_47/Ss/2008            ..............................................t..........................g..a... [ 64000]
KP055815.1_strain_BA71           ..............................................t..........................g..a... [ 64000]
MH025920.1_strain_R35            ...................................................................g............ [ 64000]
MH025919.1_strain_N10            ...................................................................g............ [ 64000]
MH025918.1_strain_R25            ...................................................................g............ [ 64000]
MH025917.1_strain_R7             ...................................................................g............ [ 64000]
MH025916.1_strain_R8             ...................................................................g............ [ 64000]
```

```
FR682468.2_Georgia_2007/1        gtttttacgtattctttaaaggtccgataggcttcttgaatacaggtaggctcaccggtataatttccatgttcatcttc [ 64080]
ASF_IND_20_CAD543                ........r....................................................................... [ 64080]
LR899193.1_Germany_2020/1        ................................................................................ [ 64080]
MN393477.1_Wuhan_2019            ................................................................................ [ 64080]
MW396979.1_Timor-Leste/2019/1    ................................................................................ [ 64080]
MK940252.1_CN/2019/InnerMongolia ................................................................................ [ 64080]
MK128995.1_China/2018/AnhuiXCGQ  ................................................................................ [ 64080]
MH681419.1_ASFV/POL/2015         ................................................................................ [ 64080]
LS478113.1_Estonia_2014          ................................................................................ [ 64080]
KP843857.1_Odintsovo_02/14       ................................................................................ [ 64080]
FN557520.1_strain_E75            ........a............................................g..t...........g........... [ 64080]
AY261366.1_Warthog               ................................................................................ [ 64080]
AY261365.1_Warmbaths             ................................................................................ [ 64080]
AY261364.1_Tengani_62            ........a............................................g..t....................... [ 64080]
AY261363.1_Pretoriuskop/96/4     ................................................................................ [ 64080]
AY261362.1_Mkuzi_1979            ........a............................................g..t...........g........... [ 64080]
AY261361.1_Malawi_Lil-20/1_1983  .......................t........................................................ [ 64080]
AY261360.1_Kenya_1950            .......................tt......................g.....t..............g........... [ 64080]
AM712240.1_OURT_88/3             ........a............................................g..t...........g........... [ 64080]
AM712239.1_Benin_97/1            ........a............................................g..t...........g........... [ 64080]
KM111295.1_Ken06.Bus             .......................tt......................g.....t..............g........... [ 64080]
KM111294.1_Ken05/Tk1             .......................tt......................g.....t..............g........... [ 64080]
KM262844.1_strain_L60            ........a............................................g..t...........g........... [ 64080]
KM102979.1_26544/OG10_Italy      ........a............................................g..t...........g........... [ 64080]
KM262845.1_strain_NHV            ........a............................................g..t...........g........... [ 64080]
KX354450.1_47/Ss/2008            ........a............................................g..t...........g........... [ 64080]
KP055815.1_strain_BA71           ........a............................................g..t...........g........... [ 64080]
MH025920.1_strain_R35            .......................tt......................g.....t..............g........... [ 64080]
MH025919.1_strain_N10            .......................tt......................g.....t..............g........... [ 64080]
MH025918.1_strain_R25            .......................tt......................g.....t..............g........... [ 64080]
MH025917.1_strain_R7             .......................tt......................g.....t..............g........... [ 64080]
MH025916.1_strain_R8             .......................tt......................g.....t..............g........... [ 64080]

FR682468.2_Georgia_2007/1        ctttaaaaagccattaaccctgtcctttctccacttaagattgtgctttccaaaaatgcgatcaagatcttgcgcctgct [ 64160]
ASF_IND_20_CAD543                .......................................................................cg....... [ 64160]
LR899193.1_Germany_2020/1        ................................................................................ [ 64160]
MN393477.1_Wuhan_2019            ................................................................................ [ 64160]
MW396979.1_Timor-Leste/2019/1    ................................................................................ [ 64160]
MK940252.1_CN/2019/InnerMongolia ................................................................................ [ 64160]
MK128995.1_China/2018/AnhuiXCGQ  ................................................................................ [ 64160]
MH681419.1_ASFV/POL/2015         ................................................................................ [ 64160]
LS478113.1_Estonia_2014          ................................................................................ [ 64160]
KP843857.1_Odintsovo_02/14       ................................................................................ [ 64160]
```

```
FN557520.1_strain_E75          .........................................................a..g..........cg....... [ 64160]
AY261366.1_Warthog             .........................................................a...................... [ 64160]
AY261365.1_Warmbaths           ..........................................a..............a...................... [ 64160]
AY261364.1_Tengani_62          ........................t................................a...................... [ 64160]
AY261363.1_Pretoriuskop/96/4   ....................g....................................a...................... [ 64160]
AY261362.1_Mkuzi_1979          ........................t.................a..................................... [ 64160]
AY261361.1_Malawi_Lil-20/1_1983 ........................t................................a.............ct....... [ 64160]
AY261360.1_Kenya_1950          ............g...t....a..t................................a.............t........ [ 64160]
AM712240.1_OURT_88/3           .........................................................a..g..........cg....... [ 64160]
AM712239.1_Benin_97/1          .........................................................a..g..........cg....... [ 64160]
KM111295.1_Ken06.Bus           ............g........a..t................................a.............c........ [ 64160]
KM111294.1_Ken05/Tk1           ............g...t....a..t................................a.............c........ [ 64160]
KM262844.1_strain_L60          .........................................................a..g..........cg....... [ 64160]
KM102979.1_26544/OG10_Italy    .........................................................a..g..........cg....... [ 64160]
KM262845.1_strain_NHV          .........................................................a..g..........cg....... [ 64160]
KX354450.1_47/Ss/2008          .........................................................a..g..........cg....... [ 64160]
KP055815.1_strain_BA71         .........................................................a..g..........cg....... [ 64160]
MH025920.1_strain_R35          ............g........a..t...................a............a.............c........ [ 64160]
MH025919.1_strain_N10          ............g........a..t...................a............a.............c........ [ 64160]
MH025918.1_strain_R25          ............g........a..t...................a............a.............c........ [ 64160]
MH025917.1_strain_R7           ............g........a..t...................a............a.............c........ [ 64160]
MH025916.1_strain_R8           ............g........a..t...................a............a.............c........ [ 64160]

FR682468.2_Georgia_2007/1      ggggtggaatcataaatccctttttaggtcgaagctttttattttttccatagcttcggccatcgcgttgcgaaacagtg [ 64240]
ASF_IND_20_CAD543              .......r........................................................................ [ 64240]
LR899193.1_Germany_2020/1      ................................................................................ [ 64240]
MN393477.1_Wuhan_2019          ................................................................................ [ 64240]
MW396979.1_Timor-Leste/2019/1  ................................................................................ [ 64240]
MK940252.1_CN/2019/InnerMongolia ................................................................................ [ 64240]
MK128995.1_China/2018/AnhuiXCGQ ................................................................................ [ 64240]
MH681419.1_ASFV/POL/2015       ................................................................................ [ 64240]
LS478113.1_Estonia_2014        ................................................................................ [ 64240]
KP843857.1_Odintsovo_02/14     ................................................................................ [ 64240]
FN557520.1_strain_E75          .......g........c............................................................... [ 64240]
AY261366.1_Warthog             ................c...........c................................................... [ 64240]
AY261365.1_Warmbaths           ................c...........c................................................... [ 64240]
AY261364.1_Tengani_62          ................c...........................................................g... [ 64240]
AY261363.1_Pretoriuskop/96/4   ................c...........c................................................... [ 64240]
AY261362.1_Mkuzi_1979          ................c...........c................................................... [ 64240]
AY261361.1_Malawi_Lil-20/1_1983 ................c................................g.............................. [ 64240]
AY261360.1_Kenya_1950          ................c.....c...............c.g........g..............a............... [ 64240]
AM712240.1_OURT_88/3           .......g........c............................................................... [ 64240]
AM712239.1_Benin_97/1          .......g........c............................................................... [ 64240]
KM111295.1_Ken06.Bus           ................c..t..c..........................g.............................. [ 64240]
```

```
KM111294.1_Ken05/Tk1            ................c.....c...............c.g........g..............a............... [ 64240]
KM262844.1_strain_L60           .......g........c............................................................... [ 64240]
KM102979.1_26544/OG10_Italy     .......g........c............................................................... [ 64240]
KM262845.1_strain_NHV           .......g........c............................................................... [ 64240]
KX354450.1_47/Ss/2008           .......g........c............................................................... [ 64240]
KP055815.1_strain_BA71          .......g........c............................................................... [ 64240]
MH025920.1_strain_R35           ................c..t..c..........................g.............................. [ 64240]
MH025919.1_strain_N10           ................c..t..c..........................g.............................. [ 64240]
MH025918.1_strain_R25           ................c..t..c..........................g.............................. [ 64240]
MH025917.1_strain_R7            ................c..t..c..........................g.............................. [ 64240]
MH025916.1_strain_R8            ................c..t..c..........................g.............................. [ 64240]

FR682468.2_Georgia_2007/1       gttaggacgcctgatagtctttccatgggcgtcgcatctaatcctatccatccaccctgatgaatatcaatggcaacaag [ 64320]
ASF_IND_20_CAD543               ................................................................................ [ 64320]
LR899193.1_Germany_2020/1       ................................................................................ [ 64320]
MN393477.1_Wuhan_2019           ................................................................................ [ 64320]
MW396979.1_Timor-Leste/2019/1   ................................................................................ [ 64320]
MK940252.1_CN/2019/InnerMongolia ................................................................................ [ 64320]
MK128995.1_China/2018/AnhuiXCGQ ................................................................................ [ 64320]
MH681419.1_ASFV/POL/2015        ................................................................................ [ 64320]
LS478113.1_Estonia_2014         ................................................................................ [ 64320]
KP843857.1_Odintsovo_02/14      ................................................................................ [ 64320]
FN557520.1_strain_E75           ...........c..c..c.................g..c......................................... [ 64320]
AY261366.1_Warthog              ..........................................................................g..... [ 64320]
AY261365.1_Warmbaths            ..........................................................................g..... [ 64320]
AY261364.1_Tengani_62           ..........................................................................g..... [ 64320]
AY261363.1_Pretoriuskop/96/4    ..........................................................................g..... [ 64320]
AY261362.1_Mkuzi_1979           ..........................................................................g..... [ 64320]
AY261361.1_Malawi_Lil-20/1_1983 ...........c..................................................g................. [ 64320]
AY261360.1_Kenya_1950           ........a..c.........................................g.......................... [ 64320]
AM712240.1_OURT_88/3            ...........c..c..c.................g..c......................................... [ 64320]
AM712239.1_Benin_97/1           ...........c..c..c.................g..c......................................... [ 64320]
KM111295.1_Ken06.Bus            ........a..c.........................................g.......................... [ 64320]
KM111294.1_Ken05/Tk1            ........a..c.........................................g.......................... [ 64320]
KM262844.1_strain_L60           ...........c..c..c.................g..c......................................... [ 64320]
KM102979.1_26544/OG10_Italy     ...........c..c..c.................g..c......................................... [ 64320]
KM262845.1_strain_NHV           ...........c..c..c.................g..c......................................... [ 64320]
KX354450.1_47/Ss/2008           ...........c..c..c.................g..c......................................... [ 64320]
KP055815.1_strain_BA71          ...........c..c..c.................g..c......................................... [ 64320]
MH025920.1_strain_R35           ........a..c.........................................g.......................... [ 64320]
MH025919.1_strain_N10           ........a..c.........................................g.......................... [ 64320]
MH025918.1_strain_R25           ........a..c.........................................g.......................... [ 64320]
MH025917.1_strain_R7            ........a..c.........................................g.......................... [ 64320]
MH025916.1_strain_R8            ........a..c.........................................g.......................... [ 64320]
```

```
FR682468.2_Georgia_2007/1        ctctcctttattttgggcaagccaagtttccaagaatgccatgctttcttcccagggataaggcccgccaacaccacggg [ 64400]
ASF_IND_20_CAD543                ................................................................................ [ 64400]
LR899193.1_Germany_2020/1        ................................................................................ [ 64400]
MN393477.1_Wuhan_2019            ................................................................................ [ 64400]
MW396979.1_Timor-Leste/2019/1    ................................................................................ [ 64400]
MK940252.1_CN/2019/InnerMongolia ................................................................................ [ 64400]
MK128995.1_China/2018/AnhuiXCGQ  ................................................................................ [ 64400]
MH681419.1_ASFV/POL/2015         ................................................................................ [ 64400]
LS478113.1_Estonia_2014          ................................................................................ [ 64400]
KP843857.1_Odintsovo_02/14       ................................................................................ [ 64400]
FN557520.1_strain_E75            ................................................................................ [ 64400]
AY261366.1_Warthog               ................................................................................ [ 64400]
AY261365.1_Warmbaths             ................................................................................ [ 64400]
AY261364.1_Tengani_62            ........................c....................................................... [ 64400]
AY261363.1_Pretoriuskop/96/4     ................................................................................ [ 64400]
AY261362.1_Mkuzi_1979            ................................................................................ [ 64400]
AY261361.1_Malawi_Lil-20/1_1983  ................................................................................ [ 64400]
AY261360.1_Kenya_1950            ................................................................................ [ 64400]
AM712240.1_OURT_88/3             ................................................................................ [ 64400]
AM712239.1_Benin_97/1            ................................................................................ [ 64400]
KM111295.1_Ken06.Bus             ........................g................................g...................... [ 64400]
KM111294.1_Ken05/Tk1             ................................................................................ [ 64400]
KM262844.1_strain_L60            ................................................................................ [ 64400]
KM102979.1_26544/OG10_Italy      ................................................................................ [ 64400]
KM262845.1_strain_NHV            ................................................................................ [ 64400]
KX354450.1_47/Ss/2008            ................................................................................ [ 64400]
KP055815.1_strain_BA71           ................................................................................ [ 64400]
MH025920.1_strain_R35            ........................g................................g...................... [ 64400]
MH025919.1_strain_N10            ........................g................................g...................... [ 64400]
MH025918.1_strain_R25            ........................g................................g...................... [ 64400]
MH025917.1_strain_R7             ........................g................................g...................... [ 64400]
MH025916.1_strain_R8             ........................g................................g...................... [ 64400]

FR682468.2_Georgia_2007/1        ttgtccaatcttgcaaggactccaggtccgacacctggtaaggctctaaagaagacggttccttgtttttgtactgcaaa [ 64480]
ASF_IND_20_CAD543                ................................................................................ [ 64480]
LR899193.1_Germany_2020/1        ................................................................................ [ 64480]
MN393477.1_Wuhan_2019            ................................................................................ [ 64480]
MW396979.1_Timor-Leste/2019/1    ................................................................................ [ 64480]
MK940252.1_CN/2019/InnerMongolia ................................................................................ [ 64480]
MK128995.1_China/2018/AnhuiXCGQ  ................................................................................ [ 64480]
MH681419.1_ASFV/POL/2015         ................................................................................ [ 64480]
LS478113.1_Estonia_2014          ................................................................................ [ 64480]
KP843857.1_Odintsovo_02/14       ................................................................................ [ 64480]
```

```
FN557520.1_strain_E75            ................................................................................ [ 64480]
AY261366.1_Warthog               ................................................................................ [ 64480]
AY261365.1_Warmbaths             ................................................................................ [ 64480]
AY261364.1_Tengani_62            ................................................................................ [ 64480]
AY261363.1_Pretoriuskop/96/4     ................................................................................ [ 64480]
AY261362.1_Mkuzi_1979            .......................................................t........................ [ 64480]
AY261361.1_Malawi_Lil-20/1_1983  ...................t...................................t..............a......... [ 64480]
AY261360.1_Kenya_1950            ...................t.........a.........................t.....a.................. [ 64480]
AM712240.1_OURT_88/3             ................................................................................ [ 64480]
AM712239.1_Benin_97/1            ................................................................................ [ 64480]
KM111295.1_Ken06.Bus             ...................t..........t........................t.....a.................. [ 64480]
KM111294.1_Ken05/Tk1             ...................t.........a.........................t.....a.................. [ 64480]
KM262844.1_strain_L60            ................................................................................ [ 64480]
KM102979.1_26544/OG10_Italy      ................................................................................ [ 64480]
KM262845.1_strain_NHV            ................................................................................ [ 64480]
KX354450.1_47/Ss/2008            ................................................................................ [ 64480]
KP055815.1_strain_BA71           ................................................................................ [ 64480]
MH025920.1_strain_R35            ...................t..........t........................t.....a.................. [ 64480]
MH025919.1_strain_N10            ...................t..........t........................t.....a.................. [ 64480]
MH025918.1_strain_R25            ...................t..........t........................t.....a.................. [ 64480]
MH025917.1_strain_R7             ...................t..........t........................t.....a.................. [ 64480]
MH025916.1_strain_R8             ...................t..........t........................t.....a.................. [ 64480]

FR682468.2_Georgia_2007/1        taagatttaatgacccatttataccatgtgtcgaaccgcagcgtggcgcctccaaagtgaaagccgtcgttgattttagg [ 64560]
ASF_IND_20_CAD543                ................................................................................ [ 64560]
LR899193.1_Germany_2020/1        ................................................................................ [ 64560]
MN393477.1_Wuhan_2019            ................................................................................ [ 64560]
MW396979.1_Timor-Leste/2019/1    ................................................................................ [ 64560]
MK940252.1_CN/2019/InnerMongolia ................................................................................ [ 64560]
MK128995.1_China/2018/AnhuiXCGQ  ................................................................................ [ 64560]
MH681419.1_ASFV/POL/2015         ................................................................................ [ 64560]
LS478113.1_Estonia_2014          ................................................................................ [ 64560]
KP843857.1_Odintsovo_02/14       ................................................................................ [ 64560]
FN557520.1_strain_E75            ..g............................................................................. [ 64560]
AY261366.1_Warthog               ..g............................................................................. [ 64560]
AY261365.1_Warmbaths             ..g............................................................................. [ 64560]
AY261364.1_Tengani_62            ..g............................................................................. [ 64560]
AY261363.1_Pretoriuskop/96/4     ..g............................................................................. [ 64560]
AY261362.1_Mkuzi_1979            ..g............................................................................. [ 64560]
AY261361.1_Malawi_Lil-20/1_1983  ..g.............................a............................................... [ 64560]
AY261360.1_Kenya_1950            ..g.............................a............................................... [ 64560]
AM712240.1_OURT_88/3             ..g............................................................................. [ 64560]
AM712239.1_Benin_97/1            ..g............................................................................. [ 64560]
KM111295.1_Ken06.Bus             ..g.............................a............................................... [ 64560]
```

```
KM111294.1_Ken05/Tk1           ..g.............................a............................................... [ 64560]
KM262844.1_strain_L60          ..g............................................................................. [ 64560]
KM102979.1_26544/OG10_Italy    ..g............................................................................. [ 64560]
KM262845.1_strain_NHV          ..g............................................................................. [ 64560]
KX354450.1_47/Ss/2008          ..g............................................................................. [ 64560]
KP055815.1_strain_BA71         ..g............................................................................. [ 64560]
MH025920.1_strain_R35          ..g.............................a............................................... [ 64560]
MH025919.1_strain_N10          ..g.............................a............................................... [ 64560]
MH025918.1_strain_R25          ..g.............................a............................................... [ 64560]
MH025917.1_strain_R7           ..g.............................a............................................... [ 64560]
MH025916.1_strain_R8           ..g.............................a............................................... [ 64560]

FR682468.2_Georgia_2007/1      atatctgcaacatatttcaaccgtacgtttgagttctgcaaaagcggccttccaaggaagtctttcgctgcgggtaagac [ 64640]
ASF_IND_20_CAD543              ..............................................................................r. [ 64640]
LR899193.1_Germany_2020/1      ................................................................................ [ 64640]
MN393477.1_Wuhan_2019          ................................................................................ [ 64640]
MW396979.1_Timor-Leste/2019/1  ................................................................................ [ 64640]
MK940252.1_CN/2019/InnerMongolia ................................................................................ [ 64640]
MK128995.1_China/2018/AnhuiXCGQ ................................................................................ [ 64640]
MH681419.1_ASFV/POL/2015       ................................................................................ [ 64640]
LS478113.1_Estonia_2014        ................................................................................ [ 64640]
KP843857.1_Odintsovo_02/14     ................................................................................ [ 64640]
FN557520.1_strain_E75          ..........................................g....................c..............g. [ 64640]
AY261366.1_Warthog             ..........................................g..a........g.......................g. [ 64640]
AY261365.1_Warmbaths           ..........................................g..a.................................. [ 64640]
AY261364.1_Tengani_62          ..........................................g..a........g......................... [ 64640]
AY261363.1_Pretoriuskop/96/4   ..........................................g..a........g......................... [ 64640]
AY261362.1_Mkuzi_1979          ..........................................g..................................... [ 64640]
AY261361.1_Malawi_Lil-20/1_1983 .....g.......................................................................... [ 64640]
AY261360.1_Kenya_1950          .....g....................................g..................................... [ 64640]
AM712240.1_OURT_88/3           ..........................................g....................c..............g. [ 64640]
AM712239.1_Benin_97/1          ..........................................g....................c..............g. [ 64640]
KM111295.1_Ken06.Bus           .........g................................g..................................... [ 64640]
KM111294.1_Ken05/Tk1           .....g....................................g..................................... [ 64640]
KM262844.1_strain_L60          ..........................................g....................c..............g. [ 64640]
KM102979.1_26544/OG10_Italy    ..........................................g....................c..............g. [ 64640]
KM262845.1_strain_NHV          ..........................................g....................c..............g. [ 64640]
KX354450.1_47/Ss/2008          ..........................................g....................c..............g. [ 64640]
KP055815.1_strain_BA71         ..........................................g....................c..............g. [ 64640]
MH025920.1_strain_R35          .........g................................g..................................... [ 64640]
MH025919.1_strain_N10          .........g................................g..................................... [ 64640]
MH025918.1_strain_R25          .........g................................g..................................... [ 64640]
MH025917.1_strain_R7           .........g................................g..................................... [ 64640]
MH025916.1_strain_R8           .........g................................g..................................... [ 64640]
```

FR682468.2_Georgia_2007/1 ggtctattttgccctgcgtgccatagcgtatggcatgtcgtgccaattgcaacaattctgacaccgatccgtgggccccg [ 64720]
ASF_IND_20_CAD543 ................................................................................ [ 64720]
LR899193.1_Germany_2020/1 ................................................................................ [ 64720]
MN393477.1_Wuhan_2019 ................................................................................ [ 64720]
MW396979.1_Timor-Leste/2019/1 ................................................................................ [ 64720]
MK940252.1_CN/2019/InnerMongolia ................................................................................ [ 64720]
MK128995.1_China/2018/AnhuiXCGQ ................................................................................ [ 64720]
MH681419.1_ASFV/POL/2015 ................................................................................ [ 64720]
LS478113.1_Estonia_2014 ................................................................................ [ 64720]
KP843857.1_Odintsovo_02/14 ................................................................................ [ 64720]
FN557520.1_strain_E75 ...................a..............g..a.......................................... [ 64720]
AY261366.1_Warthog ................t............................................................... [ 64720]
AY261365.1_Warmbaths ................................................................................ [ 64720]
AY261364.1_Tengani_62 ................................................................................ [ 64720]
AY261363.1_Pretoriuskop/96/4 ................................................................................ [ 64720]
AY261362.1_Mkuzi_1979 ................................................................................ [ 64720]
AY261361.1_Malawi_Lil-20/1_1983 ....g...........t....................a........g................................. [ 64720]
AY261360.1_Kenya_1950 ................t.....g..............a......................c................... [ 64720]
AM712240.1_OURT_88/3 ...................a..............g..a.......................................... [ 64720]
AM712239.1_Benin_97/1 ...................a..............g..a.......................................... [ 64720]
KM111295.1_Ken06.Bus ................t.....g..............a......................c...t............... [ 64720]
KM111294.1_Ken05/Tk1 ................t.....g..............a......................c..................a [ 64720]
KM262844.1_strain_L60 ...................a..............g..a.......................................... [ 64720]
KM102979.1_26544/OG10_Italy ...................a..............g..a.......................................... [ 64720]
KM262845.1_strain_NHV ...................a..............g..a.......................................... [ 64720]
KX354450.1_47/Ss/2008 ...................a..............g..a.......................................... [ 64720]
KP055815.1_strain_BA71 ...................a..............g..a.......................................... [ 64720]
MH025920.1_strain_R35 ................t.....g..............a......................c...t............... [ 64720]
MH025919.1_strain_N10 ................t.....g..............a......................c...t............... [ 64720]
MH025918.1_strain_R25 ................t.....g..............a......................c...t............... [ 64720]
MH025917.1_strain_R7 ................t.....g..............a......................c...t............... [ 64720]
MH025916.1_strain_R8 ................t.....g..............a......................c...t............... [ 64720]

FR682468.2_Georgia_2007/1 atccagtttatcggataggcaacctccgaagggtttaaaagatgctcgtaaaagcgtggatcttcagatgccaaggcgtc [ 64800]
ASF_IND_20_CAD543 ................................................................................ [ 64800]
LR899193.1_Germany_2020/1 ................................................................................ [ 64800]
MN393477.1_Wuhan_2019 ................................................................................ [ 64800]
MW396979.1_Timor-Leste/2019/1 ................................................................................ [ 64800]
MK940252.1_CN/2019/InnerMongolia ................................................................................ [ 64800]
MK128995.1_China/2018/AnhuiXCGQ ................................................................................ [ 64800]
MH681419.1_ASFV/POL/2015 ................................................................................ [ 64800]
LS478113.1_Estonia_2014 ................................................................................ [ 64800]
KP843857.1_Odintsovo_02/14 ................................................................................ [ 64800]

```
FN557520.1_strain_E75          ..........................t.........................................c........... [ 64800]
AY261366.1_Warthog             .................................................................g.............. [ 64800]
AY261365.1_Warmbaths           .....................................................a...........g.............. [ 64800]
AY261364.1_Tengani_62          ................................................................................ [ 64800]
AY261363.1_Pretoriuskop/96/4   ................................................................................ [ 64800]
AY261362.1_Mkuzi_1979          ....................................................................c........... [ 64800]
AY261361.1_Malawi_Lil-20/1_1983 .................................................................g.............. [ 64800]
AY261360.1_Kenya_1950          .................................................................g.............. [ 64800]
AM712240.1_OURT_88/3           ..........................t.........................................c........... [ 64800]
AM712239.1_Benin_97/1          ..........................t.........................................c........... [ 64800]
KM111295.1_Ken06.Bus           ................................a................................g.............. [ 64800]
KM111294.1_Ken05/Tk1           .................................................................g.............. [ 64800]
KM262844.1_strain_L60          ..........................t.........................................c........... [ 64800]
KM102979.1_26544/OG10_Italy    ..........................t.........................................c........... [ 64800]
KM262845.1_strain_NHV          ..........................t.........................................c........... [ 64800]
KX354450.1_47/Ss/2008          ..........................t.........................................c........... [ 64800]
KP055815.1_strain_BA71         ..........................t.........................................c........... [ 64800]
MH025920.1_strain_R35          ................................a................................g.............. [ 64800]
MH025919.1_strain_N10          ................................a................................g.............. [ 64800]
MH025918.1_strain_R25          ................................a................................g.............. [ 64800]
MH025917.1_strain_R7           ................................a................................g.............. [ 64800]
MH025916.1_strain_R8           ................................a................................g.............. [ 64800]

FR682468.2_Georgia_2007/1      tgcaaaggggataatgctagaaaacctgtctagacatacgttttctgtgtttacttctaaaggtagaaaaatggttgcgt [ 64880]
ASF_IND_20_CAD543              ................................................................................ [ 64880]
LR899193.1_Germany_2020/1      ................................................................................ [ 64880]
MN393477.1_Wuhan_2019          ................................................................................ [ 64880]
MW396979.1_Timor-Leste/2019/1  ................................................................................ [ 64880]
MK940252.1_CN/2019/InnerMongolia ................................................................................ [ 64880]
MK128995.1_China/2018/AnhuiXCGQ ................................................................................ [ 64880]
MH681419.1_ASFV/POL/2015       ................................................................................ [ 64880]
LS478113.1_Estonia_2014        ................................................................................ [ 64880]
KP843857.1_Odintsovo_02/14     ................................................................................ [ 64880]
FN557520.1_strain_E75          ......a........a.................g.............................................. [ 64880]
AY261366.1_Warthog             ...............a...................................c............................ [ 64880]
AY261365.1_Warmbaths           ...............a...................................c............................ [ 64880]
AY261364.1_Tengani_62          ...............a................................................................ [ 64880]
AY261363.1_Pretoriuskop/96/4   ...............a...................................c............................ [ 64880]
AY261362.1_Mkuzi_1979          ......a........a.................g.............................................. [ 64880]
AY261361.1_Malawi_Lil-20/1_1983 .........a.....a..g......t..................................g...........a....... [ 64880]
AY261360.1_Kenya_1950          ...............a..g......t.......g.............................................. [ 64880]
AM712240.1_OURT_88/3           ......a........a.................g.............................................. [ 64880]
AM712239.1_Benin_97/1          ......a........a.................g.............................................. [ 64880]
KM111295.1_Ken06.Bus           ...............a..g......t...................................................... [ 64880]
```

```
KM111294.1_Ken05/Tk1             ...............a..g......t.......g.............................................. [ 64880]
KM262844.1_strain_L60            ......a........a.................g.............................................. [ 64880]
KM102979.1_26544/OG10_Italy      ......a........a.................g.............................................. [ 64880]
KM262845.1_strain_NHV            ......a........a.................g.............................................. [ 64880]
KX354450.1_47/Ss/2008            ......a........a.................g.............................................. [ 64880]
KP055815.1_strain_BA71           ......a........a.................g.............................................. [ 64880]
MH025920.1_strain_R35            ...............a..g......t...................................................... [ 64880]
MH025919.1_strain_N10            ...............a..g......t...................................................... [ 64880]
MH025918.1_strain_R25            ...............a..g......t...................................................... [ 64880]
MH025917.1_strain_R7             ...............a..g......t...................................................... [ 64880]
MH025916.1_strain_R8             ...............a..g......t...................................................... [ 64880]

FR682468.2_Georgia_2007/1        gaggcttttgaacctgcttgttcagcggtctgcatatgctttgaataatgtctctaggactatgtcgcggcgctgcaaaa [ 64960]
ASF_IND_20_CAD543                ................................................................................ [ 64960]
LR899193.1_Germany_2020/1        ................................................................................ [ 64960]
MN393477.1_Wuhan_2019            ................................................................................ [ 64960]
MW396979.1_Timor-Leste/2019/1    ................................................................................ [ 64960]
MK940252.1_CN/2019/InnerMongolia ................................................................................ [ 64960]
MK128995.1_China/2018/AnhuiXCGQ  ................................................................................ [ 64960]
MH681419.1_ASFV/POL/2015         ................................................................................ [ 64960]
LS478113.1_Estonia_2014          ................................................................................ [ 64960]
KP843857.1_Odintsovo_02/14       ................................................................................ [ 64960]
FN557520.1_strain_E75            ................................................................a............... [ 64960]
AY261366.1_Warthog               ................................................................................ [ 64960]
AY261365.1_Warmbaths             ................................................................................ [ 64960]
AY261364.1_Tengani_62            ................................................................................ [ 64960]
AY261363.1_Pretoriuskop/96/4     .....................................a.......................................... [ 64960]
AY261362.1_Mkuzi_1979            ................................................................................ [ 64960]
AY261361.1_Malawi_Lil-20/1_1983  ............................c................................................... [ 64960]
AY261360.1_Kenya_1950            ............................c..............t.................................... [ 64960]
AM712240.1_OURT_88/3             ................................................................a............... [ 64960]
AM712239.1_Benin_97/1            ................................................................a............... [ 64960]
KM111295.1_Ken06.Bus             ............................c..............t.................................... [ 64960]
KM111294.1_Ken05/Tk1             ............................c..............t.................................... [ 64960]
KM262844.1_strain_L60            ................................................................a............... [ 64960]
KM102979.1_26544/OG10_Italy      ................................................................a............... [ 64960]
KM262845.1_strain_NHV            ................................................................a............... [ 64960]
KX354450.1_47/Ss/2008            ................................................................a............... [ 64960]
KP055815.1_strain_BA71           ................................................................a............... [ 64960]
MH025920.1_strain_R35            ............................c..............t.................................... [ 64960]
MH025919.1_strain_N10            ............................c..............t.................................... [ 64960]
MH025918.1_strain_R25            ............................c..............t.................................... [ 64960]
MH025917.1_strain_R7             ............................c..............t.................................... [ 64960]
MH025916.1_strain_R8             ............................c..............t.................................... [ 64960]
```

```
FR682468.2_Georgia_2007/1        aataccgcgtttagttctggaacctctacgccctcttgaaagagtcgacagtttaataaaataacgggttcctttgagga [ 65040]
ASF_IND_20_CAD543                ................................................................................ [ 65040]
LR899193.1_Germany_2020/1        ................................................................................ [ 65040]
MN393477.1_Wuhan_2019            ................................................................................ [ 65040]
MW396979.1_Timor-Leste/2019/1    ................................................................................ [ 65040]
MK940252.1_CN/2019/InnerMongolia ................................................................................ [ 65040]
MK128995.1_China/2018/AnhuiXCGQ  ................................................................................ [ 65040]
MH681419.1_ASFV/POL/2015         ................................................................................ [ 65040]
LS478113.1_Estonia_2014          ................................................................................ [ 65040]
KP843857.1_Odintsovo_02/14       ................................................................................ [ 65040]
FN557520.1_strain_E75            .................c.............................................................. [ 65040]
AY261366.1_Warthog               ................................................................................ [ 65040]
AY261365.1_Warmbaths             ................................................................................ [ 65040]
AY261364.1_Tengani_62            ....................g........................................................... [ 65040]
AY261363.1_Pretoriuskop/96/4     ....................g........................................................... [ 65040]
AY261362.1_Mkuzi_1979            ................................................................................ [ 65040]
AY261361.1_Malawi_Lil-20/1_1983  ........a...........t........c.................g................................ [ 65040]
AY261360.1_Kenya_1950            ....................t........c.................g................................ [ 65040]
AM712240.1_OURT_88/3             .................c.............................................................. [ 65040]
AM712239.1_Benin_97/1            .................c.............................................................. [ 65040]
KM111295.1_Ken06.Bus             .....t..............t........................................................... [ 65040]
KM111294.1_Ken05/Tk1             ....................t........c.................g................................ [ 65040]
KM262844.1_strain_L60            .................c.............................................................. [ 65040]
KM102979.1_26544/OG10_Italy      .................c.............................................................. [ 65040]
KM262845.1_strain_NHV            .................c.............................................................. [ 65040]
KX354450.1_47/Ss/2008            .................c.............................................................. [ 65040]
KP055815.1_strain_BA71           .................c.............................................................. [ 65040]
MH025920.1_strain_R35            .....t..............t........................................................... [ 65040]
MH025919.1_strain_N10            .....t..............t........................................................... [ 65040]
MH025918.1_strain_R25            .....t..............t........................................................... [ 65040]
MH025917.1_strain_R7             .....t..............t........................................................... [ 65040]
MH025916.1_strain_R8             .....t..............t........................................................... [ 65040]

FR682468.2_Georgia_2007/1        acaaaattctgtaaatgttttgaggataacctgtcgcggcagggttgagtgagctatcagggcatagacccccttggtcta [ 65120]
ASF_IND_20_CAD543                ................................................................................ [ 65120]
LR899193.1_Germany_2020/1        ................................................................................ [ 65120]
MN393477.1_Wuhan_2019            ................................................................................ [ 65120]
MW396979.1_Timor-Leste/2019/1    ................................................................................ [ 65120]
MK940252.1_CN/2019/InnerMongolia ................................................................................ [ 65120]
MK128995.1_China/2018/AnhuiXCGQ  ................................................................................ [ 65120]
MH681419.1_ASFV/POL/2015         ................................................................................ [ 65120]
LS478113.1_Estonia_2014          ................................................................................ [ 65120]
KP843857.1_Odintsovo_02/14       ................................................................................ [ 65120]
```

```
FN557520.1_strain_E75            ................................................................................ [ 65120]
AY261366.1_Warthog               ........................................................................c....... [ 65120]
AY261365.1_Warmbaths             ..................................................................a.....c....... [ 65120]
AY261364.1_Tengani_62            ........................................................................c....... [ 65120]
AY261363.1_Pretoriuskop/96/4     ........................................................................c....... [ 65120]
AY261362.1_Mkuzi_1979            ................................................................................ [ 65120]
AY261361.1_Malawi_Lil-20/1_1983  g.............................t..c.......................................a...... [ 65120]
AY261360.1_Kenya_1950            g..............................................................g.........a....c. [ 65120]
AM712240.1_OURT_88/3             ................................................................................ [ 65120]
AM712239.1_Benin_97/1            ................................................................................ [ 65120]
KM111295.1_Ken06.Bus             g........................................................................a....c. [ 65120]
KM111294.1_Ken05/Tk1             g..............................................................g.........a....c. [ 65120]
KM262844.1_strain_L60            ................................................................................ [ 65120]
KM102979.1_26544/OG10_Italy      ................................................................................ [ 65120]
KM262845.1_strain_NHV            ................................................................................ [ 65120]
KX354450.1_47/Ss/2008            ................................................................................ [ 65120]
KP055815.1_strain_BA71           ................................................................................ [ 65120]
MH025920.1_strain_R35            g........................................................................a....c. [ 65120]
MH025919.1_strain_N10            g........................................................................a....c. [ 65120]
MH025918.1_strain_R25            g........................................................................a....c. [ 65120]
MH025917.1_strain_R7             g........................................................................a....c. [ 65120]
MH025916.1_strain_R8             g........................................................................a....c. [ 65120]

FR682468.2_Georgia_2007/1        ccaacgccgcgtatagctccttggcctgtttaatatcacgggtaaataccagcattttaggagccggtatattggttttt [ 65200]
ASF_IND_20_CAD543                ................................................................................ [ 65200]
LR899193.1_Germany_2020/1        ................................................................................ [ 65200]
MN393477.1_Wuhan_2019            ................................................................................ [ 65200]
MW396979.1_Timor-Leste/2019/1    ................................................................................ [ 65200]
MK940252.1_CN/2019/InnerMongolia ................................................................................ [ 65200]
MK128995.1_China/2018/AnhuiXCGQ  ................................................................................ [ 65200]
MH681419.1_ASFV/POL/2015         ................................................................................ [ 65200]
LS478113.1_Estonia_2014          ................................................................................ [ 65200]
KP843857.1_Odintsovo_02/14       ................................................................................ [ 65200]
FN557520.1_strain_E75            .....t..............c......................g.................................... [ 65200]
AY261366.1_Warthog               .....t.........................g..................................t............. [ 65200]
AY261365.1_Warmbaths             .....t............................................................t............. [ 65200]
AY261364.1_Tengani_62            .....a............................................................t............. [ 65200]
AY261363.1_Pretoriuskop/96/4     .....t............................................................t............. [ 65200]
AY261362.1_Mkuzi_1979            .....t.....................................g.................................... [ 65200]
AY261361.1_Malawi_Lil-20/1_1983  .....t.....................................g.................c.a..t............. [ 65200]
AY261360.1_Kenya_1950            ...g.t.....................................g.................c.a.ct..c.......... [ 65200]
AM712240.1_OURT_88/3             .....t..............c......................g.................................... [ 65200]
AM712239.1_Benin_97/1            .....t..............c......................g.................................... [ 65200]
KM111295.1_Ken06.Bus             ...g.t.....................................g.................c.a.ct..c.......... [ 65200]
```

```
KM111294.1_Ken05/Tk1            ...g.t.....................................g.................c.a.ct..c.......... [ 65200]
KM262844.1_strain_L60           .....t..............c......................g.................................... [ 65200]
KM102979.1_26544/OG10_Italy     .....t..............c......................g.................................... [ 65200]
KM262845.1_strain_NHV           .....t..............c......................g.................................... [ 65200]
KX354450.1_47/Ss/2008           .....t..............c......................g.................................... [ 65200]
KP055815.1_strain_BA71          .....t..............c......................g.................................... [ 65200]
MH025920.1_strain_R35           ...g.t.....................................g.................c.a.ct..c.......... [ 65200]
MH025919.1_strain_N10           ...g.t.....................................g.................c.a.ct..c.......... [ 65200]
MH025918.1_strain_R25           ...g.t.....................................g.................c.a.ct..c.......... [ 65200]
MH025917.1_strain_R7            ...g.t.....................................g.................c.a.ct..c.......... [ 65200]
MH025916.1_strain_R8            ...g.t.....................................g.................c.a.ct..c.......... [ 65200]

FR682468.2_Georgia_2007/1       aaataggctaaggccattataatttgctttactatgatctgtttcgtggtctcctctttggtactcggttggtgggccaa [ 65280]
ASF_IND_20_CAD543               ................................................................................ [ 65280]
LR899193.1_Germany_2020/1       ................................................................................ [ 65280]
MN393477.1_Wuhan_2019           ................................................................................ [ 65280]
MW396979.1_Timor-Leste/2019/1   ................................................................................ [ 65280]
MK940252.1_CN/2019/InnerMongolia ................................................................................ [ 65280]
MK128995.1_China/2018/AnhuiXCGQ ................................................................................ [ 65280]
MH681419.1_ASFV/POL/2015        ................................................................................ [ 65280]
LS478113.1_Estonia_2014         ................................................................................ [ 65280]
KP843857.1_Odintsovo_02/14      ................................................................................ [ 65280]
FN557520.1_strain_E75           .........................................................c..t.............a..... [ 65280]
AY261366.1_Warthog              ....g........................................................................... [ 65280]
AY261365.1_Warmbaths            ....g........................................................................... [ 65280]
AY261364.1_Tengani_62           ....g..........................................c................................ [ 65280]
AY261363.1_Pretoriuskop/96/4    ....g........................................................................... [ 65280]
AY261362.1_Mkuzi_1979           .........................................................c..t.............a..... [ 65280]
AY261361.1_Malawi_Lil-20/1_1983 ...........a..............t........t..............t......c..t.....a.c........... [ 65280]
AY261360.1_Kenya_1950           ..........................t.....g..t.....................c..t.....a............. [ 65280]
AM712240.1_OURT_88/3            .........................................................c..t.............a..... [ 65280]
AM712239.1_Benin_97/1           .........................................................c..t.............a..... [ 65280]
KM111295.1_Ken06.Bus            ................................g..t.............c.......c..t...cg.............. [ 65280]
KM111294.1_Ken05/Tk1            ..........................t.....g..t.....................c..t....ta............. [ 65280]
KM262844.1_strain_L60           .........................................................c..t.............a..... [ 65280]
KM102979.1_26544/OG10_Italy     .........................................................c..t.............a..... [ 65280]
KM262845.1_strain_NHV           .........................................................c..t.............a..... [ 65280]
KX354450.1_47/Ss/2008           .........................................................c..t.............a..... [ 65280]
KP055815.1_strain_BA71          .........................................................c..t.............a..... [ 65280]
MH025920.1_strain_R35           ................................g..t.............c.......c..t...cg.............. [ 65280]
MH025919.1_strain_N10           ................................g..t.............c.......c..t...cg.............. [ 65280]
MH025918.1_strain_R25           ................................g..t.............c.......c..t...cg.............. [ 65280]
MH025917.1_strain_R7            ................................g..t.............c.......c..t...cg.............. [ 65280]
MH025916.1_strain_R8            ................................g..t.............c.......c..t...cg.............. [ 65280]
```

```
FR682468.2_Georgia_2007/1        tttaggcgcggctaccatctgcaattcaaaatcatttacatagccggcctctatgccttctcgcagatagtagcgaaagg [ 65360]
ASF_IND_20_CAD543                ................................................................................ [ 65360]
LR899193.1_Germany_2020/1        ................................................................................ [ 65360]
MN393477.1_Wuhan_2019            ................................................................................ [ 65360]
MW396979.1_Timor-Leste/2019/1    ................................................................................ [ 65360]
MK940252.1_CN/2019/InnerMongolia ................................................................................ [ 65360]
MK128995.1_China/2018/AnhuiXCGQ  ................................................................................ [ 65360]
MH681419.1_ASFV/POL/2015         ................................................................................ [ 65360]
LS478113.1_Estonia_2014          ................................................................................ [ 65360]
KP843857.1_Odintsovo_02/14       ................................................................................ [ 65360]
FN557520.1_strain_E75            ...g..............t..t.......................................................... [ 65360]
AY261366.1_Warthog               ................................................................................ [ 65360]
AY261365.1_Warmbaths             ................................................................................ [ 65360]
AY261364.1_Tengani_62            ................................................................................ [ 65360]
AY261363.1_Pretoriuskop/96/4     ...g............................................................................ [ 65360]
AY261362.1_Mkuzi_1979            ...g...................g........................................................ [ 65360]
AY261361.1_Malawi_Lil-20/1_1983  ...g........c.................g.......................a......................... [ 65360]
AY261360.1_Kenya_1950            ...g........c.................g................................................. [ 65360]
AM712240.1_OURT_88/3             ...g..............t..t.......................................................... [ 65360]
AM712239.1_Benin_97/1            ...g..............t..t.......................................................... [ 65360]
KM111295.1_Ken06.Bus             ...g........c.................g................................g................ [ 65360]
KM111294.1_Ken05/Tk1             ...g........c.................g................................................. [ 65360]
KM262844.1_strain_L60            ...g..............t..t.......................................................... [ 65360]
KM102979.1_26544/OG10_Italy      ...g..............t..t.......................................................... [ 65360]
KM262845.1_strain_NHV            ...g..............t..t.......................................................... [ 65360]
KX354450.1_47/Ss/2008            ...g..............t..t.......................................................... [ 65360]
KP055815.1_strain_BA71           ...g..............t..t.......................................................... [ 65360]
MH025920.1_strain_R35            ...g........c.................g................................g................ [ 65360]
MH025919.1_strain_N10            ...g........c.................g................................g................ [ 65360]
MH025918.1_strain_R25            ...g........c.................g................................g................ [ 65360]
MH025917.1_strain_R7             ...g........c.................g................................g................ [ 65360]
MH025916.1_strain_R8             ...g........c.................g................................g................ [ 65360]

FR682468.2_Georgia_2007/1        caacgccgccaaaaagttcacgatttttcatggaaagcggggtgtcgtacctgggcgttgccgttaaaaaaagtcggtgc [ 65440]
ASF_IND_20_CAD543                ................................................................................ [ 65440]
LR899193.1_Germany_2020/1        ................................................................................ [ 65440]
MN393477.1_Wuhan_2019            ................................................................................ [ 65440]
MW396979.1_Timor-Leste/2019/1    ................................................................................ [ 65440]
MK940252.1_CN/2019/InnerMongolia ................................................................................ [ 65440]
MK128995.1_China/2018/AnhuiXCGQ  ................................................................................ [ 65440]
MH681419.1_ASFV/POL/2015         ................................................................................ [ 65440]
LS478113.1_Estonia_2014          ................................................................................ [ 65440]
KP843857.1_Odintsovo_02/14       ................................................................................ [ 65440]
```

```
FN557520.1_strain_E75             ................................................................................ [ 65440]
AY261366.1_Warthog                ................................................................................ [ 65440]
AY261365.1_Warmbaths              ................................................................................ [ 65440]
AY261364.1_Tengani_62             ................................................................................ [ 65440]
AY261363.1_Pretoriuskop/96/4      ................................................................................ [ 65440]
AY261362.1_Mkuzi_1979             ................................................................................ [ 65440]
AY261361.1_Malawi_Lil-20/1_1983   ................g.............................a................................. [ 65440]
AY261360.1_Kenya_1950             ......................g......................................................... [ 65440]
AM712240.1_OURT_88/3              ................................................................................ [ 65440]
AM712239.1_Benin_97/1             ................................................................................ [ 65440]
KM111295.1_Ken06.Bus              ................g.............................a................................. [ 65440]
KM111294.1_Ken05/Tk1              ................................................................................ [ 65440]
KM262844.1_strain_L60             ................................................................................ [ 65440]
KM102979.1_26544/OG10_Italy       ................................................................................ [ 65440]
KM262845.1_strain_NHV             ................................................................................ [ 65440]
KX354450.1_47/Ss/2008             ................................................................................ [ 65440]
KP055815.1_strain_BA71            ................................................................................ [ 65440]
MH025920.1_strain_R35             ................g.............................a................................. [ 65440]
MH025919.1_strain_N10             ................g.............................a................................. [ 65440]
MH025918.1_strain_R25             ................g.............................a................................. [ 65440]
MH025917.1_strain_R7              ................g.............................a................................. [ 65440]
MH025916.1_strain_R8              ................g.............................a................................. [ 65440]

FR682468.2_Georgia_2007/1         ccttttttaaagttgagcaacacgtgggtaaagggccgtgtctcccattcgccgcaaatccggtgacattcatcgctaat [ 65520]
ASF_IND_20_CAD543                 ................................................................................ [ 65520]
LR899193.1_Germany_2020/1         ................................................................................ [ 65520]
MN393477.1_Wuhan_2019             ................................................................................ [ 65520]
MW396979.1_Timor-Leste/2019/1     ................................................................................ [ 65520]
MK940252.1_CN/2019/InnerMongolia  ................................................................................ [ 65520]
MK128995.1_China/2018/AnhuiXCGQ   ................................................................................ [ 65520]
MH681419.1_ASFV/POL/2015          ................................................................................ [ 65520]
LS478113.1_Estonia_2014           ................................................................................ [ 65520]
KP843857.1_Odintsovo_02/14        ................................................................................ [ 65520]
FN557520.1_strain_E75             .......................a..........................a............................. [ 65520]
AY261366.1_Warthog                .......................a.....g.................................................. [ 65520]
AY261365.1_Warmbaths              .......................a........................................................ [ 65520]
AY261364.1_Tengani_62             ................................................................................ [ 65520]
AY261363.1_Pretoriuskop/96/4      .......................a.....g.................................................. [ 65520]
AY261362.1_Mkuzi_1979             .......................a..........................a............................. [ 65520]
AY261361.1_Malawi_Lil-20/1_1983   ..c....................a.....g..a......a..........a............................. [ 65520]
AY261360.1_Kenya_1950             ..............t........a.....g.................................................. [ 65520]
AM712240.1_OURT_88/3              .......................a..........................a............................. [ 65520]
AM712239.1_Benin_97/1             .......................a..........................a............................. [ 65520]
KM111295.1_Ken06.Bus              ..............t........a.....g.................................................. [ 65520]
```

```
KM111294.1_Ken05/Tk1          .......................a.....g.................................................. [ 65520]
KM262844.1_strain_L60         .......................a..........................a............................. [ 65520]
KM102979.1_26544/OG10_Italy   .......................a..........................a............................. [ 65520]
KM262845.1_strain_NHV         .......................a..........................a............................. [ 65520]
KX354450.1_47/Ss/2008         .......................a..........................a............................. [ 65520]
KP055815.1_strain_BA71        .......................a..........................a............................. [ 65520]
MH025920.1_strain_R35         ..............t........a.....g....a............................................. [ 65520]
MH025919.1_strain_N10         ..............t........a.....g....a............................................. [ 65520]
MH025918.1_strain_R25         ..............t........a.....g....a............................................. [ 65520]
MH025917.1_strain_R7          ..............t........a.....g....a............................................. [ 65520]
MH025916.1_strain_R8          ..............t........a.....g....a............................................. [ 65520]

FR682468.2_Georgia_2007/1     aataagatcgaaatcatccaccagtagcgtggaggattggtaggtggcaatcacaagaagagaaggggcctcccgtatcc [ 65600]
ASF_IND_20_CAD543             ................................................................................ [ 65600]
LR899193.1_Germany_2020/1     ................................................................................ [ 65600]
MN393477.1_Wuhan_2019         ................................................................................ [ 65600]
MW396979.1_Timor-Leste/2019/1 ................................................................................ [ 65600]
MK940252.1_CN/2019/InnerMongolia ................................................................................ [ 65600]
MK128995.1_China/2018/AnhuiXCGQ ................................................................................ [ 65600]
MH681419.1_ASFV/POL/2015      ................................................................................ [ 65600]
LS478113.1_Estonia_2014       ................................................................................ [ 65600]
KP843857.1_Odintsovo_02/14    ................................................................................ [ 65600]
FN557520.1_strain_E75         ................................................................................ [ 65600]
AY261366.1_Warthog            .....................t...............................................t.......... [ 65600]
AY261365.1_Warmbaths          ................................................................................ [ 65600]
AY261364.1_Tengani_62         ................................................................................ [ 65600]
AY261363.1_Pretoriuskop/96/4  ................................................................................ [ 65600]
AY261362.1_Mkuzi_1979         ................................................................................ [ 65600]
AY261361.1_Malawi_Lil-20/1_1983 ...................................................a.............a.......t...... [ 65600]
AY261360.1_Kenya_1950         ...................................................a.............a.......t...... [ 65600]
AM712240.1_OURT_88/3          ................................................................................ [ 65600]
AM712239.1_Benin_97/1         ................................................................................ [ 65600]
KM111295.1_Ken06.Bus          .......................a...........................a.............a.......t...... [ 65600]
KM111294.1_Ken05/Tk1          ...................................................a.............a.......t...... [ 65600]
KM262844.1_strain_L60         ................................................................................ [ 65600]
KM102979.1_26544/OG10_Italy   ................................................................................ [ 65600]
KM262845.1_strain_NHV         ................................................................................ [ 65600]
KX354450.1_47/Ss/2008         ................................................................................ [ 65600]
KP055815.1_strain_BA71        ................................................................................ [ 65600]
MH025920.1_strain_R35         .......................a...........................a.............a.......t...... [ 65600]
MH025919.1_strain_N10         .......................a...........................a.............a.......t...... [ 65600]
MH025918.1_strain_R25         .......................a...........................a.............a.......t...... [ 65600]
MH025917.1_strain_R7          .......................a...........................a.............a.......t...... [ 65600]
MH025916.1_strain_R8          .......................a...........................a.............a.......t...... [ 65600]
```

```
FR682468.2_Georgia_2007/1        gttttgcaataaagacaggattggtggtcatttctatattgtcgtgatttagcacaatgcgggtctggtcagaccccaca [ 65680]
ASF_IND_20_CAD543                ................................................................................ [ 65680]
LR899193.1_Germany_2020/1        ................................................................................ [ 65680]
MN393477.1_Wuhan_2019            ................................................................................ [ 65680]
MW396979.1_Timor-Leste/2019/1    ................................................................................ [ 65680]
MK940252.1_CN/2019/InnerMongolia ................................................................................ [ 65680]
MK128995.1_China/2018/AnhuiXCGQ  ................................................................................ [ 65680]
MH681419.1_ASFV/POL/2015         ................................................................................ [ 65680]
LS478113.1_Estonia_2014          ................................................................................ [ 65680]
KP843857.1_Odintsovo_02/14       ................................................................................ [ 65680]
FN557520.1_strain_E75            ................................................................................ [ 65680]
AY261366.1_Warthog               ......................................................................g......... [ 65680]
AY261365.1_Warmbaths             ......................................................................g......... [ 65680]
AY261364.1_Tengani_62            .......................................................g..............g......... [ 65680]
AY261363.1_Pretoriuskop/96/4     ......................................................................g......... [ 65680]
AY261362.1_Mkuzi_1979            ................................................................................ [ 65680]
AY261361.1_Malawi_Lil-20/1_1983  ......................................................................g......... [ 65680]
AY261360.1_Kenya_1950            ....ca......t......g.....a............g...............................g......... [ 65680]
AM712240.1_OURT_88/3             ................................................................................ [ 65680]
AM712239.1_Benin_97/1            ................................................................................ [ 65680]
KM111295.1_Ken06.Bus             ....ca......t......g.....a............g...............................g......... [ 65680]
KM111294.1_Ken05/Tk1             ....ca......t......g.....a............g...............................g......... [ 65680]
KM262844.1_strain_L60            ................................................................................ [ 65680]
KM102979.1_26544/OG10_Italy      ................................................................................ [ 65680]
KM262845.1_strain_NHV            ................................................................................ [ 65680]
KX354450.1_47/Ss/2008            ................................................................................ [ 65680]
KP055815.1_strain_BA71           ................................................................................ [ 65680]
MH025920.1_strain_R35            ....ca......t......g.....a............g...............................g......... [ 65680]
MH025919.1_strain_N10            ....ca......t......g.....a............g...............................g......... [ 65680]
MH025918.1_strain_R25            ....ca......t......g.....a............g...............................g......... [ 65680]
MH025917.1_strain_R7             ....ca......t......g.....a............g...............................g......... [ 65680]
MH025916.1_strain_R8             ....ca......t......g.....a............g...............................g......... [ 65680]

FR682468.2_Georgia_2007/1        agcaaaacgttcttcaaagaaattccatactgatagagtttttccagagtctgccgtagtagggacaggcccggcaccag [ 65760]
ASF_IND_20_CAD543                ................................................................................ [ 65760]
LR899193.1_Germany_2020/1        ................................................................................ [ 65760]
MN393477.1_Wuhan_2019            ................................................................................ [ 65760]
MW396979.1_Timor-Leste/2019/1    ................................................................................ [ 65760]
MK940252.1_CN/2019/InnerMongolia ................................................................................ [ 65760]
MK128995.1_China/2018/AnhuiXCGQ  ................................................................................ [ 65760]
MH681419.1_ASFV/POL/2015         ................................................................................ [ 65760]
LS478113.1_Estonia_2014          ................................................................................ [ 65760]
KP843857.1_Odintsovo_02/14       ................................................................................ [ 65760]
```

```
FN557520.1_strain_E75            ................................................................................ [ 65760]
AY261366.1_Warthog               .................g.............................................................. [ 65760]
AY261365.1_Warmbaths             ..t..............g.............................................................. [ 65760]
AY261364.1_Tengani_62            .................g.............................................................. [ 65760]
AY261363.1_Pretoriuskop/96/4     .................g.............................................................. [ 65760]
AY261362.1_Mkuzi_1979            .......t........................................................................ [ 65760]
AY261361.1_Malawi_Lil-20/1_1983  .....c...........g............a....c.............................t.............. [ 65760]
AY261360.1_Kenya_1950            ....t............g............a................g.................t.............. [ 65760]
AM712240.1_OURT_88/3             ................................................................................ [ 65760]
AM712239.1_Benin_97/1            ................................................................................ [ 65760]
KM111295.1_Ken06.Bus             ....t..t.........g............a................g.................t.............. [ 65760]
KM111294.1_Ken05/Tk1             ....t............g............a................g.................t.............. [ 65760]
KM262844.1_strain_L60            ................................................................................ [ 65760]
KM102979.1_26544/OG10_Italy      ................................................................................ [ 65760]
KM262845.1_strain_NHV            ................................................................................ [ 65760]
KX354450.1_47/Ss/2008            ................................................................................ [ 65760]
KP055815.1_strain_BA71           ................................................................................ [ 65760]
MH025920.1_strain_R35            ....t..t.........g............a................g.................t.............. [ 65760]
MH025919.1_strain_N10            ....t..t.........g............a................g.................t.............. [ 65760]
MH025918.1_strain_R25            ....t..t.........g............a................g.................t.............. [ 65760]
MH025917.1_strain_R7             ....t..t.........g............a................g.................t.............. [ 65760]
MH025916.1_strain_R8             ....t..t.........g............a................g.................t.............. [ 65760]

FR682468.2_Georgia_2007/1        gtacaaaacttttccttgaagataattggagaggataagataggcgacgcgagttttgccgcatcggcaggccatctgca [ 65840]
ASF_IND_20_CAD543                ................................................................................ [ 65840]
LR899193.1_Germany_2020/1        ................................................................................ [ 65840]
MN393477.1_Wuhan_2019            ................................................................................ [ 65840]
MW396979.1_Timor-Leste/2019/1    ................................................................................ [ 65840]
MK940252.1_CN/2019/InnerMongolia ................................................................................ [ 65840]
MK128995.1_China/2018/AnhuiXCGQ  ................................................................................ [ 65840]
MH681419.1_ASFV/POL/2015         ................................................................................ [ 65840]
LS478113.1_Estonia_2014          ................................................................................ [ 65840]
KP843857.1_Odintsovo_02/14       ................................................................................ [ 65840]
FN557520.1_strain_E75            ...............................................................g................ [ 65840]
AY261366.1_Warthog               ................................................................................ [ 65840]
AY261365.1_Warmbaths             .................................a.............................................. [ 65840]
AY261364.1_Tengani_62            ................................................................................ [ 65840]
AY261363.1_Pretoriuskop/96/4     ................................................................................ [ 65840]
AY261362.1_Mkuzi_1979            ...............................................................g................ [ 65840]
AY261361.1_Malawi_Lil-20/1_1983  .................................a.................g.................a.......... [ 65840]
AY261360.1_Kenya_1950            .................................a.................g............................ [ 65840]
AM712240.1_OURT_88/3             ...............................................................g................ [ 65840]
AM712239.1_Benin_97/1            ...............................................................g................ [ 65840]
KM111295.1_Ken06.Bus             ........t........................a.................g............................ [ 65840]
```

```
KM111294.1_Ken05/Tk1           ...........................a.....a.................g............................ [ 65840]
KM262844.1_strain_L60          ...............................................................g................ [ 65840]
KM102979.1_26544/OG10_Italy    ...............................................................g................ [ 65840]
KM262845.1_strain_NHV          ...............................................................g................ [ 65840]
KX354450.1_47/Ss/2008          ...............................................................g................ [ 65840]
KP055815.1_strain_BA71         ...............................................................g................ [ 65840]
MH025920.1_strain_R35          ........t........................a.................g............................ [ 65840]
MH025919.1_strain_N10          ........t........................a.................g............................ [ 65840]
MH025918.1_strain_R25          ........t........................a.................g............................ [ 65840]
MH025917.1_strain_R7           ........t........................a.................g............................ [ 65840]
MH025916.1_strain_R8           ........t........................a.................g............................ [ 65840]

FR682468.2_Georgia_2007/1      gaatggccctcccacttcgccgcagctcctgatagcccatattggccgcctccttctgataaagtcgatcctcgattgca [ 65920]
ASF_IND_20_CAD543              ................................................................................ [ 65920]
LR899193.1_Germany_2020/1      ................................................................................ [ 65920]
MN393477.1_Wuhan_2019          ................................................................................ [ 65920]
MW396979.1_Timor-Leste/2019/1  ................................................................................ [ 65920]
MK940252.1_CN/2019/InnerMongolia ................................................................................ [ 65920]
MK128995.1_China/2018/AnhuiXCGQ ................................................................................ [ 65920]
MH681419.1_ASFV/POL/2015       ................................................................................ [ 65920]
LS478113.1_Estonia_2014        ................................................................................ [ 65920]
KP843857.1_Odintsovo_02/14     ................................................................................ [ 65920]
FN557520.1_strain_E75          ..........t.....................................................c............... [ 65920]
AY261366.1_Warthog             ..........t..................................................................... [ 65920]
AY261365.1_Warmbaths           ..........t..................................................................... [ 65920]
AY261364.1_Tengani_62          ................................................................................ [ 65920]
AY261363.1_Pretoriuskop/96/4   ..........t..................................................................... [ 65920]
AY261362.1_Mkuzi_1979          ..........t..................................................................... [ 65920]
AY261361.1_Malawi_Lil-20/1_1983 ..........t........tt....................................................a..g... [ 65920]
AY261360.1_Kenya_1950          ..........t..g......t................................c...................a..g.t. [ 65920]
AM712240.1_OURT_88/3           ..........t.....................................................c............... [ 65920]
AM712239.1_Benin_97/1          ..........t.....................................................c............... [ 65920]
KM111295.1_Ken06.Bus           ....a..t..t..g......t................................c...................a..g.t. [ 65920]
KM111294.1_Ken05/Tk1           ..........t..g......t................................c...................a..g.t. [ 65920]
KM262844.1_strain_L60          ..........t.....................................................c............... [ 65920]
KM102979.1_26544/OG10_Italy    ..........t.....................................................c............... [ 65920]
KM262845.1_strain_NHV          ..........t.....................................................c............... [ 65920]
KX354450.1_47/Ss/2008          ..........t.....................................................c............... [ 65920]
KP055815.1_strain_BA71         ..........t.....................................................c............... [ 65920]
MH025920.1_strain_R35          ....a.tt..t..g......t................................c...................a..g.t. [ 65920]
MH025919.1_strain_N10          ....a.tt..t..g......t................................c...................a..g.t. [ 65920]
MH025918.1_strain_R25          ....a.tt..t..g......t................................c...................a..g.t. [ 65920]
MH025917.1_strain_R7           ....a.tt..t..g......t................................c...................a..g.t. [ 65920]
MH025916.1_strain_R8           ....a.tt..t..g......t................................c...................a..g.t. [ 65920]
```

```
FR682468.2_Georgia_2007/1        gtccgtgtctcatctgtagaaaaaaataatacgtcatctgcgaaatgttc---atcttccacaggagttatcaccaggtg [ 66000]
ASF_IND_20_CAD543                ..................................................ttc........................... [ 66000]
LR899193.1_Germany_2020/1        ..................................................---........................... [ 66000]
MN393477.1_Wuhan_2019            ..................................................---........................... [ 66000]
MW396979.1_Timor-Leste/2019/1    ..................................................---........................... [ 66000]
MK940252.1_CN/2019/InnerMongolia ..................................................---........................... [ 66000]
MK128995.1_China/2018/AnhuiXCGQ  ..................................................---........................... [ 66000]
MH681419.1_ASFV/POL/2015         ..................................................---........................... [ 66000]
LS478113.1_Estonia_2014          ........................................t.........---........................... [ 66000]
KP843857.1_Odintsovo_02/14       ..................................................---........................... [ 66000]
FN557520.1_strain_E75            ..................................................ttc........................... [ 66000]
AY261366.1_Warthog               ..............c...................................ttc........................... [ 66000]
AY261365.1_Warmbaths             ..............c...................................ttc........................... [ 66000]
AY261364.1_Tengani_62            ..................................................ttc........................... [ 66000]
AY261363.1_Pretoriuskop/96/4     ..............c...................................ttc........................... [ 66000]
AY261362.1_Mkuzi_1979            ..................................................ttc........................... [ 66000]
AY261361.1_Malawi_Lil-20/1_1983  ...............................................a..ttcg.....t.....ga.g..t....a... [ 66000]
AY261360.1_Kenya_1950            ...........tc................................c.a..ttc......t.....g.............. [ 66000]
AM712240.1_OURT_88/3             ..................................................ttc........................... [ 66000]
AM712239.1_Benin_97/1            ..................................................ttc........................... [ 66000]
KM111295.1_Ken06.Bus             ...........tc..................................a..ttcg.....t.....g.............. [ 66000]
KM111294.1_Ken05/Tk1             ...........tc................................c.a..ttc......t.....g.............. [ 66000]
KM262844.1_strain_L60            ..................................................ttc........................... [ 66000]
KM102979.1_26544/OG10_Italy      ..................................................ttc........................... [ 66000]
KM262845.1_strain_NHV            ..................................................ttc........................... [ 66000]
KX354450.1_47/Ss/2008            ..................................................ttc........................... [ 66000]
KP055815.1_strain_BA71           ..................................................ttc........................... [ 66000]
MH025920.1_strain_R35            ...........tc..................................a..ttcg.....t.....g.............. [ 66000]
MH025919.1_strain_N10            ...........tc..................................a..ttcg.....t.....g.............. [ 66000]
MH025918.1_strain_R25            ...........tc..................................a..ttcg.....t.....g.............. [ 66000]
MH025917.1_strain_R7             ...........tc..................................a..ttcg.....t.....g.............. [ 66000]
MH025916.1_strain_R8             ...........tc..................................a..ttcg.....t.....g.............. [ 66000]

FR682468.2_Georgia_2007/1        tctcagtttctccttgcttatcagcggatcagagggcaaagatggctcaaccactatcgtggaatcattcatctcatagg [ 66080]
ASF_IND_20_CAD543                ................................................................................ [ 66080]
LR899193.1_Germany_2020/1        ................................................................................ [ 66080]
MN393477.1_Wuhan_2019            ................................................................................ [ 66080]
MW396979.1_Timor-Leste/2019/1    ................................................................................ [ 66080]
MK940252.1_CN/2019/InnerMongolia ................................................................................ [ 66080]
MK128995.1_China/2018/AnhuiXCGQ  ................................................................................ [ 66080]
MH681419.1_ASFV/POL/2015         ................................................................................ [ 66080]
LS478113.1_Estonia_2014          ................................................................................ [ 66080]
KP843857.1_Odintsovo_02/14       ................................................................................ [ 66080]
```

```
FN557520.1_strain_E75            ..............................................................................a. [ 66080]
AY261366.1_Warthog               ................................................................................ [ 66080]
AY261365.1_Warmbaths             ................................................................................ [ 66080]
AY261364.1_Tengani_62            ................................................................................ [ 66080]
AY261363.1_Pretoriuskop/96/4     ................................................................................ [ 66080]
AY261362.1_Mkuzi_1979            ..............................................................................a. [ 66080]
AY261361.1_Malawi_Lil-20/1_1983  .............................................t.................................. [ 66080]
AY261360.1_Kenya_1950            ...t.................................g.......t.................................. [ 66080]
AM712240.1_OURT_88/3             ..............................................................................a. [ 66080]
AM712239.1_Benin_97/1            ..............................................................................a. [ 66080]
KM111295.1_Ken06.Bus             ...t.................................g.......t.................................. [ 66080]
KM111294.1_Ken05/Tk1             ...t.................................g.......t.................................. [ 66080]
KM262844.1_strain_L60            ..............................................................................a. [ 66080]
KM102979.1_26544/OG10_Italy      ..............................................................................a. [ 66080]
KM262845.1_strain_NHV            ..............................................................................a. [ 66080]
KX354450.1_47/Ss/2008            ..............................................................................a. [ 66080]
KP055815.1_strain_BA71           ..............................................................................a. [ 66080]
MH025920.1_strain_R35            ...t.................................g.......t.................................. [ 66080]
MH025919.1_strain_N10            ...t.................................g.......t.................................. [ 66080]
MH025918.1_strain_R25            ...t.................................g.......t.................................. [ 66080]
MH025917.1_strain_R7             ...t.................................g.......t.................................. [ 66080]
MH025916.1_strain_R8             ...t.................................g.......t.................................. [ 66080]

FR682468.2_Georgia_2007/1        cgggagaatcacacaaagtatagcttatgtccagacagtttgcaacatcctcagccaattgttttattttttcgggtaaa [ 66160]
ASF_IND_20_CAD543                ................................................................................ [ 66160]
LR899193.1_Germany_2020/1        ................................................................................ [ 66160]
MN393477.1_Wuhan_2019            ................................................................................ [ 66160]
MW396979.1_Timor-Leste/2019/1    ................................................................................ [ 66160]
MK940252.1_CN/2019/InnerMongolia ................................................................................ [ 66160]
MK128995.1_China/2018/AnhuiXCGQ  ................................................................................ [ 66160]
MH681419.1_ASFV/POL/2015         ................................................................................ [ 66160]
LS478113.1_Estonia_2014          ................................................................................ [ 66160]
KP843857.1_Odintsovo_02/14       ................................................................................ [ 66160]
FN557520.1_strain_E75            t............................................................................... [ 66160]
AY261366.1_Warthog               ................................................................................ [ 66160]
AY261365.1_Warmbaths             ................................................................................ [ 66160]
AY261364.1_Tengani_62            ................................................................................ [ 66160]
AY261363.1_Pretoriuskop/96/4     .................................a.............................................. [ 66160]
AY261362.1_Mkuzi_1979            t............................................................................... [ 66160]
AY261361.1_Malawi_Lil-20/1_1983  t.................................g..t...................................c...... [ 66160]
AY261360.1_Kenya_1950            t.....c.......................t............g.............................a...... [ 66160]
AM712240.1_OURT_88/3             t............................................................................... [ 66160]
AM712239.1_Benin_97/1            t............................................................................... [ 66160]
KM111295.1_Ken06.Bus             t.....c....................................g.............................a...... [ 66160]
```

```
KM111294.1_Ken05/Tk1            t.....c.......................t............g.............................a...... [ 66160]
KM262844.1_strain_L60           t............................................................................... [ 66160]
KM102979.1_26544/OG10_Italy     t............................................................................... [ 66160]
KM262845.1_strain_NHV           t............................................................................... [ 66160]
KX354450.1_47/Ss/2008           t............................................................................... [ 66160]
KP055815.1_strain_BA71          t............................................................................... [ 66160]
MH025920.1_strain_R35           t.....c....................................g.............................a...... [ 66160]
MH025919.1_strain_N10           t.....c....................................g.............................a...... [ 66160]
MH025918.1_strain_R25           t.....c....................................g.............................a...... [ 66160]
MH025917.1_strain_R7            t.....c....................................g.............................a...... [ 66160]
MH025916.1_strain_R8            t.....c....................................g.............................a...... [ 66160]

FR682468.2_Georgia_2007/1       agacatacgagttctttgtttttgacgcgaaaaaactgtgcacaatataacacccctgcttcaattttttgcgcatcctt [ 66240]
ASF_IND_20_CAD543               ................................................................................ [ 66240]
LR899193.1_Germany_2020/1       ................................................................................ [ 66240]
MN393477.1_Wuhan_2019           ................................................................................ [ 66240]
MW396979.1_Timor-Leste/2019/1   ................................................................................ [ 66240]
MK940252.1_CN/2019/InnerMongolia ................................................................................ [ 66240]
MK128995.1_China/2018/AnhuiXCGQ ................................................................................ [ 66240]
MH681419.1_ASFV/POL/2015        ................................................................................ [ 66240]
LS478113.1_Estonia_2014         ................................................................................ [ 66240]
KP843857.1_Odintsovo_02/14      ................................................................................ [ 66240]
FN557520.1_strain_E75           .........................................g.....c................................ [ 66240]
AY261366.1_Warthog              ................................................................................ [ 66240]
AY261365.1_Warmbaths            .......g.............................................t.......................... [ 66240]
AY261364.1_Tengani_62           ................................................................................ [ 66240]
AY261363.1_Pretoriuskop/96/4    ................................................................................ [ 66240]
AY261362.1_Mkuzi_1979           .........................................g.....c................................ [ 66240]
AY261361.1_Malawi_Lil-20/1_1983 ..g....g..................t....................c.....t.................t........ [ 66240]
AY261360.1_Kenya_1950           ..g....g..................t............................................t........ [ 66240]
AM712240.1_OURT_88/3            .........................................g.....c................................ [ 66240]
AM712239.1_Benin_97/1           .........................................g.....c................................ [ 66240]
KM111295.1_Ken06.Bus            ..g....g..................t............................................t........ [ 66240]
KM111294.1_Ken05/Tk1            ..g....g..................t............................................t........ [ 66240]
KM262844.1_strain_L60           .........................................g.....c................................ [ 66240]
KM102979.1_26544/OG10_Italy     .........................................g.....c................................ [ 66240]
KM262845.1_strain_NHV           .........................................g.....c................................ [ 66240]
KX354450.1_47/Ss/2008           .........................................g.....c................................ [ 66240]
KP055815.1_strain_BA71          .........................................g.....c................................ [ 66240]
MH025920.1_strain_R35           ..g....g..................t............................................t........ [ 66240]
MH025919.1_strain_N10           ..g....g..................t............................................t........ [ 66240]
MH025918.1_strain_R25           ..g....g..................t............................................t........ [ 66240]
MH025917.1_strain_R7            ..g....g..................t............................................t........ [ 66240]
MH025916.1_strain_R8            ..g....g..................t............................................t........ [ 66240]
```

```
FR682468.2_Georgia_2007/1        ctttgtagatgtttccaatgtgaaacaatacttccattcatccgtaaaacaggttgtataagatccatcatgaagcctag [ 66320]
ASF_IND_20_CAD543                ................................................................................ [ 66320]
LR899193.1_Germany_2020/1        ................................................................................ [ 66320]
MN393477.1_Wuhan_2019            ................................................................................ [ 66320]
MW396979.1_Timor-Leste/2019/1    ................................................................................ [ 66320]
MK940252.1_CN/2019/InnerMongolia ................................................................................ [ 66320]
MK128995.1_China/2018/AnhuiXCGQ  ................................................................................ [ 66320]
MH681419.1_ASFV/POL/2015         ................................................................................ [ 66320]
LS478113.1_Estonia_2014          ................................................................................ [ 66320]
KP843857.1_Odintsovo_02/14       ................................................................................ [ 66320]
FN557520.1_strain_E75            ........................................................................g....... [ 66320]
AY261366.1_Warthog               ......................................................g......................... [ 66320]
AY261365.1_Warmbaths             ...............................c...................a..c......................... [ 66320]
AY261364.1_Tengani_62            ................................................................................ [ 66320]
AY261363.1_Pretoriuskop/96/4     ......................................................g......................... [ 66320]
AY261362.1_Mkuzi_1979            ..................c.....................................................g....... [ 66320]
AY261361.1_Malawi_Lil-20/1_1983  ........g............a....................t.......................g............. [ 66320]
AY261360.1_Kenya_1950            ........gc...........a........tc................................................ [ 66320]
AM712240.1_OURT_88/3             ........................................................................g....... [ 66320]
AM712239.1_Benin_97/1            ........................................................................g....... [ 66320]
KM111295.1_Ken06.Bus             ........gc.....t.....a.........c..........................................a..... [ 66320]
KM111294.1_Ken05/Tk1             ........gc...........a........tc................................................ [ 66320]
KM262844.1_strain_L60            ........................................................................g....... [ 66320]
KM102979.1_26544/OG10_Italy      ........................................................................g....... [ 66320]
KM262845.1_strain_NHV            ........................................................................g....... [ 66320]
KX354450.1_47/Ss/2008            ........................................................................g....... [ 66320]
KP055815.1_strain_BA71           ........................................................................g....... [ 66320]
MH025920.1_strain_R35            ........gc.....t.....a.........c..........................................a..... [ 66320]
MH025919.1_strain_N10            ........gc.....t.....a.........c..........................................a..... [ 66320]
MH025918.1_strain_R25            ........gc.....t.....a.........c..........................................a..... [ 66320]
MH025917.1_strain_R7             ........gc.....t.....a.........c..........................................a..... [ 66320]
MH025916.1_strain_R8             ........gc.....t.....a.........c..........................................a..... [ 66320]

FR682468.2_Georgia_2007/1        cggccaagtttcctgtgtgcccaactttatgtaaggattgggcctccagccagggatgaaccgccacgtaaaatcctgcg [ 66400]
ASF_IND_20_CAD543                ................................................................................ [ 66400]
LR899193.1_Germany_2020/1        ................................................................................ [ 66400]
MN393477.1_Wuhan_2019            ................................................................................ [ 66400]
MW396979.1_Timor-Leste/2019/1    ................................................................................ [ 66400]
MK940252.1_CN/2019/InnerMongolia ................................................................................ [ 66400]
MK128995.1_China/2018/AnhuiXCGQ  ................................................................................ [ 66400]
MH681419.1_ASFV/POL/2015         ................................................................................ [ 66400]
LS478113.1_Estonia_2014          ................................................................................ [ 66400]
KP843857.1_Odintsovo_02/14       ................................................................................ [ 66400]
```

```
FN557520.1_strain_E75         ................................................................................ [ 66400]
AY261366.1_Warthog            ................................................................................ [ 66400]
AY261365.1_Warmbaths          ..............................................t................................. [ 66400]
AY261364.1_Tengani_62         ................................................................................ [ 66400]
AY261363.1_Pretoriuskop/96/4  ................................................................................ [ 66400]
AY261362.1_Mkuzi_1979         ................................................................................ [ 66400]
AY261361.1_Malawi_Lil-20/1_1983 ..........................................t..............................c..c... [ 66400]
AY261360.1_Kenya_1950         ..................................a.....a.t.....................t........c...... [ 66400]
AM712240.1_OURT_88/3          ................................................................................ [ 66400]
AM712239.1_Benin_97/1         ................................................................................ [ 66400]
KM111295.1_Ken06.Bus          ...............................c..a.....a.t.....................t........c...... [ 66400]
KM111294.1_Ken05/Tk1          ..................................a.....a.t.....................t........c...... [ 66400]
KM262844.1_strain_L60         ................................................................................ [ 66400]
KM102979.1_26544/OG10_Italy   ................................................................................ [ 66400]
KM262845.1_strain_NHV         ................................................................................ [ 66400]
KX354450.1_47/Ss/2008         ................................................................................ [ 66400]
KP055815.1_strain_BA71        ................................................................................ [ 66400]
MH025920.1_strain_R35         ...............................c..a.....a.t.....................t........c...... [ 66400]
MH025919.1_strain_N10         ...............................c..a.....a.t.....................t........c...... [ 66400]
MH025918.1_strain_R25         ...............................c..a.....a.t.....................t........c...... [ 66400]
MH025917.1_strain_R7          ...............................c..a.....a.t.....................t........c...... [ 66400]
MH025916.1_strain_R8          ...............................c..a.....a.t.....................t........c...... [ 66400]

FR682468.2_Georgia_2007/1     cacatgctatatcaaattgcagtttcttaataactgtacacaggatctgaaaaacatgtgattacaaaatttagataaga [ 66480]
ASF_IND_20_CAD543             ................................................................................ [ 66480]
LR899193.1_Germany_2020/1     ................................................................................ [ 66480]
MN393477.1_Wuhan_2019         ................................................................................ [ 66480]
MW396979.1_Timor-Leste/2019/1 ................................................................................ [ 66480]
MK940252.1_CN/2019/InnerMongolia ................................................................................ [ 66480]
MK128995.1_China/2018/AnhuiXCGQ ................................................................................ [ 66480]
MH681419.1_ASFV/POL/2015      ................................................................................ [ 66480]
LS478113.1_Estonia_2014       ................................................................................ [ 66480]
KP843857.1_Odintsovo_02/14    ................................................................................ [ 66480]
FN557520.1_strain_E75         ................................................................................ [ 66480]
AY261366.1_Warthog            ......t......................................................................... [ 66480]
AY261365.1_Warmbaths          ................................................................................ [ 66480]
AY261364.1_Tengani_62         ................................................................................ [ 66480]
AY261363.1_Pretoriuskop/96/4  ................................................................................ [ 66480]
AY261362.1_Mkuzi_1979         ................................................................................ [ 66480]
AY261361.1_Malawi_Lil-20/1_1983 t...............................................t............................... [ 66480]
AY261360.1_Kenya_1950         .........g.......................a............t.t............................... [ 66480]
AM712240.1_OURT_88/3          ................................................................................ [ 66480]
AM712239.1_Benin_97/1         ................................................................................ [ 66480]
KM111295.1_Ken06.Bus          .........g.......................a............t.t............................... [ 66480]
```

```
KM111294.1_Ken05/Tk1           .........g.......................a..............t............................... [ 66480]
KM262844.1_strain_L60          ................................................................................ [ 66480]
KM102979.1_26544/OG10_Italy    ................................................................................ [ 66480]
KM262845.1_strain_NHV          ................................................................................ [ 66480]
KX354450.1_47/Ss/2008          ................................................................................ [ 66480]
KP055815.1_strain_BA71         ................................................................................ [ 66480]
MH025920.1_strain_R35          .........g.......................a............t.t............................... [ 66480]
MH025919.1_strain_N10          .........g.......................a............t.t............................... [ 66480]
MH025918.1_strain_R25          .........g.......................a............t.t............................... [ 66480]
MH025917.1_strain_R7           .........g.......................a............t.t............................... [ 66480]
MH025916.1_strain_R8           .........g.......................a............t.t............................... [ 66480]

FR682468.2_Georgia_2007/1      aatatttaatattaaaaatcacagaatacatgtcactgtgtagagagaaagccaaaaactcctcttgaccgccgtgggaa [ 66560]
ASF_IND_20_CAD543              ......................r......................................................... [ 66560]
LR899193.1_Germany_2020/1      ................................................................................ [ 66560]
MN393477.1_Wuhan_2019          ................................................................................ [ 66560]
MW396979.1_Timor-Leste/2019/1  ................................................................................ [ 66560]
MK940252.1_CN/2019/InnerMongolia ................................................................................ [ 66560]
MK128995.1_China/2018/AnhuiXCGQ ................................................................................ [ 66560]
MH681419.1_ASFV/POL/2015       ................................................................................ [ 66560]
LS478113.1_Estonia_2014        ................................................................................ [ 66560]
KP843857.1_Odintsovo_02/14     ................................................................................ [ 66560]
FN557520.1_strain_E75          ......................g......................................................... [ 66560]
AY261366.1_Warthog             ......................g.....t................................................... [ 66560]
AY261365.1_Warmbaths           ......................g.....t.......................t........................... [ 66560]
AY261364.1_Tengani_62          ......................g.....a................................................... [ 66560]
AY261363.1_Pretoriuskop/96/4   ......................g.....t.............a..................................... [ 66560]
AY261362.1_Mkuzi_1979          ......................g......................................................... [ 66560]
AY261361.1_Malawi_Lil-20/1_1983 ......................g.....a................................................... [ 66560]
AY261360.1_Kenya_1950          ......................g.....a................................................... [ 66560]
AM712240.1_OURT_88/3           ......................g......................................................... [ 66560]
AM712239.1_Benin_97/1          ......................g......................................................... [ 66560]
KM111295.1_Ken06.Bus           ......................g.....a................................................... [ 66560]
KM111294.1_Ken05/Tk1           ......................g.....a................................................... [ 66560]
KM262844.1_strain_L60          ......................g......................................................... [ 66560]
KM102979.1_26544/OG10_Italy    ......................g......................................................... [ 66560]
KM262845.1_strain_NHV          ......................g......................................................... [ 66560]
KX354450.1_47/Ss/2008          ......................g......................................................... [ 66560]
KP055815.1_strain_BA71         ......................g......................................................... [ 66560]
MH025920.1_strain_R35          ......................g.....a................................................... [ 66560]
MH025919.1_strain_N10          ......................g.....a................................................... [ 66560]
MH025918.1_strain_R25          ......................g.....a................................................... [ 66560]
MH025917.1_strain_R7           ......................g.....a................................................... [ 66560]
MH025916.1_strain_R8           ......................g.....a................................................... [ 66560]
```

```
FR682468.2_Georgia_2007/1        atcatccagggtagtaggttgtgtttcataaagttgtatgccgtagtgatcaccgtggactccagatggttattggcatc [ 66640]
ASF_IND_20_CAD543                .............................r.................................................. [ 66640]
LR899193.1_Germany_2020/1        ................................................................................ [ 66640]
MN393477.1_Wuhan_2019            ................................................................................ [ 66640]
MW396979.1_Timor-Leste/2019/1    ................................................................................ [ 66640]
MK940252.1_CN/2019/InnerMongolia ................................................................................ [ 66640]
MK128995.1_China/2018/AnhuiXCGQ  ................................................................................ [ 66640]
MH681419.1_ASFV/POL/2015         ................................................................................ [ 66640]
LS478113.1_Estonia_2014          ................................................................................ [ 66640]
KP843857.1_Odintsovo_02/14       ................................................................................ [ 66640]
FN557520.1_strain_E75            .............................g...............................................g.. [ 66640]
AY261366.1_Warthog               .............................g.................................................. [ 66640]
AY261365.1_Warmbaths             .............................g.................................................. [ 66640]
AY261364.1_Tengani_62            .............................g.................................................. [ 66640]
AY261363.1_Pretoriuskop/96/4     .............................g.................................................. [ 66640]
AY261362.1_Mkuzi_1979            .............................g.................................................. [ 66640]
AY261361.1_Malawi_Lil-20/1_1983  ........t....................g.................................................. [ 66640]
AY261360.1_Kenya_1950            .............................g.........................................g........ [ 66640]
AM712240.1_OURT_88/3             .............................g...............................................g.. [ 66640]
AM712239.1_Benin_97/1            .............................g...............................................g.. [ 66640]
KM111295.1_Ken06.Bus             .............................g.................................................. [ 66640]
KM111294.1_Ken05/Tk1             .............................g.........................................g........ [ 66640]
KM262844.1_strain_L60            .............................g...............................................g.. [ 66640]
KM102979.1_26544/OG10_Italy      .............................g...............................................g.. [ 66640]
KM262845.1_strain_NHV            .............................g...............................................g.. [ 66640]
KX354450.1_47/Ss/2008            .............................g...............................................g.. [ 66640]
KP055815.1_strain_BA71           .............................g...............................................g.. [ 66640]
MH025920.1_strain_R35            .............................g.................................................. [ 66640]
MH025919.1_strain_N10            .............................g.................................................. [ 66640]
MH025918.1_strain_R25            .............................g.................................................. [ 66640]
MH025917.1_strain_R7             .............................g.................................................. [ 66640]
MH025916.1_strain_R8             .............................g.................................................. [ 66640]

FR682468.2_Georgia_2007/1        tttgcaatactttgccatcttggcagaaaagacgataaatccacaaattctaccccagttgataagatccttaaacagct [ 66720]
ASF_IND_20_CAD543                ................................................................................ [ 66720]
LR899193.1_Germany_2020/1        ................................................................................ [ 66720]
MN393477.1_Wuhan_2019            ................................................................................ [ 66720]
MW396979.1_Timor-Leste/2019/1    ................................................................................ [ 66720]
MK940252.1_CN/2019/InnerMongolia ................................................................................ [ 66720]
MK128995.1_China/2018/AnhuiXCGQ  ................................................................................ [ 66720]
MH681419.1_ASFV/POL/2015         ................................................................................ [ 66720]
LS478113.1_Estonia_2014          ................................................................................ [ 66720]
KP843857.1_Odintsovo_02/14       ................................................................................ [ 66720]
```

```
FN557520.1_strain_E75            ................................................................................ [ 66720]
AY261366.1_Warthog               .............................................................................a.. [ 66720]
AY261365.1_Warmbaths             .............................................................................a.. [ 66720]
AY261364.1_Tengani_62            ................................................c............................a.. [ 66720]
AY261363.1_Pretoriuskop/96/4     .............................................................................a.. [ 66720]
AY261362.1_Mkuzi_1979            ................................................................................ [ 66720]
AY261361.1_Malawi_Lil-20/1_1983  ...................c.........................................................a.. [ 66720]
AY261360.1_Kenya_1950            ......g............c.........................................................a.. [ 66720]
AM712240.1_OURT_88/3             ................................................................................ [ 66720]
AM712239.1_Benin_97/1            ................................................................................ [ 66720]
KM111295.1_Ken06.Bus             ......g............c.........................................................a.. [ 66720]
KM111294.1_Ken05/Tk1             ......g............c.........................................................a.. [ 66720]
KM262844.1_strain_L60            ................................................................................ [ 66720]
KM102979.1_26544/OG10_Italy      ................................................................................ [ 66720]
KM262845.1_strain_NHV            ................................................................................ [ 66720]
KX354450.1_47/Ss/2008            ................................................................................ [ 66720]
KP055815.1_strain_BA71           ................................................................................ [ 66720]
MH025920.1_strain_R35            ......g............c.........................................................a.. [ 66720]
MH025919.1_strain_N10            ......g............c.........................................................a.. [ 66720]
MH025918.1_strain_R25            ......g............c.........................................................a.. [ 66720]
MH025917.1_strain_R7             ......g............c.........................................................a.. [ 66720]
MH025916.1_strain_R8             ......g............c.........................................................a.. [ 66720]

FR682468.2_Georgia_2007/1        cagtcacaaccccagtaaactgggttttaatttcttgaacactcgtaagagaaaaggtaattgtaacctgtttgttcaaa [ 66800]
ASF_IND_20_CAD543                .....................................r.......................................... [ 66800]
LR899193.1_Germany_2020/1        ................................................................................ [ 66800]
MN393477.1_Wuhan_2019            ................................................................................ [ 66800]
MW396979.1_Timor-Leste/2019/1    ................................................................................ [ 66800]
MK940252.1_CN/2019/InnerMongolia ................................................................................ [ 66800]
MK128995.1_China/2018/AnhuiXCGQ  ................................................................................ [ 66800]
MH681419.1_ASFV/POL/2015         ................................................................................ [ 66800]
LS478113.1_Estonia_2014          ................................................................................ [ 66800]
KP843857.1_Odintsovo_02/14       ................................................................................ [ 66800]
FN557520.1_strain_E75            .....................................g.......................................... [ 66800]
AY261366.1_Warthog               .....................................gg....t.................................... [ 66800]
AY261365.1_Warmbaths             .....................................gg....t.................................... [ 66800]
AY261364.1_Tengani_62            .g...................................gg....t.................................... [ 66800]
AY261363.1_Pretoriuskop/96/4     .............g.......................gg....t.................................... [ 66800]
AY261362.1_Mkuzi_1979            .............g.......................g.......................................... [ 66800]
AY261361.1_Malawi_Lil-20/1_1983  .....................................gg..t...................g.................. [ 66800]
AY261360.1_Kenya_1950            ...........t.........................gg...............................c......... [ 66800]
AM712240.1_OURT_88/3             .....................................g.......................................... [ 66800]
AM712239.1_Benin_97/1            .....................................g........................a................. [ 66800]
KM111295.1_Ken06.Bus             ...........t.........................gg...............................c......... [ 66800]
```

```
KM111294.1_Ken05/Tk1              ...........t.........................gg...............................c......... [ 66800]
KM262844.1_strain_L60             .....................................g.......................................... [ 66800]
KM102979.1_26544/OG10_Italy       .....................................g.......................................... [ 66800]
KM262845.1_strain_NHV             .....................................g.......................................... [ 66800]
KX354450.1_47/Ss/2008             .....................................g.......................................... [ 66800]
KP055815.1_strain_BA71            .....................................g.......................................... [ 66800]
MH025920.1_strain_R35             ...........t.........................gg...............................c......... [ 66800]
MH025919.1_strain_N10             ...........t.........................gg...............................c......... [ 66800]
MH025918.1_strain_R25             ...........t.........................gg...............................c......... [ 66800]
MH025917.1_strain_R7              ...........t.........................gg...............................c......... [ 66800]
MH025916.1_strain_R8              ...........t.........................gg...............................c......... [ 66800]

FR682468.2_Georgia_2007/1         cactcatcataataggttaaaattttttttatttgttgttgatatgggctaagctcatgctctgaaatatcattaatgta [ 66880]
ASF_IND_20_CAD543                 ................................................................................ [ 66880]
LR899193.1_Germany_2020/1         ................................................................................ [ 66880]
MN393477.1_Wuhan_2019             ................................................................................ [ 66880]
MW396979.1_Timor-Leste/2019/1     ................................................................................ [ 66880]
MK940252.1_CN/2019/InnerMongolia  ................................................................................ [ 66880]
MK128995.1_China/2018/AnhuiXCGQ   ................................................................................ [ 66880]
MH681419.1_ASFV/POL/2015          ................................................................................ [ 66880]
LS478113.1_Estonia_2014           ................................................................................ [ 66880]
KP843857.1_Odintsovo_02/14        .........................-...................................................... [ 66880]
FN557520.1_strain_E75             ................................................................................ [ 66880]
AY261366.1_Warthog                .....................................................t.......................... [ 66880]
AY261365.1_Warmbaths              ......................................c...........t.at.......................... [ 66880]
AY261364.1_Tengani_62             .....................................................t.......................... [ 66880]
AY261363.1_Pretoriuskop/96/4      .....................................................t.......................... [ 66880]
AY261362.1_Mkuzi_1979             ................................................................................ [ 66880]
AY261361.1_Malawi_Lil-20/1_1983   ..t..g................................c..g..c........t....................c..... [ 66880]
AY261360.1_Kenya_1950             ..a...................................c..g..c........t....................c..... [ 66880]
AM712240.1_OURT_88/3              ................................................................................ [ 66880]
AM712239.1_Benin_97/1             ................................................................................ [ 66880]
KM111295.1_Ken06.Bus              ..a...................................c..g..c........t....................c..... [ 66880]
KM111294.1_Ken05/Tk1              ..a...................................c..g..c........t....................c..... [ 66880]
KM262844.1_strain_L60             ................................................................................ [ 66880]
KM102979.1_26544/OG10_Italy       ................................................................................ [ 66880]
KM262845.1_strain_NHV             ................................................................................ [ 66880]
KX354450.1_47/Ss/2008             ................................................................................ [ 66880]
KP055815.1_strain_BA71            ................................................................................ [ 66880]
MH025920.1_strain_R35             ..a...................................c..g..c........t....................c..... [ 66880]
MH025919.1_strain_N10             ..a...................................c..g..c........t....................c..... [ 66880]
MH025918.1_strain_R25             ..a...................................c..g..c........t....................c..... [ 66880]
MH025917.1_strain_R7              ..a...................................c..g..c........t....................c..... [ 66880]
MH025916.1_strain_R8              ..a...................................c..g..c........t....................c..... [ 66880]
```

```
FR682468.2_Georgia_2007/1        atatttaatatatcccactagtatttcattaatgatattatgatatattaactcttctccctccatagcggcaccctata [ 66960]
ASF_IND_20_CAD543                ................................................................................ [ 66960]
LR899193.1_Germany_2020/1        ................................................................................ [ 66960]
MN393477.1_Wuhan_2019            ................................................................................ [ 66960]
MW396979.1_Timor-Leste/2019/1    ................................................................................ [ 66960]
MK940252.1_CN/2019/InnerMongolia ................................................................................ [ 66960]
MK128995.1_China/2018/AnhuiXCGQ  ................................................................................ [ 66960]
MH681419.1_ASFV/POL/2015         ................................................................................ [ 66960]
LS478113.1_Estonia_2014          ................................................................................ [ 66960]
KP843857.1_Odintsovo_02/14       ................................................................................ [ 66960]
FN557520.1_strain_E75            ............a................................................................... [ 66960]
AY261366.1_Warthog               .........g.....t.........................................c...................... [ 66960]
AY261365.1_Warmbaths             .........g.....t................................................................ [ 66960]
AY261364.1_Tengani_62            .........g.....t................................................................ [ 66960]
AY261363.1_Pretoriuskop/96/4     .........g.....t.........................................c...................... [ 66960]
AY261362.1_Mkuzi_1979            ................................................................................ [ 66960]
AY261361.1_Malawi_Lil-20/1_1983  ............c.....g...............................g............................. [ 66960]
AY261360.1_Kenya_1950            ..................g.............................g.g............................. [ 66960]
AM712240.1_OURT_88/3             ............a................................................................... [ 66960]
AM712239.1_Benin_97/1            ............a................................................................... [ 66960]
KM111295.1_Ken06.Bus             ........cg........g.............................g.g............................. [ 66960]
KM111294.1_Ken05/Tk1             ..................g.............................g.g............................. [ 66960]
KM262844.1_strain_L60            ............a................................................................... [ 66960]
KM102979.1_26544/OG10_Italy      ............a................................................................... [ 66960]
KM262845.1_strain_NHV            ............a................................................................... [ 66960]
KX354450.1_47/Ss/2008            ............a................................................................... [ 66960]
KP055815.1_strain_BA71           ............a................................................................... [ 66960]
MH025920.1_strain_R35            ........cg........g.............................g.g............................. [ 66960]
MH025919.1_strain_N10            ........cg........g.............................g.g............................. [ 66960]
MH025918.1_strain_R25            ........cg........g.............................g.g............................. [ 66960]
MH025917.1_strain_R7             ........cg........g.............................g.g............................. [ 66960]
MH025916.1_strain_R8             ........cg........g.............................g.g............................. [ 66960]

FR682468.2_Georgia_2007/1        tttttttatttaggtttcaatgttatcacaattgcgata-------caattgtgata-------caattgtg-----aca [ 67040]
ASF_IND_20_CAD543                .......................................-------...........-------........-----.t. [ 67040]
LR899193.1_Germany_2020/1        .......................................-------...........-------........-----... [ 67040]
MN393477.1_Wuhan_2019            .......................................-------...........-------........-----... [ 67040]
MW396979.1_Timor-Leste/2019/1    .......................................-------...........-------........-----... [ 67040]
MK940252.1_CN/2019/InnerMongolia .......................................-------...........-------........-----... [ 67040]
MK128995.1_China/2018/AnhuiXCGQ  .......................................-------...........-------........-----... [ 67040]
MH681419.1_ASFV/POL/2015         .......................................-------...........-------........-----... [ 67040]
LS478113.1_Estonia_2014          .......................................-------...........-------........-----... [ 67040]
KP843857.1_Odintsovo_02/14       .......................................-------...........-------........-----... [ 67040]
```

```
FN557520.1_strain_E75              .......................................-------...........-------........-----.t. [ 67040]
AY261366.1_Warthog                 ..................................t..c.-------.t.......c.-------........-----... [ 67040]
AY261365.1_Warmbaths               .............................t....t..c.-------.........c.-------........-----..g [ 67040]
AY261364.1_Tengani_62              ..................................t..c.-------...------------------------------- [ 67040]
AY261363.1_Pretoriuskop/96/4       .............................t....t..c.-------.t.......c.-------...c....-------- [ 67040]
AY261362.1_Mkuzi_1979              ...................................------------------....-------........-----.t. [ 67040]
AY261361.1_Malawi_Lil-20/1_1983    .............................g....t.cc.-------.......agc.-------........tgtat.t. [ 67040]
AY261360.1_Kenya_1950              ........................g......a.at.tg.tatacac...a....tg.tatacac...a....tg-at.tg [ 67040]
AM712240.1_OURT_88/3               .......................................-------...........-------........-----.t. [ 67040]
AM712239.1_Benin_97/1              .......................................-------...........-------........-----.t. [ 67040]
KM111295.1_Ken06.Bus               ........................gg........t.gc.-------------------------...a.a..tg-at.t. [ 67040]
KM111294.1_Ken05/Tk1               ........................gc........t.gc.-------........tc.-------...a.a..tg-at.t. [ 67040]
KM262844.1_strain_L60              .......................................-------...........-------........-----.t. [ 67040]
KM102979.1_26544/OG10_Italy        .......................................-------...........-------........-----.t. [ 67040]
KM262845.1_strain_NHV              .......................................-------...........-------........-----.t. [ 67040]
KX354450.1_47/Ss/2008              .......................................-------...........-------........-----.t. [ 67040]
KP055815.1_strain_BA71             .......................................-------...........-------........-----.t. [ 67040]
MH025920.1_strain_R35              ........................gg........t.gc.-------------------------...a.a..tg-at.t. [ 67040]
MH025919.1_strain_N10              ........................gg........t.gc.-------------------------...a.a..tg-at.t. [ 67040]
MH025918.1_strain_R25              ........................gg........t.gc.-------------------------...a.a..tg-at.t. [ 67040]
MH025917.1_strain_R7               ........................gg........t.gc.-------------------------...a.a..tg-at.t. [ 67040]
MH025916.1_strain_R8               ........................gg........t.gc.-------------------------...a.a..tg-at.t. [ 67040]

FR682468.2_Georgia_2007/1          caa-------------------------------ctgtgttgtatacaacaaatgttaggccacgtatagcaacctatat [ 67120]
ASF_IND_20_CAD543                  ...t-----tgtgacacg---------------a.............................................. [ 67120]
LR899193.1_Germany_2020/1          ...-------------------------------.............................................. [ 67120]
MN393477.1_Wuhan_2019              ...-------------------------------.............................................. [ 67120]
MW396979.1_Timor-Leste/2019/1      ...-------------------------------.............................................. [ 67120]
MK940252.1_CN/2019/InnerMongolia   ...-------------------------------.............................................. [ 67120]
MK128995.1_China/2018/AnhuiXCGQ    ...-------------------------------.............................................. [ 67120]
MH681419.1_ASFV/POL/2015           ...-------------------------------.............................................. [ 67120]
LS478113.1_Estonia_2014            ...-------------------------------.............................................. [ 67120]
KP843857.1_Odintsovo_02/14         ...-------------------------------.............................................. [ 67120]
FN557520.1_strain_E75              ...t-----tgtgacacg---------------a.............................................. [ 67120]
AY261366.1_Warthog                 ...------------------------------------c........................................ [ 67120]
AY261365.1_Warmbaths               ...t-----tgtgacaca---------------g.............................................. [ 67120]
AY261364.1_Tengani_62              ----------------------------------.............................................. [ 67120]
AY261363.1_Pretoriuskop/96/4       ---------------------------------------......................................... [ 67120]
AY261362.1_Mkuzi_1979              ...t-----tgcgacacg---------------a.............................................. [ 67120]
AY261361.1_Malawi_Lil-20/1_1983    ...t-----gttgttatatatacaatgttatatataa....a...................................... [ 67120]
AY261360.1_Kenya_1950              ...caaatgtgtgatatg----caaca------aa...a.ga...g.............a.................... [ 67120]
AM712240.1_OURT_88/3               ...t-----tgtgacacg---------------a.............................................. [ 67120]
AM712239.1_Benin_97/1              ...t-----tgtgacacg---------------a.............................................. [ 67120]
KM111295.1_Ken06.Bus               ...caaatgtgtaatata----caaca------aa.....ga............................t......... [ 67120]
```

```
KM111294.1_Ken05/Tk1             ...caaatatgtgatata----caaca------aa.....ga...g.................................. [ 67120]
KM262844.1_strain_L60            ...t-----tgtgacacg---------------a.............................................. [ 67120]
KM102979.1_26544/OG10_Italy      ...t-----tgtgacacg---------------a.............................................. [ 67120]
KM262845.1_strain_NHV            ...t-----tgtgacacg---------------a.............................................. [ 67120]
KX354450.1_47/Ss/2008            ...t-----tgtgacacg---------------a.............................................. [ 67120]
KP055815.1_strain_BA71           ...t-----tgtgacacg---------------a.............................................. [ 67120]
MH025920.1_strain_R35            ...caaatgtgtaatata----caaca------aa.....ga............................t......... [ 67120]
MH025919.1_strain_N10            ...caaatgtgtaatata----caaca------aa.....ga............................t......... [ 67120]
MH025918.1_strain_R25            ...caaatgtgtaatata----caaca------aa.....ga............................t......... [ 67120]
MH025917.1_strain_R7             ...caaatgtgtaatata----caaca------aa.....ga............................t......... [ 67120]
MH025916.1_strain_R8             ...caaatgtgtaatata----caaca------aa.....ga............................t......... [ 67120]

FR682468.2_Georgia_2007/1        gttaagaaatatttttat-cccaaca---------ttagttggaaacgagcagccgca---------------------- [ 67200]
ASF_IND_20_CAD543                ..................-.......---------.......................---------------------- [ 67200]
LR899193.1_Germany_2020/1        ..................-.......---------.......................---------------------- [ 67200]
MN393477.1_Wuhan_2019            ..................-.......---------.......................---------------------- [ 67200]
MW396979.1_Timor-Leste/2019/1    ..................-.......---------.......................---------------------- [ 67200]
MK940252.1_CN/2019/InnerMongolia ..................-.......---------.......................---------------------- [ 67200]
MK128995.1_China/2018/AnhuiXCGQ  ..................-.......---------.......................---------------------- [ 67200]
MH681419.1_ASFV/POL/2015         ..................-.......---------.......................---------------------- [ 67200]
LS478113.1_Estonia_2014          ..................-.......---------.......................---------------------- [ 67200]
KP843857.1_Odintsovo_02/14       ..................-.......---------.......................---------------------- [ 67200]
FN557520.1_strain_E75            ..................-.......---------..g....................---------------------- [ 67200]
AY261366.1_Warthog               ..................-.......---------..g....t.t...........t.aac------------------- [ 67200]
AY261365.1_Warmbaths             ..................-.......---------..g....t.............t.aac------------------- [ 67200]
AY261364.1_Tengani_62            ..................-.......--------t..g....t............at.aacaagt--------------- [ 67200]
AY261363.1_Pretoriuskop/96/4     ..................-.......---------..g....t.............t.aac------------------- [ 67200]
AY261362.1_Mkuzi_1979            ..................-.......---------..g....................---------------------- [ 67200]
AY261361.1_Malawi_Lil-20/1_1983  ..................a.......ttggtttct..g....t....a...g..t.t.aatgagtggctgtaaatgaaca [ 67200]
AY261360.1_Kenya_1950            ..................a.......ttggtttcc-.g.c.at...t-......tat.aat-agcagctgtatatgagca [ 67200]
AM712240.1_OURT_88/3             ..................-.......---------..g....................---------------------- [ 67200]
AM712239.1_Benin_97/1            ..................-.......---------..g....................---------------------- [ 67200]
KM111295.1_Ken06.Bus             .....a............a.......ttggtttcc..g.c..t..t.-......t.t.aat------------------- [ 67200]
KM111294.1_Ken05/Tk1             ..................a.......ttggtttcc..g.c.at...t-......tat.aat-agcagctgtaaacgagca [ 67200]
KM262844.1_strain_L60            ..................-.......---------..g....................---------------------- [ 67200]
KM102979.1_26544/OG10_Italy      ..................-.......---------..g....................---------------------- [ 67200]
KM262845.1_strain_NHV            ..................-.......---------..g....................---------------------- [ 67200]
KX354450.1_47/Ss/2008            ..................-.......---------..g....................---------------------- [ 67200]
KP055815.1_strain_BA71           ..................-.......---------..g....................---------------------- [ 67200]
MH025920.1_strain_R35            .....a............a.......ttggtttcc..g.c..t..t.-......t.t.aat------------------- [ 67200]
MH025919.1_strain_N10            .....a............a.......ttggtttcc..g.c..t..t.-......t.t.aat------------------- [ 67200]
MH025918.1_strain_R25            .....a............a.......ttggtttcc..g.c..t..t.-......t.t.aat------------------- [ 67200]
MH025917.1_strain_R7             .....a............a.......ttggtttcc..g.c..t..t.-......t.t.aat------------------- [ 67200]
MH025916.1_strain_R8             .....a............a.......ttggtttcc..g.c..t..t.-......t.t.aat------------------- [ 67200]
```

```
FR682468.2_Georgia_2007/1        --------------------------------aagaagtc---atttaaaataagccatttaaagatttagaatttatat [ 67280]
ASF_IND_20_CAD543                --------------------------------........---..................................... [ 67280]
LR899193.1_Germany_2020/1        --------------------------------........---..................................... [ 67280]
MN393477.1_Wuhan_2019            --------------------------------........---..................................... [ 67280]
MW396979.1_Timor-Leste/2019/1    --------------------------------........---..................................... [ 67280]
MK940252.1_CN/2019/InnerMongolia --------------------------------........---..................................... [ 67280]
MK128995.1_China/2018/AnhuiXCGQ  --------------------------------........---..................................... [ 67280]
MH681419.1_ASFV/POL/2015         --------------------------------........---..................................... [ 67280]
LS478113.1_Estonia_2014          --------------------------------........---..................................... [ 67280]
KP843857.1_Odintsovo_02/14       --------------------------------........---..................................... [ 67280]
FN557520.1_strain_E75            --------------------------------........---..................................... [ 67280]
AY261366.1_Warthog               ----------------------------aagc..ag....---..................................... [ 67280]
AY261365.1_Warmbaths             ----------------------------aagc..ag....---...........................a......... [ 67280]
AY261364.1_Tengani_62            ---------------------------aaaga........---..................................... [ 67280]
AY261363.1_Pretoriuskop/96/4     ----------------------------aagc..ag....---...........................a......... [ 67280]
AY261362.1_Mkuzi_1979            --------------------------------........---..................................... [ 67280]
AY261361.1_Malawi_Lil-20/1_1983  gct----gtaaacgagcagctgtaaacgagca........---....-......................a......... [ 67280]
AY261360.1_Kenya_1950            gctgttggtatatgagcagctgt-----atacc..c..ct-gt..acc..ca..att............ga..-.c.... [ 67280]
AM712240.1_OURT_88/3             --------------------------------........---.-................................... [ 67280]
AM712239.1_Benin_97/1            --------------------------------........---..................................... [ 67280]
KM111295.1_Ken06.Bus             --------------------------------c..c..ct-gtg.gcc..ca..att............ga..-.c.... [ 67280]
KM111294.1_Ken05/Tk1             gct----gtaaacgagcagctgt-----atacc..c..ctggt..acc..ca..att............ga..-.c.... [ 67280]
KM262844.1_strain_L60            --------------------------------........---..................................... [ 67280]
KM102979.1_26544/OG10_Italy      --------------------------------........---..................................... [ 67280]
KM262845.1_strain_NHV            --------------------------------........---.-................................... [ 67280]
KX354450.1_47/Ss/2008            --------------------------------........---..................................... [ 67280]
KP055815.1_strain_BA71           --------------------------------........---..................................... [ 67280]
MH025920.1_strain_R35            --------------------------------c..c..ct-gtg.gcc..ca..att............ga..-.c.... [ 67280]
MH025919.1_strain_N10            --------------------------------c..c..ct-gtg.gcc..ca..att............ga..-.c.... [ 67280]
MH025918.1_strain_R25            --------------------------------c..c..ct-gtg.gcc..ca..att............ga..-.c.... [ 67280]
MH025917.1_strain_R7             --------------------------------c..c..ct-gtg.gcc..ca..att............ga..-.c.... [ 67280]
MH025916.1_strain_R8             --------------------------------c..c..ct-gtg.gcc..ca..att............ga..-.c.... [ 67280]

FR682468.2_Georgia_2007/1        gtatacaactgt-acaatggaagcagttcttaccaaactcgaccaggaggaaaaaaaggctctccaaaattttcatcgtt [ 67360]
ASF_IND_20_CAD543                ............-................................................................... [ 67360]
LR899193.1_Germany_2020/1        ............-................................................................... [ 67360]
MN393477.1_Wuhan_2019            ............-................................................................... [ 67360]
MW396979.1_Timor-Leste/2019/1    ............-................................................................... [ 67360]
MK940252.1_CN/2019/InnerMongolia ............-................................................................... [ 67360]
MK128995.1_China/2018/AnhuiXCGQ  ............-................................................................... [ 67360]
MH681419.1_ASFV/POL/2015         ............-................................................................... [ 67360]
LS478113.1_Estonia_2014          ............-................................................................... [ 67360]
KP843857.1_Odintsovo_02/14       ............-................................................................... [ 67360]
```

```
FN557520.1_strain_E75            ............-................................................................... [ 67360]
AY261366.1_Warthog               ............-..........................t................g..........g............ [ 67360]
AY261365.1_Warmbaths             ............-..........................t..........................c..c...t...... [ 67360]
AY261364.1_Tengani_62            ............-..........................t................ga.........g............ [ 67360]
AY261363.1_Pretoriuskop/96/4     ............-..........................t................g..........g............ [ 67360]
AY261362.1_Mkuzi_1979            ............-................................................................... [ 67360]
AY261361.1_Malawi_Lil-20/1_1983  .....t......-.............c......t.....t................g..........gcc.a.t...... [ 67360]
AY261360.1_Kenya_1950            ..g..t....a.-............a.............t........t..............t....cc...t...aa. [ 67360]
AM712240.1_OURT_88/3             ............-................................................................... [ 67360]
AM712239.1_Benin_97/1            ............-................................................................... [ 67360]
KM111295.1_Ken06.Bus             ..g..t....a.a............a.............t........t..............t...gcc...t...aa. [ 67360]
KM111294.1_Ken05/Tk1             ..g..t....a.-............a.............t........t..............t....cc...t...aa. [ 67360]
KM262844.1_strain_L60            ............-................................................................... [ 67360]
KM102979.1_26544/OG10_Italy      ............-................................................................... [ 67360]
KM262845.1_strain_NHV            ............-................................................................... [ 67360]
KX354450.1_47/Ss/2008            ............-................................................................... [ 67360]
KP055815.1_strain_BA71           ............-................................................................... [ 67360]
MH025920.1_strain_R35            ..g..t....a.a............a.............t........t..............t...gcc...t...aa. [ 67360]
MH025919.1_strain_N10            ..g..t....a.a............a.............t........t..............t...gcc...t...aa. [ 67360]
MH025918.1_strain_R25            ..g..t....a.a............a.............t........t..............t...gcc...t...aa. [ 67360]
MH025917.1_strain_R7             ..g..t....a.a............a.............t........t..............t...gcc...t...aa. [ 67360]
MH025916.1_strain_R8             ..g..t....a.a............a.............t........t..............t...gcc...t...aa. [ 67360]

FR682468.2_Georgia_2007/1        gtgcttgggaagaaactaaaaatattataaacgattttcttgaaatccctgaggaacgatgcacctataaattcaactca [ 67440]
ASF_IND_20_CAD543                ................................................................................ [ 67440]
LR899193.1_Germany_2020/1        ................................................................................ [ 67440]
MN393477.1_Wuhan_2019            ................................................................................ [ 67440]
MW396979.1_Timor-Leste/2019/1    ................................................................................ [ 67440]
MK940252.1_CN/2019/InnerMongolia ................................................................................ [ 67440]
MK128995.1_China/2018/AnhuiXCGQ  ................................................................................ [ 67440]
MH681419.1_ASFV/POL/2015         ................................................................................ [ 67440]
LS478113.1_Estonia_2014          ................................................................................ [ 67440]
KP843857.1_Odintsovo_02/14       ................................................................................ [ 67440]
FN557520.1_strain_E75            ................................................................................ [ 67440]
AY261366.1_Warthog               ............................................................................tc.. [ 67440]
AY261365.1_Warmbaths             ....a........................................................................... [ 67440]
AY261364.1_Tengani_62            ......................c......................c...............t...............c.. [ 67440]
AY261363.1_Pretoriuskop/96/4     ....a..................g.....................................................c.. [ 67440]
AY261362.1_Mkuzi_1979            ................................................................................ [ 67440]
AY261361.1_Malawi_Lil-20/1_1983  .c..a..................................................................c........ [ 67440]
AY261360.1_Kenya_1950            .c..g................g.......g..............g.........tt..t...g................. [ 67440]
AM712240.1_OURT_88/3             ................................................................................ [ 67440]
AM712239.1_Benin_97/1            ................................................................................ [ 67440]
KM111295.1_Ken06.Bus             ....g................g.......g..............gc.......act..t...g................. [ 67440]
```

```
KM111294.1_Ken05/Tk1            .c..g................g.......g..............gc........tt..t...g................. [ 67440]
KM262844.1_strain_L60           ................................................................................ [ 67440]
KM102979.1_26544/OG10_Italy     ................................................................................ [ 67440]
KM262845.1_strain_NHV           ................................................................................ [ 67440]
KX354450.1_47/Ss/2008           ................................................................................ [ 67440]
KP055815.1_strain_BA71          ................................................................................ [ 67440]
MH025920.1_strain_R35           ....g................g.......g..............gc.......act..t...g................. [ 67440]
MH025919.1_strain_N10           ....g................g.......g..............gc.......act..t...g................. [ 67440]
MH025918.1_strain_R25           ....g................g.......g..............gc.......act..t...g................. [ 67440]
MH025917.1_strain_R7            ....g................g.......g..............gc.......act..t...g................. [ 67440]
MH025916.1_strain_R8            ....g................g.......g..............gc.......act..t...g................. [ 67440]

FR682468.2_Georgia_2007/1       tacacaaaaaaaatggagcttttatttacccctgaattccacaccgcctggcatgaagttcctgagtgcagagagttcat [ 67520]
ASF_IND_20_CAD543               ................................................................................ [ 67520]
LR899193.1_Germany_2020/1       ................................................................................ [ 67520]
MN393477.1_Wuhan_2019           ................................................................................ [ 67520]
MW396979.1_Timor-Leste/2019/1   ................................................................................ [ 67520]
MK940252.1_CN/2019/InnerMongolia ................................................................................ [ 67520]
MK128995.1_China/2018/AnhuiXCGQ ................................................................................ [ 67520]
MH681419.1_ASFV/POL/2015        ................................................................................ [ 67520]
LS478113.1_Estonia_2014         ................................................................................ [ 67520]
KP843857.1_Odintsovo_02/14      ................................................................................ [ 67520]
FN557520.1_strain_E75           ................................c............................................... [ 67520]
AY261366.1_Warthog              .......................g........c...........t........g.......................... [ 67520]
AY261365.1_Warmbaths            .......................g.....................a.......g.......................... [ 67520]
AY261364.1_Tengani_62           .......................g........c...........ta.......g.......................... [ 67520]
AY261363.1_Pretoriuskop/96/4    .......................g.............................g.......................... [ 67520]
AY261362.1_Mkuzi_1979           ................................c............................................... [ 67520]
AY261361.1_Malawi_Lil-20/1_1983 .......................g........c....................a......tt.................. [ 67520]
AY261360.1_Kenya_1950           c......................g.....t..c....c..............cg.....c..........a......t.. [ 67520]
AM712240.1_OURT_88/3            ................................c............................................... [ 67520]
AM712239.1_Benin_97/1           ................................c............................................... [ 67520]
KM111295.1_Ken06.Bus            .g.....g...............g.....t..c...................cg.....c........t.a......... [ 67520]
KM111294.1_Ken05/Tk1            .......................g.....t..c...................cg.....c..........a......... [ 67520]
KM262844.1_strain_L60           ................................c............................................... [ 67520]
KM102979.1_26544/OG10_Italy     ................................c............................................... [ 67520]
KM262845.1_strain_NHV           ................................c............................................... [ 67520]
KX354450.1_47/Ss/2008           ................................c............................................... [ 67520]
KP055815.1_strain_BA71          ................................c............................................... [ 67520]
MH025920.1_strain_R35           .g.....g...............g.....t..c...................cg.....c........t.a......... [ 67520]
MH025919.1_strain_N10           .g.....g...............g.....t..c...................cg.....c........t.a......... [ 67520]
MH025918.1_strain_R25           .g.....g...............g.....t..c...................cg.....c........t.a......... [ 67520]
MH025917.1_strain_R7            .g.....g...............g.....t..c...................cg.....c........t.a......... [ 67520]
MH025916.1_strain_R8            .g.....g...............g.....t..c...................cg.....c........t.a......... [ 67520]
```

```
FR682468.2_Georgia_2007/1        attaaactttttgagactcatttcgggacatcgagtggtattaaaaggccctacatttgtttttacaaaagagatcaaga [ 67600]
ASF_IND_20_CAD543                ................................................................................ [ 67600]
LR899193.1_Germany_2020/1        ................................................................................ [ 67600]
MN393477.1_Wuhan_2019            ................................................................................ [ 67600]
MW396979.1_Timor-Leste/2019/1    ................................................................................ [ 67600]
MK940252.1_CN/2019/InnerMongolia ................................................................................ [ 67600]
MK128995.1_China/2018/AnhuiXCGQ  ................................................................................ [ 67600]
MH681419.1_ASFV/POL/2015         ................................................................................ [ 67600]
LS478113.1_Estonia_2014          ................................................................................ [ 67600]
KP843857.1_Odintsovo_02/14       ................................................................................ [ 67600]
FN557520.1_strain_E75            ..........................................................................c..... [ 67600]
AY261366.1_Warthog               ........................a............................t.a....a............g...... [ 67600]
AY261365.1_Warmbaths             .....................................................t......a................... [ 67600]
AY261364.1_Tengani_62            .....................................................t.a....a................... [ 67600]
AY261363.1_Pretoriuskop/96/4     .......................................................a....a................... [ 67600]
AY261362.1_Mkuzi_1979            ..........................................................................c..... [ 67600]
AY261361.1_Malawi_Lil-20/1_1983  .a....................a.a..g...........gc...........g..a..................c..... [ 67600]
AY261360.1_Kenya_1950            ........a..........g.g..........................at..t..a..c...............ct.... [ 67600]
AM712240.1_OURT_88/3             ..........................................................................c..... [ 67600]
AM712239.1_Benin_97/1            ..........................................................................c..... [ 67600]
KM111295.1_Ken06.Bus             ........a..........g.g...........................t...a....c...............ct.... [ 67600]
KM111294.1_Ken05/Tk1             ........a..........g.g..........................at.....a..c...............ct.... [ 67600]
KM262844.1_strain_L60            ..........................................................................c..... [ 67600]
KM102979.1_26544/OG10_Italy      ..........................................................................c..... [ 67600]
KM262845.1_strain_NHV            ..........................................................................c..... [ 67600]
KX354450.1_47/Ss/2008            ..........................................................................c..... [ 67600]
KP055815.1_strain_BA71           ..........................................................................c..... [ 67600]
MH025920.1_strain_R35            ........a..........g.g...........................t...a....c...............ct.... [ 67600]
MH025919.1_strain_N10            ........a..........g.g...........................t...a....c...............ct.... [ 67600]
MH025918.1_strain_R25            ........a..........g.g...........................t...a....c...............ct.... [ 67600]
MH025917.1_strain_R7             ........a..........g.g...........................t...a....c...............ct.... [ 67600]
MH025916.1_strain_R8             ........a..........g.g...........................t...a....c...............ct.... [ 67600]

FR682468.2_Georgia_2007/1        atctgggcattcctagtaccatcaatgttgactttcaggccaacattgaaaatatggatgatctacagaagggaaatctc [ 67680]
ASF_IND_20_CAD543                ................................................................................ [ 67680]
LR899193.1_Germany_2020/1        ................................................................................ [ 67680]
MN393477.1_Wuhan_2019            ................................................................................ [ 67680]
MW396979.1_Timor-Leste/2019/1    ................................................................................ [ 67680]
MK940252.1_CN/2019/InnerMongolia ................................................................................ [ 67680]
MK128995.1_China/2018/AnhuiXCGQ  ................................................................................ [ 67680]
MH681419.1_ASFV/POL/2015         ................................................................................ [ 67680]
LS478113.1_Estonia_2014          ................................................................................ [ 67680]
KP843857.1_Odintsovo_02/14       ................................................................................ [ 67680]
```

```
FN557520.1_strain_E75            .c.............................................................................. [ 67680]
AY261366.1_Warthog               .c.............................................................................. [ 67680]
AY261365.1_Warmbaths             .c..........................................................................c... [ 67680]
AY261364.1_Tengani_62            .c.............................................................................. [ 67680]
AY261363.1_Pretoriuskop/96/4     .c.............................................................................. [ 67680]
AY261362.1_Mkuzi_1979            .c.............................................................................. [ 67680]
AY261361.1_Malawi_Lil-20/1_1983  .c...............................................g...........c.....a..a........t [ 67680]
AY261360.1_Kenya_1950            .c...............................................g...........c.................t [ 67680]
AM712240.1_OURT_88/3             .c.............................................................................. [ 67680]
AM712239.1_Benin_97/1            .c.............................................................................. [ 67680]
KM111295.1_Ken06.Bus             .c.....t.........................................g...........c.................t [ 67680]
KM111294.1_Ken05/Tk1             .c...............................................g...........c.................t [ 67680]
KM262844.1_strain_L60            .c.............................................................................. [ 67680]
KM102979.1_26544/OG10_Italy      .c.............................................................................. [ 67680]
KM262845.1_strain_NHV            .c.............................................................................. [ 67680]
KX354450.1_47/Ss/2008            .c.............................................................................. [ 67680]
KP055815.1_strain_BA71           .c.............................................................................. [ 67680]
MH025920.1_strain_R35            .c.....t.........................................g...........c.................t [ 67680]
MH025919.1_strain_N10            .c.....t.........................................g...........c.................t [ 67680]
MH025918.1_strain_R25            .c.....t.........................................g...........c.................t [ 67680]
MH025917.1_strain_R7             .c.....t.........................................g...........c.................t [ 67680]
MH025916.1_strain_R8             .c.....t.........................................g...........c.................t [ 67680]

FR682468.2_Georgia_2007/1        atcggcaagatgaatatcaaagaaggctaaataaaac------------------------------------------- [ 67760]
ASF_IND_20_CAD543                .....................................------------------------------------------- [ 67760]
LR899193.1_Germany_2020/1        .....................................------------------------------------------- [ 67760]
MN393477.1_Wuhan_2019            .....................................------------------------------------------- [ 67760]
MW396979.1_Timor-Leste/2019/1    .....................................------------------------------------------- [ 67760]
MK940252.1_CN/2019/InnerMongolia .....................................------------------------------------------- [ 67760]
MK128995.1_China/2018/AnhuiXCGQ  .....................................------------------------------------------- [ 67760]
MH681419.1_ASFV/POL/2015         .....................................------------------------------------------- [ 67760]
LS478113.1_Estonia_2014          .....................................------------------------------------------- [ 67760]
KP843857.1_Odintsovo_02/14       .....................................------------------------------------------- [ 67760]
FN557520.1_strain_E75            .....................................------------------------------------------- [ 67760]
AY261366.1_Warthog               ........a...............a.t..........------------------------------------------- [ 67760]
AY261365.1_Warmbaths             ........a...............a.t..........------------------------------------------- [ 67760]
AY261364.1_Tengani_62            .....................................------------------------------------------- [ 67760]
AY261363.1_Pretoriuskop/96/4     ........................a.t..........------------------------------------------- [ 67760]
AY261362.1_Mkuzi_1979            .....................................------------------------------------------- [ 67760]
AY261361.1_Malawi_Lil-20/1_1983  ........a...............aat.....g...taaattttaaa------acacgctgacatacatc---------- [ 67760]
AY261360.1_Kenya_1950            ..t.....a........t......a.t..g..g...taaattttaaataaaacgagtggcatcaaaaact---------- [ 67760]
AM712240.1_OURT_88/3             .....................................------------------------------------------- [ 67760]
AM712239.1_Benin_97/1            .....................................------------------------------------------- [ 67760]
KM111295.1_Ken06.Bus             ..t.....a........t......a.t.....g...taaattttaaatataacgagtggcatcaaaaattaagtaaaatc [ 67760]
```

```
KM111294.1_Ken05/Tk1             ..t.....a........t......a.t..g..g...taaattttaaataaaacgagtggcatcaaaaact---------- [ 67760]
KM262844.1_strain_L60            .....................................------------------------------------------- [ 67760]
KM102979.1_26544/OG10_Italy      .....................................------------------------------------------- [ 67760]
KM262845.1_strain_NHV            .....................................------------------------------------------- [ 67760]
KX354450.1_47/Ss/2008            .....................................------------------------------------------- [ 67760]
KP055815.1_strain_BA71           .....................................------------------------------------------- [ 67760]
MH025920.1_strain_R35            ..t.....a........t......a.t.....g...taaattttaaatataacgagtggcatcaaaaattaagtaaaatc [ 67760]
MH025919.1_strain_N10            ..t.....a........t......a.t.....g...taaattttaaatataacgagtggcatcaaaaattaagtaaaatc [ 67760]
MH025918.1_strain_R25            ..t.....a........t......a.t.....g...taaattttaaatataacgagtggcatcaaaaattaagtaaaatc [ 67760]
MH025917.1_strain_R7             ..t.....a........t......a.t.....g...taaattttaaatataacgagtggcatcaaaaattaagtaaaatc [ 67760]
MH025916.1_strain_R8             ..t.....a........t......a.t.....g...taaattttaaatataacgagtggcatcaaaaattaagtaaaatc [ 67760]

FR682468.2_Georgia_2007/1        -------aactaacatcaaaaaacattaaaggctatgttgtggacgatgcctttgtctcaatagtttcgaggtcatccaa [ 67840]
ASF_IND_20_CAD543                -------......................................................................... [ 67840]
LR899193.1_Germany_2020/1        -------......................................................................... [ 67840]
MN393477.1_Wuhan_2019            -------......................................................................... [ 67840]
MW396979.1_Timor-Leste/2019/1    -------......................................................................... [ 67840]
MK940252.1_CN/2019/InnerMongolia -------......................................................................... [ 67840]
MK128995.1_China/2018/AnhuiXCGQ  -------......................................................................... [ 67840]
MH681419.1_ASFV/POL/2015         -------......................................................................... [ 67840]
LS478113.1_Estonia_2014          -------......................................................................... [ 67840]
KP843857.1_Odintsovo_02/14       -------......................................................................... [ 67840]
FN557520.1_strain_E75            -------......................................................................... [ 67840]
AY261366.1_Warthog               -------....g..................a.......g.ac..t.tg................................ [ 67840]
AY261365.1_Warmbaths             -------....g..................a.......g.ac..t.tg.......................a........ [ 67840]
AY261364.1_Tengani_62            -------................t......-.......---....................................... [ 67840]
AY261363.1_Pretoriuskop/96/4     -------....g..................a.......g.ac..t.tg............................t... [ 67840]
AY261362.1_Mkuzi_1979            -------......................................................................... [ 67840]
AY261361.1_Malawi_Lil-20/1_1983  ------a...a..a.c....t..a......a.......---........tt..c.....g.....c.....a....t... [ 67840]
AY261360.1_Kenya_1950            aagtaaa.t.a..a.....gt..a......aa......---........................c..a..a........ [ 67840]
AM712240.1_OURT_88/3             -------......................................................................... [ 67840]
AM712239.1_Benin_97/1            -------......................................................................... [ 67840]
KM111295.1_Ken06.Bus             aagtaaa.t.a..a.....gt..a......aa......---........................c..a..a....t... [ 67840]
KM111294.1_Ken05/Tk1             aagtaaa.t.a..a.....gt..a......aa......---........................c..a..a........ [ 67840]
KM262844.1_strain_L60            -------......................................................................... [ 67840]
KM102979.1_26544/OG10_Italy      -------......................................................................... [ 67840]
KM262845.1_strain_NHV            -------......................................................................... [ 67840]
KX354450.1_47/Ss/2008            -------......................................................................... [ 67840]
KP055815.1_strain_BA71           -------......................................................................... [ 67840]
MH025920.1_strain_R35            aagtaaa.t.a..a.....gt..a......aa......---........................c..a..a....t... [ 67840]
MH025919.1_strain_N10            aagtaaa.t.a..a.....gt..a......aa......---........................c..a..a....tt.. [ 67840]
MH025918.1_strain_R25            aagtaaa.t.a..a.....gt..a......aa......---........................c..a..a....t... [ 67840]
MH025917.1_strain_R7             aagtaaa.t.a..a.....gt..a......aa......---........................c..a..a....t... [ 67840]
MH025916.1_strain_R8             aagtaaa.t.a..a.....gt..a......aa......---........................c..a..a....t... [ 67840]
```

```
FR682468.2_Georgia_2007/1        taactcatgtaacgtaaaaaagttggtccatttttttgaaaacattaaaagacgttcgtcttcataaataaaaaagtcat [ 67920]
ASF_IND_20_CAD543                ................................................................................ [ 67920]
LR899193.1_Germany_2020/1        ................................................................................ [ 67920]
MN393477.1_Wuhan_2019            ................................................................................ [ 67920]
MW396979.1_Timor-Leste/2019/1    ................................................................................ [ 67920]
MK940252.1_CN/2019/InnerMongolia ................................................................................ [ 67920]
MK128995.1_China/2018/AnhuiXCGQ  ................................................................................ [ 67920]
MH681419.1_ASFV/POL/2015         ................................................................................ [ 67920]
LS478113.1_Estonia_2014          ................................................................................ [ 67920]
KP843857.1_Odintsovo_02/14       ................................................................................ [ 67920]
FN557520.1_strain_E75            ................................................................................ [ 67920]
AY261366.1_Warthog               ......g................................................g........................ [ 67920]
AY261365.1_Warmbaths             ......g................................................g........................ [ 67920]
AY261364.1_Tengani_62            .......................................................g........................ [ 67920]
AY261363.1_Pretoriuskop/96/4     ......g................................................g........................ [ 67920]
AY261362.1_Mkuzi_1979            ................................................................................ [ 67920]
AY261361.1_Malawi_Lil-20/1_1983  ............t............................t............aa.......g................ [ 67920]
AY261360.1_Kenya_1950            ...t...........g................................................................ [ 67920]
AM712240.1_OURT_88/3             ................................................................................ [ 67920]
AM712239.1_Benin_97/1            ................................................................................ [ 67920]
KM111295.1_Ken06.Bus             ...t............................................................................ [ 67920]
KM111294.1_Ken05/Tk1             ...t...........g................................................................ [ 67920]
KM262844.1_strain_L60            ................................................................................ [ 67920]
KM102979.1_26544/OG10_Italy      ................................................................................ [ 67920]
KM262845.1_strain_NHV            ................................................................................ [ 67920]
KX354450.1_47/Ss/2008            ................................................................................ [ 67920]
KP055815.1_strain_BA71           ................................................................................ [ 67920]
MH025920.1_strain_R35            ...t............................................................................ [ 67920]
MH025919.1_strain_N10            ...t............................................................................ [ 67920]
MH025918.1_strain_R25            ...t............................................................................ [ 67920]
MH025917.1_strain_R7             ...t............................................................................ [ 67920]
MH025916.1_strain_R8             ...t............................................................................ [ 67920]

FR682468.2_Georgia_2007/1        tcgaaggaaaaatgatatactcaataccatagtcttgtaatattttttttaggtctctcagggtccagggatttaccagg [ 68000]
ASF_IND_20_CAD543                ................................................................................ [ 68000]
LR899193.1_Germany_2020/1        ................................................................................ [ 68000]
MN393477.1_Wuhan_2019            ................................................................................ [ 68000]
MW396979.1_Timor-Leste/2019/1    ................................................................................ [ 68000]
MK940252.1_CN/2019/InnerMongolia ................................................................................ [ 68000]
MK128995.1_China/2018/AnhuiXCGQ  ................................................................................ [ 68000]
MH681419.1_ASFV/POL/2015         ................................................................................ [ 68000]
LS478113.1_Estonia_2014          ................................................................................ [ 68000]
KP843857.1_Odintsovo_02/14       ................................................................................ [ 68000]
```

FN557520.1_strain_E75 ................................................................................ [ 68000]
AY261366.1_Warthog .............a.................................................................. [ 68000]
AY261365.1_Warmbaths .............a.................................................................. [ 68000]
AY261364.1_Tengani_62 ....g........................................................................... [ 68000]
AY261363.1_Pretoriuskop/96/4 .............a.................................................................. [ 68000]
AY261362.1_Mkuzi_1979 ............................g................................................... [ 68000]
AY261361.1_Malawi_Lil-20/1_1983 .t........................................................t.................t... [ 68000]
AY261360.1_Kenya_1950 ...............................a.....c.............a......t..a..........g...t... [ 68000]
AM712240.1_OURT_88/3 ................................................................................ [ 68000]
AM712239.1_Benin_97/1 ................................................................................ [ 68000]
KM111295.1_Ken06.Bus ...............................a.....c.............a......t..a..........g...t... [ 68000]
KM111294.1_Ken05/Tk1 ...............................a.....c.............a......t.aa..........g...t... [ 68000]
KM262844.1_strain_L60 ................................................................................ [ 68000]
KM102979.1_26544/OG10_Italy ................................................................................ [ 68000]
KM262845.1_strain_NHV ................................................................................ [ 68000]
KX354450.1_47/Ss/2008 ................................................................................ [ 68000]
KP055815.1_strain_BA71 ................................................................................ [ 68000]
MH025920.1_strain_R35 ...............................a.....c.............a......t..a..........g...t... [ 68000]
MH025919.1_strain_N10 ...............................a.....c.............a......t..a..........g...t... [ 68000]
MH025918.1_strain_R25 ...............................a.....c.............a......t..a..........g...t... [ 68000]
MH025917.1_strain_R7 ...............................a.....c.............a......t..a..........g...t... [ 68000]
MH025916.1_strain_R8 ...............................a.....c.............a......t..a..........g...t... [ 68000]

FR682468.2_Georgia_2007/1 cttctacgcgaagtgagcatcataaaaatatctaatattttttgcgccataagccagcgcggattctcattggcccacaa [ 68080]
ASF_IND_20_CAD543 ................................................................................ [ 68080]
LR899193.1_Germany_2020/1 ................................................................................ [ 68080]
MN393477.1_Wuhan_2019 ................................................................................ [ 68080]
MW396979.1_Timor-Leste/2019/1 ................................................................................ [ 68080]
MK940252.1_CN/2019/InnerMongolia ................................................................................ [ 68080]
MK128995.1_China/2018/AnhuiXCGQ ................................................................................ [ 68080]
MH681419.1_ASFV/POL/2015 ................................................................................ [ 68080]
LS478113.1_Estonia_2014 ................................................................................ [ 68080]
KP843857.1_Odintsovo_02/14 ................................................................................ [ 68080]
FN557520.1_strain_E75 ..................................................g............................. [ 68080]
AY261366.1_Warthog ..................................................g..............t.............. [ 68080]
AY261365.1_Warmbaths ..................................................g............................. [ 68080]
AY261364.1_Tengani_62 ..................................................g...........g................. [ 68080]
AY261363.1_Pretoriuskop/96/4 ..............c...................................g............................. [ 68080]
AY261362.1_Mkuzi_1979 ................................................................................ [ 68080]
AY261361.1_Malawi_Lil-20/1_1983 ..............a...................................g...........g................. [ 68080]
AY261360.1_Kenya_1950 ..................................................g.....a....................... [ 68080]
AM712240.1_OURT_88/3 ..................................................g............................. [ 68080]
AM712239.1_Benin_97/1 ..................................................g............................. [ 68080]
KM111295.1_Ken06.Bus ..................................................g.....a....................... [ 68080]

```
KM111294.1_Ken05/Tk1             ..................................................g.....a....................... [ 68080]
KM262844.1_strain_L60            ..................................................g............................. [ 68080]
KM102979.1_26544/OG10_Italy      ..................................................g............................. [ 68080]
KM262845.1_strain_NHV            ..................................................g............................. [ 68080]
KX354450.1_47/Ss/2008            ..................................................g............................. [ 68080]
KP055815.1_strain_BA71           ..................................................g............................. [ 68080]
MH025920.1_strain_R35            ..................................................g.....a....................... [ 68080]
MH025919.1_strain_N10            ..................................................g.....a....................... [ 68080]
MH025918.1_strain_R25            ..................................................g.....a....................... [ 68080]
MH025917.1_strain_R7             ..................................................g.....a....................... [ 68080]
MH025916.1_strain_R8             ..................................................g.....a....................... [ 68080]

FR682468.2_Georgia_2007/1        atcaacaataattctcttatcaaccgtgagcattcctacttgattcgaagaaatgattagatgcccagcagtccacccca [ 68160]
ASF_IND_20_CAD543                ................................................................................ [ 68160]
LR899193.1_Germany_2020/1        ................................................................................ [ 68160]
MN393477.1_Wuhan_2019            ................................................................................ [ 68160]
MW396979.1_Timor-Leste/2019/1    ................................................................................ [ 68160]
MK940252.1_CN/2019/InnerMongolia ................................................................................ [ 68160]
MK128995.1_China/2018/AnhuiXCGQ  ................................................................................ [ 68160]
MH681419.1_ASFV/POL/2015         ................................................................................ [ 68160]
LS478113.1_Estonia_2014          ................................................................................ [ 68160]
KP843857.1_Odintsovo_02/14       ................................................................................ [ 68160]
FN557520.1_strain_E75            ..................................................................g..g.......... [ 68160]
AY261366.1_Warthog               ................................................g....................g.......... [ 68160]
AY261365.1_Warmbaths             ................................................g.................g..g.......... [ 68160]
AY261364.1_Tengani_62            ................................................g....................g.......... [ 68160]
AY261363.1_Pretoriuskop/96/4     ................................................g.................g..g.......... [ 68160]
AY261362.1_Mkuzi_1979            .............................................t..g.................g..g.......... [ 68160]
AY261361.1_Malawi_Lil-20/1_1983  .....t.........t..g..........................t..g....................g.......... [ 68160]
AY261360.1_Kenya_1950            .....t............g.............................g....................g.......... [ 68160]
AM712240.1_OURT_88/3             ..................................................................g..g.......... [ 68160]
AM712239.1_Benin_97/1            ..................................................................g..g.......... [ 68160]
KM111295.1_Ken06.Bus             .....t............g..........................t..g....................g.......... [ 68160]
KM111294.1_Ken05/Tk1             .....t............g.............................g....................g.......... [ 68160]
KM262844.1_strain_L60            ..................................................................g..g.......... [ 68160]
KM102979.1_26544/OG10_Italy      ..................................................................g..g.......... [ 68160]
KM262845.1_strain_NHV            ..................................................................g..g.......... [ 68160]
KX354450.1_47/Ss/2008            ..................................................................g..g.......... [ 68160]
KP055815.1_strain_BA71           ..................................................................g..g.......... [ 68160]
MH025920.1_strain_R35            .....t............g..........................t..g....................g.......... [ 68160]
MH025919.1_strain_N10            .....t............g..........................t..g....................g.......... [ 68160]
MH025918.1_strain_R25            .....t............g..........................t..g....................g.......... [ 68160]
MH025917.1_strain_R7             .....t............g..........................t..g....................g.......... [ 68160]
MH025916.1_strain_R8             .....t............g..........................t..g....................g.......... [ 68160]
```

```
FR682468.2_Georgia_2007/1        tgagtagataacgcagcgttgtagaaatgtcacatatggaaggcattcctccacaacatgaacccaaattaggatgcgtg [ 68240]
ASF_IND_20_CAD543                ................................................................................ [ 68240]
LR899193.1_Germany_2020/1        ................................................................................ [ 68240]
MN393477.1_Wuhan_2019            ................................................................................ [ 68240]
MW396979.1_Timor-Leste/2019/1    ................................................................................ [ 68240]
MK940252.1_CN/2019/InnerMongolia ................................................................................ [ 68240]
MK128995.1_China/2018/AnhuiXCGQ  ................................................................................ [ 68240]
MH681419.1_ASFV/POL/2015         ................................................................................ [ 68240]
LS478113.1_Estonia_2014          ................................................................................ [ 68240]
KP843857.1_Odintsovo_02/14       ................................................................................ [ 68240]
FN557520.1_strain_E75            .a.............................................................................. [ 68240]
AY261366.1_Warthog               ..............................................c...........c..................... [ 68240]
AY261365.1_Warmbaths             ............................a.................c...........c..................... [ 68240]
AY261364.1_Tengani_62            ............................a.................c...........c..................... [ 68240]
AY261363.1_Pretoriuskop/96/4     ............................a.................c...........c..................... [ 68240]
AY261362.1_Mkuzi_1979            ................................................................................ [ 68240]
AY261361.1_Malawi_Lil-20/1_1983  .t..g.........................................c.....g.....c...........g......... [ 68240]
AY261360.1_Kenya_1950            .t..g................................a........c..c..g.....c..................... [ 68240]
AM712240.1_OURT_88/3             .a.............................................................................. [ 68240]
AM712239.1_Benin_97/1            .a.............................................................................. [ 68240]
KM111295.1_Ken06.Bus             .t..g................................a........c.....g.....c..................... [ 68240]
KM111294.1_Ken05/Tk1             .t..g................................a........c.....g.....c..................... [ 68240]
KM262844.1_strain_L60            .a.............................................................................. [ 68240]
KM102979.1_26544/OG10_Italy      .a.............................................................................. [ 68240]
KM262845.1_strain_NHV            .a.............................................................................. [ 68240]
KX354450.1_47/Ss/2008            .a.............................................................................. [ 68240]
KP055815.1_strain_BA71           .a.............................................................................. [ 68240]
MH025920.1_strain_R35            .t..g................................a........c.....g.....c..................... [ 68240]
MH025919.1_strain_N10            .t..g................................a........c.....g.....c..................... [ 68240]
MH025918.1_strain_R25            .t..g................................a........c.....g.....c..................... [ 68240]
MH025917.1_strain_R7             .t..g................................a........c.....g.....c..................... [ 68240]
MH025916.1_strain_R8             .t..g................................a........c.....g.....c..................... [ 68240]

FR682468.2_Georgia_2007/1        tgaaacacaaacatagcaggcttgttggccaccctgctataaatatcagcaggcatcatagcctcgctgccaaaataaat [ 68320]
ASF_IND_20_CAD543                ................................................................................ [ 68320]
LR899193.1_Germany_2020/1        ................................................................................ [ 68320]
MN393477.1_Wuhan_2019            ................................................................................ [ 68320]
MW396979.1_Timor-Leste/2019/1    ................................................................................ [ 68320]
MK940252.1_CN/2019/InnerMongolia ................................................................................ [ 68320]
MK128995.1_China/2018/AnhuiXCGQ  ................................................................................ [ 68320]
MH681419.1_ASFV/POL/2015         ................................................................................ [ 68320]
LS478113.1_Estonia_2014          ................................................................................ [ 68320]
KP843857.1_Odintsovo_02/14       ................................................................................ [ 68320]
```

```
FN557520.1_strain_E75            ................................................................................ [ 68320]
AY261366.1_Warthog               ........g...................................c................................... [ 68320]
AY261365.1_Warmbaths             ........g..................................cc................................... [ 68320]
AY261364.1_Tengani_62            ............................................c................................... [ 68320]
AY261363.1_Pretoriuskop/96/4     ........g...................................c................................... [ 68320]
AY261362.1_Mkuzi_1979            ................................................................................ [ 68320]
AY261361.1_Malawi_Lil-20/1_1983  ........g.....g.....t.......................c........t.......................... [ 68320]
AY261360.1_Kenya_1950            ........g...................................c........t...........a........g..... [ 68320]
AM712240.1_OURT_88/3             ................................................................................ [ 68320]
AM712239.1_Benin_97/1            ...............................................................................c [ 68320]
KM111295.1_Ken06.Bus             ........g...................................c........t...........a........g..... [ 68320]
KM111294.1_Ken05/Tk1             ........g...................................c........t...........a........g..... [ 68320]
KM262844.1_strain_L60            ................................................................................ [ 68320]
KM102979.1_26544/OG10_Italy      ................................................................................ [ 68320]
KM262845.1_strain_NHV            ................................................................................ [ 68320]
KX354450.1_47/Ss/2008            ................................................................................ [ 68320]
KP055815.1_strain_BA71           ................................................................................ [ 68320]
MH025920.1_strain_R35            ........g...................................c........t...........a........g..... [ 68320]
MH025919.1_strain_N10            ........g...................................c........t...........a........g..... [ 68320]
MH025918.1_strain_R25            ........g...................................c........t...........a........g..... [ 68320]
MH025917.1_strain_R7             ........g...................................c........t...........a........g..... [ 68320]
MH025916.1_strain_R8             ........g...................................c........t...........a........g..... [ 68320]

FR682468.2_Georgia_2007/1        gttctctcctgccctataggggcttggaatgatttccactatctcgggtacaccgtttatcatattaatgcggccgcacc [ 68400]
ASF_IND_20_CAD543                ................................................................................ [ 68400]
LR899193.1_Germany_2020/1        ................................................................................ [ 68400]
MN393477.1_Wuhan_2019            ................................................................................ [ 68400]
MW396979.1_Timor-Leste/2019/1    ................................................................................ [ 68400]
MK940252.1_CN/2019/InnerMongolia ................................................................................ [ 68400]
MK128995.1_China/2018/AnhuiXCGQ  ................................................................................ [ 68400]
MH681419.1_ASFV/POL/2015         ................................................................................ [ 68400]
LS478113.1_Estonia_2014          ................................................................................ [ 68400]
KP843857.1_Odintsovo_02/14       ................................................................................ [ 68400]
FN557520.1_strain_E75            ................................................................................ [ 68400]
AY261366.1_Warthog               ...........................................................c.................... [ 68400]
AY261365.1_Warmbaths             ............t..............................................c.................... [ 68400]
AY261364.1_Tengani_62            ..........a..................................................................... [ 68400]
AY261363.1_Pretoriuskop/96/4     ...........................................................c.................... [ 68400]
AY261362.1_Mkuzi_1979            ................................................................................ [ 68400]
AY261361.1_Malawi_Lil-20/1_1983  .............t.................................................................. [ 68400]
AY261360.1_Kenya_1950            .............t.................................................................. [ 68400]
AM712240.1_OURT_88/3             ................................................................................ [ 68400]
AM712239.1_Benin_97/1            ................................................................................ [ 68400]
KM111295.1_Ken06.Bus             .............t.....................................g............................ [ 68400]
```

```
KM111294.1_Ken05/Tk1           .............t..................................................................  [ 68400]
KM262844.1_strain_L60          ................................................................................  [ 68400]
KM102979.1_26544/OG10_Italy    ................................................................................  [ 68400]
KM262845.1_strain_NHV          ................................................................................  [ 68400]
KX354450.1_47/Ss/2008          ................................................................................  [ 68400]
KP055815.1_strain_BA71         ................................................................................  [ 68400]
MH025920.1_strain_R35          .............t.....................................g............................  [ 68400]
MH025919.1_strain_N10          .............t.....................................g............................  [ 68400]
MH025918.1_strain_R25          .............t.....................................g............................  [ 68400]
MH025917.1_strain_R7           .............t.....................................g............................  [ 68400]
MH025916.1_strain_R8           .............t.....................................g............................  [ 68400]

FR682468.2_Georgia_2007/1      attcacggtcatcgtccaaaaattttttgatggcaccccgaacattgtcccagttaagcaacagagtattcacaatctca  [ 68480]
ASF_IND_20_CAD543              ....................................................r...........................  [ 68480]
LR899193.1_Germany_2020/1      ................................................................................  [ 68480]
MN393477.1_Wuhan_2019          ................................................................................  [ 68480]
MW396979.1_Timor-Leste/2019/1  ................................................................................  [ 68480]
MK940252.1_CN/2019/InnerMongolia ..............................................................................  [ 68480]
MK128995.1_China/2018/AnhuiXCGQ ...............................................................................  [ 68480]
MH681419.1_ASFV/POL/2015       ................................................................................  [ 68480]
LS478113.1_Estonia_2014        ................................................................................  [ 68480]
KP843857.1_Odintsovo_02/14     ................................................................................  [ 68480]
FN557520.1_strain_E75          ....................................................a..............g............  [ 68480]
AY261366.1_Warthog             ...................................g.......g........a.......................t...  [ 68480]
AY261365.1_Warmbaths           .......a..g..a.t...................g.......g........a.......................t...  [ 68480]
AY261364.1_Tengani_62          ............................a.......................a...........................  [ 68480]
AY261363.1_Pretoriuskop/96/4   .......a..g..a.t...................g.......g........a...........................  [ 68480]
AY261362.1_Mkuzi_1979          ....................................................a...........................  [ 68480]
AY261361.1_Malawi_Lil-20/1_1983 .......a.....a.t...................g................a...........c...........t...  [ 68480]
AY261360.1_Kenya_1950          .......a.....a.t..............g.........g..g........a...........................  [ 68480]
AM712240.1_OURT_88/3           ....................................................a..............g............  [ 68480]
AM712239.1_Benin_97/1          ....................................................a..............g............  [ 68480]
KM111295.1_Ken06.Bus           .......a.....c.t..............g.........g..g........a.................t.........  [ 68480]
KM111294.1_Ken05/Tk1           .......a.....a.t..............g.........g..g........a...........................  [ 68480]
KM262844.1_strain_L60          ....................................................a..............g............  [ 68480]
KM102979.1_26544/OG10_Italy    ....................................................a..............g............  [ 68480]
KM262845.1_strain_NHV          ....................................................a..............g............  [ 68480]
KX354450.1_47/Ss/2008          ....................................................a..............g............  [ 68480]
KP055815.1_strain_BA71         ....................................................a..............g............  [ 68480]
MH025920.1_strain_R35          .......a.....c.t..............g.........g..g........a.................t.........  [ 68480]
MH025919.1_strain_N10          .......a.....c.t..............g.........g..g........a.................t.........  [ 68480]
MH025918.1_strain_R25          .......a.....c.t..............g.........g..g........a.................t.........  [ 68480]
MH025917.1_strain_R7           .......a.....c.t..............g.........g..g........a.................t.........  [ 68480]
MH025916.1_strain_R8           .......a.....c.t..............g.........g..g........a.................t.........  [ 68480]
```

```
FR682468.2_Georgia_2007/1        ttacgctccgcccagtattccttaaaacttcttttagacttgctgagctgttcccaggattcgaactcagtccaatgttt [ 68560]
ASF_IND_20_CAD543                .......................r........................................................ [ 68560]
LR899193.1_Germany_2020/1        ................................................................................ [ 68560]
MN393477.1_Wuhan_2019            ................................................................................ [ 68560]
MW396979.1_Timor-Leste/2019/1    ................................................................................ [ 68560]
MK940252.1_CN/2019/InnerMongolia ................................................................................ [ 68560]
MK128995.1_China/2018/AnhuiXCGQ  ................................................................................ [ 68560]
MH681419.1_ASFV/POL/2015         ................................................................................ [ 68560]
LS478113.1_Estonia_2014          ................................................................................ [ 68560]
KP843857.1_Odintsovo_02/14       .............................................................................-.. [ 68560]
FN557520.1_strain_E75            .......................g............................................g........... [ 68560]
AY261366.1_Warthog               ..........t............g............................................g........... [ 68560]
AY261365.1_Warmbaths             ..........t............g............................................g........... [ 68560]
AY261364.1_Tengani_62            ..........t......c.....g............................................g........... [ 68560]
AY261363.1_Pretoriuskop/96/4     ..........t............g............................................g........... [ 68560]
AY261362.1_Mkuzi_1979            .....t.................g............................................g........... [ 68560]
AY261361.1_Malawi_Lil-20/1_1983  ..........t............g..g..c................................a.....g........... [ 68560]
AY261360.1_Kenya_1950            ..c....................g....................a.................a.....g........... [ 68560]
AM712240.1_OURT_88/3             .......................g............................................g........... [ 68560]
AM712239.1_Benin_97/1            .......................g............................................g........... [ 68560]
KM111295.1_Ken06.Bus             ..c.......t......c...c.g....................a.................a.....g........... [ 68560]
KM111294.1_Ken05/Tk1             ..c..................c.g....................a.................a.....g........... [ 68560]
KM262844.1_strain_L60            .......................g............................................g........... [ 68560]
KM102979.1_26544/OG10_Italy      ....................................................................g........... [ 68560]
KM262845.1_strain_NHV            .......................g............................................g........... [ 68560]
KX354450.1_47/Ss/2008            ....................................................................g........... [ 68560]
KP055815.1_strain_BA71           .......................g............................................g........... [ 68560]
MH025920.1_strain_R35            ..c.......t......c...c.g....................a.................a.....g........... [ 68560]
MH025919.1_strain_N10            ..c.......t......c...c.g....................a.................a.....g........... [ 68560]
MH025918.1_strain_R25            ..c.......t......c...c.g....................a.................a.....g........... [ 68560]
MH025917.1_strain_R7             ..c.......t......c...c.g....................a.................a.....g........... [ 68560]
MH025916.1_strain_R8             ..c.......t......c...c.g....................a.................a.....g........... [ 68560]

FR682468.2_Georgia_2007/1        ttttcttttggggaagacttccctttttgaaacattttttgcggctccaccatc------tacactatgattttccaaaa [ 68640]
ASF_IND_20_CAD543                ......................................................------.................... [ 68640]
LR899193.1_Germany_2020/1        ......................................................------.................... [ 68640]
MN393477.1_Wuhan_2019            ......................................................------.................... [ 68640]
MW396979.1_Timor-Leste/2019/1    ......................................................------.................... [ 68640]
MK940252.1_CN/2019/InnerMongolia ......................................................------.................... [ 68640]
MK128995.1_China/2018/AnhuiXCGQ  ......................................................------.................... [ 68640]
MH681419.1_ASFV/POL/2015         ......................................................------.................... [ 68640]
LS478113.1_Estonia_2014          ......................................................------.................... [ 68640]
KP843857.1_Odintsovo_02/14       ......................................................------.................... [ 68640]
```

```
FN557520.1_strain_E75          ......................................................------.....c.............. [ 68640]
AY261366.1_Warthog             ...............................g......................------.................... [ 68640]
AY261365.1_Warmbaths           ...............................g......................------.................g.. [ 68640]
AY261364.1_Tengani_62          ................................................g.....------.....c.............. [ 68640]
AY261363.1_Pretoriuskop/96/4   ...............................g................g.....------.................... [ 68640]
AY261362.1_Mkuzi_1979          ......................................................------.....c.............. [ 68640]
AY261361.1_Malawi_Lil-20/1_1983 ..............g......a................................tacgtt........t........g.. [ 68640]
AY261360.1_Kenya_1950          .....................a................................------........cc.......... [ 68640]
AM712240.1_OURT_88/3           ......................................................------.....c.............. [ 68640]
AM712239.1_Benin_97/1          ......................................................------.....c.............. [ 68640]
KM111295.1_Ken06.Bus           .....................a................................------........cc.......... [ 68640]
KM111294.1_Ken05/Tk1           .....................a................................------........cc.......... [ 68640]
KM262844.1_strain_L60          ......................................................------.....c.............. [ 68640]
KM102979.1_26544/OG10_Italy    ......................................................------.....c.............. [ 68640]
KM262845.1_strain_NHV          ......................................................------.....c.............. [ 68640]
KX354450.1_47/Ss/2008          ......................................................------.....c.............. [ 68640]
KP055815.1_strain_BA71         ......................................................------.....c.............. [ 68640]
MH025920.1_strain_R35          .....................a................................------........cc.......... [ 68640]
MH025919.1_strain_N10          .....................a................................------........cc.......... [ 68640]
MH025918.1_strain_R25          .....................a................................------........cc.......... [ 68640]
MH025917.1_strain_R7           .....................a................................------........cc.......... [ 68640]
MH025916.1_strain_R8           .....................a................................------........cc.......... [ 68640]

FR682468.2_Georgia_2007/1      taatctccttcatcgtttgagttatatgggcattgctaagcaccttagtggtaacctgtttacctatgtgatttagcaga [ 68720]
ASF_IND_20_CAD543              ................................................................................ [ 68720]
LR899193.1_Germany_2020/1      ................................................................................ [ 68720]
MN393477.1_Wuhan_2019          ................................................................................ [ 68720]
MW396979.1_Timor-Leste/2019/1  ................................................................................ [ 68720]
MK940252.1_CN/2019/InnerMongolia ................................................................................ [ 68720]
MK128995.1_China/2018/AnhuiXCGQ ................................................................................ [ 68720]
MH681419.1_ASFV/POL/2015       ................................................................................ [ 68720]
LS478113.1_Estonia_2014        ................................................................................ [ 68720]
KP843857.1_Odintsovo_02/14     ................................................................................ [ 68720]
FN557520.1_strain_E75          ................................................................................ [ 68720]
AY261366.1_Warthog             .............g.............................t.............................c...... [ 68720]
AY261365.1_Warmbaths           .............a.............................t.................................... [ 68720]
AY261364.1_Tengani_62          .............a.....g.......................t.............................c...... [ 68720]
AY261363.1_Pretoriuskop/96/4   .............g.............................t.............................c...... [ 68720]
AY261362.1_Mkuzi_1979          ................................................................................ [ 68720]
AY261361.1_Malawi_Lil-20/1_1983 .............a.............................t..........t.......................a. [ 68720]
AY261360.1_Kenya_1950          .............at...................a........t..........t..................c...... [ 68720]
AM712240.1_OURT_88/3           ................................................................................ [ 68720]
AM712239.1_Benin_97/1          ................................................................................ [ 68720]
KM111295.1_Ken06.Bus           .............at...................a........t..........t............a.....c...... [ 68720]
```

```
KM111294.1_Ken05/Tk1             ............at....................a........t..........t............a.....c...... [ 68720]
KM262844.1_strain_L60            ................................................................................ [ 68720]
KM102979.1_26544/OG10_Italy      ................................................................................ [ 68720]
KM262845.1_strain_NHV            ................................................................................ [ 68720]
KX354450.1_47/Ss/2008            ................................................................................ [ 68720]
KP055815.1_strain_BA71           ................................................................................ [ 68720]
MH025920.1_strain_R35            ............at....................a........t..........t............a.....c...... [ 68720]
MH025919.1_strain_N10            ............at....................at.......t..........t............a.....c...... [ 68720]
MH025918.1_strain_R25            ............at....................a........t..........t............a.....c...... [ 68720]
MH025917.1_strain_R7             ............at....................a........t..........t............a.....c...... [ 68720]
MH025916.1_strain_R8             ............at....................a........t..........t............a.....c...... [ 68720]

FR682468.2_Georgia_2007/1        aaaccaagtttgtccatttgtgtctcaaccatttattcttaacaaaaca----aaaaaaaattaaaaatcatcgtcgttt [ 68800]
ASF_IND_20_CAD543                ................................................-----........................... [ 68800]
LR899193.1_Germany_2020/1        .................................................----........................... [ 68800]
MN393477.1_Wuhan_2019            .................................................----........................... [ 68800]
MW396979.1_Timor-Leste/2019/1    .................................................----........................... [ 68800]
MK940252.1_CN/2019/InnerMongolia .................................................----........................... [ 68800]
MK128995.1_China/2018/AnhuiXCGQ  .................................................----........................... [ 68800]
MH681419.1_ASFV/POL/2015         .................................................----........................... [ 68800]
LS478113.1_Estonia_2014          .................................................----........................... [ 68800]
KP843857.1_Odintsovo_02/14       ................................................-----........................... [ 68800]
FN557520.1_strain_E75            ................................................-----........................... [ 68800]
AY261366.1_Warthog               ................................................-----........................... [ 68800]
AY261365.1_Warmbaths             ................................................-----........................... [ 68800]
AY261364.1_Tengani_62            .....g....................................t.....-----........................... [ 68800]
AY261363.1_Pretoriuskop/96/4     ................................................-----........................... [ 68800]
AY261362.1_Mkuzi_1979            ................................................-----........................... [ 68800]
AY261361.1_Malawi_Lil-20/1_1983  ...........a.....................................aaac.........................c. [ 68800]
AY261360.1_Kenya_1950            ...........a....................................-----........................... [ 68800]
AM712240.1_OURT_88/3             ................................................-----........................... [ 68800]
AM712239.1_Benin_97/1            ................................................-----........................... [ 68800]
KM111295.1_Ken06.Bus             ...........a....................................-----........................... [ 68800]
KM111294.1_Ken05/Tk1             ...........a....................................-----........................... [ 68800]
KM262844.1_strain_L60            ................................................-----........................... [ 68800]
KM102979.1_26544/OG10_Italy      ................................................-----........................... [ 68800]
KM262845.1_strain_NHV            ................................................-----........................... [ 68800]
KX354450.1_47/Ss/2008            ................................................-----........................... [ 68800]
KP055815.1_strain_BA71           ................................................-----........................... [ 68800]
MH025920.1_strain_R35            ...........a.....................................----........................... [ 68800]
MH025919.1_strain_N10            ...........a....................................-----........................... [ 68800]
MH025918.1_strain_R25            ...........a.....................................----........................... [ 68800]
MH025917.1_strain_R7             ...........a.....................................----........................... [ 68800]
MH025916.1_strain_R8             ...........a.....................................----........................... [ 68800]
```

```
FR682468.2_Georgia_2007/1          aaaaagagtttgaaggcaaacgcatcatccttaacacagttctgatactgcgtaggtcttaactcgaaaaagttggtttt [ 68880]
ASF_IND_20_CAD543                  .................................................r.............................. [ 68880]
LR899193.1_Germany_2020/1          ................................................................................ [ 68880]
MN393477.1_Wuhan_2019              ................................................................................ [ 68880]
MW396979.1_Timor-Leste/2019/1      ................................................................................ [ 68880]
MK940252.1_CN/2019/InnerMongolia   ................................................................................ [ 68880]
MK128995.1_China/2018/AnhuiXCGQ    ................................................................................ [ 68880]
MH681419.1_ASFV/POL/2015           ................................................................................ [ 68880]
LS478113.1_Estonia_2014            ................................................................................ [ 68880]
KP843857.1_Odintsovo_02/14         ................................................................................ [ 68880]
FN557520.1_strain_E75              ................................................................................ [ 68880]
AY261366.1_Warthog                 .................g.......................t................................a..... [ 68880]
AY261365.1_Warmbaths               ................................................................................ [ 68880]
AY261364.1_Tengani_62              ..................................................t............................. [ 68880]
AY261363.1_Pretoriuskop/96/4       .................g........................................................a..... [ 68880]
AY261362.1_Mkuzi_1979              ................................................................................ [ 68880]
AY261361.1_Malawi_Lil-20/1_1983    ....................t........................................................... [ 68880]
AY261360.1_Kenya_1950              ................................................................................ [ 68880]
AM712240.1_OURT_88/3               ................................................................................ [ 68880]
AM712239.1_Benin_97/1              ................................................................................ [ 68880]
KM111295.1_Ken06.Bus               ................................................................................ [ 68880]
KM111294.1_Ken05/Tk1               ................................................................................ [ 68880]
KM262844.1_strain_L60              ................................................................................ [ 68880]
KM102979.1_26544/OG10_Italy        ................................................................................ [ 68880]
KM262845.1_strain_NHV              ................................................................................ [ 68880]
KX354450.1_47/Ss/2008              ................................................................................ [ 68880]
KP055815.1_strain_BA71             ................................................................................ [ 68880]
MH025920.1_strain_R35              ..................................................t............................. [ 68880]
MH025919.1_strain_N10              ..................................................t............................. [ 68880]
MH025918.1_strain_R25              ..................................................t............................. [ 68880]
MH025917.1_strain_R7               ..................................................t............................. [ 68880]
MH025916.1_strain_R8               ..................................................t............................. [ 68880]

FR682468.2_Georgia_2007/1          ttctacttcattaagaaagaatttagtcatctgaggaaaagggtttcccaccttataaatgcttttgcactgcatcatga [ 68960]
ASF_IND_20_CAD543                  ...................................................y............................ [ 68960]
LR899193.1_Germany_2020/1          ................................................................................ [ 68960]
MN393477.1_Wuhan_2019              ................................................................................ [ 68960]
MW396979.1_Timor-Leste/2019/1      ................................................................................ [ 68960]
MK940252.1_CN/2019/InnerMongolia   ................................................................................ [ 68960]
MK128995.1_China/2018/AnhuiXCGQ    ................................................................................ [ 68960]
MH681419.1_ASFV/POL/2015           ................................................................................ [ 68960]
LS478113.1_Estonia_2014            ................................................................................ [ 68960]
KP843857.1_Odintsovo_02/14         ................................................................................ [ 68960]
```

```
FN557520.1_strain_E75             ...................................................t........a................... [ 68960]
AY261366.1_Warthog                ................................................................................ [ 68960]
AY261365.1_Warmbaths              ................................................................................ [ 68960]
AY261364.1_Tengani_62             ................................................................................ [ 68960]
AY261363.1_Pretoriuskop/96/4      ................................................................................ [ 68960]
AY261362.1_Mkuzi_1979             ................................................................................ [ 68960]
AY261361.1_Malawi_Lil-20/1_1983   .............................................c.....a........a................... [ 68960]
AY261360.1_Kenya_1950             ..............................................t.............a.............g..... [ 68960]
AM712240.1_OURT_88/3              ...................................................t........a................... [ 68960]
AM712239.1_Benin_97/1             ...................................................t........a................... [ 68960]
KM111295.1_Ken06.Bus              ..............................................t.............a.............a..... [ 68960]
KM111294.1_Ken05/Tk1              ..............................................t.............a.............g..... [ 68960]
KM262844.1_strain_L60             ...................................................t........a................... [ 68960]
KM102979.1_26544/OG10_Italy       ...................................................t........a................... [ 68960]
KM262845.1_strain_NHV             ...................................................t........a................... [ 68960]
KX354450.1_47/Ss/2008             ...................................................t........a................... [ 68960]
KP055815.1_strain_BA71            ...................................................t........a................... [ 68960]
MH025920.1_strain_R35             ..............................................t.............a.............g..... [ 68960]
MH025919.1_strain_N10             ..............................................t.............a.............g..... [ 68960]
MH025918.1_strain_R25             ..............................................t.............a.............g..... [ 68960]
MH025917.1_strain_R7              ..............................................t.............a.............g..... [ 68960]
MH025916.1_strain_R8              ..............................................t.............a.............g..... [ 68960]

FR682468.2_Georgia_2007/1         agcacaaattatctgtaaagtagcgtatatattgaaatagcatttcttttgaaaaaccgggaactcttcctcttgccttg [ 69040]
ASF_IND_20_CAD543                 .....................................y.......................................... [ 69040]
LR899193.1_Germany_2020/1         ................................................................................ [ 69040]
MN393477.1_Wuhan_2019             ................................................................................ [ 69040]
MW396979.1_Timor-Leste/2019/1     ................................................................................ [ 69040]
MK940252.1_CN/2019/InnerMongolia  ................................................................................ [ 69040]
MK128995.1_China/2018/AnhuiXCGQ   ................................................................................ [ 69040]
MH681419.1_ASFV/POL/2015          ................................................................................ [ 69040]
LS478113.1_Estonia_2014           ................................................................................ [ 69040]
KP843857.1_Odintsovo_02/14        ................................................................................ [ 69040]
FN557520.1_strain_E75             .....................................c.......................................... [ 69040]
AY261366.1_Warthog                ................................................................................ [ 69040]
AY261365.1_Warmbaths              ................................................................................ [ 69040]
AY261364.1_Tengani_62             ................................................................................ [ 69040]
AY261363.1_Pretoriuskop/96/4      ................................................................................ [ 69040]
AY261362.1_Mkuzi_1979             ................................................................................ [ 69040]
AY261361.1_Malawi_Lil-20/1_1983   .....................................c....................a...........c......... [ 69040]
AY261360.1_Kenya_1950             ......g......a..................................................g.....c......... [ 69040]
AM712240.1_OURT_88/3              .....................................c.......................................... [ 69040]
AM712239.1_Benin_97/1             .....................................c.......................................... [ 69040]
KM111295.1_Ken06.Bus              ......g...............................................................c......... [ 69040]
```

```
KM111294.1_Ken05/Tk1            ......g......a........................................................c......... [ 69040]
KM262844.1_strain_L60           .....................................c.......................................... [ 69040]
KM102979.1_26544/OG10_Italy     .....................................c.......................................... [ 69040]
KM262845.1_strain_NHV           .....................................c.......................................... [ 69040]
KX354450.1_47/Ss/2008           .....................................c.......................................... [ 69040]
KP055815.1_strain_BA71          .....................................c.......................................... [ 69040]
MH025920.1_strain_R35           ......g...............................................................c......... [ 69040]
MH025919.1_strain_N10           ......g...............................................................c......... [ 69040]
MH025918.1_strain_R25           ......g...............................................................c......... [ 69040]
MH025917.1_strain_R7            ......g...............................................................c......... [ 69040]
MH025916.1_strain_R8            ......g...............................................................c......... [ 69040]

FR682468.2_Georgia_2007/1       tcaaaggcatagttaataaactcatccaccaactccacagcctccttcaaaattttgtgaatgatcttttcctcgggaat [ 69120]
ASF_IND_20_CAD543               ................................................................................ [ 69120]
LR899193.1_Germany_2020/1       ................................................................................ [ 69120]
MN393477.1_Wuhan_2019           ................................................................................ [ 69120]
MW396979.1_Timor-Leste/2019/1   ................................................................................ [ 69120]
MK940252.1_CN/2019/InnerMongolia ................................................................................ [ 69120]
MK128995.1_China/2018/AnhuiXCGQ ................................................................................ [ 69120]
MH681419.1_ASFV/POL/2015        ................................................................................ [ 69120]
LS478113.1_Estonia_2014         ................................................................................ [ 69120]
KP843857.1_Odintsovo_02/14      ................................................................................ [ 69120]
FN557520.1_strain_E75           .................g.............................................................. [ 69120]
AY261366.1_Warthog              ............c....g........................................................a..... [ 69120]
AY261365.1_Warmbaths            ............c....g........................................................a..... [ 69120]
AY261364.1_Tengani_62           .................g...........................................................g.. [ 69120]
AY261363.1_Pretoriuskop/96/4    .................g................................................c.......a..g.. [ 69120]
AY261362.1_Mkuzi_1979           .................g.............................................................. [ 69120]
AY261361.1_Malawi_Lil-20/1_1983 .................g........t....gt......................................t........ [ 69120]
AY261360.1_Kenya_1950           ............c....g........t....gt.....g..........................t...........g.. [ 69120]
AM712240.1_OURT_88/3            .................g.............................................................. [ 69120]
AM712239.1_Benin_97/1           .................g.............................................................. [ 69120]
KM111295.1_Ken06.Bus            ............c....g.............gt.....g..........g..............c......t........ [ 69120]
KM111294.1_Ken05/Tk1            ............c.............t....gt.....g..........................t...........g.. [ 69120]
KM262844.1_strain_L60           .................g.............................................................. [ 69120]
KM102979.1_26544/OG10_Italy     .................g.............................................................. [ 69120]
KM262845.1_strain_NHV           .................g.............................................................. [ 69120]
KX354450.1_47/Ss/2008           .................g.............................................................. [ 69120]
KP055815.1_strain_BA71          .................g.............................................................. [ 69120]
MH025920.1_strain_R35           ............c....g.............gt.....g..........g..............c......t........ [ 69120]
MH025919.1_strain_N10           ............c....g.............gt.....g..........g..............c......t........ [ 69120]
MH025918.1_strain_R25           ............c....g.............gt.....g..........g..............c......t........ [ 69120]
MH025917.1_strain_R7            ............c....g.............gt.....g..........g..............c......t........ [ 69120]
MH025916.1_strain_R8            ............c....g.............gt.....g..........g..............c......t........ [ 69120]
```

```
FR682468.2_Georgia_2007/1        gttatacacgtaatttgagataagaaaacacgcaaaactacagtgcatcccttcatcacgtgagataaactcattatagc [ 69200]
ASF_IND_20_CAD543                ................................................................................ [ 69200]
LR899193.1_Germany_2020/1        ................................................................................ [ 69200]
MN393477.1_Wuhan_2019            ................................................................................ [ 69200]
MW396979.1_Timor-Leste/2019/1    ................................................................................ [ 69200]
MK940252.1_CN/2019/InnerMongolia ................................................................................ [ 69200]
MK128995.1_China/2018/AnhuiXCGQ  ................................................................................ [ 69200]
MH681419.1_ASFV/POL/2015         ................................................................................ [ 69200]
LS478113.1_Estonia_2014          ................................................................................ [ 69200]
KP843857.1_Odintsovo_02/14       ................................................................................ [ 69200]
FN557520.1_strain_E75            ...............c...................................c.....g.....a................ [ 69200]
AY261366.1_Warthog               ..............................t.....g..........cg..c.....g.....a................ [ 69200]
AY261365.1_Warmbaths             ..............................t.....g..........cg..c.....g.....a................ [ 69200]
AY261364.1_Tengani_62            ...............c...................................c.....g...................... [ 69200]
AY261363.1_Pretoriuskop/96/4     ..............................t.....g..........c...c.....g.....a................ [ 69200]
AY261362.1_Mkuzi_1979            ...............c...................................c.....g.....a................ [ 69200]
AY261361.1_Malawi_Lil-20/1_1983  .........a.....c..............t.....g.....a.....t..c.....g...................... [ 69200]
AY261360.1_Kenya_1950            .....t..ga..........................g.....a........c.....g.....a................ [ 69200]
AM712240.1_OURT_88/3             ...............c...................................c.....g.....a................ [ 69200]
AM712239.1_Benin_97/1            ...............c...................................c.....g.....a................ [ 69200]
KM111295.1_Ken06.Bus             .........a..........c...............g........t.ct..c.....g...................... [ 69200]
KM111294.1_Ken05/Tk1             ........ga.....c....................g.....a........c.....g.....a................ [ 69200]
KM262844.1_strain_L60            ...............c...................................c.....g.....a................ [ 69200]
KM102979.1_26544/OG10_Italy      ...............c...................................c.....g.....a................ [ 69200]
KM262845.1_strain_NHV            ...............c...................................c.....g.....a................ [ 69200]
KX354450.1_47/Ss/2008            ...............c...................................c.....g.....a................ [ 69200]
KP055815.1_strain_BA71           ...............c...................................c.....g.....a................ [ 69200]
MH025920.1_strain_R35            .........a..........c...............g........t.ct..c.....g...................... [ 69200]
MH025919.1_strain_N10            .........a..........c...............g........t.ct..c.....g...................... [ 69200]
MH025918.1_strain_R25            .........a..........c...............g........t.ct..c.....g...................... [ 69200]
MH025917.1_strain_R7             .........a..........c...............g........t.ct..c.....g...................... [ 69200]
MH025916.1_strain_R8             .........a..........c...............g........t.ct..c.....g...................... [ 69200]

FR682468.2_Georgia_2007/1        ttacaagccccggcataatattctgttccttaagaaactggatcgccacaaagtggttttgaaataaaatgccttctacg [ 69280]
ASF_IND_20_CAD543                ................................................................................ [ 69280]
LR899193.1_Germany_2020/1        ................................................................................ [ 69280]
MN393477.1_Wuhan_2019            ................................................................................ [ 69280]
MW396979.1_Timor-Leste/2019/1    ................................................................................ [ 69280]
MK940252.1_CN/2019/InnerMongolia ................................................................................ [ 69280]
MK128995.1_China/2018/AnhuiXCGQ  ................................................................................ [ 69280]
MH681419.1_ASFV/POL/2015         ................................................................................ [ 69280]
LS478113.1_Estonia_2014          ................................................................................ [ 69280]
KP843857.1_Odintsovo_02/14       ................................................................................ [ 69280]
```

```
FN557520.1_strain_E75          ................................................................................ [ 69280]
AY261366.1_Warthog             .......t...........g.....................................................c..c... [ 69280]
AY261365.1_Warmbaths           .......t...........g.....................................................c..c... [ 69280]
AY261364.1_Tengani_62          ................................................................................ [ 69280]
AY261363.1_Pretoriuskop/96/4   .......t...........g.....................................................c..c... [ 69280]
AY261362.1_Mkuzi_1979          ................................................................................ [ 69280]
AY261361.1_Malawi_Lil-20/1_1983 ......................t........g.........................................c..g... [ 69280]
AY261360.1_Kenya_1950          ......................t..................................................c..c... [ 69280]
AM712240.1_OURT_88/3           ................................................................................ [ 69280]
AM712239.1_Benin_97/1          ................................................................................ [ 69280]
KM111295.1_Ken06.Bus           ......................t..................................................c..c... [ 69280]
KM111294.1_Ken05/Tk1           ......................t..................................................c..c... [ 69280]
KM262844.1_strain_L60          ................................................................................ [ 69280]
KM102979.1_26544/OG10_Italy    ................................................................................ [ 69280]
KM262845.1_strain_NHV          ................................................................................ [ 69280]
KX354450.1_47/Ss/2008          ................................................................................ [ 69280]
KP055815.1_strain_BA71         ................................................................................ [ 69280]
MH025920.1_strain_R35          ......................t..................................................c..c... [ 69280]
MH025919.1_strain_N10          ......................t..................................................c..c... [ 69280]
MH025918.1_strain_R25          ......................t..................................................c..c... [ 69280]
MH025917.1_strain_R7           ......................t..................................................c..c... [ 69280]
MH025916.1_strain_R8           ......................t..................................................c..c... [ 69280]

FR682468.2_Georgia_2007/1      gcggcgaagcccaccagccgctcacctagagtgttcctgtcggggtccatccactgccgcacccactgcgccattttttt [ 69360]
ASF_IND_20_CAD543              ................................................................................ [ 69360]
LR899193.1_Germany_2020/1      ................................................................................ [ 69360]
MN393477.1_Wuhan_2019          ................................................................................ [ 69360]
MW396979.1_Timor-Leste/2019/1  ................................................................................ [ 69360]
MK940252.1_CN/2019/InnerMongolia ................................................................................ [ 69360]
MK128995.1_China/2018/AnhuiXCGQ ................................................................................ [ 69360]
MH681419.1_ASFV/POL/2015       ................................................................................ [ 69360]
LS478113.1_Estonia_2014        ................................................................................ [ 69360]
KP843857.1_Odintsovo_02/14     ................................................................................ [ 69360]
FN557520.1_strain_E75          ..........................c..................................................... [ 69360]
AY261366.1_Warthog             ..............a..t.....g..c...c.....t..gt....................................... [ 69360]
AY261365.1_Warmbaths           ................a...................t..gt...........................a........... [ 69360]
AY261364.1_Tengani_62          ....................................t...t....................................... [ 69360]
AY261363.1_Pretoriuskop/96/4   ..............a..t.....g..c...c.....t..gt...........................a........... [ 69360]
AY261362.1_Mkuzi_1979          ................................................................................ [ 69360]
AY261361.1_Malawi_Lil-20/1_1983 ........a.....a..t.....g..c...c........g............................a........... [ 69360]
AY261360.1_Kenya_1950          .....a........a..t..t..g..c...c.....t..a.................t.......t..a.........c. [ 69360]
AM712240.1_OURT_88/3           ..........................c..................................................... [ 69360]
AM712239.1_Benin_97/1          ..........................c..................................................... [ 69360]
KM111295.1_Ken06.Bus           ..............a..t.....g..c...c....tt.ag................t...........a..t........ [ 69360]
```

```
KM111294.1_Ken05/Tk1            .....a........a..t..t..g..c...c.....t..a.................t.......t..a.........c. [ 69360]
KM262844.1_strain_L60           ..........................c..................................................... [ 69360]
KM102979.1_26544/OG10_Italy     ..........................c..................................................... [ 69360]
KM262845.1_strain_NHV           ..........................c..................................................... [ 69360]
KX354450.1_47/Ss/2008           ..........................c..................................................... [ 69360]
KP055815.1_strain_BA71          ..........................c..................................................... [ 69360]
MH025920.1_strain_R35           ..............a..t.....g..c...c....tt.ag................t...........a..t........ [ 69360]
MH025919.1_strain_N10           ..............a..t.....g..c...c....tt.ag................t...........a..t........ [ 69360]
MH025918.1_strain_R25           ..............a..t.....g..c...c....tt.ag................t...........a..t........ [ 69360]
MH025917.1_strain_R7            ..............a..t.....g..c...c....tt.ag................t...........a..t........ [ 69360]
MH025916.1_strain_R8            ..............a..t.....g..c...c....tt.ag................t...........a..t........ [ 69360]

FR682468.2_Georgia_2007/1       tatgatagggtgtttttcaatgccgctaaagatgcgctgttgttccttctcatccgggatcagcgtttttacctgtattg [ 69440]
ASF_IND_20_CAD543               ................................................................................ [ 69440]
LR899193.1_Germany_2020/1       ................................................................................ [ 69440]
MN393477.1_Wuhan_2019           ................................................................................ [ 69440]
MW396979.1_Timor-Leste/2019/1   ................................................................................ [ 69440]
MK940252.1_CN/2019/InnerMongolia ................................................................................ [ 69440]
MK128995.1_China/2018/AnhuiXCGQ ................................................................................ [ 69440]
MH681419.1_ASFV/POL/2015        ................................................................................ [ 69440]
LS478113.1_Estonia_2014         ................................................................................ [ 69440]
KP843857.1_Odintsovo_02/14      ................................................................................ [ 69440]
FN557520.1_strain_E75           ................................................................................ [ 69440]
AY261366.1_Warthog              a........a.................g.................................................... [ 69440]
AY261365.1_Warmbaths            .........a.................g....................t............................... [ 69440]
AY261364.1_Tengani_62           .........a.................g....................t............................... [ 69440]
AY261363.1_Pretoriuskop/96/4    .........a.................g.................................................... [ 69440]
AY261362.1_Mkuzi_1979           ................................................................................ [ 69440]
AY261361.1_Malawi_Lil-20/1_1983 .........a...........a.....g..........................t.................t....... [ 69440]
AY261360.1_Kenya_1950           .........a...........a.......................................................... [ 69440]
AM712240.1_OURT_88/3            ................................................................................ [ 69440]
AM712239.1_Benin_97/1           ................................................................................ [ 69440]
KM111295.1_Ken06.Bus            .........a...........a.....g....................t.....t................t........ [ 69440]
KM111294.1_Ken05/Tk1            .........a...........a..................................................t....... [ 69440]
KM262844.1_strain_L60           ................................................................................ [ 69440]
KM102979.1_26544/OG10_Italy     ................................................................................ [ 69440]
KM262845.1_strain_NHV           ................................................................................ [ 69440]
KX354450.1_47/Ss/2008           ................................................................................ [ 69440]
KP055815.1_strain_BA71          ................................................................................ [ 69440]
MH025920.1_strain_R35           .........a...........a.....g....................t.....t................t........ [ 69440]
MH025919.1_strain_N10           .........a...........a.....g....................t.....t................t........ [ 69440]
MH025918.1_strain_R25           .........a...........a.....g....................t.....t................t........ [ 69440]
MH025917.1_strain_R7            .........a...........a.....g....................t.....t................t........ [ 69440]
MH025916.1_strain_R8            .........a...........a.....g....................t.....t................t........ [ 69440]
```

```
FR682468.2_Georgia_2007/1        agtaggcttcgctatgaacgcactcttgggcagcctgcattgtataaaagtataacacttcctttactttaatttcgcgc [ 69520]
ASF_IND_20_CAD543                ..........r........r............................................................ [ 69520]
LR899193.1_Germany_2020/1        ................................................................................ [ 69520]
MN393477.1_Wuhan_2019            ................................................................................ [ 69520]
MW396979.1_Timor-Leste/2019/1    ................................................................................ [ 69520]
MK940252.1_CN/2019/InnerMongolia ................................................................................ [ 69520]
MK128995.1_China/2018/AnhuiXCGQ  ................................................................................ [ 69520]
MH681419.1_ASFV/POL/2015         ................................................................................ [ 69520]
LS478113.1_Estonia_2014          ................................................................................ [ 69520]
KP843857.1_Odintsovo_02/14       ................................................................................ [ 69520]
FN557520.1_strain_E75            ..........a........a............................................................ [ 69520]
AY261366.1_Warthog               ......................................................g......................... [ 69520]
AY261365.1_Warmbaths             ......................................................g......................... [ 69520]
AY261364.1_Tengani_62            ......................t......................................................... [ 69520]
AY261363.1_Pretoriuskop/96/4     ......................................................g......................... [ 69520]
AY261362.1_Mkuzi_1979            ..........a..................................................................... [ 69520]
AY261361.1_Malawi_Lil-20/1_1983  ................................................................................ [ 69520]
AY261360.1_Kenya_1950            .......c.....g........................................g......................... [ 69520]
AM712240.1_OURT_88/3             ..........a........a............................................................ [ 69520]
AM712239.1_Benin_97/1            ..........a........a............................................................ [ 69520]
KM111295.1_Ken06.Bus             .......c..............................................g......................... [ 69520]
KM111294.1_Ken05/Tk1             .......c.....g........................................g......................... [ 69520]
KM262844.1_strain_L60            ..........a........a............................................................ [ 69520]
KM102979.1_26544/OG10_Italy      ..........a........a............................................................ [ 69520]
KM262845.1_strain_NHV            ..........a........a............................................................ [ 69520]
KX354450.1_47/Ss/2008            ..........a........a............................................................ [ 69520]
KP055815.1_strain_BA71           ..........a........a............................................................ [ 69520]
MH025920.1_strain_R35            .......c..............................................g......................... [ 69520]
MH025919.1_strain_N10            .......c..............................................g......................... [ 69520]
MH025918.1_strain_R25            .......c..............................................g......................... [ 69520]
MH025917.1_strain_R7             .......c..............................................g......................... [ 69520]
MH025916.1_strain_R8             .......c..............................................g......................... [ 69520]

FR682468.2_Georgia_2007/1        ataaaattggttaaaaggttttcgataacaatttcgtcggcaacaacaaagaaggctaaaatttgtttataaaattcgcg [ 69600]
ASF_IND_20_CAD543                ................................................................................ [ 69600]
LR899193.1_Germany_2020/1        ................................................................................ [ 69600]
MN393477.1_Wuhan_2019            ................................................................................ [ 69600]
MW396979.1_Timor-Leste/2019/1    ................................................................................ [ 69600]
MK940252.1_CN/2019/InnerMongolia ................................................................................ [ 69600]
MK128995.1_China/2018/AnhuiXCGQ  ................................................................................ [ 69600]
MH681419.1_ASFV/POL/2015         ................................................................................ [ 69600]
LS478113.1_Estonia_2014          ................................................................................ [ 69600]
KP843857.1_Odintsovo_02/14       ................................................................................ [ 69600]
```

```
FN557520.1_strain_E75          ................................................................................ [ 69600]
AY261366.1_Warthog             ................................................................................ [ 69600]
AY261365.1_Warmbaths           ................................................................................ [ 69600]
AY261364.1_Tengani_62          ........a....................c.................................................. [ 69600]
AY261363.1_Pretoriuskop/96/4   ................................................................................ [ 69600]
AY261362.1_Mkuzi_1979          ................................................................................ [ 69600]
AY261361.1_Malawi_Lil-20/1_1983 ..........................c..................................................... [ 69600]
AY261360.1_Kenya_1950          .....g......................................g..............g.................... [ 69600]
AM712240.1_OURT_88/3           ................................................................................ [ 69600]
AM712239.1_Benin_97/1          ................................................................................ [ 69600]
KM111295.1_Ken06.Bus           .....g......................................g................................... [ 69600]
KM111294.1_Ken05/Tk1           .....g......................................g..............g.................a.. [ 69600]
KM262844.1_strain_L60          ................................................................................ [ 69600]
KM102979.1_26544/OG10_Italy    ................................................................................ [ 69600]
KM262845.1_strain_NHV          ................................................................................ [ 69600]
KX354450.1_47/Ss/2008          ................................................................................ [ 69600]
KP055815.1_strain_BA71         ................................................................................ [ 69600]
MH025920.1_strain_R35          .....g......................................g................................... [ 69600]
MH025919.1_strain_N10          .....g......................................g................................... [ 69600]
MH025918.1_strain_R25          .....g......................................g................................... [ 69600]
MH025917.1_strain_R7           .....g......................................g................................... [ 69600]
MH025916.1_strain_R8           .....g......................................g................................... [ 69600]

FR682468.2_Georgia_2007/1      ctgtggctttggcatggcttcccaatcatcaatgtccttacacatgtccacctcctgcgccgtccacgtcaaactttcta [ 69680]
ASF_IND_20_CAD543              ................................................................................ [ 69680]
LR899193.1_Germany_2020/1      ................................................................................ [ 69680]
MN393477.1_Wuhan_2019          ................................................................................ [ 69680]
MW396979.1_Timor-Leste/2019/1  ................................................................................ [ 69680]
MK940252.1_CN/2019/InnerMongolia ................................................................................ [ 69680]
MK128995.1_China/2018/AnhuiXCGQ ................................................................................ [ 69680]
MH681419.1_ASFV/POL/2015       ................................................................................ [ 69680]
LS478113.1_Estonia_2014        ................................................................................ [ 69680]
KP843857.1_Odintsovo_02/14     ................................................................................ [ 69680]
FN557520.1_strain_E75          ................................................t............................... [ 69680]
AY261366.1_Warthog             ..................c............................................................. [ 69680]
AY261365.1_Warmbaths           ..................c............................................................. [ 69680]
AY261364.1_Tengani_62          ..................c............................................................. [ 69680]
AY261363.1_Pretoriuskop/96/4   .....a............c............................................................. [ 69680]
AY261362.1_Mkuzi_1979          ...........a....................................t............................... [ 69680]
AY261361.1_Malawi_Lil-20/1_1983 ...c.....c........c....................................................t........ [ 69680]
AY261360.1_Kenya_1950          ...c..t........c..............t................................................. [ 69680]
AM712240.1_OURT_88/3           ................................................t............................... [ 69680]
AM712239.1_Benin_97/1          ................................................t............................... [ 69680]
KM111295.1_Ken06.Bus           ...c..............c.....g.....c................................................. [ 69680]
```

```
KM111294.1_Ken05/Tk1            ...c..t........c..............t................................................. [ 69680]
KM262844.1_strain_L60           ................................................t............................... [ 69680]
KM102979.1_26544/OG10_Italy     ................................................t............................... [ 69680]
KM262845.1_strain_NHV           ................................................t............................... [ 69680]
KX354450.1_47/Ss/2008           ................................................t............................... [ 69680]
KP055815.1_strain_BA71          ................................................t............................... [ 69680]
MH025920.1_strain_R35           ...c..............c.....g.....c................................................. [ 69680]
MH025919.1_strain_N10           ...c..............c.....g.....c................................................. [ 69680]
MH025918.1_strain_R25           ...c..............c.....g.....c................................................. [ 69680]
MH025917.1_strain_R7            ...c..............c.....g.....c................................................. [ 69680]
MH025916.1_strain_R8            ...c..............c.....g.....c................................................. [ 69680]

FR682468.2_Georgia_2007/1       attttttataccagttccaacattcggggtgctgaataggaaaaatagtgaaacgttgggaattttcaattagtaattcc [ 69760]
ASF_IND_20_CAD543               ................................................................................ [ 69760]
LR899193.1_Germany_2020/1       ................................................................................ [ 69760]
MN393477.1_Wuhan_2019           ................................................................................ [ 69760]
MW396979.1_Timor-Leste/2019/1   ................................................................................ [ 69760]
MK940252.1_CN/2019/InnerMongolia ................................................................................ [ 69760]
MK128995.1_China/2018/AnhuiXCGQ ................................................................................ [ 69760]
MH681419.1_ASFV/POL/2015        ................................................................................ [ 69760]
LS478113.1_Estonia_2014         ................................................................................ [ 69760]
KP843857.1_Odintsovo_02/14      ................................................................................ [ 69760]
FN557520.1_strain_E75           ................................................................................ [ 69760]
AY261366.1_Warthog              .......g.................t...................................................... [ 69760]
AY261365.1_Warmbaths            .......g.................t.............................g........................ [ 69760]
AY261364.1_Tengani_62           .......g.................t...................................................... [ 69760]
AY261363.1_Pretoriuskop/96/4    .......g...............................................g........................ [ 69760]
AY261362.1_Mkuzi_1979           ................................................................................ [ 69760]
AY261361.1_Malawi_Lil-20/1_1983 .......g.................t...................................................... [ 69760]
AY261360.1_Kenya_1950           .......g.................t.....................................c................ [ 69760]
AM712240.1_OURT_88/3            ................................................................................ [ 69760]
AM712239.1_Benin_97/1           ................................................................................ [ 69760]
KM111295.1_Ken06.Bus            .......g.................t...................................................... [ 69760]
KM111294.1_Ken05/Tk1            .......g.................t.....................................c................ [ 69760]
KM262844.1_strain_L60           ................................................................................ [ 69760]
KM102979.1_26544/OG10_Italy     ................................................................................ [ 69760]
KM262845.1_strain_NHV           ................................................................................ [ 69760]
KX354450.1_47/Ss/2008           ................................................................................ [ 69760]
KP055815.1_strain_BA71          ................................................................................ [ 69760]
MH025920.1_strain_R35           .......g.................t...................................................... [ 69760]
MH025919.1_strain_N10           .......g.................t...................................................... [ 69760]
MH025918.1_strain_R25           .......g.................t...................................................... [ 69760]
MH025917.1_strain_R7            .......g.................t...................................................... [ 69760]
MH025916.1_strain_R8            .......g.................t...................................................... [ 69760]
```

```
FR682468.2_Georgia_2007/1        tccatatttgaaataaatattaacatcttcaaatttattggctgccatgg---agacgttttttattg---agacgttgg [ 69840]
ASF_IND_20_CAD543                ..................................................---...............---......... [ 69840]
LR899193.1_Germany_2020/1        ..................................................---...............---......... [ 69840]
MN393477.1_Wuhan_2019            ..................................................---...............---......... [ 69840]
MW396979.1_Timor-Leste/2019/1    ..................................................---...............---......... [ 69840]
MK940252.1_CN/2019/InnerMongolia ..................................................---...............---......... [ 69840]
MK128995.1_China/2018/AnhuiXCGQ  ..................................................---...............---......... [ 69840]
MH681419.1_ASFV/POL/2015         ..................................................---...............---......... [ 69840]
LS478113.1_Estonia_2014          ..................................................---...............---......... [ 69840]
KP843857.1_Odintsovo_02/14       ..................................................---...............---......... [ 69840]
FN557520.1_strain_E75            ..................................................---...............---......... [ 69840]
AY261366.1_Warthog               .......................t..........................---.........g.....---......... [ 69840]
AY261365.1_Warmbaths             .......................t..........................---.........g...c.---......... [ 69840]
AY261364.1_Tengani_62            ..................................................---..t............---......... [ 69840]
AY261363.1_Pretoriuskop/96/4     .......................t..........................---...............---......... [ 69840]
AY261362.1_Mkuzi_1979            ..................................................---...............---......... [ 69840]
AY261361.1_Malawi_Lil-20/1_1983  .......................t..........................---...ac........c.tta...a..... [ 69840]
AY261360.1_Kenya_1950            .......................t................a.........---...ac..........tta...a..... [ 69840]
AM712240.1_OURT_88/3             ..................................................---...............---......... [ 69840]
AM712239.1_Benin_97/1            ..................................................---...............---......... [ 69840]
KM111295.1_Ken06.Bus             .......................t.....................g....aga...a.c...c.....ttg......... [ 69840]
KM111294.1_Ken05/Tk1             .......................t................a.........---...ac..........tta...a..... [ 69840]
KM262844.1_strain_L60            ..................................................---...............---......... [ 69840]
KM102979.1_26544/OG10_Italy      ..................................................---...............---......... [ 69840]
KM262845.1_strain_NHV            ..................................................---...............---......... [ 69840]
KX354450.1_47/Ss/2008            ..................................................---...............---......... [ 69840]
KP055815.1_strain_BA71           ..................................................---...............---......... [ 69840]
MH025920.1_strain_R35            .......................t.....................g....aga...a.c...c.....ttg......... [ 69840]
MH025919.1_strain_N10            .......................t.....................g....aga...a.c...c.....ttg......... [ 69840]
MH025918.1_strain_R25            .......................t.....................g....aga...a.c...c.....ttg......... [ 69840]
MH025917.1_strain_R7             .......................t.....................g....aga...a.c...c.....ttg......... [ 69840]
MH025916.1_strain_R8             .......................t.....................g....aga...a.c...c.....ttg......... [ 69840]

FR682468.2_Georgia_2007/1        catctgatgtgtatggaaaggcgttaaatgttgatttagatagactatcgcaggcgcaggttaaatatacccttcaagag [ 69920]
ASF_IND_20_CAD543                ................................................................................ [ 69920]
LR899193.1_Germany_2020/1        ................................................................................ [ 69920]
MN393477.1_Wuhan_2019            ................................................................................ [ 69920]
MW396979.1_Timor-Leste/2019/1    ................................................................................ [ 69920]
MK940252.1_CN/2019/InnerMongolia ................................................................................ [ 69920]
MK128995.1_China/2018/AnhuiXCGQ  ................................................................................ [ 69920]
MH681419.1_ASFV/POL/2015         ................................................................................ [ 69920]
LS478113.1_Estonia_2014          ................................................................................ [ 69920]
KP843857.1_Odintsovo_02/14       ................................................................................ [ 69920]
```

```
FN557520.1_strain_E75            .........................g...................................................... [ 69920]
AY261366.1_Warthog               ........ac...c.............................................a.c........t......... [ 69920]
AY261365.1_Warmbaths             ...........................................................a.................... [ 69920]
AY261364.1_Tengani_62            ........ac...c.............................................a..........t......... [ 69920]
AY261363.1_Pretoriuskop/96/4     .............c.................c...........................a..........t......... [ 69920]
AY261362.1_Mkuzi_1979            ................................................................................ [ 69920]
AY261361.1_Malawi_Lil-20/1_1983  ......c.ac...c........................a...a....cta..a....taaa.c.tc...g........g. [ 69920]
AY261360.1_Kenya_1950            ........ac......................................ta.........aa.........t......... [ 69920]
AM712240.1_OURT_88/3             .........................g...................................................... [ 69920]
AM712239.1_Benin_97/1            .........................g...................................................... [ 69920]
KM111295.1_Ken06.Bus             ......c.a....c.............................ga...ta.........aa.........t......... [ 69920]
KM111294.1_Ken05/Tk1             ........ac.........a............................ta..a......aa.........t......... [ 69920]
KM262844.1_strain_L60            .........................g...................................................... [ 69920]
KM102979.1_26544/OG10_Italy      .........................g...................................................... [ 69920]
KM262845.1_strain_NHV            .........................g...................................................... [ 69920]
KX354450.1_47/Ss/2008            .........................g...................................................... [ 69920]
KP055815.1_strain_BA71           .........................g...................................................... [ 69920]
MH025920.1_strain_R35            ......c.ac...c.............................ga...ta.........aa.........t......... [ 69920]
MH025919.1_strain_N10            ......c.ac...c.............................ga...ta.........aa.........t......... [ 69920]
MH025918.1_strain_R25            ......c.ac...c.............................ga...ta.........aa.........t......... [ 69920]
MH025917.1_strain_R7             ......c.ac...c.............................ga...ta.........aa.........t......... [ 69920]
MH025916.1_strain_R8             ......c.ac...c.............................ga...ta.........aa.........t......... [ 69920]

FR682468.2_Georgia_2007/1        cttatttcctactgcagcgctctaaccattttacattatgactattcaacccttgcggcgcgtctttcggtgtaccagct [ 70000]
ASF_IND_20_CAD543                ................................................................................ [ 70000]
LR899193.1_Germany_2020/1        ................................................................................ [ 70000]
MN393477.1_Wuhan_2019            ................................................................................ [ 70000]
MW396979.1_Timor-Leste/2019/1    ................................................................................ [ 70000]
MK940252.1_CN/2019/InnerMongolia ................................................................................ [ 70000]
MK128995.1_China/2018/AnhuiXCGQ  ................................................................................ [ 70000]
MH681419.1_ASFV/POL/2015         ................................................................................ [ 70000]
LS478113.1_Estonia_2014          ................................................................................ [ 70000]
KP843857.1_Odintsovo_02/14       ................................................................................ [ 70000]
FN557520.1_strain_E75            ..............................................................a................. [ 70000]
AY261366.1_Warthog               ..........................g.......................t........a..a................. [ 70000]
AY261365.1_Warmbaths             ..............................................................a................. [ 70000]
AY261364.1_Tengani_62            ................................................................................ [ 70000]
AY261363.1_Pretoriuskop/96/4     ..............................................................a................. [ 70000]
AY261362.1_Mkuzi_1979            ..............................................................a................. [ 70000]
AY261361.1_Malawi_Lil-20/1_1983  ..........................t.................................................ta.. [ 70000]
AY261360.1_Kenya_1950            ..........................t..a...........t.................a................ta.. [ 70000]
AM712240.1_OURT_88/3             ..............................................................a................. [ 70000]
AM712239.1_Benin_97/1            ..............................................................a................. [ 70000]
KM111295.1_Ken06.Bus             .............................a..........................a.....a.............ta.. [ 70000]
```

```
KM111294.1_Ken05/Tk1             .............................a.............................a................ta.. [ 70000]
KM262844.1_strain_L60            ..............................................................a................. [ 70000]
KM102979.1_26544/OG10_Italy      ..............................................................a................. [ 70000]
KM262845.1_strain_NHV            ..............................................................a................. [ 70000]
KX354450.1_47/Ss/2008            ..............................................................a................. [ 70000]
KP055815.1_strain_BA71           ..............................................................a................. [ 70000]
MH025920.1_strain_R35            .............................a..........................a.....a.............ta.. [ 70000]
MH025919.1_strain_N10            .............................a..........................a.....a.............ta.. [ 70000]
MH025918.1_strain_R25            .............................a..........................a.....a.............ta.. [ 70000]
MH025917.1_strain_R7             .............................a..........................a.....a.............ta.. [ 70000]
MH025916.1_strain_R8             .............................a..........................a.....a.............ta.. [ 70000]

FR682468.2_Georgia_2007/1        gcaccagtcaacggcctcctccttctcaaaggcggtgaggctgcaggccgcacaatcctgctcacgcctgtccccccagt [ 70080]
ASF_IND_20_CAD543                ................................................................................ [ 70080]
LR899193.1_Germany_2020/1        ................................................................................ [ 70080]
MN393477.1_Wuhan_2019            ................................................................................ [ 70080]
MW396979.1_Timor-Leste/2019/1    ................................................................................ [ 70080]
MK940252.1_CN/2019/InnerMongolia ................................................................................ [ 70080]
MK128995.1_China/2018/AnhuiXCGQ  ................................................................................ [ 70080]
MH681419.1_ASFV/POL/2015         ................................................................................ [ 70080]
LS478113.1_Estonia_2014          ................................................................................ [ 70080]
KP843857.1_Odintsovo_02/14       ................................................................................ [ 70080]
FN557520.1_strain_E75            ................................................................................ [ 70080]
AY261366.1_Warthog               ................................................................................ [ 70080]
AY261365.1_Warmbaths             ................................................................................ [ 70080]
AY261364.1_Tengani_62            ................................................................................ [ 70080]
AY261363.1_Pretoriuskop/96/4     ................................................................................ [ 70080]
AY261362.1_Mkuzi_1979            ..............................................................................c. [ 70080]
AY261361.1_Malawi_Lil-20/1_1983  ........................t...........a..c................................tt....t. [ 70080]
AY261360.1_Kenya_1950            ............................g.a.....a..c..a.............................t....... [ 70080]
AM712240.1_OURT_88/3             .......................................a........................................ [ 70080]
AM712239.1_Benin_97/1            ................................................................................ [ 70080]
KM111295.1_Ken06.Bus             ...t........................g.......a..c................................t....... [ 70080]
KM111294.1_Ken05/Tk1             ...t........................g.a.....a..c..a.............................t....... [ 70080]
KM262844.1_strain_L60            ................................................................................ [ 70080]
KM102979.1_26544/OG10_Italy      ................................................................................ [ 70080]
KM262845.1_strain_NHV            .......................................a........................................ [ 70080]
KX354450.1_47/Ss/2008            ................................................................................ [ 70080]
KP055815.1_strain_BA71           ................................................................................ [ 70080]
MH025920.1_strain_R35            ...t........................g.......a..c................................t....... [ 70080]
MH025919.1_strain_N10            ...t........................g.......a..c................................t....... [ 70080]
MH025918.1_strain_R25            ...t........................g.......a..c................................t....... [ 70080]
MH025917.1_strain_R7             ...t........................g.......a..c................................t....... [ 70080]
MH025916.1_strain_R8             ...t........................g.......a..c................................t....... [ 70080]
```

```
FR682468.2_Georgia_2007/1        ttgtggacgtcgtttacaagtacaaagccatttttgacagctacattgactatagcagagattacaagctgtccctcctg [ 70160]
ASF_IND_20_CAD543                ................................................................................ [ 70160]
LR899193.1_Germany_2020/1        ................................................................................ [ 70160]
MN393477.1_Wuhan_2019            ................................................................................ [ 70160]
MW396979.1_Timor-Leste/2019/1    ................................................................................ [ 70160]
MK940252.1_CN/2019/InnerMongolia ................................................................................ [ 70160]
MK128995.1_China/2018/AnhuiXCGQ  ................................................................................ [ 70160]
MH681419.1_ASFV/POL/2015         ................................................................................ [ 70160]
LS478113.1_Estonia_2014          ................................................................................ [ 70160]
KP843857.1_Odintsovo_02/14       ................................................................................ [ 70160]
FN557520.1_strain_E75            ......................................................a......................... [ 70160]
AY261366.1_Warthog               ......................................................a......................... [ 70160]
AY261365.1_Warmbaths             ......................................................a......................... [ 70160]
AY261364.1_Tengani_62            .............c.................................................................. [ 70160]
AY261363.1_Pretoriuskop/96/4     ................................................................................ [ 70160]
AY261362.1_Mkuzi_1979            ......................................................a......................... [ 70160]
AY261361.1_Malawi_Lil-20/1_1983  ....a........c........t......................................................... [ 70160]
AY261360.1_Kenya_1950            .............c........t...................................g...........aa........ [ 70160]
AM712240.1_OURT_88/3             ......................................................a......................... [ 70160]
AM712239.1_Benin_97/1            ......................................................a......................... [ 70160]
KM111295.1_Ken06.Bus             ....a........c........t..............................................aa........ [ 70160]
KM111294.1_Ken05/Tk1             .............c........t..............................................aa........ [ 70160]
KM262844.1_strain_L60            ......................................................a......................... [ 70160]
KM102979.1_26544/OG10_Italy      ......................................................a......................... [ 70160]
KM262845.1_strain_NHV            ......................................................a......................... [ 70160]
KX354450.1_47/Ss/2008            ......................................................a......................... [ 70160]
KP055815.1_strain_BA71           ......................................................a......................... [ 70160]
MH025920.1_strain_R35            ....a........c........t..............................................aa........ [ 70160]
MH025919.1_strain_N10            ....a........c........t..............................................aa........ [ 70160]
MH025918.1_strain_R25            ....a........c........t..............................................aa........ [ 70160]
MH025917.1_strain_R7             ....a........c........t..............................................aa........ [ 70160]
MH025916.1_strain_R8             ....a........c........t..............................................aa........ [ 70160]

FR682468.2_Georgia_2007/1        gggatagaaaccatgaaaaattcttatttgttaaaaaataaagatggggtcatcatggaacgcccgcaggatgcttatat [ 70240]
ASF_IND_20_CAD543                .......................y........................................................ [ 70240]
LR899193.1_Germany_2020/1        ................................................................................ [ 70240]
MN393477.1_Wuhan_2019            ................................................................................ [ 70240]
MW396979.1_Timor-Leste/2019/1    ................................................................................ [ 70240]
MK940252.1_CN/2019/InnerMongolia ................................................................................ [ 70240]
MK128995.1_China/2018/AnhuiXCGQ  ................................................................................ [ 70240]
MH681419.1_ASFV/POL/2015         ................................................................................ [ 70240]
LS478113.1_Estonia_2014          ................................................................................ [ 70240]
KP843857.1_Odintsovo_02/14       ................................................................................ [ 70240]
```

```
FN557520.1_strain_E75          .......................c...............................................c..c..... [ 70240]
AY261366.1_Warthog             ...........t...........c.......................c.......................c..c..... [ 70240]
AY261365.1_Warmbaths           .......................c...............................................c..c..... [ 70240]
AY261364.1_Tengani_62          ...............................................c.......................c..c..... [ 70240]
AY261363.1_Pretoriuskop/96/4   .......................c.......................c.......................c..c..... [ 70240]
AY261362.1_Mkuzi_1979          .......................c...............................................c........ [ 70240]
AY261361.1_Malawi_Lil-20/1_1983 .......................c..c..a.................c..............t........c..c..... [ 70240]
AY261360.1_Kenya_1950          ...........t...........c..c....................c.......................c........ [ 70240]
AM712240.1_OURT_88/3           .......................c...............................................c..c..... [ 70240]
AM712239.1_Benin_97/1          .......................c...............................................c..c..... [ 70240]
KM111295.1_Ken06.Bus           ..a.......................c....................c.......................c........ [ 70240]
KM111294.1_Ken05/Tk1           ...........t...........c..c....................c.......................c........ [ 70240]
KM262844.1_strain_L60          .......................c...............................................c..c..... [ 70240]
KM102979.1_26544/OG10_Italy    .......................c...............................................c..c..... [ 70240]
KM262845.1_strain_NHV          .......................c...............................................c..c..... [ 70240]
KX354450.1_47/Ss/2008          .......................c...............................................c..c..... [ 70240]
KP055815.1_strain_BA71         .......................c...............................................c..c..... [ 70240]
MH025920.1_strain_R35          ..a.......................c....................t.......................c........ [ 70240]
MH025919.1_strain_N10          ..a.......................c....................t.......................c........ [ 70240]
MH025918.1_strain_R25          ..a.......................c....................t.......................c........ [ 70240]
MH025917.1_strain_R7           ..a.......................c....................t.......................c........ [ 70240]
MH025916.1_strain_R8           ..a.......................c....................t.......................c........ [ 70240]

FR682468.2_Georgia_2007/1      gcgggttgccatcatgatctatgggatgggaagagtggtcaatatgaaaatgattctgctaacctatgacctgctttccc [ 70320]
ASF_IND_20_CAD543              ........................r....................a..g............................... [ 70320]
LR899193.1_Germany_2020/1      ................................................................................ [ 70320]
MN393477.1_Wuhan_2019          ................................................................................ [ 70320]
MW396979.1_Timor-Leste/2019/1  ................................................................................ [ 70320]
MK940252.1_CN/2019/InnerMongolia ................................................................................ [ 70320]
MK128995.1_China/2018/AnhuiXCGQ ................................................................................ [ 70320]
MH681419.1_ASFV/POL/2015       ................................................................................ [ 70320]
LS478113.1_Estonia_2014        ................................................................................ [ 70320]
KP843857.1_Odintsovo_02/14     ................................................................................ [ 70320]
FN557520.1_strain_E75          ...c..c.....t...........a.......a...a..t.....a..g...........g................... [ 70320]
AY261366.1_Warthog             ...c..c.................a.......a............a................................t. [ 70320]
AY261365.1_Warmbaths           ...c..c.................a.......a............a.................................. [ 70320]
AY261364.1_Tengani_62          ...c..c.................a.......a...a........a................................t. [ 70320]
AY261363.1_Pretoriuskop/96/4   ...c..c.................a.......a............a.................................. [ 70320]
AY261362.1_Mkuzi_1979          ........................a.......a...a..t.....a..g...........g................... [ 70320]
AY261361.1_Malawi_Lil-20/1_1983 ...c..c.....t......c................a..t........g...........g................... [ 70320]
AY261360.1_Kenya_1950          ...c..c.....t.....t.............................g...........g...........a....... [ 70320]
AM712240.1_OURT_88/3           ...c..c.....t...........a.......a...a..t.....a..g...........g................... [ 70320]
AM712239.1_Benin_97/1          ...c..c.....t...........a.......a...a..t.....a..g...........g................... [ 70320]
KM111295.1_Ken06.Bus           ...c..c.....t.....tc............................g...........g................... [ 70320]
```

```
KM111294.1_Ken05/Tk1             ...c..c.....t.....tc............................g...........g...........a....... [ 70320]
KM262844.1_strain_L60            ...c..c.....t...........a.......a...a..t.....a..g...........g................... [ 70320]
KM102979.1_26544/OG10_Italy      ...c..c.....t...........a.......a...a..t.....a..g...........g................... [ 70320]
KM262845.1_strain_NHV            ...c..c.....t...........a.......a...a..t.....a..g...........g................... [ 70320]
KX354450.1_47/Ss/2008            ...c..c.....t...........a.......a...a..t.....a..g...........g................... [ 70320]
KP055815.1_strain_BA71           ...c..c.....t...........a.......a...a..t.....a..g...........g................... [ 70320]
MH025920.1_strain_R35            ...c..c.....t.....tc............................g...........g................... [ 70320]
MH025919.1_strain_N10            ...c..c.....t.....tc............................g...........g................... [ 70320]
MH025918.1_strain_R25            ...c..c.....t.....tc............................g...........g................... [ 70320]
MH025917.1_strain_R7             ...c..c.....t.....tc............................g...........g................... [ 70320]
MH025916.1_strain_R8             ...c..c.....t.....tc............................g...........g................... [ 70320]

FR682468.2_Georgia_2007/1        agcacgtcatcacacacgcgtcgcccaccatgttcaatgcaggcaccaaaaagccacaactctccagctgtttcctgcta [ 70400]
ASF_IND_20_CAD543                ......................................................................c..t...... [ 70400]
LR899193.1_Germany_2020/1        ................................................................................ [ 70400]
MN393477.1_Wuhan_2019            ................................................................................ [ 70400]
MW396979.1_Timor-Leste/2019/1    ................................................................................ [ 70400]
MK940252.1_CN/2019/InnerMongolia ................................................................................ [ 70400]
MK128995.1_China/2018/AnhuiXCGQ  ................................................................................ [ 70400]
MH681419.1_ASFV/POL/2015         ................................................................................ [ 70400]
LS478113.1_Estonia_2014          ................................................................................ [ 70400]
KP843857.1_Odintsovo_02/14       ................................................................................ [ 70400]
FN557520.1_strain_E75            g......................................................g..g..t........c..t...... [ 70400]
AY261366.1_Warthog               .....a.................................................g..g..............t..a... [ 70400]
AY261365.1_Warmbaths             .....a.................................................g..g..............t..a... [ 70400]
AY261364.1_Tengani_62            ................................................................................ [ 70400]
AY261363.1_Pretoriuskop/96/4     g....a.................................................g..g..................... [ 70400]
AY261362.1_Mkuzi_1979            g.................................t....................g..g..t........c..t...... [ 70400]
AY261361.1_Malawi_Lil-20/1_1983  g...............t......................................g..g.....t..t............ [ 70400]
AY261360.1_Kenya_1950            g...............t..a....................g..............g..g..t........c......... [ 70400]
AM712240.1_OURT_88/3             g......................................................g..g..t........c..t...... [ 70400]
AM712239.1_Benin_97/1            g......................................................g..g..t........c..t...... [ 70400]
KM111295.1_Ken06.Bus             g...............t..a....................g..............g..g..t........c......... [ 70400]
KM111294.1_Ken05/Tk1             g...............t..a....................g..............g..g..t........c......... [ 70400]
KM262844.1_strain_L60            g......................................................g..g..t........c..t...... [ 70400]
KM102979.1_26544/OG10_Italy      g......................................................g..g..t........c..t...... [ 70400]
KM262845.1_strain_NHV            g......................................................g..g..t........c..t...... [ 70400]
KX354450.1_47/Ss/2008            g......................................................g..g..t........c..t...... [ 70400]
KP055815.1_strain_BA71           g......................................................g..g..t........c..t...... [ 70400]
MH025920.1_strain_R35            g...............t..a....................g..............g..g..t........c......... [ 70400]
MH025919.1_strain_N10            g...............t..a....................g..............g..g..t........c......... [ 70400]
MH025918.1_strain_R25            g...............t..a....................g..............g..g..t........c......... [ 70400]
MH025917.1_strain_R7             g...............t..a....................g..............g..g..t........c......... [ 70400]
MH025916.1_strain_R8             g...............t..a....................g..............g..g..t........c......... [ 70400]
```

```
FR682468.2_Georgia_2007/1         aatgtaaatgataatttagaaaatttatatgatatggtcaaaacggccggcatcatttcaggcggcggcggtggaatagg [ 70480]
ASF_IND_20_CAD543                 ................................................................................ [ 70480]
LR899193.1_Germany_2020/1         ................................................................................ [ 70480]
MN393477.1_Wuhan_2019             ................................................................................ [ 70480]
MW396979.1_Timor-Leste/2019/1     ................................................................................ [ 70480]
MK940252.1_CN/2019/InnerMongolia  ................................................................................ [ 70480]
MK128995.1_China/2018/AnhuiXCGQ   ................................................................................ [ 70480]
MH681419.1_ASFV/POL/2015          ................................................................................ [ 70480]
LS478113.1_Estonia_2014           ................................................................................ [ 70480]
KP843857.1_Odintsovo_02/14        ................................................................................ [ 70480]
FN557520.1_strain_E75             ....................g........c.................................................. [ 70480]
AY261366.1_Warthog                .............................c.........................................c........ [ 70480]
AY261365.1_Warmbaths              .............................c.........................................c........ [ 70480]
AY261364.1_Tengani_62             ................................................................................ [ 70480]
AY261363.1_Pretoriuskop/96/4      .............................c.........................................c........ [ 70480]
AY261362.1_Mkuzi_1979             ....................g........c.................................................. [ 70480]
AY261361.1_Malawi_Lil-20/1_1983   .............................c.........................................c........ [ 70480]
AY261360.1_Kenya_1950             ........c....................c........a................................c........ [ 70480]
AM712240.1_OURT_88/3              ....................g........c.................................................. [ 70480]
AM712239.1_Benin_97/1             ....................g........c.................................................. [ 70480]
KM111295.1_Ken06.Bus              ........c....................c........a................................c........ [ 70480]
KM111294.1_Ken05/Tk1              ........c....................c........a................................c........ [ 70480]
KM262844.1_strain_L60             ....................g........c.................................................. [ 70480]
KM102979.1_26544/OG10_Italy       ....................g........c.................................................. [ 70480]
KM262845.1_strain_NHV             ....................g........c.................................................. [ 70480]
KX354450.1_47/Ss/2008             ....................g........c.................................................. [ 70480]
KP055815.1_strain_BA71            ....................g........c.................................................. [ 70480]
MH025920.1_strain_R35             ........c....................c........a................................c........ [ 70480]
MH025919.1_strain_N10             ........c....................c........a................................c........ [ 70480]
MH025918.1_strain_R25             ........c....................c........a................................c........ [ 70480]
MH025917.1_strain_R7              ........c....................c........a................................c........ [ 70480]
MH025916.1_strain_R8              ........c....................c........a................................c........ [ 70480]

FR682468.2_Georgia_2007/1         gctgtgcttgtcaggaatacgggcaaagaatagttttatttctggtagtggtcttaaaagtaacggcatacagaattata [ 70560]
ASF_IND_20_CAD543                 ...........................a.....n.............................................. [ 70560]
LR899193.1_Germany_2020/1         ................................................................................ [ 70560]
MN393477.1_Wuhan_2019             ................................................................................ [ 70560]
MW396979.1_Timor-Leste/2019/1     ................................................................................ [ 70560]
MK940252.1_CN/2019/InnerMongolia  ................................................................................ [ 70560]
MK128995.1_China/2018/AnhuiXCGQ   ................................................................................ [ 70560]
MH681419.1_ASFV/POL/2015          ................................................................................ [ 70560]
LS478113.1_Estonia_2014           ................................................................................ [ 70560]
KP843857.1_Odintsovo_02/14        ................................................................................ [ 70560]
```

```
FN557520.1_strain_E75           .....................c.....a.....c...........c..........g...c........t..a..c.... [ 70560]
AY261366.1_Warthog              tt....t..a...........c.....a.....c...........c..........g...c........t..a..c.... [ 70560]
AY261365.1_Warmbaths            tt.......a...........c.....a.....c...........c..........g...c........t..a....... [ 70560]
AY261364.1_Tengani_62           ................................................................................ [ 70560]
AY261363.1_Pretoriuskop/96/4    tt...................c.....a.................c..........g...c........t..a....... [ 70560]
AY261362.1_Mkuzi_1979           .....................c.....a.....c...........c..........g...c........t..a..c.... [ 70560]
AY261361.1_Malawi_Lil-20/1_1983 ......t....................a.....c.....c..c.............g.........t.....a....... [ 70560]
AY261360.1_Kenya_1950           ...........................a.....c.....c..c.....c.......g...............a....... [ 70560]
AM712240.1_OURT_88/3            .....................c.....a.....c...........c..........g...c........t..a..c.... [ 70560]
AM712239.1_Benin_97/1           .....................c.....a.....c...........c..........g...c........t..a..c.... [ 70560]
KM111295.1_Ken06.Bus            ...........................a.....c.....c..c.....c.......g...............a....... [ 70560]
KM111294.1_Ken05/Tk1            ............g..............a.....c.....c..c.....c.......g...............a....... [ 70560]
KM262844.1_strain_L60           .....................c.....a.....c...........c..........g...c........t..a..c.... [ 70560]
KM102979.1_26544/OG10_Italy     .....................c.....a.....c.........a.c..........g...c........t..a..c.... [ 70560]
KM262845.1_strain_NHV           .....................c.....a.....c...........c..........g...c........t..a..c.... [ 70560]
KX354450.1_47/Ss/2008           .....................c.....a.....c.........a.c..........g...c........t..a..c.... [ 70560]
KP055815.1_strain_BA71          .....................c.....a.....c...........c..........g...c........t..a..c.... [ 70560]
MH025920.1_strain_R35           ...........................a.....c.....c..c.....c.......g...............a....... [ 70560]
MH025919.1_strain_N10           ...........................a.....c.....c..c.....c.......g...............a....... [ 70560]
MH025918.1_strain_R25           ...........................a.....c.....c..c.....c.......g...............a....... [ 70560]
MH025917.1_strain_R7            ...........................a.....c.....c..c.....c.......g...............a....... [ 70560]
MH025916.1_strain_R8            ...........................a.....c.....c..c.....c.......g...............a....... [ 70560]

FR682468.2_Georgia_2007/1       ttgtgctgcaaaatgcttcacaatgctacgcgaaccagggaggcctacgtcccggagcctacgccgtctacttagagctg [ 70640]
ASF_IND_20_CAD543               .................................................c..t........................... [ 70640]
LR899193.1_Germany_2020/1       ................................................................................ [ 70640]
MN393477.1_Wuhan_2019           ................................................................................ [ 70640]
MW396979.1_Timor-Leste/2019/1   ................................................................................ [ 70640]
MK940252.1_CN/2019/InnerMongolia ................................................................................ [ 70640]
MK128995.1_China/2018/AnhuiXCGQ ................................................................................ [ 70640]
MH681419.1_ASFV/POL/2015        ................................................................................ [ 70640]
LS478113.1_Estonia_2014         ................................................................................ [ 70640]
KP843857.1_Odintsovo_02/14      ................................................................................ [ 70640]
FN557520.1_strain_E75           ..a....a........c................................c..t..g.....t..g...........a..a [ 70640]
AY261366.1_Warthog              ..a....a........c................................c..g..g........a...........a..a [ 70640]
AY261365.1_Warmbaths            ..a....a........c................................c..t..g.....t..g...........a..a [ 70640]
AY261364.1_Tengani_62           ................................................................................ [ 70640]
AY261363.1_Pretoriuskop/96/4    ..a....a........c...........................t....c..............a...........a..a [ 70640]
AY261362.1_Mkuzi_1979           ..a....a........c................................c..t..g.....t..g...........a..a [ 70640]
AY261361.1_Malawi_Lil-20/1_1983 ....tt.a..g.................t..c..t..............c.....g........a............... [ 70640]
AY261360.1_Kenya_1950           ....tt.a..g....................c........g........c..............a......c........ [ 70640]
AM712240.1_OURT_88/3            ..a....a........c................................c..t..g.....t..g...........a..a [ 70640]
AM712239.1_Benin_97/1           ..a....a........c................................c..t..g.....t..g...........a..a [ 70640]
KM111295.1_Ken06.Bus            ....tt.a..g....................c........g........c..............a......c........ [ 70640]
```

```
KM111294.1_Ken05/Tk1             ....tt.a..g....................c........g........c..............a......c........ [ 70640]
KM262844.1_strain_L60            ..a....a........c................................c..t..g.....t..g...........a..a [ 70640]
KM102979.1_26544/OG10_Italy      ..a....a........c................................c..t..g.....t..g...........a..a [ 70640]
KM262845.1_strain_NHV            ..a....a........c................................c..t..g.....t..g...........a..a [ 70640]
KX354450.1_47/Ss/2008            ..a....a........c................................c..t..g.....t..g...........a..a [ 70640]
KP055815.1_strain_BA71           ..a....a........c................................c..t..g.....t..g...........a..a [ 70640]
MH025920.1_strain_R35            ....tt.a..g....................c........g........c..............a......c........ [ 70640]
MH025919.1_strain_N10            ....tt.a..g....................c........g........c..............a......c........ [ 70640]
MH025918.1_strain_R25            ....tt.a..g....................c........g........c..............a......c........ [ 70640]
MH025917.1_strain_R7             ....tt.a..g....................c........g........c..............a......c........ [ 70640]
MH025916.1_strain_R8             ....tt.a..g....................c........g........c..............a......c........ [ 70640]

FR682468.2_Georgia_2007/1        tggcaccaagacatctttacatttttacaaatgcctcgcctaaaaggacaaatggctgaacaacggcttaatgcccctaa [ 70720]
ASF_IND_20_CAD543                ..............................................................ga................ [ 70720]
LR899193.1_Germany_2020/1        ................................................................................ [ 70720]
MN393477.1_Wuhan_2019            ................................................................................ [ 70720]
MW396979.1_Timor-Leste/2019/1    ................................................................................ [ 70720]
MK940252.1_CN/2019/InnerMongolia ................................................................................ [ 70720]
MK128995.1_China/2018/AnhuiXCGQ  ................................................................................ [ 70720]
MH681419.1_ASFV/POL/2015         ................................................................................ [ 70720]
LS478113.1_Estonia_2014          ................................................................................ [ 70720]
KP843857.1_Odintsovo_02/14       ................................................................................ [ 70720]
FN557520.1_strain_E75            ........................c.g..g.....c...t.g....................ga.............c.. [ 70720]
AY261366.1_Warthog               .....t..................c.gg.......c...t.g....................ga.............c.. [ 70720]
AY261365.1_Warmbaths             .....t..................c.gg.......c...t.g....................ga.............c.. [ 70720]
AY261364.1_Tengani_62            ..............................................................g................. [ 70720]
AY261363.1_Pretoriuskop/96/4     ..............t.........c.gg.......c...t.g....................ga.............c.. [ 70720]
AY261362.1_Mkuzi_1979            ........................c.g........c...t.g....................ga.............c.. [ 70720]
AY261361.1_Malawi_Lil-20/1_1983  ........................c..........c...t.g....................ga..........t..... [ 70720]
AY261360.1_Kenya_1950            .....t.............................c..........................ga.............c.. [ 70720]
AM712240.1_OURT_88/3             ........................c.g..g.....c...t.g....................ga.............c.. [ 70720]
AM712239.1_Benin_97/1            ........................c.g..g.....c...t.g....................ga.............c.. [ 70720]
KM111295.1_Ken06.Bus             ...................................c..........................ga.............c.. [ 70720]
KM111294.1_Ken05/Tk1             .....t.............................c..........................ga.............c.. [ 70720]
KM262844.1_strain_L60            ........................c.g..g.....c...t.g....................ga.............c.. [ 70720]
KM102979.1_26544/OG10_Italy      ........................c.g..g.....c...t.g....................ga.............c.. [ 70720]
KM262845.1_strain_NHV            ........................c.g..g.....c...t.g....................ga.............c.. [ 70720]
KX354450.1_47/Ss/2008            ........................c.g..g.....c...t.g....................ga.............c.. [ 70720]
KP055815.1_strain_BA71           ........................c.g..g.....c...t.g....................ga.............c.. [ 70720]
MH025920.1_strain_R35            ...................................c..........................ga.............c.. [ 70720]
MH025919.1_strain_N10            ...................................c..........................ga.............c.. [ 70720]
MH025918.1_strain_R25            ...................................c..........................ga.............c.. [ 70720]
MH025917.1_strain_R7             ...................................c..........................ga.............c.. [ 70720]
MH025916.1_strain_R8             ...................................c..........................ga.............c.. [ 70720]
```

```
FR682468.2_Georgia_2007/1        tctcaagtacggcctatgggtccccgacctattcatggaaatacttgaagaccaaatacacaacagaggcgacggcaaat [ 70800]
ASF_IND_20_CAD543                ................................................................................ [ 70800]
LR899193.1_Germany_2020/1        ................................................................................ [ 70800]
MN393477.1_Wuhan_2019            ................................................................................ [ 70800]
MW396979.1_Timor-Leste/2019/1    ................................................................................ [ 70800]
MK940252.1_CN/2019/InnerMongolia ................................................................................ [ 70800]
MK128995.1_China/2018/AnhuiXCGQ  ................................................................................ [ 70800]
MH681419.1_ASFV/POL/2015         ................................................................................ [ 70800]
LS478113.1_Estonia_2014          ................................................................................ [ 70800]
KP843857.1_Odintsovo_02/14       ................................................................................ [ 70800]
FN557520.1_strain_E75            ............t....................t..........................t................cc. [ 70800]
AY261366.1_Warthog               ............t....................t...........................................g.. [ 70800]
AY261365.1_Warmbaths             ............t....................t..........................t................g.. [ 70800]
AY261364.1_Tengani_62            ................................................................................ [ 70800]
AY261363.1_Pretoriuskop/96/4     ............t...........t........t...........................................g.. [ 70800]
AY261362.1_Mkuzi_1979            ............tt...................t..........................t................cc. [ 70800]
AY261361.1_Malawi_Lil-20/1_1983  ........................t........t........t..................................c.. [ 70800]
AY261360.1_Kenya_1950            .....................t...........t...........c...............g...............c.. [ 70800]
AM712240.1_OURT_88/3             ............t....................t..........................t................cc. [ 70800]
AM712239.1_Benin_97/1            ............t....................t..........................t................cc. [ 70800]
KM111295.1_Ken06.Bus             .....................t...........t...........c...............g...............c.. [ 70800]
KM111294.1_Ken05/Tk1             .....................t...........t...........c...............g...............c.. [ 70800]
KM262844.1_strain_L60            ............t....................t..........................t................cc. [ 70800]
KM102979.1_26544/OG10_Italy      ............t....................t..........................t................cc. [ 70800]
KM262845.1_strain_NHV            ............t....................t..........................t................cc. [ 70800]
KX354450.1_47/Ss/2008            ............t....................t..........................t................cc. [ 70800]
KP055815.1_strain_BA71           ............t....................t..........................t................cc. [ 70800]
MH025920.1_strain_R35            .....................t...........t...........c...............g...............c.. [ 70800]
MH025919.1_strain_N10            .....................t...........t...........c...............g...............c.. [ 70800]
MH025918.1_strain_R25            .....................t...........t...........c...............g...............c.. [ 70800]
MH025917.1_strain_R7             .....................t...........t...........c...............g...............c.. [ 70800]
MH025916.1_strain_R8             .....................t...........t...........c...............g...............c.. [ 70800]

FR682468.2_Georgia_2007/1        ggtacctcttttcgccggatcaggcccccaatctacataaggtctttgatttggaacggtcgcagcacgaaaacgcacac [ 70880]
ASF_IND_20_CAD543                ................................................................................ [ 70880]
LR899193.1_Germany_2020/1        ................................................................................ [ 70880]
MN393477.1_Wuhan_2019            ................................................................................ [ 70880]
MW396979.1_Timor-Leste/2019/1    ................................................................................ [ 70880]
MK940252.1_CN/2019/InnerMongolia ................................................................................ [ 70880]
MK128995.1_China/2018/AnhuiXCGQ  ................................................................................ [ 70880]
MH681419.1_ASFV/POL/2015         ................................................................................ [ 70880]
LS478113.1_Estonia_2014          ................................................................................ [ 70880]
KP843857.1_Odintsovo_02/14       ................................................................................ [ 70880]
```

```
FN557520.1_strain_E75            .............a.............................t.................a.................t [ 70880]
AY261366.1_Warthog               .............a.................................................................. [ 70880]
AY261365.1_Warmbaths             .............a..a............................................a.................. [ 70880]
AY261364.1_Tengani_62            ................................................................................ [ 70880]
AY261363.1_Pretoriuskop/96/4     .............a..........................a....................................... [ 70880]
AY261362.1_Mkuzi_1979            ....t........a..a..........................t.................a.................t [ 70880]
AY261361.1_Malawi_Lil-20/1_1983  .............................................................a...........t.....t [ 70880]
AY261360.1_Kenya_1950            ............................t.............................a....gc...a....t.....t [ 70880]
AM712240.1_OURT_88/3             .............a.............................t.................a.................t [ 70880]
AM712239.1_Benin_97/1            .............a.............................t.................a.................t [ 70880]
KM111295.1_Ken06.Bus             ............................t...........a.................a....gct..a....t.....t [ 70880]
KM111294.1_Ken05/Tk1             ......................a.....t.............................a....gc...a....t.....t [ 70880]
KM262844.1_strain_L60            .............a.............................t.................a.................t [ 70880]
KM102979.1_26544/OG10_Italy      .............a.............................t.................a.................t [ 70880]
KM262845.1_strain_NHV            .............a.............................t.................a.................t [ 70880]
KX354450.1_47/Ss/2008            .............a.............................t.................a.................t [ 70880]
KP055815.1_strain_BA71           .............a.............................t.................a.................t [ 70880]
MH025920.1_strain_R35            ............................t...........a.................a....gct..a....t.....t [ 70880]
MH025919.1_strain_N10            ............................t...........a.................a....gct..a....t.....t [ 70880]
MH025918.1_strain_R25            ............................t...........a.................a....gct..a....t.....t [ 70880]
MH025917.1_strain_R7             ............................t...........a.................a....gct..a....t.....t [ 70880]
MH025916.1_strain_R8             ............................t...........a.................a....gct..a....t.....t [ 70880]

FR682468.2_Georgia_2007/1        cgcgaatttaaaaagctttactatcagtatgttgctgaaaaaaggtacaccggcgtcacaacggccaaagagattatcaa [ 70960]
ASF_IND_20_CAD543                ................................................................................ [ 70960]
LR899193.1_Germany_2020/1        ................................................................................ [ 70960]
MN393477.1_Wuhan_2019            ................................................................................ [ 70960]
MW396979.1_Timor-Leste/2019/1    ................................................................................ [ 70960]
MK940252.1_CN/2019/InnerMongolia ................................................................................ [ 70960]
MK128995.1_China/2018/AnhuiXCGQ  ................................................................................ [ 70960]
MH681419.1_ASFV/POL/2015         ................................................................................ [ 70960]
LS478113.1_Estonia_2014          ................................................................................ [ 70960]
KP843857.1_Odintsovo_02/14       ................................................................................ [ 70960]
FN557520.1_strain_E75            .....................................................t..t..g........g..a..c..t.. [ 70960]
AY261366.1_Warthog               ..t...............................................t............................. [ 70960]
AY261365.1_Warmbaths             ..................................................t............................. [ 70960]
AY261364.1_Tengani_62            ................................................................................ [ 70960]
AY261363.1_Pretoriuskop/96/4     ..................................................t............................. [ 70960]
AY261362.1_Mkuzi_1979            .....................................................t..t..g........g..a..c..t.. [ 70960]
AY261361.1_Malawi_Lil-20/1_1983  ..............a.....t........................................................... [ 70960]
AY261360.1_Kenya_1950            ..........g.........t..............c.....g.....t................................ [ 70960]
AM712240.1_OURT_88/3             .....................................................t..t..g........g..a..c..t.. [ 70960]
AM712239.1_Benin_97/1            .....................................................t..t..g........g..a..c..t.. [ 70960]
KM111295.1_Ken06.Bus             ..........g.........t..............c.....g.....t................................ [ 70960]
```

```
KM111294.1_Ken05/Tk1              ..........g.........t..............c.....g.....t................................ [ 70960]
KM262844.1_strain_L60             .....................................................t..t..g........g..a..c..t.. [ 70960]
KM102979.1_26544/OG10_Italy       .....................................................t..t..g........g..a..c..t.. [ 70960]
KM262845.1_strain_NHV             .....................................................t..t..g........g..a..c..t.. [ 70960]
KX354450.1_47/Ss/2008             .....................................................t..t..g........g..a..c..t.. [ 70960]
KP055815.1_strain_BA71            .....................................................t..t..g........g..a..c..t.. [ 70960]
MH025920.1_strain_R35             ..........g.........t..............c.....g.....t................................ [ 70960]
MH025919.1_strain_N10             ..........g.........t..............c.....g.....t................................ [ 70960]
MH025918.1_strain_R25             ..........g.........t..............c.....g.....t................................ [ 70960]
MH025917.1_strain_R7              ..........g.........t..............c.....g.....t................................ [ 70960]
MH025916.1_strain_R8              ..........g.........t..............c.....g.....t................................ [ 70960]

FR682468.2_Georgia_2007/1         agagtggttcaaaacagttgttcaagtagggaatccctatatcgggtttaaagatgccataaatcgtaaaagtaatcttt [ 71040]
ASF_IND_20_CAD543                 ...................r.y..r..g..a..c.............................................. [ 71040]
LR899193.1_Germany_2020/1         ................................................................................ [ 71040]
MN393477.1_Wuhan_2019             ................................................................................ [ 71040]
MW396979.1_Timor-Leste/2019/1     ................................................................................ [ 71040]
MK940252.1_CN/2019/InnerMongolia  ................................................................................ [ 71040]
MK128995.1_China/2018/AnhuiXCGQ   ................................................................................ [ 71040]
MH681419.1_ASFV/POL/2015          ................................................................................ [ 71040]
LS478113.1_Estonia_2014           ................................................................................ [ 71040]
KP843857.1_Odintsovo_02/14        ................................................................................ [ 71040]
FN557520.1_strain_E75             ...................a.c..g..g..a..c...........a.................................. [ 71040]
AY261366.1_Warthog                ..c...............ca.c..g..g..a..c...........a.................................. [ 71040]
AY261365.1_Warmbaths              ..c...............ca.c..g.....a..c...........a.................................. [ 71040]
AY261364.1_Tengani_62             ...a............................................................................ [ 71040]
AY261363.1_Pretoriuskop/96/4      ..c...............ca.c.....g..a..c...........a.................................. [ 71040]
AY261362.1_Mkuzi_1979             ...................a.c..g..g..a..c.....c.....a.................................. [ 71040]
AY261361.1_Malawi_Lil-20/1_1983   ..................c.....g..g..............t........g.....g..............c..c.... [ 71040]
AY261360.1_Kenya_1950             .........t........ca....g..g.......................g.....g..c...........c..c.... [ 71040]
AM712240.1_OURT_88/3              ...................a.c..g..g..a..c...........a.................................. [ 71040]
AM712239.1_Benin_97/1             ...................a.c..g..g..a..c...........a.................................. [ 71040]
KM111295.1_Ken06.Bus              ................c.c.....g..g.......................g.....g..c...........c..c.... [ 71040]
KM111294.1_Ken05/Tk1              .........t........ca....g..g.......................g.....g..c...........c..c.... [ 71040]
KM262844.1_strain_L60             ...................a.c..g..g..a..c...........a.................................. [ 71040]
KM102979.1_26544/OG10_Italy       ...................a.c..g..g..a..c...........a.................................. [ 71040]
KM262845.1_strain_NHV             ...................a.c..g..g..a..c...........a.................................. [ 71040]
KX354450.1_47/Ss/2008             ...................a.c..g..g..a..c...........a.................................. [ 71040]
KP055815.1_strain_BA71            ...................a.c..g..g..a..c...........a.................................. [ 71040]
MH025920.1_strain_R35             ................c.c.....g..g.......................g.....g..c...........c..c.... [ 71040]
MH025919.1_strain_N10             ................c.c.....g..g.......................g.....g..c...........c..c.... [ 71040]
MH025918.1_strain_R25             ................c.c.....g..g.......................g.....g..c...........c..c.... [ 71040]
MH025917.1_strain_R7              ................c.c.....g..g.......................g.....g..c...........c..c.... [ 71040]
MH025916.1_strain_R8              ................c.c.....g..g.......................g.....g..c...........c..c.... [ 71040]
```

```
FR682468.2_Georgia_2007/1        cacatgtaggcactatcacgaactccaatctttgtattgaagtcacaatcccctgctgggagggtgataaggctgaacaa [ 71120]
ASF_IND_20_CAD543                ................................................................................ [ 71120]
LR899193.1_Germany_2020/1        ................................................................................ [ 71120]
MN393477.1_Wuhan_2019            ................................................................................ [ 71120]
MW396979.1_Timor-Leste/2019/1    ................................................................................ [ 71120]
MK940252.1_CN/2019/InnerMongolia ................................................................................ [ 71120]
MK128995.1_China/2018/AnhuiXCGQ  ................................................................................ [ 71120]
MH681419.1_ASFV/POL/2015         ................................................................................ [ 71120]
LS478113.1_Estonia_2014          ................................................................................ [ 71120]
KP843857.1_Odintsovo_02/14       ................................................................................ [ 71120]
FN557520.1_strain_E75            ................................................................................ [ 71120]
AY261366.1_Warthog               .................................................t.....t........................ [ 71120]
AY261365.1_Warmbaths             ................................................................................ [ 71120]
AY261364.1_Tengani_62            ................................................................................ [ 71120]
AY261363.1_Pretoriuskop/96/4     .........................................................................c...... [ 71120]
AY261362.1_Mkuzi_1979            .........................................................................c...... [ 71120]
AY261361.1_Malawi_Lil-20/1_1983  .c...........c.....t.....t.....c........ga.t.....t..............ca..g....c..g..g [ 71120]
AY261360.1_Kenya_1950            ....c..g.....c.....t.....t..c..c........ga.......t..............cag.g....c..g..g [ 71120]
AM712240.1_OURT_88/3             ................................................................................ [ 71120]
AM712239.1_Benin_97/1            .............................................t.................................. [ 71120]
KM111295.1_Ken06.Bus             ....c..g.....c.....t.....t..c..c........ga.......t..............cag.g....c..g..g [ 71120]
KM111294.1_Ken05/Tk1             ....c..g.....c.....t.....t..c..c........ga.......t..............cag.g....c..g..g [ 71120]
KM262844.1_strain_L60            ................................................................................ [ 71120]
KM102979.1_26544/OG10_Italy      ................................................................................ [ 71120]
KM262845.1_strain_NHV            ................................................................................ [ 71120]
KX354450.1_47/Ss/2008            ................................................................................ [ 71120]
KP055815.1_strain_BA71           ................................................................................ [ 71120]
MH025920.1_strain_R35            ....c..g.....c.....t.....t..c..c........ga.......t..............cag.g....c..g..g [ 71120]
MH025919.1_strain_N10            ....c..g.....c.....t.....t..c..c........ga.......t..............cag.g....c..g..g [ 71120]
MH025918.1_strain_R25            ....c..g.....c.....t.....t..c..c........ga.......t..............cag.g....c..g..g [ 71120]
MH025917.1_strain_R7             ....c..g.....c.....t.....t..c..c........ga.......t..............cag.g....c..g..g [ 71120]
MH025916.1_strain_R8             ....c..g.....c.....t.....t..c..c........ga.......t..............cag.g....c..g..g [ 71120]

FR682468.2_Georgia_2007/1        ggtgtttgtaatctggccgcagtaaatctagccgcctttatacgtgaaaatggctacgactaccgtgggctcatagaagc [ 71200]
ASF_IND_20_CAD543                ................................................................................ [ 71200]
LR899193.1_Germany_2020/1        ................................................................................ [ 71200]
MN393477.1_Wuhan_2019            ................................................................................ [ 71200]
MW396979.1_Timor-Leste/2019/1    ................................................................................ [ 71200]
MK940252.1_CN/2019/InnerMongolia ................................................................................ [ 71200]
MK128995.1_China/2018/AnhuiXCGQ  ................................................................................ [ 71200]
MH681419.1_ASFV/POL/2015         ................................................................................ [ 71200]
LS478113.1_Estonia_2014          ................................................................................ [ 71200]
KP843857.1_Odintsovo_02/14       ................................................................................ [ 71200]
```

```
FN557520.1_strain_E75          ..c........c........g...................................t....................... [ 71200]
AY261366.1_Warthog             ....................g.....c.....t............................................... [ 71200]
AY261365.1_Warmbaths           ...........c........g.....c..................................................... [ 71200]
AY261364.1_Tengani_62          ................................................................................ [ 71200]
AY261363.1_Pretoriuskop/96/4   ...........c........g.....ct............................t....................... [ 71200]
AY261362.1_Mkuzi_1979          ..c........c........g...................................t.....t..............g.. [ 71200]
AY261361.1_Malawi_Lil-20/1_1983 ..g.....c.....t.....g........g...................g.a.t........t.....a........g.. [ 71200]
AY261360.1_Kenya_1950          ..g.....c..c..t.....g........g..............c.....ca....t..t..t..............g.. [ 71200]
AM712240.1_OURT_88/3           ..c........c........g...................................t....................... [ 71200]
AM712239.1_Benin_97/1          ..c........c........g...................................t....................... [ 71200]
KM111295.1_Ken06.Bus           ..g.....c..c..t.....g........g....................ca....t..t..t..c...........g.. [ 71200]
KM111294.1_Ken05/Tk1           ..g.....c..c..t.....g........g..............c.....ca....t..t..t..............g.. [ 71200]
KM262844.1_strain_L60          ..c........c........g...................................t....................... [ 71200]
KM102979.1_26544/OG10_Italy    ..c........c........g...................................t.................g..... [ 71200]
KM262845.1_strain_NHV          ..c........c........g...................................t....................... [ 71200]
KX354450.1_47/Ss/2008          ..c........c........g...................................t.................g..... [ 71200]
KP055815.1_strain_BA71         ..c........c........g...................................t....................... [ 71200]
MH025920.1_strain_R35          ..g.....c..c..t.....g........g....................ca....t..t..t..c...........g.. [ 71200]
MH025919.1_strain_N10          ..g.....c..c..t.....g........g....................ca....t..t..t..c...........g.. [ 71200]
MH025918.1_strain_R25          ..g.....c..c..t.....g........g....................ca....t..t..t..c...........g.. [ 71200]
MH025917.1_strain_R7           ..g.....c..c..t.....g........g....................ca....t..t..t..c...........g.. [ 71200]
MH025916.1_strain_R8           ..g.....c..c..t.....g........g....................ca....t..t..t..c...........g.. [ 71200]

FR682468.2_Georgia_2007/1      atcaggcaatgtcacagaaaatttagataatattatagataatggctactaccccacagaagccacgcggagaagcaata [ 71280]
ASF_IND_20_CAD543              ................................................................................ [ 71280]
LR899193.1_Germany_2020/1      ................................................................................ [ 71280]
MN393477.1_Wuhan_2019          ................................................................................ [ 71280]
MW396979.1_Timor-Leste/2019/1  ................................................................................ [ 71280]
MK940252.1_CN/2019/InnerMongolia ................................................................................ [ 71280]
MK128995.1_China/2018/AnhuiXCGQ ................................................................................ [ 71280]
MH681419.1_ASFV/POL/2015       ................................................................................ [ 71280]
LS478113.1_Estonia_2014        ................................................................................ [ 71280]
KP843857.1_Odintsovo_02/14     ................................................................................ [ 71280]
FN557520.1_strain_E75          .........c...................................................................... [ 71280]
AY261366.1_Warthog             ......................................................t......................... [ 71280]
AY261365.1_Warmbaths           ................................................................................ [ 71280]
AY261364.1_Tengani_62          ................................................................................ [ 71280]
AY261363.1_Pretoriuskop/96/4   ......................................................t......................... [ 71280]
AY261362.1_Mkuzi_1979          .........c...................................................................... [ 71280]
AY261361.1_Malawi_Lil-20/1_1983 .g............................c.................t..t.....g..g.....c..t.......... [ 71280]
AY261360.1_Kenya_1950          .g.......c................................c..............g..g.....c..t.......... [ 71280]
AM712240.1_OURT_88/3           .........c...................................................................... [ 71280]
AM712239.1_Benin_97/1          .........c...................................................................... [ 71280]
KM111295.1_Ken06.Bus           .g.......c................................c..............g..g.....c..t.......... [ 71280]
```

```
KM111294.1_Ken05/Tk1              .g.......c................................c..............g..g.....c..t.......... [ 71280]
KM262844.1_strain_L60             .........c...................................................................... [ 71280]
KM102979.1_26544/OG10_Italy       .........c...................................................................... [ 71280]
KM262845.1_strain_NHV             .........c...................................................................... [ 71280]
KX354450.1_47/Ss/2008             .........c...................................................................... [ 71280]
KP055815.1_strain_BA71            .........c...................................................................... [ 71280]
MH025920.1_strain_R35             .g.......c................................c..............g..g.....c..t.......... [ 71280]
MH025919.1_strain_N10             .g.......c................................c..............g..g.....c..t.......... [ 71280]
MH025918.1_strain_R25             .g.......c................................c..............g..g.....c..t.......... [ 71280]
MH025917.1_strain_R7              .g.......c................................c..............g..g.....c..t.......... [ 71280]
MH025916.1_strain_R8              .g.......c................................c..............g..g.....c..t.......... [ 71280]

FR682468.2_Georgia_2007/1         tgcgtcaccgacctattggcatcggggtctttggcctagccgacgtgtttgcgtctttaaaaatgaaatttggttcaccc [ 71360]
ASF_IND_20_CAD543                 ................................................................................ [ 71360]
LR899193.1_Germany_2020/1         ................................................................................ [ 71360]
MN393477.1_Wuhan_2019             ................................................................................ [ 71360]
MW396979.1_Timor-Leste/2019/1     ................................................................................ [ 71360]
MK940252.1_CN/2019/InnerMongolia  ................................................................................ [ 71360]
MK128995.1_China/2018/AnhuiXCGQ   ................................................................................ [ 71360]
MH681419.1_ASFV/POL/2015          ................................................................................ [ 71360]
LS478113.1_Estonia_2014           ................................................................................ [ 71360]
KP843857.1_Odintsovo_02/14        ................................................................................ [ 71360]
FN557520.1_strain_E75             ................................................................................ [ 71360]
AY261366.1_Warthog                ....g...............................................a...........a............... [ 71360]
AY261365.1_Warmbaths              ....g........................................................................... [ 71360]
AY261364.1_Tengani_62             ................................................................................ [ 71360]
AY261363.1_Pretoriuskop/96/4      ....g...............................................a...........a............... [ 71360]
AY261362.1_Mkuzi_1979             ................................................................................ [ 71360]
AY261361.1_Malawi_Lil-20/1_1983   ....g........................................................................... [ 71360]
AY261360.1_Kenya_1950             ....g........c.......................g....................t..................... [ 71360]
AM712240.1_OURT_88/3              ................................................................................ [ 71360]
AM712239.1_Benin_97/1             ................................................................................ [ 71360]
KM111295.1_Ken06.Bus              ....g........c.......................g....................t..................... [ 71360]
KM111294.1_Ken05/Tk1              ....g........c.......................g....................t..................... [ 71360]
KM262844.1_strain_L60             ................................................................................ [ 71360]
KM102979.1_26544/OG10_Italy       ................................................................................ [ 71360]
KM262845.1_strain_NHV             ................................................................................ [ 71360]
KX354450.1_47/Ss/2008             ................................................................................ [ 71360]
KP055815.1_strain_BA71            ................................................................................ [ 71360]
MH025920.1_strain_R35             ....g........c.......................g....................t..................... [ 71360]
MH025919.1_strain_N10             ....g........c.......................g....................t..................... [ 71360]
MH025918.1_strain_R25             ....g........c.......................g....................t..................... [ 71360]
MH025917.1_strain_R7              ....g........c.......................g....................t..................... [ 71360]
MH025916.1_strain_R8              ....g........c.......................g....................t..................... [ 71360]
```

```
FR682468.2_Georgia_2007/1        gaggccattgccatggatgaggccatccatgcggccctatactacggggccatgcgacgatccatagaacttgcaaaaga [ 71440]
ASF_IND_20_CAD543                ................................................................................ [ 71440]
LR899193.1_Germany_2020/1        ................................................................................ [ 71440]
MN393477.1_Wuhan_2019            ................................................................................ [ 71440]
MW396979.1_Timor-Leste/2019/1    ................................................................................ [ 71440]
MK940252.1_CN/2019/InnerMongolia ................................................................................ [ 71440]
MK128995.1_China/2018/AnhuiXCGQ  ................................................................................ [ 71440]
MH681419.1_ASFV/POL/2015         ................................................................................ [ 71440]
LS478113.1_Estonia_2014          ................................................................................ [ 71440]
KP843857.1_Odintsovo_02/14       ................................................................................ [ 71440]
FN557520.1_strain_E75            ................................................................................ [ 71440]
AY261366.1_Warthog               .......................t..t........t...........t........g.....t..............g.. [ 71440]
AY261365.1_Warmbaths             ...................................t..c........t........g.....t..............g.. [ 71440]
AY261364.1_Tengani_62            ................................................................................ [ 71440]
AY261363.1_Pretoriuskop/96/4     .......................t..t........t...........t........g.....t..............g.. [ 71440]
AY261362.1_Mkuzi_1979            ...............................................t..............t................. [ 71440]
AY261361.1_Malawi_Lil-20/1_1983  ...........g..a..c.................t...................................c........ [ 71440]
AY261360.1_Kenya_1950            ................................a..............c...............g................ [ 71440]
AM712240.1_OURT_88/3             ................................................................................ [ 71440]
AM712239.1_Benin_97/1            ................................................................................ [ 71440]
KM111295.1_Ken06.Bus             ................................a..............c...............g.t.............. [ 71440]
KM111294.1_Ken05/Tk1             ................................a..............c...............g................ [ 71440]
KM262844.1_strain_L60            ................................................................................ [ 71440]
KM102979.1_26544/OG10_Italy      ................................................................................ [ 71440]
KM262845.1_strain_NHV            ................................................................................ [ 71440]
KX354450.1_47/Ss/2008            ................................................................................ [ 71440]
KP055815.1_strain_BA71           ................................................................................ [ 71440]
MH025920.1_strain_R35            ................................a..............c...............g.t.............. [ 71440]
MH025919.1_strain_N10            ................................a..............c...............g.t.............. [ 71440]
MH025918.1_strain_R25            ................................a..............c...............g.t.............. [ 71440]
MH025917.1_strain_R7             ................................a..............c...............g.t.............. [ 71440]
MH025916.1_strain_R8             ................................a..............c...............g.t.............. [ 71440]

FR682468.2_Georgia_2007/1        aaaaggaagtcatcccagctttccggggtctgcggcctcaaagggtctactgcagcccgacctatgggttcgctgtggtg [ 71520]
ASF_IND_20_CAD543                ................................................................................ [ 71520]
LR899193.1_Germany_2020/1        ................................................................................ [ 71520]
MN393477.1_Wuhan_2019            ................................................................................ [ 71520]
MW396979.1_Timor-Leste/2019/1    ................................................................................ [ 71520]
MK940252.1_CN/2019/InnerMongolia ................................................................................ [ 71520]
MK128995.1_China/2018/AnhuiXCGQ  ................................................................................ [ 71520]
MH681419.1_ASFV/POL/2015         ................................................................................ [ 71520]
LS478113.1_Estonia_2014          ................................................................................ [ 71520]
KP843857.1_Odintsovo_02/14       ................................................................................ [ 71520]
```

```
FN557520.1_strain_E75          ......................................................a....................c.... [ 71520]
AY261366.1_Warthog             .........c........t........a..a..............c..g.....a..t...................... [ 71520]
AY261365.1_Warmbaths           ...........................a..a..............c..g..........................c.... [ 71520]
AY261364.1_Tengani_62          ................................................................................ [ 71520]
AY261363.1_Pretoriuskop/96/4   .........c........t.....a..a..a..............c..g..a..a..t...................... [ 71520]
AY261362.1_Mkuzi_1979          .............................................c..g.....a....................c.... [ 71520]
AY261361.1_Malawi_Lil-20/1_1983 ......g..........................a..t........c..gt.a..a......................... [ 71520]
AY261360.1_Kenya_1950          ............c.....t.......................a..c..gt....a....................c.a.. [ 71520]
AM712240.1_OURT_88/3           ......................................................a....................c.... [ 71520]
AM712239.1_Benin_97/1          ......................................................a....................c.... [ 71520]
KM111295.1_Ken06.Bus           ............c.....t.......................a..c..gt....a....................c.a.. [ 71520]
KM111294.1_Ken05/Tk1           ............c.....t.......................a..c..gt....a....................c.a.. [ 71520]
KM262844.1_strain_L60          ......................................................a....................c.... [ 71520]
KM102979.1_26544/OG10_Italy    ......................................................a....................c.... [ 71520]
KM262845.1_strain_NHV          ......................................................a....................c.... [ 71520]
KX354450.1_47/Ss/2008          ......................................................a....................c.... [ 71520]
KP055815.1_strain_BA71         ......................................................a....................c.... [ 71520]
MH025920.1_strain_R35          ............c.....t.......................a..c..gt....a....................c.a.. [ 71520]
MH025919.1_strain_N10          ............c.....t.......................a..c..gt....a....................c.a.. [ 71520]
MH025918.1_strain_R25          ............c.....t.......................a..c..gt....a....................c.a.. [ 71520]
MH025917.1_strain_R7           ............c.....t.......................a..c..gt....a....................c.a.. [ 71520]
MH025916.1_strain_R8           ............c.....t.......................a..c..gt....a....................c.a.. [ 71520]

FR682468.2_Georgia_2007/1      atttagtttcctcctgggaagaacgcgtggcacagacgacgcagggtgtgttgacgccgaaaaggtggtcgcagctacgc [ 71600]
ASF_IND_20_CAD543              .....ag......................................................................... [ 71600]
LR899193.1_Germany_2020/1      ................................................................................ [ 71600]
MN393477.1_Wuhan_2019          ................................................................................ [ 71600]
MW396979.1_Timor-Leste/2019/1  ................................................................................ [ 71600]
MK940252.1_CN/2019/InnerMongolia ................................................................................ [ 71600]
MK128995.1_China/2018/AnhuiXCGQ ................................................................................ [ 71600]
MH681419.1_ASFV/POL/2015       ................................................................................ [ 71600]
LS478113.1_Estonia_2014        ................................................................................ [ 71600]
KP843857.1_Odintsovo_02/14     ................................................................................ [ 71600]
FN557520.1_strain_E75          .....ag.............................................a....g......c....g...a.....g [ 71600]
AY261366.1_Warthog             .....a..c....t.......................a........a.....a....g......c...cg...a.....g [ 71600]
AY261365.1_Warmbaths           .....a...............................a........aa.a..a....g......c....g...a.....g [ 71600]
AY261364.1_Tengani_62          ..........................................t..................................... [ 71600]
AY261363.1_Pretoriuskop/96/4   .....a..c....t.......................a........a.....a....g......c...cg...a.....g [ 71600]
AY261362.1_Mkuzi_1979          .....a..............................................a....g......c....g...a.....g [ 71600]
AY261361.1_Malawi_Lil-20/1_1983 .....a..c...........a.c........g.....a.........................a.....g...a.....g [ 71600]
AY261360.1_Kenya_1950          .........t...........................a....................a....a.....g.........g [ 71600]
AM712240.1_OURT_88/3           .....ag.............................................a....g......c....g...a.....g [ 71600]
AM712239.1_Benin_97/1          .....ag.............................................a....g......c....g...a.....g [ 71600]
KM111295.1_Ken06.Bus           .........t...........................a....................a....a.....g.........g [ 71600]
```

```
KM111294.1_Ken05/Tk1             .........t...........................a....................a....a.....g.........g [ 71600]
KM262844.1_strain_L60            .....ag.............................................a....g......c....g...a.....g [ 71600]
KM102979.1_26544/OG10_Italy      .....ag.............................................a....g......c....g...a.....g [ 71600]
KM262845.1_strain_NHV            .....ag.............................................a....g......c....g...a.....g [ 71600]
KX354450.1_47/Ss/2008            .....ag.............................................a....g......c....g...a.....g [ 71600]
KP055815.1_strain_BA71           .....ag.............................................a....g......c....g...a.....g [ 71600]
MH025920.1_strain_R35            .........t...........................a....................a....a.....g.........g [ 71600]
MH025919.1_strain_N10            .........t...........................a....................a....a.....g.........g [ 71600]
MH025918.1_strain_R25            .........t...........................a....................a....a.....g.........g [ 71600]
MH025917.1_strain_R7             .........t...........................a....................a....a.....g.........g [ 71600]
MH025916.1_strain_R8             .........t...........................a....................a....a.....g.........g [ 71600]

FR682468.2_Georgia_2007/1        ctggcggctatgcagggacttcgaaatggatatgtcacagctcttatgcccaccgcaacctcctcaaattctacaggaaa [ 71680]
ASF_IND_20_CAD543                ................................................................................ [ 71680]
LR899193.1_Germany_2020/1        ................................................................................ [ 71680]
MN393477.1_Wuhan_2019            ................................................................................ [ 71680]
MW396979.1_Timor-Leste/2019/1    ................................................................................ [ 71680]
MK940252.1_CN/2019/InnerMongolia ................................................................................ [ 71680]
MK128995.1_China/2018/AnhuiXCGQ  ................................................................................ [ 71680]
MH681419.1_ASFV/POL/2015         ................................................................................ [ 71680]
LS478113.1_Estonia_2014          ................................................................................ [ 71680]
KP843857.1_Odintsovo_02/14       ................................................................................ [ 71680]
FN557520.1_strain_E75            ........c.........g..............c....g..c...................................... [ 71680]
AY261366.1_Warthog               ........g..a..a..cg..............c.......c..c................................g.. [ 71680]
AY261365.1_Warmbaths             ........g.....a..cg..............c.......c..c................................... [ 71680]
AY261364.1_Tengani_62            ................................................................................ [ 71680]
AY261363.1_Pretoriuskop/96/4     ........g..a..a..cg..............c.......c..c................................g.. [ 71680]
AY261362.1_Mkuzi_1979            ........c.........g..............c.......c..c................................... [ 71680]
AY261361.1_Malawi_Lil-20/1_1983  ........g..a......g.g............c.t....g...................................c.. [ 71680]
AY261360.1_Kenya_1950            ........g.........g.c............c....g..c................................c..c.. [ 71680]
AM712240.1_OURT_88/3             ........c.........g..............c....g..c...................................... [ 71680]
AM712239.1_Benin_97/1            ........c.........g..............c....g..c...................................... [ 71680]
KM111295.1_Ken06.Bus             ........g.........g..............c....g..c................................c..c.. [ 71680]
KM111294.1_Ken05/Tk1             ........g.........g.c............c....g..c................................c..c.. [ 71680]
KM262844.1_strain_L60            ........c.........g..............c....g..c...................................... [ 71680]
KM102979.1_26544/OG10_Italy      ........c.........g..............c....g..c...................................... [ 71680]
KM262845.1_strain_NHV            ........c.........g..............c....g..c...................................... [ 71680]
KX354450.1_47/Ss/2008            ........c.........g..............c....g..c...................................... [ 71680]
KP055815.1_strain_BA71           ........c.........g..............c....g..c...................................... [ 71680]
MH025920.1_strain_R35            ........g.........g..............c....g..c................................c..c.. [ 71680]
MH025919.1_strain_N10            ........g.........g..............c....g..c................................c..c.. [ 71680]
MH025918.1_strain_R25            ........g.........g..............c....g..c................................c..c.. [ 71680]
MH025917.1_strain_R7             ........g.........g..............c....g..c................................c..c.. [ 71680]
MH025916.1_strain_R8             ........g.........g..............c....g..c................................c..c.. [ 71680]
```

```
FR682468.2_Georgia_2007/1        aaacgaatgttttgagccctttacatccaatctatatacacgtagaacgttaagcggggagtttattgttttaaataagt [ 71760]
ASF_IND_20_CAD543                ................................................................................ [ 71760]
LR899193.1_Germany_2020/1        ................................................................................ [ 71760]
MN393477.1_Wuhan_2019            ................................................................................ [ 71760]
MW396979.1_Timor-Leste/2019/1    ................................................................................ [ 71760]
MK940252.1_CN/2019/InnerMongolia ................................................................................ [ 71760]
MK128995.1_China/2018/AnhuiXCGQ  ................................................................................ [ 71760]
MH681419.1_ASFV/POL/2015         ................................................................................ [ 71760]
LS478113.1_Estonia_2014          ................................................................................ [ 71760]
KP843857.1_Odintsovo_02/14       ................................................................................ [ 71760]
FN557520.1_strain_E75            ........................c....................................................... [ 71760]
AY261366.1_Warthog               ...............a................................a.....t.....a................... [ 71760]
AY261365.1_Warmbaths             ................................................................................ [ 71760]
AY261364.1_Tengani_62            ................................................................................ [ 71760]
AY261363.1_Pretoriuskop/96/4     .........c......................................a.....t.....a................... [ 71760]
AY261362.1_Mkuzi_1979            ................................................................................ [ 71760]
AY261361.1_Malawi_Lil-20/1_1983  ........................g..t....................a...........a......a............ [ 71760]
AY261360.1_Kenya_1950            .........c.....................t................a...........a................... [ 71760]
AM712240.1_OURT_88/3             ........................c....................................................... [ 71760]
AM712239.1_Benin_97/1            ........................c....................................................... [ 71760]
KM111295.1_Ken06.Bus             .........c.....................t................a...........a................... [ 71760]
KM111294.1_Ken05/Tk1             .........c.....................t................a........a..a................... [ 71760]
KM262844.1_strain_L60            ........................c....................................................... [ 71760]
KM102979.1_26544/OG10_Italy      ........................c....................................................... [ 71760]
KM262845.1_strain_NHV            ........................c....................................................... [ 71760]
KX354450.1_47/Ss/2008            ........................c....................................................... [ 71760]
KP055815.1_strain_BA71           ........................c....................................................... [ 71760]
MH025920.1_strain_R35            .........c.....................t................a...........a................... [ 71760]
MH025919.1_strain_N10            .........c.....................t................a...........a................... [ 71760]
MH025918.1_strain_R25            .........c.....................t................a...........a................... [ 71760]
MH025917.1_strain_R7             .........c.....................t................a...........a................... [ 71760]
MH025916.1_strain_R8             .........c.....................t................a...........a................... [ 71760]

FR682468.2_Georgia_2007/1        atttaatagacgatttaaaagaaattaatctttggacagaagccattcaacagcagctactaaatgcgggaggtagcatt [ 71840]
ASF_IND_20_CAD543                ...................................................................r............ [ 71840]
LR899193.1_Germany_2020/1        ................................................................................ [ 71840]
MN393477.1_Wuhan_2019            ................................................................................ [ 71840]
MW396979.1_Timor-Leste/2019/1    ................................................................................ [ 71840]
MK940252.1_CN/2019/InnerMongolia ................................................................................ [ 71840]
MK128995.1_China/2018/AnhuiXCGQ  ................................................................................ [ 71840]
MH681419.1_ASFV/POL/2015         ................................................................................ [ 71840]
LS478113.1_Estonia_2014          ................................................................................ [ 71840]
KP843857.1_Odintsovo_02/14       ................................................................................ [ 71840]
```

```
FN557520.1_strain_E75            ..........................g........................................a............ [ 71840]
AY261366.1_Warthog               ...................................................t...............a............ [ 71840]
AY261365.1_Warmbaths             ..........t....................c...................t............................ [ 71840]
AY261364.1_Tengani_62            ................................................................................ [ 71840]
AY261363.1_Pretoriuskop/96/4     ...................................................t...............a............ [ 71840]
AY261362.1_Mkuzi_1979            ..........t....................c...................t............................ [ 71840]
AY261361.1_Malawi_Lil-20/1_1983  .......g..t......g.................t......a...........................g.....t... [ 71840]
AY261360.1_Kenya_1950            ..........t.............................g.............................g......... [ 71840]
AM712240.1_OURT_88/3             ..........................g........................................a............ [ 71840]
AM712239.1_Benin_97/1            ..........................g........................................a............ [ 71840]
KM111295.1_Ken06.Bus             .........................g..............g.............................g......... [ 71840]
KM111294.1_Ken05/Tk1             ..........t.............................g.............................g......... [ 71840]
KM262844.1_strain_L60            ..........................g........................................a............ [ 71840]
KM102979.1_26544/OG10_Italy      ..........................g........................................a............ [ 71840]
KM262845.1_strain_NHV            ..........................g........................................a............ [ 71840]
KX354450.1_47/Ss/2008            ..........................g........................................a............ [ 71840]
KP055815.1_strain_BA71           ..........................g........................................a............ [ 71840]
MH025920.1_strain_R35            .........................g..............g.............................g......... [ 71840]
MH025919.1_strain_N10            .........................g..............g.............................g......... [ 71840]
MH025918.1_strain_R25            .........................g..............g.............................g......... [ 71840]
MH025917.1_strain_R7             .........................g..............g.............................g......... [ 71840]
MH025916.1_strain_R8             .........................g..............g.............................g......... [ 71840]

FR682468.2_Georgia_2007/1        cagcacattttggatataccggccgagatccgcgatcggtataaaacctccagggaaatgaatcaaaaaaattttaacaaa [ 71920]
ASF_IND_20_CAD543                ..................................................y............................. [ 71920]
LR899193.1_Germany_2020/1        ................................................................................ [ 71920]
MN393477.1_Wuhan_2019            ................................................................................ [ 71920]
MW396979.1_Timor-Leste/2019/1    ................................................................................ [ 71920]
MK940252.1_CN/2019/InnerMongolia ................................................................................ [ 71920]
MK128995.1_China/2018/AnhuiXCGQ  ................................................................................ [ 71920]
MH681419.1_ASFV/POL/2015         ................................................................................ [ 71920]
LS478113.1_Estonia_2014          ................................................................................ [ 71920]
KP843857.1_Odintsovo_02/14       ................................................................................ [ 71920]
FN557520.1_strain_E75            ................................t.................t............................. [ 71920]
AY261366.1_Warthog               ..a.............................t..............t..............c................. [ 71920]
AY261365.1_Warmbaths             ..........................a...................................c................. [ 71920]
AY261364.1_Tengani_62            ................................................................................ [ 71920]
AY261363.1_Pretoriuskop/96/4     ..a.............................t..............t..............c................. [ 71920]
AY261362.1_Mkuzi_1979            ..........................a...................................c................. [ 71920]
AY261361.1_Malawi_Lil-20/1_1983  .........c.a....................t..a..a...........tc.a...........g.............. [ 71920]
AY261360.1_Kenya_1950            .........c......................t..a..a...........t.a........c..g.............. [ 71920]
AM712240.1_OURT_88/3             ................................t.................t............................. [ 71920]
AM712239.1_Benin_97/1            ................................t.................t............................. [ 71920]
KM111295.1_Ken06.Bus             .........c......................t..a..a...........t.a........c..g.............. [ 71920]
```

```
KM111294.1_Ken05/Tk1           .........c......................t..a..a...........t..a........c..g.............. [ 71920]
KM262844.1_strain_L60          ................................t.................t............................. [ 71920]
KM102979.1_26544/OG10_Italy    ................................t.................t............................. [ 71920]
KM262845.1_strain_NHV          ................................t.................t............................. [ 71920]
KX354450.1_47/Ss/2008          ................................t.................t............................. [ 71920]
KP055815.1_strain_BA71         ................................t.................t............................. [ 71920]
MH025920.1_strain_R35          .........c......................t..a..a...........t..a........c..g.............. [ 71920]
MH025919.1_strain_N10          .........c......................t..a..a...........t..a........c..g.............. [ 71920]
MH025918.1_strain_R25          .........c......................t..a..a...........t..a........c..g.............. [ 71920]
MH025917.1_strain_R7           .........c......................t..a..a...........t..a........c..g.............. [ 71920]
MH025916.1_strain_R8           .........c......................t..a..a...........t..a........c..g.............. [ 71920]

FR682468.2_Georgia_2007/1      acacgcggccgcacgaaacccctttgtatcccaaagtatgtccttgaactattacttttatgaacctgaactaagccagg [ 72000]
ASF_IND_20_CAD543              ................................................................................ [ 72000]
LR899193.1_Germany_2020/1      ................................................................................ [ 72000]
MN393477.1_Wuhan_2019          ................................................................................ [ 72000]
MW396979.1_Timor-Leste/2019/1  ................................................................................ [ 72000]
MK940252.1_CN/2019/InnerMongolia ................................................................................ [ 72000]
MK128995.1_China/2018/AnhuiXCGQ ................................................................................ [ 72000]
MH681419.1_ASFV/POL/2015       ................................................................................ [ 72000]
LS478113.1_Estonia_2014        ................................................................................ [ 72000]
KP843857.1_Odintsovo_02/14     ................................................................................ [ 72000]
FN557520.1_strain_E75          ...........................g..............t........c............................ [ 72000]
AY261366.1_Warthog             ...................................................c........c..g......t......... [ 72000]
AY261365.1_Warmbaths           ...................................................c........c..g......t.......a. [ 72000]
AY261364.1_Tengani_62          ................................................................................ [ 72000]
AY261363.1_Pretoriuskop/96/4   ...................................................c........c..g......t......... [ 72000]
AY261362.1_Mkuzi_1979          ...................................................c........c..g......t.......a. [ 72000]
AY261361.1_Malawi_Lil-20/1_1983 ......a...........t.....c..g........c.....t........c..t..c............t......... [ 72000]
AY261360.1_Kenya_1950          ..................t.....c..g........c.....tc.......c.....c............t.......a. [ 72000]
AM712240.1_OURT_88/3           ...........................g..............t........c............................ [ 72000]
AM712239.1_Benin_97/1          ...........................g..............t........c............................ [ 72000]
KM111295.1_Ken06.Bus           .........t.....c........c..g........c.....tc.......c.....c............t.......a. [ 72000]
KM111294.1_Ken05/Tk1           ..................t........g........c.....tc.......c.....c............t.......a. [ 72000]
KM262844.1_strain_L60          ...........................g..............t........c............................ [ 72000]
KM102979.1_26544/OG10_Italy    ...........................g..............t........c............................ [ 72000]
KM262845.1_strain_NHV          ...........................g..............t........c............................ [ 72000]
KX354450.1_47/Ss/2008          ...........................g..............t........c............................ [ 72000]
KP055815.1_strain_BA71         ...........................g..............t........c............................ [ 72000]
MH025920.1_strain_R35          .........t.....c........c..g........c.....tc.......c.....c............t.......a. [ 72000]
MH025919.1_strain_N10          .........t.....c........c..g........c.....tc.......c.....c............t.......a. [ 72000]
MH025918.1_strain_R25          .........t.....c........c..g........c.....tc.......c.....c............t.......a. [ 72000]
MH025917.1_strain_R7           .........t.....c........c..g........c.....tc.......c.....c............t.......a. [ 72000]
MH025916.1_strain_R8           .........t.....c........c..g........c.....tc.......c.....c............t.......a. [ 72000]
```

```
FR682468.2_Georgia_2007/1        tacttacagtgctcgtcctaggctggaaaaaaggtttaactaccggttcctattactgtcattttagccctggagcgggt [ 72080]
ASF_IND_20_CAD543                ................................................................................ [ 72080]
LR899193.1_Germany_2020/1        ................................................................................ [ 72080]
MN393477.1_Wuhan_2019            ................................................................................ [ 72080]
MW396979.1_Timor-Leste/2019/1    ................................................................................ [ 72080]
MK940252.1_CN/2019/InnerMongolia ................................................................................ [ 72080]
MK128995.1_China/2018/AnhuiXCGQ  ................................................................................ [ 72080]
MH681419.1_ASFV/POL/2015         ................................................................................ [ 72080]
LS478113.1_Estonia_2014          ................................................................................ [ 72080]
KP843857.1_Odintsovo_02/14       ................................................................................ [ 72080]
FN557520.1_strain_E75            .......g...........................c....c....................................... [ 72080]
AY261366.1_Warthog               .......g.....t..g.....t........g..c.....c..t..c................................. [ 72080]
AY261365.1_Warmbaths             .......g.....t..g.....t........g..c........t..c................................. [ 72080]
AY261364.1_Tengani_62            .......g..a..................................................................... [ 72080]
AY261363.1_Pretoriuskop/96/4     .......g.....t..g.....t........g..c.....c..t..c................................. [ 72080]
AY261362.1_Mkuzi_1979            .......g.....t..g.....t........g..c........t..c................................. [ 72080]
AY261361.1_Malawi_Lil-20/1_1983  .............g.....................c....c.....c...........c..c.....t..c......... [ 72080]
AY261360.1_Kenya_1950            .g.............................g...c....c.....c...........c..c........c......... [ 72080]
AM712240.1_OURT_88/3             .......g...........................c....c....................................... [ 72080]
AM712239.1_Benin_97/1            .......g...........................c....c....................................... [ 72080]
KM111295.1_Ken06.Bus             ...............................g...c....c.....c...........c..c........c.....a... [ 72080]
KM111294.1_Ken05/Tk1             .g.............................g...c....c.....c...........c..c........c......... [ 72080]
KM262844.1_strain_L60            .......g...........................c....c....................................... [ 72080]
KM102979.1_26544/OG10_Italy      .......g...........................c....c....................................... [ 72080]
KM262845.1_strain_NHV            .......g...........................c....c....................................... [ 72080]
KX354450.1_47/Ss/2008            .......g...........................c....c....................................... [ 72080]
KP055815.1_strain_BA71           .......g...........................c....c....................................... [ 72080]
MH025920.1_strain_R35            ...............................g...c....c.....c...........c..c........c.....a... [ 72080]
MH025919.1_strain_N10            ...............................g...c....c.....c...........c..c........c.....a... [ 72080]
MH025918.1_strain_R25            ...............................g...c....c.....c...........c..c........c.....a... [ 72080]
MH025917.1_strain_R7             ...............................g...c....c.....c...........c..c........c.....a... [ 72080]
MH025916.1_strain_R8             ...............................g...c....c.....c...........c..c........c.....a... [ 72080]

FR682468.2_Georgia_2007/1        acccaaaaaaagattataagaaactctgagaaagcgtgtaatgcggactgcgaggcgtgtcttctgtaggtgtctcgcgg [ 72160]
ASF_IND_20_CAD543                ................................................................................ [ 72160]
LR899193.1_Germany_2020/1        ................................................................................ [ 72160]
MN393477.1_Wuhan_2019            ................................................................................ [ 72160]
MW396979.1_Timor-Leste/2019/1    ................................................................................ [ 72160]
MK940252.1_CN/2019/InnerMongolia ................................................................................ [ 72160]
MK128995.1_China/2018/AnhuiXCGQ  ................................................................................ [ 72160]
MH681419.1_ASFV/POL/2015         ................................................................................ [ 72160]
LS478113.1_Estonia_2014          ................................................................................ [ 72160]
KP843857.1_Odintsovo_02/14       ................................................................................ [ 72160]
```

```
FN557520.1_strain_E75          ......................................................................a......... [ 72160]
AY261366.1_Warthog             ......................................................................a......... [ 72160]
AY261365.1_Warmbaths           ......................................................................a......... [ 72160]
AY261364.1_Tengani_62          ................................................................................ [ 72160]
AY261363.1_Pretoriuskop/96/4   ......................................................................a......... [ 72160]
AY261362.1_Mkuzi_1979          ......................................................................a......... [ 72160]
AY261361.1_Malawi_Lil-20/1_1983 .....g................................................................a......... [ 72160]
AY261360.1_Kenya_1950          ........................................g.............................a......... [ 72160]
AM712240.1_OURT_88/3           ......................................................................a......... [ 72160]
AM712239.1_Benin_97/1          ......................................................................a......... [ 72160]
KM111295.1_Ken06.Bus           ........................................g.............................a......... [ 72160]
KM111294.1_Ken05/Tk1           ........................................g.............................a......... [ 72160]
KM262844.1_strain_L60          ......................................................................a......... [ 72160]
KM102979.1_26544/OG10_Italy    ......................................................................a......... [ 72160]
KM262845.1_strain_NHV          ......................................................................a......... [ 72160]
KX354450.1_47/Ss/2008          ......................................................................a......... [ 72160]
KP055815.1_strain_BA71         ......................................................................a......... [ 72160]
MH025920.1_strain_R35          ........................................g.............................a......... [ 72160]
MH025919.1_strain_N10          ........................................g.............................a......... [ 72160]
MH025918.1_strain_R25          ........................................g.............................a......... [ 72160]
MH025917.1_strain_R7           ........................................g.............................a......... [ 72160]
MH025916.1_strain_R8           ........................................g.............................a......... [ 72160]

FR682468.2_Georgia_2007/1      taaaagagcagcggggaccatatggtaaaccccaacaagaggataatgaataaaaaaagtaaacaggcatccattagttc [ 72240]
ASF_IND_20_CAD543              ................................................................................ [ 72240]
LR899193.1_Germany_2020/1      ................................................................................ [ 72240]
MN393477.1_Wuhan_2019          ................................................................................ [ 72240]
MW396979.1_Timor-Leste/2019/1  ................................................................................ [ 72240]
MK940252.1_CN/2019/InnerMongolia ................................................................................ [ 72240]
MK128995.1_China/2018/AnhuiXCGQ ................................................................................ [ 72240]
MH681419.1_ASFV/POL/2015       ................................................................................ [ 72240]
LS478113.1_Estonia_2014        ................................................................................ [ 72240]
KP843857.1_Odintsovo_02/14     ................................................................................ [ 72240]
FN557520.1_strain_E75          .........................c.........t............................................ [ 72240]
AY261366.1_Warthog             .........................c...................................................... [ 72240]
AY261365.1_Warmbaths           .........................c...................................................... [ 72240]
AY261364.1_Tengani_62          .........................c...................................................... [ 72240]
AY261363.1_Pretoriuskop/96/4   ..............a..........c.........t............................................ [ 72240]
AY261362.1_Mkuzi_1979          .........................c...................................................... [ 72240]
AY261361.1_Malawi_Lil-20/1_1983 ...........t.............c........g............a................................ [ 72240]
AY261360.1_Kenya_1950          ....g....................c........g............................................. [ 72240]
AM712240.1_OURT_88/3           .........................c.........t............................................ [ 72240]
AM712239.1_Benin_97/1          .........................c.........t............................................ [ 72240]
KM111295.1_Ken06.Bus           .....a...................c........g............................................. [ 72240]
```

```
KM111294.1_Ken05/Tk1             .....a...................c........g............................................. [ 72240]
KM262844.1_strain_L60            .........................c........t............................................. [ 72240]
KM102979.1_26544/OG10_Italy      .........................c........t............................................. [ 72240]
KM262845.1_strain_NHV            .........................c........t............................................. [ 72240]
KX354450.1_47/Ss/2008            .........................c........t............................................. [ 72240]
KP055815.1_strain_BA71           .........................c........t............................................. [ 72240]
MH025920.1_strain_R35            .....a...................c........g............................................. [ 72240]
MH025919.1_strain_N10            .....a...................c........g............................................. [ 72240]
MH025918.1_strain_R25            .....a...................c........g............................................. [ 72240]
MH025917.1_strain_R7             .....a...................c........g............................................. [ 72240]
MH025916.1_strain_R8             .....a...................c........g............................................. [ 72240]

FR682468.2_Georgia_2007/1        catattaaa-tttttttttcttctatataatggaatattttgttgcggtagacaatgaaacctccttgggggtttttact [ 72320]
ASF_IND_20_CAD543                .........-...................................................................... [ 72320]
LR899193.1_Germany_2020/1        .........-...................................................................... [ 72320]
MN393477.1_Wuhan_2019            .........-...................................................................... [ 72320]
MW396979.1_Timor-Leste/2019/1    .........-...................................................................... [ 72320]
MK940252.1_CN/2019/InnerMongolia .........-...................................................................... [ 72320]
MK128995.1_China/2018/AnhuiXCGQ  .........-...................................................................... [ 72320]
MH681419.1_ASFV/POL/2015         .........-...................................................................... [ 72320]
LS478113.1_Estonia_2014          .........-...................................................................... [ 72320]
KP843857.1_Odintsovo_02/14       .........--..................................................................... [ 72320]
FN557520.1_strain_E75            .........-....................................................c................. [ 72320]
AY261366.1_Warthog               .........-...................................................................... [ 72320]
AY261365.1_Warmbaths             .........-...................................................................... [ 72320]
AY261364.1_Tengani_62            .........t...................................................................... [ 72320]
AY261363.1_Pretoriuskop/96/4     .........-...................................................................... [ 72320]
AY261362.1_Mkuzi_1979            .........-....................................................c................. [ 72320]
AY261361.1_Malawi_Lil-20/1_1983  .........-...................................................................... [ 72320]
AY261360.1_Kenya_1950            .........-........c..........................................tc................. [ 72320]
AM712240.1_OURT_88/3             .........-....................................................c................. [ 72320]
AM712239.1_Benin_97/1            .........-....................................................c................. [ 72320]
KM111295.1_Ken06.Bus             .........-....................................................c................. [ 72320]
KM111294.1_Ken05/Tk1             .........-........c..........................................tc................. [ 72320]
KM262844.1_strain_L60            .........-....................................................c................. [ 72320]
KM102979.1_26544/OG10_Italy      .........-....................................................c................. [ 72320]
KM262845.1_strain_NHV            .........-....................................................c................. [ 72320]
KX354450.1_47/Ss/2008            .........-....................................................c................. [ 72320]
KP055815.1_strain_BA71           .........-....................................................c................. [ 72320]
MH025920.1_strain_R35            .........-....................................................c................. [ 72320]
MH025919.1_strain_N10            .........-....................................................c................. [ 72320]
MH025918.1_strain_R25            .........-....................................................c................. [ 72320]
MH025917.1_strain_R7             .........-....................................................c................. [ 72320]
MH025916.1_strain_R8             .........-....................................................c................. [ 72320]
```

| | | |
|---|---|---|
| FR682468.2_Georgia_2007/1 | tctatagagcaatgtgaagaaacgatgaaacaataccccggcctccattatgtcgtttttaagtatatgtgtccggcgga | [ 72400] |
| ASF_IND_20_CAD543 | ...................................................................ct........... | [ 72400] |
| LR899193.1_Germany_2020/1 | ................................................................................ | [ 72400] |
| MN393477.1_Wuhan_2019 | ................................................................................ | [ 72400] |
| MW396979.1_Timor-Leste/2019/1 | ................................................................................ | [ 72400] |
| MK940252.1_CN/2019/InnerMongolia | ................................................................................ | [ 72400] |
| MK128995.1_China/2018/AnhuiXCGQ | ................................................................................ | [ 72400] |
| MH681419.1_ASFV/POL/2015 | ................................................................................ | [ 72400] |
| LS478113.1_Estonia_2014 | ................................................................................ | [ 72400] |
| KP843857.1_Odintsovo_02/14 | ................................................................................ | [ 72400] |
| FN557520.1_strain_E75 | ...................................................................ct........... | [ 72400] |
| AY261366.1_Warthog | ...................................t..t............................ct........... | [ 72400] |
| AY261365.1_Warmbaths | .....g................................t.a............g.............ct........... | [ 72400] |
| AY261364.1_Tengani_62 | ................................................................................ | [ 72400] |
| AY261363.1_Pretoriuskop/96/4 | ...................................t..t............................ct........... | [ 72400] |
| AY261362.1_Mkuzi_1979 | .....g...............................................g.............ct........... | [ 72400] |
| AY261361.1_Malawi_Lil-20/1_1983 | .....g....................................................................a..... | [ 72400] |
| AY261360.1_Kenya_1950 | ..c..g........c.......................t...........c................c............ | [ 72400] |
| AM712240.1_OURT_88/3 | ...................................................................ct........... | [ 72400] |
| AM712239.1_Benin_97/1 | ...................................................................ct........... | [ 72400] |
| KM111295.1_Ken06.Bus | ..c..g........c..g................................c................c............ | [ 72400] |
| KM111294.1_Ken05/Tk1 | ..c..g........c...................................c................c............ | [ 72400] |
| KM262844.1_strain_L60 | ...................................................................ct........... | [ 72400] |
| KM102979.1_26544/OG10_Italy | ...................................................................ct........... | [ 72400] |
| KM262845.1_strain_NHV | ...................................................................ct........... | [ 72400] |
| KX354450.1_47/Ss/2008 | ...................................................................ct........... | [ 72400] |
| KP055815.1_strain_BA71 | ...................................................................ct........... | [ 72400] |
| MH025920.1_strain_R35 | ..c..g........c..g................................c................c............ | [ 72400] |
| MH025919.1_strain_N10 | ..c..g........c..g................................c................c............ | [ 72400] |
| MH025918.1_strain_R25 | ..c..g........c..g................................c................c............ | [ 72400] |
| MH025917.1_strain_R7 | ..c..g........c..g................................c................c............ | [ 72400] |
| MH025916.1_strain_R8 | ..c..g........c..g................................c................c............ | [ 72400] |

| | | |
|---|---|---|
| FR682468.2_Georgia_2007/1 | tgcagaaaatacagatgttgtatatttaatacctcgttaaccttgcatacccccatgtttgtagaccactgtccaaatc | [ 72480] |
| ASF_IND_20_CAD543 | ................................................................................ | [ 72480] |
| LR899193.1_Germany_2020/1 | ................................................................................ | [ 72480] |
| MN393477.1_Wuhan_2019 | ................................................................................ | [ 72480] |
| MW396979.1_Timor-Leste/2019/1 | ................................................................................ | [ 72480] |
| MK940252.1_CN/2019/InnerMongolia | ................................................................................ | [ 72480] |
| MK128995.1_China/2018/AnhuiXCGQ | ................................................................................ | [ 72480] |
| MH681419.1_ASFV/POL/2015 | ................................................................................ | [ 72480] |
| LS478113.1_Estonia_2014 | ................................................................................ | [ 72480] |
| KP843857.1_Odintsovo_02/14 | ................................................................................ | [ 72480] |

```
FN557520.1_strain_E75           ................................................c.......................c....... [ 72480]
AY261366.1_Warthog              ....................................a...........c............................... [ 72480]
AY261365.1_Warmbaths            ................................................c............................... [ 72480]
AY261364.1_Tengani_62           ................................................c............................... [ 72480]
AY261363.1_Pretoriuskop/96/4    ................................................c............................... [ 72480]
AY261362.1_Mkuzi_1979           ................................................c............................... [ 72480]
AY261361.1_Malawi_Lil-20/1_1983 c...............................................c.............................c. [ 72480]
AY261360.1_Kenya_1950           c........c......................................c.......................c.....c. [ 72480]
AM712240.1_OURT_88/3            ................................................c.......................c....... [ 72480]
AM712239.1_Benin_97/1           ................................................c.......................c....... [ 72480]
KM111295.1_Ken06.Bus            c...............................................c.......................c..g..c. [ 72480]
KM111294.1_Ken05/Tk1            c........c.......a..............................c.............................c. [ 72480]
KM262844.1_strain_L60           ................................................c.......................c....... [ 72480]
KM102979.1_26544/OG10_Italy     ................................................c.......................c....... [ 72480]
KM262845.1_strain_NHV           ................................................c.......................c....... [ 72480]
KX354450.1_47/Ss/2008           ................................................c.......................c....... [ 72480]
KP055815.1_strain_BA71          ................................................c.......................c....... [ 72480]
MH025920.1_strain_R35           c...............................................c.......................c..g..c. [ 72480]
MH025919.1_strain_N10           c...............................................c.......................c..g..c. [ 72480]
MH025918.1_strain_R25           c...............................................c.......................c..g..c. [ 72480]
MH025917.1_strain_R7            c...............................................c.......................c..g..c. [ 72480]
MH025916.1_strain_R8            c...............................................c.......................c..g..c. [ 72480]

FR682468.2_Georgia_2007/1       gtaccaaacaagcacgacacgtattgaaaaaaataaacttagtgttcgaggaagagtctattgaaaattggaaggtttca [ 72560]
ASF_IND_20_CAD543               ................................................................................ [ 72560]
LR899193.1_Germany_2020/1       ................................................................................ [ 72560]
MN393477.1_Wuhan_2019           ................................................................................ [ 72560]
MW396979.1_Timor-Leste/2019/1   ................................................................................ [ 72560]
MK940252.1_CN/2019/InnerMongolia ................................................................................ [ 72560]
MK128995.1_China/2018/AnhuiXCGQ ................................................................................ [ 72560]
MH681419.1_ASFV/POL/2015        ................................................................................ [ 72560]
LS478113.1_Estonia_2014         ................................................................................ [ 72560]
KP843857.1_Odintsovo_02/14      ................................................................................ [ 72560]
FN557520.1_strain_E75           ................................................................................ [ 72560]
AY261366.1_Warthog              ....t....................a...................................................... [ 72560]
AY261365.1_Warmbaths            ....t........................................................................... [ 72560]
AY261364.1_Tengani_62           ................................................................................ [ 72560]
AY261363.1_Pretoriuskop/96/4    ....t........................................................................... [ 72560]
AY261362.1_Mkuzi_1979           ....t........................................................................... [ 72560]
AY261361.1_Malawi_Lil-20/1_1983 ....t..............t............................................................ [ 72560]
AY261360.1_Kenya_1950           ....t..............t..............................................c............. [ 72560]
AM712240.1_OURT_88/3            ................................................................................ [ 72560]
AM712239.1_Benin_97/1           ................................................................................ [ 72560]
KM111295.1_Ken06.Bus            ....t..............t.............................a.....a..........c............. [ 72560]
```

```
KM111294.1_Ken05/Tk1           ....t..............t.......................a......................c............. [ 72560]
KM262844.1_strain_L60          ................................................................................ [ 72560]
KM102979.1_26544/OG10_Italy    ................................................................................ [ 72560]
KM262845.1_strain_NHV          ................................................................................ [ 72560]
KX354450.1_47/Ss/2008          ................................................................................ [ 72560]
KP055815.1_strain_BA71         ................................................................................ [ 72560]
MH025920.1_strain_R35          ....t..............t.............................a.....a..........c............. [ 72560]
MH025919.1_strain_N10          ....t..............t.............................a.....a..........c............. [ 72560]
MH025918.1_strain_R25          ....t..............t.............................a.....a..........c............. [ 72560]
MH025917.1_strain_R7           ....t..............t.............................a.....a..........c............. [ 72560]
MH025916.1_strain_R8           ....t..............t.............................a.....a..........c............. [ 72560]

FR682468.2_Georgia_2007/1      gtaaatactgtgttcccccatgttcacaacagattatctgcgccgaaactttccatcgacgaggctaatgaagccgtaga [ 72640]
ASF_IND_20_CAD543              ...........r.................................................................... [ 72640]
LR899193.1_Germany_2020/1      ................................................................................ [ 72640]
MN393477.1_Wuhan_2019          ................................................................................ [ 72640]
MW396979.1_Timor-Leste/2019/1  ................................................................................ [ 72640]
MK940252.1_CN/2019/InnerMongolia ................................................................................ [ 72640]
MK128995.1_China/2018/AnhuiXCGQ ................................................................................ [ 72640]
MH681419.1_ASFV/POL/2015       ................................................................................ [ 72640]
LS478113.1_Estonia_2014        ................................................................................ [ 72640]
KP843857.1_Odintsovo_02/14     ................................................................................ [ 72640]
FN557520.1_strain_E75          ...........a....................................t..........t.....c.............. [ 72640]
AY261366.1_Warthog             ............................................a................................... [ 72640]
AY261365.1_Warmbaths           ................................................................................ [ 72640]
AY261364.1_Tengani_62          ................................................................................ [ 72640]
AY261363.1_Pretoriuskop/96/4   ............................................a........g.......................... [ 72640]
AY261362.1_Mkuzi_1979          ................................................t..........t.....c.............. [ 72640]
AY261361.1_Malawi_Lil-20/1_1983 .................t...........t...c..a......................t.....c.............. [ 72640]
AY261360.1_Kenya_1950          .................t..c........t...................................c.............. [ 72640]
AM712240.1_OURT_88/3           ...........a....................................t..........t.....c.............. [ 72640]
AM712239.1_Benin_97/1          ...........a....................................t..........t.....c.............. [ 72640]
KM111295.1_Ken06.Bus           .................t..c........t...................................c.............. [ 72640]
KM111294.1_Ken05/Tk1           .....c...........t..c........t...................................c........t..... [ 72640]
KM262844.1_strain_L60          ...........a....................................t..........t.....c.............. [ 72640]
KM102979.1_26544/OG10_Italy    ...........a....................................t..........t.....c.............. [ 72640]
KM262845.1_strain_NHV          ...........a....................................t..........t.....c.............. [ 72640]
KX354450.1_47/Ss/2008          ...........a....................................t..........t.....c.............. [ 72640]
KP055815.1_strain_BA71         ...........a....................................t..........t.....c.............. [ 72640]
MH025920.1_strain_R35          .................t..c........t...................................c.............. [ 72640]
MH025919.1_strain_N10          .................t..c........t...................................c.............. [ 72640]
MH025918.1_strain_R25          .................t..c........t...................................c.............. [ 72640]
MH025917.1_strain_R7           .................t..c........t...................................c.............. [ 72640]
MH025916.1_strain_R8           .................t..c........t...................................c.............. [ 72640]
```

```
FR682468.2_Georgia_2007/1        aaagtttttgatacaagcaggacgactcatgtctctgtaaatgtctct-------------------------------- [ 72720]
ASF_IND_20_CAD543                ...............................................--------------------------------- [ 72720]
LR899193.1_Germany_2020/1        ................................................-------------------------------- [ 72720]
MN393477.1_Wuhan_2019            ................................................-------------------------------- [ 72720]
MW396979.1_Timor-Leste/2019/1    ................................................-------------------------------- [ 72720]
MK940252.1_CN/2019/InnerMongolia ................................................-------------------------------- [ 72720]
MK128995.1_China/2018/AnhuiXCGQ  ................................................-------------------------------- [ 72720]
MH681419.1_ASFV/POL/2015         ................................................-------------------------------- [ 72720]
LS478113.1_Estonia_2014          ................................................-------------------------------- [ 72720]
KP843857.1_Odintsovo_02/14       ................................................-------------------------------- [ 72720]
FN557520.1_strain_E75            .....................g.........................--------------------------------- [ 72720]
AY261366.1_Warthog               ........................................g......--------------------------------- [ 72720]
AY261365.1_Warmbaths             ........................................g.-------------------------------------- [ 72720]
AY261364.1_Tengani_62            ..g.............................................-------------------------------- [ 72720]
AY261363.1_Pretoriuskop/96/4     ...a...........................................--------------------------------- [ 72720]
AY261362.1_Mkuzi_1979            .....................g..................g......--------------------------------- [ 72720]
AY261361.1_Malawi_Lil-20/1_1983  ................................................-------------------------------- [ 72720]
AY261360.1_Kenya_1950            ...............g.....g...................a....tc-------------------------------- [ 72720]
AM712240.1_OURT_88/3             .....................g.........................--------------------------------- [ 72720]
AM712239.1_Benin_97/1            .....................g.........................--------------------------------- [ 72720]
KM111295.1_Ken06.Bus             ...............g.....g..............a....a.....c-------------------------------- [ 72720]
KM111294.1_Ken05/Tk1             ...............g.....g...................a....tctcctttattgaagaagtctcttcccttgtcgg [ 72720]
KM262844.1_strain_L60            .....................g.........................--------------------------------- [ 72720]
KM102979.1_26544/OG10_Italy      .....................g.........................--------------------------------- [ 72720]
KM262845.1_strain_NHV            .....................g.........................--------------------------------- [ 72720]
KX354450.1_47/Ss/2008            .....................g.........................--------------------------------- [ 72720]
KP055815.1_strain_BA71           .....................g.........................--------------------------------- [ 72720]
MH025920.1_strain_R35            ...............g.....g..............a....a..g..c-------------------------------- [ 72720]
MH025919.1_strain_N10            ...............g.....g..............a....a..g..c-------------------------------- [ 72720]
MH025918.1_strain_R25            ...............g.....g..............a....a..g..c-------------------------------- [ 72720]
MH025917.1_strain_R7             ...............g.....g..............a....a..g..c-------------------------------- [ 72720]
MH025916.1_strain_R8             ...............g.....g..............a....a..g..c-------------------------------- [ 72720]

FR682468.2_Georgia_2007/1        ----------tcctttatgggtgacgtctct------------------------------------------------- [ 72800]
ASF_IND_20_CAD543                ------------...................gt----------------------------------------------- [ 72800]
LR899193.1_Germany_2020/1        ----------.....................------------------------------------------------- [ 72800]
MN393477.1_Wuhan_2019            ----------.....................------------------------------------------------- [ 72800]
MW396979.1_Timor-Leste/2019/1    ----------.....................------------------------------------------------- [ 72800]
MK940252.1_CN/2019/InnerMongolia ----------.....................------------------------------------------------- [ 72800]
MK128995.1_China/2018/AnhuiXCGQ  ----------.....................------------------------------------------------- [ 72800]
MH681419.1_ASFV/POL/2015         ----------.....................------------------------------------------------- [ 72800]
LS478113.1_Estonia_2014          ----------.....................------------------------------------------------- [ 72800]
KP843857.1_Odintsovo_02/14       ----------.....................------------------------------------------------- [ 72800]
```

```
FN557520.1_strain_E75             ------------.........c..t......gt----------------------------------------------- [ 72800]
AY261366.1_Warthog                ------------...................------------------------------------------------- [ 72800]
AY261365.1_Warmbaths              -------------------------..t...------------------------------------------------- [ 72800]
AY261364.1_Tengani_62             ----------.....................------------------------------------------------- [ 72800]
AY261363.1_Pretoriuskop/96/4      ------------...................------------------------------------------------- [ 72800]
AY261362.1_Mkuzi_1979             ------------............t......gt----------------------------------------------- [ 72800]
AY261361.1_Malawi_Lil-20/1_1983   ----------........t.aa..a......tcctttattgagcaagtctcttcttttatcggagaaatctcttccttta [ 72800]
AY261360.1_Kenya_1950             ----------........c.aa..a......tc----------------------------------------------- [ 72800]
AM712240.1_OURT_88/3              ------------.........c..t......gt----------------------------------------------- [ 72800]
AM712239.1_Benin_97/1             ------------.........c..t......gt----------------------------------------------- [ 72800]
KM111295.1_Ken06.Bus              ----------.tt.....c.aa..a......tc----------------------------------------------- [ 72800]
KM111294.1_Ken05/Tk1              agaagactct........c.aa..a......tc----------------------------------------------- [ 72800]
KM262844.1_strain_L60             ------------.........c..t......gt----------------------------------------------- [ 72800]
KM102979.1_26544/OG10_Italy       ------------.........c..t......gt----------------------------------------------- [ 72800]
KM262845.1_strain_NHV             ------------.........c..t......gt----------------------------------------------- [ 72800]
KX354450.1_47/Ss/2008             ------------.........c..t......gt----------------------------------------------- [ 72800]
KP055815.1_strain_BA71            ------------.........c..t......gt----------------------------------------------- [ 72800]
MH025920.1_strain_R35             ----------.tt.....c.aa..a......tc----------------------------------------------- [ 72800]
MH025919.1_strain_N10             ----------.tt.....c.aa..a......tc----------------------------------------------- [ 72800]
MH025918.1_strain_R25             ----------.tt.....c.aa..a......tc----------------------------------------------- [ 72800]
MH025917.1_strain_R7              ----------.tt.....c.aa..a......tc----------------------------------------------- [ 72800]
MH025916.1_strain_R8              ----------.tt.....c.aa..a......tc----------------------------------------------- [ 72800]

FR682468.2_Georgia_2007/1         -------------------------------------------------------------------------------- [ 72880]
ASF_IND_20_CAD543                 -------------------------------------------------------------------------------- [ 72880]
LR899193.1_Germany_2020/1         -------------------------------------------------------------------------------- [ 72880]
MN393477.1_Wuhan_2019             -------------------------------------------------------------------------------- [ 72880]
MW396979.1_Timor-Leste/2019/1     -------------------------------------------------------------------------------- [ 72880]
MK940252.1_CN/2019/InnerMongolia  -------------------------------------------------------------------------------- [ 72880]
MK128995.1_China/2018/AnhuiXCGQ   -------------------------------------------------------------------------------- [ 72880]
MH681419.1_ASFV/POL/2015          -------------------------------------------------------------------------------- [ 72880]
LS478113.1_Estonia_2014           -------------------------------------------------------------------------------- [ 72880]
KP843857.1_Odintsovo_02/14        -------------------------------------------------------------------------------- [ 72880]
FN557520.1_strain_E75             -------------------------------------------------------------------------------- [ 72880]
AY261366.1_Warthog                -------------------------------------------------------------------------------- [ 72880]
AY261365.1_Warmbaths              -------------------------------------------------------------------------------- [ 72880]
AY261364.1_Tengani_62             -------------------------------------------------------------------------------- [ 72880]
AY261363.1_Pretoriuskop/96/4      -------------------------------------------------------------------------------- [ 72880]
AY261362.1_Mkuzi_1979             -------------------------------------------------------------------------------- [ 72880]
AY261361.1_Malawi_Lil-20/1_1983   ttgagcaagtctcttcttttatcggagaaatctcttcctttattgagcaagtctcttcctttgttgaataagtctcttcc [ 72880]
AY261360.1_Kenya_1950             -------------------------------------------------------------------------------- [ 72880]
AM712240.1_OURT_88/3              -------------------------------------------------------------------------------- [ 72880]
AM712239.1_Benin_97/1             -------------------------------------------------------------------------------- [ 72880]
KM111295.1_Ken06.Bus              -------------------------------------------------------------------------------- [ 72880]
```

```
KM111294.1_Ken05/Tk1              -------------------------------------------------------------------------------- [ 72880]
KM262844.1_strain_L60             -------------------------------------------------------------------------------- [ 72880]
KM102979.1_26544/OG10_Italy       -------------------------------------------------------------------------------- [ 72880]
KM262845.1_strain_NHV             -------------------------------------------------------------------------------- [ 72880]
KX354450.1_47/Ss/2008             -------------------------------------------------------------------------------- [ 72880]
KP055815.1_strain_BA71            -------------------------------------------------------------------------------- [ 72880]
MH025920.1_strain_R35             -------------------------------------------------------------------------------- [ 72880]
MH025919.1_strain_N10             -------------------------------------------------------------------------------- [ 72880]
MH025918.1_strain_R25             -------------------------------------------------------------------------------- [ 72880]
MH025917.1_strain_R7              -------------------------------------------------------------------------------- [ 72880]
MH025916.1_strain_R8              -------------------------------------------------------------------------------- [ 72880]

FR682468.2_Georgia_2007/1         ---------------------------------------tcctttgccgaggaagtctctgttatgggcaagaggtttga [ 72960]
ASF_IND_20_CAD543                 -----------------------------aagtgtctcc......r.y................................ [ 72960]
LR899193.1_Germany_2020/1         ---------------------------------------......................................... [ 72960]
MN393477.1_Wuhan_2019             ---------------------------------------......................................... [ 72960]
MW396979.1_Timor-Leste/2019/1     ---------------------------------------......................................... [ 72960]
MK940252.1_CN/2019/InnerMongolia  ---------------------------------------......................................... [ 72960]
MK128995.1_China/2018/AnhuiXCGQ   ---------------------------------------......................................... [ 72960]
MH681419.1_ASFV/POL/2015          ---------------------------------------......................................... [ 72960]
LS478113.1_Estonia_2014           ---------------------------------------......................................... [ 72960]
KP843857.1_Odintsovo_02/14        ---------------------------------------......................................... [ 72960]
FN557520.1_strain_E75             -----------------------------aagtgtctcc......a.t................................ [ 72960]
AY261366.1_Warthog                ---------------------------------------......a.................................. [ 72960]
AY261365.1_Warmbaths              ---------------------------------------......a.................................. [ 72960]
AY261364.1_Tengani_62             ---------------------------------------......................................... [ 72960]
AY261363.1_Pretoriuskop/96/4      ---------------------------------------......a.................................. [ 72960]
AY261362.1_Mkuzi_1979             -----------------------------aagtgtctcc......a.t....................t........... [ 72960]
AY261361.1_Malawi_Lil-20/1_1983   tttgttgaataagtctcttcctttgtcgaagaaatctct.......t...a..c.a........................ [ 72960]
AY261360.1_Kenya_1950             --------------------ccttgtcggagaagactct.......t....a.g..............t..a........ [ 72960]
AM712240.1_OURT_88/3              -----------------------------aagtgtctcc......a.t................................ [ 72960]
AM712239.1_Benin_97/1             -----------------------------aagtgtctcc......a.t................................ [ 72960]
KM111295.1_Ken06.Bus              --------------------ctttatcggaaaagtctct......att....................t........... [ 72960]
KM111294.1_Ken05/Tk1              --------------------ttttgtcggagaagtctct.......t....a.g..............t..a........ [ 72960]
KM262844.1_strain_L60             -----------------------------aagtgtctcc......a.t................................ [ 72960]
KM102979.1_26544/OG10_Italy       -----------------------------aagtgtctcc......a.t................................ [ 72960]
KM262845.1_strain_NHV             -----------------------------aagtgtctcc......a.t................................ [ 72960]
KX354450.1_47/Ss/2008             -----------------------------aagtgtctcc......a.t................................ [ 72960]
KP055815.1_strain_BA71            -----------------------------aagtgtctcc......a.t................................ [ 72960]
MH025920.1_strain_R35             --------------------ctttatcggaaaagtctct......att....................t........... [ 72960]
MH025919.1_strain_N10             --------------------ctttatcggaaaagtctct......att....................t........... [ 72960]
MH025918.1_strain_R25             --------------------ctttatcggaaaagtctct......att....................t........... [ 72960]
MH025917.1_strain_R7              --------------------ctttatcggaaaagtctct......att....................t........... [ 72960]
MH025916.1_strain_R8              --------------------ctttatcggaaaagtctct......att....................t........... [ 72960]
```

```
FR682468.2_Georgia_2007/1        aacaacgcaaggactctgcttaatctgctgtctcacaaagggaatcaaactacctgctttcgtatttttaatgtagtaat [ 73040]
ASF_IND_20_CAD543                ................................................................................ [ 73040]
LR899193.1_Germany_2020/1        ................................................................................ [ 73040]
MN393477.1_Wuhan_2019            ................................................................................ [ 73040]
MW396979.1_Timor-Leste/2019/1    ................................................................................ [ 73040]
MK940252.1_CN/2019/InnerMongolia ................................................................................ [ 73040]
MK128995.1_China/2018/AnhuiXCGQ  ................................................................................ [ 73040]
MH681419.1_ASFV/POL/2015         ................................................................................ [ 73040]
LS478113.1_Estonia_2014          ................................................................................ [ 73040]
KP843857.1_Odintsovo_02/14       ................................................................................ [ 73040]
FN557520.1_strain_E75            .........g..g................................................................... [ 73040]
AY261366.1_Warthog               .........g..g....................t.....a........................................ [ 73040]
AY261365.1_Warmbaths             .........g..g....................t.............................................. [ 73040]
AY261364.1_Tengani_62            ................................................................................ [ 73040]
AY261363.1_Pretoriuskop/96/4     .........g..g....................t.............................................. [ 73040]
AY261362.1_Mkuzi_1979            .........g..g................................................................... [ 73040]
AY261361.1_Malawi_Lil-20/1_1983  .........g..g............g...............................c...................... [ 73040]
AY261360.1_Kenya_1950            .........g.................t..c..t....................c..c..............a....... [ 73040]
AM712240.1_OURT_88/3             .........g..g................................................................... [ 73040]
AM712239.1_Benin_97/1            .........g..g...............................................t................... [ 73040]
KM111295.1_Ken06.Bus             .........g..g.........................................c..c...................... [ 73040]
KM111294.1_Ken05/Tk1             .........g..g............g............................c..c.....g................ [ 73040]
KM262844.1_strain_L60            .........g..g................................................................... [ 73040]
KM102979.1_26544/OG10_Italy      .........g..g................................................................... [ 73040]
KM262845.1_strain_NHV            .........g..g................................................................... [ 73040]
KX354450.1_47/Ss/2008            .........g..g................................................................... [ 73040]
KP055815.1_strain_BA71           .........g..g................................................................... [ 73040]
MH025920.1_strain_R35            .........g..g.........................................c..c...................... [ 73040]
MH025919.1_strain_N10            .........g..g.........................................c..c...................... [ 73040]
MH025918.1_strain_R25            .........g..g.........................................c..c...................... [ 73040]
MH025917.1_strain_R7             .........g..g.........................................c..c...................... [ 73040]
MH025916.1_strain_R8             .........g..g.........................................c..c...................... [ 73040]

FR682468.2_Georgia_2007/1        tacccttgttgtgatgaattttaagaccatagcgtagtcccagtactttattaatgaattttaaaattgtttgagggtcc [ 73120]
ASF_IND_20_CAD543                .........................................................................g..a... [ 73120]
LR899193.1_Germany_2020/1        ................................................................................ [ 73120]
MN393477.1_Wuhan_2019            ................................................................................ [ 73120]
MW396979.1_Timor-Leste/2019/1    ................................................................................ [ 73120]
MK940252.1_CN/2019/InnerMongolia ................................................................................ [ 73120]
MK128995.1_China/2018/AnhuiXCGQ  ................................................................................ [ 73120]
MH681419.1_ASFV/POL/2015         ................................................................................ [ 73120]
LS478113.1_Estonia_2014          ................................................................................ [ 73120]
KP843857.1_Odintsovo_02/14       ................................................................................ [ 73120]
```

```
FN557520.1_strain_E75            ..........a..g.......................a.....c.............................g..a... [ 73120]
AY261366.1_Warthog               .............g.............................c...................................t [ 73120]
AY261365.1_Warmbaths             .............g.............................c.................................... [ 73120]
AY261364.1_Tengani_62            ................................................................................ [ 73120]
AY261363.1_Pretoriuskop/96/4     .............g.............................c.................................... [ 73120]
AY261362.1_Mkuzi_1979            ..........a..g.......................a.....c.............................g..a... [ 73120]
AY261361.1_Malawi_Lil-20/1_1983  .............g...........g.................c.............................g..a... [ 73120]
AY261360.1_Kenya_1950            .............g......................t.....ac.............................g...... [ 73120]
AM712240.1_OURT_88/3             ..........a..g.......................a.....c.............................g..a... [ 73120]
AM712239.1_Benin_97/1            ..........a..g.......................a.....c.............................g..a... [ 73120]
KM111295.1_Ken06.Bus             .............g......................t.....ac.............................g..a... [ 73120]
KM111294.1_Ken05/Tk1             .............g......................t.....ac.............................g...... [ 73120]
KM262844.1_strain_L60            ..........a..g.......................a.....c.............................g..a... [ 73120]
KM102979.1_26544/OG10_Italy      ..........a..g.......................a.....c.............................g..a... [ 73120]
KM262845.1_strain_NHV            ..........a..g.......................a.....c.............................g..a... [ 73120]
KX354450.1_47/Ss/2008            ..........a..g.......................a.....c.............................g..a... [ 73120]
KP055815.1_strain_BA71           ..........a..g.......................a.....c.............................g..a... [ 73120]
MH025920.1_strain_R35            .............g......................t.....ac.............................g..a... [ 73120]
MH025919.1_strain_N10            .............g......................t.....ac.............................g..a... [ 73120]
MH025918.1_strain_R25            .............g......................t.....ac.............................g..a... [ 73120]
MH025917.1_strain_R7             .............g......................t.....ac.............................g..a... [ 73120]
MH025916.1_strain_R8             .............g......................t.....ac.............................g..a... [ 73120]

FR682468.2_Georgia_2007/1        gttttattgggctttttaagcttaaactcaaagctgatcgcgcttaaatcatactgaacaaattcatcaacgagtttcgt [ 73200]
ASF_IND_20_CAD543                ................................................................................ [ 73200]
LR899193.1_Germany_2020/1        ................................................................................ [ 73200]
MN393477.1_Wuhan_2019            ................................................................................ [ 73200]
MW396979.1_Timor-Leste/2019/1    ................................................................................ [ 73200]
MK940252.1_CN/2019/InnerMongolia ................................................................................ [ 73200]
MK128995.1_China/2018/AnhuiXCGQ  ................................................................................ [ 73200]
MH681419.1_ASFV/POL/2015         ................................................................................ [ 73200]
LS478113.1_Estonia_2014          ................................................................................ [ 73200]
KP843857.1_Odintsovo_02/14       ................................................................................ [ 73200]
FN557520.1_strain_E75            ......c.a............................c.......................................t.. [ 73200]
AY261366.1_Warthog               .....g..a............................c.........................................c [ 73200]
AY261365.1_Warmbaths             .....g..a............................c.........................................c [ 73200]
AY261364.1_Tengani_62            ................................................................................ [ 73200]
AY261363.1_Pretoriuskop/96/4     ........a............................c...............t.........................c [ 73200]
AY261362.1_Mkuzi_1979            ........a............................c.......................................t.. [ 73200]
AY261361.1_Malawi_Lil-20/1_1983  a....g..a.................t..........c.......................................... [ 73200]
AY261360.1_Kenya_1950            a.......a..........................a.ct......................................... [ 73200]
AM712240.1_OURT_88/3             ......c.a............................c.......................................t.. [ 73200]
AM712239.1_Benin_97/1            ......c.a............................c.......................................t.. [ 73200]
KM111295.1_Ken06.Bus             a.......a..........................a.ct............................t............ [ 73200]
```

```
KM111294.1_Ken05/Tk1             a.......a..........................a.ct......................................... [ 73200]
KM262844.1_strain_L60            ......c.a............................c.......................................t.. [ 73200]
KM102979.1_26544/OG10_Italy      ......c.a............................c.......................................t.. [ 73200]
KM262845.1_strain_NHV            ......c.a............................c.......................................t.. [ 73200]
KX354450.1_47/Ss/2008            ......c.a............................c.......................................t.. [ 73200]
KP055815.1_strain_BA71           ......c.a............................c.......................................t.. [ 73200]
MH025920.1_strain_R35            a.......a..........................a.ct............................t............ [ 73200]
MH025919.1_strain_N10            a.......a..........................a.ct............................t............ [ 73200]
MH025918.1_strain_R25            a.......a..........................a.ct............................t............ [ 73200]
MH025917.1_strain_R7             a.......a..........................a.ct............................t............ [ 73200]
MH025916.1_strain_R8             a.......a..........................a.ct............................t............ [ 73200]

FR682468.2_Georgia_2007/1        cattaattgttcattggtcaatatattagggtcctgaacgcatttaaagccgcacttagttaatagcataatagcgtaca [ 73280]
ASF_IND_20_CAD543                ................................................................................ [ 73280]
LR899193.1_Germany_2020/1        ................................................................................ [ 73280]
MN393477.1_Wuhan_2019            ................................................................................ [ 73280]
MW396979.1_Timor-Leste/2019/1    ................................................................................ [ 73280]
MK940252.1_CN/2019/InnerMongolia ................................................................................ [ 73280]
MK128995.1_China/2018/AnhuiXCGQ  ................................................................................ [ 73280]
MH681419.1_ASFV/POL/2015         ................................................................................ [ 73280]
LS478113.1_Estonia_2014          ................................................................................ [ 73280]
KP843857.1_Odintsovo_02/14       ................................................................................ [ 73280]
FN557520.1_strain_E75            ...c...........................................................c........g....... [ 73280]
AY261366.1_Warthog               ...c...........................................................c........g....... [ 73280]
AY261365.1_Warmbaths             ...c.....................................................t.....c........g....... [ 73280]
AY261364.1_Tengani_62            ...............................................................c........g....... [ 73280]
AY261363.1_Pretoriuskop/96/4     ...c...........................................................c................ [ 73280]
AY261362.1_Mkuzi_1979            ...c...........................................................c........g....... [ 73280]
AY261361.1_Malawi_Lil-20/1_1983  ...c...........a.................t.................a...........c........c....... [ 73280]
AY261360.1_Kenya_1950            ...c.....c.........................................a...........c........g....... [ 73280]
AM712240.1_OURT_88/3             ...c...........................................................c........g....... [ 73280]
AM712239.1_Benin_97/1            ...c...........................................................c........g....... [ 73280]
KM111295.1_Ken06.Bus             ...c.....c..........................g..........................c...........a.... [ 73280]
KM111294.1_Ken05/Tk1             ...c.....c.........................................a...........c........g....... [ 73280]
KM262844.1_strain_L60            ...c...........................................................c........g....... [ 73280]
KM102979.1_26544/OG10_Italy      ...c...........................................................c........g....... [ 73280]
KM262845.1_strain_NHV            ...c...........................................................c........g....... [ 73280]
KX354450.1_47/Ss/2008            ...c...........................................................c........g....... [ 73280]
KP055815.1_strain_BA71           ...c...........................................................c........g....... [ 73280]
MH025920.1_strain_R35            ...c.....c..........................g..........................c...........a.... [ 73280]
MH025919.1_strain_N10            ...c.....c..........................g..........................c...........a.... [ 73280]
MH025918.1_strain_R25            ...c.....c..........................g..........................c...........a.... [ 73280]
MH025917.1_strain_R7             ...c.....c..........................g..........................c...........a.... [ 73280]
MH025916.1_strain_R8             ...c.....c..........................g..........................c...........a.... [ 73280]
```

```
FR682468.2_Georgia_2007/1        tatgagattgaaaactataattaaattgtagatcatgatgctctgcgtgttgcatggcccattgatgaaagtttaattcc [ 73360]
ASF_IND_20_CAD543                ................................................................................ [ 73360]
LR899193.1_Germany_2020/1        ................................................................................ [ 73360]
MN393477.1_Wuhan_2019            ................................................................................ [ 73360]
MW396979.1_Timor-Leste/2019/1    ................................................................................ [ 73360]
MK940252.1_CN/2019/InnerMongolia ................................................................................ [ 73360]
MK128995.1_China/2018/AnhuiXCGQ  ................................................................................ [ 73360]
MH681419.1_ASFV/POL/2015         ................................................................................ [ 73360]
LS478113.1_Estonia_2014          ................................................................................ [ 73360]
KP843857.1_Odintsovo_02/14       ................................................................................ [ 73360]
FN557520.1_strain_E75            ....g........g..............c.................a....................g............ [ 73360]
AY261366.1_Warthog               ....g........g.........t....c.................a................................. [ 73360]
AY261365.1_Warmbaths             ....g........g.........t....c.................a....................g............ [ 73360]
AY261364.1_Tengani_62            .............g.................................................................. [ 73360]
AY261363.1_Pretoriuskop/96/4     ....g........g.........t....c..g........t.....a....................g............ [ 73360]
AY261362.1_Mkuzi_1979            ....g........g..............c.................a....................g............ [ 73360]
AY261361.1_Malawi_Lil-20/1_1983  ....g........g..............c.................a........a........g..g........c... [ 73360]
AY261360.1_Kenya_1950            ....g........g..............c.................a.................g..g............ [ 73360]
AM712240.1_OURT_88/3             ....g........g..............c.................a....................g............ [ 73360]
AM712239.1_Benin_97/1            ....g........g..............c.................a....................g............ [ 73360]
KM111295.1_Ken06.Bus             ....g........g..............c.................a.................g..g............ [ 73360]
KM111294.1_Ken05/Tk1             ....g........g..............c.................a.................g..g............ [ 73360]
KM262844.1_strain_L60            ....g........g..............c.................a....................g............ [ 73360]
KM102979.1_26544/OG10_Italy      ....g........g..............c.................a....................g............ [ 73360]
KM262845.1_strain_NHV            ....g........g..............c.................a....................g............ [ 73360]
KX354450.1_47/Ss/2008            ....g........g..............c.................a....................g............ [ 73360]
KP055815.1_strain_BA71           ....g........g..............c.................a....................g............ [ 73360]
MH025920.1_strain_R35            ....g........g..............c.................a.................g..g............ [ 73360]
MH025919.1_strain_N10            ....g........g..............c.................a.................g..g............ [ 73360]
MH025918.1_strain_R25            ....g........g..............c.................a.................g..g............ [ 73360]
MH025917.1_strain_R7             ....g........g..............c.................a.................g..g............ [ 73360]
MH025916.1_strain_R8             ....g........g..............c.................a.................g..g............ [ 73360]

FR682468.2_Georgia_2007/1        tgagtttgtaacatagtgagcgactcgtatactgtctttccgcggcttatttggacacggccagtatagttctgttttgt [ 73440]
ASF_IND_20_CAD543                ................................................................................ [ 73440]
LR899193.1_Germany_2020/1        ................................................................................ [ 73440]
MN393477.1_Wuhan_2019            ................................................................................ [ 73440]
MW396979.1_Timor-Leste/2019/1    ................................................................................ [ 73440]
MK940252.1_CN/2019/InnerMongolia ................................................................................ [ 73440]
MK128995.1_China/2018/AnhuiXCGQ  ................................................................................ [ 73440]
MH681419.1_ASFV/POL/2015         ................................................................................ [ 73440]
LS478113.1_Estonia_2014          ................................................................................ [ 73440]
KP843857.1_Odintsovo_02/14       ................................................................................ [ 73440]
```

```
FN557520.1_strain_E75            .......................t...........t.............................g.............. [ 73440]
AY261366.1_Warthog               .......................t...........t.....a....................g..g.............. [ 73440]
AY261365.1_Warmbaths             ...........g.....a.....t...........t.....a....................g..g.............. [ 73440]
AY261364.1_Tengani_62            ................................................................................ [ 73440]
AY261363.1_Pretoriuskop/96/4     .......................t...........t.....a....................g..g.............. [ 73440]
AY261362.1_Mkuzi_1979            .......................t...........t.............................g.............. [ 73440]
AY261361.1_Malawi_Lil-20/1_1983  .................a.....t...........t.................a.....a.....g.............. [ 73440]
AY261360.1_Kenya_1950            .................a.....t....g......t.................a........c..g.............. [ 73440]
AM712240.1_OURT_88/3             .......................t...........t.............................g.............. [ 73440]
AM712239.1_Benin_97/1            .......................t...........t.............................g.............. [ 73440]
KM111295.1_Ken06.Bus             .................a.....t....g..t...t.................a.....a..c..g.............. [ 73440]
KM111294.1_Ken05/Tk1             .................a.....t....g......t.................a........c..g.............. [ 73440]
KM262844.1_strain_L60            .......................t...........t.............................g.............. [ 73440]
KM102979.1_26544/OG10_Italy      .......................t...........t.............................g.............. [ 73440]
KM262845.1_strain_NHV            .......................t...........t.............................g.............. [ 73440]
KX354450.1_47/Ss/2008            .......................t...........t.............................g.............. [ 73440]
KP055815.1_strain_BA71           .......................t...........t.............................g.............. [ 73440]
MH025920.1_strain_R35            .................a.....t....g..t...t.................a.....a..c..g.............. [ 73440]
MH025919.1_strain_N10            .................a.....t....g..t...t.................a.....a..c..g.............. [ 73440]
MH025918.1_strain_R25            .................a.....t....g..t...t.................a.....a..c..g.............. [ 73440]
MH025917.1_strain_R7             .................a.....t....g..t...t.................a.....a..c..g.............. [ 73440]
MH025916.1_strain_R8             .................a.....t....g..t...t.................a.....a..c..g.............. [ 73440]

FR682468.2_Georgia_2007/1        cataaaactattgtattgttcaacaaatttgggagtaattttatgaccgtgccatgcataaaattcgagtagtttatact [ 73520]
ASF_IND_20_CAD543                .................................r.............................................. [ 73520]
LR899193.1_Germany_2020/1        ................................................................................ [ 73520]
MN393477.1_Wuhan_2019            ................................................................................ [ 73520]
MW396979.1_Timor-Leste/2019/1    ................................................................................ [ 73520]
MK940252.1_CN/2019/InnerMongolia ................................................................................ [ 73520]
MK128995.1_China/2018/AnhuiXCGQ  ................................................................................ [ 73520]
MH681419.1_ASFV/POL/2015         ................................................................................ [ 73520]
LS478113.1_Estonia_2014          ................................................................................ [ 73520]
KP843857.1_Odintsovo_02/14       ................................................................................ [ 73520]
FN557520.1_strain_E75            .................................g............................................t. [ 73520]
AY261366.1_Warthog               ............a................................................................... [ 73520]
AY261365.1_Warmbaths             ................................................................................ [ 73520]
AY261364.1_Tengani_62            ................................................................................ [ 73520]
AY261363.1_Pretoriuskop/96/4     ............a................................................................... [ 73520]
AY261362.1_Mkuzi_1979            .................................g.............................c................ [ 73520]
AY261361.1_Malawi_Lil-20/1_1983  ...g........a....................g..........t................................... [ 73520]
AY261360.1_Kenya_1950            ............a....................g..........t................................... [ 73520]
AM712240.1_OURT_88/3             .................................g............................................t. [ 73520]
AM712239.1_Benin_97/1            .................................g............................................t. [ 73520]
KM111295.1_Ken06.Bus             ............a....................g..........t................................... [ 73520]
```

```
KM111294.1_Ken05/Tk1            ............a....................g.............................................. [ 73520]
KM262844.1_strain_L60           .................................g............................................t. [ 73520]
KM102979.1_26544/OG10_Italy     .................................g............................................t. [ 73520]
KM262845.1_strain_NHV           .................................g............................................t. [ 73520]
KX354450.1_47/Ss/2008           .................................g............................................t. [ 73520]
KP055815.1_strain_BA71          .................................g............................................t. [ 73520]
MH025920.1_strain_R35           ............a....................g..........t................................... [ 73520]
MH025919.1_strain_N10           ............a....................g..........t................................... [ 73520]
MH025918.1_strain_R25           ............a....................g..........t................................... [ 73520]
MH025917.1_strain_R7            ............a....................g..........t................................... [ 73520]
MH025916.1_strain_R8            ............a....................g..........t................................... [ 73520]

FR682468.2_Georgia_2007/1       tttcatacgcaaataggtcttgctggtctactgtgatgccttcctttaagttttgtttaatttgtaaagctttattggca [ 73600]
ASF_IND_20_CAD543               ................................................................................ [ 73600]
LR899193.1_Germany_2020/1       ................................................................................ [ 73600]
MN393477.1_Wuhan_2019           ................................................................................ [ 73600]
MW396979.1_Timor-Leste/2019/1   ................................................................................ [ 73600]
MK940252.1_CN/2019/InnerMongolia ................................................................................ [ 73600]
MK128995.1_China/2018/AnhuiXCGQ ................................................................................ [ 73600]
MH681419.1_ASFV/POL/2015        ................................................................................ [ 73600]
LS478113.1_Estonia_2014         ................................................................................ [ 73600]
KP843857.1_Odintsovo_02/14      ................................................................................ [ 73600]
FN557520.1_strain_E75           ................................................................................ [ 73600]
AY261366.1_Warthog              ..............................................................a................. [ 73600]
AY261365.1_Warmbaths            ..............................................................a..............a.. [ 73600]
AY261364.1_Tengani_62           ................................................................................ [ 73600]
AY261363.1_Pretoriuskop/96/4    ..............................................................a................. [ 73600]
AY261362.1_Mkuzi_1979           ................................................................................ [ 73600]
AY261361.1_Malawi_Lil-20/1_1983 ....................c........................................................... [ 73600]
AY261360.1_Kenya_1950           ................a.................a........t..................a..........g...... [ 73600]
AM712240.1_OURT_88/3            ................................................................................ [ 73600]
AM712239.1_Benin_97/1           ................................................................................ [ 73600]
KM111295.1_Ken06.Bus            ..........g..c.t................................................c............... [ 73600]
KM111294.1_Ken05/Tk1            ................a.................a........t..................a................. [ 73600]
KM262844.1_strain_L60           ................................................................................ [ 73600]
KM102979.1_26544/OG10_Italy     ................................................................................ [ 73600]
KM262845.1_strain_NHV           ................................................................................ [ 73600]
KX354450.1_47/Ss/2008           ................................................................................ [ 73600]
KP055815.1_strain_BA71          ................................................................................ [ 73600]
MH025920.1_strain_R35           ..........g..c.t................................................c............... [ 73600]
MH025919.1_strain_N10           ..........g..c.t................................................c............... [ 73600]
MH025918.1_strain_R25           ..........g..c.t................................................c............... [ 73600]
MH025917.1_strain_R7            ..........g..c.t................................................c............... [ 73600]
MH025916.1_strain_R8            ..........g..c.t................................................c............... [ 73600]
```

```
FR682468.2_Georgia_2007/1        tcaatggtttcagccgaggcaatgtttacatagtcctggtgtttaatttccattttaatgcttgtatattgtttgactgt [ 73680]
ASF_IND_20_CAD543                ................................................................................ [ 73680]
LR899193.1_Germany_2020/1        ................................................................................ [ 73680]
MN393477.1_Wuhan_2019            ................................................................................ [ 73680]
MW396979.1_Timor-Leste/2019/1    ................................................................................ [ 73680]
MK940252.1_CN/2019/InnerMongolia ................................................................................ [ 73680]
MK128995.1_China/2018/AnhuiXCGQ  ................................................................................ [ 73680]
MH681419.1_ASFV/POL/2015         ................................................................................ [ 73680]
LS478113.1_Estonia_2014          ................................................................................ [ 73680]
KP843857.1_Odintsovo_02/14       ................................................................................ [ 73680]
FN557520.1_strain_E75            .........................................c..........................c........... [ 73680]
AY261366.1_Warthog               ..............t.................a........c..........................c........... [ 73680]
AY261365.1_Warmbaths             .........................................c..........................c........... [ 73680]
AY261364.1_Tengani_62            ....................................................................c........... [ 73680]
AY261363.1_Pretoriuskop/96/4     ..............t.................a........c..........................c........... [ 73680]
AY261362.1_Mkuzi_1979            .........................................c........t.................c........... [ 73680]
AY261361.1_Malawi_Lil-20/1_1983  .....c................c...c................................a........c........g.. [ 73680]
AY261360.1_Kenya_1950            .....c.................a..g.....a........c.................a........c........c.. [ 73680]
AM712240.1_OURT_88/3             .........................................c..........................c........... [ 73680]
AM712239.1_Benin_97/1            .........................................c..........................c........... [ 73680]
KM111295.1_Ken06.Bus             .....a....................g.....a........c.................a........c........c.. [ 73680]
KM111294.1_Ken05/Tk1             .....c.................a..g.....a........c.................a........c........c.. [ 73680]
KM262844.1_strain_L60            .........................................c..........................c........... [ 73680]
KM102979.1_26544/OG10_Italy      .........................................c..........................c........... [ 73680]
KM262845.1_strain_NHV            .........................................c..........................c........... [ 73680]
KX354450.1_47/Ss/2008            .........................................c..........................c........... [ 73680]
KP055815.1_strain_BA71           .........................................c..........................c........... [ 73680]
MH025920.1_strain_R35            .....a....................g.....a........c.................a........c........c.. [ 73680]
MH025919.1_strain_N10            .....a....................g.....a........c.................a........c........c.. [ 73680]
MH025918.1_strain_R25            .....a....................g.....a........c.................a........c........c.. [ 73680]
MH025917.1_strain_R7             .....a....................g.....a........c.................a........c........c.. [ 73680]
MH025916.1_strain_R8             .....a....................g.....a........c.................a........c........c.. [ 73680]

FR682468.2_Georgia_2007/1        ctccagcttttcacccgtcagtataaacaccttagcgccggtgtcggcgatctggttaataaatcgggttataaagtgat [ 73760]
ASF_IND_20_CAD543                ................................................................................ [ 73760]
LR899193.1_Germany_2020/1        ................................................................................ [ 73760]
MN393477.1_Wuhan_2019            ................................................................................ [ 73760]
MW396979.1_Timor-Leste/2019/1    ................................................................................ [ 73760]
MK940252.1_CN/2019/InnerMongolia ................................................................................ [ 73760]
MK128995.1_China/2018/AnhuiXCGQ  ................................................................................ [ 73760]
MH681419.1_ASFV/POL/2015         ................................................................................ [ 73760]
LS478113.1_Estonia_2014          ................................................................................ [ 73760]
KP843857.1_Odintsovo_02/14       ................................................................................ [ 73760]
```

```
FN557520.1_strain_E75          ..................................................c...............t............. [ 73760]
AY261366.1_Warthog             ..................................................................t............. [ 73760]
AY261365.1_Warmbaths           ..............................t...................................t...........g. [ 73760]
AY261364.1_Tengani_62          ................................................................................ [ 73760]
AY261363.1_Pretoriuskop/96/4   ..................................................................t............. [ 73760]
AY261362.1_Mkuzi_1979          ..................................................c...a...........t............. [ 73760]
AY261361.1_Malawi_Lil-20/1_1983 ......t...........................................c...............t............. [ 73760]
AY261360.1_Kenya_1950          ...t..............................................c............a..t............. [ 73760]
AM712240.1_OURT_88/3           ..................................................c...............t............. [ 73760]
AM712239.1_Benin_97/1          ..................................................c...............t............. [ 73760]
KM111295.1_Ken06.Bus           ......t.............a........................a....ct..............t............. [ 73760]
KM111294.1_Ken05/Tk1           ...t..............................................c...............t............. [ 73760]
KM262844.1_strain_L60          ..................................................c...............t............. [ 73760]
KM102979.1_26544/OG10_Italy    ..................................................c...............t............. [ 73760]
KM262845.1_strain_NHV          ..................................................c...............t............. [ 73760]
KX354450.1_47/Ss/2008          ..................................................c...............t............. [ 73760]
KP055815.1_strain_BA71         ..................................................c...............t............. [ 73760]
MH025920.1_strain_R35          ......t.............a........................a....ct..............t............. [ 73760]
MH025919.1_strain_N10          ......t.............a........................a....ct..............t............. [ 73760]
MH025918.1_strain_R25          ......t.............a........................a....ct..............t............. [ 73760]
MH025917.1_strain_R7           ......t.............a........................a....ct..............t............. [ 73760]
MH025916.1_strain_R8           ......t.............a........................a....ct..............t............. [ 73760]

FR682468.2_Georgia_2007/1      tttttgatagatgttgtatccgcattgtttcgagccatagatggtagtatggagttttataatatatcggcctacctgtt [ 73840]
ASF_IND_20_CAD543              ................................................................................ [ 73840]
LR899193.1_Germany_2020/1      ................................................................................ [ 73840]
MN393477.1_Wuhan_2019          ................................................................................ [ 73840]
MW396979.1_Timor-Leste/2019/1  ................................................................................ [ 73840]
MK940252.1_CN/2019/InnerMongolia ................................................................................ [ 73840]
MK128995.1_China/2018/AnhuiXCGQ ................................................................................ [ 73840]
MH681419.1_ASFV/POL/2015       ................................................................................ [ 73840]
LS478113.1_Estonia_2014        ................................................................................ [ 73840]
KP843857.1_Odintsovo_02/14     ................................................................................ [ 73840]
FN557520.1_strain_E75          ..................................................................c............. [ 73840]
AY261366.1_Warthog             ................................................................................ [ 73840]
AY261365.1_Warmbaths           ................................................................................ [ 73840]
AY261364.1_Tengani_62          ................................................................................ [ 73840]
AY261363.1_Pretoriuskop/96/4   ................................................................................ [ 73840]
AY261362.1_Mkuzi_1979          ................................................................................ [ 73840]
AY261361.1_Malawi_Lil-20/1_1983 .................................................c.................t............ [ 73840]
AY261360.1_Kenya_1950          ...............................a.......a.........c.............................. [ 73840]
AM712240.1_OURT_88/3           ..................................................................c............. [ 73840]
AM712239.1_Benin_97/1          ..................................................................c............. [ 73840]
KM111295.1_Ken06.Bus           .........a...c.................a.......a.........c.............................. [ 73840]
```

```
KM111294.1_Ken05/Tk1             ...............................a.......a.........c.............................. [ 73840]
KM262844.1_strain_L60            ..................................................................c............. [ 73840]
KM102979.1_26544/OG10_Italy      ..................................................................c............. [ 73840]
KM262845.1_strain_NHV            ..................................................................c............. [ 73840]
KX354450.1_47/Ss/2008            ..................................................................c............. [ 73840]
KP055815.1_strain_BA71           ..................................................................c............. [ 73840]
MH025920.1_strain_R35            .........a...c.................a.......a.........c.............................. [ 73840]
MH025919.1_strain_N10            .........a...c.................a.......a.........c.............................. [ 73840]
MH025918.1_strain_R25            .........a...c.................a.......a.........c.............................. [ 73840]
MH025917.1_strain_R7             .........a...c.................a.......a.........c.............................. [ 73840]
MH025916.1_strain_R8             .........a...c.................a.......a.........c.............................. [ 73840]

FR682468.2_Georgia_2007/1        tccttactatacgtgaaggaaagctggtgattgcttatggtctgaaaaagggtgtcacgtttttgtaacgtaaacatttc [ 73920]
ASF_IND_20_CAD543                .................................y.............................................. [ 73920]
LR899193.1_Germany_2020/1        ................................................................................ [ 73920]
MN393477.1_Wuhan_2019            ................................................................................ [ 73920]
MW396979.1_Timor-Leste/2019/1    ................................................................................ [ 73920]
MK940252.1_CN/2019/InnerMongolia ................................................................................ [ 73920]
MK128995.1_China/2018/AnhuiXCGQ  ................................................................................ [ 73920]
MH681419.1_ASFV/POL/2015         ................................................................................ [ 73920]
LS478113.1_Estonia_2014          ................................................................................ [ 73920]
KP843857.1_Odintsovo_02/14       ................................................................................ [ 73920]
FN557520.1_strain_E75            .................................t.............g.......................g........ [ 73920]
AY261366.1_Warthog               ...............................................g................................ [ 73920]
AY261365.1_Warmbaths             ......t........................................g.......................g........ [ 73920]
AY261364.1_Tengani_62            ...............................................g................................ [ 73920]
AY261363.1_Pretoriuskop/96/4     ......t........................................g.......................g........ [ 73920]
AY261362.1_Mkuzi_1979            ...............................................g.......................g........ [ 73920]
AY261361.1_Malawi_Lil-20/1_1983  .......a........................a..............g..a..a..g..............g........ [ 73920]
AY261360.1_Kenya_1950            ................................at....c........g.......................g...g.... [ 73920]
AM712240.1_OURT_88/3             .................................t.............g.......................g........ [ 73920]
AM712239.1_Benin_97/1            .................................t.............g.......................g........ [ 73920]
KM111295.1_Ken06.Bus             ................................at....c........g....................t..g...g.... [ 73920]
KM111294.1_Ken05/Tk1             ................................at....c........g.......................g...g.... [ 73920]
KM262844.1_strain_L60            .................................t.............g.......................g........ [ 73920]
KM102979.1_26544/OG10_Italy      .................................t.............g.......................g........ [ 73920]
KM262845.1_strain_NHV            .................................t.............g.......................g........ [ 73920]
KX354450.1_47/Ss/2008            .................................t.............g.......................g........ [ 73920]
KP055815.1_strain_BA71           .................................t.............g.......................g........ [ 73920]
MH025920.1_strain_R35            ................................at....c........g....................t..g...g.... [ 73920]
MH025919.1_strain_N10            ................................at....c........g....................t..g...g.... [ 73920]
MH025918.1_strain_R25            ................................at....c........g....................t..g...g.... [ 73920]
MH025917.1_strain_R7             ................................at....c........g....................t..g...g.... [ 73920]
MH025916.1_strain_R8             ................................at....c........g....................t..g...g.... [ 73920]
```

```
FR682468.2_Georgia_2007/1        aatgtcttcgatggtttctggatagtaattttgtttcccctgtaagcagattttataacacttactttttaattcacgca [ 74000]
ASF_IND_20_CAD543                ................................................................................ [ 74000]
LR899193.1_Germany_2020/1        ................................................................................ [ 74000]
MN393477.1_Wuhan_2019            ................................................................................ [ 74000]
MW396979.1_Timor-Leste/2019/1    ................................................................................ [ 74000]
MK940252.1_CN/2019/InnerMongolia ................................................................................ [ 74000]
MK128995.1_China/2018/AnhuiXCGQ  ................................................................................ [ 74000]
MH681419.1_ASFV/POL/2015         ................................................................................ [ 74000]
LS478113.1_Estonia_2014          ................................................................................ [ 74000]
KP843857.1_Odintsovo_02/14       ................................................................................ [ 74000]
FN557520.1_strain_E75            .....................g.......................................................... [ 74000]
AY261366.1_Warthog               .........a............a.............t.......................t................... [ 74000]
AY261365.1_Warmbaths             .........a............a.............t..t....................t................... [ 74000]
AY261364.1_Tengani_62            .........a............a.....................................t................... [ 74000]
AY261363.1_Pretoriuskop/96/4     .........a............a.............t.......................t................... [ 74000]
AY261362.1_Mkuzi_1979            .........a...................................................................... [ 74000]
AY261361.1_Malawi_Lil-20/1_1983  .........a........c...a.....................g...............t.....c............. [ 74000]
AY261360.1_Kenya_1950            .........a........c...a.............t........a............gtt.....c............. [ 74000]
AM712240.1_OURT_88/3             .....................g.......................................................... [ 74000]
AM712239.1_Benin_97/1            .....................g.......................................................... [ 74000]
KM111295.1_Ken06.Bus             .........a..t.....c...a.............t........a..............t.....c............. [ 74000]
KM111294.1_Ken05/Tk1             .........a........c...a.............t........a............g.t.....c............. [ 74000]
KM262844.1_strain_L60            .....................g.......................................................... [ 74000]
KM102979.1_26544/OG10_Italy      .....................g.......................................................... [ 74000]
KM262845.1_strain_NHV            .....................g.......................................................... [ 74000]
KX354450.1_47/Ss/2008            .....................g.......................................................... [ 74000]
KP055815.1_strain_BA71           .....................g.......................................................... [ 74000]
MH025920.1_strain_R35            .........a..t.....c...a.............t........a..............t.....c............. [ 74000]
MH025919.1_strain_N10            .........a..t.....c...a.............t........a..............t.....c............. [ 74000]
MH025918.1_strain_R25            .........a..t.....c...a.............t........a..............t.....c............. [ 74000]
MH025917.1_strain_R7             .........a..t.....c...a.............t........a..............t.....c............. [ 74000]
MH025916.1_strain_R8             .........a..t.....c...a.............t........a..............t.....c............. [ 74000]

FR682468.2_Georgia_2007/1        cgcggcccaacatttggcaacatgtttctacgtcacacgacatattgttaaaaaagccgtataaaacatcaaatctctta [ 74080]
ASF_IND_20_CAD543                ................................................................................ [ 74080]
LR899193.1_Germany_2020/1        ................................................................................ [ 74080]
MN393477.1_Wuhan_2019            ................................................................................ [ 74080]
MW396979.1_Timor-Leste/2019/1    ................................................................................ [ 74080]
MK940252.1_CN/2019/InnerMongolia ................................................................................ [ 74080]
MK128995.1_China/2018/AnhuiXCGQ  ................................................................................ [ 74080]
MH681419.1_ASFV/POL/2015         ................................................................................ [ 74080]
LS478113.1_Estonia_2014          ................................................................................ [ 74080]
KP843857.1_Odintsovo_02/14       ................................................................................ [ 74080]
```

```
FN557520.1_strain_E75             ................................................................................ [ 74080]
AY261366.1_Warthog                ................................................................................ [ 74080]
AY261365.1_Warmbaths              ...............................a................................................ [ 74080]
AY261364.1_Tengani_62             ................a............................................................... [ 74080]
AY261363.1_Pretoriuskop/96/4      ................................................................................ [ 74080]
AY261362.1_Mkuzi_1979             ................................................................................ [ 74080]
AY261361.1_Malawi_Lil-20/1_1983   .....................................t.......................................... [ 74080]
AY261360.1_Kenya_1950             .........................................................................c...... [ 74080]
AM712240.1_OURT_88/3              ................................................................................ [ 74080]
AM712239.1_Benin_97/1             ................................................................................ [ 74080]
KM111295.1_Ken06.Bus              ................a........................................................c...... [ 74080]
KM111294.1_Ken05/Tk1              .........................................................................c...... [ 74080]
KM262844.1_strain_L60             ................................................................................ [ 74080]
KM102979.1_26544/OG10_Italy       ................................................................................ [ 74080]
KM262845.1_strain_NHV             ................................................................................ [ 74080]
KX354450.1_47/Ss/2008             ................................................................................ [ 74080]
KP055815.1_strain_BA71            ................................................................................ [ 74080]
MH025920.1_strain_R35             ................a........................................................c...... [ 74080]
MH025919.1_strain_N10             ................a........................................................c...... [ 74080]
MH025918.1_strain_R25             ................a........................................................c...... [ 74080]
MH025917.1_strain_R7              ................a........................................................c...... [ 74080]
MH025916.1_strain_R8              ................a........................................................c...... [ 74080]

FR682468.2_Georgia_2007/1         tcttcgtatgaaacacccgctgaaatcgtgggcgtatagataaggatatcaacgagcccccaataatacgatacattatt [ 74160]
ASF_IND_20_CAD543                 ................................................................................ [ 74160]
LR899193.1_Germany_2020/1         ................................................................................ [ 74160]
MN393477.1_Wuhan_2019             ................................................................................ [ 74160]
MW396979.1_Timor-Leste/2019/1     ................................................................................ [ 74160]
MK940252.1_CN/2019/InnerMongolia  ................................................................................ [ 74160]
MK128995.1_China/2018/AnhuiXCGQ   ................................................................................ [ 74160]
MH681419.1_ASFV/POL/2015          ................................................................................ [ 74160]
LS478113.1_Estonia_2014           ................................................................................ [ 74160]
KP843857.1_Odintsovo_02/14        ................................................................................ [ 74160]
FN557520.1_strain_E75             ........c...........c........................................................... [ 74160]
AY261366.1_Warthog                ....................................................................t..c........ [ 74160]
AY261365.1_Warmbaths              ....................................................................t..c........ [ 74160]
AY261364.1_Tengani_62             ................................t...................................g........... [ 74160]
AY261363.1_Pretoriuskop/96/4      ....................................................................t..c........ [ 74160]
AY261362.1_Mkuzi_1979             ....................................................................t..c........ [ 74160]
AY261361.1_Malawi_Lil-20/1_1983   ....................................................t...............g........... [ 74160]
AY261360.1_Kenya_1950             ........c.............................a.....a.......t...t...........g.....g..... [ 74160]
AM712240.1_OURT_88/3              ........c...........c........................................................... [ 74160]
AM712239.1_Benin_97/1             ........c...........c........................................................... [ 74160]
KM111295.1_Ken06.Bus              .....a..c...........................................t...t...........g.....g..... [ 74160]
```

```
KM111294.1_Ken05/Tk1           ........c...........................................t...t...........g.....g..... [ 74160]
KM262844.1_strain_L60          ........c...........c........................................................... [ 74160]
KM102979.1_26544/OG10_Italy    ........c...........c........................................................... [ 74160]
KM262845.1_strain_NHV          ........c...........c........................................................... [ 74160]
KX354450.1_47/Ss/2008          ........c...........c........................................................... [ 74160]
KP055815.1_strain_BA71         ........c...........c........................................................... [ 74160]
MH025920.1_strain_R35          ........c...........................................t...t...........g.....g..... [ 74160]
MH025919.1_strain_N10          ........c...........................................t...t...........g.....g..... [ 74160]
MH025918.1_strain_R25          ........c...........................................t...t...........g.....g..... [ 74160]
MH025917.1_strain_R7           ........c...........................................t...t...........g.....g..... [ 74160]
MH025916.1_strain_R8           ........c...........................................t...t...........g.....g..... [ 74160]

FR682468.2_Georgia_2007/1      aaaatgggattcccgttcatgagcagtgcttttagaactataaaacccaatttttttttccggaaactttttttggataa [ 74240]
ASF_IND_20_CAD543              ................................................................................ [ 74240]
LR899193.1_Germany_2020/1      ................................................................................ [ 74240]
MN393477.1_Wuhan_2019          ................................................................................ [ 74240]
MW396979.1_Timor-Leste/2019/1  ................................................................................ [ 74240]
MK940252.1_CN/2019/InnerMongolia ................................................................................ [ 74240]
MK128995.1_China/2018/AnhuiXCGQ ................................................................................ [ 74240]
MH681419.1_ASFV/POL/2015       ................................................................................ [ 74240]
LS478113.1_Estonia_2014        ................................................................................ [ 74240]
KP843857.1_Odintsovo_02/14     ................................................................................ [ 74240]
FN557520.1_strain_E75          ................................................g............................... [ 74240]
AY261366.1_Warthog             ................................................g............................... [ 74240]
AY261365.1_Warmbaths           ................................................g..c............................ [ 74240]
AY261364.1_Tengani_62          ............t.....t.............................g............................... [ 74240]
AY261363.1_Pretoriuskop/96/4   ................................................g............................... [ 74240]
AY261362.1_Mkuzi_1979          ................................................g............................... [ 74240]
AY261361.1_Malawi_Lil-20/1_1983 ...g......................a.....................g..c...........g...........a.... [ 74240]
AY261360.1_Kenya_1950          ....................................g...........g..c....................c..a.... [ 74240]
AM712240.1_OURT_88/3           ................................................g............................... [ 74240]
AM712239.1_Benin_97/1          ................................................g............................... [ 74240]
KM111295.1_Ken06.Bus           .........................c......................g..c...........g........c..a.... [ 74240]
KM111294.1_Ken05/Tk1           ............t..............a........g...........g..c....................c..a.... [ 74240]
KM262844.1_strain_L60          ................................................g............................... [ 74240]
KM102979.1_26544/OG10_Italy    .............................................t..g............................... [ 74240]
KM262845.1_strain_NHV          ................................................g............................... [ 74240]
KX354450.1_47/Ss/2008          .............................................t..g............................... [ 74240]
KP055815.1_strain_BA71         ................................................g............................... [ 74240]
MH025920.1_strain_R35          .........................c......................g..c...........g........c..a.... [ 74240]
MH025919.1_strain_N10          .........................c......................g..c...........g........c..a.... [ 74240]
MH025918.1_strain_R25          .........................c......................g..c...........g........c..a.... [ 74240]
MH025917.1_strain_R7           .........................c......................g..c...........g........c..a.... [ 74240]
MH025916.1_strain_R8           .........................c......................g..c...........g........c..a.... [ 74240]
```

```
FR682468.2_Georgia_2007/1        atgattgcaacagccgggcctccattaatgaatttgtagggataacaatttttttgtcttctagcaaatcctttaaaagg [ 74320]
ASF_IND_20_CAD543                ................................................................................ [ 74320]
LR899193.1_Germany_2020/1        ................................................................................ [ 74320]
MN393477.1_Wuhan_2019            ................................................................................ [ 74320]
MW396979.1_Timor-Leste/2019/1    ................................................................................ [ 74320]
MK940252.1_CN/2019/InnerMongolia ................................................................................ [ 74320]
MK128995.1_China/2018/AnhuiXCGQ  ................................................................................ [ 74320]
MH681419.1_ASFV/POL/2015         ................................................................................ [ 74320]
LS478113.1_Estonia_2014          ................................................................................ [ 74320]
KP843857.1_Odintsovo_02/14       ................................................................................ [ 74320]
FN557520.1_strain_E75            ................................................................................ [ 74320]
AY261366.1_Warthog               ................................................................................ [ 74320]
AY261365.1_Warmbaths             ................................................................................ [ 74320]
AY261364.1_Tengani_62            .............a.............................................................g.... [ 74320]
AY261363.1_Pretoriuskop/96/4     ................................................................................ [ 74320]
AY261362.1_Mkuzi_1979            ................................................................................ [ 74320]
AY261361.1_Malawi_Lil-20/1_1983  .c.t......t..a..........................a.....g................................a [ 74320]
AY261360.1_Kenya_1950            ...t.........a..........................a....................g.................a [ 74320]
AM712240.1_OURT_88/3             ................................................................................ [ 74320]
AM712239.1_Benin_97/1            ................................................................................ [ 74320]
KM111295.1_Ken06.Bus             .c.....t.....a..........................a......................................a [ 74320]
KM111294.1_Ken05/Tk1             .......t.....a..........................a.....g................................. [ 74320]
KM262844.1_strain_L60            ................................................................................ [ 74320]
KM102979.1_26544/OG10_Italy      ................................................................................ [ 74320]
KM262845.1_strain_NHV            ................................................................................ [ 74320]
KX354450.1_47/Ss/2008            ................................................................................ [ 74320]
KP055815.1_strain_BA71           ................................................................................ [ 74320]
MH025920.1_strain_R35            .c.....t.....a..........................a......................................a [ 74320]
MH025919.1_strain_N10            .c.....t.....a..........................a......................................a [ 74320]
MH025918.1_strain_R25            .c.....t.....a..........................a......................................a [ 74320]
MH025917.1_strain_R7             .c.....t.....a..........................a......................................a [ 74320]
MH025916.1_strain_R8             .c.....t.....a..........................a......................................a [ 74320]

FR682468.2_Georgia_2007/1        ttatttaaccaagtttctcgtgaagaggtaaaataatacgtgtcatgctgggcccttttatattgattccagtgaaagaa [ 74400]
ASF_IND_20_CAD543                ................................................................................ [ 74400]
LR899193.1_Germany_2020/1        ................................................................................ [ 74400]
MN393477.1_Wuhan_2019            ................................................................................ [ 74400]
MW396979.1_Timor-Leste/2019/1    ................................................................................ [ 74400]
MK940252.1_CN/2019/InnerMongolia ................................................................................ [ 74400]
MK128995.1_China/2018/AnhuiXCGQ  ................................................................................ [ 74400]
MH681419.1_ASFV/POL/2015         ................................................................................ [ 74400]
LS478113.1_Estonia_2014          ................................................................................ [ 74400]
KP843857.1_Odintsovo_02/14       ................................................................................ [ 74400]
```

```
FN557520.1_strain_E75            .............................g.................................................. [ 74400]
AY261366.1_Warthog               .............................g.........a........................................ [ 74400]
AY261365.1_Warmbaths             .............................g.........a........................................ [ 74400]
AY261364.1_Tengani_62            .............................g.........a.a.....t..............c................. [ 74400]
AY261363.1_Pretoriuskop/96/4     .............................g.........a........................................ [ 74400]
AY261362.1_Mkuzi_1979            .............................g.........a........................................ [ 74400]
AY261361.1_Malawi_Lil-20/1_1983  ............a..........................a....g.t.......t...g..................... [ 74400]
AY261360.1_Kenya_1950            ............a................g.........a......t.......t...gg..c........a........ [ 74400]
AM712240.1_OURT_88/3             .............................g.................................................. [ 74400]
AM712239.1_Benin_97/1            .............................g.................................................. [ 74400]
KM111295.1_Ken06.Bus             ............a................g.........a......t.......t...gg..c........a........ [ 74400]
KM111294.1_Ken05/Tk1             ............a.......c........g.........a......t.......t...gg.................... [ 74400]
KM262844.1_strain_L60            .............................g.................................................. [ 74400]
KM102979.1_26544/OG10_Italy      .............................g.................................................. [ 74400]
KM262845.1_strain_NHV            .............................g.................................................. [ 74400]
KX354450.1_47/Ss/2008            .............................g.................................................. [ 74400]
KP055815.1_strain_BA71           .............................g.................................................. [ 74400]
MH025920.1_strain_R35            ............a................g.........a......t.......t...gg..c........a........ [ 74400]
MH025919.1_strain_N10            ............a................g.........a......t.......t...gg..c........a........ [ 74400]
MH025918.1_strain_R25            ............a................g.........a......t.......t...gg..c........a........ [ 74400]
MH025917.1_strain_R7             ............a................g.........a......t.......t...gg..c........a........ [ 74400]
MH025916.1_strain_R8             ............a................g.........a......t.......t...gg..c........a........ [ 74400]

FR682468.2_Georgia_2007/1        gatagggacatccccgcgaaaacgctgtagaatattatacgttcgatttcctaggtttgcgtccaagcatataacataat [ 74480]
ASF_IND_20_CAD543                ................................................................................ [ 74480]
LR899193.1_Germany_2020/1        ................................................................................ [ 74480]
MN393477.1_Wuhan_2019            ................................................................................ [ 74480]
MW396979.1_Timor-Leste/2019/1    ................................................................................ [ 74480]
MK940252.1_CN/2019/InnerMongolia ................................................................................ [ 74480]
MK128995.1_China/2018/AnhuiXCGQ  ................................................................................ [ 74480]
MH681419.1_ASFV/POL/2015         ................................................................................ [ 74480]
LS478113.1_Estonia_2014          ................................................................................ [ 74480]
KP843857.1_Odintsovo_02/14       ................................................................................ [ 74480]
FN557520.1_strain_E75            ......a......................................................................... [ 74480]
AY261366.1_Warthog               ......a......................................................................... [ 74480]
AY261365.1_Warmbaths             .............................a.................................................. [ 74480]
AY261364.1_Tengani_62            ................................................................................ [ 74480]
AY261363.1_Pretoriuskop/96/4     ................................................................................ [ 74480]
AY261362.1_Mkuzi_1979            ................................................................................ [ 74480]
AY261361.1_Malawi_Lil-20/1_1983  ............t..a............................................a................g.. [ 74480]
AY261360.1_Kenya_1950            ......t........a........t...................................a................g.. [ 74480]
AM712240.1_OURT_88/3             ......a......................................................................... [ 74480]
AM712239.1_Benin_97/1            ......a......................................................................... [ 74480]
KM111295.1_Ken06.Bus             ......t........a........t...................................a................... [ 74480]
```

```
KM111294.1_Ken05/Tk1              ........................t...................................a................g.. [ 74480]
KM262844.1_strain_L60             ......a......................................................................... [ 74480]
KM102979.1_26544/OG10_Italy       ......a......................................................................... [ 74480]
KM262845.1_strain_NHV             ......a......................................................................... [ 74480]
KX354450.1_47/Ss/2008             ......a......................................................................... [ 74480]
KP055815.1_strain_BA71            ......a......................................................................... [ 74480]
MH025920.1_strain_R35             ......t........a........t...................................a................... [ 74480]
MH025919.1_strain_N10             ......t........a........t...................................a................... [ 74480]
MH025918.1_strain_R25             ......t........a........t...................................a................... [ 74480]
MH025917.1_strain_R7              ......t........a........t...................................a................... [ 74480]
MH025916.1_strain_R8              ......t........a........t...................................a................... [ 74480]

FR682468.2_Georgia_2007/1         ttgccgtttcgagcatccacatgaaaatggcaaaagagggagcaaagtatttgtgcaggccgctattgaattgattaaaa [ 74560]
ASF_IND_20_CAD543                 ................................................................................ [ 74560]
LR899193.1_Germany_2020/1         ................................................................................ [ 74560]
MN393477.1_Wuhan_2019             ................................................................................ [ 74560]
MW396979.1_Timor-Leste/2019/1     ................................................................................ [ 74560]
MK940252.1_CN/2019/InnerMongolia  ................................................................................ [ 74560]
MK128995.1_China/2018/AnhuiXCGQ   ................................................................................ [ 74560]
MH681419.1_ASFV/POL/2015          ................................................................................ [ 74560]
LS478113.1_Estonia_2014           ................................................................................ [ 74560]
KP843857.1_Odintsovo_02/14        ................................................................................ [ 74560]
FN557520.1_strain_E75             ................................................................................ [ 74560]
AY261366.1_Warthog                ......................a......................................................... [ 74560]
AY261365.1_Warmbaths              .......c..............a......................................................... [ 74560]
AY261364.1_Tengani_62             ................................................................................ [ 74560]
AY261363.1_Pretoriuskop/96/4      ......................a......................................................... [ 74560]
AY261362.1_Mkuzi_1979             ................................................................................ [ 74560]
AY261361.1_Malawi_Lil-20/1_1983   ....t..c...........................................................a............ [ 74560]
AY261360.1_Kenya_1950             ..........a.................a....................c.....t........................ [ 74560]
AM712240.1_OURT_88/3              ................................................................................ [ 74560]
AM712239.1_Benin_97/1             ................................................................................ [ 74560]
KM111295.1_Ken06.Bus              ..........a.................a......................................a...........g [ 74560]
KM111294.1_Ken05/Tk1              .......c...............................................t........................ [ 74560]
KM262844.1_strain_L60             ................................................................................ [ 74560]
KM102979.1_26544/OG10_Italy       ................................................................................ [ 74560]
KM262845.1_strain_NHV             ................................................................................ [ 74560]
KX354450.1_47/Ss/2008             ................................................................................ [ 74560]
KP055815.1_strain_BA71            ................................................................................ [ 74560]
MH025920.1_strain_R35             ..........a.................a......................................a...........g [ 74560]
MH025919.1_strain_N10             ..........a.................a......................................a...........g [ 74560]
MH025918.1_strain_R25             ..........a.................a......................................a...........g [ 74560]
MH025917.1_strain_R7              ..........a.................a......................................a...........g [ 74560]
MH025916.1_strain_R8              ..........a.................a......................................a...........g [ 74560]
```

```
FR682468.2_Georgia_2007/1        atcgattctacctcatccaaaataagtaggtctacaggctcggctgtggaggttagccggaaaagtgattctacctgaat [ 74640]
ASF_IND_20_CAD543                ................................................................................ [ 74640]
LR899193.1_Germany_2020/1        ................................................................................ [ 74640]
MN393477.1_Wuhan_2019            ................................................................................ [ 74640]
MW396979.1_Timor-Leste/2019/1    ................................................................................ [ 74640]
MK940252.1_CN/2019/InnerMongolia ................................................................................ [ 74640]
MK128995.1_China/2018/AnhuiXCGQ  ................................................................................ [ 74640]
MH681419.1_ASFV/POL/2015         ................................................................................ [ 74640]
LS478113.1_Estonia_2014          ................................................................................ [ 74640]
KP843857.1_Odintsovo_02/14       ................................................................................ [ 74640]
FN557520.1_strain_E75            ...........t....................c......................................c.......c [ 74640]
AY261366.1_Warthog               ................................c............................................... [ 74640]
AY261365.1_Warmbaths             ................................c............................................... [ 74640]
AY261364.1_Tengani_62            ................................c............................................... [ 74640]
AY261363.1_Pretoriuskop/96/4     ................................c............................................... [ 74640]
AY261362.1_Mkuzi_1979            ................................c............................................... [ 74640]
AY261361.1_Malawi_Lil-20/1_1983  ................................c........a.....a........g....................... [ 74640]
AY261360.1_Kenya_1950            .....c....................c.....c..g....................g....................... [ 74640]
AM712240.1_OURT_88/3             ...........t....................c......................................c.......c [ 74640]
AM712239.1_Benin_97/1            ...........t....................c......................................c.......c [ 74640]
KM111295.1_Ken06.Bus             ..........................c.....c..g.....a..............g.................t..... [ 74640]
KM111294.1_Ken05/Tk1             .....c....................c.....c..g..............a.....g....................... [ 74640]
KM262844.1_strain_L60            ...........t....................c......................................c.......c [ 74640]
KM102979.1_26544/OG10_Italy      ...........t....................c......................................c.......c [ 74640]
KM262845.1_strain_NHV            ...........t....................c......................................c.......c [ 74640]
KX354450.1_47/Ss/2008            ...........t....................c......................................c.......c [ 74640]
KP055815.1_strain_BA71           ...........t....................c......................................c.......c [ 74640]
MH025920.1_strain_R35            ..........................c.....c..g.....a..............g.................t..... [ 74640]
MH025919.1_strain_N10            ..........................c.....c..g.....a..............g.................t..... [ 74640]
MH025918.1_strain_R25            ..........................c.....c..g.....a..............g.................t..... [ 74640]
MH025917.1_strain_R7             ..........................c.....c..g.....a..............g.................t..... [ 74640]
MH025916.1_strain_R8             ..........................c.....c..g.....a..............g.................t..... [ 74640]

FR682468.2_Georgia_2007/1        gatgactctttcgtagctgtccaaatctccagttacttcgctgtacaatgtgaaattcggtagccgggattgtatatttt [ 74720]
ASF_IND_20_CAD543                ................................................................................ [ 74720]
LR899193.1_Germany_2020/1        ................................................................................ [ 74720]
MN393477.1_Wuhan_2019            ................................................................................ [ 74720]
MW396979.1_Timor-Leste/2019/1    ................................................................................ [ 74720]
MK940252.1_CN/2019/InnerMongolia ................................................................................ [ 74720]
MK128995.1_China/2018/AnhuiXCGQ  ................................................................................ [ 74720]
MH681419.1_ASFV/POL/2015         ................................................................................ [ 74720]
LS478113.1_Estonia_2014          ................................................................................ [ 74720]
KP843857.1_Odintsovo_02/14       ................................................................................ [ 74720]
```

```
FN557520.1_strain_E75          .......................................a.....t...........t...........c.......... [ 74720]
AY261366.1_Warthog             ............a........................................................c.......... [ 74720]
AY261365.1_Warmbaths           .....................................................................c.......... [ 74720]
AY261364.1_Tengani_62          a.................a..t..g..............a..a..t.......................c.......... [ 74720]
AY261363.1_Pretoriuskop/96/4   ............a........................................................c.......... [ 74720]
AY261362.1_Mkuzi_1979          .......................................a.....t.......................c.......... [ 74720]
AY261361.1_Malawi_Lil-20/1_1983 ............a...........g...................................c.a...a.t........... [ 74720]
AY261360.1_Kenya_1950          ........................g..........................a.....t..c.....a.t........... [ 74720]
AM712240.1_OURT_88/3           .......................................a.....t...........t...........c.......... [ 74720]
AM712239.1_Benin_97/1          .......................................a.....t...........t...........c.......... [ 74720]
KM111295.1_Ken06.Bus           ........................g................................t..c.......t........... [ 74720]
KM111294.1_Ken05/Tk1           ........................g..........................a.....t..c.......t........... [ 74720]
KM262844.1_strain_L60          .......................................a.....t...........t...........c.......... [ 74720]
KM102979.1_26544/OG10_Italy    .......................................a.....t...........t...........c.......... [ 74720]
KM262845.1_strain_NHV          .......................................a.....t...........t...........c.......... [ 74720]
KX354450.1_47/Ss/2008          .......................................a.....t...........t...........c.......... [ 74720]
KP055815.1_strain_BA71         .......................................a.....t...........t...........c.......... [ 74720]
MH025920.1_strain_R35          ........................g................................t.ac.......t........... [ 74720]
MH025919.1_strain_N10          ........................g................................t..c.......t........... [ 74720]
MH025918.1_strain_R25          ........................g................................t.ac.......t........... [ 74720]
MH025917.1_strain_R7           ........................g................................t.ac.......t........... [ 74720]
MH025916.1_strain_R8           ........................g................................t.ac.......t........... [ 74720]

FR682468.2_Georgia_2007/1      ttgagaagatctgtcgaaacgtcacaaaccgtatggtttgttgttttgaaatagaattattgccgtagtatttttgcaaa [ 74800]
ASF_IND_20_CAD543              ................................................................................ [ 74800]
LR899193.1_Germany_2020/1      ................................................................................ [ 74800]
MN393477.1_Wuhan_2019          ................................................................................ [ 74800]
MW396979.1_Timor-Leste/2019/1  ................................................................................ [ 74800]
MK940252.1_CN/2019/InnerMongolia ................................................................................ [ 74800]
MK128995.1_China/2018/AnhuiXCGQ ................................................................................ [ 74800]
MH681419.1_ASFV/POL/2015       ................................................................................ [ 74800]
LS478113.1_Estonia_2014        ................................................................................ [ 74800]
KP843857.1_Odintsovo_02/14     ................................................................................ [ 74800]
FN557520.1_strain_E75          ...................t............................................................ [ 74800]
AY261366.1_Warthog             ...................t.....g...................................a.................. [ 74800]
AY261365.1_Warmbaths           ...................t.....g...................................a..............t... [ 74800]
AY261364.1_Tengani_62          ...................t......................................g..................... [ 74800]
AY261363.1_Pretoriuskop/96/4   ...................t.....g......................................a............... [ 74800]
AY261362.1_Mkuzi_1979          ........................................c....................................... [ 74800]
AY261361.1_Malawi_Lil-20/1_1983 .g.......................................................c...a..a............... [ 74800]
AY261360.1_Kenya_1950          .g........t.............................c................c...a..a............... [ 74800]
AM712240.1_OURT_88/3           ...................t............................................................ [ 74800]
AM712239.1_Benin_97/1          ...................t............................................................ [ 74800]
KM111295.1_Ken06.Bus           .g........t.............................c................c...a..a............... [ 74800]
```

```
KM111294.1_Ken05/Tk1              .g........t.............................c................c...a..a............... [ 74800]
KM262844.1_strain_L60             ...................t............................................................ [ 74800]
KM102979.1_26544/OG10_Italy       ...................t............................................................ [ 74800]
KM262845.1_strain_NHV             ...................t............................................................ [ 74800]
KX354450.1_47/Ss/2008             ...................t............................................................ [ 74800]
KP055815.1_strain_BA71            ...................t............................................................ [ 74800]
MH025920.1_strain_R35             .g........t.............................c................c...a..a............... [ 74800]
MH025919.1_strain_N10             .g........t.............................c................c...a..a............... [ 74800]
MH025918.1_strain_R25             .g........t.............................c................c...a..a............... [ 74800]
MH025917.1_strain_R7              .g........t.............................c................c...a..a............... [ 74800]
MH025916.1_strain_R8              .g........t.............................c................c...a..a............... [ 74800]

FR682468.2_Georgia_2007/1         tagttgcgcagttggacggttttacctattttcatttgagcctttacaacaagcgtagggactcgttcatattctcgcat [ 74880]
ASF_IND_20_CAD543                 ................................................................................ [ 74880]
LR899193.1_Germany_2020/1         ................................................................................ [ 74880]
MN393477.1_Wuhan_2019             ................................................................................ [ 74880]
MW396979.1_Timor-Leste/2019/1     ................................................................................ [ 74880]
MK940252.1_CN/2019/InnerMongolia  ................................................................................ [ 74880]
MK128995.1_China/2018/AnhuiXCGQ   ................................................................................ [ 74880]
MH681419.1_ASFV/POL/2015          ................................................................................ [ 74880]
LS478113.1_Estonia_2014           ................................................................................ [ 74880]
KP843857.1_Odintsovo_02/14        ................................................................................ [ 74880]
FN557520.1_strain_E75             ................................................................................ [ 74880]
AY261366.1_Warthog                ................t......................................................c..c..... [ 74880]
AY261365.1_Warmbaths              .....a........a.t......................................................c..c..... [ 74880]
AY261364.1_Tengani_62             ................t......g...............................................c..c..... [ 74880]
AY261363.1_Pretoriuskop/96/4      ................t......................................................c..c..... [ 74880]
AY261362.1_Mkuzi_1979             ................................................................................ [ 74880]
AY261361.1_Malawi_Lil-20/1_1983   ................t......g..........................g........c..c........c....t... [ 74880]
AY261360.1_Kenya_1950             ................t......g................................g.....c........c........ [ 74880]
AM712240.1_OURT_88/3              ................................................................................ [ 74880]
AM712239.1_Benin_97/1             ................................................................................ [ 74880]
KM111295.1_Ken06.Bus              ................t......g................................g.....c........c........ [ 74880]
KM111294.1_Ken05/Tk1              ................t......g................................g.....c........c........ [ 74880]
KM262844.1_strain_L60             ................................................................................ [ 74880]
KM102979.1_26544/OG10_Italy       ................................................................................ [ 74880]
KM262845.1_strain_NHV             ................................................................................ [ 74880]
KX354450.1_47/Ss/2008             ................................................................................ [ 74880]
KP055815.1_strain_BA71            ................................................................................ [ 74880]
MH025920.1_strain_R35             ................t......g................................g.....c........c........ [ 74880]
MH025919.1_strain_N10             ................t......g................................g.....c........c........ [ 74880]
MH025918.1_strain_R25             ................t......g................................g.....c........c........ [ 74880]
MH025917.1_strain_R7              ................t......g................................g.....c........c........ [ 74880]
MH025916.1_strain_R8              ................t......g................................g.....c........c........ [ 74880]
```

```
FR682468.2_Georgia_2007/1        actactttcatcatagatgtgtttttgagtatcaggcagttcttcaaagagaatggactcatgaacctctatgctctttg [ 74960]
ASF_IND_20_CAD543                ................................................................................ [ 74960]
LR899193.1_Germany_2020/1        ................................................................................ [ 74960]
MN393477.1_Wuhan_2019            ................................................................................ [ 74960]
MW396979.1_Timor-Leste/2019/1    ................................................................................ [ 74960]
MK940252.1_CN/2019/InnerMongolia ................................................................................ [ 74960]
MK128995.1_China/2018/AnhuiXCGQ  ................................................................................ [ 74960]
MH681419.1_ASFV/POL/2015         ................................................................................ [ 74960]
LS478113.1_Estonia_2014          ................................................................................ [ 74960]
KP843857.1_Odintsovo_02/14       ................................................................................ [ 74960]
FN557520.1_strain_E75            ...............................................................g................ [ 74960]
AY261366.1_Warthog               ...............a..a..........g.................................g................ [ 74960]
AY261365.1_Warmbaths             .............................g.................................................. [ 74960]
AY261364.1_Tengani_62            ...............................................................g................ [ 74960]
AY261363.1_Pretoriuskop/96/4     ...............a..a..........g.................................g................ [ 74960]
AY261362.1_Mkuzi_1979            .............................g.................................................. [ 74960]
AY261361.1_Malawi_Lil-20/1_1983  ............g..............g...................................g.....g.......... [ 74960]
AY261360.1_Kenya_1950            ...............................................................g................ [ 74960]
AM712240.1_OURT_88/3             ...............................................................g................ [ 74960]
AM712239.1_Benin_97/1            ...............................................................g................ [ 74960]
KM111295.1_Ken06.Bus             ...............................................................g................ [ 74960]
KM111294.1_Ken05/Tk1             ...............................................................g................ [ 74960]
KM262844.1_strain_L60            ...............................................................g................ [ 74960]
KM102979.1_26544/OG10_Italy      ...............................................................g................ [ 74960]
KM262845.1_strain_NHV            ...............................................................g................ [ 74960]
KX354450.1_47/Ss/2008            ...............................................................g................ [ 74960]
KP055815.1_strain_BA71           ...............................................................g................ [ 74960]
MH025920.1_strain_R35            ...............................................................g................ [ 74960]
MH025919.1_strain_N10            ...............................................................g................ [ 74960]
MH025918.1_strain_R25            ...............................................................g................ [ 74960]
MH025917.1_strain_R7             ...............................................................g................ [ 74960]
MH025916.1_strain_R8             ...............................................................g................ [ 74960]

FR682468.2_Georgia_2007/1        tcatcacttggtccacatatgtttccacaaaattatttgtgccggaaaggctgcccatgagaaggctatgtttattgtca [ 75040]
ASF_IND_20_CAD543                ................................................................................ [ 75040]
LR899193.1_Germany_2020/1        ................................................................................ [ 75040]
MN393477.1_Wuhan_2019            ................................................................................ [ 75040]
MW396979.1_Timor-Leste/2019/1    ................................................................................ [ 75040]
MK940252.1_CN/2019/InnerMongolia ................................................................................ [ 75040]
MK128995.1_China/2018/AnhuiXCGQ  ................................................................................ [ 75040]
MH681419.1_ASFV/POL/2015         ................................................................................ [ 75040]
LS478113.1_Estonia_2014          ................................................................................ [ 75040]
KP843857.1_Odintsovo_02/14       ................................................................................ [ 75040]
```

```
FN557520.1_strain_E75          ...................................g....a....................................... [ 75040]
AY261366.1_Warthog             ................g.......................a....................................... [ 75040]
AY261365.1_Warmbaths           ................g.......................a....................................... [ 75040]
AY261364.1_Tengani_62          .............t..........................a.........t.............a............... [ 75040]
AY261363.1_Pretoriuskop/96/4   ................g.......................a....................................... [ 75040]
AY261362.1_Mkuzi_1979          ................g.......................a....................................... [ 75040]
AY261361.1_Malawi_Lil-20/1_1983 a.........t.............................a..a........a...........a............... [ 75040]
AY261360.1_Kenya_1950          ...................ga.............gg....a....................................... [ 75040]
AM712240.1_OURT_88/3           ...................................g....a....................................... [ 75040]
AM712239.1_Benin_97/1          ...................................g....a....................................... [ 75040]
KM111295.1_Ken06.Bus           ..........c........g..............gg....a....................................... [ 75040]
KM111294.1_Ken05/Tk1           ...................ga.............gg....a....................................... [ 75040]
KM262844.1_strain_L60          ...................................g....a....................................... [ 75040]
KM102979.1_26544/OG10_Italy    ...................................g....a....................................... [ 75040]
KM262845.1_strain_NHV          ...................................g....a....................................... [ 75040]
KX354450.1_47/Ss/2008          ...................................g....a....................................... [ 75040]
KP055815.1_strain_BA71         ...................................g....a....................................... [ 75040]
MH025920.1_strain_R35          ..........c........g..............gg....a....................................... [ 75040]
MH025919.1_strain_N10          ..........c........g..............gg....a....................................... [ 75040]
MH025918.1_strain_R25          ..........c........g..............gg....a....................................... [ 75040]
MH025917.1_strain_R7           ..........c........g..............gg....a....................................... [ 75040]
MH025916.1_strain_R8           ..........c........g..............gg....a....................................... [ 75040]

FR682468.2_Georgia_2007/1      tggcgacagtgttgatacactttgtttcccgtgactcttaaaattagggtattgtccttatcatgcatacgcttacatat [ 75120]
ASF_IND_20_CAD543              ................................................................................ [ 75120]
LR899193.1_Germany_2020/1      ................................................................................ [ 75120]
MN393477.1_Wuhan_2019          ................................................................................ [ 75120]
MW396979.1_Timor-Leste/2019/1  ................................................................................ [ 75120]
MK940252.1_CN/2019/InnerMongolia ................................................................................ [ 75120]
MK128995.1_China/2018/AnhuiXCGQ ................................................................................ [ 75120]
MH681419.1_ASFV/POL/2015       ................................................................................ [ 75120]
LS478113.1_Estonia_2014        ................................................................................ [ 75120]
KP843857.1_Odintsovo_02/14     ................................................................................ [ 75120]
FN557520.1_strain_E75          .....................................................a.......................... [ 75120]
AY261366.1_Warthog             ........a....................g...............................................a.. [ 75120]
AY261365.1_Warmbaths           ........a.....................................................................a.. [ 75120]
AY261364.1_Tengani_62          ........................................g..................g.................a.. [ 75120]
AY261363.1_Pretoriuskop/96/4   ........a....................g...............................................a.. [ 75120]
AY261362.1_Mkuzi_1979          ........a.....................................................................a.. [ 75120]
AY261361.1_Malawi_Lil-20/1_1983 ........a..............a.......................a.....a.....g.................a.. [ 75120]
AY261360.1_Kenya_1950          ........a..............a.......c.....................a........g..............a.. [ 75120]
AM712240.1_OURT_88/3           .....................................................a.......................... [ 75120]
AM712239.1_Benin_97/1          .....................................................a.......................... [ 75120]
KM111295.1_Ken06.Bus           ........a..............a.......c..............................g..............a.. [ 75120]
```

```
KM111294.1_Ken05/Tk1             ........a..............a.......c.....................a........g..............a.. [ 75120]
KM262844.1_strain_L60            .....................................................a.......................... [ 75120]
KM102979.1_26544/OG10_Italy      .....................................................a.......................... [ 75120]
KM262845.1_strain_NHV            .....................................................a.......................... [ 75120]
KX354450.1_47/Ss/2008            .....................................................a.......................... [ 75120]
KP055815.1_strain_BA71           .....................................................a.......................... [ 75120]
MH025920.1_strain_R35            ........a..............a.......c..............................g..............a.. [ 75120]
MH025919.1_strain_N10            ........a..............a.......c..............................g..............a.. [ 75120]
MH025918.1_strain_R25            ........a..............a.......c..............................g..............a.. [ 75120]
MH025917.1_strain_R7             ........a..............a.......c..............................g..............a.. [ 75120]
MH025916.1_strain_R8             ........a..............a.......c..............................g..............a.. [ 75120]

FR682468.2_Georgia_2007/1        ttcgcagtaacttggacttgtacgtttaaacaatactaaatttttatgaacacggaggaagcaatgatttttacatagtg [ 75200]
ASF_IND_20_CAD543                ................................................................................ [ 75200]
LR899193.1_Germany_2020/1        ................................................................................ [ 75200]
MN393477.1_Wuhan_2019            ................................................................................ [ 75200]
MW396979.1_Timor-Leste/2019/1    ................................................................................ [ 75200]
MK940252.1_CN/2019/InnerMongolia ................................................................................ [ 75200]
MK128995.1_China/2018/AnhuiXCGQ  ................................................................................ [ 75200]
MH681419.1_ASFV/POL/2015         ................................................................................ [ 75200]
LS478113.1_Estonia_2014          ................................................................................ [ 75200]
KP843857.1_Odintsovo_02/14       ................................................................................ [ 75200]
FN557520.1_strain_E75            ................................................................................ [ 75200]
AY261366.1_Warthog               ................................................................................ [ 75200]
AY261365.1_Warmbaths             ................................................................................ [ 75200]
AY261364.1_Tengani_62            ................................................................................ [ 75200]
AY261363.1_Pretoriuskop/96/4     ................................................................................ [ 75200]
AY261362.1_Mkuzi_1979            ................................................................................ [ 75200]
AY261361.1_Malawi_Lil-20/1_1983  ...............................................a................................ [ 75200]
AY261360.1_Kenya_1950            ...a............................................g........a..a................aca [ 75200]
AM712240.1_OURT_88/3             ................................................................................ [ 75200]
AM712239.1_Benin_97/1            ................................................................................ [ 75200]
KM111295.1_Ken06.Bus             ...a............................................g........a..a................aca [ 75200]
KM111294.1_Ken05/Tk1             ...a............................................g........a..a................aca [ 75200]
KM262844.1_strain_L60            ................................................................................ [ 75200]
KM102979.1_26544/OG10_Italy      ................................................................................ [ 75200]
KM262845.1_strain_NHV            ................................................................................ [ 75200]
KX354450.1_47/Ss/2008            ................................................................................ [ 75200]
KP055815.1_strain_BA71           ................................................................................ [ 75200]
MH025920.1_strain_R35            ...a............................................g........a..a................aca [ 75200]
MH025919.1_strain_N10            ...a............................................g........a..a................aca [ 75200]
MH025918.1_strain_R25            ...a............................................g........a..a................aca [ 75200]
MH025917.1_strain_R7             ...a............................................g........a..a................aca [ 75200]
MH025916.1_strain_R8             ...a............................................g........a..a................aca [ 75200]
```

```
FR682468.2_Georgia_2007/1        ttcctgcaaattttaatacctcttcaagttcactttgttggatagtatcgcaggaactcggtgttgtttcttttacattt [ 75280]
ASF_IND_20_CAD543                ................................................................................ [ 75280]
LR899193.1_Germany_2020/1        ................................................................................ [ 75280]
MN393477.1_Wuhan_2019            ................................................................................ [ 75280]
MW396979.1_Timor-Leste/2019/1    ................................................................................ [ 75280]
MK940252.1_CN/2019/InnerMongolia ................................................................................ [ 75280]
MK128995.1_China/2018/AnhuiXCGQ  ................................................................................ [ 75280]
MH681419.1_ASFV/POL/2015         ....................a........................................................... [ 75280]
LS478113.1_Estonia_2014          ................................................................................ [ 75280]
KP843857.1_Odintsovo_02/14       ................................................................................ [ 75280]
FN557520.1_strain_E75            ................................................................................ [ 75280]
AY261366.1_Warthog               .......g................................a...................a..c................ [ 75280]
AY261365.1_Warmbaths             .......g.......................................................c................ [ 75280]
AY261364.1_Tengani_62            .......g........................................................................ [ 75280]
AY261363.1_Pretoriuskop/96/4     .......g................................a....................................... [ 75280]
AY261362.1_Mkuzi_1979            ................................................................................ [ 75280]
AY261361.1_Malawi_Lil-20/1_1983  ................c..t.........................a............t.a...c............... [ 75280]
AY261360.1_Kenya_1950            c......g........c..t............t.......a..g.a...a......t..---..ca............g. [ 75280]
AM712240.1_OURT_88/3             ................................................................................ [ 75280]
AM712239.1_Benin_97/1            ................................................................................ [ 75280]
KM111295.1_Ken06.Bus             c......g........c..t............t..........g.a...a......t..---..c.............g. [ 75280]
KM111294.1_Ken05/Tk1             c......g........c..t............t.......a..g.a...a......t..---..ca............g. [ 75280]
KM262844.1_strain_L60            ................................................................................ [ 75280]
KM102979.1_26544/OG10_Italy      ................................................................................ [ 75280]
KM262845.1_strain_NHV            ................................................................................ [ 75280]
KX354450.1_47/Ss/2008            ................................................................................ [ 75280]
KP055815.1_strain_BA71           ................................................................................ [ 75280]
MH025920.1_strain_R35            c......g........c..t............t..........g.a...a......t..---..c.............g. [ 75280]
MH025919.1_strain_N10            c......g........c..t............t..........g.a...a......t..---..c.............g. [ 75280]
MH025918.1_strain_R25            c......g........c..t............t..........g.a...a......t..---..c.............g. [ 75280]
MH025917.1_strain_R7             c......g........c..t............t..........g.a...a......t..---..c.............g. [ 75280]
MH025916.1_strain_R8             c......g........c..t............t..........g.a...a......t..---..c.............g. [ 75280]

FR682468.2_Georgia_2007/1        gtgaagatacaaggtaaacacgtcgtttcaaagggggttgctataagggtatcactctttttcgtggttgtactggtctc [ 75360]
ASF_IND_20_CAD543                ................................................................................ [ 75360]
LR899193.1_Germany_2020/1        ................................................................................ [ 75360]
MN393477.1_Wuhan_2019            ................................................................................ [ 75360]
MW396979.1_Timor-Leste/2019/1    ................................................................................ [ 75360]
MK940252.1_CN/2019/InnerMongolia ................................................................................ [ 75360]
MK128995.1_China/2018/AnhuiXCGQ  ................................................................................ [ 75360]
MH681419.1_ASFV/POL/2015         ................................................................................ [ 75360]
LS478113.1_Estonia_2014          ................................................................................ [ 75360]
KP843857.1_Odintsovo_02/14       ................................................................................ [ 75360]
```

```
FN557520.1_strain_E75              ............................................g................................... [ 75360]
AY261366.1_Warthog                 ................................................................................ [ 75360]
AY261365.1_Warmbaths               ................................................................................ [ 75360]
AY261364.1_Tengani_62              ................................................................................ [ 75360]
AY261363.1_Pretoriuskop/96/4       .......................................................................g........ [ 75360]
AY261362.1_Mkuzi_1979              ................................................................................ [ 75360]
AY261361.1_Malawi_Lil-20/1_1983    ..a.g..................t................a...g..a....................ca.......... [ 75360]
AY261360.1_Kenya_1950              ....g...................................t...g..a.....g..............ca.......... [ 75360]
AM712240.1_OURT_88/3               ............................................g................................... [ 75360]
AM712239.1_Benin_97/1              ............................................g................................... [ 75360]
KM111295.1_Ken06.Bus               ....g...................................t...g..a.....g..............ca.......... [ 75360]
KM111294.1_Ken05/Tk1               ....g...................................t...g..a.....g..............ca.......... [ 75360]
KM262844.1_strain_L60              ............................................g................................... [ 75360]
KM102979.1_26544/OG10_Italy        ............................................g................................... [ 75360]
KM262845.1_strain_NHV              ............................................g................................... [ 75360]
KX354450.1_47/Ss/2008              ............................................g................................... [ 75360]
KP055815.1_strain_BA71             ............................................g................................... [ 75360]
MH025920.1_strain_R35              ....g...................................t...g..a.....g..............ca.......... [ 75360]
MH025919.1_strain_N10              ....g...................................t...g..a.....g..............ca.......... [ 75360]
MH025918.1_strain_R25              ....g...................................t...g..a.....g..............ca.......... [ 75360]
MH025917.1_strain_R7               ....g...................................t...g..a.....g..............ca.......... [ 75360]
MH025916.1_strain_R8               ....g...................................t...g..a.....g..............ca.......... [ 75360]

FR682468.2_Georgia_2007/1          aaacacctctgcaagctcctcattaaacattttaacacgcatgctaccttttttatgagaccctatgatgcgaaaatttt [ 75440]
ASF_IND_20_CAD543                  ..................y............................................................. [ 75440]
LR899193.1_Germany_2020/1          ................................................................................ [ 75440]
MN393477.1_Wuhan_2019              ................................................................................ [ 75440]
MW396979.1_Timor-Leste/2019/1      ................................................................................ [ 75440]
MK940252.1_CN/2019/InnerMongolia   ................................................................................ [ 75440]
MK128995.1_China/2018/AnhuiXCGQ    ................................................................................ [ 75440]
MH681419.1_ASFV/POL/2015           ................................................................................ [ 75440]
LS478113.1_Estonia_2014            ................................................................................ [ 75440]
KP843857.1_Odintsovo_02/14         ................................................................................ [ 75440]
FN557520.1_strain_E75              ..................t............................................................. [ 75440]
AY261366.1_Warthog                 ......a......................................................................... [ 75440]
AY261365.1_Warmbaths               ......a......................................................................... [ 75440]
AY261364.1_Tengani_62              ................................................................................ [ 75440]
AY261363.1_Pretoriuskop/96/4       ......a......................................................................... [ 75440]
AY261362.1_Mkuzi_1979              ......a...........t............................................................. [ 75440]
AY261361.1_Malawi_Lil-20/1_1983    ......a..............g......................................t................... [ 75440]
AY261360.1_Kenya_1950              ......a...........t..g......................................t.................c. [ 75440]
AM712240.1_OURT_88/3               ..................t............................................................. [ 75440]
AM712239.1_Benin_97/1              ..................t............................................................. [ 75440]
KM111295.1_Ken06.Bus               ......a...........t..g......................................t.................c. [ 75440]
```

```
KM111294.1_Ken05/Tk1            ......a...........t..g......................................t.................c. [ 75440]
KM262844.1_strain_L60           ..................t............................................................. [ 75440]
KM102979.1_26544/OG10_Italy     ..................t............................................................. [ 75440]
KM262845.1_strain_NHV           ..................t............................................................. [ 75440]
KX354450.1_47/Ss/2008           ..................t............................................................. [ 75440]
KP055815.1_strain_BA71          ..................t............................................................. [ 75440]
MH025920.1_strain_R35           ......a...........t..g......................................t.................c. [ 75440]
MH025919.1_strain_N10           ......a...........t..g......................................t.................c. [ 75440]
MH025918.1_strain_R25           ......a...........t..g......................................t.................c. [ 75440]
MH025917.1_strain_R7            ......a...........t..g......................................t.................c. [ 75440]
MH025916.1_strain_R8            ......a...........t..g......................................t.................c. [ 75440]

FR682468.2_Georgia_2007/1       gaatacttttgttgacctgggggtcaacaaaaggataaacgtgtttgggaagattttctaacactttggatgtaaagact [ 75520]
ASF_IND_20_CAD543               ....r........................................................................r.. [ 75520]
LR899193.1_Germany_2020/1       ................................................................................ [ 75520]
MN393477.1_Wuhan_2019           ................................................................................ [ 75520]
MW396979.1_Timor-Leste/2019/1   ................................................................................ [ 75520]
MK940252.1_CN/2019/InnerMongolia ................................................................................ [ 75520]
MK128995.1_China/2018/AnhuiXCGQ ................................................................................ [ 75520]
MH681419.1_ASFV/POL/2015        ................................................................................ [ 75520]
LS478113.1_Estonia_2014         ................................................................................ [ 75520]
KP843857.1_Odintsovo_02/14      ................................................................................ [ 75520]
FN557520.1_strain_E75           ....g..............a.........................................................g.. [ 75520]
AY261366.1_Warthog              ...............................g........a....................................g.. [ 75520]
AY261365.1_Warmbaths            ...............................g.............................................g.. [ 75520]
AY261364.1_Tengani_62           ...............................g..............................................a. [ 75520]
AY261363.1_Pretoriuskop/96/4    .............a.................g........a....................................... [ 75520]
AY261362.1_Mkuzi_1979           ....g..............a.........................................................g.. [ 75520]
AY261361.1_Malawi_Lil-20/1_1983 ..........a....................g.......t.....................t...............g.. [ 75520]
AY261360.1_Kenya_1950           ....g..........................g......................................c......g.. [ 75520]
AM712240.1_OURT_88/3            ....g..............a.........................................................g.. [ 75520]
AM712239.1_Benin_97/1           ....g..............a.........................................................g.. [ 75520]
KM111295.1_Ken06.Bus            ....g..........................g......................................c......g.. [ 75520]
KM111294.1_Ken05/Tk1            ....g..........................g......................................c......g.. [ 75520]
KM262844.1_strain_L60           ....g..............a.........................................................g.. [ 75520]
KM102979.1_26544/OG10_Italy     ....g..............a.........................................................g.. [ 75520]
KM262845.1_strain_NHV           ....g..............a.........................................................g.. [ 75520]
KX354450.1_47/Ss/2008           ....g..............a.........................................................g.. [ 75520]
KP055815.1_strain_BA71          ....g..............a.........................................................g.. [ 75520]
MH025920.1_strain_R35           ....g..........................g......................................c......g.. [ 75520]
MH025919.1_strain_N10           ....g..........................g......................................c......g.. [ 75520]
MH025918.1_strain_R25           ....g..........................g......................................c......g.. [ 75520]
MH025917.1_strain_R7            ....g..........................g......................................c......g.. [ 75520]
MH025916.1_strain_R8            ....g..........................g......................................c......g.. [ 75520]
```

```
FR682468.2_Georgia_2007/1        ttggcctcattattgtttaatactgagtatgtataaagtatgatatgaaaggagtatttaagttctcgctttttatttaa [ 75600]
ASF_IND_20_CAD543                ................................................................................ [ 75600]
LR899193.1_Germany_2020/1        ................................................................................ [ 75600]
MN393477.1_Wuhan_2019            ................................................................................ [ 75600]
MW396979.1_Timor-Leste/2019/1    ................................................................................ [ 75600]
MK940252.1_CN/2019/InnerMongolia ................................................................................ [ 75600]
MK128995.1_China/2018/AnhuiXCGQ  ................................................................................ [ 75600]
MH681419.1_ASFV/POL/2015         ................................................................................ [ 75600]
LS478113.1_Estonia_2014          ................................................................................ [ 75600]
KP843857.1_Odintsovo_02/14       ................................................................................ [ 75600]
FN557520.1_strain_E75            ................................................................................ [ 75600]
AY261366.1_Warthog               ...........g..a.............gc.................................................. [ 75600]
AY261365.1_Warmbaths             ...........g................gc.................................................. [ 75600]
AY261364.1_Tengani_62            .............................c.................................................. [ 75600]
AY261363.1_Pretoriuskop/96/4     ............................gc.................................................. [ 75600]
AY261362.1_Mkuzi_1979            ................................................................................ [ 75600]
AY261361.1_Malawi_Lil-20/1_1983  ...........................................................g........t........... [ 75600]
AY261360.1_Kenya_1950            ...........g.................c.............................g........t........... [ 75600]
AM712240.1_OURT_88/3             ................................................................................ [ 75600]
AM712239.1_Benin_97/1            ................................................................................ [ 75600]
KM111295.1_Ken06.Bus             ...........g.................c.............................g........t........... [ 75600]
KM111294.1_Ken05/Tk1             ...........g.................c.............................g........t........... [ 75600]
KM262844.1_strain_L60            ................................................................................ [ 75600]
KM102979.1_26544/OG10_Italy      ................................................................................ [ 75600]
KM262845.1_strain_NHV            ................................................................................ [ 75600]
KX354450.1_47/Ss/2008            ................................................................................ [ 75600]
KP055815.1_strain_BA71           ................................................................................ [ 75600]
MH025920.1_strain_R35            ...........g.................c.............................g........t........... [ 75600]
MH025919.1_strain_N10            ...........g.................c.............................g........t........... [ 75600]
MH025918.1_strain_R25            ...........g.................c.............................g........t........... [ 75600]
MH025917.1_strain_R7             ...........g.................c.............................g........t........... [ 75600]
MH025916.1_strain_R8             ...........g.................c.............................g........t........... [ 75600]

FR682468.2_Georgia_2007/1        tccgatagaatctgttagcaaaatttgttcacgcgttagattgatgttataaggtaaagaatatgtctcgtaaaatacat [ 75680]
ASF_IND_20_CAD543                ................................................................................ [ 75680]
LR899193.1_Germany_2020/1        ................................................................................ [ 75680]
MN393477.1_Wuhan_2019            ................................................................................ [ 75680]
MW396979.1_Timor-Leste/2019/1    ................................................................................ [ 75680]
MK940252.1_CN/2019/InnerMongolia ................................................................................ [ 75680]
MK128995.1_China/2018/AnhuiXCGQ  ................................................................................ [ 75680]
MH681419.1_ASFV/POL/2015         ................................................................................ [ 75680]
LS478113.1_Estonia_2014          ................................................................................ [ 75680]
KP843857.1_Odintsovo_02/14       ................................................................................ [ 75680]
```

```
FN557520.1_strain_E75            c............................................................................... [ 75680]
AY261366.1_Warthog               c......................................g.......................c..............g. [ 75680]
AY261365.1_Warmbaths             c......................................g.......................c................ [ 75680]
AY261364.1_Tengani_62            c............................................................................... [ 75680]
AY261363.1_Pretoriuskop/96/4     c......................................g.......................c..............g. [ 75680]
AY261362.1_Mkuzi_1979            c............................................................................... [ 75680]
AY261361.1_Malawi_Lil-20/1_1983  c................a....................ag..a..................................... [ 75680]
AY261360.1_Kenya_1950            c.................t....................ag..a....................c................ [ 75680]
AM712240.1_OURT_88/3             c............................................................................... [ 75680]
AM712239.1_Benin_97/1            c............................................................................... [ 75680]
KM111295.1_Ken06.Bus             c..a.c............t....................g..a....................c................ [ 75680]
KM111294.1_Ken05/Tk1             c.................t....................g..a....................c................ [ 75680]
KM262844.1_strain_L60            c............................................................................... [ 75680]
KM102979.1_26544/OG10_Italy      c............................................................................... [ 75680]
KM262845.1_strain_NHV            c............................................................................... [ 75680]
KX354450.1_47/Ss/2008            c............................................................................... [ 75680]
KP055815.1_strain_BA71           c............................................................................... [ 75680]
MH025920.1_strain_R35            c....c............t....................g..a....................c................ [ 75680]
MH025919.1_strain_N10            c....c............t....................g..a....................c................ [ 75680]
MH025918.1_strain_R25            c....c............t....................g..a....................c................ [ 75680]
MH025917.1_strain_R7             c....c............t....................g..a....................c................ [ 75680]
MH025916.1_strain_R8             c....c............t....................g..a....................c................ [ 75680]

FR682468.2_Georgia_2007/1        ccatgatgacgttaattatcatgtcaaggatgtcatagacattgtcttc------gacattatcattgtcatcaacattg [ 75760]
ASF_IND_20_CAD543                .......r.........................................gtcata....c.g....c.....tg...... [ 75760]
LR899193.1_Germany_2020/1        .................................................------......................... [ 75760]
MN393477.1_Wuhan_2019            .................................................------......................... [ 75760]
MW396979.1_Timor-Leste/2019/1    .................................................------......................... [ 75760]
MK940252.1_CN/2019/InnerMongolia .................................................------......................... [ 75760]
MK128995.1_China/2018/AnhuiXCGQ  .................................................------......................... [ 75760]
MH681419.1_ASFV/POL/2015         .................................................------......................... [ 75760]
LS478113.1_Estonia_2014          .................................................------......................... [ 75760]
KP843857.1_Odintsovo_02/14       .................................................------......................... [ 75760]
FN557520.1_strain_E75            .......a......................................a..gtcata....c.g....c.....tg...... [ 75760]
AY261366.1_Warthog               ..............................................a..gtcatt......g....c.a...tct...c. [ 75760]
AY261365.1_Warmbaths             ..............................................a..gttatt......g......a...tgt.g.c. [ 75760]
AY261364.1_Tengani_62            ..............................................a.tgacatt.t...cga.........t....... [ 75760]
AY261363.1_Pretoriuskop/96/4     ..............................................a..gtcatt....a.g....c.a...tct...c. [ 75760]
AY261362.1_Mkuzi_1979            .......a......................................a..gtcatt......g..g.c.....tg------ [ 75760]
AY261361.1_Malawi_Lil-20/1_1983  .......a.....t...................t............a..------......g....c.a...tg------ [ 75760]
AY261360.1_Kenya_1950            ................g..g........t....t.........a..a..attgt-------------------------- [ 75760]
AM712240.1_OURT_88/3             .t.....a......................................a..gtcata....c.g....c.....tg...... [ 75760]
AM712239.1_Benin_97/1            .......a......................................a..gtcata....c.g....c.....tg...... [ 75760]
KM111295.1_Ken06.Bus             ................g..g........t....tg........a..a..atcat-------------------------- [ 75760]
```

```
KM111294.1_Ken05/Tk1             ................g..g........t....t.........a..a..attgt-------------------------- [ 75760]
KM262844.1_strain_L60            .......a......................................a..gtcata....c.g....c.....tg...... [ 75760]
KM102979.1_26544/OG10_Italy      .......a......................................a..gtcata....c.g....c.....tg...... [ 75760]
KM262845.1_strain_NHV            .t.....a......................................a..gtcata....c.g....c.....tg...... [ 75760]
KX354450.1_47/Ss/2008            .......a......................................a..gtcata....c.g....c.....tg...... [ 75760]
KP055815.1_strain_BA71           .......a......................................a..gtcata....c.g....c.....tg...... [ 75760]
MH025920.1_strain_R35            ................g..g........t....tg........a..a..atcat-------------------------- [ 75760]
MH025919.1_strain_N10            ................g..g........t....tg........a..a..atcat-------------------------- [ 75760]
MH025918.1_strain_R25            ................g..g........t....tg........a..a..atcat-------------------------- [ 75760]
MH025917.1_strain_R7             ................g..g........t....tg........a..a..atcat-------------------------- [ 75760]
MH025916.1_strain_R8             ................g..g........t....tg........a..a..atcat-------------------------- [ 75760]

FR682468.2_Georgia_2007/1        t------------catcagagtatgacttatttaccggaaagtcgatgtcaaattttaagcgctgaggcaaaaacccaaa [ 75840]
ASF_IND_20_CAD543                .cattat------.........................g..a...................................... [ 75840]
LR899193.1_Germany_2020/1        .------------................................................................... [ 75840]
MN393477.1_Wuhan_2019            .------------................................................................... [ 75840]
MW396979.1_Timor-Leste/2019/1    .------------................................................................... [ 75840]
MK940252.1_CN/2019/InnerMongolia .------------................................................................... [ 75840]
MK128995.1_China/2018/AnhuiXCGQ  .------------................................................................... [ 75840]
MH681419.1_ASFV/POL/2015         .------------................................................................... [ 75840]
LS478113.1_Estonia_2014          .------------................................................................... [ 75840]
KP843857.1_Odintsovo_02/14       .------------................................................................... [ 75840]
FN557520.1_strain_E75            .catcgtcattga.........................g..a................................t..... [ 75840]
AY261366.1_Warthog               acattgt------...t...a.................g......................................... [ 75840]
AY261365.1_Warmbaths             .cattgt------.......a.................g......................................... [ 75840]
AY261364.1_Tengani_62            .cattaacattgt.......a......a..........g....................a..............t..... [ 75840]
AY261363.1_Pretoriuskop/96/4     acattgt------...t...a.................g......................................... [ 75840]
AY261362.1_Mkuzi_1979            ------------t.......a.................g......................................... [ 75840]
AY261361.1_Malawi_Lil-20/1_1983  ------------t...t...a.................g...................................t..... [ 75840]
AY261360.1_Kenya_1950            ----------ctt.t.....a.................g...........g.................a.....t..... [ 75840]
AM712240.1_OURT_88/3             .cattat------.........................g..a................................t..... [ 75840]
AM712239.1_Benin_97/1            .cattat------.........................g..a................................t..... [ 75840]
KM111295.1_Ken06.Bus             ----------ctt.t.....a.................g...........g.......................t..... [ 75840]
KM111294.1_Ken05/Tk1             ----------ctt.t.....a.................g...........g.................a.....t..... [ 75840]
KM262844.1_strain_L60            .cattat------.........................g..a................................t..... [ 75840]
KM102979.1_26544/OG10_Italy      .cattat------.........................g..a................................t..... [ 75840]
KM262845.1_strain_NHV            .cattat------.........................g..a................................t..... [ 75840]
KX354450.1_47/Ss/2008            .cattat------.........................g..a................................t..... [ 75840]
KP055815.1_strain_BA71           .cattat------.........................g..a................................t..... [ 75840]
MH025920.1_strain_R35            ----------ctt.t...a.a.................g...........g.......................t..... [ 75840]
MH025919.1_strain_N10            ----------ctt.t...a.a.................g...........g.......................t..... [ 75840]
MH025918.1_strain_R25            ----------ctt.t...a.a.................g...........g.......................t..... [ 75840]
MH025917.1_strain_R7             ----------ctt.t...a.a.................g...........g.......................t..... [ 75840]
MH025916.1_strain_R8             ----------ctt.t...a.a.................g...........g.......................t..... [ 75840]
```

```
FR682468.2_Georgia_2007/1        taccacttcgtggaaacacttctgctcaaagggctgagccgcctcccactcccaaaagtcatcacgacttgaaaaaactc [ 75920]
ASF_IND_20_CAD543                ..............................................................................y. [ 75920]
LR899193.1_Germany_2020/1        ................................................................................ [ 75920]
MN393477.1_Wuhan_2019            ................................................................................ [ 75920]
MW396979.1_Timor-Leste/2019/1    ................................................................................ [ 75920]
MK940252.1_CN/2019/InnerMongolia ................................................................................ [ 75920]
MK128995.1_China/2018/AnhuiXCGQ  ................................................................................ [ 75920]
MH681419.1_ASFV/POL/2015         ................................................................................ [ 75920]
LS478113.1_Estonia_2014          ................................................................................ [ 75920]
KP843857.1_Odintsovo_02/14       ................................................................................ [ 75920]
FN557520.1_strain_E75            ...................................................................t..........c. [ 75920]
AY261366.1_Warthog               c.................................................................g............. [ 75920]
AY261365.1_Warmbaths             c..t...........................................................t..g............. [ 75920]
AY261364.1_Tengani_62            ...t..........................................................................c. [ 75920]
AY261363.1_Pretoriuskop/96/4     c.................................................................g............. [ 75920]
AY261362.1_Mkuzi_1979            c..t..............................................................g............. [ 75920]
AY261361.1_Malawi_Lil-20/1_1983  c..t..........................a........t..t.......t............g..g..c.......... [ 75920]
AY261360.1_Kenya_1950            c..t.................................t...................a........g............. [ 75920]
AM712240.1_OURT_88/3             ...................................................................t..........c. [ 75920]
AM712239.1_Benin_97/1            ...................................................................t..........c. [ 75920]
KM111295.1_Ken06.Bus             c........a..a............................................a........g............. [ 75920]
KM111294.1_Ken05/Tk1             c..t.................................t...................a........g............. [ 75920]
KM262844.1_strain_L60            ...................................................................t..........c. [ 75920]
KM102979.1_26544/OG10_Italy      ...................................................................t..........c. [ 75920]
KM262845.1_strain_NHV            ...................................................................t..........c. [ 75920]
KX354450.1_47/Ss/2008            ...................................................................t..........c. [ 75920]
KP055815.1_strain_BA71           ...................................................................t..........c. [ 75920]
MH025920.1_strain_R35            c........a..a............................................a........g............. [ 75920]
MH025919.1_strain_N10            c........a..a............................................a........g............. [ 75920]
MH025918.1_strain_R25            c........a..a............................................a........g............. [ 75920]
MH025917.1_strain_R7             c........a..a............................................a........g............. [ 75920]
MH025916.1_strain_R8             c........a..a............................................a........g............. [ 75920]

FR682468.2_Georgia_2007/1        taaaaagattattatattcatctcgcaccacgaagtgattctttaaggtttcgagagaatatttatcctctacggcttct [ 76000]
ASF_IND_20_CAD543                ................................................................................ [ 76000]
LR899193.1_Germany_2020/1        ................................................................................ [ 76000]
MN393477.1_Wuhan_2019            ................................................................................ [ 76000]
MW396979.1_Timor-Leste/2019/1    ................................................................................ [ 76000]
MK940252.1_CN/2019/InnerMongolia ................................................................................ [ 76000]
MK128995.1_China/2018/AnhuiXCGQ  ................................................................................ [ 76000]
MH681419.1_ASFV/POL/2015         ................................................................................ [ 76000]
LS478113.1_Estonia_2014          ................................................................................ [ 76000]
KP843857.1_Odintsovo_02/14       ................................................................................ [ 76000]
```

```
FN557520.1_strain_E75            ................................................................................ [ 76000]
AY261366.1_Warthog               ................................................................................ [ 76000]
AY261365.1_Warmbaths             ................................................................................ [ 76000]
AY261364.1_Tengani_62            ................................................................................ [ 76000]
AY261363.1_Pretoriuskop/96/4     ................................................................................ [ 76000]
AY261362.1_Mkuzi_1979            ................................................................................ [ 76000]
AY261361.1_Malawi_Lil-20/1_1983  ......................a..t.t...a........t.....a................................. [ 76000]
AY261360.1_Kenya_1950            ..........g....................a........t.....a....t...............t............ [ 76000]
AM712240.1_OURT_88/3             ................................................................................ [ 76000]
AM712239.1_Benin_97/1            ................................................................................ [ 76000]
KM111295.1_Ken06.Bus             ..........g....................a........t..c..a....t...............t............ [ 76000]
KM111294.1_Ken05/Tk1             ..........g....................a........t.....a....t...............t............ [ 76000]
KM262844.1_strain_L60            ................................................................................ [ 76000]
KM102979.1_26544/OG10_Italy      ................................................................................ [ 76000]
KM262845.1_strain_NHV            ................................................................................ [ 76000]
KX354450.1_47/Ss/2008            ................................................................................ [ 76000]
KP055815.1_strain_BA71           ................................................................................ [ 76000]
MH025920.1_strain_R35            ..........g....................a........t..c..a....t...............t............ [ 76000]
MH025919.1_strain_N10            ..........g....................a........t..c..a....t...............t............ [ 76000]
MH025918.1_strain_R25            ..........g....................a........t..c..a....t...............t............ [ 76000]
MH025917.1_strain_R7             ..........g....................a........t..c..a....t...............t............ [ 76000]
MH025916.1_strain_R8             ..........g....................a........t..c..a....t...............t............ [ 76000]

FR682468.2_Georgia_2007/1        ccttgggagttacagcgaagaaacttgaatgtttcttgcattttgatattta-aaattaaatcaattatgatgcggccgc [ 76080]
ASF_IND_20_CAD543                ................................................w...-......................t.--- [ 76080]
LR899193.1_Germany_2020/1        ....................................................-........................... [ 76080]
MN393477.1_Wuhan_2019            ....................................................-........................... [ 76080]
MW396979.1_Timor-Leste/2019/1    ....................................................-........................... [ 76080]
MK940252.1_CN/2019/InnerMongolia ....................................................-........................... [ 76080]
MK128995.1_China/2018/AnhuiXCGQ  ....................................................-........................... [ 76080]
MH681419.1_ASFV/POL/2015         ....................................................-........................... [ 76080]
LS478113.1_Estonia_2014          ....................................................-........................... [ 76080]
KP843857.1_Odintsovo_02/14       ....................................................-........................... [ 76080]
FN557520.1_strain_E75            ................................................a...-..................--------- [ 76080]
AY261366.1_Warthog               ................................................a...-......................----- [ 76080]
AY261365.1_Warmbaths             ................................................a...-......................----- [ 76080]
AY261364.1_Tengani_62            ................................................a...-........................... [ 76080]
AY261363.1_Pretoriuskop/96/4     ................................................a...-......................----- [ 76080]
AY261362.1_Mkuzi_1979            ..........................a.....................a...-........................... [ 76080]
AY261361.1_Malawi_Lil-20/1_1983  ............g...............................a...ac..c.g...g................----- [ 76080]
AY261360.1_Kenya_1950            .............................c..............a..ta...c.g...g.................tg.a [ 76080]
AM712240.1_OURT_88/3             ................................................a...-..................--------- [ 76080]
AM712239.1_Benin_97/1            ................................................a...-..................--------- [ 76080]
KM111295.1_Ken06.Bus             ..........................a..a..............a..ta...c.g......................g.a [ 76080]
```

```
KM111294.1_Ken05/Tk1           .............................c..............a..ta...c.g...g.................tg.a [ 76080]
KM262844.1_strain_L60          ................................................a...-..................--------- [ 76080]
KM102979.1_26544/OG10_Italy    ................................................a...-..................--------- [ 76080]
KM262845.1_strain_NHV          ................................................a...-..................--------- [ 76080]
KX354450.1_47/Ss/2008          ................................................a...-..................--------- [ 76080]
KP055815.1_strain_BA71         ................................................a...-..................--------- [ 76080]
MH025920.1_strain_R35          ..........................a..a..............a..ta...c.g......................g.a [ 76080]
MH025919.1_strain_N10          ..........................a..a..............a..ta...c.g......................g.a [ 76080]
MH025918.1_strain_R25          ..........................a..a..............a..ta...c.g......................g.a [ 76080]
MH025917.1_strain_R7           ..........................a..a..............a..ta...c.g......................g.a [ 76080]
MH025916.1_strain_R8           ..........................a..a..............a..ta...c.g......................g.a [ 76080]

FR682468.2_Georgia_2007/1      taa------------tgcggcggttgacgcggccg-cgccgctgacgcagccatc------------atacataaagcgg [ 76160]
ASF_IND_20_CAD543              --------------------------------nnn-n..................------------............. [ 76160]
LR899193.1_Germany_2020/1      ...------------....................-...................------------............. [ 76160]
MN393477.1_Wuhan_2019          ...------------....................-...................------------............. [ 76160]
MW396979.1_Timor-Leste/2019/1  ...------------....................-...................------------............. [ 76160]
MK940252.1_CN/2019/InnerMongolia ...------------....................-...................------------............. [ 76160]
MK128995.1_China/2018/AnhuiXCGQ ...------------....................-...................------------............. [ 76160]
MH681419.1_ASFV/POL/2015       ...------------....................-...................------------............. [ 76160]
LS478113.1_Estonia_2014        ...------------....................-...................------------............. [ 76160]
KP843857.1_Odintsovo_02/14     ...------------....................-...................------------............. [ 76160]
FN557520.1_strain_E75          ---------------....t.a..g------------...............g..------------...a......... [ 76160]
AY261366.1_Warthog             -------------------------------------..g.t....a.g...gc-------------------------- [ 76160]
AY261365.1_Warmbaths           -------------------------------------...............g..------------............. [ 76160]
AY261364.1_Tengani_62          ...tacggcggctaac...................cgt..a.......g...g..------------............. [ 76160]
AY261363.1_Pretoriuskop/96/4   -------------------------------------..g.t....a.g...gc.tcagtggccgtt............. [ 76160]
AY261362.1_Mkuzi_1979          ...cgcgg------c....t.a..g------------..------.......g..------------...a......... [ 76160]
AY261361.1_Malawi_Lil-20/1_1983 -------------------------------------------------...g..-----agc----------------- [ 76160]
AY261360.1_Kenya_1950          c--------------------------------------------...c....ct------------...---------- [ 76160]
AM712240.1_OURT_88/3           ---------------....t.a..g------------...............g..------------...a......... [ 76160]
AM712239.1_Benin_97/1          ---------------....t.a..g------------...............g..------------...a......... [ 76160]
KM111295.1_Ken06.Bus           c--------------------------------------------...c....--------------------------- [ 76160]
KM111294.1_Ken05/Tk1           c--------------------------------------------...c....ct------------...tg.------- [ 76160]
KM262844.1_strain_L60          ---------------....t.a..g------------...............g..------------...a......... [ 76160]
KM102979.1_26544/OG10_Italy    ---------------....t.a..g------------...............g..------------...a......... [ 76160]
KM262845.1_strain_NHV          ---------------....t.a..g------------...............g..------------...a......... [ 76160]
KX354450.1_47/Ss/2008          ---------------....t.a..g------------...............g..------------...a......... [ 76160]
KP055815.1_strain_BA71         ---------------....t.a..g------------...............g..------------...a......... [ 76160]
MH025920.1_strain_R35          c--------------------------------------------...c....--------------------------- [ 76160]
MH025919.1_strain_N10          c--------------------------------------------...c....--------------------------- [ 76160]
MH025918.1_strain_R25          c--------------------------------------------...c....--------------------------- [ 76160]
MH025917.1_strain_R7           c--------------------------------------------...c....--------------------------- [ 76160]
MH025916.1_strain_R8           c--------------------------------------------...c....--------------------------- [ 76160]
```

```
FR682468.2_Georgia_2007/1        catggccgttttataacgac-----------tagtcggccgttatatgacgaactatataaaaatgaattcttttaatta [ 76240]
ASF_IND_20_CAD543                .--.................-----------.....................r.y......................... [ 76240]
LR899193.1_Germany_2020/1        ....................-----------................................................. [ 76240]
MN393477.1_Wuhan_2019            ....................-----------................................................. [ 76240]
MW396979.1_Timor-Leste/2019/1    ....................-----------................................................. [ 76240]
MK940252.1_CN/2019/InnerMongolia ....................-----------................................................. [ 76240]
MK128995.1_China/2018/AnhuiXCGQ  ....................-----------................................................. [ 76240]
MH681419.1_ASFV/POL/2015         ....................-----------................................................. [ 76240]
LS478113.1_Estonia_2014          ....................-----------................................................. [ 76240]
KP843857.1_Odintsovo_02/14       ....................-----------................................................. [ 76240]
FN557520.1_strain_E75            .--......a..........-----------....t.....c....g.....g.c......................... [ 76240]
AY261366.1_Warthog               -------------------.-----------.cagt.....c..c....t....c......................... [ 76240]
AY261365.1_Warmbaths             ....................-----------..........c..c.........c......................... [ 76240]
AY261364.1_Tengani_62            .--a................-----------.........a..t..........c......................... [ 76240]
AY261363.1_Pretoriuskop/96/4     ....................-----------..........c..c.........c......................... [ 76240]
AY261362.1_Mkuzi_1979            .--......a..........-----------....t.....c..........g.c......................... [ 76240]
AY261361.1_Malawi_Lil-20/1_1983  ---c...a......g...g.cgtcggccgcg.c.g.ca...c.t....t.....cc........................ [ 76240]
AY261360.1_Kenya_1950            --------------------------------t........c.g--------.tc......................... [ 76240]
AM712240.1_OURT_88/3             .--......a..........-----------....t.....c....g.....g.c......................... [ 76240]
AM712239.1_Benin_97/1            .--......a..........-----------....t.....c....g.....g.c......................... [ 76240]
KM111295.1_Ken06.Bus             ----------------------------------.a...g.c.----------tc......................... [ 76240]
KM111294.1_Ken05/Tk1             ---c.g.cgc....gt..g.---------cgcta.a.....c.g--------.tc......................... [ 76240]
KM262844.1_strain_L60            .--......a..........-----------....t.....c....g.....g.c......................... [ 76240]
KM102979.1_26544/OG10_Italy      .--......a..........-----------....t.....c....g.....g.c......................... [ 76240]
KM262845.1_strain_NHV            .--......a..........-----------....t.....c....g.....g.c......................... [ 76240]
KX354450.1_47/Ss/2008            .--......a..........-----------....t.....c....g.....g.c......................... [ 76240]
KP055815.1_strain_BA71           .--......a..........-----------....t.....c....g.....g.c......................... [ 76240]
MH025920.1_strain_R35            ----------------------------------.a...g.c.----------tc......................... [ 76240]
MH025919.1_strain_N10            ----------------------------------.a...g.c.----------tc......................... [ 76240]
MH025918.1_strain_R25            ----------------------------------.a...g.c.----------tc......................... [ 76240]
MH025917.1_strain_R7             ----------------------------------.a...g.c.----------tc......................... [ 76240]
MH025916.1_strain_R8             ----------------------------------.a...g.c.----------tc......................... [ 76240]

FR682468.2_Georgia_2007/1        gagttaagtattgttgat--tgtataatccatcatggttgagccacgcgaacagttttttcaagatctgctttcagcagt [ 76320]
ASF_IND_20_CAD543                ..................--............y............................................... [ 76320]
LR899193.1_Germany_2020/1        ..................--............................................................ [ 76320]
MN393477.1_Wuhan_2019            ..................--............................................................ [ 76320]
MW396979.1_Timor-Leste/2019/1    ..................--............................................................ [ 76320]
MK940252.1_CN/2019/InnerMongolia ..................--............................................................ [ 76320]
MK128995.1_China/2018/AnhuiXCGQ  ..................--............................................................ [ 76320]
MH681419.1_ASFV/POL/2015         ..................--............................................................ [ 76320]
LS478113.1_Estonia_2014          ..................--............................................................ [ 76320]
KP843857.1_Odintsovo_02/14       ..................--............................................................ [ 76320]
```

```
FN557520.1_strain_E75           ..................--........t...t................................c.............. [ 76320]
AY261366.1_Warthog              ......c...........--............................................................ [ 76320]
AY261365.1_Warmbaths            ......c...........--...........c................................................ [ 76320]
AY261364.1_Tengani_62           ..................--............................................................ [ 76320]
AY261363.1_Pretoriuskop/96/4    ......c...........--............................................................ [ 76320]
AY261362.1_Mkuzi_1979           ..................--........................g....................c.............. [ 76320]
AY261361.1_Malawi_Lil-20/1_1983 ..................--.......c.a.................................................. [ 76320]
AY261360.1_Kenya_1950           ......g...........ta.a.......a.................................................. [ 76320]
AM712240.1_OURT_88/3            ..................--........t...t................................c.............. [ 76320]
AM712239.1_Benin_97/1           ..................--........t...t................................c.............. [ 76320]
KM111295.1_Ken06.Bus            ......g...........--.a.......a.................................................. [ 76320]
KM111294.1_Ken05/Tk1            ......g...........ta.a.......a.................................................. [ 76320]
KM262844.1_strain_L60           ..................--........t...t................................c.............. [ 76320]
KM102979.1_26544/OG10_Italy     ..................--........t...t................................c.............. [ 76320]
KM262845.1_strain_NHV           ..................--........t...t................................c.............. [ 76320]
KX354450.1_47/Ss/2008           ..................--........t...t................................c.............. [ 76320]
KP055815.1_strain_BA71          ..................--........t...t................................c.............. [ 76320]
MH025920.1_strain_R35           ......g...........--.a.......a.................................................. [ 76320]
MH025919.1_strain_N10           ......g...........--.a.......a.................................................. [ 76320]
MH025918.1_strain_R25           ......g...........--.a.......a.................................................. [ 76320]
MH025917.1_strain_R7            ......g...........--.a.......a.................................................. [ 76320]
MH025916.1_strain_R8            ......g...........--.a.......a.................................................. [ 76320]

FR682468.2_Georgia_2007/1       ggatcaacaaatggacactgtaaaaaatgacataaaagacattatgaaagaaaaaacgtcttttatggtatcattcgaaa [ 76400]
ASF_IND_20_CAD543               .........................................................r...........r.......... [ 76400]
LR899193.1_Germany_2020/1       ................................................................................ [ 76400]
MN393477.1_Wuhan_2019           ................................................................................ [ 76400]
MW396979.1_Timor-Leste/2019/1   ................................................................................ [ 76400]
MK940252.1_CN/2019/InnerMongolia ................................................................................ [ 76400]
MK128995.1_China/2018/AnhuiXCGQ ................................................................................ [ 76400]
MH681419.1_ASFV/POL/2015        ................................................................................ [ 76400]
LS478113.1_Estonia_2014         ................................................................................ [ 76400]
KP843857.1_Odintsovo_02/14      ................................................................................ [ 76400]
FN557520.1_strain_E75           ..........................................c..............a...........g.......... [ 76400]
AY261366.1_Warthog              a.........................................c..............a...................... [ 76400]
AY261365.1_Warmbaths            .............a............................c..............a...................... [ 76400]
AY261364.1_Tengani_62           .........................................................a...................... [ 76400]
AY261363.1_Pretoriuskop/96/4    ..........................................c..............a...................... [ 76400]
AY261362.1_Mkuzi_1979           .............a............................c..............a...................... [ 76400]
AY261361.1_Malawi_Lil-20/1_1983 ....a.....................................c..............a..........cg.......... [ 76400]
AY261360.1_Kenya_1950           ....a.........................t....t....g.c..............a......c.t............. [ 76400]
AM712240.1_OURT_88/3            ..........................................c..............a...........g.......... [ 76400]
AM712239.1_Benin_97/1           ..........................................c..............a...........g.......... [ 76400]
KM111295.1_Ken06.Bus            ....a.........................t....t....g.c..............a......c.t............. [ 76400]
```

```
KM111294.1_Ken05/Tk1           ....a.........................t....t....g.c..............a......c.t............. [ 76400]
KM262844.1_strain_L60          ..........................................c..............a...........g.......... [ 76400]
KM102979.1_26544/OG10_Italy    ..........................................c..............a...........g.......... [ 76400]
KM262845.1_strain_NHV          ..........................................c..............a...........g.......... [ 76400]
KX354450.1_47/Ss/2008          ..........................................c..............a...........g.......... [ 76400]
KP055815.1_strain_BA71         ..........................................c..............a...........g.......... [ 76400]
MH025920.1_strain_R35          ....a.........................t....t....g.c..............a......c.t............. [ 76400]
MH025919.1_strain_N10          ....a.........................t....t....g.c..............a......c.t............. [ 76400]
MH025918.1_strain_R25          ....a.........................t....t....g.c..............a......c.t............. [ 76400]
MH025917.1_strain_R7           ....a.........................t....t....g.c..............a......c.t............. [ 76400]
MH025916.1_strain_R8           ....a.........................t....t....g.c..............a......c.t............. [ 76400]

FR682468.2_Georgia_2007/1      actttatagaacgttacgataccatggaaaaaaatattcaagaccttcagaataagtacgaagaaatggcggccaacctt [ 76480]
ASF_IND_20_CAD543              ................................................................................ [ 76480]
LR899193.1_Germany_2020/1      ................................................................................ [ 76480]
MN393477.1_Wuhan_2019          ................................................................................ [ 76480]
MW396979.1_Timor-Leste/2019/1  ................................................................................ [ 76480]
MK940252.1_CN/2019/InnerMongolia ................................................................................ [ 76480]
MK128995.1_China/2018/AnhuiXCGQ ................................................................................ [ 76480]
MH681419.1_ASFV/POL/2015       ................................................................................ [ 76480]
LS478113.1_Estonia_2014        ................................................................................ [ 76480]
KP843857.1_Odintsovo_02/14     ................................................................................ [ 76480]
FN557520.1_strain_E75          ................................................................................ [ 76480]
AY261366.1_Warthog             ............................................................................t... [ 76480]
AY261365.1_Warmbaths           ............................................................................t... [ 76480]
AY261364.1_Tengani_62          ................................................................................ [ 76480]
AY261363.1_Pretoriuskop/96/4   ...........................................t................................t... [ 76480]
AY261362.1_Mkuzi_1979          ............................................................................t... [ 76480]
AY261361.1_Malawi_Lil-20/1_1983 .................................................a....................cat....... [ 76480]
AY261360.1_Kenya_1950          .......g.....g.............................t...........................aa...t... [ 76480]
AM712240.1_OURT_88/3           ................................................................................ [ 76480]
AM712239.1_Benin_97/1          ................................................................................ [ 76480]
KM111295.1_Ken06.Bus           .......g.....g.........................................................aa...t... [ 76480]
KM111294.1_Ken05/Tk1           .......g.....g.............................t...........................aa...t... [ 76480]
KM262844.1_strain_L60          ................................................................................ [ 76480]
KM102979.1_26544/OG10_Italy    ................................................................................ [ 76480]
KM262845.1_strain_NHV          ................................................................................ [ 76480]
KX354450.1_47/Ss/2008          ................................................................................ [ 76480]
KP055815.1_strain_BA71         ................................................................................ [ 76480]
MH025920.1_strain_R35          .......g.....g.........................................................aa...t... [ 76480]
MH025919.1_strain_N10          .......g.....g.........................................................aa...t... [ 76480]
MH025918.1_strain_R25          .......g.....g.........................................................aa...t... [ 76480]
MH025917.1_strain_R7           .......g.....g.........................................................aa...t... [ 76480]
MH025916.1_strain_R8           .......g.....g.........................................................aa...t... [ 76480]
```

```
FR682468.2_Georgia_2007/1        atgaccgtcatgacggatacaaaaattcagcttggagccattatcgcccaacttgagattctaatgataaatggcactcc [ 76560]
ASF_IND_20_CAD543                ................................................................................ [ 76560]
LR899193.1_Germany_2020/1        ..............a................................................................. [ 76560]
MN393477.1_Wuhan_2019            ................................................................................ [ 76560]
MW396979.1_Timor-Leste/2019/1    ................................................................................ [ 76560]
MK940252.1_CN/2019/InnerMongolia ................................................................................ [ 76560]
MK128995.1_China/2018/AnhuiXCGQ  ................................................................................ [ 76560]
MH681419.1_ASFV/POL/2015         ................................................................................ [ 76560]
LS478113.1_Estonia_2014          ................................................................................ [ 76560]
KP843857.1_Odintsovo_02/14       ................................................................................ [ 76560]
FN557520.1_strain_E75            ..............................................................g................. [ 76560]
AY261366.1_Warthog               ........t................................c...................................... [ 76560]
AY261365.1_Warmbaths             .........................................c...................................... [ 76560]
AY261364.1_Tengani_62            .........................................................................a...... [ 76560]
AY261363.1_Pretoriuskop/96/4     .........................................c...................................... [ 76560]
AY261362.1_Mkuzi_1979            .........................................c...................................... [ 76560]
AY261361.1_Malawi_Lil-20/1_1983  gcag.t..t......................................t..............g................. [ 76560]
AY261360.1_Kenya_1950            g.tg........g.a.....g.......................t..t............a.....g....c........ [ 76560]
AM712240.1_OURT_88/3             ..............................................................g................. [ 76560]
AM712239.1_Benin_97/1            ..............................................................g................. [ 76560]
KM111295.1_Ken06.Bus             g.tg....t...g.a.....g.......................t..t............a.....g....c........ [ 76560]
KM111294.1_Ken05/Tk1             g.tg........g.a.....g.......................t..t............a.....g....c........ [ 76560]
KM262844.1_strain_L60            ..............................................................g................. [ 76560]
KM102979.1_26544/OG10_Italy      ..............................................................g................. [ 76560]
KM262845.1_strain_NHV            ..............................................................g................. [ 76560]
KX354450.1_47/Ss/2008            ..............................................................g................. [ 76560]
KP055815.1_strain_BA71           ..............................................................g................. [ 76560]
MH025920.1_strain_R35            g.tg....t...g.a.....g.......................t..t............a.....g....c........ [ 76560]
MH025919.1_strain_N10            g.tg....t...g.a.....g.......................t..t............a.....g....c........ [ 76560]
MH025918.1_strain_R25            g.tg....t...g.a.....g.......................t..t............a.....g....c........ [ 76560]
MH025917.1_strain_R7             g.tg....t...g.a.....g.......................t..t............a.....g....c........ [ 76560]
MH025916.1_strain_R8             g.tg....t...g.a.....g.......................t..t............a.....g....c........ [ 76560]

FR682468.2_Georgia_2007/1        acttccggcaaaaaagacaacaattaaggaggctatgcccttaccttcatcaaacacgaataatgaacaaacg------- [ 76640]
ASF_IND_20_CAD543                ........................................y................................------- [ 76640]
LR899193.1_Germany_2020/1        .........................................................................------- [ 76640]
MN393477.1_Wuhan_2019            .........................................................................------- [ 76640]
MW396979.1_Timor-Leste/2019/1    .........................................................................------- [ 76640]
MK940252.1_CN/2019/InnerMongolia .........................................................................------- [ 76640]
MK128995.1_China/2018/AnhuiXCGQ  .........................................................................------- [ 76640]
MH681419.1_ASFV/POL/2015         .........................................................................------- [ 76640]
LS478113.1_Estonia_2014          .........................................................................------- [ 76640]
KP843857.1_Odintsovo_02/14       .........................................................................------- [ 76640]
```

```
FN557520.1_strain_E75            ...............a.....g..................c...................c.....t......------- [ 76640]
AY261366.1_Warthog               .....................g...........c.......................................------- [ 76640]
AY261365.1_Warmbaths             .....................g...........c.......................................------- [ 76640]
AY261364.1_Tengani_62            .....................g........t............................g......t......------- [ 76640]
AY261363.1_Pretoriuskop/96/4     .....................g...........c.............t.........................------- [ 76640]
AY261362.1_Mkuzi_1979            .....................g...........c.......................................------- [ 76640]
AY261361.1_Malawi_Lil-20/1_1983  ......a........a.....g.............c..................................t..atgtcta [ 76640]
AY261360.1_Kenya_1950            .....................g..g.....t....cat.......ac..c.....c.t............t..gtgttta [ 76640]
AM712240.1_OURT_88/3             ...............a.....g..................c...................c.....t......------- [ 76640]
AM712239.1_Benin_97/1            ...............a.....g..................c...................c.....t......------- [ 76640]
KM111295.1_Ken06.Bus             .....................g..g.....ta.c.cat.......a.........c.t............t..gtgttta [ 76640]
KM111294.1_Ken05/Tk1             .....................g..g.....t....cat.......ac..c.....c.t............t..gtgttta [ 76640]
KM262844.1_strain_L60            ...............a.....g..................c...................c.....t......------- [ 76640]
KM102979.1_26544/OG10_Italy      ...............a.....g..................c...................c.....t......------- [ 76640]
KM262845.1_strain_NHV            ...............a.....g..................c...................c.....t......------- [ 76640]
KX354450.1_47/Ss/2008            ...............a.....g..................c...................c.....t......------- [ 76640]
KP055815.1_strain_BA71           ...............a.....g..................c...................c.....t......------- [ 76640]
MH025920.1_strain_R35            .....................g..g.....ta.c.cat.......a.........c.t............t..gtgttta [ 76640]
MH025919.1_strain_N10            .....................g..g.....ta.c.cat.......a.........c.t............t..gtgttta [ 76640]
MH025918.1_strain_R25            .....................g..g.....ta.c.cat.......a.........c.t............t..gtgttta [ 76640]
MH025917.1_strain_R7             .....................g..g.....ta.c.cat.......a.........c.t............t..gtgttta [ 76640]
MH025916.1_strain_R8             .....................g..g.....ta.c.cat.......a.........c.t............t..gtgttta [ 76640]

FR682468.2_Georgia_2007/1        -----agtcctcccgcctcaggcaaaacaagtgaaacacctaaaaaaaatcccacgaatgcgatgttcttcacgcgtagc [ 76720]
ASF_IND_20_CAD543                -----........................................................................... [ 76720]
LR899193.1_Germany_2020/1        -----........................................................................... [ 76720]
MN393477.1_Wuhan_2019            -----........................................................................... [ 76720]
MW396979.1_Timor-Leste/2019/1    -----........................................................................... [ 76720]
MK940252.1_CN/2019/InnerMongolia -----........................................................................... [ 76720]
MK128995.1_China/2018/AnhuiXCGQ  -----........................................................................... [ 76720]
MH681419.1_ASFV/POL/2015         -----........................................................................... [ 76720]
LS478113.1_Estonia_2014          -----........................................................................... [ 76720]
KP843857.1_Odintsovo_02/14       -----........................................................................... [ 76720]
FN557520.1_strain_E75            -----........................................................a.................. [ 76720]
AY261366.1_Warthog               -----.........a................................................................. [ 76720]
AY261365.1_Warmbaths             -----............c.............................................................. [ 76720]
AY261364.1_Tengani_62            -----...........g..........t.................................................... [ 76720]
AY261363.1_Pretoriuskop/96/4     -----........................................................................... [ 76720]
AY261362.1_Mkuzi_1979            -----............c.............................................................. [ 76720]
AY261361.1_Malawi_Lil-20/1_1983  cctat...t..t..att.....................gtc.............................t......... [ 76720]
AY261360.1_Kenya_1950            ccaatg..t.....a.t..........t.g........gtc..........t............................ [ 76720]
AM712240.1_OURT_88/3             -----........................................................a.................. [ 76720]
AM712239.1_Benin_97/1            -----........................................................a.................. [ 76720]
KM111295.1_Ken06.Bus             ccaatg..t....ta.t.....................g.c...........t........................... [ 76720]
```

```
KM111294.1_Ken05/Tk1             ccaatg..t.....a............t.g........gtc....................................... [ 76720]
KM262844.1_strain_L60            -----........................................................a.................. [ 76720]
KM102979.1_26544/OG10_Italy      -----........................................................a.................. [ 76720]
KM262845.1_strain_NHV            -----........................................................a.................. [ 76720]
KX354450.1_47/Ss/2008            -----........................................................a.................. [ 76720]
KP055815.1_strain_BA71           -----........................................................a.................. [ 76720]
MH025920.1_strain_R35            ccaatg..t....ta.t.....................g.c...........t........................... [ 76720]
MH025919.1_strain_N10            ccaatg..t....ta.t.....................g.c...........t........................... [ 76720]
MH025918.1_strain_R25            ccaatg..t....ta.t.....................g.c...........t........................... [ 76720]
MH025917.1_strain_R7             ccaatg..t....ta.t.....................g.c...........t........................... [ 76720]
MH025916.1_strain_R8             ccaatg..t....ta.t.....................g.c...........t........................... [ 76720]

FR682468.2_Georgia_2007/1        gaatgggcatcctcgaatacttttcgagaaaagtttttaacaccagaaattcaagccatattggatgagcagtttgcaaa [ 76800]
ASF_IND_20_CAD543                ................................................................................ [ 76800]
LR899193.1_Germany_2020/1        ................................................................................ [ 76800]
MN393477.1_Wuhan_2019            ................................................................................ [ 76800]
MW396979.1_Timor-Leste/2019/1    ................................................................................ [ 76800]
MK940252.1_CN/2019/InnerMongolia ................................................................................ [ 76800]
MK128995.1_China/2018/AnhuiXCGQ  ................................................................................ [ 76800]
MH681419.1_ASFV/POL/2015         ................................................................................ [ 76800]
LS478113.1_Estonia_2014          ................................................................................ [ 76800]
KP843857.1_Odintsovo_02/14       ................................................................................ [ 76800]
FN557520.1_strain_E75            .................a...................................g.......................... [ 76800]
AY261366.1_Warthog               ...............g................................................................ [ 76800]
AY261365.1_Warmbaths             ...............gt.......................................t....................... [ 76800]
AY261364.1_Tengani_62            ....................c........................................................... [ 76800]
AY261363.1_Pretoriuskop/96/4     ..............a................................................................. [ 76800]
AY261362.1_Mkuzi_1979            ...............gt.......................................t....................... [ 76800]
AY261361.1_Malawi_Lil-20/1_1983  ...........t..t..agag.......................t................................... [ 76800]
AY261360.1_Kenya_1950            ..............tg.ac.a......c.................................................... [ 76800]
AM712240.1_OURT_88/3             .................a...................................g.......................... [ 76800]
AM712239.1_Benin_97/1            .................a...................................g.......................... [ 76800]
KM111295.1_Ken06.Bus             ..............tg.ag.a......c.................................................... [ 76800]
KM111294.1_Ken05/Tk1             ........t.....tg.ag.a......c.................................................... [ 76800]
KM262844.1_strain_L60            .................a...................................g.......................... [ 76800]
KM102979.1_26544/OG10_Italy      .................a...................................g.......................... [ 76800]
KM262845.1_strain_NHV            .................a...................................g.......................... [ 76800]
KX354450.1_47/Ss/2008            .................a...................................g.......................... [ 76800]
KP055815.1_strain_BA71           .................a...................................g.......................... [ 76800]
MH025920.1_strain_R35            ..............tg.ag.a......c.................................................... [ 76800]
MH025919.1_strain_N10            ..............tg.ag.a......c.................................................... [ 76800]
MH025918.1_strain_R25            ..............tg.ag.a......c.................................................... [ 76800]
MH025917.1_strain_R7             ..............tg.ag.a......c.................................................... [ 76800]
MH025916.1_strain_R8             ..............tg.ag.a......c.................................................... [ 76800]
```

```
FR682468.2_Georgia_2007/1        caagaccgggatcgaaagattgcatgccgagggtctttacatgtggagaacccaattctctgacgaacagaagaaaatgg [ 76880]
ASF_IND_20_CAD543                ...............................................a................................ [ 76880]
LR899193.1_Germany_2020/1        ................................................................................ [ 76880]
MN393477.1_Wuhan_2019            ................................................................................ [ 76880]
MW396979.1_Timor-Leste/2019/1    ................................................................................ [ 76880]
MK940252.1_CN/2019/InnerMongolia ................................................................................ [ 76880]
MK128995.1_China/2018/AnhuiXCGQ  ................................................................................ [ 76880]
MH681419.1_ASFV/POL/2015         ................................................................................ [ 76880]
LS478113.1_Estonia_2014          ................................................................................ [ 76880]
KP843857.1_Odintsovo_02/14       ................................................................................ [ 76880]
FN557520.1_strain_E75            ................................................................................ [ 76880]
AY261366.1_Warthog               ..................................................g............................. [ 76880]
AY261365.1_Warmbaths             ..................................................g............................. [ 76880]
AY261364.1_Tengani_62            ................................................................................ [ 76880]
AY261363.1_Pretoriuskop/96/4     ................................................................................ [ 76880]
AY261362.1_Mkuzi_1979            ..................................................g............................. [ 76880]
AY261361.1_Malawi_Lil-20/1_1983  ...a..t..a............................................g......................... [ 76880]
AY261360.1_Kenya_1950            .........t.................t.....c....................g..t...................... [ 76880]
AM712240.1_OURT_88/3             ................................................................................ [ 76880]
AM712239.1_Benin_97/1            ................................................................................ [ 76880]
KM111295.1_Ken06.Bus             t........t.................t.....c....................g..t..c................... [ 76880]
KM111294.1_Ken05/Tk1             .........t.................t.....c....................g..t....................a. [ 76880]
KM262844.1_strain_L60            ................................................................................ [ 76880]
KM102979.1_26544/OG10_Italy      ................................................................................ [ 76880]
KM262845.1_strain_NHV            ................................................................................ [ 76880]
KX354450.1_47/Ss/2008            ................................................................................ [ 76880]
KP055815.1_strain_BA71           ................................................................................ [ 76880]
MH025920.1_strain_R35            t........t.................t.....c....................g..t..c................... [ 76880]
MH025919.1_strain_N10            t........t.................t.....c....................g..t..c................... [ 76880]
MH025918.1_strain_R25            t........t.................t.....c....................g..t..c................... [ 76880]
MH025917.1_strain_R7             t........t.................t.....c....................g..t..c................... [ 76880]
MH025916.1_strain_R8             t........t.................t.....c....................g..t..c................... [ 76880]

FR682468.2_Georgia_2007/1        tcaaagagatgatgaagaagt-aatatttttggtaaaaatatttttatcaaaatttttttaccaataataaaaaatatt [ 76960]
ASF_IND_20_CAD543                .....................-.......................................................... [ 76960]
LR899193.1_Germany_2020/1        .....................-.......................................................... [ 76960]
MN393477.1_Wuhan_2019            .....................-.......................................................... [ 76960]
MW396979.1_Timor-Leste/2019/1    .....................-.......................................................... [ 76960]
MK940252.1_CN/2019/InnerMongolia .....................-.......................................................... [ 76960]
MK128995.1_China/2018/AnhuiXCGQ  .....................-.......................................................... [ 76960]
MH681419.1_ASFV/POL/2015         .....................-.......................................................... [ 76960]
LS478113.1_Estonia_2014          .....................-.......................................................... [ 76960]
KP843857.1_Odintsovo_02/14       .....................-.......................................................... [ 76960]
```

```
FN557520.1_strain_E75            .....................-................--ta............-......................... [ 76960]
AY261366.1_Warthog               .....................-.........................c................................ [ 76960]
AY261365.1_Warmbaths             .....................-.........................c................................ [ 76960]
AY261364.1_Tengani_62            .....................-.......................................t................-- [ 76960]
AY261363.1_Pretoriuskop/96/4     .....................-...................-.....c................................ [ 76960]
AY261362.1_Mkuzi_1979            .....................-.........................c................................ [ 76960]
AY261361.1_Malawi_Lil-20/1_1983  .....................g.........acc............ggt...ta........t.....t........... [ 76960]
AY261360.1_Kenya_1950            .....................-.......g.acc............ggt.c..a.......t...............t.. [ 76960]
AM712240.1_OURT_88/3             .....................-........................................................-. [ 76960]
AM712239.1_Benin_97/1            .....................-.......................................................... [ 76960]
KM111295.1_Ken06.Bus             .....................-.........at.......--------------c......................... [ 76960]
KM111294.1_Ken05/Tk1             .....................-.........ac.......--------------c..c...................t.. [ 76960]
KM262844.1_strain_L60            .....................-.......................................................... [ 76960]
KM102979.1_26544/OG10_Italy      .....................-.......................................................... [ 76960]
KM262845.1_strain_NHV            .....................-........................................................-. [ 76960]
KX354450.1_47/Ss/2008            .....................-.......................................................... [ 76960]
KP055815.1_strain_BA71           .....................-.......................................................... [ 76960]
MH025920.1_strain_R35            .....................-.........at.......--------------c......................... [ 76960]
MH025919.1_strain_N10            .....................-.........at.......--------------c......................... [ 76960]
MH025918.1_strain_R25            .....................-.........at.......--------------c......................... [ 76960]
MH025917.1_strain_R7             .....................-.........at.......--------------c......................... [ 76960]
MH025916.1_strain_R8             .....................-.........at.......--------------c......................... [ 76960]

FR682468.2_Georgia_2007/1        ttttacttttttttcttcataatatacatagaatgcctacaaaagctggcacaaaaagtaccgcaaataaaaaaacaacg [ 77040]
ASF_IND_20_CAD543                ................................................................................ [ 77040]
LR899193.1_Germany_2020/1        ................................................................................ [ 77040]
MN393477.1_Wuhan_2019            ................................................................................ [ 77040]
MW396979.1_Timor-Leste/2019/1    ................................................................................ [ 77040]
MK940252.1_CN/2019/InnerMongolia ................................................................................ [ 77040]
MK128995.1_China/2018/AnhuiXCGQ  ................................................................................ [ 77040]
MH681419.1_ASFV/POL/2015         ................................................................................ [ 77040]
LS478113.1_Estonia_2014          ................................................................................ [ 77040]
KP843857.1_Odintsovo_02/14       ...--ac......................................................................... [ 77040]
FN557520.1_strain_E75            ................................................................................ [ 77040]
AY261366.1_Warthog               ................................................................................ [ 77040]
AY261365.1_Warmbaths             ................................................................................ [ 77040]
AY261364.1_Tengani_62            ------.......................................................................... [ 77040]
AY261363.1_Pretoriuskop/96/4     ................................................................................ [ 77040]
AY261362.1_Mkuzi_1979            ................................................................................ [ 77040]
AY261361.1_Malawi_Lil-20/1_1983  ..............................................c..................c.............. [ 77040]
AY261360.1_Kenya_1950            ............................................................................g..c [ 77040]
AM712240.1_OURT_88/3             ................................................................................ [ 77040]
AM712239.1_Benin_97/1            ................................................................................ [ 77040]
KM111295.1_Ken06.Bus             ............................................................................g..c [ 77040]
```

```
KM111294.1_Ken05/Tk1             ......-.....................................................................g..c [ 77040]
KM262844.1_strain_L60            ................................................................................ [ 77040]
KM102979.1_26544/OG10_Italy      ................................................................................ [ 77040]
KM262845.1_strain_NHV            ................................................................................ [ 77040]
KX354450.1_47/Ss/2008            ................................................................................ [ 77040]
KP055815.1_strain_BA71           ................................................................................ [ 77040]
MH025920.1_strain_R35            ............................................................................g..c [ 77040]
MH025919.1_strain_N10            ............................................................................g..c [ 77040]
MH025918.1_strain_R25            ............................................................................g..c [ 77040]
MH025917.1_strain_R7             ............................................................................g..c [ 77040]
MH025916.1_strain_R8             ............................................................................g..c [ 77040]

FR682468.2_Georgia_2007/1        ---aagggctcctccaaatctggttcttccagaggccacaccggcaaaacccatgcttcttcgtccatgcattccgggat [ 77120]
ASF_IND_20_CAD543                ---.......................y..................................................... [ 77120]
LR899193.1_Germany_2020/1        ---............................................................................. [ 77120]
MN393477.1_Wuhan_2019            ---............................................................................. [ 77120]
MW396979.1_Timor-Leste/2019/1    ---............................................................................. [ 77120]
MK940252.1_CN/2019/InnerMongolia ---............................................................................. [ 77120]
MK128995.1_China/2018/AnhuiXCGQ  ---............................................................................. [ 77120]
MH681419.1_ASFV/POL/2015         ---............................................................................. [ 77120]
LS478113.1_Estonia_2014          ---............................................................................. [ 77120]
KP843857.1_Odintsovo_02/14       ---............................................................................. [ 77120]
FN557520.1_strain_E75            ---.......................c..................................................... [ 77120]
AY261366.1_Warthog               ---.....t..................g.............................c..c................... [ 77120]
AY261365.1_Warmbaths             ---.....t..................c.............................c..c................... [ 77120]
AY261364.1_Tengani_62            ---.....t..................g.............................c..c.a................. [ 77120]
AY261363.1_Pretoriuskop/96/4     ---.....t..................c.............................c..c................... [ 77120]
AY261362.1_Mkuzi_1979            ---.....t................................................c..c................... [ 77120]
AY261361.1_Malawi_Lil-20/1_1983  acc.....gc...........---------------.c..aa..t..........---..a..g..c.a...cag..... [ 77120]
AY261360.1_Kenya_1950            ---.....gc.t...............g...a.......................---..a..g..c.....cat..... [ 77120]
AM712240.1_OURT_88/3             ---.......................c..................................................... [ 77120]
AM712239.1_Benin_97/1            ---.......................c..................................................... [ 77120]
KM111295.1_Ken06.Bus             ---.....gc.................g...a...................t...---..a..g..c.....caa..... [ 77120]
KM111294.1_Ken05/Tk1             ---.....gc.t...............g...a.......................---..a..g..c.....caa..a.. [ 77120]
KM262844.1_strain_L60            ---.......................c..................................................... [ 77120]
KM102979.1_26544/OG10_Italy      ---.......................c..................................................... [ 77120]
KM262845.1_strain_NHV            ---.......................c..................................................... [ 77120]
KX354450.1_47/Ss/2008            ---.......................c..................................................... [ 77120]
KP055815.1_strain_BA71           ---.......................c..................................................... [ 77120]
MH025920.1_strain_R35            ---.....gc.................g...a.......................---..a..g..c.....caa..... [ 77120]
MH025919.1_strain_N10            ---.....gc.................g...a.......................---..a..g..c.....caa..... [ 77120]
MH025918.1_strain_R25            ---.....gc.................g...a.......................---..a..g..c.....caa..... [ 77120]
MH025917.1_strain_R7             ---.....gc.................g...a.......................---..a..g..c.....caa..... [ 77120]
MH025916.1_strain_R8             ---.....gc.................g...a.......................---..a..g..c.....caa..... [ 77120]
```

```
FR682468.2_Georgia_2007/1        gctctataaagatatggtaaatattgctagatctagaggcattccgatttaccagaatggatcgcgtcttactaaaagtg [ 77200]
ASF_IND_20_CAD543                ................................................................................ [ 77200]
LR899193.1_Germany_2020/1        ................................................................................ [ 77200]
MN393477.1_Wuhan_2019            ................................................................................ [ 77200]
MW396979.1_Timor-Leste/2019/1    ................................................................................ [ 77200]
MK940252.1_CN/2019/InnerMongolia ................................................................................ [ 77200]
MK128995.1_China/2018/AnhuiXCGQ  ................................................................................ [ 77200]
MH681419.1_ASFV/POL/2015         ................................................................................ [ 77200]
LS478113.1_Estonia_2014          ................................................................................ [ 77200]
KP843857.1_Odintsovo_02/14       ................................................................................ [ 77200]
FN557520.1_strain_E75            ................................................................................ [ 77200]
AY261366.1_Warthog               ...................................a...............t.........a.................. [ 77200]
AY261365.1_Warmbaths             ...................................a.........................a.................. [ 77200]
AY261364.1_Tengani_62            ...................................a...............t.........a.a................ [ 77200]
AY261363.1_Pretoriuskop/96/4     ...........................c.......a.........................a.................. [ 77200]
AY261362.1_Mkuzi_1979            ................................................................................ [ 77200]
AY261361.1_Malawi_Lil-20/1_1983  ......................g....c.a.....ag..........................a........c....... [ 77200]
AY261360.1_Kenya_1950            .....................c.......a.....a...t.......................a................ [ 77200]
AM712240.1_OURT_88/3             ................................................................................ [ 77200]
AM712239.1_Benin_97/1            ................................................................................ [ 77200]
KM111295.1_Ken06.Bus             .....................c.....c.a.....ag..........................a................ [ 77200]
KM111294.1_Ken05/Tk1             .....................c.......a.....a...t.......................a................ [ 77200]
KM262844.1_strain_L60            ................................................................................ [ 77200]
KM102979.1_26544/OG10_Italy      ................................................................................ [ 77200]
KM262845.1_strain_NHV            ................................................................................ [ 77200]
KX354450.1_47/Ss/2008            ................................................................................ [ 77200]
KP055815.1_strain_BA71           ................................................................................ [ 77200]
MH025920.1_strain_R35            .....................c.....c.a.....ag..........................a................ [ 77200]
MH025919.1_strain_N10            .....................c.....c.a.....ag..........................a................ [ 77200]
MH025918.1_strain_R25            .....................c.....c.a.....ag..........................a................ [ 77200]
MH025917.1_strain_R7             .....................c.....c.a.....ag..........................a................ [ 77200]
MH025916.1_strain_R8             .....................c.....c.a.....ag..........................a................ [ 77200]

FR682468.2_Georgia_2007/1        aattggagaaaaaaattaaacggtcaaaatgaatataatcaggaaacttaagcctggaacaattagccttgtgctgggac [ 77280]
ASF_IND_20_CAD543                ................................................................................ [ 77280]
LR899193.1_Germany_2020/1        ................................................................................ [ 77280]
MN393477.1_Wuhan_2019            ................................................................................ [ 77280]
MW396979.1_Timor-Leste/2019/1    ................................................................................ [ 77280]
MK940252.1_CN/2019/InnerMongolia ................................................................................ [ 77280]
MK128995.1_China/2018/AnhuiXCGQ  ................................................................................ [ 77280]
MH681419.1_ASFV/POL/2015         ................................................................................ [ 77280]
LS478113.1_Estonia_2014          ................................................................................ [ 77280]
KP843857.1_Odintsovo_02/14       ................................................................................ [ 77280]
```

```
FN557520.1_strain_E75           .......................c...............t.....g.................................. [ 77280]
AY261366.1_Warthog              .......................................t.....g.....a............................ [ 77280]
AY261365.1_Warmbaths            ..................g..........................g.....a............................ [ 77280]
AY261364.1_Tengani_62           .............................................g.................................. [ 77280]
AY261363.1_Pretoriuskop/96/4    ..................g..........................g.....a............................ [ 77280]
AY261362.1_Mkuzi_1979           .......................................t.....g.................................. [ 77280]
AY261361.1_Malawi_Lil-20/1_1983 ..................g..........................g.................................. [ 77280]
AY261360.1_Kenya_1950           ................................g.t.......a..g.................................. [ 77280]
AM712240.1_OURT_88/3            .......................................t.....g.................................. [ 77280]
AM712239.1_Benin_97/1           .......................................t.....g.................................. [ 77280]
KM111295.1_Ken06.Bus            ................................g.c.......a..g.................................. [ 77280]
KM111294.1_Ken05/Tk1            ................................g.t.......a..g.................................. [ 77280]
KM262844.1_strain_L60           .......................................t.....g.................................. [ 77280]
KM102979.1_26544/OG10_Italy     .......................................t.....g.................................. [ 77280]
KM262845.1_strain_NHV           .......................................t.....g.................................. [ 77280]
KX354450.1_47/Ss/2008           .......................................t.....g.................................. [ 77280]
KP055815.1_strain_BA71          .......................................t.....g.................................. [ 77280]
MH025920.1_strain_R35           ................................g.c.......a..g.................................. [ 77280]
MH025919.1_strain_N10           ................................g.c.......a..g.................................. [ 77280]
MH025918.1_strain_R25           ................................g.c.......a..g.................................. [ 77280]
MH025917.1_strain_R7            ................................g.c.......a..g.................................. [ 77280]
MH025916.1_strain_R8            ................................g.c.......a..g.................................. [ 77280]

FR682468.2_Georgia_2007/1       ccatgtttgccggcaaaactacgtttcttattcattgcatttacatgctcgaacgtttggaaaaaaaagtagtcttcata [ 77360]
ASF_IND_20_CAD543               ................................................................................ [ 77360]
LR899193.1_Germany_2020/1       ................................................................................ [ 77360]
MN393477.1_Wuhan_2019           ................................................................................ [ 77360]
MW396979.1_Timor-Leste/2019/1   ................................................................................ [ 77360]
MK940252.1_CN/2019/InnerMongolia ................................................................................ [ 77360]
MK128995.1_China/2018/AnhuiXCGQ ................................................................................ [ 77360]
MH681419.1_ASFV/POL/2015        ................................................................................ [ 77360]
LS478113.1_Estonia_2014         ................................................................................ [ 77360]
KP843857.1_Odintsovo_02/14      ................................................................................ [ 77360]
FN557520.1_strain_E75           ................................................................................ [ 77360]
AY261366.1_Warthog              ......................................................a......................... [ 77360]
AY261365.1_Warmbaths            ................................................................................ [ 77360]
AY261364.1_Tengani_62           ........................................................................c....... [ 77360]
AY261363.1_Pretoriuskop/96/4    ................................................................................ [ 77360]
AY261362.1_Mkuzi_1979           ................................................................................ [ 77360]
AY261361.1_Malawi_Lil-20/1_1983 ................................................................................ [ 77360]
AY261360.1_Kenya_1950           .t.......................................a................a..................... [ 77360]
AM712240.1_OURT_88/3            ................................................................................ [ 77360]
AM712239.1_Benin_97/1           ................................................................................ [ 77360]
KM111295.1_Ken06.Bus            .t........t........g.....................a................a..................... [ 77360]
```

```
KM111294.1_Ken05/Tk1          .........................................a...c............a..................... [ 77360]
KM262844.1_strain_L60         ................................................................................ [ 77360]
KM102979.1_26544/OG10_Italy   ................................................................................ [ 77360]
KM262845.1_strain_NHV         ................................................................................ [ 77360]
KX354450.1_47/Ss/2008         ................................................................................ [ 77360]
KP055815.1_strain_BA71        ................................................................................ [ 77360]
MH025920.1_strain_R35         .t........t........g.....................a................a..................... [ 77360]
MH025919.1_strain_N10         .t........t........g.....................a................a..................... [ 77360]
MH025918.1_strain_R25         .t........t........g.....................a................a..................... [ 77360]
MH025917.1_strain_R7          .t........t........g.....................a................a..................... [ 77360]
MH025916.1_strain_R8          .t........t........g.....................a................a..................... [ 77360]

FR682468.2_Georgia_2007/1     aaatctaccaaaaacacccgagacaaaactattaaaacacactccggtatacagctacgacccaaacaatgtaaaatcat [ 77440]
ASF_IND_20_CAD543             .................................................................r.............. [ 77440]
LR899193.1_Germany_2020/1     ................................................................................ [ 77440]
MN393477.1_Wuhan_2019         ................................................................................ [ 77440]
MW396979.1_Timor-Leste/2019/1 ................................................................................ [ 77440]
MK940252.1_CN/2019/InnerMongolia ................................................................................ [ 77440]
MK128995.1_China/2018/AnhuiXCGQ ................................................................................ [ 77440]
MH681419.1_ASFV/POL/2015      ................................................................................ [ 77440]
LS478113.1_Estonia_2014       ................................................................................ [ 77440]
KP843857.1_Odintsovo_02/14    ................................................................................ [ 77440]
FN557520.1_strain_E75         .................................................................g.............. [ 77440]
AY261366.1_Warthog            .................................................................g..t........... [ 77440]
AY261365.1_Warmbaths          ........a...................................................t....g.............. [ 77440]
AY261364.1_Tengani_62         .................................................................g.............. [ 77440]
AY261363.1_Pretoriuskop/96/4  ........a........t...............................................g...........t.. [ 77440]
AY261362.1_Mkuzi_1979         .................................................................g.............. [ 77440]
AY261361.1_Malawi_Lil-20/1_1983 ........a.................................................a.t....g......g....... [ 77440]
AY261360.1_Kenya_1950         ........a..............t.....g...c.........................t.....t.............. [ 77440]
AM712240.1_OURT_88/3          .................................................................g.............. [ 77440]
AM712239.1_Benin_97/1         .................................................................g.............. [ 77440]
KM111295.1_Ken06.Bus          ........a....................g...c...............................t...........t.. [ 77440]
KM111294.1_Ken05/Tk1          ........a..............t.....g...c.........................t.....t.............. [ 77440]
KM262844.1_strain_L60         .................................................................g.............. [ 77440]
KM102979.1_26544/OG10_Italy   .................................................................g.............. [ 77440]
KM262845.1_strain_NHV         .................................................................g.............. [ 77440]
KX354450.1_47/Ss/2008         .................................................................g.............. [ 77440]
KP055815.1_strain_BA71        .................................................................g.............. [ 77440]
MH025920.1_strain_R35         ........a....................g...c...............................t...........t.. [ 77440]
MH025919.1_strain_N10         ........a....................g...c...............................t...........t.. [ 77440]
MH025918.1_strain_R25         ........a....................g...c...............................t...........t.. [ 77440]
MH025917.1_strain_R7          ........a....................g...c...............................t...........t.. [ 77440]
MH025916.1_strain_R8          ........a....................g...c...............................t...........t.. [ 77440]
```

| | | |
|---|---|---|
| FR682468.2_Georgia_2007/1 | agaaagcacacagttatctgacgtgggttctctcaccgatatccatgcagttgtcgtagatgaagcgcatttttttgacg | [ 77520] |
| ASF_IND_20_CAD543 | ................................................................................ | [ 77520] |
| LR899193.1_Germany_2020/1 | ................................................................................ | [ 77520] |
| MN393477.1_Wuhan_2019 | ................................................................................ | [ 77520] |
| MW396979.1_Timor-Leste/2019/1 | ................................................................................ | [ 77520] |
| MK940252.1_CN/2019/InnerMongolia | ................................................................................ | [ 77520] |
| MK128995.1_China/2018/AnhuiXCGQ | ................................................................................ | [ 77520] |
| MH681419.1_ASFV/POL/2015 | ................................................................................ | [ 77520] |
| LS478113.1_Estonia_2014 | ................................................................................ | [ 77520] |
| KP843857.1_Odintsovo_02/14 | ................................................................................ | [ 77520] |
| FN557520.1_strain_E75 | ...........................a.................................................... | [ 77520] |
| AY261366.1_Warthog | ...........................a...........................a........................ | [ 77520] |
| AY261365.1_Warmbaths | ...........................a...........................a........................ | [ 77520] |
| AY261364.1_Tengani_62 | ...........................a...........................a........................ | [ 77520] |
| AY261363.1_Pretoriuskop/96/4 | ...........................a...........................a........................ | [ 77520] |
| AY261362.1_Mkuzi_1979 | ...........................a.................................................... | [ 77520] |
| AY261361.1_Malawi_Lil-20/1_1983 | ............a..............a.....a....................ta..........a............. | [ 77520] |
| AY261360.1_Kenya_1950 | .......g.....c.......t.....a.....a.......c............ga........................ | [ 77520] |
| AM712240.1_OURT_88/3 | ...........................a.................................................... | [ 77520] |
| AM712239.1_Benin_97/1 | ...........................a.................................................... | [ 77520] |
| KM111295.1_Ken06.Bus | .............c.......t.....a.....a.......c............ga........................ | [ 77520] |
| KM111294.1_Ken05/Tk1 | .............c.......t.....a..a..a.......c............ga........................ | [ 77520] |
| KM262844.1_strain_L60 | ...........................a.................................................... | [ 77520] |
| KM102979.1_26544/OG10_Italy | ...........................a.................................................... | [ 77520] |
| KM262845.1_strain_NHV | ...........................a.................................................... | [ 77520] |
| KX354450.1_47/Ss/2008 | ...........................a.................................................... | [ 77520] |
| KP055815.1_strain_BA71 | ...........................a.................................................... | [ 77520] |
| MH025920.1_strain_R35 | .............c.......t.....a.....a.......c............ga........................ | [ 77520] |
| MH025919.1_strain_N10 | .............c.......t.....a.....a.......c............ga........................ | [ 77520] |
| MH025918.1_strain_R25 | .............c.......t.....a.....a.......c............ga........................ | [ 77520] |
| MH025917.1_strain_R7 | .............c.......t.....a.....a.......c............ga........................ | [ 77520] |
| MH025916.1_strain_R8 | .............c.......t.....a.....a.......c............ga........................ | [ 77520] |

| | | |
|---|---|---|
| FR682468.2_Georgia_2007/1 | atttaatcacatgccgcacttgggcagaggaagaaaaaattattattcttgcgggactcaatgcttccttcgagcagaaa | [ 77600] |
| ASF_IND_20_CAD543 | ................................................................................ | [ 77600] |
| LR899193.1_Germany_2020/1 | ................................................................................ | [ 77600] |
| MN393477.1_Wuhan_2019 | ................................................................................ | [ 77600] |
| MW396979.1_Timor-Leste/2019/1 | ................................................................................ | [ 77600] |
| MK940252.1_CN/2019/InnerMongolia | ................................................................................ | [ 77600] |
| MK128995.1_China/2018/AnhuiXCGQ | ................................................................................ | [ 77600] |
| MH681419.1_ASFV/POL/2015 | ................................................................................ | [ 77600] |
| LS478113.1_Estonia_2014 | ................................................................................ | [ 77600] |
| KP843857.1_Odintsovo_02/14 | ................................................................................ | [ 77600] |

```
FN557520.1_strain_E75           .......a.a.........c............................................................ [ 77600]
AY261366.1_Warthog              .......a.a.........c........t................................................... [ 77600]
AY261365.1_Warmbaths            .......a.a.........c........t................................................... [ 77600]
AY261364.1_Tengani_62           .......a.a.........c........t...............................................a... [ 77600]
AY261363.1_Pretoriuskop/96/4    .......a.a.........c........t................................................... [ 77600]
AY261362.1_Mkuzi_1979           .......a.a.........c............................................................ [ 77600]
AY261361.1_Malawi_Lil-20/1_1983 .......a.a.......g.c........t................................................... [ 77600]
AY261360.1_Kenya_1950           .......a.g.........c........t...a............................................... [ 77600]
AM712240.1_OURT_88/3            .......a.a.........c............................................................ [ 77600]
AM712239.1_Benin_97/1           .......a.a.........c............................................................ [ 77600]
KM111295.1_Ken06.Bus            .......a.g.........c........t...a............................................... [ 77600]
KM111294.1_Ken05/Tk1            .......a.g.........c........t...a............................................... [ 77600]
KM262844.1_strain_L60           .......a.a.........c............................................................ [ 77600]
KM102979.1_26544/OG10_Italy     .......a.a.........c............................................................ [ 77600]
KM262845.1_strain_NHV           .......a.a.........c............................................................ [ 77600]
KX354450.1_47/Ss/2008           .......a.a.........c............................................................ [ 77600]
KP055815.1_strain_BA71          .......a.a.........c............................................................ [ 77600]
MH025920.1_strain_R35           .......a.g.........c........t...a............................................... [ 77600]
MH025919.1_strain_N10           .......a.g.........c........t...a............................................... [ 77600]
MH025918.1_strain_R25           .......a.g.........c........t...a............................................... [ 77600]
MH025917.1_strain_R7            .......a.g.........c........t...a............................................... [ 77600]
MH025916.1_strain_R8            .......a.g.........c........t...a............................................... [ 77600]

FR682468.2_Georgia_2007/1       atgtttccgcccatcgttcgtatttttccttactgcagctgggttaagtatattggccgcacctgtatgaaatgtaacca [ 77680]
ASF_IND_20_CAD543               ................................................................................ [ 77680]
LR899193.1_Germany_2020/1       ................................................................................ [ 77680]
MN393477.1_Wuhan_2019           ................................................................................ [ 77680]
MW396979.1_Timor-Leste/2019/1   ................................................................................ [ 77680]
MK940252.1_CN/2019/InnerMongolia ................................................................................ [ 77680]
MK128995.1_China/2018/AnhuiXCGQ ................................................................................ [ 77680]
MH681419.1_ASFV/POL/2015        ................................................................................ [ 77680]
LS478113.1_Estonia_2014         ................................................................................ [ 77680]
KP843857.1_Odintsovo_02/14      ................................................................................ [ 77680]
FN557520.1_strain_E75           ................................................................................ [ 77680]
AY261366.1_Warthog              ...............................................................................g [ 77680]
AY261365.1_Warmbaths            ...............................................................................g [ 77680]
AY261364.1_Tengani_62           ............................................g..................................g [ 77680]
AY261363.1_Pretoriuskop/96/4    ........at.....................................................................g [ 77680]
AY261362.1_Mkuzi_1979           ................................................................................ [ 77680]
AY261361.1_Malawi_Lil-20/1_1983 .......a...........a......................a....................................g [ 77680]
AY261360.1_Kenya_1950           .......a.............................a..................t......................g [ 77680]
AM712240.1_OURT_88/3            ................................................................................ [ 77680]
AM712239.1_Benin_97/1           ................................................................................ [ 77680]
KM111295.1_Ken06.Bus            .......a.............................a..................t......................g [ 77680]
```

```
KM111294.1_Ken05/Tk1           .......a.............................a..................t......................g [ 77680]
KM262844.1_strain_L60          ................................................................................ [ 77680]
KM102979.1_26544/OG10_Italy    ................................................................................ [ 77680]
KM262845.1_strain_NHV          ................................................................................ [ 77680]
KX354450.1_47/Ss/2008          ................................................................................ [ 77680]
KP055815.1_strain_BA71         ................................................................................ [ 77680]
MH025920.1_strain_R35          .......a.............................a..................t......................g [ 77680]
MH025919.1_strain_N10          .......a.............................a..................t......................g [ 77680]
MH025918.1_strain_R25          .......a.............................a..................t......................g [ 77680]
MH025917.1_strain_R7           .......a.............................a..................t......................g [ 77680]
MH025916.1_strain_R8           .......a.............................a..................t......................g [ 77680]

FR682468.2_Georgia_2007/1      acataatgcatgctttaatgtgcgtaagaacgcagacaagacgcttatccttgcgggaggaagtgaactgtacgtaacat [ 77760]
ASF_IND_20_CAD543              ................................................................................ [ 77760]
LR899193.1_Germany_2020/1      ................................................................................ [ 77760]
MN393477.1_Wuhan_2019          ................................................................................ [ 77760]
MW396979.1_Timor-Leste/2019/1  ................................................................................ [ 77760]
MK940252.1_CN/2019/InnerMongolia ................................................................................ [ 77760]
MK128995.1_China/2018/AnhuiXCGQ ................................................................................ [ 77760]
MH681419.1_ASFV/POL/2015       ................................................................................ [ 77760]
LS478113.1_Estonia_2014        ................................................................................ [ 77760]
KP843857.1_Odintsovo_02/14     ................................................................................ [ 77760]
FN557520.1_strain_E75          ................................................................................ [ 77760]
AY261366.1_Warthog             ................................................................................ [ 77760]
AY261365.1_Warmbaths           ................................................................................ [ 77760]
AY261364.1_Tengani_62          ........................................g...................g................... [ 77760]
AY261363.1_Pretoriuskop/96/4   ................................................t............................... [ 77760]
AY261362.1_Mkuzi_1979          ................................................................................ [ 77760]
AY261361.1_Malawi_Lil-20/1_1983 .................................g..........................g..c................ [ 77760]
AY261360.1_Kenya_1950          .................................g..t.......................g........a...a...... [ 77760]
AM712240.1_OURT_88/3           ................................................................................ [ 77760]
AM712239.1_Benin_97/1          ................................................................................ [ 77760]
KM111295.1_Ken06.Bus           .................................g......t...................g........a.......... [ 77760]
KM111294.1_Ken05/Tk1           .................................g..t.......................g........a...a...... [ 77760]
KM262844.1_strain_L60          ................................................................................ [ 77760]
KM102979.1_26544/OG10_Italy    ................................................................................ [ 77760]
KM262845.1_strain_NHV          ................................................................................ [ 77760]
KX354450.1_47/Ss/2008          ................................................................................ [ 77760]
KP055815.1_strain_BA71         ................................................................................ [ 77760]
MH025920.1_strain_R35          .................................g......t...................g........a.......... [ 77760]
MH025919.1_strain_N10          .................................g......t...................g........a.......... [ 77760]
MH025918.1_strain_R25          .................................g......t...................g........a.......... [ 77760]
MH025917.1_strain_R7           .................................g......t...................g........a.......... [ 77760]
MH025916.1_strain_R8           .................................g......t...................g........a.......... [ 77760]
```

```
FR682468.2_Georgia_2007/1         gttgtaacaactgtct-aaaaaatacatttattaag-------------------------------------------- [ 77840]
ASF_IND_20_CAD543                 ................-...................-------------------------------------------- [ 77840]
LR899193.1_Germany_2020/1         ................-...................-------------------------------------------- [ 77840]
MN393477.1_Wuhan_2019             ................-...................-------------------------------------------- [ 77840]
MW396979.1_Timor-Leste/2019/1     ................-...................-------------------------------------------- [ 77840]
MK940252.1_CN/2019/InnerMongolia  ................-...................-------------------------------------------- [ 77840]
MK128995.1_China/2018/AnhuiXCGQ   ................-...................-------------------------------------------- [ 77840]
MH681419.1_ASFV/POL/2015          ................-...................-------------------------------------------- [ 77840]
LS478113.1_Estonia_2014           ................-...................-------------------------------------------- [ 77840]
KP843857.1_Odintsovo_02/14        ................-...................-------------------------------------------- [ 77840]
FN557520.1_strain_E75             ................-...................-------------------------------------------- [ 77840]
AY261366.1_Warthog                ................-...................-------------------------------------------- [ 77840]
AY261365.1_Warmbaths              ................-...................-------------------------------------------- [ 77840]
AY261364.1_Tengani_62             ................-.........t.........-------------------------------------------- [ 77840]
AY261363.1_Pretoriuskop/96/4      ................a...................-------------------------------------------- [ 77840]
AY261362.1_Mkuzi_1979             ................-...................-------------------------------------------- [ 77840]
AY261361.1_Malawi_Lil-20/1_1983   .c.....t........a......c.a..g......a-------------------------------------------- [ 77840]
AY261360.1_Kenya_1950             .c.....t........a........a.........atagttgcaacctattaagtagttgcaacctattaagtagtttgc [ 77840]
AM712240.1_OURT_88/3              ................-...................-------------------------------------------- [ 77840]
AM712239.1_Benin_97/1             ................-...................-------------------------------------------- [ 77840]
KM111295.1_Ken06.Bus              .c.....t........a........a.........a-------------------------------------------- [ 77840]
KM111294.1_Ken05/Tk1              .c.....t........a........a.........atggttgcaacctattaag-------------------------- [ 77840]
KM262844.1_strain_L60             ................-...................-------------------------------------------- [ 77840]
KM102979.1_26544/OG10_Italy       ................-...................-------------------------------------------- [ 77840]
KM262845.1_strain_NHV             ................-...................-------------------------------------------- [ 77840]
KX354450.1_47/Ss/2008             ................-...................-------------------------------------------- [ 77840]
KP055815.1_strain_BA71            ................-...................-------------------------------------------- [ 77840]
MH025920.1_strain_R35             .c.....t........a........a.........a-------------------------------------------- [ 77840]
MH025919.1_strain_N10             .c.....t........a........a.........a-------------------------------------------- [ 77840]
MH025918.1_strain_R25             .c.....t........a........a.........a-------------------------------------------- [ 77840]
MH025917.1_strain_R7              .c.....t........a........a.........a-------------------------------------------- [ 77840]
MH025916.1_strain_R8              .c.....t........a........a.........a-------------------------------------------- [ 77840]

FR682468.2_Georgia_2007/1         -----------------------------------------------cagttgcaacctattaaatattaaaaatcttat [ 77920]
ASF_IND_20_CAD543                 -----------------------------------------------................................. [ 77920]
LR899193.1_Germany_2020/1         -----------------------------------------------................................. [ 77920]
MN393477.1_Wuhan_2019             -----------------------------------------------................................. [ 77920]
MW396979.1_Timor-Leste/2019/1     -----------------------------------------------................................. [ 77920]
MK940252.1_CN/2019/InnerMongolia  -----------------------------------------------................................. [ 77920]
MK128995.1_China/2018/AnhuiXCGQ   -----------------------------------------------................................. [ 77920]
MH681419.1_ASFV/POL/2015          -----------------------------------------------................................. [ 77920]
LS478113.1_Estonia_2014           -----------------------------------------------................................. [ 77920]
KP843857.1_Odintsovo_02/14        -----------------------------------------------................................. [ 77920]
```

```
FN557520.1_strain_E75              -----------------------------------------------................................. [ 77920]
AY261366.1_Warthog                 -----------------------------------------------................................. [ 77920]
AY261365.1_Warmbaths               -----------------------------------------------................................. [ 77920]
AY261364.1_Tengani_62              -----------------------------------------------................................. [ 77920]
AY261363.1_Pretoriuskop/96/4       -----------------------------------------------................................. [ 77920]
AY261362.1_Mkuzi_1979              -----------------------------------------------................................. [ 77920]
AY261361.1_Malawi_Lil-20/1_1983    -----------------------------------------------.ca......................t....... [ 77920]
AY261360.1_Kenya_1950              aacctattaaatggttgcgacctattaaatggttgcgacctattaagt......g............a....t..a.... [ 77920]
AM712240.1_OURT_88/3               -----------------------------------------------................................. [ 77920]
AM712239.1_Benin_97/1              -----------------------------------------------................................. [ 77920]
KM111295.1_Ken06.Bus               -----------------------------------------------tg.....g............a....t..a.... [ 77920]
KM111294.1_Ken05/Tk1               -----------------------------------------------t......g............a....t..a.... [ 77920]
KM262844.1_strain_L60              -----------------------------------------------................................. [ 77920]
KM102979.1_26544/OG10_Italy        -----------------------------------------------................................. [ 77920]
KM262845.1_strain_NHV              -----------------------------------------------................................. [ 77920]
KX354450.1_47/Ss/2008              -----------------------------------------------................................. [ 77920]
KP055815.1_strain_BA71             -----------------------------------------------................................. [ 77920]
MH025920.1_strain_R35              -----------------------------------------------tg.....g............a....t..a.... [ 77920]
MH025919.1_strain_N10              -----------------------------------------------tg.....g............a....t..a.... [ 77920]
MH025918.1_strain_R25              -----------------------------------------------tg.....g............a....t..a.... [ 77920]
MH025917.1_strain_R7               -----------------------------------------------tg.....g............a....t..a.... [ 77920]
MH025916.1_strain_R8               -----------------------------------------------tg.....g............a....t..a.... [ 77920]

FR682468.2_Georgia_2007/1          acaataatggatcattatcttaaaaaattacaagatatttatacgaagctcgagggtcatccctttctttttagcccgtc [ 78000]
ASF_IND_20_CAD543                  ................................................................................ [ 78000]
LR899193.1_Germany_2020/1          ................................................................................ [ 78000]
MN393477.1_Wuhan_2019              ................................................................................ [ 78000]
MW396979.1_Timor-Leste/2019/1      ................................................................................ [ 78000]
MK940252.1_CN/2019/InnerMongolia   ................................................................................ [ 78000]
MK128995.1_China/2018/AnhuiXCGQ    ................................................................................ [ 78000]
MH681419.1_ASFV/POL/2015           ................................................................................ [ 78000]
LS478113.1_Estonia_2014            ................................................................................ [ 78000]
KP843857.1_Odintsovo_02/14         ................................................................................ [ 78000]
FN557520.1_strain_E75              ........................................................c....................... [ 78000]
AY261366.1_Warthog                 ........................................................c....................... [ 78000]
AY261365.1_Warmbaths               ........................................................c....................... [ 78000]
AY261364.1_Tengani_62              ................................................................................ [ 78000]
AY261363.1_Pretoriuskop/96/4       ...................................a....................c....................... [ 78000]
AY261362.1_Mkuzi_1979              ........................................................c....................... [ 78000]
AY261361.1_Malawi_Lil-20/1_1983    ..t...........................g.g..........aa.....t.......................t..... [ 78000]
AY261360.1_Kenya_1950              ..t.............................g..........a.......t........c..............t..... [ 78000]
AM712240.1_OURT_88/3               ........................................................c....................... [ 78000]
AM712239.1_Benin_97/1              ........................................................c....................... [ 78000]
KM111295.1_Ken06.Bus               ..t.............................g..........a.......t........c..............t..... [ 78000]
```

| | | |
|---|---|---|
| KM111294.1_Ken05/Tk1 | ..t.............................g..........a......t........c..............t..... | [ 78000] |
| KM262844.1_strain_L60 | ........................................................c....................... | [ 78000] |
| KM102979.1_26544/OG10_Italy | ........................................................c....................... | [ 78000] |
| KM262845.1_strain_NHV | ........................................................c....................... | [ 78000] |
| KX354450.1_47/Ss/2008 | ........................................................c....................... | [ 78000] |
| KP055815.1_strain_BA71 | ........................................................c....................... | [ 78000] |
| MH025920.1_strain_R35 | ..t.............................g..........a......t........c..............t..... | [ 78000] |
| MH025919.1_strain_N10 | ..t.............................g..........a......t........c..............t..... | [ 78000] |
| MH025918.1_strain_R25 | ..t.............................g..........a......t........c..............t..... | [ 78000] |
| MH025917.1_strain_R7 | ..t.............................g..........a......t........c..............t..... | [ 78000] |
| MH025916.1_strain_R8 | ..t.............................g..........a......t........c..............t..... | [ 78000] |

| | | |
|---|---|---|
| FR682468.2_Georgia_2007/1 | gaaaaccaatgaaaaagagtttattactctgctaaaccaggccttggcctcaacgcagctttaccgcagcatacaacagc | [ 78080] |
| ASF_IND_20_CAD543 | ................................................................................ | [ 78080] |
| LR899193.1_Germany_2020/1 | ................................................................................ | [ 78080] |
| MN393477.1_Wuhan_2019 | ................................................................................ | [ 78080] |
| MW396979.1_Timor-Leste/2019/1 | ................................................................................ | [ 78080] |
| MK940252.1_CN/2019/InnerMongolia | ................................................................................ | [ 78080] |
| MK128995.1_China/2018/AnhuiXCGQ | ................................................................................ | [ 78080] |
| MH681419.1_ASFV/POL/2015 | ................................................................................ | [ 78080] |
| LS478113.1_Estonia_2014 | ................................................................................ | [ 78080] |
| KP843857.1_Odintsovo_02/14 | ................................................................................ | [ 78080] |
| FN557520.1_strain_E75 | ................................................................................ | [ 78080] |
| AY261366.1_Warthog | ................................................................................ | [ 78080] |
| AY261365.1_Warmbaths | ................................................................................ | [ 78080] |
| AY261364.1_Tengani_62 | ................................................................................ | [ 78080] |
| AY261363.1_Pretoriuskop/96/4 | ...............................................................t................ | [ 78080] |
| AY261362.1_Mkuzi_1979 | ................................................................................ | [ 78080] |
| AY261361.1_Malawi_Lil-20/1_1983 | ...............................t.......a.......................t................ | [ 78080] |
| AY261360.1_Kenya_1950 | .......................................a...........g............................ | [ 78080] |
| AM712240.1_OURT_88/3 | ................................................................................ | [ 78080] |
| AM712239.1_Benin_97/1 | ................................................................................ | [ 78080] |
| KM111295.1_Ken06.Bus | .......................................a...........g............................ | [ 78080] |
| KM111294.1_Ken05/Tk1 | .......................................a...........g............................ | [ 78080] |
| KM262844.1_strain_L60 | ................................................................................ | [ 78080] |
| KM102979.1_26544/OG10_Italy | ................................................................................ | [ 78080] |
| KM262845.1_strain_NHV | ................................................................................ | [ 78080] |
| KX354450.1_47/Ss/2008 | ................................................................................ | [ 78080] |
| KP055815.1_strain_BA71 | ................................................................................ | [ 78080] |
| MH025920.1_strain_R35 | .......................................a...........g............................ | [ 78080] |
| MH025919.1_strain_N10 | .......................................a...........g............................ | [ 78080] |
| MH025918.1_strain_R25 | .......................................a...........g............................ | [ 78080] |
| MH025917.1_strain_R7 | .......................................a...........g............................ | [ 78080] |
| MH025916.1_strain_R8 | .......................................a...........g............................ | [ 78080] |

```
FR682468.2_Georgia_2007/1        tgtttttaacgatgtataagctagatcccattgggtttattaactatattaaaacgagtaaacaagagtatttatgcctg [ 78160]
ASF_IND_20_CAD543                ................................................................................ [ 78160]
LR899193.1_Germany_2020/1        ................................................................................ [ 78160]
MN393477.1_Wuhan_2019            ................................................................................ [ 78160]
MW396979.1_Timor-Leste/2019/1    ................................................................................ [ 78160]
MK940252.1_CN/2019/InnerMongolia ................................................................................ [ 78160]
MK128995.1_China/2018/AnhuiXCGQ  ................................................................................ [ 78160]
MH681419.1_ASFV/POL/2015         ................................................................................ [ 78160]
LS478113.1_Estonia_2014          ................................................................................ [ 78160]
KP843857.1_Odintsovo_02/14       ................................................................................ [ 78160]
FN557520.1_strain_E75            ................................................................................ [ 78160]
AY261366.1_Warthog               ................................................................................ [ 78160]
AY261365.1_Warmbaths             ............................t..a................................................ [ 78160]
AY261364.1_Tengani_62            ................................................................................ [ 78160]
AY261363.1_Pretoriuskop/96/4     ................................................................................ [ 78160]
AY261362.1_Mkuzi_1979            ............................t..a................................................ [ 78160]
AY261361.1_Malawi_Lil-20/1_1983  ................................................................................ [ 78160]
AY261360.1_Kenya_1950            ......................................g..............g......................t... [ 78160]
AM712240.1_OURT_88/3             ................................................................................ [ 78160]
AM712239.1_Benin_97/1            ................................................................................ [ 78160]
KM111295.1_Ken06.Bus             ......................................g..............g......................t... [ 78160]
KM111294.1_Ken05/Tk1             ......................................g..............g......................t... [ 78160]
KM262844.1_strain_L60            ................................................................................ [ 78160]
KM102979.1_26544/OG10_Italy      ................................................................................ [ 78160]
KM262845.1_strain_NHV            ................................................................................ [ 78160]
KX354450.1_47/Ss/2008            ................................................................................ [ 78160]
KP055815.1_strain_BA71           ................................................................................ [ 78160]
MH025920.1_strain_R35            ......................................g..............g......................t... [ 78160]
MH025919.1_strain_N10            ......................................g..............g......................t... [ 78160]
MH025918.1_strain_R25            ......................................g..............g......................t... [ 78160]
MH025917.1_strain_R7             ......................................g..............g......................t... [ 78160]
MH025916.1_strain_R8             ......................................g..............g......................t... [ 78160]

FR682468.2_Georgia_2007/1        ttaattaatcctaaactcgttactaagtttttaaaaataacgagctttaaaatttacattaatttcaggctgaaaacttt [ 78240]
ASF_IND_20_CAD543                ..r............................................................................. [ 78240]
LR899193.1_Germany_2020/1        ................................................................................ [ 78240]
MN393477.1_Wuhan_2019            ................................................................................ [ 78240]
MW396979.1_Timor-Leste/2019/1    ................................................................................ [ 78240]
MK940252.1_CN/2019/InnerMongolia ................................................................................ [ 78240]
MK128995.1_China/2018/AnhuiXCGQ  ................................................................................ [ 78240]
MH681419.1_ASFV/POL/2015         ................................................................................ [ 78240]
LS478113.1_Estonia_2014          ................................................................................ [ 78240]
KP843857.1_Odintsovo_02/14       ................................................................................ [ 78240]
```

```
FN557520.1_strain_E75           ..g.................................................................a........... [ 78240]
AY261366.1_Warthog              ..g.................c...............................................a........... [ 78240]
AY261365.1_Warmbaths            ..g.................................................................a........... [ 78240]
AY261364.1_Tengani_62           ..g.................................................................a........... [ 78240]
AY261363.1_Pretoriuskop/96/4    ..g.................c...............................................a........... [ 78240]
AY261362.1_Mkuzi_1979           ..g.................................................................a........... [ 78240]
AY261361.1_Malawi_Lil-20/1_1983 ..g..c...........a.....g....................t................................... [ 78240]
AY261360.1_Kenya_1950           ..g..............a..c............................................t.............. [ 78240]
AM712240.1_OURT_88/3            ..g.................................................................a........... [ 78240]
AM712239.1_Benin_97/1           ..g.................................................................a........... [ 78240]
KM111295.1_Ken06.Bus            ..g..............a..c..................................................a........ [ 78240]
KM111294.1_Ken05/Tk1            ..g..............a..c............................................t.............. [ 78240]
KM262844.1_strain_L60           ..g.................................................................a........... [ 78240]
KM102979.1_26544/OG10_Italy     ..g.................................................................a........... [ 78240]
KM262845.1_strain_NHV           ..g.................................................................a........... [ 78240]
KX354450.1_47/Ss/2008           ..g.................................................................a........... [ 78240]
KP055815.1_strain_BA71          ..g.................................................................a........... [ 78240]
MH025920.1_strain_R35           ..g..............a..c..................................................a........ [ 78240]
MH025919.1_strain_N10           ..g..............a..c..................................................a........ [ 78240]
MH025918.1_strain_R25           ..g..............a..c..................................................a........ [ 78240]
MH025917.1_strain_R7            ..g..............a..c..................................................a........ [ 78240]
MH025916.1_strain_R8            ..g..............a..c..................................................a........ [ 78240]

FR682468.2_Georgia_2007/1       ttatataagtcctaataagtataataatttttacaccgctccctctgaagaaaagactaaccatcttctaaaagaagaaa [ 78320]
ASF_IND_20_CAD543               ....................................y.............................y............. [ 78320]
LR899193.1_Germany_2020/1       ................................................................................ [ 78320]
MN393477.1_Wuhan_2019           ................................................................................ [ 78320]
MW396979.1_Timor-Leste/2019/1   ................................................................................ [ 78320]
MK940252.1_CN/2019/InnerMongolia ................................................................................ [ 78320]
MK128995.1_China/2018/AnhuiXCGQ ................................................................................ [ 78320]
MH681419.1_ASFV/POL/2015        ................................................................................ [ 78320]
LS478113.1_Estonia_2014         ................................................................................ [ 78320]
KP843857.1_Odintsovo_02/14      ................................................................................ [ 78320]
FN557520.1_strain_E75           ....................................t.............................c............. [ 78320]
AY261366.1_Warthog              .........................................................c........c............. [ 78320]
AY261365.1_Warmbaths            .........................................................c........c............. [ 78320]
AY261364.1_Tengani_62           ..................................................................c............. [ 78320]
AY261363.1_Pretoriuskop/96/4    .........................................................c........c............. [ 78320]
AY261362.1_Mkuzi_1979           .........................................................c........c............. [ 78320]
AY261361.1_Malawi_Lil-20/1_1983 ..................a....................................g.c..t.....c............. [ 78320]
AY261360.1_Kenya_1950           ...................................t...................g.c..t.....c............. [ 78320]
AM712240.1_OURT_88/3            ....................................t.............................c............. [ 78320]
AM712239.1_Benin_97/1           ....................................t.............................c............. [ 78320]
KM111295.1_Ken06.Bus            c..................................t...................g.c..t.....c............. [ 78320]
```

```
KM111294.1_Ken05/Tk1             ...................................t...................g.c..t.....c............. [ 78320]
KM262844.1_strain_L60            ....................................t.............................c............. [ 78320]
KM102979.1_26544/OG10_Italy      ....................................t.............................c............. [ 78320]
KM262845.1_strain_NHV            ....................................t.............................c............. [ 78320]
KX354450.1_47/Ss/2008            ....................................t.............................c............. [ 78320]
KP055815.1_strain_BA71           ....................................t.............................c............. [ 78320]
MH025920.1_strain_R35            c..................................t...................g.c..t.....c............. [ 78320]
MH025919.1_strain_N10            c..................................t...................g.c..t.....c............. [ 78320]
MH025918.1_strain_R25            c..................................t...................g.c..t.....c............. [ 78320]
MH025917.1_strain_R7             c..................................t...................g.c..t.....c............. [ 78320]
MH025916.1_strain_R8             c..................................t...................g.c..t.....c............. [ 78320]

FR682468.2_Georgia_2007/1        aaacttgggcaaagattgttgaaga---aggaggagaagaatcctaagtc-gcttacatttttttttgctatttttatag [ 78400]
ASF_IND_20_CAD543                .........................---......................-..................c.-........ [ 78400]
LR899193.1_Germany_2020/1        .........................---..............a.......-............................. [ 78400]
MN393477.1_Wuhan_2019            .........................---......................-............................. [ 78400]
MW396979.1_Timor-Leste/2019/1    .........................---......................-............................. [ 78400]
MK940252.1_CN/2019/InnerMongolia .........................---......................-............................. [ 78400]
MK128995.1_China/2018/AnhuiXCGQ  .........................---......................-............................. [ 78400]
MH681419.1_ASFV/POL/2015         .........................---......................-............................. [ 78400]
LS478113.1_Estonia_2014          .........................---......................-............................. [ 78400]
KP843857.1_Odintsovo_02/14       .........................---......................-............................. [ 78400]
FN557520.1_strain_E75            .........................---......................-..................c.-........ [ 78400]
AY261366.1_Warthog               .........................---......................-............................. [ 78400]
AY261365.1_Warmbaths             .........................agg......................-.....t....................... [ 78400]
AY261364.1_Tengani_62            .........................---......................-............................. [ 78400]
AY261363.1_Pretoriuskop/96/4     .........................---......................-............................. [ 78400]
AY261362.1_Mkuzi_1979            .........................agg......................-............................. [ 78400]
AY261361.1_Malawi_Lil-20/1_1983  ....c....................---...................aa.-a.................c.......... [ 78400]
AY261360.1_Kenya_1950            ....c....................---....................a.-a............................ [ 78400]
AM712240.1_OURT_88/3             .........................---......................-..................c.-........ [ 78400]
AM712239.1_Benin_97/1            .........................---......................-..................c.-........ [ 78400]
KM111295.1_Ken06.Bus             ....c....................---....................a.aa............................ [ 78400]
KM111294.1_Ken05/Tk1             ....c....................---....................a.-a............................ [ 78400]
KM262844.1_strain_L60            .........................---......................-..................c.-........ [ 78400]
KM102979.1_26544/OG10_Italy      .........................---.....a................-..................c.-........ [ 78400]
KM262845.1_strain_NHV            .........................---......................-..................c.-........ [ 78400]
KX354450.1_47/Ss/2008            .........................---.....a................-..................c.-........ [ 78400]
KP055815.1_strain_BA71           .........................---......................-..................c.-........ [ 78400]
MH025920.1_strain_R35            ....c....................---....................a.aa............................ [ 78400]
MH025919.1_strain_N10            ....c....................---....................a.aa............................ [ 78400]
MH025918.1_strain_R25            ....c....................---....................a.aa............................ [ 78400]
MH025917.1_strain_R7             ....c....................---....................a.aa............................ [ 78400]
MH025916.1_strain_R8             ....c....................---....................a.aa............................ [ 78400]
```

```
FR682468.2_Georgia_2007/1        a---atgtacacgcatgttgatgttgtcggaatagctgaagcctcagcggccctctacgtgcaaaaagatagggatcgct [ 78480]
ASF_IND_20_CAD543                .---............................................................................ [ 78480]
LR899193.1_Germany_2020/1        .---............................................................................ [ 78480]
MN393477.1_Wuhan_2019            .---............................................................................ [ 78480]
MW396979.1_Timor-Leste/2019/1    .---............................................................................ [ 78480]
MK940252.1_CN/2019/InnerMongolia .---............................................................................ [ 78480]
MK128995.1_China/2018/AnhuiXCGQ  .---............................................................................ [ 78480]
MH681419.1_ASFV/POL/2015         .---............................................................................ [ 78480]
LS478113.1_Estonia_2014          .---............................................................................ [ 78480]
KP843857.1_Odintsovo_02/14       .---............................................................................ [ 78480]
FN557520.1_strain_E75            .---............................................................................ [ 78480]
AY261366.1_Warthog               .---.............................t....................t......................... [ 78480]
AY261365.1_Warmbaths             .---.............................t....................t......................... [ 78480]
AY261364.1_Tengani_62            .---............................................................................ [ 78480]
AY261363.1_Pretoriuskop/96/4     .---.............................t....................t......................... [ 78480]
AY261362.1_Mkuzi_1979            .---..................................................t......................... [ 78480]
AY261361.1_Malawi_Lil-20/1_1983  .---............................................................................ [ 78480]
AY261360.1_Kenya_1950            gaat.....t.................g..g.....g........g........t......................... [ 78480]
AM712240.1_OURT_88/3             .---............................................................................ [ 78480]
AM712239.1_Benin_97/1            .---............................................................................ [ 78480]
KM111295.1_Ken06.Bus             gaat.....t.................g..g.....g........g........t......................... [ 78480]
KM111294.1_Ken05/Tk1             gaat.....t.................g..g.....g........g........t......................... [ 78480]
KM262844.1_strain_L60            .---............................................................................ [ 78480]
KM102979.1_26544/OG10_Italy      .---............................................................................ [ 78480]
KM262845.1_strain_NHV            .---............................................................................ [ 78480]
KX354450.1_47/Ss/2008            .---............................................................................ [ 78480]
KP055815.1_strain_BA71           .---............................................................................ [ 78480]
MH025920.1_strain_R35            gaat.....t.................g..g.....g........g........t......................... [ 78480]
MH025919.1_strain_N10            gaat.....t.................g..g.....g........g........t......................... [ 78480]
MH025918.1_strain_R25            gaat.....t.................g..g.....g........g........t......................... [ 78480]
MH025917.1_strain_R7             gaat.....t.................g..g.....g........g........t......................... [ 78480]
MH025916.1_strain_R8             gaat.....t.................g..g.....g........g........t......................... [ 78480]

FR682468.2_Georgia_2007/1        acttagacgtgctaacaaccattgaaaactttatttaccaacacaaatgcatcataacaggggaaagcgcccacctactc [ 78560]
ASF_IND_20_CAD543                ................................................................................ [ 78560]
LR899193.1_Germany_2020/1        ................................................................................ [ 78560]
MN393477.1_Wuhan_2019            ................................................................................ [ 78560]
MW396979.1_Timor-Leste/2019/1    ................................................................................ [ 78560]
MK940252.1_CN/2019/InnerMongolia ................................................................................ [ 78560]
MK128995.1_China/2018/AnhuiXCGQ  ................................................................................ [ 78560]
MH681419.1_ASFV/POL/2015         ................................................................................ [ 78560]
LS478113.1_Estonia_2014          ................................................................................ [ 78560]
KP843857.1_Odintsovo_02/14       ................................................................................ [ 78560]
```

```
FN557520.1_strain_E75          ....g.................c......................................................... [ 78560]
AY261366.1_Warthog             ...cg........................................................................... [ 78560]
AY261365.1_Warmbaths           ...cg........................................................................... [ 78560]
AY261364.1_Tengani_62          ....g.................c...................................................t..... [ 78560]
AY261363.1_Pretoriuskop/96/4   ...cg........................................................................... [ 78560]
AY261362.1_Mkuzi_1979          ....g..............t..c......................................................... [ 78560]
AY261361.1_Malawi_Lil-20/1_1983 ....g...........t.....c..............................t.......................... [ 78560]
AY261360.1_Kenya_1950          ....g...........tg....c......................................a..............c... [ 78560]
AM712240.1_OURT_88/3           ....g.................c......................................................... [ 78560]
AM712239.1_Benin_97/1          ....g.................c......................................................... [ 78560]
KM111295.1_Ken06.Bus           ....g...........tg....c.....................................................c... [ 78560]
KM111294.1_Ken05/Tk1           ....g...........tg....c......................................a..............c... [ 78560]
KM262844.1_strain_L60          ....g.................c......................................................... [ 78560]
KM102979.1_26544/OG10_Italy    ....g.................c......................................................... [ 78560]
KM262845.1_strain_NHV          ....g.................c......................................................... [ 78560]
KX354450.1_47/Ss/2008          ....g.................c......................................................... [ 78560]
KP055815.1_strain_BA71         ....g.................c......................................................... [ 78560]
MH025920.1_strain_R35          ....g...........tg....c.....................................................c... [ 78560]
MH025919.1_strain_N10          ....g...........tg....c.....................................................c... [ 78560]
MH025918.1_strain_R25          ....g...........tg....c.....................................................c... [ 78560]
MH025917.1_strain_R7           ....g...........tg....c.....................................................c... [ 78560]
MH025916.1_strain_R8           ....g...........tg....c.....................................................c... [ 78560]

FR682468.2_Georgia_2007/1      tttttaaaaaaaaatatttatctttacgaattttactccaacaatgtggcggagcacagcaaggctttggcgaccctgct [ 78640]
ASF_IND_20_CAD543              ................................................................................ [ 78640]
LR899193.1_Germany_2020/1      ................................................................................ [ 78640]
MN393477.1_Wuhan_2019          ................................................................................ [ 78640]
MW396979.1_Timor-Leste/2019/1  ................................................................................ [ 78640]
MK940252.1_CN/2019/InnerMongolia ................................................................................ [ 78640]
MK128995.1_China/2018/AnhuiXCGQ ................................................................................ [ 78640]
MH681419.1_ASFV/POL/2015       ................................................................................ [ 78640]
LS478113.1_Estonia_2014        ................................................................................ [ 78640]
KP843857.1_Odintsovo_02/14     ................................................................................ [ 78640]
FN557520.1_strain_E75          ............-................................................................... [ 78640]
AY261366.1_Warthog             ................................................................................ [ 78640]
AY261365.1_Warmbaths           ................................................................................ [ 78640]
AY261364.1_Tengani_62          ................................................................................ [ 78640]
AY261363.1_Pretoriuskop/96/4   ................................................................................ [ 78640]
AY261362.1_Mkuzi_1979          .......................................g........................................ [ 78640]
AY261361.1_Malawi_Lil-20/1_1983 .....g.....g.g.........................g....c................................... [ 78640]
AY261360.1_Kenya_1950          .....g.......g.................g........................t..t.................... [ 78640]
AM712240.1_OURT_88/3           ................................................................................ [ 78640]
AM712239.1_Benin_97/1          ................................................................................ [ 78640]
KM111295.1_Ken06.Bus           .....g.......g.................g........................t..t.................... [ 78640]
```

```
KM111294.1_Ken05/Tk1            .....g.......g.................g.................t......t..t.................... [ 78640]
KM262844.1_strain_L60           ................................................................................ [ 78640]
KM102979.1_26544/OG10_Italy     ................................................................................ [ 78640]
KM262845.1_strain_NHV           ................................................................................ [ 78640]
KX354450.1_47/Ss/2008           ................................................................................ [ 78640]
KP055815.1_strain_BA71          ................................................................................ [ 78640]
MH025920.1_strain_R35           .....g.........................g........................t..t.................... [ 78640]
MH025919.1_strain_N10           .....g.........................g........................t..t.................... [ 78640]
MH025918.1_strain_R25           .....g.........................g........................t..t.................... [ 78640]
MH025917.1_strain_R7            .....g.........................g........................t..t.................... [ 78640]
MH025916.1_strain_R8            .....g.........................g........................t..t.................... [ 78640]

FR682468.2_Georgia_2007/1       ttataaacttgatccggaatacctcactcgttacacagtactcattaccaaaattcccaaccattggtatgtgattaacg [ 78720]
ASF_IND_20_CAD543               ................................................................................ [ 78720]
LR899193.1_Germany_2020/1       ................................................................................ [ 78720]
MN393477.1_Wuhan_2019           ................................................................................ [ 78720]
MW396979.1_Timor-Leste/2019/1   ................................................................................ [ 78720]
MK940252.1_CN/2019/InnerMongolia ................................................................................ [ 78720]
MK128995.1_China/2018/AnhuiXCGQ ................................................................................ [ 78720]
MH681419.1_ASFV/POL/2015        ................................................................................ [ 78720]
LS478113.1_Estonia_2014         ................................................................................ [ 78720]
KP843857.1_Odintsovo_02/14      ................................................................................ [ 78720]
FN557520.1_strain_E75           ...c.........................................c.....g............................ [ 78720]
AY261366.1_Warthog              ...c.........................................c.....g............................ [ 78720]
AY261365.1_Warmbaths            ...c.........................................c.....g............................ [ 78720]
AY261364.1_Tengani_62           ...c......................................t..c.....g............................ [ 78720]
AY261363.1_Pretoriuskop/96/4    ...c.........................................c.....g............................ [ 78720]
AY261362.1_Mkuzi_1979           ......g......................................c.....g............................ [ 78720]
AY261361.1_Malawi_Lil-20/1_1983 ......t........c..........................g........g....................t....... [ 78720]
AY261360.1_Kenya_1950           ...................................................gg.c........a.c......t....... [ 78720]
AM712240.1_OURT_88/3            ...c.........................................c.....g............................ [ 78720]
AM712239.1_Benin_97/1           ...c.........................................c.....g............................ [ 78720]
KM111295.1_Ken06.Bus            ...................................................gg.c........a.c......t....... [ 78720]
KM111294.1_Ken05/Tk1            ............c......................................gg.c........a.c......t....... [ 78720]
KM262844.1_strain_L60           ...c.........................................c.....g............................ [ 78720]
KM102979.1_26544/OG10_Italy     ...c.........................................c.....g............................ [ 78720]
KM262845.1_strain_NHV           ...c.........................................c.....g............................ [ 78720]
KX354450.1_47/Ss/2008           ...c.........................................c.....g............................ [ 78720]
KP055815.1_strain_BA71          ...c.........................................c.....g............................ [ 78720]
MH025920.1_strain_R35           ...................................................gg.c........a.c......t....... [ 78720]
MH025919.1_strain_N10           ...................................................gg.c........a.c......t....... [ 78720]
MH025918.1_strain_R25           ...................................................gg.c........a.c......t....... [ 78720]
MH025917.1_strain_R7            ...................................................gg.c........a.c......t....... [ 78720]
MH025916.1_strain_R8            ...................................................gg.c........a.c......t....... [ 78720]
```

```
FR682468.2_Georgia_2007/1        tagatcagcgagaatttgtgcgcctatatgccatcccggcagttaaacaacacttaccgattcccattttacccttctat [ 78800]
ASF_IND_20_CAD543                ................................................................................ [ 78800]
LR899193.1_Germany_2020/1        ................................................................................ [ 78800]
MN393477.1_Wuhan_2019            ................................................................................ [ 78800]
MW396979.1_Timor-Leste/2019/1    ................................................................................ [ 78800]
MK940252.1_CN/2019/InnerMongolia ................................................................................ [ 78800]
MK128995.1_China/2018/AnhuiXCGQ  ................................................................................ [ 78800]
MH681419.1_ASFV/POL/2015         ................................................................................ [ 78800]
LS478113.1_Estonia_2014          ................................................................................ [ 78800]
KP843857.1_Odintsovo_02/14       ................................................................................ [ 78800]
FN557520.1_strain_E75            .t................................t............................................. [ 78800]
AY261366.1_Warthog               ..................................t............................................. [ 78800]
AY261365.1_Warmbaths             ..................................t............................................. [ 78800]
AY261364.1_Tengani_62            .g................................t............................................. [ 78800]
AY261363.1_Pretoriuskop/96/4     ..................................t............................................. [ 78800]
AY261362.1_Mkuzi_1979            ................................................................................ [ 78800]
AY261361.1_Malawi_Lil-20/1_1983  ..........g.......................t..........................................c.. [ 78800]
AY261360.1_Kenya_1950            ..................................t..a.........g..........a..................c.. [ 78800]
AM712240.1_OURT_88/3             .t................................t............................................. [ 78800]
AM712239.1_Benin_97/1            .t................................t............................................. [ 78800]
KM111295.1_Ken06.Bus             ..................................t..a.........g..........a..................c.. [ 78800]
KM111294.1_Ken05/Tk1             ..................................t..a.........g..........a...........g......c.. [ 78800]
KM262844.1_strain_L60            .t................................t............................................. [ 78800]
KM102979.1_26544/OG10_Italy      .t................................t............................................. [ 78800]
KM262845.1_strain_NHV            .t................................t............................................. [ 78800]
KX354450.1_47/Ss/2008            .t................................t............................................. [ 78800]
KP055815.1_strain_BA71           .t................................t............................................. [ 78800]
MH025920.1_strain_R35            ..................................t..a.........g..........a..................c.. [ 78800]
MH025919.1_strain_N10            ..................................t..a.........g..........a..................c.. [ 78800]
MH025918.1_strain_R25            ..................................t..a.........g..........a..................c.. [ 78800]
MH025917.1_strain_R7             ..................................t..a.........g..........a..................c.. [ 78800]
MH025916.1_strain_R8             ..................................t..a.........g..........a..................c.. [ 78800]

FR682468.2_Georgia_2007/1        tgcaccagcgcactcacccagcaagaattgttttgtttaggacctgaactgcagttaatacaaatatattccaagctctg [ 78880]
ASF_IND_20_CAD543                ................................................................................ [ 78880]
LR899193.1_Germany_2020/1        ................................................................................ [ 78880]
MN393477.1_Wuhan_2019            ................................................................................ [ 78880]
MW396979.1_Timor-Leste/2019/1    ................................................................................ [ 78880]
MK940252.1_CN/2019/InnerMongolia ................................................................................ [ 78880]
MK128995.1_China/2018/AnhuiXCGQ  ................................................................................ [ 78880]
MH681419.1_ASFV/POL/2015         ................................................................................ [ 78880]
LS478113.1_Estonia_2014          ................................................................................ [ 78880]
KP843857.1_Odintsovo_02/14       ................................................................................ [ 78880]
```

```
FN557520.1_strain_E75             ....................c........................................................... [ 78880]
AY261366.1_Warthog                ................................................................................ [ 78880]
AY261365.1_Warmbaths              ................................................................................ [ 78880]
AY261364.1_Tengani_62             ................................................................................ [ 78880]
AY261363.1_Pretoriuskop/96/4      ................................................................................ [ 78880]
AY261362.1_Mkuzi_1979             ................................................................................ [ 78880]
AY261361.1_Malawi_Lil-20/1_1983   ..........................c.................c................................... [ 78880]
AY261360.1_Kenya_1950             ...........t.....t.........c................c.....a............................. [ 78880]
AM712240.1_OURT_88/3              ....................c........................................................... [ 78880]
AM712239.1_Benin_97/1             ....................c........................................................... [ 78880]
KM111295.1_Ken06.Bus              ...........t.....t.........c................c.....a............................. [ 78880]
KM111294.1_Ken05/Tk1              ...........t.....t.........c................c.....a............................. [ 78880]
KM262844.1_strain_L60             ....................c........................................................... [ 78880]
KM102979.1_26544/OG10_Italy       ....................c........................................................... [ 78880]
KM262845.1_strain_NHV             ....................c........................................................... [ 78880]
KX354450.1_47/Ss/2008             ....................c........................................................... [ 78880]
KP055815.1_strain_BA71            ....................c........................................................... [ 78880]
MH025920.1_strain_R35             ...........t.....t.........c................c.....a............................. [ 78880]
MH025919.1_strain_N10             ...........t.....t.........c................c.....a............................. [ 78880]
MH025918.1_strain_R25             ...........t.....t.........c................c.....a............................. [ 78880]
MH025917.1_strain_R7              ...........t.....t.........c................c.....a............................. [ 78880]
MH025916.1_strain_R8              ...........t.....t.........c................c.....a............................. [ 78880]

FR682468.2_Georgia_2007/1         taaccccaactttgtcgaggaatggcctacgttgctcgactacgaaaaaagcatgcggatgttatttttagaacagtttc [ 78960]
ASF_IND_20_CAD543                 ................................................................................ [ 78960]
LR899193.1_Germany_2020/1         ................................................................................ [ 78960]
MN393477.1_Wuhan_2019             ................................................................................ [ 78960]
MW396979.1_Timor-Leste/2019/1     ................................................................................ [ 78960]
MK940252.1_CN/2019/InnerMongolia  ................................................................................ [ 78960]
MK128995.1_China/2018/AnhuiXCGQ   ................................................................................ [ 78960]
MH681419.1_ASFV/POL/2015          ................................................................................ [ 78960]
LS478113.1_Estonia_2014           ................................................................................ [ 78960]
KP843857.1_Odintsovo_02/14        ................................................................................ [ 78960]
FN557520.1_strain_E75             ................................................................................ [ 78960]
AY261366.1_Warthog                ...........................................................c.................... [ 78960]
AY261365.1_Warmbaths              ................................................................................ [ 78960]
AY261364.1_Tengani_62             ................................................................................ [ 78960]
AY261363.1_Pretoriuskop/96/4      ...........................................................c.................... [ 78960]
AY261362.1_Mkuzi_1979             ................................................................................ [ 78960]
AY261361.1_Malawi_Lil-20/1_1983   ...........................c.....a................a........c.................... [ 78960]
AY261360.1_Kenya_1950             ..................................................a........c.c.................t [ 78960]
AM712240.1_OURT_88/3              ................................................................................ [ 78960]
AM712239.1_Benin_97/1             ................................................................................ [ 78960]
KM111295.1_Ken06.Bus              ..................................................a........c.c.................t [ 78960]
```

```
KM111294.1_Ken05/Tk1          ..................................................a........c.c.................t [ 78960]
KM262844.1_strain_L60         ................................................................................ [ 78960]
KM102979.1_26544/OG10_Italy   ................................................................................ [ 78960]
KM262845.1_strain_NHV         ................................................................................ [ 78960]
KX354450.1_47/Ss/2008         ................................................................................ [ 78960]
KP055815.1_strain_BA71        ................................................................................ [ 78960]
MH025920.1_strain_R35         ..................................................a........c.c.................t [ 78960]
MH025919.1_strain_N10         ..................................................a........c.c.................t [ 78960]
MH025918.1_strain_R25         ..................................................a........c.c.................t [ 78960]
MH025917.1_strain_R7          ..................................................a........c.c.................t [ 78960]
MH025916.1_strain_R8          ..................................................a........c.c.................t [ 78960]

FR682468.2_Georgia_2007/1     cgcaaagattggaaatgacgggcgggaagaaggaggagaaggaaaagcatgaaagtatcattaaaaaaataatactagaa [ 79040]
ASF_IND_20_CAD543             ................ag..................---......................................... [ 79040]
LR899193.1_Germany_2020/1     ................................................................................ [ 79040]
MN393477.1_Wuhan_2019         ................................................................................ [ 79040]
MW396979.1_Timor-Leste/2019/1 ................................................................................ [ 79040]
MK940252.1_CN/2019/InnerMongolia ................................................................................ [ 79040]
MK128995.1_China/2018/AnhuiXCGQ ................................................................................ [ 79040]
MH681419.1_ASFV/POL/2015      ................................................................................ [ 79040]
LS478113.1_Estonia_2014       ................................................................................ [ 79040]
KP843857.1_Odintsovo_02/14    ................................................................................ [ 79040]
FN557520.1_strain_E75         ...g............ag..................---......................................... [ 79040]
AY261366.1_Warthog            ......................................g......................................... [ 79040]
AY261365.1_Warmbaths          ..............................---.....g......................................... [ 79040]
AY261364.1_Tengani_62         ......................t.............---......................................... [ 79040]
AY261363.1_Pretoriuskop/96/4  ......................................g......................................... [ 79040]
AY261362.1_Mkuzi_1979         ....................................---......................................... [ 79040]
AY261361.1_Malawi_Lil-20/1_1983 ..g...a........a...a........a......a..g..........c.............................. [ 79040]
AY261360.1_Kenya_1950         ......................................g..........ca.......................t..... [ 79040]
AM712240.1_OURT_88/3          ...g............ag..................---......................................... [ 79040]
AM712239.1_Benin_97/1         ...g............ag..................---......................................... [ 79040]
KM111295.1_Ken06.Bus          ...............a.........------.......g.t.....a..ca...a...t..............t...... [ 79040]
KM111294.1_Ken05/Tk1          ...............a.........------.......g.a........ca.......................t..... [ 79040]
KM262844.1_strain_L60         ...g............ag..................---......................................... [ 79040]
KM102979.1_26544/OG10_Italy   ...g............ag..................---......................................... [ 79040]
KM262845.1_strain_NHV         ...g............ag..................---......................................... [ 79040]
KX354450.1_47/Ss/2008         ...g............ag..................---......................................... [ 79040]
KP055815.1_strain_BA71        ...g............ag..................---......................................... [ 79040]
MH025920.1_strain_R35         ...............a.........------.......g.t.....a..ca...a...t..............t...... [ 79040]
MH025919.1_strain_N10         ...............a.........------.......g.t.....a..ca...a...t..............t...... [ 79040]
MH025918.1_strain_R25         ...............a.........------.......g.t.....a..ca...a...t..............t...... [ 79040]
MH025917.1_strain_R7          ...............a.........------.......g.t.....a..ca...a...t..............t...... [ 79040]
MH025916.1_strain_R8          ...............a.........------.......g.t.....a..ca...a...t..............t...... [ 79040]
```

```
FR682468.2_Georgia_2007/1        atggtctctacccgtcagcgaatcgttgttgggggttacatacaaaaaaacctgtacaaccatgtactcaagaatagaaa [ 79120]
ASF_IND_20_CAD543                ................................r............................................... [ 79120]
LR899193.1_Germany_2020/1        ................................................................................ [ 79120]
MN393477.1_Wuhan_2019            ................................................................................ [ 79120]
MW396979.1_Timor-Leste/2019/1    ................................................................................ [ 79120]
MK940252.1_CN/2019/InnerMongolia ................................................................................ [ 79120]
MK128995.1_China/2018/AnhuiXCGQ  ................................................................................ [ 79120]
MH681419.1_ASFV/POL/2015         ................................................................................ [ 79120]
LS478113.1_Estonia_2014          ................................................................................ [ 79120]
KP843857.1_Odintsovo_02/14       ................................................................................ [ 79120]
FN557520.1_strain_E75            ................................a............................................... [ 79120]
AY261366.1_Warthog               .....t.......................................................................... [ 79120]
AY261365.1_Warmbaths             .....t..............c.............................t............................. [ 79120]
AY261364.1_Tengani_62            .....t.......................................................................... [ 79120]
AY261363.1_Pretoriuskop/96/4     .....t.......................................................................... [ 79120]
AY261362.1_Mkuzi_1979            .....t.......................................................................... [ 79120]
AY261361.1_Malawi_Lil-20/1_1983  .....t.......a....................................t..t..t..t.....ca....a........ [ 79120]
AY261360.1_Kenya_1950            .....t.....t..................................................c..cg....a........ [ 79120]
AM712240.1_OURT_88/3             ................................a............................................... [ 79120]
AM712239.1_Benin_97/1            ................................a...........................................a... [ 79120]
KM111295.1_Ken06.Bus             .....t.....t..................................................c..cg....a........ [ 79120]
KM111294.1_Ken05/Tk1             .....t.....t..................................................c..cg....a........ [ 79120]
KM262844.1_strain_L60            ................................a............................................... [ 79120]
KM102979.1_26544/OG10_Italy      ................................a...........................................a... [ 79120]
KM262845.1_strain_NHV            ................................a............................................... [ 79120]
KX354450.1_47/Ss/2008            ................................a...........................................a... [ 79120]
KP055815.1_strain_BA71           ................................a............................................... [ 79120]
MH025920.1_strain_R35            .....t.....t..................................................c..cg....a........ [ 79120]
MH025919.1_strain_N10            .....t.....t..................................................c..cg....a........ [ 79120]
MH025918.1_strain_R25            .....t.....t..................................................c..cg....a........ [ 79120]
MH025917.1_strain_R7             .....t.....t..................................................c..cg....a........ [ 79120]
MH025916.1_strain_R8             .....t.....t..................................................c..cg....a........ [ 79120]

FR682468.2_Georgia_2007/1        tcgtttacagcttattacgagcttaaatatttatgaagaaaaagatatcatccagcaattttgtgattcaaatggactga [ 79200]
ASF_IND_20_CAD543                ................................................................................ [ 79200]
LR899193.1_Germany_2020/1        ................................................................................ [ 79200]
MN393477.1_Wuhan_2019            ................................................................................ [ 79200]
MW396979.1_Timor-Leste/2019/1    ................................................................................ [ 79200]
MK940252.1_CN/2019/InnerMongolia ................................................................................ [ 79200]
MK128995.1_China/2018/AnhuiXCGQ  ................................................................................ [ 79200]
MH681419.1_ASFV/POL/2015         ................................................................................ [ 79200]
LS478113.1_Estonia_2014          ................................................................................ [ 79200]
KP843857.1_Odintsovo_02/14       ................................................................................ [ 79200]
```

```
FN557520.1_strain_E75             .........................g...................................................... [ 79200]
AY261366.1_Warthog                ................................................................................ [ 79200]
AY261365.1_Warmbaths              .........................g...................................................... [ 79200]
AY261364.1_Tengani_62             ................................................................................ [ 79200]
AY261363.1_Pretoriuskop/96/4      ................................................................................ [ 79200]
AY261362.1_Mkuzi_1979             .........................g...................................................... [ 79200]
AY261361.1_Malawi_Lil-20/1_1983   .....................t..........................a............................... [ 79200]
AY261360.1_Kenya_1950             .....................t.......................g.................................. [ 79200]
AM712240.1_OURT_88/3              .........................g...................................................... [ 79200]
AM712239.1_Benin_97/1             .........................g...................................................... [ 79200]
KM111295.1_Ken06.Bus              .....................t.......................g.................................. [ 79200]
KM111294.1_Ken05/Tk1              .....................t.......................g.................................. [ 79200]
KM262844.1_strain_L60             .........................g...................................................... [ 79200]
KM102979.1_26544/OG10_Italy       .........................g...................................................... [ 79200]
KM262845.1_strain_NHV             .........................g...................................................... [ 79200]
KX354450.1_47/Ss/2008             .........................g...................................................... [ 79200]
KP055815.1_strain_BA71            .........................g...................................................... [ 79200]
MH025920.1_strain_R35             .....................t.......................g.................................. [ 79200]
MH025919.1_strain_N10             .....................t.......................g.................................. [ 79200]
MH025918.1_strain_R25             .....................t.......................g.................................. [ 79200]
MH025917.1_strain_R7              .....................t.......................g.................................. [ 79200]
MH025916.1_strain_R8              .....................t.......................g.................................. [ 79200]

FR682468.2_Georgia_2007/1         agatcaaaatacgtatcaacaatccgctcttgcctacaaatccggaattacggcgtttgactatttattttaatcataat [ 79280]
ASF_IND_20_CAD543                 ................................................................................ [ 79280]
LR899193.1_Germany_2020/1         ................................................................................ [ 79280]
MN393477.1_Wuhan_2019             ................................................................................ [ 79280]
MW396979.1_Timor-Leste/2019/1     ................................................................................ [ 79280]
MK940252.1_CN/2019/InnerMongolia  ................................................................................ [ 79280]
MK128995.1_China/2018/AnhuiXCGQ   ................................................................................ [ 79280]
MH681419.1_ASFV/POL/2015          ................................................................................ [ 79280]
LS478113.1_Estonia_2014           ................................................................................ [ 79280]
KP843857.1_Odintsovo_02/14        ................................................................................ [ 79280]
FN557520.1_strain_E75             ................................................................................ [ 79280]
AY261366.1_Warthog                ..........................................................................a...c. [ 79280]
AY261365.1_Warmbaths              ..........................................................................a----- [ 79280]
AY261364.1_Tengani_62             ................................................................................ [ 79280]
AY261363.1_Pretoriuskop/96/4      ..............................................g.....a.....................a...c. [ 79280]
AY261362.1_Mkuzi_1979             ................................................................................ [ 79280]
AY261361.1_Malawi_Lil-20/1_1983   .a..t......t..g..............c.............a.................c............a...-- [ 79280]
AY261360.1_Kenya_1950             ...........................................a...........c..a............------... [ 79280]
AM712240.1_OURT_88/3              ................................................................................ [ 79280]
AM712239.1_Benin_97/1             ................................................................................ [ 79280]
KM111295.1_Ken06.Bus              ...........................................a...........c..a...........---------. [ 79280]
```

```
KM111294.1_Ken05/Tk1             ...........................................a...........c..a..............---.... [ 79280]
KM262844.1_strain_L60            ................................................................................ [ 79280]
KM102979.1_26544/OG10_Italy      ................................................................................ [ 79280]
KM262845.1_strain_NHV            ................................................................................ [ 79280]
KX354450.1_47/Ss/2008            ................................................................................ [ 79280]
KP055815.1_strain_BA71           ................................................................................ [ 79280]
MH025920.1_strain_R35            ...........................................a...........c..a...........---------. [ 79280]
MH025919.1_strain_N10            ...........................................a...........c..a...........---------. [ 79280]
MH025918.1_strain_R25            ...........................................a...........c..a...........---------. [ 79280]
MH025917.1_strain_R7             ...........................................a...........c..a...........---------. [ 79280]
MH025916.1_strain_R8             ...........................................a...........c..a...........---------. [ 79280]

FR682468.2_Georgia_2007/1        aatgatgatgatcagtcatatctaatagtagatatgtacaacacgggaagctatgagctagtgcctacaaatcagataaa [ 79360]
ASF_IND_20_CAD543                ................................................................................ [ 79360]
LR899193.1_Germany_2020/1        ................................................................................ [ 79360]
MN393477.1_Wuhan_2019            ................................................................................ [ 79360]
MW396979.1_Timor-Leste/2019/1    ................................................................................ [ 79360]
MK940252.1_CN/2019/InnerMongolia ................................................................................ [ 79360]
MK128995.1_China/2018/AnhuiXCGQ  ................................................................................ [ 79360]
MH681419.1_ASFV/POL/2015         ................................................................................ [ 79360]
LS478113.1_Estonia_2014          ................................................................................ [ 79360]
KP843857.1_Odintsovo_02/14       ................................................................................ [ 79360]
FN557520.1_strain_E75            ................................................................................ [ 79360]
AY261366.1_Warthog               ................................................................................ [ 79360]
AY261365.1_Warmbaths             ----............................................................................ [ 79360]
AY261364.1_Tengani_62            ...............a............................................a.a............g.... [ 79360]
AY261363.1_Pretoriuskop/96/4     ................................................................................ [ 79360]
AY261362.1_Mkuzi_1979            ................................................................................ [ 79360]
AY261361.1_Malawi_Lil-20/1_1983  ----.......c..............t..................................................... [ 79360]
AY261360.1_Kenya_1950            ...a.......c..............t..........................c.....g.....c..c...g..g.... [ 79360]
AM712240.1_OURT_88/3             ................................................................................ [ 79360]
AM712239.1_Benin_97/1            ................................................................................ [ 79360]
KM111295.1_Ken06.Bus             ...a.......c..............t..........................c..a..g.....c..c......g.... [ 79360]
KM111294.1_Ken05/Tk1             ...........c..............t..........................c.....g.....c..c......g.... [ 79360]
KM262844.1_strain_L60            ................................................................................ [ 79360]
KM102979.1_26544/OG10_Italy      ................................................................................ [ 79360]
KM262845.1_strain_NHV            ................................................................................ [ 79360]
KX354450.1_47/Ss/2008            ................................................................................ [ 79360]
KP055815.1_strain_BA71           ................................................................................ [ 79360]
MH025920.1_strain_R35            ...a.......c..............t..........................c..a..g.....c..c......g.... [ 79360]
MH025919.1_strain_N10            ...a.......c..............t..........................c..a..g.....c..c......g.... [ 79360]
MH025918.1_strain_R25            ...a.......c..............t..........................c..a..g.....c..c......g.... [ 79360]
MH025917.1_strain_R7             ...a.......c..............t..........................c..a..g.....c..c......g.... [ 79360]
MH025916.1_strain_R8             ...a.......c..............t..........................c..a..g.....c..c......g.... [ 79360]
```

```
FR682468.2_Georgia_2007/1        cacgcttgatggcagctttttaataggaacacccttcgtgcaagcgcgatttttgttggtagagatctgggtgcttatgc [ 79440]
ASF_IND_20_CAD543                ................................................................................ [ 79440]
LR899193.1_Germany_2020/1        ................................................................................ [ 79440]
MN393477.1_Wuhan_2019            ................................................................................ [ 79440]
MW396979.1_Timor-Leste/2019/1    ................................................................................ [ 79440]
MK940252.1_CN/2019/InnerMongolia ................................................................................ [ 79440]
MK128995.1_China/2018/AnhuiXCGQ  ................................................................................ [ 79440]
MH681419.1_ASFV/POL/2015         ................................................................................ [ 79440]
LS478113.1_Estonia_2014          ................................................................................ [ 79440]
KP843857.1_Odintsovo_02/14       ................................................................................ [ 79440]
FN557520.1_strain_E75            ...............t................................................................ [ 79440]
AY261366.1_Warthog               ............t.............................g..................................... [ 79440]
AY261365.1_Warmbaths             ............g................................................................... [ 79440]
AY261364.1_Tengani_62            ..........................................g..................................... [ 79440]
AY261363.1_Pretoriuskop/96/4     ................................................................................ [ 79440]
AY261362.1_Mkuzi_1979            ................................................................................ [ 79440]
AY261361.1_Malawi_Lil-20/1_1983  ..........aa...t..............g..t........g.....g............................... [ 79440]
AY261360.1_Kenya_1950            ...a...........t.................t........g.....g............................... [ 79440]
AM712240.1_OURT_88/3             ...............t................................................................ [ 79440]
AM712239.1_Benin_97/1            ...............t................................................................ [ 79440]
KM111295.1_Ken06.Bus             t..a.............................t........g.....g............................... [ 79440]
KM111294.1_Ken05/Tk1             ...a...........t.................t........g.....g............................... [ 79440]
KM262844.1_strain_L60            ...............t................................................................ [ 79440]
KM102979.1_26544/OG10_Italy      ...............t................................................................ [ 79440]
KM262845.1_strain_NHV            ...............t................................................................ [ 79440]
KX354450.1_47/Ss/2008            ...............t................................................................ [ 79440]
KP055815.1_strain_BA71           ...............t................................................................ [ 79440]
MH025920.1_strain_R35            t..a.............................t........g.....g............................... [ 79440]
MH025919.1_strain_N10            t..a.............................t........g.....g............................... [ 79440]
MH025918.1_strain_R25            t..a.............................t........g.....g............................... [ 79440]
MH025917.1_strain_R7             t..a.............................t........g.....g............................... [ 79440]
MH025916.1_strain_R8             t..a.............................t........g.....g............................... [ 79440]

FR682468.2_Georgia_2007/1        ttattgcgcagcaaactaaaaaggacaccaaaaaaataatacaattttttataaatcaatatgaaatgcttatgaatagt [ 79520]
ASF_IND_20_CAD543                ................................................................................ [ 79520]
LR899193.1_Germany_2020/1        ................................................................................ [ 79520]
MN393477.1_Wuhan_2019            ................................................................................ [ 79520]
MW396979.1_Timor-Leste/2019/1    ................................................................................ [ 79520]
MK940252.1_CN/2019/InnerMongolia ................................................................................ [ 79520]
MK128995.1_China/2018/AnhuiXCGQ  ................................................................................ [ 79520]
MH681419.1_ASFV/POL/2015         ................................................................................ [ 79520]
LS478113.1_Estonia_2014          ................................................................................ [ 79520]
KP843857.1_Odintsovo_02/14       .............................-.................................................. [ 79520]
```

```
FN557520.1_strain_E75            ................................................................................ [ 79520]
AY261366.1_Warthog               ................g............................................................... [ 79520]
AY261365.1_Warmbaths             ................................................................................ [ 79520]
AY261364.1_Tengani_62            .............g..c............................................................... [ 79520]
AY261363.1_Pretoriuskop/96/4     ................g............................................................... [ 79520]
AY261362.1_Mkuzi_1979            ................................................................................ [ 79520]
AY261361.1_Malawi_Lil-20/1_1983  ......................a..t........................................c............c [ 79520]
AY261360.1_Kenya_1950            .........................t..............c..............c..........c..........g.. [ 79520]
AM712240.1_OURT_88/3             ................................................................................ [ 79520]
AM712239.1_Benin_97/1            ................................................................................ [ 79520]
KM111295.1_Ken06.Bus             ........................................c..t...........c.....c....c..........g.. [ 79520]
KM111294.1_Ken05/Tk1             .........................t..............c..............c..........c..........g.. [ 79520]
KM262844.1_strain_L60            ................................................................................ [ 79520]
KM102979.1_26544/OG10_Italy      ................................................................................ [ 79520]
KM262845.1_strain_NHV            ................................................................................ [ 79520]
KX354450.1_47/Ss/2008            ................................................................................ [ 79520]
KP055815.1_strain_BA71           ................................................................................ [ 79520]
MH025920.1_strain_R35            ........................................c..t...........c.....c....c..........g.. [ 79520]
MH025919.1_strain_N10            ........................................c..t...........c.....c....c..........g.. [ 79520]
MH025918.1_strain_R25            ........................................c..t...........c.....c....c..........g.. [ 79520]
MH025917.1_strain_R7             ........................................c..t...........c.....c....c..........g.. [ 79520]
MH025916.1_strain_R8             ........................................c..t...........c.....c....c..........g.. [ 79520]

FR682468.2_Georgia_2007/1        ccttggcccagtatggaggccctttttccctcaagcagtaaaagatatttaggcaactatgtagaccctaacgcgctcat [ 79600]
ASF_IND_20_CAD543                ................................................................................ [ 79600]
LR899193.1_Germany_2020/1        ................................................................................ [ 79600]
MN393477.1_Wuhan_2019            ................................................................................ [ 79600]
MW396979.1_Timor-Leste/2019/1    ................................................................................ [ 79600]
MK940252.1_CN/2019/InnerMongolia ................................................................................ [ 79600]
MK128995.1_China/2018/AnhuiXCGQ  ................................................................................ [ 79600]
MH681419.1_ASFV/POL/2015         ................................................................................ [ 79600]
LS478113.1_Estonia_2014          ................................................................................ [ 79600]
KP843857.1_Odintsovo_02/14       ................................................................................ [ 79600]
FN557520.1_strain_E75            ................................................................................ [ 79600]
AY261366.1_Warthog               ........t..c.................................................................... [ 79600]
AY261365.1_Warmbaths             ...........c.................................................................... [ 79600]
AY261364.1_Tengani_62            ...........c.................................................................... [ 79600]
AY261363.1_Pretoriuskop/96/4     ...........c.................................................................... [ 79600]
AY261362.1_Mkuzi_1979            ................................................................................ [ 79600]
AY261361.1_Malawi_Lil-20/1_1983  ........t.....a.........................................t....................... [ 79600]
AY261360.1_Kenya_1950            ........t..c.....a..............g........g...................................... [ 79600]
AM712240.1_OURT_88/3             ................................................................................ [ 79600]
AM712239.1_Benin_97/1            ................................................................................ [ 79600]
KM111295.1_Ken06.Bus             ........t..c....................g........c.............................t........ [ 79600]
```

```
KM111294.1_Ken05/Tk1            ........t..c.....a..............g........g...................................... [ 79600]
KM262844.1_strain_L60           ................................................................................ [ 79600]
KM102979.1_26544/OG10_Italy     ................................................................................ [ 79600]
KM262845.1_strain_NHV           ................................................................................ [ 79600]
KX354450.1_47/Ss/2008           ................................................................................ [ 79600]
KP055815.1_strain_BA71          ................................................................................ [ 79600]
MH025920.1_strain_R35           ........t..c....................g........c.............................t........ [ 79600]
MH025919.1_strain_N10           ........t..c....................g........c.............................t........ [ 79600]
MH025918.1_strain_R25           ........t..c....................g........c.............................t........ [ 79600]
MH025917.1_strain_R7            ........t..c....................g........c.............................t........ [ 79600]
MH025916.1_strain_R8            ........t..c....................g........c.............................t........ [ 79600]

FR682468.2_Georgia_2007/1       aaagtgggcacaactcaaattaaaaagaataccgcctttttatcctggaaagccggatgaagaatcatgtta-agccgat [ 79680]
ASF_IND_20_CAD543               ........................................................................-....... [ 79680]
LR899193.1_Germany_2020/1       ........................................................................-....... [ 79680]
MN393477.1_Wuhan_2019           ........................................................................-....... [ 79680]
MW396979.1_Timor-Leste/2019/1   ........................................................................-....... [ 79680]
MK940252.1_CN/2019/InnerMongolia ........................................................................-....... [ 79680]
MK128995.1_China/2018/AnhuiXCGQ ........................................................................-....... [ 79680]
MH681419.1_ASFV/POL/2015        ........................................................................-....... [ 79680]
LS478113.1_Estonia_2014         ........................................................................-....... [ 79680]
KP843857.1_Odintsovo_02/14      ........................................................................-....... [ 79680]
FN557520.1_strain_E75           .............................................c..........................-....... [ 79680]
AY261366.1_Warthog              .............................................c...............------------------- [ 79680]
AY261365.1_Warmbaths            .............................................c..........................-...t... [ 79680]
AY261364.1_Tengani_62           .............................................c.........................--------- [ 79680]
AY261363.1_Pretoriuskop/96/4    .............................................c..........................-...t.g. [ 79680]
AY261362.1_Mkuzi_1979           .............................................c..........................-.....g. [ 79680]
AY261361.1_Malawi_Lil-20/1_1983 ...............a..............t...................g.....................-...ta.. [ 79680]
AY261360.1_Kenya_1950           ...a.....t.....a..............t....t......c..c....g............ggag.aa.gt.tattta [ 79680]
AM712240.1_OURT_88/3            .............................................c..........................-....... [ 79680]
AM712239.1_Benin_97/1           .............................................c..........................-....... [ 79680]
KM111295.1_Ken06.Bus            .........t.....a..............t....t......c..c....g.............ga.gag..---at.ta [ 79680]
KM111294.1_Ken05/Tk1            ...a.....t.....a..............t....t......c..c....g............ggag.aa.g-------- [ 79680]
KM262844.1_strain_L60           .............................................c..........................-....... [ 79680]
KM102979.1_26544/OG10_Italy     .............................................c..........................-....... [ 79680]
KM262845.1_strain_NHV           .............................................c..........................-....... [ 79680]
KX354450.1_47/Ss/2008           .............................................c..........................-....... [ 79680]
KP055815.1_strain_BA71          .............................................c..........................-....... [ 79680]
MH025920.1_strain_R35           .........t.....a..............t....t......c..c....g.............ga.gag..---at.ta [ 79680]
MH025919.1_strain_N10           .........t.....a..............t....t......c..c....g.............ga.gag..---at.ta [ 79680]
MH025918.1_strain_R25           .........t.....a..............t....t......c..c....g.............ga.gag..---at.ta [ 79680]
MH025917.1_strain_R7            .........t.....a..............t....t......c..c....g.............ga.gag..---at.ta [ 79680]
MH025916.1_strain_R8            .........t.....a..............t....t......c..c....g.............ga.gag..---at.ta [ 79680]
```

```
FR682468.2_Georgia_2007/1        taaaaaatcatgttaagctggttgaaaaatcatgttaagctggttgaaaaactcttggtgaaagcacggatgtaatatta [ 79760]
ASF_IND_20_CAD543                .r............----------------------............................................ [ 79760]
LR899193.1_Germany_2020/1        ................................................................................ [ 79760]
MN393477.1_Wuhan_2019            ................................................................................ [ 79760]
MW396979.1_Timor-Leste/2019/1    ................................................................................ [ 79760]
MK940252.1_CN/2019/InnerMongolia ................................................................................ [ 79760]
MK128995.1_China/2018/AnhuiXCGQ  ................................................................................ [ 79760]
MH681419.1_ASFV/POL/2015         ................................................................................ [ 79760]
LS478113.1_Estonia_2014          ................................................................................ [ 79760]
KP843857.1_Odintsovo_02/14       ................................................................................ [ 79760]
FN557520.1_strain_E75            .g............----------------------....c........t.............................. [ 79760]
AY261366.1_Warthog               ----..............g.....-...............c........t.............................. [ 79760]
AY261365.1_Warmbaths             .--g.......a......g.....-...............c........t.............................. [ 79760]
AY261364.1_Tengani_62            ------------------------------------....c........t.............................. [ 79760]
AY261363.1_Pretoriuskop/96/4     .g................g.....-...............c........t.............................. [ 79760]
AY261362.1_Mkuzi_1979            .g............----------------------....c........t.............................. [ 79760]
AY261361.1_Malawi_Lil-20/1_1983  .g-................aa...-............c..g..c.....t..........................g... [ 79760]
AY261360.1_Kenya_1950            .gt.t.t.t..a.------------.t.t.t..a......c..c.....t.............................. [ 79760]
AM712240.1_OURT_88/3             .g............----------------------....c........t.............................. [ 79760]
AM712239.1_Benin_97/1            .g............----------------------....c........t.............................. [ 79760]
KM111295.1_Ken06.Bus             ..t.t.t.t.aa.------------.t.--t..a......c..c.....t..............t............... [ 79760]
KM111294.1_Ken05/Tk1             --t.t.t.t..a.------------.t.t.t..a......c..c.....t.............................. [ 79760]
KM262844.1_strain_L60            .g............----------------------....c........t.............................. [ 79760]
KM102979.1_26544/OG10_Italy      .g............----------------------....c........t.............................. [ 79760]
KM262845.1_strain_NHV            .g............----------------------....c........t.............................. [ 79760]
KX354450.1_47/Ss/2008            .g............----------------------....c........t.............................. [ 79760]
KP055815.1_strain_BA71           .g............----------------------....c........t.............................. [ 79760]
MH025920.1_strain_R35            ..t.t.t.t.aa.------------.t.--t..a......c..c.....t..............t............... [ 79760]
MH025919.1_strain_N10            ..t.t.t.t.aa.------------.t.--t..a......c..c.....t..............t............... [ 79760]
MH025918.1_strain_R25            ..t.t.t.t.aa.------------.t.--t..a......c..c.....t..............t............... [ 79760]
MH025917.1_strain_R7             ..t.t.t.t.aa.------------.t.--t..a......c..c.....t..............t............... [ 79760]
MH025916.1_strain_R8             ..t.t.t.t.aa.------------.t.--t..a......c..c.....t..............t............... [ 79760]

FR682468.2_Georgia_2007/1        acattggccgctcgcatttcgtgttgaaatacgatggaagagcgacggctatctaccatgccgatatcggcctggacatc [ 79840]
ASF_IND_20_CAD543                ................................................................................ [ 79840]
LR899193.1_Germany_2020/1        ................................................................................ [ 79840]
MN393477.1_Wuhan_2019            ................................................................................ [ 79840]
MW396979.1_Timor-Leste/2019/1    ................................................................................ [ 79840]
MK940252.1_CN/2019/InnerMongolia ................................................................................ [ 79840]
MK128995.1_China/2018/AnhuiXCGQ  ................................................................................ [ 79840]
MH681419.1_ASFV/POL/2015         ................................................................................ [ 79840]
LS478113.1_Estonia_2014          ................................................................................ [ 79840]
KP843857.1_Odintsovo_02/14       ................................................................................ [ 79840]
```

```
FN557520.1_strain_E75              ...........c..........................g......................................... [ 79840]
AY261366.1_Warthog                 ...........c..........................g..............c.......................... [ 79840]
AY261365.1_Warmbaths               ...........c..........................g..............c.......................... [ 79840]
AY261364.1_Tengani_62              ...........c..........................g..............c.................t........ [ 79840]
AY261363.1_Pretoriuskop/96/4       ........t..c..........................g..............c.......................... [ 79840]
AY261362.1_Mkuzi_1979              .....................................................c.......................... [ 79840]
AY261361.1_Malawi_Lil-20/1_1983    ..g......................................a...........c.......t.................. [ 79840]
AY261360.1_Kenya_1950              ..g...................................g..a...........c.......t.................. [ 79840]
AM712240.1_OURT_88/3               ...........c..........................g......................................... [ 79840]
AM712239.1_Benin_97/1              ...........c..........................g......................................... [ 79840]
KM111295.1_Ken06.Bus               ..g...................................g..a...........c.......t.................. [ 79840]
KM111294.1_Ken05/Tk1               ..g...................................g..a...........c.......t.................. [ 79840]
KM262844.1_strain_L60              ...........c..........................g......................................... [ 79840]
KM102979.1_26544/OG10_Italy        ...........ct.........................g......................................... [ 79840]
KM262845.1_strain_NHV              ...........c..........................g......................................... [ 79840]
KX354450.1_47/Ss/2008              ...........ct.........................g......................................... [ 79840]
KP055815.1_strain_BA71             ...........c..........................g......................................... [ 79840]
MH025920.1_strain_R35              ..g...................................g..a...........c.......t.................. [ 79840]
MH025919.1_strain_N10              ..g...................................g..a...........c.......t.................. [ 79840]
MH025918.1_strain_R25              ..g...................................g..a...........c.......t.................. [ 79840]
MH025917.1_strain_R7               ..g...................................g..a...........c.......t.................. [ 79840]
MH025916.1_strain_R8               ..g...................................g..a...........c.......t.................. [ 79840]

FR682468.2_Georgia_2007/1          acagttcatgcacttgtagatgggatgactcgcgttatagatggcaggctcgccacagtttctacagatgtaggagatgc [ 79920]
ASF_IND_20_CAD543                  ................................................................................ [ 79920]
LR899193.1_Germany_2020/1          ................................................................................ [ 79920]
MN393477.1_Wuhan_2019              ................................................................................ [ 79920]
MW396979.1_Timor-Leste/2019/1      ................................................................................ [ 79920]
MK940252.1_CN/2019/InnerMongolia   ................................................................................ [ 79920]
MK128995.1_China/2018/AnhuiXCGQ    ................................................................................ [ 79920]
MH681419.1_ASFV/POL/2015           ................................................................................ [ 79920]
LS478113.1_Estonia_2014            ................................................................................ [ 79920]
KP843857.1_Odintsovo_02/14         ................................................................................ [ 79920]
FN557520.1_strain_E75              ................................................................................ [ 79920]
AY261366.1_Warthog                 ....................................................................c........... [ 79920]
AY261365.1_Warmbaths               ................................................................................ [ 79920]
AY261364.1_Tengani_62              .............................................g.................................. [ 79920]
AY261363.1_Pretoriuskop/96/4       .................................a..................................c.........a. [ 79920]
AY261362.1_Mkuzi_1979              ................................................................................ [ 79920]
AY261361.1_Malawi_Lil-20/1_1983    ................................................................................ [ 79920]
AY261360.1_Kenya_1950              g............................................................................... [ 79920]
AM712240.1_OURT_88/3               ................................................................................ [ 79920]
AM712239.1_Benin_97/1              ................................................................................ [ 79920]
KM111295.1_Ken06.Bus               g............................................................................... [ 79920]
```

```
KM111294.1_Ken05/Tk1              g............................................................................... [ 79920]
KM262844.1_strain_L60             ................................................................................ [ 79920]
KM102979.1_26544/OG10_Italy       ................................................................................ [ 79920]
KM262845.1_strain_NHV             ................................................................................ [ 79920]
KX354450.1_47/Ss/2008             ................................................................................ [ 79920]
KP055815.1_strain_BA71            ................................................................................ [ 79920]
MH025920.1_strain_R35             g............................................................................... [ 79920]
MH025919.1_strain_N10             g............................................................................... [ 79920]
MH025918.1_strain_R25             g............................................................................... [ 79920]
MH025917.1_strain_R7              g............................................................................... [ 79920]
MH025916.1_strain_R8              g............................................................................... [ 79920]

FR682468.2_Georgia_2007/1         agccatccgagtcgtcgtgcgatttttctatgatggtttgcatggcgccctgcgccgtaagcacccaatgctccatttct [ 80000]
ASF_IND_20_CAD543                 ................................................................................ [ 80000]
LR899193.1_Germany_2020/1         ................................................................................ [ 80000]
MN393477.1_Wuhan_2019             ................................................................................ [ 80000]
MW396979.1_Timor-Leste/2019/1     ................................................................................ [ 80000]
MK940252.1_CN/2019/InnerMongolia  ................................................................................ [ 80000]
MK128995.1_China/2018/AnhuiXCGQ   ................................................................................ [ 80000]
MH681419.1_ASFV/POL/2015          ................................................................................ [ 80000]
LS478113.1_Estonia_2014           ................................................................................ [ 80000]
KP843857.1_Odintsovo_02/14        ................................................................................ [ 80000]
FN557520.1_strain_E75             ................................................................................ [ 80000]
AY261366.1_Warthog                ................................................................................ [ 80000]
AY261365.1_Warmbaths              ....g........................................................................... [ 80000]
AY261364.1_Tengani_62             ..............................................a................................. [ 80000]
AY261363.1_Pretoriuskop/96/4      ................................................................................ [ 80000]
AY261362.1_Mkuzi_1979             ................................................................................ [ 80000]
AY261361.1_Malawi_Lil-20/1_1983   .............a................................a.............................c... [ 80000]
AY261360.1_Kenya_1950             .............a................................a.............................c... [ 80000]
AM712240.1_OURT_88/3              ................................................................................ [ 80000]
AM712239.1_Benin_97/1             ................................................................................ [ 80000]
KM111295.1_Ken06.Bus              .............a..............................................................c... [ 80000]
KM111294.1_Ken05/Tk1              .............a..............................................................c... [ 80000]
KM262844.1_strain_L60             ................................................................................ [ 80000]
KM102979.1_26544/OG10_Italy       ................................................................................ [ 80000]
KM262845.1_strain_NHV             ................................................................................ [ 80000]
KX354450.1_47/Ss/2008             ................................................................................ [ 80000]
KP055815.1_strain_BA71            ................................................................................ [ 80000]
MH025920.1_strain_R35             .............a..............................................................c... [ 80000]
MH025919.1_strain_N10             .............a..............................................................c... [ 80000]
MH025918.1_strain_R25             .............a..............................................................c... [ 80000]
MH025917.1_strain_R7              .............a..............................................................c... [ 80000]
MH025916.1_strain_R8              .............a..............................................................c... [ 80000]
```

```
FR682468.2_Georgia_2007/1        cccagacgaagacctccgtgcgatcgtttgccgtccaacggctggcctgtgagggcatccgtgggcccatagcttgcaac [ 80080]
ASF_IND_20_CAD543                ................................................................................ [ 80080]
LR899193.1_Germany_2020/1        ................................................................................ [ 80080]
MN393477.1_Wuhan_2019            ................................................................................ [ 80080]
MW396979.1_Timor-Leste/2019/1    ................................................................................ [ 80080]
MK940252.1_CN/2019/InnerMongolia ................................................................................ [ 80080]
MK128995.1_China/2018/AnhuiXCGQ  ................................................................................ [ 80080]
MH681419.1_ASFV/POL/2015         ................................................................................ [ 80080]
LS478113.1_Estonia_2014          ................................................................................ [ 80080]
KP843857.1_Odintsovo_02/14       ................................................................................ [ 80080]
FN557520.1_strain_E75            ................................................................................ [ 80080]
AY261366.1_Warthog               ................................................................................ [ 80080]
AY261365.1_Warmbaths             .................................................................t.............. [ 80080]
AY261364.1_Tengani_62            ..........................c..................................................... [ 80080]
AY261363.1_Pretoriuskop/96/4     ................................................................................ [ 80080]
AY261362.1_Mkuzi_1979            ................................................................................ [ 80080]
AY261361.1_Malawi_Lil-20/1_1983  ...................................t............................................ [ 80080]
AY261360.1_Kenya_1950            ......................................t..t...................................... [ 80080]
AM712240.1_OURT_88/3             ................................................................................ [ 80080]
AM712239.1_Benin_97/1            ................................................................................ [ 80080]
KM111295.1_Ken06.Bus             .....g.......................................................................... [ 80080]
KM111294.1_Ken05/Tk1             ................................................................................ [ 80080]
KM262844.1_strain_L60            ................................................................................ [ 80080]
KM102979.1_26544/OG10_Italy      ................................................................................ [ 80080]
KM262845.1_strain_NHV            ................................................................................ [ 80080]
KX354450.1_47/Ss/2008            ................................................................................ [ 80080]
KP055815.1_strain_BA71           ................................................................................ [ 80080]
MH025920.1_strain_R35            .....g.......................................................................... [ 80080]
MH025919.1_strain_N10            .....g.......................................................................... [ 80080]
MH025918.1_strain_R25            .....g.......................................................................... [ 80080]
MH025917.1_strain_R7             .....g.......................................................................... [ 80080]
MH025916.1_strain_R8             .....g.......................................................................... [ 80080]

FR682468.2_Georgia_2007/1        ggcgtatcggtcatccagcacaaatttttgcaggcgctggtgataggtcggtcctatgaagatggccgcatcaaagtact [ 80160]
ASF_IND_20_CAD543                ................................................................................ [ 80160]
LR899193.1_Germany_2020/1        ................................................................................ [ 80160]
MN393477.1_Wuhan_2019            ................................................................................ [ 80160]
MW396979.1_Timor-Leste/2019/1    ................................................................................ [ 80160]
MK940252.1_CN/2019/InnerMongolia ................................................................................ [ 80160]
MK128995.1_China/2018/AnhuiXCGQ  ................................................................................ [ 80160]
MH681419.1_ASFV/POL/2015         ................................................................................ [ 80160]
LS478113.1_Estonia_2014          ................................................................................ [ 80160]
KP843857.1_Odintsovo_02/14       ................................................................................ [ 80160]
```

```
FN557520.1_strain_E75           ................................................................................ [ 80160]
AY261366.1_Warthog              ................................................................................ [ 80160]
AY261365.1_Warmbaths            ......c.........................................t.................t........a.... [ 80160]
AY261364.1_Tengani_62           ................................................................................ [ 80160]
AY261363.1_Pretoriuskop/96/4    .................a.........................................................a.... [ 80160]
AY261362.1_Mkuzi_1979           ................................................................................ [ 80160]
AY261361.1_Malawi_Lil-20/1_1983 ........................c.......................t.............................t. [ 80160]
AY261360.1_Kenya_1950           ......c..a................................g.....t............................... [ 80160]
AM712240.1_OURT_88/3            ................................................................................ [ 80160]
AM712239.1_Benin_97/1           ................................................................................ [ 80160]
KM111295.1_Ken06.Bus            ......c..a................................g.....t....................g.......... [ 80160]
KM111294.1_Ken05/Tk1            ......c..a................................g.....t............................... [ 80160]
KM262844.1_strain_L60           ................................................................................ [ 80160]
KM102979.1_26544/OG10_Italy     ................................................................................ [ 80160]
KM262845.1_strain_NHV           ................................................................................ [ 80160]
KX354450.1_47/Ss/2008           ................................................................................ [ 80160]
KP055815.1_strain_BA71          ................................................................................ [ 80160]
MH025920.1_strain_R35           ......c..a................................g.....t....................g.......... [ 80160]
MH025919.1_strain_N10           ......c..a................................g.....t....................g.......... [ 80160]
MH025918.1_strain_R25           ......c..a................................g.....t....................g.......... [ 80160]
MH025917.1_strain_R7            ......c..a................................g.....t....................g.......... [ 80160]
MH025916.1_strain_R8            ......c..a................................g.....t....................g.......... [ 80160]

FR682468.2_Georgia_2007/1       cgccggtctggccgttgaacattttttggcatccattgaagcgtagaccttcttgcgccagtctttctgaaagaagctgc [ 80240]
ASF_IND_20_CAD543               ................................................................................ [ 80240]
LR899193.1_Germany_2020/1       ................................................................................ [ 80240]
MN393477.1_Wuhan_2019           ................................................................................ [ 80240]
MW396979.1_Timor-Leste/2019/1   ................................................................................ [ 80240]
MK940252.1_CN/2019/InnerMongolia ................................................................................ [ 80240]
MK128995.1_China/2018/AnhuiXCGQ ................................................................................ [ 80240]
MH681419.1_ASFV/POL/2015        ................................................................................ [ 80240]
LS478113.1_Estonia_2014         ................................................................................ [ 80240]
KP843857.1_Odintsovo_02/14      ................................................................................ [ 80240]
FN557520.1_strain_E75           ................................................................................ [ 80240]
AY261366.1_Warthog              ................................................................................ [ 80240]
AY261365.1_Warmbaths            ................................................................................ [ 80240]
AY261364.1_Tengani_62           ....a........................................................................... [ 80240]
AY261363.1_Pretoriuskop/96/4    .............a.................................................................. [ 80240]
AY261362.1_Mkuzi_1979           ................................................................................ [ 80240]
AY261361.1_Malawi_Lil-20/1_1983 ................a..............................................................t [ 80240]
AY261360.1_Kenya_1950           ................a...........a................................................... [ 80240]
AM712240.1_OURT_88/3            ................................................................................ [ 80240]
AM712239.1_Benin_97/1           ..................................................................t............. [ 80240]
KM111295.1_Ken06.Bus            .a...........................................a.................................. [ 80240]
```

```
KM111294.1_Ken05/Tk1            .a.............................................................................. [ 80240]
KM262844.1_strain_L60           ................................................................................ [ 80240]
KM102979.1_26544/OG10_Italy     ................................................................................ [ 80240]
KM262845.1_strain_NHV           ................................................................................ [ 80240]
KX354450.1_47/Ss/2008           ................................................................................ [ 80240]
KP055815.1_strain_BA71          ................................................................................ [ 80240]
MH025920.1_strain_R35           .a...........................................a.................................. [ 80240]
MH025919.1_strain_N10           .a...........................................a.................................. [ 80240]
MH025918.1_strain_R25           .a...........................................a.................................. [ 80240]
MH025917.1_strain_R7            .a...........................................a.................................. [ 80240]
MH025916.1_strain_R8            .a...........................................a.................................. [ 80240]

FR682468.2_Georgia_2007/1       acattaataggcaggaatgcggtgccgtctgttaccaccccctgtagggcatttgctagaccaaccgtggtttctatcat [ 80320]
ASF_IND_20_CAD543               ................................................................................ [ 80320]
LR899193.1_Germany_2020/1       ................................................................................ [ 80320]
MN393477.1_Wuhan_2019           ................................................................................ [ 80320]
MW396979.1_Timor-Leste/2019/1   ................................................................................ [ 80320]
MK940252.1_CN/2019/InnerMongolia ................................................................................ [ 80320]
MK128995.1_China/2018/AnhuiXCGQ ................................................................................ [ 80320]
MH681419.1_ASFV/POL/2015        ................................................................................ [ 80320]
LS478113.1_Estonia_2014         ................................................................................ [ 80320]
KP843857.1_Odintsovo_02/14      ................................................................................ [ 80320]
FN557520.1_strain_E75           ..g............................................................................. [ 80320]
AY261366.1_Warthog              ..g.................a........a.................................................. [ 80320]
AY261365.1_Warmbaths            ..g............................................................................. [ 80320]
AY261364.1_Tengani_62           ..g............................................a.................t..c........... [ 80320]
AY261363.1_Pretoriuskop/96/4    ..g...................................t........a................................ [ 80320]
AY261362.1_Mkuzi_1979           ..g............................................................................. [ 80320]
AY261361.1_Malawi_Lil-20/1_1983 ........g...........................................................a........... [ 80320]
AY261360.1_Kenya_1950           .....g..g.............................t........c....................c........... [ 80320]
AM712240.1_OURT_88/3            ..g............................................................................. [ 80320]
AM712239.1_Benin_97/1           ..g............................................................................. [ 80320]
KM111295.1_Ken06.Bus            ........g.....a....................t..t........a....................c........... [ 80320]
KM111294.1_Ken05/Tk1            .....g..g.............................t........c....................c........... [ 80320]
KM262844.1_strain_L60           ..g............................................................................. [ 80320]
KM102979.1_26544/OG10_Italy     ..g............................................................................. [ 80320]
KM262845.1_strain_NHV           ..g............................................................................. [ 80320]
KX354450.1_47/Ss/2008           ..g............................................................................. [ 80320]
KP055815.1_strain_BA71          ..g............................................................................. [ 80320]
MH025920.1_strain_R35           ........g.....a....................t..t........a....................c........... [ 80320]
MH025919.1_strain_N10           ........g.....a....................t..t........a....................c........... [ 80320]
MH025918.1_strain_R25           ........g.....a....................t..t........a....................c........... [ 80320]
MH025917.1_strain_R7            ........g.....a....................t..t........a....................c........... [ 80320]
MH025916.1_strain_R8            ........g.....a....................t..t........a....................c........... [ 80320]
```

```
FR682468.2_Georgia_2007/1        ttgaccgttggtcattcgggagggatgtgagtgggggtttacaatgaggtcgggctgcaatccgtcctctgtgaagggca [ 80400]
ASF_IND_20_CAD543                ................................................................................ [ 80400]
LR899193.1_Germany_2020/1        ................................................................................ [ 80400]
MN393477.1_Wuhan_2019            ................................................................................ [ 80400]
MW396979.1_Timor-Leste/2019/1    ................................................................................ [ 80400]
MK940252.1_CN/2019/InnerMongolia ................................................................................ [ 80400]
MK128995.1_China/2018/AnhuiXCGQ  ................................................................................ [ 80400]
MH681419.1_ASFV/POL/2015         ................................................................................ [ 80400]
LS478113.1_Estonia_2014          ................................................................................ [ 80400]
KP843857.1_Odintsovo_02/14       ................................................................................ [ 80400]
FN557520.1_strain_E75            ................................................a..........g.................... [ 80400]
AY261366.1_Warthog               ...........................................................g.................... [ 80400]
AY261365.1_Warmbaths             ...........................................................gg................... [ 80400]
AY261364.1_Tengani_62            ...........................................................g.................... [ 80400]
AY261363.1_Pretoriuskop/96/4     ......a....................................................g.................... [ 80400]
AY261362.1_Mkuzi_1979            ................................................a..........g.................... [ 80400]
AY261361.1_Malawi_Lil-20/1_1983  ..............................a..t..a......................g.................... [ 80400]
AY261360.1_Kenya_1950            ......a.......................a..t.........................g...a................ [ 80400]
AM712240.1_OURT_88/3             ................................................a..........g.................... [ 80400]
AM712239.1_Benin_97/1            ................................................a..........g.................... [ 80400]
KM111295.1_Ken06.Bus             ......a.......................a..t.........................g...............a.... [ 80400]
KM111294.1_Ken05/Tk1             ......a.......................a..t.........................g...............a.... [ 80400]
KM262844.1_strain_L60            ................................................a..........g.................... [ 80400]
KM102979.1_26544/OG10_Italy      ................................................a..........g.................... [ 80400]
KM262845.1_strain_NHV            ................................................a..........g.................... [ 80400]
KX354450.1_47/Ss/2008            ................................................a..........g.................... [ 80400]
KP055815.1_strain_BA71           ................................................a..........g.................... [ 80400]
MH025920.1_strain_R35            ......a.......................a..t.........................g...............a.... [ 80400]
MH025919.1_strain_N10            ......a.......................a..t.........................g...............a.... [ 80400]
MH025918.1_strain_R25            ......a.......................a..t.........................g...............a.... [ 80400]
MH025917.1_strain_R7             ......a.......................a..t.........................g...............a.... [ 80400]
MH025916.1_strain_R8             ......a.......................a..t.........................g...............a.... [ 80400]

FR682468.2_Georgia_2007/1        tgtctgaagtgggcagggccagcgccgcaatgcccttgttcccgctgcgagaactcattttgtcgcctatattgagattt [ 80480]
ASF_IND_20_CAD543                .......r........................................................................ [ 80480]
LR899193.1_Germany_2020/1        ................................................................................ [ 80480]
MN393477.1_Wuhan_2019            ................................................................................ [ 80480]
MW396979.1_Timor-Leste/2019/1    ................................................................................ [ 80480]
MK940252.1_CN/2019/InnerMongolia ................................................................................ [ 80480]
MK128995.1_China/2018/AnhuiXCGQ  ................................................................................ [ 80480]
MH681419.1_ASFV/POL/2015         ................................................................................ [ 80480]
LS478113.1_Estonia_2014          ................................................................................ [ 80480]
KP843857.1_Odintsovo_02/14       ................................................................................ [ 80480]
```

```
FN557520.1_strain_E75             .......g..................................................c..................... [ 80480]
AY261366.1_Warthog                .................................................g........c..................... [ 80480]
AY261365.1_Warmbaths              .................................................g........c..................... [ 80480]
AY261364.1_Tengani_62             .................................................g.............................. [ 80480]
AY261363.1_Pretoriuskop/96/4      .................................................g........c..................... [ 80480]
AY261362.1_Mkuzi_1979             ................................................................................ [ 80480]
AY261361.1_Malawi_Lil-20/1_1983   .......c...........t......................................c........a............ [ 80480]
AY261360.1_Kenya_1950             .a.....c.................t.................a.a............c........a............ [ 80480]
AM712240.1_OURT_88/3              .......g..................................................c..................... [ 80480]
AM712239.1_Benin_97/1             .......g..................................................c..................... [ 80480]
KM111295.1_Ken06.Bus              .a.....g.......a..........................................c.....a..a............ [ 80480]
KM111294.1_Ken05/Tk1              .a.....g.......a...........................a.a............c........a............ [ 80480]
KM262844.1_strain_L60             .......g..................................................c..................... [ 80480]
KM102979.1_26544/OG10_Italy       .......g..................................................c..................... [ 80480]
KM262845.1_strain_NHV             .......g..................................................c..................... [ 80480]
KX354450.1_47/Ss/2008             .......g..................................................c..................... [ 80480]
KP055815.1_strain_BA71            .......g..................................................c..................... [ 80480]
MH025920.1_strain_R35             .a.....g.......a..........................................c.....a..a............ [ 80480]
MH025919.1_strain_N10             .a.....g.......a..........................................c.....a..a............ [ 80480]
MH025918.1_strain_R25             .a.....g.......a..........................................c.....a..a............ [ 80480]
MH025917.1_strain_R7              .a.....g.......a..........................................c.....a..a............ [ 80480]
MH025916.1_strain_R8              .a.....g.......a..........................................c.....a..a............ [ 80480]

FR682468.2_Georgia_2007/1         ctttcatagcgcaggcgcatgaggccaaagatctcgtcattaggcccatggggacgcatcacagcatccacgacggccgg [ 80560]
ASF_IND_20_CAD543                 ................................................................................ [ 80560]
LR899193.1_Germany_2020/1         ................................................................................ [ 80560]
MN393477.1_Wuhan_2019             ................................................................................ [ 80560]
MW396979.1_Timor-Leste/2019/1     ................................................................................ [ 80560]
MK940252.1_CN/2019/InnerMongolia  ................................................................................ [ 80560]
MK128995.1_China/2018/AnhuiXCGQ   ................................................................................ [ 80560]
MH681419.1_ASFV/POL/2015          ................................................................................ [ 80560]
LS478113.1_Estonia_2014           ................................................................................ [ 80560]
KP843857.1_Odintsovo_02/14        ................................................................................ [ 80560]
FN557520.1_strain_E75             .........................................g...................................... [ 80560]
AY261366.1_Warthog                ..........................g.................t................................... [ 80560]
AY261365.1_Warmbaths              ..........................g........a.....g..t................................... [ 80560]
AY261364.1_Tengani_62             ..........................g..................................................... [ 80560]
AY261363.1_Pretoriuskop/96/4      ..........................g..................................................... [ 80560]
AY261362.1_Mkuzi_1979             .........................................g...................................... [ 80560]
AY261361.1_Malawi_Lil-20/1_1983   ...........t..............g..............g..t.......................t.......... [ 80560]
AY261360.1_Kenya_1950             ..........................g..............g..t.......................t.......... [ 80560]
AM712240.1_OURT_88/3              .........................................g...................................... [ 80560]
AM712239.1_Benin_97/1             .........................................g...................................... [ 80560]
KM111295.1_Ken06.Bus              ..............a...........g..............g..t.......................t.......... [ 80560]
```

```
KM111294.1_Ken05/Tk1            ..........................g..............g..t.......................t........... [ 80560]
KM262844.1_strain_L60           .........................................g...................................... [ 80560]
KM102979.1_26544/OG10_Italy     .........................................g...................................... [ 80560]
KM262845.1_strain_NHV           .........................................g...................................... [ 80560]
KX354450.1_47/Ss/2008           .........................................g...................................... [ 80560]
KP055815.1_strain_BA71          .........................................g...................................... [ 80560]
MH025920.1_strain_R35           ..............a...........g..............g..t.......................t........... [ 80560]
MH025919.1_strain_N10           ..............a...........g..............g..t.......................t........... [ 80560]
MH025918.1_strain_R25           ..............a...........g..............g..t.......................t........... [ 80560]
MH025917.1_strain_R7            ..............a...........g..............g..t.......................t........... [ 80560]
MH025916.1_strain_R8            ..............a...........g..............g..t.......................t........... [ 80560]

FR682468.2_Georgia_2007/1       ctcatcgaagccgtacatgacagaccggtcgatgtatttgttgagttcgtctttttcgccccgtattttggccacttttc [ 80640]
ASF_IND_20_CAD543               ................................................................................ [ 80640]
LR899193.1_Germany_2020/1       ................................................................................ [ 80640]
MN393477.1_Wuhan_2019           ................................................................................ [ 80640]
MW396979.1_Timor-Leste/2019/1   ................................................................................ [ 80640]
MK940252.1_CN/2019/InnerMongolia ................................................................................ [ 80640]
MK128995.1_China/2018/AnhuiXCGQ ................................................................................ [ 80640]
MH681419.1_ASFV/POL/2015        ................................................................................ [ 80640]
LS478113.1_Estonia_2014         ................................................................................ [ 80640]
KP843857.1_Odintsovo_02/14      ................................................................................ [ 80640]
FN557520.1_strain_E75           ................................................................................ [ 80640]
AY261366.1_Warthog              ................................................a............................... [ 80640]
AY261365.1_Warmbaths            ................................................................................ [ 80640]
AY261364.1_Tengani_62           ................................................................................ [ 80640]
AY261363.1_Pretoriuskop/96/4    ................................................................................ [ 80640]
AY261362.1_Mkuzi_1979           ................................................................................ [ 80640]
AY261361.1_Malawi_Lil-20/1_1983 ..................a........a..............a..........................a.......... [ 80640]
AY261360.1_Kenya_1950           .......................................a.............................a.......... [ 80640]
AM712240.1_OURT_88/3            ................................................................................ [ 80640]
AM712239.1_Benin_97/1           ................................................................................ [ 80640]
KM111295.1_Ken06.Bus            ...........................a.................................................... [ 80640]
KM111294.1_Ken05/Tk1            ...........................a.................................................... [ 80640]
KM262844.1_strain_L60           ................................................................................ [ 80640]
KM102979.1_26544/OG10_Italy     ................................................................................ [ 80640]
KM262845.1_strain_NHV           ................................................................................ [ 80640]
KX354450.1_47/Ss/2008           ................................................................................ [ 80640]
KP055815.1_strain_BA71          ................................................................................ [ 80640]
MH025920.1_strain_R35           ...........................a.................................................... [ 80640]
MH025919.1_strain_N10           ...........................a.................................................... [ 80640]
MH025918.1_strain_R25           ...........................a.................................................... [ 80640]
MH025917.1_strain_R7            ...........................a.................................................... [ 80640]
MH025916.1_strain_R8            ...........................a.................................................... [ 80640]
```

```
FR682468.2_Georgia_2007/1        ctataatgatgtcgcccttttttgaccaccgttcctacgggcacgaatccatctacaagcttttcgtaattagcaccaggc [ 80720]
ASF_IND_20_CAD543                ...........................................a..c.............................c... [ 80720]
LR899193.1_Germany_2020/1        ................................................................................ [ 80720]
MN393477.1_Wuhan_2019            ................................................................................ [ 80720]
MW396979.1_Timor-Leste/2019/1    ................................................................................ [ 80720]
MK940252.1_CN/2019/InnerMongolia ................................................................................ [ 80720]
MK128995.1_China/2018/AnhuiXCGQ  ................................................................................ [ 80720]
MH681419.1_ASFV/POL/2015         ................................................................................ [ 80720]
LS478113.1_Estonia_2014          ................................................................................ [ 80720]
KP843857.1_Odintsovo_02/14       ................................................................................ [ 80720]
FN557520.1_strain_E75            ..................................c........a..c.............................c... [ 80720]
AY261366.1_Warthog               ....g.......................t.....c.....t..a..c................................. [ 80720]
AY261365.1_Warmbaths             ....g.......................t.....c.....t..a..c................................. [ 80720]
AY261364.1_Tengani_62            ....g...........t...........t.....c........a..c.......................g.....t... [ 80720]
AY261363.1_Pretoriuskop/96/4     ....g........a..............t.....c........a..c................................. [ 80720]
AY261362.1_Mkuzi_1979            ..................................c........a..c.............................c... [ 80720]
AY261361.1_Malawi_Lil-20/1_1983  ................t.................c.....t.......................a.....g......... [ 80720]
AY261360.1_Kenya_1950            .........................t........c........a....................a.....g.....g... [ 80720]
AM712240.1_OURT_88/3             ..................................c........a..c.............................c... [ 80720]
AM712239.1_Benin_97/1            ..................................c........a..c.............................c... [ 80720]
KM111295.1_Ken06.Bus             ....g...........t...............................................a.....g.....g... [ 80720]
KM111294.1_Ken05/Tk1             ....g...........t...............................................a.....g.....g... [ 80720]
KM262844.1_strain_L60            ..................................c........a..c.............................c... [ 80720]
KM102979.1_26544/OG10_Italy      ..................................c........a..c.............................c... [ 80720]
KM262845.1_strain_NHV            ..................................c........a..c.............................c... [ 80720]
KX354450.1_47/Ss/2008            ..................................c........a..c.............................c... [ 80720]
KP055815.1_strain_BA71           ..................................c........a..c.............................c... [ 80720]
MH025920.1_strain_R35            ....g...........t...............................................a.....g.....g... [ 80720]
MH025919.1_strain_N10            ....g...........t...............................................a.....g.....g... [ 80720]
MH025918.1_strain_R25            ....g...........t...............................................a.....g.....g... [ 80720]
MH025917.1_strain_R7             ....g...........t...............................................a.....g.....g... [ 80720]
MH025916.1_strain_R8             ....g...........t...............................................a.....g.....g... [ 80720]
```

```
FR682468.2_Georgia_2007/1        ttaagatttttggtgattaaagggtcgggcttcccaaacgactctatatcgctttctaattctactttttcttctcggta [ 80800]
ASF_IND_20_CAD543                ................................................................................ [ 80800]
LR899193.1_Germany_2020/1        ................................................................................ [ 80800]
MN393477.1_Wuhan_2019            ................................................................................ [ 80800]
MW396979.1_Timor-Leste/2019/1    ................................................................................ [ 80800]
MK940252.1_CN/2019/InnerMongolia ................................................................................ [ 80800]
MK128995.1_China/2018/AnhuiXCGQ  ................................................................................ [ 80800]
MH681419.1_ASFV/POL/2015         ................................................................................ [ 80800]
LS478113.1_Estonia_2014          ................................................................................ [ 80800]
KP843857.1_Odintsovo_02/14       ................................................................................ [ 80800]
```

```
FN557520.1_strain_E75            ................................................................................ [ 80800]
AY261366.1_Warthog               ........................................................c........c.............. [ 80800]
AY261365.1_Warmbaths             ....................g............................................c.............. [ 80800]
AY261364.1_Tengani_62            ........................................................c........c.............. [ 80800]
AY261363.1_Pretoriuskop/96/4     .................................................................c.............. [ 80800]
AY261362.1_Mkuzi_1979            ................................................................................ [ 80800]
AY261361.1_Malawi_Lil-20/1_1983  .....g...........c..g..a........t........t.....g................................ [ 80800]
AY261360.1_Kenya_1950            ....................g..a........t..............g................................ [ 80800]
AM712240.1_OURT_88/3             ................................................................................ [ 80800]
AM712239.1_Benin_97/1            ................................................................................ [ 80800]
KM111295.1_Ken06.Bus             ....................g..a........t..............g................................ [ 80800]
KM111294.1_Ken05/Tk1             ....................g..a........t..............g................................ [ 80800]
KM262844.1_strain_L60            ................................................................................ [ 80800]
KM102979.1_26544/OG10_Italy      ................................................................................ [ 80800]
KM262845.1_strain_NHV            ................................................................................ [ 80800]
KX354450.1_47/Ss/2008            ................................................................................ [ 80800]
KP055815.1_strain_BA71           ................................................................................ [ 80800]
MH025920.1_strain_R35            ....................g..a........t..............g................................ [ 80800]
MH025919.1_strain_N10            ....................g..a........t..............g................................ [ 80800]
MH025918.1_strain_R25            ....................g..a........t..............g................................ [ 80800]
MH025917.1_strain_R7             ....................g..a........t..............g................................ [ 80800]
MH025916.1_strain_R8             ....................g..a........t..............g................................ [ 80800]

FR682468.2_Georgia_2007/1        gaaggtgccggcaaagccgcccctgtcaataaaggactgcgacacgatcacagagtcctcctgattgtagccgccgtaga [ 80880]
ASF_IND_20_CAD543                ................................................................................ [ 80880]
LR899193.1_Germany_2020/1        ................................................................................ [ 80880]
MN393477.1_Wuhan_2019            ................................................................................ [ 80880]
MW396979.1_Timor-Leste/2019/1    ................................................................................ [ 80880]
MK940252.1_CN/2019/InnerMongolia ................................................................................ [ 80880]
MK128995.1_China/2018/AnhuiXCGQ  ................................................................................ [ 80880]
MH681419.1_ASFV/POL/2015         ................................................................................ [ 80880]
LS478113.1_Estonia_2014          ................................................................................ [ 80880]
KP843857.1_Odintsovo_02/14       ................................................................................ [ 80880]
FN557520.1_strain_E75            ................................................................................ [ 80880]
AY261366.1_Warthog               ..........................................t..a.................................. [ 80880]
AY261365.1_Warmbaths             ........................a..............t.....a.................................. [ 80880]
AY261364.1_Tengani_62            ..................a.................t.....t..a.................................. [ 80880]
AY261363.1_Pretoriuskop/96/4     ..........................................t..a.................................. [ 80880]
AY261362.1_Mkuzi_1979            ................................................................................ [ 80880]
AY261361.1_Malawi_Lil-20/1_1983  .......................................a.................t.................a.... [ 80880]
AY261360.1_Kenya_1950            .......................................t.............................a.......... [ 80880]
AM712240.1_OURT_88/3             ................................................................................ [ 80880]
AM712239.1_Benin_97/1            ................................................................................ [ 80880]
KM111295.1_Ken06.Bus             .......................................t.............................a.......... [ 80880]
```

```
KM111294.1_Ken05/Tk1          .......................................t.............................a.......... [ 80880]
KM262844.1_strain_L60         ................................................................................ [ 80880]
KM102979.1_26544/OG10_Italy   ................................................................................ [ 80880]
KM262845.1_strain_NHV         ................................................................................ [ 80880]
KX354450.1_47/Ss/2008         ................................................................................ [ 80880]
KP055815.1_strain_BA71        ................................................................................ [ 80880]
MH025920.1_strain_R35         .......................................t.............................a.......... [ 80880]
MH025919.1_strain_N10         .......................................t.............................a.......... [ 80880]
MH025918.1_strain_R25         .......................................t.............................a.......... [ 80880]
MH025917.1_strain_R7          .......................................t.............................a.......... [ 80880]
MH025916.1_strain_R8          .......................................t.............................a.......... [ 80880]

FR682468.2_Georgia_2007/1        tcatataagccacaatggtattaagcccgttgggtatgacatagttatgtgctatggtctttacaagcggcatttcattg [ 80960]
ASF_IND_20_CAD543                ................................................................................ [ 80960]
LR899193.1_Germany_2020/1        ................................................................................ [ 80960]
MN393477.1_Wuhan_2019            ................................................................................ [ 80960]
MW396979.1_Timor-Leste/2019/1    ................................................................................ [ 80960]
MK940252.1_CN/2019/InnerMongolia ................................................................................ [ 80960]
MK128995.1_China/2018/AnhuiXCGQ  ................................................................................ [ 80960]
MH681419.1_ASFV/POL/2015         ................................................................................ [ 80960]
LS478113.1_Estonia_2014          ................................................................................ [ 80960]
KP843857.1_Odintsovo_02/14       ................................................................................ [ 80960]
FN557520.1_strain_E75            .......g........................................................................ [ 80960]
AY261366.1_Warthog               .......g................................g....................................... [ 80960]
AY261365.1_Warmbaths             .......g........................................................................ [ 80960]
AY261364.1_Tengani_62            .......g....................a................................................... [ 80960]
AY261363.1_Pretoriuskop/96/4     .......g................................g....................................... [ 80960]
AY261362.1_Mkuzi_1979            .......g...................................a..............t..................... [ 80960]
AY261361.1_Malawi_Lil-20/1_1983  .......g.................t...................................................... [ 80960]
AY261360.1_Kenya_1950            .......g.................t.....a.................c.............................. [ 80960]
AM712240.1_OURT_88/3             .......g........................................................................ [ 80960]
AM712239.1_Benin_97/1            .......g........................................................................ [ 80960]
KM111295.1_Ken06.Bus             .......g..............t........a..a............................................. [ 80960]
KM111294.1_Ken05/Tk1             .......g.................t.......................c.............................. [ 80960]
KM262844.1_strain_L60            .......g........................................................................ [ 80960]
KM102979.1_26544/OG10_Italy      .......g........................................................................ [ 80960]
KM262845.1_strain_NHV            .......g........................................................................ [ 80960]
KX354450.1_47/Ss/2008            .......g........................................................................ [ 80960]
KP055815.1_strain_BA71           .......g........................................................................ [ 80960]
MH025920.1_strain_R35            .......g..............t........a..a............................................. [ 80960]
MH025919.1_strain_N10            .......g..............t........a..a............................................. [ 80960]
MH025918.1_strain_R25            .......g..............t........a..a............................................. [ 80960]
MH025917.1_strain_R7             .......g..............t........a..a............................................. [ 80960]
MH025916.1_strain_R8             .......g..............t........a..a............................................. [ 80960]
```

```
FR682468.2_Georgia_2007/1        taaaactggaagaagcggttcatgtcgacacgatatggccagctaaagcaataccagccccccgtttgccggccttggtt [ 81040]
ASF_IND_20_CAD543                .................................................................y....kr..y..r.. [ 81040]
LR899193.1_Germany_2020/1        ................................................................................ [ 81040]
MN393477.1_Wuhan_2019            ................................................................................ [ 81040]
MW396979.1_Timor-Leste/2019/1    ................................................................................ [ 81040]
MK940252.1_CN/2019/InnerMongolia ................................................................................ [ 81040]
MK128995.1_China/2018/AnhuiXCGQ  ................................................................................ [ 81040]
MH681419.1_ASFV/POL/2015         ................................................................................ [ 81040]
LS478113.1_Estonia_2014          ................................................................................ [ 81040]
KP843857.1_Odintsovo_02/14       ................................................................................ [ 81040]
FN557520.1_strain_E75            ..............a...........t.................g.....g.....a........c....ta..c..a.. [ 81040]
AY261366.1_Warthog               .......................a..t................................t.................... [ 81040]
AY261365.1_Warmbaths             ..............a..a........t.............................a....................... [ 81040]
AY261364.1_Tengani_62            ..........................t..................................................... [ 81040]
AY261363.1_Pretoriuskop/96/4     .......................a..t................................t.................... [ 81040]
AY261362.1_Mkuzi_1979            .......................a..t.........................................t........... [ 81040]
AY261361.1_Malawi_Lil-20/1_1983  ..........................t..............a..............a.....a................. [ 81040]
AY261360.1_Kenya_1950            .......................a..t..............a....................a.....t..a........ [ 81040]
AM712240.1_OURT_88/3             ..............a...........t.................g.....g.....a........c....ta..c..a.. [ 81040]
AM712239.1_Benin_97/1            ..............a...........t.................g.....g.....a........c....ta..c..a.. [ 81040]
KM111295.1_Ken06.Bus             .......................a..t..............a....................g.....t........... [ 81040]
KM111294.1_Ken05/Tk1             .......................a..t..............a.................t..g.....t........... [ 81040]
KM262844.1_strain_L60            ..............a...........t.................g.....g.....a........c....ta..c..a.. [ 81040]
KM102979.1_26544/OG10_Italy      ..............a...........t.................g.....g.....a........c....ta..c..a.. [ 81040]
KM262845.1_strain_NHV            ..............a...........t.................g.....g.....a........c....ta..c..a.. [ 81040]
KX354450.1_47/Ss/2008            ..............a...........t.................g.....g.....a........c....ta..c..a.. [ 81040]
KP055815.1_strain_BA71           ..............a...........t.................g.....g.....a........c....ta..c..a.. [ 81040]
MH025920.1_strain_R35            .......................a..t..............a....................g.....t........... [ 81040]
MH025919.1_strain_N10            .......................a..t..............a....................g.....t........... [ 81040]
MH025918.1_strain_R25            .......................a..t..............a....................g.....t........... [ 81040]
MH025917.1_strain_R7             .......................a..t..............a....................g.....t........... [ 81040]
MH025916.1_strain_R8             .......................a..t..............a....................g.....t........... [ 81040]

FR682468.2_Georgia_2007/1        tgtttcataggtaacacgcgcaggttgggtacagtttgcgtagggggacactagggcggcaaggcccaaaatagcttggg [ 81120]
ASF_IND_20_CAD543                ................................................................................ [ 81120]
LR899193.1_Germany_2020/1        ................................................................................ [ 81120]
MN393477.1_Wuhan_2019            ................................................................................ [ 81120]
MW396979.1_Timor-Leste/2019/1    ................................................................................ [ 81120]
MK940252.1_CN/2019/InnerMongolia ................................................................................ [ 81120]
MK128995.1_China/2018/AnhuiXCGQ  ................................................................................ [ 81120]
MH681419.1_ASFV/POL/2015         ................................................................................ [ 81120]
LS478113.1_Estonia_2014          ................................................................................ [ 81120]
KP843857.1_Odintsovo_02/14       ................................................................................ [ 81120]
```

```
FN557520.1_strain_E75             .........a........a............................................................. [ 81120]
AY261366.1_Warthog                ...............t........................................................g....... [ 81120]
AY261365.1_Warmbaths              ...............t...............................................a........g....... [ 81120]
AY261364.1_Tengani_62             ...............t.................a.............................................. [ 81120]
AY261363.1_Pretoriuskop/96/4      ...............t................................................................ [ 81120]
AY261362.1_Mkuzi_1979             ..................a........a..g................................................. [ 81120]
AY261361.1_Malawi_Lil-20/1_1983   ...............t.............................a.....g....................g.....a. [ 81120]
AY261360.1_Kenya_1950             .........a........t.......................a..a.....g....................g.a...a. [ 81120]
AM712240.1_OURT_88/3              .........a........a............................................................. [ 81120]
AM712239.1_Benin_97/1             .........a........a............................................................. [ 81120]
KM111295.1_Ken06.Bus              ..............................g..............a..g..g....................g.....a. [ 81120]
KM111294.1_Ken05/Tk1              ..............................g..............a.....g....................g.a...a. [ 81120]
KM262844.1_strain_L60             .........a........a............................................................. [ 81120]
KM102979.1_26544/OG10_Italy       .........a........a............................................................. [ 81120]
KM262845.1_strain_NHV             .........a........a............................................................. [ 81120]
KX354450.1_47/Ss/2008             .........a........a............................................................. [ 81120]
KP055815.1_strain_BA71            .........a........a............................................................. [ 81120]
MH025920.1_strain_R35             ..............................g..............a..g..g....................g.....a. [ 81120]
MH025919.1_strain_N10             ..............................g..............a..g..g....................g.....a. [ 81120]
MH025918.1_strain_R25             ..............................g..............a..g..g....................g.....a. [ 81120]
MH025917.1_strain_R7              ..............................g..............a..g..g....................g.....a. [ 81120]
MH025916.1_strain_R8              ..............................g..............a..g..g....................g.....a. [ 81120]

FR682468.2_Georgia_2007/1         gcacgtccacgtgtgtgaaacgacgcgttacatcatgtttatgtttgcgtagctcgatgatggagaaggcaacaagacag [ 81200]
ASF_IND_20_CAD543                 ................................................................................ [ 81200]
LR899193.1_Germany_2020/1         ................................................................................ [ 81200]
MN393477.1_Wuhan_2019             ................................................................................ [ 81200]
MW396979.1_Timor-Leste/2019/1     ................................................................................ [ 81200]
MK940252.1_CN/2019/InnerMongolia  ................................................................................ [ 81200]
MK128995.1_China/2018/AnhuiXCGQ   ................................................................................ [ 81200]
MH681419.1_ASFV/POL/2015          ................................................................................ [ 81200]
LS478113.1_Estonia_2014           ................................................................................ [ 81200]
KP843857.1_Odintsovo_02/14        ................................................................................ [ 81200]
FN557520.1_strain_E75             .........................t..............g..c.................a.................. [ 81200]
AY261366.1_Warthog                ........................................g..c.......................a..g......... [ 81200]
AY261365.1_Warmbaths              ........................................g.ac.....c......g...................g... [ 81200]
AY261364.1_Tengani_62             .........................t..............g..c...................................a [ 81200]
AY261363.1_Pretoriuskop/96/4      ........................................g..c.................................... [ 81200]
AY261362.1_Mkuzi_1979             .........................t..............g..c.................................... [ 81200]
AY261361.1_Malawi_Lil-20/1_1983   .t.....t.....c.....g.....t........g.....g.....a..c..t..ag.a..c.....a..g.....g... [ 81200]
AY261360.1_Kenya_1950             .......t...........g..g...........g.....g.....a..c..t..ag.a........a..g........a [ 81200]
AM712240.1_OURT_88/3              .........................t..............g..c.................a.................. [ 81200]
AM712239.1_Benin_97/1             .........................t..............g..c.................a.................. [ 81200]
KM111295.1_Ken06.Bus              .t.....t.....c.....g........c.....g.....g..c.....c..t..ag.a...........g........a [ 81200]
```

```
KM111294.1_Ken05/Tk1           .......t.....c.....g........c.....g.....g..c.....c..t..ag.a...........g........a [ 81200]
KM262844.1_strain_L60          .........................t..............g..c.................a.................. [ 81200]
KM102979.1_26544/OG10_Italy    .........................t..............g..c.................a.................. [ 81200]
KM262845.1_strain_NHV          .........................t..............g..c.................a.................. [ 81200]
KX354450.1_47/Ss/2008          .........................t..............g..c.................a.................. [ 81200]
KP055815.1_strain_BA71         .........................t..............g..c.................a.................. [ 81200]
MH025920.1_strain_R35          .t.....t.....c.....g........c.....g.....g..c.....c..t..ag.a...........g........a [ 81200]
MH025919.1_strain_N10          .t.....t.....c.....g........c.....g.....g..c.....c..t..ag.a...........g........a [ 81200]
MH025918.1_strain_R25          .t.....t.....c.....g........c.....g.....g..c.....c..t..ag.a...........g........a [ 81200]
MH025917.1_strain_R7           .t.....t.....c.....g........c.....g.....g..c.....c..t..ag.a...........g........a [ 81200]
MH025916.1_strain_R8           .t.....t.....c.....g........c.....g.....g..c.....c..t..ag.a...........g........a [ 81200]

FR682468.2_Georgia_2007/1      ttttccgcctcctcgggggtaatgaactcacagatgccctgtgctacgagatcttcaagtgtaagcgttccggctaaaat [ 81280]
ASF_IND_20_CAD543              ................................................................................ [ 81280]
LR899193.1_Germany_2020/1      ................................................................................ [ 81280]
MN393477.1_Wuhan_2019          ................................................................................ [ 81280]
MW396979.1_Timor-Leste/2019/1  ................................................................................ [ 81280]
MK940252.1_CN/2019/InnerMongolia ................................................................................ [ 81280]
MK128995.1_China/2018/AnhuiXCGQ ................................................................................ [ 81280]
MH681419.1_ASFV/POL/2015       ................................................................................ [ 81280]
LS478113.1_Estonia_2014        ................................................................................ [ 81280]
KP843857.1_Odintsovo_02/14     ................................................................................ [ 81280]
FN557520.1_strain_E75          ........t..t..............t..............c.......................t.............. [ 81280]
AY261366.1_Warthog             .........................................c...................................... [ 81280]
AY261365.1_Warmbaths           .....t..t.....a..a.......................c...................................... [ 81280]
AY261364.1_Tengani_62          ........t..t.....a.......................c.................a.................... [ 81280]
AY261363.1_Pretoriuskop/96/4   .........................................c...................................... [ 81280]
AY261362.1_Mkuzi_1979          ........t..t..............t..............c...................................... [ 81280]
AY261361.1_Malawi_Lil-20/1_1983 ................................a........c...................................... [ 81280]
AY261360.1_Kenya_1950          .....t........a..........................c...................................... [ 81280]
AM712240.1_OURT_88/3           ........t..t..............t..............c.......................t.............. [ 81280]
AM712239.1_Benin_97/1          ........t..t..............t..............c.......................t.............. [ 81280]
KM111295.1_Ken06.Bus           .....t........a..........................c...................................... [ 81280]
KM111294.1_Ken05/Tk1           .....t........a..........................c...................................... [ 81280]
KM262844.1_strain_L60          ........t..t..............t..............c.......................t.............. [ 81280]
KM102979.1_26544/OG10_Italy    ........t..t..............t..............c.......................t.............. [ 81280]
KM262845.1_strain_NHV          ........t..t..............t..............c.......................t.............. [ 81280]
KX354450.1_47/Ss/2008          ........t..t..............t..............c.......................t.............. [ 81280]
KP055815.1_strain_BA71         ........t..t..............t..............c.......................t.............. [ 81280]
MH025920.1_strain_R35          .....t........a..........................c...................................... [ 81280]
MH025919.1_strain_N10          .....t........a..........................c...................................... [ 81280]
MH025918.1_strain_R25          .....t........a..........................c...................................... [ 81280]
MH025917.1_strain_R7           .....t........a..........................c...................................... [ 81280]
MH025916.1_strain_R8           .....t........a..........................c...................................... [ 81280]
```

```
FR682468.2_Georgia_2007/1         gtcttttgccatttgaggcgtaaatcgcgtattttgaatgaaagggattttatgtttttcccagtctttatcgccttttt [ 81360]
ASF_IND_20_CAD543                 ................................................................................ [ 81360]
LR899193.1_Germany_2020/1         ................................................................................ [ 81360]
MN393477.1_Wuhan_2019             ................................................................................ [ 81360]
MW396979.1_Timor-Leste/2019/1     ................................................................................ [ 81360]
MK940252.1_CN/2019/InnerMongolia  ................................................................................ [ 81360]
MK128995.1_China/2018/AnhuiXCGQ   ................................................................................ [ 81360]
MH681419.1_ASFV/POL/2015          ................................................................................ [ 81360]
LS478113.1_Estonia_2014           ................................................................................ [ 81360]
KP843857.1_Odintsovo_02/14        ................................................................................ [ 81360]
FN557520.1_strain_E75             .................a.............................................................. [ 81360]
AY261366.1_Warthog                .................a....................c......................................... [ 81360]
AY261365.1_Warmbaths              a................a.............................................................. [ 81360]
AY261364.1_Tengani_62             .................a....................ca........................................ [ 81360]
AY261363.1_Pretoriuskop/96/4      .................a....................c......................................... [ 81360]
AY261362.1_Mkuzi_1979             .................a.............................................................. [ 81360]
AY261361.1_Malawi_Lil-20/1_1983   a................a...........................t.................................. [ 81360]
AY261360.1_Kenya_1950             ag...............a....................c.....at.........................t........ [ 81360]
AM712240.1_OURT_88/3              .................a.............................................................. [ 81360]
AM712239.1_Benin_97/1             .................a.............................................................. [ 81360]
KM111295.1_Ken06.Bus              ag...............a....................c.....at.................................. [ 81360]
KM111294.1_Ken05/Tk1              ag...............a....................c.....at.........................t........ [ 81360]
KM262844.1_strain_L60             .................a.............................................................. [ 81360]
KM102979.1_26544/OG10_Italy       .................a.............................................................. [ 81360]
KM262845.1_strain_NHV             .................a.............................................................. [ 81360]
KX354450.1_47/Ss/2008             .................a.............................................................. [ 81360]
KP055815.1_strain_BA71            .................a.............................................................. [ 81360]
MH025920.1_strain_R35             ag...............a....................c.....at.................................. [ 81360]
MH025919.1_strain_N10             ag...............a....................c.....at.................................. [ 81360]
MH025918.1_strain_R25             ag...............a....................c.....at.................................. [ 81360]
MH025917.1_strain_R7              ag...............a....................c.....at.................................. [ 81360]
MH025916.1_strain_R8              ag...............a....................c.....at.................................. [ 81360]

FR682468.2_Georgia_2007/1         ttctggcctctgcggccttgtagcaggcttgattgtatttttcaatattattatctacaatgagtagggggcgggtcagc [ 81440]
ASF_IND_20_CAD543                 ................................................................................ [ 81440]
LR899193.1_Germany_2020/1         ................................................................................ [ 81440]
MN393477.1_Wuhan_2019             ................................................................................ [ 81440]
MW396979.1_Timor-Leste/2019/1     ................................................................................ [ 81440]
MK940252.1_CN/2019/InnerMongolia  ................................................................................ [ 81440]
MK128995.1_China/2018/AnhuiXCGQ   ................................................................................ [ 81440]
MH681419.1_ASFV/POL/2015          ................................................................................ [ 81440]
LS478113.1_Estonia_2014           ................................................................................ [ 81440]
KP843857.1_Odintsovo_02/14        ................................................................................ [ 81440]
```

```
FN557520.1_strain_E75          ......................a.........................................c............... [ 81440]
AY261366.1_Warthog             .......t........................................................c..a............ [ 81440]
AY261365.1_Warmbaths           ................................................................c............... [ 81440]
AY261364.1_Tengani_62          .......t..............a.........................................c............... [ 81440]
AY261363.1_Pretoriuskop/96/4   .......t.....a........a.........................................c..............t [ 81440]
AY261362.1_Mkuzi_1979          .......t........................................................c............... [ 81440]
AY261361.1_Malawi_Lil-20/1_1983 .............a..............c.t...a...................................c......... [ 81440]
AY261360.1_Kenya_1950          .............a...........a..c.c...a............g................c..a............ [ 81440]
AM712240.1_OURT_88/3           ......................a.........................................c............... [ 81440]
AM712239.1_Benin_97/1          ......................a.........................................c............... [ 81440]
KM111295.1_Ken06.Bus           .............a...........a..c.c...a............g................c..a............ [ 81440]
KM111294.1_Ken05/Tk1           .............a...........a..c.c...a.............................c..a........t... [ 81440]
KM262844.1_strain_L60          ......................a.........................................c............... [ 81440]
KM102979.1_26544/OG10_Italy    ......................a.........................................c............... [ 81440]
KM262845.1_strain_NHV          ......................a.........................................c............... [ 81440]
KX354450.1_47/Ss/2008          ......................a.........................................c............... [ 81440]
KP055815.1_strain_BA71         ......................a.........................................c............... [ 81440]
MH025920.1_strain_R35          .............a...........a..c.c...a............g................c..a............ [ 81440]
MH025919.1_strain_N10          .............a...........a..c.c...a............g................c..a............ [ 81440]
MH025918.1_strain_R25          .............a...........a..c.c...a............g................c..a............ [ 81440]
MH025917.1_strain_R7           .............a...........a..c.c...a............g................c..a............ [ 81440]
MH025916.1_strain_R8           .............a...........a..c.c...a............g................c..a............ [ 81440]

FR682468.2_Georgia_2007/1      ctaccgacgtccaaccaaaattctacttcgtctaccatgctatcccagtagatggtggtatggggatgcacaaccttgcc [ 81520]
ASF_IND_20_CAD543              ................................................................................ [ 81520]
LR899193.1_Germany_2020/1      ................................................................................ [ 81520]
MN393477.1_Wuhan_2019          ................................................................................ [ 81520]
MW396979.1_Timor-Leste/2019/1  ................................................................................ [ 81520]
MK940252.1_CN/2019/InnerMongolia ................................................................................ [ 81520]
MK128995.1_China/2018/AnhuiXCGQ ................................................................................ [ 81520]
MH681419.1_ASFV/POL/2015       ................................................................................ [ 81520]
LS478113.1_Estonia_2014        ................................................................................ [ 81520]
KP843857.1_Odintsovo_02/14     ................................................................................ [ 81520]
FN557520.1_strain_E75          .................g.......................................................t...... [ 81520]
AY261366.1_Warthog             ..g......................................................................t...... [ 81520]
AY261365.1_Warmbaths           ..g..............g.............................................................. [ 81520]
AY261364.1_Tengani_62          ..g..a...........g...............................................g.......t...... [ 81520]
AY261363.1_Pretoriuskop/96/4   ..g.....a................................................................t...... [ 81520]
AY261362.1_Mkuzi_1979          ..g..a...........g.......................................................t...... [ 81520]
AY261361.1_Malawi_Lil-20/1_1983 ..g..............g...........a.......................a.................g.t...... [ 81520]
AY261360.1_Kenya_1950          ..g..............g...........a......................................t..g.t...... [ 81520]
AM712240.1_OURT_88/3           .................g.......................................................t...... [ 81520]
AM712239.1_Benin_97/1          .................g.......................................................t...... [ 81520]
KM111295.1_Ken06.Bus           ..g..............g...........a......................................t..g.t...... [ 81520]
```

```
KM111294.1_Ken05/Tk1             ..g..............g...........a......................................t..g.t...... [ 81520]
KM262844.1_strain_L60            .................g.......................................................t...... [ 81520]
KM102979.1_26544/OG10_Italy      .................g.......................................................t...... [ 81520]
KM262845.1_strain_NHV            .................g.......................................................t...... [ 81520]
KX354450.1_47/Ss/2008            .................g.......................................................t...... [ 81520]
KP055815.1_strain_BA71           .................g.......................................................t...... [ 81520]
MH025920.1_strain_R35            ..g..............g...........a......................................t..g.t...... [ 81520]
MH025919.1_strain_N10            ..g..............g...........a......................................t..g.t...... [ 81520]
MH025918.1_strain_R25            ..g..............g...........a......................................t..g.t...... [ 81520]
MH025917.1_strain_R7             ..g..............g...........a......................................t..g.t...... [ 81520]
MH025916.1_strain_R8             ..g..............g...........a......................................t..g.t...... [ 81520]

FR682468.2_Georgia_2007/1        ctcacggcgaagcattctataccgctgagcaagctcaaaggcattggtgcagcagccgatccattctccgttgataaata [ 81600]
ASF_IND_20_CAD543                ................................................................................ [ 81600]
LR899193.1_Germany_2020/1        ................................................................................ [ 81600]
MN393477.1_Wuhan_2019            ................................................................................ [ 81600]
MW396979.1_Timor-Leste/2019/1    ................................................................................ [ 81600]
MK940252.1_CN/2019/InnerMongolia ................................................................................ [ 81600]
MK128995.1_China/2018/AnhuiXCGQ  ................................................................................ [ 81600]
MH681419.1_ASFV/POL/2015         ................................................................................ [ 81600]
LS478113.1_Estonia_2014          ................................................................................ [ 81600]
KP843857.1_Odintsovo_02/14       ................................................................................ [ 81600]
FN557520.1_strain_E75            ...........a..............................g..............a...................... [ 81600]
AY261366.1_Warthog               ................................................................................ [ 81600]
AY261365.1_Warmbaths             ................................................................................ [ 81600]
AY261364.1_Tengani_62            ..........................................g..................................... [ 81600]
AY261363.1_Pretoriuskop/96/4     ................................................................................ [ 81600]
AY261362.1_Mkuzi_1979            ..........................................g..............a...................... [ 81600]
AY261361.1_Malawi_Lil-20/1_1983  ..........................................g........a..t..a.....c................ [ 81600]
AY261360.1_Kenya_1950            ..........................................g....................c................ [ 81600]
AM712240.1_OURT_88/3             ...........a..............................g..............a...................... [ 81600]
AM712239.1_Benin_97/1            ...........a..............................g..............a...................... [ 81600]
KM111295.1_Ken06.Bus             t..........a..............................g.....a..............c................ [ 81600]
KM111294.1_Ken05/Tk1             .......................................a.......................c..............c. [ 81600]
KM262844.1_strain_L60            ...........a..............................g..............a...................... [ 81600]
KM102979.1_26544/OG10_Italy      ...........a..............................g..............a...................... [ 81600]
KM262845.1_strain_NHV            ...........a..............................g..............a...................... [ 81600]
KX354450.1_47/Ss/2008            ...........a..............................g..............a...................... [ 81600]
KP055815.1_strain_BA71           ...........a..............................g..............a...................... [ 81600]
MH025920.1_strain_R35            t..........a..............................g.....a..............c................ [ 81600]
MH025919.1_strain_N10            t..........a..............................g.....a..............c................ [ 81600]
MH025918.1_strain_R25            t..........a..............................g.....a..............c................ [ 81600]
MH025917.1_strain_R7             t..........a..............................g.....a..............c................ [ 81600]
MH025916.1_strain_R8             t..........a..............................g.....a..............c................ [ 81600]
```

```
FR682468.2_Georgia_2007/1        cgcgcgctaggccctttcgtacaatgtccttgttggaaacatcggctaactgttgaatggccggatctgatagaaggcgt [ 81680]
ASF_IND_20_CAD543                ................................................................................ [ 81680]
LR899193.1_Germany_2020/1        ................................................................................ [ 81680]
MN393477.1_Wuhan_2019            ................................................................................ [ 81680]
MW396979.1_Timor-Leste/2019/1    ................................................................................ [ 81680]
MK940252.1_CN/2019/InnerMongolia ................................................................................ [ 81680]
MK128995.1_China/2018/AnhuiXCGQ  ................................................................................ [ 81680]
MH681419.1_ASFV/POL/2015         ................................................................................ [ 81680]
LS478113.1_Estonia_2014          ................................................................................ [ 81680]
KP843857.1_Odintsovo_02/14       ................................................................................ [ 81680]
FN557520.1_strain_E75            ................................................................................ [ 81680]
AY261366.1_Warthog               .a.............................................................................. [ 81680]
AY261365.1_Warmbaths             ................................................................................ [ 81680]
AY261364.1_Tengani_62            ................................................................................ [ 81680]
AY261363.1_Pretoriuskop/96/4     .a.............................................................................. [ 81680]
AY261362.1_Mkuzi_1979            ....t........................................................................... [ 81680]
AY261361.1_Malawi_Lil-20/1_1983  .a........a.......................a............................................. [ 81680]
AY261360.1_Kenya_1950            ....t.....a.......................a..........................a.................. [ 81680]
AM712240.1_OURT_88/3             ................................................................................ [ 81680]
AM712239.1_Benin_97/1            ................................................................................ [ 81680]
KM111295.1_Ken06.Bus             ....t.....a.......................a......................................g...... [ 81680]
KM111294.1_Ken05/Tk1             ....t.....a.......................a............................................. [ 81680]
KM262844.1_strain_L60            ................................................................................ [ 81680]
KM102979.1_26544/OG10_Italy      ................................................................................ [ 81680]
KM262845.1_strain_NHV            ................................................................................ [ 81680]
KX354450.1_47/Ss/2008            ................................................................................ [ 81680]
KP055815.1_strain_BA71           ................................................................................ [ 81680]
MH025920.1_strain_R35            a...t.....a.......................a......................................g...... [ 81680]
MH025919.1_strain_N10            a...t.....a.......................a......................................g...... [ 81680]
MH025918.1_strain_R25            a...t.....a.......................a......................................g...... [ 81680]
MH025917.1_strain_R7             a...t.....a.......................a......................................g...... [ 81680]
MH025916.1_strain_R8             a...t.....a.......................a......................................g...... [ 81680]

FR682468.2_Georgia_2007/1        tgttttaacgaaagtacttctccggcggtgcagacattggcagtgatggctaactgtttagacatgcctactttttcacc [ 81760]
ASF_IND_20_CAD543                ................................................................................ [ 81760]
LR899193.1_Germany_2020/1        ................................................................................ [ 81760]
MN393477.1_Wuhan_2019            ................................................................................ [ 81760]
MW396979.1_Timor-Leste/2019/1    ................................................................................ [ 81760]
MK940252.1_CN/2019/InnerMongolia ................................................................................ [ 81760]
MK128995.1_China/2018/AnhuiXCGQ  ................................................................................ [ 81760]
MH681419.1_ASFV/POL/2015         ............................................a................................... [ 81760]
LS478113.1_Estonia_2014          ................................................................................ [ 81760]
KP843857.1_Odintsovo_02/14       ................................................................................ [ 81760]
```

```
FN557520.1_strain_E75          .......................a.....a...........g..a....................a.............. [ 81760]
AY261366.1_Warthog             ................................................................................ [ 81760]
AY261365.1_Warmbaths           ................................................................................ [ 81760]
AY261364.1_Tengani_62          .........................................g..a................................... [ 81760]
AY261363.1_Pretoriuskop/96/4   ................................................................................ [ 81760]
AY261362.1_Mkuzi_1979          .......................a.....a...........g..a..............g.................... [ 81760]
AY261361.1_Malawi_Lil-20/1_1983 .................c..c.....a..............g...........t.......................... [ 81760]
AY261360.1_Kenya_1950          .................c..c.....a..............g..a..............g.................g.. [ 81760]
AM712240.1_OURT_88/3           .......................a.....a...........g..a....................a.............. [ 81760]
AM712239.1_Benin_97/1          .......................a.....a...........g..a....................a.............. [ 81760]
KM111295.1_Ken06.Bus           ....................c.....a..............g...........t.....g.................g.. [ 81760]
KM111294.1_Ken05/Tk1           .................c..c.....a........g.....g...........t.....g.................g.. [ 81760]
KM262844.1_strain_L60          .......................a.....a...........g..a....................a.............. [ 81760]
KM102979.1_26544/OG10_Italy    .......................a.....a...........g..a....................a.............. [ 81760]
KM262845.1_strain_NHV          .......................a.....a...........g..a....................a.............. [ 81760]
KX354450.1_47/Ss/2008          .......................a.....a...........g..a....................a.............. [ 81760]
KP055815.1_strain_BA71         .......................a.....a...........g..a....................a.............. [ 81760]
MH025920.1_strain_R35          .............a......c.....a..............g...........t.....g.................g.. [ 81760]
MH025919.1_strain_N10          .............a......c.....a..............g...........t.....g.................g.. [ 81760]
MH025918.1_strain_R25          .............a......c.....a..............g...........t.....g.................g.. [ 81760]
MH025917.1_strain_R7           .............a......c.....a..............g...........t.....g.................g.. [ 81760]
MH025916.1_strain_R8           .............a......c.....a..............g...........t.....g.................g.. [ 81760]

FR682468.2_Georgia_2007/1      agtatcggctgactgggctacgcagatgtatccaggataggatgcgtgcacgcgacgcatcatgtcagccctttctgttt [ 81840]
ASF_IND_20_CAD543              ................................................................................ [ 81840]
LR899193.1_Germany_2020/1      ................................................................................ [ 81840]
MN393477.1_Wuhan_2019          ................................................................................ [ 81840]
MW396979.1_Timor-Leste/2019/1  ................................................................................ [ 81840]
MK940252.1_CN/2019/InnerMongolia ................................................................................ [ 81840]
MK128995.1_China/2018/AnhuiXCGQ ................................................................................ [ 81840]
MH681419.1_ASFV/POL/2015       ................................................................................ [ 81840]
LS478113.1_Estonia_2014        ................................................................................ [ 81840]
KP843857.1_Odintsovo_02/14     ................................................................................ [ 81840]
FN557520.1_strain_E75          .......................................a........................................ [ 81840]
AY261366.1_Warthog             .........g.................a.................................................... [ 81840]
AY261365.1_Warmbaths           ................................................................................ [ 81840]
AY261364.1_Tengani_62          ...........................a...................................a................ [ 81840]
AY261363.1_Pretoriuskop/96/4   .........g.................a.................................................... [ 81840]
AY261362.1_Mkuzi_1979          ......a................................a........................................ [ 81840]
AY261361.1_Malawi_Lil-20/1_1983 .................................t..g........a.......................t.....c.... [ 81840]
AY261360.1_Kenya_1950          .................................t..g........a.......................t.....c.... [ 81840]
AM712240.1_OURT_88/3           .......................................a........................................ [ 81840]
AM712239.1_Benin_97/1          .......................................a........................................ [ 81840]
KM111295.1_Ken06.Bus           .................................t..g........a.......................t.....c.... [ 81840]
```

```
KM111294.1_Ken05/Tk1           .................................t..g........a.......................t.....c.... [ 81840]
KM262844.1_strain_L60          .......................................a........................................ [ 81840]
KM102979.1_26544/OG10_Italy    .......................................a........................................ [ 81840]
KM262845.1_strain_NHV          .......................................a........................................ [ 81840]
KX354450.1_47/Ss/2008          .......................................a........................................ [ 81840]
KP055815.1_strain_BA71         .......................................a........................................ [ 81840]
MH025920.1_strain_R35          .................................t..g........a.......................t.....c.... [ 81840]
MH025919.1_strain_N10          .................................t..g........a.......................t.....c.... [ 81840]
MH025918.1_strain_R25          .................................t..g........a.......................t.....c.... [ 81840]
MH025917.1_strain_R7           .................................t..g........a.......................t.....c.... [ 81840]
MH025916.1_strain_R8           .................................t..g........a.......................t.....c.... [ 81840]

FR682468.2_Georgia_2007/1      gtttggatgcgttggtggtgttatgagtatttaccgtacgcaatgctgaaatggtatttaataaattttttctttccaaa [ 81920]
ASF_IND_20_CAD543              ................................................................................ [ 81920]
LR899193.1_Germany_2020/1      ................................................................................ [ 81920]
MN393477.1_Wuhan_2019          ................................................................................ [ 81920]
MW396979.1_Timor-Leste/2019/1  ................................................................................ [ 81920]
MK940252.1_CN/2019/InnerMongolia ................................................................................ [ 81920]
MK128995.1_China/2018/AnhuiXCGQ ................................................................................ [ 81920]
MH681419.1_ASFV/POL/2015       ................................................................................ [ 81920]
LS478113.1_Estonia_2014        ................................................................................ [ 81920]
KP843857.1_Odintsovo_02/14     ................................................................................ [ 81920]
FN557520.1_strain_E75          ...........................................c........c........................... [ 81920]
AY261366.1_Warthog             ................................................................................ [ 81920]
AY261365.1_Warmbaths           ................................................................................ [ 81920]
AY261364.1_Tengani_62          ...........................................c........c..g........................ [ 81920]
AY261363.1_Pretoriuskop/96/4   ................................................................................ [ 81920]
AY261362.1_Mkuzi_1979          ...........................................c........c........................... [ 81920]
AY261361.1_Malawi_Lil-20/1_1983 ....................c....g.....g...........c.................................... [ 81920]
AY261360.1_Kenya_1950          ....................c....g.................c........a........c.................g [ 81920]
AM712240.1_OURT_88/3           ...........................................c........c........................... [ 81920]
AM712239.1_Benin_97/1          ...........................................c........c........................... [ 81920]
KM111295.1_Ken06.Bus           ....................c....g.................c........a........c.................g [ 81920]
KM111294.1_Ken05/Tk1           .............a......c...................t..c........a........c.................g [ 81920]
KM262844.1_strain_L60          ...........................................c........c........................... [ 81920]
KM102979.1_26544/OG10_Italy    ...........................................c........c........................... [ 81920]
KM262845.1_strain_NHV          ...........................................c........c........................... [ 81920]
KX354450.1_47/Ss/2008          ...........................................c........c........................... [ 81920]
KP055815.1_strain_BA71         ...........................................c........c........................... [ 81920]
MH025920.1_strain_R35          ....................c....g.................c........a........c.................g [ 81920]
MH025919.1_strain_N10          ....................c....g.................c........a........c.................g [ 81920]
MH025918.1_strain_R25          ....................c....g.................c........a........c.................g [ 81920]
MH025917.1_strain_R7           ....................c....g.................c........a........c.................g [ 81920]
MH025916.1_strain_R8           ....................c....g.................c........a........c.................g [ 81920]
```

```
FR682468.2_Georgia_2007/1        ctttgagtagatactctgtttacaatggggcgctgtcgcaccatgatggttttatttcctgaaatgatagactgttccat [ 82000]
ASF_IND_20_CAD543                ................................................................................ [ 82000]
LR899193.1_Germany_2020/1        ................................................................................ [ 82000]
MN393477.1_Wuhan_2019            ................................................................................ [ 82000]
MW396979.1_Timor-Leste/2019/1    ................................................................................ [ 82000]
MK940252.1_CN/2019/InnerMongolia ................................................................................ [ 82000]
MK128995.1_China/2018/AnhuiXCGQ  ................................................................................ [ 82000]
MH681419.1_ASFV/POL/2015         ................................................................................ [ 82000]
LS478113.1_Estonia_2014          ................................................................................ [ 82000]
KP843857.1_Odintsovo_02/14       ................................................................................ [ 82000]
FN557520.1_strain_E75            ................................................................................ [ 82000]
AY261366.1_Warthog               .......................................................................t........ [ 82000]
AY261365.1_Warmbaths             ................................................................................ [ 82000]
AY261364.1_Tengani_62            ...............................................a.......................t........ [ 82000]
AY261363.1_Pretoriuskop/96/4     .......................................................................t........ [ 82000]
AY261362.1_Mkuzi_1979            ................................................................................ [ 82000]
AY261361.1_Malawi_Lil-20/1_1983  .....g....................c........c............................................ [ 82000]
AY261360.1_Kenya_1950            .....g....................c........c..........................g................. [ 82000]
AM712240.1_OURT_88/3             ................................................................................ [ 82000]
AM712239.1_Benin_97/1            ................................................................................ [ 82000]
KM111295.1_Ken06.Bus             .....g....................a........c............................................ [ 82000]
KM111294.1_Ken05/Tk1             .....g....................c........c..........................g................. [ 82000]
KM262844.1_strain_L60            ................................................................................ [ 82000]
KM102979.1_26544/OG10_Italy      ................................................................................ [ 82000]
KM262845.1_strain_NHV            ................................................................................ [ 82000]
KX354450.1_47/Ss/2008            ................................................................................ [ 82000]
KP055815.1_strain_BA71           ................................................................................ [ 82000]
MH025920.1_strain_R35            .....g....................a........c............................................ [ 82000]
MH025919.1_strain_N10            .....g....................a........c............................................ [ 82000]
MH025918.1_strain_R25            .....g....................a........c............................................ [ 82000]
MH025917.1_strain_R7             .....g....................a........c............................................ [ 82000]
MH025916.1_strain_R8             .....g....................a........c............................................ [ 82000]

FR682468.2_Georgia_2007/1        actgcgattaagatcggaggcggtattttttgataaagcggcagaaaatgcctcgataatgtttcgctgagtaagctcct [ 82080]
ASF_IND_20_CAD543                ................................................................................ [ 82080]
LR899193.1_Germany_2020/1        ................................................................................ [ 82080]
MN393477.1_Wuhan_2019            ................................................................................ [ 82080]
MW396979.1_Timor-Leste/2019/1    ................................................................................ [ 82080]
MK940252.1_CN/2019/InnerMongolia ................................................................................ [ 82080]
MK128995.1_China/2018/AnhuiXCGQ  ................................................................................ [ 82080]
MH681419.1_ASFV/POL/2015         ................................................................................ [ 82080]
LS478113.1_Estonia_2014          ................................................................................ [ 82080]
KP843857.1_Odintsovo_02/14       ................................................................................ [ 82080]
```

```
FN557520.1_strain_E75           .........g...........................................................g..t....... [ 82080]
AY261366.1_Warthog              .........g..............g................................g...........g..g....... [ 82080]
AY261365.1_Warmbaths            .........g..............g................................g...........g..g....... [ 82080]
AY261364.1_Tengani_62           .........................................................g...........g..g....... [ 82080]
AY261363.1_Pretoriuskop/96/4    .........g..............g................................g...........g..g....... [ 82080]
AY261362.1_Mkuzi_1979           .........g...........................................................g.......... [ 82080]
AY261361.1_Malawi_Lil-20/1_1983 ......g..g.....a......c.........................c..t..............t..g..t..a.... [ 82080]
AY261360.1_Kenya_1950           ......g..g........a.....g.......................c....................g..g....... [ 82080]
AM712240.1_OURT_88/3            .........g...........................................................g..t....... [ 82080]
AM712239.1_Benin_97/1           .........g...........................................................g..t....... [ 82080]
KM111295.1_Ken06.Bus            ......g..g..............g.......................c....................g..c....... [ 82080]
KM111294.1_Ken05/Tk1            ......g..g..............g.......................c....................g..g....... [ 82080]
KM262844.1_strain_L60           .........g...........................................................g..t....... [ 82080]
KM102979.1_26544/OG10_Italy     .........g...........................................................g..t....... [ 82080]
KM262845.1_strain_NHV           .........g...........................................................g..t....... [ 82080]
KX354450.1_47/Ss/2008           .........g...........................................................g..t....... [ 82080]
KP055815.1_strain_BA71          .........g...........................................................g..t....... [ 82080]
MH025920.1_strain_R35           ......g..g..............g.......................c....................g..c....... [ 82080]
MH025919.1_strain_N10           ......g..g..............g.......................c....................g..c....... [ 82080]
MH025918.1_strain_R25           ......g..g..............g.......................c....................g..c....... [ 82080]
MH025917.1_strain_R7            ......g..g..............g.......................c....................g..c....... [ 82080]
MH025916.1_strain_R8            ......g..g..............g.......................c....................g..c....... [ 82080]

FR682468.2_Georgia_2007/1       caaaggctgtttgtttaagaagttctttgaacccattgatgatgggtgctatcacggaagtattaaaaatagccttaaag [ 82160]
ASF_IND_20_CAD543               ................................................................................ [ 82160]
LR899193.1_Germany_2020/1       ................................................................................ [ 82160]
MN393477.1_Wuhan_2019           ................................................................................ [ 82160]
MW396979.1_Timor-Leste/2019/1   ................................................................................ [ 82160]
MK940252.1_CN/2019/InnerMongolia ................................................................................ [ 82160]
MK128995.1_China/2018/AnhuiXCGQ ................................................................................ [ 82160]
MH681419.1_ASFV/POL/2015        ................................................................................ [ 82160]
LS478113.1_Estonia_2014         ................................................................................ [ 82160]
KP843857.1_Odintsovo_02/14      ................................................................................ [ 82160]
FN557520.1_strain_E75           ....a.....................................................t..................... [ 82160]
AY261366.1_Warthog              .....................................a....................t..g.....g............ [ 82160]
AY261365.1_Warmbaths            ..............................................c...........t..................... [ 82160]
AY261364.1_Tengani_62           ..........................................................t....................a [ 82160]
AY261363.1_Pretoriuskop/96/4    .....................................a....................t..g.....g............ [ 82160]
AY261362.1_Mkuzi_1979           ..........................................................t..................... [ 82160]
AY261361.1_Malawi_Lil-20/1_1983 ........................................a..a........a.....t..g.....g............ [ 82160]
AY261360.1_Kenya_1950           ............................a.......................a.....t..g.....g.....t.....a [ 82160]
AM712240.1_OURT_88/3            ....a.....................................................t..................... [ 82160]
AM712239.1_Benin_97/1           ....a.....................................................t..................... [ 82160]
KM111295.1_Ken06.Bus            ....................................................a.....t..g.....g.....t.....a [ 82160]
```

```
KM111294.1_Ken05/Tk1             ............................a.......................a.....t..g.....g.....t.....a [ 82160]
KM262844.1_strain_L60            ....a.....................................................t..................... [ 82160]
KM102979.1_26544/OG10_Italy      ....a.....................................................t..................... [ 82160]
KM262845.1_strain_NHV            ....a.....................................................t..................... [ 82160]
KX354450.1_47/Ss/2008            ....a.....................................................t..................... [ 82160]
KP055815.1_strain_BA71           ....a.....................................................t..................... [ 82160]
MH025920.1_strain_R35            ....................................................a.....t..g.....g.....t.....a [ 82160]
MH025919.1_strain_N10            ....................................................a.....t..g.....g.....t.....a [ 82160]
MH025918.1_strain_R25            ....................................................a.....t..g.....g.....t.....a [ 82160]
MH025917.1_strain_R7             ....................................................a.....t..g.....g.....t.....a [ 82160]
MH025916.1_strain_R8             ....................................................a.....t..g.....g.....t.....a [ 82160]

FR682468.2_Georgia_2007/1        gccttggcgagtgagacccctgagccgtgcacccgcttggtgcggtagctatcacggtccgtgggtggaaacacattcat [ 82240]
ASF_IND_20_CAD543                ................................................................................ [ 82240]
LR899193.1_Germany_2020/1        ................................................................................ [ 82240]
MN393477.1_Wuhan_2019            ................................................................................ [ 82240]
MW396979.1_Timor-Leste/2019/1    ................................................................................ [ 82240]
MK940252.1_CN/2019/InnerMongolia ................................................................................ [ 82240]
MK128995.1_China/2018/AnhuiXCGQ  ................................................................................ [ 82240]
MH681419.1_ASFV/POL/2015         ................................................................................ [ 82240]
LS478113.1_Estonia_2014          ................................................................................ [ 82240]
KP843857.1_Odintsovo_02/14       ................................................................................ [ 82240]
FN557520.1_strain_E75            .....a..a..c.................................................................... [ 82240]
AY261366.1_Warthog               ...........c.................................................................... [ 82240]
AY261365.1_Warmbaths             ...........c.................................................................... [ 82240]
AY261364.1_Tengani_62            ...........c.................t..t...........a..a................................ [ 82240]
AY261363.1_Pretoriuskop/96/4     ...........c.................................................................... [ 82240]
AY261362.1_Mkuzi_1979            .....a..a..c..t.................t............................................... [ 82240]
AY261361.1_Malawi_Lil-20/1_1983  ........a..c.................t.............................t.................... [ 82240]
AY261360.1_Kenya_1950            ..t.....a..c..a........a.............................g.....t.....c.....t..g..... [ 82240]
AM712240.1_OURT_88/3             .....a..a..c.................................................................... [ 82240]
AM712239.1_Benin_97/1            .....a..a..c.................................................................... [ 82240]
KM111295.1_Ken06.Bus             ..t.....a..c..a......................................g...........c.....t..g..... [ 82240]
KM111294.1_Ken05/Tk1             ..t.....a..c..a......................................g.....t.....c.....t..g..... [ 82240]
KM262844.1_strain_L60            .....a..a..c.................................................................... [ 82240]
KM102979.1_26544/OG10_Italy      .....a..a..c.................................................................... [ 82240]
KM262845.1_strain_NHV            .....a..a..c.................................................................... [ 82240]
KX354450.1_47/Ss/2008            .....a..a..c.................................................................... [ 82240]
KP055815.1_strain_BA71           .....a..a..c.................................................................... [ 82240]
MH025920.1_strain_R35            ..t.....a..c..a......................................g...........c.....t..g..... [ 82240]
MH025919.1_strain_N10            ..t.....a..c..a......................................g...........c.....t..g..... [ 82240]
MH025918.1_strain_R25            ..t.....a..c..a......................................g...........c.....t..g..... [ 82240]
MH025917.1_strain_R7             ..t.....a..c..a......................................g...........c.....t..g..... [ 82240]
MH025916.1_strain_R8             ..t.....a..c..a......................................g...........c.....t..g..... [ 82240]
```

| | | |
|---|---|---|
| FR682468.2_Georgia_2007/1 | aatgacaagaagtattttatgaataagcaggcctaaaaagcgcagctttcgtacacgtgtatctgcggtttggcccatgt | [ 82320] |
| ASF_IND_20_CAD543 | ................................................................................ | [ 82320] |
| LR899193.1_Germany_2020/1 | ................................................................................ | [ 82320] |
| MN393477.1_Wuhan_2019 | ................................................................................ | [ 82320] |
| MW396979.1_Timor-Leste/2019/1 | ................................................................................ | [ 82320] |
| MK940252.1_CN/2019/InnerMongolia | ................................................................................ | [ 82320] |
| MK128995.1_China/2018/AnhuiXCGQ | ................................................................................ | [ 82320] |
| MH681419.1_ASFV/POL/2015 | ................................................................................ | [ 82320] |
| LS478113.1_Estonia_2014 | ................................................................................ | [ 82320] |
| KP843857.1_Odintsovo_02/14 | ................................................................................ | [ 82320] |
| FN557520.1_strain_E75 | .............................................t.................................. | [ 82320] |
| AY261366.1_Warthog | ................................................................................ | [ 82320] |
| AY261365.1_Warmbaths | ................................................................................ | [ 82320] |
| AY261364.1_Tengani_62 | ...........................t.................................................... | [ 82320] |
| AY261363.1_Pretoriuskop/96/4 | ................................................................................ | [ 82320] |
| AY261362.1_Mkuzi_1979 | .....................t........a................................................. | [ 82320] |
| AY261361.1_Malawi_Lil-20/1_1983 | .......................................c.................c....................a. | [ 82320] |
| AY261360.1_Kenya_1950 | ........................t....................................................... | [ 82320] |
| AM712240.1_OURT_88/3 | .............................................t.................................. | [ 82320] |
| AM712239.1_Benin_97/1 | .......................................a.....t.................................. | [ 82320] |
| KM111295.1_Ken06.Bus | ................................................................................ | [ 82320] |
| KM111294.1_Ken05/Tk1 | ........................t....................................................... | [ 82320] |
| KM262844.1_strain_L60 | .............................................t.................................. | [ 82320] |
| KM102979.1_26544/OG10_Italy | .............................................t.................................. | [ 82320] |
| KM262845.1_strain_NHV | .............................................t.................................. | [ 82320] |
| KX354450.1_47/Ss/2008 | .............................................t.................................. | [ 82320] |
| KP055815.1_strain_BA71 | .............................................t.................................. | [ 82320] |
| MH025920.1_strain_R35 | ................................................................................ | [ 82320] |
| MH025919.1_strain_N10 | ................................................................................ | [ 82320] |
| MH025918.1_strain_R25 | ................................................................................ | [ 82320] |
| MH025917.1_strain_R7 | ................................................................................ | [ 82320] |
| MH025916.1_strain_R8 | ................................................................................ | [ 82320] |

| | | |
|---|---|---|
| FR682468.2_Georgia_2007/1 | gtggcagcaatattttgtctaaaatagtaagttgtctttcatttaagtattgtaccgcattttcatcgcttttgtaagca | [ 82400] |
| ASF_IND_20_CAD543 | ................................................................................ | [ 82400] |
| LR899193.1_Germany_2020/1 | ................................................................................ | [ 82400] |
| MN393477.1_Wuhan_2019 | ................................................................................ | [ 82400] |
| MW396979.1_Timor-Leste/2019/1 | ................................................................................ | [ 82400] |
| MK940252.1_CN/2019/InnerMongolia | ................................................................................ | [ 82400] |
| MK128995.1_China/2018/AnhuiXCGQ | ................................................................................ | [ 82400] |
| MH681419.1_ASFV/POL/2015 | ................................................................................ | [ 82400] |
| LS478113.1_Estonia_2014 | ................................................................................ | [ 82400] |
| KP843857.1_Odintsovo_02/14 | ................................................................................ | [ 82400] |

```
FN557520.1_strain_E75            ......a....................................c.................................... [ 82400]
AY261366.1_Warthog               ................................................................................ [ 82400]
AY261365.1_Warmbaths             ................................................................................ [ 82400]
AY261364.1_Tengani_62            ........................................g....................a...........a...... [ 82400]
AY261363.1_Pretoriuskop/96/4     ................................................................................ [ 82400]
AY261362.1_Mkuzi_1979            ...........................................c.................................... [ 82400]
AY261361.1_Malawi_Lil-20/1_1983  .........................g.................c.......................a............ [ 82400]
AY261360.1_Kenya_1950            ...............................c...........c..............g..................... [ 82400]
AM712240.1_OURT_88/3             ......a....................................c.................................... [ 82400]
AM712239.1_Benin_97/1            ......a....................................c.................................... [ 82400]
KM111295.1_Ken06.Bus             ...............................c...........c..............g..................... [ 82400]
KM111294.1_Ken05/Tk1             ...............................c..........................g..................... [ 82400]
KM262844.1_strain_L60            ......a....................................c.................................... [ 82400]
KM102979.1_26544/OG10_Italy      ......a....................................c.................................... [ 82400]
KM262845.1_strain_NHV            ......a....................................c.................................... [ 82400]
KX354450.1_47/Ss/2008            ......a....................................c.................................... [ 82400]
KP055815.1_strain_BA71           ......a....................................c.................................... [ 82400]
MH025920.1_strain_R35            ...............................c...........c..............g..................... [ 82400]
MH025919.1_strain_N10            ...............................c...........c..............g..................... [ 82400]
MH025918.1_strain_R25            ...............................c...........c..............g..................... [ 82400]
MH025917.1_strain_R7             ...............................c...........c..............g..................... [ 82400]
MH025916.1_strain_R8             ...............................c...........c..............g..................... [ 82400]

FR682468.2_Georgia_2007/1        gatgggtttgagacaaatttggaaaccttctcggataaaaactggataatttttctcggttcagctcgtgttggaccgg [ 82480]
ASF_IND_20_CAD543                ................................................................................ [ 82480]
LR899193.1_Germany_2020/1        ................................................................................ [ 82480]
MN393477.1_Wuhan_2019            ................................................................................ [ 82480]
MW396979.1_Timor-Leste/2019/1    ................................................................................ [ 82480]
MK940252.1_CN/2019/InnerMongolia ................................................................................ [ 82480]
MK128995.1_China/2018/AnhuiXCGQ  ................................................................................ [ 82480]
MH681419.1_ASFV/POL/2015         ................................................................................ [ 82480]
LS478113.1_Estonia_2014          ................................................................................ [ 82480]
KP843857.1_Odintsovo_02/14       ................................................................................ [ 82480]
FN557520.1_strain_E75            ................................a............................................... [ 82480]
AY261366.1_Warthog               ................................................................................ [ 82480]
AY261365.1_Warmbaths             ...........................................................a.................... [ 82480]
AY261364.1_Tengani_62            ..c.............................a.............................t.a............... [ 82480]
AY261363.1_Pretoriuskop/96/4     ................................................................................ [ 82480]
AY261362.1_Mkuzi_1979            ....................a...........a............................................... [ 82480]
AY261361.1_Malawi_Lil-20/1_1983  .............................t..a.............................t...........a..... [ 82480]
AY261360.1_Kenya_1950            ...........a........a........................................................... [ 82480]
AM712240.1_OURT_88/3             ................................a............................................... [ 82480]
AM712239.1_Benin_97/1            ................................a............................................... [ 82480]
KM111295.1_Ken06.Bus             ..c.....g...........a...................................c....................... [ 82480]
```

```
KM111294.1_Ken05/Tk1            ...........a........a...................................c....................... [ 82480]
KM262844.1_strain_L60           ................................a............................................... [ 82480]
KM102979.1_26544/OG10_Italy     ................................a............................................... [ 82480]
KM262845.1_strain_NHV           ................................a............................................... [ 82480]
KX354450.1_47/Ss/2008           ................................a............................................... [ 82480]
KP055815.1_strain_BA71          ................................a............................................... [ 82480]
MH025920.1_strain_R35           ..c.....g...........a...................................c....................... [ 82480]
MH025919.1_strain_N10           ..c.....g...........a...................................c....................... [ 82480]
MH025918.1_strain_R25           ..c.....g...........a...................................c....................... [ 82480]
MH025917.1_strain_R7            ..c.....g...........a...................................c....................... [ 82480]
MH025916.1_strain_R8            ..c.....g...........a...................................c....................... [ 82480]

FR682468.2_Georgia_2007/1       ttgaaatatggggtctaaaacatgaatggatttttccagaatttctatcatgaaggtattcacaagggagttggattcta [ 82560]
ASF_IND_20_CAD543               ................................................................................ [ 82560]
LR899193.1_Germany_2020/1       ................................................................................ [ 82560]
MN393477.1_Wuhan_2019           ................................................................................ [ 82560]
MW396979.1_Timor-Leste/2019/1   ................................................................................ [ 82560]
MK940252.1_CN/2019/InnerMongolia ................................................................................ [ 82560]
MK128995.1_China/2018/AnhuiXCGQ ................................................................................ [ 82560]
MH681419.1_ASFV/POL/2015        ................................................................................ [ 82560]
LS478113.1_Estonia_2014         ................................................................................ [ 82560]
KP843857.1_Odintsovo_02/14      ................................................................................ [ 82560]
FN557520.1_strain_E75           ................................................................g.............c. [ 82560]
AY261366.1_Warthog              ................................................................................ [ 82560]
AY261365.1_Warmbaths            .........................................................g...................... [ 82560]
AY261364.1_Tengani_62           ......................................................a.....t...g............... [ 82560]
AY261363.1_Pretoriuskop/96/4    ................................................................................ [ 82560]
AY261362.1_Mkuzi_1979           ................................................................g.............c. [ 82560]
AY261361.1_Malawi_Lil-20/1_1983 ................g.....................a.....................t...g.......a....... [ 82560]
AY261360.1_Kenya_1950           ................g.....................ag...........a..a.....t...g............... [ 82560]
AM712240.1_OURT_88/3            ................................................................g.............c. [ 82560]
AM712239.1_Benin_97/1           ................................................................g.............c. [ 82560]
KM111295.1_Ken06.Bus            c........a......g.....................ag.......c............t...g............... [ 82560]
KM111294.1_Ken05/Tk1            ................g.....................ag...........a..a.....t...g............... [ 82560]
KM262844.1_strain_L60           ................................................................g.............c. [ 82560]
KM102979.1_26544/OG10_Italy     ................................................................g.............c. [ 82560]
KM262845.1_strain_NHV           ................................................................g.............c. [ 82560]
KX354450.1_47/Ss/2008           ................................................................g.............c. [ 82560]
KP055815.1_strain_BA71          ................................................................g.............c. [ 82560]
MH025920.1_strain_R35           c........a......g.....................ag.......c............t...g............... [ 82560]
MH025919.1_strain_N10           c........a......g.....................ag.......c............t...g............... [ 82560]
MH025918.1_strain_R25           c........a......g.....................ag.......c............t...g............... [ 82560]
MH025917.1_strain_R7            c........a......g.....................ag.......c............t...g............... [ 82560]
MH025916.1_strain_R8            c........a......g.....................ag.......c............t...g............... [ 82560]
```

```
FR682468.2_Georgia_2007/1        gatcaaataccacttgctcaatgatgctgtcatcgcctgtcattccaaacatgcgaaagatgagataccaaggtatgcga [ 82640]
ASF_IND_20_CAD543                ................................................................................ [ 82640]
LR899193.1_Germany_2020/1        ................................................................................ [ 82640]
MN393477.1_Wuhan_2019            ................................................................................ [ 82640]
MW396979.1_Timor-Leste/2019/1    ................................................................................ [ 82640]
MK940252.1_CN/2019/InnerMongolia ................................................................................ [ 82640]
MK128995.1_China/2018/AnhuiXCGQ  ................................................................................ [ 82640]
MH681419.1_ASFV/POL/2015         ................................................................................ [ 82640]
LS478113.1_Estonia_2014          ................................................................................ [ 82640]
KP843857.1_Odintsovo_02/14       ................................................................................ [ 82640]
FN557520.1_strain_E75            ....g........................................................................... [ 82640]
AY261366.1_Warthog               ..........t................................c.................................... [ 82640]
AY261365.1_Warmbaths             ...........................................c.................................... [ 82640]
AY261364.1_Tengani_62            ....g..........................................................................g [ 82640]
AY261363.1_Pretoriuskop/96/4     ....g...............................................a........................... [ 82640]
AY261362.1_Mkuzi_1979            ....g........................................................................... [ 82640]
AY261361.1_Malawi_Lil-20/1_1983  ..........t.......................a........c........a.......................t... [ 82640]
AY261360.1_Kenya_1950            ....g......................................c....................g.....g......... [ 82640]
AM712240.1_OURT_88/3             ....g........................................................................... [ 82640]
AM712239.1_Benin_97/1            ....g........................................................................... [ 82640]
KM111295.1_Ken06.Bus             ....g......................................c....................g.....g......... [ 82640]
KM111294.1_Ken05/Tk1             ....g......................................c....................g.....g......... [ 82640]
KM262844.1_strain_L60            ....g........................................................................... [ 82640]
KM102979.1_26544/OG10_Italy      ....g........................................................................... [ 82640]
KM262845.1_strain_NHV            ....g........................................................................... [ 82640]
KX354450.1_47/Ss/2008            ....g........................................................................... [ 82640]
KP055815.1_strain_BA71           ....g........................................................................... [ 82640]
MH025920.1_strain_R35            ....g......................................c....................g.....g......... [ 82640]
MH025919.1_strain_N10            ....g......................................c....................g.....g......... [ 82640]
MH025918.1_strain_R25            ....g......................................c....................g.....g......... [ 82640]
MH025917.1_strain_R7             ....g......................................c....................g.....g......... [ 82640]
MH025916.1_strain_R8             ....g......................................c....................g.....g......... [ 82640]

FR682468.2_Georgia_2007/1        agttttgagaacttggtgctattgatttcaatggtaatggcgccggtggtcatgtagcgtataataatttgagagctatt [ 82720]
ASF_IND_20_CAD543                .........................................r..r................................... [ 82720]
LR899193.1_Germany_2020/1        ................................................................................ [ 82720]
MN393477.1_Wuhan_2019            ................................................................................ [ 82720]
MW396979.1_Timor-Leste/2019/1    ................................................................................ [ 82720]
MK940252.1_CN/2019/InnerMongolia ................................................................................ [ 82720]
MK128995.1_China/2018/AnhuiXCGQ  ................................................................................ [ 82720]
MH681419.1_ASFV/POL/2015         ................................................................................ [ 82720]
LS478113.1_Estonia_2014          ................................................................................ [ 82720]
KP843857.1_Odintsovo_02/14       ................................................................................ [ 82720]
```

```
FN557520.1_strain_E75           .........................................a..a..a................................ [ 82720]
AY261366.1_Warthog              ................................................................................ [ 82720]
AY261365.1_Warmbaths            ................................................................................ [ 82720]
AY261364.1_Tengani_62           .......................t........................................................ [ 82720]
AY261363.1_Pretoriuskop/96/4    ................................................................................ [ 82720]
AY261362.1_Mkuzi_1979           ............................................a..a........a....................... [ 82720]
AY261361.1_Malawi_Lil-20/1_1983 ................................c.....a..a.....a..........................a..... [ 82720]
AY261360.1_Kenya_1950           ..c...................................a..........................g.............. [ 82720]
AM712240.1_OURT_88/3            .........................................a..a..a................................ [ 82720]
AM712239.1_Benin_97/1           .........................................a..a..a................................ [ 82720]
KM111295.1_Ken06.Bus            ..c...................................a........a.................g.............. [ 82720]
KM111294.1_Ken05/Tk1            ..c....................a..............a..a.......................g.............. [ 82720]
KM262844.1_strain_L60           .........................................a..a..a................................ [ 82720]
KM102979.1_26544/OG10_Italy     .........................................a..a..a................................ [ 82720]
KM262845.1_strain_NHV           .........................................a..a..a................................ [ 82720]
KX354450.1_47/Ss/2008           .........................................a..a..a................................ [ 82720]
KP055815.1_strain_BA71          .........................................a..a..a................................ [ 82720]
MH025920.1_strain_R35           ..c...................................a........a.................g.............. [ 82720]
MH025919.1_strain_N10           ..c...................................a........a.................g.............. [ 82720]
MH025918.1_strain_R25           ..c...................................a........a.................g.............. [ 82720]
MH025917.1_strain_R7            ..c...................................a........a.................g.............. [ 82720]
MH025916.1_strain_R8            ..c...................................a........a.................g.............. [ 82720]

FR682468.2_Georgia_2007/1       ttcgaaggcacctcccggttgggagataaactcgccgcgaatgatttcattattcccttgttgcatggtatggtaatgga [ 82800]
ASF_IND_20_CAD543               ................................................................................ [ 82800]
LR899193.1_Germany_2020/1       ................................................................................ [ 82800]
MN393477.1_Wuhan_2019           ................................................................................ [ 82800]
MW396979.1_Timor-Leste/2019/1   ................................................................................ [ 82800]
MK940252.1_CN/2019/InnerMongolia ............................................................................... [ 82800]
MK128995.1_China/2018/AnhuiXCGQ ................................................................................ [ 82800]
MH681419.1_ASFV/POL/2015        ................................................................................ [ 82800]
LS478113.1_Estonia_2014         ................................................................................ [ 82800]
KP843857.1_Odintsovo_02/14      ................................................................................ [ 82800]
FN557520.1_strain_E75           ...............t.....a.......................................................... [ 82800]
AY261366.1_Warthog              ...............t.................a.............................................. [ 82800]
AY261365.1_Warmbaths            ...............t......................................t...........a............. [ 82800]
AY261364.1_Tengani_62           ...............t...........g.................................................... [ 82800]
AY261363.1_Pretoriuskop/96/4    ...............t................................................................ [ 82800]
AY261362.1_Mkuzi_1979           ...............t.....a.......................................................... [ 82800]
AY261361.1_Malawi_Lil-20/1_1983 ...a...........t..c............................................................. [ 82800]
AY261360.1_Kenya_1950           ...a...........t...................................g..t......................... [ 82800]
AM712240.1_OURT_88/3            ...............t.....a.......................................................... [ 82800]
AM712239.1_Benin_97/1           ...............t.....a.......................................................... [ 82800]
KM111295.1_Ken06.Bus            ...a...........t.....c.............................g..t.....c................... [ 82800]
```

```
KM111294.1_Ken05/Tk1              ...a...........t...................................g..t......................... [ 82800]
KM262844.1_strain_L60             ...............t.....a.......................................................... [ 82800]
KM102979.1_26544/OG10_Italy       ...............t.....a.......................................................... [ 82800]
KM262845.1_strain_NHV             ...............t.....a.......................................................... [ 82800]
KX354450.1_47/Ss/2008             ...............t.....a..................................................a....... [ 82800]
KP055815.1_strain_BA71            ...............t.....a.......................................................... [ 82800]
MH025920.1_strain_R35             ...a...........t.....c.............................g..t.....c................... [ 82800]
MH025919.1_strain_N10             ...a...........t.....c.............................g..t.....c................... [ 82800]
MH025918.1_strain_R25             ...a...........t.....c.............................g..t.....c................... [ 82800]
MH025917.1_strain_R7              ...a...........t.....c.............................g..t.....c................... [ 82800]
MH025916.1_strain_R8              ...a...........t.....c.............................g..t.....c................... [ 82800]

FR682468.2_Georgia_2007/1         tgtgaagcgtgttaaagcggatgttttctaagaggtctacgacccattccccgcctcgggctataaagtagccgccgggt [ 82880]
ASF_IND_20_CAD543                 ................................................................................ [ 82880]
LR899193.1_Germany_2020/1         ................................................................................ [ 82880]
MN393477.1_Wuhan_2019             ................................................................................ [ 82880]
MW396979.1_Timor-Leste/2019/1     ................................................................................ [ 82880]
MK940252.1_CN/2019/InnerMongolia  ................................................................................ [ 82880]
MK128995.1_China/2018/AnhuiXCGQ   ................................................................................ [ 82880]
MH681419.1_ASFV/POL/2015          ................................................................................ [ 82880]
LS478113.1_Estonia_2014           ................................................................................ [ 82880]
KP843857.1_Odintsovo_02/14        ................................................................................ [ 82880]
FN557520.1_strain_E75             ................................................................................ [ 82880]
AY261366.1_Warthog                ............................c..a............................................a... [ 82880]
AY261365.1_Warmbaths              ......................a.....c..a................................................ [ 82880]
AY261364.1_Tengani_62             ...............................a............................................a... [ 82880]
AY261363.1_Pretoriuskop/96/4      ............................c..a................................................ [ 82880]
AY261362.1_Mkuzi_1979             ....................................................a........................... [ 82880]
AY261361.1_Malawi_Lil-20/1_1983   ..............................ga....................a....................a...... [ 82880]
AY261360.1_Kenya_1950             ............................c..a.....c..............c.................a......... [ 82880]
AM712240.1_OURT_88/3              ................................................................................ [ 82880]
AM712239.1_Benin_97/1             ................................................................................ [ 82880]
KM111295.1_Ken06.Bus              ............................c..a.....c..............c.................a......... [ 82880]
KM111294.1_Ken05/Tk1              ............................c..a.....c..............c.................a......... [ 82880]
KM262844.1_strain_L60             ................................................................................ [ 82880]
KM102979.1_26544/OG10_Italy       ................................................................................ [ 82880]
KM262845.1_strain_NHV             ................................................................................ [ 82880]
KX354450.1_47/Ss/2008             ................................................................................ [ 82880]
KP055815.1_strain_BA71            ................................................................................ [ 82880]
MH025920.1_strain_R35             ............................c..a.....c..............c.................a......... [ 82880]
MH025919.1_strain_N10             ............................c..a.....c..............c.................a......... [ 82880]
MH025918.1_strain_R25             ............................c..a.....c..............c.................a......... [ 82880]
MH025917.1_strain_R7              ............................c..a.....c..............c.................a......... [ 82880]
MH025916.1_strain_R8              ............................c..a.....c..............c.................a......... [ 82880]
```

```
FR682468.2_Georgia_2007/1        tcattagggtcttctcctatttctttttttgcggtttttgataggtgatgagtgtggcagcggttgctgccccgcatgat [ 82960]
ASF_IND_20_CAD543                ................................................................................ [ 82960]
LR899193.1_Germany_2020/1        ................................................................................ [ 82960]
MN393477.1_Wuhan_2019            ................................................................................ [ 82960]
MW396979.1_Timor-Leste/2019/1    ................................................................................ [ 82960]
MK940252.1_CN/2019/InnerMongolia ................................................................................ [ 82960]
MK128995.1_China/2018/AnhuiXCGQ  ................................................................................ [ 82960]
MH681419.1_ASFV/POL/2015         ................................................................................ [ 82960]
LS478113.1_Estonia_2014          ................................................................................ [ 82960]
KP843857.1_Odintsovo_02/14       ................................................................................ [ 82960]
FN557520.1_strain_E75            ................................................................................ [ 82960]
AY261366.1_Warthog               ................................a..........................c.................... [ 82960]
AY261365.1_Warmbaths             ................................a............................................... [ 82960]
AY261364.1_Tengani_62            ..................................................g............................. [ 82960]
AY261363.1_Pretoriuskop/96/4     ..................................................g..a.......................... [ 82960]
AY261362.1_Mkuzi_1979            ................................................................................ [ 82960]
AY261361.1_Malawi_Lil-20/1_1983  ................................a.....g..........ag........a.................... [ 82960]
AY261360.1_Kenya_1950            ..g.............................a.....g.......c...c...........a................. [ 82960]
AM712240.1_OURT_88/3             ................................................................................ [ 82960]
AM712239.1_Benin_97/1            ................................................................................ [ 82960]
KM111295.1_Ken06.Bus             ................................a.....g.......c...c...........a..a.............. [ 82960]
KM111294.1_Ken05/Tk1             ................................a.....g.......c...c...........a........t........ [ 82960]
KM262844.1_strain_L60            ................................................................................ [ 82960]
KM102979.1_26544/OG10_Italy      ................................................................................ [ 82960]
KM262845.1_strain_NHV            ................................................................................ [ 82960]
KX354450.1_47/Ss/2008            ................................................................................ [ 82960]
KP055815.1_strain_BA71           ................................................................................ [ 82960]
MH025920.1_strain_R35            ................................a.....g.......c...c...........a..a.............. [ 82960]
MH025919.1_strain_N10            ................................a.....g.......c...c...........a..a.............. [ 82960]
MH025918.1_strain_R25            ................................a.....g.......c...c...........a..a.............. [ 82960]
MH025917.1_strain_R7             ................................a.....g.......c...c...........a..a.............. [ 82960]
MH025916.1_strain_R8             ................................a.....g.......c...c...........a..a.............. [ 82960]

FR682468.2_Georgia_2007/1        gggaaatgtagatacctgaaaaggaggaatacttgctcgttttacctcctgccgaccattgctgtagtgcgccgttaaaa [ 83040]
ASF_IND_20_CAD543                ................................................................................ [ 83040]
LR899193.1_Germany_2020/1        ................................................................................ [ 83040]
MN393477.1_Wuhan_2019            ................................................................................ [ 83040]
MW396979.1_Timor-Leste/2019/1    ................................................................................ [ 83040]
MK940252.1_CN/2019/InnerMongolia ................................................................................ [ 83040]
MK128995.1_China/2018/AnhuiXCGQ  ................................................................................ [ 83040]
MH681419.1_ASFV/POL/2015         ................................................................................ [ 83040]
LS478113.1_Estonia_2014          ................................................................................ [ 83040]
KP843857.1_Odintsovo_02/14       ................................................................................ [ 83040]
```

```
FN557520.1_strain_E75            .........g..c........t.......................................................... [ 83040]
AY261366.1_Warthog               ................................................................................ [ 83040]
AY261365.1_Warmbaths             ................................................................................ [ 83040]
AY261364.1_Tengani_62            ............c........t.......................................................... [ 83040]
AY261363.1_Pretoriuskop/96/4     .........g..c......................................t............................ [ 83040]
AY261362.1_Mkuzi_1979            .........g..c........t.......................................................... [ 83040]
AY261361.1_Malawi_Lil-20/1_1983  ............c........g........g................................................. [ 83040]
AY261360.1_Kenya_1950            ............c........g........g...........c...........g......................... [ 83040]
AM712240.1_OURT_88/3             .........g..c........t.......................................................... [ 83040]
AM712239.1_Benin_97/1            .........g..c........t.......................................................... [ 83040]
KM111295.1_Ken06.Bus             ..............................g...........c...........g........a..a............. [ 83040]
KM111294.1_Ken05/Tk1             ............c........g........g...........c...........g......................... [ 83040]
KM262844.1_strain_L60            .........g..c........t.......................................................... [ 83040]
KM102979.1_26544/OG10_Italy      .........g..c........t.......................................................... [ 83040]
KM262845.1_strain_NHV            .........g..c........t.......................................................... [ 83040]
KX354450.1_47/Ss/2008            .........g..c........t.......................................................... [ 83040]
KP055815.1_strain_BA71           .........g..c........t.......................................................... [ 83040]
MH025920.1_strain_R35            ..............................g...........c...........g........a..a............. [ 83040]
MH025919.1_strain_N10            ..............................g...........c...........g........a..a............. [ 83040]
MH025918.1_strain_R25            ..............................g...........c...........g........a..a............. [ 83040]
MH025917.1_strain_R7             ..............................g...........c...........g........a..a............. [ 83040]
MH025916.1_strain_R8             ..............................g...........c...........g........a..a............. [ 83040]

FR682468.2_Georgia_2007/1        taacctcggcggctagattaaccgggcccgaataggaaaggccacacaggcgtgccttattgggtagtaaatttatcttg [ 83120]
ASF_IND_20_CAD543                ................................................................................ [ 83120]
LR899193.1_Germany_2020/1        ................................................................................ [ 83120]
MN393477.1_Wuhan_2019            ................................................................................ [ 83120]
MW396979.1_Timor-Leste/2019/1    ................................................................................ [ 83120]
MK940252.1_CN/2019/InnerMongolia ................................................................................ [ 83120]
MK128995.1_China/2018/AnhuiXCGQ  ................................................................................ [ 83120]
MH681419.1_ASFV/POL/2015         ................................................................................ [ 83120]
LS478113.1_Estonia_2014          ................................................................................ [ 83120]
KP843857.1_Odintsovo_02/14       ................................................................................ [ 83120]
FN557520.1_strain_E75            ......................a......................................a.................. [ 83120]
AY261366.1_Warthog               ......................a..a...................................................... [ 83120]
AY261365.1_Warmbaths             .......a.....c........a......................................................... [ 83120]
AY261364.1_Tengani_62            .....................................c.......................................... [ 83120]
AY261363.1_Pretoriuskop/96/4     .............c........a......................................................... [ 83120]
AY261362.1_Mkuzi_1979            ......................a......................................a.................. [ 83120]
AY261361.1_Malawi_Lil-20/1_1983  ....t..a...........c..a..........................a..............c.a............. [ 83120]
AY261360.1_Kenya_1950            .......a.....c..t.....a......................t...............a..c............... [ 83120]
AM712240.1_OURT_88/3             ......................a......................................a.................. [ 83120]
AM712239.1_Benin_97/1            ......................a......................................a.................. [ 83120]
KM111295.1_Ken06.Bus             .............c..t.....a.....t...........a.......................c............... [ 83120]
```

```
KM111294.1_Ken05/Tk1            .......a.....c..t.....a.........................................c............... [ 83120]
KM262844.1_strain_L60           ......................a......................................a.................. [ 83120]
KM102979.1_26544/OG10_Italy     ......................a......................................a.................. [ 83120]
KM262845.1_strain_NHV           ......................a......................................a.................. [ 83120]
KX354450.1_47/Ss/2008           ......................a......................................a.................. [ 83120]
KP055815.1_strain_BA71          ......................a......................................a.................. [ 83120]
MH025920.1_strain_R35           .............c..t.....a.....t...........a.......................c............... [ 83120]
MH025919.1_strain_N10           .............c..t.....a.....t...........a.......................c............... [ 83120]
MH025918.1_strain_R25           .............c..t.....a.....t...........a.......................c............... [ 83120]
MH025917.1_strain_R7            .............c..t.....a.....t...........a.......................c............... [ 83120]
MH025916.1_strain_R8            .............c..t.....a.....t...........a.......................c............... [ 83120]

FR682468.2_Georgia_2007/1       tttccctgtgaatagtttcgatgttgcgggcgttcaatgttcacatctgtaaagttaaattggatctgaactgattcccg [ 83200]
ASF_IND_20_CAD543               ................................................................................ [ 83200]
LR899193.1_Germany_2020/1       ................................................................................ [ 83200]
MN393477.1_Wuhan_2019           ................................................................................ [ 83200]
MW396979.1_Timor-Leste/2019/1   ................................................................................ [ 83200]
MK940252.1_CN/2019/InnerMongolia ................................................................................ [ 83200]
MK128995.1_China/2018/AnhuiXCGQ ................................................................................ [ 83200]
MH681419.1_ASFV/POL/2015        ................................................................................ [ 83200]
LS478113.1_Estonia_2014         ................................................................................ [ 83200]
KP843857.1_Odintsovo_02/14      ................................................................................ [ 83200]
FN557520.1_strain_E75           .......................................................................c........ [ 83200]
AY261366.1_Warthog              .......................................................................c.....t.. [ 83200]
AY261365.1_Warmbaths            .......................................................................c........ [ 83200]
AY261364.1_Tengani_62           .......................................................................c........ [ 83200]
AY261363.1_Pretoriuskop/96/4    .......................................................................c........ [ 83200]
AY261362.1_Mkuzi_1979           .......................................................................c........ [ 83200]
AY261361.1_Malawi_Lil-20/1_1983 ..........................t............................................g.....t.. [ 83200]
AY261360.1_Kenya_1950           ....................g..........................g.......................a........ [ 83200]
AM712240.1_OURT_88/3            .......................................................................c........ [ 83200]
AM712239.1_Benin_97/1           .......................................................................c........ [ 83200]
KM111295.1_Ken06.Bus            ....................g..........................g.......................a........ [ 83200]
KM111294.1_Ken05/Tk1            ....................g..........................g.......................a........ [ 83200]
KM262844.1_strain_L60           .......................................................................c........ [ 83200]
KM102979.1_26544/OG10_Italy     .......................................................................c........ [ 83200]
KM262845.1_strain_NHV           .......................................................................c........ [ 83200]
KX354450.1_47/Ss/2008           .......................................................................c........ [ 83200]
KP055815.1_strain_BA71          .......................................................................c........ [ 83200]
MH025920.1_strain_R35           ....................g..........................g.......................a........ [ 83200]
MH025919.1_strain_N10           ....................g..........................g.......................a........ [ 83200]
MH025918.1_strain_R25           ....................g..........................g.......................a........ [ 83200]
MH025917.1_strain_R7            ....................g..........................g.......................a........ [ 83200]
MH025916.1_strain_R8            ....................g..........................g.......................a........ [ 83200]
```

```
FR682468.2_Georgia_2007/1        aagcttatctatttcagtatggtcgcgttggtctttataagtaatatccacgttaaacatttgttttacaatttgcggaa [ 83280]
ASF_IND_20_CAD543                ................................................................................ [ 83280]
LR899193.1_Germany_2020/1        ................................................................................ [ 83280]
MN393477.1_Wuhan_2019            ................................................................................ [ 83280]
MW396979.1_Timor-Leste/2019/1    ................................................................................ [ 83280]
MK940252.1_CN/2019/InnerMongolia ................................................................................ [ 83280]
MK128995.1_China/2018/AnhuiXCGQ  ................................................................................ [ 83280]
MH681419.1_ASFV/POL/2015         ................................................................................ [ 83280]
LS478113.1_Estonia_2014          ................................................................................ [ 83280]
KP843857.1_Odintsovo_02/14       ................................................................................ [ 83280]
FN557520.1_strain_E75            ..............................................................................g. [ 83280]
AY261366.1_Warthog               ..............................................................................g. [ 83280]
AY261365.1_Warmbaths             ..............................................................................g. [ 83280]
AY261364.1_Tengani_62            ............................................................c.................g. [ 83280]
AY261363.1_Pretoriuskop/96/4     .......................................c......................................g. [ 83280]
AY261362.1_Mkuzi_1979            ................................................................................ [ 83280]
AY261361.1_Malawi_Lil-20/1_1983  ...t.............................c.....g................................c..t.... [ 83280]
AY261360.1_Kenya_1950            .................................c.....g........t.....g......................... [ 83280]
AM712240.1_OURT_88/3             ..............................................................................g. [ 83280]
AM712239.1_Benin_97/1            ..............................................................................g. [ 83280]
KM111295.1_Ken06.Bus             .................................c.....g........t.....g......................... [ 83280]
KM111294.1_Ken05/Tk1             .................................c.....g........t.....g......................... [ 83280]
KM262844.1_strain_L60            ..............................................................................g. [ 83280]
KM102979.1_26544/OG10_Italy      ..............................................................................g. [ 83280]
KM262845.1_strain_NHV            ..............................................................................g. [ 83280]
KX354450.1_47/Ss/2008            ..............................................................................g. [ 83280]
KP055815.1_strain_BA71           ..............................................................................g. [ 83280]
MH025920.1_strain_R35            .................................c.....g........t.....g......................... [ 83280]
MH025919.1_strain_N10            .................................c.....g........t.....g......................... [ 83280]
MH025918.1_strain_R25            .................................c.....g........t.....g......................... [ 83280]
MH025917.1_strain_R7             .................................c.....g........t.....g......................... [ 83280]
MH025916.1_strain_R8             .................................c.....g........t.....g......................... [ 83280]

FR682468.2_Georgia_2007/1        ttccattgtccataagatcgtcgaagcttttgatgttataccctatcaatcctgtagagtttactgcagcggagataaag [ 83360]
ASF_IND_20_CAD543                ................................................................................ [ 83360]
LR899193.1_Germany_2020/1        ................................................................................ [ 83360]
MN393477.1_Wuhan_2019            ................................................................................ [ 83360]
MW396979.1_Timor-Leste/2019/1    ................................................................................ [ 83360]
MK940252.1_CN/2019/InnerMongolia ................................................................................ [ 83360]
MK128995.1_China/2018/AnhuiXCGQ  ................................................................................ [ 83360]
MH681419.1_ASFV/POL/2015         ................................................................................ [ 83360]
LS478113.1_Estonia_2014          ................................................................................ [ 83360]
KP843857.1_Odintsovo_02/14       ................................................................................ [ 83360]
```

```
FN557520.1_strain_E75            ................................................................................ [ 83360]
AY261366.1_Warthog               ....................................................g........................... [ 83360]
AY261365.1_Warmbaths             ........................................t........c.............................. [ 83360]
AY261364.1_Tengani_62            .......a..........................a..............c.............................. [ 83360]
AY261363.1_Pretoriuskop/96/4     ........................................t...........g........................... [ 83360]
AY261362.1_Mkuzi_1979            ...................................................................g............ [ 83360]
AY261361.1_Malawi_Lil-20/1_1983  ................g.....a.................t...........g..............g...........a [ 83360]
AY261360.1_Kenya_1950            ...................a....................t........c..a..............g............ [ 83360]
AM712240.1_OURT_88/3             ................................................................................ [ 83360]
AM712239.1_Benin_97/1            ................................................................................ [ 83360]
KM111295.1_Ken06.Bus             ...................a....................t........c..a..............g............ [ 83360]
KM111294.1_Ken05/Tk1             ...................a....................t........c..a..............g............ [ 83360]
KM262844.1_strain_L60            ................................................................................ [ 83360]
KM102979.1_26544/OG10_Italy      ................................................................................ [ 83360]
KM262845.1_strain_NHV            ................................................................................ [ 83360]
KX354450.1_47/Ss/2008            ................................................................................ [ 83360]
KP055815.1_strain_BA71           ................................................................................ [ 83360]
MH025920.1_strain_R35            ...................a....................t........c..a..............g............ [ 83360]
MH025919.1_strain_N10            ...................a....................t........c..a..............g............ [ 83360]
MH025918.1_strain_R25            ...................a....................t........c..a..............g............ [ 83360]
MH025917.1_strain_R7             ...................a....................t........c..a..............g............ [ 83360]
MH025916.1_strain_R8             ...................a....................t........c..a..............g............ [ 83360]

FR682468.2_Georgia_2007/1        ctcagcatatcagcctctgtaagctcctcattatccacggtttcaatggggccgtaggttatttgcggccgcaagggttc [ 83440]
ASF_IND_20_CAD543                ................................................................................ [ 83440]
LR899193.1_Germany_2020/1        ................................................................................ [ 83440]
MN393477.1_Wuhan_2019            ................................................................................ [ 83440]
MW396979.1_Timor-Leste/2019/1    ................................................................................ [ 83440]
MK940252.1_CN/2019/InnerMongolia ................................................................................ [ 83440]
MK128995.1_China/2018/AnhuiXCGQ  ................................................................................ [ 83440]
MH681419.1_ASFV/POL/2015         ................................................................................ [ 83440]
LS478113.1_Estonia_2014          ................................................................................ [ 83440]
KP843857.1_Odintsovo_02/14       ................................................................................ [ 83440]
FN557520.1_strain_E75            ................................................................................ [ 83440]
AY261366.1_Warthog               ................................................................................ [ 83440]
AY261365.1_Warmbaths             ................................................................................ [ 83440]
AY261364.1_Tengani_62            ................................................................................ [ 83440]
AY261363.1_Pretoriuskop/96/4     ................................................................................ [ 83440]
AY261362.1_Mkuzi_1979            ................................................................................ [ 83440]
AY261361.1_Malawi_Lil-20/1_1983  ....a.............................t.....cc........c....................t........ [ 83440]
AY261360.1_Kenya_1950            ................................................................................ [ 83440]
AM712240.1_OURT_88/3             ................................................................................ [ 83440]
AM712239.1_Benin_97/1            ................................................................................ [ 83440]
KM111295.1_Ken06.Bus             ................................................................................ [ 83440]
```

```
KM111294.1_Ken05/Tk1              ................................................................................ [ 83440]
KM262844.1_strain_L60             ................................................................................ [ 83440]
KM102979.1_26544/OG10_Italy       ................................................................................ [ 83440]
KM262845.1_strain_NHV             ................................................................................ [ 83440]
KX354450.1_47/Ss/2008             ................................................................................ [ 83440]
KP055815.1_strain_BA71            ................................................................................ [ 83440]
MH025920.1_strain_R35             ................................................................................ [ 83440]
MH025919.1_strain_N10             ................................................................................ [ 83440]
MH025918.1_strain_R25             ................................................................................ [ 83440]
MH025917.1_strain_R7              ................................................................................ [ 83440]
MH025916.1_strain_R8              ................................................................................ [ 83440]

FR682468.2_Georgia_2007/1         catgattatgaagtactacattaatattcagttattctttaaaataaatctttatttataaatcttatttataatataag [ 83520]
ASF_IND_20_CAD543                 ................................................................................ [ 83520]
LR899193.1_Germany_2020/1         ................................................................................ [ 83520]
MN393477.1_Wuhan_2019             ................................................................................ [ 83520]
MW396979.1_Timor-Leste/2019/1     ................................................................................ [ 83520]
MK940252.1_CN/2019/InnerMongolia  ................................................................................ [ 83520]
MK128995.1_China/2018/AnhuiXCGQ   ................................................................................ [ 83520]
MH681419.1_ASFV/POL/2015          ................................................................................ [ 83520]
LS478113.1_Estonia_2014           ................................................................................ [ 83520]
KP843857.1_Odintsovo_02/14        ................................................................................ [ 83520]
FN557520.1_strain_E75             ...a............................................................................ [ 83520]
AY261366.1_Warthog                ................................................................................ [ 83520]
AY261365.1_Warmbaths              ................................................................................ [ 83520]
AY261364.1_Tengani_62             ................................................................................ [ 83520]
AY261363.1_Pretoriuskop/96/4      ................................................................................ [ 83520]
AY261362.1_Mkuzi_1979             ................................................................................ [ 83520]
AY261361.1_Malawi_Lil-20/1_1983   ................................................................................ [ 83520]
AY261360.1_Kenya_1950             ..........t..................................................................... [ 83520]
AM712240.1_OURT_88/3              ...a............................................................................ [ 83520]
AM712239.1_Benin_97/1             ...a............................................................................ [ 83520]
KM111295.1_Ken06.Bus              ................................................................................ [ 83520]
KM111294.1_Ken05/Tk1              ..........t..................................................................... [ 83520]
KM262844.1_strain_L60             ...a............................................................................ [ 83520]
KM102979.1_26544/OG10_Italy       ...a............................................................................ [ 83520]
KM262845.1_strain_NHV             ...a............................................................................ [ 83520]
KX354450.1_47/Ss/2008             ...a............................................................................ [ 83520]
KP055815.1_strain_BA71            ...a............................................................................ [ 83520]
MH025920.1_strain_R35             ................................................................................ [ 83520]
MH025919.1_strain_N10             ................................................................................ [ 83520]
MH025918.1_strain_R25             ................................................................................ [ 83520]
MH025917.1_strain_R7              ................................................................................ [ 83520]
MH025916.1_strain_R8              ................................................................................ [ 83520]
```

FR682468.2_Georgia_2007/1 aatgccttatgcaagagacatcacaaagtttattacggcaacggaaccagaggtgggtcttcccctgttggcgctgcagc [ 83600]
ASF_IND_20_CAD543 ..........k..................................................................... [ 83600]
LR899193.1_Germany_2020/1 ................................................................................ [ 83600]
MN393477.1_Wuhan_2019 ................................................................................ [ 83600]
MW396979.1_Timor-Leste/2019/1 ................................................................................ [ 83600]
MK940252.1_CN/2019/InnerMongolia ................................................................................ [ 83600]
MK128995.1_China/2018/AnhuiXCGQ ................................................................................ [ 83600]
MH681419.1_ASFV/POL/2015 ................................................................................ [ 83600]
LS478113.1_Estonia_2014 ................................................................................ [ 83600]
KP843857.1_Odintsovo_02/14 ................................................................................ [ 83600]
FN557520.1_strain_E75 ..........t.......t..............c..........................c.................a. [ 83600]
AY261366.1_Warthog ..........t......................c.......................a..c.................a. [ 83600]
AY261365.1_Warmbaths ..........t.......t..............c..........................c.................a. [ 83600]
AY261364.1_Tengani_62 ................................................................................ [ 83600]
AY261363.1_Pretoriuskop/96/4 ................................................................................ [ 83600]
AY261362.1_Mkuzi_1979 ..........t.......t..............c..........................c.................a. [ 83600]
AY261361.1_Malawi_Lil-20/1_1983 ..........t..........t...........c.......................c..c................... [ 83600]
AY261360.1_Kenya_1950 ..........t..........t...........c..........................c............t...... [ 83600]
AM712240.1_OURT_88/3 ..........t.......t..............c..........................c.................a. [ 83600]
AM712239.1_Benin_97/1 ..........t.......t..............c..........................c.................a. [ 83600]
KM111295.1_Ken06.Bus ..........t..........t...........c.............................................. [ 83600]
KM111294.1_Ken05/Tk1 ..........t..........t...........c..........................c................... [ 83600]
KM262844.1_strain_L60 ..........t.......t..............c..........................c.................a. [ 83600]
KM102979.1_26544/OG10_Italy ..........t.......t..............c..........................c.................a. [ 83600]
KM262845.1_strain_NHV ..........t.......t..............c..........................c.................a. [ 83600]
KX354450.1_47/Ss/2008 ..........t.......t..............c..........................c.................a. [ 83600]
KP055815.1_strain_BA71 ..........t.......t..............c..........................c.................a. [ 83600]
MH025920.1_strain_R35 ..........t..........t...........c.............................................. [ 83600]
MH025919.1_strain_N10 ..........t..........t...........c.............................................. [ 83600]
MH025918.1_strain_R25 ..........t..........t...........c.............................................. [ 83600]
MH025917.1_strain_R7 ..........t..........t...........c.............................................. [ 83600]
MH025916.1_strain_R8 ..........t..........t...........c.............................................. [ 83600]

FR682468.2_Georgia_2007/1 gctccaaatccatcataggggttattcttcttgtaataagtttgttatttattttcattggcattattatattatcagtg [ 83680]
ASF_IND_20_CAD543 ................................................................................ [ 83680]
LR899193.1_Germany_2020/1 ................................................................................ [ 83680]
MN393477.1_Wuhan_2019 ................................................................................ [ 83680]
MW396979.1_Timor-Leste/2019/1 ................................................................................ [ 83680]
MK940252.1_CN/2019/InnerMongolia ................................................................................ [ 83680]
MK128995.1_China/2018/AnhuiXCGQ ................................................................................ [ 83680]
MH681419.1_ASFV/POL/2015 ................................................................................ [ 83680]
LS478113.1_Estonia_2014 ................................................................................ [ 83680]
KP843857.1_Odintsovo_02/14 ................................................................................ [ 83680]

```
FN557520.1_strain_E75            a......g...g...........................................................c....g... [ 83680]
AY261366.1_Warthog               a......g.................a............t................g...............c..g..... [ 83680]
AY261365.1_Warmbaths             a......g...g...........................................g...............c..g....a [ 83680]
AY261364.1_Tengani_62            ..........t..................................................................agt [ 83680]
AY261363.1_Pretoriuskop/96/4     .......................................................................c....g... [ 83680]
AY261362.1_Mkuzi_1979            a......g...g.......aa....a.............................g...............c..g..... [ 83680]
AY261361.1_Malawi_Lil-20/1_1983  .......g.................c.............................................c....g... [ 83680]
AY261360.1_Kenya_1950            .......g...g........a....c.......................a.....................c....g... [ 83680]
AM712240.1_OURT_88/3             a......g...g...........................................................c....g... [ 83680]
AM712239.1_Benin_97/1            a......g...g...........................................................c....g... [ 83680]
KM111295.1_Ken06.Bus             .......g.................c.......................a.....................c....g... [ 83680]
KM111294.1_Ken05/Tk1             .......g...g.............c.......................a.....................c....g... [ 83680]
KM262844.1_strain_L60            a......g...g...........................................................c....g... [ 83680]
KM102979.1_26544/OG10_Italy      a......g...g...........................................................c....g... [ 83680]
KM262845.1_strain_NHV            a......g...g...........................................................c....g... [ 83680]
KX354450.1_47/Ss/2008            a......g...g...........................................................c....g... [ 83680]
KP055815.1_strain_BA71           a......g...g...........................................................c....g... [ 83680]
MH025920.1_strain_R35            .......g.................c.......................a.....................c....g... [ 83680]
MH025919.1_strain_N10            .......g.................c.......................a.....................c....g... [ 83680]
MH025918.1_strain_R25            .......g.................c.......................a.....................c....g... [ 83680]
MH025917.1_strain_R7             .......g.................c.......................a.....................c....g... [ 83680]
MH025916.1_strain_R8             .......g.................c.......................a.....................c....g... [ 83680]

FR682468.2_Georgia_2007/1        agtagtggtcataccacagcagcctctatatttatcgtattgagtcttatcctaggtggcggtggtttttttcttattta [ 83760]
ASF_IND_20_CAD543                ................................................................................ [ 83760]
LR899193.1_Germany_2020/1        ................................................................................ [ 83760]
MN393477.1_Wuhan_2019            ................................................................................ [ 83760]
MW396979.1_Timor-Leste/2019/1    ................................................................................ [ 83760]
MK940252.1_CN/2019/InnerMongolia ................................................................................ [ 83760]
MK128995.1_China/2018/AnhuiXCGQ  ................................................................................ [ 83760]
MH681419.1_ASFV/POL/2015         ................................................................................ [ 83760]
LS478113.1_Estonia_2014          ................................................................................ [ 83760]
KP843857.1_Odintsovo_02/14       ................................................................................ [ 83760]
FN557520.1_strain_E75            ................................................................................ [ 83760]
AY261366.1_Warthog               g..g---....cg.........gt.........g.....c........................................ [ 83760]
AY261365.1_Warmbaths             g...---....c......a...g..........g..a..c........................................ [ 83760]
AY261364.1_Tengani_62            ...g---......................................................................... [ 83760]
AY261363.1_Pretoriuskop/96/4     ....---..t....t..ca...g..........g.....c........................................ [ 83760]
AY261362.1_Mkuzi_1979            g...---....c......a...g..........g..a..c........................................ [ 83760]
AY261361.1_Malawi_Lil-20/1_1983  ...t---c.t.......ga...g..........g.....c................c....................... [ 83760]
AY261360.1_Kenya_1950            ..---.a..........c....g....g....gg.............................................. [ 83760]
AM712240.1_OURT_88/3             ......a......................................................................... [ 83760]
AM712239.1_Benin_97/1            ................................................................................ [ 83760]
KM111295.1_Ken06.Bus             ......a..........c....gt........gg.t...c....c................................... [ 83760]
```

```
KM111294.1_Ken05/Tk1             g.---.......g.........g....g....gg.............................................. [ 83760]
KM262844.1_strain_L60            ................................................................................ [ 83760]
KM102979.1_26544/OG10_Italy      ................................................................................ [ 83760]
KM262845.1_strain_NHV            ......a......................................................................... [ 83760]
KX354450.1_47/Ss/2008            ................................................................................ [ 83760]
KP055815.1_strain_BA71           ................................................................................ [ 83760]
MH025920.1_strain_R35            ......a..........c....gt........gg.t...c....c................................... [ 83760]
MH025919.1_strain_N10            ......a..........c....gt........gg.t...c....c................................... [ 83760]
MH025918.1_strain_R25            ......a..........c....gt........gg.t...c....c................................... [ 83760]
MH025917.1_strain_R7             ......a..........c....gt........gg.t...c....c................................... [ 83760]
MH025916.1_strain_R8             ......a..........c....gt........gg.t...c....c................................... [ 83760]

FR682468.2_Georgia_2007/1        taaagataattcttaacccacataaaatttgaaaaaatatagagtaagaaaatgtccaattactattattactatggcgg [ 83840]
ASF_IND_20_CAD543                .........................................a...................................... [ 83840]
LR899193.1_Germany_2020/1        ................................................................................ [ 83840]
MN393477.1_Wuhan_2019            ................................................................................ [ 83840]
MW396979.1_Timor-Leste/2019/1    ................................................................................ [ 83840]
MK940252.1_CN/2019/InnerMongolia ................................................................................ [ 83840]
MK128995.1_China/2018/AnhuiXCGQ  ................................................................................ [ 83840]
MH681419.1_ASFV/POL/2015         ................................................................................ [ 83840]
LS478113.1_Estonia_2014          ................................................................................ [ 83840]
KP843857.1_Odintsovo_02/14       ................................................................................ [ 83840]
FN557520.1_strain_E75            .........................................a...................................... [ 83840]
AY261366.1_Warthog               ................................................................................ [ 83840]
AY261365.1_Warmbaths             ................................................................................ [ 83840]
AY261364.1_Tengani_62            ................................................................................ [ 83840]
AY261363.1_Pretoriuskop/96/4     ....................................t........................................... [ 83840]
AY261362.1_Mkuzi_1979            .........................................a...................................... [ 83840]
AY261361.1_Malawi_Lil-20/1_1983  ..................t............................................................. [ 83840]
AY261360.1_Kenya_1950            ....................................g........................................... [ 83840]
AM712240.1_OURT_88/3             .........................................a...................................... [ 83840]
AM712239.1_Benin_97/1            .........................................a...................................... [ 83840]
KM111295.1_Ken06.Bus             ....................................g........................................... [ 83840]
KM111294.1_Ken05/Tk1             ....................................g........................................... [ 83840]
KM262844.1_strain_L60            .........................................a...................................... [ 83840]
KM102979.1_26544/OG10_Italy      .........................................a...................................... [ 83840]
KM262845.1_strain_NHV            .........................................a...................................... [ 83840]
KX354450.1_47/Ss/2008            .........................................a...................................... [ 83840]
KP055815.1_strain_BA71           .........................................a...................................... [ 83840]
MH025920.1_strain_R35            ....................................g........................................... [ 83840]
MH025919.1_strain_N10            ....................................g........................................... [ 83840]
MH025918.1_strain_R25            ....................................g........................................... [ 83840]
MH025917.1_strain_R7             ....................................g........................................... [ 83840]
MH025916.1_strain_R8             ....................................g........................................... [ 83840]
```

```
FR682468.2_Georgia_2007/1        ggggagatatgattggttaaaaacagtagaacccactaatttttaaaaatcgggttgccttaccaggcacacccattac [ 83920]
ASF_IND_20_CAD543                ................................................................................ [ 83920]
LR899193.1_Germany_2020/1        ................................................................................ [ 83920]
MN393477.1_Wuhan_2019            ................................................................................ [ 83920]
MW396979.1_Timor-Leste/2019/1    ................................................................................ [ 83920]
MK940252.1_CN/2019/InnerMongolia ................................................................................ [ 83920]
MK128995.1_China/2018/AnhuiXCGQ  ................................................................................ [ 83920]
MH681419.1_ASFV/POL/2015         ................................................................................ [ 83920]
LS478113.1_Estonia_2014          ................................................................................ [ 83920]
KP843857.1_Odintsovo_02/14       ................................................................................ [ 83920]
FN557520.1_strain_E75            ................................................................................ [ 83920]
AY261366.1_Warthog               ................................................................................ [ 83920]
AY261365.1_Warmbaths             ................................................................................ [ 83920]
AY261364.1_Tengani_62            ................................................................................ [ 83920]
AY261363.1_Pretoriuskop/96/4     ................................................................................ [ 83920]
AY261362.1_Mkuzi_1979            ...................................................t............................ [ 83920]
AY261361.1_Malawi_Lil-20/1_1983  a..a........c................................................................... [ 83920]
AY261360.1_Kenya_1950            ...a........c...........g....................................................... [ 83920]
AM712240.1_OURT_88/3             ................................................................................ [ 83920]
AM712239.1_Benin_97/1            ................................................................................ [ 83920]
KM111295.1_Ken06.Bus             ...a........c...........g....................................................... [ 83920]
KM111294.1_Ken05/Tk1             ...a........c...........g....................................................... [ 83920]
KM262844.1_strain_L60            ................................................................................ [ 83920]
KM102979.1_26544/OG10_Italy      ................................................................................ [ 83920]
KM262845.1_strain_NHV            ................................................................................ [ 83920]
KX354450.1_47/Ss/2008            ................................................................................ [ 83920]
KP055815.1_strain_BA71           ................................................................................ [ 83920]
MH025920.1_strain_R35            ...a........c...........g....................................................... [ 83920]
MH025919.1_strain_N10            ...a........c...........g....................................................... [ 83920]
MH025918.1_strain_R25            ...a........c...........g....................................................... [ 83920]
MH025917.1_strain_R7             ...a........c...........g....................................................... [ 83920]
MH025916.1_strain_R8             ...a........c...........g....................................................... [ 83920]

FR682468.2_Georgia_2007/1        atcttcaacatcaggca---actactcccccatctatcttagaaaaatttaaacgagcagacattcttcttaatgaggtg [ 84000]
ASF_IND_20_CAD543                .c..c............act............................................................ [ 84000]
LR899193.1_Germany_2020/1        .................---............................................................ [ 84000]
MN393477.1_Wuhan_2019            .................---............................................................ [ 84000]
MW396979.1_Timor-Leste/2019/1    .................---............................................................ [ 84000]
MK940252.1_CN/2019/InnerMongolia .................---............................................................ [ 84000]
MK128995.1_China/2018/AnhuiXCGQ  .................---............................................................ [ 84000]
MH681419.1_ASFV/POL/2015         .................---............................................................ [ 84000]
LS478113.1_Estonia_2014          .................---............................................................ [ 84000]
KP843857.1_Odintsovo_02/14       .................---............................................................ [ 84000]
```

```
FN557520.1_strain_E75          .c..c............act..................c......................t.................. [ 84000]
AY261366.1_Warthog             .c..c.........a..---............................................................ [ 84000]
AY261365.1_Warmbaths           .c..c............---............................................................ [ 84000]
AY261364.1_Tengani_62          .................---...........................................................a [ 84000]
AY261363.1_Pretoriuskop/96/4   .c..c............---............................................................ [ 84000]
AY261362.1_Mkuzi_1979          .c..c............act............................................................ [ 84000]
AY261361.1_Malawi_Lil-20/1_1983 .c..c............---........t.........c......................g.................t [ 84000]
AY261360.1_Kenya_1950          .c.....g.........---..g...............c............................c...........t [ 84000]
AM712240.1_OURT_88/3           .c..c............act..................c......................t.................. [ 84000]
AM712239.1_Benin_97/1          .c..c............act..................c......................t.................. [ 84000]
KM111295.1_Ken06.Bus           .c..c..........g.---..g..c......g.....c..................t...t.....c...........t [ 84000]
KM111294.1_Ken05/Tk1           .c.....g.........---..g...............c............................c...........t [ 84000]
KM262844.1_strain_L60          .c..c............act..................c......................t.................. [ 84000]
KM102979.1_26544/OG10_Italy    .c..c............act..................c......................t.................. [ 84000]
KM262845.1_strain_NHV          .c..c............act..................c......................t.................. [ 84000]
KX354450.1_47/Ss/2008          .c..c............act..................c......................t.................. [ 84000]
KP055815.1_strain_BA71         .c..c............act..................c......................t.................. [ 84000]
MH025920.1_strain_R35          .c..c..........g.---..g..c......g.....c..................t...t.....c...........t [ 84000]
MH025919.1_strain_N10          .c..c..........g.---..g..c......g.....c..................t...t.....c...........t [ 84000]
MH025918.1_strain_R25          .c..c..........g.---..g..c......g.....c..................t...t.....c...........t [ 84000]
MH025917.1_strain_R7           .c..c..........g.---..g..c......g.....c..................t...t.....c...........t [ 84000]
MH025916.1_strain_R8           .c..c..........g.---..g..c......g.....c..................t...t.....c...........t [ 84000]

FR682468.2_Georgia_2007/1      aaggccgaaatggacccactcatgttacaaccagaaaccgaaaaaaaactattccagatattgagtagtattgatatgtt [ 84080]
ASF_IND_20_CAD543              ................................................................................ [ 84080]
LR899193.1_Germany_2020/1      ................................................................................ [ 84080]
MN393477.1_Wuhan_2019          ................................................................................ [ 84080]
MW396979.1_Timor-Leste/2019/1  ................................................................................ [ 84080]
MK940252.1_CN/2019/InnerMongolia ................................................................................ [ 84080]
MK128995.1_China/2018/AnhuiXCGQ ................................................................................ [ 84080]
MH681419.1_ASFV/POL/2015       ................................................................................ [ 84080]
LS478113.1_Estonia_2014        ................................................................................ [ 84080]
KP843857.1_Odintsovo_02/14     ........................................-....................................... [ 84080]
FN557520.1_strain_E75          ................................................t...a..........g................ [ 84080]
AY261366.1_Warthog             ................................................t............................... [ 84080]
AY261365.1_Warmbaths           ..............t.................................t............................... [ 84080]
AY261364.1_Tengani_62          ..............t.................................t........g...................... [ 84080]
AY261363.1_Pretoriuskop/96/4   ................................................t............................... [ 84080]
AY261362.1_Mkuzi_1979          .................c..............................t....t.........g................ [ 84080]
AY261361.1_Malawi_Lil-20/1_1983 .....t..............................................at..a......g.c..c..c........ [ 84080]
AY261360.1_Kenya_1950          ..at.............c.................g................a....g.....g.c..c..c........ [ 84080]
AM712240.1_OURT_88/3           ................................................t...a..........g................ [ 84080]
AM712239.1_Benin_97/1          ................................................t...a.........ag................ [ 84080]
KM111295.1_Ken06.Bus           ..at.............c.................g............t...at.........g.c..c..c........ [ 84080]
```

```
KM111294.1_Ken05/Tk1             ..at.............c.................g................a....g.....g.c..c..c........ [ 84080]
KM262844.1_strain_L60            ................................................t...a.........ag................ [ 84080]
KM102979.1_26544/OG10_Italy      ................................................t...a.........ag................ [ 84080]
KM262845.1_strain_NHV            ................................................t...a..........g................ [ 84080]
KX354450.1_47/Ss/2008            ................................................t...a.........ag................ [ 84080]
KP055815.1_strain_BA71           ................................................t...a..........g................ [ 84080]
MH025920.1_strain_R35            ..at.............c.................g............t...at.........g.c..c..c........ [ 84080]
MH025919.1_strain_N10            ..at.............c.................g............t...at.........g.c..c..c........ [ 84080]
MH025918.1_strain_R25            ..at.............c.................g............t...at.........g.c..c..c........ [ 84080]
MH025917.1_strain_R7             ..at.............c.................g............t...at.........g.c..c..c........ [ 84080]
MH025916.1_strain_R8             ..at.............c.................g............t...at.........g.c..c..c........ [ 84080]

FR682468.2_Georgia_2007/1        caaaggtctgcgaaaaaaagtagaattcacgtacaatgctcaaattgttacgaatgcttggcttaaaatgtatgagctgc [ 84160]
ASF_IND_20_CAD543                ................................................................................ [ 84160]
LR899193.1_Germany_2020/1        ................................................................................ [ 84160]
MN393477.1_Wuhan_2019            ................................................................................ [ 84160]
MW396979.1_Timor-Leste/2019/1    ................................................................................ [ 84160]
MK940252.1_CN/2019/InnerMongolia ................................................................................ [ 84160]
MK128995.1_China/2018/AnhuiXCGQ  ................................................................................ [ 84160]
MH681419.1_ASFV/POL/2015         ................................................................................ [ 84160]
LS478113.1_Estonia_2014          ................................................................................ [ 84160]
KP843857.1_Odintsovo_02/14       ................................................................................ [ 84160]
FN557520.1_strain_E75            ...........................t.................................................... [ 84160]
AY261366.1_Warthog               ......c....................t.................................................... [ 84160]
AY261365.1_Warmbaths             ...........................t...............................................a.... [ 84160]
AY261364.1_Tengani_62            ...........................t.................................................... [ 84160]
AY261363.1_Pretoriuskop/96/4     ................................................................................ [ 84160]
AY261362.1_Mkuzi_1979            ...........................t.................................................... [ 84160]
AY261361.1_Malawi_Lil-20/1_1983  ......a......................................................................... [ 84160]
AY261360.1_Kenya_1950            ......a..............g....a..................................................... [ 84160]
AM712240.1_OURT_88/3             ...........................t.................................................... [ 84160]
AM712239.1_Benin_97/1            ...........................t.................................................... [ 84160]
KM111295.1_Ken06.Bus             ......a..............g.....t..a................................................. [ 84160]
KM111294.1_Ken05/Tk1             ......a..............g.....t.......................a............................ [ 84160]
KM262844.1_strain_L60            ...........................t.................................................... [ 84160]
KM102979.1_26544/OG10_Italy      ...........................t.................................................... [ 84160]
KM262845.1_strain_NHV            ...........................t.................................................... [ 84160]
KX354450.1_47/Ss/2008            ...........................t.................................................... [ 84160]
KP055815.1_strain_BA71           ...........................t.................................................... [ 84160]
MH025920.1_strain_R35            ......a..............g.....t.................................................... [ 84160]
MH025919.1_strain_N10            ......a..............g.....t.................................................... [ 84160]
MH025918.1_strain_R25            ......a..............g.....t.................................................... [ 84160]
MH025917.1_strain_R7             ......a..............g.....t.................................................... [ 84160]
MH025916.1_strain_R8             ......a..............g.....t.................................................... [ 84160]
```

```
FR682468.2_Georgia_2007/1        taaataccatgaattttaataatacatctcaggcattttgcaattgtgagcttccaggagggtttataagtgcaattaac [ 84240]
ASF_IND_20_CAD543                ...............................................................................y [ 84240]
LR899193.1_Germany_2020/1        ................................................................................ [ 84240]
MN393477.1_Wuhan_2019            ................................................................................ [ 84240]
MW396979.1_Timor-Leste/2019/1    ................................................................................ [ 84240]
MK940252.1_CN/2019/InnerMongolia ................................................................................ [ 84240]
MK128995.1_China/2018/AnhuiXCGQ  ................................................................................ [ 84240]
MH681419.1_ASFV/POL/2015         ................................................................................ [ 84240]
LS478113.1_Estonia_2014          ................................................................................ [ 84240]
KP843857.1_Odintsovo_02/14       ................................................................................ [ 84240]
FN557520.1_strain_E75            ...............................................................................t [ 84240]
AY261366.1_Warthog               .............................................................a.................. [ 84240]
AY261365.1_Warmbaths             ................................................................................ [ 84240]
AY261364.1_Tengani_62            ................................................................................ [ 84240]
AY261363.1_Pretoriuskop/96/4     ................................................................................ [ 84240]
AY261362.1_Mkuzi_1979            ................................................................................ [ 84240]
AY261361.1_Malawi_Lil-20/1_1983  .......................................................g.....a.................. [ 84240]
AY261360.1_Kenya_1950            ......................................................................c......... [ 84240]
AM712240.1_OURT_88/3             ...............................................................................t [ 84240]
AM712239.1_Benin_97/1            ...............................................................................t [ 84240]
KM111295.1_Ken06.Bus             ......................................................................c......... [ 84240]
KM111294.1_Ken05/Tk1             .......................................................g........................ [ 84240]
KM262844.1_strain_L60            ...............................................................................t [ 84240]
KM102979.1_26544/OG10_Italy      ...............................................................................t [ 84240]
KM262845.1_strain_NHV            ...............................................................................t [ 84240]
KX354450.1_47/Ss/2008            ...............................................................................t [ 84240]
KP055815.1_strain_BA71           ...............................................................................t [ 84240]
MH025920.1_strain_R35            ......................................................................c......... [ 84240]
MH025919.1_strain_N10            ......................................................................c......... [ 84240]
MH025918.1_strain_R25            ......................................................................c......... [ 84240]
MH025917.1_strain_R7             ......................................................................c......... [ 84240]
MH025916.1_strain_R8             ......................................................................c......... [ 84240]

FR682468.2_Georgia_2007/1        cattttaattatacaatgatgcattaccctacttttaactgggtagcttcctccctttaccccagttcggaaacagatgc [ 84320]
ASF_IND_20_CAD543                ...............................................y................................ [ 84320]
LR899193.1_Germany_2020/1        ................................................................................ [ 84320]
MN393477.1_Wuhan_2019            ................................................................................ [ 84320]
MW396979.1_Timor-Leste/2019/1    ................................................................................ [ 84320]
MK940252.1_CN/2019/InnerMongolia ................................................................................ [ 84320]
MK128995.1_China/2018/AnhuiXCGQ  ................................................................................ [ 84320]
MH681419.1_ASFV/POL/2015         ................................................................................ [ 84320]
LS478113.1_Estonia_2014          ................................................................................ [ 84320]
KP843857.1_Odintsovo_02/14       ................................................................................ [ 84320]
```

```
FN557520.1_strain_E75           ...............................................c................................ [ 84320]
AY261366.1_Warthog              ...............................................c................................ [ 84320]
AY261365.1_Warmbaths            ...............................................c................................ [ 84320]
AY261364.1_Tengani_62           ...............................................c................................ [ 84320]
AY261363.1_Pretoriuskop/96/4    ...............................................c................................ [ 84320]
AY261362.1_Mkuzi_1979           ......................................t........c................................ [ 84320]
AY261361.1_Malawi_Lil-20/1_1983 .........c.....................gc..............c................................ [ 84320]
AY261360.1_Kenya_1950           .........c......................c..............c....................a........... [ 84320]
AM712240.1_OURT_88/3            ...............................................c................................ [ 84320]
AM712239.1_Benin_97/1           ...............................................c................................ [ 84320]
KM111295.1_Ken06.Bus            .........c......................c..............c....................a........... [ 84320]
KM111294.1_Ken05/Tk1            .........c.....................................c................................ [ 84320]
KM262844.1_strain_L60           ...............................................c................................ [ 84320]
KM102979.1_26544/OG10_Italy     ...............................................c................................ [ 84320]
KM262845.1_strain_NHV           ...............................................c................................ [ 84320]
KX354450.1_47/Ss/2008           ...............................................c................................ [ 84320]
KP055815.1_strain_BA71          ...............................................c................................ [ 84320]
MH025920.1_strain_R35           .........c......................c..............c....................a........... [ 84320]
MH025919.1_strain_N10           .........c......................c..............c....................a........... [ 84320]
MH025918.1_strain_R25           .........c......................c..............c....................a........... [ 84320]
MH025917.1_strain_R7            .........c......................c..............c....................a........... [ 84320]
MH025916.1_strain_R8            .........c......................c..............c....................a........... [ 84320]

FR682468.2_Georgia_2007/1       cctggaagatcactatggtctttatcagtgcaatccggataactggttgatgcaatctcctttactgaaaaaaaatatag [ 84400]
ASF_IND_20_CAD543               ..................................................................r.........r.r. [ 84400]
LR899193.1_Germany_2020/1       ................................................................................ [ 84400]
MN393477.1_Wuhan_2019           ................................................................................ [ 84400]
MW396979.1_Timor-Leste/2019/1   ................................................................................ [ 84400]
MK940252.1_CN/2019/InnerMongolia ................................................................................ [ 84400]
MK128995.1_China/2018/AnhuiXCGQ ................................................................................ [ 84400]
MH681419.1_ASFV/POL/2015        ................................................................................ [ 84400]
LS478113.1_Estonia_2014         ................................................................................ [ 84400]
KP843857.1_Odintsovo_02/14      ................................................................................ [ 84400]
FN557520.1_strain_E75           .........c....................................c...................a.........g.g. [ 84400]
AY261366.1_Warthog              .........c....................................c...............................g. [ 84400]
AY261365.1_Warmbaths            .t.......c....................................................................g. [ 84400]
AY261364.1_Tengani_62           t..a........................................................a.................g. [ 84400]
AY261363.1_Pretoriuskop/96/4    .........c....................................................................g. [ 84400]
AY261362.1_Mkuzi_1979           ..............................................c...............................g. [ 84400]
AY261361.1_Malawi_Lil-20/1_1983 t........c........a...........................c.c.............................g. [ 84400]
AY261360.1_Kenya_1950           .t.......c....................................c.c.....g.......................g. [ 84400]
AM712240.1_OURT_88/3            .........c....................................c...................a.........g.g. [ 84400]
AM712239.1_Benin_97/1           .........c....................................c...................a.........g.g. [ 84400]
KM111295.1_Ken06.Bus            .t.......c....................................c.c.....g.......................g. [ 84400]
```

```
KM111294.1_Ken05/Tk1             t........c....................................c.c.................a...........g. [ 84400]
KM262844.1_strain_L60            .........c....................................c...................a.........g.g. [ 84400]
KM102979.1_26544/OG10_Italy      .........c....................................c...................a.........g.g. [ 84400]
KM262845.1_strain_NHV            .........c....................................c...................a.........g.g. [ 84400]
KX354450.1_47/Ss/2008            .........c....................................c...................a.........g.g. [ 84400]
KP055815.1_strain_BA71           .........c....................................c...................a.........g.g. [ 84400]
MH025920.1_strain_R35            .t.......c....................................c.c.....g.......................g. [ 84400]
MH025919.1_strain_N10            .t.......c....................................c.c.....g.......................g. [ 84400]
MH025918.1_strain_R25            .t.......c....................................c.c.....g.......................g. [ 84400]
MH025917.1_strain_R7             .t.......c....................................c.c.....g.......................g. [ 84400]
MH025916.1_strain_R8             .t.......c....................................c.c.....g.......................g. [ 84400]

FR682468.2_Georgia_2007/1        attataataacggggacgtaaccatcgctagcaatgtaaaaaacctagcgcttagagccacacaaaggctgacgcccatc [ 84480]
ASF_IND_20_CAD543                ................................................................................ [ 84480]
LR899193.1_Germany_2020/1        ................................................................................ [ 84480]
MN393477.1_Wuhan_2019            ................................................................................ [ 84480]
MW396979.1_Timor-Leste/2019/1    ................................................................................ [ 84480]
MK940252.1_CN/2019/InnerMongolia ................................................................................ [ 84480]
MK128995.1_China/2018/AnhuiXCGQ  ................................................................................ [ 84480]
MH681419.1_ASFV/POL/2015         ................................................................................ [ 84480]
LS478113.1_Estonia_2014          ................................................................................ [ 84480]
KP843857.1_Odintsovo_02/14       ................................................................................ [ 84480]
FN557520.1_strain_E75            ........g................t.................t....................g............... [ 84480]
AY261366.1_Warthog               ..............................................g.................g.....a..t...... [ 84480]
AY261365.1_Warmbaths             ...................g..........................g..a........t.....g............... [ 84480]
AY261364.1_Tengani_62            ...................g.............................a..............g........t.....t [ 84480]
AY261363.1_Pretoriuskop/96/4     ...................g..........................g.................g.....a..t...... [ 84480]
AY261362.1_Mkuzi_1979            ...................g.....t.................t..g.................g............... [ 84480]
AY261361.1_Malawi_Lil-20/1_1983  ..........t........g.............................a.....g..t.....g........t...... [ 84480]
AY261360.1_Kenya_1950            ..........t........g.....a.................t..g.......a.........g.a......t...... [ 84480]
AM712240.1_OURT_88/3             ........g................t.................t....................g............... [ 84480]
AM712239.1_Benin_97/1            ........g................t.................t....................g............... [ 84480]
KM111295.1_Ken06.Bus             ...................g..t..a.................t..g.......a.........g.a......t...... [ 84480]
KM111294.1_Ken05/Tk1             ...................g.....a......................................g.a.t....t...... [ 84480]
KM262844.1_strain_L60            ........g................t.................t....................g............... [ 84480]
KM102979.1_26544/OG10_Italy      ........g................t.................t....................g............... [ 84480]
KM262845.1_strain_NHV            ........g................t.................t....................g............... [ 84480]
KX354450.1_47/Ss/2008            ........g................t.................t....................g............... [ 84480]
KP055815.1_strain_BA71           ........g................t.................t....................g............... [ 84480]
MH025920.1_strain_R35            ...................g..t..a.................t..g.......a.........g.a......t...... [ 84480]
MH025919.1_strain_N10            ...................g..t..a.................t..g.......a.........g.a......t...... [ 84480]
MH025918.1_strain_R25            ...................g..t..a.................t..g.......a.........g.a......t...... [ 84480]
MH025917.1_strain_R7             ...................g..t..a.................t..g.......a.........g.a......t...... [ 84480]
MH025916.1_strain_R8             ...................g..t..a.................t..g.......a.........g.a......t...... [ 84480]
```

```
FR682468.2_Georgia_2007/1        catctatatacggctgatgggggtattaatgtaggacatgactacaataaacaggaagaattaaatcttaagcttcactt [ 84560]
ASF_IND_20_CAD543                ...................................r............................................ [ 84560]
LR899193.1_Germany_2020/1        ................................................................................ [ 84560]
MN393477.1_Wuhan_2019            ................................................................................ [ 84560]
MW396979.1_Timor-Leste/2019/1    ................................................................................ [ 84560]
MK940252.1_CN/2019/InnerMongolia ................................................................................ [ 84560]
MK128995.1_China/2018/AnhuiXCGQ  ................................................................................ [ 84560]
MH681419.1_ASFV/POL/2015         ................................................................................ [ 84560]
LS478113.1_Estonia_2014          ................................................................................ [ 84560]
KP843857.1_Odintsovo_02/14       ................................................................................ [ 84560]
FN557520.1_strain_E75            ...........c.....c..a..............g............................................ [ 84560]
AY261366.1_Warthog               ....................a................................a.......................t.. [ 84560]
AY261365.1_Warmbaths             .....................................................a.......................... [ 84560]
AY261364.1_Tengani_62            .....................................................a.....g.................t.. [ 84560]
AY261363.1_Pretoriuskop/96/4     ...........c........a................................a.......................... [ 84560]
AY261362.1_Mkuzi_1979            ................................................................................ [ 84560]
AY261361.1_Malawi_Lil-20/1_1983  ........c..c........a................................a.....g.................... [ 84560]
AY261360.1_Kenya_1950            ....................a................................a.....g.................... [ 84560]
AM712240.1_OURT_88/3             ...........c.....c..a..............g............................................ [ 84560]
AM712239.1_Benin_97/1            ...........c.....c..a..............g............................................ [ 84560]
KM111295.1_Ken06.Bus             ....................a.......................t........a.....g.................... [ 84560]
KM111294.1_Ken05/Tk1             ....................a................................a.....g.................... [ 84560]
KM262844.1_strain_L60            ...........c.....c..a..............g............................................ [ 84560]
KM102979.1_26544/OG10_Italy      ...........c.....c..a..............g............................................ [ 84560]
KM262845.1_strain_NHV            ...........c.....c..a..............g............................................ [ 84560]
KX354450.1_47/Ss/2008            ...........c.....c..a..............g............................................ [ 84560]
KP055815.1_strain_BA71           ...........c.....c..a..............g............................................ [ 84560]
MH025920.1_strain_R35            ....................a.......................t........a.....g.................... [ 84560]
MH025919.1_strain_N10            ....................a.......................t........a.....g.................... [ 84560]
MH025918.1_strain_R25            ....................a.......................t........a.....g.................... [ 84560]
MH025917.1_strain_R7             ....................a.......................t........a.....g.................... [ 84560]
MH025916.1_strain_R8             ....................a.......................t........a.....g.................... [ 84560]

FR682468.2_Georgia_2007/1        tggtcaagcccttacgggtttgttgagtcttagcaaaggcggaaacatgatactcaaacactataccttaaatcatgcat [ 84640]
ASF_IND_20_CAD543                ................................................................................ [ 84640]
LR899193.1_Germany_2020/1        ................................................................................ [ 84640]
MN393477.1_Wuhan_2019            ................................................................................ [ 84640]
MW396979.1_Timor-Leste/2019/1    ................................................................................ [ 84640]
MK940252.1_CN/2019/InnerMongolia ................................................................................ [ 84640]
MK128995.1_China/2018/AnhuiXCGQ  ................................................................................ [ 84640]
MH681419.1_ASFV/POL/2015         ................................................................................ [ 84640]
LS478113.1_Estonia_2014          ................................................................................ [ 84640]
KP843857.1_Odintsovo_02/14       ................................................................................ [ 84640]
```

```
FN557520.1_strain_E75            .........t.....a...........c.................................................... [ 84640]
AY261366.1_Warthog               .........t.....a................................................................ [ 84640]
AY261365.1_Warmbaths             .........t.....a................................................................ [ 84640]
AY261364.1_Tengani_62            .........t.....a...a............................................................ [ 84640]
AY261363.1_Pretoriuskop/96/4     .........t.....a..a............................................................. [ 84640]
AY261362.1_Mkuzi_1979            .........t.....a................................................................ [ 84640]
AY261361.1_Malawi_Lil-20/1_1983  c........t.................c.................................................... [ 84640]
AY261360.1_Kenya_1950            c........t..........................g........................................... [ 84640]
AM712240.1_OURT_88/3             .........t.....a...........c.................................................... [ 84640]
AM712239.1_Benin_97/1            .........t.....a...........c.................................................... [ 84640]
KM111295.1_Ken06.Bus             c........t..........................g........................................... [ 84640]
KM111294.1_Ken05/Tk1             c........t.................c...........t..g..................................... [ 84640]
KM262844.1_strain_L60            .........t.....a...........c.................................................... [ 84640]
KM102979.1_26544/OG10_Italy      .........t.....a...........c.................................................... [ 84640]
KM262845.1_strain_NHV            .........t.....a...........c.................................................... [ 84640]
KX354450.1_47/Ss/2008            .........t.....a...........c.................................................... [ 84640]
KP055815.1_strain_BA71           .........t.....a...........c.................................................... [ 84640]
MH025920.1_strain_R35            c........t..........................g........................................... [ 84640]
MH025919.1_strain_N10            c........t..........................g........................................... [ 84640]
MH025918.1_strain_R25            c........t..........................g........................................... [ 84640]
MH025917.1_strain_R7             c........t..........................g........................................... [ 84640]
MH025916.1_strain_R8             c........t..........................g........................................... [ 84640]

FR682468.2_Georgia_2007/1        ttactctttctttaatatgtgtattttctcacttttttgaggaactatacattaccaaacctacctcctctcggcccaca [ 84720]
ASF_IND_20_CAD543                ................................................................................ [ 84720]
LR899193.1_Germany_2020/1        ................................................................................ [ 84720]
MN393477.1_Wuhan_2019            ................................................................................ [ 84720]
MW396979.1_Timor-Leste/2019/1    ................................................................................ [ 84720]
MK940252.1_CN/2019/InnerMongolia ................................................................................ [ 84720]
MK128995.1_China/2018/AnhuiXCGQ  ................................................................................ [ 84720]
MH681419.1_ASFV/POL/2015         ................................................................................ [ 84720]
LS478113.1_Estonia_2014          ................................................................................ [ 84720]
KP843857.1_Odintsovo_02/14       ................................................................................ [ 84720]
FN557520.1_strain_E75            ...................c...........................................................g [ 84720]
AY261366.1_Warthog               ...................c...........................................................g [ 84720]
AY261365.1_Warmbaths             .......................................................................a.......g [ 84720]
AY261364.1_Tengani_62            ...............................................................................g [ 84720]
AY261363.1_Pretoriuskop/96/4     ...................c...........................................................g [ 84720]
AY261362.1_Mkuzi_1979            ...............................................................................g [ 84720]
AY261361.1_Malawi_Lil-20/1_1983  ...................c...........................................................g [ 84720]
AY261360.1_Kenya_1950            ...................c...............................................t.....a...t.g [ 84720]
AM712240.1_OURT_88/3             ...................c...........................................................g [ 84720]
AM712239.1_Benin_97/1            ...................c...........................................................g [ 84720]
KM111295.1_Ken06.Bus             ...................c...............................................t.....a...t.g [ 84720]
```

```
KM111294.1_Ken05/Tk1            ...................c...........................................................g [ 84720]
KM262844.1_strain_L60           ...................c...........................................................g [ 84720]
KM102979.1_26544/OG10_Italy     ...................c...........................................................g [ 84720]
KM262845.1_strain_NHV           ...................c...........................................................g [ 84720]
KX354450.1_47/Ss/2008           ...................c...........................................................g [ 84720]
KP055815.1_strain_BA71          ...................c...........................................................g [ 84720]
MH025920.1_strain_R35           ...................c...............................................t.....a...t.g [ 84720]
MH025919.1_strain_N10           ...................c...............................................t.....a...t.g [ 84720]
MH025918.1_strain_R25           ...................c...............................................t.....a...t.g [ 84720]
MH025917.1_strain_R7            ...................c...............................................t.....a...t.g [ 84720]
MH025916.1_strain_R8            ...................c...............................................t.....a...t.g [ 84720]

FR682468.2_Georgia_2007/1       aactctgaaacctatattgtgggtaaaaacagattacgcttatttacccccaaggaagaacaagtccttctaaaacggct [ 84800]
ASF_IND_20_CAD543               ...............................................y...............r.m...y.r........ [ 84800]
LR899193.1_Germany_2020/1       ................................................................................ [ 84800]
MN393477.1_Wuhan_2019           ................................................................................ [ 84800]
MW396979.1_Timor-Leste/2019/1   ................................................................................ [ 84800]
MK940252.1_CN/2019/InnerMongolia ................................................................................ [ 84800]
MK128995.1_China/2018/AnhuiXCGQ ................................................................................ [ 84800]
MH681419.1_ASFV/POL/2015        ................................................................................ [ 84800]
LS478113.1_Estonia_2014         ................................................................................ [ 84800]
KP843857.1_Odintsovo_02/14      ................................................................................ [ 84800]
FN557520.1_strain_E75           ................................gc.....c.......t...............a.a...t.g.....a.. [ 84800]
AY261366.1_Warthog              .................................................................t..c........... [ 84800]
AY261365.1_Warmbaths            .................................................................t..c..g.....a.. [ 84800]
AY261364.1_Tengani_62           ......................................t.............................c..g........ [ 84800]
AY261363.1_Pretoriuskop/96/4    .......................................c.........................t..c..g........ [ 84800]
AY261362.1_Mkuzi_1979           .............................t.......................................t.......... [ 84800]
AY261361.1_Malawi_Lil-20/1_1983 ...........a...........................c.......t.....................t.......... [ 84800]
AY261360.1_Kenya_1950           ...........a.................t.........c.......t....................gt.......... [ 84800]
AM712240.1_OURT_88/3            ................................gc.....c.......t...............a.a...t.g.....a.. [ 84800]
AM712239.1_Benin_97/1           ................................gc.....c.......t...............a.a...t.g.....a.. [ 84800]
KM111295.1_Ken06.Bus            ...........a.................t.........c............................g........... [ 84800]
KM111294.1_Ken05/Tk1            ...........a...........a.....t.........c.......t....................g........... [ 84800]
KM262844.1_strain_L60           ................................gc.....c.......t...............a.a...t.g.....a.. [ 84800]
KM102979.1_26544/OG10_Italy     ................................gc.....c.......t...............a.a...t.g.....a.. [ 84800]
KM262845.1_strain_NHV           ................................gc.....c.......t...............a.a...t.g.....a.. [ 84800]
KX354450.1_47/Ss/2008           ................................gc.....c.......t...............a.a...t.g.....a.. [ 84800]
KP055815.1_strain_BA71          ................................gc.....c.......t...............a.a...t.g.....a.. [ 84800]
MH025920.1_strain_R35           ...........a.................t.........c............................g........... [ 84800]
MH025919.1_strain_N10           ...........a.................t.........c............................g........... [ 84800]
MH025918.1_strain_R25           ...........a.................t.........c............................g........... [ 84800]
MH025917.1_strain_R7            ...........a.................t.........c............................g........... [ 84800]
MH025916.1_strain_R8            ...........a.................t.........c............................g........... [ 84800]
```

```
FR682468.2_Georgia_2007/1        agaatttttaatgatacgcccctcgtagacctaagtctttaccaaaatttacttgaaagcgtttactttgccgtagaaa [ 84880]
ASF_IND_20_CAD543                .............................................................r.................. [ 84880]
LR899193.1_Germany_2020/1        ................................................................................ [ 84880]
MN393477.1_Wuhan_2019            ................................................................................ [ 84880]
MW396979.1_Timor-Leste/2019/1    ................................................................................ [ 84880]
MK940252.1_CN/2019/InnerMongolia ................................................................................ [ 84880]
MK128995.1_China/2018/AnhuiXCGQ  ................................................................................ [ 84880]
MH681419.1_ASFV/POL/2015         ................................................................................ [ 84880]
LS478113.1_Estonia_2014          ................................................................................ [ 84880]
KP843857.1_Odintsovo_02/14       ................................................................................ [ 84880]
FN557520.1_strain_E75            .............................................................a.................. [ 84880]
AY261366.1_Warthog               ................................................................................ [ 84880]
AY261365.1_Warmbaths             ......c......................................................................... [ 84880]
AY261364.1_Tengani_62            ........................t....................................a.................. [ 84880]
AY261363.1_Pretoriuskop/96/4     ................................................................................ [ 84880]
AY261362.1_Mkuzi_1979            .............................................................a.................. [ 84880]
AY261361.1_Malawi_Lil-20/1_1983  .....................t................................a......................... [ 84880]
AY261360.1_Kenya_1950            ........................a....................................a.................. [ 84880]
AM712240.1_OURT_88/3             .............................................................a.................. [ 84880]
AM712239.1_Benin_97/1            .............................................................a.................. [ 84880]
KM111295.1_Ken06.Bus             ........................a..........a.........................a.................. [ 84880]
KM111294.1_Ken05/Tk1             ........................a..........a............................................ [ 84880]
KM262844.1_strain_L60            .............................................................a.................. [ 84880]
KM102979.1_26544/OG10_Italy      .............................................................a.................. [ 84880]
KM262845.1_strain_NHV            .............................................................a.................. [ 84880]
KX354450.1_47/Ss/2008            .............................................................a.................. [ 84880]
KP055815.1_strain_BA71           .............................................................a.................. [ 84880]
MH025920.1_strain_R35            ........................a..........a.........................a.................. [ 84880]
MH025919.1_strain_N10            ........................a..........a.........................a.................. [ 84880]
MH025918.1_strain_R25            ........................a..........a.........................a.................. [ 84880]
MH025917.1_strain_R7             ........................a..........a.........................a.................. [ 84880]
MH025916.1_strain_R8             ........................a..........a.........................a.................. [ 84880]

FR682468.2_Georgia_2007/1        caatacatctaaaacaacaaatagaatttctaaacttcggaatgaaatgttatcgacatttttataacaagattaaacta [ 84960]
ASF_IND_20_CAD543                ................r........r...................................................... [ 84960]
LR899193.1_Germany_2020/1        ................................................................................ [ 84960]
MN393477.1_Wuhan_2019            ................................................................................ [ 84960]
MW396979.1_Timor-Leste/2019/1    ................................................................................ [ 84960]
MK940252.1_CN/2019/InnerMongolia ................................................................................ [ 84960]
MK128995.1_China/2018/AnhuiXCGQ  ................................................................................ [ 84960]
MH681419.1_ASFV/POL/2015         ................................................................................ [ 84960]
LS478113.1_Estonia_2014          ................................................................................ [ 84960]
KP843857.1_Odintsovo_02/14       ................................................................................ [ 84960]
```

```
FN557520.1_strain_E75            ................g........g..........................c........................... [ 84960]
AY261366.1_Warthog               ....................................................c........................... [ 84960]
AY261365.1_Warmbaths             ................g........g..........................c........................... [ 84960]
AY261364.1_Tengani_62            .........................g...t......................c........................... [ 84960]
AY261363.1_Pretoriuskop/96/4     ....................................................c........................... [ 84960]
AY261362.1_Mkuzi_1979            ................g........g...................................................... [ 84960]
AY261361.1_Malawi_Lil-20/1_1983  .........................g..c................................................... [ 84960]
AY261360.1_Kenya_1950            .........................g...........t..............c........................... [ 84960]
AM712240.1_OURT_88/3             ................g........g..........................c........................... [ 84960]
AM712239.1_Benin_97/1            ................g........g..........................c........................... [ 84960]
KM111295.1_Ken06.Bus             .........................g...........t..............c........................... [ 84960]
KM111294.1_Ken05/Tk1             .........................g...........t..............c........................... [ 84960]
KM262844.1_strain_L60            ................g........g..........................c........................... [ 84960]
KM102979.1_26544/OG10_Italy      ................g........g..........................c........................... [ 84960]
KM262845.1_strain_NHV            ................g........g..........................c........................... [ 84960]
KX354450.1_47/Ss/2008            ................g........g..........................c........................... [ 84960]
KP055815.1_strain_BA71           ................g........g..........................c........................... [ 84960]
MH025920.1_strain_R35            .........................g...........t..............c........................... [ 84960]
MH025919.1_strain_N10            .........................g...........t..............c........................... [ 84960]
MH025918.1_strain_R25            .........................g...........t..............c........................... [ 84960]
MH025917.1_strain_R7             .........................g...........t..............c........................... [ 84960]
MH025916.1_strain_R8             .........................g...........t..............c........................... [ 84960]

FR682468.2_Georgia_2007/1        cttaacgattatttagctccgaaaaaaaagatttttcaggataggtggcgtgtgcttaataagctttatgttcttgaaaa [ 85040]
ASF_IND_20_CAD543                ................................................................................ [ 85040]
LR899193.1_Germany_2020/1        ................................................................................ [ 85040]
MN393477.1_Wuhan_2019            ................................................................................ [ 85040]
MW396979.1_Timor-Leste/2019/1    ................................................................................ [ 85040]
MK940252.1_CN/2019/InnerMongolia ................................................................................ [ 85040]
MK128995.1_China/2018/AnhuiXCGQ  ................................................................................ [ 85040]
MH681419.1_ASFV/POL/2015         ................................................................................ [ 85040]
LS478113.1_Estonia_2014          ................................................................................ [ 85040]
KP843857.1_Odintsovo_02/14       ................................................................................ [ 85040]
FN557520.1_strain_E75            ........a....................................................................... [ 85040]
AY261366.1_Warthog               .....t.......................................................................... [ 85040]
AY261365.1_Warmbaths             .....t.......................................................................... [ 85040]
AY261364.1_Tengani_62            .................c............................................a................. [ 85040]
AY261363.1_Pretoriuskop/96/4     ................................................................................ [ 85040]
AY261362.1_Mkuzi_1979            ................................................................................ [ 85040]
AY261361.1_Malawi_Lil-20/1_1983  ........a.............................a......................................... [ 85040]
AY261360.1_Kenya_1950            .....t..a........................................................c.............. [ 85040]
AM712240.1_OURT_88/3             ........a....................................................................... [ 85040]
AM712239.1_Benin_97/1            ........a....................................................................... [ 85040]
KM111295.1_Ken06.Bus             ........a...............................................c........c.............. [ 85040]
```

```
KM111294.1_Ken05/Tk1             ........a................................................c.......c.............. [ 85040]
KM262844.1_strain_L60            ........a....................................................................... [ 85040]
KM102979.1_26544/OG10_Italy      ........a....................................................................... [ 85040]
KM262845.1_strain_NHV            ........a....................................................................... [ 85040]
KX354450.1_47/Ss/2008            ........a....................................................................... [ 85040]
KP055815.1_strain_BA71           ........a....................................................................... [ 85040]
MH025920.1_strain_R35            ........a................................................c.......c.............. [ 85040]
MH025919.1_strain_N10            ........a................................................c.......c.............. [ 85040]
MH025918.1_strain_R25            ........a................................................c.......c.............. [ 85040]
MH025917.1_strain_R7             ........a................................................c.......c.............. [ 85040]
MH025916.1_strain_R8             ........a................................................c.......c.............. [ 85040]

FR682468.2_Georgia_2007/1        aaagcataaacttaagctttg---tgcctcctagggatctgttgcttaa---------------tttaacagatgcaatc [ 85120]
ASF_IND_20_CAD543                .....................---....c.tc.............c...---------------c............... [ 85120]
LR899193.1_Germany_2020/1        .....................---.........................---------------................ [ 85120]
MN393477.1_Wuhan_2019            .....................---.........................---------------................ [ 85120]
MW396979.1_Timor-Leste/2019/1    .....................---.........................---------------................ [ 85120]
MK940252.1_CN/2019/InnerMongolia .....................---.........................---------------................ [ 85120]
MK128995.1_China/2018/AnhuiXCGQ  .....................---.........................---------------................ [ 85120]
MH681419.1_ASFV/POL/2015         .....................---.........................---------------................ [ 85120]
LS478113.1_Estonia_2014          .....................---.........................---------------................ [ 85120]
KP843857.1_Odintsovo_02/14       .....................---.........................---------------................ [ 85120]
FN557520.1_strain_E75            .....................---....c.tc.............c...cttaacagatgcaatc............... [ 85120]
AY261366.1_Warthog               .....................tgc.......c.................---------------...........t..a. [ 85120]
AY261365.1_Warmbaths             .....................tgc.......c.............c...---------------c........c.t..a. [ 85120]
AY261364.1_Tengani_62            .....................---.......c.............c...-----------------c......ca...a. [ 85120]
AY261363.1_Pretoriuskop/96/4     .....................tgc.......c.............c...---------------c........c.t..a- [ 85120]
AY261362.1_Mkuzi_1979            .....................---.......c.................---------------................ [ 85120]
AY261361.1_Malawi_Lil-20/1_1983  .....................tgc.c..cttc.a..g........c...---------------c.........------ [ 85120]
AY261360.1_Kenya_1950            ...a...........a.....tac.t...ttc....g........c...---------------.......a....g... [ 85120]
AM712240.1_OURT_88/3             .....................---....c.tc.............c...---------------c............... [ 85120]
AM712239.1_Benin_97/1            .....................---....c.tc.............c...---------------c............... [ 85120]
KM111295.1_Ken06.Bus             .....................tac.t...ttc....g.......tc...------------------------------. [ 85120]
KM111294.1_Ken05/Tk1             .....................tac.t...ttc....g........c...---------------.......a....g... [ 85120]
KM262844.1_strain_L60            .....................---....c.tc.............c...---------------c............... [ 85120]
KM102979.1_26544/OG10_Italy      .....................---....c.tc.............c...---------------c............... [ 85120]
KM262845.1_strain_NHV            .....................---....c.tc.............c...---------------c............... [ 85120]
KX354450.1_47/Ss/2008            .....................---....c.tc.............c...---------------c............... [ 85120]
KP055815.1_strain_BA71           .....................---....c.tc.............c...---------------c............... [ 85120]
MH025920.1_strain_R35            .....................tac.t...ttc....g.......tc...------------------------------. [ 85120]
MH025919.1_strain_N10            .....................tac.t...ttc....g.......tc...------------------------------. [ 85120]
MH025918.1_strain_R25            .....................tac.t...ttc....g.......tc...------------------------------. [ 85120]
MH025917.1_strain_R7             .....................tac.t...ttc....g.......tc...------------------------------. [ 85120]
MH025916.1_strain_R8             .....................tac.t...ttc....g.......tc...------------------------------. [ 85120]
```

```
FR682468.2_Georgia_2007/1        ttaac------------------------------agatgtaaactaaaaagtgtgttcatacaaggattgtatttatg- [ 85200]
ASF_IND_20_CAD543                .....agatg---------------caatcttaac............................................- [ 85200]
LR899193.1_Germany_2020/1        .....------------------------------............................................- [ 85200]
MN393477.1_Wuhan_2019            .....------------------------------............................................- [ 85200]
MW396979.1_Timor-Leste/2019/1    .....------------------------------............................................- [ 85200]
MK940252.1_CN/2019/InnerMongolia .....------------------------------............................................- [ 85200]
MK128995.1_China/2018/AnhuiXCGQ  .....------------------------------............................................- [ 85200]
MH681419.1_ASFV/POL/2015         .....------------------------------............................................- [ 85200]
LS478113.1_Estonia_2014          .....------------------------------............................................- [ 85200]
KP843857.1_Odintsovo_02/14       .....------------------------------............................................- [ 85200]
FN557520.1_strain_E75            .....agatg---------------caatcttaac...c........................................- [ 85200]
AY261366.1_Warthog               .....agaca---------------caatcttaac.....c......................................- [ 85200]
AY261365.1_Warmbaths             ....tagacataaacttaatagacgcaatcttaac.....c......................................- [ 85200]
AY261364.1_Tengani_62            .....------------------------------.....c..t...................................- [ 85200]
AY261363.1_Pretoriuskop/96/4     -----------------------------cttaac.....c......................................- [ 85200]
AY261362.1_Mkuzi_1979            .....agatg---------------taatcttaac............................................- [ 85200]
AY261361.1_Malawi_Lil-20/1_1983  -----------------------tgcaaacttaac.....c......................................- [ 85200]
AY261360.1_Kenya_1950            .....aaatg---------------------tggcca.aagct....................................a [ 85200]
AM712240.1_OURT_88/3             .....agatg---------------caatcttaac...c........................................- [ 85200]
AM712239.1_Benin_97/1            .....agatg---------------caatcttaac...c........................................- [ 85200]
KM111295.1_Ken06.Bus             .....agatg---------------------tggcct.aagct..................t.................- [ 85200]
KM111294.1_Ken05/Tk1             .....aaatg---------------------tggcca.aagct....................................- [ 85200]
KM262844.1_strain_L60            .....agatg---------------caatcttaac...c........................................- [ 85200]
KM102979.1_26544/OG10_Italy      .....agatg---------------caatcttaac...c........................................- [ 85200]
KM262845.1_strain_NHV            .....agatg---------------caatcttaac...c........................................- [ 85200]
KX354450.1_47/Ss/2008            .....agatg---------------caatcttaac...c........................................- [ 85200]
KP055815.1_strain_BA71           .....agatg---------------caatcttaac...c........................................- [ 85200]
MH025920.1_strain_R35            .....agatg---------------------tggcct.aagct..................t.................- [ 85200]
MH025919.1_strain_N10            .....agatg---------------------tggcct.aagct..................t.................- [ 85200]
MH025918.1_strain_R25            .....agatg---------------------tggcct.aagct..................t.................- [ 85200]
MH025917.1_strain_R7             .....agatg---------------------tggcct.aagct..................t.................- [ 85200]
MH025916.1_strain_R8             .....agatg---------------------tggcct.aagct..................t.................- [ 85200]

FR682468.2_Georgia_2007/1        ---------aatatttattaacatataaggttgtg--------atgtaacactgtataac-ctatataact--------- [ 85280]
ASF_IND_20_CAD543                ---------..........................atggtgc-.................-..........--------- [ 85280]
LR899193.1_Germany_2020/1        ---------..........................--------.................-..........--------- [ 85280]
MN393477.1_Wuhan_2019            ---------..........................--------.................-..........--------- [ 85280]
MW396979.1_Timor-Leste/2019/1    ---------..........................--------.................-..........--------- [ 85280]
MK940252.1_CN/2019/InnerMongolia ---------..........................--------.................-..........--------- [ 85280]
MK128995.1_China/2018/AnhuiXCGQ  ---------..........................--------.................-..........--------- [ 85280]
MH681419.1_ASFV/POL/2015         ---------..........................--------.................-..........--------- [ 85280]
LS478113.1_Estonia_2014          ---------..........................--------.................-..........--------- [ 85280]
KP843857.1_Odintsovo_02/14       ---------..........................--------.................-..........--------- [ 85280]
```

```
FN557520.1_strain_E75              ---------..........................atggtgc-.................-..........--------- [ 85280]
AY261366.1_Warthog                 ---------.............--...........atggtgc-.................-....--------------- [ 85280]
AY261365.1_Warmbaths               ---------.............--...........atggtgc-.................-....--------------- [ 85280]
AY261364.1_Tengani_62              ---------.............--...........atggtgc-g................-....--------------- [ 85280]
AY261363.1_Pretoriuskop/96/4       ---------.............--...........atggtgc-.................-...c--------------- [ 85280]
AY261362.1_Mkuzi_1979              ---------.............--...........atagtgc-g................-..........--------- [ 85280]
AY261361.1_Malawi_Lil-20/1_1983    ---------.............--...........atggtgctg.............gg.agc..g....---------- [ 85280]
AY261360.1_Kenya_1950              atatttatt.............--...........atggtgctg.......g...g.g.ta..g.g..taaccg----ta [ 85280]
AM712240.1_OURT_88/3               ---------..........................atggtgc-.................-..........--------- [ 85280]
AM712239.1_Benin_97/1              ---------..........................atggtgc-.................-..........--------- [ 85280]
KM111295.1_Ken06.Bus               ---------.............--.....a.....atagtgctg.......g........a..g......atagggttta [ 85280]
KM111294.1_Ken05/Tk1               ---------.............--...........atagtgctg.....t.g...g....a.cg.g.gtaacactgtata [ 85280]
KM262844.1_strain_L60              ---------..........................atggtgc-.................-..........--------- [ 85280]
KM102979.1_26544/OG10_Italy        ---------............t.............atggtgc-.................-..........--------- [ 85280]
KM262845.1_strain_NHV              ---------..........................atggtgc-.................-..........--------- [ 85280]
KX354450.1_47/Ss/2008              ---------............t.............atggtgc-.................-..........--------- [ 85280]
KP055815.1_strain_BA71             ---------..........................atggtgc-.................-..........--------- [ 85280]
MH025920.1_strain_R35              ---------.............--.....a.....atagtgctg.......g........a..g......atagggttta [ 85280]
MH025919.1_strain_N10              ---------.............--.....a.....atagtgctg.......g........a..g......atagggttta [ 85280]
MH025918.1_strain_R25              ---------.............--.....a.....atagtgctg.......g........a..g......atagggttta [ 85280]
MH025917.1_strain_R7               ---------.............--.....a.....atagtgctg.......g........a..g......atagggttta [ 85280]
MH025916.1_strain_R8               ---------.............--.....a.....atagtgctg.......g........a..g......atagggttta [ 85280]

FR682468.2_Georgia_2007/1          ------------acac--tatgaagcacggcgtataataatttatattgaacacgatgttgactcatttatttgcaaaca [ 85360]
ASF_IND_20_CAD543                  ------------....--.............................................................. [ 85360]
LR899193.1_Germany_2020/1          ------------....--.............................................................. [ 85360]
MN393477.1_Wuhan_2019              ------------....--.............................................................. [ 85360]
MW396979.1_Timor-Leste/2019/1      ------------....--.............................................................. [ 85360]
MK940252.1_CN/2019/InnerMongolia   ------------....--.............................................................. [ 85360]
MK128995.1_China/2018/AnhuiXCGQ    ------------....--.............................................................. [ 85360]
MH681419.1_ASFV/POL/2015           ------------....--.............................................................. [ 85360]
LS478113.1_Estonia_2014            ------------....--.............................................................. [ 85360]
KP843857.1_Odintsovo_02/14         ------------....--.............................................................. [ 85360]
FN557520.1_strain_E75              ------------....--...........a.................................................. [ 85360]
AY261366.1_Warthog                 --------------..--...........-----.............................................. [ 85360]
AY261365.1_Warmbaths               --------------..--.............a...........................c.................... [ 85360]
AY261364.1_Tengani_62              --------------..--.............a.................................g.............. [ 85360]
AY261363.1_Pretoriuskop/96/4       --------------..--...........a.a..............a..g...t.....c..................t. [ 85360]
AY261362.1_Mkuzi_1979              ------------....--...........a.................................................. [ 85360]
AY261361.1_Malawi_Lil-20/1_1983    --------------..--....tga......a..............a..g...t.....c....a............... [ 85360]
AY261360.1_Kenya_1950              taactgtgtata....--....tg.......a..............a..g..........a...a............... [ 85360]
AM712240.1_OURT_88/3               ------------....--...........a.................................................. [ 85360]
AM712239.1_Benin_97/1              ------------....--...........a.................................................. [ 85360]
KM111295.1_Ken06.Bus               acaatacgtatat...--....tg.......a.................g..g...........a............... [ 85360]
```

```
KM111294.1_Ken05/Tk1             acaacgtatata...tag.g.tt.a..atata.................g..g...........a............... [ 85360]
KM262844.1_strain_L60            ------------....--...........a.................................................. [ 85360]
KM102979.1_26544/OG10_Italy      ------------....--...........a.................................................. [ 85360]
KM262845.1_strain_NHV            ------------....--...........a.................................................. [ 85360]
KX354450.1_47/Ss/2008            ------------....--...........a.................................................. [ 85360]
KP055815.1_strain_BA71           ------------....--...........a.................................................. [ 85360]
MH025920.1_strain_R35            acaatacgtatat...--....tg.......a.................g..g...........a............... [ 85360]
MH025919.1_strain_N10            acaatacgtatat...--....tg.......a.................g..g...........a............... [ 85360]
MH025918.1_strain_R25            acaatacgtatat...--....tg.......a.................g..g...........a............... [ 85360]
MH025917.1_strain_R7             acaatacgtatat...--....tg.......a.................g..g...........a............... [ 85360]
MH025916.1_strain_R8             acaatacgtatat...--....tg.......a.................g..g...........a............... [ 85360]

FR682468.2_Georgia_2007/1        aatatttgtttgca------------------------------------------------------------------ [ 85440]
ASF_IND_20_CAD543                ..............------------------------------------------------------------------ [ 85440]
LR899193.1_Germany_2020/1        ..............------------------------------------------------------------------ [ 85440]
MN393477.1_Wuhan_2019            ..............------------------------------------------------------------------ [ 85440]
MW396979.1_Timor-Leste/2019/1    ..............------------------------------------------------------------------ [ 85440]
MK940252.1_CN/2019/InnerMongolia ..............------------------------------------------------------------------ [ 85440]
MK128995.1_China/2018/AnhuiXCGQ  ..............------------------------------------------------------------------ [ 85440]
MH681419.1_ASFV/POL/2015         ..............------------------------------------------------------------------ [ 85440]
LS478113.1_Estonia_2014          ..............------------------------------------------------------------------ [ 85440]
KP843857.1_Odintsovo_02/14       ..............------------------------------------------------------------------ [ 85440]
FN557520.1_strain_E75            ..............------------------------------------------------------------------ [ 85440]
AY261366.1_Warthog               ..............------------------------------------------------------------------ [ 85440]
AY261365.1_Warmbaths             ..............------------------------------------------------------------------ [ 85440]
AY261364.1_Tengani_62            ..............------------------------------------------------------------------ [ 85440]
AY261363.1_Pretoriuskop/96/4     ..............atcgcagatgtctgca-gcatttactaatatgttgtcaattaatttatttgca------------- [ 85440]
AY261362.1_Mkuzi_1979            ............t.------------------------------------------------------------------ [ 85440]
AY261361.1_Malawi_Lil-20/1_1983  ..............aatgcagacgtctgcatgcatttattaatatgttctcaactaatttatttgc-------------- [ 85440]
AY261360.1_Kenya_1950            ..............aacgcagacgtctacacgcatttactaatatgttgtcaactaatttatttacaaacaagatgtttg [ 85440]
AM712240.1_OURT_88/3             ..............------------------------------------------------------------------ [ 85440]
AM712239.1_Benin_97/1            ..............------------------------------------------------------------------ [ 85440]
KM111295.1_Ken06.Bus             ...g....--------------------cacgcatttactaatatgttgacaa--------------------------- [ 85440]
KM111294.1_Ken05/Tk1             ...g....--------------------catgcatttactaatatgttgttaactagtatgttgacaactagtata---- [ 85440]
KM262844.1_strain_L60            ..............------------------------------------------------------------------ [ 85440]
KM102979.1_26544/OG10_Italy      ..............------------------------------------------------------------------ [ 85440]
KM262845.1_strain_NHV            ..............------------------------------------------------------------------ [ 85440]
KX354450.1_47/Ss/2008            ..............------------------------------------------------------------------ [ 85440]
KP055815.1_strain_BA71           ..............------------------------------------------------------------------ [ 85440]
MH025920.1_strain_R35            ...g....--------------------cacgcatttactaatatgttgacaa--------------------------- [ 85440]
MH025919.1_strain_N10            ...g....--------------------cacgcatttactaatatgttgacaa--------------------------- [ 85440]
MH025918.1_strain_R25            ...g....--------------------cacgcatttactaatatgttgacaa--------------------------- [ 85440]
MH025917.1_strain_R7             ...g....--------------------cacgcatttactaatatgttgacaa--------------------------- [ 85440]
MH025916.1_strain_R8             ...g....--------------------cacgcatttactaatatgttgacaa--------------------------- [ 85440]
```

```
FR682468.2_Georgia_2007/1        -------------------------------------------agacgtttgcat-----gcatttactaatatgttgtt [ 85520]
ASF_IND_20_CAD543                -------------------------------------------............-----.................... [ 85520]
LR899193.1_Germany_2020/1        -------------------------------------------............-----.................... [ 85520]
MN393477.1_Wuhan_2019            -------------------------------------------............-----.................... [ 85520]
MW396979.1_Timor-Leste/2019/1    -------------------------------------------............-----.................... [ 85520]
MK940252.1_CN/2019/InnerMongolia -------------------------------------------............-----.................... [ 85520]
MK128995.1_China/2018/AnhuiXCGQ  -------------------------------------------............-----.................... [ 85520]
MH681419.1_ASFV/POL/2015         -------------------------------------------............-----.................... [ 85520]
LS478113.1_Estonia_2014          -------------------------------------------............-----.................... [ 85520]
KP843857.1_Odintsovo_02/14       -------------------------------------------............-----.................... [ 85520]
FN557520.1_strain_E75            -------------------------------------------............-----.................... [ 85520]
AY261366.1_Warthog               -------------------------------------------............-----.................... [ 85520]
AY261365.1_Warmbaths             -------------------------------------------............-----.................... [ 85520]
AY261364.1_Tengani_62            ------------------------------------------------------.-----.................... [ 85520]
AY261363.1_Pretoriuskop/96/4     --------------------------------------aacgc...t.......c-----.................... [ 85520]
AY261362.1_Mkuzi_1979            -------------------------------------------............-----.................... [ 85520]
AY261361.1_Malawi_Lil-20/1_1983  -------------------------------------------.a....c.a.g.ttgca.................... [ 85520]
AY261360.1_Kenya_1950            tttacaaacgtttacgtttgcaaacgcagacgtttgcaaacgc...........c-----.................a.. [ 85520]
AM712240.1_OURT_88/3             -------------------------------------------............-----.................... [ 85520]
AM712239.1_Benin_97/1            -------------------------------------------............-----.................... [ 85520]
KM111295.1_Ken06.Bus             -------------------------------------------------------------------..g........ac [ 85520]
KM111294.1_Ken05/Tk1             -----------------ttgttaactaatttatttgcaaacat.a.-.......c-----............g...c.ac [ 85520]
KM262844.1_strain_L60            -------------------------------------------............-----.................... [ 85520]
KM102979.1_26544/OG10_Italy      -------------------------------------------............-----.................... [ 85520]
KM262845.1_strain_NHV            -------------------------------------------............-----.................... [ 85520]
KX354450.1_47/Ss/2008            -------------------------------------------............-----.................... [ 85520]
KP055815.1_strain_BA71           -------------------------------------------............-----.................... [ 85520]
MH025920.1_strain_R35            -------------------------------------------------------------------..g........ac [ 85520]
MH025919.1_strain_N10            -------------------------------------------------------------------..g........ac [ 85520]
MH025918.1_strain_R25            -------------------------------------------------------------------..g........ac [ 85520]
MH025917.1_strain_R7             -------------------------------------------------------------------..g........ac [ 85520]
MH025916.1_strain_R8             -------------------------------------------------------------------..g........ac [ 85520]

FR682468.2_Georgia_2007/1        gactagtttatttgcaaac-tagatgtttgattgcaaactagatgtttgcacgtatttatttgaactaatatacactcct [ 85600]
ASF_IND_20_CAD543                .....r.............-..........-------------------..y............................ [ 85600]
LR899193.1_Germany_2020/1        ...................-............................................................ [ 85600]
MN393477.1_Wuhan_2019            ...................-............................................................ [ 85600]
MW396979.1_Timor-Leste/2019/1    ...................-............................................................ [ 85600]
MK940252.1_CN/2019/InnerMongolia ...................-............................................................ [ 85600]
MK128995.1_China/2018/AnhuiXCGQ  ...................-............................................................ [ 85600]
MH681419.1_ASFV/POL/2015         ...................-............................................................ [ 85600]
LS478113.1_Estonia_2014          ...................-............................................................ [ 85600]
KP843857.1_Odintsovo_02/14       ...................-............................................................ [ 85600]
```

```
FN557520.1_strain_E75          .....a.............-..........-------------------..t............................ [ 85600]
AY261366.1_Warthog             .....a.............-c.........-------------------............................... [ 85600]
AY261365.1_Warmbaths           .....a.............-c.........-------------------.............a................. [ 85600]
AY261364.1_Tengani_62          ...................-............................................................ [ 85600]
AY261363.1_Pretoriuskop/96/4   .....a.............-..........-------------------............................... [ 85600]
AY261362.1_Mkuzi_1979          ...................-..........-------------------..t..........a................. [ 85600]
AY261361.1_Malawi_Lil-20/1_1983 a....a...g...a.....-..........-------------------t.............................. [ 85600]
AY261360.1_Kenya_1950          ...................-..........-------------------............................... [ 85600]
AM712240.1_OURT_88/3           .....a.............-..........-------------------..t............................ [ 85600]
AM712239.1_Benin_97/1          .....a.............-..........-------------------..t......................t..... [ 85600]
KM111295.1_Ken06.Bus           a....a.............a.g........-------------------............................... [ 85600]
KM111294.1_Ken05/Tk1           a....a.............a.gat......-------------------............................... [ 85600]
KM262844.1_strain_L60          .....a.............-..........-------------------..t......................t..... [ 85600]
KM102979.1_26544/OG10_Italy    .....a.............-..........-------------------..t......................t..... [ 85600]
KM262845.1_strain_NHV          .....a.............-..........-------------------..t............................ [ 85600]
KX354450.1_47/Ss/2008          .....a.............-..........-------------------..t......................t..... [ 85600]
KP055815.1_strain_BA71         .....a.............-..........-------------------..t............................ [ 85600]
MH025920.1_strain_R35          a....a.............a.g........-------------------............................... [ 85600]
MH025919.1_strain_N10          a....a.............a.g........-------------------............................... [ 85600]
MH025918.1_strain_R25          a....a.............a.g........-------------------............................... [ 85600]
MH025917.1_strain_R7           a....a.............a.g........-------------------............................... [ 85600]
MH025916.1_strain_R8           a....a.............a.g........-------------------............................... [ 85600]

FR682468.2_Georgia_2007/1      tgttttatttgttatatacacagcatacataagtgtatattgtttacacttatgtttataactcgacgtaataacatttt [ 85680]
ASF_IND_20_CAD543              ..........r..................................................................... [ 85680]
LR899193.1_Germany_2020/1      ................................................................................ [ 85680]
MN393477.1_Wuhan_2019          ................................................................................ [ 85680]
MW396979.1_Timor-Leste/2019/1  ................................................................................ [ 85680]
MK940252.1_CN/2019/InnerMongolia ................................................................................ [ 85680]
MK128995.1_China/2018/AnhuiXCGQ ................................................................................ [ 85680]
MH681419.1_ASFV/POL/2015       ................................................................................ [ 85680]
LS478113.1_Estonia_2014        ................................................................................ [ 85680]
KP843857.1_Odintsovo_02/14     ................................................................................ [ 85680]
FN557520.1_strain_E75          ..........a..............................a...................................... [ 85680]
AY261366.1_Warthog             .......c.............g..................a..a.................................... [ 85680]
AY261365.1_Warmbaths           .......c.............g...................a...................................... [ 85680]
AY261364.1_Tengani_62          .......c...............t.................a...................................... [ 85680]
AY261363.1_Pretoriuskop/96/4   .......c..................g..............a...................................... [ 85680]
AY261362.1_Mkuzi_1979          .......................t.................a...................................... [ 85680]
AY261361.1_Malawi_Lil-20/1_1983 .......c................g.g..........c.c.-----------.a.g..c.t................... [ 85680]
AY261360.1_Kenya_1950          .......c..................g..........c.c.---------.....ac....................gc- [ 85680]
AM712240.1_OURT_88/3           ..........a..............................a...................................... [ 85680]
AM712239.1_Benin_97/1          ..........a..............................a...................................... [ 85680]
KM111295.1_Ken06.Bus           .......c.........-------------------------------------------.................ac- [ 85680]
```

```
KM111294.1_Ken05/Tk1           .......c.........-------------------------------------------...................- [ 85680]
KM262844.1_strain_L60          ..........a..............................a...................................... [ 85680]
KM102979.1_26544/OG10_Italy    ..........a..............................a...................................... [ 85680]
KM262845.1_strain_NHV          ..........a..............................a...................................... [ 85680]
KX354450.1_47/Ss/2008          ..........a..............................a...................................... [ 85680]
KP055815.1_strain_BA71         ..........a..............................a...................................... [ 85680]
MH025920.1_strain_R35          .......c.........-------------------------------------------.................ac- [ 85680]
MH025919.1_strain_N10          .......c.........-------------------------------------------.................ac- [ 85680]
MH025918.1_strain_R25          .......c.........-------------------------------------------.................ac- [ 85680]
MH025917.1_strain_R7           .......c.........-------------------------------------------.................ac- [ 85680]
MH025916.1_strain_R8           .......c.........-------------------------------------------.................ac- [ 85680]

FR682468.2_Georgia_2007/1      acacgct-ttttttttgcaaatcttaataatattgtatgataaatcaaacaatgtcttatat---atgtggtttattatt [ 85760]
ASF_IND_20_CAD543              .......-......................................................---.....r......... [ 85760]
LR899193.1_Germany_2020/1      .......-......................................................---............... [ 85760]
MN393477.1_Wuhan_2019          .......-......................................................---............... [ 85760]
MW396979.1_Timor-Leste/2019/1  .......-......................................................---............... [ 85760]
MK940252.1_CN/2019/InnerMongolia .......-......................................................---............... [ 85760]
MK128995.1_China/2018/AnhuiXCGQ .......-......................................................---............... [ 85760]
MH681419.1_ASFV/POL/2015       .......-......................................................---............... [ 85760]
LS478113.1_Estonia_2014        .......t......................................................---............... [ 85760]
KP843857.1_Odintsovo_02/14     .......-......................................................---............... [ 85760]
FN557520.1_strain_E75          .......-.............c.................................g......---.....a......... [ 85760]
AY261366.1_Warthog             .......-............tc.................................g......---............... [ 85760]
AY261365.1_Warmbaths           .......-.............c................t................g......---............... [ 85760]
AY261364.1_Tengani_62          .......-.............c.................................g......---............... [ 85760]
AY261363.1_Pretoriuskop/96/4   ......--............tc.................................g......---............... [ 85760]
AY261362.1_Mkuzi_1979          .......-.............c................t................g......---.....a......... [ 85760]
AY261361.1_Malawi_Lil-20/1_1983 ...t...-.............c..........c..c..-.............a..g......---.........t.ag.. [ 85760]
AY261360.1_Kenya_1950          ------.-..............................t...c..t......a..g......---..........c.... [ 85760]
AM712240.1_OURT_88/3           .......-.............c.................................g......---.....a......... [ 85760]
AM712239.1_Benin_97/1          .......-.............c.................................g......---.....a......... [ 85760]
KM111295.1_Ken06.Bus           ..---gc-........a....a................t...----..t...a..g......ata....a.....c.... [ 85760]
KM111294.1_Ken05/Tk1           .......-........a..t.a.............c..t...----..t...a..g......att....a.....c.... [ 85760]
KM262844.1_strain_L60          .......-.............c.................................g......---.....a......... [ 85760]
KM102979.1_26544/OG10_Italy    .......-.............c.................................g......---.....a......... [ 85760]
KM262845.1_strain_NHV          .......-.............c.................................g......---.....a......... [ 85760]
KX354450.1_47/Ss/2008          .......-.............c.................................g......---.....a......... [ 85760]
KP055815.1_strain_BA71         .......-.............c.................................g......---.....a......... [ 85760]
MH025920.1_strain_R35          ..---gc-........a....a................t...----..t...a..g......ata....a.....c.... [ 85760]
MH025919.1_strain_N10          ..---gc-........a....a................t...----..t...a..g......ata....a.....c.... [ 85760]
MH025918.1_strain_R25          ..---gc-........a....a................t...----..t...a..g......ata....a.....c.... [ 85760]
MH025917.1_strain_R7           ..---gc-........a....a................t...----..t...a..g......ata....a.....c.... [ 85760]
MH025916.1_strain_R8           ..---gc-........a....a................t...----..t...a..g......ata....a.....c.... [ 85760]
```

```
FR682468.2_Georgia_2007/1        ttaggcgccgcaagatg---tactccattctcattgcatgcttggtgttattactctgtctagttatatatgtcggtcat [ 85840]
ASF_IND_20_CAD543                .................---............................................................ [ 85840]
LR899193.1_Germany_2020/1        .................---............................................................ [ 85840]
MN393477.1_Wuhan_2019            .................---............................................................ [ 85840]
MW396979.1_Timor-Leste/2019/1    .................---............................................................ [ 85840]
MK940252.1_CN/2019/InnerMongolia .................---............................................................ [ 85840]
MK128995.1_China/2018/AnhuiXCGQ  .................---............................................................ [ 85840]
MH681419.1_ASFV/POL/2015         .................---............................................................ [ 85840]
LS478113.1_Estonia_2014          .................---............................................................ [ 85840]
KP843857.1_Odintsovo_02/14       .................---............................................................ [ 85840]
FN557520.1_strain_E75            .................---..t........................c................................ [ 85840]
AY261366.1_Warthog               .............t...---..........................cc.......t......a................. [ 85840]
AY261365.1_Warmbaths             .............t...---.......................................t.................... [ 85840]
AY261364.1_Tengani_62            .............t...---..........................cc.......t...t.................... [ 85840]
AY261363.1_Pretoriuskop/96/4     .............t...---..........................cc.......t......a................. [ 85840]
AY261362.1_Mkuzi_1979            .................---..t........................c................................ [ 85840]
AY261361.1_Malawi_Lil-20/1_1983  ........g....c...tac.....a...a.t...............c................................ [ 85840]
AY261360.1_Kenya_1950            .............c...---.....a...a.t..............cc.......................t........ [ 85840]
AM712240.1_OURT_88/3             .................---..t........................c................................ [ 85840]
AM712239.1_Benin_97/1            .................---..t........................c................................ [ 85840]
KM111295.1_Ken06.Bus             ---.at...a...c...---.....a...a.t..............cc..............a........t........ [ 85840]
KM111294.1_Ken05/Tk1             ---..........c...---.....a...a.t..............cc..............a........t........ [ 85840]
KM262844.1_strain_L60            .................---..t........................c................................ [ 85840]
KM102979.1_26544/OG10_Italy      .................---..t........................c................................ [ 85840]
KM262845.1_strain_NHV            .................---..t........................c................................ [ 85840]
KX354450.1_47/Ss/2008            .................---..t........................c................................ [ 85840]
KP055815.1_strain_BA71           .................---..t........................c................................ [ 85840]
MH025920.1_strain_R35            ---.at...a...c...---.....a...a.t..............cc..............a........t........ [ 85840]
MH025919.1_strain_N10            ---.at...a...c...---.....a...a.t..............cc..............a........t........ [ 85840]
MH025918.1_strain_R25            ---.at...a...c...---.....a...a.t..............cc..............a........t........ [ 85840]
MH025917.1_strain_R7             ---.at...a...c...---.....a...a.t..............cc..............a........t........ [ 85840]
MH025916.1_strain_R8             ---.at...a...c...---.....a...a.t..............cc..............a........t........ [ 85840]

FR682468.2_Georgia_2007/1        cgtgccgatcatgcacgaaaatatttagaaggaatgtggcatggagatccggtttttctaaaacagtcggggctacaatc [ 85920]
ASF_IND_20_CAD543                ................................................................................ [ 85920]
LR899193.1_Germany_2020/1        ................................................................................ [ 85920]
MN393477.1_Wuhan_2019            ................................................................................ [ 85920]
MW396979.1_Timor-Leste/2019/1    ................................................................................ [ 85920]
MK940252.1_CN/2019/InnerMongolia ................................................................................ [ 85920]
MK128995.1_China/2018/AnhuiXCGQ  ................................................................................ [ 85920]
MH681419.1_ASFV/POL/2015         ................................................................................ [ 85920]
LS478113.1_Estonia_2014          ................................................................................ [ 85920]
KP843857.1_Odintsovo_02/14       ................................................................................ [ 85920]
```

```
FN557520.1_strain_E75          ................................................................................ [ 85920]
AY261366.1_Warthog             .......................................................................a........ [ 85920]
AY261365.1_Warmbaths           .......................................................................a........ [ 85920]
AY261364.1_Tengani_62          .......................................................................a........ [ 85920]
AY261363.1_Pretoriuskop/96/4   .......................................................................a........ [ 85920]
AY261362.1_Mkuzi_1979          ................................................................................ [ 85920]
AY261361.1_Malawi_Lil-20/1_1983 ................t................................................a.............. [ 85920]
AY261360.1_Kenya_1950          ................t.................................a..............a..a........... [ 85920]
AM712240.1_OURT_88/3           ................................................................................ [ 85920]
AM712239.1_Benin_97/1          ................................................................................ [ 85920]
KM111295.1_Ken06.Bus           ................t.......................t.........a..............a..a........... [ 85920]
KM111294.1_Ken05/Tk1           ................t.................................a..............a..a........... [ 85920]
KM262844.1_strain_L60          ................................................................................ [ 85920]
KM102979.1_26544/OG10_Italy    ................................................................................ [ 85920]
KM262845.1_strain_NHV          ................................................................................ [ 85920]
KX354450.1_47/Ss/2008          ................................................................................ [ 85920]
KP055815.1_strain_BA71         ................................................................................ [ 85920]
MH025920.1_strain_R35          ................t.......................t.........a..............a..a........... [ 85920]
MH025919.1_strain_N10          ................t.......................t.........a..............a..a........... [ 85920]
MH025918.1_strain_R25          ................t.......................t.........a..............a..a........... [ 85920]
MH025917.1_strain_R7           ................t.......................t.........a..............a..a........... [ 85920]
MH025916.1_strain_R8           ................t.......................t.........a..............a..a........... [ 85920]

FR682468.2_Georgia_2007/1      cttttatctctacatacaacctgaccatacatgtttttttagcattgtgaataaaaatggtgaaaagctgatggaaacca [ 86000]
ASF_IND_20_CAD543              ................................................................................ [ 86000]
LR899193.1_Germany_2020/1      ................................................................................ [ 86000]
MN393477.1_Wuhan_2019          ................................................................................ [ 86000]
MW396979.1_Timor-Leste/2019/1  ................................................................................ [ 86000]
MK940252.1_CN/2019/InnerMongolia ................................................................................ [ 86000]
MK128995.1_China/2018/AnhuiXCGQ ................................................................................ [ 86000]
MH681419.1_ASFV/POL/2015       ................................................................................ [ 86000]
LS478113.1_Estonia_2014        ................................................................................ [ 86000]
KP843857.1_Odintsovo_02/14     ................................................................................ [ 86000]
FN557520.1_strain_E75          ............t..........g...c.................................................... [ 86000]
AY261366.1_Warthog             ............t..............................g.................................... [ 86000]
AY261365.1_Warmbaths           ................................................................................ [ 86000]
AY261364.1_Tengani_62          ......................a......................................................t.. [ 86000]
AY261363.1_Pretoriuskop/96/4   ............t..............................g.................................... [ 86000]
AY261362.1_Mkuzi_1979          ................................................................................ [ 86000]
AY261361.1_Malawi_Lil-20/1_1983 ............t..........g...c.................................................... [ 86000]
AY261360.1_Kenya_1950          a....................c.g...c...............g.............a...........t.......... [ 86000]
AM712240.1_OURT_88/3           ............t..........g...c.................................................... [ 86000]
AM712239.1_Benin_97/1          ............t..........g...c.................................................... [ 86000]
KM111295.1_Ken06.Bus           a....................c.g...c..............tg.............a...........t.......... [ 86000]
```

```
KM111294.1_Ken05/Tk1            a....................c.g...c...............g.............a...........t.......... [ 86000]
KM262844.1_strain_L60           ............t..........g...c.................................................... [ 86000]
KM102979.1_26544/OG10_Italy     ............t..........g...c.................................................... [ 86000]
KM262845.1_strain_NHV           ............t..........g...c.................................................... [ 86000]
KX354450.1_47/Ss/2008           ............t..........g...c.................................................... [ 86000]
KP055815.1_strain_BA71          ............t..........g...c.................................................... [ 86000]
MH025920.1_strain_R35           a....................c.g...c..............tg.............a...........t.......... [ 86000]
MH025919.1_strain_N10           a....................c.g...c..............tg.............a...........t.......... [ 86000]
MH025918.1_strain_R25           a....................c.g...c..............tg.............a...........t.......... [ 86000]
MH025917.1_strain_R7            a....................c.g...c..............tg.............a...........t.......... [ 86000]
MH025916.1_strain_R8            a....................c.g...c..............tg.............a...........t.......... [ 86000]

FR682468.2_Georgia_2007/1       aaataccttgtacgataacaaataaaatatatatgttttttaaacctatttttgaatttcatgttgtgatggaagacata [ 86080]
ASF_IND_20_CAD543               ................................................................................ [ 86080]
LR899193.1_Germany_2020/1       ................................................................................ [ 86080]
MN393477.1_Wuhan_2019           ................................................................................ [ 86080]
MW396979.1_Timor-Leste/2019/1   ................................................................................ [ 86080]
MK940252.1_CN/2019/InnerMongolia ................................................................................ [ 86080]
MK128995.1_China/2018/AnhuiXCGQ ................................................................................ [ 86080]
MH681419.1_ASFV/POL/2015        ................................................................................ [ 86080]
LS478113.1_Estonia_2014         ................................................................................ [ 86080]
KP843857.1_Odintsovo_02/14      ................................................................................ [ 86080]
FN557520.1_strain_E75           ................................................................................ [ 86080]
AY261366.1_Warthog              ...................................................................a............ [ 86080]
AY261365.1_Warmbaths            .................................................................aca............ [ 86080]
AY261364.1_Tengani_62           .............a................................................................c. [ 86080]
AY261363.1_Pretoriuskop/96/4    ................................................................................ [ 86080]
AY261362.1_Mkuzi_1979           ................................................................................ [ 86080]
AY261361.1_Malawi_Lil-20/1_1983 ........................................................................c.c...c. [ 86080]
AY261360.1_Kenya_1950           ......a.................................................................c.c...c. [ 86080]
AM712240.1_OURT_88/3            ................................................................................ [ 86080]
AM712239.1_Benin_97/1           ................................................................................ [ 86080]
KM111295.1_Ken06.Bus            ......a..........................................................a........c...c. [ 86080]
KM111294.1_Ken05/Tk1            ......a..........................................................a.a......c...c. [ 86080]
KM262844.1_strain_L60           ................................................................................ [ 86080]
KM102979.1_26544/OG10_Italy     ................................................................................ [ 86080]
KM262845.1_strain_NHV           ................................................................................ [ 86080]
KX354450.1_47/Ss/2008           ................................................................................ [ 86080]
KP055815.1_strain_BA71          ................................................................................ [ 86080]
MH025920.1_strain_R35           ......a..........................................................a........c...c. [ 86080]
MH025919.1_strain_N10           ......a..........................................................a........c...c. [ 86080]
MH025918.1_strain_R25           ......a..........................................................a........c...c. [ 86080]
MH025917.1_strain_R7            ......a..........................................................a........c...c. [ 86080]
MH025916.1_strain_R8            ......a..........................................................a........c...c. [ 86080]
```

```
FR682468.2_Georgia_2007/1        catagctacttccctaagcagtttaactttctgttagatagtacagaaggtaaacttattttagaaaacaatcacgttat [ 86160]
ASF_IND_20_CAD543                .........................................cg..................................... [ 86160]
LR899193.1_Germany_2020/1        ................................................................................ [ 86160]
MN393477.1_Wuhan_2019            ................................................................................ [ 86160]
MW396979.1_Timor-Leste/2019/1    ................................................................................ [ 86160]
MK940252.1_CN/2019/InnerMongolia ................................................................................ [ 86160]
MK128995.1_China/2018/AnhuiXCGQ  ................................................................................ [ 86160]
MH681419.1_ASFV/POL/2015         ................................................................................ [ 86160]
LS478113.1_Estonia_2014          ................................................................................ [ 86160]
KP843857.1_Odintsovo_02/14       ................................................................................ [ 86160]
FN557520.1_strain_E75            .....a.....a..a.......................c..cg..................................... [ 86160]
AY261366.1_Warthog               ......................................c..cg..................................... [ 86160]
AY261365.1_Warmbaths             .....a.....a.....a....................c..c...........g.......................... [ 86160]
AY261364.1_Tengani_62            ......................................c..cg..........g.......................... [ 86160]
AY261363.1_Pretoriuskop/96/4     ......................................c..c...........g.......................... [ 86160]
AY261362.1_Mkuzi_1979            .....a.....a..........................c..cg..................................... [ 86160]
AY261361.1_Malawi_Lil-20/1_1983  t..........t.................................................................... [ 86160]
AY261360.1_Kenya_1950            ...........t.................................................................... [ 86160]
AM712240.1_OURT_88/3             .....a.....a..a.......................c..cg..................................... [ 86160]
AM712239.1_Benin_97/1            .....a.....a..a.......................c..cg..................................... [ 86160]
KM111295.1_Ken06.Bus             .....t.....t.............g............c..c...................................... [ 86160]
KM111294.1_Ken05/Tk1             ...........t..........................c..c..........................t........... [ 86160]
KM262844.1_strain_L60            .....a.....a..a.......................c..cg..................................... [ 86160]
KM102979.1_26544/OG10_Italy      .....a.....a..a.......................c..cg..................................... [ 86160]
KM262845.1_strain_NHV            .....a.....a..a.......................c..cg..................................... [ 86160]
KX354450.1_47/Ss/2008            .....a.....a..a.......................c..cg..................................... [ 86160]
KP055815.1_strain_BA71           .....a.....a..a.......................c..cg..................................... [ 86160]
MH025920.1_strain_R35            .....t.....t.............g............c..c...................................... [ 86160]
MH025919.1_strain_N10            .....t.....t.............g............c..c...................................... [ 86160]
MH025918.1_strain_R25            .....t.....t.............g............c..c...................................... [ 86160]
MH025917.1_strain_R7             .....t.....t.............g............c..c...................................... [ 86160]
MH025916.1_strain_R8             .....t.....t.............g............c..c...................................... [ 86160]

FR682468.2_Georgia_2007/1        ttatgctgtattgtataaggataatttcgccaccgcactaggaaaaacggttgaaaaatatataacacaaaattaatcat [ 86240]
ASF_IND_20_CAD543                ................................................................................ [ 86240]
LR899193.1_Germany_2020/1        ................................................................................ [ 86240]
MN393477.1_Wuhan_2019            ................................................................................ [ 86240]
MW396979.1_Timor-Leste/2019/1    ................................................................................ [ 86240]
MK940252.1_CN/2019/InnerMongolia ................................................................................ [ 86240]
MK128995.1_China/2018/AnhuiXCGQ  ................................................................................ [ 86240]
MH681419.1_ASFV/POL/2015         ................................................................................ [ 86240]
LS478113.1_Estonia_2014          ................................................................................ [ 86240]
KP843857.1_Odintsovo_02/14       ................................................................................ [ 86240]
```

```
FN557520.1_strain_E75            ........................c....................................................... [ 86240]
AY261366.1_Warthog               .....................c..c............t..........t..ca..............t............ [ 86240]
AY261365.1_Warmbaths             .....................c..c............t..............a..............a............ [ 86240]
AY261364.1_Tengani_62            ........................c....................................................... [ 86240]
AY261363.1_Pretoriuskop/96/4     ........................c........t...t..............a..............t............ [ 86240]
AY261362.1_Mkuzi_1979            ........................c....................................................... [ 86240]
AY261361.1_Malawi_Lil-20/1_1983  ........................c............t..........t...a..............t............ [ 86240]
AY261360.1_Kenya_1950            ......g.................c.......................t..................t.g---.c.c... [ 86240]
AM712240.1_OURT_88/3             ........................c....................................................... [ 86240]
AM712239.1_Benin_97/1            ........................c....................................................... [ 86240]
KM111295.1_Ken06.Bus             ........................c............t.........tca.ac.........a....t............ [ 86240]
KM111294.1_Ken05/Tk1             ........................c.......................ca.ac........................... [ 86240]
KM262844.1_strain_L60            ........................c....................................................... [ 86240]
KM102979.1_26544/OG10_Italy      ........................c....................................................... [ 86240]
KM262845.1_strain_NHV            ........................c....................................................... [ 86240]
KX354450.1_47/Ss/2008            ........................c....................................................... [ 86240]
KP055815.1_strain_BA71           ........................c....................................................... [ 86240]
MH025920.1_strain_R35            ........................c............t.........tca.ac.........a....t............ [ 86240]
MH025919.1_strain_N10            ........................c............t.........tca.ac.........a....t............ [ 86240]
MH025918.1_strain_R25            ........................c............t.........tca.ac.........a....t............ [ 86240]
MH025917.1_strain_R7             ........................c............t.........tca.ac.........a....t............ [ 86240]
MH025916.1_strain_R8             ........................c............t.........tca.ac.........a....t............ [ 86240]

FR682468.2_Georgia_2007/1        gttttctaacaaaaagtacatcggtcttatcaataagaaggagggtttgaa---aaaaaaa------------------- [ 86320]
ASF_IND_20_CAD543                ..a..t...s.....a..............nn.......-----c.g....---.......tattannnnnnnnnagtac [ 86320]
LR899193.1_Germany_2020/1        ...................................................---.......------------------- [ 86320]
MN393477.1_Wuhan_2019            ...................................................---.......------------------- [ 86320]
MW396979.1_Timor-Leste/2019/1    ...................................................---.......------------------- [ 86320]
MK940252.1_CN/2019/InnerMongolia ...................................................---.......------------------- [ 86320]
MK128995.1_China/2018/AnhuiXCGQ  ...................................................---.......------------------- [ 86320]
MH681419.1_ASFV/POL/2015         ...................................................---.......------------------- [ 86320]
LS478113.1_Estonia_2014          ...................................................---.......------------------- [ 86320]
KP843857.1_Odintsovo_02/14       ...................................................---.......------------------- [ 86320]
FN557520.1_strain_E75            ..a..t...g.....a..............tg.......-----c.g....---.......tattagatgattctagtac [ 86320]
AY261366.1_Warthog               .........t...........t.................-----.gag.g.ttg.......aaatagatgattatagtat [ 86320]
AY261365.1_Warmbaths             ..a..t...g....................tg.......-----c.g....---.......tattagatgattgtactac [ 86320]
AY261364.1_Tengani_62            ..................tccga........-----------------------g......aaatggatgacttgatgac [ 86320]
AY261363.1_Pretoriuskop/96/4     ..a..t...g....................tg.......-----c.g....---.......tattagatgattgtactac [ 86320]
AY261362.1_Mkuzi_1979            ..a..t...g.....a..............tg.......-----c.g....---.......tattagatgattctagtac [ 86320]
AY261361.1_Malawi_Lil-20/1_1983  ..............................tg.c...t.-----c.g....---.......tattagatgattctagtac [ 86320]
AY261360.1_Kenya_1950            .....tg...........tccga........g-----------------------......aaatggatgatttgatgac [ 86320]
AM712240.1_OURT_88/3             ..a..t...g.....a..............tg.......-----c.g..-.---.......tattagatgattctagtac [ 86320]
AM712239.1_Benin_97/1            ..a..t...g.....a..............tg.......-----c.g....---.......tattagatgattctagtac [ 86320]
KM111295.1_Ken06.Bus             ..a......g....................tg.c..a.g-----c.g....---.......tattagatgatcctactac [ 86320]
```

```
KM111294.1_Ken05/Tk1             ..a......g....................tg.c.....-----c.g....---.......tattagatgatgctactac [ 86320]
KM262844.1_strain_L60            ..a..t...g.....a..............tg.......-----c.g....---.......tattagatgattctagtac [ 86320]
KM102979.1_26544/OG10_Italy      ..a..t...g.....a..............tg.......-----c.g....---.......tattagatgattctagtac [ 86320]
KM262845.1_strain_NHV            ..a..t...g.....a..............tg.......-----c.g..-.---.......tattagatgattctagtac [ 86320]
KX354450.1_47/Ss/2008            ..a..t...g.....a..............tg.......-----c.g....---.......tattagatgattctagtac [ 86320]
KP055815.1_strain_BA71           ..a..t...g.....a..............tg.......-----c.g....---.......tattagatgattctagtac [ 86320]
MH025920.1_strain_R35            ..a......g....................tg.c..a.g-----c.g....---.......tattagatgatcctactac [ 86320]
MH025919.1_strain_N10            ..a......g....................tg.c..a.g-----c.g....---.......tattagatgatcctactac [ 86320]
MH025918.1_strain_R25            ..a......g....................tg.c..a.g-----c.g....---.......tattagatgatcctactac [ 86320]
MH025917.1_strain_R7             ..a......g....................tg.c..a.g-----c.g....---.......tattagatgatcctactac [ 86320]
MH025916.1_strain_R8             ..a......g....................tg.c..a.g-----c.g....---.......tattagatgatcctactac [ 86320]

FR682468.2_Georgia_2007/1        -atagatgattatagtatattaataattggaatattaattggaactaacatcttaagccttattataaatataataggag [ 86400]
ASF_IND_20_CAD543                a...a.aat..g.tac....---.........................t..ga...ct......tat...t.c...ttct [ 86400]
LR899193.1_Germany_2020/1        -............................................................................... [ 86400]
MN393477.1_Wuhan_2019            -............................................................................... [ 86400]
MW396979.1_Timor-Leste/2019/1    -............................................................................... [ 86400]
MK940252.1_CN/2019/InnerMongolia -............................................................................... [ 86400]
MK128995.1_China/2018/AnhuiXCGQ  -............................................................................... [ 86400]
MH681419.1_ASFV/POL/2015         -............................................................................... [ 86400]
LS478113.1_Estonia_2014          -............................................................................... [ 86400]
KP843857.1_Odintsovo_02/14       -............................................................................... [ 86400]
FN557520.1_strain_E75            a...a.aat..g.tac....---.............g...........t..ga...ct......tat...t.c...ttct [ 86400]
AY261366.1_Warthog               at..a------------------......................................................... [ 86400]
AY261365.1_Warmbaths             a...a.aat..g.tac....---.............g...........t..ga...ct......tat...t.c...tttt [ 86400]
AY261364.1_Tengani_62            tt..a.gtt..g.tac..ca---.t....c...c..g..catg.....t..t..t......g..g.g...---...t.g. [ 86400]
AY261363.1_Pretoriuskop/96/4     a...a.aat..g.tac....---.............g...........t..ga...ct......tat...t.c...tttt [ 86400]
AY261362.1_Mkuzi_1979            a...a.aat..g.tac....---.............g...........t..ga...ct......tat...t.c...ttct [ 86400]
AY261361.1_Malawi_Lil-20/1_1983  a...a.aat..g.tac....---...........c.g...........t..ga...ct......tat...t.c...tttt [ 86400]
AY261360.1_Kenya_1950            tt..a.gtt..g.tact..a---....cac.t.t.....cat...c..t..t..t......g-ctat..at--...t.g. [ 86400]
AM712240.1_OURT_88/3             a...a.aat..g.tac....---.............g...........t..ga...ct......tat...t.c...ttct [ 86400]
AM712239.1_Benin_97/1            a...a.aat..g.tac....---.............g...........t..ga...ct......tat...t.c...ttct [ 86400]
KM111295.1_Ken06.Bus             a...a.aat..g.tac....---.............g...........t..ta...ct.....ctat...t.c...tttt [ 86400]
KM111294.1_Ken05/Tk1             a...a.aat..g.tac....---.............g...........t..ta...ct.....ctat...t.c...tttt [ 86400]
KM262844.1_strain_L60            a...a.aat..g.tac....---.............g...........t..ga...ct......tat...t.c...ttct [ 86400]
KM102979.1_26544/OG10_Italy      a...a.aat..g.tac....---.............g...........t..ga...ct......tat...t.c...ttct [ 86400]
KM262845.1_strain_NHV            a...a.aat..g.tac....---.............g...........t..ga...ct......tat...t.c...ttct [ 86400]
KX354450.1_47/Ss/2008            a...a.aat..g.tac....---.............g...........t..ga...ct......tat...t.c...ttct [ 86400]
KP055815.1_strain_BA71           a...a.aat..g.tac....---.............g...........t..ga...ct......tat...t.c...ttct [ 86400]
MH025920.1_strain_R35            a...a.aat..g.tac....---.............g...........t..ta...ct.....ctat...t.c...tttt [ 86400]
MH025919.1_strain_N10            a...a.aat..g.tac....---.............g...........t..ta...ct.....ctat...t.c...tttt [ 86400]
MH025918.1_strain_R25            a...a.aat..g.tac....---.............g...........t..ta...ct.....ctat...t.c...tttt [ 86400]
MH025917.1_strain_R7             a...a.aat..g.tac....---.............g...........t..ta...ct.....ctat...t.c...tttt [ 86400]
MH025916.1_strain_R8             a...a.aat..g.tac....---.............g...........t..ta...ct.....ctat...t.c...tttt [ 86400]
```

```
FR682468.2_Georgia_2007/1        agattaataaaccaatatgttaccaaaatgatgataagatattttattgccctaaagattgggttggatataataatgtt [ 86480]
ASF_IND_20_CAD543                g.gannnnnnnnnnnn.............r.......a..r.....y..y.....w.....................r.. [ 86480]
LR899193.1_Germany_2020/1        ................................................................................ [ 86480]
MN393477.1_Wuhan_2019            ................................................................................ [ 86480]
MW396979.1_Timor-Leste/2019/1    ................................................................................ [ 86480]
MK940252.1_CN/2019/InnerMongolia ................................................................................ [ 86480]
MK128995.1_China/2018/AnhuiXCGQ  ................................................................................ [ 86480]
MH681419.1_ASFV/POL/2015         ................................................................................ [ 86480]
LS478113.1_Estonia_2014          ................................................................................ [ 86480]
KP843857.1_Odintsovo_02/14       ................................................................................ [ 86480]
FN557520.1_strain_E75            g.ga....t.tat..a........g....a.......a..g.....c..t.....t.....................a.. [ 86480]
AY261366.1_Warthog               .............................a.................................................. [ 86480]
AY261365.1_Warmbaths             g.ga.c..t.tatg.c....a..---..aa.a........g.....c..t.............................. [ 86480]
AY261364.1_Tengani_62            g.ggggg.g.t.g.ca..c...t---------..g..tg..........t............................ca [ 86480]
AY261363.1_Pretoriuskop/96/4     g.ga.c..t.tatg.c....a..---..aa.a........g.....c..t.............................. [ 86480]
AY261362.1_Mkuzi_1979            g.ga....t.tat..a........g....a.......a..g.....c..t.....t........................ [ 86480]
AY261361.1_Malawi_Lil-20/1_1983  g.gag...t.tat..c....a.......a---.......cg.....c..t.............................. [ 86480]
AY261360.1_Kenya_1950            g.ggggg.g.tatg...ga.cga....g.tg...a..t........c..t.................c..........cg [ 86480]
AM712240.1_OURT_88/3             g.ga....t.tat..a........g....a.......a..g.....c..t.....t.....................a.. [ 86480]
AM712239.1_Benin_97/1            g.ga....t.tat..a........g....a.......a..g.....c..t.....t.....................a.. [ 86480]
KM111295.1_Ken06.Bus             g.gag...t.tat..a....at..g...............g.....c................................g [ 86480]
KM111294.1_Ken05/Tk1             g.gag...t.tat..a....at..................g.....c..t.................g..........cg [ 86480]
KM262844.1_strain_L60            g.ga....t.tat..a........g....a.......a..g.....c..t.....t.....................a.. [ 86480]
KM102979.1_26544/OG10_Italy      g.ga....t.tat..a........g....a.......a..g.....c..t.....t.....................a.. [ 86480]
KM262845.1_strain_NHV            g.ga....t.tat..a........g....a.......a..g.....c..t.....t.....................a.. [ 86480]
KX354450.1_47/Ss/2008            g.ga....t.tat..a........g....a.......a..g.....c..t.....t.....................a.. [ 86480]
KP055815.1_strain_BA71           g.ga....t.tat..a........g....a.......a..g.....c..t.....t.....................a.. [ 86480]
MH025920.1_strain_R35            g.gag...t.tat..a....at..g...............g.....c................................g [ 86480]
MH025919.1_strain_N10            g.gag...t.tat..a....at..g...............g.....c................................g [ 86480]
MH025918.1_strain_R25            g.gag...t.tat..a....at..g...............g.....c................................g [ 86480]
MH025917.1_strain_R7             g.gag...t.tat..a....at..g...............g.....c................................g [ 86480]
MH025916.1_strain_R8             g.gag...t.tat..a....at..g...............g.....c................................g [ 86480]

FR682468.2_Georgia_2007/1        tgttattattttggcaatgaagaa------aaaaat------tataataatgcaagtaattattgtaagcaatta---aa [ 86560]
ASF_IND_20_CAD543                .....c......a.nnnnnnnnnn------nnnnnnnnnnnnnnnnnnnnnn.t.......t.....ga..nnnn---nn [ 86560]
LR899193.1_Germany_2020/1        ........................------......------.................................---.. [ 86560]
MN393477.1_Wuhan_2019            ........................------......------.................................---.. [ 86560]
MW396979.1_Timor-Leste/2019/1    ........................------......------.................................---.. [ 86560]
MK940252.1_CN/2019/InnerMongolia ........................------......------.................................---.. [ 86560]
MK128995.1_China/2018/AnhuiXCGQ  ........................------......------.................................---.. [ 86560]
MH681419.1_ASFV/POL/2015         ........................------......------.................................---.. [ 86560]
LS478113.1_Estonia_2014          ........................------......------.................................---.. [ 86560]
KP843857.1_Odintsovo_02/14       ........................------......------.................................---.. [ 86560]
```

```
FN557520.1_strain_E75          .....c......a.t....gtagtttttct......------....cagc...t.......t.....ga......---.. [ 86560]
AY261366.1_Warthog             ............a.t.....ta.---tggc..t...------....ca.c...gga....a.a.....a......aat.. [ 86560]
AY261365.1_Warmbaths           ............a.t.....tagttttact..g...------....cat............c......aa..a..---.. [ 86560]
AY261364.1_Tengani_62          ............a.t.....tatg------......------....cag.a.................aa.t...tat.. [ 86560]
AY261363.1_Pretoriuskop/96/4   ............aat.....tag------t..g...------....ca.c...t.c.....c......a......---.. [ 86560]
AY261362.1_Mkuzi_1979          .....c......a.t....gtagttttttt......------....cag....t.a............aa..a..---.. [ 86560]
AY261361.1_Malawi_Lil-20/1_1983 ....................t...------......------.................................---.. [ 86560]
AY261360.1_Kenya_1950          ............a.t...a.ca.t------......------....cag......a............aa.t.c.cat.. [ 86560]
AM712240.1_OURT_88/3           .....c......a.t....gtagtttttct......------....cagc...t.......t.....ga......---.. [ 86560]
AM712239.1_Benin_97/1          .....c......a.t....gtagtttttct......------....cagc...t.......t.....ga......---.. [ 86560]
KM111295.1_Ken06.Bus           ............a.t..c..ta.tg---ga......------....tg.............t......ta.---.aaa.. [ 86560]
KM111294.1_Ken05/Tk1           ............a.t...a.ca.t------......------....cag......a............aa.t.c.cat.. [ 86560]
KM262844.1_strain_L60          .....c......a.t....gtagtttttct......------....cagc...t.......t.....ga......---.. [ 86560]
KM102979.1_26544/OG10_Italy    .....c......a.t....gtagtttttct......------....cagc...t.......t.....ga......---.. [ 86560]
KM262845.1_strain_NHV          .....c......a.t....gtagtttttct......------....cagc...t.......t.....ga......---.. [ 86560]
KX354450.1_47/Ss/2008          .....c......a.t....gtagtttttct......------....cagc...t.......t.....ga......---.. [ 86560]
KP055815.1_strain_BA71         .....c......a.t....gtagtttttct......------....cagc...t.......t.....ga......---.. [ 86560]
MH025920.1_strain_R35          ............a.t..c..ta.tg---ga......------....tg.............t......ta.---.aaa.. [ 86560]
MH025919.1_strain_N10          ............a.t..c..ta.tg---ga......------....tg.............t......ta.---.aaa.. [ 86560]
MH025918.1_strain_R25          ............a.t..c..ta.tg---ga......------....tg.............t......ta.---.aaa.. [ 86560]
MH025917.1_strain_R7           ............a.t..c..ta.tg---ga......------....tg.............t......ta.---.aaa.. [ 86560]
MH025916.1_strain_R8           ............a.t..c..ta.tg---ga......------....tg.............t......ta.---.aaa.. [ 86560]

FR682468.2_Georgia_2007/1      tagtacgcttactaataataatactattttagtaaatcttactaaaacattaaatcttactaaaacatataatcacgaat [ 86640]
ASF_IND_20_CAD543              nnnnnnnnnnnnn......g......a....------------------..nnnnn.a.......t.......a.tc..n [ 86640]
LR899193.1_Germany_2020/1      ................................................................................ [ 86640]
MN393477.1_Wuhan_2019          ................................................................................ [ 86640]
MW396979.1_Timor-Leste/2019/1  ................................................................................ [ 86640]
MK940252.1_CN/2019/InnerMongolia ................................................................................ [ 86640]
MK128995.1_China/2018/AnhuiXCGQ ................................................................................ [ 86640]
MH681419.1_ASFV/POL/2015       ................................................................................ [ 86640]
LS478113.1_Estonia_2014        ................................................................................ [ 86640]
KP843857.1_Odintsovo_02/14     ................................................................................ [ 86640]
FN557520.1_strain_E75          .g....a...g........g......a....------------------........a.......t.......a.tc... [ 86640]
AY261366.1_Warthog             ......a..gg........ct....ga....t..........g.gtttc...............tt....c....tc.t. [ 86640]
AY261365.1_Warmbaths           ......a...g........g......a....------------------c.........t....gt.......a.t.t.. [ 86640]
AY261364.1_Tengani_62          ......aa..gt..............a.a....................................t.........t...a [ 86640]
AY261363.1_Pretoriuskop/96/4   ......a...g........g......a....------------------...............gt....c.c..t..ta [ 86640]
AY261362.1_Mkuzi_1979          ......t...g........g......a....------------------....g.t.a.......t...c...a.tc... [ 86640]
AY261361.1_Malawi_Lil-20/1_1983 ..........................a..................................................... [ 86640]
AY261360.1_Kenya_1950          ......a...g........g......aa...t..............tt..........t.....tt....t.ca.t..t. [ 86640]
AM712240.1_OURT_88/3           .g....a...g........g......a....------------------........a.......t.......a.tc... [ 86640]
AM712239.1_Benin_97/1          .g....a...g........g......a....------------------........a.......t.......a.tc... [ 86640]
KM111295.1_Ken06.Bus           ...c......g.........cg....a....................t.......t....c...ca.........t.... [ 86640]
```

```
KM111294.1_Ken05/Tk1            ......a...g........g......aa...t..............tt..........t.....tt....t.ca.t..t. [ 86640]
KM262844.1_strain_L60           .g....a...g........g......a....------------------........a.......t.......a.tc... [ 86640]
KM102979.1_26544/OG10_Italy     .g....a...g........g......a....------------------........a.......t.......a.tc... [ 86640]
KM262845.1_strain_NHV           .g....a...g........g......a....------------------........a.......t.......a.tc... [ 86640]
KX354450.1_47/Ss/2008           .g....a...g........g......a....------------------........a.......t.......a.tc... [ 86640]
KP055815.1_strain_BA71          .g....a...g........g......a....------------------........a.......t.......a.tc... [ 86640]
MH025920.1_strain_R35           ...c......g.........cg....a....................t.......t....c...ca.........t.... [ 86640]
MH025919.1_strain_N10           ...c......g.........cg....a....................t.......t....c...ca.........t.... [ 86640]
MH025918.1_strain_R25           ...c......g.........cg....a....................t.......t....c...ca.........t.... [ 86640]
MH025917.1_strain_R7            ...c......g.........cg....a....................t.......t....c...ca.........t.... [ 86640]
MH025916.1_strain_R8            ...c......g.........cg....a....................t.......t....c...ca.........t.... [ 86640]

FR682468.2_Georgia_2007/1       ctaattattgggttaattattctttaattaaaaatgagtcagtactattacgtgatagtggatattaca------a---a [ 86720]
ASF_IND_20_CAD543               nnnnnnnnnnnnnnnnnnnnnnnnnnnnnnnnnnnnnnnnnnnnnnnnnnnnnnnnnnnnnnnnn..t.----------- [ 86720]
LR899193.1_Germany_2020/1       .....................................................................------.---. [ 86720]
MN393477.1_Wuhan_2019           .....................................................................aaaaac.---. [ 86720]
MW396979.1_Timor-Leste/2019/1   .....................................................................------.---. [ 86720]
MK940252.1_CN/2019/InnerMongolia .....................................................................------.---. [ 86720]
MK128995.1_China/2018/AnhuiXCGQ .....................................................................------.---. [ 86720]
MH681419.1_ASFV/POL/2015        .....................................................................------.---. [ 86720]
LS478113.1_Estonia_2014         .....................................................................------.---. [ 86720]
KP843857.1_Odintsovo_02/14      .....................................................................------.---. [ 86720]
FN557520.1_strain_E75           ...tg...........ca..------------------a.g...a.........g.ga.aata.a..t.---gtc.---. [ 86720]
AY261366.1_Warthog              ..c........................a...t...t.t.......c....at.......aa....a.tttaaataggaa. [ 86720]
AY261365.1_Warmbaths            ..g.g........------------------t...a.ta.....ac........g.ga.cata.a....taaata.gca. [ 86720]
AY261364.1_Tengani_62           g..............................t.........t..ac......g..c...aat.t.cctt---cttcagg. [ 86720]
AY261363.1_Pretoriuskop/96/4    aatta...........c..........a.g.c...ttt...t..tc.......a.....---------.catatg.aaa. [ 86720]
AY261362.1_Mkuzi_1979           ...tg...........ca..------------------a.....gc........g.ga.a.tgta..tg---cac.aag. [ 86720]
AY261361.1_Malawi_Lil-20/1_1983 ....................................................ca.............t.---------a. [ 86720]
AY261360.1_Kenya_1950           ...ca...........a.........cc.......a.ag.....ac.......a.c...acc..ca.at---at---gat [ 86720]
AM712240.1_OURT_88/3            ...tg...........ca..------------------a.g...a.........g.ga.aata.a..t.---gtc.---. [ 86720]
AM712239.1_Benin_97/1           ...tg...........ca..------------------a.g...a.........g.ga.aata.a..t.---gtc.---. [ 86720]
KM111295.1_Ken06.Bus            .act.......................g...t...at....t..gc.....aga.....aat.t.cctt---cttcagg. [ 86720]
KM111294.1_Ken05/Tk1            ...ca...........a.........cc.......a.ag.....ac.......a.c...acc..ca.at---at---gat [ 86720]
KM262844.1_strain_L60           ...tg...........ca..------------------a.g...a.........g.ga.aata.a..t.---gtc.---. [ 86720]
KM102979.1_26544/OG10_Italy     ...tg...........ca..------------------a.g...a.........g.ga.aata.a..t.---gtc.---. [ 86720]
KM262845.1_strain_NHV           ...tg...........ca..------------------a.g...a.........g.ga.aata.a..t.---gtc.---. [ 86720]
KX354450.1_47/Ss/2008           ...tg...........ca..------------------a.g...a.........g.ga.aata.a..t.---gtc.---. [ 86720]
KP055815.1_strain_BA71          ...tg...........ca..------------------a.g...a.........g.ga.aata.a..t.---gtc.---. [ 86720]
MH025920.1_strain_R35           .act.......................g...t...at....t..gc.....aga.....aat.t.cctt---cttcagg. [ 86720]
MH025919.1_strain_N10           .act.......................g...t...at....t..gc.....aga.....aat.t.cctt---cttcagg. [ 86720]
MH025918.1_strain_R25           .act.......................g...t...at....t..gc.....aga.....aat.t.cctt---cttcagg. [ 86720]
MH025917.1_strain_R7            .act.......................g...t...at....t..gc.....aga.....aat.t.cctt---cttcagg. [ 86720]
MH025916.1_strain_R8            .act.......................g...t...at....t..gc.....aga.....aat.t.cctt---cttcagg. [ 86720]
```

```
FR682468.2_Georgia_2007/1        aaacaaaaacatgtaagtttattatatatttgtagtaaata--atatt---tttaattacttaaaatttttatatataag [ 86800]
ASF_IND_20_CAD543                -------------c.ga........t.......g.......--..ga.aaa..a..ca.ta.gt.........gag..gt [ 86800]
LR899193.1_Germany_2020/1        .........................................--.....---............................. [ 86800]
MN393477.1_Wuhan_2019            .........................................--.....---............................. [ 86800]
MW396979.1_Timor-Leste/2019/1    .........................................--.....---............................. [ 86800]
MK940252.1_CN/2019/InnerMongolia .........................................--.....---............................. [ 86800]
MK128995.1_China/2018/AnhuiXCGQ  .........................................--.....---............................. [ 86800]
MH681419.1_ASFV/POL/2015         .........................................--.....---............................. [ 86800]
LS478113.1_Estonia_2014          .........................................--.....---............................. [ 86800]
KP843857.1_Odintsovo_02/14       .........................................--.....---............................. [ 86800]
FN557520.1_strain_E75            ...gtt..ct..ac.ga........t.......g.......--..ga.aaa..a..ca.ta.gt.........gag..gt [ 86800]
AY261366.1_Warthog               ...agtc.ct..ac.ga....c...t......c..c.....----.a.act..g.....ta---t.g..c..c.ag.t.a [ 86800]
AY261365.1_Warmbaths             ..ta.ttgta..ac.gaa..t....t...........g...ggc..actaa....ga..ta---g.a....ta.ag...a [ 86800]
AY261364.1_Tengani_62            ....gt.c.t..a..ga............a.......t...----.g.taa..ag....ta-gg..gag.....a..t.c [ 86800]
AY261363.1_Pretoriuskop/96/4     .g.tct...t..t.gcca.......t......c........----.a.aat----.a..ta---g....g....a...g. [ 86800]
AY261362.1_Mkuzi_1979            ...gtt..ta..tc.ga.c......t...............--.--a.aggcc....a.a..gtt..a.aa..t.t..gt [ 86800]
AY261361.1_Malawi_Lil-20/1_1983  .........................................-----a.att............................. [ 86800]
AY261360.1_Kenya_1950            .g.gtt...t..ac.gaaac...t.t............c..-a.c.aatatc.a.t...tg--------------..t.a [ 86800]
AM712240.1_OURT_88/3             ...gtt..ct..ac.ga........t.......g.......--..ga.aaa..a..ca.ta.gt.........gag..gt [ 86800]
AM712239.1_Benin_97/1            ...gtt..ct..ac.ga........t.......g.......--..ga.aaa..a..ca.ta.gt.........gag..gt [ 86800]
KM111295.1_Ken06.Bus             .g..gt...t..a.ggac...........a.......t....-ag.ta--at..ag....tg--------------gg..a [ 86800]
KM111294.1_Ken05/Tk1             .g.gtt...tc.ac.gaaac...t.t............c..-a.c.aatatc.a.t...tg--------------..t.a [ 86800]
KM262844.1_strain_L60            ...gtt..ct..ac.ga........t.......g.......--..ga.aaa..a..ca.ta.gt.........gag..gt [ 86800]
KM102979.1_26544/OG10_Italy      ...gtt..ct..ac.ga........t.......g.......--..ga.aaa..a..ca.ta.gt.........gag..gt [ 86800]
KM262845.1_strain_NHV            ...gtt..ct..ac.ga........t.......g.......--..ga.aaa..a..ca.ta.gt.........gag..gt [ 86800]
KX354450.1_47/Ss/2008            ...gtt..ct..ac.ga........t.......g.......--..ga.aaa..a..ca.ta.gt.........gag..gt [ 86800]
KP055815.1_strain_BA71           ...gtt..ct..ac.ga........t.......g.......--..ga.aaa..a..ca.ta.gt.........gag..gt [ 86800]
MH025920.1_strain_R35            .g..gt...t..a.ggac...........a.....c.t....-ag.ta--at..ag....tg--------------gg..a [ 86800]
MH025919.1_strain_N10            .g..gt...t..a.ggac...........a.....c.t....-ag.ta--at..ag....tg--------------gg..a [ 86800]
MH025918.1_strain_R25            .g..gt...t..a.ggac...........a.....c.t....-ag.ta--at..ag....tg--------------gg..a [ 86800]
MH025917.1_strain_R7             .g..gt...t..a.ggac...........a.....c.t....-ag.ta--at..ag....tg--------------gg..a [ 86800]
MH025916.1_strain_R8             .g..gt...t..a.ggac...........a.....c.t....-ag.ta--at..ag....tg--------------gg..a [ 86800]

FR682468.2_Georgia_2007/1        tttttgatac------------------------------ta-tattataaaacatatgttcataaa-----atgataat [ 86880]
ASF_IND_20_CAD543                aaaaaa...t------------------------------..t...............a.gtg..t.aaatg..a..... [ 86880]
LR899193.1_Germany_2020/1        ..........------------------------------..-........................-----........ [ 86880]
MN393477.1_Wuhan_2019            ..........------------------------------..-........................-----........ [ 86880]
MW396979.1_Timor-Leste/2019/1    ..........------------------------------..-........................-----........ [ 86880]
MK940252.1_CN/2019/InnerMongolia ..........------------------------------..-........................-----........ [ 86880]
MK128995.1_China/2018/AnhuiXCGQ  ..........------------------------------..-........................-----........ [ 86880]
MH681419.1_ASFV/POL/2015         ..........------------------------------..-........................-----........ [ 86880]
LS478113.1_Estonia_2014          ..........------------------------------..-........................-----........ [ 86880]
KP843857.1_Odintsovo_02/14       ..........------------------------------..-........................-----........ [ 86880]
```

```
FN557520.1_strain_E75            aaaaaa...t------------------------------..t...............a.gtg..t.aaatg..a..... [ 86880]
AY261366.1_Warthog               --------------------------------tattttag..t.................--g..t.aaatg..a..t.a [ 86880]
AY261365.1_Warmbaths             aaaa---.ttacacat----------atgttttataataa..t..a..............--g....aaatg..a..... [ 86880]
AY261364.1_Tengani_62            aaaaaat.tt-tatac--------ata---gttttaaaaa..t..........t.....a--g..t.aaatg..a....a [ 86880]
AY261363.1_Pretoriuskop/96/4     aaaaaat.ttataca-----------------tatgttta..a..a....tg.t--------------aatgt.c....c [ 86880]
AY261362.1_Mkuzi_1979            .aaaaa..ttataccc--------atatgttttatgataa..t.................gt.....tgatg..a..t.. [ 86880]
AY261361.1_Malawi_Lil-20/1_1983  ------------------------------tttttgatac..-.................--gc.t.aaatg..a..... [ 86880]
AY261360.1_Kenya_1950            aaaaaat.taaatcat----------------------at..t...a.a.c....c...c--...t.aaatg..a....a [ 86880]
AM712240.1_OURT_88/3             aaaaaa...t------------------------------..t...............a.gtg..t.aaatg..a..... [ 86880]
AM712239.1_Benin_97/1            aaaaaa...t------------------------------..t...............a.gtg..t.aaatg..a..... [ 86880]
KM111295.1_Ken06.Bus             aga.aa..tacaatattttcatacata---gttttaaaaa..t..........t.....a--g..t.aaatg..a....a [ 86880]
KM111294.1_Ken05/Tk1             aaaaaat.taaatcat----------------------at..t...a.a.c....c...c--...t.aaatg..a....a [ 86880]
KM262844.1_strain_L60            aaaaaa...t------------------------------..t...............a.gtg..t.aaatg..a..... [ 86880]
KM102979.1_26544/OG10_Italy      aaaaaa...t------------------------------..t...............a.gtg..t.aaatg..a..... [ 86880]
KM262845.1_strain_NHV            aaaaaa...t------------------------------..t...............a.gtg..t.aaatg..a..... [ 86880]
KX354450.1_47/Ss/2008            aaaaaa...t------------------------------..t...............a.gtg..t.aaatg..a..... [ 86880]
KP055815.1_strain_BA71           aaaaaa...t------------------------------..t...............a.gtg..t.aaatg..a..... [ 86880]
MH025920.1_strain_R35            aga.aa..tacaatattttcatacata---gttttaaaaa..t..........t.....a--g..t.aaatg..a....a [ 86880]
MH025919.1_strain_N10            aga.aa..tacaatattttcatacata---gttttaaaaa..t..........t.....a--g..t.aaatg..a....a [ 86880]
MH025918.1_strain_R25            aga.aa..tacaatattttcatacata---gttttaaaaa..t..........t.....a--g..t.aaatg..a....a [ 86880]
MH025917.1_strain_R7             aga.aa..tacaatattttcatacata---gttttaaaaa..t..........t.....a--g..t.aaatg..a....a [ 86880]
MH025916.1_strain_R8             aga.aa..tacaatattttcatacata---gttttaaaaa..t..........t.....a--g..t.aaatg..a....a [ 86880]

FR682468.2_Georgia_2007/1        acttatttttttaatattttctaacatagtttta---------------------------------------------a [ 86960]
ASF_IND_20_CAD543                .g.............g.g..ta..a.........------------------------------------------aac. [ 86960]
LR899193.1_Germany_2020/1        ..................................---------------------------------------------. [ 86960]
MN393477.1_Wuhan_2019            ..................................---------------------------------------------. [ 86960]
MW396979.1_Timor-Leste/2019/1    ..................................---------------------------------------------. [ 86960]
MK940252.1_CN/2019/InnerMongolia ..................................---------------------------------------------. [ 86960]
MK128995.1_China/2018/AnhuiXCGQ  ..................................---------------------------------------------. [ 86960]
MH681419.1_ASFV/POL/2015         ..................................---------------------------------------------. [ 86960]
LS478113.1_Estonia_2014          ..................................---------------------------------------------. [ 86960]
KP843857.1_Odintsovo_02/14       ..................................---------------------------------------------. [ 86960]
FN557520.1_strain_E75            .g...-.........g.g..ta..a.........------------------------------------------aac. [ 86960]
AY261366.1_Warthog               ...............t.g..tc..a.........ag---------------------------------------tatt. [ 86960]
AY261365.1_Warmbaths             c................a..ta..ac.g..c...aa---------------------------------------taat. [ 86960]
AY261364.1_Tengani_62            gg...........g.t..c.t...a..g......------------------------------------------agt. [ 86960]
AY261363.1_Pretoriuskop/96/4     ..............gt.a.at...t.........ag---------------------------------------taat. [ 86960]
AY261362.1_Mkuzi_1979            .................g..ta..a.........aa---------------------------------------taat. [ 86960]
AY261361.1_Malawi_Lil-20/1_1983  ................a..c..............---------------------------------------------. [ 86960]
AY261360.1_Kenya_1950            ...............t.g..t...a.........agtattgataataaaacaaaatttaatgaaacattaattttagat. [ 86960]
AM712240.1_OURT_88/3             .g.............g.g..ta..a.........------------------------------------------aac. [ 86960]
AM712239.1_Benin_97/1            .g.............g.g..ta..a.........------------------------------------------aac. [ 86960]
KM111295.1_Ken06.Bus             g................ca.t...a..ga.....gg-------------------------------------------- [ 86960]
```

```
KM111294.1_Ken05/Tk1             ...............t.g..t...a.........agtattgataataaaacaaaatttaatgaaacattaattttagat. [ 86960]
KM262844.1_strain_L60            .g.............g.g..ta..a.........------------------------------------------aac. [ 86960]
KM102979.1_26544/OG10_Italy      .g.............g.g..ta..a.........------------------------------------------aac. [ 86960]
KM262845.1_strain_NHV            .g.............g.g..ta..a.........------------------------------------------aac. [ 86960]
KX354450.1_47/Ss/2008            .g.............g.g..ta..a.........------------------------------------------aac. [ 86960]
KP055815.1_strain_BA71           .g.............g.g..ta..a.........------------------------------------------aac. [ 86960]
MH025920.1_strain_R35            g................ca.t...a..ga.....gg-------------------------------------------- [ 86960]
MH025919.1_strain_N10            g................ca.t...a..ga.....gg-------------------------------------------- [ 86960]
MH025918.1_strain_R25            g................ca.t...a..ga.....gg-------------------------------------------- [ 86960]
MH025917.1_strain_R7             g................ca.t...a..ga.....gg-------------------------------------------- [ 86960]
MH025916.1_strain_R8             g................ca.t...a..ga.....gg-------------------------------------------- [ 86960]

FR682468.2_Georgia_2007/1        gtattgattattgggtt---agttttaataaaacaataattttagatagtaatattactaatgataataat--------- [ 87040]
ASF_IND_20_CAD543                a...nnnnnnnnnnnnn---nn...a...c....tg..t.....a...a..t.t....a.t.nnnnnnnnn--------- [ 87040]
LR899193.1_Germany_2020/1        .................---...................................................--------- [ 87040]
MN393477.1_Wuhan_2019            .................---...................................................--------- [ 87040]
MW396979.1_Timor-Leste/2019/1    .................---...................................................--------- [ 87040]
MK940252.1_CN/2019/InnerMongolia .................---...................................................--------- [ 87040]
MK128995.1_China/2018/AnhuiXCGQ  .................---...................................................--------- [ 87040]
MH681419.1_ASFV/POL/2015         .................---...................................................--------- [ 87040]
LS478113.1_Estonia_2014          .................---...................................................--------- [ 87040]
KP843857.1_Odintsovo_02/14       .................---...................................................--------- [ 87040]
FN557520.1_strain_E75            a....ataata...ag.---.c...a...c....tg..t.....a...a..t.t....a.t.a..g...ca--------- [ 87040]
AY261366.1_Warthog               a.ta.tgggt.a..ta.---------...g.t...g...c....a..........c.a..gc..g.cag.---------- [ 87040]
AY261365.1_Warmbaths             a....at.ata...ca.cct.c...a...g.......tt.....a..gt..ca.c..taggga...t..g---------- [ 87040]
AY261364.1_Tengani_62            a....ataata...ag.---.c...a.....c......ta....a...t....ta.t.ag..ag....tta--------- [ 87040]
AY261363.1_Pretoriuskop/96/4     a.ta.tgggc.a.ac..---------..cg.........c....a.......c.............c....--------- [ 87040]
AY261362.1_Mkuzi_1979            a....ataata...ca.---.c...a...g.......tt.....a...a..g......att.acgg.....--------- [ 87040]
AY261361.1_Malawi_Lil-20/1_1983  .................---...................................................--------- [ 87040]
AY261360.1_Kenya_1950            a...............a---.ag......g.t.......c....................g...---------------- [ 87040]
AM712240.1_OURT_88/3             a....ataata...ag.---.c...a...c....tg..-.....a...a..t.t....a.t.a..g...ca--------- [ 87040]
AM712239.1_Benin_97/1            a....ataata...ag.---.c...a...c....tg..t.....a...a..t.t....a.t.a..g...ca--------- [ 87040]
KM111295.1_Ken06.Bus             -................---...c.......t....................c.......t.a......t.actaatact [ 87040]
KM111294.1_Ken05/Tk1             a...............a---.ag......g.t.......c....................g...---------------- [ 87040]
KM262844.1_strain_L60            a....ataata...ag.---.c...a...c....tg..t.....a...a..t.t....a.t.a..g...ca--------- [ 87040]
KM102979.1_26544/OG10_Italy      a....ataata...ag.---.c...a...c....tg..t.....a...a..t.t....a.t.a..g...ca--------- [ 87040]
KM262845.1_strain_NHV            a....ataata...ag.---.c...a...c....tg..-.....a...a..t.t....a.t.a..g...ca--------- [ 87040]
KX354450.1_47/Ss/2008            a....ataata...ag.---.c...a...c....tg..t.....a...a..t.t....a.t.a..g...ca--------- [ 87040]
KP055815.1_strain_BA71           a....ataata...ag.---.c...a...c....tg..t.....a...a..t.t....a.t.a..g...ca--------- [ 87040]
MH025920.1_strain_R35            -................---...c.......t....................c.......t.a......t.actaatact [ 87040]
MH025919.1_strain_N10            -................---...c.......t....................c.......t.a......t.actaatact [ 87040]
MH025918.1_strain_R25            -................---...c.......t....................c.......t.a......t.actaatact [ 87040]
MH025917.1_strain_R7             -................---...c.......t....................c.......t.a......t.actaatact [ 87040]
MH025916.1_strain_R8             -................---...c.......t....................c.......t.a......t.actaatact [ 87040]
```

```
FR682468.2_Georgia_2007/1        --------------gatat---aaatggagtatcatgga------------attttttaataattcttttaatacacta [ 87120]
ASF_IND_20_CAD543                --------------nnnnn---nnnnnnnnnnnnnnnnn------------nnnnnnnnnnnnnnnnnnnnnn.att.t [ 87120]
LR899193.1_Germany_2020/1        --------------.....---.................------------............................ [ 87120]
MN393477.1_Wuhan_2019            --------------.....---.................------------............................ [ 87120]
MW396979.1_Timor-Leste/2019/1    --------------.....---.................------------............................ [ 87120]
MK940252.1_CN/2019/InnerMongolia --------------.....---.................------------............................ [ 87120]
MK128995.1_China/2018/AnhuiXCGQ  --------------.....---.................------------............................ [ 87120]
MH681419.1_ASFV/POL/2015         --------------.....---.................------------............................ [ 87120]
LS478113.1_Estonia_2014          --------------.....---.................------------............................ [ 87120]
KP843857.1_Odintsovo_02/14       --------------.....---.................------------............................ [ 87120]
FN557520.1_strain_E75            --------------t..gg---tgg.ct.t.t.ggaat.------------ca.a..a.g....aa.cg..g..att.t [ 87120]
AY261366.1_Warthog               ----------------------.gg.at.t.t.ggaat--------------....a......a.........ta.t [ 87120]
AY261365.1_Warmbaths             ----------------------tgg.ct.t.t.ggaat.------------ca.a..a......aa.cg..g.ga...t [ 87120]
AY261364.1_Tengani_62            --------------t...c---tgg.ct.t.t.ggaat.------------.a.a..a......a..cg.....att.t [ 87120]
AY261363.1_Pretoriuskop/96/4     --------------a..gaatt.gg.at.t.t.ggaattcttataataatac..a..a......a.a.......ata.t [ 87120]
AY261362.1_Mkuzi_1979            --------------ct.gg---tgg.ct.t.t.ggaat.------------ca.a..a......aa.ca..g..tc..t [ 87120]
AY261361.1_Malawi_Lil-20/1_1983  --------------.....aaatgga.t.tc..ggaatt---------------.cc..........c.......... [ 87120]
AY261360.1_Kenya_1950            --------------a...ga--.gc..t.tct.ggaatttt---------------.a......a.......ct...t [ 87120]
AM712240.1_OURT_88/3             --------------t..gg---tgg.ct.t.t.ggaat.------------ca.a..a.g....aa.cg..g..att.t [ 87120]
AM712239.1_Benin_97/1            --------------t..gg---tgg.ct.t.t.ggaat.------------ca.a..a.g....aa.cg..g..att.t [ 87120]
KM111295.1_Ken06.Bus             actaataatcctacat..aa--tgg.at.t.t.ggaat.tttataata---..ag..a......a.......tt...t [ 87120]
KM111294.1_Ken05/Tk1             --------------a...ga--.gc..t.tct.ggaatttt---------------.a......a.......ct...t [ 87120]
KM262844.1_strain_L60            --------------t..gg---tgg.ct.t.t.ggaat.------------ca.a..a.g....aa.cg..g..att.t [ 87120]
KM102979.1_26544/OG10_Italy      --------------t..gg---tgg.ct.t.t.ggaat.------------ca.a..a.g....aa.cg..g..att.t [ 87120]
KM262845.1_strain_NHV            --------------t..gg---tgg.ct.t.t.ggaat.------------ca.a..a.g....aa.cg..g..att.t [ 87120]
KX354450.1_47/Ss/2008            --------------t..gg---tgg.ct.t.t.ggaat.------------ca.a..a.g....aa.cg..g..att.t [ 87120]
KP055815.1_strain_BA71           --------------t..gg---tgg.ct.t.t.ggaat.------------ca.a..a.g....aa.cg..g..att.t [ 87120]
MH025920.1_strain_R35            actaataatcctacat..aa--tgg.at.t.t.ggaat.tttataata---..ag..a......a.......tt...t [ 87120]
MH025919.1_strain_N10            actaataatcctacat..aa--tgg.at.t.t.ggaat.tttataata---..ag..a......a.......tt...t [ 87120]
MH025918.1_strain_R25            actaataatcctacat..aa--tgg.at.t.t.ggaat.tttataata---..ag..a......a.......tt...t [ 87120]
MH025917.1_strain_R7             actaataatcctacat..aa--tgg.at.t.t.ggaat.tttataata---..ag..a......a.......tt...t [ 87120]
MH025916.1_strain_R8             actaataatcctacat..aa--tgg.at.t.t.ggaat.tttataata---..ag..a......a.......tt...t [ 87120]

FR682468.2_Georgia_2007/1        gctacatgtggaaaagcaggtaac---ttttgtgaatg---ttc---taat----------------------------- [ 87200]
ASF_IND_20_CAD543                a..tat.......t......a..t---.a....tnnnn---nnn---nnn.----------------------------- [ 87200]
LR899193.1_Germany_2020/1        ........................---..........---...---....----------------------------- [ 87200]
MN393477.1_Wuhan_2019            ........................---..........---...---....----------------------------- [ 87200]
MW396979.1_Timor-Leste/2019/1    ........................---..........---...---....----------------------------- [ 87200]
MK940252.1_CN/2019/InnerMongolia ........................---..........---...---....----------------------------- [ 87200]
MK128995.1_China/2018/AnhuiXCGQ  ........................---..........---...---....----------------------------- [ 87200]
MH681419.1_ASFV/POL/2015         ........................---..........---...---....----------------------------- [ 87200]
LS478113.1_Estonia_2014          ........................---..........---...---....----------------------------- [ 87200]
KP843857.1_Odintsovo_02/14       ........................---..........---...---....----------------------------- [ 87200]
```

```
FN557520.1_strain_E75            a..tat.......t......a..t---.a....tc...---..g---.gg.----------------------------- [ 87200]
AY261366.1_Warthog               ..c............aa.aa...t---g..........---...---....----------------------------- [ 87200]
AY261365.1_Warmbaths             a...ag..c....t.tataa...t---.a....tc...---.ga---..g.----------------------------- [ 87200]
AY261364.1_Tengani_62            a....t....t..c.t..aa..gtagc.a....ac...---.aa---aggc----------------------------- [ 87200]
AY261363.1_Pretoriuskop/96/4     ....t..........aa......tt---..........---.aa---..t.aattataatacatctatatctaatacatc [ 87200]
AY261362.1_Mkuzi_1979            a...gt..c....t.tatcaa..t---.a....tc...cac.aa---....----------------------------- [ 87200]
AY261361.1_Malawi_Lil-20/1_1983  ........................---...........---c..---....----------------------------- [ 87200]
AY261360.1_Kenya_1950            ....t.............a.....---......agt..---...---....----------------------------- [ 87200]
AM712240.1_OURT_88/3             a..tat.......t......a..t---.a....tc...---..g---.gg.----------------------------- [ 87200]
AM712239.1_Benin_97/1            a..tat.......t......a..t---.a....tc...---..g---.gg.----------------------------- [ 87200]
KM111295.1_Ken06.Bus             a.c........g..ta..ta...t---a.a...tct..---...taa....----------------------------- [ 87200]
KM111294.1_Ken05/Tk1             ..................a.....---......agt..---...---....----------------------------- [ 87200]
KM262844.1_strain_L60            a..tat.......t......a..t---.a....tc...---..g---.gg.----------------------------- [ 87200]
KM102979.1_26544/OG10_Italy      a..tat.......t......a..t---.a....tc...---..g---.gg.----------------------------- [ 87200]
KM262845.1_strain_NHV            a..tat.......t......a..t---.a....tc...---..g---.gg.----------------------------- [ 87200]
KX354450.1_47/Ss/2008            a..tat.......t......a..t---.a....tc...---..g---.gg.----------------------------- [ 87200]
KP055815.1_strain_BA71           a..tat.......t......a..t---.a....tc...---..g---.gg.----------------------------- [ 87200]
MH025920.1_strain_R35            a.c........g..ta..ta...t---a.a...tct..---...taa....----------------------------- [ 87200]
MH025919.1_strain_N10            a.c........g..ta..ta...t---a.a...tct..---...taa....----------------------------- [ 87200]
MH025918.1_strain_R25            a.c........g..ta..ta...t---a.a...tct..---...taa....----------------------------- [ 87200]
MH025917.1_strain_R7             a.c........g..ta..ta...t---a.a...tct..---...taa....----------------------------- [ 87200]
MH025916.1_strain_R8             a.c........g..ta..ta...t---a.a...tct..---...taa....----------------------------- [ 87200]

FR682468.2_Georgia_2007/1        ----tatagtacatcaatatataat------ataacaaataattgtagcttaactatttttcctcataat---gatgtat [ 87280]
ASF_IND_20_CAD543                ----c...ac.t....t.g...nnn------nnnnnnnnnnnnnnnnnnnnnnnnnnnnnnn..a.c...aca..aa... [ 87280]
LR899193.1_Germany_2020/1        ----.....................------.......................................---....... [ 87280]
MN393477.1_Wuhan_2019            ----.....................------.......................................---....... [ 87280]
MW396979.1_Timor-Leste/2019/1    ----.....................------.......................................---....... [ 87280]
MK940252.1_CN/2019/InnerMongolia ----.....................------.......................................---....... [ 87280]
MK128995.1_China/2018/AnhuiXCGQ  ----.....................------.......................................---....... [ 87280]
MH681419.1_ASFV/POL/2015         ----.....................------.......................................---....... [ 87280]
LS478113.1_Estonia_2014          ----.....................------.......................................---....... [ 87280]
KP843857.1_Odintsovo_02/14       ----.....................------.......................................---....... [ 87280]
FN557520.1_strain_E75            ----c...ac.t....t.g......------.c...............t....t..........a.c...aca..aa... [ 87280]
AY261366.1_Warthog               ----...ga..a...ct........------.................t.....a........aa.....acaa.aa... [ 87280]
AY261365.1_Warmbaths             ----ct..a.......t.g......------..---............t...gt.........aa.....acaa.aa... [ 87280]
AY261364.1_Tengani_62            ----c...ac......t.g......------.................t....tc........aa.....acaa.aa... [ 87280]
AY261363.1_Pretoriuskop/96/4     tata.c..a................------g................t.....g.......t.t..g..gata..a... [ 87280]
AY261362.1_Mkuzi_1979            ----.....c......t........------...............g.t....ta........aa.....acaa.aag.. [ 87280]
AY261361.1_Malawi_Lil-20/1_1983  ----............c........------...g..c......................c.........---....... [ 87280]
AY261360.1_Kenya_1950            ----............t.t.t....------..t..............t............ttaatag.cgagacgt... [ 87280]
AM712240.1_OURT_88/3             ----c...ac.t....t.g......------.c...............t....t..........a.c...aca..aa... [ 87280]
AM712239.1_Benin_97/1            ----c...ac.t....t.g......------.c...............t....t..........a.c...aca..aa... [ 87280]
KM111295.1_Ken06.Bus             ----....a....at.t...t....tataat.cc..............t....t.....c.......g..gaaa.aa.t. [ 87280]
```

```
KM111294.1_Ken05/Tk1             ----............t.t.t....------..t..............t............ttaatag.cgagacgt... [ 87280]
KM262844.1_strain_L60            ----c...ac.t....t.g......------.c...............t....t..........a.c...aca..aa... [ 87280]
KM102979.1_26544/OG10_Italy      ----c...ac.t....t.g......------.c...............t....t..........a.c...aca..aa... [ 87280]
KM262845.1_strain_NHV            ----c...ac.t....t.g......------.c...............t....t..........a.c...aca..aa... [ 87280]
KX354450.1_47/Ss/2008            ----c...ac.t....t.g......------.c...............t....t..........a.c...aca..aa... [ 87280]
KP055815.1_strain_BA71           ----c...ac.t....t.g......------.c...............t....t..........a.c...aca..aa... [ 87280]
MH025920.1_strain_R35            ----....a....at.t...t....tataac.cc..............t....t.....c.......g..gaaa.aa.t. [ 87280]
MH025919.1_strain_N10            ----....a....at.t...t....tataac.cc..............t....t.....c.......g..gaaa.aa.t. [ 87280]
MH025918.1_strain_R25            ----....a....at.t...t....tataac.cc..............t....t.....c.......g..gaaa.aa.t. [ 87280]
MH025917.1_strain_R7             ----....a....at.t...t....tataac.cc..............t....t.....c.......g..gaaa.aa.t. [ 87280]
MH025916.1_strain_R8             ----....a....at.t...t....tataac.cc..............t....t.....c.......g..gaaa.aa.t. [ 87280]

FR682468.2_Georgia_2007/1        ttgatacaacatatcaagtagtatg---gaatcaaataattaattatacaataaaattattaacacctgctactccccca [ 87360]
ASF_IND_20_CAD543                ..a...g....nnnnnnnnnn...attt.g.ca...a.............g..nnnnnnnnnnnnnnnnnnnnnnn.... [ 87360]
LR899193.1_Germany_2020/1        .........................---.................................................... [ 87360]
MN393477.1_Wuhan_2019            .........................---.................................................... [ 87360]
MW396979.1_Timor-Leste/2019/1    .........................---.................................................... [ 87360]
MK940252.1_CN/2019/InnerMongolia .........................---.................................................... [ 87360]
MK128995.1_China/2018/AnhuiXCGQ  .........................---.................................................... [ 87360]
MH681419.1_ASFV/POL/2015         .........................---.................................................... [ 87360]
LS478113.1_Estonia_2014          .........................---.................................................... [ 87360]
KP843857.1_Odintsovo_02/14       .........................---.................................................... [ 87360]
FN557520.1_strain_E75            ..a...g.......g..t......attt.g.ca...a.............g.....c......a.t...t.ga.t..... [ 87360]
AY261366.1_Warthog               ..a..............t......attcacgca.t.g.........c........t..g...tt.....t...at.t... [ 87360]
AY261365.1_Warmbaths             ..a......t....g..t..a...attcagg.a.tca..c..................gc..ga.t..a.......t... [ 87360]
AY261364.1_Tengani_62            ..a...........g..t......attta...a.t.a................c..a.gg..ca.....t.ga...t... [ 87360]
AY261363.1_Pretoriuskop/96/4     ..a.---..........t......atca....t.c.a..................c......tt.....t....t.t... [ 87360]
AY261362.1_Mkuzi_1979            ..a.c.....g...g..t......attta...a.t.gt.c...a...........c......gt.....t......t... [ 87360]
AY261361.1_Malawi_Lil-20/1_1983  ...g....c................---.........................................t.......... [ 87360]
AY261360.1_Kenya_1950            ..a..............a......attca.c.a.t..............a.....c.......t.....t......t... [ 87360]
AM712240.1_OURT_88/3             ..a...g.......g..t......attt.g.ca...a.............g.....c......a.t...t.ga.t..... [ 87360]
AM712239.1_Benin_97/1            ..a...g.......g..t......attt.g.ca...a.............g.....c......a.t...t.ga.t..... [ 87360]
KM111295.1_Ken06.Bus             .......g.tg.t...ga..a...attta.c.a.t.a................cg..gg...ca.....t.ga....... [ 87360]
KM111294.1_Ken05/Tk1             ..a..............a......attca.c.a.t..............a.....c.......t.....t......t... [ 87360]
KM262844.1_strain_L60            ..a...g.......g..t......attt.g.ca...a.............g.....c......a.t...t.ga.t..... [ 87360]
KM102979.1_26544/OG10_Italy      ..a...g.......g..t......attt.g.ca...a.............g.....c......a.t...t.ga.t..... [ 87360]
KM262845.1_strain_NHV            ..a...g.......g..t......attt.g.ca...a.............g.....c......a.t...t.ga.t..... [ 87360]
KX354450.1_47/Ss/2008            ..a...g.......g..t......attt.g.ca...a.............g.....c......a.t...t.ga.t..... [ 87360]
KP055815.1_strain_BA71           ..a...g.......g..t......attt.g.ca...a.............g.....c......a.t...t.ga.t..... [ 87360]
MH025920.1_strain_R35            .........tg.t...ga..a...attta.c.a.t.a................cg..gg...ca.....t.ga....... [ 87360]
MH025919.1_strain_N10            .......g.tg.t...ga..a...attta.c.a.t.a................cg..gg...ca.....t.ga....... [ 87360]
MH025918.1_strain_R25            .........tg.t...ga..a...attta.c.a.t.a................cg..gg...ca.....t.ga....... [ 87360]
MH025917.1_strain_R7             .........tg.t...ga..a...attta.c.a.t.a................cg..gg...ca.....t.ga....... [ 87360]
MH025916.1_strain_R8             .........tg.t...ga..a...attta.c.a.t.a................cg..gg...ca.....t.ga....... [ 87360]
```

```
FR682468.2_Georgia_2007/1        aatatcacatataattgtac---taattttttaataacatgtaaaaaaaataatggaacaaacactaatatatatttaaa [ 87440]
ASF_IND_20_CAD543                .c...t..............---........................n..............tgt..............t [ 87440]
LR899193.1_Germany_2020/1        ....................---......................................................... [ 87440]
MN393477.1_Wuhan_2019            ....................---......................................................... [ 87440]
MW396979.1_Timor-Leste/2019/1    ....................---......................................................... [ 87440]
MK940252.1_CN/2019/InnerMongolia ....................---......................................................... [ 87440]
MK128995.1_China/2018/AnhuiXCGQ  ....................---......................................................... [ 87440]
MH681419.1_ASFV/POL/2015         ....................---......................................................... [ 87440]
LS478113.1_Estonia_2014          ....................---......................................................... [ 87440]
KP843857.1_Odintsovo_02/14       ....................---...................-..................................... [ 87440]
FN557520.1_strain_E75            .c...t..............---.....c..................t........g.....tgt..............t [ 87440]
AY261366.1_Warthog               .t...t..............---.c.a.c.............g......c............t.t.c..t...tc..... [ 87440]
AY261365.1_Warmbaths             .....t..............---..ca.c.g.........................g.....t..........t.....c [ 87440]
AY261364.1_Tengani_62            .t...t.t.......ga...taa.....c.............g................g..t...g.a..c......t. [ 87440]
AY261363.1_Pretoriuskop/96/4     .....t..............---...c.c.............g........g....g..............g.t....tc [ 87440]
AY261362.1_Mkuzi_1979            .....t..............---..ca.c.g....................g....c.....t........g.t...... [ 87440]
AY261361.1_Malawi_Lil-20/1_1983  .....t..............---.......................................t....t.........c.. [ 87440]
AY261360.1_Kenya_1950            .t...tt...........g.---.....g..ct....at..........g......t.....t........t.t....tc [ 87440]
AM712240.1_OURT_88/3             .c...t..............---.....c..................t........g.....tgt..............t [ 87440]
AM712239.1_Benin_97/1            .c...t..............---.....c..................t........g.....tgt..............t [ 87440]
KM111295.1_Ken06.Bus             .....tt...t....ga..g---.....c........a....g...................t...g.a..c......t. [ 87440]
KM111294.1_Ken05/Tk1             .t...tt.............---.....g..ct....at..........g......t.....t........t.t....tc [ 87440]
KM262844.1_strain_L60            .c...t..............---.....c..................t........g.....tgt..............t [ 87440]
KM102979.1_26544/OG10_Italy      .c...t..............---.....c..................t........g.....tgt..............t [ 87440]
KM262845.1_strain_NHV            .c...t..............---.....c..................t........g.....tgt..............t [ 87440]
KX354450.1_47/Ss/2008            .c...t..............---.....c..................t........g.....tgt..............t [ 87440]
KP055815.1_strain_BA71           .c...t..............---.....c..................t........g.....tgt..............t [ 87440]
MH025920.1_strain_R35            .....tt...t....ga..g---.....c........a....g...................t...g.a..c......t. [ 87440]
MH025919.1_strain_N10            .....tt...t....ga..g---.....c........a....g...................t...g.a..c......t. [ 87440]
MH025918.1_strain_R25            .....tt...t....ga..g---.....c........a....g...................t...g.a..c......t. [ 87440]
MH025917.1_strain_R7             .....tt...t....ga..g---.....c........a....g...................t...g.a..c......t. [ 87440]
MH025916.1_strain_R8             .....tt...t....ga..g---.....c........a....g...................t...g.a..c......t. [ 87440]

FR682468.2_Georgia_2007/1        tataaatgatacttttgttaaatatactaatgaaagtatacttgaatataactggaataatagtaac---attaacaatt [ 87520]
ASF_IND_20_CAD543                ...t..ca....aa...nnnnnnnnnnnnnnnnnnnnnnnn..a.t...t.t......gg..a---------...t.... [ 87520]
LR899193.1_Germany_2020/1        ...................................................................---.......... [ 87520]
MN393477.1_Wuhan_2019            ...................................................................---.......... [ 87520]
MW396979.1_Timor-Leste/2019/1    ...................................................................---.......... [ 87520]
MK940252.1_CN/2019/InnerMongolia ...................................................................---.......... [ 87520]
MK128995.1_China/2018/AnhuiXCGQ  ...................................................................---.......... [ 87520]
MH681419.1_ASFV/POL/2015         ...................................................................---.......... [ 87520]
LS478113.1_Estonia_2014          ...................................................................---.......... [ 87520]
KP843857.1_Odintsovo_02/14       ...................................................................---.......... [ 87520]
```

```
FN557520.1_strain_E75            ...t..ca....aa.......tg.........g.ga...c...a.t...t.t......gg.---------.a...t.... [ 87520]
AY261366.1_Warthog               .t..........aa..aa.g..........ca.....t.t..ca.t..ct.t.......g....g.gtta.a...t.t.. [ 87520]
AY261365.1_Warmbaths             .......a....ga.......t.c..........ga.g.ta..t.c..ct.t.......ga...g.att---...t..c. [ 87520]
AY261364.1_Tengani_62            .t..........a...a....ta............a..gta..a....ct.t......tg...cg.atta.acc.t...a [ 87520]
AY261363.1_Pretoriuskop/96/4     .......a...taacca....tc........c..ga...t...act..ct.t............g.gtt---...t.... [ 87520]
AY261362.1_Mkuzi_1979            a...........aa.c......ag..g..........g.tt.......ctct.......g..c.g.ttta.a........ [ 87520]
AY261361.1_Malawi_Lil-20/1_1983  .............aa....................................................tt---........ [ 87520]
AY261360.1_Kenya_1950            ............aa......c..................t...a.t...g.t.at..ctg.....tttta.a...t.... [ 87520]
AM712240.1_OURT_88/3             ...t..ca....aa.......tg.........g.ga...c...a.t...t.t......gg.---------.a...t.... [ 87520]
AM712239.1_Benin_97/1            ...t..ca....aa.......tg.........g.ga...c...a.t...t.t......gg.---------.a...t.... [ 87520]
KM111295.1_Ken06.Bus             .t..........a...a....ta............a.gat...a.....t.t......tg....g.gtta.act.t...a [ 87520]
KM111294.1_Ken05/Tk1             ............aa......c..................t...a.t...g.t.at..ctg.....tttta.a...t.... [ 87520]
KM262844.1_strain_L60            .------------------------------------------------------------------------------- [ 87520]
KM102979.1_26544/OG10_Italy      ...t..ca....aa.......tg.........g.ga...c...a.t...t.t......gg.---------.a...t.... [ 87520]
KM262845.1_strain_NHV            ...t..ca....aa.......tg.........g.ga...c...a.t...t.t......gg.---------.a...t.... [ 87520]
KX354450.1_47/Ss/2008            ...t..ca....aa.......tg.........g.ga...c...a.t...t.t......gg.---------.a...t.... [ 87520]
KP055815.1_strain_BA71           ...t..ca....aa.......tg.........g.ga...c...a.t...t.t......gg.---------.a...t.... [ 87520]
MH025920.1_strain_R35            .t..........a...a....ta............a.gat...a.....t.t......tg....g.gtta.act.t...a [ 87520]
MH025919.1_strain_N10            .t..........a...a....ta............a.gat...a.....t.t......tg....g.gtta.act.t...a [ 87520]
MH025918.1_strain_R25            .t..........a...a....ta............a.gat...a.....t.t......tg....g.gtta.act.t...a [ 87520]
MH025917.1_strain_R7             .t..........a...a....ta............a.gat...a.....t.t......tg....g.gtta.act.t...a [ 87520]
MH025916.1_strain_R8             .t..........a...a....ta............a.gat...a.....t.t......tg....g.gtta.act.t...a [ 87520]

FR682468.2_Georgia_2007/1        ttacagctacatgtataattaataatacaattagtacatctaatgaaacaacacttataaattgtacttat---ttaaca [ 87600]
ASF_IND_20_CAD543                ...................................t...tg.........ga.aa...........nnnnn---nn..t. [ 87600]
LR899193.1_Germany_2020/1        .......................................................................---...... [ 87600]
MN393477.1_Wuhan_2019            .......................................................................---...... [ 87600]
MW396979.1_Timor-Leste/2019/1    .......................................................................---...... [ 87600]
MK940252.1_CN/2019/InnerMongolia .......................................................................---...... [ 87600]
MK128995.1_China/2018/AnhuiXCGQ  .......................................................................---...... [ 87600]
MH681419.1_ASFV/POL/2015         .......................................................................---...... [ 87600]
LS478113.1_Estonia_2014          .......................................................................---...... [ 87600]
KP843857.1_Odintsovo_02/14       .......................................................................---...... [ 87600]
FN557520.1_strain_E75            ................g..................t...tg.........ga.aa.............a..---cc..t. [ 87600]
AY261366.1_Warthog               ..ctt...........t............c...a.t..g.a...act....a.g....c.....c...a.c---cc.tt. [ 87600]
AY261365.1_Warmbaths             ................g................a.t....a...at.t..ga.a.....g.c......a..aca...tt. [ 87600]
AY261364.1_Tengani_62            .....a..........t............c...a.t.gg.a...acc....a.g..........c...a..---c..ttg [ 87600]
AY261363.1_Pretoriuskop/96/4     ................g............c...a.t..g.a...acc....a.g....c.....c...a..---cc.tt. [ 87600]
AY261362.1_Mkuzi_1979            ....g....t.......................a.t....a.....c...ca.ac....g.a...c.ca..---a..tt. [ 87600]
AY261361.1_Malawi_Lil-20/1_1983  .................................a.t....a.....t...ca.ac....g.c..c.tca..---ac.tt. [ 87600]
AY261360.1_Kenya_1950            ......t...........................c.....a...act...ga.t.......c..c...a..---a..tt. [ 87600]
AM712240.1_OURT_88/3             ................g..................t...tg.........ga.aa.............a..---cc..t. [ 87600]
AM712239.1_Benin_97/1            ................g..................t...tg.........ga.aa.............a..---cc..t. [ 87600]
KM111295.1_Ken06.Bus             ................t............c...a.t.gg.a...acc....a.g..........c...a..---c..tt. [ 87600]
```

```
KM111294.1_Ken05/Tk1              ......t.................................a...act...ga.t.......c..c...a..---a..tt. [ 87600]
KM262844.1_strain_L60             -----------------..................t...tg.........ga.aa.............a..---cc..t. [ 87600]
KM102979.1_26544/OG10_Italy       ................g..................t...tg.........ga.aa.............a..---cc..t. [ 87600]
KM262845.1_strain_NHV             ................g..................t...tg.........ga.aa.............a..---cc..t. [ 87600]
KX354450.1_47/Ss/2008             ................g..................t...tg.........ga.aa.............a..---cc..t. [ 87600]
KP055815.1_strain_BA71            ................g..................t...tg.........ga.aa.............a..---cc..t. [ 87600]
MH025920.1_strain_R35             ................t............c...a.t.gg.a...acc....a.g..........c...a..---c..tt. [ 87600]
MH025919.1_strain_N10             ................t............c...a.t.gg.a...acc....a.g..........c...a..---c..tt. [ 87600]
MH025918.1_strain_R25             ................t............c...a.t.gg.a...acc....a.g..........c...a..---c..tt. [ 87600]
MH025917.1_strain_R7              ................t............c...a.t.gg.a...acc....a.g..........c...a..---c..tt. [ 87600]
MH025916.1_strain_R8              ................t............c...a.t.gg.a...acc....a.g..........c...a..---c..tt. [ 87600]

FR682468.2_Georgia_2007/1         ttgtcatctaactattt---------------ttatactttttttaaattatattatattccattaagcatcataattgg [ 87680]
ASF_IND_20_CAD543                 ..a------..a...ca---------------aa..ta...a.cc.c.....t......cat.a.tttt..tnnnnnnnn [ 87680]
LR899193.1_Germany_2020/1         .................---------------................................................ [ 87680]
MN393477.1_Wuhan_2019             .................---------------................................................ [ 87680]
MW396979.1_Timor-Leste/2019/1     .................---------------................................................ [ 87680]
MK940252.1_CN/2019/InnerMongolia  .................---------------................................................ [ 87680]
MK128995.1_China/2018/AnhuiXCGQ   .................---------------................................................ [ 87680]
MH681419.1_ASFV/POL/2015          .................---------------................................................ [ 87680]
LS478113.1_Estonia_2014           .................---------------................................................ [ 87680]
KP843857.1_Odintsovo_02/14        .................---------------................................................ [ 87680]
FN557520.1_strain_E75             ..a------..a...ca---------------aa..ta...a.cc.c.....t......cat.a.tttt..tg.g.g... [ 87680]
AY261366.1_Warthog                ..a------..a.c..accaaaattattttct.g.a.a.a..ca..c.....t......gat.a.tttt..tg..tc... [ 87680]
AY261365.1_Warmbaths              ..a------...ag..atttagacttttttcaagtag.cgg..a..---.g.t......gat.a.tttt..tg...c... [ 87680]
AY261364.1_Tengani_62             ..a------..a.c.gaccaaaattattttct.a.a.a.a..ca..c.....t......cat.a.tttt..tg.ggg... [ 87680]
AY261363.1_Pretoriuskop/96/4      ..a------..t.c..accaaaattattttct.g.a.a.a..ca..c.....t......cat.a.tttt..tg.g.g... [ 87680]
AY261362.1_Mkuzi_1979             ..a------...a...atttagac--ttttttca.gtaac.ag...c..-g.t......gat.a.tttt..tg...c... [ 87680]
AY261361.1_Malawi_Lil-20/1_1983   ..a------.g.ag..atttagacttttttcaagtag.cag..a..---.g.t......gat.a.tttt..tgc..c... [ 87680]
AY261360.1_Kenya_1950             ..a------.gtag..atctagatttttttcaagta...ag..a..---...t......gat.a.tttt..tg...c... [ 87680]
AM712240.1_OURT_88/3              ..a------..a...ca---------------aa..ta...a.cc.c.....t......cat.a.tttt..tg.g.g... [ 87680]
AM712239.1_Benin_97/1             ..a------..a...ca---------------aa..ta...a.cc.c.....t......cat.a.tttt..tg.g.g... [ 87680]
KM111295.1_Ken06.Bus              ..a------..a.c.gaccaaaattattttct.a.a.aca..ca..c.....t......cat.a.tttt..tg...c... [ 87680]
KM111294.1_Ken05/Tk1              ..a------.gtag..atctagatttttttcaagta...ag..a..---...t......gat.a.tttt..tg...c... [ 87680]
KM262844.1_strain_L60             ..a------..a...ca---------------aa..ta...a.cc.c.....t......cat.a.tttt..tg.g.g... [ 87680]
KM102979.1_26544/OG10_Italy       ..a------..a...ca---------------aa..ta...a.cc.c.....t......cat.a.tttt..tg.g.g... [ 87680]
KM262845.1_strain_NHV             ..a------..a...ca---------------aa..ta...a.cc.c.....t......cat.a.tttt..tg.g.g... [ 87680]
KX354450.1_47/Ss/2008             ..a------..a...ca---------------aa..ta...a.cc.c.....t......cat.a.tttt..tg.g.g... [ 87680]
KP055815.1_strain_BA71            ..a------..a...ca---------------aa..ta...a.cc.c.....t......cat.a.tttt..tg.g.g... [ 87680]
MH025920.1_strain_R35             ..a------..a.c.gaccaaaattattttct.a.a.aca..ca..c.....t......cat.a.tttt..tg...c... [ 87680]
MH025919.1_strain_N10             ..a------..a.c.gaccaaaattattttct.a.a.aca..ca..c.....t......cat.a.tttt..tg...c... [ 87680]
MH025918.1_strain_R25             ..a------..a.c.gaccaaaattattttct.a.a.aca..ca..c.....t......cat.a.tttt..tg...c... [ 87680]
MH025917.1_strain_R7              ..a------..a.c.gaccaaaattattttct.a.a.aca..ca..c.....t......cat.a.tttt..tg...c... [ 87680]
MH025916.1_strain_R8              ..a------..a.c.gaccaaaattattttct.a.a.aca..ca..c.....t......cat.a.tttt..tg...c... [ 87680]
```

```
FR682468.2_Georgia_2007/1       gataacaataagtattcttcttatatccatcataacttttttatctttac---gaaaaagaaaaaaacatgttgaagaaa [ 87760]
ASF_IND_20_CAD543               nnnn.t....g...---..t....t..a......t..nnnnnnnnnnn..gaa........................... [ 87760]
LR899193.1_Germany_2020/1       ..................................................---........................... [ 87760]
MN393477.1_Wuhan_2019           ..................................................---........................... [ 87760]
MW396979.1_Timor-Leste/2019/1   ..................................................---........................... [ 87760]
MK940252.1_CN/2019/InnerMongolia ..................................................---........................... [ 87760]
MK128995.1_China/2018/AnhuiXCGQ ..................................................---........................... [ 87760]
MH681419.1_ASFV/POL/2015        ..................................................---........................... [ 87760]
LS478113.1_Estonia_2014         ..................................................---........................... [ 87760]
KP843857.1_Odintsovo_02/14      ..................................................---........................... [ 87760]
FN557520.1_strain_E75           at...t....g...---..t....t..a......t..g.a......a...gaa........................... [ 87760]
AY261366.1_Warthog              a.............---..t.......a......................---........................... [ 87760]
AY261365.1_Warmbaths            a......g......---..t....t..a......................---........................... [ 87760]
AY261364.1_Tengani_62           a.c.tt......c.---..a.......a......................---........................... [ 87760]
AY261363.1_Pretoriuskop/96/4    at...t.gc.....---..t....t..a..t...................---........................... [ 87760]
AY261362.1_Mkuzi_1979           a....t.gc.....---..t....t..a......................---........................... [ 87760]
AY261361.1_Malawi_Lil-20/1_1983 a....t.gc.....---..t.......a......................---........................... [ 87760]
AY261360.1_Kenya_1950           a......g......---......................c....t.a...---.......---................. [ 87760]
AM712240.1_OURT_88/3            at...t....g...---..t....t..a......t..g.a......a...gaa........................... [ 87760]
AM712239.1_Benin_97/1           at...t....g...---..t....t..a......t..g.a......a...gaa........................... [ 87760]
KM111295.1_Ken06.Bus            .....t........---..t.....g.a......................---........................... [ 87760]
KM111294.1_Ken05/Tk1            a......g......---......................c....t.a...---.......---................. [ 87760]
KM262844.1_strain_L60           at...t....g...---..t....t..a......t..g.a......a...gaa........................... [ 87760]
KM102979.1_26544/OG10_Italy     at...t....g...---..t....t..a......t..g.a......a...gaa........................... [ 87760]
KM262845.1_strain_NHV           at...t....g...---..t....t..a......t..g.a......a...gaa........................... [ 87760]
KX354450.1_47/Ss/2008           at...t....g...---..t....t..a......t..g.a......a...gaa........................... [ 87760]
KP055815.1_strain_BA71          at...t....g...---..t....t..a......t..g.a......a...gaa........................... [ 87760]
MH025920.1_strain_R35           .....t........---..t.....g.a......................---........................... [ 87760]
MH025919.1_strain_N10           .....t........---..t.....g.a......................---........................... [ 87760]
MH025918.1_strain_R25           .....t........---..t.....g.a......................---........................... [ 87760]
MH025917.1_strain_R7            .....t........---..t.....g.a......................---........................... [ 87760]
MH025916.1_strain_R8            .....t........---..t.....g.a......................---........................... [ 87760]

FR682468.2_Georgia_2007/1       tagaaagtccaccac---ctgaatctaatgaa---gaagaacaatgtcagcatgatgacaccacttccatacatgaacca [ 87840]
ASF_IND_20_CAD543               ...............cct..............---.....tatg.c.---.............................. [ 87840]
LR899193.1_Germany_2020/1       ...............---..............---............................................. [ 87840]
MN393477.1_Wuhan_2019           ...............---..............---............................................. [ 87840]
MW396979.1_Timor-Leste/2019/1   ...............---..............---............................................. [ 87840]
MK940252.1_CN/2019/InnerMongolia ...............---..............---............................................. [ 87840]
MK128995.1_China/2018/AnhuiXCGQ ...............---..............---............................................. [ 87840]
MH681419.1_ASFV/POL/2015        ...............---..............---............................................. [ 87840]
LS478113.1_Estonia_2014         ...............---..............---............................................. [ 87840]
KP843857.1_Odintsovo_02/14      ...............---..............---............................................. [ 87840]
```

```
FN557520.1_strain_E75              ...............cct..............---.....tatt.c.---..c........................... [ 87840]
AY261366.1_Warthog                 ....g..........---..............---................................t............ [ 87840]
AY261365.1_Warmbaths               .........t.....---..............---......tgtcag..tg............................. [ 87840]
AY261364.1_Tengani_62              ...............cct..............---........t..............t..................... [ 87840]
AY261363.1_Pretoriuskop/96/4       ...............---..............---............................................. [ 87840]
AY261362.1_Mkuzi_1979              ...............cct..............---.....tatt.c.---..c........................... [ 87840]
AY261361.1_Malawi_Lil-20/1_1983    ...........t.g.cat..............---............................................g [ 87840]
AY261360.1_Kenya_1950              .............g.---..............---.........caatgt...c.............t...........g [ 87840]
AM712240.1_OURT_88/3               ...............cct..............---.....tatt.c.---..c........................... [ 87840]
AM712239.1_Benin_97/1              ...............cct..............---.....tatt.c.---..c........................... [ 87840]
KM111295.1_Ken06.Bus               ...........t.g.---..............gag.........caa..t...c.......t.................g [ 87840]
KM111294.1_Ken05/Tk1               .............g.---..............---.........caatgt...c.............t...........g [ 87840]
KM262844.1_strain_L60              ...............cct..............---.....tatt.c.---..c........................... [ 87840]
KM102979.1_26544/OG10_Italy        ...............cct..............---.....tatt.c.---..c........................... [ 87840]
KM262845.1_strain_NHV              ...............cct..............---.....tatt.c.---..c........................... [ 87840]
KX354450.1_47/Ss/2008              ...............cct..............---.....tatt.c.---..c........................... [ 87840]
KP055815.1_strain_BA71             ...............cct..............---.....tatt.c.---..c........................... [ 87840]
MH025920.1_strain_R35              ...........t.g.---..............gag.........caa..t...c.......t.................g [ 87840]
MH025919.1_strain_N10              ...........t.g.---..............gag.........caa..t...c.......t.................g [ 87840]
MH025918.1_strain_R25              ...........t.g.---..............gag.........caa..t...c.......t.................g [ 87840]
MH025917.1_strain_R7               ...........t.g.---..............gag.........caa..t...c.......t.................g [ 87840]
MH025916.1_strain_R8               ...........t.g.---..............gag.........caa..t...c.......t.................g [ 87840]

FR682468.2_Georgia_2007/1          tctcccagagaaccattacttcctaagccttacagtcgttatcagtataatacacctatttactacatgcgtccctcaac [ 87920]
ASF_IND_20_CAD543                  ................................................................................ [ 87920]
LR899193.1_Germany_2020/1          ................................................................................ [ 87920]
MN393477.1_Wuhan_2019              ................................................................................ [ 87920]
MW396979.1_Timor-Leste/2019/1      ................................................................................ [ 87920]
MK940252.1_CN/2019/InnerMongolia   ................................................................................ [ 87920]
MK128995.1_China/2018/AnhuiXCGQ    ................................................................................ [ 87920]
MH681419.1_ASFV/POL/2015           ................................................................................ [ 87920]
LS478113.1_Estonia_2014            ................................................................................ [ 87920]
KP843857.1_Odintsovo_02/14         ................................................................................ [ 87920]
FN557520.1_strain_E75              ................................................................................ [ 87920]
AY261366.1_Warthog                 ..........................a..................................................... [ 87920]
AY261365.1_Warmbaths               ..c.......................a..................................................... [ 87920]
AY261364.1_Tengani_62              ................................................................................ [ 87920]
AY261363.1_Pretoriuskop/96/4       ..........................a.................a................................... [ 87920]
AY261362.1_Mkuzi_1979              ................................................................................ [ 87920]
AY261361.1_Malawi_Lil-20/1_1983    ......................................a..........................t..........t... [ 87920]
AY261360.1_Kenya_1950              ............................................a................................... [ 87920]
AM712240.1_OURT_88/3               ..-............................................................................. [ 87920]
AM712239.1_Benin_97/1              ................................................................................ [ 87920]
KM111295.1_Ken06.Bus               ..............................................................t................. [ 87920]
```

```
KM111294.1_Ken05/Tk1              ............................................a................................... [ 87920]
KM262844.1_strain_L60             ................................................................................ [ 87920]
KM102979.1_26544/OG10_Italy       ................................................................................ [ 87920]
KM262845.1_strain_NHV             ..-............................................................................. [ 87920]
KX354450.1_47/Ss/2008             ................................................................................ [ 87920]
KP055815.1_strain_BA71            .........a...................................................................... [ 87920]
MH025920.1_strain_R35             ..............................................................t................. [ 87920]
MH025919.1_strain_N10             ..............................................................t................. [ 87920]
MH025918.1_strain_R25             ..............................................................t................. [ 87920]
MH025917.1_strain_R7              ..............................................................t................. [ 87920]
MH025916.1_strain_R8              ..............................................................t................. [ 87920]

FR682468.2_Georgia_2007/1         a---caaccactcaacccatttcccttacctaaaccgtgtcctccacccaaaccatgtccgccac--------------- [ 88000]
ASF_IND_20_CAD543                 .---.....................y.......................................ccaagccatgcccgc [ 88000]
LR899193.1_Germany_2020/1         .---.............................................................--------------- [ 88000]
MN393477.1_Wuhan_2019             .---.............................................................--------------- [ 88000]
MW396979.1_Timor-Leste/2019/1     .---.............................................................--------------- [ 88000]
MK940252.1_CN/2019/InnerMongolia  .---.............................................................--------------- [ 88000]
MK128995.1_China/2018/AnhuiXCGQ   .---.............................................................--------------- [ 88000]
MH681419.1_ASFV/POL/2015          .---.............................................................--------------- [ 88000]
LS478113.1_Estonia_2014           .---.............................................................--------------- [ 88000]
KP843857.1_Odintsovo_02/14        .---.............................................................--------------- [ 88000]
FN557520.1_strain_E75             .---.....................c..........a..c..g.....t...........t....ccaagccatgcccgc [ 88000]
AY261366.1_Warthog                .---.....................c..........a..c..g.....t.....g------------------tgtcctc [ 88000]
AY261365.1_Warmbaths              .---.....................c..........a..c..a.....t.....g..........ccaaaccatgtcctc [ 88000]
AY261364.1_Tengani_62             .---.....................c..........a...........t.....g....--------------------- [ 88000]
AY261363.1_Pretoriuskop/96/4      .---.....................c.......c..a...........t.....g.....t....ccaaaccatgcccgc [ 88000]
AY261362.1_Mkuzi_1979             .---.....................c..........a..c..g.....t...------------------ccatgcccgc [ 88000]
AY261361.1_Malawi_Lil-20/1_1983   .---...........t....c....c....c....t..................g....--------------------- [ 88000]
AY261360.1_Kenya_1950             .caa....t.t.t..at...a.t.t.....c.....a.....c.....t..........--------------------- [ 88000]
AM712240.1_OURT_88/3              .---.....................c..........a..c..g.....t...........t....ccaagccatgcccgc [ 88000]
AM712239.1_Benin_97/1             .---.....................c..........a..c..g.....t...........t....ccaagccatgcccgc [ 88000]
KM111295.1_Ken06.Bus              .caa....t.t.t..at...a.t.t.....c.....a.....c.....t..........--------------------- [ 88000]
KM111294.1_Ken05/Tk1              .caa....t.t.t..at...a.t.t.....c.....a.....c.....t..........--------------------- [ 88000]
KM262844.1_strain_L60             .---.....................c..........a..c..g.....t...........t....ccaagccatgcccgc [ 88000]
KM102979.1_26544/OG10_Italy       .---.....................c..........a..c..g.....t...........t....ccaagccatgcccgc [ 88000]
KM262845.1_strain_NHV             .---.....................c..........a..c..g.....t...........t....ccaagccatgcccgc [ 88000]
KX354450.1_47/Ss/2008             .---.....................c..........a..c..g.....t...........t....ccaagccatgcccgc [ 88000]
KP055815.1_strain_BA71            .---.....................c..........a..c..g.....t...........t....ccaagccatgcccgc [ 88000]
MH025920.1_strain_R35             .caa....t.t.t..at...a.t.t.....c.....a.....c.....t..........--------------------- [ 88000]
MH025919.1_strain_N10             .caa....t.t.t..at...a.t.t.....c.....a.....c.....t..........--------------------- [ 88000]
MH025918.1_strain_R25             .caa....t.t.t..at...a.t.t.....c.....a.....c.....t..........--------------------- [ 88000]
MH025917.1_strain_R7              .caa....t.t.t..at...a.t.t.....c.....a.....c.....t..........--------------------- [ 88000]
MH025916.1_strain_R8              .caa....t.t.t..at...a.t.t.....c.....a.....c.....t..........--------------------- [ 88000]
```

```
FR682468.2_Georgia_2007/1        -------------------------------------------------------------------------------- [ 88080]
ASF_IND_20_CAD543                cacccaaaccatgtcctnnnnnnnnnnnnnnnnnnnnnnnnnnnnnnnnnnnnnnnnnnnnnnnnnnnnnnnnncctaaa [ 88080]
LR899193.1_Germany_2020/1        -------------------------------------------------------------------------------- [ 88080]
MN393477.1_Wuhan_2019            -------------------------------------------------------------------------------- [ 88080]
MW396979.1_Timor-Leste/2019/1    -------------------------------------------------------------------------------- [ 88080]
MK940252.1_CN/2019/InnerMongolia -------------------------------------------------------------------------------- [ 88080]
MK128995.1_China/2018/AnhuiXCGQ  -------------------------------------------------------------------------------- [ 88080]
MH681419.1_ASFV/POL/2015         -------------------------------------------------------------------------------- [ 88080]
LS478113.1_Estonia_2014          -------------------------------------------------------------------------------- [ 88080]
KP843857.1_Odintsovo_02/14       -------------------------------------------------------------------------------- [ 88080]
FN557520.1_strain_E75            cacccaaaccatgtcctccacctaaaccgtgtt------------------------------------ctccacccaaa [ 88080]
AY261366.1_Warthog               cacccaaaccatgcc------------------------------------------------------cgccacccaaa [ 88080]
AY261365.1_Warmbaths             cacccaaaccatgcc------------------------------------------------------cgccacccaaa [ 88080]
AY261364.1_Tengani_62            ---------------------------------------------------------------------ctccacccaaa [ 88080]
AY261363.1_Pretoriuskop/96/4     cacctaaaccatgtc------------------------------------------------------ccccacccaaa [ 88080]
AY261362.1_Mkuzi_1979            cacccaaaccatgtcctccacctaaaccgtgtc------------------------------------ctccacccaaa [ 88080]
AY261361.1_Malawi_Lil-20/1_1983  ---------------------------------------------------------------------ctccacccaaa [ 88080]
AY261360.1_Kenya_1950            ---------------------------------------------------------------------ctccacctaaa [ 88080]
AM712240.1_OURT_88/3             cacccaaacc-tgtcctccacctaaaccgtgtt------------------------------------ctccacccaaa [ 88080]
AM712239.1_Benin_97/1            cacccaaaccatgtcctccacctaaaccgtgtt------------------------------------ctccacccaaa [ 88080]
KM111295.1_Ken06.Bus             ---------------------------------------------------------------------ctccacctaaa [ 88080]
KM111294.1_Ken05/Tk1             ---------------------------------------------------------------------ctccacccaaa [ 88080]
KM262844.1_strain_L60            cacccaaaccatgtcctccacctaaaccgtgttctccacccaaaccgtgtcgtccacctaaaccatgtcctc-------- [ 88080]
KM102979.1_26544/OG10_Italy      cacccaaaccatgtcctccacctaaaccgtgtt------------------------------------ctccacccaaa [ 88080]
KM262845.1_strain_NHV            cacccaaacc-tgtcctccacctaaaccgtgtt------------------------------------ctccacccaaa [ 88080]
KX354450.1_47/Ss/2008            cacccaaaccatgtcctccacctaaaccgtgtt------------------------------------ctccacccaaa [ 88080]
KP055815.1_strain_BA71           cacccaaaccatgtcctccacctaaaccgtgtt------------------------------------ctccacccaaa [ 88080]
MH025920.1_strain_R35            ---------------------------------------------------------------------ctccacctaaa [ 88080]
MH025919.1_strain_N10            ---------------------------------------------------------------------ctccacctaaa [ 88080]
MH025918.1_strain_R25            ---------------------------------------------------------------------ctccacctaaa [ 88080]
MH025917.1_strain_R7             ---------------------------------------------------------------------ctccacctaaa [ 88080]
MH025916.1_strain_R8             ---------------------------------------------------------------------ctccacctaaa [ 88080]

FR682468.2_Georgia_2007/1        -------------------------------ccaaaccatgtcctccaccta------------------aaccatgtcc [ 88160]
ASF_IND_20_CAD543                ccgtgtcctccacccaaaccatgtccgccac.................t.c.------------------.......... [ 88160]
LR899193.1_Germany_2020/1        -------------------------------.....................------------------.......... [ 88160]
MN393477.1_Wuhan_2019            -------------------------------.....................------------------.......... [ 88160]
MW396979.1_Timor-Leste/2019/1    -------------------------------.....................------------------.......... [ 88160]
MK940252.1_CN/2019/InnerMongolia -------------------------------.....................------------------.......... [ 88160]
MK128995.1_China/2018/AnhuiXCGQ  -------------------------------.....................------------------.......... [ 88160]
MH681419.1_ASFV/POL/2015         -------------------------------.....................------------------.......... [ 88160]
LS478113.1_Estonia_2014          -------------------------------.....................------------------.......... [ 88160]
KP843857.1_Odintsovo_02/14       -------------------------------.....................------------------.......... [ 88160]
```

```
FN557520.1_strain_E75          ccgtgtcgtccacctaaaccatgtcctccac.t...................------------------.......... [ 88160]
AY261366.1_Warthog             ccatgcccgccacctaaaccatgtcctccac..........c..g.......------------------.......... [ 88160]
AY261365.1_Warmbaths           ccatgcccgccacctaaaccatgtcctccac.tg..t.....t..t.....g------------------..t.....t. [ 88160]
AY261364.1_Tengani_62          ccatgtcctccacctaaaccgtgtcctccac.t.....g.....c.......------------------.......... [ 88160]
AY261363.1_Pretoriuskop/96/4   ccatgcccgccacctaaaccgtgtcctccac.............a.......------------------........t. [ 88160]
AY261362.1_Mkuzi_1979          ccgtgtcgtccacctaaaccatgtcctccac.t........c..a.......------------------.......... [ 88160]
AY261361.1_Malawi_Lil-20/1_1983 ccgtgtcctccacccaaaccgtgtcctccac.......g......t..t..g------------------..t.....t. [ 88160]
AY261360.1_Kenya_1950          ccatgtcccccacctaaaccatgtcctccat....................gaaccatattctccaccca.......... [ 88160]
AM712240.1_OURT_88/3           ccgtgtcgtccacctaaaccatgtcctccac.t...................------------------.......... [ 88160]
AM712239.1_Benin_97/1          ccgtgtcgtccacctaaaccatgtcctccac.t...................------------------.......... [ 88160]
KM111295.1_Ken06.Bus           ccatgtcccccacccaaaccatgtccctcac.ag..t...a...ct.....g---------------------------- [ 88160]
KM111294.1_Ken05/Tk1           ccatgtcccccacccaaaccatgtcctccac.................t.c.---------------------------- [ 88160]
KM262844.1_strain_L60          ----------------------------cac.t...................------------------.......... [ 88160]
KM102979.1_26544/OG10_Italy    ccgtgtcgtccacctaaaccatgtcctccac.t...................------------------.......... [ 88160]
KM262845.1_strain_NHV          ccgtgtcgtccacctaaaccatgtcctccac.t...................------------------.......... [ 88160]
KX354450.1_47/Ss/2008          ccgtgtcgtccacctaaaccatgtcctccac.t...................------------------.......... [ 88160]
KP055815.1_strain_BA71         ccgtgtcgtccacctaaaccatgtcctccac.t...................------------------.......... [ 88160]
MH025920.1_strain_R35          ccatgtcccccacccaaaccatgtccctcac.tg..t...a...ct.....g---------------------------- [ 88160]
MH025919.1_strain_N10          ccatgtcccccacccaaaccatgtccctcac.tg..t...a...ct.....g---------------------------- [ 88160]
MH025918.1_strain_R25          ccatgtcccccacccaaaccatgtccctcac.tg..t...a...ct.....g---------------------------- [ 88160]
MH025917.1_strain_R7           ccatgtcccccacccaaaccatgtccctcac.tg..t...a...ct.....g---------------------------- [ 88160]
MH025916.1_strain_R8           ccatgtcccccacccaaaccatgtccctcac.tg..t...a...ct.....g---------------------------- [ 88160]

FR682468.2_Georgia_2007/1      ttcagctgaatcct------------------------------------attctccacccaaaccactacctagtatcc [ 88240]
ASF_IND_20_CAD543              ..............------------------------------------.............................. [ 88240]
LR899193.1_Germany_2020/1      ..............------------------------------------.............................. [ 88240]
MN393477.1_Wuhan_2019          ..............------------------------------------.............................. [ 88240]
MW396979.1_Timor-Leste/2019/1  ..............------------------------------------.............................. [ 88240]
MK940252.1_CN/2019/InnerMongolia ..............------------------------------------.............................. [ 88240]
MK128995.1_China/2018/AnhuiXCGQ ..............------------------------------------.............................. [ 88240]
MH681419.1_ASFV/POL/2015       ..............------------------------------------.............................. [ 88240]
LS478113.1_Estonia_2014        ..............------------------------------------.............................. [ 88240]
KP843857.1_Odintsovo_02/14     ..............------------------------------------.............................. [ 88240]
FN557520.1_strain_E75          .c..c..a..c.a.gtcctccatccaaaccatgtccttcacctgaatcct.............................. [ 88240]
AY261366.1_Warthog             .c..c.......a.------------------------------------.............................. [ 88240]
AY261365.1_Warmbaths           ....c.......a.------------------------------------.............................. [ 88240]
AY261364.1_Tengani_62          ....c........c------------------------------------.............................. [ 88240]
AY261363.1_Pretoriuskop/96/4   ....c.......a.------------------------------------.............................. [ 88240]
AY261362.1_Mkuzi_1979          ....c.....a.a.------------------------------------........................a..... [ 88240]
AY261361.1_Malawi_Lil-20/1_1983 .c..c.......a.------------------------------------......t.................a..... [ 88240]
AY261360.1_Kenya_1950          .c..c.ca..c.a.------------------------------------..c..---------t............... [ 88240]
AM712240.1_OURT_88/3           .c..c..a..c.a.gtcctccatccaaaccatgtccttcacctgaatcct.............................. [ 88240]
AM712239.1_Benin_97/1          .c..c..a..c.a.gtcctccatccaaaccatgtccttcacctgaatcct.............................. [ 88240]
KM111295.1_Ken06.Bus           --------..c.a.------------------------------------..c..---------.t........a..... [ 88240]
```

```
KM111294.1_Ken05/Tk1           --------..c.a.------------------------------------..c..---------t............... [ 88240]
KM262844.1_strain_L60          .c..c..a..c.a.gtcctccatccaaaccatgtccttcacctgaatcct.............................. [ 88240]
KM102979.1_26544/OG10_Italy    .c..c..a..c.a.gtcctccatccaaaccatgtccttcacctgaatcct.............................. [ 88240]
KM262845.1_strain_NHV          .c..c..a..c.a.gtcctccatccaaaccatgtccttcacctgaatcct.............................. [ 88240]
KX354450.1_47/Ss/2008          .c..c..a..c.a.gtcctccacctaaaccatgtccttcacctgaatcct.............................. [ 88240]
KP055815.1_strain_BA71         .c..c..a..c.a.gtcctccatccaaaccatgtccttcacctgaatcct.............................. [ 88240]
MH025920.1_strain_R35          --------..c.a.------------------------------------..c..---------.t........a..... [ 88240]
MH025919.1_strain_N10          --------..c.a.------------------------------------..c..---------.t........a..... [ 88240]
MH025918.1_strain_R25          --------..c.a.------------------------------------..c..---------.t........a..... [ 88240]
MH025917.1_strain_R7           --------..c.a.------------------------------------..c..---------.t........a..... [ 88240]
MH025916.1_strain_R8           --------..c.a.------------------------------------..c..---------.t........a..... [ 88240]

FR682468.2_Georgia_2007/1      cgctactacccaatatcccgccattatctacccaaaatatttcgcttattcacgtagatagaat---------------- [ 88320]
ASF_IND_20_CAD543              ....................................................y...........tatttaataaatatca [ 88320]
LR899193.1_Germany_2020/1      ................................................................---------------- [ 88320]
MN393477.1_Wuhan_2019          ................................................................---------------- [ 88320]
MW396979.1_Timor-Leste/2019/1  ................................................................---------------- [ 88320]
MK940252.1_CN/2019/InnerMongolia ................................................................---------------- [ 88320]
MK128995.1_China/2018/AnhuiXCGQ ................................................................---------------- [ 88320]
MH681419.1_ASFV/POL/2015       ................................................................---------------- [ 88320]
LS478113.1_Estonia_2014        ................................................................---------------- [ 88320]
KP843857.1_Odintsovo_02/14     ................................................................---------------- [ 88320]
FN557520.1_strain_E75          ..t............................a....................t...........tatttaataaatatca [ 88320]
AY261366.1_Warthog             ....g..........................a....................t...........tatttaataaaaatca [ 88320]
AY261365.1_Warmbaths           ....g..........................a....................t...........tatttaataaaaatca [ 88320]
AY261364.1_Tengani_62          ....g...........t..c...........a....................t...........tatttaataaatacca [ 88320]
AY261363.1_Pretoriuskop/96/4   ...................a...........a...........a........t.........------------------ [ 88320]
AY261362.1_Mkuzi_1979          ....g..........................a.............................g..tatttaataaaaatca [ 88320]
AY261361.1_Malawi_Lil-20/1_1983 ...............................a........c.......................tatttaatatgtacta [ 88320]
AY261360.1_Kenya_1950          ......c....g...................a................................tatttaataaaaacca [ 88320]
AM712240.1_OURT_88/3           ..t............................a....................t...........tatttaataaatatca [ 88320]
AM712239.1_Benin_97/1          ..t............................a....................t...........tatttaataaatatca [ 88320]
KM111295.1_Ken06.Bus           .a....c........................a................................tatttaatatgtacca [ 88320]
KM111294.1_Ken05/Tk1           ......c....g...................a................................tatttaataaaaacca [ 88320]
KM262844.1_strain_L60          ..t............................a....................t...........tatttaataaatatca [ 88320]
KM102979.1_26544/OG10_Italy    ..t............................a....................t...........tatttaataaatatca [ 88320]
KM262845.1_strain_NHV          ..t............................a....................t...........tatttaataaatatca [ 88320]
KX354450.1_47/Ss/2008          ..t............................a....................t...........tatttaataaatatca [ 88320]
KP055815.1_strain_BA71         ..t............................a....................t...........tatttaataaatatca [ 88320]
MH025920.1_strain_R35          .a....c........................a................................tatttaatatgtacca [ 88320]
MH025919.1_strain_N10          .a....c........................a................................tatttaatatgtacca [ 88320]
MH025918.1_strain_R25          .a....c........................a................................tatttaatatgtacca [ 88320]
MH025917.1_strain_R7           .a....c........................a................................tatttaatatgtacca [ 88320]
MH025916.1_strain_R8           .a....c........................a................................tatttaatatgtacca [ 88320]
```

```
FR682468.2_Georgia_2007/1        -----------------------------tatttaatatgtactatatattaattatttaacctttcaagctggtcttca [ 88400]
ASF_IND_20_CAD543                tgtat---ttat------taaataccacg..........r.....y..r..m............................ [ 88400]
LR899193.1_Germany_2020/1        -----------------------------................................................... [ 88400]
MN393477.1_Wuhan_2019            -----------------------------................................................... [ 88400]
MW396979.1_Timor-Leste/2019/1    -----------------------------................................................... [ 88400]
MK940252.1_CN/2019/InnerMongolia -----------------------------................................................... [ 88400]
MK128995.1_China/2018/AnhuiXCGQ  -----------------------------................................................... [ 88400]
MH681419.1_ASFV/POL/2015         -----------------------------................................................... [ 88400]
LS478113.1_Estonia_2014          -----------------------------................................................... [ 88400]
KP843857.1_Odintsovo_02/14       -----------------------------................................................... [ 88400]
FN557520.1_strain_E75            tgtat---ttat------taaataccacg..........a.....c..g..c............................ [ 88400]
AY261366.1_Warthog               tgtat---ttat------taaataccacg..........a.....cg.g..c.........t............c..... [ 88400]
AY261365.1_Warmbaths             tgtatttattat------taaataccacg..........a.....cg.g..c.........t.................. [ 88400]
AY261364.1_Tengani_62            t---------------------------g......t...a...c.cg.g..c..........t................. [ 88400]
AY261363.1_Pretoriuskop/96/4     ----------------------------------------------------.........t...t............. [ 88400]
AY261362.1_Mkuzi_1979            t---------------------------g.....c....a.....cg.g..c.........t.................. [ 88400]
AY261361.1_Malawi_Lil-20/1_1983  tatat---------------------------------------..g....c..................tc........ [ 88400]
AY261360.1_Kenya_1950            tgtattt---------------------------------------------.a.................c........ [ 88400]
AM712240.1_OURT_88/3             tgtat---ttat------taaataccacg..........a.....c..g..c............................ [ 88400]
AM712239.1_Benin_97/1            tgtat---ttat------taaataccacg..........a.....c..g..c............................ [ 88400]
KM111295.1_Ken06.Bus             tatgtaccgtatgtaccatatgtactata.g.acc....a........g.a.....................a.c..... [ 88400]
KM111294.1_Ken05/Tk1             tgtattt---------------------------.....a.....cg.g..c...................c........ [ 88400]
KM262844.1_strain_L60            tgtat---ttat------taaataccacg..........a.....c..g..c............................ [ 88400]
KM102979.1_26544/OG10_Italy      tgtat---ttat------taaataccacg..........a.....c..g..c............................ [ 88400]
KM262845.1_strain_NHV            tgtat---ttat------taaataccacg..........a.....c..g..c............................ [ 88400]
KX354450.1_47/Ss/2008            tgtat---ttat------taaataccacg..........a.....c..g..c............................ [ 88400]
KP055815.1_strain_BA71           tgtat---ttat------taaataccacg..........a.....c..g..c............................ [ 88400]
MH025920.1_strain_R35            tatgtaccgtatgtaccatatgtactata.g.acc....a........g.a.....................a.c..... [ 88400]
MH025919.1_strain_N10            tatgtaccgtatgtaccatatgtactata.g.acc....a........g.a.....................a.c..... [ 88400]
MH025918.1_strain_R25            tatgtaccgtatgtaccatatgtactata.g.acc....a........g.a.....................a.c..... [ 88400]
MH025917.1_strain_R7             tatgtaccgtatgtaccatatgtactata.g.acc....a........g.a.....................a.c..... [ 88400]
MH025916.1_strain_R8             tatgtaccgtatgtaccatatgtactata.g.acc....a........g.a.....................a.c..... [ 88400]

FR682468.2_Georgia_2007/1        tttaaatttaaaatccactaat---aaaatgtattttctagtagcagatcatcgagaacatcatgtgattccttttctta [ 88480]
ASF_IND_20_CAD543                ......................aaa....................................................... [ 88480]
LR899193.1_Germany_2020/1        ......................---....................................................... [ 88480]
MN393477.1_Wuhan_2019            ......................---....................................................... [ 88480]
MW396979.1_Timor-Leste/2019/1    ......................---....................................................... [ 88480]
MK940252.1_CN/2019/InnerMongolia ......................---....................................................... [ 88480]
MK128995.1_China/2018/AnhuiXCGQ  ......................---....................................................... [ 88480]
MH681419.1_ASFV/POL/2015         ......................---....................................................... [ 88480]
LS478113.1_Estonia_2014          ......................---....................................................... [ 88480]
KP843857.1_Odintsovo_02/14       ......................---....................................................... [ 88480]
```

```
FN557520.1_strain_E75          ......................aaa.................g..g.................................. [ 88480]
AY261366.1_Warthog             ......................-aa....................g.................................. [ 88480]
AY261365.1_Warmbaths           ......................-aa....................................................... [ 88480]
AY261364.1_Tengani_62          ......................-aa....................................................... [ 88480]
AY261363.1_Pretoriuskop/96/4   ......................-aa..............g................................a....... [ 88480]
AY261362.1_Mkuzi_1979          ......................-aa....................................................... [ 88480]
AY261361.1_Malawi_Lil-20/1_1983 ......................-aa....................................................... [ 88480]
AY261360.1_Kenya_1950          ........c.............aaa-..........c........g...........g....................c. [ 88480]
AM712240.1_OURT_88/3           ......................aaa.................g..g.................................. [ 88480]
AM712239.1_Benin_97/1          ......................aaa.................g..g.................................. [ 88480]
KM111295.1_Ken06.Bus           ...........g..........aaa-.............g.....g...........g....................c. [ 88480]
KM111294.1_Ken05/Tk1           ......................aaa....................g...........g....................c. [ 88480]
KM262844.1_strain_L60          ......................aaa.................g..g.................................. [ 88480]
KM102979.1_26544/OG10_Italy    ......................aaa.................g..g.................................. [ 88480]
KM262845.1_strain_NHV          ......................aaa.................g..g.................................. [ 88480]
KX354450.1_47/Ss/2008          ......................aaa.................g..g.................................. [ 88480]
KP055815.1_strain_BA71         ......................aaa.................g..g.................................. [ 88480]
MH025920.1_strain_R35          ...........g..........aaa-.............g.....g...........g....................c. [ 88480]
MH025919.1_strain_N10          ...........g..........aaa-.............g.....g...........g....................c. [ 88480]
MH025918.1_strain_R25          ...........g..........aaa-.............g.....g...........g....................c. [ 88480]
MH025917.1_strain_R7           ...........g..........aaa-.............g.....g...........g....................c. [ 88480]
MH025916.1_strain_R8           ...........g..........aaa-.............g.....g...........g....................c. [ 88480]

FR682468.2_Georgia_2007/1      aaaccgatttccatcacatgcatcaaaatcctatacaaa---aaaatcaagctctcctagaaatcaaacagctttttact [ 88560]
ASF_IND_20_CAD543              ......................w..n.....a..gttt.cgc....g................................. [ 88560]
LR899193.1_Germany_2020/1      .......................................---...................................... [ 88560]
MN393477.1_Wuhan_2019          .......................................---...................................... [ 88560]
MW396979.1_Timor-Leste/2019/1  .......................................---...................................... [ 88560]
MK940252.1_CN/2019/InnerMongolia .......................................---...................................... [ 88560]
MK128995.1_China/2018/AnhuiXCGQ .......................................---...................................... [ 88560]
MH681419.1_ASFV/POL/2015       .......................................---...................................... [ 88560]
LS478113.1_Estonia_2014        .......................................---...................................... [ 88560]
KP843857.1_Odintsovo_02/14     .......................................---...................................... [ 88560]
FN557520.1_strain_E75          ..........t...g.t.....a..t.....a..gttt.cgc....g...........g........g...........c [ 88560]
AY261366.1_Warthog             ..........t............................---.........................g...........c [ 88560]
AY261365.1_Warmbaths           ..........t............................---...........t.............g...........c [ 88560]
AY261364.1_Tengani_62          ..........t.........t..................---...................................... [ 88560]
AY261363.1_Pretoriuskop/96/4   ..........t............................---...........t.............g...........c [ 88560]
AY261362.1_Mkuzi_1979          ..........t............................---................t........g...........c [ 88560]
AY261361.1_Malawi_Lil-20/1_1983 ..........t............................---.........................g...........c [ 88560]
AY261360.1_Kenya_1950          ........c.t..g..t..at............cc....---....a...at..c...g........g...........c [ 88560]
AM712240.1_OURT_88/3           ..........t...g.t.....a..t.....a..gttt.cgc....g...........g........g...........c [ 88560]
AM712239.1_Benin_97/1          ..........t...g.t.....a..t.....a..gttt.cgc....g...........g........g...........c [ 88560]
KM111295.1_Ken06.Bus           ....t...c.t.....t..at............cc....---....a....t..c............g...........c [ 88560]
```

```
KM111294.1_Ken05/Tk1             ........c.t.....t..at............cc....---....a...at..c...g........g...........c [ 88560]
KM262844.1_strain_L60            ..........t...g.t.....a..t.....a..gttt.cgc....g...........g........g...........c [ 88560]
KM102979.1_26544/OG10_Italy      ..........t...g.t.....a..t.....a..gttt.cgc....g...........g........g...........c [ 88560]
KM262845.1_strain_NHV            ..........t...g.t.....a..t.....a..gttt.cgc....g...........g........g...........c [ 88560]
KX354450.1_47/Ss/2008            ..........t...g.t.....a..t.....a..gttt.cgc....g...........g........g...........c [ 88560]
KP055815.1_strain_BA71           ..........t...g.t.....a..t.....a..gttt.cgc....g...........g........g...........c [ 88560]
MH025920.1_strain_R35            ....t...c.t.....t..at............cc....---....a....t..c............g...........c [ 88560]
MH025919.1_strain_N10            ....t...c.t.....t..at............cc....---....a....t..c............g...........c [ 88560]
MH025918.1_strain_R25            ....t...c.t.....t..at............cc....---....a....t..c............g...........c [ 88560]
MH025917.1_strain_R7             ....t...c.t.....t..at............cc....---....a....t..c............g...........c [ 88560]
MH025916.1_strain_R8             ....t...c.t.....t..at............cc....---....a....t..c............g...........c [ 88560]

FR682468.2_Georgia_2007/1        ggagattatctcatctgcaaaagcccttctaccattctggcctgtattgaacgaaaaacctacaaagactttgcggcttc [ 88640]
ASF_IND_20_CAD543                ...........................a.c..t............................................... [ 88640]
LR899193.1_Germany_2020/1        ................................................................................ [ 88640]
MN393477.1_Wuhan_2019            ................................................................................ [ 88640]
MW396979.1_Timor-Leste/2019/1    ................................................................................ [ 88640]
MK940252.1_CN/2019/InnerMongolia ................................................................................ [ 88640]
MK128995.1_China/2018/AnhuiXCGQ  ................................................................................ [ 88640]
MH681419.1_ASFV/POL/2015         ................................................................................ [ 88640]
LS478113.1_Estonia_2014          ................................................................................ [ 88640]
KP843857.1_Odintsovo_02/14       ................................................................................ [ 88640]
FN557520.1_strain_E75            ...........................a.c..t............................................... [ 88640]
AY261366.1_Warthog               .............................c..a.........................................a..... [ 88640]
AY261365.1_Warmbaths             .............................c..g.........................................a..... [ 88640]
AY261364.1_Tengani_62            ................................................................................ [ 88640]
AY261363.1_Pretoriuskop/96/4     .......................t.....c..g.........................................a..... [ 88640]
AY261362.1_Mkuzi_1979            .............................c..g...........c................................... [ 88640]
AY261361.1_Malawi_Lil-20/1_1983  ..............a........t.....c..a.........................................c..... [ 88640]
AY261360.1_Kenya_1950            ..............a..............c..g........t................................c..... [ 88640]
AM712240.1_OURT_88/3             ...........................a.c..t............................................... [ 88640]
AM712239.1_Benin_97/1            ...........................a.c..t............................................... [ 88640]
KM111295.1_Ken06.Bus             ..............a..............c..g.........................................c..... [ 88640]
KM111294.1_Ken05/Tk1             ...........t..a.................g.........................................c..... [ 88640]
KM262844.1_strain_L60            ...........................a.c..t............................................... [ 88640]
KM102979.1_26544/OG10_Italy      ...........................a.c..t............................................... [ 88640]
KM262845.1_strain_NHV            ...........................a.c..t............................................... [ 88640]
KX354450.1_47/Ss/2008            ...........................a.c..t............................................... [ 88640]
KP055815.1_strain_BA71           ...........................a.c..t............................................... [ 88640]
MH025920.1_strain_R35            ..............a..............c..g.........................................c..... [ 88640]
MH025919.1_strain_N10            ..............a..............c..g.........................................c..... [ 88640]
MH025918.1_strain_R25            ..............a..............c..g.........................................c..... [ 88640]
MH025917.1_strain_R7             ..............a..............c..g.........................................c..... [ 88640]
MH025916.1_strain_R8             ..............a..............c..g.........................................c..... [ 88640]
```

```
FR682468.2_Georgia_2007/1        tttgaaagatggacgttataaaaatcgccaaaaaatgctgtcgctgcgagaacaaaccaactgtcaactttatttttttg [ 88720]
ASF_IND_20_CAD543                ................................................................................ [ 88720]
LR899193.1_Germany_2020/1        ................................................................................ [ 88720]
MN393477.1_Wuhan_2019            ................................................................................ [ 88720]
MW396979.1_Timor-Leste/2019/1    ................................................................................ [ 88720]
MK940252.1_CN/2019/InnerMongolia ................................................................................ [ 88720]
MK128995.1_China/2018/AnhuiXCGQ  ................................................................................ [ 88720]
MH681419.1_ASFV/POL/2015         ................................................................................ [ 88720]
LS478113.1_Estonia_2014          ................................................................................ [ 88720]
KP843857.1_Odintsovo_02/14       ................................................................................ [ 88720]
FN557520.1_strain_E75            .......................................a........................................ [ 88720]
AY261366.1_Warthog               ................................................................................ [ 88720]
AY261365.1_Warmbaths             ................................................................................ [ 88720]
AY261364.1_Tengani_62            .......................................a........................................ [ 88720]
AY261363.1_Pretoriuskop/96/4     ................................................................................ [ 88720]
AY261362.1_Mkuzi_1979            .......................................a........................................ [ 88720]
AY261361.1_Malawi_Lil-20/1_1983  ...................c.....................t...a..............a................... [ 88720]
AY261360.1_Kenya_1950            ...............c............................................a................... [ 88720]
AM712240.1_OURT_88/3             .......................................a........................................ [ 88720]
AM712239.1_Benin_97/1            .......................................a........................................ [ 88720]
KM111295.1_Ken06.Bus             ...............c.............................a..............a................... [ 88720]
KM111294.1_Ken05/Tk1             ...............c............................................a................... [ 88720]
KM262844.1_strain_L60            .......................................a........................................ [ 88720]
KM102979.1_26544/OG10_Italy      .......................................a........................................ [ 88720]
KM262845.1_strain_NHV            .......................................a........................................ [ 88720]
KX354450.1_47/Ss/2008            .......................................a........................................ [ 88720]
KP055815.1_strain_BA71           .......................................a........................................ [ 88720]
MH025920.1_strain_R35            ...............c.............................a..............a................... [ 88720]
MH025919.1_strain_N10            ...............c.............................a..............a................... [ 88720]
MH025918.1_strain_R25            ...............c.............................a..............a................... [ 88720]
MH025917.1_strain_R7             ...............c.............................a..............a................... [ 88720]
MH025916.1_strain_R8             ...............c.............................a..............a................... [ 88720]

FR682468.2_Georgia_2007/1        tagaaggcccggcatttcctaaccctcaaaaaaaaattaatcacgttgcctatgcaagcattattactgctatgacgcat [ 88800]
ASF_IND_20_CAD543                ................y............................................................... [ 88800]
LR899193.1_Germany_2020/1        ................................................................................ [ 88800]
MN393477.1_Wuhan_2019            ................................................................................ [ 88800]
MW396979.1_Timor-Leste/2019/1    ................................................................................ [ 88800]
MK940252.1_CN/2019/InnerMongolia ................................................................................ [ 88800]
MK128995.1_China/2018/AnhuiXCGQ  ................................................................................ [ 88800]
MH681419.1_ASFV/POL/2015         ................................................................................ [ 88800]
LS478113.1_Estonia_2014          ................................................................................ [ 88800]
KP843857.1_Odintsovo_02/14       ...........................-.................................................... [ 88800]
```

```
FN557520.1_strain_E75           ................c............................................................... [ 88800]
AY261366.1_Warthog              ................................................................................ [ 88800]
AY261365.1_Warmbaths            ................................................................................ [ 88800]
AY261364.1_Tengani_62           ................c..................................................g............ [ 88800]
AY261363.1_Pretoriuskop/96/4    ................................................................................ [ 88800]
AY261362.1_Mkuzi_1979           ..........................................................t..................... [ 88800]
AY261361.1_Malawi_Lil-20/1_1983 ........................................c..t.................................... [ 88800]
AY261360.1_Kenya_1950           ...................................................................g............ [ 88800]
AM712240.1_OURT_88/3            ................c............................................................... [ 88800]
AM712239.1_Benin_97/1           ................c............................................................... [ 88800]
KM111295.1_Ken06.Bus            ...................................................................g............ [ 88800]
KM111294.1_Ken05/Tk1            .......................................................g...........g............ [ 88800]
KM262844.1_strain_L60           ................c............................................................... [ 88800]
KM102979.1_26544/OG10_Italy     ................c............................................................... [ 88800]
KM262845.1_strain_NHV           ................c............................................................... [ 88800]
KX354450.1_47/Ss/2008           ................c............................................................... [ 88800]
KP055815.1_strain_BA71          ................c............................................................... [ 88800]
MH025920.1_strain_R35           ...................................................................g............ [ 88800]
MH025919.1_strain_N10           ...................................................................g............ [ 88800]
MH025918.1_strain_R25           ...................................................................g............ [ 88800]
MH025917.1_strain_R7            ...................................................................g............ [ 88800]
MH025916.1_strain_R8            ...................................................................g............ [ 88800]

FR682468.2_Georgia_2007/1       cttatggttagagatcatatttttgtcattcaaacgaaaaatgaggcccacagttcccaaaagcttgtgcagctttttta [ 88880]
ASF_IND_20_CAD543               ................................................................................ [ 88880]
LR899193.1_Germany_2020/1       ................................................................................ [ 88880]
MN393477.1_Wuhan_2019           ................................................................................ [ 88880]
MW396979.1_Timor-Leste/2019/1   ................................................................................ [ 88880]
MK940252.1_CN/2019/InnerMongolia ................................................................................ [ 88880]
MK128995.1_China/2018/AnhuiXCGQ ................................................................................ [ 88880]
MH681419.1_ASFV/POL/2015        ................................................................................ [ 88880]
LS478113.1_Estonia_2014         ................................................................................ [ 88880]
KP843857.1_Odintsovo_02/14      ................................................................................ [ 88880]
FN557520.1_strain_E75           ................................................................................ [ 88880]
AY261366.1_Warthog              ....................g........................................................... [ 88880]
AY261365.1_Warmbaths            ....................g........................................................... [ 88880]
AY261364.1_Tengani_62           ...............t................................................................ [ 88880]
AY261363.1_Pretoriuskop/96/4    ....................g........................................................... [ 88880]
AY261362.1_Mkuzi_1979           ................................................................................ [ 88880]
AY261361.1_Malawi_Lil-20/1_1983 ........................a.........................................a............. [ 88880]
AY261360.1_Kenya_1950           ......................................g..............c........a..g.............. [ 88880]
AM712240.1_OURT_88/3            ................................................................................ [ 88880]
AM712239.1_Benin_97/1           ................................................................................ [ 88880]
KM111295.1_Ken06.Bus            ......................................g..............c........a..g.............. [ 88880]
```

```
KM111294.1_Ken05/Tk1             ......................................g..............c........a..g.............. [ 88880]
KM262844.1_strain_L60            ................................................................................ [ 88880]
KM102979.1_26544/OG10_Italy      ................................................................................ [ 88880]
KM262845.1_strain_NHV            ................................................................................ [ 88880]
KX354450.1_47/Ss/2008            ................................................................................ [ 88880]
KP055815.1_strain_BA71           ................................................................................ [ 88880]
MH025920.1_strain_R35            ......................................g..............c........a..g.............. [ 88880]
MH025919.1_strain_N10            ......................................g..............c........a..g.............. [ 88880]
MH025918.1_strain_R25            ......................................g..............c........a..g.............. [ 88880]
MH025917.1_strain_R7             ......................................g..............c........a..g.............. [ 88880]
MH025916.1_strain_R8             ......................................g..............c........a..g.............. [ 88880]

FR682468.2_Georgia_2007/1        tgccttttctaaggaaatggtgtgcgtcgttcccacctccctcacccccacggatgaagagctatgcatcaagctatggt [ 88960]
ASF_IND_20_CAD543                ................................................................................ [ 88960]
LR899193.1_Germany_2020/1        ................................................................................ [ 88960]
MN393477.1_Wuhan_2019            ................................................................................ [ 88960]
MW396979.1_Timor-Leste/2019/1    ................................................................................ [ 88960]
MK940252.1_CN/2019/InnerMongolia ................................................................................ [ 88960]
MK128995.1_China/2018/AnhuiXCGQ  ................................................................................ [ 88960]
MH681419.1_ASFV/POL/2015         ................................................................................ [ 88960]
LS478113.1_Estonia_2014          ................................................................................ [ 88960]
KP843857.1_Odintsovo_02/14       ................................................................................ [ 88960]
FN557520.1_strain_E75            ................................................................................ [ 88960]
AY261366.1_Warthog               ................................................................................ [ 88960]
AY261365.1_Warmbaths             ................................................................................ [ 88960]
AY261364.1_Tengani_62            ................................................................................ [ 88960]
AY261363.1_Pretoriuskop/96/4     ................................................................................ [ 88960]
AY261362.1_Mkuzi_1979            ................................................................................ [ 88960]
AY261361.1_Malawi_Lil-20/1_1983  .......................................t....................a............t...... [ 88960]
AY261360.1_Kenya_1950            ................................................................................ [ 88960]
AM712240.1_OURT_88/3             ................................................................................ [ 88960]
AM712239.1_Benin_97/1            ................................................................................ [ 88960]
KM111295.1_Ken06.Bus             ..................................g.....................................a....... [ 88960]
KM111294.1_Ken05/Tk1             .........................a..............................................a....... [ 88960]
KM262844.1_strain_L60            ................................................................................ [ 88960]
KM102979.1_26544/OG10_Italy      ................................................................................ [ 88960]
KM262845.1_strain_NHV            ................................................................................ [ 88960]
KX354450.1_47/Ss/2008            ................................................................................ [ 88960]
KP055815.1_strain_BA71           ................................................................................ [ 88960]
MH025920.1_strain_R35            ..................................g.....................................a....... [ 88960]
MH025919.1_strain_N10            ..................................g.....................................a....... [ 88960]
MH025918.1_strain_R25            ..................................g.....................................a....... [ 88960]
MH025917.1_strain_R7             ..................................g.....................................a....... [ 88960]
MH025916.1_strain_R8             ..................................g.....................................a....... [ 88960]
```

```
FR682468.2_Georgia_2007/1        cttctctttctggtatttcaggcgtgataggtaaaatcttggcaaacacttgttccgtagctcatttggttcatggaaag [ 89040]
ASF_IND_20_CAD543                .......................................................................ag....... [ 89040]
LR899193.1_Germany_2020/1        ................................................................................ [ 89040]
MN393477.1_Wuhan_2019            ................................................................................ [ 89040]
MW396979.1_Timor-Leste/2019/1    ................................................................................ [ 89040]
MK940252.1_CN/2019/InnerMongolia ................................................................................ [ 89040]
MK128995.1_China/2018/AnhuiXCGQ  ................................................................................ [ 89040]
MH681419.1_ASFV/POL/2015         ................................................................................ [ 89040]
LS478113.1_Estonia_2014          ................................................................................ [ 89040]
KP843857.1_Odintsovo_02/14       ................................................................................ [ 89040]
FN557520.1_strain_E75            ..............................................t........................ag....... [ 89040]
AY261366.1_Warthog               .......................................................................ag....... [ 89040]
AY261365.1_Warmbaths             ......................................................................aag....... [ 89040]
AY261364.1_Tengani_62            .....................................t.................................ag...g... [ 89040]
AY261363.1_Pretoriuskop/96/4     .......................................................................ag....... [ 89040]
AY261362.1_Mkuzi_1979            ..............................................t........................ag......t [ 89040]
AY261361.1_Malawi_Lil-20/1_1983  .c....................................a................a...............ag...g... [ 89040]
AY261360.1_Kenya_1950            .......................................................a...............ag...gc.. [ 89040]
AM712240.1_OURT_88/3             ..............................................t........................ag....... [ 89040]
AM712239.1_Benin_97/1            ..............................................t........................ag....... [ 89040]
KM111295.1_Ken06.Bus             .......................................................a...............ag...gc.. [ 89040]
KM111294.1_Ken05/Tk1             .......................................................a...............ag...gc.. [ 89040]
KM262844.1_strain_L60            ..............................................t........................ag....... [ 89040]
KM102979.1_26544/OG10_Italy      ..............................................t........................ag....... [ 89040]
KM262845.1_strain_NHV            ..............................................t........................ag....... [ 89040]
KX354450.1_47/Ss/2008            ..............................................t........................ag....... [ 89040]
KP055815.1_strain_BA71           ..............................................t........................ag....... [ 89040]
MH025920.1_strain_R35            .......................................................a...............ag...gc.. [ 89040]
MH025919.1_strain_N10            .......................................................a...............ag...gc.. [ 89040]
MH025918.1_strain_R25            .......................................................a...............ag...gc.. [ 89040]
MH025917.1_strain_R7             .......................................................a...............ag...gc.. [ 89040]
MH025916.1_strain_R8             .......................................................a...............ag...gc.. [ 89040]

FR682468.2_Georgia_2007/1        ctttcatcgcagaatattgatcagttaaaaactccctccaaccgaccattccccaaaaaagtaaaacgtatgcttataag [ 89120]
ASF_IND_20_CAD543                ................................................................................ [ 89120]
LR899193.1_Germany_2020/1        ................................................................................ [ 89120]
MN393477.1_Wuhan_2019            ................................................................................ [ 89120]
MW396979.1_Timor-Leste/2019/1    ................................................................................ [ 89120]
MK940252.1_CN/2019/InnerMongolia ................................................................................ [ 89120]
MK128995.1_China/2018/AnhuiXCGQ  ................................................................................ [ 89120]
MH681419.1_ASFV/POL/2015         ................................................................................ [ 89120]
LS478113.1_Estonia_2014          ................................................................................ [ 89120]
KP843857.1_Odintsovo_02/14       ................................................................................ [ 89120]
```

```
FN557520.1_strain_E75          .........................................t..g................................... [ 89120]
AY261366.1_Warthog             ...c.....................................t..g................................... [ 89120]
AY261365.1_Warmbaths           ...c............................c..t........g................................... [ 89120]
AY261364.1_Tengani_62          ................................c...........g................................... [ 89120]
AY261363.1_Pretoriuskop/96/4   ...c........................................g................................... [ 89120]
AY261362.1_Mkuzi_1979          ............................................g................................... [ 89120]
AY261361.1_Malawi_Lil-20/1_1983 ...c............................c..t........g..................................a [ 89120]
AY261360.1_Kenya_1950          ...c..c.........................c...........g................................... [ 89120]
AM712240.1_OURT_88/3           .........................................t..g................................... [ 89120]
AM712239.1_Benin_97/1          .........................................t..g................................... [ 89120]
KM111295.1_Ken06.Bus           ...c..c.........................c..t........g................................... [ 89120]
KM111294.1_Ken05/Tk1           ...c..c.........................c..a........g................................... [ 89120]
KM262844.1_strain_L60          .........................................t..g................................... [ 89120]
KM102979.1_26544/OG10_Italy    .........................................t..g................................... [ 89120]
KM262845.1_strain_NHV          .........................................t..g................................... [ 89120]
KX354450.1_47/Ss/2008          .........................................t..g................................... [ 89120]
KP055815.1_strain_BA71         .........................................t..g................................... [ 89120]
MH025920.1_strain_R35          ...c..c.........................c..t........g................................... [ 89120]
MH025919.1_strain_N10          ...c..c.........................c..t........g................................... [ 89120]
MH025918.1_strain_R25          ...c..c.........................c..t........g................................... [ 89120]
MH025917.1_strain_R7           ...c..c.........................c..t........g................................... [ 89120]
MH025916.1_strain_R8           ...c..c.........................c..t........g................................... [ 89120]

FR682468.2_Georgia_2007/1      cattagcaaaggaaataaggagttagaaataaaattgctctcgggggttcccaatatcgggaaaaaattagctgccgaaa [ 89200]
ASF_IND_20_CAD543              ................................................................................ [ 89200]
LR899193.1_Germany_2020/1      ................................................................................ [ 89200]
MN393477.1_Wuhan_2019          ................................................................................ [ 89200]
MW396979.1_Timor-Leste/2019/1  ................................................................................ [ 89200]
MK940252.1_CN/2019/InnerMongolia ................................................................................ [ 89200]
MK128995.1_China/2018/AnhuiXCGQ ................................................................................ [ 89200]
MH681419.1_ASFV/POL/2015       ................................................................................ [ 89200]
LS478113.1_Estonia_2014        ................................................................................ [ 89200]
KP843857.1_Odintsovo_02/14     ................................................................................ [ 89200]
FN557520.1_strain_E75          ..................................c............................................. [ 89200]
AY261366.1_Warthog             ................................................................................ [ 89200]
AY261365.1_Warmbaths           ..................a............................................................. [ 89200]
AY261364.1_Tengani_62          ......t............................................................c............ [ 89200]
AY261363.1_Pretoriuskop/96/4   ......t......................................................................... [ 89200]
AY261362.1_Mkuzi_1979          ......t......................................................................... [ 89200]
AY261361.1_Malawi_Lil-20/1_1983 t.....t.........................................c........t...................... [ 89200]
AY261360.1_Kenya_1950          .....................a..............a....................t...................... [ 89200]
AM712240.1_OURT_88/3           ..................................c............................................. [ 89200]
AM712239.1_Benin_97/1          ..................................c............................................. [ 89200]
KM111295.1_Ken06.Bus           .........................................................t...................... [ 89200]
```

```
KM111294.1_Ken05/Tk1            .....................a..............a....................t...................... [ 89200]
KM262844.1_strain_L60           ..................................c............................................. [ 89200]
KM102979.1_26544/OG10_Italy     ..................................c............................................. [ 89200]
KM262845.1_strain_NHV           ..................................c............................................. [ 89200]
KX354450.1_47/Ss/2008           ..................................c............................................. [ 89200]
KP055815.1_strain_BA71          ..................................c............................................. [ 89200]
MH025920.1_strain_R35           .........................................................t...................... [ 89200]
MH025919.1_strain_N10           .........................................................t...................... [ 89200]
MH025918.1_strain_R25           .........................................................t...................... [ 89200]
MH025917.1_strain_R7            .........................................................t...................... [ 89200]
MH025916.1_strain_R8            .........................................................t...................... [ 89200]

FR682468.2_Georgia_2007/1       ttttaaaagatcatgcgcttcttttttttctaaatcagcccgtagaatgcttggcaaatatacaaatcgttcaaaaaacc [ 89280]
ASF_IND_20_CAD543               ...................................................................y............ [ 89280]
LR899193.1_Germany_2020/1       ................................................................................ [ 89280]
MN393477.1_Wuhan_2019           ................................................................................ [ 89280]
MW396979.1_Timor-Leste/2019/1   ................................................................................ [ 89280]
MK940252.1_CN/2019/InnerMongolia ................................................................................ [ 89280]
MK128995.1_China/2018/AnhuiXCGQ ................................................................................ [ 89280]
MH681419.1_ASFV/POL/2015        ................................................................................ [ 89280]
LS478113.1_Estonia_2014         ................................................................................ [ 89280]
KP843857.1_Odintsovo_02/14      ................................................................................ [ 89280]
FN557520.1_strain_E75           ...................................................................t............ [ 89280]
AY261366.1_Warthog              .....................................................................c.......... [ 89280]
AY261365.1_Warmbaths            .....................................................................c.......... [ 89280]
AY261364.1_Tengani_62           .....................................................................c.......... [ 89280]
AY261363.1_Pretoriuskop/96/4    .............................................................t.......c.......... [ 89280]
AY261362.1_Mkuzi_1979           ...................................................................t............ [ 89280]
AY261361.1_Malawi_Lil-20/1_1983 ..................................................c.................a.c......... [ 89280]
AY261360.1_Kenya_1950           ..........c......................................t...................cc......... [ 89280]
AM712240.1_OURT_88/3            ...................................................................t............ [ 89280]
AM712239.1_Benin_97/1           ...................................................................t............ [ 89280]
KM111295.1_Ken06.Bus            ..........c.....................................a....................cc......... [ 89280]
KM111294.1_Ken05/Tk1            ..........c...................................t..t...................cc......... [ 89280]
KM262844.1_strain_L60           ...................................................................t............ [ 89280]
KM102979.1_26544/OG10_Italy     ...................................................................t............ [ 89280]
KM262845.1_strain_NHV           ...................................................................t............ [ 89280]
KX354450.1_47/Ss/2008           ...................................................................t............ [ 89280]
KP055815.1_strain_BA71          ...................................................................t............ [ 89280]
MH025920.1_strain_R35           ..........c.....................................a....................cc......... [ 89280]
MH025919.1_strain_N10           ..........c.....................................a....................cc......... [ 89280]
MH025918.1_strain_R25           ..........c.....................................a....................cc......... [ 89280]
MH025917.1_strain_R7            ..........c.....................................a....................cc......... [ 89280]
MH025916.1_strain_R8            ..........c.....................................a....................cc......... [ 89280]
```

```
FR682468.2_Georgia_2007/1        cgtacgattaagttgggaatgaagcgagccgaagcgattcattattttttaaactggtgtggctctgcccatgtaaccga [ 89360]
ASF_IND_20_CAD543                ................................................................................ [ 89360]
LR899193.1_Germany_2020/1        ................................................................................ [ 89360]
MN393477.1_Wuhan_2019            ................................................................................ [ 89360]
MW396979.1_Timor-Leste/2019/1    ................................................................................ [ 89360]
MK940252.1_CN/2019/InnerMongolia ................................................................................ [ 89360]
MK128995.1_China/2018/AnhuiXCGQ  ................................................................................ [ 89360]
MH681419.1_ASFV/POL/2015         ................................................................................ [ 89360]
LS478113.1_Estonia_2014          ................................................................................ [ 89360]
KP843857.1_Odintsovo_02/14       ................................................................................ [ 89360]
FN557520.1_strain_E75            ..............................................................g................. [ 89360]
AY261366.1_Warthog               ..............................................................g..............a.t [ 89360]
AY261365.1_Warmbaths             ..............................................................g................. [ 89360]
AY261364.1_Tengani_62            ............c.................................................a................. [ 89360]
AY261363.1_Pretoriuskop/96/4     ............c.................................................g..............a.t [ 89360]
AY261362.1_Mkuzi_1979            ....................a.........a...............................g................. [ 89360]
AY261361.1_Malawi_Lil-20/1_1983  ......................................c.......................g.......g......a.t [ 89360]
AY261360.1_Kenya_1950            ..............................................................g.......g......a.t [ 89360]
AM712240.1_OURT_88/3             ..............................................................g................. [ 89360]
AM712239.1_Benin_97/1            ..............................................................g................. [ 89360]
KM111295.1_Ken06.Bus             ..............................................................g.......g......a.t [ 89360]
KM111294.1_Ken05/Tk1             ..............................................................g.......g......a.t [ 89360]
KM262844.1_strain_L60            ..............................................................g................. [ 89360]
KM102979.1_26544/OG10_Italy      ..............................................................g................. [ 89360]
KM262845.1_strain_NHV            ..............................................................g................. [ 89360]
KX354450.1_47/Ss/2008            ..............................................................g................. [ 89360]
KP055815.1_strain_BA71           ..............................................................g................. [ 89360]
MH025920.1_strain_R35            ..............................................................g.......g......a.t [ 89360]
MH025919.1_strain_N10            ..............................................................g.......g......a.t [ 89360]
MH025918.1_strain_R25            ..............................................................g.......g......a.t [ 89360]
MH025917.1_strain_R7             ..............................................................g.......g......a.t [ 89360]
MH025916.1_strain_R8             ..............................................................g.......g......a.t [ 89360]

FR682468.2_Georgia_2007/1        tgatagccaaaatatcacagaggcgtcgcggtcca---caatgcaggtcgcgacgcagtccgccgcaatacagc------ [ 89440]
ASF_IND_20_CAD543                ...............................c..------------------------------..........------ [ 89440]
LR899193.1_Germany_2020/1        ...................................---....................................------ [ 89440]
MN393477.1_Wuhan_2019            ...................................---....................................------ [ 89440]
MW396979.1_Timor-Leste/2019/1    ...................................---....................................------ [ 89440]
MK940252.1_CN/2019/InnerMongolia ...................................---....................................------ [ 89440]
MK128995.1_China/2018/AnhuiXCGQ  ...................................---....................................------ [ 89440]
MH681419.1_ASFV/POL/2015         ...................................---....................................------ [ 89440]
LS478113.1_Estonia_2014          ...................................---....................................------ [ 89440]
KP843857.1_Odintsovo_02/14       ...................................---....................................------ [ 89440]
```

```
FN557520.1_strain_E75          ...............................c..------------------------------....c.....------ [ 89440]
AY261366.1_Warthog             g..............................c...---.................---------------....------ [ 89440]
AY261365.1_Warmbaths           ...................a...........c...---.................---------------....------ [ 89440]
AY261364.1_Tengani_62          ...............................c...---...........---------------..g.cg....------ [ 89440]
AY261363.1_Pretoriuskop/96/4   g..................a...........c...---....................ct.att....c.....------ [ 89440]
AY261362.1_Mkuzi_1979          ..............................ac..------------------------------....c.....------ [ 89440]
AY261361.1_Malawi_Lil-20/1_1983 g.....t.................a......c.t.---t.......cc...------------ta....g....------ [ 89440]
AY261360.1_Kenya_1950          g.....................a.a..a...c..gcca....a....c.a.------------...g.cg....------ [ 89440]
AM712240.1_OURT_88/3           ...............................c..------------------------------....c.....------ [ 89440]
AM712239.1_Benin_97/1          ...............................c..------------------------------....c.....------ [ 89440]
KM111295.1_Ken06.Bus           g.......................a..a...c..gcca....a....c.a.------------...g.cg..ag------ [ 89440]
KM111294.1_Ken05/Tk1           g.....................a.a..a...c..gcca....a....c.a.------------...g.cg..agccgttg [ 89440]
KM262844.1_strain_L60          ...............................c..------------------------------....c.....------ [ 89440]
KM102979.1_26544/OG10_Italy    ...............................c..------------------------------....c.....------ [ 89440]
KM262845.1_strain_NHV          ...............................c..------------------------------....c.....------ [ 89440]
KX354450.1_47/Ss/2008          ......................a........c..------------------------------....c.....------ [ 89440]
KP055815.1_strain_BA71         ...............................c..------------------------------....c.....------ [ 89440]
MH025920.1_strain_R35          g.......................a..a...c..gcca....a....c.a.------------...g.cg..ag------ [ 89440]
MH025919.1_strain_N10          g.......................a..a...c..gcca....a....c.a.------------...g.cg..ag------ [ 89440]
MH025918.1_strain_R25          g.......................a..a...c..gcca....a....c.a.------------...g.cg..ag------ [ 89440]
MH025917.1_strain_R7           g.......................a..a...c..gcca....a....c.a.------------...g.cg..ag------ [ 89440]
MH025916.1_strain_R8           g.......................a..a...c..gcca....a....c.a.------------...g.cg..ag------ [ 89440]

FR682468.2_Georgia_2007/1      ---------ccgctgcaacgcagccattgcacg----------------------------------------------- [ 89520]
ASF_IND_20_CAD543              ---------........................----------------------------------------------- [ 89520]
LR899193.1_Germany_2020/1      ---------........................----------------------------------------------- [ 89520]
MN393477.1_Wuhan_2019          ---------........................----------------------------------------------- [ 89520]
MW396979.1_Timor-Leste/2019/1  ---------........................----------------------------------------------- [ 89520]
MK940252.1_CN/2019/InnerMongolia ---------........................----------------------------------------------- [ 89520]
MK128995.1_China/2018/AnhuiXCGQ ---------........................----------------------------------------------- [ 89520]
MH681419.1_ASFV/POL/2015       ---------........................----------------------------------------------- [ 89520]
LS478113.1_Estonia_2014        ---------........................----------------------------------------------- [ 89520]
KP843857.1_Odintsovo_02/14     ---------........................----------------------------------------------- [ 89520]
FN557520.1_strain_E75          ---------.t......................----------------------------------------------- [ 89520]
AY261366.1_Warthog             ---------.......................a----------------------------------------------- [ 89520]
AY261365.1_Warmbaths           ---------........................----------------------------------------------- [ 89520]
AY261364.1_Tengani_62          ---------....c...................----------------------------------------------- [ 89520]
AY261363.1_Pretoriuskop/96/4   ---------........................----------------------------------------------- [ 89520]
AY261362.1_Mkuzi_1979          ---------........................----------------------------------------------- [ 89520]
AY261361.1_Malawi_Lil-20/1_1983 ---------t....a.................a----------------------------------------------- [ 89520]
AY261360.1_Kenya_1950          ---------.......g................aaatatcaaatg----------------------------------- [ 89520]
AM712240.1_OURT_88/3           ---------.t......................----------------------------------------------- [ 89520]
AM712239.1_Benin_97/1          ---------.t......................----------------------------------------------- [ 89520]
KM111295.1_Ken06.Bus           ---------........................aagcatcaaatgatgcatcaaatgatgcatcaaatgaagcatcaaat [ 89520]
```

```
KM111294.1_Ken05/Tk1             caacgcagc..............t.........aaatatcaaatgaagcatcaaatg----------------------- [ 89520]
KM262844.1_strain_L60            ---------.t......................----------------------------------------------- [ 89520]
KM102979.1_26544/OG10_Italy      ---------.t......................----------------------------------------------- [ 89520]
KM262845.1_strain_NHV            ---------.t......................----------------------------------------------- [ 89520]
KX354450.1_47/Ss/2008            ---------.t.................a....----------------------------------------------- [ 89520]
KP055815.1_strain_BA71           ---------.t......................----------------------------------------------- [ 89520]
MH025920.1_strain_R35            ---------........................----------------------------------------------- [ 89520]
MH025919.1_strain_N10            ---------........................----------------------------------------------- [ 89520]
MH025918.1_strain_R25            ---------........................----------------------------------------------- [ 89520]
MH025917.1_strain_R7             ---------........................----------------------------------------------- [ 89520]
MH025916.1_strain_R8             ---------........................----------------------------------------------- [ 89520]

FR682468.2_Georgia_2007/1        -aagtatcagatgatgca---------tcatcagatg---------------------------cttcatcacccgtagg [ 89600]
ASF_IND_20_CAD543                -.................acatctaatg.........---------------------------................ [ 89600]
LR899193.1_Germany_2020/1        -.................---------..........---------------------------................ [ 89600]
MN393477.1_Wuhan_2019            -.................---------..........---------------------------................ [ 89600]
MW396979.1_Timor-Leste/2019/1    -.................---------..........---------------------------................ [ 89600]
MK940252.1_CN/2019/InnerMongolia -.................---------..........---------------------------................ [ 89600]
MK128995.1_China/2018/AnhuiXCGQ  -.................---------..........---------------------------................ [ 89600]
MH681419.1_ASFV/POL/2015         -.................---------..........---------------------------................ [ 89600]
LS478113.1_Estonia_2014          -.................---------..........---------------------------................ [ 89600]
KP843857.1_Odintsovo_02/14       -.................---------..........---------------------------................ [ 89600]
FN557520.1_strain_E75            -.................acatctaatg........a---------------------------.a..c......a.... [ 89600]
AY261366.1_Warthog               -.................t---------.........---------------------------.c.........ac... [ 89600]
AY261365.1_Warmbaths             -.................tcatctaatg.........---------------------------...........ac... [ 89600]
AY261364.1_Tengani_62            -..............a..tcatctaatg........a---------------------------.a..c......ac... [ 89600]
AY261363.1_Pretoriuskop/96/4     -.c...............t---------.........---------------------------.c.........ac... [ 89600]
AY261362.1_Mkuzi_1979            -.................acatctaatg........a---------------------------.a..c......a.... [ 89600]
AY261361.1_Malawi_Lil-20/1_1983  -.............a...tcagagaatg.....c...atgcatcagagaatgcatcagataaagta........tac... [ 89600]
AY261360.1_Kenya_1950            -...c..t.a....a...tcaaatgatg.....a...---------------------------aagtca.t.a.ac... [ 89600]
AM712240.1_OURT_88/3             -.................acatctaatg........a---------------------------.a..c......a.... [ 89600]
AM712239.1_Benin_97/1            -.................acatctaatg........a---------------------------.a..c......a.... [ 89600]
KM111295.1_Ken06.Bus             a...c....a....a...tcaaataaag.....a..a---------------------------aagtca.t.a.ac... [ 89600]
KM111294.1_Ken05/Tk1             -.t.c....a........acagataaag.....a...---------------------------aagtca.t.a.ac... [ 89600]
KM262844.1_strain_L60            -.................acatctaatg........a---------------------------.a..c......a.... [ 89600]
KM102979.1_26544/OG10_Italy      -.................acatctaatg........a---------------------------.a..c......a.... [ 89600]
KM262845.1_strain_NHV            -.................acatctaatg........a---------------------------.a..c......a.... [ 89600]
KX354450.1_47/Ss/2008            -.................acatctaatg........a---------------------------.a..c......a.... [ 89600]
KP055815.1_strain_BA71           -.................acatctaatg........a---------------------------.a..c......a.... [ 89600]
MH025920.1_strain_R35            -...c....a....a...tcaaataaag.....a..a---------------------------aagtca.t.a.ac... [ 89600]
MH025919.1_strain_N10            -...c....a....a...tcaaataaag.....a..a---------------------------aagtca.t.a.ac... [ 89600]
MH025918.1_strain_R25            -...c....a....a...tcaaataaag.....a..a---------------------------aagtca.t.a.ac... [ 89600]
MH025917.1_strain_R7             -...c....a....a...tcaaataaag.....a..a---------------------------aagtca.t.a.ac... [ 89600]
MH025916.1_strain_R8             -...c....a....a...tcaaataaag.....a..a---------------------------aagtca.t.a.ac... [ 89600]
```

```
FR682468.2_Georgia_2007/1        gtatcaaacattatcta------------aagaaatgttattgaacacagcctgatgtt-aataattcactacatctaaa [ 89680]
ASF_IND_20_CAD543                .................------------..............................-.................... [ 89680]
LR899193.1_Germany_2020/1        .................------------..............................-.................... [ 89680]
MN393477.1_Wuhan_2019            .................------------..............................-.................... [ 89680]
MW396979.1_Timor-Leste/2019/1    .................------------..............................-.................... [ 89680]
MK940252.1_CN/2019/InnerMongolia .................------------..............................-.................... [ 89680]
MK128995.1_China/2018/AnhuiXCGQ  .................------------..............................-.................... [ 89680]
MH681419.1_ASFV/POL/2015         .................------------..............................-.................... [ 89680]
LS478113.1_Estonia_2014          .................------------..............................-.................... [ 89680]
KP843857.1_Odintsovo_02/14       .................------------..............................-.................... [ 89680]
FN557520.1_strain_E75            .c...............------------..............................-...t................ [ 89680]
AY261366.1_Warthog               .c...............------------.........c...........t.c......-...t................ [ 89680]
AY261365.1_Warmbaths             .c...............------------........cc....................-...t................ [ 89680]
AY261364.1_Tengani_62            .c...............------------........cc....................-...t................ [ 89680]
AY261363.1_Pretoriuskop/96/4     .c...............------------.........c....................-...t................ [ 89680]
AY261362.1_Mkuzi_1979            .c...............------------..............................-...t................ [ 89680]
AY261361.1_Malawi_Lil-20/1_1983  .c........a......aatatatatcta.......a.c............----..c.-..atga...a..t.a..... [ 89680]
AY261360.1_Kenya_1950            .c............t..--------tcta.......a.c..................c.a...t..c..a.........- [ 89680]
AM712240.1_OURT_88/3             .c...............------------..............................-...t................ [ 89680]
AM712239.1_Benin_97/1            .c...............------------..............................-...t................ [ 89680]
KM111295.1_Ken06.Bus             .c........----t..--------tcta.......a.c..................c.a...t..c..a.......... [ 89680]
KM111294.1_Ken05/Tk1             .c............t..--------tcta.......a.c..................c.a...t..c..a.........- [ 89680]
KM262844.1_strain_L60            .c...............------------..............................-...t................ [ 89680]
KM102979.1_26544/OG10_Italy      .c...............------------..............................-...t................ [ 89680]
KM262845.1_strain_NHV            .c...............------------..............................-...t................ [ 89680]
KX354450.1_47/Ss/2008            .c...............------------..............................-...t................ [ 89680]
KP055815.1_strain_BA71           .c...............------------..............................-...t................ [ 89680]
MH025920.1_strain_R35            .c........----t..--------tcta.......a.c.......g..........c.a...t..c..a.......... [ 89680]
MH025919.1_strain_N10            .c........----t..--------tcta.......a.c.......g..........c.a...t..c..a.......... [ 89680]
MH025918.1_strain_R25            .c........----t..--------tcta.......a.c.......g..........c.a...t..c..a.......... [ 89680]
MH025917.1_strain_R7             .c........----t..--------tcta.......a.c.......g..........c.a...t..c..a.......... [ 89680]
MH025916.1_strain_R8             .c........----t..--------tcta.......a.c.......g..........c.a...t..c..a.......... [ 89680]

FR682468.2_Georgia_2007/1        gaaatgttaacctcgatactaaaaagtcattgaacacaactactggggcgctaagttgtccaacacatctaaagaaatgt [ 89760]
ASF_IND_20_CAD543                ....................w.....ct.................................................... [ 89760]
LR899193.1_Germany_2020/1        ................................................................................ [ 89760]
MN393477.1_Wuhan_2019            ................................................................................ [ 89760]
MW396979.1_Timor-Leste/2019/1    ................................................................................ [ 89760]
MK940252.1_CN/2019/InnerMongolia ................................................................................ [ 89760]
MK128995.1_China/2018/AnhuiXCGQ  ................................................................................ [ 89760]
MH681419.1_ASFV/POL/2015         ................................................................................ [ 89760]
LS478113.1_Estonia_2014          ................................................................................ [ 89760]
KP843857.1_Odintsovo_02/14       ................................................................................ [ 89760]
```

```
FN557520.1_strain_E75           ....................t.....ct.............................a...................... [ 89760]
AY261366.1_Warthog              .........................act.................................................... [ 89760]
AY261365.1_Warmbaths            ..........t.........-....ac....................................t................ [ 89760]
AY261364.1_Tengani_62           ..........................c..................................................... [ 89760]
AY261363.1_Pretoriuskop/96/4    ..........................ct...................................t................ [ 89760]
AY261362.1_Mkuzi_1979           ..........................ct...................................t................ [ 89760]
AY261361.1_Malawi_Lil-20/1_1983 -.c.......t..............c.........................c.....a.....t................ [ 89760]
AY261360.1_Kenya_1950           -..tg......aa.--ct.......c.....................................a................ [ 89760]
AM712240.1_OURT_88/3            ....................t.....ct.............................a...................... [ 89760]
AM712239.1_Benin_97/1           ....................t.....ct.............................a...................... [ 89760]
KM111295.1_Ken06.Bus            -..tg.....taa.--ct.......c.....................................a................ [ 89760]
KM111294.1_Ken05/Tk1            -..tg......aa.--ct.......c.....................................a................ [ 89760]
KM262844.1_strain_L60           ....................t.....ct.............................a...................... [ 89760]
KM102979.1_26544/OG10_Italy     ....................t.....ct.............................a...................... [ 89760]
KM262845.1_strain_NHV           ....................t.....ct.............................a...................... [ 89760]
KX354450.1_47/Ss/2008           ....................t.....ct.............................a...................... [ 89760]
KP055815.1_strain_BA71          ....................t.....ct.............................a...................... [ 89760]
MH025920.1_strain_R35           -..tg.....taa.--ct.......c.....................................a................ [ 89760]
MH025919.1_strain_N10           -..tg.....taa.--ct.......c.....................................a................ [ 89760]
MH025918.1_strain_R25           -..tg.....taa.--ct.......c.....................................a................ [ 89760]
MH025917.1_strain_R7            -..tg.....taa.--ct.......c.....................................a................ [ 89760]
MH025916.1_strain_R8            -..tg.....taa.--ct.......c.....................................a................ [ 89760]

FR682468.2_Georgia_2007/1       caacatcctcgatgctaaaagggtcatcgagccggtcaataatgtcttccccaaaaagtccgggagaactgtaggccgag [ 89840]
ASF_IND_20_CAD543               ................................................................................ [ 89840]
LR899193.1_Germany_2020/1       ................................................................................ [ 89840]
MN393477.1_Wuhan_2019           ................................................................................ [ 89840]
MW396979.1_Timor-Leste/2019/1   ................................................................................ [ 89840]
MK940252.1_CN/2019/InnerMongolia ................................................................................ [ 89840]
MK128995.1_China/2018/AnhuiXCGQ ................................................................................ [ 89840]
MH681419.1_ASFV/POL/2015        ................................................................................ [ 89840]
LS478113.1_Estonia_2014         ................................................................................ [ 89840]
KP843857.1_Odintsovo_02/14      ................................................................................ [ 89840]
FN557520.1_strain_E75           .................................................g.............................. [ 89840]
AY261366.1_Warthog              ................................................................................ [ 89840]
AY261365.1_Warmbaths            ................................................................................ [ 89840]
AY261364.1_Tengani_62           ................................................................................ [ 89840]
AY261363.1_Pretoriuskop/96/4    ................................................................................ [ 89840]
AY261362.1_Mkuzi_1979           ................................................................................ [ 89840]
AY261361.1_Malawi_Lil-20/1_1983 .c..............................t............................................... [ 89840]
AY261360.1_Kenya_1950           ..........................................................c........g............ [ 89840]
AM712240.1_OURT_88/3            .................................................g.............................. [ 89840]
AM712239.1_Benin_97/1           .................................................g.............................. [ 89840]
KM111295.1_Ken06.Bus            .............a............................................c........g.....a...... [ 89840]
```

```
KM111294.1_Ken05/Tk1             ..................................................c........g............ [ 89840]
KM262844.1_strain_L60            .................................................g.............................. [ 89840]
KM102979.1_26544/OG10_Italy      .................................................g.............................. [ 89840]
KM262845.1_strain_NHV            .................................................g.............................. [ 89840]
KX354450.1_47/Ss/2008            .................................................g.............................. [ 89840]
KP055815.1_strain_BA71           .................................................g.............................. [ 89840]
MH025920.1_strain_R35            .............a............................................c........g.....a...... [ 89840]
MH025919.1_strain_N10            .............a............................................c........g.....a...... [ 89840]
MH025918.1_strain_R25            .............a............................................c........g.....a...... [ 89840]
MH025917.1_strain_R7             .............a............................................c........g.....a...... [ 89840]
MH025916.1_strain_R8             .............a............................................c........g.....a...... [ 89840]

FR682468.2_Georgia_2007/1        atgtcgtccatggagctatcttccccagagcacacaaagtcctctccaaaaatcataaagttaaatgcaccgggcttact [ 89920]
ASF_IND_20_CAD543                ................................................................................ [ 89920]
LR899193.1_Germany_2020/1        ................................................................................ [ 89920]
MN393477.1_Wuhan_2019            ................................................................................ [ 89920]
MW396979.1_Timor-Leste/2019/1    ................................................................................ [ 89920]
MK940252.1_CN/2019/InnerMongolia ................................................................................ [ 89920]
MK128995.1_China/2018/AnhuiXCGQ  ................................................................................ [ 89920]
MH681419.1_ASFV/POL/2015         ................................................................................ [ 89920]
LS478113.1_Estonia_2014          ................................................................................ [ 89920]
KP843857.1_Odintsovo_02/14       ................................................................................ [ 89920]
FN557520.1_strain_E75            .....a......................................c................................... [ 89920]
AY261366.1_Warthog               .....a......................................c................................... [ 89920]
AY261365.1_Warmbaths             .....a......................................c................................... [ 89920]
AY261364.1_Tengani_62            ................................................................................ [ 89920]
AY261363.1_Pretoriuskop/96/4     .....a......................................c................................... [ 89920]
AY261362.1_Mkuzi_1979            .....a......................................c................................... [ 89920]
AY261361.1_Malawi_Lil-20/1_1983  .....a.......................................................................... [ 89920]
AY261360.1_Kenya_1950            .....a......................................c................................... [ 89920]
AM712240.1_OURT_88/3             .....a......................................c................................... [ 89920]
AM712239.1_Benin_97/1            .....a......................................c................................... [ 89920]
KM111295.1_Ken06.Bus             .....a......................................c................................... [ 89920]
KM111294.1_Ken05/Tk1             .....a......................................c.......................g........... [ 89920]
KM262844.1_strain_L60            .....a......................................c................................... [ 89920]
KM102979.1_26544/OG10_Italy      .....a......................................c................................... [ 89920]
KM262845.1_strain_NHV            .....a......................................c................................... [ 89920]
KX354450.1_47/Ss/2008            .....a......................................c................................... [ 89920]
KP055815.1_strain_BA71           ....aa......................................c................................... [ 89920]
MH025920.1_strain_R35            .....a......................................c................................... [ 89920]
MH025919.1_strain_N10            .....a......................................c................................... [ 89920]
MH025918.1_strain_R25            .....a......................................c................................... [ 89920]
MH025917.1_strain_R7             .....a......................................c................................... [ 89920]
MH025916.1_strain_R8             .....a......................................c................................... [ 89920]
```

```
FR682468.2_Georgia_2007/1        taacagcttttcgctttgaataatagtgttgagttctgtcagcgcaaactctctcacaatattcacaacccaggagggct [ 90000]
ASF_IND_20_CAD543                ................................................................................ [ 90000]
LR899193.1_Germany_2020/1        ................................................................................ [ 90000]
MN393477.1_Wuhan_2019            ................................................................................ [ 90000]
MW396979.1_Timor-Leste/2019/1    ................................................................................ [ 90000]
MK940252.1_CN/2019/InnerMongolia ................................................................................ [ 90000]
MK128995.1_China/2018/AnhuiXCGQ  ................................................................................ [ 90000]
MH681419.1_ASFV/POL/2015         ................................................................................ [ 90000]
LS478113.1_Estonia_2014          ................................................................................ [ 90000]
KP843857.1_Odintsovo_02/14       ................................................................................ [ 90000]
FN557520.1_strain_E75            .................................c..............t..........................a.... [ 90000]
AY261366.1_Warthog               .................................c.........................................a.... [ 90000]
AY261365.1_Warmbaths             .................................c.........................................a.... [ 90000]
AY261364.1_Tengani_62            .................................c.............................................. [ 90000]
AY261363.1_Pretoriuskop/96/4     .................................c.........................................a.... [ 90000]
AY261362.1_Mkuzi_1979            .................................c..............t..........................a.... [ 90000]
AY261361.1_Malawi_Lil-20/1_1983  .................................c....................t.t..................a.... [ 90000]
AY261360.1_Kenya_1950            ...........................a.....c.........................................a.... [ 90000]
AM712240.1_OURT_88/3             .................................c..............t..........................a.... [ 90000]
AM712239.1_Benin_97/1            .................................c..............t..........................a.... [ 90000]
KM111295.1_Ken06.Bus             ...........................a.....c.........................................a.... [ 90000]
KM111294.1_Ken05/Tk1             ...........................a.....c.........................................a.... [ 90000]
KM262844.1_strain_L60            .................................c..............t..........................a.... [ 90000]
KM102979.1_26544/OG10_Italy      .................................c..............t..........................a.... [ 90000]
KM262845.1_strain_NHV            .................................c..............t..........................a.... [ 90000]
KX354450.1_47/Ss/2008            .................................c..............t..........................a.... [ 90000]
KP055815.1_strain_BA71           .................................c..............t..........................a.... [ 90000]
MH025920.1_strain_R35            ...........................a.....c.........................................a.... [ 90000]
MH025919.1_strain_N10            ...........................a.....c.........a...............................a.... [ 90000]
MH025918.1_strain_R25            ...........................a.....c.........................................a.... [ 90000]
MH025917.1_strain_R7             ...........................a.....c.........................................a.... [ 90000]
MH025916.1_strain_R8             ...........................a.....c.........................................a.... [ 90000]

FR682468.2_Georgia_2007/1        ctttaatttcatacagcgttaagaaacttatacataaaaattctatagagtaaagcaaggcgctggcaggatctgttacc [ 90080]
ASF_IND_20_CAD543                ..............................y................................................. [ 90080]
LR899193.1_Germany_2020/1        ................................................................................ [ 90080]
MN393477.1_Wuhan_2019            ................................................................................ [ 90080]
MW396979.1_Timor-Leste/2019/1    ................................................................................ [ 90080]
MK940252.1_CN/2019/InnerMongolia ................................................................................ [ 90080]
MK128995.1_China/2018/AnhuiXCGQ  ................................................................................ [ 90080]
MH681419.1_ASFV/POL/2015         ................................................................................ [ 90080]
LS478113.1_Estonia_2014          ................................................................................ [ 90080]
KP843857.1_Odintsovo_02/14       ................................................................................ [ 90080]
```

```
FN557520.1_strain_E75           ..............................c................................................. [ 90080]
AY261366.1_Warthog              .......g........................................................................ [ 90080]
AY261365.1_Warmbaths            .........................................................g...................... [ 90080]
AY261364.1_Tengani_62           ................................................................................ [ 90080]
AY261363.1_Pretoriuskop/96/4    ................................................................................ [ 90080]
AY261362.1_Mkuzi_1979           ..............................c................................................. [ 90080]
AY261361.1_Malawi_Lil-20/1_1983 ....................................g............a........t..................... [ 90080]
AY261360.1_Kenya_1950           ....................................................g........................... [ 90080]
AM712240.1_OURT_88/3            ..............................c................................................. [ 90080]
AM712239.1_Benin_97/1           ..............................c................................................. [ 90080]
KM111295.1_Ken06.Bus            ....................................................g........................... [ 90080]
KM111294.1_Ken05/Tk1            ................................................................................ [ 90080]
KM262844.1_strain_L60           ..............................c................................................. [ 90080]
KM102979.1_26544/OG10_Italy     ..............................c................................................. [ 90080]
KM262845.1_strain_NHV           ..............................c................................................. [ 90080]
KX354450.1_47/Ss/2008           ..............................c................................................. [ 90080]
KP055815.1_strain_BA71          ..............................c................................................. [ 90080]
MH025920.1_strain_R35           ....................................................g........................... [ 90080]
MH025919.1_strain_N10           ....................................................g........................... [ 90080]
MH025918.1_strain_R25           ....................................................g........................... [ 90080]
MH025917.1_strain_R7            ....................................................g........................... [ 90080]
MH025916.1_strain_R8            ....................................................g........................... [ 90080]

FR682468.2_Georgia_2007/1       cgtaggtgtttaaatgtagtgtgatattcattcacaacgttaggcagcaccttttccaaatcctccttttcctcgtacga [ 90160]
ASF_IND_20_CAD543               ................................................................................ [ 90160]
LR899193.1_Germany_2020/1       ................................................................................ [ 90160]
MN393477.1_Wuhan_2019           ................................................................................ [ 90160]
MW396979.1_Timor-Leste/2019/1   ................................................................................ [ 90160]
MK940252.1_CN/2019/InnerMongolia ................................................................................ [ 90160]
MK128995.1_China/2018/AnhuiXCGQ ................................................................................ [ 90160]
MH681419.1_ASFV/POL/2015        ................................................................................ [ 90160]
LS478113.1_Estonia_2014         ................................................................................ [ 90160]
KP843857.1_Odintsovo_02/14      ................................................................................ [ 90160]
FN557520.1_strain_E75           ...................................g..............t.......................a..... [ 90160]
AY261366.1_Warthog              ...................................g..............t............................. [ 90160]
AY261365.1_Warmbaths            ...................................g..............t............................. [ 90160]
AY261364.1_Tengani_62           ..................................................t............................. [ 90160]
AY261363.1_Pretoriuskop/96/4    ...................................g..............t............................. [ 90160]
AY261362.1_Mkuzi_1979           ...................................g..............t.......................a..... [ 90160]
AY261361.1_Malawi_Lil-20/1_1983 .....t..........................t..g..............t............................. [ 90160]
AY261360.1_Kenya_1950           .....t........a....................g..............t........g.................... [ 90160]
AM712240.1_OURT_88/3            ...................................g..............t.......................a..... [ 90160]
AM712239.1_Benin_97/1           ...................................g..............t.......................a..... [ 90160]
KM111295.1_Ken06.Bus            .....t........a....................g...........t..t..........t.................. [ 90160]
```

```
KM111294.1_Ken05/Tk1             .....t........a....................g..............t........g.................... [ 90160]
KM262844.1_strain_L60            ...................................g..............t.......................a..... [ 90160]
KM102979.1_26544/OG10_Italy      ...................................g..............t.......................a..... [ 90160]
KM262845.1_strain_NHV            ...................................g..............t.......................a..... [ 90160]
KX354450.1_47/Ss/2008            ...................................g..............t.......................a..... [ 90160]
KP055815.1_strain_BA71           ...................................g..............t.......................a..... [ 90160]
MH025920.1_strain_R35            .....t........a....................g...........t..t............................. [ 90160]
MH025919.1_strain_N10            .....t........a....................g...........t..t............................. [ 90160]
MH025918.1_strain_R25            .....t........a....................g...........t..t............................. [ 90160]
MH025917.1_strain_R7             .....t........a....................g...........t..t............................. [ 90160]
MH025916.1_strain_R8             .....t........a....................g...........t..t............................. [ 90160]

FR682468.2_Georgia_2007/1        caggtgctttacaagcctttcaacatgtataggaggcttgttaaatgtactaacgtgccgcaaacagttataattatata [ 90240]
ASF_IND_20_CAD543                ................................................................................ [ 90240]
LR899193.1_Germany_2020/1        ................................................................................ [ 90240]
MN393477.1_Wuhan_2019            ................................................................................ [ 90240]
MW396979.1_Timor-Leste/2019/1    ................................................................................ [ 90240]
MK940252.1_CN/2019/InnerMongolia ................................................................................ [ 90240]
MK128995.1_China/2018/AnhuiXCGQ  ................................................................................ [ 90240]
MH681419.1_ASFV/POL/2015         ................................................................................ [ 90240]
LS478113.1_Estonia_2014          ................................................................................ [ 90240]
KP843857.1_Odintsovo_02/14       ................................................................................ [ 90240]
FN557520.1_strain_E75            ...a.......................c..........................a......................... [ 90240]
AY261366.1_Warthog               ...........................c..........................a......................... [ 90240]
AY261365.1_Warmbaths             ...........................c..........................a......................... [ 90240]
AY261364.1_Tengani_62            ...........................c..........................a......................... [ 90240]
AY261363.1_Pretoriuskop/96/4     ...........................c..........................a......................... [ 90240]
AY261362.1_Mkuzi_1979            ...a.......................c..........................a......................... [ 90240]
AY261361.1_Malawi_Lil-20/1_1983  .......................t..ac........t........g.......t.......................... [ 90240]
AY261360.1_Kenya_1950            .......................t...c........t.....g..g.......t.......................... [ 90240]
AM712240.1_OURT_88/3             ...a.......................c..........................a......................... [ 90240]
AM712239.1_Benin_97/1            ...a.......................c..........................a......................... [ 90240]
KM111295.1_Ken06.Bus             .......................t...c........t.....g..g.......t.......................... [ 90240]
KM111294.1_Ken05/Tk1             .......................t...c........t.....g..g.......t.......................... [ 90240]
KM262844.1_strain_L60            ...a.......................c..........................a......................... [ 90240]
KM102979.1_26544/OG10_Italy      ...a.......................c..........................a......................... [ 90240]
KM262845.1_strain_NHV            ...a.......................c..........................a......................... [ 90240]
KX354450.1_47/Ss/2008            ...a.......................c..........................a......................... [ 90240]
KP055815.1_strain_BA71           ...a.......................c..........................a......................... [ 90240]
MH025920.1_strain_R35            .......................t...c........t.....g..g.......t.......................... [ 90240]
MH025919.1_strain_N10            .......................t...c........t.....g..g.......t.......................... [ 90240]
MH025918.1_strain_R25            .......................t...c........t.....g..g.......t.......................... [ 90240]
MH025917.1_strain_R7             .......................t...c........t.....g..g.......t.......................... [ 90240]
MH025916.1_strain_R8             .......................t...c........t.....g..g.......t.......................... [ 90240]
```

```
FR682468.2_Georgia_2007/1        agaaaatacgtacggcagagtcgaccgccatgagccttggatcatccattgaggtaggtggtggcggggcaccctggcct [ 90320]
ASF_IND_20_CAD543                ................................................................................ [ 90320]
LR899193.1_Germany_2020/1        ................................................................................ [ 90320]
MN393477.1_Wuhan_2019            ................................................................................ [ 90320]
MW396979.1_Timor-Leste/2019/1    ................................................................................ [ 90320]
MK940252.1_CN/2019/InnerMongolia ................................................................................ [ 90320]
MK128995.1_China/2018/AnhuiXCGQ  ................................................................................ [ 90320]
MH681419.1_ASFV/POL/2015         ................................................................................ [ 90320]
LS478113.1_Estonia_2014          ................................................................................ [ 90320]
KP843857.1_Odintsovo_02/14       ................................................................................ [ 90320]
FN557520.1_strain_E75            ................................................................................ [ 90320]
AY261366.1_Warthog               ................................................................................ [ 90320]
AY261365.1_Warmbaths             ................................................................................ [ 90320]
AY261364.1_Tengani_62            ................................................................................ [ 90320]
AY261363.1_Pretoriuskop/96/4     ...........................................g.................................... [ 90320]
AY261362.1_Mkuzi_1979            ................................................................................ [ 90320]
AY261361.1_Malawi_Lil-20/1_1983  ......................a..t........t..a.......................c..g............... [ 90320]
AY261360.1_Kenya_1950            .........................t...........a.......................c..a............... [ 90320]
AM712240.1_OURT_88/3             ................................................................................ [ 90320]
AM712239.1_Benin_97/1            ................................................................................ [ 90320]
KM111295.1_Ken06.Bus             .........................t...........a.......................c..a............... [ 90320]
KM111294.1_Ken05/Tk1             .........................t...........a.......................c..a............... [ 90320]
KM262844.1_strain_L60            ................................................................................ [ 90320]
KM102979.1_26544/OG10_Italy      ................................................................................ [ 90320]
KM262845.1_strain_NHV            ................................................................................ [ 90320]
KX354450.1_47/Ss/2008            ................................................................................ [ 90320]
KP055815.1_strain_BA71           ................................................................................ [ 90320]
MH025920.1_strain_R35            .........................t...........a.......................c..a............... [ 90320]
MH025919.1_strain_N10            .........................t...........a.......................c..a............... [ 90320]
MH025918.1_strain_R25            .........................t...........a.......................c..a............... [ 90320]
MH025917.1_strain_R7             .........................t...........a.......................c..a............... [ 90320]
MH025916.1_strain_R8             .........................t...........a.......................c..a............... [ 90320]

FR682468.2_Georgia_2007/1        tccctgatgtctgcgtaggagcgcccctccatggcccctatggcctctatcacagcaggactgatatccaaaatcttggc [ 90400]
ASF_IND_20_CAD543                ................................................................................ [ 90400]
LR899193.1_Germany_2020/1        ................................................................................ [ 90400]
MN393477.1_Wuhan_2019            ................................................................................ [ 90400]
MW396979.1_Timor-Leste/2019/1    ................................................................................ [ 90400]
MK940252.1_CN/2019/InnerMongolia ................................................................................ [ 90400]
MK128995.1_China/2018/AnhuiXCGQ  ................................................................................ [ 90400]
MH681419.1_ASFV/POL/2015         ................................................................................ [ 90400]
LS478113.1_Estonia_2014          ................................................................................ [ 90400]
KP843857.1_Odintsovo_02/14       ................................................................................ [ 90400]
```

```
FN557520.1_strain_E75             ...................................t............................................ [ 90400]
AY261366.1_Warthog                ...................................t..............t............................. [ 90400]
AY261365.1_Warmbaths              ...................................t............................................ [ 90400]
AY261364.1_Tengani_62             ...................................t..............t.......................t..... [ 90400]
AY261363.1_Pretoriuskop/96/4      ...................................t............................................ [ 90400]
AY261362.1_Mkuzi_1979             ................................................................................ [ 90400]
AY261361.1_Malawi_Lil-20/1_1983   ...t...............................t.................g.......................... [ 90400]
AY261360.1_Kenya_1950             ...t...................t...........t.............c.............................. [ 90400]
AM712240.1_OURT_88/3              ...................................t............................................ [ 90400]
AM712239.1_Benin_97/1             ...................................t............................................ [ 90400]
KM111295.1_Ken06.Bus              ...t...................t...........t.............c.............................. [ 90400]
KM111294.1_Ken05/Tk1              ...t................a..t........a..t.............c........................t..a.. [ 90400]
KM262844.1_strain_L60             ...................................t............................................ [ 90400]
KM102979.1_26544/OG10_Italy       ...................................t............................................ [ 90400]
KM262845.1_strain_NHV             ...................................t............................................ [ 90400]
KX354450.1_47/Ss/2008             ...................................t............................................ [ 90400]
KP055815.1_strain_BA71            ...................................t............................................ [ 90400]
MH025920.1_strain_R35             ...t...................t...........t.............c.............................. [ 90400]
MH025919.1_strain_N10             ...t...................t...........t.............c.............................. [ 90400]
MH025918.1_strain_R25             ...t...................t...........t.............c.............................. [ 90400]
MH025917.1_strain_R7              ...t...................t...........t.............c.............................. [ 90400]
MH025916.1_strain_R8              ...t...................t...........t.............c.............................. [ 90400]

FR682468.2_Georgia_2007/1         cgtcttgattatttttccgtaatcgaaagtccatggctcctgtggaggcttgggttgtgtttcggtggagggcgtggtca [ 90480]
ASF_IND_20_CAD543                 ................................................................................ [ 90480]
LR899193.1_Germany_2020/1         ................................................................................ [ 90480]
MN393477.1_Wuhan_2019             ................................................................................ [ 90480]
MW396979.1_Timor-Leste/2019/1     ................................................................................ [ 90480]
MK940252.1_CN/2019/InnerMongolia  ................................................................................ [ 90480]
MK128995.1_China/2018/AnhuiXCGQ   ................................................................................ [ 90480]
MH681419.1_ASFV/POL/2015          ................................................................................ [ 90480]
LS478113.1_Estonia_2014           ................................................................................ [ 90480]
KP843857.1_Odintsovo_02/14        ................................................................................ [ 90480]
FN557520.1_strain_E75             ...........................g....................................c............... [ 90480]
AY261366.1_Warthog                ...........................g.............................................a...... [ 90480]
AY261365.1_Warmbaths              ...........................g.................................................... [ 90480]
AY261364.1_Tengani_62             ...........................g.................................................... [ 90480]
AY261363.1_Pretoriuskop/96/4      ...........................g.................................................... [ 90480]
AY261362.1_Mkuzi_1979             ...........................g.................................................... [ 90480]
AY261361.1_Malawi_Lil-20/1_1983   ...........................g........t......t......c..............c...a...ac..ca. [ 90480]
AY261360.1_Kenya_1950             ..................a.................t.t....t........................g....ac..ca. [ 90480]
AM712240.1_OURT_88/3              ...........................g....................................c............... [ 90480]
AM712239.1_Benin_97/1             ...........................g....................................c............... [ 90480]
KM111295.1_Ken06.Bus              ..................a.................t.t....t........................g....ac..cag [ 90480]
```

```
KM111294.1_Ken05/Tk1            ..................a.................t.t....t..............a......gt...a..ac..ca. [ 90480]
KM262844.1_strain_L60           ...........................g....................................c............... [ 90480]
KM102979.1_26544/OG10_Italy     ...........................g....................................c............... [ 90480]
KM262845.1_strain_NHV           ...........................g....................................c............... [ 90480]
KX354450.1_47/Ss/2008           ...........................g....................................c............... [ 90480]
KP055815.1_strain_BA71          ...........................g....................................c............... [ 90480]
MH025920.1_strain_R35           ..................a.................t.t....t........................g....ac..cag [ 90480]
MH025919.1_strain_N10           ..................a.................t.t....t........................g....ac..cag [ 90480]
MH025918.1_strain_R25           ..................a.................t.t....t........................g....ac..cag [ 90480]
MH025917.1_strain_R7            ..................a.................t.t....t........................g....ac..cag [ 90480]
MH025916.1_strain_R8            ..................a.................t.t....t........................g....ac..cag [ 90480]

FR682468.2_Georgia_2007/1       tatctttctttatttgaatagaacggatcgacatcttttccttatcgtactggtctttataattattataatagtcatga [ 90560]
ASF_IND_20_CAD543               ....................................................r........................... [ 90560]
LR899193.1_Germany_2020/1       ................................................................................ [ 90560]
MN393477.1_Wuhan_2019           ................................................................................ [ 90560]
MW396979.1_Timor-Leste/2019/1   ................................................................................ [ 90560]
MK940252.1_CN/2019/InnerMongolia ................................................................................ [ 90560]
MK128995.1_China/2018/AnhuiXCGQ ................................................................................ [ 90560]
MH681419.1_ASFV/POL/2015        ................................................................................ [ 90560]
LS478113.1_Estonia_2014         ................................................................................ [ 90560]
KP843857.1_Odintsovo_02/14      ................................................................................ [ 90560]
FN557520.1_strain_E75           ....................................................a........................... [ 90560]
AY261366.1_Warthog              ....................................................a........................... [ 90560]
AY261365.1_Warmbaths            ....................................................a........................... [ 90560]
AY261364.1_Tengani_62           .................................................t.............................. [ 90560]
AY261363.1_Pretoriuskop/96/4    ....................................................a........................... [ 90560]
AY261362.1_Mkuzi_1979           ....................................................a........................... [ 90560]
AY261361.1_Malawi_Lil-20/1_1983 ..........c............t.a..........................a........g.................. [ 90560]
AY261360.1_Kenya_1950           ...................t...t.a.................c........a........................... [ 90560]
AM712240.1_OURT_88/3            ....................................................a........................... [ 90560]
AM712239.1_Benin_97/1           ....................................................a........................... [ 90560]
KM111295.1_Ken06.Bus            ...................t...t.a.................c........a........................... [ 90560]
KM111294.1_Ken05/Tk1            ...................t...taa.................c........a........................... [ 90560]
KM262844.1_strain_L60           ....................................................a........................... [ 90560]
KM102979.1_26544/OG10_Italy     ....................................................a........................... [ 90560]
KM262845.1_strain_NHV           ....................................................a........................... [ 90560]
KX354450.1_47/Ss/2008           ....................................................a........................... [ 90560]
KP055815.1_strain_BA71          ....................................................a........................... [ 90560]
MH025920.1_strain_R35           ...................t...t.a.................c........a........................... [ 90560]
MH025919.1_strain_N10           ...................t...t.a.................c........a........................... [ 90560]
MH025918.1_strain_R25           ...................t...t.a.................c........a........................... [ 90560]
MH025917.1_strain_R7            ...................t...t.a.................c........a........................... [ 90560]
MH025916.1_strain_R8            ...................t...t.a.................c........a........................... [ 90560]
```

```
FR682468.2_Georgia_2007/1         actaattcgggttgagaaagatgatcgtatataatataggtaaaaagtccgcacttgacacattttttatcctggaagtc [ 90640]
ASF_IND_20_CAD543                 ................................................................................ [ 90640]
LR899193.1_Germany_2020/1         ................................................................................ [ 90640]
MN393477.1_Wuhan_2019             ................................................................................ [ 90640]
MW396979.1_Timor-Leste/2019/1     ................................................................................ [ 90640]
MK940252.1_CN/2019/InnerMongolia  ................................................................................ [ 90640]
MK128995.1_China/2018/AnhuiXCGQ   ................................................................................ [ 90640]
MH681419.1_ASFV/POL/2015          ................................................................................ [ 90640]
LS478113.1_Estonia_2014           ................................................................................ [ 90640]
KP843857.1_Odintsovo_02/14        ................................................................................ [ 90640]
FN557520.1_strain_E75             ................................................................................ [ 90640]
AY261366.1_Warthog                ................................................................................ [ 90640]
AY261365.1_Warmbaths              ................................................................................ [ 90640]
AY261364.1_Tengani_62             ................................................................................ [ 90640]
AY261363.1_Pretoriuskop/96/4      ................................................................................ [ 90640]
AY261362.1_Mkuzi_1979             ................................................................................ [ 90640]
AY261361.1_Malawi_Lil-20/1_1983   .......................g.....................................................a.. [ 90640]
AY261360.1_Kenya_1950             ...........c..............................................t..................... [ 90640]
AM712240.1_OURT_88/3              ................................................................................ [ 90640]
AM712239.1_Benin_97/1             ................................................................................ [ 90640]
KM111295.1_Ken06.Bus              ..........................................................t..................... [ 90640]
KM111294.1_Ken05/Tk1              ...........c..............................................t..................... [ 90640]
KM262844.1_strain_L60             ................................................................................ [ 90640]
KM102979.1_26544/OG10_Italy       ................................................................................ [ 90640]
KM262845.1_strain_NHV             ................................................................................ [ 90640]
KX354450.1_47/Ss/2008             ................................................................................ [ 90640]
KP055815.1_strain_BA71            ................................................................................ [ 90640]
MH025920.1_strain_R35             ..........................................................t..................... [ 90640]
MH025919.1_strain_N10             ..........................................................t..................... [ 90640]
MH025918.1_strain_R25             ..........................................................t..................... [ 90640]
MH025917.1_strain_R7              ..........................................................t..................... [ 90640]
MH025916.1_strain_R8              ..........................................................t..................... [ 90640]

FR682468.2_Georgia_2007/1         gtgtaatcctcccttggggcagcgtgactcgtagaaggcataaaaggtgttaaattctaagctcgcctttagggctgttt [ 90720]
ASF_IND_20_CAD543                 ................................................................................ [ 90720]
LR899193.1_Germany_2020/1         ................................................................................ [ 90720]
MN393477.1_Wuhan_2019             ................................................................................ [ 90720]
MW396979.1_Timor-Leste/2019/1     ................................................................................ [ 90720]
MK940252.1_CN/2019/InnerMongolia  ................................................................................ [ 90720]
MK128995.1_China/2018/AnhuiXCGQ   ................................................................................ [ 90720]
MH681419.1_ASFV/POL/2015          ................................................................................ [ 90720]
LS478113.1_Estonia_2014           ................................................................................ [ 90720]
KP843857.1_Odintsovo_02/14        ................................................................................ [ 90720]
```

```
FN557520.1_strain_E75          .........c...................................................................... [ 90720]
AY261366.1_Warthog             ................................................................................ [ 90720]
AY261365.1_Warmbaths           ..............................a................................................. [ 90720]
AY261364.1_Tengani_62          .........a...........a.....t....................a............................... [ 90720]
AY261363.1_Pretoriuskop/96/4   ................................................a............................... [ 90720]
AY261362.1_Mkuzi_1979          .........c...................................................................... [ 90720]
AY261361.1_Malawi_Lil-20/1_1983 .........a.....c.....................................................c.......... [ 90720]
AY261360.1_Kenya_1950          .........a.....c...........t.................................................... [ 90720]
AM712240.1_OURT_88/3           .........c...................................................................... [ 90720]
AM712239.1_Benin_97/1          .........c...................................................................... [ 90720]
KM111295.1_Ken06.Bus           .........g.....c................................................................ [ 90720]
KM111294.1_Ken05/Tk1           .........g.....c................................................................ [ 90720]
KM262844.1_strain_L60          .........c...................................................................... [ 90720]
KM102979.1_26544/OG10_Italy    .........c...................................................................... [ 90720]
KM262845.1_strain_NHV          .........c...................................................................... [ 90720]
KX354450.1_47/Ss/2008          .........c...................................................................... [ 90720]
KP055815.1_strain_BA71         .........c...................................................................... [ 90720]
MH025920.1_strain_R35          .........g.....c................................................................ [ 90720]
MH025919.1_strain_N10          .........g.....c................................................................ [ 90720]
MH025918.1_strain_R25          .........g.....c................................................................ [ 90720]
MH025917.1_strain_R7           .........g.....c................................................................ [ 90720]
MH025916.1_strain_R8           .........g.....c................................................................ [ 90720]

FR682468.2_Georgia_2007/1      ggaccttttttatgtttaattgccccacctcatgttgtagcacgtggcatacagaacagcgtagatcggcaagtgcataa [ 90800]
ASF_IND_20_CAD543              ................................................................................ [ 90800]
LR899193.1_Germany_2020/1      ................................................................................ [ 90800]
MN393477.1_Wuhan_2019          ................................................................................ [ 90800]
MW396979.1_Timor-Leste/2019/1  ................................................................................ [ 90800]
MK940252.1_CN/2019/InnerMongolia ................................................................................ [ 90800]
MK128995.1_China/2018/AnhuiXCGQ ................................................................................ [ 90800]
MH681419.1_ASFV/POL/2015       ................................................................................ [ 90800]
LS478113.1_Estonia_2014        ................................................................................ [ 90800]
KP843857.1_Odintsovo_02/14     ................................................................................ [ 90800]
FN557520.1_strain_E75          .........................................................................c...... [ 90800]
AY261366.1_Warthog             .........................................................................c..g... [ 90800]
AY261365.1_Warmbaths           .........................................................................c...... [ 90800]
AY261364.1_Tengani_62          ......................t..............a......................................g... [ 90800]
AY261363.1_Pretoriuskop/96/4   .........................t.........................................a.....c...... [ 90800]
AY261362.1_Mkuzi_1979          .........................................................................c...... [ 90800]
AY261361.1_Malawi_Lil-20/1_1983 .........................................................................c...... [ 90800]
AY261360.1_Kenya_1950          .........................................................................a...... [ 90800]
AM712240.1_OURT_88/3           .........................................................................c...... [ 90800]
AM712239.1_Benin_97/1          .........................................................................c...... [ 90800]
KM111295.1_Ken06.Bus           ......................t..................................................a...... [ 90800]
```

```
KM111294.1_Ken05/Tk1           ......................t..................................................a...... [ 90800]
KM262844.1_strain_L60          .........................................................................c...... [ 90800]
KM102979.1_26544/OG10_Italy    .............a...........................................................c...... [ 90800]
KM262845.1_strain_NHV          .........................................................................c...... [ 90800]
KX354450.1_47/Ss/2008          .............a...........................................................c...... [ 90800]
KP055815.1_strain_BA71         .........................................................................c...... [ 90800]
MH025920.1_strain_R35          ......................t..................................................a...... [ 90800]
MH025919.1_strain_N10          ......................t..................................................a...... [ 90800]
MH025918.1_strain_R25          ......................t..................................................a...... [ 90800]
MH025917.1_strain_R7           ......................t..................................................a...... [ 90800]
MH025916.1_strain_R8           ......................t..................................................a...... [ 90800]

FR682468.2_Georgia_2007/1      tggttgtcaattttttttatgacgtctttgcgtgttacttcaatctcggcgggtttctgcgaactgtctacggccttgta [ 90880]
ASF_IND_20_CAD543              ................................................................................ [ 90880]
LR899193.1_Germany_2020/1      ................................................................................ [ 90880]
MN393477.1_Wuhan_2019          ................................................................................ [ 90880]
MW396979.1_Timor-Leste/2019/1  ................................................................................ [ 90880]
MK940252.1_CN/2019/InnerMongolia ................................................................................ [ 90880]
MK128995.1_China/2018/AnhuiXCGQ ................................................................................ [ 90880]
MH681419.1_ASFV/POL/2015       ................................................................................ [ 90880]
LS478113.1_Estonia_2014        ................................................................................ [ 90880]
KP843857.1_Odintsovo_02/14     ................................................................................ [ 90880]
FN557520.1_strain_E75          ....................................................................g........... [ 90880]
AY261366.1_Warthog             .................................................t........................t..... [ 90880]
AY261365.1_Warmbaths           ................................................................................ [ 90880]
AY261364.1_Tengani_62          ................................................................................ [ 90880]
AY261363.1_Pretoriuskop/96/4   ....c.....................................................................t..... [ 90880]
AY261362.1_Mkuzi_1979          ....................................................................g........... [ 90880]
AY261361.1_Malawi_Lil-20/1_1983 ..................................a............c......................t......... [ 90880]
AY261360.1_Kenya_1950          ................................c.a............c......................t......... [ 90880]
AM712240.1_OURT_88/3           ....................................................................g........... [ 90880]
AM712239.1_Benin_97/1          ....................................................................g........... [ 90880]
KM111295.1_Ken06.Bus           ...................................g...........c....................c.t......... [ 90880]
KM111294.1_Ken05/Tk1           ...................................g...........c......................t......... [ 90880]
KM262844.1_strain_L60          ....................................................................g........... [ 90880]
KM102979.1_26544/OG10_Italy    ....................................................................g........... [ 90880]
KM262845.1_strain_NHV          ....................................................................g........... [ 90880]
KX354450.1_47/Ss/2008          ....................................................................g........... [ 90880]
KP055815.1_strain_BA71         ....................................................................g........... [ 90880]
MH025920.1_strain_R35          ...................................g...........c....................c.t......... [ 90880]
MH025919.1_strain_N10          ...................................g...........c....................c.t......... [ 90880]
MH025918.1_strain_R25          ...................................g...........c....................c.t......... [ 90880]
MH025917.1_strain_R7           ...................................g...........c....................c.t......... [ 90880]
MH025916.1_strain_R8           ...................................g...........c....................c.t......... [ 90880]
```

```
FR682468.2_Georgia_2007/1        aacgtaaatggtccacttatgaggaagccccctttcatcgtatagggttgaaatgggaagccttttatactcaaacagcc [ 90960]
ASF_IND_20_CAD543                ................................................................................ [ 90960]
LR899193.1_Germany_2020/1        ................................................................................ [ 90960]
MN393477.1_Wuhan_2019            ................................................................................ [ 90960]
MW396979.1_Timor-Leste/2019/1    ................................................................................ [ 90960]
MK940252.1_CN/2019/InnerMongolia ................................................................................ [ 90960]
MK128995.1_China/2018/AnhuiXCGQ  ................................................................................ [ 90960]
MH681419.1_ASFV/POL/2015         ................................................................................ [ 90960]
LS478113.1_Estonia_2014          ................................................................................ [ 90960]
KP843857.1_Odintsovo_02/14       ................................................................................ [ 90960]
FN557520.1_strain_E75            .......................................a........................................ [ 90960]
AY261366.1_Warthog               ................................................................................ [ 90960]
AY261365.1_Warmbaths             ................................................................................ [ 90960]
AY261364.1_Tengani_62            ..............................g........a........................................ [ 90960]
AY261363.1_Pretoriuskop/96/4     ............................................a................................... [ 90960]
AY261362.1_Mkuzi_1979            ................................................................................ [ 90960]
AY261361.1_Malawi_Lil-20/1_1983  c..a...........................t....g..a.....................t.................. [ 90960]
AY261360.1_Kenya_1950            ...a....................c......t.............................t.................. [ 90960]
AM712240.1_OURT_88/3             .......................................a........................................ [ 90960]
AM712239.1_Benin_97/1            .......................................a........................................ [ 90960]
KM111295.1_Ken06.Bus             ...a....................c......t.............................t.................. [ 90960]
KM111294.1_Ken05/Tk1             ........................c......t.......a.....................t.................. [ 90960]
KM262844.1_strain_L60            .......................................a........................................ [ 90960]
KM102979.1_26544/OG10_Italy      .......................................a........................................ [ 90960]
KM262845.1_strain_NHV            .......................................a........................................ [ 90960]
KX354450.1_47/Ss/2008            .......................................a........................................ [ 90960]
KP055815.1_strain_BA71           .......................................a........................................ [ 90960]
MH025920.1_strain_R35            ...a....................c......t.............................t.................. [ 90960]
MH025919.1_strain_N10            ...a....................c......t.............................t.................. [ 90960]
MH025918.1_strain_R25            ...a....................c......t.............................t.................. [ 90960]
MH025917.1_strain_R7             ...a....................c......t.............................t.................. [ 90960]
MH025916.1_strain_R8             ...a....................c......t.............................t.................. [ 90960]

FR682468.2_Georgia_2007/1        gagtccgttggtcggctcttcctgtgttaggatcaaatatgttataaaatccttgctgagcaagcagggccttttgctcg [ 91040]
ASF_IND_20_CAD543                ................................................................................ [ 91040]
LR899193.1_Germany_2020/1        ................................................................................ [ 91040]
MN393477.1_Wuhan_2019            ................................................................................ [ 91040]
MW396979.1_Timor-Leste/2019/1    ................................................................................ [ 91040]
MK940252.1_CN/2019/InnerMongolia ................................................................................ [ 91040]
MK128995.1_China/2018/AnhuiXCGQ  ................................................................................ [ 91040]
MH681419.1_ASFV/POL/2015         ................................................................................ [ 91040]
LS478113.1_Estonia_2014          ................................................................................ [ 91040]
KP843857.1_Odintsovo_02/14       ................................................................................ [ 91040]
```

```
FN557520.1_strain_E75             ................................................................................ [ 91040]
AY261366.1_Warthog                ................c............................................................... [ 91040]
AY261365.1_Warmbaths              ..........a.....c...........................................................t..a [ 91040]
AY261364.1_Tengani_62             ................c..............................................................a [ 91040]
AY261363.1_Pretoriuskop/96/4      ................c.......................a..........................a............ [ 91040]
AY261362.1_Mkuzi_1979             ............................................................................t..a [ 91040]
AY261361.1_Malawi_Lil-20/1_1983   ....t.....a.....c...........g.....................................aa...c.......a [ 91040]
AY261360.1_Kenya_1950             .g........a.....ct................................................a............a [ 91040]
AM712240.1_OURT_88/3              ................................................................................ [ 91040]
AM712239.1_Benin_97/1             ................................................................................ [ 91040]
KM111295.1_Ken06.Bus              .g........a.....ct................................................a............a [ 91040]
KM111294.1_Ken05/Tk1              .g........a.....ct................................................a............a [ 91040]
KM262844.1_strain_L60             ................................................................................ [ 91040]
KM102979.1_26544/OG10_Italy       ................................................................................ [ 91040]
KM262845.1_strain_NHV             ................................................................................ [ 91040]
KX354450.1_47/Ss/2008             ................................................................................ [ 91040]
KP055815.1_strain_BA71            ................................................................................ [ 91040]
MH025920.1_strain_R35             .g........a.....ct................................................a............a [ 91040]
MH025919.1_strain_N10             .g........a.....ct................................................a............a [ 91040]
MH025918.1_strain_R25             .g........a.....ct................................................a............a [ 91040]
MH025917.1_strain_R7              .g........a.....ct................................................a............a [ 91040]
MH025916.1_strain_R8              .g........a.....ct................................................a............a [ 91040]

FR682468.2_Georgia_2007/1         ccataagcattttcgtacgttttgaattctgcaagttcggagttaaaattaggtgcattttgtaaatacttaagaaataa [ 91120]
ASF_IND_20_CAD543                 ................................................................................ [ 91120]
LR899193.1_Germany_2020/1         ................................................................................ [ 91120]
MN393477.1_Wuhan_2019             ................................................................................ [ 91120]
MW396979.1_Timor-Leste/2019/1     ................................................................................ [ 91120]
MK940252.1_CN/2019/InnerMongolia  ................................................................................ [ 91120]
MK128995.1_China/2018/AnhuiXCGQ   ................................................................................ [ 91120]
MH681419.1_ASFV/POL/2015          ................................................................................ [ 91120]
LS478113.1_Estonia_2014           ................................................................................ [ 91120]
KP843857.1_Odintsovo_02/14        ................................................................................ [ 91120]
FN557520.1_strain_E75             ................................................................................ [ 91120]
AY261366.1_Warthog                ................................................................................ [ 91120]
AY261365.1_Warmbaths              ............................................................c................... [ 91120]
AY261364.1_Tengani_62             ................................................................................ [ 91120]
AY261363.1_Pretoriuskop/96/4      ................................................................................ [ 91120]
AY261362.1_Mkuzi_1979             ............................................................c................... [ 91120]
AY261361.1_Malawi_Lil-20/1_1983   ..............a..tt..........c.........ag....................................... [ 91120]
AY261360.1_Kenya_1950             .................tt.c........c.......................a.......................... [ 91120]
AM712240.1_OURT_88/3              ................................................................................ [ 91120]
AM712239.1_Benin_97/1             ................................................................................ [ 91120]
KM111295.1_Ken06.Bus              .................tt.c........c.......................a.......................... [ 91120]
```

```
KM111294.1_Ken05/Tk1             .................tt.c........c.......................a.......................... [ 91120]
KM262844.1_strain_L60            ................................................................................ [ 91120]
KM102979.1_26544/OG10_Italy      ................................................................................ [ 91120]
KM262845.1_strain_NHV            ................................................................................ [ 91120]
KX354450.1_47/Ss/2008            ................................................................................ [ 91120]
KP055815.1_strain_BA71           ................................................................................ [ 91120]
MH025920.1_strain_R35            .................tt.c........c.......................a.......................... [ 91120]
MH025919.1_strain_N10            .................tt.c........c.......................a.......................... [ 91120]
MH025918.1_strain_R25            .................tt.c........c.......................a.......................... [ 91120]
MH025917.1_strain_R7             .................tt.c........c.......................a.......................... [ 91120]
MH025916.1_strain_R8             .................tt.c........c.......................a.......................... [ 91120]

FR682468.2_Georgia_2007/1        ttcataggctctaaggtaaatgagagttgaggttttttcctcatcccg---tcctccccaccacacccgcaggctttctt [ 91200]
ASF_IND_20_CAD543                ................................................---............................. [ 91200]
LR899193.1_Germany_2020/1        ................................................---............................. [ 91200]
MN393477.1_Wuhan_2019            ................................................---............................. [ 91200]
MW396979.1_Timor-Leste/2019/1    ................................................---............................. [ 91200]
MK940252.1_CN/2019/InnerMongolia ................................................---............................. [ 91200]
MK128995.1_China/2018/AnhuiXCGQ  ................................................---............................. [ 91200]
MH681419.1_ASFV/POL/2015         ................................................---............................. [ 91200]
LS478113.1_Estonia_2014          ................................................---............................. [ 91200]
KP843857.1_Odintsovo_02/14       ................................................---............................. [ 91200]
FN557520.1_strain_E75            ................................................---............................. [ 91200]
AY261366.1_Warthog               ................................................---............................. [ 91200]
AY261365.1_Warmbaths             ..................t..a..........................---............................. [ 91200]
AY261364.1_Tengani_62            ........................c.......................---............................. [ 91200]
AY261363.1_Pretoriuskop/96/4     ..................g..a..........................---............................. [ 91200]
AY261362.1_Mkuzi_1979            ..................t..a..........................---............................. [ 91200]
AY261361.1_Malawi_Lil-20/1_1983  ..........t.............c......a..........g.....---............................. [ 91200]
AY261360.1_Kenya_1950            ........................c......a..........g.....---............................. [ 91200]
AM712240.1_OURT_88/3             ................................................---............................. [ 91200]
AM712239.1_Benin_97/1            ................................................---............................. [ 91200]
KM111295.1_Ken06.Bus             ...............................a..........g.....---............................. [ 91200]
KM111294.1_Ken05/Tk1             ........................c......a................tcc............................. [ 91200]
KM262844.1_strain_L60            ................................................---............................. [ 91200]
KM102979.1_26544/OG10_Italy      ................................................---............................. [ 91200]
KM262845.1_strain_NHV            ................................................---............................. [ 91200]
KX354450.1_47/Ss/2008            ................................................---............................. [ 91200]
KP055815.1_strain_BA71           ................................................---............................. [ 91200]
MH025920.1_strain_R35            ...............................a..........g.....---............................. [ 91200]
MH025919.1_strain_N10            ...............................a..........g.....---............................. [ 91200]
MH025918.1_strain_R25            ...............................a..........g.....---............................. [ 91200]
MH025917.1_strain_R7             ...............................a..........g.....---............................. [ 91200]
MH025916.1_strain_R8             ...............................a..........g.....---............................. [ 91200]
```

```
FR682468.2_Georgia_2007/1        cttgaaaatagatgtcattcagacgcgtcaactgcgtaaaatcaggccgatatttagaggtataaattttatcataaaat [ 91280]
ASF_IND_20_CAD543                ................................................................................ [ 91280]
LR899193.1_Germany_2020/1        ................................................................................ [ 91280]
MN393477.1_Wuhan_2019            ................................................................................ [ 91280]
MW396979.1_Timor-Leste/2019/1    ................................................................................ [ 91280]
MK940252.1_CN/2019/InnerMongolia ................................................................................ [ 91280]
MK128995.1_China/2018/AnhuiXCGQ  ................................................................................ [ 91280]
MH681419.1_ASFV/POL/2015         ................................................................................ [ 91280]
LS478113.1_Estonia_2014          ................................................................................ [ 91280]
KP843857.1_Odintsovo_02/14       ................................................................................ [ 91280]
FN557520.1_strain_E75            ...........................c.................................................... [ 91280]
AY261366.1_Warthog               ................c..........c..g................................................. [ 91280]
AY261365.1_Warmbaths             ................c..........c.................................................... [ 91280]
AY261364.1_Tengani_62            ................c........t.c............................a....................... [ 91280]
AY261363.1_Pretoriuskop/96/4     ................c..........c.................................................... [ 91280]
AY261362.1_Mkuzi_1979            ................c..........c.................................................... [ 91280]
AY261361.1_Malawi_Lil-20/1_1983  ................g..........c............................a....................... [ 91280]
AY261360.1_Kenya_1950            ............c..............c...........................g........................ [ 91280]
AM712240.1_OURT_88/3             ...........................c.................................................... [ 91280]
AM712239.1_Benin_97/1            ...........................c.................................................... [ 91280]
KM111295.1_Ken06.Bus             ............c..............c...........................g........................ [ 91280]
KM111294.1_Ken05/Tk1             ...........................c...........................g........................ [ 91280]
KM262844.1_strain_L60            ...........................c.................................................... [ 91280]
KM102979.1_26544/OG10_Italy      ...........................c.................................................... [ 91280]
KM262845.1_strain_NHV            ...........................c.................................................... [ 91280]
KX354450.1_47/Ss/2008            ...........................c.................................................... [ 91280]
KP055815.1_strain_BA71           ...........................c.................................................... [ 91280]
MH025920.1_strain_R35            ............c..............c...........................g........................ [ 91280]
MH025919.1_strain_N10            ............c..............c...........................g........................ [ 91280]
MH025918.1_strain_R25            ............c..............c...........................g........................ [ 91280]
MH025917.1_strain_R7             ............c..............c...........................g........................ [ 91280]
MH025916.1_strain_R8             ............c..............c...........................g........................ [ 91280]

FR682468.2_Georgia_2007/1        tctttttgcgataatagctcggccggggtacgtcctatcacggttttaaactcatattcagcctccttgggagtccgtgg [ 91360]
ASF_IND_20_CAD543                ................................................................................ [ 91360]
LR899193.1_Germany_2020/1        ................................................................................ [ 91360]
MN393477.1_Wuhan_2019            ................................................................................ [ 91360]
MW396979.1_Timor-Leste/2019/1    ................................................................................ [ 91360]
MK940252.1_CN/2019/InnerMongolia ................................................................................ [ 91360]
MK128995.1_China/2018/AnhuiXCGQ  ................................................................................ [ 91360]
MH681419.1_ASFV/POL/2015         ................................................................................ [ 91360]
LS478113.1_Estonia_2014          ................................................................................ [ 91360]
KP843857.1_Odintsovo_02/14       ................................................................................ [ 91360]
```

```
FN557520.1_strain_E75          ................................................................................ [ 91360]
AY261366.1_Warthog             ..........................a...................................t................. [ 91360]
AY261365.1_Warmbaths           ..........................a..................................................c.. [ 91360]
AY261364.1_Tengani_62          ................................................................................ [ 91360]
AY261363.1_Pretoriuskop/96/4   ..........................a..................................................... [ 91360]
AY261362.1_Mkuzi_1979          ..........................a..................................................... [ 91360]
AY261361.1_Malawi_Lil-20/1_1983 ......................t................................................g...t.... [ 91360]
AY261360.1_Kenya_1950          .......................................................................g.....c.. [ 91360]
AM712240.1_OURT_88/3           ................................................................................ [ 91360]
AM712239.1_Benin_97/1          ................................................................................ [ 91360]
KM111295.1_Ken06.Bus           .......................................................................g........ [ 91360]
KM111294.1_Ken05/Tk1           ....................a..................................................g.....c.. [ 91360]
KM262844.1_strain_L60          ................................................................................ [ 91360]
KM102979.1_26544/OG10_Italy    ................................................................................ [ 91360]
KM262845.1_strain_NHV          ................................................................................ [ 91360]
KX354450.1_47/Ss/2008          ................................................................................ [ 91360]
KP055815.1_strain_BA71         ................................................................................ [ 91360]
MH025920.1_strain_R35          .......................................................................g........ [ 91360]
MH025919.1_strain_N10          .......................................................................g........ [ 91360]
MH025918.1_strain_R25          .......................................................................g........ [ 91360]
MH025917.1_strain_R7           .......................................................................g........ [ 91360]
MH025916.1_strain_R8           .......................................................................g........ [ 91360]

FR682468.2_Georgia_2007/1      tttgtgcatagggatgctgccgtcaatacgggccactgtggcagcataatcatacatggggtccagcagaatctctgtca [ 91440]
ASF_IND_20_CAD543              ....................................w........................................... [ 91440]
LR899193.1_Germany_2020/1      ................................................................................ [ 91440]
MN393477.1_Wuhan_2019          ................................................................................ [ 91440]
MW396979.1_Timor-Leste/2019/1  ................................................................................ [ 91440]
MK940252.1_CN/2019/InnerMongolia ................................................................................ [ 91440]
MK128995.1_China/2018/AnhuiXCGQ ................................................................................ [ 91440]
MH681419.1_ASFV/POL/2015       ................................................................................ [ 91440]
LS478113.1_Estonia_2014        ................................................................................ [ 91440]
KP843857.1_Odintsovo_02/14     ................................................................................ [ 91440]
FN557520.1_strain_E75          c..a................................a........................................... [ 91440]
AY261366.1_Warthog             c......................................a........................................ [ 91440]
AY261365.1_Warmbaths           c............................................................................... [ 91440]
AY261364.1_Tengani_62          c............................................................................... [ 91440]
AY261363.1_Pretoriuskop/96/4   c....a................................................t.a......t................ [ 91440]
AY261362.1_Mkuzi_1979          c............................................................................... [ 91440]
AY261361.1_Malawi_Lil-20/1_1983 c...................................g...................a...............t....... [ 91440]
AY261360.1_Kenya_1950          c...................................g...................a......t....a...t....... [ 91440]
AM712240.1_OURT_88/3           c..a................................a........................................... [ 91440]
AM712239.1_Benin_97/1          c..a................................a........................................... [ 91440]
KM111295.1_Ken06.Bus           c...................................g..a............................a........... [ 91440]
```

```
KM111294.1_Ken05/Tk1           c...................................g..a............................a........... [ 91440]
KM262844.1_strain_L60          c..a................................a........................................... [ 91440]
KM102979.1_26544/OG10_Italy    c..a................................a........................................... [ 91440]
KM262845.1_strain_NHV          c..a................................a........................................... [ 91440]
KX354450.1_47/Ss/2008          c..a................................a........................................... [ 91440]
KP055815.1_strain_BA71         c..a................................a........................................... [ 91440]
MH025920.1_strain_R35          c...................................g..a............................a........... [ 91440]
MH025919.1_strain_N10          c...................................g..a............................a........... [ 91440]
MH025918.1_strain_R25          c...................................g..a............................a........... [ 91440]
MH025917.1_strain_R7           c...................................g..a............................a........... [ 91440]
MH025916.1_strain_R8           c...................................g..a............................a........... [ 91440]

FR682468.2_Georgia_2007/1      aaagtaccttggtgtcgtcctgcacgctaagcccttgtagcccattttggtggataatttttttgaaagcctcccgaaaa [ 91520]
ASF_IND_20_CAD543              ................................................................................ [ 91520]
LR899193.1_Germany_2020/1      ................................................................................ [ 91520]
MN393477.1_Wuhan_2019          ................................................................................ [ 91520]
MW396979.1_Timor-Leste/2019/1  ................................................................................ [ 91520]
MK940252.1_CN/2019/InnerMongolia ................................................................................ [ 91520]
MK128995.1_China/2018/AnhuiXCGQ ................................................................................ [ 91520]
MH681419.1_ASFV/POL/2015       ................................................................................ [ 91520]
LS478113.1_Estonia_2014        ................................................................................ [ 91520]
KP843857.1_Odintsovo_02/14     ................................................................................ [ 91520]
FN557520.1_strain_E75          ................................................................................ [ 91520]
AY261366.1_Warthog             ........................................t..g..c................................. [ 91520]
AY261365.1_Warmbaths           .......t..............t......................................................... [ 91520]
AY261364.1_Tengani_62          ................a.....t......................................................... [ 91520]
AY261363.1_Pretoriuskop/96/4   ................a.....t......................................................... [ 91520]
AY261362.1_Mkuzi_1979          ........................................t..g..c................................. [ 91520]
AY261361.1_Malawi_Lil-20/1_1983 .............a..........................t..g..c................................. [ 91520]
AY261360.1_Kenya_1950          .....................................c.....g..c..........................t...... [ 91520]
AM712240.1_OURT_88/3           ................................................................................ [ 91520]
AM712239.1_Benin_97/1          ................................................................................ [ 91520]
KM111295.1_Ken06.Bus           ........................................t..g..c..........................t...... [ 91520]
KM111294.1_Ken05/Tk1           .....................................c.....g..c..........................t...... [ 91520]
KM262844.1_strain_L60          ................................................................................ [ 91520]
KM102979.1_26544/OG10_Italy    ................................................................................ [ 91520]
KM262845.1_strain_NHV          ................................................................................ [ 91520]
KX354450.1_47/Ss/2008          ................................................................................ [ 91520]
KP055815.1_strain_BA71         ................................................................................ [ 91520]
MH025920.1_strain_R35          ........................................t..g..c..........................t...... [ 91520]
MH025919.1_strain_N10          ........................................t..g..c..........................t...... [ 91520]
MH025918.1_strain_R25          ........................................t..g..c..........................t...... [ 91520]
MH025917.1_strain_R7           ........................................t..g..c..........................t...... [ 91520]
MH025916.1_strain_R8           ........................................t..g..c..........................t...... [ 91520]
```

```
FR682468.2_Georgia_2007/1        ttattagcaatccactgatccgtaatctcagatagctgatttattataccgctatattgctgcatcattttctccaaaag [ 91600]
ASF_IND_20_CAD543                ........................................................................y........ [ 91600]
LR899193.1_Germany_2020/1        ................................................................................ [ 91600]
MN393477.1_Wuhan_2019            ................................................................................ [ 91600]
MW396979.1_Timor-Leste/2019/1    ................................................................................ [ 91600]
MK940252.1_CN/2019/InnerMongolia ................................................................................ [ 91600]
MK128995.1_China/2018/AnhuiXCGQ  ................................................................................ [ 91600]
MH681419.1_ASFV/POL/2015         ................................................................................ [ 91600]
LS478113.1_Estonia_2014          ................................................................................ [ 91600]
KP843857.1_Odintsovo_02/14       ................................................................................ [ 91600]
FN557520.1_strain_E75            ........................................................................t........ [ 91600]
AY261366.1_Warthog               ................................................................................ [ 91600]
AY261365.1_Warmbaths             ................................................................................ [ 91600]
AY261364.1_Tengani_62            ................................................................................ [ 91600]
AY261363.1_Pretoriuskop/96/4     ................................................................................ [ 91600]
AY261362.1_Mkuzi_1979            ........................................................................t........ [ 91600]
AY261361.1_Malawi_Lil-20/1_1983  ..........................t..........................g.........g.......t........ [ 91600]
AY261360.1_Kenya_1950            .....g...............................................g.........g.......t........ [ 91600]
AM712240.1_OURT_88/3             ........................................................................t........ [ 91600]
AM712239.1_Benin_97/1            ........................................................................t........ [ 91600]
KM111295.1_Ken06.Bus             .....g...............................................g...........a.g...t........ [ 91600]
KM111294.1_Ken05/Tk1             .....g...............................................g.........g.......t........ [ 91600]
KM262844.1_strain_L60            ........................................................................t........ [ 91600]
KM102979.1_26544/OG10_Italy      ........................................................................t........ [ 91600]
KM262845.1_strain_NHV            ........................................................................t........ [ 91600]
KX354450.1_47/Ss/2008            ........................................................................t........ [ 91600]
KP055815.1_strain_BA71           ........................................................................t........ [ 91600]
MH025920.1_strain_R35            .....g...............................................g...........a.g...t........ [ 91600]
MH025919.1_strain_N10            .....g...............................................g...........a.g...t........ [ 91600]
MH025918.1_strain_R25            .....g...............................................g...........a.g...t........ [ 91600]
MH025917.1_strain_R7             .....g...............................................g...........a.g...t........ [ 91600]
MH025916.1_strain_R8             .....g...............................................g...........a.g...t........ [ 91600]

FR682468.2_Georgia_2007/1        aaaggtcacgtatgcattcaaagagctatccgccttcattccatgaatggtaatcgtaagaaattctttatttttttgcg [ 91680]
ASF_IND_20_CAD543                ................................................................................ [ 91680]
LR899193.1_Germany_2020/1        ................................................................................ [ 91680]
MN393477.1_Wuhan_2019            ................................................................................ [ 91680]
MW396979.1_Timor-Leste/2019/1    ................................................................................ [ 91680]
MK940252.1_CN/2019/InnerMongolia ................................................................................ [ 91680]
MK128995.1_China/2018/AnhuiXCGQ  ................................................................................ [ 91680]
MH681419.1_ASFV/POL/2015         ................................................................................ [ 91680]
LS478113.1_Estonia_2014          ................................................................................ [ 91680]
KP843857.1_Odintsovo_02/14       ................................................................................ [ 91680]
```

```
FN557520.1_strain_E75              .....................c.......................................................... [ 91680]
AY261366.1_Warthog                 ................................................................................ [ 91680]
AY261365.1_Warmbaths               ......................................................t......................... [ 91680]
AY261364.1_Tengani_62              g.....g......................................................................... [ 91680]
AY261363.1_Pretoriuskop/96/4       ................................................................................ [ 91680]
AY261362.1_Mkuzi_1979              .....................c.......................................................... [ 91680]
AY261361.1_Malawi_Lil-20/1_1983    .........a...................................................................... [ 91680]
AY261360.1_Kenya_1950              ...............g................................................................ [ 91680]
AM712240.1_OURT_88/3               .....................c.......................................................... [ 91680]
AM712239.1_Benin_97/1              .....................c.......................................................... [ 91680]
KM111295.1_Ken06.Bus               ...............g................................................................ [ 91680]
KM111294.1_Ken05/Tk1               ...............g................................................................ [ 91680]
KM262844.1_strain_L60              .....................c.......................................................... [ 91680]
KM102979.1_26544/OG10_Italy        .....................c.......................................................... [ 91680]
KM262845.1_strain_NHV              .....................c.......................................................... [ 91680]
KX354450.1_47/Ss/2008              .....................c.......................................................... [ 91680]
KP055815.1_strain_BA71             .....................c.......................................................... [ 91680]
MH025920.1_strain_R35              ...............g................................................................ [ 91680]
MH025919.1_strain_N10              ...............g................................................................ [ 91680]
MH025918.1_strain_R25              ...............g................................................................ [ 91680]
MH025917.1_strain_R7               ...............g................................................................ [ 91680]
MH025916.1_strain_R8               ...............g................................................................ [ 91680]

FR682468.2_Georgia_2007/1          agctataaatgagattcaaaatataggcatagatgtagatcacagcatacagctgcgttaaaggatcgtaatcctcttcc [ 91760]
ASF_IND_20_CAD543                  ............................................................................y... [ 91760]
LR899193.1_Germany_2020/1          ................................................................................ [ 91760]
MN393477.1_Wuhan_2019              ................................................................................ [ 91760]
MW396979.1_Timor-Leste/2019/1      ................................................................................ [ 91760]
MK940252.1_CN/2019/InnerMongolia   ................................................................................ [ 91760]
MK128995.1_China/2018/AnhuiXCGQ    ................................................................................ [ 91760]
MH681419.1_ASFV/POL/2015           ................................................................................ [ 91760]
LS478113.1_Estonia_2014            ................................................................................ [ 91760]
KP843857.1_Odintsovo_02/14         ................................................................................ [ 91760]
FN557520.1_strain_E75              ............................................................................c... [ 91760]
AY261366.1_Warthog                 ............................................................................c... [ 91760]
AY261365.1_Warmbaths               ............................................................................c... [ 91760]
AY261364.1_Tengani_62              ............................................................................c... [ 91760]
AY261363.1_Pretoriuskop/96/4       ............................................................................c... [ 91760]
AY261362.1_Mkuzi_1979              ............................................................................c... [ 91760]
AY261361.1_Malawi_Lil-20/1_1983    ....g..................................................a....................c..t [ 91760]
AY261360.1_Kenya_1950              ........................................g..............a....................c... [ 91760]
AM712240.1_OURT_88/3               ............................................................................c... [ 91760]
AM712239.1_Benin_97/1              ............................................................................c... [ 91760]
KM111295.1_Ken06.Bus               ..........c................................g...........a....g...............c... [ 91760]
```

```
KM111294.1_Ken05/Tk1            .......................................................g....................c... [ 91760]
KM262844.1_strain_L60           ............................................................................c... [ 91760]
KM102979.1_26544/OG10_Italy     ............................................................................c... [ 91760]
KM262845.1_strain_NHV           ............................................................................c... [ 91760]
KX354450.1_47/Ss/2008           ............................................................................c... [ 91760]
KP055815.1_strain_BA71          ............................................................................c... [ 91760]
MH025920.1_strain_R35           ..........c................................g...........a....................c... [ 91760]
MH025919.1_strain_N10           ..........c................................g...........a....................c... [ 91760]
MH025918.1_strain_R25           ..........c................................g...........a....................c... [ 91760]
MH025917.1_strain_R7            ..........c................................g...........a....................c... [ 91760]
MH025916.1_strain_R8            ..........c................................g...........a....................c... [ 91760]

FR682468.2_Georgia_2007/1       tttttaatattttcgatgctatacacgagcggcaggcagacatttacggctatattggcaaactgtttcacgtctacaag [ 91840]
ASF_IND_20_CAD543               ................................................................................ [ 91840]
LR899193.1_Germany_2020/1       ................................................................................ [ 91840]
MN393477.1_Wuhan_2019           ................................................................................ [ 91840]
MW396979.1_Timor-Leste/2019/1   ................................................................................ [ 91840]
MK940252.1_CN/2019/InnerMongolia ................................................................................ [ 91840]
MK128995.1_China/2018/AnhuiXCGQ ................................................................................ [ 91840]
MH681419.1_ASFV/POL/2015        ................................................................................ [ 91840]
LS478113.1_Estonia_2014         ................................................................................ [ 91840]
KP843857.1_Odintsovo_02/14      ................................................................................ [ 91840]
FN557520.1_strain_E75           ..........................................................................c..... [ 91840]
AY261366.1_Warthog              ..........................................................................c..... [ 91840]
AY261365.1_Warmbaths            ..........................................................................c..... [ 91840]
AY261364.1_Tengani_62           .......................................................................a..c..... [ 91840]
AY261363.1_Pretoriuskop/96/4    ..........................................................................c..... [ 91840]
AY261362.1_Mkuzi_1979           ................................t...................................t........... [ 91840]
AY261361.1_Malawi_Lil-20/1_1983 ..................t....t.t............a......................................... [ 91840]
AY261360.1_Kenya_1950           ..............a........t.t...................................................... [ 91840]
AM712240.1_OURT_88/3            ..........................................................................c..... [ 91840]
AM712239.1_Benin_97/1           ..........................................................................c..... [ 91840]
KM111295.1_Ken06.Bus            ..............a........t.t...................................................... [ 91840]
KM111294.1_Ken05/Tk1            .......................t.t...................................................... [ 91840]
KM262844.1_strain_L60           ..........................................................................c..... [ 91840]
KM102979.1_26544/OG10_Italy     ..........................................................................c..... [ 91840]
KM262845.1_strain_NHV           ..........................................................................c..... [ 91840]
KX354450.1_47/Ss/2008           ..........................................................................c..... [ 91840]
KP055815.1_strain_BA71          ..........................................................................c..... [ 91840]
MH025920.1_strain_R35           ..............a...t....t.t...................................................... [ 91840]
MH025919.1_strain_N10           ..............a........t.t...................................................... [ 91840]
MH025918.1_strain_R25           ..............a...t....t.t...................................................... [ 91840]
MH025917.1_strain_R7            ..............a...t....t.t...................................................... [ 91840]
MH025916.1_strain_R8            ..............a...t....t.t...................................................... [ 91840]
```

```
FR682468.2_Georgia_2007/1        ctttccaaagtggataaacgtgcaggccttcatggtttcctgccaaataaaaacacggagcttactattaagatcgccga [ 91920]
ASF_IND_20_CAD543                ................................................................................ [ 91920]
LR899193.1_Germany_2020/1        ................................................................................ [ 91920]
MN393477.1_Wuhan_2019            ................................................................................ [ 91920]
MW396979.1_Timor-Leste/2019/1    ................................................................................ [ 91920]
MK940252.1_CN/2019/InnerMongolia ................................................................................ [ 91920]
MK128995.1_China/2018/AnhuiXCGQ  ................................................................................ [ 91920]
MH681419.1_ASFV/POL/2015         ................................................................................ [ 91920]
LS478113.1_Estonia_2014          ................................................................................ [ 91920]
KP843857.1_Odintsovo_02/14       ................................................................................ [ 91920]
FN557520.1_strain_E75            .....................................................t.......................... [ 91920]
AY261366.1_Warthog               ...c...................................t.............t.......................... [ 91920]
AY261365.1_Warmbaths             .....................................................t...a...................... [ 91920]
AY261364.1_Tengani_62            ..................t..................................t.......................... [ 91920]
AY261363.1_Pretoriuskop/96/4     .................................a...................t...a...................... [ 91920]
AY261362.1_Mkuzi_1979            .....................................................t.......................... [ 91920]
AY261361.1_Malawi_Lil-20/1_1983  ...c........a..........................t.............t.......................... [ 91920]
AY261360.1_Kenya_1950            ...c.................a...............................t.......................... [ 91920]
AM712240.1_OURT_88/3             .....................................................t.......................... [ 91920]
AM712239.1_Benin_97/1            .....................................................t.......................... [ 91920]
KM111295.1_Ken06.Bus             ...c.................a...............................t.......................... [ 91920]
KM111294.1_Ken05/Tk1             ...c.................a...............................t.......................... [ 91920]
KM262844.1_strain_L60            .....................................................t.......................... [ 91920]
KM102979.1_26544/OG10_Italy      .....................................................t.......................... [ 91920]
KM262845.1_strain_NHV            .....................................................t.......................... [ 91920]
KX354450.1_47/Ss/2008            .....................................................t.......................... [ 91920]
KP055815.1_strain_BA71           .....................................................t.......................... [ 91920]
MH025920.1_strain_R35            ...c.................a...............................t.......................... [ 91920]
MH025919.1_strain_N10            ...c.................a.....t.........................t.......................... [ 91920]
MH025918.1_strain_R25            ...c.................a...............................t.......................... [ 91920]
MH025917.1_strain_R7             ...c.................a...............................t.......................... [ 91920]
MH025916.1_strain_R8             ...c.................a...............................t.......................... [ 91920]

FR682468.2_Georgia_2007/1        tgatgcccacatctgccgtacgatcctcttgaataaaatgggccagctcttcgccacaaattttgcaaaagtaggagtaa [ 92000]
ASF_IND_20_CAD543                ................................................................................ [ 92000]
LR899193.1_Germany_2020/1        ................................................................................ [ 92000]
MN393477.1_Wuhan_2019            ................................................................................ [ 92000]
MW396979.1_Timor-Leste/2019/1    ................................................................................ [ 92000]
MK940252.1_CN/2019/InnerMongolia ................................................................................ [ 92000]
MK128995.1_China/2018/AnhuiXCGQ  ................................................................................ [ 92000]
MH681419.1_ASFV/POL/2015         ................................................................................ [ 92000]
LS478113.1_Estonia_2014          ................................................................................ [ 92000]
KP843857.1_Odintsovo_02/14       ................................................................................ [ 92000]
```

```
FN557520.1_strain_E75           ...c............................................................................ [ 92000]
AY261366.1_Warthog              ...c............................................................................ [ 92000]
AY261365.1_Warmbaths            ...c..................g......................................................... [ 92000]
AY261364.1_Tengani_62           ...c............................................................................ [ 92000]
AY261363.1_Pretoriuskop/96/4    ...c............................................................................ [ 92000]
AY261362.1_Mkuzi_1979           ...c............................................................................ [ 92000]
AY261361.1_Malawi_Lil-20/1_1983 ...c.............................c.............................................. [ 92000]
AY261360.1_Kenya_1950           ...c......................................................t....................t [ 92000]
AM712240.1_OURT_88/3            ...c............................................................................ [ 92000]
AM712239.1_Benin_97/1           ...c............................................................................ [ 92000]
KM111295.1_Ken06.Bus            ...c..............................g.......................t....................t [ 92000]
KM111294.1_Ken05/Tk1            ...c......................................................t....................t [ 92000]
KM262844.1_strain_L60           ...c............................................................................ [ 92000]
KM102979.1_26544/OG10_Italy     ...c............................................................................ [ 92000]
KM262845.1_strain_NHV           ...c............................................................................ [ 92000]
KX354450.1_47/Ss/2008           ...c............................................................................ [ 92000]
KP055815.1_strain_BA71          ...c............................................................................ [ 92000]
MH025920.1_strain_R35           ...c..............................g..................t....t....................t [ 92000]
MH025919.1_strain_N10           ...c..............................g..................t....t....................t [ 92000]
MH025918.1_strain_R25           ...c..............................g..................t....t....................t [ 92000]
MH025917.1_strain_R7            ...c..............................g..................t....t....................t [ 92000]
MH025916.1_strain_R8            ...c..............................g..................t....t....................t [ 92000]

FR682468.2_Georgia_2007/1       ataagcccctggttgttttctttctccttgtttattcctgaaaatttcattagcttggttcgcatggtgtcgtaggacgc [ 92080]
ASF_IND_20_CAD543               ................................................................................ [ 92080]
LR899193.1_Germany_2020/1       ................................................................................ [ 92080]
MN393477.1_Wuhan_2019           ................................................................................ [ 92080]
MW396979.1_Timor-Leste/2019/1   ................................................................................ [ 92080]
MK940252.1_CN/2019/InnerMongolia ................................................................................ [ 92080]
MK128995.1_China/2018/AnhuiXCGQ ................................................................................ [ 92080]
MH681419.1_ASFV/POL/2015        ................................................................................ [ 92080]
LS478113.1_Estonia_2014         ................................................................................ [ 92080]
KP843857.1_Odintsovo_02/14      ................................................................................ [ 92080]
FN557520.1_strain_E75           ................................................................................ [ 92080]
AY261366.1_Warthog              ................................................................................ [ 92080]
AY261365.1_Warmbaths            .............................................................................t.. [ 92080]
AY261364.1_Tengani_62           ..g....................t........................................................ [ 92080]
AY261363.1_Pretoriuskop/96/4    .......................t........................................................ [ 92080]
AY261362.1_Mkuzi_1979           ................................................................................ [ 92080]
AY261361.1_Malawi_Lil-20/1_1983 ....................................................a........................... [ 92080]
AY261360.1_Kenya_1950           ................................................................................ [ 92080]
AM712240.1_OURT_88/3            ................................................................................ [ 92080]
AM712239.1_Benin_97/1           ................................................................................ [ 92080]
KM111295.1_Ken06.Bus            ................................................................................ [ 92080]
```

```
KM111294.1_Ken05/Tk1               ................................................................................ [ 92080]
KM262844.1_strain_L60              ................................................................................ [ 92080]
KM102979.1_26544/OG10_Italy        ................................................................................ [ 92080]
KM262845.1_strain_NHV              ................................................................................ [ 92080]
KX354450.1_47/Ss/2008              ................................................................................ [ 92080]
KP055815.1_strain_BA71             ................................................................................ [ 92080]
MH025920.1_strain_R35              ................................................................................ [ 92080]
MH025919.1_strain_N10              ................................................................................ [ 92080]
MH025918.1_strain_R25              ................................................................................ [ 92080]
MH025917.1_strain_R7               ................................................................................ [ 92080]
MH025916.1_strain_R8               ................................................................................ [ 92080]

FR682468.2_Georgia_2007/1          ttctgccgcttgaagctgtataagcatgtccacatggggacaaagcagcttaaacccgcaggctttgcatagattccaat [ 92160]
ASF_IND_20_CAD543                  ................................................................................ [ 92160]
LR899193.1_Germany_2020/1          ................................................................................ [ 92160]
MN393477.1_Wuhan_2019              ................................................................................ [ 92160]
MW396979.1_Timor-Leste/2019/1      ................................................................................ [ 92160]
MK940252.1_CN/2019/InnerMongolia   ................................................................................ [ 92160]
MK128995.1_China/2018/AnhuiXCGQ    ................................................................................ [ 92160]
MH681419.1_ASFV/POL/2015           ................................................................................ [ 92160]
LS478113.1_Estonia_2014            ................................................................................ [ 92160]
KP843857.1_Odintsovo_02/14         ................................................................................ [ 92160]
FN557520.1_strain_E75              .......................................g........................................ [ 92160]
AY261366.1_Warthog                 .......................................g.......a................................ [ 92160]
AY261365.1_Warmbaths               .......................................g........................................ [ 92160]
AY261364.1_Tengani_62              .......................................g........t............................... [ 92160]
AY261363.1_Pretoriuskop/96/4       .......................................g........................................ [ 92160]
AY261362.1_Mkuzi_1979              .......................................g........................................ [ 92160]
AY261361.1_Malawi_Lil-20/1_1983    ....................................a..................................a........ [ 92160]
AY261360.1_Kenya_1950              ....................................a........................................... [ 92160]
AM712240.1_OURT_88/3               .......................................g........................................ [ 92160]
AM712239.1_Benin_97/1              .......................................g........................................ [ 92160]
KM111295.1_Ken06.Bus               ....................................a.................t...........a.....g....... [ 92160]
KM111294.1_Ken05/Tk1               ....................................a........................................... [ 92160]
KM262844.1_strain_L60              .......................................g........................................ [ 92160]
KM102979.1_26544/OG10_Italy        .......................................g........................................ [ 92160]
KM262845.1_strain_NHV              .......................................g........................................ [ 92160]
KX354450.1_47/Ss/2008              .......................................g........................................ [ 92160]
KP055815.1_strain_BA71             .......................................g........................................ [ 92160]
MH025920.1_strain_R35              ....................................a.................t...........a.....g....... [ 92160]
MH025919.1_strain_N10              ....................................a.................t...........a.....g....... [ 92160]
MH025918.1_strain_R25              ....................................a.................t...........a.....g....... [ 92160]
MH025917.1_strain_R7               ....................................a.................t...........a.....g....... [ 92160]
MH025916.1_strain_R8               ....................................a.................t...........a.....g....... [ 92160]
```

```
FR682468.2_Georgia_2007/1        tggtggtattgtttttttccttgtagagtacacgaatactttctaatacttttaataactccgcgtattgaagacccgaa [ 92240]
ASF_IND_20_CAD543                ................................................................................ [ 92240]
LR899193.1_Germany_2020/1        ................................................................................ [ 92240]
MN393477.1_Wuhan_2019            ................................................................................ [ 92240]
MW396979.1_Timor-Leste/2019/1    ................................................................................ [ 92240]
MK940252.1_CN/2019/InnerMongolia ................................................................................ [ 92240]
MK128995.1_China/2018/AnhuiXCGQ  ................................................................................ [ 92240]
MH681419.1_ASFV/POL/2015         ................................................................................ [ 92240]
LS478113.1_Estonia_2014          ................................................................................ [ 92240]
KP843857.1_Odintsovo_02/14       ................................................................................ [ 92240]
FN557520.1_strain_E75            ................................................................................ [ 92240]
AY261366.1_Warthog               ........................................c...............g.t..a...........g...... [ 92240]
AY261365.1_Warmbaths             ........................................................g...a...........g...... [ 92240]
AY261364.1_Tengani_62            ........................................................g...a...........g...... [ 92240]
AY261363.1_Pretoriuskop/96/4     ........................................c...............g.t..a..a........g...... [ 92240]
AY261362.1_Mkuzi_1979            ............................................................................t... [ 92240]
AY261361.1_Malawi_Lil-20/1_1983  ............................c...........................g.t..a.....c............ [ 92240]
AY261360.1_Kenya_1950            ............................c...........................g.t..a.....c.....g...... [ 92240]
AM712240.1_OURT_88/3             ................................................................................ [ 92240]
AM712239.1_Benin_97/1            ................................................................................ [ 92240]
KM111295.1_Ken06.Bus             ............................c...........................g.t..a.....c.....g...... [ 92240]
KM111294.1_Ken05/Tk1             ............................c...........................g.t..a.....c.....g...... [ 92240]
KM262844.1_strain_L60            ................................................................................ [ 92240]
KM102979.1_26544/OG10_Italy      ................................................................................ [ 92240]
KM262845.1_strain_NHV            ................................................................................ [ 92240]
KX354450.1_47/Ss/2008            ................................................................................ [ 92240]
KP055815.1_strain_BA71           ................................................................................ [ 92240]
MH025920.1_strain_R35            ............................c...........................g.t..a.....c.....g...... [ 92240]
MH025919.1_strain_N10            ............................c...........................g.t..a.....c.....g...... [ 92240]
MH025918.1_strain_R25            ............................c...........................g.t..a.....c.....g...... [ 92240]
MH025917.1_strain_R7             ............................c...........................g.t..a.....c.....g...... [ 92240]
MH025916.1_strain_R8             ............................c...........................g.t..a.....c.....g...... [ 92240]

FR682468.2_Georgia_2007/1        cgcaactgttttaccagcttgagatgagcacatgcatttttttcttggagttcccactgttttttaatgtttaggtattc [ 92320]
ASF_IND_20_CAD543                ................................................................................ [ 92320]
LR899193.1_Germany_2020/1        ................................................................................ [ 92320]
MN393477.1_Wuhan_2019            ................................................................................ [ 92320]
MW396979.1_Timor-Leste/2019/1    ................................................................................ [ 92320]
MK940252.1_CN/2019/InnerMongolia ................................................................................ [ 92320]
MK128995.1_China/2018/AnhuiXCGQ  ................................................................................ [ 92320]
MH681419.1_ASFV/POL/2015         ................................................................................ [ 92320]
LS478113.1_Estonia_2014          ................................................................................ [ 92320]
KP843857.1_Odintsovo_02/14       ................................................................................ [ 92320]
```

```
FN557520.1_strain_E75           ...................................g............................................ [ 92320]
AY261366.1_Warthog              ...................................g............................................ [ 92320]
AY261365.1_Warmbaths            ................a..................g............................................ [ 92320]
AY261364.1_Tengani_62           ...................................g............................................ [ 92320]
AY261363.1_Pretoriuskop/96/4    ................a..................g............................................ [ 92320]
AY261362.1_Mkuzi_1979           ................a..............................a................................ [ 92320]
AY261361.1_Malawi_Lil-20/1_1983 ...................................g..............c............................. [ 92320]
AY261360.1_Kenya_1950           ...................................g............................................ [ 92320]
AM712240.1_OURT_88/3            ...................................g............................................ [ 92320]
AM712239.1_Benin_97/1           ...................................g............................................ [ 92320]
KM111295.1_Ken06.Bus            ...................................g............................................ [ 92320]
KM111294.1_Ken05/Tk1            ...................................g............................................ [ 92320]
KM262844.1_strain_L60           ...................................g............................................ [ 92320]
KM102979.1_26544/OG10_Italy     ...................................g............................................ [ 92320]
KM262845.1_strain_NHV           ...................................g............................................ [ 92320]
KX354450.1_47/Ss/2008           ...................................g............................................ [ 92320]
KP055815.1_strain_BA71          ...................................g............................................ [ 92320]
MH025920.1_strain_R35           ...................................g............................................ [ 92320]
MH025919.1_strain_N10           ...................................g............................................ [ 92320]
MH025918.1_strain_R25           ...................................g............................................ [ 92320]
MH025917.1_strain_R7            ...................................g............................................ [ 92320]
MH025916.1_strain_R8            ...................................g............................................ [ 92320]

FR682468.2_Georgia_2007/1       tgttgtaataagttctgcctcctgtttcccacaggctttaatgacttcttgaaggatgctgttagggtcatccactttac [ 92400]
ASF_IND_20_CAD543               ................................................................................ [ 92400]
LR899193.1_Germany_2020/1       ................................................................................ [ 92400]
MN393477.1_Wuhan_2019           ................................................................................ [ 92400]
MW396979.1_Timor-Leste/2019/1   ................................................................................ [ 92400]
MK940252.1_CN/2019/InnerMongolia ................................................................................ [ 92400]
MK128995.1_China/2018/AnhuiXCGQ ................................................................................ [ 92400]
MH681419.1_ASFV/POL/2015        ................................................................................ [ 92400]
LS478113.1_Estonia_2014         ................................................................................ [ 92400]
KP843857.1_Odintsovo_02/14      ................................................................................ [ 92400]
FN557520.1_strain_E75           ................................................................................ [ 92400]
AY261366.1_Warthog              .....................................................................g.....c.... [ 92400]
AY261365.1_Warmbaths            ...........................................................................c.... [ 92400]
AY261364.1_Tengani_62           ......................................................a....................c.... [ 92400]
AY261363.1_Pretoriuskop/96/4    .....................................................................g.....c.... [ 92400]
AY261362.1_Mkuzi_1979           ................................................................................ [ 92400]
AY261361.1_Malawi_Lil-20/1_1983 ..............................g................................g..a........c.... [ 92400]
AY261360.1_Kenya_1950           ............c..c..............g.......................a......c.............c..g. [ 92400]
AM712240.1_OURT_88/3            ................................................................................ [ 92400]
AM712239.1_Benin_97/1           ................................................................................ [ 92400]
KM111295.1_Ken06.Bus            ............c..c..............g.......................a......c.............c..g. [ 92400]
```

```
KM111294.1_Ken05/Tk1             ............c..c..............g.......................a......c.............c..g. [ 92400]
KM262844.1_strain_L60            ................................................................................ [ 92400]
KM102979.1_26544/OG10_Italy      ................................................................................ [ 92400]
KM262845.1_strain_NHV            ................................................................................ [ 92400]
KX354450.1_47/Ss/2008            ................................................................................ [ 92400]
KP055815.1_strain_BA71           ................................................................................ [ 92400]
MH025920.1_strain_R35            ............c..c..............g.......................a......c.............c..g. [ 92400]
MH025919.1_strain_N10            ............c..c..............g.......................a......c.............c..g. [ 92400]
MH025918.1_strain_R25            ............c..c..............g.......................a......c.............c..g. [ 92400]
MH025917.1_strain_R7             ............c..c..............g.......................a......c.............c..g. [ 92400]
MH025916.1_strain_R8             ............c..c..............g.......................a......c.............c..g. [ 92400]

FR682468.2_Georgia_2007/1        cctccattgtaagaatttcacgtatagcatccgactgcaccctacctattttttcttccataattttaaaatactgtctc [ 92480]
ASF_IND_20_CAD543                ................................................................................ [ 92480]
LR899193.1_Germany_2020/1        ................................................................................ [ 92480]
MN393477.1_Wuhan_2019            ................................................................................ [ 92480]
MW396979.1_Timor-Leste/2019/1    ................................................................................ [ 92480]
MK940252.1_CN/2019/InnerMongolia ................................................................................ [ 92480]
MK128995.1_China/2018/AnhuiXCGQ  ................................................................................ [ 92480]
MH681419.1_ASFV/POL/2015         ................................................................................ [ 92480]
LS478113.1_Estonia_2014          ................................................................................ [ 92480]
KP843857.1_Odintsovo_02/14       ................................................................................ [ 92480]
FN557520.1_strain_E75            ......................................................t......................... [ 92480]
AY261366.1_Warthog               .......................................................c........................ [ 92480]
AY261365.1_Warmbaths             .......................................................c........................ [ 92480]
AY261364.1_Tengani_62            .......................................................c........................ [ 92480]
AY261363.1_Pretoriuskop/96/4     .......................................................c........................ [ 92480]
AY261362.1_Mkuzi_1979            ................................................................................ [ 92480]
AY261361.1_Malawi_Lil-20/1_1983  .t..........................g..........................c........................ [ 92480]
AY261360.1_Kenya_1950            .t.........................t...........................c........................ [ 92480]
AM712240.1_OURT_88/3             ................................................................................ [ 92480]
AM712239.1_Benin_97/1            ......................................................t......................... [ 92480]
KM111295.1_Ken06.Bus             .t.....................................................c.......................t [ 92480]
KM111294.1_Ken05/Tk1             .t..........................g..........................c........................ [ 92480]
KM262844.1_strain_L60            ................................................................................ [ 92480]
KM102979.1_26544/OG10_Italy      ......................................................t......................... [ 92480]
KM262845.1_strain_NHV            ................................................................................ [ 92480]
KX354450.1_47/Ss/2008            ......................................................t......................... [ 92480]
KP055815.1_strain_BA71           ................................................................................ [ 92480]
MH025920.1_strain_R35            .t.....................................................c.......................t [ 92480]
MH025919.1_strain_N10            .t.....................................................c.......................t [ 92480]
MH025918.1_strain_R25            .t.....................................................c.......................t [ 92480]
MH025917.1_strain_R7             .t.....................................................c.......................t [ 92480]
MH025916.1_strain_R8             .t.....................................................c.......................t [ 92480]
```

```
FR682468.2_Georgia_2007/1        gcctgggtaatgacctctgtgagcttcatgtccacctgctgcagaatcatttgctccttttcacgctgttcagcatgttg [ 92560]
ASF_IND_20_CAD543                ................................................................................ [ 92560]
LR899193.1_Germany_2020/1        ................................................................................ [ 92560]
MN393477.1_Wuhan_2019            ................................................................................ [ 92560]
MW396979.1_Timor-Leste/2019/1    ................................................................................ [ 92560]
MK940252.1_CN/2019/InnerMongolia ................................................................................ [ 92560]
MK128995.1_China/2018/AnhuiXCGQ  ................................................................................ [ 92560]
MH681419.1_ASFV/POL/2015         ................................................................................ [ 92560]
LS478113.1_Estonia_2014          ................................................................................ [ 92560]
KP843857.1_Odintsovo_02/14       ................................................................................ [ 92560]
FN557520.1_strain_E75            .................c.............................................................. [ 92560]
AY261366.1_Warthog               .........g.......ca............................................................. [ 92560]
AY261365.1_Warmbaths             .........g.......ca............................................................. [ 92560]
AY261364.1_Tengani_62            .........g.......ca...........................................g................. [ 92560]
AY261363.1_Pretoriuskop/96/4     .........g.......ca............................................................. [ 92560]
AY261362.1_Mkuzi_1979            .................c...............................a.............................. [ 92560]
AY261361.1_Malawi_Lil-20/1_1983  .........g.......c...........a.............a.........t.......................... [ 92560]
AY261360.1_Kenya_1950            .....a...g.......c...................................t..t....................... [ 92560]
AM712240.1_OURT_88/3             .................c.............................................................. [ 92560]
AM712239.1_Benin_97/1            .................c.............................................................. [ 92560]
KM111295.1_Ken06.Bus             .....a...g.......c.............................................................. [ 92560]
KM111294.1_Ken05/Tk1             .....a...g.......c...................................t..t....................... [ 92560]
KM262844.1_strain_L60            .................c.............................................................. [ 92560]
KM102979.1_26544/OG10_Italy      .................c.............................................................. [ 92560]
KM262845.1_strain_NHV            .................c.............................................................. [ 92560]
KX354450.1_47/Ss/2008            .................c.............................................................. [ 92560]
KP055815.1_strain_BA71           .................c.............................................................. [ 92560]
MH025920.1_strain_R35            .....a...g.a.....c.............................................................. [ 92560]
MH025919.1_strain_N10            .....a...g.......c.............................................................. [ 92560]
MH025918.1_strain_R25            .....a...g.a.....c.............................................................. [ 92560]
MH025917.1_strain_R7             .....a...g.a.....c.............................................................. [ 92560]
MH025916.1_strain_R8             .....a...g.a.....c.............................................................. [ 92560]

FR682468.2_Georgia_2007/1        taaaaacttttgttctacagggttccaaagcacctccaaatagcctgctctatataggtcataaagcaagggcatgtatc [ 92640]
ASF_IND_20_CAD543                ................................................................................ [ 92640]
LR899193.1_Germany_2020/1        ................................................................................ [ 92640]
MN393477.1_Wuhan_2019            ................................................................................ [ 92640]
MW396979.1_Timor-Leste/2019/1    ................................................................................ [ 92640]
MK940252.1_CN/2019/InnerMongolia ................................................................................ [ 92640]
MK128995.1_China/2018/AnhuiXCGQ  ................................................................................ [ 92640]
MH681419.1_ASFV/POL/2015         ................................................................................ [ 92640]
LS478113.1_Estonia_2014          ................................................................................ [ 92640]
KP843857.1_Odintsovo_02/14       ................................................................................ [ 92640]
```

```
FN557520.1_strain_E75            ................................................................................ [ 92640]
AY261366.1_Warthog               c..........................g.....t...........c.................................. [ 92640]
AY261365.1_Warmbaths             c..........................g.....t...........c.................................. [ 92640]
AY261364.1_Tengani_62            c..........................g.....t.............................................. [ 92640]
AY261363.1_Pretoriuskop/96/4     c..........................g.....t...........c.................................. [ 92640]
AY261362.1_Mkuzi_1979            ................................................................................ [ 92640]
AY261361.1_Malawi_Lil-20/1_1983  ......t....................g..t..............c.................................. [ 92640]
AY261360.1_Kenya_1950            ..................g........g.................................................... [ 92640]
AM712240.1_OURT_88/3             ................................................................................ [ 92640]
AM712239.1_Benin_97/1            ................................................................................ [ 92640]
KM111295.1_Ken06.Bus             ..................g........g.................................................... [ 92640]
KM111294.1_Ken05/Tk1             ..................g........g.................................................... [ 92640]
KM262844.1_strain_L60            ................................................................................ [ 92640]
KM102979.1_26544/OG10_Italy      ................................................................................ [ 92640]
KM262845.1_strain_NHV            ................................................................................ [ 92640]
KX354450.1_47/Ss/2008            ................................................................................ [ 92640]
KP055815.1_strain_BA71           ................................................................................ [ 92640]
MH025920.1_strain_R35            ..................g........g.................................................... [ 92640]
MH025919.1_strain_N10            ..................g........g.................................................... [ 92640]
MH025918.1_strain_R25            ..................g........g.................................................... [ 92640]
MH025917.1_strain_R7             ..................g........g.................................................... [ 92640]
MH025916.1_strain_R8             ..................g........g.................................................... [ 92640]

FR682468.2_Georgia_2007/1        ccgatgtaaaaaccggggacaccgagtacatcgtagacaactcttttaaaaaaaatatcacgcgcttaatgtt---ctcc [ 92720]
ASF_IND_20_CAD543                .........................................................................---.... [ 92720]
LR899193.1_Germany_2020/1        .........................................................................---.... [ 92720]
MN393477.1_Wuhan_2019            .........................................................................---.... [ 92720]
MW396979.1_Timor-Leste/2019/1    .........................................................................---.... [ 92720]
MK940252.1_CN/2019/InnerMongolia .........................................................................---.... [ 92720]
MK128995.1_China/2018/AnhuiXCGQ  .........................................................................---.... [ 92720]
MH681419.1_ASFV/POL/2015         .........................................................................---.... [ 92720]
LS478113.1_Estonia_2014          .........................................................................---.... [ 92720]
KP843857.1_Odintsovo_02/14       .........................................................................---.... [ 92720]
FN557520.1_strain_E75            .........................................................................---.... [ 92720]
AY261366.1_Warthog               ..........c...........t..................................................---.... [ 92720]
AY261365.1_Warmbaths             ..........c..t.....................................................g..a..---.... [ 92720]
AY261364.1_Tengani_62            ..........c...........g............................................g.....---.... [ 92720]
AY261363.1_Pretoriuskop/96/4     ..........c...........t..................................................---.... [ 92720]
AY261362.1_Mkuzi_1979            .........................................................................---.... [ 92720]
AY261361.1_Malawi_Lil-20/1_1983  ..........c......a.............g..g................................g.....---.... [ 92720]
AY261360.1_Kenya_1950            ..........c......a.............a..g........c.......................g.....ctc.... [ 92720]
AM712240.1_OURT_88/3             .........................................................................---.... [ 92720]
AM712239.1_Benin_97/1            .........................................................................---.... [ 92720]
KM111295.1_Ken06.Bus             ..........c......a.t...........a..g........c.......................g.....ctc.... [ 92720]
```

```
KM111294.1_Ken05/Tk1               ..........c..t...a.............a..g........c.......................g.....---.... [ 92720]
KM262844.1_strain_L60              .........................................................................---.... [ 92720]
KM102979.1_26544/OG10_Italy        .........................................................................---.... [ 92720]
KM262845.1_strain_NHV              .........................................................................---.... [ 92720]
KX354450.1_47/Ss/2008              .........................................................................---.... [ 92720]
KP055815.1_strain_BA71             .........................................................................---.... [ 92720]
MH025920.1_strain_R35              ..........c......a.t...........a..g........c.......................g.....ctc.... [ 92720]
MH025919.1_strain_N10              ..........c......a.t...........a..g........c.......................g.....ctc.... [ 92720]
MH025918.1_strain_R25              ..........c......a.t...........a..g........c.......................g.....ctc.... [ 92720]
MH025917.1_strain_R7               ..........c......a.t...........a..g........c.......................g.....ctc.... [ 92720]
MH025916.1_strain_R8               ..........c......a.t...........a..g........c.......................g.....ctc.... [ 92720]

FR682468.2_Georgia_2007/1          tccggttcaatctcctcggtttcaacgatattagatatatgactgccctgatcctcacggtctagctttcggtgtaccat [ 92800]
ASF_IND_20_CAD543                  ................................................................................ [ 92800]
LR899193.1_Germany_2020/1          ................................................................................ [ 92800]
MN393477.1_Wuhan_2019              ................................................................................ [ 92800]
MW396979.1_Timor-Leste/2019/1      ................................................................................ [ 92800]
MK940252.1_CN/2019/InnerMongolia   ................................................................................ [ 92800]
MK128995.1_China/2018/AnhuiXCGQ    ................................................................................ [ 92800]
MH681419.1_ASFV/POL/2015           ................................................................................ [ 92800]
LS478113.1_Estonia_2014            ................................................................................ [ 92800]
KP843857.1_Odintsovo_02/14         ................................................................................ [ 92800]
FN557520.1_strain_E75              ................................................................................ [ 92800]
AY261366.1_Warthog                 ................................................................................ [ 92800]
AY261365.1_Warmbaths               .t........................a..................................................... [ 92800]
AY261364.1_Tengani_62              .................a..............................c............................... [ 92800]
AY261363.1_Pretoriuskop/96/4       ................................................................................ [ 92800]
AY261362.1_Mkuzi_1979              ................................................................................ [ 92800]
AY261361.1_Malawi_Lil-20/1_1983    ........g................t..c.c..................................t.............. [ 92800]
AY261360.1_Kenya_1950              ..........c......a.......t....c..........g..........t........................... [ 92800]
AM712240.1_OURT_88/3               ................................................................................ [ 92800]
AM712239.1_Benin_97/1              ................................................................................ [ 92800]
KM111295.1_Ken06.Bus               .........................t....c..........g...................................... [ 92800]
KM111294.1_Ken05/Tk1               .........................t....c..........g...................................... [ 92800]
KM262844.1_strain_L60              ................................................................................ [ 92800]
KM102979.1_26544/OG10_Italy        ................................................................................ [ 92800]
KM262845.1_strain_NHV              ................................................................................ [ 92800]
KX354450.1_47/Ss/2008              ................................................................................ [ 92800]
KP055815.1_strain_BA71             ................................................................................ [ 92800]
MH025920.1_strain_R35              .........................t....c..........g...................................... [ 92800]
MH025919.1_strain_N10              .........................t....c..........g...................................... [ 92800]
MH025918.1_strain_R25              .........................t....c..........g...................................... [ 92800]
MH025917.1_strain_R7               .........................t....c..........g...................................... [ 92800]
MH025916.1_strain_R8               .........................t....c..........g...................................... [ 92800]
```

```
FR682468.2_Georgia_2007/1        ctcctctgctagccgattaatgagccagctatgcccgccgctccgcaaaaacttataaagttcgatatactggtgcgtaa [ 92880]
ASF_IND_20_CAD543                ................................................................................ [ 92880]
LR899193.1_Germany_2020/1        ................................................................................ [ 92880]
MN393477.1_Wuhan_2019            ................................................................................ [ 92880]
MW396979.1_Timor-Leste/2019/1    ................................................................................ [ 92880]
MK940252.1_CN/2019/InnerMongolia ................................................................................ [ 92880]
MK128995.1_China/2018/AnhuiXCGQ  ................................................................................ [ 92880]
MH681419.1_ASFV/POL/2015         ................................................................................ [ 92880]
LS478113.1_Estonia_2014          ................................................................................ [ 92880]
KP843857.1_Odintsovo_02/14       ................................................................................ [ 92880]
FN557520.1_strain_E75            ................................................................................ [ 92880]
AY261366.1_Warthog               ................................................................................ [ 92880]
AY261365.1_Warmbaths             ................................................................................ [ 92880]
AY261364.1_Tengani_62            .......................................a...................................t.... [ 92880]
AY261363.1_Pretoriuskop/96/4     ................................................................................ [ 92880]
AY261362.1_Mkuzi_1979            ................................................................................ [ 92880]
AY261361.1_Malawi_Lil-20/1_1983  ..................g.......................a..................................... [ 92880]
AY261360.1_Kenya_1950            ......c...........g.................c.....t........t............................ [ 92880]
AM712240.1_OURT_88/3             ................................................................................ [ 92880]
AM712239.1_Benin_97/1            ................................................................................ [ 92880]
KM111295.1_Ken06.Bus             ......c....a......g.......................t........t............................ [ 92880]
KM111294.1_Ken05/Tk1             ......c...........g................................t...........a..g............. [ 92880]
KM262844.1_strain_L60            ................................................................................ [ 92880]
KM102979.1_26544/OG10_Italy      ................................................................................ [ 92880]
KM262845.1_strain_NHV            ................................................................................ [ 92880]
KX354450.1_47/Ss/2008            ................................................................................ [ 92880]
KP055815.1_strain_BA71           ................................................................................ [ 92880]
MH025920.1_strain_R35            ......c....a......g.......................t........t............................ [ 92880]
MH025919.1_strain_N10            ......c....a......g.......................t........t............................ [ 92880]
MH025918.1_strain_R25            ......c....a......g.......................t........t............................ [ 92880]
MH025917.1_strain_R7             ......c....a......g.......................t........t............................ [ 92880]
MH025916.1_strain_R8             ......c....a......g.......................t........t............................ [ 92880]

FR682468.2_Georgia_2007/1        actggatgatgttttccttggtggttacgacaaccccttctccgttttttttccaggtttcttgatccacgcatttcata [ 92960]
ASF_IND_20_CAD543                ................................................................................ [ 92960]
LR899193.1_Germany_2020/1        ................................................................................ [ 92960]
MN393477.1_Wuhan_2019            ................................................................................ [ 92960]
MW396979.1_Timor-Leste/2019/1    ................................................................................ [ 92960]
MK940252.1_CN/2019/InnerMongolia ................................................................................ [ 92960]
MK128995.1_China/2018/AnhuiXCGQ  ................................................................................ [ 92960]
MH681419.1_ASFV/POL/2015         ................................................................................ [ 92960]
LS478113.1_Estonia_2014          ................................................................................ [ 92960]
KP843857.1_Odintsovo_02/14       ................................................................................ [ 92960]
```

```
FN557520.1_strain_E75             ........................................c.......................g............... [ 92960]
AY261366.1_Warthog                ........................................c.......................g............... [ 92960]
AY261365.1_Warmbaths              .................................t......c.......................g............... [ 92960]
AY261364.1_Tengani_62             ...................a....................c.......................g............... [ 92960]
AY261363.1_Pretoriuskop/96/4      ........................................c.......................g............... [ 92960]
AY261362.1_Mkuzi_1979             ........................................c.......................g............... [ 92960]
AY261361.1_Malawi_Lil-20/1_1983   .................................t......c..a....................g............... [ 92960]
AY261360.1_Kenya_1950             ....t.......................c...........c..a.................................... [ 92960]
AM712240.1_OURT_88/3              ........................................c.......................g............... [ 92960]
AM712239.1_Benin_97/1             ........................................c.......................g............... [ 92960]
KM111295.1_Ken06.Bus              ....t.......................c...........c..a............c....c..g............... [ 92960]
KM111294.1_Ken05/Tk1              ............................c...........c..a.................................... [ 92960]
KM262844.1_strain_L60             ........................................c.......................g............... [ 92960]
KM102979.1_26544/OG10_Italy       ........................................c.......................g............... [ 92960]
KM262845.1_strain_NHV             ........................................c.......................g............... [ 92960]
KX354450.1_47/Ss/2008             ........................................c.......................g............... [ 92960]
KP055815.1_strain_BA71            ........................................c.......................g............... [ 92960]
MH025920.1_strain_R35             ....t.......................c...........c..a............c....c..g............... [ 92960]
MH025919.1_strain_N10             ....t.......................c...........c..a............c....c..g............... [ 92960]
MH025918.1_strain_R25             ....t.......................c...........c..a............c....c..g............... [ 92960]
MH025917.1_strain_R7              ....t.......................c...........c..a............c....c..g............... [ 92960]
MH025916.1_strain_R8              ....t.......................c...........c..a............c....c..g............... [ 92960]

FR682468.2_Georgia_2007/1         aatactcgaataaaattggtcaaattggctcctgaggcgacgtagcccaaggtttcaggcgagaaggagcctatctcagc [ 93040]
ASF_IND_20_CAD543                 ................................................................................ [ 93040]
LR899193.1_Germany_2020/1         ................................................................................ [ 93040]
MN393477.1_Wuhan_2019             ................................................................................ [ 93040]
MW396979.1_Timor-Leste/2019/1     ................................................................................ [ 93040]
MK940252.1_CN/2019/InnerMongolia  ..........................................................................t..... [ 93040]
MK128995.1_China/2018/AnhuiXCGQ   ................................................................................ [ 93040]
MH681419.1_ASFV/POL/2015          ................................................................................ [ 93040]
LS478113.1_Estonia_2014           ................................................................................ [ 93040]
KP843857.1_Odintsovo_02/14        ................................................................................ [ 93040]
FN557520.1_strain_E75             ................................................................................ [ 93040]
AY261366.1_Warthog                ................................................................................ [ 93040]
AY261365.1_Warmbaths              ................................................................................ [ 93040]
AY261364.1_Tengani_62             ............................................a...........g.....a................. [ 93040]
AY261363.1_Pretoriuskop/96/4      ................................................................................ [ 93040]
AY261362.1_Mkuzi_1979             ................................................................................ [ 93040]
AY261361.1_Malawi_Lil-20/1_1983   ..........................a...........a.....................a.............t..... [ 93040]
AY261360.1_Kenya_1950             ................................................................................ [ 93040]
AM712240.1_OURT_88/3              ................................................................................ [ 93040]
AM712239.1_Benin_97/1             ................................................................................ [ 93040]
KM111295.1_Ken06.Bus              ................................................................................ [ 93040]
```

```
KM111294.1_Ken05/Tk1             ................................................................................ [ 93040]
KM262844.1_strain_L60            ................................................................................ [ 93040]
KM102979.1_26544/OG10_Italy      ................................................................................ [ 93040]
KM262845.1_strain_NHV            ................................................................................ [ 93040]
KX354450.1_47/Ss/2008            ................................................................................ [ 93040]
KP055815.1_strain_BA71           ................................................................................ [ 93040]
MH025920.1_strain_R35            ................................................................................ [ 93040]
MH025919.1_strain_N10            ................................................................................ [ 93040]
MH025918.1_strain_R25            ................................................................................ [ 93040]
MH025917.1_strain_R7             ................................................................................ [ 93040]
MH025916.1_strain_R8             ................................................................................ [ 93040]

FR682468.2_Georgia_2007/1        catacgcataaaacactgcggggaaaaagtttttagccgcaacttaagtccatagatttcaatgggggcttctgcgggaa [ 93120]
ASF_IND_20_CAD543                ............................s................................................... [ 93120]
LR899193.1_Germany_2020/1        ................................................................................ [ 93120]
MN393477.1_Wuhan_2019            ................................................................................ [ 93120]
MW396979.1_Timor-Leste/2019/1    ................................................................................ [ 93120]
MK940252.1_CN/2019/InnerMongolia .......................................................................t........ [ 93120]
MK128995.1_China/2018/AnhuiXCGQ  ................................................................................ [ 93120]
MH681419.1_ASFV/POL/2015         ................................................................................ [ 93120]
LS478113.1_Estonia_2014          ................................................................................ [ 93120]
KP843857.1_Odintsovo_02/14       ................................................................................ [ 93120]
FN557520.1_strain_E75            ............g...............c................................................... [ 93120]
AY261366.1_Warthog               ............g.....................................................c............. [ 93120]
AY261365.1_Warmbaths             ............g.....................................................c............. [ 93120]
AY261364.1_Tengani_62            ............g.....................................................t............. [ 93120]
AY261363.1_Pretoriuskop/96/4     ............g.....................................................c............. [ 93120]
AY261362.1_Mkuzi_1979            ............g................................................................... [ 93120]
AY261361.1_Malawi_Lil-20/1_1983  ............g..t.............................g....................ca....c....... [ 93120]
AY261360.1_Kenya_1950            ............g..t....................t........g....................ca............ [ 93120]
AM712240.1_OURT_88/3             ............g...............c................................................... [ 93120]
AM712239.1_Benin_97/1            ............g...............c................................................... [ 93120]
KM111295.1_Ken06.Bus             ............g..t....................t.............................ta............ [ 93120]
KM111294.1_Ken05/Tk1             ............g..t....................t........g........a...........ca............ [ 93120]
KM262844.1_strain_L60            ............g...............c................................................... [ 93120]
KM102979.1_26544/OG10_Italy      ............g...............c..................................c................ [ 93120]
KM262845.1_strain_NHV            ............g...............c................................................... [ 93120]
KX354450.1_47/Ss/2008            ............g...............c................................................... [ 93120]
KP055815.1_strain_BA71           ............g...............c................................................... [ 93120]
MH025920.1_strain_R35            ............g..t....................t.............................ca............ [ 93120]
MH025919.1_strain_N10            ............g..t....................t.............................ca............ [ 93120]
MH025918.1_strain_R25            ............g..t....................t.............................ca............ [ 93120]
MH025917.1_strain_R7             ............g..t....................t.............................ca............ [ 93120]
MH025916.1_strain_R8             ............g..t....................t.............................ca............ [ 93120]
```

```
FR682468.2_Georgia_2007/1        cggccaggtgcgtcccattaattaaaaaaatttctttgcgtgtgctagggcgaacacgtaattcctttttttttttcactc [ 93200]
ASF_IND_20_CAD543                ............................................................................r... [ 93200]
LR899193.1_Germany_2020/1        ................................................................................ [ 93200]
MN393477.1_Wuhan_2019            ................................................................................ [ 93200]
MW396979.1_Timor-Leste/2019/1    ................................................................................ [ 93200]
MK940252.1_CN/2019/InnerMongolia ................................................................................ [ 93200]
MK128995.1_China/2018/AnhuiXCGQ  ................................................................................ [ 93200]
MH681419.1_ASFV/POL/2015         ................................................................................ [ 93200]
LS478113.1_Estonia_2014          ................................................................................ [ 93200]
KP843857.1_Odintsovo_02/14       .................................................................-.............. [ 93200]
FN557520.1_strain_E75            .................................................................-..........g... [ 93200]
AY261366.1_Warthog               ................................................................................ [ 93200]
AY261365.1_Warmbaths             ................................................................................ [ 93200]
AY261364.1_Tengani_62            ................................................................................ [ 93200]
AY261363.1_Pretoriuskop/96/4     ................................................................................ [ 93200]
AY261362.1_Mkuzi_1979            ................................................................................ [ 93200]
AY261361.1_Malawi_Lil-20/1_1983  .......a.............................t.......................................... [ 93200]
AY261360.1_Kenya_1950            ....t................................t...........a.............................. [ 93200]
AM712240.1_OURT_88/3             ............................................................................g... [ 93200]
AM712239.1_Benin_97/1            ............................................................................g... [ 93200]
KM111295.1_Ken06.Bus             .....................................t..c........a.............................. [ 93200]
KM111294.1_Ken05/Tk1             ....t................................t...........a.............................. [ 93200]
KM262844.1_strain_L60            .................................................................-..........g... [ 93200]
KM102979.1_26544/OG10_Italy      ............................................................................g... [ 93200]
KM262845.1_strain_NHV            ............................................................................g... [ 93200]
KX354450.1_47/Ss/2008            ............................................................................g... [ 93200]
KP055815.1_strain_BA71           ............................................................................g... [ 93200]
MH025920.1_strain_R35            .....................................t..c........a.............................. [ 93200]
MH025919.1_strain_N10            .....................................t..c........a.............................. [ 93200]
MH025918.1_strain_R25            .....................................t..c........a.............................. [ 93200]
MH025917.1_strain_R7             .....................................t..c........a.............................. [ 93200]
MH025916.1_strain_R8             .....................................t..c........a.............................. [ 93200]

FR682468.2_Georgia_2007/1        acgatggggaccacatcggggtctaccagcagttgacgtatgtaggcctctatgggcatggatagatcgggcagctttga [ 93280]
ASF_IND_20_CAD543                ................................................................................ [ 93280]
LR899193.1_Germany_2020/1        ................................................................................ [ 93280]
MN393477.1_Wuhan_2019            ................................................................................ [ 93280]
MW396979.1_Timor-Leste/2019/1    ................................................................................ [ 93280]
MK940252.1_CN/2019/InnerMongolia ................................................................................ [ 93280]
MK128995.1_China/2018/AnhuiXCGQ  ................................................................................ [ 93280]
MH681419.1_ASFV/POL/2015         ................................................................................ [ 93280]
LS478113.1_Estonia_2014          ................................................................................ [ 93280]
KP843857.1_Odintsovo_02/14       ................................................................................ [ 93280]
```

```
FN557520.1_strain_E75            ..........................t.......................c............................. [ 93280]
AY261366.1_Warthog               ........a..........a..............................c............................. [ 93280]
AY261365.1_Warmbaths             ..a......................t........................c............................. [ 93280]
AY261364.1_Tengani_62            ..................................................c............................. [ 93280]
AY261363.1_Pretoriuskop/96/4     ........a..........a..............................c............................. [ 93280]
AY261362.1_Mkuzi_1979            ..................................................c............................. [ 93280]
AY261361.1_Malawi_Lil-20/1_1983  ..........t......a..a.t...........................c................t.........c.. [ 93280]
AY261360.1_Kenya_1950            ......................tc..............a...........c........a.................... [ 93280]
AM712240.1_OURT_88/3             ..........................t.......................c............................. [ 93280]
AM712239.1_Benin_97/1            ..........................t.......................c............................. [ 93280]
KM111295.1_Ken06.Bus             ......................tc..............a...........c........a.......t............ [ 93280]
KM111294.1_Ken05/Tk1             .....a................tc..............a...........c........a.......t............ [ 93280]
KM262844.1_strain_L60            ..........................t.......................c............................. [ 93280]
KM102979.1_26544/OG10_Italy      ..........................t.......................c............................. [ 93280]
KM262845.1_strain_NHV            ..........................t.......................c............................. [ 93280]
KX354450.1_47/Ss/2008            ..........................t.......................c............................. [ 93280]
KP055815.1_strain_BA71           ..........................t.......................c............................. [ 93280]
MH025920.1_strain_R35            ......................tc..............a...........c........a.......t............ [ 93280]
MH025919.1_strain_N10            ......................tc..............a...........c........a.......t............ [ 93280]
MH025918.1_strain_R25            ......................tc..............a...........c........a.......t............ [ 93280]
MH025917.1_strain_R7             ......................tc..............a...........c........a.......t............ [ 93280]
MH025916.1_strain_R8             ......................tc..............a...........c........a.......t............ [ 93280]

FR682468.2_Georgia_2007/1        ctgctcggcgcgaacatggttcacaaaatcttttagagtgaaaagaaagtctattaaacgtatgttttttatatcattag [ 93360]
ASF_IND_20_CAD543                ................................................................................ [ 93360]
LR899193.1_Germany_2020/1        ................................................................................ [ 93360]
MN393477.1_Wuhan_2019            ................................................................................ [ 93360]
MW396979.1_Timor-Leste/2019/1    ................................................................................ [ 93360]
MK940252.1_CN/2019/InnerMongolia ................................................................................ [ 93360]
MK128995.1_China/2018/AnhuiXCGQ  ................................................................................ [ 93360]
MH681419.1_ASFV/POL/2015         ................................................................................ [ 93360]
LS478113.1_Estonia_2014          ................................................................................ [ 93360]
KP843857.1_Odintsovo_02/14       ................................................................................ [ 93360]
FN557520.1_strain_E75            ................................................................................ [ 93360]
AY261366.1_Warthog               ................................................................................ [ 93360]
AY261365.1_Warmbaths             ................................................................................ [ 93360]
AY261364.1_Tengani_62            ................................................................................ [ 93360]
AY261363.1_Pretoriuskop/96/4     ................................................................................ [ 93360]
AY261362.1_Mkuzi_1979            ................................................................................ [ 93360]
AY261361.1_Malawi_Lil-20/1_1983  ....................................c........................................... [ 93360]
AY261360.1_Kenya_1950            ...t................................t.a...g..................................... [ 93360]
AM712240.1_OURT_88/3             ................................................................................ [ 93360]
AM712239.1_Benin_97/1            ................................................................................ [ 93360]
KM111295.1_Ken06.Bus             ..................a.................t........................................... [ 93360]
```

| | | |
|---|---|---|
| KM111294.1_Ken05/Tk1 | ...t................................t.a......................................... | [ 93360] |
| KM262844.1_strain_L60 | ................................................................................ | [ 93360] |
| KM102979.1_26544/OG10_Italy | ................................................................................ | [ 93360] |
| KM262845.1_strain_NHV | ................................................................................ | [ 93360] |
| KX354450.1_47/Ss/2008 | ................................................................................ | [ 93360] |
| KP055815.1_strain_BA71 | ................................................................................ | [ 93360] |
| MH025920.1_strain_R35 | ..................a.................t........................................... | [ 93360] |
| MH025919.1_strain_N10 | ..................a.................t........................................... | [ 93360] |
| MH025918.1_strain_R25 | ..................a.................t........................................... | [ 93360] |
| MH025917.1_strain_R7 | ..................a.................t........................................... | [ 93360] |
| MH025916.1_strain_R8 | ..................a.................t........................................... | [ 93360] |

| | | |
|---|---|---|
| FR682468.2_Georgia_2007/1 | accctttaagggtagagtagatttcatccactagtgcctcgatttcctcattattgagcgataagatatctgtgccacgg | [ 93440] |
| ASF_IND_20_CAD543 | ................................................................................ | [ 93440] |
| LR899193.1_Germany_2020/1 | ................................................................................ | [ 93440] |
| MN393477.1_Wuhan_2019 | ................................................................................ | [ 93440] |
| MW396979.1_Timor-Leste/2019/1 | ................................................................................ | [ 93440] |
| MK940252.1_CN/2019/InnerMongolia | ................................................................................ | [ 93440] |
| MK128995.1_China/2018/AnhuiXCGQ | ................................................................................ | [ 93440] |
| MH681419.1_ASFV/POL/2015 | ................................................................................ | [ 93440] |
| LS478113.1_Estonia_2014 | ................................................................................ | [ 93440] |
| KP843857.1_Odintsovo_02/14 | ................................................................................ | [ 93440] |
| FN557520.1_strain_E75 | ................................................................................ | [ 93440] |
| AY261366.1_Warthog | .......g.....g.................................................................. | [ 93440] |
| AY261365.1_Warmbaths | .............g.................................................................. | [ 93440] |
| AY261364.1_Tengani_62 | .............g.................................................................. | [ 93440] |
| AY261363.1_Pretoriuskop/96/4 | .......g.....g.................................................................. | [ 93440] |
| AY261362.1_Mkuzi_1979 | ................................................................................ | [ 93440] |
| AY261361.1_Malawi_Lil-20/1_1983 | .................cga......................................t.....a............... | [ 93440] |
| AY261360.1_Kenya_1950 | .a..............a..a...........c.a..................g........................... | [ 93440] |
| AM712240.1_OURT_88/3 | ................................................................................ | [ 93440] |
| AM712239.1_Benin_97/1 | ................................................................................ | [ 93440] |
| KM111295.1_Ken06.Bus | .a..............a..a...........c.a.............................................. | [ 93440] |
| KM111294.1_Ken05/Tk1 | .a..............a..a...........c.a.............................................. | [ 93440] |
| KM262844.1_strain_L60 | ................................................................................ | [ 93440] |
| KM102979.1_26544/OG10_Italy | ................................................................................ | [ 93440] |
| KM262845.1_strain_NHV | ................................................................................ | [ 93440] |
| KX354450.1_47/Ss/2008 | ................................................................................ | [ 93440] |
| KP055815.1_strain_BA71 | ................................................................................ | [ 93440] |
| MH025920.1_strain_R35 | .a..............a..a...........c.a.............................................. | [ 93440] |
| MH025919.1_strain_N10 | .a..............a..a...........c.a..........................................t... | [ 93440] |
| MH025918.1_strain_R25 | .a..............a..a...........c.a.............................................. | [ 93440] |
| MH025917.1_strain_R7 | .a..............a..a...........c.a.............................................. | [ 93440] |
| MH025916.1_strain_R8 | .a..............a..a...........c.a.............................................. | [ 93440] |

```
FR682468.2_Georgia_2007/1        tggactatttgcgcgatcgtaattacttcctccattagatagaaactgaatattatatttaaa--ataaatacaaaatgt [ 93520]
ASF_IND_20_CAD543                ...............................................................--............... [ 93520]
LR899193.1_Germany_2020/1        ...............................................................--............... [ 93520]
MN393477.1_Wuhan_2019            ...............................................................--............... [ 93520]
MW396979.1_Timor-Leste/2019/1    ...............................................................--............... [ 93520]
MK940252.1_CN/2019/InnerMongolia ...............................................................--............... [ 93520]
MK128995.1_China/2018/AnhuiXCGQ  ...............................................................--............... [ 93520]
MH681419.1_ASFV/POL/2015         ...............................................................--............... [ 93520]
LS478113.1_Estonia_2014          ...............................................................--............... [ 93520]
KP843857.1_Odintsovo_02/14       ...............................................................--............... [ 93520]
FN557520.1_strain_E75            ...............................................................--............... [ 93520]
AY261366.1_Warthog               ...............................................................--............... [ 93520]
AY261365.1_Warmbaths             ...............................................................--............... [ 93520]
AY261364.1_Tengani_62            ...............................................................--............... [ 93520]
AY261363.1_Pretoriuskop/96/4     ...............................................................--............... [ 93520]
AY261362.1_Mkuzi_1979            .....a.........................................................--............... [ 93520]
AY261361.1_Malawi_Lil-20/1_1983  .......................................g.......................--............... [ 93520]
AY261360.1_Kenya_1950            ..............a........c............................c..........at............... [ 93520]
AM712240.1_OURT_88/3             ...............................................................--............... [ 93520]
AM712239.1_Benin_97/1            ...............................................................--............... [ 93520]
KM111295.1_Ken06.Bus             ..............a........c.....t......................c..........at............... [ 93520]
KM111294.1_Ken05/Tk1             ..............a........c............................c..........at............... [ 93520]
KM262844.1_strain_L60            ...............................................................--............... [ 93520]
KM102979.1_26544/OG10_Italy      ...............................................................--............... [ 93520]
KM262845.1_strain_NHV            ...............................................................--............... [ 93520]
KX354450.1_47/Ss/2008            ...............................................................--............... [ 93520]
KP055815.1_strain_BA71           ...............................................................--............... [ 93520]
MH025920.1_strain_R35            ..............a........c.....t......................c..........at............... [ 93520]
MH025919.1_strain_N10            ..............a........c.....t......................c..........at............... [ 93520]
MH025918.1_strain_R25            ..............a........c.....t......................c..........at............... [ 93520]
MH025917.1_strain_R7             ..............a........c.....t......................c..........at............... [ 93520]
MH025916.1_strain_R8             ..............a........c.....t......................c..........at............... [ 93520]

FR682468.2_Georgia_2007/1        caaatgaaagttttcccgaaacgttggaaaacttactttcaatgttacagaccaaacagcaaaacgcaattcagtcagag [ 93600]
ASF_IND_20_CAD543                ................................................................................ [ 93600]
LR899193.1_Germany_2020/1        ................................................................................ [ 93600]
MN393477.1_Wuhan_2019            ................................................................................ [ 93600]
MW396979.1_Timor-Leste/2019/1    ................................................................................ [ 93600]
MK940252.1_CN/2019/InnerMongolia ................................................................................ [ 93600]
MK128995.1_China/2018/AnhuiXCGQ  ................................................................................ [ 93600]
MH681419.1_ASFV/POL/2015         ................................................................................ [ 93600]
LS478113.1_Estonia_2014          ................................................................................ [ 93600]
KP843857.1_Odintsovo_02/14       ................................................................................ [ 93600]
```

```
FN557520.1_strain_E75            ................................c.......g.c..................................... [ 93600]
AY261366.1_Warthog               ........................................g.c..................................... [ 93600]
AY261365.1_Warmbaths             ........................................g.c..................................... [ 93600]
AY261364.1_Tengani_62            ...............................t........g.c.c......................g............ [ 93600]
AY261363.1_Pretoriuskop/96/4     ........................................g.c..................................... [ 93600]
AY261362.1_Mkuzi_1979            ........................................gtc..................................... [ 93600]
AY261361.1_Malawi_Lil-20/1_1983  ......................c.................g.c........................g...........a [ 93600]
AY261360.1_Kenya_1950            ........................................g.c..................................... [ 93600]
AM712240.1_OURT_88/3             ................................c.......g.c..................................... [ 93600]
AM712239.1_Benin_97/1            ................................c.......g.c..................................... [ 93600]
KM111295.1_Ken06.Bus             ........................................g.c.c................................... [ 93600]
KM111294.1_Ken05/Tk1             ........................................g.c..................................... [ 93600]
KM262844.1_strain_L60            ................................c.......g.c..................................... [ 93600]
KM102979.1_26544/OG10_Italy      ................................c.......g.c..................................... [ 93600]
KM262845.1_strain_NHV            ................................c.......g.c..................................... [ 93600]
KX354450.1_47/Ss/2008            ................................c.......g.c..................................... [ 93600]
KP055815.1_strain_BA71           ................................c.......g.c..................................... [ 93600]
MH025920.1_strain_R35            ........................................g.c.c................................... [ 93600]
MH025919.1_strain_N10            ........................................g.c.c................................... [ 93600]
MH025918.1_strain_R25            ........................................g.c.c................................... [ 93600]
MH025917.1_strain_R7             ........................................g.c.c................................... [ 93600]
MH025916.1_strain_R8             ........................................g.c.c................................... [ 93600]

FR682468.2_Georgia_2007/1        gtgattgaatggctgcacagcttttgtgaaacctttcacttaaaaatacactgccataaacagtttattcctagcgggga [ 93680]
ASF_IND_20_CAD543                ................................................................................ [ 93680]
LR899193.1_Germany_2020/1        ................................................................................ [ 93680]
MN393477.1_Wuhan_2019            ................................................................................ [ 93680]
MW396979.1_Timor-Leste/2019/1    ................................................................................ [ 93680]
MK940252.1_CN/2019/InnerMongolia ................................................................................ [ 93680]
MK128995.1_China/2018/AnhuiXCGQ  ................................................................................ [ 93680]
MH681419.1_ASFV/POL/2015         ................................................................................ [ 93680]
LS478113.1_Estonia_2014          ................................................................................ [ 93680]
KP843857.1_Odintsovo_02/14       ...........................................................................-.... [ 93680]
FN557520.1_strain_E75            ................................................................................ [ 93680]
AY261366.1_Warthog               ..a................a............................................................ [ 93680]
AY261365.1_Warmbaths             ................................................................................ [ 93680]
AY261364.1_Tengani_62            ................................................................................ [ 93680]
AY261363.1_Pretoriuskop/96/4     ..a................a............................................................ [ 93680]
AY261362.1_Mkuzi_1979            ................................................................................ [ 93680]
AY261361.1_Malawi_Lil-20/1_1983  ..........................c...........t......................................... [ 93680]
AY261360.1_Kenya_1950            .....ca...............................t......................................... [ 93680]
AM712240.1_OURT_88/3             ................................................................................ [ 93680]
AM712239.1_Benin_97/1            ................................................................................ [ 93680]
KM111295.1_Ken06.Bus             .....ca...............................t.......................t................. [ 93680]
```

```
KM111294.1_Ken05/Tk1             .....ca...............................t......................................... [ 93680]
KM262844.1_strain_L60            ................................................................................ [ 93680]
KM102979.1_26544/OG10_Italy      ................................................................................ [ 93680]
KM262845.1_strain_NHV            ................................................................................ [ 93680]
KX354450.1_47/Ss/2008            ................................................................................ [ 93680]
KP055815.1_strain_BA71           ................................................................................ [ 93680]
MH025920.1_strain_R35            .....ca...............................t......................................... [ 93680]
MH025919.1_strain_N10            .....ca...............................t......................................... [ 93680]
MH025918.1_strain_R25            .....ca...............................t......................................... [ 93680]
MH025917.1_strain_R7             .....ca...............................t......................................... [ 93680]
MH025916.1_strain_R8             .....ca...............................t......................................... [ 93680]

FR682468.2_Georgia_2007/1        aaaaaaacgagctaaaatacccgctcaagaaacacagggaaacacgcagccctcccaccatgtgtaccgggttgttctct [ 93760]
ASF_IND_20_CAD543                .....................y.....r....y............................................... [ 93760]
LR899193.1_Germany_2020/1        ................................................................................ [ 93760]
MN393477.1_Wuhan_2019            ................................................................................ [ 93760]
MW396979.1_Timor-Leste/2019/1    ................................................................................ [ 93760]
MK940252.1_CN/2019/InnerMongolia ................................................................................ [ 93760]
MK128995.1_China/2018/AnhuiXCGQ  ................................................................................ [ 93760]
MH681419.1_ASFV/POL/2015         ................................................................................ [ 93760]
LS478113.1_Estonia_2014          ................................................................................ [ 93760]
KP843857.1_Odintsovo_02/14       ................................................................................ [ 93760]
FN557520.1_strain_E75            .....................t.....g....t...............................c.............t. [ 93760]
AY261366.1_Warthog               ...........................g....................................c.....a......... [ 93760]
AY261365.1_Warmbaths             ...........................g....................................c.............t. [ 93760]
AY261364.1_Tengani_62            .........g.................g....................................c............... [ 93760]
AY261363.1_Pretoriuskop/96/4     ...........................g....................................c.....a......... [ 93760]
AY261362.1_Mkuzi_1979            ...........................g....................................c.............t. [ 93760]
AY261361.1_Malawi_Lil-20/1_1983  .......t.g............a....g.....g....a......a.....------...c...c.g............. [ 93760]
AY261360.1_Kenya_1950            .......t.gc.c........ta....g.....g....a......a......c...........c............... [ 93760]
AM712240.1_OURT_88/3             .....................t.....g....t...............................c.............t. [ 93760]
AM712239.1_Benin_97/1            .....................t.....g....t...............................c.............t. [ 93760]
KM111295.1_Ken06.Bus             .......t.gc.c........ta....g.....g....a......a......c...........c............... [ 93760]
KM111294.1_Ken05/Tk1             .......t.gc.c........ta....g.....g....a......a......c...........c............... [ 93760]
KM262844.1_strain_L60            .....................t.....g....t...............................c.............t. [ 93760]
KM102979.1_26544/OG10_Italy      .....................t.....g....t................t..............c.............t. [ 93760]
KM262845.1_strain_NHV            .....................t.....g....t...............................c.............t. [ 93760]
KX354450.1_47/Ss/2008            .....................t.....g....t................t..............c.............t. [ 93760]
KP055815.1_strain_BA71           .....................t.....g....t...............................c.............t. [ 93760]
MH025920.1_strain_R35            .......t.gc.c........ta....g.....g....a......a......c...........c............... [ 93760]
MH025919.1_strain_N10            .......t.gc.c........ta....g.....g....a......a......c...........c............... [ 93760]
MH025918.1_strain_R25            .......t.gc.c........ta....g.....g....a......a......c...........c............... [ 93760]
MH025917.1_strain_R7             .......t.gc.c........ta....g.....g....a......a......c...........c............... [ 93760]
MH025916.1_strain_R8             .......t.gc.c........ta....g.....g....a......a......c...........c............... [ 93760]
```

```
FR682468.2_Georgia_2007/1        ccagagcacagccagtcaaagcacaggaatctctgctaacaaccatgtgcaacggactggtgctagatgcaaacacatgg [ 93840]
ASF_IND_20_CAD543                ................................................................................ [ 93840]
LR899193.1_Germany_2020/1        ................................................................................ [ 93840]
MN393477.1_Wuhan_2019            ................................................................................ [ 93840]
MW396979.1_Timor-Leste/2019/1    ................................................................................ [ 93840]
MK940252.1_CN/2019/InnerMongolia ................................................................................ [ 93840]
MK128995.1_China/2018/AnhuiXCGQ  ................................................................................ [ 93840]
MH681419.1_ASFV/POL/2015         ................................................................................ [ 93840]
LS478113.1_Estonia_2014          ................................................................................ [ 93840]
KP843857.1_Odintsovo_02/14       ................................................................................ [ 93840]
FN557520.1_strain_E75            .............g........g......................................................... [ 93840]
AY261366.1_Warthog               .............g....................a............................................. [ 93840]
AY261365.1_Warmbaths             .............g.................................................................. [ 93840]
AY261364.1_Tengani_62            .............g........g......................................at................. [ 93840]
AY261363.1_Pretoriuskop/96/4     .............g.................................................................. [ 93840]
AY261362.1_Mkuzi_1979            .............g.................................................................. [ 93840]
AY261361.1_Malawi_Lil-20/1_1983  .............gc....g.tg...........a.....c.....................t................. [ 93840]
AY261360.1_Kenya_1950            ..............c......tg.......................................t................. [ 93840]
AM712240.1_OURT_88/3             .............g........g......................................................... [ 93840]
AM712239.1_Benin_97/1            .............g........g......................................................... [ 93840]
KM111295.1_Ken06.Bus             .............gc.......g.......................................t................. [ 93840]
KM111294.1_Ken05/Tk1             ..............c.......g.......................................t................. [ 93840]
KM262844.1_strain_L60            .............g........g......................................................... [ 93840]
KM102979.1_26544/OG10_Italy      .............g.......tg......................................................... [ 93840]
KM262845.1_strain_NHV            .............g........g......................................................... [ 93840]
KX354450.1_47/Ss/2008            .............g.......tg......................................................... [ 93840]
KP055815.1_strain_BA71           .............g........g......................................................... [ 93840]
MH025920.1_strain_R35            .............gc.......g.......................................t................. [ 93840]
MH025919.1_strain_N10            .............gc.......g.......................................t................. [ 93840]
MH025918.1_strain_R25            .............gc.......g.......................................t................. [ 93840]
MH025917.1_strain_R7             .............gc.......g.......................................t................. [ 93840]
MH025916.1_strain_R8             .............gc.......g.......................................t................. [ 93840]

FR682468.2_Georgia_2007/1        acatgcctagccattcctccgcctgcgccctttcaacaggcgacccgccaggtccaacacttttaccgtaacaatttcta [ 93920]
ASF_IND_20_CAD543                .....y.......................................................................... [ 93920]
LR899193.1_Germany_2020/1        ................................................................................ [ 93920]
MN393477.1_Wuhan_2019            ................................................................................ [ 93920]
MW396979.1_Timor-Leste/2019/1    ................................................................................ [ 93920]
MK940252.1_CN/2019/InnerMongolia ................................................................................ [ 93920]
MK128995.1_China/2018/AnhuiXCGQ  ................................................................................ [ 93920]
MH681419.1_ASFV/POL/2015         ................................................................................ [ 93920]
LS478113.1_Estonia_2014          ................................................................................ [ 93920]
KP843857.1_Odintsovo_02/14       ................................................................................ [ 93920]
```

FN557520.1_strain_E75 .....t.........................................t..a............................. [ 93920]
AY261366.1_Warthog ................................................................................ [ 93920]
AY261365.1_Warmbaths ................................................................................ [ 93920]
AY261364.1_Tengani_62 ............................................................c................... [ 93920]
AY261363.1_Pretoriuskop/96/4 ................................................................................ [ 93920]
AY261362.1_Mkuzi_1979 ................................................................................ [ 93920]
AY261361.1_Malawi_Lil-20/1_1983 ........g...g.............a.............t...t.....aa.........a......g..t..c..t.. [ 93920]
AY261360.1_Kenya_1950 ........g...g.......a...................t...t.....a..........ac...........a..... [ 93920]
AM712240.1_OURT_88/3 .....t.........................................t..a............................. [ 93920]
AM712239.1_Benin_97/1 .....t.........................................t..a............................. [ 93920]
KM111295.1_Ken06.Bus ........g...g.......a...................t...t.....a..g.......ac...........a..... [ 93920]
KM111294.1_Ken05/Tk1 ........g...........a...................t...t.....a..........a.........t..c..... [ 93920]
KM262844.1_strain_L60 .....t.........................................t..a............................. [ 93920]
KM102979.1_26544/OG10_Italy .....t.........................................t..a............................. [ 93920]
KM262845.1_strain_NHV .....t.........................................t..a............................. [ 93920]
KX354450.1_47/Ss/2008 .....t.........................................t..a............................. [ 93920]
KP055815.1_strain_BA71 .....t.........................................t..a............................. [ 93920]
MH025920.1_strain_R35 ........g...g.......a...................t...t.....a..g.......ac...........a..... [ 93920]
MH025919.1_strain_N10 ........g...g.......a...................t...t.....a..g.......ac...........a..... [ 93920]
MH025918.1_strain_R25 ........g...g.......a...................t...t.....a..g.......ac...........a..... [ 93920]
MH025917.1_strain_R7 ........g...g.......a...................t...t.....a..g.......ac...........a..... [ 93920]
MH025916.1_strain_R8 ........g...g.......a...................t...t.....a..g.......ac...........a..... [ 93920]

FR682468.2_Georgia_2007/1 cgaagtggttcccatccaggatggcacccttctcacaatctaccactgggatgaccctgaatatggcccctcctggtgcc [ 94000]
ASF_IND_20_CAD543 ...........................................y................gc.................. [ 94000]
LR899193.1_Germany_2020/1 ................................................................................ [ 94000]
MN393477.1_Wuhan_2019 ................................................................................ [ 94000]
MW396979.1_Timor-Leste/2019/1 ................................................................................ [ 94000]
MK940252.1_CN/2019/InnerMongolia ................................................................................ [ 94000]
MK128995.1_China/2018/AnhuiXCGQ ................................................................................ [ 94000]
MH681419.1_ASFV/POL/2015 ................................................................................ [ 94000]
LS478113.1_Estonia_2014 ................................................................................ [ 94000]
KP843857.1_Odintsovo_02/14 ................................................................................ [ 94000]
FN557520.1_strain_E75 ...........................................t.............c..gc.................. [ 94000]
AY261366.1_Warthog ...........................................t.............c...c.................. [ 94000]
AY261365.1_Warmbaths ...........................................t.................c..........t....... [ 94000]
AY261364.1_Tengani_62 ...........................................t.............c...c.................. [ 94000]
AY261363.1_Pretoriuskop/96/4 ...........................................t.............c...c.................. [ 94000]
AY261362.1_Mkuzi_1979 ...........................................t.............c..gc.................. [ 94000]
AY261361.1_Malawi_Lil-20/1_1983 t..........................t...............t................gc.................t [ 94000]
AY261360.1_Kenya_1950 ........c.g................t...............t..........t.....gc.................. [ 94000]
AM712240.1_OURT_88/3 ...........................................t.............c..gc.................. [ 94000]
AM712239.1_Benin_97/1 ...........................................t.............c..gc.................. [ 94000]
KM111295.1_Ken06.Bus ........c.g....t...........t...............t..........t......c.................. [ 94000]

```
KM111294.1_Ken05/Tk1             t.....a........t...........................t................gc..........t......t [ 94000]
KM262844.1_strain_L60            ...........................................t.............c..gc.................. [ 94000]
KM102979.1_26544/OG10_Italy      ...........................................t.............c..gc.................. [ 94000]
KM262845.1_strain_NHV            ...........................................t.............c..gc.................. [ 94000]
KX354450.1_47/Ss/2008            ...........................................t.............c..gc.................. [ 94000]
KP055815.1_strain_BA71           ...........................................t.............c..gc.................. [ 94000]
MH025920.1_strain_R35            ........c.g....t...........t...............t..........t......c.................. [ 94000]
MH025919.1_strain_N10            ........c.g....t...........t...............t..........t......c.................. [ 94000]
MH025918.1_strain_R25            ........c.g....t...........t...............t..........t......c.................. [ 94000]
MH025917.1_strain_R7             ........c.g....t...........t...............t..........t......c.................. [ 94000]
MH025916.1_strain_R8             ........c.g....t...........t...............t..........t......c.................. [ 94000]

FR682468.2_Georgia_2007/1        tagcaagtacccacggatatgatgtgagtaactactgttggataggcgacaaaaccttcgccgagcttgtatacgaattg [ 94080]
ASF_IND_20_CAD543                ................................................................................ [ 94080]
LR899193.1_Germany_2020/1        ................................................................................ [ 94080]
MN393477.1_Wuhan_2019            ................................................................................ [ 94080]
MW396979.1_Timor-Leste/2019/1    ................................................................................ [ 94080]
MK940252.1_CN/2019/InnerMongolia ................................................................................ [ 94080]
MK128995.1_China/2018/AnhuiXCGQ  ................................................................................ [ 94080]
MH681419.1_ASFV/POL/2015         ................................................................................ [ 94080]
LS478113.1_Estonia_2014          ................................................................................ [ 94080]
KP843857.1_Odintsovo_02/14       ................................................................................ [ 94080]
FN557520.1_strain_E75            ................................................................................ [ 94080]
AY261366.1_Warthog               .............t..............c................................................... [ 94080]
AY261365.1_Warmbaths             ............................c................................................... [ 94080]
AY261364.1_Tengani_62            ..........t..................................................................... [ 94080]
AY261363.1_Pretoriuskop/96/4     .............t.................................................................. [ 94080]
AY261362.1_Mkuzi_1979            ............................c....................t.............................. [ 94080]
AY261361.1_Malawi_Lil-20/1_1983  .............t..............c................................................... [ 94080]
AY261360.1_Kenya_1950            .......c.....t...........a..c.............................t..............t...... [ 94080]
AM712240.1_OURT_88/3             ................................................................................ [ 94080]
AM712239.1_Benin_97/1            ................................................................................ [ 94080]
KM111295.1_Ken06.Bus             .......c.....t...........a..c............................................t...... [ 94080]
KM111294.1_Ken05/Tk1             ............................c..t................................................ [ 94080]
KM262844.1_strain_L60            ................................................................................ [ 94080]
KM102979.1_26544/OG10_Italy      ................................................................................ [ 94080]
KM262845.1_strain_NHV            ................................................................................ [ 94080]
KX354450.1_47/Ss/2008            ................................................................................ [ 94080]
KP055815.1_strain_BA71           ................................................................................ [ 94080]
MH025920.1_strain_R35            .......c.....t...........a..c............................................t...... [ 94080]
MH025919.1_strain_N10            .......c.....t...........a..c............................................t...... [ 94080]
MH025918.1_strain_R25            .......c.....t...........a..c............................................t...... [ 94080]
MH025917.1_strain_R7             .......c.....t...........a..c............................................t...... [ 94080]
MH025916.1_strain_R8             .......c.....t...........a..c............................................t...... [ 94080]
```

```
FR682468.2_Georgia_2007/1        ctgcagcagcactctacctgcgacgtcaccctggaaaaaaataaaacgcggggaacgcgtcttttctttgataacttaaa [ 94160]
ASF_IND_20_CAD543                ................................................................................ [ 94160]
LR899193.1_Germany_2020/1        ................................................................................ [ 94160]
MN393477.1_Wuhan_2019            ................................................................................ [ 94160]
MW396979.1_Timor-Leste/2019/1    ................................................................................ [ 94160]
MK940252.1_CN/2019/InnerMongolia ................................................................................ [ 94160]
MK128995.1_China/2018/AnhuiXCGQ  ................................................................................ [ 94160]
MH681419.1_ASFV/POL/2015         ................................................................................ [ 94160]
LS478113.1_Estonia_2014          ................................................................................ [ 94160]
KP843857.1_Odintsovo_02/14       ................................................................................ [ 94160]
FN557520.1_strain_E75            ..a.....................................................a............a.c........ [ 94160]
AY261366.1_Warthog               t....................................................................a.c........ [ 94160]
AY261365.1_Warmbaths             .....................................................................a.c........ [ 94160]
AY261364.1_Tengani_62            ..................................................a........g.........a.c........ [ 94160]
AY261363.1_Pretoriuskop/96/4     t..........t.........................................................a.c........ [ 94160]
AY261362.1_Mkuzi_1979            .....................................................................a.c........ [ 94160]
AY261361.1_Malawi_Lil-20/1_1983  ..............c..........................ac....a...........g.........a.c........ [ 94160]
AY261360.1_Kenya_1950            ..............c......a...................c.................g.........a.cg.g..... [ 94160]
AM712240.1_OURT_88/3             ..a.....................................................a............a.c........ [ 94160]
AM712239.1_Benin_97/1            ..a.....................................................a............a.c........ [ 94160]
KM111295.1_Ken06.Bus             ..............c......a...................c.................g.........a.cg.g..... [ 94160]
KM111294.1_Ken05/Tk1             ..............c...........g..t....................a........g.........a.c........ [ 94160]
KM262844.1_strain_L60            ..a.....................................................a............a.c........ [ 94160]
KM102979.1_26544/OG10_Italy      ..a.....................................................a............a.c........ [ 94160]
KM262845.1_strain_NHV            ..a.....................................................a............a.c........ [ 94160]
KX354450.1_47/Ss/2008            ..a.....................................................a............a.c........ [ 94160]
KP055815.1_strain_BA71           ..a.....................................................a............a.c........ [ 94160]
MH025920.1_strain_R35            ..............c......a...................c.................g.........a.cg.g..... [ 94160]
MH025919.1_strain_N10            ..............c......a...................c.................g.........a.cg.g..... [ 94160]
MH025918.1_strain_R25            ..............c......a...................c.................g.........a.cg.g..... [ 94160]
MH025917.1_strain_R7             ..............c......a...................c.................g.........a.cg.g..... [ 94160]
MH025916.1_strain_R8             ..............c......a...................c.................g.........a.cg.g..... [ 94160]

FR682468.2_Georgia_2007/1        tcccgattactgctatacgattggaatccggcaccataatttacagccgctcatctatgaccctcaaaatatttgggcga [ 94240]
ASF_IND_20_CAD543                ................................................................................ [ 94240]
LR899193.1_Germany_2020/1        ................................................................................ [ 94240]
MN393477.1_Wuhan_2019            ................................................................................ [ 94240]
MW396979.1_Timor-Leste/2019/1    ................................................................................ [ 94240]
MK940252.1_CN/2019/InnerMongolia ................................................................................ [ 94240]
MK128995.1_China/2018/AnhuiXCGQ  ................................................................................ [ 94240]
MH681419.1_ASFV/POL/2015         ................................................................................ [ 94240]
LS478113.1_Estonia_2014          ................................................................................ [ 94240]
KP843857.1_Odintsovo_02/14       ................................................................................ [ 94240]
```

```
FN557520.1_strain_E75          ...........................t.......................t............................ [ 94240]
AY261366.1_Warthog             ...........................t...................................a................ [ 94240]
AY261365.1_Warmbaths           ...........................t...................................a................ [ 94240]
AY261364.1_Tengani_62          ......c....................t.................................................... [ 94240]
AY261363.1_Pretoriuskop/96/4   ...........................t...................................a................ [ 94240]
AY261362.1_Mkuzi_1979          ...........................t.......................t............................ [ 94240]
AY261361.1_Malawi_Lil-20/1_1983 ..gt..c.......................c........cc....a..a........c...................... [ 94240]
AY261360.1_Kenya_1950          ..gt........t.................c........cc....a..a..t...c....t................... [ 94240]
AM712240.1_OURT_88/3           ...........................t.......................t............................ [ 94240]
AM712239.1_Benin_97/1          ...........................t.......................t............................ [ 94240]
KM111295.1_Ken06.Bus           ..gt........t.................c........c.....a..a..t...c........................ [ 94240]
KM111294.1_Ken05/Tk1           ..gt..c.....t..............t..c..t.....cc....a..c............................... [ 94240]
KM262844.1_strain_L60          ...........................t.......................t............................ [ 94240]
KM102979.1_26544/OG10_Italy    ...........................t.......................t............................ [ 94240]
KM262845.1_strain_NHV          ...........................t.......................t............................ [ 94240]
KX354450.1_47/Ss/2008          ...........................t.......................t............................ [ 94240]
KP055815.1_strain_BA71         ...........................t.......................t............................ [ 94240]
MH025920.1_strain_R35          ..gt........t.................c........c.....a..a..t...c........................ [ 94240]
MH025919.1_strain_N10          ..gt........t.................c........c.....a..a..t...c........................ [ 94240]
MH025918.1_strain_R25          ..gt........t.................c........c.....a..a..t...c........................ [ 94240]
MH025917.1_strain_R7           ..gt........t.................c........c.....a..a..t...c........................ [ 94240]
MH025916.1_strain_R8           ..gt........t.................c........c.....a..a..t...c........................ [ 94240]

FR682468.2_Georgia_2007/1      ttcaatctacaaacctaaaaacgcttaaaacggtatatccagaatactacggctatataggcattccaggaattcagagt [ 94320]
ASF_IND_20_CAD543              ................................................................................ [ 94320]
LR899193.1_Germany_2020/1      ................................................................................ [ 94320]
MN393477.1_Wuhan_2019          ................................................................................ [ 94320]
MW396979.1_Timor-Leste/2019/1  ................................................................................ [ 94320]
MK940252.1_CN/2019/InnerMongolia ................................................................................ [ 94320]
MK128995.1_China/2018/AnhuiXCGQ ................................................................................ [ 94320]
MH681419.1_ASFV/POL/2015       ................................................................................ [ 94320]
LS478113.1_Estonia_2014        ................................................................................ [ 94320]
KP843857.1_Odintsovo_02/14     ................................................................................ [ 94320]
FN557520.1_strain_E75          ................................................................................ [ 94320]
AY261366.1_Warthog             ..................................g............................................. [ 94320]
AY261365.1_Warmbaths           ..................................g...........t................................. [ 94320]
AY261364.1_Tengani_62          ..................................g............................................. [ 94320]
AY261363.1_Pretoriuskop/96/4   ..................................g............................................. [ 94320]
AY261362.1_Mkuzi_1979          ................................................................................ [ 94320]
AY261361.1_Malawi_Lil-20/1_1983 .....a.......tt...................g............c........g..........g........a... [ 94320]
AY261360.1_Kenya_1950          ....g..g.....tt...................g.....................g....................... [ 94320]
AM712240.1_OURT_88/3           ................................................................................ [ 94320]
AM712239.1_Benin_97/1          ................................................................................ [ 94320]
KM111295.1_Ken06.Bus           ....g..g.....tt...................g.....................g....................... [ 94320]
```

```
KM111294.1_Ken05/Tk1               .....a.c.....tt...................g............................................. [ 94320]
KM262844.1_strain_L60              ................................................................................ [ 94320]
KM102979.1_26544/OG10_Italy        ................................................................................ [ 94320]
KM262845.1_strain_NHV              ................................................................................ [ 94320]
KX354450.1_47/Ss/2008              ................................................................................ [ 94320]
KP055815.1_strain_BA71             ................................................................................ [ 94320]
MH025920.1_strain_R35              ....g..g.....tt...................g.....................g....................... [ 94320]
MH025919.1_strain_N10              ....g..g.....tt...................g.....................g....................... [ 94320]
MH025918.1_strain_R25              ....g..g.....tt...................g.....................g....................... [ 94320]
MH025917.1_strain_R7               ....g..g.....tt...................g.....................g....................... [ 94320]
MH025916.1_strain_R8               ....g..g.....tt...................g.....................g....................... [ 94320]

FR682468.2_Georgia_2007/1          caagttcctgagcttccccagtatgatttaccttatctaatacgatcttataaaactgctatgaatcaagccaaaaatgc [ 94400]
ASF_IND_20_CAD543                  ................................................................................ [ 94400]
LR899193.1_Germany_2020/1          ................................................................................ [ 94400]
MN393477.1_Wuhan_2019              ................................................................................ [ 94400]
MW396979.1_Timor-Leste/2019/1      ................................................................................ [ 94400]
MK940252.1_CN/2019/InnerMongolia   ................................................................................ [ 94400]
MK128995.1_China/2018/AnhuiXCGQ    ................................................................................ [ 94400]
MH681419.1_ASFV/POL/2015           ................................................................................ [ 94400]
LS478113.1_Estonia_2014            ................................................................................ [ 94400]
KP843857.1_Odintsovo_02/14         ................................................................................ [ 94400]
FN557520.1_strain_E75              ...............................................c.....g.......................... [ 94400]
AY261366.1_Warthog                 .....................................................g.....c.................... [ 94400]
AY261365.1_Warmbaths               ...........................................................c.................c.. [ 94400]
AY261364.1_Tengani_62              ...............................................a.....g.......................... [ 94400]
AY261363.1_Pretoriuskop/96/4       .....................................................g.....c.................... [ 94400]
AY261362.1_Mkuzi_1979              ...............................................c................................ [ 94400]
AY261361.1_Malawi_Lil-20/1_1983    ........c.................a....................c...........c.................... [ 94400]
AY261360.1_Kenya_1950              ........c.............t...ac..............a....c....g......c.................... [ 94400]
AM712240.1_OURT_88/3               ...............................................c.....g.......................... [ 94400]
AM712239.1_Benin_97/1              ...............................................c.....g.......................... [ 94400]
KM111295.1_Ken06.Bus               ........c.............t....c..............a....c....g...c....................... [ 94400]
KM111294.1_Ken05/Tk1               ........c.................a............c.......c...........c.................... [ 94400]
KM262844.1_strain_L60              ...............................................c.....g.......................... [ 94400]
KM102979.1_26544/OG10_Italy        ...............................................c.....g.......................... [ 94400]
KM262845.1_strain_NHV              ...............................................c.....g.......................... [ 94400]
KX354450.1_47/Ss/2008              ...............................................c.....g.......................... [ 94400]
KP055815.1_strain_BA71             ...............................................c.....g.......................... [ 94400]
MH025920.1_strain_R35              ........c.............t....c..............a....c....g...c....................... [ 94400]
MH025919.1_strain_N10              ........c.............t....c..............a....c....g...c....................... [ 94400]
MH025918.1_strain_R25              ........c.............t....c..............a....c....g...c....................... [ 94400]
MH025917.1_strain_R7               ........c.............t....c..............a....c....g...c....................... [ 94400]
MH025916.1_strain_R8               ........c.............t....c..............a....c....g...c....................... [ 94400]
```

```
FR682468.2_Georgia_2007/1        tataaaaaatggcaaaaaagacaagggatactttaattatggctatttactcatttcgcgagcgcctgccattactaaaa [ 94480]
ASF_IND_20_CAD543                ................................................................................ [ 94480]
LR899193.1_Germany_2020/1        ................................................................................ [ 94480]
MN393477.1_Wuhan_2019            ................................................................................ [ 94480]
MW396979.1_Timor-Leste/2019/1    ................................................................................ [ 94480]
MK940252.1_CN/2019/InnerMongolia ................................................................................ [ 94480]
MK128995.1_China/2018/AnhuiXCGQ  ................................................................................ [ 94480]
MH681419.1_ASFV/POL/2015         ................................................................................ [ 94480]
LS478113.1_Estonia_2014          ................................................................................ [ 94480]
KP843857.1_Odintsovo_02/14       ................................................................................ [ 94480]
FN557520.1_strain_E75            ..........................a..................................................... [ 94480]
AY261366.1_Warthog               .......................................c........................................ [ 94480]
AY261365.1_Warmbaths             ..................g....................c........................................ [ 94480]
AY261364.1_Tengani_62            ..................g....................c........................................ [ 94480]
AY261363.1_Pretoriuskop/96/4     .......................................c........................................ [ 94480]
AY261362.1_Mkuzi_1979            ..........................a..................................................... [ 94480]
AY261361.1_Malawi_Lil-20/1_1983  ..................g....................c............g.............a.....c....... [ 94480]
AY261360.1_Kenya_1950            ..................g..g....ag...........c.......................a................ [ 94480]
AM712240.1_OURT_88/3             ..........................a..................................................... [ 94480]
AM712239.1_Benin_97/1            ..........................a..................................................... [ 94480]
KM111295.1_Ken06.Bus             ..................g..g....ag...........c........................................ [ 94480]
KM111294.1_Ken05/Tk1             ..................g....................c.......................a................ [ 94480]
KM262844.1_strain_L60            ..........................a..................................................... [ 94480]
KM102979.1_26544/OG10_Italy      ..........................a..................................................... [ 94480]
KM262845.1_strain_NHV            ..........................a..................................................... [ 94480]
KX354450.1_47/Ss/2008            ..........................a..................................................... [ 94480]
KP055815.1_strain_BA71           ..........................a..................................................... [ 94480]
MH025920.1_strain_R35            ..................g..g....ag...........c........................................ [ 94480]
MH025919.1_strain_N10            ..................g..g....ag...........c........................................ [ 94480]
MH025918.1_strain_R25            ..................g..g....ag...........c........................................ [ 94480]
MH025917.1_strain_R7             ..................g..g....ag...........c........................................ [ 94480]
MH025916.1_strain_R8             ..................g..g....ag...........c........................................ [ 94480]

FR682468.2_Georgia_2007/1        gtacttctaatgttttgttaaaatcgcctctgctggtatttttacaaaaaagtgtgtaccagaaaaaacacaatatctct [ 94560]
ASF_IND_20_CAD543                ................................................................................ [ 94560]
LR899193.1_Germany_2020/1        ................................................................................ [ 94560]
MN393477.1_Wuhan_2019            ................................................................................ [ 94560]
MW396979.1_Timor-Leste/2019/1    ................................................................................ [ 94560]
MK940252.1_CN/2019/InnerMongolia ................................................................................ [ 94560]
MK128995.1_China/2018/AnhuiXCGQ  ................................................................................ [ 94560]
MH681419.1_ASFV/POL/2015         ................................................................................ [ 94560]
LS478113.1_Estonia_2014          ................................................................................ [ 94560]
KP843857.1_Odintsovo_02/14       ................................................................................ [ 94560]
```

```
FN557520.1_strain_E75            ................................................................................ [ 94560]
AY261366.1_Warthog               ................................................................................ [ 94560]
AY261365.1_Warmbaths             ..............c................................................................. [ 94560]
AY261364.1_Tengani_62            ................................t....g.......................................... [ 94560]
AY261363.1_Pretoriuskop/96/4     ................................................................................ [ 94560]
AY261362.1_Mkuzi_1979            ................................................................................ [ 94560]
AY261361.1_Malawi_Lil-20/1_1983  ...t...c.c...........................g.............................g......c..a.. [ 94560]
AY261360.1_Kenya_1950            ...t...c.t...............a..ct.......g.............................gt........... [ 94560]
AM712240.1_OURT_88/3             ................................................................................ [ 94560]
AM712239.1_Benin_97/1            ................................................................................ [ 94560]
KM111295.1_Ken06.Bus             ...t...c.t...............a...t.......g.............................gt........... [ 94560]
KM111294.1_Ken05/Tk1             ...t...c.t...............a..ct.......g.............................gt........... [ 94560]
KM262844.1_strain_L60            ................................................................................ [ 94560]
KM102979.1_26544/OG10_Italy      ................................................................................ [ 94560]
KM262845.1_strain_NHV            ................................................................................ [ 94560]
KX354450.1_47/Ss/2008            ................................................................................ [ 94560]
KP055815.1_strain_BA71           ................................................................................ [ 94560]
MH025920.1_strain_R35            ...t...c.t...............a...t.......g.............................gt........... [ 94560]
MH025919.1_strain_N10            ...t...c.t...............a...t.......g.............................gt........... [ 94560]
MH025918.1_strain_R25            ...t...c.t...............a...t.......g.............................gt........... [ 94560]
MH025917.1_strain_R7             ...t...c.t...............a...t.......g.............................gt........... [ 94560]
MH025916.1_strain_R8             ...t...c.t...............a...t.......g.............................gt........... [ 94560]

FR682468.2_Georgia_2007/1        aacagccagcgactagaatttattatactgcaaaactacttgatgcagcattttcgagatcatttcattgctctatttcc [ 94640]
ASF_IND_20_CAD543                .....................................................................nnnnnnnnnn. [ 94640]
LR899193.1_Germany_2020/1        ................................................................................ [ 94640]
MN393477.1_Wuhan_2019            ................................................................................ [ 94640]
MW396979.1_Timor-Leste/2019/1    ................................................................................ [ 94640]
MK940252.1_CN/2019/InnerMongolia ................................................................................ [ 94640]
MK128995.1_China/2018/AnhuiXCGQ  ................................................................................ [ 94640]
MH681419.1_ASFV/POL/2015         ................................................................................ [ 94640]
LS478113.1_Estonia_2014          ................................................................................ [ 94640]
KP843857.1_Odintsovo_02/14       ................................................................................ [ 94640]
FN557520.1_strain_E75            ...........g.................................................................... [ 94640]
AY261366.1_Warthog               .................................................................t.............. [ 94640]
AY261365.1_Warmbaths             .....t.......................................................................... [ 94640]
AY261364.1_Tengani_62            .................g.............................................................. [ 94640]
AY261363.1_Pretoriuskop/96/4     ................................................................................ [ 94640]
AY261362.1_Mkuzi_1979            ................................................................................ [ 94640]
AY261361.1_Malawi_Lil-20/1_1983  .g.....t.........g..........................................a.c................. [ 94640]
AY261360.1_Kenya_1950            .g.....t.........g...........a..............................a.c................. [ 94640]
AM712240.1_OURT_88/3             ...........g.................................................................... [ 94640]
AM712239.1_Benin_97/1            ...........g.................................................................... [ 94640]
KM111295.1_Ken06.Bus             .g.....t.........g...........a.................a............a.c................. [ 94640]
```

```
KM111294.1_Ken05/Tk1            .g.....t.........g...........a..............................a.c................. [ 94640]
KM262844.1_strain_L60           ...........g.................................................................... [ 94640]
KM102979.1_26544/OG10_Italy     ...........g.................................................................... [ 94640]
KM262845.1_strain_NHV           ...........g.................................................................... [ 94640]
KX354450.1_47/Ss/2008           ...........g.................................................................... [ 94640]
KP055815.1_strain_BA71          ...........g.................................................................... [ 94640]
MH025920.1_strain_R35           .g.....t.........g...........a.................a............a.c................. [ 94640]
MH025919.1_strain_N10           .g.....t.........g...........a.................a............a.c................. [ 94640]
MH025918.1_strain_R25           .g.....t.........g...........a.................a............a.c................. [ 94640]
MH025917.1_strain_R7            .g.....t.........g...........a.................a............a.c................. [ 94640]
MH025916.1_strain_R8            .g.....t.........g...........a.................a............a.c................. [ 94640]

FR682468.2_Georgia_2007/1       gcagtacatatcctattatacgaaataccaaaacatgttgaatatgattatccatagtattgcaactaaagataaagatc [ 94720]
ASF_IND_20_CAD543               ..................................................................c............. [ 94720]
LR899193.1_Germany_2020/1       ................................................................................ [ 94720]
MN393477.1_Wuhan_2019           ................................................................................ [ 94720]
MW396979.1_Timor-Leste/2019/1   ................................................................................ [ 94720]
MK940252.1_CN/2019/InnerMongolia ................................................................................ [ 94720]
MK128995.1_China/2018/AnhuiXCGQ ................................................................................ [ 94720]
MH681419.1_ASFV/POL/2015        ................................................................................ [ 94720]
LS478113.1_Estonia_2014         ................................................................................ [ 94720]
KP843857.1_Odintsovo_02/14      ................................................................................ [ 94720]
FN557520.1_strain_E75           ..................................................................c............. [ 94720]
AY261366.1_Warthog              ....................................................a.............c............. [ 94720]
AY261365.1_Warmbaths            ...................................................ta.............c............. [ 94720]
AY261364.1_Tengani_62           ..........................................c.......................c............. [ 94720]
AY261363.1_Pretoriuskop/96/4    ....................................................a.............c............. [ 94720]
AY261362.1_Mkuzi_1979           ..................................................................c............. [ 94720]
AY261361.1_Malawi_Lil-20/1_1983 a..................................................t.............ta.......t..... [ 94720]
AY261360.1_Kenya_1950           ....................t..............................t.............ta..g.......... [ 94720]
AM712240.1_OURT_88/3            ..................................................................c............. [ 94720]
AM712239.1_Benin_97/1           ..................................................................c............. [ 94720]
KM111295.1_Ken06.Bus            .........g..........t..............................t.............ta..g.......... [ 94720]
KM111294.1_Ken05/Tk1            .........g..........t..............................t.............ta..g.......... [ 94720]
KM262844.1_strain_L60           ..................................................................c............. [ 94720]
KM102979.1_26544/OG10_Italy     ..................................................................c............. [ 94720]
KM262845.1_strain_NHV           ..................................................................c............. [ 94720]
KX354450.1_47/Ss/2008           ..................................................................c............. [ 94720]
KP055815.1_strain_BA71          ..................................................................c............. [ 94720]
MH025920.1_strain_R35           .........g..........t..............................t.............ta..g.......... [ 94720]
MH025919.1_strain_N10           .........g..........t..............................t.............ta..g.......... [ 94720]
MH025918.1_strain_R25           .........g..........t..............................t.............ta..g.......... [ 94720]
MH025917.1_strain_R7            .........g..........t..............................t.............ta..g.......... [ 94720]
MH025916.1_strain_R8            .........g..........t..............................t.............ta..g.......... [ 94720]
```

```
FR682468.2_Georgia_2007/1        atccctttgcaggagccgtggtaaaaaaagtgttggaagatattgaaaacgccgaaaacattattgatcatacaaccatt [ 94800]
ASF_IND_20_CAD543                ................................................................................ [ 94800]
LR899193.1_Germany_2020/1        ................................................................................ [ 94800]
MN393477.1_Wuhan_2019            ................................................................................ [ 94800]
MW396979.1_Timor-Leste/2019/1    ................................................................................ [ 94800]
MK940252.1_CN/2019/InnerMongolia ................................................................................ [ 94800]
MK128995.1_China/2018/AnhuiXCGQ  ................................................................................ [ 94800]
MH681419.1_ASFV/POL/2015         ................................................................................ [ 94800]
LS478113.1_Estonia_2014          ................................................................................ [ 94800]
KP843857.1_Odintsovo_02/14       ................................................................................ [ 94800]
FN557520.1_strain_E75            ...............................a................................................ [ 94800]
AY261366.1_Warthog               ................................c.........................t..................... [ 94800]
AY261365.1_Warmbaths             ................................c.........................t.................t... [ 94800]
AY261364.1_Tengani_62            ................t...............c...................t.....t.................t... [ 94800]
AY261363.1_Pretoriuskop/96/4     ................................c.........................t..................... [ 94800]
AY261362.1_Mkuzi_1979            ................................c.........................t.................t... [ 94800]
AY261361.1_Malawi_Lil-20/1_1983  ....t.........a................a........c....................................... [ 94800]
AY261360.1_Kenya_1950            ............................g....................t.................c........t... [ 94800]
AM712240.1_OURT_88/3             ...............................a................................................ [ 94800]
AM712239.1_Benin_97/1            ...............................a................................................ [ 94800]
KM111295.1_Ken06.Bus             ............................g...........c....................................... [ 94800]
KM111294.1_Ken05/Tk1             ............................g....................t..t........................... [ 94800]
KM262844.1_strain_L60            ...............................a................................................ [ 94800]
KM102979.1_26544/OG10_Italy      ...............................a................................................ [ 94800]
KM262845.1_strain_NHV            ...............................a................................................ [ 94800]
KX354450.1_47/Ss/2008            ...............................a................................................ [ 94800]
KP055815.1_strain_BA71           ...............................a................................................ [ 94800]
MH025920.1_strain_R35            ............................g...........c........t.............................. [ 94800]
MH025919.1_strain_N10            ............................g...........c........t.............................. [ 94800]
MH025918.1_strain_R25            ............................g...........c........t.............................. [ 94800]
MH025917.1_strain_R7             ............................g...........c........t.............................. [ 94800]
MH025916.1_strain_R8             ............................g...........c........t.............................. [ 94800]

FR682468.2_Georgia_2007/1        caaaactatgcccatcaaagcaagtacgccatgctttacttgtcaattatttcccattttttaatctaatacggccaaagc [ 94880]
ASF_IND_20_CAD543                ................................................................s............... [ 94880]
LR899193.1_Germany_2020/1        ................................................................................ [ 94880]
MN393477.1_Wuhan_2019            ................................................................................ [ 94880]
MW396979.1_Timor-Leste/2019/1    ................................................................................ [ 94880]
MK940252.1_CN/2019/InnerMongolia ................................................................................ [ 94880]
MK128995.1_China/2018/AnhuiXCGQ  ................................................................................ [ 94880]
MH681419.1_ASFV/POL/2015         ................................................................................ [ 94880]
LS478113.1_Estonia_2014          ................................................................................ [ 94880]
KP843857.1_Odintsovo_02/14       ................................................................................ [ 94880]
```

```
FN557520.1_strain_E75          .............................t............................................t..... [ 94880]
AY261366.1_Warthog             ..........t..................................................................... [ 94880]
AY261365.1_Warmbaths           ..........t...............................................................t..... [ 94880]
AY261364.1_Tengani_62          ..........t....................................................................t [ 94880]
AY261363.1_Pretoriuskop/96/4   ..........t..................................................................... [ 94880]
AY261362.1_Mkuzi_1979          ..........t...............................................................t..... [ 94880]
AY261361.1_Malawi_Lil-20/1_1983 ..........t...............t..............a.......................c...------..... [ 94880]
AY261360.1_Kenya_1950          ...........gt.c.....t..................c...........a............gc....-----..gc. [ 94880]
AM712240.1_OURT_88/3           .............................t............................................t..... [ 94880]
AM712239.1_Benin_97/1          .............................t............................................t..... [ 94880]
KM111295.1_Ken06.Bus           ...........gt.c....................................a.............c....-----..gt. [ 94880]
KM111294.1_Ken05/Tk1           ...........gt.c.....t..................c...........a............gc....-----...c. [ 94880]
KM262844.1_strain_L60          .............................t............................................t..... [ 94880]
KM102979.1_26544/OG10_Italy    .............................t............................................t..... [ 94880]
KM262845.1_strain_NHV          .............................t............................................t..... [ 94880]
KX354450.1_47/Ss/2008          .............................t............................................t..... [ 94880]
KP055815.1_strain_BA71         .............................t............................................t..... [ 94880]
MH025920.1_strain_R35          ...........gt.c....................................a.............c....-----..gt. [ 94880]
MH025919.1_strain_N10          ...........gt.c....................................a.............c....-----..gt. [ 94880]
MH025918.1_strain_R25          ...........gt.c....................................a.............c....-----..gt. [ 94880]
MH025917.1_strain_R7           ...........gt.c....................................a.............c....-----..gt. [ 94880]
MH025916.1_strain_R8           ...........gt.c....................................a.............c....-----..gt. [ 94880]

FR682468.2_Georgia_2007/1      cgcgggttttttaataaactaacatttaaaaaaactgttttattaaaaattataatacttttat--tatatatggaacat [ 94960]
ASF_IND_20_CAD543              y.....-..........................g-.............................ta.............. [ 94960]
LR899193.1_Germany_2020/1      ................................................................--.............. [ 94960]
MN393477.1_Wuhan_2019          ................................................................--.............. [ 94960]
MW396979.1_Timor-Leste/2019/1  ................................................................--.............. [ 94960]
MK940252.1_CN/2019/InnerMongolia ................................................................--.............. [ 94960]
MK128995.1_China/2018/AnhuiXCGQ ................................................................--.............. [ 94960]
MH681419.1_ASFV/POL/2015       ................................................................--.............. [ 94960]
LS478113.1_Estonia_2014        ................................................................--.............. [ 94960]
KP843857.1_Odintsovo_02/14     ................................................................--.............. [ 94960]
FN557520.1_strain_E75          t.....-....................-......g.............................ta.............. [ 94960]
AY261366.1_Warthog             ......--...................-......g.........................a...--.............. [ 94960]
AY261365.1_Warmbaths           ......--...................-......g.........................a...--.............. [ 94960]
AY261364.1_Tengani_62          .a.........................-......g.............................--.............. [ 94960]
AY261363.1_Pretoriuskop/96/4   ......--...................-......g.........................a...--.............. [ 94960]
AY261362.1_Mkuzi_1979          .a....-...........................g.........................a...--.............. [ 94960]
AY261361.1_Malawi_Lil-20/1_1983 ..t.ca-.........----.......-......g....a...................c....--.............. [ 94960]
AY261360.1_Kenya_1950          t...----...........a.......-......g........................c....--.............. [ 94960]
AM712240.1_OURT_88/3           t.....-....................-......g.............................ta.............. [ 94960]
AM712239.1_Benin_97/1          t.....-....................-......g.............................ta.............. [ 94960]
KM111295.1_Ken06.Bus           t...----...........a.......-......g........................c....--.............. [ 94960]
```

```
KM111294.1_Ken05/Tk1             t...----...........a.......-......g........................c....--.............. [ 94960]
KM262844.1_strain_L60            t.....-....................-......g.............................ta.............. [ 94960]
KM102979.1_26544/OG10_Italy      t.....-....................-......g.............................ta.............. [ 94960]
KM262845.1_strain_NHV            t.....-....................-......g.............................ta.............. [ 94960]
KX354450.1_47/Ss/2008            t.....-....................-......g.............................ta.............. [ 94960]
KP055815.1_strain_BA71           t.....-....................-......g.............................ta.............. [ 94960]
MH025920.1_strain_R35            t...----...........a.......-......g........................c....--.............. [ 94960]
MH025919.1_strain_N10            t...----...........a.......-......g........................c....--.............. [ 94960]
MH025918.1_strain_R25            t...----...........a.......-......g........................c....--.............. [ 94960]
MH025917.1_strain_R7             t...----...........a.......-......g........................c....--.............. [ 94960]
MH025916.1_strain_R8             t...----...........a.......-......g........................c....--.............. [ 94960]

FR682468.2_Georgia_2007/1        ccatctacaaactatactcccgaacagcaacacgaaaaattaaaacattatgttttaatccctaaacacctttggtctta [ 95040]
ASF_IND_20_CAD543                ................................................................................ [ 95040]
LR899193.1_Germany_2020/1        ................................................................................ [ 95040]
MN393477.1_Wuhan_2019            ................................................................................ [ 95040]
MW396979.1_Timor-Leste/2019/1    ................................................................................ [ 95040]
MK940252.1_CN/2019/InnerMongolia ................................................................................ [ 95040]
MK128995.1_China/2018/AnhuiXCGQ  ................................................................................ [ 95040]
MH681419.1_ASFV/POL/2015         ................................................................................ [ 95040]
LS478113.1_Estonia_2014          ................................................................................ [ 95040]
KP843857.1_Odintsovo_02/14       ................................................................................ [ 95040]
FN557520.1_strain_E75            ........g....................................t....c........a.................... [ 95040]
AY261366.1_Warthog               ...................t...g.....................a....c............................. [ 95040]
AY261365.1_Warmbaths             .......................g..........................c............................. [ 95040]
AY261364.1_Tengani_62            .......................g..........................c............................. [ 95040]
AY261363.1_Pretoriuskop/96/4     ...................t...g.....................a....c............................. [ 95040]
AY261362.1_Mkuzi_1979            ..................................................c........a.................... [ 95040]
AY261361.1_Malawi_Lil-20/1_1983  ........c..t.....a................................c..........................c.. [ 95040]
AY261360.1_Kenya_1950            ........c........c................................ca..........c.g...........g... [ 95040]
AM712240.1_OURT_88/3             ........g....................................t....c........a.................... [ 95040]
AM712239.1_Benin_97/1            ........g....................................t....c........a.................... [ 95040]
KM111295.1_Ken06.Bus             ........c........c................................ca..........c.............g... [ 95040]
KM111294.1_Ken05/Tk1             ........c........c................................ca..........c.g...........g... [ 95040]
KM262844.1_strain_L60            ........g....................................t....c........a.................... [ 95040]
KM102979.1_26544/OG10_Italy      ........g....................................t....c........a.................... [ 95040]
KM262845.1_strain_NHV            ........g....................................t....c........a.................... [ 95040]
KX354450.1_47/Ss/2008            ........g....................................t....c........a.................... [ 95040]
KP055815.1_strain_BA71           ........g....................................t....c........a.................... [ 95040]
MH025920.1_strain_R35            ........c........c................................ca..........c.............g... [ 95040]
MH025919.1_strain_N10            ........c........c................................ca..........c.............g... [ 95040]
MH025918.1_strain_R25            ........c........c................................ca..........c.............g... [ 95040]
MH025917.1_strain_R7             ........c........c................................ca..........c.............g... [ 95040]
MH025916.1_strain_R8             ........c........c................................ca..........c.............g... [ 95040]
```

```
FR682468.2_Georgia_2007/1        tattaaatacggaacgcatgtccggtactacaccacacaaaatgttttccgagtcggtggctttgtgcttcaaaatccct [ 95120]
ASF_IND_20_CAD543                ................................................................................ [ 95120]
LR899193.1_Germany_2020/1        ................................................................................ [ 95120]
MN393477.1_Wuhan_2019            ................................................................................ [ 95120]
MW396979.1_Timor-Leste/2019/1    ................................................................................ [ 95120]
MK940252.1_CN/2019/InnerMongolia ................................................................................ [ 95120]
MK128995.1_China/2018/AnhuiXCGQ  ................................................................................ [ 95120]
MH681419.1_ASFV/POL/2015         ................................................................................ [ 95120]
LS478113.1_Estonia_2014          ................................................................................ [ 95120]
KP843857.1_Odintsovo_02/14       ................................................................................ [ 95120]
FN557520.1_strain_E75            ............g.......g........................................................... [ 95120]
AY261366.1_Warthog               ............g..c...................a............................................ [ 95120]
AY261365.1_Warmbaths             ............g..c...................a............................................ [ 95120]
AY261364.1_Tengani_62            ............g..a................................................................ [ 95120]
AY261363.1_Pretoriuskop/96/4     ............g..c...................a............................................ [ 95120]
AY261362.1_Mkuzi_1979            ............g.......g........................................................... [ 95120]
AY261361.1_Malawi_Lil-20/1_1983  ............t................................c.....t..g...........a............. [ 95120]
AY261360.1_Kenya_1950            ............t..............t.................c.....g............................ [ 95120]
AM712240.1_OURT_88/3             ............g.......g........................................................... [ 95120]
AM712239.1_Benin_97/1            ............g.......g........................................................... [ 95120]
KM111295.1_Ken06.Bus             ............t..............t.................c.....g............................ [ 95120]
KM111294.1_Ken05/Tk1             ............t..............t.................c.....g............................ [ 95120]
KM262844.1_strain_L60            ............g.......g........................................................... [ 95120]
KM102979.1_26544/OG10_Italy      ............g.......g........................................................... [ 95120]
KM262845.1_strain_NHV            ............g.......g........................................................... [ 95120]
KX354450.1_47/Ss/2008            ............g.......g........................................................... [ 95120]
KP055815.1_strain_BA71           ............g.......g........................................................... [ 95120]
MH025920.1_strain_R35            ............t..............t.................c.....g............................ [ 95120]
MH025919.1_strain_N10            ............t..............t...............a.c.....g............................ [ 95120]
MH025918.1_strain_R25            ............t..............t.................c.....g............................ [ 95120]
MH025917.1_strain_R7             ............t..............t.................c.....g............................ [ 95120]
MH025916.1_strain_R8             ............t..............t.................c.....g............................ [ 95120]

FR682468.2_Georgia_2007/1        acgaagccgttataaaaaatgaggtaaaaacagcaataagactgcaaaatagttttaacacaaaagcgaaagggcatgta [ 95200]
ASF_IND_20_CAD543                ................................................................................ [ 95200]
LR899193.1_Germany_2020/1        ................................................................................ [ 95200]
MN393477.1_Wuhan_2019            ................................................................................ [ 95200]
MW396979.1_Timor-Leste/2019/1    ................................................................................ [ 95200]
MK940252.1_CN/2019/InnerMongolia ................................................................................ [ 95200]
MK128995.1_China/2018/AnhuiXCGQ  ................................................................................ [ 95200]
MH681419.1_ASFV/POL/2015         ................................................................................ [ 95200]
LS478113.1_Estonia_2014          ................................................................................ [ 95200]
KP843857.1_Odintsovo_02/14       ................................................................................ [ 95200]
```

```
FN557520.1_strain_E75            .........................................................................a...... [ 95200]
AY261366.1_Warthog               ...........g.................................................................... [ 95200]
AY261365.1_Warmbaths             ...........g.................................................................... [ 95200]
AY261364.1_Tengani_62            ...........g.................................................................... [ 95200]
AY261363.1_Pretoriuskop/96/4     ...........g.................................................................... [ 95200]
AY261362.1_Mkuzi_1979            ................................................................................ [ 95200]
AY261361.1_Malawi_Lil-20/1_1983  ..........c...................tg.........t................................t..... [ 95200]
AY261360.1_Kenya_1950            ...............................g................................................ [ 95200]
AM712240.1_OURT_88/3             .........................................................................a...... [ 95200]
AM712239.1_Benin_97/1            .........................................................................a...... [ 95200]
KM111295.1_Ken06.Bus             ...............................g...................a............................ [ 95200]
KM111294.1_Ken05/Tk1             ...............................g................................................ [ 95200]
KM262844.1_strain_L60            .........................................................................a...... [ 95200]
KM102979.1_26544/OG10_Italy      .........................................................................a...... [ 95200]
KM262845.1_strain_NHV            .........................................................................a...... [ 95200]
KX354450.1_47/Ss/2008            .........................................................................a...... [ 95200]
KP055815.1_strain_BA71           .........................................................................a...... [ 95200]
MH025920.1_strain_R35            ...............................g...................a............................ [ 95200]
MH025919.1_strain_N10            ...............................g...................a............................ [ 95200]
MH025918.1_strain_R25            ...............................g...................a............................ [ 95200]
MH025917.1_strain_R7             ...............................g...................a............................ [ 95200]
MH025916.1_strain_R8             ...............................g...................a............................ [ 95200]

FR682468.2_Georgia_2007/1        acgtgggccgtcccatatgataatattagcaagctatatgccaaaccagatgcaattatgcttaccatacaagaaaatgt [ 95280]
ASF_IND_20_CAD543                ................................................................................ [ 95280]
LR899193.1_Germany_2020/1        ................................................................................ [ 95280]
MN393477.1_Wuhan_2019            ................................................................................ [ 95280]
MW396979.1_Timor-Leste/2019/1    ................................................................................ [ 95280]
MK940252.1_CN/2019/InnerMongolia ................................................................................ [ 95280]
MK128995.1_China/2018/AnhuiXCGQ  ................................................................................ [ 95280]
MH681419.1_ASFV/POL/2015         ................................................................................ [ 95280]
LS478113.1_Estonia_2014          ................................................................................ [ 95280]
KP843857.1_Odintsovo_02/14       ................................................................................ [ 95280]
FN557520.1_strain_E75            ................................................................................ [ 95280]
AY261366.1_Warthog               ............t........g.......................................................... [ 95280]
AY261365.1_Warmbaths             ............t........g.......................................................... [ 95280]
AY261364.1_Tengani_62            ..a.........t........g.......................................................... [ 95280]
AY261363.1_Pretoriuskop/96/4     ............t........g.......................................................... [ 95280]
AY261362.1_Mkuzi_1979            ................................................................................ [ 95280]
AY261361.1_Malawi_Lil-20/1_1983  ......a.....g.....a..g..................................c....................... [ 95280]
AY261360.1_Kenya_1950            ......a.......c......g.........................g................................ [ 95280]
AM712240.1_OURT_88/3             ................................................................................ [ 95280]
AM712239.1_Benin_97/1            ................................................................................ [ 95280]
KM111295.1_Ken06.Bus             ......a.ta....c......g.........................g................................ [ 95280]
```

```
KM111294.1_Ken05/Tk1            ......a.......c......g.........................g................................ [ 95280]
KM262844.1_strain_L60           ................................................................................ [ 95280]
KM102979.1_26544/OG10_Italy     ................................................................................ [ 95280]
KM262845.1_strain_NHV           ................................................................................ [ 95280]
KX354450.1_47/Ss/2008           ................................................................................ [ 95280]
KP055815.1_strain_BA71          ................................................................................ [ 95280]
MH025920.1_strain_R35           ......a.t.....c......g.........................g................................ [ 95280]
MH025919.1_strain_N10           ......a.t.....c......g.........................g................................ [ 95280]
MH025918.1_strain_R25           ......a.t.....c......g.........................g................................ [ 95280]
MH025917.1_strain_R7            ......a.t.....c......g.........................g................................ [ 95280]
MH025916.1_strain_R8            ......a.t.....c......g.........................g................................ [ 95280]

FR682468.2_Georgia_2007/1       tgaaaaagctcttcatgctttaaaccaaaacgtactgacgctcgcatcaaaaatacgttaaatataa--tttttgtagag [ 95360]
ASF_IND_20_CAD543               ...................................................................---.......... [ 95360]
LR899193.1_Germany_2020/1       ...................................................................--........... [ 95360]
MN393477.1_Wuhan_2019           ...................................................................--........... [ 95360]
MW396979.1_Timor-Leste/2019/1   ...................................................................--........... [ 95360]
MK940252.1_CN/2019/InnerMongolia ...................................................................--........... [ 95360]
MK128995.1_China/2018/AnhuiXCGQ ...................................................................--........... [ 95360]
MH681419.1_ASFV/POL/2015        ...................................................................--........... [ 95360]
LS478113.1_Estonia_2014         ...................................................................--........... [ 95360]
KP843857.1_Odintsovo_02/14      ...................................................................--........... [ 95360]
FN557520.1_strain_E75           ...................................................................---.......... [ 95360]
AY261366.1_Warthog              .............................................................c.t...--.c......... [ 95360]
AY261365.1_Warmbaths            ..................................t......................------t...--.c......... [ 95360]
AY261364.1_Tengani_62           ..............................t..............................c.t...--.c......... [ 95360]
AY261363.1_Pretoriuskop/96/4    .............................................................c.t...--.c......... [ 95360]
AY261362.1_Mkuzi_1979           ...................................................................--........... [ 95360]
AY261361.1_Malawi_Lil-20/1_1983 ..........................................t..................c.cc..--.c........t [ 95360]
AY261360.1_Kenya_1950           ..................c.......................t..gg..............c.tc..tc..........c [ 95360]
AM712240.1_OURT_88/3            ...................................................................---.......... [ 95360]
AM712239.1_Benin_97/1           ...................................................................---.......... [ 95360]
KM111295.1_Ken06.Bus            ..................c.......................t..gg..............c.cc..--..........c [ 95360]
KM111294.1_Ken05/Tk1            ..................c.......................t..gg..............c.cc..--.c...a....c [ 95360]
KM262844.1_strain_L60           ...................................................................---.......... [ 95360]
KM102979.1_26544/OG10_Italy     ...................................................................---.......... [ 95360]
KM262845.1_strain_NHV           ...................................................................---.......... [ 95360]
KX354450.1_47/Ss/2008           ...................................................................---.......... [ 95360]
KP055815.1_strain_BA71          ...................................................................---.......... [ 95360]
MH025920.1_strain_R35           ..................c.......................t..gg..............c.cc..--..........c [ 95360]
MH025919.1_strain_N10           ..................c.......................t..gg..............c.cc..--..........c [ 95360]
MH025918.1_strain_R25           ..................c.......................t..gg..............c.cc..--..........c [ 95360]
MH025917.1_strain_R7            ..................c.......................t..gg..............c.cc..--..........c [ 95360]
MH025916.1_strain_R8            ..................c.......................t..gg..............c.cc..--..........c [ 95360]
```

```
FR682468.2_Georgia_2007/1        gat----aaaaagctattttagctaaa---------------------------------aaataattcatatacgttat [ 95440]
ASF_IND_20_CAD543                ...----....................---------------------------------.................... [ 95440]
LR899193.1_Germany_2020/1        ...----....................---------------------------------.................... [ 95440]
MN393477.1_Wuhan_2019            ...----....................---------------------------------.................... [ 95440]
MW396979.1_Timor-Leste/2019/1    ...----....................---------------------------------.................... [ 95440]
MK940252.1_CN/2019/InnerMongolia ...----....................---------------------------------.................... [ 95440]
MK128995.1_China/2018/AnhuiXCGQ  ...----....................---------------------------------.................... [ 95440]
MH681419.1_ASFV/POL/2015         ...----....................---------------------------------.................... [ 95440]
LS478113.1_Estonia_2014          ...----....................---------------------------------.................... [ 95440]
KP843857.1_Odintsovo_02/14       ...----....................---------------------------------.................... [ 95440]
FN557520.1_strain_E75            ...----....................---------------------------------.................... [ 95440]
AY261366.1_Warthog               ...----.......g............---------------------------------.................... [ 95440]
AY261365.1_Warmbaths             ...----.......g............---------------------------------.................... [ 95440]
AY261364.1_Tengani_62            ...aaaa......ta............---------------------------------.................... [ 95440]
AY261363.1_Pretoriuskop/96/4     ...-----......g............---------------------------------.................... [ 95440]
AY261362.1_Mkuzi_1979            ...----....................---------------------------------.................... [ 95440]
AY261361.1_Malawi_Lil-20/1_1983  ...----......tc........c...---------------------------------........--..a....... [ 95440]
AY261360.1_Kenya_1950            ...----......ta.....g..c...-----ttttagccaaaaaagcaattttagccaa...a..a.a...a....... [ 95440]
AM712240.1_OURT_88/3             ...----....................---------------------------------.................... [ 95440]
AM712239.1_Benin_97/1            ...----....................---------------------------------.................... [ 95440]
KM111295.1_Ken06.Bus             ...----......ta.....g......tttgtttttggctaaaa------tttagcc--...a..a.a...a........ [ 95440]
KM111294.1_Ken05/Tk1             ...----......ta.....g..c...-----ttttagccaaaaaagcaattttagctaa...a..a.a...a....... [ 95440]
KM262844.1_strain_L60            ...----....................---------------------------------.................... [ 95440]
KM102979.1_26544/OG10_Italy      ...----....................---------------------------------.................... [ 95440]
KM262845.1_strain_NHV            ...----....................---------------------------------.................... [ 95440]
KX354450.1_47/Ss/2008            ...----....................---------------------------------.................... [ 95440]
KP055815.1_strain_BA71           ...----....................---------------------------------.................... [ 95440]
MH025920.1_strain_R35            ...----......ta.....g......tttgtttttggctaaaa------tttagcc--...a..a.a...a........ [ 95440]
MH025919.1_strain_N10            ...----......ta.....g......tttgtttttggctaaaa------tttagcc--...a..a.a...a........ [ 95440]
MH025918.1_strain_R25            ...----......ta.....g......tttgtttttggctaaaa------tttagcc--...a..a.a...a........ [ 95440]
MH025917.1_strain_R7             ...----......ta.....g......tttgtttttggctaaaa------tttagcc--...a..a.a...a........ [ 95440]
MH025916.1_strain_R8             ...----......ta.....g......tttgtttttggctaaaa------tttagcc--...a..a.a...a........ [ 95440]

FR682468.2_Georgia_2007/1        gcagaggaagaacggtggctttcaaattcagattgcatccacgtagaccgtagcgtttttttgcttctggtttatatcg [ 95520]
ASF_IND_20_CAD543                ................................................................................ [ 95520]
LR899193.1_Germany_2020/1        ................................................................................ [ 95520]
MN393477.1_Wuhan_2019            ................................................................................ [ 95520]
MW396979.1_Timor-Leste/2019/1    ................................................................................ [ 95520]
MK940252.1_CN/2019/InnerMongolia ................................................................................ [ 95520]
MK128995.1_China/2018/AnhuiXCGQ  ................................................................................ [ 95520]
MH681419.1_ASFV/POL/2015         ................................................................................ [ 95520]
LS478113.1_Estonia_2014          ................................................................................ [ 95520]
KP843857.1_Odintsovo_02/14       ................................................................................ [ 95520]
```

```
FN557520.1_strain_E75            .....a.......................................................................... [ 95520]
AY261366.1_Warthog               ................................................................................ [ 95520]
AY261365.1_Warmbaths             ................................................................................ [ 95520]
AY261364.1_Tengani_62            ................................................................................ [ 95520]
AY261363.1_Pretoriuskop/96/4     ................................................................................ [ 95520]
AY261362.1_Mkuzi_1979            ................................................................................ [ 95520]
AY261361.1_Malawi_Lil-20/1_1983  ..g...............................t......t..........................c........... [ 95520]
AY261360.1_Kenya_1950            ..g.....g.................a.......t............................................. [ 95520]
AM712240.1_OURT_88/3             .....a.......................................................................... [ 95520]
AM712239.1_Benin_97/1            .....a.......................................................................... [ 95520]
KM111295.1_Ken06.Bus             ..g.......................a.......t............................................. [ 95520]
KM111294.1_Ken05/Tk1             ..g.....g.................a.......t............................................. [ 95520]
KM262844.1_strain_L60            .....a.......................................................................... [ 95520]
KM102979.1_26544/OG10_Italy      .....a.......................................................................... [ 95520]
KM262845.1_strain_NHV            .....a.......................................................................... [ 95520]
KX354450.1_47/Ss/2008            .....a.......................................................................... [ 95520]
KP055815.1_strain_BA71           .....a.......................................................................... [ 95520]
MH025920.1_strain_R35            ..g.......................a.......t............................................. [ 95520]
MH025919.1_strain_N10            ..g.......................a.......t............................................. [ 95520]
MH025918.1_strain_R25            ..g.......................a.......t............................................. [ 95520]
MH025917.1_strain_R7             ..g.......................a.......t............................................. [ 95520]
MH025916.1_strain_R8             ..g.......................a.......t............................................. [ 95520]

FR682468.2_Georgia_2007/1        taaaccgtaataaacatcatcatttgtatccgttggatctttttcccactccggataaaaaatcggttttcttttttttt [ 95600]
ASF_IND_20_CAD543                ........................n....................................................... [ 95600]
LR899193.1_Germany_2020/1        ................................................................................ [ 95600]
MN393477.1_Wuhan_2019            ................................................................................ [ 95600]
MW396979.1_Timor-Leste/2019/1    ................................................................................ [ 95600]
MK940252.1_CN/2019/InnerMongolia ................................................................................ [ 95600]
MK128995.1_China/2018/AnhuiXCGQ  ................................................................................ [ 95600]
MH681419.1_ASFV/POL/2015         ................................................................................ [ 95600]
LS478113.1_Estonia_2014          ................................................................................ [ 95600]
KP843857.1_Odintsovo_02/14       .......................................................................-........ [ 95600]
FN557520.1_strain_E75            ..............................................................c................. [ 95600]
AY261366.1_Warthog               ..............................t...............................c................. [ 95600]
AY261365.1_Warmbaths             ..............................t...............................c...............a. [ 95600]
AY261364.1_Tengani_62            ..............................t...............................c................. [ 95600]
AY261363.1_Pretoriuskop/96/4     ..............................t...............................c................. [ 95600]
AY261362.1_Mkuzi_1979            ..............................................................c................. [ 95600]
AY261361.1_Malawi_Lil-20/1_1983  ................g.............t.................t.............c...............a. [ 95600]
AY261360.1_Kenya_1950            ...c.......................g..t...............................ct................ [ 95600]
AM712240.1_OURT_88/3             ..............................................................c................. [ 95600]
AM712239.1_Benin_97/1            ..............................................................c................. [ 95600]
KM111295.1_Ken06.Bus             ...........................g..t...............................ct................ [ 95600]
```

```
KM111294.1_Ken05/Tk1           ...........................g..t...............................ct................ [ 95600]
KM262844.1_strain_L60          ..............................................................c................. [ 95600]
KM102979.1_26544/OG10_Italy    ..............................................................c................. [ 95600]
KM262845.1_strain_NHV          ..............................................................c................. [ 95600]
KX354450.1_47/Ss/2008          ..............................................................c................. [ 95600]
KP055815.1_strain_BA71         ..............................................................c................. [ 95600]
MH025920.1_strain_R35          ...........................g..t...............................ct................ [ 95600]
MH025919.1_strain_N10          ...........................g..t...............................ct................ [ 95600]
MH025918.1_strain_R25          ...........................g..t...............................ct................ [ 95600]
MH025917.1_strain_R7           ...........................g..t...............................ct................ [ 95600]
MH025916.1_strain_R8           ...........................g..t...............................ct................ [ 95600]

FR682468.2_Georgia_2007/1      ggtcgttttttgcagtaagctgtaaattaagggaatatagcttatcgaaaagttgttcctgatccatataaatagcagca [ 95680]
ASF_IND_20_CAD543              ................................................................................ [ 95680]
LR899193.1_Germany_2020/1      ................................................................................ [ 95680]
MN393477.1_Wuhan_2019          ................................................................................ [ 95680]
MW396979.1_Timor-Leste/2019/1  ................................................................................ [ 95680]
MK940252.1_CN/2019/InnerMongolia ................................................................................ [ 95680]
MK128995.1_China/2018/AnhuiXCGQ ................................................................................ [ 95680]
MH681419.1_ASFV/POL/2015       ................................................................................ [ 95680]
LS478113.1_Estonia_2014        ................................................................................ [ 95680]
KP843857.1_Odintsovo_02/14     ................................................................................ [ 95680]
FN557520.1_strain_E75          ................................................................................ [ 95680]
AY261366.1_Warthog             ..........................................................t..............---.... [ 95680]
AY261365.1_Warmbaths           ..........................................................t..............---.... [ 95680]
AY261364.1_Tengani_62          .....g....................................................t...............a..... [ 95680]
AY261363.1_Pretoriuskop/96/4   ..........................................................t..............---.... [ 95680]
AY261362.1_Mkuzi_1979          ................................................................................ [ 95680]
AY261361.1_Malawi_Lil-20/1_1983 ......................c...................................t..............---.... [ 95680]
AY261360.1_Kenya_1950          ....a..........c.........................c................t...........c..---.... [ 95680]
AM712240.1_OURT_88/3           ................................................................................ [ 95680]
AM712239.1_Benin_97/1          ................................................................................ [ 95680]
KM111295.1_Ken06.Bus           ....a..........c.........................c................t...........c..---..t. [ 95680]
KM111294.1_Ken05/Tk1           ....a........g.c.........................c................t...........c..---.... [ 95680]
KM262844.1_strain_L60          ................................................................................ [ 95680]
KM102979.1_26544/OG10_Italy    ................................................................................ [ 95680]
KM262845.1_strain_NHV          ................................................................................ [ 95680]
KX354450.1_47/Ss/2008          ................................................................................ [ 95680]
KP055815.1_strain_BA71         ................................................................................ [ 95680]
MH025920.1_strain_R35          ....a..........c.........................c................t...........c..---..t. [ 95680]
MH025919.1_strain_N10          ....a..........c.........................c................t...........c..---..t. [ 95680]
MH025918.1_strain_R25          ....a..........c.........................c................t...........c..---..t. [ 95680]
MH025917.1_strain_R7           ....a..........c.........................c................t...........c..---..t. [ 95680]
MH025916.1_strain_R8           ....a..........c.........................c................t...........c..---..t. [ 95680]
```

```
FR682468.2_Georgia_2007/1        tatattaaaaaaaaat-aaaaaaagacgcttcaacgagtcagtaccactgcttgccaacgatttacgttggttggtgcat [ 95760]
ASF_IND_20_CAD543                ................-............................................................... [ 95760]
LR899193.1_Germany_2020/1        ................-............................................................... [ 95760]
MN393477.1_Wuhan_2019            ................-............................................................... [ 95760]
MW396979.1_Timor-Leste/2019/1    ................-............................................................... [ 95760]
MK940252.1_CN/2019/InnerMongolia ................-............................................................... [ 95760]
MK128995.1_China/2018/AnhuiXCGQ  ................-............................................................... [ 95760]
MH681419.1_ASFV/POL/2015         ................-............................................................... [ 95760]
LS478113.1_Estonia_2014          ................-............................................................... [ 95760]
KP843857.1_Odintsovo_02/14       ......-.........-............................................................... [ 95760]
FN557520.1_strain_E75            ................-............................................................... [ 95760]
AY261366.1_Warthog               ................-............................................................... [ 95760]
AY261365.1_Warmbaths             ................-............................................................... [ 95760]
AY261364.1_Tengani_62            ................-....................................a.......................... [ 95760]
AY261363.1_Pretoriuskop/96/4     ................-............................................................... [ 95760]
AY261362.1_Mkuzi_1979            ................-............................................................... [ 95760]
AY261361.1_Malawi_Lil-20/1_1983  .....--.........-....................................a.....-..............a..... [ 95760]
AY261360.1_Kenya_1950            ................a..............................................c................ [ 95760]
AM712240.1_OURT_88/3             ................-............................................................... [ 95760]
AM712239.1_Benin_97/1            ................-............................................................... [ 95760]
KM111295.1_Ken06.Bus             ................a.............................................gc............a... [ 95760]
KM111294.1_Ken05/Tk1             ................a..............................................c................ [ 95760]
KM262844.1_strain_L60            ................-............................................................... [ 95760]
KM102979.1_26544/OG10_Italy      ................-............................................................... [ 95760]
KM262845.1_strain_NHV            ................-............................................................... [ 95760]
KX354450.1_47/Ss/2008            ................-............................................................... [ 95760]
KP055815.1_strain_BA71           ................-............................................................... [ 95760]
MH025920.1_strain_R35            ................a.............................................gc............a... [ 95760]
MH025919.1_strain_N10            ................-.............................................gc............a... [ 95760]
MH025918.1_strain_R25            ................a.............................................gc............a... [ 95760]
MH025917.1_strain_R7             ................a.............................................gc............a... [ 95760]
MH025916.1_strain_R8             ................a.............................................gc............a... [ 95760]

FR682468.2_Georgia_2007/1        tatggtgatatagtaatgagtgcctgcac-aagtgcttgcacaagtgcctgcacaa------------gtgcttgcacaa [ 95840]
ASF_IND_20_CAD543                .............................-..........................------------............ [ 95840]
LR899193.1_Germany_2020/1        .............................-..........................------------............ [ 95840]
MN393477.1_Wuhan_2019            .............................-..........................------------............ [ 95840]
MW396979.1_Timor-Leste/2019/1    .............................-..........................------------............ [ 95840]
MK940252.1_CN/2019/InnerMongolia .............................-..........................------------............ [ 95840]
MK128995.1_China/2018/AnhuiXCGQ  .............................-..........................------------............ [ 95840]
MH681419.1_ASFV/POL/2015         .............................-..........................------------............ [ 95840]
LS478113.1_Estonia_2014          .............................-..........................------------............ [ 95840]
KP843857.1_Odintsovo_02/14       .............................-..........................------------............ [ 95840]
```

```
FN557520.1_strain_E75             .............................-.............g....t......g------------............ [ 95840]
AY261366.1_Warthog                .......................t.....-..............------------------------............ [ 95840]
AY261365.1_Warmbaths              ...................------------------------------------------------------------- [ 95840]
AY261364.1_Tengani_62             ...................------------------------------------------------------------- [ 95840]
AY261363.1_Pretoriuskop/96/4      ...................------------------------------------------------------------- [ 95840]
AY261362.1_Mkuzi_1979             .............................-..------------------------------------............ [ 95840]
AY261361.1_Malawi_Lil-20/1_1983   .t..a............ata...aa....------------------------------ttt------...........- [ 95840]
AY261360.1_Kenya_1950             .t..a.............ta..gaa...t.t............g------------------------............ [ 95840]
AM712240.1_OURT_88/3              .............................-.............g....t......g------------............ [ 95840]
AM712239.1_Benin_97/1             .............................-.............g....t......ggtgcttgcacaa............ [ 95840]
KM111295.1_Ken06.Bus              .t................ca-------------------------------------------------..t.......g [ 95840]
KM111294.1_Ken05/Tk1              .t..a.............ca..tt.....-.............g------------------------...........g [ 95840]
KM262844.1_strain_L60             .............................-.............g....t......g------------............ [ 95840]
KM102979.1_26544/OG10_Italy       .............................-.............g....t......g------------............ [ 95840]
KM262845.1_strain_NHV             .............................-.............g....t......g------------............ [ 95840]
KX354450.1_47/Ss/2008             .............................-.............g....t......g------------............ [ 95840]
KP055815.1_strain_BA71            .............................-.............g....t......g------------............ [ 95840]
MH025920.1_strain_R35             .t................ca-------------------------------------------------..t.......g [ 95840]
MH025919.1_strain_N10             .t................ca-------------------------------------------------..t.......g [ 95840]
MH025918.1_strain_R25             .t................ca-------------------------------------------------..t.......g [ 95840]
MH025917.1_strain_R7              .t................ca-------------------------------------------------..t.......g [ 95840]
MH025916.1_strain_R8              .t................ca-------------------------------------------------..t.......g [ 95840]

FR682468.2_Georgia_2007/1         gtgcttgcacaagtgcttacacaagtgcttgcacaagtgcctgtacacattactgcatcgccaaagcacctgcaatgcct [ 95920]
ASF_IND_20_CAD543                 ........................................y....................................... [ 95920]
LR899193.1_Germany_2020/1         ................................................................................ [ 95920]
MN393477.1_Wuhan_2019             ................................................................................ [ 95920]
MW396979.1_Timor-Leste/2019/1     ................................................................................ [ 95920]
MK940252.1_CN/2019/InnerMongolia  ................................................................................ [ 95920]
MK128995.1_China/2018/AnhuiXCGQ   ................................................................................ [ 95920]
MH681419.1_ASFV/POL/2015          ................................................................................ [ 95920]
LS478113.1_Estonia_2014           ................................................................................ [ 95920]
KP843857.1_Odintsovo_02/14        ................................................................................ [ 95920]
FN557520.1_strain_E75             ..................g.........c...........t....................................... [ 95920]
AY261366.1_Warthog                ................c.g.........c................................................... [ 95920]
AY261365.1_Warmbaths              ................c.g............................................................. [ 95920]
AY261364.1_Tengani_62             ------------......g........................c.................................... [ 95920]
AY261363.1_Pretoriuskop/96/4      ......a...........g.........c................................................... [ 95920]
AY261362.1_Mkuzi_1979             ....c........g.................................................................. [ 95920]
AY261361.1_Malawi_Lil-20/1_1983   ..................g....g................t....................................... [ 95920]
AY261360.1_Kenya_1950             ..................g.......................ac.................................... [ 95920]
AM712240.1_OURT_88/3              ..................g.........c...........t....................................... [ 95920]
AM712239.1_Benin_97/1             ..................g.........c...........t....................................... [ 95920]
KM111295.1_Ken06.Bus              ..................g....g....aa............ac.................................... [ 95920]
```

```
KM111294.1_Ken05/Tk1              ...........g......g....g...........g......ac.................................... [ 95920]
KM262844.1_strain_L60             ..................g.........c...........t....................................... [ 95920]
KM102979.1_26544/OG10_Italy       ..................g.........c...........t....................................... [ 95920]
KM262845.1_strain_NHV             ..................g.........c...........t....................................... [ 95920]
KX354450.1_47/Ss/2008             ..................g.........c...........t....................................... [ 95920]
KP055815.1_strain_BA71            ..................g.........c...........t....................................... [ 95920]
MH025920.1_strain_R35             ..................g....g....aa............ac.................................... [ 95920]
MH025919.1_strain_N10             ..................g....g....aa............ac.................................... [ 95920]
MH025918.1_strain_R25             ..................g....g....aa............ac.................................... [ 95920]
MH025917.1_strain_R7              ..................g....g....aa............ac.................................... [ 95920]
MH025916.1_strain_R8              ..................g....g....aa............ac.................................... [ 95920]

FR682468.2_Georgia_2007/1         acttcctcaacagagtacgataactaaatgcttttaagcaccgcttgcgtcgatgtgtccttcggggcaatcgggttcaa [ 96000]
ASF_IND_20_CAD543                 ................................................................................ [ 96000]
LR899193.1_Germany_2020/1         ................................................................................ [ 96000]
MN393477.1_Wuhan_2019             ................................................................................ [ 96000]
MW396979.1_Timor-Leste/2019/1     ................................................................................ [ 96000]
MK940252.1_CN/2019/InnerMongolia  ................................................................................ [ 96000]
MK128995.1_China/2018/AnhuiXCGQ   ................................................................................ [ 96000]
MH681419.1_ASFV/POL/2015          ................................................................................ [ 96000]
LS478113.1_Estonia_2014           ................................................................................ [ 96000]
KP843857.1_Odintsovo_02/14        ................................................................................ [ 96000]
FN557520.1_strain_E75             ................................................................................ [ 96000]
AY261366.1_Warthog                ................................................................................ [ 96000]
AY261365.1_Warmbaths              ................................................................................ [ 96000]
AY261364.1_Tengani_62             ................................................................................ [ 96000]
AY261363.1_Pretoriuskop/96/4      ................................................................................ [ 96000]
AY261362.1_Mkuzi_1979             ................................................................................ [ 96000]
AY261361.1_Malawi_Lil-20/1_1983   ................................................................................ [ 96000]
AY261360.1_Kenya_1950             ................................................................................ [ 96000]
AM712240.1_OURT_88/3              ................................................................................ [ 96000]
AM712239.1_Benin_97/1             ................................................................................ [ 96000]
KM111295.1_Ken06.Bus              ...................................................a............................ [ 96000]
KM111294.1_Ken05/Tk1              ................................................................................ [ 96000]
KM262844.1_strain_L60             ................................................................................ [ 96000]
KM102979.1_26544/OG10_Italy       ................................................................................ [ 96000]
KM262845.1_strain_NHV             ................................................................................ [ 96000]
KX354450.1_47/Ss/2008             ................................................................................ [ 96000]
KP055815.1_strain_BA71            ................................................................................ [ 96000]
MH025920.1_strain_R35             ................................................................................ [ 96000]
MH025919.1_strain_N10             ................................................................................ [ 96000]
MH025918.1_strain_R25             ................................................................................ [ 96000]
MH025917.1_strain_R7              ................................................................................ [ 96000]
MH025916.1_strain_R8              ................................................................................ [ 96000]
```

```
FR682468.2_Georgia_2007/1        ttggatccaatattattagtcataattacctaatacttattcaattttatc------ttttttaccttgtaaga------ [ 96080]
ASF_IND_20_CAD543                ...................................................------................m------ [ 96080]
LR899193.1_Germany_2020/1        ...................................................------.................------ [ 96080]
MN393477.1_Wuhan_2019            ...................................................------.................------ [ 96080]
MW396979.1_Timor-Leste/2019/1    ...................................................------.................------ [ 96080]
MK940252.1_CN/2019/InnerMongolia ...................................................------.................------ [ 96080]
MK128995.1_China/2018/AnhuiXCGQ  ...................................................------.................------ [ 96080]
MH681419.1_ASFV/POL/2015         ...................................................------.................------ [ 96080]
LS478113.1_Estonia_2014          ...................................................------.................------ [ 96080]
KP843857.1_Odintsovo_02/14       ...................................................------.................------ [ 96080]
FN557520.1_strain_E75            ............................t......................------................c------ [ 96080]
AY261366.1_Warthog               ...................................................------................c------ [ 96080]
AY261365.1_Warmbaths             ...................................................------.........c......c------ [ 96080]
AY261364.1_Tengani_62            ..................................c................------................c------ [ 96080]
AY261363.1_Pretoriuskop/96/4     ...................................................-------...............c------ [ 96080]
AY261362.1_Mkuzi_1979            ............................t......................------.........c......c------ [ 96080]
AY261361.1_Malawi_Lil-20/1_1983  ...................................tg......g......tttatct......ga..g.....c------ [ 96080]
AY261360.1_Kenya_1950            ....................................g.............tttatct......g...ca....ctagctt [ 96080]
AM712240.1_OURT_88/3             ............................t......................------................c------ [ 96080]
AM712239.1_Benin_97/1            ............................t......................------................c------ [ 96080]
KM111295.1_Ken06.Bus             ....................................g.............tttatct......gt..c...----tgctt [ 96080]
KM111294.1_Ken05/Tk1             ....................................a.............tttatct......g...ca....ctagctt [ 96080]
KM262844.1_strain_L60            ............................t......................------................c------ [ 96080]
KM102979.1_26544/OG10_Italy      ............................t......................------................c------ [ 96080]
KM262845.1_strain_NHV            ............................t......................------................c------ [ 96080]
KX354450.1_47/Ss/2008            ............................t......................------................c------ [ 96080]
KP055815.1_strain_BA71           ............................t......................------................c------ [ 96080]
MH025920.1_strain_R35            ....................................g.............tttatct......gt..c...----tgctt [ 96080]
MH025919.1_strain_N10            ....................................g.............tttatct......gt..c...----tgctt [ 96080]
MH025918.1_strain_R25            ....................................g.............tttatct......gt..c...----tgctt [ 96080]
MH025917.1_strain_R7             ....................................g.............tttatct......gt..c...----tgctt [ 96080]
MH025916.1_strain_R8             ....................................g.............tttatct......gt..c...----tgctt [ 96080]

FR682468.2_Georgia_2007/1        --------tttaaacagcgttttagct-tgtttaaagca----------acgtttaaaacaagctaaaatgctgtttaaa [ 96160]
ASF_IND_20_CAD543                --------.....------------------------------------------------------------------- [ 96160]
LR899193.1_Germany_2020/1        --------...................-...........----------............................... [ 96160]
MN393477.1_Wuhan_2019            --------...................-...........----------............................... [ 96160]
MW396979.1_Timor-Leste/2019/1    --------...................-...........----------............................... [ 96160]
MK940252.1_CN/2019/InnerMongolia --------...................-...........----------............................... [ 96160]
MK128995.1_China/2018/AnhuiXCGQ  --------...................-...........----------............................... [ 96160]
MH681419.1_ASFV/POL/2015         --------...................-...........----------............................... [ 96160]
LS478113.1_Estonia_2014          --------...................-...........----------............................... [ 96160]
KP843857.1_Odintsovo_02/14       --------...................-...........----------............................... [ 96160]
```

```
FN557520.1_strain_E75           --------.......gtt.....--------------------------------------------------------- [ 96160]
AY261366.1_Warthog              --------........tt.....---------------------------------.............c.......... [ 96160]
AY261365.1_Warmbaths            --------......tgtt.....---------------------------------.............c.......... [ 96160]
AY261364.1_Tengani_62           --------.......gtt.....---------------------------------.............c.......... [ 96160]
AY261363.1_Pretoriuskop/96/4    --------......tgat.....---------------------------------.....t.......c.......... [ 96160]
AY261362.1_Mkuzi_1979           --------......tgtt......aac-aagc....a.g----------ct.........----------.t.....t.. [ 96160]
AY261361.1_Malawi_Lil-20/1_1983 --------..a..cttatt.------------------------------------.....c.t.....c.t...-.... [ 96160]
AY261360.1_Kenya_1950           aaacttat..a....gtt......aact........a.gttgttttaaatt....-...t.tatc....c.t.....t-- [ 96160]
AM712240.1_OURT_88/3            --------.......gtt.....--------------------------------------------------------- [ 96160]
AM712239.1_Benin_97/1           --------.......gtt.....--------------------------------------------------------- [ 96160]
KM111295.1_Ken06.Bus            aaacttat..a.....tt......aac----------------------tt....-...t.t.tc....c.t.....t.. [ 96160]
KM111294.1_Ken05/Tk1            aaacttat..a....gtt......aac----------------------tt....-...t.tatc....c.t.....t-- [ 96160]
KM262844.1_strain_L60           --------.......gtt.....--------------------------------------------------------- [ 96160]
KM102979.1_26544/OG10_Italy     --------.......gtt.....--------------------------------------------------------- [ 96160]
KM262845.1_strain_NHV           --------.......gtt.....--------------------------------------------------------- [ 96160]
KX354450.1_47/Ss/2008           --------.......gtt.....--------------------------------------------------------- [ 96160]
KP055815.1_strain_BA71          --------.......gtt.....--------------------------------------------------------- [ 96160]
MH025920.1_strain_R35           aaacttat..a.....tt......aac----------------------tt....-...t.t.tc....c.t.....t.. [ 96160]
MH025919.1_strain_N10           aaacttat..a.....tt......aac----------------------tt....-...t.t.tc....c.t.....t.. [ 96160]
MH025918.1_strain_R25           aaacttat..a.....tt......aac----------------------tt....-...t.t.tc....c.t.....t.. [ 96160]
MH025917.1_strain_R7            aaacttat..a.....tt......aac----------------------tt....-...t.t.tc....c.t.....t.. [ 96160]
MH025916.1_strain_R8            aaacttat..a.....tt......aac----------------------tt....-...t.t.tc....c.t.....t.. [ 96160]

FR682468.2_Georgia_2007/1       acaacgtttt------------aaacaagttaaaacaaataagcttataaatataccatgacaaaattagcccaatggat [ 96240]
ASF_IND_20_CAD543               -.gtt.....------------.......................................................... [ 96240]
LR899193.1_Germany_2020/1       ..........------------.......................................................... [ 96240]
MN393477.1_Wuhan_2019           ..........------------.......................................................... [ 96240]
MW396979.1_Timor-Leste/2019/1   ..........------------.......................................................... [ 96240]
MK940252.1_CN/2019/InnerMongolia ..........------------.......................................................... [ 96240]
MK128995.1_China/2018/AnhuiXCGQ ..........------------.......................................................... [ 96240]
MH681419.1_ASFV/POL/2015        ..........------------.......................................................... [ 96240]
LS478113.1_Estonia_2014         ..........------------.......................................................... [ 96240]
KP843857.1_Odintsovo_02/14      ..........------------.......................................................... [ 96240]
FN557520.1_strain_E75           ----------------------.......................................................... [ 96240]
AY261366.1_Warthog              ..gtt.....------------.......................................................... [ 96240]
AY261365.1_Warmbaths            ..gtt.....aaacgttgttta.......................................................... [ 96240]
AY261364.1_Tengani_62           ..........------------.......................................................... [ 96240]
AY261363.1_Pretoriuskop/96/4    ..........------------.......................................................... [ 96240]
AY261362.1_Mkuzi_1979           ..gtt....a------------.......................................................... [ 96240]
AY261361.1_Malawi_Lil-20/1_1983 .tgtt.....------------.......................................................... [ 96240]
AY261360.1_Kenya_1950           ----------------------.................................................t........ [ 96240]
AM712240.1_OURT_88/3            ----------------------.......................................................... [ 96240]
AM712239.1_Benin_97/1           ----------------------.......................................................... [ 96240]
KM111295.1_Ken06.Bus            ....t....a------------.................................................t........ [ 96240]
```

```
KM111294.1_Ken05/Tk1           ----------------------.................................................t........ [ 96240]
KM262844.1_strain_L60          ----------------------.......................................................... [ 96240]
KM102979.1_26544/OG10_Italy    ----------------------.......................................................... [ 96240]
KM262845.1_strain_NHV          ----------------------.......................................................... [ 96240]
KX354450.1_47/Ss/2008          ----------------------.......................................................... [ 96240]
KP055815.1_strain_BA71         ----------------------.......................................................... [ 96240]
MH025920.1_strain_R35          ....t....a------------.................................................t........ [ 96240]
MH025919.1_strain_N10          ....t....a------------.................................................t........ [ 96240]
MH025918.1_strain_R25          ....t....a------------.................................................t........ [ 96240]
MH025917.1_strain_R7           ....t....a------------.................................................t........ [ 96240]
MH025916.1_strain_R8           ....t....a------------.................................................t........ [ 96240]

FR682468.2_Georgia_2007/1      gtttgagcagtatgtcaaagatttaaacctaaaaaatcgagggtccccctcgttccgcaaatggctcacattgcaaccct [ 96320]
ASF_IND_20_CAD543              ................................................................................ [ 96320]
LR899193.1_Germany_2020/1      ................................................................................ [ 96320]
MN393477.1_Wuhan_2019          ................................................................................ [ 96320]
MW396979.1_Timor-Leste/2019/1  ................................................................................ [ 96320]
MK940252.1_CN/2019/InnerMongolia ................................................................................ [ 96320]
MK128995.1_China/2018/AnhuiXCGQ ................................................................................ [ 96320]
MH681419.1_ASFV/POL/2015       ................................................................................ [ 96320]
LS478113.1_Estonia_2014        ................................................................................ [ 96320]
KP843857.1_Odintsovo_02/14     ................................................................................ [ 96320]
FN557520.1_strain_E75          .......................................g........................................ [ 96320]
AY261366.1_Warthog             .................................g........a..................................... [ 96320]
AY261365.1_Warmbaths           .................................g........a.............................a....... [ 96320]
AY261364.1_Tengani_62          ...........................t..............a..t.................................. [ 96320]
AY261363.1_Pretoriuskop/96/4   .................................g........a..................................... [ 96320]
AY261362.1_Mkuzi_1979          ................................................................................ [ 96320]
AY261361.1_Malawi_Lil-20/1_1983 .................................g.....g.....t.....a............................ [ 96320]
AY261360.1_Kenya_1950          .................................t........a........a....................a....... [ 96320]
AM712240.1_OURT_88/3           .......................................g........................................ [ 96320]
AM712239.1_Benin_97/1          .......................................g........................................ [ 96320]
KM111295.1_Ken06.Bus           .................................t........a........a............................ [ 96320]
KM111294.1_Ken05/Tk1           ............................t....t........a........a....................a....... [ 96320]
KM262844.1_strain_L60          .......................................g........................................ [ 96320]
KM102979.1_26544/OG10_Italy    .......................................g........................................ [ 96320]
KM262845.1_strain_NHV          .......................................g........................................ [ 96320]
KX354450.1_47/Ss/2008          .......................................g........................................ [ 96320]
KP055815.1_strain_BA71         .......................................g........................................ [ 96320]
MH025920.1_strain_R35          .................................t........a........a............................ [ 96320]
MH025919.1_strain_N10          .................................t........a........a............................ [ 96320]
MH025918.1_strain_R25          .................................t........a........a............................ [ 96320]
MH025917.1_strain_R7           .................................t........a........a............................ [ 96320]
MH025916.1_strain_R8           .................................t........a........a............................ [ 96320]
```

FR682468.2_Georgia_2007/1 cactgctgcgctattcgggtgtgatgcgtgctaacgcctttgacatcctaaaatatggctatcctatgcagcagtcaggt [ 96400]
ASF_IND_20_CAD543 ................................................................................ [ 96400]
LR899193.1_Germany_2020/1 ................................................................................ [ 96400]
MN393477.1_Wuhan_2019 ................................................................................ [ 96400]
MW396979.1_Timor-Leste/2019/1 ................................................................................ [ 96400]
MK940252.1_CN/2019/InnerMongolia ................................................................................ [ 96400]
MK128995.1_China/2018/AnhuiXCGQ ................................................................................ [ 96400]
MH681419.1_ASFV/POL/2015 ................................................................................ [ 96400]
LS478113.1_Estonia_2014 ................................................................................ [ 96400]
KP843857.1_Odintsovo_02/14 ................................................................................ [ 96400]
FN557520.1_strain_E75 ................................................................................ [ 96400]
AY261366.1_Warthog ..................................t..t......................................c... [ 96400]
AY261365.1_Warmbaths ..................................t.........................................c... [ 96400]
AY261364.1_Tengani_62 ..................................t............................................. [ 96400]
AY261363.1_Pretoriuskop/96/4 ..................................t..t......................................c... [ 96400]
AY261362.1_Mkuzi_1979 ................................................................................ [ 96400]
AY261361.1_Malawi_Lil-20/1_1983 ..................................t............................................. [ 96400]
AY261360.1_Kenya_1950 .g..............a.............................t................................. [ 96400]
AM712240.1_OURT_88/3 ................................................................................ [ 96400]
AM712239.1_Benin_97/1 ................................................................................ [ 96400]
KM111295.1_Ken06.Bus ..............................................t.............................c... [ 96400]
KM111294.1_Ken05/Tk1 ................a.............................t................................. [ 96400]
KM262844.1_strain_L60 ................................................................................ [ 96400]
KM102979.1_26544/OG10_Italy ................................................................................ [ 96400]
KM262845.1_strain_NHV ................................................................................ [ 96400]
KX354450.1_47/Ss/2008 ................................................................................ [ 96400]
KP055815.1_strain_BA71 ................................................................................ [ 96400]
MH025920.1_strain_R35 ..............................................t.............................c... [ 96400]
MH025919.1_strain_N10 ..............................................t.............................c... [ 96400]
MH025918.1_strain_R25 ..............................................t.............................c... [ 96400]
MH025917.1_strain_R7 ..............................................t.............................c... [ 96400]
MH025916.1_strain_R8 ..............................................t.............................c... [ 96400]

FR682468.2_Georgia_2007/1 tatacggttgctacgcttgaaatccactttaaaaatattaggtcttcctttgccaacatttactggaaccgtgatagcga [ 96480]
ASF_IND_20_CAD543 ................................................................................ [ 96480]
LR899193.1_Germany_2020/1 ................................................................................ [ 96480]
MN393477.1_Wuhan_2019 ................................................................................ [ 96480]
MW396979.1_Timor-Leste/2019/1 ................................................................................ [ 96480]
MK940252.1_CN/2019/InnerMongolia ................................................................................ [ 96480]
MK128995.1_China/2018/AnhuiXCGQ ................................................................................ [ 96480]
MH681419.1_ASFV/POL/2015 ................................................................................ [ 96480]
LS478113.1_Estonia_2014 ................................................................................ [ 96480]
KP843857.1_Odintsovo_02/14 ................................................................................ [ 96480]

```
FN557520.1_strain_E75             ................................................................................ [ 96480]
AY261366.1_Warthog                ..........................t..............a...................................... [ 96480]
AY261365.1_Warmbaths              ..........................t..............a...................................... [ 96480]
AY261364.1_Tengani_62             ..........................t..............a...................................... [ 96480]
AY261363.1_Pretoriuskop/96/4      ..........................t..............a...................................... [ 96480]
AY261362.1_Mkuzi_1979             ................................................................................ [ 96480]
AY261361.1_Malawi_Lil-20/1_1983   .....c....................t..............a...................................... [ 96480]
AY261360.1_Kenya_1950             ...........c..............t..............a...................................... [ 96480]
AM712240.1_OURT_88/3              ................................................................................ [ 96480]
AM712239.1_Benin_97/1             ................................................................................ [ 96480]
KM111295.1_Ken06.Bus              ..........................t..............a...................................... [ 96480]
KM111294.1_Ken05/Tk1              ...........c..............t..............a...................................... [ 96480]
KM262844.1_strain_L60             ................................................................................ [ 96480]
KM102979.1_26544/OG10_Italy       ................................................................................ [ 96480]
KM262845.1_strain_NHV             ................................................................................ [ 96480]
KX354450.1_47/Ss/2008             ................................................................................ [ 96480]
KP055815.1_strain_BA71            ................................................................................ [ 96480]
MH025920.1_strain_R35             ..........................t..............a...................................a.. [ 96480]
MH025919.1_strain_N10             ..........................t..............a...................................a.. [ 96480]
MH025918.1_strain_R25             ..........................t..............a...................................a.. [ 96480]
MH025917.1_strain_R7              ..........................t..............a...................................a.. [ 96480]
MH025916.1_strain_R8              ..........................t..............a...................................a.. [ 96480]

FR682468.2_Georgia_2007/1         ggagcctgagtacgtctgctgttgtgccacctatcaatcgcacgatggcgaataccggtatcgatttgtttggtaccaac [ 96560]
ASF_IND_20_CAD543                 ................................................................................ [ 96560]
LR899193.1_Germany_2020/1         ................................................................................ [ 96560]
MN393477.1_Wuhan_2019             ................................................................................ [ 96560]
MW396979.1_Timor-Leste/2019/1     ................................................................................ [ 96560]
MK940252.1_CN/2019/InnerMongolia  ................................................................................ [ 96560]
MK128995.1_China/2018/AnhuiXCGQ   ................................................................................ [ 96560]
MH681419.1_ASFV/POL/2015          ................................................................................ [ 96560]
LS478113.1_Estonia_2014           ................................................................................ [ 96560]
KP843857.1_Odintsovo_02/14        ................................................................................ [ 96560]
FN557520.1_strain_E75             .......................................a........................................ [ 96560]
AY261366.1_Warthog                ............t........c................................t..............c.......... [ 96560]
AY261365.1_Warmbaths              .....................c.............................g.................c.......... [ 96560]
AY261364.1_Tengani_62             .....................c...............................................c.......... [ 96560]
AY261363.1_Pretoriuskop/96/4      .....................c................................t..............c.......... [ 96560]
AY261362.1_Mkuzi_1979             ................................................................................ [ 96560]
AY261361.1_Malawi_Lil-20/1_1983   .........a................................t.....t............................... [ 96560]
AY261360.1_Kenya_1950             .........a.......................................t............................... [ 96560]
AM712240.1_OURT_88/3              .......................................a........................................ [ 96560]
AM712239.1_Benin_97/1             .......................................a........................................ [ 96560]
KM111295.1_Ken06.Bus              .........a.......................................t............................... [ 96560]
```

```
KM111294.1_Ken05/Tk1            .........a......................................t............................... [ 96560]
KM262844.1_strain_L60           .......................................a........................................ [ 96560]
KM102979.1_26544/OG10_Italy     .......................................a........................................ [ 96560]
KM262845.1_strain_NHV           .......................................a........................................ [ 96560]
KX354450.1_47/Ss/2008           .......................................a........................................ [ 96560]
KP055815.1_strain_BA71          .......................................a........................................ [ 96560]
MH025920.1_strain_R35           .........a......................................t............................... [ 96560]
MH025919.1_strain_N10           .........a......................................t............................... [ 96560]
MH025918.1_strain_R25           .........a......................................t............................... [ 96560]
MH025917.1_strain_R7            .........a......................................t............................... [ 96560]
MH025916.1_strain_R8            .........a......................................t............................... [ 96560]

FR682468.2_Georgia_2007/1       ccttcatagaggcttataatgccatagaggcggccctggatcccctggaaaccattatcctgaacctcattgcggcacga [ 96640]
ASF_IND_20_CAD543               ................................................................................ [ 96640]
LR899193.1_Germany_2020/1       ................................................................................ [ 96640]
MN393477.1_Wuhan_2019           ................................................................................ [ 96640]
MW396979.1_Timor-Leste/2019/1   ................................................................................ [ 96640]
MK940252.1_CN/2019/InnerMongolia ................................................................................ [ 96640]
MK128995.1_China/2018/AnhuiXCGQ ................................................................................ [ 96640]
MH681419.1_ASFV/POL/2015        ................................................................................ [ 96640]
LS478113.1_Estonia_2014         ................................................................................ [ 96640]
KP843857.1_Odintsovo_02/14      ................................................................................ [ 96640]
FN557520.1_strain_E75           ................................................................................ [ 96640]
AY261366.1_Warthog              ....t................................a........a................................. [ 96640]
AY261365.1_Warmbaths            .....................................a........a................................. [ 96640]
AY261364.1_Tengani_62           .....................................a........a................................. [ 96640]
AY261363.1_Pretoriuskop/96/4    .....................................a........a................................. [ 96640]
AY261362.1_Mkuzi_1979           ................................................................................ [ 96640]
AY261361.1_Malawi_Lil-20/1_1983 .............................a....t..a...t..t.......t..............t............ [ 96640]
AY261360.1_Kenya_1950           .......c..............t......a.a..t..a..c..............c...t.................... [ 96640]
AM712240.1_OURT_88/3            ................................................................................ [ 96640]
AM712239.1_Benin_97/1           ................................................................................ [ 96640]
KM111295.1_Ken06.Bus            .......c..............t......ata..t..a.....t..a................................. [ 96640]
KM111294.1_Ken05/Tk1            .......c..............t......a.a..t..a..c..............c........................ [ 96640]
KM262844.1_strain_L60           ................................................................................ [ 96640]
KM102979.1_26544/OG10_Italy     ................................................................................ [ 96640]
KM262845.1_strain_NHV           ................................................................................ [ 96640]
KX354450.1_47/Ss/2008           ................................................................................ [ 96640]
KP055815.1_strain_BA71          ................................................................................ [ 96640]
MH025920.1_strain_R35           .......c..............t......ata..t..a.....t.................................... [ 96640]
MH025919.1_strain_N10           .......c..............t......ata..t..a.....t.................................... [ 96640]
MH025918.1_strain_R25           .......c..............t......ata..t..a.....t.................................... [ 96640]
MH025917.1_strain_R7            .......c..............t......ata..t..a.....t.................................... [ 96640]
MH025916.1_strain_R8            .......c..............t......ata..t..a.....t.................................... [ 96640]
```

```
FR682468.2_Georgia_2007/1        gatctagacttcgttgttcacatatttccttataataagggccatgaagactatttggcctccacgcaacttattctcaa [ 96720]
ASF_IND_20_CAD543                ................................................................................ [ 96720]
LR899193.1_Germany_2020/1        ................................................................................ [ 96720]
MN393477.1_Wuhan_2019            ................................................................................ [ 96720]
MW396979.1_Timor-Leste/2019/1    ................................................................................ [ 96720]
MK940252.1_CN/2019/InnerMongolia ................................................................................ [ 96720]
MK128995.1_China/2018/AnhuiXCGQ  ................................................................................ [ 96720]
MH681419.1_ASFV/POL/2015         ................................................................................ [ 96720]
LS478113.1_Estonia_2014          ................................................................................ [ 96720]
KP843857.1_Odintsovo_02/14       ................................................................................ [ 96720]
FN557520.1_strain_E75            ................................................................................ [ 96720]
AY261366.1_Warthog               ................................................................................ [ 96720]
AY261365.1_Warmbaths             ................................................................................ [ 96720]
AY261364.1_Tengani_62            ................................................................................ [ 96720]
AY261363.1_Pretoriuskop/96/4     ................................................................................ [ 96720]
AY261362.1_Mkuzi_1979            ........t....................................................................... [ 96720]
AY261361.1_Malawi_Lil-20/1_1983  .......................................................................c........ [ 96720]
AY261360.1_Kenya_1950            ...........t.............................a...........cc................c........ [ 96720]
AM712240.1_OURT_88/3             ................................................................................ [ 96720]
AM712239.1_Benin_97/1            ................................................................................ [ 96720]
KM111295.1_Ken06.Bus             ...........t.............................a...........cc................c........ [ 96720]
KM111294.1_Ken05/Tk1             ...........t.............................a...........cc................c........ [ 96720]
KM262844.1_strain_L60            ................................................................................ [ 96720]
KM102979.1_26544/OG10_Italy      ................................................................................ [ 96720]
KM262845.1_strain_NHV            ................................................................................ [ 96720]
KX354450.1_47/Ss/2008            ................................................................................ [ 96720]
KP055815.1_strain_BA71           ................................................................................ [ 96720]
MH025920.1_strain_R35            ...........t.............................at..........cc................c........ [ 96720]
MH025919.1_strain_N10            ...........t.............................a...........cc................c........ [ 96720]
MH025918.1_strain_R25            ...........t.............................at..........cc................c........ [ 96720]
MH025917.1_strain_R7             ...........t.............................at..........cc................c........ [ 96720]
MH025916.1_strain_R8             ...........t.............................at..........cc................c........ [ 96720]

FR682468.2_Georgia_2007/1        aatctttattgcgacgcttttaatggacattttaagaattaaagacaacacgttggacgttcacttaaattccgactata [ 96800]
ASF_IND_20_CAD543                ................................................................................ [ 96800]
LR899193.1_Germany_2020/1        ................................................................................ [ 96800]
MN393477.1_Wuhan_2019            ................................................................................ [ 96800]
MW396979.1_Timor-Leste/2019/1    ................................................................................ [ 96800]
MK940252.1_CN/2019/InnerMongolia ................................................................................ [ 96800]
MK128995.1_China/2018/AnhuiXCGQ  ................................................................................ [ 96800]
MH681419.1_ASFV/POL/2015         ................................................................................ [ 96800]
LS478113.1_Estonia_2014          ................................................................................ [ 96800]
KP843857.1_Odintsovo_02/14       ................................................................................ [ 96800]
```

```
FN557520.1_strain_E75           ................................................................................ [ 96800]
AY261366.1_Warthog              ................................................................................ [ 96800]
AY261365.1_Warmbaths            ................................................................................ [ 96800]
AY261364.1_Tengani_62           .........................................................t...................... [ 96800]
AY261363.1_Pretoriuskop/96/4    ................................................................................ [ 96800]
AY261362.1_Mkuzi_1979           ................................................................................ [ 96800]
AY261361.1_Malawi_Lil-20/1_1983 ................................................................................ [ 96800]
AY261360.1_Kenya_1950           ................................................t............................... [ 96800]
AM712240.1_OURT_88/3            ................................................................................ [ 96800]
AM712239.1_Benin_97/1           ................................................................................ [ 96800]
KM111295.1_Ken06.Bus            ............a...................................t............................... [ 96800]
KM111294.1_Ken05/Tk1            ................................................t............................... [ 96800]
KM262844.1_strain_L60           ................................................................................ [ 96800]
KM102979.1_26544/OG10_Italy     ................................................................................ [ 96800]
KM262845.1_strain_NHV           ................................................................................ [ 96800]
KX354450.1_47/Ss/2008           ................................................................................ [ 96800]
KP055815.1_strain_BA71          ................................................................................ [ 96800]
MH025920.1_strain_R35           ............a...................................t............................... [ 96800]
MH025919.1_strain_N10           ............a...................................t............................... [ 96800]
MH025918.1_strain_R25           ............a...................................t............................... [ 96800]
MH025917.1_strain_R7            ............a...................................t............................... [ 96800]
MH025916.1_strain_R8            ............a...................................t............................... [ 96800]

FR682468.2_Georgia_2007/1       ttattgtgatggagcggctttggcctcacataaaggatgccatagaacacttttttgaagcccataaggacttactaggg [ 96880]
ASF_IND_20_CAD543               ................................................................................ [ 96880]
LR899193.1_Germany_2020/1       ................................................................................ [ 96880]
MN393477.1_Wuhan_2019           ................................................................................ [ 96880]
MW396979.1_Timor-Leste/2019/1   ................................................................................ [ 96880]
MK940252.1_CN/2019/InnerMongolia ................................................................................ [ 96880]
MK128995.1_China/2018/AnhuiXCGQ ................................................................................ [ 96880]
MH681419.1_ASFV/POL/2015        ................................................................................ [ 96880]
LS478113.1_Estonia_2014         ................................................................................ [ 96880]
KP843857.1_Odintsovo_02/14      ................................................................................ [ 96880]
FN557520.1_strain_E75           ................................................................................ [ 96880]
AY261366.1_Warthog              ............................................................................g... [ 96880]
AY261365.1_Warmbaths            ......................................................................t.....g... [ 96880]
AY261364.1_Tengani_62           ............................................................................g... [ 96880]
AY261363.1_Pretoriuskop/96/4    ......................................................................t.....g... [ 96880]
AY261362.1_Mkuzi_1979           ................................................................................ [ 96880]
AY261361.1_Malawi_Lil-20/1_1983 ............................t.....a..............g..........................g... [ 96880]
AY261360.1_Kenya_1950           ...................c..............a.........................................g... [ 96880]
AM712240.1_OURT_88/3            ................................................................................ [ 96880]
AM712239.1_Benin_97/1           ................................................................................ [ 96880]
KM111295.1_Ken06.Bus            ............................t.....a.........................................g..a [ 96880]
```

```
KM111294.1_Ken05/Tk1              ...................c..............a.........................................g... [ 96880]
KM262844.1_strain_L60             ................................................................................ [ 96880]
KM102979.1_26544/OG10_Italy       ................................................................................ [ 96880]
KM262845.1_strain_NHV             ................................................................................ [ 96880]
KX354450.1_47/Ss/2008             ................................................................................ [ 96880]
KP055815.1_strain_BA71            ................................................................................ [ 96880]
MH025920.1_strain_R35             ............................t.....a.........................................g..a [ 96880]
MH025919.1_strain_N10             ............................t.....a.........................................g..a [ 96880]
MH025918.1_strain_R25             ............................t.....a.........................................g..a [ 96880]
MH025917.1_strain_R7              ............................t.....a.........................................g..a [ 96880]
MH025916.1_strain_R8              ............................t.....a.........................................g..a [ 96880]

FR682468.2_Georgia_2007/1         tacttaattgcctttcgcaatggggggaactttgcaggaagtcttagaccctcctgtgggcaaaagattgttcccctaac [ 96960]
ASF_IND_20_CAD543                 ................................................................................ [ 96960]
LR899193.1_Germany_2020/1         ................................................................................ [ 96960]
MN393477.1_Wuhan_2019             ................................................................................ [ 96960]
MW396979.1_Timor-Leste/2019/1     ................................................................................ [ 96960]
MK940252.1_CN/2019/InnerMongolia  ................................................................................ [ 96960]
MK128995.1_China/2018/AnhuiXCGQ   ................................................................................ [ 96960]
MH681419.1_ASFV/POL/2015          ................................................................................ [ 96960]
LS478113.1_Estonia_2014           ................................................................................ [ 96960]
KP843857.1_Odintsovo_02/14        ................................................................................ [ 96960]
FN557520.1_strain_E75             ................................................................................ [ 96960]
AY261366.1_Warthog                ...c................c........t.................................................. [ 96960]
AY261365.1_Warmbaths              ...c................c........................................................... [ 96960]
AY261364.1_Tengani_62             ...c................c.....a..t.................................................. [ 96960]
AY261363.1_Pretoriuskop/96/4      ...c................c.....a..t.................................................. [ 96960]
AY261362.1_Mkuzi_1979             ................................................................................ [ 96960]
AY261361.1_Malawi_Lil-20/1_1983   .......................t..a..t.................................................. [ 96960]
AY261360.1_Kenya_1950             ...c...................t..a..................................................... [ 96960]
AM712240.1_OURT_88/3              ................................................................................ [ 96960]
AM712239.1_Benin_97/1             ................................................................................ [ 96960]
KM111295.1_Ken06.Bus              ...c...................t..a..................................................... [ 96960]
KM111294.1_Ken05/Tk1              ...c...................t..a..................................................... [ 96960]
KM262844.1_strain_L60             ................................................................................ [ 96960]
KM102979.1_26544/OG10_Italy       ................................................................................ [ 96960]
KM262845.1_strain_NHV             ................................................................................ [ 96960]
KX354450.1_47/Ss/2008             ................................................................................ [ 96960]
KP055815.1_strain_BA71            ................................................................................ [ 96960]
MH025920.1_strain_R35             ...c...................t..a..................................................... [ 96960]
MH025919.1_strain_N10             ...c...................t..a..................................................... [ 96960]
MH025918.1_strain_R25             ...c...................t..a..................................................... [ 96960]
MH025917.1_strain_R7              ...c...................t..a..................................................... [ 96960]
MH025916.1_strain_R8              ...c...................t..a..................................................... [ 96960]
```

```
FR682468.2_Georgia_2007/1        gattcgagaggtcctacaaatgaatgatattaatttagccgtatggcgggaggtgtttattatgcaggaatgttccgact [ 97040]
ASF_IND_20_CAD543                ................................................................................ [ 97040]
LR899193.1_Germany_2020/1        ................................................................................ [ 97040]
MN393477.1_Wuhan_2019            .....................................................g.......................... [ 97040]
MW396979.1_Timor-Leste/2019/1    ................................................................................ [ 97040]
MK940252.1_CN/2019/InnerMongolia ................................................................................ [ 97040]
MK128995.1_China/2018/AnhuiXCGQ  ................................................................................ [ 97040]
MH681419.1_ASFV/POL/2015         ................................................................................ [ 97040]
LS478113.1_Estonia_2014          ................................................................................ [ 97040]
KP843857.1_Odintsovo_02/14       ................................................................................ [ 97040]
FN557520.1_strain_E75            ................................................................................ [ 97040]
AY261366.1_Warthog               ...........c..............................g..................................... [ 97040]
AY261365.1_Warmbaths             ...........c.................................................................... [ 97040]
AY261364.1_Tengani_62            ...........c......................c........................................t.... [ 97040]
AY261363.1_Pretoriuskop/96/4     ...........c.................................................................... [ 97040]
AY261362.1_Mkuzi_1979            ................................................................................ [ 97040]
AY261361.1_Malawi_Lil-20/1_1983  ...........c......g..t....................g..................................... [ 97040]
AY261360.1_Kenya_1950            ......t....c.t.......t.......................................................... [ 97040]
AM712240.1_OURT_88/3             ................................................................................ [ 97040]
AM712239.1_Benin_97/1            ................................................................................ [ 97040]
KM111295.1_Ken06.Bus             .........a.c.........t.......................................................... [ 97040]
KM111294.1_Ken05/Tk1             ......t....c.........t.......................................................... [ 97040]
KM262844.1_strain_L60            ................................................................................ [ 97040]
KM102979.1_26544/OG10_Italy      ................................................................................ [ 97040]
KM262845.1_strain_NHV            ................................................................................ [ 97040]
KX354450.1_47/Ss/2008            ................................................................................ [ 97040]
KP055815.1_strain_BA71           ................................................................................ [ 97040]
MH025920.1_strain_R35            .........a.c.........t.......................................................... [ 97040]
MH025919.1_strain_N10            .........a.c.........t.......................................................... [ 97040]
MH025918.1_strain_R25            .........a.c.........t.......................................................... [ 97040]
MH025917.1_strain_R7             .........a.c.........t.......................................................... [ 97040]
MH025916.1_strain_R8             .........a.c.........t.......................................................... [ 97040]

FR682468.2_Georgia_2007/1        tagtcatcaatgggatagcgccctgtttccccatttttaacacgtggacgtatttgcaaggtattaaccagatttttttt [ 97120]
ASF_IND_20_CAD543                ................................................................................ [ 97120]
LR899193.1_Germany_2020/1        ................................................................................ [ 97120]
MN393477.1_Wuhan_2019            ................................................................................ [ 97120]
MW396979.1_Timor-Leste/2019/1    ................................................................................ [ 97120]
MK940252.1_CN/2019/InnerMongolia ................................................................................ [ 97120]
MK128995.1_China/2018/AnhuiXCGQ  ................................................................................ [ 97120]
MH681419.1_ASFV/POL/2015         ................................................................................ [ 97120]
LS478113.1_Estonia_2014          ................................................................................ [ 97120]
KP843857.1_Odintsovo_02/14       ................................................................................ [ 97120]
```

```
FN557520.1_strain_E75              .............................................................c.................. [ 97120]
AY261366.1_Warthog                 .............................................................c.................. [ 97120]
AY261365.1_Warmbaths               .............................................................c.................. [ 97120]
AY261364.1_Tengani_62              ..........................................................g..c.................. [ 97120]
AY261363.1_Pretoriuskop/96/4       ...........................................t..............g..c.................. [ 97120]
AY261362.1_Mkuzi_1979              .............................................................c.................. [ 97120]
AY261361.1_Malawi_Lil-20/1_1983    ...............................t.............................c.................. [ 97120]
AY261360.1_Kenya_1950              .............a.................t...........t..............g..c.................. [ 97120]
AM712240.1_OURT_88/3               .............................................................c.................. [ 97120]
AM712239.1_Benin_97/1              .............................................................c.................. [ 97120]
KM111295.1_Ken06.Bus               .............a.....a...........t........t..t..............g..c.................. [ 97120]
KM111294.1_Ken05/Tk1               .............a........g........t...........t..............g..c.................. [ 97120]
KM262844.1_strain_L60              .............................................................c.................. [ 97120]
KM102979.1_26544/OG10_Italy        .................a...........................................c.................. [ 97120]
KM262845.1_strain_NHV              .............................................................c.................. [ 97120]
KX354450.1_47/Ss/2008              .................a...........................................c.................. [ 97120]
KP055815.1_strain_BA71             .............................................................c.................. [ 97120]
MH025920.1_strain_R35              .............a.....a...........t........t..t..............g..c.................. [ 97120]
MH025919.1_strain_N10              .............a.....a...........t........t..t..............g..c.................. [ 97120]
MH025918.1_strain_R25              .............a.....a...........t........t..t..............g..c.................. [ 97120]
MH025917.1_strain_R7               .............a.....a...........t........t..t..............g..c.................. [ 97120]
MH025916.1_strain_R8               .............a.....a...........t........t..t..............g..c.................. [ 97120]

FR682468.2_Georgia_2007/1          gaaaacacgtctttgcaggagaaatttaaaaaagattttattgcccgagagctttccaaagaaattatcaagggccaaaa [ 97200]
ASF_IND_20_CAD543                  ................................................................................ [ 97200]
LR899193.1_Germany_2020/1          ................................................................................ [ 97200]
MN393477.1_Wuhan_2019              ................................................................................ [ 97200]
MW396979.1_Timor-Leste/2019/1      ................................................................................ [ 97200]
MK940252.1_CN/2019/InnerMongolia   ................................................................................ [ 97200]
MK128995.1_China/2018/AnhuiXCGQ    ................................................................................ [ 97200]
MH681419.1_ASFV/POL/2015           ................................................................................ [ 97200]
LS478113.1_Estonia_2014            ................................................................................ [ 97200]
KP843857.1_Odintsovo_02/14         ................................................................................ [ 97200]
FN557520.1_strain_E75              ................................................................................ [ 97200]
AY261366.1_Warthog                 ................................................................................ [ 97200]
AY261365.1_Warmbaths               ....................a............................................c.............. [ 97200]
AY261364.1_Tengani_62              ....................a............................................c.............. [ 97200]
AY261363.1_Pretoriuskop/96/4       .........................................c.......................c.............. [ 97200]
AY261362.1_Mkuzi_1979              ....................a........................................................... [ 97200]
AY261361.1_Malawi_Lil-20/1_1983    .....t..........................g..............g..............g........a........ [ 97200]
AY261360.1_Kenya_1950              .....t..........................g..............g..............g........a........ [ 97200]
AM712240.1_OURT_88/3               ................................................................................ [ 97200]
AM712239.1_Benin_97/1              ................................................................................ [ 97200]
KM111295.1_Ken06.Bus               .....t..........................g..............g..............g........a........ [ 97200]
```

```
KM111294.1_Ken05/Tk1           .....t..........................g..............g..............g........a........ [ 97200]
KM262844.1_strain_L60          ................................................................................ [ 97200]
KM102979.1_26544/OG10_Italy    ................................................................................ [ 97200]
KM262845.1_strain_NHV          ................................................................................ [ 97200]
KX354450.1_47/Ss/2008          ................................................................................ [ 97200]
KP055815.1_strain_BA71         ................................................................................ [ 97200]
MH025920.1_strain_R35          .....t..........................g..............g..............g........a........ [ 97200]
MH025919.1_strain_N10          .....t..........................g..............g..............g........a........ [ 97200]
MH025918.1_strain_R25          .....t..........................g..............g..............g........a........ [ 97200]
MH025917.1_strain_R7           .....t..........................g..............g..............g........a........ [ 97200]
MH025916.1_strain_R8           .....t..........................g..............g..............g........a........ [ 97200]

FR682468.2_Georgia_2007/1      aacgttgaatgacaaggagtttaaaaagttaagcctacatcaaatccagtacatggaatcctttctacttatgtcggatg [ 97280]
ASF_IND_20_CAD543              ................................................................................ [ 97280]
LR899193.1_Germany_2020/1      ................................................................................ [ 97280]
MN393477.1_Wuhan_2019          ................................................................................ [ 97280]
MW396979.1_Timor-Leste/2019/1  ................................................................................ [ 97280]
MK940252.1_CN/2019/InnerMongolia ................................................................................ [ 97280]
MK128995.1_China/2018/AnhuiXCGQ ................................................................................ [ 97280]
MH681419.1_ASFV/POL/2015       ................................................................................ [ 97280]
LS478113.1_Estonia_2014        ................................................................................ [ 97280]
KP843857.1_Odintsovo_02/14     ................................................................................ [ 97280]
FN557520.1_strain_E75          .................................t..............................t..........a.... [ 97280]
AY261366.1_Warthog             ................................................................................ [ 97280]
AY261365.1_Warmbaths           .............................................t.....t............................ [ 97280]
AY261364.1_Tengani_62          ................................................................................ [ 97280]
AY261363.1_Pretoriuskop/96/4   ...a.........................................t.....t............................ [ 97280]
AY261362.1_Mkuzi_1979          .............................................t.....t............................ [ 97280]
AY261361.1_Malawi_Lil-20/1_1983 ..tt..........ta..................t..........t.....t.......................a.... [ 97280]
AY261360.1_Kenya_1950          ..at..a........a.......................c.....t.....t............................ [ 97280]
AM712240.1_OURT_88/3           .................................t..............................t..........a.... [ 97280]
AM712239.1_Benin_97/1          .................................t..............................t..........a.... [ 97280]
KM111295.1_Ken06.Bus           ..at..a........a.......................c.....t.....t............................ [ 97280]
KM111294.1_Ken05/Tk1           ..at..a........a.....................t.c.....t.....t............................ [ 97280]
KM262844.1_strain_L60          .................................t..............................t..........a.... [ 97280]
KM102979.1_26544/OG10_Italy    .................................t..............................t..........a.... [ 97280]
KM262845.1_strain_NHV          .................................t..............................t..........a.... [ 97280]
KX354450.1_47/Ss/2008          .................................t..............................t..........a.... [ 97280]
KP055815.1_strain_BA71         .................................t..............................t..........a.... [ 97280]
MH025920.1_strain_R35          ..at..a........a.......................c.....t.....t............................ [ 97280]
MH025919.1_strain_N10          ..at..a........a.......................c.....t.....t............................ [ 97280]
MH025918.1_strain_R25          ..at..a........a.......................c.....t.....t............................ [ 97280]
MH025917.1_strain_R7           ..at..a........a.......................c.....t.....t............................ [ 97280]
MH025916.1_strain_R8           ..at..a........a.......................c.....t.....t............................ [ 97280]
```

```
FR682468.2_Georgia_2007/1        ttgccattatgattaccacagagtatgttggctatacccttcaatccctgccgggtattatttcgcgatccagctattta [ 97360]
ASF_IND_20_CAD543                ................................................................................ [ 97360]
LR899193.1_Germany_2020/1        ................................................................................ [ 97360]
MN393477.1_Wuhan_2019            ................................................................................ [ 97360]
MW396979.1_Timor-Leste/2019/1    ................................................................................ [ 97360]
MK940252.1_CN/2019/InnerMongolia ................................................................................ [ 97360]
MK128995.1_China/2018/AnhuiXCGQ  ................................................................................ [ 97360]
MH681419.1_ASFV/POL/2015         ................................................................................ [ 97360]
LS478113.1_Estonia_2014          ................................................................................ [ 97360]
KP843857.1_Odintsovo_02/14       ................................................................................ [ 97360]
FN557520.1_strain_E75            .........................c........c............................................. [ 97360]
AY261366.1_Warthog               ......................a......................................................... [ 97360]
AY261365.1_Warmbaths             ................................................................................ [ 97360]
AY261364.1_Tengani_62            ................................................................................ [ 97360]
AY261363.1_Pretoriuskop/96/4     .....................................t.......................................... [ 97360]
AY261362.1_Mkuzi_1979            ................................................................................ [ 97360]
AY261361.1_Malawi_Lil-20/1_1983  .........................c.......................a.............................. [ 97360]
AY261360.1_Kenya_1950            ..................................c..t.........t.a...........c...............g.. [ 97360]
AM712240.1_OURT_88/3             .........................c........c............................................. [ 97360]
AM712239.1_Benin_97/1            .........................c........c............................................. [ 97360]
KM111295.1_Ken06.Bus             ..................................c..t.........t.a...........c..............cg.. [ 97360]
KM111294.1_Ken05/Tk1             ..................................c..t.........t.a...........c...............g.. [ 97360]
KM262844.1_strain_L60            .........................c........c............................................. [ 97360]
KM102979.1_26544/OG10_Italy      .........................c........c............................................. [ 97360]
KM262845.1_strain_NHV            .........................c........c............................................. [ 97360]
KX354450.1_47/Ss/2008            .........................c........c............................................. [ 97360]
KP055815.1_strain_BA71           .........................c........c............................................. [ 97360]
MH025920.1_strain_R35            ..................................c..t.........t.a...........c..............cg.. [ 97360]
MH025919.1_strain_N10            ..................................c..t.........t.a...........c..............cg.. [ 97360]
MH025918.1_strain_R25            ..................................c..t.........t.a...........c..............cg.. [ 97360]
MH025917.1_strain_R7             ..................................c..t.........t.a...........c..............cg.. [ 97360]
MH025916.1_strain_R8             ..................................c..t.........t.a...........c..............cg.. [ 97360]

FR682468.2_Georgia_2007/1        tcccccatcgtgaaaaacattttgatggacgaagactcttttatgtccctactatttgacctatgctatggcgcctacgt [ 97440]
ASF_IND_20_CAD543                ................................................................................ [ 97440]
LR899193.1_Germany_2020/1        ................................................................................ [ 97440]
MN393477.1_Wuhan_2019            ................................................................................ [ 97440]
MW396979.1_Timor-Leste/2019/1    ................................................................................ [ 97440]
MK940252.1_CN/2019/InnerMongolia ................................................................................ [ 97440]
MK128995.1_China/2018/AnhuiXCGQ  ................................................................................ [ 97440]
MH681419.1_ASFV/POL/2015         ................................................................................ [ 97440]
LS478113.1_Estonia_2014          ................................................................................ [ 97440]
KP843857.1_Odintsovo_02/14       ................................................................................ [ 97440]
```

```
FN557520.1_strain_E75            ................................................................................ [ 97440]
AY261366.1_Warthog               .............................t.........................................t........ [ 97440]
AY261365.1_Warmbaths             ................................................................................ [ 97440]
AY261364.1_Tengani_62            ...............................................t.................t.............. [ 97440]
AY261363.1_Pretoriuskop/96/4     ................................................................................ [ 97440]
AY261362.1_Mkuzi_1979            ................................................................................ [ 97440]
AY261361.1_Malawi_Lil-20/1_1983  ..t..............t...........t..g.............................g................. [ 97440]
AY261360.1_Kenya_1950            .........................c.........t...............t....c.....g................. [ 97440]
AM712240.1_OURT_88/3             ................................................................................ [ 97440]
AM712239.1_Benin_97/1            ................................................................................ [ 97440]
KM111295.1_Ken06.Bus             .........................c.........t...............t....c.....g................. [ 97440]
KM111294.1_Ken05/Tk1             ........t................c.........t...............t....c.....g................. [ 97440]
KM262844.1_strain_L60            ................................................................................ [ 97440]
KM102979.1_26544/OG10_Italy      ................................................................................ [ 97440]
KM262845.1_strain_NHV            ................................................................................ [ 97440]
KX354450.1_47/Ss/2008            ................................................................................ [ 97440]
KP055815.1_strain_BA71           ................................................................................ [ 97440]
MH025920.1_strain_R35            .........................c.........t...............t....c.....g................. [ 97440]
MH025919.1_strain_N10            .........................c.........t...............t....c.....g................. [ 97440]
MH025918.1_strain_R25            .........................c.........t...............t....c.....g................. [ 97440]
MH025917.1_strain_R7             .........................c.........t...............t....c.....g................. [ 97440]
MH025916.1_strain_R8             .........................c.........t...............t....c.....g................. [ 97440]

FR682468.2_Georgia_2007/1        gttgcataaaaaagaaaatgtgattcacgcggatttgcacctgaataacatgacctactaccatttcaacccaaccagtt [ 97520]
ASF_IND_20_CAD543                ................................................................................ [ 97520]
LR899193.1_Germany_2020/1        ................................................................................ [ 97520]
MN393477.1_Wuhan_2019            ................................................................................ [ 97520]
MW396979.1_Timor-Leste/2019/1    ................................................................................ [ 97520]
MK940252.1_CN/2019/InnerMongolia ................................................................................ [ 97520]
MK128995.1_China/2018/AnhuiXCGQ  ................................................................................ [ 97520]
MH681419.1_ASFV/POL/2015         ................................................................................ [ 97520]
LS478113.1_Estonia_2014          ................................................................................ [ 97520]
KP843857.1_Odintsovo_02/14       ................................................................................ [ 97520]
FN557520.1_strain_E75            ...a.........................................c..t............................... [ 97520]
AY261366.1_Warthog               .c.c.................a....................a..c.................................. [ 97520]
AY261365.1_Warmbaths             .c.c.................a.......................c.................................. [ 97520]
AY261364.1_Tengani_62            .c...........................................c.................................. [ 97520]
AY261363.1_Pretoriuskop/96/4     .c.c.................a.......................c.................................. [ 97520]
AY261362.1_Mkuzi_1979            .............................................c..t............................... [ 97520]
AY261361.1_Malawi_Lil-20/1_1983  ...........................t..a..............c....................t............. [ 97520]
AY261360.1_Kenya_1950            .c................c..a.......................c.................................. [ 97520]
AM712240.1_OURT_88/3             .............................................c..t............................... [ 97520]
AM712239.1_Benin_97/1            ...a.........................................c..t............................... [ 97520]
KM111295.1_Ken06.Bus             .c................c..........................c.................................. [ 97520]
```

```
KM111294.1_Ken05/Tk1            .c................c..........................c.................................. [ 97520]
KM262844.1_strain_L60           .............................................c..t............................... [ 97520]
KM102979.1_26544/OG10_Italy     ...a.........................................c..t............................... [ 97520]
KM262845.1_strain_NHV           .............................................c..t............................... [ 97520]
KX354450.1_47/Ss/2008           ...a.........................................c..t............................... [ 97520]
KP055815.1_strain_BA71          .............................................c..t............................... [ 97520]
MH025920.1_strain_R35           .c................c..........................c.................................. [ 97520]
MH025919.1_strain_N10           .c................c..........................c.................................. [ 97520]
MH025918.1_strain_R25           .c................c..........................c.................................. [ 97520]
MH025917.1_strain_R7            .c................c..........................c.................................. [ 97520]
MH025916.1_strain_R8            .c................c..........................c.................................. [ 97520]

FR682468.2_Georgia_2007/1       ttacagatcgcaacaaaccaggaaaatacaccttaaaggtcaagaatcctgtgattgcctttataaccgggcccaaagtc [ 97600]
ASF_IND_20_CAD543               ................................................................................ [ 97600]
LR899193.1_Germany_2020/1       ................................................................................ [ 97600]
MN393477.1_Wuhan_2019           ................................................................................ [ 97600]
MW396979.1_Timor-Leste/2019/1   ................................................................................ [ 97600]
MK940252.1_CN/2019/InnerMongolia ................................................................................ [ 97600]
MK128995.1_China/2018/AnhuiXCGQ ................................................................................ [ 97600]
MH681419.1_ASFV/POL/2015        ................................................................................ [ 97600]
LS478113.1_Estonia_2014         ................................................................................ [ 97600]
KP843857.1_Odintsovo_02/14      ................................................................................ [ 97600]
FN557520.1_strain_E75           ................................................................................ [ 97600]
AY261366.1_Warthog              ................................................................................ [ 97600]
AY261365.1_Warmbaths            ................................................................................ [ 97600]
AY261364.1_Tengani_62           ..........................................................t..................... [ 97600]
AY261363.1_Pretoriuskop/96/4    ................................................................................ [ 97600]
AY261362.1_Mkuzi_1979           ................................................................................ [ 97600]
AY261361.1_Malawi_Lil-20/1_1983 .c..........................t..............t.....c.............................. [ 97600]
AY261360.1_Kenya_1950           .c..............g.....c....................t..c..c.................g............ [ 97600]
AM712240.1_OURT_88/3            ................................................................................ [ 97600]
AM712239.1_Benin_97/1           ................................................................................ [ 97600]
KM111295.1_Ken06.Bus            .c..............g.....c....................t.....c.................g............ [ 97600]
KM111294.1_Ken05/Tk1            .c..............g.....c....................t..c..c.................g............ [ 97600]
KM262844.1_strain_L60           ................................................................................ [ 97600]
KM102979.1_26544/OG10_Italy     ................................................................................ [ 97600]
KM262845.1_strain_NHV           ................................................................................ [ 97600]
KX354450.1_47/Ss/2008           ................................................................................ [ 97600]
KP055815.1_strain_BA71          ................................................................................ [ 97600]
MH025920.1_strain_R35           .c..............g.....c....................t.....c.................g............ [ 97600]
MH025919.1_strain_N10           .c..............g.....c....................t.....c.................g............ [ 97600]
MH025918.1_strain_R25           .c..............g.....c....................t.....c.................g............ [ 97600]
MH025917.1_strain_R7            .c..............g.....c....................t.....c.................g............ [ 97600]
MH025916.1_strain_R8            .c..............g.....c....................t.....c.................g............ [ 97600]
```

```
FR682468.2_Georgia_2007/1        gaaaccgaaacgtacgtgttcaagcacatagatgggttcggctgcatcattgactttagcagagccattatggggccaaa [ 97680]
ASF_IND_20_CAD543                ................................................................................ [ 97680]
LR899193.1_Germany_2020/1        ................................................................................ [ 97680]
MN393477.1_Wuhan_2019            ................................................................................ [ 97680]
MW396979.1_Timor-Leste/2019/1    ................................................................................ [ 97680]
MK940252.1_CN/2019/InnerMongolia ................................................................................ [ 97680]
MK128995.1_China/2018/AnhuiXCGQ  ................................................................................ [ 97680]
MH681419.1_ASFV/POL/2015         ................................................................................ [ 97680]
LS478113.1_Estonia_2014          ................................................................................ [ 97680]
KP843857.1_Odintsovo_02/14       ................................................................................ [ 97680]
FN557520.1_strain_E75            .............................................................................t.. [ 97680]
AY261366.1_Warthog               ...........a.................................................................... [ 97680]
AY261365.1_Warmbaths             ............................................t..t................................ [ 97680]
AY261364.1_Tengani_62            ...............................................t.................t.............. [ 97680]
AY261363.1_Pretoriuskop/96/4     ................................................................................ [ 97680]
AY261362.1_Mkuzi_1979            .............................................................................t.. [ 97680]
AY261361.1_Malawi_Lil-20/1_1983  .............................................................................t.. [ 97680]
AY261360.1_Kenya_1950            ..........................t..................................................c.. [ 97680]
AM712240.1_OURT_88/3             .............................................................................t.. [ 97680]
AM712239.1_Benin_97/1            .............................................................................t.. [ 97680]
KM111295.1_Ken06.Bus             .............................................................................c.. [ 97680]
KM111294.1_Ken05/Tk1             ..........................t..................................................c.. [ 97680]
KM262844.1_strain_L60            .............................................................................t.. [ 97680]
KM102979.1_26544/OG10_Italy      .............................................................................t.. [ 97680]
KM262845.1_strain_NHV            .............................................................................t.. [ 97680]
KX354450.1_47/Ss/2008            .............................................................................t.. [ 97680]
KP055815.1_strain_BA71           .............................................................................t.. [ 97680]
MH025920.1_strain_R35            .............................................................................c.. [ 97680]
MH025919.1_strain_N10            .............................................................................c.. [ 97680]
MH025918.1_strain_R25            .............................................................................c.. [ 97680]
MH025917.1_strain_R7             .............................................................................c.. [ 97680]
MH025916.1_strain_R8             .............................................................................c.. [ 97680]

FR682468.2_Georgia_2007/1        ccatgcaatcaagcttgagcggcagtacggcctcgcttttgtaaacaccttttaccgcaatcaaagtgagcatattttaa [ 97760]
ASF_IND_20_CAD543                ................................................................................ [ 97760]
LR899193.1_Germany_2020/1        ................................................................................ [ 97760]
MN393477.1_Wuhan_2019            ................................................................................ [ 97760]
MW396979.1_Timor-Leste/2019/1    ................................................................................ [ 97760]
MK940252.1_CN/2019/InnerMongolia ................................................................................ [ 97760]
MK128995.1_China/2018/AnhuiXCGQ  ................................................................................ [ 97760]
MH681419.1_ASFV/POL/2015         ................................................................................ [ 97760]
LS478113.1_Estonia_2014          ................................................................................ [ 97760]
KP843857.1_Odintsovo_02/14       ................................................................................ [ 97760]
```

```
FN557520.1_strain_E75          .........t.................................................................c.... [ 97760]
AY261366.1_Warthog             .........t.......................t.............................................. [ 97760]
AY261365.1_Warmbaths           .........t.......................t.............................................. [ 97760]
AY261364.1_Tengani_62          .........t.......................t.............................................. [ 97760]
AY261363.1_Pretoriuskop/96/4   .........t.......................t.............................................. [ 97760]
AY261362.1_Mkuzi_1979          .........t...................................t.................................. [ 97760]
AY261361.1_Malawi_Lil-20/1_1983 .........t..........................c........................................... [ 97760]
AY261360.1_Kenya_1950          .........t..........................c........................................... [ 97760]
AM712240.1_OURT_88/3           .........t.................................................................c.... [ 97760]
AM712239.1_Benin_97/1          .........t.................................................................c.... [ 97760]
KM111295.1_Ken06.Bus           .........t.................t........c........................................... [ 97760]
KM111294.1_Ken05/Tk1           .........t.................t........c........................................... [ 97760]
KM262844.1_strain_L60          .........t.................................................................c.... [ 97760]
KM102979.1_26544/OG10_Italy    .........t.................................................................c.... [ 97760]
KM262845.1_strain_NHV          .........t.................................................................c.... [ 97760]
KX354450.1_47/Ss/2008          .........t.................................................................c.... [ 97760]
KP055815.1_strain_BA71         .........t.................................................................c.... [ 97760]
MH025920.1_strain_R35          .........t.................t........c........................................... [ 97760]
MH025919.1_strain_N10          .........t.................t........c........................................... [ 97760]
MH025918.1_strain_R25          .........t.................t........c........................................... [ 97760]
MH025917.1_strain_R7           .........t.................t........c........................................... [ 97760]
MH025916.1_strain_R8           .........t.................t........c........................................... [ 97760]

FR682468.2_Georgia_2007/1      aggtattacggtactattttcctgaaatgctaaccaatcgcgaaaacgaaatacagggggtgattttatcaaactttaat [ 97840]
ASF_IND_20_CAD543              ...................y.........y.................................................. [ 97840]
LR899193.1_Germany_2020/1      ................................................................................ [ 97840]
MN393477.1_Wuhan_2019          ................................................................................ [ 97840]
MW396979.1_Timor-Leste/2019/1  ................................................................................ [ 97840]
MK940252.1_CN/2019/InnerMongolia ................................................................................ [ 97840]
MK128995.1_China/2018/AnhuiXCGQ ................................................................................ [ 97840]
MH681419.1_ASFV/POL/2015       ................................................................................ [ 97840]
LS478113.1_Estonia_2014        ................................................................................ [ 97840]
KP843857.1_Odintsovo_02/14     ................................................................................ [ 97840]
FN557520.1_strain_E75          ...................c.........t.................................................. [ 97840]
AY261366.1_Warthog             ..........c.t......c..............a...........t................................. [ 97840]
AY261365.1_Warmbaths           ..........c.t......c..............a............................................. [ 97840]
AY261364.1_Tengani_62          ..........c........c..............a............................................. [ 97840]
AY261363.1_Pretoriuskop/96/4   ..........c.t......c..............a............................................. [ 97840]
AY261362.1_Mkuzi_1979          ...................c............................................................ [ 97840]
AY261361.1_Malawi_Lil-20/1_1983 ...................c..............a...........t................................. [ 97840]
AY261360.1_Kenya_1950          .......g...........c..............a...........t................................. [ 97840]
AM712240.1_OURT_88/3           ...................c.........t.................................................. [ 97840]
AM712239.1_Benin_97/1          ...................c.........t.................................................. [ 97840]
KM111295.1_Ken06.Bus           .......g...........c..............a...........t................................. [ 97840]
```

```
KM111294.1_Ken05/Tk1            .......g...........c..............a...........t................................. [ 97840]
KM262844.1_strain_L60           ...................c.........t.................................................. [ 97840]
KM102979.1_26544/OG10_Italy     ...................c.........t.................................................. [ 97840]
KM262845.1_strain_NHV           ...................c.........t.................................................. [ 97840]
KX354450.1_47/Ss/2008           ...................c.........t.................................................. [ 97840]
KP055815.1_strain_BA71          ...................c.........t.................................................. [ 97840]
MH025920.1_strain_R35           .......g...........c..............a...........t................................. [ 97840]
MH025919.1_strain_N10           .......g...........c..............a...........t................................. [ 97840]
MH025918.1_strain_R25           .......g...........c..............a...........t................................. [ 97840]
MH025917.1_strain_R7            .......g...........c..............a...........t................................. [ 97840]
MH025916.1_strain_R8            .......g...........c..............a...........t................................. [ 97840]

FR682468.2_Georgia_2007/1       ttctttttcaatagcattactgccattgatttttacgccattgctagaaacctacgtagtatgctttctttggactattt [ 97920]
ASF_IND_20_CAD543               ................................................................................ [ 97920]
LR899193.1_Germany_2020/1       ................................................................................ [ 97920]
MN393477.1_Wuhan_2019           ................................................................................ [ 97920]
MW396979.1_Timor-Leste/2019/1   ................................................................................ [ 97920]
MK940252.1_CN/2019/InnerMongolia ................................................................................ [ 97920]
MK128995.1_China/2018/AnhuiXCGQ ................................................................................ [ 97920]
MH681419.1_ASFV/POL/2015        ................................................................................ [ 97920]
LS478113.1_Estonia_2014         ................................................................................ [ 97920]
KP843857.1_Odintsovo_02/14      ................................................................................ [ 97920]
FN557520.1_strain_E75           ................................................................................ [ 97920]
AY261366.1_Warthog              ................................................................................ [ 97920]
AY261365.1_Warmbaths            ................................................................................ [ 97920]
AY261364.1_Tengani_62           ...................................................t............................ [ 97920]
AY261363.1_Pretoriuskop/96/4    .....................................................g.......................... [ 97920]
AY261362.1_Mkuzi_1979           ................................................................................ [ 97920]
AY261361.1_Malawi_Lil-20/1_1983 .......................t...............................................a........ [ 97920]
AY261360.1_Kenya_1950           ..................................................t............................. [ 97920]
AM712240.1_OURT_88/3            ................................................................................ [ 97920]
AM712239.1_Benin_97/1           ................................................................................ [ 97920]
KM111295.1_Ken06.Bus            .......................................................................a........ [ 97920]
KM111294.1_Ken05/Tk1            ...................................t..............t............................. [ 97920]
KM262844.1_strain_L60           ................................................................................ [ 97920]
KM102979.1_26544/OG10_Italy     ................................................................................ [ 97920]
KM262845.1_strain_NHV           ................................................................................ [ 97920]
KX354450.1_47/Ss/2008           ................................................................................ [ 97920]
KP055815.1_strain_BA71          ................................................................................ [ 97920]
MH025920.1_strain_R35           .......................................................................a........ [ 97920]
MH025919.1_strain_N10           .......................................................................a........ [ 97920]
MH025918.1_strain_R25           .......................................................................a........ [ 97920]
MH025917.1_strain_R7            .......................................................................a........ [ 97920]
MH025916.1_strain_R8            .......................................................................a........ [ 97920]
```

```
FR682468.2_Georgia_2007/1        acacacctctgaggtgaaacgaaacgtagaaatttcgcaaacatttttggatacatgtcaatttttggaggaaaaggccg [ 98000]
ASF_IND_20_CAD543                ................................................................................ [ 98000]
LR899193.1_Germany_2020/1        ................................................................................ [ 98000]
MN393477.1_Wuhan_2019            ................................................................................ [ 98000]
MW396979.1_Timor-Leste/2019/1    ................................................................................ [ 98000]
MK940252.1_CN/2019/InnerMongolia ................................................................................ [ 98000]
MK128995.1_China/2018/AnhuiXCGQ  ................................................................................ [ 98000]
MH681419.1_ASFV/POL/2015         ................................................................................ [ 98000]
LS478113.1_Estonia_2014          ................................................................................ [ 98000]
KP843857.1_Odintsovo_02/14       ................................................................................ [ 98000]
FN557520.1_strain_E75            ..............................g................................................. [ 98000]
AY261366.1_Warthog               ..............................g................................................. [ 98000]
AY261365.1_Warmbaths             ..............................g................................................. [ 98000]
AY261364.1_Tengani_62            ...............a..............g.....a...........a............................... [ 98000]
AY261363.1_Pretoriuskop/96/4     ...............a..............g................................................. [ 98000]
AY261362.1_Mkuzi_1979            ..............................g................................................. [ 98000]
AY261361.1_Malawi_Lil-20/1_1983  ............a.................g...........c...........g......................... [ 98000]
AY261360.1_Kenya_1950            ............a.................g...........c..................................... [ 98000]
AM712240.1_OURT_88/3             ..............................g................................................. [ 98000]
AM712239.1_Benin_97/1            ..............................g................................................. [ 98000]
KM111295.1_Ken06.Bus             ............a.................g.......................g......................... [ 98000]
KM111294.1_Ken05/Tk1             ............a.................g...........c...........g......................... [ 98000]
KM262844.1_strain_L60            ..............................g................................................. [ 98000]
KM102979.1_26544/OG10_Italy      ..............................g................................................. [ 98000]
KM262845.1_strain_NHV            ..............................g................................................. [ 98000]
KX354450.1_47/Ss/2008            ..............................g................................................. [ 98000]
KP055815.1_strain_BA71           ..............................g................................................. [ 98000]
MH025920.1_strain_R35            ............a.................g.......................g......................... [ 98000]
MH025919.1_strain_N10            ............a.................g.......................g......................... [ 98000]
MH025918.1_strain_R25            ............a.................g.......................g......................... [ 98000]
MH025917.1_strain_R7             ............a.................g.......................g......................... [ 98000]
MH025916.1_strain_R8             ............a.................g.......................g......................... [ 98000]

FR682468.2_Georgia_2007/1        tggaatttttgtttaaaaatcttcatactgtcttatctggcaagccggtcgaaaaaacggccggggatgtgcttttaccc [ 98080]
ASF_IND_20_CAD543                .................................................t.............................. [ 98080]
LR899193.1_Germany_2020/1        ................................................................................ [ 98080]
MN393477.1_Wuhan_2019            ................................................................................ [ 98080]
MW396979.1_Timor-Leste/2019/1    ................................................................................ [ 98080]
MK940252.1_CN/2019/InnerMongolia ................................................................................ [ 98080]
MK128995.1_China/2018/AnhuiXCGQ  ................................................................................ [ 98080]
MH681419.1_ASFV/POL/2015         ................................................................................ [ 98080]
LS478113.1_Estonia_2014          ................................................................................ [ 98080]
KP843857.1_Odintsovo_02/14       ................................................................................ [ 98080]
```

```
FN557520.1_strain_E75             .................................................t.............................. [ 98080]
AY261366.1_Warthog                ................................................................................ [ 98080]
AY261365.1_Warmbaths              ...............................t................................................ [ 98080]
AY261364.1_Tengani_62             ................................................................................ [ 98080]
AY261363.1_Pretoriuskop/96/4      ................................................................................ [ 98080]
AY261362.1_Mkuzi_1979             ................................................................................ [ 98080]
AY261361.1_Malawi_Lil-20/1_1983   ................................................................................ [ 98080]
AY261360.1_Kenya_1950             ................................a...........................................g..t [ 98080]
AM712240.1_OURT_88/3              .................................................t.............................. [ 98080]
AM712239.1_Benin_97/1             .................................................t.............................. [ 98080]
KM111295.1_Ken06.Bus              ................................a...........................................g..t [ 98080]
KM111294.1_Ken05/Tk1              ................................a...........................................g..t [ 98080]
KM262844.1_strain_L60             .................................................t.............................. [ 98080]
KM102979.1_26544/OG10_Italy       .................................................t.............................. [ 98080]
KM262845.1_strain_NHV             .................................................t.............................. [ 98080]
KX354450.1_47/Ss/2008             .................................................t.............................. [ 98080]
KP055815.1_strain_BA71            .................................................t.............................. [ 98080]
MH025920.1_strain_R35             ................................a...........................................g..t [ 98080]
MH025919.1_strain_N10             ................................a...........................................g..t [ 98080]
MH025918.1_strain_R25             ................................a...........................................g..t [ 98080]
MH025917.1_strain_R7              ................................a...........................................g..t [ 98080]
MH025916.1_strain_R8              ................................a...........................................g..t [ 98080]

FR682468.2_Georgia_2007/1         atcgtatttaaaaaattttatacccaaatattcctaaaaatatattacggtcttttaccgtaatagatgtatacaatta [ 98160]
ASF_IND_20_CAD543                 ................................................................................ [ 98160]
LR899193.1_Germany_2020/1         ................................................................................ [ 98160]
MN393477.1_Wuhan_2019             ................................................................................ [ 98160]
MW396979.1_Timor-Leste/2019/1     ................................................................................ [ 98160]
MK940252.1_CN/2019/InnerMongolia  ................................................................................ [ 98160]
MK128995.1_China/2018/AnhuiXCGQ   ................................................................................ [ 98160]
MH681419.1_ASFV/POL/2015          ................................................................................ [ 98160]
LS478113.1_Estonia_2014           ................................................................................ [ 98160]
KP843857.1_Odintsovo_02/14        ................................................................................ [ 98160]
FN557520.1_strain_E75             ................................................................................ [ 98160]
AY261366.1_Warthog                ................................................................................ [ 98160]
AY261365.1_Warmbaths              ................................................................................ [ 98160]
AY261364.1_Tengani_62             ................................................................................ [ 98160]
AY261363.1_Pretoriuskop/96/4      ................................................................................ [ 98160]
AY261362.1_Mkuzi_1979             ................................................................................ [ 98160]
AY261361.1_Malawi_Lil-20/1_1983   ................................................................................ [ 98160]
AY261360.1_Kenya_1950             ................................................................................ [ 98160]
AM712240.1_OURT_88/3              ................................................................................ [ 98160]
AM712239.1_Benin_97/1             ................................................................................ [ 98160]
KM111295.1_Ken06.Bus              ...........................................................t.................... [ 98160]
```

```
KM111294.1_Ken05/Tk1             ................................................................................ [ 98160]
KM262844.1_strain_L60            ................................................................................ [ 98160]
KM102979.1_26544/OG10_Italy      ................................................................................ [ 98160]
KM262845.1_strain_NHV            ................................................................................ [ 98160]
KX354450.1_47/Ss/2008            ................................................................................ [ 98160]
KP055815.1_strain_BA71           ................................................................................ [ 98160]
MH025920.1_strain_R35            ...........................................................t.................... [ 98160]
MH025919.1_strain_N10            ...........................................................t.................... [ 98160]
MH025918.1_strain_R25            ...........................................................t.................... [ 98160]
MH025917.1_strain_R7             ...........................................................t.................... [ 98160]
MH025916.1_strain_R8             ...........................................................t.................... [ 98160]

FR682468.2_Georgia_2007/1        taataatataaagcgttattctgggaaagctatacaaacgtttccaccctgggctcaaaccaaagaaatcttgacgcacg [ 98240]
ASF_IND_20_CAD543                ................................................................................ [ 98240]
LR899193.1_Germany_2020/1        ................................................................................ [ 98240]
MN393477.1_Wuhan_2019            ................................................................................ [ 98240]
MW396979.1_Timor-Leste/2019/1    ................................................................................ [ 98240]
MK940252.1_CN/2019/InnerMongolia ................................................................................ [ 98240]
MK128995.1_China/2018/AnhuiXCGQ  ................................................................................ [ 98240]
MH681419.1_ASFV/POL/2015         ................................................................................ [ 98240]
LS478113.1_Estonia_2014          ................................................................................ [ 98240]
KP843857.1_Odintsovo_02/14       ................................................................................ [ 98240]
FN557520.1_strain_E75            .....................c..c....................................................... [ 98240]
AY261366.1_Warthog               ................................................................................ [ 98240]
AY261365.1_Warmbaths             .....................c.......................................................... [ 98240]
AY261364.1_Tengani_62            c....................c.......................................................... [ 98240]
AY261363.1_Pretoriuskop/96/4     ................................................................................ [ 98240]
AY261362.1_Mkuzi_1979            .....................c.......................................................... [ 98240]
AY261361.1_Malawi_Lil-20/1_1983  .....................c.......................................................... [ 98240]
AY261360.1_Kenya_1950            ..........................................c..................................... [ 98240]
AM712240.1_OURT_88/3             .....................c..c....................................................... [ 98240]
AM712239.1_Benin_97/1            .....................c..c....................................................... [ 98240]
KM111295.1_Ken06.Bus             ..........................................c..................................... [ 98240]
KM111294.1_Ken05/Tk1             ..........................................c..................................... [ 98240]
KM262844.1_strain_L60            .....................c..c....................................................... [ 98240]
KM102979.1_26544/OG10_Italy      .....................c..c....................................................... [ 98240]
KM262845.1_strain_NHV            .....................c..c....................................................... [ 98240]
KX354450.1_47/Ss/2008            .....................c..c....................................................... [ 98240]
KP055815.1_strain_BA71           .....................c..c....................................................... [ 98240]
MH025920.1_strain_R35            ..........................................c..................................... [ 98240]
MH025919.1_strain_N10            ..........................................c..................................... [ 98240]
MH025918.1_strain_R25            ..........................................c..................................... [ 98240]
MH025917.1_strain_R7             ..........................................c..................................... [ 98240]
MH025916.1_strain_R8             ..........................................c..................................... [ 98240]
```

```
FR682468.2_Georgia_2007/1        ccgagggtcgtacatttgaagatatttttcctagaggagaattagtttttaaaaaggcttacgcagaaaacaaccatttg [ 98320]
ASF_IND_20_CAD543                ................................................................................ [ 98320]
LR899193.1_Germany_2020/1        ................................................................................ [ 98320]
MN393477.1_Wuhan_2019            ................................................................................ [ 98320]
MW396979.1_Timor-Leste/2019/1    ................................................................................ [ 98320]
MK940252.1_CN/2019/InnerMongolia ................................................................................ [ 98320]
MK128995.1_China/2018/AnhuiXCGQ  ................................................................................ [ 98320]
MH681419.1_ASFV/POL/2015         ................................................................................ [ 98320]
LS478113.1_Estonia_2014          ................................................................................ [ 98320]
KP843857.1_Odintsovo_02/14       ................................................................................ [ 98320]
FN557520.1_strain_E75            ................................................................................ [ 98320]
AY261366.1_Warthog               ................................................................................ [ 98320]
AY261365.1_Warmbaths             ................................................................................ [ 98320]
AY261364.1_Tengani_62            ................................................................................ [ 98320]
AY261363.1_Pretoriuskop/96/4     ................................................................................ [ 98320]
AY261362.1_Mkuzi_1979            ............................................................................-... [ 98320]
AY261361.1_Malawi_Lil-20/1_1983  ..........................................................................t..... [ 98320]
AY261360.1_Kenya_1950            ..........................................................................t..... [ 98320]
AM712240.1_OURT_88/3             ................................................................................ [ 98320]
AM712239.1_Benin_97/1            ................................................................................ [ 98320]
KM111295.1_Ken06.Bus             ..........................................................................t..... [ 98320]
KM111294.1_Ken05/Tk1             ..........................................................................t..... [ 98320]
KM262844.1_strain_L60            ................................................................................ [ 98320]
KM102979.1_26544/OG10_Italy      ................................................................................ [ 98320]
KM262845.1_strain_NHV            ................................................................................ [ 98320]
KX354450.1_47/Ss/2008            ................................................................................ [ 98320]
KP055815.1_strain_BA71           ................................................................................ [ 98320]
MH025920.1_strain_R35            ..........................................................................t..... [ 98320]
MH025919.1_strain_N10            ..........................................................................t..... [ 98320]
MH025918.1_strain_R25            ..........................................................................t..... [ 98320]
MH025917.1_strain_R7             ..........................................................................t..... [ 98320]
MH025916.1_strain_R8             ..........................................................................t..... [ 98320]

FR682468.2_Georgia_2007/1        gacaaaattttacagcgtattcgtgagcagcttgctaatgaaaatttgtaaggcttgcagttcttgtatggtcagaacct [ 98400]
ASF_IND_20_CAD543                ................................................................................ [ 98400]
LR899193.1_Germany_2020/1        ................................................................................ [ 98400]
MN393477.1_Wuhan_2019            ................................................................................ [ 98400]
MW396979.1_Timor-Leste/2019/1    ................................................................................ [ 98400]
MK940252.1_CN/2019/InnerMongolia ................................................................................ [ 98400]
MK128995.1_China/2018/AnhuiXCGQ  ................................................................................ [ 98400]
MH681419.1_ASFV/POL/2015         ................................................................................ [ 98400]
LS478113.1_Estonia_2014          ................................................................................ [ 98400]
KP843857.1_Odintsovo_02/14       ................................................................................ [ 98400]
```

```
FN557520.1_strain_E75           ................................................................................ [ 98400]
AY261366.1_Warthog              .......................c........................................................ [ 98400]
AY261365.1_Warmbaths            .......................c........................................................ [ 98400]
AY261364.1_Tengani_62           .......................c.................................t...................... [ 98400]
AY261363.1_Pretoriuskop/96/4    .......................c........................................................ [ 98400]
AY261362.1_Mkuzi_1979           .......................c........................................................ [ 98400]
AY261361.1_Malawi_Lil-20/1_1983 .......................c........................................................ [ 98400]
AY261360.1_Kenya_1950           ..t....................c................................................a....... [ 98400]
AM712240.1_OURT_88/3            ................................................................................ [ 98400]
AM712239.1_Benin_97/1           ................................................................................ [ 98400]
KM111295.1_Ken06.Bus            ..t....................c................................................a....... [ 98400]
KM111294.1_Ken05/Tk1            ..t....................c................................................a....... [ 98400]
KM262844.1_strain_L60           ................................................................................ [ 98400]
KM102979.1_26544/OG10_Italy     ................................................................................ [ 98400]
KM262845.1_strain_NHV           ................................................................................ [ 98400]
KX354450.1_47/Ss/2008           ................................................................................ [ 98400]
KP055815.1_strain_BA71          ................................................................................ [ 98400]
MH025920.1_strain_R35           ..t....................c................................................a....... [ 98400]
MH025919.1_strain_N10           ..t....................c................................................a....... [ 98400]
MH025918.1_strain_R25           ..t....................c................................................a....... [ 98400]
MH025917.1_strain_R7            ..t....................c................................................a....... [ 98400]
MH025916.1_strain_R8            ..t....................c................................................a....... [ 98400]

FR682468.2_Georgia_2007/1       atgtcgatggaaacattatttttcgctgcagctgcggcgaaagcgttcaaggggatagtcagaacttgctcgtctctagc [ 98480]
ASF_IND_20_CAD543               ................................................................................ [ 98480]
LR899193.1_Germany_2020/1       ................................................................................ [ 98480]
MN393477.1_Wuhan_2019           ................................................................................ [ 98480]
MW396979.1_Timor-Leste/2019/1   ................................................................................ [ 98480]
MK940252.1_CN/2019/InnerMongolia ................................................................................ [ 98480]
MK128995.1_China/2018/AnhuiXCGQ ................................................................................ [ 98480]
MH681419.1_ASFV/POL/2015        ................................................................................ [ 98480]
LS478113.1_Estonia_2014         ................................................................................ [ 98480]
KP843857.1_Odintsovo_02/14      ................................................................................ [ 98480]
FN557520.1_strain_E75           .............................................................a.................. [ 98480]
AY261366.1_Warthog              ..........c....................t.......................c........................ [ 98480]
AY261365.1_Warmbaths            ..........c............................................c........................ [ 98480]
AY261364.1_Tengani_62           ..........c................................t...........c........................ [ 98480]
AY261363.1_Pretoriuskop/96/4    ..........c............................................c........................ [ 98480]
AY261362.1_Mkuzi_1979           .............................................................a.................. [ 98480]
AY261361.1_Malawi_Lil-20/1_1983 ..........c..................................................................... [ 98480]
AY261360.1_Kenya_1950           ..........c............................................c........................ [ 98480]
AM712240.1_OURT_88/3            .............................................................a.................. [ 98480]
AM712239.1_Benin_97/1           .............................................................a.................. [ 98480]
KM111295.1_Ken06.Bus            ..........c............................................c........................ [ 98480]
```

```
KM111294.1_Ken05/Tk1            ..........c........................................................a............ [ 98480]
KM262844.1_strain_L60           .............................................................a.................. [ 98480]
KM102979.1_26544/OG10_Italy     .............................................................a.................. [ 98480]
KM262845.1_strain_NHV           .............................................................a.................. [ 98480]
KX354450.1_47/Ss/2008           .............................................................a.................. [ 98480]
KP055815.1_strain_BA71          .............................................................a.................. [ 98480]
MH025920.1_strain_R35           ..........c............................................c........................ [ 98480]
MH025919.1_strain_N10           ..........c............................................c........................ [ 98480]
MH025918.1_strain_R25           ..........c............................................c........................ [ 98480]
MH025917.1_strain_R7            ..........c............................................c........................ [ 98480]
MH025916.1_strain_R8            ..........c............................................c........................ [ 98480]

FR682468.2_Georgia_2007/1       aaggtgtaccacaccggggaaatggaagataagtacaagatttttattaaaaatgcaccctttgaccccacgaattgcca [ 98560]
ASF_IND_20_CAD543               ................................................................................ [ 98560]
LR899193.1_Germany_2020/1       ................................................................................ [ 98560]
MN393477.1_Wuhan_2019           ................................................................................ [ 98560]
MW396979.1_Timor-Leste/2019/1   ................................................................................ [ 98560]
MK940252.1_CN/2019/InnerMongolia ................................................................................ [ 98560]
MK128995.1_China/2018/AnhuiXCGQ ................................................................................ [ 98560]
MH681419.1_ASFV/POL/2015        ................................................................................ [ 98560]
LS478113.1_Estonia_2014         ................................................................................ [ 98560]
KP843857.1_Odintsovo_02/14      ................................................................................ [ 98560]
FN557520.1_strain_E75           ................................................................................ [ 98560]
AY261366.1_Warthog              ................................................................................ [ 98560]
AY261365.1_Warmbaths            ................................................................................ [ 98560]
AY261364.1_Tengani_62           ................................................................................ [ 98560]
AY261363.1_Pretoriuskop/96/4    ................................................................................ [ 98560]
AY261362.1_Mkuzi_1979           ................................................................................ [ 98560]
AY261361.1_Malawi_Lil-20/1_1983 ................................................................................ [ 98560]
AY261360.1_Kenya_1950           .....a..................................................g..............a........ [ 98560]
AM712240.1_OURT_88/3            ................................................................................ [ 98560]
AM712239.1_Benin_97/1           ................................................................................ [ 98560]
KM111295.1_Ken06.Bus            ..a.....t.....t................................................................. [ 98560]
KM111294.1_Ken05/Tk1            .....a..................................................g....................... [ 98560]
KM262844.1_strain_L60           ................................................................................ [ 98560]
KM102979.1_26544/OG10_Italy     .............................................................................t.. [ 98560]
KM262845.1_strain_NHV           ................................................................................ [ 98560]
KX354450.1_47/Ss/2008           .............................................................................t.. [ 98560]
KP055815.1_strain_BA71          ................................................................................ [ 98560]
MH025920.1_strain_R35           ..a.....t.....t................................................................. [ 98560]
MH025919.1_strain_N10           ..a.....t.....t................................................................. [ 98560]
MH025918.1_strain_R25           ..a.....t.....t................................................................. [ 98560]
MH025917.1_strain_R7            ..a.....t.....t................................................................. [ 98560]
MH025916.1_strain_R8            ..a.....t.....t................................................................. [ 98560]
```

```
FR682468.2_Georgia_2007/1        aataaaaaaggattgcccaaattgtcatttagactatttgacacaaatctgtattggaagccaaaaaatcattatattgg [ 98640]
ASF_IND_20_CAD543                ................................................................................ [ 98640]
LR899193.1_Germany_2020/1        ................................................................................ [ 98640]
MN393477.1_Wuhan_2019            ................................................................................ [ 98640]
MW396979.1_Timor-Leste/2019/1    ................................................................................ [ 98640]
MK940252.1_CN/2019/InnerMongolia ................................................................................ [ 98640]
MK128995.1_China/2018/AnhuiXCGQ  ................................................................................ [ 98640]
MH681419.1_ASFV/POL/2015         ................................................................................ [ 98640]
LS478113.1_Estonia_2014          ................................................................................ [ 98640]
KP843857.1_Odintsovo_02/14       ................................................................................ [ 98640]
FN557520.1_strain_E75            ..................g............................................................. [ 98640]
AY261366.1_Warthog               .....................................c.............c............................ [ 98640]
AY261365.1_Warmbaths             ...................................................c............................ [ 98640]
AY261364.1_Tengani_62            .....................................c.............c............................ [ 98640]
AY261363.1_Pretoriuskop/96/4     .....................................c.............c............................ [ 98640]
AY261362.1_Mkuzi_1979            ..................g............................................................. [ 98640]
AY261361.1_Malawi_Lil-20/1_1983  ........................c.................g........c.................t......c... [ 98640]
AY261360.1_Kenya_1950            ..........................................g........c.................t......c... [ 98640]
AM712240.1_OURT_88/3             ..................g............................................................. [ 98640]
AM712239.1_Benin_97/1            ..................g............................................................. [ 98640]
KM111295.1_Ken06.Bus             ..........................................g........c.................t......c... [ 98640]
KM111294.1_Ken05/Tk1             .......................................a..g........c.................t......c... [ 98640]
KM262844.1_strain_L60            ..................g............................................................. [ 98640]
KM102979.1_26544/OG10_Italy      ..................g............................................................. [ 98640]
KM262845.1_strain_NHV            ..................g............................................................. [ 98640]
KX354450.1_47/Ss/2008            ..................g............................................................. [ 98640]
KP055815.1_strain_BA71           ..................g............................................................. [ 98640]
MH025920.1_strain_R35            ..........................................g........c.................t......c... [ 98640]
MH025919.1_strain_N10            ..........................................g........c.................t......c... [ 98640]
MH025918.1_strain_R25            ..........................................g........c.................t......c... [ 98640]
MH025917.1_strain_R7             ..........................................g........c.................t......c... [ 98640]
MH025916.1_strain_R8             ..........................................g........c.................t......c... [ 98640]

FR682468.2_Georgia_2007/1        tgtgccgctgtggctatatgagcaacagaggataa---accatatcatcccaccgaattatgacattcctttaaaaccgt [ 98720]
ASF_IND_20_CAD543                ...................................---.......................................... [ 98720]
LR899193.1_Germany_2020/1        ...................................---.......................................... [ 98720]
MN393477.1_Wuhan_2019            ...................................---.......................................... [ 98720]
MW396979.1_Timor-Leste/2019/1    ...................................---.......................................... [ 98720]
MK940252.1_CN/2019/InnerMongolia ...................................---.......................................... [ 98720]
MK128995.1_China/2018/AnhuiXCGQ  ...................................---.......................................... [ 98720]
MH681419.1_ASFV/POL/2015         ...................................---.......................................... [ 98720]
LS478113.1_Estonia_2014          ...................................---.......................................... [ 98720]
KP843857.1_Odintsovo_02/14       ...................................---.......................................... [ 98720]
```

```
FN557520.1_strain_E75            ...................................---..................t..g.................... [ 98720]
AY261366.1_Warthog               ....t.............c................---..................................g....... [ 98720]
AY261365.1_Warmbaths             ....t.............c................acc..................................g....... [ 98720]
AY261364.1_Tengani_62            ....t.............c................---..................................g....... [ 98720]
AY261363.1_Pretoriuskop/96/4     ....t.............c................---..................................g....... [ 98720]
AY261362.1_Mkuzi_1979            ...................................---.......................................... [ 98720]
AY261361.1_Malawi_Lil-20/1_1983  ....t.............ca...............---...--...........a.................g....... [ 98720]
AY261360.1_Kenya_1950            ....t...........c.ca........g......---...--.c......g......c.............g....... [ 98720]
AM712240.1_OURT_88/3             ...................................---..................t..g.................... [ 98720]
AM712239.1_Benin_97/1            ...................................---..................t..g.................... [ 98720]
KM111295.1_Ken06.Bus             ....t...........c.ca........g......---...--.c......g......c.............g....... [ 98720]
KM111294.1_Ken05/Tk1             ....t...........c.ca........g......---...--.c......g......c.............g....... [ 98720]
KM262844.1_strain_L60            ...................................---..................t..g.................... [ 98720]
KM102979.1_26544/OG10_Italy      ...................................---..................t..g.................... [ 98720]
KM262845.1_strain_NHV            ...................................---..................t..g.................... [ 98720]
KX354450.1_47/Ss/2008            ...................................---..................t..g.................... [ 98720]
KP055815.1_strain_BA71           ...................................---..................t..g.................... [ 98720]
MH025920.1_strain_R35            ....t...........c.ca........g......---...--.c......g......c.............g....... [ 98720]
MH025919.1_strain_N10            ....t...........c.ca........g......---...--.c......g......c.............g....... [ 98720]
MH025918.1_strain_R25            ....t...........c.ca........g......---...--.c......g......c.............g....... [ 98720]
MH025917.1_strain_R7             ....t...........c.ca........g......---...--.c......g......c.............g....... [ 98720]
MH025916.1_strain_R8             ....t...........c.ca........g......---...--.c......g......c.............g....... [ 98720]

FR682468.2_Georgia_2007/1        -ccgcctaaatagttttcacacctttggtggcagactattttataaaaagtaatgttggttcatgaagataa--agtgtg [ 98800]
ASF_IND_20_CAD543                -................................................a......................--...... [ 98800]
LR899193.1_Germany_2020/1        -.......................................................................--...... [ 98800]
MN393477.1_Wuhan_2019            -.......................................................................--...... [ 98800]
MW396979.1_Timor-Leste/2019/1    -.......................................................................--...... [ 98800]
MK940252.1_CN/2019/InnerMongolia -.......................................................................--...... [ 98800]
MK128995.1_China/2018/AnhuiXCGQ  -.......................................................................--...... [ 98800]
MH681419.1_ASFV/POL/2015         -.......................................................................--...... [ 98800]
LS478113.1_Estonia_2014          -.......................................................................--...... [ 98800]
KP843857.1_Odintsovo_02/14       -.......................................................................--...... [ 98800]
FN557520.1_strain_E75            -................................................a......................--...... [ 98800]
AY261366.1_Warthog               -................................................a......................--...... [ 98800]
AY261365.1_Warmbaths             -................................................a......................--...... [ 98800]
AY261364.1_Tengani_62            -................................................a......................--...... [ 98800]
AY261363.1_Pretoriuskop/96/4     -................t...............................a......................--...... [ 98800]
AY261362.1_Mkuzi_1979            -................................................a......................--...... [ 98800]
AY261361.1_Malawi_Lil-20/1_1983  -.........--..................a..a..--...........a......................at...... [ 98800]
AY261360.1_Kenya_1950            c..a..........c........c.........a..a............a....a.................at...... [ 98800]
AM712240.1_OURT_88/3             -................................................a......................--...... [ 98800]
AM712239.1_Benin_97/1            -................................................a......................--...... [ 98800]
KM111295.1_Ken06.Bus             c..a..........c........c.........a..a............a....a.................at...... [ 98800]
```

```
KM111294.1_Ken05/Tk1            c..a..........c........c.........a..a............a....a.................at...... [ 98800]
KM262844.1_strain_L60           -................................................a......................--...... [ 98800]
KM102979.1_26544/OG10_Italy     -................................................a......................--...... [ 98800]
KM262845.1_strain_NHV           -................................................a......................--...... [ 98800]
KX354450.1_47/Ss/2008           -................................................a......................--...... [ 98800]
KP055815.1_strain_BA71          -................................................a......................--...... [ 98800]
MH025920.1_strain_R35           c..a..........c........c.........a..a............a....a.................at...... [ 98800]
MH025919.1_strain_N10           c..a..........c........c.........a..a............a....a.................at...... [ 98800]
MH025918.1_strain_R25           c..a..........c........c.........a..a............a....a.................at...... [ 98800]
MH025917.1_strain_R7            c..a..........c........c.........a..a............a....a.................at...... [ 98800]
MH025916.1_strain_R8            c..a..........c........c.........a..a............a....a.................at...... [ 98800]

FR682468.2_Georgia_2007/1       ccaaagaaacttttataaacaaatgattaatgtaggtgctagtcgtgtgtactt-aaacagggtattctatagccaagta [ 98880]
ASF_IND_20_CAD543               ......................................................-..........--------------- [ 98880]
LR899193.1_Germany_2020/1       ......................................................-......................... [ 98880]
MN393477.1_Wuhan_2019           ......................................................-......................... [ 98880]
MW396979.1_Timor-Leste/2019/1   ......................................................-......................... [ 98880]
MK940252.1_CN/2019/InnerMongolia ......................................................-......................... [ 98880]
MK128995.1_China/2018/AnhuiXCGQ ......................................................-......................... [ 98880]
MH681419.1_ASFV/POL/2015        ......................................................-......................... [ 98880]
LS478113.1_Estonia_2014         ......................................................-......................... [ 98880]
KP843857.1_Odintsovo_02/14      ......................................................-......................... [ 98880]
FN557520.1_strain_E75           ......................................................-..........--------------- [ 98880]
AY261366.1_Warthog              ...................t..............-...................-......................... [ 98880]
AY261365.1_Warmbaths            ...................t..............-...................-......................... [ 98880]
AY261364.1_Tengani_62           ...................t..............-...................-......................... [ 98880]
AY261363.1_Pretoriuskop/96/4    ...................t..............-...................-......................... [ 98880]
AY261362.1_Mkuzi_1979           ......................................................-..........--------------- [ 98880]
AY261361.1_Malawi_Lil-20/1_1983 ...................t..t.tg.........a..................a..........--------------- [ 98880]
AY261360.1_Kenya_1950           ..............-----t....tg........-...................-..............g.......... [ 98880]
AM712240.1_OURT_88/3            ......................................................-..........--------------- [ 98880]
AM712239.1_Benin_97/1           ......................................................-..........--------------- [ 98880]
KM111295.1_Ken06.Bus            ..............-----t....tg........-...................-..............g..a.------ [ 98880]
KM111294.1_Ken05/Tk1            ..............-----t....tg........-...................-..............g.......... [ 98880]
KM262844.1_strain_L60           ......................................................-..........--------------- [ 98880]
KM102979.1_26544/OG10_Italy     ......................................................-..........--------------- [ 98880]
KM262845.1_strain_NHV           ......................................................-..........--------------- [ 98880]
KX354450.1_47/Ss/2008           ......................................................-..........--------------- [ 98880]
KP055815.1_strain_BA71          ......................................................-..........--------------- [ 98880]
MH025920.1_strain_R35           ..............-----t....tg........-...................-..............g..a.------ [ 98880]
MH025919.1_strain_N10           ..............-----t....tg........-...................-..............g..a.------ [ 98880]
MH025918.1_strain_R25           ..............-----t....tg........-...................-..............g..a.------ [ 98880]
MH025917.1_strain_R7            ..............-----t....tg........-...................-..............g..a.------ [ 98880]
MH025916.1_strain_R8            ..............-----t....tg........-...................-..............g..a.------ [ 98880]
```

```
FR682468.2_Georgia_2007/1        ttttctatagccaagtattttcta---------------------------------tagccagtattagtcaagtattt [ 98960]
ASF_IND_20_CAD543                --......................---------------------------------....................... [ 98960]
LR899193.1_Germany_2020/1        ........................---------------------------------....................... [ 98960]
MN393477.1_Wuhan_2019            ........................---------------------------------....................... [ 98960]
MW396979.1_Timor-Leste/2019/1    ........................---------------------------------....................... [ 98960]
MK940252.1_CN/2019/InnerMongolia ........................---------------------------------....................... [ 98960]
MK128995.1_China/2018/AnhuiXCGQ  ........................---------------------------------....................... [ 98960]
MH681419.1_ASFV/POL/2015         ........................---------------------------------....................... [ 98960]
LS478113.1_Estonia_2014          ........................---------------------------------....................... [ 98960]
KP843857.1_Odintsovo_02/14       ........................---------------------------------....................... [ 98960]
FN557520.1_strain_E75            --......................---------------------------------..................c.... [ 98960]
AY261366.1_Warthog               ...........t............---------------------------------....................c.. [ 98960]
AY261365.1_Warmbaths             ...-----------------....---------------------------------....................... [ 98960]
AY261364.1_Tengani_62            ........................tagctaagtattttctgtagccagtattttctg....................... [ 98960]
AY261363.1_Pretoriuskop/96/4     ...........t............---------------------------------....................... [ 98960]
AY261362.1_Mkuzi_1979            --......................---------------------------------.........------------.. [ 98960]
AY261361.1_Malawi_Lil-20/1_1983  --.....................g---------------------------------.............c...a..... [ 98960]
AY261360.1_Kenya_1950            .......................g-------------------------------------------.........g... [ 98960]
AM712240.1_OURT_88/3             --......................---------------------------------..................c.... [ 98960]
AM712239.1_Benin_97/1            --......................---------------------------------..................c.... [ 98960]
KM111295.1_Ken06.Bus             -----------------------------------------------------------------------.....g... [ 98960]
KM111294.1_Ken05/Tk1             .......................g-------------------------------------------.........g... [ 98960]
KM262844.1_strain_L60            --......................---------------------------------..................c.... [ 98960]
KM102979.1_26544/OG10_Italy      --......................---------------------------------..................c.... [ 98960]
KM262845.1_strain_NHV            --......................---------------------------------..................c.... [ 98960]
KX354450.1_47/Ss/2008            --......................---------------------------------..................c.... [ 98960]
KP055815.1_strain_BA71           --......................---------------------------------..................c.... [ 98960]
MH025920.1_strain_R35            -----------------------------------------------------------------------.....g... [ 98960]
MH025919.1_strain_N10            -----------------------------------------------------------------------.....g... [ 98960]
MH025918.1_strain_R25            -----------------------------------------------------------------------.....g... [ 98960]
MH025917.1_strain_R7             -----------------------------------------------------------------------.....g... [ 98960]
MH025916.1_strain_R8             -----------------------------------------------------------------------.....g... [ 98960]

FR682468.2_Georgia_2007/1        agatgtcag------------------------------ggtatttttatagccagta---------------tttttct [ 99040]
ASF_IND_20_CAD543                .........------------------------------...................---------------....... [ 99040]
LR899193.1_Germany_2020/1        .........------------------------------...................---------------....... [ 99040]
MN393477.1_Wuhan_2019            .........------------------------------...................---------------....... [ 99040]
MW396979.1_Timor-Leste/2019/1    .........------------------------------...................---------------....... [ 99040]
MK940252.1_CN/2019/InnerMongolia .........------------------------------...................---------------....... [ 99040]
MK128995.1_China/2018/AnhuiXCGQ  .........------------------------------...................---------------....... [ 99040]
MH681419.1_ASFV/POL/2015         .........------------------------------...................---------------....... [ 99040]
LS478113.1_Estonia_2014          .........------------------------------...................---------------....... [ 99040]
KP843857.1_Odintsovo_02/14       .........------------------------------...................---------------....... [ 99040]
```

```
FN557520.1_strain_E75            .........-------------------------------..................---------------....... [ 99040]
AY261366.1_Warthog               ........--------------------------------..................---------------....... [ 99040]
AY261365.1_Warmbaths             ........--------------------------------..................---------------....... [ 99040]
AY261364.1_Tengani_62            .........-------------------------------..................---------------....... [ 99040]
AY261363.1_Pretoriuskop/96/4     ........--------------------------------..................---------------....... [ 99040]
AY261362.1_Mkuzi_1979            .........-------------------------------..................---------------....... [ 99040]
AY261361.1_Malawi_Lil-20/1_1983  ........aatat---------------tttagatgttaa..................--------------t....... [ 99040]
AY261360.1_Kenya_1950            ......agtaggtttttcataactagtatttttataacca..................tttttatagccagta....... [ 99040]
AM712240.1_OURT_88/3             .........-------------------------------..................---------------....... [ 99040]
AM712239.1_Benin_97/1            .........-------------------------------..................---------------....... [ 99040]
KM111295.1_Ken06.Bus             ......agtaggttttttatagccagtattttcataacca..................---------------....... [ 99040]
KM111294.1_Ken05/Tk1             ......agtagg----------------ttttcataacca..................---------------....... [ 99040]
KM262844.1_strain_L60            .........-------------------------------..................---------------....... [ 99040]
KM102979.1_26544/OG10_Italy      .........-------------------------------..................---------------....... [ 99040]
KM262845.1_strain_NHV            .........-------------------------------..................---------------....... [ 99040]
KX354450.1_47/Ss/2008            .........-------------------------------..................---------------....... [ 99040]
KP055815.1_strain_BA71           .........-------------------------------..................---------------....... [ 99040]
MH025920.1_strain_R35            ......agtaggttttttatagccagtattttcataacca..................----------------...... [ 99040]
MH025919.1_strain_N10            ......agtaggttttttatagccagtattttcataacca..................----------------...... [ 99040]
MH025918.1_strain_R25            ......agtaggttttttatagccagtattttcataacca..................----------------...... [ 99040]
MH025917.1_strain_R7             ......agtaggttttttatagccagtattttcataacca..................----------------...... [ 99040]
MH025916.1_strain_R8             ......agtaggttttttatagccagtattttcataacca..................----------------...... [ 99040]

FR682468.2_Georgia_2007/1        ---atatgtacaaactattccagtaaacatatgtgtgttctttattgagcag-catcat--------------------- [ 99120]
ASF_IND_20_CAD543                ata.................................................-......--------------------- [ 99120]
LR899193.1_Germany_2020/1        ---.................................................-......--------------------- [ 99120]
MN393477.1_Wuhan_2019            ---.................................................-......--------------------- [ 99120]
MW396979.1_Timor-Leste/2019/1    ---.................................................-......--------------------- [ 99120]
MK940252.1_CN/2019/InnerMongolia ---.................................................-......--------------------- [ 99120]
MK128995.1_China/2018/AnhuiXCGQ  ---.................................................-......--------------------- [ 99120]
MH681419.1_ASFV/POL/2015         ---.................................................-......--------------------- [ 99120]
LS478113.1_Estonia_2014          ---.................................................-......--------------------- [ 99120]
KP843857.1_Odintsovo_02/14       ---.................................................-......--------------------- [ 99120]
FN557520.1_strain_E75            ata.................................................-......--------------------- [ 99120]
AY261366.1_Warthog               ata........................-...--.................g.tt.....--------------------- [ 99120]
AY261365.1_Warmbaths             ata........................-......................g.tt.....--------------------- [ 99120]
AY261364.1_Tengani_62            atc...............................................g.tt.....--------------------- [ 99120]
AY261363.1_Pretoriuskop/96/4     ---........................-......................g.tt.....ggcatt--------------- [ 99120]
AY261362.1_Mkuzi_1979            ---.................................................-......--------------------- [ 99120]
AY261361.1_Malawi_Lil-20/1_1983  ata..........ca............a.c.........t...g.c....g.tt.....ggcatt--------------- [ 99120]
AY261360.1_Kenya_1950            ata...........a.......a....a.c.--..........gc.....gctt..a..ggcattaaaagaatgagtcta [ 99120]
AM712240.1_OURT_88/3             ata.................................................-......--------------------- [ 99120]
AM712239.1_Benin_97/1            ata.................................................-......--------------------- [ 99120]
KM111295.1_Ken06.Bus             ata...........a.......a....a.c.............gc.....g.tt..a..ggcatt--------------- [ 99120]
```

```
KM111294.1_Ken05/Tk1             ata...........a.......a....a.c.--..........gc.....gctt..a..ggcattaaaagaatgagtcta [ 99120]
KM262844.1_strain_L60            ata.................................................-......--------------------- [ 99120]
KM102979.1_26544/OG10_Italy      ata.................................................-......--------------------- [ 99120]
KM262845.1_strain_NHV            ata.................................................-......--------------------- [ 99120]
KX354450.1_47/Ss/2008            ata.................................................-......--------------------- [ 99120]
KP055815.1_strain_BA71           ata.................................................-......--------------------- [ 99120]
MH025920.1_strain_R35            ata...........a.......a....a.c.............gc.....g.tt..a..ggcatt--------------- [ 99120]
MH025919.1_strain_N10            ata...........a.......a....a.c.............gc.....g.tt..a..ggcatt--------------- [ 99120]
MH025918.1_strain_R25            ata...........a.......a....a.c.............gc.....g.tt..a..ggcatt--------------- [ 99120]
MH025917.1_strain_R7             ata...........a.......a....a.c.............gc.....g.tt..a..ggcatt--------------- [ 99120]
MH025916.1_strain_R8             ata...........a.......a....a.c.............gc.....g.tt..a..ggcatt--------------- [ 99120]

FR682468.2_Georgia_2007/1        -------------------------------------------------------------------------------- [ 99200]
ASF_IND_20_CAD543                -------------------------------------------------------------------------------- [ 99200]
LR899193.1_Germany_2020/1        -------------------------------------------------------------------------------- [ 99200]
MN393477.1_Wuhan_2019            -------------------------------------------------------------------------------- [ 99200]
MW396979.1_Timor-Leste/2019/1    -------------------------------------------------------------------------------- [ 99200]
MK940252.1_CN/2019/InnerMongolia -------------------------------------------------------------------------------- [ 99200]
MK128995.1_China/2018/AnhuiXCGQ  -------------------------------------------------------------------------------- [ 99200]
MH681419.1_ASFV/POL/2015         -------------------------------------------------------------------------------- [ 99200]
LS478113.1_Estonia_2014          -------------------------------------------------------------------------------- [ 99200]
KP843857.1_Odintsovo_02/14       -------------------------------------------------------------------------------- [ 99200]
FN557520.1_strain_E75            -------------------------------------------------------------------------------- [ 99200]
AY261366.1_Warthog               -------------------------------------------------------------------------------- [ 99200]
AY261365.1_Warmbaths             -------------------------------------------------------------------------------- [ 99200]
AY261364.1_Tengani_62            -------------------------------------------------------------------------------- [ 99200]
AY261363.1_Pretoriuskop/96/4     ----------------------aaaaaa---------------------------------------------------- [ 99200]
AY261362.1_Mkuzi_1979            -------------------------------------------------------------------------------- [ 99200]
AY261361.1_Malawi_Lil-20/1_1983  ----------------------aaaaaagcgagtctaataaaccgc---------------------------------- [ 99200]
AY261360.1_Kenya_1950            ataagacgctttaat-ggcattaaaacaaagagtctaataagacgctttaatgggcattaaaagaatgagtctaataagc [ 99200]
AM712240.1_OURT_88/3             -------------------------------------------------------------------------------- [ 99200]
AM712239.1_Benin_97/1            -------------------------------------------------------------------------------- [ 99200]
KM111295.1_Ken06.Bus             ----------------------aaaacaatgagtctaataagacgctttaatggacattaaaacaatgagtttaataaac [ 99200]
KM111294.1_Ken05/Tk1             ataagccgaactaatgggcattaaaacaatgagtctaataagctgatctaatgggcattaaaacaatgagtctaataagc [ 99200]
KM262844.1_strain_L60            -------------------------------------------------------------------------------- [ 99200]
KM102979.1_26544/OG10_Italy      -------------------------------------------------------------------------------- [ 99200]
KM262845.1_strain_NHV            -------------------------------------------------------------------------------- [ 99200]
KX354450.1_47/Ss/2008            -------------------------------------------------------------------------------- [ 99200]
KP055815.1_strain_BA71           -------------------------------------------------------------------------------- [ 99200]
MH025920.1_strain_R35            ----------------------aaaacaatgagtctaataagacgctttaatgggcattaaaacaatgagtttaataaat [ 99200]
MH025919.1_strain_N10            ----------------------aaaacaatgagtctaataagacgctttaatgggcattaaaacaatgagtttaataaat [ 99200]
MH025918.1_strain_R25            ----------------------aaaacaatgagtctaataagacgctttaatgggcattaaaacaatgagtttaataaat [ 99200]
MH025917.1_strain_R7             ----------------------aaaacaatgagtctaataagacgctttaatgggcattaaaacaatgagtttaataaat [ 99200]
MH025916.1_strain_R8             ----------------------aaaacaatgagtctaataagacgctttaatgggcattaaaacaatgagtttaataaat [ 99200]
```

```
FR682468.2_Georgia_2007/1        -------------------------------------------------------------------------------- [ 99280]
ASF_IND_20_CAD543                -------------------------------------------------------------------------------- [ 99280]
LR899193.1_Germany_2020/1        -------------------------------------------------------------------------------- [ 99280]
MN393477.1_Wuhan_2019            -------------------------------------------------------------------------------- [ 99280]
MW396979.1_Timor-Leste/2019/1    -------------------------------------------------------------------------------- [ 99280]
MK940252.1_CN/2019/InnerMongolia -------------------------------------------------------------------------------- [ 99280]
MK128995.1_China/2018/AnhuiXCGQ  -------------------------------------------------------------------------------- [ 99280]
MH681419.1_ASFV/POL/2015         -------------------------------------------------------------------------------- [ 99280]
LS478113.1_Estonia_2014          -------------------------------------------------------------------------------- [ 99280]
KP843857.1_Odintsovo_02/14       -------------------------------------------------------------------------------- [ 99280]
FN557520.1_strain_E75            -------------------------------------------------------------------------------- [ 99280]
AY261366.1_Warthog               -------------------------------------------------------------------------------- [ 99280]
AY261365.1_Warmbaths             -------------------------------------------------------------------------------- [ 99280]
AY261364.1_Tengani_62            -------------------------------------------------------------------------------- [ 99280]
AY261363.1_Pretoriuskop/96/4     ------------------------------------------------------------cgaatttattagaccgctct [ 99280]
AY261362.1_Mkuzi_1979            -------------------------------------------------------------------------------- [ 99280]
AY261361.1_Malawi_Lil-20/1_1983  ---------------------------------------------agggcattaaaaaagcgagtctaataaaccgcag- [ 99280]
AY261360.1_Kenya_1950            tgatctaatgggcattaaaacaatgagtctaataagacgctttaatgagcattaaaagaatgagtctaataagctgatct [ 99280]
AM712240.1_OURT_88/3             -------------------------------------------------------------------------------- [ 99280]
AM712239.1_Benin_97/1            -------------------------------------------------------------------------------- [ 99280]
KM111295.1_Ken06.Bus             tgatctaatgggcattaaaagaatgagtctaataagacgctttaatgggtattaaaagaatgaatctaataagacgcttt [ 99280]
KM111294.1_Ken05/Tk1             tgatctaatgggcattaaaacaatgagtctaataagctgatctaatgggcattataacaaagagtctaataagacgcttt [ 99280]
KM262844.1_strain_L60            -------------------------------------------------------------------------------- [ 99280]
KM102979.1_26544/OG10_Italy      -------------------------------------------------------------------------------- [ 99280]
KM262845.1_strain_NHV            -------------------------------------------------------------------------------- [ 99280]
KX354450.1_47/Ss/2008            -------------------------------------------------------------------------------- [ 99280]
KP055815.1_strain_BA71           -------------------------------------------------------------------------------- [ 99280]
MH025920.1_strain_R35            tgatctaatgggcattaaaagaatgagtctaataagacgctttaatgggtattaaaagaatgaatctaataagacgcttt [ 99280]
MH025919.1_strain_N10            tgatctaatgggcattaaaagaatgagtctaataagacgctttaatgggtattaaaagaatgaatctaataagacgcttt [ 99280]
MH025918.1_strain_R25            tgatctaatgggcattaaaagaatgagtctaataagacgctttaatgggtattaaaagaatgaatctaataagacgcttt [ 99280]
MH025917.1_strain_R7             tgatctaatgggcattaaaagaatgagtctaataagacgctttaatgggtattaaaagaatgaatctaataagacgcttt [ 99280]
MH025916.1_strain_R8             tgatctaatgggcattaaaagaatgagtctaataagacgctttaatgggtattaaaagaatgaatctaataagacgcttt [ 99280]

FR682468.2_Georgia_2007/1        ----ggcatt---------------------------------------aacaagtttattaaactgc--tctaatgggc [ 99360]
ASF_IND_20_CAD543                ----......---------------------------------------...................--.......... [ 99360]
LR899193.1_Germany_2020/1        ----......---------------------------------------...................--.......... [ 99360]
MN393477.1_Wuhan_2019            ----......---------------------------------------...................--.......... [ 99360]
MW396979.1_Timor-Leste/2019/1    ----......---------------------------------------...................--.......... [ 99360]
MK940252.1_CN/2019/InnerMongolia ----......---------------------------------------...................--.......... [ 99360]
MK128995.1_China/2018/AnhuiXCGQ  ----......---------------------------------------...................--.......... [ 99360]
MH681419.1_ASFV/POL/2015         ----......---------------------------------------...................--.......... [ 99360]
LS478113.1_Estonia_2014          ----......---------------------------------------...................--.......... [ 99360]
KP843857.1_Odintsovo_02/14       ----......---------------------------------------...................--.......... [ 99360]
```

```
FN557520.1_strain_E75             ----......---------------------------------------................c..--.......... [ 99360]
AY261366.1_Warthog                ----......aaaa---------------------------------ac...g.a.......g..c..--...c...... [ 99360]
AY261365.1_Warmbaths              ----......aaaa---------------------------------ac...g.a.......g..c..--...c...... [ 99360]
AY261364.1_Tengani_62             ----......aaaa---------------------------------ac...g.a.......g..c..--...c...... [ 99360]
AY261363.1_Pretoriuskop/96/4      catg......aaaa---------------------------------ac...g.a.......g..c..--...c...... [ 99360]
AY261362.1_Mkuzi_1979             ----......---------------------------------------...................--.......... [ 99360]
AY261361.1_Malawi_Lil-20/1_1983   ----......aaaa---------------------------------a-.g.....c..a.....c..ag.......... [ 99360]
AY261360.1_Kenya_1950             aatg......aaa----------------------------------ac..tg...c..a...gac..--.t........ [ 99360]
AM712240.1_OURT_88/3              ----......---------------------------------------................c..--.......... [ 99360]
AM712239.1_Benin_97/1             ----......---------------------------------------................c..--.......... [ 99360]
KM111295.1_Ken06.Bus              aatg..t...aaaagaatgaatctaataagctgatctaatgggcattac..tg...c..a...gac..--.t........ [ 99360]
KM111294.1_Ken05/Tk1              aatg......aaa----------------------------------ac..ag...c..a...gac..--.t........ [ 99360]
KM262844.1_strain_L60             ----......---------------------------------------................c..--.......... [ 99360]
KM102979.1_26544/OG10_Italy       ----......---------------------------------------................c..--.......... [ 99360]
KM262845.1_strain_NHV             ----......---------------------------------------................c..--.......... [ 99360]
KX354450.1_47/Ss/2008             ----......---------------------------------------................c..--.......... [ 99360]
KP055815.1_strain_BA71            ----......---------------------------------------................c..--.......... [ 99360]
MH025920.1_strain_R35             aatg..t...aaaagaatgaatctaataagctgatctaatgggcattac..tg...c..a...gac..--.t........ [ 99360]
MH025919.1_strain_N10             aatg..t...aaaagaatgaatctaataagctgatctaatgggcattac..tg...c..a...gac..--.t........ [ 99360]
MH025918.1_strain_R25             aatg..t...aaaagaatgaatctaataagctgatctaatgggcattac..tg...c..a...gac..--.t........ [ 99360]
MH025917.1_strain_R7              aatg..t...aaaagaatgaatctaataagctgatctaatgggcattac..tg...c..a...gac..--.t........ [ 99360]
MH025916.1_strain_R8              aatg..t...aaaagaatgaatctaataagctgatctaatgggcattac..tg...c..a...gac..--.t........ [ 99360]

FR682468.2_Georgia_2007/1         attaa--atgacaa-ctcggtgcttagcaaaagtgcctataccttttaacaattagggccgggaggcattcccagctttt [ 99440]
ASF_IND_20_CAD543                 .....--.......-................................................................. [ 99440]
LR899193.1_Germany_2020/1         .....--.......-................................................................. [ 99440]
MN393477.1_Wuhan_2019             .....--.......-................................................................. [ 99440]
MW396979.1_Timor-Leste/2019/1     .....--.......-................................................................. [ 99440]
MK940252.1_CN/2019/InnerMongolia  .....--.......-................................................................. [ 99440]
MK128995.1_China/2018/AnhuiXCGQ   .....--.......-................................................................. [ 99440]
MH681419.1_ASFV/POL/2015          .....--.......-................................................................. [ 99440]
LS478113.1_Estonia_2014           .....--.......-................................................................. [ 99440]
KP843857.1_Odintsovo_02/14        .....--.......-................................................................. [ 99440]
FN557520.1_strain_E75             .....--.......-..................g...........c..g............................... [ 99440]
AY261366.1_Warthog                .....--..a....c.c............................c..g..g........a................... [ 99440]
AY261365.1_Warmbaths              ..------.a....c.c............................c..g..g......................a..... [ 99440]
AY261364.1_Tengani_62             .....--.......-.c......................c.....c..g..g............................ [ 99440]
AY261363.1_Pretoriuskop/96/4      .....--..a....c.c............................c..g..g............................ [ 99440]
AY261362.1_Mkuzi_1979             .....--.......-..................g...........c..g............................... [ 99440]
AY261361.1_Malawi_Lil-20/1_1983   .....ag..tgt..-.c.................a.t.......cc..g..g......a...............at.... [ 99440]
AY261360.1_Kenya_1950             .....ag..ag...--cg...a....a......g.t---..a..cc..g..g.....a...................... [ 99440]
AM712240.1_OURT_88/3              .....--.......-..................g...........c..g............................... [ 99440]
AM712239.1_Benin_97/1             .....--.......-..................g...........c..g............................... [ 99440]
KM111295.1_Ken06.Bus              .....ag..ag...--.g...a....a......g.tt-......cc..g..g.....a...................... [ 99440]
```

```
KM111294.1_Ken05/Tk1            .....ag..ag...--cg...a....a......g.t---..a..cc..g..g.....a...........c.......... [ 99440]
KM262844.1_strain_L60           .....--.......-..................g...........c..g............................... [ 99440]
KM102979.1_26544/OG10_Italy     .....--.......-..................g...........c..g............................... [ 99440]
KM262845.1_strain_NHV           .....--.......-..................g...........c..g............................... [ 99440]
KX354450.1_47/Ss/2008           .....--.......-..................g...........c..g............................... [ 99440]
KP055815.1_strain_BA71          .....--.......-..................g...........c..g............................... [ 99440]
MH025920.1_strain_R35           .....ag..ag...--.g...a....a......g.tt-......cc..g..g.....a...................... [ 99440]
MH025919.1_strain_N10           .....ag..ag...--.g...a....a......g.tt-......cc..g..g.....a...................... [ 99440]
MH025918.1_strain_R25           .....ag..ag...--.g...a....a......g.tt-......cc..g..g.....a...................... [ 99440]
MH025917.1_strain_R7            .....ag..ag...--.g...a....a......g.tt-......cc..g..g.....a...................... [ 99440]
MH025916.1_strain_R8            .....ag..ag...--.g...a....a......g.tt-......cc..g..g.....a...................... [ 99440]

FR682468.2_Georgia_2007/1       ttctataatcagccatacagtacccctgagcctcatacacgggaataaggtccttccattccttgttgggatcggcgggc [ 99520]
ASF_IND_20_CAD543               ................................................................................ [ 99520]
LR899193.1_Germany_2020/1       ................................................................................ [ 99520]
MN393477.1_Wuhan_2019           ................................................................................ [ 99520]
MW396979.1_Timor-Leste/2019/1   ................................................................................ [ 99520]
MK940252.1_CN/2019/InnerMongolia ................................................................................ [ 99520]
MK128995.1_China/2018/AnhuiXCGQ ................................................................................ [ 99520]
MH681419.1_ASFV/POL/2015        ................................................................................ [ 99520]
LS478113.1_Estonia_2014         ................................................................................ [ 99520]
KP843857.1_Odintsovo_02/14      ................................................................................ [ 99520]
FN557520.1_strain_E75           .......................................t......g................................. [ 99520]
AY261366.1_Warthog              .................................................c.............................. [ 99520]
AY261365.1_Warmbaths            ................................................................................ [ 99520]
AY261364.1_Tengani_62           .........................t.............t........................................ [ 99520]
AY261363.1_Pretoriuskop/96/4    .................................................c...........t.................. [ 99520]
AY261362.1_Mkuzi_1979           ................................................................................ [ 99520]
AY261361.1_Malawi_Lil-20/1_1983 .................................................c..........................a... [ 99520]
AY261360.1_Kenya_1950           ..........g.............................a........a.............................. [ 99520]
AM712240.1_OURT_88/3            .......................................t......g................................. [ 99520]
AM712239.1_Benin_97/1           .......................................t......g................................. [ 99520]
KM111295.1_Ken06.Bus            ..........g.............................a........a.............................. [ 99520]
KM111294.1_Ken05/Tk1            ..........g.............................a........a.................a............ [ 99520]
KM262844.1_strain_L60           .......................................t......g................................. [ 99520]
KM102979.1_26544/OG10_Italy     .......................................t......g................................. [ 99520]
KM262845.1_strain_NHV           .......................................t......g................................. [ 99520]
KX354450.1_47/Ss/2008           .......................................t......g................................. [ 99520]
KP055815.1_strain_BA71          .......................................t......g................................. [ 99520]
MH025920.1_strain_R35           ..........g.............................a........a.............................. [ 99520]
MH025919.1_strain_N10           ..........g.............................a........a.............................. [ 99520]
MH025918.1_strain_R25           ..........g.............................a........a.............................. [ 99520]
MH025917.1_strain_R7            ..........g.............................a........a.............................. [ 99520]
MH025916.1_strain_R8            ..........g.............................a........a.............................. [ 99520]
```

```
FR682468.2_Georgia_2007/1        cagctctcaaatgaggtgtgaatgtaagggtcctgttctttttccttaatgaagcgtttaatctccatttgatgttgttt [ 99600]
ASF_IND_20_CAD543                ................................................................................ [ 99600]
LR899193.1_Germany_2020/1        ................................................................................ [ 99600]
MN393477.1_Wuhan_2019            ................................................................................ [ 99600]
MW396979.1_Timor-Leste/2019/1    ................................................................................ [ 99600]
MK940252.1_CN/2019/InnerMongolia ................................................................................ [ 99600]
MK128995.1_China/2018/AnhuiXCGQ  ................................................................................ [ 99600]
MH681419.1_ASFV/POL/2015         ................................................................................ [ 99600]
LS478113.1_Estonia_2014          ................................................................................ [ 99600]
KP843857.1_Odintsovo_02/14       ................................................................................ [ 99600]
FN557520.1_strain_E75            ................................................................................ [ 99600]
AY261366.1_Warthog               ................................................................................ [ 99600]
AY261365.1_Warmbaths             ................................................................................ [ 99600]
AY261364.1_Tengani_62            ................................................................................ [ 99600]
AY261363.1_Pretoriuskop/96/4     ................................................................................ [ 99600]
AY261362.1_Mkuzi_1979            ................................................................................ [ 99600]
AY261361.1_Malawi_Lil-20/1_1983  ....t........................................................................... [ 99600]
AY261360.1_Kenya_1950            ...........................................................................a.... [ 99600]
AM712240.1_OURT_88/3             ................................................................................ [ 99600]
AM712239.1_Benin_97/1            ................................................................................ [ 99600]
KM111295.1_Ken06.Bus             ...........................................................................a.... [ 99600]
KM111294.1_Ken05/Tk1             ...........................................................................a.... [ 99600]
KM262844.1_strain_L60            ................................................................................ [ 99600]
KM102979.1_26544/OG10_Italy      ................................................................................ [ 99600]
KM262845.1_strain_NHV            ................................................................................ [ 99600]
KX354450.1_47/Ss/2008            ................................................................................ [ 99600]
KP055815.1_strain_BA71           ................................................................................ [ 99600]
MH025920.1_strain_R35            ...........................................................................a.... [ 99600]
MH025919.1_strain_N10            ...........................................................................a.... [ 99600]
MH025918.1_strain_R25            ...........................................................................a.... [ 99600]
MH025917.1_strain_R7             ...........................................................................a.... [ 99600]
MH025916.1_strain_R8             ...........................................................................a.... [ 99600]

FR682468.2_Georgia_2007/1        acttttttgtttgcggcggagcgtgttccgcaccaatacgtaaaaaataccaagaatcacacataaaagaattattaaaa [ 99680]
ASF_IND_20_CAD543                ................................................................................ [ 99680]
LR899193.1_Germany_2020/1        ................................................................................ [ 99680]
MN393477.1_Wuhan_2019            ................................................................................ [ 99680]
MW396979.1_Timor-Leste/2019/1    ................................................................................ [ 99680]
MK940252.1_CN/2019/InnerMongolia ................................................................................ [ 99680]
MK128995.1_China/2018/AnhuiXCGQ  ................................................................................ [ 99680]
MH681419.1_ASFV/POL/2015         ................................................................................ [ 99680]
LS478113.1_Estonia_2014          ................................................................................ [ 99680]
KP843857.1_Odintsovo_02/14       ................................................................................ [ 99680]
```

```
FN557520.1_strain_E75          ............................................................g................... [ 99680]
AY261366.1_Warthog             .a.............................................................................. [ 99680]
AY261365.1_Warmbaths           .a............................t................................................. [ 99680]
AY261364.1_Tengani_62          ................................................................................ [ 99680]
AY261363.1_Pretoriuskop/96/4   .a..........................................................g................... [ 99680]
AY261362.1_Mkuzi_1979          .a.............................................................................. [ 99680]
AY261361.1_Malawi_Lil-20/1_1983 .........................................................a.t.................... [ 99680]
AY261360.1_Kenya_1950          .t.......................................................g..g................... [ 99680]
AM712240.1_OURT_88/3           ............................................................g................... [ 99680]
AM712239.1_Benin_97/1          ............................................................g................... [ 99680]
KM111295.1_Ken06.Bus           .t.......................................................g..g................... [ 99680]
KM111294.1_Ken05/Tk1           .t.......................................................g..g................... [ 99680]
KM262844.1_strain_L60          ............................................................g................... [ 99680]
KM102979.1_26544/OG10_Italy    ............................................................g................... [ 99680]
KM262845.1_strain_NHV          ............................................................g................... [ 99680]
KX354450.1_47/Ss/2008          ............................................................g................... [ 99680]
KP055815.1_strain_BA71         ............................................................g................... [ 99680]
MH025920.1_strain_R35          .t.......................................................g..g................... [ 99680]
MH025919.1_strain_N10          .t.......................................................g..g................... [ 99680]
MH025918.1_strain_R25          .t.......................................................g..g................... [ 99680]
MH025917.1_strain_R7           .t.......................................................g..g................... [ 99680]
MH025916.1_strain_R8           .t.......................................................g..g................... [ 99680]

FR682468.2_Georgia_2007/1      aaaatatcatcatcgcggggtttaaaaaacgatcccatgcaacaggaatcgttcttaaaaccttgtctggcagggctgta [ 99760]
ASF_IND_20_CAD543              ................................................................................ [ 99760]
LR899193.1_Germany_2020/1      ................................................................................ [ 99760]
MN393477.1_Wuhan_2019          ................................................................................ [ 99760]
MW396979.1_Timor-Leste/2019/1  ................................................................................ [ 99760]
MK940252.1_CN/2019/InnerMongolia ................................................................................ [ 99760]
MK128995.1_China/2018/AnhuiXCGQ ................................................................................ [ 99760]
MH681419.1_ASFV/POL/2015       ................................................................................ [ 99760]
LS478113.1_Estonia_2014        ................................................................................ [ 99760]
KP843857.1_Odintsovo_02/14     ................................................................................ [ 99760]
FN557520.1_strain_E75          ................................................................................ [ 99760]
AY261366.1_Warthog             ..........................................t...g.......................t......... [ 99760]
AY261365.1_Warmbaths           ..........................................t...g.......................t......... [ 99760]
AY261364.1_Tengani_62          ..............................................g.......................t......... [ 99760]
AY261363.1_Pretoriuskop/96/4   ..........................................t...g.......................t......... [ 99760]
AY261362.1_Mkuzi_1979          ..........................................t...g.......................t......... [ 99760]
AY261361.1_Malawi_Lil-20/1_1983 .............t........................a.......g.............................c... [ 99760]
AY261360.1_Kenya_1950          ..........g..a................................g..........................a..c... [ 99760]
AM712240.1_OURT_88/3           ................................................................................ [ 99760]
AM712239.1_Benin_97/1          ................................................................................ [ 99760]
KM111295.1_Ken06.Bus           ..........g..a................................g..........................a..c... [ 99760]
```

```
KM111294.1_Ken05/Tk1             ..........g..a................................g.............................c... [ 99760]
KM262844.1_strain_L60            ................................................................................ [ 99760]
KM102979.1_26544/OG10_Italy      ................................................................................ [ 99760]
KM262845.1_strain_NHV            ................................................................................ [ 99760]
KX354450.1_47/Ss/2008            ................................................................................ [ 99760]
KP055815.1_strain_BA71           ................................................................................ [ 99760]
MH025920.1_strain_R35            ..........g..a................................g..........................a..c... [ 99760]
MH025919.1_strain_N10            ..........g..a................................g..........................a..c... [ 99760]
MH025918.1_strain_R25            ..........g..a................................g..........................a..c... [ 99760]
MH025917.1_strain_R7             ..........g..a................................g..........................a..c... [ 99760]
MH025916.1_strain_R8             ..........g..a................................g..........................a..c... [ 99760]

FR682468.2_Georgia_2007/1        aacatgaagtctcctcctataatcggggtgggactgtagcctaacagttcaaggtcctgtcgttctagatacttattggc [ 99840]
ASF_IND_20_CAD543                ................................................................................ [ 99840]
LR899193.1_Germany_2020/1        ................................................................................ [ 99840]
MN393477.1_Wuhan_2019            ................................................................................ [ 99840]
MW396979.1_Timor-Leste/2019/1    ................................................................................ [ 99840]
MK940252.1_CN/2019/InnerMongolia ................................................................................ [ 99840]
MK128995.1_China/2018/AnhuiXCGQ  ................................................................................ [ 99840]
MH681419.1_ASFV/POL/2015         ................................................................................ [ 99840]
LS478113.1_Estonia_2014          ................................................................................ [ 99840]
KP843857.1_Odintsovo_02/14       ................................................................................ [ 99840]
FN557520.1_strain_E75            ................................g............................................... [ 99840]
AY261366.1_Warthog               ................................g...................................g........... [ 99840]
AY261365.1_Warmbaths             ................................g...................................g........... [ 99840]
AY261364.1_Tengani_62            .....t.g..................a.........................................g........... [ 99840]
AY261363.1_Pretoriuskop/96/4     ................................g...................................g........... [ 99840]
AY261362.1_Mkuzi_1979            ................................g...................................g........... [ 99840]
AY261361.1_Malawi_Lil-20/1_1983  .....t.ga.............g..ac.....g...........t..c...........g................ga.. [ 99840]
AY261360.1_Kenya_1950            .....t.g............g.g...c.....g...........t..c...........g.................a.. [ 99840]
AM712240.1_OURT_88/3             ................................g............................................... [ 99840]
AM712239.1_Benin_97/1            ................................g............................................... [ 99840]
KM111295.1_Ken06.Bus             .....t.g............g.g...c.....g...........t..c.....a.....g.................a.. [ 99840]
KM111294.1_Ken05/Tk1             .....t.g............g.g...c.....g...........t..c...........g.................a.. [ 99840]
KM262844.1_strain_L60            ................................g............................................... [ 99840]
KM102979.1_26544/OG10_Italy      ................................g............................................... [ 99840]
KM262845.1_strain_NHV            ................................g............................................... [ 99840]
KX354450.1_47/Ss/2008            ................................g............................................... [ 99840]
KP055815.1_strain_BA71           ................................g............................................... [ 99840]
MH025920.1_strain_R35            .....t.g............g.g...c.....g...........t..c.....a.....g.................a.. [ 99840]
MH025919.1_strain_N10            .....t.g............g.g...c.....g...........t..c.....a.....g.................a.. [ 99840]
MH025918.1_strain_R25            .....t.g............g.g...c.....g...........t..c.....a.....g.................a.. [ 99840]
MH025917.1_strain_R7             .....t.g............g.g...c.....g...........t..c.....a.....g.................a.. [ 99840]
MH025916.1_strain_R8             .....t.g............g.g...c.....g...........t..c.....a.....g.................a.. [ 99840]
```

```
FR682468.2_Georgia_2007/1        gaactgcccaccctttgcccccgtttttttattaatcaagcagcgctgcattttccaccattctaaatcttcaggagaaa [ 99920]
ASF_IND_20_CAD543                ................................................................................ [ 99920]
LR899193.1_Germany_2020/1        ................................................................................ [ 99920]
MN393477.1_Wuhan_2019            ................................................................................ [ 99920]
MW396979.1_Timor-Leste/2019/1    ................................................................................ [ 99920]
MK940252.1_CN/2019/InnerMongolia ................................................................................ [ 99920]
MK128995.1_China/2018/AnhuiXCGQ  ................................................................................ [ 99920]
MH681419.1_ASFV/POL/2015         ................................................................................ [ 99920]
LS478113.1_Estonia_2014          ................................................................................ [ 99920]
KP843857.1_Odintsovo_02/14       ................................................................................ [ 99920]
FN557520.1_strain_E75            ................................................................................ [ 99920]
AY261366.1_Warthog               ................................................................................ [ 99920]
AY261365.1_Warmbaths             ................................................................................ [ 99920]
AY261364.1_Tengani_62            ................................................................................ [ 99920]
AY261363.1_Pretoriuskop/96/4     ................................................................................ [ 99920]
AY261362.1_Mkuzi_1979            ................................................................................ [ 99920]
AY261361.1_Malawi_Lil-20/1_1983  a..t..t..g..............g........t.............................c.c.............. [ 99920]
AY261360.1_Kenya_1950            a..t..t..g..............g....................................................... [ 99920]
AM712240.1_OURT_88/3             ................................................................................ [ 99920]
AM712239.1_Benin_97/1            ................................................................................ [ 99920]
KM111295.1_Ken06.Bus             a..t..t..g.....c........g....................................................... [ 99920]
KM111294.1_Ken05/Tk1             a..t..t..g...........t..g....................................................... [ 99920]
KM262844.1_strain_L60            ................................................................................ [ 99920]
KM102979.1_26544/OG10_Italy      ................................................................................ [ 99920]
KM262845.1_strain_NHV            ................................................................................ [ 99920]
KX354450.1_47/Ss/2008            ................................................................................ [ 99920]
KP055815.1_strain_BA71           ................................................................................ [ 99920]
MH025920.1_strain_R35            a..t..t..g.....c........g....................................................... [ 99920]
MH025919.1_strain_N10            a..t..t..g.....c........g....................................................... [ 99920]
MH025918.1_strain_R25            a..t..t..g.....c........g....................................................... [ 99920]
MH025917.1_strain_R7             a..t..t..g.....c........g....................................................... [ 99920]
MH025916.1_strain_R8             a..t..t..g.....c........g....................................................... [ 99920]

FR682468.2_Georgia_2007/1        gctcaatgccatatatcaactttaacgttattgcatctttttcaatatccttatcaatttggctgagcttttgagcttta [100000]
ASF_IND_20_CAD543                ...................y............................................................ [100000]
LR899193.1_Germany_2020/1        ................................................................................ [100000]
MN393477.1_Wuhan_2019            ................................................................................ [100000]
MW396979.1_Timor-Leste/2019/1    ................................................................................ [100000]
MK940252.1_CN/2019/InnerMongolia ................................................................................ [100000]
MK128995.1_China/2018/AnhuiXCGQ  ................................................................................ [100000]
MH681419.1_ASFV/POL/2015         ................................................................................ [100000]
LS478113.1_Estonia_2014          ................................................................................ [100000]
KP843857.1_Odintsovo_02/14       ................................................................................ [100000]
```

```
FN557520.1_strain_E75              ...................t............................................................ [100000]
AY261366.1_Warthog                 ...............a......................................t............t............ [100000]
AY261365.1_Warmbaths               ...............a......................................t............t............ [100000]
AY261364.1_Tengani_62              ...............a..................g...................t.......t................. [100000]
AY261363.1_Pretoriuskop/96/4       ...............a......................................t.......t................. [100000]
AY261362.1_Mkuzi_1979              ...............a......................................t............t............ [100000]
AY261361.1_Malawi_Lil-20/1_1983    ....cg.c..g.......g...............g..............t....t......atc................ [100000]
AY261360.1_Kenya_1950              ....g...............c....a..g.....g..............t....t.......tc.............c.. [100000]
AM712240.1_OURT_88/3               ...................t............................................................ [100000]
AM712239.1_Benin_97/1              ...................t............................................................ [100000]
KM111295.1_Ken06.Bus               .........................a..g.....g..............t....t.......tc.............c.. [100000]
KM111294.1_Ken05/Tk1               ....g...............c....a..g.....g..............t....t.......tc.............c.. [100000]
KM262844.1_strain_L60              ...................t............................................................ [100000]
KM102979.1_26544/OG10_Italy        ...................t............................................................ [100000]
KM262845.1_strain_NHV              ...................t............................................................ [100000]
KX354450.1_47/Ss/2008              ...................t............................................................ [100000]
KP055815.1_strain_BA71             ...................t............................................................ [100000]
MH025920.1_strain_R35              .........................a..g.....g..............t....t.......tc.............c.. [100000]
MH025919.1_strain_N10              .........................a..g.....g..............t....t.......tc.............c.. [100000]
MH025918.1_strain_R25              .........................a..g.....g..............t....t.......tc.............c.. [100000]
MH025917.1_strain_R7               .........................a..g.....g..............t....t.......tc.............c.. [100000]
MH025916.1_strain_R8               .........................a..g.....g..............t....t.......tc.............c.. [100000]

FR682468.2_Georgia_2007/1          agcgggtctagtgtgtacttccatttaaacttagtgtcctgtagtttggctacatgaaatacggaacatttcggcggggc [100080]
ASF_IND_20_CAD543                  ................................................................................ [100080]
LR899193.1_Germany_2020/1          ................................................................................ [100080]
MN393477.1_Wuhan_2019              ................................................................................ [100080]
MW396979.1_Timor-Leste/2019/1      ................................................................................ [100080]
MK940252.1_CN/2019/InnerMongolia   ................................................................................ [100080]
MK128995.1_China/2018/AnhuiXCGQ    ................................................................................ [100080]
MH681419.1_ASFV/POL/2015           ................................................................................ [100080]
LS478113.1_Estonia_2014            ................................................................................ [100080]
KP843857.1_Odintsovo_02/14         ................................................................................ [100080]
FN557520.1_strain_E75              ..........................................................................t..... [100080]
AY261366.1_Warthog                 ..........a.............................................g.............c......... [100080]
AY261365.1_Warmbaths               ..g.....................................................g.............c......... [100080]
AY261364.1_Tengani_62              ..........t.............................................g.............c......... [100080]
AY261363.1_Pretoriuskop/96/4       ..g.....................................................g.............c......... [100080]
AY261362.1_Mkuzi_1979              ..g.....................................................g.............c......... [100080]
AY261361.1_Malawi_Lil-20/1_1983    .....a..............................................a.................c......c.. [100080]
AY261360.1_Kenya_1950              ..t..a........a.......................................................c...t..... [100080]
AM712240.1_OURT_88/3               .......................................................................t..t..... [100080]
AM712239.1_Benin_97/1              ..........................................................................t..... [100080]
KM111295.1_Ken06.Bus               ..t..a........a.......................................................c...t..... [100080]
```

KM111294.1_Ken05/Tk1 ..t..a........a.......................................................c...t..... [100080]
KM262844.1_strain_L60 ..........................................................................t..... [100080]
KM102979.1_26544/OG10_Italy ..........................................................................t..... [100080]
KM262845.1_strain_NHV .......................................................................t..t..... [100080]
KX354450.1_47/Ss/2008 ..........................................................................t..... [100080]
KP055815.1_strain_BA71 ..........................................................................t..... [100080]
MH025920.1_strain_R35 ..t..a........a.......................................................c...t..... [100080]
MH025919.1_strain_N10 ..t..a........a.......................................................c...t..... [100080]
MH025918.1_strain_R25 ..t..a........a.......................................................c...t..... [100080]
MH025917.1_strain_R7 ..t..a........a.......................................................c...t..... [100080]
MH025916.1_strain_R8 ..t..a........a.......................................................c...t..... [100080]

FR682468.2_Georgia_2007/1 ctttgtgacgcccttacactgcggaagtttatcattaggacaggcgcatagatgagactgcgccacagcatcgcgaacta [100160]
ASF_IND_20_CAD543 ................................................................................ [100160]
LR899193.1_Germany_2020/1 ................................................................................ [100160]
MN393477.1_Wuhan_2019 ................................................................................ [100160]
MW396979.1_Timor-Leste/2019/1 ................................................................................ [100160]
MK940252.1_CN/2019/InnerMongolia ................................................................................ [100160]
MK128995.1_China/2018/AnhuiXCGQ ................................................................................ [100160]
MH681419.1_ASFV/POL/2015 ................................................................................ [100160]
LS478113.1_Estonia_2014 ................................................................................ [100160]
KP843857.1_Odintsovo_02/14 ................................................................................ [100160]
FN557520.1_strain_E75 ................................................................................ [100160]
AY261366.1_Warthog ........................................................................a....... [100160]
AY261365.1_Warmbaths ........................................................................a....... [100160]
AY261364.1_Tengani_62 ................................................................................ [100160]
AY261363.1_Pretoriuskop/96/4 ................................................................................ [100160]
AY261362.1_Mkuzi_1979 ................................................................................ [100160]
AY261361.1_Malawi_Lil-20/1_1983 ............t........t...................................t...................... [100160]
AY261360.1_Kenya_1950 .....................t...........g........t......................t.............. [100160]
AM712240.1_OURT_88/3 ................................................................................ [100160]
AM712239.1_Benin_97/1 ................................................................................ [100160]
KM111295.1_Ken06.Bus .....................t...........g........t..................................... [100160]
KM111294.1_Ken05/Tk1 .....................t...........g........t......................t.............. [100160]
KM262844.1_strain_L60 ................................................................................ [100160]
KM102979.1_26544/OG10_Italy ................................................................................ [100160]
KM262845.1_strain_NHV ................................................................................ [100160]
KX354450.1_47/Ss/2008 ................................................................................ [100160]
KP055815.1_strain_BA71 ................................................................................ [100160]
MH025920.1_strain_R35 .....................t...........g........t..................................... [100160]
MH025919.1_strain_N10 .....................t...........g........t..................................... [100160]
MH025918.1_strain_R25 .....................t...........g........t..................................... [100160]
MH025917.1_strain_R7 .....................t...........g........t..................................... [100160]
MH025916.1_strain_R8 .....................t...........g........t..................................... [100160]

```
FR682468.2_Georgia_2007/1        catcgcagacggagtacattttcctcctatgttaaacaataaatttttttcatagctgaaatttgtgggcctatcttttc [100240]
ASF_IND_20_CAD543                ................................................................................ [100240]
LR899193.1_Germany_2020/1        ................................................................................ [100240]
MN393477.1_Wuhan_2019            ................................................................................ [100240]
MW396979.1_Timor-Leste/2019/1    ................................................................................ [100240]
MK940252.1_CN/2019/InnerMongolia ................................................................................ [100240]
MK128995.1_China/2018/AnhuiXCGQ  ................................................................................ [100240]
MH681419.1_ASFV/POL/2015         ................................................................................ [100240]
LS478113.1_Estonia_2014          ................................................................................ [100240]
KP843857.1_Odintsovo_02/14       ................................................................................ [100240]
FN557520.1_strain_E75            ................................................................................ [100240]
AY261366.1_Warthog               .....................................................................tt.c....... [100240]
AY261365.1_Warmbaths             .....................................................................tt.c....... [100240]
AY261364.1_Tengani_62            .....................................................................tt.c....... [100240]
AY261363.1_Pretoriuskop/96/4     .......................................................t.............tt.c....... [100240]
AY261362.1_Mkuzi_1979            ................................................................................ [100240]
AY261361.1_Malawi_Lil-20/1_1983  .......a.....a.................................................................. [100240]
AY261360.1_Kenya_1950            .......ag.t..................................................................... [100240]
AM712240.1_OURT_88/3             .........................................g...................................... [100240]
AM712239.1_Benin_97/1            ................................................................................ [100240]
KM111295.1_Ken06.Bus             .......ag.t..................................................................... [100240]
KM111294.1_Ken05/Tk1             .......ag.t..................................................................... [100240]
KM262844.1_strain_L60            ................................................................................ [100240]
KM102979.1_26544/OG10_Italy      ................................................................................ [100240]
KM262845.1_strain_NHV            .........................................g...................................... [100240]
KX354450.1_47/Ss/2008            ................................................................................ [100240]
KP055815.1_strain_BA71           ................................................................................ [100240]
MH025920.1_strain_R35            .......ag.t..................................................................... [100240]
MH025919.1_strain_N10            .......ag.t..................................................................... [100240]
MH025918.1_strain_R25            .......ag.t..................................................................... [100240]
MH025917.1_strain_R7             .......ag.t..................................................................... [100240]
MH025916.1_strain_R8             .......ag.t..................................................................... [100240]

FR682468.2_Georgia_2007/1        ccttgcccggataataattataagggagtgttgaaacatctgggagagaattgcttaaaaaatgggtttttgggaggggt [100320]
ASF_IND_20_CAD543                ................................................................................ [100320]
LR899193.1_Germany_2020/1        ................................................................................ [100320]
MN393477.1_Wuhan_2019            ................................................................................ [100320]
MW396979.1_Timor-Leste/2019/1    ................................................................................ [100320]
MK940252.1_CN/2019/InnerMongolia ................................................................................ [100320]
MK128995.1_China/2018/AnhuiXCGQ  ................................................................................ [100320]
MH681419.1_ASFV/POL/2015         ................................................................................ [100320]
LS478113.1_Estonia_2014          ................................................................................ [100320]
KP843857.1_Odintsovo_02/14       ................................................................................ [100320]
```

```
FN557520.1_strain_E75           .....................................g.......................................... [100320]
AY261366.1_Warthog              ....................................tg..c................................c.....c [100320]
AY261365.1_Warmbaths            .....................................g.......................................... [100320]
AY261364.1_Tengani_62           .....................................g......................................a... [100320]
AY261363.1_Pretoriuskop/96/4    .........................t..........tg...................................c.....c [100320]
AY261362.1_Mkuzi_1979           .....................................g.......................................... [100320]
AY261361.1_Malawi_Lil-20/1_1983 .........................t..c.......tg.....a...a............................a... [100320]
AY261360.1_Kenya_1950           .........................t..........tg..c......a.........................a...... [100320]
AM712240.1_OURT_88/3            .....................................g.......................................... [100320]
AM712239.1_Benin_97/1           .....................................g.......................................... [100320]
KM111295.1_Ken06.Bus            .........................t..........tg..c......a................................ [100320]
KM111294.1_Ken05/Tk1            .........................t..........tg..c......a................................ [100320]
KM262844.1_strain_L60           .....................................g.......................................... [100320]
KM102979.1_26544/OG10_Italy     .....................................g.......................................... [100320]
KM262845.1_strain_NHV           .....................................g.......................................... [100320]
KX354450.1_47/Ss/2008           .....................................g.......................................... [100320]
KP055815.1_strain_BA71          .....................................g.......................................... [100320]
MH025920.1_strain_R35           .........................t..........tg..c......a................................ [100320]
MH025919.1_strain_N10           .........................t..........tg..c......a................................ [100320]
MH025918.1_strain_R25           .........................t..........tg..c......a................................ [100320]
MH025917.1_strain_R7            .........................t..........tg..c......a................................ [100320]
MH025916.1_strain_R8            .........................t..........tg..c......a................................ [100320]

FR682468.2_Georgia_2007/1       aactgcgactgttgtacgtcgttggccagggagattctatatgccgggctaaaggtgcaacgttcctgtgaacaacttag [100400]
ASF_IND_20_CAD543               ................................................................................ [100400]
LR899193.1_Germany_2020/1       ................................................................................ [100400]
MN393477.1_Wuhan_2019           ................................................................................ [100400]
MW396979.1_Timor-Leste/2019/1   ................................................................................ [100400]
MK940252.1_CN/2019/InnerMongolia ................................................................................ [100400]
MK128995.1_China/2018/AnhuiXCGQ ................................................................................ [100400]
MH681419.1_ASFV/POL/2015        ................................................................................ [100400]
LS478113.1_Estonia_2014         ................................................................................ [100400]
KP843857.1_Odintsovo_02/14      ................................................................................ [100400]
FN557520.1_strain_E75           .......................a........................................................ [100400]
AY261366.1_Warthog              ................................................................................ [100400]
AY261365.1_Warmbaths            .................a.............................................................. [100400]
AY261364.1_Tengani_62           ..........................................................................g..... [100400]
AY261363.1_Pretoriuskop/96/4    ..........................t...............................................g..... [100400]
AY261362.1_Mkuzi_1979           .......................a........................................................ [100400]
AY261361.1_Malawi_Lil-20/1_1983 ..........................t.................g..a..........................g....a [100400]
AY261360.1_Kenya_1950           .....t..........................a..............a..............g...........c..... [100400]
AM712240.1_OURT_88/3            .......................a........................................................ [100400]
AM712239.1_Benin_97/1           .......................a........................................................ [100400]
KM111295.1_Ken06.Bus            ................................a..............a..............g...........c..... [100400]
```

```
KM111294.1_Ken05/Tk1            ...............................................a..............g...........c..... [100400]
KM262844.1_strain_L60           .......................a........................................................ [100400]
KM102979.1_26544/OG10_Italy     .......................a........................................................ [100400]
KM262845.1_strain_NHV           .......................a........................................................ [100400]
KX354450.1_47/Ss/2008           .......................a........................................................ [100400]
KP055815.1_strain_BA71          .......................a........................................................ [100400]
MH025920.1_strain_R35           ................................a..............a..............g...........c..... [100400]
MH025919.1_strain_N10           ................................a..............a..............g...........c..... [100400]
MH025918.1_strain_R25           ................................a..............a..............g...........c..... [100400]
MH025917.1_strain_R7            ................................a..............a..............g...........c..... [100400]
MH025916.1_strain_R8            ................................a..............a..............g...........c..... [100400]

FR682468.2_Georgia_2007/1       tacgcgcgttgttaatacaaatggactggtattagcaaacctcgtaaactcttccggacttgtttgtttttgtatgatgt [100480]
ASF_IND_20_CAD543               ................................................................................ [100480]
LR899193.1_Germany_2020/1       ................................................................................ [100480]
MN393477.1_Wuhan_2019           ................................................................................ [100480]
MW396979.1_Timor-Leste/2019/1   ................................................................................ [100480]
MK940252.1_CN/2019/InnerMongolia ................................................................................ [100480]
MK128995.1_China/2018/AnhuiXCGQ ................................................................................ [100480]
MH681419.1_ASFV/POL/2015        ................................................................................ [100480]
LS478113.1_Estonia_2014         ................................................................................ [100480]
KP843857.1_Odintsovo_02/14      ................................................................................ [100480]
FN557520.1_strain_E75           ................................................................................ [100480]
AY261366.1_Warthog              .........c..............g....................................................... [100480]
AY261365.1_Warmbaths            ................................................................................ [100480]
AY261364.1_Tengani_62           .........c..............g........g......t.................t..................... [100480]
AY261363.1_Pretoriuskop/96/4    .........c...................................................................... [100480]
AY261362.1_Mkuzi_1979           ..........................................t..................................... [100480]
AY261361.1_Malawi_Lil-20/1_1983 .........c..............g........g....................t....................c.... [100480]
AY261360.1_Kenya_1950           c........c...........c...........g......t..a...............................t.... [100480]
AM712240.1_OURT_88/3            ................................................................................ [100480]
AM712239.1_Benin_97/1           ................................................................................ [100480]
KM111295.1_Ken06.Bus            .........c...........c..g..a.....g......t..a...............................t.... [100480]
KM111294.1_Ken05/Tk1            .........c...........c...........g......t..a...............................t.... [100480]
KM262844.1_strain_L60           ................................................................................ [100480]
KM102979.1_26544/OG10_Italy     ................................................................................ [100480]
KM262845.1_strain_NHV           ................................................................................ [100480]
KX354450.1_47/Ss/2008           ................................................................................ [100480]
KP055815.1_strain_BA71          ................................................................................ [100480]
MH025920.1_strain_R35           .........c...........c..g..a.....g......t..a...............................t.... [100480]
MH025919.1_strain_N10           .........c...........c..g..a.....g......t..a...............................t.... [100480]
MH025918.1_strain_R25           .........c...........c..g..a.....g......t..a...............................t.... [100480]
MH025917.1_strain_R7            .........c...........c..g..a.....g......t..a...............................t.... [100480]
MH025916.1_strain_R8            .........c...........c..g..a.....g......t..a...............................t.... [100480]
```

```
FR682468.2_Georgia_2007/1        ttagcagggagtctgccttttcgagaatccaaagcgtcgcattgtagtaaaataaaaatagcgacttatcggcaggcgtt [100560]
ASF_IND_20_CAD543                ................................................................................ [100560]
LR899193.1_Germany_2020/1        ................................................................................ [100560]
MN393477.1_Wuhan_2019            ................................................................................ [100560]
MW396979.1_Timor-Leste/2019/1    ................................................................................ [100560]
MK940252.1_CN/2019/InnerMongolia ................................................................................ [100560]
MK128995.1_China/2018/AnhuiXCGQ  ................................................................................ [100560]
MH681419.1_ASFV/POL/2015         ................................................................................ [100560]
LS478113.1_Estonia_2014          ................................................................................ [100560]
KP843857.1_Odintsovo_02/14       ................................................................................ [100560]
FN557520.1_strain_E75            ................................................................................ [100560]
AY261366.1_Warthog               ................................................................................ [100560]
AY261365.1_Warmbaths             ................................................................................ [100560]
AY261364.1_Tengani_62            ................................................................................ [100560]
AY261363.1_Pretoriuskop/96/4     .......a..............t....................a.......................g............ [100560]
AY261362.1_Mkuzi_1979            ................................................................................ [100560]
AY261361.1_Malawi_Lil-20/1_1983  ............................................................................g... [100560]
AY261360.1_Kenya_1950            ...............................t...............................c............g... [100560]
AM712240.1_OURT_88/3             ................................................................................ [100560]
AM712239.1_Benin_97/1            ................................................................................ [100560]
KM111295.1_Ken06.Bus             ...............................................................c........t...ga.. [100560]
KM111294.1_Ken05/Tk1             ...at...................................g......................c............ga.. [100560]
KM262844.1_strain_L60            ................................................................................ [100560]
KM102979.1_26544/OG10_Italy      ................................................................................ [100560]
KM262845.1_strain_NHV            ................................................................................ [100560]
KX354450.1_47/Ss/2008            ................................................................................ [100560]
KP055815.1_strain_BA71           ................................................................................ [100560]
MH025920.1_strain_R35            ...............................................................c........t...ga.. [100560]
MH025919.1_strain_N10            ...............................................................c........t...ga.. [100560]
MH025918.1_strain_R25            ...............................................................c........t...ga.. [100560]
MH025917.1_strain_R7             ...............................................................c........t...ga.. [100560]
MH025916.1_strain_R8             ...............................................................c........t...ga.. [100560]

FR682468.2_Georgia_2007/1        gcaaaagcgccgtatagaaaataaagcagtaagtactggggagacaccacaataaggttatcttgaatgatagatatcgc [100640]
ASF_IND_20_CAD543                ................................................................................ [100640]
LR899193.1_Germany_2020/1        ................................................................................ [100640]
MN393477.1_Wuhan_2019            ................................................................................ [100640]
MW396979.1_Timor-Leste/2019/1    ................................................................................ [100640]
MK940252.1_CN/2019/InnerMongolia ................................................................................ [100640]
MK128995.1_China/2018/AnhuiXCGQ  ................................................................................ [100640]
MH681419.1_ASFV/POL/2015         ................................................................................ [100640]
LS478113.1_Estonia_2014          ................................................................................ [100640]
KP843857.1_Odintsovo_02/14       ................................................................................ [100640]
```

```
FN557520.1_strain_E75            ..............................................................c................. [100640]
AY261366.1_Warthog               ................................................................................ [100640]
AY261365.1_Warmbaths             ................................................................................ [100640]
AY261364.1_Tengani_62            ................................................................................ [100640]
AY261363.1_Pretoriuskop/96/4     ..........................t.....a...........t...........a...........t........... [100640]
AY261362.1_Mkuzi_1979            ................................................................................ [100640]
AY261361.1_Malawi_Lil-20/1_1983  ................................a......................t............c........... [100640]
AY261360.1_Kenya_1950            ................................a......................t............c........... [100640]
AM712240.1_OURT_88/3             ..............................................................c................. [100640]
AM712239.1_Benin_97/1            ..............................................................c................. [100640]
KM111295.1_Ken06.Bus             ................................a....................c.ta...........a........... [100640]
KM111294.1_Ken05/Tk1             ................................a......................t............c........... [100640]
KM262844.1_strain_L60            ..............................................................c................. [100640]
KM102979.1_26544/OG10_Italy      ..............................................................c................. [100640]
KM262845.1_strain_NHV            ..............................................................c................. [100640]
KX354450.1_47/Ss/2008            ..............................................................c................. [100640]
KP055815.1_strain_BA71           ..............................................................c................. [100640]
MH025920.1_strain_R35            ................................a....................c.ta...........a........... [100640]
MH025919.1_strain_N10            ................................a....................c.ta...........a........... [100640]
MH025918.1_strain_R25            ................................a....................c.ta...........a........... [100640]
MH025917.1_strain_R7             ................................a....................c.ta...........a........... [100640]
MH025916.1_strain_R8             ................................a....................c.ta...........a........... [100640]

FR682468.2_Georgia_2007/1        tagctctttaaacatagtgctaaaaaaatgtatgtcgttcgtcttgaatatagggggactatagtccatgtagggctcac [100720]
ASF_IND_20_CAD543                ...............r................................................................ [100720]
LR899193.1_Germany_2020/1        ................................................................................ [100720]
MN393477.1_Wuhan_2019            ................................................................................ [100720]
MW396979.1_Timor-Leste/2019/1    ................................................................................ [100720]
MK940252.1_CN/2019/InnerMongolia ................................................................................ [100720]
MK128995.1_China/2018/AnhuiXCGQ  ................................................................................ [100720]
MH681419.1_ASFV/POL/2015         ................................................................................ [100720]
LS478113.1_Estonia_2014          ................................................................................ [100720]
KP843857.1_Odintsovo_02/14       ................................................................................ [100720]
FN557520.1_strain_E75            ...............g................................................................ [100720]
AY261366.1_Warthog               ................................................................................ [100720]
AY261365.1_Warmbaths             ................................................................................ [100720]
AY261364.1_Tengani_62            ..................a..................................................a.......... [100720]
AY261363.1_Pretoriuskop/96/4     ....................................a........................................... [100720]
AY261362.1_Mkuzi_1979            ................................................................................ [100720]
AY261361.1_Malawi_Lil-20/1_1983  ....................................t..t..............a.................a....... [100720]
AY261360.1_Kenya_1950            ....................................t.................a..g..............a....... [100720]
AM712240.1_OURT_88/3             ...............g................................................................ [100720]
AM712239.1_Benin_97/1            ...............g................................................................ [100720]
KM111295.1_Ken06.Bus             ...............c....................t.................a..............a.......... [100720]
```

```
KM111294.1_Ken05/Tk1             ....................................t.................a..g..............a....... [100720]
KM262844.1_strain_L60            ...............g................................................................ [100720]
KM102979.1_26544/OG10_Italy      ...............g................................................................ [100720]
KM262845.1_strain_NHV            ...............g................................................................ [100720]
KX354450.1_47/Ss/2008            ...............g................................................................ [100720]
KP055815.1_strain_BA71           ...............g................................................................ [100720]
MH025920.1_strain_R35            ...............c...................tt.................a......................... [100720]
MH025919.1_strain_N10            ...............c...................tt.................a......................... [100720]
MH025918.1_strain_R25            ...............c...................tt.................a......................... [100720]
MH025917.1_strain_R7             ...............c...................tt.................a......................... [100720]
MH025916.1_strain_R8             ...............c...................tt.................a......................... [100720]

FR682468.2_Georgia_2007/1        atatctcagtcaggtgaaggcccatttcttttatgacttcttccgggttgtacgtcgctaacaccagcgcgggataggct [100800]
ASF_IND_20_CAD543                ................................................................................ [100800]
LR899193.1_Germany_2020/1        ................................................................................ [100800]
MN393477.1_Wuhan_2019            ................................................................................ [100800]
MW396979.1_Timor-Leste/2019/1    ................................................................................ [100800]
MK940252.1_CN/2019/InnerMongolia ................................................................................ [100800]
MK128995.1_China/2018/AnhuiXCGQ  ................................................................................ [100800]
MH681419.1_ASFV/POL/2015         ................................................................................ [100800]
LS478113.1_Estonia_2014          ................................................................................ [100800]
KP843857.1_Odintsovo_02/14       ................................................................................ [100800]
FN557520.1_strain_E75            ................................................................................ [100800]
AY261366.1_Warthog               .............a..g...........c.........c......................................... [100800]
AY261365.1_Warmbaths             .............ac.g...........c.........c......................................... [100800]
AY261364.1_Tengani_62            ................g...........c.........c......................................... [100800]
AY261363.1_Pretoriuskop/96/4     ................g.....................c......................................... [100800]
AY261362.1_Mkuzi_1979            .............ac.g............................................................... [100800]
AY261361.1_Malawi_Lil-20/1_1983  ................g.......................c..a..............c..........a.......... [100800]
AY261360.1_Kenya_1950            ................g...................t...c..t.........................a...g...... [100800]
AM712240.1_OURT_88/3             ................................................................................ [100800]
AM712239.1_Benin_97/1            ................................................................................ [100800]
KM111295.1_Ken06.Bus             .............a......................t.......................g........a...g...... [100800]
KM111294.1_Ken05/Tk1             ................g...................t...c..t.........................a...g...... [100800]
KM262844.1_strain_L60            ................................................................................ [100800]
KM102979.1_26544/OG10_Italy      ................................................................................ [100800]
KM262845.1_strain_NHV            ................................................................................ [100800]
KX354450.1_47/Ss/2008            ................................................................................ [100800]
KP055815.1_strain_BA71           ................................................................................ [100800]
MH025920.1_strain_R35            .............a......................t.......................g........a...g...... [100800]
MH025919.1_strain_N10            .............a......................t.......................g........a...g...... [100800]
MH025918.1_strain_R25            .............a......................t.......................g........a...g...... [100800]
MH025917.1_strain_R7             .............a......................t.......................g........a...g...... [100800]
MH025916.1_strain_R8             .............a......................t.......................g........a...g...... [100800]
```

```
FR682468.2_Georgia_2007/1        ttgggcatatccacggtaagtgttatgtttttatcattcttatggtaggagtaagatggttgtggaaattctgttttcca [100880]
ASF_IND_20_CAD543                ................................................................................ [100880]
LR899193.1_Germany_2020/1        ................................................................................ [100880]
MN393477.1_Wuhan_2019            ................................................................................ [100880]
MW396979.1_Timor-Leste/2019/1    ................................................................................ [100880]
MK940252.1_CN/2019/InnerMongolia ................................................................................ [100880]
MK128995.1_China/2018/AnhuiXCGQ  ................................................................................ [100880]
MH681419.1_ASFV/POL/2015         ................................................................................ [100880]
LS478113.1_Estonia_2014          ................................................................................ [100880]
KP843857.1_Odintsovo_02/14       ................................................................................ [100880]
FN557520.1_strain_E75            ................................................................................ [100880]
AY261366.1_Warthog               c..........t.t.................................................................. [100880]
AY261365.1_Warmbaths             c..........t.................................................................... [100880]
AY261364.1_Tengani_62            c..........t.................................c.................................. [100880]
AY261363.1_Pretoriuskop/96/4     c..........t............................................c....................... [100880]
AY261362.1_Mkuzi_1979            ................................................................................ [100880]
AY261361.1_Malawi_Lil-20/1_1983  c..........t.................................................................... [100880]
AY261360.1_Kenya_1950            .....................................................t.......................... [100880]
AM712240.1_OURT_88/3             ................................................................................ [100880]
AM712239.1_Benin_97/1            ................................................................................ [100880]
KM111295.1_Ken06.Bus             ......................................................................t......... [100880]
KM111294.1_Ken05/Tk1             .....................................................t.......................... [100880]
KM262844.1_strain_L60            ................................................................................ [100880]
KM102979.1_26544/OG10_Italy      ................................................................................ [100880]
KM262845.1_strain_NHV            ................................................................................ [100880]
KX354450.1_47/Ss/2008            ................................................................................ [100880]
KP055815.1_strain_BA71           ................................................................................ [100880]
MH025920.1_strain_R35            ......................................................................t......... [100880]
MH025919.1_strain_N10            ......................................................................t......... [100880]
MH025918.1_strain_R25            ......................................................................t......... [100880]
MH025917.1_strain_R7             ......................................................................t......... [100880]
MH025916.1_strain_R8             ......................................................................t......... [100880]

FR682468.2_Georgia_2007/1        ctccgggactttgcaggtaattctcagctcatttagagtctggtacaggagggcgtatgccgcaaagccgtgtatggcca [100960]
ASF_IND_20_CAD543                ................................................................................ [100960]
LR899193.1_Germany_2020/1        ................................................................................ [100960]
MN393477.1_Wuhan_2019            ................................................................................ [100960]
MW396979.1_Timor-Leste/2019/1    ................................................................................ [100960]
MK940252.1_CN/2019/InnerMongolia ................................................................................ [100960]
MK128995.1_China/2018/AnhuiXCGQ  ................................................................................ [100960]
MH681419.1_ASFV/POL/2015         ................................................................................ [100960]
LS478113.1_Estonia_2014          ................................................................................ [100960]
KP843857.1_Odintsovo_02/14       ................................................................................ [100960]
```

```
FN557520.1_strain_E75           .........c.................t.......a............................................ [100960]
AY261366.1_Warthog              ......a..c..........................c...........a..a............................ [100960]
AY261365.1_Warmbaths            .........c..........................c........................................... [100960]
AY261364.1_Tengani_62           .........c..........................c................t.......................... [100960]
AY261363.1_Pretoriuskop/96/4    .........c.................t........c........................................... [100960]
AY261362.1_Mkuzi_1979           t.....a..c...................................................................... [100960]
AY261361.1_Malawi_Lil-20/1_1983 .......g.c..a......................a............................................ [100960]
AY261360.1_Kenya_1950           t........c........c........t........c................................a.......... [100960]
AM712240.1_OURT_88/3            .........c.................t.......a............................................ [100960]
AM712239.1_Benin_97/1           .........c.................t.......a............................................ [100960]
KM111295.1_Ken06.Bus            t........c........c.................c........................................... [100960]
KM111294.1_Ken05/Tk1            t........c........c........t........c................................a.......... [100960]
KM262844.1_strain_L60           .........c.................t.......a............................................ [100960]
KM102979.1_26544/OG10_Italy     .........c.................t.......a............................................ [100960]
KM262845.1_strain_NHV           .........c.................t.......a............................................ [100960]
KX354450.1_47/Ss/2008           .........c.................t.......a............................................ [100960]
KP055815.1_strain_BA71          .........c.................t.......a............................................ [100960]
MH025920.1_strain_R35           t........c........c.................c........................................... [100960]
MH025919.1_strain_N10           t........c........c.................c........................................... [100960]
MH025918.1_strain_R25           t........c........c.................c........................................... [100960]
MH025917.1_strain_R7            t........c........c.................c........................................... [100960]
MH025916.1_strain_R8            t........c........c.................c........................................... [100960]

FR682468.2_Georgia_2007/1       cttgtttaaagggaattgaaaacgttttactttcgtatgtcgacttcacaggaacaacgggaatggggtaatatttttct [101040]
ASF_IND_20_CAD543               ................................................................................ [101040]
LR899193.1_Germany_2020/1       ................................................................................ [101040]
MN393477.1_Wuhan_2019           ................................................................................ [101040]
MW396979.1_Timor-Leste/2019/1   ................................................................................ [101040]
MK940252.1_CN/2019/InnerMongolia ................................................................................ [101040]
MK128995.1_China/2018/AnhuiXCGQ ................................................................................ [101040]
MH681419.1_ASFV/POL/2015        ................................................................................ [101040]
LS478113.1_Estonia_2014         ................................................................................ [101040]
KP843857.1_Odintsovo_02/14      ................................................................................ [101040]
FN557520.1_strain_E75           ......................................a......................................... [101040]
AY261366.1_Warthog              ................................................................................ [101040]
AY261365.1_Warmbaths            ..................................a............................................. [101040]
AY261364.1_Tengani_62           ................................................................................ [101040]
AY261363.1_Pretoriuskop/96/4    ................................................................................ [101040]
AY261362.1_Mkuzi_1979           ......................t............................................a............ [101040]
AY261361.1_Malawi_Lil-20/1_1983 .c.............c......t...................t........................a............ [101040]
AY261360.1_Kenya_1950           .cc..........................t.......g..............g........................... [101040]
AM712240.1_OURT_88/3            ......................................a......................................... [101040]
AM712239.1_Benin_97/1           ......................................a......................................... [101040]
KM111295.1_Ken06.Bus            .cc..........................t.......g..............g........................... [101040]
```

```
KM111294.1_Ken05/Tk1           .cc...................t......t.......g..............g........................... [101040]
KM262844.1_strain_L60          ......................................a......................................... [101040]
KM102979.1_26544/OG10_Italy    ......................................a......................................... [101040]
KM262845.1_strain_NHV          ......................................a......................................... [101040]
KX354450.1_47/Ss/2008          ......................................a......................................... [101040]
KP055815.1_strain_BA71         ......................................a......................................... [101040]
MH025920.1_strain_R35          .cc..........................t.......g..............g........................... [101040]
MH025919.1_strain_N10          .cc..........................t.......g..............g........................... [101040]
MH025918.1_strain_R25          .cc..........................t.......g..............g........................... [101040]
MH025917.1_strain_R7           .cc..........................t.......g..............g........................... [101040]
MH025916.1_strain_R8           .cc..........................t.......g..............g........................... [101040]

FR682468.2_Georgia_2007/1      atgaggttataccgctgcaaatcctttttaaacctgctaaaaacatcttcccttggtgggttatcaaaaggaaagcaaaa [101120]
ASF_IND_20_CAD543              ................................................................................ [101120]
LR899193.1_Germany_2020/1      ................................................................................ [101120]
MN393477.1_Wuhan_2019          ................................................................................ [101120]
MW396979.1_Timor-Leste/2019/1  ................................................................................ [101120]
MK940252.1_CN/2019/InnerMongolia ................................................................................ [101120]
MK128995.1_China/2018/AnhuiXCGQ ................................................................................ [101120]
MH681419.1_ASFV/POL/2015       ................................................................................ [101120]
LS478113.1_Estonia_2014        ................................................................................ [101120]
KP843857.1_Odintsovo_02/14     ................................................................................ [101120]
FN557520.1_strain_E75          ................................................................................ [101120]
AY261366.1_Warthog             ...........................................................a........g........... [101120]
AY261365.1_Warmbaths           ....................................................................g........... [101120]
AY261364.1_Tengani_62          ....................................................................g........... [101120]
AY261363.1_Pretoriuskop/96/4   ....................................................................g........... [101120]
AY261362.1_Mkuzi_1979          ..................................................t............................. [101120]
AY261361.1_Malawi_Lil-20/1_1983 .....a..g.......................t...................................g........... [101120]
AY261360.1_Kenya_1950          ................................t...................................g........... [101120]
AM712240.1_OURT_88/3           ................................................................................ [101120]
AM712239.1_Benin_97/1          ................................................................................ [101120]
KM111295.1_Ken06.Bus           ................................t...................................g.....a..... [101120]
KM111294.1_Ken05/Tk1           ................................t...................................g........... [101120]
KM262844.1_strain_L60          ................................................................................ [101120]
KM102979.1_26544/OG10_Italy    ................................................................................ [101120]
KM262845.1_strain_NHV          ................................................................................ [101120]
KX354450.1_47/Ss/2008          ................................................................................ [101120]
KP055815.1_strain_BA71         ................................................................................ [101120]
MH025920.1_strain_R35          ................................t...................................g.....a..... [101120]
MH025919.1_strain_N10          ................................t...................................g.....a..... [101120]
MH025918.1_strain_R25          ................................t...................................g.....a..... [101120]
MH025917.1_strain_R7           ................................t...................................g.....a..... [101120]
MH025916.1_strain_R8           ................................t...................................g.....a..... [101120]
```

```
FR682468.2_Georgia_2007/1        tgctaggtgtagcccggcccgctggtaatcggggtgaatgattttaaggtttttatacgttaatgtgggtatggtgttaa [101200]
ASF_IND_20_CAD543                ................................................................................ [101200]
LR899193.1_Germany_2020/1        ................................................................................ [101200]
MN393477.1_Wuhan_2019            ................................................................................ [101200]
MW396979.1_Timor-Leste/2019/1    ................................................................................ [101200]
MK940252.1_CN/2019/InnerMongolia ................................................................................ [101200]
MK128995.1_China/2018/AnhuiXCGQ  ................................................................................ [101200]
MH681419.1_ASFV/POL/2015         ................................................................................ [101200]
LS478113.1_Estonia_2014          ................................................................................ [101200]
KP843857.1_Odintsovo_02/14       ................................................................................ [101200]
FN557520.1_strain_E75            ................................................................................ [101200]
AY261366.1_Warthog               .............................................................................c.. [101200]
AY261365.1_Warmbaths             .............................................................................c.. [101200]
AY261364.1_Tengani_62            ................................................a.......................a....... [101200]
AY261363.1_Pretoriuskop/96/4     .............................................................................c.. [101200]
AY261362.1_Mkuzi_1979            ...............a................................................................ [101200]
AY261361.1_Malawi_Lil-20/1_1983  ........................a........a.....a.....t...............................c.. [101200]
AY261360.1_Kenya_1950            c........c...................................t...............................c.. [101200]
AM712240.1_OURT_88/3             ................................................................................ [101200]
AM712239.1_Benin_97/1            ................................................................................ [101200]
KM111295.1_Ken06.Bus             c........c...................................t...............................c.. [101200]
KM111294.1_Ken05/Tk1             c........c...................................t...............................c.. [101200]
KM262844.1_strain_L60            ................................................................................ [101200]
KM102979.1_26544/OG10_Italy      ................................................................................ [101200]
KM262845.1_strain_NHV            ................................................................................ [101200]
KX354450.1_47/Ss/2008            ................................................................................ [101200]
KP055815.1_strain_BA71           ................................................................................ [101200]
MH025920.1_strain_R35            c........c...................................t...............................c.. [101200]
MH025919.1_strain_N10            c........c...................................t...............................c.. [101200]
MH025918.1_strain_R25            c........c...................................t...............................c.. [101200]
MH025917.1_strain_R7             c........c...................................t...............................c.. [101200]
MH025916.1_strain_R8             c........c...................................t...............................c.. [101200]

FR682468.2_Georgia_2007/1        agatattggggggcatatatgaaagatcagcaacccacacaaagtccgtgcgcacccgcatggtctgcacatggatggcg [101280]
ASF_IND_20_CAD543                ................................................................................ [101280]
LR899193.1_Germany_2020/1        ................................................................................ [101280]
MN393477.1_Wuhan_2019            ................................................................................ [101280]
MW396979.1_Timor-Leste/2019/1    ................................................................................ [101280]
MK940252.1_CN/2019/InnerMongolia ................................................................................ [101280]
MK128995.1_China/2018/AnhuiXCGQ  ................................................................................ [101280]
MH681419.1_ASFV/POL/2015         ................................................................................ [101280]
LS478113.1_Estonia_2014          ................................................................................ [101280]
KP843857.1_Odintsovo_02/14       ................................................................................ [101280]
```

```
FN557520.1_strain_E75             .........................g...................................................... [101280]
AY261366.1_Warthog                .a.......................g...................................................... [101280]
AY261365.1_Warmbaths              .a.......................g...................................................... [101280]
AY261364.1_Tengani_62             .........................g...................................................... [101280]
AY261363.1_Pretoriuskop/96/4      .........................g...................................................... [101280]
AY261362.1_Mkuzi_1979             .........................g...................................................... [101280]
AY261361.1_Malawi_Lil-20/1_1983   .......a.............................g.........................................c [101280]
AY261360.1_Kenya_1950             .......a.a......g........g..g..g.....g......................................t..c [101280]
AM712240.1_OURT_88/3              .........................g...................................................... [101280]
AM712239.1_Benin_97/1             .........................g...................................................... [101280]
KM111295.1_Ken06.Bus              .......a.a......g........g..g..g.....g......................................t..c [101280]
KM111294.1_Ken05/Tk1              .......a.a......g........g..g..g.....g......................................t..c [101280]
KM262844.1_strain_L60             .........................g...................................................... [101280]
KM102979.1_26544/OG10_Italy       .........................g...................................................... [101280]
KM262845.1_strain_NHV             .........................g...................................................... [101280]
KX354450.1_47/Ss/2008             .........................g...................................................... [101280]
KP055815.1_strain_BA71            .........................g...................................................... [101280]
MH025920.1_strain_R35             .......a.a......g........g..g..g.....g......................................t..c [101280]
MH025919.1_strain_N10             .......a.a......g........g..g..g.....g......................................t..c [101280]
MH025918.1_strain_R25             .......a.a......g........g..g..g.....g......................................t..c [101280]
MH025917.1_strain_R7              .......a.a......g........g..g..g.....g......................................t..c [101280]
MH025916.1_strain_R8              .......a.a......g........g..g..g.....g......................................t..c [101280]

FR682468.2_Georgia_2007/1         cgcaccgtgcccacctgcttgaagcccttttcatacaaaatgtcagcaagttcgtaggcgtcctcaacgtggttggggga [101360]
ASF_IND_20_CAD543                 ................................................................................ [101360]
LR899193.1_Germany_2020/1         ................................................................................ [101360]
MN393477.1_Wuhan_2019             ................................................................................ [101360]
MW396979.1_Timor-Leste/2019/1     ................................................................................ [101360]
MK940252.1_CN/2019/InnerMongolia  ................................................................................ [101360]
MK128995.1_China/2018/AnhuiXCGQ   ................................................................................ [101360]
MH681419.1_ASFV/POL/2015          ................................................................................ [101360]
LS478113.1_Estonia_2014           ................................................................................ [101360]
KP843857.1_Odintsovo_02/14        ................................................................................ [101360]
FN557520.1_strain_E75             ...........................................................a.................... [101360]
AY261366.1_Warthog                .......................c........................................................ [101360]
AY261365.1_Warmbaths              .......................c......................................t................. [101360]
AY261364.1_Tengani_62             .......................c..t..................................................... [101360]
AY261363.1_Pretoriuskop/96/4      .......................c............................................a........... [101360]
AY261362.1_Mkuzi_1979             ................................................................................ [101360]
AY261361.1_Malawi_Lil-20/1_1983   .......................c.....................................................a.. [101360]
AY261360.1_Kenya_1950             .......................c................g...g................................... [101360]
AM712240.1_OURT_88/3              ...........................................................a.................... [101360]
AM712239.1_Benin_97/1             ...........................................................a.................... [101360]
KM111295.1_Ken06.Bus              .......................c................g...g................................... [101360]
```

| | | |
|---|---|---|
| KM111294.1_Ken05/Tk1 | .......................c................g...g................................... | [101360] |
| KM262844.1_strain_L60 | ...........................................................a.................... | [101360] |
| KM102979.1_26544/OG10_Italy | ...........................................................a.................... | [101360] |
| KM262845.1_strain_NHV | ...........................................................a.................... | [101360] |
| KX354450.1_47/Ss/2008 | ...........................................................a.................... | [101360] |
| KP055815.1_strain_BA71 | ...........................................................a.................... | [101360] |
| MH025920.1_strain_R35 | .......................c................g...g................................... | [101360] |
| MH025919.1_strain_N10 | .......................c................g...g................................... | [101360] |
| MH025918.1_strain_R25 | .......................c................g...g................................... | [101360] |
| MH025917.1_strain_R7 | .......................c................g...g................................... | [101360] |
| MH025916.1_strain_R8 | .......................c................g...g................................... | [101360] |

| | | |
|---|---|---|
| FR682468.2_Georgia_2007/1 | aaacatatcaaagtcgggtctttctccctcgggataaattgagctgcctttaagatgcagggcataatcaatggcaatcc | [101440] |
| ASF_IND_20_CAD543 | ................................................................................ | [101440] |
| LR899193.1_Germany_2020/1 | ................................................................................ | [101440] |
| MN393477.1_Wuhan_2019 | ................................................................................ | [101440] |
| MW396979.1_Timor-Leste/2019/1 | ................................................................................ | [101440] |
| MK940252.1_CN/2019/InnerMongolia | ................................................................................ | [101440] |
| MK128995.1_China/2018/AnhuiXCGQ | ................................................................................ | [101440] |
| MH681419.1_ASFV/POL/2015 | ................................................................................ | [101440] |
| LS478113.1_Estonia_2014 | ................................................................................ | [101440] |
| KP843857.1_Odintsovo_02/14 | ................................................................................ | [101440] |
| FN557520.1_strain_E75 | .......................................c........................................ | [101440] |
| AY261366.1_Warthog | .......................................c...................................t.... | [101440] |
| AY261365.1_Warmbaths | .......................................c...................................t.... | [101440] |
| AY261364.1_Tengani_62 | .......................................c........................................ | [101440] |
| AY261363.1_Pretoriuskop/96/4 | .......................................c........................................ | [101440] |
| AY261362.1_Mkuzi_1979 | .......................................c........................................ | [101440] |
| AY261361.1_Malawi_Lil-20/1_1983 | .........g..............c..............c....................a.................g. | [101440] |
| AY261360.1_Kenya_1950 | .........g..............c..............c........................................ | [101440] |
| AM712240.1_OURT_88/3 | .......................................c........................................ | [101440] |
| AM712239.1_Benin_97/1 | .......................................c........................................ | [101440] |
| KM111295.1_Ken06.Bus | .........g..............c..............c........c.......................a....... | [101440] |
| KM111294.1_Ken05/Tk1 | .........g..a...........c..............c........................................ | [101440] |
| KM262844.1_strain_L60 | .......................................c........................................ | [101440] |
| KM102979.1_26544/OG10_Italy | .......................................c........................................ | [101440] |
| KM262845.1_strain_NHV | .......................................c........................................ | [101440] |
| KX354450.1_47/Ss/2008 | .......................................c........................................ | [101440] |
| KP055815.1_strain_BA71 | .......................................c........................................ | [101440] |
| MH025920.1_strain_R35 | .........g..............c..............c........c.......................a....... | [101440] |
| MH025919.1_strain_N10 | .........g..............c..............c........c.......................a....... | [101440] |
| MH025918.1_strain_R25 | .........g..............c..............c........c.......................a....... | [101440] |
| MH025917.1_strain_R7 | .........g..............c..............c........c.......................a....... | [101440] |
| MH025916.1_strain_R8 | .........g..............c..............c........c.......................a....... | [101440] |

FR682468.2_Georgia_2007/1 ccccgtacaaaataagctttttctttatgataaattcgcggaccacctccaaagccgcctcaatctccacggcatttgcc [101520]
ASF_IND_20_CAD543 ................................................................................ [101520]
LR899193.1_Germany_2020/1 ................................................................................ [101520]
MN393477.1_Wuhan_2019 ................................................................................ [101520]
MW396979.1_Timor-Leste/2019/1 ................................................................................ [101520]
MK940252.1_CN/2019/InnerMongolia ................................................................................ [101520]
MK128995.1_China/2018/AnhuiXCGQ ................................................................................ [101520]
MH681419.1_ASFV/POL/2015 ................................................................................ [101520]
LS478113.1_Estonia_2014 ................................................................................ [101520]
KP843857.1_Odintsovo_02/14 ................................................................................ [101520]
FN557520.1_strain_E75 ................................................................................ [101520]
AY261366.1_Warthog .............g.................................................................. [101520]
AY261365.1_Warmbaths .............g.................................................................. [101520]
AY261364.1_Tengani_62 .............g....................c........t........................g........... [101520]
AY261363.1_Pretoriuskop/96/4 .............g.........................................t........................ [101520]
AY261362.1_Mkuzi_1979 .............g........t......................................................... [101520]
AY261361.1_Malawi_Lil-20/1_1983 ....a...................................t........t.g............................ [101520]
AY261360.1_Kenya_1950 ....a........g.....................................g................g........... [101520]
AM712240.1_OURT_88/3 ................................................................................ [101520]
AM712239.1_Benin_97/1 ................................................................................ [101520]
KM111295.1_Ken06.Bus ....a........g.....................................g................g........... [101520]
KM111294.1_Ken05/Tk1 ....a........g.....................................g................g........... [101520]
KM262844.1_strain_L60 ................................................................................ [101520]
KM102979.1_26544/OG10_Italy ................................................................................ [101520]
KM262845.1_strain_NHV ................................................................................ [101520]
KX354450.1_47/Ss/2008 ................................................................................ [101520]
KP055815.1_strain_BA71 ................................................................................ [101520]
MH025920.1_strain_R35 ....a........g.....................................g................g........... [101520]
MH025919.1_strain_N10 ....a........g.....................................g................g........... [101520]
MH025918.1_strain_R25 ....a........g.....................................g................g........... [101520]
MH025917.1_strain_R7 ....a........g.....................................g................g........... [101520]
MH025916.1_strain_R8 ....a........g.....................................g................g........... [101520]

FR682468.2_Georgia_2007/1 tcacgtttttgagcaatgagccggtacttagaaacattaaaatcagtctttagtaaagacgtcataaatagtgtttaata [101600]
ASF_IND_20_CAD543 ................................................................................ [101600]
LR899193.1_Germany_2020/1 ................................................................................ [101600]
MN393477.1_Wuhan_2019 ................................................................................ [101600]
MW396979.1_Timor-Leste/2019/1 ................................................................................ [101600]
MK940252.1_CN/2019/InnerMongolia ................................................................................ [101600]
MK128995.1_China/2018/AnhuiXCGQ ................................................................................ [101600]
MH681419.1_ASFV/POL/2015 ................................................................................ [101600]
LS478113.1_Estonia_2014 ................................................................................ [101600]
KP843857.1_Odintsovo_02/14 ................................................................................ [101600]

| | |
|---|---|
| FN557520.1_strain_E75 | .....c.......................................................a.................. [101600] |
| AY261366.1_Warthog | ..........................t...................g..............a.................. [101600] |
| AY261365.1_Warmbaths | ..........................t...................g..............a.................. [101600] |
| AY261364.1_Tengani_62 | .............................................................a.................. [101600] |
| AY261363.1_Pretoriuskop/96/4 | ..........................t..................................a.................. [101600] |
| AY261362.1_Mkuzi_1979 | .............................................................a.................. [101600] |
| AY261361.1_Malawi_Lil-20/1_1983 | .....................t....t........................g.........a.................. [101600] |
| AY261360.1_Kenya_1950 | .....................t....t....g............g......g.........a........c......... [101600] |
| AM712240.1_OURT_88/3 | .....c.......................................................a.................. [101600] |
| AM712239.1_Benin_97/1 | .....c.......................................................a.................. [101600] |
| KM111295.1_Ken06.Bus | ..........................t....g............g......g.........a........c......... [101600] |
| KM111294.1_Ken05/Tk1 | ..........................t....g............g......g.........a........c......... [101600] |
| KM262844.1_strain_L60 | .....c.......................................................a.................. [101600] |
| KM102979.1_26544/OG10_Italy | .....c.................a.....................................a.................. [101600] |
| KM262845.1_strain_NHV | .....c.......................................................a.................. [101600] |
| KX354450.1_47/Ss/2008 | .....c.................a.....................................a.................. [101600] |
| KP055815.1_strain_BA71 | .....c.......................................................a.................. [101600] |
| MH025920.1_strain_R35 | ..........................t....g............g......g.........a........c......... [101600] |
| MH025919.1_strain_N10 | ..........................t....g............g......g.........a........c......... [101600] |
| MH025918.1_strain_R25 | ..........................t....g............g......g.........a........c......... [101600] |
| MH025917.1_strain_R7 | ..........................t....g............g......g.........a........c......... [101600] |
| MH025916.1_strain_R8 | ..........................t....g............g......g.........a........c......... [101600] |

| | |
|---|---|
| FR682468.2_Georgia_2007/1 | tatattaaaggtttgaataaaatactaaatagtaaaaatggatgccctattaaaggaaatagaaaagttatcgcagc--- [101680] |
| ASF_IND_20_CAD543 | .............................................................................--- [101680] |
| LR899193.1_Germany_2020/1 | .............................................................................--- [101680] |
| MN393477.1_Wuhan_2019 | .............................................................................--- [101680] |
| MW396979.1_Timor-Leste/2019/1 | .............................................................................--- [101680] |
| MK940252.1_CN/2019/InnerMongolia | .............................................................................--- [101680] |
| MK128995.1_China/2018/AnhuiXCGQ | .............................................................................--- [101680] |
| MH681419.1_ASFV/POL/2015 | .............................................................................--- [101680] |
| LS478113.1_Estonia_2014 | .............................................................................--- [101680] |
| KP843857.1_Odintsovo_02/14 | .............................................................................--- [101680] |
| FN557520.1_strain_E75 | .............................................................................--- [101680] |
| AY261366.1_Warthog | .............................................................................--- [101680] |
| AY261365.1_Warmbaths | .............................................................................--- [101680] |
| AY261364.1_Tengani_62 | .............................................................................--- [101680] |
| AY261363.1_Pretoriuskop/96/4 | .............................................................................--- [101680] |
| AY261362.1_Mkuzi_1979 | .............................................................................--- [101680] |
| AY261361.1_Malawi_Lil-20/1_1983 | .................................-........c...t..............................--- [101680] |
| AY261360.1_Kenya_1950 | ............................................................c................cat [101680] |
| AM712240.1_OURT_88/3 | .............................................................................--- [101680] |
| AM712239.1_Benin_97/1 | .............................................................................--- [101680] |
| KM111295.1_Ken06.Bus | ............................................................c................cat [101680] |

```
KM111294.1_Ken05/Tk1              ............................................................c................cat [101680]
KM262844.1_strain_L60             .............................................................................--- [101680]
KM102979.1_26544/OG10_Italy       .............................................................................--- [101680]
KM262845.1_strain_NHV             .............................................................................--- [101680]
KX354450.1_47/Ss/2008             .............................................................................--- [101680]
KP055815.1_strain_BA71            .............................................................................--- [101680]
MH025920.1_strain_R35             ............................................................c................cat [101680]
MH025919.1_strain_N10             ............................................................c................cat [101680]
MH025918.1_strain_R25             ............................................................c................cat [101680]
MH025917.1_strain_R7              ............................................................c................cat [101680]
MH025916.1_strain_R8              ............................................................c................cat [101680]

FR682468.2_Georgia_2007/1         catccttgcagaaagaaaacaatgatgtatgcgatctctgttttatgcaaatgaaaaaaatttctaactatcagctttta [101760]
ASF_IND_20_CAD543                 ................................................................................ [101760]
LR899193.1_Germany_2020/1         ................................................................................ [101760]
MN393477.1_Wuhan_2019             ................................................................................ [101760]
MW396979.1_Timor-Leste/2019/1     ................................................................................ [101760]
MK940252.1_CN/2019/InnerMongolia  ................................................................................ [101760]
MK128995.1_China/2018/AnhuiXCGQ   ................................................................................ [101760]
MH681419.1_ASFV/POL/2015          ................................................................................ [101760]
LS478113.1_Estonia_2014           ................................................................................ [101760]
KP843857.1_Odintsovo_02/14        ................................................................................ [101760]
FN557520.1_strain_E75             ................................................................................ [101760]
AY261366.1_Warthog                ................................................................................ [101760]
AY261365.1_Warmbaths              ................................................................................ [101760]
AY261364.1_Tengani_62             ..........a..................................................................... [101760]
AY261363.1_Pretoriuskop/96/4      ................................................................................ [101760]
AY261362.1_Mkuzi_1979             ................................................................................ [101760]
AY261361.1_Malawi_Lil-20/1_1983   ............................g................................................... [101760]
AY261360.1_Kenya_1950             ..........a..................................................................... [101760]
AM712240.1_OURT_88/3              ................................................................................ [101760]
AM712239.1_Benin_97/1             ................................................................................ [101760]
KM111295.1_Ken06.Bus              ..........a..................................................................... [101760]
KM111294.1_Ken05/Tk1              ..........a..................................................................... [101760]
KM262844.1_strain_L60             ................................................................................ [101760]
KM102979.1_26544/OG10_Italy       ................................................................................ [101760]
KM262845.1_strain_NHV             ................................................................................ [101760]
KX354450.1_47/Ss/2008             ................................................................................ [101760]
KP055815.1_strain_BA71            ................................................................................ [101760]
MH025920.1_strain_R35             ..........a..................................................................... [101760]
MH025919.1_strain_N10             ..........a..................................................................... [101760]
MH025918.1_strain_R25             ..........a..................................................................... [101760]
MH025917.1_strain_R7              ..........a..................................................................... [101760]
MH025916.1_strain_R8              ..........a..................................................................... [101760]
```

```
FR682468.2_Georgia_2007/1        tgcgaagagtgcggtcagctgaaggactggtttgaacctgaatataatgaaaaattcacggtatattctcgtctaaagat [101840]
ASF_IND_20_CAD543                ................................................................................ [101840]
LR899193.1_Germany_2020/1        ................................................................................ [101840]
MN393477.1_Wuhan_2019            ................................................................................ [101840]
MW396979.1_Timor-Leste/2019/1    ................................................................................ [101840]
MK940252.1_CN/2019/InnerMongolia ................................................................................ [101840]
MK128995.1_China/2018/AnhuiXCGQ  ................................................................................ [101840]
MH681419.1_ASFV/POL/2015         ................................................................................ [101840]
LS478113.1_Estonia_2014          ................................................................................ [101840]
KP843857.1_Odintsovo_02/14       ................................................................................ [101840]
FN557520.1_strain_E75            ................................................................................ [101840]
AY261366.1_Warthog               ................................................................................ [101840]
AY261365.1_Warmbaths             ................................................................................ [101840]
AY261364.1_Tengani_62            .........................................................t...................... [101840]
AY261363.1_Pretoriuskop/96/4     ................................................................................ [101840]
AY261362.1_Mkuzi_1979            ................................................................................ [101840]
AY261361.1_Malawi_Lil-20/1_1983  ...............................................c.............................a.. [101840]
AY261360.1_Kenya_1950            .................a.......................t...................................... [101840]
AM712240.1_OURT_88/3             ................................................................................ [101840]
AM712239.1_Benin_97/1            ................................................................................ [101840]
KM111295.1_Ken06.Bus             .................a.......................t..............................t....... [101840]
KM111294.1_Ken05/Tk1             .................a.......................t...................................... [101840]
KM262844.1_strain_L60            ................................................................................ [101840]
KM102979.1_26544/OG10_Italy      ................................................................................ [101840]
KM262845.1_strain_NHV            ................................................................................ [101840]
KX354450.1_47/Ss/2008            ................................................................................ [101840]
KP055815.1_strain_BA71           ................................................................................ [101840]
MH025920.1_strain_R35            .................a.......................t..............................t....... [101840]
MH025919.1_strain_N10            .................a.......................t..............................t....... [101840]
MH025918.1_strain_R25            .................a.......................t..............................t....... [101840]
MH025917.1_strain_R7             .................a.......................t..............................t....... [101840]
MH025916.1_strain_R8             .................a.......................t..............................t....... [101840]

FR682468.2_Georgia_2007/1        cgtgggtgccaatagttcctatcaccagcgcgatttggacaaggccaactcaagtgactatagctccttgcaatttcatc [101920]
ASF_IND_20_CAD543                ..................................................................t............. [101920]
LR899193.1_Germany_2020/1        ................................................................................ [101920]
MN393477.1_Wuhan_2019            ................................................................................ [101920]
MW396979.1_Timor-Leste/2019/1    ................................................................................ [101920]
MK940252.1_CN/2019/InnerMongolia ................................................................................ [101920]
MK128995.1_China/2018/AnhuiXCGQ  ................................................................................ [101920]
MH681419.1_ASFV/POL/2015         ................................................................................ [101920]
LS478113.1_Estonia_2014          ................................................................................ [101920]
KP843857.1_Odintsovo_02/14       ................................................................................ [101920]
```

```
FN557520.1_strain_E75              ...............c..................................................t............. [101920]
AY261366.1_Warthog                 ................................................................................ [101920]
AY261365.1_Warmbaths               t............................................................................... [101920]
AY261364.1_Tengani_62              ......................................................c...........t............. [101920]
AY261363.1_Pretoriuskop/96/4       ................................................................................ [101920]
AY261362.1_Mkuzi_1979              .........t........................................................t............. [101920]
AY261361.1_Malawi_Lil-20/1_1983    t.................................................................t............. [101920]
AY261360.1_Kenya_1950              .........t......................................t.................t............. [101920]
AM712240.1_OURT_88/3               ...............c..................................................t............. [101920]
AM712239.1_Benin_97/1              ...............c..................................................t............. [101920]
KM111295.1_Ken06.Bus               ..................................................................t............. [101920]
KM111294.1_Ken05/Tk1               .........t......................................t.................t............. [101920]
KM262844.1_strain_L60              ...............c..................................................t............. [101920]
KM102979.1_26544/OG10_Italy        ...............c..................................................t............. [101920]
KM262845.1_strain_NHV              ...............c..................................................t............. [101920]
KX354450.1_47/Ss/2008              ...............c..................................................t............. [101920]
KP055815.1_strain_BA71             ...............c..................................................t............. [101920]
MH025920.1_strain_R35              ..................................................................t............. [101920]
MH025919.1_strain_N10              ..................................................................t............. [101920]
MH025918.1_strain_R25              ..................................................................t............. [101920]
MH025917.1_strain_R7               ..................................................................t............. [101920]
MH025916.1_strain_R8               ..................................................................t............. [101920]

FR682468.2_Georgia_2007/1          acattttagaggagctcaaatccctaaatgttaagtatatggatgcggggcaaaagccctttcctattcaggtgttaaaa [102000]
ASF_IND_20_CAD543                  ................................................................................ [102000]
LR899193.1_Germany_2020/1          ................................................................................ [102000]
MN393477.1_Wuhan_2019              ................................................................................ [102000]
MW396979.1_Timor-Leste/2019/1      ................................................................................ [102000]
MK940252.1_CN/2019/InnerMongolia   ................................................................................ [102000]
MK128995.1_China/2018/AnhuiXCGQ    ................................................................................ [102000]
MH681419.1_ASFV/POL/2015           ................................................................................ [102000]
LS478113.1_Estonia_2014            ................................................................................ [102000]
KP843857.1_Odintsovo_02/14         ................................................................................ [102000]
FN557520.1_strain_E75              ...............................................................................g [102000]
AY261366.1_Warthog                 ....................a................c...........a.....a.................a.....g [102000]
AY261365.1_Warmbaths               ....................a................c.................a.................a.....g [102000]
AY261364.1_Tengani_62              .....................................c...................................a.....g [102000]
AY261363.1_Pretoriuskop/96/4       .....................................c.................a..............a..a.....g [102000]
AY261362.1_Mkuzi_1979              ...............................................................................g [102000]
AY261361.1_Malawi_Lil-20/1_1983    ..............................................a..a.....a........c........c.....g [102000]
AY261360.1_Kenya_1950              ................................................................c........a.....g [102000]
AM712240.1_OURT_88/3               ...............................................................................g [102000]
AM712239.1_Benin_97/1              ...............................................................................g [102000]
KM111295.1_Ken06.Bus               ..............................................a.................c........a.....g [102000]
```

```
KM111294.1_Ken05/Tk1             ................................................................c........a.....g [102000]
KM262844.1_strain_L60            ...............................................................................g [102000]
KM102979.1_26544/OG10_Italy      ...............................................................................g [102000]
KM262845.1_strain_NHV            ...............................................................................g [102000]
KX354450.1_47/Ss/2008            ...............................................................................g [102000]
KP055815.1_strain_BA71           ...............................................................................g [102000]
MH025920.1_strain_R35            ..............................................a.................c........a.....g [102000]
MH025919.1_strain_N10            ..............................................a.................c........a.....g [102000]
MH025918.1_strain_R25            ..............................................a.................c........a.....g [102000]
MH025917.1_strain_R7             ..............................................a.................c........a.....g [102000]
MH025916.1_strain_R8             ..............................................a.................c........a.....g [102000]

FR682468.2_Georgia_2007/1        gaaactgctcacagttataaccaagtacaacaacatcgggtcatacgcagcattacaaagcttcagatcttagccagtat [102080]
ASF_IND_20_CAD543                ................................................................................ [102080]
LR899193.1_Germany_2020/1        ................................................................................ [102080]
MN393477.1_Wuhan_2019            ................................................................................ [102080]
MW396979.1_Timor-Leste/2019/1    ................................................................................ [102080]
MK940252.1_CN/2019/InnerMongolia ................................................................................ [102080]
MK128995.1_China/2018/AnhuiXCGQ  ................................................................................ [102080]
MH681419.1_ASFV/POL/2015         ................................................................................ [102080]
LS478113.1_Estonia_2014          ................................................................................ [102080]
KP843857.1_Odintsovo_02/14       ................................................................................ [102080]
FN557520.1_strain_E75            .....a.................g........................................................ [102080]
AY261366.1_Warthog               .......................g.............................................c.......... [102080]
AY261365.1_Warmbaths             ...........t...........g.............................................c.......... [102080]
AY261364.1_Tengani_62            .......................g.............................................c.......... [102080]
AY261363.1_Pretoriuskop/96/4     ...........t...........g............................................tc.......... [102080]
AY261362.1_Mkuzi_1979            .......................g........................................................ [102080]
AY261361.1_Malawi_Lil-20/1_1983  .......................g..................................................t..... [102080]
AY261360.1_Kenya_1950            ....................t..g.....................................................c.. [102080]
AM712240.1_OURT_88/3             .......................g........................................................ [102080]
AM712239.1_Benin_97/1            .......................g........................................................ [102080]
KM111295.1_Ken06.Bus             ....................t..g.....................................................c.. [102080]
KM111294.1_Ken05/Tk1             ....................t..g.....................................................c.. [102080]
KM262844.1_strain_L60            .......................g........................................................ [102080]
KM102979.1_26544/OG10_Italy      .......................g........................................................ [102080]
KM262845.1_strain_NHV            .......................g........................................................ [102080]
KX354450.1_47/Ss/2008            .......................g........................................................ [102080]
KP055815.1_strain_BA71           .......................g........................................................ [102080]
MH025920.1_strain_R35            ....................t..g.....................................................c.. [102080]
MH025919.1_strain_N10            ....................t..g.....................................................c.. [102080]
MH025918.1_strain_R25            ....................t..g.....................................................c.. [102080]
MH025917.1_strain_R7             ....................t..g.....................................................c.. [102080]
MH025916.1_strain_R8             ....................t..g.....................................................c.. [102080]
```

```
FR682468.2_Georgia_2007/1        tctacgtagcatttgtttaaaattaaacattgcttgtacggtggcagacgccgcgaggtttactcaacttaataccaaag [102160]
ASF_IND_20_CAD543                ................................................................................ [102160]
LR899193.1_Germany_2020/1        ................................................................................ [102160]
MN393477.1_Wuhan_2019            ................................................................................ [102160]
MW396979.1_Timor-Leste/2019/1    ................................................................................ [102160]
MK940252.1_CN/2019/InnerMongolia ................................................................................ [102160]
MK128995.1_China/2018/AnhuiXCGQ  ................................................................................ [102160]
MH681419.1_ASFV/POL/2015         ................................................................................ [102160]
LS478113.1_Estonia_2014          ................................................................................ [102160]
KP843857.1_Odintsovo_02/14       ................................................................................ [102160]
FN557520.1_strain_E75            .t.............................................................................. [102160]
AY261366.1_Warthog               ................................................................................ [102160]
AY261365.1_Warmbaths             ................................................................................ [102160]
AY261364.1_Tengani_62            ................................................................................ [102160]
AY261363.1_Pretoriuskop/96/4     ................................................................................ [102160]
AY261362.1_Mkuzi_1979            .t.......................................................a...................... [102160]
AY261361.1_Malawi_Lil-20/1_1983  .........t................................a..............a...................... [102160]
AY261360.1_Kenya_1950            c........t..................................................c.....g............. [102160]
AM712240.1_OURT_88/3             .t.............................................................................. [102160]
AM712239.1_Benin_97/1            .t.............................................................................. [102160]
KM111295.1_Ken06.Bus             c........t......................................t.....a.....c.....g............. [102160]
KM111294.1_Ken05/Tk1             c........t..................................................c.....g............. [102160]
KM262844.1_strain_L60            .t.............................................................................. [102160]
KM102979.1_26544/OG10_Italy      .t.............................................................................. [102160]
KM262845.1_strain_NHV            .t.............................................................................. [102160]
KX354450.1_47/Ss/2008            .t.............................................................................. [102160]
KP055815.1_strain_BA71           .t.............................................................................. [102160]
MH025920.1_strain_R35            c........t......................................t.....a.....c.....g............. [102160]
MH025919.1_strain_N10            c........t......................................t.....a.....c.....g............. [102160]
MH025918.1_strain_R25            c........t......................................t.....a.....c.....g............. [102160]
MH025917.1_strain_R7             c........t......................................t.....a.....c.....g............. [102160]
MH025916.1_strain_R8             c........t......................................t.....a.....c.....g............. [102160]

FR682468.2_Georgia_2007/1        ggatctcaaggggcatggatcttctgcgctccctatttgtagacaataaaattactttaaacgttgatttaaaccctata [102240]
ASF_IND_20_CAD543                ................................................................................ [102240]
LR899193.1_Germany_2020/1        ................................................................................ [102240]
MN393477.1_Wuhan_2019            ................................................................................ [102240]
MW396979.1_Timor-Leste/2019/1    ................................................................................ [102240]
MK940252.1_CN/2019/InnerMongolia ................................................................................ [102240]
MK128995.1_China/2018/AnhuiXCGQ  ................................................................................ [102240]
MH681419.1_ASFV/POL/2015         ................................................................................ [102240]
LS478113.1_Estonia_2014          ................................................................................ [102240]
KP843857.1_Odintsovo_02/14       ................................................................................ [102240]
```

| | | |
|---|---|---|
| FN557520.1_strain_E75 | ................................................................................ | [102240] |
| AY261366.1_Warthog | ................................................................................ | [102240] |
| AY261365.1_Warmbaths | ................................................................................ | [102240] |
| AY261364.1_Tengani_62 | ................................................................................ | [102240] |
| AY261363.1_Pretoriuskop/96/4 | ................................................................................ | [102240] |
| AY261362.1_Mkuzi_1979 | ................................................................................ | [102240] |
| AY261361.1_Malawi_Lil-20/1_1983 | ....t..g.aa.........................................c..c........................ | [102240] |
| AY261360.1_Kenya_1950 | .......................t...............a............c..c........................ | [102240] |
| AM712240.1_OURT_88/3 | ................................................................................ | [102240] |
| AM712239.1_Benin_97/1 | ................................................................................ | [102240] |
| KM111295.1_Ken06.Bus | .......................t...............a............c..c........................ | [102240] |
| KM111294.1_Ken05/Tk1 | .......................t...............a............c..c........................ | [102240] |
| KM262844.1_strain_L60 | ................................................................................ | [102240] |
| KM102979.1_26544/OG10_Italy | ................................................................................ | [102240] |
| KM262845.1_strain_NHV | ................................................................................ | [102240] |
| KX354450.1_47/Ss/2008 | .............................................................t.................. | [102240] |
| KP055815.1_strain_BA71 | ................................................................................ | [102240] |
| MH025920.1_strain_R35 | .......................t...............a............c..c........................ | [102240] |
| MH025919.1_strain_N10 | .......................t...............a............c..c........................ | [102240] |
| MH025918.1_strain_R25 | .......................t...............a............c..c........................ | [102240] |
| MH025917.1_strain_R7 | .......................t...............a............c..c........................ | [102240] |
| MH025916.1_strain_R8 | .......................t...............a............c..c........................ | [102240] |
| FR682468.2_Georgia_2007/1 | gacagctttattaatagtacctacagtgccttacaaattaaacaaatccaccaagaactgcaggaggaaaatgtttataa | [102320] |
| ASF_IND_20_CAD543 | ................................................................................ | [102320] |
| LR899193.1_Germany_2020/1 | ................................................................................ | [102320] |
| MN393477.1_Wuhan_2019 | ................................................................................ | [102320] |
| MW396979.1_Timor-Leste/2019/1 | ................................................................................ | [102320] |
| MK940252.1_CN/2019/InnerMongolia | ................................................................................ | [102320] |
| MK128995.1_China/2018/AnhuiXCGQ | ................................................................................ | [102320] |
| MH681419.1_ASFV/POL/2015 | ................................................................................ | [102320] |
| LS478113.1_Estonia_2014 | ................................................................................ | [102320] |
| KP843857.1_Odintsovo_02/14 | ................................................................................ | [102320] |
| FN557520.1_strain_E75 | ..........................c..................................................... | [102320] |
| AY261366.1_Warthog | .................c.......ac....................t................................ | [102320] |
| AY261365.1_Warmbaths | .................c.......ac....................t................................ | [102320] |
| AY261364.1_Tengani_62 | ..........................c....................t................................ | [102320] |
| AY261363.1_Pretoriuskop/96/4 | .................c.......ac....................t................................ | [102320] |
| AY261362.1_Mkuzi_1979 | ..........................c..................................................... | [102320] |
| AY261361.1_Malawi_Lil-20/1_1983 | ..........................c...c................t..t.......................a..... | [102320] |
| AY261360.1_Kenya_1950 | ....................t.....c...c...................t............................. | [102320] |
| AM712240.1_OURT_88/3 | ..........................c..................................................... | [102320] |
| AM712239.1_Benin_97/1 | ..........................c..................................................... | [102320] |
| KM111295.1_Ken06.Bus | .....t...........c........c...c...................t..............a.............. | [102320] |

```
KM111294.1_Ken05/Tk1             ..........................c...c...................t............................. [102320]
KM262844.1_strain_L60            ..........................c..................................................... [102320]
KM102979.1_26544/OG10_Italy      ..........................c..................................................... [102320]
KM262845.1_strain_NHV            ..........................c..................................................... [102320]
KX354450.1_47/Ss/2008            ..........................c..................................................... [102320]
KP055815.1_strain_BA71           ..........................c..................................................... [102320]
MH025920.1_strain_R35            .....t...........c........c...c...................t..............a.............. [102320]
MH025919.1_strain_N10            .....t...........c........c...c...................t..............a.............. [102320]
MH025918.1_strain_R25            .....t...........c........c...c...................t..............a.............. [102320]
MH025917.1_strain_R7             .....t...........c........c...c...................t..............a.............. [102320]
MH025916.1_strain_R8             .....t...........c........c...c...................t..............a.............. [102320]

FR682468.2_Georgia_2007/1        tttaaaagaaattgttaagagctttatattatacgcggatgagaagaacatcggcgtcgatcttaacaggagaaccgttg [102400]
ASF_IND_20_CAD543                ................................................................................ [102400]
LR899193.1_Germany_2020/1        ................................................................................ [102400]
MN393477.1_Wuhan_2019            ................................................................................ [102400]
MW396979.1_Timor-Leste/2019/1    ................................................................................ [102400]
MK940252.1_CN/2019/InnerMongolia ................................................................................ [102400]
MK128995.1_China/2018/AnhuiXCGQ  ................................................................................ [102400]
MH681419.1_ASFV/POL/2015         ................................................................................ [102400]
LS478113.1_Estonia_2014          ................................................................................ [102400]
KP843857.1_Odintsovo_02/14       ................................................................................ [102400]
FN557520.1_strain_E75            ........................................................................g....... [102400]
AY261366.1_Warthog               ........................................................................g....... [102400]
AY261365.1_Warmbaths             ........................................................................g....... [102400]
AY261364.1_Tengani_62            ........................................................................g....... [102400]
AY261363.1_Pretoriuskop/96/4     ........................................................................g....... [102400]
AY261362.1_Mkuzi_1979            ................................................................................ [102400]
AY261361.1_Malawi_Lil-20/1_1983  ...........................t..............a..a.....t...........g.....a..g....... [102400]
AY261360.1_Kenya_1950            ...............g.....t.....tg......................t....................g..t.... [102400]
AM712240.1_OURT_88/3             ........................................................................g....... [102400]
AM712239.1_Benin_97/1            ........................................................................g....... [102400]
KM111295.1_Ken06.Bus             ...............g.....t.....tg...........................................g....... [102400]
KM111294.1_Ken05/Tk1             ...............g.....t.....tg......................t....................g..t.... [102400]
KM262844.1_strain_L60            ........................................................................g....... [102400]
KM102979.1_26544/OG10_Italy      ........................................................................g....... [102400]
KM262845.1_strain_NHV            ........................................................................g....... [102400]
KX354450.1_47/Ss/2008            ........................................................................g....... [102400]
KP055815.1_strain_BA71           ........................................................................g....... [102400]
MH025920.1_strain_R35            ...............g.....t.....tg...........................................g....... [102400]
MH025919.1_strain_N10            ...............g.....t.....tg...........................................g....... [102400]
MH025918.1_strain_R25            ...............g.....t.....tg...........................................g....... [102400]
MH025917.1_strain_R7             ...............g.....t.....tg...........................................g....... [102400]
MH025916.1_strain_R8             ...............g.....t.....tg...........................................g....... [102400]
```

```
FR682468.2_Georgia_2007/1        tgattgctacgatgtataatgttttacgccgtgcctactaccccatagaaattgatacggtggtgtatcaatgtaaaata [102480]
ASF_IND_20_CAD543                ................................................................................ [102480]
LR899193.1_Germany_2020/1        ................................................................................ [102480]
MN393477.1_Wuhan_2019            ................................................................................ [102480]
MW396979.1_Timor-Leste/2019/1    ................................................................................ [102480]
MK940252.1_CN/2019/InnerMongolia ................................................................................ [102480]
MK128995.1_China/2018/AnhuiXCGQ  ................................................................................ [102480]
MH681419.1_ASFV/POL/2015         ................................................................................ [102480]
LS478113.1_Estonia_2014          ................................................................................ [102480]
KP843857.1_Odintsovo_02/14       ................................................................................ [102480]
FN557520.1_strain_E75            ..................................t............................................. [102480]
AY261366.1_Warthog               ..................................t.............................a............... [102480]
AY261365.1_Warmbaths             ..................................t............................................. [102480]
AY261364.1_Tengani_62            ...................................................................c..g......... [102480]
AY261363.1_Pretoriuskop/96/4     .......c..........................t............................................. [102480]
AY261362.1_Mkuzi_1979            ..................................t............................................. [102480]
AY261361.1_Malawi_Lil-20/1_1983  ...................c.........................................a.................. [102480]
AY261360.1_Kenya_1950            .......c........c..c.....g..........................c..............c............ [102480]
AM712240.1_OURT_88/3             ..................................t............................................. [102480]
AM712239.1_Benin_97/1            ..................................t............................................. [102480]
KM111295.1_Ken06.Bus             .......c........c..c.....g..........................c..............c..g......... [102480]
KM111294.1_Ken05/Tk1             .......c........c..c.....g..........................c..............c............ [102480]
KM262844.1_strain_L60            ..................................t............................................. [102480]
KM102979.1_26544/OG10_Italy      ..................................t............................................. [102480]
KM262845.1_strain_NHV            ..................................t............................................. [102480]
KX354450.1_47/Ss/2008            ..................................t............................................. [102480]
KP055815.1_strain_BA71           ..................................t............................................. [102480]
MH025920.1_strain_R35            .......c........c..c.....g..........................c..............c..g......... [102480]
MH025919.1_strain_N10            .......c........c..c.....g..........................c..............c..g......... [102480]
MH025918.1_strain_R25            .......c........c..c.....g..........................c..............c..g......... [102480]
MH025917.1_strain_R7             .......c........c..c.....g..........................c..............c..g......... [102480]
MH025916.1_strain_R8             .......c........c..c.....g..........................c..............c..g......... [102480]

FR682468.2_Georgia_2007/1        cgaaaaaatacaattacacgtgctcttaaaatgtatgaggattactactcccactttaagtctctttatgagcagtatca [102560]
ASF_IND_20_CAD543                ................................................................................ [102560]
LR899193.1_Germany_2020/1        ................................................................................ [102560]
MN393477.1_Wuhan_2019            ................................................................................ [102560]
MW396979.1_Timor-Leste/2019/1    ................................................................................ [102560]
MK940252.1_CN/2019/InnerMongolia ................................................................................ [102560]
MK128995.1_China/2018/AnhuiXCGQ  ................................................................................ [102560]
MH681419.1_ASFV/POL/2015         ................................................................................ [102560]
LS478113.1_Estonia_2014          ................................................................................ [102560]
KP843857.1_Odintsovo_02/14       ................................................................................ [102560]
```

```
FN557520.1_strain_E75              ................................................................................ [102560]
AY261366.1_Warthog                 ................................................................................ [102560]
AY261365.1_Warmbaths               ................................................................................ [102560]
AY261364.1_Tengani_62              ...............................................t................................ [102560]
AY261363.1_Pretoriuskop/96/4       ................................................................................ [102560]
AY261362.1_Mkuzi_1979              ................................................................................ [102560]
AY261361.1_Malawi_Lil-20/1_1983    ..................................................t..t.................a........ [102560]
AY261360.1_Kenya_1950              .........................................c........t............................. [102560]
AM712240.1_OURT_88/3               ................................................................................ [102560]
AM712239.1_Benin_97/1              ................................................................................ [102560]
KM111295.1_Ken06.Bus               .........................................c........t............................. [102560]
KM111294.1_Ken05/Tk1               .........................................c........t............................. [102560]
KM262844.1_strain_L60              ................................................................................ [102560]
KM102979.1_26544/OG10_Italy        ................................................................................ [102560]
KM262845.1_strain_NHV              ................................................................................ [102560]
KX354450.1_47/Ss/2008              ................................................................................ [102560]
KP055815.1_strain_BA71             ................................................................................ [102560]
MH025920.1_strain_R35              .........................................c........t............................. [102560]
MH025919.1_strain_N10              .........................................c........t............................. [102560]
MH025918.1_strain_R25              .........................................c........t............................. [102560]
MH025917.1_strain_R7               .........................................c........t............................. [102560]
MH025916.1_strain_R8               .........................................c........t............................. [102560]

FR682468.2_Georgia_2007/1          tttaaacgcggcaaaaaaattaatttaaactaaacgtttaaactaaatgtttaaactaaac------------------- [102640]
ASF_IND_20_CAD543                  .............................................................gtt-------------taa [102640]
LR899193.1_Germany_2020/1          .............................................................------------------- [102640]
MN393477.1_Wuhan_2019              .............................................................------------------- [102640]
MW396979.1_Timor-Leste/2019/1      .............................................................------------------- [102640]
MK940252.1_CN/2019/InnerMongolia   .............................................................------------------- [102640]
MK128995.1_China/2018/AnhuiXCGQ    .............................................................------------------- [102640]
MH681419.1_ASFV/POL/2015           .............................................................------------------- [102640]
LS478113.1_Estonia_2014            .............................................................------------------- [102640]
KP843857.1_Odintsovo_02/14         .............................................................------------------- [102640]
FN557520.1_strain_E75              ...............................................c..---------------------------taa [102640]
AY261366.1_Warthog                 ..............................--......----------------------------------------aa [102640]
AY261365.1_Warmbaths               ..............................--................-------------gtt-------------taa [102640]
AY261364.1_Tengani_62              ..............................--...............ca........----------------------- [102640]
AY261363.1_Pretoriuskop/96/4       ..............................--...............c.g..........agtt-------------taa [102640]
AY261362.1_Mkuzi_1979              .................................g.............c...----------------------------- [102640]
AY261361.1_Malawi_Lil-20/1_1983    c..g.........................t-----.........------------------tt---------------- [102640]
AY261360.1_Kenya_1950              g..............................-tta............-..........t.-gtt-------------taa [102640]
AM712240.1_OURT_88/3               ...............................................c.............gtttaaactaaacgtttaa [102640]
AM712239.1_Benin_97/1              ...............................................c..---------------------------taa [102640]
KM111295.1_Ken06.Bus               g........a.................g...-..a..c.........-..c.........-gtt-------------taa [102640]
```

```
KM111294.1_Ken05/Tk1             g..............................-tta.-------------------------------------------- [102640]
KM262844.1_strain_L60            ...............................................c...--------------------------taa [102640]
KM102979.1_26544/OG10_Italy      ...............................................c...--------------------------taa [102640]
KM262845.1_strain_NHV            ...............................................c.............gtttaaactaaacgtttaa [102640]
KX354450.1_47/Ss/2008            ...............................................c...--------------------------taa [102640]
KP055815.1_strain_BA71           ...............................................c...--------------------------taa [102640]
MH025920.1_strain_R35            g........a.................g...-..a..c.........-..c.........-gtt-------------taa [102640]
MH025919.1_strain_N10            g........a.................g...-..a..c.........-..c.........-gtt-------------taa [102640]
MH025918.1_strain_R25            g........a.................g...-..a..c.........-..c.........-gtt-------------taa [102640]
MH025917.1_strain_R7             g........a.................g...-..a..c.........-..c.........-gtt-------------taa [102640]
MH025916.1_strain_R8             g........a.................g...-..a..c.........-..c.........-gtt-------------taa [102640]

FR682468.2_Georgia_2007/1        -----------gttaaaactaaacatttcgactaaagtttaaaacctagtctaacagcgggatgcccatttccctggggt [102720]
ASF_IND_20_CAD543                actaaa----a..................................................................... [102720]
LR899193.1_Germany_2020/1        -----------..................................................................... [102720]
MN393477.1_Wuhan_2019            -----------..................................................................... [102720]
MW396979.1_Timor-Leste/2019/1    -----------..................................................................... [102720]
MK940252.1_CN/2019/InnerMongolia -----------..................................................................... [102720]
MK128995.1_China/2018/AnhuiXCGQ  -----------..................................................................... [102720]
MH681419.1_ASFV/POL/2015         -----------..................................................................... [102720]
LS478113.1_Estonia_2014          -----------..................................................................... [102720]
KP843857.1_Odintsovo_02/14       -----------..................................................................... [102720]
FN557520.1_strain_E75            actaaa----a..................................................................... [102720]
AY261366.1_Warthog               actaaa----a.................a......................c............................ [102720]
AY261365.1_Warmbaths             actaaa----a...t.............a......................c............................ [102720]
AY261364.1_Tengani_62            -----------------------------------................c..........................a. [102720]
AY261363.1_Pretoriuskop/96/4     actaaa----a.................a......................c............................ [102720]
AY261362.1_Mkuzi_1979            ---------------................................................................. [102720]
AY261361.1_Malawi_Lil-20/1_1983  -----------------------tg...aa...tt.........t...a..c........a........c.......... [102720]
AY261360.1_Kenya_1950            actagacttta..ct......g..t..---.g.ctgaac....ctt.....c....at..a.............g..... [102720]
AM712240.1_OURT_88/3             actaaa----a..................................................................... [102720]
AM712239.1_Benin_97/1            actaaa----a..................................................................... [102720]
KM111295.1_Ken06.Bus             ac------------------------------.tt..ac....gtt.....c....at..a.............g..... [102720]
KM111294.1_Ken05/Tk1             --------------------------------.ct.aac....ctt.....c....at..a.............g..... [102720]
KM262844.1_strain_L60            actaaa----a..................................................................... [102720]
KM102979.1_26544/OG10_Italy      actaaa----a..................................................................... [102720]
KM262845.1_strain_NHV            actaaa----a..................................................................... [102720]
KX354450.1_47/Ss/2008            actaaa----a..................................................................... [102720]
KP055815.1_strain_BA71           actaaa----a..................................................................... [102720]
MH025920.1_strain_R35            ac------------------------------.tt..ac....gtt.....c....at..a.............g..... [102720]
MH025919.1_strain_N10            ac------------------------------.tt..ac....gtt.....c....at..a.............g..... [102720]
MH025918.1_strain_R25            ac------------------------------.tt..ac....gtt.....c....at..a.............g..... [102720]
MH025917.1_strain_R7             ac------------------------------.tt..ac....gtt.....c....at..a.............g..... [102720]
MH025916.1_strain_R8             ac------------------------------.tt..ac....gtt.....c....at..a.............g..... [102720]
```

```
FR682468.2_Georgia_2007/1        tccatatttcaacaattttttgaccttcgggtgttaccttgatgcagcgcatgacgagcagtggaattttcctattaaag [102800]
ASF_IND_20_CAD543                ................................................................................ [102800]
LR899193.1_Germany_2020/1        ................................................................................ [102800]
MN393477.1_Wuhan_2019            ................................................................................ [102800]
MW396979.1_Timor-Leste/2019/1    ................................................................................ [102800]
MK940252.1_CN/2019/InnerMongolia ................................................................................ [102800]
MK128995.1_China/2018/AnhuiXCGQ  ................................................................................ [102800]
MH681419.1_ASFV/POL/2015         ................................................................................ [102800]
LS478113.1_Estonia_2014          ................................................................................ [102800]
KP843857.1_Odintsovo_02/14       ................................................................................ [102800]
FN557520.1_strain_E75            ............................-................................................... [102800]
AY261366.1_Warthog               .........................g...................................................... [102800]
AY261365.1_Warmbaths             ......................................................................t......... [102800]
AY261364.1_Tengani_62            ............................................................a.........t......... [102800]
AY261363.1_Pretoriuskop/96/4     .........................g.................a.................................... [102800]
AY261362.1_Mkuzi_1979            ................................................................................ [102800]
AY261361.1_Malawi_Lil-20/1_1983  ......................t........c................................................ [102800]
AY261360.1_Kenya_1950            .........................a..........................a....................g...... [102800]
AM712240.1_OURT_88/3             ................................................................................ [102800]
AM712239.1_Benin_97/1            ................................................................................ [102800]
KM111295.1_Ken06.Bus             .........................a..........................a....................g...... [102800]
KM111294.1_Ken05/Tk1             .........................a..........................a....................g...... [102800]
KM262844.1_strain_L60            ................................................................................ [102800]
KM102979.1_26544/OG10_Italy      ................................................................................ [102800]
KM262845.1_strain_NHV            ................................................................................ [102800]
KX354450.1_47/Ss/2008            ................................................................................ [102800]
KP055815.1_strain_BA71           ................................................................................ [102800]
MH025920.1_strain_R35            .........................a..........................a....................g...... [102800]
MH025919.1_strain_N10            .........................a..........................a....................g...... [102800]
MH025918.1_strain_R25            .........................a..........................a....................g...... [102800]
MH025917.1_strain_R7             .........................a..........................a....................g...... [102800]
MH025916.1_strain_R8             .........................a..........................a....................g...... [102800]

FR682468.2_Georgia_2007/1        agttcttgcttagctatatcaataggactgctatatttttttttaagcattgtagatccattaattgccaattgttgcgc [102880]
ASF_IND_20_CAD543                ................................................................................ [102880]
LR899193.1_Germany_2020/1        ................................................................................ [102880]
MN393477.1_Wuhan_2019            ................................................................................ [102880]
MW396979.1_Timor-Leste/2019/1    ................................................................................ [102880]
MK940252.1_CN/2019/InnerMongolia ................................................................................ [102880]
MK128995.1_China/2018/AnhuiXCGQ  ................................................................................ [102880]
MH681419.1_ASFV/POL/2015         ................................................................................ [102880]
LS478113.1_Estonia_2014          ................................................................................ [102880]
KP843857.1_Odintsovo_02/14       ................................................................................ [102880]
```

```
FN557520.1_strain_E75           ........................................................c..............c........ [102880]
AY261366.1_Warthog              .................................................................c....g......... [102880]
AY261365.1_Warmbaths            ......................................................................g......... [102880]
AY261364.1_Tengani_62           ...........................................................g..........g......... [102880]
AY261363.1_Pretoriuskop/96/4    ......................................................................g......... [102880]
AY261362.1_Mkuzi_1979           ........................................................c..............c........ [102880]
AY261361.1_Malawi_Lil-20/1_1983 ........................................................a....................... [102880]
AY261360.1_Kenya_1950           ..............................................................g..............t.. [102880]
AM712240.1_OURT_88/3            ........................................................c..............c........ [102880]
AM712239.1_Benin_97/1           ........................................................c..............c........ [102880]
KM111295.1_Ken06.Bus            ..............................................................g..............t.. [102880]
KM111294.1_Ken05/Tk1            ..............................................................g..............t.. [102880]
KM262844.1_strain_L60           ........................................................c..............c........ [102880]
KM102979.1_26544/OG10_Italy     ........................................................c..............c........ [102880]
KM262845.1_strain_NHV           ........................................................c..............c........ [102880]
KX354450.1_47/Ss/2008           ........................................................c..............c........ [102880]
KP055815.1_strain_BA71          ........................................................c..............c........ [102880]
MH025920.1_strain_R35           ..............................................................g..............t.. [102880]
MH025919.1_strain_N10           ..............................................................g..............t.. [102880]
MH025918.1_strain_R25           ..............................................................g..............t.. [102880]
MH025917.1_strain_R7            ..............................................................g..............t.. [102880]
MH025916.1_strain_R8            ..............................................................g..............t.. [102880]

FR682468.2_Georgia_2007/1       tctaacggcgaccaaccttgtggcctcaaaggtggttaaaacgttggaggtaatgcgctcgttatcgggtataatgacca [102960]
ASF_IND_20_CAD543               ................................................................................ [102960]
LR899193.1_Germany_2020/1       ................................................................................ [102960]
MN393477.1_Wuhan_2019           ................................................................................ [102960]
MW396979.1_Timor-Leste/2019/1   ................................................................................ [102960]
MK940252.1_CN/2019/InnerMongolia ................................................................................ [102960]
MK128995.1_China/2018/AnhuiXCGQ ................................................................................ [102960]
MH681419.1_ASFV/POL/2015        ................................................................................ [102960]
LS478113.1_Estonia_2014         ................................................................................ [102960]
KP843857.1_Odintsovo_02/14      ................................................................................ [102960]
FN557520.1_strain_E75           ................................................................................ [102960]
AY261366.1_Warthog              ................................................................................ [102960]
AY261365.1_Warmbaths            ...........................g.................................................... [102960]
AY261364.1_Tengani_62           ................................................................................ [102960]
AY261363.1_Pretoriuskop/96/4    ...........................g.................................................... [102960]
AY261362.1_Mkuzi_1979           ................................................................................ [102960]
AY261361.1_Malawi_Lil-20/1_1983 ........................t....................................................... [102960]
AY261360.1_Kenya_1950           ........................t........c........................................c..... [102960]
AM712240.1_OURT_88/3            ............................g................................................... [102960]
AM712239.1_Benin_97/1           ................................................................................ [102960]
KM111295.1_Ken06.Bus            ........................t........c........................................c..... [102960]
```

```
KM111294.1_Ken05/Tk1             ........................t........c........................................c..... [102960]
KM262844.1_strain_L60            ................................................................................ [102960]
KM102979.1_26544/OG10_Italy      ................................................................................ [102960]
KM262845.1_strain_NHV            ................................................................................ [102960]
KX354450.1_47/Ss/2008            ................................................................................ [102960]
KP055815.1_strain_BA71           ................................................................................ [102960]
MH025920.1_strain_R35            ........................t........c........................................c..... [102960]
MH025919.1_strain_N10            ........................t........c........................................c..... [102960]
MH025918.1_strain_R25            ........................t........c........................................c..... [102960]
MH025917.1_strain_R7             ........................t........c........................................c..... [102960]
MH025916.1_strain_R8             ........................t........c........................................c..... [102960]

FR682468.2_Georgia_2007/1        atgtttgcgacgaggcctgcacaaagccctcgcagatggacggagactccacgatctcgtccttgtcctcggac------ [103040]
ASF_IND_20_CAD543                ..........................................................................------ [103040]
LR899193.1_Germany_2020/1        ..........................................................................------ [103040]
MN393477.1_Wuhan_2019            ..........................................................................------ [103040]
MW396979.1_Timor-Leste/2019/1    ..........................................................................------ [103040]
MK940252.1_CN/2019/InnerMongolia ..........................................................................------ [103040]
MK128995.1_China/2018/AnhuiXCGQ  ..........................................................................------ [103040]
MH681419.1_ASFV/POL/2015         ..........................................................................------ [103040]
LS478113.1_Estonia_2014          ..........................................................................------ [103040]
KP843857.1_Odintsovo_02/14       ..........................................................................------ [103040]
FN557520.1_strain_E75            ..........................................................................------ [103040]
AY261366.1_Warthog               ...............................a..........................a...............------ [103040]
AY261365.1_Warmbaths             ..........................................................................------ [103040]
AY261364.1_Tengani_62            ...............................a..........................a..t............------ [103040]
AY261363.1_Pretoriuskop/96/4     ....................................................a........t............------ [103040]
AY261362.1_Mkuzi_1979            ..............................................................c...........------ [103040]
AY261361.1_Malawi_Lil-20/1_1983  ...............................a..............t...........................ttatcc [103040]
AY261360.1_Kenya_1950            ...............................a..............t....................t......------ [103040]
AM712240.1_OURT_88/3             ..........................................................................------ [103040]
AM712239.1_Benin_97/1            ..........................................................................------ [103040]
KM111295.1_Ken06.Bus             ...............................a..............t....................t......------ [103040]
KM111294.1_Ken05/Tk1             ...............................a..............t...........a........t......------ [103040]
KM262844.1_strain_L60            ..........................................................................------ [103040]
KM102979.1_26544/OG10_Italy      ..........................................................................------ [103040]
KM262845.1_strain_NHV            ..........................................................................------ [103040]
KX354450.1_47/Ss/2008            ..........................................................................------ [103040]
KP055815.1_strain_BA71           ..........................................................................------ [103040]
MH025920.1_strain_R35            ...............................a..............t....................t......------ [103040]
MH025919.1_strain_N10            ...............................a..............t....................t......------ [103040]
MH025918.1_strain_R25            ...............................a..............t....................t......------ [103040]
MH025917.1_strain_R7             ...............................a..............t....................t......------ [103040]
MH025916.1_strain_R8             ...............................a..............t....................t......------ [103040]
```

```
FR682468.2_Georgia_2007/1        ------tcctcctcactgtcgacgaggttctcctcttccgtttccacatattcctccacgaggtcatccatgataagatc [103120]
ASF_IND_20_CAD543                ------.......................................................................... [103120]
LR899193.1_Germany_2020/1        ------.......................................................................... [103120]
MN393477.1_Wuhan_2019            ------.......................................................................... [103120]
MW396979.1_Timor-Leste/2019/1    ------.......................................................................... [103120]
MK940252.1_CN/2019/InnerMongolia ------.......................................................................... [103120]
MK128995.1_China/2018/AnhuiXCGQ  ------.......................................................................... [103120]
MH681419.1_ASFV/POL/2015         ------.......................................................................... [103120]
LS478113.1_Estonia_2014          ------.......................................................................... [103120]
KP843857.1_Odintsovo_02/14       ------.......................................................................... [103120]
FN557520.1_strain_E75            ------...................a..................................................t... [103120]
AY261366.1_Warthog               ------...................a..................................................c... [103120]
AY261365.1_Warmbaths             ------...................a..................................................c... [103120]
AY261364.1_Tengani_62            ------.....t.............a..................................................c... [103120]
AY261363.1_Pretoriuskop/96/4     ------...................a..................................................c... [103120]
AY261362.1_Mkuzi_1979            ------...................a...t..............................................c... [103120]
AY261361.1_Malawi_Lil-20/1_1983  tcggac...................aa..t.....c........................................c... [103120]
AY261360.1_Kenya_1950            ------...................aa........c.......................................ct... [103120]
AM712240.1_OURT_88/3             ------...................a............................................c.....t... [103120]
AM712239.1_Benin_97/1            ------...................a..................................................t... [103120]
KM111295.1_Ken06.Bus             ------...................aa........c.......................................ct... [103120]
KM111294.1_Ken05/Tk1             ------...................aa........c.......................................ct... [103120]
KM262844.1_strain_L60            ------...................a..................................................t... [103120]
KM102979.1_26544/OG10_Italy      ------...................a..................................................t... [103120]
KM262845.1_strain_NHV            ------...................a............................................c.....t... [103120]
KX354450.1_47/Ss/2008            ------...................a..................................................t... [103120]
KP055815.1_strain_BA71           ------...................a..................................................t... [103120]
MH025920.1_strain_R35            ------...................aa........c.......................................ct... [103120]
MH025919.1_strain_N10            ------...................aa........c.......................................ct... [103120]
MH025918.1_strain_R25            ------...................aa........c.......................................ct... [103120]
MH025917.1_strain_R7             ------...................aa........c.......................................ct... [103120]
MH025916.1_strain_R8             ------...................aa........c.......................................ct... [103120]

FR682468.2_Georgia_2007/1        ctcgttgtcattatcagccatattacactgttatcaaatgtactgtttaatacgcaaatggatttactacgttttaattg [103200]
ASF_IND_20_CAD543                ................................................................................ [103200]
LR899193.1_Germany_2020/1        ................................................................................ [103200]
MN393477.1_Wuhan_2019            ................................................................................ [103200]
MW396979.1_Timor-Leste/2019/1    ................................................................................ [103200]
MK940252.1_CN/2019/InnerMongolia ................................................................................ [103200]
MK128995.1_China/2018/AnhuiXCGQ  ................................................................................ [103200]
MH681419.1_ASFV/POL/2015         ................................................................................ [103200]
LS478113.1_Estonia_2014          ................................................................................ [103200]
KP843857.1_Odintsovo_02/14       ................................................................................ [103200]
```

```
FN557520.1_strain_E75            .....................g.......................................................... [103200]
AY261366.1_Warthog               ................................................................................ [103200]
AY261365.1_Warmbaths             ................................................................................ [103200]
AY261364.1_Tengani_62            ...............g................................................................ [103200]
AY261363.1_Pretoriuskop/96/4     ................................................................................ [103200]
AY261362.1_Mkuzi_1979            ................................................................................ [103200]
AY261361.1_Malawi_Lil-20/1_1983  ...............................................................a.....t.......... [103200]
AY261360.1_Kenya_1950            ...a..................................................t........a................ [103200]
AM712240.1_OURT_88/3             .....................g.......................................................... [103200]
AM712239.1_Benin_97/1            .....................g.......................................................... [103200]
KM111295.1_Ken06.Bus             t.....................................................t........a................ [103200]
KM111294.1_Ken05/Tk1             ...a..................................................t........a................ [103200]
KM262844.1_strain_L60            .....................g.......................................................... [103200]
KM102979.1_26544/OG10_Italy      .....................g.......................................................... [103200]
KM262845.1_strain_NHV            .....................g.......................................................... [103200]
KX354450.1_47/Ss/2008            .....................g.......................................................... [103200]
KP055815.1_strain_BA71           .....................g.......................................................... [103200]
MH025920.1_strain_R35            t.....................................................t........a................ [103200]
MH025919.1_strain_N10            t.....................................................t........a................ [103200]
MH025918.1_strain_R25            t.....................................................t........a................ [103200]
MH025917.1_strain_R7             t.....................................................t........a................ [103200]
MH025916.1_strain_R8             t.....................................................t........a................ [103200]

FR682468.2_Georgia_2007/1        tatgtcttcatgtgcaggctctagtggaaagtaattttctcacaatttttggcaccgttacacttgtgcccacaaaaacc [103280]
ASF_IND_20_CAD543                ................................................................................ [103280]
LR899193.1_Germany_2020/1        ................................................................................ [103280]
MN393477.1_Wuhan_2019            ................................................................................ [103280]
MW396979.1_Timor-Leste/2019/1    ................................................................................ [103280]
MK940252.1_CN/2019/InnerMongolia ................................................................................ [103280]
MK128995.1_China/2018/AnhuiXCGQ  ................................................................................ [103280]
MH681419.1_ASFV/POL/2015         ................................................................................ [103280]
LS478113.1_Estonia_2014          ................................................................................ [103280]
KP843857.1_Odintsovo_02/14       ................................................................................ [103280]
FN557520.1_strain_E75            ............................t................................................... [103280]
AY261366.1_Warthog               ...........a................t...................................c............... [103280]
AY261365.1_Warmbaths             ...........a................t...................................c............... [103280]
AY261364.1_Tengani_62            ...........a................t...................................c............... [103280]
AY261363.1_Pretoriuskop/96/4     .t.........a................t...................................c..............t [103280]
AY261362.1_Mkuzi_1979            ............................t................................................... [103280]
AY261361.1_Malawi_Lil-20/1_1983  ...........a.......c........c.................c...........g.....c..a............ [103280]
AY261360.1_Kenya_1950            ...........a.---aa.c.cgac...t.a...g..c....................c.....g..............g [103280]
AM712240.1_OURT_88/3             ............................t................................................... [103280]
AM712239.1_Benin_97/1            ............................t................................................... [103280]
KM111295.1_Ken06.Bus             ...........a.---aa.c.cgac...t.a...g..c..t.................c.....c..............g [103280]
```

```
KM111294.1_Ken05/Tk1            ...........a.---aa.c.cgac...t.a...g..c....................c...t.g..............g [103280]
KM262844.1_strain_L60           ............................t................................................... [103280]
KM102979.1_26544/OG10_Italy     ............................t................................................... [103280]
KM262845.1_strain_NHV           ............................t................................................... [103280]
KX354450.1_47/Ss/2008           ............................t................................................... [103280]
KP055815.1_strain_BA71          ............................t................................................... [103280]
MH025920.1_strain_R35           ...........a.---aa.c.cgac...t.a...g..c..t.................c.....c..............g [103280]
MH025919.1_strain_N10           ...........a.---aa.c.cgac...t.a...g..c..t.................c.....c..............g [103280]
MH025918.1_strain_R25           ...........a.---aa.c.cgac...t.a...g..c..t.................c.....c..............g [103280]
MH025917.1_strain_R7            ...........a.---aa.c.cgac...t.a...g..c..t.................c.....c..............g [103280]
MH025916.1_strain_R8            ...........a.---aa.c.cgac...t.a...g..c..t.................c.....c..............g [103280]

FR682468.2_Georgia_2007/1       cgcgatttttttattttatattacttttggaagtacgagtttaaccagtcgctttcaaaccttatgcgtctatctcgcca [103360]
ASF_IND_20_CAD543               ................................................................................ [103360]
LR899193.1_Germany_2020/1       ................................................................................ [103360]
MN393477.1_Wuhan_2019           ................................................................................ [103360]
MW396979.1_Timor-Leste/2019/1   ................................................................................ [103360]
MK940252.1_CN/2019/InnerMongolia ................................................................................ [103360]
MK128995.1_China/2018/AnhuiXCGQ ................................................................................ [103360]
MH681419.1_ASFV/POL/2015        ................................................................................ [103360]
LS478113.1_Estonia_2014         ................................................................................ [103360]
KP843857.1_Odintsovo_02/14      ................................................................................ [103360]
FN557520.1_strain_E75           ................................................................................ [103360]
AY261366.1_Warthog              ..........................c..................................................... [103360]
AY261365.1_Warmbaths            ..........................c..................................................... [103360]
AY261364.1_Tengani_62           ..........................c..................................................... [103360]
AY261363.1_Pretoriuskop/96/4    ..........................c..................................................... [103360]
AY261362.1_Mkuzi_1979           ................................................................................ [103360]
AY261361.1_Malawi_Lil-20/1_1983 ..........................c.............................g....................a.. [103360]
AY261360.1_Kenya_1950           ..t.......................c...............g..........g.t............c........g.. [103360]
AM712240.1_OURT_88/3            ................................................................................ [103360]
AM712239.1_Benin_97/1           ................................................................................ [103360]
KM111295.1_Ken06.Bus            ..t.......................c...............g..........g.t............c........g.. [103360]
KM111294.1_Ken05/Tk1            ..t.......................c...............g..........g.t............c........g.. [103360]
KM262844.1_strain_L60           ................................................................................ [103360]
KM102979.1_26544/OG10_Italy     ................................................................................ [103360]
KM262845.1_strain_NHV           ................................................................................ [103360]
KX354450.1_47/Ss/2008           ................................................................................ [103360]
KP055815.1_strain_BA71          ................................................................................ [103360]
MH025920.1_strain_R35           ..t.......................c...............g..........g.t............c........g.. [103360]
MH025919.1_strain_N10           ..t.......................c...............g..........g.t............c........g.. [103360]
MH025918.1_strain_R25           ..t.......................c...............g..........g.t............c........g.. [103360]
MH025917.1_strain_R7            ..t.......................c...............g..........g.t............c........g.. [103360]
MH025916.1_strain_R8            ..t.......................c...............g..........g.t............c........g.. [103360]
```

```
FR682468.2_Georgia_2007/1        aaaaacgctcacagcggtgttggatattacctttaaaaaaataacattaatttttaccacagagggcgtattgcgtatgg [103440]
ASF_IND_20_CAD543                ................................................................................ [103440]
LR899193.1_Germany_2020/1        ................................................................................ [103440]
MN393477.1_Wuhan_2019            ................................................................................ [103440]
MW396979.1_Timor-Leste/2019/1    ................................................................................ [103440]
MK940252.1_CN/2019/InnerMongolia ................................................................................ [103440]
MK128995.1_China/2018/AnhuiXCGQ  ................................................................................ [103440]
MH681419.1_ASFV/POL/2015         ................................................................................ [103440]
LS478113.1_Estonia_2014          ................................................................................ [103440]
KP843857.1_Odintsovo_02/14       ................................................................................ [103440]
FN557520.1_strain_E75            ................................................................................ [103440]
AY261366.1_Warthog               .............................................................a.......g..t..c.... [103440]
AY261365.1_Warmbaths             .............................................................a.......g..t..c.... [103440]
AY261364.1_Tengani_62            .....................................................................g..t..c.... [103440]
AY261363.1_Pretoriuskop/96/4     .............................................................a.......g..t..c.... [103440]
AY261362.1_Mkuzi_1979            ................................................................................ [103440]
AY261361.1_Malawi_Lil-20/1_1983  .....................a.......................................a....t..g..t..c.... [103440]
AY261360.1_Kenya_1950            ..........................c.................t................a.......g..t..c.... [103440]
AM712240.1_OURT_88/3             ................................................................................ [103440]
AM712239.1_Benin_97/1            ................................................................................ [103440]
KM111295.1_Ken06.Bus             ............g.............c.................t................a..........t..c.... [103440]
KM111294.1_Ken05/Tk1             ......a...................c.................t................a.......g..t..c.... [103440]
KM262844.1_strain_L60            ................................................................................ [103440]
KM102979.1_26544/OG10_Italy      ................................................................................ [103440]
KM262845.1_strain_NHV            ................................................................................ [103440]
KX354450.1_47/Ss/2008            ................................................................................ [103440]
KP055815.1_strain_BA71           ................................................................................ [103440]
MH025920.1_strain_R35            ............g.............c.................t................a..........t..c.... [103440]
MH025919.1_strain_N10            ............g.............c.................t................a..........t..c.... [103440]
MH025918.1_strain_R25            ............g.............c.................t................a..........t..c.... [103440]
MH025917.1_strain_R7             ............g.............c.................t................a..........t..c.... [103440]
MH025916.1_strain_R8             ............g.............c.................t................a..........t..c.... [103440]

FR682468.2_Georgia_2007/1        attctacgaataagccaggcgtgccactcgatatagacccccagttcattgaccttgatagtattttaatggaactggat [103520]
ASF_IND_20_CAD543                ................................................................................ [103520]
LR899193.1_Germany_2020/1        ................................................................................ [103520]
MN393477.1_Wuhan_2019            ................................................................................ [103520]
MW396979.1_Timor-Leste/2019/1    ................................................................................ [103520]
MK940252.1_CN/2019/InnerMongolia ................................................................................ [103520]
MK128995.1_China/2018/AnhuiXCGQ  ................................................................................ [103520]
MH681419.1_ASFV/POL/2015         ................................................................................ [103520]
LS478113.1_Estonia_2014          ................................................................................ [103520]
KP843857.1_Odintsovo_02/14       ................................................................................ [103520]
```

```
FN557520.1_strain_E75             ................................................................................ [103520]
AY261366.1_Warthog                ..c..........a............................t.................................a... [103520]
AY261365.1_Warmbaths              ..c..........a.......a....................t.................................a... [103520]
AY261364.1_Tengani_62             ..c..........a.................................................................. [103520]
AY261363.1_Pretoriuskop/96/4      ..c..........a............................t.................................a... [103520]
AY261362.1_Mkuzi_1979             ................................................................................ [103520]
AY261361.1_Malawi_Lil-20/1_1983   ..ca...a.............c...............................................c.......... [103520]
AY261360.1_Kenya_1950             ...a..aa........g....c....a.............t....................................... [103520]
AM712240.1_OURT_88/3              ................................................................................ [103520]
AM712239.1_Benin_97/1             ................................................................................ [103520]
KM111295.1_Ken06.Bus              ...a..aa........g....c....a.............t....................................... [103520]
KM111294.1_Ken05/Tk1              ...a..aa........g....c....a.............t....................................... [103520]
KM262844.1_strain_L60             ................................................................................ [103520]
KM102979.1_26544/OG10_Italy       ................................................................................ [103520]
KM262845.1_strain_NHV             ................................................................................ [103520]
KX354450.1_47/Ss/2008             ................................................................................ [103520]
KP055815.1_strain_BA71            ................................................................................ [103520]
MH025920.1_strain_R35             ...a..aa........g....c....a.............t....................................... [103520]
MH025919.1_strain_N10             ...a..aa........g....c....a.............t....................................... [103520]
MH025918.1_strain_R25             ...a..aa........g....c....a.............t....................................... [103520]
MH025917.1_strain_R7              ...a..aa........g....c....a.............t....................................... [103520]
MH025916.1_strain_R8              ...a..aa........g....c....a.............t....................................... [103520]

FR682468.2_Georgia_2007/1         cattaggacc-tctcccgcccatttaaatttttagtttctacaataataaaatgcgcgaggaatcatgggaagaccacga [103600]
ASF_IND_20_CAD543                 ..........-..................................................................... [103600]
LR899193.1_Germany_2020/1         ..........-..................................................................... [103600]
MN393477.1_Wuhan_2019             ..........-..................................................................... [103600]
MW396979.1_Timor-Leste/2019/1     ..........-..................................................................... [103600]
MK940252.1_CN/2019/InnerMongolia  ..........-..................................................................... [103600]
MK128995.1_China/2018/AnhuiXCGQ   ..........-..................................................................... [103600]
MH681419.1_ASFV/POL/2015          ..........-..................................................................... [103600]
LS478113.1_Estonia_2014           ..........-..................................................................... [103600]
KP843857.1_Odintsovo_02/14        ..........-..................................................................... [103600]
FN557520.1_strain_E75             ..........-..................................................................... [103600]
AY261366.1_Warthog                ..........-..-.................................................................. [103600]
AY261365.1_Warmbaths              ..........-..-.................................................................. [103600]
AY261364.1_Tengani_62             ..........-..................................................................... [103600]
AY261363.1_Pretoriuskop/96/4      ..........-..-.................................................................. [103600]
AY261362.1_Mkuzi_1979             ..........-..................................................................... [103600]
AY261361.1_Malawi_Lil-20/1_1983   .....a....-.--..................c........t................................a..t.. [103600]
AY261360.1_Kenya_1950             ..........-c.....................g........................................a..t.. [103600]
AM712240.1_OURT_88/3              ..........-..................................................................... [103600]
AM712239.1_Benin_97/1             ..........-..................................................................... [103600]
KM111295.1_Ken06.Bus              ..........tc.....................g....t...................................a..t.. [103600]
```

```
KM111294.1_Ken05/Tk1            ..........-c.....................g........................................a..t.. [103600]
KM262844.1_strain_L60           ..........-..................................................................... [103600]
KM102979.1_26544/OG10_Italy     ..........-..................................................................... [103600]
KM262845.1_strain_NHV           ..........-..................................................................... [103600]
KX354450.1_47/Ss/2008           ..........-..................................................................... [103600]
KP055815.1_strain_BA71          ..........-..................................................................... [103600]
MH025920.1_strain_R35           ..........tc.....................g....t...................................a..t.. [103600]
MH025919.1_strain_N10           ..........tc.....................g....t...................................a..t.. [103600]
MH025918.1_strain_R25           ..........tc.....................g....t...................................a..t.. [103600]
MH025917.1_strain_R7            ..........tc.....................g....t...................................a..t.. [103600]
MH025916.1_strain_R8            ..........tc.....................g....t...................................a..t.. [103600]

FR682468.2_Georgia_2007/1       taccattcagctcaccgctcagcgcaaatacctcgccgaggtgcaagctctagagacccttttgactcgagagctttcag [103680]
ASF_IND_20_CAD543               ................................................................................ [103680]
LR899193.1_Germany_2020/1       ................................................................................ [103680]
MN393477.1_Wuhan_2019           ................................................................................ [103680]
MW396979.1_Timor-Leste/2019/1   ................................................................................ [103680]
MK940252.1_CN/2019/InnerMongolia ................................................................................ [103680]
MK128995.1_China/2018/AnhuiXCGQ ................................................................................ [103680]
MH681419.1_ASFV/POL/2015        ................................................................................ [103680]
LS478113.1_Estonia_2014         ................................................................................ [103680]
KP843857.1_Odintsovo_02/14      ................................................................................ [103680]
FN557520.1_strain_E75           ..............................t................................................. [103680]
AY261366.1_Warthog              ..............................t.......................a......................... [103680]
AY261365.1_Warmbaths            ..............................t.......................a......................... [103680]
AY261364.1_Tengani_62           ..............................t...........a.....ct.............................. [103680]
AY261363.1_Pretoriuskop/96/4    ..............................t................................................. [103680]
AY261362.1_Mkuzi_1979           ..............................t................................................. [103680]
AY261361.1_Malawi_Lil-20/1_1983 ..............................t...........a..................................... [103680]
AY261360.1_Kenya_1950           ..............................t...........a.....................g.............c. [103680]
AM712240.1_OURT_88/3            ..............................t................................................. [103680]
AM712239.1_Benin_97/1           ..............................t................................................. [103680]
KM111295.1_Ken06.Bus            ..............................t...........a.....................g.............c. [103680]
KM111294.1_Ken05/Tk1            ..............................t...a.......a...................................c. [103680]
KM262844.1_strain_L60           ..............................t................................................. [103680]
KM102979.1_26544/OG10_Italy     ..............................t................................................. [103680]
KM262845.1_strain_NHV           ..............................t................................................. [103680]
KX354450.1_47/Ss/2008           ..............................t................................................. [103680]
KP055815.1_strain_BA71          ..............................t................................................. [103680]
MH025920.1_strain_R35           ..............................t...........a.....................g.............c. [103680]
MH025919.1_strain_N10           ..............................t...........a.....................g.............c. [103680]
MH025918.1_strain_R25           ..............................t...........a.....................g.............c. [103680]
MH025917.1_strain_R7            ..............................t...........a.....................g.............c. [103680]
MH025916.1_strain_R8            ..............................t...........a.....................g.............c. [103680]
```

```
FR682468.2_Georgia_2007/1        tctttctcacagagccaggcagcaaaaaaacaaatattattaatagaatcacaggaaaaacctacgcacttcccagcaca [103760]
ASF_IND_20_CAD543                ................................................................................ [103760]
LR899193.1_Germany_2020/1        ................................................................................ [103760]
MN393477.1_Wuhan_2019            ................................................................................ [103760]
MW396979.1_Timor-Leste/2019/1    ................................................................................ [103760]
MK940252.1_CN/2019/InnerMongolia ................................................................................ [103760]
MK128995.1_China/2018/AnhuiXCGQ  ................................................................................ [103760]
MH681419.1_ASFV/POL/2015         ................................................................................ [103760]
LS478113.1_Estonia_2014          ................................................................................ [103760]
KP843857.1_Odintsovo_02/14       ................................................................................ [103760]
FN557520.1_strain_E75            c............................................................................... [103760]
AY261366.1_Warthog               ................................................................t............... [103760]
AY261365.1_Warmbaths             ................................................................................ [103760]
AY261364.1_Tengani_62            c.........g..................................................................... [103760]
AY261363.1_Pretoriuskop/96/4     .................................................t.............................. [103760]
AY261362.1_Mkuzi_1979            c............................................................................... [103760]
AY261361.1_Malawi_Lil-20/1_1983  .......t..c................................c.................................... [103760]
AY261360.1_Kenya_1950            ................................................................................ [103760]
AM712240.1_OURT_88/3             c............................................................................... [103760]
AM712239.1_Benin_97/1            c............................................................................... [103760]
KM111295.1_Ken06.Bus             ................................................................................ [103760]
KM111294.1_Ken05/Tk1             ...........................................c..g..............t.................. [103760]
KM262844.1_strain_L60            c............................................................................... [103760]
KM102979.1_26544/OG10_Italy      c............................................................................... [103760]
KM262845.1_strain_NHV            c............................................................................... [103760]
KX354450.1_47/Ss/2008            c............................................................................... [103760]
KP055815.1_strain_BA71           c............................................................................... [103760]
MH025920.1_strain_R35            ................................................................................ [103760]
MH025919.1_strain_N10            ................................................................................ [103760]
MH025918.1_strain_R25            ................................................................................ [103760]
MH025917.1_strain_R7             ................................................................................ [103760]
MH025916.1_strain_R8             ................................................................................ [103760]

FR682468.2_Georgia_2007/1        gagctactaagactctacgagcatctcgagcaatgtcgcaagcaaggcgccctcatgtatttttggaaagacaggggac [103840]
ASF_IND_20_CAD543                ................................................................................ [103840]
LR899193.1_Germany_2020/1        ................................................................................ [103840]
MN393477.1_Wuhan_2019            ................................................................................ [103840]
MW396979.1_Timor-Leste/2019/1    ................................................................................ [103840]
MK940252.1_CN/2019/InnerMongolia ................................................................................ [103840]
MK128995.1_China/2018/AnhuiXCGQ  ................................................................................ [103840]
MH681419.1_ASFV/POL/2015         ................................................................................ [103840]
LS478113.1_Estonia_2014          ................................................................................ [103840]
KP843857.1_Odintsovo_02/14       ................................................................................ [103840]
```

```
FN557520.1_strain_E75          ................................................................................ [103840]
AY261366.1_Warthog             ..........................t..................................................... [103840]
AY261365.1_Warmbaths           ..........................t..................................................... [103840]
AY261364.1_Tengani_62          ................................................................................ [103840]
AY261363.1_Pretoriuskop/96/4   ....................a.....t..................................................... [103840]
AY261362.1_Mkuzi_1979          ................................................................................ [103840]
AY261361.1_Malawi_Lil-20/1_1983 ..........................t..................................................... [103840]
AY261360.1_Kenya_1950          ..a.......................t..................................................... [103840]
AM712240.1_OURT_88/3           ................................................................................ [103840]
AM712239.1_Benin_97/1          ................................................................................ [103840]
KM111295.1_Ken06.Bus           ..a.........t.............t..a.................................................. [103840]
KM111294.1_Ken05/Tk1           ..............t...........t..................................................... [103840]
KM262844.1_strain_L60          ................................................................................ [103840]
KM102979.1_26544/OG10_Italy    ................................................................................ [103840]
KM262845.1_strain_NHV          ................................................................................ [103840]
KX354450.1_47/Ss/2008          ................................................................................ [103840]
KP055815.1_strain_BA71         ................................................................................ [103840]
MH025920.1_strain_R35          ..a.........t.............t..a.................................................. [103840]
MH025919.1_strain_N10          ..a.........t.............t..a.................................................. [103840]
MH025918.1_strain_R25          ..a.........t.............t..a.................................................. [103840]
MH025917.1_strain_R7           ..a.........t.............t..a.................................................. [103840]
MH025916.1_strain_R8           ..a.........t.............t..a.................................................. [103840]

FR682468.2_Georgia_2007/1      ctactcgggtctcatgttggactatgaccttaaactcaatacaaatgctgttcccccgctggaaccccccgcgctatcac [103920]
ASF_IND_20_CAD543              ........................................................y....................... [103920]
LR899193.1_Germany_2020/1      ................................................................................ [103920]
MN393477.1_Wuhan_2019          ................................................................................ [103920]
MW396979.1_Timor-Leste/2019/1  ................................................................................ [103920]
MK940252.1_CN/2019/InnerMongolia ................................................................................ [103920]
MK128995.1_China/2018/AnhuiXCGQ ................................................................................ [103920]
MH681419.1_ASFV/POL/2015       ................................................................................ [103920]
LS478113.1_Estonia_2014        ................................................................................ [103920]
KP843857.1_Odintsovo_02/14     ................................................................................ [103920]
FN557520.1_strain_E75          ................................................................-............... [103920]
AY261366.1_Warthog             ...t..............................................c............................. [103920]
AY261365.1_Warmbaths           ..................................................c............................. [103920]
AY261364.1_Tengani_62          ..................................................cc...............t............ [103920]
AY261363.1_Pretoriuskop/96/4   ..................................................cc............................ [103920]
AY261362.1_Mkuzi_1979          ................................................................................ [103920]
AY261361.1_Malawi_Lil-20/1_1983 ................................................c.c.............t....t.......... [103920]
AY261360.1_Kenya_1950          ................c.............a.................c.cc...t........t..t...t......g. [103920]
AM712240.1_OURT_88/3           ................................................................................ [103920]
AM712239.1_Benin_97/1          ................................................................................ [103920]
KM111295.1_Ken06.Bus           ................c..........t..a.................c.cc...t........t..t...t......g. [103920]
```

```
KM111294.1_Ken05/Tk1             ...............................................t..c.............t....t.......... [103920]
KM262844.1_strain_L60            ................................................................................ [103920]
KM102979.1_26544/OG10_Italy      ................................................................................ [103920]
KM262845.1_strain_NHV            ................................................................................ [103920]
KX354450.1_47/Ss/2008            ................................................................................ [103920]
KP055815.1_strain_BA71           ................................................................................ [103920]
MH025920.1_strain_R35            ................c..........t..a.................c.cc...t........t..t...t......g. [103920]
MH025919.1_strain_N10            ................c..........t..a.................c.cc...t........t..t...t......g. [103920]
MH025918.1_strain_R25            ................c..........t..a.................c.cc...t........t..t...t......g. [103920]
MH025917.1_strain_R7             ................c..........t..a.................c.cc...t........t..t...t......g. [103920]
MH025916.1_strain_R8             ................c..........t..a.................c.cc...t........t..t...t......g. [103920]

FR682468.2_Georgia_2007/1        ggctttgccatcgaatatttgtgcatataaaaaacagcagtgtgctgcctgagggcagccataaaatccacttctttttt [104000]
ASF_IND_20_CAD543                ................................................................................ [104000]
LR899193.1_Germany_2020/1        ................................................................................ [104000]
MN393477.1_Wuhan_2019            ................................................................................ [104000]
MW396979.1_Timor-Leste/2019/1    ................................................................................ [104000]
MK940252.1_CN/2019/InnerMongolia ................................................................................ [104000]
MK128995.1_China/2018/AnhuiXCGQ  ................................................................................ [104000]
MH681419.1_ASFV/POL/2015         ................................................................................ [104000]
LS478113.1_Estonia_2014          ................................................................................ [104000]
KP843857.1_Odintsovo_02/14       ................................................................................ [104000]
FN557520.1_strain_E75            ......................a......................................................... [104000]
AY261366.1_Warthog               ..............................................a................................. [104000]
AY261365.1_Warmbaths             ..............................................a................................. [104000]
AY261364.1_Tengani_62            ..............................................a................................. [104000]
AY261363.1_Pretoriuskop/96/4     ..............................................a................................. [104000]
AY261362.1_Mkuzi_1979            ..............................................a................................. [104000]
AY261361.1_Malawi_Lil-20/1_1983  ....................a...............ct........a..c....................t......... [104000]
AY261360.1_Kenya_1950            ..........c..........................t........a..c.............................. [104000]
AM712240.1_OURT_88/3             ......................a......................................................... [104000]
AM712239.1_Benin_97/1            ......................a......................................................... [104000]
KM111295.1_Ken06.Bus             ..........c..........................t........a..c.............................. [104000]
KM111294.1_Ken05/Tk1             ..........c..........................t........a..c.............................. [104000]
KM262844.1_strain_L60            ......................a......................................................... [104000]
KM102979.1_26544/OG10_Italy      ......................a......................................................... [104000]
KM262845.1_strain_NHV            ......................a......................................................... [104000]
KX354450.1_47/Ss/2008            ......................a......................................................... [104000]
KP055815.1_strain_BA71           ......................a......................................................... [104000]
MH025920.1_strain_R35            ..........c..........................t........a..c.............................. [104000]
MH025919.1_strain_N10            ..........c..........................t........a..c.............................. [104000]
MH025918.1_strain_R25            ..........c..........................t........a..c.............................. [104000]
MH025917.1_strain_R7             ..........c..........................t........a..c.............................. [104000]
MH025916.1_strain_R8             ..........c..........................t........a..c.............................. [104000]
```

```
FR682468.2_Georgia_2007/1        acattaaaacctgaagtggttcagggcaaatatgggttccatgtgctcattcctggtctcaagctggcggcttctaccaa [104080]
ASF_IND_20_CAD543                ................................................................................ [104080]
LR899193.1_Germany_2020/1        ................................................................................ [104080]
MN393477.1_Wuhan_2019            ................................................................................ [104080]
MW396979.1_Timor-Leste/2019/1    ................................................................................ [104080]
MK940252.1_CN/2019/InnerMongolia ................................................................................ [104080]
MK128995.1_China/2018/AnhuiXCGQ  ................................................................................ [104080]
MH681419.1_ASFV/POL/2015         ................................................................................ [104080]
LS478113.1_Estonia_2014          ................................................................................ [104080]
KP843857.1_Odintsovo_02/14       ................................................................................ [104080]
FN557520.1_strain_E75            ....................c........................................................... [104080]
AY261366.1_Warthog               ....................c........................................................... [104080]
AY261365.1_Warmbaths             ....................c........................................................... [104080]
AY261364.1_Tengani_62            .......................................................................c........ [104080]
AY261363.1_Pretoriuskop/96/4     ....................c..................................................c........ [104080]
AY261362.1_Mkuzi_1979            ....................c........................................................... [104080]
AY261361.1_Malawi_Lil-20/1_1983  ....................................................................a........... [104080]
AY261360.1_Kenya_1950            ................c...c..........................................a....a..c........ [104080]
AM712240.1_OURT_88/3             ....................c........................................................... [104080]
AM712239.1_Benin_97/1            ....................c........................................................... [104080]
KM111295.1_Ken06.Bus             ................c...c..........................................a....a..c........ [104080]
KM111294.1_Ken05/Tk1             ................c...c..........................................a....a..c........ [104080]
KM262844.1_strain_L60            ....................c........................................................... [104080]
KM102979.1_26544/OG10_Italy      ....................c........................................................... [104080]
KM262845.1_strain_NHV            ....................c........................................................... [104080]
KX354450.1_47/Ss/2008            ....................c........................................................... [104080]
KP055815.1_strain_BA71           ....................c........................................................... [104080]
MH025920.1_strain_R35            ................c...c..........................................a....a..c........ [104080]
MH025919.1_strain_N10            ................c...c..........................................a....a..c........ [104080]
MH025918.1_strain_R25            ................c...c..........................................a....a..c........ [104080]
MH025917.1_strain_R7             ................c...c..........................................a....a..c........ [104080]
MH025916.1_strain_R8             ................c...c..........................................a....a..c........ [104080]

FR682468.2_Georgia_2007/1        aaaaagcattataggatccctacagcacgatgccaccgtacaaaaaattctacacgagcagggcgttacaaatcctgagt [104160]
ASF_IND_20_CAD543                ................................................................................ [104160]
LR899193.1_Germany_2020/1        ................................................................................ [104160]
MN393477.1_Wuhan_2019            ................................................................................ [104160]
MW396979.1_Timor-Leste/2019/1    ................................................................................ [104160]
MK940252.1_CN/2019/InnerMongolia ................................................................................ [104160]
MK128995.1_China/2018/AnhuiXCGQ  ................................................................................ [104160]
MH681419.1_ASFV/POL/2015         ................................................................................ [104160]
LS478113.1_Estonia_2014          ................................................................................ [104160]
KP843857.1_Odintsovo_02/14       ................................................................................ [104160]
```

FN557520.1_strain_E75 .........c...............................................a.........g............ [104160]
AY261366.1_Warthog ...................................................................g............ [104160]
AY261365.1_Warmbaths ...................................................................g............ [104160]
AY261364.1_Tengani_62 ...................................................................g............ [104160]
AY261363.1_Pretoriuskop/96/4 ...........................t.......................................g............ [104160]
AY261362.1_Mkuzi_1979 .........................................................a.........g............ [104160]
AY261361.1_Malawi_Lil-20/1_1983 ......t.......c......g......................................a...........c..c.... [104160]
AY261360.1_Kenya_1950 ..............c....t.g.................g...........................g....c....... [104160]
AM712240.1_OURT_88/3 .........c...............................................a.........g............ [104160]
AM712239.1_Benin_97/1 .........c...............................................a.........g............ [104160]
KM111295.1_Ken06.Bus ..............c....t...............................................g....c....... [104160]
KM111294.1_Ken05/Tk1 ..............c.........a..............g................................c....... [104160]
KM262844.1_strain_L60 .........c...............................................a.........g............ [104160]
KM102979.1_26544/OG10_Italy .........c...............................................a.........g............ [104160]
KM262845.1_strain_NHV .........c...............................................a.........g............ [104160]
KX354450.1_47/Ss/2008 .........c...............................................a.........g............ [104160]
KP055815.1_strain_BA71 .........c...............................................a.........g............ [104160]
MH025920.1_strain_R35 ..............c....t...............................................g....c....... [104160]
MH025919.1_strain_N10 ..............c....t...............................................g....c....... [104160]
MH025918.1_strain_R25 ..............c....t...............................................g....c....... [104160]
MH025917.1_strain_R7 ..............c....t...............................................g....c....... [104160]
MH025916.1_strain_R8 ..............c....t...............................................g....c....... [104160]

FR682468.2_Georgia_2007/1 cctgtctggacccccactccgcctccgttccctcgctcctctacggctcctccaaactaaaccacaagccctaccaactg [104240]
ASF_IND_20_CAD543 .........................y...................................................... [104240]
LR899193.1_Germany_2020/1 ................................................................................ [104240]
MN393477.1_Wuhan_2019 ................................................................................ [104240]
MW396979.1_Timor-Leste/2019/1 ................................................................................ [104240]
MK940252.1_CN/2019/InnerMongolia ................................................................................ [104240]
MK128995.1_China/2018/AnhuiXCGQ ................................................................................ [104240]
MH681419.1_ASFV/POL/2015 ................................................................................ [104240]
LS478113.1_Estonia_2014 ................................................................................ [104240]
KP843857.1_Odintsovo_02/14 ................................................................................ [104240]
FN557520.1_strain_E75 .........................t...................................................... [104240]
AY261366.1_Warthog ..........t........g............................................t..............a [104240]
AY261365.1_Warmbaths ..........t........g...........................................................a [104240]
AY261364.1_Tengani_62 .........................t.........................................a...........a [104240]
AY261363.1_Pretoriuskop/96/4 ..........t..t.....g............................................t..............a [104240]
AY261362.1_Mkuzi_1979 .........................t...................................................... [104240]
AY261361.1_Malawi_Lil-20/1_1983 ........................................................t......................a [104240]
AY261360.1_Kenya_1950 ........................................................t.g....................a [104240]
AM712240.1_OURT_88/3 .........................t...................................................... [104240]
AM712239.1_Benin_97/1 .........................t...................................................... [104240]
KM111295.1_Ken06.Bus ........................................................t.g.......g............a [104240]

```
KM111294.1_Ken05/Tk1           ........................................................t.g....................a [104240]
KM262844.1_strain_L60          .........................t...................................................... [104240]
KM102979.1_26544/OG10_Italy    .........................t...................................................... [104240]
KM262845.1_strain_NHV          .........................t...................................................... [104240]
KX354450.1_47/Ss/2008          .........................t...................................................... [104240]
KP055815.1_strain_BA71         .........................t...................................................... [104240]
MH025920.1_strain_R35          ........................................................t.g.......g............a [104240]
MH025919.1_strain_N10          ........................................................t.g.......g............a [104240]
MH025918.1_strain_R25          ........................................................t.g.......g............a [104240]
MH025917.1_strain_R7           ........................................................t.g.......g............a [104240]
MH025916.1_strain_R8           ........................................................t.g.......g............a [104240]

FR682468.2_Georgia_2007/1      aaaaccggctttgagttagtctttgatagctctgatcccgactacattcccattcatcaaataaaaaatttagaatctta [104320]
ASF_IND_20_CAD543              .....................................................................w.......... [104320]
LR899193.1_Germany_2020/1      ................................................................................ [104320]
MN393477.1_Wuhan_2019          ................................................................................ [104320]
MW396979.1_Timor-Leste/2019/1  ................................................................................ [104320]
MK940252.1_CN/2019/InnerMongolia ................................................................................ [104320]
MK128995.1_China/2018/AnhuiXCGQ ................................................................................ [104320]
MH681419.1_ASFV/POL/2015       ................................................................................ [104320]
LS478113.1_Estonia_2014        ................................................................................ [104320]
KP843857.1_Odintsovo_02/14     ................................................................................ [104320]
FN557520.1_strain_E75          .....................................................................a.......... [104320]
AY261366.1_Warthog             .....................................................................a.......... [104320]
AY261365.1_Warmbaths           .....................................................................a.......... [104320]
AY261364.1_Tengani_62          ....t......................................................t.........a.......... [104320]
AY261363.1_Pretoriuskop/96/4   .....................................................................a.......... [104320]
AY261362.1_Mkuzi_1979          .....................................................................a.......... [104320]
AY261361.1_Malawi_Lil-20/1_1983 ......................................t..............................a.....g.... [104320]
AY261360.1_Kenya_1950          ......................................t........c....................ca.c........ [104320]
AM712240.1_OURT_88/3           .....................................................................a.......... [104320]
AM712239.1_Benin_97/1          .....................................................................a.......... [104320]
KM111295.1_Ken06.Bus           ......................................t........c.....................a.c........ [104320]
KM111294.1_Ken05/Tk1           ......................................t........c....................ca.c........ [104320]
KM262844.1_strain_L60          .....................................................................a.......... [104320]
KM102979.1_26544/OG10_Italy    .....................................................................a.......... [104320]
KM262845.1_strain_NHV          .....................................................................a.......... [104320]
KX354450.1_47/Ss/2008          .....................................................................a.......... [104320]
KP055815.1_strain_BA71         .....................................................................a.......... [104320]
MH025920.1_strain_R35          ......................................t........c.....................a.c........ [104320]
MH025919.1_strain_N10          ......................................t........c.....................a.c........ [104320]
MH025918.1_strain_R25          ......................................t........c.....................a.c........ [104320]
MH025917.1_strain_R7           ......................................t........c.....................a.c........ [104320]
MH025916.1_strain_R8           ......................................t........c.....................a.c........ [104320]
```

```
FR682468.2_Georgia_2007/1        taatttagtttctgagttgagccttacgaatgaacagggaagccttgtaagacctgtctattgcgcggcagacattgccg [104400]
ASF_IND_20_CAD543                ................................................................................ [104400]
LR899193.1_Germany_2020/1        ................................................................................ [104400]
MN393477.1_Wuhan_2019            ................................................................................ [104400]
MW396979.1_Timor-Leste/2019/1    ................................................................................ [104400]
MK940252.1_CN/2019/InnerMongolia ................................................................................ [104400]
MK128995.1_China/2018/AnhuiXCGQ  ................................................................................ [104400]
MH681419.1_ASFV/POL/2015         ................................................................................ [104400]
LS478113.1_Estonia_2014          ................................................................................ [104400]
KP843857.1_Odintsovo_02/14       ................................................................................ [104400]
FN557520.1_strain_E75            ....................................a..............g............................ [104400]
AY261366.1_Warthog               ................................................................................ [104400]
AY261365.1_Warmbaths             ................................................................................ [104400]
AY261364.1_Tengani_62            ................................................................................ [104400]
AY261363.1_Pretoriuskop/96/4     .........................................................t...................... [104400]
AY261362.1_Mkuzi_1979            ....................................a..............g............................ [104400]
AY261361.1_Malawi_Lil-20/1_1983  ..................................................a.........c....a.............. [104400]
AY261360.1_Kenya_1950            ............................................................c................... [104400]
AM712240.1_OURT_88/3             ....................................a..............g............................ [104400]
AM712239.1_Benin_97/1            ....................................a..............g............................ [104400]
KM111295.1_Ken06.Bus             ............c...............................................c................... [104400]
KM111294.1_Ken05/Tk1             ............................................................c................... [104400]
KM262844.1_strain_L60            ....................................a..............g............................ [104400]
KM102979.1_26544/OG10_Italy      ....................................a..............g............................ [104400]
KM262845.1_strain_NHV            ....................................a..............g............................ [104400]
KX354450.1_47/Ss/2008            ....................................a..............g............................ [104400]
KP055815.1_strain_BA71           ....................................a..............g............................ [104400]
MH025920.1_strain_R35            ............c...............................................c................... [104400]
MH025919.1_strain_N10            ............c...............................................c................... [104400]
MH025918.1_strain_R25            ............c...............................................c................... [104400]
MH025917.1_strain_R7             ............c...............................................c................... [104400]
MH025916.1_strain_R8             ............c...............................................c................... [104400]

FR682468.2_Georgia_2007/1        ctgagaaggaggaagagatcccgaccgaggatcactcgctctccatattaatgctacatgatcccgaagcccggtattta [104480]
ASF_IND_20_CAD543                .............r.................................................................. [104480]
LR899193.1_Germany_2020/1        ................................................................................ [104480]
MN393477.1_Wuhan_2019            ................................................................................ [104480]
MW396979.1_Timor-Leste/2019/1    ................................................................................ [104480]
MK940252.1_CN/2019/InnerMongolia ................................................................................ [104480]
MK128995.1_China/2018/AnhuiXCGQ  ................................................................................ [104480]
MH681419.1_ASFV/POL/2015         ................................................................................ [104480]
LS478113.1_Estonia_2014          ................................................................................ [104480]
KP843857.1_Odintsovo_02/14       ................................................................................ [104480]
```

```
FN557520.1_strain_E75           .............g..............c..c.....a................................t......... [104480]
AY261366.1_Warthog              .............g........................................................t......... [104480]
AY261365.1_Warmbaths            .............g.................................................................. [104480]
AY261364.1_Tengani_62           .............g.................................................................. [104480]
AY261363.1_Pretoriuskop/96/4    .............g.................................................................. [104480]
AY261362.1_Mkuzi_1979           .............g..............c..c.....a................................t......... [104480]
AY261361.1_Malawi_Lil-20/1_1983 .......................gt...c........a.......................................... [104480]
AY261360.1_Kenya_1950           .......................g....t........a..t..............t.....c.................. [104480]
AM712240.1_OURT_88/3            .............g..............c..c.....a................................t......... [104480]
AM712239.1_Benin_97/1           .............g........a.....c..c.....a................................t......... [104480]
KM111295.1_Ken06.Bus            .......................g....t........a..t....................................... [104480]
KM111294.1_Ken05/Tk1            .......................g....t........a..t..............t.....c.................. [104480]
KM262844.1_strain_L60           .............g..............c..c.....a................................t......... [104480]
KM102979.1_26544/OG10_Italy     .............g..............c..c.....a................................t......... [104480]
KM262845.1_strain_NHV           .............g..............c..c.....a................................t......... [104480]
KX354450.1_47/Ss/2008           .............g..............c..c.....a................................t......... [104480]
KP055815.1_strain_BA71          .............g..............c..c.....a................................t......... [104480]
MH025920.1_strain_R35           .......................g....t........a..t....................................... [104480]
MH025919.1_strain_N10           .......................g....t........a..t....................................... [104480]
MH025918.1_strain_R25           .......................g....t........a..t....................................... [104480]
MH025917.1_strain_R7            .......................g....t........a..t....................................... [104480]
MH025916.1_strain_R8            .......................g....t........a..t....................................... [104480]

FR682468.2_Georgia_2007/1       cataaaattttaaatctgcttcctccggagtattatgtagagtaccccctatggagcaacgtcgtattcgctttggccaa [104560]
ASF_IND_20_CAD543               ................................................................................ [104560]
LR899193.1_Germany_2020/1       ................................................................................ [104560]
MN393477.1_Wuhan_2019           ................................................................................ [104560]
MW396979.1_Timor-Leste/2019/1   ................................................................................ [104560]
MK940252.1_CN/2019/InnerMongolia ................................................................................ [104560]
MK128995.1_China/2018/AnhuiXCGQ ................................................................................ [104560]
MH681419.1_ASFV/POL/2015        ................................................................................ [104560]
LS478113.1_Estonia_2014         ................................................................................ [104560]
KP843857.1_Odintsovo_02/14      ................................................................................ [104560]
FN557520.1_strain_E75           ..............c...........a...........g................................c........ [104560]
AY261366.1_Warthog              ..............c.............................t..........................cc....... [104560]
AY261365.1_Warmbaths            ..............c...........a...........g................................cc....... [104560]
AY261364.1_Tengani_62           ..............c........................................................cc....... [104560]
AY261363.1_Pretoriuskop/96/4    ..............c.......................g.............................t..cc....... [104560]
AY261362.1_Mkuzi_1979           ..............c...........a...........g................................c........ [104560]
AY261361.1_Malawi_Lil-20/1_1983 ........c.....c.............................t..............t..t........cc.a..... [104560]
AY261360.1_Kenya_1950           ..............c........c....................t...........................c....... [104560]
AM712240.1_OURT_88/3            ..............c...........a...........g................................c........ [104560]
AM712239.1_Benin_97/1           ..............c...........a...........g................................c........ [104560]
KM111295.1_Ken06.Bus            ..............c........c....................t...........................c....... [104560]
```

```
KM111294.1_Ken05/Tk1            ..............c........c....................t...........................c....... [104560]
KM262844.1_strain_L60           ..............c...........a...........g................................c........ [104560]
KM102979.1_26544/OG10_Italy     ..............c...........a...........g................................c........ [104560]
KM262845.1_strain_NHV           ..............c...........a...........g................................c........ [104560]
KX354450.1_47/Ss/2008           ..............c...........a...........g................................c........ [104560]
KP055815.1_strain_BA71          ..............c...........a...........g................................c........ [104560]
MH025920.1_strain_R35           ..............c........c....................t...........................c....... [104560]
MH025919.1_strain_N10           ..............c........c....................t...........................c....... [104560]
MH025918.1_strain_R25           ..............c........c....................t...........................c....... [104560]
MH025917.1_strain_R7            ..............c........c....................t...........................c....... [104560]
MH025916.1_strain_R8            ..............c........c....................t...........................c....... [104560]

FR682468.2_Georgia_2007/1       tacatccgctaactatcggcccctcgccgaatggttttcgcaaaaatgccctgaaaaatggaatacgggaggaaaagaga [104640]
ASF_IND_20_CAD543               ................................................y............................... [104640]
LR899193.1_Germany_2020/1       ................................................................................ [104640]
MN393477.1_Wuhan_2019           ................................................................................ [104640]
MW396979.1_Timor-Leste/2019/1   ................................................................................ [104640]
MK940252.1_CN/2019/InnerMongolia ................................................................................ [104640]
MK128995.1_China/2018/AnhuiXCGQ ................................................................................ [104640]
MH681419.1_ASFV/POL/2015        ................................................................................ [104640]
LS478113.1_Estonia_2014         ................................................................................ [104640]
KP843857.1_Odintsovo_02/14      ................................................................................ [104640]
FN557520.1_strain_E75           .........c.............................a........t............................... [104640]
AY261366.1_Warthog              ...............c.......................a........t............................... [104640]
AY261365.1_Warmbaths            .........c.....c................................t............................... [104640]
AY261364.1_Tengani_62           ................................................t............................... [104640]
AY261363.1_Pretoriuskop/96/4    ................................................t............................... [104640]
AY261362.1_Mkuzi_1979           .........c.............................a........t............................... [104640]
AY261361.1_Malawi_Lil-20/1_1983 .........c...........t..t.......................t............................... [104640]
AY261360.1_Kenya_1950           .........c........a.....t..............a........t............................... [104640]
AM712240.1_OURT_88/3            .........c.............................a........t............................... [104640]
AM712239.1_Benin_97/1           .........c.............................a........t............................... [104640]
KM111295.1_Ken06.Bus            .........c........a.....t.......................t............................... [104640]
KM111294.1_Ken05/Tk1            .........c........a.....t.......................t............................... [104640]
KM262844.1_strain_L60           .........c.............................a........t............................... [104640]
KM102979.1_26544/OG10_Italy     .........c.............................a........t............................... [104640]
KM262845.1_strain_NHV           .........c.............................a........t............................... [104640]
KX354450.1_47/Ss/2008           .........c.............................a........t............................... [104640]
KP055815.1_strain_BA71          .........c.............................a........t............................... [104640]
MH025920.1_strain_R35           .........c........a.....t.......................t............................... [104640]
MH025919.1_strain_N10           .........c........a.....t.......................t............................... [104640]
MH025918.1_strain_R25           .........c........a.....t.......................t............................... [104640]
MH025917.1_strain_R7            .........c........a.....t.......................t............................... [104640]
MH025916.1_strain_R8            .........c........a.....t.......................t............................... [104640]
```

```
FR682468.2_Georgia_2007/1        aactagaaaaactttggaatgatgcctcgcaccacactgaaaagaaaatcaccaagcggtccattatgtactgggcccac [104720]
ASF_IND_20_CAD543                ................................................................................ [104720]
LR899193.1_Germany_2020/1        ................................................................................ [104720]
MN393477.1_Wuhan_2019            ................................................................................ [104720]
MW396979.1_Timor-Leste/2019/1    ................................................................................ [104720]
MK940252.1_CN/2019/InnerMongolia ................................................................................ [104720]
MK128995.1_China/2018/AnhuiXCGQ  ................................................................................ [104720]
MH681419.1_ASFV/POL/2015         ................................................................................ [104720]
LS478113.1_Estonia_2014          ................................................................................ [104720]
KP843857.1_Odintsovo_02/14       ................................................................................ [104720]
FN557520.1_strain_E75            ..............................g.......................................t......... [104720]
AY261366.1_Warthog               ........c.....................g..................t.............................. [104720]
AY261365.1_Warmbaths             ........c.....................g................................................. [104720]
AY261364.1_Tengani_62            ........c.....................g...t.......................a..................... [104720]
AY261363.1_Pretoriuskop/96/4     ........c.....................g.....................t........................... [104720]
AY261362.1_Mkuzi_1979            ..............................g................................................. [104720]
AY261361.1_Malawi_Lil-20/1_1983  ..............................g............a.................................... [104720]
AY261360.1_Kenya_1950            ............................a..............a.................................... [104720]
AM712240.1_OURT_88/3             ..............................g.......................................t......... [104720]
AM712239.1_Benin_97/1            ..............................g.......................................t......... [104720]
KM111295.1_Ken06.Bus             ............................a.g............a.................................... [104720]
KM111294.1_Ken05/Tk1             ............................a.g............a...........a........................ [104720]
KM262844.1_strain_L60            ..............................g.......................................t......... [104720]
KM102979.1_26544/OG10_Italy      ..............................g.......................................t......... [104720]
KM262845.1_strain_NHV            ..............................g.......................................t......... [104720]
KX354450.1_47/Ss/2008            ..............................g.......................................t......... [104720]
KP055815.1_strain_BA71           ..............................g.......................................t......... [104720]
MH025920.1_strain_R35            ............................a.g............a.................................... [104720]
MH025919.1_strain_N10            ............................a.g............a.................................... [104720]
MH025918.1_strain_R25            ............................a.g............a.................................... [104720]
MH025917.1_strain_R7             ............................a.g............a.................................... [104720]
MH025916.1_strain_R8             ............................a.g............a.................................... [104720]

FR682468.2_Georgia_2007/1        aaacatgccccccagcaatacaaagaaattgtagaacaaggctacttttccattctcgctgaatatgtgtatagctataa [104800]
ASF_IND_20_CAD543                ................................................................................ [104800]
LR899193.1_Germany_2020/1        ................................................................................ [104800]
MN393477.1_Wuhan_2019            ................................................................................ [104800]
MW396979.1_Timor-Leste/2019/1    ................................................................................ [104800]
MK940252.1_CN/2019/InnerMongolia ................................................................................ [104800]
MK128995.1_China/2018/AnhuiXCGQ  ................................................................................ [104800]
MH681419.1_ASFV/POL/2015         ................................................................................ [104800]
LS478113.1_Estonia_2014          ................................................................................ [104800]
KP843857.1_Odintsovo_02/14       ................................................................................ [104800]
```

| | | |
|---|---|---|
| FN557520.1_strain_E75 | ..g.....t................................t...................................... | [104800] |
| AY261366.1_Warthog | ..g..............g.....g...........g............................................ | [104800] |
| AY261365.1_Warmbaths | ..g..............g.....g...........g............................................ | [104800] |
| AY261364.1_Tengani_62 | ..g..............g.....g...........g.............................c.............. | [104800] |
| AY261363.1_Pretoriuskop/96/4 | ..g..............g.....g...........g............................................ | [104800] |
| AY261362.1_Mkuzi_1979 | ..g..............g.....g...........g............................................ | [104800] |
| AY261361.1_Malawi_Lil-20/1_1983 | ..g..............g.....g...........g............................................ | [104800] |
| AY261360.1_Kenya_1950 | ..g..............g.....g........g..g.........................................c.. | [104800] |
| AM712240.1_OURT_88/3 | ..g.....t................................t...................................... | [104800] |
| AM712239.1_Benin_97/1 | ..g.....t................................t...................................... | [104800] |
| KM111295.1_Ken06.Bus | ..g..............g.....g........g..g....................t.................t..c.. | [104800] |
| KM111294.1_Ken05/Tk1 | ..g..............g..............g..g....................t....................c.. | [104800] |
| KM262844.1_strain_L60 | ..g.....t................................t...................................... | [104800] |
| KM102979.1_26544/OG10_Italy | ..g.....t................................t...................................... | [104800] |
| KM262845.1_strain_NHV | ..g.....t................................t...................................... | [104800] |
| KX354450.1_47/Ss/2008 | ..g.....t................................t...................................... | [104800] |
| KP055815.1_strain_BA71 | ..g.....t................................t...................................... | [104800] |
| MH025920.1_strain_R35 | ..g..............g.....g........g..g....................t.................t..c.. | [104800] |
| MH025919.1_strain_N10 | ..g..............g.....g........g..g....................t.................t..c.. | [104800] |
| MH025918.1_strain_R25 | ..g..............g.....g........g..g....................t.................t..c.. | [104800] |
| MH025917.1_strain_R7 | ..g..............g.....g........g..g....................t.................t..c.. | [104800] |
| MH025916.1_strain_R8 | ..g..............g.....g........g..g....................t.................t..c.. | [104800] |

| | | |
|---|---|---|
| FR682468.2_Georgia_2007/1 | cggcatgcttgagcactacatgatcgccaaagtcatctatgctatgatgggcaacaagtttgtagtggacgtggattcaa | [104880] |
| ASF_IND_20_CAD543 | ................................................................................ | [104880] |
| LR899193.1_Germany_2020/1 | ................................................................................ | [104880] |
| MN393477.1_Wuhan_2019 | ................................................................................ | [104880] |
| MW396979.1_Timor-Leste/2019/1 | ................................................................................ | [104880] |
| MK940252.1_CN/2019/InnerMongolia | ................................................................................ | [104880] |
| MK128995.1_China/2018/AnhuiXCGQ | ................................................................................ | [104880] |
| MH681419.1_ASFV/POL/2015 | ................................................................................ | [104880] |
| LS478113.1_Estonia_2014 | ................................................................................ | [104880] |
| KP843857.1_Odintsovo_02/14 | ................................................................................ | [104880] |
| FN557520.1_strain_E75 | ..........................................c....................g...........c.... | [104880] |
| AY261366.1_Warthog | ..........................................c....................g...........c.... | [104880] |
| AY261365.1_Warmbaths | ..........................................c....................g...........c.... | [104880] |
| AY261364.1_Tengani_62 | .....c....................................c....................g...........c.... | [104880] |
| AY261363.1_Pretoriuskop/96/4 | ......a.................t.................c....................g..a........c.... | [104880] |
| AY261362.1_Mkuzi_1979 | ..........................................c....................g...........c.... | [104880] |
| AY261361.1_Malawi_Lil-20/1_1983 | .....c..............................t.................t........g...........c.... | [104880] |
| AY261360.1_Kenya_1950 | .....c....................................c....................g...........c.... | [104880] |
| AM712240.1_OURT_88/3 | ..........................................c....................g...........c.... | [104880] |
| AM712239.1_Benin_97/1 | ..........................................c....................g...........c.... | [104880] |
| KM111295.1_Ken06.Bus | .....c....................................c....................g...........c.... | [104880] |

| | | |
|---|---|---|
| KM111294.1_Ken05/Tk1 | .....c....................................c....................g...........c.... | [104880] |
| KM262844.1_strain_L60 | ..........................................c....................g...........c.... | [104880] |
| KM102979.1_26544/OG10_Italy | ..........................................c....................g...........c.... | [104880] |
| KM262845.1_strain_NHV | ..........................................c....................g...........c.... | [104880] |
| KX354450.1_47/Ss/2008 | ..........................................c....................g...........c.... | [104880] |
| KP055815.1_strain_BA71 | ..........................................c....................g...........c.... | [104880] |
| MH025920.1_strain_R35 | .....c....................................c....................g...........c.... | [104880] |
| MH025919.1_strain_N10 | .....c....................................c....................g...........c.... | [104880] |
| MH025918.1_strain_R25 | .....c....................................c....................g...........c.... | [104880] |
| MH025917.1_strain_R7 | .....c....................................c....................g...........c.... | [104880] |
| MH025916.1_strain_R8 | .....c....................................c....................g...........c.... | [104880] |

| | | |
|---|---|---|
| FR682468.2_Georgia_2007/1 | acgggaagtacgtttggttcgaatttgtgctaccgggccagccaatgaatcagggagaaatatggaagtggcgcaaggag | [104960] |
| ASF_IND_20_CAD543 | ................................................................................ | [104960] |
| LR899193.1_Germany_2020/1 | ................................................................................ | [104960] |
| MN393477.1_Wuhan_2019 | ................................................................................ | [104960] |
| MW396979.1_Timor-Leste/2019/1 | ................................................................................ | [104960] |
| MK940252.1_CN/2019/InnerMongolia | ................................................................................ | [104960] |
| MK128995.1_China/2018/AnhuiXCGQ | ................................................................................ | [104960] |
| MH681419.1_ASFV/POL/2015 | ................................................................................ | [104960] |
| LS478113.1_Estonia_2014 | ................................................................................ | [104960] |
| KP843857.1_Odintsovo_02/14 | ................................................................................ | [104960] |
| FN557520.1_strain_E75 | ............................................................................a... | [104960] |
| AY261366.1_Warthog | ................................................................................ | [104960] |
| AY261365.1_Warmbaths | ................................................................................ | [104960] |
| AY261364.1_Tengani_62 | ................................................................................ | [104960] |
| AY261363.1_Pretoriuskop/96/4 | ................................................................................ | [104960] |
| AY261362.1_Mkuzi_1979 | ................................................................................ | [104960] |
| AY261361.1_Malawi_Lil-20/1_1983 | ...........................................g.................................... | [104960] |
| AY261360.1_Kenya_1950 | ...............................g...........g.................................... | [104960] |
| AM712240.1_OURT_88/3 | ............................................................................a... | [104960] |
| AM712239.1_Benin_97/1 | ............................................................................a... | [104960] |
| KM111295.1_Ken06.Bus | .t.............................g................................................ | [104960] |
| KM111294.1_Ken05/Tk1 | ...............................g................................................ | [104960] |
| KM262844.1_strain_L60 | ............................................................................a... | [104960] |
| KM102979.1_26544/OG10_Italy | ............................................................................a... | [104960] |
| KM262845.1_strain_NHV | ............................................................................a... | [104960] |
| KX354450.1_47/Ss/2008 | ............................................................................a... | [104960] |
| KP055815.1_strain_BA71 | ............................................................................a... | [104960] |
| MH025920.1_strain_R35 | .t.............................g................................................ | [104960] |
| MH025919.1_strain_N10 | .t.............................g................................................ | [104960] |
| MH025918.1_strain_R25 | .t.............................g................................................ | [104960] |
| MH025917.1_strain_R7 | .t.............................g................................................ | [104960] |
| MH025916.1_strain_R8 | .t.............................g................................................ | [104960] |

```
FR682468.2_Georgia_2007/1        gtaaacccggatgagctgcacatctatatttccgaaaacttttcaagggtgatggaccgaatcacggagcacatcaaata [105040]
ASF_IND_20_CAD543                ................................................................................ [105040]
LR899193.1_Germany_2020/1        ................................................................................ [105040]
MN393477.1_Wuhan_2019            ................................................................................ [105040]
MW396979.1_Timor-Leste/2019/1    ................................................................................ [105040]
MK940252.1_CN/2019/InnerMongolia ................................................................................ [105040]
MK128995.1_China/2018/AnhuiXCGQ  ................................................................................ [105040]
MH681419.1_ASFV/POL/2015         ................................................................................ [105040]
LS478113.1_Estonia_2014          ................................................................................ [105040]
KP843857.1_Odintsovo_02/14       ................................................................................ [105040]
FN557520.1_strain_E75            ................................................................................ [105040]
AY261366.1_Warthog               ................................................................................ [105040]
AY261365.1_Warmbaths             ................................................................................ [105040]
AY261364.1_Tengani_62            ........................................................t....................... [105040]
AY261363.1_Pretoriuskop/96/4     ................................................................................ [105040]
AY261362.1_Mkuzi_1979            ................................................................................ [105040]
AY261361.1_Malawi_Lil-20/1_1983  ................................................................................ [105040]
AY261360.1_Kenya_1950            ................................................................................ [105040]
AM712240.1_OURT_88/3             ................................................................................ [105040]
AM712239.1_Benin_97/1            ................................................................................ [105040]
KM111295.1_Ken06.Bus             ................................................................................ [105040]
KM111294.1_Ken05/Tk1             ................................................................................ [105040]
KM262844.1_strain_L60            ................................................................................ [105040]
KM102979.1_26544/OG10_Italy      ................................................................................ [105040]
KM262845.1_strain_NHV            ................................................................................ [105040]
KX354450.1_47/Ss/2008            ................................................................................ [105040]
KP055815.1_strain_BA71           ................................................................................ [105040]
MH025920.1_strain_R35            ................................................................................ [105040]
MH025919.1_strain_N10            ................................................................................ [105040]
MH025918.1_strain_R25            ................................................................................ [105040]
MH025917.1_strain_R7             ................................................................................ [105040]
MH025916.1_strain_R8             ................................................................................ [105040]

FR682468.2_Georgia_2007/1        ccacctcagtcaaccccatgaaagcaatattttaaattattataaaaaactattaaaagcctttgaacgctctaaaagta [105120]
ASF_IND_20_CAD543                .......................s........................................................ [105120]
LR899193.1_Germany_2020/1        ................................................................................ [105120]
MN393477.1_Wuhan_2019            ................................................................................ [105120]
MW396979.1_Timor-Leste/2019/1    ................................................................................ [105120]
MK940252.1_CN/2019/InnerMongolia ................................................................................ [105120]
MK128995.1_China/2018/AnhuiXCGQ  ................................................................................ [105120]
MH681419.1_ASFV/POL/2015         ................................................................................ [105120]
LS478113.1_Estonia_2014          ................................................................................ [105120]
KP843857.1_Odintsovo_02/14       ................................................................................ [105120]
```

```
FN557520.1_strain_E75             ............g..........c............................c........................... [105120]
AY261366.1_Warthog                ............g..........c............................c........................... [105120]
AY261365.1_Warmbaths              ............g..........c............................c........................... [105120]
AY261364.1_Tengani_62             ......t.....g..........c............................c........................... [105120]
AY261363.1_Pretoriuskop/96/4      ............g..........c...c........................c........................... [105120]
AY261362.1_Mkuzi_1979             ............g..........c............................c........................... [105120]
AY261361.1_Malawi_Lil-20/1_1983   ............g..........c............................c........................... [105120]
AY261360.1_Kenya_1950             ............g..........c............................c........................... [105120]
AM712240.1_OURT_88/3              ............g..........c............................c........................... [105120]
AM712239.1_Benin_97/1             ............g..........c............................c........................... [105120]
KM111295.1_Ken06.Bus              ............g..........c............................c........................... [105120]
KM111294.1_Ken05/Tk1              ............g..........c............................c........................... [105120]
KM262844.1_strain_L60             ............g..........c............................c........................... [105120]
KM102979.1_26544/OG10_Italy       ............g..........c............................c........................... [105120]
KM262845.1_strain_NHV             ............g..........c............................c........................... [105120]
KX354450.1_47/Ss/2008             ............g..........c............................c........................... [105120]
KP055815.1_strain_BA71            ............g..........c............................c........................... [105120]
MH025920.1_strain_R35             ............g..........c............................c........................... [105120]
MH025919.1_strain_N10             ............g..........c............................c........................... [105120]
MH025918.1_strain_R25             ............g..........c............................c........................... [105120]
MH025917.1_strain_R7              ............g..........c............................c........................... [105120]
MH025916.1_strain_R8              ............g..........c............................c........................... [105120]

FR682468.2_Georgia_2007/1         aaatctttaatgacagctttaaaaagggagttatcaggcaagctgagtttttatttcgccaaagaagctttattcaaact [105200]
ASF_IND_20_CAD543                 ................y............................................................... [105200]
LR899193.1_Germany_2020/1         ................................................................................ [105200]
MN393477.1_Wuhan_2019             ................................................................................ [105200]
MW396979.1_Timor-Leste/2019/1     ................................................................................ [105200]
MK940252.1_CN/2019/InnerMongolia  ................................................................................ [105200]
MK128995.1_China/2018/AnhuiXCGQ   ................................................................................ [105200]
MH681419.1_ASFV/POL/2015          ................................................................................ [105200]
LS478113.1_Estonia_2014           ................................................................................ [105200]
KP843857.1_Odintsovo_02/14        ................................................................................ [105200]
FN557520.1_strain_E75             ................t.................t........c.................................... [105200]
AY261366.1_Warthog                ...........................................c.......................t............ [105200]
AY261365.1_Warmbaths              ...........................................c.......................t............ [105200]
AY261364.1_Tengani_62             ................................................................................ [105200]
AY261363.1_Pretoriuskop/96/4      ...........................................c.................................... [105200]
AY261362.1_Mkuzi_1979             ...........................................c.................................... [105200]
AY261361.1_Malawi_Lil-20/1_1983   ...............................g...........c.......................t............ [105200]
AY261360.1_Kenya_1950             ................t..............g...........c.......................t............ [105200]
AM712240.1_OURT_88/3              ................t.................t........c.................................... [105200]
AM712239.1_Benin_97/1             ................t.................t........c.................................... [105200]
KM111295.1_Ken06.Bus              ........................................g..c.......................t............ [105200]
```

```
KM111294.1_Ken05/Tk1             ................t..........................c.......................t............ [105200]
KM262844.1_strain_L60            ................t.................t........c.................................... [105200]
KM102979.1_26544/OG10_Italy      ................t.................t........c.................................... [105200]
KM262845.1_strain_NHV            ................t.................t........c.................................... [105200]
KX354450.1_47/Ss/2008            ................t.................t........c.................................... [105200]
KP055815.1_strain_BA71           ................t.................t........c.................................... [105200]
MH025920.1_strain_R35            ........................................g..c.......................t............ [105200]
MH025919.1_strain_N10            ........................................g..c.......................t............ [105200]
MH025918.1_strain_R25            ........................................g..c.......................t............ [105200]
MH025917.1_strain_R7             ........................................g..c.......................t............ [105200]
MH025916.1_strain_R8             ........................................g..c.......................t............ [105200]

FR682468.2_Georgia_2007/1        ctggataccaatccccacctactgggggttggcaacggggttctctccattgagaccatcccggctaagctcattaatca [105280]
ASF_IND_20_CAD543                ................................................................................ [105280]
LR899193.1_Germany_2020/1        ................................................................................ [105280]
MN393477.1_Wuhan_2019            ................................................................................ [105280]
MW396979.1_Timor-Leste/2019/1    ................................................................................ [105280]
MK940252.1_CN/2019/InnerMongolia ................................................................................ [105280]
MK128995.1_China/2018/AnhuiXCGQ  ................................................................................ [105280]
MH681419.1_ASFV/POL/2015         ................................................................................ [105280]
LS478113.1_Estonia_2014          ................................................................................ [105280]
KP843857.1_Odintsovo_02/14       ................................................................................ [105280]
FN557520.1_strain_E75            .................................................................c.............. [105280]
AY261366.1_Warthog               .............................................................................c.. [105280]
AY261365.1_Warmbaths             .............................................................................c.. [105280]
AY261364.1_Tengani_62            ........................................................t........c...........c.. [105280]
AY261363.1_Pretoriuskop/96/4     ..........................a..................................................c.. [105280]
AY261362.1_Mkuzi_1979            ................................................................................ [105280]
AY261361.1_Malawi_Lil-20/1_1983  ....................................................................a........... [105280]
AY261360.1_Kenya_1950            ............................................t...........t....................... [105280]
AM712240.1_OURT_88/3             .................................................................c.............. [105280]
AM712239.1_Benin_97/1            .................................................................c.............. [105280]
KM111295.1_Ken06.Bus             ...............t.....t......................t................................... [105280]
KM111294.1_Ken05/Tk1             ............................................t...........t....................... [105280]
KM262844.1_strain_L60            .................................................................c.............. [105280]
KM102979.1_26544/OG10_Italy      .................................................................c.............. [105280]
KM262845.1_strain_NHV            .................................................................c.............. [105280]
KX354450.1_47/Ss/2008            .................................................................c.............. [105280]
KP055815.1_strain_BA71           .................................................................c.............. [105280]
MH025920.1_strain_R35            ...............t.....t......................t................................... [105280]
MH025919.1_strain_N10            ...............t.....t......................t................................... [105280]
MH025918.1_strain_R25            ...............t.....t......................t................................... [105280]
MH025917.1_strain_R7             ...............t.....t......................t................................... [105280]
MH025916.1_strain_R8             ...............t.....t......................t................................... [105280]
```

FR682468.2_Georgia_2007/1 ttttcacgagcatcccattcatcagtacacacacatatgttatgtgccctttaatcccgaaaacccctggacaaaactat [105360]
ASF_IND_20_CAD543 ................................................................................ [105360]
LR899193.1_Germany_2020/1 ................................................................................ [105360]
MN393477.1_Wuhan_2019 ................................................................................ [105360]
MW396979.1_Timor-Leste/2019/1 ................................................................................ [105360]
MK940252.1_CN/2019/InnerMongolia ................................................................................ [105360]
MK128995.1_China/2018/AnhuiXCGQ ................................................................................ [105360]
MH681419.1_ASFV/POL/2015 ................................................................................ [105360]
LS478113.1_Estonia_2014 ................................................................................ [105360]
KP843857.1_Odintsovo_02/14 ................................................................................ [105360]
FN557520.1_strain_E75 ................................................................................ [105360]
AY261366.1_Warthog .................................t...........c...........t...................... [105360]
AY261365.1_Warmbaths .................................t.............................t................ [105360]
AY261364.1_Tengani_62 ................................................................................ [105360]
AY261363.1_Pretoriuskop/96/4 ..........t......................t.............................t................ [105360]
AY261362.1_Mkuzi_1979 ..............................g.............................................t... [105360]
AY261361.1_Malawi_Lil-20/1_1983 ...........................t.....t..........a............g...................... [105360]
AY261360.1_Kenya_1950 ..................c..............t....................c......................... [105360]
AM712240.1_OURT_88/3 ................................g............................................... [105360]
AM712239.1_Benin_97/1 ................................................................................ [105360]
KM111295.1_Ken06.Bus ..................c..............t..........a.........c......................... [105360]
KM111294.1_Ken05/Tk1 ..................c..............t....................c......................... [105360]
KM262844.1_strain_L60 ................................................................................ [105360]
KM102979.1_26544/OG10_Italy ................................................................................ [105360]
KM262845.1_strain_NHV ................................................................................ [105360]
KX354450.1_47/Ss/2008 ................................................................................ [105360]
KP055815.1_strain_BA71 ................................................................................ [105360]
MH025920.1_strain_R35 ..................c..............t..........a.........c......................... [105360]
MH025919.1_strain_N10 ..................c..............t..........a.........c......................... [105360]
MH025918.1_strain_R25 ..................c..............t..........a.........c......................... [105360]
MH025917.1_strain_R7 ..................c..............t..........a.........c......................... [105360]
MH025916.1_strain_R8 ..................c..............t..........a.........c......................... [105360]

FR682468.2_Georgia_2007/1 tattgaatgcactccaagacatcatcccagaacttgatgctaggctgtggatcatgttctacctaagcacggccatattt [105440]
ASF_IND_20_CAD543 ................................................................................ [105440]
LR899193.1_Germany_2020/1 ................................................................................ [105440]
MN393477.1_Wuhan_2019 ................................................................................ [105440]
MW396979.1_Timor-Leste/2019/1 ................................................................................ [105440]
MK940252.1_CN/2019/InnerMongolia ................................................................................ [105440]
MK128995.1_China/2018/AnhuiXCGQ ................................................................................ [105440]
MH681419.1_ASFV/POL/2015 ................................................................................ [105440]
LS478113.1_Estonia_2014 ................................................................................ [105440]
KP843857.1_Odintsovo_02/14 ................................................................................ [105440]

```
FN557520.1_strain_E75             ................................................................g............... [105440]
AY261366.1_Warthog                ................................................................................ [105440]
AY261365.1_Warmbaths              ...................t..........................................t.g............... [105440]
AY261364.1_Tengani_62             ....a.....g...................................................t.g............... [105440]
AY261363.1_Pretoriuskop/96/4      ..............................................................t.g............... [105440]
AY261362.1_Mkuzi_1979             ....a........t.....t............................................g............... [105440]
AY261361.1_Malawi_Lil-20/1_1983   ....a...................................c.......................g............... [105440]
AY261360.1_Kenya_1950             ....a.....g.................................t...................g........a..c... [105440]
AM712240.1_OURT_88/3              ................................................................g............... [105440]
AM712239.1_Benin_97/1             ................................................................g............... [105440]
KM111295.1_Ken06.Bus              ....a...........................................................g........a..c... [105440]
KM111294.1_Ken05/Tk1              ....a.....g.................................t...................g........a..c... [105440]
KM262844.1_strain_L60             ................................................................g............... [105440]
KM102979.1_26544/OG10_Italy       ................................................................g............... [105440]
KM262845.1_strain_NHV             ................................................................g............... [105440]
KX354450.1_47/Ss/2008             ................................................................g............... [105440]
KP055815.1_strain_BA71            ................................................................g............... [105440]
MH025920.1_strain_R35             ....a...........................................................g........a..c... [105440]
MH025919.1_strain_N10             ....a...........................................................g........a..c... [105440]
MH025918.1_strain_R25             ....a...........................................................g........a..c... [105440]
MH025917.1_strain_R7              ....a...........................................................g........a..c... [105440]
MH025916.1_strain_R8              ....a...........................................................g........a..c... [105440]

FR682468.2_Georgia_2007/1         cgcggcctgaaggaggctctgatgcttttgtggcttggaggcggctgcaatggaaaaacttttctaatgcgacttgtggc [105520]
ASF_IND_20_CAD543                 .........................................y...................................... [105520]
LR899193.1_Germany_2020/1         ................................................................................ [105520]
MN393477.1_Wuhan_2019             ................................................................................ [105520]
MW396979.1_Timor-Leste/2019/1     ................................................................................ [105520]
MK940252.1_CN/2019/InnerMongolia  ................................................................................ [105520]
MK128995.1_China/2018/AnhuiXCGQ   ................................................................................ [105520]
MH681419.1_ASFV/POL/2015          ................................................................................ [105520]
LS478113.1_Estonia_2014           ................................................................................ [105520]
KP843857.1_Odintsovo_02/14        ................................................................................ [105520]
FN557520.1_strain_E75             .........................................t...................................... [105520]
AY261366.1_Warthog                .................c.........................................c.....g.............. [105520]
AY261365.1_Warmbaths              .................c.........................................c.....g.............. [105520]
AY261364.1_Tengani_62             ......t..........c...............................................g.............. [105520]
AY261363.1_Pretoriuskop/96/4      .................c...............................................g.............. [105520]
AY261362.1_Mkuzi_1979             .........................................t...................................... [105520]
AY261361.1_Malawi_Lil-20/1_1983   ......t.......a..c.........................................c.....g.............. [105520]
AY261360.1_Kenya_1950             .................c.........................................c.....g.............. [105520]
AM712240.1_OURT_88/3              .........................................t...................................... [105520]
AM712239.1_Benin_97/1             .........................................t...................................... [105520]
KM111295.1_Ken06.Bus              .................c.......................g.................c.....g.............. [105520]
```

```
KM111294.1_Ken05/Tk1              .................c.........................................c.....g.............. [105520]
KM262844.1_strain_L60             .........................................t...................................... [105520]
KM102979.1_26544/OG10_Italy       .........................................t...................................... [105520]
KM262845.1_strain_NHV             .........................................t...................................... [105520]
KX354450.1_47/Ss/2008             .........................................t...................................... [105520]
KP055815.1_strain_BA71            .........................................t...................................... [105520]
MH025920.1_strain_R35             .................c.......................g.................c.....g.............. [105520]
MH025919.1_strain_N10             .................c.......................g.................c.....g.............. [105520]
MH025918.1_strain_R25             .................c.......................g.................c.....g.............. [105520]
MH025917.1_strain_R7              .................c.......................g.................c.....g.............. [105520]
MH025916.1_strain_R8              .................c.......................g.................c.....g.............. [105520]

FR682468.2_Georgia_2007/1         catggtattgggcgatcactatgcctccaagctcaacatcagccttcttacaagctgcagagaaaccgcggaaaaaccca [105600]
ASF_IND_20_CAD543                 ................................................................................ [105600]
LR899193.1_Germany_2020/1         ................................................................................ [105600]
MN393477.1_Wuhan_2019             ................................................................................ [105600]
MW396979.1_Timor-Leste/2019/1     ................................................................................ [105600]
MK940252.1_CN/2019/InnerMongolia  ................................................................................ [105600]
MK128995.1_China/2018/AnhuiXCGQ   ................................................................................ [105600]
MH681419.1_ASFV/POL/2015          ................................................................................ [105600]
LS478113.1_Estonia_2014           ................................................................................ [105600]
KP843857.1_Odintsovo_02/14        ................................................................................ [105600]
FN557520.1_strain_E75             ................................................................................ [105600]
AY261366.1_Warthog                ......g................................t................a......g................ [105600]
AY261365.1_Warmbaths              ......g................................t................a......g................ [105600]
AY261364.1_Tengani_62             ......g................................t................a......g................ [105600]
AY261363.1_Pretoriuskop/96/4      ......g................................t................a......g................ [105600]
AY261362.1_Mkuzi_1979             ................................................................................ [105600]
AY261361.1_Malawi_Lil-20/1_1983   ......g.....t..........................t..........................g............. [105600]
AY261360.1_Kenya_1950             t.....g................................t................a.........t..a.......... [105600]
AM712240.1_OURT_88/3              ................................................................................ [105600]
AM712239.1_Benin_97/1             ................................................................................ [105600]
KM111295.1_Ken06.Bus              t.....g................................t................a.........t..a.......... [105600]
KM111294.1_Ken05/Tk1              t.....g................................t................a.........t..a.......... [105600]
KM262844.1_strain_L60             ................................................................................ [105600]
KM102979.1_26544/OG10_Italy       ................................................................................ [105600]
KM262845.1_strain_NHV             ................................................................................ [105600]
KX354450.1_47/Ss/2008             ................................................................................ [105600]
KP055815.1_strain_BA71            ................................................................................ [105600]
MH025920.1_strain_R35             t.....g................................t................a.........t..a.......... [105600]
MH025919.1_strain_N10             t.....g................................t................a.........t..a.......... [105600]
MH025918.1_strain_R25             t.....g................................t................a.........t..a.......... [105600]
MH025917.1_strain_R7              t.....g................................t................a.........t..a.......... [105600]
MH025916.1_strain_R8              t.....g................................t................a.........t..a.......... [105600]
```

```
FR682468.2_Georgia_2007/1        acagtgcctttatgcggcttaaggggcggggatatgggtactttgaggaaaccaacaaaagcgaggttctaaatacgtcg [105680]
ASF_IND_20_CAD543                ................................................................................ [105680]
LR899193.1_Germany_2020/1        ................................................................................ [105680]
MN393477.1_Wuhan_2019            ................................................................................ [105680]
MW396979.1_Timor-Leste/2019/1    ................................................................................ [105680]
MK940252.1_CN/2019/InnerMongolia ................................................................................ [105680]
MK128995.1_China/2018/AnhuiXCGQ  ................................................................................ [105680]
MH681419.1_ASFV/POL/2015         ................................................................................ [105680]
LS478113.1_Estonia_2014          ................................................................................ [105680]
KP843857.1_Odintsovo_02/14       ................................................................................ [105680]
FN557520.1_strain_E75            ..........c..................................................................... [105680]
AY261366.1_Warthog               ..........c..................................................................... [105680]
AY261365.1_Warmbaths             ..........c...............a..................................................... [105680]
AY261364.1_Tengani_62            ........c.c..................................................................... [105680]
AY261363.1_Pretoriuskop/96/4     ..........c............................................t........................ [105680]
AY261362.1_Mkuzi_1979            ..........c..................................................................... [105680]
AY261361.1_Malawi_Lil-20/1_1983  ....c.....c..................................................................... [105680]
AY261360.1_Kenya_1950            ..........c........c.....a.....g................................................ [105680]
AM712240.1_OURT_88/3             ..........c..................................................................... [105680]
AM712239.1_Benin_97/1            ..........c..................................................................... [105680]
KM111295.1_Ken06.Bus             ..........c........c................................t........t...a.............. [105680]
KM111294.1_Ken05/Tk1             ...................c.....a.....g.................................a.............. [105680]
KM262844.1_strain_L60            ..........c..................................................................... [105680]
KM102979.1_26544/OG10_Italy      ..........c..................................................................... [105680]
KM262845.1_strain_NHV            ..........c..................................................................... [105680]
KX354450.1_47/Ss/2008            ..........c..................................................................... [105680]
KP055815.1_strain_BA71           ..........c..................................................................... [105680]
MH025920.1_strain_R35            ..........c........c................................t........t...a.............. [105680]
MH025919.1_strain_N10            ..........c........c................................t........t...a.............. [105680]
MH025918.1_strain_R25            ..........c........c................................t........t...a.............. [105680]
MH025917.1_strain_R7             ..........c........c................................t........t...a.............. [105680]
MH025916.1_strain_R8             ..........c........c................................t........t...a.............. [105680]

FR682468.2_Georgia_2007/1        cggctgaaggaaatggtaaatccgggcgatgtcaccgctcgagagcttaatcaaaaacaggaaagctttcagatgacggc [105760]
ASF_IND_20_CAD543                ................................................................................ [105760]
LR899193.1_Germany_2020/1        ................................................................................ [105760]
MN393477.1_Wuhan_2019            ................................................................................ [105760]
MW396979.1_Timor-Leste/2019/1    ................................................................................ [105760]
MK940252.1_CN/2019/InnerMongolia ................................................................................ [105760]
MK128995.1_China/2018/AnhuiXCGQ  ................................................................................ [105760]
MH681419.1_ASFV/POL/2015         ................................................................................ [105760]
LS478113.1_Estonia_2014          ................................................................................ [105760]
KP843857.1_Odintsovo_02/14       ................................................................................ [105760]
```

```
FN557520.1_strain_E75             ................................................................................ [105760]
AY261366.1_Warthog                ................................................................................ [105760]
AY261365.1_Warmbaths              ..........................t..................................................... [105760]
AY261364.1_Tengani_62             .......................a........................................................ [105760]
AY261363.1_Pretoriuskop/96/4      ............................................a................................... [105760]
AY261362.1_Mkuzi_1979             ...............................................g.............................a.. [105760]
AY261361.1_Malawi_Lil-20/1_1983   ................................................................................ [105760]
AY261360.1_Kenya_1950             ................................................................................ [105760]
AM712240.1_OURT_88/3              ................................................................................ [105760]
AM712239.1_Benin_97/1             ................................................................................ [105760]
KM111295.1_Ken06.Bus              .......................................................................a........ [105760]
KM111294.1_Ken05/Tk1              ................................................................................ [105760]
KM262844.1_strain_L60             ................................................................................ [105760]
KM102979.1_26544/OG10_Italy       ................................................................................ [105760]
KM262845.1_strain_NHV             ................................................................................ [105760]
KX354450.1_47/Ss/2008             ................................................................................ [105760]
KP055815.1_strain_BA71            ................................................................................ [105760]
MH025920.1_strain_R35             .......................................................................a........ [105760]
MH025919.1_strain_N10             .......................................................................a........ [105760]
MH025918.1_strain_R25             .......................................................................a........ [105760]
MH025917.1_strain_R7              .......................................................................a........ [105760]
MH025916.1_strain_R8              .......................................................................a........ [105760]

FR682468.2_Georgia_2007/1         caccatggtcgccgcgtccaactataacttcatcattgacacgacggaccacggcacatggagaagactgcggcattatc [105840]
ASF_IND_20_CAD543                 ................................................................................ [105840]
LR899193.1_Germany_2020/1         ................................................................................ [105840]
MN393477.1_Wuhan_2019             ................................................................................ [105840]
MW396979.1_Timor-Leste/2019/1     ................................................................................ [105840]
MK940252.1_CN/2019/InnerMongolia  ................................................................................ [105840]
MK128995.1_China/2018/AnhuiXCGQ   ................................................................................ [105840]
MH681419.1_ASFV/POL/2015          ................................................................................ [105840]
LS478113.1_Estonia_2014           ................................................................................ [105840]
KP843857.1_Odintsovo_02/14        ................................................................................ [105840]
FN557520.1_strain_E75             ................................................................................ [105840]
AY261366.1_Warthog                ................................................................................ [105840]
AY261365.1_Warmbaths              ................................................................................ [105840]
AY261364.1_Tengani_62             ................................................................................ [105840]
AY261363.1_Pretoriuskop/96/4      .........t...................................................................... [105840]
AY261362.1_Mkuzi_1979             ................................................................................ [105840]
AY261361.1_Malawi_Lil-20/1_1983   .........................................................c...................... [105840]
AY261360.1_Kenya_1950             .........................................................c...................... [105840]
AM712240.1_OURT_88/3              ................................................................................ [105840]
AM712239.1_Benin_97/1             ................................................................................ [105840]
KM111295.1_Ken06.Bus              ..............................t........t........................................ [105840]
```

```
KM111294.1_Ken05/Tk1            .........................................................c...................... [105840]
KM262844.1_strain_L60           ................................................................................ [105840]
KM102979.1_26544/OG10_Italy     ................................................................................ [105840]
KM262845.1_strain_NHV           ................................................................................ [105840]
KX354450.1_47/Ss/2008           ................................................................................ [105840]
KP055815.1_strain_BA71          ................................................................................ [105840]
MH025920.1_strain_R35           ..............................t........t........................................ [105840]
MH025919.1_strain_N10           ..............................t........t........................................ [105840]
MH025918.1_strain_R25           ..............................t........t........................................ [105840]
MH025917.1_strain_R7            ..............................t........t........................................ [105840]
MH025916.1_strain_R8            ..............................t........t........................................ [105840]

FR682468.2_Georgia_2007/1       ggtcaaaggtgaaattctgccataaccccgaccccagtaacccctacgagaaaaaggaagatcctcgctttattcacgag [105920]
ASF_IND_20_CAD543               ................................................................................ [105920]
LR899193.1_Germany_2020/1       ................................................................................ [105920]
MN393477.1_Wuhan_2019           ................................................................................ [105920]
MW396979.1_Timor-Leste/2019/1   ................................................................................ [105920]
MK940252.1_CN/2019/InnerMongolia ................................................................................ [105920]
MK128995.1_China/2018/AnhuiXCGQ ................................................................................ [105920]
MH681419.1_ASFV/POL/2015        ................................................................................ [105920]
LS478113.1_Estonia_2014         ................................................................................ [105920]
KP843857.1_Odintsovo_02/14      ................................................................................ [105920]
FN557520.1_strain_E75           .t.................................g............................................ [105920]
AY261366.1_Warthog              .t.......................t..t.......a...tt.......a.............................. [105920]
AY261365.1_Warmbaths            .t.......................t..t.......a...tt.......a.............................. [105920]
AY261364.1_Tengani_62           .t.......................t..t.......a...tt.......a.............................. [105920]
AY261363.1_Pretoriuskop/96/4    .t.......................t..t.......a...tt.......a.............................. [105920]
AY261362.1_Mkuzi_1979           .t.......................t..t................................................... [105920]
AY261361.1_Malawi_Lil-20/1_1983 .t..............t........t..t..t....a....t.......a.............................. [105920]
AY261360.1_Kenya_1950           .t.......................t..t..t....a....t.......a.............................. [105920]
AM712240.1_OURT_88/3            .t.................................g............................................ [105920]
AM712239.1_Benin_97/1           .t.................................g............................................ [105920]
KM111295.1_Ken06.Bus            .t.......................t..t..t....a....t.......a.............................. [105920]
KM111294.1_Ken05/Tk1            .t.......................t..t..t....a....t.......a.............................. [105920]
KM262844.1_strain_L60           .t.................................g............................................ [105920]
KM102979.1_26544/OG10_Italy     .t.................................g............................................ [105920]
KM262845.1_strain_NHV           .t.................................g............................................ [105920]
KX354450.1_47/Ss/2008           .t.................................g............................................ [105920]
KP055815.1_strain_BA71          .t.................................g............................................ [105920]
MH025920.1_strain_R35           .t.......................t..t..t....a....t.......a.............................. [105920]
MH025919.1_strain_N10           .t.......................t..t..t....a....t.......a.............................. [105920]
MH025918.1_strain_R25           .t.......................t..t..t....a....t.......a.............................. [105920]
MH025917.1_strain_R7            .t.......................t..t..t....a....t.......a.............................. [105920]
MH025916.1_strain_R8            .t.......................t..t..t....a....t.......a.............................. [105920]
```

```
FR682468.2_Georgia_2007/1        tacatcatggatccagactgccaaaacgcattcttcagcatactcgtctatttttgggagaagctacagaaggaatacaa [106000]
ASF_IND_20_CAD543                ................n............................................................... [106000]
LR899193.1_Germany_2020/1        ................................................................................ [106000]
MN393477.1_Wuhan_2019            ................................................................................ [106000]
MW396979.1_Timor-Leste/2019/1    ................................................................................ [106000]
MK940252.1_CN/2019/InnerMongolia ................................................................................ [106000]
MK128995.1_China/2018/AnhuiXCGQ  ................................................................................ [106000]
MH681419.1_ASFV/POL/2015         ................................................................................ [106000]
LS478113.1_Estonia_2014          ................................................................................ [106000]
KP843857.1_Odintsovo_02/14       ................................................................................ [106000]
FN557520.1_strain_E75            ................................................................................ [106000]
AY261366.1_Warthog               ...............a...............................a.....c.......................... [106000]
AY261365.1_Warmbaths             ...........c...a...............................a.....c.......................... [106000]
AY261364.1_Tengani_62            ...............a...............................a.....c.......................... [106000]
AY261363.1_Pretoriuskop/96/4     ...............a...............................a.....c.......................... [106000]
AY261362.1_Mkuzi_1979            ................................................................................ [106000]
AY261361.1_Malawi_Lil-20/1_1983  ...............a...................t...........a................................ [106000]
AY261360.1_Kenya_1950            ...............a.............g.................a..............a................. [106000]
AM712240.1_OURT_88/3             ................................................................................ [106000]
AM712239.1_Benin_97/1            ................................................................................ [106000]
KM111295.1_Ken06.Bus             ...............a.............g.................a..............a................. [106000]
KM111294.1_Ken05/Tk1             ...............a.............g.................a..............a................. [106000]
KM262844.1_strain_L60            ................................................................................ [106000]
KM102979.1_26544/OG10_Italy      ................................................................................ [106000]
KM262845.1_strain_NHV            ................................................................................ [106000]
KX354450.1_47/Ss/2008            ................................................................................ [106000]
KP055815.1_strain_BA71           ................................................................................ [106000]
MH025920.1_strain_R35            ...............a.............g.................a..............a................. [106000]
MH025919.1_strain_N10            ...............a.............g.................a..............a................. [106000]
MH025918.1_strain_R25            ...............a.............g.................a..............a................. [106000]
MH025917.1_strain_R7             ...............a.............g.................a..............a................. [106000]
MH025916.1_strain_R8             ...............a.............g.................a..............a................. [106000]

FR682468.2_Georgia_2007/1        cgggcagattaaaaaagtgttttgtcccaccattgagagcgaaacggaggcgtacagaaagtcacaagatacgctacata [106080]
ASF_IND_20_CAD543                ................................................................................ [106080]
LR899193.1_Germany_2020/1        ................................................................................ [106080]
MN393477.1_Wuhan_2019            ................................................................................ [106080]
MW396979.1_Timor-Leste/2019/1    ................................................................................ [106080]
MK940252.1_CN/2019/InnerMongolia ................................................................................ [106080]
MK128995.1_China/2018/AnhuiXCGQ  ................................................................................ [106080]
MH681419.1_ASFV/POL/2015         ................................................................................ [106080]
LS478113.1_Estonia_2014          ................................................................................ [106080]
KP843857.1_Odintsovo_02/14       ................................................................................ [106080]
```

```
FN557520.1_strain_E75            ................................................................................ [106080]
AY261366.1_Warthog               ......a...........a........t..t..............a...........g...................... [106080]
AY261365.1_Warmbaths             ......a....................t.................a...........g...................... [106080]
AY261364.1_Tengani_62            ......a....................t.................a..............a................... [106080]
AY261363.1_Pretoriuskop/96/4     ......a....................t.................a...........g...................... [106080]
AY261362.1_Mkuzi_1979            ......a........................................................................ [106080]
AY261361.1_Malawi_Lil-20/1_1983  ......a....................t.................a.................................. [106080]
AY261360.1_Kenya_1950            ......a....................t.................a.....a..t......................... [106080]
AM712240.1_OURT_88/3             ................................................................................ [106080]
AM712239.1_Benin_97/1            ................................................................................ [106080]
KM111295.1_Ken06.Bus             ......a....................t.................a..............a................... [106080]
KM111294.1_Ken05/Tk1             ......a....................t.................a.....a..t......................... [106080]
KM262844.1_strain_L60            ................................................................................ [106080]
KM102979.1_26544/OG10_Italy      ................................................................................ [106080]
KM262845.1_strain_NHV            ................................................................................ [106080]
KX354450.1_47/Ss/2008            ................................................................................ [106080]
KP055815.1_strain_BA71           ................................................................................ [106080]
MH025920.1_strain_R35            ......a....................t.................a..............a................... [106080]
MH025919.1_strain_N10            ......a....................t.................a..............a................... [106080]
MH025918.1_strain_R25            ......a....................t.................a..............a................... [106080]
MH025917.1_strain_R7             ......a....................t.................a..............a................... [106080]
MH025916.1_strain_R8             ......a....................t.................a..............a................... [106080]

FR682468.2_Georgia_2007/1        ggtttatcacagaaagagtcgtggagtcgccctccgcagaaactgtgtacaacctatccgaggtcgtgacggcctacgcg [106160]
ASF_IND_20_CAD543                ................................................................................ [106160]
LR899193.1_Germany_2020/1        ................................................................................ [106160]
MN393477.1_Wuhan_2019            ................................................................................ [106160]
MW396979.1_Timor-Leste/2019/1    ................................................................................ [106160]
MK940252.1_CN/2019/InnerMongolia ................................................................................ [106160]
MK128995.1_China/2018/AnhuiXCGQ  ................................................................................ [106160]
MH681419.1_ASFV/POL/2015         ................................................................................ [106160]
LS478113.1_Estonia_2014          ................................................................................ [106160]
KP843857.1_Odintsovo_02/14       ................................................................................ [106160]
FN557520.1_strain_E75            ..................................g............................................. [106160]
AY261366.1_Warthog               .................a.............................................................a [106160]
AY261365.1_Warmbaths             .................a.............................................................a [106160]
AY261364.1_Tengani_62            ...............................................................................a [106160]
AY261363.1_Pretoriuskop/96/4     .................a.............................................................a [106160]
AY261362.1_Mkuzi_1979            ................................................................................ [106160]
AY261361.1_Malawi_Lil-20/1_1983  ..................................t................................a...........a [106160]
AY261360.1_Kenya_1950            .........................a..............g..............c.......................a [106160]
AM712240.1_OURT_88/3             ..................................g............................................. [106160]
AM712239.1_Benin_97/1            ..................................g............................................. [106160]
KM111295.1_Ken06.Bus             ...................t.....a..............g..............c.......................a [106160]
```

```
KM111294.1_Ken05/Tk1            .........................a..............g..............c.......................a [106160]
KM262844.1_strain_L60           ..................................g............................................. [106160]
KM102979.1_26544/OG10_Italy     ..................................g............................................. [106160]
KM262845.1_strain_NHV           ..................................g............................................. [106160]
KX354450.1_47/Ss/2008           ..................................g............................................. [106160]
KP055815.1_strain_BA71          ..................................g............................................. [106160]
MH025920.1_strain_R35           ...................t.....a..............g..............c.......................a [106160]
MH025919.1_strain_N10           ...................t.....a..............g..............c.......................a [106160]
MH025918.1_strain_R25           ...................t.....a..............g..............c.......................a [106160]
MH025917.1_strain_R7            ...................t.....a..............g..............c.......................a [106160]
MH025916.1_strain_R8            ...................t.....a..............g..............c.......................a [106160]

FR682468.2_Georgia_2007/1       gaatggtacaacaccaacattaacgtaaagcgccatattgccctcgagctatcccaggagttagaaaactctgtgctaga [106240]
ASF_IND_20_CAD543               ................................................................................ [106240]
LR899193.1_Germany_2020/1       ................................................................................ [106240]
MN393477.1_Wuhan_2019           ................................................................................ [106240]
MW396979.1_Timor-Leste/2019/1   ................................................................................ [106240]
MK940252.1_CN/2019/InnerMongolia ................................................................................ [106240]
MK128995.1_China/2018/AnhuiXCGQ ................................................................................ [106240]
MH681419.1_ASFV/POL/2015        ................................................................................ [106240]
LS478113.1_Estonia_2014         ................................................................................ [106240]
KP843857.1_Odintsovo_02/14      ................................................................................ [106240]
FN557520.1_strain_E75           ............g...............................t................................... [106240]
AY261366.1_Warthog              ............g.............g..................................................... [106240]
AY261365.1_Warmbaths            ............g.............g..............g...................................... [106240]
AY261364.1_Tengani_62           ............g.............g..................................................... [106240]
AY261363.1_Pretoriuskop/96/4    ............g.............g..................................................... [106240]
AY261362.1_Mkuzi_1979           ............................................t................................... [106240]
AY261361.1_Malawi_Lil-20/1_1983 ........t...........c........................................................... [106240]
AY261360.1_Kenya_1950           ........t...g.......c.......................t................................... [106240]
AM712240.1_OURT_88/3            ............g...............................t................................... [106240]
AM712239.1_Benin_97/1           ............g...............................t................................... [106240]
KM111295.1_Ken06.Bus            ........t...g.......c.......................t................................... [106240]
KM111294.1_Ken05/Tk1            ........t...g.......c.......................t................................... [106240]
KM262844.1_strain_L60           ............g...............................t................................... [106240]
KM102979.1_26544/OG10_Italy     ............g...............................t................................... [106240]
KM262845.1_strain_NHV           ............g...............................t................................... [106240]
KX354450.1_47/Ss/2008           ............g...............................t................................... [106240]
KP055815.1_strain_BA71          ............g...............................t................................... [106240]
MH025920.1_strain_R35           ........t...g.......c.......................t................................... [106240]
MH025919.1_strain_N10           ........t...g.......c.......................t................................... [106240]
MH025918.1_strain_R25           ........t...g.......c.......................t................................... [106240]
MH025917.1_strain_R7            ........t...g.......c.......................t................................... [106240]
MH025916.1_strain_R8            ........t...g.......c.......................t................................... [106240]
```

```
FR682468.2_Georgia_2007/1        aaaataccttcagtggtctcccaacaaaacgcgaattctaaagggttgccgtattttgcataaatttgaaacgctgcagc [106320]
ASF_IND_20_CAD543                ................................................................................ [106320]
LR899193.1_Germany_2020/1        ................................................................................ [106320]
MN393477.1_Wuhan_2019            ................................................................................ [106320]
MW396979.1_Timor-Leste/2019/1    ................................................................................ [106320]
MK940252.1_CN/2019/InnerMongolia ................................................................................ [106320]
MK128995.1_China/2018/AnhuiXCGQ  ................................................................................ [106320]
MH681419.1_ASFV/POL/2015         ................................................................................ [106320]
LS478113.1_Estonia_2014          ................................................................................ [106320]
KP843857.1_Odintsovo_02/14       ................................................................................ [106320]
FN557520.1_strain_E75            ................................................................................ [106320]
AY261366.1_Warthog               ................................................................................ [106320]
AY261365.1_Warmbaths             ................................................................................ [106320]
AY261364.1_Tengani_62            ................................................................................ [106320]
AY261363.1_Pretoriuskop/96/4     ................................................................................ [106320]
AY261362.1_Mkuzi_1979            ................................................................................ [106320]
AY261361.1_Malawi_Lil-20/1_1983  ..............................a................................g................ [106320]
AY261360.1_Kenya_1950            ............................................................c................... [106320]
AM712240.1_OURT_88/3             ................................................................................ [106320]
AM712239.1_Benin_97/1            ................................................................................ [106320]
KM111295.1_Ken06.Bus             ............................................................c................... [106320]
KM111294.1_Ken05/Tk1             ............................................................c................... [106320]
KM262844.1_strain_L60            ................................................................................ [106320]
KM102979.1_26544/OG10_Italy      ................................................................................ [106320]
KM262845.1_strain_NHV            ................................................................................ [106320]
KX354450.1_47/Ss/2008            ................................................................................ [106320]
KP055815.1_strain_BA71           ................................................................................ [106320]
MH025920.1_strain_R35            ............................................................c................... [106320]
MH025919.1_strain_N10            ............................................................c................... [106320]
MH025918.1_strain_R25            ............................................................c................... [106320]
MH025917.1_strain_R7             ............................................................c................... [106320]
MH025916.1_strain_R8             ............................................................c................... [106320]

FR682468.2_Georgia_2007/1        ccggcgaatcctacattggggtgtccacggccggcacactcctaaacacacccatatgcgagccaaaaaataaatggtgg [106400]
ASF_IND_20_CAD543                ................................................................................ [106400]
LR899193.1_Germany_2020/1        ................................................................................ [106400]
MN393477.1_Wuhan_2019            ................................................................................ [106400]
MW396979.1_Timor-Leste/2019/1    ................................................................................ [106400]
MK940252.1_CN/2019/InnerMongolia ................................................................................ [106400]
MK128995.1_China/2018/AnhuiXCGQ  ................................................................................ [106400]
MH681419.1_ASFV/POL/2015         ................................................................................ [106400]
LS478113.1_Estonia_2014          ................................................................................ [106400]
KP843857.1_Odintsovo_02/14       ................................................................................ [106400]
```

```
FN557520.1_strain_E75              ................................................................................ [106400]
AY261366.1_Warthog                 ...................a............................................................ [106400]
AY261365.1_Warmbaths               ................................................................................ [106400]
AY261364.1_Tengani_62              ................................................................................ [106400]
AY261363.1_Pretoriuskop/96/4       .............................a..........t....................................... [106400]
AY261362.1_Mkuzi_1979              ................................................................................ [106400]
AY261361.1_Malawi_Lil-20/1_1983    .............................................................a.................. [106400]
AY261360.1_Kenya_1950              ...................a......t..a..........t....................................... [106400]
AM712240.1_OURT_88/3               ................................................................................ [106400]
AM712239.1_Benin_97/1              ................................................................................ [106400]
KM111295.1_Ken06.Bus               ...................a......t..a..........t....................a.................. [106400]
KM111294.1_Ken05/Tk1               ...................a......t.............t....................a.................. [106400]
KM262844.1_strain_L60              ................................................................................ [106400]
KM102979.1_26544/OG10_Italy        ................................................................................ [106400]
KM262845.1_strain_NHV              ................................................................................ [106400]
KX354450.1_47/Ss/2008              ................................................................................ [106400]
KP055815.1_strain_BA71             ................................................................................ [106400]
MH025920.1_strain_R35              ...................a......t..a..........t....................a.................. [106400]
MH025919.1_strain_N10              ...................a......t..a..........t....................a.................. [106400]
MH025918.1_strain_R25              ...................a......t..a..........t....................a.................. [106400]
MH025917.1_strain_R7               ...................a......t..a..........t....................a.................. [106400]
MH025916.1_strain_R8               ...................a......t..a..........t....................a.................. [106400]

FR682468.2_Georgia_2007/1          gaatggtcccctaatccctctgcccctcctgagaaagaagcgtctgcaccaactccttagggaatatccttagaagcatg [106480]
ASF_IND_20_CAD543                  .............................................................r.................. [106480]
LR899193.1_Germany_2020/1          ................................................................................ [106480]
MN393477.1_Wuhan_2019              ................................................................................ [106480]
MW396979.1_Timor-Leste/2019/1      ................................................................................ [106480]
MK940252.1_CN/2019/InnerMongolia   ................................................................................ [106480]
MK128995.1_China/2018/AnhuiXCGQ    ................................................................................ [106480]
MH681419.1_ASFV/POL/2015           ................................................................................ [106480]
LS478113.1_Estonia_2014            ................................................................................ [106480]
KP843857.1_Odintsovo_02/14         ................................................................................ [106480]
FN557520.1_strain_E75              ........t.......t............................................a.................. [106480]
AY261366.1_Warthog                 ..............c................................................................. [106480]
AY261365.1_Warmbaths               ............................t................................................... [106480]
AY261364.1_Tengani_62              ....................................................t........................... [106480]
AY261363.1_Pretoriuskop/96/4       ................................................................................ [106480]
AY261362.1_Mkuzi_1979              ................................................................................ [106480]
AY261361.1_Malawi_Lil-20/1_1983    ..............c.....................................................t........... [106480]
AY261360.1_Kenya_1950              .............................................................a........c......... [106480]
AM712240.1_OURT_88/3               ........t.......t............................................a.................. [106480]
AM712239.1_Benin_97/1              ........t.......t............................................a.................. [106480]
KM111295.1_Ken06.Bus               ......................................................................c......... [106480]
```

```
KM111294.1_Ken05/Tk1             ............................t.........................................c......... [106480]
KM262844.1_strain_L60            ........t.......t............................................a.................. [106480]
KM102979.1_26544/OG10_Italy      ........t.......t............................................a.................. [106480]
KM262845.1_strain_NHV            ........t.......t............................................a.................. [106480]
KX354450.1_47/Ss/2008            ........t.......t............................................a.................. [106480]
KP055815.1_strain_BA71           ........t.......t............................................a.................. [106480]
MH025920.1_strain_R35            ......................................................................c......... [106480]
MH025919.1_strain_N10            ......................................................................c......... [106480]
MH025918.1_strain_R25            ......................................................................c......... [106480]
MH025917.1_strain_R7             ......................................................................c......... [106480]
MH025916.1_strain_R8             ......................................................................c......... [106480]

FR682468.2_Georgia_2007/1        tctttcggcagagccattaccggtagcaaaaaagcaacattgagtatattatatgccttagcctgctcataagcgtcctt [106560]
ASF_IND_20_CAD543                ................................................................................ [106560]
LR899193.1_Germany_2020/1        ................................................................................ [106560]
MN393477.1_Wuhan_2019            ................................................................................ [106560]
MW396979.1_Timor-Leste/2019/1    ................................................................................ [106560]
MK940252.1_CN/2019/InnerMongolia ................................................................................ [106560]
MK128995.1_China/2018/AnhuiXCGQ  ................................................................................ [106560]
MH681419.1_ASFV/POL/2015         ................................................................................ [106560]
LS478113.1_Estonia_2014          ................................................................................ [106560]
KP843857.1_Odintsovo_02/14       ................................................................................ [106560]
FN557520.1_strain_E75            ................................................................................ [106560]
AY261366.1_Warthog               ..c............................................................................. [106560]
AY261365.1_Warmbaths             ..c............................................................................. [106560]
AY261364.1_Tengani_62            ..c............................................................................. [106560]
AY261363.1_Pretoriuskop/96/4     ..c............................................................................. [106560]
AY261362.1_Mkuzi_1979            ................................................................................ [106560]
AY261361.1_Malawi_Lil-20/1_1983  ..c.............g........................................................a..... [106560]
AY261360.1_Kenya_1950            ..cc.............g...........................g.................................. [106560]
AM712240.1_OURT_88/3             ................................................................................ [106560]
AM712239.1_Benin_97/1            ................................................................................ [106560]
KM111295.1_Ken06.Bus             ..cc.............g...........................g.................................. [106560]
KM111294.1_Ken05/Tk1             ..cc.............g...........................g.................................. [106560]
KM262844.1_strain_L60            ................................................................................ [106560]
KM102979.1_26544/OG10_Italy      ................................................................................ [106560]
KM262845.1_strain_NHV            ................................................................................ [106560]
KX354450.1_47/Ss/2008            ................................................................................ [106560]
KP055815.1_strain_BA71           ................................................................................ [106560]
MH025920.1_strain_R35            ..cc.............g...........................g.................................. [106560]
MH025919.1_strain_N10            ..cc.............g...........................g.................................. [106560]
MH025918.1_strain_R25            ..cc.............g...........................g.................................. [106560]
MH025917.1_strain_R7             ..cc.............g...........................g.................................. [106560]
MH025916.1_strain_R8             ..cc.............g...........................g.................................. [106560]
```

```
FR682468.2_Georgia_2007/1        ttttttcatggtattttatgtttt------taaatatttttaattattttttaaatacgatgaacagttcgtgctccgaa [106640]
ASF_IND_20_CAD543                ........................------..t............................................... [106640]
LR899193.1_Germany_2020/1        ........................------.................................................. [106640]
MN393477.1_Wuhan_2019            ........................------.................................................. [106640]
MW396979.1_Timor-Leste/2019/1    ........................------.................................................. [106640]
MK940252.1_CN/2019/InnerMongolia ........................------.................................................. [106640]
MK128995.1_China/2018/AnhuiXCGQ  ........................------.................................................. [106640]
MH681419.1_ASFV/POL/2015         ........................------.................................................. [106640]
LS478113.1_Estonia_2014          ........................------.................................................. [106640]
KP843857.1_Odintsovo_02/14       ........................------.................................................. [106640]
FN557520.1_strain_E75            ........................------..t............................................... [106640]
AY261366.1_Warthog               ...................a....------.................................................. [106640]
AY261365.1_Warmbaths             ........................------.................................................. [106640]
AY261364.1_Tengani_62            ........................------............................................c..... [106640]
AY261363.1_Pretoriuskop/96/4     ........................------.................................................. [106640]
AY261362.1_Mkuzi_1979            ........................------..t............................................... [106640]
AY261361.1_Malawi_Lil-20/1_1983  ..............a.........tattta....a........c..............a...........a.....t..g [106640]
AY261360.1_Kenya_1950            ...................a.c.a-------t.t.t......................a....................g [106640]
AM712240.1_OURT_88/3             ........................------..t............................................... [106640]
AM712239.1_Benin_97/1            ........................------..t............................................... [106640]
KM111295.1_Ken06.Bus             ...................a.a.a---------tat................t..........................g [106640]
KM111294.1_Ken05/Tk1             ...................a.c.a-------t.t.t................t..........................g [106640]
KM262844.1_strain_L60            ........................------..t............................................... [106640]
KM102979.1_26544/OG10_Italy      ........................------..t............................................... [106640]
KM262845.1_strain_NHV            ........................------..t............................................... [106640]
KX354450.1_47/Ss/2008            ........................------..t............................................... [106640]
KP055815.1_strain_BA71           ........................------..t............................................... [106640]
MH025920.1_strain_R35            ...................a.a.at-----at.tat................t..........................g [106640]
MH025919.1_strain_N10            ...................a.a.a-------t.tat................t..........................g [106640]
MH025918.1_strain_R25            ...................a.a.at-----at.tat................t..........................g [106640]
MH025917.1_strain_R7             ...................a.a.at-----at.tat................t..........................g [106640]
MH025916.1_strain_R8             ...................a.a.at-----at.tat................t..........................g [106640]

FR682468.2_Georgia_2007/1        ggctgtttactaaaaatcggtgtgaatccgcattctttaaatatggtttcccattcggggatggtatggaaatccatgtc [106720]
ASF_IND_20_CAD543                ................................................................................ [106720]
LR899193.1_Germany_2020/1        ................................................................................ [106720]
MN393477.1_Wuhan_2019            ................................................................................ [106720]
MW396979.1_Timor-Leste/2019/1    ................................................................................ [106720]
MK940252.1_CN/2019/InnerMongolia ................................................................................ [106720]
MK128995.1_China/2018/AnhuiXCGQ  ................................................................................ [106720]
MH681419.1_ASFV/POL/2015         ................................................................................ [106720]
LS478113.1_Estonia_2014          ................................................................................ [106720]
KP843857.1_Odintsovo_02/14       ................................................................................ [106720]
```

```
FN557520.1_strain_E75           ................................................................................ [106720]
AY261366.1_Warthog              ................................................................................ [106720]
AY261365.1_Warmbaths            ................................................................................ [106720]
AY261364.1_Tengani_62           ..........................c........a...................................t........ [106720]
AY261363.1_Pretoriuskop/96/4    ................................................................................ [106720]
AY261362.1_Mkuzi_1979           ................................................................................ [106720]
AY261361.1_Malawi_Lil-20/1_1983 ......................a...c.................a........c....t..................... [106720]
AY261360.1_Kenya_1950           ........gt....c.............................a........c...ct..................... [106720]
AM712240.1_OURT_88/3            ................................................................................ [106720]
AM712239.1_Benin_97/1           ................................................................................ [106720]
KM111295.1_Ken06.Bus            .....c..gt...........................................c...ct..................... [106720]
KM111294.1_Ken05/Tk1            ........gt....c......................................c...ct..................... [106720]
KM262844.1_strain_L60           ................................................................................ [106720]
KM102979.1_26544/OG10_Italy     ................................................................................ [106720]
KM262845.1_strain_NHV           ................................................................................ [106720]
KX354450.1_47/Ss/2008           ................................................................................ [106720]
KP055815.1_strain_BA71          ................................................................................ [106720]
MH025920.1_strain_R35           .....c..gt...........................................c...ct..................... [106720]
MH025919.1_strain_N10           .....c..gt...........................................c...ct..................... [106720]
MH025918.1_strain_R25           .....c..gt...........................................c...ct..................... [106720]
MH025917.1_strain_R7            .....c..gt...........................................c...ct..................... [106720]
MH025916.1_strain_R8            .....c..gt...........................................c...ct..................... [106720]

FR682468.2_Georgia_2007/1       tctacgaatagtatggtgcccaagtgcgtcctgcaggctgtgaagccagaaggcctcctgaccttgatgaaggtcgtaca [106800]
ASF_IND_20_CAD543               ................................................................................ [106800]
LR899193.1_Germany_2020/1       ................................................................................ [106800]
MN393477.1_Wuhan_2019           ................................................................................ [106800]
MW396979.1_Timor-Leste/2019/1   ................................................................................ [106800]
MK940252.1_CN/2019/InnerMongolia ................................................................................ [106800]
MK128995.1_China/2018/AnhuiXCGQ ................................................................................ [106800]
MH681419.1_ASFV/POL/2015        ................................................................................ [106800]
LS478113.1_Estonia_2014         ................................................................................ [106800]
KP843857.1_Odintsovo_02/14      ................................................................................ [106800]
FN557520.1_strain_E75           ........................c.............................t..........t.........a.... [106800]
AY261366.1_Warthog              ........................c.....................................t................. [106800]
AY261365.1_Warmbaths            ........................c.............................t.......t................. [106800]
AY261364.1_Tengani_62           ........................c..a..........................t.......t...g........a.... [106800]
AY261363.1_Pretoriuskop/96/4    ........................c.............................t.......t................. [106800]
AY261362.1_Mkuzi_1979           ........................c.............................t..........t.........a.... [106800]
AY261361.1_Malawi_Lil-20/1_1983 .........c..g...........c........t.a.............t....t.......ta.t.............. [106800]
AY261360.1_Kenya_1950           .........c...........g..c.....t....a..................t.......t..t.....a........ [106800]
AM712240.1_OURT_88/3            ........................c.............................t..........t.........a.... [106800]
AM712239.1_Benin_97/1           ........................c.............................t..........t.........a.... [106800]
KM111295.1_Ken06.Bus            .t......................c.....t....a..................t.......t..t.....a........ [106800]
```

```
KM111294.1_Ken05/Tk1              .........c...........g..c.....t....a..................t.......t..t.....a........ [106800]
KM262844.1_strain_L60             ........................c.............................t..........t.........a.... [106800]
KM102979.1_26544/OG10_Italy       ........................c.............................t..........t.........a.... [106800]
KM262845.1_strain_NHV             ........................c.............................t..........t.........a.... [106800]
KX354450.1_47/Ss/2008             ........................c.............................t..........t.........a.... [106800]
KP055815.1_strain_BA71            ........................c.............................t..........t.........a.... [106800]
MH025920.1_strain_R35             .t......................c.....t....a..................t.......t..t.....a........ [106800]
MH025919.1_strain_N10             .t......................c.....t....a..................t.......t..t.....a........ [106800]
MH025918.1_strain_R25             .t......................c.....t....a..................t.......t..t.....a........ [106800]
MH025917.1_strain_R7              .t......................c.....t....a..................t.......t..t.....a........ [106800]
MH025916.1_strain_R8              .t......................c.....t....a..................t.......t..t.....a........ [106800]

FR682468.2_Georgia_2007/1         tgataagaaaaccatcaggtttcaacagatggtaaagcttgttaaaatcgtttatcgtaagatgatgcgccgccataggt [106880]
ASF_IND_20_CAD543                 ................................................................................ [106880]
LR899193.1_Germany_2020/1         ................................................................................ [106880]
MN393477.1_Wuhan_2019             ................................................................................ [106880]
MW396979.1_Timor-Leste/2019/1     ................................................................................ [106880]
MK940252.1_CN/2019/InnerMongolia  ................................................................................ [106880]
MK128995.1_China/2018/AnhuiXCGQ   ................................................................................ [106880]
MH681419.1_ASFV/POL/2015          ................................................................................ [106880]
LS478113.1_Estonia_2014           ................................................................................ [106880]
KP843857.1_Odintsovo_02/14        ................................................................................ [106880]
FN557520.1_strain_E75             c............................................................................... [106880]
AY261366.1_Warthog                cc.....................................................a........................ [106880]
AY261365.1_Warmbaths              cc.....................................................a........................ [106880]
AY261364.1_Tengani_62             c............................................................................... [106880]
AY261363.1_Pretoriuskop/96/4      cc.....................................................a........................ [106880]
AY261362.1_Mkuzi_1979             c................................................a.............................. [106880]
AY261361.1_Malawi_Lil-20/1_1983   ca.........................ctc...................a.....a........................ [106880]
AY261360.1_Kenya_1950             ca..........................t...........a........a.gc.......................g... [106880]
AM712240.1_OURT_88/3              c............................................................................... [106880]
AM712239.1_Benin_97/1             c............................................................................... [106880]
KM111295.1_Ken06.Bus              ca..........................t...........a........a.gc.......................g... [106880]
KM111294.1_Ken05/Tk1              ca..........................t...........a........a.gc.......................g... [106880]
KM262844.1_strain_L60             c............................................................................... [106880]
KM102979.1_26544/OG10_Italy       c............................................................................... [106880]
KM262845.1_strain_NHV             c............................................................................... [106880]
KX354450.1_47/Ss/2008             c............................................................................... [106880]
KP055815.1_strain_BA71            c............................................................................... [106880]
MH025920.1_strain_R35             ca..........................t...........a........a.gc.......................g... [106880]
MH025919.1_strain_N10             ca..........................t...........a........a.gc.......................g... [106880]
MH025918.1_strain_R25             ca..........................t...........a........a.gc.......................g... [106880]
MH025917.1_strain_R7              ca..........................t...........a........a.gc.......................g... [106880]
MH025916.1_strain_R8              ca..........................t...........a........a.gc.......................g... [106880]
```

```
FR682468.2_Georgia_2007/1          aaccctatgagctccacagagttttcatgctggacatcgtccatatcggtataaaacgtttcacagtaaatgagacgctt [106960]
ASF_IND_20_CAD543                  ................................................................................ [106960]
LR899193.1_Germany_2020/1          ................................................................................ [106960]
MN393477.1_Wuhan_2019              ................................................................................ [106960]
MW396979.1_Timor-Leste/2019/1      ................................................................................ [106960]
MK940252.1_CN/2019/InnerMongolia   ................................................................................ [106960]
MK128995.1_China/2018/AnhuiXCGQ    ................................................................................ [106960]
MH681419.1_ASFV/POL/2015           ................................................................................ [106960]
LS478113.1_Estonia_2014            ................................................................................ [106960]
KP843857.1_Odintsovo_02/14         ................................................................................ [106960]
FN557520.1_strain_E75              .................................................................a.............. [106960]
AY261366.1_Warthog                 ...............................................a........t..............c........ [106960]
AY261365.1_Warmbaths               ................................................................................ [106960]
AY261364.1_Tengani_62              ...................................g..a......................................... [106960]
AY261363.1_Pretoriuskop/96/4       .................................................................a.............. [106960]
AY261362.1_Mkuzi_1979              .g...............................................................a.............. [106960]
AY261361.1_Malawi_Lil-20/1_1983    ...................c............a.............t.........t....................... [106960]
AY261360.1_Kenya_1950              ..t....................c..c.....a.......................t.....g................. [106960]
AM712240.1_OURT_88/3               .................................................................a.............. [106960]
AM712239.1_Benin_97/1              .................................................................a.............. [106960]
KM111295.1_Ken06.Bus               ..t..............g.....c..c.....a.......................t.....g................. [106960]
KM111294.1_Ken05/Tk1               ..t....................c..c.....a.......................t.....g................. [106960]
KM262844.1_strain_L60              .................................................................a.............. [106960]
KM102979.1_26544/OG10_Italy        .................................................................a.............. [106960]
KM262845.1_strain_NHV              .................................................................a.............. [106960]
KX354450.1_47/Ss/2008              .................................................................a.............. [106960]
KP055815.1_strain_BA71             .................................................................a.............. [106960]
MH025920.1_strain_R35              ..t..............g.....c..c.....a.......................t.....g................. [106960]
MH025919.1_strain_N10              ..t..............g.....c..c.....a.......................t.....g................. [106960]
MH025918.1_strain_R25              ..t..............g.....c..c.....a.......................t.....g................. [106960]
MH025917.1_strain_R7               ..t..............g.....c..c.....a.......................t.....g................. [106960]
MH025916.1_strain_R8               ..t..............g.....c..c.....a.......................t.....g................. [106960]

FR682468.2_Georgia_2007/1          aaacgagtatcgatgacaaacatttatttccaagtaggtttgcactacgtttttaggtatatcgggaatcatgttgatta [107040]
ASF_IND_20_CAD543                  ................................................................................ [107040]
LR899193.1_Germany_2020/1          ................................................................................ [107040]
MN393477.1_Wuhan_2019              ................................................................................ [107040]
MW396979.1_Timor-Leste/2019/1      ................................................................................ [107040]
MK940252.1_CN/2019/InnerMongolia   ................................................................................ [107040]
MK128995.1_China/2018/AnhuiXCGQ    ................................................................................ [107040]
MH681419.1_ASFV/POL/2015           ................................................................................ [107040]
LS478113.1_Estonia_2014            ................................................................................ [107040]
KP843857.1_Odintsovo_02/14         ................................................................................ [107040]
```

```
FN557520.1_strain_E75             ..........t..................................c.................................. [107040]
AY261366.1_Warthog                ..........t....g......c...................a..................................... [107040]
AY261365.1_Warmbaths              ..........t....g......c.........g.........a..................................... [107040]
AY261364.1_Tengani_62             ..........t....g.................a........a..................................... [107040]
AY261363.1_Pretoriuskop/96/4      ..........t....g......c.........g.........a..................................... [107040]
AY261362.1_Mkuzi_1979             ..........t..................................c.................................. [107040]
AY261361.1_Malawi_Lil-20/1_1983   ......a...t.g.............................a..................................... [107040]
AY261360.1_Kenya_1950             ..........t...........c...................a.....a....................t.....a.... [107040]
AM712240.1_OURT_88/3              ..........t..................................c.................................. [107040]
AM712239.1_Benin_97/1             ..........t..................................c.................................. [107040]
KM111295.1_Ken06.Bus              ..........t...........c...................a..c..a..............a.....t....g..... [107040]
KM111294.1_Ken05/Tk1              ..........t...........c...................a..c..a............................... [107040]
KM262844.1_strain_L60             ..........t..................................c.................................. [107040]
KM102979.1_26544/OG10_Italy       ..........t..................................c.................................. [107040]
KM262845.1_strain_NHV             ..........t..................................c.................................. [107040]
KX354450.1_47/Ss/2008             ..........t..................................c.................................. [107040]
KP055815.1_strain_BA71            ..........t..................................c.................................. [107040]
MH025920.1_strain_R35             ..........t...........c...................a..c..a..............a.....t....g..... [107040]
MH025919.1_strain_N10             ..........t...........c...................a..c..a..............a.....t....g..... [107040]
MH025918.1_strain_R25             ..........t...........c...................a..c..a..............a.....t....g..... [107040]
MH025917.1_strain_R7              ..........t...........c...................a..c..a..............a.....t....g..... [107040]
MH025916.1_strain_R8              ..........t...........c...................a..c..a..............a.....t....g..... [107040]

FR682468.2_Georgia_2007/1         aggttgtttcgggaaacttaatcatctgactaggcttcattttcaactctttaaaggatttcccggagaagtgaaaatgg [107120]
ASF_IND_20_CAD543                 ................................................................................ [107120]
LR899193.1_Germany_2020/1         ................................................................................ [107120]
MN393477.1_Wuhan_2019             ................................................................................ [107120]
MW396979.1_Timor-Leste/2019/1     ................................................................................ [107120]
MK940252.1_CN/2019/InnerMongolia  ................................................................................ [107120]
MK128995.1_China/2018/AnhuiXCGQ   ................................................................................ [107120]
MH681419.1_ASFV/POL/2015          ................................................................................ [107120]
LS478113.1_Estonia_2014           ................................................................................ [107120]
KP843857.1_Odintsovo_02/14        ................................................................................ [107120]
FN557520.1_strain_E75             ................................................................................ [107120]
AY261366.1_Warthog                .......................g...g..c................................................. [107120]
AY261365.1_Warmbaths              .......................g...g..c................................................. [107120]
AY261364.1_Tengani_62             .......................g...g..c................................................. [107120]
AY261363.1_Pretoriuskop/96/4      .......................g...g..c................................................. [107120]
AY261362.1_Mkuzi_1979             ................................................................................ [107120]
AY261361.1_Malawi_Lil-20/1_1983   .............................................................t..a............... [107120]
AY261360.1_Kenya_1950             .......................g.....tg...t...........................................t. [107120]
AM712240.1_OURT_88/3              ................................................................................ [107120]
AM712239.1_Benin_97/1             ................................................................................ [107120]
KM111295.1_Ken06.Bus              .................g.....g.....tg...............t.................a.............ta [107120]
```

```
KM111294.1_Ken05/Tk1            ............a..........g.....tg.......................t.......................t. [107120]
KM262844.1_strain_L60           ................................................................................ [107120]
KM102979.1_26544/OG10_Italy     ................................................................................ [107120]
KM262845.1_strain_NHV           ................................................................................ [107120]
KX354450.1_47/Ss/2008           ................................................................................ [107120]
KP055815.1_strain_BA71          ................................................................................ [107120]
MH025920.1_strain_R35           .................g.....g.....tg...............t.................a.............ta [107120]
MH025919.1_strain_N10           .................g.....g.....tg...............t.................a.............ta [107120]
MH025918.1_strain_R25           .................g.....g.....tg...............t.................a.............ta [107120]
MH025917.1_strain_R7            .................g.....g.....tg...............t.................a.............ta [107120]
MH025916.1_strain_R8            .................g.....g.....tg...............t.................a.............ta [107120]

FR682468.2_Georgia_2007/1       gtctttacgtatttatgtaaaaatacctgaatgggcagagggggctcctcctcttc---gttctcgacg---cctcccaa [107200]
ASF_IND_20_CAD543               ........................................................---..........---........ [107200]
LR899193.1_Germany_2020/1       ........................................................---..........---........ [107200]
MN393477.1_Wuhan_2019           ........................................................---..........---........ [107200]
MW396979.1_Timor-Leste/2019/1   ........................................................---..........---........ [107200]
MK940252.1_CN/2019/InnerMongolia ........................................................---..........---........ [107200]
MK128995.1_China/2018/AnhuiXCGQ ........................................................---..........---........ [107200]
MH681419.1_ASFV/POL/2015        ........................................................---..........---........ [107200]
LS478113.1_Estonia_2014         ........................................................---..........---........ [107200]
KP843857.1_Odintsovo_02/14      ........................................................---..........---........ [107200]
FN557520.1_strain_E75           ...................................g....................---..c.......---........ [107200]
AY261366.1_Warthog              ........................................................---..c.......---........ [107200]
AY261365.1_Warmbaths            ........................................................---..c......a---........ [107200]
AY261364.1_Tengani_62           ........................................................---..c....g.---........ [107200]
AY261363.1_Pretoriuskop/96/4    ..................................................t.....---..c....g.---..c..... [107200]
AY261362.1_Mkuzi_1979           ........................................................ttt..c.......---........ [107200]
AY261361.1_Malawi_Lil-20/1_1983 ...........g................................t........c..ttca.c.......---........ [107200]
AY261360.1_Kenya_1950           ........................g.............g.....t...........---a.c.......cct........ [107200]
AM712240.1_OURT_88/3            ...................................g....................---..c.......---........ [107200]
AM712239.1_Benin_97/1           ...................................g....................---..c.......---........ [107200]
KM111295.1_Ken06.Bus            ........................g.............g.....t...........---..c.......cct........ [107200]
KM111294.1_Ken05/Tk1            ........................g.............g.....t...........---..c.......cct........ [107200]
KM262844.1_strain_L60           ...................................g....................---..c.......---........ [107200]
KM102979.1_26544/OG10_Italy     ...................................g....................---..c.......---........ [107200]
KM262845.1_strain_NHV           ...................................g....................---..c.......---........ [107200]
KX354450.1_47/Ss/2008           ...................................g....................---..c.......---........ [107200]
KP055815.1_strain_BA71          ...................................g....................---..c.......---........ [107200]
MH025920.1_strain_R35           ........................g.............g.....t...........---..c.......cct........ [107200]
MH025919.1_strain_N10           ........................g.............g.....t...........---..c.......cct........ [107200]
MH025918.1_strain_R25           ........................g.............g.....t...........---..c.......cct........ [107200]
MH025917.1_strain_R7            ........................g.............g.....t...........---..c.......cct........ [107200]
MH025916.1_strain_R8            ........................g.............g.....t...........---..c.......cct........ [107200]
```

```
FR682468.2_Georgia_2007/1        aatatttggaatttcctgacgtggcaaaagaaagtttatgtccacgtttacgaatccatcgaggacggacacaaagcttg [107280]
ASF_IND_20_CAD543                ................................................................................ [107280]
LR899193.1_Germany_2020/1        ................................................................................ [107280]
MN393477.1_Wuhan_2019            ................................................................................ [107280]
MW396979.1_Timor-Leste/2019/1    ................................................................................ [107280]
MK940252.1_CN/2019/InnerMongolia ................................................................................ [107280]
MK128995.1_China/2018/AnhuiXCGQ  ................................................................................ [107280]
MH681419.1_ASFV/POL/2015         ................................................................................ [107280]
LS478113.1_Estonia_2014          ................................................................................ [107280]
KP843857.1_Odintsovo_02/14       ................................................................................ [107280]
FN557520.1_strain_E75            .........................g...................................................... [107280]
AY261366.1_Warthog               ................................................................................ [107280]
AY261365.1_Warmbaths             ........................t....................................................... [107280]
AY261364.1_Tengani_62            .....................c.......................................................... [107280]
AY261363.1_Pretoriuskop/96/4     ................................................................................ [107280]
AY261362.1_Mkuzi_1979            .........................g...................................................... [107280]
AY261361.1_Malawi_Lil-20/1_1983  ............................................t................................... [107280]
AY261360.1_Kenya_1950            ..................g.............................c............................... [107280]
AM712240.1_OURT_88/3             .........................g...................................................... [107280]
AM712239.1_Benin_97/1            .........................g...................................................... [107280]
KM111295.1_Ken06.Bus             ..................g.............................c........g...................... [107280]
KM111294.1_Ken05/Tk1             ..................g.............................c............................... [107280]
KM262844.1_strain_L60            .........................g...................................................... [107280]
KM102979.1_26544/OG10_Italy      .........................g...................................................... [107280]
KM262845.1_strain_NHV            .........................g...................................................... [107280]
KX354450.1_47/Ss/2008            .........................g...................................................... [107280]
KP055815.1_strain_BA71           .........................g...................................................... [107280]
MH025920.1_strain_R35            ..................g.............................c........g...................... [107280]
MH025919.1_strain_N10            ..................g.............................c........g...................... [107280]
MH025918.1_strain_R25            ..................g.............................c........g...................... [107280]
MH025917.1_strain_R7             ..................g.............................c........g...................... [107280]
MH025916.1_strain_R8             ..................g.............................c........g...................... [107280]

FR682468.2_Georgia_2007/1        gctctaatctccattccatatactgtttagaaacgggagatagcataatcctaggcgtcacaatgcacgaagggttttta [107360]
ASF_IND_20_CAD543                ................................................................................ [107360]
LR899193.1_Germany_2020/1        ................................................................................ [107360]
MN393477.1_Wuhan_2019            ................................................................................ [107360]
MW396979.1_Timor-Leste/2019/1    ................................................................................ [107360]
MK940252.1_CN/2019/InnerMongolia ................................................................................ [107360]
MK128995.1_China/2018/AnhuiXCGQ  ................................................................................ [107360]
MH681419.1_ASFV/POL/2015         ................................................................................ [107360]
LS478113.1_Estonia_2014          ................................................................................ [107360]
KP843857.1_Odintsovo_02/14       ................................................................................ [107360]
```

```
FN557520.1_strain_E75           ..........................................................t..............a...... [107360]
AY261366.1_Warthog              .............c...................................t.............................. [107360]
AY261365.1_Warmbaths            .................................................t........t..................... [107360]
AY261364.1_Tengani_62           ................................................................................ [107360]
AY261363.1_Pretoriuskop/96/4    .............c.................................................................. [107360]
AY261362.1_Mkuzi_1979           ..........................................a...............t.....a........a...... [107360]
AY261361.1_Malawi_Lil-20/1_1983 .............c.........................cc...........g.....t..............a...... [107360]
AY261360.1_Kenya_1950           ..................................a...........c...t.......t....................g [107360]
AM712240.1_OURT_88/3            ..........................................................t..............a...... [107360]
AM712239.1_Benin_97/1           ..........................................................t..............a...... [107360]
KM111295.1_Ken06.Bus            .............c................................t...t.......t....................g [107360]
KM111294.1_Ken05/Tk1            .............c................................c...t.......t....................g [107360]
KM262844.1_strain_L60           ..........................................................t..............a...... [107360]
KM102979.1_26544/OG10_Italy     ..........................................................t..............a...... [107360]
KM262845.1_strain_NHV           ..........................................................t..............a...... [107360]
KX354450.1_47/Ss/2008           ..........................................................t..............a...... [107360]
KP055815.1_strain_BA71          ..........................................................t..............a...... [107360]
MH025920.1_strain_R35           .............c................................t...t.......t....................g [107360]
MH025919.1_strain_N10           .............c................................t...t.......t....................g [107360]
MH025918.1_strain_R25           .............c................................t...t.......t....................g [107360]
MH025917.1_strain_R7            .............c................................t...t.......t....................g [107360]
MH025916.1_strain_R8            .............c................................t...t.......t....................g [107360]

FR682468.2_Georgia_2007/1       atcaccgcatcgtggtaagaaaagtgtattccatttcttccagtataaagaagcctatgttcgtcgtagcagaaacaatt [107440]
ASF_IND_20_CAD543               ................................................................................ [107440]
LR899193.1_Germany_2020/1       ................................................................................ [107440]
MN393477.1_Wuhan_2019           ................................................................................ [107440]
MW396979.1_Timor-Leste/2019/1   ................................................................................ [107440]
MK940252.1_CN/2019/InnerMongolia ................................................................................ [107440]
MK128995.1_China/2018/AnhuiXCGQ ................................................................................ [107440]
MH681419.1_ASFV/POL/2015        ................................................................................ [107440]
LS478113.1_Estonia_2014         ................................................................................ [107440]
KP843857.1_Odintsovo_02/14      ................................................................................ [107440]
FN557520.1_strain_E75           .............................................................t.................. [107440]
AY261366.1_Warthog              ......................................g......................................... [107440]
AY261365.1_Warmbaths            ......................................g......................................... [107440]
AY261364.1_Tengani_62           ................................................................................ [107440]
AY261363.1_Pretoriuskop/96/4    ......................................g......................................... [107440]
AY261362.1_Mkuzi_1979           ................................................................................ [107440]
AY261361.1_Malawi_Lil-20/1_1983 .......t...a.................................................................... [107440]
AY261360.1_Kenya_1950           .......t...c.............................c..............t....................... [107440]
AM712240.1_OURT_88/3            .............................................................t.................. [107440]
AM712239.1_Benin_97/1           .............................................................t.................. [107440]
KM111295.1_Ken06.Bus            .......t...c.............................c..............t....................... [107440]
```

```
KM111294.1_Ken05/Tk1            .......t...c.............................c..............t....................... [107440]
KM262844.1_strain_L60           .............................................................t.................. [107440]
KM102979.1_26544/OG10_Italy     .............................................................t.................. [107440]
KM262845.1_strain_NHV           .............................................................t.................. [107440]
KX354450.1_47/Ss/2008           .............................................................t.................. [107440]
KP055815.1_strain_BA71          .............................................................t.................. [107440]
MH025920.1_strain_R35           .......t...c.............................c..............t....................... [107440]
MH025919.1_strain_N10           .......t...c.............................c..............t....................... [107440]
MH025918.1_strain_R25           .......t...c.............................c..............t....................... [107440]
MH025917.1_strain_R7            .......t...c.............................c..............t....................... [107440]
MH025916.1_strain_R8            .......t...c.............................c..............t....................... [107440]

FR682468.2_Georgia_2007/1       aaggcggtatgcctcatacatacactgtttcaaagtacaaacacgttttaaaaaggtttctgcattggcggaggccaagc [107520]
ASF_IND_20_CAD543               ................................................................................ [107520]
LR899193.1_Germany_2020/1       ................................................................................ [107520]
MN393477.1_Wuhan_2019           ................................................................................ [107520]
MW396979.1_Timor-Leste/2019/1   ................................................................................ [107520]
MK940252.1_CN/2019/InnerMongolia ................................................................................ [107520]
MK128995.1_China/2018/AnhuiXCGQ ................................................................................ [107520]
MH681419.1_ASFV/POL/2015        ................................................................................ [107520]
LS478113.1_Estonia_2014         ................................................................................ [107520]
KP843857.1_Odintsovo_02/14      ................................................................................ [107520]
FN557520.1_strain_E75           ................................................................................ [107520]
AY261366.1_Warthog              ......a..c...................................................................g.. [107520]
AY261365.1_Warmbaths            ......a..c...................................................................g.. [107520]
AY261364.1_Tengani_62           ......a..c........................a............................................. [107520]
AY261363.1_Pretoriuskop/96/4    ...a..a..c...................................................................g.. [107520]
AY261362.1_Mkuzi_1979           ................................................................................ [107520]
AY261361.1_Malawi_Lil-20/1_1983 ......a..c........a............................................g................ [107520]
AY261360.1_Kenya_1950           ......a..c........a.a.........................................................a. [107520]
AM712240.1_OURT_88/3            ................................................................................ [107520]
AM712239.1_Benin_97/1           ................................................................................ [107520]
KM111295.1_Ken06.Bus            ......a..c........a.a...................................................c.....a. [107520]
KM111294.1_Ken05/Tk1            ......a..c........a.a..............................g...........g................ [107520]
KM262844.1_strain_L60           ................................................................................ [107520]
KM102979.1_26544/OG10_Italy     ................................................................................ [107520]
KM262845.1_strain_NHV           ................................................................................ [107520]
KX354450.1_47/Ss/2008           ................................................................................ [107520]
KP055815.1_strain_BA71          ................................................................................ [107520]
MH025920.1_strain_R35           ......a..c........a.a...................................................c.....a. [107520]
MH025919.1_strain_N10           ......a..c........a.a...................................................c.....a. [107520]
MH025918.1_strain_R25           ......a..c........a.a...................................................c.....a. [107520]
MH025917.1_strain_R7            ......a..c........a.a...................................................c.....a. [107520]
MH025916.1_strain_R8            ......a..c........a.a...................................................c.....a. [107520]
```

```
FR682468.2_Georgia_2007/1        ggttttgccattggtggaaggggttcaatcctacaatggccagctcgtttaaaatatcttcgcggcgcgctaaaatctgc [107600]
ASF_IND_20_CAD543                ................................................................................ [107600]
LR899193.1_Germany_2020/1        ................................................................................ [107600]
MN393477.1_Wuhan_2019            ................................................................................ [107600]
MW396979.1_Timor-Leste/2019/1    ................................................................................ [107600]
MK940252.1_CN/2019/InnerMongolia ................................................................................ [107600]
MK128995.1_China/2018/AnhuiXCGQ  ................................................................................ [107600]
MH681419.1_ASFV/POL/2015         ................................................................................ [107600]
LS478113.1_Estonia_2014          ................................................................................ [107600]
KP843857.1_Odintsovo_02/14       ................................................................................ [107600]
FN557520.1_strain_E75            ......................................................................c......... [107600]
AY261366.1_Warthog               ..........a.................c.........................................c......... [107600]
AY261365.1_Warmbaths             ............................c...........t.............................c......... [107600]
AY261364.1_Tengani_62            .........................................................................g...... [107600]
AY261363.1_Pretoriuskop/96/4     ......................................................................c......... [107600]
AY261362.1_Mkuzi_1979            ......................................................................c......... [107600]
AY261361.1_Malawi_Lil-20/1_1983  .......................................................g..c..t..c.....c......... [107600]
AY261360.1_Kenya_1950            .........................t..........cc.....t..a..............c..c.....c......... [107600]
AM712240.1_OURT_88/3             ......................................................................c......... [107600]
AM712239.1_Benin_97/1            ......................................................................c......... [107600]
KM111295.1_Ken06.Bus             .........................t..........cc.....t..a..............c..c.....c......... [107600]
KM111294.1_Ken05/Tk1             ...................a................cc.....t..a..............c..c.....c......... [107600]
KM262844.1_strain_L60            ......................................................................c......... [107600]
KM102979.1_26544/OG10_Italy      ......................................................................c......... [107600]
KM262845.1_strain_NHV            ......................................................................c......... [107600]
KX354450.1_47/Ss/2008            ......................................................................c......... [107600]
KP055815.1_strain_BA71           ......................................................................c......... [107600]
MH025920.1_strain_R35            .........................t..........cc.....t..a..............c..c.....c......... [107600]
MH025919.1_strain_N10            .........................t..........cc.....t..a..............c..c....tc......... [107600]
MH025918.1_strain_R25            .........................t..........cc.....t..a..............c..c.....c......... [107600]
MH025917.1_strain_R7             .........................t..........cc.....t..a..............c..c.....c......... [107600]
MH025916.1_strain_R8             .........................t..........cc.....t..a..............c..c.....c......... [107600]

FR682468.2_Georgia_2007/1        accatagaagaatactttagcatttttttttcgcaccattcgcgaagatgtttagctacattattaaccttattattgat [107680]
ASF_IND_20_CAD543                ..............t................................................................. [107680]
LR899193.1_Germany_2020/1        ................................................................................ [107680]
MN393477.1_Wuhan_2019            ................................................................................ [107680]
MW396979.1_Timor-Leste/2019/1    ................................................................................ [107680]
MK940252.1_CN/2019/InnerMongolia ................................................................................ [107680]
MK128995.1_China/2018/AnhuiXCGQ  ................................................................................ [107680]
MH681419.1_ASFV/POL/2015         ................................................................................ [107680]
LS478113.1_Estonia_2014          ................................................................................ [107680]
KP843857.1_Odintsovo_02/14       ................................................................................ [107680]
```

```
FN557520.1_strain_E75             ..............t................................................................. [107680]
AY261366.1_Warthog                ..............t.........................................c....................... [107680]
AY261365.1_Warmbaths              ..............t................................................................. [107680]
AY261364.1_Tengani_62             ..............t................................................................. [107680]
AY261363.1_Pretoriuskop/96/4      ...............................................g................................ [107680]
AY261362.1_Mkuzi_1979             ..............t................................................................. [107680]
AY261361.1_Malawi_Lil-20/1_1983   ..........................................................a..................... [107680]
AY261360.1_Kenya_1950             .......c.......................................g..........a..................... [107680]
AM712240.1_OURT_88/3              ..............t................................................................. [107680]
AM712239.1_Benin_97/1             ..............t................................................................. [107680]
KM111295.1_Ken06.Bus              .......c.......................................g..........a..................... [107680]
KM111294.1_Ken05/Tk1              .......c.......................................g..........a..................... [107680]
KM262844.1_strain_L60             ..............t................................................................. [107680]
KM102979.1_26544/OG10_Italy       ..............t................................................................. [107680]
KM262845.1_strain_NHV             ..............t................................................................. [107680]
KX354450.1_47/Ss/2008             ..............t................................................................. [107680]
KP055815.1_strain_BA71            ..............t................................................................. [107680]
MH025920.1_strain_R35             .......c.......................................g..........a..................... [107680]
MH025919.1_strain_N10             .......c.......................................g..........a..................... [107680]
MH025918.1_strain_R25             .......c.......................................g..........a..................... [107680]
MH025917.1_strain_R7              .......c.......................................g..........a..................... [107680]
MH025916.1_strain_R8              .......c.......................................g..........a..................... [107680]

FR682468.2_Georgia_2007/1         aaagtatacgatggcatgttggaagccttcaaaaataaagagcccctccaaaagatcatctgccaatagaagatggatgt [107760]
ASF_IND_20_CAD543                 ................................................................................ [107760]
LR899193.1_Germany_2020/1         ................................................................................ [107760]
MN393477.1_Wuhan_2019             ................................................................................ [107760]
MW396979.1_Timor-Leste/2019/1     ................................................................................ [107760]
MK940252.1_CN/2019/InnerMongolia  ................................................................................ [107760]
MK128995.1_China/2018/AnhuiXCGQ   ................................................................................ [107760]
MH681419.1_ASFV/POL/2015          ................................................................................ [107760]
LS478113.1_Estonia_2014           ................................................................................ [107760]
KP843857.1_Odintsovo_02/14        ................................................................................ [107760]
FN557520.1_strain_E75             ................................................................................ [107760]
AY261366.1_Warthog                ........................................................................g..a.... [107760]
AY261365.1_Warmbaths              ........................................................................g....... [107760]
AY261364.1_Tengani_62             ...a................................................................a...g....... [107760]
AY261363.1_Pretoriuskop/96/4      ........................................................................g....... [107760]
AY261362.1_Mkuzi_1979             ....................................................................a........... [107760]
AY261361.1_Malawi_Lil-20/1_1983   ..........................................t.......................a.....g....... [107760]
AY261360.1_Kenya_1950             ........t..................c...........t..........................a.....g....... [107760]
AM712240.1_OURT_88/3              ................................................................................ [107760]
AM712239.1_Benin_97/1             ................................................................................ [107760]
KM111295.1_Ken06.Bus              ........t..................c...........t..........................a.....g....... [107760]
```

```
KM111294.1_Ken05/Tk1          ........t..................c...........t..........................a............. [107760]
KM262844.1_strain_L60         ................................................................................ [107760]
KM102979.1_26544/OG10_Italy   ................................................................................ [107760]
KM262845.1_strain_NHV         ................................................................................ [107760]
KX354450.1_47/Ss/2008         ................................................................................ [107760]
KP055815.1_strain_BA71        ................................................................................ [107760]
MH025920.1_strain_R35         ........t..................c...........t..........................a.....g....... [107760]
MH025919.1_strain_N10         ........t..................c...........t..........................a.....g....... [107760]
MH025918.1_strain_R25         ........t..................c...........t..........................a.....g....... [107760]
MH025917.1_strain_R7          ........t..................c...........t..........................a.....g....... [107760]
MH025916.1_strain_R8          ........t..................c...........t..........................a.....g....... [107760]

FR682468.2_Georgia_2007/1     tggtgtaagcattgtcaatattttgtagaaacggcggaatgcctgccaaaaccgcttcagcaagcatagctccgttccgt [107840]
ASF_IND_20_CAD543             ................................................................................ [107840]
LR899193.1_Germany_2020/1     ................................................................................ [107840]
MN393477.1_Wuhan_2019         ................................................................................ [107840]
MW396979.1_Timor-Leste/2019/1 ................................................................................ [107840]
MK940252.1_CN/2019/InnerMongolia ................................................................................ [107840]
MK128995.1_China/2018/AnhuiXCGQ ................................................................................ [107840]
MH681419.1_ASFV/POL/2015      ................................................................................ [107840]
LS478113.1_Estonia_2014       ................................................................................ [107840]
KP843857.1_Odintsovo_02/14    ................................................................................ [107840]
FN557520.1_strain_E75         ................................................................................ [107840]
AY261366.1_Warthog            ....a....................................................................a...... [107840]
AY261365.1_Warmbaths          ......................................................................c..a...... [107840]
AY261364.1_Tengani_62         .........................................................................a...... [107840]
AY261363.1_Pretoriuskop/96/4  .........................................................................a...... [107840]
AY261362.1_Mkuzi_1979         ................................................................................ [107840]
AY261361.1_Malawi_Lil-20/1_1983 ....a................g.....a.......................................c.....a...... [107840]
AY261360.1_Kenya_1950         ..c.a...........c..........a.............................................a.....c [107840]
AM712240.1_OURT_88/3          ................................................................................ [107840]
AM712239.1_Benin_97/1         ................................................................................ [107840]
KM111295.1_Ken06.Bus          ..c.a...........c..........a.............................................a.....c [107840]
KM111294.1_Ken05/Tk1          ..c.a...........t..........a.............................................a.....c [107840]
KM262844.1_strain_L60         ................................................................................ [107840]
KM102979.1_26544/OG10_Italy   ................................................................................ [107840]
KM262845.1_strain_NHV         ................................................................................ [107840]
KX354450.1_47/Ss/2008         ................................................................................ [107840]
KP055815.1_strain_BA71        ................................................................................ [107840]
MH025920.1_strain_R35         ..c.a...........c..........a.............................................a.....c [107840]
MH025919.1_strain_N10         ..c.a...........c..........a.............................................a.....c [107840]
MH025918.1_strain_R25         ..c.a...........c..........a.............................................a.....c [107840]
MH025917.1_strain_R7          ..c.a...........c..........a.............................................a.....c [107840]
MH025916.1_strain_R8          ..c.a...........c..........a.............................................a.....c [107840]
```

```
FR682468.2_Georgia_2007/1        tgtttactgtccaatagattcgtaagttttttgtccgcaacagacacgacggctaggatggttgcaatgtcagaaatggc [107920]
ASF_IND_20_CAD543                ................................................................................ [107920]
LR899193.1_Germany_2020/1        ................................................................................ [107920]
MN393477.1_Wuhan_2019            ................................................................................ [107920]
MW396979.1_Timor-Leste/2019/1    ................................................................................ [107920]
MK940252.1_CN/2019/InnerMongolia ................................................................................ [107920]
MK128995.1_China/2018/AnhuiXCGQ  ................................................................................ [107920]
MH681419.1_ASFV/POL/2015         ................................................................................ [107920]
LS478113.1_Estonia_2014          ................................................................................ [107920]
KP843857.1_Odintsovo_02/14       ................................................................................ [107920]
FN557520.1_strain_E75            ................................................................................ [107920]
AY261366.1_Warthog               ................................................................................ [107920]
AY261365.1_Warmbaths             ................................................................................ [107920]
AY261364.1_Tengani_62            ................................................................................ [107920]
AY261363.1_Pretoriuskop/96/4     ................................................................................ [107920]
AY261362.1_Mkuzi_1979            ...........................................................a.................... [107920]
AY261361.1_Malawi_Lil-20/1_1983  ................................a..ta.......t........c.a....a................... [107920]
AY261360.1_Kenya_1950            ...................................ta..........a.....c..a....................... [107920]
AM712240.1_OURT_88/3             ................................................................................ [107920]
AM712239.1_Benin_97/1            ................................................................................ [107920]
KM111295.1_Ken06.Bus             ...................................ta..........a................................ [107920]
KM111294.1_Ken05/Tk1             ...................................ta..........a.....c.......................... [107920]
KM262844.1_strain_L60            ................................................................................ [107920]
KM102979.1_26544/OG10_Italy      ................................................................................ [107920]
KM262845.1_strain_NHV            ................................................................................ [107920]
KX354450.1_47/Ss/2008            ................................................................................ [107920]
KP055815.1_strain_BA71           ................................................................................ [107920]
MH025920.1_strain_R35            ...................................ta..........a................................ [107920]
MH025919.1_strain_N10            ...................................ta..........a................................ [107920]
MH025918.1_strain_R25            ...................................ta..........a................................ [107920]
MH025917.1_strain_R7             ...................................ta..........a................................ [107920]
MH025916.1_strain_R8             ...................................ta..........a................................ [107920]

FR682468.2_Georgia_2007/1        ggcttgccagaaataacccgaaaagcacatgcgcgcttcttctatagataaaaacgaaaagcgagaggcaatgtctccga [108000]
ASF_IND_20_CAD543                ................................................................................ [108000]
LR899193.1_Germany_2020/1        ................................................................................ [108000]
MN393477.1_Wuhan_2019            ................................................................................ [108000]
MW396979.1_Timor-Leste/2019/1    ................................................................................ [108000]
MK940252.1_CN/2019/InnerMongolia ................................................................................ [108000]
MK128995.1_China/2018/AnhuiXCGQ  ................................................................................ [108000]
MH681419.1_ASFV/POL/2015         ................................................................................ [108000]
LS478113.1_Estonia_2014          ................................................................................ [108000]
KP843857.1_Odintsovo_02/14       ................................................................................ [108000]
```

```
FN557520.1_strain_E75             ................................................................................ [108000]
AY261366.1_Warthog                ...............c........a....................................................... [108000]
AY261365.1_Warmbaths              ...............c........a....................................................... [108000]
AY261364.1_Tengani_62             ........................a....................................................... [108000]
AY261363.1_Pretoriuskop/96/4      ...............c........a....................................................... [108000]
AY261362.1_Mkuzi_1979             ................................................................................ [108000]
AY261361.1_Malawi_Lil-20/1_1983   a.......................a............................................g........a. [108000]
AY261360.1_Kenya_1950             ........................a....................g.......................g.......... [108000]
AM712240.1_OURT_88/3              ................................................................................ [108000]
AM712239.1_Benin_97/1             ................................................................................ [108000]
KM111295.1_Ken06.Bus              ........................a............................................g.......... [108000]
KM111294.1_Ken05/Tk1              ........................a....................g.......................g.......... [108000]
KM262844.1_strain_L60             ................................................................................ [108000]
KM102979.1_26544/OG10_Italy       ................................................................................ [108000]
KM262845.1_strain_NHV             ................................................................................ [108000]
KX354450.1_47/Ss/2008             ................................................................................ [108000]
KP055815.1_strain_BA71            ................................................................................ [108000]
MH025920.1_strain_R35             ........................a............................................g.......... [108000]
MH025919.1_strain_N10             ........................a............................................g.......... [108000]
MH025918.1_strain_R25             ........................a............................................g.......... [108000]
MH025917.1_strain_R7              ........................a............................................g.......... [108000]
MH025916.1_strain_R8              ........................a............................................g.......... [108000]

FR682468.2_Georgia_2007/1         gctgcgtgagttgaagacctttttctcctctggttaaaaggcctgccacaatggcccgctcaatggctgatgccagcgca [108080]
ASF_IND_20_CAD543                 ................................................................................ [108080]
LR899193.1_Germany_2020/1         ................................................................................ [108080]
MN393477.1_Wuhan_2019             ................................................................................ [108080]
MW396979.1_Timor-Leste/2019/1     ................................................................................ [108080]
MK940252.1_CN/2019/InnerMongolia  ................................................................................ [108080]
MK128995.1_China/2018/AnhuiXCGQ   ................................................................................ [108080]
MH681419.1_ASFV/POL/2015          ................................................................................ [108080]
LS478113.1_Estonia_2014           ................................................................................ [108080]
KP843857.1_Odintsovo_02/14        ................................................................................ [108080]
FN557520.1_strain_E75             .......................................................t........................ [108080]
AY261366.1_Warthog                ................................................................................ [108080]
AY261365.1_Warmbaths              ................................................................................ [108080]
AY261364.1_Tengani_62             ................................................................................ [108080]
AY261363.1_Pretoriuskop/96/4      ................................................................................ [108080]
AY261362.1_Mkuzi_1979             ................................................................................ [108080]
AY261361.1_Malawi_Lil-20/1_1983   ................g........c..................c..........g..t........c.......a.... [108080]
AY261360.1_Kenya_1950             ............................................c..........g.......g...c............ [108080]
AM712240.1_OURT_88/3              .......................................................t........................ [108080]
AM712239.1_Benin_97/1             .......................................................t........................ [108080]
KM111295.1_Ken06.Bus              ............................................c..........g.......g...c............ [108080]
```

```
KM111294.1_Ken05/Tk1            ............................................c..........g.......g...c............ [108080]
KM262844.1_strain_L60           .......................................................t........................ [108080]
KM102979.1_26544/OG10_Italy     .......................................................t........................ [108080]
KM262845.1_strain_NHV           .......................................................t........................ [108080]
KX354450.1_47/Ss/2008           .......................................................t........................ [108080]
KP055815.1_strain_BA71          .......................................................t........................ [108080]
MH025920.1_strain_R35           ............................................c..........g.......g...c............ [108080]
MH025919.1_strain_N10           ............................................c..........g.......g...c............ [108080]
MH025918.1_strain_R25           ............................................c..........g.......g...c............ [108080]
MH025917.1_strain_R7            ............................................c..........g.......g...c............ [108080]
MH025916.1_strain_R8            ............................................c..........g.......g...c............ [108080]

FR682468.2_Georgia_2007/1       tccgtggggggaggatccagcatatcaatctcctctgccttaaacacgccttccttattttttttttaatcgtttctacgac [108160]
ASF_IND_20_CAD543               ................................................................................ [108160]
LR899193.1_Germany_2020/1       ................................................................................ [108160]
MN393477.1_Wuhan_2019           ................................................................................ [108160]
MW396979.1_Timor-Leste/2019/1   ................................................................................ [108160]
MK940252.1_CN/2019/InnerMongolia ................................................................................ [108160]
MK128995.1_China/2018/AnhuiXCGQ ................................................................................ [108160]
MH681419.1_ASFV/POL/2015        ................................................................................ [108160]
LS478113.1_Estonia_2014         ................................................................................ [108160]
KP843857.1_Odintsovo_02/14      ................................................................................ [108160]
FN557520.1_strain_E75           ................................................................................ [108160]
AY261366.1_Warthog              ................................................................................ [108160]
AY261365.1_Warmbaths            ................................................................................ [108160]
AY261364.1_Tengani_62           ......................................t......................................... [108160]
AY261363.1_Pretoriuskop/96/4    ................................................................................ [108160]
AY261362.1_Mkuzi_1979           ................................................................................ [108160]
AY261361.1_Malawi_Lil-20/1_1983 ..t..............t.a.........................................................a.. [108160]
AY261360.1_Kenya_1950           ..................................g.a.....................................c..... [108160]
AM712240.1_OURT_88/3            ................................................................................ [108160]
AM712239.1_Benin_97/1           ................................................................................ [108160]
KM111295.1_Ken06.Bus            ....................................a.....................................c..... [108160]
KM111294.1_Ken05/Tk1            ...........g......................g.a.....................................c..... [108160]
KM262844.1_strain_L60           ................................................................................ [108160]
KM102979.1_26544/OG10_Italy     ................................................................................ [108160]
KM262845.1_strain_NHV           ................................................................................ [108160]
KX354450.1_47/Ss/2008           ................................................................................ [108160]
KP055815.1_strain_BA71          ................................................................................ [108160]
MH025920.1_strain_R35           ....................................a.....................................c..... [108160]
MH025919.1_strain_N10           ....................................a.....................................c..... [108160]
MH025918.1_strain_R25           ....................................a.....................................c..... [108160]
MH025917.1_strain_R7            ....................................a.....................................c..... [108160]
MH025916.1_strain_R8            ....................................a.....................................c..... [108160]
```

```
FR682468.2_Georgia_2007/1          aatgctaagaaaaatggccccagggccttccgtaatgatttcaggatactgctgcactggtatttgctcaaagacgtgtt [108240]
ASF_IND_20_CAD543                  ................................................................................ [108240]
LR899193.1_Germany_2020/1          ................................................................................ [108240]
MN393477.1_Wuhan_2019              ................................................................................ [108240]
MW396979.1_Timor-Leste/2019/1      ................................................................................ [108240]
MK940252.1_CN/2019/InnerMongolia   ................................................................................ [108240]
MK128995.1_China/2018/AnhuiXCGQ    ................................................................................ [108240]
MH681419.1_ASFV/POL/2015           ................................................................................ [108240]
LS478113.1_Estonia_2014            ................................................................................ [108240]
KP843857.1_Odintsovo_02/14         ................................................................................ [108240]
FN557520.1_strain_E75              .......................................................g........................ [108240]
AY261366.1_Warthog                 .......................................................g........................ [108240]
AY261365.1_Warmbaths               .......................................................g........................ [108240]
AY261364.1_Tengani_62              ................................................................................ [108240]
AY261363.1_Pretoriuskop/96/4       .......................................................g........................ [108240]
AY261362.1_Mkuzi_1979              .......................................................g........................ [108240]
AY261361.1_Malawi_Lil-20/1_1983    ....t.............t................................t..tg.c...................... [108240]
AY261360.1_Kenya_1950              ....t..............................................t...g.c...................... [108240]
AM712240.1_OURT_88/3               .......................................................g........................ [108240]
AM712239.1_Benin_97/1              .......................................................g........................ [108240]
KM111295.1_Ken06.Bus               ....t..............................................t...g.c...................... [108240]
KM111294.1_Ken05/Tk1               ....t..............................................t...g.c...................... [108240]
KM262844.1_strain_L60              .......................................................g........................ [108240]
KM102979.1_26544/OG10_Italy        .......................................................g........................ [108240]
KM262845.1_strain_NHV              .......................................................g........................ [108240]
KX354450.1_47/Ss/2008              .......................................................g........................ [108240]
KP055815.1_strain_BA71             .......................................................g........................ [108240]
MH025920.1_strain_R35              ....t..............................................t...g.c...................... [108240]
MH025919.1_strain_N10              ....t..............................................t...g.c...................... [108240]
MH025918.1_strain_R25              ....t..............................................t...g.c...................... [108240]
MH025917.1_strain_R7               ....t..............................................t...g.c...................... [108240]
MH025916.1_strain_R8               ....t..............................................t...g.c...................... [108240]

FR682468.2_Georgia_2007/1          ttgtgtaaagcgggtaaaagtgcccaggaaatactctccctacacgcccctttctttgctcgatacggctttgagccgcg [108320]
ASF_IND_20_CAD543                  ................................................................................ [108320]
LR899193.1_Germany_2020/1          ................................................................................ [108320]
MN393477.1_Wuhan_2019              ................................................................................ [108320]
MW396979.1_Timor-Leste/2019/1      ................................................................................ [108320]
MK940252.1_CN/2019/InnerMongolia   ................................................................................ [108320]
MK128995.1_China/2018/AnhuiXCGQ    ................................................................................ [108320]
MH681419.1_ASFV/POL/2015           ................................................................................ [108320]
LS478113.1_Estonia_2014            ................................................................................ [108320]
KP843857.1_Odintsovo_02/14         ................................................................................ [108320]
```

```
FN557520.1_strain_E75            ...........................................g..t................................. [108320]
AY261366.1_Warthog               ...........................................g..t................................. [108320]
AY261365.1_Warmbaths             ..........t................................g..t................................. [108320]
AY261364.1_Tengani_62            ...................................................................a............ [108320]
AY261363.1_Pretoriuskop/96/4     ...........................................g..t................................. [108320]
AY261362.1_Mkuzi_1979            ...........................................g..t................................. [108320]
AY261361.1_Malawi_Lil-20/1_1983  ....a......................................g..t....................a...........a [108320]
AY261360.1_Kenya_1950            ....a......................................g..t................................. [108320]
AM712240.1_OURT_88/3             ...........................................g..t................................. [108320]
AM712239.1_Benin_97/1            ...........................................g..t................................. [108320]
KM111295.1_Ken06.Bus             ....a......................................g..t..............a.................. [108320]
KM111294.1_Ken05/Tk1             ....a......................................g..t................................. [108320]
KM262844.1_strain_L60            ...........................................g..t................................. [108320]
KM102979.1_26544/OG10_Italy      ...........................................g..t................................. [108320]
KM262845.1_strain_NHV            ...........................................g..t................................. [108320]
KX354450.1_47/Ss/2008            ...........................................g..t................................. [108320]
KP055815.1_strain_BA71           ...........................................g..t................................. [108320]
MH025920.1_strain_R35            ....a......................................g..t..............a.................. [108320]
MH025919.1_strain_N10            ....a......................................g..t..............a.................. [108320]
MH025918.1_strain_R25            ....a......................................g..t..............a.................. [108320]
MH025917.1_strain_R7             ....a......................................g..t..............a.................. [108320]
MH025916.1_strain_R8             ....a......................................g..t..............a.................. [108320]

FR682468.2_Georgia_2007/1        gggcgcgtaataagccctcccgcccattcgggatagtaggtttcaatgcttctgttccacccgggatctatgacgtactt [108400]
ASF_IND_20_CAD543                .............................a..g............................................... [108400]
LR899193.1_Germany_2020/1        ................................................................................ [108400]
MN393477.1_Wuhan_2019            ................................................................................ [108400]
MW396979.1_Timor-Leste/2019/1    ................................................................................ [108400]
MK940252.1_CN/2019/InnerMongolia ................................................................................ [108400]
MK128995.1_China/2018/AnhuiXCGQ  ................................................................................ [108400]
MH681419.1_ASFV/POL/2015         ................................................................................ [108400]
LS478113.1_Estonia_2014          ................................................................................ [108400]
KP843857.1_Odintsovo_02/14       ................................................................................ [108400]
FN557520.1_strain_E75            ...........g.................a..g............................................... [108400]
AY261366.1_Warthog               ...........g...............................................g.................... [108400]
AY261365.1_Warmbaths             ...........g..................................c................................. [108400]
AY261364.1_Tengani_62            ...........g....................g..........................g.................... [108400]
AY261363.1_Pretoriuskop/96/4     ...........g.................................................................... [108400]
AY261362.1_Mkuzi_1979            ...........g.................a..g..........................t.................... [108400]
AY261361.1_Malawi_Lil-20/1_1983  ...........c..t..............a................c...............a........a........ [108400]
AY261360.1_Kenya_1950            ...........g.................a................c...............a................. [108400]
AM712240.1_OURT_88/3             ...........g.................a..g............................................... [108400]
AM712239.1_Benin_97/1            ...........g.................a..g............................................... [108400]
KM111295.1_Ken06.Bus             ...........g..t..............a................c...............a................. [108400]
```

```
KM111294.1_Ken05/Tk1             ...........g.................a................c...............a................. [108400]
KM262844.1_strain_L60            ...........g.................a..g............................................... [108400]
KM102979.1_26544/OG10_Italy      ...........g.................a..g............................................... [108400]
KM262845.1_strain_NHV            ...........g.................a..g............................................... [108400]
KX354450.1_47/Ss/2008            ...........g.................a..g............................................... [108400]
KP055815.1_strain_BA71           ...........g.................a..g............................................... [108400]
MH025920.1_strain_R35            ...........g..t..............a................c...............a................. [108400]
MH025919.1_strain_N10            ...........g..t..............a................c...............a................. [108400]
MH025918.1_strain_R25            ...........g..t..............a................c...............a................. [108400]
MH025917.1_strain_R7             ...........g..t..............a................c...............a................. [108400]
MH025916.1_strain_R8             ...........g..t..............a................c...............a................. [108400]

FR682468.2_Georgia_2007/1        cagcgtttcaatggtaaggcccgtttccgcaacaaccgtggaaacaatgacccttcttaaaggtttttccactttagcgg [108480]
ASF_IND_20_CAD543                ................................................................................ [108480]
LR899193.1_Germany_2020/1        ................................................................................ [108480]
MN393477.1_Wuhan_2019            ................................................................................ [108480]
MW396979.1_Timor-Leste/2019/1    ................................................................................ [108480]
MK940252.1_CN/2019/InnerMongolia ................................................................................ [108480]
MK128995.1_China/2018/AnhuiXCGQ  ................................................................................ [108480]
MH681419.1_ASFV/POL/2015         ................................................................................ [108480]
LS478113.1_Estonia_2014          ................................................................................ [108480]
KP843857.1_Odintsovo_02/14       ................................................................................ [108480]
FN557520.1_strain_E75            .................................g.............................................. [108480]
AY261366.1_Warthog               ...............................................................c................ [108480]
AY261365.1_Warmbaths             ...............................................................c................ [108480]
AY261364.1_Tengani_62            .............................................g.................................. [108480]
AY261363.1_Pretoriuskop/96/4     ................................................a............................... [108480]
AY261362.1_Mkuzi_1979            ..................a............................................................. [108480]
AY261361.1_Malawi_Lil-20/1_1983  ...t........a..........................a........a..t....................c....... [108480]
AY261360.1_Kenya_1950            ................................................a............................... [108480]
AM712240.1_OURT_88/3             ................................................................................ [108480]
AM712239.1_Benin_97/1            ................................................................................ [108480]
KM111295.1_Ken06.Bus             ................................................a............................... [108480]
KM111294.1_Ken05/Tk1             ................................................a............................... [108480]
KM262844.1_strain_L60            ................................................................................ [108480]
KM102979.1_26544/OG10_Italy      ................................................................................ [108480]
KM262845.1_strain_NHV            ................................................................................ [108480]
KX354450.1_47/Ss/2008            ................................................................................ [108480]
KP055815.1_strain_BA71           ................................................................................ [108480]
MH025920.1_strain_R35            ................................................a............................... [108480]
MH025919.1_strain_N10            ................................................a............................... [108480]
MH025918.1_strain_R25            ................................................a............................... [108480]
MH025917.1_strain_R7             ................................................a............................... [108480]
MH025916.1_strain_R8             ................................................a............................... [108480]
```

```
FR682468.2_Georgia_2007/1        ttaagggatttttcacccacagattcttaatttccgctttcaggccaaggtaggcctcattttcctgcgcaatcgcctca [108560]
ASF_IND_20_CAD543                ................................................................................ [108560]
LR899193.1_Germany_2020/1        ................................................................................ [108560]
MN393477.1_Wuhan_2019            ................................................................................ [108560]
MW396979.1_Timor-Leste/2019/1    ................................................................................ [108560]
MK940252.1_CN/2019/InnerMongolia ................................................................................ [108560]
MK128995.1_China/2018/AnhuiXCGQ  ................................................................................ [108560]
MH681419.1_ASFV/POL/2015         ................................................................................ [108560]
LS478113.1_Estonia_2014          ................................................................................ [108560]
KP843857.1_Odintsovo_02/14       ................................................................................ [108560]
FN557520.1_strain_E75            ................................................................................ [108560]
AY261366.1_Warthog               ...................a....................t....................................... [108560]
AY261365.1_Warmbaths             ...................a....................t....................................... [108560]
AY261364.1_Tengani_62            ...................a....................t....................................... [108560]
AY261363.1_Pretoriuskop/96/4     ...................a....................t....................................... [108560]
AY261362.1_Mkuzi_1979            ................................................................................ [108560]
AY261361.1_Malawi_Lil-20/1_1983  ...................a....................t......................................g [108560]
AY261360.1_Kenya_1950            .......g...........a....c...............t......................................g [108560]
AM712240.1_OURT_88/3             ................................................................................ [108560]
AM712239.1_Benin_97/1            ................................................................................ [108560]
KM111295.1_Ken06.Bus             .......g...........a....c...............t........a.............................g [108560]
KM111294.1_Ken05/Tk1             .c.....g...........a....c...............t......................................g [108560]
KM262844.1_strain_L60            ................................................................................ [108560]
KM102979.1_26544/OG10_Italy      ................................................................................ [108560]
KM262845.1_strain_NHV            ................................................................................ [108560]
KX354450.1_47/Ss/2008            ................................................................................ [108560]
KP055815.1_strain_BA71           ................................................................................ [108560]
MH025920.1_strain_R35            .......g...........a....c...............t........a.............................g [108560]
MH025919.1_strain_N10            .......g...........a....c...............t........a.............................g [108560]
MH025918.1_strain_R25            .......g...........a....c...............t........a.............................g [108560]
MH025917.1_strain_R7             .......g...........a....c...............t........a.............................g [108560]
MH025916.1_strain_R8             .......g...........a....c...............t........a.............................g [108560]

FR682468.2_Georgia_2007/1        ctatcgatcggcaaaatcaacattaacggcagcttttctttggcaaggtccatatttgcattattcagcaacatcgaaag [108640]
ASF_IND_20_CAD543                ................................................................................ [108640]
LR899193.1_Germany_2020/1        ................................................................................ [108640]
MN393477.1_Wuhan_2019            ................................................................................ [108640]
MW396979.1_Timor-Leste/2019/1    ................................................................................ [108640]
MK940252.1_CN/2019/InnerMongolia ................................................................................ [108640]
MK128995.1_China/2018/AnhuiXCGQ  ................................................................................ [108640]
MH681419.1_ASFV/POL/2015         ................................................................................ [108640]
LS478113.1_Estonia_2014          ................................................................................ [108640]
KP843857.1_Odintsovo_02/14       ................................................................................ [108640]
```

```
FN557520.1_strain_E75               ................................................................................ [108640]
AY261366.1_Warthog                  .....................................................g.......................... [108640]
AY261365.1_Warmbaths                .....................................................g.............a............ [108640]
AY261364.1_Tengani_62               ................................................................................ [108640]
AY261363.1_Pretoriuskop/96/4        ...................................................................a............ [108640]
AY261362.1_Mkuzi_1979               ................................................................................ [108640]
AY261361.1_Malawi_Lil-20/1_1983     ..............................................................g.........g....... [108640]
AY261360.1_Kenya_1950               ...................g...........................a.............g.c................ [108640]
AM712240.1_OURT_88/3                ................................................................................ [108640]
AM712239.1_Benin_97/1               ................................................................................ [108640]
KM111295.1_Ken06.Bus                ...................g...........................a.............g.c................ [108640]
KM111294.1_Ken05/Tk1                ...................g...........................a.............g.c................ [108640]
KM262844.1_strain_L60               ................................................................................ [108640]
KM102979.1_26544/OG10_Italy         ................................................................................ [108640]
KM262845.1_strain_NHV               ................................................................................ [108640]
KX354450.1_47/Ss/2008               ................................................................................ [108640]
KP055815.1_strain_BA71              ................................................................................ [108640]
MH025920.1_strain_R35               ...................g...........................a.............g.c................ [108640]
MH025919.1_strain_N10               ...................g...........................a.............g.c................ [108640]
MH025918.1_strain_R25               ...................g...........................a.............g.c................ [108640]
MH025917.1_strain_R7                ...................g...........................a.............g.c................ [108640]
MH025916.1_strain_R8                ...................g...........................a.............g.c................ [108640]

FR682468.2_Georgia_2007/1           gaagcgtatttcagccataccgggcatgaaaattaaaatatctgcttccgtgggacgatcatgaatgttttctttatgaa [108720]
ASF_IND_20_CAD543                   ................................................................................ [108720]
LR899193.1_Germany_2020/1           ................................................................................ [108720]
MN393477.1_Wuhan_2019               ................................................................................ [108720]
MW396979.1_Timor-Leste/2019/1       ................................................................................ [108720]
MK940252.1_CN/2019/InnerMongolia    ................................................................................ [108720]
MK128995.1_China/2018/AnhuiXCGQ     ................................................................................ [108720]
MH681419.1_ASFV/POL/2015            ................................................................................ [108720]
LS478113.1_Estonia_2014             ................................................................................ [108720]
KP843857.1_Odintsovo_02/14          ................................................................................ [108720]
FN557520.1_strain_E75               ................................................................................ [108720]
AY261366.1_Warthog                  ................................................................................ [108720]
AY261365.1_Warmbaths                ................................................................................ [108720]
AY261364.1_Tengani_62               ................................................................................ [108720]
AY261363.1_Pretoriuskop/96/4        ................................................................................ [108720]
AY261362.1_Mkuzi_1979               ................................................................................ [108720]
AY261361.1_Malawi_Lil-20/1_1983     .............c.................................................................. [108720]
AY261360.1_Kenya_1950               .............c..............................................g................... [108720]
AM712240.1_OURT_88/3                ................................................................................ [108720]
AM712239.1_Benin_97/1               ................................................................................ [108720]
KM111295.1_Ken06.Bus                ............................................................g................... [108720]
```

```
KM111294.1_Ken05/Tk1             .............c..............................................g................... [108720]
KM262844.1_strain_L60            ................................................................................ [108720]
KM102979.1_26544/OG10_Italy      ................................................................................ [108720]
KM262845.1_strain_NHV            ................................................................................ [108720]
KX354450.1_47/Ss/2008            ................................................................................ [108720]
KP055815.1_strain_BA71           ................................................................................ [108720]
MH025920.1_strain_R35            ............................................................g................... [108720]
MH025919.1_strain_N10            ............................................................g................... [108720]
MH025918.1_strain_R25            ............................................................g................... [108720]
MH025917.1_strain_R7             ............................................................g................... [108720]
MH025916.1_strain_R8             ............................................................g................... [108720]

FR682468.2_Georgia_2007/1        tagtgagagccgtttcgcaggcggtcttaatgtagttgttggtgttatacagcggccagtgggtttccacaccgtactgt [108800]
ASF_IND_20_CAD543                ................................................................................ [108800]
LR899193.1_Germany_2020/1        ................................................................................ [108800]
MN393477.1_Wuhan_2019            ................................................................................ [108800]
MW396979.1_Timor-Leste/2019/1    ................................................................................ [108800]
MK940252.1_CN/2019/InnerMongolia ................................................................................ [108800]
MK128995.1_China/2018/AnhuiXCGQ  ................................................................................ [108800]
MH681419.1_ASFV/POL/2015         ................................................................................ [108800]
LS478113.1_Estonia_2014          ................................................................................ [108800]
KP843857.1_Odintsovo_02/14       ................................................................................ [108800]
FN557520.1_strain_E75            ................................................................................ [108800]
AY261366.1_Warthog               ................................................................................ [108800]
AY261365.1_Warmbaths             ..............................c................................................. [108800]
AY261364.1_Tengani_62            ................................................................................ [108800]
AY261363.1_Pretoriuskop/96/4     ................................................................................ [108800]
AY261362.1_Mkuzi_1979            ................................................................................ [108800]
AY261361.1_Malawi_Lil-20/1_1983  .c........................................................................a..... [108800]
AY261360.1_Kenya_1950            .c..............a.........................................................a..... [108800]
AM712240.1_OURT_88/3             ................................................................................ [108800]
AM712239.1_Benin_97/1            ................................................................................ [108800]
KM111295.1_Ken06.Bus             .c........................................................................a..... [108800]
KM111294.1_Ken05/Tk1             .c........................................................................a..... [108800]
KM262844.1_strain_L60            ................................................................................ [108800]
KM102979.1_26544/OG10_Italy      ................................................................................ [108800]
KM262845.1_strain_NHV            ................................................................................ [108800]
KX354450.1_47/Ss/2008            ................................................................................ [108800]
KP055815.1_strain_BA71           ................................................................................ [108800]
MH025920.1_strain_R35            .c........................................................................a..... [108800]
MH025919.1_strain_N10            .c........................................................................a..... [108800]
MH025918.1_strain_R25            .c........................................................................a..... [108800]
MH025917.1_strain_R7             .c........................................................................a..... [108800]
MH025916.1_strain_R8             .c........................................................................a..... [108800]
```

```
FR682468.2_Georgia_2007/1        cgtccttccaccaaaataatgttttcttttccgataccaaaataggttgagtatttatgggtatcaatggtggcggaggt [108880]
ASF_IND_20_CAD543                ................................................................................ [108880]
LR899193.1_Germany_2020/1        ................................................................................ [108880]
MN393477.1_Wuhan_2019            ................................................................................ [108880]
MW396979.1_Timor-Leste/2019/1    ................................................................................ [108880]
MK940252.1_CN/2019/InnerMongolia ................................................................................ [108880]
MK128995.1_China/2018/AnhuiXCGQ  ................................................................................ [108880]
MH681419.1_ASFV/POL/2015         ................................................................................ [108880]
LS478113.1_Estonia_2014          ................................................................................ [108880]
KP843857.1_Odintsovo_02/14       ................................................................................ [108880]
FN557520.1_strain_E75            ................................................................................ [108880]
AY261366.1_Warthog               ........................................................g..a.................... [108880]
AY261365.1_Warmbaths             ........................................................g..a.................... [108880]
AY261364.1_Tengani_62            ...........................................................a.................... [108880]
AY261363.1_Pretoriuskop/96/4     ........................................................g..a.................... [108880]
AY261362.1_Mkuzi_1979            ................................................................................ [108880]
AY261361.1_Malawi_Lil-20/1_1983  ...........t......................................a..c.......................... [108880]
AY261360.1_Kenya_1950            ..................................................a.......a..................... [108880]
AM712240.1_OURT_88/3             ................................................................................ [108880]
AM712239.1_Benin_97/1            ................................................................................ [108880]
KM111295.1_Ken06.Bus             ....................a.............................a............................. [108880]
KM111294.1_Ken05/Tk1             ..................................................a.......a..................... [108880]
KM262844.1_strain_L60            ................................................................................ [108880]
KM102979.1_26544/OG10_Italy      ................................................................................ [108880]
KM262845.1_strain_NHV            ................................................................................ [108880]
KX354450.1_47/Ss/2008            ................................................................................ [108880]
KP055815.1_strain_BA71           ................................................................................ [108880]
MH025920.1_strain_R35            ....................a.............................a............................. [108880]
MH025919.1_strain_N10            ....................a.............................a............................. [108880]
MH025918.1_strain_R25            ....................a.............................a............................. [108880]
MH025917.1_strain_R7             ....................a.............................a............................. [108880]
MH025916.1_strain_R8             ....................a.............................a............................. [108880]

FR682468.2_Georgia_2007/1        taaaattacaaagggaatacgcagcgcccctatgcttcctctttgcaacatgcgctgaagcatacttttaatatacatga [108960]
ASF_IND_20_CAD543                ................................................................................ [108960]
LR899193.1_Germany_2020/1        ................................................................................ [108960]
MN393477.1_Wuhan_2019            ................................................................................ [108960]
MW396979.1_Timor-Leste/2019/1    ................................................................................ [108960]
MK940252.1_CN/2019/InnerMongolia ................................................................................ [108960]
MK128995.1_China/2018/AnhuiXCGQ  ................................................................................ [108960]
MH681419.1_ASFV/POL/2015         ................................................................................ [108960]
LS478113.1_Estonia_2014          ................................................................................ [108960]
KP843857.1_Odintsovo_02/14       ................................................................................ [108960]
```

```
FN557520.1_strain_E75            ................................................................................ [108960]
AY261366.1_Warthog               ........................................................c....................... [108960]
AY261365.1_Warmbaths             ........................................................c....................... [108960]
AY261364.1_Tengani_62            ................................................................................ [108960]
AY261363.1_Pretoriuskop/96/4     ........................................................c....................... [108960]
AY261362.1_Mkuzi_1979            ....................................c........................................... [108960]
AY261361.1_Malawi_Lil-20/1_1983  ....................................g...................c....................... [108960]
AY261360.1_Kenya_1950            ......g...........g.....................................c....................... [108960]
AM712240.1_OURT_88/3             ................................................................................ [108960]
AM712239.1_Benin_97/1            ................................................................................ [108960]
KM111295.1_Ken06.Bus             ......g.................t...............................c....................... [108960]
KM111294.1_Ken05/Tk1             ......g...........g.....................................c....................... [108960]
KM262844.1_strain_L60            ................................................................................ [108960]
KM102979.1_26544/OG10_Italy      ................................................................................ [108960]
KM262845.1_strain_NHV            ................................................................................ [108960]
KX354450.1_47/Ss/2008            ................................................................................ [108960]
KP055815.1_strain_BA71           ................................................................................ [108960]
MH025920.1_strain_R35            ......g.................t...............................c....................... [108960]
MH025919.1_strain_N10            ......g.................t...............................c....................... [108960]
MH025918.1_strain_R25            ......g.................t...............................c....................... [108960]
MH025917.1_strain_R7             ......g.................t...............................c....................... [108960]
MH025916.1_strain_R8             ......g.................t...............................c....................... [108960]

FR682468.2_Georgia_2007/1        gcataaggtcgatgcctagggctcgctcatgggcctcatctataatcataaaggcatagcgggaagctatctcatcatcc [109040]
ASF_IND_20_CAD543                ................................................................................ [109040]
LR899193.1_Germany_2020/1        ................................................................................ [109040]
MN393477.1_Wuhan_2019            ................................................................................ [109040]
MW396979.1_Timor-Leste/2019/1    ................................................................................ [109040]
MK940252.1_CN/2019/InnerMongolia ................................................................................ [109040]
MK128995.1_China/2018/AnhuiXCGQ  ................................................................................ [109040]
MH681419.1_ASFV/POL/2015         ................................................................................ [109040]
LS478113.1_Estonia_2014          ................................................................................ [109040]
KP843857.1_Odintsovo_02/14       ................................................................................ [109040]
FN557520.1_strain_E75            ................................................................................ [109040]
AY261366.1_Warthog               ................................................................................ [109040]
AY261365.1_Warmbaths             ................................................................................ [109040]
AY261364.1_Tengani_62            ................................................................................ [109040]
AY261363.1_Pretoriuskop/96/4     ................................................................................ [109040]
AY261362.1_Mkuzi_1979            ................................................................................ [109040]
AY261361.1_Malawi_Lil-20/1_1983  ....g....................t.................g.................................... [109040]
AY261360.1_Kenya_1950            ....g......................................g...........g........................ [109040]
AM712240.1_OURT_88/3             ................................................................................ [109040]
AM712239.1_Benin_97/1            ................................................................................ [109040]
KM111295.1_Ken06.Bus             ....g......................................g...........g........................ [109040]
```

```
KM111294.1_Ken05/Tk1            ....g......................................g...........g........................ [109040]
KM262844.1_strain_L60           ................................................................................ [109040]
KM102979.1_26544/OG10_Italy     ................................................................................ [109040]
KM262845.1_strain_NHV           ................................................................................ [109040]
KX354450.1_47/Ss/2008           ................................................................................ [109040]
KP055815.1_strain_BA71          ................................................................................ [109040]
MH025920.1_strain_R35           ....g......................................g...........g........................ [109040]
MH025919.1_strain_N10           ....g......................................g...........g........................ [109040]
MH025918.1_strain_R25           ....g......................................g...........g........................ [109040]
MH025917.1_strain_R7            ....g......................................g...........g........................ [109040]
MH025916.1_strain_R8            ....g......................................g...........g........................ [109040]

FR682468.2_Georgia_2007/1       gtcattgtatgtagctgcgccaacagaacccccgcggttgcataaataaggccccgattgggtttttccgtcagaggctt [109120]
ASF_IND_20_CAD543               ................................................................................ [109120]
LR899193.1_Germany_2020/1       ................................................................................ [109120]
MN393477.1_Wuhan_2019           ................................................................................ [109120]
MW396979.1_Timor-Leste/2019/1   ................................................................................ [109120]
MK940252.1_CN/2019/InnerMongolia ................................................................................ [109120]
MK128995.1_China/2018/AnhuiXCGQ ................................................................................ [109120]
MH681419.1_ASFV/POL/2015        ................................................................................ [109120]
LS478113.1_Estonia_2014         ................................................................................ [109120]
KP843857.1_Odintsovo_02/14      ................................................................................ [109120]
FN557520.1_strain_E75           ............................................g................................... [109120]
AY261366.1_Warthog              .........................................g..g................................... [109120]
AY261365.1_Warmbaths            .........................................g..g................................... [109120]
AY261364.1_Tengani_62           .........................................g..g................................... [109120]
AY261363.1_Pretoriuskop/96/4    .........................................g..g................................... [109120]
AY261362.1_Mkuzi_1979           .........................................g..g................................... [109120]
AY261361.1_Malawi_Lil-20/1_1983 .............a........g......t.....a.....g..g................................... [109120]
AY261360.1_Kenya_1950           .............a........g..g..t...........g..g.................c................. [109120]
AM712240.1_OURT_88/3            ............................................g................................... [109120]
AM712239.1_Benin_97/1           ............................................g................................... [109120]
KM111295.1_Ken06.Bus            .............a........gt.g..t...........g..g................................... [109120]
KM111294.1_Ken05/Tk1            .............a........g..g..t...........g..g................................... [109120]
KM262844.1_strain_L60           ............................................g................................... [109120]
KM102979.1_26544/OG10_Italy     ............................................g................................... [109120]
KM262845.1_strain_NHV           ............................................g................................... [109120]
KX354450.1_47/Ss/2008           ............................................g................................... [109120]
KP055815.1_strain_BA71          ............................................g................................... [109120]
MH025920.1_strain_R35           .............a........gt.g..t...........g..g................................... [109120]
MH025919.1_strain_N10           .............a........gt.g..t...........g..g................................... [109120]
MH025918.1_strain_R25           .............a........gt.g..t...........g..g................................... [109120]
MH025917.1_strain_R7            .............a........gt.g..t...........g..g................................... [109120]
MH025916.1_strain_R8            .............a........gt.g..t...........g..g................................... [109120]
```

```
FR682468.2_Georgia_2007/1        cgtttggtagcccactgtttggcctaatatcatgtcggggtagtgggttgaggcgccgatgtctttggcgagggtcaccg [109200]
ASF_IND_20_CAD543                ................................................................................ [109200]
LR899193.1_Germany_2020/1        ................................................................................ [109200]
MN393477.1_Wuhan_2019            ................................................................................ [109200]
MW396979.1_Timor-Leste/2019/1    ................................................................................ [109200]
MK940252.1_CN/2019/InnerMongolia ................................................................................ [109200]
MK128995.1_China/2018/AnhuiXCGQ  ................................................................................ [109200]
MH681419.1_ASFV/POL/2015         ................................................................................ [109200]
LS478113.1_Estonia_2014          ................................................................................ [109200]
KP843857.1_Odintsovo_02/14       ................................................................................ [109200]
FN557520.1_strain_E75            ...c..........................................................................t. [109200]
AY261366.1_Warthog               ...c............................................................................ [109200]
AY261365.1_Warmbaths             ...c.............................a............................................g. [109200]
AY261364.1_Tengani_62            ...c............................................................................ [109200]
AY261363.1_Pretoriuskop/96/4     ...c..........................................................................g. [109200]
AY261362.1_Mkuzi_1979            ...c............................................................................ [109200]
AY261361.1_Malawi_Lil-20/1_1983  ...c...........c....................a........................................... [109200]
AY261360.1_Kenya_1950            ...c...........c...........g..................................................t. [109200]
AM712240.1_OURT_88/3             ...c..........................................................................t. [109200]
AM712239.1_Benin_97/1            ...c..........................................................................t. [109200]
KM111295.1_Ken06.Bus             ...c.......................g..................................................t. [109200]
KM111294.1_Ken05/Tk1             ...c...........c...........g.................................................... [109200]
KM262844.1_strain_L60            ...c..........................................................................t. [109200]
KM102979.1_26544/OG10_Italy      ...c..........................................................................t. [109200]
KM262845.1_strain_NHV            ...c..........................................................................t. [109200]
KX354450.1_47/Ss/2008            ...c..........................................................................t. [109200]
KP055815.1_strain_BA71           ...c..........................................................................t. [109200]
MH025920.1_strain_R35            ...c.......................g..................................................t. [109200]
MH025919.1_strain_N10            ...c.......................g..................................................t. [109200]
MH025918.1_strain_R25            ...c.......................g..................................................t. [109200]
MH025917.1_strain_R7             ...c.......................g..................................................t. [109200]
MH025916.1_strain_R8             ...c.......................g..................................................t. [109200]

FR682468.2_Georgia_2007/1        cggttaggactcttggctgggtacaaataaccgagcgtcccaagtatttttggaaagaatgcgtgttttcatttctcaga [109280]
ASF_IND_20_CAD543                ................................................................................ [109280]
LR899193.1_Germany_2020/1        ................................................................................ [109280]
MN393477.1_Wuhan_2019            ................................................................................ [109280]
MW396979.1_Timor-Leste/2019/1    ................................................................................ [109280]
MK940252.1_CN/2019/InnerMongolia ................................................................................ [109280]
MK128995.1_China/2018/AnhuiXCGQ  ................................................................................ [109280]
MH681419.1_ASFV/POL/2015         ................................................................................ [109280]
LS478113.1_Estonia_2014          ................................................................................ [109280]
KP843857.1_Odintsovo_02/14       ................................................................................ [109280]
```

```
FN557520.1_strain_E75            ................................................................................ [109280]
AY261366.1_Warthog               ...........................................a....................a............... [109280]
AY261365.1_Warmbaths             .a.........................................a.................................... [109280]
AY261364.1_Tengani_62            ...........................................a.................................... [109280]
AY261363.1_Pretoriuskop/96/4     .a.........................................a.................................... [109280]
AY261362.1_Mkuzi_1979            ................................................................................ [109280]
AY261361.1_Malawi_Lil-20/1_1983  .a...................................c.....a...................................t [109280]
AY261360.1_Kenya_1950            .....................................g.....a.................................... [109280]
AM712240.1_OURT_88/3             ................................................................................ [109280]
AM712239.1_Benin_97/1            ................................................................................ [109280]
KM111295.1_Ken06.Bus             .....................................g.....a.................................... [109280]
KM111294.1_Ken05/Tk1             .a...................................c.....a.........g....g....................t [109280]
KM262844.1_strain_L60            ................................................................................ [109280]
KM102979.1_26544/OG10_Italy      ................................................................................ [109280]
KM262845.1_strain_NHV            ................................................................................ [109280]
KX354450.1_47/Ss/2008            ................................................................................ [109280]
KP055815.1_strain_BA71           ................................................................................ [109280]
MH025920.1_strain_R35            .....................................g.....a.................................... [109280]
MH025919.1_strain_N10            .....................................g.....a.................................... [109280]
MH025918.1_strain_R25            .....................................g.....a.................................... [109280]
MH025917.1_strain_R7             .....................................g.....a.................................... [109280]
MH025916.1_strain_R8             .....................................g.....a.................................... [109280]

FR682468.2_Georgia_2007/1        attctgaacacgtgtacgggtaaggccgtggattttccggaaccagtgcgtgactttataatgagcacccggtctgcgag [109360]
ASF_IND_20_CAD543                ................................................................................ [109360]
LR899193.1_Germany_2020/1        ................................................................................ [109360]
MN393477.1_Wuhan_2019            ................................................................................ [109360]
MW396979.1_Timor-Leste/2019/1    ................................................................................ [109360]
MK940252.1_CN/2019/InnerMongolia ................................................................................ [109360]
MK128995.1_China/2018/AnhuiXCGQ  ................................................................................ [109360]
MH681419.1_ASFV/POL/2015         ................................................................................ [109360]
LS478113.1_Estonia_2014          ................................................................................ [109360]
KP843857.1_Odintsovo_02/14       ................................................................................ [109360]
FN557520.1_strain_E75            ....................c.g..................t...................................... [109360]
AY261366.1_Warthog               ......................g......a...........t...................................... [109360]
AY261365.1_Warmbaths             ......................g..................t...................................... [109360]
AY261364.1_Tengani_62            ......................g..................t...................................... [109360]
AY261363.1_Pretoriuskop/96/4     .....a................g..................t...................................... [109360]
AY261362.1_Mkuzi_1979            ....................c.g..................t...................................... [109360]
AY261361.1_Malawi_Lil-20/1_1983  ......................g..................t..g.................a................. [109360]
AY261360.1_Kenya_1950            ..............c.......g..................t..g........t.......................... [109360]
AM712240.1_OURT_88/3             ....................c.g..................t...................................... [109360]
AM712239.1_Benin_97/1            ....................c.g..................t...................................... [109360]
KM111295.1_Ken06.Bus             .................a....g..................t..g........t.......................... [109360]
```

```
KM111294.1_Ken05/Tk1           ......................g..................t..g........t.......................... [109360]
KM262844.1_strain_L60          ....................c.g..................t...................................... [109360]
KM102979.1_26544/OG10_Italy    ....................c.g..................t...................................... [109360]
KM262845.1_strain_NHV          ....................c.g..................t...................................... [109360]
KX354450.1_47/Ss/2008          ....................c.g..................t...................................... [109360]
KP055815.1_strain_BA71         ....................c.g..................t...................................... [109360]
MH025920.1_strain_R35          .................a....g..................t..g........t.......................... [109360]
MH025919.1_strain_N10          .................a....g..................t..g........t.......................... [109360]
MH025918.1_strain_R25          .................a....g..................t..g........t.......................... [109360]
MH025917.1_strain_R7           .................a....g..................t..g........t.......................... [109360]
MH025916.1_strain_R8           .................a....g..................t..g........t.......................... [109360]

FR682468.2_Georgia_2007/1      ggaggttggaatggcccctccaaactccgggagacgttgttttatccaagtgatgatgtaatgaataggaacatcattct [109440]
ASF_IND_20_CAD543              ................................................................................ [109440]
LR899193.1_Germany_2020/1      ................................................................................ [109440]
MN393477.1_Wuhan_2019          ................................................................................ [109440]
MW396979.1_Timor-Leste/2019/1  ................................................................................ [109440]
MK940252.1_CN/2019/InnerMongolia ................................................................................ [109440]
MK128995.1_China/2018/AnhuiXCGQ ................................................................................ [109440]
MH681419.1_ASFV/POL/2015       ................................................................................ [109440]
LS478113.1_Estonia_2014        ................................................................................ [109440]
KP843857.1_Odintsovo_02/14     ................................................................................ [109440]
FN557520.1_strain_E75          ................................................c............................... [109440]
AY261366.1_Warthog             ................................................................................ [109440]
AY261365.1_Warmbaths           ................................................................................ [109440]
AY261364.1_Tengani_62          a............................................................................... [109440]
AY261363.1_Pretoriuskop/96/4   ................................................................................ [109440]
AY261362.1_Mkuzi_1979          ................................................c............................... [109440]
AY261361.1_Malawi_Lil-20/1_1983 ...............t.......................................................t........ [109440]
AY261360.1_Kenya_1950          ...a.....t...ctt..c.....t.....a........................................t........ [109440]
AM712240.1_OURT_88/3           ................................................c............................... [109440]
AM712239.1_Benin_97/1          ................................................c............................... [109440]
KM111295.1_Ken06.Bus           .........t...ctt..c...........a........................................t........ [109440]
KM111294.1_Ken05/Tk1           ...............t.......................................................t........ [109440]
KM262844.1_strain_L60          ................................................c............................... [109440]
KM102979.1_26544/OG10_Italy    ................................................c............................... [109440]
KM262845.1_strain_NHV          ................................................c............................... [109440]
KX354450.1_47/Ss/2008          ................................................c............................... [109440]
KP055815.1_strain_BA71         ................................................c............................... [109440]
MH025920.1_strain_R35          .........t...ctt..c...........a........................................t........ [109440]
MH025919.1_strain_N10          .........t...ctt..c...........a........................................t........ [109440]
MH025918.1_strain_R25          .........t...ctt..c...........a........................................t........ [109440]
MH025917.1_strain_R7           .........t...ctt..c...........a........................................t........ [109440]
MH025916.1_strain_R8           .........t...ctt..c...........a........................................t........ [109440]
```

```
FR682468.2_Georgia_2007/1        tgtgctcagcgggcacgttatagagatgaccaggctccaataaagtcggttttcccatattctattgttttaaggattga [109520]
ASF_IND_20_CAD543                ................................................................................ [109520]
LR899193.1_Germany_2020/1        ................................................................................ [109520]
MN393477.1_Wuhan_2019            ................................................................................ [109520]
MW396979.1_Timor-Leste/2019/1    ................................................................................ [109520]
MK940252.1_CN/2019/InnerMongolia ................................................................................ [109520]
MK128995.1_China/2018/AnhuiXCGQ  ................................................................................ [109520]
MH681419.1_ASFV/POL/2015         ................................................................................ [109520]
LS478113.1_Estonia_2014          ................................................................................ [109520]
KP843857.1_Odintsovo_02/14       ................................................................................ [109520]
FN557520.1_strain_E75            ................................................................................ [109520]
AY261366.1_Warthog               ................................................................................ [109520]
AY261365.1_Warmbaths             ................................................................................ [109520]
AY261364.1_Tengani_62            ................................................................................ [109520]
AY261363.1_Pretoriuskop/96/4     .............t.................................................................. [109520]
AY261362.1_Mkuzi_1979            ................................................................................ [109520]
AY261361.1_Malawi_Lil-20/1_1983  .............t......................................................a........g.. [109520]
AY261360.1_Kenya_1950            .............t...............................................................g.. [109520]
AM712240.1_OURT_88/3             ................................................................................ [109520]
AM712239.1_Benin_97/1            ................................................................................ [109520]
KM111295.1_Ken06.Bus             .............t.................................................................. [109520]
KM111294.1_Ken05/Tk1             .............t.................................................................. [109520]
KM262844.1_strain_L60            ................................................................................ [109520]
KM102979.1_26544/OG10_Italy      ................................................................................ [109520]
KM262845.1_strain_NHV            ................................................................................ [109520]
KX354450.1_47/Ss/2008            ................................................................................ [109520]
KP055815.1_strain_BA71           ................................................................................ [109520]
MH025920.1_strain_R35            .............t.................................................................. [109520]
MH025919.1_strain_N10            .............t.................................................................. [109520]
MH025918.1_strain_R25            .............t.................................................................. [109520]
MH025917.1_strain_R7             .............t.................................................................. [109520]
MH025916.1_strain_R8             .............t.................................................................. [109520]

FR682468.2_Georgia_2007/1        ttgttcataaatatttttatactctgaccaagaaattatttttttattaagccggttatttacgttgttatggaacgcga [109600]
ASF_IND_20_CAD543                ................................................................................ [109600]
LR899193.1_Germany_2020/1        ................................................................................ [109600]
MN393477.1_Wuhan_2019            ................................................................................ [109600]
MW396979.1_Timor-Leste/2019/1    ................................................................................ [109600]
MK940252.1_CN/2019/InnerMongolia ................................................................................ [109600]
MK128995.1_China/2018/AnhuiXCGQ  ................................................................................ [109600]
MH681419.1_ASFV/POL/2015         ................................................................................ [109600]
LS478113.1_Estonia_2014          ................................................................................ [109600]
KP843857.1_Odintsovo_02/14       ................................................................................ [109600]
```

```
FN557520.1_strain_E75            ................................................................................ [109600]
AY261366.1_Warthog               ......................................................a...........a.....a....... [109600]
AY261365.1_Warmbaths             ......................................................a.................a....... [109600]
AY261364.1_Tengani_62            .....................t................................a.................a....... [109600]
AY261363.1_Pretoriuskop/96/4     ......................................................a.................a....... [109600]
AY261362.1_Mkuzi_1979            ................................................................................ [109600]
AY261361.1_Malawi_Lil-20/1_1983  ......................................................t........a........a....... [109600]
AY261360.1_Kenya_1950            .....................t................................t....................t.... [109600]
AM712240.1_OURT_88/3             ................................................................................ [109600]
AM712239.1_Benin_97/1            ................................................................................ [109600]
KM111295.1_Ken06.Bus             ......................................................t....................t.... [109600]
KM111294.1_Ken05/Tk1             ......................................................t....................t.... [109600]
KM262844.1_strain_L60            ................................................................................ [109600]
KM102979.1_26544/OG10_Italy      ................................................................................ [109600]
KM262845.1_strain_NHV            ................................................................................ [109600]
KX354450.1_47/Ss/2008            ................................................................................ [109600]
KP055815.1_strain_BA71           ................................................................................ [109600]
MH025920.1_strain_R35            ......................................................t....................t.... [109600]
MH025919.1_strain_N10            ......................................................t....................t.... [109600]
MH025918.1_strain_R25            ......................................................t....................t.... [109600]
MH025917.1_strain_R7             ......................................................t....................t.... [109600]
MH025916.1_strain_R8             ......................................................t....................t.... [109600]

FR682468.2_Georgia_2007/1        aggtccagtactgaaagtcctccgagttgtttaatgtcaagggattttttgtaagatacgaaaaggcgtggtgctggcac [109680]
ASF_IND_20_CAD543                ................................................................................ [109680]
LR899193.1_Germany_2020/1        ................................................................................ [109680]
MN393477.1_Wuhan_2019            ................................................................................ [109680]
MW396979.1_Timor-Leste/2019/1    ................................................................................ [109680]
MK940252.1_CN/2019/InnerMongolia ................................................................................ [109680]
MK128995.1_China/2018/AnhuiXCGQ  ................................................................................ [109680]
MH681419.1_ASFV/POL/2015         ................................................................................ [109680]
LS478113.1_Estonia_2014          ................................................................................ [109680]
KP843857.1_Odintsovo_02/14       ................................................................................ [109680]
FN557520.1_strain_E75            ............................................................................a... [109680]
AY261366.1_Warthog               ..........................................................................c..... [109680]
AY261365.1_Warmbaths             ................................................................................ [109680]
AY261364.1_Tengani_62            ................................................................................ [109680]
AY261363.1_Pretoriuskop/96/4     ..........................................................................c....t [109680]
AY261362.1_Mkuzi_1979            ................................................................................ [109680]
AY261361.1_Malawi_Lil-20/1_1983  ............................a........t..a.......................a.........c.a... [109680]
AY261360.1_Kenya_1950            ..................................c.....a..g..............t...............c.a... [109680]
AM712240.1_OURT_88/3             ............................................................................a... [109680]
AM712239.1_Benin_97/1            ............................................................................a... [109680]
KM111295.1_Ken06.Bus             ..................................c.....a..g..............t..............tc.a... [109680]
```

```
KM111294.1_Ken05/Tk1          ..................................c.....a..g..............t...............c.a... [109680]
KM262844.1_strain_L60         ............................................................................a... [109680]
KM102979.1_26544/OG10_Italy   ............................................................................a... [109680]
KM262845.1_strain_NHV         ............................................................................a... [109680]
KX354450.1_47/Ss/2008         ............................................................................a... [109680]
KP055815.1_strain_BA71        ............................................................................a... [109680]
MH025920.1_strain_R35         ..................................c.....a..g..............t..............tc.a... [109680]
MH025919.1_strain_N10         ..................................c.....a..g..............t..............tc.a... [109680]
MH025918.1_strain_R25         ..................................c.....a..g..............t..............tc.a... [109680]
MH025917.1_strain_R7          ..................................c.....a..g..............t..............tc.a... [109680]
MH025916.1_strain_R8          ..................................c.....a..g..............t..............tc.a... [109680]

FR682468.2_Georgia_2007/1     ctggtgcatggcagagactcgataaagttcagtatccattggatggcttcatatttttctttccagctaggagcgtctga [109760]
ASF_IND_20_CAD543             ................................................................................ [109760]
LR899193.1_Germany_2020/1     ................................................................................ [109760]
MN393477.1_Wuhan_2019         ................................................................................ [109760]
MW396979.1_Timor-Leste/2019/1 ................................................................................ [109760]
MK940252.1_CN/2019/InnerMongolia ................................................................................ [109760]
MK128995.1_China/2018/AnhuiXCGQ ................................................................................ [109760]
MH681419.1_ASFV/POL/2015      ................................................................................ [109760]
LS478113.1_Estonia_2014       ................................................................................ [109760]
KP843857.1_Odintsovo_02/14    ................................................................................ [109760]
FN557520.1_strain_E75         ................................................................................ [109760]
AY261366.1_Warthog            ...............................................c...................c......a..... [109760]
AY261365.1_Warmbaths          ...................................................................c............ [109760]
AY261364.1_Tengani_62         ...................................................................c.....t...... [109760]
AY261363.1_Pretoriuskop/96/4  ...................................................................c......a..... [109760]
AY261362.1_Mkuzi_1979         ................................................................................ [109760]
AY261361.1_Malawi_Lil-20/1_1983 a.............c...........a........................................c.....t...... [109760]
AY261360.1_Kenya_1950         a.............g........g....................a.....................t.......a..... [109760]
AM712240.1_OURT_88/3          ................................................................................ [109760]
AM712239.1_Benin_97/1         ................................................................................ [109760]
KM111295.1_Ken06.Bus          a.............g........g..a..g..............a.....................t............. [109760]
KM111294.1_Ken05/Tk1          a.............g........g..a.................a.....................t............. [109760]
KM262844.1_strain_L60         ................................................................................ [109760]
KM102979.1_26544/OG10_Italy   ................................................................................ [109760]
KM262845.1_strain_NHV         ................................................................................ [109760]
KX354450.1_47/Ss/2008         ................................................................................ [109760]
KP055815.1_strain_BA71        ................................................................................ [109760]
MH025920.1_strain_R35         a.............g........g..a..g..............a.....................t............. [109760]
MH025919.1_strain_N10         a.............g........g..a..g..............a.....................t............. [109760]
MH025918.1_strain_R25         a.............g........g..a..g..............a.....................t............. [109760]
MH025917.1_strain_R7          a.............g........g..a..g..............a.....................t............. [109760]
MH025916.1_strain_R8          a.............g........g..a..g..............a.....................t............. [109760]
```

```
FR682468.2_Georgia_2007/1        aaaaaagatagcatatagatgcaaggatcgccagtatttaggtccccaatgcaacatttataaccttttgaaaaatctca [109840]
ASF_IND_20_CAD543                ................................................................................ [109840]
LR899193.1_Germany_2020/1        ................................................................................ [109840]
MN393477.1_Wuhan_2019            ................................................................................ [109840]
MW396979.1_Timor-Leste/2019/1    ................................................................................ [109840]
MK940252.1_CN/2019/InnerMongolia ................................................................................ [109840]
MK128995.1_China/2018/AnhuiXCGQ  ................................................................................ [109840]
MH681419.1_ASFV/POL/2015         ................................................................................ [109840]
LS478113.1_Estonia_2014          ................................................................................ [109840]
KP843857.1_Odintsovo_02/14       ................................................................................ [109840]
FN557520.1_strain_E75            ................................................................................ [109840]
AY261366.1_Warthog               ..........................c..................................................... [109840]
AY261365.1_Warmbaths             ..........................c..................................................... [109840]
AY261364.1_Tengani_62            ..........................c..................................................... [109840]
AY261363.1_Pretoriuskop/96/4     ..........................c..................................................... [109840]
AY261362.1_Mkuzi_1979            ................................................................................ [109840]
AY261361.1_Malawi_Lil-20/1_1983  ..........................t.......a.............................t............... [109840]
AY261360.1_Kenya_1950            ............g.............c......aa............................................. [109840]
AM712240.1_OURT_88/3             ....g........................................................................... [109840]
AM712239.1_Benin_97/1            ................................................................................ [109840]
KM111295.1_Ken06.Bus             ..........................c......aa............................................. [109840]
KM111294.1_Ken05/Tk1             ............g.............c......aa............................................. [109840]
KM262844.1_strain_L60            ................................................................................ [109840]
KM102979.1_26544/OG10_Italy      ................................................................................ [109840]
KM262845.1_strain_NHV            ....g........................................................................... [109840]
KX354450.1_47/Ss/2008            ................................................................................ [109840]
KP055815.1_strain_BA71           ................................................................................ [109840]
MH025920.1_strain_R35            ..........................c......aa............................................. [109840]
MH025919.1_strain_N10            ..........................c......aa............................................. [109840]
MH025918.1_strain_R25            ..........................c......aa............................................. [109840]
MH025917.1_strain_R7             ..........................c......aa............................................. [109840]
MH025916.1_strain_R8             ..........................c......aa............................................. [109840]

FR682468.2_Georgia_2007/1        ttccatatagaggtaaatatttttttccatggagaattttttttgcactcttgaagggattgcgccacatcgtcaaatgt [109920]
ASF_IND_20_CAD543                ................................................................................ [109920]
LR899193.1_Germany_2020/1        ................................................................................ [109920]
MN393477.1_Wuhan_2019            ................................................................................ [109920]
MW396979.1_Timor-Leste/2019/1    ................................................................................ [109920]
MK940252.1_CN/2019/InnerMongolia ................................................................................ [109920]
MK128995.1_China/2018/AnhuiXCGQ  ................................................................................ [109920]
MH681419.1_ASFV/POL/2015         ................................................................................ [109920]
LS478113.1_Estonia_2014          ................................................................................ [109920]
KP843857.1_Odintsovo_02/14       ................................................................................ [109920]
```

```
FN557520.1_strain_E75           ................................................................................ [109920]
AY261366.1_Warthog              ...............t.........................................a...................... [109920]
AY261365.1_Warmbaths            ...............t...............a................................................ [109920]
AY261364.1_Tengani_62           ...............t................................................................ [109920]
AY261363.1_Pretoriuskop/96/4    ...............t................................................................ [109920]
AY261362.1_Mkuzi_1979           ................................................................................ [109920]
AY261361.1_Malawi_Lil-20/1_1983 ...............t.............t.......................g.....c.........a.......... [109920]
AY261360.1_Kenya_1950           ...............t...........t.........................c.....c.........a.......... [109920]
AM712240.1_OURT_88/3            ................................................................................ [109920]
AM712239.1_Benin_97/1           ................................................................................ [109920]
KM111295.1_Ken06.Bus            ...............t...........t.........................c.....c..a....a.......... [109920]
KM111294.1_Ken05/Tk1            ...............t...........t.........................c.....c.........a.......... [109920]
KM262844.1_strain_L60           ................................................................................ [109920]
KM102979.1_26544/OG10_Italy     ................................................................................ [109920]
KM262845.1_strain_NHV           ................................................................................ [109920]
KX354450.1_47/Ss/2008           ................................................................................ [109920]
KP055815.1_strain_BA71          ................................................................................ [109920]
MH025920.1_strain_R35           ...............t...........t.........................c.....c..a....a.......... [109920]
MH025919.1_strain_N10           ...............t...........t.........................c.....c..a....a.......... [109920]
MH025918.1_strain_R25           ...............t...........t.........................c.....c..a....a.......... [109920]
MH025917.1_strain_R7            ...............t...........t.........................c.....c..a....a.......... [109920]
MH025916.1_strain_R8            ...............t...........t.........................c.....c..a....a.......... [109920]

FR682468.2_Georgia_2007/1       tttttgttttccatgtattttggcgtaattccagccagtatctgtgtcatggtccttaatgtcatccgctaactgaaagg [110000]
ASF_IND_20_CAD543               ................................................................................ [110000]
LR899193.1_Germany_2020/1       ................................................................................ [110000]
MN393477.1_Wuhan_2019           ................................................................................ [110000]
MW396979.1_Timor-Leste/2019/1   ................................................................................ [110000]
MK940252.1_CN/2019/InnerMongolia ................................................................................ [110000]
MK128995.1_China/2018/AnhuiXCGQ ................................................................................ [110000]
MH681419.1_ASFV/POL/2015        ................................................................................ [110000]
LS478113.1_Estonia_2014         ................................................................................ [110000]
KP843857.1_Odintsovo_02/14      ................................................................................ [110000]
FN557520.1_strain_E75           ................................................................................ [110000]
AY261366.1_Warthog              ....c.........................................................................a. [110000]
AY261365.1_Warmbaths            ....c........................................................................... [110000]
AY261364.1_Tengani_62           ....c........................................................................... [110000]
AY261363.1_Pretoriuskop/96/4    ....c........................................................................... [110000]
AY261362.1_Mkuzi_1979           ................................................................................ [110000]
AY261361.1_Malawi_Lil-20/1_1983 ....c................a..........ca.............................................. [110000]
AY261360.1_Kenya_1950           ....c.........................g.g...c..........................g................ [110000]
AM712240.1_OURT_88/3            ................................................................................ [110000]
AM712239.1_Benin_97/1           ................................................................................ [110000]
KM111295.1_Ken06.Bus            ..............................g.g...c..........................g................ [110000]
```

```
KM111294.1_Ken05/Tk1           ....c.........................g.g...c..........................g................ [110000]
KM262844.1_strain_L60          ................................................................................ [110000]
KM102979.1_26544/OG10_Italy    ................................................................................ [110000]
KM262845.1_strain_NHV          ................................................................................ [110000]
KX354450.1_47/Ss/2008          ................................................................................ [110000]
KP055815.1_strain_BA71         ................................................................................ [110000]
MH025920.1_strain_R35          ....c.........................g.g...c..........................g................ [110000]
MH025919.1_strain_N10          ....c.........................g.g...c..........................g................ [110000]
MH025918.1_strain_R25          ....c.........................g.g...c..........................g................ [110000]
MH025917.1_strain_R7           ....c.........................g.g...c..........................g................ [110000]
MH025916.1_strain_R8           ....c.........................g.g...c..........................g................ [110000]

FR682468.2_Georgia_2007/1      catgtccaaaacaatgggcagccctttcaatcatcccaatgtcttcaacggatccagttcctaaaacccagcccataata [110080]
ASF_IND_20_CAD543              ................................................tc.t...t..g..c.................. [110080]
LR899193.1_Germany_2020/1      ................................................................................ [110080]
MN393477.1_Wuhan_2019          ................................................................................ [110080]
MW396979.1_Timor-Leste/2019/1  ................................................................................ [110080]
MK940252.1_CN/2019/InnerMongolia ................................................................................ [110080]
MK128995.1_China/2018/AnhuiXCGQ ................................................................................ [110080]
MH681419.1_ASFV/POL/2015       ................................................................................ [110080]
LS478113.1_Estonia_2014        ................................................................................ [110080]
KP843857.1_Odintsovo_02/14     ................................................................................ [110080]
FN557520.1_strain_E75          .............g..................................tc.t...t..g..c.................. [110080]
AY261366.1_Warthog             ...t............................................................................ [110080]
AY261365.1_Warmbaths           ...t............................................................................ [110080]
AY261364.1_Tengani_62          ...t.......................................c....tc.t...t..g..c.................. [110080]
AY261363.1_Pretoriuskop/96/4   ...t............................................tc.t...t..g..c.................. [110080]
AY261362.1_Mkuzi_1979          ................................................................................ [110080]
AY261361.1_Malawi_Lil-20/1_1983 ...t...........................a.c..............tc.t...t..g..c.................. [110080]
AY261360.1_Kenya_1950          ...t...............g...........agc..............ta.t...t..g..c.................. [110080]
AM712240.1_OURT_88/3           .............g..................................tc.t...t..g..c.................. [110080]
AM712239.1_Benin_97/1          .............g..................................tc.t...t..g..c.................. [110080]
KM111295.1_Ken06.Bus           ...t...............g...........agc..............ta.t...t..g..c.................. [110080]
KM111294.1_Ken05/Tk1           ...t...............g...........agc..............ta.t...t..g..c.................. [110080]
KM262844.1_strain_L60          .............g..................................tc.t...t..g..c.................. [110080]
KM102979.1_26544/OG10_Italy    .............g..................................tc.t...t..g..c.................. [110080]
KM262845.1_strain_NHV          .............g..................................tc.t...t..g..c.................. [110080]
KX354450.1_47/Ss/2008          .............g..................................tc.t...t..g..c.................. [110080]
KP055815.1_strain_BA71         .............g..................................tc.t...t..g..c.................. [110080]
MH025920.1_strain_R35          ...t...............g...........agc..............ta.t...t..g..c.................. [110080]
MH025919.1_strain_N10          ...t...............g...........agc..............ta.t...t..g..c.................. [110080]
MH025918.1_strain_R25          ...t...............g...........agc..............ta.t...t..g..c.................. [110080]
MH025917.1_strain_R7           ...t...............g...........agc..............ta.t...t..g..c.................. [110080]
MH025916.1_strain_R8           ...t...............g...........agc..............ta.t...t..g..c.................. [110080]
```

```
FR682468.2_Georgia_2007/1        aacgcgatcttaaaaaagggaatggttttttctggagtgtctactaactgaccggaacccgcgctgtttagagagtggct [110160]
ASF_IND_20_CAD543                ................................................................................ [110160]
LR899193.1_Germany_2020/1        ................................................................................ [110160]
MN393477.1_Wuhan_2019            ................................................................................ [110160]
MW396979.1_Timor-Leste/2019/1    ................................................................................ [110160]
MK940252.1_CN/2019/InnerMongolia ................................................................................ [110160]
MK128995.1_China/2018/AnhuiXCGQ  ................................................................................ [110160]
MH681419.1_ASFV/POL/2015         ................................................................................ [110160]
LS478113.1_Estonia_2014          ................................................................................ [110160]
KP843857.1_Odintsovo_02/14       ................................................................................ [110160]
FN557520.1_strain_E75            ................................................................................ [110160]
AY261366.1_Warthog               ...................................g............................................ [110160]
AY261365.1_Warmbaths             ...................................g............................................ [110160]
AY261364.1_Tengani_62            .......................................................................g........ [110160]
AY261363.1_Pretoriuskop/96/4     ...................................g............................................ [110160]
AY261362.1_Mkuzi_1979            ...................................g............................................ [110160]
AY261361.1_Malawi_Lil-20/1_1983  .................a.....t...................................t.................... [110160]
AY261360.1_Kenya_1950            ........t........a.....t..............a....................t.................... [110160]
AM712240.1_OURT_88/3             ................................................................................ [110160]
AM712239.1_Benin_97/1            ................................................................................ [110160]
KM111295.1_Ken06.Bus             ........t........a.....t..............a....................t.................... [110160]
KM111294.1_Ken05/Tk1             ........t........a.....t..............a....................t.................... [110160]
KM262844.1_strain_L60            ................................................................................ [110160]
KM102979.1_26544/OG10_Italy      ................................................................................ [110160]
KM262845.1_strain_NHV            ................................................................................ [110160]
KX354450.1_47/Ss/2008            ................................................................................ [110160]
KP055815.1_strain_BA71           ................................................................................ [110160]
MH025920.1_strain_R35            ........t........a.....t..............a....................t.................... [110160]
MH025919.1_strain_N10            ........t........a.....t..............a....................t.................... [110160]
MH025918.1_strain_R25            ........t........a.....t..............a....................t.................... [110160]
MH025917.1_strain_R7             ........t........a.....t..............a....................t.................... [110160]
MH025916.1_strain_R8             ........t........a.....t..............a....................t.................... [110160]

FR682468.2_Georgia_2007/1        tacaaaggtacacagcagcgctcccagttggttgggatccggaaaccttggacagtgttccttaatccagtcgatttgcc [110240]
ASF_IND_20_CAD543                ................................................................................ [110240]
LR899193.1_Germany_2020/1        ................................................................................ [110240]
MN393477.1_Wuhan_2019            ................................................................................ [110240]
MW396979.1_Timor-Leste/2019/1    ................................................................................ [110240]
MK940252.1_CN/2019/InnerMongolia ................................................................................ [110240]
MK128995.1_China/2018/AnhuiXCGQ  ................................................................................ [110240]
MH681419.1_ASFV/POL/2015         ................................................................................ [110240]
LS478113.1_Estonia_2014          ................................................................................ [110240]
KP843857.1_Odintsovo_02/14       ................................................................................ [110240]
```

```
FN557520.1_strain_E75            ................................................................................ [110240]
AY261366.1_Warthog               .................................a.................g............................ [110240]
AY261365.1_Warmbaths             .................................a.................g............................ [110240]
AY261364.1_Tengani_62            .................................a.................g............................ [110240]
AY261363.1_Pretoriuskop/96/4     .................................a.................g............................ [110240]
AY261362.1_Mkuzi_1979            ................................................................................ [110240]
AY261361.1_Malawi_Lil-20/1_1983  .................................a.................g............................ [110240]
AY261360.1_Kenya_1950            .........g.......................a..g..g...........g....t...............c.c..... [110240]
AM712240.1_OURT_88/3             ................................................................................ [110240]
AM712239.1_Benin_97/1            ................................................................................ [110240]
KM111295.1_Ken06.Bus             .........g.......................a.....g...........g....t...............c.c..... [110240]
KM111294.1_Ken05/Tk1             .........g.......................a..g..g...........g....t...............c.c..... [110240]
KM262844.1_strain_L60            ................................................................................ [110240]
KM102979.1_26544/OG10_Italy      ................................................................................ [110240]
KM262845.1_strain_NHV            ................................................................................ [110240]
KX354450.1_47/Ss/2008            ................................................................................ [110240]
KP055815.1_strain_BA71           ................................................................................ [110240]
MH025920.1_strain_R35            .........g.......................a.....g...........g....t...............c.c..... [110240]
MH025919.1_strain_N10            .........g.......................a.....g...........g....t...............c.c..... [110240]
MH025918.1_strain_R25            .........g.......................a.....g...........g....t...............c.c..... [110240]
MH025917.1_strain_R7             .........g.......................a.....g...........g....t...............c.c..... [110240]
MH025916.1_strain_R8             .........g.......................a.....g...........g....t...............c.c..... [110240]

FR682468.2_Georgia_2007/1        ggcaaatattttgaaatccttgcatggttagcgccagagcgctcatctgcgccttggctacgccaaagcgggcccacact [110320]
ASF_IND_20_CAD543                ................................................................................ [110320]
LR899193.1_Germany_2020/1        ................................................................................ [110320]
MN393477.1_Wuhan_2019            ................................................................................ [110320]
MW396979.1_Timor-Leste/2019/1    ................................................................................ [110320]
MK940252.1_CN/2019/InnerMongolia ................................................................................ [110320]
MK128995.1_China/2018/AnhuiXCGQ  ................................................................................ [110320]
MH681419.1_ASFV/POL/2015         ................................................................................ [110320]
LS478113.1_Estonia_2014          ................................................................................ [110320]
KP843857.1_Odintsovo_02/14       ................................................................................ [110320]
FN557520.1_strain_E75            .......g........................................................................ [110320]
AY261366.1_Warthog               .......g........................................................................ [110320]
AY261365.1_Warmbaths             .......g........a............................................................... [110320]
AY261364.1_Tengani_62            .......g.....................................................a.................. [110320]
AY261363.1_Pretoriuskop/96/4     .......g........................................................................ [110320]
AY261362.1_Mkuzi_1979            ................................................................................ [110320]
AY261361.1_Malawi_Lil-20/1_1983  .......g.............................g.......................t........a......... [110320]
AY261360.1_Kenya_1950            .......g.............................g.................a..............a......... [110320]
AM712240.1_OURT_88/3             .......g........................................................................ [110320]
AM712239.1_Benin_97/1            .......g........................................................................ [110320]
KM111295.1_Ken06.Bus             .a.....g.............................g...........t.....a..............a......... [110320]
```

```
KM111294.1_Ken05/Tk1             .......g.............................g.................a..............a......... [110320]
KM262844.1_strain_L60            .......g........................................................................ [110320]
KM102979.1_26544/OG10_Italy      .......g........................................................................ [110320]
KM262845.1_strain_NHV            .......g........................................................................ [110320]
KX354450.1_47/Ss/2008            .......g........................................................................ [110320]
KP055815.1_strain_BA71           .......g........................................................................ [110320]
MH025920.1_strain_R35            .a.....g.............................g...........t.....a..............a......... [110320]
MH025919.1_strain_N10            .a.....g.............................g...........t.....a..............a......... [110320]
MH025918.1_strain_R25            .a.....g.............................g...........t.....a..............a......... [110320]
MH025917.1_strain_R7             .a.....g.............................g...........t.....a..............a......... [110320]
MH025916.1_strain_R8             .a.....g.............................g...........t.....a..............a......... [110320]

FR682468.2_Georgia_2007/1        gtatctttatttcgccgcttcacatcgttgtcaaaggagggcatatcatcgataatcaaagaagctacgtgaaagtactc [110400]
ASF_IND_20_CAD543                ................................................................................ [110400]
LR899193.1_Germany_2020/1        ................................................................................ [110400]
MN393477.1_Wuhan_2019            ................................................................................ [110400]
MW396979.1_Timor-Leste/2019/1    ................................................................................ [110400]
MK940252.1_CN/2019/InnerMongolia ................................................................................ [110400]
MK128995.1_China/2018/AnhuiXCGQ  ................................................................................ [110400]
MH681419.1_ASFV/POL/2015         ................................................................................ [110400]
LS478113.1_Estonia_2014          ................................................................................ [110400]
KP843857.1_Odintsovo_02/14       ................................................................................ [110400]
FN557520.1_strain_E75            ...............................................g................................ [110400]
AY261366.1_Warthog               ...............................................g................................ [110400]
AY261365.1_Warmbaths             ..............................................................c................. [110400]
AY261364.1_Tengani_62            ..........................a....................g................................ [110400]
AY261363.1_Pretoriuskop/96/4     ...............................................g................................ [110400]
AY261362.1_Mkuzi_1979            ...............................................g................................ [110400]
AY261361.1_Malawi_Lil-20/1_1983  ....................a..........................g..a.....t........c.............. [110400]
AY261360.1_Kenya_1950            ....................a..........................g.................c.............. [110400]
AM712240.1_OURT_88/3             ...............................................g................................ [110400]
AM712239.1_Benin_97/1            ...............................................g................................ [110400]
KM111295.1_Ken06.Bus             ....................a..........................g.................c.............. [110400]
KM111294.1_Ken05/Tk1             ....................a..........................g.................c.............. [110400]
KM262844.1_strain_L60            ...............................................g................................ [110400]
KM102979.1_26544/OG10_Italy      ...............................................g................................ [110400]
KM262845.1_strain_NHV            ...............................................g................................ [110400]
KX354450.1_47/Ss/2008            ...............................................g................................ [110400]
KP055815.1_strain_BA71           ...............................................g................................ [110400]
MH025920.1_strain_R35            ....................a..........................g.................c.............. [110400]
MH025919.1_strain_N10            ....................a..........................g.................c.............. [110400]
MH025918.1_strain_R25            ....................a..........................g.................c.............. [110400]
MH025917.1_strain_R7             ....................a..........................g.................c.............. [110400]
MH025916.1_strain_R8             ....................a..........................g.................c.............. [110400]
```

```
FR682468.2_Georgia_2007/1        cgctgctagggcggcctctgccggataaataggcgccccaaaggaatgttgcaactgacaggcccgaacaatttccatca [110480]
ASF_IND_20_CAD543                ................................................................................ [110480]
LR899193.1_Germany_2020/1        ................................................................................ [110480]
MN393477.1_Wuhan_2019            ................................................................................ [110480]
MW396979.1_Timor-Leste/2019/1    ................................................................................ [110480]
MK940252.1_CN/2019/InnerMongolia ................................................................................ [110480]
MK128995.1_China/2018/AnhuiXCGQ  ................................................................................ [110480]
MH681419.1_ASFV/POL/2015         ................................................................................ [110480]
LS478113.1_Estonia_2014          ................................................................................ [110480]
KP843857.1_Odintsovo_02/14       ................................................................................ [110480]
FN557520.1_strain_E75            .a..........a................................................................... [110480]
AY261366.1_Warthog               .a.............................................................................. [110480]
AY261365.1_Warmbaths             .a.............................................................................. [110480]
AY261364.1_Tengani_62            .a.............................................................................. [110480]
AY261363.1_Pretoriuskop/96/4     .a..........a................................................................... [110480]
AY261362.1_Mkuzi_1979            ................................................................................ [110480]
AY261361.1_Malawi_Lil-20/1_1983  .a.c..c.....a................................................................... [110480]
AY261360.1_Kenya_1950            .a..........a.................g......................g.......................... [110480]
AM712240.1_OURT_88/3             .a..........a................................................................... [110480]
AM712239.1_Benin_97/1            .a..........a................................................................... [110480]
KM111295.1_Ken06.Bus             .a.c..........................g......................g.......................... [110480]
KM111294.1_Ken05/Tk1             .a.c........a.................g......................g.......................... [110480]
KM262844.1_strain_L60            .a..........a................................................................... [110480]
KM102979.1_26544/OG10_Italy      .a..........a................................................................... [110480]
KM262845.1_strain_NHV            .a..........a................................................................... [110480]
KX354450.1_47/Ss/2008            .a..........a................................................................... [110480]
KP055815.1_strain_BA71           .a..........a................................................................... [110480]
MH025920.1_strain_R35            .a.c..........................g......................g.......................... [110480]
MH025919.1_strain_N10            .a.c..........................g......................g.......................... [110480]
MH025918.1_strain_R25            .a.c..........................g......................g.......................... [110480]
MH025917.1_strain_R7             .a.c..........................g......................g.......................... [110480]
MH025916.1_strain_R8             .a.c..........................g......................g.......................... [110480]

FR682468.2_Georgia_2007/1        ggataatgggacggatatacttcccacctcttagagcgtaagagcaaggctctgttagttgtcccttaaagtccccatct [110560]
ASF_IND_20_CAD543                ................................................................................ [110560]
LR899193.1_Germany_2020/1        ................................................................................ [110560]
MN393477.1_Wuhan_2019            ................................................................................ [110560]
MW396979.1_Timor-Leste/2019/1    ................................................................................ [110560]
MK940252.1_CN/2019/InnerMongolia ................................................................................ [110560]
MK128995.1_China/2018/AnhuiXCGQ  ................................................................................ [110560]
MH681419.1_ASFV/POL/2015         ................................................................................ [110560]
LS478113.1_Estonia_2014          ................................................................................ [110560]
KP843857.1_Odintsovo_02/14       ................................................................................ [110560]
```

```
FN557520.1_strain_E75          ................................................................................ [110560]
AY261366.1_Warthog             ................................................................................ [110560]
AY261365.1_Warmbaths           .............................................................c.................. [110560]
AY261364.1_Tengani_62          ......................t......................................c.................. [110560]
AY261363.1_Pretoriuskop/96/4   .............................................................c.................. [110560]
AY261362.1_Mkuzi_1979          ..........g.....................................................t............... [110560]
AY261361.1_Malawi_Lil-20/1_1983 .................................a.............................................. [110560]
AY261360.1_Kenya_1950          ..........g.................c.....g..........................c.................. [110560]
AM712240.1_OURT_88/3           ................................................................................ [110560]
AM712239.1_Benin_97/1          ................................................................................ [110560]
KM111295.1_Ken06.Bus           ..........g.................c.....g..........................c.................. [110560]
KM111294.1_Ken05/Tk1           ..........g.................c.....g..........................c.................. [110560]
KM262844.1_strain_L60          ................................................................................ [110560]
KM102979.1_26544/OG10_Italy    ................................................................................ [110560]
KM262845.1_strain_NHV          ................................................................................ [110560]
KX354450.1_47/Ss/2008          ................................................................................ [110560]
KP055815.1_strain_BA71         ................................................................................ [110560]
MH025920.1_strain_R35          ..........g.................c.....g..........................c.................. [110560]
MH025919.1_strain_N10          ..........g.................c.....g..........................c.................. [110560]
MH025918.1_strain_R25          ..........g.................c.....g..........................c.................. [110560]
MH025917.1_strain_R7           ..........g.................c.....g..........................c.................. [110560]
MH025916.1_strain_R8           ..........g.................c.....g..........................c.................. [110560]

FR682468.2_Georgia_2007/1      tcaatagcattatttaagatggtctcaaactcttcactaaaggttttataatttttaggattcagtggatgtattccatg [110640]
ASF_IND_20_CAD543              ................................................................................ [110640]
LR899193.1_Germany_2020/1      ................................................................................ [110640]
MN393477.1_Wuhan_2019          ................................................................................ [110640]
MW396979.1_Timor-Leste/2019/1  ................................................................................ [110640]
MK940252.1_CN/2019/InnerMongolia ................................................................................ [110640]
MK128995.1_China/2018/AnhuiXCGQ ................................................................................ [110640]
MH681419.1_ASFV/POL/2015       ................................................................................ [110640]
LS478113.1_Estonia_2014        ................................................................................ [110640]
KP843857.1_Odintsovo_02/14     ................................................................................ [110640]
FN557520.1_strain_E75          ...........g......................t............................................. [110640]
AY261366.1_Warthog             ..g....................t........................................................ [110640]
AY261365.1_Warmbaths           ..g....................t........................................................ [110640]
AY261364.1_Tengani_62          ..g............................................................................. [110640]
AY261363.1_Pretoriuskop/96/4   ..g....................t...................................c.................... [110640]
AY261362.1_Mkuzi_1979          ................................................................................ [110640]
AY261361.1_Malawi_Lil-20/1_1983 .......................t...........g.......................g.................... [110640]
AY261360.1_Kenya_1950          ..g....................t...........g.......................g..t..c.....g........ [110640]
AM712240.1_OURT_88/3           ...........g......................t............................................. [110640]
AM712239.1_Benin_97/1          ...........g......................t............................................. [110640]
KM111295.1_Ken06.Bus           ..g..............a.....t...........g.......................g..t..c.....g........ [110640]
```

```
KM111294.1_Ken05/Tk1            ..g....................t...........g.......................g..t.ac.....g........ [110640]
KM262844.1_strain_L60           ...........g......................t............................................. [110640]
KM102979.1_26544/OG10_Italy     ...........g......................t............................................. [110640]
KM262845.1_strain_NHV           ...........g......................t............................................. [110640]
KX354450.1_47/Ss/2008           ...........g......................t............................................. [110640]
KP055815.1_strain_BA71          ...........g......................t............................................. [110640]
MH025920.1_strain_R35           ..g..............a.....t...........g.......................g..t..c.....g........ [110640]
MH025919.1_strain_N10           ..g..............a.....t...........g.......................g..t..c.....g........ [110640]
MH025918.1_strain_R25           ..g..............a.....t...........g.......................g..t..c.....g........ [110640]
MH025917.1_strain_R7            ..g..............a.....t...........g.......................g..t..c.....g........ [110640]
MH025916.1_strain_R8            ..g..............a.....t...........g.......................g..t..c.....g........ [110640]

FR682468.2_Georgia_2007/1       aaaaagcgcgacactacgcggtgctgtgattctaaaatacttaggtttgcgcgtataggatattaaaataataataagaa [110720]
ASF_IND_20_CAD543               ................................................................................ [110720]
LR899193.1_Germany_2020/1       ................................................................................ [110720]
MN393477.1_Wuhan_2019           ................................................................................ [110720]
MW396979.1_Timor-Leste/2019/1   ................................................................................ [110720]
MK940252.1_CN/2019/InnerMongolia ................................................................................ [110720]
MK128995.1_China/2018/AnhuiXCGQ ................................................................................ [110720]
MH681419.1_ASFV/POL/2015        ................................................................................ [110720]
LS478113.1_Estonia_2014         ................................................................................ [110720]
KP843857.1_Odintsovo_02/14      ................................................................................ [110720]
FN557520.1_strain_E75           .....................a.......................................................... [110720]
AY261366.1_Warthog              .....................a.......................................................... [110720]
AY261365.1_Warmbaths            .....................a.......................................................... [110720]
AY261364.1_Tengani_62           .........a...........a....................................................c..... [110720]
AY261363.1_Pretoriuskop/96/4    .....................a.......................................................... [110720]
AY261362.1_Mkuzi_1979           .....................a.......................................................... [110720]
AY261361.1_Malawi_Lil-20/1_1983 ........tc.....g.....a.a.........................t.............................. [110720]
AY261360.1_Kenya_1950           .........c..g........a.a.................g......c............................... [110720]
AM712240.1_OURT_88/3            .....................a.......................................................... [110720]
AM712239.1_Benin_97/1           .....................a.......................................................... [110720]
KM111295.1_Ken06.Bus            .........c..g........a.a.................g......c............................... [110720]
KM111294.1_Ken05/Tk1            .........c..g........a.a.................g......c............................... [110720]
KM262844.1_strain_L60           .....................a.......................................................... [110720]
KM102979.1_26544/OG10_Italy     .....................a.......................................................... [110720]
KM262845.1_strain_NHV           .....................a.......................................................... [110720]
KX354450.1_47/Ss/2008           .....................a.......................................................... [110720]
KP055815.1_strain_BA71          .....................a.......................................................... [110720]
MH025920.1_strain_R35           .........c..g........a.a.................g......c............................... [110720]
MH025919.1_strain_N10           .........c..g........a.a.................g......c............................... [110720]
MH025918.1_strain_R25           .........c..g........a.a.................g......c............................... [110720]
MH025917.1_strain_R7            .........c..g........a.a.................g......c............................... [110720]
MH025916.1_strain_R8            .........c..g........a.a.................g......c............................... [110720]
```

```
FR682468.2_Georgia_2007/1        ctacaatgatggagatatagatgagatgcaacatgctgagttgtctccccgcagggaatggtccttttccgcgcttgtta [110800]
ASF_IND_20_CAD543                ................................................................................ [110800]
LR899193.1_Germany_2020/1        ................................................................................ [110800]
MN393477.1_Wuhan_2019            ................................................................................ [110800]
MW396979.1_Timor-Leste/2019/1    ................................................................................ [110800]
MK940252.1_CN/2019/InnerMongolia ................................................................................ [110800]
MK128995.1_China/2018/AnhuiXCGQ  ................................................................................ [110800]
MH681419.1_ASFV/POL/2015         ................................................................................ [110800]
LS478113.1_Estonia_2014          ................................................................................ [110800]
KP843857.1_Odintsovo_02/14       ................................................................................ [110800]
FN557520.1_strain_E75            ........................................................................a..g.... [110800]
AY261366.1_Warthog               ...........................................................................g.... [110800]
AY261365.1_Warmbaths             ...........................................................................g.... [110800]
AY261364.1_Tengani_62            ...........................................................................g.... [110800]
AY261363.1_Pretoriuskop/96/4     ...........................................................................g.... [110800]
AY261362.1_Mkuzi_1979            ................................................................................ [110800]
AY261361.1_Malawi_Lil-20/1_1983  ..............................g................................................. [110800]
AY261360.1_Kenya_1950            .............................................c.................................. [110800]
AM712240.1_OURT_88/3             ........................................................................a..g.... [110800]
AM712239.1_Benin_97/1            ........................................................................a..g.... [110800]
KM111295.1_Ken06.Bus             .............................................c.................................. [110800]
KM111294.1_Ken05/Tk1             .............................................c.................................. [110800]
KM262844.1_strain_L60            ........................................................................a..g.... [110800]
KM102979.1_26544/OG10_Italy      ........................................................................a..g.... [110800]
KM262845.1_strain_NHV            ........................................................................a..g.... [110800]
KX354450.1_47/Ss/2008            ........................................................................a..g.... [110800]
KP055815.1_strain_BA71           ........................................................................a..g.... [110800]
MH025920.1_strain_R35            .............................................c.................................. [110800]
MH025919.1_strain_N10            .............................................c.................................. [110800]
MH025918.1_strain_R25            .............................................c.................................. [110800]
MH025917.1_strain_R7             .............................................c.................................. [110800]
MH025916.1_strain_R8             .............................................c.................................. [110800]

FR682468.2_Georgia_2007/1        acggtaccgaggaggcgttgaaatctttaggaaaggtgctgtctagtttggaatctccaattcctcccgtatatttaggt [110880]
ASF_IND_20_CAD543                ................................................................................ [110880]
LR899193.1_Germany_2020/1        ................................................................................ [110880]
MN393477.1_Wuhan_2019            ................................................................................ [110880]
MW396979.1_Timor-Leste/2019/1    ................................................................................ [110880]
MK940252.1_CN/2019/InnerMongolia ................................................................................ [110880]
MK128995.1_China/2018/AnhuiXCGQ  ................................................................................ [110880]
MH681419.1_ASFV/POL/2015         ................................................................................ [110880]
LS478113.1_Estonia_2014          ................................................................................ [110880]
KP843857.1_Odintsovo_02/14       ................................................................................ [110880]
```

```
FN557520.1_strain_E75              .......t........................................................................ [110880]
AY261366.1_Warthog                 .......t.......................................c...............................g [110880]
AY261365.1_Warmbaths               .......t........................................................................ [110880]
AY261364.1_Tengani_62              .......t........................................................................ [110880]
AY261363.1_Pretoriuskop/96/4       .......t.......................................................................g [110880]
AY261362.1_Mkuzi_1979              ................................................................................ [110880]
AY261361.1_Malawi_Lil-20/1_1983    .......t..............................g............................t............ [110880]
AY261360.1_Kenya_1950              .......t...........................c....a..........................t............ [110880]
AM712240.1_OURT_88/3               .......t........................................................................ [110880]
AM712239.1_Benin_97/1              .......t........................................................................ [110880]
KM111295.1_Ken06.Bus               .......t...........................c....a..........................t............ [110880]
KM111294.1_Ken05/Tk1               .......t...........................c....a..........................t............ [110880]
KM262844.1_strain_L60              .......t........................................................................ [110880]
KM102979.1_26544/OG10_Italy        .......t........................................................................ [110880]
KM262845.1_strain_NHV              .......t........................................................................ [110880]
KX354450.1_47/Ss/2008              .......t........................................................................ [110880]
KP055815.1_strain_BA71             .......t........................................................................ [110880]
MH025920.1_strain_R35              .......t...........................c....a..........................t............ [110880]
MH025919.1_strain_N10              .......t...........................c....a..........................t............ [110880]
MH025918.1_strain_R25              .......t...........................c....a..........................t............ [110880]
MH025917.1_strain_R7               .......t...........................c....a..........................t............ [110880]
MH025916.1_strain_R8               .......t...........................c....a..........................t............ [110880]

FR682468.2_Georgia_2007/1          atataattattgtgtctagaaattgtttgctttgaggtatcaaaatattcagcctgaccgctatttcttttagaataatt [110960]
ASF_IND_20_CAD543                  ..............y................................................................. [110960]
LR899193.1_Germany_2020/1          ................................................................................ [110960]
MN393477.1_Wuhan_2019              ................................................................................ [110960]
MW396979.1_Timor-Leste/2019/1      ................................................................................ [110960]
MK940252.1_CN/2019/InnerMongolia   ................................................................................ [110960]
MK128995.1_China/2018/AnhuiXCGQ    ................................................................................ [110960]
MH681419.1_ASFV/POL/2015           ................................................................................ [110960]
LS478113.1_Estonia_2014            ................................................................................ [110960]
KP843857.1_Odintsovo_02/14         ................................................................................ [110960]
FN557520.1_strain_E75              ..............c................................................................. [110960]
AY261366.1_Warthog                 ............................................................t................... [110960]
AY261365.1_Warmbaths               ............................................................t................... [110960]
AY261364.1_Tengani_62              ............................................................t................... [110960]
AY261363.1_Pretoriuskop/96/4       ............................................................t................... [110960]
AY261362.1_Mkuzi_1979              ................................................................................ [110960]
AY261361.1_Malawi_Lil-20/1_1983    ...................................t....................g..a....c............... [110960]
AY261360.1_Kenya_1950              ................................c..t....................g......................g [110960]
AM712240.1_OURT_88/3               ..............c................................................................. [110960]
AM712239.1_Benin_97/1              ..............c................................................................. [110960]
KM111295.1_Ken06.Bus               ................................c..t....................g......................g [110960]
```

```
KM111294.1_Ken05/Tk1             ................................c..t....................g......................g [110960]
KM262844.1_strain_L60            ..............c................................................................. [110960]
KM102979.1_26544/OG10_Italy      ..............c................................................................. [110960]
KM262845.1_strain_NHV            ..............c................................................................. [110960]
KX354450.1_47/Ss/2008            ..............c................................................................. [110960]
KP055815.1_strain_BA71           ..............c................................................................. [110960]
MH025920.1_strain_R35            ................................c..t....................g......................g [110960]
MH025919.1_strain_N10            ................................c..t....................g......................g [110960]
MH025918.1_strain_R25            ................................c..t....................g......................g [110960]
MH025917.1_strain_R7             ................................c..t....................g......................g [110960]
MH025916.1_strain_R8             ................................c..t....................g......................g [110960]

FR682468.2_Georgia_2007/1        cggtatagggcttgagtagttggcaatactcttaaaccggggcaccaaggtaacaatattttccatataatgggtttgat [111040]
ASF_IND_20_CAD543                ................................................................................ [111040]
LR899193.1_Germany_2020/1        ................................................................................ [111040]
MN393477.1_Wuhan_2019            ................................................................................ [111040]
MW396979.1_Timor-Leste/2019/1    ................................................................................ [111040]
MK940252.1_CN/2019/InnerMongolia ................................................................................ [111040]
MK128995.1_China/2018/AnhuiXCGQ  ................................................................................ [111040]
MH681419.1_ASFV/POL/2015         ................................................................................ [111040]
LS478113.1_Estonia_2014          ................................................................................ [111040]
KP843857.1_Odintsovo_02/14       ................................................................................ [111040]
FN557520.1_strain_E75            ................................................................................ [111040]
AY261366.1_Warthog               a............................................................................... [111040]
AY261365.1_Warmbaths             a.......................................................................a....... [111040]
AY261364.1_Tengani_62            a.......................................................................a....... [111040]
AY261363.1_Pretoriuskop/96/4     a......................................a................................a....... [111040]
AY261362.1_Mkuzi_1979            ................................................................................ [111040]
AY261361.1_Malawi_Lil-20/1_1983  a..............a..........................t.............................a....... [111040]
AY261360.1_Kenya_1950            a..............a...................................g....................a....... [111040]
AM712240.1_OURT_88/3             ................................................................................ [111040]
AM712239.1_Benin_97/1            ................................................................................ [111040]
KM111295.1_Ken06.Bus             a..............a........................................................a....... [111040]
KM111294.1_Ken05/Tk1             a..............a...................................g....................a....... [111040]
KM262844.1_strain_L60            ................................................................................ [111040]
KM102979.1_26544/OG10_Italy      ................................................................................ [111040]
KM262845.1_strain_NHV            ................................................................................ [111040]
KX354450.1_47/Ss/2008            ................................................................................ [111040]
KP055815.1_strain_BA71           ................................................................................ [111040]
MH025920.1_strain_R35            a..............a........................................................a....... [111040]
MH025919.1_strain_N10            a..............a........................................................a....... [111040]
MH025918.1_strain_R25            a..............a........................................................a....... [111040]
MH025917.1_strain_R7             a..............a........................................................a....... [111040]
MH025916.1_strain_R8             a..............a........................................................a....... [111040]
```

```
FR682468.2_Georgia_2007/1        acgctttgtttaaaaatgggcttaccggctttatgcttgttagttgtgcattgagtaccggtatgtcttctaggatttgt [111120]
ASF_IND_20_CAD543                ................................................................................ [111120]
LR899193.1_Germany_2020/1        ................................................................................ [111120]
MN393477.1_Wuhan_2019            ................................................................................ [111120]
MW396979.1_Timor-Leste/2019/1    ................................................................................ [111120]
MK940252.1_CN/2019/InnerMongolia ................................................................................ [111120]
MK128995.1_China/2018/AnhuiXCGQ  ................................................................................ [111120]
MH681419.1_ASFV/POL/2015         ................................................................................ [111120]
LS478113.1_Estonia_2014          ................................................................................ [111120]
KP843857.1_Odintsovo_02/14       ................................................................................ [111120]
FN557520.1_strain_E75            ................................................................................ [111120]
AY261366.1_Warthog               .......................................c..............................c......... [111120]
AY261365.1_Warmbaths             ..................................a....c..............................c......... [111120]
AY261364.1_Tengani_62            .t.....................................c..............................c......... [111120]
AY261363.1_Pretoriuskop/96/4     .......................................c..............................c......... [111120]
AY261362.1_Mkuzi_1979            ................................................................................ [111120]
AY261361.1_Malawi_Lil-20/1_1983  .t.................a....t...t........c..............t.a.gt......a.....c......... [111120]
AY261360.1_Kenya_1950            ................c....................c..............t...gt............c.aa...... [111120]
AM712240.1_OURT_88/3             ................................................................................ [111120]
AM712239.1_Benin_97/1            ................................................................................ [111120]
KM111295.1_Ken06.Bus             .....................................c..............t...gt............c..a....c. [111120]
KM111294.1_Ken05/Tk1             ................c....................c..............t....t............c.aa...... [111120]
KM262844.1_strain_L60            ................................................................................ [111120]
KM102979.1_26544/OG10_Italy      ................................................................................ [111120]
KM262845.1_strain_NHV            ................................................................................ [111120]
KX354450.1_47/Ss/2008            ................................................................................ [111120]
KP055815.1_strain_BA71           ................................................................................ [111120]
MH025920.1_strain_R35            .....................................c..............t...gt............c..a....c. [111120]
MH025919.1_strain_N10            .....................................c..............t...gt............c..a....c. [111120]
MH025918.1_strain_R25            .....................................c..............t...gt............c..a....c. [111120]
MH025917.1_strain_R7             .....................................c..............t...gt............c..a....c. [111120]
MH025916.1_strain_R8             .....................................c..............t...gt............c..a....c. [111120]

FR682468.2_Georgia_2007/1        ggctttatagaatgattagcaaacacagaatgtagtatattagatacttgtagcatatgtctatttgcggaaaattcctg [111200]
ASF_IND_20_CAD543                ................................................................................ [111200]
LR899193.1_Germany_2020/1        ................................................................................ [111200]
MN393477.1_Wuhan_2019            ................................................................................ [111200]
MW396979.1_Timor-Leste/2019/1    ................................................................................ [111200]
MK940252.1_CN/2019/InnerMongolia ................................................................................ [111200]
MK128995.1_China/2018/AnhuiXCGQ  ................................................................................ [111200]
MH681419.1_ASFV/POL/2015         ................................................................................ [111200]
LS478113.1_Estonia_2014          ................................................................................ [111200]
KP843857.1_Odintsovo_02/14       ................................................................................ [111200]
```

```
FN557520.1_strain_E75             ...........................................c............g....................... [111200]
AY261366.1_Warthog                ................................................................................ [111200]
AY261365.1_Warmbaths              ................................................................................ [111200]
AY261364.1_Tengani_62             ................................................................................ [111200]
AY261363.1_Pretoriuskop/96/4      ................................................................................ [111200]
AY261362.1_Mkuzi_1979             ................................................................................ [111200]
AY261361.1_Malawi_Lil-20/1_1983   ................................................................................ [111200]
AY261360.1_Kenya_1950             ................................c...................a.....................c..... [111200]
AM712240.1_OURT_88/3              ...........................................c............g....................... [111200]
AM712239.1_Benin_97/1             ...........................................c............g....................... [111200]
KM111295.1_Ken06.Bus              ................................c...................a.....................c..... [111200]
KM111294.1_Ken05/Tk1              ................................c.........................................c..... [111200]
KM262844.1_strain_L60             ...........................................c............g....................... [111200]
KM102979.1_26544/OG10_Italy       ...........................................c............g....................... [111200]
KM262845.1_strain_NHV             ...........................................c............g....................... [111200]
KX354450.1_47/Ss/2008             ...........................................c............g....................... [111200]
KP055815.1_strain_BA71            ...........................................c............g....................... [111200]
MH025920.1_strain_R35             ................................c...................a.....................c..... [111200]
MH025919.1_strain_N10             ................................c...................a.....................c..... [111200]
MH025918.1_strain_R25             ................................c...................a.....................c..... [111200]
MH025917.1_strain_R7              ................................c...................a.....................c..... [111200]
MH025916.1_strain_R8              ................................c...................a.....................c..... [111200]

FR682468.2_Georgia_2007/1         gtattctctgccgtgttgcgaatctttgggcggaaggggaccaagcatcggcacgtccgtgtaggtactggtggatttta [111280]
ASF_IND_20_CAD543                 ................................................................................ [111280]
LR899193.1_Germany_2020/1         ................................................................................ [111280]
MN393477.1_Wuhan_2019             ................................................................................ [111280]
MW396979.1_Timor-Leste/2019/1     ................................................................................ [111280]
MK940252.1_CN/2019/InnerMongolia  ................................................................................ [111280]
MK128995.1_China/2018/AnhuiXCGQ   ................................................................................ [111280]
MH681419.1_ASFV/POL/2015          ................................................................................ [111280]
LS478113.1_Estonia_2014           ................................................................................ [111280]
KP843857.1_Odintsovo_02/14        ................................................................................ [111280]
FN557520.1_strain_E75             .......................................................c........................ [111280]
AY261366.1_Warthog                .......................................................c....a................... [111280]
AY261365.1_Warmbaths              .......................................................c........................ [111280]
AY261364.1_Tengani_62             .......................................................c...............c........ [111280]
AY261363.1_Pretoriuskop/96/4      ..g.................................a........t.........c........................ [111280]
AY261362.1_Mkuzi_1979             .......................................................c........................ [111280]
AY261361.1_Malawi_Lil-20/1_1983   ...................a.........at............---..g..t.t.c.........cg.a..c........ [111280]
AY261360.1_Kenya_1950             ...................a.........at..............a..g..t...c..........g.a........... [111280]
AM712240.1_OURT_88/3              .......................................................c........................ [111280]
AM712239.1_Benin_97/1             .......................................................c........................ [111280]
KM111295.1_Ken06.Bus              ...................a.........at..............a..g..t..............g.a........... [111280]
```

```
KM111294.1_Ken05/Tk1             ...................a.........at..............a..g..t...c..........g.a........... [111280]
KM262844.1_strain_L60            .......................................................c........................ [111280]
KM102979.1_26544/OG10_Italy      .......................................................c........................ [111280]
KM262845.1_strain_NHV            .......................................................c........................ [111280]
KX354450.1_47/Ss/2008            .......................................................c........................ [111280]
KP055815.1_strain_BA71           .......................................................c........................ [111280]
MH025920.1_strain_R35            ...................a.........at..............a..g..t..............g.a........... [111280]
MH025919.1_strain_N10            ...................a.........at..............a..g..t..............g.a........... [111280]
MH025918.1_strain_R25            ...................a.........at..............a..g..t..............g.a........... [111280]
MH025917.1_strain_R7             ...................a.........at..............a..g..t..............g.a........... [111280]
MH025916.1_strain_R8             ...................a.........at..............a..g..t..............g.a........... [111280]

FR682468.2_Georgia_2007/1        tgagttcctgctctatgttcggtttgacatgtggatttcctaaaggaatacctctacctgcaatccctttttctaccgac [111360]
ASF_IND_20_CAD543                ................................................................................ [111360]
LR899193.1_Germany_2020/1        ................................................................................ [111360]
MN393477.1_Wuhan_2019            ................................................................................ [111360]
MW396979.1_Timor-Leste/2019/1    ................................................................................ [111360]
MK940252.1_CN/2019/InnerMongolia ................................................................................ [111360]
MK128995.1_China/2018/AnhuiXCGQ  ................................................................................ [111360]
MH681419.1_ASFV/POL/2015         ................................................................................ [111360]
LS478113.1_Estonia_2014          ................................................................................ [111360]
KP843857.1_Odintsovo_02/14       ................................................................................ [111360]
FN557520.1_strain_E75            ................................................................................ [111360]
AY261366.1_Warthog               .......................................................g.........t..........a... [111360]
AY261365.1_Warmbaths             .......................................................g.........t..........a... [111360]
AY261364.1_Tengani_62            .......................................................g.........t..........a... [111360]
AY261363.1_Pretoriuskop/96/4     ...........................................g...........g.........t..........a... [111360]
AY261362.1_Mkuzi_1979            ..........................................................c..................... [111360]
AY261361.1_Malawi_Lil-20/1_1983  ....c....................t..........................c..g.........a....c.....a..t [111360]
AY261360.1_Kenya_1950            ..........t.....a..............c....................c..g.....c..ta..........a... [111360]
AM712240.1_OURT_88/3             ................................................................................ [111360]
AM712239.1_Benin_97/1            ................................................................................ [111360]
KM111295.1_Ken06.Bus             ................a..............c....................c..g.....c..ta.......c..a... [111360]
KM111294.1_Ken05/Tk1             ..........t.....a..............c....................c..g.....c..ta..........a... [111360]
KM262844.1_strain_L60            ................................................................................ [111360]
KM102979.1_26544/OG10_Italy      ................................................................................ [111360]
KM262845.1_strain_NHV            ................................................................................ [111360]
KX354450.1_47/Ss/2008            ................................................................................ [111360]
KP055815.1_strain_BA71           ................................................................................ [111360]
MH025920.1_strain_R35            ................a..............c....................c..g.....c..ta.......c..a... [111360]
MH025919.1_strain_N10            ................a..............c....................c..g.....c..ta.......c..a... [111360]
MH025918.1_strain_R25            ................a..............c....................c..g.....c..ta.......c..a... [111360]
MH025917.1_strain_R7             ................a..............c....................c..g.....c..ta.......c..a... [111360]
MH025916.1_strain_R8             ................a..............c....................c..g.....c..ta.......c..a... [111360]
```

```
FR682468.2_Georgia_2007/1        gcaggtagattgtgcgctaaacacaaaatattgtacacgtctttgtgcggaatatatccgttatagtgctggcccggcat [111440]
ASF_IND_20_CAD543                ................................................................................ [111440]
LR899193.1_Germany_2020/1        ................................................................................ [111440]
MN393477.1_Wuhan_2019            ................................................................................ [111440]
MW396979.1_Timor-Leste/2019/1    ................................................................................ [111440]
MK940252.1_CN/2019/InnerMongolia ................................................................................ [111440]
MK128995.1_China/2018/AnhuiXCGQ  ................................................................................ [111440]
MH681419.1_ASFV/POL/2015         ................................................................................ [111440]
LS478113.1_Estonia_2014          ................................................................................ [111440]
KP843857.1_Odintsovo_02/14       ................................................................................ [111440]
FN557520.1_strain_E75            ................................................................................ [111440]
AY261366.1_Warthog               .......ac..........................t.......ga......................a............ [111440]
AY261365.1_Warmbaths             .......ac..................................ga......................a............ [111440]
AY261364.1_Tengani_62            .......ac..................................ga..g...................a............ [111440]
AY261363.1_Pretoriuskop/96/4     .......ac..................................ga.....................ca............ [111440]
AY261362.1_Mkuzi_1979            ................................................................................ [111440]
AY261361.1_Malawi_Lil-20/1_1983  ..g....a...............t....................a..............a.......a........a... [111440]
AY261360.1_Kenya_1950            .....c......................................a..............a.................... [111440]
AM712240.1_OURT_88/3             ................................................................................ [111440]
AM712239.1_Benin_97/1            ................................................................................ [111440]
KM111295.1_Ken06.Bus             .....c......................................a..............a.................t.. [111440]
KM111294.1_Ken05/Tk1             .....c......................................a..............a.................... [111440]
KM262844.1_strain_L60            ................................................................................ [111440]
KM102979.1_26544/OG10_Italy      ................................................................................ [111440]
KM262845.1_strain_NHV            ................................................................................ [111440]
KX354450.1_47/Ss/2008            ................................................................................ [111440]
KP055815.1_strain_BA71           ................................................................................ [111440]
MH025920.1_strain_R35            .....c......................................a..............a.................t.. [111440]
MH025919.1_strain_N10            .....c......................................a..............a.................t.. [111440]
MH025918.1_strain_R25            .....c......................................a..............a.................t.. [111440]
MH025917.1_strain_R7             .....c......................................a..............a.................t.. [111440]
MH025916.1_strain_R8             .....c......................................a..............a.................t.. [111440]

FR682468.2_Georgia_2007/1        ctgatcgccaaggtgctgctcatgcttaatggtaccctttgttctgagtttaggaagatcctcgtacgaaaaaaattttg [111520]
ASF_IND_20_CAD543                ................................................................................ [111520]
LR899193.1_Germany_2020/1        ................................................................................ [111520]
MN393477.1_Wuhan_2019            ................................................................................ [111520]
MW396979.1_Timor-Leste/2019/1    ................................................................................ [111520]
MK940252.1_CN/2019/InnerMongolia ................................................................................ [111520]
MK128995.1_China/2018/AnhuiXCGQ  ................................................................................ [111520]
MH681419.1_ASFV/POL/2015         ................................................................................ [111520]
LS478113.1_Estonia_2014          ................................................................................ [111520]
KP843857.1_Odintsovo_02/14       ................................................................................ [111520]
```

```
FN557520.1_strain_E75            ...........................................t............t....................... [111520]
AY261366.1_Warthog               t............................................................................... [111520]
AY261365.1_Warmbaths             t............................................................................... [111520]
AY261364.1_Tengani_62            ................................................................................ [111520]
AY261363.1_Pretoriuskop/96/4     t............................................................................... [111520]
AY261362.1_Mkuzi_1979            ...........................................t............t....................... [111520]
AY261361.1_Malawi_Lil-20/1_1983  ................................c..........t...................a................ [111520]
AY261360.1_Kenya_1950            ...g....................................a...............c......a...............a [111520]
AM712240.1_OURT_88/3             ...........................................t............t....................... [111520]
AM712239.1_Benin_97/1            ...........................................t............t....................... [111520]
KM111295.1_Ken06.Bus             ........................................a...............c.c....a................ [111520]
KM111294.1_Ken05/Tk1             ...g....................................a......a........c......a................ [111520]
KM262844.1_strain_L60            ...........................................t............t....................... [111520]
KM102979.1_26544/OG10_Italy      ...........................................t............t....................... [111520]
KM262845.1_strain_NHV            ...........................................t............t....................... [111520]
KX354450.1_47/Ss/2008            ...........................................t............t....................... [111520]
KP055815.1_strain_BA71           ...........................................t............t....................... [111520]
MH025920.1_strain_R35            ........................................a...............c.c....a................ [111520]
MH025919.1_strain_N10            ........................................a...............c.c....a................ [111520]
MH025918.1_strain_R25            ........................................a...............c.c....a................ [111520]
MH025917.1_strain_R7             ........................................a...............c.c....a................ [111520]
MH025916.1_strain_R8             ........................................a...............c.c....a................ [111520]

FR682468.2_Georgia_2007/1        tgtgctcgctgaacctcgtagaaggaaccgaactattttttgggttttttaaggaaggcaatgaggaaggctgggtcaga [111600]
ASF_IND_20_CAD543                ................................................................................ [111600]
LR899193.1_Germany_2020/1        ................................................................................ [111600]
MN393477.1_Wuhan_2019            ................................................................................ [111600]
MW396979.1_Timor-Leste/2019/1    ................................................................................ [111600]
MK940252.1_CN/2019/InnerMongolia ................................................................................ [111600]
MK128995.1_China/2018/AnhuiXCGQ  ................................................................................ [111600]
MH681419.1_ASFV/POL/2015         ................................................................................ [111600]
LS478113.1_Estonia_2014          ................................................................................ [111600]
KP843857.1_Odintsovo_02/14       ................................................................................ [111600]
FN557520.1_strain_E75            ......................c......................................................... [111600]
AY261366.1_Warthog               .........................................................................a...... [111600]
AY261365.1_Warmbaths             .............t.................................................................. [111600]
AY261364.1_Tengani_62            ................................................................................ [111600]
AY261363.1_Pretoriuskop/96/4     .......a..................................................t..................... [111600]
AY261362.1_Mkuzi_1979            ......................c......................................................... [111600]
AY261361.1_Malawi_Lil-20/1_1983  ..........a.......c.............a........................................a..t... [111600]
AY261360.1_Kenya_1950            ......................g..c.t....a...........................................t... [111600]
AM712240.1_OURT_88/3             ......................c......................................................... [111600]
AM712239.1_Benin_97/1            ......................c......................................................... [111600]
KM111295.1_Ken06.Bus             .......a..............g..c.t....a...........................................t... [111600]
```

```
KM111294.1_Ken05/Tk1             ......................g..c.t....a...................................aa......t... [111600]
KM262844.1_strain_L60            ......................c......................................................... [111600]
KM102979.1_26544/OG10_Italy      ......................c......................................................... [111600]
KM262845.1_strain_NHV            ......................c......................................................... [111600]
KX354450.1_47/Ss/2008            ......................c......................................................... [111600]
KP055815.1_strain_BA71           ......................c......................................................... [111600]
MH025920.1_strain_R35            .......a..............g..c.t....a...........................................t... [111600]
MH025919.1_strain_N10            .......a..............g..c.t....a...........................................t... [111600]
MH025918.1_strain_R25            .......a..............g..c.t....a...........................................t... [111600]
MH025917.1_strain_R7             .......a..............g..c.t....a...........................................t... [111600]
MH025916.1_strain_R8             .......a..............g..c.t....a...........................................t... [111600]

FR682468.2_Georgia_2007/1        caatttttctgtgtgccctttaagctagccacctgcggaaatgtttttttttccgtacgaacaacattgcgcctaattag [111680]
ASF_IND_20_CAD543                ................................................................................ [111680]
LR899193.1_Germany_2020/1        ................................................................................ [111680]
MN393477.1_Wuhan_2019            ................................................................................ [111680]
MW396979.1_Timor-Leste/2019/1    ................................................................................ [111680]
MK940252.1_CN/2019/InnerMongolia ................................................................................ [111680]
MK128995.1_China/2018/AnhuiXCGQ  ................................................................................ [111680]
MH681419.1_ASFV/POL/2015         ................................................................................ [111680]
LS478113.1_Estonia_2014          ................................................................................ [111680]
KP843857.1_Odintsovo_02/14       ............................................-................................... [111680]
FN557520.1_strain_E75            ................................................................................ [111680]
AY261366.1_Warthog               ...............t.........c..a........................t..................t....... [111680]
AY261365.1_Warmbaths             ...............t.........c..at.......................t.......................... [111680]
AY261364.1_Tengani_62            ...............t.....................................t.......................... [111680]
AY261363.1_Pretoriuskop/96/4     ...............t.........c..a........................t..................t....... [111680]
AY261362.1_Mkuzi_1979            ................................................................................ [111680]
AY261361.1_Malawi_Lil-20/1_1983  ....g.........a.t....................................t........g................. [111680]
AY261360.1_Kenya_1950            ..............a.t.......g................c...........t.......................... [111680]
AM712240.1_OURT_88/3             ................................................................................ [111680]
AM712239.1_Benin_97/1            ................................................................................ [111680]
KM111295.1_Ken06.Bus             ..............a.t.......g....t...........c...........t.......................... [111680]
KM111294.1_Ken05/Tk1             ............a.a.t.......g................c...........t.......................... [111680]
KM262844.1_strain_L60            ................................................................................ [111680]
KM102979.1_26544/OG10_Italy      ................................................................................ [111680]
KM262845.1_strain_NHV            ................................................................................ [111680]
KX354450.1_47/Ss/2008            ................................................................................ [111680]
KP055815.1_strain_BA71           ................................................................................ [111680]
MH025920.1_strain_R35            ..............a.t.......g....t...........c...........t.......................... [111680]
MH025919.1_strain_N10            ..............a.t.......g....t...........c...........t.......................... [111680]
MH025918.1_strain_R25            ..............a.t.......g....t...........c...........t.......................... [111680]
MH025917.1_strain_R7             ..............a.t.......g....t...........c...........t.......................... [111680]
MH025916.1_strain_R8             ..............a.t.......g....t...........c...........t.......................... [111680]
```

```
FR682468.2_Georgia_2007/1        gttttccgtatgggttgaaaaagcaggacgatgatttttaaaatgattaaaaagtttattttttggaatggagctgtacg [111760]
ASF_IND_20_CAD543                ................................................................................ [111760]
LR899193.1_Germany_2020/1        ................................................................................ [111760]
MN393477.1_Wuhan_2019            ................................................................................ [111760]
MW396979.1_Timor-Leste/2019/1    ................................................................................ [111760]
MK940252.1_CN/2019/InnerMongolia ................................................................................ [111760]
MK128995.1_China/2018/AnhuiXCGQ  ................................................................................ [111760]
MH681419.1_ASFV/POL/2015         ................................................................................ [111760]
LS478113.1_Estonia_2014          ................................................................................ [111760]
KP843857.1_Odintsovo_02/14       ................................................................................ [111760]
FN557520.1_strain_E75            ................................................................................ [111760]
AY261366.1_Warthog               ................................a............................................... [111760]
AY261365.1_Warmbaths             ................................a............................................... [111760]
AY261364.1_Tengani_62            ................................................................................ [111760]
AY261363.1_Pretoriuskop/96/4     ................................a............................................... [111760]
AY261362.1_Mkuzi_1979            ................................................................................ [111760]
AY261361.1_Malawi_Lil-20/1_1983  a.......................g...................t................................... [111760]
AY261360.1_Kenya_1950            ........................g..g......a......g..t..............c...........ga..a.... [111760]
AM712240.1_OURT_88/3             ................................................................................ [111760]
AM712239.1_Benin_97/1            ................................................................................ [111760]
KM111295.1_Ken06.Bus             ........................g..g......a......g..t..............cc..........ga....... [111760]
KM111294.1_Ken05/Tk1             ........................g..g......a.c....g..t..............c...........ga..a.... [111760]
KM262844.1_strain_L60            ................................................................................ [111760]
KM102979.1_26544/OG10_Italy      ................................................................................ [111760]
KM262845.1_strain_NHV            ................................................................................ [111760]
KX354450.1_47/Ss/2008            ................................................................................ [111760]
KP055815.1_strain_BA71           ................................................................................ [111760]
MH025920.1_strain_R35            ........................g..g......a......g..t..............cc..........ga....... [111760]
MH025919.1_strain_N10            ........................g..g......a......g..t..............cc..........ga....... [111760]
MH025918.1_strain_R25            ........................g..g......a......g..t..............cc..........ga....... [111760]
MH025917.1_strain_R7             ........................g..g......a......g..t..............cc..........ga....... [111760]
MH025916.1_strain_R8             ........................g..g......a......g..t..............cc..........ga....... [111760]

FR682468.2_Georgia_2007/1        gctccagatcttgcgcatcgccgtaaccaatgtttttgtgctgagggttcagcataaaagaaaagttacgtagatcactg [111840]
ASF_IND_20_CAD543                ................................................................................ [111840]
LR899193.1_Germany_2020/1        ................................................................................ [111840]
MN393477.1_Wuhan_2019            ................................................................................ [111840]
MW396979.1_Timor-Leste/2019/1    ................................................................................ [111840]
MK940252.1_CN/2019/InnerMongolia ................................................................................ [111840]
MK128995.1_China/2018/AnhuiXCGQ  ................................................................................ [111840]
MH681419.1_ASFV/POL/2015         ................................................................................ [111840]
LS478113.1_Estonia_2014          ................................................................................ [111840]
KP843857.1_Odintsovo_02/14       ................................................................................ [111840]
```

```
FN557520.1_strain_E75             ................................................................................ [111840]
AY261366.1_Warthog                ........................................................................a....... [111840]
AY261365.1_Warmbaths              ......a..................................................................g...... [111840]
AY261364.1_Tengani_62             ...............................a................................................ [111840]
AY261363.1_Pretoriuskop/96/4      .........................................................................g...... [111840]
AY261362.1_Mkuzi_1979             ................................................................................ [111840]
AY261361.1_Malawi_Lil-20/1_1983   ....t.ag...........a............................................................ [111840]
AY261360.1_Kenya_1950             ....t..g.......................a..............a................................c [111840]
AM712240.1_OURT_88/3              ................................................................................ [111840]
AM712239.1_Benin_97/1             ................................................................................ [111840]
KM111295.1_Ken06.Bus              ....t..g......................................a................................c [111840]
KM111294.1_Ken05/Tk1              ....t..g.......................a..............a................................c [111840]
KM262844.1_strain_L60             ................................................................................ [111840]
KM102979.1_26544/OG10_Italy       ................................................................................ [111840]
KM262845.1_strain_NHV             ................................................................................ [111840]
KX354450.1_47/Ss/2008             ................................................................................ [111840]
KP055815.1_strain_BA71            ................................................................................ [111840]
MH025920.1_strain_R35             ....t..g......................................a................................c [111840]
MH025919.1_strain_N10             ....t..g......................................a................................c [111840]
MH025918.1_strain_R25             ....t..g......................................a................................c [111840]
MH025917.1_strain_R7              ....t..g......................................a................................c [111840]
MH025916.1_strain_R8              ....t..g......................................a................................c [111840]

FR682468.2_Georgia_2007/1         agttgcaatcccttttcagccttttcaggactattagtgtattcattgtatacaggcgcagctccatttttgttgccgca [111920]
ASF_IND_20_CAD543                 ................................................................................ [111920]
LR899193.1_Germany_2020/1         ................................................................................ [111920]
MN393477.1_Wuhan_2019             ................................................................................ [111920]
MW396979.1_Timor-Leste/2019/1     ................................................................................ [111920]
MK940252.1_CN/2019/InnerMongolia  ................................................................................ [111920]
MK128995.1_China/2018/AnhuiXCGQ   ................................................................................ [111920]
MH681419.1_ASFV/POL/2015          ................................................................................ [111920]
LS478113.1_Estonia_2014           ................................................................................ [111920]
KP843857.1_Odintsovo_02/14        ................................................................................ [111920]
FN557520.1_strain_E75             ................................................................................ [111920]
AY261366.1_Warthog                ........................c..........g...................................a........ [111920]
AY261365.1_Warmbaths              ........................c..........g...................................a........ [111920]
AY261364.1_Tengani_62             ...................................g........g................................... [111920]
AY261363.1_Pretoriuskop/96/4      ......................g.c..........g...................................a........ [111920]
AY261362.1_Mkuzi_1979             ................................................................................ [111920]
AY261361.1_Malawi_Lil-20/1_1983   ............c...........c.g........g...ct................t.............a........ [111920]
AY261360.1_Kenya_1950             ........c...............c.g........g...c...................g...........a........ [111920]
AM712240.1_OURT_88/3              ................................................................................ [111920]
AM712239.1_Benin_97/1             ................................................................................ [111920]
KM111295.1_Ken06.Bus              ........................c.g........g...c...................g...........a........ [111920]
```

```
KM111294.1_Ken05/Tk1            ........c...............c.g........g...c................t..g...........a........ [111920]
KM262844.1_strain_L60           ................................................................................ [111920]
KM102979.1_26544/OG10_Italy     ................................................................................ [111920]
KM262845.1_strain_NHV           ................................................................................ [111920]
KX354450.1_47/Ss/2008           ................................................................................ [111920]
KP055815.1_strain_BA71          ................................................................................ [111920]
MH025920.1_strain_R35           ........................c.g........g...c...................g...........a........ [111920]
MH025919.1_strain_N10           ........................c.g........g...c...................g...........a........ [111920]
MH025918.1_strain_R25           ........................c.g........g...c...................g...........a........ [111920]
MH025917.1_strain_R7            ........................c.g........g...c...................g...........a........ [111920]
MH025916.1_strain_R8            ........................c.g........g...c...................g...........a........ [111920]

FR682468.2_Georgia_2007/1       gtaccgggaatttagtatattatcagaataccggttatgacgcggcaaatcgctttcccaaagaggtggatctgacctat [112000]
ASF_IND_20_CAD543               ................................................................................ [112000]
LR899193.1_Germany_2020/1       ................................................................................ [112000]
MN393477.1_Wuhan_2019           ................................................................................ [112000]
MW396979.1_Timor-Leste/2019/1   ................................................................................ [112000]
MK940252.1_CN/2019/InnerMongolia ................................................................................ [112000]
MK128995.1_China/2018/AnhuiXCGQ ................................................................................ [112000]
MH681419.1_ASFV/POL/2015        ................................................................................ [112000]
LS478113.1_Estonia_2014         ................................................................................ [112000]
KP843857.1_Odintsovo_02/14      ................................................................................ [112000]
FN557520.1_strain_E75           ................................................................................ [112000]
AY261366.1_Warthog              .................................a.....g........................................ [112000]
AY261365.1_Warmbaths            .................................a.....g........................................ [112000]
AY261364.1_Tengani_62           .......................................g................................c....... [112000]
AY261363.1_Pretoriuskop/96/4    .................................a.....g........................................ [112000]
AY261362.1_Mkuzi_1979           ................................................................................ [112000]
AY261361.1_Malawi_Lil-20/1_1983 ................................................g............................... [112000]
AY261360.1_Kenya_1950           ........c.....a...............t...................................c............. [112000]
AM712240.1_OURT_88/3            ................................................................................ [112000]
AM712239.1_Benin_97/1           ................................................................................ [112000]
KM111295.1_Ken06.Bus            ........c.....a...............t..............t....................c............. [112000]
KM111294.1_Ken05/Tk1            ........c.....a...............t...................................c............. [112000]
KM262844.1_strain_L60           ................................................................................ [112000]
KM102979.1_26544/OG10_Italy     ................................................................................ [112000]
KM262845.1_strain_NHV           ................................................................................ [112000]
KX354450.1_47/Ss/2008           ................................................................................ [112000]
KP055815.1_strain_BA71          ................................................................................ [112000]
MH025920.1_strain_R35           ........c.....a...............t..............t....................c............. [112000]
MH025919.1_strain_N10           ........c.....a...............t..............t....................c............. [112000]
MH025918.1_strain_R25           ........c.....a...............t..............t....................c............. [112000]
MH025917.1_strain_R7            ........c.....a...............t..............t....................c............. [112000]
MH025916.1_strain_R8            ........c.....a...............t..............t....................c............. [112000]
```

```
FR682468.2_Georgia_2007/1        aatcggctaacagctttgaagcataatcatgatacattgtatataaaagttaattattatattgagaaggcataattact [112080]
ASF_IND_20_CAD543                ................................................................................ [112080]
LR899193.1_Germany_2020/1        ................................................................................ [112080]
MN393477.1_Wuhan_2019            ................................................................................ [112080]
MW396979.1_Timor-Leste/2019/1    ................................................................................ [112080]
MK940252.1_CN/2019/InnerMongolia ................................................................................ [112080]
MK128995.1_China/2018/AnhuiXCGQ  ................................................................................ [112080]
MH681419.1_ASFV/POL/2015         ................................................................................ [112080]
LS478113.1_Estonia_2014          ................................................................................ [112080]
KP843857.1_Odintsovo_02/14       ................................................................................ [112080]
FN557520.1_strain_E75            ...............................................................................c [112080]
AY261366.1_Warthog               ...............................................................................c [112080]
AY261365.1_Warmbaths             ...............................................................................c [112080]
AY261364.1_Tengani_62            .........................c.....................................................c [112080]
AY261363.1_Pretoriuskop/96/4     ...............................................................................c [112080]
AY261362.1_Mkuzi_1979            ...............................................................................c [112080]
AY261361.1_Malawi_Lil-20/1_1983  ....c........................................................................... [112080]
AY261360.1_Kenya_1950            ....c........................................................................... [112080]
AM712240.1_OURT_88/3             ...............................................................................c [112080]
AM712239.1_Benin_97/1            ...............................................................................c [112080]
KM111295.1_Ken06.Bus             ....c...........g............................................................... [112080]
KM111294.1_Ken05/Tk1             ....c........................................................................... [112080]
KM262844.1_strain_L60            ...............................................................................c [112080]
KM102979.1_26544/OG10_Italy      ...............................................................................c [112080]
KM262845.1_strain_NHV            ...............................................................................c [112080]
KX354450.1_47/Ss/2008            ...............................................................................c [112080]
KP055815.1_strain_BA71           ...............................................................................c [112080]
MH025920.1_strain_R35            ....c...........g............................................................... [112080]
MH025919.1_strain_N10            ....c...........g............................................................... [112080]
MH025918.1_strain_R25            ....c...........g............................................................... [112080]
MH025917.1_strain_R7             ....c...........g............................................................... [112080]
MH025916.1_strain_R8             ....c...........g............................................................... [112080]

FR682468.2_Georgia_2007/1        tcttgtaggggtacaagaggctttgaatcaggcaaactgacgggttttgaat---------------cggccggctttgg [112160]
ASF_IND_20_CAD543                ....................................................---------------............. [112160]
LR899193.1_Germany_2020/1        ....................................................---------------............. [112160]
MN393477.1_Wuhan_2019            ....................................................---------------............. [112160]
MW396979.1_Timor-Leste/2019/1    ....................................................---------------............. [112160]
MK940252.1_CN/2019/InnerMongolia ....................................................---------------............. [112160]
MK128995.1_China/2018/AnhuiXCGQ  ....................................................---------------............. [112160]
MH681419.1_ASFV/POL/2015         ....................................................---------------............. [112160]
LS478113.1_Estonia_2014          ....................................................---------------............. [112160]
KP843857.1_Odintsovo_02/14       ....................................................---------------............. [112160]
```

```
FN557520.1_strain_E75              ...............................c.g.....g.c..c.......---------------............. [112160]
AY261366.1_Warthog                 ...................a...........c------------------------------------...........a [112160]
AY261365.1_Warmbaths               ...............................ctg...------------------------------t...a.......a [112160]
AY261364.1_Tengani_62              ........................a......c.g.....g.c..c.......---------------......aa....a [112160]
AY261363.1_Pretoriuskop/96/4       ...............................ctg.....g.c..........---------------.......t....a [112160]
AY261362.1_Mkuzi_1979              ...............................c.g.......c..c.......---------------............. [112160]
AY261361.1_Malawi_Lil-20/1_1983    .......a...................a.g.ctggttcag.t..........---------------.....a.t----- [112160]
AY261360.1_Kenya_1950              .......a.....t.............a.g.c.ggtt------------------------------.......t..aaa [112160]
AM712240.1_OURT_88/3               ...............................c.g.....g.c..c.......cggccggctttggac............. [112160]
AM712239.1_Benin_97/1              ...............................c.g.....g.c..c.......cggccggctttggac............. [112160]
KM111295.1_Ken06.Bus               .......a.....t.............a.g.c.ggt-------------------------------------------- [112160]
KM111294.1_Ken05/Tk1               .......a.....t.............a.g.c.ggtt------------------------------.......t..aaa [112160]
KM262844.1_strain_L60              ...............................c.g.....g.c..c.......---------------............. [112160]
KM102979.1_26544/OG10_Italy        ...............................c.g.....g.c..c.......---------------............. [112160]
KM262845.1_strain_NHV              ...............................c.g.....g.c..c.......cggccggctttggac............. [112160]
KX354450.1_47/Ss/2008              ...............................c.g.....g.c..c.......---------------............. [112160]
KP055815.1_strain_BA71             ...............................c.g.....g.c..c.......---------------............. [112160]
MH025920.1_strain_R35              .......a.....t.............a.g.c.ggt-------------------------------------------- [112160]
MH025919.1_strain_N10              .......a.....t.............a.g.c.ggt-------------------------------------------- [112160]
MH025918.1_strain_R25              .......a.....t.............a.g.c.ggt-------------------------------------------- [112160]
MH025917.1_strain_R7               .......a.....t.............a.g.c.ggt-------------------------------------------- [112160]
MH025916.1_strain_R8               .......a.....t.............a.g.c.ggt-------------------------------------------- [112160]

FR682468.2_Georgia_2007/1          accg---------------gcaggtatctttttaggttgatcttcttctagctcattagacacggatgggggagaaatag [112240]
ASF_IND_20_CAD543                  ....gccggctttggaccg............................................................. [112240]
LR899193.1_Germany_2020/1          ....---------------............................................................. [112240]
MN393477.1_Wuhan_2019              ....---------------............................................................. [112240]
MW396979.1_Timor-Leste/2019/1      ....---------------............................................................. [112240]
MK940252.1_CN/2019/InnerMongolia   ....---------------............................................................. [112240]
MK128995.1_China/2018/AnhuiXCGQ    ....---------------............................................................. [112240]
MH681419.1_ASFV/POL/2015           ....---------------............................................................. [112240]
LS478113.1_Estonia_2014            ....---------------............................................................. [112240]
KP843857.1_Odintsovo_02/14         ....---------------............................................................. [112240]
FN557520.1_strain_E75              ....gccggctttggaccg............................................................. [112240]
AY261366.1_Warthog                 .t..gccggttttggaccg..g....................c..................................... [112240]
AY261365.1_Warmbaths               .t..gccggctttgcaccg..g.c..................c..................................... [112240]
AY261364.1_Tengani_62              ....gccggttttgaaccg..g...........t........c......................c.............. [112240]
AY261363.1_Pretoriuskop/96/4       .t..gccggttttgaaccg..g.c..................c..................................... [112240]
AY261362.1_Mkuzi_1979              ....gccggttttggaccg............................................a................ [112240]
AY261361.1_Malawi_Lil-20/1_1983    -...gccagttttgaaccg..ga..............c.............t........a.t..............c.. [112240]
AY261360.1_Kenya_1950              .t..gccggttttgaaccg..g......a........c.............t....c..ga..a.............c.. [112240]
AM712240.1_OURT_88/3               ....gccggctttggaccg............................................................. [112240]
AM712239.1_Benin_97/1              ....gccggctttggaccg............................................................. [112240]
KM111295.1_Ken06.Bus               -t..gccggttttgaaccg..g.a....a........c.............t....c..ga..a.............c.. [112240]
```

```
KM111294.1_Ken05/Tk1            .t..gccggttttgaaccg.gg.a....a........c.............t....c..ga..a.............c.. [112240]
KM262844.1_strain_L60           ....gccggctttggaccg............................................................. [112240]
KM102979.1_26544/OG10_Italy     ....gccggctttggaccg............................................................. [112240]
KM262845.1_strain_NHV           ....gccggctttggaccg............................................................. [112240]
KX354450.1_47/Ss/2008           ....gccggctttggaccg............................................................. [112240]
KP055815.1_strain_BA71          ....gccggctttggaccg............................................................. [112240]
MH025920.1_strain_R35           -t..gccggttttgaaccg..g.a....a........c.............t....c..ga..a.............c.. [112240]
MH025919.1_strain_N10           -t..gccggttttgaaccg..g.a....a........c.............t....c..ga..a.............c.. [112240]
MH025918.1_strain_R25           -t..gccggttttgaaccg..g.a....a........c.............t....c..ga..a.............c.. [112240]
MH025917.1_strain_R7            -t..gccggttttgaaccg..g.a....a........c.............t....c..ga..a.............c.. [112240]
MH025916.1_strain_R8            -t..gccggttttgaaccg..g.a....a........c.............t....c..ga..a.............c.. [112240]

FR682468.2_Georgia_2007/1       gaggaataatttcatctccgcccttatatttgtcatggatagaagaaacaattacatccatgtttgatttattataaatg [112320]
ASF_IND_20_CAD543               ................................................................................ [112320]
LR899193.1_Germany_2020/1       ................................................................................ [112320]
MN393477.1_Wuhan_2019           ................................................................................ [112320]
MW396979.1_Timor-Leste/2019/1   ................................................................................ [112320]
MK940252.1_CN/2019/InnerMongolia ................................................................................ [112320]
MK128995.1_China/2018/AnhuiXCGQ ................................................................................ [112320]
MH681419.1_ASFV/POL/2015        ................................................................................ [112320]
LS478113.1_Estonia_2014         ................................................................................ [112320]
KP843857.1_Odintsovo_02/14      ................................................................................ [112320]
FN557520.1_strain_E75           ..................................g............................................. [112320]
AY261366.1_Warthog              ..................................g............................................c [112320]
AY261365.1_Warmbaths            ..................................g...........................................cc [112320]
AY261364.1_Tengani_62           ..................................g............................................. [112320]
AY261363.1_Pretoriuskop/96/4    ..................................g............................................. [112320]
AY261362.1_Mkuzi_1979           ..................................g............................................. [112320]
AY261361.1_Malawi_Lil-20/1_1983 ...................t..t.............................g..............c............ [112320]
AY261360.1_Kenya_1950           .......g...........t..t...........g.................g........................... [112320]
AM712240.1_OURT_88/3            ..................................g............................................. [112320]
AM712239.1_Benin_97/1           ..................................g............................................. [112320]
KM111295.1_Ken06.Bus            .......g...........t..t...........g.................g........................... [112320]
KM111294.1_Ken05/Tk1            .......g...........t..t...........g.................g........................... [112320]
KM262844.1_strain_L60           ..................................g............................................. [112320]
KM102979.1_26544/OG10_Italy     ..................................g............................................. [112320]
KM262845.1_strain_NHV           ..................................g............................................. [112320]
KX354450.1_47/Ss/2008           ..................................g............................................. [112320]
KP055815.1_strain_BA71          ..................................g............................................. [112320]
MH025920.1_strain_R35           .......g...........t..t...........g.................g........................... [112320]
MH025919.1_strain_N10           .......g...........t..t...........g.................g........................... [112320]
MH025918.1_strain_R25           .......g...........t..t...........g.................g........................... [112320]
MH025917.1_strain_R7            .......g...........t..t...........g.................g........................... [112320]
MH025916.1_strain_R8            .......g...........t..t...........g.................g........................... [112320]
```

```
FR682468.2_Georgia_2007/1        tcgtttaactggtgatttaaaacataataatgcaaaaataatagggctacaatgcatatatatacgtaaatagccgtctt [112400]
ASF_IND_20_CAD543                ................................................................................ [112400]
LR899193.1_Germany_2020/1        ................................................................................ [112400]
MN393477.1_Wuhan_2019            ................................................................................ [112400]
MW396979.1_Timor-Leste/2019/1    ................................................................................ [112400]
MK940252.1_CN/2019/InnerMongolia ................................................................................ [112400]
MK128995.1_China/2018/AnhuiXCGQ  ................................................................................ [112400]
MH681419.1_ASFV/POL/2015         ................................................................................ [112400]
LS478113.1_Estonia_2014          ................................................................................ [112400]
KP843857.1_Odintsovo_02/14       ................................................................................ [112400]
FN557520.1_strain_E75            ................................................................................ [112400]
AY261366.1_Warthog               .................................................................a.............. [112400]
AY261365.1_Warmbaths             ................................................................................ [112400]
AY261364.1_Tengani_62            ...........a..........................c......................................... [112400]
AY261363.1_Pretoriuskop/96/4     ................................................................................ [112400]
AY261362.1_Mkuzi_1979            ................................................................................ [112400]
AY261361.1_Malawi_Lil-20/1_1983  ..a.......................g.................t...................g............... [112400]
AY261360.1_Kenya_1950            ............................................t...................t............... [112400]
AM712240.1_OURT_88/3             ................................................................................ [112400]
AM712239.1_Benin_97/1            ................................................................................ [112400]
KM111295.1_Ken06.Bus             ............................................t........a..........ta.............. [112400]
KM111294.1_Ken05/Tk1             ............................................t...................t............... [112400]
KM262844.1_strain_L60            ................................................................................ [112400]
KM102979.1_26544/OG10_Italy      ................................................................................ [112400]
KM262845.1_strain_NHV            ................................................................................ [112400]
KX354450.1_47/Ss/2008            ................................................................................ [112400]
KP055815.1_strain_BA71           ................................................................................ [112400]
MH025920.1_strain_R35            ............................................t........a..........ta.............. [112400]
MH025919.1_strain_N10            ............................................t........a..........ta.............. [112400]
MH025918.1_strain_R25            ............................................t........a..........ta.............. [112400]
MH025917.1_strain_R7             ............................................t........a..........ta.............. [112400]
MH025916.1_strain_R8             ............................................t........a..........ta.............. [112400]

FR682468.2_Georgia_2007/1        cgtttttcgttttttatccaccggcggattacaaattgcaaaaaatacaactaataccaccgctgtaatgattaaggcca [112480]
ASF_IND_20_CAD543                ................................................................................ [112480]
LR899193.1_Germany_2020/1        ................................................................................ [112480]
MN393477.1_Wuhan_2019            ................................................................................ [112480]
MW396979.1_Timor-Leste/2019/1    ................................................................................ [112480]
MK940252.1_CN/2019/InnerMongolia ................................................................................ [112480]
MK128995.1_China/2018/AnhuiXCGQ  ................................................................................ [112480]
MH681419.1_ASFV/POL/2015         ................................................................................ [112480]
LS478113.1_Estonia_2014          ................................................................................ [112480]
KP843857.1_Odintsovo_02/14       ................................................................................ [112480]
```

```
FN557520.1_strain_E75            ................................................................................ [112480]
AY261366.1_Warthog               ...............................................t................................ [112480]
AY261365.1_Warmbaths             ................................................................................ [112480]
AY261364.1_Tengani_62            ................................................................................ [112480]
AY261363.1_Pretoriuskop/96/4     ................................................................................ [112480]
AY261362.1_Mkuzi_1979            ................................................................................ [112480]
AY261361.1_Malawi_Lil-20/1_1983  ....................................g.........................................t. [112480]
AY261360.1_Kenya_1950            ....................................g................g......t.................t. [112480]
AM712240.1_OURT_88/3             ................................................................................ [112480]
AM712239.1_Benin_97/1            ................................................................................ [112480]
KM111295.1_Ken06.Bus             ....................................g................g......t.................t. [112480]
KM111294.1_Ken05/Tk1             ....................................g................g......t.................t. [112480]
KM262844.1_strain_L60            ................................................................................ [112480]
KM102979.1_26544/OG10_Italy      ................................................................................ [112480]
KM262845.1_strain_NHV            ................................................................................ [112480]
KX354450.1_47/Ss/2008            ................................................................................ [112480]
KP055815.1_strain_BA71           ................................................................................ [112480]
MH025920.1_strain_R35            ....................................g................g......t.................t. [112480]
MH025919.1_strain_N10            ....................................g................g......t.................t. [112480]
MH025918.1_strain_R25            ....................................g................g......t.................t. [112480]
MH025917.1_strain_R7             ....................................g................g......t.................t. [112480]
MH025916.1_strain_R8             ....................................g................g......t.................t. [112480]

FR682468.2_Georgia_2007/1        caatgaaaggattttgaaaggatgttttgaacggttcgcacgtataaattttttctcctaaattattgatacccgcaata [112560]
ASF_IND_20_CAD543                ................................................................................ [112560]
LR899193.1_Germany_2020/1        ................................................................................ [112560]
MN393477.1_Wuhan_2019            ................................................................................ [112560]
MW396979.1_Timor-Leste/2019/1    ................................................................................ [112560]
MK940252.1_CN/2019/InnerMongolia ................................................................................ [112560]
MK128995.1_China/2018/AnhuiXCGQ  ................................................................................ [112560]
MH681419.1_ASFV/POL/2015         ................................................................................ [112560]
LS478113.1_Estonia_2014          ................................................................................ [112560]
KP843857.1_Odintsovo_02/14       ................................................................................ [112560]
FN557520.1_strain_E75            ...............................t................................................ [112560]
AY261366.1_Warthog               ..................................c............................................. [112560]
AY261365.1_Warmbaths             ...............................t..c............................................. [112560]
AY261364.1_Tengani_62            ...............................t..c............................................. [112560]
AY261363.1_Pretoriuskop/96/4     ..................................c............................................. [112560]
AY261362.1_Mkuzi_1979            ................g.................c............................................. [112560]
AY261361.1_Malawi_Lil-20/1_1983  ...............................g..c.............................g............... [112560]
AY261360.1_Kenya_1950            ..................................c.....t.......................g............... [112560]
AM712240.1_OURT_88/3             ...............................t................................................ [112560]
AM712239.1_Benin_97/1            ...............................t................................................ [112560]
KM111295.1_Ken06.Bus             ..................................c.....t.......................g............... [112560]
```

```
KM111294.1_Ken05/Tk1             ..................................c.....t.......................g............... [112560]
KM262844.1_strain_L60            ...............................t................................................ [112560]
KM102979.1_26544/OG10_Italy      ...............................t................................................ [112560]
KM262845.1_strain_NHV            ...............................t................................................ [112560]
KX354450.1_47/Ss/2008            ...............................t................................................ [112560]
KP055815.1_strain_BA71           ...............................t................................................ [112560]
MH025920.1_strain_R35            ..................................c.....t.......................g............... [112560]
MH025919.1_strain_N10            ..................................c.....t.......................g............... [112560]
MH025918.1_strain_R25            ..................................c.....t.......................g............... [112560]
MH025917.1_strain_R7             ..................................c.....t.......................g............... [112560]
MH025916.1_strain_R8             ..................................c.....t.......................g............... [112560]

FR682468.2_Georgia_2007/1        aaatctacattcattttatatatttataaattatgaaaaatttagagttacatctccgccggaccaatcattgctaaaat [112640]
ASF_IND_20_CAD543                ............................................r................................... [112640]
LR899193.1_Germany_2020/1        ................................................................................ [112640]
MN393477.1_Wuhan_2019            ................................................................................ [112640]
MW396979.1_Timor-Leste/2019/1    ................................................................................ [112640]
MK940252.1_CN/2019/InnerMongolia ................................................................................ [112640]
MK128995.1_China/2018/AnhuiXCGQ  ................................................................................ [112640]
MH681419.1_ASFV/POL/2015         ................................................................................ [112640]
LS478113.1_Estonia_2014          ................................................................................ [112640]
KP843857.1_Odintsovo_02/14       ................................................................................ [112640]
FN557520.1_strain_E75            ............................................a................................... [112640]
AY261366.1_Warthog               ................................................................................ [112640]
AY261365.1_Warmbaths             .........................................................a...................... [112640]
AY261364.1_Tengani_62            ................................................................................ [112640]
AY261363.1_Pretoriuskop/96/4     .........................................................a..a................... [112640]
AY261362.1_Mkuzi_1979            ................................................................................ [112640]
AY261361.1_Malawi_Lil-20/1_1983  .........................................................a...................... [112640]
AY261360.1_Kenya_1950            .........................................................a...................... [112640]
AM712240.1_OURT_88/3             ............................................a................................... [112640]
AM712239.1_Benin_97/1            ............................................a................................... [112640]
KM111295.1_Ken06.Bus             .........................................................a...................... [112640]
KM111294.1_Ken05/Tk1             .........................................................a...................... [112640]
KM262844.1_strain_L60            ............................................a................................... [112640]
KM102979.1_26544/OG10_Italy      ............................................a................................... [112640]
KM262845.1_strain_NHV            ............................................a................................... [112640]
KX354450.1_47/Ss/2008            ............................................a................................... [112640]
KP055815.1_strain_BA71           ............................................a................................... [112640]
MH025920.1_strain_R35            .........................................................a...................... [112640]
MH025919.1_strain_N10            .........................................................a...................... [112640]
MH025918.1_strain_R25            .........................................................a...................... [112640]
MH025917.1_strain_R7             .........................................................a...................... [112640]
MH025916.1_strain_R8             .........................................................a...................... [112640]
```

```
FR682468.2_Georgia_2007/1        ttgaagattcttcaaaaaggcccgactggttgaatgtcttctgctcaggtttccaaaaattttccaagaatggattttga [112720]
ASF_IND_20_CAD543                ................................................................................ [112720]
LR899193.1_Germany_2020/1        ................................................................................ [112720]
MN393477.1_Wuhan_2019            ................................................................................ [112720]
MW396979.1_Timor-Leste/2019/1    ................................................................................ [112720]
MK940252.1_CN/2019/InnerMongolia ................................................................................ [112720]
MK128995.1_China/2018/AnhuiXCGQ  ................................................................................ [112720]
MH681419.1_ASFV/POL/2015         ................................................................................ [112720]
LS478113.1_Estonia_2014          ................................................................................ [112720]
KP843857.1_Odintsovo_02/14       ................................................................................ [112720]
FN557520.1_strain_E75            ................................................................................ [112720]
AY261366.1_Warthog               ......................t....................t..............g..................... [112720]
AY261365.1_Warmbaths             ......................t....................t..............g........a............ [112720]
AY261364.1_Tengani_62            ......t...............t....................t..............g..................... [112720]
AY261363.1_Pretoriuskop/96/4     a.....................t....................t..............g..................... [112720]
AY261362.1_Mkuzi_1979            ................................................................................ [112720]
AY261361.1_Malawi_Lil-20/1_1983  c...g.....................................................g........a............ [112720]
AY261360.1_Kenya_1950            c...g.....................................................g........a............ [112720]
AM712240.1_OURT_88/3             ................................................................................ [112720]
AM712239.1_Benin_97/1            ................................................................................ [112720]
KM111295.1_Ken06.Bus             c...g.....................................................g........a............ [112720]
KM111294.1_Ken05/Tk1             c...g.....................................................g........a............ [112720]
KM262844.1_strain_L60            ................................................................................ [112720]
KM102979.1_26544/OG10_Italy      ................................................................................ [112720]
KM262845.1_strain_NHV            ................................................................................ [112720]
KX354450.1_47/Ss/2008            ................................................................................ [112720]
KP055815.1_strain_BA71           ................................................................................ [112720]
MH025920.1_strain_R35            c...g.....................................................g........a............ [112720]
MH025919.1_strain_N10            c...g.....................................................g........a............ [112720]
MH025918.1_strain_R25            c...g.....................................................g........a............ [112720]
MH025917.1_strain_R7             c...g.....................................................g........a............ [112720]
MH025916.1_strain_R8             c...g.....................................................g........a............ [112720]

FR682468.2_Georgia_2007/1        acaataggctcatct------tgatttcttcttcaaggatattttctttgata-------------------------tc [112800]
ASF_IND_20_CAD543                ...............------................................-------------------------.. [112800]
LR899193.1_Germany_2020/1        ...............------................................-------------------------.. [112800]
MN393477.1_Wuhan_2019            ...............------................................-------------------------.. [112800]
MW396979.1_Timor-Leste/2019/1    ...............------................................-------------------------.. [112800]
MK940252.1_CN/2019/InnerMongolia ...............------................................-------------------------.. [112800]
MK128995.1_China/2018/AnhuiXCGQ  ...............------................................-------------------------.. [112800]
MH681419.1_ASFV/POL/2015         ...............------................................-------------------------.. [112800]
LS478113.1_Estonia_2014          ...............------................................-------------------------.. [112800]
KP843857.1_Odintsovo_02/14       ...............------................................-------------------------.. [112800]
```

```
FN557520.1_strain_E75            ...............------..............................g..------------------------.. [112800]
AY261366.1_Warthog               .....c.........------..........................c..a...------------------------.. [112800]
AY261365.1_Warmbaths             .....c.........------..........................c..a...------------------------.. [112800]
AY261364.1_Tengani_62            .....c.........------..........................c..a...------------------------.. [112800]
AY261363.1_Pretoriuskop/96/4     .....t.........------..........................c..a...------------------------.. [112800]
AY261362.1_Mkuzi_1979            ...............------.................................------------------------.. [112800]
AY261361.1_Malawi_Lil-20/1_1983  ...............------......................c.......t..------------------------.. [112800]
AY261360.1_Kenya_1950            .....g.........tcttta..........t......a...ac.....cttc.atctcggggttatcttgatttaac.t [112800]
AM712240.1_OURT_88/3             ...............------..............................g..------------------------.. [112800]
AM712239.1_Benin_97/1            ...............------..............................g..------------------------.. [112800]
KM111295.1_Ken06.Bus             .....g.........tcttta......................ca......t..------------------------c. [112800]
KM111294.1_Ken05/Tk1             .....g.........tcttta..........t.---------------------------------------------.. [112800]
KM262844.1_strain_L60            ...............------..............................g..------------------------.. [112800]
KM102979.1_26544/OG10_Italy      ...............------..............................g..------------------------.. [112800]
KM262845.1_strain_NHV            ...............------..............................g..------------------------.. [112800]
KX354450.1_47/Ss/2008            ...............------..............................g..------------------------.. [112800]
KP055815.1_strain_BA71           ...............------..............................g..------------------------.. [112800]
MH025920.1_strain_R35            .....g.........tcttta......................ca......t..------------------------c. [112800]
MH025919.1_strain_N10            .....g.........tcttta......................ca......t..------------------------c. [112800]
MH025918.1_strain_R25            .....g.........tcttta......................ca......t..------------------------c. [112800]
MH025917.1_strain_R7             .....g.........tcttta......................ca......t..------------------------c. [112800]
MH025916.1_strain_R8             .....g.........tcttta......................ca......t..------------------------c. [112800]

FR682468.2_Georgia_2007/1        aagaaca---gcttctttaaactcaggtgtatcttgattaaa------------------ctcag---gtttatcctgat [112880]
ASF_IND_20_CAD543                .......---................................------------------.....---............ [112880]
LR899193.1_Germany_2020/1        .......---................................------------------.....---............ [112880]
MN393477.1_Wuhan_2019            .......---................................------------------.....---............ [112880]
MW396979.1_Timor-Leste/2019/1    .......---................................------------------.....---............ [112880]
MK940252.1_CN/2019/InnerMongolia .......---................................------------------.....---............ [112880]
MK128995.1_China/2018/AnhuiXCGQ  .......---................................------------------.....---............ [112880]
MH681419.1_ASFV/POL/2015         .......---................................------------------.....---............ [112880]
LS478113.1_Estonia_2014          .......---................................------------------.....---............ [112880]
KP843857.1_Odintsovo_02/14       .......---................................------------------.....---............ [112880]
FN557520.1_strain_E75            .......---................................------------------.....---............ [112880]
AY261366.1_Warthog               .......---..................t.............------------------t....---.......t.... [112880]
AY261365.1_Warmbaths             .......---..................t.............------------------t....---.......t.... [112880]
AY261364.1_Tengani_62            .......---..................t.............-------------------------------------- [112880]
AY261363.1_Pretoriuskop/96/4     .......---..................t.............ctcaagaatatcttcttc.....---..........t. [112880]
AY261362.1_Mkuzi_1979            .......---................................------------------.....---............ [112880]
AY261361.1_Malawi_Lil-20/1_1983  .....------t......gtta...a.aa..gt..ct.....------------------.....gtt............ [112880]
AY261360.1_Kenya_1950            .....t.---t......c..t...g..gt.............-------------------------------------- [112880]
AM712240.1_OURT_88/3             .......---................................------------------.....---............ [112880]
AM712239.1_Benin_97/1            .......---................................------------------.....---............ [112880]
KM111295.1_Ken06.Bus             .....t.actt...........c.g...t....c........-------------------------------------- [112880]
```

```
KM111294.1_Ken05/Tk1             .....t.actt...........c.g...t.............-------------------------------------- [112880]
KM262844.1_strain_L60            .......---................................------------------.....---............ [112880]
KM102979.1_26544/OG10_Italy      .......---................................------------------.....---............ [112880]
KM262845.1_strain_NHV            .......---................................------------------.....---............ [112880]
KX354450.1_47/Ss/2008            .......---................................------------------.....---............ [112880]
KP055815.1_strain_BA71           .......---................................------------------.....---............ [112880]
MH025920.1_strain_R35            .....t.actt...........c.g...t....c........-------------------------------------- [112880]
MH025919.1_strain_N10            .....t.actt...........c.g...t....c........-------------------------------------- [112880]
MH025918.1_strain_R25            .....t.actt...........c.g...t....c........-------------------------------------- [112880]
MH025917.1_strain_R7             .....t.actt...........c.g...t....c........-------------------------------------- [112880]
MH025916.1_strain_R8             .....t.actt...........c.g...t....c........-------------------------------------- [112880]

FR682468.2_Georgia_2007/1        caatcgcaaaaatattatcttc---ttcag---------atatatcctgtttaatcgcaagaatagtttcttc------c [112960]
ASF_IND_20_CAD543                .........r......w.....---.....---------r.................................------. [112960]
LR899193.1_Germany_2020/1        ......................---.....---------..................................------. [112960]
MN393477.1_Wuhan_2019            ......................---.....---------..................................------. [112960]
MW396979.1_Timor-Leste/2019/1    ......................---.....---------..................................------. [112960]
MK940252.1_CN/2019/InnerMongolia ......................---.....---------..................................------. [112960]
MK128995.1_China/2018/AnhuiXCGQ  ......................---.....---------..................................------. [112960]
MH681419.1_ASFV/POL/2015         ......................---.....---------..................................------. [112960]
LS478113.1_Estonia_2014          ......................---.....---------..................................------. [112960]
KP843857.1_Odintsovo_02/14       ......................---.....---------..................................------. [112960]
FN557520.1_strain_E75            .........g......t.....---.....---------g.........................t.......------. [112960]
AY261366.1_Warthog               t..a.t...g.....ct.....---c....---------ggt.......................tc......------. [112960]
AY261365.1_Warmbaths             t..a.t...g.....ct.....---c....---------g.t.......................tc......------. [112960]
AY261364.1_Tengani_62            ----.t...g.....ct.....---c....---------ggt.......................tc......------. [112960]
AY261363.1_Pretoriuskop/96/4     t........g.....ct.....---c....---------g.t.......................tc......------. [112960]
AY261362.1_Mkuzi_1979            .........g......t.....---.....---------g.................................------. [112960]
AY261361.1_Malawi_Lil-20/1_1983  t..a.t...g....gct.....---..t.aacataggttg.t...t...a..g.a.t...c............tttaat. [112960]
AY261360.1_Kenya_1950            ----.tt..g...g.ct.....aatc..g.---------ggt....t..a....a.t........tc......---aat. [112960]
AM712240.1_OURT_88/3             .........g......t.....---.....---------g.........................t.......------. [112960]
AM712239.1_Benin_97/1            .........g......t.....---.....---------g.........................t.......------. [112960]
KM111295.1_Ken06.Bus             ----.tt..g.....ct.....aatc..g.---------ggt....t..a....a.t........tc......---aat. [112960]
KM111294.1_Ken05/Tk1             ----.tt..g.....ct.....aatc..g.---------ggt....t..a..t.a.t........tc......---aat. [112960]
KM262844.1_strain_L60            .........g......t.....---.....---------g.........................t.......------. [112960]
KM102979.1_26544/OG10_Italy      .........g......t.....---.....---------g.........................t.......------. [112960]
KM262845.1_strain_NHV            .........g......t.....---.....---------g.........................t.......------. [112960]
KX354450.1_47/Ss/2008            .........g......t.....---.....---------g.........................t.......------. [112960]
KP055815.1_strain_BA71           .........g......t.....---.....---------g.........................t.......------. [112960]
MH025920.1_strain_R35            ----.tt..g.....ct.....aatc..g.---------ggt....t..a....a.t........tc......---aat. [112960]
MH025919.1_strain_N10            ----.tt..g.....ct.....aatc..g.---------ggt....t..a....a.t........tc......---aat. [112960]
MH025918.1_strain_R25            ----.tt..g.....ct.....aatc..g.---------ggt....t..a....a.t........tc......---aat. [112960]
MH025917.1_strain_R7             ----.tt..g.....ct.....aatc..g.---------ggt....t..a....a.t........tc......---aat. [112960]
MH025916.1_strain_R8             ----.tt..g.....ct.....aatc..g.---------ggt....t..a....a.t........tc......---aat. [112960]
```

```
FR682468.2_Georgia_2007/1        tcaggtttatcctgatcaatcgcaagaatattttcttctt---------------------cag---gtttatcctgacc [113040]
ASF_IND_20_CAD543                ........................................---------------------...---............. [113040]
LR899193.1_Germany_2020/1        ........................................---------------------...---............. [113040]
MN393477.1_Wuhan_2019            ........................................---------------------...---............. [113040]
MW396979.1_Timor-Leste/2019/1    ........................................---------------------...---............. [113040]
MK940252.1_CN/2019/InnerMongolia ........................................---------------------...---............. [113040]
MK128995.1_China/2018/AnhuiXCGQ  ........................................---------------------...---............. [113040]
MH681419.1_ASFV/POL/2015         ........................................---------------------...---............. [113040]
LS478113.1_Estonia_2014          ........................................---------------------...---............. [113040]
KP843857.1_Odintsovo_02/14       ........................................---------------------...---............. [113040]
FN557520.1_strain_E75            ......................................c.---------------------...---............. [113040]
AY261366.1_Warthog               .....g........t.t.....................c.---------------------...---............t [113040]
AY261365.1_Warmbaths             .....g........t.t.....................c.---------------------...---............t [113040]
AY261364.1_Tengani_62            .....g........t.t.....................c.caggtttatcccgattaaaca...---.......t...tt [113040]
AY261363.1_Pretoriuskop/96/4     .....g..........t.t...................c.caggtttatcctgtttaaaca...---............t [113040]
AY261362.1_Mkuzi_1979            ..............................g.......c.---------------------...---............. [113040]
AY261361.1_Malawi_Lil-20/1_1983  ...........t....t..a.t...c....gc........---------------taaacat..gtt...........tt [113040]
AY261360.1_Kenya_1950            ..g..g.....t....t..a.t.........c......aa----tct--------------.g.---.g.....t...tt [113040]
AM712240.1_OURT_88/3             ......................................c.---------------------...---............. [113040]
AM712239.1_Benin_97/1            ......................................c.---------------------...---............. [113040]
KM111295.1_Ken06.Bus             ..g..g.....t....t..a.t.....c...c......aa----tct--------------.g.---.g.....t...tt [113040]
KM111294.1_Ken05/Tk1             ..g..g.....t....tt.a.t.........c......aa----tct--------------tg.---.a.....t...tt [113040]
KM262844.1_strain_L60            ......................................c.---------------------...---............. [113040]
KM102979.1_26544/OG10_Italy      ......................................c.---------------------...---............. [113040]
KM262845.1_strain_NHV            ......................................c.---------------------...---............. [113040]
KX354450.1_47/Ss/2008            ......................................c.---------------------...---............. [113040]
KP055815.1_strain_BA71           ......................................c.---------------------...---............. [113040]
MH025920.1_strain_R35            ..g..g.....t....t..a.t.....c...c......aa----tct--------------.g.---.g.....t...tt [113040]
MH025919.1_strain_N10            ..g..g.....t....t..a.t.....c...c......aa----tct--------------.g.---.g.....t...tt [113040]
MH025918.1_strain_R25            ..g..g.....t....t..a.t.....c...c......aa----tct--------------.g.---.g.....t...tt [113040]
MH025917.1_strain_R7             ..g..g.....t....t..a.t.....c...c......aa----tct--------------.g.---.g.....t...tt [113040]
MH025916.1_strain_R8             ..g..g.....t....t..a.t.....c...c......aa----tct--------------.g.---.g.....t...tt [113040]

FR682468.2_Georgia_2007/1        aaactcaacaatatctttctcgctaaatccgtttttagtgtgaagctcttggttttgaagagaattatcaaaatctattt [113120]
ASF_IND_20_CAD543                ........y....................................................................... [113120]
LR899193.1_Germany_2020/1        ................................................................................ [113120]
MN393477.1_Wuhan_2019            ................................................................................ [113120]
MW396979.1_Timor-Leste/2019/1    ................................................................................ [113120]
MK940252.1_CN/2019/InnerMongolia ................................................................................ [113120]
MK128995.1_China/2018/AnhuiXCGQ  ................................................................................ [113120]
MH681419.1_ASFV/POL/2015         ................................................................................ [113120]
LS478113.1_Estonia_2014          ................................................................................ [113120]
KP843857.1_Odintsovo_02/14       ................................................................................ [113120]
```

```
FN557520.1_strain_E75             .............................................t...............................g.. [113120]
AY261366.1_Warthog                ..............................a..............t.................................. [113120]
AY261365.1_Warmbaths              ................c.............a..............t...............................g.. [113120]
AY261364.1_Tengani_62             ........t.....................a.............------...c...........g.........c.g.. [113120]
AY261363.1_Pretoriuskop/96/4      ..............................a..............t...............................g.. [113120]
AY261362.1_Mkuzi_1979             .............................................t...............................g.. [113120]
AY261361.1_Malawi_Lil-20/1_1983   .............g................a.......c............c..............g............. [113120]
AY261360.1_Kenya_1950             t.......g....gt....c.a........a......aca.t........................g............. [113120]
AM712240.1_OURT_88/3              .............................................t...............................g.. [113120]
AM712239.1_Benin_97/1             .............................................t...............................g.. [113120]
KM111295.1_Ken06.Bus              ........g....gt......a........a......aca.t........................g.........t... [113120]
KM111294.1_Ken05/Tk1              t.......g....gt...t..a........a......aca.t........................g............. [113120]
KM262844.1_strain_L60             .............................................t...............................g.. [113120]
KM102979.1_26544/OG10_Italy       .............................................t...............................g.. [113120]
KM262845.1_strain_NHV             .............................................t...............................g.. [113120]
KX354450.1_47/Ss/2008             .............................................t...............................g.. [113120]
KP055815.1_strain_BA71            .............................................t...............................g.. [113120]
MH025920.1_strain_R35             ........g....gt......a........a......aca.t........................g.........t... [113120]
MH025919.1_strain_N10             ........g....gt......a........a......aca.t........................g.........t... [113120]
MH025918.1_strain_R25             ........g....gt......a........a......aca.t........................g.........t... [113120]
MH025917.1_strain_R7              ........g....gt......a........a......aca.t........................g.........t... [113120]
MH025916.1_strain_R8              ........g....gt......a........a......aca.t........................g.........t... [113120]

FR682468.2_Georgia_2007/1         tagttgttgtcctagaccgtggcacgggatagttatctaatggtttacttactatagtcctcgaatgtggcacgggataa [113200]
ASF_IND_20_CAD543                 ................................................................................ [113200]
LR899193.1_Germany_2020/1         ................................................................................ [113200]
MN393477.1_Wuhan_2019             ................................................................................ [113200]
MW396979.1_Timor-Leste/2019/1     ................................................................................ [113200]
MK940252.1_CN/2019/InnerMongolia  ................................................................................ [113200]
MK128995.1_China/2018/AnhuiXCGQ   ................................................................................ [113200]
MH681419.1_ASFV/POL/2015          ................................................................................ [113200]
LS478113.1_Estonia_2014           ................................................................................ [113200]
KP843857.1_Odintsovo_02/14        ................................................................................ [113200]
FN557520.1_strain_E75             ..........t.....t........a............gg........................................ [113200]
AY261366.1_Warthog                ...---...................a............g..................................a...... [113200]
AY261365.1_Warmbaths              ...---...................a............g..................................a..g... [113200]
AY261364.1_Tengani_62             ...---...................a............g..................................a..g... [113200]
AY261363.1_Pretoriuskop/96/4      ...---...................a............g..................................a..g... [113200]
AY261362.1_Mkuzi_1979             .....a....t.....t........a............g......................................... [113200]
AY261361.1_Malawi_Lil-20/1_1983   ...---...a...............a............g........t................t........a.....g [113200]
AY261360.1_Kenya_1950             ...---a..a.......t....................g.........................t........a.....g [113200]
AM712240.1_OURT_88/3              ..........t.....t........a............gg........................................ [113200]
AM712239.1_Benin_97/1             ..........t.....t........a............gg........................................ [113200]
KM111295.1_Ken06.Bus              ...---...a.......t....................g.........................t........a.....g [113200]
```

```
KM111294.1_Ken05/Tk1             ...---...a.......t....................g.........................t........a.....g [113200]
KM262844.1_strain_L60            ..........t.....t........a............gg........................................ [113200]
KM102979.1_26544/OG10_Italy      ..........t.....t........a............gg........................................ [113200]
KM262845.1_strain_NHV            ..........t.....t........a............gg........................................ [113200]
KX354450.1_47/Ss/2008            ..........t.....t........a............gg........................................ [113200]
KP055815.1_strain_BA71           ..........t.....t........a............gg........................................ [113200]
MH025920.1_strain_R35            ...---...a.......t....................g.........................t........a.....g [113200]
MH025919.1_strain_N10            ...---...a.......t....................g.........................t........a.....g [113200]
MH025918.1_strain_R25            ...---...a.......t....................g.........................t........a.....g [113200]
MH025917.1_strain_R7             ...---...a.......t....................g.........................t........a.....g [113200]
MH025916.1_strain_R8             ...---...a.......t....................g.........................t........a.....g [113200]

FR682468.2_Georgia_2007/1        ttgtttggtgacttgctggttagctcttggcttgttaatagttct---tgttttctcaataattccatctct-------- [113280]
ASF_IND_20_CAD543                .............................................---........s...............-------- [113280]
LR899193.1_Germany_2020/1        .............................................---........................-------- [113280]
MN393477.1_Wuhan_2019            .............................................---........................-------- [113280]
MW396979.1_Timor-Leste/2019/1    .............................................---........................-------- [113280]
MK940252.1_CN/2019/InnerMongolia .............................................---........................-------- [113280]
MK128995.1_China/2018/AnhuiXCGQ  .............................................---........................-------- [113280]
MH681419.1_ASFV/POL/2015         .............................................---........................-------- [113280]
LS478113.1_Estonia_2014          .............................................---........................-------- [113280]
KP843857.1_Odintsovo_02/14       .............................................---........................-------- [113280]
FN557520.1_strain_E75            .............................................---........................-------- [113280]
AY261366.1_Warthog               .............................................---..................g.....-------- [113280]
AY261365.1_Warmbaths             .............................................---..................g.....-------- [113280]
AY261364.1_Tengani_62            ...............t........................a....---....cc............g.....-------- [113280]
AY261363.1_Pretoriuskop/96/4     .............................................---..................g.....-------- [113280]
AY261362.1_Mkuzi_1979            .............................................---....................a...-------- [113280]
AY261361.1_Malawi_Lil-20/1_1983  ......................c...........cc.........---....c............tg.t...tcttgttc [113280]
AY261360.1_Kenya_1950            ...c.............a..c.a...........cc.........tcc....gct..........tg.....-------- [113280]
AM712240.1_OURT_88/3             .............................................---........................-------- [113280]
AM712239.1_Benin_97/1            .............................................---........................-------- [113280]
KM111295.1_Ken06.Bus             .................a..c.a...........cc.........tcc....cc...........t......-------- [113280]
KM111294.1_Ken05/Tk1             .................a..c.a...........cc.........tcc....cc...........tg.....-------- [113280]
KM262844.1_strain_L60            .............................................---........................-------- [113280]
KM102979.1_26544/OG10_Italy      .............................................---........................-------- [113280]
KM262845.1_strain_NHV            .............................................---........................-------- [113280]
KX354450.1_47/Ss/2008            .............................................---........................-------- [113280]
KP055815.1_strain_BA71           .............................................---........................-------- [113280]
MH025920.1_strain_R35            .................a..c.a...........cc.........tcc....cc...........t......-------- [113280]
MH025919.1_strain_N10            .................a..c.a...........cc.........tcc....cc...........t......-------- [113280]
MH025918.1_strain_R25            .................a..c.a...........cc.........tcc....cc...........t......-------- [113280]
MH025917.1_strain_R7             .................a..c.a...........cc.........tcc....cc...........t......-------- [113280]
MH025916.1_strain_R8             .................a..c.a...........cc.........tcc....cc...........t......-------- [113280]
```

```
FR682468.2_Georgia_2007/1        ----actacttctttttgatccgctggtgtctctttttggtattcttcattagaaaaatgttcagagggtaatgtttcaa [113360]
ASF_IND_20_CAD543                ----............................................................................ [113360]
LR899193.1_Germany_2020/1        ----............................................................................ [113360]
MN393477.1_Wuhan_2019            ----............................................................................ [113360]
MW396979.1_Timor-Leste/2019/1    ----............................................................................ [113360]
MK940252.1_CN/2019/InnerMongolia ----............................................................................ [113360]
MK128995.1_China/2018/AnhuiXCGQ  ----............................................................................ [113360]
MH681419.1_ASFV/POL/2015         ----............................................................................ [113360]
LS478113.1_Estonia_2014          ----............................................................................ [113360]
KP843857.1_Odintsovo_02/14       ----............................................................................ [113360]
FN557520.1_strain_E75            ----....................c....................................................... [113360]
AY261366.1_Warthog               ----....................c................................g...................... [113360]
AY261365.1_Warmbaths             ----....................c.....a................................................. [113360]
AY261364.1_Tengani_62            ----....................c................................g...................... [113360]
AY261363.1_Pretoriuskop/96/4     ----....................c....................................................... [113360]
AY261362.1_Mkuzi_1979            ----....................c....................................................... [113360]
AY261361.1_Malawi_Lil-20/1_1983  tctc.a..t.a.............c.a..ct................................................. [113360]
AY261360.1_Kenya_1950            -------....g......g...ctc....c............g..................................... [113360]
AM712240.1_OURT_88/3             ----....................c....................................................... [113360]
AM712239.1_Benin_97/1            ----....................c....................................................... [113360]
KM111295.1_Ken06.Bus             -------....g......g...ctc....c............g..................................... [113360]
KM111294.1_Ken05/Tk1             -------....g......g...ctc....c............g..................................... [113360]
KM262844.1_strain_L60            ----....................c....................................................... [113360]
KM102979.1_26544/OG10_Italy      ----....................c....................................................... [113360]
KM262845.1_strain_NHV            ----....................c....................................................... [113360]
KX354450.1_47/Ss/2008            ----....................c....................................................... [113360]
KP055815.1_strain_BA71           ----....................c....................................................... [113360]
MH025920.1_strain_R35            -------....g......g...ctc....c............g..................................... [113360]
MH025919.1_strain_N10            -------....g......g...ctc....c............g..................................... [113360]
MH025918.1_strain_R25            -------....g......g...ctc....c............g..................................... [113360]
MH025917.1_strain_R7             -------....g......g...ctc....c............g..................................... [113360]
MH025916.1_strain_R8             -------....g......g...ctc....c............g..................................... [113360]

FR682468.2_Georgia_2007/1        taaactttgtgagtggatagctgctctttgatgtagaagagcgttgaatttgctgataaaggagttgaacaagtcgccgg [113440]
ASF_IND_20_CAD543                ................................................................................ [113440]
LR899193.1_Germany_2020/1        ................................................................................ [113440]
MN393477.1_Wuhan_2019            ................................................................................ [113440]
MW396979.1_Timor-Leste/2019/1    ................................................................................ [113440]
MK940252.1_CN/2019/InnerMongolia ................................................................................ [113440]
MK128995.1_China/2018/AnhuiXCGQ  ................................................................................ [113440]
MH681419.1_ASFV/POL/2015         ................................................................................ [113440]
LS478113.1_Estonia_2014          ................................................................................ [113440]
KP843857.1_Odintsovo_02/14       ................................................................................ [113440]
```

```
FN557520.1_strain_E75          ................................................................................ [113440]
AY261366.1_Warthog             ................................................................................ [113440]
AY261365.1_Warmbaths           ................................................................................ [113440]
AY261364.1_Tengani_62          ................................................................................ [113440]
AY261363.1_Pretoriuskop/96/4   ................................................................................ [113440]
AY261362.1_Mkuzi_1979          ................................................................................ [113440]
AY261361.1_Malawi_Lil-20/1_1983 .......................g.........a.....................g..c..............a..t... [113440]
AY261360.1_Kenya_1950          .......................t.t.......a...........................a...........g...... [113440]
AM712240.1_OURT_88/3           ................................................................................ [113440]
AM712239.1_Benin_97/1          ................................................................................ [113440]
KM111295.1_Ken06.Bus           .......................t.t.......a.........c.............................g...... [113440]
KM111294.1_Ken05/Tk1           .......................t.t.......a.........c.................a...........g...... [113440]
KM262844.1_strain_L60          ................................................................................ [113440]
KM102979.1_26544/OG10_Italy    ................................................................................ [113440]
KM262845.1_strain_NHV          ................................................................................ [113440]
KX354450.1_47/Ss/2008          ................................................................................ [113440]
KP055815.1_strain_BA71         ................................................................................ [113440]
MH025920.1_strain_R35          .......................t.t.......a.........c.............................g...... [113440]
MH025919.1_strain_N10          .......................t.t.......a.........c.............................g...... [113440]
MH025918.1_strain_R25          .......................t.t.......a.........c.............................g...... [113440]
MH025917.1_strain_R7           .......................t.t.......a.........c.............................g...... [113440]
MH025916.1_strain_R8           .......................t.t.......a.........c.............................g...... [113440]

FR682468.2_Georgia_2007/1      tattcactctgtcttttttcatattttttacgtagcgtggagagatctgctaagagcgacttgttttcagatgttaattc [113520]
ASF_IND_20_CAD543              ..............................................................k................. [113520]
LR899193.1_Germany_2020/1      ................................................................................ [113520]
MN393477.1_Wuhan_2019          ................................................................................ [113520]
MW396979.1_Timor-Leste/2019/1  ................................................................................ [113520]
MK940252.1_CN/2019/InnerMongolia ................................................................................ [113520]
MK128995.1_China/2018/AnhuiXCGQ ................................................................................ [113520]
MH681419.1_ASFV/POL/2015       ................................................................................ [113520]
LS478113.1_Estonia_2014        ................................................................................ [113520]
KP843857.1_Odintsovo_02/14     ................................................................................ [113520]
FN557520.1_strain_E75          ...............................................c..............t................. [113520]
AY261366.1_Warthog             ................................................................................ [113520]
AY261365.1_Warmbaths           ..............................................................t................. [113520]
AY261364.1_Tengani_62          .....................................a........................t................. [113520]
AY261363.1_Pretoriuskop/96/4   ..............................................................t................. [113520]
AY261362.1_Mkuzi_1979          ..............................................................t................. [113520]
AY261361.1_Malawi_Lil-20/1_1983 ......................................a.....t...t....a.a...t..t.............g... [113520]
AY261360.1_Kenya_1950          ............................................t........a.....t..t................. [113520]
AM712240.1_OURT_88/3           ...............................................c..............t................. [113520]
AM712239.1_Benin_97/1          ...............................................c..............t................. [113520]
KM111295.1_Ken06.Bus           ............................................t........a.....t..t................. [113520]
```

```
KM111294.1_Ken05/Tk1            ............................................t........a.....t..t................. [113520]
KM262844.1_strain_L60           ...............................................c..............t................. [113520]
KM102979.1_26544/OG10_Italy     ...............................................c..............t................. [113520]
KM262845.1_strain_NHV           ...............................................c..............t................. [113520]
KX354450.1_47/Ss/2008           ...............................................c..............t................. [113520]
KP055815.1_strain_BA71          ...............................................c..............t................. [113520]
MH025920.1_strain_R35           ............................................t........a.....t..t................. [113520]
MH025919.1_strain_N10           ............................................t........a.....t..t................. [113520]
MH025918.1_strain_R25           ............................................t........a.....t..t................. [113520]
MH025917.1_strain_R7            ............................................t........a.....t..t................. [113520]
MH025916.1_strain_R8            ............................................t........a.....t..t................. [113520]

FR682468.2_Georgia_2007/1       ttcaatttgatgaagaaggctgcgattgtatgaactaagtcttgcatacgtttcttctaattctgtctccggctccacat [113600]
ASF_IND_20_CAD543               ................................................................................ [113600]
LR899193.1_Germany_2020/1       ................................................................................ [113600]
MN393477.1_Wuhan_2019           ................................................................................ [113600]
MW396979.1_Timor-Leste/2019/1   ................................................................................ [113600]
MK940252.1_CN/2019/InnerMongolia ................................................................................ [113600]
MK128995.1_China/2018/AnhuiXCGQ ................................................................................ [113600]
MH681419.1_ASFV/POL/2015        ................................................................................ [113600]
LS478113.1_Estonia_2014         ................................................................................ [113600]
KP843857.1_Odintsovo_02/14      ................................................................................ [113600]
FN557520.1_strain_E75           ................................................................................ [113600]
AY261366.1_Warthog              ................................................................................ [113600]
AY261365.1_Warmbaths            ............................................................c................... [113600]
AY261364.1_Tengani_62           ................................................................................ [113600]
AY261363.1_Pretoriuskop/96/4    ................................................................................ [113600]
AY261362.1_Mkuzi_1979           ................................................................................ [113600]
AY261361.1_Malawi_Lil-20/1_1983 ...........a.........................................................a.......... [113600]
AY261360.1_Kenya_1950           ...........a................................t.......................ta.....t.... [113600]
AM712240.1_OURT_88/3            ................................................................................ [113600]
AM712239.1_Benin_97/1           ................................................................................ [113600]
KM111295.1_Ken06.Bus            ...g.......a..................c.............t.......................ta.....t.... [113600]
KM111294.1_Ken05/Tk1            ...........a................................t.......................ta.....t.... [113600]
KM262844.1_strain_L60           ................................................................................ [113600]
KM102979.1_26544/OG10_Italy     ................................................................................ [113600]
KM262845.1_strain_NHV           ................................................................................ [113600]
KX354450.1_47/Ss/2008           ................................................................................ [113600]
KP055815.1_strain_BA71          ................................................................................ [113600]
MH025920.1_strain_R35           ...g.......a..................c.............t.......................ta.....t.... [113600]
MH025919.1_strain_N10           ...g.......a..................c.............t.......................ta.....t.... [113600]
MH025918.1_strain_R25           ...g.......a..................c.............t.......................ta.....t.... [113600]
MH025917.1_strain_R7            ...g.......a..................c.............t.......................ta.....t.... [113600]
MH025916.1_strain_R8            ...g.......a..................c.............t.......................ta.....t.... [113600]
```

```
FR682468.2_Georgia_2007/1        aggcctgttttcgcagaaatttattgtatagttccattctttttttgagcagaaaggtaagactataatcttgcatttct [113680]
ASF_IND_20_CAD543                ................................................................................ [113680]
LR899193.1_Germany_2020/1        ................................................................................ [113680]
MN393477.1_Wuhan_2019            ................................................................................ [113680]
MW396979.1_Timor-Leste/2019/1    ................................................................................ [113680]
MK940252.1_CN/2019/InnerMongolia ................................................................................ [113680]
MK128995.1_China/2018/AnhuiXCGQ  ................................................................................ [113680]
MH681419.1_ASFV/POL/2015         ................................................................................ [113680]
LS478113.1_Estonia_2014          ................................................................................ [113680]
KP843857.1_Odintsovo_02/14       ................................................................................ [113680]
FN557520.1_strain_E75            ...............a......g......................................................... [113680]
AY261366.1_Warthog               ................................................................................ [113680]
AY261365.1_Warmbaths             ................................................................................ [113680]
AY261364.1_Tengani_62            ...................................................a............................ [113680]
AY261363.1_Pretoriuskop/96/4     ................................................................................ [113680]
AY261362.1_Mkuzi_1979            ...............a......g......................................................... [113680]
AY261361.1_Malawi_Lil-20/1_1983  ......................g.....................................................c... [113680]
AY261360.1_Kenya_1950            ......................g.....c......................a........................c... [113680]
AM712240.1_OURT_88/3             ...............a......g......................................................... [113680]
AM712239.1_Benin_97/1            ...............a......g......................................................... [113680]
KM111295.1_Ken06.Bus             ......................g.....c......................a........................c... [113680]
KM111294.1_Ken05/Tk1             ............................c......................a........................c... [113680]
KM262844.1_strain_L60            ...............a......g......................................................... [113680]
KM102979.1_26544/OG10_Italy      ...............a......g......................................................... [113680]
KM262845.1_strain_NHV            ...............a......g......................................................... [113680]
KX354450.1_47/Ss/2008            ...............a......g......................................................... [113680]
KP055815.1_strain_BA71           ...............a......g......................................................... [113680]
MH025920.1_strain_R35            ......................g.....c......................a........................c... [113680]
MH025919.1_strain_N10            ......................g.....c......................a........................c... [113680]
MH025918.1_strain_R25            ......................g.....c......................a........................c... [113680]
MH025917.1_strain_R7             ......................g.....c......................a........................c... [113680]
MH025916.1_strain_R8             ......................g.....c......................a........................c... [113680]

FR682468.2_Georgia_2007/1        ttcgtaactttatggtagttttctttccggtttttgataataaagggcagcattttttctgttgtgataaaggtgcccag [113760]
ASF_IND_20_CAD543                ................................................................................ [113760]
LR899193.1_Germany_2020/1        ................................................................................ [113760]
MN393477.1_Wuhan_2019            ................................................................................ [113760]
MW396979.1_Timor-Leste/2019/1    ................................................................................ [113760]
MK940252.1_CN/2019/InnerMongolia ................................................................................ [113760]
MK128995.1_China/2018/AnhuiXCGQ  ................................................................................ [113760]
MH681419.1_ASFV/POL/2015         ................................................................................ [113760]
LS478113.1_Estonia_2014          ................................................................................ [113760]
KP843857.1_Odintsovo_02/14       ................................................................................ [113760]
```

```
FN557520.1_strain_E75            ...............................................................................a [113760]
AY261366.1_Warthog               ................................................................................ [113760]
AY261365.1_Warmbaths             .......................................................................a........ [113760]
AY261364.1_Tengani_62            ................................................................................ [113760]
AY261363.1_Pretoriuskop/96/4     ................................................................................ [113760]
AY261362.1_Mkuzi_1979            .......................................................................a........ [113760]
AY261361.1_Malawi_Lil-20/1_1983  ..g................................a.c........................g...........a..... [113760]
AY261360.1_Kenya_1950            ..g...t.c....................a.....a.c.......................................... [113760]
AM712240.1_OURT_88/3             ...............................................................................a [113760]
AM712239.1_Benin_97/1            ...............................................................................a [113760]
KM111295.1_Ken06.Bus             ..g.....c....................a.....a.c.......................................t.. [113760]
KM111294.1_Ken05/Tk1             ..g...t.c....................a.....a.c.................................a........ [113760]
KM262844.1_strain_L60            ...............................................................................a [113760]
KM102979.1_26544/OG10_Italy      ...............................................................................a [113760]
KM262845.1_strain_NHV            ...............................................................................a [113760]
KX354450.1_47/Ss/2008            ...............................................................................a [113760]
KP055815.1_strain_BA71           ...............................................................................a [113760]
MH025920.1_strain_R35            ..g.....c....................a.....a.c.......................................t.. [113760]
MH025919.1_strain_N10            ..g.....c....................a.....a.c.......................................t.. [113760]
MH025918.1_strain_R25            ..g.....c....................a.....a.c.......................................t.. [113760]
MH025917.1_strain_R7             ..g.....c....................a.....a.c.......................................t.. [113760]
MH025916.1_strain_R8             ..g.....c....................a.....a.c.......................................t.. [113760]

FR682468.2_Georgia_2007/1        attgctaatgtagtcgcacagtagcaattccaagatagattctttcttttcaaggcttatagattggctgtattctttag [113840]
ASF_IND_20_CAD543                ................................................................................ [113840]
LR899193.1_Germany_2020/1        ................................................................................ [113840]
MN393477.1_Wuhan_2019            ................................................................................ [113840]
MW396979.1_Timor-Leste/2019/1    ................................................................................ [113840]
MK940252.1_CN/2019/InnerMongolia ................................................................................ [113840]
MK128995.1_China/2018/AnhuiXCGQ  ................................................................................ [113840]
MH681419.1_ASFV/POL/2015         ................................................................................ [113840]
LS478113.1_Estonia_2014          ................................................................................ [113840]
KP843857.1_Odintsovo_02/14       ................................................................................ [113840]
FN557520.1_strain_E75            ................................................................................ [113840]
AY261366.1_Warthog               ................................................................................ [113840]
AY261365.1_Warmbaths             ................................................................................ [113840]
AY261364.1_Tengani_62            ................................................................................ [113840]
AY261363.1_Pretoriuskop/96/4     .....................c.......................................................... [113840]
AY261362.1_Mkuzi_1979            ................................................................................ [113840]
AY261361.1_Malawi_Lil-20/1_1983  .......................................a........................................ [113840]
AY261360.1_Kenya_1950            .......................................a........................................ [113840]
AM712240.1_OURT_88/3             ................................................................................ [113840]
AM712239.1_Benin_97/1            ................................................................................ [113840]
KM111295.1_Ken06.Bus             .......................................a........................................ [113840]
```

```
KM111294.1_Ken05/Tk1              .......................................a........................................ [113840]
KM262844.1_strain_L60             ................................................................................ [113840]
KM102979.1_26544/OG10_Italy       ................................................................................ [113840]
KM262845.1_strain_NHV             ................................................................................ [113840]
KX354450.1_47/Ss/2008             ................................................................................ [113840]
KP055815.1_strain_BA71            ................................................................................ [113840]
MH025920.1_strain_R35             .......................................a........................................ [113840]
MH025919.1_strain_N10             .......................................a........................................ [113840]
MH025918.1_strain_R25             .......................................a........................................ [113840]
MH025917.1_strain_R7              .......................................a........................................ [113840]
MH025916.1_strain_R8              .......................................a........................................ [113840]

FR682468.2_Georgia_2007/1         gtatgaaagaatcaacaatcgttgttacgaagtttgaaaagtttaatgttttgctgttaatttgggtaatgttacaaaaa [113920]
ASF_IND_20_CAD543                 ...........................y.................................................... [113920]
LR899193.1_Germany_2020/1         ................................................................................ [113920]
MN393477.1_Wuhan_2019             ................................................................................ [113920]
MW396979.1_Timor-Leste/2019/1     ................................................................................ [113920]
MK940252.1_CN/2019/InnerMongolia  ................................................................................ [113920]
MK128995.1_China/2018/AnhuiXCGQ   ................................................................................ [113920]
MH681419.1_ASFV/POL/2015          ................................................................................ [113920]
LS478113.1_Estonia_2014           ................................................................................ [113920]
KP843857.1_Odintsovo_02/14        ................................................................................ [113920]
FN557520.1_strain_E75             ...........................t.................................................... [113920]
AY261366.1_Warthog                ..........g.............a......................................................g [113920]
AY261365.1_Warmbaths              ..........g.............a..............................................c.......g [113920]
AY261364.1_Tengani_62             ..........g.............a......................................................g [113920]
AY261363.1_Pretoriuskop/96/4      ..........g.............a......................................................g [113920]
AY261362.1_Mkuzi_1979             ................................................................................ [113920]
AY261361.1_Malawi_Lil-20/1_1983   ...a....................a.............................................a......... [113920]
AY261360.1_Kenya_1950             ...a....................a................c............................a......... [113920]
AM712240.1_OURT_88/3              ...........................t.................................................... [113920]
AM712239.1_Benin_97/1             ...........................t.................................................... [113920]
KM111295.1_Ken06.Bus              ...a....................a................c......................a.....a......... [113920]
KM111294.1_Ken05/Tk1              ...a....................a................c............................a......... [113920]
KM262844.1_strain_L60             ...........................t.................................................... [113920]
KM102979.1_26544/OG10_Italy       ...........................t.................................................... [113920]
KM262845.1_strain_NHV             ...........................t.................................................... [113920]
KX354450.1_47/Ss/2008             ...........................t.................................................... [113920]
KP055815.1_strain_BA71            ...........................t.................................................... [113920]
MH025920.1_strain_R35             ...a....................a................c......................a.....a......... [113920]
MH025919.1_strain_N10             ...a....................a................c......................a.....a......... [113920]
MH025918.1_strain_R25             ...a....................a................c......................a.....a......... [113920]
MH025917.1_strain_R7              ...a....................a................c......................a.....a......... [113920]
MH025916.1_strain_R8              ...a....................a................c......................a.....a......... [113920]
```

```
FR682468.2_Georgia_2007/1        tatttgtaaaaactatctagcattttttcataaagttttttattttgtttaacccctaaaatatagccctttacttgata [114000]
ASF_IND_20_CAD543                ................................................................................ [114000]
LR899193.1_Germany_2020/1        ................................................................................ [114000]
MN393477.1_Wuhan_2019            ................................................................................ [114000]
MW396979.1_Timor-Leste/2019/1    ................................................................................ [114000]
MK940252.1_CN/2019/InnerMongolia ................................................................................ [114000]
MK128995.1_China/2018/AnhuiXCGQ  ................................................................................ [114000]
MH681419.1_ASFV/POL/2015         ................................................................................ [114000]
LS478113.1_Estonia_2014          ................................................................................ [114000]
KP843857.1_Odintsovo_02/14       ................................................................................ [114000]
FN557520.1_strain_E75            ................................................................................ [114000]
AY261366.1_Warthog               ................................................................................ [114000]
AY261365.1_Warmbaths             ................................................................................ [114000]
AY261364.1_Tengani_62            ................................................................................ [114000]
AY261363.1_Pretoriuskop/96/4     ................................................................................ [114000]
AY261362.1_Mkuzi_1979            ................................................................................ [114000]
AY261361.1_Malawi_Lil-20/1_1983  ................................................................................ [114000]
AY261360.1_Kenya_1950            ................................................................................ [114000]
AM712240.1_OURT_88/3             ................................................................................ [114000]
AM712239.1_Benin_97/1            ................................................................................ [114000]
KM111295.1_Ken06.Bus             ................................................................................ [114000]
KM111294.1_Ken05/Tk1             ................................................................................ [114000]
KM262844.1_strain_L60            ................................................................................ [114000]
KM102979.1_26544/OG10_Italy      ................................................................................ [114000]
KM262845.1_strain_NHV            ................................................................................ [114000]
KX354450.1_47/Ss/2008            ................................................................................ [114000]
KP055815.1_strain_BA71           ................................................................................ [114000]
MH025920.1_strain_R35            ................................................................................ [114000]
MH025919.1_strain_N10            ................................................................................ [114000]
MH025918.1_strain_R25            ................................................................................ [114000]
MH025917.1_strain_R7             ................................................................................ [114000]
MH025916.1_strain_R8             ................................................................................ [114000]

FR682468.2_Georgia_2007/1        ctgatattccgtaacaatggaatgttttttgtatagtgcatttttgtataaaaagttataaaaaatgttgataaaatacg [114080]
ASF_IND_20_CAD543                ................................................................................ [114080]
LR899193.1_Germany_2020/1        ................................................................................ [114080]
MN393477.1_Wuhan_2019            ................................................................................ [114080]
MW396979.1_Timor-Leste/2019/1    ................................................................................ [114080]
MK940252.1_CN/2019/InnerMongolia ................................................................................ [114080]
MK128995.1_China/2018/AnhuiXCGQ  ................................................................................ [114080]
MH681419.1_ASFV/POL/2015         ................................................................................ [114080]
LS478113.1_Estonia_2014          ................................................................................ [114080]
KP843857.1_Odintsovo_02/14       ................................................................................ [114080]
```

```
FN557520.1_strain_E75            ................................................................................ [114080]
AY261366.1_Warthog               ................................................................................ [114080]
AY261365.1_Warmbaths             ................................................................................ [114080]
AY261364.1_Tengani_62            ................................................................................ [114080]
AY261363.1_Pretoriuskop/96/4     ................................................................................ [114080]
AY261362.1_Mkuzi_1979            ................................................................................ [114080]
AY261361.1_Malawi_Lil-20/1_1983  ..........a..................................................................... [114080]
AY261360.1_Kenya_1950            ..........a.................................................................t. [114080]
AM712240.1_OURT_88/3             ................................................................................ [114080]
AM712239.1_Benin_97/1            ................................................................................ [114080]
KM111295.1_Ken06.Bus             ..........a.................................................................t. [114080]
KM111294.1_Ken05/Tk1             ..........a.................................................................t. [114080]
KM262844.1_strain_L60            ................................................................................ [114080]
KM102979.1_26544/OG10_Italy      ................................................................................ [114080]
KM262845.1_strain_NHV            ................................................................................ [114080]
KX354450.1_47/Ss/2008            ................................................................................ [114080]
KP055815.1_strain_BA71           ................................................................................ [114080]
MH025920.1_strain_R35            ..........a.................................................................t. [114080]
MH025919.1_strain_N10            ..........a.................................................................t. [114080]
MH025918.1_strain_R25            ..........a.................................................................t. [114080]
MH025917.1_strain_R7             ..........a.................................................................t. [114080]
MH025916.1_strain_R8             ..........a.................................................................t. [114080]

FR682468.2_Georgia_2007/1        caccaagggtttcaaaaatacttataacgtgggattcttcctgatccattatatcatatgtaatattattttaataaaaa [114160]
ASF_IND_20_CAD543                ................................................................................ [114160]
LR899193.1_Germany_2020/1        ................................................................................ [114160]
MN393477.1_Wuhan_2019            ................................................................................ [114160]
MW396979.1_Timor-Leste/2019/1    ................................................................................ [114160]
MK940252.1_CN/2019/InnerMongolia ................................................................................ [114160]
MK128995.1_China/2018/AnhuiXCGQ  ................................................................................ [114160]
MH681419.1_ASFV/POL/2015         ................................................................................ [114160]
LS478113.1_Estonia_2014          ................................................................................ [114160]
KP843857.1_Odintsovo_02/14       ................................................................................ [114160]
FN557520.1_strain_E75            ................................................................................ [114160]
AY261366.1_Warthog               .............................c.................................................. [114160]
AY261365.1_Warmbaths             ................................................................................ [114160]
AY261364.1_Tengani_62            ................................................................................ [114160]
AY261363.1_Pretoriuskop/96/4     ................................................................................ [114160]
AY261362.1_Mkuzi_1979            ................................................................................ [114160]
AY261361.1_Malawi_Lil-20/1_1983  .g....ca.............................c..t................................g...... [114160]
AY261360.1_Kenya_1950            .g....a.................................t................................g...... [114160]
AM712240.1_OURT_88/3             ................................................................................ [114160]
AM712239.1_Benin_97/1            ................................................................................ [114160]
KM111295.1_Ken06.Bus             .g....a......g..........................t.................c..............g...... [114160]
```

```
KM111294.1_Ken05/Tk1             .g....a.................................t................................g...... [114160]
KM262844.1_strain_L60            ................................................................................ [114160]
KM102979.1_26544/OG10_Italy      ................................................................................ [114160]
KM262845.1_strain_NHV            ................................................................................ [114160]
KX354450.1_47/Ss/2008            ................................................................................ [114160]
KP055815.1_strain_BA71           ................................................................................ [114160]
MH025920.1_strain_R35            .g....a......g..........................t.................c..............g...... [114160]
MH025919.1_strain_N10            .g....a......g..........................t.................c..............g...... [114160]
MH025918.1_strain_R25            .g....a......g..........................t.................c..............g...... [114160]
MH025917.1_strain_R7             .g....a......g..........................t.................c..............g...... [114160]
MH025916.1_strain_R8             .g....a......g..........................t.................c..............g...... [114160]

FR682468.2_Georgia_2007/1        attactgacgaataacatgc-aaaaaaaatatgtttaaacttat-------------tttaagctagcacttatttaaaa [114240]
ASF_IND_20_CAD543                ....................-.......................-------------....................... [114240]
LR899193.1_Germany_2020/1        ....................-.......................-------------....................... [114240]
MN393477.1_Wuhan_2019            ....................-.......................-------------....................... [114240]
MW396979.1_Timor-Leste/2019/1    ....................-.......................-------------....................... [114240]
MK940252.1_CN/2019/InnerMongolia ....................-.......................-------------....................... [114240]
MK128995.1_China/2018/AnhuiXCGQ  ....................-.......................-------------....................... [114240]
MH681419.1_ASFV/POL/2015         ....................-.......................-------------....................... [114240]
LS478113.1_Estonia_2014          ....................-.......................-------------....................... [114240]
KP843857.1_Odintsovo_02/14       ....................-.......................-------------....................... [114240]
FN557520.1_strain_E75            .........t..........-.......................-------------....................... [114240]
AY261366.1_Warthog               ....................-............----------------------------------------------- [114240]
AY261365.1_Warmbaths             ....................-............----------------------------------------------- [114240]
AY261364.1_Tengani_62            .........t..........-............----------------------------------------------- [114240]
AY261363.1_Pretoriuskop/96/4     ....................-............----------------------------------------------- [114240]
AY261362.1_Mkuzi_1979            .........t..........a.......................-------------....................... [114240]
AY261361.1_Malawi_Lil-20/1_1983  t.......at..........-...........t....g......tttaaaacgtttg.....ata..t............ [114240]
AY261360.1_Kenya_1950            ........at....c.....-...........t....g......-------------........t..g.-.t.....t. [114240]
AM712240.1_OURT_88/3             .........t..........-.......................-------------....................... [114240]
AM712239.1_Benin_97/1            .........t..........-.......................-------------....................... [114240]
KM111295.1_Ken06.Bus             .----...at....c.....-...........t....g......-------------........t..g...t.....t. [114240]
KM111294.1_Ken05/Tk1             ........at....c.....-...........t....g......-------------........t..g...t.....t. [114240]
KM262844.1_strain_L60            .........t..........-.......................-------------....................... [114240]
KM102979.1_26544/OG10_Italy      .........t..........-.......................-------------....................... [114240]
KM262845.1_strain_NHV            .........t..........-.......................-------------....................... [114240]
KX354450.1_47/Ss/2008            .........t..........-.......................-------------....................... [114240]
KP055815.1_strain_BA71           .........t..........-.......................-------------....................... [114240]
MH025920.1_strain_R35            .----...at....c.....-...........t....g......-------------........t..g...t.....t. [114240]
MH025919.1_strain_N10            .----...at....c.....-...........t....g......-------------........t..g...t.....t. [114240]
MH025918.1_strain_R25            .----...at....c.....-...........t....g......-------------........t..g...t.....t. [114240]
MH025917.1_strain_R7             .----...at....c.....-...........t....g......-------------........t..g...t.....t. [114240]
MH025916.1_strain_R8             .----...at....c.....-...........t....g......-------------........t..g...t.....t. [114240]
```

```
FR682468.2_Georgia_2007/1        g---------------------tgttttaaacacgttttaaat---------tgtatgttaatacacttaaaaattaagc [114320]
ASF_IND_20_CAD543                .-------------------ct.....................---------..............s............. [114320]
LR899193.1_Germany_2020/1        .---------------------.....................---------............................ [114320]
MN393477.1_Wuhan_2019            .---------------------.....................---------............................ [114320]
MW396979.1_Timor-Leste/2019/1    .---------------------.....................---------............................ [114320]
MK940252.1_CN/2019/InnerMongolia .---------------------.....................---------............................ [114320]
MK128995.1_China/2018/AnhuiXCGQ  .---------------------.....................---------............................ [114320]
MH681419.1_ASFV/POL/2015         .---------------------.....................---------............................ [114320]
LS478113.1_Estonia_2014          .---------------------.....................---------............................ [114320]
KP843857.1_Odintsovo_02/14       .---------------------.....................---------............................ [114320]
FN557520.1_strain_E75            .-------------------ct.....................---------..............g............. [114320]
AY261366.1_Warthog               -------------------------..................---------..........a................g [114320]
AY261365.1_Warmbaths             -------------------------..................---------..........a................g [114320]
AY261364.1_Tengani_62            -------------------------..................---------..........a................g [114320]
AY261363.1_Pretoriuskop/96/4     -------------------------..a....c..........---------..........a...........g....g [114320]
AY261362.1_Mkuzi_1979            .-------------------ct.....................---------..............g............. [114320]
AY261361.1_Malawi_Lil-20/1_1983  aaagtactttttaaaaagcact.t...a...............gcgtttaaaaa........a................t [114320]
AY261360.1_Kenya_1950            agcg---------------------c.c........c......---------.a.t.t...--..t.............t [114320]
AM712240.1_OURT_88/3             .-------------------ct.....................---------..............g............. [114320]
AM712239.1_Benin_97/1            .-------------------ct.....................---------..............g............. [114320]
KM111295.1_Ken06.Bus             aacg---------------------c..........c......---------.a.c.t...gg................t [114320]
KM111294.1_Ken05/Tk1             agcg---------------------c.c........c......---------.a.t.t...--..t.............t [114320]
KM262844.1_strain_L60            .-------------------ct.....................---------..............g............. [114320]
KM102979.1_26544/OG10_Italy      .-------------------ct.....................---------..............g............. [114320]
KM262845.1_strain_NHV            .-------------------ct.....................---------..............g............. [114320]
KX354450.1_47/Ss/2008            .-------------------ct.....................---------..............g............. [114320]
KP055815.1_strain_BA71           .-------------------ct.....................---------..............g............. [114320]
MH025920.1_strain_R35            aacg---------------------c..........c......---------.a.c.t...gg................t [114320]
MH025919.1_strain_N10            aacg---------------------c..........c......---------.a.c.t...gg................t [114320]
MH025918.1_strain_R25            aacg---------------------c..........c......---------.a.c.t...gg................t [114320]
MH025917.1_strain_R7             aacg---------------------c..........c......---------.a.c.t...gg................t [114320]
MH025916.1_strain_R8             aacg---------------------c..........c......---------.a.c.t...gg................t [114320]

FR682468.2_Georgia_2007/1        cgaaatttgctccaataaggattacttttatcaatgaccacctctttactataaacggctttacataattttaataatgc [114400]
ASF_IND_20_CAD543                .............................................................................w.. [114400]
LR899193.1_Germany_2020/1        ................................................................................ [114400]
MN393477.1_Wuhan_2019            ................................................................................ [114400]
MW396979.1_Timor-Leste/2019/1    ................................................................................ [114400]
MK940252.1_CN/2019/InnerMongolia ................................................................................ [114400]
MK128995.1_China/2018/AnhuiXCGQ  ................................................................................ [114400]
MH681419.1_ASFV/POL/2015         ........a....................................................................... [114400]
LS478113.1_Estonia_2014          ................................................................................ [114400]
KP843857.1_Odintsovo_02/14       ................................................................................ [114400]
```

```
FN557520.1_strain_E75          .............................................................................a.. [114400]
AY261366.1_Warthog             ........a............................................g.......................a.. [114400]
AY261365.1_Warmbaths           ........a............................................g.......................a.. [114400]
AY261364.1_Tengani_62          ........a............................................g.......................a.. [114400]
AY261363.1_Pretoriuskop/96/4   ........a............................................g.......................a.. [114400]
AY261362.1_Mkuzi_1979          .............................................................................a.. [114400]
AY261361.1_Malawi_Lil-20/1_1983 ........a.....g..c.......................................................g...a.. [114400]
AY261360.1_Kenya_1950          ........a.....g........g..............g..............g.......................a.. [114400]
AM712240.1_OURT_88/3           .............................................................................a.. [114400]
AM712239.1_Benin_97/1          .............................................................................a.. [114400]
KM111295.1_Ken06.Bus           ........a.....g........g..............g..............g.......................a.. [114400]
KM111294.1_Ken05/Tk1           ........a.....g........g..............g..............g.......................a.. [114400]
KM262844.1_strain_L60          .............................................................................a.. [114400]
KM102979.1_26544/OG10_Italy    .............................................................................a.. [114400]
KM262845.1_strain_NHV          .............................................................................a.. [114400]
KX354450.1_47/Ss/2008          .............................................................................a.. [114400]
KP055815.1_strain_BA71         .............................................................................a.. [114400]
MH025920.1_strain_R35          ........a.....g........g..............g..............g.......................a.. [114400]
MH025919.1_strain_N10          ........a.....g........g..............g..............g.......................a.. [114400]
MH025918.1_strain_R25          ........a.....g........g..............g..............g.......................a.. [114400]
MH025917.1_strain_R7           ........a.....g........g..............g..............g.......................a.. [114400]
MH025916.1_strain_R8           ........a.....g........g..............g..............g.......................a.. [114400]

FR682468.2_Georgia_2007/1      tttagagccaaagctgaaggcagtgggaagcggcactgtactatggtaaaaatgttgccgatgttcatcctcgcggatgt [114480]
ASF_IND_20_CAD543              ................................................................................ [114480]
LR899193.1_Germany_2020/1      ................................................................................ [114480]
MN393477.1_Wuhan_2019          ................................................................................ [114480]
MW396979.1_Timor-Leste/2019/1  ................................................................................ [114480]
MK940252.1_CN/2019/InnerMongolia ................................................................................ [114480]
MK128995.1_China/2018/AnhuiXCGQ ................................................................................ [114480]
MH681419.1_ASFV/POL/2015       ................................................................................ [114480]
LS478113.1_Estonia_2014        ................................................................................ [114480]
KP843857.1_Odintsovo_02/14     ................................................................................ [114480]
FN557520.1_strain_E75          ...g..........................................................t................. [114480]
AY261366.1_Warthog             .....................g........................................t................. [114480]
AY261365.1_Warmbaths           ..............................................................t................. [114480]
AY261364.1_Tengani_62          ................................................................................ [114480]
AY261363.1_Pretoriuskop/96/4   ......a.......................................................t................. [114480]
AY261362.1_Mkuzi_1979          ..............................................................t................. [114480]
AY261361.1_Malawi_Lil-20/1_1983 ......a..................................................t....t................. [114480]
AY261360.1_Kenya_1950          .....................g.....c.....t..c.........................a................. [114480]
AM712240.1_OURT_88/3           ...g..........................................................t................. [114480]
AM712239.1_Benin_97/1          ...g..........................................................t................. [114480]
KM111295.1_Ken06.Bus           .....................g.....c..................................a................. [114480]
```

KM111294.1_Ken05/Tk1 ....................tg.....c.....t..c.........................a................. [114480]
KM262844.1_strain_L60 ...g..........................................................t................. [114480]
KM102979.1_26544/OG10_Italy ...g..........................................................t................. [114480]
KM262845.1_strain_NHV ...g..........................................................t................. [114480]
KX354450.1_47/Ss/2008 ...g..........................................................t................. [114480]
KP055815.1_strain_BA71 ...g..........................................................t................. [114480]
MH025920.1_strain_R35 .....................g.....c..................................a................. [114480]
MH025919.1_strain_N10 .....................g.....c..................................a................. [114480]
MH025918.1_strain_R25 .....................g.....c..................................a................. [114480]
MH025917.1_strain_R7 .....................g.....c..................................a................. [114480]
MH025916.1_strain_R8 .....................g.....c..................................a................. [114480]

FR682468.2_Georgia_2007/1 acacaagtttcctatatcctttaaacacaatatggctaatttcttccacatactccttatcctgtttggaatagcggttg [114560]
ASF_IND_20_CAD543 ................................................................................ [114560]
LR899193.1_Germany_2020/1 ................................................................................ [114560]
MN393477.1_Wuhan_2019 ................................................................................ [114560]
MW396979.1_Timor-Leste/2019/1 ................................................................................ [114560]
MK940252.1_CN/2019/InnerMongolia ................................................................................ [114560]
MK128995.1_China/2018/AnhuiXCGQ ................................................................................ [114560]
MH681419.1_ASFV/POL/2015 ................................................................................ [114560]
LS478113.1_Estonia_2014 ................................................................................ [114560]
KP843857.1_Odintsovo_02/14 ................................................................................ [114560]
FN557520.1_strain_E75 ................................................................................ [114560]
AY261366.1_Warthog .................................................g...........t.................. [114560]
AY261365.1_Warmbaths ................c................................g.............................. [114560]
AY261364.1_Tengani_62 ................c.............................t..g.............................. [114560]
AY261363.1_Pretoriuskop/96/4 ................c................................g.............................. [114560]
AY261362.1_Mkuzi_1979 ................................................................................ [114560]
AY261361.1_Malawi_Lil-20/1_1983 .........................t.....g.................g.................a............ [114560]
AY261360.1_Kenya_1950 ................c....................g...........g.............................. [114560]
AM712240.1_OURT_88/3 ................................................................................ [114560]
AM712239.1_Benin_97/1 ................................................................................ [114560]
KM111295.1_Ken06.Bus ................c....................g...........g.............................. [114560]
KM111294.1_Ken05/Tk1 ................c....................g...........g.............................. [114560]
KM262844.1_strain_L60 ................................................................................ [114560]
KM102979.1_26544/OG10_Italy ................................................................................ [114560]
KM262845.1_strain_NHV ................................................................................ [114560]
KX354450.1_47/Ss/2008 ................................................................................ [114560]
KP055815.1_strain_BA71 ................................................................................ [114560]
MH025920.1_strain_R35 ................c....................g...........g.............................. [114560]
MH025919.1_strain_N10 ................c....................g...........g.............................. [114560]
MH025918.1_strain_R25 ................c....................g...........g.............................. [114560]
MH025917.1_strain_R7 ................c....................g...........g.............................. [114560]
MH025916.1_strain_R8 ................c....................g...........g.............................. [114560]

```
FR682468.2_Georgia_2007/1        ctttgacgggaaaaattcgacatacaaatagaggcatttgtaaaaatggaaacaaatgcgtttttacgaagattggcggg [114640]
ASF_IND_20_CAD543                ................................................................................ [114640]
LR899193.1_Germany_2020/1        ................................................................................ [114640]
MN393477.1_Wuhan_2019            ................................................................................ [114640]
MW396979.1_Timor-Leste/2019/1    ................................................................................ [114640]
MK940252.1_CN/2019/InnerMongolia ................................................................................ [114640]
MK128995.1_China/2018/AnhuiXCGQ  ................................................................................ [114640]
MH681419.1_ASFV/POL/2015         ................................................................................ [114640]
LS478113.1_Estonia_2014          ................................................................................ [114640]
KP843857.1_Odintsovo_02/14       ................................................................................ [114640]
FN557520.1_strain_E75            .....t.......................................................................... [114640]
AY261366.1_Warthog               ...................................g............................................ [114640]
AY261365.1_Warmbaths             .......................g...........g.......................a.................a.. [114640]
AY261364.1_Tengani_62            .......................g......................................................a.. [114640]
AY261363.1_Pretoriuskop/96/4     .......................g...........g.......................a.................a.. [114640]
AY261362.1_Mkuzi_1979            .....t.......................................................................... [114640]
AY261361.1_Malawi_Lil-20/1_1983  ...................................g....................c..a.................... [114640]
AY261360.1_Kenya_1950            ...................................g.......................a.................... [114640]
AM712240.1_OURT_88/3             .....t.......................................................................... [114640]
AM712239.1_Benin_97/1            .....t.......................................................................... [114640]
KM111295.1_Ken06.Bus             .................t.................g.......................a.................... [114640]
KM111294.1_Ken05/Tk1             ...................................g.......................a.................... [114640]
KM262844.1_strain_L60            .....t.......................................................................... [114640]
KM102979.1_26544/OG10_Italy      .....t.......................................................................... [114640]
KM262845.1_strain_NHV            .....t.......................................................................... [114640]
KX354450.1_47/Ss/2008            .....t.......................................................................... [114640]
KP055815.1_strain_BA71           .....t.......................................................................... [114640]
MH025920.1_strain_R35            .................t.................g.......................a.................... [114640]
MH025919.1_strain_N10            .................t.................g.......................a.................... [114640]
MH025918.1_strain_R25            .................t.................g.......................a.................... [114640]
MH025917.1_strain_R7             .................t.................g.......................a.................... [114640]
MH025916.1_strain_R8             .................t.................g.......................a.................... [114640]

FR682468.2_Georgia_2007/1        taaatcggtatcatcttggcagcaaataatcatcgaaataaaacagtgacgattttggtaaaaaaacttttttaaaaattt [114720]
ASF_IND_20_CAD543                ................................................................................ [114720]
LR899193.1_Germany_2020/1        ................................................................................ [114720]
MN393477.1_Wuhan_2019            ................................................................................ [114720]
MW396979.1_Timor-Leste/2019/1    ................................................................................ [114720]
MK940252.1_CN/2019/InnerMongolia ................................................................................ [114720]
MK128995.1_China/2018/AnhuiXCGQ  ................................................................................ [114720]
MH681419.1_ASFV/POL/2015         ................................................................................ [114720]
LS478113.1_Estonia_2014          ................................................................................ [114720]
KP843857.1_Odintsovo_02/14       ................................................................................ [114720]
```

```
FN557520.1_strain_E75           .........g...................................................................... [114720]
AY261366.1_Warthog              .........g...................................................................... [114720]
AY261365.1_Warmbaths            .........g..g................................................................... [114720]
AY261364.1_Tengani_62           .........g..g................................................................... [114720]
AY261363.1_Pretoriuskop/96/4    .........g..g................................................................... [114720]
AY261362.1_Mkuzi_1979           .........g...................................................................... [114720]
AY261361.1_Malawi_Lil-20/1_1983 ......a..g...................................a.................................. [114720]
AY261360.1_Kenya_1950           .........g..g...............................................g................... [114720]
AM712240.1_OURT_88/3            .........g...................................................................... [114720]
AM712239.1_Benin_97/1           .........g...................................................................... [114720]
KM111295.1_Ken06.Bus            .........g..g................................................................... [114720]
KM111294.1_Ken05/Tk1            .........g..g...............................................g................... [114720]
KM262844.1_strain_L60           .........g...................................................................... [114720]
KM102979.1_26544/OG10_Italy     .........g...................................................................... [114720]
KM262845.1_strain_NHV           .........g...................................................................... [114720]
KX354450.1_47/Ss/2008           .........g...................................................................... [114720]
KP055815.1_strain_BA71          .........g...................................................................... [114720]
MH025920.1_strain_R35           .........g..g................................................................... [114720]
MH025919.1_strain_N10           .........g..g................................................................... [114720]
MH025918.1_strain_R25           .........g..g................................................................... [114720]
MH025917.1_strain_R7            .........g..g................................................................... [114720]
MH025916.1_strain_R8            .........g..g................................................................... [114720]

FR682468.2_Georgia_2007/1       cttttgtaaataatgggtgcagttcggccgcgcagtcgtctaatattaaaagtaaacgaggattaagattgatatagttt [114800]
ASF_IND_20_CAD543               ................................................................................ [114800]
LR899193.1_Germany_2020/1       ................................................................................ [114800]
MN393477.1_Wuhan_2019           ................................................................................ [114800]
MW396979.1_Timor-Leste/2019/1   ................................................................................ [114800]
MK940252.1_CN/2019/InnerMongolia ................................................................................ [114800]
MK128995.1_China/2018/AnhuiXCGQ ................................................................................ [114800]
MH681419.1_ASFV/POL/2015        ................................................................................ [114800]
LS478113.1_Estonia_2014         ................................................................................ [114800]
KP843857.1_Odintsovo_02/14      ................................................................................ [114800]
FN557520.1_strain_E75           ................................................................................ [114800]
AY261366.1_Warthog              .....................................a.......................................... [114800]
AY261365.1_Warmbaths            ................................................................................ [114800]
AY261364.1_Tengani_62           ............................t........a.......................................... [114800]
AY261363.1_Pretoriuskop/96/4    ................................................................................ [114800]
AY261362.1_Mkuzi_1979           ................................................................................ [114800]
AY261361.1_Malawi_Lil-20/1_1983 .........................a..g.....a..a.......................................... [114800]
AY261360.1_Kenya_1950           ..........a.g............a..g................................................... [114800]
AM712240.1_OURT_88/3            ................................................................................ [114800]
AM712239.1_Benin_97/1           ................................................................................ [114800]
KM111295.1_Ken06.Bus            ..........a.g............a..g.........................................a......... [114800]
```

```
KM111294.1_Ken05/Tk1              ..........a.g............a..g................................................... [114800]
KM262844.1_strain_L60             ................................................................................ [114800]
KM102979.1_26544/OG10_Italy       ................................................................................ [114800]
KM262845.1_strain_NHV             ................................................................................ [114800]
KX354450.1_47/Ss/2008             ................................................................................ [114800]
KP055815.1_strain_BA71            ................................................................................ [114800]
MH025920.1_strain_R35             ..........a.g............a..g.........................................a......... [114800]
MH025919.1_strain_N10             ..........a.g............a..g.........................................a......... [114800]
MH025918.1_strain_R25             ..........a.g............a..g.........................................a......... [114800]
MH025917.1_strain_R7              ..........a.g............a..g.........................................a......... [114800]
MH025916.1_strain_R8              ..........a.g............a..g.........................................a......... [114800]

FR682468.2_Georgia_2007/1         aacgtaaacttttcatcctctgtaaggcataagtttttatacatatgaatgttctgtataataattttttttaaaagttg [114880]
ASF_IND_20_CAD543                 ................................................................................ [114880]
LR899193.1_Germany_2020/1         ................................................................................ [114880]
MN393477.1_Wuhan_2019             ................................................................................ [114880]
MW396979.1_Timor-Leste/2019/1     ................................................................................ [114880]
MK940252.1_CN/2019/InnerMongolia  ................................................................................ [114880]
MK128995.1_China/2018/AnhuiXCGQ   ................................................................................ [114880]
MH681419.1_ASFV/POL/2015          ................................................................................ [114880]
LS478113.1_Estonia_2014           ................................................................................ [114880]
KP843857.1_Odintsovo_02/14        ................................................................................ [114880]
FN557520.1_strain_E75             ................................................................................ [114880]
AY261366.1_Warthog                ................t.........a..........................t.......................... [114880]
AY261365.1_Warmbaths              ..........................a..............t...................................... [114880]
AY261364.1_Tengani_62             ................................................................................ [114880]
AY261363.1_Pretoriuskop/96/4      ..........................a..................................................... [114880]
AY261362.1_Mkuzi_1979             ................................................................................ [114880]
AY261361.1_Malawi_Lil-20/1_1983   .....g..t.................a..................................................... [114880]
AY261360.1_Kenya_1950             .....g....................ag.................................................... [114880]
AM712240.1_OURT_88/3              ................................................................................ [114880]
AM712239.1_Benin_97/1             ................................................................................ [114880]
KM111295.1_Ken06.Bus              ..........................ag.................................................... [114880]
KM111294.1_Ken05/Tk1              .....g....................ag.................................................... [114880]
KM262844.1_strain_L60             ................................................................................ [114880]
KM102979.1_26544/OG10_Italy       ................................................................................ [114880]
KM262845.1_strain_NHV             ................................................................................ [114880]
KX354450.1_47/Ss/2008             ................................................................................ [114880]
KP055815.1_strain_BA71            ................................................................................ [114880]
MH025920.1_strain_R35             ..........................ag.................................................... [114880]
MH025919.1_strain_N10             ..........................ag.................................................... [114880]
MH025918.1_strain_R25             ..........................ag.................................................... [114880]
MH025917.1_strain_R7              ..........................ag.................................................... [114880]
MH025916.1_strain_R8              ..........................ag.................................................... [114880]
```

```
FR682468.2_Georgia_2007/1        ctgataaagcgatgtaatcttttcttcttttttttggtccgtttgttcagcctttaagcactccacttttgcaatatttt [114960]
ASF_IND_20_CAD543                ................................................................................ [114960]
LR899193.1_Germany_2020/1        ................................................................................ [114960]
MN393477.1_Wuhan_2019            ................................................................................ [114960]
MW396979.1_Timor-Leste/2019/1    ................................................................................ [114960]
MK940252.1_CN/2019/InnerMongolia ................................................................................ [114960]
MK128995.1_China/2018/AnhuiXCGQ  ................................................................................ [114960]
MH681419.1_ASFV/POL/2015         ................................................................................ [114960]
LS478113.1_Estonia_2014          ................................................................................ [114960]
KP843857.1_Odintsovo_02/14       ................................................................................ [114960]
FN557520.1_strain_E75            ................................................................................ [114960]
AY261366.1_Warthog               ................................................................................ [114960]
AY261365.1_Warmbaths             ................................................................................ [114960]
AY261364.1_Tengani_62            ................................................................................ [114960]
AY261363.1_Pretoriuskop/96/4     .........................................................a...................... [114960]
AY261362.1_Mkuzi_1979            ................................................................................ [114960]
AY261361.1_Malawi_Lil-20/1_1983  ................................................g............................... [114960]
AY261360.1_Kenya_1950            .........a..c................................................................... [114960]
AM712240.1_OURT_88/3             ................................................................................ [114960]
AM712239.1_Benin_97/1            ................................................................................ [114960]
KM111295.1_Ken06.Bus             .........a..c................................................................... [114960]
KM111294.1_Ken05/Tk1             .........a..c................................................................... [114960]
KM262844.1_strain_L60            ................................................................................ [114960]
KM102979.1_26544/OG10_Italy      ................................................................................ [114960]
KM262845.1_strain_NHV            ................................................................................ [114960]
KX354450.1_47/Ss/2008            ................................................................................ [114960]
KP055815.1_strain_BA71           ................................................................................ [114960]
MH025920.1_strain_R35            .........a..c................................................................... [114960]
MH025919.1_strain_N10            .........a..c................................................................... [114960]
MH025918.1_strain_R25            .........a..c................................................................... [114960]
MH025917.1_strain_R7             .........a..c................................................................... [114960]
MH025916.1_strain_R8             .........a..c................................................................... [114960]

FR682468.2_Georgia_2007/1        tgttttccttttgctgtatatcgatcggaagtttatgatacaatgtttttagcatatcgatgttgtttactcgactgtag [115040]
ASF_IND_20_CAD543                ................................................................................ [115040]
LR899193.1_Germany_2020/1        ................................................................................ [115040]
MN393477.1_Wuhan_2019            ................................................................................ [115040]
MW396979.1_Timor-Leste/2019/1    ................................................................................ [115040]
MK940252.1_CN/2019/InnerMongolia ................................................................................ [115040]
MK128995.1_China/2018/AnhuiXCGQ  ................................................................................ [115040]
MH681419.1_ASFV/POL/2015         ................................................................................ [115040]
LS478113.1_Estonia_2014          ................................................................................ [115040]
KP843857.1_Odintsovo_02/14       ................................................................................ [115040]
```

```
FN557520.1_strain_E75          ................................................................................ [115040]
AY261366.1_Warthog             .........................t...................................................... [115040]
AY261365.1_Warmbaths           .........................t...................................................... [115040]
AY261364.1_Tengani_62          ...................g.....t...................................................... [115040]
AY261363.1_Pretoriuskop/96/4   .........................t...................................................... [115040]
AY261362.1_Mkuzi_1979          ................................................................................ [115040]
AY261361.1_Malawi_Lil-20/1_1983 .a.....................................................g........a............... [115040]
AY261360.1_Kenya_1950          .........................t.............................g........................ [115040]
AM712240.1_OURT_88/3           ................................................................................ [115040]
AM712239.1_Benin_97/1          ................................................................................ [115040]
KM111295.1_Ken06.Bus           .........................t.............................g.......................a [115040]
KM111294.1_Ken05/Tk1           .........................t.............................g........................ [115040]
KM262844.1_strain_L60          ................................................................................ [115040]
KM102979.1_26544/OG10_Italy    ................................................................................ [115040]
KM262845.1_strain_NHV          ................................................................................ [115040]
KX354450.1_47/Ss/2008          ................................................................................ [115040]
KP055815.1_strain_BA71         ................................................................................ [115040]
MH025920.1_strain_R35          .........................t.............................g.......................a [115040]
MH025919.1_strain_N10          .........................t.............................g.......................a [115040]
MH025918.1_strain_R25          .........................t.............................g.......................a [115040]
MH025917.1_strain_R7           .........................t.............................g.......................a [115040]
MH025916.1_strain_R8           .........................t.............................g.......................a [115040]

FR682468.2_Georgia_2007/1      atggaggacatcatagtttgccgctgccagatggcctccaaaaagcgttcagcgcccttgttgtcattttttttttgctt [115120]
ASF_IND_20_CAD543              ................................................................................ [115120]
LR899193.1_Germany_2020/1      ................................................................................ [115120]
MN393477.1_Wuhan_2019          ................................................................................ [115120]
MW396979.1_Timor-Leste/2019/1  ................................................................................ [115120]
MK940252.1_CN/2019/InnerMongolia ................................................................................ [115120]
MK128995.1_China/2018/AnhuiXCGQ ................................................................................ [115120]
MH681419.1_ASFV/POL/2015       ................................................................................ [115120]
LS478113.1_Estonia_2014        ................................................................................ [115120]
KP843857.1_Odintsovo_02/14     ........................................................................-....... [115120]
FN557520.1_strain_E75          ................................................................................ [115120]
AY261366.1_Warthog             ..............g..............................................................t.. [115120]
AY261365.1_Warmbaths           ...............................c.....................a.......................t.. [115120]
AY261364.1_Tengani_62          .....a...............................................a.......................t.. [115120]
AY261363.1_Pretoriuskop/96/4   ..............g..............................................................t.. [115120]
AY261362.1_Mkuzi_1979          ................................................................................ [115120]
AY261361.1_Malawi_Lil-20/1_1983 ..............g................................................................. [115120]
AY261360.1_Kenya_1950          ..............g.........................................t....................... [115120]
AM712240.1_OURT_88/3           ................................................................................ [115120]
AM712239.1_Benin_97/1          ............................................a................................... [115120]
KM111295.1_Ken06.Bus           ..............g.........................................t....................... [115120]
```

```
KM111294.1_Ken05/Tk1            ..............g.......................................t..t....................... [115120]
KM262844.1_strain_L60           ................................................................................ [115120]
KM102979.1_26544/OG10_Italy     ................................................................................ [115120]
KM262845.1_strain_NHV           ................................................................................ [115120]
KX354450.1_47/Ss/2008           ................................................................................ [115120]
KP055815.1_strain_BA71          ................................................................................ [115120]
MH025920.1_strain_R35           ..............g..........................................t....................... [115120]
MH025919.1_strain_N10           ..............g..........................................t....................... [115120]
MH025918.1_strain_R25           ..............g..........................................t....................... [115120]
MH025917.1_strain_R7            ..............g..........................................t....................... [115120]
MH025916.1_strain_R8            ..............g..........................................t....................... [115120]

FR682468.2_Georgia_2007/1       atcggcgagccacaagcggtagtgtattagagttggatgtacaaaaccctcatatgaacgatttgagggttccgaggggg [115200]
ASF_IND_20_CAD543               ........................w....................................................... [115200]
LR899193.1_Germany_2020/1       ................................................................................ [115200]
MN393477.1_Wuhan_2019           ................................................................................ [115200]
MW396979.1_Timor-Leste/2019/1   ................................................................................ [115200]
MK940252.1_CN/2019/InnerMongolia ................................................................................ [115200]
MK128995.1_China/2018/AnhuiXCGQ ................................................................................ [115200]
MH681419.1_ASFV/POL/2015        ................................................................................ [115200]
LS478113.1_Estonia_2014         ................................................................................ [115200]
KP843857.1_Odintsovo_02/14      ................................................................................ [115200]
FN557520.1_strain_E75           ........................a....................................................... [115200]
AY261366.1_Warthog              ........................a.............................c........c..a..c.......... [115200]
AY261365.1_Warmbaths            ...a....................a.............................c...........a..c........a. [115200]
AY261364.1_Tengani_62           .....................a..a...........g.................c..............c........a. [115200]
AY261363.1_Pretoriuskop/96/4    ........................a...........g.................c........c..a..c.......... [115200]
AY261362.1_Mkuzi_1979           ........................a....................................................... [115200]
AY261361.1_Malawi_Lil-20/1_1983 ...a..a.....t...........a..g..............g...........c..g..............t.....a. [115200]
AY261360.1_Kenya_1950           ...a..a.................a....a...............g........c...........a..c..t.....a. [115200]
AM712240.1_OURT_88/3            ........................a....................................................... [115200]
AM712239.1_Benin_97/1           ........................a....................................................... [115200]
KM111295.1_Ken06.Bus            ...a..a.................a....a...............g........c...........a..c........a. [115200]
KM111294.1_Ken05/Tk1            ...a..a.................a....a...............g........c...........a..c..t.....a. [115200]
KM262844.1_strain_L60           ........................a....................................................... [115200]
KM102979.1_26544/OG10_Italy     ........................a....................................................... [115200]
KM262845.1_strain_NHV           ........................a....................................................... [115200]
KX354450.1_47/Ss/2008           ........................a....................................................... [115200]
KP055815.1_strain_BA71          ........................a....................................................... [115200]
MH025920.1_strain_R35           ...a..a.................a....a...............g........c...........a..c........a. [115200]
MH025919.1_strain_N10           ...a..a.................a....a...............g........c...........a..c........a. [115200]
MH025918.1_strain_R25           ...a..a.................a....a...............g........c...........a..c........a. [115200]
MH025917.1_strain_R7            ...a..a.................a....a...............g........c...........a..c........a. [115200]
MH025916.1_strain_R8            ...a..a.................a....a...............g........c...........a..c........a. [115200]
```

```
FR682468.2_Georgia_2007/1        caaccactaaaatttgttcaatatggggttgcaggattttcataatatgtttaacgtacacggttttgcctgtttttgag [115280]
ASF_IND_20_CAD543                ................................................................................ [115280]
LR899193.1_Germany_2020/1        ................................................................................ [115280]
MN393477.1_Wuhan_2019            ................................................................................ [115280]
MW396979.1_Timor-Leste/2019/1    ................................................................................ [115280]
MK940252.1_CN/2019/InnerMongolia ................................................................................ [115280]
MK128995.1_China/2018/AnhuiXCGQ  ................................................................................ [115280]
MH681419.1_ASFV/POL/2015         ................................................................................ [115280]
LS478113.1_Estonia_2014          ................................................................................ [115280]
KP843857.1_Odintsovo_02/14       ................................................................................ [115280]
FN557520.1_strain_E75            ................................................................................ [115280]
AY261366.1_Warthog               .................................a.....................a.....a.................. [115280]
AY261365.1_Warmbaths             .................................a.....................a........................ [115280]
AY261364.1_Tengani_62            .......c.........................a.....................a.....a.................. [115280]
AY261363.1_Pretoriuskop/96/4     .................................a.....................a.....a.................. [115280]
AY261362.1_Mkuzi_1979            ................................................................................ [115280]
AY261361.1_Malawi_Lil-20/1_1983  .................................a.....................a.....a.................. [115280]
AY261360.1_Kenya_1950            .................................a...........................a.................. [115280]
AM712240.1_OURT_88/3             ................................................................................ [115280]
AM712239.1_Benin_97/1            ................................................................................ [115280]
KM111295.1_Ken06.Bus             ....t..................................................a.....a.................. [115280]
KM111294.1_Ken05/Tk1             .................................a...........................a.................. [115280]
KM262844.1_strain_L60            ................................................................................ [115280]
KM102979.1_26544/OG10_Italy      ................................................................................ [115280]
KM262845.1_strain_NHV            ................................................................................ [115280]
KX354450.1_47/Ss/2008            ................................................................................ [115280]
KP055815.1_strain_BA71           ................................................................................ [115280]
MH025920.1_strain_R35            ....t..................................................a.....a.................. [115280]
MH025919.1_strain_N10            ....t..................................................a.....a.................. [115280]
MH025918.1_strain_R25            ....t..................................................a.....a.................. [115280]
MH025917.1_strain_R7             ....t..................................................a.....a.................. [115280]
MH025916.1_strain_R8             ....t..................................................a.....a.................. [115280]

FR682468.2_Georgia_2007/1        gggccatatagcacagttgttttatctataaaatgatgtgctttgaactgtagttcaggaattagcttccctgaatgggt [115360]
ASF_IND_20_CAD543                ..............................................................y................. [115360]
LR899193.1_Germany_2020/1        ................................................................................ [115360]
MN393477.1_Wuhan_2019            ................................................................................ [115360]
MW396979.1_Timor-Leste/2019/1    ................................................................................ [115360]
MK940252.1_CN/2019/InnerMongolia ................................................................................ [115360]
MK128995.1_China/2018/AnhuiXCGQ  ................................................................................ [115360]
MH681419.1_ASFV/POL/2015         ................................................................................ [115360]
LS478113.1_Estonia_2014          ................................................................................ [115360]
KP843857.1_Odintsovo_02/14       ................................................................................ [115360]
```

```
FN557520.1_strain_E75            ..............................................................c................. [115360]
AY261366.1_Warthog               ................................................................................ [115360]
AY261365.1_Warmbaths             ................................................................................ [115360]
AY261364.1_Tengani_62            ........c....................................................................... [115360]
AY261363.1_Pretoriuskop/96/4     ........c....................................................................... [115360]
AY261362.1_Mkuzi_1979            ................................................................................ [115360]
AY261361.1_Malawi_Lil-20/1_1983  ..........a...c........g..g....................t..............g..t..t........... [115360]
AY261360.1_Kenya_1950            ..............g........g..g....................t....................t......g.a.. [115360]
AM712240.1_OURT_88/3             ..............................................................c................. [115360]
AM712239.1_Benin_97/1            ..............................................................c................. [115360]
KM111295.1_Ken06.Bus             ..............g........g..g....................t....................t......g.a.. [115360]
KM111294.1_Ken05/Tk1             .......................g..g....................t....................t......g.a.. [115360]
KM262844.1_strain_L60            ..............................................................c................. [115360]
KM102979.1_26544/OG10_Italy      ..............................................................c................. [115360]
KM262845.1_strain_NHV            ..............................................................c................. [115360]
KX354450.1_47/Ss/2008            ..............................................................c................. [115360]
KP055815.1_strain_BA71           ..............................................................c................. [115360]
MH025920.1_strain_R35            ..............g........g..g....................t....................t......g.a.. [115360]
MH025919.1_strain_N10            ..............g........g..g....................t....................t......g.a.. [115360]
MH025918.1_strain_R25            ..............g........g..g....................t....................t......g.a.. [115360]
MH025917.1_strain_R7             ..............g........g..g....................t....................t......g.a.. [115360]
MH025916.1_strain_R8             ..............g........g..g....................t....................t......g.a.. [115360]

FR682468.2_Georgia_2007/1        cgttagggccatctctatattattacaattctgcttttgtatataaaatttctttttcgagtttattattattgttgacc [115440]
ASF_IND_20_CAD543                ................................................................................ [115440]
LR899193.1_Germany_2020/1        ................................................................................ [115440]
MN393477.1_Wuhan_2019            ................................................................................ [115440]
MW396979.1_Timor-Leste/2019/1    ................................................................................ [115440]
MK940252.1_CN/2019/InnerMongolia ................................................................................ [115440]
MK128995.1_China/2018/AnhuiXCGQ  ................................................................................ [115440]
MH681419.1_ASFV/POL/2015         ................................................................................ [115440]
LS478113.1_Estonia_2014          ................................................................................ [115440]
KP843857.1_Odintsovo_02/14       ................................................................................ [115440]
FN557520.1_strain_E75            ................................................................................ [115440]
AY261366.1_Warthog               ................................................................................ [115440]
AY261365.1_Warmbaths             ................................................................g..g.....a...... [115440]
AY261364.1_Tengani_62            ................................................................g..g.....a...... [115440]
AY261363.1_Pretoriuskop/96/4     ................................................................g..g.....a...... [115440]
AY261362.1_Mkuzi_1979            ................................................................................ [115440]
AY261361.1_Malawi_Lil-20/1_1983  ................................................................g............... [115440]
AY261360.1_Kenya_1950            t...............................................................g...........a... [115440]
AM712240.1_OURT_88/3             ................................................................................ [115440]
AM712239.1_Benin_97/1            ................................................................................ [115440]
KM111295.1_Ken06.Bus             t...............................................................g............... [115440]
```

```
KM111294.1_Ken05/Tk1             t...............................................................g............... [115440]
KM262844.1_strain_L60            ................................................................................ [115440]
KM102979.1_26544/OG10_Italy      ................................................................................ [115440]
KM262845.1_strain_NHV            ................................................................................ [115440]
KX354450.1_47/Ss/2008            ................................................................................ [115440]
KP055815.1_strain_BA71           ................................................................................ [115440]
MH025920.1_strain_R35            t...............................................................g............... [115440]
MH025919.1_strain_N10            t...............................................................g............... [115440]
MH025918.1_strain_R25            t...............................................................g............... [115440]
MH025917.1_strain_R7             t...............................................................g............... [115440]
MH025916.1_strain_R8             t...............................................................g............... [115440]

FR682468.2_Georgia_2007/1        cacatatctacccgtatcgtatcatcaggcacattgagcatttcaagcgcattatctaactgtttttttgtttttatcag [115520]
ASF_IND_20_CAD543                .......................................................y........................ [115520]
LR899193.1_Germany_2020/1        ................................................................................ [115520]
MN393477.1_Wuhan_2019            ................................................................................ [115520]
MW396979.1_Timor-Leste/2019/1    ................................................................................ [115520]
MK940252.1_CN/2019/InnerMongolia ................................................................................ [115520]
MK128995.1_China/2018/AnhuiXCGQ  ................................................................................ [115520]
MH681419.1_ASFV/POL/2015         ................................................................................ [115520]
LS478113.1_Estonia_2014          ................................................................................ [115520]
KP843857.1_Odintsovo_02/14       ..............................................................-................. [115520]
FN557520.1_strain_E75            .......................................................t........................ [115520]
AY261366.1_Warthog               .......................................................t........................ [115520]
AY261365.1_Warmbaths             ................................g............................................... [115520]
AY261364.1_Tengani_62            .......................................................t........................ [115520]
AY261363.1_Pretoriuskop/96/4     .......................................................t........................ [115520]
AY261362.1_Mkuzi_1979            ................................................................................ [115520]
AY261361.1_Malawi_Lil-20/1_1983  ........................................................c....................... [115520]
AY261360.1_Kenya_1950            ........c.........................................g..t.....................tc... [115520]
AM712240.1_OURT_88/3             .......................................................t........................ [115520]
AM712239.1_Benin_97/1            .......................................................t........................ [115520]
KM111295.1_Ken06.Bus             ........c............................................t.....................tc... [115520]
KM111294.1_Ken05/Tk1             ........c............................................t.....................tc... [115520]
KM262844.1_strain_L60            .......................................................t........................ [115520]
KM102979.1_26544/OG10_Italy      .......................................................t........................ [115520]
KM262845.1_strain_NHV            .......................................................t........................ [115520]
KX354450.1_47/Ss/2008            .......................................................t........................ [115520]
KP055815.1_strain_BA71           .......................................................t........................ [115520]
MH025920.1_strain_R35            ........c............................................t.....................tc... [115520]
MH025919.1_strain_N10            ........c............................................t.....................tc... [115520]
MH025918.1_strain_R25            ........c............................................t.....................tc... [115520]
MH025917.1_strain_R7             ........c............................................t.....................tc... [115520]
MH025916.1_strain_R8             ........c............................................t.....................tc... [115520]
```

```
FR682468.2_Georgia_2007/1        ctcgctttcttcatcgggggttaaattttctttactaagcagttgcttaattttttcttcgcagtcgtctataaaatcat [115600]
ASF_IND_20_CAD543                ................................................................................ [115600]
LR899193.1_Germany_2020/1        ................................................................................ [115600]
MN393477.1_Wuhan_2019            ................................................................................ [115600]
MW396979.1_Timor-Leste/2019/1    ................................................................................ [115600]
MK940252.1_CN/2019/InnerMongolia ................................................................................ [115600]
MK128995.1_China/2018/AnhuiXCGQ  ................................................................................ [115600]
MH681419.1_ASFV/POL/2015         ................................................................................ [115600]
LS478113.1_Estonia_2014          ................................................................................ [115600]
KP843857.1_Odintsovo_02/14       ................................................................................ [115600]
FN557520.1_strain_E75            ................................................................................ [115600]
AY261366.1_Warthog               ...t.............a.............................................................. [115600]
AY261365.1_Warmbaths             .................a......t....................................................... [115600]
AY261364.1_Tengani_62            .................a..............................................g............... [115600]
AY261363.1_Pretoriuskop/96/4     ..................................................................a............. [115600]
AY261362.1_Mkuzi_1979            ................................................................................ [115600]
AY261361.1_Malawi_Lil-20/1_1983  ........t........a.............................................t..a............. [115600]
AY261360.1_Kenya_1950            ..g.....t......a..........................c.....g.................a............. [115600]
AM712240.1_OURT_88/3             ................................................................................ [115600]
AM712239.1_Benin_97/1            ................................................................................ [115600]
KM111295.1_Ken06.Bus             ..g.....t.................................c.....g.................a.....g....... [115600]
KM111294.1_Ken05/Tk1             ..g.....t......a..........................c.....g.................a............. [115600]
KM262844.1_strain_L60            ................................................................................ [115600]
KM102979.1_26544/OG10_Italy      ................................................................................ [115600]
KM262845.1_strain_NHV            ................................................................................ [115600]
KX354450.1_47/Ss/2008            ................................................................................ [115600]
KP055815.1_strain_BA71           ................................................................................ [115600]
MH025920.1_strain_R35            ..g.....t.................................c.....g.................a.....g....... [115600]
MH025919.1_strain_N10            ..g.....t.................................c.....g.................a.....g....... [115600]
MH025918.1_strain_R25            ..g.....t.................................c.....g.................a.....g....... [115600]
MH025917.1_strain_R7             ..g.....t.................................c.....g.................a.....g....... [115600]
MH025916.1_strain_R8             ..g.....t.................................c.....g.................a.....g....... [115600]

FR682468.2_Georgia_2007/1        actctcgagcttttttgatatttccagatgctttttctaggttttttagctccttaaaggaaagcagtcccttaatcccg [115680]
ASF_IND_20_CAD543                ................................................................................ [115680]
LR899193.1_Germany_2020/1        ................................................................................ [115680]
MN393477.1_Wuhan_2019            ................................................................................ [115680]
MW396979.1_Timor-Leste/2019/1    ................................................................................ [115680]
MK940252.1_CN/2019/InnerMongolia ................................................................................ [115680]
MK128995.1_China/2018/AnhuiXCGQ  ................................................................................ [115680]
MH681419.1_ASFV/POL/2015         ................................................................................ [115680]
LS478113.1_Estonia_2014          ................................................................................ [115680]
KP843857.1_Odintsovo_02/14       ................................................................................ [115680]
```

```
FN557520.1_strain_E75              ................................................................................ [115680]
AY261366.1_Warthog                 ................................................a............................... [115680]
AY261365.1_Warmbaths               ..................c......g...................................................... [115680]
AY261364.1_Tengani_62              ................................................................................ [115680]
AY261363.1_Pretoriuskop/96/4       ................................................................................ [115680]
AY261362.1_Mkuzi_1979              ................................................................................ [115680]
AY261361.1_Malawi_Lil-20/1_1983    ..................c.....................a.............................t......... [115680]
AY261360.1_Kenya_1950              ........................................a....................................... [115680]
AM712240.1_OURT_88/3               ................................................................................ [115680]
AM712239.1_Benin_97/1              ................................................................................ [115680]
KM111295.1_Ken06.Bus               ..................................c.....a....................................... [115680]
KM111294.1_Ken05/Tk1               ........................................a....................................... [115680]
KM262844.1_strain_L60              ................................................................................ [115680]
KM102979.1_26544/OG10_Italy        ..........................................................a..................... [115680]
KM262845.1_strain_NHV              ................................................................................ [115680]
KX354450.1_47/Ss/2008              ..........................................................a..................... [115680]
KP055815.1_strain_BA71             ................................................................................ [115680]
MH025920.1_strain_R35              ..................................c.....a....................................... [115680]
MH025919.1_strain_N10              ..................................c.....a....................................... [115680]
MH025918.1_strain_R25              ..................................c.....a....................................... [115680]
MH025917.1_strain_R7               ..................................c.....a....................................... [115680]
MH025916.1_strain_R8               ..................................c.....a....................................... [115680]

FR682468.2_Georgia_2007/1          ctatccgtgtgaaaggttgaattatagatggagagccccggagcatccgggccagtttcttgtatattttttgctttttt [115760]
ASF_IND_20_CAD543                  ................................................................................ [115760]
LR899193.1_Germany_2020/1          ................................................................................ [115760]
MN393477.1_Wuhan_2019              ................................................................................ [115760]
MW396979.1_Timor-Leste/2019/1      ................................................................................ [115760]
MK940252.1_CN/2019/InnerMongolia   ................................................................................ [115760]
MK128995.1_China/2018/AnhuiXCGQ    ................................................................................ [115760]
MH681419.1_ASFV/POL/2015           ................................................................................ [115760]
LS478113.1_Estonia_2014            ................................................................................ [115760]
KP843857.1_Odintsovo_02/14         ................................................................................ [115760]
FN557520.1_strain_E75              .....................................................g.......................... [115760]
AY261366.1_Warthog                 ................................................................................ [115760]
AY261365.1_Warmbaths               ......................................t......................................... [115760]
AY261364.1_Tengani_62              ...........g.................................................................... [115760]
AY261363.1_Pretoriuskop/96/4       ...................................t............................................ [115760]
AY261362.1_Mkuzi_1979              ................................................................................ [115760]
AY261361.1_Malawi_Lil-20/1_1983    ..g...................................t..............g.......................... [115760]
AY261360.1_Kenya_1950              ..g..t................................t......................................... [115760]
AM712240.1_OURT_88/3               .....................................................g.......................... [115760]
AM712239.1_Benin_97/1              .....................................................g.......................... [115760]
KM111295.1_Ken06.Bus               ..g..t................................t......................................... [115760]
```

```
KM111294.1_Ken05/Tk1             ..g..t................................t......................................... [115760]
KM262844.1_strain_L60            .....................................................g.......................... [115760]
KM102979.1_26544/OG10_Italy      .....................................................g.......................... [115760]
KM262845.1_strain_NHV            .....................................................g.......................... [115760]
KX354450.1_47/Ss/2008            .....................................................g.......................... [115760]
KP055815.1_strain_BA71           .....................................................g.......................... [115760]
MH025920.1_strain_R35            ..g..t................................t......................................... [115760]
MH025919.1_strain_N10            ..g..t................................t......................................... [115760]
MH025918.1_strain_R25            ..g..t................................t......................................... [115760]
MH025917.1_strain_R7             ..g..t................................t......................................... [115760]
MH025916.1_strain_R8             ..g..t................................t......................................... [115760]

FR682468.2_Georgia_2007/1        gtggtaaatagtatttcgtaaaatctcttttcctatctttaggtcttcctcatgacggtccaaaatccgttttattattt [115840]
ASF_IND_20_CAD543                ................................................................................ [115840]
LR899193.1_Germany_2020/1        ................................................................................ [115840]
MN393477.1_Wuhan_2019            ................................................................................ [115840]
MW396979.1_Timor-Leste/2019/1    ................................................................................ [115840]
MK940252.1_CN/2019/InnerMongolia ................................................................................ [115840]
MK128995.1_China/2018/AnhuiXCGQ  ................................................................................ [115840]
MH681419.1_ASFV/POL/2015         ................................................................................ [115840]
LS478113.1_Estonia_2014          ................................................................................ [115840]
KP843857.1_Odintsovo_02/14       ................................................................................ [115840]
FN557520.1_strain_E75            .........g..........g........................................................... [115840]
AY261366.1_Warthog               .........g...................................................................... [115840]
AY261365.1_Warmbaths             .........g..........g........................................................... [115840]
AY261364.1_Tengani_62            .........g...................................................................... [115840]
AY261363.1_Pretoriuskop/96/4     .........g...................................................................... [115840]
AY261362.1_Mkuzi_1979            .........g..........g........................................................... [115840]
AY261361.1_Malawi_Lil-20/1_1983  ...a.....g................................a.......................t..........c.. [115840]
AY261360.1_Kenya_1950            .........g..........................t.....a..................................... [115840]
AM712240.1_OURT_88/3             .........g..........g........................................................... [115840]
AM712239.1_Benin_97/1            .........g..........g........................................................... [115840]
KM111295.1_Ken06.Bus             .........g..........................t.....a..................................... [115840]
KM111294.1_Ken05/Tk1             .........g....................c.....t.....a..................................... [115840]
KM262844.1_strain_L60            .........g..........g........................................................... [115840]
KM102979.1_26544/OG10_Italy      .........g..........g........................................................... [115840]
KM262845.1_strain_NHV            .........g..........g........................................................... [115840]
KX354450.1_47/Ss/2008            .........g..........g........................................................... [115840]
KP055815.1_strain_BA71           .........g..........g........................................................... [115840]
MH025920.1_strain_R35            .........g..........................t.....a..................................... [115840]
MH025919.1_strain_N10            .........g..........................t.....a..................................... [115840]
MH025918.1_strain_R25            .........g..........................t.....a..................................... [115840]
MH025917.1_strain_R7             .........g..........................t.....a..................................... [115840]
MH025916.1_strain_R8             .........g..........................t.....a..................................... [115840]
```

```
FR682468.2_Georgia_2007/1        cattattttgattaaaataattgtagcgctctctgttggccttaaagcttcccaggagtgtccagttgcctaattgaatg [115920]
ASF_IND_20_CAD543                ................................................................................ [115920]
LR899193.1_Germany_2020/1        ................................................................................ [115920]
MN393477.1_Wuhan_2019            ................................................................................ [115920]
MW396979.1_Timor-Leste/2019/1    ................................................................................ [115920]
MK940252.1_CN/2019/InnerMongolia ................................................................................ [115920]
MK128995.1_China/2018/AnhuiXCGQ  ................................................................................ [115920]
MH681419.1_ASFV/POL/2015         ................................................................................ [115920]
LS478113.1_Estonia_2014          ................................................................................ [115920]
KP843857.1_Odintsovo_02/14       ................................................................................ [115920]
FN557520.1_strain_E75            ................................................................................ [115920]
AY261366.1_Warthog               ...................g.....a...........a.......................................... [115920]
AY261365.1_Warmbaths             ...................g............................................................ [115920]
AY261364.1_Tengani_62            ...................g.....a...................................................... [115920]
AY261363.1_Pretoriuskop/96/4     ...................g...................................a........................ [115920]
AY261362.1_Mkuzi_1979            ...................g............................................................ [115920]
AY261361.1_Malawi_Lil-20/1_1983  ...................g............................................................ [115920]
AY261360.1_Kenya_1950            ...................g...................................a...........a............ [115920]
AM712240.1_OURT_88/3             ................................................................................ [115920]
AM712239.1_Benin_97/1            ................................................................................ [115920]
KM111295.1_Ken06.Bus             ...................g...................................a........................ [115920]
KM111294.1_Ken05/Tk1             ...................g...................................a........................ [115920]
KM262844.1_strain_L60            ................................................................................ [115920]
KM102979.1_26544/OG10_Italy      ................................................................................ [115920]
KM262845.1_strain_NHV            ................................................................................ [115920]
KX354450.1_47/Ss/2008            ................................................................................ [115920]
KP055815.1_strain_BA71           ................................................................................ [115920]
MH025920.1_strain_R35            ...................g...................................a........................ [115920]
MH025919.1_strain_N10            ...................g...................................a........................ [115920]
MH025918.1_strain_R25            ...................g...................................a........................ [115920]
MH025917.1_strain_R7             ...................g...................................a........................ [115920]
MH025916.1_strain_R8             ...................g...................................a........................ [115920]

FR682468.2_Georgia_2007/1        gatgaaacctctgagaaaatctggtctttatatttataataaaattcatcaaccttttgttggttg---ctgctatccac [116000]
ASF_IND_20_CAD543                ..................................................................---........... [116000]
LR899193.1_Germany_2020/1        ..................................................................---........... [116000]
MN393477.1_Wuhan_2019            ..................................................................---........... [116000]
MW396979.1_Timor-Leste/2019/1    ..................................................................---........... [116000]
MK940252.1_CN/2019/InnerMongolia ..................................................................---........... [116000]
MK128995.1_China/2018/AnhuiXCGQ  ..................................................................---........... [116000]
MH681419.1_ASFV/POL/2015         ..................................................................---........... [116000]
LS478113.1_Estonia_2014          ..................................................................---........... [116000]
KP843857.1_Odintsovo_02/14       ..................................................................---........... [116000]
```

```
FN557520.1_strain_E75               ..................................................................---........... [116000]
AY261366.1_Warthog                  ..............................................................a...---........... [116000]
AY261365.1_Warmbaths                ..................................................................---........... [116000]
AY261364.1_Tengani_62               ..................................................................---........... [116000]
AY261363.1_Pretoriuskop/96/4        ..................................................................---........... [116000]
AY261362.1_Mkuzi_1979               ..................................................................---........... [116000]
AY261361.1_Malawi_Lil-20/1_1983     ....................t.....c.........................t.............---........... [116000]
AY261360.1_Kenya_1950               ....................................................t......c......cta..a........ [116000]
AM712240.1_OURT_88/3                ..................................................................---........... [116000]
AM712239.1_Benin_97/1               ..................................................................---........... [116000]
KM111295.1_Ken06.Bus                ....................................................t......c.....acta..a........ [116000]
KM111294.1_Ken05/Tk1                ....................................................t......c......cta..a........ [116000]
KM262844.1_strain_L60               ..................................................................---........... [116000]
KM102979.1_26544/OG10_Italy         ..................................................................---........... [116000]
KM262845.1_strain_NHV               ..................................................................---........... [116000]
KX354450.1_47/Ss/2008               ..................................................................---........... [116000]
KP055815.1_strain_BA71              ..................................................................---........... [116000]
MH025920.1_strain_R35               ....................................................t......c.....acta..a........ [116000]
MH025919.1_strain_N10               ....................................................t......c.....acta..a........ [116000]
MH025918.1_strain_R25               ....................................................t......c.....acta..a........ [116000]
MH025917.1_strain_R7                ....................................................t......c.....acta..a........ [116000]
MH025916.1_strain_R8                ....................................................t......c.....acta..a........ [116000]

FR682468.2_Georgia_2007/1           cacatcataaataatgaaggcaaactctaggtcgggtttttctgggtagatgctttccgtagcggcccgcaactcttcgt [116080]
ASF_IND_20_CAD543                   ................................................................................ [116080]
LR899193.1_Germany_2020/1           ................................................................................ [116080]
MN393477.1_Wuhan_2019               ................................................................................ [116080]
MW396979.1_Timor-Leste/2019/1       ................................................................................ [116080]
MK940252.1_CN/2019/InnerMongolia    ................................................................................ [116080]
MK128995.1_China/2018/AnhuiXCGQ     ................................................................................ [116080]
MH681419.1_ASFV/POL/2015            ................................................................................ [116080]
LS478113.1_Estonia_2014             ................................................................................ [116080]
KP843857.1_Odintsovo_02/14          ................................................................................ [116080]
FN557520.1_strain_E75               ....................................c........................................... [116080]
AY261366.1_Warthog                  ...............a.............................a.................................. [116080]
AY261365.1_Warmbaths                ................................................................................ [116080]
AY261364.1_Tengani_62               ..............................a................................a................ [116080]
AY261363.1_Pretoriuskop/96/4        ................................................................................ [116080]
AY261362.1_Mkuzi_1979               ....................................c........................................... [116080]
AY261361.1_Malawi_Lil-20/1_1983     ...............a................................................................ [116080]
AY261360.1_Kenya_1950               ...............a................................................................ [116080]
AM712240.1_OURT_88/3                ....................................c........................................... [116080]
AM712239.1_Benin_97/1               ....................................c........................................... [116080]
KM111295.1_Ken06.Bus                ...............a................................................................ [116080]
```

```
KM111294.1_Ken05/Tk1             ...............a........................................................t....... [116080]
KM262844.1_strain_L60            ....................................c........................................... [116080]
KM102979.1_26544/OG10_Italy      ....................................c........................................... [116080]
KM262845.1_strain_NHV            ....................................c........................................... [116080]
KX354450.1_47/Ss/2008            ....................................c........................................... [116080]
KP055815.1_strain_BA71           ....................................c........................................... [116080]
MH025920.1_strain_R35            ...............a................................................................ [116080]
MH025919.1_strain_N10            ...............a................................................................ [116080]
MH025918.1_strain_R25            ...............a................................................................ [116080]
MH025917.1_strain_R7             ...............a................................................................ [116080]
MH025916.1_strain_R8             ...............a................................................................ [116080]

FR682468.2_Georgia_2007/1        aattatcctcaatgtaataattccacttataaaaagtatcctgaggtggaatatgctgcgaaagatatctagtaattttt [116160]
ASF_IND_20_CAD543                ................................................................................ [116160]
LR899193.1_Germany_2020/1        ................................................................................ [116160]
MN393477.1_Wuhan_2019            ................................................................................ [116160]
MW396979.1_Timor-Leste/2019/1    ................................................................................ [116160]
MK940252.1_CN/2019/InnerMongolia ................................................................................ [116160]
MK128995.1_China/2018/AnhuiXCGQ  ................................................................................ [116160]
MH681419.1_ASFV/POL/2015         ................................................................................ [116160]
LS478113.1_Estonia_2014          ................................................................................ [116160]
KP843857.1_Odintsovo_02/14       ................................................................................ [116160]
FN557520.1_strain_E75            ................................................................................ [116160]
AY261366.1_Warthog               .......t.....a......................................g..t........................ [116160]
AY261365.1_Warmbaths             ................................................................................ [116160]
AY261364.1_Tengani_62            ................................................................................ [116160]
AY261363.1_Pretoriuskop/96/4     .............a......................................g........................... [116160]
AY261362.1_Mkuzi_1979            ................................................................................ [116160]
AY261361.1_Malawi_Lil-20/1_1983  ..................................c.................g........................... [116160]
AY261360.1_Kenya_1950            ..................................c.................g........................... [116160]
AM712240.1_OURT_88/3             ................................................................................ [116160]
AM712239.1_Benin_97/1            ................................................................................ [116160]
KM111295.1_Ken06.Bus             .........................t........c.................g..t........................ [116160]
KM111294.1_Ken05/Tk1             ..................................c.................g..t........................ [116160]
KM262844.1_strain_L60            ................................................................................ [116160]
KM102979.1_26544/OG10_Italy      ................................................................................ [116160]
KM262845.1_strain_NHV            ................................................................................ [116160]
KX354450.1_47/Ss/2008            ................................................................................ [116160]
KP055815.1_strain_BA71           ................................................................................ [116160]
MH025920.1_strain_R35            .........................t........c.................g..t........................ [116160]
MH025919.1_strain_N10            .........................t........c.................g..t........................ [116160]
MH025918.1_strain_R25            .........................t........c.................g..t........................ [116160]
MH025917.1_strain_R7             .........................t........c.................g..t........................ [116160]
MH025916.1_strain_R8             .........................t........c.................g..t........................ [116160]
```

```
FR682468.2_Georgia_2007/1        gtgttaaagagaatgggtttaaacgccctcggattttcaagcatatgtttaatgctttggtgaagttctatattttgtaa [116240]
ASF_IND_20_CAD543                ................................................................................ [116240]
LR899193.1_Germany_2020/1        ................................................................................ [116240]
MN393477.1_Wuhan_2019            ................................................................................ [116240]
MW396979.1_Timor-Leste/2019/1    ................................................................................ [116240]
MK940252.1_CN/2019/InnerMongolia ................................................................................ [116240]
MK128995.1_China/2018/AnhuiXCGQ  ................................................................................ [116240]
MH681419.1_ASFV/POL/2015         ................................................................................ [116240]
LS478113.1_Estonia_2014          ................................................................................ [116240]
KP843857.1_Odintsovo_02/14       ................................................................................ [116240]
FN557520.1_strain_E75            ...........................................................a.................... [116240]
AY261366.1_Warthog               .............................t.................................................. [116240]
AY261365.1_Warmbaths             ................................................................................ [116240]
AY261364.1_Tengani_62            .....................................................a.......................... [116240]
AY261363.1_Pretoriuskop/96/4     .............................t.................................................. [116240]
AY261362.1_Mkuzi_1979            ...........................................................a.................... [116240]
AY261361.1_Malawi_Lil-20/1_1983  ..a....................t..t..t......................c........................... [116240]
AY261360.1_Kenya_1950            ..a....................t...t.t....................t............................. [116240]
AM712240.1_OURT_88/3             ...........................................................a.................... [116240]
AM712239.1_Benin_97/1            ...........................................................a.................... [116240]
KM111295.1_Ken06.Bus             ..a....................ta..t.t....................t............................. [116240]
KM111294.1_Ken05/Tk1             ..a....................t...t.t....................t............................. [116240]
KM262844.1_strain_L60            ...........................................................a.................... [116240]
KM102979.1_26544/OG10_Italy      ...........................................................a.................... [116240]
KM262845.1_strain_NHV            ...........................................................a.................... [116240]
KX354450.1_47/Ss/2008            ...........................................................a.................... [116240]
KP055815.1_strain_BA71           ...........................................................a.................... [116240]
MH025920.1_strain_R35            ..a....................ta..t.t....................t............................. [116240]
MH025919.1_strain_N10            ..a....................ta..t.t....................t............................. [116240]
MH025918.1_strain_R25            ..a....................ta..t.t....................t............................. [116240]
MH025917.1_strain_R7             ..a....................ta..t.t....................t............................. [116240]
MH025916.1_strain_R8             ..a....................ta..t.t....................t............................. [116240]

FR682468.2_Georgia_2007/1        tatgtgggctgctgccctatagccctgtggggtttgggtgattgcatcaatatcggcctgaagctcattaggcacattta [116320]
ASF_IND_20_CAD543                ........y....................................................................... [116320]
LR899193.1_Germany_2020/1        ................................................................................ [116320]
MN393477.1_Wuhan_2019            ................................................................................ [116320]
MW396979.1_Timor-Leste/2019/1    ................................................................................ [116320]
MK940252.1_CN/2019/InnerMongolia ................................................................................ [116320]
MK128995.1_China/2018/AnhuiXCGQ  ................................................................................ [116320]
MH681419.1_ASFV/POL/2015         ................................................................................ [116320]
LS478113.1_Estonia_2014          ................................................................................ [116320]
KP843857.1_Odintsovo_02/14       ................................................................................ [116320]
```

```
FN557520.1_strain_E75            .......................................a........................................ [116320]
AY261366.1_Warthog               ................................................................................ [116320]
AY261365.1_Warmbaths             ................................................................................ [116320]
AY261364.1_Tengani_62            ................................................................................ [116320]
AY261363.1_Pretoriuskop/96/4     ................................................................................ [116320]
AY261362.1_Mkuzi_1979            ...........................c...........a........................................ [116320]
AY261361.1_Malawi_Lil-20/1_1983  .....................a........a................................................. [116320]
AY261360.1_Kenya_1950            ..................g.....t...c.......a...................................t....... [116320]
AM712240.1_OURT_88/3             .......................................a........................................ [116320]
AM712239.1_Benin_97/1            .......................................a........................................ [116320]
KM111295.1_Ken06.Bus             ....................................a...................................t....... [116320]
KM111294.1_Ken05/Tk1             ........................t...........a.................a.................t....... [116320]
KM262844.1_strain_L60            .......................................a........................................ [116320]
KM102979.1_26544/OG10_Italy      .......................................a........................................ [116320]
KM262845.1_strain_NHV            .......................................a........................................ [116320]
KX354450.1_47/Ss/2008            .......................................a........................................ [116320]
KP055815.1_strain_BA71           .......................................a........................................ [116320]
MH025920.1_strain_R35            ....................................a...................................t....... [116320]
MH025919.1_strain_N10            ....................................a...................................t....... [116320]
MH025918.1_strain_R25            ....................................a...................................t....... [116320]
MH025917.1_strain_R7             ....................................a...................................t....... [116320]
MH025916.1_strain_R8             ....................................a...................................t....... [116320]

FR682468.2_Georgia_2007/1        atgttttttgcatgatgtgtaaagggatgcgctcaggatctgctaaatcggtgtattct--------------------- [116400]
ASF_IND_20_CAD543                ...........................................................--------------------- [116400]
LR899193.1_Germany_2020/1        ...........................................................--------------------- [116400]
MN393477.1_Wuhan_2019            ...........................................................--------------------- [116400]
MW396979.1_Timor-Leste/2019/1    ...........................................................--------------------- [116400]
MK940252.1_CN/2019/InnerMongolia ...........................................................--------------------- [116400]
MK128995.1_China/2018/AnhuiXCGQ  ...........................................................--------------------- [116400]
MH681419.1_ASFV/POL/2015         ...........................................................--------------------- [116400]
LS478113.1_Estonia_2014          ...........................................................--------------------- [116400]
KP843857.1_Odintsovo_02/14       ...........................................................--------------------- [116400]
FN557520.1_strain_E75            ...........................................................--------------------- [116400]
AY261366.1_Warthog               ...................................................c.......--------------------- [116400]
AY261365.1_Warmbaths             ...................................................c.......--------------------- [116400]
AY261364.1_Tengani_62            ..........................................t.........a......--------------------- [116400]
AY261363.1_Pretoriuskop/96/4     ...................................................c.......gtgtttgcacaagtgcttgca [116400]
AY261362.1_Mkuzi_1979            ...........................................................--------------------- [116400]
AY261361.1_Malawi_Lil-20/1_1983  .........................a................t................--------------------- [116400]
AY261360.1_Kenya_1950            ........................................cat................--------------------- [116400]
AM712240.1_OURT_88/3             ...........................................................--------------------- [116400]
AM712239.1_Benin_97/1            ...........................................................--------------------- [116400]
KM111295.1_Ken06.Bus             ........................................cat................--------------------- [116400]
```

```
KM111294.1_Ken05/Tk1             ........................................cat................--------------------- [116400]
KM262844.1_strain_L60            ...........................................................--------------------- [116400]
KM102979.1_26544/OG10_Italy      ...........................................................--------------------- [116400]
KM262845.1_strain_NHV            ...........................................................--------------------- [116400]
KX354450.1_47/Ss/2008            ...........................................................--------------------- [116400]
KP055815.1_strain_BA71           ...........................................................--------------------- [116400]
MH025920.1_strain_R35            ........................................cat................--------------------- [116400]
MH025919.1_strain_N10            ........................................cat................--------------------- [116400]
MH025918.1_strain_R25            ........................................cat................--------------------- [116400]
MH025917.1_strain_R7             ........................................cat................--------------------- [116400]
MH025916.1_strain_R8             ........................................cat................--------------------- [116400]

FR682468.2_Georgia_2007/1        ---------------------------------------------------gtgcttgtacaagtgc------------t [116480]
ASF_IND_20_CAD543                ---------------------------------------------------................ttgyacargtgc. [116480]
LR899193.1_Germany_2020/1        ---------------------------------------------------................------------. [116480]
MN393477.1_Wuhan_2019            ---------------------------------------------------................------------. [116480]
MW396979.1_Timor-Leste/2019/1    ---------------------------------------------------................------------. [116480]
MK940252.1_CN/2019/InnerMongolia ---------------------------------------------------................------------. [116480]
MK128995.1_China/2018/AnhuiXCGQ  ---------------------------------------------------................------------. [116480]
MH681419.1_ASFV/POL/2015         ---------------------------------------------------................------------. [116480]
LS478113.1_Estonia_2014          ---------------------------------------------------................------------. [116480]
KP843857.1_Odintsovo_02/14       ---------------------------------------------------................------------. [116480]
FN557520.1_strain_E75            ---------------------------------------------------.......c...g...tttgcacaggtgc. [116480]
AY261366.1_Warthog               ---------------------------------------------------................ttgcacaagtgc. [116480]
AY261365.1_Warmbaths             ---------------------------------------------------...t............ttgcacatgtatc [116480]
AY261364.1_Tengani_62            ---------------------------------------------------...............t------------- [116480]
AY261363.1_Pretoriuskop/96/4     caagtgcttacacaagtgtctgcgcatgtgtctacattggtatctgcacaa.......c........ttgcacaagtgc. [116480]
AY261362.1_Mkuzi_1979            ---------------------------------------------------...t...c........ttgcacaggtgc. [116480]
AY261361.1_Malawi_Lil-20/1_1983  ---------------------------------------------------...t...c.ttga..tttgcattgatgtc [116480]
AY261360.1_Kenya_1950            ---------------------------------------------------.......c...g...tctgcacaagt--- [116480]
AM712240.1_OURT_88/3             ---------------------------------------------------.......c...g...tttgcacaggtgc. [116480]
AM712239.1_Benin_97/1            ---------------------------------------------------.......c...g...tttgcacaggtgc. [116480]
KM111295.1_Ken06.Bus             ---------------------------------------------------.......c...g...tctgcacaagt--- [116480]
KM111294.1_Ken05/Tk1             ---------------------------------------------------.......c...g...tctgcacaa----- [116480]
KM262844.1_strain_L60            ---------------------------------------------------.......c...g...tttgcacaggtgc. [116480]
KM102979.1_26544/OG10_Italy      ---------------------------------------------------.......c...g...tttgcacaggtgc. [116480]
KM262845.1_strain_NHV            ---------------------------------------------------.......c...g...tttgcacaggtgc. [116480]
KX354450.1_47/Ss/2008            ---------------------------------------------------.......c...g...tttgcacaggtgc. [116480]
KP055815.1_strain_BA71           ---------------------------------------------------.......c...g...tttgcacaggtgc. [116480]
MH025920.1_strain_R35            ---------------------------------------------------.......c...g...tctgcacaagt--- [116480]
MH025919.1_strain_N10            ---------------------------------------------------.......c...g...tctgcacaagt--- [116480]
MH025918.1_strain_R25            ---------------------------------------------------.......c...g...tctgcacaagt--- [116480]
MH025917.1_strain_R7             ---------------------------------------------------.......c...g...tctgcacaagt--- [116480]
MH025916.1_strain_R8             ---------------------------------------------------.......c...g...tctgcacaagt--- [116480]
```

```
FR682468.2_Georgia_2007/1        tgcacaggtatctacattggtatctgcacacat----------------------------------------------- [116560]
ASF_IND_20_CAD543                .................................----------------------------------------------- [116560]
LR899193.1_Germany_2020/1        .................................----------------------------------------------- [116560]
MN393477.1_Wuhan_2019            .................................----------------------------------------------- [116560]
MW396979.1_Timor-Leste/2019/1    .................................----------------------------------------------- [116560]
MK940252.1_CN/2019/InnerMongolia .................................----------------------------------------------- [116560]
MK128995.1_China/2018/AnhuiXCGQ  .................................----------------------------------------------- [116560]
MH681419.1_ASFV/POL/2015         .................................----------------------------------------------- [116560]
LS478113.1_Estonia_2014          .................................----------------------------------------------- [116560]
KP843857.1_Odintsovo_02/14       .................................----------------------------------------------- [116560]
FN557520.1_strain_E75            .........g.......................------------gcttgcacaagtgtttacattggtatctgcacaag [116560]
AY261366.1_Warthog               ......a..g...t....a..----------------------------------------------------------- [116560]
AY261365.1_Warmbaths             ......a..gct.g..caa..----------------------------------------------------------- [116560]
AY261364.1_Tengani_62            -------------------------------------------------------------------------------- [116560]
AY261363.1_Pretoriuskop/96/4     ......a..gct.g..ca...----------------------------------------------------------- [116560]
AY261362.1_Mkuzi_1979            -------------------------------------------------------------------------------- [116560]
AY261361.1_Malawi_Lil-20/1_1983  ....tt...g.t.g.....a.g......ttgg.------------gtctacatttgtgcttgcattggtgcttgcattta [116560]
AY261360.1_Kenya_1950            -------------------------------------------------------------------------------- [116560]
AM712240.1_OURT_88/3             .........g.......................------------gcttgcacaagtgtttacattggtatctgcacaag [116560]
AM712239.1_Benin_97/1            .........g....................ag.gcttgcacctgtgcttgcacaagtgcttgcacaagtgcttgcacaag [116560]
KM111295.1_Ken06.Bus             -------------------------------------------------------------------------------- [116560]
KM111294.1_Ken05/Tk1             -------------------------------------------------------------------------------- [116560]
KM262844.1_strain_L60            .........g.......................------------gcttgcacaagtgtttacattggtatctgcacaag [116560]
KM102979.1_26544/OG10_Italy      .........g......................------------------------------------------------ [116560]
KM262845.1_strain_NHV            .........g.......................------------gcttgcacaagtgtttacattggtatctgcacaag [116560]
KX354450.1_47/Ss/2008            .........g......................------------------------------------------------ [116560]
KP055815.1_strain_BA71           .........g.......................------------gcttgcacaagtgtttacattggtatctgcacaag [116560]
MH025920.1_strain_R35            -------------------------------------------------------------------------------- [116560]
MH025919.1_strain_N10            -------------------------------------------------------------------------------- [116560]
MH025918.1_strain_R25            -------------------------------------------------------------------------------- [116560]
MH025917.1_strain_R7             -------------------------------------------------------------------------------- [116560]
MH025916.1_strain_R8             -------------------------------------------------------------------------------- [116560]

FR682468.2_Georgia_2007/1        -gcttgcacaggtgtctacattggtatc---------------------------------------------------- [116640]
ASF_IND_20_CAD543                -...........................---------------------------------------------------- [116640]
LR899193.1_Germany_2020/1        -...........................---------------------------------------------------- [116640]
MN393477.1_Wuhan_2019            -...........................---------------------------------------------------- [116640]
MW396979.1_Timor-Leste/2019/1    -...........................---------------------------------------------------- [116640]
MK940252.1_CN/2019/InnerMongolia -...........................---------------------------------------------------- [116640]
MK128995.1_China/2018/AnhuiXCGQ  -...........................---------------------------------------------------- [116640]
MH681419.1_ASFV/POL/2015         -...........................---------------------------------------------------- [116640]
LS478113.1_Estonia_2014          -...........................---------------------------------------------------- [116640]
KP843857.1_Odintsovo_02/14       -...........................---------------------------------------------------- [116640]
```

```
FN557520.1_strain_E75             t...........................---------------------------------------------------- [116640]
AY261366.1_Warthog                -.tc..t....c......t.........---------------------------------------------------- [116640]
AY261365.1_Warmbaths              -.........a...c..........g..---------------------------------------------------- [116640]
AY261364.1_Tengani_62             ---------------..t....a..g..---------------------------------------------------- [116640]
AY261363.1_Pretoriuskop/96/4      -.tc...g..t.................---------------------------------------------------- [116640]
AY261362.1_Mkuzi_1979             -------------------------------------------------------------------------------- [116640]
AY261361.1_Malawi_Lil-20/1_1983   t.......tt............t..gct---------------------------------------------------- [116640]
AY261360.1_Kenya_1950             -.........a..............g..---------------------------------------------------- [116640]
AM712240.1_OURT_88/3              t...........................------------------------tgcacaagtgcttgcacaggtgtctaca [116640]
AM712239.1_Benin_97/1             t...........................tgcacaagtgcttgcacctgtgcttgcacaagtgcttgcacaagtgcttgca [116640]
KM111295.1_Ken06.Bus              -....a....a..............g..---------------------------------------------------- [116640]
KM111294.1_Ken05/Tk1              -----------------------..g..---------------------------------------------------- [116640]
KM262844.1_strain_L60             t...........................---------------------------------------------------- [116640]
KM102979.1_26544/OG10_Italy       -------------------------------------------------------------------------------- [116640]
KM262845.1_strain_NHV             t...........................------------------------tgcacaagtgcttgcacaggtgtctaca [116640]
KX354450.1_47/Ss/2008             -------------------------------------------------------------------------------- [116640]
KP055815.1_strain_BA71            t...........................---------------------------------------------------- [116640]
MH025920.1_strain_R35             -....a....a..............g..---------------------------------------------------- [116640]
MH025919.1_strain_N10             -....a....a..............g..---------------------------------------------------- [116640]
MH025918.1_strain_R25             -....a....a..............g..---------------------------------------------------- [116640]
MH025917.1_strain_R7              -....a....a..............g..---------------------------------------------------- [116640]
MH025916.1_strain_R8              -....a....a..............g..---------------------------------------------------- [116640]

FR682468.2_Georgia_2007/1         --------tgcacacatgcttgcacaagtgtctacattggtatc------------------------------------ [116720]
ASF_IND_20_CAD543                 --------......ag............................t----------------------------------- [116720]
LR899193.1_Germany_2020/1         --------....................................------------------------------------ [116720]
MN393477.1_Wuhan_2019             --------....................................------------------------------------ [116720]
MW396979.1_Timor-Leste/2019/1     --------....................................------------------------------------ [116720]
MK940252.1_CN/2019/InnerMongolia  --------....................................------------------------------------ [116720]
MK128995.1_China/2018/AnhuiXCGQ   --------....................................------------------------------------ [116720]
MH681419.1_ASFV/POL/2015          --------....................................------------------------------------ [116720]
LS478113.1_Estonia_2014           --------....................................------------------------------------ [116720]
KP843857.1_Odintsovo_02/14        --------....................................------------------------------------ [116720]
FN557520.1_strain_E75             --------......ag..........g.................t----------------------------------- [116720]
AY261366.1_Warthog                --------......ag.................t....a..g..t----------------------------------- [116720]
AY261365.1_Warmbaths              --------....ttgg......------------..........t----------------------------------- [116720]
AY261364.1_Tengani_62             --------......ag..tc........................t----------------------------------- [116720]
AY261363.1_Pretoriuskop/96/4      --------......ag.a........g.................t----------------------------------- [116720]
AY261362.1_Mkuzi_1979             --------......ag..........g.................t----------------------------------- [116720]
AY261361.1_Malawi_Lil-20/1_1983   --------....ttgg.............a..............t----------------------------------- [116720]
AY261360.1_Kenya_1950             --------......gg..........g.....------------------------------------------------ [116720]
AM712240.1_OURT_88/3              ttggtatc......ag..........g.................tgcacaagtgcttgcacctgtgcttgcacaagtgct [116720]
AM712239.1_Benin_97/1             caagtgct......ag..........g.................tgcacaagtgcttgcacct----------------- [116720]
KM111295.1_Ken06.Bus              --------......ag.........................g..tgcac------------------------------- [116720]
```

```
KM111294.1_Ken05/Tk1             --------......gg.....a........ct------------------------------------------------ [116720]
KM262844.1_strain_L60            --------......ag..........g.................t----------------------------------- [116720]
KM102979.1_26544/OG10_Italy      -------------------------------------------------------------------------------- [116720]
KM262845.1_strain_NHV            ttggtatc......ag..........g.................tgcacaagtgcttgcacctgtgcttgcacaagtgct [116720]
KX354450.1_47/Ss/2008            -------------------------------------------------------------------------------- [116720]
KP055815.1_strain_BA71           --------......ag..........g.................t----------------------------------- [116720]
MH025920.1_strain_R35            --------......ag.........................g..tgcacaagtatct----------------------- [116720]
MH025919.1_strain_N10            --------......ag.........................g..tgcacaagtatct----------------------- [116720]
MH025918.1_strain_R25            --------......ag.........................g..tgcacaagtatct----------------------- [116720]
MH025917.1_strain_R7             --------......ag.........................g..tgcacaagtatct----------------------- [116720]
MH025916.1_strain_R8             --------......ag.........................g..tgcacaagtatct----------------------- [116720]

FR682468.2_Georgia_2007/1        -------------------------------------------------------------------------------- [116800]
ASF_IND_20_CAD543                -------------------------gcacaagtgcttgcacaggtgtctacattggtatctgcacaagtgcttgcacagg [116800]
LR899193.1_Germany_2020/1        -------------------------------------------------------------------------------- [116800]
MN393477.1_Wuhan_2019            -------------------------------------------------------------------------------- [116800]
MW396979.1_Timor-Leste/2019/1    -------------------------------------------------------------------------------- [116800]
MK940252.1_CN/2019/InnerMongolia -------------------------------------------------------------------------------- [116800]
MK128995.1_China/2018/AnhuiXCGQ  -------------------------------------------------------------------------------- [116800]
MH681419.1_ASFV/POL/2015         -------------------------------------------------------------------------------- [116800]
LS478113.1_Estonia_2014          -------------------------------------------------------------------------------- [116800]
KP843857.1_Odintsovo_02/14       -------------------------------------------------------------------------------- [116800]
FN557520.1_strain_E75            -------------------------------------------------------------gcacaagtgcttgcacagg [116800]
AY261366.1_Warthog               -------------------------------------------------------------------------------- [116800]
AY261365.1_Warmbaths             -------------------------------------------------------------------------------- [116800]
AY261364.1_Tengani_62            -------------------------------------------------------------------------------- [116800]
AY261363.1_Pretoriuskop/96/4     -------------------------------------------------------------------------------- [116800]
AY261362.1_Mkuzi_1979            -------------------------------------------------------------------------------- [116800]
AY261361.1_Malawi_Lil-20/1_1983  -------------gcatttatgcttgcattggtgcttgcattggtgtctacatttgtgcttgcattggtgcttgcattta [116800]
AY261360.1_Kenya_1950            -------------------------------------------------------------------------------- [116800]
AM712240.1_OURT_88/3             tgcacaagtgcttgcacaagtgcttgcacaagtgcttgcacaggtgtctacattggtatctgcacaagtgcttgcacagg [116800]
AM712239.1_Benin_97/1            -------------------gtgcttgcacaagtgcttgcacaagtgcttgcacaagtgcttgcacaagtgcttgcacagg [116800]
KM111295.1_Ken06.Bus             -----------------aagtgcttacacaagtgcttacacaagtgtctgcacaagtatctgcacaagtgcttgcacaag [116800]
KM111294.1_Ken05/Tk1             -------------------------------------------------------------------------------- [116800]
KM262844.1_strain_L60            -------------------------------------------------------------gcacaagtgcttgcacagg [116800]
KM102979.1_26544/OG10_Italy      -------------------------------------------------------------------------------- [116800]
KM262845.1_strain_NHV            tgcacaagtgcttgcacaagtgcttgcacaagtgcttgcacaggtgtctacattggtatctgcacaagtgcttgcacagg [116800]
KX354450.1_47/Ss/2008            -------------------------------------------------------------------------------- [116800]
KP055815.1_strain_BA71           -------------------------gcacaagtgcttgcacaggtgtctacattggtatctgcacaagtgcttgcacagg [116800]
MH025920.1_strain_R35            -------------gcacaagtgcttacacaagtgcttacacaagtgtctgcacaagtatctgcacaagtgcttgcacaag [116800]
MH025919.1_strain_N10            -------------gcacaagtgcttacacaagtgcttacacaagtgtctgcacaagtatctgcacaagtgcttgcacaag [116800]
MH025918.1_strain_R25            -------------gcacaagtgcttacacaagtgcttacacaagtgtctgcacaagtatctgcacaagtgcttgcacaag [116800]
MH025917.1_strain_R7             -------------gcacaagtgcttacacaagtgcttacacaagtgtctgcacaagtatctgcacaagtgcttgcacaag [116800]
MH025916.1_strain_R8             -------------gcacaagtgcttacacaagtgcttacacaagtgtctgcacaagtatctgcacaagtgcttgcacaag [116800]
```

```
FR682468.2_Georgia_2007/1        -------------------------------------------------------------------------------- [116880]
ASF_IND_20_CAD543                tgtctacattggtatctgcacaagtnnnnnnnnnt--------------------------------------------- [116880]
LR899193.1_Germany_2020/1        -------------------------------------------------------------------------------- [116880]
MN393477.1_Wuhan_2019            -------------------------------------------------------------------------------- [116880]
MW396979.1_Timor-Leste/2019/1    -------------------------------------------------------------------------------- [116880]
MK940252.1_CN/2019/InnerMongolia -------------------------------------------------------------------------------- [116880]
MK128995.1_China/2018/AnhuiXCGQ  -------------------------------------------------------------------------------- [116880]
MH681419.1_ASFV/POL/2015         -------------------------------------------------------------------------------- [116880]
LS478113.1_Estonia_2014          -------------------------------------------------------------------------------- [116880]
KP843857.1_Odintsovo_02/14       -------------------------------------------------------------------------------- [116880]
FN557520.1_strain_E75            tgtctacattggtatctgcacaagtgcttgcacctg-------------------------------------------- [116880]
AY261366.1_Warthog               -------------------------------------------------------------------------------- [116880]
AY261365.1_Warmbaths             -------------------------------------------------------------------------------- [116880]
AY261364.1_Tengani_62            -------------------------------------------------------------------------------- [116880]
AY261363.1_Pretoriuskop/96/4     -------------------------------------------------------------------------------- [116880]
AY261362.1_Mkuzi_1979            -------------------------------------------------------------------------------- [116880]
AY261361.1_Malawi_Lil-20/1_1983  tgcttgcattggtgtctacatttgtgcttgcattg--------------------------------------------- [116880]
AY261360.1_Kenya_1950            -------------------------------------------------------------------------------- [116880]
AM712240.1_OURT_88/3             tgtctacattggtatctgcacaagtgcttgcacctgtgcttgcacctgtgcttgcacaagtgcttgcacaagtgcttgca [116880]
AM712239.1_Benin_97/1            tgtctacattggtatctgcacaagtgcttgcacct--------------------------------------------- [116880]
KM111295.1_Ken06.Bus             tgtctacattggtgtctgcacaaatgtctgcacaa--------------------------------------------- [116880]
KM111294.1_Ken05/Tk1             -------------------------------------------------------------------------------- [116880]
KM262844.1_strain_L60            tgtctacattggtatctgcacaagtgcttgcacct--------------------------------------------- [116880]
KM102979.1_26544/OG10_Italy      -------------------------------------------------------------------------------- [116880]
KM262845.1_strain_NHV            tgtctacattggtatctgcacaagtgcttgcacctgtgcttgcacctgtgcttgcacaa------------gtgcttgca [116880]
KX354450.1_47/Ss/2008            -------------------------------------------------------------------------------- [116880]
KP055815.1_strain_BA71           tgtctacattggtatctgcacaagtgcttgcacct--------------------------------------------- [116880]
MH025920.1_strain_R35            tgtctacattggtgtctgcacaagtgtctgcacaa--------------------------------------------- [116880]
MH025919.1_strain_N10            tgtctacattggtgtctgcacaagtgtctgcacaa--------------------------------------------- [116880]
MH025918.1_strain_R25            tgtctacattggtgtctgcacaagtgtctgcacaa--------------------------------------------- [116880]
MH025917.1_strain_R7             tgtctacattggtgtctgcacaagtgtctgcacaa--------------------------------------------- [116880]
MH025916.1_strain_R8             tgtctacattggtgtctgcacaagtgtctgcacaa--------------------------------------------- [116880]

FR682468.2_Georgia_2007/1        -------------------------------------------------------------------------------- [116960]
ASF_IND_20_CAD543                ---------------------------------------------------------------------------gtgct [116960]
LR899193.1_Germany_2020/1        -------------------------------------------------------------------------------- [116960]
MN393477.1_Wuhan_2019            -------------------------------------------------------------------------------- [116960]
MW396979.1_Timor-Leste/2019/1    -------------------------------------------------------------------------------- [116960]
MK940252.1_CN/2019/InnerMongolia -------------------------------------------------------------------------------- [116960]
MK128995.1_China/2018/AnhuiXCGQ  -------------------------------------------------------------------------------- [116960]
MH681419.1_ASFV/POL/2015         -------------------------------------------------------------------------------- [116960]
LS478113.1_Estonia_2014          -------------------------------------------------------------------------------- [116960]
KP843857.1_Odintsovo_02/14       -------------------------------------------------------------------------------- [116960]
```

```
FN557520.1_strain_E75              ----------------------------------------------------------------tgcttgcacaagtgct [116960]
AY261366.1_Warthog                 -------------------------------------------------------------------------------- [116960]
AY261365.1_Warmbaths               -------------------------------------------------------------------------------- [116960]
AY261364.1_Tengani_62              -------------------------------------------------------------------------------- [116960]
AY261363.1_Pretoriuskop/96/4       -------------------------------------------------------------------------------- [116960]
AY261362.1_Mkuzi_1979              -------------------------------------------------------------------------------- [116960]
AY261361.1_Malawi_Lil-20/1_1983    ---------------------------------------------------------------------------gtgct [116960]
AY261360.1_Kenya_1950              -------------------------------------------------------------------------------- [116960]
AM712240.1_OURT_88/3               caagtgcttgcacaagtgcttgcacaagtgcttgcacaagtgcttgcacaagtgcttgcacaagtgcttgcacaagtgct [116960]
AM712239.1_Benin_97/1              ---------------------------------------------------------------------------gtgct [116960]
KM111295.1_Ken06.Bus               ---------------------------------------------------------------------------gtgct [116960]
KM111294.1_Ken05/Tk1               -------------------------------------------------------------------------------- [116960]
KM262844.1_strain_L60              ---------------------------------------------------------------------------gtgct [116960]
KM102979.1_26544/OG10_Italy        ----------------------------------------------------------------------------tgct [116960]
KM262845.1_strain_NHV              caagtgcttgcacaagtgcttgcacaagtgcttgcacaagtgcttgcacaagtgcttgcacaagtgcttgcacaagtgct [116960]
KX354450.1_47/Ss/2008              ----------------------------------------------------------------------------tgct [116960]
KP055815.1_strain_BA71             ---------------------------------------------------------------------------gtgct [116960]
MH025920.1_strain_R35              ---------------------------------------------------------------------------gtgct [116960]
MH025919.1_strain_N10              ---------------------------------------------------------------------------gtgct [116960]
MH025918.1_strain_R25              ---------------------------------------------------------------------------gtgct [116960]
MH025917.1_strain_R7               ---------------------------------------------------------------------------gtgct [116960]
MH025916.1_strain_R8               ---------------------------------------------------------------------------gtgct [116960]

FR682468.2_Georgia_2007/1          -------------------------------------------------------------------------------- [117040]
ASF_IND_20_CAD543                  tgcacaagtgcttnnnnnnnnnnnnnnnnnnnnnnnnnnnnnnnnnnnnnnnnnnnnnnnnnnnnnntgcttgcacaag [117040]
LR899193.1_Germany_2020/1          -------------------------------------------------------------------------------- [117040]
MN393477.1_Wuhan_2019              -------------------------------------------------------------------------------- [117040]
MW396979.1_Timor-Leste/2019/1      -------------------------------------------------------------------------------- [117040]
MK940252.1_CN/2019/InnerMongolia   -------------------------------------------------------------------------------- [117040]
MK128995.1_China/2018/AnhuiXCGQ    -------------------------------------------------------------------------------- [117040]
MH681419.1_ASFV/POL/2015           -------------------------------------------------------------------------------- [117040]
LS478113.1_Estonia_2014            -------------------------------------------------------------------------------- [117040]
KP843857.1_Odintsovo_02/14         -------------------------------------------------------------------------------- [117040]
FN557520.1_strain_E75              tgcacaagtgcttgc------------------------------------acaagtgcttgcacaagtgcttgcacagg [117040]
AY261366.1_Warthog                 -gtacagctgtctat------------------------------------attggtatctgcacaagtgcttgcacagg [117040]
AY261365.1_Warmbaths               -gcactggtgcttgc------------------------------------attggtgtctgcactagtgcttg------ [117040]
AY261364.1_Tengani_62              -gcacaagtgct--------------------------------------------------------------------- [117040]
AY261363.1_Pretoriuskop/96/4       -gcacaagtgcttgc------------------------------------attggtgtctgcactggtgcttg------ [117040]
AY261362.1_Mkuzi_1979              -gcacaagtgtctac------------------------------------attagtatttgcacaagtgcttgcacagg [117040]
AY261361.1_Malawi_Lil-20/1_1983    tgcacaagtatctac------------attggtatctgcatttatgcttgcattggtgcttgcacttgtgtctgcattgg [117040]
AY261360.1_Kenya_1950              tgtacaagtgtctac------------------------------------attggtgtctgcacaggtgcttgcacaag [117040]
AM712240.1_OURT_88/3               tgcacaagtgcttgc------------------------------------acaagtgcttgcacaagtgcttgcacagg [117040]
AM712239.1_Benin_97/1              tgcacaagtgcttgc------------------------------------acaagtgcttgcacaagtgcttgcacagg [117040]
KM111295.1_Ken06.Bus               tgcacaagtgtctac------------------------------------attggtgtc------------tgcacaag [117040]
```

```
KM111294.1_Ken05/Tk1           tgcacaagtgtctac------------------------------------attggtgtc-------------------- [117040]
KM262844.1_strain_L60          tgcacaagtgcttgc------------------------------------acaagtgcttgcacaagtgcttgcacagg [117040]
KM102979.1_26544/OG10_Italy    tgcacaagtgcttgc------------------------------------acaagtgcttgcacaagtgcttgcacagg [117040]
KM262845.1_strain_NHV          tgcacaagtgcttgc------------------------------------acaagtgcttgcacaagtgcttgcacagg [117040]
KX354450.1_47/Ss/2008          tgcacaagtgcttgc------------------------------------acaagtgcttgcacaagtgcttgcacagg [117040]
KP055815.1_strain_BA71         tgcacaagtgcttgc------------------------------------acaagtgcttgcacaagtgcttgcacagg [117040]
MH025920.1_strain_R35          tgcacaagtgtctac------------------------------------attggtgtctgcacaagtgcttgcacaag [117040]
MH025919.1_strain_N10          tgcacaagtgtctac------------------------------------attggtgtctgcacaagtgcttgcacaag [117040]
MH025918.1_strain_R25          tgcacaagtgtctac------------------------------------attggtgtctgcacaagtgcttgcacaag [117040]
MH025917.1_strain_R7           tgcacaagtgtctac------------------------------------attggtgtctgcacaagtgcttgcacaag [117040]
MH025916.1_strain_R8           tgcacaagtgtctac------------------------------------attggtgtctgcacaagtgcttgcacaag [117040]

FR682468.2_Georgia_2007/1        ----------------------------tgcacaagtatacgcactttgagcatgaagattaggatcaaacacaaaatgt [117120]
ASF_IND_20_CAD543                tgtctacattggtatctgcacaagtgct.................................................... [117120]
LR899193.1_Germany_2020/1        ----------------------------.................................................... [117120]
MN393477.1_Wuhan_2019            ----------------------------.................................................... [117120]
MW396979.1_Timor-Leste/2019/1    ----------------------------.................................................... [117120]
MK940252.1_CN/2019/InnerMongolia ----------------------------.................................................... [117120]
MK128995.1_China/2018/AnhuiXCGQ  ----------------------------.................................................... [117120]
MH681419.1_ASFV/POL/2015         ----------------------------.................................................... [117120]
LS478113.1_Estonia_2014          ----------------------------.................................................... [117120]
KP843857.1_Odintsovo_02/14       ----------------------------.................................................... [117120]
FN557520.1_strain_E75            tgtctacattggtatctgcacaagtgct.................................................... [117120]
AY261366.1_Warthog               tgtctacattggtatc------------.................................................... [117120]
AY261365.1_Warmbaths             ------cattggtgtc------------..........................................t......... [117120]
AY261364.1_Tengani_62            ----------------------------.................................................... [117120]
AY261363.1_Pretoriuskop/96/4     ------cattggtgtc------------.................................................... [117120]
AY261362.1_Mkuzi_1979            tgtctgcactagtgcttgcattggtgtc.................................................... [117120]
AY261361.1_Malawi_Lil-20/1_1983  tgcttgcacttgtgtctgcattggtgct......g............................................. [117120]
AY261360.1_Kenya_1950            tgcttgcacaggtgcttgcacaggtgct......g............................................. [117120]
AM712240.1_OURT_88/3             tgtctacattggtatctgcacaagtgct.................................................... [117120]
AM712239.1_Benin_97/1            tgtctacattggtatctgcacaagtgct.................................................... [117120]
KM111295.1_Ken06.Bus             tgcttgcacaagtgcttgcacaagtgct......g............................................. [117120]
KM111294.1_Ken05/Tk1             ----tgcacaggtgcttgcacaggtgct......g............................................. [117120]
KM262844.1_strain_L60            tgtctacattggtatctgcacaagtgct.................................................... [117120]
KM102979.1_26544/OG10_Italy      tgtctacattggtatctgcacaagtgct.................................................... [117120]
KM262845.1_strain_NHV            tgtctacattggtatctgcacaagtgct.................................................... [117120]
KX354450.1_47/Ss/2008            tgtctacattggtatctgcacaagtgct.................................................... [117120]
KP055815.1_strain_BA71           tgtctacattggtatctgcacaagtgct.................................................... [117120]
MH025920.1_strain_R35            tgcttgcacaagtgcttgcacaagtgct......g............................................. [117120]
MH025919.1_strain_N10            tgcttgcacaagtgcttgcacaagtgct......g................t............................ [117120]
MH025918.1_strain_R25            tgcttgcacaagtgcttgcacaagtgct......g............................................. [117120]
MH025917.1_strain_R7             tgcttgcacaagtgcttgcacaagtgct......g............................................. [117120]
MH025916.1_strain_R8             tgcttgcacaagtgcttgcacaagtgct......g............................................. [117120]
```

```
FR682468.2_Georgia_2007/1        tctcgtaaaaagctatcgatcgttgttttagcttccttgcttttctgcgtctgggttttgcagctatctgctatagataa [117200]
ASF_IND_20_CAD543                ................................................................................ [117200]
LR899193.1_Germany_2020/1        ................................................................................ [117200]
MN393477.1_Wuhan_2019            ................................................................................ [117200]
MW396979.1_Timor-Leste/2019/1    ................................................................................ [117200]
MK940252.1_CN/2019/InnerMongolia ................................................................................ [117200]
MK128995.1_China/2018/AnhuiXCGQ  ................................................................................ [117200]
MH681419.1_ASFV/POL/2015         ................................................................................ [117200]
LS478113.1_Estonia_2014          ................................................................................ [117200]
KP843857.1_Odintsovo_02/14       ................................................................................ [117200]
FN557520.1_strain_E75            ....................................................................c........... [117200]
AY261366.1_Warthog               ...............................................t....................c........... [117200]
AY261365.1_Warmbaths             ......................c........................t....................c........... [117200]
AY261364.1_Tengani_62            ......................c.....................t.......................c........... [117200]
AY261363.1_Pretoriuskop/96/4     ....................................................................c........... [117200]
AY261362.1_Mkuzi_1979            ......................c........................t....................c........... [117200]
AY261361.1_Malawi_Lil-20/1_1983  ..............g...................................t..............g..c.......c... [117200]
AY261360.1_Kenya_1950            ..c...........g.....................g..........t..t..a.........t.g.............. [117200]
AM712240.1_OURT_88/3             ....................................................................c........... [117200]
AM712239.1_Benin_97/1            ....................................................................c........... [117200]
KM111295.1_Ken06.Bus             ..c...........g.....t..........................t..t..a.........t.g.............. [117200]
KM111294.1_Ken05/Tk1             ..c...........g.....................g..........t..t..a.........t.g.............. [117200]
KM262844.1_strain_L60            ....................................................................c........... [117200]
KM102979.1_26544/OG10_Italy      ....................................................................c........... [117200]
KM262845.1_strain_NHV            ....................................................................c........... [117200]
KX354450.1_47/Ss/2008            ....................................................................c........... [117200]
KP055815.1_strain_BA71           ....................................................................c........... [117200]
MH025920.1_strain_R35            ..c...........g.....t..........................t..t..a.........t.g.............. [117200]
MH025919.1_strain_N10            ..c...........g.....t..........................t..t..a.........t.g.............. [117200]
MH025918.1_strain_R25            ..c...........g.....t..........................t..t..a.........t.g.............. [117200]
MH025917.1_strain_R7             ..c...........g.....t..........................t..t..a.........t.g.............. [117200]
MH025916.1_strain_R8             ..c...........g.....t..........................t..t..a.........t.g.............. [117200]

FR682468.2_Georgia_2007/1        aattgtatttactaccgattcagagggaacatcattagtttcctgtttcaaagtatcaactaacgttattagctcactga [117280]
ASF_IND_20_CAD543                ................................................................................ [117280]
LR899193.1_Germany_2020/1        ................................................................................ [117280]
MN393477.1_Wuhan_2019            ................................................................................ [117280]
MW396979.1_Timor-Leste/2019/1    ................................................................................ [117280]
MK940252.1_CN/2019/InnerMongolia ................................................................................ [117280]
MK128995.1_China/2018/AnhuiXCGQ  ................................................................................ [117280]
MH681419.1_ASFV/POL/2015         ................................................................................ [117280]
LS478113.1_Estonia_2014          ................................................................................ [117280]
KP843857.1_Odintsovo_02/14       ................................................................................ [117280]
```

```
FN557520.1_strain_E75             ................................................................................ [117280]
AY261366.1_Warthog                .......................................................................a........ [117280]
AY261365.1_Warmbaths              ................................................................................ [117280]
AY261364.1_Tengani_62             ................................................................................ [117280]
AY261363.1_Pretoriuskop/96/4      ................................................................................ [117280]
AY261362.1_Mkuzi_1979             ................................................................................ [117280]
AY261361.1_Malawi_Lil-20/1_1983   ...............t...............................................t................ [117280]
AY261360.1_Kenya_1950             ..g.................................t......................................g.... [117280]
AM712240.1_OURT_88/3              ................................................................................ [117280]
AM712239.1_Benin_97/1             ................................................................................ [117280]
KM111295.1_Ken06.Bus              ..g............................................................t...........g.... [117280]
KM111294.1_Ken05/Tk1              ..g.................................t......................................g.... [117280]
KM262844.1_strain_L60             ................................................................................ [117280]
KM102979.1_26544/OG10_Italy       ................................................................................ [117280]
KM262845.1_strain_NHV             ................................................................................ [117280]
KX354450.1_47/Ss/2008             ................................................................................ [117280]
KP055815.1_strain_BA71            ................................................................................ [117280]
MH025920.1_strain_R35             ..g............................................................t...........g.... [117280]
MH025919.1_strain_N10             ..g............................................................t...........g.... [117280]
MH025918.1_strain_R25             ..g............................................................t...........g.... [117280]
MH025917.1_strain_R7              ..g............................................................t...........g.... [117280]
MH025916.1_strain_R8              ..g............................................................t...........g.... [117280]

FR682468.2_Georgia_2007/1         gaagagttttggtcgtgtgggtaggttttgaataggaaggcatccattcctgcagagctttgaagacatatccaataaag [117360]
ASF_IND_20_CAD543                 ................................................................................ [117360]
LR899193.1_Germany_2020/1         ................................................................................ [117360]
MN393477.1_Wuhan_2019             ................................................................................ [117360]
MW396979.1_Timor-Leste/2019/1     ................................................................................ [117360]
MK940252.1_CN/2019/InnerMongolia  ................................................................................ [117360]
MK128995.1_China/2018/AnhuiXCGQ   ................................................................................ [117360]
MH681419.1_ASFV/POL/2015          ................................................................................ [117360]
LS478113.1_Estonia_2014           ................................................................................ [117360]
KP843857.1_Odintsovo_02/14        ................................................................................ [117360]
FN557520.1_strain_E75             ................................................................................ [117360]
AY261366.1_Warthog                ......................g..............g.............................g...........a [117360]
AY261365.1_Warmbaths              ......................g..............g.............................g...........a [117360]
AY261364.1_Tengani_62             ......................g..............g.........................................a [117360]
AY261363.1_Pretoriuskop/96/4      ......................g..............g.............................g...........a [117360]
AY261362.1_Mkuzi_1979             ......................g..............g.............................g...........a [117360]
AY261361.1_Malawi_Lil-20/1_1983   ......................g..............g........................................ga [117360]
AY261360.1_Kenya_1950             ....c..........................................................................a [117360]
AM712240.1_OURT_88/3              ................................................................................ [117360]
AM712239.1_Benin_97/1             ................................................................................ [117360]
KM111295.1_Ken06.Bus              ....c.............a...g..............g.............................g...........a [117360]
```

```
KM111294.1_Ken05/Tk1           ....c..........................................................................a [117360]
KM262844.1_strain_L60          ................................................................................ [117360]
KM102979.1_26544/OG10_Italy    ................................................................................ [117360]
KM262845.1_strain_NHV          ................................................................................ [117360]
KX354450.1_47/Ss/2008          ................................................................................ [117360]
KP055815.1_strain_BA71         ................................................................................ [117360]
MH025920.1_strain_R35          ....c.............a...g..............g.............................g...........a [117360]
MH025919.1_strain_N10          ....c.............a...g..............g.............................g...........a [117360]
MH025918.1_strain_R25          ....c.............a...g..............g.............................g...........a [117360]
MH025917.1_strain_R7           ....c.............a...g..............g.............................g...........a [117360]
MH025916.1_strain_R8           ....c.............a...g..............g.............................g...........a [117360]

FR682468.2_Georgia_2007/1      ctagtcattataagacgtcgaatatactgctcccgcaaatttgtaaaagagcaaaaggccaccctgctatcatttttgaa [117440]
ASF_IND_20_CAD543              ................................................................................ [117440]
LR899193.1_Germany_2020/1      ................................................................................ [117440]
MN393477.1_Wuhan_2019          ................................................................................ [117440]
MW396979.1_Timor-Leste/2019/1  ................................................................................ [117440]
MK940252.1_CN/2019/InnerMongolia ................................................................................ [117440]
MK128995.1_China/2018/AnhuiXCGQ ................................................................................ [117440]
MH681419.1_ASFV/POL/2015       ................................................................................ [117440]
LS478113.1_Estonia_2014        ................................................................................ [117440]
KP843857.1_Odintsovo_02/14     ................................................................................ [117440]
FN557520.1_strain_E75          ................................................................................ [117440]
AY261366.1_Warthog             ...............................................g................................ [117440]
AY261365.1_Warmbaths           ...............................................g................................ [117440]
AY261364.1_Tengani_62          ...............................................g................................ [117440]
AY261363.1_Pretoriuskop/96/4   ...............................................g................................ [117440]
AY261362.1_Mkuzi_1979          ...............................................g................................ [117440]
AY261361.1_Malawi_Lil-20/1_1983 ...............................................g................................ [117440]
AY261360.1_Kenya_1950          ...............................................g.............................a.. [117440]
AM712240.1_OURT_88/3           ................................................................................ [117440]
AM712239.1_Benin_97/1          ................................................................................ [117440]
KM111295.1_Ken06.Bus           ...............................................g.............................a.. [117440]
KM111294.1_Ken05/Tk1           ...............................................g.............................a.. [117440]
KM262844.1_strain_L60          ................................................................................ [117440]
KM102979.1_26544/OG10_Italy    ........................................................................c....... [117440]
KM262845.1_strain_NHV          ................................................................................ [117440]
KX354450.1_47/Ss/2008          ................................................................................ [117440]
KP055815.1_strain_BA71         ................................................................................ [117440]
MH025920.1_strain_R35          ...............................................g.............................a.. [117440]
MH025919.1_strain_N10          ...............................................g.............................a.. [117440]
MH025918.1_strain_R25          ...............................................g.............................a.. [117440]
MH025917.1_strain_R7           ...............................................g.............................a.. [117440]
MH025916.1_strain_R8           ...............................................g.............................a.. [117440]
```

```
FR682468.2_Georgia_2007/1        ctgtttgtaagggttcgtcctttggtaaagctgtttaagcgtttcttcggatatttcagtagagggatcctccaatacgt [117520]
ASF_IND_20_CAD543                ................................................................................ [117520]
LR899193.1_Germany_2020/1        ................................................................................ [117520]
MN393477.1_Wuhan_2019            ................................................................................ [117520]
MW396979.1_Timor-Leste/2019/1    ................................................................................ [117520]
MK940252.1_CN/2019/InnerMongolia ................................................................................ [117520]
MK128995.1_China/2018/AnhuiXCGQ  ................................................................................ [117520]
MH681419.1_ASFV/POL/2015         ................................................................................ [117520]
LS478113.1_Estonia_2014          ................................................................................ [117520]
KP843857.1_Odintsovo_02/14       ................................................................................ [117520]
FN557520.1_strain_E75            ........................a....................................................... [117520]
AY261366.1_Warthog               ........................a................................c..........t........... [117520]
AY261365.1_Warmbaths             ........................a................................c...................... [117520]
AY261364.1_Tengani_62            ........................a................................c...................... [117520]
AY261363.1_Pretoriuskop/96/4     ........................a................................c..........t........... [117520]
AY261362.1_Mkuzi_1979            ........................a................................c...................... [117520]
AY261361.1_Malawi_Lil-20/1_1983  ......a..g..............a.......................a.................g............. [117520]
AY261360.1_Kenya_1950            ......a.....a..t...t....a........c..........................g................... [117520]
AM712240.1_OURT_88/3             ........................a....................................................... [117520]
AM712239.1_Benin_97/1            ........................a....................................................... [117520]
KM111295.1_Ken06.Bus             ......a.....a..t..tt....a.....t..c..........................g................... [117520]
KM111294.1_Ken05/Tk1             ......a.....a..t...t....a........c..........................g................... [117520]
KM262844.1_strain_L60            ........................a....................................................... [117520]
KM102979.1_26544/OG10_Italy      ........................a....................................................... [117520]
KM262845.1_strain_NHV            ........................a....................................................... [117520]
KX354450.1_47/Ss/2008            ........................a....................................................... [117520]
KP055815.1_strain_BA71           ........................a....................................................... [117520]
MH025920.1_strain_R35            ......a.....a..t..tt....a.....t..c..........................g................... [117520]
MH025919.1_strain_N10            ......a.....a..t..tt....a.....t..c..........................g................... [117520]
MH025918.1_strain_R25            ......a.....a..t..tt....a.....t..c..........................g................... [117520]
MH025917.1_strain_R7             ......a.....a..t..tt....a.....t..c..........................g................... [117520]
MH025916.1_strain_R8             ......a.....a..t..tt....a.....t..c..........................g................... [117520]

FR682468.2_Georgia_2007/1        ttttgagaagctcatcaatattaaattctgccatatcttagagtttattatatacatattaaagctttaatataa-gggg [117600]
ASF_IND_20_CAD543                ...........................................................................-.... [117600]
LR899193.1_Germany_2020/1        ...........................................................................-.... [117600]
MN393477.1_Wuhan_2019            ...........................................................................g.... [117600]
MW396979.1_Timor-Leste/2019/1    ...........................................................................g.... [117600]
MK940252.1_CN/2019/InnerMongolia ...........................................................................g.... [117600]
MK128995.1_China/2018/AnhuiXCGQ  ...........................................................................g.... [117600]
MH681419.1_ASFV/POL/2015         ...........................................................................-.... [117600]
LS478113.1_Estonia_2014          ...........................................................................-.... [117600]
KP843857.1_Odintsovo_02/14       ...........................................................................-.... [117600]
```

```
FN557520.1_strain_E75              ...........................................................................--... [117600]
AY261366.1_Warthog                 ...........................................g...............................-.... [117600]
AY261365.1_Warmbaths               ...........................................g...............................-.... [117600]
AY261364.1_Tengani_62              ...........................................g...............................-.... [117600]
AY261363.1_Pretoriuskop/96/4       ...........................................g...............................-.... [117600]
AY261362.1_Mkuzi_1979              ...........................................................................-.... [117600]
AY261361.1_Malawi_Lil-20/1_1983    .......................................g..-................................g.... [117600]
AY261360.1_Kenya_1950              .........................c...............g-............g...................-.... [117600]
AM712240.1_OURT_88/3               ...........................................................................-.... [117600]
AM712239.1_Benin_97/1              ...........................................................................-.... [117600]
KM111295.1_Ken06.Bus               .........................................g-............g...................-.... [117600]
KM111294.1_Ken05/Tk1               .........................c...............g-............g...................-.... [117600]
KM262844.1_strain_L60              ...........................................................................-.... [117600]
KM102979.1_26544/OG10_Italy        ...........................................................................-.... [117600]
KM262845.1_strain_NHV              ...........................................................................-.... [117600]
KX354450.1_47/Ss/2008              ...........................................................................-.... [117600]
KP055815.1_strain_BA71             ...........................................................................-.... [117600]
MH025920.1_strain_R35              .........................................g-............g...................-.... [117600]
MH025919.1_strain_N10              .........................................g-............g...................-.... [117600]
MH025918.1_strain_R25              .........................................g-............g...................-.... [117600]
MH025917.1_strain_R7               .........................................g-............g...................-.... [117600]
MH025916.1_strain_R8               .........................................g-............g...................-.... [117600]

FR682468.2_Georgia_2007/1          ggtataacaatggacgaaatcatcaataaataccaagctgttgaaaaactttttaaggaaattcagcaaggattggccgc [117680]
ASF_IND_20_CAD543                  ................................................................................ [117680]
LR899193.1_Germany_2020/1          ................................................................................ [117680]
MN393477.1_Wuhan_2019              ................................................................................ [117680]
MW396979.1_Timor-Leste/2019/1      ................................................................................ [117680]
MK940252.1_CN/2019/InnerMongolia   ................................................................................ [117680]
MK128995.1_China/2018/AnhuiXCGQ    ................................................................................ [117680]
MH681419.1_ASFV/POL/2015           ................................................................................ [117680]
LS478113.1_Estonia_2014            ................................................................................ [117680]
KP843857.1_Odintsovo_02/14         ................................................................................ [117680]
FN557520.1_strain_E75              ................................................................................ [117680]
AY261366.1_Warthog                 ................................................................................ [117680]
AY261365.1_Warmbaths               .............................................................................t.. [117680]
AY261364.1_Tengani_62              .......................................a..........................g...........at [117680]
AY261363.1_Pretoriuskop/96/4       ..................................................................g...........at [117680]
AY261362.1_Mkuzi_1979              .............................................................................t.. [117680]
AY261361.1_Malawi_Lil-20/1_1983    ........................................................a.........g............. [117680]
AY261360.1_Kenya_1950              ....................t..t.......................g........a.........g..........t.. [117680]
AM712240.1_OURT_88/3               ................................................................................ [117680]
AM712239.1_Benin_97/1              ................................................................................ [117680]
KM111295.1_Ken06.Bus               ....................t..t................................a.........g..........t.. [117680]
```

```
KM111294.1_Ken05/Tk1             ....................t..t.......................g........a.........g..........t.. [117680]
KM262844.1_strain_L60            ................................................................................ [117680]
KM102979.1_26544/OG10_Italy      ................................................................................ [117680]
KM262845.1_strain_NHV            ................................................................................ [117680]
KX354450.1_47/Ss/2008            ................................................................................ [117680]
KP055815.1_strain_BA71           ................................................................................ [117680]
MH025920.1_strain_R35            ....................t..t................................a.........g..........t.. [117680]
MH025919.1_strain_N10            ....................t..t................................a.........g..........t.. [117680]
MH025918.1_strain_R25            ....................t..t................................a.........g..........t.. [117680]
MH025917.1_strain_R7             ....................t..t................................a.........g..........t.. [117680]
MH025916.1_strain_R8             ....................t..t................................a.........g..........t.. [117680]

FR682468.2_Georgia_2007/1        gtatgatcaatacaagaccttaattagtgaaatgatgcactataataatcatatcaagcaggagtattttaacttttttaa [117760]
ASF_IND_20_CAD543                ................................................................................ [117760]
LR899193.1_Germany_2020/1        ................................................................................ [117760]
MN393477.1_Wuhan_2019            ................................................................................ [117760]
MW396979.1_Timor-Leste/2019/1    ................................................................................ [117760]
MK940252.1_CN/2019/InnerMongolia ................................................................................ [117760]
MK128995.1_China/2018/AnhuiXCGQ  ................................................................................ [117760]
MH681419.1_ASFV/POL/2015         ................................................................................ [117760]
LS478113.1_Estonia_2014          ................................................................................ [117760]
KP843857.1_Odintsovo_02/14       ................................................................................ [117760]
FN557520.1_strain_E75            ................................................................................ [117760]
AY261366.1_Warthog               ................................................................................ [117760]
AY261365.1_Warmbaths             ................................................................................ [117760]
AY261364.1_Tengani_62            ...............a...............c..t.............c.....t......................... [117760]
AY261363.1_Pretoriuskop/96/4     ...............a..........a....c..t............................................. [117760]
AY261362.1_Mkuzi_1979            ................................................................................ [117760]
AY261361.1_Malawi_Lil-20/1_1983  ...............a...........c...t.at.....................................t....... [117760]
AY261360.1_Kenya_1950            ...............................t.at.....................................t....... [117760]
AM712240.1_OURT_88/3             ................................................................................ [117760]
AM712239.1_Benin_97/1            ................................................................................ [117760]
KM111295.1_Ken06.Bus             ...............................t.at.....................................t....... [117760]
KM111294.1_Ken05/Tk1             ...............................t.at.....................................t....... [117760]
KM262844.1_strain_L60            ................................................................................ [117760]
KM102979.1_26544/OG10_Italy      ................................................................................ [117760]
KM262845.1_strain_NHV            ................................................................................ [117760]
KX354450.1_47/Ss/2008            ................................................................................ [117760]
KP055815.1_strain_BA71           ................................................................................ [117760]
MH025920.1_strain_R35            ...............................t.at.....................................t....... [117760]
MH025919.1_strain_N10            ...............................t.at.....................................t....... [117760]
MH025918.1_strain_R25            ...............................t.at.....................................t....... [117760]
MH025917.1_strain_R7             ...............................t.at.....................................t....... [117760]
MH025916.1_strain_R8             ...............................t.at.....................................t....... [117760]
```

```
FR682468.2_Georgia_2007/1        tgattatttcaccttatcttattagggcgcatagcggagaaacgctgcgaaacaaagtaaataatgaaattaaacgtctt [117840]
ASF_IND_20_CAD543                ................................................................................ [117840]
LR899193.1_Germany_2020/1        ................................................................................ [117840]
MN393477.1_Wuhan_2019            ................................................................................ [117840]
MW396979.1_Timor-Leste/2019/1    ................................................................................ [117840]
MK940252.1_CN/2019/InnerMongolia ................................................................................ [117840]
MK128995.1_China/2018/AnhuiXCGQ  ................................................................................ [117840]
MH681419.1_ASFV/POL/2015         ................................................................................ [117840]
LS478113.1_Estonia_2014          ................................................................................ [117840]
KP843857.1_Odintsovo_02/14       ................................................................................ [117840]
FN557520.1_strain_E75            ................................................................................ [117840]
AY261366.1_Warthog               ................................................................................ [117840]
AY261365.1_Warmbaths             ............................a................................................... [117840]
AY261364.1_Tengani_62            ................................................................................ [117840]
AY261363.1_Pretoriuskop/96/4     ................................................................................ [117840]
AY261362.1_Mkuzi_1979            ................................................................................ [117840]
AY261361.1_Malawi_Lil-20/1_1983  ............................................t................................... [117840]
AY261360.1_Kenya_1950            ................................................................................ [117840]
AM712240.1_OURT_88/3             ................................................................................ [117840]
AM712239.1_Benin_97/1            ................................................................................ [117840]
KM111295.1_Ken06.Bus             ...........................................a.................................... [117840]
KM111294.1_Ken05/Tk1             ................................................................................ [117840]
KM262844.1_strain_L60            ................................................................................ [117840]
KM102979.1_26544/OG10_Italy      ................................................................................ [117840]
KM262845.1_strain_NHV            ................................................................................ [117840]
KX354450.1_47/Ss/2008            ................................................................................ [117840]
KP055815.1_strain_BA71           ................................................................................ [117840]
MH025920.1_strain_R35            ...........................................a.................................... [117840]
MH025919.1_strain_N10            ...........................................a.................................... [117840]
MH025918.1_strain_R25            ...........................................a.................................... [117840]
MH025917.1_strain_R7             ...........................................a.................................... [117840]
MH025916.1_strain_R8             ...........................................a.................................... [117840]

FR682468.2_Georgia_2007/1        attttggttgaaaatatcaataccaaaatatctaaaacgctggtaagtgttaattttttactacagaaaaaactttcaac [117920]
ASF_IND_20_CAD543                ................................................................................ [117920]
LR899193.1_Germany_2020/1        ................................................................................ [117920]
MN393477.1_Wuhan_2019            ................................................................................ [117920]
MW396979.1_Timor-Leste/2019/1    ................................................................................ [117920]
MK940252.1_CN/2019/InnerMongolia ................................................................................ [117920]
MK128995.1_China/2018/AnhuiXCGQ  ................................................................................ [117920]
MH681419.1_ASFV/POL/2015         ................................................................................ [117920]
LS478113.1_Estonia_2014          ................................................................................ [117920]
KP843857.1_Odintsovo_02/14       ................................................................................ [117920]
```

```
FN557520.1_strain_E75          ................................................................................ [117920]
AY261366.1_Warthog             ................................................................................ [117920]
AY261365.1_Warmbaths           ................................................................................ [117920]
AY261364.1_Tengani_62          ........c.................g..................................................... [117920]
AY261363.1_Pretoriuskop/96/4   ................................................................................ [117920]
AY261362.1_Mkuzi_1979          .................................................................a.............. [117920]
AY261361.1_Malawi_Lil-20/1_1983 ................................c.............................................gt [117920]
AY261360.1_Kenya_1950          ........c................................c....................................g. [117920]
AM712240.1_OURT_88/3           ................................................................................ [117920]
AM712239.1_Benin_97/1          ................................................................................ [117920]
KM111295.1_Ken06.Bus           ........c................................c....................................g. [117920]
KM111294.1_Ken05/Tk1           ........c.............................t..c....................................g. [117920]
KM262844.1_strain_L60          ................................................................................ [117920]
KM102979.1_26544/OG10_Italy    ................................................................................ [117920]
KM262845.1_strain_NHV          ................................................................................ [117920]
KX354450.1_47/Ss/2008          ................................................................................ [117920]
KP055815.1_strain_BA71         ................................................................................ [117920]
MH025920.1_strain_R35          ........c................................c....................................g. [117920]
MH025919.1_strain_N10          ........c................................c....................................g. [117920]
MH025918.1_strain_R25          ........c................................c....................................g. [117920]
MH025917.1_strain_R7           ........c................................c....................................g. [117920]
MH025916.1_strain_R8           ........c................................c....................................g. [117920]

FR682468.2_Georgia_2007/1      ggacggggtgaaaacgaaaaacatgtggtgcaccaataatcccatgctgcaggtaaaaacagcccacaacctttttaagc [118000]
ASF_IND_20_CAD543              ................................................................................ [118000]
LR899193.1_Germany_2020/1      ................................................................................ [118000]
MN393477.1_Wuhan_2019          ................................................................................ [118000]
MW396979.1_Timor-Leste/2019/1  ................................................................................ [118000]
MK940252.1_CN/2019/InnerMongolia ................................................................................ [118000]
MK128995.1_China/2018/AnhuiXCGQ ................................................................................ [118000]
MH681419.1_ASFV/POL/2015       ................................................................................ [118000]
LS478113.1_Estonia_2014        ................................................................................ [118000]
KP843857.1_Odintsovo_02/14     ................................................................................ [118000]
FN557520.1_strain_E75          ................................................................................ [118000]
AY261366.1_Warthog             ..............................................................................a. [118000]
AY261365.1_Warmbaths           ........................................................................c....... [118000]
AY261364.1_Tengani_62          ................................................................................ [118000]
AY261363.1_Pretoriuskop/96/4   ................................................................................ [118000]
AY261362.1_Mkuzi_1979          ................................................................................ [118000]
AY261361.1_Malawi_Lil-20/1_1983 a..t.................................................................t.......... [118000]
AY261360.1_Kenya_1950          ...t...a................................................g....................... [118000]
AM712240.1_OURT_88/3           ................................................................................ [118000]
AM712239.1_Benin_97/1          ................................................................................ [118000]
KM111295.1_Ken06.Bus           ...t...a................t...............................g.....................a. [118000]
```

```
KM111294.1_Ken05/Tk1            ...t...a................................................g....................... [118000]
KM262844.1_strain_L60           ................................................................................ [118000]
KM102979.1_26544/OG10_Italy     ................................................................................ [118000]
KM262845.1_strain_NHV           ................................................................................ [118000]
KX354450.1_47/Ss/2008           ................................................................................ [118000]
KP055815.1_strain_BA71          ................................................................................ [118000]
MH025920.1_strain_R35           ...t...a................t...............................g.....................a. [118000]
MH025919.1_strain_N10           ...t...a................t...............................g.....................a. [118000]
MH025918.1_strain_R25           ...t...a................t...............................g.....................a. [118000]
MH025917.1_strain_R7            ...t...a................t...............................g.....................a. [118000]
MH025916.1_strain_R8            ...t...a................t...............................g.....................a. [118000]

FR682468.2_Georgia_2007/1       aactatgcgacacacagtccaaaactcaatgggtacaaactttaaaatataaggaatgcaagtattgtcataccgacatg [118080]
ASF_IND_20_CAD543               .................................................y.............................. [118080]
LR899193.1_Germany_2020/1       ................................................................................ [118080]
MN393477.1_Wuhan_2019           ................................................................................ [118080]
MW396979.1_Timor-Leste/2019/1   ................................................................................ [118080]
MK940252.1_CN/2019/InnerMongolia ................................................................................ [118080]
MK128995.1_China/2018/AnhuiXCGQ ................................................................................ [118080]
MH681419.1_ASFV/POL/2015        ................................................................................ [118080]
LS478113.1_Estonia_2014         ................................................................................ [118080]
KP843857.1_Odintsovo_02/14      ................................................................................ [118080]
FN557520.1_strain_E75           ........................................c........c.............................. [118080]
AY261366.1_Warthog              ................................................................................ [118080]
AY261365.1_Warmbaths            ................................................................................ [118080]
AY261364.1_Tengani_62           ................................................................................ [118080]
AY261363.1_Pretoriuskop/96/4    ................................................................................ [118080]
AY261362.1_Mkuzi_1979           ................................................................................ [118080]
AY261361.1_Malawi_Lil-20/1_1983 ....g.......................g...............................................t... [118080]
AY261360.1_Kenya_1950           ..t..........c...................................c........t........c........t... [118080]
AM712240.1_OURT_88/3            ........................................c........c.............................. [118080]
AM712239.1_Benin_97/1           ........................................c........c.............................. [118080]
KM111295.1_Ken06.Bus            ..t..........c...................................c..........................t... [118080]
KM111294.1_Ken05/Tk1            ..t..........c...................................c........t........c........t... [118080]
KM262844.1_strain_L60           ........................................c........c.............................. [118080]
KM102979.1_26544/OG10_Italy     ........................................c........c.............................. [118080]
KM262845.1_strain_NHV           ........................................c........c.............................. [118080]
KX354450.1_47/Ss/2008           ........................................c........c.............................. [118080]
KP055815.1_strain_BA71          ........................................c........c.............................. [118080]
MH025920.1_strain_R35           ..t..........c...................................c..........................t... [118080]
MH025919.1_strain_N10           ..t..........c...................................c..........................t... [118080]
MH025918.1_strain_R25           ..t..........c...................................c..........................t... [118080]
MH025917.1_strain_R7            ..t..........c...................................c..........................t... [118080]
MH025916.1_strain_R8            ..t..........c...................................c..........................t... [118080]
```

```
FR682468.2_Georgia_2007/1        gtgtttaacaccacgcagtttgggctgcaatgtcctaactgcggttgtattcaagaattgatgggaaccatttttgatga [118160]
ASF_IND_20_CAD543                ................................................................................ [118160]
LR899193.1_Germany_2020/1        ................................................................................ [118160]
MN393477.1_Wuhan_2019            ................................................................................ [118160]
MW396979.1_Timor-Leste/2019/1    ................................................................................ [118160]
MK940252.1_CN/2019/InnerMongolia ................................................................................ [118160]
MK128995.1_China/2018/AnhuiXCGQ  ................................................................................ [118160]
MH681419.1_ASFV/POL/2015         ................................................................................ [118160]
LS478113.1_Estonia_2014          ................................................................................ [118160]
KP843857.1_Odintsovo_02/14       ................................................................................ [118160]
FN557520.1_strain_E75            ................................................................................ [118160]
AY261366.1_Warthog               ....................................................................t........... [118160]
AY261365.1_Warmbaths             ....................c......................................a.................... [118160]
AY261364.1_Tengani_62            .......................a..a.........................................t........... [118160]
AY261363.1_Pretoriuskop/96/4     ....................c...............................................t........... [118160]
AY261362.1_Mkuzi_1979            ................................................................................ [118160]
AY261361.1_Malawi_Lil-20/1_1983  ....................................................................t........... [118160]
AY261360.1_Kenya_1950            ..a.....................................................g...........t........c.. [118160]
AM712240.1_OURT_88/3             ................................................................................ [118160]
AM712239.1_Benin_97/1            ................................................................................ [118160]
KM111295.1_Ken06.Bus             ..a.....................................................g...........t........c.. [118160]
KM111294.1_Ken05/Tk1             ..a.....t....................g..........................g...........t........c.. [118160]
KM262844.1_strain_L60            ................................................................................ [118160]
KM102979.1_26544/OG10_Italy      ................................................................................ [118160]
KM262845.1_strain_NHV            ................................................................................ [118160]
KX354450.1_47/Ss/2008            ................................................................................ [118160]
KP055815.1_strain_BA71           ................................................................................ [118160]
MH025920.1_strain_R35            ..a.....................................................g...........t........c.. [118160]
MH025919.1_strain_N10            ..a.....................................................g...........t........c.. [118160]
MH025918.1_strain_R25            ..a.....................................................g...........t........c.. [118160]
MH025917.1_strain_R7             ..a.....................................................g...........t........c.. [118160]
MH025916.1_strain_R8             ..a.....................................................g...........t........c.. [118160]

FR682468.2_Georgia_2007/1        aacacatttttacaaccatgatgggcagaaagcaaagtcaggtatctttaaccctaaccgtcactatcggttttggatag [118240]
ASF_IND_20_CAD543                ................................................................................ [118240]
LR899193.1_Germany_2020/1        ................................................................................ [118240]
MN393477.1_Wuhan_2019            ................................................................................ [118240]
MW396979.1_Timor-Leste/2019/1    ................................................................................ [118240]
MK940252.1_CN/2019/InnerMongolia ................................................................................ [118240]
MK128995.1_China/2018/AnhuiXCGQ  ................................................................................ [118240]
MH681419.1_ASFV/POL/2015         ................................................................................ [118240]
LS478113.1_Estonia_2014          ................................................................................ [118240]
KP843857.1_Odintsovo_02/14       ................................................................................ [118240]
```

```
FN557520.1_strain_E75              ................................................................................ [118240]
AY261366.1_Warthog                 ...g............................................................................ [118240]
AY261365.1_Warmbaths               ...............................................................t................ [118240]
AY261364.1_Tengani_62              ...g............................................................................ [118240]
AY261363.1_Pretoriuskop/96/4       ................................................................................ [118240]
AY261362.1_Mkuzi_1979              ................................................................................ [118240]
AY261361.1_Malawi_Lil-20/1_1983    ...g.................c.....a.................................................... [118240]
AY261360.1_Kenya_1950              ...g.......................a............................................c....... [118240]
AM712240.1_OURT_88/3               ................................................................................ [118240]
AM712239.1_Benin_97/1              ................................................................................ [118240]
KM111295.1_Ken06.Bus               ...g.......................a............................................c....... [118240]
KM111294.1_Ken05/Tk1               ...g.......................a............................................c....... [118240]
KM262844.1_strain_L60              ................................................................................ [118240]
KM102979.1_26544/OG10_Italy        ................................................................................ [118240]
KM262845.1_strain_NHV              ................................................................................ [118240]
KX354450.1_47/Ss/2008              ................................................................................ [118240]
KP055815.1_strain_BA71             ................................................................................ [118240]
MH025920.1_strain_R35              ...g.......................a............................................c....... [118240]
MH025919.1_strain_N10              ...g.......................a............................................c....... [118240]
MH025918.1_strain_R25              ...g.......................a............................................c....... [118240]
MH025917.1_strain_R7               ...g.......................a............................................c....... [118240]
MH025916.1_strain_R8               ...g.......................a............................................c....... [118240]

FR682468.2_Georgia_2007/1          aacatattcttggtagaaatccagaacaagagttggggaccaaacaagatccctgcggaaccaaggtgttgcaacaacta [118320]
ASF_IND_20_CAD543                  ................................................................................ [118320]
LR899193.1_Germany_2020/1          ................................................................................ [118320]
MN393477.1_Wuhan_2019              ................................................................................ [118320]
MW396979.1_Timor-Leste/2019/1      ................................................................................ [118320]
MK940252.1_CN/2019/InnerMongolia   ................................................................................ [118320]
MK128995.1_China/2018/AnhuiXCGQ    ................................................................................ [118320]
MH681419.1_ASFV/POL/2015           ................................................................................ [118320]
LS478113.1_Estonia_2014            ................................................................................ [118320]
KP843857.1_Odintsovo_02/14         ................................................................................ [118320]
FN557520.1_strain_E75              ................................................................................ [118320]
AY261366.1_Warthog                 ....................t.g......................................................... [118320]
AY261365.1_Warmbaths               ....................................................t........................... [118320]
AY261364.1_Tengani_62              ....................t.g......................................................... [118320]
AY261363.1_Pretoriuskop/96/4       ....................t.g......................................................... [118320]
AY261362.1_Mkuzi_1979              ......................g......................................................... [118320]
AY261361.1_Malawi_Lil-20/1_1983    .............c......t..........a..a..............c.....t........................ [118320]
AY261360.1_Kenya_1950              ....................t.g................................t...........c............ [118320]
AM712240.1_OURT_88/3               ................................................................................ [118320]
AM712239.1_Benin_97/1              ................................................................................ [118320]
KM111295.1_Ken06.Bus               ....................t..................................t...........c............ [118320]
```

```
KM111294.1_Ken05/Tk1           ....................t..................................t...........c............ [118320]
KM262844.1_strain_L60          ................................................................................ [118320]
KM102979.1_26544/OG10_Italy    ................................................................................ [118320]
KM262845.1_strain_NHV          ................................................................................ [118320]
KX354450.1_47/Ss/2008          ................................................................................ [118320]
KP055815.1_strain_BA71         ................................................................................ [118320]
MH025920.1_strain_R35          ....................t..................................t...........c............ [118320]
MH025919.1_strain_N10          ....................t..................................t...........c............ [118320]
MH025918.1_strain_R25          ....................t..................................t...........c............ [118320]
MH025917.1_strain_R7           ....................t..................................t...........c............ [118320]
MH025916.1_strain_R8           ....................t..................................t...........c............ [118320]

FR682468.2_Georgia_2007/1      aaaaaaattattaagcgcgataataaatgcatcgcgcttttgacggtcgaaaatattcgaaaaatgttaaaagagataaa [118400]
ASF_IND_20_CAD543              ................................................................................ [118400]
LR899193.1_Germany_2020/1      ................................................................................ [118400]
MN393477.1_Wuhan_2019          ................................................................................ [118400]
MW396979.1_Timor-Leste/2019/1  ................................................................................ [118400]
MK940252.1_CN/2019/InnerMongolia ................................................................................ [118400]
MK128995.1_China/2018/AnhuiXCGQ ................................................................................ [118400]
MH681419.1_ASFV/POL/2015       ................................................................................ [118400]
LS478113.1_Estonia_2014        ................................................................................ [118400]
KP843857.1_Odintsovo_02/14     ................................................................................ [118400]
FN557520.1_strain_E75          ................................................................................ [118400]
AY261366.1_Warthog             ................................................................................ [118400]
AY261365.1_Warmbaths           ................................................................................ [118400]
AY261364.1_Tengani_62          ................................................................................ [118400]
AY261363.1_Pretoriuskop/96/4   .................t.............................................................. [118400]
AY261362.1_Mkuzi_1979          ................................................................................ [118400]
AY261361.1_Malawi_Lil-20/1_1983 ..........................................................................a..... [118400]
AY261360.1_Kenya_1950          ................................................................................ [118400]
AM712240.1_OURT_88/3           ................................................................................ [118400]
AM712239.1_Benin_97/1          ................................................................................ [118400]
KM111295.1_Ken06.Bus           ................................................................................ [118400]
KM111294.1_Ken05/Tk1           ................................................................................ [118400]
KM262844.1_strain_L60          ................................................................................ [118400]
KM102979.1_26544/OG10_Italy    ................................................................................ [118400]
KM262845.1_strain_NHV          ................................................................................ [118400]
KX354450.1_47/Ss/2008          ................................................................................ [118400]
KP055815.1_strain_BA71         ................................................................................ [118400]
MH025920.1_strain_R35          ................................................................................ [118400]
MH025919.1_strain_N10          ................................................................................ [118400]
MH025918.1_strain_R25          ................................................................................ [118400]
MH025917.1_strain_R7           ................................................................................ [118400]
MH025916.1_strain_R8           ................................................................................ [118400]
```

```
FR682468.2_Georgia_2007/1        ccgcacagacttaaataattgtgtttctcttatattgcgtaaacttaccggagtagggccgcctcaaatatcagagtcga [118480]
ASF_IND_20_CAD543                ................................................................................ [118480]
LR899193.1_Germany_2020/1        ................................................................................ [118480]
MN393477.1_Wuhan_2019            ................................................................................ [118480]
MW396979.1_Timor-Leste/2019/1    ................................................................................ [118480]
MK940252.1_CN/2019/InnerMongolia ................................................................................ [118480]
MK128995.1_China/2018/AnhuiXCGQ  ................................................................................ [118480]
MH681419.1_ASFV/POL/2015         ................................................................................ [118480]
LS478113.1_Estonia_2014          ................................................................................ [118480]
KP843857.1_Odintsovo_02/14       ................................................................................ [118480]
FN557520.1_strain_E75            ................................................................................ [118480]
AY261366.1_Warthog               ................................................................................ [118480]
AY261365.1_Warmbaths             ................................................................................ [118480]
AY261364.1_Tengani_62            ................................................................................ [118480]
AY261363.1_Pretoriuskop/96/4     ................................................................................ [118480]
AY261362.1_Mkuzi_1979            ................................................................................ [118480]
AY261361.1_Malawi_Lil-20/1_1983  ................................................................................ [118480]
AY261360.1_Kenya_1950            ..............................c.......................g......................... [118480]
AM712240.1_OURT_88/3             ................................................................................ [118480]
AM712239.1_Benin_97/1            ................................................................................ [118480]
KM111295.1_Ken06.Bus             ................................................................................ [118480]
KM111294.1_Ken05/Tk1             ................................................t.....g......................... [118480]
KM262844.1_strain_L60            ................................................................................ [118480]
KM102979.1_26544/OG10_Italy      ................................................................................ [118480]
KM262845.1_strain_NHV            ................................................................................ [118480]
KX354450.1_47/Ss/2008            ................................................................................ [118480]
KP055815.1_strain_BA71           ................................................................................ [118480]
MH025920.1_strain_R35            ................................................................................ [118480]
MH025919.1_strain_N10            ................................................................................ [118480]
MH025918.1_strain_R25            ................................................................................ [118480]
MH025917.1_strain_R7             ................................................................................ [118480]
MH025916.1_strain_R8             ................................................................................ [118480]

FR682468.2_Georgia_2007/1        ttttactacgaggcgaatacatatttacagaggcaattaagatacgggaaaaagtgtgtaaaaaagggcgtattaatagg [118560]
ASF_IND_20_CAD543                ................................................................................ [118560]
LR899193.1_Germany_2020/1        ................................................................................ [118560]
MN393477.1_Wuhan_2019            ................................................................................ [118560]
MW396979.1_Timor-Leste/2019/1    ................................................................................ [118560]
MK940252.1_CN/2019/InnerMongolia ................................................................................ [118560]
MK128995.1_China/2018/AnhuiXCGQ  ................................................................................ [118560]
MH681419.1_ASFV/POL/2015         ................................................................................ [118560]
LS478113.1_Estonia_2014          ................................................................................ [118560]
KP843857.1_Odintsovo_02/14       ................................................................................ [118560]
```

```
FN557520.1_strain_E75             ................................................................................ [118560]
AY261366.1_Warthog                ..c....................................................a........................ [118560]
AY261365.1_Warmbaths              .......................................................a........................ [118560]
AY261364.1_Tengani_62             ................................................................................ [118560]
AY261363.1_Pretoriuskop/96/4      ..c....................................................a........................ [118560]
AY261362.1_Mkuzi_1979             ................................................................................ [118560]
AY261361.1_Malawi_Lil-20/1_1983   .............t......................................g..a........................ [118560]
AY261360.1_Kenya_1950             .............t..g...................................g..a........................ [118560]
AM712240.1_OURT_88/3              ................................................................................ [118560]
AM712239.1_Benin_97/1             ................................................................................ [118560]
KM111295.1_Ken06.Bus              .............t..g......................................a....................c... [118560]
KM111294.1_Ken05/Tk1              .............t..g......................................a........................ [118560]
KM262844.1_strain_L60             ................................................................................ [118560]
KM102979.1_26544/OG10_Italy       ................................................................................ [118560]
KM262845.1_strain_NHV             ................................................................................ [118560]
KX354450.1_47/Ss/2008             ................................................................................ [118560]
KP055815.1_strain_BA71            ................................................................................ [118560]
MH025920.1_strain_R35             .............t..g......................................a....................c... [118560]
MH025919.1_strain_N10             .............t..g......................................a....................c... [118560]
MH025918.1_strain_R25             .............t..g......................................a....................c... [118560]
MH025917.1_strain_R7              .............t..g......................................a....................c... [118560]
MH025916.1_strain_R8              .............t..g......................................a....................c... [118560]

FR682468.2_Georgia_2007/1         aattattatccgtattatatatataaaatttttgacgccattttgcctccaaatgataccacgaatcgacgcattttaca [118640]
ASF_IND_20_CAD543                 ................................................................................ [118640]
LR899193.1_Germany_2020/1         ................................................................................ [118640]
MN393477.1_Wuhan_2019             ................................................................................ [118640]
MW396979.1_Timor-Leste/2019/1     ................................................................................ [118640]
MK940252.1_CN/2019/InnerMongolia  ................................................................................ [118640]
MK128995.1_China/2018/AnhuiXCGQ   ................................................................................ [118640]
MH681419.1_ASFV/POL/2015          ................................................................................ [118640]
LS478113.1_Estonia_2014           ................................................................................ [118640]
KP843857.1_Odintsovo_02/14        ................................................................................ [118640]
FN557520.1_strain_E75             ................................................................................ [118640]
AY261366.1_Warthog                ............................................a................................... [118640]
AY261365.1_Warmbaths              ........c..c...................................a................................ [118640]
AY261364.1_Tengani_62             ...........................................................t.................... [118640]
AY261363.1_Pretoriuskop/96/4      ........c......................................c................................ [118640]
AY261362.1_Mkuzi_1979             ................................................................................ [118640]
AY261361.1_Malawi_Lil-20/1_1983   ...........a..c....................t........a..c.......................t........ [118640]
AY261360.1_Kenya_1950             ...........a................................a..c...........t...........t........ [118640]
AM712240.1_OURT_88/3              ................................................................................ [118640]
AM712239.1_Benin_97/1             ................................................................................ [118640]
KM111295.1_Ken06.Bus              ........c..a...................................c.......................t........ [118640]
```

```
KM111294.1_Ken05/Tk1              ...........a................................a..c...........t...........t........ [118640]
KM262844.1_strain_L60             ................................................................................ [118640]
KM102979.1_26544/OG10_Italy       ................................................................................ [118640]
KM262845.1_strain_NHV             ................................................................................ [118640]
KX354450.1_47/Ss/2008             ................................................................................ [118640]
KP055815.1_strain_BA71            ................................................................................ [118640]
MH025920.1_strain_R35             ........c..a...................................c.......................t........ [118640]
MH025919.1_strain_N10             ........c..a...................................c.......................t........ [118640]
MH025918.1_strain_R25             ........c..a...................................c.......................t........ [118640]
MH025917.1_strain_R7              ........c..a...................................c.......................t........ [118640]
MH025916.1_strain_R8              ........c..a...................................c.......................t........ [118640]

FR682468.2_Georgia_2007/1         atatattcatttgcaaggaaatgatacgctagctaataatgatagtgagtgggaatctatctgtatggagctccctgaaa [118720]
ASF_IND_20_CAD543                 ......w......................................................................... [118720]
LR899193.1_Germany_2020/1         ................................................................................ [118720]
MN393477.1_Wuhan_2019             ................................................................................ [118720]
MW396979.1_Timor-Leste/2019/1     ................................................................................ [118720]
MK940252.1_CN/2019/InnerMongolia  ................................................................................ [118720]
MK128995.1_China/2018/AnhuiXCGQ   ................................................................................ [118720]
MH681419.1_ASFV/POL/2015          ................................................................................ [118720]
LS478113.1_Estonia_2014           ................................................................................ [118720]
KP843857.1_Odintsovo_02/14        ................................................................................ [118720]
FN557520.1_strain_E75             ......a......................................................................... [118720]
AY261366.1_Warthog                ........................................................................t....... [118720]
AY261365.1_Warmbaths              ............a...........................................................t....... [118720]
AY261364.1_Tengani_62             ........................................................................t....... [118720]
AY261363.1_Pretoriuskop/96/4      ........................................................................t....... [118720]
AY261362.1_Mkuzi_1979             ................................................................................ [118720]
AY261361.1_Malawi_Lil-20/1_1983   ........................................................................t....... [118720]
AY261360.1_Kenya_1950             ........................................................................t....... [118720]
AM712240.1_OURT_88/3              ......a......................................................................... [118720]
AM712239.1_Benin_97/1             ......a......................................................................... [118720]
KM111295.1_Ken06.Bus              ............a........c..................................................t....... [118720]
KM111294.1_Ken05/Tk1              ........................................................................t....... [118720]
KM262844.1_strain_L60             ......a......................................................................... [118720]
KM102979.1_26544/OG10_Italy       ......a......................................................................... [118720]
KM262845.1_strain_NHV             ......a......................................................................... [118720]
KX354450.1_47/Ss/2008             ......a......................................................................... [118720]
KP055815.1_strain_BA71            ......a......................................................................... [118720]
MH025920.1_strain_R35             ............a........c..................................................t....... [118720]
MH025919.1_strain_N10             ............a........c..................................................t....... [118720]
MH025918.1_strain_R25             ............a........c..................................................t....... [118720]
MH025917.1_strain_R7              ............a........c..................................................t....... [118720]
MH025916.1_strain_R8              ............a........c..................................................t....... [118720]
```

```
FR682468.2_Georgia_2007/1        taaaatggaagcccacagatcgaacccattgtgttcattttttttaaagatgaaga----------------------tt [118800]
ASF_IND_20_CAD543                ........................................................----------------------.. [118800]
LR899193.1_Germany_2020/1        ........................................................----------------------.. [118800]
MN393477.1_Wuhan_2019            ........................................................----------------------.. [118800]
MW396979.1_Timor-Leste/2019/1    ........................................................----------------------.. [118800]
MK940252.1_CN/2019/InnerMongolia ........................................................----------------------.. [118800]
MK128995.1_China/2018/AnhuiXCGQ  ........................................................----------------------.. [118800]
MH681419.1_ASFV/POL/2015         ........................................................----------------------.. [118800]
LS478113.1_Estonia_2014          ........................................................----------------------.. [118800]
KP843857.1_Odintsovo_02/14       ........................................................----------------------.. [118800]
FN557520.1_strain_E75            ........................................................----------------------.. [118800]
AY261366.1_Warthog               .....................................................----------------------agt.. [118800]
AY261365.1_Warmbaths             .......................................................tttttttagatattttttttagt.. [118800]
AY261364.1_Tengani_62            ........................................................----------------------.. [118800]
AY261363.1_Pretoriuskop/96/4     ...................................................a...-----------------------.. [118800]
AY261362.1_Mkuzi_1979            .....................................................c..----------------------.. [118800]
AY261361.1_Malawi_Lil-20/1_1983  ....g..........................c........................-----------------------. [118800]
AY261360.1_Kenya_1950            ....g..........................c................a..a....------------------------ [118800]
AM712240.1_OURT_88/3             ........................................................----------------------.. [118800]
AM712239.1_Benin_97/1            ........................................................----------------------.. [118800]
KM111295.1_Ken06.Bus             ....g..........................c................a..a....------------------------ [118800]
KM111294.1_Ken05/Tk1             ....g..........................c................a..a....------------------------ [118800]
KM262844.1_strain_L60            ........................................................----------------------.. [118800]
KM102979.1_26544/OG10_Italy      ........................................................----------------------.. [118800]
KM262845.1_strain_NHV            ........................................................----------------------.. [118800]
KX354450.1_47/Ss/2008            ........................................................----------------------.. [118800]
KP055815.1_strain_BA71           ........................................................----------------------.. [118800]
MH025920.1_strain_R35            ....g..........................c................a..a....------------------------ [118800]
MH025919.1_strain_N10            ....g..........................c................a..a....------------------------ [118800]
MH025918.1_strain_R25            ....g..........................c................a..a....------------------------ [118800]
MH025917.1_strain_R7             ....g..........................c................a..a....------------------------ [118800]
MH025916.1_strain_R8             ....g..........................c................a..a....------------------------ [118800]

FR682468.2_Georgia_2007/1        ttttagatga-ttttttttagtttttta-aaagacgaaaaaattttttaaaagatgaatattctt-aaaccccgcaaatt [118880]
ASF_IND_20_CAD543                ..........-.................a.........................y..........-.............. [118880]
LR899193.1_Germany_2020/1        ..........-.................-....................................-.............. [118880]
MN393477.1_Wuhan_2019            ..........-.................-....................................-.............. [118880]
MW396979.1_Timor-Leste/2019/1    ..........-.................-....................................-.............. [118880]
MK940252.1_CN/2019/InnerMongolia ..........-.................-....................................-.............. [118880]
MK128995.1_China/2018/AnhuiXCGQ  ..........-.................-....................................-.............. [118880]
MH681419.1_ASFV/POL/2015         ..........-.................-....................................-.............. [118880]
LS478113.1_Estonia_2014          ..........-.................-....................................-.............. [118880]
KP843857.1_Odintsovo_02/14       ..........-.................-....................................-.............. [118880]
```

FN557520.1_strain_E75 ..........-.................a.........................c..........-..t........... [118880]
AY261366.1_Warthog ........-.-................g-----.....................c..........-....t......... [118880]
AY261365.1_Warmbaths ........-.-................g-----.g...................c..........-.............. [118880]
AY261364.1_Tengani_62 ..........t................t---t........................cg.-..t..-...aa...t..... [118880]
AY261363.1_Pretoriuskop/96/4 ........---................g-----.....................c..........-..t........... [118880]
AY261362.1_Mkuzi_1979 ..........-.................a.........................c..........-..t........... [118880]
AY261361.1_Malawi_Lil-20/1_1983 .......----.................-.....t.cg....aa..............c...t..a...t.......t.. [118880]
AY261360.1_Kenya_1950 ------------...............t--------.tg...................atg.tc.-...--.t....ca. [118880]
AM712240.1_OURT_88/3 ..........-.................a.........................c..........-..t........... [118880]
AM712239.1_Benin_97/1 ..........-.................a.........................c..........-..t........... [118880]
KM111295.1_Ken06.Bus ------------...............t--------.tg...-...............atg.tc.-------t.t..ca. [118880]
KM111294.1_Ken05/Tk1 ------------...............t--------.tg...................atg.tc.-...--.t....ca. [118880]
KM262844.1_strain_L60 ..........-.................a.........................c..........-..t........... [118880]
KM102979.1_26544/OG10_Italy ..........-.................a.........................c..........-..t........... [118880]
KM262845.1_strain_NHV ..........-.................a.........................c..........-..t........... [118880]
KX354450.1_47/Ss/2008 ..........-.................a.........................c..........-..t........... [118880]
KP055815.1_strain_BA71 ..........-.................a.........................c..........-..t........... [118880]
MH025920.1_strain_R35 ------------...............t--------.tg...-...............atg.tc.-------t.t..ca. [118880]
MH025919.1_strain_N10 ------------...............t--------.tg...-...............atg.tc.-------t.t..ca. [118880]
MH025918.1_strain_R25 ------------...............t--------.tg...-...............atg.tc.-------t.t..ca. [118880]
MH025917.1_strain_R7 ------------...............t--------.tg...-...............atg.tc.-------t.t..ca. [118880]
MH025916.1_strain_R8 ------------...............t--------.tg...-...............atg.tc.-------t.t..ca. [118880]

FR682468.2_Georgia_2007/1 ac-ttttttttaggtactgtaacgcagcacagctgaaccgttctgaagaagaagaaagttaatagcagatgccgatacca [118960]
ASF_IND_20_CAD543 ..t............................................................................. [118960]
LR899193.1_Germany_2020/1 ..-............................................................................. [118960]
MN393477.1_Wuhan_2019 ..-............................................................................. [118960]
MW396979.1_Timor-Leste/2019/1 ..-............................................................................. [118960]
MK940252.1_CN/2019/InnerMongolia ..-............................................................................. [118960]
MK128995.1_China/2018/AnhuiXCGQ ..-............................................................................. [118960]
MH681419.1_ASFV/POL/2015 ..-............................................................................. [118960]
LS478113.1_Estonia_2014 ..-............................................................................. [118960]
KP843857.1_Odintsovo_02/14 ..--............................................................................ [118960]
FN557520.1_strain_E75 ..t.-........................................................................... [118960]
AY261366.1_Warthog ..t-............................................................................ [118960]
AY261365.1_Warmbaths ..---..................................a........................................ [118960]
AY261364.1_Tengani_62 ..---..................................a................................t....... [118960]
AY261363.1_Pretoriuskop/96/4 ..---..................................a........................................ [118960]
AY261362.1_Mkuzi_1979 ..t............................................................................. [118960]
AY261361.1_Malawi_Lil-20/1_1983 .t--...................................a................................t....... [118960]
AY261360.1_Kenya_1950 ta---..........g..........a............a..........................g............. [118960]
AM712240.1_OURT_88/3 ..t............................................................................. [118960]
AM712239.1_Benin_97/1 ..t.-........................................................................... [118960]
KM111295.1_Ken06.Bus ta---.....................a............a..........................g............. [118960]

```
KM111294.1_Ken05/Tk1             ta---.....................a............a..........................g............. [118960]
KM262844.1_strain_L60            ..t.-........................................................................... [118960]
KM102979.1_26544/OG10_Italy      ..t.-........................................................................... [118960]
KM262845.1_strain_NHV            ..t............................................................................. [118960]
KX354450.1_47/Ss/2008            ..t.-........................................................................... [118960]
KP055815.1_strain_BA71           ..t............................................................................. [118960]
MH025920.1_strain_R35            ta---.....................a............a..........................g............. [118960]
MH025919.1_strain_N10            ta---.....................a............a..........................g............. [118960]
MH025918.1_strain_R25            ta---.....................a............a..........................g............. [118960]
MH025917.1_strain_R7             ta---.....................a............a..........................g............. [118960]
MH025916.1_strain_R8             ta---.....................a............a..........................g............. [118960]

FR682468.2_Georgia_2007/1        caagatcagccgtagtgatagaccccacgtaatccgtgtcccaactaatataaaattctcttgctctggatacgttaata [119040]
ASF_IND_20_CAD543                ................................................................................ [119040]
LR899193.1_Germany_2020/1        ................................................................................ [119040]
MN393477.1_Wuhan_2019            ................................................................................ [119040]
MW396979.1_Timor-Leste/2019/1    ................................................................................ [119040]
MK940252.1_CN/2019/InnerMongolia ................................................................................ [119040]
MK128995.1_China/2018/AnhuiXCGQ  ................................................................................ [119040]
MH681419.1_ASFV/POL/2015         ................................................................................ [119040]
LS478113.1_Estonia_2014          ................................................................................ [119040]
KP843857.1_Odintsovo_02/14       ................................................................................ [119040]
FN557520.1_strain_E75            ................................................................................ [119040]
AY261366.1_Warthog               .....................................a.............................a.....a...... [119040]
AY261365.1_Warmbaths             .............g.......................a.......................................... [119040]
AY261364.1_Tengani_62            .....................................a.............................a............ [119040]
AY261363.1_Pretoriuskop/96/4     .....................................a.............................a............ [119040]
AY261362.1_Mkuzi_1979            ................................................................................ [119040]
AY261361.1_Malawi_Lil-20/1_1983  .............g..a.....t.....a................................................... [119040]
AY261360.1_Kenya_1950            .......g.....g..............a.....t..a.....g...........c........................ [119040]
AM712240.1_OURT_88/3             ................................................................................ [119040]
AM712239.1_Benin_97/1            ................................................................................ [119040]
KM111295.1_Ken06.Bus             .............g..............a.....t........g.................................... [119040]
KM111294.1_Ken05/Tk1             .......g.....g..............a.....t........g.................................... [119040]
KM262844.1_strain_L60            ................................................................................ [119040]
KM102979.1_26544/OG10_Italy      ................................................................................ [119040]
KM262845.1_strain_NHV            ................................................................................ [119040]
KX354450.1_47/Ss/2008            ................................................................................ [119040]
KP055815.1_strain_BA71           ................................................................................ [119040]
MH025920.1_strain_R35            .............g..............a.....t........g.................................... [119040]
MH025919.1_strain_N10            .............g..............a.....t........g.................................... [119040]
MH025918.1_strain_R25            .............g..............a.....t........g.................................... [119040]
MH025917.1_strain_R7             .............g..............a.....t........g.................................... [119040]
MH025916.1_strain_R8             .............g..............a.....t........g.................................... [119040]
```

```
FR682468.2_Georgia_2007/1        tgaccactgggttggtattcctcccgtggcttcaaagcaaaggtaatcatcatcgcacccggatcatcgggggttttaat [119120]
ASF_IND_20_CAD543                ................................................................................ [119120]
LR899193.1_Germany_2020/1        ................................................................................ [119120]
MN393477.1_Wuhan_2019            ................................................................................ [119120]
MW396979.1_Timor-Leste/2019/1    ................................................................................ [119120]
MK940252.1_CN/2019/InnerMongolia ................................................................................ [119120]
MK128995.1_China/2018/AnhuiXCGQ  ................................................................................ [119120]
MH681419.1_ASFV/POL/2015         ................................................................................ [119120]
LS478113.1_Estonia_2014          ................................................................................ [119120]
KP843857.1_Odintsovo_02/14       ................................................................................ [119120]
FN557520.1_strain_E75            ................................................................................ [119120]
AY261366.1_Warthog               .................................................................g.............. [119120]
AY261365.1_Warmbaths             .................................................................g.............. [119120]
AY261364.1_Tengani_62            ...................................g............................................ [119120]
AY261363.1_Pretoriuskop/96/4     .................................................................g.............. [119120]
AY261362.1_Mkuzi_1979            ................................................................................ [119120]
AY261361.1_Malawi_Lil-20/1_1983  ......................................g.................g...........a........... [119120]
AY261360.1_Kenya_1950            .............................t..........................g........g.............. [119120]
AM712240.1_OURT_88/3             ................................................................................ [119120]
AM712239.1_Benin_97/1            ................................................................................ [119120]
KM111295.1_Ken06.Bus             .....g.......................t..........................g........g.............. [119120]
KM111294.1_Ken05/Tk1             .....g.......................t..........................g........g.............. [119120]
KM262844.1_strain_L60            ................................................................................ [119120]
KM102979.1_26544/OG10_Italy      ................................................................................ [119120]
KM262845.1_strain_NHV            ................................................................................ [119120]
KX354450.1_47/Ss/2008            ................................................................................ [119120]
KP055815.1_strain_BA71           ................................................................................ [119120]
MH025920.1_strain_R35            .....g.......................t..........................g........g.............. [119120]
MH025919.1_strain_N10            .....g.......................t..........................g........g.............. [119120]
MH025918.1_strain_R25            .....g.......................t..........................g........g.............. [119120]
MH025917.1_strain_R7             .....g.......................t..........................g........g.............. [119120]
MH025916.1_strain_R8             .....g.......................t..........................g........g.............. [119120]

FR682468.2_Georgia_2007/1        cgcattgcctccgtagtggaagggtatgtaagagctgcagaactttgatggaaatttatcgataagattgataccatgag [119200]
ASF_IND_20_CAD543                ................................................................................ [119200]
LR899193.1_Germany_2020/1        ................................................................................ [119200]
MN393477.1_Wuhan_2019            ................................................................................ [119200]
MW396979.1_Timor-Leste/2019/1    ................................................................................ [119200]
MK940252.1_CN/2019/InnerMongolia ................................................................................ [119200]
MK128995.1_China/2018/AnhuiXCGQ  ................................................................................ [119200]
MH681419.1_ASFV/POL/2015         ................................................................................ [119200]
LS478113.1_Estonia_2014          ................................................................................ [119200]
KP843857.1_Odintsovo_02/14       ................................................................................ [119200]
```

```
FN557520.1_strain_E75            t.....................................................c......................... [119200]
AY261366.1_Warthog               ................................................................................ [119200]
AY261365.1_Warmbaths             ................................................................................ [119200]
AY261364.1_Tengani_62            .....................a.......................................................... [119200]
AY261363.1_Pretoriuskop/96/4     ................................................................................ [119200]
AY261362.1_Mkuzi_1979            t...c.................................................c......................... [119200]
AY261361.1_Malawi_Lil-20/1_1983  ..a.........a.........................................c......................... [119200]
AY261360.1_Kenya_1950            ..a...............a............................................................. [119200]
AM712240.1_OURT_88/3             t.....................................................c......................... [119200]
AM712239.1_Benin_97/1            t.....................................................c......................... [119200]
KM111295.1_Ken06.Bus             ..a........................................................................g.... [119200]
KM111294.1_Ken05/Tk1             ..a...............a............................................................. [119200]
KM262844.1_strain_L60            t.....................................................c......................... [119200]
KM102979.1_26544/OG10_Italy      t.....................................................c......................... [119200]
KM262845.1_strain_NHV            t.....................................................c......................... [119200]
KX354450.1_47/Ss/2008            t.....................................................c......................... [119200]
KP055815.1_strain_BA71           t.....................................................c......................... [119200]
MH025920.1_strain_R35            ..a........................................................................g.... [119200]
MH025919.1_strain_N10            ..a........................................................................g.... [119200]
MH025918.1_strain_R25            ..a........................................................................g.... [119200]
MH025917.1_strain_R7             ..a........................................................................g.... [119200]
MH025916.1_strain_R8             ..a........................................................................g.... [119200]

FR682468.2_Georgia_2007/1        cagttacggaaatgtttttaataataggtaatgtgatcggatacgtaacggggctaatatcagatatagatgaacatgcg [119280]
ASF_IND_20_CAD543                ................................................................................ [119280]
LR899193.1_Germany_2020/1        ................................................................................ [119280]
MN393477.1_Wuhan_2019            ................................................................................ [119280]
MW396979.1_Timor-Leste/2019/1    ................................................................................ [119280]
MK940252.1_CN/2019/InnerMongolia ................................................................................ [119280]
MK128995.1_China/2018/AnhuiXCGQ  ................................................................................ [119280]
MH681419.1_ASFV/POL/2015         ................................................................................ [119280]
LS478113.1_Estonia_2014          ................................................................................ [119280]
KP843857.1_Odintsovo_02/14       ................................................................................ [119280]
FN557520.1_strain_E75            .............................................................c.................. [119280]
AY261366.1_Warthog               ...............................c................t............................... [119280]
AY261365.1_Warmbaths             ...............................c................................................ [119280]
AY261364.1_Tengani_62            ..................................a...........g................................. [119280]
AY261363.1_Pretoriuskop/96/4     ...............................c................................................ [119280]
AY261362.1_Mkuzi_1979            .............................................................c.................. [119280]
AY261361.1_Malawi_Lil-20/1_1983  ...........................................a.........t.......................... [119280]
AY261360.1_Kenya_1950            ....g..........................c...........a....t....ga......t.....g...........a [119280]
AM712240.1_OURT_88/3             .............................................................c.................. [119280]
AM712239.1_Benin_97/1            .............................................................c.................. [119280]
KM111295.1_Ken06.Bus             ....a......................................a....t....g.............g...........a [119280]
```

```
KM111294.1_Ken05/Tk1              ....g..........................c...........a....t....g.......t.....g...........a [119280]
KM262844.1_strain_L60             .............................................................c.................. [119280]
KM102979.1_26544/OG10_Italy       .............................................................c.................. [119280]
KM262845.1_strain_NHV             .............................................................c.................. [119280]
KX354450.1_47/Ss/2008             .............................................................c.................. [119280]
KP055815.1_strain_BA71            .............................................................c.................. [119280]
MH025920.1_strain_R35             ....a......................................a....t....g.............g...........a [119280]
MH025919.1_strain_N10             ....a......................................a....t....g.............g...........a [119280]
MH025918.1_strain_R25             ....a......................................a....t....g.............g...........a [119280]
MH025917.1_strain_R7              ....a......................................a....t....g.............g...........a [119280]
MH025916.1_strain_R8              ....a......................................a....t....g.............g...........a [119280]

FR682468.2_Georgia_2007/1         tctggaagagctgtatctctatcctgaaagcttatctctgcgtggtgagtgggctgcataatggcgttaacaacatgtcc [119360]
ASF_IND_20_CAD543                 ................................................................................ [119360]
LR899193.1_Germany_2020/1         ................................................................................ [119360]
MN393477.1_Wuhan_2019             ................................................................................ [119360]
MW396979.1_Timor-Leste/2019/1     ................................................................................ [119360]
MK940252.1_CN/2019/InnerMongolia  ................................................................................ [119360]
MK128995.1_China/2018/AnhuiXCGQ   ................................................................................ [119360]
MH681419.1_ASFV/POL/2015          ................................................................................ [119360]
LS478113.1_Estonia_2014           ................................................................................ [119360]
KP843857.1_Odintsovo_02/14        ................................................................................ [119360]
FN557520.1_strain_E75             ..................................................a............................. [119360]
AY261366.1_Warthog                ..................................................a............................. [119360]
AY261365.1_Warmbaths              ..................................................a............................. [119360]
AY261364.1_Tengani_62             ..................................................a............................. [119360]
AY261363.1_Pretoriuskop/96/4      ..................................................a............................. [119360]
AY261362.1_Mkuzi_1979             ..................................................a............................. [119360]
AY261361.1_Malawi_Lil-20/1_1983   ...................................t..............a.......................g..... [119360]
AY261360.1_Kenya_1950             ..................................c............g..a............................. [119360]
AM712240.1_OURT_88/3              ..................................................a............................. [119360]
AM712239.1_Benin_97/1             ..................................................a............................. [119360]
KM111295.1_Ken06.Bus              ..................................c............g..a............................. [119360]
KM111294.1_Ken05/Tk1              ..................................c............g..a............................. [119360]
KM262844.1_strain_L60             ..................................................a............................. [119360]
KM102979.1_26544/OG10_Italy       ..................................................a............................. [119360]
KM262845.1_strain_NHV             ..................................................a............................. [119360]
KX354450.1_47/Ss/2008             ..................................................a............................. [119360]
KP055815.1_strain_BA71            ..................................................a............................. [119360]
MH025920.1_strain_R35             ..................................c............g..a............................. [119360]
MH025919.1_strain_N10             ..................................c............g..a............................. [119360]
MH025918.1_strain_R25             ..................................c............g..a............................. [119360]
MH025917.1_strain_R7              ..................................c............g..a............................. [119360]
MH025916.1_strain_R8              ..................................c............g..a............................. [119360]
```

```
FR682468.2_Georgia_2007/1        gaacttgtgccaatctcggtgttgatgaggattttgatcggagatgttccaggtaggttttaatcctataaacatatatt [119440]
ASF_IND_20_CAD543                ................................................................................ [119440]
LR899193.1_Germany_2020/1        ................................................................................ [119440]
MN393477.1_Wuhan_2019            ................................................................................ [119440]
MW396979.1_Timor-Leste/2019/1    ................................................................................ [119440]
MK940252.1_CN/2019/InnerMongolia ................................................................................ [119440]
MK128995.1_China/2018/AnhuiXCGQ  ................................................................................ [119440]
MH681419.1_ASFV/POL/2015         ................................................................................ [119440]
LS478113.1_Estonia_2014          ................................................................................ [119440]
KP843857.1_Odintsovo_02/14       ................................................................................ [119440]
FN557520.1_strain_E75            ................................................................................ [119440]
AY261366.1_Warthog               ................................................................................ [119440]
AY261365.1_Warmbaths             ................................................................................ [119440]
AY261364.1_Tengani_62            ................................................................................ [119440]
AY261363.1_Pretoriuskop/96/4     ................................................................................ [119440]
AY261362.1_Mkuzi_1979            .....................c.......................................................... [119440]
AY261361.1_Malawi_Lil-20/1_1983  ...................................................a............................ [119440]
AY261360.1_Kenya_1950            ....................................g.....a..a.................................. [119440]
AM712240.1_OURT_88/3             ................................................................................ [119440]
AM712239.1_Benin_97/1            ................................................................................ [119440]
KM111295.1_Ken06.Bus             ....................................g.....a..a........g......................... [119440]
KM111294.1_Ken05/Tk1             ....................................g.....a..a.................................. [119440]
KM262844.1_strain_L60            ................................................................................ [119440]
KM102979.1_26544/OG10_Italy      ................................................................................ [119440]
KM262845.1_strain_NHV            ................................................................................ [119440]
KX354450.1_47/Ss/2008            ................................................................................ [119440]
KP055815.1_strain_BA71           ................................................................................ [119440]
MH025920.1_strain_R35            ....................................g.....a..a........g......................... [119440]
MH025919.1_strain_N10            ....................................g.....a..a........g......................... [119440]
MH025918.1_strain_R25            ....................................g.....a..a........g......................... [119440]
MH025917.1_strain_R7             ....................................g.....a..a........g......................... [119440]
MH025916.1_strain_R8             ....................................g.....a..a........g......................... [119440]

FR682468.2_Georgia_2007/1        caatgggccatttaagagcagacattagtttttcatcgtggtggttattgttggtgtgggtcacctgcgttttatggaca [119520]
ASF_IND_20_CAD543                ................................................................................ [119520]
LR899193.1_Germany_2020/1        ................................................................................ [119520]
MN393477.1_Wuhan_2019            ................................................................................ [119520]
MW396979.1_Timor-Leste/2019/1    ................................................................................ [119520]
MK940252.1_CN/2019/InnerMongolia ................................................................................ [119520]
MK128995.1_China/2018/AnhuiXCGQ  ................................................................................ [119520]
MH681419.1_ASFV/POL/2015         ................................................................................ [119520]
LS478113.1_Estonia_2014          ................................................................................ [119520]
KP843857.1_Odintsovo_02/14       ................................................................................ [119520]
```

```
FN557520.1_strain_E75            ................................................................................ [119520]
AY261366.1_Warthog               .......................................................a........................ [119520]
AY261365.1_Warmbaths             ................................................................................ [119520]
AY261364.1_Tengani_62            ................................................................................ [119520]
AY261363.1_Pretoriuskop/96/4     ................................................................................ [119520]
AY261362.1_Mkuzi_1979            ...............................................................................g [119520]
AY261361.1_Malawi_Lil-20/1_1983  ........................................a..a........c........................... [119520]
AY261360.1_Kenya_1950            .g......................................a........a..c........................... [119520]
AM712240.1_OURT_88/3             ................................................................................ [119520]
AM712239.1_Benin_97/1            ................................................................................ [119520]
KM111295.1_Ken06.Bus             .g......................................a...........c........................... [119520]
KM111294.1_Ken05/Tk1             .g......................................a........a..c........................... [119520]
KM262844.1_strain_L60            ................................................................................ [119520]
KM102979.1_26544/OG10_Italy      ................................................................................ [119520]
KM262845.1_strain_NHV            ................................................................................ [119520]
KX354450.1_47/Ss/2008            ................................................................................ [119520]
KP055815.1_strain_BA71           ................................................................................ [119520]
MH025920.1_strain_R35            .g......................................a...........c........................... [119520]
MH025919.1_strain_N10            .g......................................a...........c........................... [119520]
MH025918.1_strain_R25            .g......................................a...........c........................... [119520]
MH025917.1_strain_R7             .g......................................a...........c........................... [119520]
MH025916.1_strain_R8             .g......................................a...........c........................... [119520]

FR682468.2_Georgia_2007/1        cgtatcagcgaaaagcgaacgcgttttacaaaaaggttgtgtatttcaggggttacaaacaggttattgatgtaaagttc [119600]
ASF_IND_20_CAD543                ................................................................................ [119600]
LR899193.1_Germany_2020/1        ................................................................................ [119600]
MN393477.1_Wuhan_2019            ................................................................................ [119600]
MW396979.1_Timor-Leste/2019/1    ................................................................................ [119600]
MK940252.1_CN/2019/InnerMongolia ................................................................................ [119600]
MK128995.1_China/2018/AnhuiXCGQ  ................................................................................ [119600]
MH681419.1_ASFV/POL/2015         ................................................................................ [119600]
LS478113.1_Estonia_2014          ................................................................................ [119600]
KP843857.1_Odintsovo_02/14       ................................................................................ [119600]
FN557520.1_strain_E75            ........g.....t................................................................. [119600]
AY261366.1_Warthog               ........a.....t..........................a...................................c.. [119600]
AY261365.1_Warmbaths             ........a.....t..............................................................c.. [119600]
AY261364.1_Tengani_62            ........a.....t..............................................................c.. [119600]
AY261363.1_Pretoriuskop/96/4     ........a.....t................................................................. [119600]
AY261362.1_Mkuzi_1979            ........g.....t................................................................. [119600]
AY261361.1_Malawi_Lil-20/1_1983  ........g.....c..............................................................c.. [119600]
AY261360.1_Kenya_1950            ........g.....t..............g...............................................c.. [119600]
AM712240.1_OURT_88/3             ........g.....t................................................................. [119600]
AM712239.1_Benin_97/1            ........g.....t................................................................. [119600]
KM111295.1_Ken06.Bus             ........g.....t....................a.........................................c.. [119600]
```

```
KM111294.1_Ken05/Tk1          ........g.....t..............g...............................................c.. [119600]
KM262844.1_strain_L60         ........g.....t................................................................. [119600]
KM102979.1_26544/OG10_Italy   ........g.....t................................................................. [119600]
KM262845.1_strain_NHV         ........g.....t................................................................. [119600]
KX354450.1_47/Ss/2008         ........g.....t................................................................. [119600]
KP055815.1_strain_BA71        ........g.....t................................................................. [119600]
MH025920.1_strain_R35         ........g.....t....................a.........................................c.. [119600]
MH025919.1_strain_N10         ........g.....t....................a.........................................c.. [119600]
MH025918.1_strain_R25         ........g.....t....................a.........................................c.. [119600]
MH025917.1_strain_R7          ........g.....t....................a.........................................c.. [119600]
MH025916.1_strain_R8          ........g.....t....................a.........................................c.. [119600]

FR682468.2_Georgia_2007/1     attattcgtgagcgagatttcattaatgactcctgggataaaccatggtttaaagcgtatattgcgtctactggggcgtc [119680]
ASF_IND_20_CAD543             ................................................................................ [119680]
LR899193.1_Germany_2020/1     ................................................................................ [119680]
MN393477.1_Wuhan_2019         ................................................................................ [119680]
MW396979.1_Timor-Leste/2019/1 ................................................................................ [119680]
MK940252.1_CN/2019/InnerMongolia ................................................................................ [119680]
MK128995.1_China/2018/AnhuiXCGQ ................................................................................ [119680]
MH681419.1_ASFV/POL/2015      ................................................................................ [119680]
LS478113.1_Estonia_2014       ................................................................................ [119680]
KP843857.1_Odintsovo_02/14    ................................................................................ [119680]
FN557520.1_strain_E75         ................................................................................ [119680]
AY261366.1_Warthog            ................................................................................ [119680]
AY261365.1_Warmbaths          ................................................................................ [119680]
AY261364.1_Tengani_62         ............t................................................................... [119680]
AY261363.1_Pretoriuskop/96/4  ...........................a.................................................... [119680]
AY261362.1_Mkuzi_1979         ................................................................................ [119680]
AY261361.1_Malawi_Lil-20/1_1983 ................................................................................ [119680]
AY261360.1_Kenya_1950         ......................c..................................g..g.................c. [119680]
AM712240.1_OURT_88/3          ................................................................................ [119680]
AM712239.1_Benin_97/1         ................................................................................ [119680]
KM111295.1_Ken06.Bus          ......................c..................................g..g.................c. [119680]
KM111294.1_Ken05/Tk1          ......................c..................................g..g.................c. [119680]
KM262844.1_strain_L60         ................................................................................ [119680]
KM102979.1_26544/OG10_Italy   ................................................................................ [119680]
KM262845.1_strain_NHV         ................................................................................ [119680]
KX354450.1_47/Ss/2008         ................................................................................ [119680]
KP055815.1_strain_BA71        ................................................................................ [119680]
MH025920.1_strain_R35         ......................c..................................g..g.................c. [119680]
MH025919.1_strain_N10         ......................c..................................g..g.................c. [119680]
MH025918.1_strain_R25         ......................c..................................g..g.................c. [119680]
MH025917.1_strain_R7          ......................c..................................g..g.................c. [119680]
MH025916.1_strain_R8          ......................c..................................g..g.................c. [119680]
```

```
FR682468.2_Georgia_2007/1        cagctataaaacgtgactggcgtacaaaaagtccaggaaattcattcaccaaatccttttgcgatgcaagctttatggtg [119760]
ASF_IND_20_CAD543                ................................................................................ [119760]
LR899193.1_Germany_2020/1        ................................................................................ [119760]
MN393477.1_Wuhan_2019            ................................................................................ [119760]
MW396979.1_Timor-Leste/2019/1    ................................................................................ [119760]
MK940252.1_CN/2019/InnerMongolia ................................................................................ [119760]
MK128995.1_China/2018/AnhuiXCGQ  ................................................................................ [119760]
MH681419.1_ASFV/POL/2015         ................................................................................ [119760]
LS478113.1_Estonia_2014          ................................................................................ [119760]
KP843857.1_Odintsovo_02/14       ................................................................................ [119760]
FN557520.1_strain_E75            ...g.........c..........t....................................................... [119760]
AY261366.1_Warthog               ................................................................................ [119760]
AY261365.1_Warmbaths             ................................................................................ [119760]
AY261364.1_Tengani_62            ................................................................................ [119760]
AY261363.1_Pretoriuskop/96/4     ................................................................................ [119760]
AY261362.1_Mkuzi_1979            ...g....................t....................................................... [119760]
AY261361.1_Malawi_Lil-20/1_1983  ...g..................a.................c....................................... [119760]
AY261360.1_Kenya_1950            ...g..................g.t.........g............................................. [119760]
AM712240.1_OURT_88/3             ...g.........c..........t....................................................... [119760]
AM712239.1_Benin_97/1            ...g.........c..........t....................................................... [119760]
KM111295.1_Ken06.Bus             ...g..................g.t.........g............................................. [119760]
KM111294.1_Ken05/Tk1             ...g..................g.t.........g............................................. [119760]
KM262844.1_strain_L60            ...g.........c..........t....................................................... [119760]
KM102979.1_26544/OG10_Italy      ...g.........c..........t....................................................... [119760]
KM262845.1_strain_NHV            ...g.........c..........t....................................................... [119760]
KX354450.1_47/Ss/2008            ...g.........c..........t....................................................... [119760]
KP055815.1_strain_BA71           ...g.........c..........t....................................................... [119760]
MH025920.1_strain_R35            ...g..................g.t.........g............................................. [119760]
MH025919.1_strain_N10            ...g..................g.t.........g............................................. [119760]
MH025918.1_strain_R25            ...g..................g.t.........g............................................. [119760]
MH025917.1_strain_R7             ...g..................g.t.........g............................................. [119760]
MH025916.1_strain_R8             ...g..................g.t.........g............................................. [119760]

FR682468.2_Georgia_2007/1        ataaagcgctcgccgaagggaatggatactgagggaatagcaaggttcacgttctcattaaaccaaaagcgcaacttaat [119840]
ASF_IND_20_CAD543                ................................................................................ [119840]
LR899193.1_Germany_2020/1        ................................................................................ [119840]
MN393477.1_Wuhan_2019            ................................................................................ [119840]
MW396979.1_Timor-Leste/2019/1    ................................................................................ [119840]
MK940252.1_CN/2019/InnerMongolia ................................................................................ [119840]
MK128995.1_China/2018/AnhuiXCGQ  ................................................................................ [119840]
MH681419.1_ASFV/POL/2015         ................................................................................ [119840]
LS478113.1_Estonia_2014          ................................................................................ [119840]
KP843857.1_Odintsovo_02/14       ................................................................................ [119840]
```

```
FN557520.1_strain_E75           .............................c..........................g................g...... [119840]
AY261366.1_Warthog              ........................................................g................g...... [119840]
AY261365.1_Warmbaths            ....................g........c..........................g................g...... [119840]
AY261364.1_Tengani_62           .............................c..........................g................g...... [119840]
AY261363.1_Pretoriuskop/96/4    ................................a.......................g................g...... [119840]
AY261362.1_Mkuzi_1979           .............................c..........................g................g...... [119840]
AY261361.1_Malawi_Lil-20/1_1983 .............................c...........................................g...... [119840]
AY261360.1_Kenya_1950           .............................c...........................................g...... [119840]
AM712240.1_OURT_88/3            .............................c..........................g................g...... [119840]
AM712239.1_Benin_97/1           .............................c..........................g................g...... [119840]
KM111295.1_Ken06.Bus            .............................c...........................................g...... [119840]
KM111294.1_Ken05/Tk1            .............................c...........................................g...... [119840]
KM262844.1_strain_L60           .............................c..........................g................g...... [119840]
KM102979.1_26544/OG10_Italy     .............................c..........................g................g...... [119840]
KM262845.1_strain_NHV           .............................c..........................g................g...... [119840]
KX354450.1_47/Ss/2008           .............................c..........................g................g...... [119840]
KP055815.1_strain_BA71          .............................c..........................g................g...... [119840]
MH025920.1_strain_R35           .......................a.....c...........................................g...... [119840]
MH025919.1_strain_N10           .......................a.....c...........................................g...... [119840]
MH025918.1_strain_R25           .......................a.....c...........................................g...... [119840]
MH025917.1_strain_R7            .......................a.....c...........................................g...... [119840]
MH025916.1_strain_R8            .......................a.....c...........................................g...... [119840]

FR682468.2_Georgia_2007/1       ccagagcgcaagagggggctgatagtatttaggggtttgaggtccattacagctgtaatgaacattacgtcttatgtcca [119920]
ASF_IND_20_CAD543               ................................................................................ [119920]
LR899193.1_Germany_2020/1       ................................................................................ [119920]
MN393477.1_Wuhan_2019           ................................................................................ [119920]
MW396979.1_Timor-Leste/2019/1   ................................................................................ [119920]
MK940252.1_CN/2019/InnerMongolia ................................................................................ [119920]
MK128995.1_China/2018/AnhuiXCGQ ................................................................................ [119920]
MH681419.1_ASFV/POL/2015        ................................................................................ [119920]
LS478113.1_Estonia_2014         ................................................................................ [119920]
KP843857.1_Odintsovo_02/14      ................................................................................ [119920]
FN557520.1_strain_E75           ................................................................................ [119920]
AY261366.1_Warthog              ................................................................................ [119920]
AY261365.1_Warmbaths            ................................................................................ [119920]
AY261364.1_Tengani_62           ................................................................................ [119920]
AY261363.1_Pretoriuskop/96/4    ................................................................................ [119920]
AY261362.1_Mkuzi_1979           ................................................................................ [119920]
AY261361.1_Malawi_Lil-20/1_1983 ..........................................g..............t...................... [119920]
AY261360.1_Kenya_1950           ................................................................................ [119920]
AM712240.1_OURT_88/3            ................................................................................ [119920]
AM712239.1_Benin_97/1           ................................................................................ [119920]
KM111295.1_Ken06.Bus            ................................................................................ [119920]
```

```
KM111294.1_Ken05/Tk1           ......................................................a.....g................... [119920]
KM262844.1_strain_L60          ................................................................................ [119920]
KM102979.1_26544/OG10_Italy    ................................................................................ [119920]
KM262845.1_strain_NHV          ................................................................................ [119920]
KX354450.1_47/Ss/2008          ................................................................................ [119920]
KP055815.1_strain_BA71         ................................................................................ [119920]
MH025920.1_strain_R35          ................................................................................ [119920]
MH025919.1_strain_N10          ................................................................................ [119920]
MH025918.1_strain_R25          ................................................................................ [119920]
MH025917.1_strain_R7           ................................................................................ [119920]
MH025916.1_strain_R8           ................................................................................ [119920]

FR682468.2_Georgia_2007/1      gatacgttgcgtccgtgataggagtaatatcttgtttacctgctgtttggatattgtgagagttctcgggaaaatgctgt [120000]
ASF_IND_20_CAD543              ................................................................................ [120000]
LR899193.1_Germany_2020/1      ................................................................................ [120000]
MN393477.1_Wuhan_2019          ................................................................................ [120000]
MW396979.1_Timor-Leste/2019/1  ................................................................................ [120000]
MK940252.1_CN/2019/InnerMongolia ................................................................................ [120000]
MK128995.1_China/2018/AnhuiXCGQ ................................................................................ [120000]
MH681419.1_ASFV/POL/2015       ................................................................................ [120000]
LS478113.1_Estonia_2014        ................................................................................ [120000]
KP843857.1_Odintsovo_02/14     ................................................................................ [120000]
FN557520.1_strain_E75          ................a...........................................................t... [120000]
AY261366.1_Warthog             .........................g..................................................t... [120000]
AY261365.1_Warmbaths           .........................g..................................................t... [120000]
AY261364.1_Tengani_62          .........................g.............................a........................ [120000]
AY261363.1_Pretoriuskop/96/4   .........................g..................................................t... [120000]
AY261362.1_Mkuzi_1979          ............................................................................t... [120000]
AY261361.1_Malawi_Lil-20/1_1983 .........ta.....a......................................a..g..................... [120000]
AY261360.1_Kenya_1950          ................a.....g...................................g..................... [120000]
AM712240.1_OURT_88/3           ................a...........................................................t... [120000]
AM712239.1_Benin_97/1          ................a...........................................................t... [120000]
KM111295.1_Ken06.Bus           ................a.....g...................................g..................... [120000]
KM111294.1_Ken05/Tk1           ................a.....g...................................g..................... [120000]
KM262844.1_strain_L60          ................a...........................................................t... [120000]
KM102979.1_26544/OG10_Italy    ................a...........................................................t... [120000]
KM262845.1_strain_NHV          ................a...........................................................t... [120000]
KX354450.1_47/Ss/2008          ................a...........................................................t... [120000]
KP055815.1_strain_BA71         ................a...........................................................t... [120000]
MH025920.1_strain_R35          ................a.....g...................................g..................... [120000]
MH025919.1_strain_N10          ................a.....g...................................g..................... [120000]
MH025918.1_strain_R25          ................a.....g...................................g..................... [120000]
MH025917.1_strain_R7           ................a.....g...................................g..................... [120000]
MH025916.1_strain_R8           ................a.....g...................................g..................... [120000]
```

```
FR682468.2_Georgia_2007/1        gaaagaaatttcgggttggtatggctacacgttcgctgcgtatcattttcatcggtaagaataggtttgctttggtgcgg [120080]
ASF_IND_20_CAD543                ................................................................................ [120080]
LR899193.1_Germany_2020/1        ................................................................................ [120080]
MN393477.1_Wuhan_2019            ................................................................................ [120080]
MW396979.1_Timor-Leste/2019/1    ................................................................................ [120080]
MK940252.1_CN/2019/InnerMongolia ................................................................................ [120080]
MK128995.1_China/2018/AnhuiXCGQ  ................................................................................ [120080]
MH681419.1_ASFV/POL/2015         ................................................................................ [120080]
LS478113.1_Estonia_2014          ................................................................................ [120080]
KP843857.1_Odintsovo_02/14       ................................................................................ [120080]
FN557520.1_strain_E75            .....g....................g..................................................... [120080]
AY261366.1_Warthog               ..........................g..................................................... [120080]
AY261365.1_Warmbaths             ..........................g..................................................... [120080]
AY261364.1_Tengani_62            ..............a...........g..................................................... [120080]
AY261363.1_Pretoriuskop/96/4     ..........................g..................................................... [120080]
AY261362.1_Mkuzi_1979            .....g.......................................................................... [120080]
AY261361.1_Malawi_Lil-20/1_1983  ........................g.g..t.................................................. [120080]
AY261360.1_Kenya_1950            ........................g.g..t.........................................c........ [120080]
AM712240.1_OURT_88/3             .....g....................g..................................................... [120080]
AM712239.1_Benin_97/1            .....g....................g..................................................... [120080]
KM111295.1_Ken06.Bus             ........................g.g..t.........................................c........ [120080]
KM111294.1_Ken05/Tk1             ........................g.g..t.........................................c........ [120080]
KM262844.1_strain_L60            .....g....................g..................................................... [120080]
KM102979.1_26544/OG10_Italy      .....g....................g..................................................... [120080]
KM262845.1_strain_NHV            .....g....................g..................................................... [120080]
KX354450.1_47/Ss/2008            .....g....................g..................................................... [120080]
KP055815.1_strain_BA71           .....g....................g..................................................... [120080]
MH025920.1_strain_R35            ........................g.g..t.........................................c........ [120080]
MH025919.1_strain_N10            ........................g.g..t.........................................c........ [120080]
MH025918.1_strain_R25            ........................g.g..t.........................................c........ [120080]
MH025917.1_strain_R7             ........................g.g..t.........................................c........ [120080]
MH025916.1_strain_R8             ........................g.g..t.........................................c........ [120080]

FR682468.2_Georgia_2007/1        cttgtgcaaatcatgaatgttgcataggagagggccactggttccctccaccgatacctcctggccaaccaagtgcttat [120160]
ASF_IND_20_CAD543                ................................................................................ [120160]
LR899193.1_Germany_2020/1        ................................................................................ [120160]
MN393477.1_Wuhan_2019            ................................................................................ [120160]
MW396979.1_Timor-Leste/2019/1    ................................................................................ [120160]
MK940252.1_CN/2019/InnerMongolia ................................................................................ [120160]
MK128995.1_China/2018/AnhuiXCGQ  ................................................................................ [120160]
MH681419.1_ASFV/POL/2015         ................................................................................ [120160]
LS478113.1_Estonia_2014          ................................................................................ [120160]
KP843857.1_Odintsovo_02/14       ................................................................................ [120160]
```

```
FN557520.1_strain_E75              .......................................a..........................g............. [120160]
AY261366.1_Warthog                 ..................................................................g............. [120160]
AY261365.1_Warmbaths               ................................................................................ [120160]
AY261364.1_Tengani_62              ..................................................................g............. [120160]
AY261363.1_Pretoriuskop/96/4       ................................................................................ [120160]
AY261362.1_Mkuzi_1979              .........................................................t........g............. [120160]
AY261361.1_Malawi_Lil-20/1_1983    .............................................t................................g. [120160]
AY261360.1_Kenya_1950              ........t...t....c.............................................................. [120160]
AM712240.1_OURT_88/3               .......................................a..........................g............. [120160]
AM712239.1_Benin_97/1              .......................................a..........................g............. [120160]
KM111295.1_Ken06.Bus               ........t...t....c.............................................................. [120160]
KM111294.1_Ken05/Tk1               ........t...t....c.............................................................. [120160]
KM262844.1_strain_L60              .......................................a..........................g............. [120160]
KM102979.1_26544/OG10_Italy        .......................................a..........................g............. [120160]
KM262845.1_strain_NHV              .......................................a..........................g............. [120160]
KX354450.1_47/Ss/2008              .......................................a..........................g............. [120160]
KP055815.1_strain_BA71             .......................................a..........................g............. [120160]
MH025920.1_strain_R35              ........t...t....c.............................................................. [120160]
MH025919.1_strain_N10              ........t...t....c.............................................................. [120160]
MH025918.1_strain_R25              ........t...t....c.............................................................. [120160]
MH025917.1_strain_R7               ........t...t....c.............................................................. [120160]
MH025916.1_strain_R8               ........t...t....c.............................................................. [120160]

FR682468.2_Georgia_2007/1          atccagtcattttatcccctgggatgcaaaatttgcgcacaagcgttgtgacatccgaactatattcgtctagggaattt [120240]
ASF_IND_20_CAD543                  ................................................................................ [120240]
LR899193.1_Germany_2020/1          ................................................................................ [120240]
MN393477.1_Wuhan_2019              ................................................................................ [120240]
MW396979.1_Timor-Leste/2019/1      ................................................................................ [120240]
MK940252.1_CN/2019/InnerMongolia   ................................................................................ [120240]
MK128995.1_China/2018/AnhuiXCGQ    ................................................................................ [120240]
MH681419.1_ASFV/POL/2015           ................................................................................ [120240]
LS478113.1_Estonia_2014            ................................................................................ [120240]
KP843857.1_Odintsovo_02/14         ................................................................................ [120240]
FN557520.1_strain_E75              ......................................................................c......... [120240]
AY261366.1_Warthog                 .........................a...................................................... [120240]
AY261365.1_Warmbaths               ................................................................................ [120240]
AY261364.1_Tengani_62              ................................................................................ [120240]
AY261363.1_Pretoriuskop/96/4       ...................................................................a............ [120240]
AY261362.1_Mkuzi_1979              ......................................................................c......... [120240]
AY261361.1_Malawi_Lil-20/1_1983    ................t..g..................................................c.a....... [120240]
AY261360.1_Kenya_1950              ................t.................a...................................c.....g... [120240]
AM712240.1_OURT_88/3               ......................................................................c......... [120240]
AM712239.1_Benin_97/1              ......................................................................c......... [120240]
KM111295.1_Ken06.Bus               ................t.................a..............a....................c.....g... [120240]
```

```
KM111294.1_Ken05/Tk1            ................t.................a..............a....................c.....g... [120240]
KM262844.1_strain_L60           ......................................................................c......... [120240]
KM102979.1_26544/OG10_Italy     ......................................................................c......... [120240]
KM262845.1_strain_NHV           ......................................................................c......... [120240]
KX354450.1_47/Ss/2008           ......................................................................c......... [120240]
KP055815.1_strain_BA71          ......................................................................c......... [120240]
MH025920.1_strain_R35           ................t.................a..............a....................c.....g... [120240]
MH025919.1_strain_N10           ................t.................a..............a....................c.....g... [120240]
MH025918.1_strain_R25           ................t.................a..............a....................c.....g... [120240]
MH025917.1_strain_R7            ................t.................a..............a....................c.....g... [120240]
MH025916.1_strain_R8            ................t.................a..............a....................c.....g... [120240]

FR682468.2_Georgia_2007/1       ccatttacatcgaatcttacgttttcataaagtcgttctccggggtattcgcagtagtaaaccaagtttcggtacgcatt [120320]
ASF_IND_20_CAD543               ................................................................................ [120320]
LR899193.1_Germany_2020/1       ................................................................................ [120320]
MN393477.1_Wuhan_2019           ................................................................................ [120320]
MW396979.1_Timor-Leste/2019/1   ................................................................................ [120320]
MK940252.1_CN/2019/InnerMongolia ................................................................................ [120320]
MK128995.1_China/2018/AnhuiXCGQ ................................................................................ [120320]
MH681419.1_ASFV/POL/2015        ................................................................................ [120320]
LS478113.1_Estonia_2014         ................................................................................ [120320]
KP843857.1_Odintsovo_02/14      ................................................................................ [120320]
FN557520.1_strain_E75           ................................................................................ [120320]
AY261366.1_Warthog              ................................................................................ [120320]
AY261365.1_Warmbaths            ................................................................................ [120320]
AY261364.1_Tengani_62           .................................................................a........t..... [120320]
AY261363.1_Pretoriuskop/96/4    ................................................................................ [120320]
AY261362.1_Mkuzi_1979           ................................................................................ [120320]
AY261361.1_Malawi_Lil-20/1_1983 ....................a..........................c................................ [120320]
AY261360.1_Kenya_1950           ....................a...........c............................................... [120320]
AM712240.1_OURT_88/3            ................................................................................ [120320]
AM712239.1_Benin_97/1           ................................................................................ [120320]
KM111295.1_Ken06.Bus            ....................a...........c............................................... [120320]
KM111294.1_Ken05/Tk1            ....................a...........c............................................... [120320]
KM262844.1_strain_L60           ................................................................................ [120320]
KM102979.1_26544/OG10_Italy     ................................................................................ [120320]
KM262845.1_strain_NHV           ................................................................................ [120320]
KX354450.1_47/Ss/2008           ................................................................................ [120320]
KP055815.1_strain_BA71          ................................................................................ [120320]
MH025920.1_strain_R35           ....................a...........c............................................... [120320]
MH025919.1_strain_N10           ....................a...........c............................................... [120320]
MH025918.1_strain_R25           ....................a...........c............................................... [120320]
MH025917.1_strain_R7            ....................a...........c............................................... [120320]
MH025916.1_strain_R8            ....................a...........c............................................... [120320]
```

FR682468.2_Georgia_2007/1 ctttgtgccgggtacaatgggtcttccaaaaggatctacaagcgtgtaaacggcgccctctaagggtgtttggttgtccc [120400]
ASF_IND_20_CAD543 ................................................................................ [120400]
LR899193.1_Germany_2020/1 ................................................................................ [120400]
MN393477.1_Wuhan_2019 ................................................................................ [120400]
MW396979.1_Timor-Leste/2019/1 ................................................................................ [120400]
MK940252.1_CN/2019/InnerMongolia ................................................................................ [120400]
MK128995.1_China/2018/AnhuiXCGQ ................................................................................ [120400]
MH681419.1_ASFV/POL/2015 ................................................................................ [120400]
LS478113.1_Estonia_2014 ................................................................................ [120400]
KP843857.1_Odintsovo_02/14 ................................................................................ [120400]
FN557520.1_strain_E75 ..................a...........g................................a................ [120400]
AY261366.1_Warthog ...............................................................a................ [120400]
AY261365.1_Warmbaths ...............................................................a................ [120400]
AY261364.1_Tengani_62 ...............................................................a................ [120400]
AY261363.1_Pretoriuskop/96/4 ...............................................................a................ [120400]
AY261362.1_Mkuzi_1979 ..................a...........g................................a................ [120400]
AY261361.1_Malawi_Lil-20/1_1983 ............g...........c...........c....................t.....a................ [120400]
AY261360.1_Kenya_1950 .....................c..c...........c.......................c..a................ [120400]
AM712240.1_OURT_88/3 ..................a...........g................................a................ [120400]
AM712239.1_Benin_97/1 ..................a...........g................................a................ [120400]
KM111295.1_Ken06.Bus ............c........c..c...........c.......................c..a................ [120400]
KM111294.1_Ken05/Tk1 .....................c..c...........c..........................a................ [120400]
KM262844.1_strain_L60 ..................a...........g................................a................ [120400]
KM102979.1_26544/OG10_Italy ..................a...........g...........t....................a................ [120400]
KM262845.1_strain_NHV ..................a...........g................................a................ [120400]
KX354450.1_47/Ss/2008 ..................a...........g................................a................ [120400]
KP055815.1_strain_BA71 ..................a...........g................................a................ [120400]
MH025920.1_strain_R35 ............c........c..c...........c.......................c..a................ [120400]
MH025919.1_strain_N10 ............c........c..c...........c.......................c..a................ [120400]
MH025918.1_strain_R25 ............c........c..c...........c.......................c..a................ [120400]
MH025917.1_strain_R7 ............c........c..c...........c.......................c..a................ [120400]
MH025916.1_strain_R8 ............c........c..c...........c.......................c..a................ [120400]

FR682468.2_Georgia_2007/1 agtcatatccgttgcgaggaaacgtttgaagctgcccatgggcccccatctgggacgtgccctgaatcggagcatcctgc [120480]
ASF_IND_20_CAD543 ..................................m............................................. [120480]
LR899193.1_Germany_2020/1 ................................................................................ [120480]
MN393477.1_Wuhan_2019 ................................................................................ [120480]
MW396979.1_Timor-Leste/2019/1 ................................................................................ [120480]
MK940252.1_CN/2019/InnerMongolia ................................................................................ [120480]
MK128995.1_China/2018/AnhuiXCGQ ................................................................................ [120480]
MH681419.1_ASFV/POL/2015 ................................................................................ [120480]
LS478113.1_Estonia_2014 ................................................................................ [120480]
KP843857.1_Odintsovo_02/14 ................................................................................ [120480]

| | | |
|---|---|---|
| FN557520.1_strain_E75 | ..................................a...................c......................... | [120480] |
| AY261366.1_Warthog | ................................................................................ | [120480] |
| AY261365.1_Warmbaths | .....................................g.................t........................ | [120480] |
| AY261364.1_Tengani_62 | ................................................................................ | [120480] |
| AY261363.1_Pretoriuskop/96/4 | ................................................................................ | [120480] |
| AY261362.1_Mkuzi_1979 | ..................................a...................c......................... | [120480] |
| AY261361.1_Malawi_Lil-20/1_1983 | ..................................a....................t.a...................... | [120480] |
| AY261360.1_Kenya_1950 | ..................................a....................t.a...................... | [120480] |
| AM712240.1_OURT_88/3 | ..................................a...................c......................... | [120480] |
| AM712239.1_Benin_97/1 | ..................................a...................c......................... | [120480] |
| KM111295.1_Ken06.Bus | ..................................a....................t.a...................... | [120480] |
| KM111294.1_Ken05/Tk1 | ..................................a....................t.a...................... | [120480] |
| KM262844.1_strain_L60 | ..................................a...................c......................... | [120480] |
| KM102979.1_26544/OG10_Italy | ..................................a...................c......................... | [120480] |
| KM262845.1_strain_NHV | ..................................a...................c......................... | [120480] |
| KX354450.1_47/Ss/2008 | ..................................a...................c......................... | [120480] |
| KP055815.1_strain_BA71 | ..................................a...................c......................... | [120480] |
| MH025920.1_strain_R35 | ..................................a....................t.a...................... | [120480] |
| MH025919.1_strain_N10 | ..................................a....................t.a...................... | [120480] |
| MH025918.1_strain_R25 | ..................................a....................t.a...................... | [120480] |
| MH025917.1_strain_R7 | ..................................a....................t.a...................... | [120480] |
| MH025916.1_strain_R8 | ..................................a....................t.a...................... | [120480] |

| | | |
|---|---|---|
| FR682468.2_Georgia_2007/1 | caggatgaatgacatgcacccaatatatgatggcccaccatatcatggaaaaagtctccgtactggggaataccaaaggt | [120560] |
| ASF_IND_20_CAD543 | .....y.......................................................................... | [120560] |
| LR899193.1_Germany_2020/1 | ................................................................................ | [120560] |
| MN393477.1_Wuhan_2019 | ................................................................................ | [120560] |
| MW396979.1_Timor-Leste/2019/1 | ................................................................................ | [120560] |
| MK940252.1_CN/2019/InnerMongolia | ................................................................................ | [120560] |
| MK128995.1_China/2018/AnhuiXCGQ | ................................................................................ | [120560] |
| MH681419.1_ASFV/POL/2015 | ................................................................................ | [120560] |
| LS478113.1_Estonia_2014 | ................................................................................ | [120560] |
| KP843857.1_Odintsovo_02/14 | ................................................................................ | [120560] |
| FN557520.1_strain_E75 | .....c.......................g.................................................. | [120560] |
| AY261366.1_Warthog | .....c.......................................................................... | [120560] |
| AY261365.1_Warmbaths | .....c.......................................................................... | [120560] |
| AY261364.1_Tengani_62 | .....c.......................................................................... | [120560] |
| AY261363.1_Pretoriuskop/96/4 | .....c.......................................................................... | [120560] |
| AY261362.1_Mkuzi_1979 | .....c.......................g.................................................. | [120560] |
| AY261361.1_Malawi_Lil-20/1_1983 | .....a...................c...g.....t.............................a...........a.. | [120560] |
| AY261360.1_Kenya_1950 | .....g........c..............g.....t............................................ | [120560] |
| AM712240.1_OURT_88/3 | .....c.......................g.................................................. | [120560] |
| AM712239.1_Benin_97/1 | .....c.......................g.................................................. | [120560] |
| KM111295.1_Ken06.Bus | .....g........c..g...........g.....t............................................ | [120560] |

```
KM111294.1_Ken05/Tk1             .....g...........g...........g.....t............................................ [120560]
KM262844.1_strain_L60            .....c.......................g.................................................. [120560]
KM102979.1_26544/OG10_Italy      .....c.......................g.................................................. [120560]
KM262845.1_strain_NHV            .....c.......................g.................................................. [120560]
KX354450.1_47/Ss/2008            .....c.......................g.................................................. [120560]
KP055815.1_strain_BA71           .....c.......................g.................................................. [120560]
MH025920.1_strain_R35            .....g........c..g...........g.....t............................................ [120560]
MH025919.1_strain_N10            .....g........c..g...........g.....t............................................ [120560]
MH025918.1_strain_R25            .....g........c..g...........g.....t............................................ [120560]
MH025917.1_strain_R7             .....g........c..g...........g.....t............................................ [120560]
MH025916.1_strain_R8             .....g........c..g...........g.....t............................................ [120560]

FR682468.2_Georgia_2007/1        aagcttgtttcccaaggtgggggtacccgtatgcgggcgtactttattgtattcaaaccctactggaacataaggcttaa [120640]
ASF_IND_20_CAD543                ................................................................................ [120640]
LR899193.1_Germany_2020/1        ................................................................................ [120640]
MN393477.1_Wuhan_2019            ................................................................................ [120640]
MW396979.1_Timor-Leste/2019/1    ................................................................................ [120640]
MK940252.1_CN/2019/InnerMongolia ................................................................................ [120640]
MK128995.1_China/2018/AnhuiXCGQ  ................................................................................ [120640]
MH681419.1_ASFV/POL/2015         ................................................................................ [120640]
LS478113.1_Estonia_2014          ................................................................................ [120640]
KP843857.1_Odintsovo_02/14       ................................................................................ [120640]
FN557520.1_strain_E75            ................................................................................ [120640]
AY261366.1_Warthog               ................................................................................ [120640]
AY261365.1_Warmbaths             ...............................................................c................ [120640]
AY261364.1_Tengani_62            ................................................................................ [120640]
AY261363.1_Pretoriuskop/96/4     ................................................................................ [120640]
AY261362.1_Mkuzi_1979            ................................................................................ [120640]
AY261361.1_Malawi_Lil-20/1_1983  ...t.....g.....t....................a..c......................t................. [120640]
AY261360.1_Kenya_1950            .........g.....t.................t..a.........................t...g............. [120640]
AM712240.1_OURT_88/3             ................................................................................ [120640]
AM712239.1_Benin_97/1            ................................................................................ [120640]
KM111295.1_Ken06.Bus             .........g.....t.................t..a.........................t...g..g.......... [120640]
KM111294.1_Ken05/Tk1             .........g.....t.................t..a.........................t...g............. [120640]
KM262844.1_strain_L60            ................................................................................ [120640]
KM102979.1_26544/OG10_Italy      ................................................................................ [120640]
KM262845.1_strain_NHV            ................................................................................ [120640]
KX354450.1_47/Ss/2008            ................................................................................ [120640]
KP055815.1_strain_BA71           ................................................................................ [120640]
MH025920.1_strain_R35            .........g.....t.................t..a.........................t...g..g.......... [120640]
MH025919.1_strain_N10            .........g.....t.................t..a.........................t...g..g.......... [120640]
MH025918.1_strain_R25            .........g.....t.................t..a.........................t...g..g.......... [120640]
MH025917.1_strain_R7             .........g.....t.................t..a.........................t...g..g.......... [120640]
MH025916.1_strain_R8             .........g.....t.................t..a.........................t...g..g.......... [120640]
```

```
FR682468.2_Georgia_2007/1        aatgcgcattaaaatgcaccaaatgtgtttcttcgatttgactcaaagtgggttcgggatcgggtttcccataacttttg [120720]
ASF_IND_20_CAD543                ................................................................................ [120720]
LR899193.1_Germany_2020/1        ................................................................................ [120720]
MN393477.1_Wuhan_2019            ................................................................................ [120720]
MW396979.1_Timor-Leste/2019/1    ................................................................................ [120720]
MK940252.1_CN/2019/InnerMongolia ................................................................................ [120720]
MK128995.1_China/2018/AnhuiXCGQ  ................................................................................ [120720]
MH681419.1_ASFV/POL/2015         ................................................................................ [120720]
LS478113.1_Estonia_2014          ................................................................................ [120720]
KP843857.1_Odintsovo_02/14       ................................................................................ [120720]
FN557520.1_strain_E75            ................a.........................................g..................... [120720]
AY261366.1_Warthog               ..........................................................g..................... [120720]
AY261365.1_Warmbaths             ..........................................................g..................... [120720]
AY261364.1_Tengani_62            ..........................................................g...........g......... [120720]
AY261363.1_Pretoriuskop/96/4     ..........................................................g..a.................. [120720]
AY261362.1_Mkuzi_1979            ................................................................................ [120720]
AY261361.1_Malawi_Lil-20/1_1983  ....t................t....................................g..................... [120720]
AY261360.1_Kenya_1950            ....t..g.............t....................................g..................... [120720]
AM712240.1_OURT_88/3             ................a.........................................g..................... [120720]
AM712239.1_Benin_97/1            ................a.........................................g..................... [120720]
KM111295.1_Ken06.Bus             ....t..g.............t....................................g..................... [120720]
KM111294.1_Ken05/Tk1             ....t..g.............t....................................g..................... [120720]
KM262844.1_strain_L60            ................a.........................................g..................... [120720]
KM102979.1_26544/OG10_Italy      ................a.........................................g..................... [120720]
KM262845.1_strain_NHV            ................a.........................................g..................... [120720]
KX354450.1_47/Ss/2008            ................a.........................................g..................... [120720]
KP055815.1_strain_BA71           ................a.........................................g..................... [120720]
MH025920.1_strain_R35            ....t..g.............t....................................g..................... [120720]
MH025919.1_strain_N10            ....t..g.............t....................................g..................... [120720]
MH025918.1_strain_R25            ....t..g.............t....................................g..................... [120720]
MH025917.1_strain_R7             ....t..g.............t....................................g..................... [120720]
MH025916.1_strain_R8             ....t..g.............t....................................g..................... [120720]

FR682468.2_Georgia_2007/1        ttcacatttttaatgttagagatcctgctattcagcaagtcttgggccaatataatcttgtcggccttcccatcgttagc [120800]
ASF_IND_20_CAD543                ....................r........................................................... [120800]
LR899193.1_Germany_2020/1        ................................................................................ [120800]
MN393477.1_Wuhan_2019            ................................................................................ [120800]
MW396979.1_Timor-Leste/2019/1    ................................................................................ [120800]
MK940252.1_CN/2019/InnerMongolia ................................................................................ [120800]
MK128995.1_China/2018/AnhuiXCGQ  ................................................................................ [120800]
MH681419.1_ASFV/POL/2015         ................................................................................ [120800]
LS478113.1_Estonia_2014          ................................................................................ [120800]
KP843857.1_Odintsovo_02/14       ................................................................................ [120800]
```

```
FN557520.1_strain_E75            ....................a...........a............................................... [120800]
AY261366.1_Warthog               ................................................................................ [120800]
AY261365.1_Warmbaths             ................................................................................ [120800]
AY261364.1_Tengani_62            ...........................................................a.................... [120800]
AY261363.1_Pretoriuskop/96/4     ................................................................................ [120800]
AY261362.1_Mkuzi_1979            ....................a...........a............................................... [120800]
AY261361.1_Malawi_Lil-20/1_1983  ..a.............................a.................g............................. [120800]
AY261360.1_Kenya_1950            ..a.............................a.................a..g.......................... [120800]
AM712240.1_OURT_88/3             ....................a...........a............................................... [120800]
AM712239.1_Benin_97/1            ....................a...........a............................................... [120800]
KM111295.1_Ken06.Bus             ..a.............................a.................a..g.......................... [120800]
KM111294.1_Ken05/Tk1             ..a.............................a.................a..g.......................... [120800]
KM262844.1_strain_L60            ....................a...........a............................................... [120800]
KM102979.1_26544/OG10_Italy      ....................a...........a............................................... [120800]
KM262845.1_strain_NHV            ....................a...........a............................................... [120800]
KX354450.1_47/Ss/2008            ....................a...........a............................................... [120800]
KP055815.1_strain_BA71           ....................a...........a............................................... [120800]
MH025920.1_strain_R35            ..a.............................a.................a..g.......................... [120800]
MH025919.1_strain_N10            ..a.............................a.................a..g.......................... [120800]
MH025918.1_strain_R25            ..a.............................a.................a..g.......................... [120800]
MH025917.1_strain_R7             ..a.............................a.................a..g.......................... [120800]
MH025916.1_strain_R8             ..a.............................a.................a..g.......................... [120800]

FR682468.2_Georgia_2007/1        aataagacaaaaagctcctcctgatgccatatataatgttataaaaataatttattgtttttattaaatatggcggttta [120880]
ASF_IND_20_CAD543                ................................................................................ [120880]
LR899193.1_Germany_2020/1        ................................................................................ [120880]
MN393477.1_Wuhan_2019            ................................................................................ [120880]
MW396979.1_Timor-Leste/2019/1    ................................................................................ [120880]
MK940252.1_CN/2019/InnerMongolia ................................................................................ [120880]
MK128995.1_China/2018/AnhuiXCGQ  ................................................................................ [120880]
MH681419.1_ASFV/POL/2015         ................................................................................ [120880]
LS478113.1_Estonia_2014          ................................................................................ [120880]
KP843857.1_Odintsovo_02/14       ................................................................................ [120880]
FN557520.1_strain_E75            ................................................................................ [120880]
AY261366.1_Warthog               ................................................................................ [120880]
AY261365.1_Warmbaths             ................................................................................ [120880]
AY261364.1_Tengani_62            .............................................................................g.. [120880]
AY261363.1_Pretoriuskop/96/4     ................................................................................ [120880]
AY261362.1_Mkuzi_1979            ................................................................................ [120880]
AY261361.1_Malawi_Lil-20/1_1983  ..........................................................................a..... [120880]
AY261360.1_Kenya_1950            ................................................................................ [120880]
AM712240.1_OURT_88/3             ................................................................................ [120880]
AM712239.1_Benin_97/1            ................................................................................ [120880]
KM111295.1_Ken06.Bus             ................................................................................ [120880]
```

```
KM111294.1_Ken05/Tk1            ................................................................................ [120880]
KM262844.1_strain_L60           ................................................................................ [120880]
KM102979.1_26544/OG10_Italy     ................................................................................ [120880]
KM262845.1_strain_NHV           ................................................................................ [120880]
KX354450.1_47/Ss/2008           ................................................................................ [120880]
KP055815.1_strain_BA71          ................................................................................ [120880]
MH025920.1_strain_R35           ................................................................................ [120880]
MH025919.1_strain_N10           ................................................................................ [120880]
MH025918.1_strain_R25           ................................................................................ [120880]
MH025917.1_strain_R7            ................................................................................ [120880]
MH025916.1_strain_R8            ................................................................................ [120880]

FR682468.2_Georgia_2007/1       tgcgaaggatcttgataataacaaagagttaaaccaaaaattaattaacgatcagcttaaaattattgacacgctcttgc [120960]
ASF_IND_20_CAD543               ................................................................................ [120960]
LR899193.1_Germany_2020/1       ................................................................................ [120960]
MN393477.1_Wuhan_2019           ................................................................................ [120960]
MW396979.1_Timor-Leste/2019/1   ................................................................................ [120960]
MK940252.1_CN/2019/InnerMongolia ................................................................................ [120960]
MK128995.1_China/2018/AnhuiXCGQ ................................................................................ [120960]
MH681419.1_ASFV/POL/2015        ................................................................................ [120960]
LS478113.1_Estonia_2014         ................................................................................ [120960]
KP843857.1_Odintsovo_02/14      ................................................................................ [120960]
FN557520.1_strain_E75           ................................................................................ [120960]
AY261366.1_Warthog              .........c...................................................................... [120960]
AY261365.1_Warmbaths            ................................................................................ [120960]
AY261364.1_Tengani_62           .........c...................................................................... [120960]
AY261363.1_Pretoriuskop/96/4    ................................................................................ [120960]
AY261362.1_Mkuzi_1979           ................................................................................ [120960]
AY261361.1_Malawi_Lil-20/1_1983 ......a...............................................a....................g.... [120960]
AY261360.1_Kenya_1950           .........c................................g..c.................c...........t.... [120960]
AM712240.1_OURT_88/3            ................................................................................ [120960]
AM712239.1_Benin_97/1           ................................................................................ [120960]
KM111295.1_Ken06.Bus            .........c................................g..c.............................t.... [120960]
KM111294.1_Ken05/Tk1            .........c................................g..c.................c...........t.... [120960]
KM262844.1_strain_L60           ................................................................................ [120960]
KM102979.1_26544/OG10_Italy     ................................................................................ [120960]
KM262845.1_strain_NHV           ................................................................................ [120960]
KX354450.1_47/Ss/2008           ................................................................................ [120960]
KP055815.1_strain_BA71          ................................................................................ [120960]
MH025920.1_strain_R35           .........c................................g..c.............................t.... [120960]
MH025919.1_strain_N10           .........c................................g..c.............................t.... [120960]
MH025918.1_strain_R25           .........c................................g..c.............................t.... [120960]
MH025917.1_strain_R7            .........c................................g..c.............................t.... [120960]
MH025916.1_strain_R8            .........c................................g..c.............................t.... [120960]
```

```
FR682468.2_Georgia_2007/1        tggcagaaaaaaaaaaacttttggtgtatgaactacctgccccttttgacttttcctccggcgaccctttggccagtcag [121040]
ASF_IND_20_CAD543                .r.............................................................................. [121040]
LR899193.1_Germany_2020/1        ................................................................................ [121040]
MN393477.1_Wuhan_2019            ................................................................................ [121040]
MW396979.1_Timor-Leste/2019/1    ................................................................................ [121040]
MK940252.1_CN/2019/InnerMongolia ................................................................................ [121040]
MK128995.1_China/2018/AnhuiXCGQ  ................................................................................ [121040]
MH681419.1_ASFV/POL/2015         ................................................................................ [121040]
LS478113.1_Estonia_2014          ................................................................................ [121040]
KP843857.1_Odintsovo_02/14       ......-......................................................................... [121040]
FN557520.1_strain_E75            .a..............................t.g............................................. [121040]
AY261366.1_Warthog               ................................................................................ [121040]
AY261365.1_Warmbaths             ................................................................................ [121040]
AY261364.1_Tengani_62            ................................................................................ [121040]
AY261363.1_Pretoriuskop/96/4     ................................................................................ [121040]
AY261362.1_Mkuzi_1979            .a..............................t.g............................................. [121040]
AY261361.1_Malawi_Lil-20/1_1983  ..........................c...............ac.a.................................. [121040]
AY261360.1_Kenya_1950            .a..........................c.............a.............................g....... [121040]
AM712240.1_OURT_88/3             .a..............................t.g............................................. [121040]
AM712239.1_Benin_97/1            .a..............................t.g............................................. [121040]
KM111295.1_Ken06.Bus             ............................c.............a.............................g....... [121040]
KM111294.1_Ken05/Tk1             .a..........................c.............a.............................g....... [121040]
KM262844.1_strain_L60            .a..............................t.g............................................. [121040]
KM102979.1_26544/OG10_Italy      .a..............................t.g............................................. [121040]
KM262845.1_strain_NHV            .a..............................t.g............................................. [121040]
KX354450.1_47/Ss/2008            .a..............................t.g............................................. [121040]
KP055815.1_strain_BA71           .a..............................t.g............................................. [121040]
MH025920.1_strain_R35            ............................c.............a.............................g....... [121040]
MH025919.1_strain_N10            ............................c.............a.............................g....... [121040]
MH025918.1_strain_R25            ............................c.............a.............................g....... [121040]
MH025917.1_strain_R7             ............................c.............a.............................g....... [121040]
MH025916.1_strain_R8             ............................c.............a.............................g....... [121040]

FR682468.2_Georgia_2007/1        cgcgacatatactatgccatcataaaaagcctcgaggagcgcgggtttactgtcaaaatatgtatgaaaggggatcgtgc [121120]
ASF_IND_20_CAD543                ................................................................................ [121120]
LR899193.1_Germany_2020/1        ................................................................................ [121120]
MN393477.1_Wuhan_2019            ................................................................................ [121120]
MW396979.1_Timor-Leste/2019/1    ................................................................................ [121120]
MK940252.1_CN/2019/InnerMongolia ................................................................................ [121120]
MK128995.1_China/2018/AnhuiXCGQ  ................................................................................ [121120]
MH681419.1_ASFV/POL/2015         ................................................................................ [121120]
LS478113.1_Estonia_2014          ................................................................................ [121120]
KP843857.1_Odintsovo_02/14       ................................................................................ [121120]
```

```
FN557520.1_strain_E75              ................................................................................ [121120]
AY261366.1_Warthog                 .............................t...............................................c.. [121120]
AY261365.1_Warmbaths               .............................t...............................................c.. [121120]
AY261364.1_Tengani_62              ..t..........................t..t............................................... [121120]
AY261363.1_Pretoriuskop/96/4       .............................t...............................................c.. [121120]
AY261362.1_Mkuzi_1979              ................................................................................ [121120]
AY261361.1_Malawi_Lil-20/1_1983    .............................t..............a..............t...........a.....c.. [121120]
AY261360.1_Kenya_1950              ........t....................t..............a.......................g..a..c..c.. [121120]
AM712240.1_OURT_88/3               ................................................................................ [121120]
AM712239.1_Benin_97/1              ................................................................................ [121120]
KM111295.1_Ken06.Bus               ........t....................t..............a.......................g..a..c..c.. [121120]
KM111294.1_Ken05/Tk1               ........t....................t..............a.......................g..a..c..c.. [121120]
KM262844.1_strain_L60              ................................................................................ [121120]
KM102979.1_26544/OG10_Italy        ................................................................................ [121120]
KM262845.1_strain_NHV              ................................................................................ [121120]
KX354450.1_47/Ss/2008              ................................................................................ [121120]
KP055815.1_strain_BA71             ................................................................................ [121120]
MH025920.1_strain_R35              ........t....................t..............a.......................g..a..c..c.. [121120]
MH025919.1_strain_N10              ........t....................t..............a.......................g..a..c..c.. [121120]
MH025918.1_strain_R25              ........t....................t..............a.......................g..a..c..c.. [121120]
MH025917.1_strain_R7               ........t....................t..............a.......................g..a..c..c.. [121120]
MH025916.1_strain_R8               ........t....................t..............a.......................g..a..c..c.. [121120]

FR682468.2_Georgia_2007/1          cctccttttcatcacctggaaaaaaatacaatccattgagataaacaaaaaagaagaatatctgcgcatgcacttcatac [121200]
ASF_IND_20_CAD543                  ...............y................................................................ [121200]
LR899193.1_Germany_2020/1          ................................................................................ [121200]
MN393477.1_Wuhan_2019              ................................................................................ [121200]
MW396979.1_Timor-Leste/2019/1      ................................................................................ [121200]
MK940252.1_CN/2019/InnerMongolia   ................................................................................ [121200]
MK128995.1_China/2018/AnhuiXCGQ    ................................................................................ [121200]
MH681419.1_ASFV/POL/2015           ................................................................................ [121200]
LS478113.1_Estonia_2014            ................................................................................ [121200]
KP843857.1_Odintsovo_02/14         ................................................................................ [121200]
FN557520.1_strain_E75              ...............t................................................................ [121200]
AY261366.1_Warthog                 ................................................................................ [121200]
AY261365.1_Warmbaths               ................................................................................ [121200]
AY261364.1_Tengani_62              .......................................a........................................ [121200]
AY261363.1_Pretoriuskop/96/4       ................................................................................ [121200]
AY261362.1_Mkuzi_1979              ...............t................................................................ [121200]
AY261361.1_Malawi_Lil-20/1_1983    ...t............................................................................ [121200]
AY261360.1_Kenya_1950              ...t......................................c..................................... [121200]
AM712240.1_OURT_88/3               ...............t................................................................ [121200]
AM712239.1_Benin_97/1              ...............t................................................................ [121200]
KM111295.1_Ken06.Bus               ...t......................................c..................................... [121200]
```

```
KM111294.1_Ken05/Tk1             ...t......................................c..................................... [121200]
KM262844.1_strain_L60            ...............t................................................................ [121200]
KM102979.1_26544/OG10_Italy      ...............t................................................................ [121200]
KM262845.1_strain_NHV            ...............t................................................................ [121200]
KX354450.1_47/Ss/2008            ...............t................................................................ [121200]
KP055815.1_strain_BA71           ...............t................................................................ [121200]
MH025920.1_strain_R35            ...t......................................c..................................... [121200]
MH025919.1_strain_N10            ...t......................................c..................................... [121200]
MH025918.1_strain_R25            ...t......................................c..................................... [121200]
MH025917.1_strain_R7             ...t......................................c..................................... [121200]
MH025916.1_strain_R8             ...t......................................c..................................... [121200]

FR682468.2_Georgia_2007/1        aagacgaagagaaagcattttattgtaaatttttagagtctagatgagcttttacgcaatgttgtacagtgttgtatata [121280]
ASF_IND_20_CAD543                ................................................................................ [121280]
LR899193.1_Germany_2020/1        ................................................................................ [121280]
MN393477.1_Wuhan_2019            ................................................................................ [121280]
MW396979.1_Timor-Leste/2019/1    ................................................................................ [121280]
MK940252.1_CN/2019/InnerMongolia ................................................................................ [121280]
MK128995.1_China/2018/AnhuiXCGQ  ................................................................................ [121280]
MH681419.1_ASFV/POL/2015         ................................................................................ [121280]
LS478113.1_Estonia_2014          ................................................................................ [121280]
KP843857.1_Odintsovo_02/14       ................................................................................ [121280]
FN557520.1_strain_E75            ................................................................................ [121280]
AY261366.1_Warthog               ................................................................................ [121280]
AY261365.1_Warmbaths             ................................................................................ [121280]
AY261364.1_Tengani_62            ..............t................................................................. [121280]
AY261363.1_Pretoriuskop/96/4     ................................................................................ [121280]
AY261362.1_Mkuzi_1979            ................................................................................ [121280]
AY261361.1_Malawi_Lil-20/1_1983  ....t...........................................t.........t...........a..a...... [121280]
AY261360.1_Kenya_1950            ................g.....................................................a..a...... [121280]
AM712240.1_OURT_88/3             ................................................................................ [121280]
AM712239.1_Benin_97/1            ................................................................................ [121280]
KM111295.1_Ken06.Bus             ................g.......................................t.............a..a...... [121280]
KM111294.1_Ken05/Tk1             ................g.....................................................a..a...... [121280]
KM262844.1_strain_L60            ................................................................................ [121280]
KM102979.1_26544/OG10_Italy      ................................................................................ [121280]
KM262845.1_strain_NHV            ................................................................................ [121280]
KX354450.1_47/Ss/2008            ................................................................................ [121280]
KP055815.1_strain_BA71           ................................................................................ [121280]
MH025920.1_strain_R35            ................g.....................................................a..a...... [121280]
MH025919.1_strain_N10            ................g.....................................................a..a...... [121280]
MH025918.1_strain_R25            ................g.....................................................a..a...... [121280]
MH025917.1_strain_R7             ................g.....................................................a..a...... [121280]
MH025916.1_strain_R8             ................g.....................................................a..a...... [121280]
```

```
FR682468.2_Georgia_2007/1        tgtcttgtaagcatttgttgtagagtaataagtaaaagataaataaaaatgactattaaaataaagcccaaaccattaaa [121360]
ASF_IND_20_CAD543                ................................................................................ [121360]
LR899193.1_Germany_2020/1        ................................................................................ [121360]
MN393477.1_Wuhan_2019            ................................................................................ [121360]
MW396979.1_Timor-Leste/2019/1    ................................................................................ [121360]
MK940252.1_CN/2019/InnerMongolia ................................................................................ [121360]
MK128995.1_China/2018/AnhuiXCGQ  ................................................................................ [121360]
MH681419.1_ASFV/POL/2015         ................................................................................ [121360]
LS478113.1_Estonia_2014          ................................................................................ [121360]
KP843857.1_Odintsovo_02/14       ................................................................................ [121360]
FN557520.1_strain_E75            ................................................................................ [121360]
AY261366.1_Warthog               ................................................................................ [121360]
AY261365.1_Warmbaths             ................................................................................ [121360]
AY261364.1_Tengani_62            ................................................................................ [121360]
AY261363.1_Pretoriuskop/96/4     ................................................................................ [121360]
AY261362.1_Mkuzi_1979            ................................................................................ [121360]
AY261361.1_Malawi_Lil-20/1_1983  ...............................t..................a............................. [121360]
AY261360.1_Kenya_1950            ......................a.............................................t........... [121360]
AM712240.1_OURT_88/3             ................................................................................ [121360]
AM712239.1_Benin_97/1            ................................................................................ [121360]
KM111295.1_Ken06.Bus             ......................a......................................................... [121360]
KM111294.1_Ken05/Tk1             ......................a...........................a............................. [121360]
KM262844.1_strain_L60            ................................................................................ [121360]
KM102979.1_26544/OG10_Italy      ................................................................................ [121360]
KM262845.1_strain_NHV            ................................................................................ [121360]
KX354450.1_47/Ss/2008            ................................................................................ [121360]
KP055815.1_strain_BA71           ................................................................................ [121360]
MH025920.1_strain_R35            ......................a......................................................... [121360]
MH025919.1_strain_N10            ......................a......................................................... [121360]
MH025918.1_strain_R25            ......................a......................................................... [121360]
MH025917.1_strain_R7             ......................a......................................................... [121360]
MH025916.1_strain_R8             ......................a......................................................... [121360]

FR682468.2_Georgia_2007/1        aatatttttatctgttagatttaatttaataaatggctcatggaatgtgtggtgcgccgctgcatgaggtg--------- [121440]
ASF_IND_20_CAD543                .......................................................................cggcc---- [121440]
LR899193.1_Germany_2020/1        .......................................................................--------- [121440]
MN393477.1_Wuhan_2019            .......................................................................--------- [121440]
MW396979.1_Timor-Leste/2019/1    .......................................................................--------- [121440]
MK940252.1_CN/2019/InnerMongolia .......................................................................--------- [121440]
MK128995.1_China/2018/AnhuiXCGQ  .......................................................................--------- [121440]
MH681419.1_ASFV/POL/2015         .......................................................................--------- [121440]
LS478113.1_Estonia_2014          .......................................................................--------- [121440]
KP843857.1_Odintsovo_02/14       .......................................................................--------- [121440]
```

```
FN557520.1_strain_E75             ....-..................................................................cggcc---- [121440]
AY261366.1_Warthog                ...............................................c...a.........a.........cggcc---- [121440]
AY261365.1_Warmbaths              ...............................................c...a...................cggct---- [121440]
AY261364.1_Tengani_62             ....................................t..............a...................cggccgcat [121440]
AY261363.1_Pretoriuskop/96/4      ...............................................c...a...................cggcc---- [121440]
AY261362.1_Mkuzi_1979             ...............................................c.......................--------- [121440]
AY261361.1_Malawi_Lil-20/1_1983   ...............................................c...a...a......t...g....cggccgcat [121440]
AY261360.1_Kenya_1950             .................................c.............c...a..g.t.....t..tgaa..ccgccgcat [121440]
AM712240.1_OURT_88/3              .......................................................................cggcc---- [121440]
AM712239.1_Benin_97/1             .......................................................................cggcc---- [121440]
KM111295.1_Ken06.Bus              .................................c.............c...a..g.......t..tgaa..ccgccgcat [121440]
KM111294.1_Ken05/Tk1              g................................c.............c...a..g.t.....t..tgaa..ccgccgcat [121440]
KM262844.1_strain_L60             .......................................................................cggcc---- [121440]
KM102979.1_26544/OG10_Italy       ..................................................a....................cggcc---- [121440]
KM262845.1_strain_NHV             .......................................................................cggcc---- [121440]
KX354450.1_47/Ss/2008             ..................................................a....................cggcc---- [121440]
KP055815.1_strain_BA71            .......................................................................cggcc---- [121440]
MH025920.1_strain_R35             .................................c.............c...a..g.......t..tgaa..ccgccgcat [121440]
MH025919.1_strain_N10             .................................c.............c...a..g.......t..tgaa..ccgccgcat [121440]
MH025918.1_strain_R25             .................................c.............c...a..g.......t..tgaa..ccgccgcat [121440]
MH025917.1_strain_R7              .................................c.............c...a..g.......t..tgaa..ccgccgcat [121440]
MH025916.1_strain_R8              .................................c.............c...a..g.......t..tgaa..ccgccgcat [121440]

FR682468.2_Georgia_2007/1         ---------------------tggccgcatgggatgtggtcgcataagatgtagctac---------------------- [121520]
ASF_IND_20_CAD543                 --------------------ac....................................---------------------- [121520]
LR899193.1_Germany_2020/1         ---------------------.....................................---------------------- [121520]
MN393477.1_Wuhan_2019             ---------------------.....................................---------------------- [121520]
MW396979.1_Timor-Leste/2019/1     ---------------------.....................................---------------------- [121520]
MK940252.1_CN/2019/InnerMongolia  ---------------------.....................................---------------------- [121520]
MK128995.1_China/2018/AnhuiXCGQ   ---------------------.....................................---------------------- [121520]
MH681419.1_ASFV/POL/2015          ---------------------.....................................---------------------- [121520]
LS478113.1_Estonia_2014           ---------------------.....................................---------------------- [121520]
KP843857.1_Odintsovo_02/14        ---------------------.....................................---------------------- [121520]
FN557520.1_strain_E75             --------------------ac....................................---------------------- [121520]
AY261366.1_Warthog                --------------------a....---------------....cgg...acg..cgt---------------------- [121520]
AY261365.1_Warmbaths              --------------------a.......gc.t...ac.......cgg...acg..cgt---------------------- [121520]
AY261364.1_Tengani_62             gggatacggc---------cg..............aca.c....cgg...ac...cg.---------------------- [121520]
AY261363.1_Pretoriuskop/96/4      --------------------a....---------------....cgg...acg..cgt---------------------- [121520]
AY261362.1_Mkuzi_1979             ---------------------c....................................---------------------- [121520]
AY261361.1_Malawi_Lil-20/1_1983   gagac---------------gcc........a..c.c..ct....gc....cc..cg.---------------------- [121520]
AY261360.1_Kenya_1950             gaggggctgctgcattgaatgcc........a.gg.cc.ct....tga...cc..cg.atgaggggccgctgcattgaat [121520]
AM712240.1_OURT_88/3              --------------------ac....................................---------------------- [121520]
AM712239.1_Benin_97/1             --------------------ac....................................---------------------- [121520]
KM111295.1_Ken06.Bus              gaggggctgctgcattgaatgcc..t.t...a.gg.cc..t....tga...cc..cg.atgggg---------------- [121520]
```

```
KM111294.1_Ken05/Tk1             gaggtgctgc---------------------------------------------------------------------- [121520]
KM262844.1_strain_L60            --------------------ac....................................---------------------- [121520]
KM102979.1_26544/OG10_Italy      --------------------ac....................................---------------------- [121520]
KM262845.1_strain_NHV            --------------------ac....................................---------------------- [121520]
KX354450.1_47/Ss/2008            --------------------ac....................................---------------------- [121520]
KP055815.1_strain_BA71           --------------------ac....................................---------------------- [121520]
MH025920.1_strain_R35            gaggggctgctgcattgaatgcc..t.t...a.gg.cc..t....tga...cc..cg.atgggg---------------- [121520]
MH025919.1_strain_N10            gaggggctgctgcattgaatgcc..t.t...a.gg.cc..t....tga...cc..cg.atgggg---------------- [121520]
MH025918.1_strain_R25            gaggggctgctgcattgaatgcc..t.t...a.gg.cc..t....tga...cc..cg.atgggg---------------- [121520]
MH025917.1_strain_R7             gaggggctgctgcattgaatgcc..t.t...a.gg.cc..t....tga...cc..cg.atgggg---------------- [121520]
MH025916.1_strain_R8             gaggggctgctgcattgaatgcc..t.t...a.gg.cc..t....tga...cc..cg.atgggg---------------- [121520]

FR682468.2_Georgia_2007/1        --------------------atgggatgtggc---atttgcttgcatgtaaggatcatgatgtgttgggtcttcatccca [121600]
ASF_IND_20_CAD543                --------------------............---............................................. [121600]
LR899193.1_Germany_2020/1        --------------------............---............................................. [121600]
MN393477.1_Wuhan_2019            --------------------............---............................................. [121600]
MW396979.1_Timor-Leste/2019/1    --------------------............---............................................. [121600]
MK940252.1_CN/2019/InnerMongolia --------------------............---............................................. [121600]
MK128995.1_China/2018/AnhuiXCGQ  --------------------............---............................................. [121600]
MH681419.1_ASFV/POL/2015         --------------------............---............................................. [121600]
LS478113.1_Estonia_2014          --------------------............---............................................. [121600]
KP843857.1_Odintsovo_02/14       --------------------............---............................................. [121600]
FN557520.1_strain_E75            --------------------............---.....................g....................... [121600]
AY261366.1_Warthog               --------------------...........t---.....................g....................... [121600]
AY261365.1_Warmbaths             --------------------............---..........g..........g........c.............. [121600]
AY261364.1_Tengani_62            --------------------...t..aac...cgt.....................g....................... [121600]
AY261363.1_Pretoriuskop/96/4     --------------------............---..........g..........g....................... [121600]
AY261362.1_Mkuzi_1979            --------------------............---.....................g....................... [121600]
AY261361.1_Malawi_Lil-20/1_1983  --------------------............---.......c....a..............g.....a........... [121600]
AY261360.1_Kenya_1950            gccgccgcatgaggtgctgc...aa...ca..cgc.......c.............g...------.a...a........ [121600]
AM712240.1_OURT_88/3             --------------------............---.....................g....................... [121600]
AM712239.1_Benin_97/1            --------------------............---.....................g....................... [121600]
KM111295.1_Ken06.Bus             -----------------------------c..ggc.......c.............g.....g....a...a........ [121600]
KM111294.1_Ken05/Tk1             --------------------...aa...ca..cgc.......c.............g.....g....a...a........ [121600]
KM262844.1_strain_L60            --------------------............---.....................g....................... [121600]
KM102979.1_26544/OG10_Italy      --------------------............---.....................g....................... [121600]
KM262845.1_strain_NHV            --------------------............---.....................g....................... [121600]
KX354450.1_47/Ss/2008            --------------------............---.....................g....................... [121600]
KP055815.1_strain_BA71           --------------------............---.....................g....................... [121600]
MH025920.1_strain_R35            -----------------------------c..ggc.......c.............g.....g....a...a........ [121600]
MH025919.1_strain_N10            -----------------------------c..ggc.......c.............g.....g....a...a........ [121600]
MH025918.1_strain_R25            -----------------------------c..ggc.......c.............g.....g....a...a........ [121600]
MH025917.1_strain_R7             -----------------------------c..ggc.......c.............g.....g....a...a........ [121600]
MH025916.1_strain_R8             -----------------------------c..ggc.......c.............g.....g....a...a........ [121600]
```

```
FR682468.2_Georgia_2007/1        gcaataatcgccatctttatctagctgaattgtatacccca-ttatatatcacttattatttttttttaatgtttcatga [121680]
ASF_IND_20_CAD543                .........................................-...................................... [121680]
LR899193.1_Germany_2020/1        .........................................-...................................... [121680]
MN393477.1_Wuhan_2019            .........................................-...................................... [121680]
MW396979.1_Timor-Leste/2019/1    .........................................-...................................... [121680]
MK940252.1_CN/2019/InnerMongolia .........................................-...................................... [121680]
MK128995.1_China/2018/AnhuiXCGQ  .........................................-...................................... [121680]
MH681419.1_ASFV/POL/2015         .........................................-...................................... [121680]
LS478113.1_Estonia_2014          .........................................-...................................... [121680]
KP843857.1_Odintsovo_02/14       .........................................-...................................... [121680]
FN557520.1_strain_E75            .........................................-...................................... [121680]
AY261366.1_Warthog               ........................t................-...................................... [121680]
AY261365.1_Warmbaths             ........................t................-...................................... [121680]
AY261364.1_Tengani_62            .........................................-...................................... [121680]
AY261363.1_Pretoriuskop/96/4     ........................t................-...................................... [121680]
AY261362.1_Mkuzi_1979            .........................................-...................................... [121680]
AY261361.1_Malawi_Lil-20/1_1983  .t.......................................-...................................... [121680]
AY261360.1_Kenya_1950            .t.......................................t...................................... [121680]
AM712240.1_OURT_88/3             .........................................-...................................... [121680]
AM712239.1_Benin_97/1            .........................................-...................................... [121680]
KM111295.1_Ken06.Bus             .t.......................................t...................................... [121680]
KM111294.1_Ken05/Tk1             .t.......a...............................t...................................... [121680]
KM262844.1_strain_L60            .........................................-...................................... [121680]
KM102979.1_26544/OG10_Italy      .........................................-...................................... [121680]
KM262845.1_strain_NHV            .........................................-...................................... [121680]
KX354450.1_47/Ss/2008            .........................................-...................................... [121680]
KP055815.1_strain_BA71           .........................................-...................................... [121680]
MH025920.1_strain_R35            .t.......................................t...................................... [121680]
MH025919.1_strain_N10            .t.......................................t...................................... [121680]
MH025918.1_strain_R25            .t.......................................t...................................... [121680]
MH025917.1_strain_R7             .t.......................................t...................................... [121680]
MH025916.1_strain_R8             .t.......................................t...................................... [121680]

FR682468.2_Georgia_2007/1        atttcattataggcggtgaaagggtcctcaggccccttctgtaaaagattatagagatcttcggacgctttatgtttcgt [121760]
ASF_IND_20_CAD543                ................................................................................ [121760]
LR899193.1_Germany_2020/1        ................................................................................ [121760]
MN393477.1_Wuhan_2019            ................................................................................ [121760]
MW396979.1_Timor-Leste/2019/1    ................................................................................ [121760]
MK940252.1_CN/2019/InnerMongolia ................................................................................ [121760]
MK128995.1_China/2018/AnhuiXCGQ  ................................................................................ [121760]
MH681419.1_ASFV/POL/2015         ................................................................................ [121760]
LS478113.1_Estonia_2014          ................................................................................ [121760]
KP843857.1_Odintsovo_02/14       ................................................................................ [121760]
```

```
FN557520.1_strain_E75          ................................................................................ [121760]
AY261366.1_Warthog             .................................................................t.............. [121760]
AY261365.1_Warmbaths           .................................................................t.............. [121760]
AY261364.1_Tengani_62          .................................................................t.............. [121760]
AY261363.1_Pretoriuskop/96/4   .................................................................t.............. [121760]
AY261362.1_Mkuzi_1979          ................................t............................................... [121760]
AY261361.1_Malawi_Lil-20/1_1983 .......................a......................................a..t...........t.. [121760]
AY261360.1_Kenya_1950          .......................a.............................c.....c..a..t.............. [121760]
AM712240.1_OURT_88/3           ................................................................................ [121760]
AM712239.1_Benin_97/1          ................................................................................ [121760]
KM111295.1_Ken06.Bus           .......................a.............................c.....c..a..t.............. [121760]
KM111294.1_Ken05/Tk1           .......................a.............................c.....c..a..t.............. [121760]
KM262844.1_strain_L60          ................................................................................ [121760]
KM102979.1_26544/OG10_Italy    ................................................................................ [121760]
KM262845.1_strain_NHV          ................................................................................ [121760]
KX354450.1_47/Ss/2008          ................................................................................ [121760]
KP055815.1_strain_BA71         ................................................................................ [121760]
MH025920.1_strain_R35          .......................a.............................c.....c..a..t.............. [121760]
MH025919.1_strain_N10          .......................a.............................c.....c..a..t.............. [121760]
MH025918.1_strain_R25          .......................a.............................c.....c..a..t.............. [121760]
MH025917.1_strain_R7           .......................a.............................c.....c..a..t.............. [121760]
MH025916.1_strain_R8           .......................a.............................c.....c..a..t.............. [121760]

FR682468.2_Georgia_2007/1      gcgaattaaggcgggatataacaaaagagagggccccagttccaaacaaattttacttagcgggctcatattttgcacca [121840]
ASF_IND_20_CAD543              ................................................................................ [121840]
LR899193.1_Germany_2020/1      ................................................................................ [121840]
MN393477.1_Wuhan_2019          ................................................................................ [121840]
MW396979.1_Timor-Leste/2019/1  ................................................................................ [121840]
MK940252.1_CN/2019/InnerMongolia ................................................................................ [121840]
MK128995.1_China/2018/AnhuiXCGQ ................................................................................ [121840]
MH681419.1_ASFV/POL/2015       ................................................................................ [121840]
LS478113.1_Estonia_2014        ................................................................................ [121840]
KP843857.1_Odintsovo_02/14     ................................................................................ [121840]
FN557520.1_strain_E75          ................................................................................ [121840]
AY261366.1_Warthog             ................................................................................ [121840]
AY261365.1_Warmbaths           ................................................................................ [121840]
AY261364.1_Tengani_62          ................................................................................ [121840]
AY261363.1_Pretoriuskop/96/4   ................................................................................ [121840]
AY261362.1_Mkuzi_1979          .....................t.......................................................... [121840]
AY261361.1_Malawi_Lil-20/1_1983 a.......................c.................t...................................a. [121840]
AY261360.1_Kenya_1950          a..............g.................t........t...................................a. [121840]
AM712240.1_OURT_88/3           ................................................................................ [121840]
AM712239.1_Benin_97/1          ................................................................................ [121840]
KM111295.1_Ken06.Bus           a..............g.....t....................t...................................a. [121840]
```

```
KM111294.1_Ken05/Tk1             a..............g.................t........t...................................a. [121840]
KM262844.1_strain_L60            ................................................................................ [121840]
KM102979.1_26544/OG10_Italy      ................................................................................ [121840]
KM262845.1_strain_NHV            ................................................................................ [121840]
KX354450.1_47/Ss/2008            ................................................................................ [121840]
KP055815.1_strain_BA71           ................................................................................ [121840]
MH025920.1_strain_R35            a..............g.....t....................t...................................a. [121840]
MH025919.1_strain_N10            a..............g.....t....................t...................................a. [121840]
MH025918.1_strain_R25            a..............g.....t....................t...................................a. [121840]
MH025917.1_strain_R7             a..............g.....t....................t...................................a. [121840]
MH025916.1_strain_R8             a..............g.....t....................t...................................a. [121840]

FR682468.2_Georgia_2007/1        agtttcccactacttgcgatgtttcataacgcattttaaagagctttatcataaaagtgttatgcaggccggtgtagtct [121920]
ASF_IND_20_CAD543                ................................................................................ [121920]
LR899193.1_Germany_2020/1        ................................................................................ [121920]
MN393477.1_Wuhan_2019            ................................................................................ [121920]
MW396979.1_Timor-Leste/2019/1    ................................................................................ [121920]
MK940252.1_CN/2019/InnerMongolia ................................................................................ [121920]
MK128995.1_China/2018/AnhuiXCGQ  ................................................................................ [121920]
MH681419.1_ASFV/POL/2015         ................................................................................ [121920]
LS478113.1_Estonia_2014          ................................................................................ [121920]
KP843857.1_Odintsovo_02/14       ................................................................................ [121920]
FN557520.1_strain_E75            .............................................................g.................. [121920]
AY261366.1_Warthog               ................................................................................ [121920]
AY261365.1_Warmbaths             ................................................................................ [121920]
AY261364.1_Tengani_62            g............................................................................... [121920]
AY261363.1_Pretoriuskop/96/4     ................................................................................ [121920]
AY261362.1_Mkuzi_1979            ....................................................g........g.................. [121920]
AY261361.1_Malawi_Lil-20/1_1983  g.........g..................................................................... [121920]
AY261360.1_Kenya_1950            ..........g.....................................................a.....a......... [121920]
AM712240.1_OURT_88/3             .............................................................g.................. [121920]
AM712239.1_Benin_97/1            .............................................................g.................. [121920]
KM111295.1_Ken06.Bus             ..........g.....................................................a............... [121920]
KM111294.1_Ken05/Tk1             ..........g.....................................................a.....a......... [121920]
KM262844.1_strain_L60            .............................................................g.................. [121920]
KM102979.1_26544/OG10_Italy      .............................................................g.................. [121920]
KM262845.1_strain_NHV            .............................................................g.................. [121920]
KX354450.1_47/Ss/2008            .............................................................g.................. [121920]
KP055815.1_strain_BA71           .............................................................g.................. [121920]
MH025920.1_strain_R35            ..........g.....................................................a............... [121920]
MH025919.1_strain_N10            ..........g.....................................................a............... [121920]
MH025918.1_strain_R25            ..........g.....................................................a............... [121920]
MH025917.1_strain_R7             ..........g.....................................................a............... [121920]
MH025916.1_strain_R8             ..........g.....................................................a............... [121920]
```

```
FR682468.2_Georgia_2007/1        ggcctatagttaaggaaggggatttctctggtaccgtcaaacacgatctcaagtcctctagcaagcccgatcaaaatttc [122000]
ASF_IND_20_CAD543                ................................................................................ [122000]
LR899193.1_Germany_2020/1        ................................................................................ [122000]
MN393477.1_Wuhan_2019            ................................................................................ [122000]
MW396979.1_Timor-Leste/2019/1    ................................................................................ [122000]
MK940252.1_CN/2019/InnerMongolia ................................................................................ [122000]
MK128995.1_China/2018/AnhuiXCGQ  ................................................................................ [122000]
MH681419.1_ASFV/POL/2015         ................................................................................ [122000]
LS478113.1_Estonia_2014          ................................................................................ [122000]
KP843857.1_Odintsovo_02/14       ................................................................................ [122000]
FN557520.1_strain_E75            .....................................................c...........t.............. [122000]
AY261366.1_Warthog               .....................................................c...........t.............. [122000]
AY261365.1_Warmbaths             ..............................................c......c...........t.............. [122000]
AY261364.1_Tengani_62            .....................................................c...........t.............. [122000]
AY261363.1_Pretoriuskop/96/4     ............................................a........c...........t.............. [122000]
AY261362.1_Mkuzi_1979            ................................................................................ [122000]
AY261361.1_Malawi_Lil-20/1_1983  ..............a......................................c...........t........t..... [122000]
AY261360.1_Kenya_1950            ..t...........a..............a.......................c...........t.............. [122000]
AM712240.1_OURT_88/3             .....................................................c...........t.............. [122000]
AM712239.1_Benin_97/1            .....................................................c...........t.............. [122000]
KM111295.1_Ken06.Bus             ..t...........a..............a..g.................g..c...........t.............. [122000]
KM111294.1_Ken05/Tk1             ..t...........a..............a.......................c...........t.............. [122000]
KM262844.1_strain_L60            .....................................................c...........t.............. [122000]
KM102979.1_26544/OG10_Italy      .....................................................c...........t.............. [122000]
KM262845.1_strain_NHV            .....................................................c...........t.............. [122000]
KX354450.1_47/Ss/2008            .....................................................c...........t.............. [122000]
KP055815.1_strain_BA71           .....................................................c...........t.............. [122000]
MH025920.1_strain_R35            ..t...........a..............a..g.................g..c...........t.............. [122000]
MH025919.1_strain_N10            ..t...........a..............a..g.................g..c...........t.............. [122000]
MH025918.1_strain_R25            ..t...........a..............a..g.................g..c...........t.............. [122000]
MH025917.1_strain_R7             ..t...........a..............a..g.................g..c...........t.............. [122000]
MH025916.1_strain_R8             ..t...........a..............a..g.................g..c...........t.............. [122000]

FR682468.2_Georgia_2007/1        ttcagcaatggatgagtatctaattcctacattacgaagcgtaagcatttctataacatcatctatttcctgcatagagg [122080]
ASF_IND_20_CAD543                ................................................................................ [122080]
LR899193.1_Germany_2020/1        ................................................................................ [122080]
MN393477.1_Wuhan_2019            ................................................................................ [122080]
MW396979.1_Timor-Leste/2019/1    ................................................................................ [122080]
MK940252.1_CN/2019/InnerMongolia ................................................................................ [122080]
MK128995.1_China/2018/AnhuiXCGQ  ................................................................................ [122080]
MH681419.1_ASFV/POL/2015         ................................................................................ [122080]
LS478113.1_Estonia_2014          ................................................................................ [122080]
KP843857.1_Odintsovo_02/14       ................................................................................ [122080]
```

```
FN557520.1_strain_E75              ................................................................................ [122080]
AY261366.1_Warthog                 ................................................................................ [122080]
AY261365.1_Warmbaths               ................................................................................ [122080]
AY261364.1_Tengani_62              ................................................................................ [122080]
AY261363.1_Pretoriuskop/96/4       ............c................................................................... [122080]
AY261362.1_Mkuzi_1979              ................................................................................ [122080]
AY261361.1_Malawi_Lil-20/1_1983    ............a..........................t...............g...................g.... [122080]
AY261360.1_Kenya_1950              .......................................t..............gg........................ [122080]
AM712240.1_OURT_88/3               ................................................................................ [122080]
AM712239.1_Benin_97/1              ................................................................................ [122080]
KM111295.1_Ken06.Bus               ............a.........................................gg........................ [122080]
KM111294.1_Ken05/Tk1               .......................................t..............gg........................ [122080]
KM262844.1_strain_L60              ................................................................................ [122080]
KM102979.1_26544/OG10_Italy        ................................................................................ [122080]
KM262845.1_strain_NHV              ................................................................................ [122080]
KX354450.1_47/Ss/2008              ................................................................................ [122080]
KP055815.1_strain_BA71             ................................................................................ [122080]
MH025920.1_strain_R35              ............a.........................................gg........................ [122080]
MH025919.1_strain_N10              ............a.........................................gg........................ [122080]
MH025918.1_strain_R25              ............a.........................................gg........................ [122080]
MH025917.1_strain_R7               ............a.........................................gg........................ [122080]
MH025916.1_strain_R8               ............a.........................................gg........................ [122080]

FR682468.2_Georgia_2007/1          aatctattgtaggaattttaatatcatctgtgctgatttgttcattcccaagataggtaagcagcatattaattttttct [122160]
ASF_IND_20_CAD543                  ................................................................................ [122160]
LR899193.1_Germany_2020/1          ................................................................................ [122160]
MN393477.1_Wuhan_2019              ................................................................................ [122160]
MW396979.1_Timor-Leste/2019/1      ................................................................................ [122160]
MK940252.1_CN/2019/InnerMongolia   ................................................................................ [122160]
MK128995.1_China/2018/AnhuiXCGQ    ................................................................................ [122160]
MH681419.1_ASFV/POL/2015           ................................................................................ [122160]
LS478113.1_Estonia_2014            ................................................................................ [122160]
KP843857.1_Odintsovo_02/14         ................................................................................ [122160]
FN557520.1_strain_E75              ................................................................................ [122160]
AY261366.1_Warthog                 ......................g......................................................... [122160]
AY261365.1_Warmbaths               ......................g......................................................... [122160]
AY261364.1_Tengani_62              ......................g.......................t................................. [122160]
AY261363.1_Pretoriuskop/96/4       ......................g......................................................... [122160]
AY261362.1_Mkuzi_1979              ................................................................................ [122160]
AY261361.1_Malawi_Lil-20/1_1983    ..........g...........gc......................tgt...g..........a..........c..... [122160]
AY261360.1_Kenya_1950              .......................c......................t................a................ [122160]
AM712240.1_OURT_88/3               ................................................................................ [122160]
AM712239.1_Benin_97/1              ................................................................................ [122160]
KM111295.1_Ken06.Bus               .......................c......................t..g.............a................ [122160]
```

```
KM111294.1_Ken05/Tk1            .......................c......................t................a................ [122160]
KM262844.1_strain_L60           ................................................................................ [122160]
KM102979.1_26544/OG10_Italy     ................................................................................ [122160]
KM262845.1_strain_NHV           ................................................................................ [122160]
KX354450.1_47/Ss/2008           ................................................................................ [122160]
KP055815.1_strain_BA71          ................................................................................ [122160]
MH025920.1_strain_R35           .......................c......................t..g.............a................ [122160]
MH025919.1_strain_N10           .......................c......................t..g.............a................ [122160]
MH025918.1_strain_R25           .......................c......................t..g.............a................ [122160]
MH025917.1_strain_R7            .......................c......................t..g.............a................ [122160]
MH025916.1_strain_R8            .......................c......................t..g.............a................ [122160]

FR682468.2_Georgia_2007/1       agctttactagcttagtcttacgctcataatcatgatcttttttataaaaagagttgggatcaccgttggaccgtagatg [122240]
ASF_IND_20_CAD543               ................................................................................ [122240]
LR899193.1_Germany_2020/1       ................................................................................ [122240]
MN393477.1_Wuhan_2019           ................................................................................ [122240]
MW396979.1_Timor-Leste/2019/1   ................................................................................ [122240]
MK940252.1_CN/2019/InnerMongolia ................................................................................ [122240]
MK128995.1_China/2018/AnhuiXCGQ ................................................................................ [122240]
MH681419.1_ASFV/POL/2015        ................................................................................ [122240]
LS478113.1_Estonia_2014         ................................................................................ [122240]
KP843857.1_Odintsovo_02/14      ................................................................................ [122240]
FN557520.1_strain_E75           ........c..........................g............................................ [122240]
AY261366.1_Warthog              ........c..........................t............................................ [122240]
AY261365.1_Warmbaths            ........c..........................t............................................ [122240]
AY261364.1_Tengani_62           ........c..........................t............................................ [122240]
AY261363.1_Pretoriuskop/96/4    ........c..........................t............................................ [122240]
AY261362.1_Mkuzi_1979           ........c....................................................................... [122240]
AY261361.1_Malawi_Lil-20/1_1983 ..t....................a...........t............................................ [122240]
AY261360.1_Kenya_1950           ..t....................a...........t............................................ [122240]
AM712240.1_OURT_88/3            ........c..........................g............................................ [122240]
AM712239.1_Benin_97/1           ........c..........................g............................................ [122240]
KM111295.1_Ken06.Bus            ..t....................a...........t..............g............................. [122240]
KM111294.1_Ken05/Tk1            ..t....................a...........t............................................ [122240]
KM262844.1_strain_L60           ........c..........................g............................................ [122240]
KM102979.1_26544/OG10_Italy     ........c..........................g............................................ [122240]
KM262845.1_strain_NHV           ........c..........................g............................................ [122240]
KX354450.1_47/Ss/2008           ........c..........................g............................................ [122240]
KP055815.1_strain_BA71          ........c..........................g............................................ [122240]
MH025920.1_strain_R35           ..t....................a...........t..............g............................. [122240]
MH025919.1_strain_N10           ..t....................a...........t..............g............................. [122240]
MH025918.1_strain_R25           ..t....................a...........t..............g............................. [122240]
MH025917.1_strain_R7            ..t....................a...........t..............g............................. [122240]
MH025916.1_strain_R8            ..t....................a...........t..............g............................. [122240]
```

```
FR682468.2_Georgia_2007/1        attaataaggcggtctacttgctttgtactaggtttaatacttttttcactatactcgctttcagcatagtggtttttac [122320]
ASF_IND_20_CAD543                ................................................................................ [122320]
LR899193.1_Germany_2020/1        ................................................................................ [122320]
MN393477.1_Wuhan_2019            ................................................................................ [122320]
MW396979.1_Timor-Leste/2019/1    ................................................................................ [122320]
MK940252.1_CN/2019/InnerMongolia ................................................................................ [122320]
MK128995.1_China/2018/AnhuiXCGQ  ................................................................................ [122320]
MH681419.1_ASFV/POL/2015         ................................................................................ [122320]
LS478113.1_Estonia_2014          ................................................................................ [122320]
KP843857.1_Odintsovo_02/14       ................................................................................ [122320]
FN557520.1_strain_E75            ...........................g.................................................... [122320]
AY261366.1_Warthog               ...........................g..................................................g. [122320]
AY261365.1_Warmbaths             ...........................g.................................................... [122320]
AY261364.1_Tengani_62            ...........................g.............................a...................... [122320]
AY261363.1_Pretoriuskop/96/4     ...........................g.................................................... [122320]
AY261362.1_Mkuzi_1979            ...........................g.................................................... [122320]
AY261361.1_Malawi_Lil-20/1_1983  ........................g..g............................................a....... [122320]
AY261360.1_Kenya_1950            g.....c.....a...........c..g............................................a.g..... [122320]
AM712240.1_OURT_88/3             ...........................g.................................................... [122320]
AM712239.1_Benin_97/1            ...........................g.................................................... [122320]
KM111295.1_Ken06.Bus             g.....c.................c..g...........................................ta.g..... [122320]
KM111294.1_Ken05/Tk1             g.....c.....a...........c..g............................................a.g..... [122320]
KM262844.1_strain_L60            ...........................g.................................................... [122320]
KM102979.1_26544/OG10_Italy      ...........................g.................................................... [122320]
KM262845.1_strain_NHV            ...........................g.................................................... [122320]
KX354450.1_47/Ss/2008            ...........................g.................................................... [122320]
KP055815.1_strain_BA71           ...........................g.................................................... [122320]
MH025920.1_strain_R35            g.....c.................c..g...........................................ta.g..... [122320]
MH025919.1_strain_N10            g.....c.................c..g...........................................ta.g..... [122320]
MH025918.1_strain_R25            g.....c.................c..g...........................................ta.g..... [122320]
MH025917.1_strain_R7             g.....c.................c..g...........................................ta.g..... [122320]
MH025916.1_strain_R8             g.....c.................c..g...........................................ta.g..... [122320]

FR682468.2_Georgia_2007/1        gatctcttttagaaatagctgttttttgagatgcctcagactctgcatatttttttctatgcgtagaaagagaataaccg [122400]
ASF_IND_20_CAD543                ...............w................................................................ [122400]
LR899193.1_Germany_2020/1        ................................................................................ [122400]
MN393477.1_Wuhan_2019            ................................................................................ [122400]
MW396979.1_Timor-Leste/2019/1    ................................................................................ [122400]
MK940252.1_CN/2019/InnerMongolia ................................................................................ [122400]
MK128995.1_China/2018/AnhuiXCGQ  ................................................................................ [122400]
MH681419.1_ASFV/POL/2015         ................................................................................ [122400]
LS478113.1_Estonia_2014          ................................................................................ [122400]
KP843857.1_Odintsovo_02/14       ................................................................................ [122400]
```

```
FN557520.1_strain_E75          ...............a.................................................................... [122400]
AY261366.1_Warthog             .............................................................t.................. [122400]
AY261365.1_Warmbaths           ................................................................................ [122400]
AY261364.1_Tengani_62          ..........g.....g...........................................................g... [122400]
AY261363.1_Pretoriuskop/96/4   ................................................................................ [122400]
AY261362.1_Mkuzi_1979          ................................................................................ [122400]
AY261361.1_Malawi_Lil-20/1_1983 .................a....................................................g......... [122400]
AY261360.1_Kenya_1950          .................a..........c..............c.................t........g.......t. [122400]
AM712240.1_OURT_88/3           ...............a................................................................ [122400]
AM712239.1_Benin_97/1          ...............a................................................................ [122400]
KM111295.1_Ken06.Bus           .................a...............t....................................g.......t. [122400]
KM111294.1_Ken05/Tk1           .................a.........................c.................t........g.......t. [122400]
KM262844.1_strain_L60          ...............a................................................................ [122400]
KM102979.1_26544/OG10_Italy    ...............a................................................................ [122400]
KM262845.1_strain_NHV          ...............a................................................................ [122400]
KX354450.1_47/Ss/2008          ...............a................................................................ [122400]
KP055815.1_strain_BA71         ...............a................................................................ [122400]
MH025920.1_strain_R35          .................a...............t....................................g.......t. [122400]
MH025919.1_strain_N10          .................a...............t....................................g.......t. [122400]
MH025918.1_strain_R25          .................a...............t....................................g.......t. [122400]
MH025917.1_strain_R7           .................a...............t....................................g.......t. [122400]
MH025916.1_strain_R8           .................a...............t....................................g.......t. [122400]

FR682468.2_Georgia_2007/1      ------cggtcattacgtgaactactgttgcatgcaaggcctcggcgcgtcttaccgctgcgcacactgccattgcgtat [122480]
ASF_IND_20_CAD543              ------...............................................................t..-------- [122480]
LR899193.1_Germany_2020/1      ------.......................................................................... [122480]
MN393477.1_Wuhan_2019          ------.......................................................................... [122480]
MW396979.1_Timor-Leste/2019/1  ------.......................................................................... [122480]
MK940252.1_CN/2019/InnerMongolia ------.......................................................................... [122480]
MK128995.1_China/2018/AnhuiXCGQ ------.......................................................................... [122480]
MH681419.1_ASFV/POL/2015       ------.......................................................................... [122480]
LS478113.1_Estonia_2014        ------.......................................................................... [122480]
KP843857.1_Odintsovo_02/14     ------.......................................................................... [122480]
FN557520.1_strain_E75          ------...............................................................t...c...c.c [122480]
AY261366.1_Warthog             ------.........................g....g..................t.................c...... [122480]
AY261365.1_Warmbaths           ------.........................g....g....................................ca..... [122480]
AY261364.1_Tengani_62          ------....................a....g.........................................c...... [122480]
AY261363.1_Pretoriuskop/96/4   ------.........................g....g..................t.................c...... [122480]
AY261362.1_Mkuzi_1979          ------...............t.......................................................... [122480]
AY261361.1_Malawi_Lil-20/1_1983 tgtatg.......g.............c...g....g..........t..tc.....................c..acgc [122480]
AY261360.1_Kenya_1950          tgtatg.....................c...g....g..........t..tc.....................ct..c.c [122480]
AM712240.1_OURT_88/3           ------...............................................................t...c...c.c [122480]
AM712239.1_Benin_97/1          ------...............................................................t...c...c.c [122480]
KM111295.1_Ken06.Bus           tgtatg.......gt............c...g....g..........t..tc.................t...ct..c.c [122480]
```

```
KM111294.1_Ken05/Tk1             tgtatg.....................c...g....g.............tc.....................ct..c.c [122480]
KM262844.1_strain_L60            ------...............................................................t...c...c.c [122480]
KM102979.1_26544/OG10_Italy      ------...............................................................t...c...c.c [122480]
KM262845.1_strain_NHV            ------...............................................................t...c...c.c [122480]
KX354450.1_47/Ss/2008            ------...............................................................t...c...c.c [122480]
KP055815.1_strain_BA71           ------...............................................................t...c...c.c [122480]
MH025920.1_strain_R35            tgtatg......gt.............c...g....g..........t..tc.................t...ct..c.c [122480]
MH025919.1_strain_N10            tgtatg......gt.............c...g....g..........t..tc.................t...ct..c.c [122480]
MH025918.1_strain_R25            tgtatg......gt.............c...g....g..........t..tc.................t...ct..c.c [122480]
MH025917.1_strain_R7             tgtatg......gt.............c...g....g..........t..tc.................t...ct..c.c [122480]
MH025916.1_strain_R8             tgtatg......gt.............c...g....g..........t..tc.................t...ct..c.c [122480]

FR682468.2_Georgia_2007/1        actgccatcgcgcacactgccgc---------------tgcgtatactgccattgcgtatactgccgctgcgtatgctgc [122560]
ASF_IND_20_CAD543                -------................---------------.......................................... [122560]
LR899193.1_Germany_2020/1        .......................---------------.......................................... [122560]
MN393477.1_Wuhan_2019            .......................---------------.......................................... [122560]
MW396979.1_Timor-Leste/2019/1    .......................---------------.......................................... [122560]
MK940252.1_CN/2019/InnerMongolia .......................---------------.......................................... [122560]
MK128995.1_China/2018/AnhuiXCGQ  .......................---------------.......................................... [122560]
MH681419.1_ASFV/POL/2015         .......................---------------.......................................... [122560]
LS478113.1_Estonia_2014          .......................---------------.......................................... [122560]
KP843857.1_Odintsovo_02/14       .......................---------------.......................................... [122560]
FN557520.1_strain_E75            ......gc------------------------------............................a............. [122560]
AY261366.1_Warthog               .t....gct...t.t......at---------------c..............ca....................a.... [122560]
AY261365.1_Warmbaths             ......gct...t.t......at---------------c...........................a.....c..a.... [122560]
AY261364.1_Tengani_62            ............t.t......atcgcgtatactaccgc..............c.......g..............a..a. [122560]
AY261363.1_Pretoriuskop/96/4     .t....gct...t.t......at---------------c..............ca....................a.... [122560]
AY261362.1_Mkuzi_1979            .....................attgcgtatactgccat..............c......................a.... [122560]
AY261361.1_Malawi_Lil-20/1_1983  ...........a.gt......atcgtgcatactgctgcca..c.c......gc.a.........t......a..ca..at [122560]
AY261360.1_Kenya_1950            ......g.------------------------------.t.............gt..c...........a..c..a.... [122560]
AM712240.1_OURT_88/3             ......gc------------------------------............................a............. [122560]
AM712239.1_Benin_97/1            ......gct...t.t......attgcgtatactgccat............................a............. [122560]
KM111295.1_Ken06.Bus             ........------------------------------.t.............gt---------------..c..a.... [122560]
KM111294.1_Ken05/Tk1             ........------------------------------.t.............gt..c...........a..c..a.... [122560]
KM262844.1_strain_L60            ......gc------------------------------............................a............. [122560]
KM102979.1_26544/OG10_Italy      ......gc------------------------------............................a............. [122560]
KM262845.1_strain_NHV            ......gc------------------------------............................a............. [122560]
KX354450.1_47/Ss/2008            ......gc------------------------------............................a............. [122560]
KP055815.1_strain_BA71           ......gc------------------------------............................a............. [122560]
MH025920.1_strain_R35            ........------------------------------.t.............gt---------------..c..a.... [122560]
MH025919.1_strain_N10            ........------------------------------.t.............gt---------------..c..a.... [122560]
MH025918.1_strain_R25            ........------------------------------.t.............gt---------------..c..a.... [122560]
MH025917.1_strain_R7             ........------------------------------.t.............gt---------------..c..a.... [122560]
MH025916.1_strain_R8             ........------------------------------.t.............gt---------------..c..a.... [122560]
```

```
FR682468.2_Georgia_2007/1        cgctgcgtatgctgccgctacatacacta---------------tcactacatatgctgtcagtacatacgctatcgcgg [122640]
ASF_IND_20_CAD543                .............................---------------.................................... [122640]
LR899193.1_Germany_2020/1        .............................---------------.................................... [122640]
MN393477.1_Wuhan_2019            .............................---------------.................................... [122640]
MW396979.1_Timor-Leste/2019/1    .............................---------------.................................... [122640]
MK940252.1_CN/2019/InnerMongolia .............................---------------.................................... [122640]
MK128995.1_China/2018/AnhuiXCGQ  .............................---------------.................................... [122640]
MH681419.1_ASFV/POL/2015         .............................---------------.................................... [122640]
LS478113.1_Estonia_2014          .............................---------------.................................... [122640]
KP843857.1_Odintsovo_02/14       .............................---------------.................................... [122640]
FN557520.1_strain_E75            .a........a..................---------------.................................... [122640]
AY261366.1_Warthog               .a........a..a...............---------------.................................... [122640]
AY261365.1_Warmbaths             .a........a..................---------------.................................... [122640]
AY261364.1_Tengani_62            ................a..g....t....---------------.................................... [122640]
AY261363.1_Pretoriuskop/96/4     .a........a..a...............---------------.................................... [122640]
AY261362.1_Mkuzi_1979            .a........a..................---------------.................................... [122640]
AY261361.1_Malawi_Lil-20/1_1983  .a..a.aa.c....t...gg.g..tg.c.---ctatgtattttt...tgg.tg..ct.a..gccg.gc.ta....g.... [122640]
AY261360.1_Kenya_1950            ......a..ca..ct.a.......t...g---------------...g......c......gcgg.g..a..c....t.. [122640]
AM712240.1_OURT_88/3             .a........a..................tcactacatatgctg.................................... [122640]
AM712239.1_Benin_97/1            .a........a..................---------------.................................... [122640]
KM111295.1_Ken06.Bus             ......a..ca..ct.a.......t...g---------------...g......c......gc---------------.. [122640]
KM111294.1_Ken05/Tk1             ......a..ca..ct.a.......t...g---------------...g......c......gcgg.g..a..c....... [122640]
KM262844.1_strain_L60            .a........a..................---------------.................................... [122640]
KM102979.1_26544/OG10_Italy      .a........a..................---------------.................................... [122640]
KM262845.1_strain_NHV            .a........a..................tcactacatatgctg.................................... [122640]
KX354450.1_47/Ss/2008            .a........a..................---------------.................................... [122640]
KP055815.1_strain_BA71           .a........a..................---------------.................................... [122640]
MH025920.1_strain_R35            ......a..ca..ct.a.......t...g---------------...g......c......gc---------------.. [122640]
MH025919.1_strain_N10            ......a..ca..ct.a.......t...g---------------...g......c......gc---------------.. [122640]
MH025918.1_strain_R25            ......a..ca..ct.a.......t...g---------------...g......c......gc---------------.. [122640]
MH025917.1_strain_R7             ......a..ca..ct.a.......t...g---------------...g......c......gc---------------.. [122640]
MH025916.1_strain_R8             ......a..ca..ct.a.......t...g---------------...g......c......gc---------------.. [122640]

FR682468.2_Georgia_2007/1        cgtatgccgccgtg---taccttatcgccgcccctacccgagggttttttagatataatactgtgtggggagtcaagcga [122720]
ASF_IND_20_CAD543                ..............---.........................................y...r................. [122720]
LR899193.1_Germany_2020/1        ..............---............................................................... [122720]
MN393477.1_Wuhan_2019            ..............---............................................................... [122720]
MW396979.1_Timor-Leste/2019/1    ..............---............................................................... [122720]
MK940252.1_CN/2019/InnerMongolia ..............---............................................................... [122720]
MK128995.1_China/2018/AnhuiXCGQ  ..............---............................................................... [122720]
MH681419.1_ASFV/POL/2015         ..............---............................................................... [122720]
LS478113.1_Estonia_2014          ..............---............................................................... [122720]
KP843857.1_Odintsovo_02/14       ..............---............................................................... [122720]
```

```
FN557520.1_strain_E75              ..............---...........g.............................c...a..c.............. [122720]
AY261366.1_Warthog                 ..............---...........t.............................c...a..c.c..c......... [122720]
AY261365.1_Warmbaths               ..............---...........t....................................c.............. [122720]
AY261364.1_Tengani_62              ..............---.........................................c...a..c.c..c......... [122720]
AY261363.1_Pretoriuskop/96/4       ..............---...........t.............................c...a..c.c............ [122720]
AY261362.1_Mkuzi_1979              ..............---.........................................c...a..c.............. [122720]
AY261361.1_Malawi_Lil-20/1_1983    .a..g.t.....c.---..------ta.ta....cg......................c...a..c.ca..a........ [122720]
AY261360.1_Kenya_1950              ....a...at..c.gct..t.a......t...t.c..g...a................c...a....c............ [122720]
AM712240.1_OURT_88/3               ..............---...........g.............................c...a..c.............. [122720]
AM712239.1_Benin_97/1              ..............---...........g.............................c...a..c.............. [122720]
KM111295.1_Ken06.Bus               ....a...at..c.gct..tgac.....t...t.c..g....................c...a....c............ [122720]
KM111294.1_Ken05/Tk1               ....a...at....gct..t.a......t...t.c..g...a................c...a....c............ [122720]
KM262844.1_strain_L60              ..............---...........g.............................c...a..c.............. [122720]
KM102979.1_26544/OG10_Italy        ..............---...........g.............................c...a..c.............. [122720]
KM262845.1_strain_NHV              ..............---...........g.............................c...a..c.............. [122720]
KX354450.1_47/Ss/2008              ..............---...........g.............................c...a..c.............. [122720]
KP055815.1_strain_BA71             ..............---...........g.............................c...a..c.............. [122720]
MH025920.1_strain_R35              ....a...at..c.gct..tgac.....t...t.c..g....................c...a....c............ [122720]
MH025919.1_strain_N10              ....a...at..c.gct..tgac.....t...t.c..g....................c...a....c............ [122720]
MH025918.1_strain_R25              ....a...at..c.gct..tgac.....t...t.c..g....................c...a....c............ [122720]
MH025917.1_strain_R7               ....a...at..c.gct..tgac.....t...t.c..g....................c...a....c............ [122720]
MH025916.1_strain_R8               ....a...at..c.gct..tgac.....t...t.c..g....................c...a....c............ [122720]

FR682468.2_Georgia_2007/1          aaattcagggtcattaaagttaatgcccaatgactttgccaatccattaagctcttcatcaaaatgatcggtaggaaaac [122800]
ASF_IND_20_CAD543                  ................................................................................ [122800]
LR899193.1_Germany_2020/1          ................................................................................ [122800]
MN393477.1_Wuhan_2019              ................................................................................ [122800]
MW396979.1_Timor-Leste/2019/1      ................................................................................ [122800]
MK940252.1_CN/2019/InnerMongolia   ................................................................................ [122800]
MK128995.1_China/2018/AnhuiXCGQ    ................................................................................ [122800]
MH681419.1_ASFV/POL/2015           ................................................................................ [122800]
LS478113.1_Estonia_2014            ................................................................................ [122800]
KP843857.1_Odintsovo_02/14         ................................................................................ [122800]
FN557520.1_strain_E75              .........a..................................................g................... [122800]
AY261366.1_Warthog                 ........................a...................................g................... [122800]
AY261365.1_Warmbaths               ............................................................g................... [122800]
AY261364.1_Tengani_62              ..........................................c........t........g................... [122800]
AY261363.1_Pretoriuskop/96/4       ...................................................t........g..g.....a.......... [122800]
AY261362.1_Mkuzi_1979              ............................................................g................... [122800]
AY261361.1_Malawi_Lil-20/1_1983    .........a..............a..............t...........t..c.....g...........g....... [122800]
AY261360.1_Kenya_1950              ........................a................g...g.....t..c.................g......t [122800]
AM712240.1_OURT_88/3               .........a..................................................g................... [122800]
AM712239.1_Benin_97/1              .........a..................................................g................... [122800]
KM111295.1_Ken06.Bus               ........................a................g...g.....t..c.................g......t [122800]
```

```
KM111294.1_Ken05/Tk1          ........................a................g...g.....t..c.................g......t [122800]
KM262844.1_strain_L60         .........a..................................................g................... [122800]
KM102979.1_26544/OG10_Italy   .........a..................................................g................... [122800]
KM262845.1_strain_NHV         .........a..................................................g................... [122800]
KX354450.1_47/Ss/2008         .........a..................................................g................... [122800]
KP055815.1_strain_BA71        .........a..................................................g................... [122800]
MH025920.1_strain_R35         ........................a................g...g.....t..c.................g......t [122800]
MH025919.1_strain_N10         ........................a................g...g.....t..c.................g......t [122800]
MH025918.1_strain_R25         ........................a................g...g.....t..c.................g......t [122800]
MH025917.1_strain_R7          ........................a................g...g.....t..c.................g......t [122800]
MH025916.1_strain_R8          ........................a................g...g.....t..c.................g......t [122800]

FR682468.2_Georgia_2007/1     tttgttgcttgcccatgacctgtttttcaagttcctccaaattggcttgctcatttatatggagattattcataagcgtc [122880]
ASF_IND_20_CAD543             ................................................................................ [122880]
LR899193.1_Germany_2020/1     ................................................................................ [122880]
MN393477.1_Wuhan_2019         ................................................................................ [122880]
MW396979.1_Timor-Leste/2019/1 ................................................................................ [122880]
MK940252.1_CN/2019/InnerMongolia ................................................................................ [122880]
MK128995.1_China/2018/AnhuiXCGQ ................................................................................ [122880]
MH681419.1_ASFV/POL/2015      ................................................................................ [122880]
LS478113.1_Estonia_2014       ................................................................................ [122880]
KP843857.1_Odintsovo_02/14    ................................................................................ [122880]
FN557520.1_strain_E75         .......t..........................t............................................t [122880]
AY261366.1_Warthog            ................................................................................ [122880]
AY261365.1_Warmbaths          ................................................................................ [122880]
AY261364.1_Tengani_62         ................................................................................ [122880]
AY261363.1_Pretoriuskop/96/4  ................................................................................ [122880]
AY261362.1_Mkuzi_1979         .......t..........................t............................................t [122880]
AY261361.1_Malawi_Lil-20/1_1983 ...................................................................g...........a [122880]
AY261360.1_Kenya_1950         ....c.....a........t.................t.....a...................................a [122880]
AM712240.1_OURT_88/3          .......t..........................t............................................t [122880]
AM712239.1_Benin_97/1         .......t..........................t............................................t [122880]
KM111295.1_Ken06.Bus          ....c.....a........t.................t.....a................................t..a [122880]
KM111294.1_Ken05/Tk1          ....c.....a........t.................t.....a................................t..a [122880]
KM262844.1_strain_L60         .......t..........................t............................................t [122880]
KM102979.1_26544/OG10_Italy   .......t..........................t............................................t [122880]
KM262845.1_strain_NHV         .......t..........................t............................................t [122880]
KX354450.1_47/Ss/2008         .......t..........................t............................................t [122880]
KP055815.1_strain_BA71        .......t..........................t............................................t [122880]
MH025920.1_strain_R35         ....c.....a........t.................t.....a................................t..a [122880]
MH025919.1_strain_N10         ....c.....a........t.................t.....a................................t..a [122880]
MH025918.1_strain_R25         ....c.....a........t.................t.....a................................t..a [122880]
MH025917.1_strain_R7          ....c.....a........t.................t.....a................................t..a [122880]
MH025916.1_strain_R8          ....c.....a........t.................t.....a................................t..a [122880]
```

```
FR682468.2_Georgia_2007/1        gtaattccagcaagatttgctccttctaaaaatgtggtgtcctccatcggatatactatactatttaaa-agcttttaaa [122960]
ASF_IND_20_CAD543                .....................................................................-.......... [122960]
LR899193.1_Germany_2020/1        .....................................................................-.......... [122960]
MN393477.1_Wuhan_2019            .....................................................................-.......... [122960]
MW396979.1_Timor-Leste/2019/1    .....................................................................-.......... [122960]
MK940252.1_CN/2019/InnerMongolia .....................................................................-.......... [122960]
MK128995.1_China/2018/AnhuiXCGQ  .....................................................................-.......... [122960]
MH681419.1_ASFV/POL/2015         .....................................................................-.......... [122960]
LS478113.1_Estonia_2014          .....................................................................-.......... [122960]
KP843857.1_Odintsovo_02/14       .....................................................................-.......... [122960]
FN557520.1_strain_E75            .....c...............................................................-.......... [122960]
AY261366.1_Warthog               ...............................................t.....................-.......... [122960]
AY261365.1_Warmbaths             ...................................a.................................-.......... [122960]
AY261364.1_Tengani_62            ..................................c..................................-.......... [122960]
AY261363.1_Pretoriuskop/96/4     ...............................................t.....................-.......... [122960]
AY261362.1_Mkuzi_1979            .....................................................................-.......... [122960]
AY261361.1_Malawi_Lil-20/1_1983  .............................g.......................................a.......... [122960]
AY261360.1_Kenya_1950            ......................................a........ta....................a.......... [122960]
AM712240.1_OURT_88/3             .....c...............................................................-.......... [122960]
AM712239.1_Benin_97/1            .....c...............................................................-.......... [122960]
KM111295.1_Ken06.Bus             .....c................................a........ta....................a.......... [122960]
KM111294.1_Ken05/Tk1             .....c................................a........ta....................a.......... [122960]
KM262844.1_strain_L60            .....c...............................................................-.......... [122960]
KM102979.1_26544/OG10_Italy      .....c...............................................................-.......... [122960]
KM262845.1_strain_NHV            .....c...............................................................-.......... [122960]
KX354450.1_47/Ss/2008            .....c...............................................................-.......... [122960]
KP055815.1_strain_BA71           .....c...............................................................-.......... [122960]
MH025920.1_strain_R35            .....c................................a........ta....................a.......... [122960]
MH025919.1_strain_N10            .....c................................a........ta....................a.......... [122960]
MH025918.1_strain_R25            .....c................................a........ta....................a.......... [122960]
MH025917.1_strain_R7             .....c................................a........ta....................a.......... [122960]
MH025916.1_strain_R8             .....c................................a........ta....................a.......... [122960]

FR682468.2_Georgia_2007/1        taaaaatgtgtttggaagaaatgctctcttcaagcgtgtgtagctcagatataaatgcctcctcagaaagctttccacca [123040]
ASF_IND_20_CAD543                ................................................................................ [123040]
LR899193.1_Germany_2020/1        ................................................................................ [123040]
MN393477.1_Wuhan_2019            ................................................................................ [123040]
MW396979.1_Timor-Leste/2019/1    ................................................................................ [123040]
MK940252.1_CN/2019/InnerMongolia ................................................................................ [123040]
MK128995.1_China/2018/AnhuiXCGQ  ................................................................................ [123040]
MH681419.1_ASFV/POL/2015         ................................................................................ [123040]
LS478113.1_Estonia_2014          ................................................................................ [123040]
KP843857.1_Odintsovo_02/14       ................................................................................ [123040]
```

```
FN557520.1_strain_E75          .......................................................c........................ [123040]
AY261366.1_Warthog             .........................t.............................c....................g... [123040]
AY261365.1_Warmbaths           ...............................t..........a..................................... [123040]
AY261364.1_Tengani_62          .........................t...................................................... [123040]
AY261363.1_Pretoriuskop/96/4   .........................t.............................c....................g... [123040]
AY261362.1_Mkuzi_1979          ................................................................................ [123040]
AY261361.1_Malawi_Lil-20/1_1983 .........c...............t........t................c...c..............t.....g... [123040]
AY261360.1_Kenya_1950          .........c...............t........g....................c..............t.....g... [123040]
AM712240.1_OURT_88/3           .......................................................c........................ [123040]
AM712239.1_Benin_97/1          .......................................................c........................ [123040]
KM111295.1_Ken06.Bus           .........c...............t........g................c...c..............t.....g... [123040]
KM111294.1_Ken05/Tk1           .........c...............t........g................c...c..............t.....g... [123040]
KM262844.1_strain_L60          .......................................................c........................ [123040]
KM102979.1_26544/OG10_Italy    .......................................................c........................ [123040]
KM262845.1_strain_NHV          .......................................................c........................ [123040]
KX354450.1_47/Ss/2008          .......................................................c........................ [123040]
KP055815.1_strain_BA71         .......................................................c........................ [123040]
MH025920.1_strain_R35          .........c...............t........g................c...c..............t.....g... [123040]
MH025919.1_strain_N10          .........c...............t........g................c...c..............t.....g... [123040]
MH025918.1_strain_R25          .........c...............t........g................c...c..............t.....g... [123040]
MH025917.1_strain_R7           .........c...............t........g................c...c..............t.....g... [123040]
MH025916.1_strain_R8           .........c...............t........g................c...c..............t.....g... [123040]

FR682468.2_Georgia_2007/1      tactcctttctcatcgtataggagggcgccggtttaatgtaggaaatccactgggaggtaaaaaaccggtacaacatatt [123120]
ASF_IND_20_CAD543              ................................................................................ [123120]
LR899193.1_Germany_2020/1      ................................................................................ [123120]
MN393477.1_Wuhan_2019          ................................................................................ [123120]
MW396979.1_Timor-Leste/2019/1  ................................................................................ [123120]
MK940252.1_CN/2019/InnerMongolia ................................................................................ [123120]
MK128995.1_China/2018/AnhuiXCGQ ................................................................................ [123120]
MH681419.1_ASFV/POL/2015       ................................................................................ [123120]
LS478113.1_Estonia_2014        ................................................................................ [123120]
KP843857.1_Odintsovo_02/14     ................................................................................ [123120]
FN557520.1_strain_E75          ................................c............................................... [123120]
AY261366.1_Warthog             ................................c............................................... [123120]
AY261365.1_Warmbaths           ................................c............................................... [123120]
AY261364.1_Tengani_62          .............a..................c............................................... [123120]
AY261363.1_Pretoriuskop/96/4   ................................c............................................... [123120]
AY261362.1_Mkuzi_1979          ................................................................................ [123120]
AY261361.1_Malawi_Lil-20/1_1983 .............a..........at......c....................a...........a.............. [123120]
AY261360.1_Kenya_1950          ..t..t..c...............a.......cc...................a.....g.....a.............. [123120]
AM712240.1_OURT_88/3           ................................c............................................... [123120]
AM712239.1_Benin_97/1          ................................c............................................... [123120]
KM111295.1_Ken06.Bus           ..t..t..c...............a.......cc...................a.....g.....a.............. [123120]
```

```
KM111294.1_Ken05/Tk1             ..t..t..c...............a..c....cc...................a.....g.....a.............. [123120]
KM262844.1_strain_L60            ................................c............................................... [123120]
KM102979.1_26544/OG10_Italy      ................................c............................................... [123120]
KM262845.1_strain_NHV            ................................c............................................... [123120]
KX354450.1_47/Ss/2008            ................................c............................................... [123120]
KP055815.1_strain_BA71           ................................c............................................... [123120]
MH025920.1_strain_R35            ..t..t..c...............a.......cc...................a.....g.....a.............. [123120]
MH025919.1_strain_N10            ..t..t..c...............a.......cc...................a.....g.....a.............. [123120]
MH025918.1_strain_R25            ..t..t..c...............a.......cc...................a.....g.....a.............. [123120]
MH025917.1_strain_R7             ..t..t..c...............a.......cc...................a.....g.....a.............. [123120]
MH025916.1_strain_R8             ..t..t..c...............a.......cc...................a.....g.....a.............. [123120]

FR682468.2_Georgia_2007/1        tagcagctcgcgggcctcccaccttttgggctccgtatagtgcacatcaacataagaggcggcgcatgaaaagctgcaaa [123200]
ASF_IND_20_CAD543                ................................................................................ [123200]
LR899193.1_Germany_2020/1        ................................................................................ [123200]
MN393477.1_Wuhan_2019            ................................................................................ [123200]
MW396979.1_Timor-Leste/2019/1    ................................................................................ [123200]
MK940252.1_CN/2019/InnerMongolia ................................................................................ [123200]
MK128995.1_China/2018/AnhuiXCGQ  ................................................................................ [123200]
MH681419.1_ASFV/POL/2015         ................................................................................ [123200]
LS478113.1_Estonia_2014          ................................................................................ [123200]
KP843857.1_Odintsovo_02/14       ................................................................................ [123200]
FN557520.1_strain_E75            ................................................................................ [123200]
AY261366.1_Warthog               ..a.............................................g............................... [123200]
AY261365.1_Warmbaths             .....................t.......................................................... [123200]
AY261364.1_Tengani_62            ................................................g............................... [123200]
AY261363.1_Pretoriuskop/96/4     ................................................g............................... [123200]
AY261362.1_Mkuzi_1979            ................................................................................ [123200]
AY261361.1_Malawi_Lil-20/1_1983  .....................tt...................t.a...............a..a................ [123200]
AY261360.1_Kenya_1950            ............................................g.....t.........a................... [123200]
AM712240.1_OURT_88/3             ................................................................................ [123200]
AM712239.1_Benin_97/1            ................................................................................ [123200]
KM111295.1_Ken06.Bus             ............................................a.....t.........a................... [123200]
KM111294.1_Ken05/Tk1             ............................................a.....t.........a................... [123200]
KM262844.1_strain_L60            ................................................................................ [123200]
KM102979.1_26544/OG10_Italy      ................................................................................ [123200]
KM262845.1_strain_NHV            ................................................................................ [123200]
KX354450.1_47/Ss/2008            ................................................................................ [123200]
KP055815.1_strain_BA71           ................................................................................ [123200]
MH025920.1_strain_R35            ............................................a.....t.........a................... [123200]
MH025919.1_strain_N10            ............................................a.....t.........a................... [123200]
MH025918.1_strain_R25            ............................................a.....t.........a................... [123200]
MH025917.1_strain_R7             ............................................a.....t.........a................... [123200]
MH025916.1_strain_R8             ............................................a.....t.........a................... [123200]
```

```
FR682468.2_Georgia_2007/1        agttgccgagaacgcccatctcaatctctcctcgctcattttcacgcatataggtgggcacgaattttgggacagtcttg [123280]
ASF_IND_20_CAD543                ................................................................................ [123280]
LR899193.1_Germany_2020/1        ................................................................................ [123280]
MN393477.1_Wuhan_2019            ................................................................................ [123280]
MW396979.1_Timor-Leste/2019/1    ................................................................................ [123280]
MK940252.1_CN/2019/InnerMongolia ................................................................................ [123280]
MK128995.1_China/2018/AnhuiXCGQ  ................................................................................ [123280]
MH681419.1_ASFV/POL/2015         ................................................................................ [123280]
LS478113.1_Estonia_2014          ................................................................................ [123280]
KP843857.1_Odintsovo_02/14       ................................................................................ [123280]
FN557520.1_strain_E75            ...................................................................c..a......... [123280]
AY261366.1_Warthog               ...................................................................c..a..c...... [123280]
AY261365.1_Warmbaths             ...................................................................c..a......... [123280]
AY261364.1_Tengani_62            .....................................g.............................c............ [123280]
AY261363.1_Pretoriuskop/96/4     ...................................................................c............ [123280]
AY261362.1_Mkuzi_1979            ................................................................................ [123280]
AY261361.1_Malawi_Lil-20/1_1983  ...................t...................................a..............a......... [123280]
AY261360.1_Kenya_1950            ...................................................................c............ [123280]
AM712240.1_OURT_88/3             ...................................................................c..a......... [123280]
AM712239.1_Benin_97/1            ...................................................................c..a......... [123280]
KM111295.1_Ken06.Bus             ....c..............................................................c............ [123280]
KM111294.1_Ken05/Tk1             ...................................................................c............ [123280]
KM262844.1_strain_L60            ...................................................................c..a......... [123280]
KM102979.1_26544/OG10_Italy      ...................................................................c..a......... [123280]
KM262845.1_strain_NHV            ...................................................................c..a......... [123280]
KX354450.1_47/Ss/2008            ...................................................................c..a......... [123280]
KP055815.1_strain_BA71           ...................................................................c..a......... [123280]
MH025920.1_strain_R35            ....c..............................................................c............ [123280]
MH025919.1_strain_N10            ....c..............................................................c............ [123280]
MH025918.1_strain_R25            ....c..............................................................c............ [123280]
MH025917.1_strain_R7             ....c..............................................................c............ [123280]
MH025916.1_strain_R8             ....c..............................................................c............ [123280]

FR682468.2_Georgia_2007/1        aaatagagatgacatgtccagcatttaaagctagaatgggtaacccatttggaaacagtggtgaatacggagggtagctt [123360]
ASF_IND_20_CAD543                ................................................................................ [123360]
LR899193.1_Germany_2020/1        ................................................................................ [123360]
MN393477.1_Wuhan_2019            ................................................................................ [123360]
MW396979.1_Timor-Leste/2019/1    ................................................................................ [123360]
MK940252.1_CN/2019/InnerMongolia ................................................................................ [123360]
MK128995.1_China/2018/AnhuiXCGQ  ................................................................................ [123360]
MH681419.1_ASFV/POL/2015         ................................................................................ [123360]
LS478113.1_Estonia_2014          ................................................................................ [123360]
KP843857.1_Odintsovo_02/14       ................................................................................ [123360]
```

```
FN557520.1_strain_E75            ..............c............................................a.................... [123360]
AY261366.1_Warthog               ..............c..............a.................................................. [123360]
AY261365.1_Warmbaths             ................................................................................ [123360]
AY261364.1_Tengani_62            ..............c................................................................. [123360]
AY261363.1_Pretoriuskop/96/4     ..............c................................................................. [123360]
AY261362.1_Mkuzi_1979            ..............c................................................................. [123360]
AY261361.1_Malawi_Lil-20/1_1983  .....a........c..........................................c................g..... [123360]
AY261360.1_Kenya_1950            .....a........c......................a...................c................a..... [123360]
AM712240.1_OURT_88/3             ..............c............................................a.................... [123360]
AM712239.1_Benin_97/1            ..............c............................................a.................... [123360]
KM111295.1_Ken06.Bus             .....a........c..........................................c................a..... [123360]
KM111294.1_Ken05/Tk1             .....a........c..........................................c................a..... [123360]
KM262844.1_strain_L60            ..............c............................................a.................... [123360]
KM102979.1_26544/OG10_Italy      ..............c............................................a.................... [123360]
KM262845.1_strain_NHV            ..............c............................................a.................... [123360]
KX354450.1_47/Ss/2008            ..............c............................................a.................... [123360]
KP055815.1_strain_BA71           ..............c............................................a.................... [123360]
MH025920.1_strain_R35            .....a........c..........................................c................a..... [123360]
MH025919.1_strain_N10            .....a........c..........................................c................a..... [123360]
MH025918.1_strain_R25            .....a........c..........................................c................a..... [123360]
MH025917.1_strain_R7             .....a........c..........................................c................a..... [123360]
MH025916.1_strain_R8             .....a........c..........................................c................a..... [123360]

FR682468.2_Georgia_2007/1        tttttcgacctcggcttcatcgtcattcgtatttaacgtatcggtggcagtttttttggattgcaagcattcttcaatgg [123440]
ASF_IND_20_CAD543                ................................................................................ [123440]
LR899193.1_Germany_2020/1        ................................................................................ [123440]
MN393477.1_Wuhan_2019            ................................................................................ [123440]
MW396979.1_Timor-Leste/2019/1    ................................................................................ [123440]
MK940252.1_CN/2019/InnerMongolia ................................................................................ [123440]
MK128995.1_China/2018/AnhuiXCGQ  ................................................................................ [123440]
MH681419.1_ASFV/POL/2015         ................................................................................ [123440]
LS478113.1_Estonia_2014          ................................................................................ [123440]
KP843857.1_Odintsovo_02/14       ................................................................................ [123440]
FN557520.1_strain_E75            ...........................................................t.................... [123440]
AY261366.1_Warthog               .............................................a.................t................ [123440]
AY261365.1_Warmbaths             ...........................................a...............t..................a. [123440]
AY261364.1_Tengani_62            .......................t.......................................t................ [123440]
AY261363.1_Pretoriuskop/96/4     ............a..................................................t..............a. [123440]
AY261362.1_Mkuzi_1979            ............aa.............................................t.................... [123440]
AY261361.1_Malawi_Lil-20/1_1983  ......................c.............t.........................................a. [123440]
AY261360.1_Kenya_1950            ......................c.............t.c......a.ag..........................g.... [123440]
AM712240.1_OURT_88/3             ...........................................................t.................... [123440]
AM712239.1_Benin_97/1            ...........................................................t.................... [123440]
KM111295.1_Ken06.Bus             ......................c.............t.cg.....a.tgt.........................g.... [123440]
```

```
KM111294.1_Ken05/Tk1              ......................c.............t.c......a.ag..........................g.... [123440]
KM262844.1_strain_L60             ...........................................................t.................... [123440]
KM102979.1_26544/OG10_Italy       ...........................................................t.................... [123440]
KM262845.1_strain_NHV             ...........................................................t.................... [123440]
KX354450.1_47/Ss/2008             ...........................................................t.................... [123440]
KP055815.1_strain_BA71            ...........................................................t.................... [123440]
MH025920.1_strain_R35             ......................c.............t.cg.....a.tgt................a........g.... [123440]
MH025919.1_strain_N10             ......................c.............t.cg.....a.tgt................a........g.... [123440]
MH025918.1_strain_R25             ......................c.............t.cg.....a.tgt................a........g.... [123440]
MH025917.1_strain_R7              ......................c.............t.cg.....a.tgt................a........g.... [123440]
MH025916.1_strain_R8              ......................c.............t.cg.....a.tgt................a........g.... [123440]

FR682468.2_Georgia_2007/1         taatcccggataagtataaaatattaggacaattagtttccataattttgatagttatttttatacaacatggatttaat [123520]
ASF_IND_20_CAD543                 ................................................................................ [123520]
LR899193.1_Germany_2020/1         ................................................................................ [123520]
MN393477.1_Wuhan_2019             ................................................................................ [123520]
MW396979.1_Timor-Leste/2019/1     ................................................................................ [123520]
MK940252.1_CN/2019/InnerMongolia  ................................................................................ [123520]
MK128995.1_China/2018/AnhuiXCGQ   ................................................................................ [123520]
MH681419.1_ASFV/POL/2015          ................................................................................ [123520]
LS478113.1_Estonia_2014           ................................................................................ [123520]
KP843857.1_Odintsovo_02/14        ................................................................................ [123520]
FN557520.1_strain_E75             ................................................................................ [123520]
AY261366.1_Warthog                ................................................................................ [123520]
AY261365.1_Warmbaths              ................................................................................ [123520]
AY261364.1_Tengani_62             ................................................................................ [123520]
AY261363.1_Pretoriuskop/96/4      ................................................................................ [123520]
AY261362.1_Mkuzi_1979             ................................................................................ [123520]
AY261361.1_Malawi_Lil-20/1_1983   ....t........................................................................... [123520]
AY261360.1_Kenya_1950             ....t..a.....a.................................................................. [123520]
AM712240.1_OURT_88/3              ................................................................................ [123520]
AM712239.1_Benin_97/1             ................................................................................ [123520]
KM111295.1_Ken06.Bus              ....t..a.....a.................................................................. [123520]
KM111294.1_Ken05/Tk1              ....t..a.....a.................................................................. [123520]
KM262844.1_strain_L60             ................................................................................ [123520]
KM102979.1_26544/OG10_Italy       ................................................................................ [123520]
KM262845.1_strain_NHV             ................................................................................ [123520]
KX354450.1_47/Ss/2008             ................................................................................ [123520]
KP055815.1_strain_BA71            ................................................................................ [123520]
MH025920.1_strain_R35             ....t..a.....a.................................................................. [123520]
MH025919.1_strain_N10             ....t..a.....a.................................................................. [123520]
MH025918.1_strain_R25             ....t..a.....a.................................................................. [123520]
MH025917.1_strain_R7              ....t..a.....a.................................................................. [123520]
MH025916.1_strain_R8              ....t..a.....a.................................................................. [123520]
```

```
FR682468.2_Georgia_2007/1        taaagataaatggaggacgaaacggaactgtgttttcggtcaaacaaggtgacgaggcttgaaatgtttgtctgcacata [123600]
ASF_IND_20_CAD543                ................................................................................ [123600]
LR899193.1_Germany_2020/1        ................................................................................ [123600]
MN393477.1_Wuhan_2019            ................................................................................ [123600]
MW396979.1_Timor-Leste/2019/1    ................................................................................ [123600]
MK940252.1_CN/2019/InnerMongolia ................................................................................ [123600]
MK128995.1_China/2018/AnhuiXCGQ  ................................................................................ [123600]
MH681419.1_ASFV/POL/2015         ................................................................................ [123600]
LS478113.1_Estonia_2014          ................................................................................ [123600]
KP843857.1_Odintsovo_02/14       ................................................................................ [123600]
FN557520.1_strain_E75            .............................................................................g.. [123600]
AY261366.1_Warthog               ................................................................................ [123600]
AY261365.1_Warmbaths             .....................................................a..a....................... [123600]
AY261364.1_Tengani_62            ................................................................................ [123600]
AY261363.1_Pretoriuskop/96/4     ................................................................................ [123600]
AY261362.1_Mkuzi_1979            .............................................................................g.. [123600]
AY261361.1_Malawi_Lil-20/1_1983  ..................................................t............................. [123600]
AY261360.1_Kenya_1950            ...........................t......................t....................g........ [123600]
AM712240.1_OURT_88/3             .............................................................................g.. [123600]
AM712239.1_Benin_97/1            .............................................................................g.. [123600]
KM111295.1_Ken06.Bus             ...........................t..................g...t....................g........ [123600]
KM111294.1_Ken05/Tk1             ...........................t......................t............................. [123600]
KM262844.1_strain_L60            .............................................................................g.. [123600]
KM102979.1_26544/OG10_Italy      .............................................................................g.. [123600]
KM262845.1_strain_NHV            .............................................................................g.. [123600]
KX354450.1_47/Ss/2008            .............................................................................g.. [123600]
KP055815.1_strain_BA71           .............................................................................g.. [123600]
MH025920.1_strain_R35            ...........................t..................g...t....................g........ [123600]
MH025919.1_strain_N10            ...........................t..................g...t....................g........ [123600]
MH025918.1_strain_R25            ...........................t..................g...t....................g........ [123600]
MH025917.1_strain_R7             ...........................t..................g...t....................g........ [123600]
MH025916.1_strain_R8             ...........................t..................g...t....................g........ [123600]

FR682468.2_Georgia_2007/1        cggggaaaaattaccagccttgcatgttcgcatatggagttaattaaaatgttgcaaattgctgagccggtgaaggcat [123680]
ASF_IND_20_CAD543                ................................................................................ [123680]
LR899193.1_Germany_2020/1        ................................................................................ [123680]
MN393477.1_Wuhan_2019            ................................................................................ [123680]
MW396979.1_Timor-Leste/2019/1    ................................................................................ [123680]
MK940252.1_CN/2019/InnerMongolia ................................................................................ [123680]
MK128995.1_China/2018/AnhuiXCGQ  ................................................................................ [123680]
MH681419.1_ASFV/POL/2015         ................................................................................ [123680]
LS478113.1_Estonia_2014          ................................................................................ [123680]
KP843857.1_Odintsovo_02/14       ................................................................................ [123680]
```

```
FN557520.1_strain_E75          ............................................................c..c................ [123680]
AY261366.1_Warthog             ..............g....................................a...........c..............g. [123680]
AY261365.1_Warmbaths           ..............g....................................a........c..c..............g. [123680]
AY261364.1_Tengani_62          ..............g....................................a........c..c..............g. [123680]
AY261363.1_Pretoriuskop/96/4   ..............g....................................a...........c..............g. [123680]
AY261362.1_Mkuzi_1979          ...................................................a........c..c..............g. [123680]
AY261361.1_Malawi_Lil-20/1_1983 ..............g...................................ga........c..c................ [123680]
AY261360.1_Kenya_1950          ..............g.................................gc.a........c..c..............g. [123680]
AM712240.1_OURT_88/3           ............................................................c..c................ [123680]
AM712239.1_Benin_97/1          ............................................................c..c................ [123680]
KM111295.1_Ken06.Bus           ..............g.................................gc.a........c..c..............g. [123680]
KM111294.1_Ken05/Tk1           ..............g.................................gc.a........c..c..............g. [123680]
KM262844.1_strain_L60          ............................................................c..c................ [123680]
KM102979.1_26544/OG10_Italy    ............................................................c..c................ [123680]
KM262845.1_strain_NHV          ............................................................c..c................ [123680]
KX354450.1_47/Ss/2008          ............................................................c..c................ [123680]
KP055815.1_strain_BA71         ............................................................c..c................ [123680]
MH025920.1_strain_R35          ..............g.................................gc.a........c..c..............g. [123680]
MH025919.1_strain_N10          ..............g.................................gc.a........c..c..............g. [123680]
MH025918.1_strain_R25          ..............g.................................gc.a........c..c..............g. [123680]
MH025917.1_strain_R7           ..............g.................................gc.a........c..c..............g. [123680]
MH025916.1_strain_R8           ..............g.................................gc.a........c..c..............g. [123680]

FR682468.2_Georgia_2007/1      tgaactgcaactttggccaccagtgcctaccgggctacgaatctttaataaagactccgaaaaaaactaaaaacatgttg [123760]
ASF_IND_20_CAD543              ................................................................................ [123760]
LR899193.1_Germany_2020/1      ................................................................................ [123760]
MN393477.1_Wuhan_2019          ................................................................................ [123760]
MW396979.1_Timor-Leste/2019/1  ................................................................................ [123760]
MK940252.1_CN/2019/InnerMongolia ................................................................................ [123760]
MK128995.1_China/2018/AnhuiXCGQ ................................................................................ [123760]
MH681419.1_ASFV/POL/2015       ................................................................................ [123760]
LS478113.1_Estonia_2014        ................................................................................ [123760]
KP843857.1_Odintsovo_02/14     ................................................................................ [123760]
FN557520.1_strain_E75          ................................................................................ [123760]
AY261366.1_Warthog             ................................................................................ [123760]
AY261365.1_Warmbaths           ................................................................................ [123760]
AY261364.1_Tengani_62          ................................................................................ [123760]
AY261363.1_Pretoriuskop/96/4   ................................................................................ [123760]
AY261362.1_Mkuzi_1979          .a.............................................................................. [123760]
AY261361.1_Malawi_Lil-20/1_1983 ...........................................c..g...........a.......a............. [123760]
AY261360.1_Kenya_1950          ..................................................................g............. [123760]
AM712240.1_OURT_88/3           ................................................................................ [123760]
AM712239.1_Benin_97/1          ................................................................................ [123760]
KM111295.1_Ken06.Bus           ..................................................................gc............ [123760]
```

```
KM111294.1_Ken05/Tk1              ..................................................................g............. [123760]
KM262844.1_strain_L60             ................................................................................ [123760]
KM102979.1_26544/OG10_Italy       ................................................................................ [123760]
KM262845.1_strain_NHV             ................................................................................ [123760]
KX354450.1_47/Ss/2008             ................................................................................ [123760]
KP055815.1_strain_BA71            ................................................................................ [123760]
MH025920.1_strain_R35             ..................................................................gc............ [123760]
MH025919.1_strain_N10             ..................................................................gc............ [123760]
MH025918.1_strain_R25             ..................................................................gc............ [123760]
MH025917.1_strain_R7              ..................................................................gc............ [123760]
MH025916.1_strain_R8              ..................................................................gc............ [123760]

FR682468.2_Georgia_2007/1         cgccgtccgcgcaaaacagaaggcgatgggacttgcttcaatagtgccattgaagcctccattttgtttaaggacaagat [123840]
ASF_IND_20_CAD543                 ........................................................y....................... [123840]
LR899193.1_Germany_2020/1         ................................................................................ [123840]
MN393477.1_Wuhan_2019             ................................................................................ [123840]
MW396979.1_Timor-Leste/2019/1     ................................................................................ [123840]
MK940252.1_CN/2019/InnerMongolia  ................................................................................ [123840]
MK128995.1_China/2018/AnhuiXCGQ   ................................................................................ [123840]
MH681419.1_ASFV/POL/2015          ................................................................................ [123840]
LS478113.1_Estonia_2014           ................................................................................ [123840]
KP843857.1_Odintsovo_02/14        ................................................................................ [123840]
FN557520.1_strain_E75             ................................c.......................t....................... [123840]
AY261366.1_Warthog                ...........t....................c.....t......................................... [123840]
AY261365.1_Warmbaths              .....c.....t....................c.....t......................................... [123840]
AY261364.1_Tengani_62             ...........t....................c.....t......................................... [123840]
AY261363.1_Pretoriuskop/96/4      ...........t....................c............................................... [123840]
AY261362.1_Mkuzi_1979             ................................c............................................... [123840]
AY261361.1_Malawi_Lil-20/1_1983   ..t..c.....t.................a..c..........................t...........a........ [123840]
AY261360.1_Kenya_1950             ...........t.................a..c...........c...........t..t...........a........ [123840]
AM712240.1_OURT_88/3              ................................c.......................t....................... [123840]
AM712239.1_Benin_97/1             ................................c.......................t....................... [123840]
KM111295.1_Ken06.Bus              ...........t.................a..c...........c...........t..t...........a........ [123840]
KM111294.1_Ken05/Tk1              ...........t.................a..c...........c...........t..t...........a........ [123840]
KM262844.1_strain_L60             ................................c.......................t....................... [123840]
KM102979.1_26544/OG10_Italy       ................................c.......................t....................... [123840]
KM262845.1_strain_NHV             ................................c.......................t....................... [123840]
KX354450.1_47/Ss/2008             ................................c.......................t....................... [123840]
KP055815.1_strain_BA71            ................................c.......................t....................... [123840]
MH025920.1_strain_R35             ...........t.................a..c...........c...........t..t...........a........ [123840]
MH025919.1_strain_N10             ...........t.................a..c...........c...........t..t...........a........ [123840]
MH025918.1_strain_R25             ...........t.................a..c...........c...........t..t...........a........ [123840]
MH025917.1_strain_R7              ...........t.................a..c...........c...........t..t...........a........ [123840]
MH025916.1_strain_R8              ...........t.................a..c...........c...........t..t...........a........ [123840]
```

```
FR682468.2_Georgia_2007/1        gtataaattaaaatgttttcctagtaccggggaaattcaggtcccgggcgtcatttttccggattttgaagacggaaaaa [123920]
ASF_IND_20_CAD543                ................................................................................ [123920]
LR899193.1_Germany_2020/1        ................................................................................ [123920]
MN393477.1_Wuhan_2019            ................................................................................ [123920]
MW396979.1_Timor-Leste/2019/1    ................................................................................ [123920]
MK940252.1_CN/2019/InnerMongolia ................................................................................ [123920]
MK128995.1_China/2018/AnhuiXCGQ  ................................................................................ [123920]
MH681419.1_ASFV/POL/2015         ................................................................................ [123920]
LS478113.1_Estonia_2014          ................................................................................ [123920]
KP843857.1_Odintsovo_02/14       ................................................................................ [123920]
FN557520.1_strain_E75            ................................................................................ [123920]
AY261366.1_Warthog               ...............c........................................................t....... [123920]
AY261365.1_Warmbaths             ...............c................................................................ [123920]
AY261364.1_Tengani_62            ...............c................................................................ [123920]
AY261363.1_Pretoriuskop/96/4     ...............c................................................................ [123920]
AY261362.1_Mkuzi_1979            ..........................................g.................a................... [123920]
AY261361.1_Malawi_Lil-20/1_1983  ...............c................................g............................... [123920]
AY261360.1_Kenya_1950            .......c.......c........c.....a.................g............................... [123920]
AM712240.1_OURT_88/3             ................................................................................ [123920]
AM712239.1_Benin_97/1            ................................................................................ [123920]
KM111295.1_Ken06.Bus             .......c.......c........c.....a.............................a................... [123920]
KM111294.1_Ken05/Tk1             .......c.......c........c.....a.............................a................... [123920]
KM262844.1_strain_L60            ................................................................................ [123920]
KM102979.1_26544/OG10_Italy      ................................................................................ [123920]
KM262845.1_strain_NHV            ................................................................................ [123920]
KX354450.1_47/Ss/2008            ................................................................................ [123920]
KP055815.1_strain_BA71           ................................................................................ [123920]
MH025920.1_strain_R35            .......c.......c........c.....a.............................a................... [123920]
MH025919.1_strain_N10            .......c.......c........c.....a.............................a................... [123920]
MH025918.1_strain_R25            .......c.......c........c.....a.............................a................... [123920]
MH025917.1_strain_R7             .......c.......c........c.....a.............................a................... [123920]
MH025916.1_strain_R8             .......c.......c........c.....a.............................a................... [123920]

FR682468.2_Georgia_2007/1        acattatacagcagtgggtagacttcttgcaacatcaacccattgaaaaaaaaatccagattattgaatttaaaacgatt [124000]
ASF_IND_20_CAD543                ...............................................................................c [124000]
LR899193.1_Germany_2020/1        ................................................................................ [124000]
MN393477.1_Wuhan_2019            ................................................................................ [124000]
MW396979.1_Timor-Leste/2019/1    ................................................................................ [124000]
MK940252.1_CN/2019/InnerMongolia ................................................................................ [124000]
MK128995.1_China/2018/AnhuiXCGQ  ................................................................................ [124000]
MH681419.1_ASFV/POL/2015         ................................................................................ [124000]
LS478113.1_Estonia_2014          ................................................................................ [124000]
KP843857.1_Odintsovo_02/14       ................................................................................ [124000]
```

```
FN557520.1_strain_E75              ...............................................................................c [124000]
AY261366.1_Warthog                 ..........a..................................................................... [124000]
AY261365.1_Warmbaths               ................................................................................ [124000]
AY261364.1_Tengani_62              ..........a..................................................................... [124000]
AY261363.1_Pretoriuskop/96/4       ..........a..................................................................... [124000]
AY261362.1_Mkuzi_1979              ...............................................................................c [124000]
AY261361.1_Malawi_Lil-20/1_1983    ..........a...........a..............................g..........c............... [124000]
AY261360.1_Kenya_1950              ..........a...........a.................t...........gg.a.....c.................. [124000]
AM712240.1_OURT_88/3               ...............................................................................c [124000]
AM712239.1_Benin_97/1              ...............................................................................c [124000]
KM111295.1_Ken06.Bus               ....c.....a...........a.................t...........gg.a.....c.................. [124000]
KM111294.1_Ken05/Tk1               ..........a...........a.................t...........gg.a.....c.................. [124000]
KM262844.1_strain_L60              ...............................................................................c [124000]
KM102979.1_26544/OG10_Italy        ...............................................................................c [124000]
KM262845.1_strain_NHV              ...............................................................................c [124000]
KX354450.1_47/Ss/2008              ...............................................................................c [124000]
KP055815.1_strain_BA71             ...............................................................................c [124000]
MH025920.1_strain_R35              ....c.....a...........a.................t...........gg.a.....c.................. [124000]
MH025919.1_strain_N10              ....c.....a...........a.................t...........gg.a.....c.................. [124000]
MH025918.1_strain_R25              ....c.....a...........a.................t...........gg.a.....c.................. [124000]
MH025917.1_strain_R7               ....c.....a...........a.................t...........gg.a.....c.................. [124000]
MH025916.1_strain_R8               ....c.....a...........a.................t...........gg.a.....c.................. [124000]

FR682468.2_Georgia_2007/1          atgattaattttaagtttcaaataaacccagtgtctccccgcgtcatcattcatttaaaaaaatttgcagctttgttgga [124080]
ASF_IND_20_CAD543                  ...............................nnnnnnnnnnnn...........................nnnnn..... [124080]
LR899193.1_Germany_2020/1          ................................................................................ [124080]
MN393477.1_Wuhan_2019              ................................................................................ [124080]
MW396979.1_Timor-Leste/2019/1      ................................................................................ [124080]
MK940252.1_CN/2019/InnerMongolia   ................................................................................ [124080]
MK128995.1_China/2018/AnhuiXCGQ    ................................................................................ [124080]
MH681419.1_ASFV/POL/2015           ................................................................................ [124080]
LS478113.1_Estonia_2014            ................................................................................ [124080]
KP843857.1_Odintsovo_02/14         ................................................................................ [124080]
FN557520.1_strain_E75              ................................................................................ [124080]
AY261366.1_Warthog                 ................................................................................ [124080]
AY261365.1_Warmbaths               .......................................................................c........ [124080]
AY261364.1_Tengani_62              .............................g.................................................. [124080]
AY261363.1_Pretoriuskop/96/4       ................................................................................ [124080]
AY261362.1_Mkuzi_1979              ................................................................................ [124080]
AY261361.1_Malawi_Lil-20/1_1983    .............................g.........................................c........ [124080]
AY261360.1_Kenya_1950              ...........................t.g......................................g..c.....a.. [124080]
AM712240.1_OURT_88/3               ................................................................................ [124080]
AM712239.1_Benin_97/1              ................................................................................ [124080]
KM111295.1_Ken06.Bus               ...........................t.g......................................g..c........ [124080]
```

```
KM111294.1_Ken05/Tk1            ...........................t.g......................................g..c.....a.. [124080]
KM262844.1_strain_L60           ................................................................................ [124080]
KM102979.1_26544/OG10_Italy     ................................................................................ [124080]
KM262845.1_strain_NHV           ................................................................................ [124080]
KX354450.1_47/Ss/2008           ................................................................................ [124080]
KP055815.1_strain_BA71          ................................................................................ [124080]
MH025920.1_strain_R35           ...........................t.g......................................g..c........ [124080]
MH025919.1_strain_N10           ...........................t.g......................................g..c........ [124080]
MH025918.1_strain_R25           ...........................t.g......................................g..c........ [124080]
MH025917.1_strain_R7            ...........................t.g......................................g..c........ [124080]
MH025916.1_strain_R8            ...........................t.g......................................g..c........ [124080]

FR682468.2_Georgia_2007/1       acacatccctactccatatcccatacgtgaaataaagcctccattagaagactcaaaagtatccgcaaaatttatggtca [124160]
ASF_IND_20_CAD543               ................................................................................ [124160]
LR899193.1_Germany_2020/1       ................................................................................ [124160]
MN393477.1_Wuhan_2019           ................................................................................ [124160]
MW396979.1_Timor-Leste/2019/1   ................................................................................ [124160]
MK940252.1_CN/2019/InnerMongolia ................................................................................ [124160]
MK128995.1_China/2018/AnhuiXCGQ ................................................................................ [124160]
MH681419.1_ASFV/POL/2015        ................................................................................ [124160]
LS478113.1_Estonia_2014         ................................................................................ [124160]
KP843857.1_Odintsovo_02/14      ................................................................................ [124160]
FN557520.1_strain_E75           ................................................................................ [124160]
AY261366.1_Warthog              .........................................................g...................... [124160]
AY261365.1_Warmbaths            ............................................................t................... [124160]
AY261364.1_Tengani_62           ............................................................g................... [124160]
AY261363.1_Pretoriuskop/96/4    ............................................................t................... [124160]
AY261362.1_Mkuzi_1979           ................................................................................ [124160]
AY261361.1_Malawi_Lil-20/1_1983 ...a..t..c..g...............................................g................... [124160]
AY261360.1_Kenya_1950           ...a..a..c..................................................g................... [124160]
AM712240.1_OURT_88/3            ................................................................................ [124160]
AM712239.1_Benin_97/1           ................................................................................ [124160]
KM111295.1_Ken06.Bus            ...a..a..c..................................................g................... [124160]
KM111294.1_Ken05/Tk1            ...a..a..c..................................................g................... [124160]
KM262844.1_strain_L60           ................................................................................ [124160]
KM102979.1_26544/OG10_Italy     ................................................................................ [124160]
KM262845.1_strain_NHV           ................................................................................ [124160]
KX354450.1_47/Ss/2008           ................................................................................ [124160]
KP055815.1_strain_BA71          ................................................................................ [124160]
MH025920.1_strain_R35           ...a..a..c..................................................g................... [124160]
MH025919.1_strain_N10           ...a..a..c..................................................g................... [124160]
MH025918.1_strain_R25           ...a..a..c..................................................g................... [124160]
MH025917.1_strain_R7            ...a..a..c..................................................g................... [124160]
MH025916.1_strain_R8            ...a..a..c..................................................g................... [124160]
```

```
FR682468.2_Georgia_2007/1        gtccgggaaaaaaagtacgcattaatgtttttcttaaaggtaagataaatattttaggctgcaacacaaaggaatccgcg [124240]
ASF_IND_20_CAD543                ................................................................................ [124240]
LR899193.1_Germany_2020/1        ................................................................................ [124240]
MN393477.1_Wuhan_2019            ................................................................................ [124240]
MW396979.1_Timor-Leste/2019/1    ................................................................................ [124240]
MK940252.1_CN/2019/InnerMongolia ................................................................................ [124240]
MK128995.1_China/2018/AnhuiXCGQ  ................................................................................ [124240]
MH681419.1_ASFV/POL/2015         ................................................................................ [124240]
LS478113.1_Estonia_2014          ................................................................................ [124240]
KP843857.1_Odintsovo_02/14       ................................................................................ [124240]
FN557520.1_strain_E75            ................................................................................ [124240]
AY261366.1_Warthog               ................................................................................ [124240]
AY261365.1_Warmbaths             ............................................................................t... [124240]
AY261364.1_Tengani_62            ................................................................................ [124240]
AY261363.1_Pretoriuskop/96/4     ............................................................................t... [124240]
AY261362.1_Mkuzi_1979            ................................................................................ [124240]
AY261361.1_Malawi_Lil-20/1_1983  .c......................................c....................................... [124240]
AY261360.1_Kenya_1950            ......................c............................................g............ [124240]
AM712240.1_OURT_88/3             ................................................................................ [124240]
AM712239.1_Benin_97/1            ................................................................................ [124240]
KM111295.1_Ken06.Bus             ......................c............................................g............ [124240]
KM111294.1_Ken05/Tk1             ......................c............................................g............ [124240]
KM262844.1_strain_L60            ................................................................................ [124240]
KM102979.1_26544/OG10_Italy      ................................................................................ [124240]
KM262845.1_strain_NHV            ................................................................................ [124240]
KX354450.1_47/Ss/2008            ................................................................................ [124240]
KP055815.1_strain_BA71           ................................................................................ [124240]
MH025920.1_strain_R35            ......................c............................................g............ [124240]
MH025919.1_strain_N10            ......................c............................................g............ [124240]
MH025918.1_strain_R25            ......................c............................................g............ [124240]
MH025917.1_strain_R7             ......................c............................................g............ [124240]
MH025916.1_strain_R8             ......................c............................................g............ [124240]

FR682468.2_Georgia_2007/1        gagaccatttatacgtttttgaaagatcttatcagcgtacattggcaagaaattttgtgcgtgttaccggtacccgatta [124320]
ASF_IND_20_CAD543                ................................................................................ [124320]
LR899193.1_Germany_2020/1        ................................................................................ [124320]
MN393477.1_Wuhan_2019            ................................................................................ [124320]
MW396979.1_Timor-Leste/2019/1    ................................................................................ [124320]
MK940252.1_CN/2019/InnerMongolia ................................................................................ [124320]
MK128995.1_China/2018/AnhuiXCGQ  ................................................................................ [124320]
MH681419.1_ASFV/POL/2015         ................................................................................ [124320]
LS478113.1_Estonia_2014          ................................................................................ [124320]
KP843857.1_Odintsovo_02/14       ................................................................................ [124320]
```

```
FN557520.1_strain_E75           ....t........................................................................... [124320]
AY261366.1_Warthog              ...........................................................t...........g........ [124320]
AY261365.1_Warmbaths            ................................................................................ [124320]
AY261364.1_Tengani_62           ..............a........................................................g........ [124320]
AY261363.1_Pretoriuskop/96/4    .....................................................................a.g........ [124320]
AY261362.1_Mkuzi_1979           ................................................................................ [124320]
AY261361.1_Malawi_Lil-20/1_1983 ...................................t.........................................c.. [124320]
AY261360.1_Kenya_1950           ............g................................................................... [124320]
AM712240.1_OURT_88/3            ....t........................................................................... [124320]
AM712239.1_Benin_97/1           ....t........................................................................... [124320]
KM111295.1_Ken06.Bus            ............g................................................................c.. [124320]
KM111294.1_Ken05/Tk1            ............g................................................................c.. [124320]
KM262844.1_strain_L60           ....t........................................................................... [124320]
KM102979.1_26544/OG10_Italy     ....t........................................................................... [124320]
KM262845.1_strain_NHV           ....t........................................................................... [124320]
KX354450.1_47/Ss/2008           ....t........................................................................... [124320]
KP055815.1_strain_BA71          ....t........................................................................... [124320]
MH025920.1_strain_R35           ............g................................................................c.. [124320]
MH025919.1_strain_N10           ............g................................................................c.. [124320]
MH025918.1_strain_R25           ............g................................................................c.. [124320]
MH025917.1_strain_R7            ............g................................................................c.. [124320]
MH025916.1_strain_R8            ............g................................................................c.. [124320]

FR682468.2_Georgia_2007/1       aagaatgttttcattaataaggtaatcgactatgctaaaaagaataacaagaaaaataccttgaagaactataccaaagt [124400]
ASF_IND_20_CAD543               ................................................................................ [124400]
LR899193.1_Germany_2020/1       ................................................................................ [124400]
MN393477.1_Wuhan_2019           ................................................................................ [124400]
MW396979.1_Timor-Leste/2019/1   ................................................................................ [124400]
MK940252.1_CN/2019/InnerMongolia ................................................................................ [124400]
MK128995.1_China/2018/AnhuiXCGQ ................................................................................ [124400]
MH681419.1_ASFV/POL/2015        ................................................................................ [124400]
LS478113.1_Estonia_2014         ................................................................................ [124400]
KP843857.1_Odintsovo_02/14      ................................................................................ [124400]
FN557520.1_strain_E75           ..............................g................................................. [124400]
AY261366.1_Warthog              ......a.......................g................................................. [124400]
AY261365.1_Warmbaths            ......a.......................g....................g............................ [124400]
AY261364.1_Tengani_62           ..............................g..............c.................................. [124400]
AY261363.1_Pretoriuskop/96/4    ......a.......................g................................................. [124400]
AY261362.1_Mkuzi_1979           ..............................g................................................. [124400]
AY261361.1_Malawi_Lil-20/1_1983 g.....a....................a..g..........................t..c................... [124400]
AY261360.1_Kenya_1950           g.....a....................ag.g.............................c...........g....... [124400]
AM712240.1_OURT_88/3            ..............................g................................................. [124400]
AM712239.1_Benin_97/1           ..............................g................................................. [124400]
KM111295.1_Ken06.Bus            g.....a....................ag.g.............................c...........g....... [124400]
```

```
KM111294.1_Ken05/Tk1              g.....a....................ag.g.............................c...........g....... [124400]
KM262844.1_strain_L60             ..............................g................................................. [124400]
KM102979.1_26544/OG10_Italy       ..............................g................................................. [124400]
KM262845.1_strain_NHV             ..............................g................................................. [124400]
KX354450.1_47/Ss/2008             ..............................g................................................. [124400]
KP055815.1_strain_BA71            ..............................g................................................. [124400]
MH025920.1_strain_R35             g.....a....................ag.g.............................c...........g....... [124400]
MH025919.1_strain_N10             g.....a....................ag.g.............................c...........g....... [124400]
MH025918.1_strain_R25             g.....a....................ag.g.............................c...........g....... [124400]
MH025917.1_strain_R7              g.....a....................ag.g.............................c...........g....... [124400]
MH025916.1_strain_R8              g.....a....................ag.g.............................c...........g....... [124400]

FR682468.2_Georgia_2007/1         aggtaggttttctgcatgtcacggcatggttaaaattgctaataatgtagtccacaaaagcattgctcaatacgactaaa [124480]
ASF_IND_20_CAD543                 ................................................................................ [124480]
LR899193.1_Germany_2020/1         ................................................................................ [124480]
MN393477.1_Wuhan_2019             ................................................................................ [124480]
MW396979.1_Timor-Leste/2019/1     ................................................................................ [124480]
MK940252.1_CN/2019/InnerMongolia  ................................................................................ [124480]
MK128995.1_China/2018/AnhuiXCGQ   ................................................................................ [124480]
MH681419.1_ASFV/POL/2015          ................................................................................ [124480]
LS478113.1_Estonia_2014           ................................................................................ [124480]
KP843857.1_Odintsovo_02/14        ................................................................................ [124480]
FN557520.1_strain_E75             ..........................................................g..................... [124480]
AY261366.1_Warthog                .....................................a....................g........t............ [124480]
AY261365.1_Warmbaths              .....................................a....................g........t............ [124480]
AY261364.1_Tengani_62             ..........................................................g..................... [124480]
AY261363.1_Pretoriuskop/96/4      ..........................................................g........t............ [124480]
AY261362.1_Mkuzi_1979             ..........................................................g..................... [124480]
AY261361.1_Malawi_Lil-20/1_1983   ...................a.......................g.............................c...... [124480]
AY261360.1_Kenya_1950             ...................a.......................g..............g..............c...... [124480]
AM712240.1_OURT_88/3              ..........................................................g..................... [124480]
AM712239.1_Benin_97/1             ..........................................................g..................... [124480]
KM111295.1_Ken06.Bus              ...................a.......................g..............g..............c...... [124480]
KM111294.1_Ken05/Tk1              ...................a.......................g..............g..............c...... [124480]
KM262844.1_strain_L60             ..........................................................g..................... [124480]
KM102979.1_26544/OG10_Italy       ..........................................................g..................... [124480]
KM262845.1_strain_NHV             ..........................................................g..................... [124480]
KX354450.1_47/Ss/2008             ..........................................................g..................... [124480]
KP055815.1_strain_BA71            ..........................................................g..................... [124480]
MH025920.1_strain_R35             ...................a.......................g..............g..............c...... [124480]
MH025919.1_strain_N10             ...................a.......................g..............g..............c...... [124480]
MH025918.1_strain_R25             ...................a.......................g..............g..............c...... [124480]
MH025917.1_strain_R7              ...................a.......................g..............g..............c...... [124480]
MH025916.1_strain_R8              ...................a.......................g..............g..............c...... [124480]
```

```
FR682468.2_Georgia_2007/1        aatagtaaaaaaaggataagtgctctttttatatccatatactttaaaacttattttttacactaataatttcctgcggc [124560]
ASF_IND_20_CAD543                ................................................................................ [124560]
LR899193.1_Germany_2020/1        ................................................................................ [124560]
MN393477.1_Wuhan_2019            ................................................................................ [124560]
MW396979.1_Timor-Leste/2019/1    ................................................................................ [124560]
MK940252.1_CN/2019/InnerMongolia ................................................................................ [124560]
MK128995.1_China/2018/AnhuiXCGQ  ................................................................................ [124560]
MH681419.1_ASFV/POL/2015         ................................................................................ [124560]
LS478113.1_Estonia_2014          ................................................................................ [124560]
KP843857.1_Odintsovo_02/14       ................................................................................ [124560]
FN557520.1_strain_E75            ......-......................................................................... [124560]
AY261366.1_Warthog               ................................................................................ [124560]
AY261365.1_Warmbaths             ................................................................................ [124560]
AY261364.1_Tengani_62            ................................................................................ [124560]
AY261363.1_Pretoriuskop/96/4     ................................................................................ [124560]
AY261362.1_Mkuzi_1979            ................................................................................ [124560]
AY261361.1_Malawi_Lil-20/1_1983  ..............................................................t................. [124560]
AY261360.1_Kenya_1950            .................c...........................................g.................. [124560]
AM712240.1_OURT_88/3             ................................................................................ [124560]
AM712239.1_Benin_97/1            ................................................................................ [124560]
KM111295.1_Ken06.Bus             .................c...........................................g.................. [124560]
KM111294.1_Ken05/Tk1             .................c...........................................g.................. [124560]
KM262844.1_strain_L60            ................................................................................ [124560]
KM102979.1_26544/OG10_Italy      ................................................................................ [124560]
KM262845.1_strain_NHV            ................................................................................ [124560]
KX354450.1_47/Ss/2008            ................................................................................ [124560]
KP055815.1_strain_BA71           ................................................................................ [124560]
MH025920.1_strain_R35            .................c...........................................g.................. [124560]
MH025919.1_strain_N10            .................c...........................................g.................. [124560]
MH025918.1_strain_R25            .................c...........................................g.................. [124560]
MH025917.1_strain_R7             .................c...........................................g.................. [124560]
MH025916.1_strain_R8             .................c...........................................g.................. [124560]

FR682468.2_Georgia_2007/1        cgcaatataaactgtaggtcatctataacgcccagacctgttaaaagtagagtactatgttttaagggatttaaaatatc [124640]
ASF_IND_20_CAD543                ................................................................................ [124640]
LR899193.1_Germany_2020/1        ................................................................................ [124640]
MN393477.1_Wuhan_2019            ................................................................................ [124640]
MW396979.1_Timor-Leste/2019/1    ................................................................................ [124640]
MK940252.1_CN/2019/InnerMongolia ................................................................................ [124640]
MK128995.1_China/2018/AnhuiXCGQ  ................................................................................ [124640]
MH681419.1_ASFV/POL/2015         ................................................................................ [124640]
LS478113.1_Estonia_2014          ................................................................................ [124640]
KP843857.1_Odintsovo_02/14       ................................................................................ [124640]
```

```
FN557520.1_strain_E75             ................................................................................ [124640]
AY261366.1_Warthog                ...................................c............................................ [124640]
AY261365.1_Warmbaths              ...g................................t............................................ [124640]
AY261364.1_Tengani_62             .............................a.....t.........................................g.. [124640]
AY261363.1_Pretoriuskop/96/4      ...................................t............................................ [124640]
AY261362.1_Mkuzi_1979             ................................................................................ [124640]
AY261361.1_Malawi_Lil-20/1_1983   .............................a.....t............................................ [124640]
AY261360.1_Kenya_1950             ....t..............................t.........................................g.. [124640]
AM712240.1_OURT_88/3              ................................................................................ [124640]
AM712239.1_Benin_97/1             ................................................................................ [124640]
KM111295.1_Ken06.Bus              ....t..............................t.............................a...........g.. [124640]
KM111294.1_Ken05/Tk1              ....t..............................t.........................................g.. [124640]
KM262844.1_strain_L60             ................................................................................ [124640]
KM102979.1_26544/OG10_Italy       ................................................................................ [124640]
KM262845.1_strain_NHV             ................................................................................ [124640]
KX354450.1_47/Ss/2008             ................................................................................ [124640]
KP055815.1_strain_BA71            ................................................................................ [124640]
MH025920.1_strain_R35             ....t..............................t.............................a...........g.. [124640]
MH025919.1_strain_N10             ....t..............................t.............................a...........g.. [124640]
MH025918.1_strain_R25             ....t..............................t.............................a...........g.. [124640]
MH025917.1_strain_R7              ....t..............................t.............................a...........g.. [124640]
MH025916.1_strain_R8              ....t..............................t.............................a...........g.. [124640]

FR682468.2_Georgia_2007/1         cgccgcaagaatgtgaatataattttcaaagtggtttacaggaatgcgtaagcgtttttttttgcactgcggttggttta [124720]
ASF_IND_20_CAD543                 ................................................................................ [124720]
LR899193.1_Germany_2020/1         ................................................................................ [124720]
MN393477.1_Wuhan_2019             ................................................................................ [124720]
MW396979.1_Timor-Leste/2019/1     ................................................................................ [124720]
MK940252.1_CN/2019/InnerMongolia  ................................................................................ [124720]
MK128995.1_China/2018/AnhuiXCGQ   ................................................................................ [124720]
MH681419.1_ASFV/POL/2015          ................................................................................ [124720]
LS478113.1_Estonia_2014           ................................................................................ [124720]
KP843857.1_Odintsovo_02/14        ................................................................................ [124720]
FN557520.1_strain_E75             ......g......................................................................... [124720]
AY261366.1_Warthog                ......g.......................................................c................. [124720]
AY261365.1_Warmbaths              ......g......................................................................... [124720]
AY261364.1_Tengani_62             ..............t...............................................c................. [124720]
AY261363.1_Pretoriuskop/96/4      ..............................................................c................. [124720]
AY261362.1_Mkuzi_1979             ......g...............................t......................................... [124720]
AY261361.1_Malawi_Lil-20/1_1983   ......g.......t...............a....................a..........c.........c....... [124720]
AY261360.1_Kenya_1950             ......g.......t...............a....................a..........c.........c..c.... [124720]
AM712240.1_OURT_88/3              ......g......................................................................... [124720]
AM712239.1_Benin_97/1             ......g......................................................................... [124720]
KM111295.1_Ken06.Bus              ......g.......t...............a....................a..........c.........c..c.... [124720]
```

```
KM111294.1_Ken05/Tk1           ......g.......t...............a....................a..........c.........c..c.... [124720]
KM262844.1_strain_L60          ......g.......................................................................... [124720]
KM102979.1_26544/OG10_Italy    ......g.......................................................................... [124720]
KM262845.1_strain_NHV          ......g.......................................................................... [124720]
KX354450.1_47/Ss/2008          ......g.......................................................................... [124720]
KP055815.1_strain_BA71         ......g.......................................................................... [124720]
MH025920.1_strain_R35          ......g.......t...............a....................a..........c.........c..c.... [124720]
MH025919.1_strain_N10          ......g.......t...............a....................a..........c.........c..c.... [124720]
MH025918.1_strain_R25          ......g.......t...............a....................a..........c.........c..c.... [124720]
MH025917.1_strain_R7           ......g.......t...............a....................a..........c.........c..c.... [124720]
MH025916.1_strain_R8           ......g.......t...............a....................a..........c.........c..c.... [124720]

FR682468.2_Georgia_2007/1      gggtcgaatactggcaggaggtatatatattaataagaccgcggtcgatggtttcaatatcttcatagaattcaatgcgc [124800]
ASF_IND_20_CAD543              ................................................................................ [124800]
LR899193.1_Germany_2020/1      ................................................................................ [124800]
MN393477.1_Wuhan_2019          ................................................................................ [124800]
MW396979.1_Timor-Leste/2019/1  ................................................................................ [124800]
MK940252.1_CN/2019/InnerMongolia ................................................................................ [124800]
MK128995.1_China/2018/AnhuiXCGQ ................................................................................ [124800]
MH681419.1_ASFV/POL/2015       ................................................................................ [124800]
LS478113.1_Estonia_2014        ................................................................................ [124800]
KP843857.1_Odintsovo_02/14     ................................................................................ [124800]
FN557520.1_strain_E75          ................................................................................ [124800]
AY261366.1_Warthog             ...................................................................a............ [124800]
AY261365.1_Warmbaths           ........................................a....................................... [124800]
AY261364.1_Tengani_62          ................................................................................ [124800]
AY261363.1_Pretoriuskop/96/4   .....................................t.......................................... [124800]
AY261362.1_Mkuzi_1979          ................................................................................ [124800]
AY261361.1_Malawi_Lil-20/1_1983 a.......................................a....................................... [124800]
AY261360.1_Kenya_1950          ................................................................................ [124800]
AM712240.1_OURT_88/3           ................................................................................ [124800]
AM712239.1_Benin_97/1          ................................................................................ [124800]
KM111295.1_Ken06.Bus           ................................................................................ [124800]
KM111294.1_Ken05/Tk1           ................................................................................ [124800]
KM262844.1_strain_L60          ................................................................................ [124800]
KM102979.1_26544/OG10_Italy    ................................................................................ [124800]
KM262845.1_strain_NHV          ................................................................................ [124800]
KX354450.1_47/Ss/2008          ................................................................................ [124800]
KP055815.1_strain_BA71         ................................................................................ [124800]
MH025920.1_strain_R35          ................................................................................ [124800]
MH025919.1_strain_N10          ................................................................................ [124800]
MH025918.1_strain_R25          ................................................................................ [124800]
MH025917.1_strain_R7           ................................................................................ [124800]
MH025916.1_strain_R8           ................................................................................ [124800]
```

```
FR682468.2_Georgia_2007/1        ggcgtcaaaagttttttaagatgttgacataactcatcatacgtgtaggactggaggggggaaagaagggtgtagtcaaa [124880]
ASF_IND_20_CAD543                ..........................................................................r..... [124880]
LR899193.1_Germany_2020/1        ................................................................................ [124880]
MN393477.1_Wuhan_2019            ................................................................................ [124880]
MW396979.1_Timor-Leste/2019/1    ................................................................................ [124880]
MK940252.1_CN/2019/InnerMongolia ................................................................................ [124880]
MK128995.1_China/2018/AnhuiXCGQ  ................................................................................ [124880]
MH681419.1_ASFV/POL/2015         ................................................................................ [124880]
LS478113.1_Estonia_2014          ................................................................................ [124880]
KP843857.1_Odintsovo_02/14       ................................................................................ [124880]
FN557520.1_strain_E75            .........................................................-................a..... [124880]
AY261366.1_Warthog               ..t.................g........................................................... [124880]
AY261365.1_Warmbaths             ....................g........................................................... [124880]
AY261364.1_Tengani_62            ....................g............................c.............................. [124880]
AY261363.1_Pretoriuskop/96/4     ....................g........................................................... [124880]
AY261362.1_Mkuzi_1979            ....................g........................................................... [124880]
AY261361.1_Malawi_Lil-20/1_1983  ....................g.............................t........a.........ac......... [124880]
AY261360.1_Kenya_1950            ....................g.a...........................t.....a..c.........a.......... [124880]
AM712240.1_OURT_88/3             ..........................................................................a..... [124880]
AM712239.1_Benin_97/1            ..........................................................................a..... [124880]
KM111295.1_Ken06.Bus             ....................g.a...........................t.....a..c.........a..cg...... [124880]
KM111294.1_Ken05/Tk1             ....................g.a...........................t.....a..c.........a...g...... [124880]
KM262844.1_strain_L60            ..........................................................................a..... [124880]
KM102979.1_26544/OG10_Italy      ..........................................................................a..... [124880]
KM262845.1_strain_NHV            ..........................................................................a..... [124880]
KX354450.1_47/Ss/2008            ..........................................................................a..... [124880]
KP055815.1_strain_BA71           ..........................................................................a..... [124880]
MH025920.1_strain_R35            ....................g.a...........................t.....a..c.........a..cg...... [124880]
MH025919.1_strain_N10            ....................g.a...........................t.....a..c.........a..cg...... [124880]
MH025918.1_strain_R25            ....................g.a...........................t.....a..c.........a..cg...... [124880]
MH025917.1_strain_R7             ....................g.a...........................t.....a..c.........a..cg...... [124880]
MH025916.1_strain_R8             ....................g.a...........................t.....a..c.........a..cg...... [124880]

FR682468.2_Georgia_2007/1        gttaaaaatgtttttttgaagaacctttaaagcatgttccgcgtccgtggtttccaaaatatgttttatggtatgaatgt [124960]
ASF_IND_20_CAD543                ................................................................................ [124960]
LR899193.1_Germany_2020/1        ................................................................................ [124960]
MN393477.1_Wuhan_2019            ................................................................................ [124960]
MW396979.1_Timor-Leste/2019/1    ................................................................................ [124960]
MK940252.1_CN/2019/InnerMongolia ................................................................................ [124960]
MK128995.1_China/2018/AnhuiXCGQ  ................................................................................ [124960]
MH681419.1_ASFV/POL/2015         ................................................................................ [124960]
LS478113.1_Estonia_2014          ................................................................................ [124960]
KP843857.1_Odintsovo_02/14       ................................................................................ [124960]
```

```
FN557520.1_strain_E75            ................................................................................ [124960]
AY261366.1_Warthog               ..........................................a..................................... [124960]
AY261365.1_Warmbaths             ..........................................a..........................a.......... [124960]
AY261364.1_Tengani_62            ..........................................a..................................... [124960]
AY261363.1_Pretoriuskop/96/4     ..........................................a..................................... [124960]
AY261362.1_Mkuzi_1979            ................................................................................ [124960]
AY261361.1_Malawi_Lil-20/1_1983  ........ca................................ac.........................t.......... [124960]
AY261360.1_Kenya_1950            ........ca..........................g.....ac............g............t.......... [124960]
AM712240.1_OURT_88/3             ................................................................................ [124960]
AM712239.1_Benin_97/1            ................................................................................ [124960]
KM111295.1_Ken06.Bus             ........ca..........................g.....ac............g......c.....tc......... [124960]
KM111294.1_Ken05/Tk1             ........ca..........................g.....ac............g....................... [124960]
KM262844.1_strain_L60            ................................................................................ [124960]
KM102979.1_26544/OG10_Italy      ................................................................................ [124960]
KM262845.1_strain_NHV            ................................................................................ [124960]
KX354450.1_47/Ss/2008            ................................................................................ [124960]
KP055815.1_strain_BA71           ................................................................................ [124960]
MH025920.1_strain_R35            ........ca..........................g.....ac............g......c.....tc......... [124960]
MH025919.1_strain_N10            ........ca..........................g.....ac............g......c.....tc......... [124960]
MH025918.1_strain_R25            ........ca..........................g.....ac............g......c.....tc......... [124960]
MH025917.1_strain_R7             ........ca..........................g.....ac............g......c.....tc......... [124960]
MH025916.1_strain_R8             ........ca..........................g.....ac............g......c.....tc......... [124960]

FR682468.2_Georgia_2007/1        catttaaatctacaaagtctgacagctttgtgtagaactcggtgacggaggttattttctggaaatcggtttttgaaaa [125040]
ASF_IND_20_CAD543                ................................................................................ [125040]
LR899193.1_Germany_2020/1        ................................................................................ [125040]
MN393477.1_Wuhan_2019            ................................................................................ [125040]
MW396979.1_Timor-Leste/2019/1    ................................................................................ [125040]
MK940252.1_CN/2019/InnerMongolia ................................................................................ [125040]
MK128995.1_China/2018/AnhuiXCGQ  ................................................................................ [125040]
MH681419.1_ASFV/POL/2015         ................................................................................ [125040]
LS478113.1_Estonia_2014          ................................................................................ [125040]
KP843857.1_Odintsovo_02/14       ................................................................................ [125040]
FN557520.1_strain_E75            ................................................................................ [125040]
AY261366.1_Warthog               .g...................................t.....c..c................................. [125040]
AY261365.1_Warmbaths             .g............................................a................................. [125040]
AY261364.1_Tengani_62            .........................t.................c..c................................. [125040]
AY261363.1_Pretoriuskop/96/4     ...........................................c..c................................. [125040]
AY261362.1_Mkuzi_1979            ................................................................................ [125040]
AY261361.1_Malawi_Lil-20/1_1983  .......g..a................................c..c...t............................. [125040]
AY261360.1_Kenya_1950            ..........a...........t....................c..c..at............................. [125040]
AM712240.1_OURT_88/3             ................................................................................ [125040]
AM712239.1_Benin_97/1            ................................................................................ [125040]
KM111295.1_Ken06.Bus             ..........a...........t.......................c...t............................. [125040]
```

```
KM111294.1_Ken05/Tk1            ..........a...........t....................c..c..at............................. [125040]
KM262844.1_strain_L60           ................................................................................ [125040]
KM102979.1_26544/OG10_Italy     ................................................................................ [125040]
KM262845.1_strain_NHV           ................................................................................ [125040]
KX354450.1_47/Ss/2008           ................................................................................ [125040]
KP055815.1_strain_BA71          ................................................................................ [125040]
MH025920.1_strain_R35           ..........a...........t.......................c...t............................. [125040]
MH025919.1_strain_N10           ..........a...........t.......................c...t............................. [125040]
MH025918.1_strain_R25           ..........a...........t.......................c...t............................. [125040]
MH025917.1_strain_R7            ..........a...........t.......................c...t............................. [125040]
MH025916.1_strain_R8            ..........a...........t.......................c...t............................. [125040]

FR682468.2_Georgia_2007/1       agattttcaatgtgtttgcgggttgagttgctttgcagtccatacaagacatcaaaaaattcaatcagcaaaaacttata [125120]
ASF_IND_20_CAD543               ................................................................................ [125120]
LR899193.1_Germany_2020/1       ................................................................................ [125120]
MN393477.1_Wuhan_2019           ................................................................................ [125120]
MW396979.1_Timor-Leste/2019/1   ................................................................................ [125120]
MK940252.1_CN/2019/InnerMongolia ................................................................................ [125120]
MK128995.1_China/2018/AnhuiXCGQ ................................................................................ [125120]
MH681419.1_ASFV/POL/2015        ................................................................................ [125120]
LS478113.1_Estonia_2014         ................................................................................ [125120]
KP843857.1_Odintsovo_02/14      ................................................................................ [125120]
FN557520.1_strain_E75           ................................................................................ [125120]
AY261366.1_Warthog              .........................t........................g........c.................... [125120]
AY261365.1_Warmbaths            ..................................................g........c.................... [125120]
AY261364.1_Tengani_62           ...gc....................t.................................c.......a............ [125120]
AY261363.1_Pretoriuskop/96/4    .........................t.................................c.................... [125120]
AY261362.1_Mkuzi_1979           ................................................................................ [125120]
AY261361.1_Malawi_Lil-20/1_1983 ....c....................t...................................................... [125120]
AY261360.1_Kenya_1950           ....c.......c.......a....ta........................................a............ [125120]
AM712240.1_OURT_88/3            ................................................................................ [125120]
AM712239.1_Benin_97/1           ................................................................................ [125120]
KM111295.1_Ken06.Bus            ....c....................t.........................................a............ [125120]
KM111294.1_Ken05/Tk1            ....c.......c.......a....ta........................................a............ [125120]
KM262844.1_strain_L60           ................................................................................ [125120]
KM102979.1_26544/OG10_Italy     ................................................................................ [125120]
KM262845.1_strain_NHV           ................................................................................ [125120]
KX354450.1_47/Ss/2008           ................................................................................ [125120]
KP055815.1_strain_BA71          ................................................................................ [125120]
MH025920.1_strain_R35           ....c....................t.........................................a............ [125120]
MH025919.1_strain_N10           ....c....................t.........................................a............ [125120]
MH025918.1_strain_R25           ....c....................t.........................................a............ [125120]
MH025917.1_strain_R7            ....c....................t.........................................a............ [125120]
MH025916.1_strain_R8            ....c....................t.........................................a............ [125120]
```

FR682468.2_Georgia_2007/1 caaatggttaatataaaaagctttgttggccttattctgctgaggatatggttcctctaggggatatagaatggcttggt [125200]
ASF_IND_20_CAD543 ................................................................................ [125200]
LR899193.1_Germany_2020/1 ................................................................................ [125200]
MN393477.1_Wuhan_2019 ................................................................................ [125200]
MW396979.1_Timor-Leste/2019/1 ................................................................................ [125200]
MK940252.1_CN/2019/InnerMongolia ................................................................................ [125200]
MK128995.1_China/2018/AnhuiXCGQ ................................................................................ [125200]
MH681419.1_ASFV/POL/2015 ................................................................................ [125200]
LS478113.1_Estonia_2014 ................................................................................ [125200]
KP843857.1_Odintsovo_02/14 ................................................................................ [125200]
FN557520.1_strain_E75 .........................................................c...................... [125200]
AY261366.1_Warthog .........................................................c...................... [125200]
AY261365.1_Warmbaths .........................................................c...................... [125200]
AY261364.1_Tengani_62 .........................................................c...................... [125200]
AY261363.1_Pretoriuskop/96/4 ....................................t....................c...................... [125200]
AY261362.1_Mkuzi_1979 .........................................................c...................... [125200]
AY261361.1_Malawi_Lil-20/1_1983 ...................................c.....................c..a..g................ [125200]
AY261360.1_Kenya_1950 ......a..c.........................c...a.....g...........c..........a........... [125200]
AM712240.1_OURT_88/3 .........................................................c...................... [125200]
AM712239.1_Benin_97/1 .........................................................c...................... [125200]
KM111295.1_Ken06.Bus ......a..c.........................c...a.....g...........c........c.a........... [125200]
KM111294.1_Ken05/Tk1 ......a..c.........................c...a.....g...........c..........a........... [125200]
KM262844.1_strain_L60 .........................................................c...................... [125200]
KM102979.1_26544/OG10_Italy .........................................................c...................... [125200]
KM262845.1_strain_NHV .........................................................c...................... [125200]
KX354450.1_47/Ss/2008 .........................................................c...................... [125200]
KP055815.1_strain_BA71 .........................................................c...................... [125200]
MH025920.1_strain_R35 ......a..c.........................c...a.....g...........c........c.a........... [125200]
MH025919.1_strain_N10 ......a..c.........................c...a.....g...........c........c.a........... [125200]
MH025918.1_strain_R25 ......a..c.........................c...a.....g...........c........c.a........... [125200]
MH025917.1_strain_R7 ......a..c.........................c...a.....g...........c........c.a........... [125200]
MH025916.1_strain_R8 ......a..c.........................c...a.....g...........c........c.a........... [125200]

FR682468.2_Georgia_2007/1 ctatatccctaggatcaatagtcaatgttgcgatgggaagcttttccagcgtagcgggaagagtttgggttggagcgtag [125280]
ASF_IND_20_CAD543 ................................................................................ [125280]
LR899193.1_Germany_2020/1 ................................................................................ [125280]
MN393477.1_Wuhan_2019 ................................................................................ [125280]
MW396979.1_Timor-Leste/2019/1 ................................................................................ [125280]
MK940252.1_CN/2019/InnerMongolia ................................................................................ [125280]
MK128995.1_China/2018/AnhuiXCGQ ................................................................................ [125280]
MH681419.1_ASFV/POL/2015 ................................................................................ [125280]
LS478113.1_Estonia_2014 ................................................................................ [125280]
KP843857.1_Odintsovo_02/14 ................................................................................ [125280]

```
FN557520.1_strain_E75            ................................................................................ [125280]
AY261366.1_Warthog               .................................................t.c............................ [125280]
AY261365.1_Warmbaths             ..................................................ac............................ [125280]
AY261364.1_Tengani_62            .........g...................................................................... [125280]
AY261363.1_Pretoriuskop/96/4     ...................................................c............................ [125280]
AY261362.1_Mkuzi_1979            ................................................................................ [125280]
AY261361.1_Malawi_Lil-20/1_1983  ................c..g............................................................ [125280]
AY261360.1_Kenya_1950            ...................g............................a............................... [125280]
AM712240.1_OURT_88/3             ................................................................................ [125280]
AM712239.1_Benin_97/1            ................................................................................ [125280]
KM111295.1_Ken06.Bus             ...................g............................................................ [125280]
KM111294.1_Ken05/Tk1             ...................g............................................................ [125280]
KM262844.1_strain_L60            ................................................................................ [125280]
KM102979.1_26544/OG10_Italy      ................................................................................ [125280]
KM262845.1_strain_NHV            ................................................................................ [125280]
KX354450.1_47/Ss/2008            ................................................................................ [125280]
KP055815.1_strain_BA71           ................................................................................ [125280]
MH025920.1_strain_R35            ...................g............................................................ [125280]
MH025919.1_strain_N10            ...................g............................................................ [125280]
MH025918.1_strain_R25            ...................g............................................................ [125280]
MH025917.1_strain_R7             ...................g............................................................ [125280]
MH025916.1_strain_R8             ...................g............................................................ [125280]

FR682468.2_Georgia_2007/1        taaaagtatagcccggtttttccctctgaaagaaagcccacaaattctttttttatattttgcagcaccgctgagggtac [125360]
ASF_IND_20_CAD543                ................................................................................ [125360]
LR899193.1_Germany_2020/1        ................................................................................ [125360]
MN393477.1_Wuhan_2019            ................................................................................ [125360]
MW396979.1_Timor-Leste/2019/1    ................................................................................ [125360]
MK940252.1_CN/2019/InnerMongolia ................................................................................ [125360]
MK128995.1_China/2018/AnhuiXCGQ  ................................................................................ [125360]
MH681419.1_ASFV/POL/2015         ................................................................................ [125360]
LS478113.1_Estonia_2014          ................................................................................ [125360]
KP843857.1_Odintsovo_02/14       ................................................................................ [125360]
FN557520.1_strain_E75            .....a..............................................-........................... [125360]
AY261366.1_Warthog               ..............a.....................................................t.....a..g.t [125360]
AY261365.1_Warmbaths             ..............a................................................................. [125360]
AY261364.1_Tengani_62            .................................................................g..g........... [125360]
AY261363.1_Pretoriuskop/96/4     ..............a.....................................................t.....a....t [125360]
AY261362.1_Mkuzi_1979            ................................................................................ [125360]
AY261361.1_Malawi_Lil-20/1_1983  ........................................................g...........t..........t [125360]
AY261360.1_Kenya_1950            ..............a......................................c..g..............c.......t [125360]
AM712240.1_OURT_88/3             ................................................................................ [125360]
AM712239.1_Benin_97/1            .....a.......................................................................... [125360]
KM111295.1_Ken06.Bus             ..............a......................................c..g..............c.......t [125360]
```

```
KM111294.1_Ken05/Tk1          ..............a......................................c..g..............c.......t [125360]
KM262844.1_strain_L60         ................................................................................ [125360]
KM102979.1_26544/OG10_Italy   .....a.......................................................................... [125360]
KM262845.1_strain_NHV         ................................................................................ [125360]
KX354450.1_47/Ss/2008         .....a.......................................................................... [125360]
KP055815.1_strain_BA71        ................................................................................ [125360]
MH025920.1_strain_R35         ..............a......................................c..g..............c.......t [125360]
MH025919.1_strain_N10         ..............a......................................c..g..............c.......t [125360]
MH025918.1_strain_R25         ..............a......................................c..g..............c.......t [125360]
MH025917.1_strain_R7          ..............a......................................c..g..............c.......t [125360]
MH025916.1_strain_R8          ..............a......................................c..g..............c.......t [125360]

FR682468.2_Georgia_2007/1     gatttcgtactgtttatactgtttgttgaaaagggtaataaatttccaggtttcttcaaagcttgcaatctgggtgggcc [125440]
ASF_IND_20_CAD543             ................................................................................ [125440]
LR899193.1_Germany_2020/1     ................................................................................ [125440]
MN393477.1_Wuhan_2019         ................................................................................ [125440]
MW396979.1_Timor-Leste/2019/1 ................................................................................ [125440]
MK940252.1_CN/2019/InnerMongolia ................................................................................ [125440]
MK128995.1_China/2018/AnhuiXCGQ ................................................................................ [125440]
MH681419.1_ASFV/POL/2015      ................................................................................ [125440]
LS478113.1_Estonia_2014       ................................................................................ [125440]
KP843857.1_Odintsovo_02/14    ................................................................................ [125440]
FN557520.1_strain_E75         ................................................................................ [125440]
AY261366.1_Warthog            ..................t.....................................t....................... [125440]
AY261365.1_Warmbaths          ..................t............................................................. [125440]
AY261364.1_Tengani_62         ..................t.....................................t....................... [125440]
AY261363.1_Pretoriuskop/96/4  ..................t.....................................t....................... [125440]
AY261362.1_Mkuzi_1979         ................................................................................ [125440]
AY261361.1_Malawi_Lil-20/1_1983 ......a...........t...............c...........................c......a.......... [125440]
AY261360.1_Kenya_1950         ................a.a...............cg.................................a.c........ [125440]
AM712240.1_OURT_88/3          ................................................................................ [125440]
AM712239.1_Benin_97/1         ................................................................................ [125440]
KM111295.1_Ken06.Bus          ................a.a...............cg.................................a.c........ [125440]
KM111294.1_Ken05/Tk1          ................a.a...............cg...............................g.a.c........ [125440]
KM262844.1_strain_L60         ................................................................................ [125440]
KM102979.1_26544/OG10_Italy   ................................................................................ [125440]
KM262845.1_strain_NHV         ................................................................................ [125440]
KX354450.1_47/Ss/2008         ................................................................................ [125440]
KP055815.1_strain_BA71        ................................................................................ [125440]
MH025920.1_strain_R35         ................a.a...............cg.................................a.c........ [125440]
MH025919.1_strain_N10         ................a.a...............cg.................................a.c........ [125440]
MH025918.1_strain_R25         ................a.a...............cg.................................a.c........ [125440]
MH025917.1_strain_R7          ................a.a...............cg.................................a.c........ [125440]
MH025916.1_strain_R8          ................a.a...............cg.................................a.c........ [125440]
```

```
FR682468.2_Georgia_2007/1        gcagatcaaagtcgatgggaatgtcgtcatgaatgtaggatgatagtcttataggaaaataaatagggcgatcggtgtct [125520]
ASF_IND_20_CAD543                ................................................................................ [125520]
LR899193.1_Germany_2020/1        ................................................................................ [125520]
MN393477.1_Wuhan_2019            ................................................................................ [125520]
MW396979.1_Timor-Leste/2019/1    ................................................................................ [125520]
MK940252.1_CN/2019/InnerMongolia ................................................................................ [125520]
MK128995.1_China/2018/AnhuiXCGQ  ................................................................................ [125520]
MH681419.1_ASFV/POL/2015         ................................................................................ [125520]
LS478113.1_Estonia_2014          ................................................................................ [125520]
KP843857.1_Odintsovo_02/14       ................................................................................ [125520]
FN557520.1_strain_E75            ................................................................................ [125520]
AY261366.1_Warthog               .g.............................................................................. [125520]
AY261365.1_Warmbaths             ................................................................................ [125520]
AY261364.1_Tengani_62            ............................................................................ac.. [125520]
AY261363.1_Pretoriuskop/96/4     .g.............................................................................. [125520]
AY261362.1_Mkuzi_1979            ................................................................................ [125520]
AY261361.1_Malawi_Lil-20/1_1983  .g.................................................................a.........c.. [125520]
AY261360.1_Kenya_1950            .a..........a.......................................g..............a............ [125520]
AM712240.1_OURT_88/3             ................................................................................ [125520]
AM712239.1_Benin_97/1            ................................................................................ [125520]
KM111295.1_Ken06.Bus             .a..........a...........................c...........g........................... [125520]
KM111294.1_Ken05/Tk1             .a..........a.......................................g..............a............ [125520]
KM262844.1_strain_L60            ................................................................................ [125520]
KM102979.1_26544/OG10_Italy      ................................................................................ [125520]
KM262845.1_strain_NHV            ................................................................................ [125520]
KX354450.1_47/Ss/2008            ................................................................................ [125520]
KP055815.1_strain_BA71           ................................................................................ [125520]
MH025920.1_strain_R35            .a..........a...........................c...........g........................... [125520]
MH025919.1_strain_N10            .a..........a...........................c...........g........................... [125520]
MH025918.1_strain_R25            .a..........a...........................c...........g........................... [125520]
MH025917.1_strain_R7             .a..........a...........................c...........g........................... [125520]
MH025916.1_strain_R8             .a..........a...........................c...........g........................... [125520]

FR682468.2_Georgia_2007/1        gaatcgataagtaaagcataacaaaagttatgcctgttgataagttttttaccaaccgtgtagccgggaatgtttttcac [125600]
ASF_IND_20_CAD543                ................................................................................ [125600]
LR899193.1_Germany_2020/1        ................................................................................ [125600]
MN393477.1_Wuhan_2019            ................................................................................ [125600]
MW396979.1_Timor-Leste/2019/1    ................................................................................ [125600]
MK940252.1_CN/2019/InnerMongolia ................................................................................ [125600]
MK128995.1_China/2018/AnhuiXCGQ  ................................................................................ [125600]
MH681419.1_ASFV/POL/2015         ................................................................................ [125600]
LS478113.1_Estonia_2014          ................................................................................ [125600]
KP843857.1_Odintsovo_02/14       ................................................................................ [125600]
```

```
FN557520.1_strain_E75          ................................................................................ [125600]
AY261366.1_Warthog             ................................................................................ [125600]
AY261365.1_Warmbaths           ................................................................................ [125600]
AY261364.1_Tengani_62          ........................................................a....................... [125600]
AY261363.1_Pretoriuskop/96/4   ........................................................a..a.................... [125600]
AY261362.1_Mkuzi_1979          ................................................................................ [125600]
AY261361.1_Malawi_Lil-20/1_1983 ..g................................a.......a.......gt........................... [125600]
AY261360.1_Kenya_1950          ..ga....c..........................................gt........................... [125600]
AM712240.1_OURT_88/3           ................................................................................ [125600]
AM712239.1_Benin_97/1          ................................................................................ [125600]
KM111295.1_Ken06.Bus           ..ga....c..........................................gt........................... [125600]
KM111294.1_Ken05/Tk1           ..gg....c..........................................gt........................... [125600]
KM262844.1_strain_L60          ................................................................................ [125600]
KM102979.1_26544/OG10_Italy    ................................................................................ [125600]
KM262845.1_strain_NHV          ................................................................................ [125600]
KX354450.1_47/Ss/2008          ................................................................................ [125600]
KP055815.1_strain_BA71         ................................................................................ [125600]
MH025920.1_strain_R35          ..ga....c..........................................gt........................... [125600]
MH025919.1_strain_N10          ..ga....c..........................................gt........................... [125600]
MH025918.1_strain_R25          ..ga....c..........................................gt........................... [125600]
MH025917.1_strain_R7           ..ga....c..........................................gt........................... [125600]
MH025916.1_strain_R8           ..ga....c..........................................gt........................... [125600]

FR682468.2_Georgia_2007/1      gtcatggatatcccaccagttatccttgcacataaactcgctcatagactggatgacctccatcacagggtcatcttcgg [125680]
ASF_IND_20_CAD543              .........r...................................................................... [125680]
LR899193.1_Germany_2020/1      ................................................................................ [125680]
MN393477.1_Wuhan_2019          ................................................................................ [125680]
MW396979.1_Timor-Leste/2019/1  ................................................................................ [125680]
MK940252.1_CN/2019/InnerMongolia ................................................................................ [125680]
MK128995.1_China/2018/AnhuiXCGQ ................................................................................ [125680]
MH681419.1_ASFV/POL/2015       ................................................................................ [125680]
LS478113.1_Estonia_2014        ................................................................................ [125680]
KP843857.1_Odintsovo_02/14     ................................................................................ [125680]
FN557520.1_strain_E75          ...........................a.........................c.......................... [125680]
AY261366.1_Warthog             ...........................a.................................................... [125680]
AY261365.1_Warmbaths           ...........................a.................................................... [125680]
AY261364.1_Tengani_62          ...........................a.................................................... [125680]
AY261363.1_Pretoriuskop/96/4   ...........................a.................................................... [125680]
AY261362.1_Mkuzi_1979          ...........................a.................................................... [125680]
AY261361.1_Malawi_Lil-20/1_1983 ...................cc...............t...........a............................... [125680]
AY261360.1_Kenya_1950          ...................cc......a....................a.......................c....... [125680]
AM712240.1_OURT_88/3           ...........................a.........................c.......................... [125680]
AM712239.1_Benin_97/1          ...........................a.........................c.......................... [125680]
KM111295.1_Ken06.Bus           ...................cc......a....................a.......................c..c.... [125680]
```

```
KM111294.1_Ken05/Tk1            ...................cc......a....................a.......................c....... [125680]
KM262844.1_strain_L60           ...........................a.........................c.......................... [125680]
KM102979.1_26544/OG10_Italy     ...........................a.........................c.......................... [125680]
KM262845.1_strain_NHV           ...........................a.........................c.......................... [125680]
KX354450.1_47/Ss/2008           ...........................a.........................c.......................... [125680]
KP055815.1_strain_BA71          ...........................a.........................c.......................... [125680]
MH025920.1_strain_R35           ...................cc......a....................a.......................c..c.... [125680]
MH025919.1_strain_N10           ...................cc......a....................a.......................c..c.... [125680]
MH025918.1_strain_R25           ...................cc......a....................a.......................c..c.... [125680]
MH025917.1_strain_R7            ...................cc......a....................a.......................c..c.... [125680]
MH025916.1_strain_R8            ...................cc......a....................a.......................c..c.... [125680]

FR682468.2_Georgia_2007/1       taaaaatatactgggcctcactgtttttcagaaatcttttttgctgggtgatggccattgggtagatcccttcgtccgtg [125760]
ASF_IND_20_CAD543               ................................................................................ [125760]
LR899193.1_Germany_2020/1       ................................................................................ [125760]
MN393477.1_Wuhan_2019           ................................................................................ [125760]
MW396979.1_Timor-Leste/2019/1   ................................................................................ [125760]
MK940252.1_CN/2019/InnerMongolia ................................................................................ [125760]
MK128995.1_China/2018/AnhuiXCGQ ................................................................................ [125760]
MH681419.1_ASFV/POL/2015        ................................................................................ [125760]
LS478113.1_Estonia_2014         ................................................................................ [125760]
KP843857.1_Odintsovo_02/14      ................................................................................ [125760]
FN557520.1_strain_E75           ................................................................................ [125760]
AY261366.1_Warthog              ................................................................................ [125760]
AY261365.1_Warmbaths            ................................................................................ [125760]
AY261364.1_Tengani_62           ............................t................................................... [125760]
AY261363.1_Pretoriuskop/96/4    ...................................................................t.....a...... [125760]
AY261362.1_Mkuzi_1979           ................................................................................ [125760]
AY261361.1_Malawi_Lil-20/1_1983 ......................a..........................c...........a...........a...... [125760]
AY261360.1_Kenya_1950           .................................................c........g..a...........a.t.... [125760]
AM712240.1_OURT_88/3            ................................................................................ [125760]
AM712239.1_Benin_97/1           ................................................................................ [125760]
KM111295.1_Ken06.Bus            .................................................c........g..a...........a.t.... [125760]
KM111294.1_Ken05/Tk1            .................................................c........g..a...........a.t.... [125760]
KM262844.1_strain_L60           ................................................................................ [125760]
KM102979.1_26544/OG10_Italy     ................................................................................ [125760]
KM262845.1_strain_NHV           ................................................................................ [125760]
KX354450.1_47/Ss/2008           ................................................................................ [125760]
KP055815.1_strain_BA71          ................................................................................ [125760]
MH025920.1_strain_R35           .................................................c........g..a...........a.t.... [125760]
MH025919.1_strain_N10           .................................................c........g..a...........a.t.... [125760]
MH025918.1_strain_R25           .................................................c........g..a...........a.t.... [125760]
MH025917.1_strain_R7            .................................................c........g..a...........a.t.... [125760]
MH025916.1_strain_R8            .................................................c........g..a...........a.t.... [125760]
```

```
FR682468.2_Georgia_2007/1        tcaaagataatggctatcttcttcgatgggctaagaattttttgtattgtgctgggggacacctcaaacccgatgtcgcc [125840]
ASF_IND_20_CAD543                ................................................................................ [125840]
LR899193.1_Germany_2020/1        ................................................................................ [125840]
MN393477.1_Wuhan_2019            ................................................................................ [125840]
MW396979.1_Timor-Leste/2019/1    ................................................................................ [125840]
MK940252.1_CN/2019/InnerMongolia ................................................................................ [125840]
MK128995.1_China/2018/AnhuiXCGQ  ................................................................................ [125840]
MH681419.1_ASFV/POL/2015         ................................................................................ [125840]
LS478113.1_Estonia_2014          ................................................................................ [125840]
KP843857.1_Odintsovo_02/14       ................................................................................ [125840]
FN557520.1_strain_E75            .....................................................................t.......... [125840]
AY261366.1_Warthog               .....................................................................t...a...c.. [125840]
AY261365.1_Warmbaths             .....................................................................t...a...c.. [125840]
AY261364.1_Tengani_62            .....................................................................t...a...c.. [125840]
AY261363.1_Pretoriuskop/96/4     ...................................................t.................t...a...c.. [125840]
AY261362.1_Mkuzi_1979            .....................................................................t.......... [125840]
AY261361.1_Malawi_Lil-20/1_1983  .............................a.................c..a........t.....g...t....a..c.. [125840]
AY261360.1_Kenya_1950            .................................................................g...t.a.....c.. [125840]
AM712240.1_OURT_88/3             .....................................................................t.......... [125840]
AM712239.1_Benin_97/1            .....................................................................t.......... [125840]
KM111295.1_Ken06.Bus             ........................................................a........g...t.......c.. [125840]
KM111294.1_Ken05/Tk1             .................................................................g...t.......c.. [125840]
KM262844.1_strain_L60            .....................................................................t.......... [125840]
KM102979.1_26544/OG10_Italy      .....................................................................t.......... [125840]
KM262845.1_strain_NHV            .....................................................................t.......... [125840]
KX354450.1_47/Ss/2008            .....................................................................t.......... [125840]
KP055815.1_strain_BA71           .....................................................................t.......... [125840]
MH025920.1_strain_R35            ........................................................a........g...t.......c.. [125840]
MH025919.1_strain_N10            ........................................................a........g...t.......c.. [125840]
MH025918.1_strain_R25            ........................................................a........g...t.......c.. [125840]
MH025917.1_strain_R7             ........................................................a........g...t.......c.. [125840]
MH025916.1_strain_R8             ........................................................a........g...t.......c.. [125840]

FR682468.2_Georgia_2007/1        ctgtttatctttaaaaaagacacagtgaaggtcgtagcatatggcaacaaggtccagaaagatgtcctgccatgtggtgt [125920]
ASF_IND_20_CAD543                ................................................................................ [125920]
LR899193.1_Germany_2020/1        ................................................................................ [125920]
MN393477.1_Wuhan_2019            ................................................................................ [125920]
MW396979.1_Timor-Leste/2019/1    ................................................................................ [125920]
MK940252.1_CN/2019/InnerMongolia ................................................................................ [125920]
MK128995.1_China/2018/AnhuiXCGQ  ................................................................................ [125920]
MH681419.1_ASFV/POL/2015         ................................................................................ [125920]
LS478113.1_Estonia_2014          ................................................................................ [125920]
KP843857.1_Odintsovo_02/14       ................................................................................ [125920]
```

```
FN557520.1_strain_E75           ................................................................................ [125920]
AY261366.1_Warthog              ........................a..g.................................................... [125920]
AY261365.1_Warmbaths            ................................................................................ [125920]
AY261364.1_Tengani_62           ................................................................................ [125920]
AY261363.1_Pretoriuskop/96/4    ........................a..g.................................................... [125920]
AY261362.1_Mkuzi_1979           ................................................................................ [125920]
AY261361.1_Malawi_Lil-20/1_1983 ...........................g.................................................... [125920]
AY261360.1_Kenya_1950           ...........................g.......................a......................c..... [125920]
AM712240.1_OURT_88/3            ................................................................................ [125920]
AM712239.1_Benin_97/1           ................................................................................ [125920]
KM111295.1_Ken06.Bus            ...........................g.......................a......................c..... [125920]
KM111294.1_Ken05/Tk1            ...........................g.......................a......................c..... [125920]
KM262844.1_strain_L60           ................................................................................ [125920]
KM102979.1_26544/OG10_Italy     ................................................................................ [125920]
KM262845.1_strain_NHV           ................................................................................ [125920]
KX354450.1_47/Ss/2008           ................................................................................ [125920]
KP055815.1_strain_BA71          ................................................................................ [125920]
MH025920.1_strain_R35           ...........................g.......................a......................c..... [125920]
MH025919.1_strain_N10           ...........................g.......................a......................c..... [125920]
MH025918.1_strain_R25           ...........................g.......................a......................c..... [125920]
MH025917.1_strain_R7            ...........................g.......................a......................c..... [125920]
MH025916.1_strain_R8            ...........................g.......................a......................c..... [125920]

FR682468.2_Georgia_2007/1       cccattgaagcagttggtttttttgttcaacaaaggtttgtaagataaggtttgccagctccgcgccgctggaaaacatg [126000]
ASF_IND_20_CAD543               ................................................................................ [126000]
LR899193.1_Germany_2020/1       ................................................................................ [126000]
MN393477.1_Wuhan_2019           ................................................................................ [126000]
MW396979.1_Timor-Leste/2019/1   ................................................................................ [126000]
MK940252.1_CN/2019/InnerMongolia ................................................................................ [126000]
MK128995.1_China/2018/AnhuiXCGQ ................................................................................ [126000]
MH681419.1_ASFV/POL/2015        ................................................................................ [126000]
LS478113.1_Estonia_2014         ................................................................................ [126000]
KP843857.1_Odintsovo_02/14      ................................................................................ [126000]
FN557520.1_strain_E75           ....c........................................................................... [126000]
AY261366.1_Warthog              ..........................................g.......c............................. [126000]
AY261365.1_Warmbaths            ..........................................g..................................... [126000]
AY261364.1_Tengani_62           ..........................................g..................................... [126000]
AY261363.1_Pretoriuskop/96/4    ..........................................g.......c............................. [126000]
AY261362.1_Mkuzi_1979           ................................................................................ [126000]
AY261361.1_Malawi_Lil-20/1_1983 ..................................a.......g..................t.................. [126000]
AY261360.1_Kenya_1950           ..........................................g.....a............................... [126000]
AM712240.1_OURT_88/3            ....c........................................................................... [126000]
AM712239.1_Benin_97/1           ....c........................................................................... [126000]
KM111295.1_Ken06.Bus            ..........................................g.....a............................... [126000]
```

```
KM111294.1_Ken05/Tk1              ..........................................g.....a............................... [126000]
KM262844.1_strain_L60             ....c........................................................................... [126000]
KM102979.1_26544/OG10_Italy       ....c........................................................................... [126000]
KM262845.1_strain_NHV             ....c........................................................................... [126000]
KX354450.1_47/Ss/2008             ....c........................................................................... [126000]
KP055815.1_strain_BA71            ....c........................................................................... [126000]
MH025920.1_strain_R35             ..........................................g.....a............................... [126000]
MH025919.1_strain_N10             ..........................................g.....a............................... [126000]
MH025918.1_strain_R25             ..........................................g.....a............................... [126000]
MH025917.1_strain_R7              ..........................................g.....a............................... [126000]
MH025916.1_strain_R8              ..........................................g.....a............................... [126000]

FR682468.2_Georgia_2007/1         ttgccggccccattccccaaaatatagtactgcggtgtgttggccgcctttgcaatttcaatggcaagggccttgggggc [126080]
ASF_IND_20_CAD543                 .............................................................................r.. [126080]
LR899193.1_Germany_2020/1         ................................................................................ [126080]
MN393477.1_Wuhan_2019             ................................................................................ [126080]
MW396979.1_Timor-Leste/2019/1     ................................................................................ [126080]
MK940252.1_CN/2019/InnerMongolia  ................................................................................ [126080]
MK128995.1_China/2018/AnhuiXCGQ   ................................................................................ [126080]
MH681419.1_ASFV/POL/2015          ................................................................................ [126080]
LS478113.1_Estonia_2014           ................................................................................ [126080]
KP843857.1_Odintsovo_02/14        ................................................................................ [126080]
FN557520.1_strain_E75             ...................................c.........................................a.. [126080]
AY261366.1_Warthog                ......................................a......................................... [126080]
AY261365.1_Warmbaths              ......................................a......................................... [126080]
AY261364.1_Tengani_62             ......................................a......................................... [126080]
AY261363.1_Pretoriuskop/96/4      ......................................a......................................... [126080]
AY261362.1_Mkuzi_1979             .............................................................................a.. [126080]
AY261361.1_Malawi_Lil-20/1_1983   ..............t....................c..a......................c...............a.. [126080]
AY261360.1_Kenya_1950             ...........g.......................c..a......................c...............a.. [126080]
AM712240.1_OURT_88/3              ...................................c.........................................a.. [126080]
AM712239.1_Benin_97/1             ...................................c.........................................a.. [126080]
KM111295.1_Ken06.Bus              ...........g.......................c..a......................c...............a.. [126080]
KM111294.1_Ken05/Tk1              ...........g.......................c..a......................c...............a.. [126080]
KM262844.1_strain_L60             ...................................c.........................................a.. [126080]
KM102979.1_26544/OG10_Italy       ...................................c.........................................a.. [126080]
KM262845.1_strain_NHV             ...................................c.........................................a.. [126080]
KX354450.1_47/Ss/2008             ...................................c.........................................a.. [126080]
KP055815.1_strain_BA71            ...................................c.........................................a.. [126080]
MH025920.1_strain_R35             ...........g.......................c..a......................c...............a.. [126080]
MH025919.1_strain_N10             ...........g.......................c..a......................c...............a.. [126080]
MH025918.1_strain_R25             ...........g.......................c..a......................c...............a.. [126080]
MH025917.1_strain_R7              ...........g.......................c..a......................c...............a.. [126080]
MH025916.1_strain_R8              ...........g.......................c..a......................c...............a.. [126080]
```

```
FR682468.2_Georgia_2007/1        aagatccaaaattcgagcaagggaataaaaaagcccggcattgctaattccaagcatggtttgctccacccccacaatgc [126160]
ASF_IND_20_CAD543                ................................................................................ [126160]
LR899193.1_Germany_2020/1        ................................................................................ [126160]
MN393477.1_Wuhan_2019            ................................................................................ [126160]
MW396979.1_Timor-Leste/2019/1    ................................................................................ [126160]
MK940252.1_CN/2019/InnerMongolia ................................................................................ [126160]
MK128995.1_China/2018/AnhuiXCGQ  ................................................................................ [126160]
MH681419.1_ASFV/POL/2015         ................................................................................ [126160]
LS478113.1_Estonia_2014          ................................................................................ [126160]
KP843857.1_Odintsovo_02/14       ................................................................................ [126160]
FN557520.1_strain_E75            ................................................................................ [126160]
AY261366.1_Warthog               .........................................g.....................t................ [126160]
AY261365.1_Warmbaths             ................................................................................ [126160]
AY261364.1_Tengani_62            ................................................................................ [126160]
AY261363.1_Pretoriuskop/96/4     ................................................................................ [126160]
AY261362.1_Mkuzi_1979            ................................................................................ [126160]
AY261361.1_Malawi_Lil-20/1_1983  ..................g.....g...................................c................... [126160]
AY261360.1_Kenya_1950            ...................................................g........c................... [126160]
AM712240.1_OURT_88/3             ................................................................................ [126160]
AM712239.1_Benin_97/1            ................................................................................ [126160]
KM111295.1_Ken06.Bus             ...................................................g........c................... [126160]
KM111294.1_Ken05/Tk1             ...................................................g........c................... [126160]
KM262844.1_strain_L60            ................................................................................ [126160]
KM102979.1_26544/OG10_Italy      ................................................................................ [126160]
KM262845.1_strain_NHV            ................................................................................ [126160]
KX354450.1_47/Ss/2008            ................................................................................ [126160]
KP055815.1_strain_BA71           ................................................................................ [126160]
MH025920.1_strain_R35            ...................................................g........c................... [126160]
MH025919.1_strain_N10            ...................................................g........c................... [126160]
MH025918.1_strain_R25            ...................................................g........c................... [126160]
MH025917.1_strain_R7             ...................................................g........c................... [126160]
MH025916.1_strain_R8             ...................................................g........c................... [126160]

FR682468.2_Georgia_2007/1        aaaaaatgtcgggctcttttatcgtatttaaaaacagttcatctgctatctggtggggtagaaaggcaatccggttcacc [126240]
ASF_IND_20_CAD543                ................................................................................ [126240]
LR899193.1_Germany_2020/1        ................................................................................ [126240]
MN393477.1_Wuhan_2019            ................................................................................ [126240]
MW396979.1_Timor-Leste/2019/1    ................................................................................ [126240]
MK940252.1_CN/2019/InnerMongolia ................................................................................ [126240]
MK128995.1_China/2018/AnhuiXCGQ  ................................................................................ [126240]
MH681419.1_ASFV/POL/2015         ................................................................................ [126240]
LS478113.1_Estonia_2014          ................................................................................ [126240]
KP843857.1_Odintsovo_02/14       ................................................................................ [126240]
```

```
FN557520.1_strain_E75            .................................................t.............................. [126240]
AY261366.1_Warthog               ......................t..........................t.............................. [126240]
AY261365.1_Warmbaths             ......................t......................................................... [126240]
AY261364.1_Tengani_62            .................................................t.............................. [126240]
AY261363.1_Pretoriuskop/96/4     ......................t..........................t.............................. [126240]
AY261362.1_Mkuzi_1979            .................................................t.............................. [126240]
AY261361.1_Malawi_Lil-20/1_1983  ............................gg..........g........t.............................. [126240]
AY261360.1_Kenya_1950            ............................ag...................t.............................. [126240]
AM712240.1_OURT_88/3             .................................................t.............................. [126240]
AM712239.1_Benin_97/1            .................................................t.............................. [126240]
KM111295.1_Ken06.Bus             ............................ag...................t.............................. [126240]
KM111294.1_Ken05/Tk1             ............................ag...................t.............................. [126240]
KM262844.1_strain_L60            .................................................t.............................. [126240]
KM102979.1_26544/OG10_Italy      .................................................t.............................. [126240]
KM262845.1_strain_NHV            .................................................t.............................. [126240]
KX354450.1_47/Ss/2008            .................................................t.............................. [126240]
KP055815.1_strain_BA71           .................................................t.............................. [126240]
MH025920.1_strain_R35            ............................ag...................t.............................. [126240]
MH025919.1_strain_N10            ............................ag...................t.............................. [126240]
MH025918.1_strain_R25            ............................ag...................t.............................. [126240]
MH025917.1_strain_R7             ............................ag...................t.............................. [126240]
MH025916.1_strain_R8             ............................ag...................t.............................. [126240]

FR682468.2_Georgia_2007/1        ggtattttttttccataggacaaggtatgacgcgatgtttgtgtattaagatcctccaggtcttgttctacaaacgtgtg [126320]
ASF_IND_20_CAD543                ................................................................................ [126320]
LR899193.1_Germany_2020/1        ................................................................................ [126320]
MN393477.1_Wuhan_2019            ................................................................................ [126320]
MW396979.1_Timor-Leste/2019/1    ................................................................................ [126320]
MK940252.1_CN/2019/InnerMongolia ................................................................................ [126320]
MK128995.1_China/2018/AnhuiXCGQ  ................................................................................ [126320]
MH681419.1_ASFV/POL/2015         ................................................................................ [126320]
LS478113.1_Estonia_2014          ................................................................................ [126320]
KP843857.1_Odintsovo_02/14       ................................................................................ [126320]
FN557520.1_strain_E75            ................................................................................ [126320]
AY261366.1_Warthog               ................................................................................ [126320]
AY261365.1_Warmbaths             ................................................................................ [126320]
AY261364.1_Tengani_62            ..g..................................................t....................t..... [126320]
AY261363.1_Pretoriuskop/96/4     ................................................................................ [126320]
AY261362.1_Mkuzi_1979            ................................................................................ [126320]
AY261361.1_Malawi_Lil-20/1_1983  ..........................g..................................................... [126320]
AY261360.1_Kenya_1950            ..........................g..................................................... [126320]
AM712240.1_OURT_88/3             ................................................................................ [126320]
AM712239.1_Benin_97/1            ................................................................................ [126320]
KM111295.1_Ken06.Bus             ..........................g..................................................... [126320]
```

```
KM111294.1_Ken05/Tk1            ..........................g..................................................... [126320]
KM262844.1_strain_L60           ................................................................................ [126320]
KM102979.1_26544/OG10_Italy     ................................................................................ [126320]
KM262845.1_strain_NHV           ................................................................................ [126320]
KX354450.1_47/Ss/2008           ................................................................................ [126320]
KP055815.1_strain_BA71          ................................................................................ [126320]
MH025920.1_strain_R35           ..........................g..................................................... [126320]
MH025919.1_strain_N10           ..........................g..................................................... [126320]
MH025918.1_strain_R25           ..........................g..................................................... [126320]
MH025917.1_strain_R7            ..........................g..................................................... [126320]
MH025916.1_strain_R8            ..........................g..................................................... [126320]

FR682468.2_Georgia_2007/1       cttggtgaggcaggtattgttaatatagaaccgctttgtgcccagcagggccttcgtcttttggcagcacggcagacagt [126400]
ASF_IND_20_CAD543               ................................................................................ [126400]
LR899193.1_Germany_2020/1       ................................................................................ [126400]
MN393477.1_Wuhan_2019           ................................................................................ [126400]
MW396979.1_Timor-Leste/2019/1   ................................................................................ [126400]
MK940252.1_CN/2019/InnerMongolia ................................................................................ [126400]
MK128995.1_China/2018/AnhuiXCGQ ................................................................................ [126400]
MH681419.1_ASFV/POL/2015        ................................................................................ [126400]
LS478113.1_Estonia_2014         ................................................................................ [126400]
KP843857.1_Odintsovo_02/14      ................................................................................ [126400]
FN557520.1_strain_E75           ................................................................................ [126400]
AY261366.1_Warthog              ................................................................................ [126400]
AY261365.1_Warmbaths            ................................................................................ [126400]
AY261364.1_Tengani_62           ...a..................................................t......................... [126400]
AY261363.1_Pretoriuskop/96/4    ................................................................................ [126400]
AY261362.1_Mkuzi_1979           ................................................................................ [126400]
AY261361.1_Malawi_Lil-20/1_1983 ..........................................t...........t........a..........a..... [126400]
AY261360.1_Kenya_1950           ..................a.......................t..................................... [126400]
AM712240.1_OURT_88/3            ................................................................................ [126400]
AM712239.1_Benin_97/1           ................................................................................ [126400]
KM111295.1_Ken06.Bus            ...a..............a.......................t..................................... [126400]
KM111294.1_Ken05/Tk1            ..................a.......................t..................................... [126400]
KM262844.1_strain_L60           ................................................................................ [126400]
KM102979.1_26544/OG10_Italy     ................................................................................ [126400]
KM262845.1_strain_NHV           ................................................................................ [126400]
KX354450.1_47/Ss/2008           ................................................................................ [126400]
KP055815.1_strain_BA71          ................................................................................ [126400]
MH025920.1_strain_R35           t..a..............a.......................t..................................... [126400]
MH025919.1_strain_N10           t..a..............a.......................t..................................... [126400]
MH025918.1_strain_R25           t..a..............a.......................t..................................... [126400]
MH025917.1_strain_R7            t..a..............a.......................t..................................... [126400]
MH025916.1_strain_R8            t..a..............a.......................t..................................... [126400]
```

```
FR682468.2_Georgia_2007/1        aatttagggggtggcggccttctagtaggcttagatgagggtagtcaggatgcgggcagctatagtaggcaggtaccccc [126480]
ASF_IND_20_CAD543                ................................................................................ [126480]
LR899193.1_Germany_2020/1        ................................................................................ [126480]
MN393477.1_Wuhan_2019            ................................................................................ [126480]
MW396979.1_Timor-Leste/2019/1    ................................................................................ [126480]
MK940252.1_CN/2019/InnerMongolia ................................................................................ [126480]
MK128995.1_China/2018/AnhuiXCGQ  ................................................................................ [126480]
MH681419.1_ASFV/POL/2015         ................................................................................ [126480]
LS478113.1_Estonia_2014          ................................................................................ [126480]
KP843857.1_Odintsovo_02/14       ................................................................................ [126480]
FN557520.1_strain_E75            ................................................................................ [126480]
AY261366.1_Warthog               ......a......................................................................... [126480]
AY261365.1_Warmbaths             ................................................................................ [126480]
AY261364.1_Tengani_62            ................................................................................ [126480]
AY261363.1_Pretoriuskop/96/4     ................................................................................ [126480]
AY261362.1_Mkuzi_1979            ................................................................................ [126480]
AY261361.1_Malawi_Lil-20/1_1983  ................................................................................ [126480]
AY261360.1_Kenya_1950            ................................................................................ [126480]
AM712240.1_OURT_88/3             ................................................................................ [126480]
AM712239.1_Benin_97/1            ................................................................................ [126480]
KM111295.1_Ken06.Bus             ................................................................................ [126480]
KM111294.1_Ken05/Tk1             ................................................................................ [126480]
KM262844.1_strain_L60            ................................................................................ [126480]
KM102979.1_26544/OG10_Italy      ................................................................................ [126480]
KM262845.1_strain_NHV            ................................................................................ [126480]
KX354450.1_47/Ss/2008            ................................................................................ [126480]
KP055815.1_strain_BA71           ................................................................................ [126480]
MH025920.1_strain_R35            ................................................................................ [126480]
MH025919.1_strain_N10            ................................................................................ [126480]
MH025918.1_strain_R25            ................................................................................ [126480]
MH025917.1_strain_R7             ................................................................................ [126480]
MH025916.1_strain_R8             ................................................................................ [126480]

FR682468.2_Georgia_2007/1        tccgtgaaattccaatactttactagctccttgcgcttggctggcggcatggacttcacctcggcctctgagtaaatgac [126560]
ASF_IND_20_CAD543                ................................................................................ [126560]
LR899193.1_Germany_2020/1        ................................................................................ [126560]
MN393477.1_Wuhan_2019            ................................................................................ [126560]
MW396979.1_Timor-Leste/2019/1    ................................................................................ [126560]
MK940252.1_CN/2019/InnerMongolia ................................................................................ [126560]
MK128995.1_China/2018/AnhuiXCGQ  ................................................................................ [126560]
MH681419.1_ASFV/POL/2015         ................................................................................ [126560]
LS478113.1_Estonia_2014          ................................................................................ [126560]
KP843857.1_Odintsovo_02/14       ................................................................................ [126560]
```

```
FN557520.1_strain_E75          ................................................................................ [126560]
AY261366.1_Warthog             ................................................................................ [126560]
AY261365.1_Warmbaths           ................................................................................ [126560]
AY261364.1_Tengani_62          ................................................................................ [126560]
AY261363.1_Pretoriuskop/96/4   .............................t....................a............................. [126560]
AY261362.1_Mkuzi_1979          ................................................................................ [126560]
AY261361.1_Malawi_Lil-20/1_1983 .................t...........t.....t..............a............................. [126560]
AY261360.1_Kenya_1950          ...................................t........t.....a..t.......................... [126560]
AM712240.1_OURT_88/3           ................................................................................ [126560]
AM712239.1_Benin_97/1          ................................................................................ [126560]
KM111295.1_Ken06.Bus           ...................................t.................t.......................... [126560]
KM111294.1_Ken05/Tk1           ...................................t........t.....a..t.......................... [126560]
KM262844.1_strain_L60          ................................................................................ [126560]
KM102979.1_26544/OG10_Italy    ................................................................................ [126560]
KM262845.1_strain_NHV          ................................................................................ [126560]
KX354450.1_47/Ss/2008          ................................................................................ [126560]
KP055815.1_strain_BA71         ................................................................................ [126560]
MH025920.1_strain_R35          ...................................t.................t.......................... [126560]
MH025919.1_strain_N10          ...................................t.................t.......................... [126560]
MH025918.1_strain_R25          ...................................t.................t.......................... [126560]
MH025917.1_strain_R7           ...................................t.................t.......................... [126560]
MH025916.1_strain_R8           ...................................t.................t.......................... [126560]

FR682468.2_Georgia_2007/1      gggtggccgtgggtgctggcataggacggagtaaaccgttgcctgcgtgtcgtacttgcgcaggtcatacaggtcggggt [126640]
ASF_IND_20_CAD543              ................................................................................ [126640]
LR899193.1_Germany_2020/1      ................................................................................ [126640]
MN393477.1_Wuhan_2019          ................................................................................ [126640]
MW396979.1_Timor-Leste/2019/1  ................................................................................ [126640]
MK940252.1_CN/2019/InnerMongolia ................................................................................ [126640]
MK128995.1_China/2018/AnhuiXCGQ ................................................................................ [126640]
MH681419.1_ASFV/POL/2015       ................................................................................ [126640]
LS478113.1_Estonia_2014        ................................................................................ [126640]
KP843857.1_Odintsovo_02/14     ................................................................................ [126640]
FN557520.1_strain_E75          ................................................................................ [126640]
AY261366.1_Warthog             ................................................................................ [126640]
AY261365.1_Warmbaths           ................................................................................ [126640]
AY261364.1_Tengani_62          ..................................................................g............. [126640]
AY261363.1_Pretoriuskop/96/4   ...c............................................................................ [126640]
AY261362.1_Mkuzi_1979          ................................................................................ [126640]
AY261361.1_Malawi_Lil-20/1_1983 ............a..........................................................a........ [126640]
AY261360.1_Kenya_1950          ............a..........a........................a..a...........a.......a........ [126640]
AM712240.1_OURT_88/3           ................................................................................ [126640]
AM712239.1_Benin_97/1          ................................................................................ [126640]
KM111295.1_Ken06.Bus           ............a......................................a...........a.......a........ [126640]
```

```
KM111294.1_Ken05/Tk1              ............a..........a...........................a...........a.......a........ [126640]
KM262844.1_strain_L60             ................................................................................ [126640]
KM102979.1_26544/OG10_Italy       ................................................................................ [126640]
KM262845.1_strain_NHV             ................................................................................ [126640]
KX354450.1_47/Ss/2008             ................................................................................ [126640]
KP055815.1_strain_BA71            ................................................................................ [126640]
MH025920.1_strain_R35             ............a......................................a...........a.......a........ [126640]
MH025919.1_strain_N10             ............a......................................a...........a.......a........ [126640]
MH025918.1_strain_R25             ............a......................................a...........a.......a........ [126640]
MH025917.1_strain_R7              ............a......................................a...........a.......a........ [126640]
MH025916.1_strain_R8              ............a......................................a...........a.......a........ [126640]

FR682468.2_Georgia_2007/1         cctgttcttgaagcgcacgtagctgagaggctccctttccttgttgtttatcgtgcagttgagagagtttattaaccaaa [126720]
ASF_IND_20_CAD543                 ................................................................................ [126720]
LR899193.1_Germany_2020/1         ................................................................................ [126720]
MN393477.1_Wuhan_2019             ................................................................................ [126720]
MW396979.1_Timor-Leste/2019/1     ................................................................................ [126720]
MK940252.1_CN/2019/InnerMongolia  ................................................................................ [126720]
MK128995.1_China/2018/AnhuiXCGQ   ................................................................................ [126720]
MH681419.1_ASFV/POL/2015          ................................................................................ [126720]
LS478113.1_Estonia_2014           ................................................................................ [126720]
KP843857.1_Odintsovo_02/14        ................................................................................ [126720]
FN557520.1_strain_E75             ................................................................................ [126720]
AY261366.1_Warthog                ................................................................................ [126720]
AY261365.1_Warmbaths              ......................................................................g......... [126720]
AY261364.1_Tengani_62             ...................................................................c............ [126720]
AY261363.1_Pretoriuskop/96/4      ......................................................................g......... [126720]
AY261362.1_Mkuzi_1979             ................................................................................ [126720]
AY261361.1_Malawi_Lil-20/1_1983   .......c....................a............................a............g.......t. [126720]
AY261360.1_Kenya_1950             .......c.................................................a............g.......t. [126720]
AM712240.1_OURT_88/3              ................................................................................ [126720]
AM712239.1_Benin_97/1             ................................................................................ [126720]
KM111295.1_Ken06.Bus              .......c.................................................a............g.......t. [126720]
KM111294.1_Ken05/Tk1              .......c.................................................a............g.......t. [126720]
KM262844.1_strain_L60             ................................................................................ [126720]
KM102979.1_26544/OG10_Italy       ................................................................................ [126720]
KM262845.1_strain_NHV             ................................................................................ [126720]
KX354450.1_47/Ss/2008             ................................................................................ [126720]
KP055815.1_strain_BA71            ................................................................................ [126720]
MH025920.1_strain_R35             .......c.................................................a............g.......t. [126720]
MH025919.1_strain_N10             .......c.................................................a............g.......t. [126720]
MH025918.1_strain_R25             .......c.................................................a............g.......t. [126720]
MH025917.1_strain_R7              .......c.................................................a............g.......t. [126720]
MH025916.1_strain_R8              .......c.................................................a............g.......t. [126720]
```

```
FR682468.2_Georgia_2007/1        attttgtcaggcccggtgatcaagttatctaaaaacacaaataggtaaacccaaagatagttaaactcttcctgggtaat [126800]
ASF_IND_20_CAD543                ................................................................................ [126800]
LR899193.1_Germany_2020/1        ................................................................................ [126800]
MN393477.1_Wuhan_2019            ................................................................................ [126800]
MW396979.1_Timor-Leste/2019/1    ................................................................................ [126800]
MK940252.1_CN/2019/InnerMongolia ................................................................................ [126800]
MK128995.1_China/2018/AnhuiXCGQ  ................................................................................ [126800]
MH681419.1_ASFV/POL/2015         ................................................................................ [126800]
LS478113.1_Estonia_2014          ................................................................................ [126800]
KP843857.1_Odintsovo_02/14       ................................................................................ [126800]
FN557520.1_strain_E75            ................................................................................ [126800]
AY261366.1_Warthog               ...................................t.............................t.............. [126800]
AY261365.1_Warmbaths             .....a.......................................................................... [126800]
AY261364.1_Tengani_62            .....a.......................................................................... [126800]
AY261363.1_Pretoriuskop/96/4     .....a...........................................................t.............. [126800]
AY261362.1_Mkuzi_1979            ................................................................................ [126800]
AY261361.1_Malawi_Lil-20/1_1983  ..............ac......t......c.....t.......a.................................... [126800]
AY261360.1_Kenya_1950            .................a...........c.............a.................................... [126800]
AM712240.1_OURT_88/3             ................................................................................ [126800]
AM712239.1_Benin_97/1            ................................................................................ [126800]
KM111295.1_Ken06.Bus             .................a....g......c.............a.................................... [126800]
KM111294.1_Ken05/Tk1             .................a...........c.............a.................................... [126800]
KM262844.1_strain_L60            ................................................................................ [126800]
KM102979.1_26544/OG10_Italy      ................................................................................ [126800]
KM262845.1_strain_NHV            ................................................................................ [126800]
KX354450.1_47/Ss/2008            ................................................................................ [126800]
KP055815.1_strain_BA71           ................................................................................ [126800]
MH025920.1_strain_R35            .................a....g......c.............a.................................... [126800]
MH025919.1_strain_N10            .................a....g......c.............a.................................... [126800]
MH025918.1_strain_R25            .................a....g......c.............a.................................... [126800]
MH025917.1_strain_R7             .................a....g......c.............a.................................... [126800]
MH025916.1_strain_R8             .................a....g......c.............a.................................... [126800]

FR682468.2_Georgia_2007/1        gttaaacatttctattttgatatctgtaaccctatggtagatgcgaatgttgcggccgccgtagattgtttcccaccggg [126880]
ASF_IND_20_CAD543                ................................................................................ [126880]
LR899193.1_Germany_2020/1        ................................................................................ [126880]
MN393477.1_Wuhan_2019            ................................................................................ [126880]
MW396979.1_Timor-Leste/2019/1    ................................................................................ [126880]
MK940252.1_CN/2019/InnerMongolia ................................................................................ [126880]
MK128995.1_China/2018/AnhuiXCGQ  ................................................................................ [126880]
MH681419.1_ASFV/POL/2015         ................................................................................ [126880]
LS478113.1_Estonia_2014          ................................................................................ [126880]
KP843857.1_Odintsovo_02/14       ................................................................................ [126880]
```

| | | |
|---|---|---|
| FN557520.1_strain_E75 | ................................................................................ | [126880] |
| AY261366.1_Warthog | ................................................................................ | [126880] |
| AY261365.1_Warmbaths | ................................................................................ | [126880] |
| AY261364.1_Tengani_62 | ................................................................................ | [126880] |
| AY261363.1_Pretoriuskop/96/4 | ................................................................................ | [126880] |
| AY261362.1_Mkuzi_1979 | ................................................................................ | [126880] |
| AY261361.1_Malawi_Lil-20/1_1983 | ................................................a.............................c. | [126880] |
| AY261360.1_Kenya_1950 | ........................g.......................a............................... | [126880] |
| AM712240.1_OURT_88/3 | ................................................................................ | [126880] |
| AM712239.1_Benin_97/1 | ................................................................................ | [126880] |
| KM111295.1_Ken06.Bus | ........................g..g....................a............................... | [126880] |
| KM111294.1_Ken05/Tk1 | ........................g.......................a.................c............. | [126880] |
| KM262844.1_strain_L60 | ................................................................................ | [126880] |
| KM102979.1_26544/OG10_Italy | ................................................................................ | [126880] |
| KM262845.1_strain_NHV | ................................................................................ | [126880] |
| KX354450.1_47/Ss/2008 | ................................................................................ | [126880] |
| KP055815.1_strain_BA71 | ................................................................................ | [126880] |
| MH025920.1_strain_R35 | ........................g..g....................a............................... | [126880] |
| MH025919.1_strain_N10 | ........................g..g....................a............................... | [126880] |
| MH025918.1_strain_R25 | ........................g..g....................a............................... | [126880] |
| MH025917.1_strain_R7 | ........................g..g....................a............................... | [126880] |
| MH025916.1_strain_R8 | ........................g..g....................a............................... | [126880] |

| | | |
|---|---|---|
| FR682468.2_Georgia_2007/1 | ccgcaacatttgtgtcaaagaggtacgcatacgtgttttggagcaacgcaacattgatgtccattttgcgccccggaccg | [126960] |
| ASF_IND_20_CAD543 | ................................................................................ | [126960] |
| LR899193.1_Germany_2020/1 | ................................................................................ | [126960] |
| MN393477.1_Wuhan_2019 | ................................................................................ | [126960] |
| MW396979.1_Timor-Leste/2019/1 | ................................................................................ | [126960] |
| MK940252.1_CN/2019/InnerMongolia | ................................................................................ | [126960] |
| MK128995.1_China/2018/AnhuiXCGQ | ................................................................................ | [126960] |
| MH681419.1_ASFV/POL/2015 | ................................................................................ | [126960] |
| LS478113.1_Estonia_2014 | ................................................................................ | [126960] |
| KP843857.1_Odintsovo_02/14 | ................................................................................ | [126960] |
| FN557520.1_strain_E75 | ............c................................................................... | [126960] |
| AY261366.1_Warthog | ................................................................................ | [126960] |
| AY261365.1_Warmbaths | ...................................................t............................ | [126960] |
| AY261364.1_Tengani_62 | ............................................................................g... | [126960] |
| AY261363.1_Pretoriuskop/96/4 | ................................................................................ | [126960] |
| AY261362.1_Mkuzi_1979 | ......................a......................................................... | [126960] |
| AY261361.1_Malawi_Lil-20/1_1983 | .........................g..........................g........................... | [126960] |
| AY261360.1_Kenya_1950 | .......g.................g.....t..a.................g.................g......... | [126960] |
| AM712240.1_OURT_88/3 | ............c................................................................... | [126960] |
| AM712239.1_Benin_97/1 | ............c................................................................... | [126960] |
| KM111295.1_Ken06.Bus | .......g.................g..........................g.................g..t...... | [126960] |

```
KM111294.1_Ken05/Tk1           .......g.................g.....t....................g.................g......... [126960]
KM262844.1_strain_L60          ............c................................................................... [126960]
KM102979.1_26544/OG10_Italy    ............c................................................................... [126960]
KM262845.1_strain_NHV          ............c................................................................... [126960]
KX354450.1_47/Ss/2008          ............c................................................................... [126960]
KP055815.1_strain_BA71         ............c................................................................... [126960]
MH025920.1_strain_R35          .......g.................g..........................g.................g..t...... [126960]
MH025919.1_strain_N10          .......g.................g..........................g.................g..t...... [126960]
MH025918.1_strain_R25          .......g.................g..........................g.................g..t...... [126960]
MH025917.1_strain_R7           .......g.................g..........................g.................g..t...... [126960]
MH025916.1_strain_R8           .......g.................g..........................g.................g..t...... [126960]

FR682468.2_Georgia_2007/1      gaggaaataatgatcatccgttcgatttcgtggggatcatacgaataaatcccctttttaaataaaaaattgtagacccc [127040]
ASF_IND_20_CAD543              ................................................................................ [127040]
LR899193.1_Germany_2020/1      ................................................................................ [127040]
MN393477.1_Wuhan_2019          ................................................................................ [127040]
MW396979.1_Timor-Leste/2019/1  ................................................................................ [127040]
MK940252.1_CN/2019/InnerMongolia ................................................................................ [127040]
MK128995.1_China/2018/AnhuiXCGQ ................................................................................ [127040]
MH681419.1_ASFV/POL/2015       ................................................................................ [127040]
LS478113.1_Estonia_2014        ................................................................................ [127040]
KP843857.1_Odintsovo_02/14     ................................................................................ [127040]
FN557520.1_strain_E75          ............................................................................t... [127040]
AY261366.1_Warthog             .........................................t...................................t.. [127040]
AY261365.1_Warmbaths           .............................................................................t.. [127040]
AY261364.1_Tengani_62          ................................................................................ [127040]
AY261363.1_Pretoriuskop/96/4   .............................................................................t.. [127040]
AY261362.1_Mkuzi_1979          ............................................................................t... [127040]
AY261361.1_Malawi_Lil-20/1_1983 ................................................................................ [127040]
AY261360.1_Kenya_1950          ........g...........c..................................................a........ [127040]
AM712240.1_OURT_88/3           ............................................................................t... [127040]
AM712239.1_Benin_97/1          ............................................................................t... [127040]
KM111295.1_Ken06.Bus           ....................c..................................................a........ [127040]
KM111294.1_Ken05/Tk1           ........g...........c..................................................a........ [127040]
KM262844.1_strain_L60          ............................................................................t... [127040]
KM102979.1_26544/OG10_Italy    ............................................................................t... [127040]
KM262845.1_strain_NHV          ............................................................................t... [127040]
KX354450.1_47/Ss/2008          ............................................................................t... [127040]
KP055815.1_strain_BA71         ............................................................................t... [127040]
MH025920.1_strain_R35          ....................c..................................................a........ [127040]
MH025919.1_strain_N10          ....................c..................................................a........ [127040]
MH025918.1_strain_R25          ....................c..................................................a........ [127040]
MH025917.1_strain_R7           ....................c..................................................a........ [127040]
MH025916.1_strain_R8           ....................c..................................................a........ [127040]
```

```
FR682468.2_Georgia_2007/1        ggtttgctggaggccccgcacggaaataatccctgcttgctcgtattcccgccaacgacttttgagctcggtaaatccct [127120]
ASF_IND_20_CAD543                ................................................................................ [127120]
LR899193.1_Germany_2020/1        ................................................................................ [127120]
MN393477.1_Wuhan_2019            ................................................................................ [127120]
MW396979.1_Timor-Leste/2019/1    ................................................................................ [127120]
MK940252.1_CN/2019/InnerMongolia ................................................................................ [127120]
MK128995.1_China/2018/AnhuiXCGQ  ................................................................................ [127120]
MH681419.1_ASFV/POL/2015         ................................................................................ [127120]
LS478113.1_Estonia_2014          ................................................................................ [127120]
KP843857.1_Odintsovo_02/14       ................................................................................ [127120]
FN557520.1_strain_E75            ................................................................................ [127120]
AY261366.1_Warthog               ................................................................................ [127120]
AY261365.1_Warmbaths             ..........................................a..................................... [127120]
AY261364.1_Tengani_62            .........a.................c.................................................... [127120]
AY261363.1_Pretoriuskop/96/4     ................................................................................ [127120]
AY261362.1_Mkuzi_1979            ................................................................................ [127120]
AY261361.1_Malawi_Lil-20/1_1983  t.....t.....a..................................................t...........a.... [127120]
AY261360.1_Kenya_1950            a........a....................t.....c.....a....................c................ [127120]
AM712240.1_OURT_88/3             ................................................................................ [127120]
AM712239.1_Benin_97/1            ................................................................................ [127120]
KM111295.1_Ken06.Bus             a.............................t...........a....tt..............c................ [127120]
KM111294.1_Ken05/Tk1             a.............................t...........a....................t................ [127120]
KM262844.1_strain_L60            ................................................................................ [127120]
KM102979.1_26544/OG10_Italy      ................................................................................ [127120]
KM262845.1_strain_NHV            ................................................................................ [127120]
KX354450.1_47/Ss/2008            ................................................................................ [127120]
KP055815.1_strain_BA71           ................................................................................ [127120]
MH025920.1_strain_R35            a.............................t...........a....................c................ [127120]
MH025919.1_strain_N10            a.............................t...........a....................c................ [127120]
MH025918.1_strain_R25            a.............................t...........a....................c................ [127120]
MH025917.1_strain_R7             a.............................t...........a....................c................ [127120]
MH025916.1_strain_R8             a.............................t...........a....................c................ [127120]

FR682468.2_Georgia_2007/1        tgctagaaagcgtatagggccaaaaggtggacaccgacatggagctgatagaaatttggatgtcctcgttggagggaagg [127200]
ASF_IND_20_CAD543                ................................................................................ [127200]
LR899193.1_Germany_2020/1        ................................................................................ [127200]
MN393477.1_Wuhan_2019            ................................................................................ [127200]
MW396979.1_Timor-Leste/2019/1    ................................................................................ [127200]
MK940252.1_CN/2019/InnerMongolia ................................................................................ [127200]
MK128995.1_China/2018/AnhuiXCGQ  ................................................................................ [127200]
MH681419.1_ASFV/POL/2015         ................................................................................ [127200]
LS478113.1_Estonia_2014          ................................................................................ [127200]
KP843857.1_Odintsovo_02/14       ................................................................................ [127200]
```

```
FN557520.1_strain_E75            ............................a................................................... [127200]
AY261366.1_Warthog               ............................a................................................... [127200]
AY261365.1_Warmbaths             ................................................................................ [127200]
AY261364.1_Tengani_62            ................................................................................ [127200]
AY261363.1_Pretoriuskop/96/4     .............................................................a.................. [127200]
AY261362.1_Mkuzi_1979            ................................................................................ [127200]
AY261361.1_Malawi_Lil-20/1_1983  ...............................t..t............................................. [127200]
AY261360.1_Kenya_1950            ............................a................................................... [127200]
AM712240.1_OURT_88/3             ............................a................................................... [127200]
AM712239.1_Benin_97/1            ............................a................................................... [127200]
KM111295.1_Ken06.Bus             ............................a................................................... [127200]
KM111294.1_Ken05/Tk1             ............................a................................................... [127200]
KM262844.1_strain_L60            ............................a................................................... [127200]
KM102979.1_26544/OG10_Italy      ............................a................................................... [127200]
KM262845.1_strain_NHV            ............................a................................................... [127200]
KX354450.1_47/Ss/2008            ............................a................................................... [127200]
KP055815.1_strain_BA71           ............................a................................................... [127200]
MH025920.1_strain_R35            ............................a................................................... [127200]
MH025919.1_strain_N10            ............................a................................................... [127200]
MH025918.1_strain_R25            ............................a................................................... [127200]
MH025917.1_strain_R7             ............................a................................................... [127200]
MH025916.1_strain_R8             ............................a................................................... [127200]

FR682468.2_Georgia_2007/1        ggcagactccctccacgaggaaacgcggcaggccccatatcattaattgtatgaataataggatttatgaaattatttag [127280]
ASF_IND_20_CAD543                ................................................................................ [127280]
LR899193.1_Germany_2020/1        ................................................................................ [127280]
MN393477.1_Wuhan_2019            ................................................................................ [127280]
MW396979.1_Timor-Leste/2019/1    ................................................................................ [127280]
MK940252.1_CN/2019/InnerMongolia ................................................................................ [127280]
MK128995.1_China/2018/AnhuiXCGQ  ................................................................................ [127280]
MH681419.1_ASFV/POL/2015         ................................................................................ [127280]
LS478113.1_Estonia_2014          ................................................................................ [127280]
KP843857.1_Odintsovo_02/14       ................................................................................ [127280]
FN557520.1_strain_E75            .................g.............................c................................ [127280]
AY261366.1_Warthog               .................g........a...................................g................. [127280]
AY261365.1_Warmbaths             .................g........a...................................g................a [127280]
AY261364.1_Tengani_62            .................g........a..............g....................g................. [127280]
AY261363.1_Pretoriuskop/96/4     .................g........a...................................g................. [127280]
AY261362.1_Mkuzi_1979            .................g.............................................................. [127280]
AY261361.1_Malawi_Lil-20/1_1983  .....g...........g........a.....t..........................g..g.....a........... [127280]
AY261360.1_Kenya_1950            .....g...........g........a...................................g.....a........... [127280]
AM712240.1_OURT_88/3             .................g.............................c................................ [127280]
AM712239.1_Benin_97/1            .................g.............................c................................ [127280]
KM111295.1_Ken06.Bus             .....g...........g........a...................................g.....a........... [127280]
```

```
KM111294.1_Ken05/Tk1              .....g...........g........a............a......................g.....a........... [127280]
KM262844.1_strain_L60             .................g.............................c................................ [127280]
KM102979.1_26544/OG10_Italy       .................g.............................c................................ [127280]
KM262845.1_strain_NHV             .................g.............................c................................ [127280]
KX354450.1_47/Ss/2008             .................g.............................c................................ [127280]
KP055815.1_strain_BA71            .................g.............................c................................ [127280]
MH025920.1_strain_R35             .....g...........g........a...................................g.....a........... [127280]
MH025919.1_strain_N10             .....g...........g........a...................................g.....a........... [127280]
MH025918.1_strain_R25             .....g...........g........a...................................g.....a........... [127280]
MH025917.1_strain_R7              .....g...........g........a...................................g.....a........... [127280]
MH025916.1_strain_R8              .....g...........g........a...................................g.....a........... [127280]

FR682468.2_Georgia_2007/1         ggtggacaccacggagttaaagtcgtggcgctcgttttctgaccaattgctttcgataaagtagtgcccattattttgta [127360]
ASF_IND_20_CAD543                 ................................................................................ [127360]
LR899193.1_Germany_2020/1         ................................................................................ [127360]
MN393477.1_Wuhan_2019             ................................................................................ [127360]
MW396979.1_Timor-Leste/2019/1     ................................................................................ [127360]
MK940252.1_CN/2019/InnerMongolia  ................................................................................ [127360]
MK128995.1_China/2018/AnhuiXCGQ   ................................................................................ [127360]
MH681419.1_ASFV/POL/2015          ................................................................................ [127360]
LS478113.1_Estonia_2014           ................................................................................ [127360]
KP843857.1_Odintsovo_02/14        ................................................................................ [127360]
FN557520.1_strain_E75             .....................................................................g..g....... [127360]
AY261366.1_Warthog                .................................a...................................g..g....... [127360]
AY261365.1_Warmbaths              ...a.................................................................g..g....... [127360]
AY261364.1_Tengani_62             .....................................................................g..g....... [127360]
AY261363.1_Pretoriuskop/96/4      .....................................................................g..g....... [127360]
AY261362.1_Mkuzi_1979             .....................................................................g..g....... [127360]
AY261361.1_Malawi_Lil-20/1_1983   ........................a............................................g..g....... [127360]
AY261360.1_Kenya_1950             .....................................................................g..g.....a. [127360]
AM712240.1_OURT_88/3              .....................................................................g..g....... [127360]
AM712239.1_Benin_97/1             .....................................................................g..g....... [127360]
KM111295.1_Ken06.Bus              ...a.................................................................g..g.....a. [127360]
KM111294.1_Ken05/Tk1              ..............................t......................................g..g.....a. [127360]
KM262844.1_strain_L60             .....................................................................g..g....... [127360]
KM102979.1_26544/OG10_Italy       .....................................................................g..g....... [127360]
KM262845.1_strain_NHV             .....................................................................g..g....... [127360]
KX354450.1_47/Ss/2008             .....................................................................g..g....... [127360]
KP055815.1_strain_BA71            .....................................................................g..g....... [127360]
MH025920.1_strain_R35             ...a.................................................................g..g.....a. [127360]
MH025919.1_strain_N10             ...a.................................................................g..g.....a. [127360]
MH025918.1_strain_R25             ...a.................................................................g..g.....a. [127360]
MH025917.1_strain_R7              ...a.................................................................g..g.....a. [127360]
MH025916.1_strain_R8              ...a.................................................................g..g.....a. [127360]
```

```
FR682468.2_Georgia_2007/1        tggtaagaataaaggcctttttattgataaagcgtattaaaataatagtgggtacacggaatgttttattgctgaatttt [127440]
ASF_IND_20_CAD543                ................................................................................ [127440]
LR899193.1_Germany_2020/1        ................................................................................ [127440]
MN393477.1_Wuhan_2019            ................................................................................ [127440]
MW396979.1_Timor-Leste/2019/1    ................................................................................ [127440]
MK940252.1_CN/2019/InnerMongolia ................................................................................ [127440]
MK128995.1_China/2018/AnhuiXCGQ  ................................................................................ [127440]
MH681419.1_ASFV/POL/2015         ................................................................................ [127440]
LS478113.1_Estonia_2014          ................................................................................ [127440]
KP843857.1_Odintsovo_02/14       ................................................................................ [127440]
FN557520.1_strain_E75            ................................................................................ [127440]
AY261366.1_Warthog               ...........................................g..g................................. [127440]
AY261365.1_Warmbaths             ...........................................g..g................................. [127440]
AY261364.1_Tengani_62            .......g......................................g................................. [127440]
AY261363.1_Pretoriuskop/96/4     ...........................................g..g................................. [127440]
AY261362.1_Mkuzi_1979            ................................................................................ [127440]
AY261361.1_Malawi_Lil-20/1_1983  ................................................................................ [127440]
AY261360.1_Kenya_1950            .............................t................g.....a.....a...........a......... [127440]
AM712240.1_OURT_88/3             ................................................................................ [127440]
AM712239.1_Benin_97/1            ................................................................................ [127440]
KM111295.1_Ken06.Bus             .............................t...............cg.....g.................a......... [127440]
KM111294.1_Ken05/Tk1             .............................t................g.....a.....a...........a......... [127440]
KM262844.1_strain_L60            ................................................................................ [127440]
KM102979.1_26544/OG10_Italy      ................................................................................ [127440]
KM262845.1_strain_NHV            ................................................................................ [127440]
KX354450.1_47/Ss/2008            ................................................................................ [127440]
KP055815.1_strain_BA71           ................................................................................ [127440]
MH025920.1_strain_R35            .............................t...............cg.....g.................a......... [127440]
MH025919.1_strain_N10            .............................t...............cg.....g.................a......... [127440]
MH025918.1_strain_R25            .............................t...............cg.....g.................a......... [127440]
MH025917.1_strain_R7             .............................t...............cg.....g.................a......... [127440]
MH025916.1_strain_R8             .............................t...............cg.....g.................a......... [127440]

FR682468.2_Georgia_2007/1        tcaggctccgtggaagttatgtggtgtttggaaaccacggtgggacctgttttactataaaagaacaccaccagctgagg [127520]
ASF_IND_20_CAD543                ................................................................................ [127520]
LR899193.1_Germany_2020/1        ................................................................................ [127520]
MN393477.1_Wuhan_2019            ................................................................................ [127520]
MW396979.1_Timor-Leste/2019/1    ................................................................................ [127520]
MK940252.1_CN/2019/InnerMongolia ................................................................................ [127520]
MK128995.1_China/2018/AnhuiXCGQ  ................................................................................ [127520]
MH681419.1_ASFV/POL/2015         ................................................................................ [127520]
LS478113.1_Estonia_2014          ................................................................................ [127520]
KP843857.1_Odintsovo_02/14       ................................................................................ [127520]
```

```
FN557520.1_strain_E75              ................................................................................ [127520]
AY261366.1_Warthog                 .....................................................g..............t........... [127520]
AY261365.1_Warmbaths               .....................................................g.......................... [127520]
AY261364.1_Tengani_62              .......................a........................................................ [127520]
AY261363.1_Pretoriuskop/96/4       .....................................................g.......................... [127520]
AY261362.1_Mkuzi_1979              ................................................................................ [127520]
AY261361.1_Malawi_Lil-20/1_1983    ..............g........a...........t.....a.....g................................ [127520]
AY261360.1_Kenya_1950              ........t.....g.....a..a........................................................ [127520]
AM712240.1_OURT_88/3               ................................................................................ [127520]
AM712239.1_Benin_97/1              ................................................................................ [127520]
KM111295.1_Ken06.Bus               ..............g.....a..a........................................................ [127520]
KM111294.1_Ken05/Tk1               ........t.....g.....a..a..................................................t..... [127520]
KM262844.1_strain_L60              ................................................................................ [127520]
KM102979.1_26544/OG10_Italy        ................................................................................ [127520]
KM262845.1_strain_NHV              ................................................................................ [127520]
KX354450.1_47/Ss/2008              ................................................................................ [127520]
KP055815.1_strain_BA71             ................................................................................ [127520]
MH025920.1_strain_R35              ..............g.....a..a........................................................ [127520]
MH025919.1_strain_N10              ..............g.....a..a........................................................ [127520]
MH025918.1_strain_R25              ..............g.....a..a........................................................ [127520]
MH025917.1_strain_R7               ..............g.....a..a........................................................ [127520]
MH025916.1_strain_R8               ..............g.....a..a........................................................ [127520]

FR682468.2_Georgia_2007/1          aatatcgggagtagctggaaataggtcgaaaacattgcgcacattaatttgaatatttacgaggggtgaaattttaatca [127600]
ASF_IND_20_CAD543                  ................................................................................ [127600]
LR899193.1_Germany_2020/1          ................................................................................ [127600]
MN393477.1_Wuhan_2019              ................................................................................ [127600]
MW396979.1_Timor-Leste/2019/1      ................................................................................ [127600]
MK940252.1_CN/2019/InnerMongolia   ................................................................................ [127600]
MK128995.1_China/2018/AnhuiXCGQ    ................................................................................ [127600]
MH681419.1_ASFV/POL/2015           ................................................................................ [127600]
LS478113.1_Estonia_2014            ................................................................................ [127600]
KP843857.1_Odintsovo_02/14         ................................................................................ [127600]
FN557520.1_strain_E75              ...............c.................g................................c............. [127600]
AY261366.1_Warthog                 ...............c.................g................................c............. [127600]
AY261365.1_Warmbaths               ...............c.................g................................c............. [127600]
AY261364.1_Tengani_62              ...............c.................g................................c............. [127600]
AY261363.1_Pretoriuskop/96/4       ...............c.................g.............................c..g............. [127600]
AY261362.1_Mkuzi_1979              .................................g................................c............. [127600]
AY261361.1_Malawi_Lil-20/1_1983    ............g..c.....c...........g................................c........g.... [127600]
AY261360.1_Kenya_1950              ......a.....g.....................................................c........g.... [127600]
AM712240.1_OURT_88/3               ...............c.................g................................c............. [127600]
AM712239.1_Benin_97/1              ...............c.................g................................c............. [127600]
KM111295.1_Ken06.Bus               ......a.....g.....................................................c........g.... [127600]
```

```
KM111294.1_Ken05/Tk1            ......a.....g.....................................................c........g.... [127600]
KM262844.1_strain_L60           ...............c.................g................................c............. [127600]
KM102979.1_26544/OG10_Italy     ...............c.................g................................c............. [127600]
KM262845.1_strain_NHV           ...............c.................g................................c............. [127600]
KX354450.1_47/Ss/2008           ...............c.................g................................c............. [127600]
KP055815.1_strain_BA71          ...............c.................g................................c............. [127600]
MH025920.1_strain_R35           ......a.....g.....................................................c........g.... [127600]
MH025919.1_strain_N10           ......a.....g.....................................................c........g.... [127600]
MH025918.1_strain_R25           ......a.....g.....................................................c........g.... [127600]
MH025917.1_strain_R7            ......a.....g.....................................................c........g.... [127600]
MH025916.1_strain_R8            ......a.....g.....................................................c........g.... [127600]

FR682468.2_Georgia_2007/1       ttgccgaggtgacggccaacgtgccgcgtgttagtctattcccctcgtacttggcaatgacttgttgtgctctggcatac [127680]
ASF_IND_20_CAD543               ................................................................................ [127680]
LR899193.1_Germany_2020/1       ................................................................................ [127680]
MN393477.1_Wuhan_2019           ................................................................................ [127680]
MW396979.1_Timor-Leste/2019/1   ................................................................................ [127680]
MK940252.1_CN/2019/InnerMongolia ................................................................................ [127680]
MK128995.1_China/2018/AnhuiXCGQ ................................................................................ [127680]
MH681419.1_ASFV/POL/2015        ................................................................................ [127680]
LS478113.1_Estonia_2014         ................................................................................ [127680]
KP843857.1_Odintsovo_02/14      ................................................................................ [127680]
FN557520.1_strain_E75           ................t............................................................... [127680]
AY261366.1_Warthog              ..........a..................................................................... [127680]
AY261365.1_Warmbaths            ................................................................................ [127680]
AY261364.1_Tengani_62           ................................................................................ [127680]
AY261363.1_Pretoriuskop/96/4    ...............................................................................t [127680]
AY261362.1_Mkuzi_1979           ................t............................................................... [127680]
AY261361.1_Malawi_Lil-20/1_1983 .......................................................................t....g... [127680]
AY261360.1_Kenya_1950           ..........a........................t.....................c...c.....c........g... [127680]
AM712240.1_OURT_88/3            ................t............................................................... [127680]
AM712239.1_Benin_97/1           ................t............................................................... [127680]
KM111295.1_Ken06.Bus            ..........a........................t.....................c...c.....c........g... [127680]
KM111294.1_Ken05/Tk1            ..........a........................t.....................c...c.....c........g..t [127680]
KM262844.1_strain_L60           ................t............................................................... [127680]
KM102979.1_26544/OG10_Italy     ................t............................................................... [127680]
KM262845.1_strain_NHV           ................t............................................................... [127680]
KX354450.1_47/Ss/2008           ................t............................................................... [127680]
KP055815.1_strain_BA71          ................t............................................................... [127680]
MH025920.1_strain_R35           ..........a........................t.....................c...c.....c........g... [127680]
MH025919.1_strain_N10           ..........a........................t.....................c...c.....c........g... [127680]
MH025918.1_strain_R25           ..........a........................t.....................c...c.....c........g... [127680]
MH025917.1_strain_R7            ..........a........................t.....................c...c.....c........g... [127680]
MH025916.1_strain_R8            ..........a........................t.....................c...c.....c........g... [127680]
```

| | | |
|---|---|---|
| FR682468.2_Georgia_2007/1 | gtaaagtttattagtttttgctctaggagaagcctctttttaagactggtcaaggatggagaaagagcaggatactgttt | [127760] |
| ASF_IND_20_CAD543 | ................................................................................ | [127760] |
| LR899193.1_Germany_2020/1 | ................................................................................ | [127760] |
| MN393477.1_Wuhan_2019 | ................................................................................ | [127760] |
| MW396979.1_Timor-Leste/2019/1 | ................................................................................ | [127760] |
| MK940252.1_CN/2019/InnerMongolia | ................................................................................ | [127760] |
| MK128995.1_China/2018/AnhuiXCGQ | ................................................................................ | [127760] |
| MH681419.1_ASFV/POL/2015 | ................................................................................ | [127760] |
| LS478113.1_Estonia_2014 | ................................................................................ | [127760] |
| KP843857.1_Odintsovo_02/14 | ................................................................................ | [127760] |
| FN557520.1_strain_E75 | .............................t.................................................. | [127760] |
| AY261366.1_Warthog | ............................................g................................... | [127760] |
| AY261365.1_Warmbaths | ............................................g................................... | [127760] |
| AY261364.1_Tengani_62 | ............................................g................................... | [127760] |
| AY261363.1_Pretoriuskop/96/4 | ................................................................................ | [127760] |
| AY261362.1_Mkuzi_1979 | ................................................................................ | [127760] |
| AY261361.1_Malawi_Lil-20/1_1983 | ....................t..a........................................................ | [127760] |
| AY261360.1_Kenya_1950 | ....................t..a........................................................ | [127760] |
| AM712240.1_OURT_88/3 | .............................t.................................................. | [127760] |
| AM712239.1_Benin_97/1 | ....................t........t.................................................. | [127760] |
| KM111295.1_Ken06.Bus | ....................t..a.............................a.......................... | [127760] |
| KM111294.1_Ken05/Tk1 | ....................t..a........................................................ | [127760] |
| KM262844.1_strain_L60 | .............................t.................................................. | [127760] |
| KM102979.1_26544/OG10_Italy | .............................t.................................................. | [127760] |
| KM262845.1_strain_NHV | .............................t.................................................. | [127760] |
| KX354450.1_47/Ss/2008 | .............................t.................................................. | [127760] |
| KP055815.1_strain_BA71 | .............................t.................................................. | [127760] |
| MH025920.1_strain_R35 | ....................t..a.............................a.......................... | [127760] |
| MH025919.1_strain_N10 | ....................t..a.............................a.......................... | [127760] |
| MH025918.1_strain_R25 | ....................t..a.............................a.......................... | [127760] |
| MH025917.1_strain_R7 | ....................t..a.............................a.......................... | [127760] |
| MH025916.1_strain_R8 | ....................t..a.............................a.......................... | [127760] |

| | | |
|---|---|---|
| FR682468.2_Georgia_2007/1 | ttccatttgtaagggagattgtaccaatagtttaaaggcatcgggggaaagaagaggccaatacttcataataaggccgt | [127840] |
| ASF_IND_20_CAD543 | ................................................................................ | [127840] |
| LR899193.1_Germany_2020/1 | ................................................................................ | [127840] |
| MN393477.1_Wuhan_2019 | ................................................................................ | [127840] |
| MW396979.1_Timor-Leste/2019/1 | ................................................................................ | [127840] |
| MK940252.1_CN/2019/InnerMongolia | ................................................................................ | [127840] |
| MK128995.1_China/2018/AnhuiXCGQ | ................................................................................ | [127840] |
| MH681419.1_ASFV/POL/2015 | ................................................................................ | [127840] |
| LS478113.1_Estonia_2014 | ................................................................................ | [127840] |
| KP843857.1_Odintsovo_02/14 | ................................................................................ | [127840] |

```
FN557520.1_strain_E75             ................................................................................ [127840]
AY261366.1_Warthog                .............................a.................................................. [127840]
AY261365.1_Warmbaths              .............................a.................................................. [127840]
AY261364.1_Tengani_62             ...........................................................................a.... [127840]
AY261363.1_Pretoriuskop/96/4      ................................................................................ [127840]
AY261362.1_Mkuzi_1979             ................................................................................ [127840]
AY261361.1_Malawi_Lil-20/1_1983   ..................c..g.t........................................................ [127840]
AY261360.1_Kenya_1950             ............a.....c....t..g..................................................... [127840]
AM712240.1_OURT_88/3              ................................................................................ [127840]
AM712239.1_Benin_97/1             ................................................................................ [127840]
KM111295.1_Ken06.Bus              ..................c....t..g..................................................... [127840]
KM111294.1_Ken05/Tk1              ............a.....c....t..g..................................................... [127840]
KM262844.1_strain_L60             ................................................................................ [127840]
KM102979.1_26544/OG10_Italy       ................................................................................ [127840]
KM262845.1_strain_NHV             ................................................................................ [127840]
KX354450.1_47/Ss/2008             ................................................................................ [127840]
KP055815.1_strain_BA71            ................................................................................ [127840]
MH025920.1_strain_R35             ..................c....t..g..................................................... [127840]
MH025919.1_strain_N10             ..................c....t..g..................................................... [127840]
MH025918.1_strain_R25             ..................c....t..g..................................................... [127840]
MH025917.1_strain_R7              ..................c....t..g..................................................... [127840]
MH025916.1_strain_R8              ..................c....t..g..................................................... [127840]

FR682468.2_Georgia_2007/1         aatagagtaagtcaaattggtaattatcctctatggcaatggagatttggcgccgcatgggggccactagcgtgttgagg [127920]
ASF_IND_20_CAD543                 ................................................................................ [127920]
LR899193.1_Germany_2020/1         ................................................................................ [127920]
MN393477.1_Wuhan_2019             ................................................................................ [127920]
MW396979.1_Timor-Leste/2019/1     ................................................................................ [127920]
MK940252.1_CN/2019/InnerMongolia  ................................................................................ [127920]
MK128995.1_China/2018/AnhuiXCGQ   ................................................................................ [127920]
MH681419.1_ASFV/POL/2015          ................................................................................ [127920]
LS478113.1_Estonia_2014           ................................................................................ [127920]
KP843857.1_Odintsovo_02/14        ................................................................................ [127920]
FN557520.1_strain_E75             ...............................c................................................ [127920]
AY261366.1_Warthog                ................................................................................ [127920]
AY261365.1_Warmbaths              ......................g......................................................... [127920]
AY261364.1_Tengani_62             ................................................................................ [127920]
AY261363.1_Pretoriuskop/96/4      ................................................................................ [127920]
AY261362.1_Mkuzi_1979             ................................................................................ [127920]
AY261361.1_Malawi_Lil-20/1_1983   ..................................a............................................. [127920]
AY261360.1_Kenya_1950             ..................................a................................a............ [127920]
AM712240.1_OURT_88/3              ...............................c................................................ [127920]
AM712239.1_Benin_97/1             ...............................c................................................ [127920]
KM111295.1_Ken06.Bus              ...................................................................g............ [127920]
```

```
KM111294.1_Ken05/Tk1          ..................................a................................a............ [127920]
KM262844.1_strain_L60         ...............................c................................................ [127920]
KM102979.1_26544/OG10_Italy   ...............................c................................................ [127920]
KM262845.1_strain_NHV         ...............................c................................................ [127920]
KX354450.1_47/Ss/2008         ...............................c................................................ [127920]
KP055815.1_strain_BA71        ...............................c................................................ [127920]
MH025920.1_strain_R35         ...................................................................g............ [127920]
MH025919.1_strain_N10         ...................................................................g............ [127920]
MH025918.1_strain_R25         ...................................................................g............ [127920]
MH025917.1_strain_R7          ...................................................................g............ [127920]
MH025916.1_strain_R8          ...................................................................g............ [127920]

FR682468.2_Georgia_2007/1     tctgctacaaagatgtgatgaatgttttttatgagctggaagctgtcgagcgcttccacatagagctcatctttttgact [128000]
ASF_IND_20_CAD543             ................................................................................ [128000]
LR899193.1_Germany_2020/1     ................................................................................ [128000]
MN393477.1_Wuhan_2019         ................................................................................ [128000]
MW396979.1_Timor-Leste/2019/1 ................................................................................ [128000]
MK940252.1_CN/2019/InnerMongolia ................................................................................ [128000]
MK128995.1_China/2018/AnhuiXCGQ ................................................................................ [128000]
MH681419.1_ASFV/POL/2015      ................................................................................ [128000]
LS478113.1_Estonia_2014       ................................................................................ [128000]
KP843857.1_Odintsovo_02/14    ................................................................................ [128000]
FN557520.1_strain_E75         .....c.......................................................................... [128000]
AY261366.1_Warthog            .....c.......................................................................... [128000]
AY261365.1_Warmbaths          .....c.......................................................................... [128000]
AY261364.1_Tengani_62         .....c..............................................................g..c........ [128000]
AY261363.1_Pretoriuskop/96/4  .....c.......................................................................... [128000]
AY261362.1_Mkuzi_1979         .....c.......................................................................... [128000]
AY261361.1_Malawi_Lil-20/1_1983 .....c.................a.......................a.................t.............. [128000]
AY261360.1_Kenya_1950         .....c.......................................................................... [128000]
AM712240.1_OURT_88/3          .....c.......................................................................... [128000]
AM712239.1_Benin_97/1         .....c.......................................................................... [128000]
KM111295.1_Ken06.Bus          .....c.......................................................................... [128000]
KM111294.1_Ken05/Tk1          .....c........a................................................................. [128000]
KM262844.1_strain_L60         .....c.......................................................................... [128000]
KM102979.1_26544/OG10_Italy   .....c.......................................................................... [128000]
KM262845.1_strain_NHV         .....c.......................................................................... [128000]
KX354450.1_47/Ss/2008         .....c.......................................................................... [128000]
KP055815.1_strain_BA71        .....c.......................................................................... [128000]
MH025920.1_strain_R35         .....c.......................................................................... [128000]
MH025919.1_strain_N10         .....c.......................................................................... [128000]
MH025918.1_strain_R25         .....c.......................................................................... [128000]
MH025917.1_strain_R7          .....c.......................................................................... [128000]
MH025916.1_strain_R8          .....c.......................................................................... [128000]
```

```
FR682468.2_Georgia_2007/1        ttccatagatgcgtcgatgttcaccccacccacctgttgaaactcctttttgtagtcgcgaatgtctaacgccaccccgc [128080]
ASF_IND_20_CAD543                ................................................................................ [128080]
LR899193.1_Germany_2020/1        ................................................................................ [128080]
MN393477.1_Wuhan_2019            ................................................................................ [128080]
MW396979.1_Timor-Leste/2019/1    ................................................................................ [128080]
MK940252.1_CN/2019/InnerMongolia ................................................................................ [128080]
MK128995.1_China/2018/AnhuiXCGQ  ................................................................................ [128080]
MH681419.1_ASFV/POL/2015         ................................................................................ [128080]
LS478113.1_Estonia_2014          ................................................................................ [128080]
KP843857.1_Odintsovo_02/14       ................................................................................ [128080]
FN557520.1_strain_E75            .......................g............................................g........... [128080]
AY261366.1_Warthog               .......................g...g........................................g........... [128080]
AY261365.1_Warmbaths             ............a..........g............................................g........... [128080]
AY261364.1_Tengani_62            ..................a....g............................................g........... [128080]
AY261363.1_Pretoriuskop/96/4     .......................g...g........................................g........... [128080]
AY261362.1_Mkuzi_1979            .......................g............................................g........... [128080]
AY261361.1_Malawi_Lil-20/1_1983  ......g................g.................................a..........g...g....... [128080]
AY261360.1_Kenya_1950            ......c.t............t.gg..g..t..t.............................a....g........... [128080]
AM712240.1_OURT_88/3             .......................g............................................g........... [128080]
AM712239.1_Benin_97/1            .......................g............................................g..t........ [128080]
KM111295.1_Ken06.Bus             ......c.t............t.gg........t.......................a.....a....g........... [128080]
KM111294.1_Ken05/Tk1             ......c.t............t.gg..g.....t.............................a....g........... [128080]
KM262844.1_strain_L60            .......................g............................................g........... [128080]
KM102979.1_26544/OG10_Italy      .......................g............................................gt.......... [128080]
KM262845.1_strain_NHV            .......................g............................................g........... [128080]
KX354450.1_47/Ss/2008            .......................g............................................gt.......... [128080]
KP055815.1_strain_BA71           .......................g............................................g........... [128080]
MH025920.1_strain_R35            ......c.t............t.gg........t.......................a.....a....g........... [128080]
MH025919.1_strain_N10            ......c.t............t.gg........t.......................a.....a....g........... [128080]
MH025918.1_strain_R25            ......c.t............t.gg........t.......................a.....a....g........... [128080]
MH025917.1_strain_R7             ......c.t............t.gg........t.......................a.....a....g........... [128080]
MH025916.1_strain_R8             ......c.t............t.gg........t.......................a.....a....g........... [128080]

FR682468.2_Georgia_2007/1        taccgcttaacaataggcgatacgttacctgaagcgcattgttttgaaaaaagaaaatgtgttgtctataagggggggatc [128160]
ASF_IND_20_CAD543                ................................................................................ [128160]
LR899193.1_Germany_2020/1        ................................................................................ [128160]
MN393477.1_Wuhan_2019            ................................................................................ [128160]
MW396979.1_Timor-Leste/2019/1    ................................................................................ [128160]
MK940252.1_CN/2019/InnerMongolia ................................................................................ [128160]
MK128995.1_China/2018/AnhuiXCGQ  ................................................................................ [128160]
MH681419.1_ASFV/POL/2015         ................................................................................ [128160]
LS478113.1_Estonia_2014          ................................................................................ [128160]
KP843857.1_Odintsovo_02/14       ................................................................................ [128160]
```

```
FN557520.1_strain_E75            .....................................g.......................................... [128160]
AY261366.1_Warthog               .......c.................c..........tg................................g......... [128160]
AY261365.1_Warmbaths             .......c.................c...........g.......................................... [128160]
AY261364.1_Tengani_62            ....................................tg................................g......... [128160]
AY261363.1_Pretoriuskop/96/4     .........................c..........tg................................g......... [128160]
AY261362.1_Mkuzi_1979            .....................................g.......................................... [128160]
AY261361.1_Malawi_Lil-20/1_1983  .....................................g.......................................... [128160]
AY261360.1_Kenya_1950            ............t........................g........................................c. [128160]
AM712240.1_OURT_88/3             .....................................g.......................................... [128160]
AM712239.1_Benin_97/1            .....................................g.......................................... [128160]
KM111295.1_Ken06.Bus             ............t........................g......................................a.c. [128160]
KM111294.1_Ken05/Tk1             ............t........................g........................................c. [128160]
KM262844.1_strain_L60            .....................................g.......................................... [128160]
KM102979.1_26544/OG10_Italy      .....................................g.......................................... [128160]
KM262845.1_strain_NHV            .....................................g.......................................... [128160]
KX354450.1_47/Ss/2008            .....................................g.......................................... [128160]
KP055815.1_strain_BA71           .....................................g.......................................... [128160]
MH025920.1_strain_R35            ............t........................g......................................a.c. [128160]
MH025919.1_strain_N10            ............t........................g......................................a.c. [128160]
MH025918.1_strain_R25            ............t........................g......................................a.c. [128160]
MH025917.1_strain_R7             ............t........................g......................................a.c. [128160]
MH025916.1_strain_R8             ............t........................g......................................a.c. [128160]

FR682468.2_Georgia_2007/1        cctgtggcaacgtaaattttttctcgaatgtctttaaaagtgtcttcagggaaaatactatactcgctatacatcgtctc [128240]
ASF_IND_20_CAD543                ................................................................................ [128240]
LR899193.1_Germany_2020/1        ................................................................................ [128240]
MN393477.1_Wuhan_2019            ................................................................................ [128240]
MW396979.1_Timor-Leste/2019/1    ................................................................................ [128240]
MK940252.1_CN/2019/InnerMongolia ................................................................................ [128240]
MK128995.1_China/2018/AnhuiXCGQ  ................................................................................ [128240]
MH681419.1_ASFV/POL/2015         ................................................................................ [128240]
LS478113.1_Estonia_2014          ................................................................................ [128240]
KP843857.1_Odintsovo_02/14       ................................................................................ [128240]
FN557520.1_strain_E75            ................................................................................ [128240]
AY261366.1_Warthog               .......................c........................................................ [128240]
AY261365.1_Warmbaths             .......................c........................................................ [128240]
AY261364.1_Tengani_62            ................................................................................ [128240]
AY261363.1_Pretoriuskop/96/4     .......................c........................................................ [128240]
AY261362.1_Mkuzi_1979            ................................................................................ [128240]
AY261361.1_Malawi_Lil-20/1_1983  ......................................g......................................... [128240]
AY261360.1_Kenya_1950            ..........................g...........g..a........a............................. [128240]
AM712240.1_OURT_88/3             ................................................................................ [128240]
AM712239.1_Benin_97/1            ................................................................................ [128240]
KM111295.1_Ken06.Bus             ..........................g...........g..a........a......................c...... [128240]
```

```
KM111294.1_Ken05/Tk1              ..........................g...........g..a........a............................. [128240]
KM262844.1_strain_L60             ................................................................................ [128240]
KM102979.1_26544/OG10_Italy       ................................................................................ [128240]
KM262845.1_strain_NHV             ................................................................................ [128240]
KX354450.1_47/Ss/2008             ................................................................................ [128240]
KP055815.1_strain_BA71            ................................................................................ [128240]
MH025920.1_strain_R35             ..........................g...........g..a........a......................c...... [128240]
MH025919.1_strain_N10             ..........................g...........g..a........a......................c...... [128240]
MH025918.1_strain_R25             ..........................g...........g..a........a......................c...... [128240]
MH025917.1_strain_R7              ..........................g...........g..a........a......................c...... [128240]
MH025916.1_strain_R8              ..........................g...........g..a........a......................c...... [128240]

FR682468.2_Georgia_2007/1         aatttctggcatcatcacgtttgtctcctcgccacgatcctccacaaaaagtttttcaaactcatctaaatcatcgctat [128320]
ASF_IND_20_CAD543                 ...................................................y............................ [128320]
LR899193.1_Germany_2020/1         ................................................................................ [128320]
MN393477.1_Wuhan_2019             ................................................................................ [128320]
MW396979.1_Timor-Leste/2019/1     ................................................................................ [128320]
MK940252.1_CN/2019/InnerMongolia  ................................................................................ [128320]
MK128995.1_China/2018/AnhuiXCGQ   ................................................................................ [128320]
MH681419.1_ASFV/POL/2015          ................................................................................ [128320]
LS478113.1_Estonia_2014           ................................................................................ [128320]
KP843857.1_Odintsovo_02/14        ................................................................................ [128320]
FN557520.1_strain_E75             ...................................t...............c....................g....... [128320]
AY261366.1_Warthog                ...................................t....................................g....c.. [128320]
AY261365.1_Warmbaths              ..................................tt....................................g....c.. [128320]
AY261364.1_Tengani_62             .......................c................................................g....... [128320]
AY261363.1_Pretoriuskop/96/4      ...................................t.........................................c.. [128320]
AY261362.1_Mkuzi_1979             ...................................t...............c....................g....... [128320]
AY261361.1_Malawi_Lil-20/1_1983   .......................c...........t....................................g....... [128320]
AY261360.1_Kenya_1950             ...................................t...t................................g....... [128320]
AM712240.1_OURT_88/3              ...................................t...............c....................g....... [128320]
AM712239.1_Benin_97/1             ...................................t...............c....................g....... [128320]
KM111295.1_Ken06.Bus              ...................................t....................................g....... [128320]
KM111294.1_Ken05/Tk1              ...................................t...t................................g....... [128320]
KM262844.1_strain_L60             ...................................t...............c....................g....... [128320]
KM102979.1_26544/OG10_Italy       ...................................t...............c....................g....... [128320]
KM262845.1_strain_NHV             ...................................t...............c....................g....... [128320]
KX354450.1_47/Ss/2008             ...................................t...............c....................g....... [128320]
KP055815.1_strain_BA71            ...................................t...............c....................g....... [128320]
MH025920.1_strain_R35             ...................................t....................................g....... [128320]
MH025919.1_strain_N10             ...................................t....................................g....... [128320]
MH025918.1_strain_R25             ...................................t....................................g....... [128320]
MH025917.1_strain_R7              ...................................t....................................g....... [128320]
MH025916.1_strain_R8              ...................................t....................................g....... [128320]
```

```
FR682468.2_Georgia_2007/1        ctccacccaccacgtattgggaaagctttttctcccaatcctcgccgtaaaaattttgtaaaatttctttgtccttaggg [128400]
ASF_IND_20_CAD543                ......................................................................r......... [128400]
LR899193.1_Germany_2020/1        ................................................................................ [128400]
MN393477.1_Wuhan_2019            ................................................................................ [128400]
MW396979.1_Timor-Leste/2019/1    ................................................................................ [128400]
MK940252.1_CN/2019/InnerMongolia ................................................................................ [128400]
MK128995.1_China/2018/AnhuiXCGQ  ................................................................................ [128400]
MH681419.1_ASFV/POL/2015         ................................................................................ [128400]
LS478113.1_Estonia_2014          ................................................................................ [128400]
KP843857.1_Odintsovo_02/14       ................................................................................ [128400]
FN557520.1_strain_E75            ....g.....................................t...........................a......... [128400]
AY261366.1_Warthog               ....g........................................................................... [128400]
AY261365.1_Warmbaths             ....g........................................................................... [128400]
AY261364.1_Tengani_62            ....g........................................................................... [128400]
AY261363.1_Pretoriuskop/96/4     ....g........................................................................... [128400]
AY261362.1_Mkuzi_1979            ....g.....................................t...........................a......... [128400]
AY261361.1_Malawi_Lil-20/1_1983  .c....................................................................a......... [128400]
AY261360.1_Kenya_1950            .c........g.............................t.t.................g.........a......... [128400]
AM712240.1_OURT_88/3             ....g.....................................t...........................a......... [128400]
AM712239.1_Benin_97/1            ....g.....................................t...........................a......... [128400]
KM111295.1_Ken06.Bus             .c........g........c....................t.t...........................a......... [128400]
KM111294.1_Ken05/Tk1             .c........g.............................t.t.................g.........a......... [128400]
KM262844.1_strain_L60            ....g.....................................t...........................a......... [128400]
KM102979.1_26544/OG10_Italy      ....g.....................................t...........................a......... [128400]
KM262845.1_strain_NHV            ....g.....................................t...........................a......... [128400]
KX354450.1_47/Ss/2008            ....g.....................................t...........................a......... [128400]
KP055815.1_strain_BA71           ....g.....................................t...........................a......... [128400]
MH025920.1_strain_R35            .c........g........c....................t.t...........................a......... [128400]
MH025919.1_strain_N10            .c........g........c....................t.t...........................a......... [128400]
MH025918.1_strain_R25            .c........g........c....................t.t...........................a......... [128400]
MH025917.1_strain_R7             .c........g........c....................t.t...........................a......... [128400]
MH025916.1_strain_R8             .c........g........c....................t.t...........................a......... [128400]

FR682468.2_Georgia_2007/1        gttcgctgcaggtctttgcggcaggcctgtaacacgtttgcaggaacggatcccaaaaaaataaacgtcttcgtgtactc [128480]
ASF_IND_20_CAD543                ................................................................................ [128480]
LR899193.1_Germany_2020/1        ................................................................................ [128480]
MN393477.1_Wuhan_2019            ................................................................................ [128480]
MW396979.1_Timor-Leste/2019/1    ................................................................................ [128480]
MK940252.1_CN/2019/InnerMongolia ................................................................................ [128480]
MK128995.1_China/2018/AnhuiXCGQ  ................................................................................ [128480]
MH681419.1_ASFV/POL/2015         ................................................................................ [128480]
LS478113.1_Estonia_2014          ................................................................................ [128480]
KP843857.1_Odintsovo_02/14       ................................................................................ [128480]
```

```
FN557520.1_strain_E75            ..........a........................a........c................................... [128480]
AY261366.1_Warthog               ................................................................................ [128480]
AY261365.1_Warmbaths             ................................................................................ [128480]
AY261364.1_Tengani_62            .............................c.....a............................................ [128480]
AY261363.1_Pretoriuskop/96/4     ................................................................................ [128480]
AY261362.1_Mkuzi_1979            ..........a........................a........c................................... [128480]
AY261361.1_Malawi_Lil-20/1_1983  ...................................a......................................a..... [128480]
AY261360.1_Kenya_1950            ........t....................................................................... [128480]
AM712240.1_OURT_88/3             ..........a........................a........c................................... [128480]
AM712239.1_Benin_97/1            ..........a........................a........c................................... [128480]
KM111295.1_Ken06.Bus             ........t.........t............................................................. [128480]
KM111294.1_Ken05/Tk1             ........t....................................................................... [128480]
KM262844.1_strain_L60            ..........a........................a........c................................... [128480]
KM102979.1_26544/OG10_Italy      ..........a........................a........c................................... [128480]
KM262845.1_strain_NHV            ..........a........................a........c................................... [128480]
KX354450.1_47/Ss/2008            ..........a........................a........c................................... [128480]
KP055815.1_strain_BA71           ..........a........................a........c................................... [128480]
MH025920.1_strain_R35            ........t.........t............................................................. [128480]
MH025919.1_strain_N10            ........t.........t............................................................. [128480]
MH025918.1_strain_R25            ........t.........t............................................................. [128480]
MH025917.1_strain_R7             ........t.........t............................................................. [128480]
MH025916.1_strain_R8             ........t.........t............................................................. [128480]

FR682468.2_Georgia_2007/1        attttccacaggattataaagagtaactcgtagaggatttgttaaaaagtcattttggaaatccat-------------- [128560]
ASF_IND_20_CAD543                ..................................................................-------------- [128560]
LR899193.1_Germany_2020/1        ..................................................................-------------- [128560]
MN393477.1_Wuhan_2019            ..................................................................-------------- [128560]
MW396979.1_Timor-Leste/2019/1    ..................................................................-------------- [128560]
MK940252.1_CN/2019/InnerMongolia ..................................................................-------------- [128560]
MK128995.1_China/2018/AnhuiXCGQ  ..................................................................-------------- [128560]
MH681419.1_ASFV/POL/2015         ..................................................................-------------- [128560]
LS478113.1_Estonia_2014          ..................................................................-------------- [128560]
KP843857.1_Odintsovo_02/14       ..................................................................-------------- [128560]
FN557520.1_strain_E75            .......g..........................................................-------------- [128560]
AY261366.1_Warthog               .......g..........................................................-------------- [128560]
AY261365.1_Warmbaths             .......g..........................................................-------------- [128560]
AY261364.1_Tengani_62            .......g..........................................................-------------- [128560]
AY261363.1_Pretoriuskop/96/4     .......g..........................................................-------------- [128560]
AY261362.1_Mkuzi_1979            .......g..........................................................-------------- [128560]
AY261361.1_Malawi_Lil-20/1_1983  ......tg................c.......................a..g........t.....tcatatacaggatt [128560]
AY261360.1_Kenya_1950            ......tg................c..........................g..g..a..t.....tcatatataggatt [128560]
AM712240.1_OURT_88/3             .......g..........................................................-------------- [128560]
AM712239.1_Benin_97/1            .......g..........................................................-------------- [128560]
KM111295.1_Ken06.Bus             ......tg...........................................g..g..a..t.....tcatatataagatt [128560]
```

```
KM111294.1_Ken05/Tk1            ......tg...........................................g..g..a..t.....tcatatataggatt [128560]
KM262844.1_strain_L60           .......g..........................................................-------------- [128560]
KM102979.1_26544/OG10_Italy     .......g..........................................................-------------- [128560]
KM262845.1_strain_NHV           .......g..........................................................-------------- [128560]
KX354450.1_47/Ss/2008           .......g..........................................................-------------- [128560]
KP055815.1_strain_BA71          .......g..........................................................-------------- [128560]
MH025920.1_strain_R35           ......tg...........................................g..g..a..t.....tcatatataagatt [128560]
MH025919.1_strain_N10           ......tg...........................................g..g..a..t.....tcatatataagatt [128560]
MH025918.1_strain_R25           ......tg...........................................g..g..a..t.....tcatatataagatt [128560]
MH025917.1_strain_R7            ......tg...........................................g..g..a..t.....tcatatataagatt [128560]
MH025916.1_strain_R8            ......tg...........................................g..g..a..t.....tcatatataagatt [128560]

FR682468.2_Georgia_2007/1       ----------------------------------------tatacccggtatagaaaataaaatttaaaataaaaaacgg [128640]
ASF_IND_20_CAD543               ----------------------------------------........................................ [128640]
LR899193.1_Germany_2020/1       ----------------------------------------........................................ [128640]
MN393477.1_Wuhan_2019           ----------------------------------------........................................ [128640]
MW396979.1_Timor-Leste/2019/1   ----------------------------------------........................................ [128640]
MK940252.1_CN/2019/InnerMongolia ----------------------------------------........................................ [128640]
MK128995.1_China/2018/AnhuiXCGQ ----------------------------------------........................................ [128640]
MH681419.1_ASFV/POL/2015        ----------------------------------------........................................ [128640]
LS478113.1_Estonia_2014         ----------------------------------------........................................ [128640]
KP843857.1_Odintsovo_02/14      ----------------------------------------........................................ [128640]
FN557520.1_strain_E75           ----------------------------------------........................................ [128640]
AY261366.1_Warthog              ----------------------------------------..............g......................... [128640]
AY261365.1_Warmbaths            ----------------------------------------........................................ [128640]
AY261364.1_Tengani_62           ----------------------------------------........................................ [128640]
AY261363.1_Pretoriuskop/96/4    ----------------------------------------..............g......................... [128640]
AY261362.1_Mkuzi_1979           ----------------------------------------........................................ [128640]
AY261361.1_Malawi_Lil-20/1_1983 ttgtcgtataccttatatgaaatatatctggtataggaaa......t.a...t.g......................... [128640]
AY261360.1_Kenya_1950           ctgtcgt------------------atttggtatatgaaa......t.......g......................... [128640]
AM712240.1_OURT_88/3            ----------------------------------------........................................ [128640]
AM712239.1_Benin_97/1           ----------------------------------------........................................ [128640]
KM111295.1_Ken06.Bus            ctgtcgt------------------atttggtatatgaaa......t.......g.......................a. [128640]
KM111294.1_Ken05/Tk1            ctgtcgt------------------atttggtatatgaaa......t.......g......................... [128640]
KM262844.1_strain_L60           ----------------------------------------........................................ [128640]
KM102979.1_26544/OG10_Italy     ----------------------------------------........................................ [128640]
KM262845.1_strain_NHV           ----------------------------------------........................................ [128640]
KX354450.1_47/Ss/2008           ----------------------------------------........................................ [128640]
KP055815.1_strain_BA71          ----------------------------------------........................................ [128640]
MH025920.1_strain_R35           ctgtcgt------------------atttggtatatgaaa......t.......g.......................a. [128640]
MH025919.1_strain_N10           ctgtcgt------------------atttggtatatgaaa......t.......g.......................a. [128640]
MH025918.1_strain_R25           ctgtcgt------------------atttggtatatgaaa......t.......g.......................a. [128640]
MH025917.1_strain_R7            ctgtcgt------------------atttggtatatgaaa......t.......g.......................a. [128640]
MH025916.1_strain_R8            ctgtcgt------------------atttggtatatgaaa......t.......g.......................a. [128640]
```

```
FR682468.2_Georgia_2007/1        atgatatctatcatggaccgttctgagattgttgcacgggagaacccggtgattacccaacgagttacaaatctcctaca [128720]
ASF_IND_20_CAD543                ................................................................................ [128720]
LR899193.1_Germany_2020/1        ................................................................................ [128720]
MN393477.1_Wuhan_2019            ................................................................................ [128720]
MW396979.1_Timor-Leste/2019/1    ................................................................................ [128720]
MK940252.1_CN/2019/InnerMongolia ................................................................................ [128720]
MK128995.1_China/2018/AnhuiXCGQ  ................................................................................ [128720]
MH681419.1_ASFV/POL/2015         ................................................................................ [128720]
LS478113.1_Estonia_2014          ................................................................................ [128720]
KP843857.1_Odintsovo_02/14       ................................................................................ [128720]
FN557520.1_strain_E75            ................................................................................ [128720]
AY261366.1_Warthog               ..a.............................................................t................. [128720]
AY261365.1_Warmbaths             ................................................................................ [128720]
AY261364.1_Tengani_62            ............................................t.................t................. [128720]
AY261363.1_Pretoriuskop/96/4     ..a.............................................................t................. [128720]
AY261362.1_Mkuzi_1979            ................................................................................ [128720]
AY261361.1_Malawi_Lil-20/1_1983  ..a................................g.....a..t................................g.g [128720]
AY261360.1_Kenya_1950            ..a......................................a..t.................g................g [128720]
AM712240.1_OURT_88/3             ................................................................................ [128720]
AM712239.1_Benin_97/1            ................................................................................ [128720]
KM111295.1_Ken06.Bus             ..a......................................a..t.................g................g [128720]
KM111294.1_Ken05/Tk1             ..a......................................a..t.................g................g [128720]
KM262844.1_strain_L60            ................................................................................ [128720]
KM102979.1_26544/OG10_Italy      ................................................................................ [128720]
KM262845.1_strain_NHV            ................................................................................ [128720]
KX354450.1_47/Ss/2008            ................................................................................ [128720]
KP055815.1_strain_BA71           ................................................................................ [128720]
MH025920.1_strain_R35            ..a......................................a..t.................g................g [128720]
MH025919.1_strain_N10            ..a......................................a..t.................g................g [128720]
MH025918.1_strain_R25            ..a......................................a..t.................g................g [128720]
MH025917.1_strain_R7             ..a......................................a..t.................g................g [128720]
MH025916.1_strain_R8             ..a......................................a..t.................g................g [128720]

FR682468.2_Georgia_2007/1        aaccaatgctcctctactattcatgcccattgatatccatgaagtacgatatggagcctacacacttttcatgtatggtt [128800]
ASF_IND_20_CAD543                ................................................................................ [128800]
LR899193.1_Germany_2020/1        ................................................................................ [128800]
MN393477.1_Wuhan_2019            ................................................................................ [128800]
MW396979.1_Timor-Leste/2019/1    ................................................................................ [128800]
MK940252.1_CN/2019/InnerMongolia ................................................................................ [128800]
MK128995.1_China/2018/AnhuiXCGQ  ................................................................................ [128800]
MH681419.1_ASFV/POL/2015         ................................................................................ [128800]
LS478113.1_Estonia_2014          ................................................................................ [128800]
KP843857.1_Odintsovo_02/14       ................................................................................ [128800]
```

```
FN557520.1_strain_E75            ................................................................................ [128800]
AY261366.1_Warthog               ................................................................................ [128800]
AY261365.1_Warmbaths             ................................................................................ [128800]
AY261364.1_Tengani_62            ................................................................................ [128800]
AY261363.1_Pretoriuskop/96/4     ..................................................................c............. [128800]
AY261362.1_Mkuzi_1979            ................................................................................ [128800]
AY261361.1_Malawi_Lil-20/1_1983  ..............................................................tg..c............. [128800]
AY261360.1_Kenya_1950            .....................t........................................tg..c............. [128800]
AM712240.1_OURT_88/3             ................................................................................ [128800]
AM712239.1_Benin_97/1            ................................................................................ [128800]
KM111295.1_Ken06.Bus             ..............................................................tg..c............. [128800]
KM111294.1_Ken05/Tk1             ..............................................................tg..c............. [128800]
KM262844.1_strain_L60            ................................................................................ [128800]
KM102979.1_26544/OG10_Italy      ................................................................................ [128800]
KM262845.1_strain_NHV            ................................................................................ [128800]
KX354450.1_47/Ss/2008            ................................................................................ [128800]
KP055815.1_strain_BA71           ................................................................................ [128800]
MH025920.1_strain_R35            ..............................................................tg..c............. [128800]
MH025919.1_strain_N10            ..............................................................tg..c............. [128800]
MH025918.1_strain_R25            ..............................................................tg..c............. [128800]
MH025917.1_strain_R7             ..............................................................tg..c............. [128800]
MH025916.1_strain_R8             ..............................................................tg..c............. [128800]

FR682468.2_Georgia_2007/1        ccctcgaaaacggttacaaagcagaagtaaggattgaaaacatcccagttttctttgacgtacagattgagttcaatgat [128880]
ASF_IND_20_CAD543                ................................................................................ [128880]
LR899193.1_Germany_2020/1        ................................................................................ [128880]
MN393477.1_Wuhan_2019            ................................................................................ [128880]
MW396979.1_Timor-Leste/2019/1    ................................................................................ [128880]
MK940252.1_CN/2019/InnerMongolia ................................................................................ [128880]
MK128995.1_China/2018/AnhuiXCGQ  ................................................................................ [128880]
MH681419.1_ASFV/POL/2015         ................................................................................ [128880]
LS478113.1_Estonia_2014          ................................................................................ [128880]
KP843857.1_Odintsovo_02/14       ................................................................................ [128880]
FN557520.1_strain_E75            ................................................................................ [128880]
AY261366.1_Warthog               ................................................................a.......c..gca.c [128880]
AY261365.1_Warmbaths             .............................................................g..a.......c..gca.c [128880]
AY261364.1_Tengani_62            ................................................................a.......c..gca.c [128880]
AY261363.1_Pretoriuskop/96/4     ................................................................a.......c..gca.c [128880]
AY261362.1_Mkuzi_1979            ................................................................................ [128880]
AY261361.1_Malawi_Lil-20/1_1983  ..........t.............................t..t....................a.......ctg..a.c [128880]
AY261360.1_Kenya_1950            ........................................................................ctg..a.c [128880]
AM712240.1_OURT_88/3             ................................................................................ [128880]
AM712239.1_Benin_97/1            ................................................................................ [128880]
KM111295.1_Ken06.Bus             ........................................................................ctg..a.c [128880]
```

```
KM111294.1_Ken05/Tk1              ........................................................................ctg..a.c [128880]
KM262844.1_strain_L60             ................................................................................ [128880]
KM102979.1_26544/OG10_Italy       ................................................................................ [128880]
KM262845.1_strain_NHV             ................................................................................ [128880]
KX354450.1_47/Ss/2008             ................................................................................ [128880]
KP055815.1_strain_BA71            ................................................................................ [128880]
MH025920.1_strain_R35             ........................................................................ctg..a.c [128880]
MH025919.1_strain_N10             ........................................................................ctg..a.c [128880]
MH025918.1_strain_R25             ........................................................................ctg..a.c [128880]
MH025917.1_strain_R7              ........................................................................ctg..a.c [128880]
MH025916.1_strain_R8              ........................................................................ctg..a.c [128880]

FR682468.2_Georgia_2007/1         acaaaccagctttttttaaagtcgctactgacggctgaaaatattgtgtatgaacggctggagacgctcacccagcgtcc [128960]
ASF_IND_20_CAD543                 ................................................................................ [128960]
LR899193.1_Germany_2020/1         ................................................................................ [128960]
MN393477.1_Wuhan_2019             ................................................................................ [128960]
MW396979.1_Timor-Leste/2019/1     ................................................................................ [128960]
MK940252.1_CN/2019/InnerMongolia  ................................................................................ [128960]
MK128995.1_China/2018/AnhuiXCGQ   ................................................................................ [128960]
MH681419.1_ASFV/POL/2015          ................................................................................ [128960]
LS478113.1_Estonia_2014           ................................................................................ [128960]
KP843857.1_Odintsovo_02/14        ................................................................................ [128960]
FN557520.1_strain_E75             ..............................................c................................. [128960]
AY261366.1_Warthog                ...................................c.........ac............a.................... [128960]
AY261365.1_Warmbaths              ...........................t..t....c.........ac............a.................... [128960]
AY261364.1_Tengani_62             ...................................c.........ac............a.................... [128960]
AY261363.1_Pretoriuskop/96/4      ..............................t....c.........ac............a.................... [128960]
AY261362.1_Mkuzi_1979             ..............................................c................................. [128960]
AY261361.1_Malawi_Lil-20/1_1983   .............................................ac............a...t.aa.t........... [128960]
AY261360.1_Kenya_1950             .....t........................g..............ac............a.................c.. [128960]
AM712240.1_OURT_88/3              ..............................................c................................. [128960]
AM712239.1_Benin_97/1             ..............................................c................................. [128960]
KM111295.1_Ken06.Bus              .....t....................g...g..............ac............a........g........c.. [128960]
KM111294.1_Ken05/Tk1              .....t........................g..............ac............a.................c.. [128960]
KM262844.1_strain_L60             ..............................................c................................. [128960]
KM102979.1_26544/OG10_Italy       ..............................................c................................. [128960]
KM262845.1_strain_NHV             ..............................................c................................. [128960]
KX354450.1_47/Ss/2008             ..............................................c................................. [128960]
KP055815.1_strain_BA71            ..............................................c................................. [128960]
MH025920.1_strain_R35             .....t....................g...g..............ac............a........g........c.. [128960]
MH025919.1_strain_N10             .....t....................g...g..............ac............a........g........c.. [128960]
MH025918.1_strain_R25             .....t....................g...g..............ac............a........g........c.. [128960]
MH025917.1_strain_R7              .....t....................g...g..............ac............a........g........c.. [128960]
MH025916.1_strain_R8              .....t....................g...g..............ac............a........g........c.. [128960]
```

```
FR682468.2_Georgia_2007/1        tgtaatggggtaccgcgagaaggaaaaagagtttgcaccatacattcgaatattttttaaaagcctgtatgagcgacgaa [129040]
ASF_IND_20_CAD543                ................................................................................ [129040]
LR899193.1_Germany_2020/1        ................................................................................ [129040]
MN393477.1_Wuhan_2019            ................................................................................ [129040]
MW396979.1_Timor-Leste/2019/1    ................................................................................ [129040]
MK940252.1_CN/2019/InnerMongolia ................................................................................ [129040]
MK128995.1_China/2018/AnhuiXCGQ  ................................................................................ [129040]
MH681419.1_ASFV/POL/2015         ................................................................................ [129040]
LS478113.1_Estonia_2014          ................................................................................ [129040]
KP843857.1_Odintsovo_02/14       ................................................................................ [129040]
FN557520.1_strain_E75            ................................................................................ [129040]
AY261366.1_Warthog               ..................................................................a............. [129040]
AY261365.1_Warmbaths             ................................................................................ [129040]
AY261364.1_Tengani_62            .......................................g........................................ [129040]
AY261363.1_Pretoriuskop/96/4     ................................................................................ [129040]
AY261362.1_Mkuzi_1979            ................................................................................ [129040]
AY261361.1_Malawi_Lil-20/1_1983  ............t....................c..............................t.a..c.......... [129040]
AY261360.1_Kenya_1950            a...........t.......................g..g........................t.a.....a..g.... [129040]
AM712240.1_OURT_88/3             ................................................................................ [129040]
AM712239.1_Benin_97/1            ................................................................................ [129040]
KM111295.1_Ken06.Bus             a...........t.......................g..g........................t.a........g.... [129040]
KM111294.1_Ken05/Tk1             a...........t.......................g..g........................t.a........g.... [129040]
KM262844.1_strain_L60            ................................................................................ [129040]
KM102979.1_26544/OG10_Italy      ................................................................................ [129040]
KM262845.1_strain_NHV            ................................................................................ [129040]
KX354450.1_47/Ss/2008            ................................................................................ [129040]
KP055815.1_strain_BA71           ................................................................................ [129040]
MH025920.1_strain_R35            a...........t.......................g..g........................t.a........g.... [129040]
MH025919.1_strain_N10            a...........t.......................g..g........................t.a........g.... [129040]
MH025918.1_strain_R25            a...........t.......................g..g........................t.a........g.... [129040]
MH025917.1_strain_R7             a...........t.......................g..g........................t.a........g.... [129040]
MH025916.1_strain_R8             a...........t.......................g..g........................t.a........g.... [129040]

FR682468.2_Georgia_2007/1        aagccattacttacttaaataatatgggctacaacacggccgcggacgacacaacctgttattaccgaatggtttcccga [129120]
ASF_IND_20_CAD543                ................................................................................ [129120]
LR899193.1_Germany_2020/1        ................................................................................ [129120]
MN393477.1_Wuhan_2019            ................................................................................ [129120]
MW396979.1_Timor-Leste/2019/1    ................................................................................ [129120]
MK940252.1_CN/2019/InnerMongolia ................................................................................ [129120]
MK128995.1_China/2018/AnhuiXCGQ  ................................................................................ [129120]
MH681419.1_ASFV/POL/2015         ................................................................................ [129120]
LS478113.1_Estonia_2014          ................................................................................ [129120]
KP843857.1_Odintsovo_02/14       ................................................................................ [129120]
```

```
FN557520.1_strain_E75           ................g...........t........c.......................c.................. [129120]
AY261366.1_Warthog              ....g...........g...........t................................c.................. [129120]
AY261365.1_Warmbaths            ....g...........g...........t................................c.................. [129120]
AY261364.1_Tengani_62           ....g...........g...........t................................c.................. [129120]
AY261363.1_Pretoriuskop/96/4    ....g...........g...........t................................c.................. [129120]
AY261362.1_Mkuzi_1979           ....g...........g............................................c.................. [129120]
AY261361.1_Malawi_Lil-20/1_1983 ....g.........c.g...........t................................c.................. [129120]
AY261360.1_Kenya_1950           .g..g.........c.g..c........t................................c.................. [129120]
AM712240.1_OURT_88/3            ................g...........t........c.......................c.................. [129120]
AM712239.1_Benin_97/1           ................g...........t........c.......................c.................. [129120]
KM111295.1_Ken06.Bus            .g..g.........c.g..c........t................................c.................. [129120]
KM111294.1_Ken05/Tk1            .g..g.........c.g..c........t................................c.................. [129120]
KM262844.1_strain_L60           ................g...........t........c.......................c.................. [129120]
KM102979.1_26544/OG10_Italy     ................g...........t........c.......................c.................. [129120]
KM262845.1_strain_NHV           ................g...........t........c.......................c.................. [129120]
KX354450.1_47/Ss/2008           ................g...........t........c.......................c.................. [129120]
KP055815.1_strain_BA71          ................g...........t........c.......................c.................. [129120]
MH025920.1_strain_R35           .g..g.........c.g..c........t................................c.................. [129120]
MH025919.1_strain_N10           .g..g.........c.g..c........t................................c.................. [129120]
MH025918.1_strain_R25           .g..g.........c.g..c........t................................c.................. [129120]
MH025917.1_strain_R7            .g..g.........c.g..c........t................................c.................. [129120]
MH025916.1_strain_R8            .g..g.........c.g..c........t................................c.................. [129120]

FR682468.2_Georgia_2007/1       gaattaaaactacctcttacaagttggatacagcttcagcactattcctacgagcctcgcggcttggtacacaggttttc [129200]
ASF_IND_20_CAD543               ................................................................................ [129200]
LR899193.1_Germany_2020/1       ................................................................................ [129200]
MN393477.1_Wuhan_2019           ................................................................................ [129200]
MW396979.1_Timor-Leste/2019/1   ................................................................................ [129200]
MK940252.1_CN/2019/InnerMongolia ................................................................................ [129200]
MK128995.1_China/2018/AnhuiXCGQ ................................................................................ [129200]
MH681419.1_ASFV/POL/2015        ................................................................................ [129200]
LS478113.1_Estonia_2014         ................................................................................ [129200]
KP843857.1_Odintsovo_02/14      ................................................................................ [129200]
FN557520.1_strain_E75           ..gc.......g.................................................................... [129200]
AY261366.1_Warthog              ..gc......................................................a..................... [129200]
AY261365.1_Warmbaths            ..gc..........a...........................................a..................... [129200]
AY261364.1_Tengani_62           ..gc......................................................a..................... [129200]
AY261363.1_Pretoriuskop/96/4    ..gc......................................................a..................... [129200]
AY261362.1_Mkuzi_1979           ..g........g...........c........................................................ [129200]
AY261361.1_Malawi_Lil-20/1_1983 ..g........g..........................aa................a.....tc................ [129200]
AY261360.1_Kenya_1950           ..g........g..........................a........................c................ [129200]
AM712240.1_OURT_88/3            ..gc.......g.................................................................... [129200]
AM712239.1_Benin_97/1           ..gc.......g.................................................................... [129200]
KM111295.1_Ken06.Bus            ..g........g.............................a...a....t............................. [129200]
```

```
KM111294.1_Ken05/Tk1              ..g........g..........................a........................c................ [129200]
KM262844.1_strain_L60             ..gc.......g.................................................................... [129200]
KM102979.1_26544/OG10_Italy       ..gc.......g.................................................................... [129200]
KM262845.1_strain_NHV             ..gc.......g.................................................................... [129200]
KX354450.1_47/Ss/2008             ..gc.......g.................................................................... [129200]
KP055815.1_strain_BA71            ..gc.......g.................................................................... [129200]
MH025920.1_strain_R35             ..g........g.............................a...a....t............................. [129200]
MH025919.1_strain_N10             ..g........g.............................a...a....t............................. [129200]
MH025918.1_strain_R25             ..g........g.............................a...a....t............................. [129200]
MH025917.1_strain_R7              ..g........g.............................a...a....t............................. [129200]
MH025916.1_strain_R8              ..g........g.............................a...a....t............................. [129200]

FR682468.2_Georgia_2007/1         cgtaacccccgaggatcttgtttcctatcagaatgatggccccacagaccacagcatcgttatggcctacgatatagaga [129280]
ASF_IND_20_CAD543                 ................................................................................ [129280]
LR899193.1_Germany_2020/1         ................................................................................ [129280]
MN393477.1_Wuhan_2019             ................................................................................ [129280]
MW396979.1_Timor-Leste/2019/1     ................................................................................ [129280]
MK940252.1_CN/2019/InnerMongolia  ................................................................................ [129280]
MK128995.1_China/2018/AnhuiXCGQ   ................................................................................ [129280]
MH681419.1_ASFV/POL/2015          ................................................................................ [129280]
LS478113.1_Estonia_2014           ................................................................................ [129280]
KP843857.1_Odintsovo_02/14        ................................................................................ [129280]
FN557520.1_strain_E75             ...............................g................................................ [129280]
AY261366.1_Warthog                a...........a..................g................................................ [129280]
AY261365.1_Warmbaths              a...........a..................g................................................ [129280]
AY261364.1_Tengani_62             a...........a..................g................................................ [129280]
AY261363.1_Pretoriuskop/96/4      a..............................g................................................ [129280]
AY261362.1_Mkuzi_1979             ............a..................g................................................ [129280]
AY261361.1_Malawi_Lil-20/1_1983   a...........c..................g.........t......t........t...................... [129280]
AY261360.1_Kenya_1950             ............a..................g..........t.................g................... [129280]
AM712240.1_OURT_88/3              ...............................g................................................ [129280]
AM712239.1_Benin_97/1             ...............................g................................................ [129280]
KM111295.1_Ken06.Bus              a........t..a.g................g......t.....................a.................a. [129280]
KM111294.1_Ken05/Tk1              ............a..................g..........t.................g................... [129280]
KM262844.1_strain_L60             ...............................g................................................ [129280]
KM102979.1_26544/OG10_Italy       ...............................g................................................ [129280]
KM262845.1_strain_NHV             ...............................g................................................ [129280]
KX354450.1_47/Ss/2008             ...............................g................................................ [129280]
KP055815.1_strain_BA71            ...............................g................................................ [129280]
MH025920.1_strain_R35             a........t..a.g................g......t.....................a.................a. [129280]
MH025919.1_strain_N10             a........t..a.g................g......t.....................a.................a. [129280]
MH025918.1_strain_R25             a........t..a.g................g......t.....................a.................a. [129280]
MH025917.1_strain_R7              a........t..a.g................g......t.....................a.................a. [129280]
MH025916.1_strain_R8              a........t..a.g................g......t.....................a.................a. [129280]
```

```
FR682468.2_Georgia_2007/1        cctatagccctgttaagggaaccgttccggacccaaatcaggcaaacgacgtggtgttcatgatatgcatgcgcattttt [129360]
ASF_IND_20_CAD543                ................................................................................ [129360]
LR899193.1_Germany_2020/1        ................................................................................ [129360]
MN393477.1_Wuhan_2019            ................................................................................ [129360]
MW396979.1_Timor-Leste/2019/1    ................................................................................ [129360]
MK940252.1_CN/2019/InnerMongolia ................................................................................ [129360]
MK128995.1_China/2018/AnhuiXCGQ  ................................................................................ [129360]
MH681419.1_ASFV/POL/2015         ................................................................................ [129360]
LS478113.1_Estonia_2014          ................................................................................ [129360]
KP843857.1_Odintsovo_02/14       ................................................................................ [129360]
FN557520.1_strain_E75            ................................................................................ [129360]
AY261366.1_Warthog               ...............................t................................................ [129360]
AY261365.1_Warmbaths             ................................................................................ [129360]
AY261364.1_Tengani_62            ................................................................................ [129360]
AY261363.1_Pretoriuskop/96/4     ......................t.....a................................................... [129360]
AY261362.1_Mkuzi_1979            ................................................................................ [129360]
AY261361.1_Malawi_Lil-20/1_1983  .............c.................t........................................a....... [129360]
AY261360.1_Kenya_1950            .........................c..a................................................... [129360]
AM712240.1_OURT_88/3             ................................................................................ [129360]
AM712239.1_Benin_97/1            ................................................................................ [129360]
KM111295.1_Ken06.Bus             ...........a.............c...................................................... [129360]
KM111294.1_Ken05/Tk1             .........................c..a......................................t............ [129360]
KM262844.1_strain_L60            ................................................................................ [129360]
KM102979.1_26544/OG10_Italy      ................................................................................ [129360]
KM262845.1_strain_NHV            ................................................................................ [129360]
KX354450.1_47/Ss/2008            ................................................................................ [129360]
KP055815.1_strain_BA71           ................................................................................ [129360]
MH025920.1_strain_R35            ...........a.............c...................................................... [129360]
MH025919.1_strain_N10            ...........a.............c...................................................... [129360]
MH025918.1_strain_R25            ...........a.............c...................................................... [129360]
MH025917.1_strain_R7             ...........a.............c...................................................... [129360]
MH025916.1_strain_R8             ...........a.............c...................................................... [129360]

FR682468.2_Georgia_2007/1        tggattcactccacagagcctctagcgagcacgtgcatcaccatggcaccctgcaaaaagtcctcagagtggaccaccat [129440]
ASF_IND_20_CAD543                ................................................................................ [129440]
LR899193.1_Germany_2020/1        ................................................................................ [129440]
MN393477.1_Wuhan_2019            ................................................................................ [129440]
MW396979.1_Timor-Leste/2019/1    ................................................................................ [129440]
MK940252.1_CN/2019/InnerMongolia ................................................................................ [129440]
MK128995.1_China/2018/AnhuiXCGQ  ................................................................................ [129440]
MH681419.1_ASFV/POL/2015         ................................................................................ [129440]
LS478113.1_Estonia_2014          ................................................................................ [129440]
KP843857.1_Odintsovo_02/14       ................................................................................ [129440]
```

```
FN557520.1_strain_E75          .........................................t...................................... [129440]
AY261366.1_Warthog             ........t..................................................a.................t.. [129440]
AY261365.1_Warmbaths           .............t..................c..........................a.................t.. [129440]
AY261364.1_Tengani_62          ..............g............................................a.................t.. [129440]
AY261363.1_Pretoriuskop/96/4   ...........................................................a.................t.. [129440]
AY261362.1_Mkuzi_1979          ................................................................................ [129440]
AY261361.1_Malawi_Lil-20/1_1983 ..............................................t......t.....a.................... [129440]
AY261360.1_Kenya_1950          ..........................a..........................t.....a...c................ [129440]
AM712240.1_OURT_88/3           .........................................t...................................... [129440]
AM712239.1_Benin_97/1          .........................................t...................................... [129440]
KM111295.1_Ken06.Bus           ..........................a..........................t.....a.................... [129440]
KM111294.1_Ken05/Tk1           ..........................a..t...........t...........t.....a...c................ [129440]
KM262844.1_strain_L60          .........................................t...................................... [129440]
KM102979.1_26544/OG10_Italy    .........................................t...................................... [129440]
KM262845.1_strain_NHV          .........................................t...................................... [129440]
KX354450.1_47/Ss/2008          .........................................t...................................... [129440]
KP055815.1_strain_BA71         .........................................t...................................... [129440]
MH025920.1_strain_R35          ..........................a..........................t.....a.................... [129440]
MH025919.1_strain_N10          ..........................a..........................t.....a.................... [129440]
MH025918.1_strain_R25          ..........................a..........................t.....a.................... [129440]
MH025917.1_strain_R7           ..........................a..........................t.....a.................... [129440]
MH025916.1_strain_R8           ..........................a..........................t.....a.................... [129440]

FR682468.2_Georgia_2007/1      tctatgctcctctgaaaaaaatttgttgttaagctttgctgaacagtttagccgctgggctcctgatatatgcacagggt [129520]
ASF_IND_20_CAD543              .........................y...................................................... [129520]
LR899193.1_Germany_2020/1      ................................................................................ [129520]
MN393477.1_Wuhan_2019          ................................................................................ [129520]
MW396979.1_Timor-Leste/2019/1  ................................................................................ [129520]
MK940252.1_CN/2019/InnerMongolia ................................................................................ [129520]
MK128995.1_China/2018/AnhuiXCGQ ................................................................................ [129520]
MH681419.1_ASFV/POL/2015       ................................................................................ [129520]
LS478113.1_Estonia_2014        ................................................................................ [129520]
KP843857.1_Odintsovo_02/14     ................................................................................ [129520]
FN557520.1_strain_E75          .........................c......................c............................... [129520]
AY261366.1_Warthog             c..............................................................g...........g.... [129520]
AY261365.1_Warmbaths           c..............................................................a...........g.... [129520]
AY261364.1_Tengani_62          c..............................................................a...........g.... [129520]
AY261363.1_Pretoriuskop/96/4   c..............................................................a...........g.... [129520]
AY261362.1_Mkuzi_1979          .........................c......................c............................... [129520]
AY261361.1_Malawi_Lil-20/1_1983 .g.........................................................t...a...........g.... [129520]
AY261360.1_Kenya_1950          cg....t..................c.....................................g...........g.... [129520]
AM712240.1_OURT_88/3           .........................c......................c............................... [129520]
AM712239.1_Benin_97/1          .........................c......................c............................... [129520]
KM111295.1_Ken06.Bus           cg....t..................c.....................................g...........g.... [129520]
```

```
KM111294.1_Ken05/Tk1              cg....t..................c.....................................g...........g.... [129520]
KM262844.1_strain_L60             .........................c......................c............................... [129520]
KM102979.1_26544/OG10_Italy       .........................c......................c............................... [129520]
KM262845.1_strain_NHV             .........................c......................c............................... [129520]
KX354450.1_47/Ss/2008             .........................c......................c............................... [129520]
KP055815.1_strain_BA71            .........................c......................c............................... [129520]
MH025920.1_strain_R35             cg....t..................c.....................................g...........g.... [129520]
MH025919.1_strain_N10             cg....t..................c.....................................g...........g.... [129520]
MH025918.1_strain_R25             cg....t..................c.....................................g...........g.... [129520]
MH025917.1_strain_R7              cg....t..................c.....................................g...........g.... [129520]
MH025916.1_strain_R8              cg....t..................c.....................................g...........g.... [129520]

FR682468.2_Georgia_2007/1         tcaatgattctcggtacgactggccctttatcgttgaaaaatctatgcagcacggtattctagaagaaatctttaacaaa [129600]
ASF_IND_20_CAD543                 ................................................................................ [129600]
LR899193.1_Germany_2020/1         ................................................................................ [129600]
MN393477.1_Wuhan_2019             ................................................................................ [129600]
MW396979.1_Timor-Leste/2019/1     ................................................................................ [129600]
MK940252.1_CN/2019/InnerMongolia  ................................................................................ [129600]
MK128995.1_China/2018/AnhuiXCGQ   ................................................................................ [129600]
MH681419.1_ASFV/POL/2015          ................................................................................ [129600]
LS478113.1_Estonia_2014           ................................................................................ [129600]
KP843857.1_Odintsovo_02/14        ................................................................................ [129600]
FN557520.1_strain_E75             ................................................................................ [129600]
AY261366.1_Warthog                .t..c.............................c............................................. [129600]
AY261365.1_Warmbaths              .t..c.............................c............................................. [129600]
AY261364.1_Tengani_62             .t..c.............................c............................................. [129600]
AY261363.1_Pretoriuskop/96/4      .t..c.............................c............................................. [129600]
AY261362.1_Mkuzi_1979             ................................................................................ [129600]
AY261361.1_Malawi_Lil-20/1_1983   .t................................c............................................. [129600]
AY261360.1_Kenya_1950             .t................................c........c..........a......c......g........... [129600]
AM712240.1_OURT_88/3              ................................................................................ [129600]
AM712239.1_Benin_97/1             ................................................................................ [129600]
KM111295.1_Ken06.Bus              .t................................c........c.................c......g........... [129600]
KM111294.1_Ken05/Tk1              .t...........a....................c........c.................c......g........... [129600]
KM262844.1_strain_L60             ................................................................................ [129600]
KM102979.1_26544/OG10_Italy       ................................................................................ [129600]
KM262845.1_strain_NHV             ................................................................................ [129600]
KX354450.1_47/Ss/2008             ................................................................................ [129600]
KP055815.1_strain_BA71            ................................................................................ [129600]
MH025920.1_strain_R35             .t................................c........c.................c......g........... [129600]
MH025919.1_strain_N10             .t................................c........c.................c......g........... [129600]
MH025918.1_strain_R25             .t................................c........c.................c......g........... [129600]
MH025917.1_strain_R7              .t................................c........c.................c......g........... [129600]
MH025916.1_strain_R8              .t................................c........c.................c......g........... [129600]
```

```
FR682468.2_Georgia_2007/1        atgagccttttctggcaccaaaagctggataccattctaaaatgctattacgtaaaggaaaagagagtcaaaatctcggc [129680]
ASF_IND_20_CAD543                ................................................................................ [129680]
LR899193.1_Germany_2020/1        ................................................................................ [129680]
MN393477.1_Wuhan_2019            ................................................................................ [129680]
MW396979.1_Timor-Leste/2019/1    ................................................................................ [129680]
MK940252.1_CN/2019/InnerMongolia ................................................................................ [129680]
MK128995.1_China/2018/AnhuiXCGQ  ................................................................................ [129680]
MH681419.1_ASFV/POL/2015         ................................................................................ [129680]
LS478113.1_Estonia_2014          ................................................................................ [129680]
KP843857.1_Odintsovo_02/14       ................................................................................ [129680]
FN557520.1_strain_E75            ..................................................t..g.......................... [129680]
AY261366.1_Warthog               ..................................................t..g.......................a.. [129680]
AY261365.1_Warmbaths             ..................................................t..g.......................a.. [129680]
AY261364.1_Tengani_62            ..................................................t..g.......................... [129680]
AY261363.1_Pretoriuskop/96/4     ..................................................t..g.......................a.. [129680]
AY261362.1_Mkuzi_1979            ..................................................t..g.......................... [129680]
AY261361.1_Malawi_Lil-20/1_1983  ...............a...................c..............t..g..............t........a.. [129680]
AY261360.1_Kenya_1950            ................c.................................t..g.......................... [129680]
AM712240.1_OURT_88/3             ..................................................t..g.......................... [129680]
AM712239.1_Benin_97/1            ..................................................t..g.......................... [129680]
KM111295.1_Ken06.Bus             ................c.................................t..g.......................... [129680]
KM111294.1_Ken05/Tk1             ................c.................................t..g.......................... [129680]
KM262844.1_strain_L60            ..................................................t..g.......................... [129680]
KM102979.1_26544/OG10_Italy      ..................................................t..g.......................... [129680]
KM262845.1_strain_NHV            ..................................................t..g.......................... [129680]
KX354450.1_47/Ss/2008            ..................................................t..g.......................... [129680]
KP055815.1_strain_BA71           ..................................................t..g.......................... [129680]
MH025920.1_strain_R35            ................c.................................t..g.......................... [129680]
MH025919.1_strain_N10            ................c.................................t..g.......................... [129680]
MH025918.1_strain_R25            ................c.................................t..g.......................... [129680]
MH025917.1_strain_R7             ................c.................................t..g.......................... [129680]
MH025916.1_strain_R8             ................c.................................t..g.......................... [129680]

FR682468.2_Georgia_2007/1        cgaaaaatcgatcatttcctcctttttgcatacccctggatgcctacccattgatgtccgcaacatgtgtatgcagcttt [129760]
ASF_IND_20_CAD543                ................................................................................ [129760]
LR899193.1_Germany_2020/1        ................................................................................ [129760]
MN393477.1_Wuhan_2019            ................................................................................ [129760]
MW396979.1_Timor-Leste/2019/1    ................................................................................ [129760]
MK940252.1_CN/2019/InnerMongolia ................................................................................ [129760]
MK128995.1_China/2018/AnhuiXCGQ  ................................................................................ [129760]
MH681419.1_ASFV/POL/2015         ................................................................................ [129760]
LS478113.1_Estonia_2014          ................................................................................ [129760]
KP843857.1_Odintsovo_02/14       ................................................................................ [129760]
```

```
FN557520.1_strain_E75          ................................................................................ [129760]
AY261366.1_Warthog             ................................................................................ [129760]
AY261365.1_Warmbaths           ................................................................................ [129760]
AY261364.1_Tengani_62          ................................................................................ [129760]
AY261363.1_Pretoriuskop/96/4   ....................................c........................................... [129760]
AY261362.1_Mkuzi_1979          ................................................................................ [129760]
AY261361.1_Malawi_Lil-20/1_1983 .........t...........g....................t........g........................a... [129760]
AY261360.1_Kenya_1950          .........t..............c.................t........g........t........c.......... [129760]
AM712240.1_OURT_88/3           ................................................................................ [129760]
AM712239.1_Benin_97/1          ................................................................................ [129760]
KM111295.1_Ken06.Bus           .........t...................................g.......................c.......... [129760]
KM111294.1_Ken05/Tk1           .........t...................................g.......................c.......... [129760]
KM262844.1_strain_L60          ................................................................................ [129760]
KM102979.1_26544/OG10_Italy    ................................................................................ [129760]
KM262845.1_strain_NHV          ................................................................................ [129760]
KX354450.1_47/Ss/2008          ................................................................................ [129760]
KP055815.1_strain_BA71         ................................................................................ [129760]
MH025920.1_strain_R35          .........t...................................g.......................c.......... [129760]
MH025919.1_strain_N10          .........t...................................g.......................c.......... [129760]
MH025918.1_strain_R25          .........t...................................g.......................c.......... [129760]
MH025917.1_strain_R7           .........t...................................g.......................c.......... [129760]
MH025916.1_strain_R8           .........t...................................g.......................c.......... [129760]

FR682468.2_Georgia_2007/1      accctaaagccgaaaaaacaagcttgaaagcgtttttagaaaattgtgggttagattcgaaggtagacctgccgtaccat [129840]
ASF_IND_20_CAD543              ................................................................................ [129840]
LR899193.1_Germany_2020/1      ................................................................................ [129840]
MN393477.1_Wuhan_2019          ................................................................................ [129840]
MW396979.1_Timor-Leste/2019/1  ................................................................................ [129840]
MK940252.1_CN/2019/InnerMongolia ................................................................................ [129840]
MK128995.1_China/2018/AnhuiXCGQ ................................................................................ [129840]
MH681419.1_ASFV/POL/2015       ................................................................................ [129840]
LS478113.1_Estonia_2014        ................................................................................ [129840]
KP843857.1_Odintsovo_02/14     ................................................................................ [129840]
FN557520.1_strain_E75          .......................c.a...............................................c...... [129840]
AY261366.1_Warthog             .......................c.a...................................................... [129840]
AY261365.1_Warmbaths           .......................c.a...................................................... [129840]
AY261364.1_Tengani_62          .......................c.a...................................................... [129840]
AY261363.1_Pretoriuskop/96/4   .......................c.a...................................................... [129840]
AY261362.1_Mkuzi_1979          .......................c.a...............................................c...... [129840]
AY261361.1_Malawi_Lil-20/1_1983 .......................c.a........................c............................. [129840]
AY261360.1_Kenya_1950          ......................tc.a........................c............................. [129840]
AM712240.1_OURT_88/3           .......................c.a...............................................c...... [129840]
AM712239.1_Benin_97/1          .......................c.a...............................................c...... [129840]
KM111295.1_Ken06.Bus           ......................tc.a........................c...........a................. [129840]
```

```
KM111294.1_Ken05/Tk1              ......................tc.a........................c............................. [129840]
KM262844.1_strain_L60             .......................c.a...............................................c...... [129840]
KM102979.1_26544/OG10_Italy       .......................c.a...............................................c...... [129840]
KM262845.1_strain_NHV             .......................c.a...............................................c...... [129840]
KX354450.1_47/Ss/2008             .......................c.a...............................................c...... [129840]
KP055815.1_strain_BA71            .......................c.a...............................................c...... [129840]
MH025920.1_strain_R35             ......................tc.a........................c...........a................. [129840]
MH025919.1_strain_N10             ......................tc.a........................c...........a................. [129840]
MH025918.1_strain_R25             ......................tc.a........................c...........a................. [129840]
MH025917.1_strain_R7              ......................tc.a........................c...........a................. [129840]
MH025916.1_strain_R8              ......................tc.a........................c...........a................. [129840]

FR682468.2_Georgia_2007/1         ctcatgtggaagtattatgaaacacgagacagcgaaaaaatagccgacgtggcctattactgcattatagatgcccagcg [129920]
ASF_IND_20_CAD543                 ................................y.....r..r...................................... [129920]
LR899193.1_Germany_2020/1         ................................................................................ [129920]
MN393477.1_Wuhan_2019             ................................................................................ [129920]
MW396979.1_Timor-Leste/2019/1     ................................................................................ [129920]
MK940252.1_CN/2019/InnerMongolia  ................................................................................ [129920]
MK128995.1_China/2018/AnhuiXCGQ   ................................................................................ [129920]
MH681419.1_ASFV/POL/2015          ................................................................................ [129920]
LS478113.1_Estonia_2014           ................................................................................ [129920]
KP843857.1_Odintsovo_02/14        ................................................................................ [129920]
FN557520.1_strain_E75             .......................g........t.....g..g..............c....................... [129920]
AY261366.1_Warthog                .........................................g..............c....................... [129920]
AY261365.1_Warmbaths              ................................t.....g..g..............c....................... [129920]
AY261364.1_Tengani_62             ...........................a....t.....g..g..............c....................... [129920]
AY261363.1_Pretoriuskop/96/4      .........................................g..............c....................... [129920]
AY261362.1_Mkuzi_1979             .......................g........t.....g..g..............c....................... [129920]
AY261361.1_Malawi_Lil-20/1_1983   ................................t.....g..g..............c.................t..... [129920]
AY261360.1_Kenya_1950             ................................t.....g..g..............c....................... [129920]
AM712240.1_OURT_88/3              .......................g........t.....g..g..............c....................... [129920]
AM712239.1_Benin_97/1             .......................g........t.....g..g..............c....................... [129920]
KM111295.1_Ken06.Bus              .......................g........t.....g..g..............c....................... [129920]
KM111294.1_Ken05/Tk1              ................................t.....g..g..............c....................... [129920]
KM262844.1_strain_L60             .......................g........t.....g..g..............c....................... [129920]
KM102979.1_26544/OG10_Italy       .......................g........t.....g..g..............c....................... [129920]
KM262845.1_strain_NHV             .......................g........t.....g..g..............c....................... [129920]
KX354450.1_47/Ss/2008             .......................g........t.....g..g..............c....................... [129920]
KP055815.1_strain_BA71            .......................g........t.....g..g..............c....................... [129920]
MH025920.1_strain_R35             .......................g........t.....g..g..............c....................... [129920]
MH025919.1_strain_N10             .......................g........t.....g..g..............c....................... [129920]
MH025918.1_strain_R25             .......................g........t.....g..g..............c....................... [129920]
MH025917.1_strain_R7              .......................g........t.....g..g..............c....................... [129920]
MH025916.1_strain_R8              .......................g........t.....g..g..............c....................... [129920]
```

```
FR682468.2_Georgia_2007/1        ctgtcaggaccttctggtgcgccacaatgttatccccgatcgcagagaggtaggaattctgtcatacacctcgctgtatg [130000]
ASF_IND_20_CAD543                .........................................................ct..................... [130000]
LR899193.1_Germany_2020/1        ................................................................................ [130000]
MN393477.1_Wuhan_2019            ................................................................................ [130000]
MW396979.1_Timor-Leste/2019/1    ................................................................................ [130000]
MK940252.1_CN/2019/InnerMongolia ................................................................................ [130000]
MK128995.1_China/2018/AnhuiXCGQ  ................................................................................ [130000]
MH681419.1_ASFV/POL/2015         ................................................................................ [130000]
LS478113.1_Estonia_2014          ................................................................................ [130000]
KP843857.1_Odintsovo_02/14       ................................................................................ [130000]
FN557520.1_strain_E75            .........................................................ct..............t...... [130000]
AY261366.1_Warthog               ...............a................................................................ [130000]
AY261365.1_Warmbaths             ................................................................................ [130000]
AY261364.1_Tengani_62            ...c............................................................................ [130000]
AY261363.1_Pretoriuskop/96/4     ................................................................................ [130000]
AY261362.1_Mkuzi_1979            .........................................................ct..............t...... [130000]
AY261361.1_Malawi_Lil-20/1_1983  .......................................c........a........ct..............t...... [130000]
AY261360.1_Kenya_1950            .........................................................ct..............t...... [130000]
AM712240.1_OURT_88/3             .........................................................ct..............t...... [130000]
AM712239.1_Benin_97/1            .........................................................ct..............t...... [130000]
KM111295.1_Ken06.Bus             ..................................g......................ct..............t...... [130000]
KM111294.1_Ken05/Tk1             .........................................................ct..............t...... [130000]
KM262844.1_strain_L60            .........................................................ct..............t...... [130000]
KM102979.1_26544/OG10_Italy      .........................................................ct..............t...... [130000]
KM262845.1_strain_NHV            .........................................................ct..............t...... [130000]
KX354450.1_47/Ss/2008            .........................................................ct..............t...... [130000]
KP055815.1_strain_BA71           .........................................................ct..............t...... [130000]
MH025920.1_strain_R35            ..................................g......................ct..............t...... [130000]
MH025919.1_strain_N10            ..................................g......................ct..............t...... [130000]
MH025918.1_strain_R25            ..................................g......................ct..............t...... [130000]
MH025917.1_strain_R7             ..................................g......................ct..............t...... [130000]
MH025916.1_strain_R8             ..................................g......................ct..............t...... [130000]

FR682468.2_Georgia_2007/1        actgtatctactacgcgggaggacacaaggtatgcaatatgctcattgcctatgccatccatgatgaatacggccgtatt [130080]
ASF_IND_20_CAD543                ................................................................................ [130080]
LR899193.1_Germany_2020/1        ................................................................................ [130080]
MN393477.1_Wuhan_2019            ................................................................................ [130080]
MW396979.1_Timor-Leste/2019/1    ................................................................................ [130080]
MK940252.1_CN/2019/InnerMongolia ................................................................................ [130080]
MK128995.1_China/2018/AnhuiXCGQ  ................................................................................ [130080]
MH681419.1_ASFV/POL/2015         ................................................................................ [130080]
LS478113.1_Estonia_2014          ................................................................................ [130080]
KP843857.1_Odintsovo_02/14       ................................................................................ [130080]
```

```
FN557520.1_strain_E75           .........................t........t....................t...........g............ [130080]
AY261366.1_Warthog              ................................................................................ [130080]
AY261365.1_Warmbaths            ................................................................................ [130080]
AY261364.1_Tengani_62           ......................g............................................g............ [130080]
AY261363.1_Pretoriuskop/96/4    ................................................................c............... [130080]
AY261362.1_Mkuzi_1979           .........................t........t....................t...........g............ [130080]
AY261361.1_Malawi_Lil-20/1_1983 ................a.....g................................t...........g............ [130080]
AY261360.1_Kenya_1950           ......................g....................t........c..............g........c... [130080]
AM712240.1_OURT_88/3            .........................t........t....................t...........g............ [130080]
AM712239.1_Benin_97/1           .........................t........t....................t...........g............ [130080]
KM111295.1_Ken06.Bus            ......................g....................t........c..............g........c... [130080]
KM111294.1_Ken05/Tk1            ......................g.............................c..............g........c... [130080]
KM262844.1_strain_L60           .........................t........t....................t...........g............ [130080]
KM102979.1_26544/OG10_Italy     .........................t........t....................t...........g............ [130080]
KM262845.1_strain_NHV           .........................t........t....................t...........g............ [130080]
KX354450.1_47/Ss/2008           .........................t........t....................t...........g............ [130080]
KP055815.1_strain_BA71          .........................t........t....................t...........g............ [130080]
MH025920.1_strain_R35           ......................g....................t........c..............g........c... [130080]
MH025919.1_strain_N10           ......................g....................t........c..............g........c... [130080]
MH025918.1_strain_R25           ......................g....................t........c..............g........c... [130080]
MH025917.1_strain_R7            ......................g....................t........c..............g........c... [130080]
MH025916.1_strain_R8            ......................g....................t........c..............g........c... [130080]

FR682468.2_Georgia_2007/1       gcttgcagtaccattgcccgaggtaagcgggaacacggaaaatatcccggcgcctttgtgatagaccccgttaaagggct [130160]
ASF_IND_20_CAD543               ................................................................................ [130160]
LR899193.1_Germany_2020/1       ................................................................................ [130160]
MN393477.1_Wuhan_2019           ................................................................................ [130160]
MW396979.1_Timor-Leste/2019/1   ................................................................................ [130160]
MK940252.1_CN/2019/InnerMongolia ................................................................................ [130160]
MK128995.1_China/2018/AnhuiXCGQ ................................................................................ [130160]
MH681419.1_ASFV/POL/2015        ................................................................................ [130160]
LS478113.1_Estonia_2014         ................................................................................ [130160]
KP843857.1_Odintsovo_02/14      ................................................................................ [130160]
FN557520.1_strain_E75           ........c........t..g..........................g..............t................. [130160]
AY261366.1_Warthog              .................t..g.........................................t................. [130160]
AY261365.1_Warmbaths            ....................g........................................................... [130160]
AY261364.1_Tengani_62           .................t..g.........................................t..............a.. [130160]
AY261363.1_Pretoriuskop/96/4    .................t..g.........................................t................. [130160]
AY261362.1_Mkuzi_1979           ........c........t..g..........................a.....t........t................. [130160]
AY261361.1_Malawi_Lil-20/1_1983 ........c........t..g..a......................................t......a.......... [130160]
AY261360.1_Kenya_1950           ..c.....c........t..g......................................a..t................. [130160]
AM712240.1_OURT_88/3            ........c........t..g..........................g..............t................. [130160]
AM712239.1_Benin_97/1           ........c........t..g..........................g..............t................. [130160]
KM111295.1_Ken06.Bus            ..c.....c........t..g.........................................t................. [130160]
```

```
KM111294.1_Ken05/Tk1            ..c.....c........t..g......................................c..t................. [130160]
KM262844.1_strain_L60           ........c........t..g..........................g..............t................. [130160]
KM102979.1_26544/OG10_Italy     ........c........t..g..........................g..............t................. [130160]
KM262845.1_strain_NHV           ........c........t..g..........................g..............t................. [130160]
KX354450.1_47/Ss/2008           ........c........t..g..........................g..............t................. [130160]
KP055815.1_strain_BA71          ........c........t..g..........................g..............t................. [130160]
MH025920.1_strain_R35           ..c.....c........t..g.........................................t................. [130160]
MH025919.1_strain_N10           ..c.....c........t..g.........................................t................. [130160]
MH025918.1_strain_R25           ..c.....c........t..g.........................................t................. [130160]
MH025917.1_strain_R7            ..c.....c........t..g.........................................t................. [130160]
MH025916.1_strain_R8            ..c.....c........t..g.........................................t................. [130160]

FR682468.2_Georgia_2007/1       tgaacaggataaacccaccacaggtctcgactttgcgtcgctgtacccctcactcatcatggcctacaacttttcgccag [130240]
ASF_IND_20_CAD543               ........................c.....t................................................. [130240]
LR899193.1_Germany_2020/1       ................................................................................ [130240]
MN393477.1_Wuhan_2019           ................................................................................ [130240]
MW396979.1_Timor-Leste/2019/1   ................................................................................ [130240]
MK940252.1_CN/2019/InnerMongolia ............................................................................... [130240]
MK128995.1_China/2018/AnhuiXCGQ ................................................................................ [130240]
MH681419.1_ASFV/POL/2015        ................................................................................ [130240]
LS478113.1_Estonia_2014         ................................................................................ [130240]
KP843857.1_Odintsovo_02/14      ................................................................................ [130240]
FN557520.1_strain_E75           .....................c..c.....t................................................. [130240]
AY261366.1_Warthog              ..................t............................................................. [130240]
AY261365.1_Warmbaths            ................................................................................ [130240]
AY261364.1_Tengani_62           ................................................................................ [130240]
AY261363.1_Pretoriuskop/96/4    ................................................................................ [130240]
AY261362.1_Mkuzi_1979           .....................c..c.....t................................................. [130240]
AY261361.1_Malawi_Lil-20/1_1983 ......a..............c..c....................t.....g...........t...........a.... [130240]
AY261360.1_Kenya_1950           .....................c..........................t..c............................ [130240]
AM712240.1_OURT_88/3            .....................c..c.....t................................................. [130240]
AM712239.1_Benin_97/1           .....................c..c.....t................................................. [130240]
KM111295.1_Ken06.Bus            .....................c..c.................a........c............................ [130240]
KM111294.1_Ken05/Tk1            .....................c..c..........................c............................ [130240]
KM262844.1_strain_L60           .....................c..c.....t................................................. [130240]
KM102979.1_26544/OG10_Italy     .....................c..c.....t................................................. [130240]
KM262845.1_strain_NHV           .....................c..c.....t................................................. [130240]
KX354450.1_47/Ss/2008           .....................c..c.....t................................................. [130240]
KP055815.1_strain_BA71          .....................c..c.....t................................................. [130240]
MH025920.1_strain_R35           .....................c..c.................a........c............................ [130240]
MH025919.1_strain_N10           .....................c..c.................a........c............................ [130240]
MH025918.1_strain_R25           .....................c..c.................a........c............................ [130240]
MH025917.1_strain_R7            .....................c..c.................a........c............................ [130240]
MH025916.1_strain_R8            .....................c..c.................a........c............................ [130240]
```

```
FR682468.2_Georgia_2007/1        aaaaatttgtagcctctcgggatgaggcaaatagcctcatggccaagggtgaatctcttcactacgtctcctttcacttt [130320]
ASF_IND_20_CAD543                ................................................................................ [130320]
LR899193.1_Germany_2020/1        ................................................................................ [130320]
MN393477.1_Wuhan_2019            ................................................................................ [130320]
MW396979.1_Timor-Leste/2019/1    ................................................................................ [130320]
MK940252.1_CN/2019/InnerMongolia ................................................................................ [130320]
MK128995.1_China/2018/AnhuiXCGQ  ................................................................................ [130320]
MH681419.1_ASFV/POL/2015         ................................................................................ [130320]
LS478113.1_Estonia_2014          ................................................................................ [130320]
KP843857.1_Odintsovo_02/14       ................................................................................ [130320]
FN557520.1_strain_E75            ...............................a................................................ [130320]
AY261366.1_Warthog               ......................a........c...........t.....c..g..g........................ [130320]
AY261365.1_Warmbaths             ......................a........c...........t.....c..g..g........................ [130320]
AY261364.1_Tengani_62            ......................a........c...........t.....c..g........................... [130320]
AY261363.1_Pretoriuskop/96/4     ......................a........c...........t.....c..g..g..................t..... [130320]
AY261362.1_Mkuzi_1979            ...........................................t..a..c..g..g........................ [130320]
AY261361.1_Malawi_Lil-20/1_1983  .........................a.......a.........t.....c..g.....c..................... [130320]
AY261360.1_Kenya_1950            ...........................................t.....c..g..........................c [130320]
AM712240.1_OURT_88/3             ...............................a................................................ [130320]
AM712239.1_Benin_97/1            ...............................a................................................ [130320]
KM111295.1_Ken06.Bus             ........................................a..t.....c..g..........................c [130320]
KM111294.1_Ken05/Tk1             ...........................................t.....c..g..........................c [130320]
KM262844.1_strain_L60            ...............................a................................................ [130320]
KM102979.1_26544/OG10_Italy      ...............................a................................................ [130320]
KM262845.1_strain_NHV            ...............................a................................................ [130320]
KX354450.1_47/Ss/2008            ...............................a................................................ [130320]
KP055815.1_strain_BA71           ...............................a................................................ [130320]
MH025920.1_strain_R35            ........................................a..t.....c..g..........................c [130320]
MH025919.1_strain_N10            ........................................a..t.....c..g..........................c [130320]
MH025918.1_strain_R25            ........................................a..t.....c..g..........................c [130320]
MH025917.1_strain_R7             ........................................a..t.....c..g..........................c [130320]
MH025916.1_strain_R8             ........................................a..t.....c..g..........................c [130320]

FR682468.2_Georgia_2007/1        aacaatcgtctcgtggaaggatggtttgtgcggcataataacgttcctgataaaatgggattgtacccaaaagtactcat [130400]
ASF_IND_20_CAD543                ..............................................................r................. [130400]
LR899193.1_Germany_2020/1        ................................................................................ [130400]
MN393477.1_Wuhan_2019            ................................................................................ [130400]
MW396979.1_Timor-Leste/2019/1    ................................................................................ [130400]
MK940252.1_CN/2019/InnerMongolia ................................................................................ [130400]
MK128995.1_China/2018/AnhuiXCGQ  ................................................................................ [130400]
MH681419.1_ASFV/POL/2015         ................................................................................ [130400]
LS478113.1_Estonia_2014          ................................................................................ [130400]
KP843857.1_Odintsovo_02/14       ................................................................................ [130400]
```

```
FN557520.1_strain_E75            ................................a.............................a................. [130400]
AY261366.1_Warthog               ...........a..a................................................................. [130400]
AY261365.1_Warmbaths             ...........a..a...........................a.a................................... [130400]
AY261364.1_Tengani_62            ................................a.........................................g..... [130400]
AY261363.1_Pretoriuskop/96/4     ...........a..a................................................................. [130400]
AY261362.1_Mkuzi_1979            ................................a............................................... [130400]
AY261361.1_Malawi_Lil-20/1_1983  ...........a....................a...........................c................... [130400]
AY261360.1_Kenya_1950            ..t..c.....a..a.................a............................................... [130400]
AM712240.1_OURT_88/3             ................................a.............................a................. [130400]
AM712239.1_Benin_97/1            ................................a.............................a................. [130400]
KM111295.1_Ken06.Bus             ..t..c.....a..a................................................................. [130400]
KM111294.1_Ken05/Tk1             ..t..c.....a..a.................a............................................... [130400]
KM262844.1_strain_L60            ................................a.............................a................. [130400]
KM102979.1_26544/OG10_Italy      ................................a.............................a................. [130400]
KM262845.1_strain_NHV            ................................a.............................a................. [130400]
KX354450.1_47/Ss/2008            ................................a.............................a................. [130400]
KP055815.1_strain_BA71           ................................a.............................a................. [130400]
MH025920.1_strain_R35            ..t..c.....a..a................................................................. [130400]
MH025919.1_strain_N10            ..t..c.....a..a................................................................. [130400]
MH025918.1_strain_R25            ..t..c.....a..a................................................................. [130400]
MH025917.1_strain_R7             ..t..c.....a..a................................................................. [130400]
MH025916.1_strain_R8             ..t..c.....a..a................................................................. [130400]

FR682468.2_Georgia_2007/1        cgatctacttaacaaacggaccgcccttaaacaagagcttaaaaaactaggtgagaaaaaagaatgtatccatgaatccc [130480]
ASF_IND_20_CAD543                .................................................k...................a....w..... [130480]
LR899193.1_Germany_2020/1        ................................................................................ [130480]
MN393477.1_Wuhan_2019            ................................................................................ [130480]
MW396979.1_Timor-Leste/2019/1    ................................................................................ [130480]
MK940252.1_CN/2019/InnerMongolia ................................................................................ [130480]
MK128995.1_China/2018/AnhuiXCGQ  ................................................................................ [130480]
MH681419.1_ASFV/POL/2015         ................................................................................ [130480]
LS478113.1_Estonia_2014          ................................................................................ [130480]
KP843857.1_Odintsovo_02/14       ................................................................................ [130480]
FN557520.1_strain_E75            ..................a............................................................. [130480]
AY261366.1_Warthog               ................................................................................ [130480]
AY261365.1_Warmbaths             ................................................................................ [130480]
AY261364.1_Tengani_62            ...................................................c............................ [130480]
AY261363.1_Pretoriuskop/96/4     ..............................................................................t. [130480]
AY261362.1_Mkuzi_1979            ..................a............................................................. [130480]
AY261361.1_Malawi_Lil-20/1_1983  t...........t.....a...at..............................a.....g........t.......... [130480]
AY261360.1_Kenya_1950            t...........t...........t.....g.......................a..g...................... [130480]
AM712240.1_OURT_88/3             ..................a............................................................. [130480]
AM712239.1_Benin_97/1            ..................a............................................................. [130480]
KM111295.1_Ken06.Bus             t...........t.....a.....t.............................a..g..g................... [130480]
```

```
KM111294.1_Ken05/Tk1             t...........t...........t.....g.......................a..g...........t.......... [130480]
KM262844.1_strain_L60            ..................a............................................................. [130480]
KM102979.1_26544/OG10_Italy      ..................a............................................................. [130480]
KM262845.1_strain_NHV            ..................a............................................................. [130480]
KX354450.1_47/Ss/2008            ..................a............................................................. [130480]
KP055815.1_strain_BA71           ..................a............................................................. [130480]
MH025920.1_strain_R35            t...........t.....a.....t.............................a..g..g................... [130480]
MH025919.1_strain_N10            t...........t.....a.....t.............................a..g..g................... [130480]
MH025918.1_strain_R25            t...........t.....a.....t.............................a..g..g................... [130480]
MH025917.1_strain_R7             t...........t.....a.....t.............................a..g..g................... [130480]
MH025916.1_strain_R8             t...........t.....a.....t.............................a..g..g................... [130480]

FR682468.2_Georgia_2007/1        atcctgggtttaaggaactacagtttcgccatgccatggtagacgcgaagcaaaaggcgttgaaaattttcatgaacacg [130560]
ASF_IND_20_CAD543                ................................................................................ [130560]
LR899193.1_Germany_2020/1        ................................................................................ [130560]
MN393477.1_Wuhan_2019            ................................................................................ [130560]
MW396979.1_Timor-Leste/2019/1    ................................................................................ [130560]
MK940252.1_CN/2019/InnerMongolia ................................................................................ [130560]
MK128995.1_China/2018/AnhuiXCGQ  ................................................................................ [130560]
MH681419.1_ASFV/POL/2015         ................................................................................ [130560]
LS478113.1_Estonia_2014          ................................................................................ [130560]
KP843857.1_Odintsovo_02/14       ................................................................................ [130560]
FN557520.1_strain_E75            ................................................................................ [130560]
AY261366.1_Warthog               .................................................a........a..................... [130560]
AY261365.1_Warmbaths             ................................................................................ [130560]
AY261364.1_Tengani_62            ...........................................t.....a.............................. [130560]
AY261363.1_Pretoriuskop/96/4     .................................................a...........a.................. [130560]
AY261362.1_Mkuzi_1979            ......................................................................t......... [130560]
AY261361.1_Malawi_Lil-20/1_1983  .........................c.......................a........a....................t [130560]
AY261360.1_Kenya_1950            .........................c.......................a........a....................t [130560]
AM712240.1_OURT_88/3             ................................................................................ [130560]
AM712239.1_Benin_97/1            ................................................................................ [130560]
KM111295.1_Ken06.Bus             .........................c.......................a........a....................t [130560]
KM111294.1_Ken05/Tk1             .........................c.......................a........a....................t [130560]
KM262844.1_strain_L60            ................................................................................ [130560]
KM102979.1_26544/OG10_Italy      ................................................................................ [130560]
KM262845.1_strain_NHV            ................................................................................ [130560]
KX354450.1_47/Ss/2008            ................................................................................ [130560]
KP055815.1_strain_BA71           ................................................................................ [130560]
MH025920.1_strain_R35            .........................c.......................a........a....................t [130560]
MH025919.1_strain_N10            .........................c.......................a........a....................t [130560]
MH025918.1_strain_R25            .........................c.......................a........a....................t [130560]
MH025917.1_strain_R7             .........................c.......................a........a....................t [130560]
MH025916.1_strain_R8             .........................c.......................a........a....................t [130560]
```

```
FR682468.2_Georgia_2007/1        ttttacggcgaggcaggtaacaatttgtcgcccttctttctgcttcctctagccggaggagtcaccagttcgggtcaata [130640]
ASF_IND_20_CAD543                ................................................................................ [130640]
LR899193.1_Germany_2020/1        ................................................................................ [130640]
MN393477.1_Wuhan_2019            ................................................................................ [130640]
MW396979.1_Timor-Leste/2019/1    ................................................................................ [130640]
MK940252.1_CN/2019/InnerMongolia ................................................................................ [130640]
MK128995.1_China/2018/AnhuiXCGQ  ................................................................................ [130640]
MH681419.1_ASFV/POL/2015         ................................................................................ [130640]
LS478113.1_Estonia_2014          ................................................................................ [130640]
KP843857.1_Odintsovo_02/14       ................................................................................ [130640]
FN557520.1_strain_E75            ................................................................................ [130640]
AY261366.1_Warthog               ....................................................................c........... [130640]
AY261365.1_Warmbaths             ....................................................................c........... [130640]
AY261364.1_Tengani_62            ....................................................................c........... [130640]
AY261363.1_Pretoriuskop/96/4     ....................................................................c........... [130640]
AY261362.1_Mkuzi_1979            ....................................................................c........... [130640]
AY261361.1_Malawi_Lil-20/1_1983  ........t..............c.....a.................................................. [130640]
AY261360.1_Kenya_1950            ..............g..............a.................................................. [130640]
AM712240.1_OURT_88/3             ................................................................................ [130640]
AM712239.1_Benin_97/1            ................................................................................ [130640]
KM111295.1_Ken06.Bus             ..............g.....t........a....................g............................. [130640]
KM111294.1_Ken05/Tk1             ..............g..............a.................................................. [130640]
KM262844.1_strain_L60            ................................................................................ [130640]
KM102979.1_26544/OG10_Italy      ................................................................................ [130640]
KM262845.1_strain_NHV            ................................................................................ [130640]
KX354450.1_47/Ss/2008            ................................................................................ [130640]
KP055815.1_strain_BA71           ................................................................................ [130640]
MH025920.1_strain_R35            ..............g..............a....................g............................. [130640]
MH025919.1_strain_N10            ..............g..............a....................g............................. [130640]
MH025918.1_strain_R25            ..............g..............a....................g............................. [130640]
MH025917.1_strain_R7             ..............g..............a....................g............................. [130640]
MH025916.1_strain_R8             ..............g..............a....................g............................. [130640]

FR682468.2_Georgia_2007/1        taatcttaaacttgtctataactttgttatcaataaaggttacggcatcaagtacggtgacaccgactcattatacatta [130720]
ASF_IND_20_CAD543                ................................................................................ [130720]
LR899193.1_Germany_2020/1        ................................................................................ [130720]
MN393477.1_Wuhan_2019            ................................................................................ [130720]
MW396979.1_Timor-Leste/2019/1    ................................................................................ [130720]
MK940252.1_CN/2019/InnerMongolia ................................................................................ [130720]
MK128995.1_China/2018/AnhuiXCGQ  ................................................................................ [130720]
MH681419.1_ASFV/POL/2015         ................................................................................ [130720]
LS478113.1_Estonia_2014          ................................................................................ [130720]
KP843857.1_Odintsovo_02/14       ................................................................................ [130720]
```

FN557520.1_strain_E75 ............c................................................................... [130720]
AY261366.1_Warthog .............................................a.................................. [130720]
AY261365.1_Warmbaths ................................................................................ [130720]
AY261364.1_Tengani_62 ................................................................................ [130720]
AY261363.1_Pretoriuskop/96/4 ................................................................................ [130720]
AY261362.1_Mkuzi_1979 ............c.................t................................................. [130720]
AY261361.1_Malawi_Lil-20/1_1983 ............c.................t..c...........a.................................. [130720]
AY261360.1_Kenya_1950 ............c....................c...........a...........c...........g.......... [130720]
AM712240.1_OURT_88/3 ............c................................................................... [130720]
AM712239.1_Benin_97/1 ............c................................................................... [130720]
KM111295.1_Ken06.Bus ............c....................c...........a...........c...........g.......... [130720]
KM111294.1_Ken05/Tk1 ............c....................c...........a...........c...........g.......... [130720]
KM262844.1_strain_L60 ............c................................................................... [130720]
KM102979.1_26544/OG10_Italy ............c................................................................... [130720]
KM262845.1_strain_NHV ............c................................................................... [130720]
KX354450.1_47/Ss/2008 ............c................................................................... [130720]
KP055815.1_strain_BA71 ............c................................................................... [130720]
MH025920.1_strain_R35 ............c....................c...........a...........c...................... [130720]
MH025919.1_strain_N10 ............c....................c...........a...........c...................... [130720]
MH025918.1_strain_R25 ............c....................c...........a...........c...................... [130720]
MH025917.1_strain_R7 ............c....................c...........a...........c...................... [130720]
MH025916.1_strain_R8 ............c....................c...........a...........c...................... [130720]

FR682468.2_Georgia_2007/1 catgcccagatagtctttatacagaggtaacagacgcatatttaaacagccaaaaaacgataaaacattatgagcaactc [130800]
ASF_IND_20_CAD543 ..........................................................r..................... [130800]
LR899193.1_Germany_2020/1 ................................................................................ [130800]
MN393477.1_Wuhan_2019 ................................................................................ [130800]
MW396979.1_Timor-Leste/2019/1 ................................................................................ [130800]
MK940252.1_CN/2019/InnerMongolia ................................................................................ [130800]
MK128995.1_China/2018/AnhuiXCGQ ................................................................................ [130800]
MH681419.1_ASFV/POL/2015 ................................................................................ [130800]
LS478113.1_Estonia_2014 ................................................................................ [130800]
KP843857.1_Odintsovo_02/14 ................................................................................ [130800]
FN557520.1_strain_E75 ..............................................t...........a..................... [130800]
AY261366.1_Warthog ..............................................t................................. [130800]
AY261365.1_Warmbaths ..............................................t................................. [130800]
AY261364.1_Tengani_62 ..............................................t...........a..................... [130800]
AY261363.1_Pretoriuskop/96/4 ..............................................t................................. [130800]
AY261362.1_Mkuzi_1979 ..............................................t...........a..................... [130800]
AY261361.1_Malawi_Lil-20/1_1983 ..................................g..g........t...........a..................... [130800]
AY261360.1_Kenya_1950 .......g................................c.....t..t........a..................... [130800]
AM712240.1_OURT_88/3 ..............................................t...........a..................... [130800]
AM712239.1_Benin_97/1 ..............................................t...........a..................... [130800]
KM111295.1_Ken06.Bus .......g................................c.....t..t........a..................... [130800]

KM111294.1_Ken05/Tk1 .......g................................c.....t..t........a..................... [130800]
KM262844.1_strain_L60 ..............................................t...........a..................... [130800]
KM102979.1_26544/OG10_Italy ..............................................t...........a..................... [130800]
KM262845.1_strain_NHV ..............................................t...........a..................... [130800]
KX354450.1_47/Ss/2008 ..............................................t...........a..................... [130800]
KP055815.1_strain_BA71 ..............................................t...........a..................... [130800]
MH025920.1_strain_R35 .......g................................c.....t..t........a..................... [130800]
MH025919.1_strain_N10 .......g................................c.....t..t........a..................... [130800]
MH025918.1_strain_R25 .......g................................c.....t..t........a..................... [130800]
MH025917.1_strain_R7 .......g................................c.....t..t........a..................... [130800]
MH025916.1_strain_R8 .......g................................c.....t..t........a..................... [130800]

FR682468.2_Georgia_2007/1 tgccacgaaaaagtgcttctgtctatgaaagccatgtctacactatgcgccgaggtgaatgaatacctgcgacaagataa [130880]
ASF_IND_20_CAD543 ................................................................................ [130880]
LR899193.1_Germany_2020/1 ................................................................................ [130880]
MN393477.1_Wuhan_2019 ................................................................................ [130880]
MW396979.1_Timor-Leste/2019/1 ................................................................................ [130880]
MK940252.1_CN/2019/InnerMongolia ................................................................................ [130880]
MK128995.1_China/2018/AnhuiXCGQ ................................................................................ [130880]
MH681419.1_ASFV/POL/2015 ................................................................................ [130880]
LS478113.1_Estonia_2014 ................................................................................ [130880]
KP843857.1_Odintsovo_02/14 ................................................................................ [130880]
FN557520.1_strain_E75 .......................................................................g........ [130880]
AY261366.1_Warthog .......................c...............................................g.g...... [130880]
AY261365.1_Warmbaths ..........................................t............................g........ [130880]
AY261364.1_Tengani_62 .......................c...............................................g........ [130880]
AY261363.1_Pretoriuskop/96/4 .......................................................................g........ [130880]
AY261362.1_Mkuzi_1979 .......................................................................g........ [130880]
AY261361.1_Malawi_Lil-20/1_1983 .....t.....g........a.............................t....................g........ [130880]
AY261360.1_Kenya_1950 .....t.....g...........c.....g.......................a...........t.............. [130880]
AM712240.1_OURT_88/3 .......................................................................g........ [130880]
AM712239.1_Benin_97/1 .......................................................................g........ [130880]
KM111295.1_Ken06.Bus ..t..t.....g...........c.....g....................t............................. [130880]
KM111294.1_Ken05/Tk1 .....t.....g...........c.....g..........t............a...........t.............. [130880]
KM262844.1_strain_L60 .......................................................................g........ [130880]
KM102979.1_26544/OG10_Italy .......................................................................g........ [130880]
KM262845.1_strain_NHV .......................................................................g........ [130880]
KX354450.1_47/Ss/2008 .......................................................................g........ [130880]
KP055815.1_strain_BA71 .......................................................................g........ [130880]
MH025920.1_strain_R35 ..t..t.....g...........c.....g....................t............................. [130880]
MH025919.1_strain_N10 ..t..t.....g...........c.....g....................t............................. [130880]
MH025918.1_strain_R25 ..t..t.....g...........c.....g....................t............................. [130880]
MH025917.1_strain_R7 ..t..t.....g...........c.....g....................t............................. [130880]
MH025916.1_strain_R8 ..t..t.....g...........c.....g....................t............................. [130880]

```
FR682468.2_Georgia_2007/1       tggcaccagttacctacgtatggcctacgaggaagtactctttcctgtgtgctttacaggcaagaaaaagtattatggta [130960]
ASF_IND_20_CAD543               ................................................................................ [130960]
LR899193.1_Germany_2020/1       ................................................................................ [130960]
MN393477.1_Wuhan_2019           ................................................................................ [130960]
MW396979.1_Timor-Leste/2019/1   ................................................................................ [130960]
MK940252.1_CN/2019/InnerMongolia ................................................................................ [130960]
MK128995.1_China/2018/AnhuiXCGQ ................................................................................ [130960]
MH681419.1_ASFV/POL/2015        ................................................................................ [130960]
LS478113.1_Estonia_2014         ................................................................................ [130960]
KP843857.1_Odintsovo_02/14      ................................................................................ [130960]
FN557520.1_strain_E75           ...........................................................................c.... [130960]
AY261366.1_Warthog              ..........................................c..c..t..........................c..c. [130960]
AY261365.1_Warmbaths            ..........................................c..c..t..........................c..c. [130960]
AY261364.1_Tengani_62           ........a...........................c...........t..........................c..c. [130960]
AY261363.1_Pretoriuskop/96/4    ..........................................c..c..t..........................c..c. [130960]
AY261362.1_Mkuzi_1979           ..........................................c.....t..........................c..c. [130960]
AY261361.1_Malawi_Lil-20/1_1983 ...t......................................c..c..t....................a.....c..c. [130960]
AY261360.1_Kenya_1950           ...t........t...........t.................c..c..t....................a.....c..c. [130960]
AM712240.1_OURT_88/3            ...........................................................................c.... [130960]
AM712239.1_Benin_97/1           ...........................................................................c.... [130960]
KM111295.1_Ken06.Bus            ...t........t.............................c..c..t....................a.....c..c. [130960]
KM111294.1_Ken05/Tk1            ...t........t.............................c..c..t....................a.....c..c. [130960]
KM262844.1_strain_L60           ...........................................................................c.... [130960]
KM102979.1_26544/OG10_Italy     ...........................................................................c.... [130960]
KM262845.1_strain_NHV           ...........................................................................c.... [130960]
KX354450.1_47/Ss/2008           ...........................................................................c.... [130960]
KP055815.1_strain_BA71          ...........................................................................c.... [130960]
MH025920.1_strain_R35           ...t........t.............................c..c..t....................a.....c..c. [130960]
MH025919.1_strain_N10           ...t........t.............................c..c..t....................a.....c..c. [130960]
MH025918.1_strain_R25           ...t........t.............................c..c..t....................a.....c..c. [130960]
MH025917.1_strain_R7            ...t........t.............................c..c..t....................a.....c..c. [130960]
MH025916.1_strain_R8            ...t........t.............................c..c..t....................a.....c..c. [130960]

FR682468.2_Georgia_2007/1       ttgctcatgtaaacacacccaattttaatacaaaagaattattcatccgcggaatagatatcattaagcagggtcaaaca [131040]
ASF_IND_20_CAD543               ................................................................................ [131040]
LR899193.1_Germany_2020/1       ................................................................................ [131040]
MN393477.1_Wuhan_2019           ................................................................................ [131040]
MW396979.1_Timor-Leste/2019/1   ................................................................................ [131040]
MK940252.1_CN/2019/InnerMongolia ................................................................................ [131040]
MK128995.1_China/2018/AnhuiXCGQ ................................................................................ [131040]
MH681419.1_ASFV/POL/2015        ................................................................................ [131040]
LS478113.1_Estonia_2014         ................................................................................ [131040]
KP843857.1_Odintsovo_02/14      ................................................................................ [131040]
```

| | | |
|---|---|---|
| FN557520.1_strain_E75 | ................................................................................ | [131040] |
| AY261366.1_Warthog | ................................................................................ | [131040] |
| AY261365.1_Warmbaths | ................................................................................ | [131040] |
| AY261364.1_Tengani_62 | ...................t............................................................ | [131040] |
| AY261363.1_Pretoriuskop/96/4 | ................................................................................ | [131040] |
| AY261362.1_Mkuzi_1979 | ................................................................................ | [131040] |
| AY261361.1_Malawi_Lil-20/1_1983 | ....................................................c........................... | [131040] |
| AY261360.1_Kenya_1950 | ................................................................................ | [131040] |
| AM712240.1_OURT_88/3 | ................................................................................ | [131040] |
| AM712239.1_Benin_97/1 | ................................................................................ | [131040] |
| KM111295.1_Ken06.Bus | ................................................................................ | [131040] |
| KM111294.1_Ken05/Tk1 | ................................................................................ | [131040] |
| KM262844.1_strain_L60 | ................................................................................ | [131040] |
| KM102979.1_26544/OG10_Italy | ................................................................................ | [131040] |
| KM262845.1_strain_NHV | ................................................................................ | [131040] |
| KX354450.1_47/Ss/2008 | ................................................................................ | [131040] |
| KP055815.1_strain_BA71 | ................................................................................ | [131040] |
| MH025920.1_strain_R35 | ................................................................................ | [131040] |
| MH025919.1_strain_N10 | ................................................................................ | [131040] |
| MH025918.1_strain_R25 | ................................................................................ | [131040] |
| MH025917.1_strain_R7 | ................................................................................ | [131040] |
| MH025916.1_strain_R8 | ................................................................................ | [131040] |

| | | |
|---|---|---|
| FR682468.2_Georgia_2007/1 | aaactcaccaaaacgataggaacgcgaattatggaagaatccatgaaactacgccgccctgaggaccatcgcccccctct | [131120] |
| ASF_IND_20_CAD543 | ................................................................................ | [131120] |
| LR899193.1_Germany_2020/1 | ................................................................................ | [131120] |
| MN393477.1_Wuhan_2019 | ................................................................................ | [131120] |
| MW396979.1_Timor-Leste/2019/1 | ................................................................................ | [131120] |
| MK940252.1_CN/2019/InnerMongolia | ................................................................................ | [131120] |
| MK128995.1_China/2018/AnhuiXCGQ | ................................................................................ | [131120] |
| MH681419.1_ASFV/POL/2015 | ................................................................................ | [131120] |
| LS478113.1_Estonia_2014 | ................................................................................ | [131120] |
| KP843857.1_Odintsovo_02/14 | ................................................................................ | [131120] |
| FN557520.1_strain_E75 | ....................t.............................g............................. | [131120] |
| AY261366.1_Warthog | ....................t.............................g............................. | [131120] |
| AY261365.1_Warmbaths | ....................t.............................g............................. | [131120] |
| AY261364.1_Tengani_62 | ....................t.............................g............................. | [131120] |
| AY261363.1_Pretoriuskop/96/4 | ....................t.............................g............................. | [131120] |
| AY261362.1_Mkuzi_1979 | ....................t.............................g............................. | [131120] |
| AY261361.1_Malawi_Lil-20/1_1983 | ....................t.t......c....................g.................c.....g..... | [131120] |
| AY261360.1_Kenya_1950 | ....................t........c....................g.......................g..... | [131120] |
| AM712240.1_OURT_88/3 | ....................t.............................g............................. | [131120] |
| AM712239.1_Benin_97/1 | ....................t.............................g............................. | [131120] |
| KM111295.1_Ken06.Bus | ....................t.............................g.......................g..... | [131120] |

```
KM111294.1_Ken05/Tk1              ....................t.t......c....................g.......................g..... [131120]
KM262844.1_strain_L60             ....................t.............................g............................. [131120]
KM102979.1_26544/OG10_Italy       ....................t.............................g............................. [131120]
KM262845.1_strain_NHV             ....................t.............................g............................. [131120]
KX354450.1_47/Ss/2008             ....................t.............................g............................. [131120]
KP055815.1_strain_BA71            ....................t.............................g............................. [131120]
MH025920.1_strain_R35             ....................t.............................g.......................g..... [131120]
MH025919.1_strain_N10             ....................t.............................g.......................g..... [131120]
MH025918.1_strain_R25             ....................t.............................g.......................g..... [131120]
MH025917.1_strain_R7              ....................t.............................g.......................g..... [131120]
MH025916.1_strain_R8              ....................t.............................g.......................g..... [131120]

FR682468.2_Georgia_2007/1         tattgaaatcgttaaaacggttttgaaggatgctgtggttaacatgaagcagtggaattttgaagacttcatccaaacag [131200]
ASF_IND_20_CAD543                 ................................................................................ [131200]
LR899193.1_Germany_2020/1         ................................................................................ [131200]
MN393477.1_Wuhan_2019             ................................................................................ [131200]
MW396979.1_Timor-Leste/2019/1     ................................................................................ [131200]
MK940252.1_CN/2019/InnerMongolia  ................................................................................ [131200]
MK128995.1_China/2018/AnhuiXCGQ   ................................................................................ [131200]
MH681419.1_ASFV/POL/2015          ................................................................................ [131200]
LS478113.1_Estonia_2014           ................................................................................ [131200]
KP843857.1_Odintsovo_02/14        ................................................................................ [131200]
FN557520.1_strain_E75             ................................................................................ [131200]
AY261366.1_Warthog                ................................................................................ [131200]
AY261365.1_Warmbaths              ................................................................................ [131200]
AY261364.1_Tengani_62             ................................................................................ [131200]
AY261363.1_Pretoriuskop/96/4      g............................................................................... [131200]
AY261362.1_Mkuzi_1979             ................................................................................ [131200]
AY261361.1_Malawi_Lil-20/1_1983   ........................a........g..a.......................c...........t....... [131200]
AY261360.1_Kenya_1950             ........................a.......................a...........c...........t....... [131200]
AM712240.1_OURT_88/3              ................................................................................ [131200]
AM712239.1_Benin_97/1             ................................................................................ [131200]
KM111295.1_Ken06.Bus              ........................a.......................a....................t.......... [131200]
KM111294.1_Ken05/Tk1              ..................a.....a.......................a...........c...........t....... [131200]
KM262844.1_strain_L60             ................................................................................ [131200]
KM102979.1_26544/OG10_Italy       ................................................................................ [131200]
KM262845.1_strain_NHV             ................................................................................ [131200]
KX354450.1_47/Ss/2008             ................................................................................ [131200]
KP055815.1_strain_BA71            ................................................................................ [131200]
MH025920.1_strain_R35             ........................a.......................a....................t.......... [131200]
MH025919.1_strain_N10             ........................a.......................a....................t.......... [131200]
MH025918.1_strain_R25             ........................a.......................a....................t.......... [131200]
MH025917.1_strain_R7              ........................a.......................a....................t.......... [131200]
MH025916.1_strain_R8              ........................a.......................a....................t.......... [131200]
```

```
FR682468.2_Georgia_2007/1        atgcgtggagaccggacaaagacaacaaagcagtccaaatctttatgtctcgcatgcacgctcggcgtgagcaactaaaa [131280]
ASF_IND_20_CAD543                ................................................................................ [131280]
LR899193.1_Germany_2020/1        ................................................................................ [131280]
MN393477.1_Wuhan_2019            ................................................................................ [131280]
MW396979.1_Timor-Leste/2019/1    ................................................................................ [131280]
MK940252.1_CN/2019/InnerMongolia ................................................................................ [131280]
MK128995.1_China/2018/AnhuiXCGQ  ................................................................................ [131280]
MH681419.1_ASFV/POL/2015         ................................................................................ [131280]
LS478113.1_Estonia_2014          ................................................................................ [131280]
KP843857.1_Odintsovo_02/14       ................................................................................ [131280]
FN557520.1_strain_E75            ................................................................................ [131280]
AY261366.1_Warthog               ...............................g................................a...........t... [131280]
AY261365.1_Warmbaths             ...............................g................................a...........t... [131280]
AY261364.1_Tengani_62            ................................................................................ [131280]
AY261363.1_Pretoriuskop/96/4     ...............................g................................a...........t... [131280]
AY261362.1_Mkuzi_1979            ................................................................................ [131280]
AY261361.1_Malawi_Lil-20/1_1983  .............a..t..............g..........................t..................... [131280]
AY261360.1_Kenya_1950            ...............................g................................................ [131280]
AM712240.1_OURT_88/3             ................................................................................ [131280]
AM712239.1_Benin_97/1            ................................................................................ [131280]
KM111295.1_Ken06.Bus             ................................................................................ [131280]
KM111294.1_Ken05/Tk1             ...............................g................................................ [131280]
KM262844.1_strain_L60            ................................................................................ [131280]
KM102979.1_26544/OG10_Italy      ................................................................................ [131280]
KM262845.1_strain_NHV            ................................................................................ [131280]
KX354450.1_47/Ss/2008            ..........................................................t..................... [131280]
KP055815.1_strain_BA71           ................................................................................ [131280]
MH025920.1_strain_R35            ................................................................................ [131280]
MH025919.1_strain_N10            ................................................................................ [131280]
MH025918.1_strain_R25            ................................................................................ [131280]
MH025917.1_strain_R7             ................................................................................ [131280]
MH025916.1_strain_R8             ................................................................................ [131280]

FR682468.2_Georgia_2007/1        aaacacgg---cgctgcagcatcgcaatttgctgagcccgagccgggagaacgcttctcctacgttatcgtggaaaaaca [131360]
ASF_IND_20_CAD543                ........---...........................y......................................... [131360]
LR899193.1_Germany_2020/1        ........---..................................................................... [131360]
MN393477.1_Wuhan_2019            ........---..................................................................... [131360]
MW396979.1_Timor-Leste/2019/1    ........---..................................................................... [131360]
MK940252.1_CN/2019/InnerMongolia ........---..................................................................... [131360]
MK128995.1_China/2018/AnhuiXCGQ  ........---..................................................................... [131360]
MH681419.1_ASFV/POL/2015         ........---..................................................................... [131360]
LS478113.1_Estonia_2014          ........---..................................................................... [131360]
KP843857.1_Odintsovo_02/14       ........---..................................................................... [131360]
```

FN557520.1_strain_E75 ........---...c.......................t......................................... [131360]
AY261366.1_Warthog ........---...c................................................................. [131360]
AY261365.1_Warmbaths ........---...c................................................................. [131360]
AY261364.1_Tengani_62 ........---...c................................................................. [131360]
AY261363.1_Pretoriuskop/96/4 ........---...c.......................t......................................... [131360]
AY261362.1_Mkuzi_1979 ........---...c.......................t......................................... [131360]
AY261361.1_Malawi_Lil-20/1_1983 ..g.....---...c..c.............................................................. [131360]
AY261360.1_Kenya_1950 ........cgc...c...a............................................................. [131360]
AM712240.1_OURT_88/3 ........---...c.......................t......................................... [131360]
AM712239.1_Benin_97/1 ........---...c.......................t......................................... [131360]
KM111295.1_Ken06.Bus ........cgc...c...a............................................................. [131360]
KM111294.1_Ken05/Tk1 ........cgc...c...a............................................................. [131360]
KM262844.1_strain_L60 ........---...c.......................t......................................... [131360]
KM102979.1_26544/OG10_Italy ........---...c.......................t......................................... [131360]
KM262845.1_strain_NHV ........---...c.......................t......................................... [131360]
KX354450.1_47/Ss/2008 ........---...c.......................t......................................... [131360]
KP055815.1_strain_BA71 ........---...c.......................t......................................... [131360]
MH025920.1_strain_R35 ........cgc...c...a............................................................. [131360]
MH025919.1_strain_N10 ........cgc...c...a............................................................. [131360]
MH025918.1_strain_R25 ........cgc...c...a............................................................. [131360]
MH025917.1_strain_R7 ........cgc...c...a............................................................. [131360]
MH025916.1_strain_R8 ........cgc...c...a............................................................. [131360]

FR682468.2_Georgia_2007/1 ggtacagtttgatatccagggccaccgcacagattcctccagaaagggggacaagatggaatacgtctctgaagcaaagg [131440]
ASF_IND_20_CAD543 ................................................................................ [131440]
LR899193.1_Germany_2020/1 ................................................................................ [131440]
MN393477.1_Wuhan_2019 ................................................................................ [131440]
MW396979.1_Timor-Leste/2019/1 ................................................................................ [131440]
MK940252.1_CN/2019/InnerMongolia ................................................................................ [131440]
MK128995.1_China/2018/AnhuiXCGQ ................................................................................ [131440]
MH681419.1_ASFV/POL/2015 ................................................................................ [131440]
LS478113.1_Estonia_2014 ................................................................................ [131440]
KP843857.1_Odintsovo_02/14 ................................................................................ [131440]
FN557520.1_strain_E75 a..............t................................................................ [131440]
AY261366.1_Warthog ...g...........t................................................................ [131440]
AY261365.1_Warmbaths ...g...........t................................................................ [131440]
AY261364.1_Tengani_62 ...g...........t................................................................ [131440]
AY261363.1_Pretoriuskop/96/4 ...g...........t................................................................ [131440]
AY261362.1_Mkuzi_1979 a..............t................................................................ [131440]
AY261361.1_Malawi_Lil-20/1_1983 ...g..a...........................a............................................. [131440]
AY261360.1_Kenya_1950 ...g..a...........a..............ca..a...g.....a.....a......................... [131440]
AM712240.1_OURT_88/3 a..............t................................................................ [131440]
AM712239.1_Benin_97/1 a..............t................................................................ [131440]
KM111295.1_Ken06.Bus ...g..a...........a..............ca..a..........a.....a......................... [131440]

```
KM111294.1_Ken05/Tk1            ...g..a...........a..............ca..a....g.....a.....a......................... [131440]
KM262844.1_strain_L60           a..............t................................................................ [131440]
KM102979.1_26544/OG10_Italy     a..............t................................................................ [131440]
KM262845.1_strain_NHV           a..............t................................................................ [131440]
KX354450.1_47/Ss/2008           a..............t................................................................ [131440]
KP055815.1_strain_BA71          a..............t................................................................ [131440]
MH025920.1_strain_R35           ...g..a...........a..............ca..a..........a.....a......................... [131440]
MH025919.1_strain_N10           ...g..a...........a..............ca..a..........a.....a......................... [131440]
MH025918.1_strain_R25           ...g..a...........a..............ca..a..........a.....a......................... [131440]
MH025917.1_strain_R7            ...g..a...........a..............ca..a..........a.....a......................... [131440]
MH025916.1_strain_R8            ...g..a...........a..............ca..a..........a.....a......................... [131440]

FR682468.2_Georgia_2007/1       ctaaaaatcttcctattgatatattgttttatatcaacaactatgttctaggcttgtgcgcgagattcattaatgaaaat [131520]
ASF_IND_20_CAD543               ................................................................................ [131520]
LR899193.1_Germany_2020/1       ................................................................................ [131520]
MN393477.1_Wuhan_2019           ................................................................................ [131520]
MW396979.1_Timor-Leste/2019/1   ................................................................................ [131520]
MK940252.1_CN/2019/InnerMongolia ................................................................................ [131520]
MK128995.1_China/2018/AnhuiXCGQ ................................................................................ [131520]
MH681419.1_ASFV/POL/2015        ................................................................................ [131520]
LS478113.1_Estonia_2014         ................................................................................ [131520]
KP843857.1_Odintsovo_02/14      ................................................................................ [131520]
FN557520.1_strain_E75           .............a.................................................................. [131520]
AY261366.1_Warthog              ................................................................................ [131520]
AY261365.1_Warmbaths            ...................c............................................................ [131520]
AY261364.1_Tengani_62           ................................................................................ [131520]
AY261363.1_Pretoriuskop/96/4    .............a.................................................................. [131520]
AY261362.1_Mkuzi_1979           .............a.................................................................. [131520]
AY261361.1_Malawi_Lil-20/1_1983 .............a.................................................................. [131520]
AY261360.1_Kenya_1950           .............a..c....................t.......................................... [131520]
AM712240.1_OURT_88/3            .............a.................................................................. [131520]
AM712239.1_Benin_97/1           .............a.................................................................. [131520]
KM111295.1_Ken06.Bus            .............a..c....................t...............c.......................... [131520]
KM111294.1_Ken05/Tk1            .............a..c....................t.......................................... [131520]
KM262844.1_strain_L60           .............a.................................................................. [131520]
KM102979.1_26544/OG10_Italy     .............a.................................................................. [131520]
KM262845.1_strain_NHV           .............a.................................................................. [131520]
KX354450.1_47/Ss/2008           .............a.................................................................. [131520]
KP055815.1_strain_BA71          .............a.................................................................. [131520]
MH025920.1_strain_R35           .............a..c....................t...............c.......................... [131520]
MH025919.1_strain_N10           .............a..c....................t...............c.......................... [131520]
MH025918.1_strain_R25           .............a..c....................t...............c.......................... [131520]
MH025917.1_strain_R7            .............a..c....................t...............c.......................... [131520]
MH025916.1_strain_R8            .............a..c....................t...............c.......................... [131520]
```

```
FR682468.2_Georgia_2007/1        gaagaatttcaaccccctgacaacgtcagcaataaggatgaatacgctcagcgccgagctaaatcctacctacaaaaatt [131600]
ASF_IND_20_CAD543                ................................................................................ [131600]
LR899193.1_Germany_2020/1        ................................................................................ [131600]
MN393477.1_Wuhan_2019            ................................................................................ [131600]
MW396979.1_Timor-Leste/2019/1    ................................................................................ [131600]
MK940252.1_CN/2019/InnerMongolia ................................................................................ [131600]
MK128995.1_China/2018/AnhuiXCGQ  ................................................................................ [131600]
MH681419.1_ASFV/POL/2015         ................................................................................ [131600]
LS478113.1_Estonia_2014          ................................................................................ [131600]
KP843857.1_Odintsovo_02/14       ................................................................................ [131600]
FN557520.1_strain_E75            .......................t...................................c.................... [131600]
AY261366.1_Warthog               .......................t........c............................................... [131600]
AY261365.1_Warmbaths             .......................t........c............................................... [131600]
AY261364.1_Tengani_62            ............................................t................................... [131600]
AY261363.1_Pretoriuskop/96/4     .......................t........c............................................... [131600]
AY261362.1_Mkuzi_1979            .......................t........................................................ [131600]
AY261361.1_Malawi_Lil-20/1_1983  .................c....................c.....t..c...........c.................... [131600]
AY261360.1_Kenya_1950            .....g.............g..................c.....t..c...........c...........g........ [131600]
AM712240.1_OURT_88/3             .......................t...................................c.................... [131600]
AM712239.1_Benin_97/1            .......................t...................................c.................... [131600]
KM111295.1_Ken06.Bus             .....g.....g.....c.g..................c........c...........c...........g........ [131600]
KM111294.1_Ken05/Tk1             .....g.............g..................c.....t..c...........c...........g........ [131600]
KM262844.1_strain_L60            .......................t...................................c.................... [131600]
KM102979.1_26544/OG10_Italy      .......................t...................................c.................... [131600]
KM262845.1_strain_NHV            .......................t...................................c.................... [131600]
KX354450.1_47/Ss/2008            .......................t...................................c.................... [131600]
KP055815.1_strain_BA71           .......................t...................................c.................... [131600]
MH025920.1_strain_R35            .....g.....g.....c.g..................c........c...........c...........g........ [131600]
MH025919.1_strain_N10            .....g.....g.....c.g..................c........c...........c...........g........ [131600]
MH025918.1_strain_R25            .....g.....g.....c.g..................c........c...........c...........g........ [131600]
MH025917.1_strain_R7             .....g.....g.....c.g..................c........c...........c...........g........ [131600]
MH025916.1_strain_R8             .....g.....g.....c.g..................c........c...........c...........g........ [131600]

FR682468.2_Georgia_2007/1        cgtgcaatccattcaccctaaagacaagtctgtcattaagcaaggcaatgttcatcgacagtgctacaaatacattcacc [131680]
ASF_IND_20_CAD543                ................................................................................ [131680]
LR899193.1_Germany_2020/1        ................................................................................ [131680]
MN393477.1_Wuhan_2019            ................................................................................ [131680]
MW396979.1_Timor-Leste/2019/1    ................................................................................ [131680]
MK940252.1_CN/2019/InnerMongolia ................................................................................ [131680]
MK128995.1_China/2018/AnhuiXCGQ  ................................................................................ [131680]
MH681419.1_ASFV/POL/2015         ................................................................................ [131680]
LS478113.1_Estonia_2014          ................................................................................ [131680]
KP843857.1_Odintsovo_02/14       ................................................................................ [131680]
```

```
FN557520.1_strain_E75             ...a...........................................t.........................g...... [131680]
AY261366.1_Warthog                ...............................................t.........................g...... [131680]
AY261365.1_Warmbaths              ...............................................t.........................g...... [131680]
AY261364.1_Tengani_62             ...............................................t................................ [131680]
AY261363.1_Pretoriuskop/96/4      .................................t.............t.........................g...... [131680]
AY261362.1_Mkuzi_1979             ...............................................t.........................g...... [131680]
AY261361.1_Malawi_Lil-20/1_1983   t..............t...............................t.........................g....t. [131680]
AY261360.1_Kenya_1950             ...............t...............................t.........................g....t. [131680]
AM712240.1_OURT_88/3              ...a...........................................t.........................g...... [131680]
AM712239.1_Benin_97/1             ...a...........................................t.........................g...... [131680]
KM111295.1_Ken06.Bus              ...............t...............................t.........................g....t. [131680]
KM111294.1_Ken05/Tk1              ...............t...............................t.........................g....t. [131680]
KM262844.1_strain_L60             ...a...........................................t.........................g...... [131680]
KM102979.1_26544/OG10_Italy       ...a...........................................t.........................g...... [131680]
KM262845.1_strain_NHV             ...a...........................................t.........................g...... [131680]
KX354450.1_47/Ss/2008             ...a...........................................t.........................g...... [131680]
KP055815.1_strain_BA71            ...a...........................................t.........................g...... [131680]
MH025920.1_strain_R35             ...............t...............................t.........................g....t. [131680]
MH025919.1_strain_N10             ...............t...............................t.........................g....t. [131680]
MH025918.1_strain_R25             ...............t...............................t.........................g....t. [131680]
MH025917.1_strain_R7              ...............t...............................t.........................g....t. [131680]
MH025916.1_strain_R8              ...............t...............................t.........................g....t. [131680]

FR682468.2_Georgia_2007/1         aagaaattaaaaaaaaaataggcatctttgccgacctttataaggaattttttaacaacaccacaaaccccatcgaaagc [131760]
ASF_IND_20_CAD543                 ................................................................................ [131760]
LR899193.1_Germany_2020/1         ................................................................................ [131760]
MN393477.1_Wuhan_2019             ................................................................................ [131760]
MW396979.1_Timor-Leste/2019/1     ................................................................................ [131760]
MK940252.1_CN/2019/InnerMongolia  ................................................................................ [131760]
MK128995.1_China/2018/AnhuiXCGQ   ................................................................................ [131760]
MH681419.1_ASFV/POL/2015          ................................................................................ [131760]
LS478113.1_Estonia_2014           ................................................................................ [131760]
KP843857.1_Odintsovo_02/14        ........-....................................................................... [131760]
FN557520.1_strain_E75             ................................................................................ [131760]
AY261366.1_Warthog                .........................................................................t...... [131760]
AY261365.1_Warmbaths              ................................................................................ [131760]
AY261364.1_Tengani_62             ................................................................................ [131760]
AY261363.1_Pretoriuskop/96/4      ...............................................................................t [131760]
AY261362.1_Mkuzi_1979             ................................................................................ [131760]
AY261361.1_Malawi_Lil-20/1_1983   ...............................t..t............................................. [131760]
AY261360.1_Kenya_1950             .......c..........................t........a.................................... [131760]
AM712240.1_OURT_88/3              ................................................................................ [131760]
AM712239.1_Benin_97/1             ................................................................................ [131760]
KM111295.1_Ken06.Bus              .......c..........................t........a..............t..................... [131760]
```

```
KM111294.1_Ken05/Tk1            .......c..........................t........a.................................... [131760]
KM262844.1_strain_L60           ................................................................................ [131760]
KM102979.1_26544/OG10_Italy     ................................................................................ [131760]
KM262845.1_strain_NHV           ................................................................................ [131760]
KX354450.1_47/Ss/2008           ................................................................................ [131760]
KP055815.1_strain_BA71          ................................................................................ [131760]
MH025920.1_strain_R35           .......c..........................t........a..............t..................... [131760]
MH025919.1_strain_N10           .......c..........................t........a..............t..................... [131760]
MH025918.1_strain_R25           .......c..........................t........a..............t..................... [131760]
MH025917.1_strain_R7            .......c..........................t........a..............t..................... [131760]
MH025916.1_strain_R8            .......c..........................t........a..............t..................... [131760]

FR682468.2_Georgia_2007/1       tttattcaaagcactcagtttatgatacaatactttgatggagaacaaaaagtaaaccattctatgaaaaaaatggttga [131840]
ASF_IND_20_CAD543               ................................................................................ [131840]
LR899193.1_Germany_2020/1       ................................................................................ [131840]
MN393477.1_Wuhan_2019           ................................................................................ [131840]
MW396979.1_Timor-Leste/2019/1   ................................................................................ [131840]
MK940252.1_CN/2019/InnerMongolia ................................................................................ [131840]
MK128995.1_China/2018/AnhuiXCGQ ................................................................................ [131840]
MH681419.1_ASFV/POL/2015        ................................................................................ [131840]
LS478113.1_Estonia_2014         ................................................................................ [131840]
KP843857.1_Odintsovo_02/14      ................................................................................ [131840]
FN557520.1_strain_E75           ............g...g.................c............................................. [131840]
AY261366.1_Warthog              ................................................................................ [131840]
AY261365.1_Warmbaths            ...........................................................................a.... [131840]
AY261364.1_Tengani_62           ................................................................................ [131840]
AY261363.1_Pretoriuskop/96/4    ...........................................................................a.... [131840]
AY261362.1_Mkuzi_1979           ............g.....................c............................................. [131840]
AY261361.1_Malawi_Lil-20/1_1983 ..................................c.....a......................g................ [131840]
AY261360.1_Kenya_1950           .............................t..........a.....................a................. [131840]
AM712240.1_OURT_88/3            ............g...g.................c............................................. [131840]
AM712239.1_Benin_97/1           ............g...g.................c............................................. [131840]
KM111295.1_Ken06.Bus            .............................t..........a.....................a................. [131840]
KM111294.1_Ken05/Tk1            .............................t..........a.....................a................. [131840]
KM262844.1_strain_L60           ............g...g.................c............................................. [131840]
KM102979.1_26544/OG10_Italy     ............g...g.................c............................................. [131840]
KM262845.1_strain_NHV           ............g...g.................c............................................. [131840]
KX354450.1_47/Ss/2008           ............g...g.................c............................................. [131840]
KP055815.1_strain_BA71          ............g...g.................c............................................. [131840]
MH025920.1_strain_R35           .............................t..........a.....................a................. [131840]
MH025919.1_strain_N10           .............................t..........a.....................a................. [131840]
MH025918.1_strain_R25           .............................t..........a.....................a................. [131840]
MH025917.1_strain_R7            .............................t..........a.....................a................. [131840]
MH025916.1_strain_R8            .............................t..........a.....................a................. [131840]
```

```
FR682468.2_Georgia_2007/1        acagcatgctac---ggctagtaatcgagctggtaagcccgctggtaat------------ccagccggcaatgcgctga [131920]
ASF_IND_20_CAD543                ............ttt.........g.c......................ccagccggcaac................... [131920]
LR899193.1_Germany_2020/1        ............---..................................------------................... [131920]
MN393477.1_Wuhan_2019            ............---..................................------------................... [131920]
MW396979.1_Timor-Leste/2019/1    ............---..................................------------................... [131920]
MK940252.1_CN/2019/InnerMongolia ............---..................................------------................... [131920]
MK128995.1_China/2018/AnhuiXCGQ  ............---..................................------------................... [131920]
MH681419.1_ASFV/POL/2015         ............---..................................------------................... [131920]
LS478113.1_Estonia_2014          ............---..................................------------................... [131920]
KP843857.1_Odintsovo_02/14       ............---..................................------------................... [131920]
FN557520.1_strain_E75            .....g......ttt...a.....g.cc.....................ccagctggcaac................... [131920]
AY261366.1_Warthog               ............------------.tcg.....c..ct.a.........------------................... [131920]
AY261365.1_Warmbaths             ............---..................c..c..g...a.....------------................... [131920]
AY261364.1_Tengani_62            ............---.....c............c..c..a.........------------................... [131920]
AY261363.1_Pretoriuskop/96/4     ............---..................c..c..a.........------------........t.......... [131920]
AY261362.1_Mkuzi_1979            .....g......ttt.------------.....................ccagctggcaac................... [131920]
AY261361.1_Malawi_Lil-20/1_1983  ...a........---..................c..c..a........cccagctggcaac................... [131920]
AY261360.1_Kenya_1950            g.........g.ttt...cg.c....cg..------------------------------------.............. [131920]
AM712240.1_OURT_88/3             .....g......ttt...a.....g.cc.....................ccagctggcaac................... [131920]
AM712239.1_Benin_97/1            .....g......ttt...a.....g.cc.....................ccagctggcaac................... [131920]
KM111295.1_Ken06.Bus             g....g.a....ttt...g....c.cg........c..g........c------------......a............ [131920]
KM111294.1_Ken05/Tk1             g.........g.ttt...cg.c....cg..------------------------------------.............. [131920]
KM262844.1_strain_L60            .....g......ttt...a.....g.cc.....................ccagctggcaac................... [131920]
KM102979.1_26544/OG10_Italy      .....g......ttt...a.....g.cc.....................ccagctggcaac................... [131920]
KM262845.1_strain_NHV            .....g......ttt...a.....g.cc.....................ccagctggcaac................... [131920]
KX354450.1_47/Ss/2008            .....g......ttt...a.....g.cc.....................ccagctggcaac................... [131920]
KP055815.1_strain_BA71           .....g......ttt...a.....g.cc.....................ccagctggcaac................... [131920]
MH025920.1_strain_R35            g....g.a....ttt...g....c.cg........c..g........c------------......a............ [131920]
MH025919.1_strain_N10            g....g.a....ttt...g....c.cg........c..g........c------------......a............ [131920]
MH025918.1_strain_R25            g....g.a....ttt...g....c.cg........c..g........c------------......a............ [131920]
MH025917.1_strain_R7             g....g.a....ttt...g....c.cg........c..g........c------------......a............ [131920]
MH025916.1_strain_R8             g....g.a....ttt...g....c.cg........c..g........c------------......a............ [131920]

FR682468.2_Georgia_2007/1        tgcgggctatatttacgcagctgattacggaagaaaaaaaaattgtacaagccttatacaataaggggatgcaatacac [132000]
ASF_IND_20_CAD543                ................................................................................ [132000]
LR899193.1_Germany_2020/1        ................................................................................ [132000]
MN393477.1_Wuhan_2019            ................................................................................ [132000]
MW396979.1_Timor-Leste/2019/1    ................................................................................ [132000]
MK940252.1_CN/2019/InnerMongolia ................................................................................ [132000]
MK128995.1_China/2018/AnhuiXCGQ  ................................................................................ [132000]
MH681419.1_ASFV/POL/2015         ................................................................................ [132000]
LS478113.1_Estonia_2014          ................................................................................ [132000]
KP843857.1_Odintsovo_02/14       .................................-.............................................. [132000]
```

```
FN557520.1_strain_E75              ................................................................................ [132000]
AY261366.1_Warthog                 .....................................................c.......................... [132000]
AY261365.1_Warmbaths               .....................................................c.......................... [132000]
AY261364.1_Tengani_62              ................................................................a............... [132000]
AY261363.1_Pretoriuskop/96/4       ................................................................................ [132000]
AY261362.1_Mkuzi_1979              ................................................................................ [132000]
AY261361.1_Malawi_Lil-20/1_1983    ....................t.......................................g.........c.a....... [132000]
AY261360.1_Kenya_1950              ....................t....gt...............................t...........c.a....... [132000]
AM712240.1_OURT_88/3               ................................................................................ [132000]
AM712239.1_Benin_97/1              ................................................................................ [132000]
KM111295.1_Ken06.Bus               ....................t.................................................c......... [132000]
KM111294.1_Ken05/Tk1               ....................t.................................................c.a....... [132000]
KM262844.1_strain_L60              ................................................................................ [132000]
KM102979.1_26544/OG10_Italy        ................................................................................ [132000]
KM262845.1_strain_NHV              ................................................................................ [132000]
KX354450.1_47/Ss/2008              ................................................................................ [132000]
KP055815.1_strain_BA71             ................................................................................ [132000]
MH025920.1_strain_R35              ....................t.................................................c......... [132000]
MH025919.1_strain_N10              ....................t.................................................c......... [132000]
MH025918.1_strain_R25              ....................t.................................................c......... [132000]
MH025917.1_strain_R7               ....................t.................................................c......... [132000]
MH025916.1_strain_R8               ....................t.................................................c......... [132000]

FR682468.2_Georgia_2007/1          gatcttctcacctatatcattaacaatataaattacaaaattgccacgtttcagacgaaacagatgttgacgttcgagtt [132080]
ASF_IND_20_CAD543                  ................................................................................ [132080]
LR899193.1_Germany_2020/1          ................................................................................ [132080]
MN393477.1_Wuhan_2019              ................................................................................ [132080]
MW396979.1_Timor-Leste/2019/1      ................................................................................ [132080]
MK940252.1_CN/2019/InnerMongolia   ................................................................................ [132080]
MK128995.1_China/2018/AnhuiXCGQ    ................................................................................ [132080]
MH681419.1_ASFV/POL/2015           ................................................................................ [132080]
LS478113.1_Estonia_2014            ................................................................................ [132080]
KP843857.1_Odintsovo_02/14         ................................................................................ [132080]
FN557520.1_strain_E75              ................................................................................ [132080]
AY261366.1_Warthog                 ................................c..........................g.................... [132080]
AY261365.1_Warmbaths               ...........................................................g.................... [132080]
AY261364.1_Tengani_62              ...........................................................g.................... [132080]
AY261363.1_Pretoriuskop/96/4       ...........................................................g.................... [132080]
AY261362.1_Mkuzi_1979              ................................................................................ [132080]
AY261361.1_Malawi_Lil-20/1_1983    ..c.............................c.....g.....t..............g..................c. [132080]
AY261360.1_Kenya_1950              ..c.............................c.....g....................g..................c. [132080]
AM712240.1_OURT_88/3               ................................................................................ [132080]
AM712239.1_Benin_97/1              ................................................................................ [132080]
KM111295.1_Ken06.Bus               ..c......t......................c..............a...........g..................c. [132080]
```

```
KM111294.1_Ken05/Tk1            ..c.............................c.....g.....t..............g..................c. [132080]
KM262844.1_strain_L60           ................................................................................ [132080]
KM102979.1_26544/OG10_Italy     ................................................................................ [132080]
KM262845.1_strain_NHV           ................................................................................ [132080]
KX354450.1_47/Ss/2008           ................................................................................ [132080]
KP055815.1_strain_BA71          ................................................................................ [132080]
MH025920.1_strain_R35           ..c......t......................c..............a...........g..................c. [132080]
MH025919.1_strain_N10           ..c......t......................c..............a...........g..................c. [132080]
MH025918.1_strain_R25           ..c......t......................c..............a...........g..................c. [132080]
MH025917.1_strain_R7            ..c......t......................c..............a...........g..................c. [132080]
MH025916.1_strain_R8            ..c......t......................c..............a...........g..................c. [132080]

FR682468.2_Georgia_2007/1       ttccagtactcatgtagaactgctattaaagctgaataaaacgtggcttattttggctggaattcatgtggcaaaaaaac [132160]
ASF_IND_20_CAD543               ................................................................................ [132160]
LR899193.1_Germany_2020/1       ................................................................................ [132160]
MN393477.1_Wuhan_2019           ................................................................................ [132160]
MW396979.1_Timor-Leste/2019/1   ................................................................................ [132160]
MK940252.1_CN/2019/InnerMongolia ................................................................................ [132160]
MK128995.1_China/2018/AnhuiXCGQ ................................................................................ [132160]
MH681419.1_ASFV/POL/2015        ................................................................................ [132160]
LS478113.1_Estonia_2014         ................................................................................ [132160]
KP843857.1_Odintsovo_02/14      ................................................................................ [132160]
FN557520.1_strain_E75           ...t...................................g................................g....... [132160]
AY261366.1_Warthog              .........................................................c...................... [132160]
AY261365.1_Warmbaths            .........................................................c...................... [132160]
AY261364.1_Tengani_62           .........................................................c...................... [132160]
AY261363.1_Pretoriuskop/96/4    .........................................................c...................... [132160]
AY261362.1_Mkuzi_1979           ................................................................................ [132160]
AY261361.1_Malawi_Lil-20/1_1983 ...t.............................a........a............a.......a................ [132160]
AY261360.1_Kenya_1950           ..............................a..a..c...................tc...........a.......... [132160]
AM712240.1_OURT_88/3            ...t...................................g................................g....... [132160]
AM712239.1_Benin_97/1           ...t...................................g................................g....... [132160]
KM111295.1_Ken06.Bus            ..............................a..a..c...................tg...........a.......... [132160]
KM111294.1_Ken05/Tk1            ...t..........................a..a..c...................tc...........a.......... [132160]
KM262844.1_strain_L60           ...t...................................g................................g....... [132160]
KM102979.1_26544/OG10_Italy     ...t...................................g................................g....... [132160]
KM262845.1_strain_NHV           ...t...................................g................................g....... [132160]
KX354450.1_47/Ss/2008           ...t...................................g................................g....... [132160]
KP055815.1_strain_BA71          ...t...................................g................................g....... [132160]
MH025920.1_strain_R35           ..............................a..a..c...................tg...........a.......... [132160]
MH025919.1_strain_N10           ..............................a..a..c...................tg...........a.......... [132160]
MH025918.1_strain_R25           ..............................a..a..c...................tg...........a.......... [132160]
MH025917.1_strain_R7            ..............................a..a..c...................tg...........a.......... [132160]
MH025916.1_strain_R8            ..............................a..a..c...................tg...........a.......... [132160]
```

```
FR682468.2_Georgia_2007/1        atctgcaagcttttttggattcatataacaatgaatcgccgtctagaacattcattcagcaggctatagaggaagaatgt [132240]
ASF_IND_20_CAD543                ............................y................................................... [132240]
LR899193.1_Germany_2020/1        ................................................................................ [132240]
MN393477.1_Wuhan_2019            ................................................................................ [132240]
MW396979.1_Timor-Leste/2019/1    ................................................................................ [132240]
MK940252.1_CN/2019/InnerMongolia ................................................................................ [132240]
MK128995.1_China/2018/AnhuiXCGQ  ................................................................................ [132240]
MH681419.1_ASFV/POL/2015         ................................................................................ [132240]
LS478113.1_Estonia_2014          ................................................................................ [132240]
KP843857.1_Odintsovo_02/14       ................................................................................ [132240]
FN557520.1_strain_E75            ...........c................t......c.a.......................................... [132240]
AY261366.1_Warthog               ...........c.......................c............................................ [132240]
AY261365.1_Warmbaths             ...........c.......................c............................................ [132240]
AY261364.1_Tengani_62            ...................................c.........a.................................. [132240]
AY261363.1_Pretoriuskop/96/4     ...........c.......................c.........ag................................. [132240]
AY261362.1_Mkuzi_1979            ...........c.......................c.a..a....................................... [132240]
AY261361.1_Malawi_Lil-20/1_1983  ...........c.......................c.c.......a......................a........... [132240]
AY261360.1_Kenya_1950            .......t.t.c......g..t..c..........c.........ag................................. [132240]
AM712240.1_OURT_88/3             ...........c................t......c.a.......................................... [132240]
AM712239.1_Benin_97/1            ...........c................t......c.a.......................................... [132240]
KM111295.1_Ken06.Bus             .......ta..c......g..t..c..........c.........ag..............a.................. [132240]
KM111294.1_Ken05/Tk1             .......t.t.c......g..t..c..........c.........ag................................. [132240]
KM262844.1_strain_L60            ...........c................t......c.a.......................................... [132240]
KM102979.1_26544/OG10_Italy      ...........c................t......c.a.......................................... [132240]
KM262845.1_strain_NHV            ...........c................t......c.a.......................................... [132240]
KX354450.1_47/Ss/2008            ...........c................t......c.a.......................................... [132240]
KP055815.1_strain_BA71           ...........c................t......c.a.......................................... [132240]
MH025920.1_strain_R35            .......ta..c......g..t..c..........c.........ag..............a.................. [132240]
MH025919.1_strain_N10            .......ta..c......g..t..c..........c.........ag..............a.................. [132240]
MH025918.1_strain_R25            .......ta..c......g..t..c..........c.........ag..............a.................. [132240]
MH025917.1_strain_R7             .......ta..c......g..t..c..........c.........ag..............a.................. [132240]
MH025916.1_strain_R8             .......ta..c......g..t..c..........c.........ag..............a.................. [132240]

FR682468.2_Georgia_2007/1        ggcagtattaaaccatcttgctacgactttatttcctaatacttcttaagaaactctttaaacaaggacttcgcatggtc [132320]
ASF_IND_20_CAD543                .......................................................................t..g..... [132320]
LR899193.1_Germany_2020/1        ................................................................................ [132320]
MN393477.1_Wuhan_2019            ................................................................................ [132320]
MW396979.1_Timor-Leste/2019/1    ................................................................................ [132320]
MK940252.1_CN/2019/InnerMongolia ................................................................................ [132320]
MK128995.1_China/2018/AnhuiXCGQ  ................................................................................ [132320]
MH681419.1_ASFV/POL/2015         ................................................................................ [132320]
LS478113.1_Estonia_2014          ................................................................................ [132320]
KP843857.1_Odintsovo_02/14       ................................................................................ [132320]
```

```
FN557520.1_strain_E75           .......................................................................t..g..c.. [132320]
AY261366.1_Warthog              ..........................................................................g..... [132320]
AY261365.1_Warmbaths            ..........................................................................g..... [132320]
AY261364.1_Tengani_62           ..........................................................................g..... [132320]
AY261363.1_Pretoriuskop/96/4    ..........................................................................g..... [132320]
AY261362.1_Mkuzi_1979           ..........................................................................g..... [132320]
AY261361.1_Malawi_Lil-20/1_1983 a.............................................................t..a.....t..g..c.. [132320]
AY261360.1_Kenya_1950           .........................................t.......................a...........c.. [132320]
AM712240.1_OURT_88/3            .......................................................................t..g..c.. [132320]
AM712239.1_Benin_97/1           .......................................................................t..g..c.. [132320]
KM111295.1_Ken06.Bus            .........................................t..t....................atc......g..c.. [132320]
KM111294.1_Ken05/Tk1            .........................................t..t....................atc......g..c.. [132320]
KM262844.1_strain_L60           .......................................................................t..g..c.. [132320]
KM102979.1_26544/OG10_Italy     .......................................................................t..g..c.. [132320]
KM262845.1_strain_NHV           .......................................................................t..g..c.. [132320]
KX354450.1_47/Ss/2008           .......................................................................t..g..c.. [132320]
KP055815.1_strain_BA71          .......................................................................t..g..c.. [132320]
MH025920.1_strain_R35           .........................................t..t....................atc......g..c.. [132320]
MH025919.1_strain_N10           .........................................t..t....................atc......g..c.. [132320]
MH025918.1_strain_R25           .........................................t..t....................atc......g..c.. [132320]
MH025917.1_strain_R7            .........................................t..t....................atc......g..c.. [132320]
MH025916.1_strain_R8            .........................................t..t....................atc......g..c.. [132320]

FR682468.2_Georgia_2007/1       aaaggttctaaacccatggcccttatgattcgccaaaaaagcggtttcatcaagattttctaaccctttcacggatgaag [132400]
ASF_IND_20_CAD543               ................................................................................ [132400]
LR899193.1_Germany_2020/1       ................................................................................ [132400]
MN393477.1_Wuhan_2019           ................................................................................ [132400]
MW396979.1_Timor-Leste/2019/1   ................................................................................ [132400]
MK940252.1_CN/2019/InnerMongolia ................................................................................ [132400]
MK128995.1_China/2018/AnhuiXCGQ ................................................................................ [132400]
MH681419.1_ASFV/POL/2015        ................................................................................ [132400]
LS478113.1_Estonia_2014         ................................................................................ [132400]
KP843857.1_Odintsovo_02/14      ................................................................................ [132400]
FN557520.1_strain_E75           g.....c...........a...........t................................................. [132400]
AY261366.1_Warthog              .........................................................c...................... [132400]
AY261365.1_Warmbaths            ................................................................................ [132400]
AY261364.1_Tengani_62           ................................................................................ [132400]
AY261363.1_Pretoriuskop/96/4    ................................................................................ [132400]
AY261362.1_Mkuzi_1979           ................................................................................ [132400]
AY261361.1_Malawi_Lil-20/1_1983 ......c.....................................c..................a.....t.......... [132400]
AY261360.1_Kenya_1950           ...............................................................a................ [132400]
AM712240.1_OURT_88/3            g.....c...........a...........t................................................. [132400]
AM712239.1_Benin_97/1           g.....c...........a...........t................................................. [132400]
KM111295.1_Ken06.Bus            ......ct.......................................................a.....t.......... [132400]
```

```
KM111294.1_Ken05/Tk1             ......c...................................c....................a................ [132400]
KM262844.1_strain_L60            g.....c...........a...........t................................................. [132400]
KM102979.1_26544/OG10_Italy      g.....c...........a...........t................................................. [132400]
KM262845.1_strain_NHV            g.....c...........a...........t................................................. [132400]
KX354450.1_47/Ss/2008            g.....c...........a...........t................................................. [132400]
KP055815.1_strain_BA71           g.....c...........a...........t................................................. [132400]
MH025920.1_strain_R35            ......ct.......................................................a.....t.......... [132400]
MH025919.1_strain_N10            ......ct.......................................................a.....t.......... [132400]
MH025918.1_strain_R25            ......ct.......................................................a.....t.......... [132400]
MH025917.1_strain_R7             ......ct.......................................................a.....t.......... [132400]
MH025916.1_strain_R8             ......ct.......................................................a.....t.......... [132400]

FR682468.2_Georgia_2007/1        aaataaggtgttcggcctcgtttgcccattttctatgattttttttcacctcgggttctagatctgttttctccatatac [132480]
ASF_IND_20_CAD543                ..............................................y................................. [132480]
LR899193.1_Germany_2020/1        ................................................................................ [132480]
MN393477.1_Wuhan_2019            ................................................................................ [132480]
MW396979.1_Timor-Leste/2019/1    ................................................................................ [132480]
MK940252.1_CN/2019/InnerMongolia ................................t............................................... [132480]
MK128995.1_China/2018/AnhuiXCGQ  ................................................................................ [132480]
MH681419.1_ASFV/POL/2015         ................................................................................ [132480]
LS478113.1_Estonia_2014          ................................................................................ [132480]
KP843857.1_Odintsovo_02/14       ................................................................................ [132480]
FN557520.1_strain_E75            ..............................................t....................c............ [132480]
AY261366.1_Warthog               ...................................................................c............ [132480]
AY261365.1_Warmbaths             ...................................................................c............ [132480]
AY261364.1_Tengani_62            ................................................................................ [132480]
AY261363.1_Pretoriuskop/96/4     ...................................................................c............ [132480]
AY261362.1_Mkuzi_1979            ...................................................................c............ [132480]
AY261361.1_Malawi_Lil-20/1_1983  .............a.....a..g............................................c........g... [132480]
AY261360.1_Kenya_1950            .............a.................................................................. [132480]
AM712240.1_OURT_88/3             ..............................................t....................c............ [132480]
AM712239.1_Benin_97/1            ..............................................t....................c............ [132480]
KM111295.1_Ken06.Bus             .............a.................................................................. [132480]
KM111294.1_Ken05/Tk1             .............a.................................................................. [132480]
KM262844.1_strain_L60            ..............................................t....................c............ [132480]
KM102979.1_26544/OG10_Italy      ..............................................t....................c............ [132480]
KM262845.1_strain_NHV            ..............................................t....................c............ [132480]
KX354450.1_47/Ss/2008            ..............................................t....................c............ [132480]
KP055815.1_strain_BA71           ..............................................t....................c............ [132480]
MH025920.1_strain_R35            .............a.................................................................. [132480]
MH025919.1_strain_N10            .............a.................................................................. [132480]
MH025918.1_strain_R25            .............a.................................................................. [132480]
MH025917.1_strain_R7             .............a.................................................................. [132480]
MH025916.1_strain_R8             .............a.................................................................. [132480]
```

```
FR682468.2_Georgia_2007/1        tcattgtggtcatatttttttttgggaggaggc---gtgggtggaggaatgggtggaggaagtacacccgactttcccgc [132560]
ASF_IND_20_CAD543                .........................ag......ggg............................................ [132560]
LR899193.1_Germany_2020/1        .................................---............................................ [132560]
MN393477.1_Wuhan_2019            .................................---............................................ [132560]
MW396979.1_Timor-Leste/2019/1    .................................---............................................ [132560]
MK940252.1_CN/2019/InnerMongolia .................................---............................................ [132560]
MK128995.1_China/2018/AnhuiXCGQ  .................................---............................................ [132560]
MH681419.1_ASFV/POL/2015         .................................---............................................ [132560]
LS478113.1_Estonia_2014          .................................---............................................ [132560]
KP843857.1_Odintsovo_02/14       .................................---............................................ [132560]
FN557520.1_strain_E75            .........................ag......ggg..................................g....t.... [132560]
AY261366.1_Warthog               ................................tggg..................................ga....t... [132560]
AY261365.1_Warmbaths             .................................ggg..................................g....t.... [132560]
AY261364.1_Tengani_62            .................................ggc............................................ [132560]
AY261363.1_Pretoriuskop/96/4     .................................gga............................................ [132560]
AY261362.1_Mkuzi_1979            .............................g...ggg..................................g.....t... [132560]
AY261361.1_Malawi_Lil-20/1_1983  .........................ag......ggg...............................t..g......... [132560]
AY261360.1_Kenya_1950            .................................gga..................................ga........ [132560]
AM712240.1_OURT_88/3             .........................ag......ggg..................................g....t.... [132560]
AM712239.1_Benin_97/1            .........................ag......ggg..................................g....t.... [132560]
KM111295.1_Ken06.Bus             .................................gga..................................ga........ [132560]
KM111294.1_Ken05/Tk1             .................................gga..................................ga........ [132560]
KM262844.1_strain_L60            .........................ag......ggg..................................g....t.... [132560]
KM102979.1_26544/OG10_Italy      .........................ag......ggg..................................g....t.... [132560]
KM262845.1_strain_NHV            .........................ag......ggg..................................g....t.... [132560]
KX354450.1_47/Ss/2008            .........................ag......ggg..................................g....t.... [132560]
KP055815.1_strain_BA71           .........................ag......ggg..................................g....t.... [132560]
MH025920.1_strain_R35            .................................gga..................................ga........ [132560]
MH025919.1_strain_N10            .................................gga..................................ga........ [132560]
MH025918.1_strain_R25            .................................gga..................................ga........ [132560]
MH025917.1_strain_R7             .................................gga..................................ga........ [132560]
MH025916.1_strain_R8             .................................gga..................................ga........ [132560]

FR682468.2_Georgia_2007/1        ttcaaccgttttataaaaaaatagaagcataatacaaagaataaggactatcgcaaatatgataaccagtgtcccagtcg [132640]
ASF_IND_20_CAD543                ................................................................................ [132640]
LR899193.1_Germany_2020/1        ................................................................................ [132640]
MN393477.1_Wuhan_2019            ................................................................................ [132640]
MW396979.1_Timor-Leste/2019/1    ................................................................................ [132640]
MK940252.1_CN/2019/InnerMongolia ................................................................................ [132640]
MK128995.1_China/2018/AnhuiXCGQ  ................................................................................ [132640]
MH681419.1_ASFV/POL/2015         ................................................................................ [132640]
LS478113.1_Estonia_2014          ................................................................................ [132640]
KP843857.1_Odintsovo_02/14       ................................................................................ [132640]
```

```
FN557520.1_strain_E75            ................................................................................ [132640]
AY261366.1_Warthog               ....g...c....................................................................... [132640]
AY261365.1_Warmbaths             ................................................................................ [132640]
AY261364.1_Tengani_62            ................................................................................ [132640]
AY261363.1_Pretoriuskop/96/4     ....g...................................................................g....... [132640]
AY261362.1_Mkuzi_1979            ....g...c..................................................................g.... [132640]
AY261361.1_Malawi_Lil-20/1_1983  ................................................................................ [132640]
AY261360.1_Kenya_1950            ....gt..........t.............g..............a.t...........g.................... [132640]
AM712240.1_OURT_88/3             ................................................................................ [132640]
AM712239.1_Benin_97/1            ................................................................................ [132640]
KM111295.1_Ken06.Bus             ....gt..........t............................a.t...........g.................... [132640]
KM111294.1_Ken05/Tk1             ....gt..........t.............g..............a.t...........g.................... [132640]
KM262844.1_strain_L60            ................................................................................ [132640]
KM102979.1_26544/OG10_Italy      ................................................................................ [132640]
KM262845.1_strain_NHV            ................................................................................ [132640]
KX354450.1_47/Ss/2008            ................................................................................ [132640]
KP055815.1_strain_BA71           ................................................................................ [132640]
MH025920.1_strain_R35            ....gt..........t............................a.t...........g.................... [132640]
MH025919.1_strain_N10            ....gt..........t............................a.t...........g.................... [132640]
MH025918.1_strain_R25            ....gt..........t............................a.t...........g.................... [132640]
MH025917.1_strain_R7             ....gt..........t............................a.t...........g.................... [132640]
MH025916.1_strain_R8             ....gt..........t............................a.t...........g.................... [132640]

FR682468.2_Georgia_2007/1        agggcattttgttatataagtaacgtttttttttattttttataattcgaatgaagaaccatgttgaata---------- [132720]
ASF_IND_20_CAD543                ......................................................................---------- [132720]
LR899193.1_Germany_2020/1        ......................................................................---------- [132720]
MN393477.1_Wuhan_2019            ......................................................................---------- [132720]
MW396979.1_Timor-Leste/2019/1    ......................................................................---------- [132720]
MK940252.1_CN/2019/InnerMongolia ......................................................................---------- [132720]
MK128995.1_China/2018/AnhuiXCGQ  ......................................................................---------- [132720]
MH681419.1_ASFV/POL/2015         ......................................................................---------- [132720]
LS478113.1_Estonia_2014          ......................................................................---------- [132720]
KP843857.1_Odintsovo_02/14       .........................-............................................---------- [132720]
FN557520.1_strain_E75            ......................................................................---------- [132720]
AY261366.1_Warthog               ...............................................................a......---------- [132720]
AY261365.1_Warmbaths             ..............................................................aa......---------- [132720]
AY261364.1_Tengani_62            ......................................................................---------- [132720]
AY261363.1_Pretoriuskop/96/4     ...............................................................a......---------- [132720]
AY261362.1_Mkuzi_1979            ..............................................................aa.a....---------- [132720]
AY261361.1_Malawi_Lil-20/1_1983  ....................................................---....t...a......---------- [132720]
AY261360.1_Kenya_1950            ...........................................................t...a......tgtcttttat [132720]
AM712240.1_OURT_88/3             ......................................................................---------- [132720]
AM712239.1_Benin_97/1            ......................................................................---------- [132720]
KM111295.1_Ken06.Bus             .........................-.................................t...a..g...tgtcttttat [132720]
```

```
KM111294.1_Ken05/Tk1          ...........................................................t...a......tgtcttttat [132720]
KM262844.1_strain_L60         ......................................................................---------- [132720]
KM102979.1_26544/OG10_Italy   ......................................................................---------- [132720]
KM262845.1_strain_NHV         ......................................................................---------- [132720]
KX354450.1_47/Ss/2008         ......................................................................---------- [132720]
KP055815.1_strain_BA71        ......................................................................---------- [132720]
MH025920.1_strain_R35         ...........................................................t...a..g...tgtcttttat [132720]
MH025919.1_strain_N10         ...........................................................t...a..g...tgtcttttat [132720]
MH025918.1_strain_R25         ...........................................................t...a..g...tgtcttttat [132720]
MH025917.1_strain_R7          ...........................................................t...a..g...tgtcttttat [132720]
MH025916.1_strain_R8          ...........................................................t...a..g...tgtcttttat [132720]

FR682468.2_Georgia_2007/1     -------------------------------------------------------------------------------- [132800]
ASF_IND_20_CAD543             -------------------------------------------------------------------------------- [132800]
LR899193.1_Germany_2020/1     -------------------------------------------------------------------------------- [132800]
MN393477.1_Wuhan_2019         -------------------------------------------------------------------------------- [132800]
MW396979.1_Timor-Leste/2019/1 -------------------------------------------------------------------------------- [132800]
MK940252.1_CN/2019/InnerMongolia -------------------------------------------------------------------------------- [132800]
MK128995.1_China/2018/AnhuiXCGQ -------------------------------------------------------------------------------- [132800]
MH681419.1_ASFV/POL/2015      -------------------------------------------------------------------------------- [132800]
LS478113.1_Estonia_2014       -------------------------------------------------------------------------------- [132800]
KP843857.1_Odintsovo_02/14    -------------------------------------------------------------------------------- [132800]
FN557520.1_strain_E75         -------------------------------------------------------------------------------- [132800]
AY261366.1_Warthog            -------------------------------------------------------------------------------- [132800]
AY261365.1_Warmbaths          -------------------------------------------------------------------------------- [132800]
AY261364.1_Tengani_62         -------------------------------------------------------------------------------- [132800]
AY261363.1_Pretoriuskop/96/4  -------------------------------------------------------------------------------- [132800]
AY261362.1_Mkuzi_1979         -------------------------------------------------------------------------------- [132800]
AY261361.1_Malawi_Lil-20/1_1983 -------------------------------------------------------------------------------- [132800]
AY261360.1_Kenya_1950         tcaagacatttcattatactcaaaggcatttcattatactcaaaggcatttcattatactcaaggacattttattatact [132800]
AM712240.1_OURT_88/3          -------------------------------------------------------------------------------- [132800]
AM712239.1_Benin_97/1         -------------------------------------------------------------------------------- [132800]
KM111295.1_Ken06.Bus          tcaagaca-------------------------------------------------------------tttattatact [132800]
KM111294.1_Ken05/Tk1          tcaagaca---------------------tttattatactcaaaggcattttattatactcaaggaca-tttattatact [132800]
KM262844.1_strain_L60         -------------------------------------------------------------------------------- [132800]
KM102979.1_26544/OG10_Italy   -------------------------------------------------------------------------------- [132800]
KM262845.1_strain_NHV         -------------------------------------------------------------------------------- [132800]
KX354450.1_47/Ss/2008         -------------------------------------------------------------------------------- [132800]
KP055815.1_strain_BA71        -------------------------------------------------------------------------------- [132800]
MH025920.1_strain_R35         tcaagaca-------------------------------------------------------------tttattatact [132800]
MH025919.1_strain_N10         tcaagaca-------------------------------------------------------------tttattatact [132800]
MH025918.1_strain_R25         tcaagaca-------------------------------------------------------------tttattatact [132800]
MH025917.1_strain_R7          tcaagaca-------------------------------------------------------------tttattatact [132800]
MH025916.1_strain_R8          tcaagaca-------------------------------------------------------------tttattatact [132800]
```

```
FR682468.2_Georgia_2007/1        -----------------------------------------------------------------------------gtc [132880]
ASF_IND_20_CAD543                -----------------------------------------------------------------------------... [132880]
LR899193.1_Germany_2020/1        -----------------------------------------------------------------------------... [132880]
MN393477.1_Wuhan_2019            -----------------------------------------------------------------------------... [132880]
MW396979.1_Timor-Leste/2019/1    -----------------------------------------------------------------------------... [132880]
MK940252.1_CN/2019/InnerMongolia -----------------------------------------------------------------------------... [132880]
MK128995.1_China/2018/AnhuiXCGQ  -----------------------------------------------------------------------------... [132880]
MH681419.1_ASFV/POL/2015         -----------------------------------------------------------------------------... [132880]
LS478113.1_Estonia_2014          -----------------------------------------------------------------------------... [132880]
KP843857.1_Odintsovo_02/14       -----------------------------------------------------------------------------... [132880]
FN557520.1_strain_E75            -----------------------------------------------------------------------------... [132880]
AY261366.1_Warthog               -----------------------------------------------------------------------------... [132880]
AY261365.1_Warmbaths             -----------------------------------------------------------------------------... [132880]
AY261364.1_Tengani_62            -----------------------------------------------------------------------------... [132880]
AY261363.1_Pretoriuskop/96/4     -----------------------------------------------------------------------------... [132880]
AY261362.1_Mkuzi_1979            -----------------------------------------------------------------------------... [132880]
AY261361.1_Malawi_Lil-20/1_1983  -----------------------------------------------------------------------------... [132880]
AY261360.1_Kenya_1950            caaggacatttcattatactcaaaggcatttcat---------------------------------------------- [132880]
AM712240.1_OURT_88/3             -----------------------------------------------------------------------------... [132880]
AM712239.1_Benin_97/1            -----------------------------------------------------------------------------... [132880]
KM111295.1_Ken06.Bus             caaaggcatttcatcatactcaagggcatttcat---------------------------------------------- [132880]
KM111294.1_Ken05/Tk1             caaaggcattttattatactcaaaggcatttattatacttactcaaaggcatttattatactcactcaaaggcattta.t [132880]
KM262844.1_strain_L60            -----------------------------------------------------------------------------... [132880]
KM102979.1_26544/OG10_Italy      -----------------------------------------------------------------------------... [132880]
KM262845.1_strain_NHV            -----------------------------------------------------------------------------... [132880]
KX354450.1_47/Ss/2008            -----------------------------------------------------------------------------... [132880]
KP055815.1_strain_BA71           -----------------------------------------------------------------------------... [132880]
MH025920.1_strain_R35            caaaggcatttcatcatactcaagggcatttcat---------------------------------------------- [132880]
MH025919.1_strain_N10            caaaggcatttcatcatactcaagggcatttcat---------------------------------------------- [132880]
MH025918.1_strain_R25            caaaggcatttcatcatactcaagggcatttcat---------------------------------------------- [132880]
MH025917.1_strain_R7             caaaggcatttcatcatactcaagggcatttcat---------------------------------------------- [132880]
MH025916.1_strain_R8             caaaggcatttcatcatactcaagggcatttcat---------------------------------------------- [132880]

FR682468.2_Georgia_2007/1        ttctactcaaagacattt---tgttatacggtaaatgagaatttataaaatccgaatatcactatcatactgtttatctg [132960]
ASF_IND_20_CAD543                ..........g.gtg...tta.a......................................................... [132960]
LR899193.1_Germany_2020/1        ..................---........................................................... [132960]
MN393477.1_Wuhan_2019            ..................---........................................................... [132960]
MW396979.1_Timor-Leste/2019/1    ..................---........................................................... [132960]
MK940252.1_CN/2019/InnerMongolia ..................---........................................................... [132960]
MK128995.1_China/2018/AnhuiXCGQ  ..................---........................................................... [132960]
MH681419.1_ASFV/POL/2015         ..................---........................................................... [132960]
LS478113.1_Estonia_2014          ..................---........................................................... [132960]
KP843857.1_Odintsovo_02/14       ..................---........................................................... [132960]
```

```
FN557520.1_strain_E75           .......t..g.gtg...tta.a......t..g..................t........g................... [132960]
AY261366.1_Warthog              .......t..ga.tg...tt-...............a........................................... [132960]
AY261365.1_Warmbaths            .......t..ga.tg...tt-.a.....................................g................... [132960]
AY261364.1_Tengani_62           ..........g.......---.......tt..g...a....a..................g................... [132960]
AY261363.1_Pretoriuskop/96/4    .......t..ga.tg...tt-.a......................................................... [132960]
AY261362.1_Mkuzi_1979           ..---..t..g.gt....tt-.a......t..g............................................... [132960]
AY261361.1_Malawi_Lil-20/1_1983 ........t.g.gt....---.a...a.ata.g........ac........t..g.....g................... [132960]
AY261360.1_Kenya_1950           -.a.......g.......---ca......t..g.....a..ac........t..g.....g................... [132960]
AM712240.1_OURT_88/3            .......t..g.gtg...tta.a......t..g..................t........g................... [132960]
AM712239.1_Benin_97/1           .......t..g.gtg...tta.a......t..g..................t........g................... [132960]
KM111295.1_Ken06.Bus            -.a.......g.g.....---.a......t..g...t....ac........t..g.....g................... [132960]
KM111294.1_Ken05/Tk1            a.a.........g.....----a......t..g.....a..a.........t..g.....g................... [132960]
KM262844.1_strain_L60           .......t..g.gtg...tta.a......t..g..................t........g................... [132960]
KM102979.1_26544/OG10_Italy     .......t..g.gtg...tta.a......t..g..................t........g................... [132960]
KM262845.1_strain_NHV           .......t..g.gtg...tta.a......t..g..................t........g................... [132960]
KX354450.1_47/Ss/2008           .......t..g.gtg...tta.a......t..g..................t........g................... [132960]
KP055815.1_strain_BA71          .......t..g.gtg...tta.a......t..g..................t........g................... [132960]
MH025920.1_strain_R35           -.a.......g.g.....---.a......t..g...t....ac........t..g.....g................... [132960]
MH025919.1_strain_N10           -.a.......g.g.....---.a......t..g...t....ac........t..g.....g................... [132960]
MH025918.1_strain_R25           -.a.......g.g.....---.a......t..g...t....ac........t..g.....g................... [132960]
MH025917.1_strain_R7            -.a.......g.g.....---.a......t..g...t....ac........t..g.....g................... [132960]
MH025916.1_strain_R8            -.a.......g.g.....---.a......t..g...t....ac........t..g.....g................... [132960]

FR682468.2_Georgia_2007/1       agaaggtctcactgggtcctgtgatggagaacccatactctgtaatgctggggtttataatgtggtcaggactgacaagc [133040]
ASF_IND_20_CAD543               ................................................................................ [133040]
LR899193.1_Germany_2020/1       ................................................................................ [133040]
MN393477.1_Wuhan_2019           ................................................................................ [133040]
MW396979.1_Timor-Leste/2019/1   ................................................................................ [133040]
MK940252.1_CN/2019/InnerMongolia ................................................................................ [133040]
MK128995.1_China/2018/AnhuiXCGQ ................................................................................ [133040]
MH681419.1_ASFV/POL/2015        ................................................................................ [133040]
LS478113.1_Estonia_2014         ................................................................................ [133040]
KP843857.1_Odintsovo_02/14      ................................................................................ [133040]
FN557520.1_strain_E75           ..................................g.........................................g... [133040]
AY261366.1_Warthog              .........................a..................................................g... [133040]
AY261365.1_Warmbaths            ............................................................................g... [133040]
AY261364.1_Tengani_62           ............................................................................g... [133040]
AY261363.1_Pretoriuskop/96/4    .........................a..................................................g... [133040]
AY261362.1_Mkuzi_1979           .......t..........................g.........................................g... [133040]
AY261361.1_Malawi_Lil-20/1_1983 ............................................................................g... [133040]
AY261360.1_Kenya_1950           ......................a.....a.....................................t.........g... [133040]
AM712240.1_OURT_88/3            ..................................g.........................................g... [133040]
AM712239.1_Benin_97/1           ..................................g.........................................g... [133040]
KM111295.1_Ken06.Bus            ......................a............................................g..g......... [133040]
```

```
KM111294.1_Ken05/Tk1             ......................a.....a...............................................g... [133040]
KM262844.1_strain_L60            ..................................g.........................................g... [133040]
KM102979.1_26544/OG10_Italy      ..................................g.........................................g... [133040]
KM262845.1_strain_NHV            ..................................g.........................................g... [133040]
KX354450.1_47/Ss/2008            ..................................g.........................................g... [133040]
KP055815.1_strain_BA71           ..................................g.........................................g... [133040]
MH025920.1_strain_R35            ......................a............................................g..g......... [133040]
MH025919.1_strain_N10            ......................a............................................g..g......... [133040]
MH025918.1_strain_R25            ......................a............................................g..g......... [133040]
MH025917.1_strain_R7             ......................a............................................g..g......... [133040]
MH025916.1_strain_R8             ......................a............................................g..g......... [133040]

FR682468.2_Georgia_2007/1        acatttctgaactgcgagagttctaggtttagacgcagtcgtaatagtcgctgtatatttgtaataaatattagattgcg [133120]
ASF_IND_20_CAD543                ................................................................................ [133120]
LR899193.1_Germany_2020/1        ................................................................................ [133120]
MN393477.1_Wuhan_2019            ................................................................................ [133120]
MW396979.1_Timor-Leste/2019/1    ................................................................................ [133120]
MK940252.1_CN/2019/InnerMongolia ................................................................................ [133120]
MK128995.1_China/2018/AnhuiXCGQ  ................................................................................ [133120]
MH681419.1_ASFV/POL/2015         ................................................................................ [133120]
LS478113.1_Estonia_2014          ................................................................................ [133120]
KP843857.1_Odintsovo_02/14       ................................................................................ [133120]
FN557520.1_strain_E75            ................................................................................ [133120]
AY261366.1_Warthog               ................................................................................ [133120]
AY261365.1_Warmbaths             ................................................................................ [133120]
AY261364.1_Tengani_62            ................................................................................ [133120]
AY261363.1_Pretoriuskop/96/4     ................................................................................ [133120]
AY261362.1_Mkuzi_1979            ................................................................................ [133120]
AY261361.1_Malawi_Lil-20/1_1983  .........................................c.............c........................ [133120]
AY261360.1_Kenya_1950            ........a................................c...................................... [133120]
AM712240.1_OURT_88/3             ................................................................................ [133120]
AM712239.1_Benin_97/1            ................................................................................ [133120]
KM111295.1_Ken06.Bus             ........a..t.............................c...................................... [133120]
KM111294.1_Ken05/Tk1             ........a................................c...................................... [133120]
KM262844.1_strain_L60            ................................................................................ [133120]
KM102979.1_26544/OG10_Italy      ................................................................................ [133120]
KM262845.1_strain_NHV            ................................................................................ [133120]
KX354450.1_47/Ss/2008            ................................................................................ [133120]
KP055815.1_strain_BA71           ................................................................................ [133120]
MH025920.1_strain_R35            ........a..t.............................c...................................... [133120]
MH025919.1_strain_N10            ........a..t.............................c...................................... [133120]
MH025918.1_strain_R25            ........a..t.............................c...................................... [133120]
MH025917.1_strain_R7             ........a..t.............................c...................................... [133120]
MH025916.1_strain_R8             ........a..t.............................c...................................... [133120]
```

```
FR682468.2_Georgia_2007/1        tatgaggcgagtgtcaaagcgatcctttccaatttgtactaaggtgggcttttgtattccaactcccacttgtttaacga [133200]
ASF_IND_20_CAD543                ................................................................................ [133200]
LR899193.1_Germany_2020/1        ................................................................................ [133200]
MN393477.1_Wuhan_2019            ................................................................................ [133200]
MW396979.1_Timor-Leste/2019/1    ................................................................................ [133200]
MK940252.1_CN/2019/InnerMongolia ................................................................................ [133200]
MK128995.1_China/2018/AnhuiXCGQ  ................................................................................ [133200]
MH681419.1_ASFV/POL/2015         ................................................................................ [133200]
LS478113.1_Estonia_2014          ................................................................................ [133200]
KP843857.1_Odintsovo_02/14       ................................................................................ [133200]
FN557520.1_strain_E75            ..............................g................................................. [133200]
AY261366.1_Warthog               ..............................g..a........a..................................... [133200]
AY261365.1_Warmbaths             ..............................g................................................. [133200]
AY261364.1_Tengani_62            ...a..........................g..............a.........c.............c........a. [133200]
AY261363.1_Pretoriuskop/96/4     ..............................g................................................. [133200]
AY261362.1_Mkuzi_1979            ..............................g................................................. [133200]
AY261361.1_Malawi_Lil-20/1_1983  ......a..t....................g...............................................a. [133200]
AY261360.1_Kenya_1950            ......a.......................g.......t.....ca.................g.....c.......... [133200]
AM712240.1_OURT_88/3             ..............................g................................................. [133200]
AM712239.1_Benin_97/1            ..............................g................................................. [133200]
KM111295.1_Ken06.Bus             a.....a.......................g.......t.....ca.................g.....c.......... [133200]
KM111294.1_Ken05/Tk1             ......a.......................g.......t.....ca.................g.....c.......... [133200]
KM262844.1_strain_L60            ..............................g................................................. [133200]
KM102979.1_26544/OG10_Italy      ..............................g................................................. [133200]
KM262845.1_strain_NHV            ..............................g................................................. [133200]
KX354450.1_47/Ss/2008            ..............................g................................................. [133200]
KP055815.1_strain_BA71           ..............................g................................................. [133200]
MH025920.1_strain_R35            a.....a.......................g.......t.....ca.................g.....c.......... [133200]
MH025919.1_strain_N10            a.....a.......................g.......t.....ca.................g.....c.......... [133200]
MH025918.1_strain_R25            a.....a.......................g.......t.....ca.................g.....c.......... [133200]
MH025917.1_strain_R7             a.....a.......................g.......t.....ca.................g.....c.......... [133200]
MH025916.1_strain_R8             a.....a.......................g.......t.....ca.................g.....c.......... [133200]

FR682468.2_Georgia_2007/1        tggaccagggtccttcttcccgattttgttccgtgatataggtcagcacactattttctgtatatgaggtatgatgtcgc [133280]
ASF_IND_20_CAD543                ...................................................................---nnnnnnnnnn [133280]
LR899193.1_Germany_2020/1        ................................................................................ [133280]
MN393477.1_Wuhan_2019            ................................................................................ [133280]
MW396979.1_Timor-Leste/2019/1    ................................................................................ [133280]
MK940252.1_CN/2019/InnerMongolia ................................................................................ [133280]
MK128995.1_China/2018/AnhuiXCGQ  ................................................................................ [133280]
MH681419.1_ASFV/POL/2015         ................................................................................ [133280]
LS478113.1_Estonia_2014          ................................................................................ [133280]
KP843857.1_Odintsovo_02/14       ................................................................................ [133280]
```

```
FN557520.1_strain_E75           ...........................................g..............c........aa.gg........ [133280]
AY261366.1_Warthog              ..........................................................c.g................... [133280]
AY261365.1_Warmbaths            ...........................................g..............c........aa.gg........ [133280]
AY261364.1_Tengani_62           ..........................................................c........a..gg........ [133280]
AY261363.1_Pretoriuskop/96/4    ...........................................g..............c..................... [133280]
AY261362.1_Mkuzi_1979           ...........................................g..............c........aa.gg........ [133280]
AY261361.1_Malawi_Lil-20/1_1983 ..........................................................c..................... [133280]
AY261360.1_Kenya_1950           .t..................t............c.........g..a...t.......c...........gc...a...t [133280]
AM712240.1_OURT_88/3            ...........................................g..............c........aa.gg........ [133280]
AM712239.1_Benin_97/1           ...........................................g..............c........aa.gg........ [133280]
KM111295.1_Ken06.Bus            .t..................t............c......a..g..a.....g.................gg...a...t [133280]
KM111294.1_Ken05/Tk1            .t..................t............c.........g..a...........c...........gc...a.... [133280]
KM262844.1_strain_L60           ...........................................g..............c........aa.gg........ [133280]
KM102979.1_26544/OG10_Italy     ...........................................g..............c........aa.gg........ [133280]
KM262845.1_strain_NHV           ...........................................g..............c........aa.gg........ [133280]
KX354450.1_47/Ss/2008           ...........................................g..............c........aa.gg........ [133280]
KP055815.1_strain_BA71          ...........................................g..............c........aa.gg........ [133280]
MH025920.1_strain_R35           .t..................t............c......a..g..a.....g.................gg...a...t [133280]
MH025919.1_strain_N10           .t..................t............c......a..g..a.....g.................gg...a...t [133280]
MH025918.1_strain_R25           .t..................t............c......a..g..a.....g.................gg...a...t [133280]
MH025917.1_strain_R7            .t..................t............c......a..g..a.....g.................gg...a...t [133280]
MH025916.1_strain_R8            .t..................t............c......a..g..a.....g.................gg...a...t [133280]

FR682468.2_Georgia_2007/1       atattaatacctggtgccattccaactggcggttgtgcaattcgggctgtaccgggacccaaccatcgtggagttttata [133360]
ASF_IND_20_CAD543               nnnnnnnnnnnnnnnnnnnnnnnnnnnnnnnnnnnnnnnnnnnnnnnnnnnnnnnnnnnnnnnnnnnnnnnnnnnnnnnn [133360]
LR899193.1_Germany_2020/1       ................................................................................ [133360]
MN393477.1_Wuhan_2019           ................................................................................ [133360]
MW396979.1_Timor-Leste/2019/1   ................................................................................ [133360]
MK940252.1_CN/2019/InnerMongolia ................................................................................ [133360]
MK128995.1_China/2018/AnhuiXCGQ ................................................................................ [133360]
MH681419.1_ASFV/POL/2015        ................................................................................ [133360]
LS478113.1_Estonia_2014         ................................................................................ [133360]
KP843857.1_Odintsovo_02/14      ................................................................................ [133360]
FN557520.1_strain_E75           ....ctc..---........a.g...g..t.c.c......c...tc....g.ta.............t....aca...a. [133360]
AY261366.1_Warthog              ................................................................................ [133360]
AY261365.1_Warmbaths            ....ctc..---........a.g...g..t.c.c......c...tc....g.ta.............t....aca...a. [133360]
AY261364.1_Tengani_62           ....ctcc.---........a.g...g..t.c.c......c...tc....g.ta.............t....aca...a. [133360]
AY261363.1_Pretoriuskop/96/4    ................................................................................ [133360]
AY261362.1_Mkuzi_1979           ....ctc..---........a.g...g..t.c.c......c...tc....g.ta.............t....aca...a. [133360]
AY261361.1_Malawi_Lil-20/1_1983 ................................................................................ [133360]
AY261360.1_Kenya_1950           ......cct---........att...a..tac.c......c...tc....g.tt.............t...gaca...a. [133360]
AM712240.1_OURT_88/3            ....ctc..---........a.g...g..t.c.c......c...tc....g.ta.............t....aca...a. [133360]
AM712239.1_Benin_97/1           ....ctc..---........a.g...g..t.c.c......c...tc....g.ta.............t....aca...a. [133360]
KM111295.1_Ken06.Bus            ......cct---..............a..t..g...........a.......ga..................t..c.... [133360]
```

```
KM111294.1_Ken05/Tk1           ......cct---........att...a..tac.c......c...tc....g.tt.............t...gaca...a. [133360]
KM262844.1_strain_L60          ....ctc..---........a.g...g..t.c.c......c...tc....g.ta.............t....aca...a. [133360]
KM102979.1_26544/OG10_Italy    ....ctc..---........a.g...g..t.c.c......c...tc....g.ta.............t....aca...a. [133360]
KM262845.1_strain_NHV          ....ctc..---........a.g...g..t.c.c......c...tc....g.ta.............t....aca...a. [133360]
KX354450.1_47/Ss/2008          ....ctc..---........a.g...g..t.c.c......c...tc....g.ta.............t....aca...a. [133360]
KP055815.1_strain_BA71         ....ctc..---........a.g...g..t.c.c......c...tc....g.ta.............t....aca...a. [133360]
MH025920.1_strain_R35          ......cct---..............a..t..g...........a.......ga..................t..c.... [133360]
MH025919.1_strain_N10          ......cct---..............a..t..g...........a.......ga..................t..c.... [133360]
MH025918.1_strain_R25          ......cct---..............a..t..g...........a.......ga..................t..c.... [133360]
MH025917.1_strain_R7           ......cct---..............a..t..g...........a.......ga..................t..c.... [133360]
MH025916.1_strain_R8           ......cct---..............a..t..g...........a.......ga..................t..c.... [133360]

FR682468.2_Georgia_2007/1      aacatatcgttctagcgtatttaaaaattccttaggttatttacgagtagcatgaagggtgctattaaaacaggtggat [133440]
ASF_IND_20_CAD543              nnnnnnnn....c.aa..t.c......nnnnnnnnnnnnnnnnnnnnnnnnnnnnnnnnnnnnnnnnnnnnnn.ca..ta [133440]
LR899193.1_Germany_2020/1      ................................................................................ [133440]
MN393477.1_Wuhan_2019          ................................................................................ [133440]
MW396979.1_Timor-Leste/2019/1  ................................................................................ [133440]
MK940252.1_CN/2019/InnerMongolia ................................................................................ [133440]
MK128995.1_China/2018/AnhuiXCGQ ................................................................................ [133440]
MH681419.1_ASFV/POL/2015       ................................................................................ [133440]
LS478113.1_Estonia_2014        ................................................................................ [133440]
KP843857.1_Odintsovo_02/14     ................................................................................ [133440]
FN557520.1_strain_E75          ............c.aa..t.c.......gttc....a.c....g....gtc.g.a.......tc....t..tg.ca..ta [133440]
AY261366.1_Warthog             ............c............................................a.............t........ [133440]
AY261365.1_Warmbaths           ............c.aa..t.c.......gttc....a.c..c.g....gtc.g.a.......tc....t..tg.ca..ta [133440]
AY261364.1_Tengani_62          ............c.aa..t.c.......gttc....a.c..c.g....gtc.g.a.......t.....t..tg.ca..ta [133440]
AY261363.1_Pretoriuskop/96/4   ............c.......................a..................................t........ [133440]
AY261362.1_Mkuzi_1979          ............c.aa..t.c.......gttc....a.c..c.g....gtc.g.a.......tc....t..tg.ca..ta [133440]
AY261361.1_Malawi_Lil-20/1_1983 ............c..........................................................t........ [133440]
AY261360.1_Kenya_1950          ............c.ta..t.c.......attc....a.c..c.g....atc.g.a......at...c.t.gtgata..t. [133440]
AM712240.1_OURT_88/3           ............c.aa..t.c.......gttc....a.c....g....gtc.g.a.......tc....t..tg.ca..ta [133440]
AM712239.1_Benin_97/1          ............c.aa..t.c.......gttc....a.c....g....gtc.g.a.......tc....t..tg.ca..ta [133440]
KM111295.1_Ken06.Bus           .......t............c......g......................t...............c...tg....... [133440]
KM111294.1_Ken05/Tk1           ............c.aa..t.c.......attc....a.c..c.g....atc.g.a.......t...c.t.gtgata..t. [133440]
KM262844.1_strain_L60          ............c.aa..t.c.......gttc....a.c....g....gtc.g.a.......tc....t..tg.ca..ta [133440]
KM102979.1_26544/OG10_Italy    ............c.aa..t.c.......gttc....a.c....g....gtc.g.a.......tc....t..tg.ca..ta [133440]
KM262845.1_strain_NHV          ............c.aa..t.c.......gttc....a.c....g....gtc.g.a.......tc....t..tg.ca..ta [133440]
KX354450.1_47/Ss/2008          ............c.aa..t.c.......gttc....a.c....g....gtc.g.a.......tc....t..tg.ca..ta [133440]
KP055815.1_strain_BA71         ............c.aa..t.c.......gttc....a.c....g....gtc.g.a.......tc....t..tg.ca..ta [133440]
MH025920.1_strain_R35          .......t............c......g......................t...............c...tg....... [133440]
MH025919.1_strain_N10          .......t............c......g......................t...............c...tg....... [133440]
MH025918.1_strain_R25          .......t............c......g......................t...............c...tg....... [133440]
MH025917.1_strain_R7           .......t............c......g......................t...............c...tg....... [133440]
MH025916.1_strain_R8           .......t............c......g......................t...............c...tg....... [133440]
```

```
FR682468.2_Georgia_2007/1        ggtttataaccattgtcataaaccattgcattgcttcaatatcattttgtaatgcttgacggggaggcggggcaggtaat [133520]
ASF_IND_20_CAD543                t.....a..tat.nnnnnnnnnnnnnnnnnnnnnnnnnnnnnnnnnnnnnnnnnnnnnnnnnnnnnnnnnnnnnnn.g.t. [133520]
LR899193.1_Germany_2020/1        ................................................................................ [133520]
MN393477.1_Wuhan_2019            ................................................................................ [133520]
MW396979.1_Timor-Leste/2019/1    ................................................................................ [133520]
MK940252.1_CN/2019/InnerMongolia ................................................................................ [133520]
MK128995.1_China/2018/AnhuiXCGQ  ................................................................................ [133520]
MH681419.1_ASFV/POL/2015         ................................................................................ [133520]
LS478113.1_Estonia_2014          ................................................................................ [133520]
KP843857.1_Odintsovo_02/14       ................................................................................ [133520]
FN557520.1_strain_E75            t.....a..tat.g..............ag.c......tc....g.......g.atc.........t.t..aa...g.t. [133520]
AY261366.1_Warthog               ..........t............................................g.........g.............. [133520]
AY261365.1_Warmbaths             t.....a..tat.g..............ag.c......tc....g.......g.atc.........t.t..aa...g.t. [133520]
AY261364.1_Tengani_62            t........tat.g..............ag.c......tc....g.......g.atc........tt.t..aa...g.t. [133520]
AY261363.1_Pretoriuskop/96/4     ....................................................t.a......................... [133520]
AY261362.1_Mkuzi_1979            t.....a..tat.g..............ag.c......tc....g.......g.atc.........t.t..aa...g.t. [133520]
AY261361.1_Malawi_Lil-20/1_1983  ..........t.................t.........................a......................... [133520]
AY261360.1_Kenya_1950            .a.......tat.g..............ag.c......tcg..gg...........c...tc....tca..aa...g.g. [133520]
AM712240.1_OURT_88/3             t.....a..tat.g..............ag.c......tc....g.......g.atc.........t.t..aa...g.t. [133520]
AM712239.1_Benin_97/1            t.....a..tat.g..............ag.c......tc....g.......g.atc.........t.t..aa...g.t. [133520]
KM111295.1_Ken06.Bus             ............................a.gc......................tg..........acg.........t. [133520]
KM111294.1_Ken05/Tk1             .a.......tat.g..............ag.c......tcg..gg...........c...tc....tca..aa...g.g. [133520]
KM262844.1_strain_L60            t.....a..tat.g..............ag.c......tc....g.......g.atc.........t.t..aa...g.t. [133520]
KM102979.1_26544/OG10_Italy      t.....a..tat.g..............ag.c......tc....g.......g.atc.........t.t..aa...g.t. [133520]
KM262845.1_strain_NHV            t.....a..tat.g..............ag.c......tc....g.......g.atc.........t.t..aa...g.t. [133520]
KX354450.1_47/Ss/2008            t.....a..tat.g..............ag.c......tc....g.......g.atc.........t.t..aa...g.t. [133520]
KP055815.1_strain_BA71           t.....a..tat.g..............ag.c......tc....g.......g.atc.........t.t..aa...g.t. [133520]
MH025920.1_strain_R35            ............................a.gc......................tg..........acg.........t. [133520]
MH025919.1_strain_N10            ............................a.gc......................tg..........acg.........t. [133520]
MH025918.1_strain_R25            ............................a.gc......................tg..........acg.........t. [133520]
MH025917.1_strain_R7             ............................a.gc......................tg..........acg.........t. [133520]
MH025916.1_strain_R8             ............................a.gc......................tg..........acg.........t. [133520]

FR682468.2_Georgia_2007/1        ccacgtatgttgaataaagcggttaattgtgcaccggctgtttggggcgtaatattttgtattaaatttatcatcgaatt [133600]
ASF_IND_20_CAD543                ...t..c.a..a..nnnnnnnnnnnnnnnnnnnnnnnnnnnnnnnnnnnnnnnnnnnnnnnnnnnnnnnnnnnnnnnnnn [133600]
LR899193.1_Germany_2020/1        ................................................................................ [133600]
MN393477.1_Wuhan_2019            ................................................................................ [133600]
MW396979.1_Timor-Leste/2019/1    ................................................................................ [133600]
MK940252.1_CN/2019/InnerMongolia ................................................................................ [133600]
MK128995.1_China/2018/AnhuiXCGQ  ................................................................................ [133600]
MH681419.1_ASFV/POL/2015         ................................................................................ [133600]
LS478113.1_Estonia_2014          ................................................................................ [133600]
KP843857.1_Odintsovo_02/14       ................................................................................ [133600]
```

```
FN557520.1_strain_E75            ...t..c.a..a..a.tga..cc.........t.gca.....ac..........g..........g.c..a...t..... [133600]
AY261366.1_Warthog               ....................a........................................................... [133600]
AY261365.1_Warmbaths             ...t..c.a..a..a.tga..cc.........t.gca.....gc.....................t.c..a...t..... [133600]
AY261364.1_Tengani_62            ...t..c.a..a..a.tga..cc.........t.gca.....ac.....................t.c..a...t..... [133600]
AY261363.1_Pretoriuskop/96/4     .....................a.......................................................... [133600]
AY261362.1_Mkuzi_1979            ...t..c.a..a..a.tga..cc.........t.gca.....gc..........g..........t.c..a...t..... [133600]
AY261361.1_Malawi_Lil-20/1_1983  ......c.a.....c.....a..............a.........acgt.................ga............ [133600]
AY261360.1_Kenya_1950            ...t..c.aa....a.cg.t............t.gca.....a......................c.g.....g...... [133600]
AM712240.1_OURT_88/3             ...t..c.a..a..a.tga..cc.........t.gca.....ac..........g..........g.c..a...t..... [133600]
AM712239.1_Benin_97/1            ...t..c.a..a..a.tga..cc.........t.gca.....ac..........g..........g.c..a...t..... [133600]
KM111295.1_Ken06.Bus             ......c.a..a..c.....a............g............cg......a...........gg............ [133600]
KM111294.1_Ken05/Tk1             ...t..c.aa....a.cg.t............t.gca.....a......................c.g.....a...... [133600]
KM262844.1_strain_L60            ...t..c.a..a..a.tga..cc.........t.gca.....ac..........g..........g.c..a...t..... [133600]
KM102979.1_26544/OG10_Italy      ...t..c.a..a..a.tga..cc.........t.gca.....ac..........g..........g.c..a...t..... [133600]
KM262845.1_strain_NHV            ...t..c.a..a..a.tga..cc.........t.gca.....ac..........g..........g.c..a...t..... [133600]
KX354450.1_47/Ss/2008            ...t..c.a..a..a.tga..cc.........t.gca.....ac..........g..........g.c..a...t..... [133600]
KP055815.1_strain_BA71           ...t..c.a..a..a.tga..cc.........t.gca.....ac..........g..........g.c..a...t..... [133600]
MH025920.1_strain_R35            ......c.a..a..c.....a............g............cg......a...........gg............ [133600]
MH025919.1_strain_N10            ......c.a..a..c.....a............g............cg......a...........gg............ [133600]
MH025918.1_strain_R25            ......c.a..a..c.....a............g............cg......a...........gg............ [133600]
MH025917.1_strain_R7             ......c.a..a..c.....a............g............cg......a...........gg............ [133600]
MH025916.1_strain_R8             ......c.a..a..c.....a............g............cg......a...........gg............ [133600]

FR682468.2_Georgia_2007/1        ---ggcttgcccggcatttcctataagatcgattaaattggttatttgacctcgatattgttgtacccagttttgaatgg [133680]
ASF_IND_20_CAD543                ---nnn......aat...........a..ta...nnnnnnnnnnnnnnnnnnnng.....c...gt..........gc.. [133680]
LR899193.1_Germany_2020/1        ---............................................................................. [133680]
MN393477.1_Wuhan_2019            ---............................................................................. [133680]
MW396979.1_Timor-Leste/2019/1    ---............................................................................. [133680]
MK940252.1_CN/2019/InnerMongolia ---............................................................................. [133680]
MK128995.1_China/2018/AnhuiXCGQ  ---............................................................................. [133680]
MH681419.1_ASFV/POL/2015         ---............................................................................. [133680]
LS478113.1_Estonia_2014          ---............................................................................. [133680]
KP843857.1_Odintsovo_02/14       ---............................................................................. [133680]
FN557520.1_strain_E75            ---..g......aat...........a..ta.....gg.atc..ccc..g....g.....c...gt..........gc.. [133680]
AY261366.1_Warthog               ---....................................a........................................ [133680]
AY261365.1_Warmbaths             ---..g......aat...........a..ta.....gg.atc..ccc..g....g.....c...g...........gc.. [133680]
AY261364.1_Tengani_62            ---..g......aat...........a..ta.....ga.atc..ccc..g....g.....c...g...........gc.. [133680]
AY261363.1_Pretoriuskop/96/4     ---......t..ta........................c.........................c............c.. [133680]
AY261362.1_Mkuzi_1979            ---..g......aat...........a..ta.....gg.atc..ccc..g....g.....c...g...........gc.. [133680]
AY261361.1_Malawi_Lil-20/1_1983  ---......t..........................g.c..........g..............c............c.. [133680]
AY261360.1_Kenya_1950            ---..g....tga.............a..ta.....gaaatc..ccc..g....g.....c...gt....g.....gc.. [133680]
AM712240.1_OURT_88/3             ---..g......aat...........a..ta.....gg.atc..ccc..g....g.....c...gt..........gc.. [133680]
AM712239.1_Benin_97/1            ---..g......aat...........a..ta.....gg.atc..ccc..g....g.....c...gt..........gc.. [133680]
KM111295.1_Ken06.Bus             gcc.c.......a.......................g.c.........................c..........c.c.. [133680]
```

```
KM111294.1_Ken05/Tk1            ---..g....tga.............a..ta.....gaaatc..ccc..g....g.....c...g.....g.....gc.. [133680]
KM262844.1_strain_L60           ---..g......aat...........a..ta.....gg.atc..ccc..g....g.....c...gt..........gc.. [133680]
KM102979.1_26544/OG10_Italy     ---..g......aat...........a..ta.....gg.atc..ccc..g....g.....c...gt..........gc.. [133680]
KM262845.1_strain_NHV           ---..g......aat...........a..ta.....gg.atc..ccc..g....g.....c...gt..........gc.. [133680]
KX354450.1_47/Ss/2008           ---..g......aat...........a..ta.....gg.atc..ccc..g....g.....c...gt..........gc.. [133680]
KP055815.1_strain_BA71          ---..g......aat...........a..ta.....gg.atc..ccc..g....g.....c...gt..........gc.. [133680]
MH025920.1_strain_R35           gcc.c.......a.......................g.c.........................c..........c.c.. [133680]
MH025919.1_strain_N10           gcc.c.......a.......................g.c.........................c..........c.c.. [133680]
MH025918.1_strain_R25           gcc.c.......a.......................g.c.........................c..........c.c.. [133680]
MH025917.1_strain_R7            gcc.c.......a.......................g.c.........................c..........c.c.. [133680]
MH025916.1_strain_R8            gcc.c.......a.......................g.c.........................c..........c.c.. [133680]

FR682468.2_Georgia_2007/1       cagcgatgatctcaggggttggattgttttgaatttcaggtgtttgtattagattattcacttctcttcgtgtatcttca [133760]
ASF_IND_20_CAD543               .t..a....c..............................w........nnnnnnnnnnnnnnnnnnnnnnnnnnnnnnn [133760]
LR899193.1_Germany_2020/1       ................................................................................ [133760]
MN393477.1_Wuhan_2019           ................................................................................ [133760]
MW396979.1_Timor-Leste/2019/1   ................................................................................ [133760]
MK940252.1_CN/2019/InnerMongolia ................................................................................ [133760]
MK128995.1_China/2018/AnhuiXCGQ ................................................................................ [133760]
MH681419.1_ASFV/POL/2015        ................................................................................ [133760]
LS478113.1_Estonia_2014         ................................................................................ [133760]
KP843857.1_Odintsovo_02/14      ................................................................................ [133760]
FN557520.1_strain_E75           .t..a....c..............................a........c.tg..t.ca...........ca..ct.a.. [133760]
AY261366.1_Warthog              ................................................................................ [133760]
AY261365.1_Warmbaths            .t..a....c..............................a........c.tg..t.ca...........ca..ct.a.. [133760]
AY261364.1_Tengani_62           .t..a....c..............................a.......ac.tg..t.ca...........ca..ct.a.. [133760]
AY261363.1_Pretoriuskop/96/4    .............g..a............c.................................................. [133760]
AY261362.1_Mkuzi_1979           .t..a....c..............................a........c.tg..t.ca...........ca..ct.a.. [133760]
AY261361.1_Malawi_Lil-20/1_1983 ................................................c...................a........... [133760]
AY261360.1_Kenya_1950           .t.......c......................................cc.tg..t.ca..........tca.cct.a.. [133760]
AM712240.1_OURT_88/3            .t..a....c..............................a........c.tg..t.ca...........ca..ct.a.. [133760]
AM712239.1_Benin_97/1           .t..a....c..............................a........c.tg..t.ca...........ca..ct.a.. [133760]
KM111295.1_Ken06.Bus            ..........t......a..ac...a..............c.......a........ct..................... [133760]
KM111294.1_Ken05/Tk1            .t.......c......................................cc.tg..t.ca..........tca.cct.a.. [133760]
KM262844.1_strain_L60           .t..a....c..............................a........c.tg..t.ca...........ca..ct.a.. [133760]
KM102979.1_26544/OG10_Italy     .t..a....c..............................a........c.tg..t.ca...........ca..ct.a.. [133760]
KM262845.1_strain_NHV           .t..a....c..............................a........c.tg..t.ca...........ca..ct.a.. [133760]
KX354450.1_47/Ss/2008           .t..a....c..............................a........c.tg..t.ca...........ca..ct.a.. [133760]
KP055815.1_strain_BA71          .t..a....c..............................a........c.tg..t.ca...........ca..ct.a.. [133760]
MH025920.1_strain_R35           ..........t......a..ac...a..............c.......a........ct..................... [133760]
MH025919.1_strain_N10           ..........t......a..ac...a..............c.......a........ct..................... [133760]
MH025918.1_strain_R25           ..........t......a..ac...a..............c.......a........ct..................... [133760]
MH025917.1_strain_R7            ..........t......a..ac...a..............c.......a........ct..................... [133760]
MH025916.1_strain_R8            ..........t......a..ac...a..............c.......a........ct..................... [133760]
```

```
FR682468.2_Georgia_2007/1        agctgagtcctaaatgcatttaactcgcctataatttggtttctatcaataacatttcttaaacctcgaactgtttcag- [133840]
ASF_IND_20_CAD543                nnnnnnnnnnnnnnnnnnnnnnnnnnnnnnnnnnnnnnnn..t..a......t........g.a......g.t.c.g..- [133840]
LR899193.1_Germany_2020/1        ...............................................................................- [133840]
MN393477.1_Wuhan_2019            ...............................................................................- [133840]
MW396979.1_Timor-Leste/2019/1    ...............................................................................- [133840]
MK940252.1_CN/2019/InnerMongolia ...............................................................................- [133840]
MK128995.1_China/2018/AnhuiXCGQ  ...............................................................................- [133840]
MH681419.1_ASFV/POL/2015         ...............................................................................- [133840]
LS478113.1_Estonia_2014          ...............................................................................- [133840]
KP843857.1_Odintsovo_02/14       ...............................................................................- [133840]
FN557520.1_strain_E75            .....ctct....t..t.g....a..cg......g.......t..a......t........g.a......g.t.c.g..- [133840]
AY261366.1_Warthog               ...............................................................................- [133840]
AY261365.1_Warmbaths             .....ctct....t..t.g....a..cg......g.......t..a......t........g.a......g.t.c.g..- [133840]
AY261364.1_Tengani_62            .....ctct....t..t.g....a..cg......g.......t..a......t........g.a......g.t.c.g..- [133840]
AY261363.1_Pretoriuskop/96/4     ...............................................................................- [133840]
AY261362.1_Mkuzi_1979            .....ctct....t..t.g....a..cg......g.......t..a......t........g.a......g.t.c.g..- [133840]
AY261361.1_Malawi_Lil-20/1_1983  ...............................................................................- [133840]
AY261360.1_Kenya_1950            .....ctct....t....gc...a..cg......g.......t..a.c....t........g.a......tat.c.g..g [133840]
AM712240.1_OURT_88/3             .....ctct....t..t.g....a..cg......g.......t..a......t........g.a......g.t.c.g..- [133840]
AM712239.1_Benin_97/1            .....ctct....t..t.g....a..cg......g.......t..a......t........g.a......g.t.c.g..- [133840]
KM111295.1_Ken06.Bus             ..t.....t................at...........................................c........- [133840]
KM111294.1_Ken05/Tk1             .....ctct....t....gc...a..cg......g.......t..a.c....t........g.a......gat.c.g..- [133840]
KM262844.1_strain_L60            .....ctct....t..t.g....a..cg......g.......t..a......t........g.a......g.t.c.g..- [133840]
KM102979.1_26544/OG10_Italy      .....ctct....t..t.g....a..cg......g.......t..a......t........g.a......g.t.c.g..- [133840]
KM262845.1_strain_NHV            .....ctct....t..t.g....a..cg......g.......t..a......t........g.a......g.t.c.g..- [133840]
KX354450.1_47/Ss/2008            .....ctct....t..t.g....a..cg......g.......t..a......t........g.a......g.t.c.g..- [133840]
KP055815.1_strain_BA71           .....ctct....t..t.g....a..cg......g.......t..a......t........g.a......g.t.c.g..- [133840]
MH025920.1_strain_R35            ..t.....t................at...........................................c........- [133840]
MH025919.1_strain_N10            ..t.....t................at...........................................c........- [133840]
MH025918.1_strain_R25            ..t.....t................at...........................................c........- [133840]
MH025917.1_strain_R7             ..t.....t................at...........................................c........- [133840]
MH025916.1_strain_R8             ..t.....t................at...........................................c........- [133840]

FR682468.2_Georgia_2007/1        --ccaatcgtatagtacgcacaatttcatgtaaggcctggtttatgtatattgacatgggatggccccaccgctcacgtc [133920]
ASF_IND_20_CAD543                --nnnnnnnnnnnnnnnnnnnnnnnnnnnnnnnnnnnnnnnnnnnnnnnnnnnnnnnnnnnnnnnnnnnnnnn....... [133920]
LR899193.1_Germany_2020/1        --.............................................................................. [133920]
MN393477.1_Wuhan_2019            --.............................................................................. [133920]
MW396979.1_Timor-Leste/2019/1    --.............................................................................. [133920]
MK940252.1_CN/2019/InnerMongolia --.............................................................................. [133920]
MK128995.1_China/2018/AnhuiXCGQ  --.............................................................................. [133920]
MH681419.1_ASFV/POL/2015         --.............................................................................. [133920]
LS478113.1_Estonia_2014          --.............................................................................. [133920]
KP843857.1_Odintsovo_02/14       --.............................................................................. [133920]
```

```
FN557520.1_strain_E75            --gt.t..t...g.....g....g.........a.....a..g......tc...t.a...t..c.....tt.t....... [133920]
AY261366.1_Warthog               --.............................................................................. [133920]
AY261365.1_Warmbaths             --gt.t..t...g.....g....g.........a.....a..g......tc...t.a...t..c......t.t....... [133920]
AY261364.1_Tengani_62            --gt.t..t...g.....g....g.........a.....a..g......tc...t.a...t..c......t.t....... [133920]
AY261363.1_Pretoriuskop/96/4     --.............................................................................. [133920]
AY261362.1_Mkuzi_1979            --gt.t..t...g.....g....g.........a.....a..g......tc...t.a...t..c......t.t....... [133920]
AY261361.1_Malawi_Lil-20/1_1983  --.............................................................................. [133920]
AY261360.1_Kenya_1950            tatt.t..t...g.....g....g.........a.....a..g..a..atc.....a...c..c......t......... [133920]
AM712240.1_OURT_88/3             --gt.t..t...g.....g....g.........a.....a..g......tc...t.a...t..c.....tt.t....... [133920]
AM712239.1_Benin_97/1            --gt.t..t...g.....g....g.........a.....a..g......tc...t.a...t..c.....tt.t....... [133920]
KM111295.1_Ken06.Bus             --.t........g.....g....c...g.....a.....a.......................c.....t........g. [133920]
KM111294.1_Ken05/Tk1             --gt.t..t...g.....g....g.........a.....a..g.....atc...a.a...c..c......t......... [133920]
KM262844.1_strain_L60            --gt.t..t...g.....g....g.........a.....a..g......tc...t.a...t..c.....tt.t....... [133920]
KM102979.1_26544/OG10_Italy      --gt.t..t...g.....g....g.........a.....a..g......tc...t.a...t..c.....tt.t....... [133920]
KM262845.1_strain_NHV            --gt.t..t...g.....g....g.........a.....a..g......tc...t.a...t..c.....tt.t....... [133920]
KX354450.1_47/Ss/2008            --gt.t..t...g.....g....g.........a.....a..g......tc...t.a...t..c.....tt.t....... [133920]
KP055815.1_strain_BA71           --gt.t..t...g.....g....g.........a.....a..g......tc...t.a...t..c.....tt.t....... [133920]
MH025920.1_strain_R35            --.t........g.....g....c...g.....a.....a.......................c.....t........g. [133920]
MH025919.1_strain_N10            --.t........g.....g....c...g.....a.....a.......................c.....t........g. [133920]
MH025918.1_strain_R25            --.t........g.....g....c...g.....a.....a.......................c.....t........g. [133920]
MH025917.1_strain_R7             --.t........g.....g....c...g.....a.....a.......................c.....t........g. [133920]
MH025916.1_strain_R8             --.t........g.....g....c...g.....a.....a.......................c.....t........g. [133920]

FR682468.2_Georgia_2007/1        cacgttgaatacctgcggccaaactaggacctgcctcgtcataatcaaattgtgtaggataaaggcttccaaatagcact [134000]
ASF_IND_20_CAD543                ...........................................................................r.... [134000]
LR899193.1_Germany_2020/1        ................................................................................ [134000]
MN393477.1_Wuhan_2019            ................................................................................ [134000]
MW396979.1_Timor-Leste/2019/1    ................................................................................ [134000]
MK940252.1_CN/2019/InnerMongolia ................................................................................ [134000]
MK128995.1_China/2018/AnhuiXCGQ  ................................................................................ [134000]
MH681419.1_ASFV/POL/2015         ................................................................................ [134000]
LS478113.1_Estonia_2014          ................................................................................ [134000]
KP843857.1_Odintsovo_02/14       ................................................................................ [134000]
FN557520.1_strain_E75            ........................cc..g.....t..a.....g.....c.....g..g..c..a..........a.... [134000]
AY261366.1_Warthog               ........................cc..g.....t..a.....g.....c.....g.....c..a..........a.... [134000]
AY261365.1_Warmbaths             ........................cc..g.....t..a.....g.....c.....g..g..c..a..........a.... [134000]
AY261364.1_Tengani_62            ........................cc..g.....t..a.....g.....c.....g.....c..a..........a.... [134000]
AY261363.1_Pretoriuskop/96/4     .........................g...................................................... [134000]
AY261362.1_Mkuzi_1979            ........................cc..g.....t..a.....g.....c..g..g..g..c..a..........a.... [134000]
AY261361.1_Malawi_Lil-20/1_1983  .........................g...................................................... [134000]
AY261360.1_Kenya_1950            .............c..........cc..g........a.....g.....c...........c..a.....g.....t... [134000]
AM712240.1_OURT_88/3             ........................cc..g.....t..a.....g.....c.....g..g..c..a..........a.... [134000]
AM712239.1_Benin_97/1            ........................cc..g.....t..a.....g.....c.....g..g..c..a..........a.... [134000]
KM111295.1_Ken06.Bus             .g...........c..........cc..g........a.....g.....c.....g.....c.............a.... [134000]
```

```
KM111294.1_Ken05/Tk1             .............c..........cc..g........a.....g.....c...........c..a.....g.....t... [134000]
KM262844.1_strain_L60            ........................cc..g.....t..a.....g.....c.....g..g..c..a..........a.... [134000]
KM102979.1_26544/OG10_Italy      ........................cc..g.....t..a.....g.....c.....g..g..c..a..........a.... [134000]
KM262845.1_strain_NHV            ........................cc..g.....t..a.....g.....c.....g..g..c..a..........a.... [134000]
KX354450.1_47/Ss/2008            ........................cc..g.....t..a.....g.....c.....g..g..c..a..........a.... [134000]
KP055815.1_strain_BA71           ........................cc..g.....t..a.....g.....c.....g..g..c..a..........a.... [134000]
MH025920.1_strain_R35            .g...........c..........cc..g........a.....g.....c.....g.....c.............a.... [134000]
MH025919.1_strain_N10            .g...........c..........cc..g........a.....g.....c.....g.....c.............a.... [134000]
MH025918.1_strain_R25            .g...........c..........cc..g........a.....g.....c.....g.....c.............a.... [134000]
MH025917.1_strain_R7             .g...........c..........cc..g........a.....g.....c.....g.....c.............a.... [134000]
MH025916.1_strain_R8             .g...........c..........cc..g........a.....g.....c.....g.....c.............a.... [134000]

FR682468.2_Georgia_2007/1        ttattgaaaatttggtcagaaagaaatttagggcggcccatatttagcgcgttgtcccctctaaagatgcgtgacatgta [134080]
ASF_IND_20_CAD543                ................................................................................ [134080]
LR899193.1_Germany_2020/1        ................................................................................ [134080]
MN393477.1_Wuhan_2019            ................................................................................ [134080]
MW396979.1_Timor-Leste/2019/1    ................................................................................ [134080]
MK940252.1_CN/2019/InnerMongolia ................................................................................ [134080]
MK128995.1_China/2018/AnhuiXCGQ  ................................................................................ [134080]
MH681419.1_ASFV/POL/2015         ................................................................................ [134080]
LS478113.1_Estonia_2014          ................................................................................ [134080]
KP843857.1_Odintsovo_02/14       ................................................................................ [134080]
FN557520.1_strain_E75            .....a...........g........c.....a........g..c.....a..a..t........a...........a.. [134080]
AY261366.1_Warthog               .....a...........g........c.....a...........c.....a..a..t........a.............. [134080]
AY261365.1_Warmbaths             .....a...........g........c.....a........g..c.....a..a..t........a...........a.. [134080]
AY261364.1_Tengani_62            .....a...........g........c.....a........g..c.....a..a..t........a...........a.. [134080]
AY261363.1_Pretoriuskop/96/4     .................g................................a..a.......................... [134080]
AY261362.1_Mkuzi_1979            .....a...........g........c.....a........g..c.....a..a..t........a...........a.. [134080]
AY261361.1_Malawi_Lil-20/1_1983  .................g................................a............................. [134080]
AY261360.1_Kenya_1950            .................g........c.....a........g..c.....a............................. [134080]
AM712240.1_OURT_88/3             .....a...........g........c.....a........g..c.....a..a..t........a...........a.. [134080]
AM712239.1_Benin_97/1            .....a...........g........c.....a........g..c.....a..a..t........a...........a.. [134080]
KM111295.1_Ken06.Bus             ...............a.g........c..............g..c..t..a.....t....................... [134080]
KM111294.1_Ken05/Tk1             .....a...........g........c.....a........g..c.....a............................. [134080]
KM262844.1_strain_L60            .....a...........g........c.....a........g..c.....a..a..t........a...........a.. [134080]
KM102979.1_26544/OG10_Italy      .....a...........g........c.....a........g..c.....a..a..t........a...........a.. [134080]
KM262845.1_strain_NHV            .....a...........g........c.....a........g..c.....a..a..t........a...........a.. [134080]
KX354450.1_47/Ss/2008            .....a...........g........c.....a........g..c.....a..a..t........a...........a.. [134080]
KP055815.1_strain_BA71           .....a...........g........c.....a........g..c.....a..a..t........a...........a.. [134080]
MH025920.1_strain_R35            ...............a.g........c..............g..c..t..a.....t....................... [134080]
MH025919.1_strain_N10            ...............a.g........c..............g..c..t..a.....t....................... [134080]
MH025918.1_strain_R25            ...............a.g........c..............g..c..t..a.....t....................... [134080]
MH025917.1_strain_R7             ...............a.g........c..............g..c..t..a.....t....................... [134080]
MH025916.1_strain_R8             ...............a.g........c..............g..c..t..a.....t....................... [134080]
```

```
FR682468.2_Georgia_2007/1        tccggcgttgcctttggatagtaactcattcccatattgagtaatagagaccgagacataggggtttataagaagtttta [134160]
ASF_IND_20_CAD543                ...nnnnnnnnnnnnnnnnnnnnnnnnnnnn................................................. [134160]
LR899193.1_Germany_2020/1        ................................................................................ [134160]
MN393477.1_Wuhan_2019            ................................................................................ [134160]
MW396979.1_Timor-Leste/2019/1    ................................................................................ [134160]
MK940252.1_CN/2019/InnerMongolia ................................................................................ [134160]
MK128995.1_China/2018/AnhuiXCGQ  ................................................................................ [134160]
MH681419.1_ASFV/POL/2015         ................................................................................ [134160]
LS478113.1_Estonia_2014          ................................................................................ [134160]
KP843857.1_Odintsovo_02/14       ................................................................................ [134160]
FN557520.1_strain_E75            ...a..ac....c...ctc.tc..t..g..t........g..g........t..a......................... [134160]
AY261366.1_Warthog               ...................................................t..a......................... [134160]
AY261365.1_Warmbaths             ...a..ac....c...ctc.tc..t..g..t........g..g........t..a......................... [134160]
AY261364.1_Tengani_62            ...a..ac....c...ctc.tc..t..g..t........g..g........t..a......................... [134160]
AY261363.1_Pretoriuskop/96/4     ...................................................t..a......................... [134160]
AY261362.1_Mkuzi_1979            ...a..ac....c...ctc.tc..t..g..t........g..g........t..a......................... [134160]
AY261361.1_Malawi_Lil-20/1_1983  ...............a......................................a.....a................... [134160]
AY261360.1_Kenya_1950            ............cc..a...a.........t.....c..g...........t.ta.................c....... [134160]
AM712240.1_OURT_88/3             ...a..ac....c...ctc.tc..t..g..t........g..g........t..a......................... [134160]
AM712239.1_Benin_97/1            ...a..ac....c...ctc.tc..t..g..t........g..g........t..a......................... [134160]
KM111295.1_Ken06.Bus             ......a.....cc.......c..............c..g.a.........t.ta.....a..a........g....... [134160]
KM111294.1_Ken05/Tk1             .....t......cc..a...a.........t.....c..g...........t.ta.................c....... [134160]
KM262844.1_strain_L60            ...a..ac....c...ctc.tc..t..g..t........g..g........t..a......................... [134160]
KM102979.1_26544/OG10_Italy      ...a..ac....c...ctc.tc..t..g..t........g..g........t..a......................... [134160]
KM262845.1_strain_NHV            ...a..ac....c...ctc.tc..t..g..t........g..g........t..a......................... [134160]
KX354450.1_47/Ss/2008            ...a..ac....c...ctc.tc..t..g..t........g..g........t..a......................... [134160]
KP055815.1_strain_BA71           ...a..ac....c...ctc.tc..t..g..t........g..g........t..a......................... [134160]
MH025920.1_strain_R35            ......a.....cc.......c..............c..g.a.........t.ta.....a..a........g....... [134160]
MH025919.1_strain_N10            ......a.....cc.......c..............c..g.a.........t.ta.....a..a........g....... [134160]
MH025918.1_strain_R25            ......a.....cc.......c..............c..g.a.........t.ta.....a..a........g....... [134160]
MH025917.1_strain_R7             ......a.....cc.......c..............c..g.a.........t.ta.....a..a........g....... [134160]
MH025916.1_strain_R8             ......a.....cc.......c..............c..g.a.........t.ta.....a..a........g....... [134160]

FR682468.2_Georgia_2007/1        gcataaattctcgagtatttatggggggacgattcggaatgtttaatacctctgcaacatctggttgaggagccgtggtg [134240]
ASF_IND_20_CAD543                ................................................................................ [134240]
LR899193.1_Germany_2020/1        ................................................................................ [134240]
MN393477.1_Wuhan_2019            ................................................................................ [134240]
MW396979.1_Timor-Leste/2019/1    ................................................................................ [134240]
MK940252.1_CN/2019/InnerMongolia ................................................................................ [134240]
MK128995.1_China/2018/AnhuiXCGQ  ................................................................................ [134240]
MH681419.1_ASFV/POL/2015         ................................................................................ [134240]
LS478113.1_Estonia_2014          ................................................................................ [134240]
KP843857.1_Odintsovo_02/14       ................................................................................ [134240]
```

```
FN557520.1_strain_E75             ................g..c.....................................t......c..........c.... [134240]
AY261366.1_Warthog                ................................................................c............... [134240]
AY261365.1_Warmbaths              a...............g..c..............t......................t......c..........c.... [134240]
AY261364.1_Tengani_62             ................g..c.....................................tg.....c..........c.... [134240]
AY261363.1_Pretoriuskop/96/4      ...................c.....................................tg.....c..........c.... [134240]
AY261362.1_Mkuzi_1979             ................g..c.....a...............................tg.....c..........c...a [134240]
AY261361.1_Malawi_Lil-20/1_1983   ................................................................c..........c.... [134240]
AY261360.1_Kenya_1950             ................g..c.....a.......................t.g............c..........c.... [134240]
AM712240.1_OURT_88/3              ................g..c.....................................t......c..........c.... [134240]
AM712239.1_Benin_97/1             ................g..c.....................................t......c..........c.... [134240]
KM111295.1_Ken06.Bus              a............c.....c....................a........ta.............c.........cc.... [134240]
KM111294.1_Ken05/Tk1              ................g..c.............................t.g............c..........c.... [134240]
KM262844.1_strain_L60             ................g..c.....................................t......c..........c.... [134240]
KM102979.1_26544/OG10_Italy       ................g..c.....................................t......c..........c.... [134240]
KM262845.1_strain_NHV             ................g..c.....................................t......c..........c.... [134240]
KX354450.1_47/Ss/2008             ................g..c.....................................t......c..........c.... [134240]
KP055815.1_strain_BA71            ................g..c.....................................t......c..........c.... [134240]
MH025920.1_strain_R35             a............c.....c....................a........ta.............c.........cc.... [134240]
MH025919.1_strain_N10             a............c.....c....................a........ta.............c.........cc.... [134240]
MH025918.1_strain_R25             a............c.....c....................a........ta.............c.........cc.... [134240]
MH025917.1_strain_R7              a............c.....c....................a........ta.............c.........cc.... [134240]
MH025916.1_strain_R8              a............c.....c....................a........ta.............c.........cc.... [134240]

FR682468.2_Georgia_2007/1         tccagagatcgtactttttcagccgaaatgccgtacataagacaagcaatttcttcaaaactatagtcatagttgtaaat [134320]
ASF_IND_20_CAD543                 .tt............................................................................. [134320]
LR899193.1_Germany_2020/1         ................................................................................ [134320]
MN393477.1_Wuhan_2019             ................................................................................ [134320]
MW396979.1_Timor-Leste/2019/1     ................................................................................ [134320]
MK940252.1_CN/2019/InnerMongolia  ................................................................................ [134320]
MK128995.1_China/2018/AnhuiXCGQ   ................................................................................ [134320]
MH681419.1_ASFV/POL/2015          ................................................................................ [134320]
LS478113.1_Estonia_2014           ................................................................................ [134320]
KP843857.1_Odintsovo_02/14        ................................................................................ [134320]
FN557520.1_strain_E75             .tt..c........c.....g........................................................... [134320]
AY261366.1_Warthog                .....c........c.....g........................................................... [134320]
AY261365.1_Warmbaths              .tt..c........c.....g........................................................... [134320]
AY261364.1_Tengani_62             .tt..c........c.....g........................................................... [134320]
AY261363.1_Pretoriuskop/96/4      .tt..c..........................a............................................... [134320]
AY261362.1_Mkuzi_1979             .tt..c........c................................................................. [134320]
AY261361.1_Malawi_Lil-20/1_1983   .tt..t........c.....g........................................................... [134320]
AY261360.1_Kenya_1950             .....c..a.....c.....g.......................g........c.....g........g........... [134320]
AM712240.1_OURT_88/3              .tt..c........c.....g........................................................... [134320]
AM712239.1_Benin_97/1             .tt..c........c.....g........................................................... [134320]
KM111295.1_Ken06.Bus              ..t..c..a.....c.....g......................................g........g........... [134320]
```

```
KM111294.1_Ken05/Tk1            .....c..a.....c.....g.......................g........c.....g........g........... [134320]
KM262844.1_strain_L60           .tt..c........c.....g........................................................... [134320]
KM102979.1_26544/OG10_Italy     .tt..c........c.....g........................................................... [134320]
KM262845.1_strain_NHV           .tt..c........c.....g........................................................... [134320]
KX354450.1_47/Ss/2008           .tt..c........c.....g........................................................... [134320]
KP055815.1_strain_BA71          .tt..c........c.....g........................................................... [134320]
MH025920.1_strain_R35           ..t..c..a.....c.....g......................................g........g........... [134320]
MH025919.1_strain_N10           ..t..c..a.....c.....g......................................g........g........... [134320]
MH025918.1_strain_R25           ..t..c..a.....c.....g......................................g........g........... [134320]
MH025917.1_strain_R7            ..t..c..a.....c.....g......................................g........g........... [134320]
MH025916.1_strain_R8            ..t..c..a.....c.....g......................................g........g........... [134320]

FR682468.2_Georgia_2007/1       attggcaagtggtatagatcgcatcagcgcatttacattgataggtataatattcatatcaaacaagttaaatatgcgct [134400]
ASF_IND_20_CAD543               ................................................................................ [134400]
LR899193.1_Germany_2020/1       ................................................................................ [134400]
MN393477.1_Wuhan_2019           ................................................................................ [134400]
MW396979.1_Timor-Leste/2019/1   ................................................................................ [134400]
MK940252.1_CN/2019/InnerMongolia ................................................................................ [134400]
MK128995.1_China/2018/AnhuiXCGQ ................................................................................ [134400]
MH681419.1_ASFV/POL/2015        ................................................................................ [134400]
LS478113.1_Estonia_2014         ................................................................................ [134400]
KP843857.1_Odintsovo_02/14      ................................................................................ [134400]
FN557520.1_strain_E75           ...c............................................................................ [134400]
AY261366.1_Warthog              ................................................................................ [134400]
AY261365.1_Warmbaths            ................................................................................ [134400]
AY261364.1_Tengani_62           ................................................................................ [134400]
AY261363.1_Pretoriuskop/96/4    ...c............................................................................ [134400]
AY261362.1_Mkuzi_1979           ..............................g................................................. [134400]
AY261361.1_Malawi_Lil-20/1_1983 ...........................t.................................................... [134400]
AY261360.1_Kenya_1950           ..................c........t..g..............c.................................. [134400]
AM712240.1_OURT_88/3            ...c............................................................................ [134400]
AM712239.1_Benin_97/1           ...c............................................................................ [134400]
KM111295.1_Ken06.Bus            ..................c..........................c.................................. [134400]
KM111294.1_Ken05/Tk1            ..................c........t..g..............c.................................. [134400]
KM262844.1_strain_L60           ...c............................................................................ [134400]
KM102979.1_26544/OG10_Italy     ...c............................................................................ [134400]
KM262845.1_strain_NHV           ...c............................................................................ [134400]
KX354450.1_47/Ss/2008           ...c............................................................................ [134400]
KP055815.1_strain_BA71          ...c............................................................................ [134400]
MH025920.1_strain_R35           ..................c..........................c.................................. [134400]
MH025919.1_strain_N10           ..................c..........................c.................................. [134400]
MH025918.1_strain_R25           ..................c..........................c.................................. [134400]
MH025917.1_strain_R7            ..................c..........................c.................................. [134400]
MH025916.1_strain_R8            ..................c..........................c.................................. [134400]
```

```
FR682468.2_Georgia_2007/1        cgcgctctctattagagccaagagtgcgtgtttgacctttcggcgacactattttgtgaatatgattgatttgctcctct [134480]
ASF_IND_20_CAD543                ................................................................................ [134480]
LR899193.1_Germany_2020/1        ................................................................................ [134480]
MN393477.1_Wuhan_2019            ................................................................................ [134480]
MW396979.1_Timor-Leste/2019/1    ................................................................................ [134480]
MK940252.1_CN/2019/InnerMongolia ................................................................................ [134480]
MK128995.1_China/2018/AnhuiXCGQ  ................................................................................ [134480]
MH681419.1_ASFV/POL/2015         ................................................................................ [134480]
LS478113.1_Estonia_2014          ................................................................................ [134480]
KP843857.1_Odintsovo_02/14       ................................................................................ [134480]
FN557520.1_strain_E75            ..................................g...........t................................. [134480]
AY261366.1_Warthog               ..................................g............................................. [134480]
AY261365.1_Warmbaths             ..................................g...........t................................. [134480]
AY261364.1_Tengani_62            ..................................g.....t....................................... [134480]
AY261363.1_Pretoriuskop/96/4     ..................................g............................................. [134480]
AY261362.1_Mkuzi_1979            ..................................g...........t................................. [134480]
AY261361.1_Malawi_Lil-20/1_1983  ..................................g...........t................t................ [134480]
AY261360.1_Kenya_1950            ...................t....a.........g.t.c........................t................ [134480]
AM712240.1_OURT_88/3             ..................................g...........t................................. [134480]
AM712239.1_Benin_97/1            ..................................g...........t................................. [134480]
KM111295.1_Ken06.Bus             .a.................t....a.........g.t......t..t................t................ [134480]
KM111294.1_Ken05/Tk1             ...................t....a.........g.t.c........................t................ [134480]
KM262844.1_strain_L60            ..................................g...........t................................. [134480]
KM102979.1_26544/OG10_Italy      ..................................g...........t................................. [134480]
KM262845.1_strain_NHV            ..................................g...........t................................. [134480]
KX354450.1_47/Ss/2008            ..................................g...........t................................. [134480]
KP055815.1_strain_BA71           ..................................g...........t................................. [134480]
MH025920.1_strain_R35            .a.................t....a.........g.t......t..t................t................ [134480]
MH025919.1_strain_N10            .a.................t....a.........g.t......t..t................t................ [134480]
MH025918.1_strain_R25            .a.................t....a.........g.t......t..t................t................ [134480]
MH025917.1_strain_R7             .a.................t....a.........g.t......t..t................t................ [134480]
MH025916.1_strain_R8             .a.................t....a.........g.t......t..t................t................ [134480]

FR682468.2_Georgia_2007/1        tggtaagagctttccacgaaggaaattacgtcttgcaatgttttacgaagcgaatacactgcattcatccctattcccgc [134560]
ASF_IND_20_CAD543                ................................................................................ [134560]
LR899193.1_Germany_2020/1        ................................................................................ [134560]
MN393477.1_Wuhan_2019            ................................................................................ [134560]
MW396979.1_Timor-Leste/2019/1    ................................................................................ [134560]
MK940252.1_CN/2019/InnerMongolia ................................................................................ [134560]
MK128995.1_China/2018/AnhuiXCGQ  ................................................................................ [134560]
MH681419.1_ASFV/POL/2015         ................................................................................ [134560]
LS478113.1_Estonia_2014          ................................................................................ [134560]
KP843857.1_Odintsovo_02/14       ................................................................................ [134560]
```

```
FN557520.1_strain_E75            ..........................c..................................................... [134560]
AY261366.1_Warthog               ..........................c.........................................t........... [134560]
AY261365.1_Warmbaths             ..........................a..................................................... [134560]
AY261364.1_Tengani_62            ..........................c............................................a........ [134560]
AY261363.1_Pretoriuskop/96/4     .................a........c.........................................t........... [134560]
AY261362.1_Mkuzi_1979            ..........................c..................................................... [134560]
AY261361.1_Malawi_Lil-20/1_1983  .....g....................c..........g............t............................. [134560]
AY261360.1_Kenya_1950            .....g....................c..a..c..t..g...........t..g........................t. [134560]
AM712240.1_OURT_88/3             ..........................c..................................................... [134560]
AM712239.1_Benin_97/1            ..........................c..................................................... [134560]
KM111295.1_Ken06.Bus             ..........................c..a......................................t........... [134560]
KM111294.1_Ken05/Tk1             .....g....................c..a..c..t..g...........t..g........................t. [134560]
KM262844.1_strain_L60            ..........................c..................................................... [134560]
KM102979.1_26544/OG10_Italy      ..........................c..................................................... [134560]
KM262845.1_strain_NHV            ..........................c..................................................... [134560]
KX354450.1_47/Ss/2008            ..........................c..................................................... [134560]
KP055815.1_strain_BA71           ..........................c..................................................... [134560]
MH025920.1_strain_R35            ..........................c..a......................................t........... [134560]
MH025919.1_strain_N10            ..........................c..a......................................t........... [134560]
MH025918.1_strain_R25            ..........................c..a......................................t........... [134560]
MH025917.1_strain_R7             ..........................c..a......................................t........... [134560]
MH025916.1_strain_R8             ..........................c..a......................................t........... [134560]

FR682468.2_Georgia_2007/1        tgttataatgggtttatcgtctctgttctcgctaataagattaactccaccaaaagtattttcattgtacatcatcactg [134640]
ASF_IND_20_CAD543                ................................................................................ [134640]
LR899193.1_Germany_2020/1        ................................................................................ [134640]
MN393477.1_Wuhan_2019            ................................................................................ [134640]
MW396979.1_Timor-Leste/2019/1    ................................................................................ [134640]
MK940252.1_CN/2019/InnerMongolia ................................................................................ [134640]
MK128995.1_China/2018/AnhuiXCGQ  ................................................................................ [134640]
MH681419.1_ASFV/POL/2015         ................................................................................ [134640]
LS478113.1_Estonia_2014          ................................................................................ [134640]
KP843857.1_Odintsovo_02/14       ................................................................................ [134640]
FN557520.1_strain_E75            .........a...................................................................... [134640]
AY261366.1_Warthog               ..................a.....a....................................................... [134640]
AY261365.1_Warmbaths             .........a...................................................................... [134640]
AY261364.1_Tengani_62            .........a...................................................................... [134640]
AY261363.1_Pretoriuskop/96/4     .........a...................................................................... [134640]
AY261362.1_Mkuzi_1979            .........a...................................................................... [134640]
AY261361.1_Malawi_Lil-20/1_1983  ..................a.t...a.c...a...........c.....g.....................g......... [134640]
AY261360.1_Kenya_1950            .........a..c.....a.......c...a.c...............g.....................g......... [134640]
AM712240.1_OURT_88/3             .........a...................................................................... [134640]
AM712239.1_Benin_97/1            .........a...................................................................... [134640]
KM111295.1_Ken06.Bus             ..................a...........a..............c........................g......... [134640]
```

```
KM111294.1_Ken05/Tk1             .........a..c.....a.......c...a.................g.....................g......... [134640]
KM262844.1_strain_L60            .........a...................................................................... [134640]
KM102979.1_26544/OG10_Italy      .........a...................................................................... [134640]
KM262845.1_strain_NHV            .........a...................................................................... [134640]
KX354450.1_47/Ss/2008            .........a...................................................................... [134640]
KP055815.1_strain_BA71           .........a...................................................................... [134640]
MH025920.1_strain_R35            ..................a...........a..............c........................g......... [134640]
MH025919.1_strain_N10            ..................a...........a..............c........................g......... [134640]
MH025918.1_strain_R25            ..................a...........a..............c........................g......... [134640]
MH025917.1_strain_R7             ..................a...........a..............c........................g......... [134640]
MH025916.1_strain_R8             ..................a...........a..............c........................g......... [134640]

FR682468.2_Georgia_2007/1        ttttaaaactacggatatttatgataaatcggagagcctgaatggcgtgggtataaaagtgttcaaatcgcgtgggagta [134720]
ASF_IND_20_CAD543                ................................................................................ [134720]
LR899193.1_Germany_2020/1        ................................................................................ [134720]
MN393477.1_Wuhan_2019            ................................................................................ [134720]
MW396979.1_Timor-Leste/2019/1    ................................................................................ [134720]
MK940252.1_CN/2019/InnerMongolia ................................................................................ [134720]
MK128995.1_China/2018/AnhuiXCGQ  ................................................................................ [134720]
MH681419.1_ASFV/POL/2015         ................................................................................ [134720]
LS478113.1_Estonia_2014          ................................................................................ [134720]
KP843857.1_Odintsovo_02/14       ................................................................................ [134720]
FN557520.1_strain_E75            .....................c.........a...........a..a.................g............... [134720]
AY261366.1_Warthog               ...........................................a..a................................. [134720]
AY261365.1_Warmbaths             .....................c.........a...........a.................................... [134720]
AY261364.1_Tengani_62            ...........................................a.................................... [134720]
AY261363.1_Pretoriuskop/96/4     .............a.............................a....................g............... [134720]
AY261362.1_Mkuzi_1979            ...........................................a....................g............... [134720]
AY261361.1_Malawi_Lil-20/1_1983  .....................ca.....c..............a..............a...........t..a.....g [134720]
AY261360.1_Kenya_1950            ............................c..............a......a................a...........g [134720]
AM712240.1_OURT_88/3             .....................c.........a...........a..a.................g............... [134720]
AM712239.1_Benin_97/1            .....................c.........a...........a..a.................g............... [134720]
KM111295.1_Ken06.Bus             ............................c.....................a................a............ [134720]
KM111294.1_Ken05/Tk1             ............................c..............a......a................a...........g [134720]
KM262844.1_strain_L60            .....................c.........a...........a..a.................g............... [134720]
KM102979.1_26544/OG10_Italy      .....................c.........a...........a..a.................g............... [134720]
KM262845.1_strain_NHV            .....................c.........a...........a..a.................g............... [134720]
KX354450.1_47/Ss/2008            .....................c.........a...........a..a.................g............... [134720]
KP055815.1_strain_BA71           .....................c.........a...........a..a.................g............... [134720]
MH025920.1_strain_R35            ............................c.....................a................a............ [134720]
MH025919.1_strain_N10            ............................c.....................a................a............ [134720]
MH025918.1_strain_R25            ............................c.....................a................a............ [134720]
MH025917.1_strain_R7             ............................c.....................a................a............ [134720]
MH025916.1_strain_R8             ............................c.....................a................a............ [134720]
```

```
FR682468.2_Georgia_2007/1        atttgttcgcgagcaactaccgtttcattatagtttttcatgataagctgtactccgggcatatctgagagctgtaccgg [134800]
ASF_IND_20_CAD543                ................................................................................ [134800]
LR899193.1_Germany_2020/1        ................................................................................ [134800]
MN393477.1_Wuhan_2019            ................................................................................ [134800]
MW396979.1_Timor-Leste/2019/1    ................................................................................ [134800]
MK940252.1_CN/2019/InnerMongolia ................................................................................ [134800]
MK128995.1_China/2018/AnhuiXCGQ  ................................................................................ [134800]
MH681419.1_ASFV/POL/2015         ................................................................................ [134800]
LS478113.1_Estonia_2014          ................................................................................ [134800]
KP843857.1_Odintsovo_02/14       ................................................................................ [134800]
FN557520.1_strain_E75            .........t.......c..t...........a............................................... [134800]
AY261366.1_Warthog               .................c..t..................................................t........ [134800]
AY261365.1_Warmbaths             .................c..t...........a............................................... [134800]
AY261364.1_Tengani_62            ................................a............................................... [134800]
AY261363.1_Pretoriuskop/96/4     .................c..t..........................t................................ [134800]
AY261362.1_Mkuzi_1979            .......................................................................t........ [134800]
AY261361.1_Malawi_Lil-20/1_1983  ....................a...................................a.....g.....a.....c..t.. [134800]
AY261360.1_Kenya_1950            ....................a................................c..............a........... [134800]
AM712240.1_OURT_88/3             .................c..t...........a............................................... [134800]
AM712239.1_Benin_97/1            .........t.......c..t...........a............................................... [134800]
KM111295.1_Ken06.Bus             ....................a................................c..............a..t........ [134800]
KM111294.1_Ken05/Tk1             ....................a................................c..............a........... [134800]
KM262844.1_strain_L60            .........t.......c..t...........a............................................... [134800]
KM102979.1_26544/OG10_Italy      .........t.......c..t...........a............................................... [134800]
KM262845.1_strain_NHV            .................c..t...........a............................................... [134800]
KX354450.1_47/Ss/2008            .........t.......c..t...........a............................................... [134800]
KP055815.1_strain_BA71           .................c..t...........a............................................... [134800]
MH025920.1_strain_R35            ....................a................................c..............a..t........ [134800]
MH025919.1_strain_N10            ....................a................................c..............a..t........ [134800]
MH025918.1_strain_R25            ....................a................................c..............a..t........ [134800]
MH025917.1_strain_R7             ....................a................................c..............a..t........ [134800]
MH025916.1_strain_R8             ....................a................................c..............a..t........ [134800]

FR682468.2_Georgia_2007/1        atcatttcccagtaattttcttgtgccgtatagtagtttaaactcgggggagccgctttcaaggttcgggtaaagaagag [134880]
ASF_IND_20_CAD543                ................................................................................ [134880]
LR899193.1_Germany_2020/1        ................................................................................ [134880]
MN393477.1_Wuhan_2019            ................................................................................ [134880]
MW396979.1_Timor-Leste/2019/1    ................................................................................ [134880]
MK940252.1_CN/2019/InnerMongolia ................................................................................ [134880]
MK128995.1_China/2018/AnhuiXCGQ  ................................................................................ [134880]
MH681419.1_ASFV/POL/2015         ................................................................................ [134880]
LS478113.1_Estonia_2014          ................................................................................ [134880]
KP843857.1_Odintsovo_02/14       ................................................................................ [134880]
```

```
FN557520.1_strain_E75             ..................................................................t............. [134880]
AY261366.1_Warthog                ..........................................t.......................g............. [134880]
AY261365.1_Warmbaths              ...................................a..............................t............. [134880]
AY261364.1_Tengani_62             ................................................................................ [134880]
AY261363.1_Pretoriuskop/96/4      ..................................................................t............. [134880]
AY261362.1_Mkuzi_1979             ..........................................t.......................t............. [134880]
AY261361.1_Malawi_Lil-20/1_1983   ..................................................................t............. [134880]
AY261360.1_Kenya_1950             ........................a........g.t............................................ [134880]
AM712240.1_OURT_88/3              ..................................................................t............. [134880]
AM712239.1_Benin_97/1             ..................................................................t............. [134880]
KM111295.1_Ken06.Bus              ........................a........g.t..................a......................... [134880]
KM111294.1_Ken05/Tk1              ........................a........g.t............................................ [134880]
KM262844.1_strain_L60             ..................................................................t............. [134880]
KM102979.1_26544/OG10_Italy       ..................................................................t............. [134880]
KM262845.1_strain_NHV             ..................................................................t............. [134880]
KX354450.1_47/Ss/2008             ..................................................................t............. [134880]
KP055815.1_strain_BA71            ..................................................................t............. [134880]
MH025920.1_strain_R35             ........................a........g.t..................a......................... [134880]
MH025919.1_strain_N10             ........................a........g.t..................a......................... [134880]
MH025918.1_strain_R25             ........................a........g.t..................a......................... [134880]
MH025917.1_strain_R7              ........................a........g.t..................a......................... [134880]
MH025916.1_strain_R8              ........................a........g.t..................a......................... [134880]

FR682468.2_Georgia_2007/1         gatcatatacctcattattttctattcttaggtcatgtaaataatagagcgaaagtgaaaatggcataagaggctcctta [134960]
ASF_IND_20_CAD543                 ................................................................................ [134960]
LR899193.1_Germany_2020/1         ................................................................................ [134960]
MN393477.1_Wuhan_2019             ................................................................................ [134960]
MW396979.1_Timor-Leste/2019/1     ................................................................................ [134960]
MK940252.1_CN/2019/InnerMongolia  ................................................................................ [134960]
MK128995.1_China/2018/AnhuiXCGQ   ................................................................................ [134960]
MH681419.1_ASFV/POL/2015          ................................................................................ [134960]
LS478113.1_Estonia_2014           ................................................................................ [134960]
KP843857.1_Odintsovo_02/14        ................................................................................ [134960]
FN557520.1_strain_E75             ...................c...............c.c.......................................... [134960]
AY261366.1_Warthog                ...................c...............c.c.............................g............ [134960]
AY261365.1_Warmbaths              ...................c...............c.c.......................................... [134960]
AY261364.1_Tengani_62             ...................................c............................................ [134960]
AY261363.1_Pretoriuskop/96/4      ...................c...............c.c...........t...........c.....g............ [134960]
AY261362.1_Mkuzi_1979             ...................c...............c.c.......................................... [134960]
AY261361.1_Malawi_Lil-20/1_1983   ...................c...............c.c...........t...........c.....g.....t...... [134960]
AY261360.1_Kenya_1950             ................g..c...............c.c...........t...........c.....g..g..t...... [134960]
AM712240.1_OURT_88/3              ...................c...............c.c.......................................... [134960]
AM712239.1_Benin_97/1             ...................c...............c.c.......................................... [134960]
KM111295.1_Ken06.Bus              ...................c..c............c.c...........t.................g.....t...... [134960]
```

```
KM111294.1_Ken05/Tk1             ................g..c...............c.c...........t...........c.....g.....t...... [134960]
KM262844.1_strain_L60            ...................c...............c.c.......................................... [134960]
KM102979.1_26544/OG10_Italy      ...................c...............c.c.......................................... [134960]
KM262845.1_strain_NHV            ...................c...............c.c.......................................... [134960]
KX354450.1_47/Ss/2008            ...................c...............c.c.......................................... [134960]
KP055815.1_strain_BA71           ...................c...............c.c.......................................... [134960]
MH025920.1_strain_R35            ...................c..c............c.c...........t.................g.....t...... [134960]
MH025919.1_strain_N10            ...................c..c............c.c...........t.................g.....t...... [134960]
MH025918.1_strain_R25            ...................c..c............c.c...........t.................g.....t...... [134960]
MH025917.1_strain_R7             ...................c..c............c.c...........t.................g.....t...... [134960]
MH025916.1_strain_R8             ...................c..c............c.c...........t.................g.....t...... [134960]

FR682468.2_Georgia_2007/1        ttgtaccgggacatatagttttgaatgaagtgttcttctgtttcaagatagatgggatgatcggtaagctcgtgcaggac [135040]
ASF_IND_20_CAD543                ................................................................................ [135040]
LR899193.1_Germany_2020/1        ................................................................................ [135040]
MN393477.1_Wuhan_2019            ................................................................................ [135040]
MW396979.1_Timor-Leste/2019/1    ................................................................................ [135040]
MK940252.1_CN/2019/InnerMongolia ................................................................................ [135040]
MK128995.1_China/2018/AnhuiXCGQ  ................................................................................ [135040]
MH681419.1_ASFV/POL/2015         ................................................................................ [135040]
LS478113.1_Estonia_2014          ................................................................................ [135040]
KP843857.1_Odintsovo_02/14       ................................................................................ [135040]
FN557520.1_strain_E75            ...........................................................g........t..a..t..... [135040]
AY261366.1_Warthog               ................................................................................ [135040]
AY261365.1_Warmbaths             .....t.....................................................g.................... [135040]
AY261364.1_Tengani_62            ................................................................................ [135040]
AY261363.1_Pretoriuskop/96/4     ...........................................................g........t..a..t..... [135040]
AY261362.1_Mkuzi_1979            .....t....................a................................g........t..a..t..... [135040]
AY261361.1_Malawi_Lil-20/1_1983  ..........................a.................................................aa.. [135040]
AY261360.1_Kenya_1950            ..............................................a...............a........a........ [135040]
AM712240.1_OURT_88/3             ...........................................................g........t..a..t..... [135040]
AM712239.1_Benin_97/1            ...........................................................g........t..a..t..... [135040]
KM111295.1_Ken06.Bus             ..a..t.................................................................a..t..... [135040]
KM111294.1_Ken05/Tk1             ..............................................a...............a........a........ [135040]
KM262844.1_strain_L60            ...........................................................g........t..a..t..... [135040]
KM102979.1_26544/OG10_Italy      ...........................................................g........t..a..t..... [135040]
KM262845.1_strain_NHV            ...........................................................g........t..a..t..... [135040]
KX354450.1_47/Ss/2008            ...........................................................g........t..a..t..... [135040]
KP055815.1_strain_BA71           ...........................................................g........t..a..t..... [135040]
MH025920.1_strain_R35            ..a..t.................................................................a..t..... [135040]
MH025919.1_strain_N10            ..a..t.................................................................a..t..... [135040]
MH025918.1_strain_R25            ..a..t.................................................................a..t..... [135040]
MH025917.1_strain_R7             ..a..t.................................................................a..t..... [135040]
MH025916.1_strain_R8             ..a..t.................................................................a..t..... [135040]
```

```
FR682468.2_Georgia_2007/1        ctccatggcagaatctgccagagtgtgagagcctctaatgatcccgtcgatcactgcgaccagtcgctttcgcacaacat [135120]
ASF_IND_20_CAD543                ................................................................................ [135120]
LR899193.1_Germany_2020/1        ................................................................................ [135120]
MN393477.1_Wuhan_2019            ................................................................................ [135120]
MW396979.1_Timor-Leste/2019/1    ................................................................................ [135120]
MK940252.1_CN/2019/InnerMongolia ................................................................................ [135120]
MK128995.1_China/2018/AnhuiXCGQ  ................................................................................ [135120]
MH681419.1_ASFV/POL/2015         ................................................................................ [135120]
LS478113.1_Estonia_2014          ................................................................................ [135120]
KP843857.1_Odintsovo_02/14       ................................................................................ [135120]
FN557520.1_strain_E75            t...........g..............g..................................a................. [135120]
AY261366.1_Warthog               ............g..............g....................a...........t.a..............t.. [135120]
AY261365.1_Warmbaths             t...........g..............g..................................a..............t.. [135120]
AY261364.1_Tengani_62            ..............................................................a................. [135120]
AY261363.1_Pretoriuskop/96/4     t...........g..............g....................a.............a..............t.. [135120]
AY261362.1_Mkuzi_1979            t...........g..............g....................a.............a..............t.. [135120]
AY261361.1_Malawi_Lil-20/1_1983  ............g..............g..................................a..............t.. [135120]
AY261360.1_Kenya_1950            ............g.t......g..aa................g....ta........a.t..a................. [135120]
AM712240.1_OURT_88/3             t...........g..............g..................................a................. [135120]
AM712239.1_Benin_97/1            t...........g..............g..................................a................. [135120]
KM111295.1_Ken06.Bus             ............g.t...t..g..aa.g.............cg....ta........a.t..a...t..........t.. [135120]
KM111294.1_Ken05/Tk1             ............g.t......g..aa................g....ta..t.....a.t..a................. [135120]
KM262844.1_strain_L60            t...........g..............g..................................a................. [135120]
KM102979.1_26544/OG10_Italy      t...........g..............g..................................a................. [135120]
KM262845.1_strain_NHV            t...........g..............g..................................a................. [135120]
KX354450.1_47/Ss/2008            t...........g..............g..................................a................. [135120]
KP055815.1_strain_BA71           t...........g..............g..................................a................. [135120]
MH025920.1_strain_R35            ............g.t...t..g..aa.g.............cg....ta........a.t..a...t..........t.. [135120]
MH025919.1_strain_N10            ............g.t...t..g..aa.g.............cg....ta........a.t..a...t..........t.. [135120]
MH025918.1_strain_R25            ............g.t...t..g..aa.g.............cg....ta........a.t..a...t..........t.. [135120]
MH025917.1_strain_R7             ............g.t...t..g..aa.g.............cg....ta........a.t..a...t..........t.. [135120]
MH025916.1_strain_R8             ............g.t...t..g..aa.g.............cg....ta........a.t..a...t..........t.. [135120]

FR682468.2_Georgia_2007/1        cgctcgtattattttgtgcgtctcctaggggcataagcgtaacattgggacgaaatacgccgccaattccccgcagggcc [135200]
ASF_IND_20_CAD543                ................................................................................ [135200]
LR899193.1_Germany_2020/1        ................................................................................ [135200]
MN393477.1_Wuhan_2019            ................................................................................ [135200]
MW396979.1_Timor-Leste/2019/1    ................................................................................ [135200]
MK940252.1_CN/2019/InnerMongolia ................................................................................ [135200]
MK128995.1_China/2018/AnhuiXCGQ  ................................................................................ [135200]
MH681419.1_ASFV/POL/2015         ................................................................................ [135200]
LS478113.1_Estonia_2014          ................................................................................ [135200]
KP843857.1_Odintsovo_02/14       ................................................................................ [135200]
```

```
FN557520.1_strain_E75            ................................................................................ [135200]
AY261366.1_Warthog               ................................................................................ [135200]
AY261365.1_Warmbaths             ................................................................................ [135200]
AY261364.1_Tengani_62            ................................................................................ [135200]
AY261363.1_Pretoriuskop/96/4     ...........c.................................................t........t....tc... [135200]
AY261362.1_Mkuzi_1979            ...........................................................................tc... [135200]
AY261361.1_Malawi_Lil-20/1_1983  .............................................................t........t....tt... [135200]
AY261360.1_Kenya_1950            ..t.g..g..g.....c....t......a........g....t..c...g..........tc.....a..t......... [135200]
AM712240.1_OURT_88/3             ................................................................................ [135200]
AM712239.1_Benin_97/1            ................................................................................ [135200]
KM111295.1_Ken06.Bus             ..t.g..g........c...........a........g....t..c...g..........tc.....a..t......... [135200]
KM111294.1_Ken05/Tk1             ..t.g..g..g.....c....t......a........g....t..c...g..........tc.....a..t......... [135200]
KM262844.1_strain_L60            ................................................................................ [135200]
KM102979.1_26544/OG10_Italy      ................................................................................ [135200]
KM262845.1_strain_NHV            ................................................................................ [135200]
KX354450.1_47/Ss/2008            ................................................................................ [135200]
KP055815.1_strain_BA71           ................................................................................ [135200]
MH025920.1_strain_R35            ..t.g..g....................a........g....t..c...g..........tc.....a..t......... [135200]
MH025919.1_strain_N10            ..t.g..g....................a........g....t..c...g..........tc.....a..t......... [135200]
MH025918.1_strain_R25            ..t.g..g....................a........g....t..c...g..........tc.....a..t......... [135200]
MH025917.1_strain_R7             ..t.g..g....................a........g....t..c...g..........tc.....a..t......... [135200]
MH025916.1_strain_R8             ..t.g..g....................a........g....t..c...g..........tc.....a..t......... [135200]

FR682468.2_Georgia_2007/1        gcctgaccgacggatagtcctgtcgcaggaacatt---gttattattataataaataacggaatcattattggctcccaa [135280]
ASF_IND_20_CAD543                ...................................aata...........r............................. [135280]
LR899193.1_Germany_2020/1        ...................................---.......................................... [135280]
MN393477.1_Wuhan_2019            ...................................---.......................................... [135280]
MW396979.1_Timor-Leste/2019/1    ...................................---.......................................... [135280]
MK940252.1_CN/2019/InnerMongolia ...................................---.......................................... [135280]
MK128995.1_China/2018/AnhuiXCGQ  ...................................---.......................................... [135280]
MH681419.1_ASFV/POL/2015         ...................................---.......................................... [135280]
LS478113.1_Estonia_2014          ...................................---.......................................... [135280]
KP843857.1_Odintsovo_02/14       ...................................---.......................................... [135280]
FN557520.1_strain_E75            ........................at.........aata...........g.......................c..... [135280]
AY261366.1_Warthog               .........................t........---t....................ta..............c..... [135280]
AY261365.1_Warmbaths             .................c......at.........aata...........g.......................c..... [135280]
AY261364.1_Tengani_62            ................ac.......t.........aata....g......g.......................c..... [135280]
AY261363.1_Pretoriuskop/96/4     .................c......at.........aata...........g.......................c..... [135280]
AY261362.1_Mkuzi_1979            .................c......at.........aata...........g.......................c..... [135280]
AY261361.1_Malawi_Lil-20/1_1983  ........a.t......c......at.........---......................................t... [135280]
AY261360.1_Kenya_1950            .............t.........attg........---............g.....g.ta.................t.. [135280]
AM712240.1_OURT_88/3             ........................at.........aata...........g.......................c..... [135280]
AM712239.1_Benin_97/1            ........................at.........aata...........g.......................c..... [135280]
KM111295.1_Ken06.Bus             ..........t............attg........---.a..........g.......ta.c...............t.. [135280]
```

```
KM111294.1_Ken05/Tk1             .............t.........attg........---............g.....g.ta.................t.. [135280]
KM262844.1_strain_L60            ........................at.........aata...........g.......................c..... [135280]
KM102979.1_26544/OG10_Italy      ........................at.........aata...........g.......................c..... [135280]
KM262845.1_strain_NHV            ........................at.........aata...........g.......................c..... [135280]
KX354450.1_47/Ss/2008            ........................at.........aata...........g.......................c..... [135280]
KP055815.1_strain_BA71           ........................at.........aata...........g.......................c..... [135280]
MH025920.1_strain_R35            ..........t............attg........---.a..........g.......ta.c...............t.. [135280]
MH025919.1_strain_N10            ..........t............attg........---.a..........g.......ta.c...............t.. [135280]
MH025918.1_strain_R25            ..........t............attg........---.a..........g.......ta.c...............t.. [135280]
MH025917.1_strain_R7             ..........t............attg........---.a..........g.......ta.c...............t.. [135280]
MH025916.1_strain_R8             ..........t............attg........---.a..........g.......ta.c...............t.. [135280]

FR682468.2_Georgia_2007/1        gagtgccgtcagattagggcgagctagttggacatttgtgtattgtataaattgttttagaagctctccctggctaataa [135360]
ASF_IND_20_CAD543                ...cc..a........................................r..........r......y............. [135360]
LR899193.1_Germany_2020/1        ................................................................................ [135360]
MN393477.1_Wuhan_2019            ................................................................................ [135360]
MW396979.1_Timor-Leste/2019/1    ................................................................................ [135360]
MK940252.1_CN/2019/InnerMongolia ................................................................................ [135360]
MK128995.1_China/2018/AnhuiXCGQ  ................................................................................ [135360]
MH681419.1_ASFV/POL/2015         ................................................................................ [135360]
LS478113.1_Estonia_2014          ................................................................................ [135360]
KP843857.1_Odintsovo_02/14       ................................................................................ [135360]
FN557520.1_strain_E75            ...cc..a..........a.....c.......................g..........a......c.....a....... [135360]
AY261366.1_Warthog               ...cc..a..........a.....c..................................a............a....... [135360]
AY261365.1_Warmbaths             ...cc..a..........a.....c.......................g..........a......c.....a....... [135360]
AY261364.1_Tengani_62            ...cc..a..........a.....c.......................g..........a......c........g.... [135360]
AY261363.1_Pretoriuskop/96/4     ..................a........................................a......c............. [135360]
AY261362.1_Mkuzi_1979            ...cc..a..........a.....c.......................g..........a......c.....a....... [135360]
AY261361.1_Malawi_Lil-20/1_1983  .........t........a........................................a......c.....a....... [135360]
AY261360.1_Kenya_1950            .........a........a..............t.........................a......c............. [135360]
AM712240.1_OURT_88/3             ...cc..a..........a.....c.......................g..........a......c.....a....... [135360]
AM712239.1_Benin_97/1            ...cc..a..........a.....c.......................g..........a......c.....a....... [135360]
KM111295.1_Ken06.Bus             .......a.a........a.....c........t..a......................a......c............. [135360]
KM111294.1_Ken05/Tk1             .........a........a..............t.........................a......c............. [135360]
KM262844.1_strain_L60            ...cc..a..........a.....c.......................g..........a......c.....a....... [135360]
KM102979.1_26544/OG10_Italy      ...cc..a..........a.....c.......................g..........a......c.....a....... [135360]
KM262845.1_strain_NHV            ...cc..a..........a.....c.......................g..........a......c.....a....... [135360]
KX354450.1_47/Ss/2008            ...cc..a..........a.....c.......................g..........a......c.....a....... [135360]
KP055815.1_strain_BA71           ...cc..a..........a.....c.......................g..........a......c.....a....... [135360]
MH025920.1_strain_R35            .......a.a........a.....c........t..a......................a......c............. [135360]
MH025919.1_strain_N10            .......a.a........a.....c........t..a......................a......c............. [135360]
MH025918.1_strain_R25            .......a.a........a.....c........t..a......................a......c............. [135360]
MH025917.1_strain_R7             .......a.a........a.....c........t..a......................a......c............. [135360]
MH025916.1_strain_R8             .......a.a........a.....c........t..a......................a......c............. [135360]
```

```
FR682468.2_Georgia_2007/1        gaatattaaacattttgttaaatagtggaagattggctctataattttctttaaggtaaatgggaatttctgttaaagta [135440]
ASF_IND_20_CAD543                .......................................................r..r..r.................. [135440]
LR899193.1_Germany_2020/1        ................................................................................ [135440]
MN393477.1_Wuhan_2019            ................................................................................ [135440]
MW396979.1_Timor-Leste/2019/1    ................................................................................ [135440]
MK940252.1_CN/2019/InnerMongolia ................................................................................ [135440]
MK128995.1_China/2018/AnhuiXCGQ  ................................................................................ [135440]
MH681419.1_ASFV/POL/2015         ................................................................................ [135440]
LS478113.1_Estonia_2014          ................................................................................ [135440]
KP843857.1_Odintsovo_02/14       ................................................................................ [135440]
FN557520.1_strain_E75            .......................................................a..g..a..............c..g [135440]
AY261366.1_Warthog               .............................................................a.................. [135440]
AY261365.1_Warmbaths             .......................................................a.....a.................. [135440]
AY261364.1_Tengani_62            .............................................................a.................. [135440]
AY261363.1_Pretoriuskop/96/4     .............................................................a.................. [135440]
AY261362.1_Mkuzi_1979            .......................................................a.....a.................. [135440]
AY261361.1_Malawi_Lil-20/1_1983  .............................................................a..............c..g [135440]
AY261360.1_Kenya_1950            .............................................................a..............c... [135440]
AM712240.1_OURT_88/3             .......................................................a..g..a..............c..g [135440]
AM712239.1_Benin_97/1            .......................................................a..g..a..............c..g [135440]
KM111295.1_Ken06.Bus             ....................t........................................a..............c... [135440]
KM111294.1_Ken05/Tk1             .............................................................a..............c... [135440]
KM262844.1_strain_L60            .......................................................a..g..a..............c..g [135440]
KM102979.1_26544/OG10_Italy      .......................................................a..g..a..............c..g [135440]
KM262845.1_strain_NHV            .......................................................a..g..a..............c..g [135440]
KX354450.1_47/Ss/2008            .......................................................a..g..a..............c..g [135440]
KP055815.1_strain_BA71           .......................................................a..g..a..............c..g [135440]
MH025920.1_strain_R35            ....................t........................................a..............c... [135440]
MH025919.1_strain_N10            ....................t........................................a..............c... [135440]
MH025918.1_strain_R25            ....................t........................................a..............c... [135440]
MH025917.1_strain_R7             ....................t........................................a..............c... [135440]
MH025916.1_strain_R8             ....................t........................................a..............c... [135440]

FR682468.2_Georgia_2007/1        gaaataagatgctgactcaggccctggcgattggtatccttaataagccgctgaagtataagtcccaaagacagaagaag [135520]
ASF_IND_20_CAD543                ..............................................................w.....r....r...... [135520]
LR899193.1_Germany_2020/1        ................................................................................ [135520]
MN393477.1_Wuhan_2019            ................................................................................ [135520]
MW396979.1_Timor-Leste/2019/1    ................................................................................ [135520]
MK940252.1_CN/2019/InnerMongolia ................................................................................ [135520]
MK128995.1_China/2018/AnhuiXCGQ  ................................................................................ [135520]
MH681419.1_ASFV/POL/2015         ................................................................................ [135520]
LS478113.1_Estonia_2014          ................................................................................ [135520]
KP843857.1_Odintsovo_02/14       ................................................................................ [135520]
```

```
FN557520.1_strain_E75           ...........t...........t.....g..a.......................c..g..a.....g....a...g.. [135520]
AY261366.1_Warthog              .......................t.....g.................................................. [135520]
AY261365.1_Warmbaths            .......................t.....g..a.......................c..g..a.....g........g.. [135520]
AY261364.1_Tengani_62           .......................t.....g.................g........c..g..a.....g........... [135520]
AY261363.1_Pretoriuskop/96/4    .......................t.....g.................................................. [135520]
AY261362.1_Mkuzi_1979           .......................t.....g..a.......................c..g..a.....g........g.. [135520]
AY261361.1_Malawi_Lil-20/1_1983 .......................t.....g.................................................. [135520]
AY261360.1_Kenya_1950           .......................t.....g..a...........c.a.........c..g..a.....g........... [135520]
AM712240.1_OURT_88/3            ...........t...........t.....g..a.......................c..g..a.....g....a...g.. [135520]
AM712239.1_Benin_97/1           ...........t...........t.....g..a.......................c..g..a.....g....a...g.. [135520]
KM111295.1_Ken06.Bus            .......................t.....g..............t.a.........c..g.................... [135520]
KM111294.1_Ken05/Tk1            .......................t.....g..a...........c.a.........c..g..a.....g........... [135520]
KM262844.1_strain_L60           ...........t...........t.....g..a.......................c..g..a.....g....a...g.. [135520]
KM102979.1_26544/OG10_Italy     ...........t...........t.....g..a.......................c..g..a.....g....a...g.. [135520]
KM262845.1_strain_NHV           ...........t...........t.....g..a.......................c..g..a.....g....a...g.. [135520]
KX354450.1_47/Ss/2008           ...........t...........t.....g..a.......................c..g..a.....g....a...g.. [135520]
KP055815.1_strain_BA71          ...........t...........t.....g..a.......................c..g..a.....g....a...g.. [135520]
MH025920.1_strain_R35           .......................t.....g..............t.a.........c..g.................... [135520]
MH025919.1_strain_N10           .......................t.....g..............t.a.........c..g.................... [135520]
MH025918.1_strain_R25           .......................t.....g..............t.a.........c..g.................... [135520]
MH025917.1_strain_R7            .......................t.....g..............t.a.........c..g.................... [135520]
MH025916.1_strain_R8            .......................t.....g..............t.a.........c..g.................... [135520]

FR682468.2_Georgia_2007/1       caccgactgctctgtggggtcgcctctatgaccaaagacgttgttattgcgtgctaagtcagggtgagcatatcccatct [135600]
ASF_IND_20_CAD543               ............................................................g..a....tg.......... [135600]
LR899193.1_Germany_2020/1       ................................................................................ [135600]
MN393477.1_Wuhan_2019           ................................................................................ [135600]
MW396979.1_Timor-Leste/2019/1   ................................................................................ [135600]
MK940252.1_CN/2019/InnerMongolia ................................................................................ [135600]
MK128995.1_China/2018/AnhuiXCGQ ................................................................................ [135600]
MH681419.1_ASFV/POL/2015        ................................................................................ [135600]
LS478113.1_Estonia_2014         ................................................................................ [135600]
KP843857.1_Odintsovo_02/14      ................................................................................ [135600]
FN557520.1_strain_E75           ........................c.....g......g..a......ca.......g...g..a....tg..c....g.. [135600]
AY261366.1_Warthog              .........t....................g......g.......................................... [135600]
AY261365.1_Warmbaths            .........t..............c.....g......g..a......ca.......g...g..a....tg..c....g.. [135600]
AY261364.1_Tengani_62           .........t..............c............g..a.a..................................... [135600]
AY261363.1_Pretoriuskop/96/4    .........t....................g......g.......................................... [135600]
AY261362.1_Mkuzi_1979           .........t..............c.....g......g..a......ca.......g...g..a....tg..c....g.. [135600]
AY261361.1_Malawi_Lil-20/1_1983 .........t.....a..............g......g....a..................................... [135600]
AY261360.1_Kenya_1950           ............c...........c.....g......g.ag......c........g...g.....g.....c....g.. [135600]
AM712240.1_OURT_88/3            ........................c.....g......g..a......ca.......g...g..a....tg..c....g.. [135600]
AM712239.1_Benin_97/1           ........................c.....g......g..a......ca.......g...g..a....tg..c....g.. [135600]
KM111295.1_Ken06.Bus            ..................a.....c.....g......g..a......c........g...g...........c....g.. [135600]
```

```
KM111294.1_Ken05/Tk1            ............c...........c.....g......g.ag......c........g...g.....g.....c....g.. [135600]
KM262844.1_strain_L60           ........................c.....g......g..a......ca.......g...g..a....tg..c....g.. [135600]
KM102979.1_26544/OG10_Italy     ........................c.....g......g..a......ca.......g...g..a....tg..c....g.. [135600]
KM262845.1_strain_NHV           ........................c.....g......g..a......ca.......g...g..a....tg..c....g.. [135600]
KX354450.1_47/Ss/2008           ........................c.....g......g..a......ca.......g...g..a....tg..c....g.. [135600]
KP055815.1_strain_BA71          ........................c.....g......g..a......ca.......g...g..a....tg..c....g.. [135600]
MH025920.1_strain_R35           ..................a.....c.....g......g..a......c........g...g...........c....g.. [135600]
MH025919.1_strain_N10           ..................a.....c.....g......g..a......c........g...g...........c....g.. [135600]
MH025918.1_strain_R25           ..................a.....c.....g......g..a......c........g...g...........c....g.. [135600]
MH025917.1_strain_R7            ..................a.....c.....g......g..a......c........g...g...........c....g.. [135600]
MH025916.1_strain_R8            ..................a.....c.....g......g..a......c........g...g...........c....g.. [135600]

FR682468.2_Georgia_2007/1       ccatcactgcttggctaaagttcccattagcgaatgcattaataagatttagatatatttttccgctgggagcatcataa [135680]
ASF_IND_20_CAD543               ................................................................................ [135680]
LR899193.1_Germany_2020/1       ................................................................................ [135680]
MN393477.1_Wuhan_2019           ................................................................................ [135680]
MW396979.1_Timor-Leste/2019/1   ................................................................................ [135680]
MK940252.1_CN/2019/InnerMongolia ................................................................................ [135680]
MK128995.1_China/2018/AnhuiXCGQ ................................................................................ [135680]
MH681419.1_ASFV/POL/2015        ................................................................................ [135680]
LS478113.1_Estonia_2014         ................................................................................ [135680]
KP843857.1_Odintsovo_02/14      ................................................................................ [135680]
FN557520.1_strain_E75           .......c.................g..g...........g........a.......................g...... [135680]
AY261366.1_Warthog              ............................g...........g........c.............................. [135680]
AY261365.1_Warmbaths            .......c.................g..g...........g........a.......................g...... [135680]
AY261364.1_Tengani_62           ...............................a........g........c.............................. [135680]
AY261363.1_Pretoriuskop/96/4    .................................................c.......................g...... [135680]
AY261362.1_Mkuzi_1979           .......c.................g..g...........g........a.......................g...... [135680]
AY261361.1_Malawi_Lil-20/1_1983 ................................................................................ [135680]
AY261360.1_Kenya_1950           ..........c........a..t...........g.t...g................................g...... [135680]
AM712240.1_OURT_88/3            .......c.................g..g...........g........a.......................g...... [135680]
AM712239.1_Benin_97/1           .......c.................g..g...........g........a.......................g...... [135680]
KM111295.1_Ken06.Bus            ..........c........a..t...........g.....g................................g...... [135680]
KM111294.1_Ken05/Tk1            ..........c........a..t...........g.....g................................g...... [135680]
KM262844.1_strain_L60           .......c.................g..g...........g........a.......................g...... [135680]
KM102979.1_26544/OG10_Italy     .......c.................g..g...........g........a.......................g...... [135680]
KM262845.1_strain_NHV           .......c.................g..g...........g........a.......................g...... [135680]
KX354450.1_47/Ss/2008           .......c.................g..g...........g........a.......................g...... [135680]
KP055815.1_strain_BA71          .......c.................g..g...........g........a.......................g...... [135680]
MH025920.1_strain_R35           ..........c........a..t...........g.....g................................g...... [135680]
MH025919.1_strain_N10           ..........c........a..t...........g.....g................................g...... [135680]
MH025918.1_strain_R25           ..........c........a..t...........g.....g................................g...... [135680]
MH025917.1_strain_R7            ..........c........a..t...........g.....g................................g...... [135680]
MH025916.1_strain_R8            ..........c........a..t...........g.....g................................g...... [135680]
```

```
FR682468.2_Georgia_2007/1        aatcgggtaatatatgaagctatgagctggttaaacaccatcatcatactacgattattttgaataccatagtctgatcc [135760]
ASF_IND_20_CAD543                .......................a........................................................ [135760]
LR899193.1_Germany_2020/1        ................................................................................ [135760]
MN393477.1_Wuhan_2019            ................................................................................ [135760]
MW396979.1_Timor-Leste/2019/1    ................................................................................ [135760]
MK940252.1_CN/2019/InnerMongolia ................................................................................ [135760]
MK128995.1_China/2018/AnhuiXCGQ  ................................................................................ [135760]
MH681419.1_ASFV/POL/2015         ................................................................................ [135760]
LS478113.1_Estonia_2014          ................................................................................ [135760]
KP843857.1_Odintsovo_02/14       ................................................................................ [135760]
FN557520.1_strain_E75            ..c....c...g..c..g....ca........................................................ [135760]
AY261366.1_Warthog               ..c....c......c.......ca..............................................g......... [135760]
AY261365.1_Warmbaths             ..c....c...g..c..g....ca........................................................ [135760]
AY261364.1_Tengani_62            ..c....c...g.....g....ca........g..............g............c....g.....a........ [135760]
AY261363.1_Pretoriuskop/96/4     ..c....c......c.......ca..............................................g......... [135760]
AY261362.1_Mkuzi_1979            ..c....c...g..c..g....ca........................................................ [135760]
AY261361.1_Malawi_Lil-20/1_1983  ..............c................................................................. [135760]
AY261360.1_Kenya_1950            .......c...g..c..g....c.........g..............g............c....g.............. [135760]
AM712240.1_OURT_88/3             ..c....c...g..c..g....ca........................................................ [135760]
AM712239.1_Benin_97/1            ..c....c...g..c..g....ca........................................................ [135760]
KM111295.1_Ken06.Bus             ..c....c...g..c..g....c.........g..............g............c................... [135760]
KM111294.1_Ken05/Tk1             .......c...g..c..g....c.........g..............g............c....g.............. [135760]
KM262844.1_strain_L60            ..c....c...g..c..g....ca........................................................ [135760]
KM102979.1_26544/OG10_Italy      ..c....c...g..c..g....ca........................................................ [135760]
KM262845.1_strain_NHV            ..c....c...g..c..g....ca........................................................ [135760]
KX354450.1_47/Ss/2008            ..c....c...g..c..g....ca........................................................ [135760]
KP055815.1_strain_BA71           ..c....c...g..c..g....ca........................................................ [135760]
MH025920.1_strain_R35            ..c....c...g..c..g....c.........g..............g............c................... [135760]
MH025919.1_strain_N10            ..c....c...g..c..g....c.........g..............g............c................... [135760]
MH025918.1_strain_R25            ..c....c...g..c..g....c.........g..............g............c................... [135760]
MH025917.1_strain_R7             ..c....c...g..c..g....c.........g..............g............c................... [135760]
MH025916.1_strain_R8             ..c....c...g..c..g....c.........g..............g............c................... [135760]

FR682468.2_Georgia_2007/1        gtataggcgataacgtcgaaggttgtttgcggcatcattgacattggcataggttct------gagcgctatgttgtccc [135840]
ASF_IND_20_CAD543                .........................................................------......c.......... [135840]
LR899193.1_Germany_2020/1        .........................................................------................. [135840]
MN393477.1_Wuhan_2019            .........................................................------................. [135840]
MW396979.1_Timor-Leste/2019/1    .........................................................------................. [135840]
MK940252.1_CN/2019/InnerMongolia .........................................................------................. [135840]
MK128995.1_China/2018/AnhuiXCGQ  .........................................................------................. [135840]
MH681419.1_ASFV/POL/2015         .........................................................------................. [135840]
LS478113.1_Estonia_2014          .........................................................------................. [135840]
KP843857.1_Odintsovo_02/14       .........................................................------................. [135840]
```

```
FN557520.1_strain_E75              a.........................................g..............------.g....c..a....... [135840]
AY261366.1_Warthog                 a.........................................g..............------................. [135840]
AY261365.1_Warmbaths               a.........................................g..............------.g....c..a....... [135840]
AY261364.1_Tengani_62              a.....a..................................tg..............------.g....c.......... [135840]
AY261363.1_Pretoriuskop/96/4       a.........................................g..............------................. [135840]
AY261362.1_Mkuzi_1979              a.........................................g..............------.g....c..a....... [135840]
AY261361.1_Malawi_Lil-20/1_1983    a........................................................------......a.......... [135840]
AY261360.1_Kenya_1950              a.....a..............a...........g.....a..g..........c...------ag......cagc.gt.. [135840]
AM712240.1_OURT_88/3               a.........................................g..............------.g....c..a....... [135840]
AM712239.1_Benin_97/1              a.........................................g..............------.g....c..a....... [135840]
KM111295.1_Ken06.Bus               a.....a..............a..a..........t.................c...aggtgcag......cagc.gt.. [135840]
KM111294.1_Ken05/Tk1               a.....a..............a...........g.....a..g..........c...------ag......cagc.gt.. [135840]
KM262844.1_strain_L60              a.........................................g..............------.g....c..a....... [135840]
KM102979.1_26544/OG10_Italy        a.........................................g..............------.g....c..a....... [135840]
KM262845.1_strain_NHV              a.........................................g..............------.g....c..a....... [135840]
KX354450.1_47/Ss/2008              a.........................................g..............------.g....c..a....... [135840]
KP055815.1_strain_BA71             a.........................................g..............------.g....c..a....... [135840]
MH025920.1_strain_R35              a.....a..............a..a..........t.................c...aggtgcag......cagc.gt.. [135840]
MH025919.1_strain_N10              a.....a..............a..a..........t.................c...aggtgcag......cagc.gt.. [135840]
MH025918.1_strain_R25              a.....a..............a..a..........t.................c...aggtgcag......cagc.gt.. [135840]
MH025917.1_strain_R7               a.....a..............a..a..........t.................c...aggtgcag......cagc.gt.. [135840]
MH025916.1_strain_R8               a.....a..............a..a..........t.................c...aggtgcag......cagc.gt.. [135840]

FR682468.2_Georgia_2007/1          agtagctaagagtattttcctcctgggcgttgttggtacgaataagattggagagtctaaagtctcctagtgccacctgc [135920]
ASF_IND_20_CAD543                  ................................................................................ [135920]
LR899193.1_Germany_2020/1          ................................................................................ [135920]
MN393477.1_Wuhan_2019              ................................................................................ [135920]
MW396979.1_Timor-Leste/2019/1      ................................................................................ [135920]
MK940252.1_CN/2019/InnerMongolia   ................................................................................ [135920]
MK128995.1_China/2018/AnhuiXCGQ    ................................................................................ [135920]
MH681419.1_ASFV/POL/2015           ................................................................................ [135920]
LS478113.1_Estonia_2014            ................................................................................ [135920]
KP843857.1_Odintsovo_02/14         ................................................................................ [135920]
FN557520.1_strain_E75              ............................a..........................................a....t... [135920]
AY261366.1_Warthog                 ............c.........t.....a..............c...........................a........ [135920]
AY261365.1_Warmbaths               ............................a....................a.....................a........ [135920]
AY261364.1_Tengani_62              .a.......................a..a................................................... [135920]
AY261363.1_Pretoriuskop/96/4       ............c.........t.....a..............c...........................a........ [135920]
AY261362.1_Mkuzi_1979              ............................a..........................................a........ [135920]
AY261361.1_Malawi_Lil-20/1_1983    ............................a................................................... [135920]
AY261360.1_Kenya_1950              ..a.........c..............................g...................................t [135920]
AM712240.1_OURT_88/3               ............................a..........................................a....t... [135920]
AM712239.1_Benin_97/1              ............................a..........................................a....t... [135920]
KM111295.1_Ken06.Bus               ..a.........c..................a...........g.....a.............................. [135920]
```

```
KM111294.1_Ken05/Tk1          ..a.........c.............................cg...................................t [135920]
KM262844.1_strain_L60         ............................a..........................................a....t... [135920]
KM102979.1_26544/OG10_Italy   ............................a..........................................a....t... [135920]
KM262845.1_strain_NHV         ............................a..........................................a....t... [135920]
KX354450.1_47/Ss/2008         ............................a..........................................a....t... [135920]
KP055815.1_strain_BA71        ............................a..........................................a....t... [135920]
MH025920.1_strain_R35         ..a.........c..................a...........g.....a.............................. [135920]
MH025919.1_strain_N10         ..a.........c..................a...........g.....a.............................. [135920]
MH025918.1_strain_R25         ..a.........c..................a...........g.....a.............................. [135920]
MH025917.1_strain_R7          ..a.........c..................a...........g.....a.............................. [135920]
MH025916.1_strain_R8          ..a.........c..................a...........g.....a.............................. [135920]

FR682468.2_Georgia_2007/1     tctacacgaagtccagagttattctccaaagcatcgtaaaatacgagtctactgaatactcttccgtattgttcaaagcg [136000]
ASF_IND_20_CAD543             ................................................................................ [136000]
LR899193.1_Germany_2020/1     ................................................................................ [136000]
MN393477.1_Wuhan_2019         ................................................................................ [136000]
MW396979.1_Timor-Leste/2019/1 ................................................................................ [136000]
MK940252.1_CN/2019/InnerMongolia ................................................................................ [136000]
MK128995.1_China/2018/AnhuiXCGQ ................................................................................ [136000]
MH681419.1_ASFV/POL/2015      ................................................................................ [136000]
LS478113.1_Estonia_2014       ................................................................................ [136000]
KP843857.1_Odintsovo_02/14    ................................................................................ [136000]
FN557520.1_strain_E75         ........g............................................a...........a.............. [136000]
AY261366.1_Warthog            ........g........................................................a.............. [136000]
AY261365.1_Warmbaths          ........g........................................................a.............. [136000]
AY261364.1_Tengani_62         ...........................................t..................a................. [136000]
AY261363.1_Pretoriuskop/96/4  ........g........................................................a.............. [136000]
AY261362.1_Mkuzi_1979         ........g........................................................a.............. [136000]
AY261361.1_Malawi_Lil-20/1_1983 .......................t........................t.............c..............a.. [136000]
AY261360.1_Kenya_1950         ........g...........c.caat...g................................c............t.... [136000]
AM712240.1_OURT_88/3          ........g............................................a...........a.............. [136000]
AM712239.1_Benin_97/1         ........g............................................a...........a.............. [136000]
KM111295.1_Ken06.Bus          ........g...........g..a.................c....................c............t.... [136000]
KM111294.1_Ken05/Tk1          ........g...........c.caat...g................................c............t.... [136000]
KM262844.1_strain_L60         ........g............................................a...........a.............. [136000]
KM102979.1_26544/OG10_Italy   ........g............................................a...........a.............. [136000]
KM262845.1_strain_NHV         ........g............................................a...........a.............. [136000]
KX354450.1_47/Ss/2008         ........g............................................a...........a.............. [136000]
KP055815.1_strain_BA71        ........g............................................a...........a.............. [136000]
MH025920.1_strain_R35         ........g...........g..a.................c....................c............t.... [136000]
MH025919.1_strain_N10         ........g...........g..a.................c....................c............t.... [136000]
MH025918.1_strain_R25         ........g...........g..a.................c....................c............t.... [136000]
MH025917.1_strain_R7          ........g...........g..a.................c....................c............t.... [136000]
MH025916.1_strain_R8          ........g...........g..a.................c....................c............t.... [136000]
```

```
FR682468.2_Georgia_2007/1        ttcagaggattggggattgttatttatttgaatattagccgcgtcccttctttgcgccccacctcgaagttgcagtacat [136080]
ASF_IND_20_CAD543                y.......r....................................................................... [136080]
LR899193.1_Germany_2020/1        ................................................................................ [136080]
MN393477.1_Wuhan_2019            ................................................................................ [136080]
MW396979.1_Timor-Leste/2019/1    ................................................................................ [136080]
MK940252.1_CN/2019/InnerMongolia ................................................................................ [136080]
MK128995.1_China/2018/AnhuiXCGQ  ................................................................................ [136080]
MH681419.1_ASFV/POL/2015         ................................................................................ [136080]
LS478113.1_Estonia_2014          ................................................................................ [136080]
KP843857.1_Odintsovo_02/14       ................................................................................ [136080]
FN557520.1_strain_E75            c.......g..............................t..a..t........ta........................ [136080]
AY261366.1_Warthog               c.......g..............................t..a..t........t.........t............... [136080]
AY261365.1_Warmbaths             c.......g..............................t..a..t........ta........................ [136080]
AY261364.1_Tengani_62            c.......g..............................t.....t.c....c....t.....c................ [136080]
AY261363.1_Pretoriuskop/96/4     c.......g....................................t.................................. [136080]
AY261362.1_Mkuzi_1979            c.......g..............................t..a..t........ta........................ [136080]
AY261361.1_Malawi_Lil-20/1_1983  c......................................t..a..t.................................. [136080]
AY261360.1_Kenya_1950            c.....aa...c......a.................g..t..a.tt.c.........t.....c........t....... [136080]
AM712240.1_OURT_88/3             c.......g..............................t..a..t........ta........................ [136080]
AM712239.1_Benin_97/1            c.......g..............................t..a..t........ta........................ [136080]
KM111295.1_Ken06.Bus             c.....aag..c......a....................t.....t.......................a..t....... [136080]
KM111294.1_Ken05/Tk1             c.....aag..c......a.................g..t..a.tt.c.........t.....c........t....... [136080]
KM262844.1_strain_L60            c.......g..............................t..a..t........ta........................ [136080]
KM102979.1_26544/OG10_Italy      c.......g..............................t..a..t........ta........................ [136080]
KM262845.1_strain_NHV            c.......g..............................t..a..t........ta........................ [136080]
KX354450.1_47/Ss/2008            c.......g..............................t..a..t........ta........................ [136080]
KP055815.1_strain_BA71           c.......g..............................t..a..t........ta........................ [136080]
MH025920.1_strain_R35            c.....aag..c......a....................t.....t.......................a..t....... [136080]
MH025919.1_strain_N10            c.....aag..c......a....................t.....t.......................a..t....... [136080]
MH025918.1_strain_R25            c.....aag..c......a....................t.....t.......................a..t....... [136080]
MH025917.1_strain_R7             c.....aag..c......a....................t.....t.......................a..t....... [136080]
MH025916.1_strain_R8             c.....aag..c......a....................t.....t.......................a..t....... [136080]

FR682468.2_Georgia_2007/1        tataaggctttgtaagcaaggtgtaggttttattaatgatttggttaaccccctccaggcccaattcaccgccaggaagc [136160]
ASF_IND_20_CAD543                .................................................y.............................. [136160]
LR899193.1_Germany_2020/1        ................................................................................ [136160]
MN393477.1_Wuhan_2019            ................................................................................ [136160]
MW396979.1_Timor-Leste/2019/1    ................................................................................ [136160]
MK940252.1_CN/2019/InnerMongolia ................................................................................ [136160]
MK128995.1_China/2018/AnhuiXCGQ  ................................................................................ [136160]
MH681419.1_ASFV/POL/2015         ................................................................................ [136160]
LS478113.1_Estonia_2014          ................................................................................ [136160]
KP843857.1_Odintsovo_02/14       ................................................................................ [136160]
```

```
FN557520.1_strain_E75          ....g..t............a............................t...........t.................. [136160]
AY261366.1_Warthog             ................................................................................ [136160]
AY261365.1_Warmbaths           ....g..t............a............................t.............................t [136160]
AY261364.1_Tengani_62          ................................................................................ [136160]
AY261363.1_Pretoriuskop/96/4   ................................................................................ [136160]
AY261362.1_Mkuzi_1979          ....g..t............a............................t........t....................t [136160]
AY261361.1_Malawi_Lil-20/1_1983 ................t.....a......................................................... [136160]
AY261360.1_Kenya_1950          ....................a................a...................a...................... [136160]
AM712240.1_OURT_88/3           ....g..t............a............................t...........t.................. [136160]
AM712239.1_Benin_97/1          ....g..t............a............................t...........t.................. [136160]
KM111295.1_Ken06.Bus           ....................a........................................................... [136160]
KM111294.1_Ken05/Tk1           ....................a....................................a...................... [136160]
KM262844.1_strain_L60          ....g..t............a............................t...........t.................. [136160]
KM102979.1_26544/OG10_Italy    ....g..t............a............................t...........t.................. [136160]
KM262845.1_strain_NHV          ....g..t............a............................t...........t.................. [136160]
KX354450.1_47/Ss/2008          ....g..t............a............................t...........t.................. [136160]
KP055815.1_strain_BA71         ....g..t............a............................t...........t.................. [136160]
MH025920.1_strain_R35          ....................a........................................................... [136160]
MH025919.1_strain_N10          ....................a........................................................... [136160]
MH025918.1_strain_R25          ....................a........................................................... [136160]
MH025917.1_strain_R7           ....................a........................................................... [136160]
MH025916.1_strain_R8           ....................a........................................................... [136160]

FR682468.2_Georgia_2007/1      ggccttcctccggcatcggtaggtggtttaataagtttgtcaattaaatgttcttccaaccagtaaaatgagccaggatt [136240]
ASF_IND_20_CAD543              ................................................................................ [136240]
LR899193.1_Germany_2020/1      ................................................................................ [136240]
MN393477.1_Wuhan_2019          ................................................................................ [136240]
MW396979.1_Timor-Leste/2019/1  ................................................................................ [136240]
MK940252.1_CN/2019/InnerMongolia ................................................................................ [136240]
MK128995.1_China/2018/AnhuiXCGQ ................................................................................ [136240]
MH681419.1_ASFV/POL/2015       ................................................................................ [136240]
LS478113.1_Estonia_2014        ................................................................................ [136240]
KP843857.1_Odintsovo_02/14     ................................................................................ [136240]
FN557520.1_strain_E75          ........c...........g..............c........c.............g..................... [136240]
AY261366.1_Warthog             ....................g..............c............................................ [136240]
AY261365.1_Warmbaths           ........c...........g..............c........c.............g..................... [136240]
AY261364.1_Tengani_62          .................a..g..............c..a.....c................................... [136240]
AY261363.1_Pretoriuskop/96/4   ....................g..............c........c.............g..................... [136240]
AY261362.1_Mkuzi_1979          ........c...........g..............c........c.............g..................... [136240]
AY261361.1_Malawi_Lil-20/1_1983 ..t.......ta........g..............c............................................ [136240]
AY261360.1_Kenya_1950          ....................g..............c.....g..c................................... [136240]
AM712240.1_OURT_88/3           ........c...........g..............c........c.............g..................... [136240]
AM712239.1_Benin_97/1          ........c...........g..............c........c.............g..................... [136240]
KM111295.1_Ken06.Bus           ....................g..............c........c.................a...........t..... [136240]
```

```
KM111294.1_Ken05/Tk1               ....................g..............c.....g..c................................... [136240]
KM262844.1_strain_L60              ........c...........g..............c........c.............g..................... [136240]
KM102979.1_26544/OG10_Italy        ........c.g.........g..............c........c.............g..................... [136240]
KM262845.1_strain_NHV              ........c...........g..............c........c.............g..................... [136240]
KX354450.1_47/Ss/2008              ........c...........g..............c........c.............g..................... [136240]
KP055815.1_strain_BA71             ........c...........g..............c........c.............g..................... [136240]
MH025920.1_strain_R35              ....................g..............c........c.................a...........t..... [136240]
MH025919.1_strain_N10              ....................g..............c........c.................a...........t..... [136240]
MH025918.1_strain_R25              ....................g..............c........c.................a...........t..... [136240]
MH025917.1_strain_R7               ....................g..............c........c.................a...........t..... [136240]
MH025916.1_strain_R8               ....................g..............c........c.................a...........t..... [136240]

FR682468.2_Georgia_2007/1          agatctattttcatagtattgaataatgtttttatcaatatgcgggcgtagaagatcaagaaaatacttcgtgtcggcca [136320]
ASF_IND_20_CAD543                  ................................................................................ [136320]
LR899193.1_Germany_2020/1          ................................................................................ [136320]
MN393477.1_Wuhan_2019              ................................................................................ [136320]
MW396979.1_Timor-Leste/2019/1      ................................................................................ [136320]
MK940252.1_CN/2019/InnerMongolia   ................................................................................ [136320]
MK128995.1_China/2018/AnhuiXCGQ    ................................................................................ [136320]
MH681419.1_ASFV/POL/2015           ................................................................................ [136320]
LS478113.1_Estonia_2014            ................................................................................ [136320]
KP843857.1_Odintsovo_02/14         ................................................................................ [136320]
FN557520.1_strain_E75              g...........g.............................t.............t............a.......... [136320]
AY261366.1_Warthog                 g.........................................t.............t....................... [136320]
AY261365.1_Warmbaths               g...........g.............................t.............t............a.......... [136320]
AY261364.1_Tengani_62              g..........................a......................a............................. [136320]
AY261363.1_Pretoriuskop/96/4       g...........g.............................t.............t....................... [136320]
AY261362.1_Mkuzi_1979              g...........g.............................t.............t............a.......... [136320]
AY261361.1_Malawi_Lil-20/1_1983    g............................................................................... [136320]
AY261360.1_Kenya_1950              g..........................a..........................g..............g.......... [136320]
AM712240.1_OURT_88/3               g...........g.............................t.............t............a.......... [136320]
AM712239.1_Benin_97/1              g...........g.............................t.............t............a.......... [136320]
KM111295.1_Ken06.Bus               g....................................................................g.....a.... [136320]
KM111294.1_Ken05/Tk1               g..........................a..........................g..............g.......... [136320]
KM262844.1_strain_L60              g...........g.............................t.............t............a.......... [136320]
KM102979.1_26544/OG10_Italy        g...........g.............................t.............t............a.......... [136320]
KM262845.1_strain_NHV              g...........g.............................t.............t............a.......... [136320]
KX354450.1_47/Ss/2008              g...........g.............................t.............t............a.......... [136320]
KP055815.1_strain_BA71             g...........g.............................t.............t............a.......... [136320]
MH025920.1_strain_R35              g....................................................................g.....a.... [136320]
MH025919.1_strain_N10              g....................................................................g.....a.... [136320]
MH025918.1_strain_R25              g....................................................................g.....a.... [136320]
MH025917.1_strain_R7               g....................................................................g.....a.... [136320]
MH025916.1_strain_R8               g....................................................................g.....a.... [136320]
```

```
FR682468.2_Georgia_2007/1        tcaaagaatcaattaaggaaataagacctgtaaaatctaaatgcacttgagcggtgctggtttcagggaagcgaacttga [136400]
ASF_IND_20_CAD543                ................................................................g..a............ [136400]
LR899193.1_Germany_2020/1        ................................................................................ [136400]
MN393477.1_Wuhan_2019            ................................................................................ [136400]
MW396979.1_Timor-Leste/2019/1    ................................................................................ [136400]
MK940252.1_CN/2019/InnerMongolia ................................................................................ [136400]
MK128995.1_China/2018/AnhuiXCGQ  ................................................................................ [136400]
MH681419.1_ASFV/POL/2015         ................................................................................ [136400]
LS478113.1_Estonia_2014          ................................................................................ [136400]
KP843857.1_Odintsovo_02/14       ................................................................................ [136400]
FN557520.1_strain_E75            ............................c...........g.................a.....g..a............ [136400]
AY261366.1_Warthog               .t......................................g..............a..a.....g..a............ [136400]
AY261365.1_Warmbaths             .t....................g.................g..............a..a.....g..a............ [136400]
AY261364.1_Tengani_62            ............................c...........g..t..c...........a.....g..a............ [136400]
AY261363.1_Pretoriuskop/96/4     .t......................................g..............a..a.....g..a............ [136400]
AY261362.1_Mkuzi_1979            .t......................................g..............a..a.....g..a............ [136400]
AY261361.1_Malawi_Lil-20/1_1983  ....g.......................c...........g....a................................. [136400]
AY261360.1_Kenya_1950            ........................................g..t..a...........a.....g............... [136400]
AM712240.1_OURT_88/3             ............................c...........g.................a.....g..a............ [136400]
AM712239.1_Benin_97/1            ............................c...........g.................a.....g..a............ [136400]
KM111295.1_Ken06.Bus             ..............................................a...........a.....g..a............ [136400]
KM111294.1_Ken05/Tk1             ........................................g..t..a...........a.....g............... [136400]
KM262844.1_strain_L60            ............................c...........g.................a.....g..a............ [136400]
KM102979.1_26544/OG10_Italy      ............................c...........g.................a.....g..a............ [136400]
KM262845.1_strain_NHV            ............................c...........g.................a.....g..a............ [136400]
KX354450.1_47/Ss/2008            ............................c...........g.................a.....g..a............ [136400]
KP055815.1_strain_BA71           ............................c...........g.................a.....g..a............ [136400]
MH025920.1_strain_R35            ..............................................a...........a.....g..a............ [136400]
MH025919.1_strain_N10            ..............................................a...........a.....g..a............ [136400]
MH025918.1_strain_R25            ..............................................a...........a.....g..a............ [136400]
MH025917.1_strain_R7             ..............................................a...........a.....g..a............ [136400]
MH025916.1_strain_R8             ..............................................a...........a.....g..a............ [136400]

FR682468.2_Georgia_2007/1        accattttgttaaaactggaggtcatttcgaagatattggtcaacaggagctgcatgattcgctgattatctactaaata [136480]
ASF_IND_20_CAD543                ................................................................................ [136480]
LR899193.1_Germany_2020/1        ................................................................................ [136480]
MN393477.1_Wuhan_2019            ................................................................................ [136480]
MW396979.1_Timor-Leste/2019/1    ................................................................................ [136480]
MK940252.1_CN/2019/InnerMongolia ................................................................................ [136480]
MK128995.1_China/2018/AnhuiXCGQ  ................................................................................ [136480]
MH681419.1_ASFV/POL/2015         ................................................................................ [136480]
LS478113.1_Estonia_2014          ................................................................................ [136480]
KP843857.1_Odintsovo_02/14       ................................................................................ [136480]
```

```
FN557520.1_strain_E75            .................a..................................................g.....c..... [136480]
AY261366.1_Warthog               .................a..................................................g.....c..... [136480]
AY261365.1_Warmbaths             ....................................................................g.....c..... [136480]
AY261364.1_Tengani_62            ......................................a....g........................g.....c..... [136480]
AY261363.1_Pretoriuskop/96/4     ....................................................................g.....c..... [136480]
AY261362.1_Mkuzi_1979            ....................................................................g.....c..... [136480]
AY261361.1_Malawi_Lil-20/1_1983  .t............g..a.....t..............a....g..............................c..... [136480]
AY261360.1_Kenya_1950            ....a......................................gt.......................g.....c..... [136480]
AM712240.1_OURT_88/3             .................a..................................................g.....c..... [136480]
AM712239.1_Benin_97/1            .................a..................................................g.....c..... [136480]
KM111295.1_Ken06.Bus             ....a.........g..a..............a..........g........................g.....c..... [136480]
KM111294.1_Ken05/Tk1             ....a......................................gt.......................g.....c..... [136480]
KM262844.1_strain_L60            .................a..................................................g.....c..... [136480]
KM102979.1_26544/OG10_Italy      .................a..................................................g.....c..... [136480]
KM262845.1_strain_NHV            .................a..................................................g.....c..... [136480]
KX354450.1_47/Ss/2008            .................a..................................................g.....c..... [136480]
KP055815.1_strain_BA71           .................a..................................................g.....c..... [136480]
MH025920.1_strain_R35            ....a.........g..a..............a..........g........................g.....c..... [136480]
MH025919.1_strain_N10            ....a.........g..a..............a..........g........................g.....c..... [136480]
MH025918.1_strain_R25            ....a.........g..a..............a..........g........................g.....c..... [136480]
MH025917.1_strain_R7             ....a.........g..a..............a..........g........................g.....c..... [136480]
MH025916.1_strain_R8             ....a.........g..a..............a..........g........................g.....c..... [136480]

FR682468.2_Georgia_2007/1        ccttgcggccaactcttgctccggacgaactcctccaccagcaggaatacccacatatggtacaatccaagcaaaaagag [136560]
ASF_IND_20_CAD543                .....................................................................r.......... [136560]
LR899193.1_Germany_2020/1        ................................................................................ [136560]
MN393477.1_Wuhan_2019            ................................................................................ [136560]
MW396979.1_Timor-Leste/2019/1    ................................................................................ [136560]
MK940252.1_CN/2019/InnerMongolia ................................................................................ [136560]
MK128995.1_China/2018/AnhuiXCGQ  ................................................................................ [136560]
MH681419.1_ASFV/POL/2015         ................................................................................ [136560]
LS478113.1_Estonia_2014          ................................................................................ [136560]
KP843857.1_Odintsovo_02/14       ................................................................................ [136560]
FN557520.1_strain_E75            t........t..t..........c.............................................g.......... [136560]
AY261366.1_Warthog               t........t..t..........c...........................t.................g.......... [136560]
AY261365.1_Warmbaths             t........t..t..........c.............................................g.......... [136560]
AY261364.1_Tengani_62            t........t..t..........c.........c........................c..gt.ga...ga......... [136560]
AY261363.1_Pretoriuskop/96/4     t........t..t..........c.............................................g.......... [136560]
AY261362.1_Mkuzi_1979            t........t..t..........c.............................................g.......... [136560]
AY261361.1_Malawi_Lil-20/1_1983  ......a...................................................t..................... [136560]
AY261360.1_Kenya_1950            ..................t............g.....t.............t...a..c..gt..a...g.......... [136560]
AM712240.1_OURT_88/3             t........t..t..........c.............................................g.......... [136560]
AM712239.1_Benin_97/1            t........t..t..........c.............................................g.......... [136560]
KM111295.1_Ken06.Bus             t.................t..........t............................t..t.......g.......... [136560]
```

```
KM111294.1_Ken05/Tk1              ..................t..................t.............t...a..c..gt..a...g.......... [136560]
KM262844.1_strain_L60             t........t..t..........c.............................................g.......... [136560]
KM102979.1_26544/OG10_Italy       t........t..t..........c.............................................g.......... [136560]
KM262845.1_strain_NHV             t........t..t..........c.............................................g.......... [136560]
KX354450.1_47/Ss/2008             t........t..t..........c.............................................g.......... [136560]
KP055815.1_strain_BA71            t........t..t..........c.............................................g.......... [136560]
MH025920.1_strain_R35             t.................t..........t............................t..t.......g.......... [136560]
MH025919.1_strain_N10             t.................t..........t............................t..t.......g.......... [136560]
MH025918.1_strain_R25             t.................t..........t............................t..t.......g.......... [136560]
MH025917.1_strain_R7              t.................t..........t............................t..t.......g.......... [136560]
MH025916.1_strain_R8              t.................t..........t............................t..t.......g.......... [136560]

FR682468.2_Georgia_2007/1         tttctgtggttaaatttcggtcttgggctgctgcagccgcttcggtagtgggatcagggtacaccatagaaagccgcata [136640]
ASF_IND_20_CAD543                 .y......ry.......y.........---g.........a.t...................................c. [136640]
LR899193.1_Germany_2020/1         ................................................................................ [136640]
MN393477.1_Wuhan_2019             ................................................................................ [136640]
MW396979.1_Timor-Leste/2019/1     ................................................................................ [136640]
MK940252.1_CN/2019/InnerMongolia  ................................................................................ [136640]
MK128995.1_China/2018/AnhuiXCGQ   ................................................................................ [136640]
MH681419.1_ASFV/POL/2015          ................................................................................ [136640]
LS478113.1_Estonia_2014           ................................................................................ [136640]
KP843857.1_Odintsovo_02/14        ................................................................................ [136640]
FN557520.1_strain_E75             .c......ac.......t.........---g.....t...a.t......t........a......c............c. [136640]
AY261366.1_Warthog                .c......ac.......t.........---g.....t...a.t......t........a......c...........gc. [136640]
AY261365.1_Warmbaths              .......a...................---......t............a........a......c...........gc. [136640]
AY261364.1_Tengani_62             ........ac........ta.....aa---g.....t...g..a.................t...tc......t...gc. [136640]
AY261363.1_Pretoriuskop/96/4      .c.........................---g.a..tt............................tc............. [136640]
AY261362.1_Mkuzi_1979             .......a...................---......t............a........a......c...........gc. [136640]
AY261361.1_Malawi_Lil-20/1_1983   .......a...................---g..t.tt...a........................tc......t...gc. [136640]
AY261360.1_Kenya_1950             ........ac........ta...a.a.---g...g.t...a..a..g..................tc......t...gc. [136640]
AM712240.1_OURT_88/3              .c......ac.......t.........---g.....t...a.t......t........a......c............c. [136640]
AM712239.1_Benin_97/1             .c......ac.......t.........---g.....t...a.t......t........a......c............c. [136640]
KM111295.1_Ken06.Bus              .c.....a....g....t.........---g.ag..t..ga.....c..a.......................t...gc. [136640]
KM111294.1_Ken05/Tk1              ........ac........ta...a.a.---g...g.t...a..a..g..................tc......t...gc. [136640]
KM262844.1_strain_L60             .c......ac.......t.........---g.....t...a.t......t........a......c............c. [136640]
KM102979.1_26544/OG10_Italy       .c......ac.......t.........---g.....t...a.t......t........a......c............c. [136640]
KM262845.1_strain_NHV             .c......ac.......t.........---g.....t...a.t......t........a......c............c. [136640]
KX354450.1_47/Ss/2008             .c......ac.......t.........---g.....t...a.t......t........a......c............c. [136640]
KP055815.1_strain_BA71            .c......ac.......t.........---g.....t...a.t......t........a......c............c. [136640]
MH025920.1_strain_R35             .c.....a....g....t.........---g.ag..t..ga.....c..a.......................t...gc. [136640]
MH025919.1_strain_N10             .c.....a....g....t.........---g.ag..t..ga.....c..a.......................t...gc. [136640]
MH025918.1_strain_R25             .c.....a....g....t.........---g.ag..t..ga.....c..a.......................t...gc. [136640]
MH025917.1_strain_R7              .c.....a....g....t.........---g.ag..t..ga.....c..a.......................t...gc. [136640]
MH025916.1_strain_R8              .c.....a....g....t.........---g.ag..t..ga.....c..a.......................t...gc. [136640]
```

```
FR682468.2_Georgia_2007/1        ttgatttctttaatgactaatcctggatttctaatctcagagatggccccgtgttttcttccgagccagtcaataagatt [136720]
ASF_IND_20_CAD543                .cat....c....ca..y.............................................................. [136720]
LR899193.1_Germany_2020/1        ................................................................................ [136720]
MN393477.1_Wuhan_2019            ................................................................................ [136720]
MW396979.1_Timor-Leste/2019/1    ................................................................................ [136720]
MK940252.1_CN/2019/InnerMongolia ................................................................................ [136720]
MK128995.1_China/2018/AnhuiXCGQ  ................................................................................ [136720]
MH681419.1_ASFV/POL/2015         ................................................................................ [136720]
LS478113.1_Estonia_2014          ................................................................................ [136720]
KP843857.1_Odintsovo_02/14       ................................................................................ [136720]
FN557520.1_strain_E75            .cat....c....ca..c.............................................................. [136720]
AY261366.1_Warthog               .cat....c....ca..c.............................................................. [136720]
AY261365.1_Warmbaths             .cat..g.c....ca..c.............................t................................ [136720]
AY261364.1_Tengani_62            .c.t..g.c.....a..c.............................................................. [136720]
AY261363.1_Pretoriuskop/96/4     ................................................................................ [136720]
AY261362.1_Mkuzi_1979            .cat..g.c....ca..c.............................................................. [136720]
AY261361.1_Malawi_Lil-20/1_1983  ..at..g.c....ca..c..........................a..t...........a.............g...... [136720]
AY261360.1_Kenya_1950            .c.t..g.c..............c.....................t........c.......c................. [136720]
AM712240.1_OURT_88/3             .cat....c....ca..c.............................................................. [136720]
AM712239.1_Benin_97/1            .cat....c....ca..c.............................................................. [136720]
KM111295.1_Ken06.Bus             .c.t..g.c........c.....c.....................t........c.......c................. [136720]
KM111294.1_Ken05/Tk1             .c.t..g.c........c.....c.....................t........c.......c................. [136720]
KM262844.1_strain_L60            .cat....c....ca..c.............................................................. [136720]
KM102979.1_26544/OG10_Italy      .cat....c....ca..c.............................................................. [136720]
KM262845.1_strain_NHV            .cat....c....ca..c.............................................................. [136720]
KX354450.1_47/Ss/2008            .cat....c....ca..c.............................................................. [136720]
KP055815.1_strain_BA71           .cat....c....ca..c.............................................................. [136720]
MH025920.1_strain_R35            .c.t..g.c........c.....c.....................t........c.......c................. [136720]
MH025919.1_strain_N10            .c.t..g.c........c.....c.....................t........c.......c................. [136720]
MH025918.1_strain_R25            .c.t..g.c........c.....c.....................t........c.......c................. [136720]
MH025917.1_strain_R7             .c.t..g.c........c.....c.....................t........c.......c................. [136720]
MH025916.1_strain_R8             .c.t..g.c........c.....c.....................t........c.......c................. [136720]

FR682468.2_Georgia_2007/1        ggcgcggttcacgttggcagcttgtgtctctcgtaaccattcgataatgcttttttgaatcgtatctaggtctaaacctt [136800]
ASF_IND_20_CAD543                ................................................................................ [136800]
LR899193.1_Germany_2020/1        ................................................................................ [136800]
MN393477.1_Wuhan_2019            ................................................................................ [136800]
MW396979.1_Timor-Leste/2019/1    ................................................................................ [136800]
MK940252.1_CN/2019/InnerMongolia ................................................................................ [136800]
MK128995.1_China/2018/AnhuiXCGQ  ................................................................................ [136800]
MH681419.1_ASFV/POL/2015         ................................................................................ [136800]
LS478113.1_Estonia_2014          ................................................................................ [136800]
KP843857.1_Odintsovo_02/14       ................................................................................ [136800]
```

```
FN557520.1_strain_E75            ...t...........a..g.....c..t..............a..g.................................. [136800]
AY261366.1_Warthog               ....t.............g.......................a..g.................................. [136800]
AY261365.1_Warmbaths             ...t..............g.....c..t..............a..g.......................a.......... [136800]
AY261364.1_Tengani_62            ...t...........a..g.....c..t..............a..g.................................. [136800]
AY261363.1_Pretoriuskop/96/4     ....t.............g.---...................a..g..............t................... [136800]
AY261362.1_Mkuzi_1979            ...t..............g.....c..t..............a..g.......................a.......... [136800]
AY261361.1_Malawi_Lil-20/1_1983  ...at.......a.c..gg.gc.....t..............a..g..a....................a.......... [136800]
AY261360.1_Kenya_1950            ...a..a...........g.....c..t..c..............g....................c..a......g.c. [136800]
AM712240.1_OURT_88/3             ...t...........a..g.....c..t..............a..g.................................. [136800]
AM712239.1_Benin_97/1            ...t...........a..g.....c..t..............a..g.................................. [136800]
KM111295.1_Ken06.Bus             ...at.............g..c..c...................c................a....c..a......g.c. [136800]
KM111294.1_Ken05/Tk1             ...a..a...........g.....c..t..c..............g....................c..a......g.c. [136800]
KM262844.1_strain_L60            ...t...........a..g.....c..t..............a..g.................................. [136800]
KM102979.1_26544/OG10_Italy      ...t...........a..g.....c..t..............a..g.................................. [136800]
KM262845.1_strain_NHV            ...t...........a..g.....c..t..............a..g.................................. [136800]
KX354450.1_47/Ss/2008            ...t...........a..g.....c..t..............a..g.................................. [136800]
KP055815.1_strain_BA71           ...t...........a..g.....c..t..............a..g.................................. [136800]
MH025920.1_strain_R35            ...at.............g..c..c...................c................a....c..a......g.c. [136800]
MH025919.1_strain_N10            ...ata............g..c..c...................c................a....c..a......g.c. [136800]
MH025918.1_strain_R25            ...at.............g..c..c...................c................a....c..a......g.c. [136800]
MH025917.1_strain_R7             ...at.............g..c..c...................c................a....c..a......g.c. [136800]
MH025916.1_strain_R8             ...at.............g..c..c...................c................a....c..a......g.c. [136800]

FR682468.2_Georgia_2007/1        taatgttattacgaaagttattaagaagtacgtaaatagcactcaataagttaagacctgtaataacggtttcatgaaac [136880]
ASF_IND_20_CAD543                ...........................................y.................................... [136880]
LR899193.1_Germany_2020/1        ................................................................................ [136880]
MN393477.1_Wuhan_2019            ................................................................................ [136880]
MW396979.1_Timor-Leste/2019/1    ................................................................................ [136880]
MK940252.1_CN/2019/InnerMongolia ................................................................................ [136880]
MK128995.1_China/2018/AnhuiXCGQ  ................................................................................ [136880]
MH681419.1_ASFV/POL/2015         ................................................................................ [136880]
LS478113.1_Estonia_2014          ................................................................................ [136880]
KP843857.1_Odintsovo_02/14       ................................................................................ [136880]
FN557520.1_strain_E75            ...........................................t.................................... [136880]
AY261366.1_Warthog               .........................g...................................................... [136880]
AY261365.1_Warmbaths             .........................g...................................................... [136880]
AY261364.1_Tengani_62            ...........................................t.................................... [136880]
AY261363.1_Pretoriuskop/96/4     .........................g...................................................... [136880]
AY261362.1_Mkuzi_1979            .........................g...................................................... [136880]
AY261361.1_Malawi_Lil-20/1_1983  ....a........................................................................... [136880]
AY261360.1_Kenya_1950            ....a............g.......g..c............................................g...... [136880]
AM712240.1_OURT_88/3             ...........................................t.................................... [136880]
AM712239.1_Benin_97/1            ...........................................t.................................... [136880]
KM111295.1_Ken06.Bus             ....a............g.......g..c............................................g...... [136880]
```

```
KM111294.1_Ken05/Tk1             ....a............g.......g..c............................................g...... [136880]
KM262844.1_strain_L60            ...........................................t.................................... [136880]
KM102979.1_26544/OG10_Italy      ...........................................t.................................... [136880]
KM262845.1_strain_NHV            ...........................................t.................................... [136880]
KX354450.1_47/Ss/2008            ...........................................t.................................... [136880]
KP055815.1_strain_BA71           ...........................................t.................................... [136880]
MH025920.1_strain_R35            ....a............g.......g..c............................................g...... [136880]
MH025919.1_strain_N10            ....a............g.......g..c............................................g...... [136880]
MH025918.1_strain_R25            ....a............g.......g..c............................................g...... [136880]
MH025917.1_strain_R7             ....a............g.......g..c............................................g...... [136880]
MH025916.1_strain_R8             ....a............g.......g..c............................................g...... [136880]

FR682468.2_Georgia_2007/1        agaaatattttgttaacatctgtatctgccagtgactcagagccttgaataagttttgaaacgatttgaattttatcggt [136960]
ASF_IND_20_CAD543                ................................................................................ [136960]
LR899193.1_Germany_2020/1        ................................................................................ [136960]
MN393477.1_Wuhan_2019            ................................................................................ [136960]
MW396979.1_Timor-Leste/2019/1    ................................................................................ [136960]
MK940252.1_CN/2019/InnerMongolia ................................................................................ [136960]
MK128995.1_China/2018/AnhuiXCGQ  ................................................................................ [136960]
MH681419.1_ASFV/POL/2015         ................................................................................ [136960]
LS478113.1_Estonia_2014          ................................................................................ [136960]
KP843857.1_Odintsovo_02/14       ................................................................................ [136960]
FN557520.1_strain_E75            ..............................................................a...........g..... [136960]
AY261366.1_Warthog               ................................c.............................a...........g..... [136960]
AY261365.1_Warmbaths             ................................c.............................a...........g..a.. [136960]
AY261364.1_Tengani_62            ................................................................................ [136960]
AY261363.1_Pretoriuskop/96/4     ................................c.............................a...........g..... [136960]
AY261362.1_Mkuzi_1979            .a..............................c.............................a...........g..a.. [136960]
AY261361.1_Malawi_Lil-20/1_1983  ................................c.........................................g..... [136960]
AY261360.1_Kenya_1950            ...........a..............c..t..c.........................................g..... [136960]
AM712240.1_OURT_88/3             ..............................................................a...........g..... [136960]
AM712239.1_Benin_97/1            ..............................................................a...........g..... [136960]
KM111295.1_Ken06.Bus             ..........................c..t..c...........c.............................g..... [136960]
KM111294.1_Ken05/Tk1             ...........a..............c..t..c...........c.............................g..... [136960]
KM262844.1_strain_L60            ..............................................................a...........g..... [136960]
KM102979.1_26544/OG10_Italy      ..............................................................a...........g..... [136960]
KM262845.1_strain_NHV            ..............................................................a...........g..... [136960]
KX354450.1_47/Ss/2008            ..............................................................a...........g..... [136960]
KP055815.1_strain_BA71           ..............................................................a...........g..... [136960]
MH025920.1_strain_R35            ..........................c..t..c...........c.............................g..... [136960]
MH025919.1_strain_N10            ..........................c..t..c...........c.............................g..... [136960]
MH025918.1_strain_R25            ..........................c..t..c...........c.............................g..... [136960]
MH025917.1_strain_R7             ..........................c..t..c...........c.............................g..... [136960]
MH025916.1_strain_R8             ..........................c..t..c...........c.............................g..... [136960]
```

```
FR682468.2_Georgia_2007/1        atgctcctttttgagttcattgatagcctggcgaatgagttcttggtaggaaattttgcccaattcttgttgcagactgg [137040]
ASF_IND_20_CAD543                ................................................................................ [137040]
LR899193.1_Germany_2020/1        ................................................................................ [137040]
MN393477.1_Wuhan_2019            ................................................................................ [137040]
MW396979.1_Timor-Leste/2019/1    ................................................................................ [137040]
MK940252.1_CN/2019/InnerMongolia ................................................................................ [137040]
MK128995.1_China/2018/AnhuiXCGQ  ................................................................................ [137040]
MH681419.1_ASFV/POL/2015         ................................................................................ [137040]
LS478113.1_Estonia_2014          ................................................................................ [137040]
KP843857.1_Odintsovo_02/14       ................................................................................ [137040]
FN557520.1_strain_E75            ..........................................c..........c..................t..g.... [137040]
AY261366.1_Warthog               g..g......................................c..........c............c............. [137040]
AY261365.1_Warmbaths             g..g......................................c..........c..................t..g.... [137040]
AY261364.1_Tengani_62            ................................................................................ [137040]
AY261363.1_Pretoriuskop/96/4     g..g......................................c..........c............c............. [137040]
AY261362.1_Mkuzi_1979            g..g........a.............................c..........c..................t....... [137040]
AY261361.1_Malawi_Lil-20/1_1983  ............a................................................................... [137040]
AY261360.1_Kenya_1950            g...........a........tg.g.....ct..........c..........c...t..t.................a. [137040]
AM712240.1_OURT_88/3             ..........................................c..........c..................t..g.... [137040]
AM712239.1_Benin_97/1            ..........................................c..........c..................t..g.... [137040]
KM111295.1_Ken06.Bus             g...........a........tg.g.....ct..........c..........c...t..t.................a. [137040]
KM111294.1_Ken05/Tk1             g...........a........tg.g.....ct..........c..........c...t..t.................a. [137040]
KM262844.1_strain_L60            ..........................................c..........c..................t..g.... [137040]
KM102979.1_26544/OG10_Italy      ..........................................c..........c..................t..g.... [137040]
KM262845.1_strain_NHV            ..........................................c..........c..................t..g.... [137040]
KX354450.1_47/Ss/2008            ..........................................c..........c..................t..g.... [137040]
KP055815.1_strain_BA71           ..........................................c..........c..................t..g.... [137040]
MH025920.1_strain_R35            g...........a........tg.g.....ct..........c..........c...t..t.................a. [137040]
MH025919.1_strain_N10            g...........a........tg.g.....ct..........c..........c...t..t.................a. [137040]
MH025918.1_strain_R25            g...........a........tg.g.....ct..........c..........c...t..t.................a. [137040]
MH025917.1_strain_R7             g...........a........tg.g.....ct..........c..........c...t..t.................a. [137040]
MH025916.1_strain_R8             g...........a........tg.g.....ct..........c..........c...t..t.................a. [137040]

FR682468.2_Georgia_2007/1        gatcttcaaacatctcactaagctgtttcctaaatttttgtaccaaatcccactgggagttgggctgcagcattcctgtt [137120]
ASF_IND_20_CAD543                ................................................................................ [137120]
LR899193.1_Germany_2020/1        ................................................................................ [137120]
MN393477.1_Wuhan_2019            ................................................................................ [137120]
MW396979.1_Timor-Leste/2019/1    ................................................................................ [137120]
MK940252.1_CN/2019/InnerMongolia ................................................................................ [137120]
MK128995.1_China/2018/AnhuiXCGQ  ................................................................................ [137120]
MH681419.1_ASFV/POL/2015         ................................................................................ [137120]
LS478113.1_Estonia_2014          ................................................................................ [137120]
KP843857.1_Odintsovo_02/14       ................................................................................ [137120]
```

```
FN557520.1_strain_E75           ..............................................g................................. [137120]
AY261366.1_Warthog              ................g.............................g................................. [137120]
AY261365.1_Warmbaths            ................g.............................g................................. [137120]
AY261364.1_Tengani_62           ................................................................................ [137120]
AY261363.1_Pretoriuskop/96/4    ................g.............................g.............................g... [137120]
AY261362.1_Mkuzi_1979           ................g.............................g................................. [137120]
AY261361.1_Malawi_Lil-20/1_1983 ................g.....t.......................g........a....................a... [137120]
AY261360.1_Kenya_1950           .............t................................g................................. [137120]
AM712240.1_OURT_88/3            ..............................................g................................. [137120]
AM712239.1_Benin_97/1           ..............................................g................................. [137120]
KM111295.1_Ken06.Bus            .............t................................g................................. [137120]
KM111294.1_Ken05/Tk1            .............t................................g................................. [137120]
KM262844.1_strain_L60           ..............................................g................................. [137120]
KM102979.1_26544/OG10_Italy     ..............................................g................................. [137120]
KM262845.1_strain_NHV           ..............................................g................................. [137120]
KX354450.1_47/Ss/2008           ..............................................g................................. [137120]
KP055815.1_strain_BA71          ..............................................g................................. [137120]
MH025920.1_strain_R35           .............t................................g................................. [137120]
MH025919.1_strain_N10           .............t................................g................................. [137120]
MH025918.1_strain_R25           .............t................................g................................. [137120]
MH025917.1_strain_R7            .............t................................g................................. [137120]
MH025916.1_strain_R8            .............t................................g................................. [137120]

FR682468.2_Georgia_2007/1       tggacatccacagagtctatattgtatagtgccgggcgccacttgggggtaggctgggttgaaggactaataaacctatc [137200]
ASF_IND_20_CAD543               ................................................................................ [137200]
LR899193.1_Germany_2020/1       ................................................................................ [137200]
MN393477.1_Wuhan_2019           ................................................................................ [137200]
MW396979.1_Timor-Leste/2019/1   ................................................................................ [137200]
MK940252.1_CN/2019/InnerMongolia ................................................................................ [137200]
MK128995.1_China/2018/AnhuiXCGQ ................................................................................ [137200]
MH681419.1_ASFV/POL/2015        ................................................................................ [137200]
LS478113.1_Estonia_2014         ................................................................................ [137200]
KP843857.1_Odintsovo_02/14      ................................................................................ [137200]
FN557520.1_strain_E75           .....................................................t.......................... [137200]
AY261366.1_Warthog              ..................................................g.......a..................... [137200]
AY261365.1_Warmbaths            ..................................................g.......a..................... [137200]
AY261364.1_Tengani_62           ................................................................................ [137200]
AY261363.1_Pretoriuskop/96/4    ..a.....t..g.......................a..............g.......a..................... [137200]
AY261362.1_Mkuzi_1979           .............................c.....a............c....t.......................... [137200]
AY261361.1_Malawi_Lil-20/1_1983 ..a.....t..g..a........a........t.................g.....a.ac.......c............ [137200]
AY261360.1_Kenya_1950           ...........g..a...................................g.....a.ac.......c............ [137200]
AM712240.1_OURT_88/3            .....................................................t.......................... [137200]
AM712239.1_Benin_97/1           .....................................................t.......................... [137200]
KM111295.1_Ken06.Bus            ...........g..a...................................g.....a.a........c............ [137200]
```

```
KM111294.1_Ken05/Tk1             ..a........g..a...................................g.....a.ac.......c............ [137200]
KM262844.1_strain_L60            .....................................................t.......................... [137200]
KM102979.1_26544/OG10_Italy      ..a..................................................t.......................... [137200]
KM262845.1_strain_NHV            .....................................................t.......................... [137200]
KX354450.1_47/Ss/2008            ..a..................................................t.......................... [137200]
KP055815.1_strain_BA71           .....................................................t.......................... [137200]
MH025920.1_strain_R35            ...........g..a...................................g.....a.a........c............ [137200]
MH025919.1_strain_N10            ...........g..a...................................g.....a.a........c............ [137200]
MH025918.1_strain_R25            ...........g..a...................................g.....a.a........c............ [137200]
MH025917.1_strain_R7             ...........g..a...................................g.....a.a........c............ [137200]
MH025916.1_strain_R8             ...........g..a...................................g.....a.a........c............ [137200]

FR682468.2_Georgia_2007/1        ggagggaagtaattgtgaggattgtgtatagccatcctcatcaggaagaatggagtagttggtttgattcatcattccaa [137280]
ASF_IND_20_CAD543                ................................................................................ [137280]
LR899193.1_Germany_2020/1        ................................................................................ [137280]
MN393477.1_Wuhan_2019            ................................................................................ [137280]
MW396979.1_Timor-Leste/2019/1    ................................................................................ [137280]
MK940252.1_CN/2019/InnerMongolia ....................................t........................................... [137280]
MK128995.1_China/2018/AnhuiXCGQ  ................................................................................ [137280]
MH681419.1_ASFV/POL/2015         ................................................................................ [137280]
LS478113.1_Estonia_2014          ................................................................................ [137280]
KP843857.1_Odintsovo_02/14       ................................................................................ [137280]
FN557520.1_strain_E75            ...............c.................g............................................g. [137280]
AY261366.1_Warthog               ...............c.................g.............................................. [137280]
AY261365.1_Warmbaths             ...............c.................g............................................g. [137280]
AY261364.1_Tengani_62            ..............................................................................g. [137280]
AY261363.1_Pretoriuskop/96/4     ...............c.................g............................................g. [137280]
AY261362.1_Mkuzi_1979            ...............c.........a.......g............................................g. [137280]
AY261361.1_Malawi_Lil-20/1_1983  ...............c.................g............................................g. [137280]
AY261360.1_Kenya_1950            ...............c..a..............g........t..................................... [137280]
AM712240.1_OURT_88/3             ...............c.................g............................................g. [137280]
AM712239.1_Benin_97/1            ...............c.................g............................................g. [137280]
KM111295.1_Ken06.Bus             ...............c..a..............g........t..................................... [137280]
KM111294.1_Ken05/Tk1             ...............c..a..............g........t..................................... [137280]
KM262844.1_strain_L60            ...............c.................g............................................g. [137280]
KM102979.1_26544/OG10_Italy      ...............c.................g............................................g. [137280]
KM262845.1_strain_NHV            ...............c.................g............................................g. [137280]
KX354450.1_47/Ss/2008            ...............c.................g............................................g. [137280]
KP055815.1_strain_BA71           ...............c.................g............................................g. [137280]
MH025920.1_strain_R35            ...............c..a..............g........t..................................... [137280]
MH025919.1_strain_N10            ...............c..a..............g........t..................................... [137280]
MH025918.1_strain_R25            ...............c..a..............g........t..................................... [137280]
MH025917.1_strain_R7             ...............c..a..............g........t..................................... [137280]
MH025916.1_strain_R8             ...............c..a..............g........t..................................... [137280]
```

```
FR682468.2_Georgia_2007/1        aatcattcatagttcgcgcttcctgaacaatgcgttgaaatttttcccattcggtgcgtgtaatgacaccgaatctgcgg [137360]
ASF_IND_20_CAD543                ....................................................r........................... [137360]
LR899193.1_Germany_2020/1        ................................................................................ [137360]
MN393477.1_Wuhan_2019            ................................................................................ [137360]
MW396979.1_Timor-Leste/2019/1    ................................................................................ [137360]
MK940252.1_CN/2019/InnerMongolia ................................................................................ [137360]
MK128995.1_China/2018/AnhuiXCGQ  ................................................................................ [137360]
MH681419.1_ASFV/POL/2015         ................................................................................ [137360]
LS478113.1_Estonia_2014          ................................................................................ [137360]
KP843857.1_Odintsovo_02/14       ................................................................................ [137360]
FN557520.1_strain_E75            .g..................................................a........g..a..g.....c...... [137360]
AY261366.1_Warthog               .g.............................................................................. [137360]
AY261365.1_Warmbaths             .g.............................................................................. [137360]
AY261364.1_Tengani_62            .g.............................a.............................g..a..g.....c...... [137360]
AY261363.1_Pretoriuskop/96/4     ................................................................................ [137360]
AY261362.1_Mkuzi_1979            .g.............................................................................. [137360]
AY261361.1_Malawi_Lil-20/1_1983  .g..............t....................g...........c..a....................c.....a [137360]
AY261360.1_Kenya_1950            .g........g......................................c.......................c...... [137360]
AM712240.1_OURT_88/3             .g..................................................a........g..a..g.....c...... [137360]
AM712239.1_Benin_97/1            .g..................................................a........g..a..g.....c...... [137360]
KM111295.1_Ken06.Bus             .g........g......................................c.......................c...... [137360]
KM111294.1_Ken05/Tk1             .g........g......................................c.......................c...... [137360]
KM262844.1_strain_L60            .g..................................................a........g..a..g.....c...... [137360]
KM102979.1_26544/OG10_Italy      .g..................................................a........g..a..g.....c...... [137360]
KM262845.1_strain_NHV            .g..................................................a........g..a..g.....c...... [137360]
KX354450.1_47/Ss/2008            .g..................................................a........g..a..g.....c...... [137360]
KP055815.1_strain_BA71           .g..................................................a........g..a..g.....c...... [137360]
MH025920.1_strain_R35            .g........g......................................c.......................c...... [137360]
MH025919.1_strain_N10            .g........g......................................c.......................c...... [137360]
MH025918.1_strain_R25            .g........g......................................c.......................c...... [137360]
MH025917.1_strain_R7             .g........g......................................c.......................c...... [137360]
MH025916.1_strain_R8             .g........g......................................c.......................c...... [137360]

FR682468.2_Georgia_2007/1        tttatttcatttacaaaatggataagcgctttttttggttgcttcttgttcaccatactctaagttaaagtgttggtaaat [137440]
ASF_IND_20_CAD543                ................................................................................ [137440]
LR899193.1_Germany_2020/1        ................................................................................ [137440]
MN393477.1_Wuhan_2019            ................................................................................ [137440]
MW396979.1_Timor-Leste/2019/1    ................................................................................ [137440]
MK940252.1_CN/2019/InnerMongolia ................................................................................ [137440]
MK128995.1_China/2018/AnhuiXCGQ  ................................................................................ [137440]
MH681419.1_ASFV/POL/2015         ................................................................................ [137440]
LS478113.1_Estonia_2014          ................................................................................ [137440]
KP843857.1_Odintsovo_02/14       ................................................................................ [137440]
```

FN557520.1_strain_E75 ................................................................................ [137440]
AY261366.1_Warthog .............................c....................g............................. [137440]
AY261365.1_Warmbaths ..................................................g............................. [137440]
AY261364.1_Tengani_62 ...................................a..c...........g............................. [137440]
AY261363.1_Pretoriuskop/96/4 .............................c........c......................................... [137440]
AY261362.1_Mkuzi_1979 ..................................................g............................. [137440]
AY261361.1_Malawi_Lil-20/1_1983 .................g.................a..g.............................a........... [137440]
AY261360.1_Kenya_1950 .................g...........c......acc...........g........c.................... [137440]
AM712240.1_OURT_88/3 ................................................................................ [137440]
AM712239.1_Benin_97/1 ................................................................................ [137440]
KM111295.1_Ken06.Bus .................g...........c......acc...........g........c.................... [137440]
KM111294.1_Ken05/Tk1 .................g...........c......acc...........g........c.................... [137440]
KM262844.1_strain_L60 ................................................................................ [137440]
KM102979.1_26544/OG10_Italy ................................................................................ [137440]
KM262845.1_strain_NHV ................................................................................ [137440]
KX354450.1_47/Ss/2008 ................................................................................ [137440]
KP055815.1_strain_BA71 ................................................................................ [137440]
MH025920.1_strain_R35 .................g...........c......acc...........g........c.................... [137440]
MH025919.1_strain_N10 .................g...........c......acc...........g........c.................... [137440]
MH025918.1_strain_R25 .................g...........c......acc...........g........c.................... [137440]
MH025917.1_strain_R7 .................g...........c......acc...........g........c.................... [137440]
MH025916.1_strain_R8 .................g...........c......acc...........g........c.................... [137440]

FR682468.2_Georgia_2007/1 gacgtttatttctttgataagctgacgaatttcggtttctgagtagtcaccaatgttaataagctcaataggacgcataa [137520]
ASF_IND_20_CAD543 ................................................................................ [137520]
LR899193.1_Germany_2020/1 ................................................................................ [137520]
MN393477.1_Wuhan_2019 ................................................................................ [137520]
MW396979.1_Timor-Leste/2019/1 ................................................................................ [137520]
MK940252.1_CN/2019/InnerMongolia ................................................................................ [137520]
MK128995.1_China/2018/AnhuiXCGQ ................................................................................ [137520]
MH681419.1_ASFV/POL/2015 ................................................................................ [137520]
LS478113.1_Estonia_2014 ................................................................................ [137520]
KP843857.1_Odintsovo_02/14 ................................................................................ [137520]
FN557520.1_strain_E75 ................................................................................ [137520]
AY261366.1_Warthog a..a..................................................a.................c....... [137520]
AY261365.1_Warmbaths ......................................................a.................c....... [137520]
AY261364.1_Tengani_62 ...........................t..........................a......................... [137520]
AY261363.1_Pretoriuskop/96/4 a..a..................................................a.................c....... [137520]
AY261362.1_Mkuzi_1979 ........................................................................c....... [137520]
AY261361.1_Malawi_Lil-20/1_1983 ..........................................a...........a......................... [137520]
AY261360.1_Kenya_1950 ...............a........g.............................a......................... [137520]
AM712240.1_OURT_88/3 ................................................................................ [137520]
AM712239.1_Benin_97/1 ................................................................................ [137520]
KM111295.1_Ken06.Bus ...............a........g.............................a......................... [137520]

```
KM111294.1_Ken05/Tk1              ...............a........g.............................a......................... [137520]
KM262844.1_strain_L60             ................................................................................ [137520]
KM102979.1_26544/OG10_Italy       ................................................................................ [137520]
KM262845.1_strain_NHV             ................................................................................ [137520]
KX354450.1_47/Ss/2008             ................................................................................ [137520]
KP055815.1_strain_BA71            ................................................................................ [137520]
MH025920.1_strain_R35             ...............a........g.............................a......................... [137520]
MH025919.1_strain_N10             ...............a........g.............................a......................... [137520]
MH025918.1_strain_R25             ...............a........g.............................a......................... [137520]
MH025917.1_strain_R7              ...............a........g.............................a......................... [137520]
MH025916.1_strain_R8              ...............a........g.............................a......................... [137520]

FR682468.2_Georgia_2007/1         agataatgcgaataagtcctgagaagattccttccagctcaggaagcatcgagatctgtacattttcatctctaaaggaa [137600]
ASF_IND_20_CAD543                 .........................k...................................................... [137600]
LR899193.1_Germany_2020/1         ................................................................................ [137600]
MN393477.1_Wuhan_2019             ................................................................................ [137600]
MW396979.1_Timor-Leste/2019/1     ................................................................................ [137600]
MK940252.1_CN/2019/InnerMongolia  ................................................................................ [137600]
MK128995.1_China/2018/AnhuiXCGQ   ................................................................................ [137600]
MH681419.1_ASFV/POL/2015          ................................................................................ [137600]
LS478113.1_Estonia_2014           ................................................................................ [137600]
KP843857.1_Odintsovo_02/14        ................................................................................ [137600]
FN557520.1_strain_E75             .........................t...............................................g...... [137600]
AY261366.1_Warthog                ...................c.....a.....c..t.................a....................g...... [137600]
AY261365.1_Warmbaths              .........................................................................g...... [137600]
AY261364.1_Tengani_62             .........................a...............................................g...... [137600]
AY261363.1_Pretoriuskop/96/4      .........................................................................g...... [137600]
AY261362.1_Mkuzi_1979             ...................c.....a.....c..t.................a........g.................. [137600]
AY261361.1_Malawi_Lil-20/1_1983   ...................c...........c.................................c.......g...... [137600]
AY261360.1_Kenya_1950             ......c...............a........c.....t..g..t...........t..c..............g...... [137600]
AM712240.1_OURT_88/3              .........................t...............................................g...... [137600]
AM712239.1_Benin_97/1             .........................t...............................................g...... [137600]
KM111295.1_Ken06.Bus              ......c...............a........c.....t..g..t...........t..c..............g...... [137600]
KM111294.1_Ken05/Tk1              ......c...............a........c.....t..g..t...........t..c..............g...... [137600]
KM262844.1_strain_L60             .........................t...............................................g...... [137600]
KM102979.1_26544/OG10_Italy       .........................t...............................................g...... [137600]
KM262845.1_strain_NHV             .........................t...............................................g...... [137600]
KX354450.1_47/Ss/2008             .........................t...............................................g...... [137600]
KP055815.1_strain_BA71            .........................t...............................................g...... [137600]
MH025920.1_strain_R35             ......c...............a........c.....t..g..t...........t..c..............g...... [137600]
MH025919.1_strain_N10             ......c...............a........c.....t..g..t...........t..c..............g...... [137600]
MH025918.1_strain_R25             ......c...............a........c.....t..g..t...........t..c..............g...... [137600]
MH025917.1_strain_R7              ......c...............a........c.....t..g..t...........t..c..............g...... [137600]
MH025916.1_strain_R8              ......c...............a........c.....t..g..t...........t..c..............g...... [137600]
```

```
FR682468.2_Georgia_2007/1        aacaacttttgataaaattcggcgaggcggggaaggcggaagtaaagctctgctgcctcgggaattacctcgggctctag [137680]
ASF_IND_20_CAD543                ....ry.......................................................................... [137680]
LR899193.1_Germany_2020/1        ................................................................................ [137680]
MN393477.1_Wuhan_2019            ................................................................................ [137680]
MW396979.1_Timor-Leste/2019/1    ................................................................................ [137680]
MK940252.1_CN/2019/InnerMongolia ................................................................................ [137680]
MK128995.1_China/2018/AnhuiXCGQ  ................................................................................ [137680]
MH681419.1_ASFV/POL/2015         ................................................................................ [137680]
LS478113.1_Estonia_2014          ................................................................................ [137680]
KP843857.1_Odintsovo_02/14       ................................................................................ [137680]
FN557520.1_strain_E75            ....gt...........c...........a.................t..c....................a..t..... [137680]
AY261366.1_Warthog               ....gt...........c..a..............c..............c....................a.......a [137680]
AY261365.1_Warmbaths             .................c..a.............................c............................a [137680]
AY261364.1_Tengani_62            ....gt...........c...........a....................c............................. [137680]
AY261363.1_Pretoriuskop/96/4     .................c..a..............c............................................ [137680]
AY261362.1_Mkuzi_1979            ..................................................c....................a..t..... [137680]
AY261361.1_Malawi_Lil-20/1_1983  ....gt...........c...........a....................c........a..............t..... [137680]
AY261360.1_Kenya_1950            ....gt.....g..............t.....g...............c.c.....t........c.....a........ [137680]
AM712240.1_OURT_88/3             ....gt...........c...........a.................t..c....................a..t..... [137680]
AM712239.1_Benin_97/1            ....gt...........c...........a.................t..c....................a..t..... [137680]
KM111295.1_Ken06.Bus             ....gt.....g..............t.....g...............c.c..............c.............. [137680]
KM111294.1_Ken05/Tk1             ....gt.....g..............t.....t...............c.c.....t........c.....a........ [137680]
KM262844.1_strain_L60            ....gt...........c...........a.................t..c....................a..t..... [137680]
KM102979.1_26544/OG10_Italy      ....gt...........c...........a.................t..c....................a..t..... [137680]
KM262845.1_strain_NHV            ....gt...........c...........a.................t..c....................a..t..... [137680]
KX354450.1_47/Ss/2008            ....gt...........c...........a.................t..c....................a..t..... [137680]
KP055815.1_strain_BA71           ....gt...........c...........a.................t..c....................a..t..... [137680]
MH025920.1_strain_R35            ....gt.....g..............t.....g...............c.c..............c.....a........ [137680]
MH025919.1_strain_N10            ....gt.....g..............t.....g...............c.c..............c.....a........ [137680]
MH025918.1_strain_R25            ....gt.....g..............t.....g...............c.c..............c.....a........ [137680]
MH025917.1_strain_R7             ....gt.....g..............t.....g...............c.c..............c.....a........ [137680]
MH025916.1_strain_R8             ....gt.....g..............t.....g...............c.c..............c.....a........ [137680]

FR682468.2_Georgia_2007/1        ctcatcggcaccccccaatatcatacgcgtgggtataagtttgtacacgggctcaggccgttcaaacatgtcgtaaatcc [137760]
ASF_IND_20_CAD543                ................................................................................ [137760]
LR899193.1_Germany_2020/1        ................................................................................ [137760]
MN393477.1_Wuhan_2019            ................................................................................ [137760]
MW396979.1_Timor-Leste/2019/1    ................................................................................ [137760]
MK940252.1_CN/2019/InnerMongolia ................................................................................ [137760]
MK128995.1_China/2018/AnhuiXCGQ  ................................................................................ [137760]
MH681419.1_ASFV/POL/2015         ................................................................................ [137760]
LS478113.1_Estonia_2014          ................................................................................ [137760]
KP843857.1_Odintsovo_02/14       ................................................................................ [137760]
```

```
FN557520.1_strain_E75             .........g..t....g.........t....................a....................a..a.....g. [137760]
AY261366.1_Warthog                ............t....g.........t....................a....................a........g. [137760]
AY261365.1_Warmbaths              ...g........t....g.........t....................a....................a..a.....g. [137760]
AY261364.1_Tengani_62             ............t....g........................a..t..a.....g..t...........a.......... [137760]
AY261363.1_Pretoriuskop/96/4      ............t....g........................a..t..a.....g..t...........a..a.....g. [137760]
AY261362.1_Mkuzi_1979             .........g..t....g.........t....................a....................a..a.....g. [137760]
AY261361.1_Malawi_Lil-20/1_1983   .........g..t....g...........................t.....t.................a..a..g..g. [137760]
AY261360.1_Kenya_1950             .................gg...................................g..t....................g. [137760]
AM712240.1_OURT_88/3              .........g..t....g.........t....................a....................a..a.....g. [137760]
AM712239.1_Benin_97/1             .........g..t....g.........t....................a....................a..a.....g. [137760]
KM111295.1_Ken06.Bus              ............t....gg...................................g..t....................g. [137760]
KM111294.1_Ken05/Tk1              ............t....gg...................................g..t....................g. [137760]
KM262844.1_strain_L60             .........g..t....g.........t....................a....................a..a.....g. [137760]
KM102979.1_26544/OG10_Italy       .........g..t....g.........t....................a....................a..a.....g. [137760]
KM262845.1_strain_NHV             .........g..t....g.........t....................a....................a..a.....g. [137760]
KX354450.1_47/Ss/2008             .........g..t....g.........t....................a....................a..a.....g. [137760]
KP055815.1_strain_BA71            .........g..t....g.........t....................a....................a..a.....g. [137760]
MH025920.1_strain_R35             ............t....gg...................................g..t....................g. [137760]
MH025919.1_strain_N10             ............t....gg...................................g..t....................g. [137760]
MH025918.1_strain_R25             ............t....gg...................................g..t....................g. [137760]
MH025917.1_strain_R7              ............t....gg...................................g..t....................g. [137760]
MH025916.1_strain_R8              ............t....gg...................................g..t....................g. [137760]

FR682468.2_Georgia_2007/1         ctaatacaataaaaatcttggcggccatacttttcagcatgaaggtgaagaagacgtcctcggtttcccagcgggttgat [137840]
ASF_IND_20_CAD543                 ..........r..................................................................... [137840]
LR899193.1_Germany_2020/1         ................................................................................ [137840]
MN393477.1_Wuhan_2019             ................................................................................ [137840]
MW396979.1_Timor-Leste/2019/1     ................................................................................ [137840]
MK940252.1_CN/2019/InnerMongolia  ..................................t............................................. [137840]
MK128995.1_China/2018/AnhuiXCGQ   ................................................................................ [137840]
MH681419.1_ASFV/POL/2015          ................................................................................ [137840]
LS478113.1_Estonia_2014           ................................................................................ [137840]
KP843857.1_Odintsovo_02/14        ................................................................................ [137840]
FN557520.1_strain_E75             ..........g........a............................................................ [137840]
AY261366.1_Warthog                ..........g........a............................................................ [137840]
AY261365.1_Warmbaths              ..........g........a............................................................ [137840]
AY261364.1_Tengani_62             ................................................................................ [137840]
AY261363.1_Pretoriuskop/96/4      ..........g........a............................................................ [137840]
AY261362.1_Mkuzi_1979             ..........g........a....................a....................................... [137840]
AY261361.1_Malawi_Lil-20/1_1983   ....c...........t.....a.....g...........a...........c.ta.....a............c..... [137840]
AY261360.1_Kenya_1950             ....c..............a..a...........a..........................a.................. [137840]
AM712240.1_OURT_88/3              ..........g........a............................................................ [137840]
AM712239.1_Benin_97/1             ..........g........a............................................................ [137840]
KM111295.1_Ken06.Bus              ....c..............a..a...........a..........................a........t......... [137840]
```

```
KM111294.1_Ken05/Tk1              ....c..............a..a...........a..........................a.................. [137840]
KM262844.1_strain_L60             ..........g........a............................................................ [137840]
KM102979.1_26544/OG10_Italy       ..........g........a............................................................ [137840]
KM262845.1_strain_NHV             ..........g........a............................................................ [137840]
KX354450.1_47/Ss/2008             ..........g........a............................................................ [137840]
KP055815.1_strain_BA71            ..........g........a............................................................ [137840]
MH025920.1_strain_R35             ....c..............a..a...........a..........................a........t......... [137840]
MH025919.1_strain_N10             ....c..............a..a...........a..........................a........t......... [137840]
MH025918.1_strain_R25             ....c..............a..a...........a..........................a........t......... [137840]
MH025917.1_strain_R7              ....c..............a..a...........a..........................a........t......... [137840]
MH025916.1_strain_R8              ....c..............a..a...........a..........................a........t......... [137840]

FR682468.2_Georgia_2007/1         agggcgtcgttaactctcacagtagagaggtagacccgctgagccgcttcctcggcagtctgtgcaagcgccatcctttg [137920]
ASF_IND_20_CAD543                 ...........................................................tc................... [137920]
LR899193.1_Germany_2020/1         ................................................................................ [137920]
MN393477.1_Wuhan_2019             ................................................................................ [137920]
MW396979.1_Timor-Leste/2019/1     ................................................................................ [137920]
MK940252.1_CN/2019/InnerMongolia  ................................................................................ [137920]
MK128995.1_China/2018/AnhuiXCGQ   ................................................................................ [137920]
MH681419.1_ASFV/POL/2015          ................................................................................ [137920]
LS478113.1_Estonia_2014           ................................................................................ [137920]
KP843857.1_Odintsovo_02/14        ................................................................................ [137920]
FN557520.1_strain_E75             .......................................................g...tc..........t.c...... [137920]
AY261366.1_Warthog                ......................................t........c.......g....c..........t.c...... [137920]
AY261365.1_Warmbaths              .......................................................g...tc..........t.c...... [137920]
AY261364.1_Tengani_62             .......................g.......................c................................ [137920]
AY261363.1_Pretoriuskop/96/4      ...............................................c.......g...tc..........t.c...... [137920]
AY261362.1_Mkuzi_1979             .....a.................................................g...tc..........t.c...... [137920]
AY261361.1_Malawi_Lil-20/1_1983   ........a.................a...........t........c................................ [137920]
AY261360.1_Kenya_1950             ...........g...............................a...c.....a.................t.c...... [137920]
AM712240.1_OURT_88/3              .......................................................g...tc..........t.c...... [137920]
AM712239.1_Benin_97/1             .......................................................g...tc..........t.c...... [137920]
KM111295.1_Ken06.Bus              ...........g...............................at..c.......................t.c...... [137920]
KM111294.1_Ken05/Tk1              .a.........g...............................a...c.......................t.c...... [137920]
KM262844.1_strain_L60             .......................................................g...tc..........t.c...... [137920]
KM102979.1_26544/OG10_Italy       .......................................................g...tc..........t.c...... [137920]
KM262845.1_strain_NHV             .......................................................g...tc..........t.c...... [137920]
KX354450.1_47/Ss/2008             .......................................................g...tc..........t.c...... [137920]
KP055815.1_strain_BA71            .......................................................g...tc..........t.c...... [137920]
MH025920.1_strain_R35             ...........g...............................at..c.......................t.c...... [137920]
MH025919.1_strain_N10             ...........g...............................at..c.......................t.c...... [137920]
MH025918.1_strain_R25             ...........g...............................at..c.......................t.c...... [137920]
MH025917.1_strain_R7              ...........g...............................at..c.......................t.c...... [137920]
MH025916.1_strain_R8              ...........g...............................at..c.......................t.c...... [137920]
```

```
FR682468.2_Georgia_2007/1        tcctccaatttctgattgatttagatttttaagtcccacggaaagcgcagaatgttgaagatattcaagcaaggttttat [138000]
ASF_IND_20_CAD543                ................................................................................ [138000]
LR899193.1_Germany_2020/1        ................................................................................ [138000]
MN393477.1_Wuhan_2019            ................................................................................ [138000]
MW396979.1_Timor-Leste/2019/1    ................................................................................ [138000]
MK940252.1_CN/2019/InnerMongolia ................................................................................ [138000]
MK128995.1_China/2018/AnhuiXCGQ  ................................................................................ [138000]
MH681419.1_ASFV/POL/2015         ................................................................................ [138000]
LS478113.1_Estonia_2014          ................................................................................ [138000]
KP843857.1_Odintsovo_02/14       ................................................................................ [138000]
FN557520.1_strain_E75            c...........a..............................................t.................... [138000]
AY261366.1_Warthog               c..........................................................t.................... [138000]
AY261365.1_Warmbaths             c..........................................................t.................... [138000]
AY261364.1_Tengani_62            ..................g...........g................................................. [138000]
AY261363.1_Pretoriuskop/96/4     c..........................................................t.................... [138000]
AY261362.1_Mkuzi_1979            c............................................................................... [138000]
AY261361.1_Malawi_Lil-20/1_1983  ..................g...........g................................................. [138000]
AY261360.1_Kenya_1950            ...........g..t...............g..............t..g..........t.................... [138000]
AM712240.1_OURT_88/3             c...........a..............................................t.................... [138000]
AM712239.1_Benin_97/1            c...........a..............................................t.................... [138000]
KM111295.1_Ken06.Bus             ...........g..t...............g..............t..g..........t.................... [138000]
KM111294.1_Ken05/Tk1             ...........g..t...............g..............t..g..........t.................... [138000]
KM262844.1_strain_L60            c...........a..............................................t.................... [138000]
KM102979.1_26544/OG10_Italy      c...........a..............................................t.................... [138000]
KM262845.1_strain_NHV            c...........a..............................................t.................... [138000]
KX354450.1_47/Ss/2008            c...........a..............................................t.................... [138000]
KP055815.1_strain_BA71           c...........a..............................................t.................... [138000]
MH025920.1_strain_R35            ...........g..t...............g..............t..g..........t.................... [138000]
MH025919.1_strain_N10            ...........g..t...............g..............t..g..........t.................... [138000]
MH025918.1_strain_R25            ...........g..t...............g..............t..g..........t.................... [138000]
MH025917.1_strain_R7             ...........g..t...............g..............t..g..........t.................... [138000]
MH025916.1_strain_R8             ...........g..t...............g..............t..g..........t.................... [138000]

FR682468.2_Georgia_2007/1        agatttgcaggggcgacatgggcaccatttgccgcagctcctctcccccaagcatgtccccaatccgggcaaaggcattg [138080]
ASF_IND_20_CAD543                ................................................................................ [138080]
LR899193.1_Germany_2020/1        ................................................................................ [138080]
MN393477.1_Wuhan_2019            ................................................................................ [138080]
MW396979.1_Timor-Leste/2019/1    ................................................................................ [138080]
MK940252.1_CN/2019/InnerMongolia ................................................................................ [138080]
MK128995.1_China/2018/AnhuiXCGQ  ................................................................................ [138080]
MH681419.1_ASFV/POL/2015         ................................................................................ [138080]
LS478113.1_Estonia_2014          ................................................................................ [138080]
KP843857.1_Odintsovo_02/14       ................................................................................ [138080]
```

```
FN557520.1_strain_E75           ....c....a................................................t..............a...... [138080]
AY261366.1_Warthog              ....c.....................................................t..................... [138080]
AY261365.1_Warmbaths            ....c.....................................................t..............a...... [138080]
AY261364.1_Tengani_62           ....c....................................................................a...... [138080]
AY261363.1_Pretoriuskop/96/4    ....c.....................................................t..................... [138080]
AY261362.1_Mkuzi_1979           ....c....a................................................t..................... [138080]
AY261361.1_Malawi_Lil-20/1_1983 .......a.......................t................................................ [138080]
AY261360.1_Kenya_1950           .a.......aa...............g...................t..g...........g.....a............ [138080]
AM712240.1_OURT_88/3            ....c....a................................................t..............a...... [138080]
AM712239.1_Benin_97/1           ....c....a................................................t..............a...... [138080]
KM111295.1_Ken06.Bus            .a.......aa...............g...................t..g...........g.....a............ [138080]
KM111294.1_Ken05/Tk1            .a.......aa...............g...................t..g...........g.....a............ [138080]
KM262844.1_strain_L60           ....c....a................................................t..............a...... [138080]
KM102979.1_26544/OG10_Italy     ....c....a................................................t..............a...... [138080]
KM262845.1_strain_NHV           ....c....a................................................t..............a...... [138080]
KX354450.1_47/Ss/2008           ....c....a................................................t..............a...... [138080]
KP055815.1_strain_BA71          ....c....a................................................t..............a...... [138080]
MH025920.1_strain_R35           .a.......aa...............g...................t..g...........g.....a............ [138080]
MH025919.1_strain_N10           .a.......aa...............g...................t..g...........g.....a............ [138080]
MH025918.1_strain_R25           .a.......aa...............g...................t..g...........g.....a............ [138080]
MH025917.1_strain_R7            .a.......aa...............g...................t..g...........g.....a............ [138080]
MH025916.1_strain_R8            .a.......aa...............g...................t..g...........g.....a............ [138080]

FR682468.2_Georgia_2007/1       atgatattttttaagcgcctgaaagttagaaagagagcgcccgataaggtcgcgaatgtttttagcctggcttgctctgac [138160]
ASF_IND_20_CAD543               ..........................r..................................................... [138160]
LR899193.1_Germany_2020/1       ................................................................................ [138160]
MN393477.1_Wuhan_2019           ................................................................................ [138160]
MW396979.1_Timor-Leste/2019/1   ................................................................................ [138160]
MK940252.1_CN/2019/InnerMongolia ................................................................................ [138160]
MK128995.1_China/2018/AnhuiXCGQ ................................................................................ [138160]
MH681419.1_ASFV/POL/2015        ................................................................................ [138160]
LS478113.1_Estonia_2014         ................................................................................ [138160]
KP843857.1_Odintsovo_02/14      ................................................................................ [138160]
FN557520.1_strain_E75           ..........................g........a............................................ [138160]
AY261366.1_Warthog              ..........................g........a............................................ [138160]
AY261365.1_Warmbaths            ..........................g........a............................................ [138160]
AY261364.1_Tengani_62           ..........................g..g.................................................. [138160]
AY261363.1_Pretoriuskop/96/4    ..........................g........a............................................ [138160]
AY261362.1_Mkuzi_1979           ..........................g..................................................a.. [138160]
AY261361.1_Malawi_Lil-20/1_1983 ..........................g..g.................................................. [138160]
AY261360.1_Kenya_1950           ..........................g..g................................g................. [138160]
AM712240.1_OURT_88/3            ..........................g........a............................................ [138160]
AM712239.1_Benin_97/1           ..........................g........a............................................ [138160]
KM111295.1_Ken06.Bus            ..........................g..g................................g................. [138160]
```

```
KM111294.1_Ken05/Tk1            ..........................g..g................................g................. [138160]
KM262844.1_strain_L60           ..........................g........a............................................ [138160]
KM102979.1_26544/OG10_Italy     ..........................g........a............................................ [138160]
KM262845.1_strain_NHV           ..........................g........a............................................ [138160]
KX354450.1_47/Ss/2008           ..........................g........a............................................ [138160]
KP055815.1_strain_BA71          ..........................g........a............................................ [138160]
MH025920.1_strain_R35           ..........................g..g................................g................. [138160]
MH025919.1_strain_N10           ..........................g..g................................g................. [138160]
MH025918.1_strain_R25           ..........................g..g................................g................. [138160]
MH025917.1_strain_R7            ..........................g..g................................g................. [138160]
MH025916.1_strain_R8            ..........................g..g................................g................. [138160]

FR682468.2_Georgia_2007/1       gggacggagggtaccaacgcttcggccttgttggatttcagccgcaactttttcgtagtagtggcccgcaggagcattat [138240]
ASF_IND_20_CAD543               ................................................................................ [138240]
LR899193.1_Germany_2020/1       ................................................................................ [138240]
MN393477.1_Wuhan_2019           ................................................................................ [138240]
MW396979.1_Timor-Leste/2019/1   ................................................................................ [138240]
MK940252.1_CN/2019/InnerMongolia ................................................................................ [138240]
MK128995.1_China/2018/AnhuiXCGQ ................................................................................ [138240]
MH681419.1_ASFV/POL/2015        ................................................................................ [138240]
LS478113.1_Estonia_2014         ................................................................................ [138240]
KP843857.1_Odintsovo_02/14      ................................................................................ [138240]
FN557520.1_strain_E75           ................................................................................ [138240]
AY261366.1_Warthog              ................................................................................ [138240]
AY261365.1_Warmbaths            ................................................................................ [138240]
AY261364.1_Tengani_62           ................................................................................ [138240]
AY261363.1_Pretoriuskop/96/4    .....................................................................g.......... [138240]
AY261362.1_Mkuzi_1979           a....................................................................g.......... [138240]
AY261361.1_Malawi_Lil-20/1_1983 ......................................................a......................... [138240]
AY261360.1_Kenya_1950           ...............g....cc..t..c..c................................................. [138240]
AM712240.1_OURT_88/3            ................................................................................ [138240]
AM712239.1_Benin_97/1           ................................................................................ [138240]
KM111295.1_Ken06.Bus            ...............g....cc..t..c..c................................................. [138240]
KM111294.1_Ken05/Tk1            ...............g....cc..t..c..c................................................. [138240]
KM262844.1_strain_L60           ................................................................................ [138240]
KM102979.1_26544/OG10_Italy     ................................................................................ [138240]
KM262845.1_strain_NHV           ................................................................................ [138240]
KX354450.1_47/Ss/2008           ................................................................................ [138240]
KP055815.1_strain_BA71          ................................................................................ [138240]
MH025920.1_strain_R35           ...............g....cc..t..c..c................................................. [138240]
MH025919.1_strain_N10           ...............g....cc..t..c..c................................................. [138240]
MH025918.1_strain_R25           ...............g....cc..t..c..c................................................. [138240]
MH025917.1_strain_R7            ...............g....cc..t..c..c................................................. [138240]
MH025916.1_strain_R8            ...............g....cc..t..c..c................................................. [138240]
```

```
FR682468.2_Georgia_2007/1        ccgtaaagacgttggagtcgttgcctgtggaggtgggaaaactttcaaagacttgtgcaagcgtgtcccctgttgtctcg [138320]
ASF_IND_20_CAD543                ................................................................................ [138320]
LR899193.1_Germany_2020/1        ................................................................................ [138320]
MN393477.1_Wuhan_2019            ................................................................................ [138320]
MW396979.1_Timor-Leste/2019/1    ................................................................................ [138320]
MK940252.1_CN/2019/InnerMongolia ................................................................................ [138320]
MK128995.1_China/2018/AnhuiXCGQ  ................................................................................ [138320]
MH681419.1_ASFV/POL/2015         ................................................................................ [138320]
LS478113.1_Estonia_2014          ................................................................................ [138320]
KP843857.1_Odintsovo_02/14       ................................................................................ [138320]
FN557520.1_strain_E75            ......................a....c...............................................c.... [138320]
AY261366.1_Warthog               ......................a....ca.....a................................t.......c.... [138320]
AY261365.1_Warmbaths             ......................a....c...............................................c.... [138320]
AY261364.1_Tengani_62            ...........................................................................c.... [138320]
AY261363.1_Pretoriuskop/96/4     ...........................c...............................................c.... [138320]
AY261362.1_Mkuzi_1979            ...................a.......ca..............................................c.... [138320]
AY261361.1_Malawi_Lil-20/1_1983  ...........................c..........................................g....c.... [138320]
AY261360.1_Kenya_1950            ...................a.......ca.....c.................c........a.............c...c [138320]
AM712240.1_OURT_88/3             ......................a....c...............................................c.... [138320]
AM712239.1_Benin_97/1            ......................a....c...............................................c.... [138320]
KM111295.1_Ken06.Bus             ...................a.......c......c.................c........a.............c...c [138320]
KM111294.1_Ken05/Tk1             ...................a.......ca.....c.................c........a.............c...c [138320]
KM262844.1_strain_L60            ......................a....c...............................................c.... [138320]
KM102979.1_26544/OG10_Italy      ......................a....c...............................................c.... [138320]
KM262845.1_strain_NHV            ......................a....c...............................................c.... [138320]
KX354450.1_47/Ss/2008            ......................a....c...............................................c.... [138320]
KP055815.1_strain_BA71           ......................a....c...............................................c.... [138320]
MH025920.1_strain_R35            ...................a.......c......c.................c........a.............c...c [138320]
MH025919.1_strain_N10            ...................a.......c......c.................c........a.............c...c [138320]
MH025918.1_strain_R25            ...................a.......c......c.................c........a.............c...c [138320]
MH025917.1_strain_R7             ...................a.......c......c.................c........a.............c...c [138320]
MH025916.1_strain_R8             ...................a.......c......c.................c........a.............c...c [138320]

FR682468.2_Georgia_2007/1        gtgaaccatcgtcctataatgcgcacgccatccagcatctgttggactgtttgaatagaatctatgttgtttacaaacgt [138400]
ASF_IND_20_CAD543                ................................................................................ [138400]
LR899193.1_Germany_2020/1        ................................................................................ [138400]
MN393477.1_Wuhan_2019            ................................................................................ [138400]
MW396979.1_Timor-Leste/2019/1    ................................................................................ [138400]
MK940252.1_CN/2019/InnerMongolia ................................................................................ [138400]
MK128995.1_China/2018/AnhuiXCGQ  ................................................................................ [138400]
MH681419.1_ASFV/POL/2015         ................................................................................ [138400]
LS478113.1_Estonia_2014          ................................................................................ [138400]
KP843857.1_Odintsovo_02/14       ................................................................................ [138400]
```

```
FN557520.1_strain_E75           ..................................a......c...................................... [138400]
AY261366.1_Warthog              ..a...............................a......c...................................... [138400]
AY261365.1_Warmbaths            ..................................a......c...................................... [138400]
AY261364.1_Tengani_62           ..................................a......c...................................... [138400]
AY261363.1_Pretoriuskop/96/4    ..................................a......c...................................... [138400]
AY261362.1_Mkuzi_1979           ..................................a......c...................................... [138400]
AY261361.1_Malawi_Lil-20/1_1983 ...........c..............t.......a......c..a................................t.. [138400]
AY261360.1_Kenya_1950           ..........................c.....t........c...........g.......................t.. [138400]
AM712240.1_OURT_88/3            ..................................a......c...................................... [138400]
AM712239.1_Benin_97/1           ..................................a......c...................................... [138400]
KM111295.1_Ken06.Bus            ..........................c..............c...........g.......................t.. [138400]
KM111294.1_Ken05/Tk1            ..........................c.....t........c...........g.......................t.. [138400]
KM262844.1_strain_L60           ..................................a......c...................................... [138400]
KM102979.1_26544/OG10_Italy     ..................................a......c...................................... [138400]
KM262845.1_strain_NHV           ..................................a......c...................................... [138400]
KX354450.1_47/Ss/2008           ..................................a......c...................................... [138400]
KP055815.1_strain_BA71          ..................................a......c...................................... [138400]
MH025920.1_strain_R35           ..........................c..............c...........g.......................t.. [138400]
MH025919.1_strain_N10           ..........................c..............c...........g.......................t.. [138400]
MH025918.1_strain_R25           ..........................c..............c...........g.......................t.. [138400]
MH025917.1_strain_R7            ..........................c..............c...........g.......................t.. [138400]
MH025916.1_strain_R8            ..........................c..............c...........g.......................t.. [138400]

FR682468.2_Georgia_2007/1       tttggtaatgtttttaagataaagatctagcccttccagagctcgatagaatcggcgttttacatcatactccagctcga [138480]
ASF_IND_20_CAD543               ................................................................................ [138480]
LR899193.1_Germany_2020/1       ................................................................................ [138480]
MN393477.1_Wuhan_2019           ................................................................................ [138480]
MW396979.1_Timor-Leste/2019/1   ................................................................................ [138480]
MK940252.1_CN/2019/InnerMongolia ................................................................................ [138480]
MK128995.1_China/2018/AnhuiXCGQ ................................................................................ [138480]
MH681419.1_ASFV/POL/2015        ................................................................................ [138480]
LS478113.1_Estonia_2014         ................................................................................ [138480]
KP843857.1_Odintsovo_02/14      ................................................................................ [138480]
FN557520.1_strain_E75           ........................g..........................g..............g............. [138480]
AY261366.1_Warthog              ........................g..c.......................c..............g............. [138480]
AY261365.1_Warmbaths            ........................g..........................c...........g................ [138480]
AY261364.1_Tengani_62           .........a..............g..........................c...........g........t....... [138480]
AY261363.1_Pretoriuskop/96/4    ........................g..........................c..............g............. [138480]
AY261362.1_Mkuzi_1979           ........................g..........................c..............g............. [138480]
AY261361.1_Malawi_Lil-20/1_1983 ......g.........................................a..c...........g................ [138480]
AY261360.1_Kenya_1950           ..................g.....g..c.......................c..a......................... [138480]
AM712240.1_OURT_88/3            ........................g..........................g..............g............. [138480]
AM712239.1_Benin_97/1           ........................g..........................g..............g............. [138480]
KM111295.1_Ken06.Bus            ..................g.....g..c.......................c...........g................ [138480]
```

KM111294.1_Ken05/Tk1 ..................g.....g..c.......................c..a......................... [138480]
KM262844.1_strain_L60 ........................g..........................g..............g............. [138480]
KM102979.1_26544/OG10_Italy ........................g..........................g..............g............. [138480]
KM262845.1_strain_NHV ........................g..........................g..............g............. [138480]
KX354450.1_47/Ss/2008 ........................g..........................g..............g............. [138480]
KP055815.1_strain_BA71 ........................g..........................g............................ [138480]
MH025920.1_strain_R35 ..................g.....g..c.......................c...........g................ [138480]
MH025919.1_strain_N10 ..................g.....g..c.......................c...........g................ [138480]
MH025918.1_strain_R25 ..................g.....g..c.......................c...........g................ [138480]
MH025917.1_strain_R7 ..................g.....g..c.......................c...........g................ [138480]
MH025916.1_strain_R8 ..................g.....g..c.......................c...........g................ [138480]

FR682468.2_Georgia_2007/1 tggcgcttacggttgccttccagtctacttcctgggcacctccaggatttgggcccacgtgtcctctggcaagatctaca [138560]
ASF_IND_20_CAD543 ................................................................................ [138560]
LR899193.1_Germany_2020/1 ................................................................................ [138560]
MN393477.1_Wuhan_2019 ................................................................................ [138560]
MW396979.1_Timor-Leste/2019/1 ................................................................................ [138560]
MK940252.1_CN/2019/InnerMongolia ................................................................................ [138560]
MK128995.1_China/2018/AnhuiXCGQ ................................................................................ [138560]
MH681419.1_ASFV/POL/2015 ................................................................................ [138560]
LS478113.1_Estonia_2014 ................................................................................ [138560]
KP843857.1_Odintsovo_02/14 ................................................................................ [138560]
FN557520.1_strain_E75 ..........a.c.............g...................g................................. [138560]
AY261366.1_Warthog ..........................t...................g................................. [138560]
AY261365.1_Warmbaths ..........a.c.............t...................g................................. [138560]
AY261364.1_Tengani_62 ..........................t....t..............g................................. [138560]
AY261363.1_Pretoriuskop/96/4 ..........a.c.................................g................................. [138560]
AY261362.1_Mkuzi_1979 ..........a.c.............g...................g........t........................ [138560]
AY261361.1_Malawi_Lil-20/1_1983 ....a.......c..................t.................a..a........................... [138560]
AY261360.1_Kenya_1950 .........t........................a..g........g........t.....c...........g...... [138560]
AM712240.1_OURT_88/3 ..........a.c.............g...................g................................. [138560]
AM712239.1_Benin_97/1 ..........a.c.............g...................g................................. [138560]
KM111295.1_Ken06.Bus .........ta.......................a..g........g........t.................g...... [138560]
KM111294.1_Ken05/Tk1 .........t........................a..g........g........t.................g...... [138560]
KM262844.1_strain_L60 ..........a.c.............g...................g................................. [138560]
KM102979.1_26544/OG10_Italy ..........a.c.............g...................g................................. [138560]
KM262845.1_strain_NHV ..........a.c.............g...................g................................. [138560]
KX354450.1_47/Ss/2008 ..........a.c.............g...................g................................. [138560]
KP055815.1_strain_BA71 ..........a.c.............g...................g................................. [138560]
MH025920.1_strain_R35 .........ta.......................a..g........g........t.................g...... [138560]
MH025919.1_strain_N10 .........ta.......................a..g........g........t.................g...... [138560]
MH025918.1_strain_R25 .........ta.......................a..g........g........t.................g...... [138560]
MH025917.1_strain_R7 .........ta.......................a..g........g........t.................g...... [138560]
MH025916.1_strain_R8 .........ta.......................a..g........g........t.................g...... [138560]

```
FR682468.2_Georgia_2007/1        gccggagaattaatgcgcgcatttttttccgtatccaactgcatgaggcgtcccgcaatagcatctccgagaatagtggc [138640]
ASF_IND_20_CAD543                ................................................................................ [138640]
LR899193.1_Germany_2020/1        ................................................................................ [138640]
MN393477.1_Wuhan_2019            ................................................................................ [138640]
MW396979.1_Timor-Leste/2019/1    ................................................................................ [138640]
MK940252.1_CN/2019/InnerMongolia ................................................................................ [138640]
MK128995.1_China/2018/AnhuiXCGQ  ................................................................................ [138640]
MH681419.1_ASFV/POL/2015         ................................................................................ [138640]
LS478113.1_Estonia_2014          ................................................................................ [138640]
KP843857.1_Odintsovo_02/14       ................................................................................ [138640]
FN557520.1_strain_E75            ................................................................................ [138640]
AY261366.1_Warthog               ................................................................................ [138640]
AY261365.1_Warmbaths             ................................................................................ [138640]
AY261364.1_Tengani_62            ..........................................................................g..... [138640]
AY261363.1_Pretoriuskop/96/4     ..t............................................................................. [138640]
AY261362.1_Mkuzi_1979            ................................................................................ [138640]
AY261361.1_Malawi_Lil-20/1_1983  .....g........a....................t..t...................................g..... [138640]
AY261360.1_Kenya_1950            ..............................................................g...........g..... [138640]
AM712240.1_OURT_88/3             ................................................................................ [138640]
AM712239.1_Benin_97/1            ................................................................................ [138640]
KM111295.1_Ken06.Bus             ..............................................................g...........g..... [138640]
KM111294.1_Ken05/Tk1             ..............................................................g...........g..... [138640]
KM262844.1_strain_L60            ................................................................................ [138640]
KM102979.1_26544/OG10_Italy      ................................................................................ [138640]
KM262845.1_strain_NHV            ................................................................................ [138640]
KX354450.1_47/Ss/2008            ................................................................................ [138640]
KP055815.1_strain_BA71           ................................................................................ [138640]
MH025920.1_strain_R35            ..............................................................g...........g..... [138640]
MH025919.1_strain_N10            ..............................................................g...........g..... [138640]
MH025918.1_strain_R25            ..............................................................g...........g..... [138640]
MH025917.1_strain_R7             ..............................................................g...........g..... [138640]
MH025916.1_strain_R8             ..............................................................g...........g..... [138640]

FR682468.2_Georgia_2007/1        atagttttcctcgtaggattgaaactcctgtttgttatgcgttaaattggagtaaatctgggccacataatagtaataca [138720]
ASF_IND_20_CAD543                ................................................................................ [138720]
LR899193.1_Germany_2020/1        ................................................................................ [138720]
MN393477.1_Wuhan_2019            ................................................................................ [138720]
MW396979.1_Timor-Leste/2019/1    ................................................................................ [138720]
MK940252.1_CN/2019/InnerMongolia ................................................................................ [138720]
MK128995.1_China/2018/AnhuiXCGQ  ................................................................................ [138720]
MH681419.1_ASFV/POL/2015         ................................................................................ [138720]
LS478113.1_Estonia_2014          ................................................................................ [138720]
KP843857.1_Odintsovo_02/14       ................................................................................ [138720]
```

FN557520.1_strain_E75 .....................................................................g.......... [138720]
AY261366.1_Warthog .....................................................................g.......... [138720]
AY261365.1_Warmbaths .....................................................................g.......... [138720]
AY261364.1_Tengani_62 .....................................................................g.......... [138720]
AY261363.1_Pretoriuskop/96/4 .....................................................................g.......... [138720]
AY261362.1_Mkuzi_1979 .....................................................................g.......... [138720]
AY261361.1_Malawi_Lil-20/1_1983 .....................................................................g.......... [138720]
AY261360.1_Kenya_1950 .....................................................................g.......... [138720]
AM712240.1_OURT_88/3 .....................................................................g.......... [138720]
AM712239.1_Benin_97/1 .....................................................................g.......... [138720]
KM111295.1_Ken06.Bus .....................................................................g.......... [138720]
KM111294.1_Ken05/Tk1 .....................................................................g.......... [138720]
KM262844.1_strain_L60 .....................................................................g.......... [138720]
KM102979.1_26544/OG10_Italy .....................................................................g.......... [138720]
KM262845.1_strain_NHV .....................................................................g.......... [138720]
KX354450.1_47/Ss/2008 .....................................................................g.......... [138720]
KP055815.1_strain_BA71 .....................................................................g.......... [138720]
MH025920.1_strain_R35 .....................................................................g.......... [138720]
MH025919.1_strain_N10 .....................................................................g.......... [138720]
MH025918.1_strain_R25 .....................................................................g.......... [138720]
MH025917.1_strain_R7 .....................................................................g.......... [138720]
MH025916.1_strain_R8 .....................................................................g.......... [138720]

FR682468.2_Georgia_2007/1 taaaggtgttaattgcctggttgaggtcaacctgcgatcgcgcggccttgctgagcccaagctcttcaactgttagggca [138800]
ASF_IND_20_CAD543 ....r..r........................................................................ [138800]
LR899193.1_Germany_2020/1 ................................................................................ [138800]
MN393477.1_Wuhan_2019 ................................................................................ [138800]
MW396979.1_Timor-Leste/2019/1 ................................................................................ [138800]
MK940252.1_CN/2019/InnerMongolia ................................................................................ [138800]
MK128995.1_China/2018/AnhuiXCGQ ................................................................................ [138800]
MH681419.1_ASFV/POL/2015 ................................................................................ [138800]
LS478113.1_Estonia_2014 ................................................................................ [138800]
KP843857.1_Odintsovo_02/14 ................................................................................ [138800]
FN557520.1_strain_E75 ....a..a.............................c...................................g..a... [138800]
AY261366.1_Warthog .......a.............................c.................t..............c..g...... [138800]
AY261365.1_Warmbaths ....a..a.............................c.................t.................g..a... [138800]
AY261364.1_Tengani_62 .......a..........................t..c...........a.......................g...... [138800]
AY261363.1_Pretoriuskop/96/4 ....a..a.............................c....................g...........c..g...... [138800]
AY261362.1_Mkuzi_1979 ....a..a.............................c.................t..............c..g...... [138800]
AY261361.1_Malawi_Lil-20/1_1983 .....................................c....................g..................... [138800]
AY261360.1_Kenya_1950 .....................................c....................g.....c..c..c......... [138800]
AM712240.1_OURT_88/3 ....a..a.............................c...................................g..a... [138800]
AM712239.1_Benin_97/1 ....a..a.............................c...................................g..a... [138800]
KM111295.1_Ken06.Bus ....a................................c....................g.....c..c..c......... [138800]

```
KM111294.1_Ken05/Tk1              .....................................c....................g.....c..c..c......... [138800]
KM262844.1_strain_L60             ....a..a.............................c...................................g..a... [138800]
KM102979.1_26544/OG10_Italy       ....a..a.............................c...................................g..a... [138800]
KM262845.1_strain_NHV             ....a..a.............................c...................................g..a... [138800]
KX354450.1_47/Ss/2008             ....a..a.............................c...................................g..a... [138800]
KP055815.1_strain_BA71            ....a..a.............................c...................................g..a... [138800]
MH025920.1_strain_R35             ....a................................c....................g.....c..c..c......... [138800]
MH025919.1_strain_N10             ....a................................c....................g.....c..c..c......... [138800]
MH025918.1_strain_R25             ....a................................c....................g.....c..c..c......... [138800]
MH025917.1_strain_R7              ....a................................c....................g.....c..c..c......... [138800]
MH025916.1_strain_R8              ....a................................c....................g.....c..c..c......... [138800]

FR682468.2_Georgia_2007/1         gcaccgcctacccttgtacactcgcagtcctcctcgcctccatactttttttgcacaatatcggtataaaaatcaataat [138880]
ASF_IND_20_CAD543                 ................................................................................ [138880]
LR899193.1_Germany_2020/1         ................................................................................ [138880]
MN393477.1_Wuhan_2019             ................................................................................ [138880]
MW396979.1_Timor-Leste/2019/1     ................................................................................ [138880]
MK940252.1_CN/2019/InnerMongolia  ................................................................................ [138880]
MK128995.1_China/2018/AnhuiXCGQ   ................................................................................ [138880]
MH681419.1_ASFV/POL/2015          ................................................................................ [138880]
LS478113.1_Estonia_2014           ................................................................................ [138880]
KP843857.1_Odintsovo_02/14        ................................................................................ [138880]
FN557520.1_strain_E75             .................g...............c.......g...................................... [138880]
AY261366.1_Warthog                .................g...............c.a.....g...................................... [138880]
AY261365.1_Warmbaths              .................g...............c.......g...................................... [138880]
AY261364.1_Tengani_62             .................g.............................................................. [138880]
AY261363.1_Pretoriuskop/96/4      .................g.......................g...................................... [138880]
AY261362.1_Mkuzi_1979             .................g...............c.......g...................................... [138880]
AY261361.1_Malawi_Lil-20/1_1983   .................................c.a....................g....................... [138880]
AY261360.1_Kenya_1950             ............t....................c.a..c.................g....................... [138880]
AM712240.1_OURT_88/3              .................g...............c.......g...................................... [138880]
AM712239.1_Benin_97/1             .................g...............c.......g...................................... [138880]
KM111295.1_Ken06.Bus              ............t..........a.........c.a..c.................g....................... [138880]
KM111294.1_Ken05/Tk1              ............t....................c.a..c.................g....................... [138880]
KM262844.1_strain_L60             .................g...............c.......g...................................... [138880]
KM102979.1_26544/OG10_Italy       .................g...............c.......g...................................... [138880]
KM262845.1_strain_NHV             .................g...............c.......g...................................... [138880]
KX354450.1_47/Ss/2008             .................g...............c.......g...................................... [138880]
KP055815.1_strain_BA71            .................g...............c.......g...................................... [138880]
MH025920.1_strain_R35             ............t..........a.........c.a..c.................g....................... [138880]
MH025919.1_strain_N10             ............t..........a.........c.a..c.................g....................... [138880]
MH025918.1_strain_R25             ............t..........a.........c.a..c.................g....................... [138880]
MH025917.1_strain_R7              ............t..........a.........c.a..c.................g....................... [138880]
MH025916.1_strain_R8              ............t..........a.........c.a..c.................g....................... [138880]
```

```
FR682468.2_Georgia_2007/1        ctgtagcaagcgagagcaggagtcataaagatttttaaaattagggtcggttttagatatctcctccaaaacatttttaa [138960]
ASF_IND_20_CAD543                ..........................................................tg......y............. [138960]
LR899193.1_Germany_2020/1        ................................................................................ [138960]
MN393477.1_Wuhan_2019            ................................................................................ [138960]
MW396979.1_Timor-Leste/2019/1    ................................................................................ [138960]
MK940252.1_CN/2019/InnerMongolia ................................................................................ [138960]
MK128995.1_China/2018/AnhuiXCGQ  ................................................................................ [138960]
MH681419.1_ASFV/POL/2015         ................................................................................ [138960]
LS478113.1_Estonia_2014          ................................................................................ [138960]
KP843857.1_Odintsovo_02/14       ................................................................................ [138960]
FN557520.1_strain_E75            ...c.a...........................................a........tg......t............. [138960]
AY261366.1_Warthog               ...c.at..........................................a........tg......t............. [138960]
AY261365.1_Warmbaths             ...c.at..........................................a........tg......t............. [138960]
AY261364.1_Tengani_62            ...c..............................................................t............. [138960]
AY261363.1_Pretoriuskop/96/4     ...c.a....................................................tg......t............. [138960]
AY261362.1_Mkuzi_1979            ...c.a...........................................a........tg......t............. [138960]
AY261361.1_Malawi_Lil-20/1_1983  ...c.........a...............................a.................................. [138960]
AY261360.1_Kenya_1950            ...c.........a..........................................cc.a......t...........c. [138960]
AM712240.1_OURT_88/3             ...c.a...........................................a........tg......t............. [138960]
AM712239.1_Benin_97/1            ...c.a...........................................a........tg......t............. [138960]
KM111295.1_Ken06.Bus             ...c.........a..........................................cc.a......t...........c. [138960]
KM111294.1_Ken05/Tk1             ...c.........a..........................................cc.a......t...........c. [138960]
KM262844.1_strain_L60            ...c.a...........................................a........tg......t............. [138960]
KM102979.1_26544/OG10_Italy      ...c.a...........................................a........tg......t............. [138960]
KM262845.1_strain_NHV            ...c.a...........................................a........tg......t............. [138960]
KX354450.1_47/Ss/2008            ...c.a...........................................a........tg......t............. [138960]
KP055815.1_strain_BA71           ...c.a...........................................a........tg......t............. [138960]
MH025920.1_strain_R35            ...c.........a..........................................cc.a......t...........c. [138960]
MH025919.1_strain_N10            ...c.........a..........................................cc.a......t...........c. [138960]
MH025918.1_strain_R25            ...c.........a..........................................cc.a......t...........c. [138960]
MH025917.1_strain_R7             ...c.........a..........................................cc.a......t...........c. [138960]
MH025916.1_strain_R8             ...c.........a..........................................cc.a......t...........c. [138960]

FR682468.2_Georgia_2007/1        caagcgtaagctgtgttaagaaggtttcgcgttcttctcgtgcggccgcattggtgtaaaagccgataagacttagatca [139040]
ASF_IND_20_CAD543                ................................................................................ [139040]
LR899193.1_Germany_2020/1        ................................................................................ [139040]
MN393477.1_Wuhan_2019            ................................................................................ [139040]
MW396979.1_Timor-Leste/2019/1    ................................................................................ [139040]
MK940252.1_CN/2019/InnerMongolia ................................................................................ [139040]
MK128995.1_China/2018/AnhuiXCGQ  ................................................................................ [139040]
MH681419.1_ASFV/POL/2015         ................................................................................ [139040]
LS478113.1_Estonia_2014          ................................................................................ [139040]
KP843857.1_Odintsovo_02/14       ................................................................................ [139040]
```

```
FN557520.1_strain_E75            .....a...a.........a..........................t..g.............................. [139040]
AY261366.1_Warthog               .....a...a.........a..a......................................................... [139040]
AY261365.1_Warmbaths             .....a...a.........a..a.......................t..g.............................. [139040]
AY261364.1_Tengani_62            ..............................................t..g.............................. [139040]
AY261363.1_Pretoriuskop/96/4     .....a...a.........a..........................t..g.............................. [139040]
AY261362.1_Mkuzi_1979            .....a...a.........a..........................t..g.............................. [139040]
AY261361.1_Malawi_Lil-20/1_1983  ....t.................a..............c...........g.............................. [139040]
AY261360.1_Kenya_1950            ..............a.c....................c...........g..at.a.....a........g.....g... [139040]
AM712240.1_OURT_88/3             .....a...a.........a..........................t..g.............................. [139040]
AM712239.1_Benin_97/1            .....a...a.........a..........................t..g.............................. [139040]
KM111295.1_Ken06.Bus             ..............a.c....................c...........g..at.a.....a........g.....g... [139040]
KM111294.1_Ken05/Tk1             ..............a.c....................c...........g..at.a.....a........g.....g... [139040]
KM262844.1_strain_L60            .....a...a.........a..........................t..g.............................. [139040]
KM102979.1_26544/OG10_Italy      .....a...a.........a..........................t..g.............................. [139040]
KM262845.1_strain_NHV            .....a...a.........a..........................t..g.............................. [139040]
KX354450.1_47/Ss/2008            .....a...a.........a..........................t..g.............................. [139040]
KP055815.1_strain_BA71           .....a...a.........a..........................t..g.............................. [139040]
MH025920.1_strain_R35            ..............a.c....................c...........g..at.a.....a........g.....g... [139040]
MH025919.1_strain_N10            ..............a.c....................c...........g..at.a.....a........g.....g... [139040]
MH025918.1_strain_R25            ..............a.c....................c...........g..at.a.....a........g.....g... [139040]
MH025917.1_strain_R7             ..............a.c....................c...........g..at.a.....a........g.....g... [139040]
MH025916.1_strain_R8             ..............a.c....................c...........g..at.a.....a........g.....g... [139040]

FR682468.2_Georgia_2007/1        agtgcgatggtgcccatatcattaatgcgcgaaagagcatctcgaagcctcgttatgttcggcgtcaaggcaatttcttt [139120]
ASF_IND_20_CAD543                ................................................................................ [139120]
LR899193.1_Germany_2020/1        ................................................................................ [139120]
MN393477.1_Wuhan_2019            ................................................................................ [139120]
MW396979.1_Timor-Leste/2019/1    ................................................................................ [139120]
MK940252.1_CN/2019/InnerMongolia ................................................................................ [139120]
MK128995.1_China/2018/AnhuiXCGQ  ................................................................................ [139120]
MH681419.1_ASFV/POL/2015         ................................................................................ [139120]
LS478113.1_Estonia_2014          ................................................................................ [139120]
KP843857.1_Odintsovo_02/14       ................................................................................ [139120]
FN557520.1_strain_E75            ..c............................................................................. [139120]
AY261366.1_Warthog               ..c.....................................................a....................c.. [139120]
AY261365.1_Warmbaths             ..c..............................................................t.............. [139120]
AY261364.1_Tengani_62            ..c.............................g............................................... [139120]
AY261363.1_Pretoriuskop/96/4     ..c............................................................................. [139120]
AY261362.1_Mkuzi_1979            ..c..........................................................................c.. [139120]
AY261361.1_Malawi_Lil-20/1_1983  ..c..a....................c.....g..............t................................ [139120]
AY261360.1_Kenya_1950            ..c.............................g...............t............................c.. [139120]
AM712240.1_OURT_88/3             ..c............................................................................. [139120]
AM712239.1_Benin_97/1            ..c............................................................................. [139120]
KM111295.1_Ken06.Bus             ..c.............................g...............t............................c.. [139120]
```

```
KM111294.1_Ken05/Tk1              ..c.............................g...............t............................c.. [139120]
KM262844.1_strain_L60             ..c............................................................................. [139120]
KM102979.1_26544/OG10_Italy       ..c............................................................................. [139120]
KM262845.1_strain_NHV             ..c............................................................................. [139120]
KX354450.1_47/Ss/2008             ..c............................................................................. [139120]
KP055815.1_strain_BA71            ..c............................................................................. [139120]
MH025920.1_strain_R35             ..c.............................g...............t............................c.. [139120]
MH025919.1_strain_N10             ..c.............................g...............t............................c.. [139120]
MH025918.1_strain_R25             ..c.............................g...............t............................c.. [139120]
MH025917.1_strain_R7              ..c.............................g...............t............................c.. [139120]
MH025916.1_strain_R8              ..c.............................g...............t............................c.. [139120]

FR682468.2_Georgia_2007/1         aacaagtttgatgcctatttttttcacattttccaaaaagtcgttataggcttgtgtgcttttattcaaaaattccatga [139200]
ASF_IND_20_CAD543                 ...............................................................r................ [139200]
LR899193.1_Germany_2020/1         ................................................................................ [139200]
MN393477.1_Wuhan_2019             ................................................................................ [139200]
MW396979.1_Timor-Leste/2019/1     ................................................................................ [139200]
MK940252.1_CN/2019/InnerMongolia  ................................................................................ [139200]
MK128995.1_China/2018/AnhuiXCGQ   ................................................................................ [139200]
MH681419.1_ASFV/POL/2015          ................................................................................ [139200]
LS478113.1_Estonia_2014           ................................................................................ [139200]
KP843857.1_Odintsovo_02/14        ................................................................................ [139200]
FN557520.1_strain_E75             ...............................................................g................ [139200]
AY261366.1_Warthog                ...................................................c.....a.....g................ [139200]
AY261365.1_Warmbaths              ...................................................c.....a.....g................ [139200]
AY261364.1_Tengani_62             ...................................................c...........g................ [139200]
AY261363.1_Pretoriuskop/96/4      ................................................a..c...........g................ [139200]
AY261362.1_Mkuzi_1979             ...................................................c.....a.....g................ [139200]
AY261361.1_Malawi_Lil-20/1_1983   ..........................................a........c.....a.....g................ [139200]
AY261360.1_Kenya_1950             ...g.....c.........................................c...........g................ [139200]
AM712240.1_OURT_88/3              ...............................................................g................ [139200]
AM712239.1_Benin_97/1             ...............................................................g................ [139200]
KM111295.1_Ken06.Bus              ...g.....c.........................................c...........g................ [139200]
KM111294.1_Ken05/Tk1              ...g.....c.........................................c...........g................ [139200]
KM262844.1_strain_L60             ...............................................................g................ [139200]
KM102979.1_26544/OG10_Italy       ...............................................................g................ [139200]
KM262845.1_strain_NHV             ...............................................................g................ [139200]
KX354450.1_47/Ss/2008             ...............................................................g................ [139200]
KP055815.1_strain_BA71            ...............................................................g................ [139200]
MH025920.1_strain_R35             ...g.....c.........................................c...........g................ [139200]
MH025919.1_strain_N10             ...g.....c.........................................c...........g................ [139200]
MH025918.1_strain_R25             ...g.....c.........................................c...........g................ [139200]
MH025917.1_strain_R7              ...g.....c.........................................c...........g................ [139200]
MH025916.1_strain_R8              ...g.....c.........................................c...........g................ [139200]
```

```
FR682468.2_Georgia_2007/1        ggatgtgctttctatccagtctttgcgcttcaatcctcctatctagtggcgttttctcctcatcgcccccctttttggca [139280]
ASF_IND_20_CAD543                .............................................................g.................. [139280]
LR899193.1_Germany_2020/1        ................................................................................ [139280]
MN393477.1_Wuhan_2019            ................................................................................ [139280]
MW396979.1_Timor-Leste/2019/1    ................................................................................ [139280]
MK940252.1_CN/2019/InnerMongolia ................................................................................ [139280]
MK128995.1_China/2018/AnhuiXCGQ  ................................................................................ [139280]
MH681419.1_ASFV/POL/2015         ................................................................................ [139280]
LS478113.1_Estonia_2014          ................................................................................ [139280]
KP843857.1_Odintsovo_02/14       ................................................................................ [139280]
FN557520.1_strain_E75            .............................................................g..a............... [139280]
AY261366.1_Warthog               ...................c.....t.............................t.....g..a............... [139280]
AY261365.1_Warmbaths             ...................c.........................................g..a............... [139280]
AY261364.1_Tengani_62            ...................c.......................a.................................... [139280]
AY261363.1_Pretoriuskop/96/4     .........................t.....g.............................g..a............... [139280]
AY261362.1_Mkuzi_1979            ....a..............c.........................................g..a............... [139280]
AY261361.1_Malawi_Lil-20/1_1983  ...................c..................t.................a....................... [139280]
AY261360.1_Kenya_1950            ...................c...........c.....tt....a...............c.................... [139280]
AM712240.1_OURT_88/3             .............................................................g..a............... [139280]
AM712239.1_Benin_97/1            .............................................................g..a............... [139280]
KM111295.1_Ken06.Bus             ...............................c.....tt....a.................................... [139280]
KM111294.1_Ken05/Tk1             ...................c...........c.....tt....a.................................... [139280]
KM262844.1_strain_L60            .............................................................g..a............... [139280]
KM102979.1_26544/OG10_Italy      .............................................................g..a............... [139280]
KM262845.1_strain_NHV            .............................................................g..a............... [139280]
KX354450.1_47/Ss/2008            .............................................................g..a............... [139280]
KP055815.1_strain_BA71           .............................................................g..a............... [139280]
MH025920.1_strain_R35            ...............................c.....tt....a.................................... [139280]
MH025919.1_strain_N10            ..g............................c.....tt....a.................................... [139280]
MH025918.1_strain_R25            ...............................c.....tt....a.................................... [139280]
MH025917.1_strain_R7             ...............................c.....tt....a.................................... [139280]
MH025916.1_strain_R8             ...............................c.....tt....a.................................... [139280]

FR682468.2_Georgia_2007/1        caactgttctcaaggattttgtggcgttcattaaaggtctgtcgcaacaggttcacggctttttcaaactcagcaatgtt [139360]
ASF_IND_20_CAD543                ..............a................................................................. [139360]
LR899193.1_Germany_2020/1        ................................................................................ [139360]
MN393477.1_Wuhan_2019            ................................................................................ [139360]
MW396979.1_Timor-Leste/2019/1    ................................................................................ [139360]
MK940252.1_CN/2019/InnerMongolia ................................................................................ [139360]
MK128995.1_China/2018/AnhuiXCGQ  ................................................................................ [139360]
MH681419.1_ASFV/POL/2015         ................................................................................ [139360]
LS478113.1_Estonia_2014          ................................................................................ [139360]
KP843857.1_Odintsovo_02/14       ................................................................................ [139360]
```

```
FN557520.1_strain_E75          ..............a........................................................g........ [139360]
AY261366.1_Warthog             .....a............................................a.....a..............g........ [139360]
AY261365.1_Warmbaths           ....................a..............a...........t.......................g........ [139360]
AY261364.1_Tengani_62          ....................a...................................a..............g........ [139360]
AY261363.1_Pretoriuskop/96/4   .......................................................................g........ [139360]
AY261362.1_Mkuzi_1979          ....................a..........................t.......................g........ [139360]
AY261361.1_Malawi_Lil-20/1_1983 ...t................a..........................t..a....................g........ [139360]
AY261360.1_Kenya_1950          ...t................a..........................t.......................g........ [139360]
AM712240.1_OURT_88/3           ..............a........................................................g........ [139360]
AM712239.1_Benin_97/1          ..............a........................................................g........ [139360]
KM111295.1_Ken06.Bus           ...t................a........g.................t.......................g........ [139360]
KM111294.1_Ken05/Tk1           ...t................a..........................t.......................g........ [139360]
KM262844.1_strain_L60          ..............a........................................................g........ [139360]
KM102979.1_26544/OG10_Italy    ..............a........................................................g........ [139360]
KM262845.1_strain_NHV          ..............a........................................................g........ [139360]
KX354450.1_47/Ss/2008          ..............a........................................................g........ [139360]
KP055815.1_strain_BA71         ..............a........................................................g........ [139360]
MH025920.1_strain_R35          ...t................a........g.................t.......................g........ [139360]
MH025919.1_strain_N10          ...t................a........g.................t.......................g........ [139360]
MH025918.1_strain_R25          ...t................a........g.................t.......................g........ [139360]
MH025917.1_strain_R7           ...t................a........g.................t.......................g........ [139360]
MH025916.1_strain_R8           ...t................a........g.................t.......................g........ [139360]

FR682468.2_Georgia_2007/1      ttctgcggagacaagaccactaaacctttttgaggtcaagctccttgtcaaactccgcccagtttttgctttgaaggtact [139440]
ASF_IND_20_CAD543              ................................................................................ [139440]
LR899193.1_Germany_2020/1      ................................................................................ [139440]
MN393477.1_Wuhan_2019          ................................................................................ [139440]
MW396979.1_Timor-Leste/2019/1  ................................................................................ [139440]
MK940252.1_CN/2019/InnerMongolia ................................................................................ [139440]
MK128995.1_China/2018/AnhuiXCGQ ................................................................................ [139440]
MH681419.1_ASFV/POL/2015       ................................................................................ [139440]
LS478113.1_Estonia_2014        ................................................................................ [139440]
KP843857.1_Odintsovo_02/14     ................................................................................ [139440]
FN557520.1_strain_E75          ...............g........t.................t................................a.... [139440]
AY261366.1_Warthog             ...............g........t....................................................... [139440]
AY261365.1_Warmbaths           ...............g........t....................................................... [139440]
AY261364.1_Tengani_62          ...............g........t....................................................... [139440]
AY261363.1_Pretoriuskop/96/4   ...............g........t..............................t........................ [139440]
AY261362.1_Mkuzi_1979          ...............g........t..................................................a.... [139440]
AY261361.1_Malawi_Lil-20/1_1983 ......t........g........t....................................................... [139440]
AY261360.1_Kenya_1950          ......c........g........t....................a.......................a.......... [139440]
AM712240.1_OURT_88/3           ...............g........t.................t................................a.... [139440]
AM712239.1_Benin_97/1          ...............g........t.................t................................a.... [139440]
KM111295.1_Ken06.Bus           ......t........g........t....................a.......................a.......... [139440]
```

```
KM111294.1_Ken05/Tk1            ......c........g........t....................a.......................a.......... [139440]
KM262844.1_strain_L60           ...............g........t................t.................................a.... [139440]
KM102979.1_26544/OG10_Italy     ...............g........t................t.................................a.... [139440]
KM262845.1_strain_NHV           ...............g........t................t.................................a.... [139440]
KX354450.1_47/Ss/2008           ...............g........t................t.................................a.... [139440]
KP055815.1_strain_BA71          ...............g........t................t.................................a.... [139440]
MH025920.1_strain_R35           ......t........g........t....................a.......................a.......... [139440]
MH025919.1_strain_N10           ......t........g........t....................a.......................a.......... [139440]
MH025918.1_strain_R25           ......t........g........t....................a.......................a.......... [139440]
MH025917.1_strain_R7            ......t........g........t....................a.......................a.......... [139440]
MH025916.1_strain_R8            ......t........g........t....................a.......................a.......... [139440]

FR682468.2_Georgia_2007/1       gttcaaccttgagtcctactttctggagagccttattaattttattcgcaacagacgcagcaatacctagattacaaagt [139520]
ASF_IND_20_CAD543               ................................................................................ [139520]
LR899193.1_Germany_2020/1       ................................................................................ [139520]
MN393477.1_Wuhan_2019           ................................................................................ [139520]
MW396979.1_Timor-Leste/2019/1   ................................................................................ [139520]
MK940252.1_CN/2019/InnerMongolia ................................................................................ [139520]
MK128995.1_China/2018/AnhuiXCGQ ................................................................................ [139520]
MH681419.1_ASFV/POL/2015        ................................................................................ [139520]
LS478113.1_Estonia_2014         ................................................................................ [139520]
KP843857.1_Odintsovo_02/14      ................................................................................ [139520]
FN557520.1_strain_E75           .......................................................t..g.....g............... [139520]
AY261366.1_Warthog              .......................................................t..g.....g............... [139520]
AY261365.1_Warmbaths            ..............................................t........t..g.....g............... [139520]
AY261364.1_Tengani_62           ..............................................g...........g.....g............... [139520]
AY261363.1_Pretoriuskop/96/4    .......................................................t..g.....g............... [139520]
AY261362.1_Mkuzi_1979           ...............................t..............t........t..g.....g............... [139520]
AY261361.1_Malawi_Lil-20/1_1983 ....................cg...a.............................t..g.....g............... [139520]
AY261360.1_Kenya_1950           ....................cgt...............................t...g.....g............... [139520]
AM712240.1_OURT_88/3            .......................................................t..g.....g............... [139520]
AM712239.1_Benin_97/1           .......................................................t..g.....g............... [139520]
KM111295.1_Ken06.Bus            ....................cgt..a............................t...g.....g............... [139520]
KM111294.1_Ken05/Tk1            ....................cgt...............................t...g.....g............... [139520]
KM262844.1_strain_L60           .......................................................t..g.....g............... [139520]
KM102979.1_26544/OG10_Italy     .......................................................t..g.....g............... [139520]
KM262845.1_strain_NHV           .......................................................t..g.....g............... [139520]
KX354450.1_47/Ss/2008           .......................................................t..g.....g............... [139520]
KP055815.1_strain_BA71          .......................................................t..g.....g............... [139520]
MH025920.1_strain_R35           ....................cgt..a............................t...g.....g............... [139520]
MH025919.1_strain_N10           ....................cgt..a............................t...g.....g............... [139520]
MH025918.1_strain_R25           ....................cgt..a............................t...g.....g............... [139520]
MH025917.1_strain_R7            ....................cgt..a............................t...g.....g............... [139520]
MH025916.1_strain_R8            ....................cgt..a............................t...g.....g............... [139520]
```

```
FR682468.2_Georgia_2007/1        gtgtacgaaagtacttttccaaaatttttggttcccaagacactatttgtatcatttaaaagtttaataatatccacctc [139600]
ASF_IND_20_CAD543                ................................................................................ [139600]
LR899193.1_Germany_2020/1        ................................................................................ [139600]
MN393477.1_Wuhan_2019            ................................................................................ [139600]
MW396979.1_Timor-Leste/2019/1    ................................................................................ [139600]
MK940252.1_CN/2019/InnerMongolia ................................................................................ [139600]
MK128995.1_China/2018/AnhuiXCGQ  ................................................................................ [139600]
MH681419.1_ASFV/POL/2015         ................................................................................ [139600]
LS478113.1_Estonia_2014          ................................................................................ [139600]
KP843857.1_Odintsovo_02/14       ................................................................................ [139600]
FN557520.1_strain_E75            ...........c..............................................g..................... [139600]
AY261366.1_Warthog               ...................................t......................g..................... [139600]
AY261365.1_Warmbaths             ...................................t..............g.......g...c................. [139600]
AY261364.1_Tengani_62            ..........................................................g..................... [139600]
AY261363.1_Pretoriuskop/96/4     .....t.............................t..............g.......g...c................. [139600]
AY261362.1_Mkuzi_1979            ...................................t..............g.......g...c................. [139600]
AY261361.1_Malawi_Lil-20/1_1983  ...........c..............................................g...c..............t.. [139600]
AY261360.1_Kenya_1950            ....................g..............t.........................t...............t.. [139600]
AM712240.1_OURT_88/3             ...........c..............................................g..................... [139600]
AM712239.1_Benin_97/1            ...........c..............................................g..................... [139600]
KM111295.1_Ken06.Bus             ....................g..............t.........................t...............t.. [139600]
KM111294.1_Ken05/Tk1             ....................g..............t.........................t...............t.. [139600]
KM262844.1_strain_L60            ...........c..............................................g..................... [139600]
KM102979.1_26544/OG10_Italy      ...........c..............................................g..................... [139600]
KM262845.1_strain_NHV            ...........c..............................................g..................... [139600]
KX354450.1_47/Ss/2008            ...........c..............................................g..................... [139600]
KP055815.1_strain_BA71           ...........c..............................................g..................... [139600]
MH025920.1_strain_R35            ....................g..............t.........................t...............t.. [139600]
MH025919.1_strain_N10            ....................g..............t.........................t...............t.. [139600]
MH025918.1_strain_R25            ....................g..............t.........................t...............t.. [139600]
MH025917.1_strain_R7             ....................g..............t.........................t...............t.. [139600]
MH025916.1_strain_R8             ....................g..............t.........................t...............t.. [139600]

FR682468.2_Georgia_2007/1        atccgtctgcagtttatcaagttccttttgggtgggagttaaaatattgtcaataaaattcgttaaaatgttgatttgca [139680]
ASF_IND_20_CAD543                ..............................................................................y. [139680]
LR899193.1_Germany_2020/1        ................................................................................ [139680]
MN393477.1_Wuhan_2019            ................................................................................ [139680]
MW396979.1_Timor-Leste/2019/1    ................................................................................ [139680]
MK940252.1_CN/2019/InnerMongolia ................................................................................ [139680]
MK128995.1_China/2018/AnhuiXCGQ  ................................................................................ [139680]
MH681419.1_ASFV/POL/2015         ................................................................................ [139680]
LS478113.1_Estonia_2014          ................................................................................ [139680]
KP843857.1_Odintsovo_02/14       ................................................................................ [139680]
```

```
FN557520.1_strain_E75              ..............................................................................t. [139680]
AY261366.1_Warthog                 ................................................................................ [139680]
AY261365.1_Warmbaths               ................................................................................ [139680]
AY261364.1_Tengani_62              ....................................c........................................... [139680]
AY261363.1_Pretoriuskop/96/4       ................................................................................ [139680]
AY261362.1_Mkuzi_1979              ................................................................................ [139680]
AY261361.1_Malawi_Lil-20/1_1983    ....................................c................................ac......... [139680]
AY261360.1_Kenya_1950              ........t...........................c.....g...........g.....g................... [139680]
AM712240.1_OURT_88/3               ..............................................................................t. [139680]
AM712239.1_Benin_97/1              ..............................................................................t. [139680]
KM111295.1_Ken06.Bus               ........t...........................c.....g...........g.....g................... [139680]
KM111294.1_Ken05/Tk1               ........t...........................c.....g...........g.....g................... [139680]
KM262844.1_strain_L60              ..............................................................................t. [139680]
KM102979.1_26544/OG10_Italy        ..............................................................................t. [139680]
KM262845.1_strain_NHV              ..............................................................................t. [139680]
KX354450.1_47/Ss/2008              ..............................................................................t. [139680]
KP055815.1_strain_BA71             ..............................................................................t. [139680]
MH025920.1_strain_R35              ........t...........................c.....g...........g.....g................... [139680]
MH025919.1_strain_N10              ........t...........................c.....g...........g.....g................... [139680]
MH025918.1_strain_R25              ........t...........................c.....g...........g.....g................... [139680]
MH025917.1_strain_R7               ........t...........................c.....g...........g.....g................... [139680]
MH025916.1_strain_R8               ........t...........................c.....g...........g.....g................... [139680]

FR682468.2_Georgia_2007/1          ggttttgttcatttaaaagtcgacgatatactgcttcaatcatggtgactgcattaatgacttcctcattgggggctgct [139760]
ASF_IND_20_CAD543                  ................................................................................ [139760]
LR899193.1_Germany_2020/1          ................................................................................ [139760]
MN393477.1_Wuhan_2019              ................................................................................ [139760]
MW396979.1_Timor-Leste/2019/1      ................................................................................ [139760]
MK940252.1_CN/2019/InnerMongolia   ................................................................................ [139760]
MK128995.1_China/2018/AnhuiXCGQ    ................................................................................ [139760]
MH681419.1_ASFV/POL/2015           ................................................................................ [139760]
LS478113.1_Estonia_2014            ................................................................................ [139760]
KP843857.1_Odintsovo_02/14         ................................................................................ [139760]
FN557520.1_strain_E75              ................................................................t............... [139760]
AY261366.1_Warthog                 ................................................................................ [139760]
AY261365.1_Warmbaths               ................................................................t........a...... [139760]
AY261364.1_Tengani_62              ................................................................................ [139760]
AY261363.1_Pretoriuskop/96/4       ................................................................t............... [139760]
AY261362.1_Mkuzi_1979              a...............................................................t............... [139760]
AY261361.1_Malawi_Lil-20/1_1983    .................................................c.............................. [139760]
AY261360.1_Kenya_1950              .a...............................................c.............................. [139760]
AM712240.1_OURT_88/3               ................................................................t............... [139760]
AM712239.1_Benin_97/1              ................................................................t............... [139760]
KM111295.1_Ken06.Bus               .................................................c.................g............ [139760]
```

```
KM111294.1_Ken05/Tk1             .................................................c.............................. [139760]
KM262844.1_strain_L60            ................................................................t............... [139760]
KM102979.1_26544/OG10_Italy      ................................................................t............... [139760]
KM262845.1_strain_NHV            ................................................................t............... [139760]
KX354450.1_47/Ss/2008            ................................................................t............... [139760]
KP055815.1_strain_BA71           ................................................................t............... [139760]
MH025920.1_strain_R35            .................................................c.................g............ [139760]
MH025919.1_strain_N10            .................................................c.................g............ [139760]
MH025918.1_strain_R25            .................................................c.................g............ [139760]
MH025917.1_strain_R7             .................................................c.................g............ [139760]
MH025916.1_strain_R8             .................................................c.................g............ [139760]

FR682468.2_Georgia_2007/1        ttggttacctccgtcaccatgcgctcgtgaagttgcttaatggcgtcgtttaacagcttgatattttcaagtgtattttc [139840]
ASF_IND_20_CAD543                ................................................................................ [139840]
LR899193.1_Germany_2020/1        ................................................................................ [139840]
MN393477.1_Wuhan_2019            ................................................................................ [139840]
MW396979.1_Timor-Leste/2019/1    ................................................................................ [139840]
MK940252.1_CN/2019/InnerMongolia ................................................................................ [139840]
MK128995.1_China/2018/AnhuiXCGQ  ................................................................................ [139840]
MH681419.1_ASFV/POL/2015         ................................................................................ [139840]
LS478113.1_Estonia_2014          ................................................................................ [139840]
KP843857.1_Odintsovo_02/14       ................................................................................ [139840]
FN557520.1_strain_E75            ..............................................................g................. [139840]
AY261366.1_Warthog               ..............................................................g................. [139840]
AY261365.1_Warmbaths             ...................................t........a.................g................. [139840]
AY261364.1_Tengani_62            ..........................a...................................g................. [139840]
AY261363.1_Pretoriuskop/96/4     ..............................................................g................. [139840]
AY261362.1_Mkuzi_1979            ..............................................................g................. [139840]
AY261361.1_Malawi_Lil-20/1_1983  ..............................................................g................. [139840]
AY261360.1_Kenya_1950            ................................................c.............g................. [139840]
AM712240.1_OURT_88/3             ..............................................................g................. [139840]
AM712239.1_Benin_97/1            ..............................................................g................. [139840]
KM111295.1_Ken06.Bus             ................................................c.......t.....g................. [139840]
KM111294.1_Ken05/Tk1             ................................................c.............g................. [139840]
KM262844.1_strain_L60            ..............................................................g................. [139840]
KM102979.1_26544/OG10_Italy      ..............................................................g................. [139840]
KM262845.1_strain_NHV            ..............................................................g................. [139840]
KX354450.1_47/Ss/2008            ..............................................................g................. [139840]
KP055815.1_strain_BA71           ..............................................................g................. [139840]
MH025920.1_strain_R35            ................................................c.......t.....g................. [139840]
MH025919.1_strain_N10            ................................................c.......t.....g................. [139840]
MH025918.1_strain_R25            ................................................c.......t.....g................. [139840]
MH025917.1_strain_R7             ................................................c.......t.....g................. [139840]
MH025916.1_strain_R8             ................................................c.......t.....g................. [139840]
```

```
FR682468.2_Georgia_2007/1        tatactgccgtgtacatcaagatactctgcgcgcagtccatgagttagggagttaatgtacagaactatttgtcgacata [139920]
ASF_IND_20_CAD543                ................................................................................ [139920]
LR899193.1_Germany_2020/1        ................................................................................ [139920]
MN393477.1_Wuhan_2019            ................................................................................ [139920]
MW396979.1_Timor-Leste/2019/1    ................................................................................ [139920]
MK940252.1_CN/2019/InnerMongolia ................................................................................ [139920]
MK128995.1_China/2018/AnhuiXCGQ  ................................................................................ [139920]
MH681419.1_ASFV/POL/2015         ................................................................................ [139920]
LS478113.1_Estonia_2014          ................................................................................ [139920]
KP843857.1_Odintsovo_02/14       ................................................................................ [139920]
FN557520.1_strain_E75            .........a..............t..c..........................g......................... [139920]
AY261366.1_Warthog               ........................t..c.......................a..g......................... [139920]
AY261365.1_Warmbaths             ........................t..c..........................g......................... [139920]
AY261364.1_Tengani_62            ...........................c........c.....c...........g......................... [139920]
AY261363.1_Pretoriuskop/96/4     ...........................c..............t..................................... [139920]
AY261362.1_Mkuzi_1979            ........................t..c..........................g......................... [139920]
AY261361.1_Malawi_Lil-20/1_1983  ........................t..c..............g........a.....a...................... [139920]
AY261360.1_Kenya_1950            ...........................c...........g..c...........g......................... [139920]
AM712240.1_OURT_88/3             .........a..............t..c..........................g......................... [139920]
AM712239.1_Benin_97/1            .........a..............t..c..........................g......................... [139920]
KM111295.1_Ken06.Bus             ...........................c...........g..c...........g....................g.... [139920]
KM111294.1_Ken05/Tk1             ...........................c...........g..c...........g......................... [139920]
KM262844.1_strain_L60            .........a..............t..c..........................g......................... [139920]
KM102979.1_26544/OG10_Italy      .........a..............t..c..........................g......................... [139920]
KM262845.1_strain_NHV            .........a..............t..c..........................g......................... [139920]
KX354450.1_47/Ss/2008            .........a..............t..c..........................g......................... [139920]
KP055815.1_strain_BA71           .........a..............t..c..........................g......................... [139920]
MH025920.1_strain_R35            ...........................c...........g..c...........g....................g.... [139920]
MH025919.1_strain_N10            ...........................c...........g..c...........g....................g.... [139920]
MH025918.1_strain_R25            ...........................c...........g..c...........g....................g.... [139920]
MH025917.1_strain_R7             ...........................c...........g..c...........g....................g.... [139920]
MH025916.1_strain_R8             ...........................c...........g..c...........g....................g.... [139920]

FR682468.2_Georgia_2007/1        tactggcggccccttcggtggtatctataagcttatcctgacctaaatcaataaattcctggttaatggcgtctgcaatc [140000]
ASF_IND_20_CAD543                ................................................................................ [140000]
LR899193.1_Germany_2020/1        ................................................................................ [140000]
MN393477.1_Wuhan_2019            ................................................................................ [140000]
MW396979.1_Timor-Leste/2019/1    ................................................................................ [140000]
MK940252.1_CN/2019/InnerMongolia ................................................................................ [140000]
MK128995.1_China/2018/AnhuiXCGQ  ................................................................................ [140000]
MH681419.1_ASFV/POL/2015         ................................................................................ [140000]
LS478113.1_Estonia_2014          ................................................................................ [140000]
KP843857.1_Odintsovo_02/14       ................................................................................ [140000]
```

```
FN557520.1_strain_E75          .............a..............g................................................... [140000]
AY261366.1_Warthog             ................................................................................ [140000]
AY261365.1_Warmbaths           .............a..............g................................................... [140000]
AY261364.1_Tengani_62          ............................g................................................... [140000]
AY261363.1_Pretoriuskop/96/4   ................................................................................ [140000]
AY261362.1_Mkuzi_1979          .............a..............g................................................... [140000]
AY261361.1_Malawi_Lil-20/1_1983 ............................g................................................... [140000]
AY261360.1_Kenya_1950          ............................g................................................... [140000]
AM712240.1_OURT_88/3           .............a..............g................................................... [140000]
AM712239.1_Benin_97/1          .............a..............g................................................... [140000]
KM111295.1_Ken06.Bus           ............................g................................................... [140000]
KM111294.1_Ken05/Tk1           ............................g................................................... [140000]
KM262844.1_strain_L60          .............a..............g................................................... [140000]
KM102979.1_26544/OG10_Italy    .............a..............g................................................... [140000]
KM262845.1_strain_NHV          .............a..............g................................................... [140000]
KX354450.1_47/Ss/2008          .............a..............g................................................... [140000]
KP055815.1_strain_BA71         .............a..............g................................................... [140000]
MH025920.1_strain_R35          ............................g................................................... [140000]
MH025919.1_strain_N10          ............................g................................................... [140000]
MH025918.1_strain_R25          ............................g................................................... [140000]
MH025917.1_strain_R7           ............................g................................................... [140000]
MH025916.1_strain_R8           ............................g................................................... [140000]

FR682468.2_Georgia_2007/1      attttacagacggtctcctgtttttccgcattttttacaaaggtggaaccggctcgaggatcgggcagttgttttttgat [140080]
ASF_IND_20_CAD543              ................................................................................ [140080]
LR899193.1_Germany_2020/1      ................................................................................ [140080]
MN393477.1_Wuhan_2019          ................................................................................ [140080]
MW396979.1_Timor-Leste/2019/1  ................................................................................ [140080]
MK940252.1_CN/2019/InnerMongolia ................................................................................ [140080]
MK128995.1_China/2018/AnhuiXCGQ ................................................................................ [140080]
MH681419.1_ASFV/POL/2015       ................................................................................ [140080]
LS478113.1_Estonia_2014        ................................................................................ [140080]
KP843857.1_Odintsovo_02/14     ................................................................................ [140080]
FN557520.1_strain_E75          ................................................................................ [140080]
AY261366.1_Warthog             ...........a.................................................................... [140080]
AY261365.1_Warmbaths           ................................................................................ [140080]
AY261364.1_Tengani_62          ................................................................................ [140080]
AY261363.1_Pretoriuskop/96/4   ................................................................................ [140080]
AY261362.1_Mkuzi_1979          ................................................................................ [140080]
AY261361.1_Malawi_Lil-20/1_1983 ................................................................................ [140080]
AY261360.1_Kenya_1950          ..........t..................................................................... [140080]
AM712240.1_OURT_88/3           ................................................................................ [140080]
AM712239.1_Benin_97/1          ................................................................................ [140080]
KM111295.1_Ken06.Bus           ..........t..............................a.....g................................ [140080]
```

```
KM111294.1_Ken05/Tk1             ..........t..................................................................... [140080]
KM262844.1_strain_L60            ................................................................................ [140080]
KM102979.1_26544/OG10_Italy      ................................................................................ [140080]
KM262845.1_strain_NHV            ................................................................................ [140080]
KX354450.1_47/Ss/2008            ................................................................................ [140080]
KP055815.1_strain_BA71           ................................................................................ [140080]
MH025920.1_strain_R35            ..........t..............................a.....g................................ [140080]
MH025919.1_strain_N10            ..........t..............................a.....g................................ [140080]
MH025918.1_strain_R25            ..........t..............................a.....g................................ [140080]
MH025917.1_strain_R7             ..........t..............................a.....g................................ [140080]
MH025916.1_strain_R8             ..........t..............................a.....g................................ [140080]

FR682468.2_Georgia_2007/1        atctttaagaatatcttcgatgggctgctttgtgtctactttgaaccctattttggcaatcgccctgataattccttcta [140160]
ASF_IND_20_CAD543                ................................................................................ [140160]
LR899193.1_Germany_2020/1        ................................................................................ [140160]
MN393477.1_Wuhan_2019            ................................................................................ [140160]
MW396979.1_Timor-Leste/2019/1    ................................................................................ [140160]
MK940252.1_CN/2019/InnerMongolia ................................................................................ [140160]
MK128995.1_China/2018/AnhuiXCGQ  ................................................................................ [140160]
MH681419.1_ASFV/POL/2015         ................................................................................ [140160]
LS478113.1_Estonia_2014          ................................................................................ [140160]
KP843857.1_Odintsovo_02/14       ................................................................................ [140160]
FN557520.1_strain_E75            ........................................................................c..c.... [140160]
AY261366.1_Warthog               ........................................................................c..c.... [140160]
AY261365.1_Warmbaths             ........................................................................c..c.... [140160]
AY261364.1_Tengani_62            ..................................................................a........c.... [140160]
AY261363.1_Pretoriuskop/96/4     ........................................................................c..c.... [140160]
AY261362.1_Mkuzi_1979            .............................................t.................................. [140160]
AY261361.1_Malawi_Lil-20/1_1983  ......................................t.................a.......t............... [140160]
AY261360.1_Kenya_1950            ....................................c...................a.......t.......c..c..g. [140160]
AM712240.1_OURT_88/3             ........................................................................c..c.... [140160]
AM712239.1_Benin_97/1            ........................................................................c..c.... [140160]
KM111295.1_Ken06.Bus             ....................................c...................a.......t.......c..c..g. [140160]
KM111294.1_Ken05/Tk1             ....................................c...................a.......t.......c..c..g. [140160]
KM262844.1_strain_L60            ........................................................................c..c.... [140160]
KM102979.1_26544/OG10_Italy      ........................................................................c..c.... [140160]
KM262845.1_strain_NHV            ........................................................................c..c.... [140160]
KX354450.1_47/Ss/2008            ........................................................................c..c.... [140160]
KP055815.1_strain_BA71           ........................................................................c..c.... [140160]
MH025920.1_strain_R35            ....................................c...................a.......t.......c..c..g. [140160]
MH025919.1_strain_N10            ....................................c...................a.......t.......c..c..g. [140160]
MH025918.1_strain_R25            ....................................c...................a.......t.......c..c..g. [140160]
MH025917.1_strain_R7             ....................................c...................a.......t.......c..c..g. [140160]
MH025916.1_strain_R8             ....................................c...................a.......t.......c..c..g. [140160]
```

```
FR682468.2_Georgia_2007/1        taatccgcagctttgctttactcgatacggagtctatgtgataatctttaatgtgttgtacaggatttttgtcccccccg [140240]
ASF_IND_20_CAD543                ................................................................................ [140240]
LR899193.1_Germany_2020/1        ................................................................................ [140240]
MN393477.1_Wuhan_2019            ................................................................................ [140240]
MW396979.1_Timor-Leste/2019/1    ................................................................................ [140240]
MK940252.1_CN/2019/InnerMongolia ................................................................................ [140240]
MK128995.1_China/2018/AnhuiXCGQ  ................................................................................ [140240]
MH681419.1_ASFV/POL/2015         ................................................................................ [140240]
LS478113.1_Estonia_2014          ................................................................................ [140240]
KP843857.1_Odintsovo_02/14       ................................................................................ [140240]
FN557520.1_strain_E75            ...c.............................................g.............................. [140240]
AY261366.1_Warthog               .................................................g.............................. [140240]
AY261365.1_Warmbaths             .................................................g.............................. [140240]
AY261364.1_Tengani_62            .................................................g.............................. [140240]
AY261363.1_Pretoriuskop/96/4     .....................................a...........g.............................. [140240]
AY261362.1_Mkuzi_1979            .........a...................................................................... [140240]
AY261361.1_Malawi_Lil-20/1_1983  .................................................g.......................t...... [140240]
AY261360.1_Kenya_1950            .....................................a...........g...........g.................. [140240]
AM712240.1_OURT_88/3             ...c.............................................g.............................. [140240]
AM712239.1_Benin_97/1            ...c.............................................g.............................. [140240]
KM111295.1_Ken06.Bus             .............................................................g.................. [140240]
KM111294.1_Ken05/Tk1             .....................................a...........g...........g.................. [140240]
KM262844.1_strain_L60            ...c.............................................g.............................. [140240]
KM102979.1_26544/OG10_Italy      ...c.............................................g.............................. [140240]
KM262845.1_strain_NHV            ...c.............................................g.............................. [140240]
KX354450.1_47/Ss/2008            ...c.............................................g.............................. [140240]
KP055815.1_strain_BA71           ...c.............................................g.............................. [140240]
MH025920.1_strain_R35            .............................................................g.................. [140240]
MH025919.1_strain_N10            .............................................................g.................. [140240]
MH025918.1_strain_R25            .............................................................g.................. [140240]
MH025917.1_strain_R7             .............................................................g.................. [140240]
MH025916.1_strain_R8             .............................................................g.................. [140240]

FR682468.2_Georgia_2007/1        ccattaaaatatcctccccctgaaaaaggacgagtttgtctttgtatatgatcctgtaacttcgcatatatatttgcttc [140320]
ASF_IND_20_CAD543                ................................................................................ [140320]
LR899193.1_Germany_2020/1        ................................................................................ [140320]
MN393477.1_Wuhan_2019            ................................................................................ [140320]
MW396979.1_Timor-Leste/2019/1    ................................................................................ [140320]
MK940252.1_CN/2019/InnerMongolia ................................................................................ [140320]
MK128995.1_China/2018/AnhuiXCGQ  ................................................................................ [140320]
MH681419.1_ASFV/POL/2015         ................................................................................ [140320]
LS478113.1_Estonia_2014          ................................................................................ [140320]
KP843857.1_Odintsovo_02/14       ................................................................................ [140320]
```

```
FN557520.1_strain_E75              ......................................................................a......... [140320]
AY261366.1_Warthog                 ................................................................................ [140320]
AY261365.1_Warmbaths               ................................................................................ [140320]
AY261364.1_Tengani_62              ...........a.................................................................... [140320]
AY261363.1_Pretoriuskop/96/4       ................................................................................ [140320]
AY261362.1_Mkuzi_1979              .................t.............................................................. [140320]
AY261361.1_Malawi_Lil-20/1_1983    .................t..a........................................................... [140320]
AY261360.1_Kenya_1950              .................t..a.................c..............t................a......... [140320]
AM712240.1_OURT_88/3               ......................................................................a......... [140320]
AM712239.1_Benin_97/1              ......................................................................a......... [140320]
KM111295.1_Ken06.Bus               .................t..a.......a........................t................a......... [140320]
KM111294.1_Ken05/Tk1               .................t..a.c...............c..............t................a......... [140320]
KM262844.1_strain_L60              ......................................................................a......... [140320]
KM102979.1_26544/OG10_Italy        ......................................................................a......... [140320]
KM262845.1_strain_NHV              ......................................................................a......... [140320]
KX354450.1_47/Ss/2008              ......................................................................a......... [140320]
KP055815.1_strain_BA71             ......................................................................a......... [140320]
MH025920.1_strain_R35              .................t..a.......a........................t................a......... [140320]
MH025919.1_strain_N10              .................t..a.......a........................t................a......... [140320]
MH025918.1_strain_R25              .................t..a.......a........................t................a......... [140320]
MH025917.1_strain_R7               .................t..a.......a........................t................a......... [140320]
MH025916.1_strain_R8               .................t..a.......a........................t................a......... [140320]

FR682468.2_Georgia_2007/1          tgatgaaggcagtggtctactagaggttgaagatccacggttacccattataataaaaaaaaataaagatttaaaactac [140400]
ASF_IND_20_CAD543                  ................................................................................ [140400]
LR899193.1_Germany_2020/1          ................................................................................ [140400]
MN393477.1_Wuhan_2019              ................................................................................ [140400]
MW396979.1_Timor-Leste/2019/1      ................................................................................ [140400]
MK940252.1_CN/2019/InnerMongolia   ................................................................................ [140400]
MK128995.1_China/2018/AnhuiXCGQ    ................................................................................ [140400]
MH681419.1_ASFV/POL/2015           ................................................................................ [140400]
LS478113.1_Estonia_2014            ................................................................................ [140400]
KP843857.1_Odintsovo_02/14         ......................................................-......................... [140400]
FN557520.1_strain_E75              .......a..g..................................................................... [140400]
AY261366.1_Warthog                 ..........g..................................................................... [140400]
AY261365.1_Warmbaths               ..........g..................................................................... [140400]
AY261364.1_Tengani_62              .......a..g..............a...................................................... [140400]
AY261363.1_Pretoriuskop/96/4       ..........g..................................................................... [140400]
AY261362.1_Mkuzi_1979              ..........g..................................................................... [140400]
AY261361.1_Malawi_Lil-20/1_1983    .......a..g..................................................................... [140400]
AY261360.1_Kenya_1950              .......a..g.............c....................................................... [140400]
AM712240.1_OURT_88/3               .......a..g..................................................................... [140400]
AM712239.1_Benin_97/1              .......a..g..................................................................... [140400]
KM111295.1_Ken06.Bus               .......a..g.............c....................................................... [140400]
```

```
KM111294.1_Ken05/Tk1             .......a..g............c........................................................ [140400]
KM262844.1_strain_L60            .......a..g..................................................................... [140400]
KM102979.1_26544/OG10_Italy      .......a..g..................................................................... [140400]
KM262845.1_strain_NHV            .......a..g..................................................................... [140400]
KX354450.1_47/Ss/2008            .......a..g..................................................................... [140400]
KP055815.1_strain_BA71           .......a..g..................................................................... [140400]
MH025920.1_strain_R35            .......a..g............c........................................................ [140400]
MH025919.1_strain_N10            .......a..g............c........................................................ [140400]
MH025918.1_strain_R25            .......a..g............c........................................................ [140400]
MH025917.1_strain_R7             .......a..g............c........................................................ [140400]
MH025916.1_strain_R8             .......a..g............c........................................................ [140400]

FR682468.2_Georgia_2007/1        aaatattttgctgtttataa--------------------acccaatcatataagacta-------------------ac [140480]
ASF_IND_20_CAD543                ..................r.--------------------.y.................-------------------.. [140480]
LR899193.1_Germany_2020/1        ....................--------------------...................-------------------.. [140480]
MN393477.1_Wuhan_2019            ....................--------------------...................-------------------.. [140480]
MW396979.1_Timor-Leste/2019/1    ....................--------------------...................-------------------.. [140480]
MK940252.1_CN/2019/InnerMongolia ....................--------------------...................-------------------.. [140480]
MK128995.1_China/2018/AnhuiXCGQ  ....................--------------------...................-------------------.. [140480]
MH681419.1_ASFV/POL/2015         ....................--------------------...................-------------------.. [140480]
LS478113.1_Estonia_2014          ....................--------------------...................-------------------.. [140480]
KP843857.1_Odintsovo_02/14       ....................--------------------...................-------------------.. [140480]
FN557520.1_strain_E75            ..................g.--------------------.t..............a..-------------------.. [140480]
AY261366.1_Warthog               ..................g.--------------------................a..-------------------.. [140480]
AY261365.1_Warmbaths             ..................g.--------------------................a..-------------------.. [140480]
AY261364.1_Tengani_62            ..................gc--------------------................a..-------------------t. [140480]
AY261363.1_Pretoriuskop/96/4     ..................g.--------------------................a..-------------------.. [140480]
AY261362.1_Mkuzi_1979            ....................--------------------................a..-------------------.. [140480]
AY261361.1_Malawi_Lil-20/1_1983  ..............cc..gg--------------------.t......--.gg.ctg..-------------------.. [140480]
AY261360.1_Kenya_1950            ..............cc..gggccaatcaattaggattgca.a.......at..g....gcaaaccaatcaggattgcc.. [140480]
AM712240.1_OURT_88/3             ..................g.--------------------.t..............a..-------------------.. [140480]
AM712239.1_Benin_97/1            ..................g.--------------------.t..............a..-------------------.. [140480]
KM111295.1_Ken06.Bus             ..............cc..gg---------------------a.......at..g....gcaaaccaatcaggattgcc.. [140480]
KM111294.1_Ken05/Tk1             ..............cc..gg---------------------a.......at..g....gcaaaccaatcaggattgcc.. [140480]
KM262844.1_strain_L60            ..................g.--------------------.t..............a..-------------------.. [140480]
KM102979.1_26544/OG10_Italy      ..................g.--------------------.t..............a..-------------------.. [140480]
KM262845.1_strain_NHV            ..................g.--------------------.t..............a..-------------------.. [140480]
KX354450.1_47/Ss/2008            ..................g.--------------------.t..............a..-------------------.. [140480]
KP055815.1_strain_BA71           ..................g.--------------------.t..............a..-------------------.. [140480]
MH025920.1_strain_R35            ..............cc..gg---------------------a.......at..g....gcaaaccaatcaggattgcc.. [140480]
MH025919.1_strain_N10            ..............cc..gg---------------------a.......at..g....gcaaaccaatcaggattgcc.. [140480]
MH025918.1_strain_R25            ..............cc..gg---------------------a.......at..g....gcaaaccaatcaggattgcc.. [140480]
MH025917.1_strain_R7             ..............cc..gg---------------------a.......at..g....gcaaaccaatcaggattgcc.. [140480]
MH025916.1_strain_R8             ..............cc..gg---------------------a.......at..g....gcaaaccaatcaggattgcc.. [140480]
```

```
FR682468.2_Georgia_2007/1        taaaacattaaatgtaggtgagataaaagcttattttttttttttaaaagtttaataaccatgagtcttaccacctcttttt [140560]
ASF_IND_20_CAD543                .................................-.............................................. [140560]
LR899193.1_Germany_2020/1        ................................................................................ [140560]
MN393477.1_Wuhan_2019            ................................................................................ [140560]
MW396979.1_Timor-Leste/2019/1    ................................................................................ [140560]
MK940252.1_CN/2019/InnerMongolia ................................................................................ [140560]
MK128995.1_China/2018/AnhuiXCGQ  ................................................................................ [140560]
MH681419.1_ASFV/POL/2015         ................................................................................ [140560]
LS478113.1_Estonia_2014          ................................................................................ [140560]
KP843857.1_Odintsovo_02/14       .................................-.............................................. [140560]
FN557520.1_strain_E75            ......................................................g......................... [140560]
AY261366.1_Warthog               ......t..........c....................................g.......t................. [140560]
AY261365.1_Warmbaths             ....g.t..........c....................................g.......t................. [140560]
AY261364.1_Tengani_62            ....c.................................................g......................... [140560]
AY261363.1_Pretoriuskop/96/4     ......t..........c....................................g......................... [140560]
AY261362.1_Mkuzi_1979            ....g.t..........c....................................g......................... [140560]
AY261361.1_Malawi_Lil-20/1_1983  ....c.......................a.........................g..............tt......... [140560]
AY261360.1_Kenya_1950            ...................a........ctc.......................g...............t......... [140560]
AM712240.1_OURT_88/3             ......................................................g......................... [140560]
AM712239.1_Benin_97/1            ......................................................g......................... [140560]
KM111295.1_Ken06.Bus             ..................ca........ctc.......................g...............t......... [140560]
KM111294.1_Ken05/Tk1             ...................a........ctc.......................g...............t......... [140560]
KM262844.1_strain_L60            ......................................................g......................... [140560]
KM102979.1_26544/OG10_Italy      ......................................................g......................... [140560]
KM262845.1_strain_NHV            .................................-....................g......................... [140560]
KX354450.1_47/Ss/2008            ......................................................g......................... [140560]
KP055815.1_strain_BA71           ......................................................g......................... [140560]
MH025920.1_strain_R35            ..................ca........ctc.......................g...............t......... [140560]
MH025919.1_strain_N10            ..................ca........ctc.......................g...............t......... [140560]
MH025918.1_strain_R25            ..................ca........ctc.......................g...............t......... [140560]
MH025917.1_strain_R7             ..................ca........ctc.......................g...............t......... [140560]
MH025916.1_strain_R8             ..................ca........ctc.......................g...............t......... [140560]

FR682468.2_Georgia_2007/1        cttcttcctttagaggggttccataaatggtttgaataaaattatgtgctctaataaccttgttaaaatcaggtgccttt [140640]
ASF_IND_20_CAD543                ................................................................................ [140640]
LR899193.1_Germany_2020/1        ................................................................................ [140640]
MN393477.1_Wuhan_2019            ................................................................................ [140640]
MW396979.1_Timor-Leste/2019/1    ................................................................................ [140640]
MK940252.1_CN/2019/InnerMongolia ................................................................................ [140640]
MK128995.1_China/2018/AnhuiXCGQ  ................................................................................ [140640]
MH681419.1_ASFV/POL/2015         ................................................................................ [140640]
LS478113.1_Estonia_2014          ................................................................................ [140640]
KP843857.1_Odintsovo_02/14       ................................................................................ [140640]
```

```
FN557520.1_strain_E75            ........................g...............g....................................... [140640]
AY261366.1_Warthog               ........................g...............g....................................... [140640]
AY261365.1_Warmbaths             ........................g...............g....................................... [140640]
AY261364.1_Tengani_62            .......t................g...............g....................................... [140640]
AY261363.1_Pretoriuskop/96/4     ........................g...............g....................................... [140640]
AY261362.1_Mkuzi_1979            ........................g...............g....................................... [140640]
AY261361.1_Malawi_Lil-20/1_1983  ....c..t....ag..........g...............g..............g.t...................... [140640]
AY261360.1_Kenya_1950            ....c..t........ac..........t.........t.g..............g........................ [140640]
AM712240.1_OURT_88/3             ........................g...............g....................................... [140640]
AM712239.1_Benin_97/1            ........................g...............g....................................... [140640]
KM111295.1_Ken06.Bus             ....c..t.........c..........t.........t.g..............g........................ [140640]
KM111294.1_Ken05/Tk1             ....c..t.........c..........t.........t.g..............g........................ [140640]
KM262844.1_strain_L60            ........................g...............g....................................... [140640]
KM102979.1_26544/OG10_Italy      ........................g...............g....................................... [140640]
KM262845.1_strain_NHV            ........................g...............g....................................... [140640]
KX354450.1_47/Ss/2008            ........................g...............g....................................... [140640]
KP055815.1_strain_BA71           ........................g...............g....................................... [140640]
MH025920.1_strain_R35            ....c..t.........c..........t.........t.g..............g........................ [140640]
MH025919.1_strain_N10            ....c..t.........c..........t.........t.g..............g........................ [140640]
MH025918.1_strain_R25            ....c..t.........c..........t.........t.g..............g........................ [140640]
MH025917.1_strain_R7             ....c..t.........c..........t.........t.g..............g........................ [140640]
MH025916.1_strain_R8             ....c..t.........c..........t.........t.g..............g........................ [140640]

FR682468.2_Georgia_2007/1        ccatattgttcaatatgttgcacagtcttttgtgcaagcatatacagcttggagtctttaggtacctc---cgatgaggg [140720]
ASF_IND_20_CAD543                ....................................................................---......... [140720]
LR899193.1_Germany_2020/1        ....................................................................---......... [140720]
MN393477.1_Wuhan_2019            ....................................................................---......... [140720]
MW396979.1_Timor-Leste/2019/1    ....................................................................---......... [140720]
MK940252.1_CN/2019/InnerMongolia ....................................................................---......... [140720]
MK128995.1_China/2018/AnhuiXCGQ  ....................................................................---......... [140720]
MH681419.1_ASFV/POL/2015         ....................................................................---......... [140720]
LS478113.1_Estonia_2014          ....................................................................---......... [140720]
KP843857.1_Odintsovo_02/14       ....................................................................---......... [140720]
FN557520.1_strain_E75            ...............................................................t....---t........ [140720]
AY261366.1_Warthog               ....................................................................---.....g... [140720]
AY261365.1_Warmbaths             ..............g.....................................................---......... [140720]
AY261364.1_Tengani_62            ..............g.................................g..............g.a..---t....g... [140720]
AY261363.1_Pretoriuskop/96/4     ....................................................................---......... [140720]
AY261362.1_Mkuzi_1979            ..............g............................................g........---......... [140720]
AY261361.1_Malawi_Lil-20/1_1983  .....c........g.......................tgc...........c..t...g...gt...tgtt..a..a.. [140720]
AY261360.1_Kenya_1950            .....c........g..........c.............gc...........t...c..ga..gt...---t........ [140720]
AM712240.1_OURT_88/3             ...............................................................t....---t........ [140720]
AM712239.1_Benin_97/1            ...............................................................t....---t........ [140720]
KM111295.1_Ken06.Bus             .....c........g..........c.............gc...........t...c..g...gt...---t........ [140720]
```

```
KM111294.1_Ken05/Tk1             .....c........g..........c.............gc...........t...c..g...gt...---t........ [140720]
KM262844.1_strain_L60            ...............................................................t....---t........ [140720]
KM102979.1_26544/OG10_Italy      ...............................................................t....---t........ [140720]
KM262845.1_strain_NHV            ...............................................................t....---t........ [140720]
KX354450.1_47/Ss/2008            ...............................................................t....---t........ [140720]
KP055815.1_strain_BA71           ...............................................................t....---t........ [140720]
MH025920.1_strain_R35            .....c........g..........c.............gc...........t...c..g...gt...---t........ [140720]
MH025919.1_strain_N10            .....c........g..........c.............gc...........t...c..g...gt...---t........ [140720]
MH025918.1_strain_R25            .....c........g..........c.............gc...........t...c..g...gt...---t........ [140720]
MH025917.1_strain_R7             .....c........g..........c.............gc...........t...c..g...gt...---t........ [140720]
MH025916.1_strain_R8             .....c........g..........c.............gc...........t...c..g...gt...---t........ [140720]

FR682468.2_Georgia_2007/1        ctcttgctcaaacaacgtttcaaaggaggatgtgcattcattgg---------------tttcattatcatttttttcat [140800]
ASF_IND_20_CAD543                ...............y............................---------------..................... [140800]
LR899193.1_Germany_2020/1        ............................................---------------..................... [140800]
MN393477.1_Wuhan_2019            ............................................---------------..................... [140800]
MW396979.1_Timor-Leste/2019/1    ............................................---------------..................... [140800]
MK940252.1_CN/2019/InnerMongolia ............................................---------------..................... [140800]
MK128995.1_China/2018/AnhuiXCGQ  ............................................---------------..................... [140800]
MH681419.1_ASFV/POL/2015         ............................................---------------..................... [140800]
LS478113.1_Estonia_2014          ............................................---------------..................... [140800]
KP843857.1_Odintsovo_02/14       ............................................---------------..................... [140800]
FN557520.1_strain_E75            ...............t............................---------------..................... [140800]
AY261366.1_Warthog               ...............t..........................a.---------------..................... [140800]
AY261365.1_Warmbaths             ...............t............................---------------..................... [140800]
AY261364.1_Tengani_62            ...............t............................---------------..................... [140800]
AY261363.1_Pretoriuskop/96/4     ...............t............................---------------..c.................. [140800]
AY261362.1_Mkuzi_1979            ...............t............................---------------..................... [140800]
AY261361.1_Malawi_Lil-20/1_1983  ......t.......gg........a..t.............caatttcatgatcattct......g.............. [140800]
AY261360.1_Kenya_1950            ..............g............t.....a........aatttcatggccatccgc..t..g.c....ag...... [140800]
AM712240.1_OURT_88/3             ...............t................c...........---------------..................... [140800]
AM712239.1_Benin_97/1            ...............t............................---------------..................... [140800]
KM111295.1_Ken06.Bus             ..............g............t..............a.tttcatggctatccgc..t..g.c....ag...... [140800]
KM111294.1_Ken05/Tk1             ..............g............t..............a.tttcatggccatccgc..t..g.c....ag...... [140800]
KM262844.1_strain_L60            ...............t............................---------------..................... [140800]
KM102979.1_26544/OG10_Italy      ...............t............................---------------..................... [140800]
KM262845.1_strain_NHV            ...............t................c...........---------------..................... [140800]
KX354450.1_47/Ss/2008            ...............t............................---------------..................... [140800]
KP055815.1_strain_BA71           ...............t............................---------------..................... [140800]
MH025920.1_strain_R35            ..............g............t..............a.tttcatggctatccgc..t..g.c....ag...... [140800]
MH025919.1_strain_N10            ..............g............t..............a.tttcatggctatccgc..t..g.c....ag...... [140800]
MH025918.1_strain_R25            ..............g............t..............a.tttcatggctatccgc..t..g.c....ag...... [140800]
MH025917.1_strain_R7             ..............g............t..............a.tttcatggctatccgc..t..g.c....ag...... [140800]
MH025916.1_strain_R8             ..............g............t..............a.tttcatggctatccgc..t..g.c....ag...... [140800]
```

```
FR682468.2_Georgia_2007/1        gaatgttctccgaagatgctgag------gattccgtctcctcttcaaacagcacatgcagaatcatattccattcttct [140880]
ASF_IND_20_CAD543                .....c.t...............------................................................... [140880]
LR899193.1_Germany_2020/1        .......................------................................................... [140880]
MN393477.1_Wuhan_2019            .......................------................................................... [140880]
MW396979.1_Timor-Leste/2019/1    .......................------................................................... [140880]
MK940252.1_CN/2019/InnerMongolia .......................------................................................... [140880]
MK128995.1_China/2018/AnhuiXCGQ  .......................------................................................... [140880]
MH681419.1_ASFV/POL/2015         .......................------................................................... [140880]
LS478113.1_Estonia_2014          .......................------................................................... [140880]
KP843857.1_Odintsovo_02/14       .......................------................................................... [140880]
FN557520.1_strain_E75            .....c.t.....t.........------.....t............................................. [140880]
AY261366.1_Warthog               .....c.t.....t.........------......................a............................ [140880]
AY261365.1_Warmbaths             .....c.t.....t....t....------................................................... [140880]
AY261364.1_Tengani_62            .......t.....t.........------........t.......................................... [140880]
AY261363.1_Pretoriuskop/96/4     .....c.t.....t.........------................................................... [140880]
AY261362.1_Mkuzi_1979            .....c.t.....t....t....------................................................... [140880]
AY261361.1_Malawi_Lil-20/1_1983  .....c....ta.g.........------..........................g...........g............ [140880]
AY261360.1_Kenya_1950            ..t.ac....ta.g....t...agatgtt...a............................g.................. [140880]
AM712240.1_OURT_88/3             .....c.t.....t.........------.....t............................................. [140880]
AM712239.1_Benin_97/1            .....c.t.....t.........------.....t............................................. [140880]
KM111295.1_Ken06.Bus             ..t..c....ta.g....t...agatgtt.......c........................g.................. [140880]
KM111294.1_Ken05/Tk1             ..t.ac....ta.g....t...agatgtt................................g.................. [140880]
KM262844.1_strain_L60            .....c.t.....t.........------.....t............................................. [140880]
KM102979.1_26544/OG10_Italy      .....c.t.....t.........------.....t............................................. [140880]
KM262845.1_strain_NHV            .....c.t.....t.........------.....t............................................. [140880]
KX354450.1_47/Ss/2008            .....c.t.....t.........------.....t............................................. [140880]
KP055815.1_strain_BA71           .....c.t.....t.........------.....t............................................. [140880]
MH025920.1_strain_R35            ..t..c....ta.g....t...agatgtt.......c........................g.................. [140880]
MH025919.1_strain_N10            ..t..c....ta.g....t...agatgtt.......c........................g.................. [140880]
MH025918.1_strain_R25            ..t..c....ta.g....t...agatgtt.......c........................g.................. [140880]
MH025917.1_strain_R7             ..t..c....ta.g....t...agatgtt.......c........................g.................. [140880]
MH025916.1_strain_R8             ..t..c....ta.g....t...agatgtt.......c........................g.................. [140880]

FR682468.2_Georgia_2007/1        tgagcctgatgttcagtatacccttgccctgcatatatacgagcagatttcacaatatcatacttaacagtactaagcaa [140960]
ASF_IND_20_CAD543                ..........................................................................r..... [140960]
LR899193.1_Germany_2020/1        ................................................................................ [140960]
MN393477.1_Wuhan_2019            ................................................................................ [140960]
MW396979.1_Timor-Leste/2019/1    ................................................................................ [140960]
MK940252.1_CN/2019/InnerMongolia ..............................................................a................. [140960]
MK128995.1_China/2018/AnhuiXCGQ  ................................................................................ [140960]
MH681419.1_ASFV/POL/2015         ................................................................................ [140960]
LS478113.1_Estonia_2014          ................................................................................ [140960]
KP843857.1_Odintsovo_02/14       ................................................................................ [140960]
```

```
FN557520.1_strain_E75            .......................................t..................................g..... [140960]
AY261366.1_Warthog               ...............................................c..........................g..... [140960]
AY261365.1_Warmbaths             ..........................................................................g..... [140960]
AY261364.1_Tengani_62            ..........................t...............................................g..... [140960]
AY261363.1_Pretoriuskop/96/4     ..........................................................................g..... [140960]
AY261362.1_Mkuzi_1979            ..........................................................................g..... [140960]
AY261361.1_Malawi_Lil-20/1_1983  ........t...........t..c........g....c...c......c.......................t.g..... [140960]
AY261360.1_Kenya_1950            ........c...........t..c..t........g..g......tt.c.........................g....g [140960]
AM712240.1_OURT_88/3             .......................................t..................................g..... [140960]
AM712239.1_Benin_97/1            .......................................t..................................g..... [140960]
KM111295.1_Ken06.Bus             ........c...........t..c..t........g..g......tt.c.........................g....g [140960]
KM111294.1_Ken05/Tk1             ........c...........t..c..t........g..g......tt.c.........................g....g [140960]
KM262844.1_strain_L60            .......................................t..................................g..... [140960]
KM102979.1_26544/OG10_Italy      .......................................t..................................g..... [140960]
KM262845.1_strain_NHV            .......................................t..................................g..... [140960]
KX354450.1_47/Ss/2008            .......................................t..................................g..... [140960]
KP055815.1_strain_BA71           .......................................t..................................g..... [140960]
MH025920.1_strain_R35            ........c...........t..c..t........g..g......tt.c.........................g....g [140960]
MH025919.1_strain_N10            ........c...........t..c..t........g..g......tt.c.........................g....g [140960]
MH025918.1_strain_R25            ........c...........t..c..t........g..g......tt.c.........................g....g [140960]
MH025917.1_strain_R7             ........c...........t..c..t........g..g......tt.c.........................g....g [140960]
MH025916.1_strain_R8             ........c...........t..c..t........g..g......tt.c.........................g....g [140960]

FR682468.2_Georgia_2007/1        tgtttttatagcggtcgtaacaattctaccgctattgataatctcaacagaaaaccaattatacaggctacccgcatgaa [141040]
ASF_IND_20_CAD543                ................................................................................ [141040]
LR899193.1_Germany_2020/1        ................................................................................ [141040]
MN393477.1_Wuhan_2019            ................................................................................ [141040]
MW396979.1_Timor-Leste/2019/1    ................................................................................ [141040]
MK940252.1_CN/2019/InnerMongolia ................................................................................ [141040]
MK128995.1_China/2018/AnhuiXCGQ  ................................................................................ [141040]
MH681419.1_ASFV/POL/2015         ................................................................................ [141040]
LS478113.1_Estonia_2014          ................................................................................ [141040]
KP843857.1_Odintsovo_02/14       ................................................................................ [141040]
FN557520.1_strain_E75            .................................................................a.............. [141040]
AY261366.1_Warthog               ..............................a..................................a.............. [141040]
AY261365.1_Warmbaths             ..............................a..................................a.............. [141040]
AY261364.1_Tengani_62            ........................................................................t....... [141040]
AY261363.1_Pretoriuskop/96/4     ..............................a................................................. [141040]
AY261362.1_Mkuzi_1979            ..............................a..................................a......t....... [141040]
AY261361.1_Malawi_Lil-20/1_1983  ..............................a.....a..........tg.........g.............t....... [141040]
AY261360.1_Kenya_1950            ...............t.....g...................................gg.......a.....g....... [141040]
AM712240.1_OURT_88/3             .................................................................a.............. [141040]
AM712239.1_Benin_97/1            .................................................................a.............. [141040]
KM111295.1_Ken06.Bus             ...............t.........................................gg.............g....... [141040]
```

```
KM111294.1_Ken05/Tk1            ...............t.....g...................................gg.......a.....a....... [141040]
KM262844.1_strain_L60           .................................................................a.............. [141040]
KM102979.1_26544/OG10_Italy     .................................................................a.............. [141040]
KM262845.1_strain_NHV           .................................................................a.............. [141040]
KX354450.1_47/Ss/2008           .................................................................a.............. [141040]
KP055815.1_strain_BA71          .................................................................a.............. [141040]
MH025920.1_strain_R35           ...............t.........................................gg.............g....... [141040]
MH025919.1_strain_N10           ...............t.........................................gg.............g....... [141040]
MH025918.1_strain_R25           ...............t.........................................gg.............g....... [141040]
MH025917.1_strain_R7            ...............t.........................................gg.............g....... [141040]
MH025916.1_strain_R8            ...............t.........................................gg.............g....... [141040]

FR682468.2_Georgia_2007/1       acacaacttgtgaagatgatcttaaatccgttttgaagatgacctccattttcatggatatatttaaaataaaatccatt [141120]
ASF_IND_20_CAD543               ................................................................................ [141120]
LR899193.1_Germany_2020/1       ................................................................................ [141120]
MN393477.1_Wuhan_2019           ................................................................................ [141120]
MW396979.1_Timor-Leste/2019/1   ................................................................................ [141120]
MK940252.1_CN/2019/InnerMongolia ................................................................................ [141120]
MK128995.1_China/2018/AnhuiXCGQ ................................................................................ [141120]
MH681419.1_ASFV/POL/2015        ................................................................................ [141120]
LS478113.1_Estonia_2014         ................................................................................ [141120]
KP843857.1_Odintsovo_02/14      ................................................................................ [141120]
FN557520.1_strain_E75           ................................................................................ [141120]
AY261366.1_Warthog              .............................................................................g.. [141120]
AY261365.1_Warmbaths            .............................................................................g.. [141120]
AY261364.1_Tengani_62           ................................................................................ [141120]
AY261363.1_Pretoriuskop/96/4    .............g...............................................................g.. [141120]
AY261362.1_Mkuzi_1979           ................................................................................ [141120]
AY261361.1_Malawi_Lil-20/1_1983 ....g..c.......................c..a.....c.............c......................... [141120]
AY261360.1_Kenya_1950           ....g........g....c.....................t....................................g.. [141120]
AM712240.1_OURT_88/3            ................................................................................ [141120]
AM712239.1_Benin_97/1           ................................................................................ [141120]
KM111295.1_Ken06.Bus            .............g....c.....................t....................................... [141120]
KM111294.1_Ken05/Tk1            ....g........g....c.....................t....................................... [141120]
KM262844.1_strain_L60           ................................................................................ [141120]
KM102979.1_26544/OG10_Italy     ................................................................................ [141120]
KM262845.1_strain_NHV           ................................................................................ [141120]
KX354450.1_47/Ss/2008           ................................................................................ [141120]
KP055815.1_strain_BA71          ................................................................................ [141120]
MH025920.1_strain_R35           .............g....c.....................t....................................... [141120]
MH025919.1_strain_N10           .............g....c.....................t....................................... [141120]
MH025918.1_strain_R25           .............g....c.....................t....................................... [141120]
MH025917.1_strain_R7            .............g....c.....................t....................................... [141120]
MH025916.1_strain_R8            .............g....c.....................t....................................... [141120]
```

```
FR682468.2_Georgia_2007/1        caattttaaaattataaaataataagaagatgccctctaatatgaaacagttttgcaagatttctgtatggctacagcag [141200]
ASF_IND_20_CAD543                ................................................................................ [141200]
LR899193.1_Germany_2020/1        ................................................................................ [141200]
MN393477.1_Wuhan_2019            ................................................................................ [141200]
MW396979.1_Timor-Leste/2019/1    ................................................................................ [141200]
MK940252.1_CN/2019/InnerMongolia ................................................................................ [141200]
MK128995.1_China/2018/AnhuiXCGQ  ................................................................................ [141200]
MH681419.1_ASFV/POL/2015         ................................................................................ [141200]
LS478113.1_Estonia_2014          ................................................................................ [141200]
KP843857.1_Odintsovo_02/14       ................................................................................ [141200]
FN557520.1_strain_E75            .........................................................................g...... [141200]
AY261366.1_Warthog               ................................................................................ [141200]
AY261365.1_Warmbaths             ................................................................................ [141200]
AY261364.1_Tengani_62            ................................................................................ [141200]
AY261363.1_Pretoriuskop/96/4     ................................................................................ [141200]
AY261362.1_Mkuzi_1979            ................................................................................ [141200]
AY261361.1_Malawi_Lil-20/1_1983  ........................................c..............t...............t........ [141200]
AY261360.1_Kenya_1950            ........................................c....................................... [141200]
AM712240.1_OURT_88/3             .........................................................................g...... [141200]
AM712239.1_Benin_97/1            .........................................................................g...... [141200]
KM111295.1_Ken06.Bus             ........................................c.....................................g. [141200]
KM111294.1_Ken05/Tk1             ........................................c....................................... [141200]
KM262844.1_strain_L60            .........................................................................g...... [141200]
KM102979.1_26544/OG10_Italy      .........................................................................g...... [141200]
KM262845.1_strain_NHV            .........................................................................g...... [141200]
KX354450.1_47/Ss/2008            .........................................................................g...... [141200]
KP055815.1_strain_BA71           .........................................................................g...... [141200]
MH025920.1_strain_R35            ........................................c.....................................g. [141200]
MH025919.1_strain_N10            ........................................c.....................................g. [141200]
MH025918.1_strain_R25            ........................................c.....................................g. [141200]
MH025917.1_strain_R7             ........................................c.....................................g. [141200]
MH025916.1_strain_R8             ........................................c.....................................g. [141200]

FR682468.2_Georgia_2007/1        cacgatccagatttattagaaattatcaacaacttatgtatgcttggcaatttatccgcggcaaagtacaaacacggagt [141280]
ASF_IND_20_CAD543                ................................................................................ [141280]
LR899193.1_Germany_2020/1        ................................................................................ [141280]
MN393477.1_Wuhan_2019            ................................................................................ [141280]
MW396979.1_Timor-Leste/2019/1    ................................................................................ [141280]
MK940252.1_CN/2019/InnerMongolia ................................................................................ [141280]
MK128995.1_China/2018/AnhuiXCGQ  ................................................................................ [141280]
MH681419.1_ASFV/POL/2015         ................................................................................ [141280]
LS478113.1_Estonia_2014          ................................................................................ [141280]
KP843857.1_Odintsovo_02/14       ................................................................................ [141280]
```

```
FN557520.1_strain_E75             ................................................................................ [141280]
AY261366.1_Warthog                ................................................................................ [141280]
AY261365.1_Warmbaths              ................................................................................ [141280]
AY261364.1_Tengani_62             ................................................................................ [141280]
AY261363.1_Pretoriuskop/96/4      ................................................................................ [141280]
AY261362.1_Mkuzi_1979             ................................................................................ [141280]
AY261361.1_Malawi_Lil-20/1_1983   ..t..............................c.............................................. [141280]
AY261360.1_Kenya_1950             ........g...c....................c.............................................. [141280]
AM712240.1_OURT_88/3              ................................................................................ [141280]
AM712239.1_Benin_97/1             ................................................................................ [141280]
KM111295.1_Ken06.Bus              ........g...c................t...c.............................................. [141280]
KM111294.1_Ken05/Tk1              ........g...c....................c.............................................. [141280]
KM262844.1_strain_L60             ................................................................................ [141280]
KM102979.1_26544/OG10_Italy       ................................................................................ [141280]
KM262845.1_strain_NHV             ................................................................................ [141280]
KX354450.1_47/Ss/2008             ................................................................................ [141280]
KP055815.1_strain_BA71            ................................................................................ [141280]
MH025920.1_strain_R35             ........g...c................t...c.............................................. [141280]
MH025919.1_strain_N10             ........g...c................t...c.............................................. [141280]
MH025918.1_strain_R25             ........g...c................t...c.............................................. [141280]
MH025917.1_strain_R7              ........g...c................t...c.............................................. [141280]
MH025916.1_strain_R8              ........g...c................t...c.............................................. [141280]

FR682468.2_Georgia_2007/1         taccttcatttaccccaaacaggcaaagatccgcgatgaaataaaaaaacatgcctactccaatgacccttcacaagcca [141360]
ASF_IND_20_CAD543                 ................................................................................ [141360]
LR899193.1_Germany_2020/1         ................................................................................ [141360]
MN393477.1_Wuhan_2019             ................................................................................ [141360]
MW396979.1_Timor-Leste/2019/1     ................................................................................ [141360]
MK940252.1_CN/2019/InnerMongolia  ................................................................................ [141360]
MK128995.1_China/2018/AnhuiXCGQ   ................................................................................ [141360]
MH681419.1_ASFV/POL/2015          ................................................................................ [141360]
LS478113.1_Estonia_2014           ................................................................................ [141360]
KP843857.1_Odintsovo_02/14        ................................................................................ [141360]
FN557520.1_strain_E75             ............t........................................................c.....g.... [141360]
AY261366.1_Warthog                ............t.......................c................................c.....g.... [141360]
AY261365.1_Warmbaths              ............t.......................c................................c.....g.... [141360]
AY261364.1_Tengani_62             ............t................................................................... [141360]
AY261363.1_Pretoriuskop/96/4      ............t.......................c.............................t..c.......... [141360]
AY261362.1_Mkuzi_1979             ............t........................................................c.....g.... [141360]
AY261361.1_Malawi_Lil-20/1_1983   ............t....................t..c....................t...........c..g..g.... [141360]
AY261360.1_Kenya_1950             g...........t....................t.......................t...........c.....g.... [141360]
AM712240.1_OURT_88/3              ............t........................................................c.....g.... [141360]
AM712239.1_Benin_97/1             ............t........................................................c.....g.... [141360]
KM111295.1_Ken06.Bus              g...........t.................a..t.......................t...........c.....g.... [141360]
```

```
KM111294.1_Ken05/Tk1             g...........t....................t.......................t...........c.....g.... [141360]
KM262844.1_strain_L60            ............t........................................................c.....g.... [141360]
KM102979.1_26544/OG10_Italy      ............t........................................................c.....g.... [141360]
KM262845.1_strain_NHV            ............t........................................................c.....g.... [141360]
KX354450.1_47/Ss/2008            ............t........................................................c.....g.... [141360]
KP055815.1_strain_BA71           ............t........................................................c.....g.... [141360]
MH025920.1_strain_R35            g...........t.................a..t.......................t...........c.....g.... [141360]
MH025919.1_strain_N10            g...........t.................a..t.......................t...........c.....g.... [141360]
MH025918.1_strain_R25            g...........t.................a..t.......................t...........c.....g.... [141360]
MH025917.1_strain_R7             g...........t.................a..t.......................t...........c.....g.... [141360]
MH025916.1_strain_R8             g...........t.................a..t.......................t...........c.....g.... [141360]

FR682468.2_Georgia_2007/1        taaagaccttagaatcactcatccttccattttacattcccactccagcggagttcaccggggaaatcggctcctacacc [141440]
ASF_IND_20_CAD543                ................................................................................ [141440]
LR899193.1_Germany_2020/1        ................................................................................ [141440]
MN393477.1_Wuhan_2019            ................................................................................ [141440]
MW396979.1_Timor-Leste/2019/1    ................................................................................ [141440]
MK940252.1_CN/2019/InnerMongolia ................................................................................ [141440]
MK128995.1_China/2018/AnhuiXCGQ  ................................................................................ [141440]
MH681419.1_ASFV/POL/2015         ................................................................................ [141440]
LS478113.1_Estonia_2014          ................................................................................ [141440]
KP843857.1_Odintsovo_02/14       ................................................................................ [141440]
FN557520.1_strain_E75            ................................................................................ [141440]
AY261366.1_Warthog               ...............................................................................t [141440]
AY261365.1_Warmbaths             ................................................................................ [141440]
AY261364.1_Tengani_62            .............................................................a.................. [141440]
AY261363.1_Pretoriuskop/96/4     ................................................................................ [141440]
AY261362.1_Mkuzi_1979            ................................................................................ [141440]
AY261361.1_Malawi_Lil-20/1_1983  .............g..c............................................................... [141440]
AY261360.1_Kenya_1950            ........c......................................at............................... [141440]
AM712240.1_OURT_88/3             ................................................................................ [141440]
AM712239.1_Benin_97/1            ................................................................................ [141440]
KM111295.1_Ken06.Bus             ........c..................................c...at............................... [141440]
KM111294.1_Ken05/Tk1             ........c......................................at..............................t [141440]
KM262844.1_strain_L60            ................................................................................ [141440]
KM102979.1_26544/OG10_Italy      ................................................................................ [141440]
KM262845.1_strain_NHV            ................................................................................ [141440]
KX354450.1_47/Ss/2008            ................................................................................ [141440]
KP055815.1_strain_BA71           ................................................................................ [141440]
MH025920.1_strain_R35            ........c..................................c...at............................... [141440]
MH025919.1_strain_N10            ........c..................................c...at............................... [141440]
MH025918.1_strain_R25            ........c..................................c...at............................... [141440]
MH025917.1_strain_R7             ........c..................................c...at............................... [141440]
MH025916.1_strain_R8             ........c..................................c...at............................... [141440]
```

```
FR682468.2_Georgia_2007/1        ggagtgaaattagaggttgaaaaaacggaggcgaataaagttattttaaaaaatggagaagcggtcctagtaccggcggc [141520]
ASF_IND_20_CAD543                ...............................................r................................ [141520]
LR899193.1_Germany_2020/1        ................................................................................ [141520]
MN393477.1_Wuhan_2019            ................................................................................ [141520]
MW396979.1_Timor-Leste/2019/1    ................................................................................ [141520]
MK940252.1_CN/2019/InnerMongolia ................................................................................ [141520]
MK128995.1_China/2018/AnhuiXCGQ  ................................................................................ [141520]
MH681419.1_ASFV/POL/2015         ................................................................................ [141520]
LS478113.1_Estonia_2014          ................................................................................ [141520]
KP843857.1_Odintsovo_02/14       ................................................................................ [141520]
FN557520.1_strain_E75            .................c.............................g..............a.....g..t........ [141520]
AY261366.1_Warthog               .................c............................................a.....g........... [141520]
AY261365.1_Warmbaths             .................c..................................................g........... [141520]
AY261364.1_Tengani_62            .................c............................................a.....g........... [141520]
AY261363.1_Pretoriuskop/96/4     .................c.............................g..............a................. [141520]
AY261362.1_Mkuzi_1979            .................c.............................g..............a.....g........... [141520]
AY261361.1_Malawi_Lil-20/1_1983  .................c.............................g.................t..ga.......... [141520]
AY261360.1_Kenya_1950            .................c.......a...a.................g........g.....a......a....a..... [141520]
AM712240.1_OURT_88/3             .................c.............................g..............a.....g..t........ [141520]
AM712239.1_Benin_97/1            .................c.............................g..............a.....g..t........ [141520]
KM111295.1_Ken06.Bus             .................c.......a...a............g....g........g.....a.....ga....t..... [141520]
KM111294.1_Ken05/Tk1             .................c.......a...a.................g........g.....a......a....a..... [141520]
KM262844.1_strain_L60            .................c.............................g..............a.....g..t........ [141520]
KM102979.1_26544/OG10_Italy      .................c.............................g..............a.....g..t........ [141520]
KM262845.1_strain_NHV            .................c.............................g..............a.....g..t........ [141520]
KX354450.1_47/Ss/2008            .................c.............................g..............a.....g..t........ [141520]
KP055815.1_strain_BA71           .................c.............................g..............a.....g..t........ [141520]
MH025920.1_strain_R35            .................c.......a...a............g....g........g.....a......a....a..... [141520]
MH025919.1_strain_N10            .................c.......a...a............g....g........g.....a......a....a..... [141520]
MH025918.1_strain_R25            .................c.......a...a............g....g........g.....a......a....a..... [141520]
MH025917.1_strain_R7             .................c.......a...a............g....g........g.....a......a....a..... [141520]
MH025916.1_strain_R8             .................c.......a...a............g....g........g.....a......a....a..... [141520]

FR682468.2_Georgia_2007/1        cgattttaagccctttcctgatcgccgactagcggtctggatcatggagtcaggctctatgcccctggagggtcccccct [141600]
ASF_IND_20_CAD543                ................................................................................ [141600]
LR899193.1_Germany_2020/1        ................................................................................ [141600]
MN393477.1_Wuhan_2019            ................................................................................ [141600]
MW396979.1_Timor-Leste/2019/1    ................................................................................ [141600]
MK940252.1_CN/2019/InnerMongolia ................................................................................ [141600]
MK128995.1_China/2018/AnhuiXCGQ  ................................................................................ [141600]
MH681419.1_ASFV/POL/2015         ................................................................................ [141600]
LS478113.1_Estonia_2014          ................................................................................ [141600]
KP843857.1_Odintsovo_02/14       ................................................................................ [141600]
```

```
FN557520.1_strain_E75            ................................................................................ [141600]
AY261366.1_Warthog               .........a................................t...................................t. [141600]
AY261365.1_Warmbaths             .........a................................t..................................... [141600]
AY261364.1_Tengani_62            .........a................................t....................t................ [141600]
AY261363.1_Pretoriuskop/96/4     .........a................................t..............c.....t...........t.... [141600]
AY261362.1_Mkuzi_1979            ................................................................................ [141600]
AY261361.1_Malawi_Lil-20/1_1983  ..................................................................a............. [141600]
AY261360.1_Kenya_1950            .........a.................g......................................a..a..g..t.... [141600]
AM712240.1_OURT_88/3             ................................................................................ [141600]
AM712239.1_Benin_97/1            ................................................................................ [141600]
KM111295.1_Ken06.Bus             t........a.................g......................................a..a..g..t.... [141600]
KM111294.1_Ken05/Tk1             .........a.........a.......g......................................a..a..g....... [141600]
KM262844.1_strain_L60            ................................................................................ [141600]
KM102979.1_26544/OG10_Italy      ................................................................................ [141600]
KM262845.1_strain_NHV            ................................................................................ [141600]
KX354450.1_47/Ss/2008            ................................................................................ [141600]
KP055815.1_strain_BA71           ................................................................................ [141600]
MH025920.1_strain_R35            t........a.................g......................................a..a..g..t.... [141600]
MH025919.1_strain_N10            t........a.................g......................................a..a..g..t.... [141600]
MH025918.1_strain_R25            t........a.................g......................................a..a..g..t.... [141600]
MH025917.1_strain_R7             t........a.................g......................................a..a..g..t.... [141600]
MH025916.1_strain_R8             t........a.................g......................................a..a..g..t.... [141600]

FR682468.2_Georgia_2007/1        ataagcggaaaaaggagggtggggggaatgacccgccggttcctaagcatatctcgccgtatactccgcgcacgcgtatt [141680]
ASF_IND_20_CAD543                ................................................................................ [141680]
LR899193.1_Germany_2020/1        ................................................................................ [141680]
MN393477.1_Wuhan_2019            ................................................................................ [141680]
MW396979.1_Timor-Leste/2019/1    ................................................................................ [141680]
MK940252.1_CN/2019/InnerMongolia ................................................................................ [141680]
MK128995.1_China/2018/AnhuiXCGQ  ................................................................................ [141680]
MH681419.1_ASFV/POL/2015         ................................................................................ [141680]
LS478113.1_Estonia_2014          ................................................................................ [141680]
KP843857.1_Odintsovo_02/14       ................................................................................ [141680]
FN557520.1_strain_E75            ................................................................................ [141680]
AY261366.1_Warthog               ................................................................................ [141680]
AY261365.1_Warmbaths             ................................................................................ [141680]
AY261364.1_Tengani_62            ...............................t................................................ [141680]
AY261363.1_Pretoriuskop/96/4     .........................a...................................................... [141680]
AY261362.1_Mkuzi_1979            ................................................................................ [141680]
AY261361.1_Malawi_Lil-20/1_1983  ................................................................................ [141680]
AY261360.1_Kenya_1950            ......................a..................t.a.................................... [141680]
AM712240.1_OURT_88/3             ................................................................................ [141680]
AM712239.1_Benin_97/1            ................................................................................ [141680]
KM111295.1_Ken06.Bus             .............a........a...............a..t.a.................................... [141680]
```

```
KM111294.1_Ken05/Tk1            ......................a..................t.a.................................... [141680]
KM262844.1_strain_L60           ................................................................................ [141680]
KM102979.1_26544/OG10_Italy     ................................................................................ [141680]
KM262845.1_strain_NHV           ................................................................................ [141680]
KX354450.1_47/Ss/2008           ................................................................................ [141680]
KP055815.1_strain_BA71          ................................................................................ [141680]
MH025920.1_strain_R35           .............a........a...............a..t.a.................................... [141680]
MH025919.1_strain_N10           .............a........a...............a..t.a.................................... [141680]
MH025918.1_strain_R25           .............a........a...............a..t.a.................................... [141680]
MH025917.1_strain_R7            .............a........a...............a..t.a.................................... [141680]
MH025916.1_strain_R8            .............a........a...............a..t.a.................................... [141680]

FR682468.2_Georgia_2007/1       gccattgaggtggaaaaggcctttgatgactgtatgcgtcaaaactggtgtagtgtcaataatccctatcttgccaagtc [141760]
ASF_IND_20_CAD543               ................................................................................ [141760]
LR899193.1_Germany_2020/1       ................................................................................ [141760]
MN393477.1_Wuhan_2019           ................................................................................ [141760]
MW396979.1_Timor-Leste/2019/1   ................................................................................ [141760]
MK940252.1_CN/2019/InnerMongolia ................................................................................ [141760]
MK128995.1_China/2018/AnhuiXCGQ ................................................................................ [141760]
MH681419.1_ASFV/POL/2015        ................................................................................ [141760]
LS478113.1_Estonia_2014         ................................................................................ [141760]
KP843857.1_Odintsovo_02/14      ................................................................................ [141760]
FN557520.1_strain_E75           .............................................................................a.. [141760]
AY261366.1_Warthog              ...........a.....a............................................c..............a.. [141760]
AY261365.1_Warmbaths            ...........a..................................................c..............a.. [141760]
AY261364.1_Tengani_62           ................................................................................ [141760]
AY261363.1_Pretoriuskop/96/4    ...........a..................................................c..............a.. [141760]
AY261362.1_Mkuzi_1979           .............................................................................a.. [141760]
AY261361.1_Malawi_Lil-20/1_1983 ...........a..........................c......................................a.. [141760]
AY261360.1_Kenya_1950           ...........a...........c.....a...............................................a.. [141760]
AM712240.1_OURT_88/3            .............................................................................a.. [141760]
AM712239.1_Benin_97/1           .............................................................................a.. [141760]
KM111295.1_Ken06.Bus            ...........a...........c.....a...............................................a.. [141760]
KM111294.1_Ken05/Tk1            ...........a...........c.....a...............................................a.. [141760]
KM262844.1_strain_L60           .............................................................................a.. [141760]
KM102979.1_26544/OG10_Italy     .............................................................................a.. [141760]
KM262845.1_strain_NHV           .............................................................................a.. [141760]
KX354450.1_47/Ss/2008           .............................................................................a.. [141760]
KP055815.1_strain_BA71          .............................................................................a.. [141760]
MH025920.1_strain_R35           ...........a...........c.....a...............................................a.. [141760]
MH025919.1_strain_N10           ...........a...........c.....a...............................................a.. [141760]
MH025918.1_strain_R25           ...........a...........c.....a...............................................a.. [141760]
MH025917.1_strain_R7            ...........a...........c.....a...............................................a.. [141760]
MH025916.1_strain_R8            ...........a...........c.....a...............................................a.. [141760]
```

```
FR682468.2_Georgia_2007/1        ggtctccttgctgtctttcttgtcgctcaaccatcccaccgagtttattaaggtactgccgcttatagactttgacccct [141840]
ASF_IND_20_CAD543                ................................................................................ [141840]
LR899193.1_Germany_2020/1        ................................................................................ [141840]
MN393477.1_Wuhan_2019            ................................................................................ [141840]
MW396979.1_Timor-Leste/2019/1    ................................................................................ [141840]
MK940252.1_CN/2019/InnerMongolia ................................................................................ [141840]
MK128995.1_China/2018/AnhuiXCGQ  ................................................................................ [141840]
MH681419.1_ASFV/POL/2015         ................................................................................ [141840]
LS478113.1_Estonia_2014          ................................................................................ [141840]
KP843857.1_Odintsovo_02/14       ................................................................................ [141840]
FN557520.1_strain_E75            a..t............................................................................ [141840]
AY261366.1_Warthog               ........................a....................................................... [141840]
AY261365.1_Warmbaths             ........................a..............t..a..............a.................t.... [141840]
AY261364.1_Tengani_62            a...................................t................................t.......... [141840]
AY261363.1_Pretoriuskop/96/4     ........................a....................................................... [141840]
AY261362.1_Mkuzi_1979            a..t............................................................................ [141840]
AY261361.1_Malawi_Lil-20/1_1983  a..............................................................................c [141840]
AY261360.1_Kenya_1950            ......................................................c......................... [141840]
AM712240.1_OURT_88/3             a..t............................................................................ [141840]
AM712239.1_Benin_97/1            a..t............................................................................ [141840]
KM111295.1_Ken06.Bus             ......................................................c......................... [141840]
KM111294.1_Ken05/Tk1             ......................................................c......................... [141840]
KM262844.1_strain_L60            a..t............................................................................ [141840]
KM102979.1_26544/OG10_Italy      a..t............................................................................ [141840]
KM262845.1_strain_NHV            a..t............................................................................ [141840]
KX354450.1_47/Ss/2008            a..t............................................................................ [141840]
KP055815.1_strain_BA71           a..t............................................................................ [141840]
MH025920.1_strain_R35            ......................................................c......................... [141840]
MH025919.1_strain_N10            ......................................................c......................... [141840]
MH025918.1_strain_R25            ......................................................c......................... [141840]
MH025917.1_strain_R7             ......................................................c......................... [141840]
MH025916.1_strain_R8             ......................................................c......................... [141840]

FR682468.2_Georgia_2007/1        tggtgaccttttatctacttcttgagccctataaaacgcatggggatgacttttttaattccggaaaccattttattcggc [141920]
ASF_IND_20_CAD543                ................................................................................ [141920]
LR899193.1_Germany_2020/1        ................................................................................ [141920]
MN393477.1_Wuhan_2019            ................................................................................ [141920]
MW396979.1_Timor-Leste/2019/1    ................................................................................ [141920]
MK940252.1_CN/2019/InnerMongolia ................................................................................ [141920]
MK128995.1_China/2018/AnhuiXCGQ  ................................................................................ [141920]
MH681419.1_ASFV/POL/2015         ................................................................................ [141920]
LS478113.1_Estonia_2014          ................................................................................ [141920]
KP843857.1_Odintsovo_02/14       ................................................................................ [141920]
```

```
FN557520.1_strain_E75            ...............................................................................t [141920]
AY261366.1_Warthog               .......t...................................a.................a.................. [141920]
AY261365.1_Warmbaths             .......t...................................a...................................t [141920]
AY261364.1_Tengani_62            .....................................a.....a.................................... [141920]
AY261363.1_Pretoriuskop/96/4     .......t...................................a.................a.................. [141920]
AY261362.1_Mkuzi_1979            ...............................................................................t [141920]
AY261361.1_Malawi_Lil-20/1_1983  ...........................................a.......................ag..........t [141920]
AY261360.1_Kenya_1950            ............................g..............a.......................a............ [141920]
AM712240.1_OURT_88/3             ...............................................................................t [141920]
AM712239.1_Benin_97/1            ...............................................................................t [141920]
KM111295.1_Ken06.Bus             ............................g..............a.......................a............ [141920]
KM111294.1_Ken05/Tk1             ............................g..............a.......................a............ [141920]
KM262844.1_strain_L60            ...............................................................................t [141920]
KM102979.1_26544/OG10_Italy      ...............................................................................t [141920]
KM262845.1_strain_NHV            ...............................................................................t [141920]
KX354450.1_47/Ss/2008            ...............................................................................t [141920]
KP055815.1_strain_BA71           ...............................................................................t [141920]
MH025920.1_strain_R35            ............................g..............a.......................a............ [141920]
MH025919.1_strain_N10            ............................g..............a.......................a............ [141920]
MH025918.1_strain_R25            ............................g..............a.......................a............ [141920]
MH025917.1_strain_R7             ............................g..............a.......................a............ [141920]
MH025916.1_strain_R8             ............................g..............a.......................a............ [141920]

FR682468.2_Georgia_2007/1        cctaccggatggaatggtacagatctgtatcaaagtgccatgctggagtttaaaaagttttttacccagattactcgcca [142000]
ASF_IND_20_CAD543                ...............................................r................................ [142000]
LR899193.1_Germany_2020/1        ................................................................................ [142000]
MN393477.1_Wuhan_2019            ................................................................................ [142000]
MW396979.1_Timor-Leste/2019/1    ................................................................................ [142000]
MK940252.1_CN/2019/InnerMongolia ................................................................................ [142000]
MK128995.1_China/2018/AnhuiXCGQ  ................................................................................ [142000]
MH681419.1_ASFV/POL/2015         ................................................................................ [142000]
LS478113.1_Estonia_2014          ................................................................................ [142000]
KP843857.1_Odintsovo_02/14       ................................................................................ [142000]
FN557520.1_strain_E75            ......................................t........a................................ [142000]
AY261366.1_Warthog               .................c...............................................g........c..... [142000]
AY261365.1_Warmbaths             .................c...............................................g........c..... [142000]
AY261364.1_Tengani_62            ....................g.....a....................a..........................c..... [142000]
AY261363.1_Pretoriuskop/96/4     .................c...............................................g........c..... [142000]
AY261362.1_Mkuzi_1979            ......................................t........a................................ [142000]
AY261361.1_Malawi_Lil-20/1_1983  ........t.....c..c........a....................a.................g........c..... [142000]
AY261360.1_Kenya_1950            ..c...........c..c..........................a..a.................g........c..... [142000]
AM712240.1_OURT_88/3             ......................................t........a................................ [142000]
AM712239.1_Benin_97/1            ......................................t........a................................ [142000]
KM111295.1_Ken06.Bus             ..c...........c..c..........................a..a.................g........c..... [142000]
```

```
KM111294.1_Ken05/Tk1            ..cg..........c..c..........................a..a.................g........c..... [142000]
KM262844.1_strain_L60           ......................................t........a................................ [142000]
KM102979.1_26544/OG10_Italy     ......................................t........a................................ [142000]
KM262845.1_strain_NHV           ......................................t........a................................ [142000]
KX354450.1_47/Ss/2008           ......................................t........a................................ [142000]
KP055815.1_strain_BA71          ......................................t........a................................ [142000]
MH025920.1_strain_R35           ..c...........c..c..........................a..a.................g........c..... [142000]
MH025919.1_strain_N10           ..c...........c..c..........................a..a.................g........c..... [142000]
MH025918.1_strain_R25           ..c...........c..c..........................a..a.................g........c..... [142000]
MH025917.1_strain_R7            ..c...........c..c..........................a..a.................g........c..... [142000]
MH025916.1_strain_R8            ..c...........c..c..........................a..a.................g........c..... [142000]

FR682468.2_Georgia_2007/1       aacctttatggacatagccgattcggctactaaggaggtagatgttcccatatgttactcggatcccgaaaccgtacatt [142080]
ASF_IND_20_CAD543               ................................................................................ [142080]
LR899193.1_Germany_2020/1       ................................................................................ [142080]
MN393477.1_Wuhan_2019           ................................................................................ [142080]
MW396979.1_Timor-Leste/2019/1   ................................................................................ [142080]
MK940252.1_CN/2019/InnerMongolia ................................................................................ [142080]
MK128995.1_China/2018/AnhuiXCGQ ................................................................................ [142080]
MH681419.1_ASFV/POL/2015        ................................................................................ [142080]
LS478113.1_Estonia_2014         ................................................................................ [142080]
KP843857.1_Odintsovo_02/14      ................................................................................ [142080]
FN557520.1_strain_E75           .......................................g........t........t...................... [142080]
AY261366.1_Warthog              ............t...........................................................t....... [142080]
AY261365.1_Warmbaths            ............t.........................................c.................t....... [142080]
AY261364.1_Tengani_62           ........................a....................................................... [142080]
AY261363.1_Pretoriuskop/96/4    ............t...............................................a.....t.....t....... [142080]
AY261362.1_Mkuzi_1979           .......................................g........t........t...................... [142080]
AY261361.1_Malawi_Lil-20/1_1983 ............t.........a....c................................a................... [142080]
AY261360.1_Kenya_1950           ......................a....c..c.......................c.....a.....t............. [142080]
AM712240.1_OURT_88/3            .......................................g........t........t...................... [142080]
AM712239.1_Benin_97/1           .......................................g........t........t...................... [142080]
KM111295.1_Ken06.Bus            ............t.........a....c..........................c.....a.....t............. [142080]
KM111294.1_Ken05/Tk1            ............t.........a....c..........................c.....a.....t............. [142080]
KM262844.1_strain_L60           .......................................g........t........t...................... [142080]
KM102979.1_26544/OG10_Italy     .......................................g........t........t...................... [142080]
KM262845.1_strain_NHV           .......................................g........t........t...................... [142080]
KX354450.1_47/Ss/2008           .......................................g........t........t...................... [142080]
KP055815.1_strain_BA71          .......................................g........t........t...................... [142080]
MH025920.1_strain_R35           ............t.........a....c..........................c.....a.....t............. [142080]
MH025919.1_strain_N10           ............t.........a....c..........................c.....a.....t............. [142080]
MH025918.1_strain_R25           ............t.........a....c..........................c.....a.....t............. [142080]
MH025917.1_strain_R7            ............t.........a....c..........................c.....a.....t............. [142080]
MH025916.1_strain_R8            ............t.........a....c..........................c.....a.....t............. [142080]
```

```
FR682468.2_Georgia_2007/1        cctatgccaatcacgtgcgtactgaaattttgcatcacaatgccgtcaataaggttacaacacctaacctcgtcgtgcag [142160]
ASF_IND_20_CAD543                .....r.......................................................................... [142160]
LR899193.1_Germany_2020/1        ................................................................................ [142160]
MN393477.1_Wuhan_2019            ................................................................................ [142160]
MW396979.1_Timor-Leste/2019/1    ................................................................................ [142160]
MK940252.1_CN/2019/InnerMongolia ................................................................................ [142160]
MK128995.1_China/2018/AnhuiXCGQ  ................................................................................ [142160]
MH681419.1_ASFV/POL/2015         ................................................................................ [142160]
LS478113.1_Estonia_2014          ................................................................................ [142160]
KP843857.1_Odintsovo_02/14       ................................................................................ [142160]
FN557520.1_strain_E75            .....a................................................................a......... [142160]
AY261366.1_Warthog               .....a.a......................................t.............g.........a..t...... [142160]
AY261365.1_Warmbaths             .....a.a....................................................g.........a..t...... [142160]
AY261364.1_Tengani_62            .....................................t......................g.........a......... [142160]
AY261363.1_Pretoriuskop/96/4     .....a.a....................................................g.........a..t...... [142160]
AY261362.1_Mkuzi_1979            .....a................................................................a......... [142160]
AY261361.1_Malawi_Lil-20/1_1983  .......t.....................c....c..t...atg..a..c..a.......g......t..a......... [142160]
AY261360.1_Kenya_1950            .......t.............................t..catg..a..c..a.......g.........g.....a... [142160]
AM712240.1_OURT_88/3             .....a................................................................a......... [142160]
AM712239.1_Benin_97/1            .....a......................................a.........................a......... [142160]
KM111295.1_Ken06.Bus             .......t.............................t..catg..a..c..a.......g.........g.....a... [142160]
KM111294.1_Ken05/Tk1             .......t.............................t..catg..a..c..a.......g.........g.....a... [142160]
KM262844.1_strain_L60            .....a................................................................a......... [142160]
KM102979.1_26544/OG10_Italy      .....a................................................................a......... [142160]
KM262845.1_strain_NHV            .....a................................................................a......... [142160]
KX354450.1_47/Ss/2008            .....a................................................................a......... [142160]
KP055815.1_strain_BA71           .....a................................................................a......... [142160]
MH025920.1_strain_R35            .......t.............................t..catg..a..c..a.......g.........g.....a... [142160]
MH025919.1_strain_N10            .......t.............................t..catg..a..c..a.......g.........g.....a... [142160]
MH025918.1_strain_R25            .......t.............................t..catg..a..c..a.......g.........g.....a... [142160]
MH025917.1_strain_R7             .......t.............................t..catg..a..c..a.......g.........g.....a... [142160]
MH025916.1_strain_R8             .......t.............................t..catg..a..c..a.......g.........g.....a... [142160]

FR682468.2_Georgia_2007/1        gcctataatgagctcgagcaaaccaataccatacgacattacggccctattttcccggaaagtaccatcaacgcactgcg [142240]
ASF_IND_20_CAD543                ................................................................................ [142240]
LR899193.1_Germany_2020/1        ................................................................................ [142240]
MN393477.1_Wuhan_2019            ................................................................................ [142240]
MW396979.1_Timor-Leste/2019/1    ................................................................................ [142240]
MK940252.1_CN/2019/InnerMongolia ................................................................................ [142240]
MK128995.1_China/2018/AnhuiXCGQ  ................................................................................ [142240]
MH681419.1_ASFV/POL/2015         ................................................................................ [142240]
LS478113.1_Estonia_2014          ................................................................................ [142240]
KP843857.1_Odintsovo_02/14       ................................................................................ [142240]
```

```
FN557520.1_strain_E75            ................................................................................ [142240]
AY261366.1_Warthog               ................................................................................ [142240]
AY261365.1_Warmbaths             ................................................................................ [142240]
AY261364.1_Tengani_62            ................................................................................ [142240]
AY261363.1_Pretoriuskop/96/4     ................................................................................ [142240]
AY261362.1_Mkuzi_1979            ................................................................................ [142240]
AY261361.1_Malawi_Lil-20/1_1983  ..........................c..a..............g...........a.................c..a.. [142240]
AY261360.1_Kenya_1950            .....c....................c..a..............g................................... [142240]
AM712240.1_OURT_88/3             ................................................................................ [142240]
AM712239.1_Benin_97/1            ................................................................................ [142240]
KM111295.1_Ken06.Bus             .....c....................c..a..............g................................... [142240]
KM111294.1_Ken05/Tk1             .....c.................t..c..a..............g................................... [142240]
KM262844.1_strain_L60            ................................................................................ [142240]
KM102979.1_26544/OG10_Italy      ................................................................................ [142240]
KM262845.1_strain_NHV            ................................................................................ [142240]
KX354450.1_47/Ss/2008            ................................................................................ [142240]
KP055815.1_strain_BA71           ................................................................................ [142240]
MH025920.1_strain_R35            .t...c....................c..a..............g................................... [142240]
MH025919.1_strain_N10            .t...c....................c..a..............g................................... [142240]
MH025918.1_strain_R25            .t...c....................c..a..............g................................... [142240]
MH025917.1_strain_R7             .t...c....................c..a..............g................................... [142240]
MH025916.1_strain_R8             .t...c....................c..a..............g................................... [142240]

FR682468.2_Georgia_2007/1        tttttggaaaaagctgtggcaggatgaacagcgatttgttatccacggcctgcaccgcacgttgatggatcaacccacct [142320]
ASF_IND_20_CAD543                ................................................................................ [142320]
LR899193.1_Germany_2020/1        ................................................................................ [142320]
MN393477.1_Wuhan_2019            ................................................................................ [142320]
MW396979.1_Timor-Leste/2019/1    ................................................................................ [142320]
MK940252.1_CN/2019/InnerMongolia ................................................................................ [142320]
MK128995.1_China/2018/AnhuiXCGQ  ................................................................................ [142320]
MH681419.1_ASFV/POL/2015         ................................................................................ [142320]
LS478113.1_Estonia_2014          ................................................................................ [142320]
KP843857.1_Odintsovo_02/14       ................................................................................ [142320]
FN557520.1_strain_E75            ...c......................................t..................................... [142320]
AY261366.1_Warthog               ...c.......................................................................t.... [142320]
AY261365.1_Warmbaths             ...c.........................................t.................................. [142320]
AY261364.1_Tengani_62            ................................................................................ [142320]
AY261363.1_Pretoriuskop/96/4     ...c............................................................................ [142320]
AY261362.1_Mkuzi_1979            ...c......................................t..................................... [142320]
AY261361.1_Malawi_Lil-20/1_1983  ...c.............................g.............................................. [142320]
AY261360.1_Kenya_1950            ...c.............................g...........t.................................. [142320]
AM712240.1_OURT_88/3             ...c......................................t..................................... [142320]
AM712239.1_Benin_97/1            ...c......................................t..................................... [142320]
KM111295.1_Ken06.Bus             ...c.............................g...........t.................................. [142320]
```

```
KM111294.1_Ken05/Tk1              ...c.............................g...........t.................................. [142320]
KM262844.1_strain_L60             ...c......................................t..................................... [142320]
KM102979.1_26544/OG10_Italy       ...c......................................t..................................... [142320]
KM262845.1_strain_NHV             ...c......................................t..................................... [142320]
KX354450.1_47/Ss/2008             ...c......................................t..................................... [142320]
KP055815.1_strain_BA71            ...c......................................t..................................... [142320]
MH025920.1_strain_R35             ...c.............................g...........t.................................. [142320]
MH025919.1_strain_N10             ...c.............................g...........t.................................. [142320]
MH025918.1_strain_R25             ...c.............................g...........t.................................. [142320]
MH025917.1_strain_R7              ...c.............................g...........t.................................. [142320]
MH025916.1_strain_R8              ...c.............................g...........t.................................. [142320]

FR682468.2_Georgia_2007/1         atgaaacctctgagtttgcagagatcgttagaaatttacggttttcgcgtcccggcaataactatataaacgagcttaat [142400]
ASF_IND_20_CAD543                 ................................................................................ [142400]
LR899193.1_Germany_2020/1         ................................................................................ [142400]
MN393477.1_Wuhan_2019             ................................................................................ [142400]
MW396979.1_Timor-Leste/2019/1     ................................................................................ [142400]
MK940252.1_CN/2019/InnerMongolia  ................................................................................ [142400]
MK128995.1_China/2018/AnhuiXCGQ   ................................................................................ [142400]
MH681419.1_ASFV/POL/2015          ................................................................................ [142400]
LS478113.1_Estonia_2014           ................................................................................ [142400]
KP843857.1_Odintsovo_02/14        ................................................................................ [142400]
FN557520.1_strain_E75             ................................................................................ [142400]
AY261366.1_Warthog                ........................................t.....a................................. [142400]
AY261365.1_Warmbaths              ........................................t....................................... [142400]
AY261364.1_Tengani_62             ........................................t....................................... [142400]
AY261363.1_Pretoriuskop/96/4      ........................................t....................................... [142400]
AY261362.1_Mkuzi_1979             ................................................................................ [142400]
AY261361.1_Malawi_Lil-20/1_1983   ........................................t...........t.................t......... [142400]
AY261360.1_Kenya_1950             ........................................t...........g..t..............t.....c... [142400]
AM712240.1_OURT_88/3              ................................................................................ [142400]
AM712239.1_Benin_97/1             ................................................................................ [142400]
KM111295.1_Ken06.Bus              ........................................t...................................c... [142400]
KM111294.1_Ken05/Tk1              ........................................t..............t..............t.....c... [142400]
KM262844.1_strain_L60             ................................................................................ [142400]
KM102979.1_26544/OG10_Italy       ................................................................................ [142400]
KM262845.1_strain_NHV             ................................................................................ [142400]
KX354450.1_47/Ss/2008             ................................................................................ [142400]
KP055815.1_strain_BA71            ................................................................................ [142400]
MH025920.1_strain_R35             ........................................t...................................c... [142400]
MH025919.1_strain_N10             ........................................t...................................c... [142400]
MH025918.1_strain_R25             ........................................t...................................c... [142400]
MH025917.1_strain_R7              ........................................t...................................c... [142400]
MH025916.1_strain_R8              ........................................t...................................c... [142400]
```

```
FR682468.2_Georgia_2007/1        attacaagtcccgctatgtacggcgacaagcataccaccggagatattgcgcccaatgatagatttgccatgttggtggc [142480]
ASF_IND_20_CAD543                ................................................................................ [142480]
LR899193.1_Germany_2020/1        ................................................................................ [142480]
MN393477.1_Wuhan_2019            ................................................................................ [142480]
MW396979.1_Timor-Leste/2019/1    ................................................................................ [142480]
MK940252.1_CN/2019/InnerMongolia ................................................................................ [142480]
MK128995.1_China/2018/AnhuiXCGQ  ................................................................................ [142480]
MH681419.1_ASFV/POL/2015         ................................................................................ [142480]
LS478113.1_Estonia_2014          ................................................................................ [142480]
KP843857.1_Odintsovo_02/14       ................................................................................ [142480]
FN557520.1_strain_E75            ...........t..............t..................................................... [142480]
AY261366.1_Warthog               ...........t.................................................................... [142480]
AY261365.1_Warmbaths             ................................................................................ [142480]
AY261364.1_Tengani_62            ................................................................................ [142480]
AY261363.1_Pretoriuskop/96/4     ................................................................................ [142480]
AY261362.1_Mkuzi_1979            ................................................................................ [142480]
AY261361.1_Malawi_Lil-20/1_1983  ..c.......................t..................................................... [142480]
AY261360.1_Kenya_1950            ..c..................................................g........g................. [142480]
AM712240.1_OURT_88/3             ...........t..............t..................................................... [142480]
AM712239.1_Benin_97/1            ...........t..............t..................................................... [142480]
KM111295.1_Ken06.Bus             ..c........t..................................................g................. [142480]
KM111294.1_Ken05/Tk1             ..c...........................................................g................. [142480]
KM262844.1_strain_L60            ...........t..............t..................................................... [142480]
KM102979.1_26544/OG10_Italy      ...........t..............t..................................................... [142480]
KM262845.1_strain_NHV            ...........t..............t..................................................... [142480]
KX354450.1_47/Ss/2008            ...........t..............t..................................................... [142480]
KP055815.1_strain_BA71           ...........t..............t..................................................... [142480]
MH025920.1_strain_R35            ..c........t..................................................g................. [142480]
MH025919.1_strain_N10            ..c........t..................................................g................. [142480]
MH025918.1_strain_R25            ..c........t..................................................g................. [142480]
MH025917.1_strain_R7             ..c........t..................................................g................. [142480]
MH025916.1_strain_R8             ..c........t..................................................g................. [142480]

FR682468.2_Georgia_2007/1        ctttatcaacagtactgactttttatacaccgcgattcccgaggaaaaggtaggggggaatgaaa------------ccc [142560]
ASF_IND_20_CAD543                .................................................................------------... [142560]
LR899193.1_Germany_2020/1        .................................................................------------... [142560]
MN393477.1_Wuhan_2019            .................................................................------------... [142560]
MW396979.1_Timor-Leste/2019/1    .................................................................------------... [142560]
MK940252.1_CN/2019/InnerMongolia .................................................................------------... [142560]
MK128995.1_China/2018/AnhuiXCGQ  .................................................................------------... [142560]
MH681419.1_ASFV/POL/2015         .................................................................------------... [142560]
LS478113.1_Estonia_2014          .................................................................------------... [142560]
KP843857.1_Odintsovo_02/14       .................................................................------------... [142560]
```

```
FN557520.1_strain_E75            .................................c........a......................------------... [142560]
AY261366.1_Warthog               .........................................................a.......------------... [142560]
AY261365.1_Warmbaths             .................................................................------------... [142560]
AY261364.1_Tengani_62            .........................................................a.......------------... [142560]
AY261363.1_Pretoriuskop/96/4     ...............a.................................................------------... [142560]
AY261362.1_Mkuzi_1979            .................................................................------------... [142560]
AY261361.1_Malawi_Lil-20/1_1983  ...............c..t..............a.............................c.------------... [142560]
AY261360.1_Kenya_1950            ......t.....c..............t...........g...........g.....a.....t.ctcaaaccggtt... [142560]
AM712240.1_OURT_88/3             .................................c........a......................------------... [142560]
AM712239.1_Benin_97/1            .................................c........a......................------------... [142560]
KM111295.1_Ken06.Bus             ......t.....c..............t...........a...........g.....a.....t.cacaaaccggtc... [142560]
KM111294.1_Ken05/Tk1             ......t.....c..............t...........g...........g.....a.....t.ctcaaaccggtc... [142560]
KM262844.1_strain_L60            .................................c........a......................------------... [142560]
KM102979.1_26544/OG10_Italy      .................................c........a......................------------... [142560]
KM262845.1_strain_NHV            .................................c........a......................------------... [142560]
KX354450.1_47/Ss/2008            .................................c........a......................------------... [142560]
KP055815.1_strain_BA71           .................................c........a......................------------... [142560]
MH025920.1_strain_R35            ......t.....c..............t...........a...........g.....a.....t.cacaaaccggtc... [142560]
MH025919.1_strain_N10            ......t.....c..............t...........a...........g.....a.....t.cacaaaccggtc... [142560]
MH025918.1_strain_R25            ......t.....c..............t...........a...........g.....a.....t.cacaaaccggtc... [142560]
MH025917.1_strain_R7             ......t.....c..............t...........a...........g.....a.....t.cacaaaccggtc... [142560]
MH025916.1_strain_R8             ......t.....c..............t...........a...........g.....a.....t.cacaaaccggtc... [142560]

FR682468.2_Georgia_2007/1        aaaccagtagccttacagacctagttccaacacggctacactcttttttaaatcataatctaagcaaacttaaaatctta [142640]
ASF_IND_20_CAD543                ................................................................................ [142640]
LR899193.1_Germany_2020/1        ................................................................................ [142640]
MN393477.1_Wuhan_2019            ................................................................................ [142640]
MW396979.1_Timor-Leste/2019/1    ................................................................................ [142640]
MK940252.1_CN/2019/InnerMongolia ................................................................................ [142640]
MK128995.1_China/2018/AnhuiXCGQ  ................................................................................ [142640]
MH681419.1_ASFV/POL/2015         ................................................................................ [142640]
LS478113.1_Estonia_2014          ................................................................................ [142640]
KP843857.1_Odintsovo_02/14       ................................................................................ [142640]
FN557520.1_strain_E75            ................................................................................ [142640]
AY261366.1_Warthog               ................................................................................ [142640]
AY261365.1_Warmbaths             ................................................................................ [142640]
AY261364.1_Tengani_62            ................................................................................ [142640]
AY261363.1_Pretoriuskop/96/4     .................................................g.............................. [142640]
AY261362.1_Mkuzi_1979            ................................................................................ [142640]
AY261361.1_Malawi_Lil-20/1_1983  ...................t................................c........................... [142640]
AY261360.1_Kenya_1950            ...................t................................c.................c.....a..g [142640]
AM712240.1_OURT_88/3             ................................................................................ [142640]
AM712239.1_Benin_97/1            ................................................................................ [142640]
KM111295.1_Ken06.Bus             ...................t................................c.................c.....a..g [142640]
```

```
KM111294.1_Ken05/Tk1          ...................t................................c.................c.....a..g [142640]
KM262844.1_strain_L60         ................................................................................ [142640]
KM102979.1_26544/OG10_Italy   ................................................................................ [142640]
KM262845.1_strain_NHV         ................................................................................ [142640]
KX354450.1_47/Ss/2008         ................................................................................ [142640]
KP055815.1_strain_BA71        ................................................................................ [142640]
MH025920.1_strain_R35         ...................t................................c.................c.....a..g [142640]
MH025919.1_strain_N10         ...................t................................c.................c.....a..g [142640]
MH025918.1_strain_R25         ...................t................................c.................c.....a..g [142640]
MH025917.1_strain_R7          ...................t................................c.................c.....a..g [142640]
MH025916.1_strain_R8          ...................t................................c.................c.....a..g [142640]

FR682468.2_Georgia_2007/1     aaccgcgcgcagcaaacggttagaaatattctttcaaatgattgtcttaatcaactgaaacattatgttaaacacacggg [142720]
ASF_IND_20_CAD543             ................................................................................ [142720]
LR899193.1_Germany_2020/1     ................................................................................ [142720]
MN393477.1_Wuhan_2019         ................................................................................ [142720]
MW396979.1_Timor-Leste/2019/1 ................................................................................ [142720]
MK940252.1_CN/2019/InnerMongolia ................................................................................ [142720]
MK128995.1_China/2018/AnhuiXCGQ ................................................................................ [142720]
MH681419.1_ASFV/POL/2015      ................................................................................ [142720]
LS478113.1_Estonia_2014       ................................................................................ [142720]
KP843857.1_Odintsovo_02/14    ................................................................................ [142720]
FN557520.1_strain_E75         ..........................c..................................................... [142720]
AY261366.1_Warthog            ................................................................................ [142720]
AY261365.1_Warmbaths          ..........................c..................................................... [142720]
AY261364.1_Tengani_62         .....t..................................................a....................... [142720]
AY261363.1_Pretoriuskop/96/4  ................................................................................ [142720]
AY261362.1_Mkuzi_1979         ..........................c..................................................... [142720]
AY261361.1_Malawi_Lil-20/1_1983 .....t..................................................a....................... [142720]
AY261360.1_Kenya_1950         ..t..t................a.................................a....................... [142720]
AM712240.1_OURT_88/3          ..........................c..................................................... [142720]
AM712239.1_Benin_97/1         ..........................c..................................................... [142720]
KM111295.1_Ken06.Bus          ..t..t................a.................................a....................... [142720]
KM111294.1_Ken05/Tk1          ..t..t................a.................................a....................... [142720]
KM262844.1_strain_L60         ......................a...c..................................................... [142720]
KM102979.1_26544/OG10_Italy   ..........................c..................................................... [142720]
KM262845.1_strain_NHV         ..........................c..................................................... [142720]
KX354450.1_47/Ss/2008         ..........................c..................................................... [142720]
KP055815.1_strain_BA71        ..........................c..................................................... [142720]
MH025920.1_strain_R35         ..t..t................a.................................a....................... [142720]
MH025919.1_strain_N10         ..t..t................a.................................a....................... [142720]
MH025918.1_strain_R25         ..t..t................a.................................a....................... [142720]
MH025917.1_strain_R7          ..t..t................a.................................a....................... [142720]
MH025916.1_strain_R8          ..t..t................a.................................a....................... [142720]
```

```
FR682468.2_Georgia_2007/1        aaaaaatgaaatactaaagttacttcaagaataagtatgttgatacctgtggtgtgttttacctgtgggtttcctattgg [142800]
ASF_IND_20_CAD543                ................................................................................ [142800]
LR899193.1_Germany_2020/1        ................................................................................ [142800]
MN393477.1_Wuhan_2019            ................................................................................ [142800]
MW396979.1_Timor-Leste/2019/1    ................................................................................ [142800]
MK940252.1_CN/2019/InnerMongolia ................................................................................ [142800]
MK128995.1_China/2018/AnhuiXCGQ  ................................................................................ [142800]
MH681419.1_ASFV/POL/2015         ................................................................................ [142800]
LS478113.1_Estonia_2014          ................................................................................ [142800]
KP843857.1_Odintsovo_02/14       ................................................................................ [142800]
FN557520.1_strain_E75            ..................................c............g................................ [142800]
AY261366.1_Warthog               ..................................c............................................. [142800]
AY261365.1_Warmbaths             ..................................c............................................. [142800]
AY261364.1_Tengani_62            ..................................a............................................. [142800]
AY261363.1_Pretoriuskop/96/4     ...................a..............c............................................. [142800]
AY261362.1_Mkuzi_1979            ..................................c............................................. [142800]
AY261361.1_Malawi_Lil-20/1_1983  ..............................t...a...............a............................. [142800]
AY261360.1_Kenya_1950            ..................................c............................................. [142800]
AM712240.1_OURT_88/3             ..................................c............g................................ [142800]
AM712239.1_Benin_97/1            ..................................c............g................................ [142800]
KM111295.1_Ken06.Bus             ...................a..............cc........t................................... [142800]
KM111294.1_Ken05/Tk1             ..................................cc............................................ [142800]
KM262844.1_strain_L60            ..................................c............g................................ [142800]
KM102979.1_26544/OG10_Italy      ..................................c............g................................ [142800]
KM262845.1_strain_NHV            ..................................c............g................................ [142800]
KX354450.1_47/Ss/2008            ..................................c............g................................ [142800]
KP055815.1_strain_BA71           ..................................c............g................................ [142800]
MH025920.1_strain_R35            ...................a..............cc........t................................... [142800]
MH025919.1_strain_N10            ...................a..............cc........t................................... [142800]
MH025918.1_strain_R25            ...................a..............cc........t................................... [142800]
MH025917.1_strain_R7             ...................a..............cc........t................................... [142800]
MH025916.1_strain_R8             ...................a..............cc........t................................... [142800]

FR682468.2_Georgia_2007/1        aacctacgcggcaatttttgacaaggctcgtaccgagtatattaaaaccaaaatgggcggaacattgccgcaaaatatcc [142880]
ASF_IND_20_CAD543                ................................................................................ [142880]
LR899193.1_Germany_2020/1        ................................................................................ [142880]
MN393477.1_Wuhan_2019            ................................................................................ [142880]
MW396979.1_Timor-Leste/2019/1    ................................................................................ [142880]
MK940252.1_CN/2019/InnerMongolia ................................................................................ [142880]
MK128995.1_China/2018/AnhuiXCGQ  ................................................................................ [142880]
MH681419.1_ASFV/POL/2015         ................................................................................ [142880]
LS478113.1_Estonia_2014          ................................................................................ [142880]
KP843857.1_Odintsovo_02/14       ................................................................................ [142880]
```

```
FN557520.1_strain_E75          ........................................................a.........a............. [142880]
AY261366.1_Warthog             ..................................................................a............. [142880]
AY261365.1_Warmbaths           ..................................................................a...........t. [142880]
AY261364.1_Tengani_62          ..................................................................a............. [142880]
AY261363.1_Pretoriuskop/96/4   ..................................................................a............. [142880]
AY261362.1_Mkuzi_1979          ........................................................a.........a............. [142880]
AY261361.1_Malawi_Lil-20/1_1983 ..................................................................a............. [142880]
AY261360.1_Kenya_1950          .....................t...........t................................a............. [142880]
AM712240.1_OURT_88/3           ........................................................a.........a............. [142880]
AM712239.1_Benin_97/1          ........................................................a.........a............. [142880]
KM111295.1_Ken06.Bus           .....................t............................................a............. [142880]
KM111294.1_Ken05/Tk1           .....................t............................................a............. [142880]
KM262844.1_strain_L60          ........................................................a.........a............. [142880]
KM102979.1_26544/OG10_Italy    ........................................................a.........a............. [142880]
KM262845.1_strain_NHV          ........................................................a.........a............. [142880]
KX354450.1_47/Ss/2008          ........................................................a.........a............. [142880]
KP055815.1_strain_BA71         ........................................................a.........a............. [142880]
MH025920.1_strain_R35          .....................t............................................a............. [142880]
MH025919.1_strain_N10          .....................t............................................a............. [142880]
MH025918.1_strain_R25          .....................t............................................a............. [142880]
MH025917.1_strain_R7           .....................t............................................a............. [142880]
MH025916.1_strain_R8           .....................t............................................a............. [142880]

FR682468.2_Georgia_2007/1      cattagatgcttctctccagattgagttaaaagacctcattacagctctgggaatcccaatgcgggtgtgttgtcgcact [142960]
ASF_IND_20_CAD543              ................................................................................ [142960]
LR899193.1_Germany_2020/1      ................................................................................ [142960]
MN393477.1_Wuhan_2019          ................................................................................ [142960]
MW396979.1_Timor-Leste/2019/1  ................................................................................ [142960]
MK940252.1_CN/2019/InnerMongolia ................................................................................ [142960]
MK128995.1_China/2018/AnhuiXCGQ ................................................................................ [142960]
MH681419.1_ASFV/POL/2015       ................................................................................ [142960]
LS478113.1_Estonia_2014        ................................................................................ [142960]
KP843857.1_Odintsovo_02/14     ................................................................................ [142960]
FN557520.1_strain_E75          ..................................t............................................. [142960]
AY261366.1_Warthog             ..........c..................................................................... [142960]
AY261365.1_Warmbaths           ..........c.....................................................a............... [142960]
AY261364.1_Tengani_62          ................................................................................ [142960]
AY261363.1_Pretoriuskop/96/4   ..........g..................................................................... [142960]
AY261362.1_Mkuzi_1979          ..................................t............................................. [142960]
AY261361.1_Malawi_Lil-20/1_1983 ..........c.....g..a......c..........t.....g........g.......................t... [142960]
AY261360.1_Kenya_1950          .g.................a......c................g.......................a...........g [142960]
AM712240.1_OURT_88/3           ..................................t............................................. [142960]
AM712239.1_Benin_97/1          ..................................t............................................. [142960]
KM111295.1_Ken06.Bus           ..........c........a......c................g...................................g [142960]
```

```
KM111294.1_Ken05/Tk1           ...................a......c................g...................................g [142960]
KM262844.1_strain_L60          ..................................t............................................. [142960]
KM102979.1_26544/OG10_Italy    ..................................t............................................. [142960]
KM262845.1_strain_NHV          ..................................t............................................. [142960]
KX354450.1_47/Ss/2008          ..................................t............................................. [142960]
KP055815.1_strain_BA71         t.................................t............................................. [142960]
MH025920.1_strain_R35          ..........c........a......c................g...................................g [142960]
MH025919.1_strain_N10          ..........c........a......c................g...................................g [142960]
MH025918.1_strain_R25          ..........c........a......c................g...................................g [142960]
MH025917.1_strain_R7           ..........c........a......c................g...................................g [142960]
MH025916.1_strain_R8           ..........c........a......c................g...................................g [142960]

FR682468.2_Georgia_2007/1      catttaattactacgttggattatcgtaaat-------------attattaatatctaaaattgaaaaaatatttttaat [143040]
ASF_IND_20_CAD543              ...............................-------------.................................... [143040]
LR899193.1_Germany_2020/1      ...............................-------------.................................... [143040]
MN393477.1_Wuhan_2019          ...............................-------------.................................... [143040]
MW396979.1_Timor-Leste/2019/1  ...............................-------------.................................... [143040]
MK940252.1_CN/2019/InnerMongolia ...............................-------------.................................... [143040]
MK128995.1_China/2018/AnhuiXCGQ ...............................-------------.................................... [143040]
MH681419.1_ASFV/POL/2015       ...............................-------------.................................... [143040]
LS478113.1_Estonia_2014        ...............................-------------.................................... [143040]
KP843857.1_Odintsovo_02/14     ...............................-------------.................................... [143040]
FN557520.1_strain_E75          ...............................-------------.................................... [143040]
AY261366.1_Warthog             ...............................-------------.................................... [143040]
AY261365.1_Warmbaths           ...............................-------------........c........................... [143040]
AY261364.1_Tengani_62          .............................g.-------------...........g.............t.......... [143040]
AY261363.1_Pretoriuskop/96/4   ...............................-------------.................................... [143040]
AY261362.1_Mkuzi_1979          ...............................-------------.................................... [143040]
AY261361.1_Malawi_Lil-20/1_1983 .............................g.attattaataagt...........g.............c.......... [143040]
AY261360.1_Kenya_1950          ...............................-------------........ccgg.............c.........g [143040]
AM712240.1_OURT_88/3           ...............................-------------.................................... [143040]
AM712239.1_Benin_97/1          ...............................-------------.................................... [143040]
KM111295.1_Ken06.Bus           ...............................-------------........ccgg.............c.........g [143040]
KM111294.1_Ken05/Tk1           ...............................-------------........ccgg.............c.........g [143040]
KM262844.1_strain_L60          ...............................-------------.................................... [143040]
KM102979.1_26544/OG10_Italy    ...............................-------------.................................... [143040]
KM262845.1_strain_NHV          ...............................-------------.................................... [143040]
KX354450.1_47/Ss/2008          ...............................-------------.................................... [143040]
KP055815.1_strain_BA71         ...............................-------------.................................... [143040]
MH025920.1_strain_R35          ...............................-------------........ccgg.............c.........g [143040]
MH025919.1_strain_N10          ...............................-------------........ccgg.............c.........g [143040]
MH025918.1_strain_R25          ...............................-------------........ccgg.............c.........g [143040]
MH025917.1_strain_R7           ...............................-------------........ccgg.............c.........g [143040]
MH025916.1_strain_R8           ...............................-------------........ccgg.............c.........g [143040]
```

```
FR682468.2_Georgia_2007/1        gttactagtaaaaatgactacacacatctttcacgcagatgatctcctacaagcattgcaacaagcaaaagcagaaaaaa [143120]
ASF_IND_20_CAD543                ................................................................................ [143120]
LR899193.1_Germany_2020/1        ................................................................................ [143120]
MN393477.1_Wuhan_2019            ................................................................................ [143120]
MW396979.1_Timor-Leste/2019/1    ................................................................................ [143120]
MK940252.1_CN/2019/InnerMongolia ................................................................................ [143120]
MK128995.1_China/2018/AnhuiXCGQ  ................................................................................ [143120]
MH681419.1_ASFV/POL/2015         ................................................................................ [143120]
LS478113.1_Estonia_2014          ................................................................................ [143120]
KP843857.1_Odintsovo_02/14       ................................................................................ [143120]
FN557520.1_strain_E75            ................................................................................ [143120]
AY261366.1_Warthog               ...............................a..................................g............. [143120]
AY261365.1_Warmbaths             ................................................................................ [143120]
AY261364.1_Tengani_62            ...............................a..........a..t....................g............. [143120]
AY261363.1_Pretoriuskop/96/4     ...............................a..........a.......................g............. [143120]
AY261362.1_Mkuzi_1979            ................................................................................ [143120]
AY261361.1_Malawi_Lil-20/1_1983  ......................aca......t..........c..t.................................. [143120]
AY261360.1_Kenya_1950            ......................---.................c..t.....c...c..............a......... [143120]
AM712240.1_OURT_88/3             ................................................................................ [143120]
AM712239.1_Benin_97/1            ................................................................................ [143120]
KM111295.1_Ken06.Bus             ......................---.................c..t.....c...c........................ [143120]
KM111294.1_Ken05/Tk1             ......................---.................c..t.....c...c........................ [143120]
KM262844.1_strain_L60            ................................................................................ [143120]
KM102979.1_26544/OG10_Italy      ................................................................................ [143120]
KM262845.1_strain_NHV            ................................................................................ [143120]
KX354450.1_47/Ss/2008            ................................................................................ [143120]
KP055815.1_strain_BA71           ................................................................................ [143120]
MH025920.1_strain_R35            ......................---.................c..t.....c...c........................ [143120]
MH025919.1_strain_N10            ......................---.................c..t.....c...c........................ [143120]
MH025918.1_strain_R25            ......................---.................c..t.....c...c........................ [143120]
MH025917.1_strain_R7             ......................---.................c..t.....c...c........................ [143120]
MH025916.1_strain_R8             ......................---.................c..t.....c...c........................ [143120]

FR682468.2_Georgia_2007/1        atttttcatctgtattttctttagattgggataaattacgcacagcgaagcgtaatacaacggttaaatatgttacggtc [143200]
ASF_IND_20_CAD543                .......w........................................................................ [143200]
LR899193.1_Germany_2020/1        ................................................................................ [143200]
MN393477.1_Wuhan_2019            ................................................................................ [143200]
MW396979.1_Timor-Leste/2019/1    ................................................................................ [143200]
MK940252.1_CN/2019/InnerMongolia ................................................................................ [143200]
MK128995.1_China/2018/AnhuiXCGQ  ................................................................................ [143200]
MH681419.1_ASFV/POL/2015         ................................................................................ [143200]
LS478113.1_Estonia_2014          ................................................................................ [143200]
KP843857.1_Odintsovo_02/14       ................................................................................ [143200]
```

```
FN557520.1_strain_E75          .......t........................................................................ [143200]
AY261366.1_Warthog             .......t.....................a.............................t.................... [143200]
AY261365.1_Warmbaths           .......t..........a........................................t.................... [143200]
AY261364.1_Tengani_62          .......t...a......a........................................t.................... [143200]
AY261363.1_Pretoriuskop/96/4   .......t.....................................................a.................. [143200]
AY261362.1_Mkuzi_1979          .......t........................................................................ [143200]
AY261361.1_Malawi_Lil-20/1_1983 .......t..................................t................t.................... [143200]
AY261360.1_Kenya_1950          .......c..................................t.........c......t...................t [143200]
AM712240.1_OURT_88/3           .......t........................................................................ [143200]
AM712239.1_Benin_97/1          .......t........................................................................ [143200]
KM111295.1_Ken06.Bus           .......t..................................t.........c......t...................t [143200]
KM111294.1_Ken05/Tk1           .......c..................................t.........c......t...................t [143200]
KM262844.1_strain_L60          .......t........................................................................ [143200]
KM102979.1_26544/OG10_Italy    .......t........................................................................ [143200]
KM262845.1_strain_NHV          .......t........................................................................ [143200]
KX354450.1_47/Ss/2008          .......t........................................................................ [143200]
KP055815.1_strain_BA71         .......t........................................................................ [143200]
MH025920.1_strain_R35          .......t..................................t.........c......t...................t [143200]
MH025919.1_strain_N10          .......t..................................t.........c......t...................t [143200]
MH025918.1_strain_R25          .......t..................................t.........c......t...................t [143200]
MH025917.1_strain_R7           .......t..................................t.........c......t...................t [143200]
MH025916.1_strain_R8           .......t..................................t.........c......t...................t [143200]

FR682468.2_Georgia_2007/1      aatgtcatagtaaaaggcaaaaaagctccgctaatgtttaactttcaaaatgaaaaacatgtaggaaccattcctcccag [143280]
ASF_IND_20_CAD543              ................................................................................ [143280]
LR899193.1_Germany_2020/1      ................................................................................ [143280]
MN393477.1_Wuhan_2019          ................................................................................ [143280]
MW396979.1_Timor-Leste/2019/1  ................................................................................ [143280]
MK940252.1_CN/2019/InnerMongolia ................................................................................ [143280]
MK128995.1_China/2018/AnhuiXCGQ ................................................................................ [143280]
MH681419.1_ASFV/POL/2015       ................................................................................ [143280]
LS478113.1_Estonia_2014        ................................................................................ [143280]
KP843857.1_Odintsovo_02/14     ................................................................................ [143280]
FN557520.1_strain_E75          ................................................................................ [143280]
AY261366.1_Warthog             t........a...................................................................... [143280]
AY261365.1_Warmbaths           t........a...................................................................... [143280]
AY261364.1_Tengani_62          t......c.a...................a.................................................. [143280]
AY261363.1_Pretoriuskop/96/4   .................t.............................................................. [143280]
AY261362.1_Mkuzi_1979          ................................................................................ [143280]
AY261361.1_Malawi_Lil-20/1_1983 .....a.cg...............................g.....................g................. [143280]
AY261360.1_Kenya_1950          c....a..g...........................................................a...t....... [143280]
AM712240.1_OURT_88/3           ................................................................................ [143280]
AM712239.1_Benin_97/1          ................................................................................ [143280]
KM111295.1_Ken06.Bus           c....a..g...........................................................a...t....... [143280]
```

```
KM111294.1_Ken05/Tk1             c....a..g...........................................................a...t....... [143280]
KM262844.1_strain_L60            ................................................................................ [143280]
KM102979.1_26544/OG10_Italy      ................................................................................ [143280]
KM262845.1_strain_NHV            ................................................................................ [143280]
KX354450.1_47/Ss/2008            ................................................................................ [143280]
KP055815.1_strain_BA71           ................................................................................ [143280]
MH025920.1_strain_R35            c....a..g...........................................................a...t....... [143280]
MH025919.1_strain_N10            c....a..g...........................................................a...t....... [143280]
MH025918.1_strain_R25            c....a..g...........................................................a...t....... [143280]
MH025917.1_strain_R7             c....a..g...........................................................a...t....... [143280]
MH025916.1_strain_R8             c....a..g...........................................................a...t....... [143280]

FR682468.2_Georgia_2007/1        taccgatgaagaggttatacggatgaatgctgaaaatccaaagtttttggtgaaaaaacgtgacagggatccctgtttgc [143360]
ASF_IND_20_CAD543                ................................................................................ [143360]
LR899193.1_Germany_2020/1        ................................................................................ [143360]
MN393477.1_Wuhan_2019            ................................................................................ [143360]
MW396979.1_Timor-Leste/2019/1    ................................................................................ [143360]
MK940252.1_CN/2019/InnerMongolia ................................................................................ [143360]
MK128995.1_China/2018/AnhuiXCGQ  ................................................................................ [143360]
MH681419.1_ASFV/POL/2015         ................................................................................ [143360]
LS478113.1_Estonia_2014          ................................................................................ [143360]
KP843857.1_Odintsovo_02/14       ................................................................................ [143360]
FN557520.1_strain_E75            ................................................................................ [143360]
AY261366.1_Warthog               ................................................................................ [143360]
AY261365.1_Warmbaths             ............................................................c..t................ [143360]
AY261364.1_Tengani_62            ...t............................................................................ [143360]
AY261363.1_Pretoriuskop/96/4     ................................................................................ [143360]
AY261362.1_Mkuzi_1979            ................................................................................ [143360]
AY261361.1_Malawi_Lil-20/1_1983  ........................................................g....................... [143360]
AY261360.1_Kenya_1950            ...t..............c...........................................................a. [143360]
AM712240.1_OURT_88/3             ................................................................................ [143360]
AM712239.1_Benin_97/1            ................................................................................ [143360]
KM111295.1_Ken06.Bus             ..................c...........................................................a. [143360]
KM111294.1_Ken05/Tk1             ...t..............c...........................................................a. [143360]
KM262844.1_strain_L60            ................................................................................ [143360]
KM102979.1_26544/OG10_Italy      ................................................................................ [143360]
KM262845.1_strain_NHV            ................................................................................ [143360]
KX354450.1_47/Ss/2008            ................................................................................ [143360]
KP055815.1_strain_BA71           ................................................................................ [143360]
MH025920.1_strain_R35            ..................c...........................................................a. [143360]
MH025919.1_strain_N10            ..................c...........................................................a. [143360]
MH025918.1_strain_R25            ..................c...........................................................a. [143360]
MH025917.1_strain_R7             ..................c...........................................................a. [143360]
MH025916.1_strain_R8             ..................c...........................................................a. [143360]
```

```
FR682468.2_Georgia_2007/1        agttcaacaaatacaaaatctcgccgccattggaagatgatggtctcactgttaaaaagaatgagcagggtgaagaaata [143440]
ASF_IND_20_CAD543                .............y.................................................................. [143440]
LR899193.1_Germany_2020/1        ................................................................................ [143440]
MN393477.1_Wuhan_2019            ................................................................................ [143440]
MW396979.1_Timor-Leste/2019/1    ................................................................................ [143440]
MK940252.1_CN/2019/InnerMongolia ................................................................................ [143440]
MK128995.1_China/2018/AnhuiXCGQ  ................................................................................ [143440]
MH681419.1_ASFV/POL/2015         ................................................................................ [143440]
LS478113.1_Estonia_2014          ................................................................................ [143440]
KP843857.1_Odintsovo_02/14       ................................................................................ [143440]
FN557520.1_strain_E75            .............t..............g................................................... [143440]
AY261366.1_Warthog               ............................g.................t................................. [143440]
AY261365.1_Warmbaths             ......................a.....g.................t................................. [143440]
AY261364.1_Tengani_62            ............................g.................t................................. [143440]
AY261363.1_Pretoriuskop/96/4     ............................g.................t................................. [143440]
AY261362.1_Mkuzi_1979            .............t.................a................................................ [143440]
AY261361.1_Malawi_Lil-20/1_1983  .......t..g..................c................t................................. [143440]
AY261360.1_Kenya_1950            ............................g...............t.t..............c.................t [143440]
AM712240.1_OURT_88/3             .............t..............g................................................... [143440]
AM712239.1_Benin_97/1            .............t..............g................................................... [143440]
KM111295.1_Ken06.Bus             ............................g.................t..............c.................c [143440]
KM111294.1_Ken05/Tk1             ............................g.................t..............c.................c [143440]
KM262844.1_strain_L60            .............t..............g................................................... [143440]
KM102979.1_26544/OG10_Italy      .............t..............g................................................... [143440]
KM262845.1_strain_NHV            .............t..............g................................................... [143440]
KX354450.1_47/Ss/2008            .............t..............g................................................... [143440]
KP055815.1_strain_BA71           .............t..............g................................................... [143440]
MH025920.1_strain_R35            ............................g.................t..............c.................c [143440]
MH025919.1_strain_N10            ............................g.................t..............c.................c [143440]
MH025918.1_strain_R25            ............................g.................t..............c.................c [143440]
MH025917.1_strain_R7             ............................g.................t..............c.................c [143440]
MH025916.1_strain_R8             ............................g.................t..............c.................c [143440]

FR682468.2_Georgia_2007/1        taccccggcgacgaagaaaaatctaagttgtttcaaattattgaactgttagaagaagcctttgaagacgctgtgcaaaa [143520]
ASF_IND_20_CAD543                ................................................................................ [143520]
LR899193.1_Germany_2020/1        ................................................................................ [143520]
MN393477.1_Wuhan_2019            ................................................................................ [143520]
MW396979.1_Timor-Leste/2019/1    ................................................................................ [143520]
MK940252.1_CN/2019/InnerMongolia ................................................................................ [143520]
MK128995.1_China/2018/AnhuiXCGQ  ................................................................................ [143520]
MH681419.1_ASFV/POL/2015         ................................................................................ [143520]
LS478113.1_Estonia_2014          ................................................................................ [143520]
KP843857.1_Odintsovo_02/14       ................................................................................ [143520]
```

```
FN557520.1_strain_E75            ..t.....t....................................................................... [143520]
AY261366.1_Warthog               ....................................................................t........... [143520]
AY261365.1_Warmbaths             ....................................................................t........... [143520]
AY261364.1_Tengani_62            ................................................................................ [143520]
AY261363.1_Pretoriuskop/96/4     ....................................................................t........... [143520]
AY261362.1_Mkuzi_1979            ................................................................................ [143520]
AY261361.1_Malawi_Lil-20/1_1983  ..........................................................................a..... [143520]
AY261360.1_Kenya_1950            ........t..t...............................................t...........c........ [143520]
AM712240.1_OURT_88/3             ..t.....t....................................................................... [143520]
AM712239.1_Benin_97/1            ..t.....t....................................................................... [143520]
KM111295.1_Ken06.Bus             ...........t...............................................t.................... [143520]
KM111294.1_Ken05/Tk1             ...........................................................t...........c........ [143520]
KM262844.1_strain_L60            ..t.....t....................................................................... [143520]
KM102979.1_26544/OG10_Italy      ..t.....t....................................................................... [143520]
KM262845.1_strain_NHV            ..t.....t....................................................................... [143520]
KX354450.1_47/Ss/2008            ..t.....t....................................................................... [143520]
KP055815.1_strain_BA71           ..t.....t....................................................................... [143520]
MH025920.1_strain_R35            ...........t...............................................t.................... [143520]
MH025919.1_strain_N10            ...........t...............................................t.................... [143520]
MH025918.1_strain_R25            ...........t...............................................t.................... [143520]
MH025917.1_strain_R7             ...........t...............................................t.................... [143520]
MH025916.1_strain_R8             ...........t...............................................t.................... [143520]

FR682468.2_Georgia_2007/1        aggtcctgaagccatgaaaacgaaacatgttataaaattaattcaaagaaaaatttctaatagcgcggttaaaaacgcag [143600]
ASF_IND_20_CAD543                ................................................................................ [143600]
LR899193.1_Germany_2020/1        ................................................................................ [143600]
MN393477.1_Wuhan_2019            ................................................................................ [143600]
MW396979.1_Timor-Leste/2019/1    ................................................................................ [143600]
MK940252.1_CN/2019/InnerMongolia ................................................................................ [143600]
MK128995.1_China/2018/AnhuiXCGQ  ................................................................................ [143600]
MH681419.1_ASFV/POL/2015         ................................................................................ [143600]
LS478113.1_Estonia_2014          ................................................................................ [143600]
KP843857.1_Odintsovo_02/14       ................................................................................ [143600]
FN557520.1_strain_E75            ................................................................................ [143600]
AY261366.1_Warthog               ..........a.........................................................c........... [143600]
AY261365.1_Warmbaths             ..........a.t.......................................................c......t.... [143600]
AY261364.1_Tengani_62            ..........a.........................................................c........... [143600]
AY261363.1_Pretoriuskop/96/4     ..........a.........................................................c........... [143600]
AY261362.1_Mkuzi_1979            ................................................................................ [143600]
AY261361.1_Malawi_Lil-20/1_1983  ......c...a..........a...a..a...........g..................g......aa.......t.... [143600]
AY261360.1_Kenya_1950            ..........aa.........a...a..a...........g.g................g..a...a.c......t.... [143600]
AM712240.1_OURT_88/3             ................................................................................ [143600]
AM712239.1_Benin_97/1            ................................................................................ [143600]
KM111295.1_Ken06.Bus             ..........aa.........a...a..a...........g.g................g..a...a.c......t.... [143600]
```

```
KM111294.1_Ken05/Tk1           ..........aa.........a...a..a...........g.g........g...............c..........g. [143600]
KM262844.1_strain_L60          ................................................................................ [143600]
KM102979.1_26544/OG10_Italy    ................................................................................ [143600]
KM262845.1_strain_NHV          ................................................................................ [143600]
KX354450.1_47/Ss/2008          ................................................................................ [143600]
KP055815.1_strain_BA71         ................................................................................ [143600]
MH025920.1_strain_R35          ..........aa.........a...a..a...........g.g................g..a...a.c......t.... [143600]
MH025919.1_strain_N10          ..........aa.........a...a..a...........g.g................g..a...a.c......t.... [143600]
MH025918.1_strain_R25          ..........aa.........a...a..a...........g.g................g..a...a.c......t.... [143600]
MH025917.1_strain_R7           ..........aa.........a...a..a...........g.g................g..a...a.c......t.... [143600]
MH025916.1_strain_R8           ..........aa.........a...a..a...........g.g................g..a...a.c......t.... [143600]

FR682468.2_Georgia_2007/1      acaaacctttgccgaatcctatcgcacgcattcgtattaaaatcaatcccgctacaagtatactaacaccaatattgctt [143680]
ASF_IND_20_CAD543              ................................................................................ [143680]
LR899193.1_Germany_2020/1      ................................................................................ [143680]
MN393477.1_Wuhan_2019          ................................................................................ [143680]
MW396979.1_Timor-Leste/2019/1  ................................................................................ [143680]
MK940252.1_CN/2019/InnerMongolia ................................................................................ [143680]
MK128995.1_China/2018/AnhuiXCGQ ................................................................................ [143680]
MH681419.1_ASFV/POL/2015       ................................................................................ [143680]
LS478113.1_Estonia_2014        ................................................................................ [143680]
KP843857.1_Odintsovo_02/14     ................................................................................ [143680]
FN557520.1_strain_E75          ......................................................................g......... [143680]
AY261366.1_Warthog             .t.............................................t...g.....a..................a... [143680]
AY261365.1_Warmbaths           .........................................................a...................... [143680]
AY261364.1_Tengani_62          ...............................................t.........a...................... [143680]
AY261363.1_Pretoriuskop/96/4   .t.............................................t...g.....a..................a... [143680]
AY261362.1_Mkuzi_1979          ......................................................................g......... [143680]
AY261361.1_Malawi_Lil-20/1_1983 .t.............................c.........g.t......a......c...g..gg.............. [143680]
AY261360.1_Kenya_1950          .....t................t..................g........at....ac...g..gg.....g........ [143680]
AM712240.1_OURT_88/3           ......a...............................................................g......... [143680]
AM712239.1_Benin_97/1          ......................................................................g......... [143680]
KM111295.1_Ken06.Bus           .....t.......a........t..................g........at....ac...gt.gg.............. [143680]
KM111294.1_Ken05/Tk1           .t...........a........t..................g........at....ac...g..gg.....g........ [143680]
KM262844.1_strain_L60          ......................................................................g......... [143680]
KM102979.1_26544/OG10_Italy    ......................................................................g......... [143680]
KM262845.1_strain_NHV          ......a...............................................................g......... [143680]
KX354450.1_47/Ss/2008          ......................................................................g......... [143680]
KP055815.1_strain_BA71         ......................................................................g......... [143680]
MH025920.1_strain_R35          .....t.......a........t..................g........at....ac...gt.gg.............. [143680]
MH025919.1_strain_N10          .....t.......a........t..................g........at....ac...gt.gg.............. [143680]
MH025918.1_strain_R25          .....t.......a........t..................g........at....ac...gt.gg.............. [143680]
MH025917.1_strain_R7           .....t.......a........t..................g........at....ac...gt.gg.............. [143680]
MH025916.1_strain_R8           .....t.......a........t..................g........at....ac...gt.gg.............. [143680]
```

```
FR682468.2_Georgia_2007/1        gataaaaataagcccattactttacagaatggtaaaacaagctttgaagagttaaaagatgaagacggcgttaaggccaa [143760]
ASF_IND_20_CAD543                ................................................................................ [143760]
LR899193.1_Germany_2020/1        ................................................................................ [143760]
MN393477.1_Wuhan_2019            ................................................................................ [143760]
MW396979.1_Timor-Leste/2019/1    ................................................................................ [143760]
MK940252.1_CN/2019/InnerMongolia ................................................................................ [143760]
MK128995.1_China/2018/AnhuiXCGQ  ................................................................................ [143760]
MH681419.1_ASFV/POL/2015         ................................................................................ [143760]
LS478113.1_Estonia_2014          ................................................................................ [143760]
KP843857.1_Odintsovo_02/14       ................................................................................ [143760]
FN557520.1_strain_E75            ................................................................................ [143760]
AY261366.1_Warthog               a......g............c........................................................... [143760]
AY261365.1_Warmbaths             .......g.......................................................a................ [143760]
AY261364.1_Tengani_62            a......g........................................................................ [143760]
AY261363.1_Pretoriuskop/96/4     a......g........................................................................ [143760]
AY261362.1_Mkuzi_1979            ................................................................................ [143760]
AY261361.1_Malawi_Lil-20/1_1983  ...................................g.................g...............aca........ [143760]
AY261360.1_Kenya_1950            .......g...........................g..............ac.g......a................... [143760]
AM712240.1_OURT_88/3             ................................................................................ [143760]
AM712239.1_Benin_97/1            ................................................................................ [143760]
KM111295.1_Ken06.Bus             .......g...........................g..............ac.g......a.............a..... [143760]
KM111294.1_Ken05/Tk1             .......g...........................g..............ac.g......a................... [143760]
KM262844.1_strain_L60            ................................................................................ [143760]
KM102979.1_26544/OG10_Italy      ................................................................................ [143760]
KM262845.1_strain_NHV            ................................................................................ [143760]
KX354450.1_47/Ss/2008            ................................................................................ [143760]
KP055815.1_strain_BA71           ................................................................................ [143760]
MH025920.1_strain_R35            .......g...........................g..............ac.g......a.............a..... [143760]
MH025919.1_strain_N10            .......g...........................g..............ac.g......a.............a..... [143760]
MH025918.1_strain_R25            .......g...........................g..............ac.g......a.............a..... [143760]
MH025917.1_strain_R7             .......g...........................g..............ac.g......a.............a..... [143760]
MH025916.1_strain_R8             .......g...........................g..............ac.g......a.............a..... [143760]

FR682468.2_Georgia_2007/1        tccggataatattcataagcttatagaatcgcattctatacatgatggcatcattaatgctagatctatttgcatcagca [143840]
ASF_IND_20_CAD543                ................................................................................ [143840]
LR899193.1_Germany_2020/1        ................................................................................ [143840]
MN393477.1_Wuhan_2019            ................................................................................ [143840]
MW396979.1_Timor-Leste/2019/1    ................................................................................ [143840]
MK940252.1_CN/2019/InnerMongolia ................................................................................ [143840]
MK128995.1_China/2018/AnhuiXCGQ  ................................................................................ [143840]
MH681419.1_ASFV/POL/2015         ................................................................................ [143840]
LS478113.1_Estonia_2014          ................................................................................ [143840]
KP843857.1_Odintsovo_02/14       ................................................................................ [143840]
```

```
FN557520.1_strain_E75            .......................................g........................................ [143840]
AY261366.1_Warthog               ........................t................................c...................... [143840]
AY261365.1_Warmbaths             ........................t................................c...................... [143840]
AY261364.1_Tengani_62            ........................t....................................................... [143840]
AY261363.1_Pretoriuskop/96/4     ........................t................................c...................... [143840]
AY261362.1_Mkuzi_1979            ................................................................................ [143840]
AY261361.1_Malawi_Lil-20/1_1983  ...............c........t....................................................... [143840]
AY261360.1_Kenya_1950            .........c..............t....................................................... [143840]
AM712240.1_OURT_88/3             .......................................g........................................ [143840]
AM712239.1_Benin_97/1            .......................................g........................................ [143840]
KM111295.1_Ken06.Bus             .........c..............t......a................................................ [143840]
KM111294.1_Ken05/Tk1             c........c..............t........a..c...........t..t............................ [143840]
KM262844.1_strain_L60            .......................................g........................................ [143840]
KM102979.1_26544/OG10_Italy      .......................................g........................................ [143840]
KM262845.1_strain_NHV            .......................................g........................................ [143840]
KX354450.1_47/Ss/2008            .......................................g........................................ [143840]
KP055815.1_strain_BA71           .......................................g........................................ [143840]
MH025920.1_strain_R35            .........c..............t......a................................................ [143840]
MH025919.1_strain_N10            .........c..............t......a................................................ [143840]
MH025918.1_strain_R25            .........c..............t......a................................................ [143840]
MH025917.1_strain_R7             .........c..............t......a................................................ [143840]
MH025916.1_strain_R8             .........c..............t......a................................................ [143840]

FR682468.2_Georgia_2007/1        atatgggcatttcatttccgctttgcttggaaatgggagttgtaaaagtttttgaaaaaaataatgggattgatgtgaac [143920]
ASF_IND_20_CAD543                .........................tc..................................................... [143920]
LR899193.1_Germany_2020/1        ................................................................................ [143920]
MN393477.1_Wuhan_2019            ................................................................................ [143920]
MW396979.1_Timor-Leste/2019/1    ................................................................................ [143920]
MK940252.1_CN/2019/InnerMongolia ................................................................................ [143920]
MK128995.1_China/2018/AnhuiXCGQ  ................................................................................ [143920]
MH681419.1_ASFV/POL/2015         ................................................................................ [143920]
LS478113.1_Estonia_2014          ................................................................................ [143920]
KP843857.1_Odintsovo_02/14       ................................................................................ [143920]
FN557520.1_strain_E75            .........................tc..................................................... [143920]
AY261366.1_Warthog               ..........................c....g.............................................g.. [143920]
AY261365.1_Warmbaths             ..........................c....g.................c...........................g.. [143920]
AY261364.1_Tengani_62            .c.................a......c....g.............................................g.. [143920]
AY261363.1_Pretoriuskop/96/4     ..........................c....g.......................................a.....g.. [143920]
AY261362.1_Mkuzi_1979            ..........................c..................................................... [143920]
AY261361.1_Malawi_Lil-20/1_1983  ..........................c....g.............................................g.. [143920]
AY261360.1_Kenya_1950            g.........................c....g........g....................................g.. [143920]
AM712240.1_OURT_88/3             .........................tc..................................................... [143920]
AM712239.1_Benin_97/1            .........................tc..................................................... [143920]
KM111295.1_Ken06.Bus             g.........................c....g........g....................................g.. [143920]
```

```
KM111294.1_Ken05/Tk1             g.........................c....g........g....................................g.. [143920]
KM262844.1_strain_L60            .........................tc..................................................... [143920]
KM102979.1_26544/OG10_Italy      .........................tc..................................................... [143920]
KM262845.1_strain_NHV            .........................tc..................................................... [143920]
KX354450.1_47/Ss/2008            .........................tc..................................................... [143920]
KP055815.1_strain_BA71           .........................tc..................................................... [143920]
MH025920.1_strain_R35            g.........................c....g........g....................................g.. [143920]
MH025919.1_strain_N10            g.........................c....g........g....................................g.. [143920]
MH025918.1_strain_R25            g.........................c....g........g....................................g.. [143920]
MH025917.1_strain_R7             g.........................c....g........g....................................g.. [143920]
MH025916.1_strain_R8             g.........................c....g........g....................................g.. [143920]

FR682468.2_Georgia_2007/1        tccatttatggctcagacgatatttcaactcttgttaatcagattgctattgcttaaacaat------------------ [144000]
ASF_IND_20_CAD543                ..............................................................------------------ [144000]
LR899193.1_Germany_2020/1        ..............................................................------------------ [144000]
MN393477.1_Wuhan_2019            ..............................................................------------------ [144000]
MW396979.1_Timor-Leste/2019/1    ..............................................................------------------ [144000]
MK940252.1_CN/2019/InnerMongolia ..............................................................------------------ [144000]
MK128995.1_China/2018/AnhuiXCGQ  ..............................................................------------------ [144000]
MH681419.1_ASFV/POL/2015         ..............................................................------------------ [144000]
LS478113.1_Estonia_2014          ..............................................................------------------ [144000]
KP843857.1_Odintsovo_02/14       ..............................................................------------------ [144000]
FN557520.1_strain_E75            ............................................c.................------------------ [144000]
AY261366.1_Warthog               .........a..................................................tc------------------ [144000]
AY261365.1_Warmbaths             .........a..................................................tc------------------ [144000]
AY261364.1_Tengani_62            .........a...................................................cttgttt------------ [144000]
AY261363.1_Pretoriuskop/96/4     .........a...................................................c------------------ [144000]
AY261362.1_Mkuzi_1979            ..............................................................------------------ [144000]
AY261361.1_Malawi_Lil-20/1_1983  ........................a...a....a........g..................c------------------ [144000]
AY261360.1_Kenya_1950            ............c.......g...a...a....a........g..a...............gctgtttaaacagcttgct [144000]
AM712240.1_OURT_88/3             ............................................c.................------------------ [144000]
AM712239.1_Benin_97/1            ............................................c.................------------------ [144000]
KM111295.1_Ken06.Bus             ............c...........a...a....a........g.................gcttgctt------------ [144000]
KM111294.1_Ken05/Tk1             ............c...........a...a....a........g..a..............gcttgctt------------ [144000]
KM262844.1_strain_L60            ............................................c.................------------------ [144000]
KM102979.1_26544/OG10_Italy      ............................................c.................------------------ [144000]
KM262845.1_strain_NHV            ............................................c.................------------------ [144000]
KX354450.1_47/Ss/2008            ............................................c.................------------------ [144000]
KP055815.1_strain_BA71           ............................................c.................------------------ [144000]
MH025920.1_strain_R35            ............c...........a...a....a........g.................gcttgctt------------ [144000]
MH025919.1_strain_N10            ............c...........a...a....a........g.................gcttgctt------------ [144000]
MH025918.1_strain_R25            ............c...........a...a....a........g.................gcttgctt------------ [144000]
MH025917.1_strain_R7             ............c...........a...a....a........g.................gcttgctt------------ [144000]
MH025916.1_strain_R8             ............c...........a...a....a........g.................gcttgctt------------ [144000]
```

```
FR682468.2_Georgia_2007/1        ---------------------------------------------------------------ttgctcaaaacaagctt [144080]
ASF_IND_20_CAD543                ---------------------------------------------------------------................. [144080]
LR899193.1_Germany_2020/1        ---------------------------------------------------------------................. [144080]
MN393477.1_Wuhan_2019            ---------------------------------------------------------------................. [144080]
MW396979.1_Timor-Leste/2019/1    ---------------------------------------------------------------................. [144080]
MK940252.1_CN/2019/InnerMongolia ---------------------------------------------------------------................. [144080]
MK128995.1_China/2018/AnhuiXCGQ  ---------------------------------------------------------------................. [144080]
MH681419.1_ASFV/POL/2015         ---------------------------------------------------------------................. [144080]
LS478113.1_Estonia_2014          ---------------------------------------------------------------................. [144080]
KP843857.1_Odintsovo_02/14       ---------------------------------------------------------------................. [144080]
FN557520.1_strain_E75            ---------------------------------------------------------------................. [144080]
AY261366.1_Warthog               ---------------------------------------------------------------................. [144080]
AY261365.1_Warmbaths             ---------------------------------------------------------------.....t........... [144080]
AY261364.1_Tengani_62            -----------------------------------------aaaacaacttgtttaaaacaac................. [144080]
AY261363.1_Pretoriuskop/96/4     ---------------------------------------------------------------................. [144080]
AY261362.1_Mkuzi_1979            ---------------------------------------------------------------................. [144080]
AY261361.1_Malawi_Lil-20/1_1983  ---------------------------------------------------------------...t.t........... [144080]
AY261360.1_Kenya_1950            taaacaagttgttttaaacaacttgcttaaacaagttgttttaaacaacttgttttaaacaac.....t.......a... [144080]
AM712240.1_OURT_88/3             ---------------------------------------------------------------................. [144080]
AM712239.1_Benin_97/1            ---------------------------------------------------------------................. [144080]
KM111295.1_Ken06.Bus             ----------------------------------------------------------------------......a... [144080]
KM111294.1_Ken05/Tk1             --------------------------------------------------------aaacaag...t.t.......a... [144080]
KM262844.1_strain_L60            ---------------------------------------------------------------................. [144080]
KM102979.1_26544/OG10_Italy      ---------------------------------------------------------------................. [144080]
KM262845.1_strain_NHV            ---------------------------------------------------------------................. [144080]
KX354450.1_47/Ss/2008            ---------------------------------------------------------------................. [144080]
KP055815.1_strain_BA71           ---------------------------------------------------------------................. [144080]
MH025920.1_strain_R35            ----------------------------------------------------------------------......a... [144080]
MH025919.1_strain_N10            ----------------------------------------------------------------------......a... [144080]
MH025918.1_strain_R25            ----------------------------------------------------------------------......a... [144080]
MH025917.1_strain_R7             ----------------------------------------------------------------------......a... [144080]
MH025916.1_strain_R8             ----------------------------------------------------------------------......a... [144080]

FR682468.2_Georgia_2007/1        ataaacgtttcttaggtatgcgatacgtaaatcctaattctttaataagttctttttcagt--------------agtga [144160]
ASF_IND_20_CAD543                .............................................................--------------..... [144160]
LR899193.1_Germany_2020/1        .............................................................agtgatttttcagt..... [144160]
MN393477.1_Wuhan_2019            .............................................................--------------..... [144160]
MW396979.1_Timor-Leste/2019/1    .............................................................--------------..... [144160]
MK940252.1_CN/2019/InnerMongolia .............................................................--------------..... [144160]
MK128995.1_China/2018/AnhuiXCGQ  .............................................................--------------..... [144160]
MH681419.1_ASFV/POL/2015         .............................................................--------------..... [144160]
LS478113.1_Estonia_2014          .............................................................--------------..... [144160]
KP843857.1_Odintsovo_02/14       .............................................................--------------..... [144160]
```

```
FN557520.1_strain_E75             .............................................................--------------..... [144160]
AY261366.1_Warthog                ...............................c.............................--------------..... [144160]
AY261365.1_Warmbaths              ...............................c.............................--------------..... [144160]
AY261364.1_Tengani_62             .............................................................--------------..... [144160]
AY261363.1_Pretoriuskop/96/4      ...............................c.............................--------------..... [144160]
AY261362.1_Mkuzi_1979             ...............................c.............................--------------..... [144160]
AY261361.1_Malawi_Lil-20/1_1983   ...............................c....................c........--------------...a. [144160]
AY261360.1_Kenya_1950             ..........t.......c............c....................cc.......--------------...a. [144160]
AM712240.1_OURT_88/3              .............................................................--------------..... [144160]
AM712239.1_Benin_97/1             .............................................................--------------..... [144160]
KM111295.1_Ken06.Bus              ..........t.......c............c....................cc.......--------------...a. [144160]
KM111294.1_Ken05/Tk1              ..........t.......c............c....................c........--------------...a. [144160]
KM262844.1_strain_L60             .............................................................--------------..... [144160]
KM102979.1_26544/OG10_Italy       .............................................................--------------..... [144160]
KM262845.1_strain_NHV             .............................................................--------------..... [144160]
KX354450.1_47/Ss/2008             .............................................................--------------..... [144160]
KP055815.1_strain_BA71            .............................................................--------------..... [144160]
MH025920.1_strain_R35             ..........t.......c............c....................cc.......--------------...a. [144160]
MH025919.1_strain_N10             ..........t.......c............c....................cc.......--------------...a. [144160]
MH025918.1_strain_R25             ..........t.......c............c....................cc.......--------------...a. [144160]
MH025917.1_strain_R7              ..........t.......c............c....................cc.......--------------...a. [144160]
MH025916.1_strain_R8              ..........t.......c............c....................cc.......--------------...a. [144160]

FR682468.2_Georgia_2007/1         tttttagaggtactaaagtttgatttttaaataatccatactgatttagcttataattc-tttttttttaacgcagctcg [144240]
ASF_IND_20_CAD543                 ...........................................................-.................... [144240]
LR899193.1_Germany_2020/1         ...........................................................-.................... [144240]
MN393477.1_Wuhan_2019             ...........................................................-.................... [144240]
MW396979.1_Timor-Leste/2019/1     ...........................................................-.................... [144240]
MK940252.1_CN/2019/InnerMongolia  ...........................................................-.................... [144240]
MK128995.1_China/2018/AnhuiXCGQ   ...........................................................-.................... [144240]
MH681419.1_ASFV/POL/2015          ...........................................................-.................... [144240]
LS478113.1_Estonia_2014           ...........................................................-.................... [144240]
KP843857.1_Odintsovo_02/14        ...........................................................-.................... [144240]
FN557520.1_strain_E75             ...........................................................-..............g..... [144240]
AY261366.1_Warthog                ....c......................................................-..............g..... [144240]
AY261365.1_Warmbaths              ....c...........g..........................................-..............g..... [144240]
AY261364.1_Tengani_62             ........................................t..................-.................... [144240]
AY261363.1_Pretoriuskop/96/4      ....c...........g..........................................-..............g..... [144240]
AY261362.1_Mkuzi_1979             ...........................................................-.................... [144240]
AY261361.1_Malawi_Lil-20/1_1983   ...........................................................-...........g..g..... [144240]
AY261360.1_Kenya_1950             ................t..........................g...........g...-..............g..... [144240]
AM712240.1_OURT_88/3              ...........................................................-..............g..... [144240]
AM712239.1_Benin_97/1             ...........................................................-..............g..... [144240]
KM111295.1_Ken06.Bus              ................t..........................g...........g...t..............g..... [144240]
```

```
KM111294.1_Ken05/Tk1            ...........................................g...........g...-..............g..... [144240]
KM262844.1_strain_L60           ...........................................................-..............g..... [144240]
KM102979.1_26544/OG10_Italy     ...........................................................-..............g..... [144240]
KM262845.1_strain_NHV           ...........................................................-..............g..... [144240]
KX354450.1_47/Ss/2008           ...........................................................-..............g..... [144240]
KP055815.1_strain_BA71          ...........................................................-..............g..... [144240]
MH025920.1_strain_R35           ................t..........................g...........g...-..............g..... [144240]
MH025919.1_strain_N10           ................t..........................g...........g...-..............g..... [144240]
MH025918.1_strain_R25           ................t..........................g...........g...-..............g..... [144240]
MH025917.1_strain_R7            ................t..........................g...........g...-..............g..... [144240]
MH025916.1_strain_R8            ................t..........................g...........g...-..............g..... [144240]

FR682468.2_Georgia_2007/1       aattcttattaaataagaaacgggacccgtaaaatgaagtactgcgtatggcttttcctcggctaaggccgtaaaaagat [144320]
ASF_IND_20_CAD543               ................................................................................ [144320]
LR899193.1_Germany_2020/1       ................................................................................ [144320]
MN393477.1_Wuhan_2019           ................................................................................ [144320]
MW396979.1_Timor-Leste/2019/1   ................................................................................ [144320]
MK940252.1_CN/2019/InnerMongolia ................................................................................ [144320]
MK128995.1_China/2018/AnhuiXCGQ ..................................................a............................. [144320]
MH681419.1_ASFV/POL/2015        ................................................................................ [144320]
LS478113.1_Estonia_2014         ................................................................................ [144320]
KP843857.1_Odintsovo_02/14      ................................................................................ [144320]
FN557520.1_strain_E75           ...........g..........................a..t.........t..............a............. [144320]
AY261366.1_Warthog              .........c.g............g....................................................... [144320]
AY261365.1_Warmbaths            .........c.g............g................................t...................... [144320]
AY261364.1_Tengani_62           .........c.g............c.................c..................................... [144320]
AY261363.1_Pretoriuskop/96/4    .........c.g............g....................................................... [144320]
AY261362.1_Mkuzi_1979           ...........g..........................a..t.........t..............a............. [144320]
AY261361.1_Malawi_Lil-20/1_1983 .........c.g..........................a..t...................................... [144320]
AY261360.1_Kenya_1950           ......g..c.g.........a..g.............a..t...a.......................t.......... [144320]
AM712240.1_OURT_88/3            ...........g..........................a..t.........t..............a............. [144320]
AM712239.1_Benin_97/1           ...........g..........................a..t.........t..............a............. [144320]
KM111295.1_Ken06.Bus            ......g..c.g.........a..g.............a..t...................................... [144320]
KM111294.1_Ken05/Tk1            ......g..c.g.........a..g.............a..t...a.......................t.......... [144320]
KM262844.1_strain_L60           ...........g..........................a..t.........t..............a............. [144320]
KM102979.1_26544/OG10_Italy     ...........g..........................a..t.........t..............a............. [144320]
KM262845.1_strain_NHV           ...........g..........................a..t.........t..............a............. [144320]
KX354450.1_47/Ss/2008           ...........g..........................a..t.........t..............a............. [144320]
KP055815.1_strain_BA71          ...........g..........................a..t.........t..............a............. [144320]
MH025920.1_strain_R35           ......g..c.g.........a..g.............a..t...................................... [144320]
MH025919.1_strain_N10           ......g..c.g.........a..g.............a..t...................................... [144320]
MH025918.1_strain_R25           ......g..c.g.........a..g.............a..t...................................... [144320]
MH025917.1_strain_R7            ......g..c.g.........a..g.............a..t...................................... [144320]
MH025916.1_strain_R8            ......g..c.g.........a..g.............a..t...................................... [144320]
```

```
FR682468.2_Georgia_2007/1        caagttgatatgtgttttttttccattcaataaaaagtacacactttcgttctccgcagacttttacagaaaaagaaaga [144400]
ASF_IND_20_CAD543                .....................c.......................................................... [144400]
LR899193.1_Germany_2020/1        ................................................................................ [144400]
MN393477.1_Wuhan_2019            ................................................................................ [144400]
MW396979.1_Timor-Leste/2019/1    ................................................................................ [144400]
MK940252.1_CN/2019/InnerMongolia ................................................................................ [144400]
MK128995.1_China/2018/AnhuiXCGQ  ..........................................................................a..... [144400]
MH681419.1_ASFV/POL/2015         ................................................................................ [144400]
LS478113.1_Estonia_2014          ................................................................................ [144400]
KP843857.1_Odintsovo_02/14       ................................................................................ [144400]
FN557520.1_strain_E75            ..........c..c.......c.......................................................... [144400]
AY261366.1_Warthog               ..........c..........................c............................t...........a. [144400]
AY261365.1_Warmbaths             ..........c..........................c........................................a. [144400]
AY261364.1_Tengani_62            ..........c..........................c........................................a. [144400]
AY261363.1_Pretoriuskop/96/4     ..........c..........................c........................................a. [144400]
AY261362.1_Mkuzi_1979            ..........c..c.......c.......................................................... [144400]
AY261361.1_Malawi_Lil-20/1_1983  ..........c..c.......c.......................................................... [144400]
AY261360.1_Kenya_1950            ..........c..c.......c...............c.....t.................................... [144400]
AM712240.1_OURT_88/3             ..........c..c.......c.......................................................... [144400]
AM712239.1_Benin_97/1            ..........c..c.......c.......................................................... [144400]
KM111295.1_Ken06.Bus             ..........c..c.......c...............c.....t.................................... [144400]
KM111294.1_Ken05/Tk1             ..........c..c.......c...............c.....t.....................g.............. [144400]
KM262844.1_strain_L60            ..........c..c.......c.......................................................... [144400]
KM102979.1_26544/OG10_Italy      ..........c..c.......c.......................................................... [144400]
KM262845.1_strain_NHV            ..........c..c.......c.......................................................... [144400]
KX354450.1_47/Ss/2008            ..........c..c.......c.......................................................... [144400]
KP055815.1_strain_BA71           ..........c..c.......c.......................................................... [144400]
MH025920.1_strain_R35            ..........c..c.......c...............c.....t.................................... [144400]
MH025919.1_strain_N10            ..........c..c.......c...............c.....t.................................... [144400]
MH025918.1_strain_R25            ..........c..c.......c...............c.....t.................................... [144400]
MH025917.1_strain_R7             ..........c..c.......c...............c.....t.................................... [144400]
MH025916.1_strain_R8             ..........c..c.......c...............c.....t.................................... [144400]

FR682468.2_Georgia_2007/1        tcctttatgcgaatgttgggcaggacgtgttttaaaagttttttttctggaacaataataagaagatccacgtcattaag [144480]
ASF_IND_20_CAD543                ................................................................................ [144480]
LR899193.1_Germany_2020/1        ................................................................................ [144480]
MN393477.1_Wuhan_2019            ................................................................................ [144480]
MW396979.1_Timor-Leste/2019/1    ................................................................................ [144480]
MK940252.1_CN/2019/InnerMongolia ................................................................................ [144480]
MK128995.1_China/2018/AnhuiXCGQ  ...................a............................................................ [144480]
MH681419.1_ASFV/POL/2015         ................................................................................ [144480]
LS478113.1_Estonia_2014          ................................................................................ [144480]
KP843857.1_Odintsovo_02/14       ................................................................................ [144480]
```

```
FN557520.1_strain_E75           c............................................................................... [144480]
AY261366.1_Warthog              ................................................................................ [144480]
AY261365.1_Warmbaths            ................................................................................ [144480]
AY261364.1_Tengani_62           ..............a..................................................g.............. [144480]
AY261363.1_Pretoriuskop/96/4    ................................................................................ [144480]
AY261362.1_Mkuzi_1979           c............................................................................... [144480]
AY261361.1_Malawi_Lil-20/1_1983 c...................................ca.......................................... [144480]
AY261360.1_Kenya_1950           c...........................a....................................g.............. [144480]
AM712240.1_OURT_88/3            c......................................................................a........ [144480]
AM712239.1_Benin_97/1           c............................................................................... [144480]
KM111295.1_Ken06.Bus            c..........g.....................................a...............g.............. [144480]
KM111294.1_Ken05/Tk1            c...........................a....................................g.............. [144480]
KM262844.1_strain_L60           c............................................................................... [144480]
KM102979.1_26544/OG10_Italy     c............................................................................... [144480]
KM262845.1_strain_NHV           c......................................................................a........ [144480]
KX354450.1_47/Ss/2008           c............................................................................... [144480]
KP055815.1_strain_BA71          c............................................................................... [144480]
MH025920.1_strain_R35           c..........g.....................................a...............g.............. [144480]
MH025919.1_strain_N10           c..........g.....................................a...............g.............. [144480]
MH025918.1_strain_R25           c..........g.....................................a...............g.............. [144480]
MH025917.1_strain_R7            c..........g.....................................a...............g.............. [144480]
MH025916.1_strain_R8            c..........g.....................................a...............g.............. [144480]

FR682468.2_Georgia_2007/1       cattttctcttcgcgtcttaagctaccaacagcaacgatgttttttgataaaatttttataagttgtccattatattcaa [144560]
ASF_IND_20_CAD543               ................................................................................ [144560]
LR899193.1_Germany_2020/1       ................................................................................ [144560]
MN393477.1_Wuhan_2019           ................................................................................ [144560]
MW396979.1_Timor-Leste/2019/1   ................................................................................ [144560]
MK940252.1_CN/2019/InnerMongolia ................................................................................ [144560]
MK128995.1_China/2018/AnhuiXCGQ ................................................................................ [144560]
MH681419.1_ASFV/POL/2015        ................................................................................ [144560]
LS478113.1_Estonia_2014         ................................................................................ [144560]
KP843857.1_Odintsovo_02/14      ................................................................................ [144560]
FN557520.1_strain_E75           .....................a.......................................................... [144560]
AY261366.1_Warthog              ....................................a........................................... [144560]
AY261365.1_Warmbaths            ................................................................................ [144560]
AY261364.1_Tengani_62           ...............................a.......................................g........ [144560]
AY261363.1_Pretoriuskop/96/4    ...................................t............................................ [144560]
AY261362.1_Mkuzi_1979           ................................................................................ [144560]
AY261361.1_Malawi_Lil-20/1_1983 .............................................c.................................. [144560]
AY261360.1_Kenya_1950           ........................t........c.t.......................................c.... [144560]
AM712240.1_OURT_88/3            .....................a.......................................................... [144560]
AM712239.1_Benin_97/1           .....................a.......................................................... [144560]
KM111295.1_Ken06.Bus            .................................c.t.......................................c.... [144560]
```

```
KM111294.1_Ken05/Tk1             .................................c.t.......................................c.... [144560]
KM262844.1_strain_L60            .....................a.......................................................... [144560]
KM102979.1_26544/OG10_Italy      .....................a.......................................................... [144560]
KM262845.1_strain_NHV            .....................a.......................................................... [144560]
KX354450.1_47/Ss/2008            .....................a.......................................................... [144560]
KP055815.1_strain_BA71           .....................a.......................................................... [144560]
MH025920.1_strain_R35            .................................c.t.......................................c.... [144560]
MH025919.1_strain_N10            .................................c.t.......................................c.... [144560]
MH025918.1_strain_R25            .................................c.t.......................................c.... [144560]
MH025917.1_strain_R7             .................................c.t.......................................c.... [144560]
MH025916.1_strain_R8             .................................c.t.......................................c.... [144560]

FR682468.2_Georgia_2007/1        acgcaagtcgggagcgtaagtcatttacaattttttttccttgaataagcgttaacattttatatttaatattaaaatct [144640]
ASF_IND_20_CAD543                .....................s.......................................................... [144640]
LR899193.1_Germany_2020/1        ................................................................................ [144640]
MN393477.1_Wuhan_2019            ................................................................................ [144640]
MW396979.1_Timor-Leste/2019/1    ................................................................................ [144640]
MK940252.1_CN/2019/InnerMongolia ................................................................................ [144640]
MK128995.1_China/2018/AnhuiXCGQ  ................................................................................ [144640]
MH681419.1_ASFV/POL/2015         ................................................................................ [144640]
LS478113.1_Estonia_2014          ................................................................................ [144640]
KP843857.1_Odintsovo_02/14       ................................................................................ [144640]
FN557520.1_strain_E75            .............a.......g.......................................................... [144640]
AY261366.1_Warthog               ................................................................................ [144640]
AY261365.1_Warmbaths             ................................................................................ [144640]
AY261364.1_Tengani_62            ................................................................................ [144640]
AY261363.1_Pretoriuskop/96/4     ........................................c....................................... [144640]
AY261362.1_Mkuzi_1979            .............a.......g.......................................................... [144640]
AY261361.1_Malawi_Lil-20/1_1983  .t...........a..c....a.......................................................... [144640]
AY261360.1_Kenya_1950            .............a..c...g.g.....................g................................... [144640]
AM712240.1_OURT_88/3             .............a.......g.......................................................... [144640]
AM712239.1_Benin_97/1            .............a.......g.......................................................... [144640]
KM111295.1_Ken06.Bus             ........t....a..c...g.......................g................................... [144640]
KM111294.1_Ken05/Tk1             .............a..c...g.g.....................g................................... [144640]
KM262844.1_strain_L60            .............a.......g.......................................................... [144640]
KM102979.1_26544/OG10_Italy      .............a.......g.......................................................... [144640]
KM262845.1_strain_NHV            .............a.......g.......................................................... [144640]
KX354450.1_47/Ss/2008            .............a.......g.......................................................... [144640]
KP055815.1_strain_BA71           .............a.......g.......................................................... [144640]
MH025920.1_strain_R35            .............a..c...g.......................g................................... [144640]
MH025919.1_strain_N10            .............a..c...g.......................g................................... [144640]
MH025918.1_strain_R25            .............a..c...g.......................g................................... [144640]
MH025917.1_strain_R7             .............a..c...g.......................g................................... [144640]
MH025916.1_strain_R8             .............a..c...g.......................g................................... [144640]
```

```
FR682468.2_Georgia_2007/1        tttcattttatatattatatacgcaaaatggcacttgatggttcaagtggtggaggctctaatgtagaaacattacttat [144720]
ASF_IND_20_CAD543                ................................................................................ [144720]
LR899193.1_Germany_2020/1        ................................................................................ [144720]
MN393477.1_Wuhan_2019            ................................................................................ [144720]
MW396979.1_Timor-Leste/2019/1    ................................................................................ [144720]
MK940252.1_CN/2019/InnerMongolia ................................................................................ [144720]
MK128995.1_China/2018/AnhuiXCGQ  ................................................................................ [144720]
MH681419.1_ASFV/POL/2015         ................................................................................ [144720]
LS478113.1_Estonia_2014          ................................................................................ [144720]
KP843857.1_Odintsovo_02/14       ................................................................................ [144720]
FN557520.1_strain_E75            ................................................................................ [144720]
AY261366.1_Warthog               .....................................................t.......................... [144720]
AY261365.1_Warmbaths             ................................................................................ [144720]
AY261364.1_Tengani_62            ................................................................................ [144720]
AY261363.1_Pretoriuskop/96/4     .........................t...................................................... [144720]
AY261362.1_Mkuzi_1979            .........................................a............a......................... [144720]
AY261361.1_Malawi_Lil-20/1_1983  ................................................................................ [144720]
AY261360.1_Kenya_1950            ................................................................................ [144720]
AM712240.1_OURT_88/3             ................................................................................ [144720]
AM712239.1_Benin_97/1            ................................................................................ [144720]
KM111295.1_Ken06.Bus             ................................................................................ [144720]
KM111294.1_Ken05/Tk1             ................................................................................ [144720]
KM262844.1_strain_L60            ................................................................................ [144720]
KM102979.1_26544/OG10_Italy      ................................................................................ [144720]
KM262845.1_strain_NHV            ................................................................................ [144720]
KX354450.1_47/Ss/2008            ................................................................................ [144720]
KP055815.1_strain_BA71           ................................................................................ [144720]
MH025920.1_strain_R35            ................................................................................ [144720]
MH025919.1_strain_N10            ................................................................................ [144720]
MH025918.1_strain_R25            ................................................................................ [144720]
MH025917.1_strain_R7             ................................................................................ [144720]
MH025916.1_strain_R8             ................................................................................ [144720]

FR682468.2_Georgia_2007/1        agtagcaatcattgtggttattatggcaatcatgctttactatttttggtggatgccccgc---cagcaaaaaaaatgta [144800]
ASF_IND_20_CAD543                .............................................................---................ [144800]
LR899193.1_Germany_2020/1        .............................................................---................ [144800]
MN393477.1_Wuhan_2019            .............................................................---................ [144800]
MW396979.1_Timor-Leste/2019/1    .............................................................---................ [144800]
MK940252.1_CN/2019/InnerMongolia .............................................................---................ [144800]
MK128995.1_China/2018/AnhuiXCGQ  .............................................................---................ [144800]
MH681419.1_ASFV/POL/2015         .............................................................---................ [144800]
LS478113.1_Estonia_2014          .............................................................---................ [144800]
KP843857.1_Odintsovo_02/14       .............................................................---................ [144800]
```

```
FN557520.1_strain_E75            c............................................................---................ [144800]
AY261366.1_Warthog               c............................................................---................ [144800]
AY261365.1_Warmbaths             c............................................................---................ [144800]
AY261364.1_Tengani_62            c............................................................---................ [144800]
AY261363.1_Pretoriuskop/96/4     c........................................................t...---................ [144800]
AY261362.1_Mkuzi_1979            t............................................................---................ [144800]
AY261361.1_Malawi_Lil-20/1_1983  t............................................................cag................ [144800]
AY261360.1_Kenya_1950            t.........g..........c.......................................cag................ [144800]
AM712240.1_OURT_88/3             c............................................................---................ [144800]
AM712239.1_Benin_97/1            c............................................................---................ [144800]
KM111295.1_Ken06.Bus             t.........g..........c...................................t...cag................ [144800]
KM111294.1_Ken05/Tk1             t.........g..........c.......................................cag..a............. [144800]
KM262844.1_strain_L60            c............................................................---................ [144800]
KM102979.1_26544/OG10_Italy      c............................................................---................ [144800]
KM262845.1_strain_NHV            c............................................................---................ [144800]
KX354450.1_47/Ss/2008            c............................................................---................ [144800]
KP055815.1_strain_BA71           c............................................................---................ [144800]
MH025920.1_strain_R35            t.........g..........c...................................t...cag................ [144800]
MH025919.1_strain_N10            t.........g..........c...................................t...cag................ [144800]
MH025918.1_strain_R25            t.........g..........c...................................t...cag................ [144800]
MH025917.1_strain_R7             t.........g..........c...................................t...cag................ [144800]
MH025916.1_strain_R8             t.........g..........c...................................t...cag................ [144800]

FR682468.2_Georgia_2007/1        gcaaggctgaagaatgcacatgtaataacggaagctgttccctaaaaacaagttaaaacatgcaattat----------a [144880]
ASF_IND_20_CAD543                ..........................................................m..........----------. [144880]
LR899193.1_Germany_2020/1        .....................................................................----------. [144880]
MN393477.1_Wuhan_2019            .....................................................................----------. [144880]
MW396979.1_Timor-Leste/2019/1    .....................................................................----------. [144880]
MK940252.1_CN/2019/InnerMongolia .....................................................................----------. [144880]
MK128995.1_China/2018/AnhuiXCGQ  .....................................................................----------. [144880]
MH681419.1_ASFV/POL/2015         .....................................................................----------. [144880]
LS478113.1_Estonia_2014          .....................................................................----------. [144880]
KP843857.1_Odintsovo_02/14       .....................................................................----------. [144880]
FN557520.1_strain_E75            ..........................................................a..........----------. [144880]
AY261366.1_Warthog               ................................................................t....----------- [144880]
AY261365.1_Warmbaths             ..........c.....................................................t....----------. [144880]
AY261364.1_Tengani_62            ..........c.............................t....................a..t..g.----------. [144880]
AY261363.1_Pretoriuskop/96/4     ...........................................................--------------------- [144880]
AY261362.1_Mkuzi_1979            ..........c..........................................................----------. [144880]
AY261361.1_Malawi_Lil-20/1_1983  ...........................................................g..t.t....ggatgc--at. [144880]
AY261360.1_Kenya_1950            ........................c...........................c........at.t....----------- [144880]
AM712240.1_OURT_88/3             ..........................................................a..........----------. [144880]
AM712239.1_Benin_97/1            ..........................................................a..........----------. [144880]
KM111295.1_Ken06.Bus             ..........c.............c............c..............c.........t.t....gcatacataa. [144880]
```

```
KM111294.1_Ken05/Tk1             .....c....c.................t.............................tg..t.t....gcatgc----. [144880]
KM262844.1_strain_L60            ..........................................................a..........----------. [144880]
KM102979.1_26544/OG10_Italy      ..........................................................a..........----------. [144880]
KM262845.1_strain_NHV            ..........................................................a..........----------. [144880]
KX354450.1_47/Ss/2008            ..........................................................a..........----------. [144880]
KP055815.1_strain_BA71           ..........................................................a..........----------. [144880]
MH025920.1_strain_R35            ..........c.............c............c..............c.........t.t....gcatacataa. [144880]
MH025919.1_strain_N10            ..........c.............c............c..............c.........t.t....gcatacataa. [144880]
MH025918.1_strain_R25            ..........c.............c............c..............c.........t.t....gcatacataa. [144880]
MH025917.1_strain_R7             ..........c.............c............c..............c.........t.t....gcatacataa. [144880]
MH025916.1_strain_R8             ..........c.............c............c..............c.........t.t....gcatacataa. [144880]

FR682468.2_Georgia_2007/1        tgcatgcatataaacgcatgcatataaacgcat---------acatat-aaaatgcgtaaatactatataaaaaactata [144960]
ASF_IND_20_CAD543                ..y.....-------------------------------------.g.-....cnnnn......c.c..--......... [144960]
LR899193.1_Germany_2020/1        .................................---------......-............................... [144960]
MN393477.1_Wuhan_2019            .................................---------......-............................... [144960]
MW396979.1_Timor-Leste/2019/1    .................................---------......-............................... [144960]
MK940252.1_CN/2019/InnerMongolia .................................---------......-............................... [144960]
MK128995.1_China/2018/AnhuiXCGQ  .................................---------......-............................... [144960]
MH681419.1_ASFV/POL/2015         .................................---------......-............................... [144960]
LS478113.1_Estonia_2014          .................................---------......-............................... [144960]
KP843857.1_Odintsovo_02/14       .................................---------......-............................... [144960]
FN557520.1_strain_E75            ..t..-------------------------------------g...g.-....c..........c.c..--......... [144960]
AY261366.1_Warthog               ---...........t....a.....gc.--------------g...g.-....c..a....................... [144960]
AY261365.1_Warmbaths             ..............t....a.....gc.--------------....g.-....c..a....................... [144960]
AY261364.1_Tengani_62            ......t.a..gc.t...------------------------g...g.-....ca......................... [144960]
AY261363.1_Pretoriuskop/96/4     ---...t.......t..........gc.--------------g.....-....c.......................... [144960]
AY261362.1_Mkuzi_1979            .................................gcatatgcag...g.-....c.......................... [144960]
AY261361.1_Malawi_Lil-20/1_1983  ....c....g....t...c....g....t...c---------g.....a....c.......................... [144960]
AY261360.1_Kenya_1950            ---........gcgt....--..gc.tgtaa..g--------c.....a.....a..............-.......... [144960]
AM712240.1_OURT_88/3             ..t..-------------------------------------g...g.-....c..........c.c..--......... [144960]
AM712239.1_Benin_97/1            ..t..-------------------------------------g...g.-....c..........c.c..--......... [144960]
KM111295.1_Ken06.Bus             .a...a......c.........cgc.tgtaa..g--------c.....a.....a..........t...-.......... [144960]
KM111294.1_Ken05/Tk1             .a...a.....gc......---------------------------..a....c.......................... [144960]
KM262844.1_strain_L60            ..t..-------------------------------------g...g.-....c..........c.c..--......... [144960]
KM102979.1_26544/OG10_Italy      ..t..-------------------------------------g...g.-....c..........c.c..--......... [144960]
KM262845.1_strain_NHV            ..t..-------------------------------------g...g.-....c..........c.c..--......... [144960]
KX354450.1_47/Ss/2008            ..t..-------------------------------------g...g.-....c..........c.c..--......... [144960]
KP055815.1_strain_BA71           ..t..-------------------------------------g...g.-....c..........c.c..--......... [144960]
MH025920.1_strain_R35            .a...a......c.........cgc.tgtaa..g--------c.....a.....a..........t...-.......... [144960]
MH025919.1_strain_N10            .a...a......c.........cgc.tgtaa..g--------c.....a.....a..........t...-.......... [144960]
MH025918.1_strain_R25            .a...a......c.........cgc.tgtaa..g--------c.....a.....a..........t...-.......... [144960]
MH025917.1_strain_R7             .a...a......c.........cgc.tgtaa..g--------c.....a.....a..........t...-.......... [144960]
MH025916.1_strain_R8             .a...a......c.........cgc.tgtaa..g--------c.....a.....a..........t...-.......... [144960]
```

```
FR682468.2_Georgia_2007/1        acatatcaatcaaggaatcaacacttttataattttccgtaatatatttttcatccataatgatgtcagagtacatggtc [145040]
ASF_IND_20_CAD543                ................................................................................ [145040]
LR899193.1_Germany_2020/1        ................................................................................ [145040]
MN393477.1_Wuhan_2019            ................................................................................ [145040]
MW396979.1_Timor-Leste/2019/1    ................................................................................ [145040]
MK940252.1_CN/2019/InnerMongolia ................................................................................ [145040]
MK128995.1_China/2018/AnhuiXCGQ  ................................................................................ [145040]
MH681419.1_ASFV/POL/2015         ................................................................................ [145040]
LS478113.1_Estonia_2014          ................................................................................ [145040]
KP843857.1_Odintsovo_02/14       ................................................................................ [145040]
FN557520.1_strain_E75            ....g.......t................................................................... [145040]
AY261366.1_Warthog               ............t................................................................... [145040]
AY261365.1_Warmbaths             ............t................................................................... [145040]
AY261364.1_Tengani_62            ............t.........................a......................................... [145040]
AY261363.1_Pretoriuskop/96/4     ............t................................................................... [145040]
AY261362.1_Mkuzi_1979            ............t................................................................... [145040]
AY261361.1_Malawi_Lil-20/1_1983  ............t......................................................g............ [145040]
AY261360.1_Kenya_1950            ............t.......................................g........................... [145040]
AM712240.1_OURT_88/3             ....g.......t................................................................... [145040]
AM712239.1_Benin_97/1            ....g.......t................................................................... [145040]
KM111295.1_Ken06.Bus             ............t.......................................g........................... [145040]
KM111294.1_Ken05/Tk1             ............t.......................................g........................... [145040]
KM262844.1_strain_L60            ....g.......t................................................................... [145040]
KM102979.1_26544/OG10_Italy      ....g.......t................................................................... [145040]
KM262845.1_strain_NHV            ....g.......t................................................................... [145040]
KX354450.1_47/Ss/2008            ....g.......t................................................................... [145040]
KP055815.1_strain_BA71           ....g.......t................................................................... [145040]
MH025920.1_strain_R35            ............t.......................................g........................... [145040]
MH025919.1_strain_N10            ............t.......................................g........................... [145040]
MH025918.1_strain_R25            ............t.......................................g........................... [145040]
MH025917.1_strain_R7             ............t.......................................g........................... [145040]
MH025916.1_strain_R8             ............t.......................................g........................... [145040]

FR682468.2_Georgia_2007/1        cctatgcgaggaacagagcccataagggtaggcgcggcaataccgtaaatgggattcacggcggagtcaaccgcagcatc [145120]
ASF_IND_20_CAD543                ................................................................................ [145120]
LR899193.1_Germany_2020/1        ................................................................................ [145120]
MN393477.1_Wuhan_2019            ................................................................................ [145120]
MW396979.1_Timor-Leste/2019/1    ................................................................................ [145120]
MK940252.1_CN/2019/InnerMongolia ................................................................................ [145120]
MK128995.1_China/2018/AnhuiXCGQ  ................................................................................ [145120]
MH681419.1_ASFV/POL/2015         ................................................................................ [145120]
LS478113.1_Estonia_2014          ................................................................................ [145120]
KP843857.1_Odintsovo_02/14       ................................................................................ [145120]
```

FN557520.1_strain_E75 ................................................................................ [145120]
AY261366.1_Warthog ................................................................................ [145120]
AY261365.1_Warmbaths ................................................................................ [145120]
AY261364.1_Tengani_62 ................................................................................ [145120]
AY261363.1_Pretoriuskop/96/4 ................................................................................ [145120]
AY261362.1_Mkuzi_1979 .........................................g..........................g........... [145120]
AY261361.1_Malawi_Lil-20/1_1983 .........................................c.......................a.............. [145120]
AY261360.1_Kenya_1950 .........................................g...........g................t......... [145120]
AM712240.1_OURT_88/3 ..........................a..................................................... [145120]
AM712239.1_Benin_97/1 ................................................................................ [145120]
KM111295.1_Ken06.Bus ................................g........g........a..g................t......... [145120]
KM111294.1_Ken05/Tk1 .........................................g...........g................t......... [145120]
KM262844.1_strain_L60 ................................................................................ [145120]
KM102979.1_26544/OG10_Italy ................................................................................ [145120]
KM262845.1_strain_NHV ..........................a..................................................... [145120]
KX354450.1_47/Ss/2008 ................................................................................ [145120]
KP055815.1_strain_BA71 ................................................................................ [145120]
MH025920.1_strain_R35 ................................g........g........a..g................t......... [145120]
MH025919.1_strain_N10 ................................g........g........a..g................t......... [145120]
MH025918.1_strain_R25 ................................g........g........a..g................t......... [145120]
MH025917.1_strain_R7 ................................g........g........a..g................t......... [145120]
MH025916.1_strain_R8 ................................g........g........a..g................t......... [145120]

FR682468.2_Georgia_2007/1 tgtcaagacctggactggagacgacaaggccattcgcaacaacacgttggaaggctctcttgcattaagccctgcctttt [145200]
ASF_IND_20_CAD543 ................................................................................ [145200]
LR899193.1_Germany_2020/1 ................................................................................ [145200]
MN393477.1_Wuhan_2019 ................................................................................ [145200]
MW396979.1_Timor-Leste/2019/1 ................................................................................ [145200]
MK940252.1_CN/2019/InnerMongolia ................................................................................ [145200]
MK128995.1_China/2018/AnhuiXCGQ ................................................................................ [145200]
MH681419.1_ASFV/POL/2015 ................................................................................ [145200]
LS478113.1_Estonia_2014 ................................................................................ [145200]
KP843857.1_Odintsovo_02/14 ................................................................................ [145200]
FN557520.1_strain_E75 ............a................................................................... [145200]
AY261366.1_Warthog c....................................................................t.......... [145200]
AY261365.1_Warmbaths .........................................g...................................... [145200]
AY261364.1_Tengani_62 c............................................................................... [145200]
AY261363.1_Pretoriuskop/96/4 c........t...............................g...................................... [145200]
AY261362.1_Mkuzi_1979 ............a................................................................... [145200]
AY261361.1_Malawi_Lil-20/1_1983 c............................................................................... [145200]
AY261360.1_Kenya_1950 c.....................................................t..............a.......... [145200]
AM712240.1_OURT_88/3 ............a................................................................... [145200]
AM712239.1_Benin_97/1 ............a................................................................... [145200]
KM111295.1_Ken06.Bus c.....................................................t..............t..c....... [145200]

```
KM111294.1_Ken05/Tk1             c.....................................................t..............a.......... [145200]
KM262844.1_strain_L60            ............a................................................................... [145200]
KM102979.1_26544/OG10_Italy      ............a................................................................... [145200]
KM262845.1_strain_NHV            ............a................................................................... [145200]
KX354450.1_47/Ss/2008            ............a................................................................... [145200]
KP055815.1_strain_BA71           ............a................................................................... [145200]
MH025920.1_strain_R35            c.....................................................t..............t..c....... [145200]
MH025919.1_strain_N10            c.....................................................t..............t..c....... [145200]
MH025918.1_strain_R25            c.....................................................t..............t..c....... [145200]
MH025917.1_strain_R7             c.....................................................t..............t..c....... [145200]
MH025916.1_strain_R8             c.....................................................t..............t..c....... [145200]

FR682468.2_Georgia_2007/1        ctagagaggtaacctgtcccgttcttgtcatgagatctgcgtacatgagtaaatgacgatggttgggacccttgtccccc [145280]
ASF_IND_20_CAD543                ..........r..................................................................... [145280]
LR899193.1_Germany_2020/1        ................................................................................ [145280]
MN393477.1_Wuhan_2019            ................................................................................ [145280]
MW396979.1_Timor-Leste/2019/1    ................................................................................ [145280]
MK940252.1_CN/2019/InnerMongolia ................................................................................ [145280]
MK128995.1_China/2018/AnhuiXCGQ  ................................................................................ [145280]
MH681419.1_ASFV/POL/2015         ................................................................................ [145280]
LS478113.1_Estonia_2014          ................................................................................ [145280]
KP843857.1_Odintsovo_02/14       ................................................................................ [145280]
FN557520.1_strain_E75            ..........g..................................................................... [145280]
AY261366.1_Warthog               .................................................g.g............................ [145280]
AY261365.1_Warmbaths             ...................................................g............................ [145280]
AY261364.1_Tengani_62            .................................................g.g............................ [145280]
AY261363.1_Pretoriuskop/96/4     ...................................................g............................ [145280]
AY261362.1_Mkuzi_1979            ......................................................................t......... [145280]
AY261361.1_Malawi_Lil-20/1_1983  ...................................................g............a........a...... [145280]
AY261360.1_Kenya_1950            ...a...............g..........................a..c.g............................ [145280]
AM712240.1_OURT_88/3             ..........g..................................................................... [145280]
AM712239.1_Benin_97/1            ..........g..................................................................... [145280]
KM111295.1_Ken06.Bus             ...a...............g..........................a..c.g............................ [145280]
KM111294.1_Ken05/Tk1             ...a...............g..........................a..c.g............................ [145280]
KM262844.1_strain_L60            ..........g..................................................................... [145280]
KM102979.1_26544/OG10_Italy      ..........g..................................................................... [145280]
KM262845.1_strain_NHV            ..........g..................................................................... [145280]
KX354450.1_47/Ss/2008            ..........g..................................................................... [145280]
KP055815.1_strain_BA71           ..........g..................................................................... [145280]
MH025920.1_strain_R35            ...a...............g..........................a..c.g............................ [145280]
MH025919.1_strain_N10            ...a...............g..........................a..c.g............................ [145280]
MH025918.1_strain_R25            ...a...............g..........................a..c.g............................ [145280]
MH025917.1_strain_R7             ...a...............g..........................a..c.g............................ [145280]
MH025916.1_strain_R8             ...a...............g..........................a..c.g............................ [145280]
```

```
FR682468.2_Georgia_2007/1        ataaccgttctaatttcactaataattttttgccgtgccgcttctatgccgtaaagctccatggtgtctcctatagagga [145360]
ASF_IND_20_CAD543                ................................................................................ [145360]
LR899193.1_Germany_2020/1        ................................................................................ [145360]
MN393477.1_Wuhan_2019            ................................................................................ [145360]
MW396979.1_Timor-Leste/2019/1    ................................................................................ [145360]
MK940252.1_CN/2019/InnerMongolia ................................................................................ [145360]
MK128995.1_China/2018/AnhuiXCGQ  ................................................................................ [145360]
MH681419.1_ASFV/POL/2015         ................................................................................ [145360]
LS478113.1_Estonia_2014          ................................................................................ [145360]
KP843857.1_Odintsovo_02/14       ................................................................................ [145360]
FN557520.1_strain_E75            ................................t..g..............a............................. [145360]
AY261366.1_Warthog               ......................................t...........a.......................g..... [145360]
AY261365.1_Warmbaths             ................................................................................ [145360]
AY261364.1_Tengani_62            ..........................................................................g..... [145360]
AY261363.1_Pretoriuskop/96/4     ....................g.....................................................g..... [145360]
AY261362.1_Mkuzi_1979            ................................t..g............................................ [145360]
AY261361.1_Malawi_Lil-20/1_1983  ................................t..g..............a.......................g..... [145360]
AY261360.1_Kenya_1950            ....................g...........t..c..............a............................. [145360]
AM712240.1_OURT_88/3             ................................t..g..............a............................. [145360]
AM712239.1_Benin_97/1            ................................t..g..............a............................. [145360]
KM111295.1_Ken06.Bus             ....................g...........t..c..............a............................. [145360]
KM111294.1_Ken05/Tk1             ....................g...........t..c..............a............................. [145360]
KM262844.1_strain_L60            ................................t..g..............a............................. [145360]
KM102979.1_26544/OG10_Italy      ................................t..g..............a............................. [145360]
KM262845.1_strain_NHV            ................................t..g..............a............................. [145360]
KX354450.1_47/Ss/2008            ................................t..g..............a............................. [145360]
KP055815.1_strain_BA71           ................................t..g..............a............................. [145360]
MH025920.1_strain_R35            ....................g...........t..c..............a............................. [145360]
MH025919.1_strain_N10            ....................g...........t..c..............a............................. [145360]
MH025918.1_strain_R25            ....................g...........t..c..............a............................. [145360]
MH025917.1_strain_R7             ....................g...........t..c..............a............................. [145360]
MH025916.1_strain_R8             ....................g...........t..c..............a............................. [145360]

FR682468.2_Georgia_2007/1        cgatacgatggtgtatgggtcgatgttatcatcaagcattgcgccaaaaatattagtcccgtttgttttgatggcgtaga [145440]
ASF_IND_20_CAD543                ................................................................................ [145440]
LR899193.1_Germany_2020/1        ................................................................................ [145440]
MN393477.1_Wuhan_2019            ................................................................................ [145440]
MW396979.1_Timor-Leste/2019/1    ................................................................................ [145440]
MK940252.1_CN/2019/InnerMongolia ................................................................................ [145440]
MK128995.1_China/2018/AnhuiXCGQ  ................................................................................ [145440]
MH681419.1_ASFV/POL/2015         ................................................................................ [145440]
LS478113.1_Estonia_2014          ................................................................................ [145440]
KP843857.1_Odintsovo_02/14       ................................................................................ [145440]
```

```
FN557520.1_strain_E75            ...c............................................................................ [145440]
AY261366.1_Warthog               ...c........a........a.......................................................... [145440]
AY261365.1_Warmbaths             ...c.................a.......................................................... [145440]
AY261364.1_Tengani_62            ...c............................................................................ [145440]
AY261363.1_Pretoriuskop/96/4     ...c..................................................t......................... [145440]
AY261362.1_Mkuzi_1979            ................................................................................ [145440]
AY261361.1_Malawi_Lil-20/1_1983  ...c.................a................................t.....a................... [145440]
AY261360.1_Kenya_1950            ...c.................a................................t..t..a................... [145440]
AM712240.1_OURT_88/3             ...c............................................................................ [145440]
AM712239.1_Benin_97/1            ...c............................................................................ [145440]
KM111295.1_Ken06.Bus             ...c.................a....................a...........t.....a................... [145440]
KM111294.1_Ken05/Tk1             ...c.................a................................t.....a................... [145440]
KM262844.1_strain_L60            ...c............................................................................ [145440]
KM102979.1_26544/OG10_Italy      ...c............................................................................ [145440]
KM262845.1_strain_NHV            ...c............................................................................ [145440]
KX354450.1_47/Ss/2008            ...c............................................................................ [145440]
KP055815.1_strain_BA71           ...c............................................................................ [145440]
MH025920.1_strain_R35            ...c.................a....................a...........t.....a................... [145440]
MH025919.1_strain_N10            ...c.................a....................a...........t.....a................... [145440]
MH025918.1_strain_R25            ...c.................a....................a...........t.....a................... [145440]
MH025917.1_strain_R7             ...c.................a....................a...........t.....a................... [145440]
MH025916.1_strain_R8             ...c.................a....................a...........t.....a................... [145440]

FR682468.2_Georgia_2007/1        tattgtctagtcttaccagtttcccctgggcatccacacggtggcgcataagcttaacaacattcgcatttttgatgcct [145520]
ASF_IND_20_CAD543                ...................c..t......................................................... [145520]
LR899193.1_Germany_2020/1        ................................................................................ [145520]
MN393477.1_Wuhan_2019            ................................................................................ [145520]
MW396979.1_Timor-Leste/2019/1    ................................................................................ [145520]
MK940252.1_CN/2019/InnerMongolia ................................................................................ [145520]
MK128995.1_China/2018/AnhuiXCGQ  ................................................................................ [145520]
MH681419.1_ASFV/POL/2015         ................................................................................ [145520]
LS478113.1_Estonia_2014          ................................................................................ [145520]
KP843857.1_Odintsovo_02/14       ................................................................................ [145520]
FN557520.1_strain_E75            ....a..............c..t......................................................... [145520]
AY261366.1_Warthog               ....a..............c..t......................................................... [145520]
AY261365.1_Warmbaths             ....a..............c..t....................a.................................... [145520]
AY261364.1_Tengani_62            ....a..............c..t......................................................... [145520]
AY261363.1_Pretoriuskop/96/4     ....a..............c..t.....................................................a... [145520]
AY261362.1_Mkuzi_1979            ....a..............c..t......................................................... [145520]
AY261361.1_Malawi_Lil-20/1_1983  ..............g....c................................t..........................a [145520]
AY261360.1_Kenya_1950            ......................t........................................................g [145520]
AM712240.1_OURT_88/3             ....a..............c..t......................................................... [145520]
AM712239.1_Benin_97/1            ....a..............c..t......................................................... [145520]
KM111295.1_Ken06.Bus             ...................c..t........................................................g [145520]
```

```
KM111294.1_Ken05/Tk1            ......................t.................a......................................g [145520]
KM262844.1_strain_L60           ....a..............c..t......................................................... [145520]
KM102979.1_26544/OG10_Italy     ....a..............c..t......................................................... [145520]
KM262845.1_strain_NHV           ....a..............c..t......................................................... [145520]
KX354450.1_47/Ss/2008           ....a..............c..t......................................................... [145520]
KP055815.1_strain_BA71          ....a..............c..t......................................................... [145520]
MH025920.1_strain_R35           ...................c..t........................................................g [145520]
MH025919.1_strain_N10           ...................c..t........................................................g [145520]
MH025918.1_strain_R25           ...................c..t........................................................g [145520]
MH025917.1_strain_R7            ...................c..t........................................................g [145520]
MH025916.1_strain_R8            ...................c..t........................................................g [145520]

FR682468.2_Georgia_2007/1       ggtattcctctaatcgtgctatttaatagtttatccaccacatttacggcaattttttcatccgtagccattcgggtatt [145600]
ASF_IND_20_CAD543               ................................................................................ [145600]
LR899193.1_Germany_2020/1       ................................................................................ [145600]
MN393477.1_Wuhan_2019           ................................................................................ [145600]
MW396979.1_Timor-Leste/2019/1   ................................................................................ [145600]
MK940252.1_CN/2019/InnerMongolia ................................................................................ [145600]
MK128995.1_China/2018/AnhuiXCGQ ................................................................................ [145600]
MH681419.1_ASFV/POL/2015        ................................................................................ [145600]
LS478113.1_Estonia_2014         ................................................................................ [145600]
KP843857.1_Odintsovo_02/14      ................................................................................ [145600]
FN557520.1_strain_E75           ..................................................g............................. [145600]
AY261366.1_Warthog              ..................................................g............................. [145600]
AY261365.1_Warmbaths            ..................................................g............................. [145600]
AY261364.1_Tengani_62           .......................................................................c........ [145600]
AY261363.1_Pretoriuskop/96/4    ..............t...................................g............................. [145600]
AY261362.1_Mkuzi_1979           ..................................................g............................. [145600]
AY261361.1_Malawi_Lil-20/1_1983 ............................a...........t........................g.............. [145600]
AY261360.1_Kenya_1950           ............................a...........a....................................... [145600]
AM712240.1_OURT_88/3            ..................................................g............................. [145600]
AM712239.1_Benin_97/1           ..................................................g............................. [145600]
KM111295.1_Ken06.Bus            ............................a...........a....................................... [145600]
KM111294.1_Ken05/Tk1            ............................a...........a....................................... [145600]
KM262844.1_strain_L60           ..................................................g............................. [145600]
KM102979.1_26544/OG10_Italy     ..................................................g............................. [145600]
KM262845.1_strain_NHV           ..................................................g............................. [145600]
KX354450.1_47/Ss/2008           ..................................................g............................. [145600]
KP055815.1_strain_BA71          ..................................................g............................. [145600]
MH025920.1_strain_R35           ............................a...........a....................................... [145600]
MH025919.1_strain_N10           ............................a...........a....................................... [145600]
MH025918.1_strain_R25           ............................a...........a....................................... [145600]
MH025917.1_strain_R7            ............................a...........a....................................... [145600]
MH025916.1_strain_R8            ............................a...........a....................................... [145600]
```

```
FR682468.2_Georgia_2007/1        ggtactgcgtctaaaggcgctttcccgtaggtatatgcgaataatgatgggaatccctgaggccgtgttttccacagaat [145680]
ASF_IND_20_CAD543                ................................................................................ [145680]
LR899193.1_Germany_2020/1        ................................................................................ [145680]
MN393477.1_Wuhan_2019            ................................................................................ [145680]
MW396979.1_Timor-Leste/2019/1    ................................................................................ [145680]
MK940252.1_CN/2019/InnerMongolia ................................................................................ [145680]
MK128995.1_China/2018/AnhuiXCGQ  ................................................................................ [145680]
MH681419.1_ASFV/POL/2015         ................................................................................ [145680]
LS478113.1_Estonia_2014          ................................................................................ [145680]
KP843857.1_Odintsovo_02/14       ................................................................................ [145680]
FN557520.1_strain_E75            a.............................a.......................a......................... [145680]
AY261366.1_Warthog               ......................................................a......................... [145680]
AY261365.1_Warmbaths             a.......................t.....a.......................a......................... [145680]
AY261364.1_Tengani_62            ......................................................a......................... [145680]
AY261363.1_Pretoriuskop/96/4     ......................................................a......................... [145680]
AY261362.1_Mkuzi_1979            a.............................a.......................a......................... [145680]
AY261361.1_Malawi_Lil-20/1_1983  ..............................a.....a.................g......................... [145680]
AY261360.1_Kenya_1950            ..............................a..............a........g......................... [145680]
AM712240.1_OURT_88/3             a.............................a.......................a......................... [145680]
AM712239.1_Benin_97/1            a.............................a.......................a......................... [145680]
KM111295.1_Ken06.Bus             ..............................a..............a........g......................... [145680]
KM111294.1_Ken05/Tk1             ..............................a.....a...........a.....g.....a................... [145680]
KM262844.1_strain_L60            a.............................a.......................a......................... [145680]
KM102979.1_26544/OG10_Italy      a.............................a.......................a......................... [145680]
KM262845.1_strain_NHV            a.............................a.......................a......................... [145680]
KX354450.1_47/Ss/2008            a.............................a.......................a......................... [145680]
KP055815.1_strain_BA71           a.............................a.......................a......................... [145680]
MH025920.1_strain_R35            ..............................a..............a........g......................... [145680]
MH025919.1_strain_N10            ..............................a..............a........g......................... [145680]
MH025918.1_strain_R25            ..............................a..............a........g......................... [145680]
MH025917.1_strain_R7             ..............................a..............a........g......................... [145680]
MH025916.1_strain_R8             ..............................a..............a........g......................... [145680]

FR682468.2_Georgia_2007/1        gcatgatgtaggtgttggggtgtttagctcttagactattaataatactttctagactaatgctttttaatatcatggtt [145760]
ASF_IND_20_CAD543                ................................................................................ [145760]
LR899193.1_Germany_2020/1        ................................................................................ [145760]
MN393477.1_Wuhan_2019            ................................................................................ [145760]
MW396979.1_Timor-Leste/2019/1    ................................................................................ [145760]
MK940252.1_CN/2019/InnerMongolia ................................................................................ [145760]
MK128995.1_China/2018/AnhuiXCGQ  ................................................................................ [145760]
MH681419.1_ASFV/POL/2015         ................................................................................ [145760]
LS478113.1_Estonia_2014          ................................................................................ [145760]
KP843857.1_Odintsovo_02/14       ................................................................................ [145760]
```

```
FN557520.1_strain_E75              ......................c......................................................... [145760]
AY261366.1_Warthog                 ................................................................................ [145760]
AY261365.1_Warmbaths               ................................................................................ [145760]
AY261364.1_Tengani_62              .......a......................................g................................. [145760]
AY261363.1_Pretoriuskop/96/4       ................................................................................ [145760]
AY261362.1_Mkuzi_1979              ......................c......................................................... [145760]
AY261361.1_Malawi_Lil-20/1_1983    ............aa.................................................................. [145760]
AY261360.1_Kenya_1950              .........................g..............c.c............g.............gg......... [145760]
AM712240.1_OURT_88/3               ......................c......................................................... [145760]
AM712239.1_Benin_97/1              ......................c......................................................... [145760]
KM111295.1_Ken06.Bus               .........................g..............c.c............g.............gg......... [145760]
KM111294.1_Ken05/Tk1               .............a....a.....................c.c............g.............gg......... [145760]
KM262844.1_strain_L60              ......................c......................................................... [145760]
KM102979.1_26544/OG10_Italy        ......................c......................................................... [145760]
KM262845.1_strain_NHV              ......................c......................................................... [145760]
KX354450.1_47/Ss/2008              ......................c......................................................... [145760]
KP055815.1_strain_BA71             ......................c......................................................... [145760]
MH025920.1_strain_R35              .........................g..............c.c............g.............gg......... [145760]
MH025919.1_strain_N10              .........................g..............c.c............g.............gg......... [145760]
MH025918.1_strain_R25              .........................g..............c.c............g.............gg......... [145760]
MH025917.1_strain_R7               .........................g..............c.c............g.............gg......... [145760]
MH025916.1_strain_R8               .........................g..............c.c............g.............gg......... [145760]

FR682468.2_Georgia_2007/1          gttttgtttaattccaagcggatacaccagtttgcaatatcctctgggggctgtagtagaggatggttttccagaaaatc [145840]
ASF_IND_20_CAD543                  .....r.......................................................................... [145840]
LR899193.1_Germany_2020/1          ................................................................................ [145840]
MN393477.1_Wuhan_2019              ................................................................................ [145840]
MW396979.1_Timor-Leste/2019/1      ................................................................................ [145840]
MK940252.1_CN/2019/InnerMongolia   ................................................................................ [145840]
MK128995.1_China/2018/AnhuiXCGQ    ................................................................................ [145840]
MH681419.1_ASFV/POL/2015           ................................................................................ [145840]
LS478113.1_Estonia_2014            ................................................................................ [145840]
KP843857.1_Odintsovo_02/14         ................................................................................ [145840]
FN557520.1_strain_E75              .....a......................................c................................... [145840]
AY261366.1_Warthog                 .........................................t..c................................... [145840]
AY261365.1_Warmbaths               ............................................c................................... [145840]
AY261364.1_Tengani_62              ............................................c................................... [145840]
AY261363.1_Pretoriuskop/96/4       .........................................t..c................................... [145840]
AY261362.1_Mkuzi_1979              .....a......................................c................................... [145840]
AY261361.1_Malawi_Lil-20/1_1983    .....a......................................c..............g.................... [145840]
AY261360.1_Kenya_1950              .....a......................................c................................... [145840]
AM712240.1_OURT_88/3               .....a......................................c................................... [145840]
AM712239.1_Benin_97/1              .....a......................................c................................... [145840]
KM111295.1_Ken06.Bus               .....a......................................c...........c....................... [145840]
```

```
KM111294.1_Ken05/Tk1              .....a......................................c................................... [145840]
KM262844.1_strain_L60             .....a......................................c................................... [145840]
KM102979.1_26544/OG10_Italy       .....a......................................c................................... [145840]
KM262845.1_strain_NHV             .....a......................................c................................... [145840]
KX354450.1_47/Ss/2008             .....a......................................c................................... [145840]
KP055815.1_strain_BA71            .....a......................................c................................... [145840]
MH025920.1_strain_R35             .....a......................................c...........c....................... [145840]
MH025919.1_strain_N10             .....a......................................c...........c....................... [145840]
MH025918.1_strain_R25             .....a......................................c...........c....................... [145840]
MH025917.1_strain_R7              .....a......................................c...........c....................... [145840]
MH025916.1_strain_R8              .....a......................................c...........c....................... [145840]

FR682468.2_Georgia_2007/1         cgtcatccattccacatcacttgcaaaatcggggtacatcaca-tttttttttgtgcttgaatacgtttcgtacaatagg [145920]
ASF_IND_20_CAD543                 ...........................................-.................................... [145920]
LR899193.1_Germany_2020/1         ...........................................-.................................... [145920]
MN393477.1_Wuhan_2019             ...........................................-.................................... [145920]
MW396979.1_Timor-Leste/2019/1     ...........................................-.................................... [145920]
MK940252.1_CN/2019/InnerMongolia  ...........................................-.................................... [145920]
MK128995.1_China/2018/AnhuiXCGQ   ...........................................-.................................... [145920]
MH681419.1_ASFV/POL/2015          ...........................................-.................................... [145920]
LS478113.1_Estonia_2014           ...........................................-.................................... [145920]
KP843857.1_Odintsovo_02/14        ...........................................-.................................... [145920]
FN557520.1_strain_E75             ...........................................t.................................... [145920]
AY261366.1_Warthog                ...........................................-...................................a [145920]
AY261365.1_Warmbaths              ...t.......................................-.................................... [145920]
AY261364.1_Tengani_62             ...........................................-.................................... [145920]
AY261363.1_Pretoriuskop/96/4      ...........................................-...................................a [145920]
AY261362.1_Mkuzi_1979             ...........................................-.................................... [145920]
AY261361.1_Malawi_Lil-20/1_1983   g...........t..............................-....................t...........a... [145920]
AY261360.1_Kenya_1950             ...........................................-...................................a [145920]
AM712240.1_OURT_88/3              ...........................................-.................................... [145920]
AM712239.1_Benin_97/1             ...........................................-.................................... [145920]
KM111295.1_Ken06.Bus              ...........................................-...................................a [145920]
KM111294.1_Ken05/Tk1              ...........................................-...................................a [145920]
KM262844.1_strain_L60             ...........................................-.................................... [145920]
KM102979.1_26544/OG10_Italy       ...........................................-.................................... [145920]
KM262845.1_strain_NHV             ...........................................-.................................... [145920]
KX354450.1_47/Ss/2008             ...........................................-.................................... [145920]
KP055815.1_strain_BA71            ...........................................-.................................... [145920]
MH025920.1_strain_R35             ...........................................-...................................a [145920]
MH025919.1_strain_N10             ...........................................-...................................a [145920]
MH025918.1_strain_R25             ...........................................-...................................a [145920]
MH025917.1_strain_R7              ...........................................-...................................a [145920]
MH025916.1_strain_R8              ...........................................-...................................a [145920]
```

```
FR682468.2_Georgia_2007/1        tgccactgcaatatcaaccgttcgaacgttataagctctatgctgttagcaatttcttgcgcatatgttttatttgtttc [146000]
ASF_IND_20_CAD543                ................................................................................ [146000]
LR899193.1_Germany_2020/1        ................................................................................ [146000]
MN393477.1_Wuhan_2019            ................................................................................ [146000]
MW396979.1_Timor-Leste/2019/1    ................................................................................ [146000]
MK940252.1_CN/2019/InnerMongolia ................................................................................ [146000]
MK128995.1_China/2018/AnhuiXCGQ  ................................................................................ [146000]
MH681419.1_ASFV/POL/2015         ................................................................................ [146000]
LS478113.1_Estonia_2014          ................................................................................ [146000]
KP843857.1_Odintsovo_02/14       ................................................................................ [146000]
FN557520.1_strain_E75            ..........g..................................................................... [146000]
AY261366.1_Warthog               ..........g..................................................................... [146000]
AY261365.1_Warmbaths             ..........g..................................................................... [146000]
AY261364.1_Tengani_62            ..........g..................................................................... [146000]
AY261363.1_Pretoriuskop/96/4     ........t.g...............t..................................................... [146000]
AY261362.1_Mkuzi_1979            ..........g..................................................................... [146000]
AY261361.1_Malawi_Lil-20/1_1983  .....t....g..................................................................... [146000]
AY261360.1_Kenya_1950            ..........g.....gt.....a..t....................c.......................g........ [146000]
AM712240.1_OURT_88/3             ..........g..................................................................... [146000]
AM712239.1_Benin_97/1            ..........g..................................................................... [146000]
KM111295.1_Ken06.Bus             ..........g.....gt.....a.......................c.......................g........ [146000]
KM111294.1_Ken05/Tk1             ..........g.....gt.....a.......................c.......................g........ [146000]
KM262844.1_strain_L60            ..........g..................................................................... [146000]
KM102979.1_26544/OG10_Italy      ..........g..................................................................... [146000]
KM262845.1_strain_NHV            ..........g..................................................................... [146000]
KX354450.1_47/Ss/2008            ..........g..................................................................... [146000]
KP055815.1_strain_BA71           ..........g..................................................................... [146000]
MH025920.1_strain_R35            ..........g.....gt.....a.......................c.......................g........ [146000]
MH025919.1_strain_N10            ..........g.....gt.....a.......................c.......................g........ [146000]
MH025918.1_strain_R25            ..........g.....gt.....a.......................c.......................g........ [146000]
MH025917.1_strain_R7             ..........g.....gt.....a.......................c.......................g........ [146000]
MH025916.1_strain_R8             ..........g.....gt.....a.......................c.......................g........ [146000]

FR682468.2_Georgia_2007/1        cacttccgggttctttagacgtaaaagcatttcagaggattgttcagcctctacgggcttcgcgctaaagatctcctggg [146080]
ASF_IND_20_CAD543                ................................................................................ [146080]
LR899193.1_Germany_2020/1        ................................................................................ [146080]
MN393477.1_Wuhan_2019            ................................................................................ [146080]
MW396979.1_Timor-Leste/2019/1    ................................................................................ [146080]
MK940252.1_CN/2019/InnerMongolia ................................................................................ [146080]
MK128995.1_China/2018/AnhuiXCGQ  ................................................................................ [146080]
MH681419.1_ASFV/POL/2015         ................................................................................ [146080]
LS478113.1_Estonia_2014          ................................................................................ [146080]
KP843857.1_Odintsovo_02/14       ................................................................................ [146080]
```

```
FN557520.1_strain_E75           ................................................................................ [146080]
AY261366.1_Warthog              ............................................................t................... [146080]
AY261365.1_Warmbaths            ................................................................................ [146080]
AY261364.1_Tengani_62           ................................................................................ [146080]
AY261363.1_Pretoriuskop/96/4    ...............................................................a................ [146080]
AY261362.1_Mkuzi_1979           ........................................................................t....... [146080]
AY261361.1_Malawi_Lil-20/1_1983 ......t.....t.........................................a......................... [146080]
AY261360.1_Kenya_1950           ......t.....t.........................................a......................... [146080]
AM712240.1_OURT_88/3            ................................................................................ [146080]
AM712239.1_Benin_97/1           ................................................................................ [146080]
KM111295.1_Ken06.Bus            ......t.....t.........................................a......................... [146080]
KM111294.1_Ken05/Tk1            ......t.....t.........................................a......................... [146080]
KM262844.1_strain_L60           ................................................................................ [146080]
KM102979.1_26544/OG10_Italy     ................................................................................ [146080]
KM262845.1_strain_NHV           ................................................................................ [146080]
KX354450.1_47/Ss/2008           ................................................................................ [146080]
KP055815.1_strain_BA71          ................................................................................ [146080]
MH025920.1_strain_R35           ......t.....t.........................................a......................... [146080]
MH025919.1_strain_N10           ......t.....t.........................................a......................... [146080]
MH025918.1_strain_R25           ......t.....t.........................................a......................... [146080]
MH025917.1_strain_R7            ......t.....t.........................................a......................... [146080]
MH025916.1_strain_R8            ......t.....t.........................................a......................... [146080]

FR682468.2_Georgia_2007/1       gccgcacaattcccgacttgttggttcccccggccacggaccggtggtgggagtccagcatatattgtgtcaagggctct [146160]
ASF_IND_20_CAD543               ................................................................................ [146160]
LR899193.1_Germany_2020/1       ................................................................................ [146160]
MN393477.1_Wuhan_2019           ................................................................................ [146160]
MW396979.1_Timor-Leste/2019/1   ................................................................................ [146160]
MK940252.1_CN/2019/InnerMongolia ................................................................................ [146160]
MK128995.1_China/2018/AnhuiXCGQ ................................................................................ [146160]
MH681419.1_ASFV/POL/2015        ................................................................................ [146160]
LS478113.1_Estonia_2014         ................................................................................ [146160]
KP843857.1_Odintsovo_02/14      ................................................................................ [146160]
FN557520.1_strain_E75           ...................................................................c........t... [146160]
AY261366.1_Warthog              ...............................a........g..........................a............ [146160]
AY261365.1_Warmbaths            ...............................a...................................a............ [146160]
AY261364.1_Tengani_62           ...............................a.............................g.....a............ [146160]
AY261363.1_Pretoriuskop/96/4    ...............................a.............................g.....a............ [146160]
AY261362.1_Mkuzi_1979           ...................................................................c........t... [146160]
AY261361.1_Malawi_Lil-20/1_1983 .............t........a........c.....a..g................a.........a..t......... [146160]
AY261360.1_Kenya_1950           ..........g....................c........g..........................a............ [146160]
AM712240.1_OURT_88/3            ...................................................................c........t... [146160]
AM712239.1_Benin_97/1           ...................................................................c........t... [146160]
KM111295.1_Ken06.Bus            ..........g....................c........g..........................a............ [146160]
```

```
KM111294.1_Ken05/Tk1           ..........g....................c........g..........................a............ [146160]
KM262844.1_strain_L60          ...................................................................c........t... [146160]
KM102979.1_26544/OG10_Italy    ...................................................................c........t... [146160]
KM262845.1_strain_NHV          ...................................................................c........t... [146160]
KX354450.1_47/Ss/2008          ...................................................................c........t... [146160]
KP055815.1_strain_BA71         ...................................................................c........t... [146160]
MH025920.1_strain_R35          ..........g....................c........g..........................a............ [146160]
MH025919.1_strain_N10          ..........g....................c........g..........................a............ [146160]
MH025918.1_strain_R25          ..........g....................c........g..........................a............ [146160]
MH025917.1_strain_R7           ..........g....................c........g..........................a............ [146160]
MH025916.1_strain_R8           ..........g....................c........g..........................a............ [146160]

FR682468.2_Georgia_2007/1      gatacggactgcgccgccaggattcccactgcctcaccgtagttaataagactttgagtatattgtagccttatgaggtc [146240]
ASF_IND_20_CAD543              ................................................................................ [146240]
LR899193.1_Germany_2020/1      ................................................................................ [146240]
MN393477.1_Wuhan_2019          ................................................................................ [146240]
MW396979.1_Timor-Leste/2019/1  ................................................................................ [146240]
MK940252.1_CN/2019/InnerMongolia ................................................................................ [146240]
MK128995.1_China/2018/AnhuiXCGQ ................................................................................ [146240]
MH681419.1_ASFV/POL/2015       ................................................................................ [146240]
LS478113.1_Estonia_2014        ................................................................................ [146240]
KP843857.1_Odintsovo_02/14     ................................................................................ [146240]
FN557520.1_strain_E75          .............................c.................................................. [146240]
AY261366.1_Warthog             ....................a........................................................... [146240]
AY261365.1_Warmbaths           ....................a........................................................... [146240]
AY261364.1_Tengani_62          ................................................................................ [146240]
AY261363.1_Pretoriuskop/96/4   ................................................................................ [146240]
AY261362.1_Mkuzi_1979          ................................................................................ [146240]
AY261361.1_Malawi_Lil-20/1_1983 ................................................................................ [146240]
AY261360.1_Kenya_1950          .................t.............................g........g....................... [146240]
AM712240.1_OURT_88/3           .............................c.................................................. [146240]
AM712239.1_Benin_97/1          .............................c.................................................. [146240]
KM111295.1_Ken06.Bus           ..............t..t.............................g........g....................... [146240]
KM111294.1_Ken05/Tk1           .................t.............................g........g....................... [146240]
KM262844.1_strain_L60          .............................c.................................................. [146240]
KM102979.1_26544/OG10_Italy    .............................c.................................................. [146240]
KM262845.1_strain_NHV          .............................c.................................................. [146240]
KX354450.1_47/Ss/2008          .............................c.................................................. [146240]
KP055815.1_strain_BA71         .............................c.................................................. [146240]
MH025920.1_strain_R35          ..............t..t.............................g........g....................... [146240]
MH025919.1_strain_N10          ..............t..t.............................g........g....................... [146240]
MH025918.1_strain_R25          ..............t..t.............................g........g....................... [146240]
MH025917.1_strain_R7           ..............t..t.............................g........g....................... [146240]
MH025916.1_strain_R8           ..............t..t.............................g........g....................... [146240]
```

```
FR682468.2_Georgia_2007/1        caggatggcactcatctgctcgcaggtaatgtttaatgttttaacggttgccagttcgatgcgaataagcatgcgcatca [146320]
ASF_IND_20_CAD543                ................................................................................ [146320]
LR899193.1_Germany_2020/1        ................................................................................ [146320]
MN393477.1_Wuhan_2019            ................................................................................ [146320]
MW396979.1_Timor-Leste/2019/1    ................................................................................ [146320]
MK940252.1_CN/2019/InnerMongolia ................................................................................ [146320]
MK128995.1_China/2018/AnhuiXCGQ  ................................................................................ [146320]
MH681419.1_ASFV/POL/2015         ................................................................................ [146320]
LS478113.1_Estonia_2014          ................................................................................ [146320]
KP843857.1_Odintsovo_02/14       ................................................................................ [146320]
FN557520.1_strain_E75            ................................................................................ [146320]
AY261366.1_Warthog               ................................................................................ [146320]
AY261365.1_Warmbaths             ........................................................................a....... [146320]
AY261364.1_Tengani_62            ................................................................................ [146320]
AY261363.1_Pretoriuskop/96/4     ...........................g.................................................... [146320]
AY261362.1_Mkuzi_1979            ................................................................................ [146320]
AY261361.1_Malawi_Lil-20/1_1983  ...........................t.................................................... [146320]
AY261360.1_Kenya_1950            ...........................t..............g..c...........a...........t.......... [146320]
AM712240.1_OURT_88/3             ................................................................................ [146320]
AM712239.1_Benin_97/1            ................................................................................ [146320]
KM111295.1_Ken06.Bus             ...........................t..............g..c...........a...........t.......... [146320]
KM111294.1_Ken05/Tk1             ...........................t..............g..c...........a...........t.......... [146320]
KM262844.1_strain_L60            ................................................................................ [146320]
KM102979.1_26544/OG10_Italy      ................................................................................ [146320]
KM262845.1_strain_NHV            ................................................................................ [146320]
KX354450.1_47/Ss/2008            ................................................................................ [146320]
KP055815.1_strain_BA71           ................................................................................ [146320]
MH025920.1_strain_R35            ...........................t..............g..c...........a...........t.......... [146320]
MH025919.1_strain_N10            ...........................t..............g..c...........a...........t.......... [146320]
MH025918.1_strain_R25            ...........................t..............g..c...........a...........t.......... [146320]
MH025917.1_strain_R7             ...........................t..............g..c...........a...........t.......... [146320]
MH025916.1_strain_R8             ...........................t..............g..c...........a...........t.......... [146320]

FR682468.2_Georgia_2007/1        gagaggcagcccgtttaagataaacgggtatgggcgtttgtagtcgttcctgaatgttgttaataaacacgtatggaaga [146400]
ASF_IND_20_CAD543                ................................................................................ [146400]
LR899193.1_Germany_2020/1        ................................................................................ [146400]
MN393477.1_Wuhan_2019            ................................................................................ [146400]
MW396979.1_Timor-Leste/2019/1    ................................................................................ [146400]
MK940252.1_CN/2019/InnerMongolia ................................................................................ [146400]
MK128995.1_China/2018/AnhuiXCGQ  ................................................................................ [146400]
MH681419.1_ASFV/POL/2015         ................................................................................ [146400]
LS478113.1_Estonia_2014          ................................................................................ [146400]
KP843857.1_Odintsovo_02/14       ................................................................................ [146400]
```

```
FN557520.1_strain_E75           ...c......t.............................c....................................... [146400]
AY261366.1_Warthog              .........................a.................c.................................... [146400]
AY261365.1_Warmbaths            ...........................................c.................................... [146400]
AY261364.1_Tengani_62           ...............................a...........c.................................... [146400]
AY261363.1_Pretoriuskop/96/4    ..........g................................c.................................... [146400]
AY261362.1_Mkuzi_1979           ................................................................................ [146400]
AY261361.1_Malawi_Lil-20/1_1983 ............t..................a.....c.....c..............a..................... [146400]
AY261360.1_Kenya_1950           .......g....t........................c.....c...........a........................ [146400]
AM712240.1_OURT_88/3            ...c......t.............................c....................................... [146400]
AM712239.1_Benin_97/1           ...c......t.............................c....................................... [146400]
KM111295.1_Ken06.Bus            .......g....t........................c.....c...........a........................ [146400]
KM111294.1_Ken05/Tk1            .......g....t........................c.....c...........a........................ [146400]
KM262844.1_strain_L60           ...c......t.............................c....................................... [146400]
KM102979.1_26544/OG10_Italy     ...c......t.............................c....................................... [146400]
KM262845.1_strain_NHV           ...c......t.............................c....................................... [146400]
KX354450.1_47/Ss/2008           ...c......t.............................c....................................... [146400]
KP055815.1_strain_BA71          ...c......t.............................c....................................... [146400]
MH025920.1_strain_R35           .......g....t........................c.....c...........a........................ [146400]
MH025919.1_strain_N10           .......g....t........................c.....c...........a........................ [146400]
MH025918.1_strain_R25           .......g....t........................c.....c...........a........................ [146400]
MH025917.1_strain_R7            .......g....t........................c.....c...........a........................ [146400]
MH025916.1_strain_R8            .......g....t........................c.....c...........a........................ [146400]

FR682468.2_Georgia_2007/1       tttttgcaaaacgttttgaccatcgcgtatttttgtagaatacttttttcgtcgaagggaagcacgccactggtggagct [146480]
ASF_IND_20_CAD543               ................................................................................ [146480]
LR899193.1_Germany_2020/1       ................................................................................ [146480]
MN393477.1_Wuhan_2019           ................................................................................ [146480]
MW396979.1_Timor-Leste/2019/1   ................................................................................ [146480]
MK940252.1_CN/2019/InnerMongolia ................................................................................ [146480]
MK128995.1_China/2018/AnhuiXCGQ ................................................................................ [146480]
MH681419.1_ASFV/POL/2015        ................................................................................ [146480]
LS478113.1_Estonia_2014         ................................................................................ [146480]
KP843857.1_Odintsovo_02/14      ................................................................................ [146480]
FN557520.1_strain_E75           ................................................................................ [146480]
AY261366.1_Warthog              .........................t................g..............................c...... [146480]
AY261365.1_Warmbaths            .........................t................g..............................c...... [146480]
AY261364.1_Tengani_62           .........................t.....................................t.........c...... [146480]
AY261363.1_Pretoriuskop/96/4    .........................t................g..........................g...c...... [146480]
AY261362.1_Mkuzi_1979           ................................................................................ [146480]
AY261361.1_Malawi_Lil-20/1_1983 .........................t...........................a.......................... [146480]
AY261360.1_Kenya_1950           .........................t...c.....c..............a.................g........... [146480]
AM712240.1_OURT_88/3            ................................................................................ [146480]
AM712239.1_Benin_97/1           ................................................................................ [146480]
KM111295.1_Ken06.Bus            .........................t...c.....c..............a.................g.....a..... [146480]
```

```
KM111294.1_Ken05/Tk1              .........................t...................................................... [146480]
KM262844.1_strain_L60             ................................................................................ [146480]
KM102979.1_26544/OG10_Italy       ................................................................................ [146480]
KM262845.1_strain_NHV             ................................................................................ [146480]
KX354450.1_47/Ss/2008             ................................................................................ [146480]
KP055815.1_strain_BA71            ................................................................................ [146480]
MH025920.1_strain_R35             .........................t...c.....c..............a.................g.....a..... [146480]
MH025919.1_strain_N10             .........................t...c.....c..............a.................g.....a..... [146480]
MH025918.1_strain_R25             .........................t...c.....c..............a.................g.....a..... [146480]
MH025917.1_strain_R7              .........................t...c.....c..............a.................g.....a..... [146480]
MH025916.1_strain_R8              .........................t...c.....c..............a.................g.....a..... [146480]

FR682468.2_Georgia_2007/1         cagtagaatgttttttacgatgctggccacgtttaccggcacctgtctaacatctgtaagcagctgactgaaattaaaat [146560]
ASF_IND_20_CAD543                 ................................................................................ [146560]
LR899193.1_Germany_2020/1         ................................................................................ [146560]
MN393477.1_Wuhan_2019             ................................................................................ [146560]
MW396979.1_Timor-Leste/2019/1     ................................................................................ [146560]
MK940252.1_CN/2019/InnerMongolia  ................................................................................ [146560]
MK128995.1_China/2018/AnhuiXCGQ   ................................................................................ [146560]
MH681419.1_ASFV/POL/2015          ................................................................................ [146560]
LS478113.1_Estonia_2014           ................................................................................ [146560]
KP843857.1_Odintsovo_02/14        ................................................................................ [146560]
FN557520.1_strain_E75             ..................................................................g............. [146560]
AY261366.1_Warthog                ................................................................................ [146560]
AY261365.1_Warmbaths              ................................................................................ [146560]
AY261364.1_Tengani_62             ................................................................................ [146560]
AY261363.1_Pretoriuskop/96/4      ..........................................t..................................... [146560]
AY261362.1_Mkuzi_1979             ................................................................................ [146560]
AY261361.1_Malawi_Lil-20/1_1983   g.....g...........................................t............................. [146560]
AY261360.1_Kenya_1950             ...............c................................................................ [146560]
AM712240.1_OURT_88/3              ..................................................................g............. [146560]
AM712239.1_Benin_97/1             ..................................................................g............. [146560]
KM111295.1_Ken06.Bus              ...............c................................................................ [146560]
KM111294.1_Ken05/Tk1              ......g......................................................................... [146560]
KM262844.1_strain_L60             ..................................................................g............. [146560]
KM102979.1_26544/OG10_Italy       ..................................................................g............. [146560]
KM262845.1_strain_NHV             ..................................................................g............. [146560]
KX354450.1_47/Ss/2008             ..................................................................g............. [146560]
KP055815.1_strain_BA71            ..................................................................g............. [146560]
MH025920.1_strain_R35             ...............c................................................................ [146560]
MH025919.1_strain_N10             ...............c................................................................ [146560]
MH025918.1_strain_R25             ...............c................................................................ [146560]
MH025917.1_strain_R7              ...............c................................................................ [146560]
MH025916.1_strain_R8              ...............c................................................................ [146560]
```

```
FR682468.2_Georgia_2007/1        tttcgacgtttaggaagatctgtcgatatttatctctatcctttttaaggcgtgaaaattcttcttcaaacaagggcgat [146640]
ASF_IND_20_CAD543                ................................................................................ [146640]
LR899193.1_Germany_2020/1        ................................................................................ [146640]
MN393477.1_Wuhan_2019            ................................................................................ [146640]
MW396979.1_Timor-Leste/2019/1    ................................................................................ [146640]
MK940252.1_CN/2019/InnerMongolia ................................................................................ [146640]
MK128995.1_China/2018/AnhuiXCGQ  ................................................................................ [146640]
MH681419.1_ASFV/POL/2015         ................................................................................ [146640]
LS478113.1_Estonia_2014          ................................................................................ [146640]
KP843857.1_Odintsovo_02/14       ................................................................................ [146640]
FN557520.1_strain_E75            ............................................................................g... [146640]
AY261366.1_Warthog               ................................................................................ [146640]
AY261365.1_Warmbaths             ................................................................................ [146640]
AY261364.1_Tengani_62            ................................................................................ [146640]
AY261363.1_Pretoriuskop/96/4     .....t.......................................................................... [146640]
AY261362.1_Mkuzi_1979            .....t......................................................................g... [146640]
AY261361.1_Malawi_Lil-20/1_1983  .....t..................................................................g...g..c [146640]
AY261360.1_Kenya_1950            .....t......................................................................a... [146640]
AM712240.1_OURT_88/3             ............................................................................g... [146640]
AM712239.1_Benin_97/1            ............................................................................g... [146640]
KM111295.1_Ken06.Bus             .....t......................................................................a... [146640]
KM111294.1_Ken05/Tk1             .....t......................................................................a... [146640]
KM262844.1_strain_L60            ............................................................................g... [146640]
KM102979.1_26544/OG10_Italy      ............................................................................g... [146640]
KM262845.1_strain_NHV            ............................................................................g... [146640]
KX354450.1_47/Ss/2008            ............................................................................g... [146640]
KP055815.1_strain_BA71           ............................................................................g... [146640]
MH025920.1_strain_R35            .....t......................................................................a... [146640]
MH025919.1_strain_N10            .....t......................................................................a... [146640]
MH025918.1_strain_R25            .....t......................................................................a... [146640]
MH025917.1_strain_R7             .....t......................................................................a... [146640]
MH025916.1_strain_R8             .....t......................................................................a... [146640]

FR682468.2_Georgia_2007/1        tgtatcccggtgtacttgaatttgtcttcaagttcctggtccgacagcatgatggtttcaaaccgtacggtttcaagctg [146720]
ASF_IND_20_CAD543                ................................................................................ [146720]
LR899193.1_Germany_2020/1        ................................................................................ [146720]
MN393477.1_Wuhan_2019            ................................................................................ [146720]
MW396979.1_Timor-Leste/2019/1    ................................................................................ [146720]
MK940252.1_CN/2019/InnerMongolia ................................................................................ [146720]
MK128995.1_China/2018/AnhuiXCGQ  ................................................................................ [146720]
MH681419.1_ASFV/POL/2015         ................................................................................ [146720]
LS478113.1_Estonia_2014          ................................................................................ [146720]
KP843857.1_Odintsovo_02/14       ................................................................................ [146720]
```

```
FN557520.1_strain_E75            ..........................c...............................................g..... [146720]
AY261366.1_Warthog               .....t..a....................................................................... [146720]
AY261365.1_Warmbaths             ................................................................................ [146720]
AY261364.1_Tengani_62            ................................................................................ [146720]
AY261363.1_Pretoriuskop/96/4     ................................................................................ [146720]
AY261362.1_Mkuzi_1979            ..........................c...............................................g..... [146720]
AY261361.1_Malawi_Lil-20/1_1983  ..............................................................a..c..c........t.. [146720]
AY261360.1_Kenya_1950            .............................g...................................c..t........t.. [146720]
AM712240.1_OURT_88/3             ..........................c...............................................g..... [146720]
AM712239.1_Benin_97/1            ..........................c...............................................g..... [146720]
KM111295.1_Ken06.Bus             .............................g........a..........................c..t........t.. [146720]
KM111294.1_Ken05/Tk1             .............................g...................................c..t........t.. [146720]
KM262844.1_strain_L60            ..........................c...............................................g..... [146720]
KM102979.1_26544/OG10_Italy      ..........................c...............................................g..... [146720]
KM262845.1_strain_NHV            ..........................c...............................................g..... [146720]
KX354450.1_47/Ss/2008            ..........................c...............................................g..... [146720]
KP055815.1_strain_BA71           ..........................c...............................................g..... [146720]
MH025920.1_strain_R35            .............................g........a..........................c..t........t.. [146720]
MH025919.1_strain_N10            .............................g........a..........................c..t........t.. [146720]
MH025918.1_strain_R25            .............................g........a..........................c..t........t.. [146720]
MH025917.1_strain_R7             .............................g........a..........................c..t........t.. [146720]
MH025916.1_strain_R8             .............................g........a..........................c..t........t.. [146720]

FR682468.2_Georgia_2007/1        gcgcgcatcaaggccgtcctctccgtacaactgctgcacaagacgcgtatcgatggaaacccgtcggtaataatccacaa [146800]
ASF_IND_20_CAD543                ................................................................................ [146800]
LR899193.1_Germany_2020/1        ................................................................................ [146800]
MN393477.1_Wuhan_2019            ................................................................................ [146800]
MW396979.1_Timor-Leste/2019/1    ................................................................................ [146800]
MK940252.1_CN/2019/InnerMongolia ................................................................................ [146800]
MK128995.1_China/2018/AnhuiXCGQ  ................................................................................ [146800]
MH681419.1_ASFV/POL/2015         ................................................................................ [146800]
LS478113.1_Estonia_2014          ................................................................................ [146800]
KP843857.1_Odintsovo_02/14       ................................................................................ [146800]
FN557520.1_strain_E75            ...t...........a.............................a.................................. [146800]
AY261366.1_Warthog               ...t............................................................................ [146800]
AY261365.1_Warmbaths             ................................................................................ [146800]
AY261364.1_Tengani_62            .............................................a........a......................... [146800]
AY261363.1_Pretoriuskop/96/4     ................................................................................ [146800]
AY261362.1_Mkuzi_1979            ...............a.............................a.................................. [146800]
AY261361.1_Malawi_Lil-20/1_1983  .............................................a.....a............................ [146800]
AY261360.1_Kenya_1950            .....................a........t..............a.....a............................ [146800]
AM712240.1_OURT_88/3             ...t...........a.............................a.................................. [146800]
AM712239.1_Benin_97/1            ...t...........a.............................a.................................. [146800]
KM111295.1_Ken06.Bus             .............................................a.....a............................ [146800]
```

```
KM111294.1_Ken05/Tk1              ..............................t..............a.....a............................ [146800]
KM262844.1_strain_L60             ...t...........a.............................a.................................. [146800]
KM102979.1_26544/OG10_Italy       ...t...........a.............................a.................................. [146800]
KM262845.1_strain_NHV             ...t...........a.............................a.................................. [146800]
KX354450.1_47/Ss/2008             ...t...........a.............................a.................................. [146800]
KP055815.1_strain_BA71            ...t...........a.............................a.................................. [146800]
MH025920.1_strain_R35             .............................................a.....a............................ [146800]
MH025919.1_strain_N10             .............................................a.....a............................ [146800]
MH025918.1_strain_R25             .............................................a.....a............................ [146800]
MH025917.1_strain_R7              .............................................a.....a............................ [146800]
MH025916.1_strain_R8              .............................................a.....a............................ [146800]

FR682468.2_Georgia_2007/1         tacaggattgaaggccaaagatggctttacggttggcatagcctgtggatgatgtcgataatgctttgttgatcaagtcg [146880]
ASF_IND_20_CAD543                 ................................................................................ [146880]
LR899193.1_Germany_2020/1         ................................................................................ [146880]
MN393477.1_Wuhan_2019             ................................................................................ [146880]
MW396979.1_Timor-Leste/2019/1     ................................................................................ [146880]
MK940252.1_CN/2019/InnerMongolia  ................................................................................ [146880]
MK128995.1_China/2018/AnhuiXCGQ   ................................................................................ [146880]
MH681419.1_ASFV/POL/2015          ................................................................................ [146880]
LS478113.1_Estonia_2014           ................................................................................ [146880]
KP843857.1_Odintsovo_02/14        ................................................................................ [146880]
FN557520.1_strain_E75             ................................................................................ [146880]
AY261366.1_Warthog                ..............................................a.............g................... [146880]
AY261365.1_Warmbaths              ......................a.......................a.............g................... [146880]
AY261364.1_Tengani_62             ...............................a............................g................... [146880]
AY261363.1_Pretoriuskop/96/4      ..............................................a.............g................... [146880]
AY261362.1_Mkuzi_1979             ................................................................................ [146880]
AY261361.1_Malawi_Lil-20/1_1983   ..............................................a.............g................... [146880]
AY261360.1_Kenya_1950             .......c.........................................c..........g............t...... [146880]
AM712240.1_OURT_88/3              ................................................................................ [146880]
AM712239.1_Benin_97/1             ................................................................................ [146880]
KM111295.1_Ken06.Bus              .......c.......................a..............a..c..........g................... [146880]
KM111294.1_Ken05/Tk1              .......c.........................................c..........g................... [146880]
KM262844.1_strain_L60             ................................................................................ [146880]
KM102979.1_26544/OG10_Italy       ................................................................................ [146880]
KM262845.1_strain_NHV             ................................................................................ [146880]
KX354450.1_47/Ss/2008             ................................................................................ [146880]
KP055815.1_strain_BA71            ................................................................................ [146880]
MH025920.1_strain_R35             .......c.......................a..............a..c..........g................... [146880]
MH025919.1_strain_N10             .......c.......................a..............a..c..........g................... [146880]
MH025918.1_strain_R25             .......c.......................a..............a..c..........g................... [146880]
MH025917.1_strain_R7              .......c.......................a..............a..c..........g................... [146880]
MH025916.1_strain_R8              .......c.......................a..............a..c..........g................... [146880]
```

```
FR682468.2_Georgia_2007/1        aatcttccattcatttccccaaagataaattcaggggaggtaaggcccgcaatatagctgttgcagatgaacccgtaggc [146960]
ASF_IND_20_CAD543                ................................................................................ [146960]
LR899193.1_Germany_2020/1        ................................................................................ [146960]
MN393477.1_Wuhan_2019            ................................................................................ [146960]
MW396979.1_Timor-Leste/2019/1    ................................................................................ [146960]
MK940252.1_CN/2019/InnerMongolia ................................................................................ [146960]
MK128995.1_China/2018/AnhuiXCGQ  ................................................................................ [146960]
MH681419.1_ASFV/POL/2015         ................................................................................ [146960]
LS478113.1_Estonia_2014          ................................................................................ [146960]
KP843857.1_Odintsovo_02/14       ................................................................................ [146960]
FN557520.1_strain_E75            ................................................................................ [146960]
AY261366.1_Warthog               .................................................................a..a........... [146960]
AY261365.1_Warmbaths             ................................................................................ [146960]
AY261364.1_Tengani_62            ................................................................................ [146960]
AY261363.1_Pretoriuskop/96/4     ................................................................................ [146960]
AY261362.1_Mkuzi_1979            ....................................................................a........... [146960]
AY261361.1_Malawi_Lil-20/1_1983  ..........................g..................................................... [146960]
AY261360.1_Kenya_1950            .....................................................g.......................... [146960]
AM712240.1_OURT_88/3             ................................................................................ [146960]
AM712239.1_Benin_97/1            ................................................................................ [146960]
KM111295.1_Ken06.Bus             .....................................................g.......................... [146960]
KM111294.1_Ken05/Tk1             .....................................................g.......................... [146960]
KM262844.1_strain_L60            ................................................................................ [146960]
KM102979.1_26544/OG10_Italy      ................................................................................ [146960]
KM262845.1_strain_NHV            ................................................................................ [146960]
KX354450.1_47/Ss/2008            ................................................................................ [146960]
KP055815.1_strain_BA71           ................................................................................ [146960]
MH025920.1_strain_R35            .....................................................g.......................... [146960]
MH025919.1_strain_N10            .....................................................g.......................... [146960]
MH025918.1_strain_R25            .....................................................g.......................... [146960]
MH025917.1_strain_R7             .....................................................g.......................... [146960]
MH025916.1_strain_R8             .....................................................g.......................... [146960]

FR682468.2_Georgia_2007/1        ctgcgcctccagggcaaacctggggtagtacaccagggtcctaccgaaggaaaactggggttgaatgcgttgtgtattaa [147040]
ASF_IND_20_CAD543                ................................................................................ [147040]
LR899193.1_Germany_2020/1        ................................................................................ [147040]
MN393477.1_Wuhan_2019            ................................................................................ [147040]
MW396979.1_Timor-Leste/2019/1    ................................................................................ [147040]
MK940252.1_CN/2019/InnerMongolia ................................................................................ [147040]
MK128995.1_China/2018/AnhuiXCGQ  ................................................................................ [147040]
MH681419.1_ASFV/POL/2015         ................................................................................ [147040]
LS478113.1_Estonia_2014          ................................................................................ [147040]
KP843857.1_Odintsovo_02/14       ................................................................................ [147040]
```

```
FN557520.1_strain_E75          ..........................................g..................................... [147040]
AY261366.1_Warthog             ..............................t...........g..........................c.......... [147040]
AY261365.1_Warmbaths           ..............................t...........g..........................c.......... [147040]
AY261364.1_Tengani_62          ..........................................g..................................... [147040]
AY261363.1_Pretoriuskop/96/4   ..........................................g..........................c.......... [147040]
AY261362.1_Mkuzi_1979          ..........................................g..................................... [147040]
AY261361.1_Malawi_Lil-20/1_1983 ..........................................g..................................... [147040]
AY261360.1_Kenya_1950          ..........................................g..................................... [147040]
AM712240.1_OURT_88/3           ..........................................g..................................... [147040]
AM712239.1_Benin_97/1          ..........................................g..................................... [147040]
KM111295.1_Ken06.Bus           ..........................................g..................................... [147040]
KM111294.1_Ken05/Tk1           ..........................................g..................................... [147040]
KM262844.1_strain_L60          ..........................................g..................................... [147040]
KM102979.1_26544/OG10_Italy    ..........................................g..................................... [147040]
KM262845.1_strain_NHV          ..........................................g..................................... [147040]
KX354450.1_47/Ss/2008          ..........................................g..................................... [147040]
KP055815.1_strain_BA71         ..........................................g..................................... [147040]
MH025920.1_strain_R35          ..........................................g..................................... [147040]
MH025919.1_strain_N10          ..........................................g..................................... [147040]
MH025918.1_strain_R25          ..........................................g..................................... [147040]
MH025917.1_strain_R7           ..........................................g..................................... [147040]
MH025916.1_strain_R8           ..........................................g..................................... [147040]

FR682468.2_Georgia_2007/1      tttcaatttggccgatgcccgccatgatgtgaatcatattggggtttgagcccttggcgccagtggccaccatctgaaaa [147120]
ASF_IND_20_CAD543              ................................................................................ [147120]
LR899193.1_Germany_2020/1      ................................................................................ [147120]
MN393477.1_Wuhan_2019          ................................................................................ [147120]
MW396979.1_Timor-Leste/2019/1  ................................................................................ [147120]
MK940252.1_CN/2019/InnerMongolia ................................................................................ [147120]
MK128995.1_China/2018/AnhuiXCGQ ................................................................................ [147120]
MH681419.1_ASFV/POL/2015       ................................................................................ [147120]
LS478113.1_Estonia_2014        ................................................................................ [147120]
KP843857.1_Odintsovo_02/14     ................................................................................ [147120]
FN557520.1_strain_E75          ................................................................................ [147120]
AY261366.1_Warthog             ................................................................................ [147120]
AY261365.1_Warmbaths           .............a.................................................................. [147120]
AY261364.1_Tengani_62          ................................................................................ [147120]
AY261363.1_Pretoriuskop/96/4   ................................................................................ [147120]
AY261362.1_Mkuzi_1979          ...................t................................t........................... [147120]
AY261361.1_Malawi_Lil-20/1_1983 .............................................................g.................. [147120]
AY261360.1_Kenya_1950          .............................................................c.................. [147120]
AM712240.1_OURT_88/3           ................................................................................ [147120]
AM712239.1_Benin_97/1          ................................................................................ [147120]
KM111295.1_Ken06.Bus           .............................................................c.................. [147120]
```

```
KM111294.1_Ken05/Tk1             .............................................................c.................. [147120]
KM262844.1_strain_L60            ................................................................................ [147120]
KM102979.1_26544/OG10_Italy      ................................................................................ [147120]
KM262845.1_strain_NHV            ................................................................................ [147120]
KX354450.1_47/Ss/2008            ................................................................................ [147120]
KP055815.1_strain_BA71           ................................................................................ [147120]
MH025920.1_strain_R35            .............................................................c.................. [147120]
MH025919.1_strain_N10            .............................................................c.................. [147120]
MH025918.1_strain_R25            .............................................................c.................. [147120]
MH025917.1_strain_R7             .............................................................c.................. [147120]
MH025916.1_strain_R8             .............................................................c.................. [147120]

FR682468.2_Georgia_2007/1        agcccattggtttccggattaatggaattcataatcggctttaaaattctatcgggaaatttaagcgcattcagctgcaa [147200]
ASF_IND_20_CAD543                ................................................................................ [147200]
LR899193.1_Germany_2020/1        ................................................................................ [147200]
MN393477.1_Wuhan_2019            ................................................................................ [147200]
MW396979.1_Timor-Leste/2019/1    ................................................................................ [147200]
MK940252.1_CN/2019/InnerMongolia ................................................................................ [147200]
MK128995.1_China/2018/AnhuiXCGQ  ................................................................................ [147200]
MH681419.1_ASFV/POL/2015         ................................................................................ [147200]
LS478113.1_Estonia_2014          ................................................................................ [147200]
KP843857.1_Odintsovo_02/14       ................................................................................ [147200]
FN557520.1_strain_E75            ...............................................c...........c.................... [147200]
AY261366.1_Warthog               ...............................................c................................ [147200]
AY261365.1_Warmbaths             ...............................................c................................ [147200]
AY261364.1_Tengani_62            ...............................................c.............................t.. [147200]
AY261363.1_Pretoriuskop/96/4     ...............................................c................................ [147200]
AY261362.1_Mkuzi_1979            ...............................................c................................ [147200]
AY261361.1_Malawi_Lil-20/1_1983  ........t......................................c................................ [147200]
AY261360.1_Kenya_1950            ........t......................................c................................ [147200]
AM712240.1_OURT_88/3             ...............................................c...........c.................... [147200]
AM712239.1_Benin_97/1            ...............................................c...........c.................... [147200]
KM111295.1_Ken06.Bus             ........t......................................c................................ [147200]
KM111294.1_Ken05/Tk1             ........t......................................c........g....................... [147200]
KM262844.1_strain_L60            ...............................................c...........c.................... [147200]
KM102979.1_26544/OG10_Italy      ...............................................c...........c.................... [147200]
KM262845.1_strain_NHV            ...............................................c...........c.................... [147200]
KX354450.1_47/Ss/2008            ...............................................c...........c.................... [147200]
KP055815.1_strain_BA71           ...............................................c...........c.................... [147200]
MH025920.1_strain_R35            ........t......................................c................................ [147200]
MH025919.1_strain_N10            ........t......................................c................................ [147200]
MH025918.1_strain_R25            ........t......................................c................................ [147200]
MH025917.1_strain_R7             ........t......................................c................................ [147200]
MH025916.1_strain_R8             ........t......................................c................................ [147200]
```

```
FR682468.2_Georgia_2007/1        tttttcgtagaagtcatgcgttgtcaggcctataggcggcatgatgtctccatgaagcagccggttgtttatttcctccg [147280]
ASF_IND_20_CAD543                ................................................................................ [147280]
LR899193.1_Germany_2020/1        ................................................................................ [147280]
MN393477.1_Wuhan_2019            ................................................................................ [147280]
MW396979.1_Timor-Leste/2019/1    ................................................................................ [147280]
MK940252.1_CN/2019/InnerMongolia ................................................................................ [147280]
MK128995.1_China/2018/AnhuiXCGQ  ................................................................................ [147280]
MH681419.1_ASFV/POL/2015         ................................................................................ [147280]
LS478113.1_Estonia_2014          ................................................................................ [147280]
KP843857.1_Odintsovo_02/14       ................................................................................ [147280]
FN557520.1_strain_E75            ................................................................................ [147280]
AY261366.1_Warthog               ................................................................................ [147280]
AY261365.1_Warmbaths             ................................................................................ [147280]
AY261364.1_Tengani_62            ................................................................................ [147280]
AY261363.1_Pretoriuskop/96/4     ................................................................................ [147280]
AY261362.1_Mkuzi_1979            ................................................................................ [147280]
AY261361.1_Malawi_Lil-20/1_1983  ................................................................................ [147280]
AY261360.1_Kenya_1950            ................................................................................ [147280]
AM712240.1_OURT_88/3             ................................................................................ [147280]
AM712239.1_Benin_97/1            ................................................................................ [147280]
KM111295.1_Ken06.Bus             ................................................................................ [147280]
KM111294.1_Ken05/Tk1             ................................................................................ [147280]
KM262844.1_strain_L60            ................................................................................ [147280]
KM102979.1_26544/OG10_Italy      ................................................................................ [147280]
KM262845.1_strain_NHV            ................................................................................ [147280]
KX354450.1_47/Ss/2008            ................................................................................ [147280]
KP055815.1_strain_BA71           ................................................................................ [147280]
MH025920.1_strain_R35            ................................................................................ [147280]
MH025919.1_strain_N10            ................................................................................ [147280]
MH025918.1_strain_R25            ................................................................................ [147280]
MH025917.1_strain_R7             ................................................................................ [147280]
MH025916.1_strain_R8             ................................................................................ [147280]

FR682468.2_Georgia_2007/1        actcaagcagcagttcattgataatttcttggacctcctgatgtgcctccggggttaggagcatgtcggccgtggacact [147360]
ASF_IND_20_CAD543                ................................................................................ [147360]
LR899193.1_Germany_2020/1        ................................................................................ [147360]
MN393477.1_Wuhan_2019            ................................................................................ [147360]
MW396979.1_Timor-Leste/2019/1    ................................................................................ [147360]
MK940252.1_CN/2019/InnerMongolia ................................................................................ [147360]
MK128995.1_China/2018/AnhuiXCGQ  ................................................................................ [147360]
MH681419.1_ASFV/POL/2015         ................................................................................ [147360]
LS478113.1_Estonia_2014          ................................................................................ [147360]
KP843857.1_Odintsovo_02/14       ................................................................................ [147360]
```

```
FN557520.1_strain_E75              ................................................................................ [147360]
AY261366.1_Warthog                 ...........................................c.............a...................... [147360]
AY261365.1_Warmbaths               ...........................................c.............a...................... [147360]
AY261364.1_Tengani_62              ...........................................c..t..........a...................... [147360]
AY261363.1_Pretoriuskop/96/4       ...........................................c.............a...................... [147360]
AY261362.1_Mkuzi_1979              ................................................................................ [147360]
AY261361.1_Malawi_Lil-20/1_1983    .t................................t........c.............a......a..............c [147360]
AY261360.1_Kenya_1950              ...........................................c.............aa....................c [147360]
AM712240.1_OURT_88/3               ................................................................................ [147360]
AM712239.1_Benin_97/1              ................................................................................ [147360]
KM111295.1_Ken06.Bus               .........................................................aa....................c [147360]
KM111294.1_Ken05/Tk1               ...........................................c.............a.........a...........c [147360]
KM262844.1_strain_L60              ................................................................................ [147360]
KM102979.1_26544/OG10_Italy        ................................................................................ [147360]
KM262845.1_strain_NHV              ................................................................................ [147360]
KX354450.1_47/Ss/2008              ................................................................................ [147360]
KP055815.1_strain_BA71             ................................................................................ [147360]
MH025920.1_strain_R35              .........................................................aa....................c [147360]
MH025919.1_strain_N10              .........................................................aa....................c [147360]
MH025918.1_strain_R25              .........................................................aa....................c [147360]
MH025917.1_strain_R7               .........................................................aa....................c [147360]
MH025916.1_strain_R8               .........................................................aa....................c [147360]

FR682468.2_Georgia_2007/1          gtgaatccggcgttgcgcacgtagtttagggcgagctgctgggtcgcaaatatcattttcaaggcctgctgcggcccata [147440]
ASF_IND_20_CAD543                  ................................................................................ [147440]
LR899193.1_Germany_2020/1          ................................................................................ [147440]
MN393477.1_Wuhan_2019              ................................................................................ [147440]
MW396979.1_Timor-Leste/2019/1      ................................................................................ [147440]
MK940252.1_CN/2019/InnerMongolia   ................................................................................ [147440]
MK128995.1_China/2018/AnhuiXCGQ    ................................................................................ [147440]
MH681419.1_ASFV/POL/2015           ................................................................................ [147440]
LS478113.1_Estonia_2014            ................................................................................ [147440]
KP843857.1_Odintsovo_02/14         ................................................................................ [147440]
FN557520.1_strain_E75              ..........................c..................................................... [147440]
AY261366.1_Warthog                 ..........................c..................................................... [147440]
AY261365.1_Warmbaths               ..........................c..................................................... [147440]
AY261364.1_Tengani_62              ..........................c..a..a............................................... [147440]
AY261363.1_Pretoriuskop/96/4       ....................a.....c..................................................... [147440]
AY261362.1_Mkuzi_1979              ..........................c..................................................... [147440]
AY261361.1_Malawi_Lil-20/1_1983    .................t........c..................................................... [147440]
AY261360.1_Kenya_1950              ..........................c..................................ga...........t..... [147440]
AM712240.1_OURT_88/3               ..........................c..................................................... [147440]
AM712239.1_Benin_97/1              ..........................c..................................................... [147440]
KM111295.1_Ken06.Bus               ..........................c..................................ga...........t..... [147440]
```

```
KM111294.1_Ken05/Tk1            ...........a.....g........c..................................ga...........t..... [147440]
KM262844.1_strain_L60           ..........................c..................................................... [147440]
KM102979.1_26544/OG10_Italy     ..........................c..................................................... [147440]
KM262845.1_strain_NHV           ..........................c..................................................... [147440]
KX354450.1_47/Ss/2008           ..........................c..................................................... [147440]
KP055815.1_strain_BA71          ..........................c..................................................... [147440]
MH025920.1_strain_R35           ..........................c..................................ga...........t..... [147440]
MH025919.1_strain_N10           ..........................c..................................ga...........t..... [147440]
MH025918.1_strain_R25           ..........................c..................................ga...........t..... [147440]
MH025917.1_strain_R7            ..........................c..................................ga...........t..... [147440]
MH025916.1_strain_R8            ..........................c..................................ga...........t..... [147440]

FR682468.2_Georgia_2007/1       cctacgcgaaataaggtgatagattccaccggaggaacccgctccgacggcctttttgtcaaggacgccttcaatgagtt [147520]
ASF_IND_20_CAD543               ................................................................................ [147520]
LR899193.1_Germany_2020/1       ................................................................................ [147520]
MN393477.1_Wuhan_2019           ................................................................................ [147520]
MW396979.1_Timor-Leste/2019/1   ................................................................................ [147520]
MK940252.1_CN/2019/InnerMongolia ................................................................................ [147520]
MK128995.1_China/2018/AnhuiXCGQ ................................................................................ [147520]
MH681419.1_ASFV/POL/2015        ................................................................................ [147520]
LS478113.1_Estonia_2014         ................................................................................ [147520]
KP843857.1_Odintsovo_02/14      ................................................................................ [147520]
FN557520.1_strain_E75           t...................................g........................................... [147520]
AY261366.1_Warthog              ....................................g........................................... [147520]
AY261365.1_Warmbaths            t...................................g........................................... [147520]
AY261364.1_Tengani_62           ....................................g........................................... [147520]
AY261363.1_Pretoriuskop/96/4    ....................................g........................................... [147520]
AY261362.1_Mkuzi_1979           ....................................g........................................... [147520]
AY261361.1_Malawi_Lil-20/1_1983 .....................a.....g........g..t.....a..t............................... [147520]
AY261360.1_Kenya_1950           .....................a.....g........g..t..c.....t............................... [147520]
AM712240.1_OURT_88/3            t...................................g........................................... [147520]
AM712239.1_Benin_97/1           t...................................g........................................... [147520]
KM111295.1_Ken06.Bus            ...........................g........g..t..c.....t............................... [147520]
KM111294.1_Ken05/Tk1            t....................a.....g........g..t..c.....t..t.......................c.... [147520]
KM262844.1_strain_L60           t...................................g........................................... [147520]
KM102979.1_26544/OG10_Italy     t...................................g........................................... [147520]
KM262845.1_strain_NHV           t...................................g........................................... [147520]
KX354450.1_47/Ss/2008           t...................................g........................................... [147520]
KP055815.1_strain_BA71          t...................................g........................................... [147520]
MH025920.1_strain_R35           ...........................g........g..t..c.....t............................... [147520]
MH025919.1_strain_N10           ...........................g........g..t..c.....t............................... [147520]
MH025918.1_strain_R25           ...........................g........g..t..c.....t............................... [147520]
MH025917.1_strain_R7            ...........................g........g..t..c.....t............................... [147520]
MH025916.1_strain_R8            ...........................g........g..t..c.....t............................... [147520]
```

```
FR682468.2_Georgia_2007/1        cgccgttgcgtatttgtgtagagatgtcctgcttgttataatgcatgtagggtgcatacacttctgagtaccatgtgggg [147600]
ASF_IND_20_CAD543                ................................................................................ [147600]
LR899193.1_Germany_2020/1        ................................................................................ [147600]
MN393477.1_Wuhan_2019            ................................................................................ [147600]
MW396979.1_Timor-Leste/2019/1    ................................................................................ [147600]
MK940252.1_CN/2019/InnerMongolia ................................................................................ [147600]
MK128995.1_China/2018/AnhuiXCGQ  ................................................................................ [147600]
MH681419.1_ASFV/POL/2015         ................................................................................ [147600]
LS478113.1_Estonia_2014          ................................................................................ [147600]
KP843857.1_Odintsovo_02/14       ................................................................................ [147600]
FN557520.1_strain_E75            ....................................................c........................... [147600]
AY261366.1_Warthog               .a..................................................c........................... [147600]
AY261365.1_Warmbaths             .a..................................................c........................... [147600]
AY261364.1_Tengani_62            .a................................a.................c.....t..................... [147600]
AY261363.1_Pretoriuskop/96/4     .a..................................................c........................... [147600]
AY261362.1_Mkuzi_1979            ....................................................c........................... [147600]
AY261361.1_Malawi_Lil-20/1_1983  .a..............c..............t....................c........c.................a [147600]
AY261360.1_Kenya_1950            .a..............c.....a........t..a..............c..c........................... [147600]
AM712240.1_OURT_88/3             ....................................................c........................... [147600]
AM712239.1_Benin_97/1            ....................................................c........................... [147600]
KM111295.1_Ken06.Bus             .a..............c.....a........t..a..............c..c........................... [147600]
KM111294.1_Ken05/Tk1             .a..............c.....a........t..a..............c..c........................... [147600]
KM262844.1_strain_L60            ....................................................c........................... [147600]
KM102979.1_26544/OG10_Italy      ....................................................c........................... [147600]
KM262845.1_strain_NHV            ....................................................c........................... [147600]
KX354450.1_47/Ss/2008            ....................................................c........................... [147600]
KP055815.1_strain_BA71           ....................................................c........................... [147600]
MH025920.1_strain_R35            .a.....a........c.....a........t..a..............c..c........................... [147600]
MH025919.1_strain_N10            .a..............c.....a........t..a..............c..c........................... [147600]
MH025918.1_strain_R25            .a.....a........c.....a........t..a..............c..c........................... [147600]
MH025917.1_strain_R7             .a.....a........c.....a........t..a..............c..c........................... [147600]
MH025916.1_strain_R8             .a.....a........c.....a........t..a..............c..c........................... [147600]

FR682468.2_Georgia_2007/1        gctcgttgataattgatgggggtctgcctcagtagcatagatacaaccgatttgccatccagcaggtcagttggggagta [147680]
ASF_IND_20_CAD543                ................................................................................ [147680]
LR899193.1_Germany_2020/1        ................................................................................ [147680]
MN393477.1_Wuhan_2019            ................................................................................ [147680]
MW396979.1_Timor-Leste/2019/1    ................................................................................ [147680]
MK940252.1_CN/2019/InnerMongolia ................................................................................ [147680]
MK128995.1_China/2018/AnhuiXCGQ  ................................................................................ [147680]
MH681419.1_ASFV/POL/2015         ................................................................................ [147680]
LS478113.1_Estonia_2014          ................................................................................ [147680]
KP843857.1_Odintsovo_02/14       ................................................................................ [147680]
```

```
FN557520.1_strain_E75            ...........................t.................................................... [147680]
AY261366.1_Warthog               ................................................................a............... [147680]
AY261365.1_Warmbaths             ..........................t.................g................................... [147680]
AY261364.1_Tengani_62            ...........................t...............................t.................... [147680]
AY261363.1_Pretoriuskop/96/4     ................................................................................ [147680]
AY261362.1_Mkuzi_1979            ....................a......t.................................................... [147680]
AY261361.1_Malawi_Lil-20/1_1983  .....c.....g.....a.........t.................................................... [147680]
AY261360.1_Kenya_1950            ........g..g...............t....................a...................c........... [147680]
AM712240.1_OURT_88/3             ...........................t.................................................... [147680]
AM712239.1_Benin_97/1            ...........................t.................................................... [147680]
KM111295.1_Ken06.Bus             ........g..g...............t....................a..........t........c........... [147680]
KM111294.1_Ken05/Tk1             ........g..g...............t..................................t.....c........... [147680]
KM262844.1_strain_L60            ...........................t.................................................... [147680]
KM102979.1_26544/OG10_Italy      ...........................t.................................................... [147680]
KM262845.1_strain_NHV            ...........................t.................................................... [147680]
KX354450.1_47/Ss/2008            ...........................t.................................................... [147680]
KP055815.1_strain_BA71           ...........................t.................................................... [147680]
MH025920.1_strain_R35            ........g..g...............t....................a..........t........c........... [147680]
MH025919.1_strain_N10            ........g..g...............t....................a..........t........c........... [147680]
MH025918.1_strain_R25            ........g..g...............t....................a..........t........c........... [147680]
MH025917.1_strain_R7             ........g..g...............t....................a..........t........c........... [147680]
MH025916.1_strain_R8             ........g..g...............t....................a..........t........c........... [147680]

FR682468.2_Georgia_2007/1        gttggcaaaacaaggtgggtcggtttgggttgtttgaaacaaccccatggcgtgcagcttgttcatcacatttttcccca [147760]
ASF_IND_20_CAD543                ................................................................................ [147760]
LR899193.1_Germany_2020/1        ................................................................................ [147760]
MN393477.1_Wuhan_2019            ................................................................................ [147760]
MW396979.1_Timor-Leste/2019/1    ................................................................................ [147760]
MK940252.1_CN/2019/InnerMongolia ................................................................................ [147760]
MK128995.1_China/2018/AnhuiXCGQ  ................................................................................ [147760]
MH681419.1_ASFV/POL/2015         ................................................................................ [147760]
LS478113.1_Estonia_2014          ................................................................................ [147760]
KP843857.1_Odintsovo_02/14       ................................................................................ [147760]
FN557520.1_strain_E75            .....................c....................t..............t...................... [147760]
AY261366.1_Warthog               .........................................................t...................... [147760]
AY261365.1_Warmbaths             ...a.................a.....a.................................................... [147760]
AY261364.1_Tengani_62            ................................................................................ [147760]
AY261363.1_Pretoriuskop/96/4     ...a.................a..........................a........t...................... [147760]
AY261362.1_Mkuzi_1979            .....................c....................t..............t...................... [147760]
AY261361.1_Malawi_Lil-20/1_1983  ...............c.....c.................t.................t...................... [147760]
AY261360.1_Kenya_1950            ............g........a.................t........a........t.................t.... [147760]
AM712240.1_OURT_88/3             .....................c....................t..............t...................... [147760]
AM712239.1_Benin_97/1            .....................c....................t..............t...................... [147760]
KM111295.1_Ken06.Bus             ............g........a.................t........a........t...................... [147760]
```

```
KM111294.1_Ken05/Tk1              ............g........a.................t........a........t.................t.... [147760]
KM262844.1_strain_L60             .....................c....................t..............t...................... [147760]
KM102979.1_26544/OG10_Italy       .....................c....................t..............t...................... [147760]
KM262845.1_strain_NHV             .....................c....................t..............t...................... [147760]
KX354450.1_47/Ss/2008             .....................c....................t..............t...................... [147760]
KP055815.1_strain_BA71            .....................c....................t..............t...................... [147760]
MH025920.1_strain_R35             ............g........a.................t........a........t...................... [147760]
MH025919.1_strain_N10             ............g........a.................t........a........t...................... [147760]
MH025918.1_strain_R25             ............g........a.................t........a........t...................... [147760]
MH025917.1_strain_R7              ............g........a.................t........a........t...................... [147760]
MH025916.1_strain_R8              ............g........a.................t........a........t...................... [147760]

FR682468.2_Georgia_2007/1         tgggggtgttcgtgcgtgtaagcaaaaagcttcccaccgtggagtcctgcacctgcccattaacgggacccgagctcttt [147840]
ASF_IND_20_CAD543                 ................................................................................ [147840]
LR899193.1_Germany_2020/1         ................................................................................ [147840]
MN393477.1_Wuhan_2019             ................................................................................ [147840]
MW396979.1_Timor-Leste/2019/1     ................................................................................ [147840]
MK940252.1_CN/2019/InnerMongolia  ................................................................................ [147840]
MK128995.1_China/2018/AnhuiXCGQ   ................................................................................ [147840]
MH681419.1_ASFV/POL/2015          ................................................................................ [147840]
LS478113.1_Estonia_2014           ................................................................................ [147840]
KP843857.1_Odintsovo_02/14        ................................................................................ [147840]
FN557520.1_strain_E75             .....................................t.....a.............................a.....c [147840]
AY261366.1_Warthog                .....................................t.........................................c [147840]
AY261365.1_Warmbaths              .....................................t.........................................c [147840]
AY261364.1_Tengani_62             .....................................t.........................................c [147840]
AY261363.1_Pretoriuskop/96/4      .....................................t...........t.......................a..t... [147840]
AY261362.1_Mkuzi_1979             .....................................t.....a...................................c [147840]
AY261361.1_Malawi_Lil-20/1_1983   ..........t..........................t.........................................c [147840]
AY261360.1_Kenya_1950             ............................a........g..........................a..............c [147840]
AM712240.1_OURT_88/3              .....................................t.....a.............................a.....c [147840]
AM712239.1_Benin_97/1             .....................................t.....a.............................a.....c [147840]
KM111295.1_Ken06.Bus              ............................a........g..........................a..............c [147840]
KM111294.1_Ken05/Tk1              ............................a........g..........................a..............c [147840]
KM262844.1_strain_L60             .....................................t.....a.............................a.....c [147840]
KM102979.1_26544/OG10_Italy       .............a.......................t.....a.............................a.....c [147840]
KM262845.1_strain_NHV             .....................................t.....a.............................a.....c [147840]
KX354450.1_47/Ss/2008             .............a.......................t.....a.............................a.....c [147840]
KP055815.1_strain_BA71            .....................................t.....a.............................a.....c [147840]
MH025920.1_strain_R35             ............................a........g..........................a..............c [147840]
MH025919.1_strain_N10             ............................a........g..........................a..............c [147840]
MH025918.1_strain_R25             ............................a........g..........................a..............c [147840]
MH025917.1_strain_R7              ............................a........g..........................a..............c [147840]
MH025916.1_strain_R8              ............................a........g..........................a..............c [147840]
```

```
FR682468.2_Georgia_2007/1        gtggaaatgaaccagtttcgcacagaacaaagtagttcggcctcaacgcggctcatgacgctccagggaacccagagatt [147920]
ASF_IND_20_CAD543                ................................................................................ [147920]
LR899193.1_Germany_2020/1        ................................................................................ [147920]
MN393477.1_Wuhan_2019            ................................................................................ [147920]
MW396979.1_Timor-Leste/2019/1    ................................................................................ [147920]
MK940252.1_CN/2019/InnerMongolia ................................................................................ [147920]
MK128995.1_China/2018/AnhuiXCGQ  ................................................................................ [147920]
MH681419.1_ASFV/POL/2015         ................................................................................ [147920]
LS478113.1_Estonia_2014          ................................................................................ [147920]
KP843857.1_Odintsovo_02/14       ................................................................................ [147920]
FN557520.1_strain_E75            ................................................................................ [147920]
AY261366.1_Warthog               ........a....................................................................... [147920]
AY261365.1_Warmbaths             ................................................................................ [147920]
AY261364.1_Tengani_62            ............................................g................................... [147920]
AY261363.1_Pretoriuskop/96/4     ........a....................................................................... [147920]
AY261362.1_Mkuzi_1979            ................................................................................ [147920]
AY261361.1_Malawi_Lil-20/1_1983  .......................g........................................................ [147920]
AY261360.1_Kenya_1950            .......................g........................................................ [147920]
AM712240.1_OURT_88/3             ................................................................................ [147920]
AM712239.1_Benin_97/1            ................................................................................ [147920]
KM111295.1_Ken06.Bus             .......................g........................................................ [147920]
KM111294.1_Ken05/Tk1             .......................g........................................................ [147920]
KM262844.1_strain_L60            ................................................................................ [147920]
KM102979.1_26544/OG10_Italy      ................................................................................ [147920]
KM262845.1_strain_NHV            ................................................................................ [147920]
KX354450.1_47/Ss/2008            ................................................................................ [147920]
KP055815.1_strain_BA71           ................................................................................ [147920]
MH025920.1_strain_R35            .......................g........................................................ [147920]
MH025919.1_strain_N10            .......................g........................................................ [147920]
MH025918.1_strain_R25            .......................g........................................................ [147920]
MH025917.1_strain_R7             .......................g........................................................ [147920]
MH025916.1_strain_R8             .......................g........................................................ [147920]

FR682468.2_Georgia_2007/1        catctgatccccgtcaaagtccgcattataccaggcacatgcgctgacattcatttgaaacgtagaaatttttgggtttt [148000]
ASF_IND_20_CAD543                ................................................................................ [148000]
LR899193.1_Germany_2020/1        ................................................................................ [148000]
MN393477.1_Wuhan_2019            ................................................................................ [148000]
MW396979.1_Timor-Leste/2019/1    ................................................................................ [148000]
MK940252.1_CN/2019/InnerMongolia ................................................................................ [148000]
MK128995.1_China/2018/AnhuiXCGQ  ................................................................................ [148000]
MH681419.1_ASFV/POL/2015         ................................................................................ [148000]
LS478113.1_Estonia_2014          ................................................................................ [148000]
KP843857.1_Odintsovo_02/14       ................................................................................ [148000]
```

```
FN557520.1_strain_E75             ............a................................................................... [148000]
AY261366.1_Warthog                ............a................................................................... [148000]
AY261365.1_Warmbaths              ............a................................................................... [148000]
AY261364.1_Tengani_62             ............a..........................g........................................ [148000]
AY261363.1_Pretoriuskop/96/4      ............a................................................................... [148000]
AY261362.1_Mkuzi_1979             ............a................................................................... [148000]
AY261361.1_Malawi_Lil-20/1_1983   ............a................................................................... [148000]
AY261360.1_Kenya_1950             ......g................................c........................................ [148000]
AM712240.1_OURT_88/3              ............a................................................................... [148000]
AM712239.1_Benin_97/1             ............a................................................................... [148000]
KM111295.1_Ken06.Bus              ......g................................c........................................ [148000]
KM111294.1_Ken05/Tk1              ............a................................................................... [148000]
KM262844.1_strain_L60             ............a................................................................... [148000]
KM102979.1_26544/OG10_Italy       ............a................................................................... [148000]
KM262845.1_strain_NHV             ............a................................................................... [148000]
KX354450.1_47/Ss/2008             ............a................................................................... [148000]
KP055815.1_strain_BA71            ............a................................................................... [148000]
MH025920.1_strain_R35             ......g................................c........................................ [148000]
MH025919.1_strain_N10             ......g................................c........................................ [148000]
MH025918.1_strain_R25             ......g................................c........................................ [148000]
MH025917.1_strain_R7              ......g................................c........................................ [148000]
MH025916.1_strain_R8              ......g................................c........................................ [148000]

FR682468.2_Georgia_2007/1         caagaacgacaatccggtgaacccctatgctgcttcgttcgagagaaggctggcgattaaaaaacgcgacgtcgccagtg [148080]
ASF_IND_20_CAD543                 ................................................................................ [148080]
LR899193.1_Germany_2020/1         ................................................................................ [148080]
MN393477.1_Wuhan_2019             ................................................................................ [148080]
MW396979.1_Timor-Leste/2019/1     ................................................................................ [148080]
MK940252.1_CN/2019/InnerMongolia  ................................................................................ [148080]
MK128995.1_China/2018/AnhuiXCGQ   ................................................................................ [148080]
MH681419.1_ASFV/POL/2015          ................................................................................ [148080]
LS478113.1_Estonia_2014           ................................................................................ [148080]
KP843857.1_Odintsovo_02/14        ................................................................................ [148080]
FN557520.1_strain_E75             ................................................................................ [148080]
AY261366.1_Warthog                ...............................................................................a [148080]
AY261365.1_Warmbaths              ................................................................................ [148080]
AY261364.1_Tengani_62             ..............................................g................................. [148080]
AY261363.1_Pretoriuskop/96/4      ............................................................................g..a [148080]
AY261362.1_Mkuzi_1979             ................................................................................ [148080]
AY261361.1_Malawi_Lil-20/1_1983   ....g........t........t.............t..............................a............ [148080]
AY261360.1_Kenya_1950             ....g..c..............t.............t.......................................g... [148080]
AM712240.1_OURT_88/3              ................................................................................ [148080]
AM712239.1_Benin_97/1             ................................................................................ [148080]
KM111295.1_Ken06.Bus              ....g..c..............t.............t.......................................g... [148080]
```

```
KM111294.1_Ken05/Tk1             ....g...............................t....................................a...... [148080]
KM262844.1_strain_L60            ................................................................................ [148080]
KM102979.1_26544/OG10_Italy      ................................................................................ [148080]
KM262845.1_strain_NHV            ................................................................................ [148080]
KX354450.1_47/Ss/2008            ................................................................................ [148080]
KP055815.1_strain_BA71           ................................................................................ [148080]
MH025920.1_strain_R35            ....g..c..............t.............t.......................................g... [148080]
MH025919.1_strain_N10            ....g..c..............t.............t.......................................g... [148080]
MH025918.1_strain_R25            ....g..c..............t.............t.......................................g... [148080]
MH025917.1_strain_R7             ....g..c..............t.............t.......................................g... [148080]
MH025916.1_strain_R8             ....g..c..............t.............t.......................................g... [148080]

FR682468.2_Georgia_2007/1        acgacgtcacggtaaaggatgtctcctacctccagcctaaagtcttgtttgagaccctcaatgtcgtgaacggattgtgt [148160]
ASF_IND_20_CAD543                ................................................................................ [148160]
LR899193.1_Germany_2020/1        ................................................................................ [148160]
MN393477.1_Wuhan_2019            ................................................................................ [148160]
MW396979.1_Timor-Leste/2019/1    ................................................................................ [148160]
MK940252.1_CN/2019/InnerMongolia ................................................................................ [148160]
MK128995.1_China/2018/AnhuiXCGQ  ................................................................................ [148160]
MH681419.1_ASFV/POL/2015         ................................................................................ [148160]
LS478113.1_Estonia_2014          ................................................................................ [148160]
KP843857.1_Odintsovo_02/14       ................................................................................ [148160]
FN557520.1_strain_E75            .....a........................................................a................. [148160]
AY261366.1_Warthog               ...................................t............................................ [148160]
AY261365.1_Warmbaths             ...................................t............................................ [148160]
AY261364.1_Tengani_62            ...................................t.................g.......................... [148160]
AY261363.1_Pretoriuskop/96/4     ..a................................t.................g.......................... [148160]
AY261362.1_Mkuzi_1979            .....a.............................t..........................a................. [148160]
AY261361.1_Malawi_Lil-20/1_1983  ....................a..............t............................................ [148160]
AY261360.1_Kenya_1950            .....a...........a........c........t............................................ [148160]
AM712240.1_OURT_88/3             .....a........................................................a................. [148160]
AM712239.1_Benin_97/1            .....a........................................................a................. [148160]
KM111295.1_Ken06.Bus             .....a...........a........c..................................................... [148160]
KM111294.1_Ken05/Tk1             ....................a..............t..........................a........a........ [148160]
KM262844.1_strain_L60            .....a........................................................a................. [148160]
KM102979.1_26544/OG10_Italy      .....a........................................................a................. [148160]
KM262845.1_strain_NHV            .....a........................................................a................. [148160]
KX354450.1_47/Ss/2008            .....a........................................................a................. [148160]
KP055815.1_strain_BA71           .....a........................................................a................. [148160]
MH025920.1_strain_R35            .....a...........a........c..................................................... [148160]
MH025919.1_strain_N10            .....a...........a........c..................................................... [148160]
MH025918.1_strain_R25            .....a...........a........c..................................................... [148160]
MH025917.1_strain_R7             .....a...........a........c..................................................... [148160]
MH025916.1_strain_R8             .....a...........a........c..................................................... [148160]
```

```
FR682468.2_Georgia_2007/1        tatttgcttatacactcttgaacaaccagggtactggcgctttccatttaaaaaatagggcattaatctattaatattat [148240]
ASF_IND_20_CAD543                ................................................................................ [148240]
LR899193.1_Germany_2020/1        ................................................................................ [148240]
MN393477.1_Wuhan_2019            ................................................................................ [148240]
MW396979.1_Timor-Leste/2019/1    ................................................................................ [148240]
MK940252.1_CN/2019/InnerMongolia ................................................................................ [148240]
MK128995.1_China/2018/AnhuiXCGQ  ................................................................................ [148240]
MH681419.1_ASFV/POL/2015         ................................................................................ [148240]
LS478113.1_Estonia_2014          ................................................................................ [148240]
KP843857.1_Odintsovo_02/14       ................................................................................ [148240]
FN557520.1_strain_E75            ................................................................................ [148240]
AY261366.1_Warthog               ......t......................................g.................................. [148240]
AY261365.1_Warmbaths             ......t......................................g.................................. [148240]
AY261364.1_Tengani_62            ......t......................................................................... [148240]
AY261363.1_Pretoriuskop/96/4     ......t......................................................................... [148240]
AY261362.1_Mkuzi_1979            ................................................................................ [148240]
AY261361.1_Malawi_Lil-20/1_1983  ...c..t.................t.....a...................................c........g.... [148240]
AY261360.1_Kenya_1950            ...c..t.................t............................................g.....g.... [148240]
AM712240.1_OURT_88/3             ................................................................................ [148240]
AM712239.1_Benin_97/1            ................................................................................ [148240]
KM111295.1_Ken06.Bus             ...c..t.................t..................................................g.... [148240]
KM111294.1_Ken05/Tk1             ...c..t.................t.........................................c........g.... [148240]
KM262844.1_strain_L60            ................................................................................ [148240]
KM102979.1_26544/OG10_Italy      ................................................................................ [148240]
KM262845.1_strain_NHV            ................................................................................ [148240]
KX354450.1_47/Ss/2008            ................................................................................ [148240]
KP055815.1_strain_BA71           ................................................................................ [148240]
MH025920.1_strain_R35            ...c..t.................t..................................................g.... [148240]
MH025919.1_strain_N10            ...c..t.................t..................................................g.... [148240]
MH025918.1_strain_R25            ...c..t.................t..................................................g.... [148240]
MH025917.1_strain_R7             ...c..t.................t..................................................g.... [148240]
MH025916.1_strain_R8             ...c..t.................t..................................................g.... [148240]

FR682468.2_Georgia_2007/1        aatgttgcactgtttccgcaacttgcagcgttcgtgcaaaggaaatgggatagccaacctcgtccaggtgaaggtctgag [148320]
ASF_IND_20_CAD543                ................................................................................ [148320]
LR899193.1_Germany_2020/1        ................................................................................ [148320]
MN393477.1_Wuhan_2019            ................................................................................ [148320]
MW396979.1_Timor-Leste/2019/1    ................................................................................ [148320]
MK940252.1_CN/2019/InnerMongolia ................................................................................ [148320]
MK128995.1_China/2018/AnhuiXCGQ  ................................................................................ [148320]
MH681419.1_ASFV/POL/2015         ................................................................................ [148320]
LS478113.1_Estonia_2014          ................................................................................ [148320]
KP843857.1_Odintsovo_02/14       ................................................................................ [148320]
```

```
FN557520.1_strain_E75            ................................................................................ [148320]
AY261366.1_Warthog               ................................................................................ [148320]
AY261365.1_Warmbaths             ................................................................................ [148320]
AY261364.1_Tengani_62            ...................g....................................................a....... [148320]
AY261363.1_Pretoriuskop/96/4     ...................g............................................................ [148320]
AY261362.1_Mkuzi_1979            ................................................................................ [148320]
AY261361.1_Malawi_Lil-20/1_1983  ..........c..............................................................a...... [148320]
AY261360.1_Kenya_1950            ....c.....c....................a...............................................a [148320]
AM712240.1_OURT_88/3             ................................................................................ [148320]
AM712239.1_Benin_97/1            ................................................................................ [148320]
KM111295.1_Ken06.Bus             ....c.....c....................a...............................................a [148320]
KM111294.1_Ken05/Tk1             ....c.....c........................................................a............ [148320]
KM262844.1_strain_L60            ................................................................................ [148320]
KM102979.1_26544/OG10_Italy      ................................................................................ [148320]
KM262845.1_strain_NHV            ................................................................................ [148320]
KX354450.1_47/Ss/2008            ................................................................................ [148320]
KP055815.1_strain_BA71           ................................................................................ [148320]
MH025920.1_strain_R35            ....c.....c....................a...............................................a [148320]
MH025919.1_strain_N10            ....c.....c....................a...............................................a [148320]
MH025918.1_strain_R25            ....c.....c....................a...............................................a [148320]
MH025917.1_strain_R7             ....c.....c....................a...............................................a [148320]
MH025916.1_strain_R8             ....c.....c....................a...............................................a [148320]

FR682468.2_Georgia_2007/1        ttcccgcagatggtggaccggctgatcgaccatacctggctgcccagtagggatttacgaattcttccctccttgcgagg [148400]
ASF_IND_20_CAD543                ................................................................................ [148400]
LR899193.1_Germany_2020/1        ................................................................................ [148400]
MN393477.1_Wuhan_2019            ................................................................................ [148400]
MW396979.1_Timor-Leste/2019/1    ................................................................................ [148400]
MK940252.1_CN/2019/InnerMongolia ................................................................................ [148400]
MK128995.1_China/2018/AnhuiXCGQ  ................................................................................ [148400]
MH681419.1_ASFV/POL/2015         ................................................................................ [148400]
LS478113.1_Estonia_2014          ................................................................................ [148400]
KP843857.1_Odintsovo_02/14       ................................................................................ [148400]
FN557520.1_strain_E75            ................................................................................ [148400]
AY261366.1_Warthog               ................................................................................ [148400]
AY261365.1_Warmbaths             ................................................................................ [148400]
AY261364.1_Tengani_62            ........................................................g....................... [148400]
AY261363.1_Pretoriuskop/96/4     ................................................................................ [148400]
AY261362.1_Mkuzi_1979            ................................................................................ [148400]
AY261361.1_Malawi_Lil-20/1_1983  ..t............................................................................. [148400]
AY261360.1_Kenya_1950            ..t..............t......................................g....................... [148400]
AM712240.1_OURT_88/3             ................................................................................ [148400]
AM712239.1_Benin_97/1            ................................................................................ [148400]
KM111295.1_Ken06.Bus             ..t..............t......................................g....................... [148400]
```

```
KM111294.1_Ken05/Tk1               ..t.....................................................g.................a..... [148400]
KM262844.1_strain_L60              ................................................................................ [148400]
KM102979.1_26544/OG10_Italy        ................................................................................ [148400]
KM262845.1_strain_NHV              ................................................................................ [148400]
KX354450.1_47/Ss/2008              ................................................................................ [148400]
KP055815.1_strain_BA71             ................................................................................ [148400]
MH025920.1_strain_R35              ..t..............t......................................g....................... [148400]
MH025919.1_strain_N10              ..t..............t......................................g....................... [148400]
MH025918.1_strain_R25              ..t..............t......................................g....................... [148400]
MH025917.1_strain_R7               ..t..............t......................................g....................... [148400]
MH025916.1_strain_R8               ..t..............t......................................g....................... [148400]

FR682468.2_Georgia_2007/1          aagtcttcgcatgatggagggagcagggcgtgcccccatgacgatcccacgctttcccgtgcctccctgggttgcggtgg [148480]
ASF_IND_20_CAD543                  ................................................................................ [148480]
LR899193.1_Germany_2020/1          ................................................................................ [148480]
MN393477.1_Wuhan_2019              ................................................................................ [148480]
MW396979.1_Timor-Leste/2019/1      ................................................................................ [148480]
MK940252.1_CN/2019/InnerMongolia   ................................................................................ [148480]
MK128995.1_China/2018/AnhuiXCGQ    ................................................................................ [148480]
MH681419.1_ASFV/POL/2015           ................................................................................ [148480]
LS478113.1_Estonia_2014            ................................................................................ [148480]
KP843857.1_Odintsovo_02/14         ................................................................................ [148480]
FN557520.1_strain_E75              ...............a........g.....g................................................. [148480]
AY261366.1_Warthog                 ........................g..a..a..........................t...........a.......... [148480]
AY261365.1_Warmbaths               ........................g.....g..........................t...................... [148480]
AY261364.1_Tengani_62              ........................g.....g..........................t...................... [148480]
AY261363.1_Pretoriuskop/96/4       ...........................a..a..........................t...................... [148480]
AY261362.1_Mkuzi_1979              ...............a........g.....g................................................. [148480]
AY261361.1_Malawi_Lil-20/1_1983    ........................g.....g..........................t...................... [148480]
AY261360.1_Kenya_1950              ........................g.....g..........................t...................... [148480]
AM712240.1_OURT_88/3               ...............a........g.....g................................................. [148480]
AM712239.1_Benin_97/1              ...............a........g.....g................................................. [148480]
KM111295.1_Ken06.Bus               ........................g.....g..........................t...................... [148480]
KM111294.1_Ken05/Tk1               ..............................g..............t...........t...................... [148480]
KM262844.1_strain_L60              ...............a........g.....g................................................. [148480]
KM102979.1_26544/OG10_Italy        ...............a........g.....g................................................. [148480]
KM262845.1_strain_NHV              ...............a........g.....g................................................. [148480]
KX354450.1_47/Ss/2008              ...............a........g.....g................................................. [148480]
KP055815.1_strain_BA71             ...............a........g.....g................................................. [148480]
MH025920.1_strain_R35              ........................g.....g..........................t...................... [148480]
MH025919.1_strain_N10              ........................g.....g..........................t...................... [148480]
MH025918.1_strain_R25              ........................g.....g..........................t...................... [148480]
MH025917.1_strain_R7               ........................g.....g..........................t...................... [148480]
MH025916.1_strain_R8               ........................g.....g..........................t...................... [148480]
```

```
FR682468.2_Georgia_2007/1        tggaaacggaatccaacaaaaagttatagtaaagttgctgtatggtttgcaaattgcggtcaatatttaaaggtattttt [148560]
ASF_IND_20_CAD543                ................................................................................ [148560]
LR899193.1_Germany_2020/1        ................................................................................ [148560]
MN393477.1_Wuhan_2019            ................................................................................ [148560]
MW396979.1_Timor-Leste/2019/1    ................................................................................ [148560]
MK940252.1_CN/2019/InnerMongolia ................................................................................ [148560]
MK128995.1_China/2018/AnhuiXCGQ  ................................................................................ [148560]
MH681419.1_ASFV/POL/2015         ................................................................................ [148560]
LS478113.1_Estonia_2014          ................................................................................ [148560]
KP843857.1_Odintsovo_02/14       ................................................................................ [148560]
FN557520.1_strain_E75            ................................................................................ [148560]
AY261366.1_Warthog               ................................................................................ [148560]
AY261365.1_Warmbaths             .................................................t.............................. [148560]
AY261364.1_Tengani_62            ................................................................................ [148560]
AY261363.1_Pretoriuskop/96/4     ................................................................................ [148560]
AY261362.1_Mkuzi_1979            ................................................................................ [148560]
AY261361.1_Malawi_Lil-20/1_1983  ...............................................................c................ [148560]
AY261360.1_Kenya_1950            ...............g........................g......................c................ [148560]
AM712240.1_OURT_88/3             ................................................................................ [148560]
AM712239.1_Benin_97/1            ................................................................................ [148560]
KM111295.1_Ken06.Bus             ...............g...............................................c................ [148560]
KM111294.1_Ken05/Tk1             ...............g........................g......................c................ [148560]
KM262844.1_strain_L60            ................................................................................ [148560]
KM102979.1_26544/OG10_Italy      ................................................................................ [148560]
KM262845.1_strain_NHV            ................................................................................ [148560]
KX354450.1_47/Ss/2008            ................................................................................ [148560]
KP055815.1_strain_BA71           ................................................................................ [148560]
MH025920.1_strain_R35            ...............g...............................................c................ [148560]
MH025919.1_strain_N10            ...............g...............................................c................ [148560]
MH025918.1_strain_R25            ...............g...............................................c................ [148560]
MH025917.1_strain_R7             ...............g...............................................c................ [148560]
MH025916.1_strain_R8             ...............g...............................................c................ [148560]

FR682468.2_Georgia_2007/1        tggccgcgcacgatttgtaggtccttcgggatcagcagattctttcgaaccagatactgaatcacgttgttaatgtcgtg [148640]
ASF_IND_20_CAD543                ................................................................................ [148640]
LR899193.1_Germany_2020/1        ................................................................................ [148640]
MN393477.1_Wuhan_2019            ................................................................................ [148640]
MW396979.1_Timor-Leste/2019/1    ................................................................................ [148640]
MK940252.1_CN/2019/InnerMongolia ................................................................................ [148640]
MK128995.1_China/2018/AnhuiXCGQ  ................................................................................ [148640]
MH681419.1_ASFV/POL/2015         ................................................................................ [148640]
LS478113.1_Estonia_2014          ................................................................................ [148640]
KP843857.1_Odintsovo_02/14       ................................................................................ [148640]
```

```
FN557520.1_strain_E75              ...........................................................g........a........... [148640]
AY261366.1_Warthog                 .................g.......................t..........................a........... [148640]
AY261365.1_Warmbaths               .............................a......................................a........... [148640]
AY261364.1_Tengani_62              ................................................................................ [148640]
AY261363.1_Pretoriuskop/96/4       .................g...................a..............................a........... [148640]
AY261362.1_Mkuzi_1979              ...........................................................g........a........... [148640]
AY261361.1_Malawi_Lil-20/1_1983    ...........a........................................................a........... [148640]
AY261360.1_Kenya_1950              .........................................t........t.t...............a........... [148640]
AM712240.1_OURT_88/3               ...........................................................g........a........... [148640]
AM712239.1_Benin_97/1              ...........................................................g........a........... [148640]
KM111295.1_Ken06.Bus               .........................................t........t.t...............a........a.. [148640]
KM111294.1_Ken05/Tk1               .........................................t........t.t...............a........... [148640]
KM262844.1_strain_L60              ...........................................................g........a........... [148640]
KM102979.1_26544/OG10_Italy        ...........................................................g........a........... [148640]
KM262845.1_strain_NHV              ...........................................................g........a........... [148640]
KX354450.1_47/Ss/2008              ...........................................................g........a........... [148640]
KP055815.1_strain_BA71             ...........................................................g........a........... [148640]
MH025920.1_strain_R35              .........................................t........t.t...............a........a.. [148640]
MH025919.1_strain_N10              .........................................t........t.t...............a........a.. [148640]
MH025918.1_strain_R25              .........................................t........t.t...............a........a.. [148640]
MH025917.1_strain_R7               .........................................t........t.t...............a........a.. [148640]
MH025916.1_strain_R8               .........................................t........t.t...............a........a.. [148640]

FR682468.2_Georgia_2007/1          aaagctttgggggcctgacccgattcccaatctgatgccaggtcgtatgctgatggggggatctgaatggccttaagca [148720]
ASF_IND_20_CAD543                  ................................................................................ [148720]
LR899193.1_Germany_2020/1          ................................................................................ [148720]
MN393477.1_Wuhan_2019              ................................................................................ [148720]
MW396979.1_Timor-Leste/2019/1      ................................................................................ [148720]
MK940252.1_CN/2019/InnerMongolia   ................................................................................ [148720]
MK128995.1_China/2018/AnhuiXCGQ    ................................................................................ [148720]
MH681419.1_ASFV/POL/2015           ................................................................................ [148720]
LS478113.1_Estonia_2014            ................................................................................ [148720]
KP843857.1_Odintsovo_02/14         ................................................................................ [148720]
FN557520.1_strain_E75              ................................................................................ [148720]
AY261366.1_Warthog                 ..................t......................................a...................... [148720]
AY261365.1_Warmbaths               ..................t.................a........................................... [148720]
AY261364.1_Tengani_62              ................................................................................ [148720]
AY261363.1_Pretoriuskop/96/4       ................................................................................ [148720]
AY261362.1_Mkuzi_1979              ................................................................................ [148720]
AY261361.1_Malawi_Lil-20/1_1983    ....................................a.....g................................c.... [148720]
AY261360.1_Kenya_1950              ..........................................g................................c.... [148720]
AM712240.1_OURT_88/3               ................................................................................ [148720]
AM712239.1_Benin_97/1              ................................................................................ [148720]
KM111295.1_Ken06.Bus               .........c................................g........a.......................c.... [148720]
```

```
KM111294.1_Ken05/Tk1            ..........................................g................................c.... [148720]
KM262844.1_strain_L60           ................................................................................ [148720]
KM102979.1_26544/OG10_Italy     ................................................................................ [148720]
KM262845.1_strain_NHV           ................................................................................ [148720]
KX354450.1_47/Ss/2008           ................................................................................ [148720]
KP055815.1_strain_BA71          ................................................................................ [148720]
MH025920.1_strain_R35           .........c................................g........a.......................c.... [148720]
MH025919.1_strain_N10           .........c................................g........a.......................c.... [148720]
MH025918.1_strain_R25           .........c................................g........a.......................c.... [148720]
MH025917.1_strain_R7            .........c................................g........a.......................c.... [148720]
MH025916.1_strain_R8            .........c................................g........a.......................c.... [148720]

FR682468.2_Georgia_2007/1       caagtttttcgggatgggagtttttacttcgccccagttttacaacggtgtcgtaggttacgcgcgaaaaaatctctctg [148800]
ASF_IND_20_CAD543               ................................................................................ [148800]
LR899193.1_Germany_2020/1       ................................................................................ [148800]
MN393477.1_Wuhan_2019           ................................................................................ [148800]
MW396979.1_Timor-Leste/2019/1   ................................................................................ [148800]
MK940252.1_CN/2019/InnerMongolia ................................................................................ [148800]
MK128995.1_China/2018/AnhuiXCGQ ................................................................................ [148800]
MH681419.1_ASFV/POL/2015        ................................................................................ [148800]
LS478113.1_Estonia_2014         ................................................................................ [148800]
KP843857.1_Odintsovo_02/14      ................................................................................ [148800]
FN557520.1_strain_E75           ................................................................................ [148800]
AY261366.1_Warthog              ......................................................................g......... [148800]
AY261365.1_Warmbaths            ......................................................................g......... [148800]
AY261364.1_Tengani_62           .................................................a....................g......... [148800]
AY261363.1_Pretoriuskop/96/4    ......................................................................g......... [148800]
AY261362.1_Mkuzi_1979           ......................................................................g......... [148800]
AY261361.1_Malawi_Lil-20/1_1983 ..............................................a.......................g......... [148800]
AY261360.1_Kenya_1950           ...............................t................................t..............a [148800]
AM712240.1_OURT_88/3            ................................................................................ [148800]
AM712239.1_Benin_97/1           ................................................................................ [148800]
KM111295.1_Ken06.Bus            .............g..................................................t............... [148800]
KM111294.1_Ken05/Tk1            ...............................t................................t..............a [148800]
KM262844.1_strain_L60           ................................................................................ [148800]
KM102979.1_26544/OG10_Italy     ................................................................................ [148800]
KM262845.1_strain_NHV           ................................................................................ [148800]
KX354450.1_47/Ss/2008           ................................................................................ [148800]
KP055815.1_strain_BA71          ................................................................................ [148800]
MH025920.1_strain_R35           .............g..................................................t............... [148800]
MH025919.1_strain_N10           .............g..................................................t............... [148800]
MH025918.1_strain_R25           .............g..................................................t............... [148800]
MH025917.1_strain_R7            .............g..................................................t............... [148800]
MH025916.1_strain_R8            .............g..................................................t............... [148800]
```

```
FR682468.2_Georgia_2007/1        atgatctgcgggtacagtttgtcaatcttgccctgctgatccgcccaaaaggtaaaataatcttccgagtccttaacaat [148880]
ASF_IND_20_CAD543                ..........................................r..................................... [148880]
LR899193.1_Germany_2020/1        ................................................................................ [148880]
MN393477.1_Wuhan_2019            ................................................................................ [148880]
MW396979.1_Timor-Leste/2019/1    ................................................................................ [148880]
MK940252.1_CN/2019/InnerMongolia ................................................................................ [148880]
MK128995.1_China/2018/AnhuiXCGQ  ................................................................................ [148880]
MH681419.1_ASFV/POL/2015         ................................................................................ [148880]
LS478113.1_Estonia_2014          ................................................................................ [148880]
KP843857.1_Odintsovo_02/14       ................................................................................ [148880]
FN557520.1_strain_E75            ..........................................a..................................... [148880]
AY261366.1_Warthog               ..a..t....................................a.................................t... [148880]
AY261365.1_Warmbaths             ..a..t...................................ta.................................t... [148880]
AY261364.1_Tengani_62            ..a..t...................................ta.................................t... [148880]
AY261363.1_Pretoriuskop/96/4     ..a..t...................................ta.................................t... [148880]
AY261362.1_Mkuzi_1979            ..a..t...................................ta.................................t... [148880]
AY261361.1_Malawi_Lil-20/1_1983  ................a.........................a................g.....a..a.......t... [148880]
AY261360.1_Kenya_1950            ..............t..........................ta................g................t... [148880]
AM712240.1_OURT_88/3             ..........................................a..................................... [148880]
AM712239.1_Benin_97/1            ..........................................a..................................... [148880]
KM111295.1_Ken06.Bus             ..............t..........................ta................g................t... [148880]
KM111294.1_Ken05/Tk1             ..............t..........................ta................g................t... [148880]
KM262844.1_strain_L60            ..........................................a..................................... [148880]
KM102979.1_26544/OG10_Italy      ..........................................a..................................... [148880]
KM262845.1_strain_NHV            ..........................................a..................................... [148880]
KX354450.1_47/Ss/2008            ..........................................a..................................... [148880]
KP055815.1_strain_BA71           ..........................................a..................................... [148880]
MH025920.1_strain_R35            ..............t..........................ta................g................t... [148880]
MH025919.1_strain_N10            ..............t..........................ta................g................t... [148880]
MH025918.1_strain_R25            ..............t..........................ta................g................t... [148880]
MH025917.1_strain_R7             ..............t..........................ta................g................t... [148880]
MH025916.1_strain_R8             ..............t..........................ta................g................t... [148880]

FR682468.2_Georgia_2007/1        tttggggtgtactgccttacagacgtagcactgctttccttcggtttggcttgaagccgcttcaataagacgcttaggcc [148960]
ASF_IND_20_CAD543                ................................................................................ [148960]
LR899193.1_Germany_2020/1        ................................................................................ [148960]
MN393477.1_Wuhan_2019            ................................................................................ [148960]
MW396979.1_Timor-Leste/2019/1    ................................................................................ [148960]
MK940252.1_CN/2019/InnerMongolia ................................................................................ [148960]
MK128995.1_China/2018/AnhuiXCGQ  ................................................................................ [148960]
MH681419.1_ASFV/POL/2015         ................................................................................ [148960]
LS478113.1_Estonia_2014          ................................................................................ [148960]
KP843857.1_Odintsovo_02/14       ................................................................................ [148960]
```

```
FN557520.1_strain_E75             ................................................................................ [148960]
AY261366.1_Warthog                .............................................c.................................. [148960]
AY261365.1_Warmbaths              .........................................................g...................... [148960]
AY261364.1_Tengani_62             ..........................................a..c...........g...........c.......... [148960]
AY261363.1_Pretoriuskop/96/4      ..............t................................................................. [148960]
AY261362.1_Mkuzi_1979             .........................................................g...................... [148960]
AY261361.1_Malawi_Lil-20/1_1983   c..a..a....t.a..........a.........................c..............c.............. [148960]
AY261360.1_Kenya_1950             c..a..a....tga..................................................gc.............. [148960]
AM712240.1_OURT_88/3              ................................................................................ [148960]
AM712239.1_Benin_97/1             ................................................................................ [148960]
KM111295.1_Ken06.Bus              c..a..a....taa..................................................gc.............. [148960]
KM111294.1_Ken05/Tk1              c..a..a....tga..................................................gc...t.......... [148960]
KM262844.1_strain_L60             ................................................................................ [148960]
KM102979.1_26544/OG10_Italy       ................................................................................ [148960]
KM262845.1_strain_NHV             ................................................................................ [148960]
KX354450.1_47/Ss/2008             ................................................................................ [148960]
KP055815.1_strain_BA71            ................................................................................ [148960]
MH025920.1_strain_R35             c..a..a....taa..................................................gc.............. [148960]
MH025919.1_strain_N10             c..a..a....taa..................................................gc.............. [148960]
MH025918.1_strain_R25             c..a..a....taa..................................................gc.............. [148960]
MH025917.1_strain_R7              c..a..a....taa..................................................gc.............. [148960]
MH025916.1_strain_R8              c..a..a....taa..................................................gc.............. [148960]

FR682468.2_Georgia_2007/1         taataaggtgctcgtacctctttaggtcaacgatgggagccccgcagttgagacatataacccttaaccatcgtcgtatt [149040]
ASF_IND_20_CAD543                 ................................................................................ [149040]
LR899193.1_Germany_2020/1         ................................................................................ [149040]
MN393477.1_Wuhan_2019             ................................................................................ [149040]
MW396979.1_Timor-Leste/2019/1     ................................................................................ [149040]
MK940252.1_CN/2019/InnerMongolia  ................................................................................ [149040]
MK128995.1_China/2018/AnhuiXCGQ   ................................................................................ [149040]
MH681419.1_ASFV/POL/2015          ................................................................................ [149040]
LS478113.1_Estonia_2014           ................................................................................ [149040]
KP843857.1_Odintsovo_02/14        ................................................................................ [149040]
FN557520.1_strain_E75             .......a.......................a....................g........................... [149040]
AY261366.1_Warthog                ....................................................g........................... [149040]
AY261365.1_Warmbaths              ..........t.............a...........................g........................... [149040]
AY261364.1_Tengani_62             .............a..t..t................................g........................... [149040]
AY261363.1_Pretoriuskop/96/4      ....................................................g........................... [149040]
AY261362.1_Mkuzi_1979             .......a............................................g........................... [149040]
AY261361.1_Malawi_Lil-20/1_1983   ....................................................g........t.................. [149040]
AY261360.1_Kenya_1950             ............................t..a..............a.....g.............g............. [149040]
AM712240.1_OURT_88/3              .......a.......................a....................g........................... [149040]
AM712239.1_Benin_97/1             .......a.......................a....................g........................... [149040]
KM111295.1_Ken06.Bus              ...............................a..............a.....g.............g............. [149040]
```

```
KM111294.1_Ken05/Tk1             ............................t..a..............a.....g.............g............. [149040]
KM262844.1_strain_L60            .......a.......................a....................g........................... [149040]
KM102979.1_26544/OG10_Italy      .......a.......................a....................g........................... [149040]
KM262845.1_strain_NHV            .......a.......................a....................g........................... [149040]
KX354450.1_47/Ss/2008            .......a.......................a....................g........................... [149040]
KP055815.1_strain_BA71           .......a.......................a....................g........................... [149040]
MH025920.1_strain_R35            ...............................a..............a.....g.............g............. [149040]
MH025919.1_strain_N10            ...............................a..............a.....g.............g............. [149040]
MH025918.1_strain_R25            ...............................a..............a.....g.............g............. [149040]
MH025917.1_strain_R7             ...............................a..............a.....g.............g............. [149040]
MH025916.1_strain_R8             ...............................a..............a.....g.............g............. [149040]

FR682468.2_Georgia_2007/1        tcggcgatgaagagcggctgaagcaccggagcatgcatctgcagtatcccagggtgtcccatacattgcttgcgctggtg [149120]
ASF_IND_20_CAD543                ................................................................................ [149120]
LR899193.1_Germany_2020/1        ................................................................................ [149120]
MN393477.1_Wuhan_2019            ................................................................................ [149120]
MW396979.1_Timor-Leste/2019/1    ................................................................................ [149120]
MK940252.1_CN/2019/InnerMongolia ................................................................................ [149120]
MK128995.1_China/2018/AnhuiXCGQ  ................................................................................ [149120]
MH681419.1_ASFV/POL/2015         ................................................................................ [149120]
LS478113.1_Estonia_2014          ................................................................................ [149120]
KP843857.1_Odintsovo_02/14       ................................................................................ [149120]
FN557520.1_strain_E75            ................................................................................ [149120]
AY261366.1_Warthog               ..........................t...................................g................. [149120]
AY261365.1_Warmbaths             ................................................................................ [149120]
AY261364.1_Tengani_62            ................................................................................ [149120]
AY261363.1_Pretoriuskop/96/4     ................................................................................ [149120]
AY261362.1_Mkuzi_1979            ................................................................................ [149120]
AY261361.1_Malawi_Lil-20/1_1983  .................t.....a.......................t..t............................. [149120]
AY261360.1_Kenya_1950            .......................a..........................t.....c.....g..c.............. [149120]
AM712240.1_OURT_88/3             ................................................................................ [149120]
AM712239.1_Benin_97/1            ................................................................................ [149120]
KM111295.1_Ken06.Bus             .......................a..........................t.....c.....g..c.............. [149120]
KM111294.1_Ken05/Tk1             .......................a..........................t.....c.....g..c.............. [149120]
KM262844.1_strain_L60            ................................................................................ [149120]
KM102979.1_26544/OG10_Italy      ................................................................................ [149120]
KM262845.1_strain_NHV            ................................................................................ [149120]
KX354450.1_47/Ss/2008            ................................................................................ [149120]
KP055815.1_strain_BA71           ................................................................................ [149120]
MH025920.1_strain_R35            .......................a..........................t.....c.....g..c.............. [149120]
MH025919.1_strain_N10            .......................a..........................t.....c.....g..c.............. [149120]
MH025918.1_strain_R25            .......................a..........................t.....c.....g..c.............. [149120]
MH025917.1_strain_R7             .......................a..........................t.....c.....g..c.............. [149120]
MH025916.1_strain_R8             .......................a..........................t.....c.....g..c.............. [149120]
```

```
FR682468.2_Georgia_2007/1        tgagcaagtgatgcatttataatggtgatcggtggttcccattcgcgcatcatagatacccccttcggcgggaagggtgc [149200]
ASF_IND_20_CAD543                ................................................................................ [149200]
LR899193.1_Germany_2020/1        ................................................................................ [149200]
MN393477.1_Wuhan_2019            ................................................................................ [149200]
MW396979.1_Timor-Leste/2019/1    ................................................................................ [149200]
MK940252.1_CN/2019/InnerMongolia ................................................................................ [149200]
MK128995.1_China/2018/AnhuiXCGQ  ................................................................................ [149200]
MH681419.1_ASFV/POL/2015         ................................................................................ [149200]
LS478113.1_Estonia_2014          ................................................................................ [149200]
KP843857.1_Odintsovo_02/14       ................................................................................ [149200]
FN557520.1_strain_E75            ...............................................................c................ [149200]
AY261366.1_Warthog               ........................a................................g.....c................ [149200]
AY261365.1_Warmbaths             ........................a................................g.....c................ [149200]
AY261364.1_Tengani_62            ........................a....................t...........g.....c................ [149200]
AY261363.1_Pretoriuskop/96/4     ........................a................................g.....c................ [149200]
AY261362.1_Mkuzi_1979            ........................a......................................c................ [149200]
AY261361.1_Malawi_Lil-20/1_1983  ......g...............................................a........c........t.....a. [149200]
AY261360.1_Kenya_1950            ......g......................................t.................c.....a..t.....a. [149200]
AM712240.1_OURT_88/3             ...............................................................c................ [149200]
AM712239.1_Benin_97/1            ...............................................................c................ [149200]
KM111295.1_Ken06.Bus             ......g......................................t..g.....a........c........t.....a. [149200]
KM111294.1_Ken05/Tk1             ......g......................................t.................c.....a..t.....a. [149200]
KM262844.1_strain_L60            ...............................................................c................ [149200]
KM102979.1_26544/OG10_Italy      ...............................................................c................ [149200]
KM262845.1_strain_NHV            ...............................................................c................ [149200]
KX354450.1_47/Ss/2008            ...............................................................c................ [149200]
KP055815.1_strain_BA71           ...............................................................c................ [149200]
MH025920.1_strain_R35            ......g......................................t..g.....a........c........t.....a. [149200]
MH025919.1_strain_N10            ......g......................................t..g.....a........c........t.....a. [149200]
MH025918.1_strain_R25            ......g......................................t..g.....a........c........t.....a. [149200]
MH025917.1_strain_R7             ......g......................................t..g.....a........c........t.....a. [149200]
MH025916.1_strain_R8             ......g......................................t..g.....a........c........t.....a. [149200]

FR682468.2_Georgia_2007/1        cctcaaataaattagaaatggtaacctccataacgccttgcctcttatgatcattgtcaccggcaatattgaactgaacg [149280]
ASF_IND_20_CAD543                ................................................................................ [149280]
LR899193.1_Germany_2020/1        ................................................................................ [149280]
MN393477.1_Wuhan_2019            ................................................................................ [149280]
MW396979.1_Timor-Leste/2019/1    ................................................................................ [149280]
MK940252.1_CN/2019/InnerMongolia ................................................................................ [149280]
MK128995.1_China/2018/AnhuiXCGQ  ................................................................................ [149280]
MH681419.1_ASFV/POL/2015         ................................................................................ [149280]
LS478113.1_Estonia_2014          ................................................................................ [149280]
KP843857.1_Odintsovo_02/14       ................................................................................ [149280]
```

```
FN557520.1_strain_E75            .................................................g.............................. [149280]
AY261366.1_Warthog               .............g..............................................................t... [149280]
AY261365.1_Warmbaths             .............g..............................................................t... [149280]
AY261364.1_Tengani_62            .............g.................................................................. [149280]
AY261363.1_Pretoriuskop/96/4     .............g..............................................................t... [149280]
AY261362.1_Mkuzi_1979            .................................................g.............................. [149280]
AY261361.1_Malawi_Lil-20/1_1983  ..........g..g..g..........................t.....g..........................t... [149280]
AY261360.1_Kenya_1950            .............g.............................t................................t... [149280]
AM712240.1_OURT_88/3             .................................................g.............................. [149280]
AM712239.1_Benin_97/1            .................................................g.............................. [149280]
KM111295.1_Ken06.Bus             .............g.............................t................................t... [149280]
KM111294.1_Ken05/Tk1             .............g.............................t................................t... [149280]
KM262844.1_strain_L60            .................................................g.............................. [149280]
KM102979.1_26544/OG10_Italy      .................................................g.............................. [149280]
KM262845.1_strain_NHV            .................................................g.............................. [149280]
KX354450.1_47/Ss/2008            .................................................g.............................. [149280]
KP055815.1_strain_BA71           .................................................g.............................. [149280]
MH025920.1_strain_R35            .............g.............................t................................t... [149280]
MH025919.1_strain_N10            .............g.............................t.................a..............t... [149280]
MH025918.1_strain_R25            .............g.............................t................................t... [149280]
MH025917.1_strain_R7             .............g.............................t................................t... [149280]
MH025916.1_strain_R8             .............g.............................t................................t... [149280]

FR682468.2_Georgia_2007/1        gcggctatttcggcatatccagcctccatatttttgctaaatacataataaaacttcaaatgtt-aaaaaaaaataaca- [149360]
ASF_IND_20_CAD543                ................................................................a..............- [149360]
LR899193.1_Germany_2020/1        ................................................................-..............- [149360]
MN393477.1_Wuhan_2019            ................................................................-..............- [149360]
MW396979.1_Timor-Leste/2019/1    ................................................................-..............- [149360]
MK940252.1_CN/2019/InnerMongolia ................................................................-..............- [149360]
MK128995.1_China/2018/AnhuiXCGQ  ................................................................-..............- [149360]
MH681419.1_ASFV/POL/2015         ................................................................-..............- [149360]
LS478113.1_Estonia_2014          ................................................................-..............- [149360]
KP843857.1_Odintsovo_02/14       ................................................................--.............- [149360]
FN557520.1_strain_E75            ................................................................a..............- [149360]
AY261366.1_Warthog               ..a........a....................................................a..............- [149360]
AY261365.1_Warmbaths             ..a.............................................................a..............- [149360]
AY261364.1_Tengani_62            ................................................................--.............- [149360]
AY261363.1_Pretoriuskop/96/4     ..a.............................................................a.........g....a [149360]
AY261362.1_Mkuzi_1979            ................................................................a..............- [149360]
AY261361.1_Malawi_Lil-20/1_1983  ....................g............c..............................a.............c- [149360]
AY261360.1_Kenya_1950            ..............g..................c..............................a.............t- [149360]
AM712240.1_OURT_88/3             ................................................................a..............- [149360]
AM712239.1_Benin_97/1            ................................................................a..............- [149360]
KM111295.1_Ken06.Bus             ..............g..................c..............................a.............t- [149360]
```

```
KM111294.1_Ken05/Tk1             ..............g..................c..............................-.............t- [149360]
KM262844.1_strain_L60            ................................................................a..............- [149360]
KM102979.1_26544/OG10_Italy      ................................................................a..............- [149360]
KM262845.1_strain_NHV            ................................................................a..............- [149360]
KX354450.1_47/Ss/2008            ................................................................a..............- [149360]
KP055815.1_strain_BA71           ................................................................a..............- [149360]
MH025920.1_strain_R35            ..............g..................c..............................a.............t- [149360]
MH025919.1_strain_N10            ..............g..................c..............................a.............t- [149360]
MH025918.1_strain_R25            ..............g..................c..............................a.............t- [149360]
MH025917.1_strain_R7             ..............g..................c..............................a.............t- [149360]
MH025916.1_strain_R8             ..............g..................c..............................a.............t- [149360]

FR682468.2_Georgia_2007/1        -----------tcggttggcata--tttttttgttaaaaccaagtgttaaatgatttctaaaacatttatcggttcacga [149440]
ASF_IND_20_CAD543                -----------............---.............y...r.....................y.............. [149440]
LR899193.1_Germany_2020/1        -----------............--........................................c.............. [149440]
MN393477.1_Wuhan_2019            -----------............--........................................c.............. [149440]
MW396979.1_Timor-Leste/2019/1    -----------............--........................................c.............. [149440]
MK940252.1_CN/2019/InnerMongolia -----------............--........................................c.............. [149440]
MK128995.1_China/2018/AnhuiXCGQ  -----------............--........................................c.............. [149440]
MH681419.1_ASFV/POL/2015         -----------............--....................................................... [149440]
LS478113.1_Estonia_2014          -----------............--........................................c.............. [149440]
KP843857.1_Odintsovo_02/14       -----------............---.......................................c.............. [149440]
FN557520.1_strain_E75            -----------............---.............t...a.................................... [149440]
AY261366.1_Warthog               -attctcttca...ag.......ta..................a....................g............... [149440]
AY261365.1_Warmbaths             -attctctaca............ta..................a....................g............... [149440]
AY261364.1_Tengani_62            -----------...ag....t..--..................a....................g.....t......... [149440]
AY261363.1_Pretoriuskop/96/4     tattctctaca....g.......t-..................a....................g............... [149440]
AY261362.1_Mkuzi_1979            -----------............---.............t...a.................................... [149440]
AY261361.1_Malawi_Lil-20/1_1983  -attctctaacc..a........ta............----..a........a...........g..............g [149440]
AY261360.1_Kenya_1950            -attctccac----------------------------.t..gt........a...........g.....t......... [149440]
AM712240.1_OURT_88/3             -----------............---.............t...a.................................... [149440]
AM712239.1_Benin_97/1            -----------............---.............t...a.................................... [149440]
KM111295.1_Ken06.Bus             -attctccgc----------------------------.t..gt........a........................... [149440]
KM111294.1_Ken05/Tk1             -attctccac----------------------------.t..gt........a...........g............... [149440]
KM262844.1_strain_L60            -----------............---.............t...a.................................... [149440]
KM102979.1_26544/OG10_Italy      -----------............---.............t...a.................................... [149440]
KM262845.1_strain_NHV            -----------............---.............t...a.................................... [149440]
KX354450.1_47/Ss/2008            -----------............---.............t...a.................................... [149440]
KP055815.1_strain_BA71           -----------............---.............t...a.................................... [149440]
MH025920.1_strain_R35            -attctccgc----------------------------.t..gt........a........................... [149440]
MH025919.1_strain_N10            -attctccgc----------------------------.t..gt........a........................... [149440]
MH025918.1_strain_R25            -attctccgc----------------------------.t..gt........a........................... [149440]
MH025917.1_strain_R7             -attctccgc----------------------------.t..gt........a........................... [149440]
MH025916.1_strain_R8             -attctccgc----------------------------.t..gt........a........................... [149440]
```

```
FR682468.2_Georgia_2007/1        aaacctaccgcacgggcctgaagaggaatgccagttttgggggaaagctcggcatattccacggtaagctcttttccata [149520]
ASF_IND_20_CAD543                ................................................................................ [149520]
LR899193.1_Germany_2020/1        ................................................................................ [149520]
MN393477.1_Wuhan_2019            ................................................................................ [149520]
MW396979.1_Timor-Leste/2019/1    ................................................................................ [149520]
MK940252.1_CN/2019/InnerMongolia ................................................................................ [149520]
MK128995.1_China/2018/AnhuiXCGQ  ................................................................................ [149520]
MH681419.1_ASFV/POL/2015         ................................................................................ [149520]
LS478113.1_Estonia_2014          ................................................................................ [149520]
KP843857.1_Odintsovo_02/14       ................................................................................ [149520]
FN557520.1_strain_E75            ................................................................................ [149520]
AY261366.1_Warthog               .......................c........g.............................a........c........ [149520]
AY261365.1_Warmbaths             .......................c.................a....................a........c........ [149520]
AY261364.1_Tengani_62            .......................g......................................a........c........ [149520]
AY261363.1_Pretoriuskop/96/4     .......................c......................................a........c........ [149520]
AY261362.1_Mkuzi_1979            ................................................................................ [149520]
AY261361.1_Malawi_Lil-20/1_1983  ...................................c.....a....................a........c........ [149520]
AY261360.1_Kenya_1950            ................................g.............................a........c........ [149520]
AM712240.1_OURT_88/3             ................................................................................ [149520]
AM712239.1_Benin_97/1            ................................................................................ [149520]
KM111295.1_Ken06.Bus             ................................g.......a.....................a........c........ [149520]
KM111294.1_Ken05/Tk1             .................t..............g.....a.......................a........c........ [149520]
KM262844.1_strain_L60            ................................................................................ [149520]
KM102979.1_26544/OG10_Italy      ................................................................................ [149520]
KM262845.1_strain_NHV            ................................................................................ [149520]
KX354450.1_47/Ss/2008            ................................................................................ [149520]
KP055815.1_strain_BA71           ................................................................................ [149520]
MH025920.1_strain_R35            ................................g.......a.....................a........c........ [149520]
MH025919.1_strain_N10            ................................g.......a.....................a........c........ [149520]
MH025918.1_strain_R25            ................................g.......a.....................a........c........ [149520]
MH025917.1_strain_R7             ................................g.......a.....................a........c........ [149520]
MH025916.1_strain_R8             ................................g.......a.....................a........c........ [149520]

FR682468.2_Georgia_2007/1        aagatgtttttaaataaggcgggcgtgagtttttgaaaaagagcataacgatccgcgtacgtcaaatgcttaggagtga [149600]
ASF_IND_20_CAD543                ................................................................................ [149600]
LR899193.1_Germany_2020/1        ................................................................................ [149600]
MN393477.1_Wuhan_2019            ................................................................................ [149600]
MW396979.1_Timor-Leste/2019/1    ................................................................................ [149600]
MK940252.1_CN/2019/InnerMongolia ................................................................................ [149600]
MK128995.1_China/2018/AnhuiXCGQ  ................................................................................ [149600]
MH681419.1_ASFV/POL/2015         ................................................................................ [149600]
LS478113.1_Estonia_2014          ................................................................................ [149600]
KP843857.1_Odintsovo_02/14       ................................................................................ [149600]
```

```
FN557520.1_strain_E75             ..............................................................................c. [149600]
AY261366.1_Warthog                ......................................................a.......................c. [149600]
AY261365.1_Warmbaths              ......................................................a.......................c. [149600]
AY261364.1_Tengani_62             .................................a....................a..a....................c. [149600]
AY261363.1_Pretoriuskop/96/4      ......................................................a.......................c. [149600]
AY261362.1_Mkuzi_1979             ..............................................................................c. [149600]
AY261361.1_Malawi_Lil-20/1_1983   ......................................................a..............t........c. [149600]
AY261360.1_Kenya_1950             .....................a........................................................c. [149600]
AM712240.1_OURT_88/3              ..............................................................................c. [149600]
AM712239.1_Benin_97/1             ..............................................................................c. [149600]
KM111295.1_Ken06.Bus              .....................a........................................................c. [149600]
KM111294.1_Ken05/Tk1              .....................a........................................................c. [149600]
KM262844.1_strain_L60             ..............................................................................c. [149600]
KM102979.1_26544/OG10_Italy       ..............................................................................c. [149600]
KM262845.1_strain_NHV             ..............................................................................c. [149600]
KX354450.1_47/Ss/2008             ..............................................................................c. [149600]
KP055815.1_strain_BA71            ..............................................................................c. [149600]
MH025920.1_strain_R35             .....................a........................................................c. [149600]
MH025919.1_strain_N10             .....................a........................................................c. [149600]
MH025918.1_strain_R25             .....................a........................................................c. [149600]
MH025917.1_strain_R7              .....................a........................................................c. [149600]
MH025916.1_strain_R8              .....................a........................................................c. [149600]

FR682468.2_Georgia_2007/1         ctacaaaccgctttttgtttggcaattcgcaaacccataaaatggcgcctaagtcctttcccttttttccctgagtatag [149680]
ASF_IND_20_CAD543                 ................................................................................ [149680]
LR899193.1_Germany_2020/1         ................................................................................ [149680]
MN393477.1_Wuhan_2019             ................................................................................ [149680]
MW396979.1_Timor-Leste/2019/1     ................................................................................ [149680]
MK940252.1_CN/2019/InnerMongolia  ................................................................................ [149680]
MK128995.1_China/2018/AnhuiXCGQ   ................................................................................ [149680]
MH681419.1_ASFV/POL/2015          ................................................................................ [149680]
LS478113.1_Estonia_2014           ................................................................................ [149680]
KP843857.1_Odintsovo_02/14        ................................................................................ [149680]
FN557520.1_strain_E75             ........................g...a................................................... [149680]
AY261366.1_Warthog                .c......................g........................c.............................. [149680]
AY261365.1_Warmbaths              .c......................g....................................................... [149680]
AY261364.1_Tengani_62             .c......................g....................................................... [149680]
AY261363.1_Pretoriuskop/96/4      .c......................g........................c.............................. [149680]
AY261362.1_Mkuzi_1979             ........................g...a................................................... [149680]
AY261361.1_Malawi_Lil-20/1_1983   ......................t.g...a..............a..............c..................... [149680]
AY261360.1_Kenya_1950             .......a................g.................................c..................... [149680]
AM712240.1_OURT_88/3              ........................g...a................................................... [149680]
AM712239.1_Benin_97/1             ........................g...a................................................... [149680]
KM111295.1_Ken06.Bus              .......a................g.................................c..................... [149680]
```

```
KM111294.1_Ken05/Tk1             .......a................g.................................c..................... [149680]
KM262844.1_strain_L60            ........................g...a................................................... [149680]
KM102979.1_26544/OG10_Italy      ........................g...a................................................... [149680]
KM262845.1_strain_NHV            ........................g...a................................................... [149680]
KX354450.1_47/Ss/2008            ........................g...a................................................... [149680]
KP055815.1_strain_BA71           ........................g...a................................................... [149680]
MH025920.1_strain_R35            .......a................g.................................c..................... [149680]
MH025919.1_strain_N10            .......a................g.................................c..................... [149680]
MH025918.1_strain_R25            .......a................g.................................c..................... [149680]
MH025917.1_strain_R7             .......a................g.................................c..................... [149680]
MH025916.1_strain_R8             .......a................g.................................c..................... [149680]

FR682468.2_Georgia_2007/1        tccactaaaataaattcagcgtctagcagcggtttcagcttggcaagatgcgctgagtggtagttgttgtatcccggctc [149760]
ASF_IND_20_CAD543                ................................................................................ [149760]
LR899193.1_Germany_2020/1        ................................................................................ [149760]
MN393477.1_Wuhan_2019            ................................................................................ [149760]
MW396979.1_Timor-Leste/2019/1    ................................................................................ [149760]
MK940252.1_CN/2019/InnerMongolia ................................................................................ [149760]
MK128995.1_China/2018/AnhuiXCGQ  ................................................................................ [149760]
MH681419.1_ASFV/POL/2015         ................................................................................ [149760]
LS478113.1_Estonia_2014          ................................................................................ [149760]
KP843857.1_Odintsovo_02/14       ................................................................................ [149760]
FN557520.1_strain_E75            .....c.......................................................................... [149760]
AY261366.1_Warthog               .....c.......................................................................... [149760]
AY261365.1_Warmbaths             ................................................................................ [149760]
AY261364.1_Tengani_62            .....c.......................................................................... [149760]
AY261363.1_Pretoriuskop/96/4     .....c.......................................................................... [149760]
AY261362.1_Mkuzi_1979            .....c.......................................................................... [149760]
AY261361.1_Malawi_Lil-20/1_1983  ..t..c..............a.......a.....................g.g........................t.. [149760]
AY261360.1_Kenya_1950            .....c..............a........t....................g.g...a..a.................... [149760]
AM712240.1_OURT_88/3             .....c.......................................................................... [149760]
AM712239.1_Benin_97/1            .....c.......................................................................... [149760]
KM111295.1_Ken06.Bus             .....c..............a........t....................g.g...a..a.................... [149760]
KM111294.1_Ken05/Tk1             .....c..............a........t....................g.g...a..a.................... [149760]
KM262844.1_strain_L60            .....c.......................................................................... [149760]
KM102979.1_26544/OG10_Italy      .....c.......................................................................... [149760]
KM262845.1_strain_NHV            .....c.......................................................................... [149760]
KX354450.1_47/Ss/2008            .....c.......................................................................... [149760]
KP055815.1_strain_BA71           .....c.......................................................................... [149760]
MH025920.1_strain_R35            .....c..............a........t....................g.g...a..a.................... [149760]
MH025919.1_strain_N10            .....c..............a........t....................g.g...a..a.................... [149760]
MH025918.1_strain_R25            .....c..............a........t....................g.g...a..a.................... [149760]
MH025917.1_strain_R7             .....c..............a........t....................g.g...a..a.................... [149760]
MH025916.1_strain_R8             .....c..............a........t....................g.g...a..a.................... [149760]
```

```
FR682468.2_Georgia_2007/1        atagggcccattggcattgcgtacgatggctccctcgtagccctccttaataaactgcgccttaagcctaagggcctcat [149840]
ASF_IND_20_CAD543                ................................................................................ [149840]
LR899193.1_Germany_2020/1        ................................................................................ [149840]
MN393477.1_Wuhan_2019            ................................................................................ [149840]
MW396979.1_Timor-Leste/2019/1    ................................................................................ [149840]
MK940252.1_CN/2019/InnerMongolia ................................................................................ [149840]
MK128995.1_China/2018/AnhuiXCGQ  ................................................................................ [149840]
MH681419.1_ASFV/POL/2015         ................................................................................ [149840]
LS478113.1_Estonia_2014          ................................................................................ [149840]
KP843857.1_Odintsovo_02/14       ................................................................................ [149840]
FN557520.1_strain_E75            ................................................................................ [149840]
AY261366.1_Warthog               ................................................................................ [149840]
AY261365.1_Warmbaths             ................................................................................ [149840]
AY261364.1_Tengani_62            ................................................................................ [149840]
AY261363.1_Pretoriuskop/96/4     ............aa.................................................................. [149840]
AY261362.1_Mkuzi_1979            ................................................................................ [149840]
AY261361.1_Malawi_Lil-20/1_1983  .........................................................t.t......t............. [149840]
AY261360.1_Kenya_1950            ...........................................................tt..c................ [149840]
AM712240.1_OURT_88/3             ................................................................................ [149840]
AM712239.1_Benin_97/1            ................................................................................ [149840]
KM111295.1_Ken06.Bus             ...........................................................tt................... [149840]
KM111294.1_Ken05/Tk1             ...........................................................tt................... [149840]
KM262844.1_strain_L60            ................................................................................ [149840]
KM102979.1_26544/OG10_Italy      ................................................................................ [149840]
KM262845.1_strain_NHV            ................................................................................ [149840]
KX354450.1_47/Ss/2008            ................................................................................ [149840]
KP055815.1_strain_BA71           ................................................................................ [149840]
MH025920.1_strain_R35            ...........................................................tt................... [149840]
MH025919.1_strain_N10            ...........................................................tt................... [149840]
MH025918.1_strain_R25            ...........................................................tt................... [149840]
MH025917.1_strain_R7             ...........................................................tt................... [149840]
MH025916.1_strain_R8             ...........................................................tt................... [149840]

FR682468.2_Georgia_2007/1        ccacattcttcacgctaaaattttcaacttggtggataaaggtaagatcttccttctgtttaaaaatatttgttaatagc [149920]
ASF_IND_20_CAD543                ............................................................................y... [149920]
LR899193.1_Germany_2020/1        ................................................................................ [149920]
MN393477.1_Wuhan_2019            ................................................................................ [149920]
MW396979.1_Timor-Leste/2019/1    ................................................................................ [149920]
MK940252.1_CN/2019/InnerMongolia ................................................................................ [149920]
MK128995.1_China/2018/AnhuiXCGQ  ................................................................................ [149920]
MH681419.1_ASFV/POL/2015         ................................................................................ [149920]
LS478113.1_Estonia_2014          ................................................................................ [149920]
KP843857.1_Odintsovo_02/14       ................................................................................ [149920]
```

```
FN557520.1_strain_E75            ............................................................................c... [149920]
AY261366.1_Warthog               ..t....t....................................................................c... [149920]
AY261365.1_Warmbaths             ..t................................................g........................c... [149920]
AY261364.1_Tengani_62            ..t.........................................................................c... [149920]
AY261363.1_Pretoriuskop/96/4     ..t.....................................a...................................c... [149920]
AY261362.1_Mkuzi_1979            ............................................................................c... [149920]
AY261361.1_Malawi_Lil-20/1_1983  ..t.........................................................................g... [149920]
AY261360.1_Kenya_1950            ..t...c...a.t............................a..........t.......................c..t [149920]
AM712240.1_OURT_88/3             ............................................................................c... [149920]
AM712239.1_Benin_97/1            ............................................................................c... [149920]
KM111295.1_Ken06.Bus             ..t...c...a.t..................a.............a......t..t....................c..t [149920]
KM111294.1_Ken05/Tk1             ..t...c...a.t............................a..........t.......................c..t [149920]
KM262844.1_strain_L60            ............................................................................c... [149920]
KM102979.1_26544/OG10_Italy      ............................................................................c... [149920]
KM262845.1_strain_NHV            ............................................................................c... [149920]
KX354450.1_47/Ss/2008            ............................................................................c... [149920]
KP055815.1_strain_BA71           ............................................................................c... [149920]
MH025920.1_strain_R35            ..t...c...a.t..................a.............a......t..t....................c..t [149920]
MH025919.1_strain_N10            ..t...c...a.t..................a.............a......t..t....................c..t [149920]
MH025918.1_strain_R25            ..t...c...a.t..................a.............a......t..t....................c..t [149920]
MH025917.1_strain_R7             ..t...c...a.t..................a.............a......t..t....................c..t [149920]
MH025916.1_strain_R8             ..t...c...a.t..................a.............a......t..t....................c..t [149920]

FR682468.2_Georgia_2007/1        tgttgtctcttgttggaaggcatttgaagctgatcactccaaaaacagtcaaacacgtaaaagtgcagctcggaggaatc [150000]
ASF_IND_20_CAD543                ..y.........................................r................................... [150000]
LR899193.1_Germany_2020/1        ................................................................................ [150000]
MN393477.1_Wuhan_2019            ................................................................................ [150000]
MW396979.1_Timor-Leste/2019/1    ................................................................................ [150000]
MK940252.1_CN/2019/InnerMongolia ................................................................................ [150000]
MK128995.1_China/2018/AnhuiXCGQ  ................................................................................ [150000]
MH681419.1_ASFV/POL/2015         ................................................................................ [150000]
LS478113.1_Estonia_2014          ................................................................................ [150000]
KP843857.1_Odintsovo_02/14       ................................................................................ [150000]
FN557520.1_strain_E75            ..c.........................................g................................... [150000]
AY261366.1_Warthog               ..c.........................................g................................... [150000]
AY261365.1_Warmbaths             ..c.........................................g................................... [150000]
AY261364.1_Tengani_62            ............................................g................................... [150000]
AY261363.1_Pretoriuskop/96/4     ..c.........................................g................................... [150000]
AY261362.1_Mkuzi_1979            ..c.........................................g................................... [150000]
AY261361.1_Malawi_Lil-20/1_1983  ..c.........................................g................................... [150000]
AY261360.1_Kenya_1950            ..............a.............................g........t.............a............ [150000]
AM712240.1_OURT_88/3             ..c.........................................g................................... [150000]
AM712239.1_Benin_97/1            ..c.........................................g................................... [150000]
KM111295.1_Ken06.Bus             ..............a.............................g................................... [150000]
```

```
KM111294.1_Ken05/Tk1            ..............a.............................g........t.............a............ [150000]
KM262844.1_strain_L60           ..c.........................................g................................... [150000]
KM102979.1_26544/OG10_Italy     ..c.........................................g................................... [150000]
KM262845.1_strain_NHV           ..c.........................................g................................... [150000]
KX354450.1_47/Ss/2008           ..c.........................................g................................... [150000]
KP055815.1_strain_BA71          ..c.........................................g................................... [150000]
MH025920.1_strain_R35           ..............a.............................g................................... [150000]
MH025919.1_strain_N10           ..............a.............................g................................... [150000]
MH025918.1_strain_R25           ..............a.............................g................................... [150000]
MH025917.1_strain_R7            ..............a.............................g................................... [150000]
MH025916.1_strain_R8            ..............a.............................g................................... [150000]

FR682468.2_Georgia_2007/1       tgtcttcgcattcgcctgccccgcgatccattgcagaggtttgcggtgtaaataaagctcaccatccaaatatactctca [150080]
ASF_IND_20_CAD543               ................................................................................ [150080]
LR899193.1_Germany_2020/1       ................................................................................ [150080]
MN393477.1_Wuhan_2019           ................................................................................ [150080]
MW396979.1_Timor-Leste/2019/1   ................................................................................ [150080]
MK940252.1_CN/2019/InnerMongolia ................................................................................ [150080]
MK128995.1_China/2018/AnhuiXCGQ ................................................................................ [150080]
MH681419.1_ASFV/POL/2015        ................................................................................ [150080]
LS478113.1_Estonia_2014         ................................................................................ [150080]
KP843857.1_Odintsovo_02/14      ................................................................................ [150080]
FN557520.1_strain_E75           ............................................................g................... [150080]
AY261366.1_Warthog              ............................................................g................... [150080]
AY261365.1_Warmbaths            .........g..................................................g................... [150080]
AY261364.1_Tengani_62           .........g..g...............................................g................... [150080]
AY261363.1_Pretoriuskop/96/4    .........g........t..t......................................g................... [150080]
AY261362.1_Mkuzi_1979           .....................t......................................g................... [150080]
AY261361.1_Malawi_Lil-20/1_1983 ..c...t..........................t.a............c...........g................... [150080]
AY261360.1_Kenya_1950           .ac...t..g........t.......................at....c...........g..............c.... [150080]
AM712240.1_OURT_88/3            ............................................................g................... [150080]
AM712239.1_Benin_97/1           ............................................................g................... [150080]
KM111295.1_Ken06.Bus            ..c...t..g..g.....t.......................at....c...........g..............c.... [150080]
KM111294.1_Ken05/Tk1            ..c...t..g........t.......................at....c...........g..............c.... [150080]
KM262844.1_strain_L60           ............................................................g................... [150080]
KM102979.1_26544/OG10_Italy     ............................................................g................... [150080]
KM262845.1_strain_NHV           ............................................................g................... [150080]
KX354450.1_47/Ss/2008           ............................................................g................... [150080]
KP055815.1_strain_BA71          ............................................................g................... [150080]
MH025920.1_strain_R35           ..c...t..g..g.....t.......................at....c...........g..............c.... [150080]
MH025919.1_strain_N10           ..c...t..g..g.....t.......................at....c...........g..............c.... [150080]
MH025918.1_strain_R25           ..c...t..g..g.....t.......................at....c...........g..............c.... [150080]
MH025917.1_strain_R7            ..c...t..g..g.....t.......................at....c...........g..............c.... [150080]
MH025916.1_strain_R8            ..c...t..g..g.....t.......................at....c...........g..............c.... [150080]
```

```
FR682468.2_Georgia_2007/1        cgtctataaataaataaagctgtttgagctcttttttaatattgtcaagacctaaaaattcctttttcgtgcgcgaatac [150160]
ASF_IND_20_CAD543                ................................................................................ [150160]
LR899193.1_Germany_2020/1        ................................................................................ [150160]
MN393477.1_Wuhan_2019            ................................................................................ [150160]
MW396979.1_Timor-Leste/2019/1    ................................................................................ [150160]
MK940252.1_CN/2019/InnerMongolia ................................................................................ [150160]
MK128995.1_China/2018/AnhuiXCGQ  ................................................................................ [150160]
MH681419.1_ASFV/POL/2015         ................................................................................ [150160]
LS478113.1_Estonia_2014          ................................................................................ [150160]
KP843857.1_Odintsovo_02/14       ................................................................................ [150160]
FN557520.1_strain_E75            ..................................................................c............. [150160]
AY261366.1_Warthog               .........................a........................................c............t [150160]
AY261365.1_Warmbaths             .a.......................a........................................c............. [150160]
AY261364.1_Tengani_62            .........................a.................a......................c............. [150160]
AY261363.1_Pretoriuskop/96/4     .........................a..t..............a......................c............. [150160]
AY261362.1_Mkuzi_1979            ..................................................................c............. [150160]
AY261361.1_Malawi_Lil-20/1_1983  .........................a..........................c.............c............. [150160]
AY261360.1_Kenya_1950            .........................a..........................c.............c............. [150160]
AM712240.1_OURT_88/3             ..................................................................c............. [150160]
AM712239.1_Benin_97/1            ..................................................................c............. [150160]
KM111295.1_Ken06.Bus             .........................a..........................c.............c............. [150160]
KM111294.1_Ken05/Tk1             .........................a..........................c.............c............. [150160]
KM262844.1_strain_L60            ..................................................................c............. [150160]
KM102979.1_26544/OG10_Italy      ..................................................................c............. [150160]
KM262845.1_strain_NHV            ..................................................................c............. [150160]
KX354450.1_47/Ss/2008            ..................................................................c............. [150160]
KP055815.1_strain_BA71           ..................................................................c............. [150160]
MH025920.1_strain_R35            .........................a..........................c.............c............. [150160]
MH025919.1_strain_N10            .........................a..........................c.............c............. [150160]
MH025918.1_strain_R25            .........................a..........................c.............c............. [150160]
MH025917.1_strain_R7             .........................a..........................c.............c............. [150160]
MH025916.1_strain_R8             .........................a..........................c.............c............. [150160]

FR682468.2_Georgia_2007/1        aagagaatgctaccatcgccctgctggcaggccacagctcgaacgccattacgcttgcgctgcacgatgggatctgtttc [150240]
ASF_IND_20_CAD543                ..........ag...............................y.................................... [150240]
LR899193.1_Germany_2020/1        ................................................................................ [150240]
MN393477.1_Wuhan_2019            ................................................................................ [150240]
MW396979.1_Timor-Leste/2019/1    ................................................................................ [150240]
MK940252.1_CN/2019/InnerMongolia ................................................................................ [150240]
MK128995.1_China/2018/AnhuiXCGQ  ................................................................................ [150240]
MH681419.1_ASFV/POL/2015         ................................................................................ [150240]
LS478113.1_Estonia_2014          ................................................................................ [150240]
KP843857.1_Odintsovo_02/14       ................................................................................ [150240]
```

```
FN557520.1_strain_E75             ..........ag......................................g............................. [150240]
AY261366.1_Warthog                .g.........t......................................g.....a....................... [150240]
AY261365.1_Warmbaths              ...........t........t..t..........................g.....a....................... [150240]
AY261364.1_Tengani_62             .g.......t........................................g.....a....................... [150240]
AY261363.1_Pretoriuskop/96/4      ...........t.....a................................g.....a....................... [150240]
AY261362.1_Mkuzi_1979             ....................t..t..........................g.....a....................... [150240]
AY261361.1_Malawi_Lil-20/1_1983   .g.........t......................................g..............a.............. [150240]
AY261360.1_Kenya_1950             ...........t......................................g............................. [150240]
AM712240.1_OURT_88/3              ..........ag......................................g............................. [150240]
AM712239.1_Benin_97/1             ..........ag......................................g............................. [150240]
KM111295.1_Ken06.Bus              .g......a..t......................................g............................. [150240]
KM111294.1_Ken05/Tk1              .g......a..t..............a.......................g......................t...... [150240]
KM262844.1_strain_L60             ..........ag......................................g............................. [150240]
KM102979.1_26544/OG10_Italy       ..........ag......................................g............................. [150240]
KM262845.1_strain_NHV             ..........ag......................................g............................. [150240]
KX354450.1_47/Ss/2008             ..........ag......................................g............................. [150240]
KP055815.1_strain_BA71            ..........ag......................................g............................. [150240]
MH025920.1_strain_R35             .g......a..t......................................g............................. [150240]
MH025919.1_strain_N10             .g......a..t......................................g............................. [150240]
MH025918.1_strain_R25             .g......a..t......................................g............................. [150240]
MH025917.1_strain_R7              .g......a..t......................................g............................. [150240]
MH025916.1_strain_R8              .g......a..t......................................g............................. [150240]

FR682468.2_Georgia_2007/1         ttcttcaaaaaatgtcttaggaattatattaaaatattttaccagcatagggggataattcctctatttgtgtgggctc [150320]
ASF_IND_20_CAD543                 ................................................................................ [150320]
LR899193.1_Germany_2020/1         ................................................................................ [150320]
MN393477.1_Wuhan_2019             ................................................................................ [150320]
MW396979.1_Timor-Leste/2019/1     ................................................................................ [150320]
MK940252.1_CN/2019/InnerMongolia  ................................................................................ [150320]
MK128995.1_China/2018/AnhuiXCGQ   ................................................................................ [150320]
MH681419.1_ASFV/POL/2015          ................................................................................ [150320]
LS478113.1_Estonia_2014           ................................................................................ [150320]
KP843857.1_Odintsovo_02/14        ................................................................................ [150320]
FN557520.1_strain_E75             .........................................................c...................... [150320]
AY261366.1_Warthog                .........................................................c......g.......a....... [150320]
AY261365.1_Warmbaths              ...c.....................................................c...................... [150320]
AY261364.1_Tengani_62             ............................................a...g........t...................... [150320]
AY261363.1_Pretoriuskop/96/4      .........................................................c....................c. [150320]
AY261362.1_Mkuzi_1979             ...c.....................................................c...................... [150320]
AY261361.1_Malawi_Lil-20/1_1983   ...........................g................a............c...................... [150320]
AY261360.1_Kenya_1950             g.......................g..........g...............a.....c...................... [150320]
AM712240.1_OURT_88/3              .........................................................c...................... [150320]
AM712239.1_Benin_97/1             .........................................................c...................... [150320]
KM111295.1_Ken06.Bus              g.......................g..........g...............a.....c......t............... [150320]
```

```
KM111294.1_Ken05/Tk1            g.......................g..........g...............a.....c...................... [150320]
KM262844.1_strain_L60           .........................................................c...................... [150320]
KM102979.1_26544/OG10_Italy     .........................................................c...................... [150320]
KM262845.1_strain_NHV           .........................................................c...................... [150320]
KX354450.1_47/Ss/2008           .........................................................c...................... [150320]
KP055815.1_strain_BA71          .........................................................c...................... [150320]
MH025920.1_strain_R35           g.......................g..........g...............a.....c......t............... [150320]
MH025919.1_strain_N10           g.......................g..........g...............a.....c......t............... [150320]
MH025918.1_strain_R25           g.......................g..........g...............a.....c......t............... [150320]
MH025917.1_strain_R7            g.......................g..........g...............a.....c......t............... [150320]
MH025916.1_strain_R8            g.......................g..........g...............a.....c......t............... [150320]

FR682468.2_Georgia_2007/1       cccgcttttgtctggcatggcgattatatttactaagggcgtccttgaatgcctgatggactaccgttgtggcatttttt [150400]
ASF_IND_20_CAD543               ................................................................................ [150400]
LR899193.1_Germany_2020/1       ................................................................................ [150400]
MN393477.1_Wuhan_2019           ................................................................................ [150400]
MW396979.1_Timor-Leste/2019/1   ................................................................................ [150400]
MK940252.1_CN/2019/InnerMongolia ................................................................................ [150400]
MK128995.1_China/2018/AnhuiXCGQ ................................................................................ [150400]
MH681419.1_ASFV/POL/2015        ................................................................................ [150400]
LS478113.1_Estonia_2014         ................................................................................ [150400]
KP843857.1_Odintsovo_02/14      ................................................................................ [150400]
FN557520.1_strain_E75           ........................................a..............g........................ [150400]
AY261366.1_Warthog              ...................a....................a..............g........................ [150400]
AY261365.1_Warmbaths            ........................................a..............g........................ [150400]
AY261364.1_Tengani_62           ...........................................t...........g........................ [150400]
AY261363.1_Pretoriuskop/96/4    ........................................a..............g.....a..t............... [150400]
AY261362.1_Mkuzi_1979           ........................................a..............g........................ [150400]
AY261361.1_Malawi_Lil-20/1_1983 ........................................a..............g........t............... [150400]
AY261360.1_Kenya_1950           .......c....................c........a..a..............g.....c..g............... [150400]
AM712240.1_OURT_88/3            ........................................a..............g........................ [150400]
AM712239.1_Benin_97/1           ........................................a..............g........................ [150400]
KM111295.1_Ken06.Bus            .......c....................c...........a..............g.....c..t............... [150400]
KM111294.1_Ken05/Tk1            .......c....................c........a..a..............g.....c..g............... [150400]
KM262844.1_strain_L60           ........................................a..............g........................ [150400]
KM102979.1_26544/OG10_Italy     ........................................a..............g........................ [150400]
KM262845.1_strain_NHV           ........................................a..............g........................ [150400]
KX354450.1_47/Ss/2008           ........................................a..............g........................ [150400]
KP055815.1_strain_BA71          ........................................a..............g........................ [150400]
MH025920.1_strain_R35           .......c....................c...........a..............g.....c..t............... [150400]
MH025919.1_strain_N10           .......c....................c...........a..............g.....c..t............... [150400]
MH025918.1_strain_R25           .......c....................c...........a..............g.....c..t............... [150400]
MH025917.1_strain_R7            .......c....................c...........a..............g.....c..t............... [150400]
MH025916.1_strain_R8            .......c....................c...........a..............g.....c..t............... [150400]
```

```
FR682468.2_Georgia_2007/1        ttacccaagttttttccctcggtaacacgtgtcatttttgatatccgcaccgccccttcttccacaaaaaattttgtgaa [150480]
ASF_IND_20_CAD543                ...............................................................r................ [150480]
LR899193.1_Germany_2020/1        ................................................................................ [150480]
MN393477.1_Wuhan_2019            ................................................................................ [150480]
MW396979.1_Timor-Leste/2019/1    ................................................................................ [150480]
MK940252.1_CN/2019/InnerMongolia ................................................................................ [150480]
MK128995.1_China/2018/AnhuiXCGQ  ................................................................................ [150480]
MH681419.1_ASFV/POL/2015         ................................................................................ [150480]
LS478113.1_Estonia_2014          ................................................................................ [150480]
KP843857.1_Odintsovo_02/14       ................................................................................ [150480]
FN557520.1_strain_E75            ................................................................................ [150480]
AY261366.1_Warthog               ....................................................tt..c....................... [150480]
AY261365.1_Warmbaths             .........................................a..........tt..c....................... [150480]
AY261364.1_Tengani_62            .....................................................t..c....................... [150480]
AY261363.1_Pretoriuskop/96/4     .........................t...............a..........tt..c....................... [150480]
AY261362.1_Mkuzi_1979            .........................................a..........tt..c....................... [150480]
AY261361.1_Malawi_Lil-20/1_1983  .....t.......................g..tg.......a..............c....................... [150480]
AY261360.1_Kenya_1950            .............................g...g.......a..............c........g.............. [150480]
AM712240.1_OURT_88/3             ................................................................................ [150480]
AM712239.1_Benin_97/1            ................................................................................ [150480]
KM111295.1_Ken06.Bus             .............................g...g.......a.......t...t..c........g.............. [150480]
KM111294.1_Ken05/Tk1             .............................g...g.......a...........t..c........g.............. [150480]
KM262844.1_strain_L60            ................................................................................ [150480]
KM102979.1_26544/OG10_Italy      ................................................................................ [150480]
KM262845.1_strain_NHV            ................................................................................ [150480]
KX354450.1_47/Ss/2008            ................................................................................ [150480]
KP055815.1_strain_BA71           ................................................................................ [150480]
MH025920.1_strain_R35            .............................g...g.......a.......t...t..c........g.............. [150480]
MH025919.1_strain_N10            .............................g...g.......a.......t...t..c........g.............. [150480]
MH025918.1_strain_R25            .............................g...g.......a.......t...t..c........g.............. [150480]
MH025917.1_strain_R7             .............................g...g.......a.......t...t..c........g.............. [150480]
MH025916.1_strain_R8             .............................g...g.......a.......t...t..c........g.............. [150480]

FR682468.2_Georgia_2007/1        aatttcagcaacggcgtcttttacatctgtggaaaacatctcatctgtgatgggaatgatcgtgttgtgctgcaccactt [150560]
ASF_IND_20_CAD543                ..............................................................................nn [150560]
LR899193.1_Germany_2020/1        ................................................................................ [150560]
MN393477.1_Wuhan_2019            ................................................................................ [150560]
MW396979.1_Timor-Leste/2019/1    ................................................................................ [150560]
MK940252.1_CN/2019/InnerMongolia ................................................................................ [150560]
MK128995.1_China/2018/AnhuiXCGQ  ................................................................................ [150560]
MH681419.1_ASFV/POL/2015         ................................................................................ [150560]
LS478113.1_Estonia_2014          ................................................................................ [150560]
KP843857.1_Odintsovo_02/14       ................................................................................ [150560]
```

```
FN557520.1_strain_E75               ............................................................t................... [150560]
AY261366.1_Warthog                  .............................................c.................................. [150560]
AY261365.1_Warmbaths                .............................................c.................................. [150560]
AY261364.1_Tengani_62               .............................................c..........c.................t..... [150560]
AY261363.1_Pretoriuskop/96/4        ...............a.............................c.................................. [150560]
AY261362.1_Mkuzi_1979               .............................................c.................................. [150560]
AY261361.1_Malawi_Lil-20/1_1983     ......g......................................c..a.......g.................t..... [150560]
AY261360.1_Kenya_1950               ......g....................c.................c..a.......g............a....t..... [150560]
AM712240.1_OURT_88/3                ............................................................t................... [150560]
AM712239.1_Benin_97/1               ............................................................t................... [150560]
KM111295.1_Ken06.Bus                ......g....................c.................c..a.......g.................t..... [150560]
KM111294.1_Ken05/Tk1                ......g....................c.................c..a.......g............a....t..... [150560]
KM262844.1_strain_L60               ............................................................t................... [150560]
KM102979.1_26544/OG10_Italy         ............................................................t................... [150560]
KM262845.1_strain_NHV               ............................................................t................... [150560]
KX354450.1_47/Ss/2008               ............................................................t................... [150560]
KP055815.1_strain_BA71              ............................................................t................... [150560]
MH025920.1_strain_R35               ......g....................c.................c..a.......g.................t..... [150560]
MH025919.1_strain_N10               ......g....................c.................c..a.......g.................t..... [150560]
MH025918.1_strain_R25               ......g....................c.................c..a.......g.................t..... [150560]
MH025917.1_strain_R7                ......g....................c.................c..a.......g.................t..... [150560]
MH025916.1_strain_R8                ......g....................c.................c..a.......g.................t..... [150560]

FR682468.2_Georgia_2007/1           gcacacaaataatccatgaggccttttttccgcttttcgtttcagactcaatcggaggaaaacaaaaaatgttgtttgaa [150640]
ASF_IND_20_CAD543                   nnnnn........................................................................... [150640]
LR899193.1_Germany_2020/1           ................................................................................ [150640]
MN393477.1_Wuhan_2019               ................................................................................ [150640]
MW396979.1_Timor-Leste/2019/1       ................................................................................ [150640]
MK940252.1_CN/2019/InnerMongolia    ................................................................................ [150640]
MK128995.1_China/2018/AnhuiXCGQ     ................................................................................ [150640]
MH681419.1_ASFV/POL/2015            ................................................................................ [150640]
LS478113.1_Estonia_2014             ................................................................................ [150640]
KP843857.1_Odintsovo_02/14          ................................................................................ [150640]
FN557520.1_strain_E75               ................................................................................ [150640]
AY261366.1_Warthog                  ....g.....g..........................a................................a......... [150640]
AY261365.1_Warmbaths                ....g.....g..........................a................................a......... [150640]
AY261364.1_Tengani_62               ....g.....g..........................a....................g...........a......... [150640]
AY261363.1_Pretoriuskop/96/4        ....g.....g..........................a................................a......... [150640]
AY261362.1_Mkuzi_1979               ..............................................t................................. [150640]
AY261361.1_Malawi_Lil-20/1_1983     ....g.....g...........t........at....a..c...........t................ca......... [150640]
AY261360.1_Kenya_1950               ....g.....g...........t..............a...............................ca......... [150640]
AM712240.1_OURT_88/3                ................................................................................ [150640]
AM712239.1_Benin_97/1               ................................................................................ [150640]
KM111295.1_Ken06.Bus                ....g.....g...........t..............a...............................ca......... [150640]
```

```
KM111294.1_Ken05/Tk1             ....g.....g..........tt..............a...............................ca......... [150640]
KM262844.1_strain_L60            ................................................................................ [150640]
KM102979.1_26544/OG10_Italy      ................................................................................ [150640]
KM262845.1_strain_NHV            ................................................................................ [150640]
KX354450.1_47/Ss/2008            ................................................................................ [150640]
KP055815.1_strain_BA71           ................................................................................ [150640]
MH025920.1_strain_R35            ....g.....g...........t..............a...............................ca......... [150640]
MH025919.1_strain_N10            ....g.....g...........t..............a...............................ca......... [150640]
MH025918.1_strain_R25            ....g.....g...........t..............a...............................ca......... [150640]
MH025917.1_strain_R7             ....g.....g...........t..............a...............................ca......... [150640]
MH025916.1_strain_R8             ....g.....g...........t..............a...............................ca......... [150640]

FR682468.2_Georgia_2007/1        tattgcccaggaaattgatttagcatgg-ttttaacaataaaataagcctatcaat-ttttttataatttgaatagttat [150720]
ASF_IND_20_CAD543                ............................-...........................-....................... [150720]
LR899193.1_Germany_2020/1        ............................-...........................-....................... [150720]
MN393477.1_Wuhan_2019            ............................-...........................-....................... [150720]
MW396979.1_Timor-Leste/2019/1    ............................-...........................-....................... [150720]
MK940252.1_CN/2019/InnerMongolia ............................-...........................-....................... [150720]
MK128995.1_China/2018/AnhuiXCGQ  ............................-...........................-....................... [150720]
MH681419.1_ASFV/POL/2015         ............................-...........................-....................... [150720]
LS478113.1_Estonia_2014          ............................-...........................-....................... [150720]
KP843857.1_Odintsovo_02/14       ............................-...........................-....................... [150720]
FN557520.1_strain_E75            ............................-...........................-....................... [150720]
AY261366.1_Warthog               c.a...........a.............--.........g..g....g........--.........a............ [150720]
AY261365.1_Warmbaths             c.a...........a.............--.........g..g..c.g........t..........a............ [150720]
AY261364.1_Tengani_62            c................g..........--.........g..g....g........-..........a............ [150720]
AY261363.1_Pretoriuskop/96/4     c.a.....g.....a.............--.........g..g....g........-..........a............ [150720]
AY261362.1_Mkuzi_1979            ............................-...........................-....................... [150720]
AY261361.1_Malawi_Lil-20/1_1983  c.................c........at......t...t..ga...g........-..........a............ [150720]
AY261360.1_Kenya_1950            agc.........................t..........t..ga...g........-..........a............ [150720]
AM712240.1_OURT_88/3             ............................-...........................-....................... [150720]
AM712239.1_Benin_97/1            ............................-...........................-....................... [150720]
KM111295.1_Ken06.Bus             agc.........................t..........t..ga...g........-..........a............ [150720]
KM111294.1_Ken05/Tk1             agc.........................t..........t..ga...g........-..........a............ [150720]
KM262844.1_strain_L60            ............................-...........................-....................... [150720]
KM102979.1_26544/OG10_Italy      ............................-...........................-....................... [150720]
KM262845.1_strain_NHV            ............................-...........................-....................... [150720]
KX354450.1_47/Ss/2008            ............................-...........................-....................... [150720]
KP055815.1_strain_BA71           ............................-...........................-....................... [150720]
MH025920.1_strain_R35            agc.........................t..........t..ga...g........-..........a............ [150720]
MH025919.1_strain_N10            agc.........................t..........t..ga...g........-..........a............ [150720]
MH025918.1_strain_R25            agc.........................t..........t..ga...g........-..........a............ [150720]
MH025917.1_strain_R7             agc.........................t..........t..ga...g........-..........a............ [150720]
MH025916.1_strain_R8             agc.........................t..........t..ga...g........-..........a............ [150720]
```

```
FR682468.2_Georgia_2007/1        tccaaattcaatatggcttctttagataatttagtggcacgatatcagaggtgctttaatgaccagtctcttaaaaatag [150800]
ASF_IND_20_CAD543                ................................................................................ [150800]
LR899193.1_Germany_2020/1        ................................................................................ [150800]
MN393477.1_Wuhan_2019            ................................................................................ [150800]
MW396979.1_Timor-Leste/2019/1    ................................................................................ [150800]
MK940252.1_CN/2019/InnerMongolia ................................................................................ [150800]
MK128995.1_China/2018/AnhuiXCGQ  ................................................................................ [150800]
MH681419.1_ASFV/POL/2015         ................................................................................ [150800]
LS478113.1_Estonia_2014          ................................................................................ [150800]
KP843857.1_Odintsovo_02/14       ................................................................................ [150800]
FN557520.1_strain_E75            ................................................................................ [150800]
AY261366.1_Warthog               ................................................................................ [150800]
AY261365.1_Warmbaths             .....................................................t.................c........ [150800]
AY261364.1_Tengani_62            ................................................................................ [150800]
AY261363.1_Pretoriuskop/96/4     .......................................................................c........ [150800]
AY261362.1_Mkuzi_1979            ................................................................................ [150800]
AY261361.1_Malawi_Lil-20/1_1983  .....g...........c........g...........c..g...................................... [150800]
AY261360.1_Kenya_1950            .......c..................g..................................................c.. [150800]
AM712240.1_OURT_88/3             ................................................................................ [150800]
AM712239.1_Benin_97/1            ................................................................................ [150800]
KM111295.1_Ken06.Bus             .t.....c..................g..................................................c.. [150800]
KM111294.1_Ken05/Tk1             ..........................g..................................................c.. [150800]
KM262844.1_strain_L60            ................................................................................ [150800]
KM102979.1_26544/OG10_Italy      ................................................................................ [150800]
KM262845.1_strain_NHV            ................................................................................ [150800]
KX354450.1_47/Ss/2008            ................................................................................ [150800]
KP055815.1_strain_BA71           ................................................................................ [150800]
MH025920.1_strain_R35            .t.....c..................g..................................................c.. [150800]
MH025919.1_strain_N10            .t.....c..................g..................................................c.. [150800]
MH025918.1_strain_R25            .t.....c..................g..................................................c.. [150800]
MH025917.1_strain_R7             .t.....c..................g..................................................c.. [150800]
MH025916.1_strain_R8             .t.....c..................g..................................................c.. [150800]

FR682468.2_Georgia_2007/1        tactattgaacttgaaatacgttttcaacagataaatttttattattcaaaaccgtatatgaggcacttgtggcacaag [150880]
ASF_IND_20_CAD543                ................................................................................ [150880]
LR899193.1_Germany_2020/1        ................................................................................ [150880]
MN393477.1_Wuhan_2019            ................................................................................ [150880]
MW396979.1_Timor-Leste/2019/1    ................................................................................ [150880]
MK940252.1_CN/2019/InnerMongolia ................................................................................ [150880]
MK128995.1_China/2018/AnhuiXCGQ  ................................................................................ [150880]
MH681419.1_ASFV/POL/2015         ................................................................................ [150880]
LS478113.1_Estonia_2014          ................................................................................ [150880]
KP843857.1_Odintsovo_02/14       ................................................................................ [150880]
```

```
FN557520.1_strain_E75              ................................................................................ [150880]
AY261366.1_Warthog                 ................................................................................ [150880]
AY261365.1_Warmbaths               ................................................................................ [150880]
AY261364.1_Tengani_62              ................................................................................ [150880]
AY261363.1_Pretoriuskop/96/4       ................................................................................ [150880]
AY261362.1_Mkuzi_1979              ................................................................................ [150880]
AY261361.1_Malawi_Lil-20/1_1983    ..............................c................................a................ [150880]
AY261360.1_Kenya_1950              ..............................c............................................g.... [150880]
AM712240.1_OURT_88/3               ................................................................................ [150880]
AM712239.1_Benin_97/1              ................................................................................ [150880]
KM111295.1_Ken06.Bus               ..............................c............................................g.... [150880]
KM111294.1_Ken05/Tk1               ..............................c............................................g.... [150880]
KM262844.1_strain_L60              ................................................................................ [150880]
KM102979.1_26544/OG10_Italy        ................................................................................ [150880]
KM262845.1_strain_NHV              ................................................................................ [150880]
KX354450.1_47/Ss/2008              ................................................................................ [150880]
KP055815.1_strain_BA71             ................................................................................ [150880]
MH025920.1_strain_R35              ..............................c............................................g.... [150880]
MH025919.1_strain_N10              ..............................c............................................g.... [150880]
MH025918.1_strain_R25              ..............................c............................................g.... [150880]
MH025917.1_strain_R7               ..............................c............................................g.... [150880]
MH025916.1_strain_R8               ..............................c............................................g.... [150880]

FR682468.2_Georgia_2007/1          agatccctagcaccatctcccacagcatccgctgcatcaaaaaagttcaccatgaaaaccactgccgggaaaaaattttg [150960]
ASF_IND_20_CAD543                  ................................................................................ [150960]
LR899193.1_Germany_2020/1          ................................................................................ [150960]
MN393477.1_Wuhan_2019              ................................................................................ [150960]
MW396979.1_Timor-Leste/2019/1      ................................................................................ [150960]
MK940252.1_CN/2019/InnerMongolia   ................................................................................ [150960]
MK128995.1_China/2018/AnhuiXCGQ    ................................................................................ [150960]
MH681419.1_ASFV/POL/2015           ................................................................................ [150960]
LS478113.1_Estonia_2014            ................................................................................ [150960]
KP843857.1_Odintsovo_02/14         ................................................................................ [150960]
FN557520.1_strain_E75              ................................................................................ [150960]
AY261366.1_Warthog                 ....t........................................................................... [150960]
AY261365.1_Warmbaths               ...................t...........t................................................ [150960]
AY261364.1_Tengani_62              ................................................................................ [150960]
AY261363.1_Pretoriuskop/96/4       ...............................t................................................ [150960]
AY261362.1_Mkuzi_1979              ................................................................................ [150960]
AY261361.1_Malawi_Lil-20/1_1983    ....t..............t.........................................t.....a............ [150960]
AY261360.1_Kenya_1950              .......c...................................g.......................a............ [150960]
AM712240.1_OURT_88/3               ................................................................................ [150960]
AM712239.1_Benin_97/1              ................................................................................ [150960]
KM111295.1_Ken06.Bus               .......c...................................g.......................a............ [150960]
```

```
KM111294.1_Ken05/Tk1             .......c...................................g.......................a............ [150960]
KM262844.1_strain_L60            ................................................................................ [150960]
KM102979.1_26544/OG10_Italy      ................................................................................ [150960]
KM262845.1_strain_NHV            ................................................................................ [150960]
KX354450.1_47/Ss/2008            ................................................................................ [150960]
KP055815.1_strain_BA71           ................................................................................ [150960]
MH025920.1_strain_R35            .......c...................................g.......................a............ [150960]
MH025919.1_strain_N10            .......c...................................g.......................a............ [150960]
MH025918.1_strain_R25            .......c...................................g.......................a............ [150960]
MH025917.1_strain_R7             .......c...................................g.......................a............ [150960]
MH025916.1_strain_R8             .......c...................................g.......................a............ [150960]

FR682468.2_Georgia_2007/1        ccgtcggaaaatctttacttcaaaaaacagcctctcatgttttttaagttttcagagcctgcatctctgggctgtaaggt [151040]
ASF_IND_20_CAD543                ................................................................................ [151040]
LR899193.1_Germany_2020/1        ................................................................................ [151040]
MN393477.1_Wuhan_2019            ................................................................................ [151040]
MW396979.1_Timor-Leste/2019/1    ................................................................................ [151040]
MK940252.1_CN/2019/InnerMongolia ................................................................................ [151040]
MK128995.1_China/2018/AnhuiXCGQ  ................................................................................ [151040]
MH681419.1_ASFV/POL/2015         ................................................................................ [151040]
LS478113.1_Estonia_2014          ................................................................................ [151040]
KP843857.1_Odintsovo_02/14       ................................................................................ [151040]
FN557520.1_strain_E75            ................................................................................ [151040]
AY261366.1_Warthog               ........g....................................................................... [151040]
AY261365.1_Warmbaths             ........g....................................................................... [151040]
AY261364.1_Tengani_62            ........g.......................c............................................... [151040]
AY261363.1_Pretoriuskop/96/4     ........g...t................................................................... [151040]
AY261362.1_Mkuzi_1979            ................................................................................ [151040]
AY261361.1_Malawi_Lil-20/1_1983  ........g....................a...................................c.............. [151040]
AY261360.1_Kenya_1950            ........t...t........................................g.......................... [151040]
AM712240.1_OURT_88/3             ................................................................................ [151040]
AM712239.1_Benin_97/1            ................................................................................ [151040]
KM111295.1_Ken06.Bus             ........t...t.................t......................g.......................... [151040]
KM111294.1_Ken05/Tk1             ........t...t........................................g.......................... [151040]
KM262844.1_strain_L60            ................................................................................ [151040]
KM102979.1_26544/OG10_Italy      ................................................................................ [151040]
KM262845.1_strain_NHV            ................................................................................ [151040]
KX354450.1_47/Ss/2008            ................................................................................ [151040]
KP055815.1_strain_BA71           ................................................................................ [151040]
MH025920.1_strain_R35            ........t...t.................t......................g.......................... [151040]
MH025919.1_strain_N10            ........t...t.................t......................g.......................... [151040]
MH025918.1_strain_R25            ........t...t.................t......................g.......................... [151040]
MH025917.1_strain_R7             ........t...t.................t......................g.......................... [151040]
MH025916.1_strain_R8             ........t...t.................t......................g.......................... [151040]
```

FR682468.2_Georgia_2007/1 ctcgctggccatcgagcagcccattcgtaaatttatcttggactcctccattctcgttcggctcaaaaatcgtacgacct [151120]
ASF_IND_20_CAD543 ................................................................................ [151120]
LR899193.1_Germany_2020/1 ................................................................................ [151120]
MN393477.1_Wuhan_2019 ................................................................................ [151120]
MW396979.1_Timor-Leste/2019/1 ................................................................................ [151120]
MK940252.1_CN/2019/InnerMongolia ................................................................................ [151120]
MK128995.1_China/2018/AnhuiXCGQ ................................................................................ [151120]
MH681419.1_ASFV/POL/2015 ................................................................................ [151120]
LS478113.1_Estonia_2014 ................................................................................ [151120]
KP843857.1_Odintsovo_02/14 ................................................................................ [151120]
FN557520.1_strain_E75 .................................................g.............................. [151120]
AY261366.1_Warthog ................................................tg.............................. [151120]
AY261365.1_Warmbaths .................................................g.............................. [151120]
AY261364.1_Tengani_62 .................................................g.............................. [151120]
AY261363.1_Pretoriuskop/96/4 .................................................g.............................. [151120]
AY261362.1_Mkuzi_1979 .................................................g.............................. [151120]
AY261361.1_Malawi_Lil-20/1_1983 .................................................g.............................. [151120]
AY261360.1_Kenya_1950 ........................c........................g.............................. [151120]
AM712240.1_OURT_88/3 .................................................g.............................. [151120]
AM712239.1_Benin_97/1 .................................................g.............................. [151120]
KM111295.1_Ken06.Bus .................................................g.............................. [151120]
KM111294.1_Ken05/Tk1 .................................................g.............................. [151120]
KM262844.1_strain_L60 .................................................g.............................. [151120]
KM102979.1_26544/OG10_Italy .................................................g.............................. [151120]
KM262845.1_strain_NHV .................................................g.............................. [151120]
KX354450.1_47/Ss/2008 .................................................g.............................. [151120]
KP055815.1_strain_BA71 .................................................g.............................. [151120]
MH025920.1_strain_R35 .................................................g.............................. [151120]
MH025919.1_strain_N10 .................................................g.............................. [151120]
MH025918.1_strain_R25 .................................................g.............................. [151120]
MH025917.1_strain_R7 .................................................g.............................. [151120]
MH025916.1_strain_R8 .................................................g.............................. [151120]

FR682468.2_Georgia_2007/1 ttcgggtatctgaactttggaaaatagagcttaccattgtaaagcagctgatgggaagcgaggtctctgcaaaacttgcc [151200]
ASF_IND_20_CAD543 ................................................................................ [151200]
LR899193.1_Germany_2020/1 ................................................................................ [151200]
MN393477.1_Wuhan_2019 ................................................................................ [151200]
MW396979.1_Timor-Leste/2019/1 ................................................................................ [151200]
MK940252.1_CN/2019/InnerMongolia ................................................................................ [151200]
MK128995.1_China/2018/AnhuiXCGQ ................................................................................ [151200]
MH681419.1_ASFV/POL/2015 ................................................................................ [151200]
LS478113.1_Estonia_2014 ................................................................................ [151200]
KP843857.1_Odintsovo_02/14 ................................................................................ [151200]

```
FN557520.1_strain_E75          ................................................................................ [151200]
AY261366.1_Warthog             .............................................................................a.. [151200]
AY261365.1_Warmbaths           .............................................................................a.. [151200]
AY261364.1_Tengani_62          ........................................g....................................a.. [151200]
AY261363.1_Pretoriuskop/96/4   .c...........................................................................a.. [151200]
AY261362.1_Mkuzi_1979          ................................................................................ [151200]
AY261361.1_Malawi_Lil-20/1_1983 ...a.a...g..t..................................t.............................a.. [151200]
AY261360.1_Kenya_1950          ...a.a...g..c......................g....g................................g...a.. [151200]
AM712240.1_OURT_88/3           ................................................................................ [151200]
AM712239.1_Benin_97/1          ................................................................................ [151200]
KM111295.1_Ken06.Bus           ...a.a..................................g................................g...a.. [151200]
KM111294.1_Ken05/Tk1           ...a.a.............................g....g................................g...a.. [151200]
KM262844.1_strain_L60          ................................................................................ [151200]
KM102979.1_26544/OG10_Italy    ................................................................................ [151200]
KM262845.1_strain_NHV          ................................................................................ [151200]
KX354450.1_47/Ss/2008          ................................................................................ [151200]
KP055815.1_strain_BA71         ................................................................................ [151200]
MH025920.1_strain_R35          ...a.a..................................g................................g...a.. [151200]
MH025919.1_strain_N10          ...a.a..................................g................................g...a.. [151200]
MH025918.1_strain_R25          ...a.a..................................g................................g...a.. [151200]
MH025917.1_strain_R7           ...a.a..................................g................................g...a.. [151200]
MH025916.1_strain_R8           ...a.a..................................g................................g...a.. [151200]

FR682468.2_Georgia_2007/1      gctttcaaaacgcttctgtttgacaccccagagcaacaaacgacaaaaaatatgatgacgttaataaacccagatgacga [151280]
ASF_IND_20_CAD543              ................................................................................ [151280]
LR899193.1_Germany_2020/1      ................................................................................ [151280]
MN393477.1_Wuhan_2019          ................................................................................ [151280]
MW396979.1_Timor-Leste/2019/1  ................................................................................ [151280]
MK940252.1_CN/2019/InnerMongolia ................................................................................ [151280]
MK128995.1_China/2018/AnhuiXCGQ ................................................................................ [151280]
MH681419.1_ASFV/POL/2015       ................................................................................ [151280]
LS478113.1_Estonia_2014        ................................................................................ [151280]
KP843857.1_Odintsovo_02/14     ................................................................................ [151280]
FN557520.1_strain_E75          ................................................................................ [151280]
AY261366.1_Warthog             ..c......................................................t...................... [151280]
AY261365.1_Warmbaths           ..c.......................t..............................t...................... [151280]
AY261364.1_Tengani_62          a.c............................................................................. [151280]
AY261363.1_Pretoriuskop/96/4   ..c...........c..........................................t...................... [151280]
AY261362.1_Mkuzi_1979          ................................................................................ [151280]
AY261361.1_Malawi_Lil-20/1_1983 ..c.............................a.........g..................................... [151280]
AY261360.1_Kenya_1950          ..c..............a.....t.....g............g.................c..........g........ [151280]
AM712240.1_OURT_88/3           ............................................................................g... [151280]
AM712239.1_Benin_97/1          ................................................................................ [151280]
KM111295.1_Ken06.Bus           ..c..............a...........g............g............................g........ [151280]
```

```
KM111294.1_Ken05/Tk1            ..c..............a.....t.....g............g.................c..........g........ [151280]
KM262844.1_strain_L60           ................................................................................ [151280]
KM102979.1_26544/OG10_Italy     ................................................................................ [151280]
KM262845.1_strain_NHV           ............................................................................g... [151280]
KX354450.1_47/Ss/2008           ................................................................................ [151280]
KP055815.1_strain_BA71          ................................................................................ [151280]
MH025920.1_strain_R35           ..c..............a...........g............g............................g........ [151280]
MH025919.1_strain_N10           ..c..............a...........g............g............................g........ [151280]
MH025918.1_strain_R25           ..c..............a...........g............g............................g........ [151280]
MH025917.1_strain_R7            ..c..............a...........g............g............................g........ [151280]
MH025916.1_strain_R8            ..c..............a...........g............g............................g........ [151280]

FR682468.2_Georgia_2007/1       atatctttacgaaatagaaatagagtatacaggaaagcccgaatccctaacggcggcagatgttataaaaattaaaaaca [151360]
ASF_IND_20_CAD543               ................................................................................ [151360]
LR899193.1_Germany_2020/1       ................................................................................ [151360]
MN393477.1_Wuhan_2019           ................................................................................ [151360]
MW396979.1_Timor-Leste/2019/1   ................................................................................ [151360]
MK940252.1_CN/2019/InnerMongolia ................................................................................ [151360]
MK128995.1_China/2018/AnhuiXCGQ ................................................................................ [151360]
MH681419.1_ASFV/POL/2015        ................................................................................ [151360]
LS478113.1_Estonia_2014         ................................................................................ [151360]
KP843857.1_Odintsovo_02/14      ................................................................................ [151360]
FN557520.1_strain_E75           ................................................................................ [151360]
AY261366.1_Warthog              .............................................................a.................. [151360]
AY261365.1_Warmbaths            .............................................................a.................. [151360]
AY261364.1_Tengani_62           ................................................................................ [151360]
AY261363.1_Pretoriuskop/96/4    .........t...................................................................... [151360]
AY261362.1_Mkuzi_1979           ................................................................................ [151360]
AY261361.1_Malawi_Lil-20/1_1983 ..........................................g..t.....a..a......................... [151360]
AY261360.1_Kenya_1950           .............................................t.................................. [151360]
AM712240.1_OURT_88/3            ................................................................................ [151360]
AM712239.1_Benin_97/1           ................................................................................ [151360]
KM111295.1_Ken06.Bus            .............................................t.................................. [151360]
KM111294.1_Ken05/Tk1            .............................................t.................................. [151360]
KM262844.1_strain_L60           ................................................................................ [151360]
KM102979.1_26544/OG10_Italy     ................................................................................ [151360]
KM262845.1_strain_NHV           ................................................................................ [151360]
KX354450.1_47/Ss/2008           ................................................................................ [151360]
KP055815.1_strain_BA71          ................................................................................ [151360]
MH025920.1_strain_R35           .............................................t.................................. [151360]
MH025919.1_strain_N10           .............................................t.................................. [151360]
MH025918.1_strain_R25           .............................................t.................................. [151360]
MH025917.1_strain_R7            .............................................t.................................. [151360]
MH025916.1_strain_R8            .............................................t.................................. [151360]
```

FR682468.2_Georgia_2007/1 cggtgttgacacttatttctccaaaccatttaatgctaacagcctaccaccaggccattgaattcattgcctcccatata [151440]
ASF_IND_20_CAD543 ................................................................................ [151440]
LR899193.1_Germany_2020/1 ................................................................................ [151440]
MN393477.1_Wuhan_2019 ................................................................................ [151440]
MW396979.1_Timor-Leste/2019/1 ................................................................................ [151440]
MK940252.1_CN/2019/InnerMongolia ................................................................................ [151440]
MK128995.1_China/2018/AnhuiXCGQ ................................................................................ [151440]
MH681419.1_ASFV/POL/2015 ................................................................................ [151440]
LS478113.1_Estonia_2014 ................................................................................ [151440]
KP843857.1_Odintsovo_02/14 ................................................................................ [151440]
FN557520.1_strain_E75 ......................g......................................................... [151440]
AY261366.1_Warthog ..........g...........g......................................................... [151440]
AY261365.1_Warmbaths ..........g...........g......................................................... [151440]
AY261364.1_Tengani_62 ..........g...........g......................................................... [151440]
AY261363.1_Pretoriuskop/96/4 ......................g......................................................... [151440]
AY261362.1_Mkuzi_1979 ......................g......................................................... [151440]
AY261361.1_Malawi_Lil-20/1_1983 ......................g...............................................a......... [151440]
AY261360.1_Kenya_1950 ......................g......................................g........a......... [151440]
AM712240.1_OURT_88/3 ......................g......................................................... [151440]
AM712239.1_Benin_97/1 ......................g......................................................... [151440]
KM111295.1_Ken06.Bus ......................g......................................g........a......... [151440]
KM111294.1_Ken05/Tk1 ......................g......................................g........a......... [151440]
KM262844.1_strain_L60 ......................g......................................................... [151440]
KM102979.1_26544/OG10_Italy ......................g......................................................... [151440]
KM262845.1_strain_NHV ......................g......................................................... [151440]
KX354450.1_47/Ss/2008 ......................g......................................................... [151440]
KP055815.1_strain_BA71 ......................g......................................................... [151440]
MH025920.1_strain_R35 ......................g......................................g........a......... [151440]
MH025919.1_strain_N10 ......................g......................................g........a......... [151440]
MH025918.1_strain_R25 ......................g......................................g........a......... [151440]
MH025917.1_strain_R7 ......................g......................................g........a......... [151440]
MH025916.1_strain_R8 ......................g......................................g........a......... [151440]

FR682468.2_Georgia_2007/1 ctgtcctcagaaatccttcttgctcgtattaagagcgggaagtgggggcttaaacgcctcctcccccaggtgaaatccat [151520]
ASF_IND_20_CAD543 ................................................................................ [151520]
LR899193.1_Germany_2020/1 ................................................................................ [151520]
MN393477.1_Wuhan_2019 ................................................................................ [151520]
MW396979.1_Timor-Leste/2019/1 ................................................................................ [151520]
MK940252.1_CN/2019/InnerMongolia ................................................................................ [151520]
MK128995.1_China/2018/AnhuiXCGQ ................................................................................ [151520]
MH681419.1_ASFV/POL/2015 ................................................................................ [151520]
LS478113.1_Estonia_2014 ................................................................................ [151520]
KP843857.1_Odintsovo_02/14 ................................................................................ [151520]

```
FN557520.1_strain_E75             ................................................................................ [151520]
AY261366.1_Warthog                .....t.................a......................................t................. [151520]
AY261365.1_Warmbaths              .....t.................a......................................t................. [151520]
AY261364.1_Tengani_62             .......................a......................................t................. [151520]
AY261363.1_Pretoriuskop/96/4      .......................a......................................t................. [151520]
AY261362.1_Mkuzi_1979             ................................................................................ [151520]
AY261361.1_Malawi_Lil-20/1_1983   ..............a.......................a......................................... [151520]
AY261360.1_Kenya_1950             ..............a....................t..a.................t....................... [151520]
AM712240.1_OURT_88/3              ...................................................................g............ [151520]
AM712239.1_Benin_97/1             ................................................................................ [151520]
KM111295.1_Ken06.Bus              ..............a....................t..a......................................... [151520]
KM111294.1_Ken05/Tk1              ..............a.......................a.................t...........a..a........ [151520]
KM262844.1_strain_L60             ................................................................................ [151520]
KM102979.1_26544/OG10_Italy       ................................................................................ [151520]
KM262845.1_strain_NHV             ...................................................................g............ [151520]
KX354450.1_47/Ss/2008             ................................................................................ [151520]
KP055815.1_strain_BA71            ....t........................................................................... [151520]
MH025920.1_strain_R35             ..............a....................t..a......................................... [151520]
MH025919.1_strain_N10             ..............a....................t..a......................................... [151520]
MH025918.1_strain_R25             ..............a....................t..a......................................... [151520]
MH025917.1_strain_R7              ..............a....................t..a......................................... [151520]
MH025916.1_strain_R8              ..............a....................t..a......................................... [151520]

FR682468.2_Georgia_2007/1         gaccaaagcggattacatgaaattttatccgcccgttggctactatgtaacggacaaagcagatggaattagaggcatcg [151600]
ASF_IND_20_CAD543                 ................................................................................ [151600]
LR899193.1_Germany_2020/1         ................................................................................ [151600]
MN393477.1_Wuhan_2019             ................................................................................ [151600]
MW396979.1_Timor-Leste/2019/1     ................................................................................ [151600]
MK940252.1_CN/2019/InnerMongolia  ................................................................................ [151600]
MK128995.1_China/2018/AnhuiXCGQ   ................................................................................ [151600]
MH681419.1_ASFV/POL/2015          ................................................................................ [151600]
LS478113.1_Estonia_2014           ................................................................................ [151600]
KP843857.1_Odintsovo_02/14        ................................................................................ [151600]
FN557520.1_strain_E75             ................................................................................ [151600]
AY261366.1_Warthog                .................................t.....t......a................................. [151600]
AY261365.1_Warmbaths              .........a.......................t.....t......a................................. [151600]
AY261364.1_Tengani_62             .................................t.............................................. [151600]
AY261363.1_Pretoriuskop/96/4      .................................t.....t......a................................. [151600]
AY261362.1_Mkuzi_1979             ................................................................................ [151600]
AY261361.1_Malawi_Lil-20/1_1983   ..................................................................g............. [151600]
AY261360.1_Kenya_1950             ............c.................c...........t.....g............................... [151600]
AM712240.1_OURT_88/3              ................................................................................ [151600]
AM712239.1_Benin_97/1             ................................................................................ [151600]
KM111295.1_Ken06.Bus              ............c.................c................................................. [151600]
```

```
KM111294.1_Ken05/Tk1             ............c.................c...........t.....g............................... [151600]
KM262844.1_strain_L60            ................................................................................ [151600]
KM102979.1_26544/OG10_Italy      ................................................................................ [151600]
KM262845.1_strain_NHV            ................................................................................ [151600]
KX354450.1_47/Ss/2008            ................................................................................ [151600]
KP055815.1_strain_BA71           ................................................................................ [151600]
MH025920.1_strain_R35            ............c.................c................................................. [151600]
MH025919.1_strain_N10            ............c.................c................................................. [151600]
MH025918.1_strain_R25            ............c.................c................................................. [151600]
MH025917.1_strain_R7             ............c.................c................................................. [151600]
MH025916.1_strain_R8             ............c.................c................................................. [151600]

FR682468.2_Georgia_2007/1        ccgtcattcaggacacgcaaatttatgtggttgcagaccagttatacagcctaggtaccaccggcattgaaccccttaaa [151680]
ASF_IND_20_CAD543                ................................................................................ [151680]
LR899193.1_Germany_2020/1        ................................................................................ [151680]
MN393477.1_Wuhan_2019            ................................................................................ [151680]
MW396979.1_Timor-Leste/2019/1    ................................................................................ [151680]
MK940252.1_CN/2019/InnerMongolia ................................................................................ [151680]
MK128995.1_China/2018/AnhuiXCGQ  ................................................................................ [151680]
MH681419.1_ASFV/POL/2015         ................................................................................ [151680]
LS478113.1_Estonia_2014          ................................................................................ [151680]
KP843857.1_Odintsovo_02/14       ................................................................................ [151680]
FN557520.1_strain_E75            .........................................c...................................... [151680]
AY261366.1_Warthog               ....t....................................c.t.................................... [151680]
AY261365.1_Warmbaths             .........................................c.t.................................... [151680]
AY261364.1_Tengani_62            .......c.................................c.t.................................... [151680]
AY261363.1_Pretoriuskop/96/4     ................a........................c.t.................................... [151680]
AY261362.1_Mkuzi_1979            .........................................c...................................... [151680]
AY261361.1_Malawi_Lil-20/1_1983  ............................c............c...................................... [151680]
AY261360.1_Kenya_1950            ......................g..................c.............c.......................g [151680]
AM712240.1_OURT_88/3             .........................................c...................................... [151680]
AM712239.1_Benin_97/1            .........................................c...................................... [151680]
KM111295.1_Ken06.Bus             .....g................g..................c....t........c.......................g [151680]
KM111294.1_Ken05/Tk1             ......................g..................c.............c..............g........g [151680]
KM262844.1_strain_L60            .........................................c...................................... [151680]
KM102979.1_26544/OG10_Italy      .........................................c...................................... [151680]
KM262845.1_strain_NHV            .........................................c...................................... [151680]
KX354450.1_47/Ss/2008            .........................................c...................................... [151680]
KP055815.1_strain_BA71           .........................................c...................................... [151680]
MH025920.1_strain_R35            .....g................g..................c....t........c.......................g [151680]
MH025919.1_strain_N10            .....g................g..................c....t........c.......................g [151680]
MH025918.1_strain_R25            .....g................g..................c....t........c.......................g [151680]
MH025917.1_strain_R7             .....g................g..................c....t........c.......................g [151680]
MH025916.1_strain_R8             .....g................g..................c....t........c.......................g [151680]
```

```
FR682468.2_Georgia_2007/1        ccaaccattttggacggtgaatttatgcctgaaaaaaaagaattttatgggtttgacgtcatcatgtatgagggcaatct [151760]
ASF_IND_20_CAD543                ................................................................................ [151760]
LR899193.1_Germany_2020/1        ................................................................................ [151760]
MN393477.1_Wuhan_2019            ................................................................................ [151760]
MW396979.1_Timor-Leste/2019/1    ................................................................................ [151760]
MK940252.1_CN/2019/InnerMongolia ................................................................................ [151760]
MK128995.1_China/2018/AnhuiXCGQ  ................................................................................ [151760]
MH681419.1_ASFV/POL/2015         ................................................................................ [151760]
LS478113.1_Estonia_2014          ................................................................................ [151760]
KP843857.1_Odintsovo_02/14       ...............................-................................................ [151760]
FN557520.1_strain_E75            ..............t................................................................. [151760]
AY261366.1_Warthog               ........................................................t..............a........ [151760]
AY261365.1_Warmbaths             ........................................................t....................... [151760]
AY261364.1_Tengani_62            ................................................................................ [151760]
AY261363.1_Pretoriuskop/96/4     ........................................................t....................... [151760]
AY261362.1_Mkuzi_1979            ................................................................................ [151760]
AY261361.1_Malawi_Lil-20/1_1983  ..............t.........................................t..t..t...........t..... [151760]
AY261360.1_Kenya_1950            .................c.................g....................t..t.................... [151760]
AM712240.1_OURT_88/3             ..............t................................................................. [151760]
AM712239.1_Benin_97/1            ..............t................................................................. [151760]
KM111295.1_Ken06.Bus             .................c.................g....................t..t.................... [151760]
KM111294.1_Ken05/Tk1             .................c.................g....................t..t.................... [151760]
KM262844.1_strain_L60            ..............t................................................................. [151760]
KM102979.1_26544/OG10_Italy      ..............t................................................................. [151760]
KM262845.1_strain_NHV            ..............t................................................................. [151760]
KX354450.1_47/Ss/2008            ..............t................................................................. [151760]
KP055815.1_strain_BA71           ..............t................................................................. [151760]
MH025920.1_strain_R35            .................c.................g....................t..t.................... [151760]
MH025919.1_strain_N10            .................c.................g....................t..t.................... [151760]
MH025918.1_strain_R25            .................c.................g....................t..t.................... [151760]
MH025917.1_strain_R7             .................c.................g....................t..t.................... [151760]
MH025916.1_strain_R8             .................c.................g....................t..t.................... [151760]

FR682468.2_Georgia_2007/1        attgacgcaacaggggtttgaaacaagaattgagtctttaagcaagggcattaaagtcttacaagcgtttaacataaaag [151840]
ASF_IND_20_CAD543                ................................................................................ [151840]
LR899193.1_Germany_2020/1        ................................................................................ [151840]
MN393477.1_Wuhan_2019            ................................................................................ [151840]
MW396979.1_Timor-Leste/2019/1    ................................................................................ [151840]
MK940252.1_CN/2019/InnerMongolia ................................................................................ [151840]
MK128995.1_China/2018/AnhuiXCGQ  ................................................................................ [151840]
MH681419.1_ASFV/POL/2015         ................................................................................ [151840]
LS478113.1_Estonia_2014          ................................................................................ [151840]
KP843857.1_Odintsovo_02/14       ................................................................................ [151840]
```

```
FN557520.1_strain_E75            g............................................................................... [151840]
AY261366.1_Warthog               ................................................................................ [151840]
AY261365.1_Warmbaths             ........................................................c.........a............. [151840]
AY261364.1_Tengani_62            g........................................................t...................... [151840]
AY261363.1_Pretoriuskop/96/4     ..................................a............................................. [151840]
AY261362.1_Mkuzi_1979            g............................................................................... [151840]
AY261361.1_Malawi_Lil-20/1_1983  g...........a..........t.................a...................................... [151840]
AY261360.1_Kenya_1950            g.............................c...g......a........................a............. [151840]
AM712240.1_OURT_88/3             g............................................................................... [151840]
AM712239.1_Benin_97/1            g............................................................................... [151840]
KM111295.1_Ken06.Bus             g.............................c...g......a...................................... [151840]
KM111294.1_Ken05/Tk1             g.............................c...g......a........................a............. [151840]
KM262844.1_strain_L60            g............................................................................... [151840]
KM102979.1_26544/OG10_Italy      g............................................................................... [151840]
KM262845.1_strain_NHV            g............................................................................... [151840]
KX354450.1_47/Ss/2008            g............................................................................... [151840]
KP055815.1_strain_BA71           g............................................................................... [151840]
MH025920.1_strain_R35            g.............................c...g......a...................................... [151840]
MH025919.1_strain_N10            g.............................c...g......a...................................... [151840]
MH025918.1_strain_R25            g.............................c...g......a...................................... [151840]
MH025917.1_strain_R7             g.............................c...g......a...................................... [151840]
MH025916.1_strain_R8             g.............................c...g......a...................................... [151840]

FR682468.2_Georgia_2007/1        cagaaatgaagccctttatttcgctaacaagtgcagatcccaacgtgctcctcaaaaactttgaaagcatttttaagaaa [151920]
ASF_IND_20_CAD543                ................................................................................ [151920]
LR899193.1_Germany_2020/1        ................................................................................ [151920]
MN393477.1_Wuhan_2019            ................................................................................ [151920]
MW396979.1_Timor-Leste/2019/1    ................................................................................ [151920]
MK940252.1_CN/2019/InnerMongolia ................................................................................ [151920]
MK128995.1_China/2018/AnhuiXCGQ  ................................................................................ [151920]
MH681419.1_ASFV/POL/2015         ................................................................................ [151920]
LS478113.1_Estonia_2014          ................................................................................ [151920]
KP843857.1_Odintsovo_02/14       ................................................................................ [151920]
FN557520.1_strain_E75            ................................................................................ [151920]
AY261366.1_Warthog               ................................................................................ [151920]
AY261365.1_Warmbaths             .................................................g.................t............ [151920]
AY261364.1_Tengani_62            ...............................c................................................ [151920]
AY261363.1_Pretoriuskop/96/4     ................................................................................ [151920]
AY261362.1_Mkuzi_1979            ................................................................................ [151920]
AY261361.1_Malawi_Lil-20/1_1983  .................................................t..................g........... [151920]
AY261360.1_Kenya_1950            .................................................t..................g........... [151920]
AM712240.1_OURT_88/3             ................................................................................ [151920]
AM712239.1_Benin_97/1            ................................................................................ [151920]
KM111295.1_Ken06.Bus             .................................................t..................g........... [151920]
```

```
KM111294.1_Ken05/Tk1               .................................................t..................g........... [151920]
KM262844.1_strain_L60              ................................................................................ [151920]
KM102979.1_26544/OG10_Italy        ................................................................................ [151920]
KM262845.1_strain_NHV              ................................................................................ [151920]
KX354450.1_47/Ss/2008              ................................................................................ [151920]
KP055815.1_strain_BA71             ................................................................................ [151920]
MH025920.1_strain_R35              .................................................t..................g........... [151920]
MH025919.1_strain_N10              .................................................t..................g........... [151920]
MH025918.1_strain_R25              .................................................t..................g........... [151920]
MH025917.1_strain_R7               .................................................t..................g........... [151920]
MH025916.1_strain_R8               .................................................t..................g........... [151920]

FR682468.2_Georgia_2007/1          aaaactcgcccatattctattgatggcatcattttagtagaacctggcaattcttatctaaatacaaacacctttaagtg [152000]
ASF_IND_20_CAD543                  ....................................................................y..y........ [152000]
LR899193.1_Germany_2020/1          ................................................................................ [152000]
MN393477.1_Wuhan_2019              ................................................................................ [152000]
MW396979.1_Timor-Leste/2019/1      ................................................................................ [152000]
MK940252.1_CN/2019/InnerMongolia   ................................................................................ [152000]
MK128995.1_China/2018/AnhuiXCGQ    ................................................................................ [152000]
MH681419.1_ASFV/POL/2015           ................................................................................ [152000]
LS478113.1_Estonia_2014            ................................................................................ [152000]
KP843857.1_Odintsovo_02/14         ................................................................................ [152000]
FN557520.1_strain_E75              ....................................................................t..t........ [152000]
AY261366.1_Warthog                 ...........g...........................................................t........ [152000]
AY261365.1_Warmbaths               ...........g...........................................................t........ [152000]
AY261364.1_Tengani_62              ........t..g................................c.................c................. [152000]
AY261363.1_Pretoriuskop/96/4       ...........g...........................................................t........ [152000]
AY261362.1_Mkuzi_1979              ....................................................................t..t........ [152000]
AY261361.1_Malawi_Lil-20/1_1983    ...............g...........c...........................................t........ [152000]
AY261360.1_Kenya_1950              .................c.....................................................g........ [152000]
AM712240.1_OURT_88/3               ....................................................................t..t........ [152000]
AM712239.1_Benin_97/1              ....................................................................t..t........ [152000]
KM111295.1_Ken06.Bus               .................c..................................................t..g........ [152000]
KM111294.1_Ken05/Tk1               ...............g.......................................................g........ [152000]
KM262844.1_strain_L60              ....................................................................t..t........ [152000]
KM102979.1_26544/OG10_Italy        ....................................................................t..t........ [152000]
KM262845.1_strain_NHV              ....................................................................t..t........ [152000]
KX354450.1_47/Ss/2008              ....................................................................t..t........ [152000]
KP055815.1_strain_BA71             ....................................................................t..t........ [152000]
MH025920.1_strain_R35              .................c..................................................t..g........ [152000]
MH025919.1_strain_N10              .................c..................................................t..g........ [152000]
MH025918.1_strain_R25              .................c..................................................t..g........ [152000]
MH025917.1_strain_R7               .................c..................................................t..g........ [152000]
MH025916.1_strain_R8               .................c..................................................t..g........ [152000]
```

```
FR682468.2_Georgia_2007/1        gaagcccacctgggataacacattagactttttggtgcgaaaatgtccggagagtttaaacgtaccagagtacgcgccca [152080]
ASF_IND_20_CAD543                ................................................................................ [152080]
LR899193.1_Germany_2020/1        ................................................................................ [152080]
MN393477.1_Wuhan_2019            ................................................................................ [152080]
MW396979.1_Timor-Leste/2019/1    ................................................................................ [152080]
MK940252.1_CN/2019/InnerMongolia ................................................................................ [152080]
MK128995.1_China/2018/AnhuiXCGQ  ................................................................................ [152080]
MH681419.1_ASFV/POL/2015         ................................................................................ [152080]
LS478113.1_Estonia_2014          ................................................................................ [152080]
KP843857.1_Odintsovo_02/14       ................................................................................ [152080]
FN557520.1_strain_E75            ...............................................................g................ [152080]
AY261366.1_Warthog               ................................................a............................... [152080]
AY261365.1_Warmbaths             ................................................a............................... [152080]
AY261364.1_Tengani_62            ................................................a............................... [152080]
AY261363.1_Pretoriuskop/96/4     ................................................a............................... [152080]
AY261362.1_Mkuzi_1979            ...............................................................g................ [152080]
AY261361.1_Malawi_Lil-20/1_1983  .........................................................g...................... [152080]
AY261360.1_Kenya_1950            ............................................................t................... [152080]
AM712240.1_OURT_88/3             ...............................................................g................ [152080]
AM712239.1_Benin_97/1            ...............................................................g................ [152080]
KM111295.1_Ken06.Bus             .............................................c.....a.a......t................... [152080]
KM111294.1_Ken05/Tk1             ............................................................t................... [152080]
KM262844.1_strain_L60            ...............................................................g................ [152080]
KM102979.1_26544/OG10_Italy      ...............................................................g................ [152080]
KM262845.1_strain_NHV            ...............................................................g................ [152080]
KX354450.1_47/Ss/2008            ...............................................................g................ [152080]
KP055815.1_strain_BA71           ...............................................................g................ [152080]
MH025920.1_strain_R35            .............................................c.....a.a......t................... [152080]
MH025919.1_strain_N10            .............................................c.....a.a......t................... [152080]
MH025918.1_strain_R25            .............................................c.....a.a......t................... [152080]
MH025917.1_strain_R7             .............................................c.....a.a......t................... [152080]
MH025916.1_strain_R8             .............................................c.....a.a......t................... [152080]

FR682468.2_Georgia_2007/1        aaaaagggttttccctgcatctactatttgtaggcatctccggagagcttttaaaaaattagcgctaaattggtgtcca [152160]
ASF_IND_20_CAD543                ................................................................................ [152160]
LR899193.1_Germany_2020/1        ................................................................................ [152160]
MN393477.1_Wuhan_2019            ................................................................................ [152160]
MW396979.1_Timor-Leste/2019/1    ................................................................................ [152160]
MK940252.1_CN/2019/InnerMongolia ................................................................................ [152160]
MK128995.1_China/2018/AnhuiXCGQ  ................................................................................ [152160]
MH681419.1_ASFV/POL/2015         ................................................................................ [152160]
LS478113.1_Estonia_2014          ................................................................................ [152160]
KP843857.1_Odintsovo_02/14       ................................................................................ [152160]
```

```
FN557520.1_strain_E75          ................................................................................ [152160]
AY261366.1_Warthog             .......................................................g........................ [152160]
AY261365.1_Warmbaths           .......................................................g........................ [152160]
AY261364.1_Tengani_62          .....................................t.................g.......................g [152160]
AY261363.1_Pretoriuskop/96/4   .....................................t..............c..g........................ [152160]
AY261362.1_Mkuzi_1979          ........................................t....................................... [152160]
AY261361.1_Malawi_Lil-20/1_1983 .................t...........................................g.................. [152160]
AY261360.1_Kenya_1950          .......c.....g.......................t.....g.................................... [152160]
AM712240.1_OURT_88/3           ................................................................................ [152160]
AM712239.1_Benin_97/1          ................................................................................ [152160]
KM111295.1_Ken06.Bus           .......c.....g....................t..t.....g................................c... [152160]
KM111294.1_Ken05/Tk1           .......c.....g.........t..........t..t.....g.....................t.............. [152160]
KM262844.1_strain_L60          ................................................................................ [152160]
KM102979.1_26544/OG10_Italy    ................................................................................ [152160]
KM262845.1_strain_NHV          ................................................................................ [152160]
KX354450.1_47/Ss/2008          ................................................................................ [152160]
KP055815.1_strain_BA71         ................................................................................ [152160]
MH025920.1_strain_R35          .......c.....g....................t..t.....g................................c... [152160]
MH025919.1_strain_N10          .......c.....g....................t..t.....g................................c... [152160]
MH025918.1_strain_R25          .......c.....g....................t..t.....g................................c... [152160]
MH025917.1_strain_R7           .......c.....g....................t..t.....g................................c... [152160]
MH025916.1_strain_R8           .......c.....g....................t..t.....g................................c... [152160]

FR682468.2_Georgia_2007/1      ggatatacgaaactattccccgttacacagcgcaaccaaaactactttccagtacagttccagccatcggattttccatt [152240]
ASF_IND_20_CAD543              ..............................................................................y. [152240]
LR899193.1_Germany_2020/1      ................................................................................ [152240]
MN393477.1_Wuhan_2019          ................................................................................ [152240]
MW396979.1_Timor-Leste/2019/1  ................................................................................ [152240]
MK940252.1_CN/2019/InnerMongolia ................................................................................ [152240]
MK128995.1_China/2018/AnhuiXCGQ ................................................................................ [152240]
MH681419.1_ASFV/POL/2015       ................................................................................ [152240]
LS478113.1_Estonia_2014        ................................................................................ [152240]
KP843857.1_Odintsovo_02/14     ................................................................................ [152240]
FN557520.1_strain_E75          ...........................................................t..................c. [152240]
AY261366.1_Warthog             .....c........g................................................................. [152240]
AY261365.1_Warmbaths           ..c..c.....................................................t.................... [152240]
AY261364.1_Tengani_62          .....c.......................................................................... [152240]
AY261363.1_Pretoriuskop/96/4   .....c.......................................................................... [152240]
AY261362.1_Mkuzi_1979          ...........................................................t.................... [152240]
AY261361.1_Malawi_Lil-20/1_1983 .....c..a...............................................a....................... [152240]
AY261360.1_Kenya_1950          .....c..a.....g...................................g.....a...........c........... [152240]
AM712240.1_OURT_88/3           ...........................................................t..................c. [152240]
AM712239.1_Benin_97/1          ...........................................................t..................c. [152240]
KM111295.1_Ken06.Bus           .....c..a.....g...................................g.....a...........c........... [152240]
```

```
KM111294.1_Ken05/Tk1             .....c..a.....g...................................g.....a.....a................. [152240]
KM262844.1_strain_L60            ...........................................................t..................c. [152240]
KM102979.1_26544/OG10_Italy      ...........................................................t..................c. [152240]
KM262845.1_strain_NHV            ...........................................................t..................c. [152240]
KX354450.1_47/Ss/2008            ...........................................................t..................c. [152240]
KP055815.1_strain_BA71           ...........................................................t..................c. [152240]
MH025920.1_strain_R35            .....c..a.....g...................................g.....a...........c........... [152240]
MH025919.1_strain_N10            .....c..a.....g...................................g.....a...........c........... [152240]
MH025918.1_strain_R25            .....c..a.....g...................................g.....a...........c........... [152240]
MH025917.1_strain_R7             .....c..a.....g...................................g.....a...........c........... [152240]
MH025916.1_strain_R8             .....c..a.....g...................................g.....a...........c........... [152240]

FR682468.2_Georgia_2007/1        ggcatttctttattaccacccagatacctcgtcattttctaatatagatggaaaggtccttgaaatgcgttgtcttaaga [152320]
ASF_IND_20_CAD543                ......................................................nnn....................... [152320]
LR899193.1_Germany_2020/1        ................................................................................ [152320]
MN393477.1_Wuhan_2019            ................................................................................ [152320]
MW396979.1_Timor-Leste/2019/1    ................................................................................ [152320]
MK940252.1_CN/2019/InnerMongolia ................................................................................ [152320]
MK128995.1_China/2018/AnhuiXCGQ  ................................................................................ [152320]
MH681419.1_ASFV/POL/2015         ................................................................................ [152320]
LS478113.1_Estonia_2014          ................................................................................ [152320]
KP843857.1_Odintsovo_02/14       ................................................................................ [152320]
FN557520.1_strain_E75            ................................................................................ [152320]
AY261366.1_Warthog               ........................................g.............a......................... [152320]
AY261365.1_Warmbaths             ..............................a.........g....................................... [152320]
AY261364.1_Tengani_62            ........................................g.....................................a. [152320]
AY261363.1_Pretoriuskop/96/4     ........................................g....................................... [152320]
AY261362.1_Mkuzi_1979            ................................................................................ [152320]
AY261361.1_Malawi_Lil-20/1_1983  ...g.......................g..a........cg.............a......................... [152320]
AY261360.1_Kenya_1950            ...........................g.....g.....cg.......c........t...................... [152320]
AM712240.1_OURT_88/3             ................................................................................ [152320]
AM712239.1_Benin_97/1            ................................................................................ [152320]
KM111295.1_Ken06.Bus             ...........................g.....g.....cg.......c........t...................... [152320]
KM111294.1_Ken05/Tk1             ...g.......................g..a..g.............................................. [152320]
KM262844.1_strain_L60            ................................................................................ [152320]
KM102979.1_26544/OG10_Italy      ................................................................................ [152320]
KM262845.1_strain_NHV            ................................................................................ [152320]
KX354450.1_47/Ss/2008            ................................................................................ [152320]
KP055815.1_strain_BA71           ................................................................................ [152320]
MH025920.1_strain_R35            ...........................g.....g.....cg.......c........t...................... [152320]
MH025919.1_strain_N10            ...........................g.....g.....cg.......c........t...................... [152320]
MH025918.1_strain_R25            ...........................g.....g.....cg.......c........t...................... [152320]
MH025917.1_strain_R7             ...........................g.....g.....cg.......c........t...................... [152320]
MH025916.1_strain_R8             ...........................g.....g.....cg.......c........t...................... [152320]
```

```
FR682468.2_Georgia_2007/1        gagaaatcaatcacgtcagctgggaaattgtaaaaatccgggaggataggcagcaggatcttaaaaccggcgggtatttt [152400]
ASF_IND_20_CAD543                ................................................................................ [152400]
LR899193.1_Germany_2020/1        ................................................................................ [152400]
MN393477.1_Wuhan_2019            ................................................................................ [152400]
MW396979.1_Timor-Leste/2019/1    ................................................................................ [152400]
MK940252.1_CN/2019/InnerMongolia ................................................................................ [152400]
MK128995.1_China/2018/AnhuiXCGQ  ................................................................................ [152400]
MH681419.1_ASFV/POL/2015         ................................................................................ [152400]
LS478113.1_Estonia_2014          ................................................................................ [152400]
KP843857.1_Odintsovo_02/14       ................................................................................ [152400]
FN557520.1_strain_E75            .......t...t.......a............................................................ [152400]
AY261366.1_Warthog               ...........................................a........a..a........................ [152400]
AY261365.1_Warmbaths             .............ta................................................................. [152400]
AY261364.1_Tengani_62            ..............a..........................................................a...... [152400]
AY261363.1_Pretoriuskop/96/4     ................................................................................ [152400]
AY261362.1_Mkuzi_1979            .......t...t.......a............................................................ [152400]
AY261361.1_Malawi_Lil-20/1_1983  .....g.............................................................a.....a...... [152400]
AY261360.1_Kenya_1950            .....g.....t.......................................................a.....c...... [152400]
AM712240.1_OURT_88/3             .......t...t.......a............................................................ [152400]
AM712239.1_Benin_97/1            .......t...t.......a............................................................ [152400]
KM111295.1_Ken06.Bus             .....g.....t.......................................................a...a.c...... [152400]
KM111294.1_Ken05/Tk1             .....g.............t....................a..........................a.....c...... [152400]
KM262844.1_strain_L60            .......t...t.......a............................................................ [152400]
KM102979.1_26544/OG10_Italy      .......t...t.......a............................................................ [152400]
KM262845.1_strain_NHV            .......t...t.......a............................................................ [152400]
KX354450.1_47/Ss/2008            .......t...t.......a............................................................ [152400]
KP055815.1_strain_BA71           .......t...t.......a............................................................ [152400]
MH025920.1_strain_R35            .....g.....t.......................................................a...a.c...... [152400]
MH025919.1_strain_N10            .....g.....t.......................................................a...a.c...... [152400]
MH025918.1_strain_R25            .....g.....t.......................................................a...a.c...... [152400]
MH025917.1_strain_R7             .....g.....t.......................................................a...a.c...... [152400]
MH025916.1_strain_R8             .....g.....t.......................................................a...a.c...... [152400]

FR682468.2_Georgia_2007/1        ggcaatgatttcaaaacagccgaactcacatggcttaactatatggatcccttttcctttgaggagctggcaaagggccc [152480]
ASF_IND_20_CAD543                ................................................................................ [152480]
LR899193.1_Germany_2020/1        ................................................................................ [152480]
MN393477.1_Wuhan_2019            ................................................................................ [152480]
MW396979.1_Timor-Leste/2019/1    ................................................................................ [152480]
MK940252.1_CN/2019/InnerMongolia ................................................................................ [152480]
MK128995.1_China/2018/AnhuiXCGQ  ................................................................................ [152480]
MH681419.1_ASFV/POL/2015         ................................................................................ [152480]
LS478113.1_Estonia_2014          ................................................................................ [152480]
KP843857.1_Odintsovo_02/14       ................................................................................ [152480]
```

```
FN557520.1_strain_E75            .................................................................a.............. [152480]
AY261366.1_Warthog               ................................................................................ [152480]
AY261365.1_Warmbaths             ..t............................................................................. [152480]
AY261364.1_Tengani_62            ...........t.....g............................................a................. [152480]
AY261363.1_Pretoriuskop/96/4     ..........................t..................................................... [152480]
AY261362.1_Mkuzi_1979            ..................................................t............................. [152480]
AY261361.1_Malawi_Lil-20/1_1983  .................g..t.........................................a..a.............. [152480]
AY261360.1_Kenya_1950            .................g.........................................c.....a.....g........ [152480]
AM712240.1_OURT_88/3             .................................................................a.............. [152480]
AM712239.1_Benin_97/1            .................................................................a.............. [152480]
KM111295.1_Ken06.Bus             .................g.........................................c.....a.....g........ [152480]
KM111294.1_Ken05/Tk1             .................g.........................................c.....a.....g........ [152480]
KM262844.1_strain_L60            .................................................................a.............. [152480]
KM102979.1_26544/OG10_Italy      .................................................................a.............. [152480]
KM262845.1_strain_NHV            .................................................................a.............. [152480]
KX354450.1_47/Ss/2008            .................................................................a.............. [152480]
KP055815.1_strain_BA71           .................................................................a.............. [152480]
MH025920.1_strain_R35            .................g.........................................c.....a.....g........ [152480]
MH025919.1_strain_N10            .................g.........................................c.....a.....g........ [152480]
MH025918.1_strain_R25            .................g.........................................c.....a.....g........ [152480]
MH025917.1_strain_R7             .................g.........................................c.....a.....g........ [152480]
MH025916.1_strain_R8             .................g.........................................c.....a.....g........ [152480]

FR682468.2_Georgia_2007/1        ttctggaatgtacttcgccggtgccaaaaccggcatataccgcgctcaaacagcacttatttcctttattaaacaagaaa [152560]
ASF_IND_20_CAD543                ................................................................................ [152560]
LR899193.1_Germany_2020/1        ................................................................................ [152560]
MN393477.1_Wuhan_2019            ................................................................................ [152560]
MW396979.1_Timor-Leste/2019/1    ................................................................................ [152560]
MK940252.1_CN/2019/InnerMongolia ................................................................................ [152560]
MK128995.1_China/2018/AnhuiXCGQ  ................................................................................ [152560]
MH681419.1_ASFV/POL/2015         ................................................................................ [152560]
LS478113.1_Estonia_2014          ................................................................................ [152560]
KP843857.1_Odintsovo_02/14       ................................................................................ [152560]
FN557520.1_strain_E75            ................................................................................ [152560]
AY261366.1_Warthog               ........................................................................g....... [152560]
AY261365.1_Warmbaths             ........................................................................g....... [152560]
AY261364.1_Tengani_62            ...............t........t....................................................... [152560]
AY261363.1_Pretoriuskop/96/4     c.......................................................................g....... [152560]
AY261362.1_Mkuzi_1979            ................................................................................ [152560]
AY261361.1_Malawi_Lil-20/1_1983  c.......................t....................................................... [152560]
AY261360.1_Kenya_1950            c.......................t.................t........g............................ [152560]
AM712240.1_OURT_88/3             ................................................................................ [152560]
AM712239.1_Benin_97/1            ................................................................................ [152560]
KM111295.1_Ken06.Bus             ........................t.................t........g............................ [152560]
```

```
KM111294.1_Ken05/Tk1             c.......................t.....t...........t........g............................ [152560]
KM262844.1_strain_L60            ................................................................................ [152560]
KM102979.1_26544/OG10_Italy      ................................................................................ [152560]
KM262845.1_strain_NHV            ................................................................................ [152560]
KX354450.1_47/Ss/2008            ................................................................................ [152560]
KP055815.1_strain_BA71           ................................................................................ [152560]
MH025920.1_strain_R35            ........................t.................t........g............................ [152560]
MH025919.1_strain_N10            ........................t.................t........g............................ [152560]
MH025918.1_strain_R25            ........................t.................t........g............................ [152560]
MH025917.1_strain_R7             ........................t.................t........g............................ [152560]
MH025916.1_strain_R8             ........................t.................t........g............................ [152560]

FR682468.2_Georgia_2007/1        tcatccaaaaaataagtcaccaatcctgggttatcgatcttggaataggaaaagggcaggacctaggacgttacctggac [152640]
ASF_IND_20_CAD543                ................................................................................ [152640]
LR899193.1_Germany_2020/1        ................................................................................ [152640]
MN393477.1_Wuhan_2019            ................................................................................ [152640]
MW396979.1_Timor-Leste/2019/1    ................................................................................ [152640]
MK940252.1_CN/2019/InnerMongolia ................................................................................ [152640]
MK128995.1_China/2018/AnhuiXCGQ  ................................................................................ [152640]
MH681419.1_ASFV/POL/2015         ................................................................................ [152640]
LS478113.1_Estonia_2014          ................................................................................ [152640]
KP843857.1_Odintsovo_02/14       ................................................................................ [152640]
FN557520.1_strain_E75            ................................................................................ [152640]
AY261366.1_Warthog               ....................................................g.....................t..... [152640]
AY261365.1_Warmbaths             ....................................................g.................c......... [152640]
AY261364.1_Tengani_62            ....................................................g.................c......... [152640]
AY261363.1_Pretoriuskop/96/4     ....................................................g.....................t..... [152640]
AY261362.1_Mkuzi_1979            ................................................................................ [152640]
AY261361.1_Malawi_Lil-20/1_1983  ....................................................g..............c...........t [152640]
AY261360.1_Kenya_1950            ...............................g....................g..............g........a..t [152640]
AM712240.1_OURT_88/3             ................................................................................ [152640]
AM712239.1_Benin_97/1            ................................................................................ [152640]
KM111295.1_Ken06.Bus             ....t..........................g....................g..............g........a..t [152640]
KM111294.1_Ken05/Tk1             ...............................g....................g..a.....t.....t........a..t [152640]
KM262844.1_strain_L60            ................................................................................ [152640]
KM102979.1_26544/OG10_Italy      ................................................................................ [152640]
KM262845.1_strain_NHV            ................................................................................ [152640]
KX354450.1_47/Ss/2008            ................................................................................ [152640]
KP055815.1_strain_BA71           ..............................................................................g. [152640]
MH025920.1_strain_R35            ....t..........................g....................g..............g........a..t [152640]
MH025919.1_strain_N10            ....t..........................g....................g..............g........a..t [152640]
MH025918.1_strain_R25            ....t..........................g....................g..............g........a..t [152640]
MH025917.1_strain_R7             ....t..........................g....................g..............g........a..t [152640]
MH025916.1_strain_R8             ....t..........................g....................g..............g........a..t [152640]
```

```
FR682468.2_Georgia_2007/1        gcagggataaggcatcttgttgggatcgataaggatcaaaccgcgcttgcggagcttgtttatcgaaaattttcgcatgc [152720]
ASF_IND_20_CAD543                ................................................................................ [152720]
LR899193.1_Germany_2020/1        ................................................................................ [152720]
MN393477.1_Wuhan_2019            ................................................................................ [152720]
MW396979.1_Timor-Leste/2019/1    ................................................................................ [152720]
MK940252.1_CN/2019/InnerMongolia ................................................................................ [152720]
MK128995.1_China/2018/AnhuiXCGQ  ................................................................................ [152720]
MH681419.1_ASFV/POL/2015         ................................................................................ [152720]
LS478113.1_Estonia_2014          ................................................................................ [152720]
KP843857.1_Odintsovo_02/14       ................................................................................ [152720]
FN557520.1_strain_E75            ......g................a...........c...........c................................ [152720]
AY261366.1_Warthog               ......g..................................t.....c....................g........... [152720]
AY261365.1_Warmbaths             ......g........................................c..a.................g........... [152720]
AY261364.1_Tengani_62            ........c......................................c................................ [152720]
AY261363.1_Pretoriuskop/96/4     ......g........................................c....................g........... [152720]
AY261362.1_Mkuzi_1979            ......g................a...........c...........c................................ [152720]
AY261361.1_Malawi_Lil-20/1_1983  ...................................c.....t.....c.........a...................... [152720]
AY261360.1_Kenya_1950            ...............................................c.........a..........g........... [152720]
AM712240.1_OURT_88/3             ......g................a...........c...........c................................ [152720]
AM712239.1_Benin_97/1            ......g................a...........c...........c................................ [152720]
KM111295.1_Ken06.Bus             ...............................................c.........a..........g........... [152720]
KM111294.1_Ken05/Tk1             .........................................t.....c.........a..........g........... [152720]
KM262844.1_strain_L60            ......g................a...........c...........c................................ [152720]
KM102979.1_26544/OG10_Italy      ......g................a...........c...........c................................ [152720]
KM262845.1_strain_NHV            ......g................a...........c...........c................................ [152720]
KX354450.1_47/Ss/2008            ......g................a...........c...........c................................ [152720]
KP055815.1_strain_BA71           ......g................a...........c...........c................................ [152720]
MH025920.1_strain_R35            ...............................................c.........a..........g........... [152720]
MH025919.1_strain_N10            ...............................................c.........a..........g........... [152720]
MH025918.1_strain_R25            ...............................................c.........a..........g........... [152720]
MH025917.1_strain_R7             ...............................................c.........a..........g........... [152720]
MH025916.1_strain_R8             ...............................................c.........a..........g........... [152720]

FR682468.2_Georgia_2007/1        tacgacccgacagcacaagcacgctaccaacatttacgtgttgcatcaagacctcgcagagcctgcgaaagaaatcagcg [152800]
ASF_IND_20_CAD543                ................................................................................ [152800]
LR899193.1_Germany_2020/1        ................................................................................ [152800]
MN393477.1_Wuhan_2019            ................................................................................ [152800]
MW396979.1_Timor-Leste/2019/1    ................................................................................ [152800]
MK940252.1_CN/2019/InnerMongolia ................................................................................ [152800]
MK128995.1_China/2018/AnhuiXCGQ  ................................................................................ [152800]
MH681419.1_ASFV/POL/2015         ................................................................................ [152800]
LS478113.1_Estonia_2014          ................................................................................ [152800]
KP843857.1_Odintsovo_02/14       ................................................................................ [152800]
```

```
FN557520.1_strain_E75            ...a............................................................................ [152800]
AY261366.1_Warthog               ...a................................t................................g.......... [152800]
AY261365.1_Warmbaths             ...a................................t.................t..............g.......... [152800]
AY261364.1_Tengani_62            ...a................................t................................g.......... [152800]
AY261363.1_Pretoriuskop/96/4     ...a................................t................................g.......... [152800]
AY261362.1_Mkuzi_1979            ...a.................................................................g.......... [152800]
AY261361.1_Malawi_Lil-20/1_1983  ...a................................t.............................t..g..g..t.... [152800]
AY261360.1_Kenya_1950            ....................................t........c...........c........t..g.......... [152800]
AM712240.1_OURT_88/3             ...a.................................................................g.......... [152800]
AM712239.1_Benin_97/1            ...a............................................................................ [152800]
KM111295.1_Ken06.Bus             ....................................t....................c........t..g.......... [152800]
KM111294.1_Ken05/Tk1             ....................................t.............................t..g..g..t.... [152800]
KM262844.1_strain_L60            ...a.................................................................g.......... [152800]
KM102979.1_26544/OG10_Italy      ...a............................................................................ [152800]
KM262845.1_strain_NHV            ...a.................................................................g.......... [152800]
KX354450.1_47/Ss/2008            ...a............................................................................ [152800]
KP055815.1_strain_BA71           ...a.................................................................g.......... [152800]
MH025920.1_strain_R35            ....................................t....................c........t..g.......... [152800]
MH025919.1_strain_N10            ....................................t....................c........t..g.......... [152800]
MH025918.1_strain_R25            ....................................t....................c........t..g.......... [152800]
MH025917.1_strain_R7             ....................................t....................c........t..g.......... [152800]
MH025916.1_strain_R8             ....................................t....................c........t..g.......... [152800]

FR682468.2_Georgia_2007/1        aaaaggtacaccaaatttacgggtttcccaaggagggagcttcttccattgttagcaacctgtttattcactatcttatg [152880]
ASF_IND_20_CAD543                ................................................................................ [152880]
LR899193.1_Germany_2020/1        ................................................................................ [152880]
MN393477.1_Wuhan_2019            ................................................................................ [152880]
MW396979.1_Timor-Leste/2019/1    ................................................................................ [152880]
MK940252.1_CN/2019/InnerMongolia ................................................................................ [152880]
MK128995.1_China/2018/AnhuiXCGQ  ................................................................................ [152880]
MH681419.1_ASFV/POL/2015         ................................................................................ [152880]
LS478113.1_Estonia_2014          ................................................................................ [152880]
KP843857.1_Odintsovo_02/14       ................................................................................ [152880]
FN557520.1_strain_E75            ................................................................................ [152880]
AY261366.1_Warthog               ................................................................................ [152880]
AY261365.1_Warmbaths             ................................................................................ [152880]
AY261364.1_Tengani_62            ................................................................................ [152880]
AY261363.1_Pretoriuskop/96/4     ................................................................................ [152880]
AY261362.1_Mkuzi_1979            ................................................................................ [152880]
AY261361.1_Malawi_Lil-20/1_1983  ....a...................................c....................................... [152880]
AY261360.1_Kenya_1950            ....a...................................c.......................c............... [152880]
AM712240.1_OURT_88/3             ................................................................................ [152880]
AM712239.1_Benin_97/1            ................................................................................ [152880]
KM111295.1_Ken06.Bus             ....a...................................c.......................c............... [152880]
```

```
KM111294.1_Ken05/Tk1             ....a...................................c....................................... [152880]
KM262844.1_strain_L60            ................................................................................ [152880]
KM102979.1_26544/OG10_Italy      ................................................................................ [152880]
KM262845.1_strain_NHV            ................................................................................ [152880]
KX354450.1_47/Ss/2008            ................................................................................ [152880]
KP055815.1_strain_BA71           ................................................................................ [152880]
MH025920.1_strain_R35            ....a.....a.............................c.......................c............... [152880]
MH025919.1_strain_N10            ....a.....a.............................c.......................c............... [152880]
MH025918.1_strain_R25            ....a.....a.............................c.......................c............... [152880]
MH025917.1_strain_R7             ....a.....a.............................c.......................c............... [152880]
MH025916.1_strain_R8             ....a.....a.............................c.......................c............... [152880]

FR682468.2_Georgia_2007/1        aaaaacacgcagcaggtggaaaacctggccgttctgtgccataagcttcttcagccggggggaatggtgtggtttaccac [152960]
ASF_IND_20_CAD543                ................................................................................ [152960]
LR899193.1_Germany_2020/1        ................................................................................ [152960]
MN393477.1_Wuhan_2019            ................................................................................ [152960]
MW396979.1_Timor-Leste/2019/1    ................................................................................ [152960]
MK940252.1_CN/2019/InnerMongolia ................................................................................ [152960]
MK128995.1_China/2018/AnhuiXCGQ  ................................................................................ [152960]
MH681419.1_ASFV/POL/2015         ................................................................................ [152960]
LS478113.1_Estonia_2014          ................................................................................ [152960]
KP843857.1_Odintsovo_02/14       ................................................................................ [152960]
FN557520.1_strain_E75            ........................................................-....................... [152960]
AY261366.1_Warthog               ................................................................................ [152960]
AY261365.1_Warmbaths             ................................................................................ [152960]
AY261364.1_Tengani_62            ................................................................................ [152960]
AY261363.1_Pretoriuskop/96/4     ................................................................................ [152960]
AY261362.1_Mkuzi_1979            ................................................................................ [152960]
AY261361.1_Malawi_Lil-20/1_1983  ......t.....................................a................................... [152960]
AY261360.1_Kenya_1950            ...tc.t.....................................a................................... [152960]
AM712240.1_OURT_88/3             ................................................................................ [152960]
AM712239.1_Benin_97/1            ................................................................................ [152960]
KM111295.1_Ken06.Bus             ...tc.t...................a.................a................................... [152960]
KM111294.1_Ken05/Tk1             ...tc.t.a...................................a................................... [152960]
KM262844.1_strain_L60            ................................................................................ [152960]
KM102979.1_26544/OG10_Italy      ................................................................................ [152960]
KM262845.1_strain_NHV            ................................................................................ [152960]
KX354450.1_47/Ss/2008            ................................................................................ [152960]
KP055815.1_strain_BA71           ................................................................................ [152960]
MH025920.1_strain_R35            ...tc.t...................a.................a................................... [152960]
MH025919.1_strain_N10            ...tc.t...................a.................a................................... [152960]
MH025918.1_strain_R25            ...tc.t...................a.................a................................... [152960]
MH025917.1_strain_R7             ...tc.t...................a.................a................................... [152960]
MH025916.1_strain_R8             ...tc.t...................a.................a................................... [152960]
```

```
FR682468.2_Georgia_2007/1        catgttgggagaacaggtcttagaattacttcatgaaaatagaatagagctcaatgaagtatgggaggctcgtgaaaacg [153040]
ASF_IND_20_CAD543                ................................................................................ [153040]
LR899193.1_Germany_2020/1        ................................................................................ [153040]
MN393477.1_Wuhan_2019            ................................................................................ [153040]
MW396979.1_Timor-Leste/2019/1    ................................................................................ [153040]
MK940252.1_CN/2019/InnerMongolia ................................................................................ [153040]
MK128995.1_China/2018/AnhuiXCGQ  ................................................................................ [153040]
MH681419.1_ASFV/POL/2015         ................................................................................ [153040]
LS478113.1_Estonia_2014          ................................................................................ [153040]
KP843857.1_Odintsovo_02/14       ................................................................................ [153040]
FN557520.1_strain_E75            .....................................................................a.......... [153040]
AY261366.1_Warthog               ....c.........g......................................................a..c....... [153040]
AY261365.1_Warmbaths             ....c.........g......................................................a.......... [153040]
AY261364.1_Tengani_62            ....c.........g......................................................a.......... [153040]
AY261363.1_Pretoriuskop/96/4     ....c.........g...................................................a..a.......... [153040]
AY261362.1_Mkuzi_1979            .....................................................................a.......... [153040]
AY261361.1_Malawi_Lil-20/1_1983  ....c.........g...t................................t..c..............g..a.....t. [153040]
AY261360.1_Kenya_1950            ....c.........g...........................gg.........................a..c.....t. [153040]
AM712240.1_OURT_88/3             .....................................................................a.......... [153040]
AM712239.1_Benin_97/1            .....................................................................a.......... [153040]
KM111295.1_Ken06.Bus             t...c.........g...........................g..........................g..c.....t. [153040]
KM111294.1_Ken05/Tk1             ....c.........g...........................gg.........................a..c.....t. [153040]
KM262844.1_strain_L60            .....................................................................a.......... [153040]
KM102979.1_26544/OG10_Italy      .....................................................................a.......... [153040]
KM262845.1_strain_NHV            .....................................................................a.......... [153040]
KX354450.1_47/Ss/2008            .....................................................................a.......... [153040]
KP055815.1_strain_BA71           .....................................................................a.......... [153040]
MH025920.1_strain_R35            t...c.........g...........................g..........................g..c.....t. [153040]
MH025919.1_strain_N10            t...c.........g...........................g..........................g..c.....t. [153040]
MH025918.1_strain_R25            t...c.........g...........................g..........................g..c.....t. [153040]
MH025917.1_strain_R7             t...c.........g...........................g..........................g..c.....t. [153040]
MH025916.1_strain_R8             t...c.........g...........................g..........................g..c.....t. [153040]

FR682468.2_Georgia_2007/1        aagtggtcaaatttgctattaaacgtctctttaaagaggatatattacaggaaactgggcaagaaattggagtcctgtta [153120]
ASF_IND_20_CAD543                ................................................................................ [153120]
LR899193.1_Germany_2020/1        ................................................................................ [153120]
MN393477.1_Wuhan_2019            ................................................................................ [153120]
MW396979.1_Timor-Leste/2019/1    ................................................................................ [153120]
MK940252.1_CN/2019/InnerMongolia ................................................................................ [153120]
MK128995.1_China/2018/AnhuiXCGQ  ................................................................................ [153120]
MH681419.1_ASFV/POL/2015         ................................................................................ [153120]
LS478113.1_Estonia_2014          ................................................................................ [153120]
KP843857.1_Odintsovo_02/14       ................................................................................ [153120]
```

FN557520.1_strain_E75 .g.............................................................................. [153120]
AY261366.1_Warthog .g...................................a....................................t..... [153120]
AY261365.1_Warmbaths .g.............................................................................. [153120]
AY261364.1_Tengani_62 .g.............................................................................. [153120]
AY261363.1_Pretoriuskop/96/4 .g.........................................................................t..... [153120]
AY261362.1_Mkuzi_1979 .g.............................................................................. [153120]
AY261361.1_Malawi_Lil-20/1_1983 .g..............a..............c....................g........................... [153120]
AY261360.1_Kenya_1950 .g..a..t..g.....c........c...............g.cc..........g.......................g [153120]
AM712240.1_OURT_88/3 .g.............................................................................. [153120]
AM712239.1_Benin_97/1 .g.............................................................................. [153120]
KM111295.1_Ken06.Bus .g..a..t.................c...............g.cc..........g....................a..g [153120]
KM111294.1_Ken05/Tk1 .g..a..t........c........c...............g.cc..........g.................t.....g [153120]
KM262844.1_strain_L60 .g.............................................................................. [153120]
KM102979.1_26544/OG10_Italy .g.............................................................................. [153120]
KM262845.1_strain_NHV .g.............................................................................. [153120]
KX354450.1_47/Ss/2008 .g.............................................................................. [153120]
KP055815.1_strain_BA71 .g.............................................................................. [153120]
MH025920.1_strain_R35 .g..a..t.................c...............g.cc..........g....................a..g [153120]
MH025919.1_strain_N10 .g..a..t.................c...............g.cc..........g....................a..g [153120]
MH025918.1_strain_R25 .g..a..t.................c...............g.cc..........g....................a..g [153120]
MH025917.1_strain_R7 .g..a..t.................c...............g.cc..........g....................a..g [153120]
MH025916.1_strain_R8 .g..a..t.................c...............g.cc..........g....................a..g [153120]

FR682468.2_Georgia_2007/1 cccttcagcaatggcgacttctacaatgaatatcttgtgaacacagcgtttttaattaaaatatttaaacatcacggctt [153200]
ASF_IND_20_CAD543 ................................................................................ [153200]
LR899193.1_Germany_2020/1 ................................................................................ [153200]
MN393477.1_Wuhan_2019 ................................................................................ [153200]
MW396979.1_Timor-Leste/2019/1 ................................................................................ [153200]
MK940252.1_CN/2019/InnerMongolia ................................................................................ [153200]
MK128995.1_China/2018/AnhuiXCGQ ................................................................................ [153200]
MH681419.1_ASFV/POL/2015 ................................................................................ [153200]
LS478113.1_Estonia_2014 ................................................................................ [153200]
KP843857.1_Odintsovo_02/14 ................................................................................ [153200]
FN557520.1_strain_E75 ................................................................................ [153200]
AY261366.1_Warthog .............................g..............g................................a.. [153200]
AY261365.1_Warmbaths ...............................................a.............................g.. [153200]
AY261364.1_Tengani_62 .....t.............a.........................................................g.. [153200]
AY261363.1_Pretoriuskop/96/4 .............................g..............g................................a.. [153200]
AY261362.1_Mkuzi_1979 ................................................................................ [153200]
AY261361.1_Malawi_Lil-20/1_1983 .......................t....................g................................... [153200]
AY261360.1_Kenya_1950 ..........................c..................................................... [153200]
AM712240.1_OURT_88/3 ................................................................................ [153200]
AM712239.1_Benin_97/1 ................................................................................ [153200]
KM111295.1_Ken06.Bus ..........................c..................................................... [153200]

```
KM111294.1_Ken05/Tk1           ..........................c..................................................... [153200]
KM262844.1_strain_L60          ................................................................................ [153200]
KM102979.1_26544/OG10_Italy    ................................................................................ [153200]
KM262845.1_strain_NHV          ................................................................................ [153200]
KX354450.1_47/Ss/2008          ................................................................................ [153200]
KP055815.1_strain_BA71         ................................................................................ [153200]
MH025920.1_strain_R35          ..........................c..................................................... [153200]
MH025919.1_strain_N10          ..........................c..................................................... [153200]
MH025918.1_strain_R25          ..........................c..................................................... [153200]
MH025917.1_strain_R7           ..........................c..................................................... [153200]
MH025916.1_strain_R8           ..........................c..................................................... [153200]

FR682468.2_Georgia_2007/1      ttccctagttcaaaagcagtcctttaaggactggattccagaatttcaaaactttagtaaaagtttgtataaaattctta [153280]
ASF_IND_20_CAD543              ................................................................................ [153280]
LR899193.1_Germany_2020/1      ................................................................................ [153280]
MN393477.1_Wuhan_2019          ................................................................................ [153280]
MW396979.1_Timor-Leste/2019/1  ................................................................................ [153280]
MK940252.1_CN/2019/InnerMongolia ................................................................................ [153280]
MK128995.1_China/2018/AnhuiXCGQ ................................................................................ [153280]
MH681419.1_ASFV/POL/2015       ................................................................................ [153280]
LS478113.1_Estonia_2014        ................................................................................ [153280]
KP843857.1_Odintsovo_02/14     ................................................................................ [153280]
FN557520.1_strain_E75          ................................................................................ [153280]
AY261366.1_Warthog             ......g..c...................................................................... [153280]
AY261365.1_Warmbaths           ......g..c...................................................................... [153280]
AY261364.1_Tengani_62          ......g..c.................a.................................................... [153280]
AY261363.1_Pretoriuskop/96/4   ......g..c...................................................................... [153280]
AY261362.1_Mkuzi_1979          ................................................................................ [153280]
AY261361.1_Malawi_Lil-20/1_1983 ..............t...................................c...........c...a............. [153280]
AY261360.1_Kenya_1950          ......g.......t...................................c...............a............. [153280]
AM712240.1_OURT_88/3           ................................................................................ [153280]
AM712239.1_Benin_97/1          ................................................................................ [153280]
KM111295.1_Ken06.Bus           ......g.......t............a..a...................c...............a............. [153280]
KM111294.1_Ken05/Tk1           ....t.........t............a......................c...............a............. [153280]
KM262844.1_strain_L60          ................................................................................ [153280]
KM102979.1_26544/OG10_Italy    ........................a....................................................... [153280]
KM262845.1_strain_NHV          ................................................................................ [153280]
KX354450.1_47/Ss/2008          ........................a....................................................... [153280]
KP055815.1_strain_BA71         ................................................................................ [153280]
MH025920.1_strain_R35          ......g.......t......t.....a..a...................c...............a............. [153280]
MH025919.1_strain_N10          ......g.......t......t.....a..a...................c...............a............. [153280]
MH025918.1_strain_R25          ......g.......t......t.....a..a...................c...............a............. [153280]
MH025917.1_strain_R7           ......g.......t......t.....a..a...................c...............a............. [153280]
MH025916.1_strain_R8           ......g.......t......t.....a..a...................c...............a............. [153280]
```

```
FR682468.2_Georgia_2007/1        cagaagccgataaaacttggacaagcctttttgggtttatttgtctgcgcaaaaattaaatattttttcataagaagtac [153360]
ASF_IND_20_CAD543                ................................................................................ [153360]
LR899193.1_Germany_2020/1        ................................................................................ [153360]
MN393477.1_Wuhan_2019            ................................................................................ [153360]
MW396979.1_Timor-Leste/2019/1    ................................................................................ [153360]
MK940252.1_CN/2019/InnerMongolia ................................................................................ [153360]
MK128995.1_China/2018/AnhuiXCGQ  ................................................................................ [153360]
MH681419.1_ASFV/POL/2015         ................................................................................ [153360]
LS478113.1_Estonia_2014          ................................................................................ [153360]
KP843857.1_Odintsovo_02/14       ................................................................................ [153360]
FN557520.1_strain_E75            ................................................................................ [153360]
AY261366.1_Warthog               ................................................................................ [153360]
AY261365.1_Warmbaths             ................................................................................ [153360]
AY261364.1_Tengani_62            ....g..........................................t................................ [153360]
AY261363.1_Pretoriuskop/96/4     ................................................................................ [153360]
AY261362.1_Mkuzi_1979            ................................................................................ [153360]
AY261361.1_Malawi_Lil-20/1_1983  ...........................................c.................t...........c...... [153360]
AY261360.1_Kenya_1950            ...........................................c................-t...........c...... [153360]
AM712240.1_OURT_88/3             ................................................................................ [153360]
AM712239.1_Benin_97/1            ................................................................................ [153360]
KM111295.1_Ken06.Bus             ....g......................................c.................t...........c...... [153360]
KM111294.1_Ken05/Tk1             ...........................................c................-t...........c...... [153360]
KM262844.1_strain_L60            ................................................................................ [153360]
KM102979.1_26544/OG10_Italy      ................................................................................ [153360]
KM262845.1_strain_NHV            ................................................................................ [153360]
KX354450.1_47/Ss/2008            ................................................................................ [153360]
KP055815.1_strain_BA71           ................................................................................ [153360]
MH025920.1_strain_R35            ....g......................................c.................t...........c...... [153360]
MH025919.1_strain_N10            ....g......................................c.................t...........c...... [153360]
MH025918.1_strain_R25            ....g......................................c.................t...........c...... [153360]
MH025917.1_strain_R7             ....g......................................c.................t...........c...... [153360]
MH025916.1_strain_R8             ....g......................................c.................t...........c...... [153360]

FR682468.2_Georgia_2007/1        taccc-aggttttaaagaaatagctaaaaatatcatatggatactgccatgcagcttaaaacgtctattggtttaattac [153440]
ASF_IND_20_CAD543                .....-.......................................................................... [153440]
LR899193.1_Germany_2020/1        .....-.......................................................................... [153440]
MN393477.1_Wuhan_2019            .....-.......................................................................... [153440]
MW396979.1_Timor-Leste/2019/1    .....-.......................................................................... [153440]
MK940252.1_CN/2019/InnerMongolia .....-.......................................................................... [153440]
MK128995.1_China/2018/AnhuiXCGQ  .....-.......................................................................... [153440]
MH681419.1_ASFV/POL/2015         .....-.......................................................................... [153440]
LS478113.1_Estonia_2014          .....-.......................................................................... [153440]
KP843857.1_Odintsovo_02/14       .....-g......................................................................... [153440]
```

```
FN557520.1_strain_E75          .....-.......................................................................... [153440]
AY261366.1_Warthog             .....-.......................................................................... [153440]
AY261365.1_Warmbaths           .....-.......................................................................... [153440]
AY261364.1_Tengani_62          ..-----.................................................c....................... [153440]
AY261363.1_Pretoriuskop/96/4   .....-.......................................................................... [153440]
AY261362.1_Mkuzi_1979          .....-.......................................................................... [153440]
AY261361.1_Malawi_Lil-20/1_1983 ..t..-................t....g................-...........c..............a........ [153440]
AY261360.1_Kenya_1950          ..t..t................t.c..g....g.......................g.....a........g........ [153440]
AM712240.1_OURT_88/3           .....-.......................................................................... [153440]
AM712239.1_Benin_97/1          .....-.......................................................................... [153440]
KM111295.1_Ken06.Bus           ..t..-................t.c..g....g.......................g.....a................. [153440]
KM111294.1_Ken05/Tk1           ..t.t-................t.c..g....g.......................g.....a................. [153440]
KM262844.1_strain_L60          .....-.......................................................................... [153440]
KM102979.1_26544/OG10_Italy    .....-.......................................................................... [153440]
KM262845.1_strain_NHV          .....-.......................................................................... [153440]
KX354450.1_47/Ss/2008          .....-.......................................................................... [153440]
KP055815.1_strain_BA71         .....-.......................................................................... [153440]
MH025920.1_strain_R35          ..t..-................t.c..g....g.......................g.....a................. [153440]
MH025919.1_strain_N10          ..t..-................t.c..g....g.......................g.....a................. [153440]
MH025918.1_strain_R25          ..t..-................t.c..g....g.......................g.....a................. [153440]
MH025917.1_strain_R7           ..t..-................t.c..g....g.......................g.....a................. [153440]
MH025916.1_strain_R8           ..t..-................t.c..g....g.......................g.....a................. [153440]

FR682468.2_Georgia_2007/1      atgtcgtatgaacacccaaaataaccaaatagaaactattctggttcaaaaacgttacagccttgctttttcagaattta [153520]
ASF_IND_20_CAD543              ................................................................................ [153520]
LR899193.1_Germany_2020/1      ................................................................................ [153520]
MN393477.1_Wuhan_2019          ................................................................................ [153520]
MW396979.1_Timor-Leste/2019/1  ................................................................................ [153520]
MK940252.1_CN/2019/InnerMongolia ................................................................................ [153520]
MK128995.1_China/2018/AnhuiXCGQ ................................................................................ [153520]
MH681419.1_ASFV/POL/2015       ................................................................................ [153520]
LS478113.1_Estonia_2014        ................................................................................ [153520]
KP843857.1_Odintsovo_02/14     ................................................................................ [153520]
FN557520.1_strain_E75          t............................................................................... [153520]
AY261366.1_Warthog             t...................................g........................................... [153520]
AY261365.1_Warmbaths           t............................................................................... [153520]
AY261364.1_Tengani_62          t............................................................................... [153520]
AY261363.1_Pretoriuskop/96/4   t............................................................................... [153520]
AY261362.1_Mkuzi_1979          t............................................................................... [153520]
AY261361.1_Malawi_Lil-20/1_1983 t...................g...t...........c........................................... [153520]
AY261360.1_Kenya_1950          t...................gc..............c........................................... [153520]
AM712240.1_OURT_88/3           t............................................................................... [153520]
AM712239.1_Benin_97/1          t............................................................................... [153520]
KM111295.1_Ken06.Bus           t...................gc..............c........................................... [153520]
```

```
KM111294.1_Ken05/Tk1            t...................g...............c........................................... [153520]
KM262844.1_strain_L60           t............................................................................... [153520]
KM102979.1_26544/OG10_Italy     t............................................................................... [153520]
KM262845.1_strain_NHV           t............................................................................... [153520]
KX354450.1_47/Ss/2008           t............................................................................... [153520]
KP055815.1_strain_BA71          t............................................................................... [153520]
MH025920.1_strain_R35           t...................gc..............c........................................... [153520]
MH025919.1_strain_N10           t...................gc..............c........................................... [153520]
MH025918.1_strain_R25           t...................gc..............c........................................... [153520]
MH025917.1_strain_R7            t...................gc..............c........................................... [153520]
MH025916.1_strain_R8            t...................gc..............c........................................... [153520]

FR682468.2_Georgia_2007/1       ttcattgtcattactctataaatgctaatcaaggtcatctgattaaaatgtttaataacatgacaattaatgaacgactg [153600]
ASF_IND_20_CAD543               ................................................................................ [153600]
LR899193.1_Germany_2020/1       ................................................................................ [153600]
MN393477.1_Wuhan_2019           ................................................................................ [153600]
MW396979.1_Timor-Leste/2019/1   ................................................................................ [153600]
MK940252.1_CN/2019/InnerMongolia ................................................................................ [153600]
MK128995.1_China/2018/AnhuiXCGQ ................................................................................ [153600]
MH681419.1_ASFV/POL/2015        ................................................................................ [153600]
LS478113.1_Estonia_2014         ................................................................................ [153600]
KP843857.1_Odintsovo_02/14      ................................................................................ [153600]
FN557520.1_strain_E75           ................g............................................................... [153600]
AY261366.1_Warthog              ................................................................................ [153600]
AY261365.1_Warmbaths            ................................................................................ [153600]
AY261364.1_Tengani_62           ................................................................................ [153600]
AY261363.1_Pretoriuskop/96/4    ................................................................g............... [153600]
AY261362.1_Mkuzi_1979           ................g............................................................... [153600]
AY261361.1_Malawi_Lil-20/1_1983 .......c...............t.......ca............................................... [153600]
AY261360.1_Kenya_1950           .......................t.......ca...................c........................t.. [153600]
AM712240.1_OURT_88/3            ................g............................................................... [153600]
AM712239.1_Benin_97/1           ................g............................................................... [153600]
KM111295.1_Ken06.Bus            .......................t.......ca...................c........................t.. [153600]
KM111294.1_Ken05/Tk1            .......................t.......ca...................c........................t.. [153600]
KM262844.1_strain_L60           ................g............................................................... [153600]
KM102979.1_26544/OG10_Italy     ................g............................................................... [153600]
KM262845.1_strain_NHV           ................g............................................................... [153600]
KX354450.1_47/Ss/2008           ................g............................................................... [153600]
KP055815.1_strain_BA71          ................g............................................................... [153600]
MH025920.1_strain_R35           .......................t.......ca...................c........................t.. [153600]
MH025919.1_strain_N10           .......................t.......ca...................c........................t.. [153600]
MH025918.1_strain_R25           .......................t.......ca...................c........................t.. [153600]
MH025917.1_strain_R7            .......................t.......ca...................c........................t.. [153600]
MH025916.1_strain_R8            .......................t.......ca...................c........................t.. [153600]
```

```
FR682468.2_Georgia_2007/1        cttgtcaaaacactggattttgaccgcatgtggtatcatatttggattgaaactccagtctacgaactataccacaaaaa [153680]
ASF_IND_20_CAD543                ................................................................................ [153680]
LR899193.1_Germany_2020/1        ................................................................................ [153680]
MN393477.1_Wuhan_2019            ................................................................................ [153680]
MW396979.1_Timor-Leste/2019/1    ................................................................................ [153680]
MK940252.1_CN/2019/InnerMongolia ................................................................................ [153680]
MK128995.1_China/2018/AnhuiXCGQ  ................................................................................ [153680]
MH681419.1_ASFV/POL/2015         ................................................................................ [153680]
LS478113.1_Estonia_2014          ................................................................................ [153680]
KP843857.1_Odintsovo_02/14       ................................................................................ [153680]
FN557520.1_strain_E75            ................................................................................ [153680]
AY261366.1_Warthog               ...........................................................a.................... [153680]
AY261365.1_Warmbaths             ...........................................................a.................... [153680]
AY261364.1_Tengani_62            ...a...........................................c...........a.................... [153680]
AY261363.1_Pretoriuskop/96/4     ...........................................................a.................... [153680]
AY261362.1_Mkuzi_1979            ................................................................................ [153680]
AY261361.1_Malawi_Lil-20/1_1983  ...a........t..........t.......................c...........a.................... [153680]
AY261360.1_Kenya_1950            ...a.t......t..................................c...........a..t................. [153680]
AM712240.1_OURT_88/3             ................................................................................ [153680]
AM712239.1_Benin_97/1            ................................................................................ [153680]
KM111295.1_Ken06.Bus             ...a........t..................................c...........a..t................. [153680]
KM111294.1_Ken05/Tk1             ...a........t..................................c...........a..t...t............. [153680]
KM262844.1_strain_L60            ................................................................................ [153680]
KM102979.1_26544/OG10_Italy      ................................................................................ [153680]
KM262845.1_strain_NHV            ................................................................................ [153680]
KX354450.1_47/Ss/2008            ................................................................................ [153680]
KP055815.1_strain_BA71           ................................................................................ [153680]
MH025920.1_strain_R35            ...a........t..................................c...........a..t................. [153680]
MH025919.1_strain_N10            ...a........t..................................c...........a..t................. [153680]
MH025918.1_strain_R25            ...a........t..................................c...........a..t................. [153680]
MH025917.1_strain_R7             ...a........t..................................c...........a..t................. [153680]
MH025916.1_strain_R8             ...a........t..................................c...........a..t................. [153680]

FR682468.2_Georgia_2007/1        ataccaaaaatttaggaaaaattggcttctcccggataatgggaaaaagcttatttcattaatcaaccaagcaaagggct [153760]
ASF_IND_20_CAD543                .................................r.............................................. [153760]
LR899193.1_Germany_2020/1        ................................................................................ [153760]
MN393477.1_Wuhan_2019            ................................................................................ [153760]
MW396979.1_Timor-Leste/2019/1    ................................................................................ [153760]
MK940252.1_CN/2019/InnerMongolia ................................................................................ [153760]
MK128995.1_China/2018/AnhuiXCGQ  ................................................................................ [153760]
MH681419.1_ASFV/POL/2015         ................................................................................ [153760]
LS478113.1_Estonia_2014          ................................................................................ [153760]
KP843857.1_Odintsovo_02/14       ................................................................................ [153760]
```

```
FN557520.1_strain_E75             .................................a.............................................. [153760]
AY261366.1_Warthog                .......................................c........................................ [153760]
AY261365.1_Warmbaths              ..............................t................................................. [153760]
AY261364.1_Tengani_62             .......................................c........................................ [153760]
AY261363.1_Pretoriuskop/96/4      .................................a.....c....................g................... [153760]
AY261362.1_Mkuzi_1979             .................................a.............................................. [153760]
AY261361.1_Malawi_Lil-20/1_1983   ..............a............ga................................................... [153760]
AY261360.1_Kenya_1950             ..............aa............a................................................... [153760]
AM712240.1_OURT_88/3              .................................a.............................................. [153760]
AM712239.1_Benin_97/1             .................................a.............................................. [153760]
KM111295.1_Ken06.Bus              ..............aa............a................................................... [153760]
KM111294.1_Ken05/Tk1              ..............aa............a....a.............................................. [153760]
KM262844.1_strain_L60             .................................a.............................................. [153760]
KM102979.1_26544/OG10_Italy       .................................a.............................................. [153760]
KM262845.1_strain_NHV             .................................a.............................................. [153760]
KX354450.1_47/Ss/2008             .................................a.............................................. [153760]
KP055815.1_strain_BA71            .................................a.............................................. [153760]
MH025920.1_strain_R35             ..............aa............a................................................... [153760]
MH025919.1_strain_N10             ..............aa............a................................................... [153760]
MH025918.1_strain_R25             ..............aa............a................................................... [153760]
MH025917.1_strain_R7              ..............aa............a................................................... [153760]
MH025916.1_strain_R8              ..............aa............a................................................... [153760]

FR682468.2_Georgia_2007/1         caggaacacttctatgggaaatccctaagggtaagccgaaggaagacgagtcggaccttacctgtgccatacgggagttt [153840]
ASF_IND_20_CAD543                 ................................................................................ [153840]
LR899193.1_Germany_2020/1         ................................................................................ [153840]
MN393477.1_Wuhan_2019             ................................................................................ [153840]
MW396979.1_Timor-Leste/2019/1     ................................................................................ [153840]
MK940252.1_CN/2019/InnerMongolia  ................................................................................ [153840]
MK128995.1_China/2018/AnhuiXCGQ   ................................................................................ [153840]
MH681419.1_ASFV/POL/2015          ................................................................................ [153840]
LS478113.1_Estonia_2014           ................................................................................ [153840]
KP843857.1_Odintsovo_02/14        ................................................................................ [153840]
FN557520.1_strain_E75             ......................t.......................t................................. [153840]
AY261366.1_Warthog                ................................................................................ [153840]
AY261365.1_Warmbaths              ................................................................................ [153840]
AY261364.1_Tengani_62             .......c........................................................................ [153840]
AY261363.1_Pretoriuskop/96/4      ................................................................................ [153840]
AY261362.1_Mkuzi_1979             .......c..............t.......................t................................. [153840]
AY261361.1_Malawi_Lil-20/1_1983   ...........t................................a..............g.................... [153840]
AY261360.1_Kenya_1950             .......g...t................................a..............g.................... [153840]
AM712240.1_OURT_88/3              ......................t.......................t................................. [153840]
AM712239.1_Benin_97/1             ......................t.......................t................................. [153840]
KM111295.1_Ken06.Bus              ...........t................................a..............g.............a...... [153840]
```

```
KM111294.1_Ken05/Tk1            ...........t................................a..............g.................... [153840]
KM262844.1_strain_L60           ......................t.......................t................................. [153840]
KM102979.1_26544/OG10_Italy     ......................t.......................t................................. [153840]
KM262845.1_strain_NHV           ......................t.......................t................................. [153840]
KX354450.1_47/Ss/2008           ......................t.......................t................................. [153840]
KP055815.1_strain_BA71          ......................t.......................t................................. [153840]
MH025920.1_strain_R35           ...........t................................a..............g.............a...... [153840]
MH025919.1_strain_N10           ...........t................................a..............g.............a...... [153840]
MH025918.1_strain_R25           ...........t................................a..............g.............a...... [153840]
MH025917.1_strain_R7            ...........t................................a..............g.............a...... [153840]
MH025916.1_strain_R8            ...........t................................a..............g.............a...... [153840]

FR682468.2_Georgia_2007/1       gaagaagaaaccgggattacccgcgaatattaccagattctcccagagtttaaaaaatctatgtcatactttgacggtaa [153920]
ASF_IND_20_CAD543               ................................................................................ [153920]
LR899193.1_Germany_2020/1       ................................................................................ [153920]
MN393477.1_Wuhan_2019           ................................................................................ [153920]
MW396979.1_Timor-Leste/2019/1   ................................................................................ [153920]
MK940252.1_CN/2019/InnerMongolia ................................................................................ [153920]
MK128995.1_China/2018/AnhuiXCGQ ................................................................................ [153920]
MH681419.1_ASFV/POL/2015        ................................................................................ [153920]
LS478113.1_Estonia_2014         ................................................................................ [153920]
KP843857.1_Odintsovo_02/14      ................................................................................ [153920]
FN557520.1_strain_E75           ................................................................................ [153920]
AY261366.1_Warthog              ................................................................................ [153920]
AY261365.1_Warmbaths            ...........t.................................................................... [153920]
AY261364.1_Tengani_62           ..............a................................................................. [153920]
AY261363.1_Pretoriuskop/96/4    ................................................................................ [153920]
AY261362.1_Mkuzi_1979           ................................................................................ [153920]
AY261361.1_Malawi_Lil-20/1_1983 ..............a...g........g..............................................g..... [153920]
AY261360.1_Kenya_1950           ..g........t..a...g........g..................ca..........................g..... [153920]
AM712240.1_OURT_88/3            ................................................................................ [153920]
AM712239.1_Benin_97/1           ................................................................................ [153920]
KM111295.1_Ken06.Bus            ..g........t..a...g........g..................ca..........................g..... [153920]
KM111294.1_Ken05/Tk1            ..g........t..a...g........g..................ca..........................g..... [153920]
KM262844.1_strain_L60           ................................................................................ [153920]
KM102979.1_26544/OG10_Italy     ................................................................................ [153920]
KM262845.1_strain_NHV           ................................................................................ [153920]
KX354450.1_47/Ss/2008           ................................................................................ [153920]
KP055815.1_strain_BA71          ................................................................................ [153920]
MH025920.1_strain_R35           ..g........t..a...g........g..................ca..........................g..... [153920]
MH025919.1_strain_N10           ..g........t..a...g........g..................ca..........................g..... [153920]
MH025918.1_strain_R25           ..g........t..a...g........g..................ca..........................g..... [153920]
MH025917.1_strain_R7            ..g........t..a...g........g..................ca..........................g..... [153920]
MH025916.1_strain_R8            ..g........t..a...g........g..................ca..........................g..... [153920]
```

```
FR682468.2_Georgia_2007/1        aacagaatataagcatatctacttccttgcaatgttatgtaagtcgttggaggaacccaatatgaatctttctttacaat [154000]
ASF_IND_20_CAD543                ................................................................................ [154000]
LR899193.1_Germany_2020/1        ................................................................................ [154000]
MN393477.1_Wuhan_2019            ................................................................................ [154000]
MW396979.1_Timor-Leste/2019/1    ................................................................................ [154000]
MK940252.1_CN/2019/InnerMongolia ................................................................................ [154000]
MK128995.1_China/2018/AnhuiXCGQ  ................................................................................ [154000]
MH681419.1_ASFV/POL/2015         ................................................................................ [154000]
LS478113.1_Estonia_2014          ................................................................................ [154000]
KP843857.1_Odintsovo_02/14       ................................................................................ [154000]
FN557520.1_strain_E75            ................................................................................ [154000]
AY261366.1_Warthog               ...............c................................................................ [154000]
AY261365.1_Warmbaths             ................................................................................ [154000]
AY261364.1_Tengani_62            .............................................t.................................. [154000]
AY261363.1_Pretoriuskop/96/4     .............................................a.................................. [154000]
AY261362.1_Mkuzi_1979            ................................................................................ [154000]
AY261361.1_Malawi_Lil-20/1_1983  ..................a............g.c........a..t.................................. [154000]
AY261360.1_Kenya_1950            ..................a............g.............t.................................. [154000]
AM712240.1_OURT_88/3             ................................................................................ [154000]
AM712239.1_Benin_97/1            ................................................................................ [154000]
KM111295.1_Ken06.Bus             ..................a............g.............t.................................. [154000]
KM111294.1_Ken05/Tk1             ..................a............g.............t.................................. [154000]
KM262844.1_strain_L60            ................................................................................ [154000]
KM102979.1_26544/OG10_Italy      ................................................................................ [154000]
KM262845.1_strain_NHV            ................................................................................ [154000]
KX354450.1_47/Ss/2008            ................................................................................ [154000]
KP055815.1_strain_BA71           ................................................................................ [154000]
MH025920.1_strain_R35            ..................a............g.............t.................................. [154000]
MH025919.1_strain_N10            ..................a............g.............t.................................. [154000]
MH025918.1_strain_R25            ..................a............g.............t.................................. [154000]
MH025917.1_strain_R7             ..................a............g.............t.................................. [154000]
MH025916.1_strain_R8             ..................a............g.............t.................................. [154000]

FR682468.2_Georgia_2007/1        acgaaaaccgaattgccgaaatttctaaaatttcttggcaaaatatggaggctgtacgttttattagcaaacgccagtca [154080]
ASF_IND_20_CAD543                ................................................................................ [154080]
LR899193.1_Germany_2020/1        ................................................................................ [154080]
MN393477.1_Wuhan_2019            ................................................................................ [154080]
MW396979.1_Timor-Leste/2019/1    ................................................................................ [154080]
MK940252.1_CN/2019/InnerMongolia ................................................................................ [154080]
MK128995.1_China/2018/AnhuiXCGQ  ................................................................................ [154080]
MH681419.1_ASFV/POL/2015         ................................................................................ [154080]
LS478113.1_Estonia_2014          ................................................................................ [154080]
KP843857.1_Odintsovo_02/14       ................................................................................ [154080]
```

```
FN557520.1_strain_E75           .......t..g..................................................................... [154080]
AY261366.1_Warthog              ..........g..................................................................... [154080]
AY261365.1_Warmbaths            ..........g.................................................................a... [154080]
AY261364.1_Tengani_62           ..........g..................................................................... [154080]
AY261363.1_Pretoriuskop/96/4    ..........g.................................................................a... [154080]
AY261362.1_Mkuzi_1979           .......t..g..................................................................... [154080]
AY261361.1_Malawi_Lil-20/1_1983 ......c......................................................................... [154080]
AY261360.1_Kenya_1950           ......c.................................................................a....... [154080]
AM712240.1_OURT_88/3            .......t..g..................................................................... [154080]
AM712239.1_Benin_97/1           .......t..g..................................................................... [154080]
KM111295.1_Ken06.Bus            ......c......................................................................... [154080]
KM111294.1_Ken05/Tk1            ......c.................................................................a....... [154080]
KM262844.1_strain_L60           .......t..g..................................................................... [154080]
KM102979.1_26544/OG10_Italy     .......t..g..................................................................... [154080]
KM262845.1_strain_NHV           .......t..g..................................................................... [154080]
KX354450.1_47/Ss/2008           .......t..g..................................................................... [154080]
KP055815.1_strain_BA71          .......t..g..................................................................... [154080]
MH025920.1_strain_R35           ......c......................................................................... [154080]
MH025919.1_strain_N10           ......c......................................................................... [154080]
MH025918.1_strain_R25           ......c......................................................................... [154080]
MH025917.1_strain_R7            ......c......................................................................... [154080]
MH025916.1_strain_R8            ......c......................................................................... [154080]

FR682468.2_Georgia_2007/1       ttaaacctggagcctatcatcgggcctgcatttaattttattaaaaactatttacgatacaa-------------gcact [154160]
ASF_IND_20_CAD543               ..............................................................-------------..... [154160]
LR899193.1_Germany_2020/1       ..............................................................-------------..... [154160]
MN393477.1_Wuhan_2019           ..............................................................-------------..... [154160]
MW396979.1_Timor-Leste/2019/1   ..............................................................-------------..... [154160]
MK940252.1_CN/2019/InnerMongolia ..............................................................-------------..... [154160]
MK128995.1_China/2018/AnhuiXCGQ ..............................................................-------------..... [154160]
MH681419.1_ASFV/POL/2015        ..............................................................-------------..... [154160]
LS478113.1_Estonia_2014         ..............................................................-------------..... [154160]
KP843857.1_Odintsovo_02/14      ..............................................................-------------..... [154160]
FN557520.1_strain_E75           ..t........................................................t..-------------..... [154160]
AY261366.1_Warthog              ...........................................................t..-------------..... [154160]
AY261365.1_Warmbaths            .................t.........................................t..-------------..... [154160]
AY261364.1_Tengani_62           ..............................................................-------------..... [154160]
AY261363.1_Pretoriuskop/96/4    .................t.........................................t..-------------..... [154160]
AY261362.1_Mkuzi_1979           ..............................................................gcactaggatgcc..... [154160]
AY261361.1_Malawi_Lil-20/1_1983 ..............c............................................t..-------------...t. [154160]
AY261360.1_Kenya_1950           ..g...........c...............................................-------------a.... [154160]
AM712240.1_OURT_88/3            ..t........................................................t..-------------..... [154160]
AM712239.1_Benin_97/1           ..t........................................................t..-------------..... [154160]
KM111295.1_Ken06.Bus            ..g...........c...............................................-------------a.... [154160]
```

```
KM111294.1_Ken05/Tk1             ..g...........c...............................................-------------a.... [154160]
KM262844.1_strain_L60            ..t........................................................t..-------------..... [154160]
KM102979.1_26544/OG10_Italy      ..t........................................................t..-------------..... [154160]
KM262845.1_strain_NHV            ..t........................................................t..-------------..... [154160]
KX354450.1_47/Ss/2008            ..t........................................................t..-------------..... [154160]
KP055815.1_strain_BA71           ..t........................................................t..-------------..... [154160]
MH025920.1_strain_R35            ..g...........c...............................................-------------a.... [154160]
MH025919.1_strain_N10            ..g...........c...............................................-------------a.... [154160]
MH025918.1_strain_R25            ..g...........c...............................................-------------a.... [154160]
MH025917.1_strain_R7             ..g...........c...............................................-------------a.... [154160]
MH025916.1_strain_R8             ..g...........c...............................................-------------a.... [154160]

FR682468.2_Georgia_2007/1        agg------------atgccgcatta--------------aaatgccaca-taaggtaatacactaggaatgtcgcac-- [154240]
ASF_IND_20_CAD543                ...------------...........--------------..........-...........................-- [154240]
LR899193.1_Germany_2020/1        ...------------...........--------------..........-...........................-- [154240]
MN393477.1_Wuhan_2019            ...------------...........--------------..........-...........................-- [154240]
MW396979.1_Timor-Leste/2019/1    ...------------...........--------------..........-...........................-- [154240]
MK940252.1_CN/2019/InnerMongolia ...------------...........--------------..........-...........................-- [154240]
MK128995.1_China/2018/AnhuiXCGQ  ...------------...........--------------..........-...........................-- [154240]
MH681419.1_ASFV/POL/2015         ...------------...........--------------..........-...........................-- [154240]
LS478113.1_Estonia_2014          ...------------...........--------------..........-...........................-- [154240]
KP843857.1_Odintsovo_02/14       ...------------...........--------------..........-...........................-- [154240]
FN557520.1_strain_E75            ...------------...........--------------.......g..-.....................c.....-- [154240]
AY261366.1_Warthog               ...------------........c..--------------gg.....g..t..gaa.gccg.......-...c.....-- [154240]
AY261365.1_Warmbaths             ...------------.....a..c..--------------gg........-c........g...........c.....-- [154240]
AY261364.1_Tengani_62            ...------------........c..--------------gg.....g..---------------------------.-- [154240]
AY261363.1_Pretoriuskop/96/4     ...------------...........--------------g......g..-------------.........c.....-- [154240]
AY261362.1_Mkuzi_1979            ...------------...........--------------.......g..-....a................c.....-- [154240]
AY261361.1_Malawi_Lil-20/1_1983  ...----------------------------------------....g..-ctg...gccg....g..--..c....tat [154240]
AY261360.1_Kenya_1950            ...----------------------------------------....g..------------ct........c....tat [154240]
AM712240.1_OURT_88/3             ...------------...........--------------.......g..-.....................c.....-- [154240]
AM712239.1_Benin_97/1            ...------------...........--------------.......g..-.....................c.....-- [154240]
KM111295.1_Ken06.Bus             ...aatgcacttagg........c..ggtgccgcatttagg......g..------------tt........c.a..tat [154240]
KM111294.1_Ken05/Tk1             ...----------------------------------------....g..------------ct........c....tat [154240]
KM262844.1_strain_L60            ...------------...........--------------.......g..-.....................c.....-- [154240]
KM102979.1_26544/OG10_Italy      ...------------...........--------------.......g..-.....................c.....-- [154240]
KM262845.1_strain_NHV            ...------------...........--------------.......g..-.....................c.....-- [154240]
KX354450.1_47/Ss/2008            ...------------...........--------------.......g..-.....................c.....-- [154240]
KP055815.1_strain_BA71           ...------------...........--------------.......g..-.....................c.....-- [154240]
MH025920.1_strain_R35            ...aatgcacttagg........c..ggtgccgcatttagg......g..------------tt........c.a..tat [154240]
MH025919.1_strain_N10            ...aatgcacttagg........c..ggtgccgcatttagg......g..------------tt........c.a..tat [154240]
MH025918.1_strain_R25            ...aatgcacttagg........c..ggtgccgcatttagg......g..------------tt........c.a..tat [154240]
MH025917.1_strain_R7             ...aatgcacttagg........c..ggtgccgcatttagg......g..------------tt........c.a..tat [154240]
MH025916.1_strain_R8             ...aatgcacttagg........c..ggtgccgcatttagg......g..------------tt........c.a..tat [154240]
```

```
FR682468.2_Georgia_2007/1        ----acgcacaagaatacaacgtcgccggagatttattatctagtacacgttttatg-tatgtacaatccgccttcattt [154320]
ASF_IND_20_CAD543                ----.....................................................-...................... [154320]
LR899193.1_Germany_2020/1        ----.....................................................-...................... [154320]
MN393477.1_Wuhan_2019            ----.....................................................-...................... [154320]
MW396979.1_Timor-Leste/2019/1    ----.....................................................-...................... [154320]
MK940252.1_CN/2019/InnerMongolia ----.....................................................-...................... [154320]
MK128995.1_China/2018/AnhuiXCGQ  ----.....................................................-...................... [154320]
MH681419.1_ASFV/POL/2015         ----.....................................................-...................... [154320]
LS478113.1_Estonia_2014          ----.....................................................-...................... [154320]
KP843857.1_Odintsovo_02/14       ----.....................................................-...................... [154320]
FN557520.1_strain_E75            ----................t....................................-...................... [154320]
AY261366.1_Warthog               ----.........-----.........................t.....a.......-...................... [154320]
AY261365.1_Warmbaths             ----.........-----.........................t.....a.......-...................... [154320]
AY261364.1_Tengani_62            ----.......................................t.....a.......-...................... [154320]
AY261363.1_Pretoriuskop/96/4     ----.........-----..t......................t.....a.......-...................... [154320]
AY261362.1_Mkuzi_1979            ----.....................................................-...................... [154320]
AY261361.1_Malawi_Lil-20/1_1983  gatg...t.t....g........t..t.............t..-..t..........t.........c............ [154320]
AY261360.1_Kenya_1950            tatg..........g......a..............a..c...-.....a-......-.......-.............. [154320]
AM712240.1_OURT_88/3             ----................t....................................-...................... [154320]
AM712239.1_Benin_97/1            ----................t....................................-...................... [154320]
KM111295.1_Ken06.Bus             tatg..........g......a..............a..c...-......-......-.......-.............. [154320]
KM111294.1_Ken05/Tk1             tatg..........g......a..............a..c...-......-......-.......-.............. [154320]
KM262844.1_strain_L60            ----................t....................................-...................... [154320]
KM102979.1_26544/OG10_Italy      ----................t....................................-...................... [154320]
KM262845.1_strain_NHV            ----................t....................................-...................... [154320]
KX354450.1_47/Ss/2008            ----................t....................................-...................... [154320]
KP055815.1_strain_BA71           ----................t....................................-...................... [154320]
MH025920.1_strain_R35            tatg..........g......a..............a..c...-......-......-.......-.............. [154320]
MH025919.1_strain_N10            tatg..........g......a..............a..c...-......-......-.......-.............. [154320]
MH025918.1_strain_R25            tatg..........g......a..............a..c...-......-......-.......-.............. [154320]
MH025917.1_strain_R7             tatg..........g......a..............a..c...-......-......-.......-.............. [154320]
MH025916.1_strain_R8             tatg..........g......a..............a..c...-......-......-.......-.............. [154320]

FR682468.2_Georgia_2007/1        aatatattgagcggatgtac-tatgtatttattttaacaaa-aaacatta-tttttttt-------------taatcttc [154400]
ASF_IND_20_CAD543                ....................-....................a........-........-------------........ [154400]
LR899193.1_Germany_2020/1        ....................-....................-........-........-------------........ [154400]
MN393477.1_Wuhan_2019            ....................-....................-........-........-------------........ [154400]
MW396979.1_Timor-Leste/2019/1    ....................-....................-........-........-------------........ [154400]
MK940252.1_CN/2019/InnerMongolia ....................-....................-........-........-------------........ [154400]
MK128995.1_China/2018/AnhuiXCGQ  ....................-....................-........-........-------------........ [154400]
MH681419.1_ASFV/POL/2015         ....................-....................-........-........-------------........ [154400]
LS478113.1_Estonia_2014          ....................-....................-........-........-------------........ [154400]
KP843857.1_Odintsovo_02/14       ....................-....................-........--.......-------------........ [154400]
```

```
FN557520.1_strain_E75          ....................-....................a........-........-------------........ [154400]
AY261366.1_Warthog             ....................-..................---........-........-------------........ [154400]
AY261365.1_Warmbaths           ....................-..................---........-........-------------........ [154400]
AY261364.1_Tengani_62          ....................-....................a........-........-------------........ [154400]
AY261363.1_Pretoriuskop/96/4   ....................-..................---........-........-------------........ [154400]
AY261362.1_Mkuzi_1979          ....................-....................a........-........-------------........ [154400]
AY261361.1_Malawi_Lil-20/1_1983 .........t..........t....c..........g....a...t.a..-........gctggtattttaa.ct..... [154400]
AY261360.1_Kenya_1950          .........t.t........-g..............g....a...t.----........---ggatttttaa.cg..a.. [154400]
AM712240.1_OURT_88/3           ....................-....................a........-........-------------........ [154400]
AM712239.1_Benin_97/1          ....................-....................a........-........-------------........ [154400]
KM111295.1_Ken06.Bus           .........t.t........-g...................a...t....-........---tgtattttac.cg..a.. [154400]
KM111294.1_Ken05/Tk1           .........t.t........-g..............g....a...t..---........---ggatttttaa.cg..a.. [154400]
KM262844.1_strain_L60          ....................-....................a........-........-------------........ [154400]
KM102979.1_26544/OG10_Italy    ....................-....................a........-........-------------........ [154400]
KM262845.1_strain_NHV          ....................-....................a........-........-------------........ [154400]
KX354450.1_47/Ss/2008          ....................-....................a........-........-------------........ [154400]
KP055815.1_strain_BA71         ....................-....................a........-........-------------........ [154400]
MH025920.1_strain_R35          .........t.t........-g...................a...t....-........---tgtattttac.cg..a.. [154400]
MH025919.1_strain_N10          .........t.t........-g...................a...t....t........---tgtattttac.cg..a.. [154400]
MH025918.1_strain_R25          .........t.t........-g...................a...t....-........---tgtattttac.cg..a.. [154400]
MH025917.1_strain_R7           .........t.t........-g...................a...t....-........---tgtattttac.cg..a.. [154400]
MH025916.1_strain_R8           .........t.t........-g...................a...t....-........---tgtattttac.cg..a.. [154400]

FR682468.2_Georgia_2007/1      atcatctgtttttataaactcagtaatatcaaaagtagcttgtggggtttcagagggttcaccttgg------------- [154480]
ASF_IND_20_CAD543              .......r...........................................................------------- [154480]
LR899193.1_Germany_2020/1      ...................................................................------------- [154480]
MN393477.1_Wuhan_2019          ...................................................................------------- [154480]
MW396979.1_Timor-Leste/2019/1  ...................................................................------------- [154480]
MK940252.1_CN/2019/InnerMongolia ...................................................................------------- [154480]
MK128995.1_China/2018/AnhuiXCGQ ...................................................................------------- [154480]
MH681419.1_ASFV/POL/2015       ...................................................................------------- [154480]
LS478113.1_Estonia_2014        ...................................................................------------- [154480]
KP843857.1_Odintsovo_02/14     ...................................................................------------- [154480]
FN557520.1_strain_E75          .......a...........................................................------------- [154480]
AY261366.1_Warthog             ...c...............................................................------------- [154480]
AY261365.1_Warmbaths           ...c................................................a..............------------- [154480]
AY261364.1_Tengani_62          ...c...............................................................tt----------- [154480]
AY261363.1_Pretoriuskop/96/4   ...c...............................................................------------- [154480]
AY261362.1_Mkuzi_1979          ...c...............................................................------------- [154480]
AY261361.1_Malawi_Lil-20/1_1983 ...c......................................c...t.......t......tg...attaattttgtggt [154480]
AY261360.1_Kenya_1950          c..c................................g..........c..t...t.a....ta...attatctctgtgga [154480]
AM712240.1_OURT_88/3           .......a...........................................................------------- [154480]
AM712239.1_Benin_97/1          .......a...........................................................------------- [154480]
KM111295.1_Ken06.Bus           c..c...a.......................................c..t...t.a....ta...attatctctgtgaa [154480]
```

```
KM111294.1_Ken05/Tk1            c..c...........................................c..t...t.a....ta...attatctctgtgga [154480]
KM262844.1_strain_L60           .......a...........................................................------------- [154480]
KM102979.1_26544/OG10_Italy     .......a...........................................................------------- [154480]
KM262845.1_strain_NHV           .......a...........................................................------------- [154480]
KX354450.1_47/Ss/2008           .......a...........................................................------------- [154480]
KP055815.1_strain_BA71          .......a...........................................................------------- [154480]
MH025920.1_strain_R35           c..c...a.......................................c..t...t.a....ta...attatctctgtgaa [154480]
MH025919.1_strain_N10           c..c...a.......................................c..t...t.a....ta...attatctctgtgaa [154480]
MH025918.1_strain_R25           c..c...a.......................................c..t...t.a....ta...attatctctgtgaa [154480]
MH025917.1_strain_R7            c..c...a.......................................c..t...t.a....ta...attatctctgtgaa [154480]
MH025916.1_strain_R8            c..c...a.......................................c..t...t.a....ta...attatctctgtgaa [154480]

FR682468.2_Georgia_2007/1       --------------------------------------------------------------ttatcctccgtgaggata [154560]
ASF_IND_20_CAD543               --------------------------------------------------------------.................. [154560]
LR899193.1_Germany_2020/1       --------------------------------------------------------------.................. [154560]
MN393477.1_Wuhan_2019           --------------------------------------------------------------.................. [154560]
MW396979.1_Timor-Leste/2019/1   --------------------------------------------------------------.................. [154560]
MK940252.1_CN/2019/InnerMongolia --------------------------------------------------------------.................. [154560]
MK128995.1_China/2018/AnhuiXCGQ --------------------------------------------------------------.................. [154560]
MH681419.1_ASFV/POL/2015        --------------------------------------------------------------.................. [154560]
LS478113.1_Estonia_2014         --------------------------------------------------------------.................. [154560]
KP843857.1_Odintsovo_02/14      --------------------------------------------------------------.................. [154560]
FN557520.1_strain_E75           --------------------------------------------------------------.................. [154560]
AY261366.1_Warthog              --------------------------------------------------------------.................. [154560]
AY261365.1_Warmbaths            --------------------------------------------------------------...........a...... [154560]
AY261364.1_Tengani_62           -------------------------------------------------------------a..............a... [154560]
AY261363.1_Pretoriuskop/96/4    --------------------------------------------------------------.................. [154560]
AY261362.1_Mkuzi_1979           --------------------------------------------------------------.................. [154560]
AY261361.1_Malawi_Lil-20/1_1983 ggttaacctcgtggtggttaatcttgtgatggttaacctcatg---------------atgg...a....atga.a.t.. [154560]
AY261360.1_Kenya_1950           gattaatctccat---------------------------------------------gaag...a....atg.g..t.. [154560]
AM712240.1_OURT_88/3            --------------------------------------------------------------.................. [154560]
AM712239.1_Benin_97/1           --------------------------------------------------------------.................. [154560]
KM111295.1_Ken06.Bus            ggttaatctccatgaggttaacctcatgagggttaacttcatgagggttaacttcatggaag...a....atg.g.at.. [154560]
KM111294.1_Ken05/Tk1            ggttaatctccatgaggttaacctcatgagggttaacttcata---------------gaag...a....atg.g..t.. [154560]
KM262844.1_strain_L60           --------------------------------------------------------------.................. [154560]
KM102979.1_26544/OG10_Italy     --------------------------------------------------------------.................. [154560]
KM262845.1_strain_NHV           --------------------------------------------------------------.................. [154560]
KX354450.1_47/Ss/2008           --------------------------------------------------------------.................. [154560]
KP055815.1_strain_BA71          --------------------------------------------------------------.................. [154560]
MH025920.1_strain_R35           ggttaatctccatgaggttaacctcatgagggttaacttcatgagggttaacttcatggaag...a....atg.g.at.. [154560]
MH025919.1_strain_N10           ggttaatctccatgaggttaacctcatgagggttaacttcatgagggttaacttcatggaag...a....atg.g.at.. [154560]
MH025918.1_strain_R25           ggttaatctccatgaggttaacctcatgagggttaacttcatgagggttaacttcatggaag...a....atg.g.at.. [154560]
MH025917.1_strain_R7            ggttaatctccatgaggttaacctcatgagggttaacttcatgagggttaacttcatggaag...a....atg.g.at.. [154560]
MH025916.1_strain_R8            ggttaatctccatgaggttaacctcatgagggttaacttcatgagggttaacttcatggaag...a....atg.g.at.. [154560]
```

```
FR682468.2_Georgia_2007/1        acatgttcttcaggttcgtcgtcactggagaacccatcatttaattcctcttcactcaacatctgtaaaaaatcttccaa [154640]
ASF_IND_20_CAD543                ................................................................................ [154640]
LR899193.1_Germany_2020/1        ................................................................................ [154640]
MN393477.1_Wuhan_2019            ................................................................................ [154640]
MW396979.1_Timor-Leste/2019/1    ................................................................................ [154640]
MK940252.1_CN/2019/InnerMongolia ................................................................................ [154640]
MK128995.1_China/2018/AnhuiXCGQ  ................................................................................ [154640]
MH681419.1_ASFV/POL/2015         ................................................................................ [154640]
LS478113.1_Estonia_2014          ................................................................................ [154640]
KP843857.1_Odintsovo_02/14       ................................................................................ [154640]
FN557520.1_strain_E75            ................................................................................ [154640]
AY261366.1_Warthog               ...........g........a.................g...........c...t......................... [154640]
AY261365.1_Warmbaths             ...........g........a........a........g...........c...t......................... [154640]
AY261364.1_Tengani_62            ...........g........a.................g...........c...t......................... [154640]
AY261363.1_Pretoriuskop/96/4     ...........g........a.................g...........c...t......................... [154640]
AY261362.1_Mkuzi_1979            ...................................g..g...............t......................... [154640]
AY261361.1_Malawi_Lil-20/1_1983  ...c.......g........a.....a......t....g...........a...t.......t...........c..... [154640]
AY261360.1_Kenya_1950            ...c.......g.....a..a.....a.......t...g.......c...a...t...t..................... [154640]
AM712240.1_OURT_88/3             ................................................................................ [154640]
AM712239.1_Benin_97/1            ................................................................................ [154640]
KM111295.1_Ken06.Bus             ...c.......g.....a..a.....a.......t...g.......c...a...t...t..................... [154640]
KM111294.1_Ken05/Tk1             ...c.......g.....a..a.....a.......t...g.......c...a...t...t..................... [154640]
KM262844.1_strain_L60            ................................................................................ [154640]
KM102979.1_26544/OG10_Italy      ................................................................................ [154640]
KM262845.1_strain_NHV            ................................................................................ [154640]
KX354450.1_47/Ss/2008            ................................................................................ [154640]
KP055815.1_strain_BA71           ................................................................................ [154640]
MH025920.1_strain_R35            ...c.......g.....a..a.....a.......t...g.......c...a...t...t..................... [154640]
MH025919.1_strain_N10            ...c.......g.....a..a.....a.......t...g.......c...a...t...t..................... [154640]
MH025918.1_strain_R25            ...c.......g.....a..a.....a.......t...g.......c...a...t...t..................... [154640]
MH025917.1_strain_R7             ...c.......g.....a..a.....a.......t...g.......c...a...t...t..................... [154640]
MH025916.1_strain_R8             ...c.......g.....a..a.....a.......t...g.......c...a...t...t..................... [154640]

FR682468.2_Georgia_2007/1        gctttcgctatcgttaaaatcctcatcatccataagaataatggtaccttcctcatcgtttcctccttgtttcgtgtcta [154720]
ASF_IND_20_CAD543                ................................................................................ [154720]
LR899193.1_Germany_2020/1        ................................................................................ [154720]
MN393477.1_Wuhan_2019            ................................................................................ [154720]
MW396979.1_Timor-Leste/2019/1    ................................................................................ [154720]
MK940252.1_CN/2019/InnerMongolia ................................................................................ [154720]
MK128995.1_China/2018/AnhuiXCGQ  ................................................................................ [154720]
MH681419.1_ASFV/POL/2015         ................................................................................ [154720]
LS478113.1_Estonia_2014          ................................................................................ [154720]
KP843857.1_Odintsovo_02/14       ................................................................................ [154720]
```

| | | |
|---|---|---|
| FN557520.1_strain_E75 | ................................................................................ | [154720] |
| AY261366.1_Warthog | ................................................................................ | [154720] |
| AY261365.1_Warmbaths | ................................................................................ | [154720] |
| AY261364.1_Tengani_62 | ................................................................................ | [154720] |
| AY261363.1_Pretoriuskop/96/4 | .............................................g.................................. | [154720] |
| AY261362.1_Mkuzi_1979 | ................................................................................ | [154720] |
| AY261361.1_Malawi_Lil-20/1_1983 | ...................................t...............................c............ | [154720] |
| AY261360.1_Kenya_1950 | a...........a......................t.........g.................................. | [154720] |
| AM712240.1_OURT_88/3 | ................................................................................ | [154720] |
| AM712239.1_Benin_97/1 | ................................................................................ | [154720] |
| KM111295.1_Ken06.Bus | a...........a......................t.........g.................................. | [154720] |
| KM111294.1_Ken05/Tk1 | a...........a......................t.........g.................................. | [154720] |
| KM262844.1_strain_L60 | ................................................................................ | [154720] |
| KM102979.1_26544/OG10_Italy | ................................................................................ | [154720] |
| KM262845.1_strain_NHV | ................................................................................ | [154720] |
| KX354450.1_47/Ss/2008 | ................................................................................ | [154720] |
| KP055815.1_strain_BA71 | ................................................................................ | [154720] |
| MH025920.1_strain_R35 | a...........a......................t.........g.................................. | [154720] |
| MH025919.1_strain_N10 | a...........a......................t.........g.................................. | [154720] |
| MH025918.1_strain_R25 | a...........a......................t.........g.................................. | [154720] |
| MH025917.1_strain_R7 | a...........a......................t.........g.................................. | [154720] |
| MH025916.1_strain_R8 | a...........a......................t.........g.................................. | [154720] |

| | | |
|---|---|---|
| FR682468.2_Georgia_2007/1 | aataggcctgcatggcatttgcaaaagtatcaaaataggctgagtcagattgctgttccaaaatatggccttgcgtatta | [154800] |
| ASF_IND_20_CAD543 | ................................................................................ | [154800] |
| LR899193.1_Germany_2020/1 | ................................................................................ | [154800] |
| MN393477.1_Wuhan_2019 | ................................................................................ | [154800] |
| MW396979.1_Timor-Leste/2019/1 | ................................................................................ | [154800] |
| MK940252.1_CN/2019/InnerMongolia | ................................................................................ | [154800] |
| MK128995.1_China/2018/AnhuiXCGQ | ................................................................................ | [154800] |
| MH681419.1_ASFV/POL/2015 | ................................................................................ | [154800] |
| LS478113.1_Estonia_2014 | ................................................................................ | [154800] |
| KP843857.1_Odintsovo_02/14 | ................................................................................ | [154800] |
| FN557520.1_strain_E75 | .......................................g........................................ | [154800] |
| AY261366.1_Warthog | .......t...............................g........................................ | [154800] |
| AY261365.1_Warmbaths | .......................................g........................................ | [154800] |
| AY261364.1_Tengani_62 | .......................................g.................................t...... | [154800] |
| AY261363.1_Pretoriuskop/96/4 | .......................................g........................................ | [154800] |
| AY261362.1_Mkuzi_1979 | .......................................g........................................ | [154800] |
| AY261361.1_Malawi_Lil-20/1_1983 | .......................................g............t..........g................ | [154800] |
| AY261360.1_Kenya_1950 | t................c.c..............g....t..g.........t....................a...... | [154800] |
| AM712240.1_OURT_88/3 | .......................................g........................................ | [154800] |
| AM712239.1_Benin_97/1 | .......................................g........................................ | [154800] |
| KM111295.1_Ken06.Bus | t..................c..............g....t..ga........t..c.................a...... | [154800] |

```
KM111294.1_Ken05/Tk1             t................c.c..............g....t..g.........t....................a...... [154800]
KM262844.1_strain_L60            .......................................g........................................ [154800]
KM102979.1_26544/OG10_Italy      .......................................g........................................ [154800]
KM262845.1_strain_NHV            .......................................g........................................ [154800]
KX354450.1_47/Ss/2008            .......................................g........................................ [154800]
KP055815.1_strain_BA71           .......................................g........................................ [154800]
MH025920.1_strain_R35            t..................c..............g....t..ga........t..c.................a...... [154800]
MH025919.1_strain_N10            t..................c..............g....t..ga........t..c.................a...... [154800]
MH025918.1_strain_R25            t..................c..............g....t..ga........t..c.................a...... [154800]
MH025917.1_strain_R7             t..................c..............g....t..ga........t..c.................a...... [154800]
MH025916.1_strain_R8             t..................c..............g....t..ga........t..c.................a...... [154800]

FR682468.2_Georgia_2007/1        aatgtggttgcatcgttgttaaatgcttgcaaatacagtaagggatttatatccattattattaagc-aaaaaaaattta [154880]
ASF_IND_20_CAD543                ...................................................................-............ [154880]
LR899193.1_Germany_2020/1        ...................................................................-............ [154880]
MN393477.1_Wuhan_2019            ...................................................................-............ [154880]
MW396979.1_Timor-Leste/2019/1    ...................................................................-............ [154880]
MK940252.1_CN/2019/InnerMongolia ...................................................................-............ [154880]
MK128995.1_China/2018/AnhuiXCGQ  ...................................................................-............ [154880]
MH681419.1_ASFV/POL/2015         ...................................................................-............ [154880]
LS478113.1_Estonia_2014          ...................................................................-............ [154880]
KP843857.1_Odintsovo_02/14       ...................................................................-............ [154880]
FN557520.1_strain_E75            ...................................................................a............ [154880]
AY261366.1_Warthog               ...................................t...............................a............ [154880]
AY261365.1_Warmbaths             ...................................t...............................a............ [154880]
AY261364.1_Tengani_62            ...................................t...............................-............ [154880]
AY261363.1_Pretoriuskop/96/4     ...................................t...............................a............ [154880]
AY261362.1_Mkuzi_1979            ...................................................................-............ [154880]
AY261361.1_Malawi_Lil-20/1_1983  ...................................t...............................-............ [154880]
AY261360.1_Kenya_1950            ...................................t.............a.................-............ [154880]
AM712240.1_OURT_88/3             ...................................................................-............ [154880]
AM712239.1_Benin_97/1            ...................................................................a............ [154880]
KM111295.1_Ken06.Bus             ..............a..a.................t.............a.................a............ [154880]
KM111294.1_Ken05/Tk1             ......a............................t.............a.................-............ [154880]
KM262844.1_strain_L60            ...................................................................-............ [154880]
KM102979.1_26544/OG10_Italy      ...................................................................a............ [154880]
KM262845.1_strain_NHV            ...................................................................-............ [154880]
KX354450.1_47/Ss/2008            ...................................................................a............ [154880]
KP055815.1_strain_BA71           ...................................................................-............ [154880]
MH025920.1_strain_R35            ..............a..a.................t.............a.................a............ [154880]
MH025919.1_strain_N10            ..............a..a.................t.............a.................a............ [154880]
MH025918.1_strain_R25            ..............a..a.................t.............a.................a............ [154880]
MH025917.1_strain_R7             ..............a..a.................t.............a.................a............ [154880]
MH025916.1_strain_R8             ..............a..a.................t.............a.................a............ [154880]
```

```
FR682468.2_Georgia_2007/1        aattatttttcgaccgatgttaggtaaa--------------------------------------attaaacaattgct [154960]
ASF_IND_20_CAD543                ............................--------------------------------------.............. [154960]
LR899193.1_Germany_2020/1        ............................--------------------------------------.............. [154960]
MN393477.1_Wuhan_2019            ............................--------------------------------------.............. [154960]
MW396979.1_Timor-Leste/2019/1    ............................--------------------------------------.............. [154960]
MK940252.1_CN/2019/InnerMongolia ............................--------------------------------------.............. [154960]
MK128995.1_China/2018/AnhuiXCGQ  ............................--------------------------------------.............. [154960]
MH681419.1_ASFV/POL/2015         ............................--------------------------------------.............. [154960]
LS478113.1_Estonia_2014          ............................--------------------------------------.............. [154960]
KP843857.1_Odintsovo_02/14       ............................--------------------------------------.............. [154960]
FN557520.1_strain_E75            ............................--------------------------------------.............. [154960]
AY261366.1_Warthog               ............................--------------------------------------.....g.------- [154960]
AY261365.1_Warmbaths             ............................--------------------------------------.....g.------- [154960]
AY261364.1_Tengani_62            ..........t.................--------------------------------------.....g.------- [154960]
AY261363.1_Pretoriuskop/96/4     ............................--------------------------------------.....g-------- [154960]
AY261362.1_Mkuzi_1979            .....................t..c...--------------------------------------.....g........ [154960]
AY261361.1_Malawi_Lil-20/1_1983  ...........a................--------------------------------------.............. [154960]
AY261360.1_Kenya_1950            ...........agt..............ttaaacacctgttgtgattttaaacacctgtttgtgatt....g..cc..t. [154960]
AM712240.1_OURT_88/3             ............................--------------------------------------.............. [154960]
AM712239.1_Benin_97/1            ............................--------------------------------------.............. [154960]
KM111295.1_Ken06.Bus             ...........ag.............c.---------------------------------------....g...c..t. [154960]
KM111294.1_Ken05/Tk1             ...........agt...........g..---------------------------------------.......cc..t. [154960]
KM262844.1_strain_L60            ............................--------------------------------------.............. [154960]
KM102979.1_26544/OG10_Italy      ............................--------------------------------------.............. [154960]
KM262845.1_strain_NHV            ............................--------------------------------------.............. [154960]
KX354450.1_47/Ss/2008            ............................--------------------------------------.............. [154960]
KP055815.1_strain_BA71           ............................--------------------------------------.............. [154960]
MH025920.1_strain_R35            ...........ag.............c.---------------------------------------....g...c..t. [154960]
MH025919.1_strain_N10            ...........ag.............c.---------------------------------------....g...c..t. [154960]
MH025918.1_strain_R25            ...........ag.............c.---------------------------------------....g...c..t. [154960]
MH025917.1_strain_R7             ...........ag.............c.---------------------------------------....g...c..t. [154960]
MH025916.1_strain_R8             ...........ag.............c.---------------------------------------....g...c..t. [154960]

FR682468.2_Georgia_2007/1        atag--------gtgttaagcaatg------------------------------------------------------- [155040]
ASF_IND_20_CAD543                ....--------.............------------------------------------------------------- [155040]
LR899193.1_Germany_2020/1        ....--------.............------------------------------------------------------- [155040]
MN393477.1_Wuhan_2019            ....--------.............------------------------------------------------------- [155040]
MW396979.1_Timor-Leste/2019/1    ....--------.............------------------------------------------------------- [155040]
MK940252.1_CN/2019/InnerMongolia ....--------.............------------------------------------------------------- [155040]
MK128995.1_China/2018/AnhuiXCGQ  ....--------.............------------------------------------------------------- [155040]
MH681419.1_ASFV/POL/2015         ....--------.............------------------------------------------------------- [155040]
LS478113.1_Estonia_2014          ....--------.............------------------------------------------------------- [155040]
KP843857.1_Odintsovo_02/14       ....--------.............------------------------------------------------------- [155040]
```

```
FN557520.1_strain_E75            ....--------.............------------------------------------------------------- [155040]
AY261366.1_Warthog               ..t.--------c.a..........------------------------------------------------------- [155040]
AY261365.1_Warmbaths             ..t.--------c.a..........c--------------------------------------------------tatt [155040]
AY261364.1_Tengani_62            ..t.--------c.a......g...ct--taatagcaat-----------------------------------actatt [155040]
AY261363.1_Pretoriuskop/96/4     ..t.--------c.a..........------------------------------------------------------- [155040]
AY261362.1_Mkuzi_1979            ....--------.............------------------------------------------------------- [155040]
AY261361.1_Malawi_Lil-20/1_1983  g.gacggccaac............tgttgtgacggtcaa-----------------------------------cgtgtt [155040]
AY261360.1_Kenya_1950            tgt.--------a.t........ctgt--ttgtgattttaaacacctgtttgtgattttaagcaattgctgatagatgtt [155040]
AM712240.1_OURT_88/3             ....--------.............------------------------------------------------------- [155040]
AM712239.1_Benin_97/1            ....--------.............------------------------------------------------------- [155040]
KM111295.1_Ken06.Bus             tgt.--------a.t.....t...tgc--tgata-------------------gatgttaagtaa-tatttattgatgtt [155040]
KM111294.1_Ken05/Tk1             tgt.--------a.t.........tgc--tgata----------------------------------------gatgtt [155040]
KM262844.1_strain_L60            ....--------.............------------------------------------------------------- [155040]
KM102979.1_26544/OG10_Italy      ....--------.............------------------------------------------------------- [155040]
KM262845.1_strain_NHV            ....--------.............------------------------------------------------------- [155040]
KX354450.1_47/Ss/2008            ....--------.............------------------------------------------------------- [155040]
KP055815.1_strain_BA71           ....--------.............------------------------------------------------------- [155040]
MH025920.1_strain_R35            tgt.--------a.t.....t...tgc--tgata-------------------gatgttaagtaa-tatttattgatgtt [155040]
MH025919.1_strain_N10            tgt.--------a.t.....t...tgc--tgata-------------------gatgttaagtaa-tatttattgatgtt [155040]
MH025918.1_strain_R25            tgt.--------a.t.....t...tgc--tgata-------------------gatgttaagtaa-tatttattgatgtt [155040]
MH025917.1_strain_R7             tgt.--------a.t.....t...tgc--tgata-------------------gatgttaagtaa-tatttattgatgtt [155040]
MH025916.1_strain_R8             tgt.--------a.t.....t...tgc--tgata-------------------gatgttaagtaa-tatttattgatgtt [155040]

FR682468.2_Georgia_2007/1        --------tttattgattttaagtactcaacaaccatgatgtaaatactatacagcacttttggatttttaatcaaatcc [155120]
ASF_IND_20_CAD543                --------........................................................................ [155120]
LR899193.1_Germany_2020/1        --------........................................................................ [155120]
MN393477.1_Wuhan_2019            --------........................................................................ [155120]
MW396979.1_Timor-Leste/2019/1    --------........................................................................ [155120]
MK940252.1_CN/2019/InnerMongolia --------........................................................................ [155120]
MK128995.1_China/2018/AnhuiXCGQ  --------........................................................................ [155120]
MH681419.1_ASFV/POL/2015         --------........................................................................ [155120]
LS478113.1_Estonia_2014          --------........................................................................ [155120]
KP843857.1_Odintsovo_02/14       --------........................................................................ [155120]
FN557520.1_strain_E75            --------........................................................................ [155120]
AY261366.1_Warthog               --------..c....................g................................................ [155120]
AY261365.1_Warmbaths             aagcaata..c....................g................................................ [155120]
AY261364.1_Tengani_62            aagcaata..c...........a........g................................................ [155120]
AY261363.1_Pretoriuskop/96/4     --------..c....................g................................................ [155120]
AY261362.1_Mkuzi_1979            --------........................................................................ [155120]
AY261361.1_Malawi_Lil-20/1_1983  aagcaatg..c...........a........g..t.a.t...............a......................... [155120]
AY261360.1_Kenya_1950            aagtaata..............a.......................g............................g.... [155120]
AM712240.1_OURT_88/3             --------........................................................................ [155120]
AM712239.1_Benin_97/1            --------........................................................................ [155120]
KM111295.1_Ken06.Bus             aagtaata..............a.......................g............................g.... [155120]
```

```
KM111294.1_Ken05/Tk1             aagtaata..............a.......................g............................g.... [155120]
KM262844.1_strain_L60            --------........................................................................ [155120]
KM102979.1_26544/OG10_Italy      --------........................................................................ [155120]
KM262845.1_strain_NHV            --------........................................................................ [155120]
KX354450.1_47/Ss/2008            --------........................................................................ [155120]
KP055815.1_strain_BA71           --------........................................................................ [155120]
MH025920.1_strain_R35            aagtaata..c...........a.......................g............................g.... [155120]
MH025919.1_strain_N10            aagtaata..c...........a.......................g............................g.... [155120]
MH025918.1_strain_R25            aagtaata..c...........a.......................g............................g.... [155120]
MH025917.1_strain_R7             aagtaata..c...........a.......................g............................g.... [155120]
MH025916.1_strain_R8             aagtaata..c...........a.......................g............................g.... [155120]

FR682468.2_Georgia_2007/1        agattaatactaacttcttttgtgatacagttcgtaataatagtatcctgctcatcgttttgtaagatttcttttaatat [155200]
ASF_IND_20_CAD543                ................................................................................ [155200]
LR899193.1_Germany_2020/1        ................................................................................ [155200]
MN393477.1_Wuhan_2019            ................................................................................ [155200]
MW396979.1_Timor-Leste/2019/1    ................................................................................ [155200]
MK940252.1_CN/2019/InnerMongolia ................................................................................ [155200]
MK128995.1_China/2018/AnhuiXCGQ  ................................................................................ [155200]
MH681419.1_ASFV/POL/2015         ................................................................................ [155200]
LS478113.1_Estonia_2014          ................................................................................ [155200]
KP843857.1_Odintsovo_02/14       ................................................................................ [155200]
FN557520.1_strain_E75            ................................................................................ [155200]
AY261366.1_Warthog               .........................................g.....t..t............................. [155200]
AY261365.1_Warmbaths             .........................................g.....t..t............................. [155200]
AY261364.1_Tengani_62            .........................................g.....t..t............................. [155200]
AY261363.1_Pretoriuskop/96/4     .........................................g.....t..t............................. [155200]
AY261362.1_Mkuzi_1979            ................................................................................ [155200]
AY261361.1_Malawi_Lil-20/1_1983  ...............................................t.......t............c........... [155200]
AY261360.1_Kenya_1950            ................................t........g..............a....................... [155200]
AM712240.1_OURT_88/3             ................................................................................ [155200]
AM712239.1_Benin_97/1            ................................................................................ [155200]
KM111295.1_Ken06.Bus             ........g.......................t........g.......................a.............. [155200]
KM111294.1_Ken05/Tk1             ................................t........g..............a....................... [155200]
KM262844.1_strain_L60            ................................................................................ [155200]
KM102979.1_26544/OG10_Italy      ................................................................................ [155200]
KM262845.1_strain_NHV            ................................................................................ [155200]
KX354450.1_47/Ss/2008            ................................................................................ [155200]
KP055815.1_strain_BA71           ................................................................................ [155200]
MH025920.1_strain_R35            ........g.......................t........g.......................a.............. [155200]
MH025919.1_strain_N10            ........g.......................t........g.......................a.............. [155200]
MH025918.1_strain_R25            ........g.......................t........g.......................a.............. [155200]
MH025917.1_strain_R7             ........g.......................t........g.......................a.............. [155200]
MH025916.1_strain_R8             ........g.......................t........g.......................a.............. [155200]
```

```
FR682468.2_Georgia_2007/1        atttttttttaccgggatactaagcaattgattattttcttttaaaaactccttttgatattcaatcgtcttattcattg [155280]
ASF_IND_20_CAD543                ................................................................................ [155280]
LR899193.1_Germany_2020/1        ................................................................................ [155280]
MN393477.1_Wuhan_2019            ................................................................................ [155280]
MW396979.1_Timor-Leste/2019/1    ................................................................................ [155280]
MK940252.1_CN/2019/InnerMongolia ................................................................................ [155280]
MK128995.1_China/2018/AnhuiXCGQ  ................................................................................ [155280]
MH681419.1_ASFV/POL/2015         ................................................................................ [155280]
LS478113.1_Estonia_2014          ................................................................................ [155280]
KP843857.1_Odintsovo_02/14       .-.............................................................................. [155280]
FN557520.1_strain_E75            ................................................................................ [155280]
AY261366.1_Warthog               .......................................c...........t............................ [155280]
AY261365.1_Warmbaths             ...........t.......................................t............................ [155280]
AY261364.1_Tengani_62            ...........t............t....................................................... [155280]
AY261363.1_Pretoriuskop/96/4     ...............a...................................t............................ [155280]
AY261362.1_Mkuzi_1979            ................................................................................ [155280]
AY261361.1_Malawi_Lil-20/1_1983  .c.................t...............................t............................ [155280]
AY261360.1_Kenya_1950            .........g.t.......t.........................g.................................. [155280]
AM712240.1_OURT_88/3             ................................................................................ [155280]
AM712239.1_Benin_97/1            ................................................................................ [155280]
KM111295.1_Ken06.Bus             .........g.t.................................g.....t.................t.......... [155280]
KM111294.1_Ken05/Tk1             .........g.t.................................g.................................. [155280]
KM262844.1_strain_L60            ................................................................................ [155280]
KM102979.1_26544/OG10_Italy      ................................................................................ [155280]
KM262845.1_strain_NHV            ................................................................................ [155280]
KX354450.1_47/Ss/2008            ................................................................................ [155280]
KP055815.1_strain_BA71           ................................................................................ [155280]
MH025920.1_strain_R35            .........g.t.................................g.....t.................t.......... [155280]
MH025919.1_strain_N10            .........g.t.................................g.....t.................t.......... [155280]
MH025918.1_strain_R25            .........g.t.................................g.....t.................t.......... [155280]
MH025917.1_strain_R7             .........g.t.................................g.....t.................t.......... [155280]
MH025916.1_strain_R8             .........g.t.................................g.....t.................t.......... [155280]

FR682468.2_Georgia_2007/1        aatatttgtatataactataattacaaatgttcaatgaattgttattca------------------------------- [155360]
ASF_IND_20_CAD543                .................................................------------------------------- [155360]
LR899193.1_Germany_2020/1        .................................................------------------------------- [155360]
MN393477.1_Wuhan_2019            .................................................------------------------------- [155360]
MW396979.1_Timor-Leste/2019/1    .................................................------------------------------- [155360]
MK940252.1_CN/2019/InnerMongolia .................................................------------------------------- [155360]
MK128995.1_China/2018/AnhuiXCGQ  .................................................------------------------------- [155360]
MH681419.1_ASFV/POL/2015         .................................................------------------------------- [155360]
LS478113.1_Estonia_2014          .................................................------------------------------- [155360]
KP843857.1_Odintsovo_02/14       .................................................------------------------------- [155360]
```

```
FN557520.1_strain_E75            .................................................------------------------------- [155360]
AY261366.1_Warthog               ........................................a...c....------------------------------- [155360]
AY261365.1_Warmbaths             ........................................a...c....------------------------------- [155360]
AY261364.1_Tengani_62            ........................................a...c....------------------------------- [155360]
AY261363.1_Pretoriuskop/96/4     ........................................a...c....------------------------------- [155360]
AY261362.1_Mkuzi_1979            .................................................------------------------------- [155360]
AY261361.1_Malawi_Lil-20/1_1983  .................................................ttaaattactattcattaaatagttatttat [155360]
AY261360.1_Kenya_1950            ................................t.......a.-------------------------------------- [155360]
AM712240.1_OURT_88/3             .................................................------------------------------- [155360]
AM712239.1_Benin_97/1            .................................................------------------------------- [155360]
KM111295.1_Ken06.Bus             ................................t.......a.-------------------------------------- [155360]
KM111294.1_Ken05/Tk1             ................................t.......a.-------------------------------------- [155360]
KM262844.1_strain_L60            .................................................------------------------------- [155360]
KM102979.1_26544/OG10_Italy      .................................................------------------------------- [155360]
KM262845.1_strain_NHV            .................................................------------------------------- [155360]
KX354450.1_47/Ss/2008            .................................................------------------------------- [155360]
KP055815.1_strain_BA71           .................................................------------------------------- [155360]
MH025920.1_strain_R35            ................................t.......a.-------------------------------------- [155360]
MH025919.1_strain_N10            ................................t.......a.-------------------------------------- [155360]
MH025918.1_strain_R25            ................................t.......a.-------------------------------------- [155360]
MH025917.1_strain_R7             ................................t.......a.-------------------------------------- [155360]
MH025916.1_strain_R8             ................................t.......a.-------------------------------------- [155360]

FR682468.2_Georgia_2007/1        -------------------------------------------------------------------------------- [155440]
ASF_IND_20_CAD543                -------------------------------------------------------------------------------- [155440]
LR899193.1_Germany_2020/1        -------------------------------------------------------------------------------- [155440]
MN393477.1_Wuhan_2019            -------------------------------------------------------------------------------- [155440]
MW396979.1_Timor-Leste/2019/1    -------------------------------------------------------------------------------- [155440]
MK940252.1_CN/2019/InnerMongolia -------------------------------------------------------------------------------- [155440]
MK128995.1_China/2018/AnhuiXCGQ  -------------------------------------------------------------------------------- [155440]
MH681419.1_ASFV/POL/2015         -------------------------------------------------------------------------------- [155440]
LS478113.1_Estonia_2014          -------------------------------------------------------------------------------- [155440]
KP843857.1_Odintsovo_02/14       -------------------------------------------------------------------------------- [155440]
FN557520.1_strain_E75            -------------------------------------------------------------------------------- [155440]
AY261366.1_Warthog               -------------------------------------------------------------------------------- [155440]
AY261365.1_Warmbaths             -------------------------------------------------------------------------------- [155440]
AY261364.1_Tengani_62            -------------------------------------------------------------------------------- [155440]
AY261363.1_Pretoriuskop/96/4     -------------------------------------------------------------------------------- [155440]
AY261362.1_Mkuzi_1979            -------------------------------------------------------------------------------- [155440]
AY261361.1_Malawi_Lil-20/1_1983  taaatgatttattaaaatgttattcaatgaattgttattcaatgaattgttattcaatgaattattattcaattaacata [155440]
AY261360.1_Kenya_1950            -------------------------------------------------------------------------------- [155440]
AM712240.1_OURT_88/3             -------------------------------------------------------------------------------- [155440]
AM712239.1_Benin_97/1            -------------------------------------------------------------------------------- [155440]
KM111295.1_Ken06.Bus             -------------------------------------------------------------------------------- [155440]
```

```
KM111294.1_Ken05/Tk1             -------------------------------------------------------------------------------- [155440]
KM262844.1_strain_L60            -------------------------------------------------------------------------------- [155440]
KM102979.1_26544/OG10_Italy      -------------------------------------------------------------------------------- [155440]
KM262845.1_strain_NHV            -------------------------------------------------------------------------------- [155440]
KX354450.1_47/Ss/2008            -------------------------------------------------------------------------------- [155440]
KP055815.1_strain_BA71           -------------------------------------------------------------------------------- [155440]
MH025920.1_strain_R35            -------------------------------------------------------------------------------- [155440]
MH025919.1_strain_N10            -------------------------------------------------------------------------------- [155440]
MH025918.1_strain_R25            -------------------------------------------------------------------------------- [155440]
MH025917.1_strain_R7             -------------------------------------------------------------------------------- [155440]
MH025916.1_strain_R8             -------------------------------------------------------------------------------- [155440]

FR682468.2_Georgia_2007/1        -tgtcgggagatggctatttaaaaatcatgtcctatttttctttgctcaataagcatccaaatattttcatggcgtttta [155520]
ASF_IND_20_CAD543                -............................................................................... [155520]
LR899193.1_Germany_2020/1        -............................................................................... [155520]
MN393477.1_Wuhan_2019            -............................................................................... [155520]
MW396979.1_Timor-Leste/2019/1    -............................................................................... [155520]
MK940252.1_CN/2019/InnerMongolia -............................................................................... [155520]
MK128995.1_China/2018/AnhuiXCGQ  -............................................................................... [155520]
MH681419.1_ASFV/POL/2015         -............................................................................... [155520]
LS478113.1_Estonia_2014          -............................................................................... [155520]
KP843857.1_Odintsovo_02/14       -............................................................................... [155520]
FN557520.1_strain_E75            -............................................................................... [155520]
AY261366.1_Warthog               -..........c....g.......................................................a....... [155520]
AY261365.1_Warmbaths             -..........c....g.......................................................a....... [155520]
AY261364.1_Tengani_62            -.......g..c....g......c.....a.t........................................a....... [155520]
AY261363.1_Pretoriuskop/96/4     -..........c....g.......................................................a....... [155520]
AY261362.1_Mkuzi_1979            -............................................................................... [155520]
AY261361.1_Malawi_Lil-20/1_1983  g.a....c...c....g......-.ct.....a.......t...............................a....... [155520]
AY261360.1_Kenya_1950            -....a.c..-.....g......------a..a.......................................a....... [155520]
AM712240.1_OURT_88/3             -............................................................................... [155520]
AM712239.1_Benin_97/1            -............................................................................... [155520]
KM111295.1_Ken06.Bus             -....a.c......t.g......------a..a.......................................a....... [155520]
KM111294.1_Ken05/Tk1             -....a.c..-.....g......------a..a.......................................a....... [155520]
KM262844.1_strain_L60            -............................................................................... [155520]
KM102979.1_26544/OG10_Italy      -............................................................................... [155520]
KM262845.1_strain_NHV            -............................................................................... [155520]
KX354450.1_47/Ss/2008            -............................................................................... [155520]
KP055815.1_strain_BA71           -............................................................................... [155520]
MH025920.1_strain_R35            -....a.c......t.g......------a..a.......................................a....... [155520]
MH025919.1_strain_N10            -....a.c......t.g......------a..a.......................................a....... [155520]
MH025918.1_strain_R25            -....a.c......t.g......------a..a.......................................a....... [155520]
MH025917.1_strain_R7             -....a.c......t.g......------a..a.......................................a....... [155520]
MH025916.1_strain_R8             -....a.c......t.g......------a..a.......................................a....... [155520]
```

```
FR682468.2_Georgia_2007/1        ttaattgttcattattgaacgtatcacaaagatcatttataaattgcagatagtttattatttctttcaagagagtaaca [155600]
ASF_IND_20_CAD543                .....................................y.......................................... [155600]
LR899193.1_Germany_2020/1        ................................................................................ [155600]
MN393477.1_Wuhan_2019            ................................................................................ [155600]
MW396979.1_Timor-Leste/2019/1    ................................................................................ [155600]
MK940252.1_CN/2019/InnerMongolia ................................................................................ [155600]
MK128995.1_China/2018/AnhuiXCGQ  ................................................................................ [155600]
MH681419.1_ASFV/POL/2015         ................................................................................ [155600]
LS478113.1_Estonia_2014          ................................................................................ [155600]
KP843857.1_Odintsovo_02/14       ................................................................................ [155600]
FN557520.1_strain_E75            .....................................c.......................................... [155600]
AY261366.1_Warthog               ...................t.................c..g.......a............................... [155600]
AY261365.1_Warmbaths             .....................................c.......................................... [155600]
AY261364.1_Tengani_62            ................................................................................ [155600]
AY261363.1_Pretoriuskop/96/4     .....................................c..g.......a............................... [155600]
AY261362.1_Mkuzi_1979            ...................t.................c.......................................... [155600]
AY261361.1_Malawi_Lil-20/1_1983  ....c.....g..........................c.......................................... [155600]
AY261360.1_Kenya_1950            ....c.....g..........................c.......................................... [155600]
AM712240.1_OURT_88/3             .....................................c.......................................... [155600]
AM712239.1_Benin_97/1            .....................................c.......................................... [155600]
KM111295.1_Ken06.Bus             ....c.....g..........................c........................................g. [155600]
KM111294.1_Ken05/Tk1             ....c.....g..........................c.......................................... [155600]
KM262844.1_strain_L60            .....................................c.......................................... [155600]
KM102979.1_26544/OG10_Italy      .....................................c.......................................... [155600]
KM262845.1_strain_NHV            .....................................c.......................................... [155600]
KX354450.1_47/Ss/2008            .....................................c.......................................... [155600]
KP055815.1_strain_BA71           .....................................c.......................................... [155600]
MH025920.1_strain_R35            ....c.....g..........................c........................................g. [155600]
MH025919.1_strain_N10            ....c.....g..........................c........................................g. [155600]
MH025918.1_strain_R25            ....c.....g..........................c........................................g. [155600]
MH025917.1_strain_R7             ....c.....g..........................c........................................g. [155600]
MH025916.1_strain_R8             ....c.....g..........................c........................................g. [155600]

FR682468.2_Georgia_2007/1        aacattacttcagcagaacatataataggtaattcagtggcgttaaaagaattttgatcttgttgatacgccaatggcga [155680]
ASF_IND_20_CAD543                ................................................................................ [155680]
LR899193.1_Germany_2020/1        ................................................................................ [155680]
MN393477.1_Wuhan_2019            ................................................................................ [155680]
MW396979.1_Timor-Leste/2019/1    ................................................................................ [155680]
MK940252.1_CN/2019/InnerMongolia ................................................................................ [155680]
MK128995.1_China/2018/AnhuiXCGQ  ................................................................................ [155680]
MH681419.1_ASFV/POL/2015         ................................................................................ [155680]
LS478113.1_Estonia_2014          ................................................................................ [155680]
KP843857.1_Odintsovo_02/14       ................................................................................ [155680]
```

```
FN557520.1_strain_E75             ................................................................................ [155680]
AY261366.1_Warthog                ................................c...................................t........... [155680]
AY261365.1_Warmbaths              ................................c...................................t........... [155680]
AY261364.1_Tengani_62             ....................................................................t........... [155680]
AY261363.1_Pretoriuskop/96/4      ....................................................................t........... [155680]
AY261362.1_Mkuzi_1979             ...................................t............................................ [155680]
AY261361.1_Malawi_Lil-20/1_1983   ........................................t...........................t........t.. [155680]
AY261360.1_Kenya_1950             ..........................g...........c.ta.....c.....g.t............t........... [155680]
AM712240.1_OURT_88/3              ................................................................................ [155680]
AM712239.1_Benin_97/1             ................................................................................ [155680]
KM111295.1_Ken06.Bus              ..........................g........g..c.ta.....c.....g.t............t........... [155680]
KM111294.1_Ken05/Tk1              ..........................g...........c.ta.....c.....g.t............t........... [155680]
KM262844.1_strain_L60             ................................................................................ [155680]
KM102979.1_26544/OG10_Italy       ................................................................................ [155680]
KM262845.1_strain_NHV             ................................................................................ [155680]
KX354450.1_47/Ss/2008             ................................................................................ [155680]
KP055815.1_strain_BA71            ................................................................................ [155680]
MH025920.1_strain_R35             ..........................g........g..c.ta.....c.....g.t............t........... [155680]
MH025919.1_strain_N10             ..........................g........g..c.ta.....c.....g.t............t........... [155680]
MH025918.1_strain_R25             ..........................g........g..c.ta.....c.....g.t............t........... [155680]
MH025917.1_strain_R7              ..........................g........g..c.ta.....c.....g.t............t........... [155680]
MH025916.1_strain_R8              ..........................g........g..c.ta.....c.....g.t............t........... [155680]

FR682468.2_Georgia_2007/1         ggacttaaggagatttgggggtcttgcccaaaaccctaggctgctgttcttgttttttagggcgtcataaagaaatgaaa [155760]
ASF_IND_20_CAD543                 ................................................................................ [155760]
LR899193.1_Germany_2020/1         ................................................................................ [155760]
MN393477.1_Wuhan_2019             ................................................................................ [155760]
MW396979.1_Timor-Leste/2019/1     ................................................................................ [155760]
MK940252.1_CN/2019/InnerMongolia  ................................................................................ [155760]
MK128995.1_China/2018/AnhuiXCGQ   ................................................................................ [155760]
MH681419.1_ASFV/POL/2015          ................................................................................ [155760]
LS478113.1_Estonia_2014           ................................................................................ [155760]
KP843857.1_Odintsovo_02/14        ................................................................................ [155760]
FN557520.1_strain_E75             ...............c..............................c................................. [155760]
AY261366.1_Warthog                c..............c..............................c................................. [155760]
AY261365.1_Warmbaths              c..............c..............................c................................. [155760]
AY261364.1_Tengani_62             c..............c..............................c................................. [155760]
AY261363.1_Pretoriuskop/96/4      c.......a......c..............................c................................. [155760]
AY261362.1_Mkuzi_1979             ...............c..............................c................................. [155760]
AY261361.1_Malawi_Lil-20/1_1983   ........aa....cc..............................c.t..a..............g............. [155760]
AY261360.1_Kenya_1950             a.......a.....cc..........................a...c....a............................ [155760]
AM712240.1_OURT_88/3              ...............c..............................c................................. [155760]
AM712239.1_Benin_97/1             ...............c..............................c................................. [155760]
KM111295.1_Ken06.Bus              a.......aa....cc..........................a...c....a............................ [155760]
```

```
KM111294.1_Ken05/Tk1            a.......a.....cc..........................a...c....a.............t.............. [155760]
KM262844.1_strain_L60           ...............c..............................c................................. [155760]
KM102979.1_26544/OG10_Italy     ...............c..............................c................................. [155760]
KM262845.1_strain_NHV           ...............c..............................c................................. [155760]
KX354450.1_47/Ss/2008           ...............c..............................c................................. [155760]
KP055815.1_strain_BA71          ...............c..............................c................................. [155760]
MH025920.1_strain_R35           a.......aa....cc..........................a...c....a............................ [155760]
MH025919.1_strain_N10           a.......aa....cc..........................a...c....a............................ [155760]
MH025918.1_strain_R25           a.......aa....cc..........................a...c....a............................ [155760]
MH025917.1_strain_R7            a.......aa....cc..........................a...c....a............................ [155760]
MH025916.1_strain_R8            a.......aa....cc..........................a...c....a............................ [155760]

FR682468.2_Georgia_2007/1       gcacattgcaaggcttaagccgcgacatctccttccccttgggccctttccatatttttagatctaagatctcatccgag [155840]
ASF_IND_20_CAD543               ................................................................................ [155840]
LR899193.1_Germany_2020/1       ................................................................................ [155840]
MN393477.1_Wuhan_2019           ................................................................................ [155840]
MW396979.1_Timor-Leste/2019/1   ................................................................................ [155840]
MK940252.1_CN/2019/InnerMongolia ................................................................................ [155840]
MK128995.1_China/2018/AnhuiXCGQ ................................................................................ [155840]
MH681419.1_ASFV/POL/2015        ................................................................................ [155840]
LS478113.1_Estonia_2014         ................................................................................ [155840]
KP843857.1_Odintsovo_02/14      ................................................................................ [155840]
FN557520.1_strain_E75           ................................................................................ [155840]
AY261366.1_Warthog              .............................................................g....g............. [155840]
AY261365.1_Warmbaths            .............................................................g....g...t......... [155840]
AY261364.1_Tengani_62           ..................................................................g............. [155840]
AY261363.1_Pretoriuskop/96/4    ...........................................t.................g....g............. [155840]
AY261362.1_Mkuzi_1979           ................................................................................ [155840]
AY261361.1_Malawi_Lil-20/1_1983 ..........c.............c.........................................g.........a... [155840]
AY261360.1_Kenya_1950           ..........c................c..............................a.......g...t.....g... [155840]
AM712240.1_OURT_88/3            ................................................................................ [155840]
AM712239.1_Benin_97/1           ................................................................................ [155840]
KM111295.1_Ken06.Bus            ..........c................c..............................a.......g...t.....g... [155840]
KM111294.1_Ken05/Tk1            ..........t................c..............................a.......g...t.....g... [155840]
KM262844.1_strain_L60           ................................................................................ [155840]
KM102979.1_26544/OG10_Italy     ................................................................................ [155840]
KM262845.1_strain_NHV           ................................................................................ [155840]
KX354450.1_47/Ss/2008           ................................................................................ [155840]
KP055815.1_strain_BA71          ................................................................................ [155840]
MH025920.1_strain_R35           ..........c................c..............................a.......g...t.....g... [155840]
MH025919.1_strain_N10           ..........c................c..............................a.......g...t.....g... [155840]
MH025918.1_strain_R25           ..........c................c..............................a.......g...t.....g... [155840]
MH025917.1_strain_R7            ..........c................c..............................a.......g...t.....g... [155840]
MH025916.1_strain_R8            ..........c................c..............................a.......g...t.....g... [155840]
```

```
FR682468.2_Georgia_2007/1        cttatagagtaggtatagtaaagtttttcaaaaaagcatatctgcttgaagtcttttttagaacgactttcaagaagcat [155920]
ASF_IND_20_CAD543                ................................................................................ [155920]
LR899193.1_Germany_2020/1        ................................................................................ [155920]
MN393477.1_Wuhan_2019            ................................................................................ [155920]
MW396979.1_Timor-Leste/2019/1    ................................................................................ [155920]
MK940252.1_CN/2019/InnerMongolia ................................................................................ [155920]
MK128995.1_China/2018/AnhuiXCGQ  ................................................................................ [155920]
MH681419.1_ASFV/POL/2015         ................................................................................ [155920]
LS478113.1_Estonia_2014          ................................................................................ [155920]
KP843857.1_Odintsovo_02/14       ................................................................................ [155920]
FN557520.1_strain_E75            ................................................................................ [155920]
AY261366.1_Warthog               ................................................................................ [155920]
AY261365.1_Warmbaths             ..................................................a............................. [155920]
AY261364.1_Tengani_62            ...................................a........t..a..a............................. [155920]
AY261363.1_Pretoriuskop/96/4     ................................................................................ [155920]
AY261362.1_Mkuzi_1979            ................................................................................ [155920]
AY261361.1_Malawi_Lil-20/1_1983  t.................................................a............................. [155920]
AY261360.1_Kenya_1950            .......g..............................g..a........a.................a.........g. [155920]
AM712240.1_OURT_88/3             ................................................................................ [155920]
AM712239.1_Benin_97/1            ................................................................................ [155920]
KM111295.1_Ken06.Bus             .......g..............................g..a........a.................a.........g. [155920]
KM111294.1_Ken05/Tk1             .......g..............................g..a........a.t...............a.........g. [155920]
KM262844.1_strain_L60            ................................................................................ [155920]
KM102979.1_26544/OG10_Italy      ................................................................................ [155920]
KM262845.1_strain_NHV            ................................................................................ [155920]
KX354450.1_47/Ss/2008            ................................................................................ [155920]
KP055815.1_strain_BA71           ................................................................................ [155920]
MH025920.1_strain_R35            .......g..............................g..a........a.................a.........g. [155920]
MH025919.1_strain_N10            .......g..............................g..a........a.................a.........g. [155920]
MH025918.1_strain_R25            .......g..............................g..a........a.................a.........g. [155920]
MH025917.1_strain_R7             .......g..............................g..a........a.................a.........g. [155920]
MH025916.1_strain_R8             .......g..............................g..a........a.................a.........g. [155920]

FR682468.2_Georgia_2007/1        ttctataatgttaacaagttttgttaggtttaaggcctgttcctgtgtaagctcctcttgcacgtgatagactgaaaaag [156000]
ASF_IND_20_CAD543                ................................................................................ [156000]
LR899193.1_Germany_2020/1        ................................................................................ [156000]
MN393477.1_Wuhan_2019            ................................................................................ [156000]
MW396979.1_Timor-Leste/2019/1    ................................................................................ [156000]
MK940252.1_CN/2019/InnerMongolia ................................................................................ [156000]
MK128995.1_China/2018/AnhuiXCGQ  ................................................................................ [156000]
MH681419.1_ASFV/POL/2015         ................................................................................ [156000]
LS478113.1_Estonia_2014          ................................................................................ [156000]
KP843857.1_Odintsovo_02/14       ................................................................................ [156000]
```

```
FN557520.1_strain_E75          ................................................................................ [156000]
AY261366.1_Warthog             ................................................................................ [156000]
AY261365.1_Warmbaths           ................................................................................ [156000]
AY261364.1_Tengani_62          ................................................................................ [156000]
AY261363.1_Pretoriuskop/96/4   ................................................................................ [156000]
AY261362.1_Mkuzi_1979          ................................................................................ [156000]
AY261361.1_Malawi_Lil-20/1_1983 ................................g...t........c........t.....t.................g. [156000]
AY261360.1_Kenya_1950          ........ca...................................c........t.......................g. [156000]
AM712240.1_OURT_88/3           ................................................................................ [156000]
AM712239.1_Benin_97/1          ................................................................................ [156000]
KM111295.1_Ken06.Bus           ........ca...................................c........t.......................g. [156000]
KM111294.1_Ken05/Tk1           ........ca...................................c........t.......................g. [156000]
KM262844.1_strain_L60          ................................................................................ [156000]
KM102979.1_26544/OG10_Italy    ................................................................................ [156000]
KM262845.1_strain_NHV          ................................................................................ [156000]
KX354450.1_47/Ss/2008          ................................................................................ [156000]
KP055815.1_strain_BA71         ................................................................................ [156000]
MH025920.1_strain_R35          ........ca...................................c........t.......................g. [156000]
MH025919.1_strain_N10          ........ca...................................c........t.......................g. [156000]
MH025918.1_strain_R25          ........ca...................................c........t.......................g. [156000]
MH025917.1_strain_R7           ........ca...................................c........t.......................g. [156000]
MH025916.1_strain_R8           ........ca...................................c........t.......................g. [156000]

FR682468.2_Georgia_2007/1      tgtgcttaggaatgaaaatactccccgtggcactggcctgttgtctgccaggtatatagtacacgctgctgttagcaagc [156080]
ASF_IND_20_CAD543              ................................................................................ [156080]
LR899193.1_Germany_2020/1      ................................................................................ [156080]
MN393477.1_Wuhan_2019          ................................................................................ [156080]
MW396979.1_Timor-Leste/2019/1  ................................................................................ [156080]
MK940252.1_CN/2019/InnerMongolia ................................................................................ [156080]
MK128995.1_China/2018/AnhuiXCGQ ................................................................................ [156080]
MH681419.1_ASFV/POL/2015       ................................................................................ [156080]
LS478113.1_Estonia_2014        ................................................................................ [156080]
KP843857.1_Odintsovo_02/14     ................................................................................ [156080]
FN557520.1_strain_E75          ................................................................................ [156080]
AY261366.1_Warthog             ......................................................................a......... [156080]
AY261365.1_Warmbaths           ................................................................................ [156080]
AY261364.1_Tengani_62          ......................................................................a......... [156080]
AY261363.1_Pretoriuskop/96/4   ................................................................................ [156080]
AY261362.1_Mkuzi_1979          ................................................................................ [156080]
AY261361.1_Malawi_Lil-20/1_1983 ................g.....t............................................a..a......... [156080]
AY261360.1_Kenya_1950          .a..............g....................t.............................a..a......... [156080]
AM712240.1_OURT_88/3           ................................................................................ [156080]
AM712239.1_Benin_97/1          ................................................................................ [156080]
KM111295.1_Ken06.Bus           .a..............g..................................................a..a......... [156080]
```

```
KM111294.1_Ken05/Tk1            .a..............g...t.................t..............................a..a......... [156080]
KM262844.1_strain_L60           ................................................................................ [156080]
KM102979.1_26544/OG10_Italy     ................................................................................ [156080]
KM262845.1_strain_NHV           ................................................................................ [156080]
KX354450.1_47/Ss/2008           ................................................................................ [156080]
KP055815.1_strain_BA71          ................................................................................ [156080]
MH025920.1_strain_R35           .a..............g..................................................a..a......... [156080]
MH025919.1_strain_N10           .a..............g..................................................a..a......... [156080]
MH025918.1_strain_R25           .a..............g..................................................a..a......... [156080]
MH025917.1_strain_R7            .a..............g..................................................a..a......... [156080]
MH025916.1_strain_R8            .a..............g..................................................a..a......... [156080]

FR682468.2_Georgia_2007/1       tgtaccggcacaatttgccccacttctgcaacattattttgcgattcggacgagggtatgacaatagttacgggttcagt [156160]
ASF_IND_20_CAD543               ................................................................................ [156160]
LR899193.1_Germany_2020/1       ................................................................................ [156160]
MN393477.1_Wuhan_2019           ................................................................................ [156160]
MW396979.1_Timor-Leste/2019/1   ................................................................................ [156160]
MK940252.1_CN/2019/InnerMongolia ................................................................................ [156160]
MK128995.1_China/2018/AnhuiXCGQ ................................................................................ [156160]
MH681419.1_ASFV/POL/2015        ................................................................................ [156160]
LS478113.1_Estonia_2014         ................................................................................ [156160]
KP843857.1_Odintsovo_02/14      ................................................................................ [156160]
FN557520.1_strain_E75           ....................t........................................................... [156160]
AY261366.1_Warthog              .................................................................c.............. [156160]
AY261365.1_Warmbaths            .................t...............................................c.............. [156160]
AY261364.1_Tengani_62           .......................c.......t.....................a...........c.............. [156160]
AY261363.1_Pretoriuskop/96/4    .................................................................c.............. [156160]
AY261362.1_Mkuzi_1979           ................................................................................ [156160]
AY261361.1_Malawi_Lil-20/1_1983 .......................c.............................a.....t.................... [156160]
AY261360.1_Kenya_1950           .................t.....c.................t........t..a.....t.................c.. [156160]
AM712240.1_OURT_88/3            ....................t........................................................... [156160]
AM712239.1_Benin_97/1           ....................t........................................................... [156160]
KM111295.1_Ken06.Bus            .................t.....c.................t...........a.....t.................c.. [156160]
KM111294.1_Ken05/Tk1            .................t.....c.................t...........a.....t.................c.. [156160]
KM262844.1_strain_L60           ....................t........................................................... [156160]
KM102979.1_26544/OG10_Italy     ....................t........................................................... [156160]
KM262845.1_strain_NHV           ....................t........................................................... [156160]
KX354450.1_47/Ss/2008           ....................t........................................................... [156160]
KP055815.1_strain_BA71          ....................t........................................................... [156160]
MH025920.1_strain_R35           .................t.....c.................t...........a.....t.................c.. [156160]
MH025919.1_strain_N10           .................t.....c.................t...........a.....t.................c.. [156160]
MH025918.1_strain_R25           .................t.....c.................t...........a.....t.................c.. [156160]
MH025917.1_strain_R7            .................t.....c.................t...........a.....t.................c.. [156160]
MH025916.1_strain_R8            .................t.....c.................t...........a.....t.................c.. [156160]
```

```
FR682468.2_Georgia_2007/1        caataggctttcgccgagaataatattactgtcatttttaataattttaacggccgctattaaatcaaaggcatttaagt [156240]
ASF_IND_20_CAD543                ................................................................................ [156240]
LR899193.1_Germany_2020/1        ................................................................................ [156240]
MN393477.1_Wuhan_2019            ................................................................................ [156240]
MW396979.1_Timor-Leste/2019/1    ................................................................................ [156240]
MK940252.1_CN/2019/InnerMongolia ................................................................................ [156240]
MK128995.1_China/2018/AnhuiXCGQ  ................................................................................ [156240]
MH681419.1_ASFV/POL/2015         ................................................................................ [156240]
LS478113.1_Estonia_2014          ................................................................................ [156240]
KP843857.1_Odintsovo_02/14       ................................................................................ [156240]
FN557520.1_strain_E75            ................................................................................ [156240]
AY261366.1_Warthog               ................................................................................ [156240]
AY261365.1_Warmbaths             ............a......................................a...c........................ [156240]
AY261364.1_Tengani_62            ................................................................................ [156240]
AY261363.1_Pretoriuskop/96/4     ................................................................................ [156240]
AY261362.1_Mkuzi_1979            ................................................................................ [156240]
AY261361.1_Malawi_Lil-20/1_1983  ...............t.................g.............................................c [156240]
AY261360.1_Kenya_1950            ...c...........t............t....g........g..................................... [156240]
AM712240.1_OURT_88/3             ................................................................................ [156240]
AM712239.1_Benin_97/1            ................................................................................ [156240]
KM111295.1_Ken06.Bus             ...c...........t............t....g........g..................................... [156240]
KM111294.1_Ken05/Tk1             ...c...........t............t....g........g..................................... [156240]
KM262844.1_strain_L60            ................................................................................ [156240]
KM102979.1_26544/OG10_Italy      ................................................................................ [156240]
KM262845.1_strain_NHV            ................................................................................ [156240]
KX354450.1_47/Ss/2008            ................................................................................ [156240]
KP055815.1_strain_BA71           ................................................................................ [156240]
MH025920.1_strain_R35            ...c...........t............t....g........g..................................... [156240]
MH025919.1_strain_N10            ...c...........t............t....g........g..................................... [156240]
MH025918.1_strain_R25            ...c...........t............t....g........g..................................... [156240]
MH025917.1_strain_R7             ...c...........t............t....g........g..................................... [156240]
MH025916.1_strain_R8             ...c...........t............t....g........g..................................... [156240]

FR682468.2_Georgia_2007/1        aagaaacaacagcagaaaatcttacatgcatatatcctcttccgctattattcgtacgcataataaaacaaggggagcgt [156320]
ASF_IND_20_CAD543                ................................................................................ [156320]
LR899193.1_Germany_2020/1        ................................................................................ [156320]
MN393477.1_Wuhan_2019            ................................................................................ [156320]
MW396979.1_Timor-Leste/2019/1    ................................................................................ [156320]
MK940252.1_CN/2019/InnerMongolia ................................................................................ [156320]
MK128995.1_China/2018/AnhuiXCGQ  ................................................................................ [156320]
MH681419.1_ASFV/POL/2015         ................................................................................ [156320]
LS478113.1_Estonia_2014          ................................................................................ [156320]
KP843857.1_Odintsovo_02/14       ................................................................................ [156320]
```

```
FN557520.1_strain_E75            ................................................................................ [156320]
AY261366.1_Warthog               .........g...................................................................... [156320]
AY261365.1_Warmbaths             .........g..........................................t........................... [156320]
AY261364.1_Tengani_62            .........g...........................c.......................t.................. [156320]
AY261363.1_Pretoriuskop/96/4     .........g...........................c.......................................... [156320]
AY261362.1_Mkuzi_1979            ................................................................................ [156320]
AY261361.1_Malawi_Lil-20/1_1983  .........g...........................c...................................t...... [156320]
AY261360.1_Kenya_1950            .........g...............g...........c..............g........................... [156320]
AM712240.1_OURT_88/3             ................................................................................ [156320]
AM712239.1_Benin_97/1            ................................................................................ [156320]
KM111295.1_Ken06.Bus             .........g...............g...........c..............g........................... [156320]
KM111294.1_Ken05/Tk1             .........g...............g...........c..............g........................... [156320]
KM262844.1_strain_L60            ................................................................................ [156320]
KM102979.1_26544/OG10_Italy      ................................................................................ [156320]
KM262845.1_strain_NHV            ................................................................................ [156320]
KX354450.1_47/Ss/2008            ................................................................................ [156320]
KP055815.1_strain_BA71           ................................................................................ [156320]
MH025920.1_strain_R35            .........g...............g...........c..............g........................... [156320]
MH025919.1_strain_N10            .........g...............g...........c..............g........................... [156320]
MH025918.1_strain_R25            .........g...............g...........c..............g........................... [156320]
MH025917.1_strain_R7             .........g...............g...........c..............g........................... [156320]
MH025916.1_strain_R8             .........g...............g...........c..............g........................... [156320]

FR682468.2_Georgia_2007/1        tgtataacgccagtaatattaagaataaaactgtttttgaaacacttacccacataaatgttttcaagctccttcaaaag [156400]
ASF_IND_20_CAD543                ................................................................................ [156400]
LR899193.1_Germany_2020/1        ................................................................................ [156400]
MN393477.1_Wuhan_2019            ................................................................................ [156400]
MW396979.1_Timor-Leste/2019/1    ................................................................................ [156400]
MK940252.1_CN/2019/InnerMongolia ................................................................................ [156400]
MK128995.1_China/2018/AnhuiXCGQ  ................................................................................ [156400]
MH681419.1_ASFV/POL/2015         ................................................................................ [156400]
LS478113.1_Estonia_2014          ................................................................................ [156400]
KP843857.1_Odintsovo_02/14       ................................................................................ [156400]
FN557520.1_strain_E75            ................................................................................ [156400]
AY261366.1_Warthog               .......t........................................................................ [156400]
AY261365.1_Warmbaths             .......t........................................................................ [156400]
AY261364.1_Tengani_62            ................................................................................ [156400]
AY261363.1_Pretoriuskop/96/4     .....t.t........................................................................ [156400]
AY261362.1_Mkuzi_1979            ................................................................................ [156400]
AY261361.1_Malawi_Lil-20/1_1983  ..................................................t............................. [156400]
AY261360.1_Kenya_1950            .....g..a.........c...t...........................t............................. [156400]
AM712240.1_OURT_88/3             ................................................................................ [156400]
AM712239.1_Benin_97/1            ........a....................................................................... [156400]
KM111295.1_Ken06.Bus             .....g..a.........c...t...........................t............................. [156400]
```

KM111294.1_Ken05/Tk1 .....g..a.........c...t...........................t............................. [156400]
KM262844.1_strain_L60 ................................................................................ [156400]
KM102979.1_26544/OG10_Italy ................................................................................ [156400]
KM262845.1_strain_NHV ................................................................................ [156400]
KX354450.1_47/Ss/2008 ................................................................................ [156400]
KP055815.1_strain_BA71 ................................................................................ [156400]
MH025920.1_strain_R35 .....g..a.........c...t...........................t............................. [156400]
MH025919.1_strain_N10 .....g..a.........c...t...........................t............................. [156400]
MH025918.1_strain_R25 .....g..a.........c...t...........................t............................. [156400]
MH025917.1_strain_R7 .....g..a.........c...t...........................t............................. [156400]
MH025916.1_strain_R8 .....g..a.........c...t...........................t............................. [156400]

FR682468.2_Georgia_2007/1 atgagcctccacatttgtacaaaaattggtaggatcatcaatattcaacgttgtctcaaaaattttttggtcgatcatat [156480]
ASF_IND_20_CAD543 ................................................................................ [156480]
LR899193.1_Germany_2020/1 ................................................................................ [156480]
MN393477.1_Wuhan_2019 ................................................................................ [156480]
MW396979.1_Timor-Leste/2019/1 ................................................................................ [156480]
MK940252.1_CN/2019/InnerMongolia ................................................................................ [156480]
MK128995.1_China/2018/AnhuiXCGQ ................................................................................ [156480]
MH681419.1_ASFV/POL/2015 ................................................................................ [156480]
LS478113.1_Estonia_2014 ................................................................................ [156480]
KP843857.1_Odintsovo_02/14 ................................................................................ [156480]
FN557520.1_strain_E75 ................................................................................ [156480]
AY261366.1_Warthog ............g.......................g.....g..................................... [156480]
AY261365.1_Warmbaths ............g.......................g.....g..................................... [156480]
AY261364.1_Tengani_62 ....................................g..g........................................ [156480]
AY261363.1_Pretoriuskop/96/4 ............g.......................g.....g..................................... [156480]
AY261362.1_Mkuzi_1979 ................................................................................ [156480]
AY261361.1_Malawi_Lil-20/1_1983 ...........................a.c........................t.....t................... [156480]
AY261360.1_Kenya_1950 ...........................a.c......................a........................... [156480]
AM712240.1_OURT_88/3 ................................................................................ [156480]
AM712239.1_Benin_97/1 ................................................................................ [156480]
KM111295.1_Ken06.Bus ...........................a.c......................a........................... [156480]
KM111294.1_Ken05/Tk1 ...........................a.c......................a........................... [156480]
KM262844.1_strain_L60 ................................................................................ [156480]
KM102979.1_26544/OG10_Italy ................................................................................ [156480]
KM262845.1_strain_NHV ................................................................................ [156480]
KX354450.1_47/Ss/2008 ................................................................................ [156480]
KP055815.1_strain_BA71 ................................................................................ [156480]
MH025920.1_strain_R35 ...........................a.c......................a........................... [156480]
MH025919.1_strain_N10 ...........................a.c......................a........................... [156480]
MH025918.1_strain_R25 ...........................a.c......................a........................... [156480]
MH025917.1_strain_R7 ...........................a.c......................a........................... [156480]
MH025916.1_strain_R8 ...........................a.c......................a........................... [156480]

```
FR682468.2_Georgia_2007/1        ctataatatattctgtctatttcaatttaaataatatacgaataaat-aacgagattatt-------------------- [156560]
ASF_IND_20_CAD543                ...............................................a............-------------------- [156560]
LR899193.1_Germany_2020/1        ...............................................-............-------------------- [156560]
MN393477.1_Wuhan_2019            ...............................................-............-------------------- [156560]
MW396979.1_Timor-Leste/2019/1    ...............................................-............-------------------- [156560]
MK940252.1_CN/2019/InnerMongolia ...............................................-............-------------------- [156560]
MK128995.1_China/2018/AnhuiXCGQ  ...............................................-............-------------------- [156560]
MH681419.1_ASFV/POL/2015         ...............................................-............-------------------- [156560]
LS478113.1_Estonia_2014          ...............................................-............-------------------- [156560]
KP843857.1_Odintsovo_02/14       ...............................................-............-------------------- [156560]
FN557520.1_strain_E75            ...............................................a............-------------------- [156560]
AY261366.1_Warthog               .......................................c.......a.......------------------------- [156560]
AY261365.1_Warmbaths             ...............................................a.......------------------------- [156560]
AY261364.1_Tengani_62            ...............................................a............aaataaatgagattgttaaa [156560]
AY261363.1_Pretoriuskop/96/4     .......................................c.......a.......------------------------- [156560]
AY261362.1_Mkuzi_1979            ...............................................a............-------------------- [156560]
AY261361.1_Malawi_Lil-20/1_1983  ...............a...............................a.g..........-------------------- [156560]
AY261360.1_Kenya_1950            ....................................g.t......cc--------------------------------- [156560]
AM712240.1_OURT_88/3             ...............................................a............-------------------- [156560]
AM712239.1_Benin_97/1            ...............................................a............-------------------- [156560]
KM111295.1_Ken06.Bus             ....................................g.t......cc--------------------------------- [156560]
KM111294.1_Ken05/Tk1             ....................................g.t......cc--------------------------------- [156560]
KM262844.1_strain_L60            ...............................................a............-------------------- [156560]
KM102979.1_26544/OG10_Italy      ...............................................a............-------------------- [156560]
KM262845.1_strain_NHV            ...............................................a............-------------------- [156560]
KX354450.1_47/Ss/2008            ...............................................a............-------------------- [156560]
KP055815.1_strain_BA71           ...............................................a............-------------------- [156560]
MH025920.1_strain_R35            ....................................g.t......cc--------------------------------- [156560]
MH025919.1_strain_N10            ....................................g.t......cc--------------------------------- [156560]
MH025918.1_strain_R25            ....................................g.t......cc--------------------------------- [156560]
MH025917.1_strain_R7             ....................................g.t......cc--------------------------------- [156560]
MH025916.1_strain_R8             ....................................g.t......cc--------------------------------- [156560]

FR682468.2_Georgia_2007/1        ---------ttattaaataagca-----------------atggtgtatacactttgtatttac---tttgagatatac- [156640]
ASF_IND_20_CAD543                ---------..............-----------------........................---............- [156640]
LR899193.1_Germany_2020/1        ---------..............-----------------........................---............- [156640]
MN393477.1_Wuhan_2019            ---------..............-----------------........................---............- [156640]
MW396979.1_Timor-Leste/2019/1    ---------..............-----------------........................---............- [156640]
MK940252.1_CN/2019/InnerMongolia ---------..............-----------------........................---............- [156640]
MK128995.1_China/2018/AnhuiXCGQ  ---------..............-----------------........................---............- [156640]
MH681419.1_ASFV/POL/2015         ---------..............-----------------........................---............- [156640]
LS478113.1_Estonia_2014          ---------..............-----------------........................---............- [156640]
KP843857.1_Odintsovo_02/14       ---------..............-----------------........................---............- [156640]
```

```
FN557520.1_strain_E75            ---------.............------------------........................---............- [156640]
AY261366.1_Warthog               ---------...........a.gagattgttaaataagca..a.g....g.g......g.a...act..atgt...c..- [156640]
AY261365.1_Warmbaths             ---------..g..........---------------------------------------------------------- [156640]
AY261364.1_Tengani_62            taaacgaga..g........a.gagattgttaaataagca..a.g....g.g......g.a...a--.aaagtg...g.a [156640]
AY261363.1_Pretoriuskop/96/4     ---------..g..........------------------..a.g....g.g......g.a...act..gtgt......- [156640]
AY261362.1_Mkuzi_1979            ---------.............------------------........................---............- [156640]
AY261361.1_Malawi_Lil-20/1_1983  ---------.a..a..g..ta..----------------c.aa......-...gaa..gca...a--caaa.tg.....a [156640]
AY261360.1_Kenya_1950            ---------------------------------------------------------.g....tt--------------- [156640]
AM712240.1_OURT_88/3             ---------.............------------------........................---............- [156640]
AM712239.1_Benin_97/1            ---------.............------------------........................---............- [156640]
KM111295.1_Ken06.Bus             --------------------------------------------------------t.g.....t--------------- [156640]
KM111294.1_Ken05/Tk1             ---------------------------------------------------------.g....tt--------------- [156640]
KM262844.1_strain_L60            ---------.............------------------........................---............- [156640]
KM102979.1_26544/OG10_Italy      ---------.............------------------........................---............- [156640]
KM262845.1_strain_NHV            ---------.............------------------........................---............- [156640]
KX354450.1_47/Ss/2008            ---------.............------------------........................---............- [156640]
KP055815.1_strain_BA71           ---------.............------------------........................---............- [156640]
MH025920.1_strain_R35            --------------------------------------------------------t.g.....t--------------- [156640]
MH025919.1_strain_N10            --------------------------------------------------------t.g.....t--------------- [156640]
MH025918.1_strain_R25            --------------------------------------------------------t.g.....t--------------- [156640]
MH025917.1_strain_R7             --------------------------------------------------------t.g.....t--------------- [156640]
MH025916.1_strain_R8             --------------------------------------------------------t.g.....t--------------- [156640]

FR682468.2_Georgia_2007/1        --tttgtgtatcacaacgtgccctaagatgtgtgcacaagtgacggcattttgtcgttaaaaaggtaaaaccagcggatt [156720]
ASF_IND_20_CAD543                --.............................................................................. [156720]
LR899193.1_Germany_2020/1        --.............................................................................. [156720]
MN393477.1_Wuhan_2019            --.............................................................................. [156720]
MW396979.1_Timor-Leste/2019/1    --.............................................................................. [156720]
MK940252.1_CN/2019/InnerMongolia --.............................................................................. [156720]
MK128995.1_China/2018/AnhuiXCGQ  --.............................................................................. [156720]
MH681419.1_ASFV/POL/2015         --.............................................................................. [156720]
LS478113.1_Estonia_2014          --.............................................................................. [156720]
KP843857.1_Odintsovo_02/14       --.............................................................................. [156720]
FN557520.1_strain_E75            --.............................................................................. [156720]
AY261366.1_Warthog               --...a.........--...t...................g....................................... [156720]
AY261365.1_Warmbaths             ------------.t.--...t.t.................g....................................... [156720]
AY261364.1_Tengani_62            ct...a..........a...t...................c....................................... [156720]
AY261363.1_Pretoriuskop/96/4     --...a.........--...t...................g....................................... [156720]
AY261362.1_Mkuzi_1979            --.............................................................................. [156720]
AY261361.1_Malawi_Lil-20/1_1983  c-...a.........taa.a............................................................ [156720]
AY261360.1_Kenya_1950            --.a.tc.........a.................t.....g....................................... [156720]
AM712240.1_OURT_88/3             --.............................................................................. [156720]
AM712239.1_Benin_97/1            --.............................................................................. [156720]
KM111295.1_Ken06.Bus             --.a.tc.........a.................t.....g....................................... [156720]
```

```
KM111294.1_Ken05/Tk1            --.a.tc.........a.................t.....g....................................... [156720]
KM262844.1_strain_L60           --.............................................................................. [156720]
KM102979.1_26544/OG10_Italy     --.............................................................................. [156720]
KM262845.1_strain_NHV           --.............................................................................. [156720]
KX354450.1_47/Ss/2008           --.............................................................................. [156720]
KP055815.1_strain_BA71          --.............................................................................. [156720]
MH025920.1_strain_R35           --.a.tc.........a.................t.....g....................................... [156720]
MH025919.1_strain_N10           --.a.tc.........a.................t.....g....................................... [156720]
MH025918.1_strain_R25           --.a.tc.........a.................t.....g....................................... [156720]
MH025917.1_strain_R7            --.a.tc.........a.................t.....g....................................... [156720]
MH025916.1_strain_R8            --.a.tc.........a.................t.....g....................................... [156720]

FR682468.2_Georgia_2007/1       ccatcctgcattccatttggttgattacgagcctccatttctttttgcaaaaggttattgcgaatgagtaagcagagctt [156800]
ASF_IND_20_CAD543               ................................................................................ [156800]
LR899193.1_Germany_2020/1       ................................................................................ [156800]
MN393477.1_Wuhan_2019           ................................................................................ [156800]
MW396979.1_Timor-Leste/2019/1   ................................................................................ [156800]
MK940252.1_CN/2019/InnerMongolia ................................................................................ [156800]
MK128995.1_China/2018/AnhuiXCGQ ................................................................................ [156800]
MH681419.1_ASFV/POL/2015        ................................................................................ [156800]
LS478113.1_Estonia_2014         ................................................................................ [156800]
KP843857.1_Odintsovo_02/14      ................................................................................ [156800]
FN557520.1_strain_E75           ................................................................................ [156800]
AY261366.1_Warthog              ......................................c......................................... [156800]
AY261365.1_Warmbaths            ......................................c......................................... [156800]
AY261364.1_Tengani_62           ......................................c......................................... [156800]
AY261363.1_Pretoriuskop/96/4    ......................................c......................................... [156800]
AY261362.1_Mkuzi_1979           ................................................................................ [156800]
AY261361.1_Malawi_Lil-20/1_1983 ........................................................g....................... [156800]
AY261360.1_Kenya_1950           ................................t.....................g.g........a.............. [156800]
AM712240.1_OURT_88/3            ................................................................................ [156800]
AM712239.1_Benin_97/1           ................................................................................ [156800]
KM111295.1_Ken06.Bus            ...........c....................t.....................g.g....................... [156800]
KM111294.1_Ken05/Tk1            ................................t.....................g.g........a.............. [156800]
KM262844.1_strain_L60           ................................................................................ [156800]
KM102979.1_26544/OG10_Italy     ................................................................................ [156800]
KM262845.1_strain_NHV           ................................................................................ [156800]
KX354450.1_47/Ss/2008           ................................................................................ [156800]
KP055815.1_strain_BA71          ................................................................................ [156800]
MH025920.1_strain_R35           ...........c....................t.....................g.g....................... [156800]
MH025919.1_strain_N10           ...........c....................t.....................g.g....................... [156800]
MH025918.1_strain_R25           ...........c....................t.....................g.g....................... [156800]
MH025917.1_strain_R7            ...........c....................t.....................g.g....................... [156800]
MH025916.1_strain_R8            ...........c....................t.....................g.g....................... [156800]
```

| | | |
|---|---|---|
| FR682468.2_Georgia_2007/1 | gatggcactaatctttgtaaggtttaaacttatgcccaattggtcagcaatttttttgttgctcctcccgtccgcgtgttt | [156880] |
| ASF_IND_20_CAD543 | ................................................................................ | [156880] |
| LR899193.1_Germany_2020/1 | ................................................................................ | [156880] |
| MN393477.1_Wuhan_2019 | ................................................................................ | [156880] |
| MW396979.1_Timor-Leste/2019/1 | ................................................................................ | [156880] |
| MK940252.1_CN/2019/InnerMongolia | ................................................................................ | [156880] |
| MK128995.1_China/2018/AnhuiXCGQ | ................................................................................ | [156880] |
| MH681419.1_ASFV/POL/2015 | ................................................................................ | [156880] |
| LS478113.1_Estonia_2014 | ................................................................................ | [156880] |
| KP843857.1_Odintsovo_02/14 | ................................................................................ | [156880] |
| FN557520.1_strain_E75 | ................................................................................ | [156880] |
| AY261366.1_Warthog | ............................................t................................... | [156880] |
| AY261365.1_Warmbaths | ............................................t................................... | [156880] |
| AY261364.1_Tengani_62 | ............................................t..............................g.... | [156880] |
| AY261363.1_Pretoriuskop/96/4 | ............................................t................................... | [156880] |
| AY261362.1_Mkuzi_1979 | ................................................................................ | [156880] |
| AY261361.1_Malawi_Lil-20/1_1983 | a.............................g.............t.....................t............. | [156880] |
| AY261360.1_Kenya_1950 | a...........t........a........a................................................. | [156880] |
| AM712240.1_OURT_88/3 | ................................................................................ | [156880] |
| AM712239.1_Benin_97/1 | ................................................................................ | [156880] |
| KM111295.1_Ken06.Bus | a...........t........a........g................................................. | [156880] |
| KM111294.1_Ken05/Tk1 | a...........t........a........g................................................. | [156880] |
| KM262844.1_strain_L60 | ................................................................................ | [156880] |
| KM102979.1_26544/OG10_Italy | ................................................................................ | [156880] |
| KM262845.1_strain_NHV | ................................................................................ | [156880] |
| KX354450.1_47/Ss/2008 | ................................................................................ | [156880] |
| KP055815.1_strain_BA71 | ................................................................................ | [156880] |
| MH025920.1_strain_R35 | a...........t........a........g................................................. | [156880] |
| MH025919.1_strain_N10 | a...........t........a........g................................................. | [156880] |
| MH025918.1_strain_R25 | a...........t........a........g................................................. | [156880] |
| MH025917.1_strain_R7 | a...........t........a........g................................................. | [156880] |
| MH025916.1_strain_R8 | a...........t........a........g................................................. | [156880] |

| | | |
|---|---|---|
| FR682468.2_Georgia_2007/1 | cgcatacggctccccggtttagcatgcgaatatcagtaatctcattctttttttaaaacctggataggtgggcggatttta | [156960] |
| ASF_IND_20_CAD543 | ................................................................................ | [156960] |
| LR899193.1_Germany_2020/1 | ................................................................................ | [156960] |
| MN393477.1_Wuhan_2019 | ................................................................................ | [156960] |
| MW396979.1_Timor-Leste/2019/1 | ................................................................................ | [156960] |
| MK940252.1_CN/2019/InnerMongolia | ................................................................................ | [156960] |
| MK128995.1_China/2018/AnhuiXCGQ | ................................................................................ | [156960] |
| MH681419.1_ASFV/POL/2015 | ................................................................................ | [156960] |
| LS478113.1_Estonia_2014 | ................................................................................ | [156960] |
| KP843857.1_Odintsovo_02/14 | ................................................................................ | [156960] |

```
FN557520.1_strain_E75            ...........................................g.................................... [156960]
AY261366.1_Warthog               .a.............................................................................. [156960]
AY261365.1_Warmbaths             ................................................................................ [156960]
AY261364.1_Tengani_62            ................................................................................ [156960]
AY261363.1_Pretoriuskop/96/4     ................................................................................ [156960]
AY261362.1_Mkuzi_1979            ...........................................g.................................... [156960]
AY261361.1_Malawi_Lil-20/1_1983  ......t....................................g.................t...........a...... [156960]
AY261360.1_Kenya_1950            ................a............................................t..g........a...... [156960]
AM712240.1_OURT_88/3             ...........................................g.................................... [156960]
AM712239.1_Benin_97/1            ...........................................g.................................... [156960]
KM111295.1_Ken06.Bus             ...............ta............................................t..g........a...... [156960]
KM111294.1_Ken05/Tk1             ................a..........................g.................t..g........a...... [156960]
KM262844.1_strain_L60            ...........................................g.................................... [156960]
KM102979.1_26544/OG10_Italy      ...........................................g.................................... [156960]
KM262845.1_strain_NHV            ...........................................g.................................... [156960]
KX354450.1_47/Ss/2008            ...........................................g.................................... [156960]
KP055815.1_strain_BA71           ...........................................g.................................... [156960]
MH025920.1_strain_R35            ...............ta............................................t..g........a...... [156960]
MH025919.1_strain_N10            ...............ta............................................t..g........a...... [156960]
MH025918.1_strain_R25            ...............ta............................................t..g........a...... [156960]
MH025917.1_strain_R7             ...............ta............................................t..g........a...... [156960]
MH025916.1_strain_R8             ...............ta............................................t..g........a...... [156960]

FR682468.2_Georgia_2007/1        aatttaagggcctttcccttgctttccatatagcctatgacgatgtcgttttcttttcgtttaacattaatattaagcat [157040]
ASF_IND_20_CAD543                ................................................................................ [157040]
LR899193.1_Germany_2020/1        ................................................................................ [157040]
MN393477.1_Wuhan_2019            ................................................................................ [157040]
MW396979.1_Timor-Leste/2019/1    ................................................................................ [157040]
MK940252.1_CN/2019/InnerMongolia ................................................................................ [157040]
MK128995.1_China/2018/AnhuiXCGQ  ................................................................................ [157040]
MH681419.1_ASFV/POL/2015         ................................................................................ [157040]
LS478113.1_Estonia_2014          ................................................................................ [157040]
KP843857.1_Odintsovo_02/14       ................................................................................ [157040]
FN557520.1_strain_E75            .........................................a...................................... [157040]
AY261366.1_Warthog               ..............................................................g................. [157040]
AY261365.1_Warmbaths             ..............................................................g................. [157040]
AY261364.1_Tengani_62            ................................................................................ [157040]
AY261363.1_Pretoriuskop/96/4     ..............................................................g................. [157040]
AY261362.1_Mkuzi_1979            ................................................................................ [157040]
AY261361.1_Malawi_Lil-20/1_1983  ..............................................................g................. [157040]
AY261360.1_Kenya_1950            ..........tt..................................................g................. [157040]
AM712240.1_OURT_88/3             .........................................a...................................... [157040]
AM712239.1_Benin_97/1            .........................................a...................................... [157040]
KM111295.1_Ken06.Bus             ..........tt..................................................g................. [157040]
```

```
KM111294.1_Ken05/Tk1            ..........tt..................................................g................. [157040]
KM262844.1_strain_L60           .........................................a...................................... [157040]
KM102979.1_26544/OG10_Italy     .........................................a...................................... [157040]
KM262845.1_strain_NHV           .........................................a...................................... [157040]
KX354450.1_47/Ss/2008           .........................................a...................................... [157040]
KP055815.1_strain_BA71          .........................................a...................................... [157040]
MH025920.1_strain_R35           ..........tt..................................................g................. [157040]
MH025919.1_strain_N10           ..........tt..................................................g................. [157040]
MH025918.1_strain_R25           ..........tt..................................................g................. [157040]
MH025917.1_strain_R7            ..........tt..................................................g................. [157040]
MH025916.1_strain_R8            ..........tt..................................................g................. [157040]

FR682468.2_Georgia_2007/1       ataaagcggaatttcatgccaggttttatcttctcgcgaggtaataagtcgcacggagtcctccgtggcatagcccacta [157120]
ASF_IND_20_CAD543               ................................................................................ [157120]
LR899193.1_Germany_2020/1       ................................................................................ [157120]
MN393477.1_Wuhan_2019           ................................................................................ [157120]
MW396979.1_Timor-Leste/2019/1   ................................................................................ [157120]
MK940252.1_CN/2019/InnerMongolia ................................................................................ [157120]
MK128995.1_China/2018/AnhuiXCGQ ................................................................................ [157120]
MH681419.1_ASFV/POL/2015        ................................................................................ [157120]
LS478113.1_Estonia_2014         ................................................................................ [157120]
KP843857.1_Odintsovo_02/14      ................................................................................ [157120]
FN557520.1_strain_E75           ...g..........................................................................c. [157120]
AY261366.1_Warthog              .................................a.............................................. [157120]
AY261365.1_Warmbaths            .................................a.............................................. [157120]
AY261364.1_Tengani_62           .................................a.............................................. [157120]
AY261363.1_Pretoriuskop/96/4    .................................a.............................................. [157120]
AY261362.1_Mkuzi_1979           ........................................................................a....... [157120]
AY261361.1_Malawi_Lil-20/1_1983 ................................ta..t..a....................t...........a....... [157120]
AY261360.1_Kenya_1950           ................................ga..t........................................... [157120]
AM712240.1_OURT_88/3            ...g..........................................................................c. [157120]
AM712239.1_Benin_97/1           ...g..........................................................................c. [157120]
KM111295.1_Ken06.Bus            .....................a..........ga..t.......................t................... [157120]
KM111294.1_Ken05/Tk1            ................................ga..t.......................t................... [157120]
KM262844.1_strain_L60           ...g..........................................................................c. [157120]
KM102979.1_26544/OG10_Italy     ...g..........................................................................c. [157120]
KM262845.1_strain_NHV           ...g..........................................................................c. [157120]
KX354450.1_47/Ss/2008           ...g..........................................................................c. [157120]
KP055815.1_strain_BA71          ...g..........................................................................c. [157120]
MH025920.1_strain_R35           .....................a..........ga..t.......................t................... [157120]
MH025919.1_strain_N10           .....................a..........ga..t.......................t................... [157120]
MH025918.1_strain_R25           .....................a..........ga..t.......................t................... [157120]
MH025917.1_strain_R7            .....................a..........ga..t.......................t................... [157120]
MH025916.1_strain_R8            .....................a..........ga..t.......................t................... [157120]
```

```
FR682468.2_Georgia_2007/1        gagtgttgtcatccccaggcacgtggcttataattttaaaaatgtccggaaatggctgaatatctttttttgaaaaagcg [157200]
ASF_IND_20_CAD543                ................................................................................ [157200]
LR899193.1_Germany_2020/1        ................................................................................ [157200]
MN393477.1_Wuhan_2019            ................................................................................ [157200]
MW396979.1_Timor-Leste/2019/1    ................................................................................ [157200]
MK940252.1_CN/2019/InnerMongolia ................................................................................ [157200]
MK128995.1_China/2018/AnhuiXCGQ  ................................................................................ [157200]
MH681419.1_ASFV/POL/2015         ................................................................................ [157200]
LS478113.1_Estonia_2014          ................................................................................ [157200]
KP843857.1_Odintsovo_02/14       ................................................................................ [157200]
FN557520.1_strain_E75            ................................................................................ [157200]
AY261366.1_Warthog               ..............................................t........t........................ [157200]
AY261365.1_Warmbaths             ..............................................t........t........................ [157200]
AY261364.1_Tengani_62            ..............................................t........t........................ [157200]
AY261363.1_Pretoriuskop/96/4     ................g.............................t........t........................ [157200]
AY261362.1_Mkuzi_1979            ................................................................................ [157200]
AY261361.1_Malawi_Lil-20/1_1983  .........................a....................t................................. [157200]
AY261360.1_Kenya_1950            ......ca.....t.............................a..t................................. [157200]
AM712240.1_OURT_88/3             ................................................................................ [157200]
AM712239.1_Benin_97/1            ................................................................................ [157200]
KM111295.1_Ken06.Bus             ......ca...................................a..t................................. [157200]
KM111294.1_Ken05/Tk1             ......ca.....t.............................a..t................................. [157200]
KM262844.1_strain_L60            ................................................................................ [157200]
KM102979.1_26544/OG10_Italy      ................................................................................ [157200]
KM262845.1_strain_NHV            ................................................................................ [157200]
KX354450.1_47/Ss/2008            ................................................................................ [157200]
KP055815.1_strain_BA71           ................................................................................ [157200]
MH025920.1_strain_R35            ......ca...................................a..t................................. [157200]
MH025919.1_strain_N10            ......ca...................................a..t................................. [157200]
MH025918.1_strain_R25            ......ca...................................a..t................................. [157200]
MH025917.1_strain_R7             ......ca...................................a..t................................. [157200]
MH025916.1_strain_R8             ......ca...................................a..t................................. [157200]

FR682468.2_Georgia_2007/1        atgaaaaactttttataaacctcgacaagggcccccatacctgcaagattatctataataagtgcttctagcatcgtata [157280]
ASF_IND_20_CAD543                ................................................................................ [157280]
LR899193.1_Germany_2020/1        ................................................................................ [157280]
MN393477.1_Wuhan_2019            ................................................................................ [157280]
MW396979.1_Timor-Leste/2019/1    ................................................................................ [157280]
MK940252.1_CN/2019/InnerMongolia ................................................................................ [157280]
MK128995.1_China/2018/AnhuiXCGQ  ................................................................................ [157280]
MH681419.1_ASFV/POL/2015         ................................................................................ [157280]
LS478113.1_Estonia_2014          ................................................................................ [157280]
KP843857.1_Odintsovo_02/14       ................................................................................ [157280]
```

FN557520.1_strain_E75 ................................................................................ [157280]
AY261366.1_Warthog ................................................................................ [157280]
AY261365.1_Warmbaths ......................................................................a......... [157280]
AY261364.1_Tengani_62 ................................t.........a..................................... [157280]
AY261363.1_Pretoriuskop/96/4 ................................................................................ [157280]
AY261362.1_Mkuzi_1979 ................................................................................ [157280]
AY261361.1_Malawi_Lil-20/1_1983 ..............................................................c...........t..... [157280]
AY261360.1_Kenya_1950 .............................a................................c...........t..... [157280]
AM712240.1_OURT_88/3 ................................................................................ [157280]
AM712239.1_Benin_97/1 ................................................................................ [157280]
KM111295.1_Ken06.Bus .........................t..a................................c...........t..... [157280]
KM111294.1_Ken05/Tk1 .............................a................................c...........t..... [157280]
KM262844.1_strain_L60 ................................................................................ [157280]
KM102979.1_26544/OG10_Italy ................................................................................ [157280]
KM262845.1_strain_NHV ................................................................................ [157280]
KX354450.1_47/Ss/2008 ................................................................................ [157280]
KP055815.1_strain_BA71 ................................................................................ [157280]
MH025920.1_strain_R35 .........................t..a................................c...........t..... [157280]
MH025919.1_strain_N10 .........................t..a................................c...........t..... [157280]
MH025918.1_strain_R25 .........................t..a................................c...........t..... [157280]
MH025917.1_strain_R7 .........................t..a................................c...........t..... [157280]
MH025916.1_strain_R8 .........................t..a................................c...........t..... [157280]

FR682468.2_Georgia_2007/1 gtgaaatgaagcggggtagtggatgagtacctgctccattggctcatcctgaaaatccttctgaaacttttcatacaata [157360]
ASF_IND_20_CAD543 ................................................................................ [157360]
LR899193.1_Germany_2020/1 ................................................................................ [157360]
MN393477.1_Wuhan_2019 ................................................................................ [157360]
MW396979.1_Timor-Leste/2019/1 ................................................................................ [157360]
MK940252.1_CN/2019/InnerMongolia ................................................................................ [157360]
MK128995.1_China/2018/AnhuiXCGQ ................................................................................ [157360]
MH681419.1_ASFV/POL/2015 ................................................................................ [157360]
LS478113.1_Estonia_2014 ................................................................................ [157360]
KP843857.1_Odintsovo_02/14 ................................................................................ [157360]
FN557520.1_strain_E75 ...............a..a............................................................. [157360]
AY261366.1_Warthog ...............a.........................................t...................g.. [157360]
AY261365.1_Warmbaths ...............a.........................................t...................g.. [157360]
AY261364.1_Tengani_62 ...............a.............................................................g.. [157360]
AY261363.1_Pretoriuskop/96/4 ...............a.............................................................g.. [157360]
AY261362.1_Mkuzi_1979 ...............a..a............................................................. [157360]
AY261361.1_Malawi_Lil-20/1_1983 a.....c........a........a...........t........................................gg. [157360]
AY261360.1_Kenya_1950 ......c........a................................t..........................t.gg. [157360]
AM712240.1_OURT_88/3 ...............a..a............................................................. [157360]
AM712239.1_Benin_97/1 ...............a..a............................................................. [157360]
KM111295.1_Ken06.Bus ......c........a........a........t..t....................t....................g. [157360]

```
KM111294.1_Ken05/Tk1             ......c........a................................t..........................t.gg. [157360]
KM262844.1_strain_L60            ...............a..a............................................................. [157360]
KM102979.1_26544/OG10_Italy      ...............a..a............................................................. [157360]
KM262845.1_strain_NHV            ...............a..a............................................................. [157360]
KX354450.1_47/Ss/2008            ...............a..a............................................................. [157360]
KP055815.1_strain_BA71           ...............a..a............................................................. [157360]
MH025920.1_strain_R35            ......c........a........a........t..t....................t....................g. [157360]
MH025919.1_strain_N10            ......c........a........a........t..t....................t....................g. [157360]
MH025918.1_strain_R25            ......c........a........a........t..t....................t....................g. [157360]
MH025917.1_strain_R7             ......c........a........a........t..t....................t....................g. [157360]
MH025916.1_strain_R8             ......c........a........a........t..t....................t....................g. [157360]

FR682468.2_Georgia_2007/1        cttgaaagggttctttggtctgcgagtgttcgaggtatttggtaatacggatgctgtgcatcgcgggaggctgaaaatcc [157440]
ASF_IND_20_CAD543                ................................................................................ [157440]
LR899193.1_Germany_2020/1        ................................................................................ [157440]
MN393477.1_Wuhan_2019            ................................................................................ [157440]
MW396979.1_Timor-Leste/2019/1    ................................................................................ [157440]
MK940252.1_CN/2019/InnerMongolia ................................................................................ [157440]
MK128995.1_China/2018/AnhuiXCGQ  ................................................................................ [157440]
MH681419.1_ASFV/POL/2015         ................................................................................ [157440]
LS478113.1_Estonia_2014          ................................................................................ [157440]
KP843857.1_Odintsovo_02/14       ................................................................................ [157440]
FN557520.1_strain_E75            ....................................................................a........... [157440]
AY261366.1_Warthog               .......a...................................g..................at................ [157440]
AY261365.1_Warmbaths             .......a...................................g...................t................ [157440]
AY261364.1_Tengani_62            .....................................c.....g...................t................ [157440]
AY261363.1_Pretoriuskop/96/4     .......a...................................g...................t................ [157440]
AY261362.1_Mkuzi_1979            ................................................................................ [157440]
AY261361.1_Malawi_Lil-20/1_1983  .......t........a..............a...........g.......c...........t................ [157440]
AY261360.1_Kenya_1950            ....g..a...................................g...................t................ [157440]
AM712240.1_OURT_88/3             ....................................................................a........... [157440]
AM712239.1_Benin_97/1            ....................................................................a........... [157440]
KM111295.1_Ken06.Bus             ....g..t..........................a........g..................at................ [157440]
KM111294.1_Ken05/Tk1             ....g..a...................................g...................t................ [157440]
KM262844.1_strain_L60            ....................................................................a........... [157440]
KM102979.1_26544/OG10_Italy      ....................................................................a........... [157440]
KM262845.1_strain_NHV            ....................................................................a........... [157440]
KX354450.1_47/Ss/2008            ....................................................................a........... [157440]
KP055815.1_strain_BA71           ....................................................................a........... [157440]
MH025920.1_strain_R35            ....g..t..........................a........g..................at................ [157440]
MH025919.1_strain_N10            ....g..t..........................a........g..................at................ [157440]
MH025918.1_strain_R25            ....g..t..........................a........g..................at................ [157440]
MH025917.1_strain_R7             ....g..t..........................a........g..................at................ [157440]
MH025916.1_strain_R8             ....g..t..........................a........g..................at................ [157440]
```

```
FR682468.2_Georgia_2007/1        cgaatatatgtttcaatatctaataccggttcctttttatggttaagcaccgcagcgacgtacaaatgctcaggctttgc [157520]
ASF_IND_20_CAD543                ................................................................................ [157520]
LR899193.1_Germany_2020/1        ................................................................................ [157520]
MN393477.1_Wuhan_2019            ................................................................................ [157520]
MW396979.1_Timor-Leste/2019/1    ................................................................................ [157520]
MK940252.1_CN/2019/InnerMongolia ................................................................................ [157520]
MK128995.1_China/2018/AnhuiXCGQ  ................................................................................ [157520]
MH681419.1_ASFV/POL/2015         ................................................................................ [157520]
LS478113.1_Estonia_2014          ................................................................................ [157520]
KP843857.1_Odintsovo_02/14       ................................................................................ [157520]
FN557520.1_strain_E75            ................................................................................ [157520]
AY261366.1_Warthog               .................g........t..................................................... [157520]
AY261365.1_Warmbaths             ..........................t..................................................... [157520]
AY261364.1_Tengani_62            ..........................t..................................................... [157520]
AY261363.1_Pretoriuskop/96/4     ..........................t..................................................... [157520]
AY261362.1_Mkuzi_1979            ................................................................................ [157520]
AY261361.1_Malawi_Lil-20/1_1983  ........c...........c..c..t..............a................t..................... [157520]
AY261360.1_Kenya_1950            ........c.....g........c................a.................t..................... [157520]
AM712240.1_OURT_88/3             ................................................................................ [157520]
AM712239.1_Benin_97/1            ................................................................................ [157520]
KM111295.1_Ken06.Bus             ........c.....g........c..................................t..................... [157520]
KM111294.1_Ken05/Tk1             ........c.....g........c..................................t..................... [157520]
KM262844.1_strain_L60            ................................................................................ [157520]
KM102979.1_26544/OG10_Italy      ................................................................................ [157520]
KM262845.1_strain_NHV            ................................................................................ [157520]
KX354450.1_47/Ss/2008            ................................................................................ [157520]
KP055815.1_strain_BA71           ................................................................................ [157520]
MH025920.1_strain_R35            ........c.....g........c..................................t..................... [157520]
MH025919.1_strain_N10            ........c.....g........c..................................t..................... [157520]
MH025918.1_strain_R25            ........c.....g........c..................................t..................... [157520]
MH025917.1_strain_R7             ........c.....g........c..................................t..................... [157520]
MH025916.1_strain_R8             ........c.....g........c..................................t..................... [157520]

FR682468.2_Georgia_2007/1        cggcacatgcataatggtgcaaagacgattctgtatccataattccttgcactggttttttgagtagcatagagaaatga [157600]
ASF_IND_20_CAD543                ................................................................................ [157600]
LR899193.1_Germany_2020/1        ................................................................................ [157600]
MN393477.1_Wuhan_2019            ................................................................................ [157600]
MW396979.1_Timor-Leste/2019/1    ................................................................................ [157600]
MK940252.1_CN/2019/InnerMongolia ................................................................................ [157600]
MK128995.1_China/2018/AnhuiXCGQ  ................................................................................ [157600]
MH681419.1_ASFV/POL/2015         ................................................................................ [157600]
LS478113.1_Estonia_2014          ................................................................................ [157600]
KP843857.1_Odintsovo_02/14       ................................................................................ [157600]
```

```
FN557520.1_strain_E75             ................................................................................ [157600]
AY261366.1_Warthog                .............................................t.................................. [157600]
AY261365.1_Warmbaths              ..............................t..............t.................................. [157600]
AY261364.1_Tengani_62             .............................................t.................................. [157600]
AY261363.1_Pretoriuskop/96/4      ..............................t..............t.................................. [157600]
AY261362.1_Mkuzi_1979             ................................................................................ [157600]
AY261361.1_Malawi_Lil-20/1_1983   .....................................a.......t..a.....a........a..a..c..g....... [157600]
AY261360.1_Kenya_1950             ....................................a........t.....t..a..............c..g....c.. [157600]
AM712240.1_OURT_88/3              ................................................................................ [157600]
AM712239.1_Benin_97/1             ................................................................................ [157600]
KM111295.1_Ken06.Bus              ....................................a........t........a..............c..g....... [157600]
KM111294.1_Ken05/Tk1              ....................................a........t.....t..a..............c..g....c.. [157600]
KM262844.1_strain_L60             ................................................................................ [157600]
KM102979.1_26544/OG10_Italy       ................................................................................ [157600]
KM262845.1_strain_NHV             ................................................................................ [157600]
KX354450.1_47/Ss/2008             ................................................................................ [157600]
KP055815.1_strain_BA71            ................................................................................ [157600]
MH025920.1_strain_R35             ....................................a........t........a..............c..g....... [157600]
MH025919.1_strain_N10             ....................................a........t........a..............c..g....... [157600]
MH025918.1_strain_R25             ....................................a........t........a..............c..g....... [157600]
MH025917.1_strain_R7              ....................................a........t........a..............c..g....... [157600]
MH025916.1_strain_R8              ....................................a........t........a..............c..g....... [157600]

FR682468.2_Georgia_2007/1         gcgccagcgcgaagttgtcctctgagaagagtttattatcgatggtaattccctgtatgagcttgggagtggaaacagcc [157680]
ASF_IND_20_CAD543                 ................................................................................ [157680]
LR899193.1_Germany_2020/1         ................................................................................ [157680]
MN393477.1_Wuhan_2019             ................................................................................ [157680]
MW396979.1_Timor-Leste/2019/1     ................................................................................ [157680]
MK940252.1_CN/2019/InnerMongolia  ................................................................................ [157680]
MK128995.1_China/2018/AnhuiXCGQ   ................................................................................ [157680]
MH681419.1_ASFV/POL/2015          ................................................................................ [157680]
LS478113.1_Estonia_2014           ................................................................................ [157680]
KP843857.1_Odintsovo_02/14        ................................................................................ [157680]
FN557520.1_strain_E75             ...................................................................c........t... [157680]
AY261366.1_Warthog                ...................................................................c........t... [157680]
AY261365.1_Warmbaths              ...................................................................c........t... [157680]
AY261364.1_Tengani_62             ...................................................................c........t... [157680]
AY261363.1_Pretoriuskop/96/4      ...................................................................c........t... [157680]
AY261362.1_Mkuzi_1979             ...................................................................c........t... [157680]
AY261361.1_Malawi_Lil-20/1_1983   .a...........................................................t.....c............ [157680]
AY261360.1_Kenya_1950             .a.................................................................c............ [157680]
AM712240.1_OURT_88/3              ...................................................................c........t... [157680]
AM712239.1_Benin_97/1             ...................................................................c........t... [157680]
KM111295.1_Ken06.Bus              .a.................................................................c............ [157680]
```

```
KM111294.1_Ken05/Tk1            .a.................................................................c............ [157680]
KM262844.1_strain_L60           ...................................................................c........t... [157680]
KM102979.1_26544/OG10_Italy     ...................................................................c........t... [157680]
KM262845.1_strain_NHV           ...................................................................c........t... [157680]
KX354450.1_47/Ss/2008           ...................................................................c........t... [157680]
KP055815.1_strain_BA71          ...................................................................c........t... [157680]
MH025920.1_strain_R35           .a.................................................................c............ [157680]
MH025919.1_strain_N10           .a.................................................................c............ [157680]
MH025918.1_strain_R25           .a.................................................................c............ [157680]
MH025917.1_strain_R7            .a.................................................................c............ [157680]
MH025916.1_strain_R8            .a.................................................................c............ [157680]

FR682468.2_Georgia_2007/1       ttccatagctcggagtacgtccacacggggcgtgccataaacaaagatataataatattagaaattgtttttacctcttg [157760]
ASF_IND_20_CAD543               ................................................................................ [157760]
LR899193.1_Germany_2020/1       ................................................................................ [157760]
MN393477.1_Wuhan_2019           ................................................................................ [157760]
MW396979.1_Timor-Leste/2019/1   ................................................................................ [157760]
MK940252.1_CN/2019/InnerMongolia ................................................................................ [157760]
MK128995.1_China/2018/AnhuiXCGQ ................................................................................ [157760]
MH681419.1_ASFV/POL/2015        ................................................................................ [157760]
LS478113.1_Estonia_2014         ................................................................................ [157760]
KP843857.1_Odintsovo_02/14      ................................................................................ [157760]
FN557520.1_strain_E75           ................................................................................ [157760]
AY261366.1_Warthog              .................................................................c.............. [157760]
AY261365.1_Warmbaths            .................................................................c.............. [157760]
AY261364.1_Tengani_62           .c...........................a.................a................................ [157760]
AY261363.1_Pretoriuskop/96/4    ................................................................................ [157760]
AY261362.1_Mkuzi_1979           .............................a.................................................. [157760]
AY261361.1_Malawi_Lil-20/1_1983 .......a...a...................................g................................ [157760]
AY261360.1_Kenya_1950           ..........................c..................................................c.. [157760]
AM712240.1_OURT_88/3            ................................................................................ [157760]
AM712239.1_Benin_97/1           ................................................................................ [157760]
KM111295.1_Ken06.Bus            .............g............c..................................................c.. [157760]
KM111294.1_Ken05/Tk1            ..........................c..................................................c.. [157760]
KM262844.1_strain_L60           ................................................................................ [157760]
KM102979.1_26544/OG10_Italy     ................................................................................ [157760]
KM262845.1_strain_NHV           ................................................................................ [157760]
KX354450.1_47/Ss/2008           ................................................................................ [157760]
KP055815.1_strain_BA71          ................................................................................ [157760]
MH025920.1_strain_R35           .............g............c..................................................c.. [157760]
MH025919.1_strain_N10           .............g............c..................................................c.. [157760]
MH025918.1_strain_R25           .............g............c..................................................c.. [157760]
MH025917.1_strain_R7            .............g............c..................................................c.. [157760]
MH025916.1_strain_R8            .............g............c..................................................c.. [157760]
```

```
FR682468.2_Georgia_2007/1        ctccccgtatccataggcctcaaaggtattgaggacggtggctccgacgtttgccggcgtgatggatggactaaggggca [157840]
ASF_IND_20_CAD543                ................................................................................ [157840]
LR899193.1_Germany_2020/1        ................................................................................ [157840]
MN393477.1_Wuhan_2019            ................................................................................ [157840]
MW396979.1_Timor-Leste/2019/1    ................................................................................ [157840]
MK940252.1_CN/2019/InnerMongolia ................................................................................ [157840]
MK128995.1_China/2018/AnhuiXCGQ  ................................................................................ [157840]
MH681419.1_ASFV/POL/2015         ................................................................................ [157840]
LS478113.1_Estonia_2014          ................................................................................ [157840]
KP843857.1_Odintsovo_02/14       ................................................................................ [157840]
FN557520.1_strain_E75            ................................................................................ [157840]
AY261366.1_Warthog               ...................................................g............................ [157840]
AY261365.1_Warmbaths             ...............a...................................g............................ [157840]
AY261364.1_Tengani_62            ...t............................................................................ [157840]
AY261363.1_Pretoriuskop/96/4     ...............a...................................g............................ [157840]
AY261362.1_Mkuzi_1979            ................................................................................ [157840]
AY261361.1_Malawi_Lil-20/1_1983  ................................................................................ [157840]
AY261360.1_Kenya_1950            ...t........g..c................................................................ [157840]
AM712240.1_OURT_88/3             ................................................................................ [157840]
AM712239.1_Benin_97/1            ................................................................................ [157840]
KM111295.1_Ken06.Bus             ...t....................a..............................................c........ [157840]
KM111294.1_Ken05/Tk1             ...t........g..c................................................................ [157840]
KM262844.1_strain_L60            ................................................................................ [157840]
KM102979.1_26544/OG10_Italy      ................................................................................ [157840]
KM262845.1_strain_NHV            ................................................................................ [157840]
KX354450.1_47/Ss/2008            ................................................................................ [157840]
KP055815.1_strain_BA71           ................................................................................ [157840]
MH025920.1_strain_R35            ...t....................a..............................................c........ [157840]
MH025919.1_strain_N10            ...t....................a..............................................c........ [157840]
MH025918.1_strain_R25            ...t....................a..............................................c........ [157840]
MH025917.1_strain_R7             ...t....................a..............................................c........ [157840]
MH025916.1_strain_R8             ...t....................a..............................................c........ [157840]

FR682468.2_Georgia_2007/1        gactttccaacataggcttatcaatcttaatctggttggtgaacccatcaatggcgtgctttcgcagcgccttatccccc [157920]
ASF_IND_20_CAD543                ................................................................................ [157920]
LR899193.1_Germany_2020/1        ................................................................................ [157920]
MN393477.1_Wuhan_2019            ................................................................................ [157920]
MW396979.1_Timor-Leste/2019/1    ................................................................................ [157920]
MK940252.1_CN/2019/InnerMongolia ................................................................................ [157920]
MK128995.1_China/2018/AnhuiXCGQ  ................................................................................ [157920]
MH681419.1_ASFV/POL/2015         ................................................................................ [157920]
LS478113.1_Estonia_2014          ................................................................................ [157920]
KP843857.1_Odintsovo_02/14       ................................................................................ [157920]
```

```
FN557520.1_strain_E75            .........g...................................................................... [157920]
AY261366.1_Warthog               .........g...........................t.......................................... [157920]
AY261365.1_Warmbaths             .........g...........................t.......................................... [157920]
AY261364.1_Tengani_62            .....................................t.......................................... [157920]
AY261363.1_Pretoriuskop/96/4     .........g...........................t.......................................... [157920]
AY261362.1_Mkuzi_1979            .........g...................................................................... [157920]
AY261361.1_Malawi_Lil-20/1_1983  .........g.....................t.....t.......................................... [157920]
AY261360.1_Kenya_1950            .........g............g..............t...........g.............................. [157920]
AM712240.1_OURT_88/3             .........g...................................................................... [157920]
AM712239.1_Benin_97/1            .........g...................................................................... [157920]
KM111295.1_Ken06.Bus             .........g............g..............t...........g.............................. [157920]
KM111294.1_Ken05/Tk1             .........g............g..............t...........g.............................. [157920]
KM262844.1_strain_L60            .........g...................................................................... [157920]
KM102979.1_26544/OG10_Italy      .........g...................................................................... [157920]
KM262845.1_strain_NHV            .........g...................................................................... [157920]
KX354450.1_47/Ss/2008            .........g...................................................................... [157920]
KP055815.1_strain_BA71           .........g...................................................................... [157920]
MH025920.1_strain_R35            .........g............g..............t...........g.............................. [157920]
MH025919.1_strain_N10            .........g............g..............t...........g.............................. [157920]
MH025918.1_strain_R25            .........g............g..............t...........g.............................. [157920]
MH025917.1_strain_R7             .........g............g..............t...........g.............................. [157920]
MH025916.1_strain_R8             .........g............g..............t...........g.............................. [157920]

FR682468.2_Georgia_2007/1        tcctgtattaaaatgtattcttttaatttttgtgcgtacttagcgagctctggccctccatcgggtgttgtcgatacgta [158000]
ASF_IND_20_CAD543                ................................................................................ [158000]
LR899193.1_Germany_2020/1        ................................................................................ [158000]
MN393477.1_Wuhan_2019            ................................................................................ [158000]
MW396979.1_Timor-Leste/2019/1    ................................................................................ [158000]
MK940252.1_CN/2019/InnerMongolia ................................................................................ [158000]
MK128995.1_China/2018/AnhuiXCGQ  ................................................................................ [158000]
MH681419.1_ASFV/POL/2015         ................................................................................ [158000]
LS478113.1_Estonia_2014          ................................................................................ [158000]
KP843857.1_Odintsovo_02/14       ................................................................................ [158000]
FN557520.1_strain_E75            ............................................a................................... [158000]
AY261366.1_Warthog               ..............................................................a..............a.. [158000]
AY261365.1_Warmbaths             ..............................................................a................. [158000]
AY261364.1_Tengani_62            ..............a................................................................. [158000]
AY261363.1_Pretoriuskop/96/4     .........................................g...................................a.. [158000]
AY261362.1_Mkuzi_1979            ............................................a................................... [158000]
AY261361.1_Malawi_Lil-20/1_1983  .........................................g.....t.......................t........ [158000]
AY261360.1_Kenya_1950            .........................................g.....t........c..g...........t..c..... [158000]
AM712240.1_OURT_88/3             ............................................a................................... [158000]
AM712239.1_Benin_97/1            ............................................a................................... [158000]
KM111295.1_Ken06.Bus             ...................................a.....g.....t........c..............t........ [158000]
```

```
KM111294.1_Ken05/Tk1              ...............................................t........c..g...........t..c..... [158000]
KM262844.1_strain_L60             ............................................a................................... [158000]
KM102979.1_26544/OG10_Italy       ............................................a................................... [158000]
KM262845.1_strain_NHV             ............................................a................................... [158000]
KX354450.1_47/Ss/2008             ............................................a................................... [158000]
KP055815.1_strain_BA71            ............................................a................................... [158000]
MH025920.1_strain_R35             ...................................a.....g.....t........c..............t........ [158000]
MH025919.1_strain_N10             ...................................a.....g.....t........c..............t........ [158000]
MH025918.1_strain_R25             ...................................a.....g.....t........c..............t........ [158000]
MH025917.1_strain_R7              ...................................a.....g.....t........c..............t........ [158000]
MH025916.1_strain_R8              ...................................a.....g.....t........c..............t........ [158000]

FR682468.2_Georgia_2007/1         caaataaattgtcacgttgcgctcactgggggggagctccatgtgtgaattttttcgcaccaccctcccaaatacctgaa [158080]
ASF_IND_20_CAD543                 ..............................................................y................. [158080]
LR899193.1_Germany_2020/1         ................................................................................ [158080]
MN393477.1_Wuhan_2019             ................................................................................ [158080]
MW396979.1_Timor-Leste/2019/1     ................................................................................ [158080]
MK940252.1_CN/2019/InnerMongolia  ................................................................................ [158080]
MK128995.1_China/2018/AnhuiXCGQ   ................................................................................ [158080]
MH681419.1_ASFV/POL/2015          ................................................................................ [158080]
LS478113.1_Estonia_2014           ................................................................................ [158080]
KP843857.1_Odintsovo_02/14        ................................................................................ [158080]
FN557520.1_strain_E75             ......g..........................a.............................................. [158080]
AY261366.1_Warthog                ................................................................................ [158080]
AY261365.1_Warmbaths              .................................a.............................................. [158080]
AY261364.1_Tengani_62             ............................................................t................... [158080]
AY261363.1_Pretoriuskop/96/4      .................................a...........c.................................. [158080]
AY261362.1_Mkuzi_1979             ................................................................................ [158080]
AY261361.1_Malawi_Lil-20/1_1983   .........................t.......a..........................t..t.g...g.....t.... [158080]
AY261360.1_Kenya_1950             ...g.....g...............t.......a...............................g.............. [158080]
AM712240.1_OURT_88/3              ......g..........................a.............................................. [158080]
AM712239.1_Benin_97/1             ......g..........................a.............................................. [158080]
KM111295.1_Ken06.Bus              ...g.....g...............t.......a...............................g.............. [158080]
KM111294.1_Ken05/Tk1              ...g.....g...............t.......a...............................g.............. [158080]
KM262844.1_strain_L60             ......g..........................a.............................................. [158080]
KM102979.1_26544/OG10_Italy       ......g..........................a.............................................. [158080]
KM262845.1_strain_NHV             ......g..........................a.............................................. [158080]
KX354450.1_47/Ss/2008             ......g..........................a.............................................. [158080]
KP055815.1_strain_BA71            ......g..........................a.............................................. [158080]
MH025920.1_strain_R35             ...g.....g...............t.......a...............................g.............. [158080]
MH025919.1_strain_N10             ...g.....g...............t.......a...............................g.............. [158080]
MH025918.1_strain_R25             ...g.....g...............t.......a...............................g.............. [158080]
MH025917.1_strain_R7              ...g.....g...............t.......a...............................g.............. [158080]
MH025916.1_strain_R8              ...g.....g...............t.......a...............................g.............. [158080]
```

```
FR682468.2_Georgia_2007/1        taagccggggaatatcaaggggcaatgacataatcatctcgtaccgcacggcctgaaagttcaaaccctccacaatcacc [158160]
ASF_IND_20_CAD543                ................................................................................ [158160]
LR899193.1_Germany_2020/1        ................................................................................ [158160]
MN393477.1_Wuhan_2019            ................................................................................ [158160]
MW396979.1_Timor-Leste/2019/1    ................................................................................ [158160]
MK940252.1_CN/2019/InnerMongolia ................................................................................ [158160]
MK128995.1_China/2018/AnhuiXCGQ  ................................................................................ [158160]
MH681419.1_ASFV/POL/2015         ................................................................................ [158160]
LS478113.1_Estonia_2014          ................................................................................ [158160]
KP843857.1_Odintsovo_02/14       ................................................................................ [158160]
FN557520.1_strain_E75            ................................................................................ [158160]
AY261366.1_Warthog               ................................................................................ [158160]
AY261365.1_Warmbaths             ................................................................................ [158160]
AY261364.1_Tengani_62            ................................................................................ [158160]
AY261363.1_Pretoriuskop/96/4     ................................................................................ [158160]
AY261362.1_Mkuzi_1979            ................................................................................ [158160]
AY261361.1_Malawi_Lil-20/1_1983  ....................................................t........t........t......... [158160]
AY261360.1_Kenya_1950            .........................a..........................t........................... [158160]
AM712240.1_OURT_88/3             ................................................................................ [158160]
AM712239.1_Benin_97/1            ................................................................................ [158160]
KM111295.1_Ken06.Bus             .........................a..........................t........................... [158160]
KM111294.1_Ken05/Tk1             .........................a..........................t........................... [158160]
KM262844.1_strain_L60            ................................................................................ [158160]
KM102979.1_26544/OG10_Italy      ................................................................................ [158160]
KM262845.1_strain_NHV            ................................................................................ [158160]
KX354450.1_47/Ss/2008            ................................................................................ [158160]
KP055815.1_strain_BA71           ................................................................................ [158160]
MH025920.1_strain_R35            .........................a..........................t........................... [158160]
MH025919.1_strain_N10            .........................a..........................t........................... [158160]
MH025918.1_strain_R25            .........................a..........................t........................... [158160]
MH025917.1_strain_R7             .........................a..........................t........................... [158160]
MH025916.1_strain_R8             .........................a..........................t........................... [158160]

FR682468.2_Georgia_2007/1        ttggacccgatgagaatacgcagctggtggccttccaggttggacgaggcgttaaaaagagccaggcttcgttcgcgtac [158240]
ASF_IND_20_CAD543                ................................................................................ [158240]
LR899193.1_Germany_2020/1        ................................................................................ [158240]
MN393477.1_Wuhan_2019            ................................................................................ [158240]
MW396979.1_Timor-Leste/2019/1    ................................................................................ [158240]
MK940252.1_CN/2019/InnerMongolia ................................................................................ [158240]
MK128995.1_China/2018/AnhuiXCGQ  ................................................................................ [158240]
MH681419.1_ASFV/POL/2015         ................................................................................ [158240]
LS478113.1_Estonia_2014          ................................................................................ [158240]
KP843857.1_Odintsovo_02/14       ................................................................................ [158240]
```

```
FN557520.1_strain_E75             ................................................................................ [158240]
AY261366.1_Warthog                ................................................................................ [158240]
AY261365.1_Warmbaths              ................................................................................ [158240]
AY261364.1_Tengani_62             ................................................................................ [158240]
AY261363.1_Pretoriuskop/96/4      ................................................................................ [158240]
AY261362.1_Mkuzi_1979             ................................................................................ [158240]
AY261361.1_Malawi_Lil-20/1_1983   .............................a.....t..a...........a............................. [158240]
AY261360.1_Kenya_1950             ...................................t..a...........a............................. [158240]
AM712240.1_OURT_88/3              ................................................................................ [158240]
AM712239.1_Benin_97/1             ................................................................................ [158240]
KM111295.1_Ken06.Bus              ...................................t..a...........a.......................a..... [158240]
KM111294.1_Ken05/Tk1              ...................................t..a...........a............................. [158240]
KM262844.1_strain_L60             ................................................................................ [158240]
KM102979.1_26544/OG10_Italy       ................................................................................ [158240]
KM262845.1_strain_NHV             ................................................................................ [158240]
KX354450.1_47/Ss/2008             ................................................................................ [158240]
KP055815.1_strain_BA71            ................................................................................ [158240]
MH025920.1_strain_R35             ...................................t..a...........a.......................a..... [158240]
MH025919.1_strain_N10             ...................................t..a...........a.......................a..... [158240]
MH025918.1_strain_R25             ...................................t..a...........a.......................a..... [158240]
MH025917.1_strain_R7              ...................................t..a...........a.......................a..... [158240]
MH025916.1_strain_R8              ...................................t..a...........a.......................a..... [158240]

FR682468.2_Georgia_2007/1         agcgggctctatttcgctgtgcagaatggtgaaccgtactggaataaactgatggtcgctatgt---gtgtgctcatcgc [158320]
ASF_IND_20_CAD543                 ................................................................---............. [158320]
LR899193.1_Germany_2020/1         ................................................................---............. [158320]
MN393477.1_Wuhan_2019             ................................................................---............. [158320]
MW396979.1_Timor-Leste/2019/1     ................................................................---............. [158320]
MK940252.1_CN/2019/InnerMongolia  ................................................................---............. [158320]
MK128995.1_China/2018/AnhuiXCGQ   ................................................................---............. [158320]
MH681419.1_ASFV/POL/2015          ................................................................---............. [158320]
LS478113.1_Estonia_2014           ................................................................---............. [158320]
KP843857.1_Odintsovo_02/14        ................................................................---............. [158320]
FN557520.1_strain_E75             ................................................................---............. [158320]
AY261366.1_Warthog                ................................................................---.....g....... [158320]
AY261365.1_Warmbaths              ................................................................---.....g....... [158320]
AY261364.1_Tengani_62             ............................................c...................---............. [158320]
AY261363.1_Pretoriuskop/96/4      ................................................................---.....g....... [158320]
AY261362.1_Mkuzi_1979             ................................................................---............. [158320]
AY261361.1_Malawi_Lil-20/1_1983   ................................................................---..a..a....... [158320]
AY261360.1_Kenya_1950             ...a.......................a.........g.............g............gtg.....g....... [158320]
AM712240.1_OURT_88/3              ................................................................---............. [158320]
AM712239.1_Benin_97/1             ................................................................---............. [158320]
KM111295.1_Ken06.Bus              ...a.................................g.............g............gtg.....g....... [158320]
```

```
KM111294.1_Ken05/Tk1           ...a.......................a.........g.............g............gtg.....g....... [158320]
KM262844.1_strain_L60          ................................................................---............. [158320]
KM102979.1_26544/OG10_Italy    ................................................................---............. [158320]
KM262845.1_strain_NHV          ................................................................---............. [158320]
KX354450.1_47/Ss/2008          ................................................................---............. [158320]
KP055815.1_strain_BA71         ................................................................---............. [158320]
MH025920.1_strain_R35          ...a.................................g.............g............gtg.....g....... [158320]
MH025919.1_strain_N10          ...a.................................g.............g............gtg.....g....... [158320]
MH025918.1_strain_R25          ...a.................................g.............g............gtg.....g....... [158320]
MH025917.1_strain_R7           ...a.................................g.............g............gtg.....g....... [158320]
MH025916.1_strain_R8           ...a.................................g.............g............gtg.....g....... [158320]

FR682468.2_Georgia_2007/1        gaatcgcggcgcagatggagcagcgggtcgttcccacaggggacgaaacttcatttaaaatgccattactttgtaaaatt [158400]
ASF_IND_20_CAD543                ................................................................................ [158400]
LR899193.1_Germany_2020/1        ................................................................................ [158400]
MN393477.1_Wuhan_2019            ................................................................................ [158400]
MW396979.1_Timor-Leste/2019/1    ................................................................................ [158400]
MK940252.1_CN/2019/InnerMongolia ................................................................................ [158400]
MK128995.1_China/2018/AnhuiXCGQ  ................................................................................ [158400]
MH681419.1_ASFV/POL/2015         ................................................................................ [158400]
LS478113.1_Estonia_2014          ................................................................................ [158400]
KP843857.1_Odintsovo_02/14       ................................................................................ [158400]
FN557520.1_strain_E75            ................................................................................ [158400]
AY261366.1_Warthog               .......a.................a...................................................... [158400]
AY261365.1_Warmbaths             .........................a...................................................... [158400]
AY261364.1_Tengani_62            .........................a...................................................... [158400]
AY261363.1_Pretoriuskop/96/4     .........................a...........g.......................................... [158400]
AY261362.1_Mkuzi_1979            ................................................................................ [158400]
AY261361.1_Malawi_Lil-20/1_1983  ....a........a...........a....................t........c.....a.................. [158400]
AY261360.1_Kenya_1950            ....a.......................a........c...........c..g..c.....a.................. [158400]
AM712240.1_OURT_88/3             ................................................................................ [158400]
AM712239.1_Benin_97/1            ................................................................................ [158400]
KM111295.1_Ken06.Bus             ....a.......................a....................c.....c.....a.................. [158400]
KM111294.1_Ken05/Tk1             ....a.......................a....................c..g..c.....a.................. [158400]
KM262844.1_strain_L60            ................................................................................ [158400]
KM102979.1_26544/OG10_Italy      ................................................................................ [158400]
KM262845.1_strain_NHV            ................................................................................ [158400]
KX354450.1_47/Ss/2008            ................................................................................ [158400]
KP055815.1_strain_BA71           ................................................................................ [158400]
MH025920.1_strain_R35            ....a.......................a....................c.....c.....a.................. [158400]
MH025919.1_strain_N10            ....a.......................a....................c.....c.....a.................. [158400]
MH025918.1_strain_R25            ....a.......................a....................c.....c.....a.................. [158400]
MH025917.1_strain_R7             ....a.......................a....................c.....c.....a.................. [158400]
MH025916.1_strain_R8             ....a.......................a....................c.....c.....a.................. [158400]
```

```
FR682468.2_Georgia_2007/1        tcttgcaagataagaacccccgacatgcggacccgattgtggtaaattaaaatttttcccccggccttgccgaataatgga [158480]
ASF_IND_20_CAD543                ................................................................................ [158480]
LR899193.1_Germany_2020/1        ................................................................................ [158480]
MN393477.1_Wuhan_2019            ................................................................................ [158480]
MW396979.1_Timor-Leste/2019/1    ................................................................................ [158480]
MK940252.1_CN/2019/InnerMongolia ................................................................................ [158480]
MK128995.1_China/2018/AnhuiXCGQ  ................................................................................ [158480]
MH681419.1_ASFV/POL/2015         ................................................................................ [158480]
LS478113.1_Estonia_2014          ................................................................................ [158480]
KP843857.1_Odintsovo_02/14       ................................................................................ [158480]
FN557520.1_strain_E75            .......g........................................................................ [158480]
AY261366.1_Warthog               .......g........................................................................ [158480]
AY261365.1_Warmbaths             .......g.....................................................................a.. [158480]
AY261364.1_Tengani_62            .......g........................................................................ [158480]
AY261363.1_Pretoriuskop/96/4     .......g............t........................................................... [158480]
AY261362.1_Mkuzi_1979            .......g........................................................................ [158480]
AY261361.1_Malawi_Lil-20/1_1983  .......g......g....................................................c............ [158480]
AY261360.1_Kenya_1950            .......g......g................................................................. [158480]
AM712240.1_OURT_88/3             .......g........................................................................ [158480]
AM712239.1_Benin_97/1            .......g........................................................................ [158480]
KM111295.1_Ken06.Bus             .......g......g................................................................. [158480]
KM111294.1_Ken05/Tk1             .......g......g................................................................. [158480]
KM262844.1_strain_L60            .......g........................................................................ [158480]
KM102979.1_26544/OG10_Italy      .......g........................................................................ [158480]
KM262845.1_strain_NHV            .......g........................................................................ [158480]
KX354450.1_47/Ss/2008            .......g........................................................................ [158480]
KP055815.1_strain_BA71           .......g........................................................................ [158480]
MH025920.1_strain_R35            .......g......g................................................................. [158480]
MH025919.1_strain_N10            .......g......g................................................................. [158480]
MH025918.1_strain_R25            .......g......g................................................................. [158480]
MH025917.1_strain_R7             .......g......g................................................................. [158480]
MH025916.1_strain_R8             .......g......g................................................................. [158480]

FR682468.2_Georgia_2007/1        aagaatgtctttcatcatttgagtgtattttccgctataaaaggccaatcccgagatgtgcgttggtggctgcagcgaca [158560]
ASF_IND_20_CAD543                ................................................................................ [158560]
LR899193.1_Germany_2020/1        ................................................................................ [158560]
MN393477.1_Wuhan_2019            ................................................................................ [158560]
MW396979.1_Timor-Leste/2019/1    ................................................................................ [158560]
MK940252.1_CN/2019/InnerMongolia ................................................................................ [158560]
MK128995.1_China/2018/AnhuiXCGQ  ................................................................................ [158560]
MH681419.1_ASFV/POL/2015         ................................................................................ [158560]
LS478113.1_Estonia_2014          ................................................................................ [158560]
KP843857.1_Odintsovo_02/14       ................................................................................ [158560]
```

```
FN557520.1_strain_E75            ................................................................................ [158560]
AY261366.1_Warthog               .........................................................a...a................t. [158560]
AY261365.1_Warmbaths             .........................................................a...................... [158560]
AY261364.1_Tengani_62            .........................................................a...a....c............. [158560]
AY261363.1_Pretoriuskop/96/4     .........................................................a........c............. [158560]
AY261362.1_Mkuzi_1979            ................................................................................ [158560]
AY261361.1_Malawi_Lil-20/1_1983  .................................a....................a..a....................t. [158560]
AY261360.1_Kenya_1950            .................................a..............c........a....................t. [158560]
AM712240.1_OURT_88/3             ................................................................................ [158560]
AM712239.1_Benin_97/1            ................................................................................ [158560]
KM111295.1_Ken06.Bus             .................................a.......................a.....a..............t. [158560]
KM111294.1_Ken05/Tk1             .................................a.......................a....................t. [158560]
KM262844.1_strain_L60            ................................................................................ [158560]
KM102979.1_26544/OG10_Italy      ................................................................................ [158560]
KM262845.1_strain_NHV            ................................................................................ [158560]
KX354450.1_47/Ss/2008            ................................................................................ [158560]
KP055815.1_strain_BA71           ................................................................................ [158560]
MH025920.1_strain_R35            .................................a.......................a.....a..............t. [158560]
MH025919.1_strain_N10            .................................a.......................a.....a..............t. [158560]
MH025918.1_strain_R25            .................................a.......................a.....a..............t. [158560]
MH025917.1_strain_R7             .................................a.......................a.....a..............t. [158560]
MH025916.1_strain_R8             .................................a.......................a.....a..............t. [158560]

FR682468.2_Georgia_2007/1        aaaagctgccactcacattaaagggggctctacgcgaaggctcaataatctgtaccccgttttccagaagccagtctgtg [158640]
ASF_IND_20_CAD543                ................................................................................ [158640]
LR899193.1_Germany_2020/1        ................................................................................ [158640]
MN393477.1_Wuhan_2019            ................................................................................ [158640]
MW396979.1_Timor-Leste/2019/1    ................................................................................ [158640]
MK940252.1_CN/2019/InnerMongolia ................................................................................ [158640]
MK128995.1_China/2018/AnhuiXCGQ  ................................................................................ [158640]
MH681419.1_ASFV/POL/2015         ................................................................................ [158640]
LS478113.1_Estonia_2014          ................................................................................ [158640]
KP843857.1_Odintsovo_02/14       ................................................................................ [158640]
FN557520.1_strain_E75            ................................................................................ [158640]
AY261366.1_Warthog               .................................................t.............................. [158640]
AY261365.1_Warmbaths             ...................................................................g............ [158640]
AY261364.1_Tengani_62            ................................................................................ [158640]
AY261363.1_Pretoriuskop/96/4     ................................................................................ [158640]
AY261362.1_Mkuzi_1979            .........................a...................................................... [158640]
AY261361.1_Malawi_Lil-20/1_1983  .........................c.t..............................a..................... [158640]
AY261360.1_Kenya_1950            g..........................t.....................t........a..................... [158640]
AM712240.1_OURT_88/3             ................................................................................ [158640]
AM712239.1_Benin_97/1            ................................................................................ [158640]
KM111295.1_Ken06.Bus             g..........................t.....................t........a..................... [158640]
```

```
KM111294.1_Ken05/Tk1             g..........................t.....................t........a..................... [158640]
KM262844.1_strain_L60            ................................................................................ [158640]
KM102979.1_26544/OG10_Italy      ................................................................................ [158640]
KM262845.1_strain_NHV            ................................................................................ [158640]
KX354450.1_47/Ss/2008            ................................................................................ [158640]
KP055815.1_strain_BA71           ................................................................................ [158640]
MH025920.1_strain_R35            g..........................t.....................t........a..................... [158640]
MH025919.1_strain_N10            g..........................t.....................t........a..................... [158640]
MH025918.1_strain_R25            g..........................t.....................t........a..................... [158640]
MH025917.1_strain_R7             g..........................t.....................t........a..................... [158640]
MH025916.1_strain_R8             g..........................t.....................t........a..................... [158640]

FR682468.2_Georgia_2007/1        cttgccatagaaagggcggtgggggtttccgtcgagttaaacaggccgtaagccttgggttccgtttgttttgaaaattt [158720]
ASF_IND_20_CAD543                ................................................................................ [158720]
LR899193.1_Germany_2020/1        ................................................................................ [158720]
MN393477.1_Wuhan_2019            ................................................................................ [158720]
MW396979.1_Timor-Leste/2019/1    ................................................................................ [158720]
MK940252.1_CN/2019/InnerMongolia ................................................................................ [158720]
MK128995.1_China/2018/AnhuiXCGQ  ................................................................................ [158720]
MH681419.1_ASFV/POL/2015         ................................................................................ [158720]
LS478113.1_Estonia_2014          ................................................................................ [158720]
KP843857.1_Odintsovo_02/14       ................................................................................ [158720]
FN557520.1_strain_E75            .................a.............................................................. [158720]
AY261366.1_Warthog               .......................a........................................................ [158720]
AY261365.1_Warmbaths             .................a.....a........................................................ [158720]
AY261364.1_Tengani_62            .................a..a.........................................t................. [158720]
AY261363.1_Pretoriuskop/96/4     ......................................g......................................... [158720]
AY261362.1_Mkuzi_1979            ....................a........................................................... [158720]
AY261361.1_Malawi_Lil-20/1_1983  ...................c...a........a............................................... [158720]
AY261360.1_Kenya_1950            ........g..........c..................g...........c............................. [158720]
AM712240.1_OURT_88/3             .................a.............................................................. [158720]
AM712239.1_Benin_97/1            .................a.............................................................. [158720]
KM111295.1_Ken06.Bus             ........g..........c..................g...........c............................. [158720]
KM111294.1_Ken05/Tk1             ........g..........c..................g...........c............................. [158720]
KM262844.1_strain_L60            .................a.............................................................. [158720]
KM102979.1_26544/OG10_Italy      .................a.............................................................. [158720]
KM262845.1_strain_NHV            .................a.............................................................. [158720]
KX354450.1_47/Ss/2008            .................a.............................................................. [158720]
KP055815.1_strain_BA71           .................a.............................................................. [158720]
MH025920.1_strain_R35            ........g..........c..................g...........c............................. [158720]
MH025919.1_strain_N10            ........g..........c..................g...........c............................. [158720]
MH025918.1_strain_R25            ........g..........c..................g...........c............................. [158720]
MH025917.1_strain_R7             ........g..........c..................g...........c............................. [158720]
MH025916.1_strain_R8             ........g..........c..................g...........c............................. [158720]
```

```
FR682468.2_Georgia_2007/1        tgggttgggaaacaccatgtcataaatgctgtacgcattactcgagattttagggtcagggcccagctgtttaagcgttt [158800]
ASF_IND_20_CAD543                ................................................................................ [158800]
LR899193.1_Germany_2020/1        ................................................................................ [158800]
MN393477.1_Wuhan_2019            ................................................................................ [158800]
MW396979.1_Timor-Leste/2019/1    ................................................................................ [158800]
MK940252.1_CN/2019/InnerMongolia ................................................................................ [158800]
MK128995.1_China/2018/AnhuiXCGQ  ................................................................................ [158800]
MH681419.1_ASFV/POL/2015         ................................................................................ [158800]
LS478113.1_Estonia_2014          ................................................................................ [158800]
KP843857.1_Odintsovo_02/14       ................................................................................ [158800]
FN557520.1_strain_E75            ................................................................................ [158800]
AY261366.1_Warthog               ...............................................................t................ [158800]
AY261365.1_Warmbaths             ................................................................................ [158800]
AY261364.1_Tengani_62            ................................................................................ [158800]
AY261363.1_Pretoriuskop/96/4     ................................................................................ [158800]
AY261362.1_Mkuzi_1979            ................................................................................ [158800]
AY261361.1_Malawi_Lil-20/1_1983  ...a........t..........................g.........................a.............. [158800]
AY261360.1_Kenya_1950            .......................................g........................................ [158800]
AM712240.1_OURT_88/3             ................................................................................ [158800]
AM712239.1_Benin_97/1            ................................................................................ [158800]
KM111295.1_Ken06.Bus             .............g.........................g........................................ [158800]
KM111294.1_Ken05/Tk1             .......................................g........................................ [158800]
KM262844.1_strain_L60            ................................................................................ [158800]
KM102979.1_26544/OG10_Italy      ................................................................................ [158800]
KM262845.1_strain_NHV            ................................................................................ [158800]
KX354450.1_47/Ss/2008            ................................................................................ [158800]
KP055815.1_strain_BA71           ................................................................................ [158800]
MH025920.1_strain_R35            .............g.........................g........................................ [158800]
MH025919.1_strain_N10            .............g.........................g........................................ [158800]
MH025918.1_strain_R25            .............g.........................g........................................ [158800]
MH025917.1_strain_R7             .............g.........................g........................................ [158800]
MH025916.1_strain_R8             .............g.........................g........................................ [158800]

FR682468.2_Georgia_2007/1        caagctgatactcagacatggggcattcgatgaaatgtaagtacggcaatgtttcgtctttataggacaacatctttccg [158880]
ASF_IND_20_CAD543                ................................................................................ [158880]
LR899193.1_Germany_2020/1        ................................................................................ [158880]
MN393477.1_Wuhan_2019            ................................................................................ [158880]
MW396979.1_Timor-Leste/2019/1    ................................................................................ [158880]
MK940252.1_CN/2019/InnerMongolia ................................................................................ [158880]
MK128995.1_China/2018/AnhuiXCGQ  ................................................................................ [158880]
MH681419.1_ASFV/POL/2015         ................................................................................ [158880]
LS478113.1_Estonia_2014          ................................................................................ [158880]
KP843857.1_Odintsovo_02/14       ................................................................................ [158880]
```

```
FN557520.1_strain_E75           ................................................................................ [158880]
AY261366.1_Warthog              .................................................ct..................g.......... [158880]
AY261365.1_Warmbaths            .................................................ct..................g.......... [158880]
AY261364.1_Tengani_62           ......................a..........................c......c....................... [158880]
AY261363.1_Pretoriuskop/96/4    .................................................ct..................g.......... [158880]
AY261362.1_Mkuzi_1979           ................................................................................ [158880]
AY261361.1_Malawi_Lil-20/1_1983 ..........a.t....................................ct........c...................a [158880]
AY261360.1_Kenya_1950           ..........a.......................g..c...........ca......t.c.................... [158880]
AM712240.1_OURT_88/3            ................................................................................ [158880]
AM712239.1_Benin_97/1           ................................................................................ [158880]
KM111295.1_Ken06.Bus            ..........a......................................ca........c....a............... [158880]
KM111294.1_Ken05/Tk1            ..........a.......................g..c...........ca........c....a............... [158880]
KM262844.1_strain_L60           ................................................................................ [158880]
KM102979.1_26544/OG10_Italy     ................................................................................ [158880]
KM262845.1_strain_NHV           ................................................................................ [158880]
KX354450.1_47/Ss/2008           ................................................................................ [158880]
KP055815.1_strain_BA71          ................................................................................ [158880]
MH025920.1_strain_R35           ..........a......................................ca........c....a............... [158880]
MH025919.1_strain_N10           ..........a......................................ca........c....a............... [158880]
MH025918.1_strain_R25           ..........a......................................ca........c....a............... [158880]
MH025917.1_strain_R7            ..........a......................................ca........c....a............... [158880]
MH025916.1_strain_R8            ..........a......................................ca........c....a............... [158880]

FR682468.2_Georgia_2007/1       gcaaatattctttcggggtaaaaattggtgttggtatccaacaaaaaagatacccttccggtgctcagtctttccacaag [158960]
ASF_IND_20_CAD543               ..............r..r.........................................k.................... [158960]
LR899193.1_Germany_2020/1       ................................................................................ [158960]
MN393477.1_Wuhan_2019           ................................................................................ [158960]
MW396979.1_Timor-Leste/2019/1   ................................................................................ [158960]
MK940252.1_CN/2019/InnerMongolia ................................................................................ [158960]
MK128995.1_China/2018/AnhuiXCGQ ................................................................................ [158960]
MH681419.1_ASFV/POL/2015        ................................................................................ [158960]
LS478113.1_Estonia_2014         ................................................................................ [158960]
KP843857.1_Odintsovo_02/14      ................................................................................ [158960]
FN557520.1_strain_E75           ..............a..a...............................................g.............. [158960]
AY261366.1_Warthog              ..............a................................................................. [158960]
AY261365.1_Warmbaths            ..............a................................................................. [158960]
AY261364.1_Tengani_62           ..............a................................................................. [158960]
AY261363.1_Pretoriuskop/96/4    ..............a................................................................. [158960]
AY261362.1_Mkuzi_1979           ..............a................................................................. [158960]
AY261361.1_Malawi_Lil-20/1_1983 ..............a................................g.....t......a................... [158960]
AY261360.1_Kenya_1950           .....g........a......t.........................g................................ [158960]
AM712240.1_OURT_88/3            ..............a..a...............................................g.............. [158960]
AM712239.1_Benin_97/1           ..............a..a...............................................g.............. [158960]
KM111295.1_Ken06.Bus            .....g........a......t.........................g................................ [158960]
```

```
KM111294.1_Ken05/Tk1            .....g........a......t.........................g................................ [158960]
KM262844.1_strain_L60           ..............a..a...............................................g.............. [158960]
KM102979.1_26544/OG10_Italy     ..............a..a...............................................g.............. [158960]
KM262845.1_strain_NHV           ..............a..a...............................................g.............. [158960]
KX354450.1_47/Ss/2008           ..............a..a...............................................g.............. [158960]
KP055815.1_strain_BA71          ..............a..a...............................................g.............. [158960]
MH025920.1_strain_R35           .....g........a......t.........................g................................ [158960]
MH025919.1_strain_N10           .....g........a......t.........................g................................ [158960]
MH025918.1_strain_R25           .....g........a......t.........................g................................ [158960]
MH025917.1_strain_R7            .....g........a......t.........................g................................ [158960]
MH025916.1_strain_R8            .....g........a......t.........................g................................ [158960]

FR682468.2_Georgia_2007/1       agctagggcgtcctttttccatttaacggaatgcccactgctgtcaaacagttgctggcgctggaggggctggccgttgg [159040]
ASF_IND_20_CAD543               ................................................................................ [159040]
LR899193.1_Germany_2020/1       ................................................................................ [159040]
MN393477.1_Wuhan_2019           ................................................................................ [159040]
MW396979.1_Timor-Leste/2019/1   ................................................................................ [159040]
MK940252.1_CN/2019/InnerMongolia ................................................................................ [159040]
MK128995.1_China/2018/AnhuiXCGQ ................................................................................ [159040]
MH681419.1_ASFV/POL/2015        ................................................................................ [159040]
LS478113.1_Estonia_2014         ................................................................................ [159040]
KP843857.1_Odintsovo_02/14      ................................................................................ [159040]
FN557520.1_strain_E75           ...c............................................................................ [159040]
AY261366.1_Warthog              ................................................................................ [159040]
AY261365.1_Warmbaths            ................................................................................ [159040]
AY261364.1_Tengani_62           ................................................................................ [159040]
AY261363.1_Pretoriuskop/96/4    ................................................................................ [159040]
AY261362.1_Mkuzi_1979           ...c............................................................................ [159040]
AY261361.1_Malawi_Lil-20/1_1983 ........................t...........g...t.......t............................... [159040]
AY261360.1_Kenya_1950           ...c.....a..............t...........g..at.........a............................. [159040]
AM712240.1_OURT_88/3            ...c............................................................................ [159040]
AM712239.1_Benin_97/1           ...c............................................................................ [159040]
KM111295.1_Ken06.Bus            ...c.....a..............t..a...........at.......t............................... [159040]
KM111294.1_Ken05/Tk1            ...c.....a..............t..............at.........a............................. [159040]
KM262844.1_strain_L60           ...c............................................................................ [159040]
KM102979.1_26544/OG10_Italy     ...c............................................................................ [159040]
KM262845.1_strain_NHV           ...c............................................................................ [159040]
KX354450.1_47/Ss/2008           ...c............................................................................ [159040]
KP055815.1_strain_BA71          ...c............................................................................ [159040]
MH025920.1_strain_R35           ...c.....a..............t..a...........at.......t............................... [159040]
MH025919.1_strain_N10           ...c.....a..............t..a...........at.......t............................... [159040]
MH025918.1_strain_R25           ...c.....a..............t..a...........at.......t............................... [159040]
MH025917.1_strain_R7            ...c.....a..............t..a...........at.......t............................... [159040]
MH025916.1_strain_R8            ...c.....a..............t..a...........at.......t............................... [159040]
```

```
FR682468.2_Georgia_2007/1        gcagctcatgccgcggaaccaaaaggtttaacaggtcgacgtattccatgacactcccggttacgggcgttgccgacatg [159120]
ASF_IND_20_CAD543                ................................................................................ [159120]
LR899193.1_Germany_2020/1        ................................................................................ [159120]
MN393477.1_Wuhan_2019            ................................................................................ [159120]
MW396979.1_Timor-Leste/2019/1    ................................................................................ [159120]
MK940252.1_CN/2019/InnerMongolia ................................................................................ [159120]
MK128995.1_China/2018/AnhuiXCGQ  ................................................................................ [159120]
MH681419.1_ASFV/POL/2015         ................................................................................ [159120]
LS478113.1_Estonia_2014          ................................................................................ [159120]
KP843857.1_Odintsovo_02/14       ................................................................................ [159120]
FN557520.1_strain_E75            ................................................................................ [159120]
AY261366.1_Warthog               ................................................................................ [159120]
AY261365.1_Warmbaths             ................................................................................ [159120]
AY261364.1_Tengani_62            .................................................a.............................. [159120]
AY261363.1_Pretoriuskop/96/4     ................................................................................ [159120]
AY261362.1_Mkuzi_1979            ................................................................................ [159120]
AY261361.1_Malawi_Lil-20/1_1983  ................................................................................ [159120]
AY261360.1_Kenya_1950            ...........................................c........g........................... [159120]
AM712240.1_OURT_88/3             ................................................................................ [159120]
AM712239.1_Benin_97/1            ................................................................................ [159120]
KM111295.1_Ken06.Bus             ...........................................c........g........................... [159120]
KM111294.1_Ken05/Tk1             ...........................................c........g........................... [159120]
KM262844.1_strain_L60            ................................................................................ [159120]
KM102979.1_26544/OG10_Italy      ................................................................................ [159120]
KM262845.1_strain_NHV            ................................................................................ [159120]
KX354450.1_47/Ss/2008            ................................................................................ [159120]
KP055815.1_strain_BA71           ................................................................................ [159120]
MH025920.1_strain_R35            ...........................................c........g........................... [159120]
MH025919.1_strain_N10            ...........................................c........g........................... [159120]
MH025918.1_strain_R25            ...........................................c........g........................... [159120]
MH025917.1_strain_R7             ...........................................c........g........................... [159120]
MH025916.1_strain_R8             ...........................................c........g........................... [159120]

FR682468.2_Georgia_2007/1        aagacggccctgggggcctggtgaggtggaaaggcatccaggacatactgtaaagcgatgccataattatttcgttcctg [159200]
ASF_IND_20_CAD543                ................................................................................ [159200]
LR899193.1_Germany_2020/1        ................................................................................ [159200]
MN393477.1_Wuhan_2019            ................................................................................ [159200]
MW396979.1_Timor-Leste/2019/1    ................................................................................ [159200]
MK940252.1_CN/2019/InnerMongolia ................................................................................ [159200]
MK128995.1_China/2018/AnhuiXCGQ  ................................................................................ [159200]
MH681419.1_ASFV/POL/2015         ................................................................................ [159200]
LS478113.1_Estonia_2014          ................................................................................ [159200]
KP843857.1_Odintsovo_02/14       ................................................................................ [159200]
```

```
FN557520.1_strain_E75            ................................................................................ [159200]
AY261366.1_Warthog               .......................g.........................................g.............. [159200]
AY261365.1_Warmbaths             .......................g.........................................g.............. [159200]
AY261364.1_Tengani_62            .......................g..c......................................g...........t.. [159200]
AY261363.1_Pretoriuskop/96/4     .......................g..............t..........................g.............. [159200]
AY261362.1_Mkuzi_1979            .......................g........................................................ [159200]
AY261361.1_Malawi_Lil-20/1_1983  ..a....................g.................a.................t........g........... [159200]
AY261360.1_Kenya_1950            ..........g............g.............................g..a..t.................... [159200]
AM712240.1_OURT_88/3             ................................................................................ [159200]
AM712239.1_Benin_97/1            ................................................................................ [159200]
KM111295.1_Ken06.Bus             ..........g............g.............................g..a..t.................... [159200]
KM111294.1_Ken05/Tk1             ..........g............g.............................g..a..t.................... [159200]
KM262844.1_strain_L60            ................................................................................ [159200]
KM102979.1_26544/OG10_Italy      ................................................................................ [159200]
KM262845.1_strain_NHV            ................................................................................ [159200]
KX354450.1_47/Ss/2008            ................................................................................ [159200]
KP055815.1_strain_BA71           ................................................................................ [159200]
MH025920.1_strain_R35            ..........g............g.............................g..a..t.................... [159200]
MH025919.1_strain_N10            ..........g............g.............................g..a..t.................... [159200]
MH025918.1_strain_R25            ..........g............g.............................g..a..t.................... [159200]
MH025917.1_strain_R7             ..........g............g.............................g..a..t.................... [159200]
MH025916.1_strain_R8             ..........g............g.............................g..a..t.................... [159200]

FR682468.2_Georgia_2007/1        gatattgtacacgttgtgtatttcatccgcaatgagcagtcctcccctaagttgctccatgattttttgattcacccgga [159280]
ASF_IND_20_CAD543                ................................................................................ [159280]
LR899193.1_Germany_2020/1        ................................................................................ [159280]
MN393477.1_Wuhan_2019            ................................................................................ [159280]
MW396979.1_Timor-Leste/2019/1    ................................................................................ [159280]
MK940252.1_CN/2019/InnerMongolia ................................................................................ [159280]
MK128995.1_China/2018/AnhuiXCGQ  ................................................................................ [159280]
MH681419.1_ASFV/POL/2015         ................................................................................ [159280]
LS478113.1_Estonia_2014          ................................................................................ [159280]
KP843857.1_Odintsovo_02/14       ................................................................................ [159280]
FN557520.1_strain_E75            ...................................................c............................ [159280]
AY261366.1_Warthog               ...................................................c............................ [159280]
AY261365.1_Warmbaths             ............a......................................c............................ [159280]
AY261364.1_Tengani_62            ...g...............................................c............................ [159280]
AY261363.1_Pretoriuskop/96/4     ..........................................c........c............................ [159280]
AY261362.1_Mkuzi_1979            ...................................................c............................ [159280]
AY261361.1_Malawi_Lil-20/1_1983  a..g......................................c..t.....c.......................a..a. [159280]
AY261360.1_Kenya_1950            a.........................................c........c........t..............a.... [159280]
AM712240.1_OURT_88/3             ...................................................c............................ [159280]
AM712239.1_Benin_97/1            ...................................................c............................ [159280]
KM111295.1_Ken06.Bus             a.............................g...........c........c........t...........g..a..a. [159280]
```

```
KM111294.1_Ken05/Tk1             a.........................................c........c........t..............a.... [159280]
KM262844.1_strain_L60            ...................................................c............................ [159280]
KM102979.1_26544/OG10_Italy      ...................................................c............................ [159280]
KM262845.1_strain_NHV            ...................................................c............................ [159280]
KX354450.1_47/Ss/2008            ...................................................c............................ [159280]
KP055815.1_strain_BA71           ...................................................c............................ [159280]
MH025920.1_strain_R35            a.............................g...........c........c........t...........g..a..a. [159280]
MH025919.1_strain_N10            a.............................g...........c........c........t...........g..a..a. [159280]
MH025918.1_strain_R25            a.............................g...........c........c........t...........g..a..a. [159280]
MH025917.1_strain_R7             a.............................g...........c........c........t...........g..a..a. [159280]
MH025916.1_strain_R8             a.............................g...........c........c........t...........g..a..a. [159280]

FR682468.2_Georgia_2007/1        tgaggccgtttgtctcggcctcgctaattttttgcacgaactgagatatatcgttctcattcaatgtatcttctgcttcg [159360]
ASF_IND_20_CAD543                ................................................................................ [159360]
LR899193.1_Germany_2020/1        ................................................................................ [159360]
MN393477.1_Wuhan_2019            ................................................................................ [159360]
MW396979.1_Timor-Leste/2019/1    ................................................................................ [159360]
MK940252.1_CN/2019/InnerMongolia ................................................................................ [159360]
MK128995.1_China/2018/AnhuiXCGQ  ................................................................................ [159360]
MH681419.1_ASFV/POL/2015         ................................................................................ [159360]
LS478113.1_Estonia_2014          ................................................................................ [159360]
KP843857.1_Odintsovo_02/14       ................................................................................ [159360]
FN557520.1_strain_E75            .........................g...................................................... [159360]
AY261366.1_Warthog               .........................g...................................................... [159360]
AY261365.1_Warmbaths             .........................g...................................................... [159360]
AY261364.1_Tengani_62            .........................g.....................................t................ [159360]
AY261363.1_Pretoriuskop/96/4     .........................g...................................................... [159360]
AY261362.1_Mkuzi_1979            .........................g...................................................... [159360]
AY261361.1_Malawi_Lil-20/1_1983  .......a.................g.....................................g................ [159360]
AY261360.1_Kenya_1950            ............at...........g...................................................... [159360]
AM712240.1_OURT_88/3             .........................g...................................................... [159360]
AM712239.1_Benin_97/1            .........................g...................................................... [159360]
KM111295.1_Ken06.Bus             .c..........at...........g.................................................t.... [159360]
KM111294.1_Ken05/Tk1             ............at...........g...................................................... [159360]
KM262844.1_strain_L60            .........................g...................................................... [159360]
KM102979.1_26544/OG10_Italy      .........................g...................................................... [159360]
KM262845.1_strain_NHV            .........................g...................................................... [159360]
KX354450.1_47/Ss/2008            .........................g...................................................... [159360]
KP055815.1_strain_BA71           .........................g...................................................... [159360]
MH025920.1_strain_R35            .c..........at...........g.................................................t.... [159360]
MH025919.1_strain_N10            .c..........at...........g.................................................t.... [159360]
MH025918.1_strain_R25            .c..........at...........g.................................................t.... [159360]
MH025917.1_strain_R7             .c..........at...........g.................................................t.... [159360]
MH025916.1_strain_R8             .c..........at...........g.................................................t.... [159360]
```

```
FR682468.2_Georgia_2007/1        tcagaacgatgaaacagagaaagcacatcaaagtttttctcttcacccttactcgtaatattgaaaagcttggatgcaaa [159440]
ASF_IND_20_CAD543                ................................................................................ [159440]
LR899193.1_Germany_2020/1        ................................................................................ [159440]
MN393477.1_Wuhan_2019            ................................................................................ [159440]
MW396979.1_Timor-Leste/2019/1    ................................................................................ [159440]
MK940252.1_CN/2019/InnerMongolia ................................................................................ [159440]
MK128995.1_China/2018/AnhuiXCGQ  ................................................................................ [159440]
MH681419.1_ASFV/POL/2015         ................................................................................ [159440]
LS478113.1_Estonia_2014          ................................................................................ [159440]
KP843857.1_Odintsovo_02/14       ................................................................................ [159440]
FN557520.1_strain_E75            ................................................................................ [159440]
AY261366.1_Warthog               ................................................................................ [159440]
AY261365.1_Warmbaths             ................................................................................ [159440]
AY261364.1_Tengani_62            ..............................................................a................. [159440]
AY261363.1_Pretoriuskop/96/4     ................................................................................ [159440]
AY261362.1_Mkuzi_1979            ....................................................................t........... [159440]
AY261361.1_Malawi_Lil-20/1_1983  ................a...............a......................................a........ [159440]
AY261360.1_Kenya_1950            .......................................................................a........ [159440]
AM712240.1_OURT_88/3             ................................................................................ [159440]
AM712239.1_Benin_97/1            ................................................................................ [159440]
KM111295.1_Ken06.Bus             ................................................................................ [159440]
KM111294.1_Ken05/Tk1             ..............t........................................................a........ [159440]
KM262844.1_strain_L60            ................................................................................ [159440]
KM102979.1_26544/OG10_Italy      ................................................................................ [159440]
KM262845.1_strain_NHV            ................................................................................ [159440]
KX354450.1_47/Ss/2008            ................................................................................ [159440]
KP055815.1_strain_BA71           ................................................................................ [159440]
MH025920.1_strain_R35            ................................................................................ [159440]
MH025919.1_strain_N10            ................................................................................ [159440]
MH025918.1_strain_R25            ................................................................................ [159440]
MH025917.1_strain_R7             ................................................................................ [159440]
MH025916.1_strain_R8             ................................................................................ [159440]

FR682468.2_Georgia_2007/1        ttccttatagccgtaaaactgaaaaaagcctccgcggtttctatcggttaaacggcgctttaacgtactaacgaacccat [159520]
ASF_IND_20_CAD543                ................................................................................ [159520]
LR899193.1_Germany_2020/1        ................................................................................ [159520]
MN393477.1_Wuhan_2019            ................................................................................ [159520]
MW396979.1_Timor-Leste/2019/1    ................................................................................ [159520]
MK940252.1_CN/2019/InnerMongolia ................................................................................ [159520]
MK128995.1_China/2018/AnhuiXCGQ  ................................................................................ [159520]
MH681419.1_ASFV/POL/2015         ................................................................................ [159520]
LS478113.1_Estonia_2014          ................................................................................ [159520]
KP843857.1_Odintsovo_02/14       ................................................................................ [159520]
```

```
FN557520.1_strain_E75            .......................................................................... [159520]
```

```
KM111294.1_Ken05/Tk1            ................a........................................................a...... [159600]
KM262844.1_strain_L60           ................a............................................................... [159600]
KM102979.1_26544/OG10_Italy     ................a............................................................... [159600]
KM262845.1_strain_NHV           ................a............................................................... [159600]
KX354450.1_47/Ss/2008           ................a............................................................... [159600]
KP055815.1_strain_BA71          ................a............................................................... [159600]
MH025920.1_strain_R35           ................a........................................................a...... [159600]
MH025919.1_strain_N10           ................a........................................................a...... [159600]
MH025918.1_strain_R25           ................a........................................................a...... [159600]
MH025917.1_strain_R7            ................a........................................................a...... [159600]
MH025916.1_strain_R8            ................a........................................................a...... [159600]

FR682468.2_Georgia_2007/1       ttgtaagaaacaaatcccagctcaggacgtcttagcatttctgtttgaatgatggcgcgtgtaaagcctaccacaaaaat [159680]
ASF_IND_20_CAD543               ................................................................................ [159680]
LR899193.1_Germany_2020/1       ................................................................................ [159680]
MN393477.1_Wuhan_2019           ................................................................................ [159680]
MW396979.1_Timor-Leste/2019/1   ................................................................................ [159680]
MK940252.1_CN/2019/InnerMongolia ................................................................................ [159680]
MK128995.1_China/2018/AnhuiXCGQ ................................................................................ [159680]
MH681419.1_ASFV/POL/2015        ................................................................................ [159680]
LS478113.1_Estonia_2014         ................................................................................ [159680]
KP843857.1_Odintsovo_02/14      ................................................................................ [159680]
FN557520.1_strain_E75           ................................................................................ [159680]
AY261366.1_Warthog              ................................................................................ [159680]
AY261365.1_Warmbaths            ................................................................................ [159680]
AY261364.1_Tengani_62           ..................................a............................................. [159680]
AY261363.1_Pretoriuskop/96/4    ................................................................................ [159680]
AY261362.1_Mkuzi_1979           .................................................................c.............. [159680]
AY261361.1_Malawi_Lil-20/1_1983 .....................................................a.t........................ [159680]
AY261360.1_Kenya_1950           ..........................................................................g..... [159680]
AM712240.1_OURT_88/3            ................................................................................ [159680]
AM712239.1_Benin_97/1           ................................................................................ [159680]
KM111295.1_Ken06.Bus            ..........................................................................g..... [159680]
KM111294.1_Ken05/Tk1            ..........................................................................g..... [159680]
KM262844.1_strain_L60           ................................................................................ [159680]
KM102979.1_26544/OG10_Italy     ................................................................................ [159680]
KM262845.1_strain_NHV           ................................................................................ [159680]
KX354450.1_47/Ss/2008           ................................................................................ [159680]
KP055815.1_strain_BA71          ................................................................................ [159680]
MH025920.1_strain_R35           ..........................................................................g..... [159680]
MH025919.1_strain_N10           ..........................................................................g..... [159680]
MH025918.1_strain_R25           ..........................................................................g..... [159680]
MH025917.1_strain_R7            ..........................................................................g..... [159680]
MH025916.1_strain_R8            ..........................................................................g..... [159680]
```

```
FR682468.2_Georgia_2007/1        ccagggcgcattttcaataaaattcatgtagtggttcataaattgacgcgcgatggcaatcgcggcaatgctttttcccg [159760]
ASF_IND_20_CAD543                ................................................................................ [159760]
LR899193.1_Germany_2020/1        ................................................................................ [159760]
MN393477.1_Wuhan_2019            ................................................................................ [159760]
MW396979.1_Timor-Leste/2019/1    ................................................................................ [159760]
MK940252.1_CN/2019/InnerMongolia ................................................................................ [159760]
MK128995.1_China/2018/AnhuiXCGQ  ................................................................................ [159760]
MH681419.1_ASFV/POL/2015         ................................................................................ [159760]
LS478113.1_Estonia_2014          ................................................................................ [159760]
KP843857.1_Odintsovo_02/14       ................................................................................ [159760]
FN557520.1_strain_E75            .......................................g.....g.....a............................ [159760]
AY261366.1_Warthog               .......................................g.....g.................................. [159760]
AY261365.1_Warmbaths             .......................................g.....g.................................. [159760]
AY261364.1_Tengani_62            .......................................g.....g.................................. [159760]
AY261363.1_Pretoriuskop/96/4     .......................................g.....g.................................. [159760]
AY261362.1_Mkuzi_1979            .......................................g.....g.................................. [159760]
AY261361.1_Malawi_Lil-20/1_1983  ...a.....g...................................................................... [159760]
AY261360.1_Kenya_1950            ...a.......................a...........g..........................g............. [159760]
AM712240.1_OURT_88/3             .......................................g.....g.....a............................ [159760]
AM712239.1_Benin_97/1            .......................................g.....g.....a............................ [159760]
KM111295.1_Ken06.Bus             ...a.......................a...........g..........................g............. [159760]
KM111294.1_Ken05/Tk1             ...a.......................a...........g..........................g............. [159760]
KM262844.1_strain_L60            .......................................g.....g.....a............................ [159760]
KM102979.1_26544/OG10_Italy      .......................................g.....g.....a............................ [159760]
KM262845.1_strain_NHV            .......................................g.....g.....a............................ [159760]
KX354450.1_47/Ss/2008            .......................................g.....g.....a............................ [159760]
KP055815.1_strain_BA71           .......................................g.....g.....a............................ [159760]
MH025920.1_strain_R35            ...a.......................a...........g..........................g............. [159760]
MH025919.1_strain_N10            ...a.......................a...........g..........................g............. [159760]
MH025918.1_strain_R25            ...a.......................a...........g..........................g............. [159760]
MH025917.1_strain_R7             ...a.......................a...........g..........................g............. [159760]
MH025916.1_strain_R8             ...a.......................a...........g..........................g............. [159760]

FR682468.2_Georgia_2007/1        tcccggtctgccagtttaataaaagacgcgagtagggcgtgttgggattttgaaagttttggacgaaaagctgggcatta [159840]
ASF_IND_20_CAD543                ................................................................................ [159840]
LR899193.1_Germany_2020/1        ................................................................................ [159840]
MN393477.1_Wuhan_2019            ................................................................................ [159840]
MW396979.1_Timor-Leste/2019/1    ................................................................................ [159840]
MK940252.1_CN/2019/InnerMongolia ................................................................................ [159840]
MK128995.1_China/2018/AnhuiXCGQ  ................................................................................ [159840]
MH681419.1_ASFV/POL/2015         ................................................................................ [159840]
LS478113.1_Estonia_2014          ................................................................................ [159840]
KP843857.1_Odintsovo_02/14       ................................................................................ [159840]
```

```
FN557520.1_strain_E75           .....................................t........g................................. [159840]
AY261366.1_Warthog              ..............................................g................................. [159840]
AY261365.1_Warmbaths            ..............................................g................................. [159840]
AY261364.1_Tengani_62           ..............................................g................................. [159840]
AY261363.1_Pretoriuskop/96/4    ..............................................g................................. [159840]
AY261362.1_Mkuzi_1979           .....................................t........g................................. [159840]
AY261361.1_Malawi_Lil-20/1_1983 .......................................................a........................ [159840]
AY261360.1_Kenya_1950           .....................................t........g..........................a...... [159840]
AM712240.1_OURT_88/3            .....................................t........g................................. [159840]
AM712239.1_Benin_97/1           .....................................t........g................................. [159840]
KM111295.1_Ken06.Bus            ..............................................g........a.................a...... [159840]
KM111294.1_Ken05/Tk1            .....................................t........g..........................a...... [159840]
KM262844.1_strain_L60           .....................................t........g................................. [159840]
KM102979.1_26544/OG10_Italy     .....................................t........g................................. [159840]
KM262845.1_strain_NHV           .....................................t........g................................. [159840]
KX354450.1_47/Ss/2008           .....................................t........g................................. [159840]
KP055815.1_strain_BA71          .....................................t........g................................. [159840]
MH025920.1_strain_R35           ..............................................g........a.................a...... [159840]
MH025919.1_strain_N10           .....................................t........g........a.................a...... [159840]
MH025918.1_strain_R25           ..............................................g........a.................a...... [159840]
MH025917.1_strain_R7            ..............................................g........a.................a...... [159840]
MH025916.1_strain_R8            ..............................................g........a.................a...... [159840]

FR682468.2_Georgia_2007/1       tgcaattggagacccttgatggaaggaaagggcgacgcgtaggggtcacacggaaaaaacgctcgcccccccttctcgca [159920]
ASF_IND_20_CAD543               ................................................................................ [159920]
LR899193.1_Germany_2020/1       ................................................................................ [159920]
MN393477.1_Wuhan_2019           ................................................................................ [159920]
MW396979.1_Timor-Leste/2019/1   ................................................................................ [159920]
MK940252.1_CN/2019/InnerMongolia ................................................................................ [159920]
MK128995.1_China/2018/AnhuiXCGQ ................................................................................ [159920]
MH681419.1_ASFV/POL/2015        ................................................................................ [159920]
LS478113.1_Estonia_2014         ................................................................................ [159920]
KP843857.1_Odintsovo_02/14      ................................................................................ [159920]
FN557520.1_strain_E75           ...................................................................-............ [159920]
AY261366.1_Warthog              .................................................................t.............. [159920]
AY261365.1_Warmbaths            ................................................................................ [159920]
AY261364.1_Tengani_62           ......................................a...........g.......................t..... [159920]
AY261363.1_Pretoriuskop/96/4    ................................................................................ [159920]
AY261362.1_Mkuzi_1979           ................................................................................ [159920]
AY261361.1_Malawi_Lil-20/1_1983 .................a................................g..............t.............. [159920]
AY261360.1_Kenya_1950           .....c.....g.....a................................g..g...........t.............. [159920]
AM712240.1_OURT_88/3            ................................................................................ [159920]
AM712239.1_Benin_97/1           ................................................................................ [159920]
KM111295.1_Ken06.Bus            .....c.....g.....a.............................g..g..g...........t.............. [159920]
```

KM111294.1_Ken05/Tk1 .....c.....g.....a................................g..g...........t.............. [159920]
KM262844.1_strain_L60 ................................................................................ [159920]
KM102979.1_26544/OG10_Italy ................................................................................ [159920]
KM262845.1_strain_NHV ................................................................................ [159920]
KX354450.1_47/Ss/2008 ................................................................................ [159920]
KP055815.1_strain_BA71 ................................................................................ [159920]
MH025920.1_strain_R35 .....c.....g.....a.............................g..g..g...........t.............. [159920]
MH025919.1_strain_N10 .....c.....g.....a.............................g..g..g...........t.............. [159920]
MH025918.1_strain_R25 .....c.....g.....a.............................g..g..g...........t.............. [159920]
MH025917.1_strain_R7 .....c.....g.....a.............................g..g..g...........t.............. [159920]
MH025916.1_strain_R8 .....c.....g.....a.............................g..g..g...........t.............. [159920]

FR682468.2_Georgia_2007/1 gccagg-cccaccgatctggacaaaatgagcccgcagatcacgaatgagctctttttggtcgacaggaggggaaatcaac [160000]
ASF_IND_20_CAD543 ......-......................................................................... [160000]
LR899193.1_Germany_2020/1 ......-......................................................................... [160000]
MN393477.1_Wuhan_2019 ......-......................................................................... [160000]
MW396979.1_Timor-Leste/2019/1 ......-......................................................................... [160000]
MK940252.1_CN/2019/InnerMongolia ......-......................................................................... [160000]
MK128995.1_China/2018/AnhuiXCGQ ......-......................................................................... [160000]
MH681419.1_ASFV/POL/2015 ......-......................................................................... [160000]
LS478113.1_Estonia_2014 ......-......................................................................... [160000]
KP843857.1_Odintsovo_02/14 ......-......................................................................... [160000]
FN557520.1_strain_E75 ......-......................................................................... [160000]
AY261366.1_Warthog ......-......................................................................... [160000]
AY261365.1_Warmbaths ......-......................................................................... [160000]
AY261364.1_Tengani_62 ......-...t..........t........t................................................. [160000]
AY261363.1_Pretoriuskop/96/4 ......-......................................................................... [160000]
AY261362.1_Mkuzi_1979 ......-......................................................................... [160000]
AY261361.1_Malawi_Lil-20/1_1983 ......c......a...........................t.........t..............a............. [160000]
AY261360.1_Kenya_1950 ...g..-......a................a..........t.t.......t............................ [160000]
AM712240.1_OURT_88/3 ......-......................................................................... [160000]
AM712239.1_Benin_97/1 ......-......................................................................... [160000]
KM111295.1_Ken06.Bus ...g..-......a................a..........t.........t............................ [160000]
KM111294.1_Ken05/Tk1 ...g..-......a................a..........t.t.......t............................ [160000]
KM262844.1_strain_L60 ......-......................................................................... [160000]
KM102979.1_26544/OG10_Italy ......-......................................................................... [160000]
KM262845.1_strain_NHV ......-......................................................................... [160000]
KX354450.1_47/Ss/2008 ......-......................................................................... [160000]
KP055815.1_strain_BA71 ......-......................................................................... [160000]
MH025920.1_strain_R35 ...g..-......a................a..........t.........t............................ [160000]
MH025919.1_strain_N10 ...g..-......a................a..........t.........t............................ [160000]
MH025918.1_strain_R25 ...g..-......a................a..........t.........t............................ [160000]
MH025917.1_strain_R7 ...g..-......a................a..........t.........t............................ [160000]
MH025916.1_strain_R8 ...g..-......a................a..........t.........t............................ [160000]

```
FR682468.2_Georgia_2007/1        gatttaaactcctttcttcgcgccaactgctgcaaaaagtctgcggcatccaattcgggatacgccatattatcataaaa [160080]
ASF_IND_20_CAD543                ................................................................................ [160080]
LR899193.1_Germany_2020/1        ................................................................................ [160080]
MN393477.1_Wuhan_2019            ................................................................................ [160080]
MW396979.1_Timor-Leste/2019/1    ................................................................................ [160080]
MK940252.1_CN/2019/InnerMongolia ................................................................................ [160080]
MK128995.1_China/2018/AnhuiXCGQ  ................................................................................ [160080]
MH681419.1_ASFV/POL/2015         ................................................................................ [160080]
LS478113.1_Estonia_2014          ................................................................................ [160080]
KP843857.1_Odintsovo_02/14       ............................................................................-... [160080]
FN557520.1_strain_E75            ..c............................................................................. [160080]
AY261366.1_Warthog               ..c............................................................................. [160080]
AY261365.1_Warmbaths             ..c............................................................................. [160080]
AY261364.1_Tengani_62            ..c............................................................................. [160080]
AY261363.1_Pretoriuskop/96/4     ..c..........................t.................................................. [160080]
AY261362.1_Mkuzi_1979            ..c............................................................................. [160080]
AY261361.1_Malawi_Lil-20/1_1983  ..c......................................c.................g.................... [160080]
AY261360.1_Kenya_1950            ..c.......................t..................................................... [160080]
AM712240.1_OURT_88/3             ..c............................................................................. [160080]
AM712239.1_Benin_97/1            ..c............................................................................. [160080]
KM111295.1_Ken06.Bus             ..c.......................t.....................a............................... [160080]
KM111294.1_Ken05/Tk1             ..c......................................c...................................... [160080]
KM262844.1_strain_L60            ..c............................................................................. [160080]
KM102979.1_26544/OG10_Italy      ..c............................................................................. [160080]
KM262845.1_strain_NHV            ..c............................................................................. [160080]
KX354450.1_47/Ss/2008            ..c............................................................................. [160080]
KP055815.1_strain_BA71           ..c............................................................................. [160080]
MH025920.1_strain_R35            ..c.......................t.....................a............................... [160080]
MH025919.1_strain_N10            ..c.......................t.....................a............................... [160080]
MH025918.1_strain_R25            ..c.......................t.....................a............................... [160080]
MH025917.1_strain_R7             ..c.......................t.....................a............................... [160080]
MH025916.1_strain_R8             ..c.......................t.....................a............................... [160080]

FR682468.2_Georgia_2007/1        aaaataaacc-tttttatgaaaacttttatgtgattctgtattgcaattgttttttatgaatactgtaaataagcgtatc [160160]
ASF_IND_20_CAD543                ..........-..................................................................... [160160]
LR899193.1_Germany_2020/1        ..........-..................................................................... [160160]
MN393477.1_Wuhan_2019            ..........-..................................................................... [160160]
MW396979.1_Timor-Leste/2019/1    ..........-..................................................................... [160160]
MK940252.1_CN/2019/InnerMongolia ..........-..................................................................... [160160]
MK128995.1_China/2018/AnhuiXCGQ  ..........-..................................................................... [160160]
MH681419.1_ASFV/POL/2015         ..........-..................................................................... [160160]
LS478113.1_Estonia_2014          ..........-..................................................................... [160160]
KP843857.1_Odintsovo_02/14       ..........-..................................................................... [160160]
```

```
FN557520.1_strain_E75           ..........-..................................................................... [160160]
AY261366.1_Warthog              ..........-................................................g.................... [160160]
AY261365.1_Warmbaths            ..........-................................................g.................... [160160]
AY261364.1_Tengani_62           ..........-............g........................................................ [160160]
AY261363.1_Pretoriuskop/96/4    ..........-................................................g.......g............ [160160]
AY261362.1_Mkuzi_1979           ..........-..................................................................... [160160]
AY261361.1_Malawi_Lil-20/1_1983 ..........t..................c..t.......................g..........g........c... [160160]
AY261360.1_Kenya_1950           ..........-.......a....g.....c..t..................................g............ [160160]
AM712240.1_OURT_88/3            ..........-..................................................................... [160160]
AM712239.1_Benin_97/1           ..........-..................................................................... [160160]
KM111295.1_Ken06.Bus            ..........-............g.....c.at.................c................g............ [160160]
KM111294.1_Ken05/Tk1            ..........-.......a....g.....c..t..................................g............ [160160]
KM262844.1_strain_L60           ..........-..................................................................... [160160]
KM102979.1_26544/OG10_Italy     ..........-..................................................................... [160160]
KM262845.1_strain_NHV           ..........-..................................................................... [160160]
KX354450.1_47/Ss/2008           ..........-..................................................................... [160160]
KP055815.1_strain_BA71          ..........-..................................................................... [160160]
MH025920.1_strain_R35           ..........-............g.....c.at.................c................g............ [160160]
MH025919.1_strain_N10           ..........-............g.....c.at.................c................g............ [160160]
MH025918.1_strain_R25           ..........-............g.....c.at.................c................g............ [160160]
MH025917.1_strain_R7            ..........-............g.....c.at.................c................g............ [160160]
MH025916.1_strain_R8            ..........-............g.....c.at.................c................g............ [160160]

FR682468.2_Georgia_2007/1       aacttgtttttctaacgaagaggcgttattctttttttctggatataaaataataataagtataataattaagactaaac [160240]
ASF_IND_20_CAD543               ................................................................................ [160240]
LR899193.1_Germany_2020/1       ................................................................................ [160240]
MN393477.1_Wuhan_2019           ................................................................................ [160240]
MW396979.1_Timor-Leste/2019/1   ................................................................................ [160240]
MK940252.1_CN/2019/InnerMongolia ................................................................................ [160240]
MK128995.1_China/2018/AnhuiXCGQ ................................................................................ [160240]
MH681419.1_ASFV/POL/2015        ................................................................................ [160240]
LS478113.1_Estonia_2014         ................................................................................ [160240]
KP843857.1_Odintsovo_02/14      ................................................................................ [160240]
FN557520.1_strain_E75           ................................................g............................... [160240]
AY261366.1_Warthog              ........................a.......................g............................... [160240]
AY261365.1_Warmbaths            ........................ag......................g...........c...........a....... [160240]
AY261364.1_Tengani_62           ........................a.......................g............................... [160240]
AY261363.1_Pretoriuskop/96/4    ................................................g...........c................... [160240]
AY261362.1_Mkuzi_1979           ........................a....................................................... [160240]
AY261361.1_Malawi_Lil-20/1_1983 ............cg......g...a.......................g...........c................... [160240]
AY261360.1_Kenya_1950           c.....g.....cg......g..ta.................g.....g............................... [160240]
AM712240.1_OURT_88/3            ................................................g............................... [160240]
AM712239.1_Benin_97/1           ................................................g............................... [160240]
KM111295.1_Ken06.Bus            c.....g.....cg......g..ta.................g.....g............................... [160240]
```

```
KM111294.1_Ken05/Tk1            c.....g.....cg......g..ta..c..............g.....g............................... [160240]
KM262844.1_strain_L60           ................................................g............................... [160240]
KM102979.1_26544/OG10_Italy     ................................................g............................... [160240]
KM262845.1_strain_NHV           ................................................g............................... [160240]
KX354450.1_47/Ss/2008           ................................................g............................... [160240]
KP055815.1_strain_BA71          ................................................g............................... [160240]
MH025920.1_strain_R35           c.....g.....cg......g..ta.................g.....g............................... [160240]
MH025919.1_strain_N10           c.....g.....cg......g..ta.................g.....g............................... [160240]
MH025918.1_strain_R25           c.....g.....cg......g..ta.................g.....g............................... [160240]
MH025917.1_strain_R7            c.....g.....cg......g..ta.................g.....g............................... [160240]
MH025916.1_strain_R8            c.....g.....cg......g..ta.................g.....g............................... [160240]

FR682468.2_Georgia_2007/1       agcaggcaatcactatcaaactcatattatacttacttttttat-aaaaagtattatatcttatgaatgcgcaagttcag [160320]
ASF_IND_20_CAD543               ............................................-................................... [160320]
LR899193.1_Germany_2020/1       ............................................-................................... [160320]
MN393477.1_Wuhan_2019           ............................................-................................... [160320]
MW396979.1_Timor-Leste/2019/1   ............................................-................................... [160320]
MK940252.1_CN/2019/InnerMongolia ............................................-................................... [160320]
MK128995.1_China/2018/AnhuiXCGQ ............................................-................................... [160320]
MH681419.1_ASFV/POL/2015        ............................................-................................... [160320]
LS478113.1_Estonia_2014         ............................................-................................... [160320]
KP843857.1_Odintsovo_02/14      ............................................-................................... [160320]
FN557520.1_strain_E75           ............................................-................................... [160320]
AY261366.1_Warthog              ............................................-................................... [160320]
AY261365.1_Warmbaths            ............................................-................................... [160320]
AY261364.1_Tengani_62           ............................................-................................... [160320]
AY261363.1_Pretoriuskop/96/4    ............................................-................................... [160320]
AY261362.1_Mkuzi_1979           ............................................-................................... [160320]
AY261361.1_Malawi_Lil-20/1_1983 ..........a.................................a........................ta......... [160320]
AY261360.1_Kenya_1950           .a........g.................................-................................... [160320]
AM712240.1_OURT_88/3            ............................................-................................... [160320]
AM712239.1_Benin_97/1           ............................................-................................... [160320]
KM111295.1_Ken06.Bus            ..........g.................................-................................... [160320]
KM111294.1_Ken05/Tk1            ..........g.................................-................................... [160320]
KM262844.1_strain_L60           ............................................-................................... [160320]
KM102979.1_26544/OG10_Italy     ............................................-................................... [160320]
KM262845.1_strain_NHV           ............................................-................................... [160320]
KX354450.1_47/Ss/2008           ............................................-................................... [160320]
KP055815.1_strain_BA71          ............................................-................................... [160320]
MH025920.1_strain_R35           ..........g.................................-................................... [160320]
MH025919.1_strain_N10           ..........g.................................-................................... [160320]
MH025918.1_strain_R25           ..........g.................................-................................... [160320]
MH025917.1_strain_R7            ..........g.................................-................................... [160320]
MH025916.1_strain_R8            ..........g.................................-................................... [160320]
```

```
FR682468.2_Georgia_2007/1        ctaattgttcgtcgcttggaatgtgggactgcagggaggtggagtttttcctttttctaaagaataccgggaaatggtgg [160400]
ASF_IND_20_CAD543                ................................................................................ [160400]
LR899193.1_Germany_2020/1        ................................................................................ [160400]
MN393477.1_Wuhan_2019            ................................................................................ [160400]
MW396979.1_Timor-Leste/2019/1    ................................................................................ [160400]
MK940252.1_CN/2019/InnerMongolia ................................................................................ [160400]
MK128995.1_China/2018/AnhuiXCGQ  ................................................................................ [160400]
MH681419.1_ASFV/POL/2015         ................................................................................ [160400]
LS478113.1_Estonia_2014          ................................................................................ [160400]
KP843857.1_Odintsovo_02/14       ................................................................................ [160400]
FN557520.1_strain_E75            ...................................................................t............ [160400]
AY261366.1_Warthog               ...g............................t............................................... [160400]
AY261365.1_Warmbaths             .........................a......t............................................... [160400]
AY261364.1_Tengani_62            ............................t................................................... [160400]
AY261363.1_Pretoriuskop/96/4     ...g.....................a......t....................c.......................... [160400]
AY261362.1_Mkuzi_1979            ................................................................................ [160400]
AY261361.1_Malawi_Lil-20/1_1983  ...........................c....t...............................a..t............ [160400]
AY261360.1_Kenya_1950            .........................---....t...t...c..........................t............ [160400]
AM712240.1_OURT_88/3             ...................................................................t............ [160400]
AM712239.1_Benin_97/1            ...................................................................t............ [160400]
KM111295.1_Ken06.Bus             .........................---....t..................................t............ [160400]
KM111294.1_Ken05/Tk1             .........................---....t...t...c..........................t............ [160400]
KM262844.1_strain_L60            ...................................................................t............ [160400]
KM102979.1_26544/OG10_Italy      ...................................................................t............ [160400]
KM262845.1_strain_NHV            ...................................................................t............ [160400]
KX354450.1_47/Ss/2008            ...................................................................t............ [160400]
KP055815.1_strain_BA71           ...................................................................t............ [160400]
MH025920.1_strain_R35            ..........a..............---....t..................................t............ [160400]
MH025919.1_strain_N10            ..........a..............---....t..................................t............ [160400]
MH025918.1_strain_R25            ..........a..............---....t..................................t............ [160400]
MH025917.1_strain_R7             ..........a..............---....t..................................t............ [160400]
MH025916.1_strain_R8             ..........a..............---....t..................................t............ [160400]

FR682468.2_Georgia_2007/1        tgaggctcaggttgttgtacatagtagctaggaggag---gtttaggtatgctcgacttgcagtcaatagtccggttata [160480]
ASF_IND_20_CAD543                .....................................---........................................ [160480]
LR899193.1_Germany_2020/1        .....................................---........................................ [160480]
MN393477.1_Wuhan_2019            .....................................---........................................ [160480]
MW396979.1_Timor-Leste/2019/1    .....................................---........................................ [160480]
MK940252.1_CN/2019/InnerMongolia .....................................---........................................ [160480]
MK128995.1_China/2018/AnhuiXCGQ  .....................................---........................................ [160480]
MH681419.1_ASFV/POL/2015         .....................................---........................................ [160480]
LS478113.1_Estonia_2014          .....................................---........................................ [160480]
KP843857.1_Odintsovo_02/14       .....................................---........................................ [160480]
```

```
FN557520.1_strain_E75            .....................................---.......c...................c..a......... [160480]
AY261366.1_Warthog               .....................................---.......c...................c..c......... [160480]
AY261365.1_Warmbaths             ................................t....---.......c...................c..c......... [160480]
AY261364.1_Tengani_62            .....................................---.......c...................c..a......... [160480]
AY261363.1_Pretoriuskop/96/4     .....................................---.......c...t....g..........c..c......... [160480]
AY261362.1_Mkuzi_1979            .....................................---........................................ [160480]
AY261361.1_Malawi_Lil-20/1_1983  ..........a........t.g.a.............---..c....c...t...............c..at.....g.. [160480]
AY261360.1_Kenya_1950            ........g...................c........gaa.......c...g...............c..a......g.. [160480]
AM712240.1_OURT_88/3             .....................................---.......c...................c..a......... [160480]
AM712239.1_Benin_97/1            .....................................---.......c...................c..a......... [160480]
KM111295.1_Ken06.Bus             ........g...................c........gaa.......c...g...............c..a......g.. [160480]
KM111294.1_Ken05/Tk1             ........g............................gaa.......c...g...............c..a......g.. [160480]
KM262844.1_strain_L60            .....................................---.......c...................c..a......... [160480]
KM102979.1_26544/OG10_Italy      .....................................---.......c...................c..a......... [160480]
KM262845.1_strain_NHV            .....................................---.......c...................c..a......... [160480]
KX354450.1_47/Ss/2008            .....................................---.......c...................c..a......... [160480]
KP055815.1_strain_BA71           .....................................---.......c...................c..a......... [160480]
MH025920.1_strain_R35            ........g...................c........gaa.......c...g...............c..a......g.. [160480]
MH025919.1_strain_N10            ........g...................c........gaa.......c...g...............c..a......g.. [160480]
MH025918.1_strain_R25            ........g...................c........gaa.......c...g...............c..a......g.. [160480]
MH025917.1_strain_R7             ........g...................c........gaa.......c...g...............c..a......g.. [160480]
MH025916.1_strain_R8             ........g...................c........gaa.......c...g...............c..a......g.. [160480]

FR682468.2_Georgia_2007/1        gtaaacgatggcaacgatgataagaataataatgagcaaaatcaaaatgcccaggagaatcgcagttgttccgggatatt [160560]
ASF_IND_20_CAD543                ................................................................................ [160560]
LR899193.1_Germany_2020/1        ................................................................................ [160560]
MN393477.1_Wuhan_2019            ................................................................................ [160560]
MW396979.1_Timor-Leste/2019/1    ................................................................................ [160560]
MK940252.1_CN/2019/InnerMongolia ................................................................................ [160560]
MK128995.1_China/2018/AnhuiXCGQ  ................................................................................ [160560]
MH681419.1_ASFV/POL/2015         ................................................................................ [160560]
LS478113.1_Estonia_2014          ................................................................................ [160560]
KP843857.1_Odintsovo_02/14       ................................................................................ [160560]
FN557520.1_strain_E75            ................................c..................t..a........................c [160560]
AY261366.1_Warthog               ................................c............................................... [160560]
AY261365.1_Warmbaths             ................................c............................................... [160560]
AY261364.1_Tengani_62            ................................c............................................... [160560]
AY261363.1_Pretoriuskop/96/4     ................................c............................................... [160560]
AY261362.1_Mkuzi_1979            ................................c..............................................c [160560]
AY261361.1_Malawi_Lil-20/1_1983  ............c...........g.......cc......................t..........a.........g.c [160560]
AY261360.1_Kenya_1950            .................c..............cc.................t.a..tg.........a.........g.c [160560]
AM712240.1_OURT_88/3             ................................c..................t..a........................c [160560]
AM712239.1_Benin_97/1            ................................c..................t..a........................c [160560]
KM111295.1_Ken06.Bus             .................c..............cc.................t...tg.........a..........c. [160560]
```

```
KM111294.1_Ken05/Tk1            .................c..............cc.................t.a..tg.........a..........c. [160560]
KM262844.1_strain_L60           ................................c..................t..a........................c [160560]
KM102979.1_26544/OG10_Italy     ................................c..................t..a........................c [160560]
KM262845.1_strain_NHV           ................................c..................t..a........................c [160560]
KX354450.1_47/Ss/2008           ................................c..................t..a........................c [160560]
KP055815.1_strain_BA71          ................................c..................t..a........................c [160560]
MH025920.1_strain_R35           .................c..............cc.................t....tg.........a..........c. [160560]
MH025919.1_strain_N10           .................c..............cc.................t....tg.........a..........c. [160560]
MH025918.1_strain_R25           .................c..............cc.................t....tg.........a..........c. [160560]
MH025917.1_strain_R7            .................c..............cc.................t....tg.........a..........c. [160560]
MH025916.1_strain_R8            .................c..............cc.................t....tg.........a..........c. [160560]

FR682468.2_Georgia_2007/1       tggcgattgtatgggctaaaaggccttgggtgctttgtttaattccctcgcgggttgacaggttatgagaaagcagtgga [160640]
ASF_IND_20_CAD543               ................................................................................ [160640]
LR899193.1_Germany_2020/1       ................................................................................ [160640]
MN393477.1_Wuhan_2019           ................................................................................ [160640]
MW396979.1_Timor-Leste/2019/1   ................................................................................ [160640]
MK940252.1_CN/2019/InnerMongolia ................................................................................ [160640]
MK128995.1_China/2018/AnhuiXCGQ ................................................................................ [160640]
MH681419.1_ASFV/POL/2015        ................................................................................ [160640]
LS478113.1_Estonia_2014         ................................................................................ [160640]
KP843857.1_Odintsovo_02/14      ................................................................................ [160640]
FN557520.1_strain_E75           .......c....................a................................................... [160640]
AY261366.1_Warthog              .............................................................a.................. [160640]
AY261365.1_Warmbaths            .............................................................a.................. [160640]
AY261364.1_Tengani_62           .......c........................................................................ [160640]
AY261363.1_Pretoriuskop/96/4    ................................................................................ [160640]
AY261362.1_Mkuzi_1979           .......c....................a................................................... [160640]
AY261361.1_Malawi_Lil-20/1_1983 .......c........................t......................a.....a.................. [160640]
AY261360.1_Kenya_1950           .......c.............a.......................................a.................. [160640]
AM712240.1_OURT_88/3            .......c....................a................................................... [160640]
AM712239.1_Benin_97/1           .......c....................a................................................... [160640]
KM111295.1_Ken06.Bus            .ca....c.............a.......................................a.................. [160640]
KM111294.1_Ken05/Tk1            .ca....c.............a.......................................a.................. [160640]
KM262844.1_strain_L60           .......c....................a................................................... [160640]
KM102979.1_26544/OG10_Italy     .......c....................a................................................... [160640]
KM262845.1_strain_NHV           .......c....................a................................................... [160640]
KX354450.1_47/Ss/2008           .......c....................a................................................... [160640]
KP055815.1_strain_BA71          .......c....................a................................................... [160640]
MH025920.1_strain_R35           .ca....c.............a.......................................a.................. [160640]
MH025919.1_strain_N10           .ca....c.............a.......................................a.................. [160640]
MH025918.1_strain_R25           .ca....c.............a.......................................a.................. [160640]
MH025917.1_strain_R7            .ca....c.............a.......................................a.................. [160640]
MH025916.1_strain_R8            .ca....c.............a.......................................a.................. [160640]
```

```
FR682468.2_Georgia_2007/1        gacgtttcagtgtccatttattacaattgaacagttatattaatctcaaataaaatataacacaaaattaattatggcca [160720]
ASF_IND_20_CAD543                ................................................................................ [160720]
LR899193.1_Germany_2020/1        ................................................................................ [160720]
MN393477.1_Wuhan_2019            ................................................................................ [160720]
MW396979.1_Timor-Leste/2019/1    ................................................................................ [160720]
MK940252.1_CN/2019/InnerMongolia ................................................................................ [160720]
MK128995.1_China/2018/AnhuiXCGQ  ................................................................................ [160720]
MH681419.1_ASFV/POL/2015         ................................................................................ [160720]
LS478113.1_Estonia_2014          ................................................................................ [160720]
KP843857.1_Odintsovo_02/14       ................................................................................ [160720]
FN557520.1_strain_E75            ................................................................................ [160720]
AY261366.1_Warthog               ..a............................................................................. [160720]
AY261365.1_Warmbaths             ..a............................................................................. [160720]
AY261364.1_Tengani_62            ..a............................................................................. [160720]
AY261363.1_Pretoriuskop/96/4     ..a............................................................................. [160720]
AY261362.1_Mkuzi_1979            ................................................................................ [160720]
AY261361.1_Malawi_Lil-20/1_1983  ................................................................................ [160720]
AY261360.1_Kenya_1950            ..a............................................................................. [160720]
AM712240.1_OURT_88/3             ................................................................................ [160720]
AM712239.1_Benin_97/1            ................................................................................ [160720]
KM111295.1_Ken06.Bus             ..a............................................................................. [160720]
KM111294.1_Ken05/Tk1             ..a............................................................................. [160720]
KM262844.1_strain_L60            ................................................................................ [160720]
KM102979.1_26544/OG10_Italy      ................................................................................ [160720]
KM262845.1_strain_NHV            ................................................................................ [160720]
KX354450.1_47/Ss/2008            ................................................................................ [160720]
KP055815.1_strain_BA71           ................................................................................ [160720]
MH025920.1_strain_R35            ..a............................................................................. [160720]
MH025919.1_strain_N10            ..a............................................................................. [160720]
MH025918.1_strain_R25            ..a............................................................................. [160720]
MH025917.1_strain_R7             ..a............................................................................. [160720]
MH025916.1_strain_R8             ..a............................................................................. [160720]

FR682468.2_Georgia_2007/1        tgcaaaagttatttacgtatatttacgagtttattgaatatcgtaagatggtgctgttggaagaaaaggtaccatatgat [160800]
ASF_IND_20_CAD543                ................................................................................ [160800]
LR899193.1_Germany_2020/1        ................................................................................ [160800]
MN393477.1_Wuhan_2019            ................................................................................ [160800]
MW396979.1_Timor-Leste/2019/1    ................................................................................ [160800]
MK940252.1_CN/2019/InnerMongolia ................................................................................ [160800]
MK128995.1_China/2018/AnhuiXCGQ  ................................................................................ [160800]
MH681419.1_ASFV/POL/2015         ................................................................................ [160800]
LS478113.1_Estonia_2014          ................................................................................ [160800]
KP843857.1_Odintsovo_02/14       ................................................................................ [160800]
```

| | | |
|---|---|---|
| FN557520.1_strain_E75 | ............................a................................................... | [160800] |
| AY261366.1_Warthog | ................................................................................ | [160800] |
| AY261365.1_Warmbaths | ................................................................................ | [160800] |
| AY261364.1_Tengani_62 | ...................................................................a...........c | [160800] |
| AY261363.1_Pretoriuskop/96/4 | ...................................a...........................................c | [160800] |
| AY261362.1_Mkuzi_1979 | ............................a................................................... | [160800] |
| AY261361.1_Malawi_Lil-20/1_1983 | ...................................................................a...........c | [160800] |
| AY261360.1_Kenya_1950 | ............................a..c...................................a...........c | [160800] |
| AM712240.1_OURT_88/3 | ............................a................................................... | [160800] |
| AM712239.1_Benin_97/1 | ............................a................................................... | [160800] |
| KM111295.1_Ken06.Bus | ............................a..c...................................a...........c | [160800] |
| KM111294.1_Ken05/Tk1 | ............................a..c...................................a...........c | [160800] |
| KM262844.1_strain_L60 | ............................a................................................... | [160800] |
| KM102979.1_26544/OG10_Italy | ............................a................................................... | [160800] |
| KM262845.1_strain_NHV | ............................a................................................... | [160800] |
| KX354450.1_47/Ss/2008 | ............................a................................................... | [160800] |
| KP055815.1_strain_BA71 | ............................a................................................... | [160800] |
| MH025920.1_strain_R35 | ............................a..c...................................a...........c | [160800] |
| MH025919.1_strain_N10 | ............................a..c...................................a...........c | [160800] |
| MH025918.1_strain_R25 | ............................a..c...................................a...........c | [160800] |
| MH025917.1_strain_R7 | ............................a..c...................................a...........c | [160800] |
| MH025916.1_strain_R8 | ............................a..c...................................a...........c | [160800] |

| | | |
|---|---|---|
| FR682468.2_Georgia_2007/1 | aagtttgttcaaatggtacttaatacaggattttttcgtattaacgcggagacgctgaatcacggaatcgtatccgtgtt | [160880] |
| ASF_IND_20_CAD543 | ................................................................................ | [160880] |
| LR899193.1_Germany_2020/1 | ................................................................................ | [160880] |
| MN393477.1_Wuhan_2019 | ................................................................................ | [160880] |
| MW396979.1_Timor-Leste/2019/1 | ................................................................................ | [160880] |
| MK940252.1_CN/2019/InnerMongolia | ................................................................................ | [160880] |
| MK128995.1_China/2018/AnhuiXCGQ | ................................................................................ | [160880] |
| MH681419.1_ASFV/POL/2015 | ................................................................................ | [160880] |
| LS478113.1_Estonia_2014 | ................................................................................ | [160880] |
| KP843857.1_Odintsovo_02/14 | ................................................................................ | [160880] |
| FN557520.1_strain_E75 | .......................................................................g........ | [160880] |
| AY261366.1_Warthog | .......................................................................g........ | [160880] |
| AY261365.1_Warmbaths | .......................................................................g........ | [160880] |
| AY261364.1_Tengani_62 | .......................c.......................a...........c...........g........ | [160880] |
| AY261363.1_Pretoriuskop/96/4 | ............................................t..........................g........ | [160880] |
| AY261362.1_Mkuzi_1979 | .......................................................................g........ | [160880] |
| AY261361.1_Malawi_Lil-20/1_1983 | ..................t....................................................g........ | [160880] |
| AY261360.1_Kenya_1950 | ...............a..t....................................................g........ | [160880] |
| AM712240.1_OURT_88/3 | .......................................................................g........ | [160880] |
| AM712239.1_Benin_97/1 | .......................................................................g........ | [160880] |
| KM111295.1_Ken06.Bus | .g.............a..t.......c............................................g........ | [160880] |

```
KM111294.1_Ken05/Tk1            ...............a..t....................................................g........ [160880]
KM262844.1_strain_L60           .......................................................................g........ [160880]
KM102979.1_26544/OG10_Italy     .......................................................................g........ [160880]
KM262845.1_strain_NHV           .......................................................................g........ [160880]
KX354450.1_47/Ss/2008           .......................................................................g........ [160880]
KP055815.1_strain_BA71          .......................................................................g........ [160880]
MH025920.1_strain_R35           .g.............a..t.......t............................................g........ [160880]
MH025919.1_strain_N10           .g.............a..t.......c............................................g........ [160880]
MH025918.1_strain_R25           .g.............a..t.......t............................................g........ [160880]
MH025917.1_strain_R7            .g.............a..t.......t............................................g........ [160880]
MH025916.1_strain_R8            .g.............a..t.......t............................................g........ [160880]

FR682468.2_Georgia_2007/1       tatctttggagcaaatggcaagtacgttcaccacggaggcgacatgagaacgcttttaacgaatacgcttaatgaaaaaa [160960]
ASF_IND_20_CAD543               ................................................................................ [160960]
LR899193.1_Germany_2020/1       ................................................................................ [160960]
MN393477.1_Wuhan_2019           ................................................................................ [160960]
MW396979.1_Timor-Leste/2019/1   ................................................................................ [160960]
MK940252.1_CN/2019/InnerMongolia ................................................................................ [160960]
MK128995.1_China/2018/AnhuiXCGQ ................................................................................ [160960]
MH681419.1_ASFV/POL/2015        ................................................................................ [160960]
LS478113.1_Estonia_2014         ................................................................................ [160960]
KP843857.1_Odintsovo_02/14      ................................................................................ [160960]
FN557520.1_strain_E75           ................................................................................ [160960]
AY261366.1_Warthog              ................................................................................ [160960]
AY261365.1_Warmbaths            ................................................................................ [160960]
AY261364.1_Tengani_62           ........................t....................................................... [160960]
AY261363.1_Pretoriuskop/96/4    ................................................................................ [160960]
AY261362.1_Mkuzi_1979           ................................................................................ [160960]
AY261361.1_Malawi_Lil-20/1_1983 ...............c...................................c....................c....... [160960]
AY261360.1_Kenya_1950           ...............c...........c................................a...g.a..g.......... [160960]
AM712240.1_OURT_88/3            ................................................................................ [160960]
AM712239.1_Benin_97/1           ................................................................................ [160960]
KM111295.1_Ken06.Bus            ...............c...........c................................a...g....g.......... [160960]
KM111294.1_Ken05/Tk1            ...............c...........c................................a...g.a..g.......... [160960]
KM262844.1_strain_L60           ................................................................................ [160960]
KM102979.1_26544/OG10_Italy     ................................................................................ [160960]
KM262845.1_strain_NHV           ................................................................................ [160960]
KX354450.1_47/Ss/2008           ................................................................................ [160960]
KP055815.1_strain_BA71          ................................................................................ [160960]
MH025920.1_strain_R35           ...............c...........c................................a...g....g.......... [160960]
MH025919.1_strain_N10           ...............c...........c................................a...g....g.......... [160960]
MH025918.1_strain_R25           ...............c...........c................................a...g....g.......... [160960]
MH025917.1_strain_R7            ...............c...........c................................a...g....g.......... [160960]
MH025916.1_strain_R8            ...............c...........c................................a...g....g.......... [160960]
```

```
FR682468.2_Georgia_2007/1        aacattatgaagaattaattttaatcgttgataagcccgttttaagcaaaaaaaaatatttagatataatcgtcgagcag [161040]
ASF_IND_20_CAD543                ................................................................................ [161040]
LR899193.1_Germany_2020/1        ................................................................................ [161040]
MN393477.1_Wuhan_2019            ................................................................................ [161040]
MW396979.1_Timor-Leste/2019/1    ................................................................................ [161040]
MK940252.1_CN/2019/InnerMongolia ................................................................................ [161040]
MK128995.1_China/2018/AnhuiXCGQ  ................................................................................ [161040]
MH681419.1_ASFV/POL/2015         ................................................................................ [161040]
LS478113.1_Estonia_2014          ................................................................................ [161040]
KP843857.1_Odintsovo_02/14       ................................................................................ [161040]
FN557520.1_strain_E75            ................................................................................ [161040]
AY261366.1_Warthog               ...............................c................................................ [161040]
AY261365.1_Warmbaths             ...............................c.....g.......................................... [161040]
AY261364.1_Tengani_62            ...............................c................................................ [161040]
AY261363.1_Pretoriuskop/96/4     ...............................c................................................ [161040]
AY261362.1_Mkuzi_1979            ..................................a............................................. [161040]
AY261361.1_Malawi_Lil-20/1_1983  ....a..........................c..a..ta.....g......................t............ [161040]
AY261360.1_Kenya_1950            ....a........g.................c......a.....g................g.....t............ [161040]
AM712240.1_OURT_88/3             ................................................................................ [161040]
AM712239.1_Benin_97/1            ................................................................................ [161040]
KM111295.1_Ken06.Bus             ....a..........................c......a.....g................g.....t............ [161040]
KM111294.1_Ken05/Tk1             ....a..........................c..a...a.....g................g.....t............ [161040]
KM262844.1_strain_L60            ................................................................................ [161040]
KM102979.1_26544/OG10_Italy      ................................................................................ [161040]
KM262845.1_strain_NHV            ................................................................................ [161040]
KX354450.1_47/Ss/2008            ................................................................................ [161040]
KP055815.1_strain_BA71           ................................................................................ [161040]
MH025920.1_strain_R35            ....a..........................c......a.....g................g.....t............ [161040]
MH025919.1_strain_N10            ....a..........................c......a.....g................g.....t............ [161040]
MH025918.1_strain_R25            ....a..........................c......a.....g................g.....t............ [161040]
MH025917.1_strain_R7             ....a..........................c......a.....g................g.....t............ [161040]
MH025916.1_strain_R8             ....a..........................c......a.....g................g.....t............ [161040]

FR682468.2_Georgia_2007/1        cgcgctgcaaatcccacgattgtaataaacatatatccctaccacctgttctgcattaacattcccaaggtgagtgccat [161120]
ASF_IND_20_CAD543                ................................................................................ [161120]
LR899193.1_Germany_2020/1        ................................................................................ [161120]
MN393477.1_Wuhan_2019            ................................................................................ [161120]
MW396979.1_Timor-Leste/2019/1    ................................................................................ [161120]
MK940252.1_CN/2019/InnerMongolia ................................................................................ [161120]
MK128995.1_China/2018/AnhuiXCGQ  ................................................................................ [161120]
MH681419.1_ASFV/POL/2015         ................................................................................ [161120]
LS478113.1_Estonia_2014          ................................................................................ [161120]
KP843857.1_Odintsovo_02/14       ................................................................................ [161120]
```

```
FN557520.1_strain_E75             ..g............................................................................. [161120]
AY261366.1_Warthog                ................................................................................ [161120]
AY261365.1_Warmbaths              ................................................................................ [161120]
AY261364.1_Tengani_62             ..g............................................................................. [161120]
AY261363.1_Pretoriuskop/96/4      ..g............................................................................. [161120]
AY261362.1_Mkuzi_1979             ..g............................................................................. [161120]
AY261361.1_Malawi_Lil-20/1_1983   ...........c......g..........................t.......t.....t..............c..... [161120]
AY261360.1_Kenya_1950             ...........c......g..........................t............................c..... [161120]
AM712240.1_OURT_88/3              ..g............................................................................. [161120]
AM712239.1_Benin_97/1             ..g............................................................................. [161120]
KM111295.1_Ken06.Bus              .....c.....c......g..........................t............................c..... [161120]
KM111294.1_Ken05/Tk1              ...........c......g..........................t............................c..... [161120]
KM262844.1_strain_L60             ..g............................................................................. [161120]
KM102979.1_26544/OG10_Italy       ..g............................................................t................ [161120]
KM262845.1_strain_NHV             ..g............................................................................. [161120]
KX354450.1_47/Ss/2008             ..g............................................................t................ [161120]
KP055815.1_strain_BA71            ..g............................................................................. [161120]
MH025920.1_strain_R35             .....c.....c......g..........................t............................c..... [161120]
MH025919.1_strain_N10             .....c.....c......g..........................t............................c..... [161120]
MH025918.1_strain_R25             .....c.....c......g..........................t............................c..... [161120]
MH025917.1_strain_R7              .....c.....c......g..........................t............................c..... [161120]
MH025916.1_strain_R8              .....c.....c......g..........................t............................c..... [161120]

FR682468.2_Georgia_2007/1         tcctaaacataaactaattactcaggaggaggcgcaggagtttttaggtcgcgaatatctgcaaccgcaggacctcatgc [161200]
ASF_IND_20_CAD543                 ................................................................................ [161200]
LR899193.1_Germany_2020/1         ................................................................................ [161200]
MN393477.1_Wuhan_2019             ................................................................................ [161200]
MW396979.1_Timor-Leste/2019/1     ................................................................................ [161200]
MK940252.1_CN/2019/InnerMongolia  ................................................................................ [161200]
MK128995.1_China/2018/AnhuiXCGQ   ................................................................................ [161200]
MH681419.1_ASFV/POL/2015          ................................................................................ [161200]
LS478113.1_Estonia_2014           ................................................................................ [161200]
KP843857.1_Odintsovo_02/14        ................................................................................ [161200]
FN557520.1_strain_E75             ...............................................................g................ [161200]
AY261366.1_Warthog                ............g........c.......................g.................g................ [161200]
AY261365.1_Warmbaths              ............g........c.......................g.................g................ [161200]
AY261364.1_Tengani_62             ............g........c.......................g..c..............g................ [161200]
AY261363.1_Pretoriuskop/96/4      .....gg.....g........c.......................g..c..............g................ [161200]
AY261362.1_Mkuzi_1979             ...............................................................g........t....... [161200]
AY261361.1_Malawi_Lil-20/1_1983   ............g..t.....c.................a.....g.................g................ [161200]
AY261360.1_Kenya_1950             .....gg.....g..t.....c..............a.c......g.....t...........g................ [161200]
AM712240.1_OURT_88/3              ...............................................................g................ [161200]
AM712239.1_Benin_97/1             ...............................................................g................ [161200]
KM111295.1_Ken06.Bus              .....gg.....g..t.....c..............a.c......g.................g................ [161200]
```

```
KM111294.1_Ken05/Tk1           .....gg.....g..t.....c..............a.c......g.....t...........g................ [161200]
KM262844.1_strain_L60          ...............................................................g................ [161200]
KM102979.1_26544/OG10_Italy    ...............................................................g................ [161200]
KM262845.1_strain_NHV          ...............................................................g................ [161200]
KX354450.1_47/Ss/2008          ...............................................................g................ [161200]
KP055815.1_strain_BA71         ...............................................................g................ [161200]
MH025920.1_strain_R35          .....gg.....g..t.....c..............a.c......g.................g................ [161200]
MH025919.1_strain_N10          .....gg.....g..t.....c..............a.c......g.................g................ [161200]
MH025918.1_strain_R25          .....gg.....g..t.....c..............a.c......g.................g................ [161200]
MH025917.1_strain_R7           .....gg.....g..t.....c..............a.c......g.................g................ [161200]
MH025916.1_strain_R8           .....gg.....g..t.....c..............a.c......g.................g................ [161200]

FR682468.2_Georgia_2007/1      aaattagcgcgtcagaccccccggtggtctggctgggaggaagaccgggagactttgtgcaaattgagcggccctcagag [161280]
ASF_IND_20_CAD543              ................................................................................ [161280]
LR899193.1_Germany_2020/1      ................................................................................ [161280]
MN393477.1_Wuhan_2019          ................................................................................ [161280]
MW396979.1_Timor-Leste/2019/1  ................................................................................ [161280]
MK940252.1_CN/2019/InnerMongolia ................................................................................ [161280]
MK128995.1_China/2018/AnhuiXCGQ ................................................................................ [161280]
MH681419.1_ASFV/POL/2015       ................................................................................ [161280]
LS478113.1_Estonia_2014        ................................................................................ [161280]
KP843857.1_Odintsovo_02/14     ................................................................................ [161280]
FN557520.1_strain_E75          ................................................................................ [161280]
AY261366.1_Warthog             ................t...........t................................................... [161280]
AY261365.1_Warmbaths           ................t............................................................... [161280]
AY261364.1_Tengani_62          ................................................................................ [161280]
AY261363.1_Pretoriuskop/96/4   ................t............................................................... [161280]
AY261362.1_Mkuzi_1979          ................................................................................ [161280]
AY261361.1_Malawi_Lil-20/1_1983 ....c..t........................................................................ [161280]
AY261360.1_Kenya_1950          .......t.....g.................................................................. [161280]
AM712240.1_OURT_88/3           ................................................................................ [161280]
AM712239.1_Benin_97/1          ................................................................................ [161280]
KM111295.1_Ken06.Bus           .......t.....g.....t............................................................ [161280]
KM111294.1_Ken05/Tk1           .......t.....g.................................................................. [161280]
KM262844.1_strain_L60          ................................................................................ [161280]
KM102979.1_26544/OG10_Italy    ................................................................................ [161280]
KM262845.1_strain_NHV          ................................................................................ [161280]
KX354450.1_47/Ss/2008          ................................................................................ [161280]
KP055815.1_strain_BA71         ................................................................................ [161280]
MH025920.1_strain_R35          .......t.....g.....t............................................................ [161280]
MH025919.1_strain_N10          .......t.....g.....t............................................................ [161280]
MH025918.1_strain_R25          .......t.....g.....t............................................................ [161280]
MH025917.1_strain_R7           .......t.....g.....t............................................................ [161280]
MH025916.1_strain_R8           .......t.....g.....t............................................................ [161280]
```

```
FR682468.2_Georgia_2007/1        acagctatgcacgctgttgttatccgctttatcaccaagtccaaaatttgagtcc------------------------- [161360]
ASF_IND_20_CAD543                .......................................................------------------------- [161360]
LR899193.1_Germany_2020/1        .......................................................------------------------- [161360]
MN393477.1_Wuhan_2019            .......................................................------------------------- [161360]
MW396979.1_Timor-Leste/2019/1    .......................................................------------------------- [161360]
MK940252.1_CN/2019/InnerMongolia .......................................................------------------------- [161360]
MK128995.1_China/2018/AnhuiXCGQ  .......................................................------------------------- [161360]
MH681419.1_ASFV/POL/2015         .......................................................------------------------- [161360]
LS478113.1_Estonia_2014          .......................................................------------------------- [161360]
KP843857.1_Odintsovo_02/14       .......................................................------------------------- [161360]
FN557520.1_strain_E75            .......................................................------------------------- [161360]
AY261366.1_Warthog               .....c.................................................------------------------- [161360]
AY261365.1_Warmbaths             .....c.................................................------------------------- [161360]
AY261364.1_Tengani_62            .....c....................t.........--------------.....------------------------- [161360]
AY261363.1_Pretoriuskop/96/4     .....c.................................................------------------------- [161360]
AY261362.1_Mkuzi_1979            .......................................................------------------------- [161360]
AY261361.1_Malawi_Lil-20/1_1983  .....c.......................c..t.....a..t............tagtgtttaaaaagctaatgtattta [161360]
AY261360.1_Kenya_1950            .....c........c..c..........ac........a........a......tagtgcttaaag-------------- [161360]
AM712240.1_OURT_88/3             .......................................................------------------------- [161360]
AM712239.1_Benin_97/1            .......................................................------------------------- [161360]
KM111295.1_Ken06.Bus             .....c......................ac........a................a------------------------ [161360]
KM111294.1_Ken05/Tk1             .....c............a.........ac........a................a------------------------ [161360]
KM262844.1_strain_L60            .......................................................------------------------- [161360]
KM102979.1_26544/OG10_Italy      .......................................................------------------------- [161360]
KM262845.1_strain_NHV            .......................................................------------------------- [161360]
KX354450.1_47/Ss/2008            .......................................................------------------------- [161360]
KP055815.1_strain_BA71           .......................................................------------------------- [161360]
MH025920.1_strain_R35            .....c......................ac........a................a------------------------ [161360]
MH025919.1_strain_N10            .....c.......t..............ac........a................a------------------------ [161360]
MH025918.1_strain_R25            .....c......................ac........a................a------------------------ [161360]
MH025917.1_strain_R7             .....c......................ac........a................a------------------------ [161360]
MH025916.1_strain_R8             .....c......................ac........a................a------------------------ [161360]

FR682468.2_Georgia_2007/1        -------------------------------------------------------------------------------- [161440]
ASF_IND_20_CAD543                -------------------------------------------------------------------------------- [161440]
LR899193.1_Germany_2020/1        -------------------------------------------------------------------------------- [161440]
MN393477.1_Wuhan_2019            -------------------------------------------------------------------------------- [161440]
MW396979.1_Timor-Leste/2019/1    -------------------------------------------------------------------------------- [161440]
MK940252.1_CN/2019/InnerMongolia -------------------------------------------------------------------------------- [161440]
MK128995.1_China/2018/AnhuiXCGQ  -------------------------------------------------------------------------------- [161440]
MH681419.1_ASFV/POL/2015         -------------------------------------------------------------------------------- [161440]
LS478113.1_Estonia_2014          -------------------------------------------------------------------------------- [161440]
KP843857.1_Odintsovo_02/14       -------------------------------------------------------------------------------- [161440]
```

```
FN557520.1_strain_E75            -------------------------------------------------------------------------------- [161440]
AY261366.1_Warthog               -------------------------------------------------------------------------------- [161440]
AY261365.1_Warmbaths             -------------------------------------------------------------------------------- [161440]
AY261364.1_Tengani_62            -------------------------------------------------------------------------------- [161440]
AY261363.1_Pretoriuskop/96/4     -------------------------------------------------------------------------------- [161440]
AY261362.1_Mkuzi_1979            -------------------------------------------------------------------------------- [161440]
AY261361.1_Malawi_Lil-20/1_1983  aagctaatgtgtttaaagctaatgtgtttaaaactataatgtgtttaaagctataatgtgtttaaagctaatgtgtttaa [161440]
AY261360.1_Kenya_1950            ----------------------------------------------------ataatgtgtttaaaaataatgtgtttaa [161440]
AM712240.1_OURT_88/3             -------------------------------------------------------------------------------- [161440]
AM712239.1_Benin_97/1            -------------------------------------------------------------------------------- [161440]
KM111295.1_Ken06.Bus             -------------------------------------------------------------------------------- [161440]
KM111294.1_Ken05/Tk1             -------------------------------------------------------------------------------- [161440]
KM262844.1_strain_L60            -------------------------------------------------------------------------------- [161440]
KM102979.1_26544/OG10_Italy      -------------------------------------------------------------------------------- [161440]
KM262845.1_strain_NHV            -------------------------------------------------------------------------------- [161440]
KX354450.1_47/Ss/2008            -------------------------------------------------------------------------------- [161440]
KP055815.1_strain_BA71           -------------------------------------------------------------------------------- [161440]
MH025920.1_strain_R35            -------------------------------------------------------------------------------- [161440]
MH025919.1_strain_N10            -------------------------------------------------------------------------------- [161440]
MH025918.1_strain_R25            -------------------------------------------------------------------------------- [161440]
MH025917.1_strain_R7             -------------------------------------------------------------------------------- [161440]
MH025916.1_strain_R8             -------------------------------------------------------------------------------- [161440]

FR682468.2_Georgia_2007/1        ---------------------cgtgtttaaagatgacagacagctaagtaagcatatctgtaaaattgtcgatgtcctct [161520]
ASF_IND_20_CAD543                ---------------------........................................................... [161520]
LR899193.1_Germany_2020/1        ---------------------........................................................... [161520]
MN393477.1_Wuhan_2019            ---------------------........................................................... [161520]
MW396979.1_Timor-Leste/2019/1    ---------------------........................................................... [161520]
MK940252.1_CN/2019/InnerMongolia ---------------------........................................................... [161520]
MK128995.1_China/2018/AnhuiXCGQ  ---------------------........................................................... [161520]
MH681419.1_ASFV/POL/2015         ---------------------........................................................... [161520]
LS478113.1_Estonia_2014          ---------------------........................................................... [161520]
KP843857.1_Odintsovo_02/14       ---------------------........................................................... [161520]
FN557520.1_strain_E75            ---------------------........................................................... [161520]
AY261366.1_Warthog               ---------------------..................c........................................ [161520]
AY261365.1_Warmbaths             ---------------------..................c........................................ [161520]
AY261364.1_Tengani_62            ---------------------..................c........................................ [161520]
AY261363.1_Pretoriuskop/96/4     ---------------------t.................c...........................a............ [161520]
AY261362.1_Mkuzi_1979            ---------------------........................................................... [161520]
AY261361.1_Malawi_Lil-20/1_1983  agatgacgtgtttaaagataa.a...........a....c.g...............t........ca............ [161520]
AY261360.1_Kenya_1950            agataacgtgtttaaagataa..................c.ga..............t........ca........t... [161520]
AM712240.1_OURT_88/3             ---------------------........................................................... [161520]
AM712239.1_Benin_97/1            ---------------------........................................................... [161520]
KM111295.1_Ken06.Bus             -------gtgtttaaatataat.................ctg...............t........ca............ [161520]
```

```
KM111294.1_Ken05/Tk1              -------gtgtttaaagataat.................c.ga..............t........ca........t... [161520]
KM262844.1_strain_L60             ---------------------........................................................... [161520]
KM102979.1_26544/OG10_Italy       ---------------------........................................................... [161520]
KM262845.1_strain_NHV             ---------------------........................................................... [161520]
KX354450.1_47/Ss/2008             ---------------------........................................................... [161520]
KP055815.1_strain_BA71            ---------------------........................................................... [161520]
MH025920.1_strain_R35             -------gtgtttaaagataat.................ctg...............t........ca............ [161520]
MH025919.1_strain_N10             -------gtgtttaaagataat.................ctg...............t........ca............ [161520]
MH025918.1_strain_R25             -------gtgtttaaagataat.................ctg...............t........ca............ [161520]
MH025917.1_strain_R7              -------gtgtttaaagataat.................ctg...............t........ca............ [161520]
MH025916.1_strain_R8              -------gtgtttaaagataat.................ctg...............t........ca............ [161520]

FR682468.2_Georgia_2007/1         gtggatagagcgctttcctctgagcagcaaattttttcatacatctccatgggggatggcgaggctttaatagtatgtag [161600]
ASF_IND_20_CAD543                 ................................................................................ [161600]
LR899193.1_Germany_2020/1         ................................................................................ [161600]
MN393477.1_Wuhan_2019             ................................................................................ [161600]
MW396979.1_Timor-Leste/2019/1     ................................................................................ [161600]
MK940252.1_CN/2019/InnerMongolia  ................................................................................ [161600]
MK128995.1_China/2018/AnhuiXCGQ   ................................................................................ [161600]
MH681419.1_ASFV/POL/2015          ................................................................................ [161600]
LS478113.1_Estonia_2014           ................................................................................ [161600]
KP843857.1_Odintsovo_02/14        ................................................................................ [161600]
FN557520.1_strain_E75             .............................g............g..........a.......................... [161600]
AY261366.1_Warthog                ..........g..............................ag..........a.......................... [161600]
AY261365.1_Warmbaths              ..........g..............................ag..........a..........t............... [161600]
AY261364.1_Tengani_62             ..a....a..g...............................g..........a..........t.........t..... [161600]
AY261363.1_Pretoriuskop/96/4      ..........g..................g...........ag..........a.......................... [161600]
AY261362.1_Mkuzi_1979             .............................g............g..........a.......................... [161600]
AY261361.1_Malawi_Lil-20/1_1983   ..a......ag......t.t.....................ggc....................tc....g......... [161600]
AY261360.1_Kenya_1950             ..........g......t.t..t......g...........agc.........a.....t....t..g............ [161600]
AM712240.1_OURT_88/3              .............................g............g..........a.......................... [161600]
AM712239.1_Benin_97/1             .............................g............g..........a.......................... [161600]
KM111295.1_Ken06.Bus              .................t.t..t......g..c........agc.........a.....t....t..g............ [161600]
KM111294.1_Ken05/Tk1              ..........g......t.t..t......g...........agct........a.....t....t..g............ [161600]
KM262844.1_strain_L60             .............................g............g..........a.......................... [161600]
KM102979.1_26544/OG10_Italy       .............................g............g..........a.......................... [161600]
KM262845.1_strain_NHV             .............................g............g..........a.......................... [161600]
KX354450.1_47/Ss/2008             .............................g............g..........a.......................... [161600]
KP055815.1_strain_BA71            .............................g............g..........a.......................... [161600]
MH025920.1_strain_R35             .................t.t..t......g..c........agc.........a.....t....t..g............ [161600]
MH025919.1_strain_N10             .................t.t..t......g..c........agc.........a.....t....t..g............ [161600]
MH025918.1_strain_R25             .................t.t..t......g..c........agc.........a.....t....t..g............ [161600]
MH025917.1_strain_R7              .................t.t..t......g..c........agc.........a.....t....t..g............ [161600]
MH025916.1_strain_R8              .................t.t..t......g..c........agc.........a.....t....t..g............ [161600]
```

```
FR682468.2_Georgia_2007/1        gtcacgtaagaactgttgtatgatgggatatttgtcttttaaaaactggggatgtttcataactggaattatttgaaaga [161680]
ASF_IND_20_CAD543                ................................................................................ [161680]
LR899193.1_Germany_2020/1        ................................................................................ [161680]
MN393477.1_Wuhan_2019            ................................................................................ [161680]
MW396979.1_Timor-Leste/2019/1    ................................................................................ [161680]
MK940252.1_CN/2019/InnerMongolia ................................................................................ [161680]
MK128995.1_China/2018/AnhuiXCGQ  ................................................................................ [161680]
MH681419.1_ASFV/POL/2015         ................................................................................ [161680]
LS478113.1_Estonia_2014          ................................................................................ [161680]
KP843857.1_Odintsovo_02/14       ................................................................................ [161680]
FN557520.1_strain_E75            ................................................................................ [161680]
AY261366.1_Warthog               ........................a....................................................... [161680]
AY261365.1_Warmbaths             ................................................................................ [161680]
AY261364.1_Tengani_62            ................................................................................ [161680]
AY261363.1_Pretoriuskop/96/4     ........................a....................................................... [161680]
AY261362.1_Mkuzi_1979            ................................................................................ [161680]
AY261361.1_Malawi_Lil-20/1_1983  ....t....a.................................................................g.... [161680]
AY261360.1_Kenya_1950            ....t....a...........a..............c......................................g.... [161680]
AM712240.1_OURT_88/3             ................................................................................ [161680]
AM712239.1_Benin_97/1            ................................................................................ [161680]
KM111295.1_Ken06.Bus             ....t....a...........a..............c......................................g.... [161680]
KM111294.1_Ken05/Tk1             ....t....a...........a.....g........c......................................g.... [161680]
KM262844.1_strain_L60            ................................................................................ [161680]
KM102979.1_26544/OG10_Italy      ................................................................................ [161680]
KM262845.1_strain_NHV            ................................................................................ [161680]
KX354450.1_47/Ss/2008            ................................................................................ [161680]
KP055815.1_strain_BA71           ................................................................................ [161680]
MH025920.1_strain_R35            ....t....a...........a..............c......................................g.... [161680]
MH025919.1_strain_N10            ....t....a...........a..............c......................................g.... [161680]
MH025918.1_strain_R25            ....t....a...........a..............c......................................g.... [161680]
MH025917.1_strain_R7             ....t....a...........a..............c......................................g.... [161680]
MH025916.1_strain_R8             ....t....a...........a..............c......................................g.... [161680]

FR682468.2_Georgia_2007/1        taaagaccttccatccaaagtagccaaccacatttggcatttcgggacacgcggtttcataaggcatagaatagtgaata [161760]
ASF_IND_20_CAD543                ................................................................................ [161760]
LR899193.1_Germany_2020/1        ................................................................................ [161760]
MN393477.1_Wuhan_2019            ................................................................................ [161760]
MW396979.1_Timor-Leste/2019/1    ................................................................................ [161760]
MK940252.1_CN/2019/InnerMongolia ................................................................................ [161760]
MK128995.1_China/2018/AnhuiXCGQ  ................................................................................ [161760]
MH681419.1_ASFV/POL/2015         ................................................................................ [161760]
LS478113.1_Estonia_2014          ................................................................................ [161760]
KP843857.1_Odintsovo_02/14       ................................................................................ [161760]
```

```
FN557520.1_strain_E75            ................................................................................ [161760]
AY261366.1_Warthog               ................................................................................ [161760]
AY261365.1_Warmbaths             ................................................................................ [161760]
AY261364.1_Tengani_62            ...........................................a........a........................... [161760]
AY261363.1_Pretoriuskop/96/4     ................................................................................ [161760]
AY261362.1_Mkuzi_1979            ................................................................................ [161760]
AY261361.1_Malawi_Lil-20/1_1983  ....................................a...............a.....g..g.................c [161760]
AY261360.1_Kenya_1950            ....................................a......a.................g...g.............c [161760]
AM712240.1_OURT_88/3             ................................................................................ [161760]
AM712239.1_Benin_97/1            ................................................................................ [161760]
KM111295.1_Ken06.Bus             ....................................a......a.................g.................c [161760]
KM111294.1_Ken05/Tk1             ....................................a......a.................g.....g...........c [161760]
KM262844.1_strain_L60            ................................................................................ [161760]
KM102979.1_26544/OG10_Italy      ................................................................................ [161760]
KM262845.1_strain_NHV            ................................................................................ [161760]
KX354450.1_47/Ss/2008            ................................................................................ [161760]
KP055815.1_strain_BA71           ................................................................................ [161760]
MH025920.1_strain_R35            ....................................a......a.................g.................c [161760]
MH025919.1_strain_N10            ....................................a......a.................g.................c [161760]
MH025918.1_strain_R25            ....................................a......a.................g.................c [161760]
MH025917.1_strain_R7             ....................................a......a.................g.................c [161760]
MH025916.1_strain_R8             ....................................a......a.................g.................c [161760]
```

```
FR682468.2_Georgia_2007/1        gtgtactgatctttttgatacagcgtttcaagtagttggcgaaatgtttccgcgtcgagcgtgccaaaatcttgaggagc [161840]
ASF_IND_20_CAD543                ................................................................................ [161840]
LR899193.1_Germany_2020/1        ................................................................................ [161840]
MN393477.1_Wuhan_2019            ................................................................................ [161840]
MW396979.1_Timor-Leste/2019/1    ................................................................................ [161840]
MK940252.1_CN/2019/InnerMongolia ................................................................................ [161840]
MK128995.1_China/2018/AnhuiXCGQ  ................................................................................ [161840]
MH681419.1_ASFV/POL/2015         ................................................................................ [161840]
LS478113.1_Estonia_2014          ................................................................................ [161840]
KP843857.1_Odintsovo_02/14       ................................................................................ [161840]
FN557520.1_strain_E75            ................................................................................ [161840]
AY261366.1_Warthog               ...........c.....g.........................................t.................... [161840]
AY261365.1_Warmbaths             .................g......................................................a....t.. [161840]
AY261364.1_Tengani_62            .................g.....................t........................................ [161840]
AY261363.1_Pretoriuskop/96/4     ...........c.....g.............................................................. [161840]
AY261362.1_Mkuzi_1979            ................................................................................ [161840]
AY261361.1_Malawi_Lil-20/1_1983  ..a.......t......g...........t..c......t......c........t...............c........ [161840]
AY261360.1_Kenya_1950            .................g....a......t.ag.a...at...............t...............c........ [161840]
AM712240.1_OURT_88/3             ................................................................................ [161840]
AM712239.1_Benin_97/1            ................................................................................ [161840]
KM111295.1_Ken06.Bus             .................g....a......t.ag.a...at...............t...............c........ [161840]
```

```
KM111294.1_Ken05/Tk1           .................g....a......t.ag.a...at...............t...............c........ [161840]
KM262844.1_strain_L60          ................................................................................ [161840]
KM102979.1_26544/OG10_Italy    ................................................................................ [161840]
KM262845.1_strain_NHV          ................................................................................ [161840]
KX354450.1_47/Ss/2008          ................................................................................ [161840]
KP055815.1_strain_BA71         ................................................................................ [161840]
MH025920.1_strain_R35          .................g....a......t.ag.a...at...............t...............c........ [161840]
MH025919.1_strain_N10          .................g....a......t.ag.a...at...............t...............c........ [161840]
MH025918.1_strain_R25          .................g....a......t.ag.a...at...............t...............c........ [161840]
MH025917.1_strain_R7           .................g....a......t.ag.a...at...............t...............c........ [161840]
MH025916.1_strain_R8           .................g....a......t.ag.a...at...............t...............c........ [161840]

FR682468.2_Georgia_2007/1      ctcggtgtgctcctgtgtagagcagatcgtgatgattccccaggcaagcgggagcatggactctggagggtggatatccg [161920]
ASF_IND_20_CAD543              ....................................................nnnnnnnnnnnnnnn............. [161920]
LR899193.1_Germany_2020/1      ................................................................................ [161920]
MN393477.1_Wuhan_2019          ................................................................................ [161920]
MW396979.1_Timor-Leste/2019/1  ................................................................................ [161920]
MK940252.1_CN/2019/InnerMongolia ................................................................................ [161920]
MK128995.1_China/2018/AnhuiXCGQ ................................................................................ [161920]
MH681419.1_ASFV/POL/2015       ................................................................................ [161920]
LS478113.1_Estonia_2014        ................................................................................ [161920]
KP843857.1_Odintsovo_02/14     ................................................................................ [161920]
FN557520.1_strain_E75          .........t...................................................................... [161920]
AY261366.1_Warthog             ........................a.....................................................aa [161920]
AY261365.1_Warmbaths           ..................g...........................................................aa [161920]
AY261364.1_Tengani_62          ............t.......................................g................a........aa [161920]
AY261363.1_Pretoriuskop/96/4   ..................g...........................................................aa [161920]
AY261362.1_Mkuzi_1979          ................................................................................ [161920]
AY261361.1_Malawi_Lil-20/1_1983 ..................g.............................a.............................a. [161920]
AY261360.1_Kenya_1950          ........a.........g..............a...................................a........a. [161920]
AM712240.1_OURT_88/3           ................................................................................ [161920]
AM712239.1_Benin_97/1          .........t...................................................................... [161920]
KM111295.1_Ken06.Bus           ........a.........g..................................................a..a.....a. [161920]
KM111294.1_Ken05/Tk1           ........a.........g..................................................a........a. [161920]
KM262844.1_strain_L60          ................................................................................ [161920]
KM102979.1_26544/OG10_Italy    .........t...................................................................... [161920]
KM262845.1_strain_NHV          ................................................................................ [161920]
KX354450.1_47/Ss/2008          .........t...................................................................... [161920]
KP055815.1_strain_BA71         ................................................................................ [161920]
MH025920.1_strain_R35          ........a.........g..................................................a..a.....a. [161920]
MH025919.1_strain_N10          ........a.........g..................................................a..a.....a. [161920]
MH025918.1_strain_R25          ........a.........g..................................................a..a.....a. [161920]
MH025917.1_strain_R7           ........a.........g..................................................a..a.....a. [161920]
MH025916.1_strain_R8           ........a.........g..................................................a..a.....a. [161920]
```

```
FR682468.2_Georgia_2007/1        tattggtctcattattcgatcccagctgatgaatgccgcacacgcgaaacatggcctcgacgtagatgcccatagagata [162000]
ASF_IND_20_CAD543                .........y...................................................................... [162000]
LR899193.1_Germany_2020/1        ................................................................................ [162000]
MN393477.1_Wuhan_2019            ................................................................................ [162000]
MW396979.1_Timor-Leste/2019/1    ................................................................................ [162000]
MK940252.1_CN/2019/InnerMongolia ................................................................................ [162000]
MK128995.1_China/2018/AnhuiXCGQ  ................................................................................ [162000]
MH681419.1_ASFV/POL/2015         ................................................................................ [162000]
LS478113.1_Estonia_2014          ................................................................................ [162000]
KP843857.1_Odintsovo_02/14       ................................................................................ [162000]
FN557520.1_strain_E75            ................................................................................ [162000]
AY261366.1_Warthog               ..........g......c.............................................................. [162000]
AY261365.1_Warmbaths             .................c......a....................................................... [162000]
AY261364.1_Tengani_62            ....a............c.............................................................. [162000]
AY261363.1_Pretoriuskop/96/4     ...........c.....c.............................................................. [162000]
AY261362.1_Mkuzi_1979            ................................................................................ [162000]
AY261361.1_Malawi_Lil-20/1_1983  ............g....c..................t......a.................a.................. [162000]
AY261360.1_Kenya_1950            .................c......................t....................a......g....g..a... [162000]
AM712240.1_OURT_88/3             ................................................................................ [162000]
AM712239.1_Benin_97/1            ................................................................................ [162000]
KM111295.1_Ken06.Bus             .................c......................t....................a......g....g..a... [162000]
KM111294.1_Ken05/Tk1             .................c......................t....................a......g....g..a... [162000]
KM262844.1_strain_L60            ................................................................................ [162000]
KM102979.1_26544/OG10_Italy      ................................................................................ [162000]
KM262845.1_strain_NHV            ................................................................................ [162000]
KX354450.1_47/Ss/2008            ................................................................................ [162000]
KP055815.1_strain_BA71           ................................................................................ [162000]
MH025920.1_strain_R35            .................c......................t....................a......g....g..a... [162000]
MH025919.1_strain_N10            .................c......................t....................a......g....g..a... [162000]
MH025918.1_strain_R25            .................c......................t....................a......g....g..a... [162000]
MH025917.1_strain_R7             .................c......................t....................a......g....g..a... [162000]
MH025916.1_strain_R8             .................c......................t....................a......g....g..a... [162000]

FR682468.2_Georgia_2007/1        ggcggcgaaagggcaagaccggattgtatttgcggcatatagtaggagggcaccgagttttttatttttcggttgaatgg [162080]
ASF_IND_20_CAD543                ................................................................................ [162080]
LR899193.1_Germany_2020/1        ................................................................................ [162080]
MN393477.1_Wuhan_2019            ................................................................................ [162080]
MW396979.1_Timor-Leste/2019/1    ................................................................................ [162080]
MK940252.1_CN/2019/InnerMongolia ................................................................................ [162080]
MK128995.1_China/2018/AnhuiXCGQ  ................................................................................ [162080]
MH681419.1_ASFV/POL/2015         ................................................................................ [162080]
LS478113.1_Estonia_2014          ................................................................................ [162080]
KP843857.1_Odintsovo_02/14       ................................................................................ [162080]
```

```
FN557520.1_strain_E75             .....t...........g.............................................................. [162080]
AY261366.1_Warthog                .....g...........g.........................................................t.... [162080]
AY261365.1_Warmbaths              .....g...........g..........................a..............................t.... [162080]
AY261364.1_Tengani_62             .....t...........g.......................a...................................... [162080]
AY261363.1_Pretoriuskop/96/4      .....g...........g..........................a..............................t.... [162080]
AY261362.1_Mkuzi_1979             .....t...........g.............................................................. [162080]
AY261361.1_Malawi_Lil-20/1_1983   .....t...........g..a...........g......................................a..a..g.. [162080]
AY261360.1_Kenya_1950             .....t...........g..a..c........g....c......a..........................a..a..... [162080]
AM712240.1_OURT_88/3              .....t...........g.............................................................. [162080]
AM712239.1_Benin_97/1             .....t...........g.............................................................. [162080]
KM111295.1_Ken06.Bus              .....t...........g..a..c........g....c.................................a..a..... [162080]
KM111294.1_Ken05/Tk1              .....t...........g..a..c........g....c......a..........................a..a..... [162080]
KM262844.1_strain_L60             .....t...........g.............................................................. [162080]
KM102979.1_26544/OG10_Italy       .....t...........g.............................................................. [162080]
KM262845.1_strain_NHV             .....t...........g.............................................................. [162080]
KX354450.1_47/Ss/2008             .....t...........g.............................................................. [162080]
KP055815.1_strain_BA71            .....t...........g.............................................................. [162080]
MH025920.1_strain_R35             .....t...........g..a..c........g....c.................................a..a..... [162080]
MH025919.1_strain_N10             .....t...........g..a..c........g....c.................................a..a..... [162080]
MH025918.1_strain_R25             .....t...........g..a..c........g....c.................................a..a..... [162080]
MH025917.1_strain_R7              .....t...........g..a..c........g....c.................................a..a..... [162080]
MH025916.1_strain_R8              .....t...........g..a..c........g....c.................................a..a..... [162080]

FR682468.2_Georgia_2007/1         ggactttatttctaccagcacggggatgcgtttcgtggcctcatagcgtacgttgttaaaaattgttttgatttcccagg [162160]
ASF_IND_20_CAD543                 ................................................................................ [162160]
LR899193.1_Germany_2020/1         ................................................................................ [162160]
MN393477.1_Wuhan_2019             ................................................................................ [162160]
MW396979.1_Timor-Leste/2019/1     ................................................................................ [162160]
MK940252.1_CN/2019/InnerMongolia  ................................................................................ [162160]
MK128995.1_China/2018/AnhuiXCGQ   ................................................................................ [162160]
MH681419.1_ASFV/POL/2015          ................................................................................ [162160]
LS478113.1_Estonia_2014           ................................................................................ [162160]
KP843857.1_Odintsovo_02/14        ................................................................................ [162160]
FN557520.1_strain_E75             ................................................................................ [162160]
AY261366.1_Warthog                ...t...................................t.......a...a............................ [162160]
AY261365.1_Warmbaths              ...t...................................t.......a...a............................ [162160]
AY261364.1_Tengani_62             ................................................................................ [162160]
AY261363.1_Pretoriuskop/96/4      ...t...................................t.......a...a............................ [162160]
AY261362.1_Mkuzi_1979             ......................................t......................................... [162160]
AY261361.1_Malawi_Lil-20/1_1983   a................at.................a.tt.......a...a.c.......................... [162160]
AY261360.1_Kenya_1950             .................a....................tt.......a...a.c......g................... [162160]
AM712240.1_OURT_88/3              ................................................................................ [162160]
AM712239.1_Benin_97/1             ................................................................................ [162160]
KM111295.1_Ken06.Bus              .................a....................tt.......a...a.c.......................... [162160]
```

```
KM111294.1_Ken05/Tk1           .................a....................tt.....a.a...a.c......g................... [162160]
KM262844.1_strain_L60          ................................................................................ [162160]
KM102979.1_26544/OG10_Italy    ................................................................................ [162160]
KM262845.1_strain_NHV          ................................................................................ [162160]
KX354450.1_47/Ss/2008          ................................................................................ [162160]
KP055815.1_strain_BA71         ................................................................................ [162160]
MH025920.1_strain_R35          .................a....................tt.......a...a.c.......................... [162160]
MH025919.1_strain_N10          .................a....................tt.......a...a.c.......................... [162160]
MH025918.1_strain_R25          .................a....................tt.......a...a.c.......................... [162160]
MH025917.1_strain_R7           .................a....................tt.......a...a.c.......................... [162160]
MH025916.1_strain_R8           .................a....................tt.......a...a.c.......................... [162160]

FR682468.2_Georgia_2007/1      actgttg---agtgtatcccagcgttaggtgacaaaacccatcggggctattactatgtccggggtatcccaaataggtc [162240]
ASF_IND_20_CAD543              .......---...................................................................... [162240]
LR899193.1_Germany_2020/1      .......---...................................................................... [162240]
MN393477.1_Wuhan_2019          .......---...................................................................... [162240]
MW396979.1_Timor-Leste/2019/1  .......---...................................................................... [162240]
MK940252.1_CN/2019/InnerMongolia .......---...................................................................... [162240]
MK128995.1_China/2018/AnhuiXCGQ .......---...................................................................... [162240]
MH681419.1_ASFV/POL/2015       .......---...................................................................... [162240]
LS478113.1_Estonia_2014        .......---...................................................................... [162240]
KP843857.1_Odintsovo_02/14     .......---...................................................................... [162240]
FN557520.1_strain_E75          .......---...................................................................... [162240]
AY261366.1_Warthog             .......---...........a........a...........................c....................t [162240]
AY261365.1_Warmbaths           .......agt....................a...........................c....................t [162240]
AY261364.1_Tengani_62          .......---....................................a...........c....................t [162240]
AY261363.1_Pretoriuskop/96/4   .......---....................a......t....................c....................t [162240]
AY261362.1_Mkuzi_1979          .......---...................................................................... [162240]
AY261361.1_Malawi_Lil-20/1_1983 .......---................................................c....................t [162240]
AY261360.1_Kenya_1950          .......---............g.......a......t..g..a........g.....c..................... [162240]
AM712240.1_OURT_88/3           .......---...................................................................... [162240]
AM712239.1_Benin_97/1          .......---...................................................................... [162240]
KM111295.1_Ken06.Bus           .......---............g.......ag.....t..g..a........g.....c..................... [162240]
KM111294.1_Ken05/Tk1           .......---............g.......a......t..g..a........g.....c..................... [162240]
KM262844.1_strain_L60          .......---...................................................................... [162240]
KM102979.1_26544/OG10_Italy    .......---...................................................................... [162240]
KM262845.1_strain_NHV          .......---...................................................................... [162240]
KX354450.1_47/Ss/2008          .......---...................................................................... [162240]
KP055815.1_strain_BA71         .......---...................................................................... [162240]
MH025920.1_strain_R35          .......---............g.......ag.....t..g..a........g.....c..................... [162240]
MH025919.1_strain_N10          .......---............g.......ag.....t..g..a........g.....c..................... [162240]
MH025918.1_strain_R25          .......---............g.......ag.....t..g..a........g.....c..................... [162240]
MH025917.1_strain_R7           .......---............g.......ag.....t..g..a........g.....c..................... [162240]
MH025916.1_strain_R8           .......---............g.......ag.....t..g..a........g.....c..................... [162240]
```

```
FR682468.2_Georgia_2007/1        ccatcaatatgaatattgtcacctatgacggtggtttggcagaacaactcaagcagatctttactaacacgctcaaaaag [162320]
ASF_IND_20_CAD543                ................................................................................ [162320]
LR899193.1_Germany_2020/1        ................................................................................ [162320]
MN393477.1_Wuhan_2019            ................................................................................ [162320]
MW396979.1_Timor-Leste/2019/1    ................................................................................ [162320]
MK940252.1_CN/2019/InnerMongolia ................................................................................ [162320]
MK128995.1_China/2018/AnhuiXCGQ  ................................................................................ [162320]
MH681419.1_ASFV/POL/2015         ................................................................................ [162320]
LS478113.1_Estonia_2014          ................................................................................ [162320]
KP843857.1_Odintsovo_02/14       ................................................................................ [162320]
FN557520.1_strain_E75            ....................................................................t........... [162320]
AY261366.1_Warthog               ....................c..........................t.....t.......................... [162320]
AY261365.1_Warmbaths             ................................................................................ [162320]
AY261364.1_Tengani_62            .................a.......................a...................................... [162320]
AY261363.1_Pretoriuskop/96/4     ....................c........................................................... [162320]
AY261362.1_Mkuzi_1979            ................................................................................ [162320]
AY261361.1_Malawi_Lil-20/1_1983  ...........g..............t............................a........................ [162320]
AY261360.1_Kenya_1950            ...........g..............t..........................t...........c.............. [162320]
AM712240.1_OURT_88/3             ....................................................................t........... [162320]
AM712239.1_Benin_97/1            ....................................................................t........... [162320]
KM111295.1_Ken06.Bus             ...........g..............t......................................c.............. [162320]
KM111294.1_Ken05/Tk1             ...........g..............t..........................t...........c.............. [162320]
KM262844.1_strain_L60            ....................................................................t........... [162320]
KM102979.1_26544/OG10_Italy      ....................................................................t........... [162320]
KM262845.1_strain_NHV            ....................................................................t........... [162320]
KX354450.1_47/Ss/2008            ....................................................................t........... [162320]
KP055815.1_strain_BA71           ....................................................................t........... [162320]
MH025920.1_strain_R35            ...........g..............t......................................c.............. [162320]
MH025919.1_strain_N10            ...........g..............t......................................c.............. [162320]
MH025918.1_strain_R25            ...........g..............t......................................c.............. [162320]
MH025917.1_strain_R7             ...........g..............t......................................c.............. [162320]
MH025916.1_strain_R8             ...........g..............t......................................c.............. [162320]

FR682468.2_Georgia_2007/1        ggttccccagctacaagcagcgcggttcaaattcttcttaaaaagatttgctttttccgccaaggttatataatagcttt [162400]
ASF_IND_20_CAD543                ................................................................r............... [162400]
LR899193.1_Germany_2020/1        ................................................................................ [162400]
MN393477.1_Wuhan_2019            ................................................................................ [162400]
MW396979.1_Timor-Leste/2019/1    ................................................................................ [162400]
MK940252.1_CN/2019/InnerMongolia ................................................................................ [162400]
MK128995.1_China/2018/AnhuiXCGQ  ................................................................................ [162400]
MH681419.1_ASFV/POL/2015         ................................................................................ [162400]
LS478113.1_Estonia_2014          ................................................................................ [162400]
KP843857.1_Odintsovo_02/14       ................................................................................ [162400]
```

```
FN557520.1_strain_E75           ................................................................a............... [162400]
AY261366.1_Warthog              ................................................................................ [162400]
AY261365.1_Warmbaths            ................................................................................ [162400]
AY261364.1_Tengani_62           ................................................................................ [162400]
AY261363.1_Pretoriuskop/96/4    ................................................................................ [162400]
AY261362.1_Mkuzi_1979           ................................................................................ [162400]
AY261361.1_Malawi_Lil-20/1_1983 ..................t...t..................................t..t................... [162400]
AY261360.1_Kenya_1950           ..........t.......t...t....t...............................................a.... [162400]
AM712240.1_OURT_88/3            ................................................................a............... [162400]
AM712239.1_Benin_97/1           ................................................................a............... [162400]
KM111295.1_Ken06.Bus            ..........t.......t...t....t...............................................a.... [162400]
KM111294.1_Ken05/Tk1            ..........t.......t...t....t...............................................a.... [162400]
KM262844.1_strain_L60           ................................................................a............... [162400]
KM102979.1_26544/OG10_Italy     ................................................................a............... [162400]
KM262845.1_strain_NHV           ................................................................a............... [162400]
KX354450.1_47/Ss/2008           ................................................................a............... [162400]
KP055815.1_strain_BA71          ................................................................a............... [162400]
MH025920.1_strain_R35           ..........t.......t...t....t...............................................a.... [162400]
MH025919.1_strain_N10           ..........t.......t...t....t...............................................a.... [162400]
MH025918.1_strain_R25           ..........t.......t...t....t...............................................a.... [162400]
MH025917.1_strain_R7            ..........t.......t...t....t...............................................a.... [162400]
MH025916.1_strain_R8            ..........t.......t...t....t...............................................a.... [162400]

FR682468.2_Georgia_2007/1       tgtaagggtttaaacctaaaacgctggcaaggtcagagccacccacctgagtgcgacgaatagcatgccaggcatcggag [162480]
ASF_IND_20_CAD543               ................................................................................ [162480]
LR899193.1_Germany_2020/1       ................................................................................ [162480]
MN393477.1_Wuhan_2019           ................................................................................ [162480]
MW396979.1_Timor-Leste/2019/1   ................................................................................ [162480]
MK940252.1_CN/2019/InnerMongolia ................................................................................ [162480]
MK128995.1_China/2018/AnhuiXCGQ ................................................................................ [162480]
MH681419.1_ASFV/POL/2015        ................................................................................ [162480]
LS478113.1_Estonia_2014         ................................................................................ [162480]
KP843857.1_Odintsovo_02/14      ................................................................................ [162480]
FN557520.1_strain_E75           .........................a...................................................... [162480]
AY261366.1_Warthog              .........................a..............c..................gc................... [162480]
AY261365.1_Warmbaths            .....................t...a..............c..................gc..................a [162480]
AY261364.1_Tengani_62           .........................a..............c............t......c................... [162480]
AY261363.1_Pretoriuskop/96/4    .....................t...a..............c............t......c................... [162480]
AY261362.1_Mkuzi_1979           .........................a...................................................... [162480]
AY261361.1_Malawi_Lil-20/1_1983 ....g................t...a.................t...............g...............t.... [162480]
AY261360.1_Kenya_1950           .........................a.................................g.g.............t.... [162480]
AM712240.1_OURT_88/3            .........................a...................................................... [162480]
AM712239.1_Benin_97/1           .........................a...................................................... [162480]
KM111295.1_Ken06.Bus            .........................a.................................g.g.............t.... [162480]
```

```
KM111294.1_Ken05/Tk1              .a.......................a.................................g.g.............t.... [162480]
KM262844.1_strain_L60             .........................a...................................................... [162480]
KM102979.1_26544/OG10_Italy       .........................a...................................................... [162480]
KM262845.1_strain_NHV             .........................a...................................................... [162480]
KX354450.1_47/Ss/2008             .........................a...................................................... [162480]
KP055815.1_strain_BA71            .........................a...................................................... [162480]
MH025920.1_strain_R35             ......................a..a.................................g.g.............t.... [162480]
MH025919.1_strain_N10             ......................a..a.................................g.g.............t.... [162480]
MH025918.1_strain_R25             ......................a..a.................................g.g.............t.... [162480]
MH025917.1_strain_R7              ......................a..a.................................g.g.............t.... [162480]
MH025916.1_strain_R8              ......................a..a.................................g.g.............t.... [162480]

FR682468.2_Georgia_2007/1         cgctgctgaggagagtctttaaacaggcgtacaaaggtttccattatacttgttttaacaggaattcaatataaaaagtc [162560]
ASF_IND_20_CAD543                 ................................................................................ [162560]
LR899193.1_Germany_2020/1         ................................................................................ [162560]
MN393477.1_Wuhan_2019             ................................................................................ [162560]
MW396979.1_Timor-Leste/2019/1     ................................................................................ [162560]
MK940252.1_CN/2019/InnerMongolia  ................................................................................ [162560]
MK128995.1_China/2018/AnhuiXCGQ   ................................................................................ [162560]
MH681419.1_ASFV/POL/2015          ................................................................................ [162560]
LS478113.1_Estonia_2014           ................................................................................ [162560]
KP843857.1_Odintsovo_02/14        ................................................................................ [162560]
FN557520.1_strain_E75             ................................................................................ [162560]
AY261366.1_Warthog                ............................a................................................... [162560]
AY261365.1_Warmbaths              ............................a................................................... [162560]
AY261364.1_Tengani_62             ................................................................................ [162560]
AY261363.1_Pretoriuskop/96/4      .........................a..a................................................... [162560]
AY261362.1_Mkuzi_1979             ..........................................................t..................... [162560]
AY261361.1_Malawi_Lil-20/1_1983   ................................................t............................... [162560]
AY261360.1_Kenya_1950             .............................c.................................................. [162560]
AM712240.1_OURT_88/3              ................................................................................ [162560]
AM712239.1_Benin_97/1             ................................................................................ [162560]
KM111295.1_Ken06.Bus              .............................c.................................................. [162560]
KM111294.1_Ken05/Tk1              .............................c.................................................. [162560]
KM262844.1_strain_L60             ................................................................................ [162560]
KM102979.1_26544/OG10_Italy       ................................................................................ [162560]
KM262845.1_strain_NHV             ................................................................................ [162560]
KX354450.1_47/Ss/2008             ................................................................................ [162560]
KP055815.1_strain_BA71            ................................................................................ [162560]
MH025920.1_strain_R35             .............................c.................................................. [162560]
MH025919.1_strain_N10             .............................c.................................................. [162560]
MH025918.1_strain_R25             .............................c.................................................. [162560]
MH025917.1_strain_R7              .............................c.................................................. [162560]
MH025916.1_strain_R8              .............................c.................................................. [162560]
```

```
FR682468.2_Georgia_2007/1        aacacagtttgcaatttttccaatctcaagatatagccatacatttttttttccaattggcgaatatgtttaagctcatg [162640]
ASF_IND_20_CAD543                ................................................................................ [162640]
LR899193.1_Germany_2020/1        ................................................................................ [162640]
MN393477.1_Wuhan_2019            ................................................................................ [162640]
MW396979.1_Timor-Leste/2019/1    ................................................................................ [162640]
MK940252.1_CN/2019/InnerMongolia ................................................................................ [162640]
MK128995.1_China/2018/AnhuiXCGQ  ................................................................................ [162640]
MH681419.1_ASFV/POL/2015         ................................................................................ [162640]
LS478113.1_Estonia_2014          ................................................................................ [162640]
KP843857.1_Odintsovo_02/14       ................................................................................ [162640]
FN557520.1_strain_E75            ................................................................................ [162640]
AY261366.1_Warthog               ..a..................................................t.....a.................... [162640]
AY261365.1_Warmbaths             ..a...................g..............................t.......................... [162640]
AY261364.1_Tengani_62            ..a..................................................t.......................... [162640]
AY261363.1_Pretoriuskop/96/4     ..a..................................................t.....a.................... [162640]
AY261362.1_Mkuzi_1979            ................................................................................ [162640]
AY261361.1_Malawi_Lil-20/1_1983  ..t..................................................t..........c.a............. [162640]
AY261360.1_Kenya_1950            ....t................................................t............a............. [162640]
AM712240.1_OURT_88/3             ................................................................................ [162640]
AM712239.1_Benin_97/1            ................................................................................ [162640]
KM111295.1_Ken06.Bus             ....t................................................t............a............. [162640]
KM111294.1_Ken05/Tk1             ....t................................................t............a............. [162640]
KM262844.1_strain_L60            ................................................................................ [162640]
KM102979.1_26544/OG10_Italy      ................................................................................ [162640]
KM262845.1_strain_NHV            ................................................................................ [162640]
KX354450.1_47/Ss/2008            ................................................................................ [162640]
KP055815.1_strain_BA71           ................................................................................ [162640]
MH025920.1_strain_R35            ....t................................................t............a............. [162640]
MH025919.1_strain_N10            ....t................................................t............a............. [162640]
MH025918.1_strain_R25            ....t................................................t............a............. [162640]
MH025917.1_strain_R7             ....t................................................t............a............. [162640]
MH025916.1_strain_R8             ....t................................................t............a............. [162640]

FR682468.2_Georgia_2007/1        tgtttcaatattagcatccggaaatttaaatgcataaagatgttcaaaggcctgatttatacacgtatcaaaggatctgt [162720]
ASF_IND_20_CAD543                ................................................................................ [162720]
LR899193.1_Germany_2020/1        ................................................................................ [162720]
MN393477.1_Wuhan_2019            ................................................................................ [162720]
MW396979.1_Timor-Leste/2019/1    ................................................................................ [162720]
MK940252.1_CN/2019/InnerMongolia ................................................................................ [162720]
MK128995.1_China/2018/AnhuiXCGQ  ................................................................................ [162720]
MH681419.1_ASFV/POL/2015         ................................................................................ [162720]
LS478113.1_Estonia_2014          ................................................................................ [162720]
KP843857.1_Odintsovo_02/14       ................................................................................ [162720]
```

```
FN557520.1_strain_E75           ........................................................................c...t... [162720]
AY261366.1_Warthog              .......................................c..........a.....................t....... [162720]
AY261365.1_Warmbaths            ..................................................a.....................t....... [162720]
AY261364.1_Tengani_62           .........................................a.....g..a............t........t....... [162720]
AY261363.1_Pretoriuskop/96/4    ..................................................a.....................t....... [162720]
AY261362.1_Mkuzi_1979           ........................................................................c...t... [162720]
AY261361.1_Malawi_Lil-20/1_1983 ......t.......................g..........t......t.a..........g.t........t..c.... [162720]
AY261360.1_Kenya_1950           g.....t.......................g..........t...g..a.a..........g.a.......gc...t... [162720]
AM712240.1_OURT_88/3            ........................................................................c...t... [162720]
AM712239.1_Benin_97/1           ........................................................................c...t... [162720]
KM111295.1_Ken06.Bus            g.....t.......................g..........t..tg..a.a..........g.a.......gt...t... [162720]
KM111294.1_Ken05/Tk1            g.....t.......................g..........t...g..a.a..........g.a.......gt...t... [162720]
KM262844.1_strain_L60           ........................................................................c...t... [162720]
KM102979.1_26544/OG10_Italy     ........................................................................c...t... [162720]
KM262845.1_strain_NHV           ........................................................................c...t... [162720]
KX354450.1_47/Ss/2008           ........................................................................c...t... [162720]
KP055815.1_strain_BA71          ........................................................................c...t... [162720]
MH025920.1_strain_R35           g.....t.......................g..........t..tg..a.a..........g.a.......gt...t... [162720]
MH025919.1_strain_N10           g.....t.......................g..........t..tg..a.a..........g.a.......gt...t... [162720]
MH025918.1_strain_R25           g.....t.......................g..........t..tg..a.a..........g.a.......gt...t... [162720]
MH025917.1_strain_R7            g.....t.......................g..........t..tg..a.a..........g.a.......gt...t... [162720]
MH025916.1_strain_R8            g.....t.......................g..........t..tg..a.a..........g.a.......gt...t... [162720]

FR682468.2_Georgia_2007/1       ggtatgttattagcttcagcatgtgtgccagatcttcaagatggtctaaatttatacggttttccacgtggtggatcatg [162800]
ASF_IND_20_CAD543               ................................................................................ [162800]
LR899193.1_Germany_2020/1       ................................................................................ [162800]
MN393477.1_Wuhan_2019           ................................................................................ [162800]
MW396979.1_Timor-Leste/2019/1   ................................................................................ [162800]
MK940252.1_CN/2019/InnerMongolia ................................................................................ [162800]
MK128995.1_China/2018/AnhuiXCGQ ................................................................................ [162800]
MH681419.1_ASFV/POL/2015        ................................................................................ [162800]
LS478113.1_Estonia_2014         ................................................................................ [162800]
KP843857.1_Odintsovo_02/14      ................................................................................ [162800]
FN557520.1_strain_E75           ................................................................................ [162800]
AY261366.1_Warthog              ...............................................................................a [162800]
AY261365.1_Warmbaths            ...............................................................................a [162800]
AY261364.1_Tengani_62           ...............................................................................a [162800]
AY261363.1_Pretoriuskop/96/4    ...............................................................................a [162800]
AY261362.1_Mkuzi_1979           ................................................................................ [162800]
AY261361.1_Malawi_Lil-20/1_1983 ..........a.................................c................................... [162800]
AY261360.1_Kenya_1950           ............................................c................................... [162800]
AM712240.1_OURT_88/3            ................................................................................ [162800]
AM712239.1_Benin_97/1           ................................................................................ [162800]
KM111295.1_Ken06.Bus            ......................a.....................c................................... [162800]
```

```
KM111294.1_Ken05/Tk1             ............................................c..................................a [162800]
KM262844.1_strain_L60            ................................................................................ [162800]
KM102979.1_26544/OG10_Italy      ................................................................................ [162800]
KM262845.1_strain_NHV            ................................................................................ [162800]
KX354450.1_47/Ss/2008            ................................................................................ [162800]
KP055815.1_strain_BA71           ................................................................................ [162800]
MH025920.1_strain_R35            ......................a.....................c................................... [162800]
MH025919.1_strain_N10            ......................a.....................c................................... [162800]
MH025918.1_strain_R25            ......................a.....................c................................... [162800]
MH025917.1_strain_R7             ......................a.....................c................................... [162800]
MH025916.1_strain_R8             ......................a.....................c................................... [162800]

FR682468.2_Georgia_2007/1        tctgccacatcttgagcccccatccaggggatcacaaggtactcccccttaaagatgattcgtcgtttttttaaaaaatc [162880]
ASF_IND_20_CAD543                ................................................................................ [162880]
LR899193.1_Germany_2020/1        ................................................................................ [162880]
MN393477.1_Wuhan_2019            ................................................................................ [162880]
MW396979.1_Timor-Leste/2019/1    ................................................................................ [162880]
MK940252.1_CN/2019/InnerMongolia ................................................................................ [162880]
MK128995.1_China/2018/AnhuiXCGQ  ................................................................................ [162880]
MH681419.1_ASFV/POL/2015         ................................................................................ [162880]
LS478113.1_Estonia_2014          ................................................................................ [162880]
KP843857.1_Odintsovo_02/14       ................................................................................ [162880]
FN557520.1_strain_E75            ................................................................................ [162880]
AY261366.1_Warthog               .................t........................................c..................... [162880]
AY261365.1_Warmbaths             .............................a.................................................. [162880]
AY261364.1_Tengani_62            .................t........................................c..................... [162880]
AY261363.1_Pretoriuskop/96/4     .............................a.................................................. [162880]
AY261362.1_Mkuzi_1979            ................................................................................ [162880]
AY261361.1_Malawi_Lil-20/1_1983  ........t........t.......................tc..t.................t................ [162880]
AY261360.1_Kenya_1950            ........g........t........................c..t...................c..g........... [162880]
AM712240.1_OURT_88/3             ................................................................................ [162880]
AM712239.1_Benin_97/1            ................................................................................ [162880]
KM111295.1_Ken06.Bus             ..c.....g........t........................c..t...................c..gc.......... [162880]
KM111294.1_Ken05/Tk1             ........g........t........................c..t...................c..g........... [162880]
KM262844.1_strain_L60            ................................................................................ [162880]
KM102979.1_26544/OG10_Italy      ................................................................................ [162880]
KM262845.1_strain_NHV            ................................................................................ [162880]
KX354450.1_47/Ss/2008            ................................................................................ [162880]
KP055815.1_strain_BA71           ................................................................................ [162880]
MH025920.1_strain_R35            ..c.....g........t........................c..t...................c..gc.......... [162880]
MH025919.1_strain_N10            ..c.....g........t........................c..t...................c..gc.......... [162880]
MH025918.1_strain_R25            ..c.....g........t........................c..t...................c..gc.......... [162880]
MH025917.1_strain_R7             ..c.....g........t........................c..t...................c..gc.......... [162880]
MH025916.1_strain_R8             ..c.....g........t........................c..t...................c..gc.......... [162880]
```

```
FR682468.2_Georgia_2007/1        atgaaaacgttttaaagcttcaagaaaggggcagttgggctttgaccccaaaatgctgacgacgatatcctcgggcatga [162960]
ASF_IND_20_CAD543                ...............................nnnnnnnn......................................... [162960]
LR899193.1_Germany_2020/1        ................................................................................ [162960]
MN393477.1_Wuhan_2019            ................................................................................ [162960]
MW396979.1_Timor-Leste/2019/1    ................................................................................ [162960]
MK940252.1_CN/2019/InnerMongolia ................................................................................ [162960]
MK128995.1_China/2018/AnhuiXCGQ  ................................................................................ [162960]
MH681419.1_ASFV/POL/2015         ................................................................................ [162960]
LS478113.1_Estonia_2014          ................................................................................ [162960]
KP843857.1_Odintsovo_02/14       ................................................................................ [162960]
FN557520.1_strain_E75            ................................................................................ [162960]
AY261366.1_Warthog               ..............................a....................g............................ [162960]
AY261365.1_Warmbaths             ..............................a....................g............................ [162960]
AY261364.1_Tengani_62            ..............................a..............................................c.. [162960]
AY261363.1_Pretoriuskop/96/4     ..............................a....................g............................ [162960]
AY261362.1_Mkuzi_1979            ................................................................................ [162960]
AY261361.1_Malawi_Lil-20/1_1983  ..............................a..a...........t....g.........a....c.............. [162960]
AY261360.1_Kenya_1950            .............................................t................................t. [162960]
AM712240.1_OURT_88/3             ................................................................................ [162960]
AM712239.1_Benin_97/1            ................................................................................ [162960]
KM111295.1_Ken06.Bus             .............................................t................................t. [162960]
KM111294.1_Ken05/Tk1             .............................................t................................t. [162960]
KM262844.1_strain_L60            ................................................................................ [162960]
KM102979.1_26544/OG10_Italy      ................................................................................ [162960]
KM262845.1_strain_NHV            ................................................................................ [162960]
KX354450.1_47/Ss/2008            ................................................................................ [162960]
KP055815.1_strain_BA71           ................................................................................ [162960]
MH025920.1_strain_R35            .............................................t................................t. [162960]
MH025919.1_strain_N10            .............................................t................................t. [162960]
MH025918.1_strain_R25            .............................................t................................t. [162960]
MH025917.1_strain_R7             .............................................t................................t. [162960]
MH025916.1_strain_R8             .............................................t................................t. [162960]

FR682468.2_Georgia_2007/1        tgtattcgcagtgaggatagtagtttacggactctaattcagcggcccgccgttttatttcgtatcttgcccagttattc [163040]
ASF_IND_20_CAD543                ................................................................................ [163040]
LR899193.1_Germany_2020/1        ................................................................................ [163040]
MN393477.1_Wuhan_2019            ................................................................................ [163040]
MW396979.1_Timor-Leste/2019/1    ................................................................................ [163040]
MK940252.1_CN/2019/InnerMongolia ................................................................................ [163040]
MK128995.1_China/2018/AnhuiXCGQ  ................................................................................ [163040]
MH681419.1_ASFV/POL/2015         ................................................................................ [163040]
LS478113.1_Estonia_2014          ................................................................................ [163040]
KP843857.1_Odintsovo_02/14       ................................................................................ [163040]
```

```
FN557520.1_strain_E75           ................................................................................ [163040]
AY261366.1_Warthog              c............................................................................... [163040]
AY261365.1_Warmbaths            c............................................................................... [163040]
AY261364.1_Tengani_62           .................................................................t.............. [163040]
AY261363.1_Pretoriuskop/96/4    c................................................................t.............. [163040]
AY261362.1_Mkuzi_1979           ...............a................................................................ [163040]
AY261361.1_Malawi_Lil-20/1_1983 ..........a................................a.................................... [163040]
AY261360.1_Kenya_1950           ...........c................a.....c........a.....................t.............. [163040]
AM712240.1_OURT_88/3            ................................................................................ [163040]
AM712239.1_Benin_97/1           ................................................................................ [163040]
KM111295.1_Ken06.Bus            ...........c................a.....c..............................t.............. [163040]
KM111294.1_Ken05/Tk1            ...........c................a.....c........a.....................t.............. [163040]
KM262844.1_strain_L60           ................................................................................ [163040]
KM102979.1_26544/OG10_Italy     ................................................................................ [163040]
KM262845.1_strain_NHV           ................................................................................ [163040]
KX354450.1_47/Ss/2008           ................................................................................ [163040]
KP055815.1_strain_BA71          ................................................................................ [163040]
MH025920.1_strain_R35           ...........c................a.....c..............................t.............. [163040]
MH025919.1_strain_N10           ...........c................a.....c..............................t.............. [163040]
MH025918.1_strain_R25           ...........c................a.....c..............................t.............. [163040]
MH025917.1_strain_R7            ...........c................a.....c..............................t.............. [163040]
MH025916.1_strain_R8            ...........c................a.....c..............................t.............. [163040]

FR682468.2_Georgia_2007/1       agagagtactccacgcctccgaccacaacagacatcctatctattaaaaaataacaat-aaaaaccttatgaaatctatg [163120]
ASF_IND_20_CAD543               ..........................................................-..................... [163120]
LR899193.1_Germany_2020/1       ..........................................................-..................... [163120]
MN393477.1_Wuhan_2019           ..........................................................-..................... [163120]
MW396979.1_Timor-Leste/2019/1   ..........................................................-..................... [163120]
MK940252.1_CN/2019/InnerMongolia ..........................................................-..................... [163120]
MK128995.1_China/2018/AnhuiXCGQ ..........................................................-..................... [163120]
MH681419.1_ASFV/POL/2015        ..........................................................-..................... [163120]
LS478113.1_Estonia_2014         ..........................................................-..................... [163120]
KP843857.1_Odintsovo_02/14      ..........................................................-..................... [163120]
FN557520.1_strain_E75           ..........................................................-..................... [163120]
AY261366.1_Warthog              ..........................................................-..................... [163120]
AY261365.1_Warmbaths            ..........................................................-..................... [163120]
AY261364.1_Tengani_62           ..........................................................-............g........ [163120]
AY261363.1_Pretoriuskop/96/4    ........t.................................................-..................... [163120]
AY261362.1_Mkuzi_1979           ..........................................................-..................... [163120]
AY261361.1_Malawi_Lil-20/1_1983 .................c..................----..................-..................... [163120]
AY261360.1_Kenya_1950           .................c..................----..................a..................... [163120]
AM712240.1_OURT_88/3            ..........................................................-..................... [163120]
AM712239.1_Benin_97/1           ..........................................................-..................... [163120]
KM111295.1_Ken06.Bus            .................c..................----..................a..................... [163120]
```

```
KM111294.1_Ken05/Tk1             .................c..................----..................-..................... [163120]
KM262844.1_strain_L60            ..........................................................-..................... [163120]
KM102979.1_26544/OG10_Italy      ..........................................................-..................... [163120]
KM262845.1_strain_NHV            ..........................................................-..................... [163120]
KX354450.1_47/Ss/2008            ..........................................................-..................... [163120]
KP055815.1_strain_BA71           ..........................................................-..................... [163120]
MH025920.1_strain_R35            .................c..................----..................a..................... [163120]
MH025919.1_strain_N10            .................c..................----..................a..................... [163120]
MH025918.1_strain_R25            .................c..................----..................a..................... [163120]
MH025917.1_strain_R7             .................c..................----..................a..................... [163120]
MH025916.1_strain_R8             .................c..................----..................a..................... [163120]

FR682468.2_Georgia_2007/1        tatagtggccgct-aaaatgtctatattagaaaaaattacgtcaagtccctctgaatgcgcagagcatcttacaaacaaa [163200]
ASF_IND_20_CAD543                .............-.................................................................. [163200]
LR899193.1_Germany_2020/1        .............-.................................................................. [163200]
MN393477.1_Wuhan_2019            .............-.................................................................. [163200]
MW396979.1_Timor-Leste/2019/1    .............-.................................................................. [163200]
MK940252.1_CN/2019/InnerMongolia .............-.................................................................. [163200]
MK128995.1_China/2018/AnhuiXCGQ  .............-.................................................................. [163200]
MH681419.1_ASFV/POL/2015         .............-.................................................................. [163200]
LS478113.1_Estonia_2014          .............-.................................................................. [163200]
KP843857.1_Odintsovo_02/14       .............-.................................................................. [163200]
FN557520.1_strain_E75            .............-.....................................................c............ [163200]
AY261366.1_Warthog               .............-.....................................................c............ [163200]
AY261365.1_Warmbaths             .............-.......................c.............................c............ [163200]
AY261364.1_Tengani_62            .............-.....................................................c............ [163200]
AY261363.1_Pretoriuskop/96/4     .............-.....................................................c............ [163200]
AY261362.1_Mkuzi_1979            .............-.....................................................c............ [163200]
AY261361.1_Malawi_Lil-20/1_1983  .............-..t..................................................ca........... [163200]
AY261360.1_Kenya_1950            .............-..t.....c...........g....................g...........ca....t...... [163200]
AM712240.1_OURT_88/3             .............-.....................................................c............ [163200]
AM712239.1_Benin_97/1            .............-.....................................................c............ [163200]
KM111295.1_Ken06.Bus             .............a..t.....c...........g....................g...........ca....tg..... [163200]
KM111294.1_Ken05/Tk1             .............-..t.....c...........g....................g...........ca....t...... [163200]
KM262844.1_strain_L60            .............-.....................................................c............ [163200]
KM102979.1_26544/OG10_Italy      .............-.....................................................c............ [163200]
KM262845.1_strain_NHV            .............-.....................................................c............ [163200]
KX354450.1_47/Ss/2008            .............-.....................................................c............ [163200]
KP055815.1_strain_BA71           .............-.....................................................c............ [163200]
MH025920.1_strain_R35            .............a..t.....c...........g....................g...........ca....tg..... [163200]
MH025919.1_strain_N10            .............a..t.....c...........g....................g...........ca....tg..... [163200]
MH025918.1_strain_R25            .............a..t.....c...........g....................g...........ca....tg..... [163200]
MH025917.1_strain_R7             .............a..t.....c...........g....................g...........ca....tg..... [163200]
MH025916.1_strain_R8             .............a..t.....c...........g....................g...........ca....tg..... [163200]
```

```
FR682468.2_Georgia_2007/1        gatagctgtttaagtaaaaaaatacaaaaagagctcacctctttttttggaaaaaaaagagacactcggttgcgattcgga [163280]
ASF_IND_20_CAD543                ................................................................................ [163280]
LR899193.1_Germany_2020/1        ................................................................................ [163280]
MN393477.1_Wuhan_2019            ................................................................................ [163280]
MW396979.1_Timor-Leste/2019/1    ................................................................................ [163280]
MK940252.1_CN/2019/InnerMongolia ................................................................................ [163280]
MK128995.1_China/2018/AnhuiXCGQ  ................................................................................ [163280]
MH681419.1_ASFV/POL/2015         ................................................................................ [163280]
LS478113.1_Estonia_2014          ................................................................................ [163280]
KP843857.1_Odintsovo_02/14       ................................................................................ [163280]
FN557520.1_strain_E75            ................................................................................ [163280]
AY261366.1_Warthog               ................................................c...................c........... [163280]
AY261365.1_Warmbaths             ................................................c...................c........... [163280]
AY261364.1_Tengani_62            ................................................c...................c........... [163280]
AY261363.1_Pretoriuskop/96/4     ................................................c...................c........... [163280]
AY261362.1_Mkuzi_1979            ................................................................................ [163280]
AY261361.1_Malawi_Lil-20/1_1983  ................................................c.........................c..... [163280]
AY261360.1_Kenya_1950            ................................................c...................g.....c..a.. [163280]
AM712240.1_OURT_88/3             ................................................................................ [163280]
AM712239.1_Benin_97/1            ................................................................................ [163280]
KM111295.1_Ken06.Bus             t...............................................c...................g.....c..a.. [163280]
KM111294.1_Ken05/Tk1             ................................................c...................g.....c..a.. [163280]
KM262844.1_strain_L60            ................................................................................ [163280]
KM102979.1_26544/OG10_Italy      ................................................................................ [163280]
KM262845.1_strain_NHV            ................................................................................ [163280]
KX354450.1_47/Ss/2008            ................................................................................ [163280]
KP055815.1_strain_BA71           ................................................................................ [163280]
MH025920.1_strain_R35            t...............................................c...................g.....c..a.. [163280]
MH025919.1_strain_N10            t...............................................c...................g.....c..a.. [163280]
MH025918.1_strain_R25            t...............................................c...................g.....c..a.. [163280]
MH025917.1_strain_R7             t...............................................c...................g.....c..a.. [163280]
MH025916.1_strain_R8             t...............................................c...................g.....c..a.. [163280]

FR682468.2_Georgia_2007/1        gtcctgcgtaattacccaccccgccgtgaaggcctatgcgcaacaaaagggactggacctctccaaagaactggagactc [163360]
ASF_IND_20_CAD543                ................................................................................ [163360]
LR899193.1_Germany_2020/1        ................................................................................ [163360]
MN393477.1_Wuhan_2019            ................................................................................ [163360]
MW396979.1_Timor-Leste/2019/1    .............m...m.............................................................. [163360]
MK940252.1_CN/2019/InnerMongolia ................................................................................ [163360]
MK128995.1_China/2018/AnhuiXCGQ  ................................................................................ [163360]
MH681419.1_ASFV/POL/2015         ................................................................................ [163360]
LS478113.1_Estonia_2014          ................................................................................ [163360]
KP843857.1_Odintsovo_02/14       ................................................................................ [163360]
```

```
FN557520.1_strain_E75            ................................................................................ [163360]
AY261366.1_Warthog               ................................................................................ [163360]
AY261365.1_Warmbaths             ................................................................................ [163360]
AY261364.1_Tengani_62            .................................t......................................a....... [163360]
AY261363.1_Pretoriuskop/96/4     ................................................................................ [163360]
AY261362.1_Mkuzi_1979            .......................................a........................................ [163360]
AY261361.1_Malawi_Lil-20/1_1983  ...t.....c.....t....................c..............t..a......................... [163360]
AY261360.1_Kenya_1950            .........g..........................c........................................... [163360]
AM712240.1_OURT_88/3             ................................................................................ [163360]
AM712239.1_Benin_97/1            ................................................................................ [163360]
KM111295.1_Ken06.Bus             .........g..........................c........................................... [163360]
KM111294.1_Ken05/Tk1             .........g..........................c........................................... [163360]
KM262844.1_strain_L60            ................................................................................ [163360]
KM102979.1_26544/OG10_Italy      ................................................................................ [163360]
KM262845.1_strain_NHV            ................................................................................ [163360]
KX354450.1_47/Ss/2008            ................................................................................ [163360]
KP055815.1_strain_BA71           ................................................................................ [163360]
MH025920.1_strain_R35            .........g..........................c........................................... [163360]
MH025919.1_strain_N10            .........g..........................c........................................... [163360]
MH025918.1_strain_R25            .........g..........................c........................................... [163360]
MH025917.1_strain_R7             .........g..........................c........................................... [163360]
MH025916.1_strain_R8             .........g..........................c........................................... [163360]

FR682468.2_Georgia_2007/1        ggtttaaagcgccaggacccagaaacaacacgggtcttcttacaaacttcaatattgatgaaacgctgcagaggtgggcc [163440]
ASF_IND_20_CAD543                ................................................................................ [163440]
LR899193.1_Germany_2020/1        ................................................................................ [163440]
MN393477.1_Wuhan_2019            ................................................................................ [163440]
MW396979.1_Timor-Leste/2019/1    ................................................................................ [163440]
MK940252.1_CN/2019/InnerMongolia ................................................................................ [163440]
MK128995.1_China/2018/AnhuiXCGQ  ................................................................................ [163440]
MH681419.1_ASFV/POL/2015         ................................................................................ [163440]
LS478113.1_Estonia_2014          ................................................................................ [163440]
KP843857.1_Odintsovo_02/14       ................................................................................ [163440]
FN557520.1_strain_E75            .......................................................c........................ [163440]
AY261366.1_Warthog               .......................................................c........................ [163440]
AY261365.1_Warmbaths             .......................................................c........................ [163440]
AY261364.1_Tengani_62            ................................................................................ [163440]
AY261363.1_Pretoriuskop/96/4     .......................................................c........................ [163440]
AY261362.1_Mkuzi_1979            .......................................................c........................ [163440]
AY261361.1_Malawi_Lil-20/1_1983  ..................................c....................c.................a...... [163440]
AY261360.1_Kenya_1950            .......g........t.................c....................c........................ [163440]
AM712240.1_OURT_88/3             .......................................................c........................ [163440]
AM712239.1_Benin_97/1            .......................................................c........................ [163440]
KM111295.1_Ken06.Bus             .......g........t.................c....................c........................ [163440]
```

```
KM111294.1_Ken05/Tk1             .......g........t.................c....................c........................ [163440]
KM262844.1_strain_L60            .......................................................c........................ [163440]
KM102979.1_26544/OG10_Italy      .......................................................c........................ [163440]
KM262845.1_strain_NHV            .......................................................c........................ [163440]
KX354450.1_47/Ss/2008            .......................................................c........................ [163440]
KP055815.1_strain_BA71           .......................................................c........................ [163440]
MH025920.1_strain_R35            .......g........t.................c....................c........................ [163440]
MH025919.1_strain_N10            .......g........t.................c....................c........................ [163440]
MH025918.1_strain_R25            .......g........t.................c....................c........................ [163440]
MH025917.1_strain_R7             .......g........t.................c....................c........................ [163440]
MH025916.1_strain_R8             .......g........t.................c....................c........................ [163440]

FR682468.2_Georgia_2007/1        ataaaatacaccaagtttttcaactgtcctttttccatgatggactttgagagggtccattataaatttaatcaagtgga [163520]
ASF_IND_20_CAD543                ................................................................................ [163520]
LR899193.1_Germany_2020/1        ................................................................................ [163520]
MN393477.1_Wuhan_2019            ................................................................................ [163520]
MW396979.1_Timor-Leste/2019/1    ................................................................................ [163520]
MK940252.1_CN/2019/InnerMongolia ................................................................................ [163520]
MK128995.1_China/2018/AnhuiXCGQ  ................................................................................ [163520]
MH681419.1_ASFV/POL/2015         ................................................................................ [163520]
LS478113.1_Estonia_2014          ................................................................................ [163520]
KP843857.1_Odintsovo_02/14       ................................................................................ [163520]
FN557520.1_strain_E75            .....g.......................................................................... [163520]
AY261366.1_Warthog               .....g.....................................................c.................... [163520]
AY261365.1_Warmbaths             .....g.........................................c...........c.................... [163520]
AY261364.1_Tengani_62            .....g.....................................................c.................... [163520]
AY261363.1_Pretoriuskop/96/4     .....g.........................................c...........c.................... [163520]
AY261362.1_Mkuzi_1979            .....g..................................................t....................... [163520]
AY261361.1_Malawi_Lil-20/1_1983  .....g...............................................ca......................... [163520]
AY261360.1_Kenya_1950            .....g................................................a....c.................... [163520]
AM712240.1_OURT_88/3             .....g.......................................................................... [163520]
AM712239.1_Benin_97/1            .....g.......................................................................... [163520]
KM111295.1_Ken06.Bus             .....g................................................a....c.................... [163520]
KM111294.1_Ken05/Tk1             .....g................................................a....c.................... [163520]
KM262844.1_strain_L60            .....g.......................................................................... [163520]
KM102979.1_26544/OG10_Italy      .....g.......................................................................... [163520]
KM262845.1_strain_NHV            .....g.......................................................................... [163520]
KX354450.1_47/Ss/2008            .....g.......................................................................... [163520]
KP055815.1_strain_BA71           .....g.......................................................................... [163520]
MH025920.1_strain_R35            .....g................................................a....c.................... [163520]
MH025919.1_strain_N10            .....g................................................a....c.................... [163520]
MH025918.1_strain_R25            .....g................................................a....c.................... [163520]
MH025917.1_strain_R7             .....g................................................a....c.................... [163520]
MH025916.1_strain_R8             .....g................................................a....c.................... [163520]
```

```
FR682468.2_Georgia_2007/1        tatggtaaaggtatataagggagaagagctacaatatgtagaaggcaaagtggtcaagcgtccttgtaacaccttcggat [163600]
ASF_IND_20_CAD543                ................................................................................ [163600]
LR899193.1_Germany_2020/1        ................................................................................ [163600]
MN393477.1_Wuhan_2019            ................................................................................ [163600]
MW396979.1_Timor-Leste/2019/1    ................................................................................ [163600]
MK940252.1_CN/2019/InnerMongolia ................................................................................ [163600]
MK128995.1_China/2018/AnhuiXCGQ  ................................................................................ [163600]
MH681419.1_ASFV/POL/2015         ................................................................................ [163600]
LS478113.1_Estonia_2014          ................................................................................ [163600]
KP843857.1_Odintsovo_02/14       ................................................................................ [163600]
FN557520.1_strain_E75            .....................g.......................................................... [163600]
AY261366.1_Warthog               .....................g..g....................................................... [163600]
AY261365.1_Warmbaths             .....................g..g....................................................... [163600]
AY261364.1_Tengani_62            .....................g..g....................................................... [163600]
AY261363.1_Pretoriuskop/96/4     .....................g..g....................................................... [163600]
AY261362.1_Mkuzi_1979            .....................g.....................................................t.... [163600]
AY261361.1_Malawi_Lil-20/1_1983  .....c...............g..g...........c.............c...g......................... [163600]
AY261360.1_Kenya_1950            .....................g..g...........c..g..........c...g......................... [163600]
AM712240.1_OURT_88/3             .....................g.......................................................... [163600]
AM712239.1_Benin_97/1            .....................g.......................................................... [163600]
KM111295.1_Ken06.Bus             .....................g..g...........c..g..........c...g......................... [163600]
KM111294.1_Ken05/Tk1             .....................g..g...........c..g..........c...g......................... [163600]
KM262844.1_strain_L60            .....................g.......................................................... [163600]
KM102979.1_26544/OG10_Italy      .....................g.......................................................... [163600]
KM262845.1_strain_NHV            .....................g.......................................................... [163600]
KX354450.1_47/Ss/2008            .....................g.......................................................... [163600]
KP055815.1_strain_BA71           .....................g.......................................................... [163600]
MH025920.1_strain_R35            .....................g.gg...........c..g..........c...g......................... [163600]
MH025919.1_strain_N10            .....................g.gg...........c..g..........c...g......................... [163600]
MH025918.1_strain_R25            .....................g.gg...........c..g..........c...g......................... [163600]
MH025917.1_strain_R7             .....................g.gg...........c..g..........c...g......................... [163600]
MH025916.1_strain_R8             .....................g.gg...........c..g..........c...g......................... [163600]

FR682468.2_Georgia_2007/1        gcgttttaaacacggacttttcaacgggcactggaaaacactgggtagccatctttgtggatatgcggggcgactgctgg [163680]
ASF_IND_20_CAD543                ................................................................................ [163680]
LR899193.1_Germany_2020/1        ................................................................................ [163680]
MN393477.1_Wuhan_2019            ................................................................................ [163680]
MW396979.1_Timor-Leste/2019/1    ................................................................................ [163680]
MK940252.1_CN/2019/InnerMongolia ................................................................................ [163680]
MK128995.1_China/2018/AnhuiXCGQ  ................................................................................ [163680]
MH681419.1_ASFV/POL/2015         ................................................................................ [163680]
LS478113.1_Estonia_2014          ................................................................................ [163680]
KP843857.1_Odintsovo_02/14       ................................................................................ [163680]
```

```
FN557520.1_strain_E75            ................................................................................ [163680]
AY261366.1_Warthog               .....................................g..t....................................... [163680]
AY261365.1_Warmbaths             .....................................g..t....................................... [163680]
AY261364.1_Tengani_62            .............a....a..................g..t....................................... [163680]
AY261363.1_Pretoriuskop/96/4     .....................................g..t....................................... [163680]
AY261362.1_Mkuzi_1979            ................................................................................ [163680]
AY261361.1_Malawi_Lil-20/1_1983  .........................a..tc.a.....g..t....................................... [163680]
AY261360.1_Kenya_1950            ............................t..g.....g..t.....t................................. [163680]
AM712240.1_OURT_88/3             ................................................................................ [163680]
AM712239.1_Benin_97/1            ................................................................................ [163680]
KM111295.1_Ken06.Bus             .........................a..t..g.....g..t.....t................................. [163680]
KM111294.1_Ken05/Tk1             .........................a..t..g.....g..t.....t................................. [163680]
KM262844.1_strain_L60            ................................................................................ [163680]
KM102979.1_26544/OG10_Italy      ................................................................................ [163680]
KM262845.1_strain_NHV            ................................................................................ [163680]
KX354450.1_47/Ss/2008            ................................................................................ [163680]
KP055815.1_strain_BA71           ................................................................................ [163680]
MH025920.1_strain_R35            .........................a..t..g.....g..t.....t................................. [163680]
MH025919.1_strain_N10            .........................a..t..g.....g..t.....t................................. [163680]
MH025918.1_strain_R25            .........................a..t..g.....g..t.....t................................. [163680]
MH025917.1_strain_R7             .........................a..t..g.....g..t.....t................................. [163680]
MH025916.1_strain_R8             .........................a..t..g.....g..t.....t................................. [163680]

FR682468.2_Georgia_2007/1        agcatcgaatattttaattcgacgggaaattctcctccaggtcccgttattcgttggatggaacgggtcaaacagcagct [163760]
ASF_IND_20_CAD543                ................................................................................ [163760]
LR899193.1_Germany_2020/1        ................................................................................ [163760]
MN393477.1_Wuhan_2019            ................................................................................ [163760]
MW396979.1_Timor-Leste/2019/1    ................................................................................ [163760]
MK940252.1_CN/2019/InnerMongolia ................................................................................ [163760]
MK128995.1_China/2018/AnhuiXCGQ  ................................................................................ [163760]
MH681419.1_ASFV/POL/2015         ................................................................................ [163760]
LS478113.1_Estonia_2014          ................................................................................ [163760]
KP843857.1_Odintsovo_02/14       ................................................................................ [163760]
FN557520.1_strain_E75            .....................g..........c..c-................c.......................... [163760]
AY261366.1_Warthog               .....................g..........c....................c.......................... [163760]
AY261365.1_Warmbaths             .....................g..........c..c.................c.......................... [163760]
AY261364.1_Tengani_62            ....................ag..........c..g.................c.......................... [163760]
AY261363.1_Pretoriuskop/96/4     .....................g........g.c..c..g..............c.......................... [163760]
AY261362.1_Mkuzi_1979            .....................g..........c..c.................c.......................... [163760]
AY261361.1_Malawi_Lil-20/1_1983  .....t...............g......................g........c........g................. [163760]
AY261360.1_Kenya_1950            .....t...............g..........c...........g........c.......................... [163760]
AM712240.1_OURT_88/3             .....................g..........c..c.................c.......................... [163760]
AM712239.1_Benin_97/1            .....................g..........c..c.................c.......................... [163760]
KM111295.1_Ken06.Bus             .....t...............g....c.....c...........g........c.......................... [163760]
```

```
KM111294.1_Ken05/Tk1          .....t...............g..........c...........g........c.......................... [163760]
KM262844.1_strain_L60         .....................g..........c..c.................c.......................... [163760]
KM102979.1_26544/OG10_Italy   .....................g..........c..c.................c.......................... [163760]
KM262845.1_strain_NHV         .....................g..........c..c.................c.......................... [163760]
KX354450.1_47/Ss/2008         .....................g..........c..c.................c.......................... [163760]
KP055815.1_strain_BA71        .....................g..........c..c.................c.......................... [163760]
MH025920.1_strain_R35         .....t...............g....c.....c...........g........c.......................... [163760]
MH025919.1_strain_N10         .....t...............g....c.....c...........g........c.......................... [163760]
MH025918.1_strain_R25         .....t...............g....c.....c...........g........c.......................... [163760]
MH025917.1_strain_R7          .....t...............g....c.....c...........g........c.......................... [163760]
MH025916.1_strain_R8          .....t...............g....c.....c...........g........c.......................... [163760]

FR682468.2_Georgia_2007/1     attaaaaatacaccacaccgtgaaaacgcttgcagttaccaacattcgtcaccaacggtcgcagaccgagtgcggcccct [163840]
ASF_IND_20_CAD543             ................................................................................ [163840]
LR899193.1_Germany_2020/1     ................................................................................ [163840]
MN393477.1_Wuhan_2019         ................................................................................ [163840]
MW396979.1_Timor-Leste/2019/1 ................................................................................ [163840]
MK940252.1_CN/2019/InnerMongolia ................................................................................ [163840]
MK128995.1_China/2018/AnhuiXCGQ ................................................................................ [163840]
MH681419.1_ASFV/POL/2015      ................................................................................ [163840]
LS478113.1_Estonia_2014       ................................................................................ [163840]
KP843857.1_Odintsovo_02/14    ................................................................................ [163840]
FN557520.1_strain_E75         .................................g....................g......................... [163840]
AY261366.1_Warthog            .................................g....................g......................... [163840]
AY261365.1_Warmbaths          .................................g....................g......................... [163840]
AY261364.1_Tengani_62         .................................g....................g......................... [163840]
AY261363.1_Pretoriuskop/96/4  .................................g..............c.....g......................... [163840]
AY261362.1_Mkuzi_1979         .................................g....................g......................... [163840]
AY261361.1_Malawi_Lil-20/1_1983 .................................g....................g......................... [163840]
AY261360.1_Kenya_1950         .....................t...........g..g.................g......................... [163840]
AM712240.1_OURT_88/3          .................................g....................g......................... [163840]
AM712239.1_Benin_97/1         .................................g....................g......................... [163840]
KM111295.1_Ken06.Bus          .....................t...........g..g..t..t...........g......................... [163840]
KM111294.1_Ken05/Tk1          ...................a.t........g..g..g.................g......................... [163840]
KM262844.1_strain_L60         .................................g....................g......................... [163840]
KM102979.1_26544/OG10_Italy   .................................g....................g......................... [163840]
KM262845.1_strain_NHV         .................................g....................g......................... [163840]
KX354450.1_47/Ss/2008         .................................g....................g......................... [163840]
KP055815.1_strain_BA71        .................................g....................g......................... [163840]
MH025920.1_strain_R35         .....................t..........tg..g..t..t...........g......................... [163840]
MH025919.1_strain_N10         .....................t..........tg..g..t..t...........g......................... [163840]
MH025918.1_strain_R25         .....................t..........tg..g..t..t...........g......................... [163840]
MH025917.1_strain_R7          .....................t..........tg..g..t..t...........g......................... [163840]
MH025916.1_strain_R8          .....................t..........tg..g..t..t...........g......................... [163840]
```

```
FR682468.2_Georgia_2007/1        acagcctgttttacatcagggcacgcctcgacaacgtgtcatacgcccattttatatccgctaggattaccgacgaagac [163920]
ASF_IND_20_CAD543                ................................................................................ [163920]
LR899193.1_Germany_2020/1        ................................................................................ [163920]
MN393477.1_Wuhan_2019            ................................................................................ [163920]
MW396979.1_Timor-Leste/2019/1    ................................................................................ [163920]
MK940252.1_CN/2019/InnerMongolia ................................................................................ [163920]
MK128995.1_China/2018/AnhuiXCGQ  ................................................................................ [163920]
MH681419.1_ASFV/POL/2015         ................................................................................ [163920]
LS478113.1_Estonia_2014          ................................................................................ [163920]
KP843857.1_Odintsovo_02/14       ................................................................................ [163920]
FN557520.1_strain_E75            ................................................................................ [163920]
AY261366.1_Warthog               ............................t................................................... [163920]
AY261365.1_Warmbaths             ................................................................................ [163920]
AY261364.1_Tengani_62            ................................................................................ [163920]
AY261363.1_Pretoriuskop/96/4     .....t.......................................................................... [163920]
AY261362.1_Mkuzi_1979            ................................................................................ [163920]
AY261361.1_Malawi_Lil-20/1_1983  .t.................a........................a.............ta...a.............a.t [163920]
AY261360.1_Kenya_1950            .t...............c..........................a.............ta...................a [163920]
AM712240.1_OURT_88/3             ................................................................................ [163920]
AM712239.1_Benin_97/1            ................................................................................ [163920]
KM111295.1_Ken06.Bus             .t...............c.a........................a.............ta.................... [163920]
KM111294.1_Ken05/Tk1             .t...............c..........................a.............ta...................a [163920]
KM262844.1_strain_L60            ................................................................................ [163920]
KM102979.1_26544/OG10_Italy      ................................................................................ [163920]
KM262845.1_strain_NHV            ................................................................................ [163920]
KX354450.1_47/Ss/2008            ................................................................................ [163920]
KP055815.1_strain_BA71           ................................................................................ [163920]
MH025920.1_strain_R35            .t...............c.a........................a.............ta.................... [163920]
MH025919.1_strain_N10            .t...............c.a........................a.............ta.................... [163920]
MH025918.1_strain_R25            .t...............c.a........................a.............ta.................... [163920]
MH025917.1_strain_R7             .t...............c.a........................a.............ta.................... [163920]
MH025916.1_strain_R8             .t...............c.a........................a.............ta.................... [163920]

FR682468.2_Georgia_2007/1        atgtataagtttagaacccatctgtttcgcatcgcataaactaataaagtttgaattctttataggaataaaaatggaag [164000]
ASF_IND_20_CAD543                ................................................................................ [164000]
LR899193.1_Germany_2020/1        ................................................................................ [164000]
MN393477.1_Wuhan_2019            ................................................................................ [164000]
MW396979.1_Timor-Leste/2019/1    ................................................................................ [164000]
MK940252.1_CN/2019/InnerMongolia ................................................................................ [164000]
MK128995.1_China/2018/AnhuiXCGQ  ................................................................................ [164000]
MH681419.1_ASFV/POL/2015         ................................................................................ [164000]
LS478113.1_Estonia_2014          ................................................................................ [164000]
KP843857.1_Odintsovo_02/14       ................................................................................ [164000]
```

FN557520.1_strain_E75 ................................................................................ [164000]
AY261366.1_Warthog .................................................a.............................. [164000]
AY261365.1_Warmbaths ................................................................................ [164000]
AY261364.1_Tengani_62 ................................................................................ [164000]
AY261363.1_Pretoriuskop/96/4 ................................................................................ [164000]
AY261362.1_Mkuzi_1979 ................................................................................ [164000]
AY261361.1_Malawi_Lil-20/1_1983 ..................................................................t............. [164000]
AY261360.1_Kenya_1950 .......................a.........................a.......t.....................a [164000]
AM712240.1_OURT_88/3 ................................................................................ [164000]
AM712239.1_Benin_97/1 ................................................................................ [164000]
KM111295.1_Ken06.Bus .................................................a.............................a [164000]
KM111294.1_Ken05/Tk1 .......................a.........................a.......t.....................a [164000]
KM262844.1_strain_L60 ................................................................................ [164000]
KM102979.1_26544/OG10_Italy ................................................................................ [164000]
KM262845.1_strain_NHV ................................................................................ [164000]
KX354450.1_47/Ss/2008 ................................................................................ [164000]
KP055815.1_strain_BA71 ................................................................................ [164000]
MH025920.1_strain_R35 .................................................a.............................a [164000]
MH025919.1_strain_N10 .................................................a.............................a [164000]
MH025918.1_strain_R25 .................................................a.............................a [164000]
MH025917.1_strain_R7 .................................................a.............................a [164000]
MH025916.1_strain_R8 .................................................a.............................a [164000]

FR682468.2_Georgia_2007/1 cgtttgaaatcagcgatttcaaagagcatgcgaagaaaaaaagcatgtgggctggcgccctcaacaaagtcactatttcg [164080]
ASF_IND_20_CAD543 ................................................................................ [164080]
LR899193.1_Germany_2020/1 ................................................................................ [164080]
MN393477.1_Wuhan_2019 ................................................................................ [164080]
MW396979.1_Timor-Leste/2019/1 ................................................................................ [164080]
MK940252.1_CN/2019/InnerMongolia ................................................................................ [164080]
MK128995.1_China/2018/AnhuiXCGQ ................................................................................ [164080]
MH681419.1_ASFV/POL/2015 ................................................................................ [164080]
LS478113.1_Estonia_2014 ................................................................................ [164080]
KP843857.1_Odintsovo_02/14 ................................................................................ [164080]
FN557520.1_strain_E75 ................................................................................ [164080]
AY261366.1_Warthog ................................................................................ [164080]
AY261365.1_Warmbaths ................................................................................ [164080]
AY261364.1_Tengani_62 ................................................................................ [164080]
AY261363.1_Pretoriuskop/96/4 ................................................................................ [164080]
AY261362.1_Mkuzi_1979 ................................................................................ [164080]
AY261361.1_Malawi_Lil-20/1_1983 .......................................................t........................ [164080]
AY261360.1_Kenya_1950 .......................................................t........................ [164080]
AM712240.1_OURT_88/3 ................................................................................ [164080]
AM712239.1_Benin_97/1 ................................................................................ [164080]
KM111295.1_Ken06.Bus .............t.........................................t........................ [164080]

```
KM111294.1_Ken05/Tk1                ...................t...................................t........................ [164080]
KM262844.1_strain_L60               ................................................................................ [164080]
KM102979.1_26544/OG10_Italy         ................................................................................ [164080]
KM262845.1_strain_NHV               ................................................................................ [164080]
KX354450.1_47/Ss/2008               ................................................................................ [164080]
KP055815.1_strain_BA71              ................................................................................ [164080]
MH025920.1_strain_R35               .............t.........................................t........................ [164080]
MH025919.1_strain_N10               .............t.........................................t........................ [164080]
MH025918.1_strain_R25               .............t.........................................t........................ [164080]
MH025917.1_strain_R7                .............t.........................................t........................ [164080]
MH025916.1_strain_R8                .............t.........................................t........................ [164080]

FR682468.2_Georgia_2007/1           ggtcttatgggggtctttaccgaagatgaggaccttatggcgttacccattcacagagaccactgccccgctttgttaaa [164160]
ASF_IND_20_CAD543                   .................................................................y.............. [164160]
LR899193.1_Germany_2020/1           ................................................................................ [164160]
MN393477.1_Wuhan_2019               ................................................................................ [164160]
MW396979.1_Timor-Leste/2019/1       ................................................................................ [164160]
MK940252.1_CN/2019/InnerMongolia    ................................................................................ [164160]
MK128995.1_China/2018/AnhuiXCGQ     ................................................................................ [164160]
MH681419.1_ASFV/POL/2015            ................................................................................ [164160]
LS478113.1_Estonia_2014             ................................................................................ [164160]
KP843857.1_Odintsovo_02/14          ................................................................................ [164160]
FN557520.1_strain_E75               .................................................................t.............. [164160]
AY261366.1_Warthog                  .........................................a.....t................................ [164160]
AY261365.1_Warmbaths                .........................................a...................................... [164160]
AY261364.1_Tengani_62               ................................................................................ [164160]
AY261363.1_Pretoriuskop/96/4        .........................................a.......................t.............. [164160]
AY261362.1_Mkuzi_1979               .................................................................t.............. [164160]
AY261361.1_Malawi_Lil-20/1_1983     ....................a.....................c................t.....t.............. [164160]
AY261360.1_Kenya_1950               ....................a.....................c......................t.............. [164160]
AM712240.1_OURT_88/3                .................................................................t.............. [164160]
AM712239.1_Benin_97/1               .................................................................t.............. [164160]
KM111295.1_Ken06.Bus                ....................a.....................c..................................... [164160]
KM111294.1_Ken05/Tk1                ....................a.....................c.....g................t.............. [164160]
KM262844.1_strain_L60               .................................................................t.............. [164160]
KM102979.1_26544/OG10_Italy         .................................................................t.............. [164160]
KM262845.1_strain_NHV               .................................................................t.............. [164160]
KX354450.1_47/Ss/2008               .................................................................t.............. [164160]
KP055815.1_strain_BA71              .................................................................t.............. [164160]
MH025920.1_strain_R35               ....................a.....................c..................................... [164160]
MH025919.1_strain_N10               ....................a.....................c..................................... [164160]
MH025918.1_strain_R25               ....................a.....................c..................................... [164160]
MH025917.1_strain_R7                ....................a.....................c..................................... [164160]
MH025916.1_strain_R8                ....................a.....................c..................................... [164160]
```

```
FR682468.2_Georgia_2007/1        aatttttgacgagatcatcgtaaatgccacggatcatgaaagagcttgccataacaaaacaaaaaaggtaacttacatta [164240]
ASF_IND_20_CAD543                ........................................................................y....... [164240]
LR899193.1_Germany_2020/1        ................................................................................ [164240]
MN393477.1_Wuhan_2019            ................................................................................ [164240]
MW396979.1_Timor-Leste/2019/1    ................................................................................ [164240]
MK940252.1_CN/2019/InnerMongolia ................................................................................ [164240]
MK128995.1_China/2018/AnhuiXCGQ  ................................................................................ [164240]
MH681419.1_ASFV/POL/2015         ................................................................................ [164240]
LS478113.1_Estonia_2014          ................................................................................ [164240]
KP843857.1_Odintsovo_02/14       ................................................................................ [164240]
FN557520.1_strain_E75            .............c.......................................g..................c.....c. [164240]
AY261366.1_Warthog               .....................................................g..................c.....c. [164240]
AY261365.1_Warmbaths             .............c.......................................g....g...................c. [164240]
AY261364.1_Tengani_62            .............g.............g....................t....g..................c.....c. [164240]
AY261363.1_Pretoriuskop/96/4     .....................................................g........................c. [164240]
AY261362.1_Mkuzi_1979            .............c....a..................................g..................c.....c. [164240]
AY261361.1_Malawi_Lil-20/1_1983  ...c.........c.......................................g............a.....c.....c. [164240]
AY261360.1_Kenya_1950            .............c..........c.........................................a.....c..t..c. [164240]
AM712240.1_OURT_88/3             .............c.......................................g..................c.....c. [164240]
AM712239.1_Benin_97/1            .............c.......................................g..................c.....c. [164240]
KM111295.1_Ken06.Bus             .............c..........c.........................................a.....c.....c. [164240]
KM111294.1_Ken05/Tk1             .............c..........c.........................................a.....c..t..c. [164240]
KM262844.1_strain_L60            .............c.......................................g..................c.....c. [164240]
KM102979.1_26544/OG10_Italy      .............c.......................................g..................c.....c. [164240]
KM262845.1_strain_NHV            .............c.......................................g..................c.....c. [164240]
KX354450.1_47/Ss/2008            .............c.......................................g..................c.....c. [164240]
KP055815.1_strain_BA71           .............c.......................................g..................c.....c. [164240]
MH025920.1_strain_R35            .............c..........c.........................................a.....c.....c. [164240]
MH025919.1_strain_N10            .............c..........c.........................................a.....c.....c. [164240]
MH025918.1_strain_R25            .............c..........c.........................................a.....c.....c. [164240]
MH025917.1_strain_R7             .............c..........c.........................................a.....c.....c. [164240]
MH025916.1_strain_R8             .............c..........c.........................................a.....c.....c. [164240]

FR682468.2_Georgia_2007/1        aaatttcgtttgataaaggtgtgttttcttgcgaaaacgatggcccgggaatccccattgcaaagcatgagcaagccagt [164320]
ASF_IND_20_CAD543                .........................................................................r...... [164320]
LR899193.1_Germany_2020/1        ................................................................................ [164320]
MN393477.1_Wuhan_2019            ................................................................................ [164320]
MW396979.1_Timor-Leste/2019/1    ................................................................................ [164320]
MK940252.1_CN/2019/InnerMongolia ................................................................................ [164320]
MK128995.1_China/2018/AnhuiXCGQ  ................................................................................ [164320]
MH681419.1_ASFV/POL/2015         ................................................................................ [164320]
LS478113.1_Estonia_2014          ................................................................................ [164320]
KP843857.1_Odintsovo_02/14       ................................................................................ [164320]
```

```
FN557520.1_strain_E75          ...................c.....................................................g...... [164320]
AY261366.1_Warthog             ...........a....................................................a........g...... [164320]
AY261365.1_Warmbaths           ...........................................................a.............g...... [164320]
AY261364.1_Tengani_62          ...........a.............................................................g...... [164320]
AY261363.1_Pretoriuskop/96/4   .........................................................................g...... [164320]
AY261362.1_Mkuzi_1979          ...................c.....................................................g...... [164320]
AY261361.1_Malawi_Lil-20/1_1983 ......................a...g..............................................g...... [164320]
AY261360.1_Kenya_1950          ....................................................t.......t............g.....c [164320]
AM712240.1_OURT_88/3           ...................c.....................................................g...... [164320]
AM712239.1_Benin_97/1          ...................c.....................................................g...... [164320]
KM111295.1_Ken06.Bus           ...............................t............................t............g.....c [164320]
KM111294.1_Ken05/Tk1           ....................................................t.......t............g.....c [164320]
KM262844.1_strain_L60          ...................c.....................................................g...... [164320]
KM102979.1_26544/OG10_Italy    ...................c.....................................................g...... [164320]
KM262845.1_strain_NHV          ...................c.....................................................g...... [164320]
KX354450.1_47/Ss/2008          ...................c.....................................................g...... [164320]
KP055815.1_strain_BA71         ...................c.....................................................g...... [164320]
MH025920.1_strain_R35          ...............................t............................t............g.....c [164320]
MH025919.1_strain_N10          ...............................t............................t............g.....c [164320]
MH025918.1_strain_R25          ...............................t............................t............g.....c [164320]
MH025917.1_strain_R7           ...............................t............................t............g.....c [164320]
MH025916.1_strain_R8           ...............................t............................t............g.....c [164320]

FR682468.2_Georgia_2007/1      cttatcgccaagcgcgatgtgtatgttcccgaggtggcttcatgtcactttttagccggaacgaacatcaataaggccaa [164400]
ASF_IND_20_CAD543              ............................................ctt....c............................ [164400]
LR899193.1_Germany_2020/1      ................................................................................ [164400]
MN393477.1_Wuhan_2019          ................................................................................ [164400]
MW396979.1_Timor-Leste/2019/1  ................................................................................ [164400]
MK940252.1_CN/2019/InnerMongolia ................................................................................ [164400]
MK128995.1_China/2018/AnhuiXCGQ ................................................................................ [164400]
MH681419.1_ASFV/POL/2015       ................................................................................ [164400]
LS478113.1_Estonia_2014        ................................................................................ [164400]
KP843857.1_Odintsovo_02/14     ................................................................................ [164400]
FN557520.1_strain_E75          ............................................ctt....c............................ [164400]
AY261366.1_Warthog             ..............g.............................c......c............................ [164400]
AY261365.1_Warmbaths           ..............g.............................ct.t...c.......g.................... [164400]
AY261364.1_Tengani_62          ..............g.............................c......c.......g.................... [164400]
AY261363.1_Pretoriuskop/96/4   ..............g.............................c......c............................ [164400]
AY261362.1_Mkuzi_1979          ............................................ctt....c............................ [164400]
AY261361.1_Malawi_Lil-20/1_1983 ..............g.............................c......c....t....................... [164400]
AY261360.1_Kenya_1950          ........a.................c.................ct.t...c..........a................. [164400]
AM712240.1_OURT_88/3           ............................................ctt....c............................ [164400]
AM712239.1_Benin_97/1          ............................................ctt....c............................ [164400]
KM111295.1_Ken06.Bus           ........a.................c........a........ct.t...c..........a..............t.. [164400]
```

```
KM111294.1_Ken05/Tk1          ........a.................c.................ct.t...c..........a................. [164400]
KM262844.1_strain_L60         ............................................ctt....c............................ [164400]
KM102979.1_26544/OG10_Italy   ............................................ctt....c............................ [164400]
KM262845.1_strain_NHV         ............................................ctt....c............................ [164400]
KX354450.1_47/Ss/2008         ............................................ctt....c............................ [164400]
KP055815.1_strain_BA71        ............................................ctt....c............................ [164400]
MH025920.1_strain_R35         ........a.................c........a........ct.t...c..........a..............t.. [164400]
MH025919.1_strain_N10         ........a.................c........a........ct.t...c..........a..............t.. [164400]
MH025918.1_strain_R25         ........a.................c........a........ct.t...c..........a..............t.. [164400]
MH025917.1_strain_R7          ........a.................c........a........ct.t...c..........a..............t.. [164400]
MH025916.1_strain_R8          ........a.................c........a........ct.t...c..........a..............t.. [164400]

FR682468.2_Georgia_2007/1     ggactgtatcaaggggggaaccaacggcgtcgggctgaagctcgccatggtgcattcgcagtgggccattcttaccaccg [164480]
ASF_IND_20_CAD543             ................................................................................ [164480]
LR899193.1_Germany_2020/1     ................................................................................ [164480]
MN393477.1_Wuhan_2019         ................................................................................ [164480]
MW396979.1_Timor-Leste/2019/1 ................................................................................ [164480]
MK940252.1_CN/2019/InnerMongolia ................................................................................ [164480]
MK128995.1_China/2018/AnhuiXCGQ ................................................................................ [164480]
MH681419.1_ASFV/POL/2015      ................................................................................ [164480]
LS478113.1_Estonia_2014       ................................................................................ [164480]
KP843857.1_Odintsovo_02/14    ................................................................................ [164480]
FN557520.1_strain_E75         ................................................................................ [164480]
AY261366.1_Warthog            ................................................................................ [164480]
AY261365.1_Warmbaths          .................................a.............................................. [164480]
AY261364.1_Tengani_62         ................................................................................ [164480]
AY261363.1_Pretoriuskop/96/4  ................................................................................ [164480]
AY261362.1_Mkuzi_1979         ................................................................................ [164480]
AY261361.1_Malawi_Lil-20/1_1983 .........................................................a...................... [164480]
AY261360.1_Kenya_1950         ..................g...........t..........................a...................... [164480]
AM712240.1_OURT_88/3          ................................................................................ [164480]
AM712239.1_Benin_97/1         ................................................................................ [164480]
KM111295.1_Ken06.Bus          ..................g...........t..........................a...................... [164480]
KM111294.1_Ken05/Tk1          ..................g...........t..........................a...................... [164480]
KM262844.1_strain_L60         ................................................................................ [164480]
KM102979.1_26544/OG10_Italy   ................................................................................ [164480]
KM262845.1_strain_NHV         ................................................................................ [164480]
KX354450.1_47/Ss/2008         ................................................................................ [164480]
KP055815.1_strain_BA71        ................................................................................ [164480]
MH025920.1_strain_R35         ..................g...........t..........................a...................... [164480]
MH025919.1_strain_N10         ..................g...........t..........................a...................... [164480]
MH025918.1_strain_R25         ..................g...........t..........................a...................... [164480]
MH025917.1_strain_R7          ..................g...........t..........................a...................... [164480]
MH025916.1_strain_R8          ..................g...........t..........................a...................... [164480]
```

```
FR682468.2_Georgia_2007/1        ccgacggcgcgcaaaagtatgttcaacatatcaaccaacgcctagatatcattgagcctcctaccattacaccctccagg [164560]
ASF_IND_20_CAD543                ................................................................................ [164560]
LR899193.1_Germany_2020/1        ................................................................................ [164560]
MN393477.1_Wuhan_2019            ................................................................................ [164560]
MW396979.1_Timor-Leste/2019/1    ................................................................................ [164560]
MK940252.1_CN/2019/InnerMongolia ................................................................................ [164560]
MK128995.1_China/2018/AnhuiXCGQ  ................................................................................ [164560]
MH681419.1_ASFV/POL/2015         ................................................................................ [164560]
LS478113.1_Estonia_2014          ................................................................................ [164560]
KP843857.1_Odintsovo_02/14       ................................................................................ [164560]
FN557520.1_strain_E75            ............................a........g.......................................... [164560]
AY261366.1_Warthog               .....................................g......................................t... [164560]
AY261365.1_Warmbaths             .....................................g.......................................... [164560]
AY261364.1_Tengani_62            .....................................g.......................................... [164560]
AY261363.1_Pretoriuskop/96/4     .....................................g.......................................... [164560]
AY261362.1_Mkuzi_1979            .....................................g.......................................... [164560]
AY261361.1_Malawi_Lil-20/1_1983  ...............................a.....g.....g....................g............... [164560]
AY261360.1_Kenya_1950            ...............................a.....g..........a.........a....................a [164560]
AM712240.1_OURT_88/3             ............................a........g.......................................... [164560]
AM712239.1_Benin_97/1            ............................a........g.......................................... [164560]
KM111295.1_Ken06.Bus             ........t......................a.....g..........a.........a....................a [164560]
KM111294.1_Ken05/Tk1             ...............................a.....g..........a.........a....................a [164560]
KM262844.1_strain_L60            ............................a........g.......................................... [164560]
KM102979.1_26544/OG10_Italy      ............................a........g.......................................... [164560]
KM262845.1_strain_NHV            ............................a........g.......................................... [164560]
KX354450.1_47/Ss/2008            ............................a........g.......................................... [164560]
KP055815.1_strain_BA71           ............................a........g.......................................... [164560]
MH025920.1_strain_R35            ........t......................a.....g..........a.........a....................a [164560]
MH025919.1_strain_N10            ........t......................a.....g..........a.........a....................a [164560]
MH025918.1_strain_R25            ........t......................a.....g..........a.........a....................a [164560]
MH025917.1_strain_R7             ........t......................a.....g..........a.........a....................a [164560]
MH025916.1_strain_R8             ........t......................a.....g..........a.........a....................a [164560]

FR682468.2_Georgia_2007/1        gaaatgtttacacgtatcgagctcatgcccgtataccaggaactagggtacgcggagcctctgtctgaaacagagcaggc [164640]
ASF_IND_20_CAD543                ................................................................................ [164640]
LR899193.1_Germany_2020/1        ................................................................................ [164640]
MN393477.1_Wuhan_2019            ................................................................................ [164640]
MW396979.1_Timor-Leste/2019/1    ................................................................................ [164640]
MK940252.1_CN/2019/InnerMongolia ................................................................................ [164640]
MK128995.1_China/2018/AnhuiXCGQ  ................................................................................ [164640]
MH681419.1_ASFV/POL/2015         ................................................................................ [164640]
LS478113.1_Estonia_2014          ................................................................................ [164640]
KP843857.1_Odintsovo_02/14       ................................................................................ [164640]
```

```
FN557520.1_strain_E75             .......................................................................g.....a.. [164640]
AY261366.1_Warthog                .......................................................................g.....a.. [164640]
AY261365.1_Warmbaths              .......................................................................g........ [164640]
AY261364.1_Tengani_62             ........................t..............................................tg........ [164640]
AY261363.1_Pretoriuskop/96/4      .......................................................................g........ [164640]
AY261362.1_Mkuzi_1979             .......................................................................g.....a.. [164640]
AY261361.1_Malawi_Lil-20/1_1983   ..................................................t...c................g.....a.. [164640]
AY261360.1_Kenya_1950             ..................................................t...c................g.....a.. [164640]
AM712240.1_OURT_88/3              .......................................................................g.....a.. [164640]
AM712239.1_Benin_97/1             .......................................................................g.....a.. [164640]
KM111295.1_Ken06.Bus              ..................................................t..ac................g.....a.. [164640]
KM111294.1_Ken05/Tk1              .............................t....................t...c................g.....a.. [164640]
KM262844.1_strain_L60             .......................................................................g.....a.. [164640]
KM102979.1_26544/OG10_Italy       .......................................................................g.....a.. [164640]
KM262845.1_strain_NHV             .......................................................................g.....a.. [164640]
KX354450.1_47/Ss/2008             .......................................................................g.....a.. [164640]
KP055815.1_strain_BA71            .......................................................................g.....a.. [164640]
MH025920.1_strain_R35             ..................................................t..ac................g.....a.. [164640]
MH025919.1_strain_N10             ..................................................t..ac................g.....a.. [164640]
MH025918.1_strain_R25             ..................................................t..ac................g.....a.. [164640]
MH025917.1_strain_R7              ..................................................t..ac................g.....a.. [164640]
MH025916.1_strain_R8              ..................................................t..ac................g.....a.. [164640]

FR682468.2_Georgia_2007/1         ggatctttccgcctggatttaccttcgcgcctgccaatgcgcggcctacgtgggaaaaggcaccaccatttattacaatg [164720]
ASF_IND_20_CAD543                 ................................................................................ [164720]
LR899193.1_Germany_2020/1         ................................................................................ [164720]
MN393477.1_Wuhan_2019             ................................................................................ [164720]
MW396979.1_Timor-Leste/2019/1     ................................................................................ [164720]
MK940252.1_CN/2019/InnerMongolia  ................................................................................ [164720]
MK128995.1_China/2018/AnhuiXCGQ   ................................................................................ [164720]
MH681419.1_ASFV/POL/2015          ................................................................................ [164720]
LS478113.1_Estonia_2014           ................................................................................ [164720]
KP843857.1_Odintsovo_02/14        ................................................................................ [164720]
FN557520.1_strain_E75             a....................t.......................................................... [164720]
AY261366.1_Warthog                a..............................................................t................ [164720]
AY261365.1_Warmbaths              a............................................................................... [164720]
AY261364.1_Tengani_62             ................................................................................ [164720]
AY261363.1_Pretoriuskop/96/4      a............................................................................... [164720]
AY261362.1_Mkuzi_1979             a............................................................................... [164720]
AY261361.1_Malawi_Lil-20/1_1983   a.................................................................t............. [164720]
AY261360.1_Kenya_1950             a..c............................................................................ [164720]
AM712240.1_OURT_88/3              a....................t.......................................................... [164720]
AM712239.1_Benin_97/1             a....................t.......................................................... [164720]
KM111295.1_Ken06.Bus              a..c............................................................................ [164720]
```

```
KM111294.1_Ken05/Tk1            a..c............................................................................ [164720]
KM262844.1_strain_L60           a....................t.......................................................... [164720]
KM102979.1_26544/OG10_Italy     a....................t.......................................................... [164720]
KM262845.1_strain_NHV           a....................t.......................................................... [164720]
KX354450.1_47/Ss/2008           a....................t.......................................................... [164720]
KP055815.1_strain_BA71          a....................t.......................................................... [164720]
MH025920.1_strain_R35           a..c............................................................................ [164720]
MH025919.1_strain_N10           a..c............................................................................ [164720]
MH025918.1_strain_R25           a..c............................................................................ [164720]
MH025917.1_strain_R7            a..c............................................................................ [164720]
MH025916.1_strain_R8            a..c............................................................................ [164720]

FR682468.2_Georgia_2007/1       ataagccttgccgcacgggctctgtgatggcgctagccaaaatgtacaccctgttgagcgcgcctaatagcacgatacat [164800]
ASF_IND_20_CAD543               ................................................................................ [164800]
LR899193.1_Germany_2020/1       ................................................................................ [164800]
MN393477.1_Wuhan_2019           ................................................................................ [164800]
MW396979.1_Timor-Leste/2019/1   ................................................................................ [164800]
MK940252.1_CN/2019/InnerMongolia ................................................................................ [164800]
MK128995.1_China/2018/AnhuiXCGQ ................................................................................ [164800]
MH681419.1_ASFV/POL/2015        ................................................................................ [164800]
LS478113.1_Estonia_2014         ................................................................................ [164800]
KP843857.1_Odintsovo_02/14      ................................................................................ [164800]
FN557520.1_strain_E75           ..................................g............................................. [164800]
AY261366.1_Warthog              .a................................g.................a...............g........... [164800]
AY261365.1_Warmbaths            ..................................g............................................. [164800]
AY261364.1_Tengani_62           ..................................g............................................. [164800]
AY261363.1_Pretoriuskop/96/4    ..................................g...............t............................. [164800]
AY261362.1_Mkuzi_1979           ..................................g............................................. [164800]
AY261361.1_Malawi_Lil-20/1_1983 ..........taa.........c..a........g.......................t..................... [164800]
AY261360.1_Kenya_1950           ...........a.....a....c..a........g..t..............ag...ct.....c........t...t.. [164800]
AM712240.1_OURT_88/3            ..................................g............................................. [164800]
AM712239.1_Benin_97/1           ..................................g..............t.............................. [164800]
KM111295.1_Ken06.Bus            ...........a.....a....c..a........g.................ag...ct.....c........c...t.. [164800]
KM111294.1_Ken05/Tk1            ...........a.....a....c..a........g..t..............ag...ct.....c........t...t.. [164800]
KM262844.1_strain_L60           ..................................g............................................. [164800]
KM102979.1_26544/OG10_Italy     ..................................g............................................. [164800]
KM262845.1_strain_NHV           ..................................g............................................. [164800]
KX354450.1_47/Ss/2008           ..................................g............................................. [164800]
KP055815.1_strain_BA71          ..................................g............................................. [164800]
MH025920.1_strain_R35           ...........a.....a....c..a........g.................ag...ct.....c........c...t.. [164800]
MH025919.1_strain_N10           ...........a.....a....c..a........g.................ag...ct.....c........c...t.. [164800]
MH025918.1_strain_R25           ...........a.....a....c..a........g.................ag...ct.....c........c...t.. [164800]
MH025917.1_strain_R7            ...........a.....a....c..a........g.................ag...ct.....c........c...t.. [164800]
MH025916.1_strain_R8            ...........a.....a....c..a........g.................ag...ct.....c........c...t.. [164800]
```

```
FR682468.2_Georgia_2007/1        acggcgaccattaaggccgacgcaaagccctatagcctgcaccccctgcaggttgcggcggtcgtgtcccccaagtttaa [164880]
ASF_IND_20_CAD543                ................................................................................ [164880]
LR899193.1_Germany_2020/1        ................................................................................ [164880]
MN393477.1_Wuhan_2019            ................................................................................ [164880]
MW396979.1_Timor-Leste/2019/1    ................................................................................ [164880]
MK940252.1_CN/2019/InnerMongolia ................................................................................ [164880]
MK128995.1_China/2018/AnhuiXCGQ  ................................................................................ [164880]
MH681419.1_ASFV/POL/2015         ................................................................................ [164880]
LS478113.1_Estonia_2014          ................................................................................ [164880]
KP843857.1_Odintsovo_02/14       ................................................................................ [164880]
FN557520.1_strain_E75            ..........................a.................t................................... [164880]
AY261366.1_Warthog               ........t...................................t................................... [164880]
AY261365.1_Warmbaths             ....................t.......................t................................... [164880]
AY261364.1_Tengani_62            ......................................a.....t................................... [164880]
AY261363.1_Pretoriuskop/96/4     ............................................t................................... [164880]
AY261362.1_Mkuzi_1979            ..........................a.................t................................... [164880]
AY261361.1_Malawi_Lil-20/1_1983  ..aa..g.............t.....a.....c..t........t................................... [164880]
AY261360.1_Kenya_1950            ...a.c...........t..............c................................c.............. [164880]
AM712240.1_OURT_88/3             ..........................a.................t................................... [164880]
AM712239.1_Benin_97/1            ..........................a.................t................................... [164880]
KM111295.1_Ken06.Bus             .....c...........t...............................................c.............. [164880]
KM111294.1_Ken05/Tk1             ...a.c...........t..............c................................c.............. [164880]
KM262844.1_strain_L60            ..........................a.................t................................... [164880]
KM102979.1_26544/OG10_Italy      ..........................a.................t................................... [164880]
KM262845.1_strain_NHV            ..........................a.................t................................... [164880]
KX354450.1_47/Ss/2008            ..........................a.................t................................... [164880]
KP055815.1_strain_BA71           ..........................a.................t................................... [164880]
MH025920.1_strain_R35            .....c...........t...............................................c.............. [164880]
MH025919.1_strain_N10            .....c...........t...............................................c.............. [164880]
MH025918.1_strain_R25            .....c...........t...............................................c.............. [164880]
MH025917.1_strain_R7             .....c...........t...............................................c.............. [164880]
MH025916.1_strain_R8             .....c...........t...............................................c.............. [164880]

FR682468.2_Georgia_2007/1        aaaatttgaacacgtgtccgttatcaacggggtaaattgcgtaaaaggagaacatgtcacctttttgaaaaagactatta [164960]
ASF_IND_20_CAD543                .......................................y........................................ [164960]
LR899193.1_Germany_2020/1        ................................................................................ [164960]
MN393477.1_Wuhan_2019            ................................................................................ [164960]
MW396979.1_Timor-Leste/2019/1    ................................................................................ [164960]
MK940252.1_CN/2019/InnerMongolia ................................................................................ [164960]
MK128995.1_China/2018/AnhuiXCGQ  ................................................................................ [164960]
MH681419.1_ASFV/POL/2015         ................................................................................ [164960]
LS478113.1_Estonia_2014          ................................................................................ [164960]
KP843857.1_Odintsovo_02/14       ................................................................................ [164960]
```

```
FN557520.1_strain_E75           ...................a...................t.................t.................c.... [164960]
AY261366.1_Warthog              ...................a............................................................ [164960]
AY261365.1_Warmbaths            ...................a............................................................ [164960]
AY261364.1_Tengani_62           ...................a..........a........t........................................ [164960]
AY261363.1_Pretoriuskop/96/4    .......................................t........................................ [164960]
AY261362.1_Mkuzi_1979           ...................a...................t.................t...................... [164960]
AY261361.1_Malawi_Lil-20/1_1983 ...................a...................t........................................ [164960]
AY261360.1_Kenya_1950           ...................a.............c.....t.....g...........................g.c.... [164960]
AM712240.1_OURT_88/3            ...................a...................t.................t.................c.... [164960]
AM712239.1_Benin_97/1           ...................a...................t.................t.................c.... [164960]
KM111295.1_Ken06.Bus            ...................a...................t.....g...........................g.c.... [164960]
KM111294.1_Ken05/Tk1            ...................a...................t.....g...........................g.c.... [164960]
KM262844.1_strain_L60           ...................a...................t.................t.................c.... [164960]
KM102979.1_26544/OG10_Italy     ...................a...................t.................t.................c.... [164960]
KM262845.1_strain_NHV           ...................a...................t.................t.................c.... [164960]
KX354450.1_47/Ss/2008           ...................a...................t.................t.................c.... [164960]
KP055815.1_strain_BA71          ...................a...................t.................t.................c.... [164960]
MH025920.1_strain_R35           ...................a...................t.....g...........................g.c.... [164960]
MH025919.1_strain_N10           ...................a...................t.....g...........................g.c.... [164960]
MH025918.1_strain_R25           ...................a...................t.....g...........................g.c.... [164960]
MH025917.1_strain_R7            ...................a...................t.....g...........................g.c.... [164960]
MH025916.1_strain_R8            ...................a...................t.....g...........................g.c.... [164960]

FR682468.2_Georgia_2007/1       atgaaatggtcgttaaaaaatttcaacaaacgattaaagataaaaaccgcaaaacaacattacgagacagctgttcaaac [165040]
ASF_IND_20_CAD543               ................................................................................ [165040]
LR899193.1_Germany_2020/1       ................................................................................ [165040]
MN393477.1_Wuhan_2019           ................................................................................ [165040]
MW396979.1_Timor-Leste/2019/1   ................................................................................ [165040]
MK940252.1_CN/2019/InnerMongolia ................................................................................ [165040]
MK128995.1_China/2018/AnhuiXCGQ ................................................................................ [165040]
MH681419.1_ASFV/POL/2015        ................................................................................ [165040]
LS478113.1_Estonia_2014         ................................................................................ [165040]
KP843857.1_Odintsovo_02/14      ................................................................................ [165040]
FN557520.1_strain_E75           ...........a................g...................................t............... [165040]
AY261366.1_Warthog              .......a....................g................................................... [165040]
AY261365.1_Warmbaths            ............................g................................................... [165040]
AY261364.1_Tengani_62           ............................g.................t................................. [165040]
AY261363.1_Pretoriuskop/96/4    ............................g................................................... [165040]
AY261362.1_Mkuzi_1979           ...........a................g...................................t............... [165040]
AY261361.1_Malawi_Lil-20/1_1983 ............................g...................................t............... [165040]
AY261360.1_Kenya_1950           ............................g...g...............................c............... [165040]
AM712240.1_OURT_88/3            ...........a................g...................................t............... [165040]
AM712239.1_Benin_97/1           ...........a................g...................................t............... [165040]
KM111295.1_Ken06.Bus            ............................g...................................c............... [165040]
```

```
KM111294.1_Ken05/Tk1            ............................g...g...............................c............... [165040]
KM262844.1_strain_L60           ...........a................g...................................t............... [165040]
KM102979.1_26544/OG10_Italy     ...........a................g...................................t............... [165040]
KM262845.1_strain_NHV           ...........a................g...................................t............... [165040]
KX354450.1_47/Ss/2008           ...........a................g...................................t............... [165040]
KP055815.1_strain_BA71          ...........a................g...................................t............... [165040]
MH025920.1_strain_R35           ............................g...................................c............... [165040]
MH025919.1_strain_N10           ............................g...................................c............... [165040]
MH025918.1_strain_R25           ............................g...................................c............... [165040]
MH025917.1_strain_R7            ............................g...................................c............... [165040]
MH025916.1_strain_R8            ............................g...................................c............... [165040]

FR682468.2_Georgia_2007/1       atctttatcgttatagtgggttccattccaggaatagaatggaccggccagcggaaggatgaacttagcatcgcggaaaa [165120]
ASF_IND_20_CAD543               ................................................................................ [165120]
LR899193.1_Germany_2020/1       ................................................................................ [165120]
MN393477.1_Wuhan_2019           ................................................................................ [165120]
MW396979.1_Timor-Leste/2019/1   ................................................................................ [165120]
MK940252.1_CN/2019/InnerMongolia ................................................................................ [165120]
MK128995.1_China/2018/AnhuiXCGQ ................................................................................ [165120]
MH681419.1_ASFV/POL/2015        ................................................................................ [165120]
LS478113.1_Estonia_2014         ................................................................................ [165120]
KP843857.1_Odintsovo_02/14      ................................................................................ [165120]
FN557520.1_strain_E75           ......g.........................c.........................................a..... [165120]
AY261366.1_Warthog              ..........................................................................a..... [165120]
AY261365.1_Warmbaths            ................................................................................ [165120]
AY261364.1_Tengani_62           .................................................................a........a..... [165120]
AY261363.1_Pretoriuskop/96/4    ..........................................................................a..... [165120]
AY261362.1_Mkuzi_1979           ......g.........................c.........................................a..... [165120]
AY261361.1_Malawi_Lil-20/1_1983 ......g.......c.................c.........................................a..... [165120]
AY261360.1_Kenya_1950           ......g.......c.................c................................a........a..... [165120]
AM712240.1_OURT_88/3            ......g.........................c.........................................a..... [165120]
AM712239.1_Benin_97/1           ......g.........................c.........................................a..... [165120]
KM111295.1_Ken06.Bus            ......g.......c.................c................................a........a..... [165120]
KM111294.1_Ken05/Tk1            ......g.......c.................c................................a........a..... [165120]
KM262844.1_strain_L60           ......g.........................c.........................................a..... [165120]
KM102979.1_26544/OG10_Italy     ......g.........................c.........................................a..... [165120]
KM262845.1_strain_NHV           ......g.........................c.........................................a..... [165120]
KX354450.1_47/Ss/2008           ......g.........................c.........................................a..... [165120]
KP055815.1_strain_BA71          ......g.........................c.........................................a..... [165120]
MH025920.1_strain_R35           ......g.......c.................c................................a........a..... [165120]
MH025919.1_strain_N10           ......g.......c.................c................................a........a..... [165120]
MH025918.1_strain_R25           ......g.......c.................c................................a........a..... [165120]
MH025917.1_strain_R7            ......g.......c.................c................................a........a..... [165120]
MH025916.1_strain_R8            ......g.......c.................c................................a........a..... [165120]
```

```
FR682468.2_Georgia_2007/1        tgtttttaaaacgcattactccattccttctagttttttaacaagtatgacaaagtctatcgtggatattcttctgcaat [165200]
ASF_IND_20_CAD543                ................................................................................ [165200]
LR899193.1_Germany_2020/1        ................................................................................ [165200]
MN393477.1_Wuhan_2019            ................................................................................ [165200]
MW396979.1_Timor-Leste/2019/1    ................................................................................ [165200]
MK940252.1_CN/2019/InnerMongolia ................................................................................ [165200]
MK128995.1_China/2018/AnhuiXCGQ  ................................................................................ [165200]
MH681419.1_ASFV/POL/2015         ................................................................................ [165200]
LS478113.1_Estonia_2014          ................................................................................ [165200]
KP843857.1_Odintsovo_02/14       ................................................................................ [165200]
FN557520.1_strain_E75            ........................c....................c.......g.......................... [165200]
AY261366.1_Warthog               .............................................c.................................. [165200]
AY261365.1_Warmbaths             .............................................c.................................. [165200]
AY261364.1_Tengani_62            ........................c....................c.................................. [165200]
AY261363.1_Pretoriuskop/96/4     .............................................c.................................. [165200]
AY261362.1_Mkuzi_1979            ........................c....................c.......g.......................... [165200]
AY261361.1_Malawi_Lil-20/1_1983  ........................c...................ac.......ga......................... [165200]
AY261360.1_Kenya_1950            c.......................c....................c.......ga......................... [165200]
AM712240.1_OURT_88/3             ........................c....................c.......g.......................... [165200]
AM712239.1_Benin_97/1            ........................c....................c.......g.......................... [165200]
KM111295.1_Ken06.Bus             c.......................c....................c.......ga......................... [165200]
KM111294.1_Ken05/Tk1             c.......................c....................c.......ga......................... [165200]
KM262844.1_strain_L60            ........................c....................c.......g.......................... [165200]
KM102979.1_26544/OG10_Italy      ........................c....................c.......g.......................... [165200]
KM262845.1_strain_NHV            ........................c....................c.......g.......................... [165200]
KX354450.1_47/Ss/2008            ........................c....................c.......g.......................... [165200]
KP055815.1_strain_BA71           ........................c....................c.......g.......................... [165200]
MH025920.1_strain_R35            c.......................c....................c.......ga......................... [165200]
MH025919.1_strain_N10            c.......................c....................c.......ga......................... [165200]
MH025918.1_strain_R25            c.......................c....................c.......ga......................... [165200]
MH025917.1_strain_R7             c.......................c....................c.......ga......................... [165200]
MH025916.1_strain_R8             c.......................c....................c.......ga......................... [165200]

FR682468.2_Georgia_2007/1        ccatttctaaaaaagataaccataaacaggtcgacgtagacaaatatacgcgtgcccgcaatgcgggaggaaaaagggcg [165280]
ASF_IND_20_CAD543                ................................................................................ [165280]
LR899193.1_Germany_2020/1        ................................................................................ [165280]
MN393477.1_Wuhan_2019            ................................................................................ [165280]
MW396979.1_Timor-Leste/2019/1    ................................................................................ [165280]
MK940252.1_CN/2019/InnerMongolia ................................................................................ [165280]
MK128995.1_China/2018/AnhuiXCGQ  ................................................................................ [165280]
MH681419.1_ASFV/POL/2015         ................................................................................ [165280]
LS478113.1_Estonia_2014          ................................................................................ [165280]
KP843857.1_Odintsovo_02/14       ................................................................................ [165280]
```

```
FN557520.1_strain_E75            ......................................................................g......... [165280]
AY261366.1_Warthog               .......................................................t........................ [165280]
AY261365.1_Warmbaths             .......................................................t........................ [165280]
AY261364.1_Tengani_62            .......................................................t........a..........a.... [165280]
AY261363.1_Pretoriuskop/96/4     .......................................................t........................ [165280]
AY261362.1_Mkuzi_1979            ................................................................................ [165280]
AY261361.1_Malawi_Lil-20/1_1983  ...........................................................................a.... [165280]
AY261360.1_Kenya_1950            .............................a.....................................t.......a.... [165280]
AM712240.1_OURT_88/3             ......................................................................g......... [165280]
AM712239.1_Benin_97/1            ......................................................................g......... [165280]
KM111295.1_Ken06.Bus             .............................a.....................................t.......a.... [165280]
KM111294.1_Ken05/Tk1             .............................a.....................................t.......a.... [165280]
KM262844.1_strain_L60            ......................................................................g......... [165280]
KM102979.1_26544/OG10_Italy      ......................................................................g......... [165280]
KM262845.1_strain_NHV            ......................................................................g......... [165280]
KX354450.1_47/Ss/2008            ......................................................................g......... [165280]
KP055815.1_strain_BA71           ......................................................................g......... [165280]
MH025920.1_strain_R35            .............................a.....................................t.......a.... [165280]
MH025919.1_strain_N10            .............................a.....................................t.......a.... [165280]
MH025918.1_strain_R25            .............................a.....................................t.......a.... [165280]
MH025917.1_strain_R7             .............................a.....................................t.......a.... [165280]
MH025916.1_strain_R8             .............................a.....................................t.......a.... [165280]

FR682468.2_Georgia_2007/1        caggactgcatgctactcgcggcggaaggggatagcgcactttccctgctgcgcacgggactaaccctgggaaagtccaa [165360]
ASF_IND_20_CAD543                ................................................................................ [165360]
LR899193.1_Germany_2020/1        ................................................................................ [165360]
MN393477.1_Wuhan_2019            ................................................................................ [165360]
MW396979.1_Timor-Leste/2019/1    ................................................................................ [165360]
MK940252.1_CN/2019/InnerMongolia ................................................................................ [165360]
MK128995.1_China/2018/AnhuiXCGQ  ................................................................................ [165360]
MH681419.1_ASFV/POL/2015         ................................................................................ [165360]
LS478113.1_Estonia_2014          ................................................................................ [165360]
KP843857.1_Odintsovo_02/14       ................................................................................ [165360]
FN557520.1_strain_E75            ................................................t.............g................. [165360]
AY261366.1_Warthog               .............................a........g.......................g................. [165360]
AY261365.1_Warmbaths             .............................a........g.......................g................. [165360]
AY261364.1_Tengani_62            ..............................................................g..............t.. [165360]
AY261363.1_Pretoriuskop/96/4     ......................................g.....t.................g................. [165360]
AY261362.1_Mkuzi_1979            ..............................................................g................. [165360]
AY261361.1_Malawi_Lil-20/1_1983  ......................................................g.......g...........a..... [165360]
AY261360.1_Kenya_1950            ........t.............................................g.......g................. [165360]
AM712240.1_OURT_88/3             ................................................t.............g................. [165360]
AM712239.1_Benin_97/1            ................................................t.............g................. [165360]
KM111295.1_Ken06.Bus             ..a.....t.............................................g.......g................. [165360]
```

```
KM111294.1_Ken05/Tk1             ........t.............................................g.......g................. [165360]
KM262844.1_strain_L60            ................................................t.............g................. [165360]
KM102979.1_26544/OG10_Italy      ................................................t.............g................. [165360]
KM262845.1_strain_NHV            ................................................t.............g................. [165360]
KX354450.1_47/Ss/2008            ................................................t.............g................. [165360]
KP055815.1_strain_BA71           ................................................t.............g................. [165360]
MH025920.1_strain_R35            ..a.....t.............................................g.......g................. [165360]
MH025919.1_strain_N10            ..a.....t.............................................g.......g................. [165360]
MH025918.1_strain_R25            ..a.....t.............................................g.......g................. [165360]
MH025917.1_strain_R7             ..a.....t.............................................g.......g................. [165360]
MH025916.1_strain_R8             ..a.....t.............................................g.......g................. [165360]

FR682468.2_Georgia_2007/1        cccaagcgggccctcctttgacttctgcggcatgatctccctgggaggagtcatcatgaatgcctgcaaaaaggtgacaa [165440]
ASF_IND_20_CAD543                ................................................................................ [165440]
LR899193.1_Germany_2020/1        ................................................................................ [165440]
MN393477.1_Wuhan_2019            ................................................................................ [165440]
MW396979.1_Timor-Leste/2019/1    ................................................................................ [165440]
MK940252.1_CN/2019/InnerMongolia ................................................................................ [165440]
MK128995.1_China/2018/AnhuiXCGQ  ................................................................................ [165440]
MH681419.1_ASFV/POL/2015         ................................................................................ [165440]
LS478113.1_Estonia_2014          ................................................................................ [165440]
KP843857.1_Odintsovo_02/14       ................................................................................ [165440]
FN557520.1_strain_E75            ................................................g............................... [165440]
AY261366.1_Warthog               ................................................................................ [165440]
AY261365.1_Warmbaths             ................................................................................ [165440]
AY261364.1_Tengani_62            t..................................................t..t......................... [165440]
AY261363.1_Pretoriuskop/96/4     ................................................................................ [165440]
AY261362.1_Mkuzi_1979            ................................................g............................... [165440]
AY261361.1_Malawi_Lil-20/1_1983  ................................................g............................... [165440]
AY261360.1_Kenya_1950            ............t...................................g............................... [165440]
AM712240.1_OURT_88/3             ................................................g............................... [165440]
AM712239.1_Benin_97/1            ................................................g............................... [165440]
KM111295.1_Ken06.Bus             ............t...................................g............................... [165440]
KM111294.1_Ken05/Tk1             ............t...................................g............................... [165440]
KM262844.1_strain_L60            ................................................g............................... [165440]
KM102979.1_26544/OG10_Italy      ................................................g............................... [165440]
KM262845.1_strain_NHV            ................................................g............................... [165440]
KX354450.1_47/Ss/2008            ................................................g............................... [165440]
KP055815.1_strain_BA71           ................................................g............................... [165440]
MH025920.1_strain_R35            ............t...................................g............................... [165440]
MH025919.1_strain_N10            ............t...................................g............................... [165440]
MH025918.1_strain_R25            ............t...................................g............................... [165440]
MH025917.1_strain_R7             ............t...................................g............................... [165440]
MH025916.1_strain_R8             ............t...................................g............................... [165440]
```

```
FR682468.2_Georgia_2007/1        acattacaacggactctggagaaaccattatggtgcgcaacgaacagcttaccaataataaagtgttgcagggaatcgtg [165520]
ASF_IND_20_CAD543                ................................................................................ [165520]
LR899193.1_Germany_2020/1        ................................................................................ [165520]
MN393477.1_Wuhan_2019            ................................................................................ [165520]
MW396979.1_Timor-Leste/2019/1    ................................................................................ [165520]
MK940252.1_CN/2019/InnerMongolia ................................................................................ [165520]
MK128995.1_China/2018/AnhuiXCGQ  ................................................................................ [165520]
MH681419.1_ASFV/POL/2015         ................................................................................ [165520]
LS478113.1_Estonia_2014          ................................................................................ [165520]
KP843857.1_Odintsovo_02/14       ................................................................................ [165520]
FN557520.1_strain_E75            ............................c..............nn...............................t... [165520]
AY261366.1_Warthog               ............................c................................................... [165520]
AY261365.1_Warmbaths             ............................c............................................g...... [165520]
AY261364.1_Tengani_62            ............................c.....a.................t..............a.....g.....a [165520]
AY261363.1_Pretoriuskop/96/4     ............................c............................................g..t... [165520]
AY261362.1_Mkuzi_1979            ............................c................................................... [165520]
AY261361.1_Malawi_Lil-20/1_1983  ..........a.....c...........c............................................g...... [165520]
AY261360.1_Kenya_1950            ............................c...................................a..a.....g...... [165520]
AM712240.1_OURT_88/3             ............................c...............................................t... [165520]
AM712239.1_Benin_97/1            ............................c...............................................t... [165520]
KM111295.1_Ken06.Bus             ............................c....................c..............a..a............ [165520]
KM111294.1_Ken05/Tk1             ..........a.....c...........c...........t...........t...........a..a.....g...... [165520]
KM262844.1_strain_L60            ............................c...............................................t... [165520]
KM102979.1_26544/OG10_Italy      ............................c...............................................t... [165520]
KM262845.1_strain_NHV            ............................c...............................................t... [165520]
KX354450.1_47/Ss/2008            ............................c...............................................t... [165520]
KP055815.1_strain_BA71           ............................c...............................................t... [165520]
MH025920.1_strain_R35            ............................c....................c..............a..a............ [165520]
MH025919.1_strain_N10            ............................c....................c..............a..a............ [165520]
MH025918.1_strain_R25            ............................c....................c..............a..a............ [165520]
MH025917.1_strain_R7             ............................c....................c..............a..a............ [165520]
MH025916.1_strain_R8             ............................c....................c..............a..a............ [165520]

FR682468.2_Georgia_2007/1        caggtattgggtctagacttcaactgccattacaaaacacaggaagagcgagcaaagctgagatacggctgcattgttgc [165600]
ASF_IND_20_CAD543                ................................................................................ [165600]
LR899193.1_Germany_2020/1        ................................................................................ [165600]
MN393477.1_Wuhan_2019            ................................................................................ [165600]
MW396979.1_Timor-Leste/2019/1    ................................................................................ [165600]
MK940252.1_CN/2019/InnerMongolia ................................................................................ [165600]
MK128995.1_China/2018/AnhuiXCGQ  ................................................................................ [165600]
MH681419.1_ASFV/POL/2015         ................................................................................ [165600]
LS478113.1_Estonia_2014          ................................................................................ [165600]
KP843857.1_Odintsovo_02/14       ................................................................................ [165600]
```

FN557520.1_strain_E75 ......................................g......................................... [165600]
AY261366.1_Warthog ........a.............................g..........................t.............. [165600]
AY261365.1_Warmbaths ........a.............................g..........................t.............. [165600]
AY261364.1_Tengani_62 ........a.............................g..........................t.............. [165600]
AY261363.1_Pretoriuskop/96/4 ........a.............................g..........................t.............. [165600]
AY261362.1_Mkuzi_1979 ......................................g......................................... [165600]
AY261361.1_Malawi_Lil-20/1_1983 ........a........t....................g..........................t.............. [165600]
AY261360.1_Kenya_1950 ......................................g.....................c....t.............. [165600]
AM712240.1_OURT_88/3 ......................................g......................................... [165600]
AM712239.1_Benin_97/1 ......................................g......................................... [165600]
KM111295.1_Ken06.Bus ......................................g.....................c....t.............. [165600]
KM111294.1_Ken05/Tk1 ......................................g.....................c....t.............. [165600]
KM262844.1_strain_L60 ......................................g......................................... [165600]
KM102979.1_26544/OG10_Italy ......................................g......................................... [165600]
KM262845.1_strain_NHV ......................................g......................................... [165600]
KX354450.1_47/Ss/2008 ......................................g......................................... [165600]
KP055815.1_strain_BA71 ......................................g......................................... [165600]
MH025920.1_strain_R35 ......................................g.....................c....t.............. [165600]
MH025919.1_strain_N10 ......................................g.....................c....t.............. [165600]
MH025918.1_strain_R25 ......................................g.....................c....t.............. [165600]
MH025917.1_strain_R7 ......................................g.....................c....t.............. [165600]
MH025916.1_strain_R8 ......................................g.....................c....t.............. [165600]

FR682468.2_Georgia_2007/1 gtgcgttgatcaagatctggatgggtgtggaaaaatccttggactgctgctggcctactttcacctgttttggcctcagc [165680]
ASF_IND_20_CAD543 ................................................................................ [165680]
LR899193.1_Germany_2020/1 ................................................................................ [165680]
MN393477.1_Wuhan_2019 ................................................................................ [165680]
MW396979.1_Timor-Leste/2019/1 ................................................................................ [165680]
MK940252.1_CN/2019/InnerMongolia ................................................................................ [165680]
MK128995.1_China/2018/AnhuiXCGQ ................................................................................ [165680]
MH681419.1_ASFV/POL/2015 ................................................................................ [165680]
LS478113.1_Estonia_2014 ................................................................................ [165680]
KP843857.1_Odintsovo_02/14 ................................................................................ [165680]
FN557520.1_strain_E75 ................................................................................ [165680]
AY261366.1_Warthog ....................................t........................................... [165680]
AY261365.1_Warmbaths ....................................t........................................... [165680]
AY261364.1_Tengani_62 ................t...................t........................................... [165680]
AY261363.1_Pretoriuskop/96/4 ....................................t..........................t................ [165680]
AY261362.1_Mkuzi_1979 ................................................................................ [165680]
AY261361.1_Malawi_Lil-20/1_1983 .........c...........c..............t........................................... [165680]
AY261360.1_Kenya_1950 .........c...........c..............t........................................... [165680]
AM712240.1_OURT_88/3 ................................................................................ [165680]
AM712239.1_Benin_97/1 ................................................................................ [165680]
KM111295.1_Ken06.Bus .........c...........c..a...........t........................................... [165680]

```
KM111294.1_Ken05/Tk1           .........c...........c..............t............t.............................. [165680]
KM262844.1_strain_L60          ................................................................................ [165680]
KM102979.1_26544/OG10_Italy    ................................................................................ [165680]
KM262845.1_strain_NHV          ................................................................................ [165680]
KX354450.1_47/Ss/2008          ................................................................................ [165680]
KP055815.1_strain_BA71         ................................................................................ [165680]
MH025920.1_strain_R35          .........c...........c..a...........t........................................... [165680]
MH025919.1_strain_N10          .........c...........c..a...........t........................................... [165680]
MH025918.1_strain_R25          .........c...........c..a...........t........................................... [165680]
MH025917.1_strain_R7           .........c...........c..a...........t........................................... [165680]
MH025916.1_strain_R8           .........c...........c..a...........t........................................... [165680]

FR682468.2_Georgia_2007/1      ttattatccatggtttcgtaaaacgactgcttaccccgctgatacgtgtgtatgaaaagggtaagaccatgcccgtggaa [165760]
ASF_IND_20_CAD543              ................................................................................ [165760]
LR899193.1_Germany_2020/1      ................................................................................ [165760]
MN393477.1_Wuhan_2019          ................................................................................ [165760]
MW396979.1_Timor-Leste/2019/1  ................................................................................ [165760]
MK940252.1_CN/2019/InnerMongolia ................................................................................ [165760]
MK128995.1_China/2018/AnhuiXCGQ ................................................................................ [165760]
MH681419.1_ASFV/POL/2015       ................................................................................ [165760]
LS478113.1_Estonia_2014        ................................................................................ [165760]
KP843857.1_Odintsovo_02/14     ................................................................................ [165760]
FN557520.1_strain_E75          ....................................................c........c.....t........a... [165760]
AY261366.1_Warthog             .............................................................c......g........... [165760]
AY261365.1_Warmbaths           ....................................................c........c......g........... [165760]
AY261364.1_Tengani_62          ....................................................c........c......g.......a... [165760]
AY261363.1_Pretoriuskop/96/4   .......t............................................c........c......g........... [165760]
AY261362.1_Mkuzi_1979          ................................................................................ [165760]
AY261361.1_Malawi_Lil-20/1_1983 ....................................................................g........... [165760]
AY261360.1_Kenya_1950          .....g....................t..................................c..t...g........... [165760]
AM712240.1_OURT_88/3           ....................................................c........c.....t........a... [165760]
AM712239.1_Benin_97/1          ....................................................c........c.....t........a... [165760]
KM111295.1_Ken06.Bus           .....g.......................................................c..t...g........... [165760]
KM111294.1_Ken05/Tk1           .....g.......................................................c..t...g........... [165760]
KM262844.1_strain_L60          ....................................................c........c.....t........a... [165760]
KM102979.1_26544/OG10_Italy    ....................................................c........c.....t........a... [165760]
KM262845.1_strain_NHV          ....................................................c........c.....t........a... [165760]
KX354450.1_47/Ss/2008          ....................................................c........c.....t........a... [165760]
KP055815.1_strain_BA71         ....................................................c........c.....t........a... [165760]
MH025920.1_strain_R35          .....g.......................................................c..t...g........... [165760]
MH025919.1_strain_N10          .....g......................................................ac..t...g........... [165760]
MH025918.1_strain_R25          .....g.......................................................c..t...g........... [165760]
MH025917.1_strain_R7           .....g.......................................................c..t...g........... [165760]
MH025916.1_strain_R8           .....g.......................................................c..t...g........... [165760]
```

```
FR682468.2_Georgia_2007/1        ttttactatgaacaagagtttgatgcctgggcaaaaaagcagaccagcttagccaaccataccgtaaaatattacaaggg [165840]
ASF_IND_20_CAD543                ................................................................................ [165840]
LR899193.1_Germany_2020/1        ................................................................................ [165840]
MN393477.1_Wuhan_2019            ................................................................................ [165840]
MW396979.1_Timor-Leste/2019/1    ................................................................................ [165840]
MK940252.1_CN/2019/InnerMongolia ................................................................................ [165840]
MK128995.1_China/2018/AnhuiXCGQ  ................................................................................ [165840]
MH681419.1_ASFV/POL/2015         ................................................................................ [165840]
LS478113.1_Estonia_2014          ................................................................................ [165840]
KP843857.1_Odintsovo_02/14       ................................................................................ [165840]
FN557520.1_strain_E75            ..............g.....................................t...t.....t................. [165840]
AY261366.1_Warthog               ..............g................................t................................ [165840]
AY261365.1_Warmbaths             ..............g................................t................................ [165840]
AY261364.1_Tengani_62            ..............g.........................................t.....t...........t..... [165840]
AY261363.1_Pretoriuskop/96/4     ..............g...........................................................t..... [165840]
AY261362.1_Mkuzi_1979            ..............g.........................................t.....t...........t..... [165840]
AY261361.1_Malawi_Lil-20/1_1983  ........c.................a.....g...............c....t..t.....t...........t..... [165840]
AY261360.1_Kenya_1950            ........c.................a.....g...........t...c.g.....t.....t...........t..... [165840]
AM712240.1_OURT_88/3             ..............g.....................................t...t.....t................. [165840]
AM712239.1_Benin_97/1            ..............g.....................................t...t.....t................. [165840]
KM111295.1_Ken06.Bus             ........c.................a.....g...........t...c.g..t..t.....t...........t..... [165840]
KM111294.1_Ken05/Tk1             ........c.................a.....g...........t...c.g..t..t.....t...........t..... [165840]
KM262844.1_strain_L60            ..............g.....................................t...t.....t................. [165840]
KM102979.1_26544/OG10_Italy      ..............g.....................................t...t.....t................. [165840]
KM262845.1_strain_NHV            ..............g.....................................t...t.....t................. [165840]
KX354450.1_47/Ss/2008            ..............g.....................................t...t.....t................. [165840]
KP055815.1_strain_BA71           ..............g.....................................t...t.....t................. [165840]
MH025920.1_strain_R35            ........c.................a.....g...........t...c.g..t..t.....t...........t..... [165840]
MH025919.1_strain_N10            ........c.................a.....g...........t...c.g..t..t.....t...........t..... [165840]
MH025918.1_strain_R25            ........c.................a.....g...........t...c.g..t..t.....t...........t..... [165840]
MH025917.1_strain_R7             ........c.................a.....g...........t...c.g..t..t.....t...........t..... [165840]
MH025916.1_strain_R8             ........c.................a.....g...........t...c.g..t..t.....t...........t..... [165840]

FR682468.2_Georgia_2007/1        attggcggcgcatgacacccatgaagtaaaaagcatgttcaaacattttgacaacatggtgtacacgtttaccctggatg [165920]
ASF_IND_20_CAD543                ................................................................................ [165920]
LR899193.1_Germany_2020/1        ................................................................................ [165920]
MN393477.1_Wuhan_2019            ................................................................................ [165920]
MW396979.1_Timor-Leste/2019/1    ................................................................................ [165920]
MK940252.1_CN/2019/InnerMongolia ................................................................................ [165920]
MK128995.1_China/2018/AnhuiXCGQ  ................................................................................ [165920]
MH681419.1_ASFV/POL/2015         ................................................................................ [165920]
LS478113.1_Estonia_2014          ................................................................................ [165920]
KP843857.1_Odintsovo_02/14       ................................................................................ [165920]
```

```
FN557520.1_strain_E75              ................................................................................ [165920]
AY261366.1_Warthog                 ...........................................................................a.... [165920]
AY261365.1_Warmbaths               ...........................................................................a.... [165920]
AY261364.1_Tengani_62              ................................................................................ [165920]
AY261363.1_Pretoriuskop/96/4       .......................................t........................................ [165920]
AY261362.1_Mkuzi_1979              ................................................................................ [165920]
AY261361.1_Malawi_Lil-20/1_1983    ................................................................................ [165920]
AY261360.1_Kenya_1950              ...................................................t..a..................t.a.... [165920]
AM712240.1_OURT_88/3               ................................................................................ [165920]
AM712239.1_Benin_97/1              ................................................................................ [165920]
KM111295.1_Ken06.Bus               ............c.........................................a....................a.... [165920]
KM111294.1_Ken05/Tk1               ...................................................t..a..................t.a.... [165920]
KM262844.1_strain_L60              ................................................................................ [165920]
KM102979.1_26544/OG10_Italy        ................................................................................ [165920]
KM262845.1_strain_NHV              ................................................................................ [165920]
KX354450.1_47/Ss/2008              ................................................................................ [165920]
KP055815.1_strain_BA71             ................................................................................ [165920]
MH025920.1_strain_R35              ............c.........................................a....................a.... [165920]
MH025919.1_strain_N10              ............c.........................................a....................a.... [165920]
MH025918.1_strain_R25              ............c.........................................a....................a.... [165920]
MH025917.1_strain_R7               ............c.........................................a....................a.... [165920]
MH025916.1_strain_R8               ............c.........................................a....................a.... [165920]

FR682468.2_Georgia_2007/1          actcagcaaaggagttgtttcatatttattttggcggggagtcggagttgcgaaaaagagagctttgcaccggcgtggtg [166000]
ASF_IND_20_CAD543                  ................................................................................ [166000]
LR899193.1_Germany_2020/1          ................................................................................ [166000]
MN393477.1_Wuhan_2019              ................................................................................ [166000]
MW396979.1_Timor-Leste/2019/1      ................................................................................ [166000]
MK940252.1_CN/2019/InnerMongolia   ................................................................................ [166000]
MK128995.1_China/2018/AnhuiXCGQ    ................................................................................ [166000]
MH681419.1_ASFV/POL/2015           ................................................................................ [166000]
LS478113.1_Estonia_2014            ................................................................................ [166000]
KP843857.1_Odintsovo_02/14         ................................................................................ [166000]
FN557520.1_strain_E75              ....g........................................................................... [166000]
AY261366.1_Warthog                 ..................................t............................................. [166000]
AY261365.1_Warmbaths               ...........................................a.................................... [166000]
AY261364.1_Tengani_62              ................................................................................ [166000]
AY261363.1_Pretoriuskop/96/4       ................................................................................ [166000]
AY261362.1_Mkuzi_1979              ................................................................................ [166000]
AY261361.1_Malawi_Lil-20/1_1983    ..........................................................g..................... [166000]
AY261360.1_Kenya_1950              ................................................................................ [166000]
AM712240.1_OURT_88/3               ....g........................................................................... [166000]
AM712239.1_Benin_97/1              ....g........................................................................... [166000]
KM111295.1_Ken06.Bus               ..................................t.........................................a... [166000]
```

```
KM111294.1_Ken05/Tk1             ................................................................................ [166000]
KM262844.1_strain_L60            ....g........................................................................... [166000]
KM102979.1_26544/OG10_Italy      ....g........................................................................... [166000]
KM262845.1_strain_NHV            ....g........................................................................... [166000]
KX354450.1_47/Ss/2008            ....g........................................................................... [166000]
KP055815.1_strain_BA71           ....g........................................................................... [166000]
MH025920.1_strain_R35            ..................................t.........................................a... [166000]
MH025919.1_strain_N10            ..................................t.........................................a... [166000]
MH025918.1_strain_R25            ..................................t.........................................a... [166000]
MH025917.1_strain_R7             ..................................t.........................................a... [166000]
MH025916.1_strain_R8             ..................................t.........................................a... [166000]

FR682468.2_Georgia_2007/1        ccgctcaccgaaacccagacgcagtccattcatagtgtccgacgaattccttgcagcctgcatctgcaagtagataccaa [166080]
ASF_IND_20_CAD543                ................................................................................ [166080]
LR899193.1_Germany_2020/1        ................................................................................ [166080]
MN393477.1_Wuhan_2019            ................................................................................ [166080]
MW396979.1_Timor-Leste/2019/1    ................................................................................ [166080]
MK940252.1_CN/2019/InnerMongolia ................................................................................ [166080]
MK128995.1_China/2018/AnhuiXCGQ  ................................................................................ [166080]
MH681419.1_ASFV/POL/2015         ................................................................................ [166080]
LS478113.1_Estonia_2014          ................................................................................ [166080]
KP843857.1_Odintsovo_02/14       ................................................................................ [166080]
FN557520.1_strain_E75            ........t...........................................................g........... [166080]
AY261366.1_Warthog               ....................................................................g........... [166080]
AY261365.1_Warmbaths             ....................a...............................................g........... [166080]
AY261364.1_Tengani_62            ...............................................................t....g........... [166080]
AY261363.1_Pretoriuskop/96/4     ........t....................c......................................g........... [166080]
AY261362.1_Mkuzi_1979            ........t...........................................................g........... [166080]
AY261361.1_Malawi_Lil-20/1_1983  ..........................t..........a.....a..................c..c..g.....c..... [166080]
AY261360.1_Kenya_1950            ..............................................................c.....g.....c..... [166080]
AM712240.1_OURT_88/3             ........t...........................................................g........... [166080]
AM712239.1_Benin_97/1            ........t...........................................................g........... [166080]
KM111295.1_Ken06.Bus             ..............................................................c.....g.....c..... [166080]
KM111294.1_Ken05/Tk1             ..............................................................c.....g.....c..... [166080]
KM262844.1_strain_L60            ........t...........................................................g........... [166080]
KM102979.1_26544/OG10_Italy      ........t...........................................................g........... [166080]
KM262845.1_strain_NHV            ........t...........................................................g........... [166080]
KX354450.1_47/Ss/2008            ........t...........................................................g........... [166080]
KP055815.1_strain_BA71           ........t...........................................................g........... [166080]
MH025920.1_strain_R35            ..............................................................c.....g.....c..... [166080]
MH025919.1_strain_N10            ..............................................................c.....g.....c..... [166080]
MH025918.1_strain_R25            ..............................................................c.....g.....c..... [166080]
MH025917.1_strain_R7             ..............................................................c.....g.....c..... [166080]
MH025916.1_strain_R8             ..............................................................c.....g.....c..... [166080]
```

FR682468.2_Georgia_2007/1 ggcttacaagctggatgccatcgagcggcagattcccaacttcttagacgggatgacgcgggcgcggcgcaaaattttag [166160]
ASF_IND_20_CAD543 ................................................................................ [166160]
LR899193.1_Germany_2020/1 ................................................................................ [166160]
MN393477.1_Wuhan_2019 ................................................................................ [166160]
MW396979.1_Timor-Leste/2019/1 ................................................................................ [166160]
MK940252.1_CN/2019/InnerMongolia ................................................................................ [166160]
MK128995.1_China/2018/AnhuiXCGQ ................................................................................ [166160]
MH681419.1_ASFV/POL/2015 ................................................................................ [166160]
LS478113.1_Estonia_2014 ................................................................................ [166160]
KP843857.1_Odintsovo_02/14 ................................................................................ [166160]
FN557520.1_strain_E75 ...................................................a............................ [166160]
AY261366.1_Warthog ................................................................................ [166160]
AY261365.1_Warmbaths ................................................................................ [166160]
AY261364.1_Tengani_62 ................................................................................ [166160]
AY261363.1_Pretoriuskop/96/4 ...c............................................................................ [166160]
AY261362.1_Mkuzi_1979 ...................................................a............................ [166160]
AY261361.1_Malawi_Lil-20/1_1983 ...c..........................a..a.......................c...................... [166160]
AY261360.1_Kenya_1950 ...c........a.................a................................................. [166160]
AM712240.1_OURT_88/3 ...................................................a............................ [166160]
AM712239.1_Benin_97/1 ...................................................a............................ [166160]
KM111295.1_Ken06.Bus ...c........a.................a.................t..............a................ [166160]
KM111294.1_Ken05/Tk1 ...c..........................a..a.......................c...................... [166160]
KM262844.1_strain_L60 ...................................................a............................ [166160]
KM102979.1_26544/OG10_Italy ...................................................a............................ [166160]
KM262845.1_strain_NHV ...................................................a............................ [166160]
KX354450.1_47/Ss/2008 ...................................................a............................ [166160]
KP055815.1_strain_BA71 ...................................................a............................ [166160]
MH025920.1_strain_R35 ...c........a.................a.................t..............a................ [166160]
MH025919.1_strain_N10 ...c........a.................a.................t..............a................ [166160]
MH025918.1_strain_R25 ...c........a.................a.................t..............a................ [166160]
MH025917.1_strain_R7 ...c........a.................a.................t..............a................ [166160]
MH025916.1_strain_R8 ...c........a.................a.................t..............a................ [166160]

FR682468.2_Georgia_2007/1 ccgggggggtgaaatgcttcgcctccaacaaccgtgaacgaaaggtttttcagttcgggggctacgttgcagatcacatg [166240]
ASF_IND_20_CAD543 ................................................................................ [166240]
LR899193.1_Germany_2020/1 ................................................................................ [166240]
MN393477.1_Wuhan_2019 ................................................................................ [166240]
MW396979.1_Timor-Leste/2019/1 ................................................................................ [166240]
MK940252.1_CN/2019/InnerMongolia ................................................................................ [166240]
MK128995.1_China/2018/AnhuiXCGQ ................................................................................ [166240]
MH681419.1_ASFV/POL/2015 ................................................................................ [166240]
LS478113.1_Estonia_2014 ................................................................................ [166240]
KP843857.1_Odintsovo_02/14 ................................................................................ [166240]

```
FN557520.1_strain_E75          ..-...................t...............................................g......... [166240]
AY261366.1_Warthog             .........................t............................................g......... [166240]
AY261365.1_Warmbaths           .......a.................t............................................g......... [166240]
AY261364.1_Tengani_62          ......................................................................g......... [166240]
AY261363.1_Pretoriuskop/96/4   .......a..............................................................g......... [166240]
AY261362.1_Mkuzi_1979          ......................t...............................................g......... [166240]
AY261361.1_Malawi_Lil-20/1_1983 ........t..............................................t..............g......... [166240]
AY261360.1_Kenya_1950          ........t.............................................................g..c...... [166240]
AM712240.1_OURT_88/3           ......................t...............................................g......... [166240]
AM712239.1_Benin_97/1          ......................t...............................................g......... [166240]
KM111295.1_Ken06.Bus           ....a...t...................t.....c.................a.................g..c...... [166240]
KM111294.1_Ken05/Tk1           ........t.............................................................g..c...... [166240]
KM262844.1_strain_L60          ......................t...............................................g......... [166240]
KM102979.1_26544/OG10_Italy    ......................t...............................................g......... [166240]
KM262845.1_strain_NHV          ......................t...............................................g......... [166240]
KX354450.1_47/Ss/2008          ......................t...............................................g......... [166240]
KP055815.1_strain_BA71         ......................t...............................................g......... [166240]
MH025920.1_strain_R35          ....a...t...................t.....c.................a.................g..c...... [166240]
MH025919.1_strain_N10          ....a...t...................t.....c.................a.................g..c...... [166240]
MH025918.1_strain_R25          ....a...t...................t.....c.................a.................g..c...... [166240]
MH025917.1_strain_R7           ....a...t...................t.....c.................a.................g..c...... [166240]
MH025916.1_strain_R8           ....a...t...................t.....c.................a.................g..c...... [166240]

FR682468.2_Georgia_2007/1      ttttatcaccatggcgacatgtcgttaaacacaagtattataaaagccgcccagtattacccaggctcctcccacctcta [166320]
ASF_IND_20_CAD543              ................................................................................ [166320]
LR899193.1_Germany_2020/1      ................................................................................ [166320]
MN393477.1_Wuhan_2019          ................................................................................ [166320]
MW396979.1_Timor-Leste/2019/1  ................................................................................ [166320]
MK940252.1_CN/2019/InnerMongolia ................................................................................ [166320]
MK128995.1_China/2018/AnhuiXCGQ ................................................................................ [166320]
MH681419.1_ASFV/POL/2015       ................................................................................ [166320]
LS478113.1_Estonia_2014        ................................................................................ [166320]
KP843857.1_Odintsovo_02/14     ................................................................................ [166320]
FN557520.1_strain_E75          ..............................................................g................. [166320]
AY261366.1_Warthog             ..............................................................g................. [166320]
AY261365.1_Warmbaths           ..............................................................g................. [166320]
AY261364.1_Tengani_62          ..............................................................g................. [166320]
AY261363.1_Pretoriuskop/96/4   ..............t...............................................g................. [166320]
AY261362.1_Mkuzi_1979          ..............................................................g................. [166320]
AY261361.1_Malawi_Lil-20/1_1983 .................t.....................................................t........ [166320]
AY261360.1_Kenya_1950          .................t....................c........t.....a..c.....g.....t.....t..... [166320]
AM712240.1_OURT_88/3           ..............................................................g................. [166320]
AM712239.1_Benin_97/1          ..............................................................g................. [166320]
KM111295.1_Ken06.Bus           .................t....................c........t.....a..c.....g.....t.....t..... [166320]
```

```
KM111294.1_Ken05/Tk1          .................t....................c........t.....a..c.....g.....t.....t..... [166320]
KM262844.1_strain_L60         ..............................................................g................. [166320]
KM102979.1_26544/OG10_Italy   ..............................................................g................. [166320]
KM262845.1_strain_NHV         ..............................................................g................. [166320]
KX354450.1_47/Ss/2008         ..............................................................g................. [166320]
KP055815.1_strain_BA71        ..............................................................g................. [166320]
MH025920.1_strain_R35         .................t....................c........t.....a..c.....g.....t.....t..... [166320]
MH025919.1_strain_N10         ...........c.....t....................c........t.....a..c.....g.....t.....t..... [166320]
MH025918.1_strain_R25         .................t....................c........t.....a..c.....g.....t.....t..... [166320]
MH025917.1_strain_R7          .................t....................c........t.....a..c.....g.....t.....t..... [166320]
MH025916.1_strain_R8          .................t....................c........t.....a..c.....g.....t.....t..... [166320]

FR682468.2_Georgia_2007/1        tccggtattcataggcataggaagttttggctccaggcacctgggaggaaaggatgcaggatccccaagatacatcagtg [166400]
ASF_IND_20_CAD543                ................................................................................ [166400]
LR899193.1_Germany_2020/1        ................................................................................ [166400]
MN393477.1_Wuhan_2019            ................................................................................ [166400]
MW396979.1_Timor-Leste/2019/1    ................................................................................ [166400]
MK940252.1_CN/2019/InnerMongolia ................................................................................ [166400]
MK128995.1_China/2018/AnhuiXCGQ  ................................................................................ [166400]
MH681419.1_ASFV/POL/2015         ................................................................................ [166400]
LS478113.1_Estonia_2014          ................................................................................ [166400]
KP843857.1_Odintsovo_02/14       ................................................................................ [166400]
FN557520.1_strain_E75            ...a....................c..c.................................................... [166400]
AY261366.1_Warthog               ...a....................c..c.................................................... [166400]
AY261365.1_Warmbaths             ...a....................c..c.................................................... [166400]
AY261364.1_Tengani_62            ...a....................c..c.................................................... [166400]
AY261363.1_Pretoriuskop/96/4     ...a....................c..c.................................................... [166400]
AY261362.1_Mkuzi_1979            ...a....................c..c.................................................... [166400]
AY261361.1_Malawi_Lil-20/1_1983  ...a....................c..c.................................................... [166400]
AY261360.1_Kenya_1950            ...t.....t..............c..............t...........a............................ [166400]
AM712240.1_OURT_88/3             ...a....................c..c.................................................... [166400]
AM712239.1_Benin_97/1            ...a....................c..c.................................................... [166400]
KM111295.1_Ken06.Bus             ...t.....t..............c..............t...........a............................ [166400]
KM111294.1_Ken05/Tk1             ...t.....t..............c..............t...........a............................ [166400]
KM262844.1_strain_L60            ...a....................c..c.................................................... [166400]
KM102979.1_26544/OG10_Italy      ...a....................c..c.................................................... [166400]
KM262845.1_strain_NHV            ...a....................c..c.................................................... [166400]
KX354450.1_47/Ss/2008            ...a....................c..c.................................................... [166400]
KP055815.1_strain_BA71           ...a....................c..c.................................................... [166400]
MH025920.1_strain_R35            ...t.....t..............c..............t...........a............................ [166400]
MH025919.1_strain_N10            ...t.....t..............c..............t...........a............................ [166400]
MH025918.1_strain_R25            ...t.....t..............c..............t...........a............................ [166400]
MH025917.1_strain_R7             ...t.....t..............c..............t...........a............................ [166400]
MH025916.1_strain_R8             ...t.....t..............c..............t...........a............................ [166400]
```

```
FR682468.2_Georgia_2007/1        tgcagcttgcgtctgaatttattaaaacaatgttccccgcggaggactcatggcttctcccctacgtctttgaggacggc [166480]
ASF_IND_20_CAD543                ................................................................................ [166480]
LR899193.1_Germany_2020/1        ................................................................................ [166480]
MN393477.1_Wuhan_2019            ................................................................................ [166480]
MW396979.1_Timor-Leste/2019/1    ................................................................................ [166480]
MK940252.1_CN/2019/InnerMongolia ................................................................................ [166480]
MK128995.1_China/2018/AnhuiXCGQ  ................................................................................ [166480]
MH681419.1_ASFV/POL/2015         ................................................................................ [166480]
LS478113.1_Estonia_2014          ................................................................................ [166480]
KP843857.1_Odintsovo_02/14       ................................................................................ [166480]
FN557520.1_strain_E75            ................................................................................ [166480]
AY261366.1_Warthog               ................................................................................ [166480]
AY261365.1_Warmbaths             ................................................................................ [166480]
AY261364.1_Tengani_62            ................................................................................ [166480]
AY261363.1_Pretoriuskop/96/4     ..................................t............................................. [166480]
AY261362.1_Mkuzi_1979            ................................................................................ [166480]
AY261361.1_Malawi_Lil-20/1_1983  .............c.......................a..a.................t..............a...... [166480]
AY261360.1_Kenya_1950            .....................................g....................t..................... [166480]
AM712240.1_OURT_88/3             ................................................................................ [166480]
AM712239.1_Benin_97/1            ................................................................................ [166480]
KM111295.1_Ken06.Bus             ......................................a............................t............ [166480]
KM111294.1_Ken05/Tk1             ....a................................g....................t..................... [166480]
KM262844.1_strain_L60            ................................................................................ [166480]
KM102979.1_26544/OG10_Italy      ................................................................................ [166480]
KM262845.1_strain_NHV            ................................................................................ [166480]
KX354450.1_47/Ss/2008            ................................................................................ [166480]
KP055815.1_strain_BA71           ................................................................................ [166480]
MH025920.1_strain_R35            ......................................a............................t............ [166480]
MH025919.1_strain_N10            ......................................a............................t............ [166480]
MH025918.1_strain_R25            ......................................a............................t............ [166480]
MH025917.1_strain_R7             ......................................a............................t............ [166480]
MH025916.1_strain_R8             ......................................a............................t............ [166480]

FR682468.2_Georgia_2007/1        cagcgggcggaaccagagtactacgtgcctgtgttgccgcttgctattatggagtacggcgccaacccatcggagggctg [166560]
ASF_IND_20_CAD543                .......................................................r........................ [166560]
LR899193.1_Germany_2020/1        ................................................................................ [166560]
MN393477.1_Wuhan_2019            ................................................................................ [166560]
MW396979.1_Timor-Leste/2019/1    ................................................................................ [166560]
MK940252.1_CN/2019/InnerMongolia ................................................................................ [166560]
MK128995.1_China/2018/AnhuiXCGQ  ................................................................................ [166560]
MH681419.1_ASFV/POL/2015         ................................................................................ [166560]
LS478113.1_Estonia_2014          ................................................................................ [166560]
KP843857.1_Odintsovo_02/14       ................................................................................ [166560]
```

```
FN557520.1_strain_E75            ................................a............................................... [166560]
AY261366.1_Warthog               ........a....................................................................... [166560]
AY261365.1_Warmbaths             ................................................................................ [166560]
AY261364.1_Tengani_62            ................................................................................ [166560]
AY261363.1_Pretoriuskop/96/4     ....................t........................................................... [166560]
AY261362.1_Mkuzi_1979            ................................a............................................... [166560]
AY261361.1_Malawi_Lil-20/1_1983  ................................a...........c................................... [166560]
AY261360.1_Kenya_1950            .................a..............a...........c................................... [166560]
AM712240.1_OURT_88/3             ................................a............................................... [166560]
AM712239.1_Benin_97/1            ................................a............................................... [166560]
KM111295.1_Ken06.Bus             ..a..............a..............a...........c................................... [166560]
KM111294.1_Ken05/Tk1             .................a..............a...........c................................... [166560]
KM262844.1_strain_L60            ................................a............................................... [166560]
KM102979.1_26544/OG10_Italy      ................................a............................................... [166560]
KM262845.1_strain_NHV            ................................a............................................... [166560]
KX354450.1_47/Ss/2008            ................................a............................................... [166560]
KP055815.1_strain_BA71           ................................a............................................... [166560]
MH025920.1_strain_R35            ..a..............a..............a...........c................................... [166560]
MH025919.1_strain_N10            ..a..............a..............a...........c................................... [166560]
MH025918.1_strain_R25            ..a..............a..............a...........c................................... [166560]
MH025917.1_strain_R7             ..a..............a..............a...........c................................... [166560]
MH025916.1_strain_R8             ..a..............a..............a...........c................................... [166560]

FR682468.2_Georgia_2007/1        gaagtacaccacttgggcccggcaactggaagacattttggccttggtgagggcctacgtcgacaaagacaacccaaaac [166640]
ASF_IND_20_CAD543                ................................................................................ [166640]
LR899193.1_Germany_2020/1        ................................................................................ [166640]
MN393477.1_Wuhan_2019            ................................................................................ [166640]
MW396979.1_Timor-Leste/2019/1    ................................................................................ [166640]
MK940252.1_CN/2019/InnerMongolia ................................................................................ [166640]
MK128995.1_China/2018/AnhuiXCGQ  ................................................................................ [166640]
MH681419.1_ASFV/POL/2015         ................................................................................ [166640]
LS478113.1_Estonia_2014          ................................................................................ [166640]
KP843857.1_Odintsovo_02/14       ................................................................................ [166640]
FN557520.1_strain_E75            ................................................................................ [166640]
AY261366.1_Warthog               .....................c.........................................t..ga............ [166640]
AY261365.1_Warmbaths             .....................c.........................................t..ga............ [166640]
AY261364.1_Tengani_62            .....................c.............................a...........t..ga............ [166640]
AY261363.1_Pretoriuskop/96/4     .....................c.........................................t..ga............ [166640]
AY261362.1_Mkuzi_1979            ................................................................................ [166640]
AY261361.1_Malawi_Lil-20/1_1983  ............c........c.........................................t..ga............ [166640]
AY261360.1_Kenya_1950            .....................c....................................a....t..ga............ [166640]
AM712240.1_OURT_88/3             ................................................................................ [166640]
AM712239.1_Benin_97/1            ................................................................................ [166640]
KM111295.1_Ken06.Bus             .....................c.........................................t..ga....t....... [166640]
```

```
KM111294.1_Ken05/Tk1             ...a.................c....................t..a..a..a...........t..ga............ [166640]
KM262844.1_strain_L60            ................................................................................ [166640]
KM102979.1_26544/OG10_Italy      ................................................................................ [166640]
KM262845.1_strain_NHV            ................................................................................ [166640]
KX354450.1_47/Ss/2008            ................................................................................ [166640]
KP055815.1_strain_BA71           ................................................................................ [166640]
MH025920.1_strain_R35            .....................c.........................................t..ga....t....... [166640]
MH025919.1_strain_N10            .....................c.........................................t..ga....t....... [166640]
MH025918.1_strain_R25            .....................c.........................................t..ga....t....... [166640]
MH025917.1_strain_R7             .....................c.........................................t..ga....t....... [166640]
MH025916.1_strain_R8             .....................c.........................................t..ga....t....... [166640]

FR682468.2_Georgia_2007/1        acgagctactgcactatgcaataaaacataagattactatactcccgctgcggccctccaattacaatttcaagggccat [166720]
ASF_IND_20_CAD543                ................................................................................ [166720]
LR899193.1_Germany_2020/1        ................................................................................ [166720]
MN393477.1_Wuhan_2019            ................................................................................ [166720]
MW396979.1_Timor-Leste/2019/1    ................................................................................ [166720]
MK940252.1_CN/2019/InnerMongolia ................................................................................ [166720]
MK128995.1_China/2018/AnhuiXCGQ  ................................................................................ [166720]
MH681419.1_ASFV/POL/2015         ................................................................................ [166720]
LS478113.1_Estonia_2014          ................................................................................ [166720]
KP843857.1_Odintsovo_02/14       ................................................................................ [166720]
FN557520.1_strain_E75            ................................................................................ [166720]
AY261366.1_Warthog               ................................................................................ [166720]
AY261365.1_Warmbaths             .......................g.c........c...g......................................... [166720]
AY261364.1_Tengani_62            ......................................................................t......... [166720]
AY261363.1_Pretoriuskop/96/4     ..........a..................................................................... [166720]
AY261362.1_Mkuzi_1979            ..........a..............................t...................................... [166720]
AY261361.1_Malawi_Lil-20/1_1983  ................c......g...g......c...g....t.....a.....t..t...........t.....t..c [166720]
AY261360.1_Kenya_1950            .......c........c......g.t............g..........a........t...........t.....t..c [166720]
AM712240.1_OURT_88/3             ................................................................................ [166720]
AM712239.1_Benin_97/1            ................................................................................ [166720]
KM111295.1_Ken06.Bus             .......c........c......g.c............g....t.....a........t...........t.....t..c [166720]
KM111294.1_Ken05/Tk1             .......c........c......g.c............g..........a........t...........t.....t..c [166720]
KM262844.1_strain_L60            ................................................................................ [166720]
KM102979.1_26544/OG10_Italy      ................................................................................ [166720]
KM262845.1_strain_NHV            ................................................................................ [166720]
KX354450.1_47/Ss/2008            ................................................................................ [166720]
KP055815.1_strain_BA71           ................................................................................ [166720]
MH025920.1_strain_R35            .......c........c......g.c............g....t.....a........t...........t.....t..c [166720]
MH025919.1_strain_N10            .......c........c......g.c............g....t.....a........t...........t.....t..c [166720]
MH025918.1_strain_R25            .......c........c......g.c............g....t.....a........t...........t.....t..c [166720]
MH025917.1_strain_R7             .......c........c......g.c............g....t.....a........t...........t.....t..c [166720]
MH025916.1_strain_R8             .......c........c......g.c............g....t.....a........t...........t.....t..c [166720]
```

```
FR682468.2_Georgia_2007/1        ttgaagcggtttggccaatactactacagctacggcacgtacgtcatctcagagcagcgaaatataattactattacgga [166800]
ASF_IND_20_CAD543                ................................................................................ [166800]
LR899193.1_Germany_2020/1        ................................................................................ [166800]
MN393477.1_Wuhan_2019            ................................................................................ [166800]
MW396979.1_Timor-Leste/2019/1    ................................................................................ [166800]
MK940252.1_CN/2019/InnerMongolia ................................................................................ [166800]
MK128995.1_China/2018/AnhuiXCGQ  ................................................................................ [166800]
MH681419.1_ASFV/POL/2015         ................................................................................ [166800]
LS478113.1_Estonia_2014          ................................................................................ [166800]
KP843857.1_Odintsovo_02/14       ................................................................................ [166800]
FN557520.1_strain_E75            ...........................................a.................................... [166800]
AY261366.1_Warthog               ............................................g................................... [166800]
AY261365.1_Warmbaths             ..............t.............................g................................... [166800]
AY261364.1_Tengani_62            ..............t...................................g............................. [166800]
AY261363.1_Pretoriuskop/96/4     ..............t................................................................. [166800]
AY261362.1_Mkuzi_1979            ................................................................................ [166800]
AY261361.1_Malawi_Lil-20/1_1983  .....a..a..c...........t..............t......g.................................. [166800]
AY261360.1_Kenya_1950            .....a..a..c..........................t......g...................g.............. [166800]
AM712240.1_OURT_88/3             ...........................................a.................................... [166800]
AM712239.1_Benin_97/1            ...........................................a.................................... [166800]
KM111295.1_Ken06.Bus             .....a..a..c..........................t......g...................g.............. [166800]
KM111294.1_Ken05/Tk1             .....a..a..c..........................t......g...................g.............. [166800]
KM262844.1_strain_L60            ...........................................a.................................... [166800]
KM102979.1_26544/OG10_Italy      ...........................................a.................................... [166800]
KM262845.1_strain_NHV            ...........................................a.................................... [166800]
KX354450.1_47/Ss/2008            ...........................................a.................................... [166800]
KP055815.1_strain_BA71           ...........................................a.................................... [166800]
MH025920.1_strain_R35            .....a..a..c..........................t......g...................g.............. [166800]
MH025919.1_strain_N10            .....a..a..c..........................t......g...................g.............. [166800]
MH025918.1_strain_R25            .....a..a..c..........................t......g...................g.............. [166800]
MH025917.1_strain_R7             .....a..a..c..........................t......g...................g.............. [166800]
MH025916.1_strain_R8             .....a..a..c..........................t......g...................g.............. [166800]

FR682468.2_Georgia_2007/1        gcttcctctgcgtgttcctacggttgcatacatcgaaagtataaaaaaatcgagtaaccgcatgacatttattgaagaaa [166880]
ASF_IND_20_CAD543                ................................................................................ [166880]
LR899193.1_Germany_2020/1        ................................................................................ [166880]
MN393477.1_Wuhan_2019            ................................................................................ [166880]
MW396979.1_Timor-Leste/2019/1    ................................................................................ [166880]
MK940252.1_CN/2019/InnerMongolia ................................................................................ [166880]
MK128995.1_China/2018/AnhuiXCGQ  ................................................................................ [166880]
MH681419.1_ASFV/POL/2015         ................................................................................ [166880]
LS478113.1_Estonia_2014          ................................................................................ [166880]
KP843857.1_Odintsovo_02/14       ................................................................................ [166880]
```

```
FN557520.1_strain_E75            ..............................t................................................. [166880]
AY261366.1_Warthog               ...........................t..t........c..........................c............. [166880]
AY261365.1_Warmbaths             ...........................t..t......................c.......................... [166880]
AY261364.1_Tengani_62            ................................................................t............... [166880]
AY261363.1_Pretoriuskop/96/4     ...........................t..t.......................c......................... [166880]
AY261362.1_Mkuzi_1979            .........................................................t...................... [166880]
AY261361.1_Malawi_Lil-20/1_1983  .........a.................t...........c........................g............... [166880]
AY261360.1_Kenya_1950            ......g....................t.....t..............................g............... [166880]
AM712240.1_OURT_88/3             ..............................t................................................. [166880]
AM712239.1_Benin_97/1            ..............................t................................................. [166880]
KM111295.1_Ken06.Bus             ...c..g....................t.....t..............................g............... [166880]
KM111294.1_Ken05/Tk1             ......g....................t.....t..............................g............... [166880]
KM262844.1_strain_L60            ..............................t................................................. [166880]
KM102979.1_26544/OG10_Italy      ..............................t................................................. [166880]
KM262845.1_strain_NHV            ..............................t................................................. [166880]
KX354450.1_47/Ss/2008            ..............................t................................................. [166880]
KP055815.1_strain_BA71           ..............................t................................................. [166880]
MH025920.1_strain_R35            ...c..g....................t.....t..............................g............... [166880]
MH025919.1_strain_N10            ...c..g....................t.....t..............................g............... [166880]
MH025918.1_strain_R25            ...c..g....................t.....t..............................g............... [166880]
MH025917.1_strain_R7             ...c..g....................t.....t..............................g............... [166880]
MH025916.1_strain_R8             ...c..g....................t.....t..............................g............... [166880]

FR682468.2_Georgia_2007/1        tcatcgactacagtagttcagaaactattgaaattctggtgaaattaaagccaaatagtcttaaccgtatcgtggaagaa [166960]
ASF_IND_20_CAD543                ................................................................................ [166960]
LR899193.1_Germany_2020/1        ................................................................................ [166960]
MN393477.1_Wuhan_2019            ................................................................................ [166960]
MW396979.1_Timor-Leste/2019/1    ................................................................................ [166960]
MK940252.1_CN/2019/InnerMongolia ................................................................................ [166960]
MK128995.1_China/2018/AnhuiXCGQ  ................................................................................ [166960]
MH681419.1_ASFV/POL/2015         ................................................................................ [166960]
LS478113.1_Estonia_2014          ................................................................................ [166960]
KP843857.1_Odintsovo_02/14       ................................................................................ [166960]
FN557520.1_strain_E75            .........................c..................c................c.................. [166960]
AY261366.1_Warthog               .........................c.....g.....a.......................c.........a........ [166960]
AY261365.1_Warmbaths             .........................c.....g..........................c..c.........a........ [166960]
AY261364.1_Tengani_62            .........................c...................................................... [166960]
AY261363.1_Pretoriuskop/96/4     .........................c.....g.............................c.........a........ [166960]
AY261362.1_Mkuzi_1979            ........................tc...................................c.................. [166960]
AY261361.1_Malawi_Lil-20/1_1983  ....t..............g.....c.................g.................c.................. [166960]
AY261360.1_Kenya_1950            ..g......................c.........t....a..gc.g...........c....g...g...a.a...... [166960]
AM712240.1_OURT_88/3             .........................c..................c................c.................. [166960]
AM712239.1_Benin_97/1            .........................c..................c................c.................. [166960]
KM111295.1_Ken06.Bus             ..g......................c.........t....a..gc.g...........c....g...g...a........ [166960]
```

```
KM111294.1_Ken05/Tk1          ..g......................c.........t....a..gc.g...........c....g...g...a.a...... [166960]
KM262844.1_strain_L60         .........................c..................c................c.................. [166960]
KM102979.1_26544/OG10_Italy   .........................c..................c................c.................. [166960]
KM262845.1_strain_NHV         .........................c..................c................c.................. [166960]
KX354450.1_47/Ss/2008         .........................c..................c................c.................. [166960]
KP055815.1_strain_BA71        .........................c..................c................c.................. [166960]
MH025920.1_strain_R35         ..g......................c.........t....a..gc.g...........c....g...g...a........ [166960]
MH025919.1_strain_N10         ..g......................c..............a..gc.g...........c....g...g...a........ [166960]
MH025918.1_strain_R25         ..g......................c.........t....a..gc.g...........c....g...g...a........ [166960]
MH025917.1_strain_R7          ..g......................c.........t....a..gc.g...........c....g...g...a........ [166960]
MH025916.1_strain_R8          ..g......................c.........t....a..gc.g...........c....g...g...a........ [166960]

FR682468.2_Georgia_2007/1     tttaaggagactgaagagcaagattccatagaaaattttctgcgcctgcgcaattgtttacattcacatctaaactttgt [167040]
ASF_IND_20_CAD543             ................................................................................ [167040]
LR899193.1_Germany_2020/1     ................................................................................ [167040]
MN393477.1_Wuhan_2019         ................................................................................ [167040]
MW396979.1_Timor-Leste/2019/1 ................................................................................ [167040]
MK940252.1_CN/2019/InnerMongolia ................................................................................ [167040]
MK128995.1_China/2018/AnhuiXCGQ ................................................................................ [167040]
MH681419.1_ASFV/POL/2015      ................................................................................ [167040]
LS478113.1_Estonia_2014       ................................................................................ [167040]
KP843857.1_Odintsovo_02/14    ................................................................................ [167040]
FN557520.1_strain_E75         .................................................................g.............. [167040]
AY261366.1_Warthog            ......tgc..a..g..................................................g.............. [167040]
AY261365.1_Warmbaths          ......tgc..a..g..................................................g.............. [167040]
AY261364.1_Tengani_62         ......tgc..a..g.............................t....................g.............. [167040]
AY261363.1_Pretoriuskop/96/4  ......tgc..a..g..................................................g.............. [167040]
AY261362.1_Mkuzi_1979         .................................................................g.............. [167040]
AY261361.1_Malawi_Lil-20/1_1983 .................a.........................................g.....g..ct.......... [167040]
AY261360.1_Kenya_1950         ..............g.....ga....t........c.....a.....a...........g.....g..c........... [167040]
AM712240.1_OURT_88/3          .................................................................g.............. [167040]
AM712239.1_Benin_97/1         .................................................................g.............. [167040]
KM111295.1_Ken06.Bus          ..............g.....ga....t..............a..tt.a.................g..c........... [167040]
KM111294.1_Ken05/Tk1          ..............g.....ga....t........c.....a.....a.................g..c........... [167040]
KM262844.1_strain_L60         .................................................................g.............. [167040]
KM102979.1_26544/OG10_Italy   .................................................................g.............. [167040]
KM262845.1_strain_NHV         .................................................................g.............. [167040]
KX354450.1_47/Ss/2008         .................................................................g.............. [167040]
KP055815.1_strain_BA71        .................................................................g.............. [167040]
MH025920.1_strain_R35         ..............g.....ga....t..............a..tt.a.................g..c........... [167040]
MH025919.1_strain_N10         ..............g.....ga....t..............a..tt.a.................g..c........... [167040]
MH025918.1_strain_R25         ..............g.....ga....t..............a..tt.a.................g..c........... [167040]
MH025917.1_strain_R7          ..............g.....ga....t..............a..tt.a.................g..c........... [167040]
MH025916.1_strain_R8          ..............g.....ga....t..............a..tt.a.................g..c........... [167040]
```

```
FR682468.2_Georgia_2007/1        aaaacctaaaggtggcattatcgagtttaacacgtattatgaaattttgtatgcgtggctaccttacaggcgtgagcttt [167120]
ASF_IND_20_CAD543                ................................................................................ [167120]
LR899193.1_Germany_2020/1        ................................................................................ [167120]
MN393477.1_Wuhan_2019            ................................................................................ [167120]
MW396979.1_Timor-Leste/2019/1    ................................................................................ [167120]
MK940252.1_CN/2019/InnerMongolia ................................................................................ [167120]
MK128995.1_China/2018/AnhuiXCGQ  ................................................................................ [167120]
MH681419.1_ASFV/POL/2015         ................................................................................ [167120]
LS478113.1_Estonia_2014          ................................................................................ [167120]
KP843857.1_Odintsovo_02/14       ................................................................................ [167120]
FN557520.1_strain_E75            ...............t...............t.a..............a............................... [167120]
AY261366.1_Warthog               ...............t.....t.......................................................... [167120]
AY261365.1_Warmbaths             ...............t.....t.......................................................... [167120]
AY261364.1_Tengani_62            ...............t.....t..................................................c....... [167120]
AY261363.1_Pretoriuskop/96/4     ...............t.....t........t................................................. [167120]
AY261362.1_Mkuzi_1979            ............c..t................................a............................... [167120]
AY261361.1_Malawi_Lil-20/1_1983  ...............t...............t................a...........g................... [167120]
AY261360.1_Kenya_1950            ............a.....c............t................a..c........g..............t.... [167120]
AM712240.1_OURT_88/3             ...............t...............t.a..............a............................... [167120]
AM712239.1_Benin_97/1            ...............t...............t.a..............a............................... [167120]
KM111295.1_Ken06.Bus             ............a.....c............t.a..............a..c..a.....g..............t.... [167120]
KM111294.1_Ken05/Tk1             ............g.....c............t.a..............a..c........g................... [167120]
KM262844.1_strain_L60            ...............t...............t.a..............a............................... [167120]
KM102979.1_26544/OG10_Italy      ...............t...............t.a..............a............................... [167120]
KM262845.1_strain_NHV            ...............t...............t.a..............a............................... [167120]
KX354450.1_47/Ss/2008            ...............t...............t.a..............a............................... [167120]
KP055815.1_strain_BA71           ...............t...............t.a..............a............................... [167120]
MH025920.1_strain_R35            ............a.....c............t.a..............a..c..a.....g..............t.... [167120]
MH025919.1_strain_N10            ............a.....c............t.a..............a..c..a.....g..............t.... [167120]
MH025918.1_strain_R25            ............a.....c............t.a..............a..c..a.....g..............t.... [167120]
MH025917.1_strain_R7             ............a.....c............t.a..............a..c..a.....g..............t.... [167120]
MH025916.1_strain_R8             ............a.....c............t.a..............a..c..a.....g..............t.... [167120]

FR682468.2_Georgia_2007/1        accaaaagcgtcttatgcgtgagcacgcggtgcttaagctgcgcattatcatggaaactgctattgtacgctacatcaat [167200]
ASF_IND_20_CAD543                ................................................................................ [167200]
LR899193.1_Germany_2020/1        ................................................................................ [167200]
MN393477.1_Wuhan_2019            ................................................................................ [167200]
MW396979.1_Timor-Leste/2019/1    ................................................................................ [167200]
MK940252.1_CN/2019/InnerMongolia ................................................................................ [167200]
MK128995.1_China/2018/AnhuiXCGQ  ................................................................................ [167200]
MH681419.1_ASFV/POL/2015         ................................................................................ [167200]
LS478113.1_Estonia_2014          ................................................................................ [167200]
KP843857.1_Odintsovo_02/14       ................................................................................ [167200]
```

```
FN557520.1_strain_E75           ................................................................................ [167200]
AY261366.1_Warthog              ...................c.................a.......................................... [167200]
AY261365.1_Warmbaths            ...................c............................................................ [167200]
AY261364.1_Tengani_62           ...................c.............................t.............................. [167200]
AY261363.1_Pretoriuskop/96/4    ...................c............................................................ [167200]
AY261362.1_Mkuzi_1979           ..........................a..................................................... [167200]
AY261361.1_Malawi_Lil-20/1_1983 .......a................g.........a.......................c..c.....g............ [167200]
AY261360.1_Kenya_1950           .......a................g...c...t.a..........................c.....g............ [167200]
AM712240.1_OURT_88/3            ................................................................................ [167200]
AM712239.1_Benin_97/1           ................................................................................ [167200]
KM111295.1_Ken06.Bus            .......a................g...c...t.a..a......c................c.....g............ [167200]
KM111294.1_Ken05/Tk1            .......a................g...c...t.a..........................c.....g............ [167200]
KM262844.1_strain_L60           ................................................................................ [167200]
KM102979.1_26544/OG10_Italy     ................................................................................ [167200]
KM262845.1_strain_NHV           ................................................................................ [167200]
KX354450.1_47/Ss/2008           ................................................................................ [167200]
KP055815.1_strain_BA71          ................................................................................ [167200]
MH025920.1_strain_R35           .......a................g...c...t.a.........c................c.....g............ [167200]
MH025919.1_strain_N10           .......a................g...c...t.a.........c................c.....g............ [167200]
MH025918.1_strain_R25           .......a................g...c...t.a.........c................c.....g............ [167200]
MH025917.1_strain_R7            .......a................g...c...t.a.........c................c.....g............ [167200]
MH025916.1_strain_R8            .......a................g...c...t.a.........c................c.....g............ [167200]

FR682468.2_Georgia_2007/1       gagtctgcagagctaaatctttcccattatgaggatgaaaaggaggcaagccgcattctaagcgagcatggatttccccc [167280]
ASF_IND_20_CAD543               ................................................................................ [167280]
LR899193.1_Germany_2020/1       ................................................................................ [167280]
MN393477.1_Wuhan_2019           ................................................................................ [167280]
MW396979.1_Timor-Leste/2019/1   ................................................................................ [167280]
MK940252.1_CN/2019/InnerMongolia ................................................................................ [167280]
MK128995.1_China/2018/AnhuiXCGQ ................................................................................ [167280]
MH681419.1_ASFV/POL/2015        ................................................................................ [167280]
LS478113.1_Estonia_2014         ................................................................................ [167280]
KP843857.1_Odintsovo_02/14      ................................................................................ [167280]
FN557520.1_strain_E75           ................................................................................ [167280]
AY261366.1_Warthog              ..........................................................................c..t.. [167280]
AY261365.1_Warmbaths            ..........................................................................c..t.. [167280]
AY261364.1_Tengani_62           ................................................................................ [167280]
AY261363.1_Pretoriuskop/96/4    ..........................................................................c..t.. [167280]
AY261362.1_Mkuzi_1979           ................................................................................ [167280]
AY261361.1_Malawi_Lil-20/1_1983 .....a.....t..............c..c.................g.....t...t.............g..c..t.. [167280]
AY261360.1_Kenya_1950           .....c..g..t..............c....................gg....t...t.g........c..g..c..... [167280]
AM712240.1_OURT_88/3            ................................................................................ [167280]
AM712239.1_Benin_97/1           ................................................................................ [167280]
KM111295.1_Ken06.Bus            .....c..g..t..............c....................gg....t...t.g........c..g..c..... [167280]
```

| | | |
|---|---|---|
| KM111294.1_Ken05/Tk1 | .....c..g..t..............c....................gg....t...t.g........c..g..c..... | [167280] |
| KM262844.1_strain_L60 | ................................................................................ | [167280] |
| KM102979.1_26544/OG10_Italy | ................................................................................ | [167280] |
| KM262845.1_strain_NHV | ................................................................................ | [167280] |
| KX354450.1_47/Ss/2008 | ................................................................................ | [167280] |
| KP055815.1_strain_BA71 | ................................................................................ | [167280] |
| MH025920.1_strain_R35 | .....c..g..t..............c....................gg....t...t.g........c..g..c..... | [167280] |
| MH025919.1_strain_N10 | .....c..g..t..............c....................gg....t...t.g........c..g..c..... | [167280] |
| MH025918.1_strain_R25 | .....c..g..t..............c....................gg....t...t.g........c..g..c..... | [167280] |
| MH025917.1_strain_R7 | .....c..g..t..............c....................gg....t...t.g........c..g..c..... | [167280] |
| MH025916.1_strain_R8 | .....c..g..t..............c....................gg....t...t.g........c..g..c..... | [167280] |

| | | |
|---|---|---|
| FR682468.2_Georgia_2007/1 | gctgaaccacacgctgatcatttcccctgagtttgcctctatagaggaactcaatcaaaaagcactgcagggctgttata | [167360] |
| ASF_IND_20_CAD543 | ................................................................................ | [167360] |
| LR899193.1_Germany_2020/1 | ................................................................................ | [167360] |
| MN393477.1_Wuhan_2019 | ................................................................................ | [167360] |
| MW396979.1_Timor-Leste/2019/1 | ................................................................................ | [167360] |
| MK940252.1_CN/2019/InnerMongolia | ................................................................................ | [167360] |
| MK128995.1_China/2018/AnhuiXCGQ | ................................................................................ | [167360] |
| MH681419.1_ASFV/POL/2015 | ................................................................................ | [167360] |
| LS478113.1_Estonia_2014 | ................................................................................ | [167360] |
| KP843857.1_Odintsovo_02/14 | ................................................................................ | [167360] |
| FN557520.1_strain_E75 | ...............................................................g................ | [167360] |
| AY261366.1_Warthog | .........g..........................t.................c........g................ | [167360] |
| AY261365.1_Warmbaths | .........g............................................c........g................ | [167360] |
| AY261364.1_Tengani_62 | .........g............................................c........g................ | [167360] |
| AY261363.1_Pretoriuskop/96/4 | .........g............................................c........g................ | [167360] |
| AY261362.1_Mkuzi_1979 | ...............................................................g................ | [167360] |
| AY261361.1_Malawi_Lil-20/1_1983 | ...a.....gt.a..c..t.c...t..g...........c.......................g................ | [167360] |
| AY261360.1_Kenya_1950 | ...a.....gt....c...cc...t..g.........a.............g..c......................... | [167360] |
| AM712240.1_OURT_88/3 | ...............................................................g................ | [167360] |
| AM712239.1_Benin_97/1 | ...............................................................g................ | [167360] |
| KM111295.1_Ken06.Bus | ...a.....gt....c...cc...t..g.........a.c..............c......................... | [167360] |
| KM111294.1_Ken05/Tk1 | ...a.....gt....c...cc...t..g.........a.............g..c......................... | [167360] |
| KM262844.1_strain_L60 | ...............................................................g................ | [167360] |
| KM102979.1_26544/OG10_Italy | ...............................................................g................ | [167360] |
| KM262845.1_strain_NHV | ...............................................................g................ | [167360] |
| KX354450.1_47/Ss/2008 | ...............................................................g................ | [167360] |
| KP055815.1_strain_BA71 | ...............................................................g................ | [167360] |
| MH025920.1_strain_R35 | ...a.....gt....c...cc...t..g.........a.c..............c......................... | [167360] |
| MH025919.1_strain_N10 | ...a.....gt....c...cc...t..g.........a.c..............c......................... | [167360] |
| MH025918.1_strain_R25 | ...a.....gt....c...cc...t..g.........a.c..............c......................... | [167360] |
| MH025917.1_strain_R7 | ...a.....gt....c...cc...t..g.........a.c..............c......................... | [167360] |
| MH025916.1_strain_R8 | ...a.....gt....c...cc...t..g.........a.c..............c......................... | [167360] |

```
FR682468.2_Georgia_2007/1        cctatatactatctttgcaggctcgagaattgcttatcgcagccaaaactcgtcgggtggaaaaaata-aaaaaaatgca [167440]
ASF_IND_20_CAD543                ....................................................................-........... [167440]
LR899193.1_Germany_2020/1        ....................................................................-........... [167440]
MN393477.1_Wuhan_2019            ....................................................................-........... [167440]
MW396979.1_Timor-Leste/2019/1    ....................................................................-........... [167440]
MK940252.1_CN/2019/InnerMongolia ....................................................................-........... [167440]
MK128995.1_China/2018/AnhuiXCGQ  ....................................................................-........... [167440]
MH681419.1_ASFV/POL/2015         ....................................................................-........... [167440]
LS478113.1_Estonia_2014          ....................................................................-........... [167440]
KP843857.1_Odintsovo_02/14       ....................................................................-........... [167440]
FN557520.1_strain_E75            ....................................................................a........... [167440]
AY261366.1_Warthog               .t......................................g...........................-........... [167440]
AY261365.1_Warmbaths             ....................................................................-........... [167440]
AY261364.1_Tengani_62            .............c......................................................-........... [167440]
AY261363.1_Pretoriuskop/96/4     ....................................................................-........... [167440]
AY261362.1_Mkuzi_1979            ....................................................................-........... [167440]
AY261361.1_Malawi_Lil-20/1_1983  .......c..............c.................g..t..g.....................-........... [167440]
AY261360.1_Kenya_1950            ................a.............................g....c...............-........... [167440]
AM712240.1_OURT_88/3             ....................................................................-........... [167440]
AM712239.1_Benin_97/1            ...................................................a................-........... [167440]
KM111295.1_Ken06.Bus             ..............................................g.....................-........... [167440]
KM111294.1_Ken05/Tk1             ..............................................g....c...............-........... [167440]
KM262844.1_strain_L60            ....................................................................-........... [167440]
KM102979.1_26544/OG10_Italy      ....................................................................-........... [167440]
KM262845.1_strain_NHV            ....................................................................-........... [167440]
KX354450.1_47/Ss/2008            ....................................................................-........... [167440]
KP055815.1_strain_BA71           ....................................................................-........... [167440]
MH025920.1_strain_R35            ..............................................g.....................-........... [167440]
MH025919.1_strain_N10            ..............................................g.....................-........... [167440]
MH025918.1_strain_R25            ..............................................g.....................-........... [167440]
MH025917.1_strain_R7             ..............................................g.....................-........... [167440]
MH025916.1_strain_R8             ..............................................g.....................-........... [167440]

FR682468.2_Georgia_2007/1        agctcgtcttgataaggttgagcagcttttgcaagagtctccctttcccggcgccagcgtatggctggaggaaattgatg [167520]
ASF_IND_20_CAD543                .................................r.............................................. [167520]
LR899193.1_Germany_2020/1        ................................................................................ [167520]
MN393477.1_Wuhan_2019            ................................................................................ [167520]
MW396979.1_Timor-Leste/2019/1    ................................................................................ [167520]
MK940252.1_CN/2019/InnerMongolia ................................................................................ [167520]
MK128995.1_China/2018/AnhuiXCGQ  ................................................................................ [167520]
MH681419.1_ASFV/POL/2015         ................................................................................ [167520]
LS478113.1_Estonia_2014          ................................................................................ [167520]
KP843857.1_Odintsovo_02/14       ................................................................................ [167520]
```

```
FN557520.1_strain_E75             .................................g.............................................. [167520]
AY261366.1_Warthog                g................................g.....a........................................ [167520]
AY261365.1_Warmbaths              g................................g.............................................. [167520]
AY261364.1_Tengani_62             g....................a...........g.............................................. [167520]
AY261363.1_Pretoriuskop/96/4      g................................g.............................................. [167520]
AY261362.1_Mkuzi_1979             g................................g.............................................. [167520]
AY261361.1_Malawi_Lil-20/1_1983   ...g.............................g..........................g................... [167520]
AY261360.1_Kenya_1950             ...g.............................g..............................t.......g....... [167520]
AM712240.1_OURT_88/3              .................................g.............................................. [167520]
AM712239.1_Benin_97/1             .................................g.............................................. [167520]
KM111295.1_Ken06.Bus              ...g.............................g..............................t.......g....... [167520]
KM111294.1_Ken05/Tk1              ...g.............................g..............................t.......g....... [167520]
KM262844.1_strain_L60             .................................g.............................................. [167520]
KM102979.1_26544/OG10_Italy       .................................g.............................................. [167520]
KM262845.1_strain_NHV             .................................g.............................................. [167520]
KX354450.1_47/Ss/2008             .................................g.............................................. [167520]
KP055815.1_strain_BA71            .................................g.............................................. [167520]
MH025920.1_strain_R35             ...g.............................g..............................t.......g....... [167520]
MH025919.1_strain_N10             ...g.............................g..............................t.......g....... [167520]
MH025918.1_strain_R25             ...g.............................g..............................t.......g....... [167520]
MH025917.1_strain_R7              ...g.............................g..............................t.......g....... [167520]
MH025916.1_strain_R8              ...g.............................g..............................t.......g....... [167520]

FR682468.2_Georgia_2007/1         cggtggaaaaggctattataaaaggaagaaatactcagtggaaatttcattaaacgc-ta-ccggttt--tatgatgtcc [167600]
ASF_IND_20_CAD543                 .........................................................-..-.....-.--.......... [167600]
LR899193.1_Germany_2020/1         .........................................................-..-.......--.......... [167600]
MN393477.1_Wuhan_2019             .........................................................-..-.......--.......... [167600]
MW396979.1_Timor-Leste/2019/1     .........................................................-..-.......--.......... [167600]
MK940252.1_CN/2019/InnerMongolia  .........................................................-..-.......--.......... [167600]
MK128995.1_China/2018/AnhuiXCGQ   .........................................................-..-.......--.......... [167600]
MH681419.1_ASFV/POL/2015          .........................................................-..-.......--.......... [167600]
LS478113.1_Estonia_2014           .........................................................-..-.......--.......... [167600]
KP843857.1_Odintsovo_02/14        .........................................................-..-.......--.......... [167600]
FN557520.1_strain_E75             .........................................................-..-.....-.--.......... [167600]
AY261366.1_Warthog                .........................................................-..-.......--.......... [167600]
AY261365.1_Warmbaths              .........................................................-..-.......--.......... [167600]
AY261364.1_Tengani_62             .........................................................-..-.......--....g..... [167600]
AY261363.1_Pretoriuskop/96/4      .....................................................g...-..c.......--.......... [167600]
AY261362.1_Mkuzi_1979             .........................................................-..-.......--.......... [167600]
AY261361.1_Malawi_Lil-20/1_1983   ..........a...................gc.........................-..-.t.a..c--......c... [167600]
AY261360.1_Kenya_1950             ..........a....................c..a...................t.ga.gcta..c..ta...t...a.- [167600]
AM712240.1_OURT_88/3              .........................................................-..-.....-.--.......... [167600]
AM712239.1_Benin_97/1             .........................................................-..-.....-.--.......... [167600]
KM111295.1_Ken06.Bus              ..........a.......................a...................t..a.gcta..c..--...t...a-- [167600]
```

```
KM111294.1_Ken05/Tk1          ..........a....................c..a...................t.ga.gcta..c..--...t...a.. [167600]
KM262844.1_strain_L60         .........................................................-..-....-..--.......... [167600]
KM102979.1_26544/OG10_Italy   .........................................................-..-....-..--.......... [167600]
KM262845.1_strain_NHV         .........................................................-..-....-..--.......... [167600]
KX354450.1_47/Ss/2008         .........................................................-..-....-..--.......... [167600]
KP055815.1_strain_BA71        .........................................................-..-....-..--.......... [167600]
MH025920.1_strain_R35         ..........a.......................a...................t..a.gcta..c..--...t...a-- [167600]
MH025919.1_strain_N10         ..........a.......................a...................t..a.gcta..c..--...t...a-- [167600]
MH025918.1_strain_R25         ..........a.......................a...................t..a.gcta..c..--...t...a-- [167600]
MH025917.1_strain_R7          ..........a.......................a...................t..a.gcta..c..--...t...a-- [167600]
MH025916.1_strain_R8          ..........a.......................a...................t..a.gcta..c..--...t...a-- [167600]

FR682468.2_Georgia_2007/1     ---aataggtgttaagcaatcagttcatcaacatttttttcaagaatttgaaaagtttggataatgttctgaatactttt [167680]
ASF_IND_20_CAD543             ---.........................................r................................... [167680]
LR899193.1_Germany_2020/1     ---............................................................................. [167680]
MN393477.1_Wuhan_2019         ---............................................................................. [167680]
MW396979.1_Timor-Leste/2019/1 ---............................................................................. [167680]
MK940252.1_CN/2019/InnerMongolia ---.......................................................................... [167680]
MK128995.1_China/2018/AnhuiXCGQ ---............................................................................ [167680]
MH681419.1_ASFV/POL/2015      ---............................................................................. [167680]
LS478113.1_Estonia_2014       ---............................................................................. [167680]
KP843857.1_Odintsovo_02/14    ---............................................................................. [167680]
FN557520.1_strain_E75         ---.........................................g..........................g........ [167680]
AY261366.1_Warthog            ---.........................................g.............................g..... [167680]
AY261365.1_Warmbaths          ---.........................................g................................... [167680]
AY261364.1_Tengani_62         --a.........................................g................................... [167680]
AY261363.1_Pretoriuskop/96/4  ---.................t.......................g.............................g..... [167680]
AY261362.1_Mkuzi_1979         ---.........................................g................................... [167680]
AY261361.1_Malawi_Lil-20/1_1983 atat................t..........t............gg.......ca......................... [167680]
AY261360.1_Kenya_1950         -cat....a...........tc.........t............gg.......ta..........a.............. [167680]
AM712240.1_OURT_88/3          ---.........................................g..........................g........ [167680]
AM712239.1_Benin_97/1         ---.........................................g..........................g........ [167680]
KM111295.1_Ken06.Bus          -cat....a...........tc.........t............gg.......ta......................... [167680]
KM111294.1_Ken05/Tk1          atat....a...........tc.........t............gg.......ta..........a.............. [167680]
KM262844.1_strain_L60         ---.........................................g..........................g........ [167680]
KM102979.1_26544/OG10_Italy   ---.........................................g..........................g........ [167680]
KM262845.1_strain_NHV         ---.........................................g..........................g........ [167680]
KX354450.1_47/Ss/2008         ---.........................................g..........................g........ [167680]
KP055815.1_strain_BA71        ---.........................................g..........................g........ [167680]
MH025920.1_strain_R35         -cat....a...........tc.........t............gg.......ta......................... [167680]
MH025919.1_strain_N10         -cat....a...........tc.........t............gg.......ta......................... [167680]
MH025918.1_strain_R25         -cat....a...........tc.........t............gg.......ta......................... [167680]
MH025917.1_strain_R7          -cat....a...........tc.........t............gg.......ta......................... [167680]
MH025916.1_strain_R8          -cat....a...........tc.........t............gg.......ta......................... [167680]
```

```
FR682468.2_Georgia_2007/1        ttctaaaagagttatcaaatcttcttgtgaggccttatgaataattgttaataccatttcttgcttatggggaacacact [167760]
ASF_IND_20_CAD543                ................................................................................ [167760]
LR899193.1_Germany_2020/1        ................................................................................ [167760]
MN393477.1_Wuhan_2019            ................................................................................ [167760]
MW396979.1_Timor-Leste/2019/1    ................................................................................ [167760]
MK940252.1_CN/2019/InnerMongolia ................................................................................ [167760]
MK128995.1_China/2018/AnhuiXCGQ  ................................................................................ [167760]
MH681419.1_ASFV/POL/2015         ................................................................................ [167760]
LS478113.1_Estonia_2014          ................................................................................ [167760]
KP843857.1_Odintsovo_02/14       ................................................................................ [167760]
FN557520.1_strain_E75            ................................................................................ [167760]
AY261366.1_Warthog               ...........................g.................................................... [167760]
AY261365.1_Warmbaths             ...........................g.................................................... [167760]
AY261364.1_Tengani_62            ...........................g.................................................... [167760]
AY261363.1_Pretoriuskop/96/4     ...........................g.................................................... [167760]
AY261362.1_Mkuzi_1979            ...........................g.................................................... [167760]
AY261361.1_Malawi_Lil-20/1_1983  .........t.................c.....t.............................................. [167760]
AY261360.1_Kenya_1950            .........g.......................t...........g.....c..............g.......t..... [167760]
AM712240.1_OURT_88/3             ................................................................................ [167760]
AM712239.1_Benin_97/1            ................................................................................ [167760]
KM111295.1_Ken06.Bus             .........g.......................t...........g.....c..............g.......t..... [167760]
KM111294.1_Ken05/Tk1             .........g.......................t...........g.....c..............g.......t..... [167760]
KM262844.1_strain_L60            ................................................................................ [167760]
KM102979.1_26544/OG10_Italy      ................................................................................ [167760]
KM262845.1_strain_NHV            ................................................................................ [167760]
KX354450.1_47/Ss/2008            ................................................................................ [167760]
KP055815.1_strain_BA71           ................................................................................ [167760]
MH025920.1_strain_R35            .........g.......................t...........g.....c..............g.......t..... [167760]
MH025919.1_strain_N10            .........g.......................t...........g.....c..............g.......t..... [167760]
MH025918.1_strain_R25            .........g.......................t...........g.....c..............g.......t..... [167760]
MH025917.1_strain_R7             .........g.......................t...........g.....c..............g.......t..... [167760]
MH025916.1_strain_R8             .........g.......................t...........g.....c..............g.......t..... [167760]

FR682468.2_Georgia_2007/1        gataccccacaaagctaatatcaggaatcatttcataaatatatgtttttagcagatttccgatggtatgggtttcatct [167840]
ASF_IND_20_CAD543                .........................................y...................................... [167840]
LR899193.1_Germany_2020/1        ................................................................................ [167840]
MN393477.1_Wuhan_2019            ................................................................................ [167840]
MW396979.1_Timor-Leste/2019/1    ................................................................................ [167840]
MK940252.1_CN/2019/InnerMongolia ................................................................................ [167840]
MK128995.1_China/2018/AnhuiXCGQ  ................................................................................ [167840]
MH681419.1_ASFV/POL/2015         ................................................................................ [167840]
LS478113.1_Estonia_2014          ................................................................................ [167840]
KP843857.1_Odintsovo_02/14       ................................................................................ [167840]
```

```
FN557520.1_strain_E75            .........................................c...................a..a.....a........c [167840]
AY261366.1_Warthog               ...................................................a.........a.................. [167840]
AY261365.1_Warmbaths             ...................................................a............................ [167840]
AY261364.1_Tengani_62            .......................................................g........................ [167840]
AY261363.1_Pretoriuskop/96/4     ...................................................a............................ [167840]
AY261362.1_Mkuzi_1979            .............................................................a..a.....a........c [167840]
AY261361.1_Malawi_Lil-20/1_1983  .........................................c..................................g... [167840]
AY261360.1_Kenya_1950            ..........g........g..g..................c.............g........................ [167840]
AM712240.1_OURT_88/3             .........................................c...................a..a.....a........c [167840]
AM712239.1_Benin_97/1            .........................................c...................a..a.....a........c [167840]
KM111295.1_Ken06.Bus             ...................g..g..................c.............g.....a.................. [167840]
KM111294.1_Ken05/Tk1             ..........g........g..g..................c.............g........................ [167840]
KM262844.1_strain_L60            .........................................c...................a..a.....a........c [167840]
KM102979.1_26544/OG10_Italy      .........................................c...................a..a.....a........c [167840]
KM262845.1_strain_NHV            .........................................c...................a..a.....a........c [167840]
KX354450.1_47/Ss/2008            .........................................c...................a..a.....a........c [167840]
KP055815.1_strain_BA71           .........................................c...................a..a.....a........c [167840]
MH025920.1_strain_R35            ...................g..g..................c.............g.....a.................. [167840]
MH025919.1_strain_N10            ...................g..g..................c.............g.....a.................. [167840]
MH025918.1_strain_R25            ...................g..g..................c.............g.....a.................. [167840]
MH025917.1_strain_R7             ...................g..g..................c.............g.....a.................. [167840]
MH025916.1_strain_R8             ...................g..g..................c.............g.....a.................. [167840]

FR682468.2_Georgia_2007/1        tttatcgtgataatggcctttgttttttcctcatccatggaaaacagcacaagttccggctgcggctcttcaaagttttc [167920]
ASF_IND_20_CAD543                ................................................................................ [167920]
LR899193.1_Germany_2020/1        ................................................................................ [167920]
MN393477.1_Wuhan_2019            ................................................................................ [167920]
MW396979.1_Timor-Leste/2019/1    ................................................................................ [167920]
MK940252.1_CN/2019/InnerMongolia ................................................................................ [167920]
MK128995.1_China/2018/AnhuiXCGQ  ................................................................................ [167920]
MH681419.1_ASFV/POL/2015         ................................................................................ [167920]
LS478113.1_Estonia_2014          ................................................................................ [167920]
KP843857.1_Odintsovo_02/14       ................................................................................ [167920]
FN557520.1_strain_E75            ..............a.................g....ac......................................... [167920]
AY261366.1_Warthog               .....t......................................g................................... [167920]
AY261365.1_Warmbaths             .....t......................................g................................... [167920]
AY261364.1_Tengani_62            .....t......................................g................................... [167920]
AY261363.1_Pretoriuskop/96/4     .....t......................................g................................... [167920]
AY261362.1_Mkuzi_1979            ..............a.................g....ac......................................... [167920]
AY261361.1_Malawi_Lil-20/1_1983  ...........g..........................a......................................... [167920]
AY261360.1_Kenya_1950            ......................................a......................................... [167920]
AM712240.1_OURT_88/3             ..............a.................g....ac......................................... [167920]
AM712239.1_Benin_97/1            ..............a.................g....ac......................................... [167920]
KM111295.1_Ken06.Bus             ......................................a......................................... [167920]
```

```
KM111294.1_Ken05/Tk1              ......................................a......................................... [167920]
KM262844.1_strain_L60             ..............a.................g....ac......................................... [167920]
KM102979.1_26544/OG10_Italy       ..............a.................g....ac......................................... [167920]
KM262845.1_strain_NHV             ..............a.................g....ac......................................... [167920]
KX354450.1_47/Ss/2008             ..............a.................g....ac......................................... [167920]
KP055815.1_strain_BA71            ..............a.................g....ac......................................... [167920]
MH025920.1_strain_R35             ......................................a......................................... [167920]
MH025919.1_strain_N10             .................t....................a......................................... [167920]
MH025918.1_strain_R25             ......................................a......................................... [167920]
MH025917.1_strain_R7              ......................................a......................................... [167920]
MH025916.1_strain_R8              ......................................a......................................... [167920]

FR682468.2_Georgia_2007/1         ataaattttttgaatgctttggattcggccaataatgatccggcaggcgttttttaaatacgtgcgaacggcctggttga [168000]
ASF_IND_20_CAD543                 ................................................................................ [168000]
LR899193.1_Germany_2020/1         ................................................................................ [168000]
MN393477.1_Wuhan_2019             ................................................................................ [168000]
MW396979.1_Timor-Leste/2019/1     ................................................................................ [168000]
MK940252.1_CN/2019/InnerMongolia  ................................................................................ [168000]
MK128995.1_China/2018/AnhuiXCGQ   ................................................................................ [168000]
MH681419.1_ASFV/POL/2015          ................................................................................ [168000]
LS478113.1_Estonia_2014           ................................................................................ [168000]
KP843857.1_Odintsovo_02/14        ................................................................................ [168000]
FN557520.1_strain_E75             ...........................................................................a.... [168000]
AY261366.1_Warthog                ......................................c......................................... [168000]
AY261365.1_Warmbaths              ......................................c......................................... [168000]
AY261364.1_Tengani_62             ......................................c......................................... [168000]
AY261363.1_Pretoriuskop/96/4      ......................................c..............................a.......... [168000]
AY261362.1_Mkuzi_1979             ...........................................................................a.... [168000]
AY261361.1_Malawi_Lil-20/1_1983   ......................................c.........a...........t.............t..... [168000]
AY261360.1_Kenya_1950             ................t.....................c.....................t.....t...a......... [168000]
AM712240.1_OURT_88/3              ...........................................................................a.... [168000]
AM712239.1_Benin_97/1             ...........................................................................a.... [168000]
KM111295.1_Ken06.Bus              ................t.....................cg....................t..c..t.t........... [168000]
KM111294.1_Ken05/Tk1              ................t.....................c.....................t.....t............. [168000]
KM262844.1_strain_L60             ...........................................................................a.... [168000]
KM102979.1_26544/OG10_Italy       ...........................................................................a.... [168000]
KM262845.1_strain_NHV             ...........................................................................a.... [168000]
KX354450.1_47/Ss/2008             ...........................................................................a.... [168000]
KP055815.1_strain_BA71            ...........................................................................a.... [168000]
MH025920.1_strain_R35             ................t.....................cg....................t..c..t.t........... [168000]
MH025919.1_strain_N10             ................t.....................cg....................t..c..t.t........... [168000]
MH025918.1_strain_R25             ................t.....................cg....................t..c..t.t........... [168000]
MH025917.1_strain_R7              ................t.....................cg....................t..c..t.t........... [168000]
MH025916.1_strain_R8              ................t.....................cg....................t..c..t.t........... [168000]
```

```
FR682468.2_Georgia_2007/1        tatgtggcagcggcaccgctggaaagcaaagccccaggcggtggtgacgcgggtctgaggtcatagagctttgcttgtaa [168080]
ASF_IND_20_CAD543                ................................................................................ [168080]
LR899193.1_Germany_2020/1        ................................................................................ [168080]
MN393477.1_Wuhan_2019            ................................................................................ [168080]
MW396979.1_Timor-Leste/2019/1    ................................................................................ [168080]
MK940252.1_CN/2019/InnerMongolia ................................................................................ [168080]
MK128995.1_China/2018/AnhuiXCGQ  ................................................................................ [168080]
MH681419.1_ASFV/POL/2015         ................................................................................ [168080]
LS478113.1_Estonia_2014          ................................................................................ [168080]
KP843857.1_Odintsovo_02/14       ................................................................................ [168080]
FN557520.1_strain_E75            .........................................................................t...... [168080]
AY261366.1_Warthog               ................................................................................ [168080]
AY261365.1_Warmbaths             ................................................................................ [168080]
AY261364.1_Tengani_62            ...................................................................a............ [168080]
AY261363.1_Pretoriuskop/96/4     ................................................................................ [168080]
AY261362.1_Mkuzi_1979            ......................g.........................................g........t...... [168080]
AY261361.1_Malawi_Lil-20/1_1983  ..c......t.............................................c...........a............ [168080]
AY261360.1_Kenya_1950            .gc....................................................c....................a..t [168080]
AM712240.1_OURT_88/3             .........................................................................t...... [168080]
AM712239.1_Benin_97/1            ........................................a................................t...... [168080]
KM111295.1_Ken06.Bus             .gc......a...t.........................................c........................ [168080]
KM111294.1_Ken05/Tk1             .gc....................................................c....................a..t [168080]
KM262844.1_strain_L60            .........................................................................t...... [168080]
KM102979.1_26544/OG10_Italy      .........................................................................t...... [168080]
KM262845.1_strain_NHV            .........................................................................t...... [168080]
KX354450.1_47/Ss/2008            .........................................................................t...... [168080]
KP055815.1_strain_BA71           .........................................................................t...... [168080]
MH025920.1_strain_R35            .gc......a...t.........................................c........................ [168080]
MH025919.1_strain_N10            .gc......a...t.........................................c........................ [168080]
MH025918.1_strain_R25            .gc......a...t.........................................c........................ [168080]
MH025917.1_strain_R7             .gc......a...t.........................................c........................ [168080]
MH025916.1_strain_R8             .gc......a...t.........................................c........................ [168080]

FR682468.2_Georgia_2007/1        ccgctaagcgccatatattctttttttatccgttgggtactgttcaatgtcaaggtgggaaaaatgtgttttaacggcaag [168160]
ASF_IND_20_CAD543                ................................................................................ [168160]
LR899193.1_Germany_2020/1        ................................................................................ [168160]
MN393477.1_Wuhan_2019            ................................................................................ [168160]
MW396979.1_Timor-Leste/2019/1    ................................................................................ [168160]
MK940252.1_CN/2019/InnerMongolia ................................................................................ [168160]
MK128995.1_China/2018/AnhuiXCGQ  ................................................................................ [168160]
MH681419.1_ASFV/POL/2015         ................................................................................ [168160]
LS478113.1_Estonia_2014          ................................................................................ [168160]
KP843857.1_Odintsovo_02/14       ................................................................................ [168160]
```

```
FN557520.1_strain_E75          ..a..........................t...........c.....a..g............................. [168160]
AY261366.1_Warthog             .........................................c...................................... [168160]
AY261365.1_Warmbaths           ......................................t..c...................................... [168160]
AY261364.1_Tengani_62          .........................................c...................................... [168160]
AY261363.1_Pretoriuskop/96/4   .........................................c.....a................................ [168160]
AY261362.1_Mkuzi_1979          ..a..........................t...........c.....a................................ [168160]
AY261361.1_Malawi_Lil-20/1_1983 .........................................c.....a................................ [168160]
AY261360.1_Kenya_1950          ...................................a.....c..g................................... [168160]
AM712240.1_OURT_88/3           ..a..........................t...........c.....a..g............................. [168160]
AM712239.1_Benin_97/1          ..a..........................t...........c.....a..g............................. [168160]
KM111295.1_Ken06.Bus           ...................................a.....c..g................................... [168160]
KM111294.1_Ken05/Tk1           ...................................a.....c..g................................... [168160]
KM262844.1_strain_L60          ..a..........................t...........c.....a..g............................. [168160]
KM102979.1_26544/OG10_Italy    ..a..........................t...........c.....a..g............................. [168160]
KM262845.1_strain_NHV          ..a..........................t...........c.....a..g............................. [168160]
KX354450.1_47/Ss/2008          ..a..........................t...........c.....a..g............................. [168160]
KP055815.1_strain_BA71         ..a..........................t...........c.....a..g............................. [168160]
MH025920.1_strain_R35          ...................................a.....c..g................................... [168160]
MH025919.1_strain_N10          ...................................a.....c..g................................... [168160]
MH025918.1_strain_R25          ...................................a.....c..g................................... [168160]
MH025917.1_strain_R7           ...................................a.....c..g................................... [168160]
MH025916.1_strain_R8           ...................................a.....c..g................................... [168160]

FR682468.2_Georgia_2007/1      attaaaggcggcatgctttcgtcctatgccctttttaatatagatatcctctataatcaacgattttccgggttgtagga [168240]
ASF_IND_20_CAD543              ................................................................................ [168240]
LR899193.1_Germany_2020/1      ................................................................................ [168240]
MN393477.1_Wuhan_2019          ................................................................................ [168240]
MW396979.1_Timor-Leste/2019/1  ................................................................................ [168240]
MK940252.1_CN/2019/InnerMongolia ................................................................................ [168240]
MK128995.1_China/2018/AnhuiXCGQ ................................................................................ [168240]
MH681419.1_ASFV/POL/2015       ................................................................................ [168240]
LS478113.1_Estonia_2014        ................................................................................ [168240]
KP843857.1_Odintsovo_02/14     ................................................................................ [168240]
FN557520.1_strain_E75          ...............t................................................................ [168240]
AY261366.1_Warthog             .....................a........................................................a. [168240]
AY261365.1_Warmbaths           .....................a........................................................a. [168240]
AY261364.1_Tengani_62          .....................a......................................t................... [168240]
AY261363.1_Pretoriuskop/96/4   .....................a.........c................................................ [168240]
AY261362.1_Mkuzi_1979          ................................................................................ [168240]
AY261361.1_Malawi_Lil-20/1_1983 ...............t.....c.....................................g...............c.... [168240]
AY261360.1_Kenya_1950          ...........................................................g.........a.......... [168240]
AM712240.1_OURT_88/3           ...............t................................................................ [168240]
AM712239.1_Benin_97/1          ...............t................................................................ [168240]
KM111295.1_Ken06.Bus           ...........................................................g.........a.......... [168240]
```

```
KM111294.1_Ken05/Tk1            ...........................................................g.........a.......... [168240]
KM262844.1_strain_L60           ...............t................................................................ [168240]
KM102979.1_26544/OG10_Italy     ...............t................................................................ [168240]
KM262845.1_strain_NHV           ...............t................................................................ [168240]
KX354450.1_47/Ss/2008           ...............t................................................................ [168240]
KP055815.1_strain_BA71          ...............t................................................................ [168240]
MH025920.1_strain_R35           ...........................................................g.........a.......... [168240]
MH025919.1_strain_N10           ...........................................................g.........a.......... [168240]
MH025918.1_strain_R25           ...........................................................g.........a.......... [168240]
MH025917.1_strain_R7            ...........................................................g.........a.......... [168240]
MH025916.1_strain_R8            ...........................................................g.........a.......... [168240]

FR682468.2_Georgia_2007/1       agccaatctcaaaggtaggattaaaaatcgggtatttaagcttagggcctgccacctggatgagatcgcggctatagatg [168320]
ASF_IND_20_CAD543               .............r.................................................................. [168320]
LR899193.1_Germany_2020/1       ................................................................................ [168320]
MN393477.1_Wuhan_2019           ................................................................................ [168320]
MW396979.1_Timor-Leste/2019/1   ................................................................................ [168320]
MK940252.1_CN/2019/InnerMongolia ................................................................................ [168320]
MK128995.1_China/2018/AnhuiXCGQ ................................................................................ [168320]
MH681419.1_ASFV/POL/2015        ................................................................................ [168320]
LS478113.1_Estonia_2014         ................................................................................ [168320]
KP843857.1_Odintsovo_02/14      ................................................................................ [168320]
FN557520.1_strain_E75           .............a............................................a..................... [168320]
AY261366.1_Warthog              ........................................t.................a..c........a......... [168320]
AY261365.1_Warmbaths            ........................................t.................a..c........a......... [168320]
AY261364.1_Tengani_62           ..............................................a...........a..c........a......... [168320]
AY261363.1_Pretoriuskop/96/4    ..............................................a...........a..c.....a..a......... [168320]
AY261362.1_Mkuzi_1979           ........................................t.................a..a.................. [168320]
AY261361.1_Malawi_Lil-20/1_1983 ..........................................................a..c........ag........ [168320]
AY261360.1_Kenya_1950           .............a....................c.......................a..c.................. [168320]
AM712240.1_OURT_88/3            .............a............................................a..................... [168320]
AM712239.1_Benin_97/1           .............a............................................a..................... [168320]
KM111295.1_Ken06.Bus            .............a....................c.......................a..c........a......... [168320]
KM111294.1_Ken05/Tk1            .............a....................c.......................a..c.................. [168320]
KM262844.1_strain_L60           .............a............................................a..................... [168320]
KM102979.1_26544/OG10_Italy     .............a............................................a..................... [168320]
KM262845.1_strain_NHV           .............a............................................a..................... [168320]
KX354450.1_47/Ss/2008           .............a............................................a..................... [168320]
KP055815.1_strain_BA71          .............a............................................a..................... [168320]
MH025920.1_strain_R35           .............a....................c.......................a..c........a......... [168320]
MH025919.1_strain_N10           .............a....................c.......................a..c........a......... [168320]
MH025918.1_strain_R25           .............a....................c.......................a..c........a......... [168320]
MH025917.1_strain_R7            .............a....................c.......................a..c........a......... [168320]
MH025916.1_strain_R8            .............a....................c.......................a..c........a......... [168320]
```

```
FR682468.2_Georgia_2007/1        gttttaacctcacagctattgtttaaactccgcagagcaaataccagtgtctcgtttttcgcataaatcggaatgaaatt [168400]
ASF_IND_20_CAD543                ................................................................................ [168400]
LR899193.1_Germany_2020/1        ................................................................................ [168400]
MN393477.1_Wuhan_2019            ................................................................................ [168400]
MW396979.1_Timor-Leste/2019/1    ................................................................................ [168400]
MK940252.1_CN/2019/InnerMongolia ................................................................................ [168400]
MK128995.1_China/2018/AnhuiXCGQ  ................................................................................ [168400]
MH681419.1_ASFV/POL/2015         ................................................................................ [168400]
LS478113.1_Estonia_2014          ................................................................................ [168400]
KP843857.1_Odintsovo_02/14       ................................................................................ [168400]
FN557520.1_strain_E75            ................................................................................ [168400]
AY261366.1_Warthog               ................................................................................ [168400]
AY261365.1_Warmbaths             ................................................................................ [168400]
AY261364.1_Tengani_62            ....................a........................................................... [168400]
AY261363.1_Pretoriuskop/96/4     ................................................................................ [168400]
AY261362.1_Mkuzi_1979            ................................................................................ [168400]
AY261361.1_Malawi_Lil-20/1_1983  .............................g.................a.a........c..................... [168400]
AY261360.1_Kenya_1950            .....c.......................g...........c................cg.................... [168400]
AM712240.1_OURT_88/3             ................................................................................ [168400]
AM712239.1_Benin_97/1            ................................................................................ [168400]
KM111295.1_Ken06.Bus             .....c.......................g...........c...........a....cg.................... [168400]
KM111294.1_Ken05/Tk1             .....c.......................g...........c................cg.................... [168400]
KM262844.1_strain_L60            ................................................................................ [168400]
KM102979.1_26544/OG10_Italy      ................................................................................ [168400]
KM262845.1_strain_NHV            ................................................................................ [168400]
KX354450.1_47/Ss/2008            ................................................................................ [168400]
KP055815.1_strain_BA71           ................................................................................ [168400]
MH025920.1_strain_R35            .....c.......................g...........c...........a....cg.................... [168400]
MH025919.1_strain_N10            .....c.......................g...........c...........a....cg.................... [168400]
MH025918.1_strain_R25            .....c.......................g...........c...........a....cg.................... [168400]
MH025917.1_strain_R7             .....c.......................g...........c...........a....cg.................... [168400]
MH025916.1_strain_R8             .....c.......................g...........c...........a....cg.................... [168400]

FR682468.2_Georgia_2007/1        aatgcggtttctaataaattgttccgtcataaacaggtccgtggaatcctcgatcttatacccaccgggcttaatatcta [168480]
ASF_IND_20_CAD543                ................................................................................ [168480]
LR899193.1_Germany_2020/1        ................................................................................ [168480]
MN393477.1_Wuhan_2019            ................................................................................ [168480]
MW396979.1_Timor-Leste/2019/1    ................................................................................ [168480]
MK940252.1_CN/2019/InnerMongolia ................................................................................ [168480]
MK128995.1_China/2018/AnhuiXCGQ  ................................................................................ [168480]
MH681419.1_ASFV/POL/2015         ................................................................................ [168480]
LS478113.1_Estonia_2014          ................................................................................ [168480]
KP843857.1_Odintsovo_02/14       ................................................................................ [168480]
```

```
FN557520.1_strain_E75           ................................................................................ [168480]
AY261366.1_Warthog              ................................................................................ [168480]
AY261365.1_Warmbaths            ................................................................................ [168480]
AY261364.1_Tengani_62           ................................................................................ [168480]
AY261363.1_Pretoriuskop/96/4    ................................................................................ [168480]
AY261362.1_Mkuzi_1979           ................................................................................ [168480]
AY261361.1_Malawi_Lil-20/1_1983 .............................................g..t............................... [168480]
AY261360.1_Kenya_1950           ..............c...............................................................a. [168480]
AM712240.1_OURT_88/3            ................................................................................ [168480]
AM712239.1_Benin_97/1           ................................................................................ [168480]
KM111295.1_Ken06.Bus            ..............................................................................a. [168480]
KM111294.1_Ken05/Tk1            ..............................................................................a. [168480]
KM262844.1_strain_L60           ................................................................................ [168480]
KM102979.1_26544/OG10_Italy     ................................................................................ [168480]
KM262845.1_strain_NHV           ................................................................................ [168480]
KX354450.1_47/Ss/2008           ................................................................................ [168480]
KP055815.1_strain_BA71          ................................................................................ [168480]
MH025920.1_strain_R35           ..............................................................................a. [168480]
MH025919.1_strain_N10           ..............................................................................a. [168480]
MH025918.1_strain_R25           ..............................................................................a. [168480]
MH025917.1_strain_R7            ..............................................................................a. [168480]
MH025916.1_strain_R8            ..............................................................................a. [168480]

FR682468.2_Georgia_2007/1       gcatataattgggaatttcatcttgcaagacccgcgacaggccgtggaccgcggctctgctaatgcccttaaagtccata [168560]
ASF_IND_20_CAD543               ................................................................................ [168560]
LR899193.1_Germany_2020/1       ................................................................................ [168560]
MN393477.1_Wuhan_2019           ................................................................................ [168560]
MW396979.1_Timor-Leste/2019/1   ................................................................................ [168560]
MK940252.1_CN/2019/InnerMongolia ................................................................................ [168560]
MK128995.1_China/2018/AnhuiXCGQ ................................................................................ [168560]
MH681419.1_ASFV/POL/2015        ................................................................................ [168560]
LS478113.1_Estonia_2014         ................................................................................ [168560]
KP843857.1_Odintsovo_02/14      ................................................................................ [168560]
FN557520.1_strain_E75           ..............................................c................................. [168560]
AY261366.1_Warthog              ...........................................................t.................... [168560]
AY261365.1_Warmbaths            .........................................................................a...... [168560]
AY261364.1_Tengani_62           ..........a..................................................................... [168560]
AY261363.1_Pretoriuskop/96/4    ...........................................................t.................... [168560]
AY261362.1_Mkuzi_1979           ................................................................................ [168560]
AY261361.1_Malawi_Lil-20/1_1983 ...............................t...........................t....a............... [168560]
AY261360.1_Kenya_1950           .......g......................................a....t.......t.t...........a.....c [168560]
AM712240.1_OURT_88/3            ..............................................c................................. [168560]
AM712239.1_Benin_97/1           ..............................................c................................. [168560]
KM111295.1_Ken06.Bus            .......g.............................t.............t.......t.t....t......a.....c [168560]
```

```
KM111294.1_Ken05/Tk1              .......g......................................a....t.......t.t...........a.....c [168560]
KM262844.1_strain_L60             ..............................................c................................. [168560]
KM102979.1_26544/OG10_Italy       ..............................................c................................. [168560]
KM262845.1_strain_NHV             ..............................................c................................. [168560]
KX354450.1_47/Ss/2008             ..............................................c................................. [168560]
KP055815.1_strain_BA71            ..............................................c................................. [168560]
MH025920.1_strain_R35             .......g.............................t.............t.......t.t....t......a.....c [168560]
MH025919.1_strain_N10             .......g.............................t.............t.......t.t....t......a.....c [168560]
MH025918.1_strain_R25             .......g.............................t.............t.......t.t....t......a.....c [168560]
MH025917.1_strain_R7              .......g.............................t.............t.......t.t....t......a.....c [168560]
MH025916.1_strain_R8              .......g.............................t.............t.......t.t....t......a.....c [168560]

FR682468.2_Georgia_2007/1         acaacattgaccgggacgaggggcaactgctcctcgagctgaaatagtttttggccgcatttttaataaagaggttgga [168640]
ASF_IND_20_CAD543                 ................................................................................ [168640]
LR899193.1_Germany_2020/1         ................................................................................ [168640]
MN393477.1_Wuhan_2019             ................................................................................ [168640]
MW396979.1_Timor-Leste/2019/1     ................................................................................ [168640]
MK940252.1_CN/2019/InnerMongolia  ................................................................................ [168640]
MK128995.1_China/2018/AnhuiXCGQ   ................................................................................ [168640]
MH681419.1_ASFV/POL/2015          ................................................................................ [168640]
LS478113.1_Estonia_2014           ................................................................................ [168640]
KP843857.1_Odintsovo_02/14        ................................................................................ [168640]
FN557520.1_strain_E75             ...........t....................................c.......................g....... [168640]
AY261366.1_Warthog                ................................................c.......t...............g....... [168640]
AY261365.1_Warmbaths              ................................................c.......................g....... [168640]
AY261364.1_Tengani_62             ................................................c.......................g....... [168640]
AY261363.1_Pretoriuskop/96/4      .................a..............................c.......................g....... [168640]
AY261362.1_Mkuzi_1979             ................................................c.......................g....... [168640]
AY261361.1_Malawi_Lil-20/1_1983   ...........t..a................................c........................g.a..... [168640]
AY261360.1_Kenya_1950             .t.........t..a.........................................................g....c.. [168640]
AM712240.1_OURT_88/3              ...........t....................................c.......................g....... [168640]
AM712239.1_Benin_97/1             ...........t....................................c.......................g....... [168640]
KM111295.1_Ken06.Bus              .t.........t..a........................................................ag....c.. [168640]
KM111294.1_Ken05/Tk1              .t.........t..a.........................................................g....c.. [168640]
KM262844.1_strain_L60             ...........t....................................c.......................g....... [168640]
KM102979.1_26544/OG10_Italy       ...........t....................................c.......................g....... [168640]
KM262845.1_strain_NHV             ...........t....................................c.......................g....... [168640]
KX354450.1_47/Ss/2008             ...........t....................................c.......................g....... [168640]
KP055815.1_strain_BA71            ...........t....................................c.......................g....... [168640]
MH025920.1_strain_R35             .t.........t..a........................................................ag....c.. [168640]
MH025919.1_strain_N10             .t.........t..a........................................................ag....c.. [168640]
MH025918.1_strain_R25             .t.........t..a........................................................ag....c.. [168640]
MH025917.1_strain_R7              .t.........t..a........................................................ag....c.. [168640]
MH025916.1_strain_R8              .t.........t..a........................................................ag....c.. [168640]
```

```
FR682468.2_Georgia_2007/1        aaagtctatcaaaaacggtttgatttccacgttttggaaaattttttccatttgtattataaatatatctatatatattc [168720]
ASF_IND_20_CAD543                ....................................................................ag.......... [168720]
LR899193.1_Germany_2020/1        ................................................................................ [168720]
MN393477.1_Wuhan_2019            ................................................................................ [168720]
MW396979.1_Timor-Leste/2019/1    ................................................................................ [168720]
MK940252.1_CN/2019/InnerMongolia ................................................................................ [168720]
MK128995.1_China/2018/AnhuiXCGQ  ................................................................................ [168720]
MH681419.1_ASFV/POL/2015         ................................................................................ [168720]
LS478113.1_Estonia_2014          ................................................................................ [168720]
KP843857.1_Odintsovo_02/14       ................................................................................ [168720]
FN557520.1_strain_E75            ....................................................................ag.......... [168720]
AY261366.1_Warthog               ....................................................................t........... [168720]
AY261365.1_Warmbaths             ....................................................................t........... [168720]
AY261364.1_Tengani_62            ....................................................................t........... [168720]
AY261363.1_Pretoriuskop/96/4     ....................................................................t........... [168720]
AY261362.1_Mkuzi_1979            ....................................................................ag.......... [168720]
AY261361.1_Malawi_Lil-20/1_1983  ..............g.........c...........................................t........... [168720]
AY261360.1_Kenya_1950            ...a..........g...c.....g...........................................t........... [168720]
AM712240.1_OURT_88/3             ....................................................................ag.......... [168720]
AM712239.1_Benin_97/1            ....................................................................ag.......... [168720]
KM111295.1_Ken06.Bus             ...a..........g...c.....a...........................................t........... [168720]
KM111294.1_Ken05/Tk1             ...a..........g...c.....g...........................................t........... [168720]
KM262844.1_strain_L60            ....................................................................ag.......... [168720]
KM102979.1_26544/OG10_Italy      ....................................................................ag.......... [168720]
KM262845.1_strain_NHV            ....................................................................ag.......... [168720]
KX354450.1_47/Ss/2008            ....................................................................ag.......... [168720]
KP055815.1_strain_BA71           ....................................................................ag.......... [168720]
MH025920.1_strain_R35            ...a..........g...c.....a...........................................t........... [168720]
MH025919.1_strain_N10            ...a..........g...c.....a...........................................t........... [168720]
MH025918.1_strain_R25            ...a..........g...c.....a...........................................t........... [168720]
MH025917.1_strain_R7             ...a..........g...c.....a...........................................t........... [168720]
MH025916.1_strain_R8             ...a..........g...c.....a...........................................t........... [168720]

FR682468.2_Georgia_2007/1        aaattatggtagtttatgacttgctcgtttctttaagtaaggaatccatagatgtgctacggtttgtagaggcaaacctt [168800]
ASF_IND_20_CAD543                ................................................................................ [168800]
LR899193.1_Germany_2020/1        ................................................................................ [168800]
MN393477.1_Wuhan_2019            ................................................................................ [168800]
MW396979.1_Timor-Leste/2019/1    ................................................................................ [168800]
MK940252.1_CN/2019/InnerMongolia ................................................................................ [168800]
MK128995.1_China/2018/AnhuiXCGQ  ................................................................................ [168800]
MH681419.1_ASFV/POL/2015         ................................................................................ [168800]
LS478113.1_Estonia_2014          ................................................................................ [168800]
KP843857.1_Odintsovo_02/14       ................................................................................ [168800]
```

```
FN557520.1_strain_E75            ................................................................................ [168800]
AY261366.1_Warthog               ................................................................................ [168800]
AY261365.1_Warmbaths             ................................................................................ [168800]
AY261364.1_Tengani_62            .......................t.....................................a.................. [168800]
AY261363.1_Pretoriuskop/96/4     ................................................................................ [168800]
AY261362.1_Mkuzi_1979            ................................................................................ [168800]
AY261361.1_Malawi_Lil-20/1_1983  .........................t..............a....................a..............t... [168800]
AY261360.1_Kenya_1950            ..........t...................................a..................a.....a....t... [168800]
AM712240.1_OURT_88/3             ................................................................................ [168800]
AM712239.1_Benin_97/1            ................................................................................ [168800]
KM111295.1_Ken06.Bus             ....................c.........................a..................a.....a....t... [168800]
KM111294.1_Ken05/Tk1             ..............................................a..................a.....a....t... [168800]
KM262844.1_strain_L60            ................................................................................ [168800]
KM102979.1_26544/OG10_Italy      ................................................................................ [168800]
KM262845.1_strain_NHV            ................................................................................ [168800]
KX354450.1_47/Ss/2008            ................................................................................ [168800]
KP055815.1_strain_BA71           ................................................................................ [168800]
MH025920.1_strain_R35            ....................c.........................a..................a.....a....t... [168800]
MH025919.1_strain_N10            ....................c.........................a..................a.....a....t... [168800]
MH025918.1_strain_R25            ....................c.........................a..................a.....a....t... [168800]
MH025917.1_strain_R7             ....................c.........................a..................a.....a....t... [168800]
MH025916.1_strain_R8             ....................c.........................a..................a.....a....t... [168800]

FR682468.2_Georgia_2007/1        gcggcgtttaaccagcagtatatttttttcaatatccaaagaaaaaactcgatcacgacaccccttctcattacgccgca [168880]
ASF_IND_20_CAD543                ................................................................................ [168880]
LR899193.1_Germany_2020/1        ................................................................................ [168880]
MN393477.1_Wuhan_2019            ................................................................................ [168880]
MW396979.1_Timor-Leste/2019/1    ................................................................................ [168880]
MK940252.1_CN/2019/InnerMongolia ................................................................................ [168880]
MK128995.1_China/2018/AnhuiXCGQ  ................................................................................ [168880]
MH681419.1_ASFV/POL/2015         ................................................................................ [168880]
LS478113.1_Estonia_2014          ................................................................................ [168880]
KP843857.1_Odintsovo_02/14       ................................................................................ [168880]
FN557520.1_strain_E75            ................................................................................ [168880]
AY261366.1_Warthog               .............................t.................................................. [168880]
AY261365.1_Warmbaths             ................................................................................ [168880]
AY261364.1_Tengani_62            ................................................................................ [168880]
AY261363.1_Pretoriuskop/96/4     .............................t.................................................. [168880]
AY261362.1_Mkuzi_1979            ................................................................................ [168880]
AY261361.1_Malawi_Lil-20/1_1983  .....................................................t..a..g..t.....t........... [168880]
AY261360.1_Kenya_1950            ..................................................a....t...g.....c.....c........ [168880]
AM712240.1_OURT_88/3             ................................................................................ [168880]
AM712239.1_Benin_97/1            ................................................................................ [168880]
KM111295.1_Ken06.Bus             ..................................................a....t...g...........c........ [168880]
```

```
KM111294.1_Ken05/Tk1             ..................................................a....t...g...........c........ [168880]
KM262844.1_strain_L60            ................................................................................ [168880]
KM102979.1_26544/OG10_Italy      ................................................................................ [168880]
KM262845.1_strain_NHV            ................................................................................ [168880]
KX354450.1_47/Ss/2008            ................................................................................ [168880]
KP055815.1_strain_BA71           ................................................................................ [168880]
MH025920.1_strain_R35            ..................................................a....t...g...........c........ [168880]
MH025919.1_strain_N10            ..................................................a....t...g...........c........ [168880]
MH025918.1_strain_R25            ..................................................a....t...g...........c........ [168880]
MH025917.1_strain_R7             ..................................................a....t...g...........c........ [168880]
MH025916.1_strain_R8             ..................................................a....t...g...........c........ [168880]

FR682468.2_Georgia_2007/1        gcaggaaaaaatttcgcaaattgttgagtttttaatggatgaatataataagaacaatagaaggccctccgggccgccgc [168960]
ASF_IND_20_CAD543                ................................................................................ [168960]
LR899193.1_Germany_2020/1        ................................................................................ [168960]
MN393477.1_Wuhan_2019            ................................................................................ [168960]
MW396979.1_Timor-Leste/2019/1    ................................................................................ [168960]
MK940252.1_CN/2019/InnerMongolia ................................................................................ [168960]
MK128995.1_China/2018/AnhuiXCGQ  ................................................................................ [168960]
MH681419.1_ASFV/POL/2015         ................................................................................ [168960]
LS478113.1_Estonia_2014          ................................................................................ [168960]
KP843857.1_Odintsovo_02/14       ................................................................................ [168960]
FN557520.1_strain_E75            ................................................................................ [168960]
AY261366.1_Warthog               ...............................................................a.....t.......... [168960]
AY261365.1_Warmbaths             .....................................................................t.......... [168960]
AY261364.1_Tengani_62            ...........................c.........................g...............t.......... [168960]
AY261363.1_Pretoriuskop/96/4     ...........................c.........................................t.......... [168960]
AY261362.1_Mkuzi_1979            ...........................................................a.................... [168960]
AY261361.1_Malawi_Lil-20/1_1983  ...........................c.................................................... [168960]
AY261360.1_Kenya_1950            ........................g..........................a.g.........a.---.t.......... [168960]
AM712240.1_OURT_88/3             ................................................................................ [168960]
AM712239.1_Benin_97/1            ................................................................................ [168960]
KM111295.1_Ken06.Bus             ........................g..........................a.g.........a.---.t.......... [168960]
KM111294.1_Ken05/Tk1             ........................g..........................a.g.........a.---.t.......... [168960]
KM262844.1_strain_L60            ................................................................................ [168960]
KM102979.1_26544/OG10_Italy      ................................................................................ [168960]
KM262845.1_strain_NHV            ................................................................................ [168960]
KX354450.1_47/Ss/2008            ................................................................................ [168960]
KP055815.1_strain_BA71           ................................................................................ [168960]
MH025920.1_strain_R35            ........................g..........................a.g.........a.---.t.......... [168960]
MH025919.1_strain_N10            ........................g..........................a.g.........a.---.t.......... [168960]
MH025918.1_strain_R25            ........................g..........................a.g.........a.---.t.......... [168960]
MH025917.1_strain_R7             ........................g..........................a.g.........a.---.t.......... [168960]
MH025916.1_strain_R8             ........................g..........................a.g.........a.---.t.......... [168960]
```

```
FR682468.2_Georgia_2007/1        gtgagcagcccatgc------acccattattgccgtatcaacaatcctcggacgaacagcccatgatgccgtatcaacag [169040]
ASF_IND_20_CAD543                ...............------........................................................... [169040]
LR899193.1_Germany_2020/1        ...............------........................................................... [169040]
MN393477.1_Wuhan_2019            ...............------........................................................... [169040]
MW396979.1_Timor-Leste/2019/1    ...............------........................................................... [169040]
MK940252.1_CN/2019/InnerMongolia ...............------........................................................... [169040]
MK128995.1_China/2018/AnhuiXCGQ  ...............------........................................................... [169040]
MH681419.1_ASFV/POL/2015         ...............------........................................................... [169040]
LS478113.1_Estonia_2014          ...............------........................................................... [169040]
KP843857.1_Odintsovo_02/14       ...............------........................................................... [169040]
FN557520.1_strain_E75            ...............------........................................................... [169040]
AY261366.1_Warthog               ...............------..........................c................................ [169040]
AY261365.1_Warmbaths             ...............------..........................c................................ [169040]
AY261364.1_Tengani_62            ...............------..........................c...................a...........a [169040]
AY261363.1_Pretoriuskop/96/4     ...............------...................g......c................................ [169040]
AY261362.1_Mkuzi_1979            ...............------........................................................... [169040]
AY261361.1_Malawi_Lil-20/1_1983  ...............------......................g........t........................... [169040]
AY261360.1_Kenya_1950            ...............aggcgt...........t..........g........a........................... [169040]
AM712240.1_OURT_88/3             ...............------........................................................... [169040]
AM712239.1_Benin_97/1            ...............------........................................................... [169040]
KM111295.1_Ken06.Bus             ...............cggcgt....g.................g........a.................a......... [169040]
KM111294.1_Ken05/Tk1             ...............aggcgt...........t..........g........a........................... [169040]
KM262844.1_strain_L60            ...............------........................................................... [169040]
KM102979.1_26544/OG10_Italy      ...............------........................................................... [169040]
KM262845.1_strain_NHV            ...............------........................................................... [169040]
KX354450.1_47/Ss/2008            ...............------........................................................... [169040]
KP055815.1_strain_BA71           ...............------........................................................... [169040]
MH025920.1_strain_R35            ...............cggcgt....g.................g........a.................a......... [169040]
MH025919.1_strain_N10            ...............cggcgt....g.................g........a.................a......... [169040]
MH025918.1_strain_R25            ...............cggcgt....g.................g........a.................a......... [169040]
MH025917.1_strain_R7             ...............cggcgt....g.................g........a.................a......... [169040]
MH025916.1_strain_R8             ...............cggcgt....g.................g........a.................a......... [169040]

FR682468.2_Georgia_2007/1        cccccggggaatgatgatcagccatatgagcaaatataccataaaaaacacgcgtcgcagcaagtaaatactgaactgaa [169120]
ASF_IND_20_CAD543                ................................................................................ [169120]
LR899193.1_Germany_2020/1        ................................................................................ [169120]
MN393477.1_Wuhan_2019            ................................................................................ [169120]
MW396979.1_Timor-Leste/2019/1    ................................................................................ [169120]
MK940252.1_CN/2019/InnerMongolia ................................................................................ [169120]
MK128995.1_China/2018/AnhuiXCGQ  ................................................................................ [169120]
MH681419.1_ASFV/POL/2015         ................................................................................ [169120]
LS478113.1_Estonia_2014          ................................................................................ [169120]
KP843857.1_Odintsovo_02/14       ................................................................................ [169120]
```

```
FN557520.1_strain_E75              ................................................................................ [169120]
AY261366.1_Warthog                 ...........................................................a...................g [169120]
AY261365.1_Warmbaths               ...........................................................a...................g [169120]
AY261364.1_Tengani_62              .....c.....................................................a...................g [169120]
AY261363.1_Pretoriuskop/96/4       ...........................................................a..g................g [169120]
AY261362.1_Mkuzi_1979              ................................................................................ [169120]
AY261361.1_Malawi_Lil-20/1_1983    ........tg.c..c............................................a...................g [169120]
AY261360.1_Kenya_1950              ........tg....c............................................a..g................g [169120]
AM712240.1_OURT_88/3               ................................................................................ [169120]
AM712239.1_Benin_97/1              ................................................................................ [169120]
KM111295.1_Ken06.Bus               ........tg....c......................................c.....a...................g [169120]
KM111294.1_Ken05/Tk1               ........tg....c............................................a...................g [169120]
KM262844.1_strain_L60              ................................................................................ [169120]
KM102979.1_26544/OG10_Italy        ................................................................................ [169120]
KM262845.1_strain_NHV              ................................................................................ [169120]
KX354450.1_47/Ss/2008              ................................................................................ [169120]
KP055815.1_strain_BA71             ................................................................................ [169120]
MH025920.1_strain_R35              ........tg....c..............t.......................c.....a...................g [169120]
MH025919.1_strain_N10              ........tg....c..............t.......................c.....a...................g [169120]
MH025918.1_strain_R25              ........tg....c..............t.......................c.....a...................g [169120]
MH025917.1_strain_R7               ........tg....c..............t.......................c.....a...................g [169120]
MH025916.1_strain_R8               ........tg....c..............t.......................c.....a...................g [169120]

FR682468.2_Georgia_2007/1          cgattattatcaacatattcttgcattaggcgatgaagacaaaggtatggacagcatgttaaaacttccagaaaaggcaa [169200]
ASF_IND_20_CAD543                  ................................................................................ [169200]
LR899193.1_Germany_2020/1          ................................................................................ [169200]
MN393477.1_Wuhan_2019              ................................................................................ [169200]
MW396979.1_Timor-Leste/2019/1      ................................................................................ [169200]
MK940252.1_CN/2019/InnerMongolia   ................................................................................ [169200]
MK128995.1_China/2018/AnhuiXCGQ    ................................................................................ [169200]
MH681419.1_ASFV/POL/2015           ................................................................................ [169200]
LS478113.1_Estonia_2014            ................................................................................ [169200]
KP843857.1_Odintsovo_02/14         ................................................................................ [169200]
FN557520.1_strain_E75              ................................................................................ [169200]
AY261366.1_Warthog                 ........................................................c.................g.a... [169200]
AY261365.1_Warmbaths               ........................................................c.................g.a... [169200]
AY261364.1_Tengani_62              ........................................................c.................g..... [169200]
AY261363.1_Pretoriuskop/96/4       ........................................................c....................... [169200]
AY261362.1_Mkuzi_1979              ................................................................................ [169200]
AY261361.1_Malawi_Lil-20/1_1983    ........................t...................t................................... [169200]
AY261360.1_Kenya_1950              ........................t.........................................c.......g..... [169200]
AM712240.1_OURT_88/3               ................................................................................ [169200]
AM712239.1_Benin_97/1              ................................................................................ [169200]
KM111295.1_Ken06.Bus               ........................t.........................................c.......g..... [169200]
```

```
KM111294.1_Ken05/Tk1             ........................t.........................................c.......g..... [169200]
KM262844.1_strain_L60            ................................................................................ [169200]
KM102979.1_26544/OG10_Italy      ................................................................................ [169200]
KM262845.1_strain_NHV            ................................................................................ [169200]
KX354450.1_47/Ss/2008            ................................................................................ [169200]
KP055815.1_strain_BA71           ................................................................................ [169200]
MH025920.1_strain_R35            ........................t.........................................c.......g..... [169200]
MH025919.1_strain_N10            ........................t.........................................c.......g..... [169200]
MH025918.1_strain_R25            ........................t.........................................c.......g..... [169200]
MH025917.1_strain_R7             ........................t.........................................c.......g..... [169200]
MH025916.1_strain_R8             ........................t.........................................c.......g..... [169200]

FR682468.2_Georgia_2007/1        aaaggga---tagcgatgatgaggacgacatgttttctat-aaaaaactaacgacgtaacaattaaacaaaaaataaaaa [169280]
ASF_IND_20_CAD543                .......---..............................-....................................... [169280]
LR899193.1_Germany_2020/1        .......---..............................-....................................... [169280]
MN393477.1_Wuhan_2019            .......---..............................-....................................... [169280]
MW396979.1_Timor-Leste/2019/1    .......---..............................-....................................... [169280]
MK940252.1_CN/2019/InnerMongolia .......---..............................-....................................... [169280]
MK128995.1_China/2018/AnhuiXCGQ  .......---..............................-....................................... [169280]
MH681419.1_ASFV/POL/2015         .......---..............................-....................................... [169280]
LS478113.1_Estonia_2014          .......---..............................-....................................... [169280]
KP843857.1_Odintsovo_02/14       .......---..............................-...........................-........... [169280]
FN557520.1_strain_E75            ......g---..............................-....................................... [169280]
AY261366.1_Warthog               .......---.................g............-....................................... [169280]
AY261365.1_Warmbaths             .......---.................g............-....................................... [169280]
AY261364.1_Tengani_62            .......---................ag............-....................................... [169280]
AY261363.1_Pretoriuskop/96/4     .......---.................g............-....................................... [169280]
AY261362.1_Mkuzi_1979            .......---..............................-....................................... [169280]
AY261361.1_Malawi_Lil-20/1_1983  .......tga................ag.......c....a....................................... [169280]
AY261360.1_Kenya_1950            .......---a.....c..................c....a....................................... [169280]
AM712240.1_OURT_88/3             ......g---..............................-....................................... [169280]
AM712239.1_Benin_97/1            ......g---..............................-....................................... [169280]
KM111295.1_Ken06.Bus             .......---a.....c..................c....a....................................... [169280]
KM111294.1_Ken05/Tk1             .......---a.....c..................c....a....................................... [169280]
KM262844.1_strain_L60            ......g---..............................-....................................... [169280]
KM102979.1_26544/OG10_Italy      ......g---..............................-....................................... [169280]
KM262845.1_strain_NHV            ......g---..............................-....................................... [169280]
KX354450.1_47/Ss/2008            ......g---..............................-....................................... [169280]
KP055815.1_strain_BA71           ......g---..............................-....................................... [169280]
MH025920.1_strain_R35            .......---a.....c..................c....a....................................... [169280]
MH025919.1_strain_N10            .......---a.....c..................c....a....................................... [169280]
MH025918.1_strain_R25            .......---a.....c..................c....a....................................... [169280]
MH025917.1_strain_R7             .......---a.....c..................c....a....................................... [169280]
MH025916.1_strain_R8             .......---a.....c..................c....a....................................... [169280]
```

```
FR682468.2_Georgia_2007/1        tcattataaaatgaatcttgaatacgtccaagttgttcaaaaatttaatcaagtactcctagaacttaccaaaaaagtat [169360]
ASF_IND_20_CAD543                ................................................................................ [169360]
LR899193.1_Germany_2020/1        ................................................................................ [169360]
MN393477.1_Wuhan_2019            ................................................................................ [169360]
MW396979.1_Timor-Leste/2019/1    ................................................................................ [169360]
MK940252.1_CN/2019/InnerMongolia ................................................................................ [169360]
MK128995.1_China/2018/AnhuiXCGQ  ................................................................................ [169360]
MH681419.1_ASFV/POL/2015         ................................................................................ [169360]
LS478113.1_Estonia_2014          ................................................................................ [169360]
KP843857.1_Odintsovo_02/14       ................................................................................ [169360]
FN557520.1_strain_E75            ................................................................................ [169360]
AY261366.1_Warthog               ................................................................................ [169360]
AY261365.1_Warmbaths             ................................................................................ [169360]
AY261364.1_Tengani_62            ................................................................................ [169360]
AY261363.1_Pretoriuskop/96/4     ................................................................................ [169360]
AY261362.1_Mkuzi_1979            ................................................................................ [169360]
AY261361.1_Malawi_Lil-20/1_1983  ...........................a..c...............................................t. [169360]
AY261360.1_Kenya_1950            ...........................g..c...............................................c. [169360]
AM712240.1_OURT_88/3             ................................................................................ [169360]
AM712239.1_Benin_97/1            ................................................................................ [169360]
KM111295.1_Ken06.Bus             ...........................g..c...............................................c. [169360]
KM111294.1_Ken05/Tk1             ...........................g..c...............................................c. [169360]
KM262844.1_strain_L60            ................................................................................ [169360]
KM102979.1_26544/OG10_Italy      ................................................................................ [169360]
KM262845.1_strain_NHV            ................................................................................ [169360]
KX354450.1_47/Ss/2008            ................................................................................ [169360]
KP055815.1_strain_BA71           ................................................................................ [169360]
MH025920.1_strain_R35            ...........................g..c...............................................c. [169360]
MH025919.1_strain_N10            ...........................g..c...............................................c. [169360]
MH025918.1_strain_R25            ...........................g..c...............................................c. [169360]
MH025917.1_strain_R7             ...........................g..c...............................................c. [169360]
MH025916.1_strain_R8             ...........................g..c...............................................c. [169360]

FR682468.2_Georgia_2007/1        gtaccgttgtgggcgggagcaaacccacctattggtatcaccacattagaagggtttgctcagaatgtccatccatgccg [169440]
ASF_IND_20_CAD543                ................................................................................ [169440]
LR899193.1_Germany_2020/1        ................................................................................ [169440]
MN393477.1_Wuhan_2019            ................................................................................ [169440]
MW396979.1_Timor-Leste/2019/1    ................................................................................ [169440]
MK940252.1_CN/2019/InnerMongolia ................................................................................ [169440]
MK128995.1_China/2018/AnhuiXCGQ  ................................................................................ [169440]
MH681419.1_ASFV/POL/2015         ................................................................................ [169440]
LS478113.1_Estonia_2014          ................................................................................ [169440]
KP843857.1_Odintsovo_02/14       ................................................................................ [169440]
```

```
FN557520.1_strain_E75            ................................................................................ [169440]
AY261366.1_Warthog               ....t...........a...................................a........................... [169440]
AY261365.1_Warmbaths             ....t...........a...................................a........................... [169440]
AY261364.1_Tengani_62            ................................................................................ [169440]
AY261363.1_Pretoriuskop/96/4     ................................................................................ [169440]
AY261362.1_Mkuzi_1979            ................................................................................ [169440]
AY261361.1_Malawi_Lil-20/1_1983  ................................................................................ [169440]
AY261360.1_Kenya_1950            ..................a........................t.................................... [169440]
AM712240.1_OURT_88/3             ................................................................................ [169440]
AM712239.1_Benin_97/1            ................................................................................ [169440]
KM111295.1_Ken06.Bus             ..................a........................t.................................... [169440]
KM111294.1_Ken05/Tk1             ..................a........................t.................................... [169440]
KM262844.1_strain_L60            ................................................................................ [169440]
KM102979.1_26544/OG10_Italy      ................................................................................ [169440]
KM262845.1_strain_NHV            ................................................................................ [169440]
KX354450.1_47/Ss/2008            ................................................................................ [169440]
KP055815.1_strain_BA71           ................................................................................ [169440]
MH025920.1_strain_R35            ..................a........................t.................................... [169440]
MH025919.1_strain_N10            ..................a........................t.................................... [169440]
MH025918.1_strain_R25            ..................a........................t.................................... [169440]
MH025917.1_strain_R7             ..................a........................t.................................... [169440]
MH025916.1_strain_R8             ..................a........................t.................................... [169440]

FR682468.2_Georgia_2007/1        atgagtatgataggtccgtatctgaatgtctataaagcccaaattctaacaagggacaagaatttttttatgaatttcga [169520]
ASF_IND_20_CAD543                ................................................................................ [169520]
LR899193.1_Germany_2020/1        ................................................................................ [169520]
MN393477.1_Wuhan_2019            ................................................................................ [169520]
MW396979.1_Timor-Leste/2019/1    ................................................................................ [169520]
MK940252.1_CN/2019/InnerMongolia ................................................................................ [169520]
MK128995.1_China/2018/AnhuiXCGQ  ................................................................................ [169520]
MH681419.1_ASFV/POL/2015         ................................................................................ [169520]
LS478113.1_Estonia_2014          ................................................................................ [169520]
KP843857.1_Odintsovo_02/14       ................................................................................ [169520]
FN557520.1_strain_E75            ................................................................................ [169520]
AY261366.1_Warthog               ....................................................a........................... [169520]
AY261365.1_Warmbaths             ....................................................a........................... [169520]
AY261364.1_Tengani_62            ..........................c.........t...............a........................... [169520]
AY261363.1_Pretoriuskop/96/4     .......................a............t...............a........................... [169520]
AY261362.1_Mkuzi_1979            ................................................................................ [169520]
AY261361.1_Malawi_Lil-20/1_1983  ....c............a..c...............a.t.............aa.......................... [169520]
AY261360.1_Kenya_1950            ....................c...............a........g......a........................... [169520]
AM712240.1_OURT_88/3             ................................................................................ [169520]
AM712239.1_Benin_97/1            ................................................................................ [169520]
KM111295.1_Ken06.Bus             ....................c...............a........g......a........................... [169520]
```

```
KM111294.1_Ken05/Tk1              ....................c...............a........g......a........................... [169520]
KM262844.1_strain_L60             ................................................................................ [169520]
KM102979.1_26544/OG10_Italy       ................................................................................ [169520]
KM262845.1_strain_NHV             ................................................................................ [169520]
KX354450.1_47/Ss/2008             ................................................................................ [169520]
KP055815.1_strain_BA71            ................................................................................ [169520]
MH025920.1_strain_R35             ....................c...............a........g......a........................... [169520]
MH025919.1_strain_N10             ....................c...............a........g......a........................... [169520]
MH025918.1_strain_R25             ....................c...............a........g......a........................... [169520]
MH025917.1_strain_R7              ....................c...............a........g......a........................... [169520]
MH025916.1_strain_R8              ....................c...............a........g......a........................... [169520]

FR682468.2_Georgia_2007/1         ---tcccgcgcataatgagtacacctttatcattcaaaaactaaaagaagcagcccgaaatatgccggaagacgaattag [169600]
ASF_IND_20_CAD543                 ---............................................................................. [169600]
LR899193.1_Germany_2020/1         ---............................................................................. [169600]
MN393477.1_Wuhan_2019             ---............................................................................. [169600]
MW396979.1_Timor-Leste/2019/1     ---............................................................................. [169600]
MK940252.1_CN/2019/InnerMongolia  ---............................................................................. [169600]
MK128995.1_China/2018/AnhuiXCGQ   ---............................................................................. [169600]
MH681419.1_ASFV/POL/2015          ---............................................................................. [169600]
LS478113.1_Estonia_2014           ---............................................................................. [169600]
KP843857.1_Odintsovo_02/14        ---............................................................................. [169600]
FN557520.1_strain_E75             ---............................................................................. [169600]
AY261366.1_Warthog                ---...t......................................................................... [169600]
AY261365.1_Warmbaths              ---...t......................................................................... [169600]
AY261364.1_Tengani_62             ---...................................................t..g..............t....... [169600]
AY261363.1_Pretoriuskop/96/4      ---...t.................................................................t....... [169600]
AY261362.1_Mkuzi_1979             ---............................................................................. [169600]
AY261361.1_Malawi_Lil-20/1_1983   ---...t................................g........................t.......g....... [169600]
AY261360.1_Kenya_1950             tcc...t..t.............................g...........c........c................... [169600]
AM712240.1_OURT_88/3              ---............................................................................. [169600]
AM712239.1_Benin_97/1             ---............................................................................. [169600]
KM111295.1_Ken06.Bus              tcc...t..t.............................g....................c................... [169600]
KM111294.1_Ken05/Tk1              tcc...t..t.............................g...........c........c................... [169600]
KM262844.1_strain_L60             ---............................................................................. [169600]
KM102979.1_26544/OG10_Italy       ---............................................................................. [169600]
KM262845.1_strain_NHV             ---............................................................................. [169600]
KX354450.1_47/Ss/2008             ---............................................................................. [169600]
KP055815.1_strain_BA71            ---............................................................................. [169600]
MH025920.1_strain_R35             tcc...t..t.............................g....................c................... [169600]
MH025919.1_strain_N10             tcc...t..t.............................g....................c................... [169600]
MH025918.1_strain_R25             tcc...t..t.............................g....................c................... [169600]
MH025917.1_strain_R7              tcc...t..t.............................g....................c................... [169600]
MH025916.1_strain_R8              tcc...t..t.............................g....................c................... [169600]
```

```
FR682468.2_Georgia_2007/1        aacagtactgggtaaaacttttatttttacttaaaagctacataaaatgtaagccctttattaattaaagaattgatgca [169680]
ASF_IND_20_CAD543                .............................................................................a.- [169680]
LR899193.1_Germany_2020/1        ................................................................................ [169680]
MN393477.1_Wuhan_2019            ................................................................................ [169680]
MW396979.1_Timor-Leste/2019/1    ................................................................................ [169680]
MK940252.1_CN/2019/InnerMongolia ................................................................................ [169680]
MK128995.1_China/2018/AnhuiXCGQ  ................................................................................ [169680]
MH681419.1_ASFV/POL/2015         ................................................................................ [169680]
LS478113.1_Estonia_2014          ................................................................................ [169680]
KP843857.1_Odintsovo_02/14       ................................................................................ [169680]
FN557520.1_strain_E75            ...............................................................................- [169680]
AY261366.1_Warthog               ................................................................................ [169680]
AY261365.1_Warmbaths             ................................................................................ [169680]
AY261364.1_Tengani_62            ................................................................................ [169680]
AY261363.1_Pretoriuskop/96/4     ...............................................................................- [169680]
AY261362.1_Mkuzi_1979            ...............................................................................- [169680]
AY261361.1_Malawi_Lil-20/1_1983  ....a........................................................................... [169680]
AY261360.1_Kenya_1950            ............c........................t.....................................t.... [169680]
AM712240.1_OURT_88/3             ...............................................................................- [169680]
AM712239.1_Benin_97/1            ...............................................................................- [169680]
KM111295.1_Ken06.Bus             ............c........................t.....................................t.... [169680]
KM111294.1_Ken05/Tk1             ............c........................t.....................................t.... [169680]
KM262844.1_strain_L60            ...............................................................................- [169680]
KM102979.1_26544/OG10_Italy      ...............................................................................- [169680]
KM262845.1_strain_NHV            ...............................................................................- [169680]
KX354450.1_47/Ss/2008            ...............................................................................- [169680]
KP055815.1_strain_BA71           .............................................................................a.- [169680]
MH025920.1_strain_R35            ............c........................t.....................................t.... [169680]
MH025919.1_strain_N10            ............c........................t.....................................t.... [169680]
MH025918.1_strain_R25            ............c........................t.....................................t.... [169680]
MH025917.1_strain_R7             ............c........................t.....................................t.... [169680]
MH025916.1_strain_R8             ............c........................t.....................................t.... [169680]

FR682468.2_Georgia_2007/1        taactaataaatggccggtcgtgttaaaataaaacagaaagagctcatagactctactgtaaaaaacaaaaatgtgatga [169760]
ASF_IND_20_CAD543                ................................................................................ [169760]
LR899193.1_Germany_2020/1        ................................................................................ [169760]
MN393477.1_Wuhan_2019            ................................................................................ [169760]
MW396979.1_Timor-Leste/2019/1    ................................................................................ [169760]
MK940252.1_CN/2019/InnerMongolia ................................................................................ [169760]
MK128995.1_China/2018/AnhuiXCGQ  ................................................................................ [169760]
MH681419.1_ASFV/POL/2015         ................................................................................ [169760]
LS478113.1_Estonia_2014          ................................................................................ [169760]
KP843857.1_Odintsovo_02/14       ................................................................................ [169760]
```

```
FN557520.1_strain_E75            ................................................................................ [169760]
AY261366.1_Warthog               ................................................................................ [169760]
AY261365.1_Warmbaths             ..c............................................................................. [169760]
AY261364.1_Tengani_62            ................................................................................ [169760]
AY261363.1_Pretoriuskop/96/4     ................................................................................ [169760]
AY261362.1_Mkuzi_1979            ................................................................................ [169760]
AY261361.1_Malawi_Lil-20/1_1983  ................................................................................ [169760]
AY261360.1_Kenya_1950            ....................................a....................c...................... [169760]
AM712240.1_OURT_88/3             ................................................................................ [169760]
AM712239.1_Benin_97/1            ................................................................................ [169760]
KM111295.1_Ken06.Bus             ....................................a....................c...................... [169760]
KM111294.1_Ken05/Tk1             ....................................a....................c...................... [169760]
KM262844.1_strain_L60            ................................................................................ [169760]
KM102979.1_26544/OG10_Italy      ................................................................................ [169760]
KM262845.1_strain_NHV            ................................................................................ [169760]
KX354450.1_47/Ss/2008            ................................................................................ [169760]
KP055815.1_strain_BA71           ................................................................................ [169760]
MH025920.1_strain_R35            ....................................a....................c...................... [169760]
MH025919.1_strain_N10            ....................................a....................c...................... [169760]
MH025918.1_strain_R25            ....................................a....................c...................... [169760]
MH025917.1_strain_R7             ....................................a....................c...................... [169760]
MH025916.1_strain_R8             ....................................a....................c...................... [169760]

FR682468.2_Georgia_2007/1        atctgttccatgaaattataggctcaaaaggcaatattaattttagcgttgtctggcccaagtttaaaaaaatcaaacag [169840]
ASF_IND_20_CAD543                ................................................................................ [169840]
LR899193.1_Germany_2020/1        ................................................................................ [169840]
MN393477.1_Wuhan_2019            ................................................................................ [169840]
MW396979.1_Timor-Leste/2019/1    ................................................................................ [169840]
MK940252.1_CN/2019/InnerMongolia ................................................................................ [169840]
MK128995.1_China/2018/AnhuiXCGQ  ................................................................................ [169840]
MH681419.1_ASFV/POL/2015         ................................................................................ [169840]
LS478113.1_Estonia_2014          ................................................................................ [169840]
KP843857.1_Odintsovo_02/14       ................................................................................ [169840]
FN557520.1_strain_E75            ................................................................................ [169840]
AY261366.1_Warthog               ..............................................t................................. [169840]
AY261365.1_Warmbaths             ..............................................t..c.............................. [169840]
AY261364.1_Tengani_62            ..............................................t................................. [169840]
AY261363.1_Pretoriuskop/96/4     ..............................................t................................. [169840]
AY261362.1_Mkuzi_1979            ................................................................................ [169840]
AY261361.1_Malawi_Lil-20/1_1983  ....a..............t...........t..c...........ta................c..............a [169840]
AY261360.1_Kenya_1950            ....a.........................................t................................a [169840]
AM712240.1_OURT_88/3             ................................................................................ [169840]
AM712239.1_Benin_97/1            ................................................................................ [169840]
KM111295.1_Ken06.Bus             ....a.........................................t................................a [169840]
```

```
KM111294.1_Ken05/Tk1             ....a.........................................t................................a [169840]
KM262844.1_strain_L60            ................................................................................ [169840]
KM102979.1_26544/OG10_Italy      ................................................................................ [169840]
KM262845.1_strain_NHV            ................................................................................ [169840]
KX354450.1_47/Ss/2008            ................................................................................ [169840]
KP055815.1_strain_BA71           ................................................................................ [169840]
MH025920.1_strain_R35            ....a.........................................t................................a [169840]
MH025919.1_strain_N10            ....a.........................................t................................a [169840]
MH025918.1_strain_R25            ....a.........................................t................................a [169840]
MH025917.1_strain_R7             ....a.........................................t................................a [169840]
MH025916.1_strain_R8             ....a.........................................t................................a [169840]

FR682468.2_Georgia_2007/1        agcgtttatgactacatttccactctttctgtgctggaaaaagcaaacgttatgcaaaactttgaagctgataagaaact [169920]
ASF_IND_20_CAD543                ................................................................................ [169920]
LR899193.1_Germany_2020/1        ................................................................................ [169920]
MN393477.1_Wuhan_2019            ................................................................................ [169920]
MW396979.1_Timor-Leste/2019/1    ................................................................................ [169920]
MK940252.1_CN/2019/InnerMongolia ................................................................................ [169920]
MK128995.1_China/2018/AnhuiXCGQ  ................................................................................ [169920]
MH681419.1_ASFV/POL/2015         ................................................................................ [169920]
LS478113.1_Estonia_2014          ................................................................................ [169920]
KP843857.1_Odintsovo_02/14       ................................................................................ [169920]
FN557520.1_strain_E75            ..............................................g................................. [169920]
AY261366.1_Warthog               ..............................................g....................ac........... [169920]
AY261365.1_Warmbaths             ..............................................g....................a............ [169920]
AY261364.1_Tengani_62            ..............................................g....................a............ [169920]
AY261363.1_Pretoriuskop/96/4     ..............................................g....................a............ [169920]
AY261362.1_Mkuzi_1979            ..............................................g................................. [169920]
AY261361.1_Malawi_Lil-20/1_1983  .....g.....a..........t.......................g....................aa.........a. [169920]
AY261360.1_Kenya_1950            .....c........................................g....................a..........a. [169920]
AM712240.1_OURT_88/3             ..............................................g................................. [169920]
AM712239.1_Benin_97/1            ..............................................g................................. [169920]
KM111295.1_Ken06.Bus             .....c........................................g....................a..........a. [169920]
KM111294.1_Ken05/Tk1             .....c........................................g....................a..........a. [169920]
KM262844.1_strain_L60            ..............................................g................................. [169920]
KM102979.1_26544/OG10_Italy      ..............................................g................................. [169920]
KM262845.1_strain_NHV            ..............................................g................................. [169920]
KX354450.1_47/Ss/2008            ..............................................g................................. [169920]
KP055815.1_strain_BA71           ..............................................g................................. [169920]
MH025920.1_strain_R35            .....c........................................g....................a..........a. [169920]
MH025919.1_strain_N10            .....c........................................g....................a..........a. [169920]
MH025918.1_strain_R25            .....c........................................g....................a..........a. [169920]
MH025917.1_strain_R7             .....c........................................g....................a..........a. [169920]
MH025916.1_strain_R8             .....c........................................g....................a..........a. [169920]
```

```
FR682468.2_Georgia_2007/1        gttggaactttttgtacaaaagctgtgggctgcctatgaaggctatttcaaatatcccgagattgaaaaatatgaggtgg [170000]
ASF_IND_20_CAD543                ................................................................................ [170000]
LR899193.1_Germany_2020/1        ................................................................................ [170000]
MN393477.1_Wuhan_2019            ................................................................................ [170000]
MW396979.1_Timor-Leste/2019/1    ................................................................................ [170000]
MK940252.1_CN/2019/InnerMongolia ................................................................................ [170000]
MK128995.1_China/2018/AnhuiXCGQ  ................................................................................ [170000]
MH681419.1_ASFV/POL/2015         ................................................................................ [170000]
LS478113.1_Estonia_2014          ................................................................................ [170000]
KP843857.1_Odintsovo_02/14       ................................................................................ [170000]
FN557520.1_strain_E75            ................................................................................ [170000]
AY261366.1_Warthog               ................................................................................ [170000]
AY261365.1_Warmbaths             ................................................................................ [170000]
AY261364.1_Tengani_62            ................................................................................ [170000]
AY261363.1_Pretoriuskop/96/4     ................................................................................ [170000]
AY261362.1_Mkuzi_1979            ................................................................................ [170000]
AY261361.1_Malawi_Lil-20/1_1983  .........................................................t...................... [170000]
AY261360.1_Kenya_1950            .............................................c........c..t...................... [170000]
AM712240.1_OURT_88/3             ................................................................................ [170000]
AM712239.1_Benin_97/1            ................................................................................ [170000]
KM111295.1_Ken06.Bus             .............................................c........c..t...................... [170000]
KM111294.1_Ken05/Tk1             ......................t......................c........c..t...................... [170000]
KM262844.1_strain_L60            ................................................................................ [170000]
KM102979.1_26544/OG10_Italy      ................................................................................ [170000]
KM262845.1_strain_NHV            ................................................................................ [170000]
KX354450.1_47/Ss/2008            ................................................................................ [170000]
KP055815.1_strain_BA71           ................................................................................ [170000]
MH025920.1_strain_R35            .............................................c........c..t...................... [170000]
MH025919.1_strain_N10            .............................................c........c..t...................... [170000]
MH025918.1_strain_R25            .............................................c........c..t...................... [170000]
MH025917.1_strain_R7             .............................................c........c..t...................... [170000]
MH025916.1_strain_R8             .............................................c........c..t...................... [170000]

FR682468.2_Georgia_2007/1        aaggccaggtaaatttcaatctcgtacctcagtgcgtcctcgaaaagtttagccagttgtataggataagaatcaattca [170080]
ASF_IND_20_CAD543                ................................................................................ [170080]
LR899193.1_Germany_2020/1        ................................................................................ [170080]
MN393477.1_Wuhan_2019            ................................................................................ [170080]
MW396979.1_Timor-Leste/2019/1    ................................................................................ [170080]
MK940252.1_CN/2019/InnerMongolia ................................................................................ [170080]
MK128995.1_China/2018/AnhuiXCGQ  ................................................................................ [170080]
MH681419.1_ASFV/POL/2015         ................................................................................ [170080]
LS478113.1_Estonia_2014          ................................................................................ [170080]
KP843857.1_Odintsovo_02/14       ................................................................................ [170080]
```

```
FN557520.1_strain_E75             ................................................................................ [170080]
AY261366.1_Warthog                ............................c....at.......................a..............t...... [170080]
AY261365.1_Warmbaths              ............................c....at.a.....................a..............t...... [170080]
AY261364.1_Tengani_62             .................................at............................................. [170080]
AY261363.1_Pretoriuskop/96/4      ............................c....at.......................a..................... [170080]
AY261362.1_Mkuzi_1979             ................................................................................ [170080]
AY261361.1_Malawi_Lil-20/1_1983   .t...............................at..............................tc......t...... [170080]
AY261360.1_Kenya_1950             .....................ag........c.at..............................tc............. [170080]
AM712240.1_OURT_88/3              ................................................................................ [170080]
AM712239.1_Benin_97/1             ................................................................................ [170080]
KM111295.1_Ken06.Bus              .....................ag........cca...............................tc............. [170080]
KM111294.1_Ken05/Tk1              .....................ag........ccat..............................tc............. [170080]
KM262844.1_strain_L60             ................................................................................ [170080]
KM102979.1_26544/OG10_Italy       ................................................................................ [170080]
KM262845.1_strain_NHV             ................................................................................ [170080]
KX354450.1_47/Ss/2008             ................................................................................ [170080]
KP055815.1_strain_BA71            ................................................................................ [170080]
MH025920.1_strain_R35             .....................ag........cca...............................tc............. [170080]
MH025919.1_strain_N10             .....................ag........cca...............................tc............. [170080]
MH025918.1_strain_R25             .....................ag........cca...............................tc............. [170080]
MH025917.1_strain_R7              .....................ag........cca...............................tc............. [170080]
MH025916.1_strain_R8              .....................ag........cca...............................tc............. [170080]

FR682468.2_Georgia_2007/1         gagcttgtcacactcatcctaaacagctgtgcctttatgagtaaatataacgattatattctcaaaaaagatccctacat [170160]
ASF_IND_20_CAD543                 ....................r........................................................... [170160]
LR899193.1_Germany_2020/1         ................................................................................ [170160]
MN393477.1_Wuhan_2019             ................................................................................ [170160]
MW396979.1_Timor-Leste/2019/1     ................................................................................ [170160]
MK940252.1_CN/2019/InnerMongolia  ................................................................................ [170160]
MK128995.1_China/2018/AnhuiXCGQ   ................................................................................ [170160]
MH681419.1_ASFV/POL/2015          ................................................................................ [170160]
LS478113.1_Estonia_2014           ................................................................................ [170160]
KP843857.1_Odintsovo_02/14        ................................................................................ [170160]
FN557520.1_strain_E75             ....................g........................................................... [170160]
AY261366.1_Warthog                .......................................................................c........ [170160]
AY261365.1_Warmbaths              .......................................................................c........ [170160]
AY261364.1_Tengani_62             ..................................................t.....c..............c........ [170160]
AY261363.1_Pretoriuskop/96/4      ..............t...................................t....................c........ [170160]
AY261362.1_Mkuzi_1979             ................................................................................ [170160]
AY261361.1_Malawi_Lil-20/1_1983   ....................................t.......g...........c..............c........ [170160]
AY261360.1_Kenya_1950             ....................................t.............t.....c..............c........ [170160]
AM712240.1_OURT_88/3              ....................g........................................................... [170160]
AM712239.1_Benin_97/1             ....................g........................................................... [170160]
KM111295.1_Ken06.Bus              ....................................t.............t.....c..............c........ [170160]
```

```
KM111294.1_Ken05/Tk1             ....................................t.............t.....c..............c........ [170160]
KM262844.1_strain_L60            ....................g........................................................... [170160]
KM102979.1_26544/OG10_Italy      ....................g........................................................... [170160]
KM262845.1_strain_NHV            ....................g........................................................... [170160]
KX354450.1_47/Ss/2008            ....................g........................................................... [170160]
KP055815.1_strain_BA71           ....................g........................................................... [170160]
MH025920.1_strain_R35            ....................................t.............t.....c..............c........ [170160]
MH025919.1_strain_N10            ....................................t.............t.....c..............c........ [170160]
MH025918.1_strain_R25            ....................................t.............t.....c..............c........ [170160]
MH025917.1_strain_R7             ....................................t.............t.....c..............c........ [170160]
MH025916.1_strain_R8             ....................................t.............t.....c..............c........ [170160]

FR682468.2_Georgia_2007/1        actaaccataacccccggcctatgcttttcccccattcccaacttcgaggacctaaattttaaacatctttacaacagtg [170240]
ASF_IND_20_CAD543                ................................................................................ [170240]
LR899193.1_Germany_2020/1        ................................................................................ [170240]
MN393477.1_Wuhan_2019            ................................................................................ [170240]
MW396979.1_Timor-Leste/2019/1    ................................................................................ [170240]
MK940252.1_CN/2019/InnerMongolia ................................................................................ [170240]
MK128995.1_China/2018/AnhuiXCGQ  ................................................................................ [170240]
MH681419.1_ASFV/POL/2015         ................................................................................ [170240]
LS478113.1_Estonia_2014          ................................................................................ [170240]
KP843857.1_Odintsovo_02/14       ................................................................................ [170240]
FN557520.1_strain_E75            ................................................................................ [170240]
AY261366.1_Warthog               ........................t....................................................... [170240]
AY261365.1_Warmbaths             ..........................................................................t..... [170240]
AY261364.1_Tengani_62            ............t........g.......................................................... [170240]
AY261363.1_Pretoriuskop/96/4     ................................................................................ [170240]
AY261362.1_Mkuzi_1979            ................................................................................ [170240]
AY261361.1_Malawi_Lil-20/1_1983  .............................................t..................t............... [170240]
AY261360.1_Kenya_1950            .............................................t..................t............... [170240]
AM712240.1_OURT_88/3             ................................................................................ [170240]
AM712239.1_Benin_97/1            ................................................................................ [170240]
KM111295.1_Ken06.Bus             .............................................t..................t............... [170240]
KM111294.1_Ken05/Tk1             .............................................t..................t............... [170240]
KM262844.1_strain_L60            ................................................................................ [170240]
KM102979.1_26544/OG10_Italy      ................................................................................ [170240]
KM262845.1_strain_NHV            ................................................................................ [170240]
KX354450.1_47/Ss/2008            ................................................................................ [170240]
KP055815.1_strain_BA71           ................................................................................ [170240]
MH025920.1_strain_R35            .............................................t..................t............... [170240]
MH025919.1_strain_N10            .............................................t..................t............... [170240]
MH025918.1_strain_R25            .............................................t..................t............... [170240]
MH025917.1_strain_R7             .............................................t..................t............... [170240]
MH025916.1_strain_R8             .............................................t..................t............... [170240]
```

```
FR682468.2_Georgia_2007/1        ataaaaattctcagcatgacaaagagtttatcatgtttatattatataagctttatacggctgccctaggagtgtacaat [170320]
ASF_IND_20_CAD543                ................................................................................ [170320]
LR899193.1_Germany_2020/1        ................................................................................ [170320]
MN393477.1_Wuhan_2019            ................................................................................ [170320]
MW396979.1_Timor-Leste/2019/1    ................................................................................ [170320]
MK940252.1_CN/2019/InnerMongolia ................................................................................ [170320]
MK128995.1_China/2018/AnhuiXCGQ  ................................................................................ [170320]
MH681419.1_ASFV/POL/2015         ................................................................................ [170320]
LS478113.1_Estonia_2014          ................................................................................ [170320]
KP843857.1_Odintsovo_02/14       ................................................................................ [170320]
FN557520.1_strain_E75            ................................................................................ [170320]
AY261366.1_Warthog               .........................c...............................t...................... [170320]
AY261365.1_Warmbaths             .........................c...................................................... [170320]
AY261364.1_Tengani_62            ................c........c...................................................... [170320]
AY261363.1_Pretoriuskop/96/4     .........................c...................................................... [170320]
AY261362.1_Mkuzi_1979            ................................................................................ [170320]
AY261361.1_Malawi_Lil-20/1_1983  ......................g..c.......................................t.............. [170320]
AY261360.1_Kenya_1950            ......................g..c...................................................... [170320]
AM712240.1_OURT_88/3             ................................................................................ [170320]
AM712239.1_Benin_97/1            ................................................................................ [170320]
KM111295.1_Ken06.Bus             ......................g..c...................................................... [170320]
KM111294.1_Ken05/Tk1             ......................g..c..................................................t... [170320]
KM262844.1_strain_L60            ................................................................................ [170320]
KM102979.1_26544/OG10_Italy      ................................................................................ [170320]
KM262845.1_strain_NHV            ................................................................................ [170320]
KX354450.1_47/Ss/2008            ................................................................................ [170320]
KP055815.1_strain_BA71           ................................................................................ [170320]
MH025920.1_strain_R35            ......................g..c...................................................... [170320]
MH025919.1_strain_N10            ......................g..c...................................................... [170320]
MH025918.1_strain_R25            ......................g..c...................................................... [170320]
MH025917.1_strain_R7             ......................g..c...................................................... [170320]
MH025916.1_strain_R8             ......................g..c...................................................... [170320]

FR682468.2_Georgia_2007/1        gccatctcgattccagacatcgacgtagaagaccttgaaaatatcatcctatcctcggtgagccagattaaaaaacaaat [170400]
ASF_IND_20_CAD543                ................................................................................ [170400]
LR899193.1_Germany_2020/1        ................................................................................ [170400]
MN393477.1_Wuhan_2019            ................................................................................ [170400]
MW396979.1_Timor-Leste/2019/1    ................................................................................ [170400]
MK940252.1_CN/2019/InnerMongolia ................................................................................ [170400]
MK128995.1_China/2018/AnhuiXCGQ  ................................................................................ [170400]
MH681419.1_ASFV/POL/2015         ................................................................................ [170400]
LS478113.1_Estonia_2014          ................................................................................ [170400]
KP843857.1_Odintsovo_02/14       ................................................................................ [170400]
```

FN557520.1_strain_E75 ............................................t................................... [170400]
AY261366.1_Warthog ........c.................g..................................................... [170400]
AY261365.1_Warmbaths ..........................g..................................................... [170400]
AY261364.1_Tengani_62 ...........................................................a.................... [170400]
AY261363.1_Pretoriuskop/96/4 ..........................g..................................................... [170400]
AY261362.1_Mkuzi_1979 ............................................t................................... [170400]
AY261361.1_Malawi_Lil-20/1_1983 ........c..............t........................t.............t................. [170400]
AY261360.1_Kenya_1950 ........c....................g.....c..........................t..............g.. [170400]
AM712240.1_OURT_88/3 ............................................t................................... [170400]
AM712239.1_Benin_97/1 ............................................t................................... [170400]
KM111295.1_Ken06.Bus ........c.....c..............g.....c..........................t..............g.. [170400]
KM111294.1_Ken05/Tk1 ........c....................g.....c..........................t..............g.. [170400]
KM262844.1_strain_L60 ............................................t................................... [170400]
KM102979.1_26544/OG10_Italy ............................................t................................... [170400]
KM262845.1_strain_NHV ............................................t................................... [170400]
KX354450.1_47/Ss/2008 ............................................t................................... [170400]
KP055815.1_strain_BA71 ............................................t................................... [170400]
MH025920.1_strain_R35 ........cg....c..............g.....c..........................t..............g.. [170400]
MH025919.1_strain_N10 ........cg....c..............g.....c..........................t..............g.. [170400]
MH025918.1_strain_R25 ........cg....c..............g.....c..........................t..............g.. [170400]
MH025917.1_strain_R7 ........cg....c..............g.....c..........................t..............g.. [170400]
MH025916.1_strain_R8 ........cg....c..............g.....c..........................t..............g.. [170400]

FR682468.2_Georgia_2007/1 tccgcgctgcaaagacgccttcaacaaaattgaatcttcggtacacctgttgcgcaaaaattttaacacatattacagtg [170480]
ASF_IND_20_CAD543 ................................................................................ [170480]
LR899193.1_Germany_2020/1 ................................................................................ [170480]
MN393477.1_Wuhan_2019 ................................................................................ [170480]
MW396979.1_Timor-Leste/2019/1 ................................................................................ [170480]
MK940252.1_CN/2019/InnerMongolia ................................................................................ [170480]
MK128995.1_China/2018/AnhuiXCGQ ................................................................................ [170480]
MH681419.1_ASFV/POL/2015 ................................................................................ [170480]
LS478113.1_Estonia_2014 ................................................................................ [170480]
KP843857.1_Odintsovo_02/14 ................................................................................ [170480]
FN557520.1_strain_E75 .................................................c.............................. [170480]
AY261366.1_Warthog .................................................c.............................. [170480]
AY261365.1_Warmbaths .................................................c............................c. [170480]
AY261364.1_Tengani_62 ..........................................g......c.............................. [170480]
AY261363.1_Pretoriuskop/96/4 .................................................c.............................. [170480]
AY261362.1_Mkuzi_1979 .................................................c.............................. [170480]
AY261361.1_Malawi_Lil-20/1_1983 ..............................c.....c............c.a..........................c. [170480]
AY261360.1_Kenya_1950 ..............................c..................c...................g........c. [170480]
AM712240.1_OURT_88/3 .................................................c.............................. [170480]
AM712239.1_Benin_97/1 .................................................c.............................. [170480]
KM111295.1_Ken06.Bus ..............................c.....c............c...................g........c. [170480]

```
KM111294.1_Ken05/Tk1              ..............................c..................c...................g........c. [170480]
KM262844.1_strain_L60             .................................................c.............................. [170480]
KM102979.1_26544/OG10_Italy       .................................................c.............................. [170480]
KM262845.1_strain_NHV             .................................................c.............................. [170480]
KX354450.1_47/Ss/2008             .................................................c.............................. [170480]
KP055815.1_strain_BA71            .................................................c.............................. [170480]
MH025920.1_strain_R35             ..............................c.....c............c...................g........c. [170480]
MH025919.1_strain_N10             ..............................c.....c............c...................g........c. [170480]
MH025918.1_strain_R25             ..............................c.....c............c...................g........c. [170480]
MH025917.1_strain_R7              ..............................c.....c............c...................g........c. [170480]
MH025916.1_strain_R8              ..............................c.....c............c...................g........c. [170480]

FR682468.2_Georgia_2007/1         actatgtgggctcaggctacaacccaaccatcattatggaacagtacattaaagacatatcacaggattccaagaacata [170560]
ASF_IND_20_CAD543                 ................................................................................ [170560]
LR899193.1_Germany_2020/1         ................................................................................ [170560]
MN393477.1_Wuhan_2019             ................................................................................ [170560]
MW396979.1_Timor-Leste/2019/1     ................................................................................ [170560]
MK940252.1_CN/2019/InnerMongolia  ................................................................................ [170560]
MK128995.1_China/2018/AnhuiXCGQ   ................................................................................ [170560]
MH681419.1_ASFV/POL/2015          ................................................................................ [170560]
LS478113.1_Estonia_2014           ................................................................................ [170560]
KP843857.1_Odintsovo_02/14        ................................................................................ [170560]
FN557520.1_strain_E75             ................................................................................ [170560]
AY261366.1_Warthog                ................................................................................ [170560]
AY261365.1_Warmbaths              ................................................................................ [170560]
AY261364.1_Tengani_62             ..........g..................................................................... [170560]
AY261363.1_Pretoriuskop/96/4      ................................................................................ [170560]
AY261362.1_Mkuzi_1979             ................................................................................ [170560]
AY261361.1_Malawi_Lil-20/1_1983   .............................................................g.................. [170560]
AY261360.1_Kenya_1950             .............g.................................................................. [170560]
AM712240.1_OURT_88/3              ................................................................................ [170560]
AM712239.1_Benin_97/1             ................................................................................ [170560]
KM111295.1_Ken06.Bus              ................................................................................ [170560]
KM111294.1_Ken05/Tk1              .............g.................................................................. [170560]
KM262844.1_strain_L60             ................................................................................ [170560]
KM102979.1_26544/OG10_Italy       ................................................................................ [170560]
KM262845.1_strain_NHV             ................................................................................ [170560]
KX354450.1_47/Ss/2008             ................................................................................ [170560]
KP055815.1_strain_BA71            ................................................................................ [170560]
MH025920.1_strain_R35             ................................................................................ [170560]
MH025919.1_strain_N10             ................................................................................ [170560]
MH025918.1_strain_R25             ................................................................................ [170560]
MH025917.1_strain_R7              ................................................................................ [170560]
MH025916.1_strain_R8              ................................................................................ [170560]
```

```
FR682468.2_Georgia_2007/1        tcaccacgcatttcctaccagtttagaaccatcatcaagtattaccgcgacatgattgccaccaggcatcaaacgatgga [170640]
ASF_IND_20_CAD543                ................................................................................ [170640]
LR899193.1_Germany_2020/1        ................................................................................ [170640]
MN393477.1_Wuhan_2019            ................................................................................ [170640]
MW396979.1_Timor-Leste/2019/1    ................................................................................ [170640]
MK940252.1_CN/2019/InnerMongolia ................................................................................ [170640]
MK128995.1_China/2018/AnhuiXCGQ  ................................................................................ [170640]
MH681419.1_ASFV/POL/2015         ................................................................................ [170640]
LS478113.1_Estonia_2014          ................................................................................ [170640]
KP843857.1_Odintsovo_02/14       ................................................................................ [170640]
FN557520.1_strain_E75            ........................................................c....................... [170640]
AY261366.1_Warthog               ................................................................a............... [170640]
AY261365.1_Warmbaths             ................................................................a............... [170640]
AY261364.1_Tengani_62            ................................................................a............... [170640]
AY261363.1_Pretoriuskop/96/4     ................................................................a............... [170640]
AY261362.1_Mkuzi_1979            ................................................................................ [170640]
AY261361.1_Malawi_Lil-20/1_1983  .....g......................................t.....t.............a............... [170640]
AY261360.1_Kenya_1950            .....g....................g.....................................a............... [170640]
AM712240.1_OURT_88/3             ........................................................c....................... [170640]
AM712239.1_Benin_97/1            ........................................................c....................... [170640]
KM111295.1_Ken06.Bus             .....g....................g.....................................a............... [170640]
KM111294.1_Ken05/Tk1             .....g....................g.....................................a............... [170640]
KM262844.1_strain_L60            ........................................................c....................... [170640]
KM102979.1_26544/OG10_Italy      ........................................................c....................... [170640]
KM262845.1_strain_NHV            ........................................................c....................... [170640]
KX354450.1_47/Ss/2008            ........................................................c....................... [170640]
KP055815.1_strain_BA71           ........................................................c....................... [170640]
MH025920.1_strain_R35            .....g....................g.....................................a............... [170640]
MH025919.1_strain_N10            .....g....................g.....................................a............... [170640]
MH025918.1_strain_R25            .....g....................g.....................................a............... [170640]
MH025917.1_strain_R7             .....g....................g.....................................a............... [170640]
MH025916.1_strain_R8             .....g....................g.....................................a............... [170640]

FR682468.2_Georgia_2007/1        cccccaggtattaaacctcgtaaagcacgtcgaaaagaaattagatatgcttgatagagaaaaaaattagtatatatagt [170720]
ASF_IND_20_CAD543                ................................................................................ [170720]
LR899193.1_Germany_2020/1        ................................................................................ [170720]
MN393477.1_Wuhan_2019            ................................................................................ [170720]
MW396979.1_Timor-Leste/2019/1    ................................................................................ [170720]
MK940252.1_CN/2019/InnerMongolia ................................................................................ [170720]
MK128995.1_China/2018/AnhuiXCGQ  ................................................................................ [170720]
MH681419.1_ASFV/POL/2015         ................................................................................ [170720]
LS478113.1_Estonia_2014          ................................................................................ [170720]
KP843857.1_Odintsovo_02/14       ................................................................................ [170720]
```

```
FN557520.1_strain_E75               ................................................................................ [170720]
AY261366.1_Warthog                  t..............................................................................c [170720]
AY261365.1_Warmbaths                t............................................................................... [170720]
AY261364.1_Tengani_62               t............................................................................... [170720]
AY261363.1_Pretoriuskop/96/4        t..............................................................................c [170720]
AY261362.1_Mkuzi_1979               t............................................................................... [170720]
AY261361.1_Malawi_Lil-20/1_1983     ........................a.........................................a............. [170720]
AY261360.1_Kenya_1950               ................................................................................ [170720]
AM712240.1_OURT_88/3                ................................................................................ [170720]
AM712239.1_Benin_97/1               ................................................................................ [170720]
KM111295.1_Ken06.Bus                ................................................................................ [170720]
KM111294.1_Ken05/Tk1                ................................................................................ [170720]
KM262844.1_strain_L60               ................................................................................ [170720]
KM102979.1_26544/OG10_Italy         ................................................................................ [170720]
KM262845.1_strain_NHV               ................................................................................ [170720]
KX354450.1_47/Ss/2008               ................................................................................ [170720]
KP055815.1_strain_BA71              ................................................................................ [170720]
MH025920.1_strain_R35               ................................................................................ [170720]
MH025919.1_strain_N10               ................................................................................ [170720]
MH025918.1_strain_R25               ................................................................................ [170720]
MH025917.1_strain_R7                ................................................................................ [170720]
MH025916.1_strain_R8                ................................................................................ [170720]

FR682468.2_Georgia_2007/1           tatggtgaatctttttcctgtttttaccttaattgtgattattacaattttaattacgactcgagaactatccaccacga [170800]
ASF_IND_20_CAD543                   ................................................................................ [170800]
LR899193.1_Germany_2020/1           ................................................................................ [170800]
MN393477.1_Wuhan_2019               ................................................................................ [170800]
MW396979.1_Timor-Leste/2019/1       ................................................................................ [170800]
MK940252.1_CN/2019/InnerMongolia    ................................................................................ [170800]
MK128995.1_China/2018/AnhuiXCGQ     ................................................................................ [170800]
MH681419.1_ASFV/POL/2015            ................................................................................ [170800]
LS478113.1_Estonia_2014             ................................................................................ [170800]
KP843857.1_Odintsovo_02/14          ................................................................................ [170800]
FN557520.1_strain_E75               ................................................................................ [170800]
AY261366.1_Warthog                  .....................................................................g.......... [170800]
AY261365.1_Warmbaths                ................................................................................ [170800]
AY261364.1_Tengani_62               .....................................................................g.......... [170800]
AY261363.1_Pretoriuskop/96/4        .....................................................................g.......... [170800]
AY261362.1_Mkuzi_1979               .........................................................a...................... [170800]
AY261361.1_Malawi_Lil-20/1_1983     .....................................................................g..t....... [170800]
AY261360.1_Kenya_1950               ..................................................................gt.g..t....... [170800]
AM712240.1_OURT_88/3                ................................................................................ [170800]
AM712239.1_Benin_97/1               ................................................................................ [170800]
KM111295.1_Ken06.Bus                ..................................................................gt.g..t....... [170800]
```

```
KM111294.1_Ken05/Tk1              ..................................................................gt.g..t....... [170800]
KM262844.1_strain_L60             ................................................................................ [170800]
KM102979.1_26544/OG10_Italy       ................................................................................ [170800]
KM262845.1_strain_NHV             ................................................................................ [170800]
KX354450.1_47/Ss/2008             ................................................................................ [170800]
KP055815.1_strain_BA71            ................................................................................ [170800]
MH025920.1_strain_R35             ..................................................................gt.g..t....... [170800]
MH025919.1_strain_N10             ..................................................................gt.g..t....... [170800]
MH025918.1_strain_R25             ..................................................................gt.g..t....... [170800]
MH025917.1_strain_R7              ..................................................................gt.g..t....... [170800]
MH025916.1_strain_R8              ..................................................................gt.g..t....... [170800]

FR682468.2_Georgia_2007/1         tgcttattgtttctcttgtaacagattatattattattaatacacagtatacggaacagcagcatgaaaacaatacattt [170880]
ASF_IND_20_CAD543                 ................................................................................ [170880]
LR899193.1_Germany_2020/1         ................................................................................ [170880]
MN393477.1_Wuhan_2019             ................................................................................ [170880]
MW396979.1_Timor-Leste/2019/1     ................................................................................ [170880]
MK940252.1_CN/2019/InnerMongolia  ................................................................................ [170880]
MK128995.1_China/2018/AnhuiXCGQ   ................................................................................ [170880]
MH681419.1_ASFV/POL/2015          ................................................................................ [170880]
LS478113.1_Estonia_2014           ................................................................................ [170880]
KP843857.1_Odintsovo_02/14        ................................................................................ [170880]
FN557520.1_strain_E75             ................................................................................ [170880]
AY261366.1_Warthog                ....c........................................................................... [170880]
AY261365.1_Warmbaths              ....c........................................................................... [170880]
AY261364.1_Tengani_62             ....c........................................................................... [170880]
AY261363.1_Pretoriuskop/96/4      ....c........................................................................... [170880]
AY261362.1_Mkuzi_1979             ................................................................................ [170880]
AY261361.1_Malawi_Lil-20/1_1983   .....................................................................tg......... [170880]
AY261360.1_Kenya_1950             ....c......................................g..........---............tg..c.a.... [170880]
AM712240.1_OURT_88/3              ................................................................................ [170880]
AM712239.1_Benin_97/1             ................................................................................ [170880]
KM111295.1_Ken06.Bus              ....c......................................g...........g.............tg..c.a.... [170880]
KM111294.1_Ken05/Tk1              ....c.................................................---............tg..c.a.... [170880]
KM262844.1_strain_L60             ................................................................................ [170880]
KM102979.1_26544/OG10_Italy       ................................................................................ [170880]
KM262845.1_strain_NHV             ................................................................................ [170880]
KX354450.1_47/Ss/2008             ................................................................................ [170880]
KP055815.1_strain_BA71            ................................................................................ [170880]
MH025920.1_strain_R35             ....c......................................g...........g.............tg..c.a.... [170880]
MH025919.1_strain_N10             ....c......................................g...........g.............tg..c.a.... [170880]
MH025918.1_strain_R25             ....c......................................g...........g.............tg..c.a.... [170880]
MH025917.1_strain_R7              ....c......................................g...........g.............tg..c.a.... [170880]
MH025916.1_strain_R8              ....c......................................g...........g.............tg..c.a.... [170880]
```

```
FR682468.2_Georgia_2007/1        ttca---------tgccgcaaaaaaattcttttaacgaatcttataataaagacaaaaaatctaatatacatattcccta [170960]
ASF_IND_20_CAD543                ....---------................................................................... [170960]
LR899193.1_Germany_2020/1        ....---------................................................................... [170960]
MN393477.1_Wuhan_2019            ....---------................................................................... [170960]
MW396979.1_Timor-Leste/2019/1    ....---------................................................................... [170960]
MK940252.1_CN/2019/InnerMongolia ....---------................................................................... [170960]
MK128995.1_China/2018/AnhuiXCGQ  ....---------................................................................... [170960]
MH681419.1_ASFV/POL/2015         ....---------................................................................... [170960]
LS478113.1_Estonia_2014          ....---------................................................................... [170960]
KP843857.1_Odintsovo_02/14       ....---------................................................................... [170960]
FN557520.1_strain_E75            .c..---------......................................................c............ [170960]
AY261366.1_Warthog               .c..---------.....................g................................c............ [170960]
AY261365.1_Warmbaths             .c..---------.....................g................................c.......t.... [170960]
AY261364.1_Tengani_62            .c..---------........................................t.............c............ [170960]
AY261363.1_Pretoriuskop/96/4     .c..---------.....................g................................c............ [170960]
AY261362.1_Mkuzi_1979            .c..---------................................................................... [170960]
AY261361.1_Malawi_Lil-20/1_1983  .ca.cgcaacagt......................................................c............ [170960]
AY261360.1_Kenya_1950            .cagcgcaacaggg..t................g..........................c......c.......t.... [170960]
AM712240.1_OURT_88/3             .c..---------......................................................c............ [170960]
AM712239.1_Benin_97/1            .c..---------......................................................c............ [170960]
KM111295.1_Ken06.Bus             .cagcgcaacaggg.tt................g..........................c......c.......t.... [170960]
KM111294.1_Ken05/Tk1             .cagcgcaacaggg..t................g.t........................c......c.......t.... [170960]
KM262844.1_strain_L60            .c..---------......................................................c............ [170960]
KM102979.1_26544/OG10_Italy      .c..---------......................................................c............ [170960]
KM262845.1_strain_NHV            .c..---------......................................................c............ [170960]
KX354450.1_47/Ss/2008            .c..---------......................................................c............ [170960]
KP055815.1_strain_BA71           .c..---------......................................................c............ [170960]
MH025920.1_strain_R35            .cagcgcaacaggg.tt................g..........................c......c.......t.... [170960]
MH025919.1_strain_N10            .cagcgcaacaggg.tt................g..........................c......c.......t.... [170960]
MH025918.1_strain_R25            .cagcgcaacaggg.tt................g..........................c......c.......t.... [170960]
MH025917.1_strain_R7             .cagcgcaacaggg.tt................g..........................c......c.......t.... [170960]
MH025916.1_strain_R8             .cagcgcaacaggg.tt................g..........................c......c.......t.... [170960]

FR682468.2_Georgia_2007/1        ccagtggctggcgcctgaactgaaggaagctgagagcaagtactggtggggcaattatgatcctcatagcgagcccgttc [171040]
ASF_IND_20_CAD543                ................................................................................ [171040]
LR899193.1_Germany_2020/1        ................................................................................ [171040]
MN393477.1_Wuhan_2019            ................................................................................ [171040]
MW396979.1_Timor-Leste/2019/1    ................................................................................ [171040]
MK940252.1_CN/2019/InnerMongolia ................................................................................ [171040]
MK128995.1_China/2018/AnhuiXCGQ  ................................................................................ [171040]
MH681419.1_ASFV/POL/2015         ................................................................................ [171040]
LS478113.1_Estonia_2014          ................................................................................ [171040]
KP843857.1_Odintsovo_02/14       ................................................................................ [171040]
```

```
FN557520.1_strain_E75           .....................a.......................................................... [171040]
AY261366.1_Warthog              ................................................................................ [171040]
AY261365.1_Warmbaths            ................................................................t............... [171040]
AY261364.1_Tengani_62           ............a...................................................t.....a......... [171040]
AY261363.1_Pretoriuskop/96/4    ................................................................................ [171040]
AY261362.1_Mkuzi_1979           .....................a.......................................................... [171040]
AY261361.1_Malawi_Lil-20/1_1983 t..................................a......t.....................t............... [171040]
AY261360.1_Kenya_1950           ...................................a...................g........t.....c......... [171040]
AM712240.1_OURT_88/3            .....................a.......................................................... [171040]
AM712239.1_Benin_97/1           .....................a.......................................................... [171040]
KM111295.1_Ken06.Bus            ...................................a...................g........t.....c......... [171040]
KM111294.1_Ken05/Tk1            ...................................a...................g........t.....c......... [171040]
KM262844.1_strain_L60           .....................a.......................................................... [171040]
KM102979.1_26544/OG10_Italy     .....................a.......................................................... [171040]
KM262845.1_strain_NHV           .....................a.......................................................... [171040]
KX354450.1_47/Ss/2008           .....................a.......................................................... [171040]
KP055815.1_strain_BA71          .....................a.......................................................... [171040]
MH025920.1_strain_R35           ...................................a...................g........t.....c......... [171040]
MH025919.1_strain_N10           ...................................a...................g........t.....c......... [171040]
MH025918.1_strain_R25           ...................................a...................g........t.....c......... [171040]
MH025917.1_strain_R7            ...................................a...................g........t.....c......... [171040]
MH025916.1_strain_R8            ...................................a...................g........t.....c......... [171040]

FR682468.2_Georgia_2007/1       tcgctggcgcatcttgaatatcttcatacgtggcacgtcaccatcaaaaacattgcccaacagcacgggcttgatataaa [171120]
ASF_IND_20_CAD543               ................................................................................ [171120]
LR899193.1_Germany_2020/1       ................................................................................ [171120]
MN393477.1_Wuhan_2019           ................................................................................ [171120]
MW396979.1_Timor-Leste/2019/1   ................................................................................ [171120]
MK940252.1_CN/2019/InnerMongolia ................................................................................ [171120]
MK128995.1_China/2018/AnhuiXCGQ ................................................................................ [171120]
MH681419.1_ASFV/POL/2015        ................................................................................ [171120]
LS478113.1_Estonia_2014         ................................................................................ [171120]
KP843857.1_Odintsovo_02/14      ................................................................................ [171120]
FN557520.1_strain_E75           ..............................................................a................. [171120]
AY261366.1_Warthog              .............................c................................a................. [171120]
AY261365.1_Warmbaths            .............................c................................a................. [171120]
AY261364.1_Tengani_62           .............................c................................a................. [171120]
AY261363.1_Pretoriuskop/96/4    .............................c................................a................. [171120]
AY261362.1_Mkuzi_1979           ..............................................................a................. [171120]
AY261361.1_Malawi_Lil-20/1_1983 ........................a....c.......c..a.....................a................. [171120]
AY261360.1_Kenya_1950           .......................ca....c.......c.ca.....................a................. [171120]
AM712240.1_OURT_88/3            ..............................................................a................. [171120]
AM712239.1_Benin_97/1           ..............................................................a................. [171120]
KM111295.1_Ken06.Bus            ......a................ca....c.......c.ca.....................a................. [171120]
```

```
KM111294.1_Ken05/Tk1             ......a................ca....c.......c.ca.....................a................. [171120]
KM262844.1_strain_L60            ..............................................................a................. [171120]
KM102979.1_26544/OG10_Italy      ..............................................................a................. [171120]
KM262845.1_strain_NHV            ..............................................................a................. [171120]
KX354450.1_47/Ss/2008            ..............................................................a................. [171120]
KP055815.1_strain_BA71           ..............................................................a................. [171120]
MH025920.1_strain_R35            ......a................ca....c.......c.ca..............t......a................. [171120]
MH025919.1_strain_N10            ......a................ca....c.......c.ca..............t......a................. [171120]
MH025918.1_strain_R25            ......a................ca....c.......c.ca..............t......a................. [171120]
MH025917.1_strain_R7             ......a................ca....c.......c.ca..............t......a................. [171120]
MH025916.1_strain_R8             ......a................ca....c.......c.ca..............t......a................. [171120]

FR682468.2_Georgia_2007/1        ggtggccattgtggtctcaacatcgcatttaaataattttttgccaatttccggggcgcttaacatcgaatgtataacct [171200]
ASF_IND_20_CAD543                ................................................................................ [171200]
LR899193.1_Germany_2020/1        ................................................................................ [171200]
MN393477.1_Wuhan_2019            ................................................................................ [171200]
MW396979.1_Timor-Leste/2019/1    ................................................................................ [171200]
MK940252.1_CN/2019/InnerMongolia ................................................................................ [171200]
MK128995.1_China/2018/AnhuiXCGQ  ................................................................................ [171200]
MH681419.1_ASFV/POL/2015         ................................................................................ [171200]
LS478113.1_Estonia_2014          ................................................................................ [171200]
KP843857.1_Odintsovo_02/14       ................................................................................ [171200]
FN557520.1_strain_E75            ................................................................................ [171200]
AY261366.1_Warthog               ...................................c................ac.......................... [171200]
AY261365.1_Warmbaths             ......t.....a......................c................ac.......................... [171200]
AY261364.1_Tengani_62            ......t.....a......................c................ac.......................... [171200]
AY261363.1_Pretoriuskop/96/4     ...................................c................ac.......................... [171200]
AY261362.1_Mkuzi_1979            ..................................................g............................. [171200]
AY261361.1_Malawi_Lil-20/1_1983  a...................a.a.......................t.....ac...a...................... [171200]
AY261360.1_Kenya_1950            .............a......aca.......................g.....acac........................ [171200]
AM712240.1_OURT_88/3             ................................................................................ [171200]
AM712239.1_Benin_97/1            ................................................................................ [171200]
KM111295.1_Ken06.Bus             .............a......aca.......................g.....acac........................ [171200]
KM111294.1_Ken05/Tk1             .............a......aca.......................g......cac........................ [171200]
KM262844.1_strain_L60            ................................................................................ [171200]
KM102979.1_26544/OG10_Italy      ................................................................................ [171200]
KM262845.1_strain_NHV            ................................................................................ [171200]
KX354450.1_47/Ss/2008            ................................................................................ [171200]
KP055815.1_strain_BA71           ................................................................................ [171200]
MH025920.1_strain_R35            .............a......aca.......................g.....acac........................ [171200]
MH025919.1_strain_N10            .............a......aca.......................g.....acac........................ [171200]
MH025918.1_strain_R25            .............a......aca.......................g.....acac........................ [171200]
MH025917.1_strain_R7             .............a......aca.......................g.....acac........................ [171200]
MH025916.1_strain_R8             .............a......aca.......................g.....acac........................ [171200]
```

```
FR682468.2_Georgia_2007/1        tccccagttgcggcatcaaggagatagacctcctatgggcgcgcattaaactatttcaacattactgcgccatcggtgcc [171280]
ASF_IND_20_CAD543                ................................................................................ [171280]
LR899193.1_Germany_2020/1        ................................................................................ [171280]
MN393477.1_Wuhan_2019            ................................................................................ [171280]
MW396979.1_Timor-Leste/2019/1    ................................................................................ [171280]
MK940252.1_CN/2019/InnerMongolia ................................................................................ [171280]
MK128995.1_China/2018/AnhuiXCGQ  ................................................................................ [171280]
MH681419.1_ASFV/POL/2015         ................................................................................ [171280]
LS478113.1_Estonia_2014          ................................................................................ [171280]
KP843857.1_Odintsovo_02/14       ................................................................................ [171280]
FN557520.1_strain_E75            .t.............................................................................. [171280]
AY261366.1_Warthog               .t...g......................................................................c... [171280]
AY261365.1_Warmbaths             .t...g......................................................................c... [171280]
AY261364.1_Tengani_62            .t........t....................t........a...................................c... [171280]
AY261363.1_Pretoriuskop/96/4     .t...g......................................................................c... [171280]
AY261362.1_Mkuzi_1979            .t.............................................................................. [171280]
AY261361.1_Malawi_Lil-20/1_1983  .t...g..........a..............t........a.............................a.....c... [171280]
AY261360.1_Kenya_1950            .t...g.c........a..............t........a.............................a.....c... [171280]
AM712240.1_OURT_88/3             .t.............................................................................. [171280]
AM712239.1_Benin_97/1            .t.............................................................................. [171280]
KM111295.1_Ken06.Bus             .t...g.c........a..............t........a.............................a.....c... [171280]
KM111294.1_Ken05/Tk1             .t...g.c..t.....a..............t........a.............................a.....c... [171280]
KM262844.1_strain_L60            .t.............................................................................. [171280]
KM102979.1_26544/OG10_Italy      .t.............................................................................. [171280]
KM262845.1_strain_NHV            .t.............................................................................. [171280]
KX354450.1_47/Ss/2008            .t.............................................................................. [171280]
KP055815.1_strain_BA71           .t.............................................................................. [171280]
MH025920.1_strain_R35            .t...g.c........a..............t........a.............................a.....c... [171280]
MH025919.1_strain_N10            .t...g.c........a..............t........a.............................a.....c... [171280]
MH025918.1_strain_R25            .t...g.c........a..............t........a.............................a.....c... [171280]
MH025917.1_strain_R7             .t...g.c........a..............t........a.............................a.....c... [171280]
MH025916.1_strain_R8             .t...g.c........a..............t........a.............................a.....c... [171280]

FR682468.2_Georgia_2007/1        cgtcttttatggctggtaagtgctgacatcaggccccctgtttcagcgtggccagccatcgccgacagtctaaaaaaggg [171360]
ASF_IND_20_CAD543                ................................................................................ [171360]
LR899193.1_Germany_2020/1        ................................................................................ [171360]
MN393477.1_Wuhan_2019            ................................................................................ [171360]
MW396979.1_Timor-Leste/2019/1    ................................................................................ [171360]
MK940252.1_CN/2019/InnerMongolia ................................................................................ [171360]
MK128995.1_China/2018/AnhuiXCGQ  ................................................................................ [171360]
MH681419.1_ASFV/POL/2015         ................................................................................ [171360]
LS478113.1_Estonia_2014          ................................................................................ [171360]
KP843857.1_Odintsovo_02/14       ................................................................................ [171360]
```

```
FN557520.1_strain_E75            ..............................................................t................. [171360]
AY261366.1_Warthog               .............................................a.......g.......................... [171360]
AY261365.1_Warmbaths             ..............a.....................t........a.......g..............c........... [171360]
AY261364.1_Tengani_62            .....................................................g.......................... [171360]
AY261363.1_Pretoriuskop/96/4     .............................................a.......g..............c........... [171360]
AY261362.1_Mkuzi_1979            ................................................................................ [171360]
AY261361.1_Malawi_Lil-20/1_1983  ............t................t......t........a.a...........t.................... [171360]
AY261360.1_Kenya_1950            .................g..................t........a..........g....................... [171360]
AM712240.1_OURT_88/3             ..............................................................t................. [171360]
AM712239.1_Benin_97/1            ..............................................................t................. [171360]
KM111295.1_Ken06.Bus             .................g..................t........a..........g....................... [171360]
KM111294.1_Ken05/Tk1             .................g..................t........a..........g....................... [171360]
KM262844.1_strain_L60            ..............................................................t................. [171360]
KM102979.1_26544/OG10_Italy      ..............................................................t................. [171360]
KM262845.1_strain_NHV            ..............................................................t................. [171360]
KX354450.1_47/Ss/2008            ..............................................................t................. [171360]
KP055815.1_strain_BA71           ..............................................................t................. [171360]
MH025920.1_strain_R35            .................g..................t........a..........g....................... [171360]
MH025919.1_strain_N10            .................g..................t........a..........g....................... [171360]
MH025918.1_strain_R25            .................g..................t........a..........g....................... [171360]
MH025917.1_strain_R7             .................g..................t........a..........g....................... [171360]
MH025916.1_strain_R8             .................g..................t........a..........g....................... [171360]

FR682468.2_Georgia_2007/1        agcagatgcggtcgttattccctacccctcccgatggaacaatcttatacctaccgtcatcaaagaaatagttgtccacc [171440]
ASF_IND_20_CAD543                ................................................................................ [171440]
LR899193.1_Germany_2020/1        ................................................................................ [171440]
MN393477.1_Wuhan_2019            ................................................................................ [171440]
MW396979.1_Timor-Leste/2019/1    ................................................................................ [171440]
MK940252.1_CN/2019/InnerMongolia ................................................................................ [171440]
MK128995.1_China/2018/AnhuiXCGQ  ................................................................................ [171440]
MH681419.1_ASFV/POL/2015         ................................................................................ [171440]
LS478113.1_Estonia_2014          ................................................................................ [171440]
KP843857.1_Odintsovo_02/14       ................................................................................ [171440]
FN557520.1_strain_E75            ............g...........t....................c.................................. [171440]
AY261366.1_Warthog               ................................................................................ [171440]
AY261365.1_Warmbaths             ...........................t.................................................... [171440]
AY261364.1_Tengani_62            ................................................................................ [171440]
AY261363.1_Pretoriuskop/96/4     ................................................................................ [171440]
AY261362.1_Mkuzi_1979            ........................t....................................................... [171440]
AY261361.1_Malawi_Lil-20/1_1983  .........a..t...g....t..t.......a....................................c.......... [171440]
AY261360.1_Kenya_1950            ................g.......t........g........c..................................g.. [171440]
AM712240.1_OURT_88/3             ............g...........t....................c.................................. [171440]
AM712239.1_Benin_97/1            ............g...........t....................c.................................. [171440]
KM111295.1_Ken06.Bus             ................g.......t........g........c..................................g.. [171440]
```

```
KM111294.1_Ken05/Tk1             ................g.......t........g........c..................................g.. [171440]
KM262844.1_strain_L60            ............g...........t....................c.................................. [171440]
KM102979.1_26544/OG10_Italy      ............g...........t....................c.................................. [171440]
KM262845.1_strain_NHV            ............g...........t....................c.................................. [171440]
KX354450.1_47/Ss/2008            ............g...........t....................c.................................. [171440]
KP055815.1_strain_BA71           ............g...........t....................c.................................. [171440]
MH025920.1_strain_R35            ................g.......t........g........c..................................g.. [171440]
MH025919.1_strain_N10            ................g.......t........g........c..................................g.. [171440]
MH025918.1_strain_R25            ................g.......t........g........c..................................g.. [171440]
MH025917.1_strain_R7             ................g.......t........g........c..................................g.. [171440]
MH025916.1_strain_R8             ................g.......t........g........c..................................g.. [171440]

FR682468.2_Georgia_2007/1        aaaaaaaatgccttgtggcggtggatgcacgccaccttgatacagatacccagattgtaggggccgggatgggctgcatc [171520]
ASF_IND_20_CAD543                ................................................................................ [171520]
LR899193.1_Germany_2020/1        ................................................................................ [171520]
MN393477.1_Wuhan_2019            ................................................................................ [171520]
MW396979.1_Timor-Leste/2019/1    ................................................................................ [171520]
MK940252.1_CN/2019/InnerMongolia ................................................................................ [171520]
MK128995.1_China/2018/AnhuiXCGQ  ................................................................................ [171520]
MH681419.1_ASFV/POL/2015         ................................................................................ [171520]
LS478113.1_Estonia_2014          ................................................................................ [171520]
KP843857.1_Odintsovo_02/14       ................................................................................ [171520]
FN557520.1_strain_E75            ..........................................................g..................... [171520]
AY261366.1_Warthog               ................................................................................ [171520]
AY261365.1_Warmbaths             ................................................................................ [171520]
AY261364.1_Tengani_62            ......................a................................c..g..................... [171520]
AY261363.1_Pretoriuskop/96/4     ................................................................................ [171520]
AY261362.1_Mkuzi_1979            ..........................................................g..................... [171520]
AY261361.1_Malawi_Lil-20/1_1983  ..........t...................a................................................. [171520]
AY261360.1_Kenya_1950            ..............................a.....................a........................... [171520]
AM712240.1_OURT_88/3             ..........................................................g..................... [171520]
AM712239.1_Benin_97/1            ..........................................................g..................... [171520]
KM111295.1_Ken06.Bus             ..............................at....................a........................... [171520]
KM111294.1_Ken05/Tk1             ..............................a.....................a........................... [171520]
KM262844.1_strain_L60            ..........................................................g..................... [171520]
KM102979.1_26544/OG10_Italy      ..........................................................g..................... [171520]
KM262845.1_strain_NHV            ..........................................................g..................... [171520]
KX354450.1_47/Ss/2008            ..........................................................g..................... [171520]
KP055815.1_strain_BA71           ..........................................................g..................... [171520]
MH025920.1_strain_R35            ..............................at....................a........................... [171520]
MH025919.1_strain_N10            ..............................at....................a........................... [171520]
MH025918.1_strain_R25            ..............................at....................a........................... [171520]
MH025917.1_strain_R7             ..............................at....................a........................... [171520]
MH025916.1_strain_R8             ..............................at....................a........................... [171520]
```

```
FR682468.2_Georgia_2007/1        gtcctaaccctaaaggcccttatggtgcgcctaagtattggcaaacagcccgttaagatactgtggcccgaccttcacgg [171600]
ASF_IND_20_CAD543                ....................................r........................................... [171600]
LR899193.1_Germany_2020/1        ................................................................................ [171600]
MN393477.1_Wuhan_2019            ................................................................................ [171600]
MW396979.1_Timor-Leste/2019/1    ................................................................................ [171600]
MK940252.1_CN/2019/InnerMongolia ................................................................................ [171600]
MK128995.1_China/2018/AnhuiXCGQ  ................................................................................ [171600]
MH681419.1_ASFV/POL/2015         ................................................................................ [171600]
LS478113.1_Estonia_2014          ................................................................................ [171600]
KP843857.1_Odintsovo_02/14       ................................................................................ [171600]
FN557520.1_strain_E75            .............................t.................................................. [171600]
AY261366.1_Warthog               ...................................................a............................ [171600]
AY261365.1_Warmbaths             ...................................................a............................ [171600]
AY261364.1_Tengani_62            .................t............................................a................. [171600]
AY261363.1_Pretoriuskop/96/4     .............................t.....................a...................t........ [171600]
AY261362.1_Mkuzi_1979            .............................t.................................................. [171600]
AY261361.1_Malawi_Lil-20/1_1983  .............................t.....................a............................ [171600]
AY261360.1_Kenya_1950            .............................t........c............a............................ [171600]
AM712240.1_OURT_88/3             .............................t.................................................. [171600]
AM712239.1_Benin_97/1            .............................t.................................................. [171600]
KM111295.1_Ken06.Bus             .............................t........c............a....a....................... [171600]
KM111294.1_Ken05/Tk1             .............................t........c............a............................ [171600]
KM262844.1_strain_L60            .............................t.................................................. [171600]
KM102979.1_26544/OG10_Italy      .............................t.................................................. [171600]
KM262845.1_strain_NHV            .............................t.................................................. [171600]
KX354450.1_47/Ss/2008            .............................t.................................................. [171600]
KP055815.1_strain_BA71           .............................t.................................................. [171600]
MH025920.1_strain_R35            .............................t........c............a....a....................... [171600]
MH025919.1_strain_N10            .............................t........c............a....a....................... [171600]
MH025918.1_strain_R25            .............................t........c............a....a....................... [171600]
MH025917.1_strain_R7             .............................t........c............a....a....................... [171600]
MH025916.1_strain_R8             .............................t........c............a....a....................... [171600]

FR682468.2_Georgia_2007/1        cactgccgagggcattcctctggagggggtggaggttggctggtttttaaacgcttatgcgcataaattaaatatacgct [171680]
ASF_IND_20_CAD543                ................................................................................ [171680]
LR899193.1_Germany_2020/1        ................................................................................ [171680]
MN393477.1_Wuhan_2019            ................................................................................ [171680]
MW396979.1_Timor-Leste/2019/1    ................................................................................ [171680]
MK940252.1_CN/2019/InnerMongolia ................................................................................ [171680]
MK128995.1_China/2018/AnhuiXCGQ  ................................................................................ [171680]
MH681419.1_ASFV/POL/2015         ................................................................................ [171680]
LS478113.1_Estonia_2014          ................................................................................ [171680]
KP843857.1_Odintsovo_02/14       ................................................................................ [171680]
```

```
FN557520.1_strain_E75           ................................................................................ [171680]
AY261366.1_Warthog              ........................a.....a................................................. [171680]
AY261365.1_Warmbaths            ........................a.....a.......................c......................... [171680]
AY261364.1_Tengani_62           ........................a.....a................................................. [171680]
AY261363.1_Pretoriuskop/96/4    ........................a.....a................................................. [171680]
AY261362.1_Mkuzi_1979           ......................................................c......................... [171680]
AY261361.1_Malawi_Lil-20/1_1983 ...g....................a..a..a....................t............................ [171680]
AY261360.1_Kenya_1950           ...c..............c.....a.....a.........................................c.....t. [171680]
AM712240.1_OURT_88/3            ................................................................................ [171680]
AM712239.1_Benin_97/1           ................................................................................ [171680]
KM111295.1_Ken06.Bus            ...c..............c.....a.....a.............................a...........c.....t. [171680]
KM111294.1_Ken05/Tk1            ...c..............c.....a.....a.........................................c.....t. [171680]
KM262844.1_strain_L60           ................................................................................ [171680]
KM102979.1_26544/OG10_Italy     ................................................................................ [171680]
KM262845.1_strain_NHV           ................................................................................ [171680]
KX354450.1_47/Ss/2008           ................................................................................ [171680]
KP055815.1_strain_BA71          ................................................................................ [171680]
MH025920.1_strain_R35           ...c..............c.....a.....a.............................a...........c.....t. [171680]
MH025919.1_strain_N10           ...c..............c.....a.....a.............................a...........c.....t. [171680]
MH025918.1_strain_R25           ...c..............c.....a.....a.............................a...........c.....t. [171680]
MH025917.1_strain_R7            ...c..............c.....a.....a.............................a...........c.....t. [171680]
MH025916.1_strain_R8            ...c..............c.....a.....a.............................a...........c.....t. [171680]

FR682468.2_Georgia_2007/1       gcctaggggctgatcatattgcgcagcacttaacttaattctttatttaaaaagtccacgcatccagtg----------- [171760]
ASF_IND_20_CAD543               .....................................................................----------- [171760]
LR899193.1_Germany_2020/1       .....................................................................----------- [171760]
MN393477.1_Wuhan_2019           .....................................................................----------- [171760]
MW396979.1_Timor-Leste/2019/1   .....................................................................----------- [171760]
MK940252.1_CN/2019/InnerMongolia .....................................................................----------- [171760]
MK128995.1_China/2018/AnhuiXCGQ .....................................................................----------- [171760]
MH681419.1_ASFV/POL/2015        ........................................t............................----------- [171760]
LS478113.1_Estonia_2014         .....................................................................----------- [171760]
KP843857.1_Odintsovo_02/14      .....................................................................----------- [171760]
FN557520.1_strain_E75           ......................................................c.............a----------- [171760]
AY261366.1_Warthog              .............g..............------------.............--------------------------- [171760]
AY261365.1_Warmbaths            .............g..............------------..............c.............a----------- [171760]
AY261364.1_Tengani_62           ...............................c......................c........t....a----------- [171760]
AY261363.1_Pretoriuskop/96/4    ......................................................c.t..at..t.....----------- [171760]
AY261362.1_Mkuzi_1979           ......................................................c.............a----------- [171760]
AY261361.1_Malawi_Lil-20/1_1983 .............a...........a.......a........ac..........c...t....tttaaaagcccatgcat [171760]
AY261360.1_Kenya_1950           ........cgg...t..................a.........c..........c.------------------------ [171760]
AM712240.1_OURT_88/3            ......................................................c.............a----------- [171760]
AM712239.1_Benin_97/1           ......................................................c.............a----------- [171760]
KM111295.1_Ken06.Bus            ........cgg...t..................a.........c..........c.------------------------ [171760]
```

```
KM111294.1_Ken05/Tk1             ........cgg...t..................a.........c..........c.------------------------ [171760]
KM262844.1_strain_L60            ......................................................c.............a----------- [171760]
KM102979.1_26544/OG10_Italy      ......................................................c.............a----------- [171760]
KM262845.1_strain_NHV            ......................................................c.............a----------- [171760]
KX354450.1_47/Ss/2008            ......................................................c.............a----------- [171760]
KP055815.1_strain_BA71           ......................................................c.............a----------- [171760]
MH025920.1_strain_R35            ........cgg...t..................a.........c..........c.------------------------ [171760]
MH025919.1_strain_N10            ........cgg...t..................a.........c..........c.------------------------ [171760]
MH025918.1_strain_R25            ........cgg...t..................a.........c..........c.------------------------ [171760]
MH025917.1_strain_R7             ........cgg...t..................a.........c..........c.------------------------ [171760]
MH025916.1_strain_R8             ........cgg...t..................a.........c..........c.------------------------ [171760]

FR682468.2_Georgia_2007/1        --------------------------gcggcctaca-ttaagggcctacgcacat-aaatatacactggctagaagtacg [171840]
ASF_IND_20_CAD543                --------------------------..........-..................-......................y. [171840]
LR899193.1_Germany_2020/1        --------------------------..........-..................-........................ [171840]
MN393477.1_Wuhan_2019            --------------------------..........-..................-........................ [171840]
MW396979.1_Timor-Leste/2019/1    --------------------------..........-..................-........................ [171840]
MK940252.1_CN/2019/InnerMongolia --------------------------..........-..................-........................ [171840]
MK128995.1_China/2018/AnhuiXCGQ  --------------------------..........-..................-........................ [171840]
MH681419.1_ASFV/POL/2015         --------------------------..........-..................-........................ [171840]
LS478113.1_Estonia_2014          --------------------------..........-..................-........................ [171840]
KP843857.1_Odintsovo_02/14       --------------------------..........-..................-........................ [171840]
FN557520.1_strain_E75            --------------------------..........-..................-........g.............t. [171840]
AY261366.1_Warthog               -------------------------------------------............-........g...........c... [171840]
AY261365.1_Warmbaths             --------------------------..........-....a.............-........g...........c... [171840]
AY261364.1_Tengani_62            --------------------------..........-....a......t......-........g..a........c... [171840]
AY261363.1_Pretoriuskop/96/4     --------------------------..........t....a...t.........-........g..-........c... [171840]
AY261362.1_Mkuzi_1979            --------------------------..........-....a.............-........g...........c... [171840]
AY261361.1_Malawi_Lil-20/1_1983  tttaaaagccgacgcatgtaaattgc..ta..c.g.agc..a...t.........a........g...........c.ta [171840]
AY261360.1_Kenya_1950            ------------------------------------------------t...tg.a..cat...g.c...-.....c.-a [171840]
AM712240.1_OURT_88/3             --------------------------..........-..................-........g.............t. [171840]
AM712239.1_Benin_97/1            --------------------------..........-..................-........g.............t. [171840]
KM111295.1_Ken06.Bus             ------------------------------------------------....tg.a..cat...g.c.........cg-a [171840]
KM111294.1_Ken05/Tk1             ------------------------------------------------t...tg.a..cat...g.c...-.....c.-a [171840]
KM262844.1_strain_L60            --------------------------..........-..................-........g.............t. [171840]
KM102979.1_26544/OG10_Italy      --------------------------..........-..................-........g.............t. [171840]
KM262845.1_strain_NHV            --------------------------..........-..................-........g.............t. [171840]
KX354450.1_47/Ss/2008            --------------------------..........-..................-........g.............t. [171840]
KP055815.1_strain_BA71           --------------------------..........-..................-........g.............t. [171840]
MH025920.1_strain_R35            ------------------------------------------------....tg.a..cat...g.c.........cg-a [171840]
MH025919.1_strain_N10            ------------------------------------------------....tg.a..cat...g.c.........cg-a [171840]
MH025918.1_strain_R25            ------------------------------------------------....tg.a..cat...g.c.........cg-a [171840]
MH025917.1_strain_R7             ------------------------------------------------....tg.a..cat...g.c.........cg-a [171840]
MH025916.1_strain_R8             ------------------------------------------------....tg.a..cat...g.c.........cg-a [171840]
```

```
FR682468.2_Georgia_2007/1        ccttcatttaaaccattgaattatttatataatggctgcaaacattattgcaacaagagccgtgccaaagatggccagca [171920]
ASF_IND_20_CAD543                ................................................................................ [171920]
LR899193.1_Germany_2020/1        ................................................................................ [171920]
MN393477.1_Wuhan_2019            ................................................................................ [171920]
MW396979.1_Timor-Leste/2019/1    ................................................................................ [171920]
MK940252.1_CN/2019/InnerMongolia ................................................................................ [171920]
MK128995.1_China/2018/AnhuiXCGQ  ................................................................................ [171920]
MH681419.1_ASFV/POL/2015         ................................................................................ [171920]
LS478113.1_Estonia_2014          ................................................................................ [171920]
KP843857.1_Odintsovo_02/14       ................................................................................ [171920]
FN557520.1_strain_E75            ......................................t.......................c................. [171920]
AY261366.1_Warthog               t...................................------------..............c........c........ [171920]
AY261365.1_Warmbaths             t...................................------------..............c........c........ [171920]
AY261364.1_Tengani_62            ..............................................................c........c........ [171920]
AY261363.1_Pretoriuskop/96/4     .............................................................ac................. [171920]
AY261362.1_Mkuzi_1979            t.............................................................c........c........ [171920]
AY261361.1_Malawi_Lil-20/1_1983  ................ca........................t.............a.....c.a....a.......... [171920]
AY261360.1_Kenya_1950            ................ca........................t.....g.............c.a...........g... [171920]
AM712240.1_OURT_88/3             ......................................t.......................c................. [171920]
AM712239.1_Benin_97/1            ......................................t.......................c................. [171920]
KM111295.1_Ken06.Bus             t...............ca..............................g.............c.a...........g... [171920]
KM111294.1_Ken05/Tk1             ................ca........................t.....g.............c.a.........t.g... [171920]
KM262844.1_strain_L60            ......................................t.......................c................. [171920]
KM102979.1_26544/OG10_Italy      ......................................t.......................c................. [171920]
KM262845.1_strain_NHV            ......................................t.......................c................. [171920]
KX354450.1_47/Ss/2008            ......................................t.......................c................. [171920]
KP055815.1_strain_BA71           ......................................t.......................c................. [171920]
MH025920.1_strain_R35            t...............ca..............................g.............c.a...........g... [171920]
MH025919.1_strain_N10            t...............ca..............................g.............c.a...........g... [171920]
MH025918.1_strain_R25            t...............ca..............................g.............c.a...........g... [171920]
MH025917.1_strain_R7             t...............ca..............................g.............c.a...........g... [171920]
MH025916.1_strain_R8             t...............ca..............................g.............c.a...........g... [171920]

FR682468.2_Georgia_2007/1        aaaaagagcatcaatactgtctgctagactcccaggaaaagcgtcatgggcattatcccttttcatttgaattaaagcct [172000]
ASF_IND_20_CAD543                ...................y............................................................ [172000]
LR899193.1_Germany_2020/1        ................................................................................ [172000]
MN393477.1_Wuhan_2019            ................................................................................ [172000]
MW396979.1_Timor-Leste/2019/1    ................................................................................ [172000]
MK940252.1_CN/2019/InnerMongolia ................................................................................ [172000]
MK128995.1_China/2018/AnhuiXCGQ  ................................................................................ [172000]
MH681419.1_ASFV/POL/2015         ................................................................................ [172000]
LS478113.1_Estonia_2014          ................................................................................ [172000]
KP843857.1_Odintsovo_02/14       ................................................................................ [172000]
```

```
FN557520.1_strain_E75           ...................c............................................................ [172000]
AY261366.1_Warthog              ...................c.............................c..............g............... [172000]
AY261365.1_Warmbaths            ...................c........................g....c.............................. [172000]
AY261364.1_Tengani_62           ...................c.............................t.............................. [172000]
AY261363.1_Pretoriuskop/96/4    ...................c........................g....c..............g............... [172000]
AY261362.1_Mkuzi_1979           ...................c........t................................................... [172000]
AY261361.1_Malawi_Lil-20/1_1983 ...................c.............................c..ac.........................c [172000]
AY261360.1_Kenya_1950           .............g.....c.........a.............c.....c.....c........c..............c [172000]
AM712240.1_OURT_88/3            ...................c............................................................ [172000]
AM712239.1_Benin_97/1           ...................c............................................................ [172000]
KM111295.1_Ken06.Bus            .............g.....c..a......a...................c.....c........c..............c [172000]
KM111294.1_Ken05/Tk1            .............g.....c..a......a...................c.....c........c..............c [172000]
KM262844.1_strain_L60           ...................c............................................................ [172000]
KM102979.1_26544/OG10_Italy     ...................c............................................................ [172000]
KM262845.1_strain_NHV           ...................c............................................................ [172000]
KX354450.1_47/Ss/2008           ...................c............................................................ [172000]
KP055815.1_strain_BA71          ...................c............................................................ [172000]
MH025920.1_strain_R35           .............g.....c..a......a...................c.....c........c..............c [172000]
MH025919.1_strain_N10           .............g.....c..a......a...................c.....c........c..............c [172000]
MH025918.1_strain_R25           .............g.....c..a......a...................c.....c........c..............c [172000]
MH025917.1_strain_R7            .............g.....c..a......a...................c.....c........c..............c [172000]
MH025916.1_strain_R8            .............g.....c..a......a...................c.....c........c..............c [172000]

FR682468.2_Georgia_2007/1       tatgggcaaacaggcgcaaatatcataggagtacagggctcacttacccatgttatcaaaatgacagtatttccatttat [172080]
ASF_IND_20_CAD543               ................................................................................ [172080]
LR899193.1_Germany_2020/1       ................................................................................ [172080]
MN393477.1_Wuhan_2019           ................................................................................ [172080]
MW396979.1_Timor-Leste/2019/1   ................................................................................ [172080]
MK940252.1_CN/2019/InnerMongolia ................................................................................ [172080]
MK128995.1_China/2018/AnhuiXCGQ ................................................................................ [172080]
MH681419.1_ASFV/POL/2015        ................................................................................ [172080]
LS478113.1_Estonia_2014         ................................................................................ [172080]
KP843857.1_Odintsovo_02/14      ................................................................................ [172080]
FN557520.1_strain_E75           .........................................g...................................... [172080]
AY261366.1_Warthog              .........................................g...................................... [172080]
AY261365.1_Warmbaths            .........................................g...................................... [172080]
AY261364.1_Tengani_62           .............................c...........g........a............................. [172080]
AY261363.1_Pretoriuskop/96/4    .........................................g...................................... [172080]
AY261362.1_Mkuzi_1979           .........................................g...................................... [172080]
AY261361.1_Malawi_Lil-20/1_1983 ..............t.g.......................................t.......t............... [172080]
AY261360.1_Kenya_1950           ..............t.....c................................cg..........g..c........... [172080]
AM712240.1_OURT_88/3            .........................................g...................................... [172080]
AM712239.1_Benin_97/1           .........................................g...................................... [172080]
KM111295.1_Ken06.Bus            ..............t.....c........g.......................cg.............t........... [172080]
```

```
KM111294.1_Ken05/Tk1            ..............t.....c................................cg.............c........... [172080]
KM262844.1_strain_L60           .........................................g...................................... [172080]
KM102979.1_26544/OG10_Italy     .........................................g...................................... [172080]
KM262845.1_strain_NHV           .........................................g...................................... [172080]
KX354450.1_47/Ss/2008           .........................................g...................................... [172080]
KP055815.1_strain_BA71          .........................................g...................................... [172080]
MH025920.1_strain_R35           ..............t.....c........g.......................cg.............t........... [172080]
MH025919.1_strain_N10           ..............t.....c........g.......................cg.............t........... [172080]
MH025918.1_strain_R25           ..............t.....c........g.......................cg.............t........... [172080]
MH025917.1_strain_R7            ..............t.....c........g.......................cg.............t........... [172080]
MH025916.1_strain_R8            ..............t.....c........g.......................cg.............t........... [172080]

FR682468.2_Georgia_2007/1       gattccttttcctttacaaaaaactcatatagatgattttattggtggacgcatttatttattttttaaggaactggaca [172160]
ASF_IND_20_CAD543               ................................................................................ [172160]
LR899193.1_Germany_2020/1       ................................................................................ [172160]
MN393477.1_Wuhan_2019           ................................................................................ [172160]
MW396979.1_Timor-Leste/2019/1   ................................................................................ [172160]
MK940252.1_CN/2019/InnerMongolia ................................................................................ [172160]
MK128995.1_China/2018/AnhuiXCGQ ................................................................................ [172160]
MH681419.1_ASFV/POL/2015        ................................................................................ [172160]
LS478113.1_Estonia_2014         ................................................................................ [172160]
KP843857.1_Odintsovo_02/14      ................................................................................ [172160]
FN557520.1_strain_E75           ................................................................................ [172160]
AY261366.1_Warthog              ......c......c.........t............................g..........................g [172160]
AY261365.1_Warmbaths            .......................t............................g..........................g [172160]
AY261364.1_Tengani_62           ..............................................a.....g................a.........g [172160]
AY261363.1_Pretoriuskop/96/4    .......................t............................g..........................g [172160]
AY261362.1_Mkuzi_1979           ...............................................................................g [172160]
AY261361.1_Malawi_Lil-20/1_1983 ..........................................c.........g....................t....t. [172160]
AY261360.1_Kenya_1950           ...a......................g...c.g...............g...g.........................t. [172160]
AM712240.1_OURT_88/3            ................................................................................ [172160]
AM712239.1_Benin_97/1           ................................................................................ [172160]
KM111295.1_Ken06.Bus            ...a......................g...c.................g...g.........................t. [172160]
KM111294.1_Ken05/Tk1            ...a......................g...c.................g...g.........................t. [172160]
KM262844.1_strain_L60           ................................................................................ [172160]
KM102979.1_26544/OG10_Italy     ................................................................................ [172160]
KM262845.1_strain_NHV           ................................................................................ [172160]
KX354450.1_47/Ss/2008           ................................................................................ [172160]
KP055815.1_strain_BA71          ................................................................................ [172160]
MH025920.1_strain_R35           ...a......................g...c.................g...g.........................t. [172160]
MH025919.1_strain_N10           ...a......................g...c.................g...g.........................t. [172160]
MH025918.1_strain_R25           ...a......................g...c.................g...g.........................t. [172160]
MH025917.1_strain_R7            ...a......................g...c.................g...g.........................t. [172160]
MH025916.1_strain_R8            ...a......................g...c.................g...g.........................t. [172160]
```

```
FR682468.2_Georgia_2007/1        tgcaagcagtttctgatgtaaatggaatgcaataccacttcgagttcaaggttgttcctgtaagccccaaccaagtagag [172240]
ASF_IND_20_CAD543                ................................................................................ [172240]
LR899193.1_Germany_2020/1        ................................................................................ [172240]
MN393477.1_Wuhan_2019            ................................................................................ [172240]
MW396979.1_Timor-Leste/2019/1    ................................................................................ [172240]
MK940252.1_CN/2019/InnerMongolia ................................................................................ [172240]
MK128995.1_China/2018/AnhuiXCGQ  ................................................................................ [172240]
MH681419.1_ASFV/POL/2015         ................................................................................ [172240]
LS478113.1_Estonia_2014          ................................................................................ [172240]
KP843857.1_Odintsovo_02/14       ................................................................................ [172240]
FN557520.1_strain_E75            ................................................................................ [172240]
AY261366.1_Warthog               ........a.....................................t........c.........t.............. [172240]
AY261365.1_Warmbaths             ........a.....................................t........c.........t.............. [172240]
AY261364.1_Tengani_62            ........a...........c.........................t........c.........t.............. [172240]
AY261363.1_Pretoriuskop/96/4     ........a...............................t.....t........c.........t.............. [172240]
AY261362.1_Mkuzi_1979            ..............................................t................................. [172240]
AY261361.1_Malawi_Lil-20/1_1983  ..............................................t...........g......t.............. [172240]
AY261360.1_Kenya_1950            ........t.....................................t........c..g......t..g........... [172240]
AM712240.1_OURT_88/3             ................................................................................ [172240]
AM712239.1_Benin_97/1            ................................................................................ [172240]
KM111295.1_Ken06.Bus             ........t....c................................t........c.tg......t..g........... [172240]
KM111294.1_Ken05/Tk1             ........t.....................................t........c..g......t..g........... [172240]
KM262844.1_strain_L60            ................................................................................ [172240]
KM102979.1_26544/OG10_Italy      ................................................................................ [172240]
KM262845.1_strain_NHV            ................................................................................ [172240]
KX354450.1_47/Ss/2008            ................................................................................ [172240]
KP055815.1_strain_BA71           ................................................................................ [172240]
MH025920.1_strain_R35            ........t....c................................t........c.tg......t..g........... [172240]
MH025919.1_strain_N10            ........t....c................................t........c.tg......t..g........... [172240]
MH025918.1_strain_R25            ........t....c................................t........c.tg......t..g........... [172240]
MH025917.1_strain_R7             ........t....c................................t........c.tg......t..g........... [172240]
MH025916.1_strain_R8             ........t....c................................t........c.tg......t..g........... [172240]

FR682468.2_Georgia_2007/1        cttcttcctgtgaataataaatataaatttacatatgctataccggtagtgcaataccttaccccaatcttttatgatct [172320]
ASF_IND_20_CAD543                ................................................................................ [172320]
LR899193.1_Germany_2020/1        ................................................................................ [172320]
MN393477.1_Wuhan_2019            ................................................................................ [172320]
MW396979.1_Timor-Leste/2019/1    ................................................................................ [172320]
MK940252.1_CN/2019/InnerMongolia ................................................................................ [172320]
MK128995.1_China/2018/AnhuiXCGQ  ................................................................................ [172320]
MH681419.1_ASFV/POL/2015         ................................................................................ [172320]
LS478113.1_Estonia_2014          ................................................................................ [172320]
KP843857.1_Odintsovo_02/14       ................................................................................ [172320]
```

```
FN557520.1_strain_E75            ...............................................g................................ [172320]
AY261366.1_Warthog               ........c........c............................ag................................ [172320]
AY261365.1_Warmbaths             ........c........c............................ag.................t.............. [172320]
AY261364.1_Tengani_62            .................c............................ag................................ [172320]
AY261363.1_Pretoriuskop/96/4     ........c........c............................ag................................ [172320]
AY261362.1_Mkuzi_1979            ...............................................g................................ [172320]
AY261361.1_Malawi_Lil-20/1_1983  ............................................a.ag...........c.................... [172320]
AY261360.1_Kenya_1950            ...................c............c...........a.agc............................... [172320]
AM712240.1_OURT_88/3             ...............................................g................................ [172320]
AM712239.1_Benin_97/1            ...............................................g................................ [172320]
KM111295.1_Ken06.Bus             ...................c............c...........a.agc..........c.................... [172320]
KM111294.1_Ken05/Tk1             ...................c............c...........a.agc..........c.................... [172320]
KM262844.1_strain_L60            ...............................................g................................ [172320]
KM102979.1_26544/OG10_Italy      ...............................................g................................ [172320]
KM262845.1_strain_NHV            ...............................................g................................ [172320]
KX354450.1_47/Ss/2008            ...............................................g................................ [172320]
KP055815.1_strain_BA71           ...............................................g................................ [172320]
MH025920.1_strain_R35            ...................c............c...........a.agc..........c.................... [172320]
MH025919.1_strain_N10            ...................c............c...........a.agc..........c.................... [172320]
MH025918.1_strain_R25            ...................c............c...........a.agc..........c.................... [172320]
MH025917.1_strain_R7             ...................c............c...........a.agc..........c.................... [172320]
MH025916.1_strain_R8             ...................c............c...........a.agc..........c.................... [172320]

FR682468.2_Georgia_2007/1        ttcgggaccgctagatttcccattagatactctttcggtccatgtggatatcctctccaatcatatacagcttcctatcc [172400]
ASF_IND_20_CAD543                ..................................................k............................. [172400]
LR899193.1_Germany_2020/1        ................................................................................ [172400]
MN393477.1_Wuhan_2019            ................................................................................ [172400]
MW396979.1_Timor-Leste/2019/1    ................................................................................ [172400]
MK940252.1_CN/2019/InnerMongolia ................................................................................ [172400]
MK128995.1_China/2018/AnhuiXCGQ  ................................................................................ [172400]
MH681419.1_ASFV/POL/2015         ................................................................................ [172400]
LS478113.1_Estonia_2014          ................................................................................ [172400]
KP843857.1_Odintsovo_02/14       ................................................................................ [172400]
FN557520.1_strain_E75            ..................................................g............................. [172400]
AY261366.1_Warthog               ......g......a....t...............................g....a........................ [172400]
AY261365.1_Warmbaths             ......g......a....................................g....a........................ [172400]
AY261364.1_Tengani_62            ......c....................................a.a..c.g....a........................ [172400]
AY261363.1_Pretoriuskop/96/4     ...a.........a....................................g...ta....a................... [172400]
AY261362.1_Mkuzi_1979            ......g......a....................................g..........t.................. [172400]
AY261361.1_Malawi_Lil-20/1_1983  ...a.................g........c...................g....a.............t.......... [172400]
AY261360.1_Kenya_1950            ..............................c...........c.....c.gt...a..c..........t.......... [172400]
AM712240.1_OURT_88/3             ..................................................g............................. [172400]
AM712239.1_Benin_97/1            ..................................................g............................. [172400]
KM111295.1_Ken06.Bus             c.....gg......................c...........c.....c.gt...a.tc..........t.......... [172400]
```

```
KM111294.1_Ken05/Tk1           ......g.......................c...c.....g.c.....c.gt...a..c..........t.......... [172400]
KM262844.1_strain_L60          ..................................................g............................. [172400]
KM102979.1_26544/OG10_Italy    ..................................................g............................. [172400]
KM262845.1_strain_NHV          ..................................................g............................. [172400]
KX354450.1_47/Ss/2008          ..................................................g............................. [172400]
KP055815.1_strain_BA71         ..................................................g............................. [172400]
MH025920.1_strain_R35          c.....gg......................c...........c.....c.gt...a.tc..........t.......... [172400]
MH025919.1_strain_N10          c.....gg......................c...........c.....c.gt...a.tc..........t.......... [172400]
MH025918.1_strain_R25          c.....gg......................c...........c.....c.gt...a.tc..........t.......... [172400]
MH025917.1_strain_R7           c.....gg......................c...........c.....c.gt...a.tc..........t.......... [172400]
MH025916.1_strain_R8           c.....gg......................c...........c.....c.gt...a.tc..........t.......... [172400]

FR682468.2_Georgia_2007/1      aaaaccataacctaacaacgggtgatcgtgtttttatttctggatataaacacctgcaaacgattgaattatgtaaaaat [172480]
ASF_IND_20_CAD543              ................................................................................ [172480]
LR899193.1_Germany_2020/1      ................................................................................ [172480]
MN393477.1_Wuhan_2019          ................................................................................ [172480]
MW396979.1_Timor-Leste/2019/1  ................................................................................ [172480]
MK940252.1_CN/2019/InnerMongolia ................................................................................ [172480]
MK128995.1_China/2018/AnhuiXCGQ ................................................................................ [172480]
MH681419.1_ASFV/POL/2015       ...........................a.................................................... [172480]
LS478113.1_Estonia_2014        ................................................................................ [172480]
KP843857.1_Odintsovo_02/14     ................................................................................ [172480]
FN557520.1_strain_E75          .......c.................c...................................................... [172480]
AY261366.1_Warthog             ......................................................................g......... [172480]
AY261365.1_Warmbaths           ......................................................................g......... [172480]
AY261364.1_Tengani_62          ......................................................................g......... [172480]
AY261363.1_Pretoriuskop/96/4   ......................................................................g......... [172480]
AY261362.1_Mkuzi_1979          ...........t.................................................................... [172480]
AY261361.1_Malawi_Lil-20/1_1983 ..................a...................................c..............cg.a....... [172480]
AY261360.1_Kenya_1950          ..................t................g.......c.................c......acg......... [172480]
AM712240.1_OURT_88/3           .......c.................c...................................................... [172480]
AM712239.1_Benin_97/1          .......c.................c...................................................... [172480]
KM111295.1_Ken06.Bus           ......................c............g.......c.................c......acg......... [172480]
KM111294.1_Ken05/Tk1           ..................t.........c..............c.................c......acg......... [172480]
KM262844.1_strain_L60          .......c.................c...................................................... [172480]
KM102979.1_26544/OG10_Italy    .......c.................c...................................................... [172480]
KM262845.1_strain_NHV          .......c.................c...................................................... [172480]
KX354450.1_47/Ss/2008          .......c.................c...................................................... [172480]
KP055815.1_strain_BA71         .......c.................c...................................................... [172480]
MH025920.1_strain_R35          ......................c..c.........g.......c.................c......acg......... [172480]
MH025919.1_strain_N10          ......................c..c.........g.......c.................c......acg......... [172480]
MH025918.1_strain_R25          ......................c..c.........g.......c.................c......acg......... [172480]
MH025917.1_strain_R7           ......................c..c.........g.......c.................c......acg......... [172480]
MH025916.1_strain_R8           ......................c..c.........g.......c.................c......acg......... [172480]
```

```
FR682468.2_Georgia_2007/1        aacaagatttttatcaaaaatataccgccgctttcatccgaaaaaataaaactatatatactaaaaaatcgaatcagaat [172560]
ASF_IND_20_CAD543                ..................w............................................................. [172560]
LR899193.1_Germany_2020/1        ................................................................................ [172560]
MN393477.1_Wuhan_2019            ................................................................................ [172560]
MW396979.1_Timor-Leste/2019/1    ................................................................................ [172560]
MK940252.1_CN/2019/InnerMongolia ................................................................................ [172560]
MK128995.1_China/2018/AnhuiXCGQ  ................................................................................ [172560]
MH681419.1_ASFV/POL/2015         ................................................................................ [172560]
LS478113.1_Estonia_2014          ................................................................................ [172560]
KP843857.1_Odintsovo_02/14       ................................................................................ [172560]
FN557520.1_strain_E75            ..................t..........................................c......c........... [172560]
AY261366.1_Warthog               ..................tg.........................................c.................. [172560]
AY261365.1_Warmbaths             ..................tg.........................................c.................. [172560]
AY261364.1_Tengani_62            ..................tg..................t......................c.................. [172560]
AY261363.1_Pretoriuskop/96/4     ..................tg..................t......................c.................. [172560]
AY261362.1_Mkuzi_1979            ..................t..........................................c.................. [172560]
AY261361.1_Malawi_Lil-20/1_1983  ....c.............tg......a.......t...a...........t..........c.................. [172560]
AY261360.1_Kenya_1950            ....ta............g...................a...........t..........c............a..... [172560]
AM712240.1_OURT_88/3             ..................t..........................................c......c........... [172560]
AM712239.1_Benin_97/1            ..................t..........................................c......c........... [172560]
KM111295.1_Ken06.Bus             ....ta............g.......a........g..a...........t..........c............a..... [172560]
KM111294.1_Ken05/Tk1             ....ta............g...................a...........t..........c............a..... [172560]
KM262844.1_strain_L60            ..................t..........................................c......c........... [172560]
KM102979.1_26544/OG10_Italy      ..................t..........................................c......c........... [172560]
KM262845.1_strain_NHV            ..................t..........................................c......c........... [172560]
KX354450.1_47/Ss/2008            ..................t..........................................c......c........... [172560]
KP055815.1_strain_BA71           ..................t..........................................c......c........... [172560]
MH025920.1_strain_R35            ....ta............g.......a........g..a...........t..........c............a..... [172560]
MH025919.1_strain_N10            ....ta............g.......a........g..a...........t..........c............a..... [172560]
MH025918.1_strain_R25            ....ta............g.......a........g..a...........t..........c............a..... [172560]
MH025917.1_strain_R7             ....ta............g.......a........g..a...........t..........c............a..... [172560]
MH025916.1_strain_R8             ....ta............g.......a........g..a...........t..........c............a..... [172560]

FR682468.2_Georgia_2007/1        tccgctatactttaaatcttt-aaaaacgtctaagtaataacatttttatagtctactcctagttccgaaataggctgaa [172640]
ASF_IND_20_CAD543                .....................a.......................................................... [172640]
LR899193.1_Germany_2020/1        .....................-.......................................................... [172640]
MN393477.1_Wuhan_2019            .....................-.......................................................... [172640]
MW396979.1_Timor-Leste/2019/1    .....................-.......................................................... [172640]
MK940252.1_CN/2019/InnerMongolia .....................-.......................................................... [172640]
MK128995.1_China/2018/AnhuiXCGQ  .....................-.......................................................... [172640]
MH681419.1_ASFV/POL/2015         .....................-.......................................................... [172640]
LS478113.1_Estonia_2014          .....................-.......................................................... [172640]
KP843857.1_Odintsovo_02/14       .....................-.......................................................... [172640]
```

```
FN557520.1_strain_E75            ...a.................a.......................................................... [172640]
AY261366.1_Warthog               ...a.................a.......................................................... [172640]
AY261365.1_Warmbaths             ...a.................a.......................................................... [172640]
AY261364.1_Tengani_62            .....................a.......................................................... [172640]
AY261363.1_Pretoriuskop/96/4     ...a..............g..a.......................................................... [172640]
AY261362.1_Mkuzi_1979            ...a.................a.......................................................... [172640]
AY261361.1_Malawi_Lil-20/1_1983  ...a..t..............-.................c...........a..............t...........g. [172640]
AY261360.1_Kenya_1950            ...a.................-....g........................a...........c................ [172640]
AM712240.1_OURT_88/3             ...a.................a.......................................................... [172640]
AM712239.1_Benin_97/1            ...a.................a.......................................................... [172640]
KM111295.1_Ken06.Bus             ...a.................-....g.a......................a............................ [172640]
KM111294.1_Ken05/Tk1             ...a.................-....g........................a...........c................ [172640]
KM262844.1_strain_L60            ...a.................a.......................................................... [172640]
KM102979.1_26544/OG10_Italy      ...a.................a.......................................................... [172640]
KM262845.1_strain_NHV            ...a.................a.......................................................... [172640]
KX354450.1_47/Ss/2008            ...a.................a.......................................................... [172640]
KP055815.1_strain_BA71           ...a.................a.......................................................... [172640]
MH025920.1_strain_R35            ...a.................-....g.a......................a............................ [172640]
MH025919.1_strain_N10            ...a.................-....g.a......................a............................ [172640]
MH025918.1_strain_R25            ...a.................-....g.a......................a............................ [172640]
MH025917.1_strain_R7             ...a.................-....g.a......................a............................ [172640]
MH025916.1_strain_R8             ...a.................-....g.a......................a............................ [172640]

FR682468.2_Georgia_2007/1        tttcttttttaagtcctttaaaccaaggatgtgatacaagactcttaaaggaaagccgcttattttcattaattgttaaa [172720]
ASF_IND_20_CAD543                ................................................................................ [172720]
LR899193.1_Germany_2020/1        ................................................................................ [172720]
MN393477.1_Wuhan_2019            ................................................................................ [172720]
MW396979.1_Timor-Leste/2019/1    ................................................................................ [172720]
MK940252.1_CN/2019/InnerMongolia ................................................................................ [172720]
MK128995.1_China/2018/AnhuiXCGQ  ................................................................................ [172720]
MH681419.1_ASFV/POL/2015         ................................................................................ [172720]
LS478113.1_Estonia_2014          ................................................................................ [172720]
KP843857.1_Odintsovo_02/14       ................................................................................ [172720]
FN557520.1_strain_E75            ................................................................................ [172720]
AY261366.1_Warthog               .............c.................................................................. [172720]
AY261365.1_Warmbaths             .............c.................c..c.......................t..................... [172720]
AY261364.1_Tengani_62            .............c...................................a.............................. [172720]
AY261363.1_Pretoriuskop/96/4     .............c.................c..c............................................. [172720]
AY261362.1_Mkuzi_1979            .............c............................................t..................... [172720]
AY261361.1_Malawi_Lil-20/1_1983  .............c............................c......a.............................. [172720]
AY261360.1_Kenya_1950            .............c............................c......a.......................g...... [172720]
AM712240.1_OURT_88/3             ................................................................................ [172720]
AM712239.1_Benin_97/1            ................................................................................ [172720]
KM111295.1_Ken06.Bus             .............c............................c......a.......................g...... [172720]
```

```
KM111294.1_Ken05/Tk1             .............c............................c......a.......................g...... [172720]
KM262844.1_strain_L60            ................................................................................ [172720]
KM102979.1_26544/OG10_Italy      ................................................................................ [172720]
KM262845.1_strain_NHV            ................................................................................ [172720]
KX354450.1_47/Ss/2008            ................................................................................ [172720]
KP055815.1_strain_BA71           ................................................................................ [172720]
MH025920.1_strain_R35            .............c............................c......a.......................g...... [172720]
MH025919.1_strain_N10            .............c............................c......a.......................g...... [172720]
MH025918.1_strain_R25            .............c............................c......a.......................g...... [172720]
MH025917.1_strain_R7             .............c............................c......a.......................g...... [172720]
MH025916.1_strain_R8             .............c............................c......a.......................g...... [172720]

FR682468.2_Georgia_2007/1        cattccgtgataaactgttttcccgtctctgaaatgttctcgggaatataattttcccgtttcaggatatcatttaaata [172800]
ASF_IND_20_CAD543                .................................................................r.............. [172800]
LR899193.1_Germany_2020/1        ................................................................................ [172800]
MN393477.1_Wuhan_2019            ................................................................................ [172800]
MW396979.1_Timor-Leste/2019/1    ................................................................................ [172800]
MK940252.1_CN/2019/InnerMongolia ................................................................................ [172800]
MK128995.1_China/2018/AnhuiXCGQ  ................................................................................ [172800]
MH681419.1_ASFV/POL/2015         ................................................................................ [172800]
LS478113.1_Estonia_2014          ................................................................................ [172800]
KP843857.1_Odintsovo_02/14       ................................................................................ [172800]
FN557520.1_strain_E75            .................................................................a.............. [172800]
AY261366.1_Warthog               ..a..............c........................................c..................... [172800]
AY261365.1_Warmbaths             ..a..............c........................................c..................... [172800]
AY261364.1_Tengani_62            ..a..t...........c........................................c.....a............... [172800]
AY261363.1_Pretoriuskop/96/4     ..a..............c........................................c..................... [172800]
AY261362.1_Mkuzi_1979            .................c...............................................a.............. [172800]
AY261361.1_Malawi_Lil-20/1_1983  ..a...................................a...........g.....t.c..................... [172800]
AY261360.1_Kenya_1950            ..a..................................ga.................t.c.........g..g........ [172800]
AM712240.1_OURT_88/3             .................................................................a.............. [172800]
AM712239.1_Benin_97/1            .................................................................a.............. [172800]
KM111295.1_Ken06.Bus             ..a..................................ga.................t.c.........g..g........ [172800]
KM111294.1_Ken05/Tk1             ..a..................................ga.................t.c.........g..g........ [172800]
KM262844.1_strain_L60            .................................................................a.............. [172800]
KM102979.1_26544/OG10_Italy      .................................................................a.............. [172800]
KM262845.1_strain_NHV            .................................................................a.............. [172800]
KX354450.1_47/Ss/2008            .................................................................a.............. [172800]
KP055815.1_strain_BA71           .................................................................a.............. [172800]
MH025920.1_strain_R35            ..a..................................ga.................t.c.........g..g........ [172800]
MH025919.1_strain_N10            ..a..................................ga.................t.c.........g..g........ [172800]
MH025918.1_strain_R25            ..a..................................ga.................t.c.........g..g........ [172800]
MH025917.1_strain_R7             ..a..................................ga.................t.c.........g..g........ [172800]
MH025916.1_strain_R8             ..a..................................ga.................t.c.........g..g........ [172800]
```

```
FR682468.2_Georgia_2007/1        aaaattttctgcacgaaatctaaaaagattaaccgcgaccatacctatcgtccacacggttaaaggaagctggtagtaat [172880]
ASF_IND_20_CAD543                ...........................................................................r.... [172880]
LR899193.1_Germany_2020/1        ................................................................................ [172880]
MN393477.1_Wuhan_2019            ................................................................................ [172880]
MW396979.1_Timor-Leste/2019/1    ................................................................................ [172880]
MK940252.1_CN/2019/InnerMongolia ................................................................................ [172880]
MK128995.1_China/2018/AnhuiXCGQ  ................................................................................ [172880]
MH681419.1_ASFV/POL/2015         ................................................................................ [172880]
LS478113.1_Estonia_2014          ................................................................................ [172880]
KP843857.1_Odintsovo_02/14       ................................................................................ [172880]
FN557520.1_strain_E75            .........................................................a.................a.... [172880]
AY261366.1_Warthog               .........................................................a...................... [172880]
AY261365.1_Warmbaths             .........................................................a...................... [172880]
AY261364.1_Tengani_62            .........................................................a...................... [172880]
AY261363.1_Pretoriuskop/96/4     ...........................g.............................a...................... [172880]
AY261362.1_Mkuzi_1979            .........................................................a...................... [172880]
AY261361.1_Malawi_Lil-20/1_1983  ........................g...........c....................c....g................. [172880]
AY261360.1_Kenya_1950            ........................g...........c....................a....g................. [172880]
AM712240.1_OURT_88/3             .........................................................a.................a.... [172880]
AM712239.1_Benin_97/1            .........................................................a.................a.... [172880]
KM111295.1_Ken06.Bus             ........................g...........c....................a....g................. [172880]
KM111294.1_Ken05/Tk1             ........................g...........c....................a....g................. [172880]
KM262844.1_strain_L60            .........................................................a.................a.... [172880]
KM102979.1_26544/OG10_Italy      .........................................................a.................a.... [172880]
KM262845.1_strain_NHV            .........................................................a.................a.... [172880]
KX354450.1_47/Ss/2008            .........................................................a.................a.... [172880]
KP055815.1_strain_BA71           .........................................................a.................a.... [172880]
MH025920.1_strain_R35            ........................g...........c....................a....g................. [172880]
MH025919.1_strain_N10            ........................g...........c....................a....g................. [172880]
MH025918.1_strain_R25            ........................g...........c....................a....g................. [172880]
MH025917.1_strain_R7             ........................g...........c....................a....g................. [172880]
MH025916.1_strain_R8             ........................g...........c....................a....g................. [172880]

FR682468.2_Georgia_2007/1        aaccataataataaaattctggacacacgtattcccatgttccaaacatattatattggggacgggtttcgtctaatcta [172960]
ASF_IND_20_CAD543                ................................................................................ [172960]
LR899193.1_Germany_2020/1        ................................................................................ [172960]
MN393477.1_Wuhan_2019            ................................................................................ [172960]
MW396979.1_Timor-Leste/2019/1    ................................................................................ [172960]
MK940252.1_CN/2019/InnerMongolia ................................................................................ [172960]
MK128995.1_China/2018/AnhuiXCGQ  ................................................................................ [172960]
MH681419.1_ASFV/POL/2015         ................................................................................ [172960]
LS478113.1_Estonia_2014          ................................................................................ [172960]
KP843857.1_Odintsovo_02/14       ................................................................................ [172960]
```

```
FN557520.1_strain_E75             ................................................................................ [172960]
AY261366.1_Warthog                ................................................................................ [172960]
AY261365.1_Warmbaths              ................................................................................ [172960]
AY261364.1_Tengani_62             ................................................................................ [172960]
AY261363.1_Pretoriuskop/96/4      ................................................................................ [172960]
AY261362.1_Mkuzi_1979             ................................................................................ [172960]
AY261361.1_Malawi_Lil-20/1_1983   ......................g...................................a...t.......a.t.....g. [172960]
AY261360.1_Kenya_1950             .g....................g.......................................t..tc...a.t.....g. [172960]
AM712240.1_OURT_88/3              ................................................................................ [172960]
AM712239.1_Benin_97/1             ................................................................................ [172960]
KM111295.1_Ken06.Bus              .g....................g...................................a...t..tc...a.t....... [172960]
KM111294.1_Ken05/Tk1              .g....................g.......................................t..tc...a.t....... [172960]
KM262844.1_strain_L60             ................................................................................ [172960]
KM102979.1_26544/OG10_Italy       ................................................................................ [172960]
KM262845.1_strain_NHV             ................................................................................ [172960]
KX354450.1_47/Ss/2008             ................................................................................ [172960]
KP055815.1_strain_BA71            ................................................................................ [172960]
MH025920.1_strain_R35             .g....................g...................................a...t..tc...a.t....... [172960]
MH025919.1_strain_N10             .g....................g...................................a...t..tc...a.t....... [172960]
MH025918.1_strain_R25             .g....................g...................................a...t..tc...a.t....... [172960]
MH025917.1_strain_R7              .g....................g...................................a...t..tc...a.t....... [172960]
MH025916.1_strain_R8              .g....................g...................................a...t..tc...a.t....... [172960]

FR682468.2_Georgia_2007/1         acagcgcttccaaagtcaatgaccttaatgatcttttgatttatgtctataataaggttctcatccttaatatccccatg [173040]
ASF_IND_20_CAD543                 ................................................................................ [173040]
LR899193.1_Germany_2020/1         ................................................................................ [173040]
MN393477.1_Wuhan_2019             ................................................................................ [173040]
MW396979.1_Timor-Leste/2019/1     ................................................................................ [173040]
MK940252.1_CN/2019/InnerMongolia  ................................................................................ [173040]
MK128995.1_China/2018/AnhuiXCGQ   ................................................................................ [173040]
MH681419.1_ASFV/POL/2015          ................................................................................ [173040]
LS478113.1_Estonia_2014           ................................................................................ [173040]
KP843857.1_Odintsovo_02/14        ................................................................................ [173040]
FN557520.1_strain_E75             ................................................................................ [173040]
AY261366.1_Warthog                ..........................t...........g......................................... [173040]
AY261365.1_Warmbaths              ..........................t...........g......................................... [173040]
AY261364.1_Tengani_62             .............................a.................................................. [173040]
AY261363.1_Pretoriuskop/96/4      ..........................t...........g......................................... [173040]
AY261362.1_Mkuzi_1979             ......................................g......................................... [173040]
AY261361.1_Malawi_Lil-20/1_1983   ........................................c...a................................g.. [173040]
AY261360.1_Kenya_1950             .............................c........t..............g...........t...........g.. [173040]
AM712240.1_OURT_88/3              ................................................................................ [173040]
AM712239.1_Benin_97/1             ................................................................................ [173040]
KM111295.1_Ken06.Bus              .............................c........t..............g........g..t...........g.. [173040]
```

```
KM111294.1_Ken05/Tk1           .............................c........t..............g...........t...........g.. [173040]
KM262844.1_strain_L60          ................................................................................ [173040]
KM102979.1_26544/OG10_Italy    ................................................................................ [173040]
KM262845.1_strain_NHV          ................................................................................ [173040]
KX354450.1_47/Ss/2008          ................................................................................ [173040]
KP055815.1_strain_BA71         ................................................................................ [173040]
MH025920.1_strain_R35          .............................c........t..............g........g..t...........g.. [173040]
MH025919.1_strain_N10          .............................c........t..............g........g..t...........g.. [173040]
MH025918.1_strain_R25          .............................c........t..............g........g..t...........g.. [173040]
MH025917.1_strain_R7           .............................c........t..............g........g..t...........g.. [173040]
MH025916.1_strain_R8           .............................c........t..............g........g..t...........g.. [173040]

FR682468.2_Georgia_2007/1      gataaagcccttctcataaatgttttgtataataagaataagctggaatattatttttttggcttcggtttcctcaagtt [173120]
ASF_IND_20_CAD543              ................................................................................ [173120]
LR899193.1_Germany_2020/1      ................................................................................ [173120]
MN393477.1_Wuhan_2019          ................................................................................ [173120]
MW396979.1_Timor-Leste/2019/1  ................................................................................ [173120]
MK940252.1_CN/2019/InnerMongolia ................................................................................ [173120]
MK128995.1_China/2018/AnhuiXCGQ ................................................................................ [173120]
MH681419.1_ASFV/POL/2015       ................................................................................ [173120]
LS478113.1_Estonia_2014        ................................................................................ [173120]
KP843857.1_Odintsovo_02/14     ................................................................................ [173120]
FN557520.1_strain_E75          ................................................................................ [173120]
AY261366.1_Warthog             ...............................................................c................ [173120]
AY261365.1_Warmbaths           ................................................................................ [173120]
AY261364.1_Tengani_62          ................................................g............................... [173120]
AY261363.1_Pretoriuskop/96/4   ...............................................................c................ [173120]
AY261362.1_Mkuzi_1979          ................................................................................ [173120]
AY261361.1_Malawi_Lil-20/1_1983 .......................................c.t...a.....................t............ [173120]
AY261360.1_Kenya_1950          .....................a..........................c..................t............ [173120]
AM712240.1_OURT_88/3           ................................................................................ [173120]
AM712239.1_Benin_97/1          ................................................................................ [173120]
KM111295.1_Ken06.Bus           .....................a..........................c..................t............ [173120]
KM111294.1_Ken05/Tk1           .....................a..........................c..................t............ [173120]
KM262844.1_strain_L60          ................................................................................ [173120]
KM102979.1_26544/OG10_Italy    ................................................................................ [173120]
KM262845.1_strain_NHV          ................................................................................ [173120]
KX354450.1_47/Ss/2008          ................................................................................ [173120]
KP055815.1_strain_BA71         ................................................................................ [173120]
MH025920.1_strain_R35          .....................a..........................c..................t............ [173120]
MH025919.1_strain_N10          .....................a..........................c..................t............ [173120]
MH025918.1_strain_R25          .....................a..........................c..................t............ [173120]
MH025917.1_strain_R7           .....................a..........................c..................t............ [173120]
MH025916.1_strain_R8           .....................a..........................c..................t............ [173120]
```

```
FR682468.2_Georgia_2007/1        ttttaaagtaatgataatgaagtagatcaacactatttggaatatattctatgattagtatatgatacatagcattttcg [173200]
ASF_IND_20_CAD543                ................................................................................ [173200]
LR899193.1_Germany_2020/1        ................................................................................ [173200]
MN393477.1_Wuhan_2019            ................................................................................ [173200]
MW396979.1_Timor-Leste/2019/1    ................................................................................ [173200]
MK940252.1_CN/2019/InnerMongolia ................................................................................ [173200]
MK128995.1_China/2018/AnhuiXCGQ  ................................................................................ [173200]
MH681419.1_ASFV/POL/2015         ................................................................................ [173200]
LS478113.1_Estonia_2014          ................................................................................ [173200]
KP843857.1_Odintsovo_02/14       ................................................................................ [173200]
FN557520.1_strain_E75            ....................................................a........................... [173200]
AY261366.1_Warthog               .........................g...................................................... [173200]
AY261365.1_Warmbaths             .........................g...................................................... [173200]
AY261364.1_Tengani_62            .........................g...................................................... [173200]
AY261363.1_Pretoriuskop/96/4     .........................g...................................................... [173200]
AY261362.1_Mkuzi_1979            ........................................g...........a........................... [173200]
AY261361.1_Malawi_Lil-20/1_1983  ..............................t........................c..............g........a [173200]
AY261360.1_Kenya_1950            .......................................................c.......a......g........a [173200]
AM712240.1_OURT_88/3             ....................................................a........................... [173200]
AM712239.1_Benin_97/1            ....................................................a........................... [173200]
KM111295.1_Ken06.Bus             .......................................................c.......a......g........a [173200]
KM111294.1_Ken05/Tk1             .......................................................c.......a......g........a [173200]
KM262844.1_strain_L60            ....................................................a........................... [173200]
KM102979.1_26544/OG10_Italy      ....................................................a........................... [173200]
KM262845.1_strain_NHV            ....................................................a........................... [173200]
KX354450.1_47/Ss/2008            ....................................................a........................... [173200]
KP055815.1_strain_BA71           ....................................................a........................... [173200]
MH025920.1_strain_R35            .......................................................c.......a......g........a [173200]
MH025919.1_strain_N10            .......................................................c.......a......g........a [173200]
MH025918.1_strain_R25            .......................................................c.......a......g........a [173200]
MH025917.1_strain_R7             .......................................................c.......a......g........a [173200]
MH025916.1_strain_R8             .......................................................c.......a......g........a [173200]

FR682468.2_Georgia_2007/1        gtatattcgataagcttaataacaccgggagtatcttgcagggctttcaacacgatgacttcatttcctggaatttcttt [173280]
ASF_IND_20_CAD543                ................................................................................ [173280]
LR899193.1_Germany_2020/1        ................................................................................ [173280]
MN393477.1_Wuhan_2019            ................................................................................ [173280]
MW396979.1_Timor-Leste/2019/1    ................................................................................ [173280]
MK940252.1_CN/2019/InnerMongolia ................................................................................ [173280]
MK128995.1_China/2018/AnhuiXCGQ  ................................................................................ [173280]
MH681419.1_ASFV/POL/2015         ................................................................................ [173280]
LS478113.1_Estonia_2014          ................................................................................ [173280]
KP843857.1_Odintsovo_02/14       ................................................................................ [173280]
```

```
FN557520.1_strain_E75            ................................................................................ [173280]
AY261366.1_Warthog               ................................................................................ [173280]
AY261365.1_Warmbaths             ...............................................t....................c........... [173280]
AY261364.1_Tengani_62            ...............................................t................................ [173280]
AY261363.1_Pretoriuskop/96/4     ......................................t......................................... [173280]
AY261362.1_Mkuzi_1979            ................................................................................ [173280]
AY261361.1_Malawi_Lil-20/1_1983  ........a..............t.........................t.....c...c.................... [173280]
AY261360.1_Kenya_1950            ........a.............tt.........................t.........c.................... [173280]
AM712240.1_OURT_88/3             ................................................................................ [173280]
AM712239.1_Benin_97/1            ................................................................................ [173280]
KM111295.1_Ken06.Bus             ........a.............tt.........................t.........c.................... [173280]
KM111294.1_Ken05/Tk1             ........a.............tt.........................t.........c.................... [173280]
KM262844.1_strain_L60            ................................................................................ [173280]
KM102979.1_26544/OG10_Italy      ................................................................................ [173280]
KM262845.1_strain_NHV            ................................................................................ [173280]
KX354450.1_47/Ss/2008            ................................................................................ [173280]
KP055815.1_strain_BA71           ................................................................................ [173280]
MH025920.1_strain_R35            ........a.............tt.........................t.........c.................... [173280]
MH025919.1_strain_N10            ........a.............tt.........................t.........c.................... [173280]
MH025918.1_strain_R25            ........a.............tt.........................t.........c.................... [173280]
MH025917.1_strain_R7             ........a.............tt.........................t.........c.................... [173280]
MH025916.1_strain_R8             ........a.............tt.........................t.........c.................... [173280]

FR682468.2_Georgia_2007/1        tttagaaacgtacttaaatataatgggttgccctacttgatgacccaaaaagacgttat---ttctgccaccctcaaaca [173360]
ASF_IND_20_CAD543                ...........................................................---.................. [173360]
LR899193.1_Germany_2020/1        ...........................................................---.................. [173360]
MN393477.1_Wuhan_2019            ...........................................................---.................. [173360]
MW396979.1_Timor-Leste/2019/1    ...........................................................---.................. [173360]
MK940252.1_CN/2019/InnerMongolia ...........................................................---.................. [173360]
MK128995.1_China/2018/AnhuiXCGQ  ...........................................................---.................. [173360]
MH681419.1_ASFV/POL/2015         ...........................................................---.................. [173360]
LS478113.1_Estonia_2014          ...........................................................---.................. [173360]
KP843857.1_Odintsovo_02/14       ...........................................................---.................. [173360]
FN557520.1_strain_E75            ...........................................................---.................. [173360]
AY261366.1_Warthog               ...........................................................---....a............. [173360]
AY261365.1_Warmbaths             ...........................................................---....a............. [173360]
AY261364.1_Tengani_62            ...........................................................---....a............. [173360]
AY261363.1_Pretoriuskop/96/4     ...........................................................---....a............. [173360]
AY261362.1_Mkuzi_1979            ...........................................................---.................. [173360]
AY261361.1_Malawi_Lil-20/1_1983  ......t..a........a.c..ct..........................t.......ttc....a............. [173360]
AY261360.1_Kenya_1950            ......t...........a.c..ct.a.....t..t...t...........a.......ttc....a............. [173360]
AM712240.1_OURT_88/3             ...........................................................---.................. [173360]
AM712239.1_Benin_97/1            ...........................................................---.................. [173360]
KM111295.1_Ken06.Bus             ......t...........a.cc.ct.a.....t..t...............a.......ttc....a............. [173360]
```

```
KM111294.1_Ken05/Tk1            ......t...........a.c..ct.a.....t..t...t...........a.......ttc....a............. [173360]
KM262844.1_strain_L60           ...........................................................---.................. [173360]
KM102979.1_26544/OG10_Italy     ...........................................................---.................. [173360]
KM262845.1_strain_NHV           ...........................................................---.................. [173360]
KX354450.1_47/Ss/2008           ...........................................................---.................. [173360]
KP055815.1_strain_BA71          ...........................................................---.................. [173360]
MH025920.1_strain_R35           ......t...........a.cc.ct.a.....t..t...............a.......ttc....a............. [173360]
MH025919.1_strain_N10           ......t...........a.cc.ct.a.....t..t...............a.......ttc....a............. [173360]
MH025918.1_strain_R25           ......t...........a.cc.ct.a.....t..t...............a.......ttc....a............. [173360]
MH025917.1_strain_R7            ......t...........a.cc.ct.a.....t..t...............a.......ttc....a............. [173360]
MH025916.1_strain_R8            ......t...........a.cc.ct.a.....t..t...............a.......ttc....a............. [173360]

FR682468.2_Georgia_2007/1       tgggtctcgtcgcaatgaaatacatgtgctgcgttgtggagatcctttccacctttgctgtaggataaaacgcatattgt [173440]
ASF_IND_20_CAD543               ................................................................................ [173440]
LR899193.1_Germany_2020/1       ................................................................................ [173440]
MN393477.1_Wuhan_2019           ................................................................................ [173440]
MW396979.1_Timor-Leste/2019/1   ................................................................................ [173440]
MK940252.1_CN/2019/InnerMongolia ................................................................................ [173440]
MK128995.1_China/2018/AnhuiXCGQ ................................................................................ [173440]
MH681419.1_ASFV/POL/2015        ................................................................................ [173440]
LS478113.1_Estonia_2014         ................................................................................ [173440]
KP843857.1_Odintsovo_02/14      ................................................................................ [173440]
FN557520.1_strain_E75           ................................................................................ [173440]
AY261366.1_Warthog              .....................................................................c.......... [173440]
AY261365.1_Warmbaths            .....................................................................c.......... [173440]
AY261364.1_Tengani_62           ............................t......................................c.c.......... [173440]
AY261363.1_Pretoriuskop/96/4    .a...................................................................c.......... [173440]
AY261362.1_Mkuzi_1979           .....................................................................c.......... [173440]
AY261361.1_Malawi_Lil-20/1_1983 .a...t....t........g..........c.........................at...........ct......... [173440]
AY261360.1_Kenya_1950           .a...t....t........g...ga.....c.........a...............a............ct......... [173440]
AM712240.1_OURT_88/3            ................................................................................ [173440]
AM712239.1_Benin_97/1           ................................................................................ [173440]
KM111295.1_Ken06.Bus            .a...t.t..t.a......g...gaa....c.........a...............a............c.......... [173440]
KM111294.1_Ken05/Tk1            .a...t....t........g...ga.....c.........a...............a............ct......... [173440]
KM262844.1_strain_L60           ................................................................................ [173440]
KM102979.1_26544/OG10_Italy     ................................................................................ [173440]
KM262845.1_strain_NHV           ................................................................................ [173440]
KX354450.1_47/Ss/2008           ................................................................................ [173440]
KP055815.1_strain_BA71          ........................................a....................................... [173440]
MH025920.1_strain_R35           .a...t.t..t........g...gaa....c.........a...............a............c.......... [173440]
MH025919.1_strain_N10           .a...t.t..t........g...gaa....c.........a...............a............c.......... [173440]
MH025918.1_strain_R25           .a...t.t..t........g...gaa....c.........a...............a............c.......... [173440]
MH025917.1_strain_R7            .a...t.t..t........g...gaa....c.........a...............a............c.......... [173440]
MH025916.1_strain_R8            .a...t.t..t........g...gaa....c.........a...............a............c.......... [173440]
```

```
FR682468.2_Georgia_2007/1        gcctggggatttttaacattttttttaagctgttgttccggcctggacatgttttattagctttatatataaagggttag [173520]
ASF_IND_20_CAD543                ........gg...................................................................... [173520]
LR899193.1_Germany_2020/1        ................................................................................ [173520]
MN393477.1_Wuhan_2019            ................................................................................ [173520]
MW396979.1_Timor-Leste/2019/1    ................................................................................ [173520]
MK940252.1_CN/2019/InnerMongolia ................................................................................ [173520]
MK128995.1_China/2018/AnhuiXCGQ  ................................................................................ [173520]
MH681419.1_ASFV/POL/2015         ................................................................................ [173520]
LS478113.1_Estonia_2014          ................................................................................ [173520]
KP843857.1_Odintsovo_02/14       ................................................................................ [173520]
FN557520.1_strain_E75            ........gg...................................................................... [173520]
AY261366.1_Warthog               .....a..g....................................................................... [173520]
AY261365.1_Warmbaths             .....a..g....................................................................... [173520]
AY261364.1_Tengani_62            ........g....................................................................... [173520]
AY261363.1_Pretoriuskop/96/4     .....a..g....................................................................... [173520]
AY261362.1_Mkuzi_1979            ........g....................................................................... [173520]
AY261361.1_Malawi_Lil-20/1_1983  .....a..g..........c........a..............................a...............a.... [173520]
AY261360.1_Kenya_1950            .....a..g..........c........a..............................at..............a.... [173520]
AM712240.1_OURT_88/3             ........gg...................................................................... [173520]
AM712239.1_Benin_97/1            ....-...gg...................................................................... [173520]
KM111295.1_Ken06.Bus             .....a..g..........c........a..............................a...............a.... [173520]
KM111294.1_Ken05/Tk1             .....a..g..........c........a..............................a...............a.... [173520]
KM262844.1_strain_L60            ........gg...................................................................... [173520]
KM102979.1_26544/OG10_Italy      ........gg...................................................................... [173520]
KM262845.1_strain_NHV            ........gg...................................................................... [173520]
KX354450.1_47/Ss/2008            ....-...gg...................................................................... [173520]
KP055815.1_strain_BA71           ........gg...................................................................... [173520]
MH025920.1_strain_R35            .....a..g..........c........a..............................a...............a.... [173520]
MH025919.1_strain_N10            .....a..g..........c........a..............................a...............a.... [173520]
MH025918.1_strain_R25            .....a..g..........c........a..............................a...............a.... [173520]
MH025917.1_strain_R7             .....a..g..........c........a..............................a...............a.... [173520]
MH025916.1_strain_R8             .....a..g..........c........a..............................a...............a.... [173520]

FR682468.2_Georgia_2007/1        aaggtttaatttcaatatatgccttaatgatgggattatattcgtaaaaggtatagcctaatcctacgtctttgtttttt [173600]
ASF_IND_20_CAD543                ................................................................................ [173600]
LR899193.1_Germany_2020/1        ................................................................................ [173600]
MN393477.1_Wuhan_2019            ................................................................................ [173600]
MW396979.1_Timor-Leste/2019/1    ................................................................................ [173600]
MK940252.1_CN/2019/InnerMongolia ................................................................................ [173600]
MK128995.1_China/2018/AnhuiXCGQ  ................................................................................ [173600]
MH681419.1_ASFV/POL/2015         ................................................................................ [173600]
LS478113.1_Estonia_2014          ................................................................................ [173600]
KP843857.1_Odintsovo_02/14       ................................................................................ [173600]
```

FN557520.1_strain_E75 .......................................................................................... [173600]
AY261366.1_Warthog ...................................................................a............ [173600]
AY261365.1_Warmbaths ...................................................................t............ [173600]
AY261364.1_Tengani_62 ............................................................g.......c........... [173600]
AY261363.1_Pretoriuskop/96/4 ...................................................................a............ [173600]
AY261362.1_Mkuzi_1979 ...................................................................t............ [173600]
AY261361.1_Malawi_Lil-20/1_1983 .....................................g.....a.......................c............ [173600]
AY261360.1_Kenya_1950 .....................................g.....a.......................cc........... [173600]
AM712240.1_OURT_88/3 ................................................................................ [173600]
AM712239.1_Benin_97/1 ................................................................................ [173600]
KM111295.1_Ken06.Bus .....................................g..........................c..cc.c......... [173600]
KM111294.1_Ken05/Tk1 .....................................g.....a.......................cc........... [173600]
KM262844.1_strain_L60 ................................................................................ [173600]
KM102979.1_26544/OG10_Italy ................................................................................ [173600]
KM262845.1_strain_NHV ................................................................................ [173600]
KX354450.1_47/Ss/2008 ................................................................................ [173600]
KP055815.1_strain_BA71 ................................................................................ [173600]
MH025920.1_strain_R35 .....................................g..........................c..cc.c......... [173600]
MH025919.1_strain_N10 .....................................g..........................c..cc.c......... [173600]
MH025918.1_strain_R25 .....................................g..........................c..cc.c......... [173600]
MH025917.1_strain_R7 .....................................g..........................c..cc.c......... [173600]
MH025916.1_strain_R8 .....................................g..........................c..cc.c......... [173600]

FR682468.2_Georgia_2007/1 tggtaaaaaaactgtttgccctcgtaggatatgctataggcttttacttcggcttttacaagcggttggcagggattggg [173680]
ASF_IND_20_CAD543 ................................................................................ [173680]
LR899193.1_Germany_2020/1 ................................................................................ [173680]
MN393477.1_Wuhan_2019 ................................................................................ [173680]
MW396979.1_Timor-Leste/2019/1 ................................................................................ [173680]
MK940252.1_CN/2019/InnerMongolia ................................................................................ [173680]
MK128995.1_China/2018/AnhuiXCGQ ................................................................................ [173680]
MH681419.1_ASFV/POL/2015 ................................................................................ [173680]
LS478113.1_Estonia_2014 ................................................................................ [173680]
KP843857.1_Odintsovo_02/14 ................................................................................ [173680]
FN557520.1_strain_E75 ................................................................................ [173680]
AY261366.1_Warthog ................................................................................ [173680]
AY261365.1_Warmbaths ................................................................................ [173680]
AY261364.1_Tengani_62 ................................................................................ [173680]
AY261363.1_Pretoriuskop/96/4 ................................................................................ [173680]
AY261362.1_Mkuzi_1979 ................................................................................ [173680]
AY261361.1_Malawi_Lil-20/1_1983 ....................................................t........................... [173680]
AY261360.1_Kenya_1950 ....................................................t........................... [173680]
AM712240.1_OURT_88/3 ................................................................................ [173680]
AM712239.1_Benin_97/1 ................................................................................ [173680]
KM111295.1_Ken06.Bus ....................................................t........................a.. [173680]

```
KM111294.1_Ken05/Tk1              ....................................................t........................... [173680]
KM262844.1_strain_L60             ................................................................................ [173680]
KM102979.1_26544/OG10_Italy       ................................................................................ [173680]
KM262845.1_strain_NHV             ................................................................................ [173680]
KX354450.1_47/Ss/2008             ................................................................................ [173680]
KP055815.1_strain_BA71            ................................................................................ [173680]
MH025920.1_strain_R35             ....................................................t........................a.. [173680]
MH025919.1_strain_N10             ....................................................t........................a.. [173680]
MH025918.1_strain_R25             ....................................................t........................a.. [173680]
MH025917.1_strain_R7              ....................................................t........................a.. [173680]
MH025916.1_strain_R8              ....................................................t........................a.. [173680]

FR682468.2_Georgia_2007/1         caaacgtaaatcgcgttcaaagttttcatgaaaaagcaaagcatttgtgggctgacacatcagacagccgctttcgccat [173760]
ASF_IND_20_CAD543                 ................................................................................ [173760]
LR899193.1_Germany_2020/1         ................................................................................ [173760]
MN393477.1_Wuhan_2019             ................................................................................ [173760]
MW396979.1_Timor-Leste/2019/1     ................................................................................ [173760]
MK940252.1_CN/2019/InnerMongolia  ................................................................................ [173760]
MK128995.1_China/2018/AnhuiXCGQ   ................................................................................ [173760]
MH681419.1_ASFV/POL/2015          ................................................................................ [173760]
LS478113.1_Estonia_2014           ................................................................................ [173760]
KP843857.1_Odintsovo_02/14        ................................................................................ [173760]
FN557520.1_strain_E75             ..............................................................................g. [173760]
AY261366.1_Warthog                ..............................................................................g. [173760]
AY261365.1_Warmbaths              ..............................................................................g. [173760]
AY261364.1_Tengani_62             ..............................................................................g. [173760]
AY261363.1_Pretoriuskop/96/4      ..............................................................................g. [173760]
AY261362.1_Mkuzi_1979             ..............................................................................g. [173760]
AY261361.1_Malawi_Lil-20/1_1983   ....................................g.g........................g.....a........g. [173760]
AY261360.1_Kenya_1950             .........g............................g........................g..............g. [173760]
AM712240.1_OURT_88/3              ..............................................................................g. [173760]
AM712239.1_Benin_97/1             ..............................................................................g. [173760]
KM111295.1_Ken06.Bus              ....................................g.g........................g..............g. [173760]
KM111294.1_Ken05/Tk1              .........g..........................g.g........................g..............g. [173760]
KM262844.1_strain_L60             ..............................................................................g. [173760]
KM102979.1_26544/OG10_Italy       ..............................................................................g. [173760]
KM262845.1_strain_NHV             ..............................................................................g. [173760]
KX354450.1_47/Ss/2008             ..............................................................................g. [173760]
KP055815.1_strain_BA71            ..............................................................................g. [173760]
MH025920.1_strain_R35             ....................................g.g........................g..............g. [173760]
MH025919.1_strain_N10             ....................................g.g........................g..............g. [173760]
MH025918.1_strain_R25             ....................................g.g........................g..............g. [173760]
MH025917.1_strain_R7              ....................................g.g........................g..............g. [173760]
MH025916.1_strain_R8              ....................................g.g........................g..............g. [173760]
```

```
FR682468.2_Georgia_2007/1        tgaaggcacattcaatggccgcccttttttagtaaatcgcggaaagcagaattaagatggctcttttcaagccccctttcg [173840]
ASF_IND_20_CAD543                ................................................................................ [173840]
LR899193.1_Germany_2020/1        ................................................................................ [173840]
MN393477.1_Wuhan_2019            ................................................................................ [173840]
MW396979.1_Timor-Leste/2019/1    ................................................................................ [173840]
MK940252.1_CN/2019/InnerMongolia ................................................................................ [173840]
MK128995.1_China/2018/AnhuiXCGQ  ................................................................................ [173840]
MH681419.1_ASFV/POL/2015         ................................................................................ [173840]
LS478113.1_Estonia_2014          ................................................................................ [173840]
KP843857.1_Odintsovo_02/14       ................................................................................ [173840]
FN557520.1_strain_E75            ................................................................................ [173840]
AY261366.1_Warthog               ................................................................................ [173840]
AY261365.1_Warmbaths             ...............................................................................a [173840]
AY261364.1_Tengani_62            ................................................................................ [173840]
AY261363.1_Pretoriuskop/96/4     ................................................................................ [173840]
AY261362.1_Mkuzi_1979            ...............................................................................a [173840]
AY261361.1_Malawi_Lil-20/1_1983  ...................t.................a.......................t.................. [173840]
AY261360.1_Kenya_1950            .......g...........t.........................................t.................. [173840]
AM712240.1_OURT_88/3             ................................................................................ [173840]
AM712239.1_Benin_97/1            ................................................................................ [173840]
KM111295.1_Ken06.Bus             .......g...........t.........................................t.................. [173840]
KM111294.1_Ken05/Tk1             .......g...........t.........................................t.................. [173840]
KM262844.1_strain_L60            ................................................................................ [173840]
KM102979.1_26544/OG10_Italy      ................................................................................ [173840]
KM262845.1_strain_NHV            ................................................................................ [173840]
KX354450.1_47/Ss/2008            ................................................................................ [173840]
KP055815.1_strain_BA71           ................................................................................ [173840]
MH025920.1_strain_R35            .......g...........t.........................................t.................. [173840]
MH025919.1_strain_N10            .......g...........t.........................................t.................. [173840]
MH025918.1_strain_R25            .......g...........t.........................................t.................. [173840]
MH025917.1_strain_R7             .......g...........t.........................................t.................. [173840]
MH025916.1_strain_R8             .......g...........t.........................................t.................. [173840]

FR682468.2_Georgia_2007/1        tgaaaacgctcatcaatcgttttttgttcctgactgccttcgggaatactataaaacattttttgattagccaccgcgat [173920]
ASF_IND_20_CAD543                ...........................................................................r.... [173920]
LR899193.1_Germany_2020/1        ................................................................................ [173920]
MN393477.1_Wuhan_2019            ................................................................................ [173920]
MW396979.1_Timor-Leste/2019/1    ................................................................................ [173920]
MK940252.1_CN/2019/InnerMongolia ................................................................................ [173920]
MK128995.1_China/2018/AnhuiXCGQ  ................................................................................ [173920]
MH681419.1_ASFV/POL/2015         ................................................................................ [173920]
LS478113.1_Estonia_2014          ................................................................................ [173920]
KP843857.1_Odintsovo_02/14       ................................................................................ [173920]
```

```
FN557520.1_strain_E75           ................................................................................ [173920]
AY261366.1_Warthog              ................................................................................ [173920]
AY261365.1_Warmbaths            ...................................a............................................ [173920]
AY261364.1_Tengani_62           ....................................................................c........... [173920]
AY261363.1_Pretoriuskop/96/4    .............................................................................a.. [173920]
AY261362.1_Mkuzi_1979           ...................................a............................................ [173920]
AY261361.1_Malawi_Lil-20/1_1983 ................................................................................ [173920]
AY261360.1_Kenya_1950           ..............g................................................................. [173920]
AM712240.1_OURT_88/3            ................................................................................ [173920]
AM712239.1_Benin_97/1           ................................................................................ [173920]
KM111295.1_Ken06.Bus            ..............g................................................................. [173920]
KM111294.1_Ken05/Tk1            ..............g.....................................................c........... [173920]
KM262844.1_strain_L60           ................................................................................ [173920]
KM102979.1_26544/OG10_Italy     ................................................................................ [173920]
KM262845.1_strain_NHV           ................................................................................ [173920]
KX354450.1_47/Ss/2008           ................................................................................ [173920]
KP055815.1_strain_BA71          ................................................................................ [173920]
MH025920.1_strain_R35           ..............g................................................................. [173920]
MH025919.1_strain_N10           ..............g................................................................. [173920]
MH025918.1_strain_R25           ..............g................................................................. [173920]
MH025917.1_strain_R7            ..............g................................................................. [173920]
MH025916.1_strain_R8            ..............g................................................................. [173920]

FR682468.2_Georgia_2007/1       gtacaaaaaaggctgtacggttttctcctcgggcggtagcgcatcgtggctaccaatgcgtataatgcgcgccttcactt [174000]
ASF_IND_20_CAD543               ................................................................................ [174000]
LR899193.1_Germany_2020/1       ................................................................................ [174000]
MN393477.1_Wuhan_2019           ................................................................................ [174000]
MW396979.1_Timor-Leste/2019/1   ................................................................................ [174000]
MK940252.1_CN/2019/InnerMongolia ................................................................................ [174000]
MK128995.1_China/2018/AnhuiXCGQ ................................................................................ [174000]
MH681419.1_ASFV/POL/2015        ................................................................................ [174000]
LS478113.1_Estonia_2014         ................................................................................ [174000]
KP843857.1_Odintsovo_02/14      ................................................................................ [174000]
FN557520.1_strain_E75           ...............c................................................................ [174000]
AY261366.1_Warthog              ................................................................................ [174000]
AY261365.1_Warmbaths            ................................................................................ [174000]
AY261364.1_Tengani_62           ...................................................g............................ [174000]
AY261363.1_Pretoriuskop/96/4    .............................................a.................................. [174000]
AY261362.1_Mkuzi_1979           ................................................................................ [174000]
AY261361.1_Malawi_Lil-20/1_1983 ..............................a................................................. [174000]
AY261360.1_Kenya_1950           ..............................a................................................. [174000]
AM712240.1_OURT_88/3            ...............c................................................................ [174000]
AM712239.1_Benin_97/1           ...............c................................................................ [174000]
KM111295.1_Ken06.Bus            ..............................a.a........t...................................... [174000]
```

```
KM111294.1_Ken05/Tk1            ..............................a................................................. [174000]
KM262844.1_strain_L60           ...............c................................................................ [174000]
KM102979.1_26544/OG10_Italy     ...............c................................................................ [174000]
KM262845.1_strain_NHV           ...............c................................................................ [174000]
KX354450.1_47/Ss/2008           ...............c................................................................ [174000]
KP055815.1_strain_BA71          ...............c................................................................ [174000]
MH025920.1_strain_R35           ..............................a.a........t...................................... [174000]
MH025919.1_strain_N10           ..............................a.a........t...................................... [174000]
MH025918.1_strain_R25           ..............................a.a........t...................................... [174000]
MH025917.1_strain_R7            ..............................a.a........t...................................... [174000]
MH025916.1_strain_R8            ..............................a.a........t...................................... [174000]

FR682468.2_Georgia_2007/1       gatcctctcgggccttatcccagtacggctctaggatatgaacctgccgcccgtatttgagatccaatccctcagctcct [174080]
ASF_IND_20_CAD543               ................................................................................ [174080]
LR899193.1_Germany_2020/1       ................................................................................ [174080]
MN393477.1_Wuhan_2019           ................................................................................ [174080]
MW396979.1_Timor-Leste/2019/1   ................................................................................ [174080]
MK940252.1_CN/2019/InnerMongolia ................................................................................ [174080]
MK128995.1_China/2018/AnhuiXCGQ ................................................................................ [174080]
MH681419.1_ASFV/POL/2015        ................................................................................ [174080]
LS478113.1_Estonia_2014         ................................................................................ [174080]
KP843857.1_Odintsovo_02/14      ................................................................................ [174080]
FN557520.1_strain_E75           ...............................................................................c [174080]
AY261366.1_Warthog              ...............................c............................................c..c [174080]
AY261365.1_Warmbaths            ...............................c............................................c..c [174080]
AY261364.1_Tengani_62           .............t.................................................................c [174080]
AY261363.1_Pretoriuskop/96/4    ...............................c...........t...................................c [174080]
AY261362.1_Mkuzi_1979           ............................................................................c..c [174080]
AY261361.1_Malawi_Lil-20/1_1983 ............................................................................c..c [174080]
AY261360.1_Kenya_1950           ...............................................................................c [174080]
AM712240.1_OURT_88/3            ...............................................................................c [174080]
AM712239.1_Benin_97/1           ...............................................................................c [174080]
KM111295.1_Ken06.Bus            ..............................................g...........a....................c [174080]
KM111294.1_Ken05/Tk1            ...............................................................................c [174080]
KM262844.1_strain_L60           ...............................................................................c [174080]
KM102979.1_26544/OG10_Italy     ...............................................................................c [174080]
KM262845.1_strain_NHV           ...............................................................................c [174080]
KX354450.1_47/Ss/2008           ...............................................................................c [174080]
KP055815.1_strain_BA71          ...............................................................................c [174080]
MH025920.1_strain_R35           ..............................................g...........a....................c [174080]
MH025919.1_strain_N10           ..............................................g...........a....................c [174080]
MH025918.1_strain_R25           ..............................................g...........a....................c [174080]
MH025917.1_strain_R7            ..............................................g...........a....................c [174080]
MH025916.1_strain_R8            ..............................................g...........a....................c [174080]
```

```
FR682468.2_Georgia_2007/1        gttttagagacgagtaaaattttaataacctctccgtgtatattcagcggcgaattccaaagctgctggatcatgtcgcg [174160]
ASF_IND_20_CAD543                ................................................................................ [174160]
LR899193.1_Germany_2020/1        ................................................................................ [174160]
MN393477.1_Wuhan_2019            ................................................................................ [174160]
MW396979.1_Timor-Leste/2019/1    ................................................................................ [174160]
MK940252.1_CN/2019/InnerMongolia ................................................................................ [174160]
MK128995.1_China/2018/AnhuiXCGQ  ................................................................................ [174160]
MH681419.1_ASFV/POL/2015         ................................................................................ [174160]
LS478113.1_Estonia_2014          ................................................................................ [174160]
KP843857.1_Odintsovo_02/14       ................................................................................ [174160]
FN557520.1_strain_E75            .............................................................c.................. [174160]
AY261366.1_Warthog               ................................................................................ [174160]
AY261365.1_Warmbaths             ................................................................................ [174160]
AY261364.1_Tengani_62            ........a..........c............................................................ [174160]
AY261363.1_Pretoriuskop/96/4     ................................................................................ [174160]
AY261362.1_Mkuzi_1979            ..........................................................................a..... [174160]
AY261361.1_Malawi_Lil-20/1_1983  ..........................c......................................t.............. [174160]
AY261360.1_Kenya_1950            .............................................................................a.. [174160]
AM712240.1_OURT_88/3             .............................................................c.................. [174160]
AM712239.1_Benin_97/1            .............................................................c.................. [174160]
KM111295.1_Ken06.Bus             ..................................................g..........................a.. [174160]
KM111294.1_Ken05/Tk1             ..................................................g..........................a.. [174160]
KM262844.1_strain_L60            .............................................................c.................. [174160]
KM102979.1_26544/OG10_Italy      .............................................................c.................. [174160]
KM262845.1_strain_NHV            .............................................................c.................. [174160]
KX354450.1_47/Ss/2008            .............................................................c.................. [174160]
KP055815.1_strain_BA71           .............................................................c.................. [174160]
MH025920.1_strain_R35            ..................................................g..........................a.. [174160]
MH025919.1_strain_N10            ..................................................g..........................a.. [174160]
MH025918.1_strain_R25            ..................................................g..........................a.. [174160]
MH025917.1_strain_R7             ..................................................g..........................a.. [174160]
MH025916.1_strain_R8             ..................................................g..........................a.. [174160]

FR682468.2_Georgia_2007/1        ctctttagataaaattttccctgtaataagcgtaaatcgtgttattttggaggacaggactaacgtatgggtcggcccat [174240]
ASF_IND_20_CAD543                ................................................................................ [174240]
LR899193.1_Germany_2020/1        ................................................................................ [174240]
MN393477.1_Wuhan_2019            ................................................................................ [174240]
MW396979.1_Timor-Leste/2019/1    ................................................................................ [174240]
MK940252.1_CN/2019/InnerMongolia ................................................................................ [174240]
MK128995.1_China/2018/AnhuiXCGQ  ................................................................................ [174240]
MH681419.1_ASFV/POL/2015         ................................................................................ [174240]
LS478113.1_Estonia_2014          ................................................................................ [174240]
KP843857.1_Odintsovo_02/14       ................................................................................ [174240]
```

| | | |
|---|---|---|
| FN557520.1_strain_E75 | ................................................................................ | [174240] |
| AY261366.1_Warthog | ......g............................................................c............ | [174240] |
| AY261365.1_Warmbaths | ......g............................................................c.......t.... | [174240] |
| AY261364.1_Tengani_62 | ..............................g....................................c............ | [174240] |
| AY261363.1_Pretoriuskop/96/4 | ......g......................................................................... | [174240] |
| AY261362.1_Mkuzi_1979 | ......g...........t............................................................. | [174240] |
| AY261361.1_Malawi_Lil-20/1_1983 | ..............................a..g..........................................t... | [174240] |
| AY261360.1_Kenya_1950 | ...c..............t...........a.............................................t... | [174240] |
| AM712240.1_OURT_88/3 | ................................................................................ | [174240] |
| AM712239.1_Benin_97/1 | ................................................................................ | [174240] |
| KM111295.1_Ken06.Bus | ..................t...........a.............................................t... | [174240] |
| KM111294.1_Ken05/Tk1 | ...c..............t...........a.............................................t... | [174240] |
| KM262844.1_strain_L60 | ................................................................................ | [174240] |
| KM102979.1_26544/OG10_Italy | ................................................................................ | [174240] |
| KM262845.1_strain_NHV | ................................................................................ | [174240] |
| KX354450.1_47/Ss/2008 | ................................................................................ | [174240] |
| KP055815.1_strain_BA71 | ................................................................................ | [174240] |
| MH025920.1_strain_R35 | ..................t...........a.......................t.....................t... | [174240] |
| MH025919.1_strain_N10 | ..................t...........a.......................t.....................t... | [174240] |
| MH025918.1_strain_R25 | ..................t...........a.......................t.....................t... | [174240] |
| MH025917.1_strain_R7 | ..................t...........a.......................t.....................t... | [174240] |
| MH025916.1_strain_R8 | ..................t...........a.......................t.....................t... | [174240] |

| | | |
|---|---|---|
| FR682468.2_Georgia_2007/1 | cttccgcaaagtttttcaccataagatctttcccatccttatgaaggaggatggtgttgtgcccttcttccaatactttt | [174320] |
| ASF_IND_20_CAD543 | ................................................................................ | [174320] |
| LR899193.1_Germany_2020/1 | ................................................................................ | [174320] |
| MN393477.1_Wuhan_2019 | ................................................................................ | [174320] |
| MW396979.1_Timor-Leste/2019/1 | ................................................................................ | [174320] |
| MK940252.1_CN/2019/InnerMongolia | ................................................................................ | [174320] |
| MK128995.1_China/2018/AnhuiXCGQ | ................................................................................ | [174320] |
| MH681419.1_ASFV/POL/2015 | ................................................................................ | [174320] |
| LS478113.1_Estonia_2014 | ................................................................................ | [174320] |
| KP843857.1_Odintsovo_02/14 | ................................................................................ | [174320] |
| FN557520.1_strain_E75 | ....g........................................................................... | [174320] |
| AY261366.1_Warthog | ............................................................................c... | [174320] |
| AY261365.1_Warmbaths | ..............................................a.............................c... | [174320] |
| AY261364.1_Tengani_62 | ................................................................................ | [174320] |
| AY261363.1_Pretoriuskop/96/4 | ................................................................................ | [174320] |
| AY261362.1_Mkuzi_1979 | ....g........................................................................... | [174320] |
| AY261361.1_Malawi_Lil-20/1_1983 | ...............................t..g...................................t.g...c... | [174320] |
| AY261360.1_Kenya_1950 | ..............................................a.......c...............t.g...c... | [174320] |
| AM712240.1_OURT_88/3 | ....g........................................................................... | [174320] |
| AM712239.1_Benin_97/1 | ....g........................................................................... | [174320] |
| KM111295.1_Ken06.Bus | ......................................................c...............t.g...c... | [174320] |

```
KM111294.1_Ken05/Tk1             ......................................................c...............t.g...c... [174320]
KM262844.1_strain_L60            ....g........................................................................... [174320]
KM102979.1_26544/OG10_Italy      ....g........................................................................... [174320]
KM262845.1_strain_NHV            ....g........................................................................... [174320]
KX354450.1_47/Ss/2008            ....g........................................................................... [174320]
KP055815.1_strain_BA71           ....g........................................................................... [174320]
MH025920.1_strain_R35            ......................................................c...............t.g...c... [174320]
MH025919.1_strain_N10            ......................................................c...............t.g...c... [174320]
MH025918.1_strain_R25            ......................................................c...............t.g...c... [174320]
MH025917.1_strain_R7             ......................................................c...............t.g...c... [174320]
MH025916.1_strain_R8             ......................................................c...............t.g...c... [174320]

FR682468.2_Georgia_2007/1        aggggctgaaggcactggtagccctctatttctaaaaagcgggccacgacgtgaaggcccaattccacaaactgtgagta [174400]
ASF_IND_20_CAD543                ................................................................................ [174400]
LR899193.1_Germany_2020/1        ................................................................................ [174400]
MN393477.1_Wuhan_2019            ................................................................................ [174400]
MW396979.1_Timor-Leste/2019/1    ................................................................................ [174400]
MK940252.1_CN/2019/InnerMongolia ................................................................................ [174400]
MK128995.1_China/2018/AnhuiXCGQ  ................................................................................ [174400]
MH681419.1_ASFV/POL/2015         ................................................................................ [174400]
LS478113.1_Estonia_2014          ................................................................................ [174400]
KP843857.1_Odintsovo_02/14       ................................................................................ [174400]
FN557520.1_strain_E75            ................................................................................ [174400]
AY261366.1_Warthog               ................................................................................ [174400]
AY261365.1_Warmbaths             ................................................................................ [174400]
AY261364.1_Tengani_62            ................................................................................ [174400]
AY261363.1_Pretoriuskop/96/4     ................................................................................ [174400]
AY261362.1_Mkuzi_1979            ...................g............................................................ [174400]
AY261361.1_Malawi_Lil-20/1_1983  .................a.............................t..........................a..... [174400]
AY261360.1_Kenya_1950            .................a.......................a.....t.............g............g..... [174400]
AM712240.1_OURT_88/3             ................................................................................ [174400]
AM712239.1_Benin_97/1            ................................................................................ [174400]
KM111295.1_Ken06.Bus             .................a.............................t.............g............g..... [174400]
KM111294.1_Ken05/Tk1             .................a.......................a.....t.............g............g..... [174400]
KM262844.1_strain_L60            ................................................................................ [174400]
KM102979.1_26544/OG10_Italy      ................................................................................ [174400]
KM262845.1_strain_NHV            ................................................................................ [174400]
KX354450.1_47/Ss/2008            ................................................................................ [174400]
KP055815.1_strain_BA71           ................................................................................ [174400]
MH025920.1_strain_R35            .................a.............................t.............g............g..... [174400]
MH025919.1_strain_N10            .................a.............................t.............g............g..... [174400]
MH025918.1_strain_R25            .................a.............................t.............g............g..... [174400]
MH025917.1_strain_R7             .................a.............................t.............g............g..... [174400]
MH025916.1_strain_R8             .................a.............................t.............g............g..... [174400]
```

```
FR682468.2_Georgia_2007/1        aatgagcacagggcccggagacgttttaatattttttagcatgcgtactattttgggactagaattttctgtgaaggcct [174480]
ASF_IND_20_CAD543                ................................................................................ [174480]
LR899193.1_Germany_2020/1        ................................................................................ [174480]
MN393477.1_Wuhan_2019            ................................................................................ [174480]
MW396979.1_Timor-Leste/2019/1    ................................................................................ [174480]
MK940252.1_CN/2019/InnerMongolia ................................................................................ [174480]
MK128995.1_China/2018/AnhuiXCGQ  ................................................................................ [174480]
MH681419.1_ASFV/POL/2015         ................................................................................ [174480]
LS478113.1_Estonia_2014          ................................................................................ [174480]
KP843857.1_Odintsovo_02/14       ................................................................................ [174480]
FN557520.1_strain_E75            .............................................c.................................. [174480]
AY261366.1_Warthog               ............................................aa.................................. [174480]
AY261365.1_Warmbaths             ......................................a.....aa.................................. [174480]
AY261364.1_Tengani_62            ............................................aa.................................. [174480]
AY261363.1_Pretoriuskop/96/4     ............................................aa.................................. [174480]
AY261362.1_Mkuzi_1979            ............a................................................................... [174480]
AY261361.1_Malawi_Lil-20/1_1983  ............a........t......................a.g......................a.......... [174480]
AY261360.1_Kenya_1950            ............................................a..................gg............... [174480]
AM712240.1_OURT_88/3             .............................................c.................................. [174480]
AM712239.1_Benin_97/1            .............................................c.................................. [174480]
KM111295.1_Ken06.Bus             ......t.....a...............................a...................g............... [174480]
KM111294.1_Ken05/Tk1             ............................................a..................gg............... [174480]
KM262844.1_strain_L60            .............................................c.................................. [174480]
KM102979.1_26544/OG10_Italy      .............................................c.................................. [174480]
KM262845.1_strain_NHV            .............................................c.................................. [174480]
KX354450.1_47/Ss/2008            .............................................c.................................. [174480]
KP055815.1_strain_BA71           .............................................c.................................. [174480]
MH025920.1_strain_R35            ......t.....a...............................a...................g............... [174480]
MH025919.1_strain_N10            ......t.....a...............................a...................g............... [174480]
MH025918.1_strain_R25            ......t.....a...............................a...................g............... [174480]
MH025917.1_strain_R7             ......t.....a...............................a...................g............... [174480]
MH025916.1_strain_R8             ......t.....a...............................a...................g............... [174480]

FR682468.2_Georgia_2007/1        ctttgggcagctgctgaacagcctctgataatttttcatcctcctttactgttagcatttcggacgcgaagatgctgatc [174560]
ASF_IND_20_CAD543                ................................................................................ [174560]
LR899193.1_Germany_2020/1        ................................................................................ [174560]
MN393477.1_Wuhan_2019            ................................................................................ [174560]
MW396979.1_Timor-Leste/2019/1    ................................................................................ [174560]
MK940252.1_CN/2019/InnerMongolia ................................................................................ [174560]
MK128995.1_China/2018/AnhuiXCGQ  ................................................................................ [174560]
MH681419.1_ASFV/POL/2015         ................................................................................ [174560]
LS478113.1_Estonia_2014          ................................................................................ [174560]
KP843857.1_Odintsovo_02/14       ................................................................................ [174560]
```

```
FN557520.1_strain_E75            ................................................................................ [174560]
AY261366.1_Warthog               .................................................g.............................. [174560]
AY261365.1_Warmbaths             ................................................................................ [174560]
AY261364.1_Tengani_62            ............................c................................................... [174560]
AY261363.1_Pretoriuskop/96/4     ................................................................................ [174560]
AY261362.1_Mkuzi_1979            ................................................................................ [174560]
AY261361.1_Malawi_Lil-20/1_1983  ..............................g......g.......................................... [174560]
AY261360.1_Kenya_1950            ...........................................t....tg.............................. [174560]
AM712240.1_OURT_88/3             ................................................................................ [174560]
AM712239.1_Benin_97/1            ................................................................................ [174560]
KM111295.1_Ken06.Bus             ...........................................t....tg.............................. [174560]
KM111294.1_Ken05/Tk1             .....................t.....................t....tg.............................. [174560]
KM262844.1_strain_L60            ................................................................................ [174560]
KM102979.1_26544/OG10_Italy      ................................................................................ [174560]
KM262845.1_strain_NHV            ................................................................................ [174560]
KX354450.1_47/Ss/2008            ................................................................................ [174560]
KP055815.1_strain_BA71           ................................................................................ [174560]
MH025920.1_strain_R35            ...........................................t....tg.............................. [174560]
MH025919.1_strain_N10            ...........................................t....tg.............................. [174560]
MH025918.1_strain_R25            ...........................................t....tg.............................. [174560]
MH025917.1_strain_R7             ...........................................t....tg.............................. [174560]
MH025916.1_strain_R8             ...........................................t....tg.............................. [174560]

FR682468.2_Georgia_2007/1        atacgggaacgcacatagtaggaggagcctgactcttgctccgatcctggcaggcagagggcggcggcatttattttttc [174640]
ASF_IND_20_CAD543                ................................................................................ [174640]
LR899193.1_Germany_2020/1        ................................................................................ [174640]
MN393477.1_Wuhan_2019            ................................................................................ [174640]
MW396979.1_Timor-Leste/2019/1    ................................................................................ [174640]
MK940252.1_CN/2019/InnerMongolia ................................................................................ [174640]
MK128995.1_China/2018/AnhuiXCGQ  ................................................................................ [174640]
MH681419.1_ASFV/POL/2015         ................................................................................ [174640]
LS478113.1_Estonia_2014          ................................................................................ [174640]
KP843857.1_Odintsovo_02/14       ................................................................................ [174640]
FN557520.1_strain_E75            .......................................................................a........ [174640]
AY261366.1_Warthog               .....................................................a.......................... [174640]
AY261365.1_Warmbaths             ................................................................................ [174640]
AY261364.1_Tengani_62            ................................................................................ [174640]
AY261363.1_Pretoriuskop/96/4     ................................................................................ [174640]
AY261362.1_Mkuzi_1979            ................................................................................ [174640]
AY261361.1_Malawi_Lil-20/1_1983  ...................................................................t...c........ [174640]
AY261360.1_Kenya_1950            ..g...........................................................a....t............ [174640]
AM712240.1_OURT_88/3             .......................................................................a........ [174640]
AM712239.1_Benin_97/1            .......................................................................a........ [174640]
KM111295.1_Ken06.Bus             ..g...........................................................a....t............ [174640]
```

```
KM111294.1_Ken05/Tk1              ..g...........................................................a....t............ [174640]
KM262844.1_strain_L60             .......................................................................a........ [174640]
KM102979.1_26544/OG10_Italy       .......................................................................a........ [174640]
KM262845.1_strain_NHV             .......................................................................a........ [174640]
KX354450.1_47/Ss/2008             .......................................................................a........ [174640]
KP055815.1_strain_BA71            .......................................................................a........ [174640]
MH025920.1_strain_R35             ..g...........................................................a....t............ [174640]
MH025919.1_strain_N10             ..g...........................................................a....t............ [174640]
MH025918.1_strain_R25             ..g...........................................................a....t............ [174640]
MH025917.1_strain_R7              ..g...........................................................a....t............ [174640]
MH025916.1_strain_R8              ..g...........................................................a....t............ [174640]

FR682468.2_Georgia_2007/1         atacattcctgagctggcgtgcttttccgcgttttcaacgtctcgggccagcagatattgcctatactgctcgggtgaca [174720]
ASF_IND_20_CAD543                 ................................................................................ [174720]
LR899193.1_Germany_2020/1         ................................................................................ [174720]
MN393477.1_Wuhan_2019             ................................................................................ [174720]
MW396979.1_Timor-Leste/2019/1     ................................................................................ [174720]
MK940252.1_CN/2019/InnerMongolia  ................................................................................ [174720]
MK128995.1_China/2018/AnhuiXCGQ   ................................................................................ [174720]
MH681419.1_ASFV/POL/2015          ................................................................................ [174720]
LS478113.1_Estonia_2014           ................................................................................ [174720]
KP843857.1_Odintsovo_02/14        ................................................................................ [174720]
FN557520.1_strain_E75             ................................................................................ [174720]
AY261366.1_Warthog                ................................................................................ [174720]
AY261365.1_Warmbaths              ................................................................................ [174720]
AY261364.1_Tengani_62             ........................................................................a....... [174720]
AY261363.1_Pretoriuskop/96/4      ................................................................................ [174720]
AY261362.1_Mkuzi_1979             ................................................................................ [174720]
AY261361.1_Malawi_Lil-20/1_1983   ......g..c................................c..a....a............................. [174720]
AY261360.1_Kenya_1950             g.g.......................................c................................g.... [174720]
AM712240.1_OURT_88/3              ................................................................................ [174720]
AM712239.1_Benin_97/1             ................................................................................ [174720]
KM111295.1_Ken06.Bus              g.g.......................................c................................g.... [174720]
KM111294.1_Ken05/Tk1              g.g.......................................c..a....a......c...................... [174720]
KM262844.1_strain_L60             ................................................................................ [174720]
KM102979.1_26544/OG10_Italy       ................................................................................ [174720]
KM262845.1_strain_NHV             ................................................................................ [174720]
KX354450.1_47/Ss/2008             ................................................................................ [174720]
KP055815.1_strain_BA71            ................................................................................ [174720]
MH025920.1_strain_R35             g.g..................t....................c................................g.... [174720]
MH025919.1_strain_N10             g.g..................t....................c................................g.... [174720]
MH025918.1_strain_R25             g.g..................t....................c................................g.... [174720]
MH025917.1_strain_R7              g.g..................t....................c................................g.... [174720]
MH025916.1_strain_R8              g.g..................t....................c................................g.... [174720]
```

```
FR682468.2_Georgia_2007/1        tttcaaccttttctataataagaggaagctctgtggggaatagcttgttgagctcattctggtttccagcgtagcttatc [174800]
ASF_IND_20_CAD543                ................................................................................ [174800]
LR899193.1_Germany_2020/1        ................................................................................ [174800]
MN393477.1_Wuhan_2019            ................................................................................ [174800]
MW396979.1_Timor-Leste/2019/1    ................................................................................ [174800]
MK940252.1_CN/2019/InnerMongolia ................................................................................ [174800]
MK128995.1_China/2018/AnhuiXCGQ  ................................................................................ [174800]
MH681419.1_ASFV/POL/2015         ................................................................................ [174800]
LS478113.1_Estonia_2014          ................................................................................ [174800]
KP843857.1_Odintsovo_02/14       ................................................................................ [174800]
FN557520.1_strain_E75            ................................................................................ [174800]
AY261366.1_Warthog               ................................................................................ [174800]
AY261365.1_Warmbaths             ................................................................................ [174800]
AY261364.1_Tengani_62            ................................................................................ [174800]
AY261363.1_Pretoriuskop/96/4     ................................................................................ [174800]
AY261362.1_Mkuzi_1979            ................................................................................ [174800]
AY261361.1_Malawi_Lil-20/1_1983  ............................t.....a.................t.....t...........a......... [174800]
AY261360.1_Kenya_1950            ............................t.....a.................t.....t...........a......... [174800]
AM712240.1_OURT_88/3             ................................................................................ [174800]
AM712239.1_Benin_97/1            ................................................................................ [174800]
KM111295.1_Ken06.Bus             ............................t.....a........t........t.....t...........a......... [174800]
KM111294.1_Ken05/Tk1             ............................t.....a.................t.....t..a........a......... [174800]
KM262844.1_strain_L60            ................................................................................ [174800]
KM102979.1_26544/OG10_Italy      ................................................................................ [174800]
KM262845.1_strain_NHV            ................................................................................ [174800]
KX354450.1_47/Ss/2008            ................................................................................ [174800]
KP055815.1_strain_BA71           ................................................................................ [174800]
MH025920.1_strain_R35            ............................t.....a........t........t.....t...........a......... [174800]
MH025919.1_strain_N10            ............................t.....a........t........t.....t...........a......... [174800]
MH025918.1_strain_R25            ............................t.....a........t........t.....t...........a......... [174800]
MH025917.1_strain_R7             ............................t.....a........t........t.....t...........a......... [174800]
MH025916.1_strain_R8             ............................t.....a........t........t.....t...........a......... [174800]

FR682468.2_Georgia_2007/1        atacccactaggcggtttagtagtttgtccgcgtttaaagggctattcgttgttttattgacataagcggtgtagaatct [174880]
ASF_IND_20_CAD543                ................................................................................ [174880]
LR899193.1_Germany_2020/1        ................................................................................ [174880]
MN393477.1_Wuhan_2019            ................................................................................ [174880]
MW396979.1_Timor-Leste/2019/1    ................................................................................ [174880]
MK940252.1_CN/2019/InnerMongolia ................................................................................ [174880]
MK128995.1_China/2018/AnhuiXCGQ  ................................................................................ [174880]
MH681419.1_ASFV/POL/2015         ................................................................................ [174880]
LS478113.1_Estonia_2014          ................................................................................ [174880]
KP843857.1_Odintsovo_02/14       ................................................................................ [174880]
```

```
FN557520.1_strain_E75           ................................................................................ [174880]
AY261366.1_Warthog              ..c......................................a.................a.................... [174880]
AY261365.1_Warmbaths            ..c.............................a..........................a.....g.............. [174880]
AY261364.1_Tengani_62           ................................................................................ [174880]
AY261363.1_Pretoriuskop/96/4    ..c............................................................................. [174880]
AY261362.1_Mkuzi_1979           ................................................................................ [174880]
AY261361.1_Malawi_Lil-20/1_1983 ....................c..c.................a...................................... [174880]
AY261360.1_Kenya_1950           ....................c..............c.....a...................................... [174880]
AM712240.1_OURT_88/3            ................................................................................ [174880]
AM712239.1_Benin_97/1           ................................................................................ [174880]
KM111295.1_Ken06.Bus            ....................c....................a...................................... [174880]
KM111294.1_Ken05/Tk1            ....................c.ac...........c.....a...................................... [174880]
KM262844.1_strain_L60           ................................................................................ [174880]
KM102979.1_26544/OG10_Italy     ................................................................................ [174880]
KM262845.1_strain_NHV           ................................................................................ [174880]
KX354450.1_47/Ss/2008           ................................................................................ [174880]
KP055815.1_strain_BA71          ................................................................................ [174880]
MH025920.1_strain_R35           ....................c....................a...................................... [174880]
MH025919.1_strain_N10           ....................c....................a...................................... [174880]
MH025918.1_strain_R25           ....................c....................a...................................... [174880]
MH025917.1_strain_R7            ....................c....................a...................................... [174880]
MH025916.1_strain_R8            ....................c....................a...................................... [174880]

FR682468.2_Georgia_2007/1       ttcatagtgaagaggtaataagattcgcccgcttagcatattaaaacagggcaccatttcaaaggggtccttcgaacacg [174960]
ASF_IND_20_CAD543               ................................................................................ [174960]
LR899193.1_Germany_2020/1       ................................................................................ [174960]
MN393477.1_Wuhan_2019           ................................................................................ [174960]
MW396979.1_Timor-Leste/2019/1   ................................................................................ [174960]
MK940252.1_CN/2019/InnerMongolia ................................................................................ [174960]
MK128995.1_China/2018/AnhuiXCGQ ................................................................................ [174960]
MH681419.1_ASFV/POL/2015        ................................................................................ [174960]
LS478113.1_Estonia_2014         ................................................................................ [174960]
KP843857.1_Odintsovo_02/14      ................................................................................ [174960]
FN557520.1_strain_E75           ................................................................................ [174960]
AY261366.1_Warthog              ................................................................................ [174960]
AY261365.1_Warmbaths            ................................................................................ [174960]
AY261364.1_Tengani_62           ...........................g.......a............................................ [174960]
AY261363.1_Pretoriuskop/96/4    ................................................................................ [174960]
AY261362.1_Mkuzi_1979           ..............................a................................................. [174960]
AY261361.1_Malawi_Lil-20/1_1983 .....................a...........c.............................a................ [174960]
AY261360.1_Kenya_1950           ..................c..a.......................................................... [174960]
AM712240.1_OURT_88/3            ................................................................................ [174960]
AM712239.1_Benin_97/1           ................................................................................ [174960]
KM111295.1_Ken06.Bus            ..................c..a.........................................a................ [174960]
```

KM111294.1_Ken05/Tk1 ..................c..a.......................................................... [174960]
KM262844.1_strain_L60 ................................................................................ [174960]
KM102979.1_26544/OG10_Italy ................................................................................ [174960]
KM262845.1_strain_NHV ................................................................................ [174960]
KX354450.1_47/Ss/2008 ................................................................................ [174960]
KP055815.1_strain_BA71 ................................................................................ [174960]
MH025920.1_strain_R35 ..................c..a.........................................a................ [174960]
MH025919.1_strain_N10 ..................c..a.........................................a................ [174960]
MH025918.1_strain_R25 ..................c..a.........................................a................ [174960]
MH025917.1_strain_R7 ..................c..a.........................................a................ [174960]
MH025916.1_strain_R8 ..................c..a.........................................a................ [174960]

FR682468.2_Georgia_2007/1 gggtgcctgttaaaaacagaatacgaatatttttagcttgcataatattattgtacagctggcgggcatttgttttatca [175040]
ASF_IND_20_CAD543 ................................................................................ [175040]
LR899193.1_Germany_2020/1 ................................................................................ [175040]
MN393477.1_Wuhan_2019 ................................................................................ [175040]
MW396979.1_Timor-Leste/2019/1 ................................................................................ [175040]
MK940252.1_CN/2019/InnerMongolia ................................................................................ [175040]
MK128995.1_China/2018/AnhuiXCGQ ................................................................................ [175040]
MH681419.1_ASFV/POL/2015 ................................................................................ [175040]
LS478113.1_Estonia_2014 ................................................................................ [175040]
KP843857.1_Odintsovo_02/14 ................................................................................ [175040]
FN557520.1_strain_E75 ................................................................................ [175040]
AY261366.1_Warthog ................................................................................ [175040]
AY261365.1_Warmbaths ................................................................................ [175040]
AY261364.1_Tengani_62 .................................................g.............................. [175040]
AY261363.1_Pretoriuskop/96/4 ..........c...................................g................................. [175040]
AY261362.1_Mkuzi_1979 ............................g................................................... [175040]
AY261361.1_Malawi_Lil-20/1_1983 ....c..............g..............c............................................. [175040]
AY261360.1_Kenya_1950 ....c.............ag............................................................ [175040]
AM712240.1_OURT_88/3 ................................................................................ [175040]
AM712239.1_Benin_97/1 ................................................................................ [175040]
KM111295.1_Ken06.Bus ....c.............ag...........................................................g [175040]
KM111294.1_Ken05/Tk1 ....c.............ag............................................................ [175040]
KM262844.1_strain_L60 ................................................................................ [175040]
KM102979.1_26544/OG10_Italy ................................................................................ [175040]
KM262845.1_strain_NHV ................................................................................ [175040]
KX354450.1_47/Ss/2008 ................................................................................ [175040]
KP055815.1_strain_BA71 ................................................................................ [175040]
MH025920.1_strain_R35 ....c.............ag...........................................................g [175040]
MH025919.1_strain_N10 ....c.............ag...........................................................g [175040]
MH025918.1_strain_R25 ....c.............ag...........................................................g [175040]
MH025917.1_strain_R7 ....c.............ag...........................................................g [175040]
MH025916.1_strain_R8 ....c.............ag...........................................................g [175040]

```
FR682468.2_Georgia_2007/1        ttggcgctattgataattcctctaaagaggttgtgtgcctcgtcaacgatgagcaggcatccatttagggaccctcccgc [175120]
ASF_IND_20_CAD543                ................................................................................ [175120]
LR899193.1_Germany_2020/1        ................................................................................ [175120]
MN393477.1_Wuhan_2019            ................................................................................ [175120]
MW396979.1_Timor-Leste/2019/1    ................................................................................ [175120]
MK940252.1_CN/2019/InnerMongolia ................................................................................ [175120]
MK128995.1_China/2018/AnhuiXCGQ  ................................................................................ [175120]
MH681419.1_ASFV/POL/2015         ................................................................................ [175120]
LS478113.1_Estonia_2014          ................................................................................ [175120]
KP843857.1_Odintsovo_02/14       ................................................................................ [175120]
FN557520.1_strain_E75            ...........................................................a..g................. [175120]
AY261366.1_Warthog               ..............................................................g................. [175120]
AY261365.1_Warmbaths             ..............................................................g................. [175120]
AY261364.1_Tengani_62            ..............................................................g................. [175120]
AY261363.1_Pretoriuskop/96/4     ..............................................................g................. [175120]
AY261362.1_Mkuzi_1979            ...........................................................a..g................. [175120]
AY261361.1_Malawi_Lil-20/1_1983  .........................................a........a...........g................. [175120]
AY261360.1_Kenya_1950            ..............t.....c.........................................g....a............ [175120]
AM712240.1_OURT_88/3             ...........................................................a..g................. [175120]
AM712239.1_Benin_97/1            ...........................................................a..g................. [175120]
KM111295.1_Ken06.Bus             ..............t.......................t..a..t..a..............g....a............ [175120]
KM111294.1_Ken05/Tk1             ..............t.....c..............................................a............ [175120]
KM262844.1_strain_L60            ...........................................................a..g................. [175120]
KM102979.1_26544/OG10_Italy      ...........................................................a..g................. [175120]
KM262845.1_strain_NHV            ...........................................................a..g................. [175120]
KX354450.1_47/Ss/2008            ...........................................................a..g................. [175120]
KP055815.1_strain_BA71           ...........................................................a..g................. [175120]
MH025920.1_strain_R35            ..............t.......................t..a..t..a..............g....a............ [175120]
MH025919.1_strain_N10            ..............t.......................t..a..t..a..............g....a............ [175120]
MH025918.1_strain_R25            ..............t.......................t..a..t..a..............g....a............ [175120]
MH025917.1_strain_R7             ..............t.......................t..a..t..a..............g....a............ [175120]
MH025916.1_strain_R8             ..............t.......................t..a..t..a..............g....a............ [175120]

FR682468.2_Georgia_2007/1        ctttatgatctgctgccccatgttgtaagcgtctagggacacaaacctgaagcgccgcgagattttttgtagctctttgg [175200]
ASF_IND_20_CAD543                ................................................................................ [175200]
LR899193.1_Germany_2020/1        ................................................................................ [175200]
MN393477.1_Wuhan_2019            ................................................................................ [175200]
MW396979.1_Timor-Leste/2019/1    ................................................................................ [175200]
MK940252.1_CN/2019/InnerMongolia ................................................................................ [175200]
MK128995.1_China/2018/AnhuiXCGQ  ................................................................................ [175200]
MH681419.1_ASFV/POL/2015         ................................................................................ [175200]
LS478113.1_Estonia_2014          ................................................................................ [175200]
KP843857.1_Odintsovo_02/14       ................................................................................ [175200]
```

```
FN557520.1_strain_E75             ................................................................................ [175200]
AY261366.1_Warthog                ............................................................a................... [175200]
AY261365.1_Warmbaths              ................................................a............................... [175200]
AY261364.1_Tengani_62             ................................................................................ [175200]
AY261363.1_Pretoriuskop/96/4      ................................................................................ [175200]
AY261362.1_Mkuzi_1979             ................................................................................ [175200]
AY261361.1_Malawi_Lil-20/1_1983   ................................................a............................... [175200]
AY261360.1_Kenya_1950             ....................................a...........a.................c............. [175200]
AM712240.1_OURT_88/3              ................................................................................ [175200]
AM712239.1_Benin_97/1             ................................................................................ [175200]
KM111295.1_Ken06.Bus              ..............................a.................a.................c............. [175200]
KM111294.1_Ken05/Tk1              ................................................a.................c............. [175200]
KM262844.1_strain_L60             ................................................................................ [175200]
KM102979.1_26544/OG10_Italy       ................................................................................ [175200]
KM262845.1_strain_NHV             ................................................................................ [175200]
KX354450.1_47/Ss/2008             ................................................................................ [175200]
KP055815.1_strain_BA71            ................................................................................ [175200]
MH025920.1_strain_R35             ..............................a.................a.................c............. [175200]
MH025919.1_strain_N10             ..............................a.................a.................c............. [175200]
MH025918.1_strain_R25             ..............................a.................a.................c............. [175200]
MH025917.1_strain_R7              ..............................a.................a.................c............. [175200]
MH025916.1_strain_R8              ..............................a.................a.................c............. [175200]

FR682468.2_Georgia_2007/1         agtgatccgtcgtttccggatataaaagtttaataagctttaacaaagactgttggaagtttgagtgcaacgacttgggt [175280]
ASF_IND_20_CAD543                 ..........................................r..................................... [175280]
LR899193.1_Germany_2020/1         ................................................................................ [175280]
MN393477.1_Wuhan_2019             ................................................................................ [175280]
MW396979.1_Timor-Leste/2019/1     ................................................................................ [175280]
MK940252.1_CN/2019/InnerMongolia  ................................................................................ [175280]
MK128995.1_China/2018/AnhuiXCGQ   ................................................................................ [175280]
MH681419.1_ASFV/POL/2015          ................................................................................ [175280]
LS478113.1_Estonia_2014           ................................................................................ [175280]
KP843857.1_Odintsovo_02/14        ................................................................................ [175280]
FN557520.1_strain_E75             ..........................................g..................................... [175280]
AY261366.1_Warthog                ..........................................g..................................... [175280]
AY261365.1_Warmbaths              ..........................................g..................................... [175280]
AY261364.1_Tengani_62             ..........................................g..................................... [175280]
AY261363.1_Pretoriuskop/96/4      ..........................................g..................................... [175280]
AY261362.1_Mkuzi_1979             ..........................................g..................................... [175280]
AY261361.1_Malawi_Lil-20/1_1983   ..........................................g..............................t...... [175280]
AY261360.1_Kenya_1950             ..g.......a...............................g........................t.....t.....c [175280]
AM712240.1_OURT_88/3              ..........................................g..................................... [175280]
AM712239.1_Benin_97/1             ..........................................g..................................... [175280]
KM111295.1_Ken06.Bus              ..........a...............................g..............................t.....c [175280]
```

```
KM111294.1_Ken05/Tk1            ..........a...............................g........................t.....t.....c [175280]
KM262844.1_strain_L60           ..........................................g..................................... [175280]
KM102979.1_26544/OG10_Italy     ..........................................g..................................... [175280]
KM262845.1_strain_NHV           ..........................................g..................................... [175280]
KX354450.1_47/Ss/2008           ..........................................g..................................... [175280]
KP055815.1_strain_BA71          ..........................................g..................................... [175280]
MH025920.1_strain_R35           ..........a...............................g..............................t.....c [175280]
MH025919.1_strain_N10           ..........a...............................g..............................t.....c [175280]
MH025918.1_strain_R25           ..........a...............................g..............................t.....c [175280]
MH025917.1_strain_R7            ..........a...............................g..............................t.....c [175280]
MH025916.1_strain_R8            ..........a...............................g..............................t.....c [175280]

FR682468.2_Georgia_2007/1       gcgatcagaatcgggttgtaaatatgtgaaagtgagatggcaagcgacaggctcaaaatggttttccccatgcccatctg [175360]
ASF_IND_20_CAD543               ................................................................................ [175360]
LR899193.1_Germany_2020/1       ................................................................................ [175360]
MN393477.1_Wuhan_2019           ................................................................................ [175360]
MW396979.1_Timor-Leste/2019/1   ................................................................................ [175360]
MK940252.1_CN/2019/InnerMongolia ................................................................................ [175360]
MK128995.1_China/2018/AnhuiXCGQ ................................................................................ [175360]
MH681419.1_ASFV/POL/2015        ................................................................................ [175360]
LS478113.1_Estonia_2014         ................................................................................ [175360]
KP843857.1_Odintsovo_02/14      ................................................................................ [175360]
FN557520.1_strain_E75           ................................................................................ [175360]
AY261366.1_Warthog              ................................................................................ [175360]
AY261365.1_Warmbaths            ................................................................................ [175360]
AY261364.1_Tengani_62           ........................................................g....................... [175360]
AY261363.1_Pretoriuskop/96/4    ................................................................................ [175360]
AY261362.1_Mkuzi_1979           ....................g........................................................... [175360]
AY261361.1_Malawi_Lil-20/1_1983 .......a........................c..a..a........a................................ [175360]
AY261360.1_Kenya_1950           ..a..t.............................a..a........a.................g.............. [175360]
AM712240.1_OURT_88/3            ................................................................................ [175360]
AM712239.1_Benin_97/1           ................................................................................ [175360]
KM111295.1_Ken06.Bus            .....t...........a.................a..a........a.................g.............. [175360]
KM111294.1_Ken05/Tk1            ..a..t.............................a..a........a.................g.............. [175360]
KM262844.1_strain_L60           ................................................................................ [175360]
KM102979.1_26544/OG10_Italy     ................................................................................ [175360]
KM262845.1_strain_NHV           ................................................................................ [175360]
KX354450.1_47/Ss/2008           ................................................................................ [175360]
KP055815.1_strain_BA71          ................................................................................ [175360]
MH025920.1_strain_R35           .....t...........a.................a..a........a.................g.............. [175360]
MH025919.1_strain_N10           .....t...........a.................a..a........a.................g.............. [175360]
MH025918.1_strain_R25           .....t...........a.................a..a........a.................g.............. [175360]
MH025917.1_strain_R7            .....t...........a.................a..a........a.................g.............. [175360]
MH025916.1_strain_R8            .....t...........a.................a..a........a.................g.............. [175360]
```

```
FR682468.2_Georgia_2007/1        gtgatagatgaggaggccccgtgtgttttccccctggcctatcccaaatttaggatccgaaaaggcggtgtaaattaaaa [175440]
ASF_IND_20_CAD543                ................................................................................ [175440]
LR899193.1_Germany_2020/1        ................................................................................ [175440]
MN393477.1_Wuhan_2019            ................................................................................ [175440]
MW396979.1_Timor-Leste/2019/1    ................................................................................ [175440]
MK940252.1_CN/2019/InnerMongolia ................................................................................ [175440]
MK128995.1_China/2018/AnhuiXCGQ  ................................................................................ [175440]
MH681419.1_ASFV/POL/2015         ................................................................................ [175440]
LS478113.1_Estonia_2014          ................................................................................ [175440]
KP843857.1_Odintsovo_02/14       ................................................................................ [175440]
FN557520.1_strain_E75            ................................................................................ [175440]
AY261366.1_Warthog               ................................................................................ [175440]
AY261365.1_Warmbaths             ................................................................................ [175440]
AY261364.1_Tengani_62            ................................................................................ [175440]
AY261363.1_Pretoriuskop/96/4     ................................................................................ [175440]
AY261362.1_Mkuzi_1979            ................................................................................ [175440]
AY261361.1_Malawi_Lil-20/1_1983  a...................................................................a........... [175440]
AY261360.1_Kenya_1950            ................................................c..........................c.... [175440]
AM712240.1_OURT_88/3             ................................................................................ [175440]
AM712239.1_Benin_97/1            ................................................................................ [175440]
KM111295.1_Ken06.Bus             ................................................c............................... [175440]
KM111294.1_Ken05/Tk1             ................................................c..........................c.... [175440]
KM262844.1_strain_L60            ................................................................................ [175440]
KM102979.1_26544/OG10_Italy      ................................................................................ [175440]
KM262845.1_strain_NHV            ................................................................................ [175440]
KX354450.1_47/Ss/2008            ................................................................................ [175440]
KP055815.1_strain_BA71           ................................................................................ [175440]
MH025920.1_strain_R35            ................................................c............................... [175440]
MH025919.1_strain_N10            ................................................c............................... [175440]
MH025918.1_strain_R25            ................................................c............................... [175440]
MH025917.1_strain_R7             ................................................c............................... [175440]
MH025916.1_strain_R8             ................................................c............................... [175440]

FR682468.2_Georgia_2007/1        actggtagtatttcagggctcgtgcaaagcgggcagtgagtgaggtgtctttgctttcctgaagctctttatatttttca [175520]
ASF_IND_20_CAD543                ................................................................................ [175520]
LR899193.1_Germany_2020/1        ................................................................................ [175520]
MN393477.1_Wuhan_2019            ................................................................................ [175520]
MW396979.1_Timor-Leste/2019/1    ................................................................................ [175520]
MK940252.1_CN/2019/InnerMongolia ................................................................................ [175520]
MK128995.1_China/2018/AnhuiXCGQ  ................................................................................ [175520]
MH681419.1_ASFV/POL/2015         ................................................................................ [175520]
LS478113.1_Estonia_2014          ................................................................................ [175520]
KP843857.1_Odintsovo_02/14       ................................................................................ [175520]
```

```
FN557520.1_strain_E75             ................t...........a................................................... [175520]
AY261366.1_Warthog                ................................................................................ [175520]
AY261365.1_Warmbaths              ................................................................................ [175520]
AY261364.1_Tengani_62             ............................a................................................... [175520]
AY261363.1_Pretoriuskop/96/4      ................................................................................ [175520]
AY261362.1_Mkuzi_1979             ................t...........a................................................... [175520]
AY261361.1_Malawi_Lil-20/1_1983   ................ta..........a................................................... [175520]
AY261360.1_Kenya_1950             .t..............t................................c.......................c...... [175520]
AM712240.1_OURT_88/3              ................t...........a................................................... [175520]
AM712239.1_Benin_97/1             ................t...........a................................................... [175520]
KM111295.1_Ken06.Bus              ................t.......................g........c.......................c...... [175520]
KM111294.1_Ken05/Tk1              .t..............t................................c.............t.........c...... [175520]
KM262844.1_strain_L60             ................t...........a................................................... [175520]
KM102979.1_26544/OG10_Italy       ................t...........a................................................... [175520]
KM262845.1_strain_NHV             ................t...........a................................................... [175520]
KX354450.1_47/Ss/2008             ................t...........a................................................... [175520]
KP055815.1_strain_BA71            ................t...........a................................................... [175520]
MH025920.1_strain_R35             ................t.......................g........c.......................c...... [175520]
MH025919.1_strain_N10             ................t.......................g........c.......................c...... [175520]
MH025918.1_strain_R25             ................t.......................g........c.......................c...... [175520]
MH025917.1_strain_R7              ................t.......................g........c.......................c...... [175520]
MH025916.1_strain_R8              ................t.......................g........c.......................c...... [175520]

FR682468.2_Georgia_2007/1         tatacctcttttaggtatgcttctatttggacggggaaggaggtgttgttgtgcacgcaagacatgactcgttataagga [175600]
ASF_IND_20_CAD543                 ................................................................................ [175600]
LR899193.1_Germany_2020/1         ................................................................................ [175600]
MN393477.1_Wuhan_2019             ................................................................................ [175600]
MW396979.1_Timor-Leste/2019/1     ................................................................................ [175600]
MK940252.1_CN/2019/InnerMongolia  ................................................................................ [175600]
MK128995.1_China/2018/AnhuiXCGQ   ................................................................................ [175600]
MH681419.1_ASFV/POL/2015          ................................................................................ [175600]
LS478113.1_Estonia_2014           ................................................................................ [175600]
KP843857.1_Odintsovo_02/14        ................................................................................ [175600]
FN557520.1_strain_E75             ................................................................................ [175600]
AY261366.1_Warthog                a.......................g....................................................... [175600]
AY261365.1_Warmbaths              a.......................g....................................................... [175600]
AY261364.1_Tengani_62             ............................................a.....................g............. [175600]
AY261363.1_Pretoriuskop/96/4      a.......................g....................................................... [175600]
AY261362.1_Mkuzi_1979             ................................................................................ [175600]
AY261361.1_Malawi_Lil-20/1_1983   a.......c............................................g.......................... [175600]
AY261360.1_Kenya_1950             a....................................................g.......................... [175600]
AM712240.1_OURT_88/3              ................................................................................ [175600]
AM712239.1_Benin_97/1             ................................................................................ [175600]
KM111295.1_Ken06.Bus              a....................................................g.......................... [175600]
```

```
KM111294.1_Ken05/Tk1            a....................................................g.......................... [175600]
KM262844.1_strain_L60           ................................................................................ [175600]
KM102979.1_26544/OG10_Italy     ................................................................................ [175600]
KM262845.1_strain_NHV           ................................................................................ [175600]
KX354450.1_47/Ss/2008           ................................................................................ [175600]
KP055815.1_strain_BA71          ................................................................................ [175600]
MH025920.1_strain_R35           a....................................................g.......................... [175600]
MH025919.1_strain_N10           a....................................................g.......................... [175600]
MH025918.1_strain_R25           a....................................................g.......................... [175600]
MH025917.1_strain_R7            a....................................................g.......................... [175600]
MH025916.1_strain_R8            a....................................................g.......................... [175600]

FR682468.2_Georgia_2007/1       tcccatattaaaacttcattagaagaatagggctgctgatagctagcgctgcacttaaaaatggggtagccctttttctt [175680]
ASF_IND_20_CAD543               ................................................................................ [175680]
LR899193.1_Germany_2020/1       ................................................................................ [175680]
MN393477.1_Wuhan_2019           ................................................................................ [175680]
MW396979.1_Timor-Leste/2019/1   ................................................................................ [175680]
MK940252.1_CN/2019/InnerMongolia ................................................................................ [175680]
MK128995.1_China/2018/AnhuiXCGQ ................................................................................ [175680]
MH681419.1_ASFV/POL/2015        ................................................................................ [175680]
LS478113.1_Estonia_2014         ................................................................................ [175680]
KP843857.1_Odintsovo_02/14      ................................................................................ [175680]
FN557520.1_strain_E75           ................................................................................ [175680]
AY261366.1_Warthog              .........................................a...................................... [175680]
AY261365.1_Warmbaths            .........................................a..g................................... [175680]
AY261364.1_Tengani_62           .........................................a...................................... [175680]
AY261363.1_Pretoriuskop/96/4    .........................................a...................................... [175680]
AY261362.1_Mkuzi_1979           ................................................................................ [175680]
AY261361.1_Malawi_Lil-20/1_1983 .........................................a..g................................... [175680]
AY261360.1_Kenya_1950           .........................................a.................g.................... [175680]
AM712240.1_OURT_88/3            ................................................................................ [175680]
AM712239.1_Benin_97/1           ................................................................................ [175680]
KM111295.1_Ken06.Bus            .............................a...........a.................g.................... [175680]
KM111294.1_Ken05/Tk1            .........................................a...................................... [175680]
KM262844.1_strain_L60           ................................................................................ [175680]
KM102979.1_26544/OG10_Italy     ................................................................................ [175680]
KM262845.1_strain_NHV           ................................................................................ [175680]
KX354450.1_47/Ss/2008           ................................................................................ [175680]
KP055815.1_strain_BA71          ................................................................................ [175680]
MH025920.1_strain_R35           .............................a...........a.................g.................... [175680]
MH025919.1_strain_N10           .............................a...........a.................g.................... [175680]
MH025918.1_strain_R25           .............................a...........a.................g.................... [175680]
MH025917.1_strain_R7            .............................a...........a.................g.................... [175680]
MH025916.1_strain_R8            .............................a...........a.................g.................... [175680]
```

```
FR682468.2_Georgia_2007/1        gtaaatccggtgcctgtcgtagacctggctagaaagcgggcttagtgtatctttaatgtccacaacgatgcgtacctttt [175760]
ASF_IND_20_CAD543                ................................................................................ [175760]
LR899193.1_Germany_2020/1        ................................................................................ [175760]
MN393477.1_Wuhan_2019            ................................................................................ [175760]
MW396979.1_Timor-Leste/2019/1    ................................................................................ [175760]
MK940252.1_CN/2019/InnerMongolia ................................................................................ [175760]
MK128995.1_China/2018/AnhuiXCGQ  ................................................................................ [175760]
MH681419.1_ASFV/POL/2015         ................................................................................ [175760]
LS478113.1_Estonia_2014          ................................................................................ [175760]
KP843857.1_Odintsovo_02/14       ................................................................................ [175760]
FN557520.1_strain_E75            ...............................................................g................ [175760]
AY261366.1_Warthog               ...............................................................g................ [175760]
AY261365.1_Warmbaths             ...............................................................g................ [175760]
AY261364.1_Tengani_62            ...............................................................g................ [175760]
AY261363.1_Pretoriuskop/96/4     .............................................c.................g................ [175760]
AY261362.1_Mkuzi_1979            ...............................................................g................ [175760]
AY261361.1_Malawi_Lil-20/1_1983  ......g..........................t.............................................. [175760]
AY261360.1_Kenya_1950            ..............................g..t........c..................................... [175760]
AM712240.1_OURT_88/3             ...............................................................g................ [175760]
AM712239.1_Benin_97/1            ...............................................................g................ [175760]
KM111295.1_Ken06.Bus             ..............................g..t........c..................................... [175760]
KM111294.1_Ken05/Tk1             .................................t..........................t................... [175760]
KM262844.1_strain_L60            ...............................................................g................ [175760]
KM102979.1_26544/OG10_Italy      ...............................................................g................ [175760]
KM262845.1_strain_NHV            ...............................................................g................ [175760]
KX354450.1_47/Ss/2008            ...............................................................g................ [175760]
KP055815.1_strain_BA71           ...............................................................g................ [175760]
MH025920.1_strain_R35            ..............................g..t..t.....c..................................... [175760]
MH025919.1_strain_N10            ..............................g..t..t.....c..................................... [175760]
MH025918.1_strain_R25            ..............................g..t..t.....c..................................... [175760]
MH025917.1_strain_R7             ..............................g..t..t.....c..................................... [175760]
MH025916.1_strain_R8             ..............................g..t..t.....c..................................... [175760]

FR682468.2_Georgia_2007/1        tttcatccgatccctgccgggtaatacgtcccaagatttgctccatgttgtttctgcggggcgttgccatgatgatcgat [175840]
ASF_IND_20_CAD543                ................................................................................ [175840]
LR899193.1_Germany_2020/1        ................................................................................ [175840]
MN393477.1_Wuhan_2019            ................................................................................ [175840]
MW396979.1_Timor-Leste/2019/1    ................................................................................ [175840]
MK940252.1_CN/2019/InnerMongolia ................................................................................ [175840]
MK128995.1_China/2018/AnhuiXCGQ  ................................................................................ [175840]
MH681419.1_ASFV/POL/2015         ................................................................................ [175840]
LS478113.1_Estonia_2014          ................................................................................ [175840]
KP843857.1_Odintsovo_02/14       ................................................................................ [175840]
```

```
FN557520.1_strain_E75              ................................................................................ [175840]
AY261366.1_Warthog                 ................................................................................ [175840]
AY261365.1_Warmbaths               ................................................................................ [175840]
AY261364.1_Tengani_62              ..............................................a................................. [175840]
AY261363.1_Pretoriuskop/96/4       ................................................................................ [175840]
AY261362.1_Mkuzi_1979              ................................................................................ [175840]
AY261361.1_Malawi_Lil-20/1_1983    ..........................................................c..................... [175840]
AY261360.1_Kenya_1950              ..........c.....t...........a................................................... [175840]
AM712240.1_OURT_88/3               ................................................................................ [175840]
AM712239.1_Benin_97/1              ................................................................................ [175840]
KM111295.1_Ken06.Bus               ............................a................................................... [175840]
KM111294.1_Ken05/Tk1               ................................................................................ [175840]
KM262844.1_strain_L60              ................................................................................ [175840]
KM102979.1_26544/OG10_Italy        ................................................................................ [175840]
KM262845.1_strain_NHV              ................................................................................ [175840]
KX354450.1_47/Ss/2008              ................................................................................ [175840]
KP055815.1_strain_BA71             ................................................................................ [175840]
MH025920.1_strain_R35              ............................a................................................... [175840]
MH025919.1_strain_N10              ............................a................................................... [175840]
MH025918.1_strain_R25              ............................a................................................... [175840]
MH025917.1_strain_R7               ............................a................................................... [175840]
MH025916.1_strain_R8               ............................a................................................... [175840]

FR682468.2_Georgia_2007/1          gtcatatgcttgaaggaaatgcctctacgcccgtagccataggtcagcaagataatggaagcgctgtgtgcctgagaaag [175920]
ASF_IND_20_CAD543                  ................................................................................ [175920]
LR899193.1_Germany_2020/1          ................................................................................ [175920]
MN393477.1_Wuhan_2019              ................................................................................ [175920]
MW396979.1_Timor-Leste/2019/1      ................................................................................ [175920]
MK940252.1_CN/2019/InnerMongolia   ................................................................................ [175920]
MK128995.1_China/2018/AnhuiXCGQ    ................................................................................ [175920]
MH681419.1_ASFV/POL/2015           ................................................................................ [175920]
LS478113.1_Estonia_2014            ................................................................................ [175920]
KP843857.1_Odintsovo_02/14         ................................................................................ [175920]
FN557520.1_strain_E75              ................................................................................ [175920]
AY261366.1_Warthog                 ................................................................................ [175920]
AY261365.1_Warmbaths               ..........................c..................................................... [175920]
AY261364.1_Tengani_62              ................................................................................ [175920]
AY261363.1_Pretoriuskop/96/4       ......................................................................t......... [175920]
AY261362.1_Mkuzi_1979              ..........................c..................................................... [175920]
AY261361.1_Malawi_Lil-20/1_1983    ..................................................a............................. [175920]
AY261360.1_Kenya_1950              .....................................................g.......................g.. [175920]
AM712240.1_OURT_88/3               ................................................................................ [175920]
AM712239.1_Benin_97/1              ................................................................................ [175920]
KM111295.1_Ken06.Bus               ..............a......................................g.......................g.. [175920]
```

```
KM111294.1_Ken05/Tk1            .....................................................g.......................... [175920]
KM262844.1_strain_L60           ................................................................................ [175920]
KM102979.1_26544/OG10_Italy     ................................................................................ [175920]
KM262845.1_strain_NHV           ................................................................................ [175920]
KX354450.1_47/Ss/2008           ................................................................................ [175920]
KP055815.1_strain_BA71          ................................................................................ [175920]
MH025920.1_strain_R35           ..............a......................................g.......................g.. [175920]
MH025919.1_strain_N10           ..............a......................................g.......................g.. [175920]
MH025918.1_strain_R25           ..............a......................................g.......................g.. [175920]
MH025917.1_strain_R7            ..............a......................................g.......................g.. [175920]
MH025916.1_strain_R8            ..............a......................................g.......................g.. [175920]

FR682468.2_Georgia_2007/1       agcggtatttgaaaccccgccgcataggagcgccacctccggaacgataatttgaacatctttgaattctttggaaagcg [176000]
ASF_IND_20_CAD543               ................................................................................ [176000]
LR899193.1_Germany_2020/1       ................................................................................ [176000]
MN393477.1_Wuhan_2019           ................................................................................ [176000]
MW396979.1_Timor-Leste/2019/1   ................................................................................ [176000]
MK940252.1_CN/2019/InnerMongolia ................................................................................ [176000]
MK128995.1_China/2018/AnhuiXCGQ ................................................................................ [176000]
MH681419.1_ASFV/POL/2015        ................................................................................ [176000]
LS478113.1_Estonia_2014         ................................................................................ [176000]
KP843857.1_Odintsovo_02/14      ................................................................................ [176000]
FN557520.1_strain_E75           ...............................................c................................ [176000]
AY261366.1_Warthog              ................................................................................ [176000]
AY261365.1_Warmbaths            ................................................................................ [176000]
AY261364.1_Tengani_62           ................................................................................ [176000]
AY261363.1_Pretoriuskop/96/4    .............................................t.................................. [176000]
AY261362.1_Mkuzi_1979           ................................................................................ [176000]
AY261361.1_Malawi_Lil-20/1_1983 ...........................a.....t...................c...g...................... [176000]
AY261360.1_Kenya_1950           ...........................a..........................t................c........ [176000]
AM712240.1_OURT_88/3            ...............................................c................................ [176000]
AM712239.1_Benin_97/1           ...............................................c................................ [176000]
KM111295.1_Ken06.Bus            ...........................a..........................t................c........ [176000]
KM111294.1_Ken05/Tk1            ...........................a.....t...................c...g.............c........ [176000]
KM262844.1_strain_L60           ...............................................c................................ [176000]
KM102979.1_26544/OG10_Italy     ...............................................c................................ [176000]
KM262845.1_strain_NHV           ...............................................c................................ [176000]
KX354450.1_47/Ss/2008           ...............................................c................................ [176000]
KP055815.1_strain_BA71          ...............................................c................................ [176000]
MH025920.1_strain_R35           ...........................a..........................t................c........ [176000]
MH025919.1_strain_N10           ...........................a..........................t................c........ [176000]
MH025918.1_strain_R25           ...........................a..........................t................c........ [176000]
MH025917.1_strain_R7            ...........................a..........................t................c........ [176000]
MH025916.1_strain_R8            ...........................a..........................t................c........ [176000]
```

```
FR682468.2_Georgia_2007/1        cctgataaaaaatttctaaaagtttgcgaaattccacgaaaatgatgatgccatacggctcatcggtcccccatttgtga [176080]
ASF_IND_20_CAD543                ................................................................................ [176080]
LR899193.1_Germany_2020/1        ................................................................................ [176080]
MN393477.1_Wuhan_2019            ................................................................................ [176080]
MW396979.1_Timor-Leste/2019/1    ................................................................................ [176080]
MK940252.1_CN/2019/InnerMongolia ................................................................................ [176080]
MK128995.1_China/2018/AnhuiXCGQ  ................................................................................ [176080]
MH681419.1_ASFV/POL/2015         ................................................................................ [176080]
LS478113.1_Estonia_2014          ................................................................................ [176080]
KP843857.1_Odintsovo_02/14       ................................................................................ [176080]
FN557520.1_strain_E75            ................................................................................ [176080]
AY261366.1_Warthog               .........................a...................................................... [176080]
AY261365.1_Warmbaths             .........................a...................................................... [176080]
AY261364.1_Tengani_62            .........................a...................................................... [176080]
AY261363.1_Pretoriuskop/96/4     .........................a...........a.......................................... [176080]
AY261362.1_Mkuzi_1979            .........................a...................................................... [176080]
AY261361.1_Malawi_Lil-20/1_1983  ....................................................g..........t............t... [176080]
AY261360.1_Kenya_1950            ....................................................g........g.t............t... [176080]
AM712240.1_OURT_88/3             ................................................................................ [176080]
AM712239.1_Benin_97/1            ................................................................................ [176080]
KM111295.1_Ken06.Bus             ....................................................g..........t............t... [176080]
KM111294.1_Ken05/Tk1             ...............................c....................g........g.t............t... [176080]
KM262844.1_strain_L60            ................................................................................ [176080]
KM102979.1_26544/OG10_Italy      ................................................................................ [176080]
KM262845.1_strain_NHV            ................................................................................ [176080]
KX354450.1_47/Ss/2008            ................................................................................ [176080]
KP055815.1_strain_BA71           ................................................................................ [176080]
MH025920.1_strain_R35            ....................................................g..........t............t... [176080]
MH025919.1_strain_N10            ....................................................g..........t............t... [176080]
MH025918.1_strain_R25            ....................................................g..........t............t... [176080]
MH025917.1_strain_R7             ....................................................g..........t............t... [176080]
MH025916.1_strain_R8             ....................................................g..........t............t... [176080]

FR682468.2_Georgia_2007/1        ggctcagcggtatgcagggagtaaagccgctttgcctcatttacgacaagttgtatacgcgaaggatcttgaagtagttt [176160]
ASF_IND_20_CAD543                ................................................................................ [176160]
LR899193.1_Germany_2020/1        ................................................................................ [176160]
MN393477.1_Wuhan_2019            ................................................................................ [176160]
MW396979.1_Timor-Leste/2019/1    ................................................................................ [176160]
MK940252.1_CN/2019/InnerMongolia ................................................................................ [176160]
MK128995.1_China/2018/AnhuiXCGQ  ................................................................................ [176160]
MH681419.1_ASFV/POL/2015         ................................................................................ [176160]
LS478113.1_Estonia_2014          ................................................................................ [176160]
KP843857.1_Odintsovo_02/14       ................................................................................ [176160]
```

```
FN557520.1_strain_E75           ................................................................................ [176160]
AY261366.1_Warthog              ..t...............................................c............................. [176160]
AY261365.1_Warmbaths            ..t...............................................c............................. [176160]
AY261364.1_Tengani_62           ..t............................................................................. [176160]
AY261363.1_Pretoriuskop/96/4    ..t............................................................................. [176160]
AY261362.1_Mkuzi_1979           ..................................................c............................. [176160]
AY261361.1_Malawi_Lil-20/1_1983 ..t.....a..............c..........t.........t................................... [176160]
AY261360.1_Kenya_1950           ..t....................c..........t...g.....t...........g.................g..... [176160]
AM712240.1_OURT_88/3            ................................................................................ [176160]
AM712239.1_Benin_97/1           ................................................................................ [176160]
KM111295.1_Ken06.Bus            ..t....................c..........t...g.....t.............................g..... [176160]
KM111294.1_Ken05/Tk1            ..t....................c..........t...g.....t...........g.................g..... [176160]
KM262844.1_strain_L60           ................................................................................ [176160]
KM102979.1_26544/OG10_Italy     ................................................................................ [176160]
KM262845.1_strain_NHV           ................................................................................ [176160]
KX354450.1_47/Ss/2008           ................................................................................ [176160]
KP055815.1_strain_BA71          ................................................................................ [176160]
MH025920.1_strain_R35           ..t....................c..........t...g.....t.............................g..... [176160]
MH025919.1_strain_N10           ..t....................c..........t...g.....t.............................g..... [176160]
MH025918.1_strain_R25           ..t....................c..........t...g.....t.............................g..... [176160]
MH025917.1_strain_R7            ..t....................c..........t...g.....t.............................g..... [176160]
MH025916.1_strain_R8            ..t....................c..........t...g.....t.............................g..... [176160]

FR682468.2_Georgia_2007/1       atcaatggtggcaatggccgataccttttcattaatatacacagggctaacgaagtcaggatgtccctgatattcgattt [176240]
ASF_IND_20_CAD543               ................................................................................ [176240]
LR899193.1_Germany_2020/1       ................................................................................ [176240]
MN393477.1_Wuhan_2019           ................................................................................ [176240]
MW396979.1_Timor-Leste/2019/1   ................................................................................ [176240]
MK940252.1_CN/2019/InnerMongolia ................................................................................ [176240]
MK128995.1_China/2018/AnhuiXCGQ ................................................................................ [176240]
MH681419.1_ASFV/POL/2015        ................................................................................ [176240]
LS478113.1_Estonia_2014         ................................................................................ [176240]
KP843857.1_Odintsovo_02/14      ................................................................................ [176240]
FN557520.1_strain_E75           .....................................c.......................................... [176240]
AY261366.1_Warthog              ...............a..............g......c..........g..a............................ [176240]
AY261365.1_Warmbaths            .....................................c..........g..a............................ [176240]
AY261364.1_Tengani_62           .....................................c.......................................... [176240]
AY261363.1_Pretoriuskop/96/4    .....................................c.............a............................ [176240]
AY261362.1_Mkuzi_1979           .....................................c...................g...................... [176240]
AY261361.1_Malawi_Lil-20/1_1983 ..............................g......c.t.....ct..........g.....c................ [176240]
AY261360.1_Kenya_1950           .....................................c.......tt................c................ [176240]
AM712240.1_OURT_88/3            .....................................c.......................................... [176240]
AM712239.1_Benin_97/1           .....................................c.......................................... [176240]
KM111295.1_Ken06.Bus            ..............................g..g...c.......tt...t......g........t............. [176240]
```

```
KM111294.1_Ken05/Tk1             .....................................c.......tt................c................ [176240]
KM262844.1_strain_L60            .....................................c.......................................... [176240]
KM102979.1_26544/OG10_Italy      .....................................c.......................................... [176240]
KM262845.1_strain_NHV            .....................................c.......................................... [176240]
KX354450.1_47/Ss/2008            .....................................c.......................................... [176240]
KP055815.1_strain_BA71           .....................................c.......................................... [176240]
MH025920.1_strain_R35            ..............................g..g...c.......tt...t......g........t............. [176240]
MH025919.1_strain_N10            ..............................g..g...c.......tt...t......g........t............. [176240]
MH025918.1_strain_R25            ..............................g..g...c.......tt...t......g........t............. [176240]
MH025917.1_strain_R7             ..............................g..g...c.......tt...t......g........t............. [176240]
MH025916.1_strain_R8             ..............................g..g...c.......tt...t......g........t............. [176240]

FR682468.2_Georgia_2007/1        ccctcacgtacccggaaaaggttgtggtgggacttacagtcctctggggctgtcctagatggtgaataataatcttgtcc [176320]
ASF_IND_20_CAD543                ................................................................................ [176320]
LR899193.1_Germany_2020/1        ................................................................................ [176320]
MN393477.1_Wuhan_2019            ................................................................................ [176320]
MW396979.1_Timor-Leste/2019/1    ................................................................................ [176320]
MK940252.1_CN/2019/InnerMongolia ................................................................................ [176320]
MK128995.1_China/2018/AnhuiXCGQ  ................................................................................ [176320]
MH681419.1_ASFV/POL/2015         ................................................................................ [176320]
LS478113.1_Estonia_2014          ................................................................................ [176320]
KP843857.1_Odintsovo_02/14       ................................................................................ [176320]
FN557520.1_strain_E75            ................................................................................ [176320]
AY261366.1_Warthog               .....................................g.............................g............ [176320]
AY261365.1_Warmbaths             ................................................................................ [176320]
AY261364.1_Tengani_62            ................................................................................ [176320]
AY261363.1_Pretoriuskop/96/4     .....................................g.............................g............ [176320]
AY261362.1_Mkuzi_1979            ................................................................................ [176320]
AY261361.1_Malawi_Lil-20/1_1983  .........................a...........g....................g........g............ [176320]
AY261360.1_Kenya_1950            .........................c..............t.........................gg............ [176320]
AM712240.1_OURT_88/3             ................................................................................ [176320]
AM712239.1_Benin_97/1            ................................................................................ [176320]
KM111295.1_Ken06.Bus             .........................a...........g..t................a........gg............ [176320]
KM111294.1_Ken05/Tk1             .........................a..............t.........................gg............ [176320]
KM262844.1_strain_L60            ................................................................................ [176320]
KM102979.1_26544/OG10_Italy      ................................................................................ [176320]
KM262845.1_strain_NHV            ................................................................................ [176320]
KX354450.1_47/Ss/2008            ................................................................................ [176320]
KP055815.1_strain_BA71           ................................................................................ [176320]
MH025920.1_strain_R35            .........................a...........g..t................a........gg............ [176320]
MH025919.1_strain_N10            .........................a...a.......g..t................a........gg............ [176320]
MH025918.1_strain_R25            .........................a...........g..t................a........gg............ [176320]
MH025917.1_strain_R7             .........................a...........g..t................a........gg............ [176320]
MH025916.1_strain_R8             .........................a...........g..t................a........gg............ [176320]
```

```
FR682468.2_Georgia_2007/1        ataccatcgggccggtccaggggtgtagcggacagtcctaatatccgactaagttgtattttccaaaaaattttgtaatt [176400]
ASF_IND_20_CAD543                ................................................................................ [176400]
LR899193.1_Germany_2020/1        ................................................................................ [176400]
MN393477.1_Wuhan_2019            ................................................................................ [176400]
MW396979.1_Timor-Leste/2019/1    ................................................................................ [176400]
MK940252.1_CN/2019/InnerMongolia ................................................................................ [176400]
MK128995.1_China/2018/AnhuiXCGQ  ................................................................................ [176400]
MH681419.1_ASFV/POL/2015         ................................................................................ [176400]
LS478113.1_Estonia_2014          ................................................................................ [176400]
KP843857.1_Odintsovo_02/14       ................................................................................ [176400]
FN557520.1_strain_E75            ..........................g..................................................... [176400]
AY261366.1_Warthog               ..................................................c............................. [176400]
AY261365.1_Warmbaths             ..................................................c............................. [176400]
AY261364.1_Tengani_62            ................................................................................ [176400]
AY261363.1_Pretoriuskop/96/4     .............................a.................................................. [176400]
AY261362.1_Mkuzi_1979            ................................................................................ [176400]
AY261361.1_Malawi_Lil-20/1_1983  .................t................................c..c.......................... [176400]
AY261360.1_Kenya_1950            ..c........a..a...........g.......................c............................. [176400]
AM712240.1_OURT_88/3             ..........................g..................................................... [176400]
AM712239.1_Benin_97/1            ..........................g..................................................... [176400]
KM111295.1_Ken06.Bus             ..c........a..............g..........................c.......................... [176400]
KM111294.1_Ken05/Tk1             ..c........a..a...........g..a....................c............................. [176400]
KM262844.1_strain_L60            ..........................g..................................................... [176400]
KM102979.1_26544/OG10_Italy      ..........................g..................................................... [176400]
KM262845.1_strain_NHV            ..........................g..................................................... [176400]
KX354450.1_47/Ss/2008            ..........................g..................................................... [176400]
KP055815.1_strain_BA71           ..........................g..................................................... [176400]
MH025920.1_strain_R35            ..c........a..............g..........................c.......................... [176400]
MH025919.1_strain_N10            ..c........a..............g..........................c.......................... [176400]
MH025918.1_strain_R25            ..c........a..............g..........................c.......................... [176400]
MH025917.1_strain_R7             ..c........a..............g..........................c.......................... [176400]
MH025916.1_strain_R8             ..c........a..............g..........................c.......................... [176400]

FR682468.2_Georgia_2007/1        ctccggcgagtgtaattcatgtgcctcatctaacacgactagaccaaagggctcaaagaactgctcaggcttcttgcgca [176480]
ASF_IND_20_CAD543                ................................................................................ [176480]
LR899193.1_Germany_2020/1        ................................................................................ [176480]
MN393477.1_Wuhan_2019            ................................................................................ [176480]
MW396979.1_Timor-Leste/2019/1    ................................................................................ [176480]
MK940252.1_CN/2019/InnerMongolia ................................................................................ [176480]
MK128995.1_China/2018/AnhuiXCGQ  ................................................................................ [176480]
MH681419.1_ASFV/POL/2015         ................................................................................ [176480]
LS478113.1_Estonia_2014          ................................................................................ [176480]
KP843857.1_Odintsovo_02/14       ................................................................................ [176480]
```

```
FN557520.1_strain_E75            ................................................................................ [176480]
AY261366.1_Warthog               t............................................g.................................. [176480]
AY261365.1_Warmbaths             t............................................g.................................. [176480]
AY261364.1_Tengani_62            ....................................a....a...g.................................. [176480]
AY261363.1_Pretoriuskop/96/4     ...........................g.................g.................................. [176480]
AY261362.1_Mkuzi_1979            ................................................................................ [176480]
AY261361.1_Malawi_Lil-20/1_1983  ...........................g.................g...............gc................. [176480]
AY261360.1_Kenya_1950            ....................................a........g...............g.................. [176480]
AM712240.1_OURT_88/3             ................................................................................ [176480]
AM712239.1_Benin_97/1            ................................................................................ [176480]
KM111295.1_Ken06.Bus             ...t...................................c.....g..............tg.................. [176480]
KM111294.1_Ken05/Tk1             .....................c.......................g...............g.................. [176480]
KM262844.1_strain_L60            ................................................................................ [176480]
KM102979.1_26544/OG10_Italy      ................................................................................ [176480]
KM262845.1_strain_NHV            ................................................................................ [176480]
KX354450.1_47/Ss/2008            ................................................................................ [176480]
KP055815.1_strain_BA71           ................................................................................ [176480]
MH025920.1_strain_R35            ...t...................................c.....g..............tg.................. [176480]
MH025919.1_strain_N10            ...t...................................c.....g..............tg.................. [176480]
MH025918.1_strain_R25            ...t...................................c.....g..............tg.................. [176480]
MH025917.1_strain_R7             ...t...................................c.....g..............tg.................. [176480]
MH025916.1_strain_R8             ...t...................................c.....g..............tg.................. [176480]

FR682468.2_Georgia_2007/1        gggtattaatgattcccacgatgacgtcgtactctttgctcgtcatgtcctttttcttgcacgctgcattattgtaagca [176560]
ASF_IND_20_CAD543                ................................................................................ [176560]
LR899193.1_Germany_2020/1        ................................................................................ [176560]
MN393477.1_Wuhan_2019            ................................................................................ [176560]
MW396979.1_Timor-Leste/2019/1    ................................................................................ [176560]
MK940252.1_CN/2019/InnerMongolia ................................................................................ [176560]
MK128995.1_China/2018/AnhuiXCGQ  ................................................................................ [176560]
MH681419.1_ASFV/POL/2015         ................................................................................ [176560]
LS478113.1_Estonia_2014          ................................................................................ [176560]
KP843857.1_Odintsovo_02/14       ................................................................................ [176560]
FN557520.1_strain_E75            ................................................................................ [176560]
AY261366.1_Warthog               ...............................t.....a.......................................... [176560]
AY261365.1_Warmbaths             .....................................a.......................................... [176560]
AY261364.1_Tengani_62            .....................................a.....g.................................... [176560]
AY261363.1_Pretoriuskop/96/4     .....................................a.......................................... [176560]
AY261362.1_Mkuzi_1979            ................................................................................ [176560]
AY261361.1_Malawi_Lil-20/1_1983  a....................................a.....a.................................... [176560]
AY261360.1_Kenya_1950            a............g.......................a.....a.......................g............ [176560]
AM712240.1_OURT_88/3             ................................................................................ [176560]
AM712239.1_Benin_97/1            ................................................................................ [176560]
KM111295.1_Ken06.Bus             a............g.......................a.....a.......................g............ [176560]
```

```
KM111294.1_Ken05/Tk1             a............g.......................a.....a.......................g............ [176560]
KM262844.1_strain_L60            ................................................................................ [176560]
KM102979.1_26544/OG10_Italy      ................................................................................ [176560]
KM262845.1_strain_NHV            ................................................................................ [176560]
KX354450.1_47/Ss/2008            ................................................................................ [176560]
KP055815.1_strain_BA71           ................................................................................ [176560]
MH025920.1_strain_R35            a............g.......................a.....a.......................g............ [176560]
MH025919.1_strain_N10            a............g.......................a.....a.......................g............ [176560]
MH025918.1_strain_R25            a............g.......................a.....a.......................g............ [176560]
MH025917.1_strain_R7             a............g.......................a.....a.......................g............ [176560]
MH025916.1_strain_R8             a............g.......................a.....a.......................g............ [176560]

FR682468.2_Georgia_2007/1        gctacacgtaggtggggcaggagcaatgttagctcgtcgatccactgtatttgaatcgccttggtgggcacgatgaccag [176640]
ASF_IND_20_CAD543                ................................................................................ [176640]
LR899193.1_Germany_2020/1        ................................................................................ [176640]
MN393477.1_Wuhan_2019            ................................................................................ [176640]
MW396979.1_Timor-Leste/2019/1    ................................................................................ [176640]
MK940252.1_CN/2019/InnerMongolia ................................................................................ [176640]
MK128995.1_China/2018/AnhuiXCGQ  ................................................................................ [176640]
MH681419.1_ASFV/POL/2015         ................................................................................ [176640]
LS478113.1_Estonia_2014          ................................................................................ [176640]
KP843857.1_Odintsovo_02/14       ................................................................................ [176640]
FN557520.1_strain_E75            ................................................................................ [176640]
AY261366.1_Warthog               ................................................................................ [176640]
AY261365.1_Warmbaths             .................................................c...........................t.. [176640]
AY261364.1_Tengani_62            .............................................................................t.. [176640]
AY261363.1_Pretoriuskop/96/4     ....................a............................c.............................. [176640]
AY261362.1_Mkuzi_1979            ................................................................................ [176640]
AY261361.1_Malawi_Lil-20/1_1983  ................................................................................ [176640]
AY261360.1_Kenya_1950            ...................a.......t................t................................... [176640]
AM712240.1_OURT_88/3             ................................................................................ [176640]
AM712239.1_Benin_97/1            ................................................................................ [176640]
KM111295.1_Ken06.Bus             ...................a.......t................t................................... [176640]
KM111294.1_Ken05/Tk1             .................t.a.......t.................................................... [176640]
KM262844.1_strain_L60            ................................................................................ [176640]
KM102979.1_26544/OG10_Italy      ................................................................................ [176640]
KM262845.1_strain_NHV            ................................................................................ [176640]
KX354450.1_47/Ss/2008            ................................................................................ [176640]
KP055815.1_strain_BA71           ................................................................................ [176640]
MH025920.1_strain_R35            ...................a.......t................t................................... [176640]
MH025919.1_strain_N10            ...................a.......t................t................................... [176640]
MH025918.1_strain_R25            ...................a.......t................t................................... [176640]
MH025917.1_strain_R7             ...................a.......t................t................................... [176640]
MH025916.1_strain_R8             ...................a.......t................t................................... [176640]
```

```
FR682468.2_Georgia_2007/1        ggtagggtacaaaagtttttgaataatgctgatcgcaatacgcgttttccccaaaccggtatttagatgtaggtaaaagc [176720]
ASF_IND_20_CAD543                ................................................................................ [176720]
LR899193.1_Germany_2020/1        ................................................................................ [176720]
MN393477.1_Wuhan_2019            ................................................................................ [176720]
MW396979.1_Timor-Leste/2019/1    ................................................................................ [176720]
MK940252.1_CN/2019/InnerMongolia ................................................................................ [176720]
MK128995.1_China/2018/AnhuiXCGQ  ................................................................................ [176720]
MH681419.1_ASFV/POL/2015         ................................................................................ [176720]
LS478113.1_Estonia_2014          ................................................................................ [176720]
KP843857.1_Odintsovo_02/14       ................................................................................ [176720]
FN557520.1_strain_E75            ................................................................................ [176720]
AY261366.1_Warthog               ................................................................................ [176720]
AY261365.1_Warmbaths             ...........................a.................................................... [176720]
AY261364.1_Tengani_62            .......................................................................a........ [176720]
AY261363.1_Pretoriuskop/96/4     ................................................................................ [176720]
AY261362.1_Mkuzi_1979            ................................................................................ [176720]
AY261361.1_Malawi_Lil-20/1_1983  ....................................g........................................... [176720]
AY261360.1_Kenya_1950            ...........................a................................g........c.a........ [176720]
AM712240.1_OURT_88/3             ................................................................................ [176720]
AM712239.1_Benin_97/1            ................................................................................ [176720]
KM111295.1_Ken06.Bus             ...........................a.........................................c.a........ [176720]
KM111294.1_Ken05/Tk1             ...........................a.........................................c.a........ [176720]
KM262844.1_strain_L60            ................................................................................ [176720]
KM102979.1_26544/OG10_Italy      ................................................................................ [176720]
KM262845.1_strain_NHV            ................................................................................ [176720]
KX354450.1_47/Ss/2008            ................................................................................ [176720]
KP055815.1_strain_BA71           ................................................................................ [176720]
MH025920.1_strain_R35            ...........................a.........................................c.a........ [176720]
MH025919.1_strain_N10            ...........................a.........................................c.a........ [176720]
MH025918.1_strain_R25            ...........................a.........................................c.a........ [176720]
MH025917.1_strain_R7             ...........................a.........................................c.a........ [176720]
MH025916.1_strain_R8             ...........................a.........................................c.a........ [176720]

FR682468.2_Georgia_2007/1        gcccatagggggacaggagctttttatgaatcttatcgaccatttcttgctggtagttaaatagtggaaattctgtttca [176800]
ASF_IND_20_CAD543                ................................................................................ [176800]
LR899193.1_Germany_2020/1        ................................................................................ [176800]
MN393477.1_Wuhan_2019            ................................................................................ [176800]
MW396979.1_Timor-Leste/2019/1    ................................................................................ [176800]
MK940252.1_CN/2019/InnerMongolia ................................................................................ [176800]
MK128995.1_China/2018/AnhuiXCGQ  ................................................................................ [176800]
MH681419.1_ASFV/POL/2015         ................................................................................ [176800]
LS478113.1_Estonia_2014          ................................................................................ [176800]
KP843857.1_Odintsovo_02/14       ................................................................................ [176800]
```

```
FN557520.1_strain_E75            ................................................................................ [176800]
AY261366.1_Warthog               ................................................................................ [176800]
AY261365.1_Warmbaths             ................................................................................ [176800]
AY261364.1_Tengani_62            .t..g...............................................a........................... [176800]
AY261363.1_Pretoriuskop/96/4     ................................................................................ [176800]
AY261362.1_Mkuzi_1979            ................................................................................ [176800]
AY261361.1_Malawi_Lil-20/1_1983  ............g................................................................... [176800]
AY261360.1_Kenya_1950            ................................................................................ [176800]
AM712240.1_OURT_88/3             ................................................................................ [176800]
AM712239.1_Benin_97/1            ................................................................................ [176800]
KM111295.1_Ken06.Bus             ................................................................................ [176800]
KM111294.1_Ken05/Tk1             .t.............................................................................. [176800]
KM262844.1_strain_L60            ................................................................................ [176800]
KM102979.1_26544/OG10_Italy      ................................................................................ [176800]
KM262845.1_strain_NHV            ................................................................................ [176800]
KX354450.1_47/Ss/2008            ................................................................................ [176800]
KP055815.1_strain_BA71           ................................................................................ [176800]
MH025920.1_strain_R35            ................................................................................ [176800]
MH025919.1_strain_N10            ................................................................................ [176800]
MH025918.1_strain_R25            ................................................................................ [176800]
MH025917.1_strain_R7             ................................................................................ [176800]
MH025916.1_strain_R8             ................................................................................ [176800]

FR682468.2_Georgia_2007/1        acgcatgggagggcccgcagcgacacggggcgcgtcgtgtaaaccatgttaaacatttcaaactgcttttgcagcaatat [176880]
ASF_IND_20_CAD543                ................................................................................ [176880]
LR899193.1_Germany_2020/1        ................................................................................ [176880]
MN393477.1_Wuhan_2019            ................................................................................ [176880]
MW396979.1_Timor-Leste/2019/1    ................................................................................ [176880]
MK940252.1_CN/2019/InnerMongolia ................................................................................ [176880]
MK128995.1_China/2018/AnhuiXCGQ  ................................................................................ [176880]
MH681419.1_ASFV/POL/2015         ................................................................................ [176880]
LS478113.1_Estonia_2014          ................................................................................ [176880]
KP843857.1_Odintsovo_02/14       ................................................................................ [176880]
FN557520.1_strain_E75            .....................................................................c.......... [176880]
AY261366.1_Warthog               ..............................t...c..................................c.......... [176880]
AY261365.1_Warmbaths             ..................................c..................................c.......... [176880]
AY261364.1_Tengani_62            ...............................a...........t.................................... [176880]
AY261363.1_Pretoriuskop/96/4     ..................................c..................................c.......... [176880]
AY261362.1_Mkuzi_1979            ..................................c..................................c.......... [176880]
AY261361.1_Malawi_Lil-20/1_1983  .............t...t.....t..a..at...ct........t........................c.......... [176880]
AY261360.1_Kenya_1950            ...........a..................t...c..............................t...c.......... [176880]
AM712240.1_OURT_88/3             .....................................................................c.......... [176880]
AM712239.1_Benin_97/1            .....................................................................c.......... [176880]
KM111295.1_Ken06.Bus             ...........a.t................t...c..................................c...a.....g [176880]
```

```
KM111294.1_Ken05/Tk1             ...........a..................t...c..............................t...c.......... [176880]
KM262844.1_strain_L60            .....................................................................c.......... [176880]
KM102979.1_26544/OG10_Italy      .....................................................................c.......... [176880]
KM262845.1_strain_NHV            .....................................................................c.......... [176880]
KX354450.1_47/Ss/2008            .....................................................................c.......... [176880]
KP055815.1_strain_BA71           .....................................................................c.......... [176880]
MH025920.1_strain_R35            ...........a.t................t...c..................................c...a.....g [176880]
MH025919.1_strain_N10            ...........a.t................t...c..................................c...a.....g [176880]
MH025918.1_strain_R25            ...........a.t................t...c..................................c...a.....g [176880]
MH025917.1_strain_R7             ...........a.t................t...c..................................c...a.....g [176880]
MH025916.1_strain_R8             ...........a.t................t...c..................................c...a.....g [176880]

FR682468.2_Georgia_2007/1        gggaaaataaatgtattccccctgcagcgtgaaggcagtttcctgtcttatggctatgtgctttggctgcccgggtaatg [176960]
ASF_IND_20_CAD543                ................................................................................ [176960]
LR899193.1_Germany_2020/1        ................................................................................ [176960]
MN393477.1_Wuhan_2019            ................................................................................ [176960]
MW396979.1_Timor-Leste/2019/1    ................................................................................ [176960]
MK940252.1_CN/2019/InnerMongolia ................................................................................ [176960]
MK128995.1_China/2018/AnhuiXCGQ  ................................................................................ [176960]
MH681419.1_ASFV/POL/2015         ................................................................................ [176960]
LS478113.1_Estonia_2014          ................................................................................ [176960]
KP843857.1_Odintsovo_02/14       ................................................................................ [176960]
FN557520.1_strain_E75            ........................................................................a..c.... [176960]
AY261366.1_Warthog               ........................................................................a....... [176960]
AY261365.1_Warmbaths             ........................................................................a....... [176960]
AY261364.1_Tengani_62            ............................................................t...........a....... [176960]
AY261363.1_Pretoriuskop/96/4     ........................................................................a....... [176960]
AY261362.1_Mkuzi_1979            ........................................................................a....... [176960]
AY261361.1_Malawi_Lil-20/1_1983  ........................................................................a....... [176960]
AY261360.1_Kenya_1950            ..................t..............a..g.........................................c. [176960]
AM712240.1_OURT_88/3             ........................................................................a..c.... [176960]
AM712239.1_Benin_97/1            ........................................................................a..c.... [176960]
KM111295.1_Ken06.Bus             ..................t..............a..g.........................................c. [176960]
KM111294.1_Ken05/Tk1             ............a.....t..............a..g................................t........c. [176960]
KM262844.1_strain_L60            ........................................................................a..c.... [176960]
KM102979.1_26544/OG10_Italy      ........................................................................a..c.... [176960]
KM262845.1_strain_NHV            ........................................................................a..c.... [176960]
KX354450.1_47/Ss/2008            ........................................................................a..c.... [176960]
KP055815.1_strain_BA71           ........................................................................a..c.... [176960]
MH025920.1_strain_R35            ..................t..............a..g.........................................c. [176960]
MH025919.1_strain_N10            ..................t..............a..g.........................................c. [176960]
MH025918.1_strain_R25            ..................t..............a..g.........................................c. [176960]
MH025917.1_strain_R7             ..................t..............a..g.........................................c. [176960]
MH025916.1_strain_R8             ..................t..............a..g.........................................c. [176960]
```

```
FR682468.2_Georgia_2007/1        cccgcgccgtaacggtgagcgccttaagaacgcgcccgaaatcatgttgtaatttactttgtagcttcttataatttatt [177040]
ASF_IND_20_CAD543                ................................................................................ [177040]
LR899193.1_Germany_2020/1        ................................................................................ [177040]
MN393477.1_Wuhan_2019            ................................................................................ [177040]
MW396979.1_Timor-Leste/2019/1    ................................................................................ [177040]
MK940252.1_CN/2019/InnerMongolia ................................................................................ [177040]
MK128995.1_China/2018/AnhuiXCGQ  ................................................................................ [177040]
MH681419.1_ASFV/POL/2015         ................................................................................ [177040]
LS478113.1_Estonia_2014          ................................................................................ [177040]
KP843857.1_Odintsovo_02/14       ................................................................................ [177040]
FN557520.1_strain_E75            ..............................g................................................. [177040]
AY261366.1_Warthog               ..............................g...........t..................................... [177040]
AY261365.1_Warmbaths             ..............................g.................................c.............. [177040]
AY261364.1_Tengani_62            ..............................ga................................c.............. [177040]
AY261363.1_Pretoriuskop/96/4     ..............................g.................................c.............. [177040]
AY261362.1_Mkuzi_1979            ..............................g.................................c.............. [177040]
AY261361.1_Malawi_Lil-20/1_1983  ............t.................a..............................c.................. [177040]
AY261360.1_Kenya_1950            ............tc.........c......a...g..a.c.....................c..t............... [177040]
AM712240.1_OURT_88/3             ..............................g................................................. [177040]
AM712239.1_Benin_97/1            ..............................g................................................. [177040]
KM111295.1_Ken06.Bus             ............t..........c......a...g..a.......................c..t............... [177040]
KM111294.1_Ken05/Tk1             ............tc.........c......a...g..a.......................c..t............... [177040]
KM262844.1_strain_L60            ..............................g................................................. [177040]
KM102979.1_26544/OG10_Italy      ..............................g................................................. [177040]
KM262845.1_strain_NHV            ..............................g................................................. [177040]
KX354450.1_47/Ss/2008            ..............................g................................................. [177040]
KP055815.1_strain_BA71           ..............................g................................................. [177040]
MH025920.1_strain_R35            ............t..........c......a...g..a.......................c..t............... [177040]
MH025919.1_strain_N10            ............t..........c......a...g..a.......................c..t............... [177040]
MH025918.1_strain_R25            ............t..........c......a...g..a.......................c..t............... [177040]
MH025917.1_strain_R7             ............t..........c......a...g..a.......................c..t............... [177040]
MH025916.1_strain_R8             ............t..........c......a...g..a.......................c..t............... [177040]

FR682468.2_Georgia_2007/1        cctattccagcaaaggatataatggcctccattctcacgctggacgggttatatgcagaggttccaaaattcttaccaga [177120]
ASF_IND_20_CAD543                ................................................................................ [177120]
LR899193.1_Germany_2020/1        ................................................................................ [177120]
MN393477.1_Wuhan_2019            ................................................................................ [177120]
MW396979.1_Timor-Leste/2019/1    ................................................................................ [177120]
MK940252.1_CN/2019/InnerMongolia ................................................................................ [177120]
MK128995.1_China/2018/AnhuiXCGQ  ................................................................................ [177120]
MH681419.1_ASFV/POL/2015         ................................................................................ [177120]
LS478113.1_Estonia_2014          ................................................................................ [177120]
KP843857.1_Odintsovo_02/14       ................................................................................ [177120]
```

```
FN557520.1_strain_E75            ....................................g.a.........c............................... [177120]
AY261366.1_Warthog               ........g.......................................c............................... [177120]
AY261365.1_Warmbaths             ................................................c............................... [177120]
AY261364.1_Tengani_62            ........g.......................................c............................... [177120]
AY261363.1_Pretoriuskop/96/4     ................................................c............................... [177120]
AY261362.1_Mkuzi_1979            ................................................c............................... [177120]
AY261361.1_Malawi_Lil-20/1_1983  ........g..........a...............tg.a.........c............................... [177120]
AY261360.1_Kenya_1950            ........g.............................a.........c............................... [177120]
AM712240.1_OURT_88/3             ....................................g.a.........c............................... [177120]
AM712239.1_Benin_97/1            ....................................g.a.........c............................... [177120]
KM111295.1_Ken06.Bus             ........g.............................a.........c............................... [177120]
KM111294.1_Ken05/Tk1             ........g.............................a.........c............................... [177120]
KM262844.1_strain_L60            ....................................g.a.........c............................... [177120]
KM102979.1_26544/OG10_Italy      ....................................g.a.........c............................... [177120]
KM262845.1_strain_NHV            ....................................g.a.........c............................... [177120]
KX354450.1_47/Ss/2008            ....................................g.a.........c............................... [177120]
KP055815.1_strain_BA71           ....................................g.a.........c............................... [177120]
MH025920.1_strain_R35            ........g.............................a.........c............................... [177120]
MH025919.1_strain_N10            ........g.............................a.........c............................... [177120]
MH025918.1_strain_R25            ........g.............................a.........c............................... [177120]
MH025917.1_strain_R7             ........g.............................a.........c............................... [177120]
MH025916.1_strain_R8             ........g.............................a.........c............................... [177120]

FR682468.2_Georgia_2007/1        ggcgttacgagagggctgtgctggcaagaatcctctaagcttttatattcaacaaattttaaatttaatgggatgtgacg [177200]
ASF_IND_20_CAD543                ................................................................................ [177200]
LR899193.1_Germany_2020/1        ................................................................................ [177200]
MN393477.1_Wuhan_2019            ................................................................................ [177200]
MW396979.1_Timor-Leste/2019/1    ................................................................................ [177200]
MK940252.1_CN/2019/InnerMongolia ................................................................................ [177200]
MK128995.1_China/2018/AnhuiXCGQ  ................................................................................ [177200]
MH681419.1_ASFV/POL/2015         ................................................................................ [177200]
LS478113.1_Estonia_2014          ................................................................................ [177200]
KP843857.1_Odintsovo_02/14       ................................................................................ [177200]
FN557520.1_strain_E75            ................................................................................ [177200]
AY261366.1_Warthog               ...............g................................................................ [177200]
AY261365.1_Warmbaths             ...............a................................................................ [177200]
AY261364.1_Tengani_62            ...............g................................................................ [177200]
AY261363.1_Pretoriuskop/96/4     ...............a................................................................ [177200]
AY261362.1_Mkuzi_1979            ................................................................................ [177200]
AY261361.1_Malawi_Lil-20/1_1983  ...............g..............a...........c..................................... [177200]
AY261360.1_Kenya_1950            ...............a..c...........a................................................. [177200]
AM712240.1_OURT_88/3             ................................................................................ [177200]
AM712239.1_Benin_97/1            ................................................................................ [177200]
KM111295.1_Ken06.Bus             ...............a..c...........a................................................. [177200]
```

```
KM111294.1_Ken05/Tk1             ...............a..c...........a...............................................t. [177200]
KM262844.1_strain_L60            ................................................................................ [177200]
KM102979.1_26544/OG10_Italy      ................................................................................ [177200]
KM262845.1_strain_NHV            ................................................................................ [177200]
KX354450.1_47/Ss/2008            ................................................................................ [177200]
KP055815.1_strain_BA71           ................................................................................ [177200]
MH025920.1_strain_R35            ...............a..c...........a................................................. [177200]
MH025919.1_strain_N10            ...............a..c...........a................................................. [177200]
MH025918.1_strain_R25            ...............a..c...........a................................................. [177200]
MH025917.1_strain_R7             ...............a..c...........a................................................. [177200]
MH025916.1_strain_R8             ...............a..c...........a................................................. [177200]

FR682468.2_Georgia_2007/1        gtaacgagtaccatgttctttttaccagcagctccgaggaagcaaatactcatatgatcatggccgccgtgcgtcgccat [177280]
ASF_IND_20_CAD543                ................................................................................ [177280]
LR899193.1_Germany_2020/1        ................................................................................ [177280]
MN393477.1_Wuhan_2019            ................................................................................ [177280]
MW396979.1_Timor-Leste/2019/1    ................................................................................ [177280]
MK940252.1_CN/2019/InnerMongolia ................................................................................ [177280]
MK128995.1_China/2018/AnhuiXCGQ  ................................................................................ [177280]
MH681419.1_ASFV/POL/2015         ................................................................................ [177280]
LS478113.1_Estonia_2014          ................................................................................ [177280]
KP843857.1_Odintsovo_02/14       ................................................................................ [177280]
FN557520.1_strain_E75            ..........................g.......t............................................. [177280]
AY261366.1_Warthog               ............................................................................g... [177280]
AY261365.1_Warmbaths             ..........t..................................................................... [177280]
AY261364.1_Tengani_62            .......................................................................t........ [177280]
AY261363.1_Pretoriuskop/96/4     ................................................................................ [177280]
AY261362.1_Mkuzi_1979            ..........................g.......t............................................. [177280]
AY261361.1_Malawi_Lil-20/1_1983  ..g.........................................................c...............g... [177280]
AY261360.1_Kenya_1950            .c................................t..............g................t......c..g... [177280]
AM712240.1_OURT_88/3             ..........................g.......t............................................. [177280]
AM712239.1_Benin_97/1            ..........................g.......t............................................. [177280]
KM111295.1_Ken06.Bus             .c................................t.......................................c..g... [177280]
KM111294.1_Ken05/Tk1             .c................................t..............g.......................c..g... [177280]
KM262844.1_strain_L60            ..........................g.......t............................................. [177280]
KM102979.1_26544/OG10_Italy      ..........................g.......t............................................. [177280]
KM262845.1_strain_NHV            ..........................g.......t............................................. [177280]
KX354450.1_47/Ss/2008            ..........................g.......t............................................. [177280]
KP055815.1_strain_BA71           ..........................g.......t............................................. [177280]
MH025920.1_strain_R35            .c................................t.......................................c..g... [177280]
MH025919.1_strain_N10            .c................................t.......................................c..g... [177280]
MH025918.1_strain_R25            .c................................t.......................................c..g... [177280]
MH025917.1_strain_R7             .c................................t.......................................c..g... [177280]
MH025916.1_strain_R8             .c................................t.......................................c..g... [177280]
```

```
FR682468.2_Georgia_2007/1        ttgctgcggacgcagcaaaggcctcatgtcattatcggagcagccgagccccctagcgtcaccgaatgtgtgaaggcatt [177360]
ASF_IND_20_CAD543                ................................................................................ [177360]
LR899193.1_Germany_2020/1        ................................................................................ [177360]
MN393477.1_Wuhan_2019            ................................................................................ [177360]
MW396979.1_Timor-Leste/2019/1    ................................................................................ [177360]
MK940252.1_CN/2019/InnerMongolia ................................................................................ [177360]
MK128995.1_China/2018/AnhuiXCGQ  ................................................................................ [177360]
MH681419.1_ASFV/POL/2015         ................................................................................ [177360]
LS478113.1_Estonia_2014          ................................................................................ [177360]
KP843857.1_Odintsovo_02/14       ................................................................................ [177360]
FN557520.1_strain_E75            ....................a........................................................... [177360]
AY261366.1_Warthog               .............gc................................................................. [177360]
AY261365.1_Warmbaths             ................................................................................ [177360]
AY261364.1_Tengani_62            ..............c..............................................................g.. [177360]
AY261363.1_Pretoriuskop/96/4     .................g.........................g.............a...................... [177360]
AY261362.1_Mkuzi_1979            ....................a........................................................... [177360]
AY261361.1_Malawi_Lil-20/1_1983  .............gc........................................................a..a..... [177360]
AY261360.1_Kenya_1950            ..............a........................................................a........ [177360]
AM712240.1_OURT_88/3             ....................a........................................................... [177360]
AM712239.1_Benin_97/1            ....................a........................................................... [177360]
KM111295.1_Ken06.Bus             ...........a..a........................................................a........ [177360]
KM111294.1_Ken05/Tk1             ..............a........................................................a........ [177360]
KM262844.1_strain_L60            ....................a........................................................... [177360]
KM102979.1_26544/OG10_Italy      ....................a........................................................... [177360]
KM262845.1_strain_NHV            ....................a........................................................... [177360]
KX354450.1_47/Ss/2008            ....................a........................................................... [177360]
KP055815.1_strain_BA71           ....................a........................................................... [177360]
MH025920.1_strain_R35            ...........a..a........................................................a........ [177360]
MH025919.1_strain_N10            ...........a..a........................................................a........ [177360]
MH025918.1_strain_R25            ...........a..a........................................................a........ [177360]
MH025917.1_strain_R7             ...........a..a........................................................a........ [177360]
MH025916.1_strain_R8             ...........a..a........................................................a........ [177360]

FR682468.2_Georgia_2007/1        ggcgcaggaaaaacgctgcgtatacaccatcatcccccctaaaaaatttgaaatagatcctgttgcggtatacgatgcca [177440]
ASF_IND_20_CAD543                ................................................................................ [177440]
LR899193.1_Germany_2020/1        ................................................................................ [177440]
MN393477.1_Wuhan_2019            ................................................................................ [177440]
MW396979.1_Timor-Leste/2019/1    ................................................................................ [177440]
MK940252.1_CN/2019/InnerMongolia ................................................................................ [177440]
MK128995.1_China/2018/AnhuiXCGQ  ................................................................................ [177440]
MH681419.1_ASFV/POL/2015         ................................................................................ [177440]
LS478113.1_Estonia_2014          ................................................................................ [177440]
KP843857.1_Odintsovo_02/14       ................................................................................ [177440]
```

```
FN557520.1_strain_E75           ................................................................................ [177440]
AY261366.1_Warthog              ................................................................................ [177440]
AY261365.1_Warmbaths            ................................................................................ [177440]
AY261364.1_Tengani_62           a....................................................................g.......... [177440]
AY261363.1_Pretoriuskop/96/4    ..................t...........t................................................. [177440]
AY261362.1_Mkuzi_1979           ................................................................................ [177440]
AY261361.1_Malawi_Lil-20/1_1983 .....g...............g..t....................................................... [177440]
AY261360.1_Kenya_1950           .....................g...............................................c.......... [177440]
AM712240.1_OURT_88/3            ................................................................................ [177440]
AM712239.1_Benin_97/1           ................................................................................ [177440]
KM111295.1_Ken06.Bus            .....................g.......................................................... [177440]
KM111294.1_Ken05/Tk1            .....................g...............................................c.......... [177440]
KM262844.1_strain_L60           ................................................................................ [177440]
KM102979.1_26544/OG10_Italy     ................................................................................ [177440]
KM262845.1_strain_NHV           ................................................................................ [177440]
KX354450.1_47/Ss/2008           ................................................................................ [177440]
KP055815.1_strain_BA71          ................................................................................ [177440]
MH025920.1_strain_R35           .....................g.......................................................... [177440]
MH025919.1_strain_N10           .....................g.......................................................... [177440]
MH025918.1_strain_R25           .....................g.......................................................... [177440]
MH025917.1_strain_R7            .....................g.......................................................... [177440]
MH025916.1_strain_R8            .....................g.......................................................... [177440]

FR682468.2_Georgia_2007/1       tacaaagcaatacctgcttagcgtgcatttcaggcactaatgctgttgtcaaaacgttcaacaaactccaggacatcagc [177520]
ASF_IND_20_CAD543               ................................................................................ [177520]
LR899193.1_Germany_2020/1       ................................................................................ [177520]
MN393477.1_Wuhan_2019           ................................................................................ [177520]
MW396979.1_Timor-Leste/2019/1   ................................................................................ [177520]
MK940252.1_CN/2019/InnerMongolia ................................................................................ [177520]
MK128995.1_China/2018/AnhuiXCGQ ................................................................................ [177520]
MH681419.1_ASFV/POL/2015        ................................................................................ [177520]
LS478113.1_Estonia_2014         ................................................................................ [177520]
KP843857.1_Odintsovo_02/14      ................................................................................ [177520]
FN557520.1_strain_E75           .....................................c.......................................... [177520]
AY261366.1_Warthog              ...............................g.....c......................................t... [177520]
AY261365.1_Warmbaths            ...............................g.....c......................................t... [177520]
AY261364.1_Tengani_62           ...............................g.....c.............................t........t... [177520]
AY261363.1_Pretoriuskop/96/4    ...............................g.....c......................................t... [177520]
AY261362.1_Mkuzi_1979           ...............................g.....c......................................t... [177520]
AY261361.1_Malawi_Lil-20/1_1983 ...............................g.....c................................a.....t... [177520]
AY261360.1_Kenya_1950           ...............................g.....c...................................a..t... [177520]
AM712240.1_OURT_88/3            .....................................c.......................................... [177520]
AM712239.1_Benin_97/1           .....................................c.......................................... [177520]
KM111295.1_Ken06.Bus            ...............................g.....c...................................a..t... [177520]
```

KM111294.1_Ken05/Tk1 ...............................g.....c...................................a..t... [177520]
KM262844.1_strain_L60 .....................................c.......................................... [177520]
KM102979.1_26544/OG10_Italy .....................................c.......................................... [177520]
KM262845.1_strain_NHV .....................................c.......................................... [177520]
KX354450.1_47/Ss/2008 .....................................c.......................................... [177520]
KP055815.1_strain_BA71 .....................................c.......................................... [177520]
MH025920.1_strain_R35 ...............................g.....c...................................a..t... [177520]
MH025919.1_strain_N10 ...............................g.....c...................................a..t... [177520]
MH025918.1_strain_R25 ...............................g.....c...................................a..t... [177520]
MH025917.1_strain_R7 ...............................g.....c...................................a..t... [177520]
MH025916.1_strain_R8 ...............................g.....c...................................a..t... [177520]

FR682468.2_Georgia_2007/1 aacgtgttaaaaggtattcccctgcactcagaagtgagtgatcttgtttatcaaggatgtattcaacaaaatccgcccgc [177600]
ASF_IND_20_CAD543 ................................................................................ [177600]
LR899193.1_Germany_2020/1 ................................................................................ [177600]
MN393477.1_Wuhan_2019 ................................................................................ [177600]
MW396979.1_Timor-Leste/2019/1 ................................................................................ [177600]
MK940252.1_CN/2019/InnerMongolia ................................................................................ [177600]
MK128995.1_China/2018/AnhuiXCGQ ................................................................................ [177600]
MH681419.1_ASFV/POL/2015 ................................................................................ [177600]
LS478113.1_Estonia_2014 ................................................................................ [177600]
KP843857.1_Odintsovo_02/14 ................................................................................ [177600]
FN557520.1_strain_E75 ..a......................................g.....................a..........t..... [177600]
AY261366.1_Warthog ..a.......g..................g....................c............a................ [177600]
AY261365.1_Warmbaths ..a.......g..................g....................c............a..........a..... [177600]
AY261364.1_Tengani_62 ..a......gg..................g....................c............................. [177600]
AY261363.1_Pretoriuskop/96/4 ..a.......g..................g....................c............a................ [177600]
AY261362.1_Mkuzi_1979 ..a.......g..................g....................c............a..........a..... [177600]
AY261361.1_Malawi_Lil-20/1_1983 ..........g......a........t..g.............................c.................... [177600]
AY261360.1_Kenya_1950 ..a...c..gg..c............t..g...a........g....................a.....c.......... [177600]
AM712240.1_OURT_88/3 ..a......................................g.....................a..........t..... [177600]
AM712239.1_Benin_97/1 ..a......................................g.....................a..........t..... [177600]
KM111295.1_Ken06.Bus ..a...c..gg..c............t..g...a........g....................a.....c.......t.. [177600]
KM111294.1_Ken05/Tk1 ..a...c..gg..c............t..g...a........g....................a.....c.......... [177600]
KM262844.1_strain_L60 ..a......................................g.....................a..........t..... [177600]
KM102979.1_26544/OG10_Italy ..a......................................g.....................a..........t..... [177600]
KM262845.1_strain_NHV ..a......................................g.....................a..........t..... [177600]
KX354450.1_47/Ss/2008 ..a......................................g.....................a..........t..... [177600]
KP055815.1_strain_BA71 ..a......................................g.....................a..........t..... [177600]
MH025920.1_strain_R35 ..a...c..gg..c............t..g...a........g....................a.....c.......t.. [177600]
MH025919.1_strain_N10 ..a...c..gg..c............t..g...a........g....................a.....c.......t.. [177600]
MH025918.1_strain_R25 ..a...c..gg..c............t..g...a........g....................a.....c.......t.. [177600]
MH025917.1_strain_R7 ..a...c..gg..c............t..g...a........g....................a.....c.......t.. [177600]
MH025916.1_strain_R8 ..a...c..gg..c............t..g...a........g....................a.....c.......t.. [177600]

```
FR682468.2_Georgia_2007/1        tgatagtttttcaataaatagtctctacggcttcctgggagtcggtgttttgggaatgaagaaaaaggtcatgcaaggat [177680]
ASF_IND_20_CAD543                ................................................y...........r................... [177680]
LR899193.1_Germany_2020/1        ................................................................................ [177680]
MN393477.1_Wuhan_2019            ................................................................................ [177680]
MW396979.1_Timor-Leste/2019/1    ................................................................................ [177680]
MK940252.1_CN/2019/InnerMongolia ................................................................................ [177680]
MK128995.1_China/2018/AnhuiXCGQ  ................................................................................ [177680]
MH681419.1_ASFV/POL/2015         ................................................................................ [177680]
LS478113.1_Estonia_2014          ................................................................................ [177680]
KP843857.1_Odintsovo_02/14       ................................................................................ [177680]
FN557520.1_strain_E75            ..................................t....g........c...........a................... [177680]
AY261366.1_Warthog               ..................................t.............c............................... [177680]
AY261365.1_Warmbaths             ..................................t.............c............................... [177680]
AY261364.1_Tengani_62            ..................................t.............c............................... [177680]
AY261363.1_Pretoriuskop/96/4     .....................c............t...........a.c............................... [177680]
AY261362.1_Mkuzi_1979            ..................................t.............c............................... [177680]
AY261361.1_Malawi_Lil-20/1_1983  .............t.g...........t..t...t....g..a.....c............................ac. [177680]
AY261360.1_Kenya_1950            g..c.........t.......c.....t..............a..c..cc.......a..........c..........c [177680]
AM712240.1_OURT_88/3             ..................................t....g........c...........a................... [177680]
AM712239.1_Benin_97/1            ..................................t....g........c...........a................... [177680]
KM111295.1_Ken06.Bus             g..c.........t.......c.....t..............a..ca.cc.......a..a.......c......g.... [177680]
KM111294.1_Ken05/Tk1             g..c.........t.......c.....t..............a..c..cc.......a..........c..........c [177680]
KM262844.1_strain_L60            ..................................t....g........c...........a................... [177680]
KM102979.1_26544/OG10_Italy      ..................................t....g........c...........a................... [177680]
KM262845.1_strain_NHV            ..................................t....g........c...........a................... [177680]
KX354450.1_47/Ss/2008            ..................................t....g........c...........a................... [177680]
KP055815.1_strain_BA71           ..................................t....g........c...........a................... [177680]
MH025920.1_strain_R35            g..c.........t.......c.....t..............a..ca.cc.......a..a.......c......g.... [177680]
MH025919.1_strain_N10            g..c.........t.......c.....t..............a..ca.cc.......a..a.......c......g.... [177680]
MH025918.1_strain_R25            g..c.........t.......c.....t..............a..ca.cc.......a..a.......c......g.... [177680]
MH025917.1_strain_R7             g..c.........t.......c.....t..............a..ca.cc.......a..a.......c......g.... [177680]
MH025916.1_strain_R8             g..c.........t.......c.....t..............a..ca.cc.......a..a.......c......g.... [177680]

FR682468.2_Georgia_2007/1        tggggccgctcatttttggaggagggctgagaggcggaagccctaatatacccggaattcatgccatgtataaaacgcta [177760]
ASF_IND_20_CAD543                ................................................................................ [177760]
LR899193.1_Germany_2020/1        ................................................................................ [177760]
MN393477.1_Wuhan_2019            ................................................................................ [177760]
MW396979.1_Timor-Leste/2019/1    ................................................................................ [177760]
MK940252.1_CN/2019/InnerMongolia ................................................................................ [177760]
MK128995.1_China/2018/AnhuiXCGQ  ................................................................................ [177760]
MH681419.1_ASFV/POL/2015         ................................................................................ [177760]
LS478113.1_Estonia_2014          ................................................................................ [177760]
KP843857.1_Odintsovo_02/14       ................................................................................ [177760]
```

```
FN557520.1_strain_E75             ......................g.................................................g....... [177760]
AY261366.1_Warthog                ........................................................................g....... [177760]
AY261365.1_Warmbaths              ........................................................................g....... [177760]
AY261364.1_Tengani_62             ..........................................................a.............g....... [177760]
AY261363.1_Pretoriuskop/96/4      ........................................................................g....... [177760]
AY261362.1_Mkuzi_1979             ........................................................................g....... [177760]
AY261361.1_Malawi_Lil-20/1_1983   ................................................................t.......g...a... [177760]
AY261360.1_Kenya_1950             ....a....................a.....g...............g............................at.. [177760]
AM712240.1_OURT_88/3              ......................g.................................................g....... [177760]
AM712239.1_Benin_97/1             ......................g.................................................g....... [177760]
KM111295.1_Ken06.Bus              ....a....................a.....g...............g.................t..........at.. [177760]
KM111294.1_Ken05/Tk1              ....a....................a.....g...............g............................at.. [177760]
KM262844.1_strain_L60             ......................g.................................................g....... [177760]
KM102979.1_26544/OG10_Italy       ......................g.................................................g....... [177760]
KM262845.1_strain_NHV             ......................g.................................................g....... [177760]
KX354450.1_47/Ss/2008             ......................g.................................................g....... [177760]
KP055815.1_strain_BA71            ......................g.................................................g....... [177760]
MH025920.1_strain_R35             ....a....................a.....g...............g.................t..........at.. [177760]
MH025919.1_strain_N10             ....a....................a.....g...............g.................t..........at.. [177760]
MH025918.1_strain_R25             ....a....................a.....g...............g.................t..........at.. [177760]
MH025917.1_strain_R7              ....a....................a.....g...............g.................t..........at.. [177760]
MH025916.1_strain_R8              ....a....................a.....g...............g.................t..........at.. [177760]

FR682468.2_Georgia_2007/1         acccagcaaaggccttctatgaaaaaaataaatacaatacatacgctgttcatgaaaactttaaaaaaacatcagcatgt [177840]
ASF_IND_20_CAD543                 .................y.............................................................. [177840]
LR899193.1_Germany_2020/1         ................................................................................ [177840]
MN393477.1_Wuhan_2019             ................................................................................ [177840]
MW396979.1_Timor-Leste/2019/1     ................................................................................ [177840]
MK940252.1_CN/2019/InnerMongolia  ................................................................................ [177840]
MK128995.1_China/2018/AnhuiXCGQ   ................................................................................ [177840]
MH681419.1_ASFV/POL/2015          ................................................................................ [177840]
LS478113.1_Estonia_2014           ................................................................................ [177840]
KP843857.1_Odintsovo_02/14        ................................................................................ [177840]
FN557520.1_strain_E75             .................c.........................a.................................... [177840]
AY261366.1_Warthog                .................c.........................a..................................a. [177840]
AY261365.1_Warmbaths              .................c.........................g......t............................. [177840]
AY261364.1_Tengani_62             .................c.........................a.................................... [177840]
AY261363.1_Pretoriuskop/96/4      .................c..a......................a......t............................. [177840]
AY261362.1_Mkuzi_1979             .................c.........................a.................................... [177840]
AY261361.1_Malawi_Lil-20/1_1983   .t.........a........c......................a......t..a.....a.................... [177840]
AY261360.1_Kenya_1950             .....a........c..c..a..............tg......a..............t..................... [177840]
AM712240.1_OURT_88/3              .................c.........................a.................................... [177840]
AM712239.1_Benin_97/1             .................c.........................a.................................... [177840]
KM111295.1_Ken06.Bus              ..............c..c..a..............tg......a..............t..................... [177840]
```

```
KM111294.1_Ken05/Tk1              .....a........c..c..a..............tg......a.t............t..................... [177840]
KM262844.1_strain_L60             .................c.........................a.................................... [177840]
KM102979.1_26544/OG10_Italy       .................c.........................a.................................... [177840]
KM262845.1_strain_NHV             .................c.........................a.................................... [177840]
KX354450.1_47/Ss/2008             .................c.........................a.................................... [177840]
KP055815.1_strain_BA71            .................c.........................a.................................... [177840]
MH025920.1_strain_R35             ..............c..c..a..............tg......a.............tt...................a. [177840]
MH025919.1_strain_N10             ..............c..c..a..............tg......a.............tt...................a. [177840]
MH025918.1_strain_R25             ..............c..c..a..............tg......a.............tt...................a. [177840]
MH025917.1_strain_R7              ..............c..c..a..............tg......a.............tt...................a. [177840]
MH025916.1_strain_R8              ..............c..c..a..............tg......a.............tt...................a. [177840]

FR682468.2_Georgia_2007/1         atatctacccatagggggcgtgtctgcagaggacacgtctgcagaaaacatatctacaaaagacatgcct---------- [177920]
ASF_IND_20_CAD543                 ......................................................................---------- [177920]
LR899193.1_Germany_2020/1         ......................................................................---------- [177920]
MN393477.1_Wuhan_2019             ......................................................................---------- [177920]
MW396979.1_Timor-Leste/2019/1     ......................................................................---------- [177920]
MK940252.1_CN/2019/InnerMongolia  ......................................................................---------- [177920]
MK128995.1_China/2018/AnhuiXCGQ   ......................................................................---------- [177920]
MH681419.1_ASFV/POL/2015          ......................................................................---------- [177920]
LS478113.1_Estonia_2014           ......................................................................---------- [177920]
KP843857.1_Odintsovo_02/14        ......................................................................---------- [177920]
FN557520.1_strain_E75             g............-....g...............................................a...---------- [177920]
AY261366.1_Warthog                .........g.........................t.........g..---c...g..........a..ctctaccgacg [177920]
AY261365.1_Warmbaths              ......................------------------------------..............a..ctctaccgaca [177920]
AY261364.1_Tengani_62             ...................................t....a..a..g....g...g..........a..ctctaccgaca [177920]
AY261363.1_Pretoriuskop/96/4      ...................................t....a..a..g....g..............a..ctctactgaca [177920]
AY261362.1_Mkuzi_1979             .............................................................t....a...---------- [177920]
AY261361.1_Malawi_Lil-20/1_1983   g.........---------------------.gttt....at...gg..g.gc.ct.t..c.g.......---------- [177920]
AY261360.1_Kenya_1950             g.....g.......a....a.....t.ta.c.g..t.c.ct.ta.cgg...gc.ct.t.gt.g...a...---------- [177920]
AM712240.1_OURT_88/3              g.................g...............................................a...---------- [177920]
AM712239.1_Benin_97/1             g.................g...............................................a...---------- [177920]
KM111295.1_Ken06.Bus              g.....g.......a.------------------------------gg...gc.ct.t..t.g...a...---------- [177920]
KM111294.1_Ken05/Tk1              g.....g.......a....a.....t.ta.c.g..t.c.ct.ta.tgg...gc.ct.t..t.g...a...---------- [177920]
KM262844.1_strain_L60             g.................g...............................................a...---------- [177920]
KM102979.1_26544/OG10_Italy       g........a........g...............................................a...---------- [177920]
KM262845.1_strain_NHV             g.................g...............................................a...---------- [177920]
KX354450.1_47/Ss/2008             g.................g...............................................a...---------- [177920]
KP055815.1_strain_BA71            g.................g...............................................a...---------- [177920]
MH025920.1_strain_R35             g.....g.......a.------------------------------gg...gc.ct.t..t.g...a...---------- [177920]
MH025919.1_strain_N10             g.....g.......a.------------------------------gg...gc.ct.t..t.g...a...---------- [177920]
MH025918.1_strain_R25             g.....g.......a.------------------------------gg...gc.ct.t..t.g...a...---------- [177920]
MH025917.1_strain_R7              g.....g.......a.------------------------------gg...gc.ct.t..t.g...a...---------- [177920]
MH025916.1_strain_R8              g.....g.......a.------------------------------gg...gc.ct.t..t.g...a...---------- [177920]
```

```
FR682468.2_Georgia_2007/1        --gttgaaggcccgaagggactcccgggctatattttatttagcgttggccgtcgcgccgaggagctacaaaaaaaaatt [178000]
ASF_IND_20_CAD543                --...........................................................r..a............... [178000]
LR899193.1_Germany_2020/1        --.............................................................................. [178000]
MN393477.1_Wuhan_2019            --.............................................................................. [178000]
MW396979.1_Timor-Leste/2019/1    --.............................................................................. [178000]
MK940252.1_CN/2019/InnerMongolia --.............................................................................. [178000]
MK128995.1_China/2018/AnhuiXCGQ  --.............................................................................. [178000]
MH681419.1_ASFV/POL/2015         --.............................................................................. [178000]
LS478113.1_Estonia_2014          --.............................................................................. [178000]
KP843857.1_Odintsovo_02/14       --.............................................................................. [178000]
FN557520.1_strain_E75            --................a..............................t...........a..a............... [178000]
AY261366.1_Warthog               tgcc..tt.........................................t.a............................ [178000]
AY261365.1_Warmbaths             tgcc..tt..........a..............................t.............................. [178000]
AY261364.1_Tengani_62            tgcc..tt.........................................t.............................. [178000]
AY261363.1_Pretoriuskop/96/4     tgc...tt.........................................t.a......t..................... [178000]
AY261362.1_Mkuzi_1979            --................a..............................t...........a..a............... [178000]
AY261361.1_Malawi_Lil-20/1_1983  --.......t.......................................t.a.tc...t.....a..............a [178000]
AY261360.1_Kenya_1950            --...............a.c........t....................t...tc........................a [178000]
AM712240.1_OURT_88/3             --................a..............................t...........a..a............... [178000]
AM712239.1_Benin_97/1            --................a..............................t...........a..a............... [178000]
KM111295.1_Ken06.Bus             --...............a.c........t....................t...tc........................a [178000]
KM111294.1_Ken05/Tk1             --...............a.c........t....................t...tc........................a [178000]
KM262844.1_strain_L60            --................a..............................t...........a..a............... [178000]
KM102979.1_26544/OG10_Italy      --................a..............................t...........a..a............... [178000]
KM262845.1_strain_NHV            --................a..............................t...........a..a............... [178000]
KX354450.1_47/Ss/2008            --................a..............................t...........a..a............... [178000]
KP055815.1_strain_BA71           --................a..............................t...........a..a............... [178000]
MH025920.1_strain_R35            --...............a.c........t....................t...tc........................a [178000]
MH025919.1_strain_N10            --...............a.c........t....................t...tc........................a [178000]
MH025918.1_strain_R25            --...............a.c........t....................t...tc........................a [178000]
MH025917.1_strain_R7             --...............a.c........t....................t...tc........................a [178000]
MH025916.1_strain_R8             --...............a.c........t....................t...tc........................a [178000]

FR682468.2_Georgia_2007/1        ttcactaaatttaatataaaggttggccgtgttgttgacttacaagagatactgtttcgtatcaaaataccccaaaaata [178080]
ASF_IND_20_CAD543                ................................................................................ [178080]
LR899193.1_Germany_2020/1        ................................................................................ [178080]
MN393477.1_Wuhan_2019            ................................................................................ [178080]
MW396979.1_Timor-Leste/2019/1    ................................................................................ [178080]
MK940252.1_CN/2019/InnerMongolia ................................................................................ [178080]
MK128995.1_China/2018/AnhuiXCGQ  ................................................................................ [178080]
MH681419.1_ASFV/POL/2015         ................................................................................ [178080]
LS478113.1_Estonia_2014          ................................................................................ [178080]
KP843857.1_Odintsovo_02/14       ................................................................................ [178080]
```

```
FN557520.1_strain_E75           ..............................a................................................. [178080]
AY261366.1_Warthog              ..............................a.....................................c.....g..... [178080]
AY261365.1_Warmbaths            ..............................a.........c.................................g..... [178080]
AY261364.1_Tengani_62           ..............................a...........................................g..... [178080]
AY261363.1_Pretoriuskop/96/4    .....................a........a...........................................g..... [178080]
AY261362.1_Mkuzi_1979           ..............................a.........c....................................... [178080]
AY261361.1_Malawi_Lil-20/1_1983 ..t............g.......a......a.....a.............g..a.......................... [178080]
AY261360.1_Kenya_1950           ...............g.......a......a.....a..........ag.g...........t........t........ [178080]
AM712240.1_OURT_88/3            ..............................a................................................. [178080]
AM712239.1_Benin_97/1           ..............................a................................................. [178080]
KM111295.1_Ken06.Bus            ..t............g.......a......a.....a..........ag.g...........t........t........ [178080]
KM111294.1_Ken05/Tk1            ...............g.......a......a.....a..........ag.g...........t........t........ [178080]
KM262844.1_strain_L60           ..............................a................................................. [178080]
KM102979.1_26544/OG10_Italy     ..............................a................................................. [178080]
KM262845.1_strain_NHV           ..............................a................................................. [178080]
KX354450.1_47/Ss/2008           ..............................a................................................. [178080]
KP055815.1_strain_BA71          ..............................a......................a.......................... [178080]
MH025920.1_strain_R35           ..t............g.......a......a.....a..........ag.g...........t........t........ [178080]
MH025919.1_strain_N10           ..t............g.......a......a.....a..........ag.g...........t........t........ [178080]
MH025918.1_strain_R25           ..t............g.......a......a.....a..........ag.g...........t........t........ [178080]
MH025917.1_strain_R7            ..t............g.......a......a.....a..........ag.g...........t........t........ [178080]
MH025916.1_strain_R8            ..t............g.......a......a.....a..........ag.g...........t........t........ [178080]

FR682468.2_Georgia_2007/1       ctgggagacattattgttcatccaattaagagataatttgaccaaagaggacataaaaagagttatggttgttttgatgc [178160]
ASF_IND_20_CAD543               ...................................................y............................ [178160]
LR899193.1_Germany_2020/1       ................................................................................ [178160]
MN393477.1_Wuhan_2019           ................................................................................ [178160]
MW396979.1_Timor-Leste/2019/1   ................................................................................ [178160]
MK940252.1_CN/2019/InnerMongolia ................................................................................ [178160]
MK128995.1_China/2018/AnhuiXCGQ ................................................................................ [178160]
MH681419.1_ASFV/POL/2015        ................................................................................ [178160]
LS478113.1_Estonia_2014         ................................................................................ [178160]
KP843857.1_Odintsovo_02/14      ................................................................................ [178160]
FN557520.1_strain_E75           ...................................................t.....g...................... [178160]
AY261366.1_Warthog              ....................................c..............t.....g...................... [178160]
AY261365.1_Warmbaths            ...................................................t.....g...................... [178160]
AY261364.1_Tengani_62           ...................................................t............................ [178160]
AY261363.1_Pretoriuskop/96/4    ...................................................t......c..................... [178160]
AY261362.1_Mkuzi_1979           ................................................................................ [178160]
AY261361.1_Malawi_Lil-20/1_1983 ..................t..t............gt.c....t........t.....g.....................t [178160]
AY261360.1_Kenya_1950           ..................t..............gg..c.............t.....g..........c.a........t [178160]
AM712240.1_OURT_88/3            ...................................................t.....g...................... [178160]
AM712239.1_Benin_97/1           ...................................................t.....g...................... [178160]
KM111295.1_Ken06.Bus            ..................t..............gg..c.a..t......a.t................c.a........t [178160]
```

```
KM111294.1_Ken05/Tk1             ..................t..............gg..c.............t.....g..........c.a........t [178160]
KM262844.1_strain_L60            ...................................................t.....g...................... [178160]
KM102979.1_26544/OG10_Italy      ...................................................t.....g...................... [178160]
KM262845.1_strain_NHV            ...................................................t.....g...................... [178160]
KX354450.1_47/Ss/2008            ...................................................t.....g...................... [178160]
KP055815.1_strain_BA71           ...................................................t.....g...................... [178160]
MH025920.1_strain_R35            ..................t..............gg..c.a..t......a.t................c.a........t [178160]
MH025919.1_strain_N10            ..................t..............gg..c.a..t......a.t................c.a........t [178160]
MH025918.1_strain_R25            ..................t..............gg..c.a..t......a.t................c.a........t [178160]
MH025917.1_strain_R7             ..................t..............gg..c.a..t......a.t................c.a........t [178160]
MH025916.1_strain_R8             ..................t..............gg..c.a..t......a.t................c.a........t [178160]

FR682468.2_Georgia_2007/1        atttagataccatcactcctcgtggctctcttcctcctccgagccactcttcttctttttcttaatcgtttttgtttgtt [178240]
ASF_IND_20_CAD543                ............................................y................................... [178240]
LR899193.1_Germany_2020/1        ................................................................................ [178240]
MN393477.1_Wuhan_2019            ................................................................................ [178240]
MW396979.1_Timor-Leste/2019/1    ................................................................................ [178240]
MK940252.1_CN/2019/InnerMongolia ................................................................................ [178240]
MK128995.1_China/2018/AnhuiXCGQ  ................................................................................ [178240]
MH681419.1_ASFV/POL/2015         ................................................................................ [178240]
LS478113.1_Estonia_2014          ................................................................................ [178240]
KP843857.1_Odintsovo_02/14       ................................................................................ [178240]
FN557520.1_strain_E75            ............................................t................................... [178240]
AY261366.1_Warthog               ............................................t................................... [178240]
AY261365.1_Warmbaths             ............................................t.............................c..... [178240]
AY261364.1_Tengani_62            ................................................................................ [178240]
AY261363.1_Pretoriuskop/96/4     ............................................t................................... [178240]
AY261362.1_Mkuzi_1979            ............................................t................................... [178240]
AY261361.1_Malawi_Lil-20/1_1983  .............t.................................................................. [178240]
AY261360.1_Kenya_1950            ...........g.t..............................t................................... [178240]
AM712240.1_OURT_88/3             ............................................t................................... [178240]
AM712239.1_Benin_97/1            ............................................t................................... [178240]
KM111295.1_Ken06.Bus             ...........g.t..............................t................................... [178240]
KM111294.1_Ken05/Tk1             ...........g.t..............................t................................... [178240]
KM262844.1_strain_L60            ............................................t................................... [178240]
KM102979.1_26544/OG10_Italy      ............................................t................................... [178240]
KM262845.1_strain_NHV            ............................................t................................... [178240]
KX354450.1_47/Ss/2008            ............................................t................................... [178240]
KP055815.1_strain_BA71           ............................................t................................... [178240]
MH025920.1_strain_R35            ...........g.t..............................t................................... [178240]
MH025919.1_strain_N10            ...........g.t..............................t................................... [178240]
MH025918.1_strain_R25            ...........g.t..............................t................................... [178240]
MH025917.1_strain_R7             ...........g.t..............................t................................... [178240]
MH025916.1_strain_R8             ...........g.t..............................t................................... [178240]
```

```
FR682468.2_Georgia_2007/1        ctataataagggaaaagaactccgtgggatcttgttccccgtacaggttatctgcgaccataaggatgcttagaatggta [178320]
ASF_IND_20_CAD543                ......................y......................................................... [178320]
LR899193.1_Germany_2020/1        ................................................................................ [178320]
MN393477.1_Wuhan_2019            ................................................................................ [178320]
MW396979.1_Timor-Leste/2019/1    ................................................................................ [178320]
MK940252.1_CN/2019/InnerMongolia ................................................................................ [178320]
MK128995.1_China/2018/AnhuiXCGQ  ................................................................................ [178320]
MH681419.1_ASFV/POL/2015         ................................................................................ [178320]
LS478113.1_Estonia_2014          ................................................................................ [178320]
KP843857.1_Odintsovo_02/14       ................................................................................ [178320]
FN557520.1_strain_E75            ......................t......................................................... [178320]
AY261366.1_Warthog               ................................................................t............... [178320]
AY261365.1_Warmbaths             ................................................................t............... [178320]
AY261364.1_Tengani_62            ................................................................t............... [178320]
AY261363.1_Pretoriuskop/96/4     ................................................................t............... [178320]
AY261362.1_Mkuzi_1979            ......................t.........................................t............... [178320]
AY261361.1_Malawi_Lil-20/1_1983  ......................t.................a.......................t...t........... [178320]
AY261360.1_Kenya_1950            ......................t.......................a.................t...t....g...... [178320]
AM712240.1_OURT_88/3             ......................t......................................................... [178320]
AM712239.1_Benin_97/1            ......................t......................................................... [178320]
KM111295.1_Ken06.Bus             ......................t.......................a.................t...t....g...... [178320]
KM111294.1_Ken05/Tk1             ......................t.......................a.................t...t....g...... [178320]
KM262844.1_strain_L60            ......................t......................................................... [178320]
KM102979.1_26544/OG10_Italy      ......................t......................................................... [178320]
KM262845.1_strain_NHV            ......................t......................................................... [178320]
KX354450.1_47/Ss/2008            ......................t......................................................... [178320]
KP055815.1_strain_BA71           ......................t......................................................... [178320]
MH025920.1_strain_R35            ......................t.......................a.................t...t....g...... [178320]
MH025919.1_strain_N10            ......................t.......................a.................t...t....g...... [178320]
MH025918.1_strain_R25            ......................t.......................a.................t...t....g...... [178320]
MH025917.1_strain_R7             ......................t.......................a.................t...t....g...... [178320]
MH025916.1_strain_R8             ......................t.......................a.................t...t....g...... [178320]

FR682468.2_Georgia_2007/1        aacaggtgagaatacataagggtttgcgttttaagaaaaccctgacgttgaatcataattgaaaacaccttgcaaagccg [178400]
ASF_IND_20_CAD543                ................................................................................ [178400]
LR899193.1_Germany_2020/1        ................................................................................ [178400]
MN393477.1_Wuhan_2019            ................................................................................ [178400]
MW396979.1_Timor-Leste/2019/1    ................................................................................ [178400]
MK940252.1_CN/2019/InnerMongolia ................................................................................ [178400]
MK128995.1_China/2018/AnhuiXCGQ  ................................................................................ [178400]
MH681419.1_ASFV/POL/2015         ................................................................................ [178400]
LS478113.1_Estonia_2014          ................................................................................ [178400]
KP843857.1_Odintsovo_02/14       ................................................................................ [178400]
```

```
FN557520.1_strain_E75             ................................................................................ [178400]
AY261366.1_Warthog                ................................................................................ [178400]
AY261365.1_Warmbaths              ............................................g................................... [178400]
AY261364.1_Tengani_62             ................................................................................ [178400]
AY261363.1_Pretoriuskop/96/4      ................................................................................ [178400]
AY261362.1_Mkuzi_1979             ............................................g................................... [178400]
AY261361.1_Malawi_Lil-20/1_1983   ...........g.....................................................t.............. [178400]
AY261360.1_Kenya_1950             ...........................................................g.................... [178400]
AM712240.1_OURT_88/3              ................................................................................ [178400]
AM712239.1_Benin_97/1             ................................................................................ [178400]
KM111295.1_Ken06.Bus              ...........................................................g.................... [178400]
KM111294.1_Ken05/Tk1              ...........................................................g.................... [178400]
KM262844.1_strain_L60             ................................................................................ [178400]
KM102979.1_26544/OG10_Italy       ................................................................................ [178400]
KM262845.1_strain_NHV             ................................................................................ [178400]
KX354450.1_47/Ss/2008             ................................................................................ [178400]
KP055815.1_strain_BA71            ................................................................................ [178400]
MH025920.1_strain_R35             ...........................................................g.................... [178400]
MH025919.1_strain_N10             ...........................................................g.................... [178400]
MH025918.1_strain_R25             ...........................................................g.................... [178400]
MH025917.1_strain_R7              ...........................................................g.................... [178400]
MH025916.1_strain_R8              ...........................................................g.................... [178400]

FR682468.2_Georgia_2007/1         actcatcagttgttctgtaatggcgttaagcattttctggaatttttcttggttttcgggtgtgattttatattcatgta [178480]
ASF_IND_20_CAD543                 ................................................................................ [178480]
LR899193.1_Germany_2020/1         ................................................................................ [178480]
MN393477.1_Wuhan_2019             ................................................................................ [178480]
MW396979.1_Timor-Leste/2019/1     ................................................................................ [178480]
MK940252.1_CN/2019/InnerMongolia  ................................................................................ [178480]
MK128995.1_China/2018/AnhuiXCGQ   ................................................................................ [178480]
MH681419.1_ASFV/POL/2015          ................................................................................ [178480]
LS478113.1_Estonia_2014           ................................................................................ [178480]
KP843857.1_Odintsovo_02/14        ................................................................................ [178480]
FN557520.1_strain_E75             ................................................................................ [178480]
AY261366.1_Warthog                .....................a..a....................................................... [178480]
AY261365.1_Warmbaths              ........................a....................................................... [178480]
AY261364.1_Tengani_62             ......t.................a..................................a.................... [178480]
AY261363.1_Pretoriuskop/96/4      ........................a..................................a.................... [178480]
AY261362.1_Mkuzi_1979             ........................a....................................................... [178480]
AY261361.1_Malawi_Lil-20/1_1983   ...............c........a................................a...................... [178480]
AY261360.1_Kenya_1950             .........c...........a..................g................a..c................... [178480]
AM712240.1_OURT_88/3              ................................................................................ [178480]
AM712239.1_Benin_97/1             ................................................................................ [178480]
KM111295.1_Ken06.Bus              .........c...........a..................g................a..c................... [178480]
```

```
KM111294.1_Ken05/Tk1            .........c...........a..................g................a..c................... [178480]
KM262844.1_strain_L60           ................................................................................ [178480]
KM102979.1_26544/OG10_Italy     ................................................................................ [178480]
KM262845.1_strain_NHV           ................................................................................ [178480]
KX354450.1_47/Ss/2008           ................................................................................ [178480]
KP055815.1_strain_BA71          ................................................................................ [178480]
MH025920.1_strain_R35           .........c...........a..................g................a..c................... [178480]
MH025919.1_strain_N10           .........c...........a..................g................a..c................... [178480]
MH025918.1_strain_R25           .........c...........a..................g................a..c................... [178480]
MH025917.1_strain_R7            .........c...........a..................g................a..c................... [178480]
MH025916.1_strain_R8            .........c...........a..................g................a..c................... [178480]

FR682468.2_Georgia_2007/1        gaaagtgtttcacacctgaggagaagaatctttcctccttcgagagcccatctttgatgatgggaagttccttgatcagg [178560]
ASF_IND_20_CAD543                ................................................................................ [178560]
LR899193.1_Germany_2020/1        ................................................................................ [178560]
MN393477.1_Wuhan_2019            ................................................................................ [178560]
MW396979.1_Timor-Leste/2019/1    ................................................................................ [178560]
MK940252.1_CN/2019/InnerMongolia ................................................................................ [178560]
MK128995.1_China/2018/AnhuiXCGQ  ................................................................................ [178560]
MH681419.1_ASFV/POL/2015         ................................................................................ [178560]
LS478113.1_Estonia_2014          ................................................................................ [178560]
KP843857.1_Odintsovo_02/14       ................................................................................ [178560]
FN557520.1_strain_E75            ................................................................................ [178560]
AY261366.1_Warthog               ................................................................g............... [178560]
AY261365.1_Warmbaths             ................................................................g............... [178560]
AY261364.1_Tengani_62            ................................................................................ [178560]
AY261363.1_Pretoriuskop/96/4     ................................................................g............... [178560]
AY261362.1_Mkuzi_1979            ................................................................g............... [178560]
AY261361.1_Malawi_Lil-20/1_1983  ................................................................................ [178560]
AY261360.1_Kenya_1950            ................................................................................ [178560]
AM712240.1_OURT_88/3             ................................................................................ [178560]
AM712239.1_Benin_97/1            ................................................................................ [178560]
KM111295.1_Ken06.Bus             ................................................................................ [178560]
KM111294.1_Ken05/Tk1             ................................................................................ [178560]
KM262844.1_strain_L60            ................................................................................ [178560]
KM102979.1_26544/OG10_Italy      ................................................................................ [178560]
KM262845.1_strain_NHV            ................................................................................ [178560]
KX354450.1_47/Ss/2008            ................................................................................ [178560]
KP055815.1_strain_BA71           ................................................................................ [178560]
MH025920.1_strain_R35            ................................................................................ [178560]
MH025919.1_strain_N10            ................................................................................ [178560]
MH025918.1_strain_R25            ................................................................................ [178560]
MH025917.1_strain_R7             ................................................................................ [178560]
MH025916.1_strain_R8             ................................................................................ [178560]
```

```
FR682468.2_Georgia_2007/1        gcaaaccattcctcctcttgggcttgcggattctgaagatactgatggcagatatggtttagaatggtgcacacgtagct [178640]
ASF_IND_20_CAD543                ................................................................................ [178640]
LR899193.1_Germany_2020/1        ................................................................................ [178640]
MN393477.1_Wuhan_2019            ................................................................................ [178640]
MW396979.1_Timor-Leste/2019/1    ................................................................................ [178640]
MK940252.1_CN/2019/InnerMongolia ................................................................................ [178640]
MK128995.1_China/2018/AnhuiXCGQ  ................................................................................ [178640]
MH681419.1_ASFV/POL/2015         ................................................................................ [178640]
LS478113.1_Estonia_2014          ................................................................................ [178640]
KP843857.1_Odintsovo_02/14       ................................................................................ [178640]
FN557520.1_strain_E75            .......................c........................................................ [178640]
AY261366.1_Warthog               ................................................................................ [178640]
AY261365.1_Warmbaths             ................................................................................ [178640]
AY261364.1_Tengani_62            .............................................................a.................. [178640]
AY261363.1_Pretoriuskop/96/4     ................................................................................ [178640]
AY261362.1_Mkuzi_1979            ................................................................................ [178640]
AY261361.1_Malawi_Lil-20/1_1983  ...........t.....................................................a.............. [178640]
AY261360.1_Kenya_1950            .............................g.................................................. [178640]
AM712240.1_OURT_88/3             .......................c........................................................ [178640]
AM712239.1_Benin_97/1            .......................c........................................................ [178640]
KM111295.1_Ken06.Bus             .............................g.................................................. [178640]
KM111294.1_Ken05/Tk1             .............................g.................................................. [178640]
KM262844.1_strain_L60            .......................c........................................................ [178640]
KM102979.1_26544/OG10_Italy      .......................c........................................................ [178640]
KM262845.1_strain_NHV            .......................c........................................................ [178640]
KX354450.1_47/Ss/2008            .......................c........................................................ [178640]
KP055815.1_strain_BA71           .......................c........................................................ [178640]
MH025920.1_strain_R35            .............................g................a................................. [178640]
MH025919.1_strain_N10            .............................g................a................................. [178640]
MH025918.1_strain_R25            .............................g................a................................. [178640]
MH025917.1_strain_R7             .............................g................a................................. [178640]
MH025916.1_strain_R8             .............................g................a................................. [178640]

FR682468.2_Georgia_2007/1        aataagctctgagctgattctttggttggttttcaaatgttggcgaaagtagtttttcaccgaagtgcatgtaataaacg [178720]
ASF_IND_20_CAD543                ................................................................................ [178720]
LR899193.1_Germany_2020/1        ................................................................................ [178720]
MN393477.1_Wuhan_2019            ................................................................................ [178720]
MW396979.1_Timor-Leste/2019/1    ................................................................................ [178720]
MK940252.1_CN/2019/InnerMongolia ................................................................................ [178720]
MK128995.1_China/2018/AnhuiXCGQ  ................................................................................ [178720]
MH681419.1_ASFV/POL/2015         ................................................................................ [178720]
LS478113.1_Estonia_2014          ................................................................................ [178720]
KP843857.1_Odintsovo_02/14       ................................................................................ [178720]
```

```
FN557520.1_strain_E75            ................................................................................ [178720]
AY261366.1_Warthog               ................................................................................ [178720]
AY261365.1_Warmbaths             ................................................................................ [178720]
AY261364.1_Tengani_62            ................................................................................ [178720]
AY261363.1_Pretoriuskop/96/4     ................................................................................ [178720]
AY261362.1_Mkuzi_1979            ................................................................................ [178720]
AY261361.1_Malawi_Lil-20/1_1983  .......................................c........................................ [178720]
AY261360.1_Kenya_1950            .......................................c........................................ [178720]
AM712240.1_OURT_88/3             ................................................................................ [178720]
AM712239.1_Benin_97/1            ................................................................................ [178720]
KM111295.1_Ken06.Bus             .......................................c........................................ [178720]
KM111294.1_Ken05/Tk1             .......................................c................................g....... [178720]
KM262844.1_strain_L60            ................................................................................ [178720]
KM102979.1_26544/OG10_Italy      ................................................................................ [178720]
KM262845.1_strain_NHV            ................................................................................ [178720]
KX354450.1_47/Ss/2008            ................................................................................ [178720]
KP055815.1_strain_BA71           ................................................................................ [178720]
MH025920.1_strain_R35            .......................................c........................................ [178720]
MH025919.1_strain_N10            .......................................c........................................ [178720]
MH025918.1_strain_R25            .......................................c........................................ [178720]
MH025917.1_strain_R7             .......................................c........................................ [178720]
MH025916.1_strain_R8             .......................................c........................................ [178720]

FR682468.2_Georgia_2007/1        tcttcattttcttataatatacaacagtatgttgagtctttaatttaaaattacaaggagttttctaggtctttatgcgt [178800]
ASF_IND_20_CAD543                ................................................................................ [178800]
LR899193.1_Germany_2020/1        ................................................................................ [178800]
MN393477.1_Wuhan_2019            ................................................................................ [178800]
MW396979.1_Timor-Leste/2019/1    ................................................................................ [178800]
MK940252.1_CN/2019/InnerMongolia ................................................................................ [178800]
MK128995.1_China/2018/AnhuiXCGQ  ................................................................................ [178800]
MH681419.1_ASFV/POL/2015         ................................................................................ [178800]
LS478113.1_Estonia_2014          ................................................................................ [178800]
KP843857.1_Odintsovo_02/14       ................................................................................ [178800]
FN557520.1_strain_E75            ................................................................................ [178800]
AY261366.1_Warthog               ................................................................................ [178800]
AY261365.1_Warmbaths             ................................................................................ [178800]
AY261364.1_Tengani_62            ................................................................................ [178800]
AY261363.1_Pretoriuskop/96/4     ................................................................................ [178800]
AY261362.1_Mkuzi_1979            ................................................................................ [178800]
AY261361.1_Malawi_Lil-20/1_1983  .................................................................c...........t.. [178800]
AY261360.1_Kenya_1950            .................................................................c........g..t.. [178800]
AM712240.1_OURT_88/3             ................................................................................ [178800]
AM712239.1_Benin_97/1            ................................................................................ [178800]
KM111295.1_Ken06.Bus             .................................................................c........g..t.. [178800]
```

```
KM111294.1_Ken05/Tk1              .................................................................c........g..t.. [178800]
KM262844.1_strain_L60             ................................................................................ [178800]
KM102979.1_26544/OG10_Italy       ................................................................................ [178800]
KM262845.1_strain_NHV             ................................................................................ [178800]
KX354450.1_47/Ss/2008             ................................................................................ [178800]
KP055815.1_strain_BA71            ................................................................................ [178800]
MH025920.1_strain_R35             .................................................................c........g..t.. [178800]
MH025919.1_strain_N10             .................................................................c........g..t.. [178800]
MH025918.1_strain_R25             .................................................................c........g..t.. [178800]
MH025917.1_strain_R7              .................................................................c........g..t.. [178800]
MH025916.1_strain_R8              .................................................................c........g..t.. [178800]

FR682468.2_Georgia_2007/1         ataggtgtttctttgtcgtaaattttcaatagccgacattgtttgtgaagcagtgttctgagtagtgactgtcgtgtaag [178880]
ASF_IND_20_CAD543                 ................................................................................ [178880]
LR899193.1_Germany_2020/1         ................................................................................ [178880]
MN393477.1_Wuhan_2019             ................................................................................ [178880]
MW396979.1_Timor-Leste/2019/1     ................................................................................ [178880]
MK940252.1_CN/2019/InnerMongolia  ................................................................................ [178880]
MK128995.1_China/2018/AnhuiXCGQ   ................................................................................ [178880]
MH681419.1_ASFV/POL/2015          ................................................................................ [178880]
LS478113.1_Estonia_2014           ................................................................................ [178880]
KP843857.1_Odintsovo_02/14        ................................................................................ [178880]
FN557520.1_strain_E75             ................................................................................ [178880]
AY261366.1_Warthog                .......c.......c....g..........................................g...........a.... [178880]
AY261365.1_Warmbaths              .......c.......c....g......................................................a.... [178880]
AY261364.1_Tengani_62             .......c.......c....g........................g.............................a.... [178880]
AY261363.1_Pretoriuskop/96/4      .......c.......c....g....................................t.................a.... [178880]
AY261362.1_Mkuzi_1979             ................................................................................ [178880]
AY261361.1_Malawi_Lil-20/1_1983   ...............c....g.......tc.....g.....................t..g..c...g.......a...a [178880]
AY261360.1_Kenya_1950             .......c.......c....g.gc.....c.....g...........................c...g....t..a...a [178880]
AM712240.1_OURT_88/3              ................................................................................ [178880]
AM712239.1_Benin_97/1             ................................................................................ [178880]
KM111295.1_Ken06.Bus              .......c.......c....g.gc.....c.....g...............................g....t..a...a [178880]
KM111294.1_Ken05/Tk1              .......c.......c....g.gca....c.....g...............................g....t..a...a [178880]
KM262844.1_strain_L60             ................................................................................ [178880]
KM102979.1_26544/OG10_Italy       ................................................................................ [178880]
KM262845.1_strain_NHV             ................................................................................ [178880]
KX354450.1_47/Ss/2008             ................................................................................ [178880]
KP055815.1_strain_BA71            ................................................................................ [178880]
MH025920.1_strain_R35             .......c.......c....g.gc.....c.....g...............................g....t..a...a [178880]
MH025919.1_strain_N10             .......c.......c....g.gc.....c.....g...............................g....t..a...a [178880]
MH025918.1_strain_R25             .......c.......c....g.gc.....c.....g...............................g....t..a...a [178880]
MH025917.1_strain_R7              .......c.......c....g.gc.....c.....g...............................g....t..a...a [178880]
MH025916.1_strain_R8              .......c.......c....g.gc.....c.....g...............................g....t..a...a [178880]
```

```
FR682468.2_Georgia_2007/1        gctcagccggatgagcagg------------------------------------------------------------a [178960]
ASF_IND_20_CAD543                ..............------------------------------------------------------------------ [178960]
LR899193.1_Germany_2020/1        ...................------------------------------------------------------------. [178960]
MN393477.1_Wuhan_2019            ...................------------------------------------------------------------. [178960]
MW396979.1_Timor-Leste/2019/1    ...................------------------------------------------------------------. [178960]
MK940252.1_CN/2019/InnerMongolia ...................------------------------------------------------------------. [178960]
MK128995.1_China/2018/AnhuiXCGQ  ...................------------------------------------------------------------. [178960]
MH681419.1_ASFV/POL/2015         ...................------------------------------------------------------------. [178960]
LS478113.1_Estonia_2014          ...................------------------------------------------------------------. [178960]
KP843857.1_Odintsovo_02/14       ...................------------------------------------------------------------. [178960]
FN557520.1_strain_E75            ......t.......------------------------------------------------------------------ [178960]
AY261366.1_Warthog               ...................agcactcgcggctgcactcgca------------------------------------gct [178960]
AY261365.1_Warmbaths             ............a......agcactcgcggctgcactcgca------------------------------------gct [178960]
AY261364.1_Tengani_62            ...................agcactcgcggctgcactcgcagctgtattcgtggccgcattcgcagccgcacttgcggcc [178960]
AY261363.1_Pretoriuskop/96/4     ...................agcactcgcggctgcacttgca------------------------------------gct [178960]
AY261362.1_Mkuzi_1979            ......t.......------------------------------------------------------------------ [178960]
AY261361.1_Malawi_Lil-20/1_1983  ...................a------------------------------------------------------------ [178960]
AY261360.1_Kenya_1950            .........a.g..------------------------------------------------------------------ [178960]
AM712240.1_OURT_88/3             ......t.......------------------------------------------------------------------ [178960]
AM712239.1_Benin_97/1            ......t.......------------------------------------------------------------------ [178960]
KM111295.1_Ken06.Bus             .........a.gc.------------------------------------------------------------------ [178960]
KM111294.1_Ken05/Tk1             .........a.g..------------------------------------------------------------------ [178960]
KM262844.1_strain_L60            ......t.......------------------------------------------------------------------ [178960]
KM102979.1_26544/OG10_Italy      ......t.......------------------------------------------------------------------ [178960]
KM262845.1_strain_NHV            ......t.......------------------------------------------------------------------ [178960]
KX354450.1_47/Ss/2008            ......t.......------------------------------------------------------------------ [178960]
KP055815.1_strain_BA71           ......t.......------------------------------------------------------------------ [178960]
MH025920.1_strain_R35            .........a.gc.------------------------------------------------------------------ [178960]
MH025919.1_strain_N10            .........a.gc.------------------------------------------------------------------ [178960]
MH025918.1_strain_R25            .........a.gc.------------------------------------------------------------------ [178960]
MH025917.1_strain_R7             .........a.gc.------------------------------------------------------------------ [178960]
MH025916.1_strain_R8             .........a.gc.------------------------------------------------------------------ [178960]

FR682468.2_Georgia_2007/1        gcactcgcggccgcaggtgcg------------gccgccggcccgccagttgccatgactagtctgtccgtaactgggtt [179040]
ASF_IND_20_CAD543                .....................------------.....t......................................... [179040]
LR899193.1_Germany_2020/1        .....................------------............................................... [179040]
MN393477.1_Wuhan_2019            .....................------------............................................... [179040]
MW396979.1_Timor-Leste/2019/1    .....................------------............................................... [179040]
MK940252.1_CN/2019/InnerMongolia .....................------------............................................... [179040]
MK128995.1_China/2018/AnhuiXCGQ  .....................------------............................................... [179040]
MH681419.1_ASFV/POL/2015         .....................------------............................................... [179040]
LS478113.1_Estonia_2014          .....................------------............................................... [179040]
KP843857.1_Odintsovo_02/14       .....................------------............................................... [179040]
```

```
FN557520.1_strain_E75            .....................------------.....t......................................... [179040]
AY261366.1_Warthog               ........a......ct....------------.....t......................................... [179040]
AY261365.1_Warmbaths             ........a......ct....gccgcacttgca.....t........t........a...g.gt.....a.......tc. [179040]
AY261364.1_Tengani_62            .....t.........ct....gccgcacttgcg.....t.......................c.............---- [179040]
AY261363.1_Pretoriuskop/96/4     ........a......ct....gccgcacttacg.....t......................................... [179040]
AY261362.1_Mkuzi_1979            .....................------------.....t......................................... [179040]
AY261361.1_Malawi_Lil-20/1_1983  .....t...------------------------.....t..........c.......c..................---- [179040]
AY261360.1_Kenya_1950            .....t...------------------------.....t.....t.............t..................... [179040]
AM712240.1_OURT_88/3             .....................------------.....t......................................... [179040]
AM712239.1_Benin_97/1            .....................------------.....t......................................... [179040]
KM111295.1_Ken06.Bus             .....t...------------------------.....t.....a.............t...................c. [179040]
KM111294.1_Ken05/Tk1             .....t...------------------------.....t.....t.............t..............t...... [179040]
KM262844.1_strain_L60            .....................------------.....t......................................... [179040]
KM102979.1_26544/OG10_Italy      .....................------------.....t......................................... [179040]
KM262845.1_strain_NHV            .....................------------.....t......................................... [179040]
KX354450.1_47/Ss/2008            .....................------------.....t......................................... [179040]
KP055815.1_strain_BA71           a....................------------.....t......................................... [179040]
MH025920.1_strain_R35            .....t...------------------------.....t.....a.............t...................c. [179040]
MH025919.1_strain_N10            .....t...------------------------.....t.....a.............t...................c. [179040]
MH025918.1_strain_R25            .....t...------------------------.....t.....a.............t...................c. [179040]
MH025917.1_strain_R7             .....t...------------------------.....t.....a.............t...................c. [179040]
MH025916.1_strain_R8             .....t...------------------------.....t.....a.............t...................c. [179040]

FR682468.2_Georgia_2007/1        gtccgtaactggtttg---------------tttgttgctggtctgtttgttg---ccggtctgcccgtgactggcttgc [179120]
ASF_IND_20_CAD543                ................---------------......................---........................ [179120]
LR899193.1_Germany_2020/1        ................---------------......................---........................ [179120]
MN393477.1_Wuhan_2019            ................---------------......................---........................ [179120]
MW396979.1_Timor-Leste/2019/1    ................---------------......................---........................ [179120]
MK940252.1_CN/2019/InnerMongolia ................---------------......................---........................ [179120]
MK128995.1_China/2018/AnhuiXCGQ  ................---------------......................---........................ [179120]
MH681419.1_ASFV/POL/2015         ................---------------......................---........................ [179120]
LS478113.1_Estonia_2014          ................---------------......................---........................ [179120]
KP843857.1_Odintsovo_02/14       ................---------------......................---........................ [179120]
FN557520.1_strain_E75            ......g.........---------------......................---...................t.... [179120]
AY261366.1_Warthog               ..tt..tg.....c.a---------------.cc..aa...............---....c...........c....... [179120]
AY261365.1_Warmbaths             ..tt..tg.....c.a---------------.cc.caa..a............---........t..........t..c. [179120]
AY261364.1_Tengani_62            -----------------------------------------............---....c................... [179120]
AY261363.1_Pretoriuskop/96/4     ..tt..tg.....c.a---------------.cc..aa...............---....c................... [179120]
AY261362.1_Mkuzi_1979            ......g.........---------------......................---........................ [179120]
AY261361.1_Malawi_Lil-20/1_1983  -----------.c...---------------.....aa....ct.......aa---.t.........a............ [179120]
AY261360.1_Kenya_1950            ..t.........cc..------------------------------.c.....---.t..c...t.t..a.t........ [179120]
AM712240.1_OURT_88/3             ......g.........---------------......................---...................t.... [179120]
AM712239.1_Benin_97/1            ......g.........---------------......................---...................t.... [179120]
KM111295.1_Ken06.Bus             ..t.a.......cc..tccgcgaccgagctg..c..aa....c..a.c.....---.t..c...t......t........ [179120]
```

```
KM111294.1_Ken05/Tk1            ..t.a.......cc..tccgcgaccgagctg..c..aa....c..a..ca.g.caa.t..c...t.t..a.t........ [179120]
KM262844.1_strain_L60           ......g.........---------------......................---...................t.... [179120]
KM102979.1_26544/OG10_Italy     ......g.........---------------......................---...................t.... [179120]
KM262845.1_strain_NHV           ......g.........---------------......................---...................t.... [179120]
KX354450.1_47/Ss/2008           ......g.........---------------......................---...................t.... [179120]
KP055815.1_strain_BA71          ......g.........---------------......................---...................t.... [179120]
MH025920.1_strain_R35           ..t.a.......cc..tccgcgaccgagctg..c..aa....c..a.c.....---.t..c...t......t........ [179120]
MH025919.1_strain_N10           ..t.a.......cc..tccgcgaccgagctg..c..aa....c..a.c.....---.t..c...t......t........ [179120]
MH025918.1_strain_R25           ..t.a.......cc..tccgcgaccgagctg..c..aa....c..a.c.....---.t..c...t......t........ [179120]
MH025917.1_strain_R7            ..t.a.......cc..tccgcgaccgagctg..c..aa....c..a.c.....---.t..c...t......t........ [179120]
MH025916.1_strain_R8            ..t.a.......cc..tccgcgaccgagctg..c..aa....c..a.c.....---.t..c...t......t........ [179120]

FR682468.2_Georgia_2007/1       ctacacttgctgtagtcgctccagctggtttagaggtacctggttgtggagtgacttctacccactgctgatcttgataa [179200]
ASF_IND_20_CAD543               ...........................................................r.................... [179200]
LR899193.1_Germany_2020/1       ................................................................................ [179200]
MN393477.1_Wuhan_2019           ................................................................................ [179200]
MW396979.1_Timor-Leste/2019/1   ................................................................................ [179200]
MK940252.1_CN/2019/InnerMongolia ................................................................................ [179200]
MK128995.1_China/2018/AnhuiXCGQ ................................................................................ [179200]
MH681419.1_ASFV/POL/2015        ................................................................................ [179200]
LS478113.1_Estonia_2014         ................................................................................ [179200]
KP843857.1_Odintsovo_02/14      ................................................................................ [179200]
FN557520.1_strain_E75           ..g......................c.................................g....t............... [179200]
AY261366.1_Warthog              .........................c...................................................... [179200]
AY261365.1_Warmbaths            .------.c................c...................................................... [179200]
AY261364.1_Tengani_62           .................a.......c...................................................... [179200]
AY261363.1_Pretoriuskop/96/4    .........................c...................................................... [179200]
AY261362.1_Mkuzi_1979           ..g......................c...................................................... [179200]
AY261361.1_Malawi_Lil-20/1_1983 ..g.....c......at........c.......caa.g..................c..g.................... [179200]
AY261360.1_Kenya_1950           ....gt.gc.......g........c...........................g..c..g.................... [179200]
AM712240.1_OURT_88/3            ..g......................c.................................g....t............... [179200]
AM712239.1_Benin_97/1           ..g......................c.................................g....t............... [179200]
KM111295.1_Ken06.Bus            .....t.gc.......g........c...........................g..c..g.................... [179200]
KM111294.1_Ken05/Tk1            ....gt.gc.......g........c.....................a.....g..c..g.................... [179200]
KM262844.1_strain_L60           ..g......................c.................................g....t............... [179200]
KM102979.1_26544/OG10_Italy     ..g......................c.................................g....t............... [179200]
KM262845.1_strain_NHV           ..g......................c.................................g....t............... [179200]
KX354450.1_47/Ss/2008           ..g......................c.................................g....t............... [179200]
KP055815.1_strain_BA71          ..g......................c.................................g....t............... [179200]
MH025920.1_strain_R35           .....t.gc.......g........c...........................g..c..g.................... [179200]
MH025919.1_strain_N10           .....t.gc.......g........c...........................g..c..g.................... [179200]
MH025918.1_strain_R25           .....t.gc.......g........c...........................g..c..g.................... [179200]
MH025917.1_strain_R7            .....t.gc.......g........c...........................g..c..g.................... [179200]
MH025916.1_strain_R8            .....t.gc.......g........c...........................g..c..g.................... [179200]
```

```
FR682468.2_Georgia_2007/1        ggatttataaactgtatatcttcctcctcaata------gcagcagcttttttctttcttgaagagaatagatagattag [179280]
ASF_IND_20_CAD543                .................................gc---g......................................... [179280]
LR899193.1_Germany_2020/1        .................................------......................................... [179280]
MN393477.1_Wuhan_2019            .................................------......................................... [179280]
MW396979.1_Timor-Leste/2019/1    .................................------......................................... [179280]
MK940252.1_CN/2019/InnerMongolia .................................------......................................... [179280]
MK128995.1_China/2018/AnhuiXCGQ  .................................------......................................... [179280]
MH681419.1_ASFV/POL/2015         .................................------......................................... [179280]
LS478113.1_Estonia_2014          .................................------......................................... [179280]
KP843857.1_Odintsovo_02/14       .................................------......................................... [179280]
FN557520.1_strain_E75            .................................gc---g......................................... [179280]
AY261366.1_Warthog               .................................gc---g..........................a........a..... [179280]
AY261365.1_Warmbaths             .................................gc---g............c.............a........a..... [179280]
AY261364.1_Tengani_62            ..........................t......gc---g..........................a........a..... [179280]
AY261363.1_Pretoriuskop/96/4     .................................gc---g....g..............................a..... [179280]
AY261362.1_Mkuzi_1979            .................................gc---g..........................a........a..... [179280]
AY261361.1_Malawi_Lil-20/1_1983  .................................gc---g..........................a........a..... [179280]
AY261360.1_Kenya_1950            .................................gcgggg..........................a..c.....a..... [179280]
AM712240.1_OURT_88/3             .................................gc---g......................................... [179280]
AM712239.1_Benin_97/1            .................................gc---g......................................... [179280]
KM111295.1_Ken06.Bus             .................................gcggcg..........................a..c.....a..... [179280]
KM111294.1_Ken05/Tk1             .................................gcggcg..........................a..c.....a..... [179280]
KM262844.1_strain_L60            .................................gc---g......................................... [179280]
KM102979.1_26544/OG10_Italy      .................................gc---g......................................... [179280]
KM262845.1_strain_NHV            .................................gc---g......................................... [179280]
KX354450.1_47/Ss/2008            .................................gc---g......................................... [179280]
KP055815.1_strain_BA71           .................................gc---g......................................... [179280]
MH025920.1_strain_R35            .................................gcggcg..........................a..c.....a..... [179280]
MH025919.1_strain_N10            .................................gcggcg..........................a..c.....a..... [179280]
MH025918.1_strain_R25            .................................gcggcg..........................a..c.....a..... [179280]
MH025917.1_strain_R7             .................................gcggcg..........................a..c.....a..... [179280]
MH025916.1_strain_R8             .................................gcggcg..........................a..c.....a..... [179280]

FR682468.2_Georgia_2007/1        aacgatgataatgatgactaagaccacgatagcaatgagaatagtatacatatgtgtggagaagaagcttggtgtagtga [179360]
ASF_IND_20_CAD543                ............r.............................................................gg.... [179360]
LR899193.1_Germany_2020/1        ................................................................................ [179360]
MN393477.1_Wuhan_2019            ................................................................................ [179360]
MW396979.1_Timor-Leste/2019/1    ................................................................................ [179360]
MK940252.1_CN/2019/InnerMongolia ................................................................................ [179360]
MK128995.1_China/2018/AnhuiXCGQ  ................................................................................ [179360]
MH681419.1_ASFV/POL/2015         ................................................................................ [179360]
LS478113.1_Estonia_2014          ................................................................................ [179360]
KP843857.1_Odintsovo_02/14       ................................................................................ [179360]
```

```
FN557520.1_strain_E75            ............a.............................................................gg.... [179360]
AY261366.1_Warthog               ............a................................................................a.. [179360]
AY261365.1_Warmbaths             ............a................................................................a.. [179360]
AY261364.1_Tengani_62            ............a................................................................a.. [179360]
AY261363.1_Pretoriuskop/96/4     ............a................................................................a.. [179360]
AY261362.1_Mkuzi_1979            ............a..............................................................g.... [179360]
AY261361.1_Malawi_Lil-20/1_1983  ............a....t..................a.....................................c..g.g [179360]
AY261360.1_Kenya_1950            ............a....t.................c..........c......................c.......a.g [179360]
AM712240.1_OURT_88/3             ............a.............................................................gg.... [179360]
AM712239.1_Benin_97/1            ............a.............................................................gg.... [179360]
KM111295.1_Ken06.Bus             ............a....t.................c..........c......................c.......a.g [179360]
KM111294.1_Ken05/Tk1             ............a....t.................c..........c......................c.......a.g [179360]
KM262844.1_strain_L60            ............a.............................................................gg.... [179360]
KM102979.1_26544/OG10_Italy      ............a.............................................................gg.... [179360]
KM262845.1_strain_NHV            ............a.............................................................gg.... [179360]
KX354450.1_47/Ss/2008            ............a.............................................................gg.... [179360]
KP055815.1_strain_BA71           ............a.............................................................gg.... [179360]
MH025920.1_strain_R35            ............a....t.................c..........c......................c.......a.g [179360]
MH025919.1_strain_N10            ............a....t.................c..........c......................c.......a.g [179360]
MH025918.1_strain_R25            ............a....t.................c..........c......................c.......a.g [179360]
MH025917.1_strain_R7             ............a....t.................c..........c......................c.......a.g [179360]
MH025916.1_strain_R8             ............a....t.................c..........c......................c.......a.g [179360]

FR682468.2_Georgia_2007/1        ctggtgacaaacactcaccataatgccgcggataaaccggttgaaaaaattcagaatccatttaagatactattataaat [179440]
ASF_IND_20_CAD543                ................................................................................ [179440]
LR899193.1_Germany_2020/1        ................................................................................ [179440]
MN393477.1_Wuhan_2019            ................................................................................ [179440]
MW396979.1_Timor-Leste/2019/1    ................................................................................ [179440]
MK940252.1_CN/2019/InnerMongolia ................................................................................ [179440]
MK128995.1_China/2018/AnhuiXCGQ  ................................................................................ [179440]
MH681419.1_ASFV/POL/2015         ................................................................................ [179440]
LS478113.1_Estonia_2014          ................................................................................ [179440]
KP843857.1_Odintsovo_02/14       ................................................................................ [179440]
FN557520.1_strain_E75            ................................................................................ [179440]
AY261366.1_Warthog               ............................t................................................... [179440]
AY261365.1_Warmbaths             .........................g...................................................... [179440]
AY261364.1_Tengani_62            ................................................................................ [179440]
AY261363.1_Pretoriuskop/96/4     ................................................................................ [179440]
AY261362.1_Mkuzi_1979            ................................................................................ [179440]
AY261361.1_Malawi_Lil-20/1_1983  t..a............g............................................................... [179440]
AY261360.1_Kenya_1950            t............t..g....................g.......................................... [179440]
AM712240.1_OURT_88/3             ................................................................................ [179440]
AM712239.1_Benin_97/1            ................................................................................ [179440]
KM111295.1_Ken06.Bus             t............t..g....................a.......................................... [179440]
```

```
KM111294.1_Ken05/Tk1             t............t..g....................a.....g.................................... [179440]
KM262844.1_strain_L60            ................................................................................ [179440]
KM102979.1_26544/OG10_Italy      ................................................................................ [179440]
KM262845.1_strain_NHV            ................................................................................ [179440]
KX354450.1_47/Ss/2008            ................................................................................ [179440]
KP055815.1_strain_BA71           ................................................................................ [179440]
MH025920.1_strain_R35            t............t..g....................a.......................................... [179440]
MH025919.1_strain_N10            t............t..g....................a.......................................... [179440]
MH025918.1_strain_R25            t............t..g....................a.......................................... [179440]
MH025917.1_strain_R7             t............t..g....................a.......................................... [179440]
MH025916.1_strain_R8             t............t..g....................a.......................................... [179440]

FR682468.2_Georgia_2007/1        aatatataaaaatgttgtggcgcaatgaaattacagaatttatggaccaactttccaagtattctcaagaaatcttaaaa [179520]
ASF_IND_20_CAD543                ................................................................................ [179520]
LR899193.1_Germany_2020/1        ................................................................................ [179520]
MN393477.1_Wuhan_2019            ................................................................................ [179520]
MW396979.1_Timor-Leste/2019/1    ................................................................................ [179520]
MK940252.1_CN/2019/InnerMongolia ................................................................................ [179520]
MK128995.1_China/2018/AnhuiXCGQ  ................................................................................ [179520]
MH681419.1_ASFV/POL/2015         ................................................................................ [179520]
LS478113.1_Estonia_2014          ................................................................................ [179520]
KP843857.1_Odintsovo_02/14       ................................................................................ [179520]
FN557520.1_strain_E75            ................................................................................ [179520]
AY261366.1_Warthog               ....................................................c........................... [179520]
AY261365.1_Warmbaths             ....................................................c........................... [179520]
AY261364.1_Tengani_62            ..................................g............................................. [179520]
AY261363.1_Pretoriuskop/96/4     ....................................................c........................... [179520]
AY261362.1_Mkuzi_1979            ................................................................................ [179520]
AY261361.1_Malawi_Lil-20/1_1983  ................................................................................ [179520]
AY261360.1_Kenya_1950            ..............................................t.................c............... [179520]
AM712240.1_OURT_88/3             ................................................................................ [179520]
AM712239.1_Benin_97/1            ................................................................................ [179520]
KM111295.1_Ken06.Bus             ..............................................t................gc............... [179520]
KM111294.1_Ken05/Tk1             ..............................................t.................c............... [179520]
KM262844.1_strain_L60            ................................................................................ [179520]
KM102979.1_26544/OG10_Italy      ................................................................................ [179520]
KM262845.1_strain_NHV            ................................................................................ [179520]
KX354450.1_47/Ss/2008            ................................................................................ [179520]
KP055815.1_strain_BA71           ................................................................................ [179520]
MH025920.1_strain_R35            ..............................................t................gc............... [179520]
MH025919.1_strain_N10            ..............................................t................gc............... [179520]
MH025918.1_strain_R25            ..............................................t................gc............... [179520]
MH025917.1_strain_R7             ..............................................t................gc............... [179520]
MH025916.1_strain_R8             ..............................................t................gc............... [179520]
```

```
FR682468.2_Georgia_2007/1        acgtttaagcaattgcgtcctagtgaatataaacaatacaatgaatttttaacacaagttacaccgttgctgcaaaaaac [179600]
ASF_IND_20_CAD543                ................................................................................ [179600]
LR899193.1_Germany_2020/1        ................................................................................ [179600]
MN393477.1_Wuhan_2019            ................................................................................ [179600]
MW396979.1_Timor-Leste/2019/1    ................................................................................ [179600]
MK940252.1_CN/2019/InnerMongolia ................................................................................ [179600]
MK128995.1_China/2018/AnhuiXCGQ  ................................................................................ [179600]
MH681419.1_ASFV/POL/2015         ................................................................................ [179600]
LS478113.1_Estonia_2014          ................................................................................ [179600]
KP843857.1_Odintsovo_02/14       ................................................................................ [179600]
FN557520.1_strain_E75            ................................................................................ [179600]
AY261366.1_Warthog               ...........................................................g...........a........ [179600]
AY261365.1_Warmbaths             ...........................................................g...........a........ [179600]
AY261364.1_Tengani_62            .....c.....................................................g..g................. [179600]
AY261363.1_Pretoriuskop/96/4     ...........................................................g.................... [179600]
AY261362.1_Mkuzi_1979            ................................................................................ [179600]
AY261361.1_Malawi_Lil-20/1_1983  ...........g...................................................t.a.....a........ [179600]
AY261360.1_Kenya_1950            ...........g.......................g............................................ [179600]
AM712240.1_OURT_88/3             ................................................................................ [179600]
AM712239.1_Benin_97/1            ................................................................................ [179600]
KM111295.1_Ken06.Bus             ........a..g.......................g............................................ [179600]
KM111294.1_Ken05/Tk1             ...........g.......................g............................................ [179600]
KM262844.1_strain_L60            ................................................................................ [179600]
KM102979.1_26544/OG10_Italy      ................................................................................ [179600]
KM262845.1_strain_NHV            ................................................................................ [179600]
KX354450.1_47/Ss/2008            ................................................................................ [179600]
KP055815.1_strain_BA71           ................................................................................ [179600]
MH025920.1_strain_R35            ........a..g.......................g............................................ [179600]
MH025919.1_strain_N10            ........a..g.......................g............................................ [179600]
MH025918.1_strain_R25            ........a..g.......................g............................................ [179600]
MH025917.1_strain_R7             ........a..g.......................g............................................ [179600]
MH025916.1_strain_R8             ........a..g.......................g............................................ [179600]

FR682468.2_Georgia_2007/1        ccctgaaaaaattccagagttggttgaccatatattcaattacctagacaacgttgaaaaaatttgtgagctcctcgtga [179680]
ASF_IND_20_CAD543                ................................................................................ [179680]
LR899193.1_Germany_2020/1        ................................................................................ [179680]
MN393477.1_Wuhan_2019            ................................................................................ [179680]
MW396979.1_Timor-Leste/2019/1    ................................................................................ [179680]
MK940252.1_CN/2019/InnerMongolia ................................................................................ [179680]
MK128995.1_China/2018/AnhuiXCGQ  ................................................................................ [179680]
MH681419.1_ASFV/POL/2015         ................................................................................ [179680]
LS478113.1_Estonia_2014          ................................................................................ [179680]
KP843857.1_Odintsovo_02/14       ................................................................................ [179680]
```

```
FN557520.1_strain_E75           .t.........................................................................g.... [179680]
AY261366.1_Warthog              ......c..............a.......................................................... [179680]
AY261365.1_Warmbaths            .....................a...............g.......................................... [179680]
AY261364.1_Tengani_62           ...................c.c.......................................................... [179680]
AY261363.1_Pretoriuskop/96/4    .....................a.......................................................... [179680]
AY261362.1_Mkuzi_1979           ...........................................................................g.... [179680]
AY261361.1_Malawi_Lil-20/1_1983 .................................t.........................................g.... [179680]
AY261360.1_Kenya_1950           ...................ga............t...g..................................a..g...c [179680]
AM712240.1_OURT_88/3            .t.........................................................................g.... [179680]
AM712239.1_Benin_97/1           .t.........................................................................g.... [179680]
KM111295.1_Ken06.Bus            ...................ga............t...g..........t.......................a..g...c [179680]
KM111294.1_Ken05/Tk1            ...................ga............t...g..........t.......................a..g...c [179680]
KM262844.1_strain_L60           .t.........................................................................g.... [179680]
KM102979.1_26544/OG10_Italy     .t.........................................................................g.... [179680]
KM262845.1_strain_NHV           .t.........................................................................g.... [179680]
KX354450.1_47/Ss/2008           .t.........................................................................g.... [179680]
KP055815.1_strain_BA71          .t.........................................................................g.... [179680]
MH025920.1_strain_R35           ...................ga............t...g..........t.......................a..g...c [179680]
MH025919.1_strain_N10           ...................ga............t...g..........t.......................a..g...c [179680]
MH025918.1_strain_R25           ...................ga............t...g..........t.......................a..g...c [179680]
MH025917.1_strain_R7            ...................ga............t...g..........t.......................a..g...c [179680]
MH025916.1_strain_R8            ...................ga............t...g..........t.......................a..g...c [179680]

FR682468.2_Georgia_2007/1       atgctagctcaattattattagttcaaaaatacgagaacaagtaaaacacggaatgagcttcagctataaagccgacctc [179760]
ASF_IND_20_CAD543               ................................................................................ [179760]
LR899193.1_Germany_2020/1       ................................................................................ [179760]
MN393477.1_Wuhan_2019           ................................................................................ [179760]
MW396979.1_Timor-Leste/2019/1   ................................................................................ [179760]
MK940252.1_CN/2019/InnerMongolia ................................................................................ [179760]
MK128995.1_China/2018/AnhuiXCGQ ................................................................................ [179760]
MH681419.1_ASFV/POL/2015        ................................................................................ [179760]
LS478113.1_Estonia_2014         ................................................................................ [179760]
KP843857.1_Odintsovo_02/14      ................................................................................ [179760]
FN557520.1_strain_E75           ................................................................................ [179760]
AY261366.1_Warthog              ................................................................................ [179760]
AY261365.1_Warmbaths            ................................................................................ [179760]
AY261364.1_Tengani_62           ................................................................................ [179760]
AY261363.1_Pretoriuskop/96/4    ....................................................c........................... [179760]
AY261362.1_Mkuzi_1979           ............................................................................t... [179760]
AY261361.1_Malawi_Lil-20/1_1983 ....g........................................................................... [179760]
AY261360.1_Kenya_1950           .......................................................................a........ [179760]
AM712240.1_OURT_88/3            ................................................................................ [179760]
AM712239.1_Benin_97/1           ................................................................................ [179760]
KM111295.1_Ken06.Bus            .......................................................................a........ [179760]
```

```
KM111294.1_Ken05/Tk1             .......................................................................a........ [179760]
KM262844.1_strain_L60            ................................................................................ [179760]
KM102979.1_26544/OG10_Italy      ................................................................................ [179760]
KM262845.1_strain_NHV            ................................................................................ [179760]
KX354450.1_47/Ss/2008            ................................................................................ [179760]
KP055815.1_strain_BA71           ................................................................................ [179760]
MH025920.1_strain_R35            .......................................................................a........ [179760]
MH025919.1_strain_N10            .......................................................................a........ [179760]
MH025918.1_strain_R25            .......................................................................a........ [179760]
MH025917.1_strain_R7             .......................................................................a........ [179760]
MH025916.1_strain_R8             .......................................................................a........ [179760]

FR682468.2_Georgia_2007/1        gactccttggcggacattctctctcaaaaacagtacgtgcttatgcatctttcaaaaaatattgcggccgagtattttaa [179840]
ASF_IND_20_CAD543                ................................................................................ [179840]
LR899193.1_Germany_2020/1        ................................................................................ [179840]
MN393477.1_Wuhan_2019            ................................................................................ [179840]
MW396979.1_Timor-Leste/2019/1    ................................................................................ [179840]
MK940252.1_CN/2019/InnerMongolia ................................................................................ [179840]
MK128995.1_China/2018/AnhuiXCGQ  ................................................................................ [179840]
MH681419.1_ASFV/POL/2015         ................................................................................ [179840]
LS478113.1_Estonia_2014          ................................................................................ [179840]
KP843857.1_Odintsovo_02/14       ................................................................................ [179840]
FN557520.1_strain_E75            .................c.............................................................. [179840]
AY261366.1_Warthog               .................c...at..............................................c.......... [179840]
AY261365.1_Warmbaths             .................c...................................................c.......... [179840]
AY261364.1_Tengani_62            .................c...................................................c.......... [179840]
AY261363.1_Pretoriuskop/96/4     ..............t..c...................................................c.......... [179840]
AY261362.1_Mkuzi_1979            .................c.............................................................. [179840]
AY261361.1_Malawi_Lil-20/1_1983  ...........a...g.c...........................a................a......c.a........ [179840]
AY261360.1_Kenya_1950            .............g...c..........................t........................c.c........ [179840]
AM712240.1_OURT_88/3             .................c.............................................................. [179840]
AM712239.1_Benin_97/1            .................c.............................................................. [179840]
KM111295.1_Ken06.Bus             .............g...c..........................t........................c.c........ [179840]
KM111294.1_Ken05/Tk1             .............g...c..........................t........................c.c........ [179840]
KM262844.1_strain_L60            .................c.............................................................. [179840]
KM102979.1_26544/OG10_Italy      .................c.............................................................. [179840]
KM262845.1_strain_NHV            .................c.............................................................. [179840]
KX354450.1_47/Ss/2008            .................c.............................................................. [179840]
KP055815.1_strain_BA71           .................c.............................................................. [179840]
MH025920.1_strain_R35            .............g...c..........................t........................c.c........ [179840]
MH025919.1_strain_N10            .............g...c..........................t........................c.c........ [179840]
MH025918.1_strain_R25            .............g...c..........................t........................c.c........ [179840]
MH025917.1_strain_R7             .............g...c..........................t........................c.c........ [179840]
MH025916.1_strain_R8             .............g...c..........................t........................c.c........ [179840]
```

```
FR682468.2_Georgia_2007/1        tacgtgtttaaaccaagggaaatccaagttagatctcaaagctgcctctgtattttatagtagtcgttcccgaacggcaa [179920]
ASF_IND_20_CAD543                ................................................................................ [179920]
LR899193.1_Germany_2020/1        ................................................................................ [179920]
MN393477.1_Wuhan_2019            ................................................................................ [179920]
MW396979.1_Timor-Leste/2019/1    ................................................................................ [179920]
MK940252.1_CN/2019/InnerMongolia ................................................................................ [179920]
MK128995.1_China/2018/AnhuiXCGQ  ................................................................................ [179920]
MH681419.1_ASFV/POL/2015         ................................................................................ [179920]
LS478113.1_Estonia_2014          ................................................................................ [179920]
KP843857.1_Odintsovo_02/14       ................................................................................ [179920]
FN557520.1_strain_E75            ..................a............................................c...c............ [179920]
AY261366.1_Warthog               ...............................................................c...c............ [179920]
AY261365.1_Warmbaths             ..................a............................................c..cc....g....... [179920]
AY261364.1_Tengani_62            ..................a............................................c..cc............ [179920]
AY261363.1_Pretoriuskop/96/4     ..................a............................................c..cc............ [179920]
AY261362.1_Mkuzi_1979            ..................a............................................c...c............ [179920]
AY261361.1_Malawi_Lil-20/1_1983  ............g.............g...............g................a...c...c....g....... [179920]
AY261360.1_Kenya_1950            ............t.............g............g...........g.......a...c...c....g..c.... [179920]
AM712240.1_OURT_88/3             ..................a............................................c...c............ [179920]
AM712239.1_Benin_97/1            ..................a............................................c...c............ [179920]
KM111295.1_Ken06.Bus             ............t.............g............g...........g.......a...c...c....g..c.... [179920]
KM111294.1_Ken05/Tk1             ...a........t.............g............g...........g.......a...c...c....g..t.... [179920]
KM262844.1_strain_L60            ..................a............................................c...c............ [179920]
KM102979.1_26544/OG10_Italy      ..................a............................................c...c............ [179920]
KM262845.1_strain_NHV            ..................a............................................c...c............ [179920]
KX354450.1_47/Ss/2008            ..................a............................................c...c............ [179920]
KP055815.1_strain_BA71           ..................a............................................c...c............ [179920]
MH025920.1_strain_R35            ............t.............g............g...........g.......a...c...c....g..c.... [179920]
MH025919.1_strain_N10            ............t.............g............g...........g.......a...c...c....g..c.... [179920]
MH025918.1_strain_R25            ............t.............g............g...........g.......a...c...c....g..c.... [179920]
MH025917.1_strain_R7             ............t.............g............g...........g.......a...c...c....g..c.... [179920]
MH025916.1_strain_R8             ............t.............g............g...........g.......a...c...c....g..c.... [179920]

FR682468.2_Georgia_2007/1        gctcagcagaactctatagaaaaatgctatacgcctatggttcaccgcaggaaattaattattatactgaaaaagcccga [180000]
ASF_IND_20_CAD543                ................................................................................ [180000]
LR899193.1_Germany_2020/1        ................................................................................ [180000]
MN393477.1_Wuhan_2019            ................................................................................ [180000]
MW396979.1_Timor-Leste/2019/1    ................................................................................ [180000]
MK940252.1_CN/2019/InnerMongolia ................................................................................ [180000]
MK128995.1_China/2018/AnhuiXCGQ  ................................................................................ [180000]
MH681419.1_ASFV/POL/2015         ................................................................................ [180000]
LS478113.1_Estonia_2014          ................................................................................ [180000]
KP843857.1_Odintsovo_02/14       ................................................................................ [180000]
```

```
FN557520.1_strain_E75          ............................................................................t... [180000]
AY261366.1_Warthog             ...........................................g.................................... [180000]
AY261365.1_Warmbaths           ...........................................g.................................... [180000]
AY261364.1_Tengani_62          ...........................................g.................................... [180000]
AY261363.1_Pretoriuskop/96/4   ...........................................g.................................... [180000]
AY261362.1_Mkuzi_1979          ............................................................................t... [180000]
AY261361.1_Malawi_Lil-20/1_1983 .............a....................t.....c..g.ta..a.............................. [180000]
AY261360.1_Kenya_1950          .............a............t.............c....ta..a.................g............ [180000]
AM712240.1_OURT_88/3           ............................................................................t... [180000]
AM712239.1_Benin_97/1          ............................................................................t... [180000]
KM111295.1_Ken06.Bus           .............a..........................c..g.ta..a.................g............ [180000]
KM111294.1_Ken05/Tk1           .............a............t.............c..g.ta..a.................g............ [180000]
KM262844.1_strain_L60          ............................................................................t... [180000]
KM102979.1_26544/OG10_Italy    ............................................................................t... [180000]
KM262845.1_strain_NHV          ............................................................................t... [180000]
KX354450.1_47/Ss/2008          ............................................................................t... [180000]
KP055815.1_strain_BA71         ............................................................................t... [180000]
MH025920.1_strain_R35          .............a..........................c..g.ta..a.................g............ [180000]
MH025919.1_strain_N10          .............a..........................c..g.ta..a.................g............ [180000]
MH025918.1_strain_R25          .............a..........................c..g.ta..a.................g............ [180000]
MH025917.1_strain_R7           .............a..........................c..g.ta..a.................g............ [180000]
MH025916.1_strain_R8           .............a..........................c..g.ta..a.................g............ [180000]

FR682468.2_Georgia_2007/1      aataagacgttggatgtggaggagagcgacagcatggccatcatcgaacgaacggcccgacacaacctttcccttatgca [180080]
ASF_IND_20_CAD543              ..............y................................................................. [180080]
LR899193.1_Germany_2020/1      ................................................................................ [180080]
MN393477.1_Wuhan_2019          ................................................................................ [180080]
MW396979.1_Timor-Leste/2019/1  ................................................................................ [180080]
MK940252.1_CN/2019/InnerMongolia ................................................................................ [180080]
MK128995.1_China/2018/AnhuiXCGQ ................................................................................ [180080]
MH681419.1_ASFV/POL/2015       ................................................................................ [180080]
LS478113.1_Estonia_2014        ................................................................................ [180080]
KP843857.1_Odintsovo_02/14     ................................................................................ [180080]
FN557520.1_strain_E75          .........c....c................................................................. [180080]
AY261366.1_Warthog             .........c....c......................................a.......................... [180080]
AY261365.1_Warmbaths           .........c...............a...................................................... [180080]
AY261364.1_Tengani_62          .........c...................................................................... [180080]
AY261363.1_Pretoriuskop/96/4   .....a...c...............a...................................................... [180080]
AY261362.1_Mkuzi_1979          .........c....c................................................................. [180080]
AY261361.1_Malawi_Lil-20/1_1983 .........c................t..................................................... [180080]
AY261360.1_Kenya_1950          .........c................t.............ct..t.....g........t.................... [180080]
AM712240.1_OURT_88/3           .........c....c................................................................. [180080]
AM712239.1_Benin_97/1          .........c....c................................................................. [180080]
KM111295.1_Ken06.Bus           .........c................t........aa.g.c.........g........t.................... [180080]
```

```
KM111294.1_Ken05/Tk1          .........c.......a........t..t..........ct..t.....g........t.................... [180080]
KM262844.1_strain_L60         .........c....c................................................................. [180080]
KM102979.1_26544/OG10_Italy   .........c....c................................................................. [180080]
KM262845.1_strain_NHV         .........c....c................................................................. [180080]
KX354450.1_47/Ss/2008         .........c....c................................................................. [180080]
KP055815.1_strain_BA71        .........c....c................................................................. [180080]
MH025920.1_strain_R35         .........c................t........aa.g.c.........g........t.................... [180080]
MH025919.1_strain_N10         .........c................t........aa.g.c.........g........t.................... [180080]
MH025918.1_strain_R25         .........c................t........aa.g.c.........g........t.................... [180080]
MH025917.1_strain_R7          .........c................t........aa.g.c.........g........t.................... [180080]
MH025916.1_strain_R8          .........c................t........aa.g.c.........g........t.................... [180080]

FR682468.2_Georgia_2007/1     cccgctagaagccatggggcttacctttggggcaaccaacacggacgccgacccggaggatctgaaggacaaaacggtga [180160]
ASF_IND_20_CAD543             ................................................................................ [180160]
LR899193.1_Germany_2020/1     ................................................................................ [180160]
MN393477.1_Wuhan_2019         ................................................................................ [180160]
MW396979.1_Timor-Leste/2019/1 ................................................................................ [180160]
MK940252.1_CN/2019/InnerMongolia ................................................................................ [180160]
MK128995.1_China/2018/AnhuiXCGQ ................................................................................ [180160]
MH681419.1_ASFV/POL/2015      ................................................................................ [180160]
LS478113.1_Estonia_2014       ................................................................................ [180160]
KP843857.1_Odintsovo_02/14    ................................................................................ [180160]
FN557520.1_strain_E75         ................................................................................ [180160]
AY261366.1_Warthog            .....................a................................a......................... [180160]
AY261365.1_Warmbaths          .....................a.......................................................... [180160]
AY261364.1_Tengani_62         .....................a....................................a....a................ [180160]
AY261363.1_Pretoriuskop/96/4  .....................a.......................................................... [180160]
AY261362.1_Mkuzi_1979         ................................................................................ [180160]
AY261361.1_Malawi_Lil-20/1_1983 .........g.........................................................a............ [180160]
AY261360.1_Kenya_1950         ..................t.............................a....................t.......... [180160]
AM712240.1_OURT_88/3          ................................................................................ [180160]
AM712239.1_Benin_97/1         ................................................................................ [180160]
KM111295.1_Ken06.Bus          ..................a........................a....a....................t.......... [180160]
KM111294.1_Ken05/Tk1          ..................t.............................a....................t.......... [180160]
KM262844.1_strain_L60         ................................................................................ [180160]
KM102979.1_26544/OG10_Italy   ................................................................................ [180160]
KM262845.1_strain_NHV         ................................................................................ [180160]
KX354450.1_47/Ss/2008         ................................................................................ [180160]
KP055815.1_strain_BA71        ................................................................................ [180160]
MH025920.1_strain_R35         ..................a........................a....a....................t.......... [180160]
MH025919.1_strain_N10         ..................a........................a....a....................t.......... [180160]
MH025918.1_strain_R25         ..................a........................a....a....................t.......... [180160]
MH025917.1_strain_R7          ..................a........................a....a....................t.......... [180160]
MH025916.1_strain_R8          ..................a........................a....a....................t.......... [180160]
```

```
FR682468.2_Georgia_2007/1        taaatttaacgctcccgcaggcaacagaaagcatcacctaccatcttaaatccctaatgcagctaaaaaaagtaagtacg [180240]
ASF_IND_20_CAD543                ................................................................................ [180240]
LR899193.1_Germany_2020/1        ................................................................................ [180240]
MN393477.1_Wuhan_2019            ................................................................................ [180240]
MW396979.1_Timor-Leste/2019/1    ................................................................................ [180240]
MK940252.1_CN/2019/InnerMongolia ................................................................................ [180240]
MK128995.1_China/2018/AnhuiXCGQ  ................................................................................ [180240]
MH681419.1_ASFV/POL/2015         ................................................................................ [180240]
LS478113.1_Estonia_2014          ................................................................................ [180240]
KP843857.1_Odintsovo_02/14       ................................................................................ [180240]
FN557520.1_strain_E75            ................................................................................ [180240]
AY261366.1_Warthog               ......................g......................................................... [180240]
AY261365.1_Warmbaths             ......................g......................................................... [180240]
AY261364.1_Tengani_62            ................................................................................ [180240]
AY261363.1_Pretoriuskop/96/4     ......................g......................................................... [180240]
AY261362.1_Mkuzi_1979            ................................................................................ [180240]
AY261361.1_Malawi_Lil-20/1_1983  ...........................................c.................................... [180240]
AY261360.1_Kenya_1950            .............t..................g................t.............................. [180240]
AM712240.1_OURT_88/3             ................................................................................ [180240]
AM712239.1_Benin_97/1            ................................................................................ [180240]
KM111295.1_Ken06.Bus             .............t..................g................t.............................. [180240]
KM111294.1_Ken05/Tk1             ..........a..t..................g................t.............................. [180240]
KM262844.1_strain_L60            ................................................................................ [180240]
KM102979.1_26544/OG10_Italy      ................................................................................ [180240]
KM262845.1_strain_NHV            ................................................................................ [180240]
KX354450.1_47/Ss/2008            ................................................................................ [180240]
KP055815.1_strain_BA71           ................................................................................ [180240]
MH025920.1_strain_R35            .............t..................g................t.............................. [180240]
MH025919.1_strain_N10            .............t..................g................t.............................. [180240]
MH025918.1_strain_R25            .............t..................g................t.............................. [180240]
MH025917.1_strain_R7             .............t..................g................t.............................. [180240]
MH025916.1_strain_R8             .............t..................g................t.............................. [180240]

FR682468.2_Georgia_2007/1        gcttcaggactaaatacaaacattttgaaagcatttgataatattatttccacccctgtgaaaaaaaataaaatggcctc [180320]
ASF_IND_20_CAD543                ................................................................................ [180320]
LR899193.1_Germany_2020/1        ................................................................................ [180320]
MN393477.1_Wuhan_2019            ................................................................................ [180320]
MW396979.1_Timor-Leste/2019/1    ................................................................................ [180320]
MK940252.1_CN/2019/InnerMongolia ................................................................................ [180320]
MK128995.1_China/2018/AnhuiXCGQ  ................................................................................ [180320]
MH681419.1_ASFV/POL/2015         ................................................................................ [180320]
LS478113.1_Estonia_2014          ................................................................................ [180320]
KP843857.1_Odintsovo_02/14       ................................................................................ [180320]
```

```
FN557520.1_strain_E75             ................................................................................ [180320]
AY261366.1_Warthog                ................................g............................................... [180320]
AY261365.1_Warmbaths              ................................................................................ [180320]
AY261364.1_Tengani_62             ................................g............................................... [180320]
AY261363.1_Pretoriuskop/96/4      ................................g............................................... [180320]
AY261362.1_Mkuzi_1979             ................................c............................................... [180320]
AY261361.1_Malawi_Lil-20/1_1983   ................................g............................................... [180320]
AY261360.1_Kenya_1950             a...............................g............................................... [180320]
AM712240.1_OURT_88/3              ................................................................................ [180320]
AM712239.1_Benin_97/1             ................................................................................ [180320]
KM111295.1_Ken06.Bus              a...............................g..................g............................ [180320]
KM111294.1_Ken05/Tk1              a...............................g..................g............................ [180320]
KM262844.1_strain_L60             ................................................................................ [180320]
KM102979.1_26544/OG10_Italy       ................................................................................ [180320]
KM262845.1_strain_NHV             ................................................................................ [180320]
KX354450.1_47/Ss/2008             ................................................................................ [180320]
KP055815.1_strain_BA71            ................................................................................ [180320]
MH025920.1_strain_R35             a...............................g..................g............................ [180320]
MH025919.1_strain_N10             a...............................g..................g............................ [180320]
MH025918.1_strain_R25             a...............................g..................g............................ [180320]
MH025917.1_strain_R7              a...............................g..................g............................ [180320]
MH025916.1_strain_R8              a...............................g..................g............................ [180320]

FR682468.2_Georgia_2007/1         caagttggcgcccgggatggatgtcgtgttcactagcgataacggaaaaacattttttactaaaaacattttaagcaaaa [180400]
ASF_IND_20_CAD543                 ................................................................................ [180400]
LR899193.1_Germany_2020/1         ................................................................................ [180400]
MN393477.1_Wuhan_2019             ................................................................................ [180400]
MW396979.1_Timor-Leste/2019/1     ................................................................................ [180400]
MK940252.1_CN/2019/InnerMongolia  ................................................................................ [180400]
MK128995.1_China/2018/AnhuiXCGQ   ................................................................................ [180400]
MH681419.1_ASFV/POL/2015          ................................................................................ [180400]
LS478113.1_Estonia_2014           ................................................................................ [180400]
KP843857.1_Odintsovo_02/14        ................................................................................ [180400]
FN557520.1_strain_E75             ................................................................................ [180400]
AY261366.1_Warthog                .............................................g.................................. [180400]
AY261365.1_Warmbaths              ................................................................................ [180400]
AY261364.1_Tengani_62             ............................................................g................... [180400]
AY261363.1_Pretoriuskop/96/4      ................................................................................ [180400]
AY261362.1_Mkuzi_1979             .......a.t...................................................................... [180400]
AY261361.1_Malawi_Lil-20/1_1983   ............................................................g................... [180400]
AY261360.1_Kenya_1950             .........a.....a............................................g......g............ [180400]
AM712240.1_OURT_88/3              ................................................................................ [180400]
AM712239.1_Benin_97/1             ................................................................................ [180400]
KM111295.1_Ken06.Bus              ...............a............................................g......g............ [180400]
```

```
KM111294.1_Ken05/Tk1            ...............a............................................g......g............ [180400]
KM262844.1_strain_L60           ................................................................................ [180400]
KM102979.1_26544/OG10_Italy     ................................................................................ [180400]
KM262845.1_strain_NHV           ................................................................................ [180400]
KX354450.1_47/Ss/2008           ................................................................................ [180400]
KP055815.1_strain_BA71          ................................................................................ [180400]
MH025920.1_strain_R35           ...............a............................................g......g............ [180400]
MH025919.1_strain_N10           ...............a............................................g......g............ [180400]
MH025918.1_strain_R25           ...............a............................................g......g............ [180400]
MH025917.1_strain_R7            ...............a............................................g......g............ [180400]
MH025916.1_strain_R8            ...............a............................................g......g............ [180400]

FR682468.2_Georgia_2007/1       acatgctagcggggcccaaagagcgggtgtttgcatataataatctcattagtaatttaaataactcctgtttcatacaa [180480]
ASF_IND_20_CAD543               ................................................................y............... [180480]
LR899193.1_Germany_2020/1       ................................................................................ [180480]
MN393477.1_Wuhan_2019           ................................................................................ [180480]
MW396979.1_Timor-Leste/2019/1   ................................................................................ [180480]
MK940252.1_CN/2019/InnerMongolia ................................................................................ [180480]
MK128995.1_China/2018/AnhuiXCGQ ................................................................................ [180480]
MH681419.1_ASFV/POL/2015        ................................................................................ [180480]
LS478113.1_Estonia_2014         ................................................................................ [180480]
KP843857.1_Odintsovo_02/14      ................................................................................ [180480]
FN557520.1_strain_E75           ................................................................t............... [180480]
AY261366.1_Warthog              ................................................................................ [180480]
AY261365.1_Warmbaths            ................................................................................ [180480]
AY261364.1_Tengani_62           ................................................................................ [180480]
AY261363.1_Pretoriuskop/96/4    ................................................................................ [180480]
AY261362.1_Mkuzi_1979           ................................................................t............... [180480]
AY261361.1_Malawi_Lil-20/1_1983 ...................................................a............................ [180480]
AY261360.1_Kenya_1950           ......................................................................c..t...... [180480]
AM712240.1_OURT_88/3            ................................................................t............... [180480]
AM712239.1_Benin_97/1           ................................................................t............... [180480]
KM111295.1_Ken06.Bus            ......................................................................c......... [180480]
KM111294.1_Ken05/Tk1            ......................................................................c..t...... [180480]
KM262844.1_strain_L60           ................................................................t............... [180480]
KM102979.1_26544/OG10_Italy     ................................................................t............... [180480]
KM262845.1_strain_NHV           ................................................................t............... [180480]
KX354450.1_47/Ss/2008           ................................................................t............... [180480]
KP055815.1_strain_BA71          ................................................................t............... [180480]
MH025920.1_strain_R35           ......................................................................c......... [180480]
MH025919.1_strain_N10           ......................................................................c......... [180480]
MH025918.1_strain_R25           ......................................................................c......... [180480]
MH025917.1_strain_R7            ......................................................................c......... [180480]
MH025916.1_strain_R8            ......................................................................c......... [180480]
```

```
FR682468.2_Georgia_2007/1        aatcacaacgattttttaagacagcaggactcttggcccttctatgacgcgcacaattttaccaacaagtttttaatgca [180560]
ASF_IND_20_CAD543                ................................................................................ [180560]
LR899193.1_Germany_2020/1        ................................................................................ [180560]
MN393477.1_Wuhan_2019            ................................................................................ [180560]
MW396979.1_Timor-Leste/2019/1    ................................................................................ [180560]
MK940252.1_CN/2019/InnerMongolia ................................................................................ [180560]
MK128995.1_China/2018/AnhuiXCGQ  ................................................................................ [180560]
MH681419.1_ASFV/POL/2015         ................................................................................ [180560]
LS478113.1_Estonia_2014          ................................................................................ [180560]
KP843857.1_Odintsovo_02/14       ................................................................................ [180560]
FN557520.1_strain_E75            ................................................................................ [180560]
AY261366.1_Warthog               ................................................................................ [180560]
AY261365.1_Warmbaths             ................................................................................ [180560]
AY261364.1_Tengani_62            ................................................................................ [180560]
AY261363.1_Pretoriuskop/96/4     ................................................................................ [180560]
AY261362.1_Mkuzi_1979            ................................................................................ [180560]
AY261361.1_Malawi_Lil-20/1_1983  .......c...........a..............................c.................a........... [180560]
AY261360.1_Kenya_1950            ..c..............g................................a.................a........... [180560]
AM712240.1_OURT_88/3             ..c............................................................................. [180560]
AM712239.1_Benin_97/1            ................................................................................ [180560]
KM111295.1_Ken06.Bus             .................g................................a.................a........... [180560]
KM111294.1_Ken05/Tk1             ..c..............g................................a.................a........... [180560]
KM262844.1_strain_L60            ................................................................................ [180560]
KM102979.1_26544/OG10_Italy      ................................................................................ [180560]
KM262845.1_strain_NHV            ..c............................................................................. [180560]
KX354450.1_47/Ss/2008            ................................................................................ [180560]
KP055815.1_strain_BA71           ................................................................................ [180560]
MH025920.1_strain_R35            .................g................................a.................a........... [180560]
MH025919.1_strain_N10            .................g................................a.................a........... [180560]
MH025918.1_strain_R25            .................g................................a.................a........... [180560]
MH025917.1_strain_R7             .................g................................a.................a........... [180560]
MH025916.1_strain_R8             .................g................................a.................a........... [180560]

FR682468.2_Georgia_2007/1        gcctattttttcggggcagacccgtcctcggcttcagggagccatggaggcggcgcatgtggaaacgcatctcacggcat [180640]
ASF_IND_20_CAD543                ................................................................................ [180640]
LR899193.1_Germany_2020/1        ................................................................................ [180640]
MN393477.1_Wuhan_2019            ................................................................................ [180640]
MW396979.1_Timor-Leste/2019/1    ................................................................................ [180640]
MK940252.1_CN/2019/InnerMongolia ................................................................................ [180640]
MK128995.1_China/2018/AnhuiXCGQ  ................................................................................ [180640]
MH681419.1_ASFV/POL/2015         ................................................................................ [180640]
LS478113.1_Estonia_2014          ................................................................................ [180640]
KP843857.1_Odintsovo_02/14       ................................................................................ [180640]
```

```
FN557520.1_strain_E75           .........c............................................................t....... [180640]
AY261366.1_Warthog              ............a..............c............................................t....... [180640]
AY261365.1_Warmbaths            ............a..............c............................................t....... [180640]
AY261364.1_Tengani_62           ........................................................g...............t....... [180640]
AY261363.1_Pretoriuskop/96/4    ........................................................................t....... [180640]
AY261362.1_Mkuzi_1979           .........c..............................................................t....... [180640]
AY261361.1_Malawi_Lil-20/1_1983 ............a..a.......................g...........c....................t.....g. [180640]
AY261360.1_Kenya_1950           ............a....................c.................c.....c..a...........t....... [180640]
AM712240.1_OURT_88/3            .........c..............................................................t....... [180640]
AM712239.1_Benin_97/1           .........c..............................................................t....... [180640]
KM111295.1_Ken06.Bus            ............a....................c.................c.....c..a.......g...t....... [180640]
KM111294.1_Ken05/Tk1            ............a....................c.................c.....c..a...........t....... [180640]
KM262844.1_strain_L60           .........c..............................................................t....... [180640]
KM102979.1_26544/OG10_Italy     .........c..............................................................t....... [180640]
KM262845.1_strain_NHV           .........c..............................................................t....... [180640]
KX354450.1_47/Ss/2008           .........c..............................................................t....... [180640]
KP055815.1_strain_BA71          .........c..............................................................t....... [180640]
MH025920.1_strain_R35           ............a....................c.................c.....c..a.......g...t....... [180640]
MH025919.1_strain_N10           ............a....................c.................c.....c..a.......g...t....... [180640]
MH025918.1_strain_R25           ............a....................c.................c.....c..a.......g...t....... [180640]
MH025917.1_strain_R7            ............a....................c.................c.....c..a.......g...t....... [180640]
MH025916.1_strain_R8            ............a....................c.................c.....c..a.......g...t....... [180640]

FR682468.2_Georgia_2007/1       ttttacaaagtattcagccctctaggccacaagatccctctgttttggcttcccccaagttatctgctctaatcttgaac [180720]
ASF_IND_20_CAD543               ................................................................................ [180720]
LR899193.1_Germany_2020/1       ................................................................................ [180720]
MN393477.1_Wuhan_2019           ................................................................................ [180720]
MW396979.1_Timor-Leste/2019/1   ................................................................................ [180720]
MK940252.1_CN/2019/InnerMongolia ................................................................................ [180720]
MK128995.1_China/2018/AnhuiXCGQ ................................................................................ [180720]
MH681419.1_ASFV/POL/2015        ................................................................................ [180720]
LS478113.1_Estonia_2014         ................................................................................ [180720]
KP843857.1_Odintsovo_02/14      ................................................................................ [180720]
FN557520.1_strain_E75           ................................................................................ [180720]
AY261366.1_Warthog              ................................................................................ [180720]
AY261365.1_Warmbaths            ............................g................................................... [180720]
AY261364.1_Tengani_62           ................................................................................ [180720]
AY261363.1_Pretoriuskop/96/4    ............................g................................................... [180720]
AY261362.1_Mkuzi_1979           ................................................................................ [180720]
AY261361.1_Malawi_Lil-20/1_1983 .......................c...................g.................................... [180720]
AY261360.1_Kenya_1950           ....g...........a..t..c..................a.c.................................... [180720]
AM712240.1_OURT_88/3            ................................................................................ [180720]
AM712239.1_Benin_97/1           ................................................................................ [180720]
KM111295.1_Ken06.Bus            ....g...........a..t..c..................a...................................... [180720]
```

```
KM111294.1_Ken05/Tk1            ....g...........a..t..c..................a...................................... [180720]
KM262844.1_strain_L60           ................................................................................ [180720]
KM102979.1_26544/OG10_Italy     ................................................................................ [180720]
KM262845.1_strain_NHV           ................................................................................ [180720]
KX354450.1_47/Ss/2008           ................................................................................ [180720]
KP055815.1_strain_BA71          ................................................................................ [180720]
MH025920.1_strain_R35           ....g...........a..t..c..................a...................................... [180720]
MH025919.1_strain_N10           ....g...........a..t..c..................a...................................... [180720]
MH025918.1_strain_R25           ....g...........a..t..c..................a...................................... [180720]
MH025917.1_strain_R7            ....g...........a..t..c..................a...................................... [180720]
MH025916.1_strain_R8            ....g...........a..t..c..................a...................................... [180720]

FR682468.2_Georgia_2007/1       ta------------------------------------------------------------------------------ [180800]
ASF_IND_20_CAD543               ..------------------------------------------------------------------------------ [180800]
LR899193.1_Germany_2020/1       ..------------------------------------------------------------------------------ [180800]
MN393477.1_Wuhan_2019           ..------------------------------------------------------------------------------ [180800]
MW396979.1_Timor-Leste/2019/1   ..------------------------------------------------------------------------------ [180800]
MK940252.1_CN/2019/InnerMongolia ..------------------------------------------------------------------------------ [180800]
MK128995.1_China/2018/AnhuiXCGQ ..------------------------------------------------------------------------------ [180800]
MH681419.1_ASFV/POL/2015        ..------------------------------------------------------------------------------ [180800]
LS478113.1_Estonia_2014         ..------------------------------------------------------------------------------ [180800]
KP843857.1_Odintsovo_02/14      ..------------------------------------------------------------------------------ [180800]
FN557520.1_strain_E75           ..------------------------------------------------------------------------------ [180800]
AY261366.1_Warthog              ..------------------------------------------------------------------------------ [180800]
AY261365.1_Warmbaths            ..------------------------------------------------------------------------------ [180800]
AY261364.1_Tengani_62           ..------------------------------------------------------------------------------ [180800]
AY261363.1_Pretoriuskop/96/4    ..------------------------------------------------------------------------------ [180800]
AY261362.1_Mkuzi_1979           ..------------------------------------------------------------------------------ [180800]
AY261361.1_Malawi_Lil-20/1_1983 .------------------------------------------------------------------------------- [180800]
AY261360.1_Kenya_1950           ..aagtcttgaa----------------------------------------------------taaagtcttgaataaa [180800]
AM712240.1_OURT_88/3            ..------------------------------------------------------------------------------ [180800]
AM712239.1_Benin_97/1           ..------------------------------------------------------------------------------ [180800]
KM111295.1_Ken06.Bus            ..----------------------------------------------------------------aaatcttgaataaa [180800]
KM111294.1_Ken05/Tk1            ..aagtcttgaataaaatcttgaactaaaatcttgaataaaatcttgaaaataaaatcttgaactaaaatcttgaataaa [180800]
KM262844.1_strain_L60           ..------------------------------------------------------------------------------ [180800]
KM102979.1_26544/OG10_Italy     ..------------------------------------------------------------------------------ [180800]
KM262845.1_strain_NHV           ..------------------------------------------------------------------------------ [180800]
KX354450.1_47/Ss/2008           ..------------------------------------------------------------------------------ [180800]
KP055815.1_strain_BA71          ..------------------------------------------------------------------------------ [180800]
MH025920.1_strain_R35           ..----------------------------------------------------------------aaatcttgaataaa [180800]
MH025919.1_strain_N10           ..----------------------------------------------------------------aaatcttgaataaa [180800]
MH025918.1_strain_R25           ..----------------------------------------------------------------aaatcttgaataaa [180800]
MH025917.1_strain_R7            ..----------------------------------------------------------------aaatcttgaataaa [180800]
MH025916.1_strain_R8            ..----------------------------------------------------------------aaatcttgaataaa [180800]
```

```
FR682468.2_Georgia_2007/1        -----------aaaacag--cctttcttggacttaaatgatggtctaccagtttttgaaataacttagagaactatgaag [180880]
ASF_IND_20_CAD543                -----------.....g.ct............................................................ [180880]
LR899193.1_Germany_2020/1        -----------.......--............................................................ [180880]
MN393477.1_Wuhan_2019            -----------.......--............................................................ [180880]
MW396979.1_Timor-Leste/2019/1    -----------.......--............................................................ [180880]
MK940252.1_CN/2019/InnerMongolia -----------.......--............................................................ [180880]
MK128995.1_China/2018/AnhuiXCGQ  -----------.......--............................................................ [180880]
MH681419.1_ASFV/POL/2015         -----------.......--............................................................ [180880]
LS478113.1_Estonia_2014          -----------.......--............................................................ [180880]
KP843857.1_Odintsovo_02/14       -----------.......--............................................................ [180880]
FN557520.1_strain_E75            -----------.....g.ct..........................g................................. [180880]
AY261366.1_Warthog               -----------....g..ct.................--------.g.................--.............. [180880]
AY261365.1_Warmbaths             -----------.......ct..........................g.................--.............. [180880]
AY261364.1_Tengani_62            -----------.....g.c-...........t..............g................................. [180880]
AY261363.1_Pretoriuskop/96/4     -----------.......c-..........................g................................. [180880]
AY261362.1_Mkuzi_1979            -----------.......ct..........................g.t............................... [180880]
AY261361.1_Malawi_Lil-20/1_1983  -----------.....g.tta....t.a.att..........a..--..ga......................ggc..-- [180880]
AY261360.1_Kenya_1950            atcttgaacta.......c-tac........t............---.a..c.........c.......c.g...a.... [180880]
AM712240.1_OURT_88/3             -----------.....g.ct..........................g................................. [180880]
AM712239.1_Benin_97/1            -----------.....g.ct..........................g................................. [180880]
KM111295.1_Ken06.Bus             atcttgaacta.......c-tac........t............---.a..c.........c.......c.g...c.... [180880]
KM111294.1_Ken05/Tk1             atcttgaacta.......c-tac........t.........c..---.a..c.........c.......c.g...c.... [180880]
KM262844.1_strain_L60            -----------.....g.ct..........................g................................. [180880]
KM102979.1_26544/OG10_Italy      -----------.....g.ct..........................g................................. [180880]
KM262845.1_strain_NHV            -----------.....g.ct..........................g................................. [180880]
KX354450.1_47/Ss/2008            -----------.....g.ct..........................g................................. [180880]
KP055815.1_strain_BA71           -----------.....g.ct..........................g................................. [180880]
MH025920.1_strain_R35            atcttgaacta.......c-tac........t............---.a..c.........c.......c.g...c.... [180880]
MH025919.1_strain_N10            atcttgaacta.......c-tac........t............---.a..c.........c.......c.g...c.... [180880]
MH025918.1_strain_R25            atcttgaacta.......c-tac........t............---.a..c.........c.......c.g...c.... [180880]
MH025917.1_strain_R7             atcttgaacta.......c-tac........t............---.a..c.........c.......c.g...c.... [180880]
MH025916.1_strain_R8             atcttgaacta.......c-tac........t............---.a..c.........c.......c.g...c.... [180880]

FR682468.2_Georgia_2007/1        a-ttttcatgaaatttaaattagagatttgcaaaggttacttgcggtcattttctgttgaattaaataatt-attcgaat [180960]
ASF_IND_20_CAD543                .-..........................................................w..........-...m.... [180960]
LR899193.1_Germany_2020/1        .-.....................................................................-........ [180960]
MN393477.1_Wuhan_2019            .-.....................................................................-........ [180960]
MW396979.1_Timor-Leste/2019/1    .-.....................................................................-........ [180960]
MK940252.1_CN/2019/InnerMongolia .-.....................................................................-........ [180960]
MK128995.1_China/2018/AnhuiXCGQ  .-.....................................................................-........ [180960]
MH681419.1_ASFV/POL/2015         .-.....................................................................-........ [180960]
LS478113.1_Estonia_2014          .-.....................................................................-........ [180960]
KP843857.1_Odintsovo_02/14       .-.....................................................................-........ [180960]
```

```
FN557520.1_strain_E75              .-..........................................................t..........-...a.... [180960]
AY261366.1_Warthog                 -------........................................................g.......-........ [180960]
AY261365.1_Warmbaths               -------................................................................-...a.... [180960]
AY261364.1_Tengani_62              .t.....................................................................-........ [180960]
AY261363.1_Pretoriuskop/96/4       .-....................a...c............................................-...a.... [180960]
AY261362.1_Mkuzi_1979              .-.......................................c..................t..........-...a.... [180960]
AY261361.1_Malawi_Lil-20/1_1983    -----c.................................................................a...a.... [180960]
AY261360.1_Kenya_1950              .-..........t.....t.........................................t..........-...a.... [180960]
AM712240.1_OURT_88/3               .-..........................................................t..........-...a.... [180960]
AM712239.1_Benin_97/1              .-..........................................................t..........-...a.... [180960]
KM111295.1_Ken06.Bus               .g..........t...............................................t..........-...a.... [180960]
KM111294.1_Ken05/Tk1               .-..........t.....t.........................................t..........-...a.... [180960]
KM262844.1_strain_L60              .-..........................................................t..........-...a.... [180960]
KM102979.1_26544/OG10_Italy        .-..........................................................t..........-...a.... [180960]
KM262845.1_strain_NHV              .-..........................................................t..........-...a.... [180960]
KX354450.1_47/Ss/2008              .-..........................................................t..........-...a.... [180960]
KP055815.1_strain_BA71             .-..........................................................t..........-...a.... [180960]
MH025920.1_strain_R35              .g..........t...............................................t..........-...a.... [180960]
MH025919.1_strain_N10              .g..........t...............................................t..........-...a.... [180960]
MH025918.1_strain_R25              .g..........t...............................................t..........-...a.... [180960]
MH025917.1_strain_R7               .g..........t...............................................t..........-...a.... [180960]
MH025916.1_strain_R8               .g..........t...............................................t..........-...a.... [180960]

FR682468.2_Georgia_2007/1          agtataatgtctgaagatattcgtcgtggtcctggcagaccgccaaagaaaagggttgttcccaactttgagcgcaaggg [181040]
ASF_IND_20_CAD543                  ................................................................................ [181040]
LR899193.1_Germany_2020/1          ................................................................................ [181040]
MN393477.1_Wuhan_2019              ................................................................................ [181040]
MW396979.1_Timor-Leste/2019/1      ................................................................................ [181040]
MK940252.1_CN/2019/InnerMongolia   ................................................................................ [181040]
MK128995.1_China/2018/AnhuiXCGQ    ................................................................................ [181040]
MH681419.1_ASFV/POL/2015           ................................................................................ [181040]
LS478113.1_Estonia_2014            ................................................................................ [181040]
KP843857.1_Odintsovo_02/14         ................................................................................ [181040]
FN557520.1_strain_E75              .............................a........g......................................... [181040]
AY261366.1_Warthog                 ................................................................................ [181040]
AY261365.1_Warmbaths               ................................................................................ [181040]
AY261364.1_Tengani_62              ................................................................................ [181040]
AY261363.1_Pretoriuskop/96/4       ..........................c..................................................... [181040]
AY261362.1_Mkuzi_1979              .............................a........g......................................... [181040]
AY261361.1_Malawi_Lil-20/1_1983    ..........................c..a.................................................. [181040]
AY261360.1_Kenya_1950              ..........................c..a.................................................. [181040]
AM712240.1_OURT_88/3               .............................a........g......................................... [181040]
AM712239.1_Benin_97/1              .............................a........g......................................... [181040]
KM111295.1_Ken06.Bus               ..........................c..g.......................a.......................... [181040]
```

```
KM111294.1_Ken05/Tk1            ..........................c..a.................................................. [181040]
KM262844.1_strain_L60           .............................a........g......................................... [181040]
KM102979.1_26544/OG10_Italy     .............................a........g......................................... [181040]
KM262845.1_strain_NHV           .............................a........g......................................... [181040]
KX354450.1_47/Ss/2008           .............................a........g......................................... [181040]
KP055815.1_strain_BA71          .............................a........g......................................... [181040]
MH025920.1_strain_R35           ..........................c..g.......................a.......................... [181040]
MH025919.1_strain_N10           ..........................c..g.......................a.......................... [181040]
MH025918.1_strain_R25           ..........................c..g.......................a.......................... [181040]
MH025917.1_strain_R7            ..........................c..g.......................a.......................... [181040]
MH025916.1_strain_R8            ..........................c..g.......................a.......................... [181040]

FR682468.2_Georgia_2007/1       cattctggaaaaaccagttcggccacaaagccgtctcgagttttcctatgataacccgctgatatttaaaaatcttttta [181120]
ASF_IND_20_CAD543               ................................................................................ [181120]
LR899193.1_Germany_2020/1       ................................................................................ [181120]
MN393477.1_Wuhan_2019           ................................................................................ [181120]
MW396979.1_Timor-Leste/2019/1   ................................................................................ [181120]
MK940252.1_CN/2019/InnerMongolia ................................................................................ [181120]
MK128995.1_China/2018/AnhuiXCGQ ................................................................................ [181120]
MH681419.1_ASFV/POL/2015        ................................................................................ [181120]
LS478113.1_Estonia_2014         ................................................................................ [181120]
KP843857.1_Odintsovo_02/14      ................................................................................ [181120]
FN557520.1_strain_E75           ................................................................................ [181120]
AY261366.1_Warthog              ................................................................................ [181120]
AY261365.1_Warmbaths            ................................................................................ [181120]
AY261364.1_Tengani_62           ................................................................................ [181120]
AY261363.1_Pretoriuskop/96/4    ....t........................................................................... [181120]
AY261362.1_Mkuzi_1979           ................................................................................ [181120]
AY261361.1_Malawi_Lil-20/1_1983 ....t....g..............t................................c...................... [181120]
AY261360.1_Kenya_1950           g........g.....c................................................................ [181120]
AM712240.1_OURT_88/3            ................................................................................ [181120]
AM712239.1_Benin_97/1           ................................................................................ [181120]
KM111295.1_Ken06.Bus            g........g.....c................................................................ [181120]
KM111294.1_Ken05/Tk1            g........g.....c................................................................ [181120]
KM262844.1_strain_L60           ................................................................................ [181120]
KM102979.1_26544/OG10_Italy     ................................................................................ [181120]
KM262845.1_strain_NHV           ................................................................................ [181120]
KX354450.1_47/Ss/2008           ................................................................................ [181120]
KP055815.1_strain_BA71          ................................................................................ [181120]
MH025920.1_strain_R35           g........g.....c................................................................ [181120]
MH025919.1_strain_N10           g........g.....c................................................................ [181120]
MH025918.1_strain_R25           g........g.....c................................................................ [181120]
MH025917.1_strain_R7            g........g.....c................................................................ [181120]
MH025916.1_strain_R8            g........g.....c................................................................ [181120]
```

```
FR682468.2_Georgia_2007/1        tttactttaaaaaccttaaaagtaaaaatattttggtgcgatgtacccccaccgagattacctttttttcacgtgaccag [181200]
ASF_IND_20_CAD543                ....t........................................................................... [181200]
LR899193.1_Germany_2020/1        ................................................................................ [181200]
MN393477.1_Wuhan_2019            ................................................................................ [181200]
MW396979.1_Timor-Leste/2019/1    ................................................................................ [181200]
MK940252.1_CN/2019/InnerMongolia ................................................................................ [181200]
MK128995.1_China/2018/AnhuiXCGQ  ................................................................................ [181200]
MH681419.1_ASFV/POL/2015         ................................................................................ [181200]
LS478113.1_Estonia_2014          ................................................................................ [181200]
KP843857.1_Odintsovo_02/14       ................................................................................ [181200]
FN557520.1_strain_E75            ....t........................................................................... [181200]
AY261366.1_Warthog               ................................................................................ [181200]
AY261365.1_Warmbaths             ............t................................................................... [181200]
AY261364.1_Tengani_62            ................................................................................ [181200]
AY261363.1_Pretoriuskop/96/4     ................................................................................ [181200]
AY261362.1_Mkuzi_1979            ................................................................................ [181200]
AY261361.1_Malawi_Lil-20/1_1983  ............t................................................................... [181200]
AY261360.1_Kenya_1950            ............t................................................................... [181200]
AM712240.1_OURT_88/3             ....t........................................................................... [181200]
AM712239.1_Benin_97/1            ....t........................................................................... [181200]
KM111295.1_Ken06.Bus             ............t................................................................... [181200]
KM111294.1_Ken05/Tk1             ............t................................................................... [181200]
KM262844.1_strain_L60            ....t........................................................................... [181200]
KM102979.1_26544/OG10_Italy      ....t........................................................................... [181200]
KM262845.1_strain_NHV            ....t........................................................................... [181200]
KX354450.1_47/Ss/2008            ....t........................................................................... [181200]
KP055815.1_strain_BA71           ....t........................................................................... [181200]
MH025920.1_strain_R35            ............t................................................................... [181200]
MH025919.1_strain_N10            ............t................................................................... [181200]
MH025918.1_strain_R25            ............t................................................................... [181200]
MH025917.1_strain_R7             ............t................................................................... [181200]
MH025916.1_strain_R8             ............t................................................................... [181200]

FR682468.2_Georgia_2007/1        tcgcaggcaagctttgttattgccaccatcgacggaaaaaacgtgaaccattattacgccagtgatgtcttttggctagg [181280]
ASF_IND_20_CAD543                ................................................................................ [181280]
LR899193.1_Germany_2020/1        ................................................................................ [181280]
MN393477.1_Wuhan_2019            ................................................................................ [181280]
MW396979.1_Timor-Leste/2019/1    ................................................................................ [181280]
MK940252.1_CN/2019/InnerMongolia ................................................................................ [181280]
MK128995.1_China/2018/AnhuiXCGQ  ................................................................................ [181280]
MH681419.1_ASFV/POL/2015         ................................................................................ [181280]
LS478113.1_Estonia_2014          ................................................................................ [181280]
KP843857.1_Odintsovo_02/14       ................................................................................ [181280]
```

```
FN557520.1_strain_E75          .....a.......................................................................g.. [181280]
AY261366.1_Warthog             ................................................................................ [181280]
AY261365.1_Warmbaths           .....a.......................................................................... [181280]
AY261364.1_Tengani_62          .....a.......................................................................... [181280]
AY261363.1_Pretoriuskop/96/4   .....a.......................................................................... [181280]
AY261362.1_Mkuzi_1979          .....a.......................................................................g.. [181280]
AY261361.1_Malawi_Lil-20/1_1983 ....................c....................t..a................................... [181280]
AY261360.1_Kenya_1950          .....a.......................................................................... [181280]
AM712240.1_OURT_88/3           .....a.......................................................................g.. [181280]
AM712239.1_Benin_97/1          .....a.......................................................................g.. [181280]
KM111295.1_Ken06.Bus           ................................................................................ [181280]
KM111294.1_Ken05/Tk1           .....a.......................................................................... [181280]
KM262844.1_strain_L60          .....a.......................................................................g.. [181280]
KM102979.1_26544/OG10_Italy    .....a.......................................................................g.. [181280]
KM262845.1_strain_NHV          .....a.......................................................................g.. [181280]
KX354450.1_47/Ss/2008          .....a.......................................................................g.. [181280]
KP055815.1_strain_BA71         .....a.......................................................................g.. [181280]
MH025920.1_strain_R35          ................................................................................ [181280]
MH025919.1_strain_N10          ................................................................................ [181280]
MH025918.1_strain_R25          ................................................................................ [181280]
MH025917.1_strain_R7           ................................................................................ [181280]
MH025916.1_strain_R8           ................................................................................ [181280]

FR682468.2_Georgia_2007/1      catcaacagagagctcgttgaaaaaatgtttaacagcattgatcgctcttttttaaaaattaccatcgttcaccgctatg [181360]
ASF_IND_20_CAD543              ................................................................................ [181360]
LR899193.1_Germany_2020/1      ................................................................................ [181360]
MN393477.1_Wuhan_2019          ................................................................................ [181360]
MW396979.1_Timor-Leste/2019/1  ................................................................................ [181360]
MK940252.1_CN/2019/InnerMongolia ................................................................................ [181360]
MK128995.1_China/2018/AnhuiXCGQ ................................................................................ [181360]
MH681419.1_ASFV/POL/2015       ................................................................................ [181360]
LS478113.1_Estonia_2014        ................................................................................ [181360]
KP843857.1_Odintsovo_02/14     ................................................................................ [181360]
FN557520.1_strain_E75          ......t...................................c..................................... [181360]
AY261366.1_Warthog             ......t...................................c..................................... [181360]
AY261365.1_Warmbaths           ......t...................................c..................................... [181360]
AY261364.1_Tengani_62          ............a.............................c..................................... [181360]
AY261363.1_Pretoriuskop/96/4   ......t...................................c..................................... [181360]
AY261362.1_Mkuzi_1979          ......t...................................c..................................... [181360]
AY261361.1_Malawi_Lil-20/1_1983 ......t................................c..c..........................c.......... [181360]
AY261360.1_Kenya_1950          ......t...................................c.................c................... [181360]
AM712240.1_OURT_88/3           ......t...................................c..................................... [181360]
AM712239.1_Benin_97/1          ......t...................................c..................................... [181360]
KM111295.1_Ken06.Bus           ......t........t..........................c..................................... [181360]
```

```
KM111294.1_Ken05/Tk1            ......t...................................c.................c................... [181360]
KM262844.1_strain_L60           ......t...................................c..................................... [181360]
KM102979.1_26544/OG10_Italy     ......t...................................c..................................... [181360]
KM262845.1_strain_NHV           ......t...................................c..................................... [181360]
KX354450.1_47/Ss/2008           ......t...................................c..................................... [181360]
KP055815.1_strain_BA71          ......t...................................c..................................... [181360]
MH025920.1_strain_R35           ......t........t..........................c.................c................... [181360]
MH025919.1_strain_N10           ......t........t..........................c.................c................... [181360]
MH025918.1_strain_R25           ......t........t..........................c.................c................... [181360]
MH025917.1_strain_R7            ......t........t..........................c.................c................... [181360]
MH025916.1_strain_R8            ......t........t..........................c.................c................... [181360]

FR682468.2_Georgia_2007/1       acaagcctgaaaccctgttttttatctttacggattttgacattgacaaggagtgcacgtatcagattacggtctcggag [181440]
ASF_IND_20_CAD543               ................................................................................ [181440]
LR899193.1_Germany_2020/1       ................................................................................ [181440]
MN393477.1_Wuhan_2019           ................................................................................ [181440]
MW396979.1_Timor-Leste/2019/1   ................................................................................ [181440]
MK940252.1_CN/2019/InnerMongolia ................................................................................ [181440]
MK128995.1_China/2018/AnhuiXCGQ ................................................................................ [181440]
MH681419.1_ASFV/POL/2015        ................................................................................ [181440]
LS478113.1_Estonia_2014         ................................................................................ [181440]
KP843857.1_Odintsovo_02/14      ................................................................................ [181440]
FN557520.1_strain_E75           ................................................................................ [181440]
AY261366.1_Warthog              .................................................a.............................. [181440]
AY261365.1_Warmbaths            ...........................................c....................c............... [181440]
AY261364.1_Tengani_62           ........................................t....................................... [181440]
AY261363.1_Pretoriuskop/96/4    ...........................................c....................c............... [181440]
AY261362.1_Mkuzi_1979           ................................................................................ [181440]
AY261361.1_Malawi_Lil-20/1_1983 ..................a...................................................a......... [181440]
AY261360.1_Kenya_1950           ................................................................................ [181440]
AM712240.1_OURT_88/3            ................................................................................ [181440]
AM712239.1_Benin_97/1           ................................................................................ [181440]
KM111295.1_Ken06.Bus            ......................................................................a......... [181440]
KM111294.1_Ken05/Tk1            ................................................................................ [181440]
KM262844.1_strain_L60           ................................................................................ [181440]
KM102979.1_26544/OG10_Italy     ................................................................................ [181440]
KM262845.1_strain_NHV           ................................................................................ [181440]
KX354450.1_47/Ss/2008           ................................................................................ [181440]
KP055815.1_strain_BA71          ................................................................................ [181440]
MH025920.1_strain_R35           ......................................................................a......... [181440]
MH025919.1_strain_N10           ......................................................................a......... [181440]
MH025918.1_strain_R25           ......................................................................a......... [181440]
MH025917.1_strain_R7            ......................................................................a......... [181440]
MH025916.1_strain_R8            ......................................................................a......... [181440]
```

```
FR682468.2_Georgia_2007/1        cccgagctcgatatggaccttatcgaaatggaaaaaagcatcagtgaagaaagactcaagaactatcctctgcgctggga [181520]
ASF_IND_20_CAD543                ................................................................................ [181520]
LR899193.1_Germany_2020/1        ................................................................................ [181520]
MN393477.1_Wuhan_2019            ................................................................................ [181520]
MW396979.1_Timor-Leste/2019/1    ................................................................................ [181520]
MK940252.1_CN/2019/InnerMongolia ................................................................................ [181520]
MK128995.1_China/2018/AnhuiXCGQ  ................................................................................ [181520]
MH681419.1_ASFV/POL/2015         ................................................................................ [181520]
LS478113.1_Estonia_2014          ................................................................................ [181520]
KP843857.1_Odintsovo_02/14       ................................................................................ [181520]
FN557520.1_strain_E75            ................................................................................ [181520]
AY261366.1_Warthog               .......................t..............t....................a.........t.......... [181520]
AY261365.1_Warmbaths             .......................t...................................a..............g..... [181520]
AY261364.1_Tengani_62            .......................t.......................g...........a..............g..... [181520]
AY261363.1_Pretoriuskop/96/4     .......................t...................................a..............g..... [181520]
AY261362.1_Mkuzi_1979            ................................................................................ [181520]
AY261361.1_Malawi_Lil-20/1_1983  ........t..............t.............................g.....a..............g..... [181520]
AY261360.1_Kenya_1950            ........t..............t.............................g.....a..............g..... [181520]
AM712240.1_OURT_88/3             ................................................................................ [181520]
AM712239.1_Benin_97/1            ................................................................................ [181520]
KM111295.1_Ken06.Bus             ........t..............t.............................g.....a..t...........g..... [181520]
KM111294.1_Ken05/Tk1             ........t..............t.............................g.....a..............g..... [181520]
KM262844.1_strain_L60            ................................................................................ [181520]
KM102979.1_26544/OG10_Italy      ................................................................................ [181520]
KM262845.1_strain_NHV            ................................................................................ [181520]
KX354450.1_47/Ss/2008            ................................................................................ [181520]
KP055815.1_strain_BA71           ................................................................................ [181520]
MH025920.1_strain_R35            ........t..............t.............................g.....a..t...........g..... [181520]
MH025919.1_strain_N10            ........t..............t.............................g.....a..t...........g..... [181520]
MH025918.1_strain_R25            ........t..............t.............................g.....a..t...........g..... [181520]
MH025917.1_strain_R7             ........t..............t.............................g.....a..t...........g..... [181520]
MH025916.1_strain_R8             ........t..............t.............................g.....a..t...........g..... [181520]

FR682468.2_Georgia_2007/1        gtttacctccaagcagctcaagaaaacatttagcgacttatcaaactacaccgagctcgtgaccattgaaaaactcggcg [181600]
ASF_IND_20_CAD543                ................................................................................ [181600]
LR899193.1_Germany_2020/1        ................................................................................ [181600]
MN393477.1_Wuhan_2019            ................................................................................ [181600]
MW396979.1_Timor-Leste/2019/1    ................................................................................ [181600]
MK940252.1_CN/2019/InnerMongolia ................................................................................ [181600]
MK128995.1_China/2018/AnhuiXCGQ  ................................................................................ [181600]
MH681419.1_ASFV/POL/2015         ................................................................................ [181600]
LS478113.1_Estonia_2014          ................................................................................ [181600]
KP843857.1_Odintsovo_02/14       ................................................................................ [181600]
```

```
FN557520.1_strain_E75           ................................................................................ [181600]
AY261366.1_Warthog              ...................................................t............................ [181600]
AY261365.1_Warmbaths            ............a................................................................... [181600]
AY261364.1_Tengani_62           ................................................................................ [181600]
AY261363.1_Pretoriuskop/96/4    ................................................................................ [181600]
AY261362.1_Mkuzi_1979           ................................................................................ [181600]
AY261361.1_Malawi_Lil-20/1_1983 ..............................c....................t............................ [181600]
AY261360.1_Kenya_1950           .................................t...c.......................................... [181600]
AM712240.1_OURT_88/3            ................................................................................ [181600]
AM712239.1_Benin_97/1           ................................................................................ [181600]
KM111295.1_Ken06.Bus            .................................t.............................................. [181600]
KM111294.1_Ken05/Tk1            .................................t...c.......................................... [181600]
KM262844.1_strain_L60           ................................................................................ [181600]
KM102979.1_26544/OG10_Italy     ................................................................................ [181600]
KM262845.1_strain_NHV           ................................................................................ [181600]
KX354450.1_47/Ss/2008           ................................................................................ [181600]
KP055815.1_strain_BA71          ................................................................................ [181600]
MH025920.1_strain_R35           .................................t.............................................. [181600]
MH025919.1_strain_N10           .................................t.............................................. [181600]
MH025918.1_strain_R25           .................................t.............................................. [181600]
MH025917.1_strain_R7            .................................t.............................................. [181600]
MH025916.1_strain_R8            .................................t.............................................. [181600]

FR682468.2_Georgia_2007/1       gcgatacgccgctgcacctgtatttccaaaagtttaactccatctcataccacgagatgtataaatcttccaacaagatc [181680]
ASF_IND_20_CAD543               ................................................................................ [181680]
LR899193.1_Germany_2020/1       ................................................................................ [181680]
MN393477.1_Wuhan_2019           ................................................................................ [181680]
MW396979.1_Timor-Leste/2019/1   ................................................................................ [181680]
MK940252.1_CN/2019/InnerMongolia ................................................................................ [181680]
MK128995.1_China/2018/AnhuiXCGQ ................................................................................ [181680]
MH681419.1_ASFV/POL/2015        ................................................................................ [181680]
LS478113.1_Estonia_2014         ................................................................................ [181680]
KP843857.1_Odintsovo_02/14      ................................................................................ [181680]
FN557520.1_strain_E75           ................................................................................ [181680]
AY261366.1_Warthog              ................................................................................ [181680]
AY261365.1_Warmbaths            .............a.................................................................. [181680]
AY261364.1_Tengani_62           ................................................................................ [181680]
AY261363.1_Pretoriuskop/96/4    ................................................................................ [181680]
AY261362.1_Mkuzi_1979           ................................................................c............... [181680]
AY261361.1_Malawi_Lil-20/1_1983 ....c.....a...........................................................t......... [181680]
AY261360.1_Kenya_1950           ........................................t....................................... [181680]
AM712240.1_OURT_88/3            ................................................................................ [181680]
AM712239.1_Benin_97/1           ................................................................................ [181680]
KM111295.1_Ken06.Bus            ................................................................................ [181680]
```

```
KM111294.1_Ken05/Tk1          ..........c.............................t....................................... [181680]
KM262844.1_strain_L60         ................................................................................ [181680]
KM102979.1_26544/OG10_Italy   ................................................................................ [181680]
KM262845.1_strain_NHV         ................................................................................ [181680]
KX354450.1_47/Ss/2008         ................................................................................ [181680]
KP055815.1_strain_BA71        ................................................................................ [181680]
MH025920.1_strain_R35         ................................................................................ [181680]
MH025919.1_strain_N10         ................................................................................ [181680]
MH025918.1_strain_R25         ................................................................................ [181680]
MH025917.1_strain_R7          ................................................................................ [181680]
MH025916.1_strain_R8          ................................................................................ [181680]

FR682468.2_Georgia_2007/1     aacctgacctcgaccattcctaagtcgcaggtgttccagataaatgttaaaattgctcacatcaagtcgctggcctcggc [181760]
ASF_IND_20_CAD543             ................................................................................ [181760]
LR899193.1_Germany_2020/1     ................................................................................ [181760]
MN393477.1_Wuhan_2019         ................................................................................ [181760]
MW396979.1_Timor-Leste/2019/1 ................................................................................ [181760]
MK940252.1_CN/2019/InnerMongolia ................................................................................ [181760]
MK128995.1_China/2018/AnhuiXCGQ ................................................................................ [181760]
MH681419.1_ASFV/POL/2015      ................................................................................ [181760]
LS478113.1_Estonia_2014       ................................................................................ [181760]
KP843857.1_Odintsovo_02/14    ................................................................................ [181760]
FN557520.1_strain_E75         ................................................................................ [181760]
AY261366.1_Warthog            ................................................................................ [181760]
AY261365.1_Warmbaths          .................c..................................................a........... [181760]
AY261364.1_Tengani_62         ................................................................................ [181760]
AY261363.1_Pretoriuskop/96/4  ................................................................................ [181760]
AY261362.1_Mkuzi_1979         ................................................................................ [181760]
AY261361.1_Malawi_Lil-20/1_1983 ...........a.................................................................... [181760]
AY261360.1_Kenya_1950         ...........a.................................................................... [181760]
AM712240.1_OURT_88/3          ................................................................................ [181760]
AM712239.1_Benin_97/1         ................................................................................ [181760]
KM111295.1_Ken06.Bus          ...........a..............a........t............................................ [181760]
KM111294.1_Ken05/Tk1          ...........a.................................................................... [181760]
KM262844.1_strain_L60         ................................................................................ [181760]
KM102979.1_26544/OG10_Italy   ................................................................................ [181760]
KM262845.1_strain_NHV         ................................................................................ [181760]
KX354450.1_47/Ss/2008         ................................................................................ [181760]
KP055815.1_strain_BA71        ................................................................................ [181760]
MH025920.1_strain_R35         ...........a..............a........t............................................ [181760]
MH025919.1_strain_N10         ...........a..............a........t............................................ [181760]
MH025918.1_strain_R25         ...........a..............a........t............................................ [181760]
MH025917.1_strain_R7          ...........a..............a........t............................................ [181760]
MH025916.1_strain_R8          ...........a..............a........t............................................ [181760]
```

```
FR682468.2_Georgia_2007/1        tatggtcaccgacaagatccgcattctgtgcgaagaaaatgggaacctaatctttcaatcggaaatggatgcccttatgt [181840]
ASF_IND_20_CAD543                ................................................................................ [181840]
LR899193.1_Germany_2020/1        ................................................................................ [181840]
MN393477.1_Wuhan_2019            ................................................................................ [181840]
MW396979.1_Timor-Leste/2019/1    ................................................................................ [181840]
MK940252.1_CN/2019/InnerMongolia ................................................................................ [181840]
MK128995.1_China/2018/AnhuiXCGQ  ................................................................................ [181840]
MH681419.1_ASFV/POL/2015         ................................................................................ [181840]
LS478113.1_Estonia_2014          ................................................................................ [181840]
KP843857.1_Odintsovo_02/14       ................................................................................ [181840]
FN557520.1_strain_E75            ................................................................................ [181840]
AY261366.1_Warthog               ............................................................a................... [181840]
AY261365.1_Warmbaths             ............................................................a................... [181840]
AY261364.1_Tengani_62            ................................................................................ [181840]
AY261363.1_Pretoriuskop/96/4     ............................................................a................... [181840]
AY261362.1_Mkuzi_1979            ................................................................................ [181840]
AY261361.1_Malawi_Lil-20/1_1983  ............................................................a................... [181840]
AY261360.1_Kenya_1950            ...............a...........a....................g..........................ac.a. [181840]
AM712240.1_OURT_88/3             ................................................................................ [181840]
AM712239.1_Benin_97/1            ................................................................................ [181840]
KM111295.1_Ken06.Bus             c..............a...........a..............a.................a..............ac.a. [181840]
KM111294.1_Ken05/Tk1             ...............a...........a....................g..........................ac.a. [181840]
KM262844.1_strain_L60            ................................................................................ [181840]
KM102979.1_26544/OG10_Italy      ................................................................................ [181840]
KM262845.1_strain_NHV            ................................................................................ [181840]
KX354450.1_47/Ss/2008            ................................................................................ [181840]
KP055815.1_strain_BA71           ................................................................................ [181840]
MH025920.1_strain_R35            c..............a...........a..............a................................ac.a. [181840]
MH025919.1_strain_N10            c..............a...........a..............a................................ac.a. [181840]
MH025918.1_strain_R25            c..............a...........a..............a................................ac.a. [181840]
MH025917.1_strain_R7             c..............a...........a..............a................................ac.a. [181840]
MH025916.1_strain_R8             c..............a...........a..............a................................ac.a. [181840]

FR682468.2_Georgia_2007/1        taaatacgattaccttgaacaccacgatatagtt-cggtaacattagat------------------------------- [181920]
ASF_IND_20_CAD543                ..................................-..............------------------------------- [181920]
LR899193.1_Germany_2020/1        ..................................-..............------------------------------- [181920]
MN393477.1_Wuhan_2019            ..................................-..............------------------------------- [181920]
MW396979.1_Timor-Leste/2019/1    ..................................-..............------------------------------- [181920]
MK940252.1_CN/2019/InnerMongolia ..................................-..............------------------------------- [181920]
MK128995.1_China/2018/AnhuiXCGQ  ..................................-..............------------------------------- [181920]
MH681419.1_ASFV/POL/2015         ..................................-..............------------------------------- [181920]
LS478113.1_Estonia_2014          ..................................-..............------------------------------- [181920]
KP843857.1_Odintsovo_02/14       ..................................-..............------------------------------- [181920]
```

```
FN557520.1_strain_E75             .....................a............-..............------------------------- [181920]
AY261366.1_Warthog                .....................a..t.........c........a.....------------------------------- [181920]
AY261365.1_Warmbaths              .....................a..t.........c........a..t..------------------------------- [181920]
AY261364.1_Tengani_62             .....................g..t.........c........ac....------------------------------- [181920]
AY261363.1_Pretoriuskop/96/4      .....................a..t.........c........a.....------------------------------- [181920]
AY261362.1_Mkuzi_1979             .....................a............-........a.....------------------------------- [181920]
AY261361.1_Malawi_Lil-20/1_1983   .......c.....ta......a..t.........-t.....aga.....------------------------------- [181920]
AY261360.1_Kenya_1950             .......c.............at.t.........-.a....a.a.g...ggtcctactattaaataccattaccttgaac [181920]
AM712240.1_OURT_88/3              .....................a............-..............------------------------------- [181920]
AM712239.1_Benin_97/1             .....................a............-..............------------------------------- [181920]
KM111295.1_Ken06.Bus              .......c.............a..t.........-......a.a.g.--------------------------------- [181920]
KM111294.1_Ken05/Tk1              .......c.............a..t.........-......a.a.g.--------------------------------- [181920]
KM262844.1_strain_L60             .....................a............-..............------------------------------- [181920]
KM102979.1_26544/OG10_Italy       .....................a............-..............------------------------------- [181920]
KM262845.1_strain_NHV             .....................a............-..............------------------------------- [181920]
KX354450.1_47/Ss/2008             .....................a............-..............------------------------------- [181920]
KP055815.1_strain_BA71            .....................a............-..............------------------------------- [181920]
MH025920.1_strain_R35             .......c.............a..t.........-......a.a.g.--------------------------------- [181920]
MH025919.1_strain_N10             .......c.............a..t.........-......a.a.g.--------------------------------- [181920]
MH025918.1_strain_R25             .......c.............a..t.........-......a.a.g.--------------------------------- [181920]
MH025917.1_strain_R7              .......c.............a..t.........-......a.a.g.--------------------------------- [181920]
MH025916.1_strain_R8              .......c.............a..t.........-......a.a.g.--------------------------------- [181920]

FR682468.2_Georgia_2007/1         -----------------------------gttctaatatttagcat-------ctaaataatacgctgtagtccggtcag [182000]
ASF_IND_20_CAD543                 -----------------------------.................-------........................... [182000]
LR899193.1_Germany_2020/1         -----------------------------.................-------........................... [182000]
MN393477.1_Wuhan_2019             -----------------------------.................-------........................... [182000]
MW396979.1_Timor-Leste/2019/1     -----------------------------.................-------........................... [182000]
MK940252.1_CN/2019/InnerMongolia  -----------------------------.................-------........................... [182000]
MK128995.1_China/2018/AnhuiXCGQ   -----------------------------.................-------........................... [182000]
MH681419.1_ASFV/POL/2015          -----------------------------.................-------........................... [182000]
LS478113.1_Estonia_2014           -----------------------------.................-------........................... [182000]
KP843857.1_Odintsovo_02/14        -----------------------------.................-------........................... [182000]
FN557520.1_strain_E75             -----------------------------.................-------........................... [182000]
AY261366.1_Warthog                -----------------------------............t....-------........................... [182000]
AY261365.1_Warmbaths              -----------------------------............t....-------........................... [182000]
AY261364.1_Tengani_62             -----------------------------..........g.t....-------........................... [182000]
AY261363.1_Pretoriuskop/96/4      -----------------------------............t....-------........................... [182000]
AY261362.1_Mkuzi_1979             -----------------------------.................-------........................... [182000]
AY261361.1_Malawi_Lil-20/1_1983   -----------------------------..c......g.......catcacc.................a......... [182000]
AY261360.1_Kenya_1950             aacatgatatagttcggtaaaaatggatg..c......g.....gcc------............t....a......... [182000]
AM712240.1_OURT_88/3              -----------------------------.................-------........................... [182000]
AM712239.1_Benin_97/1             -----------------------------.................-------........................... [182000]
KM111295.1_Ken06.Bus              --------------------------------......g.....gcc------........c...t....a......... [182000]
```

```
KM111294.1_Ken05/Tk1          --------------------------atg..c......g......cc------............t....a......... [182000]
KM262844.1_strain_L60         -----------------------------.................-------........................... [182000]
KM102979.1_26544/OG10_Italy   -----------------------------.................-------........................... [182000]
KM262845.1_strain_NHV         -----------------------------.................-------........................... [182000]
KX354450.1_47/Ss/2008         -----------------------------.................-------........................... [182000]
KP055815.1_strain_BA71        -----------------------------.................-------........................... [182000]
MH025920.1_strain_R35         --------------------------------......g.....gcc------........c...t....a......... [182000]
MH025919.1_strain_N10         --------------------------------......g.....gcc------........c...t....a......... [182000]
MH025918.1_strain_R25         --------------------------------......g.....gcc------........c...t....a......... [182000]
MH025917.1_strain_R7          --------------------------------......g.....gcc------........c...t....a......... [182000]
MH025916.1_strain_R8          --------------------------------......g.....gcc------........c...t....a......... [182000]

FR682468.2_Georgia_2007/1     ggttgcgtcacagttttcccatttttttgcctcgtcggcggtggccaccgttgccctatcatttacgcccggtaagacaa [182080]
ASF_IND_20_CAD543             ................................................................................ [182080]
LR899193.1_Germany_2020/1     ................................................................................ [182080]
MN393477.1_Wuhan_2019         ................................................................................ [182080]
MW396979.1_Timor-Leste/2019/1 ................................................................................ [182080]
MK940252.1_CN/2019/InnerMongolia ................................................................................ [182080]
MK128995.1_China/2018/AnhuiXCGQ ................................................................................ [182080]
MH681419.1_ASFV/POL/2015      ................................................................................ [182080]
LS478113.1_Estonia_2014       ................................................................................ [182080]
KP843857.1_Odintsovo_02/14    ................................................................................ [182080]
FN557520.1_strain_E75         ..............................g................................................. [182080]
AY261366.1_Warthog            ..............................g............a.........................t.......... [182080]
AY261365.1_Warmbaths          .................................c....at....t........................t.......... [182080]
AY261364.1_Tengani_62         .................................c...................................t.......... [182080]
AY261363.1_Pretoriuskop/96/4  ..............................g......................................t.......... [182080]
AY261362.1_Mkuzi_1979         .................................c........a..................................... [182080]
AY261361.1_Malawi_Lil-20/1_1983 ......a..g.....c.................a.......................g...........t.......t.. [182080]
AY261360.1_Kenya_1950         a.....a.......................g..........................g.....c..c..t.......t.. [182080]
AM712240.1_OURT_88/3          ..............................g................................................. [182080]
AM712239.1_Benin_97/1         ..............................g................................................. [182080]
KM111295.1_Ken06.Bus          a.....a..........................c..........t............g.....c..c..t....g..t.. [182080]
KM111294.1_Ken05/Tk1          a.....a.......................g..........................g.....c..c..t.......t.. [182080]
KM262844.1_strain_L60         ..............................g................................................. [182080]
KM102979.1_26544/OG10_Italy   ..............................g................................................. [182080]
KM262845.1_strain_NHV         ..............................g................................................. [182080]
KX354450.1_47/Ss/2008         ..............................g................................................. [182080]
KP055815.1_strain_BA71        ..............................g................................................. [182080]
MH025920.1_strain_R35         a.....a..........................c..........t............g.....c..c..t....g..t.. [182080]
MH025919.1_strain_N10         a.....a..........................c..........t............g.....c..c..t....g..t.. [182080]
MH025918.1_strain_R25         a.....a..........................c..........t............g.....c..c..t....g..t.. [182080]
MH025917.1_strain_R7          a.....a..........................c..........t............g.....c..c..t....g..t.. [182080]
MH025916.1_strain_R8          a.....a..........................c..........t............g.....c..c..t....g..t.. [182080]
```

```
FR682468.2_Georgia_2007/1         agctaaaggcgttcagcggggcttggcaatgcccgcccagcgtgaaggagctcggaggattttgcgcatcccgaaatccc [182160]
ASF_IND_20_CAD543                 ................................................................................ [182160]
LR899193.1_Germany_2020/1         ................................................................................ [182160]
MN393477.1_Wuhan_2019             ................................................................................ [182160]
MW396979.1_Timor-Leste/2019/1     ................................................................................ [182160]
MK940252.1_CN/2019/InnerMongolia  ................................................................................ [182160]
MK128995.1_China/2018/AnhuiXCGQ   ................................................................................ [182160]
MH681419.1_ASFV/POL/2015          ................................................................................ [182160]
LS478113.1_Estonia_2014           ................................................................................ [182160]
KP843857.1_Odintsovo_02/14        ................................................................................ [182160]
FN557520.1_strain_E75             ................................................................................ [182160]
AY261366.1_Warthog                ..............................................a................................. [182160]
AY261365.1_Warmbaths              ................................................................................ [182160]
AY261364.1_Tengani_62             ................................................................................ [182160]
AY261363.1_Pretoriuskop/96/4      ................................................................................ [182160]
AY261362.1_Mkuzi_1979             ........................................t....................................... [182160]
AY261361.1_Malawi_Lil-20/1_1983   .........................a.............a........................................ [182160]
AY261360.1_Kenya_1950             ................t..a..c................a....................................a... [182160]
AM712240.1_OURT_88/3              ................................................................................ [182160]
AM712239.1_Benin_97/1             ................................................................................ [182160]
KM111295.1_Ken06.Bus              ................t..a..c..a.............a...a................................a... [182160]
KM111294.1_Ken05/Tk1              ................t..a..c................a....................................a... [182160]
KM262844.1_strain_L60             ................................................................................ [182160]
KM102979.1_26544/OG10_Italy       ................................................................................ [182160]
KM262845.1_strain_NHV             ................................................................................ [182160]
KX354450.1_47/Ss/2008             ................................................................................ [182160]
KP055815.1_strain_BA71            ................................................................................ [182160]
MH025920.1_strain_R35             ................t..a..c..a.............a...a................................a... [182160]
MH025919.1_strain_N10             ................t..a..c..a.............a...a................................a... [182160]
MH025918.1_strain_R25             ................t..a..c..a.............a...a................................a... [182160]
MH025917.1_strain_R7              ................t..a..c..a.............a...a................................a... [182160]
MH025916.1_strain_R8              ................t..a..c..a.............a...a................................a... [182160]

FR682468.2_Georgia_2007/1         ttagccatgttgtttaacacttcggttacgtcaatcgagtgaagggatcccttgggatccgtgaatgtaaagacgcagtt [182240]
ASF_IND_20_CAD543                 .........................................................................y...... [182240]
LR899193.1_Germany_2020/1         ................................................................................ [182240]
MN393477.1_Wuhan_2019             ................................................................................ [182240]
MW396979.1_Timor-Leste/2019/1     ................................................................................ [182240]
MK940252.1_CN/2019/InnerMongolia  ................................................................................ [182240]
MK128995.1_China/2018/AnhuiXCGQ   ................................................................................ [182240]
MH681419.1_ASFV/POL/2015          ................................................................................ [182240]
LS478113.1_Estonia_2014           ................................................................................ [182240]
KP843857.1_Odintsovo_02/14        ................................................................................ [182240]
```

```
FN557520.1_strain_E75             .........................................................................t...... [182240]
AY261366.1_Warthog                ............................................a............................t...... [182240]
AY261365.1_Warmbaths              ............................................a............................t...... [182240]
AY261364.1_Tengani_62             .........................................................................t...... [182240]
AY261363.1_Pretoriuskop/96/4      .........................................................................t...... [182240]
AY261362.1_Mkuzi_1979             .........................................................................t...... [182240]
AY261361.1_Malawi_Lil-20/1_1983   ........a........t..a.t......a....ct.....g..t..c................................ [182240]
AY261360.1_Kenya_1950             ..g.....a........t..a.t......a...........g..c..c........g....................... [182240]
AM712240.1_OURT_88/3              .........................................................................t...... [182240]
AM712239.1_Benin_97/1             .........................................................................t...... [182240]
KM111295.1_Ken06.Bus              ..g.....a........t..a.t......a...........g..t..c........g....................... [182240]
KM111294.1_Ken05/Tk1              ..g.....a........t..a.t......a...........g..c..c........g....................... [182240]
KM262844.1_strain_L60             .........................................................................t...... [182240]
KM102979.1_26544/OG10_Italy       .........................................................................t...... [182240]
KM262845.1_strain_NHV             .........................................................................t...... [182240]
KX354450.1_47/Ss/2008             .........................................................................t...... [182240]
KP055815.1_strain_BA71            .........................................................................t...... [182240]
MH025920.1_strain_R35             ..g.....a........t..a.t......a...........g..t..c........g....................... [182240]
MH025919.1_strain_N10             ..g.....a........t..a.t......a...........g..t..c........g....................... [182240]
MH025918.1_strain_R25             ..g.....a........t..a.t......a...........g..t..c........g....................... [182240]
MH025917.1_strain_R7              ..g.....a........t..a.t......a...........g..t..c........g....................... [182240]
MH025916.1_strain_R8              ..g.....a........t..a.t......a...........g..t..c........g....................... [182240]

FR682468.2_Georgia_2007/1         tctaaagcgcatgtatgcgatggacgattcatcgggggttttgaaggtaacagtgttccccttgctgtacttaaaggggg [182320]
ASF_IND_20_CAD543                 ................................................................................ [182320]
LR899193.1_Germany_2020/1         ................................................................................ [182320]
MN393477.1_Wuhan_2019             ................................................................................ [182320]
MW396979.1_Timor-Leste/2019/1     ................................................................................ [182320]
MK940252.1_CN/2019/InnerMongolia  ................................................................................ [182320]
MK128995.1_China/2018/AnhuiXCGQ   ................................................................................ [182320]
MH681419.1_ASFV/POL/2015          ................................................................................ [182320]
LS478113.1_Estonia_2014           ................................................................................ [182320]
KP843857.1_Odintsovo_02/14        ................................................................................ [182320]
FN557520.1_strain_E75             ................................................................................ [182320]
AY261366.1_Warthog                ..........................................a........ta.a....................a.... [182320]
AY261365.1_Warmbaths              ...................................................t..a....................a.... [182320]
AY261364.1_Tengani_62             ............a................................................................... [182320]
AY261363.1_Pretoriuskop/96/4      .................................a..a.....a........t..a..............t.....a.... [182320]
AY261362.1_Mkuzi_1979             ...g............................................................................ [182320]
AY261361.1_Malawi_Lil-20/1_1983   ct..........a......................................t..a.......................a. [182320]
AY261360.1_Kenya_1950             .t..........a..............a.......................t..a......................... [182320]
AM712240.1_OURT_88/3              ................................................................................ [182320]
AM712239.1_Benin_97/1             ................................................................................ [182320]
KM111295.1_Ken06.Bus              .t..........a..............a.......................t..a......................... [182320]
```

```
KM111294.1_Ken05/Tk1             .t..........a..............a.......................t..a......................... [182320]
KM262844.1_strain_L60            ................................................................................ [182320]
KM102979.1_26544/OG10_Italy      ................................................................................ [182320]
KM262845.1_strain_NHV            ................................................................................ [182320]
KX354450.1_47/Ss/2008            ................................................................................ [182320]
KP055815.1_strain_BA71           ................................................................................ [182320]
MH025920.1_strain_R35            .t..........a..............a.......................t..a......................... [182320]
MH025919.1_strain_N10            .t..........a..............a.......................t..a......................... [182320]
MH025918.1_strain_R25            .t..........a..............a.......................t..a......................... [182320]
MH025917.1_strain_R7             .t..........a..............a.......................t..a......................... [182320]
MH025916.1_strain_R8             .t..........a..............a.......................t..a......................... [182320]

FR682468.2_Georgia_2007/1        accatccggtaaaattataccaaatgaaagcaataataattaaaataaccaacacaatagttatagacaacacaaagtct [182400]
ASF_IND_20_CAD543                ................................................................................ [182400]
LR899193.1_Germany_2020/1        ................................................................................ [182400]
MN393477.1_Wuhan_2019            ................................................................................ [182400]
MW396979.1_Timor-Leste/2019/1    ................................................................................ [182400]
MK940252.1_CN/2019/InnerMongolia ................................................................................ [182400]
MK128995.1_China/2018/AnhuiXCGQ  ................................................................................ [182400]
MH681419.1_ASFV/POL/2015         ................................................................................ [182400]
LS478113.1_Estonia_2014          ................................................................................ [182400]
KP843857.1_Odintsovo_02/14       ................................................................................ [182400]
FN557520.1_strain_E75            ................................................................................ [182400]
AY261366.1_Warthog               ............................................................................a... [182400]
AY261365.1_Warmbaths             ............................................................................a... [182400]
AY261364.1_Tengani_62            ..........................................................g.................a... [182400]
AY261363.1_Pretoriuskop/96/4     ............................................................................a... [182400]
AY261362.1_Mkuzi_1979            ................................................................................ [182400]
AY261361.1_Malawi_Lil-20/1_1983  ..........................................................g.................a... [182400]
AY261360.1_Kenya_1950            ..........................t..................................g..............a... [182400]
AM712240.1_OURT_88/3             ................................................................................ [182400]
AM712239.1_Benin_97/1            ................................................................................ [182400]
KM111295.1_Ken06.Bus             ..........................t.....................t............g..............a... [182400]
KM111294.1_Ken05/Tk1             ..........................t............................g.....g..............a... [182400]
KM262844.1_strain_L60            ................................................................................ [182400]
KM102979.1_26544/OG10_Italy      ................................................................................ [182400]
KM262845.1_strain_NHV            ................................................................................ [182400]
KX354450.1_47/Ss/2008            ................................................................................ [182400]
KP055815.1_strain_BA71           ................................................................................ [182400]
MH025920.1_strain_R35            ..........................t.....................t............g..............a... [182400]
MH025919.1_strain_N10            ..........................t.....................t............g..............a... [182400]
MH025918.1_strain_R25            ..........................t.....................t............g..............a... [182400]
MH025917.1_strain_R7             ..........................t.....................t............g..............a... [182400]
MH025916.1_strain_R8             ..........................t.....................t............g..............a... [182400]
```

```
FR682468.2_Georgia_2007/1         gtagtgccgcccattattaaataaaaatattttagaccgccggcttaaaatttactta----ttgctcatagcttaagtc [182480]
ASF_IND_20_CAD543                 ..........................................................----.................. [182480]
LR899193.1_Germany_2020/1         ..........................................................----.................. [182480]
MN393477.1_Wuhan_2019             ..........................................................----.................. [182480]
MW396979.1_Timor-Leste/2019/1     ..........................................................----.................. [182480]
MK940252.1_CN/2019/InnerMongolia  ..........................................................----.................. [182480]
MK128995.1_China/2018/AnhuiXCGQ   ..........................................................----.................. [182480]
MH681419.1_ASFV/POL/2015          ..........................................................----.................. [182480]
LS478113.1_Estonia_2014           ..........................................................----.................. [182480]
KP843857.1_Odintsovo_02/14        ..........................................................----.................. [182480]
FN557520.1_strain_E75             ..........................................................----........g......... [182480]
AY261366.1_Warthog                .c........................................................----....-------------- [182480]
AY261365.1_Warmbaths              .c...........-..........................................c.----.a..-------------- [182480]
AY261364.1_Tengani_62             ..........................................................----...t.............. [182480]
AY261363.1_Pretoriuskop/96/4      .c........................................................----.................. [182480]
AY261362.1_Mkuzi_1979             ..........................................................----........g......... [182480]
AY261361.1_Malawi_Lil-20/1_1983   .............-........................ag..................----.................g [182480]
AY261360.1_Kenya_1950             .............-.........................g..................agtc........g.....c..t [182480]
AM712240.1_OURT_88/3              ..........................................................----........g......... [182480]
AM712239.1_Benin_97/1             ..........................................................----........g......... [182480]
KM111295.1_Ken06.Bus              .............-.........................g..................agtc........g.....c..t [182480]
KM111294.1_Ken05/Tk1              .............-.........................g..................agtc........g.....c..t [182480]
KM262844.1_strain_L60             ..........................................................----........g......... [182480]
KM102979.1_26544/OG10_Italy       ..........................................................----........g......... [182480]
KM262845.1_strain_NHV             ..........................................................----........g......... [182480]
KX354450.1_47/Ss/2008             ..........................................................----........g......... [182480]
KP055815.1_strain_BA71            ..........................................................----........g......... [182480]
MH025920.1_strain_R35             .............-.........................g..................agtc........g.....c..t [182480]
MH025919.1_strain_N10             .............-.........................g..................agtc........g.....c..t [182480]
MH025918.1_strain_R25             .............-.........................g..................agtc........g.....c..t [182480]
MH025917.1_strain_R7              .............-.........................g..................agtc........g.....c..t [182480]
MH025916.1_strain_R8              .............-.........................g..................agtc........g.....c..t [182480]

FR682468.2_Georgia_2007/1         tat------------------------tttattcatagcttaagtttattgctca--------------tggcttaagtc [182560]
ASF_IND_20_CAD543                 ...------------------------............................--------------........... [182560]
LR899193.1_Germany_2020/1         ...------------------------............................--------------........... [182560]
MN393477.1_Wuhan_2019             ...------------------------............................--------------........... [182560]
MW396979.1_Timor-Leste/2019/1     ...------------------------............................--------------........... [182560]
MK940252.1_CN/2019/InnerMongolia  ...------------------------............................--------------........... [182560]
MK128995.1_China/2018/AnhuiXCGQ   ...------------------------............................--------------........... [182560]
MH681419.1_ASFV/POL/2015          ...------------------------............................--------------........... [182560]
LS478113.1_Estonia_2014           ...------------------------............................--------------........... [182560]
KP843857.1_Odintsovo_02/14        ...------------------------............................--------------........... [182560]
```

```
FN557520.1_strain_E75              ...------------------------............................--------------.a......... [182560]
AY261366.1_Warthog                 -------------------------------------------------------------------------------- [182560]
AY261365.1_Warmbaths               -------------------------------------------------------------------------------- [182560]
AY261364.1_Tengani_62              ...g---------------------------------------------------------------------------- [182560]
AY261363.1_Pretoriuskop/96/4       ...ga------ttcatagcctaagtcta..gc.............c.g.......--------------........... [182560]
AY261362.1_Mkuzi_1979              ..--------------------------.a.........................--------------........... [182560]
AY261361.1_Malawi_Lil-20/1_1983    gc.taaatatgctgttagcttaaa---.a.gc.gt........a.a.gc..t.ggcttaaatatgctgt........a.a [182560]
AY261360.1_Kenya_1950              gc.aa-------------------------------------------------------------tca.---------- [182560]
AM712240.1_OURT_88/3               ...------------------------............................--------------.a......... [182560]
AM712239.1_Benin_97/1              ...------------------------............................--------------.a......... [182560]
KM111295.1_Ken06.Bus               ac.aa-------------------------------------------------------------tga.a.t....c.a [182560]
KM111294.1_Ken05/Tk1               gc.aa-------------------------------------------------------------tca........c.a [182560]
KM262844.1_strain_L60              ...------------------------............................--------------.a......... [182560]
KM102979.1_26544/OG10_Italy        ...------------------------............................--------------.a......... [182560]
KM262845.1_strain_NHV              ...------------------------............................--------------.a......... [182560]
KX354450.1_47/Ss/2008              ...------------------------............................--------------.a......... [182560]
KP055815.1_strain_BA71             ...------------------------............................--------------.a......... [182560]
MH025920.1_strain_R35              ac.aa-------------------------------------------------------------tga.a.t....c.a [182560]
MH025919.1_strain_N10              ac.aa-------------------------------------------------------------tga.a.t....c.a [182560]
MH025918.1_strain_R25              ac.aa-------------------------------------------------------------tga.a.t....c.a [182560]
MH025917.1_strain_R7               ac.aa-------------------------------------------------------------tga.a.t....c.a [182560]
MH025916.1_strain_R8               ac.aa-------------------------------------------------------------tga.a.t....c.a [182560]

FR682468.2_Georgia_2007/1          tattgctta-----------------------------------------------------------------tagctt [182640]
ASF_IND_20_CAD543                  .....t.c.-----------------------------------------------------------------...... [182640]
LR899193.1_Germany_2020/1          .........-----------------------------------------------------------------...... [182640]
MN393477.1_Wuhan_2019              .........-----------------------------------------------------------------...... [182640]
MW396979.1_Timor-Leste/2019/1      .........-----------------------------------------------------------------...... [182640]
MK940252.1_CN/2019/InnerMongolia   .........-----------------------------------------------------------------...... [182640]
MK128995.1_China/2018/AnhuiXCGQ    .........-----------------------------------------------------------------...... [182640]
MH681419.1_ASFV/POL/2015           .....t...-----------------------------------------------------------------...... [182640]
LS478113.1_Estonia_2014            .........-----------------------------------------------------------------...... [182640]
KP843857.1_Odintsovo_02/14         .........-----------------------------------------------------------------...... [182640]
FN557520.1_strain_E75              ..------------------------------------------------------------------------------ [182640]
AY261366.1_Warthog                 -------------------------------------------------------------------------------- [182640]
AY261365.1_Warmbaths               -------------------------------------------------------------------------------. [182640]
AY261364.1_Tengani_62              -------------------------------------------------------------------------------- [182640]
AY261363.1_Pretoriuskop/96/4       .g.....c.-----------------------------------------------------------------.g.... [182640]
AY261362.1_Mkuzi_1979              ....a..c.-----------------------------------------------------------------...... [182640]
AY261361.1_Malawi_Lil-20/1_1983    .gc..t.agcttaaatatgctgttagcttaaatatgctgttggcttaaatatgctgttggcttaaatatgctgt...... [182640]
AY261360.1_Kenya_1950              -------------------------------------------------------------------------------- [182640]
AM712240.1_OURT_88/3               ..------------------------------------------------------------------------------ [182640]
AM712239.1_Benin_97/1              ..------------------------------------------------------------------------------ [182640]
KM111295.1_Ken06.Bus               .gc-----------------------------------------------------------------------...... [182640]
```

```
KM111294.1_Ken05/Tk1             .gc-----------------------------------------------------------------------...... [182640]
KM262844.1_strain_L60            ..------------------------------------------------------------------------------ [182640]
KM102979.1_26544/OG10_Italy      ..------------------------------------------------------------------------------ [182640]
KM262845.1_strain_NHV            ..------------------------------------------------------------------------------ [182640]
KX354450.1_47/Ss/2008            ..------------------------------------------------------------------------------ [182640]
KP055815.1_strain_BA71           ..------------------------------------------------------------------------------ [182640]
MH025920.1_strain_R35            .gc-----------------------------------------------------------------------...... [182640]
MH025919.1_strain_N10            .gc-----------------------------------------------------------------------...... [182640]
MH025918.1_strain_R25            .gc-----------------------------------------------------------------------...... [182640]
MH025917.1_strain_R7             .gc-----------------------------------------------------------------------...... [182640]
MH025916.1_strain_R8             .gc-----------------------------------------------------------------------...... [182640]

FR682468.2_Georgia_2007/1        aagtctat------------tttattcatagcttaagtctattgttcatggcttaagtttgtt-gctca----------- [182720]
ASF_IND_20_CAD543                .......------------------------------------------------------..-at...----------- [182720]
LR899193.1_Germany_2020/1        ........------------...........................................-.....----------- [182720]
MN393477.1_Wuhan_2019            ........------------...........................................-.....----------- [182720]
MW396979.1_Timor-Leste/2019/1    ........------------...........................................-.....----------- [182720]
MK940252.1_CN/2019/InnerMongolia ........------------...........................................-.....----------- [182720]
MK128995.1_China/2018/AnhuiXCGQ  ........------------...........................................-.....----------- [182720]
MH681419.1_ASFV/POL/2015         ........------------..........................t................-.t.t.----------- [182720]
LS478113.1_Estonia_2014          ........------------...........................................-.....----------- [182720]
KP843857.1_Odintsovo_02/14       ........------------...........................................-.....----------- [182720]
FN557520.1_strain_E75            -----------------------------------------.................c.a..-a....----------- [182720]
AY261366.1_Warthog               -------------------------....g............ga.....a........c....-.....----------- [182720]
AY261365.1_Warmbaths             .......--------------.g.....................c....a....g...c....-.....----------- [182720]
AY261364.1_Tengani_62            -------------------------------------------a..............c....-.....----------- [182720]
AY261363.1_Pretoriuskop/96/4     g.....g--------------..gc....g..............c....a........c....-.....----------- [182720]
AY261362.1_Mkuzi_1979            ....t..--------------..g.....g....--........c....a........c.a..-a....----------- [182720]
AY261361.1_Malawi_Lil-20/1_1983  ..a.a.gctgttagcttaaa.a.gc.gt.g......a.---a..c.gt........a.a..c.-.t.agcttaaatatgc [182720]
AY261360.1_Kenya_1950            -------------------------------------------------------------------------------- [182720]
AM712240.1_OURT_88/3             -----------------------------------------.................c.a..-a....----------- [182720]
AM712239.1_Benin_97/1            -----------------------------------------.................c.a..-a....----------- [182720]
KM111295.1_Ken06.Bus             ..c.a..c----------------------------------------.a......c.a.ac.----------------- [182720]
KM111294.1_Ken05/Tk1             ..c.a.gc----------------------------------------........c.a..c.----------------- [182720]
KM262844.1_strain_L60            -----------------------------------------.................c.a..-a....----------- [182720]
KM102979.1_26544/OG10_Italy      -----------------------------------------.................c.a..-a....----------- [182720]
KM262845.1_strain_NHV            -----------------------------------------.................c.a..-a....----------- [182720]
KX354450.1_47/Ss/2008            -----------------------------------------.................c.a..-a....----------- [182720]
KP055815.1_strain_BA71           -----------------------------------------.................c.a..-a....----------- [182720]
MH025920.1_strain_R35            ..c.a..c----------------------------------------.a......c.a.ac.a...t.actatac---- [182720]
MH025919.1_strain_N10            ..c.a..c----------------------------------------.a......c.a.ac.a...t.actatac---- [182720]
MH025918.1_strain_R25            ..c.a..c----------------------------------------.a......c.a.ac.a...t.actatac---- [182720]
MH025917.1_strain_R7             ..c.a..c----------------------------------------.a......c.a.ac.a...t.actatac---- [182720]
MH025916.1_strain_R8             ..c.a..c----------------------------------------.a......c.a.ac.a...t.actatac---- [182720]
```

```
FR682468.2_Georgia_2007/1        ---tagcttaactccattactga--------------------------------------------------------- [182800]
ASF_IND_20_CAD543                ---.............---------------------------------------------------------------- [182800]
LR899193.1_Germany_2020/1        ---....................--------------------------------------------------------- [182800]
MN393477.1_Wuhan_2019            ---....................--------------------------------------------------------- [182800]
MW396979.1_Timor-Leste/2019/1    ---....................--------------------------------------------------------- [182800]
MK940252.1_CN/2019/InnerMongolia ---....................--------------------------------------------------------- [182800]
MK128995.1_China/2018/AnhuiXCGQ  ---....................--------------------------------------------------------- [182800]
MH681419.1_ASFV/POL/2015         ---....................--------------------------------------------------------- [182800]
LS478113.1_Estonia_2014          ---....................--------------------------------------------------------- [182800]
KP843857.1_Odintsovo_02/14       ---....................--------------------------------------------------------- [182800]
FN557520.1_strain_E75            ---.............---------------------------------------------------------------- [182800]
AY261366.1_Warthog               ---......g.g..t......c.--------------------------------------------------------- [182800]
AY261365.1_Warmbaths             ---.g......g..t......c.--------------------------------------------------------- [182800]
AY261364.1_Tengani_62            ---......g.g..tg..g..c.--------------------------------------------------------- [182800]
AY261363.1_Pretoriuskop/96/4     ---.g....g.g..t......c.--------------------------------------------------------- [182800]
AY261362.1_Mkuzi_1979            ---....................--------------------------------------------------------- [182800]
AY261361.1_Malawi_Lil-20/1_1983  tgt........a.atgc.gt..gcttaaatatgctgttggcttaaatatgctgttggcttaaatatgctgttggcttaaa [182800]
AY261360.1_Kenya_1950            ----g........atgc--------------------------------------------------------------- [182800]
AM712240.1_OURT_88/3             ---.............---------------------------------------------------------------- [182800]
AM712239.1_Benin_97/1            ---.............---------------------------------------------------------------- [182800]
KM111295.1_Ken06.Bus             ----.........at.c--------------------------------------------------------------- [182800]
KM111294.1_Ken05/Tk1             ----.........atgc--------------------------------------------------------------- [182800]
KM262844.1_strain_L60            ---.............---------------------------------------------------------------- [182800]
KM102979.1_26544/OG10_Italy      ---.............---------------------------------------------------------------- [182800]
KM262845.1_strain_NHV            ---.............---------------------------------------------------------------- [182800]
KX354450.1_47/Ss/2008            ---.............---------------------------------------------------------------- [182800]
KP055815.1_strain_BA71           ---.............---------------------------------------------------------------- [182800]
MH025920.1_strain_R35            ---..........at.c--------------------------------------------------------------- [182800]
MH025919.1_strain_N10            ---..........at.c--------------------------------------------------------------- [182800]
MH025918.1_strain_R25            ---..........at.c--------------------------------------------------------------- [182800]
MH025917.1_strain_R7             ---..........at.c--------------------------------------------------------------- [182800]
MH025916.1_strain_R8             ---..........at.c--------------------------------------------------------------- [182800]

FR682468.2_Georgia_2007/1        ----------------------------------------------------------------------------tagc [182880]
ASF_IND_20_CAD543                -------------------------------------------------------------------------------- [182880]
LR899193.1_Germany_2020/1        ----------------------------------------------------------------------------.... [182880]
MN393477.1_Wuhan_2019            ----------------------------------------------------------------------------.... [182880]
MW396979.1_Timor-Leste/2019/1    ----------------------------------------------------------------------------.... [182880]
MK940252.1_CN/2019/InnerMongolia ----------------------------------------------------------------------------.... [182880]
MK128995.1_China/2018/AnhuiXCGQ  ----------------------------------------------------------------------------.... [182880]
MH681419.1_ASFV/POL/2015         ----------------------------------------------------------------------------.... [182880]
LS478113.1_Estonia_2014          ----------------------------------------------------------------------------.... [182880]
KP843857.1_Odintsovo_02/14       ----------------------------------------------------------------------------.... [182880]
```

```
FN557520.1_strain_E75             -------------------------------------------------------------------------------- [182880]
AY261366.1_Warthog                ----------------------------------------------------------------------------.... [182880]
AY261365.1_Warmbaths              ----------------------------------------------------------------------------.... [182880]
AY261364.1_Tengani_62             ----------------------------------------------------------------------------.... [182880]
AY261363.1_Pretoriuskop/96/4      ----------------------------------------------------------------------------.... [182880]
AY261362.1_Mkuzi_1979             ----------------------------------------------------------------------------.... [182880]
AY261361.1_Malawi_Lil-20/1_1983   tatgctgttagcttaaatatgctgttggcttaaatatgctgttggcttaaatatgctgttagcttaaatatgctgt.... [182880]
AY261360.1_Kenya_1950             ----------------------------------------------------------------------------.... [182880]
AM712240.1_OURT_88/3              -------------------------------------------------------------------------------- [182880]
AM712239.1_Benin_97/1             -------------------------------------------------------------------------------- [182880]
KM111295.1_Ken06.Bus              ----------------------------------------------------------------------------.... [182880]
KM111294.1_Ken05/Tk1              ----------------------------------------------------------------------------.... [182880]
KM262844.1_strain_L60             -------------------------------------------------------------------------------- [182880]
KM102979.1_26544/OG10_Italy       -------------------------------------------------------------------------------- [182880]
KM262845.1_strain_NHV             -------------------------------------------------------------------------------- [182880]
KX354450.1_47/Ss/2008             -------------------------------------------------------------------------------- [182880]
KP055815.1_strain_BA71            -------------------------------------------------------------------------------- [182880]
MH025920.1_strain_R35             ----------------------------------------------------------------------------.... [182880]
MH025919.1_strain_N10             ----------------------------------------------------------------------------.... [182880]
MH025918.1_strain_R25             ----------------------------------------------------------------------------.... [182880]
MH025917.1_strain_R7              ----------------------------------------------------------------------------.... [182880]
MH025916.1_strain_R8              ----------------------------------------------------------------------------.... [182880]

FR682468.2_Georgia_2007/1         ttac--------------------tgatcatgacttaaata-----------aaaatattttgcccgcttaaaaattgtt [182960]
ASF_IND_20_CAD543                 ....--------------------.................-----------............................ [182960]
LR899193.1_Germany_2020/1         ....--------------------.................-----------............................ [182960]
MN393477.1_Wuhan_2019             ....--------------------.................-----------............................ [182960]
MW396979.1_Timor-Leste/2019/1     ....--------------------.................-----------............................ [182960]
MK940252.1_CN/2019/InnerMongolia  ....--------------------.................-----------............................ [182960]
MK128995.1_China/2018/AnhuiXCGQ   ....--------------------.................-----------............................ [182960]
MH681419.1_ASFV/POL/2015          ....--------------------.................-----------............................ [182960]
LS478113.1_Estonia_2014           ....--------------------.................-----------............................ [182960]
KP843857.1_Odintsovo_02/14        ....--------------------.................-----------............................ [182960]
FN557520.1_strain_E75             ....--------------------.................-----------............................ [182960]
AY261366.1_Warthog                ....--------------------.................-----------.......g.....g.............. [182960]
AY261365.1_Warmbaths              ....--------------------.................-----------.......g.....g.a............ [182960]
AY261364.1_Tengani_62             ....--------------------.................-----------.......g.....g.............. [182960]
AY261363.1_Pretoriuskop/96/4      ....--------------------.................-----------.......g.....g.............. [182960]
AY261362.1_Mkuzi_1979             ....--------------------.................-----------.....................g...... [182960]
AY261361.1_Malawi_Lil-20/1_1983   ...aatatgctgttggcttaaata..c.gt..g........tgctgttggctt......gc..ttg.............. [182960]
AY261360.1_Kenya_1950             ...actatgc--------------........a.-.......-----------.......g....tga............. [182960]
AM712240.1_OURT_88/3              ....--------------------.................-----------............................ [182960]
AM712239.1_Benin_97/1             ....--------------------.................-----------............................ [182960]
KM111295.1_Ken06.Bus              ..gactatgc--------------....t..a.-.......-----------.......g.....g.............. [182960]
```

```
KM111294.1_Ken05/Tk1             ...actatgc--------------.......a.-...t...-----------.......g....tga............. [182960]
KM262844.1_strain_L60            ....--------------------.................-----------............................ [182960]
KM102979.1_26544/OG10_Italy      ....--------------------.................-----------............................ [182960]
KM262845.1_strain_NHV            ....--------------------.................-----------............................ [182960]
KX354450.1_47/Ss/2008            ....--------------------.................-----------............................ [182960]
KP055815.1_strain_BA71           ....--------------------.................-----------............................ [182960]
MH025920.1_strain_R35            ..gactatgc--------------....t..a.-.......-----------.......g.....g.............. [182960]
MH025919.1_strain_N10            ..gactatgc--------------....t..a.-.......-----------.......g.....g.............. [182960]
MH025918.1_strain_R25            ..gactatgc--------------....t..a.-.......-----------.......g.....g.............. [182960]
MH025917.1_strain_R7             ..gactatgc--------------....t..a.-.......-----------.......g.....g.............. [182960]
MH025916.1_strain_R8             ..gactatgc--------------....t..a.-.......-----------.......g.....g.............. [182960]

FR682468.2_Georgia_2007/1        taggtttgaaaaaataagagatggagggggcaacttatcgtcattgtgtttacccccactg---gaagacatcaaacggt [183040]
ASF_IND_20_CAD543                .............................................................---................ [183040]
LR899193.1_Germany_2020/1        .............................................................---................ [183040]
MN393477.1_Wuhan_2019            .............................................................---................ [183040]
MW396979.1_Timor-Leste/2019/1    .............................................................---................ [183040]
MK940252.1_CN/2019/InnerMongolia .............................................................---................ [183040]
MK128995.1_China/2018/AnhuiXCGQ  .............................................................---................ [183040]
MH681419.1_ASFV/POL/2015         .............................................................---................ [183040]
LS478113.1_Estonia_2014          .............................................................---................ [183040]
KP843857.1_Odintsovo_02/14       .............................................................---................ [183040]
FN557520.1_strain_E75            .............................................................---....g...t....... [183040]
AY261366.1_Warthog               .............................................................gaa........t....... [183040]
AY261365.1_Warmbaths             .............................................................gaa........t....... [183040]
AY261364.1_Tengani_62            ..............................................g..............---........t....... [183040]
AY261363.1_Pretoriuskop/96/4     .............................................................---................ [183040]
AY261362.1_Mkuzi_1979            ..a..........................................................---................ [183040]
AY261361.1_Malawi_Lil-20/1_1983  .................................t.....t.....................---........t....... [183040]
AY261360.1_Kenya_1950            .................................t.....tc....t......g..g.....---........t....... [183040]
AM712240.1_OURT_88/3             .............................................................---....g........... [183040]
AM712239.1_Benin_97/1            .............................................................---....g........... [183040]
KM111295.1_Ken06.Bus             .................................t.....tc....c......g........---........t.....a. [183040]
KM111294.1_Ken05/Tk1             .................................t.....tc....t......g..g.....---........t....... [183040]
KM262844.1_strain_L60            .............................................................---....g........... [183040]
KM102979.1_26544/OG10_Italy      .............................................................---....g........... [183040]
KM262845.1_strain_NHV            .............................................................---....g........... [183040]
KX354450.1_47/Ss/2008            .............................................................---....g........... [183040]
KP055815.1_strain_BA71           .............................................................---....g........... [183040]
MH025920.1_strain_R35            .................................t.....tc....c......g........---........t.....a. [183040]
MH025919.1_strain_N10            .................................t.....tc....c......g........---........t.....a. [183040]
MH025918.1_strain_R25            .................................t.....tc....c......g........---........t.....a. [183040]
MH025917.1_strain_R7             .................................t.....tc....c......g........---........t.....a. [183040]
MH025916.1_strain_R8             .................................t.....tc....c......g........---........t.....a. [183040]
```

```
FR682468.2_Georgia_2007/1        aaataattataagaatcaaaatgattaatataagggttaaaaaaggatgattcatcacattaattaaaaacgtatttata [183120]
ASF_IND_20_CAD543                ................................................................................ [183120]
LR899193.1_Germany_2020/1        ................................................................................ [183120]
MN393477.1_Wuhan_2019            ................................................................................ [183120]
MW396979.1_Timor-Leste/2019/1    ................................................................................ [183120]
MK940252.1_CN/2019/InnerMongolia ................................................................................ [183120]
MK128995.1_China/2018/AnhuiXCGQ  ................................................................................ [183120]
MH681419.1_ASFV/POL/2015         ................................................................................ [183120]
LS478113.1_Estonia_2014          ................................................................................ [183120]
KP843857.1_Odintsovo_02/14       ................................................................................ [183120]
FN557520.1_strain_E75            ................................................................................ [183120]
AY261366.1_Warthog               ................................................................................ [183120]
AY261365.1_Warmbaths             ................................................................................ [183120]
AY261364.1_Tengani_62            ................................................................................ [183120]
AY261363.1_Pretoriuskop/96/4     ................................................................................ [183120]
AY261362.1_Mkuzi_1979            ................................................................................ [183120]
AY261361.1_Malawi_Lil-20/1_1983  .......c.c......t............................................................... [183120]
AY261360.1_Kenya_1950            .........c......t............................................................... [183120]
AM712240.1_OURT_88/3             ................................................................................ [183120]
AM712239.1_Benin_97/1            ................................................................................ [183120]
KM111295.1_Ken06.Bus             .........c......t..........................g.................................... [183120]
KM111294.1_Ken05/Tk1             .........c......t............................................................... [183120]
KM262844.1_strain_L60            ................................................................................ [183120]
KM102979.1_26544/OG10_Italy      ................................................................................ [183120]
KM262845.1_strain_NHV            ................................................................................ [183120]
KX354450.1_47/Ss/2008            ................................................................................ [183120]
KP055815.1_strain_BA71           ................................................................................ [183120]
MH025920.1_strain_R35            .........c......t..........................g.................................... [183120]
MH025919.1_strain_N10            .........c......t..........................g.................................... [183120]
MH025918.1_strain_R25            .........c......t..........................g.................................... [183120]
MH025917.1_strain_R7             .........c......t..........................g.................................... [183120]
MH025916.1_strain_R8             .........c......t..........................g.................................... [183120]

FR682468.2_Georgia_2007/1        acgctgttgcagttgaaattttggtataggtcggaaatattgcccgagcctccgtattctgcaatgttctgacatatggt [183200]
ASF_IND_20_CAD543                ....................................................t........................... [183200]
LR899193.1_Germany_2020/1        ................................................................................ [183200]
MN393477.1_Wuhan_2019            ................................................................................ [183200]
MW396979.1_Timor-Leste/2019/1    ................................................................................ [183200]
MK940252.1_CN/2019/InnerMongolia ................................................................................ [183200]
MK128995.1_China/2018/AnhuiXCGQ  ................................................................................ [183200]
MH681419.1_ASFV/POL/2015         ....................................................t........................... [183200]
LS478113.1_Estonia_2014          ................................................................................ [183200]
KP843857.1_Odintsovo_02/14       ................................................................................ [183200]
```

```
FN557520.1_strain_E75            ....................................................t........................... [183200]
AY261366.1_Warthog               ................................a..t.........................................a.. [183200]
AY261365.1_Warmbaths             ................................a..t.........................................a.. [183200]
AY261364.1_Tengani_62            ................................a..t.........................................a.. [183200]
AY261363.1_Pretoriuskop/96/4     .............................a.................................................. [183200]
AY261362.1_Mkuzi_1979            ..........................................t.........t........................... [183200]
AY261361.1_Malawi_Lil-20/1_1983  ........a................................a....t................................. [183200]
AY261360.1_Kenya_1950            .........................................a....t................................. [183200]
AM712240.1_OURT_88/3             ....................................................t........................... [183200]
AM712239.1_Benin_97/1            ...................................................t............................ [183200]
KM111295.1_Ken06.Bus             .........................................a....t................................. [183200]
KM111294.1_Ken05/Tk1             .........................................a....t................................. [183200]
KM262844.1_strain_L60            ....................................................t........................... [183200]
KM102979.1_26544/OG10_Italy      ................................................................................ [183200]
KM262845.1_strain_NHV            ....................................................t........................... [183200]
KX354450.1_47/Ss/2008            ................................................................................ [183200]
KP055815.1_strain_BA71           ................................................................................ [183200]
MH025920.1_strain_R35            .........................................a....t................................. [183200]
MH025919.1_strain_N10            .........................................a....t................................. [183200]
MH025918.1_strain_R25            .........................................a....t................................. [183200]
MH025917.1_strain_R7             .........................................a....t................................. [183200]
MH025916.1_strain_R8             .........................................a....t................................. [183200]

FR682468.2_Georgia_2007/1        gagtccggaggggcactgcttgttggtcaaaatatttctttgctccgttgttttataggcatttttatttccattacacg [183280]
ASF_IND_20_CAD543                ................................................................................ [183280]
LR899193.1_Germany_2020/1        ................................................................................ [183280]
MN393477.1_Wuhan_2019            ................................................................................ [183280]
MW396979.1_Timor-Leste/2019/1    ................................................................................ [183280]
MK940252.1_CN/2019/InnerMongolia ................................................................................ [183280]
MK128995.1_China/2018/AnhuiXCGQ  ................................................................................ [183280]
MH681419.1_ASFV/POL/2015         ................................................................................ [183280]
LS478113.1_Estonia_2014          ................................................................................ [183280]
KP843857.1_Odintsovo_02/14       ................................................................................ [183280]
FN557520.1_strain_E75            ................................................................................ [183280]
AY261366.1_Warthog               ...a............................................c..........................g.... [183280]
AY261365.1_Warmbaths             ............a...................................c............................... [183280]
AY261364.1_Tengani_62            .......................................a........c............................... [183280]
AY261363.1_Pretoriuskop/96/4     ................................................c..........................g.... [183280]
AY261362.1_Mkuzi_1979            ..................t............................................................. [183280]
AY261361.1_Malawi_Lil-20/1_1983  a.....a.................c..............a..t..................................... [183280]
AY261360.1_Kenya_1950            ........................c...............................................g....... [183280]
AM712240.1_OURT_88/3             ................................................................................ [183280]
AM712239.1_Benin_97/1            ................................................................................ [183280]
KM111295.1_Ken06.Bus             ........................c..............a.................t..............g....... [183280]
```

```
KM111294.1_Ken05/Tk1                ........................c............t..................................g....... [183280]
KM262844.1_strain_L60               ................................................................................ [183280]
KM102979.1_26544/OG10_Italy         ................................................................................ [183280]
KM262845.1_strain_NHV               ................................................................................ [183280]
KX354450.1_47/Ss/2008               ................................................................................ [183280]
KP055815.1_strain_BA71              .......................................a........................................ [183280]
MH025920.1_strain_R35               ........................c..............a.................t..............g....... [183280]
MH025919.1_strain_N10               ........................c..............a.................t..............g....... [183280]
MH025918.1_strain_R25               ........................c..............a.................t..............g....... [183280]
MH025917.1_strain_R7                ........................c..............a.................t..............g....... [183280]
MH025916.1_strain_R8                ........................c..............a.................t..............g....... [183280]

FR682468.2_Georgia_2007/1           gagcaaacgcacattcagcccatagggtgccggagttcacacaggcacaatactggctatacgcatactcatcctttgag [183360]
ASF_IND_20_CAD543                   ................................................................................ [183360]
LR899193.1_Germany_2020/1           ................................................................................ [183360]
MN393477.1_Wuhan_2019               ................................................................................ [183360]
MW396979.1_Timor-Leste/2019/1       ................................................................................ [183360]
MK940252.1_CN/2019/InnerMongolia    ................................................................................ [183360]
MK128995.1_China/2018/AnhuiXCGQ     ................................................................................ [183360]
MH681419.1_ASFV/POL/2015            ................................................................................ [183360]
LS478113.1_Estonia_2014             ................................................................................ [183360]
KP843857.1_Odintsovo_02/14          ................................................................................ [183360]
FN557520.1_strain_E75               ................................................................................ [183360]
AY261366.1_Warthog                  ........................a....................................................... [183360]
AY261365.1_Warmbaths                ........................a....................................................... [183360]
AY261364.1_Tengani_62               ........................a....................................................... [183360]
AY261363.1_Pretoriuskop/96/4        ........................a....................................................... [183360]
AY261362.1_Mkuzi_1979               ................................................................................ [183360]
AY261361.1_Malawi_Lil-20/1_1983     ........................................................................t....... [183360]
AY261360.1_Kenya_1950               ........................................g....................................... [183360]
AM712240.1_OURT_88/3                ................................................................................ [183360]
AM712239.1_Benin_97/1               ................................................................................ [183360]
KM111295.1_Ken06.Bus                ........................................g.............................g........a [183360]
KM111294.1_Ken05/Tk1                ........................................g....................................... [183360]
KM262844.1_strain_L60               ................................................................................ [183360]
KM102979.1_26544/OG10_Italy         .................c.............................................................. [183360]
KM262845.1_strain_NHV               ................................................................................ [183360]
KX354450.1_47/Ss/2008               .................c.............................................................. [183360]
KP055815.1_strain_BA71              ................................................................................ [183360]
MH025920.1_strain_R35               ........................................g.............................g........a [183360]
MH025919.1_strain_N10               ........................................g.............................g........a [183360]
MH025918.1_strain_R25               ........................................g.............................g........a [183360]
MH025917.1_strain_R7                ........................................g.............................g........a [183360]
MH025916.1_strain_R8                ........................................g.............................g........a [183360]
```

```
FR682468.2_Georgia_2007/1        cacaatccctgtttatcgcatatgctcccaataatattgtcatcctccgccgtttgttgatttgtatgcgagcgtaaaat [183440]
ASF_IND_20_CAD543                ................................................................................ [183440]
LR899193.1_Germany_2020/1        ................................................................................ [183440]
MN393477.1_Wuhan_2019            ................................................................................ [183440]
MW396979.1_Timor-Leste/2019/1    ................................................................................ [183440]
MK940252.1_CN/2019/InnerMongolia ................................................................................ [183440]
MK128995.1_China/2018/AnhuiXCGQ  ................................................................................ [183440]
MH681419.1_ASFV/POL/2015         ................................................................................ [183440]
LS478113.1_Estonia_2014          ................................................................................ [183440]
KP843857.1_Odintsovo_02/14       ................................................................................ [183440]
FN557520.1_strain_E75            ................................................................................ [183440]
AY261366.1_Warthog               ..................................................................c.........c... [183440]
AY261365.1_Warmbaths             ..................................................................c............. [183440]
AY261364.1_Tengani_62            ....................................g........................................... [183440]
AY261363.1_Pretoriuskop/96/4     ....................................g........................................... [183440]
AY261362.1_Mkuzi_1979            ................................................................................ [183440]
AY261361.1_Malawi_Lil-20/1_1983  ..............g.....................g.......t...............c..c.t.............. [183440]
AY261360.1_Kenya_1950            ..............g..............t......g.......t..................t.t.............. [183440]
AM712240.1_OURT_88/3             .........c...................................................................... [183440]
AM712239.1_Benin_97/1            ................................................................................ [183440]
KM111295.1_Ken06.Bus             ..............g..............t......g.......t..................t.t.........g.... [183440]
KM111294.1_Ken05/Tk1             ..............g..............t......g.......t..................t.t.............. [183440]
KM262844.1_strain_L60            ................................................................................ [183440]
KM102979.1_26544/OG10_Italy      ................................................................................ [183440]
KM262845.1_strain_NHV            .........c...................................................................... [183440]
KX354450.1_47/Ss/2008            ................................................................................ [183440]
KP055815.1_strain_BA71           ................................................................................ [183440]
MH025920.1_strain_R35            ..............g..............t......g.......t..................t.t.........g.... [183440]
MH025919.1_strain_N10            ..............g..............t......g.......t..................t.t.........g.... [183440]
MH025918.1_strain_R25            ..............g..............t......g.......t..................t.t.........g.... [183440]
MH025917.1_strain_R7             ..............g..............t......g.......t..................t.t.........g.... [183440]
MH025916.1_strain_R8             ..............g..............t......g.......t..................t.t.........g.... [183440]

FR682468.2_Georgia_2007/1        agcggcccaggccttgggctcctttttttgcagctcggaaatcgaagggcctgtacagctaaagtcgacccaaatatcat [183520]
ASF_IND_20_CAD543                ................................................................................ [183520]
LR899193.1_Germany_2020/1        ................................................................................ [183520]
MN393477.1_Wuhan_2019            ................................................................................ [183520]
MW396979.1_Timor-Leste/2019/1    ................................................................................ [183520]
MK940252.1_CN/2019/InnerMongolia ................................................................................ [183520]
MK128995.1_China/2018/AnhuiXCGQ  ................................................................................ [183520]
MH681419.1_ASFV/POL/2015         ................................................................................ [183520]
LS478113.1_Estonia_2014          ................................................................................ [183520]
KP843857.1_Odintsovo_02/14       ................................................................................ [183520]
```

```
FN557520.1_strain_E75            ................................................................................ [183520]
AY261366.1_Warthog               ..........................................t..................................... [183520]
AY261365.1_Warmbaths             ................................................................................ [183520]
AY261364.1_Tengani_62            ................................................................................ [183520]
AY261363.1_Pretoriuskop/96/4     ................................................................................ [183520]
AY261362.1_Mkuzi_1979            .........................................c...................................... [183520]
AY261361.1_Malawi_Lil-20/1_1983  ....................................a....g...................................... [183520]
AY261360.1_Kenya_1950            c...........g............................c..........................t........... [183520]
AM712240.1_OURT_88/3             ................................................................................ [183520]
AM712239.1_Benin_97/1            ................................................................................ [183520]
KM111295.1_Ken06.Bus             c...........a.........................................g......................... [183520]
KM111294.1_Ken05/Tk1             c...........a.......................................................t........... [183520]
KM262844.1_strain_L60            ................................................................................ [183520]
KM102979.1_26544/OG10_Italy      ................................................................................ [183520]
KM262845.1_strain_NHV            ................................................................................ [183520]
KX354450.1_47/Ss/2008            ................................................................................ [183520]
KP055815.1_strain_BA71           ................................................................................ [183520]
MH025920.1_strain_R35            c...........a.........................................g......................... [183520]
MH025919.1_strain_N10            c...........a.........................................g......................... [183520]
MH025918.1_strain_R25            c...........a.........................................g......................... [183520]
MH025917.1_strain_R7             c...........a.........................................g......................... [183520]
MH025916.1_strain_R8             c...........a.........................................g......................... [183520]

FR682468.2_Georgia_2007/1        tgcatttcgtggaaactggcatgcaa---gacataattgaaataattaataagtatatatcatggcaacaaattttttta [183600]
ASF_IND_20_CAD543                ..........................---................................................... [183600]
LR899193.1_Germany_2020/1        ..........................---................................................... [183600]
MN393477.1_Wuhan_2019            ..........................---................................................... [183600]
MW396979.1_Timor-Leste/2019/1    ..........................---................................................... [183600]
MK940252.1_CN/2019/InnerMongolia ..........................---................................................... [183600]
MK128995.1_China/2018/AnhuiXCGQ  ..........................---................................................... [183600]
MH681419.1_ASFV/POL/2015         ..........................---................................................... [183600]
LS478113.1_Estonia_2014          ..........................---................................................... [183600]
KP843857.1_Odintsovo_02/14       ..........................---................................................... [183600]
FN557520.1_strain_E75            ..........................---................................................... [183600]
AY261366.1_Warthog               ..........................---................................................... [183600]
AY261365.1_Warmbaths             ..........................---................................................... [183600]
AY261364.1_Tengani_62            ..........................---................................................... [183600]
AY261363.1_Pretoriuskop/96/4     ..........................---................................................... [183600]
AY261362.1_Mkuzi_1979            ..........................gac................................................... [183600]
AY261361.1_Malawi_Lil-20/1_1983  ..........c....t.........t---..........................................t........ [183600]
AY261360.1_Kenya_1950            ....c.....c.........g....g---..............t....t............................... [183600]
AM712240.1_OURT_88/3             ..........................---................................................... [183600]
AM712239.1_Benin_97/1            ..........................---................................................... [183600]
KM111295.1_Ken06.Bus             ....c.....c..............g---..............t....t............................... [183600]
```

```
KM111294.1_Ken05/Tk1              ....c.....c.........g....g---..............t....t............................... [183600]
KM262844.1_strain_L60             ..........................---................................................... [183600]
KM102979.1_26544/OG10_Italy       ..........................---................................................... [183600]
KM262845.1_strain_NHV             ..........................---................................................... [183600]
KX354450.1_47/Ss/2008             ..........................---................................................... [183600]
KP055815.1_strain_BA71            ..........................---................................................... [183600]
MH025920.1_strain_R35             ....c.....c..............g---..............t....t............................... [183600]
MH025919.1_strain_N10             ....c.....c..............g---..............t....t............................... [183600]
MH025918.1_strain_R25             ....c.....c..............g---..............t....t............................... [183600]
MH025917.1_strain_R7              ....c.....c..............g---..............t....t............................... [183600]
MH025916.1_strain_R8              ....c.....c..............g---..............t....t............................... [183600]

FR682468.2_Georgia_2007/1         ttcaacctatcaccgaagaagctgaagcatactacccaccttccgtgataacgaataaacggaaggacctgggggtagac [183680]
ASF_IND_20_CAD543                 ................................................................................ [183680]
LR899193.1_Germany_2020/1         ................................................................................ [183680]
MN393477.1_Wuhan_2019             ................................................................................ [183680]
MW396979.1_Timor-Leste/2019/1     ................................................................................ [183680]
MK940252.1_CN/2019/InnerMongolia  ................................................................................ [183680]
MK128995.1_China/2018/AnhuiXCGQ   ................................................................................ [183680]
MH681419.1_ASFV/POL/2015          ................................................................................ [183680]
LS478113.1_Estonia_2014           ................................................................................ [183680]
KP843857.1_Odintsovo_02/14        ................................................................................ [183680]
FN557520.1_strain_E75             ..............c................................................................. [183680]
AY261366.1_Warthog                .............t..............................................................g..t [183680]
AY261365.1_Warmbaths              ............................................................................g..t [183680]
AY261364.1_Tengani_62             ..........................a.................................................g..t [183680]
AY261363.1_Pretoriuskop/96/4      ............................................................................g..t [183680]
AY261362.1_Mkuzi_1979             ......................c.....................................................g..t [183680]
AY261361.1_Malawi_Lil-20/1_1983   ...........................................ta................a...........a..g..t [183680]
AY261360.1_Kenya_1950             .c........................a.......t..c.......................ac................t [183680]
AM712240.1_OURT_88/3              ..............c................................................................. [183680]
AM712239.1_Benin_97/1             ..............c................................................................. [183680]
KM111295.1_Ken06.Bus              .c........................a..........c.......................ac.............g..t [183680]
KM111294.1_Ken05/Tk1              .c........................a..........c.......................ac.............g..t [183680]
KM262844.1_strain_L60             ..............c................................................................. [183680]
KM102979.1_26544/OG10_Italy       ..............c................................................................. [183680]
KM262845.1_strain_NHV             ..............c................................................................. [183680]
KX354450.1_47/Ss/2008             ..............c................................................................. [183680]
KP055815.1_strain_BA71            ..............c................................................................. [183680]
MH025920.1_strain_R35             .c........................a..........c.......................ac.............g..t [183680]
MH025919.1_strain_N10             .c........................a..........c.......................ac.............g..t [183680]
MH025918.1_strain_R25             .c........................a..........c.......................ac.............g..t [183680]
MH025917.1_strain_R7              .c........................a..........c.......................ac.............g..t [183680]
MH025916.1_strain_R8              .c........................a..........c.......................ac.............g..t [183680]
```

```
FR682468.2_Georgia_2007/1        gtatactgttgctccgacctagtgcttcaacctggactaaatattgttcgcctgcatattaaagtagcatgcgaacacat [183760]
ASF_IND_20_CAD543                ................................................................................ [183760]
LR899193.1_Germany_2020/1        ................................................................................ [183760]
MN393477.1_Wuhan_2019            ................................................................................ [183760]
MW396979.1_Timor-Leste/2019/1    ................................................................................ [183760]
MK940252.1_CN/2019/InnerMongolia ................................................................................ [183760]
MK128995.1_China/2018/AnhuiXCGQ  ................................................................................ [183760]
MH681419.1_ASFV/POL/2015         ................................................................................ [183760]
LS478113.1_Estonia_2014          ................................................................................ [183760]
KP843857.1_Odintsovo_02/14       ................................................................................ [183760]
FN557520.1_strain_E75            ................................................................................ [183760]
AY261366.1_Warthog               ................................................................................ [183760]
AY261365.1_Warmbaths             ................................................................................ [183760]
AY261364.1_Tengani_62            ................................c........c...........a.......................... [183760]
AY261363.1_Pretoriuskop/96/4     ................................................................................ [183760]
AY261362.1_Mkuzi_1979            ................................................................................ [183760]
AY261361.1_Malawi_Lil-20/1_1983  ..............ta......................g......a...c.............................. [183760]
AY261360.1_Kenya_1950            ................................c..............................................g [183760]
AM712240.1_OURT_88/3             ................................................................................ [183760]
AM712239.1_Benin_97/1            ................................................................................ [183760]
KM111295.1_Ken06.Bus             ................................c......................................t.......a [183760]
KM111294.1_Ken05/Tk1             ................................c..............................................g [183760]
KM262844.1_strain_L60            ................................................................................ [183760]
KM102979.1_26544/OG10_Italy      ................................................................................ [183760]
KM262845.1_strain_NHV            ................................................................................ [183760]
KX354450.1_47/Ss/2008            ................................................................................ [183760]
KP055815.1_strain_BA71           ................................................................................ [183760]
MH025920.1_strain_R35            ................................c......................................t.......a [183760]
MH025919.1_strain_N10            ................................c......................................t.......a [183760]
MH025918.1_strain_R25            ................................c......................................t.......a [183760]
MH025917.1_strain_R7             ................................c......................................t.......a [183760]
MH025916.1_strain_R8             ................................c......................................t.......a [183760]

FR682468.2_Georgia_2007/1        gggcaaaaaatgcggttttaaaatcatggcgagaagcagtatgtgcacccatgaacggctgctcatccttgcaaacggaa [183840]
ASF_IND_20_CAD543                ................................................................................ [183840]
LR899193.1_Germany_2020/1        ................................................................................ [183840]
MN393477.1_Wuhan_2019            ................................................................................ [183840]
MW396979.1_Timor-Leste/2019/1    ................................................................................ [183840]
MK940252.1_CN/2019/InnerMongolia ................................................................................ [183840]
MK128995.1_China/2018/AnhuiXCGQ  ................................................................................ [183840]
MH681419.1_ASFV/POL/2015         ................................................................................ [183840]
LS478113.1_Estonia_2014          ................................................................................ [183840]
KP843857.1_Odintsovo_02/14       ................................................................................ [183840]
```

```
FN557520.1_strain_E75            ...........................................................................t.... [183840]
AY261366.1_Warthog               ..............................a..g...............t.............................. [183840]
AY261365.1_Warmbaths             .................................g...............t.............................. [183840]
AY261364.1_Tengani_62            ...............g.................g.........................................t..g. [183840]
AY261363.1_Pretoriuskop/96/4     .................................g...............t.............................. [183840]
AY261362.1_Mkuzi_1979            .................................g...............t.............................. [183840]
AY261361.1_Malawi_Lil-20/1_1983  .........g........................................................t...........g. [183840]
AY261360.1_Kenya_1950            .........g.....a.......................c..........................t.........a.c. [183840]
AM712240.1_OURT_88/3             ...........................................................................t.... [183840]
AM712239.1_Benin_97/1            ...........................................................................t.... [183840]
KM111295.1_Ken06.Bus             .......................................c..........................t.........a.c. [183840]
KM111294.1_Ken05/Tk1             .......................................c..........................t.........a.c. [183840]
KM262844.1_strain_L60            ...........................................................................t.... [183840]
KM102979.1_26544/OG10_Italy      ...........................................................................t.... [183840]
KM262845.1_strain_NHV            ...........................................................................t.... [183840]
KX354450.1_47/Ss/2008            ...........................................................................t.... [183840]
KP055815.1_strain_BA71           ...........................................................................t.... [183840]
MH025920.1_strain_R35            .......................................c..........................t.........a.c. [183840]
MH025919.1_strain_N10            .......................................c..........................t.........a.c. [183840]
MH025918.1_strain_R25            .......................................c..........................t.........a.c. [183840]
MH025917.1_strain_R7             .......................................c..........................t.........a.c. [183840]
MH025916.1_strain_R8             .......................................c..........................t.........a.c. [183840]

FR682468.2_Georgia_2007/1        ttggtttaatagacccgggttatgtgggcgagctcatgctcaagatcattaatcttggcgacaccccggtccaaatatgg [183920]
ASF_IND_20_CAD543                ................................................................................ [183920]
LR899193.1_Germany_2020/1        ................................................................................ [183920]
MN393477.1_Wuhan_2019            ................................................................................ [183920]
MW396979.1_Timor-Leste/2019/1    ................................................................................ [183920]
MK940252.1_CN/2019/InnerMongolia ................................................................................ [183920]
MK128995.1_China/2018/AnhuiXCGQ  ................................................................................ [183920]
MH681419.1_ASFV/POL/2015         ................................................................................ [183920]
LS478113.1_Estonia_2014          ................................................................................ [183920]
KP843857.1_Odintsovo_02/14       ................................................................................ [183920]
FN557520.1_strain_E75            ...................c............................................................ [183920]
AY261366.1_Warthog               ...................c........t................................................... [183920]
AY261365.1_Warmbaths             ...................c........t................................................... [183920]
AY261364.1_Tengani_62            ...................c............................................................ [183920]
AY261363.1_Pretoriuskop/96/4     ...................c............................................................ [183920]
AY261362.1_Mkuzi_1979            ...................c........t................................................... [183920]
AY261361.1_Malawi_Lil-20/1_1983  .............t.....c..c........................................t................ [183920]
AY261360.1_Kenya_1950            .............t.....c..c....................a.................t........a......... [183920]
AM712240.1_OURT_88/3             ...................c............................................................ [183920]
AM712239.1_Benin_97/1            ...................c............................................................ [183920]
KM111295.1_Ken06.Bus             .............t.....c..c....................a..........................a......... [183920]
```

```
KM111294.1_Ken05/Tk1             .............t.....c..c....................a.................t........a......... [183920]
KM262844.1_strain_L60            ...................c............................................................ [183920]
KM102979.1_26544/OG10_Italy      ...................c............................................................ [183920]
KM262845.1_strain_NHV            ...................c............................................................ [183920]
KX354450.1_47/Ss/2008            ...................c............................................................ [183920]
KP055815.1_strain_BA71           ...................c............................................................ [183920]
MH025920.1_strain_R35            .............t.....c..c....................a.................t........a......... [183920]
MH025919.1_strain_N10            .............t.....c..c....................a.................t........a......... [183920]
MH025918.1_strain_R25            .............t.....c..c....................a.................t........a......... [183920]
MH025917.1_strain_R7             .............t.....c..c....................a.................t........a......... [183920]
MH025916.1_strain_R8             .............t.....c..c....................a.................t........a......... [183920]

FR682468.2_Georgia_2007/1        gccaaagaatgtttggtgcagttggtggcccaaggtgaccatgtgcctgaccatatcaacatcctaaaaagaaaccaaat [184000]
ASF_IND_20_CAD543                ................................................................................ [184000]
LR899193.1_Germany_2020/1        ................................................................................ [184000]
MN393477.1_Wuhan_2019            ................................................................................ [184000]
MW396979.1_Timor-Leste/2019/1    ................................................................................ [184000]
MK940252.1_CN/2019/InnerMongolia ................................................................................ [184000]
MK128995.1_China/2018/AnhuiXCGQ  ................................................................................ [184000]
MH681419.1_ASFV/POL/2015         ................................................................................ [184000]
LS478113.1_Estonia_2014          ................................................................................ [184000]
KP843857.1_Odintsovo_02/14       ................................................................................ [184000]
FN557520.1_strain_E75            ................................................................................ [184000]
AY261366.1_Warthog               ...............................................................t................ [184000]
AY261365.1_Warmbaths             ................................................................................ [184000]
AY261364.1_Tengani_62            ...............................................................t................ [184000]
AY261363.1_Pretoriuskop/96/4     ...............................................................t................ [184000]
AY261362.1_Mkuzi_1979            ................................................................................ [184000]
AY261361.1_Malawi_Lil-20/1_1983  ............c.....................................t............t..........tg.... [184000]
AY261360.1_Kenya_1950            ..g..................c...................c........t..c.......................... [184000]
AM712240.1_OURT_88/3             ................................................................................ [184000]
AM712239.1_Benin_97/1            ................................................................................ [184000]
KM111295.1_Ken06.Bus             ..g......................................ca.......t...g................g........ [184000]
KM111294.1_Ken05/Tk1             ..g..................c...................c........t............................. [184000]
KM262844.1_strain_L60            ................................................................................ [184000]
KM102979.1_26544/OG10_Italy      ................................................................................ [184000]
KM262845.1_strain_NHV            ................................................................................ [184000]
KX354450.1_47/Ss/2008            ................................................................................ [184000]
KP055815.1_strain_BA71           ................................................................................ [184000]
MH025920.1_strain_R35            ..g......................................ca.......t...g................g........ [184000]
MH025919.1_strain_N10            ..g......................................ca.......t...g................g........ [184000]
MH025918.1_strain_R25            ..g......................................ca.......t...g................g........ [184000]
MH025917.1_strain_R7             ..g......................................ca.......t...g................g........ [184000]
MH025916.1_strain_R8             ..g......................................ca.......t...g................g........ [184000]
```

```
FR682468.2_Georgia_2007/1        atttccgctgtttgcgcctaccccaagaggcgagggtagatttgggagcacgggcgaggccgggattatgagaacttaat [184080]
ASF_IND_20_CAD543                ................................................................................ [184080]
LR899193.1_Germany_2020/1        ................................................................................ [184080]
MN393477.1_Wuhan_2019            ................................................................................ [184080]
MW396979.1_Timor-Leste/2019/1    ................................................................................ [184080]
MK940252.1_CN/2019/InnerMongolia ................................................................................ [184080]
MK128995.1_China/2018/AnhuiXCGQ  ................................................................................ [184080]
MH681419.1_ASFV/POL/2015         ................................................................................ [184080]
LS478113.1_Estonia_2014          ................................................................................ [184080]
KP843857.1_Odintsovo_02/14       ................................................................................ [184080]
FN557520.1_strain_E75            ................................................................................ [184080]
AY261366.1_Warthog               ................................................................................ [184080]
AY261365.1_Warmbaths             ................................................................................ [184080]
AY261364.1_Tengani_62            ................................................................................ [184080]
AY261363.1_Pretoriuskop/96/4     ................................................................................ [184080]
AY261362.1_Mkuzi_1979            ................................................................................ [184080]
AY261361.1_Malawi_Lil-20/1_1983  ........................g..............................a...-.................... [184080]
AY261360.1_Kenya_1950            .........a...........g...........a.............................................. [184080]
AM712240.1_OURT_88/3             .............................................................................g.. [184080]
AM712239.1_Benin_97/1            ................................................................................ [184080]
KM111295.1_Ken06.Bus             .........a...........g........t..a.............................................. [184080]
KM111294.1_Ken05/Tk1             .........a...........g...........a.............................................. [184080]
KM262844.1_strain_L60            ................................................................................ [184080]
KM102979.1_26544/OG10_Italy      ................................................................................ [184080]
KM262845.1_strain_NHV            .............................................................................g.. [184080]
KX354450.1_47/Ss/2008            ................................................................................ [184080]
KP055815.1_strain_BA71           ................................................................................ [184080]
MH025920.1_strain_R35            .........a...........g........t..a.............................................. [184080]
MH025919.1_strain_N10            .........a...........g........t..a.............................................. [184080]
MH025918.1_strain_R25            .........a...........g........t..a.............................................. [184080]
MH025917.1_strain_R7             .........a...........g........t..a.............................................. [184080]
MH025916.1_strain_R8             .........a...........g........t..a.............................................. [184080]

FR682468.2_Georgia_2007/1        tttatttttttcttaacataatgggaggctctacaagcaaaaattcctttaaaaatacgaccaacattatcagcaattc [184160]
ASF_IND_20_CAD543                ................................................................................ [184160]
LR899193.1_Germany_2020/1        ................................................................................ [184160]
MN393477.1_Wuhan_2019            ................................................................................ [184160]
MW396979.1_Timor-Leste/2019/1    ................................................................................ [184160]
MK940252.1_CN/2019/InnerMongolia ................................................................................ [184160]
MK128995.1_China/2018/AnhuiXCGQ  ................................................................................ [184160]
MH681419.1_ASFV/POL/2015         ................................................................................ [184160]
LS478113.1_Estonia_2014          ................................................................................ [184160]
KP843857.1_Odintsovo_02/14       ....-........................................................................... [184160]
```

```
FN557520.1_strain_E75            ................................................................................ [184160]
AY261366.1_Warthog               ..................................................c............................. [184160]
AY261365.1_Warmbaths             ..................................................c............................. [184160]
AY261364.1_Tengani_62            ................................................................................ [184160]
AY261363.1_Pretoriuskop/96/4     ..................................................c............................. [184160]
AY261362.1_Mkuzi_1979            ..................................................c............................. [184160]
AY261361.1_Malawi_Lil-20/1_1983  ..........................................t................................c.... [184160]
AY261360.1_Kenya_1950            ...........................................................................c.... [184160]
AM712240.1_OURT_88/3             ................................................................................ [184160]
AM712239.1_Benin_97/1            ................................................................................ [184160]
KM111295.1_Ken06.Bus             ...........................................................a...............c.... [184160]
KM111294.1_Ken05/Tk1             ...........................................................................c.... [184160]
KM262844.1_strain_L60            ................................................................................ [184160]
KM102979.1_26544/OG10_Italy      ................................................................................ [184160]
KM262845.1_strain_NHV            ................................................................................ [184160]
KX354450.1_47/Ss/2008            ................................................................................ [184160]
KP055815.1_strain_BA71           ................................................................................ [184160]
MH025920.1_strain_R35            ...........................................................a...............c.... [184160]
MH025919.1_strain_N10            ...........................................................a...............c.... [184160]
MH025918.1_strain_R25            ...........................................................a...............c.... [184160]
MH025917.1_strain_R7             ...........................................................a...............c.... [184160]
MH025916.1_strain_R8             ...........................................................a...............c.... [184160]

FR682468.2_Georgia_2007/1        cattttcaatcagatgcaaagttgtatttccatgttggatggcaaaaattacataggcgtattcggtgatggaaatattt [184240]
ASF_IND_20_CAD543                ....................r........................................................... [184240]
LR899193.1_Germany_2020/1        ................................................................................ [184240]
MN393477.1_Wuhan_2019            ................................................................................ [184240]
MW396979.1_Timor-Leste/2019/1    ................................................................................ [184240]
MK940252.1_CN/2019/InnerMongolia ................................................................................ [184240]
MK128995.1_China/2018/AnhuiXCGQ  ................................................................................ [184240]
MH681419.1_ASFV/POL/2015         ................................................................................ [184240]
LS478113.1_Estonia_2014          ................................................................................ [184240]
KP843857.1_Odintsovo_02/14       ................................................................................ [184240]
FN557520.1_strain_E75            ....................a........................................................... [184240]
AY261366.1_Warthog               ........................c.........c............................................. [184240]
AY261365.1_Warmbaths             ....................a........................................................... [184240]
AY261364.1_Tengani_62            .........a...................................................................... [184240]
AY261363.1_Pretoriuskop/96/4     ........................c....................................................... [184240]
AY261362.1_Mkuzi_1979            ....................a........................................................... [184240]
AY261361.1_Malawi_Lil-20/1_1983  ....................a.........t................................................a [184240]
AY261360.1_Kenya_1950            t...................a.........t.............c..................................a [184240]
AM712240.1_OURT_88/3             ....................a........................................................... [184240]
AM712239.1_Benin_97/1            ....................a........................................................... [184240]
KM111295.1_Ken06.Bus             t.........................................t....................................a [184240]
```

```
KM111294.1_Ken05/Tk1            t..............................................................................a [184240]
KM262844.1_strain_L60           ....................a........................................................... [184240]
KM102979.1_26544/OG10_Italy     ....................a........................................................... [184240]
KM262845.1_strain_NHV           ....................a........................................................... [184240]
KX354450.1_47/Ss/2008           ....................a........................................................... [184240]
KP055815.1_strain_BA71          ....................a........................................................... [184240]
MH025920.1_strain_R35           t..............................................................................a [184240]
MH025919.1_strain_N10           t..............................................................................a [184240]
MH025918.1_strain_R25           t..............................................................................a [184240]
MH025917.1_strain_R7            t..............................................................................a [184240]
MH025916.1_strain_R8            t..............................................................................a [184240]

FR682468.2_Georgia_2007/1       taaaccacgttttccaggatttaaacttatcattaaacacaagttgcgtgcaaaagcacgtaaacgaggaaaatttcatt [184320]
ASF_IND_20_CAD543               ....y........................................................................... [184320]
LR899193.1_Germany_2020/1       ................................................................................ [184320]
MN393477.1_Wuhan_2019           ................................................................................ [184320]
MW396979.1_Timor-Leste/2019/1   ................................................................................ [184320]
MK940252.1_CN/2019/InnerMongolia ................................................................................ [184320]
MK128995.1_China/2018/AnhuiXCGQ ................................................................................ [184320]
MH681419.1_ASFV/POL/2015        ................................................................................ [184320]
LS478113.1_Estonia_2014         ................................................................................ [184320]
KP843857.1_Odintsovo_02/14      ................................................................................ [184320]
FN557520.1_strain_E75           ....t...........................c.gg....t....................................... [184320]
AY261366.1_Warthog              ................a...............c.gg....t........................a.............. [184320]
AY261365.1_Warmbaths            ........................................t....................................... [184320]
AY261364.1_Tengani_62           ................................c.gg....t....................................... [184320]
AY261363.1_Pretoriuskop/96/4    ................................................................................ [184320]
AY261362.1_Mkuzi_1979           ........................................t....................................... [184320]
AY261361.1_Malawi_Lil-20/1_1983 ................a.......................t........c.................a............ [184320]
AY261360.1_Kenya_1950           ................a..................g.t..t........c..........................t... [184320]
AM712240.1_OURT_88/3            ....t...........................c.gg....t....................................... [184320]
AM712239.1_Benin_97/1           ....t...........................c.gg....t....................................... [184320]
KM111295.1_Ken06.Bus            ................a..................g....t.....t....................a............ [184320]
KM111294.1_Ken05/Tk1            ................a....................t..t.....t....................a............ [184320]
KM262844.1_strain_L60           ....t...........................c.gg....t....................................... [184320]
KM102979.1_26544/OG10_Italy     ....t...........................c.gg....t....................................... [184320]
KM262845.1_strain_NHV           ....t...........................c.gg....t....................................... [184320]
KX354450.1_47/Ss/2008           ....t...........................c.gg....t....................................... [184320]
KP055815.1_strain_BA71          ....t...........................c.gg....t....................................... [184320]
MH025920.1_strain_R35           ................a..................g....t.....t....................a............ [184320]
MH025919.1_strain_N10           ................a..................g....t.....t....................a............ [184320]
MH025918.1_strain_R25           ................a..................g....t.....t....................a............ [184320]
MH025917.1_strain_R7            ................a..................g....t.....t....................a............ [184320]
MH025916.1_strain_R8            ................a..................g....t.....t....................a............ [184320]
```

```
FR682468.2_Georgia_2007/1        acaaatctttcgaaccaaattactcaaaatttaaaagaccaagaagttgcgttaacccaatggatggacgcaggaactca [184400]
ASF_IND_20_CAD543                ...........................................................................ca... [184400]
LR899193.1_Germany_2020/1        ................................................................................ [184400]
MN393477.1_Wuhan_2019            ................................................................................ [184400]
MW396979.1_Timor-Leste/2019/1    ................................................................................ [184400]
MK940252.1_CN/2019/InnerMongolia ................................................................................ [184400]
MK128995.1_China/2018/AnhuiXCGQ  ................................................................................ [184400]
MH681419.1_ASFV/POL/2015         ................................................................................ [184400]
LS478113.1_Estonia_2014          ................................................................................ [184400]
KP843857.1_Odintsovo_02/14       ................................................................................ [184400]
FN557520.1_strain_E75            ...........................................................................ca... [184400]
AY261366.1_Warthog               ...........................................................................ca... [184400]
AY261365.1_Warmbaths             ....................a........................................................... [184400]
AY261364.1_Tengani_62            ....................a......................................................ca... [184400]
AY261363.1_Pretoriuskop/96/4     ................................................................................ [184400]
AY261362.1_Mkuzi_1979            ....................a........................................................... [184400]
AY261361.1_Malawi_Lil-20/1_1983  ...........................................................................ca... [184400]
AY261360.1_Kenya_1950            ...........................................................................ca... [184400]
AM712240.1_OURT_88/3             ...........................................................................ca... [184400]
AM712239.1_Benin_97/1            ...........................................................................ca... [184400]
KM111295.1_Ken06.Bus             ......................................t....................................ca... [184400]
KM111294.1_Ken05/Tk1             ..................................................a........................ca... [184400]
KM262844.1_strain_L60            ...........................................................................ca... [184400]
KM102979.1_26544/OG10_Italy      ...........................................................................ca... [184400]
KM262845.1_strain_NHV            ...........................................................................ca... [184400]
KX354450.1_47/Ss/2008            ...........................................................................ca... [184400]
KP055815.1_strain_BA71           ...........................................................................ca... [184400]
MH025920.1_strain_R35            ......................................t....................................ca... [184400]
MH025919.1_strain_N10            ......................................t....................................ca... [184400]
MH025918.1_strain_R25            ......................................t....................................ca... [184400]
MH025917.1_strain_R7             ......................................t....................................ca... [184400]
MH025916.1_strain_R8             ......................................t....................................ca... [184400]

FR682468.2_Georgia_2007/1        cgatcagaaaacggatatagaagaaaatataaaggtaaacttaacaaccacacttattcaaaactgcgtttcatccctgt [184480]
ASF_IND_20_CAD543                ................................................................................ [184480]
LR899193.1_Germany_2020/1        ................................................................................ [184480]
MN393477.1_Wuhan_2019            ................................................................................ [184480]
MW396979.1_Timor-Leste/2019/1    ................................................................................ [184480]
MK940252.1_CN/2019/InnerMongolia ................................................................................ [184480]
MK128995.1_China/2018/AnhuiXCGQ  ................................................................................ [184480]
MH681419.1_ASFV/POL/2015         ................................................................................ [184480]
LS478113.1_Estonia_2014          ................................................................................ [184480]
KP843857.1_Odintsovo_02/14       ................................................................................ [184480]
```

```
FN557520.1_strain_E75            ................................................................................ [184480]
AY261366.1_Warthog               ................................................................................ [184480]
AY261365.1_Warmbaths             ................................................................................ [184480]
AY261364.1_Tengani_62            ................................................................................ [184480]
AY261363.1_Pretoriuskop/96/4     ................................................................................ [184480]
AY261362.1_Mkuzi_1979            ................................................................................ [184480]
AY261361.1_Malawi_Lil-20/1_1983  ...c...................................t....a............................g...... [184480]
AY261360.1_Kenya_1950            ...c...........c............................a............................g...... [184480]
AM712240.1_OURT_88/3             ................................................................................ [184480]
AM712239.1_Benin_97/1            ................................................................................ [184480]
KM111295.1_Ken06.Bus             ...c...........c............................a............................g...... [184480]
KM111294.1_Ken05/Tk1             ...c...........c............................a............................g...... [184480]
KM262844.1_strain_L60            ................................................................................ [184480]
KM102979.1_26544/OG10_Italy      ................................................................................ [184480]
KM262845.1_strain_NHV            ................................................................................ [184480]
KX354450.1_47/Ss/2008            ................................................................................ [184480]
KP055815.1_strain_BA71           ................................................................................ [184480]
MH025920.1_strain_R35            ...c........a..c............................a.....................t......g...... [184480]
MH025919.1_strain_N10            ...c........a..c............................a.....................t......g...... [184480]
MH025918.1_strain_R25            ...c........a..c............................a.....................t......g...... [184480]
MH025917.1_strain_R7             ...c........a..c............................a.....................t......g...... [184480]
MH025916.1_strain_R8             ...c........a..c............................a.....................t......g...... [184480]

FR682468.2_Georgia_2007/1        cgggtatgaacgtgctggtggtgaaggggaatggcaacattgttgaaaacgcaactcagaagcagtcgcagcaaatcatc [184560]
ASF_IND_20_CAD543                ................................................................................ [184560]
LR899193.1_Germany_2020/1        ................................................................................ [184560]
MN393477.1_Wuhan_2019            ................................................................................ [184560]
MW396979.1_Timor-Leste/2019/1    ................................................................................ [184560]
MK940252.1_CN/2019/InnerMongolia ................................................................................ [184560]
MK128995.1_China/2018/AnhuiXCGQ  ................................................................................ [184560]
MH681419.1_ASFV/POL/2015         ................................................................................ [184560]
LS478113.1_Estonia_2014          ................................................................................ [184560]
KP843857.1_Odintsovo_02/14       ................................................................................ [184560]
FN557520.1_strain_E75            ................................................................................ [184560]
AY261366.1_Warthog               ....a........................................................................... [184560]
AY261365.1_Warmbaths             ....a........................................................................... [184560]
AY261364.1_Tengani_62            .a..g........................................................................... [184560]
AY261363.1_Pretoriuskop/96/4     ....a.....t.................a................................................... [184560]
AY261362.1_Mkuzi_1979            ................................................................................ [184560]
AY261361.1_Malawi_Lil-20/1_1983  ....a........................................................................... [184560]
AY261360.1_Kenya_1950            ....c........................................................................... [184560]
AM712240.1_OURT_88/3             ................................................................................ [184560]
AM712239.1_Benin_97/1            ................................................................................ [184560]
KM111295.1_Ken06.Bus             ....a........................................................................... [184560]
```

```
KM111294.1_Ken05/Tk1              ....a........................................................................... [184560]
KM262844.1_strain_L60             ................................................................................ [184560]
KM102979.1_26544/OG10_Italy       ................................................................................ [184560]
KM262845.1_strain_NHV             ................................................................................ [184560]
KX354450.1_47/Ss/2008             ................................................................................ [184560]
KP055815.1_strain_BA71            ................................................................................ [184560]
MH025920.1_strain_R35             ....a........................................................................... [184560]
MH025919.1_strain_N10             ....a........................................................................... [184560]
MH025918.1_strain_R25             ....a........................................................................... [184560]
MH025917.1_strain_R7              ....a........................................................................... [184560]
MH025916.1_strain_R8              ....a........................................................................... [184560]

FR682468.2_Georgia_2007/1         tctaactgcttgcaggggagcaagcaggccatagacaccacaaccggcatcactaacacggtaaatcagtactcacacta [184640]
ASF_IND_20_CAD543                 ................................................................................ [184640]
LR899193.1_Germany_2020/1         ................................................................................ [184640]
MN393477.1_Wuhan_2019             ................................................................................ [184640]
MW396979.1_Timor-Leste/2019/1     ................................................................................ [184640]
MK940252.1_CN/2019/InnerMongolia  ................................................................................ [184640]
MK128995.1_China/2018/AnhuiXCGQ   ................................................................................ [184640]
MH681419.1_ASFV/POL/2015          ................................................................................ [184640]
LS478113.1_Estonia_2014           ................................................................................ [184640]
KP843857.1_Odintsovo_02/14        ................................................................................ [184640]
FN557520.1_strain_E75             .................................................................c.............. [184640]
AY261366.1_Warthog                ..........................................................................c..... [184640]
AY261365.1_Warmbaths              ..........................................................................c..... [184640]
AY261364.1_Tengani_62             ..........................................................................c..... [184640]
AY261363.1_Pretoriuskop/96/4      ..........................................................................c..... [184640]
AY261362.1_Mkuzi_1979             .................................................................c.............. [184640]
AY261361.1_Malawi_Lil-20/1_1983   ........t........................................................c.............. [184640]
AY261360.1_Kenya_1950             ........t............................................g.......................... [184640]
AM712240.1_OURT_88/3              .................................................................c.............. [184640]
AM712239.1_Benin_97/1             .................................................................c.............. [184640]
KM111295.1_Ken06.Bus              ........t............................................g.......................... [184640]
KM111294.1_Ken05/Tk1              ........t........a...................................g.......................... [184640]
KM262844.1_strain_L60             .................................................................c.............. [184640]
KM102979.1_26544/OG10_Italy       .................................................................c.............. [184640]
KM262845.1_strain_NHV             .................................................................c.............. [184640]
KX354450.1_47/Ss/2008             .................................................................c.............. [184640]
KP055815.1_strain_BA71            .................................................................c.............. [184640]
MH025920.1_strain_R35             ........t............................................g.......................... [184640]
MH025919.1_strain_N10             ........t............................................g.......................... [184640]
MH025918.1_strain_R25             ........t............................................g.......................... [184640]
MH025917.1_strain_R7              ........t............................................g.......................... [184640]
MH025916.1_strain_R8              ........t............................................g.......................... [184640]
```

```
FR682468.2_Georgia_2007/1         cacctcaaaaaacttttttgacttcattgcagacgcaatttcggctgtttttaaaaacatcatggtcgcggctgtagtta [184720]
ASF_IND_20_CAD543                 ................................................................................ [184720]
LR899193.1_Germany_2020/1         ................................................................................ [184720]
MN393477.1_Wuhan_2019             ................................................................................ [184720]
MW396979.1_Timor-Leste/2019/1     ................................................................................ [184720]
MK940252.1_CN/2019/InnerMongolia  ................................................................................ [184720]
MK128995.1_China/2018/AnhuiXCGQ   ................................................................................ [184720]
MH681419.1_ASFV/POL/2015          ................................................................................ [184720]
LS478113.1_Estonia_2014           ................................................................................ [184720]
KP843857.1_Odintsovo_02/14        ................................................................................ [184720]
FN557520.1_strain_E75             ..................................................................t.....c....... [184720]
AY261366.1_Warthog                ........................................................................c....... [184720]
AY261365.1_Warmbaths              ....................................t...................................c....... [184720]
AY261364.1_Tengani_62             ...............c........................................................c....... [184720]
AY261363.1_Pretoriuskop/96/4      ........................................................................c....... [184720]
AY261362.1_Mkuzi_1979             ........................................................................c....... [184720]
AY261361.1_Malawi_Lil-20/1_1983   ........................................................................c..c.... [184720]
AY261360.1_Kenya_1950             .....................g..................................................c..g.... [184720]
AM712240.1_OURT_88/3              ..................................................................t.....c....... [184720]
AM712239.1_Benin_97/1             ..................................................................t.....c....... [184720]
KM111295.1_Ken06.Bus              .....................g..................................................c..g.... [184720]
KM111294.1_Ken05/Tk1              .....................g..................................................c..g.... [184720]
KM262844.1_strain_L60             ..................................................................t.....c....... [184720]
KM102979.1_26544/OG10_Italy       ..................................................................t.....c....... [184720]
KM262845.1_strain_NHV             ..................................................................t.....c....... [184720]
KX354450.1_47/Ss/2008             ..................................................................t.....c....... [184720]
KP055815.1_strain_BA71            ..................................................................t.....c....... [184720]
MH025920.1_strain_R35             .....................g..................................................c..g.... [184720]
MH025919.1_strain_N10             .....................g..................................................c..g.... [184720]
MH025918.1_strain_R25             .....................g..................................................c..g.... [184720]
MH025917.1_strain_R7              .....................g..................................................c..g.... [184720]
MH025916.1_strain_R8              .....................g..................................................c..g.... [184720]

FR682468.2_Georgia_2007/1         tcgttctaatcatcgtagggtttatagccgtcttttactttttgcattcacggcaccgccatgaggaggaagaagaagct [184800]
ASF_IND_20_CAD543                 ................................................................................ [184800]
LR899193.1_Germany_2020/1         ................................................................................ [184800]
MN393477.1_Wuhan_2019             ................................................................................ [184800]
MW396979.1_Timor-Leste/2019/1     ................................................................................ [184800]
MK940252.1_CN/2019/InnerMongolia  ................................................................................ [184800]
MK128995.1_China/2018/AnhuiXCGQ   ................................................................................ [184800]
MH681419.1_ASFV/POL/2015          ................................................................................ [184800]
LS478113.1_Estonia_2014           ................................................................................ [184800]
KP843857.1_Odintsovo_02/14        ................................................................................ [184800]
```

```
FN557520.1_strain_E75            .............................................................c.................. [184800]
AY261366.1_Warthog               ......................c......................................c.................. [184800]
AY261365.1_Warmbaths             .............................................................c.................. [184800]
AY261364.1_Tengani_62            .............................................................c.................. [184800]
AY261363.1_Pretoriuskop/96/4     ......................c......................................c.................. [184800]
AY261362.1_Mkuzi_1979            .............................................................c.................. [184800]
AY261361.1_Malawi_Lil-20/1_1983  ............................g..a...................-..................g......... [184800]
AY261360.1_Kenya_1950            .....g......................g..a...............................................g [184800]
AM712240.1_OURT_88/3             .............................................................c.................. [184800]
AM712239.1_Benin_97/1            .............................................................c.................. [184800]
KM111295.1_Ken06.Bus             .....g......................g..a.......................t..............g........a [184800]
KM111294.1_Ken05/Tk1             .....g......................g..a...............................................g [184800]
KM262844.1_strain_L60            .............................................................c.................. [184800]
KM102979.1_26544/OG10_Italy      .............................................................c.................. [184800]
KM262845.1_strain_NHV            .............................................................c.................. [184800]
KX354450.1_47/Ss/2008            .............................................................c.................. [184800]
KP055815.1_strain_BA71           .............................................................c.................. [184800]
MH025920.1_strain_R35            .....g......................g..a.......................t..............g........a [184800]
MH025919.1_strain_N10            .....g......................g..a.......................t..............g........a [184800]
MH025918.1_strain_R25            .....g......................g..a.......................t..............g........a [184800]
MH025917.1_strain_R7             .....g......................g..a.......................t..............g........a [184800]
MH025916.1_strain_R8             .....g......................g..a.......................t..............g........a [184800]

FR682468.2_Georgia_2007/1        gaaccactcataagcaacaaggtattaaaaaatgctgccgtttcgtaataatttaattaaaagt-aaaaaaaaaaggtat [184880]
ASF_IND_20_CAD543                ................................................................-............... [184880]
LR899193.1_Germany_2020/1        ................................................................-............... [184880]
MN393477.1_Wuhan_2019            ................................................................-............... [184880]
MW396979.1_Timor-Leste/2019/1    ................................................................a............... [184880]
MK940252.1_CN/2019/InnerMongolia ................................................................-............... [184880]
MK128995.1_China/2018/AnhuiXCGQ  ................................................................-............... [184880]
MH681419.1_ASFV/POL/2015         ................................................................-............... [184880]
LS478113.1_Estonia_2014          ................................................................-............... [184880]
KP843857.1_Odintsovo_02/14       ................................................................--.............. [184880]
FN557520.1_strain_E75            .....g..........................................................-....-.......... [184880]
AY261366.1_Warthog               .....g..........................................................-............... [184880]
AY261365.1_Warmbaths             .....g...........t..............................................-............... [184880]
AY261364.1_Tengani_62            .....g...........t....................t.........................-............... [184880]
AY261363.1_Pretoriuskop/96/4     .................t....................t.........................-............... [184880]
AY261362.1_Mkuzi_1979            .....g..........................................................-............... [184880]
AY261361.1_Malawi_Lil-20/1_1983  .....g...........t...t............t....---..ac..................-............... [184880]
AY261360.1_Kenya_1950            .....g.......c.tct...a.......................c..................-............... [184880]
AM712240.1_OURT_88/3             .....g..........................................................-............... [184880]
AM712239.1_Benin_97/1            .....g..........................................................-............... [184880]
KM111295.1_Ken06.Bus             .....g.......c.tct...a.......................c..................-............... [184880]
```

KM111294.1_Ken05/Tk1 .....g.......c.tct...a.......................c..................-............... [184880]
KM262844.1_strain_L60 .....g..........................................................-....-.......... [184880]
KM102979.1_26544/OG10_Italy .....g..........................................................-............... [184880]
KM262845.1_strain_NHV .....g..........................................................-............... [184880]
KX354450.1_47/Ss/2008 .....g..........................................................-............... [184880]
KP055815.1_strain_BA71 .....g..........................................................-............... [184880]
MH025920.1_strain_R35 .....g.......c.tct...a.......................c..................-............... [184880]
MH025919.1_strain_N10 .....g.......c.tct...a.......................c..................-............... [184880]
MH025918.1_strain_R25 .....g.......c.tct...a.......................c..................-............... [184880]
MH025917.1_strain_R7 .....g.......c.tct...a.......................c..................-............... [184880]
MH025916.1_strain_R8 .....g.......c.tct...a.......................c..................-............... [184880]

FR682468.2_Georgia_2007/1 tgttatagtgatggcagattttaattctccaatccagtatttgaaagaagattcgagggaccggacctctataggttctc [184960]
ASF_IND_20_CAD543 ................................................................................ [184960]
LR899193.1_Germany_2020/1 ................................................................................ [184960]
MN393477.1_Wuhan_2019 ................................................................................ [184960]
MW396979.1_Timor-Leste/2019/1 ................................................................................ [184960]
MK940252.1_CN/2019/InnerMongolia ................................................................................ [184960]
MK128995.1_China/2018/AnhuiXCGQ ................................................................................ [184960]
MH681419.1_ASFV/POL/2015 ................................................................................ [184960]
LS478113.1_Estonia_2014 ................................................................................ [184960]
KP843857.1_Odintsovo_02/14 ................................................................................ [184960]
FN557520.1_strain_E75 ................................................................................ [184960]
AY261366.1_Warthog ...............................................................................t [184960]
AY261365.1_Warmbaths ................................................................................ [184960]
AY261364.1_Tengani_62 ........................c.......................t.....t........a...............t [184960]
AY261363.1_Pretoriuskop/96/4 ...............................................................................t [184960]
AY261362.1_Mkuzi_1979 ................................................................................ [184960]
AY261361.1_Malawi_Lil-20/1_1983 ........................c.............................c........................t [184960]
AY261360.1_Kenya_1950 ...................c..........t.................t....tc.a.....aa...............t [184960]
AM712240.1_OURT_88/3 ................................................................................ [184960]
AM712239.1_Benin_97/1 ................................................................................ [184960]
KM111295.1_Ken06.Bus ...................c..........t.................t....tc.a.....aa...............t [184960]
KM111294.1_Ken05/Tk1 ...................c..........t.................t....tc.a.....aa...............t [184960]
KM262844.1_strain_L60 ................................................................................ [184960]
KM102979.1_26544/OG10_Italy ................................................................................ [184960]
KM262845.1_strain_NHV ................................................................................ [184960]
KX354450.1_47/Ss/2008 ................................................................................ [184960]
KP055815.1_strain_BA71 ................................................................................ [184960]
MH025920.1_strain_R35 ...................c..........t.................t....tc.a.....aa...............t [184960]
MH025919.1_strain_N10 ...................c..........t.................t....tc.a.....aa...............t [184960]
MH025918.1_strain_R25 ...................c..........t.................t....tc.a.....aa...............t [184960]
MH025917.1_strain_R7 ...................c..........t.................t....tc.a.....aa...............t [184960]
MH025916.1_strain_R8 ...................c..........t.................t....tc.a.....aa...............t [184960]

```
FR682468.2_Georgia_2007/1        tagaatacgatgaaaatgccgacacgatgataccgagcttcgcagcaggcttggaagagtttgaacccattcccgactat [185040]
ASF_IND_20_CAD543                ..........................................................................------ [185040]
LR899193.1_Germany_2020/1        ................................................................................ [185040]
MN393477.1_Wuhan_2019            ................................................................................ [185040]
MW396979.1_Timor-Leste/2019/1    ................................................................................ [185040]
MK940252.1_CN/2019/InnerMongolia ................................................................................ [185040]
MK128995.1_China/2018/AnhuiXCGQ  ................................................................................ [185040]
MH681419.1_ASFV/POL/2015         ................................................................................ [185040]
LS478113.1_Estonia_2014          ................................................................................ [185040]
KP843857.1_Odintsovo_02/14       ................................................................................ [185040]
FN557520.1_strain_E75            .................t..........t.............................c...............------ [185040]
AY261366.1_Warthog               .g............................................................................t. [185040]
AY261365.1_Warmbaths             .................t..........t.................................................t. [185040]
AY261364.1_Tengani_62            .g........................................................a...................t. [185040]
AY261363.1_Pretoriuskop/96/4     .................t.......a..t.............................c...............ag.cc. [185040]
AY261362.1_Mkuzi_1979            .................t..........t.............................c...............------ [185040]
AY261361.1_Malawi_Lil-20/1_1983  .g.............................................................................c [185040]
AY261360.1_Kenya_1950            .................t................c.....................a.c.a.............------ [185040]
AM712240.1_OURT_88/3             .................t..........t.............................c...............------ [185040]
AM712239.1_Benin_97/1            .................t..........t.............................c...............------ [185040]
KM111295.1_Ken06.Bus             .................t................c.....................a.c.a.............------ [185040]
KM111294.1_Ken05/Tk1             .................t................c.....................a.c.a.............------ [185040]
KM262844.1_strain_L60            .................t..........t.............................c...............------ [185040]
KM102979.1_26544/OG10_Italy      .................t..........t.............................c...............------ [185040]
KM262845.1_strain_NHV            .................t..........t.............................c...............------ [185040]
KX354450.1_47/Ss/2008            .................t..........t.............................c...............------ [185040]
KP055815.1_strain_BA71           .................t..........t.............................c...............------ [185040]
MH025920.1_strain_R35            .................t................c.....................a.c.a.............------ [185040]
MH025919.1_strain_N10            .................t................c.....................a.c.a.............------ [185040]
MH025918.1_strain_R25            .................t................c.....................a.c.a.............------ [185040]
MH025917.1_strain_R7             .................t................c.....................a.c.a.............------ [185040]
MH025916.1_strain_R8             .................t................c.....................a.c.a.............------ [185040]

FR682468.2_Georgia_2007/1        gaccctaccacatcaacttccctgtattcacaattgacccacaacatggaaaaaatcgcagaggaagaggatagtaattt [185120]
ASF_IND_20_CAD543                ag.............................................................................. [185120]
LR899193.1_Germany_2020/1        ................................................................................ [185120]
MN393477.1_Wuhan_2019            ................................................................................ [185120]
MW396979.1_Timor-Leste/2019/1    ................................................................................ [185120]
MK940252.1_CN/2019/InnerMongolia ................................................................................ [185120]
MK128995.1_China/2018/AnhuiXCGQ  ................................................................................ [185120]
MH681419.1_ASFV/POL/2015         ................................................................................ [185120]
LS478113.1_Estonia_2014          ................................................................................ [185120]
KP843857.1_Odintsovo_02/14       ................................................................................ [185120]
```

```
FN557520.1_strain_E75           ag.........................................................g.............t...... [185120]
AY261366.1_Warthog              .....a..................................................t................a...... [185120]
AY261365.1_Warmbaths            .....a.....................................................g..a..........c...... [185120]
AY261364.1_Tengani_62           ...........................................................g........t....a...... [185120]
AY261363.1_Pretoriuskop/96/4    a---.......................................................g........a....t...... [185120]
AY261362.1_Mkuzi_1979           ag.........................................................g.............t...... [185120]
AY261361.1_Malawi_Lil-20/1_1983 ..t.......g.........t..a...........a................g......g........t....t...... [185120]
AY261360.1_Kenya_1950           ag.............t....g..a..c.....c.......................t..g........t....t...... [185120]
AM712240.1_OURT_88/3            ag.........................................................g.............t...... [185120]
AM712239.1_Benin_97/1           ag.........................................................g.............t...... [185120]
KM111295.1_Ken06.Bus            ag.............t....g..a..c.....c.......................t..g........t....t...... [185120]
KM111294.1_Ken05/Tk1            ag.............t....g..a..c.....c.......................t..g........t....t...... [185120]
KM262844.1_strain_L60           ag.........................................................g.............t...... [185120]
KM102979.1_26544/OG10_Italy     ag.........................................................g.............t...... [185120]
KM262845.1_strain_NHV           ag.........................................................g.............t...... [185120]
KX354450.1_47/Ss/2008           ag.........................................................g.............t...... [185120]
KP055815.1_strain_BA71          ag.........................................................g.............t...... [185120]
MH025920.1_strain_R35           ag.............t....g..a..c.....c.......................t..g........t....t...... [185120]
MH025919.1_strain_N10           ag.............t....g..a..c.....c.......................t..g........t....t...... [185120]
MH025918.1_strain_R25           ag.............t....g..a..c.....c.......................t..g........t....t...... [185120]
MH025917.1_strain_R7            ag.............t....g..a..c.....c.......................t..g........t....t...... [185120]
MH025916.1_strain_R8            ag.............t....g..a..c.....c.......................t..g........t....t...... [185120]

FR682468.2_Georgia_2007/1       tctacacgatactagggagtttacttcactggtccccgatgaggcagac---aataaaccggaagatgacgaagaaa--- [185200]
ASF_IND_20_CAD543               .................................................---.........................--- [185200]
LR899193.1_Germany_2020/1       .................................................---.........................--- [185200]
MN393477.1_Wuhan_2019           .................................................---.........................--- [185200]
MW396979.1_Timor-Leste/2019/1   .................................................---.........................--- [185200]
MK940252.1_CN/2019/InnerMongolia .................................................---.........................--- [185200]
MK128995.1_China/2018/AnhuiXCGQ .................................................---.........................--- [185200]
MH681419.1_ASFV/POL/2015        .................................................---.........................--- [185200]
LS478113.1_Estonia_2014         .................................................---.........................--- [185200]
KP843857.1_Odintsovo_02/14      .................................................---.........................--- [185200]
FN557520.1_strain_E75           .................................................---.........................--- [185200]
AY261366.1_Warthog              ..................a..............................---.........................--- [185200]
AY261365.1_Warmbaths            ..................a..............................---.........................--- [185200]
AY261364.1_Tengani_62           ..................a...........a..t...............aat........................t--- [185200]
AY261363.1_Pretoriuskop/96/4    ..................a..............t.....a...a.....---.........................--- [185200]
AY261362.1_Mkuzi_1979           .................................................---.........................--- [185200]
AY261361.1_Malawi_Lil-20/1_1983 ..a...............a....................a...a.....---..............c..........--- [185200]
AY261360.1_Kenya_1950           ............g.....a..........................g...---.........................gcg [185200]
AM712240.1_OURT_88/3            .................................................---.........................--- [185200]
AM712239.1_Benin_97/1           .................................................---.........................--- [185200]
KM111295.1_Ken06.Bus            ............g.....a..........................g...---.........................gcg [185200]
```

```
KM111294.1_Ken05/Tk1            ............g.....a..........................g...---.........................gcg [185200]
KM262844.1_strain_L60           .................................................---.........................--- [185200]
KM102979.1_26544/OG10_Italy     .................................................---.........................--- [185200]
KM262845.1_strain_NHV           .................................................---.........................--- [185200]
KX354450.1_47/Ss/2008           .................................................---.........................--- [185200]
KP055815.1_strain_BA71          .................................................---.........................--- [185200]
MH025920.1_strain_R35           ............g.....a..........................g...---.........................gcg [185200]
MH025919.1_strain_N10           ............g.....a.........t................g...---.........................gcg [185200]
MH025918.1_strain_R25           ............g.....a..........................g...---.........................gcg [185200]
MH025917.1_strain_R7            ............g.....a..........................g...---.........................gcg [185200]
MH025916.1_strain_R8            ............g.....a..........................g...---.........................gcg [185200]

FR682468.2_Georgia_2007/1       gcggtgcaaaacctaaaaagaaaaaacatttgtttccaaaattaagctcgcataaatcgaagtaaaaattgaagcgaaaa [185280]
ASF_IND_20_CAD543               ................................................................................ [185280]
LR899193.1_Germany_2020/1       ................................................................................ [185280]
MN393477.1_Wuhan_2019           ................................................................................ [185280]
MW396979.1_Timor-Leste/2019/1   ................................................................................ [185280]
MK940252.1_CN/2019/InnerMongolia ................................................................................ [185280]
MK128995.1_China/2018/AnhuiXCGQ ................................................................................ [185280]
MH681419.1_ASFV/POL/2015        ................................................................................ [185280]
LS478113.1_Estonia_2014         ................................................................................ [185280]
KP843857.1_Odintsovo_02/14      ................................................................................ [185280]
FN557520.1_strain_E75           ................................................................................ [185280]
AY261366.1_Warthog              ................................................................................ [185280]
AY261365.1_Warmbaths            ....c........................................................................... [185280]
AY261364.1_Tengani_62           ................................................................................ [185280]
AY261363.1_Pretoriuskop/96/4    ....c........................................................................... [185280]
AY261362.1_Mkuzi_1979           ................................................................................ [185280]
AY261361.1_Malawi_Lil-20/1_1983 ....c.........................................t...........c...............t..... [185280]
AY261360.1_Kenya_1950           ....c.........................................t................................. [185280]
AM712240.1_OURT_88/3            ...........................g.................................................... [185280]
AM712239.1_Benin_97/1           ................................................................................ [185280]
KM111295.1_Ken06.Bus            ....c...............................................c....................a..-... [185280]
KM111294.1_Ken05/Tk1            ....c.........................................t................................. [185280]
KM262844.1_strain_L60           ................................................................................ [185280]
KM102979.1_26544/OG10_Italy     ................................................................................ [185280]
KM262845.1_strain_NHV           ...........................g.................................................... [185280]
KX354450.1_47/Ss/2008           ................................................................................ [185280]
KP055815.1_strain_BA71          ................................................................................ [185280]
MH025920.1_strain_R35           ....c...............................................c....................a...... [185280]
MH025919.1_strain_N10           ....c...............................................c....................a...... [185280]
MH025918.1_strain_R25           ....c...............................................c....................a...... [185280]
MH025917.1_strain_R7            ....c...............................................c....................a...... [185280]
MH025916.1_strain_R8            ....c...............................................c....................a...... [185280]
```

```
FR682468.2_Georgia_2007/1        aaagtagaaaaaaaatgtttggagcttttgtaagccaccgtttgtggtcagatagtggttgtacgaccacctgcatcaca [185360]
ASF_IND_20_CAD543                ...agtag........................................................................ [185360]
LR899193.1_Germany_2020/1        ................................................................................ [185360]
MN393477.1_Wuhan_2019            ................................................................................ [185360]
MW396979.1_Timor-Leste/2019/1    ................................................................................ [185360]
MK940252.1_CN/2019/InnerMongolia ................................................................................ [185360]
MK128995.1_China/2018/AnhuiXCGQ  ................................................................................ [185360]
MH681419.1_ASFV/POL/2015         ................................................................................ [185360]
LS478113.1_Estonia_2014          ................................................................................ [185360]
KP843857.1_Odintsovo_02/14       ................................................................................ [185360]
FN557520.1_strain_E75            ...agtag...........................................................t............ [185360]
AY261366.1_Warthog               ...agtag...........................................................t............ [185360]
AY261365.1_Warmbaths             ...agtag..................................................g..................... [185360]
AY261364.1_Tengani_62            ................................................................................ [185360]
AY261363.1_Pretoriuskop/96/4     ..........................................................g..................... [185360]
AY261362.1_Mkuzi_1979            ...agtag...........................................................t............ [185360]
AY261361.1_Malawi_Lil-20/1_1983  ....-tag...........................................................t............ [185360]
AY261360.1_Kenya_1950            ...agtag.......................g..t................................t.........g.. [185360]
AM712240.1_OURT_88/3             ...agtag...........................................................t............ [185360]
AM712239.1_Benin_97/1            ...agtag...........................................................t............ [185360]
KM111295.1_Ken06.Bus             ...aatag.......................g..t.....................a..........t.........g.. [185360]
KM111294.1_Ken05/Tk1             ...agtag.......................g..t................................t.........g.. [185360]
KM262844.1_strain_L60            ...agtag...........................................................t............ [185360]
KM102979.1_26544/OG10_Italy      ...agtag...........................................................t............ [185360]
KM262845.1_strain_NHV            ...agtag...........................................................t............ [185360]
KX354450.1_47/Ss/2008            ...agtag...........................................................t............ [185360]
KP055815.1_strain_BA71           ...agtag...........................................................t............ [185360]
MH025920.1_strain_R35            ...aatag.......................g..t.....................a..........t.........g.. [185360]
MH025919.1_strain_N10            ...aatag.......................g..t.....................a..........t.........g.. [185360]
MH025918.1_strain_R25            ...aatag.......................g..t.....................a..........t.........g.. [185360]
MH025917.1_strain_R7             ...aatag.......................g..t.....................a..........t.........g.. [185360]
MH025916.1_strain_R8             ...aatag.......................g..t.....................a..........t.........g.. [185360]

FR682468.2_Georgia_2007/1        aacagcattgctaattatgtagccttcggcgaacaaattggatttccctttaaatcagctcaggtatttattgccggccc [185440]
ASF_IND_20_CAD543                ................................................................................ [185440]
LR899193.1_Germany_2020/1        ................................................................................ [185440]
MN393477.1_Wuhan_2019            ................................................................................ [185440]
MW396979.1_Timor-Leste/2019/1    ................................................................................ [185440]
MK940252.1_CN/2019/InnerMongolia ................................................................................ [185440]
MK128995.1_China/2018/AnhuiXCGQ  ................................................................................ [185440]
MH681419.1_ASFV/POL/2015         ................................................................................ [185440]
LS478113.1_Estonia_2014          ................................................................................ [185440]
KP843857.1_Odintsovo_02/14       ................................................................................ [185440]
```

```
FN557520.1_strain_E75          ..............................................................a................. [185440]
AY261366.1_Warthog             ..............................................................a................. [185440]
AY261365.1_Warmbaths           ..............................................................a................. [185440]
AY261364.1_Tengani_62          ................................................................................ [185440]
AY261363.1_Pretoriuskop/96/4   ..............................................................a..............t.. [185440]
AY261362.1_Mkuzi_1979          ........................................................t.....a................. [185440]
AY261361.1_Malawi_Lil-20/1_1983 ..............c...........t..t........c.......................a................. [185440]
AY261360.1_Kenya_1950          ...........c..c...........t.............a...............g..c..a................. [185440]
AM712240.1_OURT_88/3           ..............................................................a................. [185440]
AM712239.1_Benin_97/1          ..............................................................a................. [185440]
KM111295.1_Ken06.Bus           ...........c..c...........t.............a...............g..c..a................. [185440]
KM111294.1_Ken05/Tk1           ...........c..c...........t.............a..................c..a................. [185440]
KM262844.1_strain_L60          ..............................................................a................. [185440]
KM102979.1_26544/OG10_Italy    ..............................................................a................. [185440]
KM262845.1_strain_NHV          ..............................................................a................. [185440]
KX354450.1_47/Ss/2008          ..............................................................a................. [185440]
KP055815.1_strain_BA71         ..............................................................a................. [185440]
MH025920.1_strain_R35          ...........c..c...........t.............a...............g..c..a................. [185440]
MH025919.1_strain_N10          ...........c..c...........t.............a...............g..c..a................. [185440]
MH025918.1_strain_R25          ...........c..c...........t.............a...............g..c..a................. [185440]
MH025917.1_strain_R7           ...........c..c...........t.............a...............g..c..a................. [185440]
MH025916.1_strain_R8           ...........c..c...........t.............a...............g..c..a................. [185440]

FR682468.2_Georgia_2007/1      tagaaaggctgtgataaatattcaggaagatgataaagttgagcttttaaagatgattgttaagcacaatctttgggttg [185520]
ASF_IND_20_CAD543              ................................................................................ [185520]
LR899193.1_Germany_2020/1      ................................................................................ [185520]
MN393477.1_Wuhan_2019          ................................................................................ [185520]
MW396979.1_Timor-Leste/2019/1  ................................................................................ [185520]
MK940252.1_CN/2019/InnerMongolia ................................................................................ [185520]
MK128995.1_China/2018/AnhuiXCGQ ................................................................................ [185520]
MH681419.1_ASFV/POL/2015       ................................................................................ [185520]
LS478113.1_Estonia_2014        ................................................................................ [185520]
KP843857.1_Odintsovo_02/14     ................................................................................ [185520]
FN557520.1_strain_E75          ................................................................................ [185520]
AY261366.1_Warthog             ................................................................................ [185520]
AY261365.1_Warmbaths           ................................................................................ [185520]
AY261364.1_Tengani_62          ................................................................................ [185520]
AY261363.1_Pretoriuskop/96/4   ................................................................................ [185520]
AY261362.1_Mkuzi_1979          .....................................................c.......................... [185520]
AY261361.1_Malawi_Lil-20/1_1983 .........................a.................................a.................... [185520]
AY261360.1_Kenya_1950          c.............................a.......c........c...........c.................... [185520]
AM712240.1_OURT_88/3           ................................................................................ [185520]
AM712239.1_Benin_97/1          ................................................................................ [185520]
KM111295.1_Ken06.Bus           ..............................a.......c........c......a....c.................... [185520]
```

```
KM111294.1_Ken05/Tk1             c.............................a.......c........c...a.......c.................... [185520]
KM262844.1_strain_L60            ................................................................................ [185520]
KM102979.1_26544/OG10_Italy      ................................................................................ [185520]
KM262845.1_strain_NHV            ................................................................................ [185520]
KX354450.1_47/Ss/2008            ................................................................................ [185520]
KP055815.1_strain_BA71           ................................................................................ [185520]
MH025920.1_strain_R35            ..............................a.......c........c......a....c.................... [185520]
MH025919.1_strain_N10            ..............................a.......c........c......a....c.................... [185520]
MH025918.1_strain_R25            ..............................a.......c........c......a....c.................... [185520]
MH025917.1_strain_R7             ..............................a.......c........c......a....c.................... [185520]
MH025916.1_strain_R8             ..............................a.......c........c......a....c.................... [185520]

FR682468.2_Georgia_2007/1        ttgctcatggaacctacttagatgtgccctggtcccgtaagagtgcgtttgttacacattttatacaacaagaactactt [185600]
ASF_IND_20_CAD543                ................................................................................ [185600]
LR899193.1_Germany_2020/1        ................................................................................ [185600]
MN393477.1_Wuhan_2019            ................................................................................ [185600]
MW396979.1_Timor-Leste/2019/1    ................................................................................ [185600]
MK940252.1_CN/2019/InnerMongolia ................................................................................ [185600]
MK128995.1_China/2018/AnhuiXCGQ  ................................................................................ [185600]
MH681419.1_ASFV/POL/2015         ................................................................................ [185600]
LS478113.1_Estonia_2014          ................................................................................ [185600]
KP843857.1_Odintsovo_02/14       ................................................................................ [185600]
FN557520.1_strain_E75            .......................................g........................................ [185600]
AY261366.1_Warthog               .......................................g........................................ [185600]
AY261365.1_Warmbaths             .......................................g..................................t..... [185600]
AY261364.1_Tengani_62            ....a..................................g........................................ [185600]
AY261363.1_Pretoriuskop/96/4     .g.....................................g........................................ [185600]
AY261362.1_Mkuzi_1979            .......................................g......a................................. [185600]
AY261361.1_Malawi_Lil-20/1_1983  .......................................g......a................................. [185600]
AY261360.1_Kenya_1950            ...................................a...g...............c...........t............ [185600]
AM712240.1_OURT_88/3             .......................................g........................................ [185600]
AM712239.1_Benin_97/1            .......................................g........................................ [185600]
KM111295.1_Ken06.Bus             ...................................a...g...............c...........t............ [185600]
KM111294.1_Ken05/Tk1             .......................................g......a................................. [185600]
KM262844.1_strain_L60            .......................................g........................................ [185600]
KM102979.1_26544/OG10_Italy      .......................................g........................................ [185600]
KM262845.1_strain_NHV            .......................................g........................................ [185600]
KX354450.1_47/Ss/2008            .......................................g........................................ [185600]
KP055815.1_strain_BA71           .......................................g........................................ [185600]
MH025920.1_strain_R35            ...................................a...g...............c...........t............ [185600]
MH025919.1_strain_N10            ...................................a...g...............c...........t............ [185600]
MH025918.1_strain_R25            ...................................a...g...............c...........t............ [185600]
MH025917.1_strain_R7             ...................................a...g...............c...........t............ [185600]
MH025916.1_strain_R8             ...................................a...g...............c...........t............ [185600]
```

```
FR682468.2_Georgia_2007/1        atatgcaaggaagtcggtattaaagggttagttttacacctaggcgctgtggagcctgaacttattatggaaggactaaa [185680]
ASF_IND_20_CAD543                ................................................................................ [185680]
LR899193.1_Germany_2020/1        ................................................................................ [185680]
MN393477.1_Wuhan_2019            ................................................................................ [185680]
MW396979.1_Timor-Leste/2019/1    ................................................................................ [185680]
MK940252.1_CN/2019/InnerMongolia ................................................................................ [185680]
MK128995.1_China/2018/AnhuiXCGQ  ................................................................................ [185680]
MH681419.1_ASFV/POL/2015         ................................................................................ [185680]
LS478113.1_Estonia_2014          ................................................................................ [185680]
KP843857.1_Odintsovo_02/14       .............................................................................-.. [185680]
FN557520.1_strain_E75            ..........................a...................tc.................cg............. [185680]
AY261366.1_Warthog               ..........................a....................c.................cg............. [185680]
AY261365.1_Warmbaths             ..........................a....................c.................cg............. [185680]
AY261364.1_Tengani_62            ..........................a..............c.....c..................g.......g..... [185680]
AY261363.1_Pretoriuskop/96/4     ..........................a....................c.................cg............. [185680]
AY261362.1_Mkuzi_1979            ..........................a....................c.................cg............. [185680]
AY261361.1_Malawi_Lil-20/1_1983  ..........................a..g........t........c..................g............. [185680]
AY261360.1_Kenya_1950            ..........................a....................c..................g............. [185680]
AM712240.1_OURT_88/3             ..........................a....................c.................cg............. [185680]
AM712239.1_Benin_97/1            ..........................a...................tc.................cg............. [185680]
KM111295.1_Ken06.Bus             ...........g..............a....................c..................g............. [185680]
KM111294.1_Ken05/Tk1             ..........................a...........t........c..................g............. [185680]
KM262844.1_strain_L60            ..........................a....................c.................cg............. [185680]
KM102979.1_26544/OG10_Italy      ..........................a...................tc.................cg............. [185680]
KM262845.1_strain_NHV            ..........................a....................c.................cg............. [185680]
KX354450.1_47/Ss/2008            ..........................a...................tc.................cg............. [185680]
KP055815.1_strain_BA71           ..........................a....................c.................cg............. [185680]
MH025920.1_strain_R35            ...........g..............a....................c..................g............. [185680]
MH025919.1_strain_N10            ...........g..............a....................c..................g............. [185680]
MH025918.1_strain_R25            ...........g..............a....................c..................g............. [185680]
MH025917.1_strain_R7             ...........g..............a....................c..................g............. [185680]
MH025916.1_strain_R8             ...........g..............a....................c..................g............. [185680]

FR682468.2_Georgia_2007/1        aaaaattaagccggttgagggggttgtcatttacctggaaaccccgcataacaaacatcatacatataaatacagtacaa [185760]
ASF_IND_20_CAD543                ................................................................................ [185760]
LR899193.1_Germany_2020/1        ................................................................................ [185760]
MN393477.1_Wuhan_2019            ................................................................................ [185760]
MW396979.1_Timor-Leste/2019/1    ................................................................................ [185760]
MK940252.1_CN/2019/InnerMongolia ................................................................................ [185760]
MK128995.1_China/2018/AnhuiXCGQ  ................................................................................ [185760]
MH681419.1_ASFV/POL/2015         ................................................................................ [185760]
LS478113.1_Estonia_2014          ................................................................................ [185760]
KP843857.1_Odintsovo_02/14       ................................................................................ [185760]
```

```
FN557520.1_strain_E75            ................................................................................ [185760]
AY261366.1_Warthog               ....................................c........................................... [185760]
AY261365.1_Warmbaths             ................................................................................ [185760]
AY261364.1_Tengani_62            ..........................................t..a................t................. [185760]
AY261363.1_Pretoriuskop/96/4     ................................................................................ [185760]
AY261362.1_Mkuzi_1979            ................................................................................ [185760]
AY261361.1_Malawi_Lil-20/1_1983  .........................a...............................c.....c................ [185760]
AY261360.1_Kenya_1950            ....................a.....................g..................g.................. [185760]
AM712240.1_OURT_88/3             ................................................................................ [185760]
AM712239.1_Benin_97/1            ................................................................................ [185760]
KM111295.1_Ken06.Bus             .................................t........g.....c............................... [185760]
KM111294.1_Ken05/Tk1             ..........................................g..................g.................. [185760]
KM262844.1_strain_L60            ................................................................................ [185760]
KM102979.1_26544/OG10_Italy      ................................................................................ [185760]
KM262845.1_strain_NHV            ................................................................................ [185760]
KX354450.1_47/Ss/2008            ................................................................................ [185760]
KP055815.1_strain_BA71           ................................................................................ [185760]
MH025920.1_strain_R35            .................................t........g.....c............................... [185760]
MH025919.1_strain_N10            .................................t........g.....c............................... [185760]
MH025918.1_strain_R25            .................................t........g.....c............................... [185760]
MH025917.1_strain_R7             .................................t........g.....c............................... [185760]
MH025916.1_strain_R8             .................................t........g.....c............................... [185760]

FR682468.2_Georgia_2007/1        ttgagcagatcaaagaattgtttttacggatacgaaataccaggttgaaacagattggtttatgcattgatacggctcac [185840]
ASF_IND_20_CAD543                ................................................................................ [185840]
LR899193.1_Germany_2020/1        ................................................................................ [185840]
MN393477.1_Wuhan_2019            ................................................................................ [185840]
MW396979.1_Timor-Leste/2019/1    ................................................................................ [185840]
MK940252.1_CN/2019/InnerMongolia ................................................................................ [185840]
MK128995.1_China/2018/AnhuiXCGQ  ................................................................................ [185840]
MH681419.1_ASFV/POL/2015         ................................................................................ [185840]
LS478113.1_Estonia_2014          ................................................................................ [185840]
KP843857.1_Odintsovo_02/14       ................................................................................ [185840]
FN557520.1_strain_E75            .g.........................................a...........c........................ [185840]
AY261366.1_Warthog               .g.................a...................................c........................ [185840]
AY261365.1_Warmbaths             .g.................a...................................c........................ [185840]
AY261364.1_Tengani_62            ...................a...................................c........................ [185840]
AY261363.1_Pretoriuskop/96/4     .g.....................................................c........................ [185840]
AY261362.1_Mkuzi_1979            .g.........................................a...........c........................ [185840]
AY261361.1_Malawi_Lil-20/1_1983  ............................t............g...................................... [185840]
AY261360.1_Kenya_1950            ...................a........t..........g....c....g..t..c....................a... [185840]
AM712240.1_OURT_88/3             .g.........................................a...........c........................ [185840]
AM712239.1_Benin_97/1            .g.........................................a...........c........................ [185840]
KM111295.1_Ken06.Bus             ...................a........t...........t...c....g..t.......................a..t [185840]
```

```
KM111294.1_Ken05/Tk1           ...................a........t..........g....c....g..t..c....................a... [185840]
KM262844.1_strain_L60          .g.........................................a...........c........................ [185840]
KM102979.1_26544/OG10_Italy    .g.........................................a...........c........................ [185840]
KM262845.1_strain_NHV          .g.........................................a...........c........................ [185840]
KX354450.1_47/Ss/2008          .g.........................................a...........c........................ [185840]
KP055815.1_strain_BA71         .g.........................................a...........c........................ [185840]
MH025920.1_strain_R35          ...................a........t..........t....c....g..t.......................a..t [185840]
MH025919.1_strain_N10          ...................a........t..........t....c....g..t.......................a..t [185840]
MH025918.1_strain_R25          ...................a........t..........t....c....g..t.......................a..t [185840]
MH025917.1_strain_R7           ...................a........t..........t....c....g..t.......................a..t [185840]
MH025916.1_strain_R8           ...................a........t..........t....c....g..t.......................a..t [185840]

FR682468.2_Georgia_2007/1      atctggtcttccggtgtcaacatctccagctataatgacgcggggcaatggctgcgctcgctggaaaacattcattccgt [185920]
ASF_IND_20_CAD543              ................................................................................ [185920]
LR899193.1_Germany_2020/1      ................................................................................ [185920]
MN393477.1_Wuhan_2019          ................................................................................ [185920]
MW396979.1_Timor-Leste/2019/1  ................................................................................ [185920]
MK940252.1_CN/2019/InnerMongolia ................................................................................ [185920]
MK128995.1_China/2018/AnhuiXCGQ ................................................................................ [185920]
MH681419.1_ASFV/POL/2015       ................................................................................ [185920]
LS478113.1_Estonia_2014        ................................................................................ [185920]
KP843857.1_Odintsovo_02/14     ................................................................................ [185920]
FN557520.1_strain_E75          ..........................................................................c..... [185920]
AY261366.1_Warthog             ..........................................................................c..... [185920]
AY261365.1_Warmbaths           ..........................................................................c..... [185920]
AY261364.1_Tengani_62          ..........................................................................c..... [185920]
AY261363.1_Pretoriuskop/96/4   ..........................................................................c..... [185920]
AY261362.1_Mkuzi_1979          ..........................................................................c..... [185920]
AY261361.1_Malawi_Lil-20/1_1983 ..t........t.....t.................cag....a...................a...g.......c..... [185920]
AY261360.1_Kenya_1950          ..t........t...............g..............................................c..... [185920]
AM712240.1_OURT_88/3           ..........................................................................c..... [185920]
AM712239.1_Benin_97/1          ..........................................................................c..... [185920]
KM111295.1_Ken06.Bus           ..t........t.....t.................g.....t................................c..... [185920]
KM111294.1_Ken05/Tk1           ..t........t.............................t................................c..... [185920]
KM262844.1_strain_L60          ..........................................................................c..... [185920]
KM102979.1_26544/OG10_Italy    ..........................................................................c..... [185920]
KM262845.1_strain_NHV          ..........................................................................c..... [185920]
KX354450.1_47/Ss/2008          ..........................................................................c..... [185920]
KP055815.1_strain_BA71         ..........................................................................c..... [185920]
MH025920.1_strain_R35          ..t........t.....t.................g.....t................................c..... [185920]
MH025919.1_strain_N10          ..t........t.....t.................g.....t................................c..... [185920]
MH025918.1_strain_R25          ..t........t.....t.................g.....t................................c..... [185920]
MH025917.1_strain_R7           ..t........t.....t.................g.....t................................c..... [185920]
MH025916.1_strain_R8           ..t........t.....t.................g.....t................................c..... [185920]
```

```
FR682468.2_Georgia_2007/1        gatcccaccaagccacattatgttccacctaaatgatgccgccacagaatgcggaagcggtatagaccgacatgcaagtc [186000]
ASF_IND_20_CAD543                ................................................................................ [186000]
LR899193.1_Germany_2020/1        ................................................................................ [186000]
MN393477.1_Wuhan_2019            ................................................................................ [186000]
MW396979.1_Timor-Leste/2019/1    ................................................................................ [186000]
MK940252.1_CN/2019/InnerMongolia ................................................................................ [186000]
MK128995.1_China/2018/AnhuiXCGQ  ................................................................................ [186000]
MH681419.1_ASFV/POL/2015         ................................................................................ [186000]
LS478113.1_Estonia_2014          ................................................................................ [186000]
KP843857.1_Odintsovo_02/14       ................................................................................ [186000]
FN557520.1_strain_E75            ................................................................................ [186000]
AY261366.1_Warthog               ................................................................................ [186000]
AY261365.1_Warmbaths             ................................................................................ [186000]
AY261364.1_Tengani_62            .....................................................................t.......... [186000]
AY261363.1_Pretoriuskop/96/4     ................................................................................ [186000]
AY261362.1_Mkuzi_1979            ................................................................................ [186000]
AY261361.1_Malawi_Lil-20/1_1983  .............................................ga.......g.....cg.................. [186000]
AY261360.1_Kenya_1950            .............................................g...........t...........g.......... [186000]
AM712240.1_OURT_88/3             ................................................................................ [186000]
AM712239.1_Benin_97/1            ................................................................................ [186000]
KM111295.1_Ken06.Bus             .............................................g...........t...........g.......... [186000]
KM111294.1_Ken05/Tk1             .............................................g...........t...........g.......... [186000]
KM262844.1_strain_L60            ................................................................................ [186000]
KM102979.1_26544/OG10_Italy      ................................................................................ [186000]
KM262845.1_strain_NHV            ................................................................................ [186000]
KX354450.1_47/Ss/2008            ................................................................................ [186000]
KP055815.1_strain_BA71           ................................................................................ [186000]
MH025920.1_strain_R35            .............................................g...........t...........g.......... [186000]
MH025919.1_strain_N10            .............................................g...........t...........g.......... [186000]
MH025918.1_strain_R25            .............................................g...........t...........g.......... [186000]
MH025917.1_strain_R7             .............................................g...........t...........g.......... [186000]
MH025916.1_strain_R8             .............................................g...........t...........g.......... [186000]

FR682468.2_Georgia_2007/1        tttttgaaggaatgatttggaaatcatatagccataaaataaagcaaagcggtttatattgttttgttgaatacgttacg [186080]
ASF_IND_20_CAD543                ............................................m................................... [186080]
LR899193.1_Germany_2020/1        ................................................................................ [186080]
MN393477.1_Wuhan_2019            ................................................................................ [186080]
MW396979.1_Timor-Leste/2019/1    ................................................................................ [186080]
MK940252.1_CN/2019/InnerMongolia ................................................................................ [186080]
MK128995.1_China/2018/AnhuiXCGQ  ................................................................................ [186080]
MH681419.1_ASFV/POL/2015         ................................................................................ [186080]
LS478113.1_Estonia_2014          ................................................................................ [186080]
KP843857.1_Odintsovo_02/14       ................................................................................ [186080]
```

```
FN557520.1_strain_E75          ............................................a.......g.....................a..... [186080]
AY261366.1_Warthog             ............................................a.......g.....................a..... [186080]
AY261365.1_Warmbaths           ............................................a.......g.....................a..... [186080]
AY261364.1_Tengani_62          ..........................................................................a..... [186080]
AY261363.1_Pretoriuskop/96/4   ..........................................................................a..... [186080]
AY261362.1_Mkuzi_1979          ............................................a.......g.....................a..... [186080]
AY261361.1_Malawi_Lil-20/1_1983 ..............................................t......c....................a..... [186080]
AY261360.1_Kenya_1950          ................a.............................t......c.g..............g...a..... [186080]
AM712240.1_OURT_88/3           ............................................a.......g.....................a..... [186080]
AM712239.1_Benin_97/1          ............................................a.......g.....................a..... [186080]
KM111295.1_Ken06.Bus           ................a.............................t......c.g..............g...a..... [186080]
KM111294.1_Ken05/Tk1           ................a.............................t......c.g..............g...a..... [186080]
KM262844.1_strain_L60          ............................................a.......g.....................a..... [186080]
KM102979.1_26544/OG10_Italy    ............................................a.......g.....................a..... [186080]
KM262845.1_strain_NHV          ............................................a.......g.....................a..... [186080]
KX354450.1_47/Ss/2008          ............................................a.......g.....................a..... [186080]
KP055815.1_strain_BA71         ............................................a.......g.....................a..... [186080]
MH025920.1_strain_R35          ................a.............................t......c.g..............g...a..... [186080]
MH025919.1_strain_N10          ................a.............................t......c.g..............g...a..... [186080]
MH025918.1_strain_R25          ................a.............................t......c.g..............g...a..... [186080]
MH025917.1_strain_R7           ................a.............................t......c.g..............g...a..... [186080]
MH025916.1_strain_R8           ................a.............................t......c.g..............g...a..... [186080]

FR682468.2_Georgia_2007/1      cgacaccagtgtccggctatattggagagaaacctcgggtcttccatgcaattacaaaccgctttaaccgcagaatttac [186160]
ASF_IND_20_CAD543              ................................................................................ [186160]
LR899193.1_Germany_2020/1      ................................................................................ [186160]
MN393477.1_Wuhan_2019          ................................................................................ [186160]
MW396979.1_Timor-Leste/2019/1  ................................................................................ [186160]
MK940252.1_CN/2019/InnerMongolia ................................................................................ [186160]
MK128995.1_China/2018/AnhuiXCGQ ................................................................................ [186160]
MH681419.1_ASFV/POL/2015       ................................................................................ [186160]
LS478113.1_Estonia_2014        ................................................................................ [186160]
KP843857.1_Odintsovo_02/14     ................................................................................ [186160]
FN557520.1_strain_E75          ..t............................................................................. [186160]
AY261366.1_Warthog             ..t............................................................................. [186160]
AY261365.1_Warmbaths           ..t............................................................................. [186160]
AY261364.1_Tengani_62          ..t............................................................................. [186160]
AY261363.1_Pretoriuskop/96/4   ................................................................................ [186160]
AY261362.1_Mkuzi_1979          ..t............................................................................. [186160]
AY261361.1_Malawi_Lil-20/1_1983 ...............................................................................a [186160]
AY261360.1_Kenya_1950          .....t.................a...........................c..........g.....ta........ca [186160]
AM712240.1_OURT_88/3           ..t............................................................................. [186160]
AM712239.1_Benin_97/1          ..t............................................................................. [186160]
KM111295.1_Ken06.Bus           .....t.................a........t...........t......c.g........g.....ta........ca [186160]
```

```
KM111294.1_Ken05/Tk1            .....t.................a...........................c..........g.....ta........ca [186160]
KM262844.1_strain_L60           ..t............................................................................. [186160]
KM102979.1_26544/OG10_Italy     ..t............................................................................. [186160]
KM262845.1_strain_NHV           ..t............................................................................. [186160]
KX354450.1_47/Ss/2008           ..t............................................................................. [186160]
KP055815.1_strain_BA71          ..t............................................................................. [186160]
MH025920.1_strain_R35           .....t.................a........t...........t......c.g........g.....ta........ca [186160]
MH025919.1_strain_N10           .....t.................a........t...........t......c.g........g.....ta........ca [186160]
MH025918.1_strain_R25           .....t.................a........t...........t......c.g........g.....ta........ca [186160]
MH025917.1_strain_R7            .....t.................a........t...........t......c.g........g.....ta........ca [186160]
MH025916.1_strain_R8            .....t.................a........t...........t......c.g........g.....ta........ca [186160]

FR682468.2_Georgia_2007/1       tacattaaaatcgttattaaaataaggatgagttttagcgaatgtcccttagttattagtgcatgcaaaaaatttctaca [186240]
ASF_IND_20_CAD543               ................................................................................ [186240]
LR899193.1_Germany_2020/1       ................................................................................ [186240]
MN393477.1_Wuhan_2019           ................................................................................ [186240]
MW396979.1_Timor-Leste/2019/1   ................................................................................ [186240]
MK940252.1_CN/2019/InnerMongolia ................................................................................ [186240]
MK128995.1_China/2018/AnhuiXCGQ ................................................................................ [186240]
MH681419.1_ASFV/POL/2015        ................................................................................ [186240]
LS478113.1_Estonia_2014         ................................................................................ [186240]
KP843857.1_Odintsovo_02/14      ................................................................................ [186240]
FN557520.1_strain_E75           ................................................................................ [186240]
AY261366.1_Warthog              ................................................................................ [186240]
AY261365.1_Warmbaths            ................................................................................ [186240]
AY261364.1_Tengani_62           ................................................................................ [186240]
AY261363.1_Pretoriuskop/96/4    ................................................................................ [186240]
AY261362.1_Mkuzi_1979           ................................................................................ [186240]
AY261361.1_Malawi_Lil-20/1_1983 ...............c..........t....................t.....c.......................... [186240]
AY261360.1_Kenya_1950           .............c............t....a......t..............c.......................... [186240]
AM712240.1_OURT_88/3            ................................................................................ [186240]
AM712239.1_Benin_97/1           ................................................................................ [186240]
KM111295.1_Ken06.Bus            .............c.......g....t....a........-..a.........c.......................... [186240]
KM111294.1_Ken05/Tk1            .............c............t....a.....................c.......................... [186240]
KM262844.1_strain_L60           ................................................................................ [186240]
KM102979.1_26544/OG10_Italy     ................................................................................ [186240]
KM262845.1_strain_NHV           ................................................................................ [186240]
KX354450.1_47/Ss/2008           ................................................................................ [186240]
KP055815.1_strain_BA71          ................................................................................ [186240]
MH025920.1_strain_R35           .............c.......g....t....a...........a.........c.......................... [186240]
MH025919.1_strain_N10           .............c.......g....t....a...........a.........c.......................... [186240]
MH025918.1_strain_R25           .............c.......g....t....a...........a.........c.......................... [186240]
MH025917.1_strain_R7            .............c.......g....t....a...........a.........c.......................... [186240]
MH025916.1_strain_R8            .............c.......g....t....a...........a.........c.......................... [186240]
```

```
FR682468.2_Georgia_2007/1        aaagcgtattacaatagagaatgaagcacttataaatgccttaataaccgctttagcgcagaccagcacgttgaatgatc [186320]
ASF_IND_20_CAD543                ................................................................................ [186320]
LR899193.1_Germany_2020/1        ................................................................................ [186320]
MN393477.1_Wuhan_2019            ................................................................................ [186320]
MW396979.1_Timor-Leste/2019/1    ................................................................................ [186320]
MK940252.1_CN/2019/InnerMongolia ................................................................................ [186320]
MK128995.1_China/2018/AnhuiXCGQ  ................................................................................ [186320]
MH681419.1_ASFV/POL/2015         ................................................................................ [186320]
LS478113.1_Estonia_2014          ................................................................................ [186320]
KP843857.1_Odintsovo_02/14       ................................................................................ [186320]
FN557520.1_strain_E75            ......c.....................................................a................... [186320]
AY261366.1_Warthog               ......c......................................................................... [186320]
AY261365.1_Warmbaths             ............t...............................................a....a.............. [186320]
AY261364.1_Tengani_62            ...........................c....................t............................... [186320]
AY261363.1_Pretoriuskop/96/4     ......c..........................................................c.............. [186320]
AY261362.1_Mkuzi_1979            ......c.....................................................a................... [186320]
AY261361.1_Malawi_Lil-20/1_1983  ......c................................t........g................a.............. [186320]
AY261360.1_Kenya_1950            g..a.......................g...........tc...................a....c............g. [186320]
AM712240.1_OURT_88/3             ......c.....................................................a................... [186320]
AM712239.1_Benin_97/1            ......c.....................................................a................... [186320]
KM111295.1_Ken06.Bus             g..a.......................g...........tc...................a....c............g. [186320]
KM111294.1_Ken05/Tk1             g..a...................................tc...................a....c............g. [186320]
KM262844.1_strain_L60            ......c.....................................................a................... [186320]
KM102979.1_26544/OG10_Italy      ......c.....................................................a................... [186320]
KM262845.1_strain_NHV            ......c.....................................................a................... [186320]
KX354450.1_47/Ss/2008            ......c.....................................................a................... [186320]
KP055815.1_strain_BA71           ......c.....................................................a................... [186320]
MH025920.1_strain_R35            g..a.......................g...........tc...................a....c............g. [186320]
MH025919.1_strain_N10            g..a.......................g...........tc...................a....c............g. [186320]
MH025918.1_strain_R25            g..a.......................g...........tc...................a....c............g. [186320]
MH025917.1_strain_R7             g..a.......................g...........tc...................a....c............g. [186320]
MH025916.1_strain_R8             g..a.......................g...........tc...................a....c............g. [186320]

FR682468.2_Georgia_2007/1        tttgtttattacctattcaaacctatttgcttagttataaaaatgcttttgagtggatacacttcgtatgtattgcaatc [186400]
ASF_IND_20_CAD543                ................................................................................ [186400]
LR899193.1_Germany_2020/1        ................................................................................ [186400]
MN393477.1_Wuhan_2019            ................................................................................ [186400]
MW396979.1_Timor-Leste/2019/1    ................................................................................ [186400]
MK940252.1_CN/2019/InnerMongolia ................................................................................ [186400]
MK128995.1_China/2018/AnhuiXCGQ  ................................................................................ [186400]
MH681419.1_ASFV/POL/2015         ................................................................................ [186400]
LS478113.1_Estonia_2014          ................................................................................ [186400]
KP843857.1_Odintsovo_02/14       ................................................................................ [186400]
```

```
FN557520.1_strain_E75          ..................................c............................................. [186400]
AY261366.1_Warthog             ..................................c............................................. [186400]
AY261365.1_Warmbaths           ..................................c............................................. [186400]
AY261364.1_Tengani_62          ..................................c............................................. [186400]
AY261363.1_Pretoriuskop/96/4   ..................................c............................................. [186400]
AY261362.1_Mkuzi_1979          ..................................c............................................. [186400]
AY261361.1_Malawi_Lil-20/1_1983 ...................g..............c...........c................................. [186400]
AY261360.1_Kenya_1950          ..........g.......................c............................................t [186400]
AM712240.1_OURT_88/3           ..................................c............................................. [186400]
AM712239.1_Benin_97/1          ..................................c............................................. [186400]
KM111295.1_Ken06.Bus           ..........g....................................................................t [186400]
KM111294.1_Ken05/Tk1           ..........g.......................c............................................t [186400]
KM262844.1_strain_L60          ..................................c............................................. [186400]
KM102979.1_26544/OG10_Italy    ..................................c............................................. [186400]
KM262845.1_strain_NHV          ..................................c............................................. [186400]
KX354450.1_47/Ss/2008          ..................................c............................................. [186400]
KP055815.1_strain_BA71         ..................................c............................................. [186400]
MH025920.1_strain_R35          ..........g....................................................................t [186400]
MH025919.1_strain_N10          ..........g....................................................................t [186400]
MH025918.1_strain_R25          ..........g....................................................................t [186400]
MH025917.1_strain_R7           ..........g....................................................................t [186400]
MH025916.1_strain_R8           ..........g....................................................................t [186400]

FR682468.2_Georgia_2007/1      accactattttggataataagtataactggaaggactgtacggtagatattaattatatttttctccatgtaacctatat [186480]
ASF_IND_20_CAD543              ..........................................................................m..... [186480]
LR899193.1_Germany_2020/1      ................................................................................ [186480]
MN393477.1_Wuhan_2019          ................................................................................ [186480]
MW396979.1_Timor-Leste/2019/1  ................................................................................ [186480]
MK940252.1_CN/2019/InnerMongolia ................................................................................ [186480]
MK128995.1_China/2018/AnhuiXCGQ ................................................................................ [186480]
MH681419.1_ASFV/POL/2015       ................................................................................ [186480]
LS478113.1_Estonia_2014        ................................................................................ [186480]
KP843857.1_Odintsovo_02/14     ................................................................................ [186480]
FN557520.1_strain_E75          ...........a..c.....................................................c.....a..... [186480]
AY261366.1_Warthog             ................g...................................................c..c........ [186480]
AY261365.1_Warmbaths           ..t........a..c.....................................................c..c........ [186480]
AY261364.1_Tengani_62          ...........a..c.....................................................c........... [186480]
AY261363.1_Pretoriuskop/96/4   ................g...................................................c..c........ [186480]
AY261362.1_Mkuzi_1979          ...........a..c.....................................................c.....a..... [186480]
AY261361.1_Malawi_Lil-20/1_1983 .....c.....a..c.g........................c..........................c........... [186480]
AY261360.1_Kenya_1950          ...........a..c...........t.....a........c.......................t..c.....a..... [186480]
AM712240.1_OURT_88/3           ...........a..c.....................................................c.....a..... [186480]
AM712239.1_Benin_97/1          ...........a..c.....................................................c.....a..... [186480]
KM111295.1_Ken06.Bus           ...........a..c...........t.....a........c.......................t..c.....a..... [186480]
```

```
KM111294.1_Ken05/Tk1              ...........a..c...........t.....a........c.......................t..c.....a..... [186480]
KM262844.1_strain_L60             ...........a..c.....................................................c.....a..... [186480]
KM102979.1_26544/OG10_Italy       ...........a..c.....................................................c.....a..... [186480]
KM262845.1_strain_NHV             ...........a..c.....................................................c.....a..... [186480]
KX354450.1_47/Ss/2008             ...........a..c.....................................................c.....a..... [186480]
KP055815.1_strain_BA71            ...........a..c.....................................................c.....a..... [186480]
MH025920.1_strain_R35             ...........a..c...........t.....a........c.......................t..c.....a..... [186480]
MH025919.1_strain_N10             ...........a..c...........t.....a........c.......................t..c.....a..... [186480]
MH025918.1_strain_R25             ...........a..c...........t.....a........c.......................t..c.....a..... [186480]
MH025917.1_strain_R7              ...........a..c...........t.....a........c.......................t..c.....a..... [186480]
MH025916.1_strain_R8              ...........a..c...........t.....a........c.......................t..c.....a..... [186480]

FR682468.2_Georgia_2007/1         ttacaatattaaaaccaaggaatacctagactactgttcttaa--actttattttttctatatttacgccaaagagaata [186560]
ASF_IND_20_CAD543                 ...........................................--...........................--g..... [186560]
LR899193.1_Germany_2020/1         ...........................................--................................... [186560]
MN393477.1_Wuhan_2019             ...........................................--................................... [186560]
MW396979.1_Timor-Leste/2019/1     ...........................................--................................... [186560]
MK940252.1_CN/2019/InnerMongolia  ...........................................--................................... [186560]
MK128995.1_China/2018/AnhuiXCGQ   ...........................................--................................... [186560]
MH681419.1_ASFV/POL/2015          ...........................................--................................... [186560]
LS478113.1_Estonia_2014           ...........................................--................................... [186560]
KP843857.1_Odintsovo_02/14        ...........................................--................................... [186560]
FN557520.1_strain_E75             ...........................................--...........................--g..... [186560]
AY261366.1_Warthog                ...........................................--........................t..--g..... [186560]
AY261365.1_Warmbaths              ...........................................--........................t..--g..... [186560]
AY261364.1_Tengani_62             ...........................................--..........................gg..a.... [186560]
AY261363.1_Pretoriuskop/96/4      .......................................a...--........................t..--g..... [186560]
AY261362.1_Mkuzi_1979             ...........................................--...........................--g..... [186560]
AY261361.1_Malawi_Lil-20/1_1983   .....cc.......................t............cct.....................at...--g..... [186560]
AY261360.1_Kenya_1950             .....cc........t...........................act.....................at...--gac..g [186560]
AM712240.1_OURT_88/3              ...........................................--...........................--g..... [186560]
AM712239.1_Benin_97/1             ...........................................--...........................--g..... [186560]
KM111295.1_Ken06.Bus              .....cc........t...........................act.....................at...--ga...g [186560]
KM111294.1_Ken05/Tk1              .....cc........t...........................act.....................at...--gac..g [186560]
KM262844.1_strain_L60             ...........................................--...........................--g..... [186560]
KM102979.1_26544/OG10_Italy       ...........................................--...........................--g..... [186560]
KM262845.1_strain_NHV             ...........................................--...........................--g..... [186560]
KX354450.1_47/Ss/2008             ...........................................--...........................--g..... [186560]
KP055815.1_strain_BA71            ...........................................--...........................--g..... [186560]
MH025920.1_strain_R35             .....cc........t...........................act.....................at...--ga...g [186560]
MH025919.1_strain_N10             .....cc........t...........................act.....................at...--ga...g [186560]
MH025918.1_strain_R25             .....cc........t...........................act.....................at...--ga...g [186560]
MH025917.1_strain_R7              .....cc........t...........................act.....................at...--ga...g [186560]
MH025916.1_strain_R8              .....cc........t...........................act.....................at...--ga...g [186560]
```

```
FR682468.2_Georgia_2007/1         tttaaa--gttttttttgaaaaaaaataatatat-gtagataaaattcagttacatgatata------------------ [186640]
ASF_IND_20_CAD543                 ......--t.................y.......-.........................-------------------- [186640]
LR899193.1_Germany_2020/1         ......--..........................-...........................------------------ [186640]
MN393477.1_Wuhan_2019             ......--..........................-...........................------------------ [186640]
MW396979.1_Timor-Leste/2019/1     ......--..........................-...........................------------------ [186640]
MK940252.1_CN/2019/InnerMongolia  ......--..........................-...........................------------------ [186640]
MK128995.1_China/2018/AnhuiXCGQ   ......--..........................-...........................------------------ [186640]
MH681419.1_ASFV/POL/2015          ......--..........................-...........................------------------ [186640]
LS478113.1_Estonia_2014           ......--..........................-...........................------------------ [186640]
KP843857.1_Odintsovo_02/14        ......--.-........t...............-...........................------------------ [186640]
FN557520.1_strain_E75             ......--t.............-...c.......-.........................-------------------- [186640]
AY261366.1_Warthog                ......g-t.......ga................g...........................agcatatgtgtaagc--- [186640]
AY261365.1_Warmbaths              ......gtt.......ca........c.......g...........................agcatatgtgtaagcata [186640]
AY261364.1_Tengani_62             ......--..................c.......-..g........................------------------ [186640]
AY261363.1_Pretoriuskop/96/4      ......att.......ca........c.......-...........................agcatatgtg-------- [186640]
AY261362.1_Mkuzi_1979             ......--a.................c.......-.........................-------------------- [186640]
AY261361.1_Malawi_Lil-20/1_1983   a.....ctt......caa........c.......-..g....-----t.....t...t....agcatatgt--------- [186640]
AY261360.1_Kenya_1950             ..-...t-t......caa........cg...ggg------......at............-------------------- [186640]
AM712240.1_OURT_88/3              ......--t.................c.......-.........................-------------------- [186640]
AM712239.1_Benin_97/1             ......--t.................c.......-.........................-------------------- [186640]
KM111295.1_Ken06.Bus              ..-...a-t........c.........g...ggg-----...c...atg...........-------------------- [186640]
KM111294.1_Ken05/Tk1              ..a..tt-t......caa........cg...ggg------......at............-------------------- [186640]
KM262844.1_strain_L60             ......--t.................c.......-.........................-------------------- [186640]
KM102979.1_26544/OG10_Italy       ......--t.................c.......-.........................-------------------- [186640]
KM262845.1_strain_NHV             ......---.................c.......-.........................-------------------- [186640]
KX354450.1_47/Ss/2008             ......--t.................c.......-.........................-------------------- [186640]
KP055815.1_strain_BA71            ......--t.................c.......-.........................-------------------- [186640]
MH025920.1_strain_R35             ..a...a-t........c.........g...ggg-----...c...atg...........-------------------- [186640]
MH025919.1_strain_N10             ..a...a-t........c.........g...ggg-----...c...atg...........-------------------- [186640]
MH025918.1_strain_R25             ..a...a-t........c.........g...ggg-----...c...atg...........-------------------- [186640]
MH025917.1_strain_R7              ..a...a-t........c.........g...ggg-----...c...atg...........-------------------- [186640]
MH025916.1_strain_R8              ..a...a-t........c.........g...ggg-----...c...atg...........-------------------- [186640]

FR682468.2_Georgia_2007/1         tgtgtaaacatgtgtggtaaacaacatatggttatgct-----------------------------------------t [186720]
ASF_IND_20_CAD543                 ----...g.......-......................-----------------------------------------. [186720]
LR899193.1_Germany_2020/1         ......................................-----------------------------------------. [186720]
MN393477.1_Wuhan_2019             ......................................-----------------------------------------. [186720]
MW396979.1_Timor-Leste/2019/1     ......................................-----------------------------------------. [186720]
MK940252.1_CN/2019/InnerMongolia  ......................................-----------------------------------------. [186720]
MK128995.1_China/2018/AnhuiXCGQ   ......................................-----------------------------------------. [186720]
MH681419.1_ASFV/POL/2015          ......................................-----------------------------------------. [186720]
LS478113.1_Estonia_2014           ......................................-----------------------------------------. [186720]
KP843857.1_Odintsovo_02/14        ......................................-----------------------------------------. [186720]
```

```
FN557520.1_strain_E75           ----...g...a...-..................c..------------------------------------------. [186720]
AY261366.1_Warthog              ---------..a...-..................c..------------------------------------------. [186720]
AY261365.1_Warmbaths            ........t..a...-..................c..------------------------------------------. [186720]
AY261364.1_Tengani_62           ...........a...-..................c..------------------------------------------. [186720]
AY261363.1_Pretoriuskop/96/4    ---........a...-..................c..------------------------------------------. [186720]
AY261362.1_Mkuzi_1979           ----...g...a...-..................c..------------------------------------------. [186720]
AY261361.1_Malawi_Lil-20/1_1983 --.a...g...a.a.-........a...c.....cat.atatcagataaatgcatatatgtaaacaaaataggttacgc. [186720]
AY261360.1_Kenya_1950           ----...g...aga.-........a.ata.....c..------------------------------------------. [186720]
AM712240.1_OURT_88/3            ----...g...a...-..................c..------------------------------------------. [186720]
AM712239.1_Benin_97/1           ----...g...a...-..................c..------------------------------------------. [186720]
KM111295.1_Ken06.Bus            ----...g...aga.-......-.a.ata.....c..------------------------------------------. [186720]
KM111294.1_Ken05/Tk1            ----...g...aga.-......-.a.ata.....c..------------------------------------------. [186720]
KM262844.1_strain_L60           ----...g...a...-..................c..------------------------------------------. [186720]
KM102979.1_26544/OG10_Italy     ----...g...a...-..................c..------------------------------------------. [186720]
KM262845.1_strain_NHV           ----...g...a...-..................c..------------------------------------------. [186720]
KX354450.1_47/Ss/2008           ----...g...a...-..................c..------------------------------------------. [186720]
KP055815.1_strain_BA71          ----...g...a...-..................c..------------------------------------------. [186720]
MH025920.1_strain_R35           ----...g...aga.-......-.a.ata.....c..------------------------------------------. [186720]
MH025919.1_strain_N10           ----...g...aga.-......-.a.ata.....c..------------------------------------------. [186720]
MH025918.1_strain_R25           ----...g...aga.-......-.a.ata.....c..------------------------------------------. [186720]
MH025917.1_strain_R7            ----...g...aga.-......-.a.ata.....c..------------------------------------------. [186720]
MH025916.1_strain_R8            ----...g...aga.-......-.a.ata.....c..------------------------------------------. [186720]

FR682468.2_Georgia_2007/1       tataagataaatgcgcat---------aatatatgtaaacaaaatatggttatgtg-ttaaat------gc--------- [186800]
ASF_IND_20_CAD543               ..................---------.w........................act-.atg------------------- [186800]
LR899193.1_Germany_2020/1       ..................---------.............................-......------..--------- [186800]
MN393477.1_Wuhan_2019           ..................---------.............................-......------..--------- [186800]
MW396979.1_Timor-Leste/2019/1   ..................---------.............................-......------..--------- [186800]
MK940252.1_CN/2019/InnerMongolia ..................---------.............................-......------..--------- [186800]
MK128995.1_China/2018/AnhuiXCGQ ..................---------.............................-......------..--------- [186800]
MH681419.1_ASFV/POL/2015        ..................---------.............................-......------..--------- [186800]
LS478113.1_Estonia_2014         ..................---------.............................-......------..--------- [186800]
KP843857.1_Odintsovo_02/14      ..................---------.............................-......------..--------- [186800]
FN557520.1_strain_E75           ...g..............---------.t........................act-.atg.g----------------- [186800]
AY261366.1_Warthog              ...g....-.......g.---------.t........................act-.atg.gataaat..--------- [186800]
AY261365.1_Warmbaths            ...g..............---------.t........................act-.atg.gataaat..--------- [186800]
AY261364.1_Tengani_62           ...g..............---------.t........................act-.atg.gataaat..--------- [186800]
AY261363.1_Pretoriuskop/96/4    ...g..............---------.t........................act-.atg.gataaat..--------- [186800]
AY261362.1_Mkuzi_1979           ...g..............---------.t........................act-.atg.gataaat..--------- [186800]
AY261361.1_Malawi_Lil-20/1_1983 ...g...........t..---------.tg.........t..c.........cactg.atg.gat---tc.--------- [186800]
AY261360.1_Kenya_1950           c..g....tc..a..t..atatgcatat.............g....-.a....actt.atg.gattcata.gtatatatg [186800]
AM712240.1_OURT_88/3            ...g..............---------.t........................act-.atg.g----------------- [186800]
AM712239.1_Benin_97/1           ...g..............---------.t........................act-.atg.g----------------- [186800]
KM111295.1_Ken06.Bus            c..g....tc..a..------------t.............g....-.a....acttcatg.gattcata.gtatat--- [186800]
```

```
KM111294.1_Ken05/Tk1             c..g....tc..a..------------t.............g....-.a....actt.atg.gattcata.gtatatatg [186800]
KM262844.1_strain_L60            ...g..............---------.t........................act-.atg.g----------------- [186800]
KM102979.1_26544/OG10_Italy      ...g..............---------.t........................act-.atg.g----------------- [186800]
KM262845.1_strain_NHV            ...g..............---------.t........................act-.atg.g----------------- [186800]
KX354450.1_47/Ss/2008            ...g..............---------.t........................act-.atg.g----------------- [186800]
KP055815.1_strain_BA71           ...g..............---------.t........................act-.atg.g----------------- [186800]
MH025920.1_strain_R35            c..g....tc..a..------------t.............g....-.a....acttcatg.gattcata.gtatat--- [186800]
MH025919.1_strain_N10            c..g....tc..a..------------t.............g....-.a....acttcatg.gattcata.gtatat--- [186800]
MH025918.1_strain_R25            c..g....tc..a..------------t.............g....-.a....acttcatg.gattcata.gtatat--- [186800]
MH025917.1_strain_R7             c..g....tc..a..------------t.............g....-.a....acttcatg.gattcata.gtatat--- [186800]
MH025916.1_strain_R8             c..g....tc..a..------------t.............g....-.a....acttcatg.gattcata.gtatat--- [186800]

FR682468.2_Georgia_2007/1        ---atataaatgtattttaacgtatatcttgtgataatggatatatg-----cattta---------------------- [186880]
ASF_IND_20_CAD543                ---.k.....................-....................-----.w....---------------------- [186880]
LR899193.1_Germany_2020/1        ---............................................-----......---------------------- [186880]
MN393477.1_Wuhan_2019            ---............................................-----......---------------------- [186880]
MW396979.1_Timor-Leste/2019/1    ---............................................-----......---------------------- [186880]
MK940252.1_CN/2019/InnerMongolia ---............................................-----......---------------------- [186880]
MK128995.1_China/2018/AnhuiXCGQ  ---............................................-----......---------------------- [186880]
MH681419.1_ASFV/POL/2015         ---............................................-----......---------------------- [186880]
LS478113.1_Estonia_2014          ---............................................-----......---------------------- [186880]
KP843857.1_Odintsovo_02/14       ---............................................-----......---------------------- [186880]
FN557520.1_strain_E75            -----.....................-....................-----.t....---------------------- [186880]
AY261366.1_Warthog               ---..................a.........................tt--ttt....---------------------- [186880]
AY261365.1_Warmbaths             ---..................a.........................-----.t....---------------------- [186880]
AY261364.1_Tengani_62            ---..................a........................a-----.t....---------------------- [186880]
AY261363.1_Pretoriuskop/96/4     ---..................a.........................-t--ttt....---------------------- [186880]
AY261362.1_Mkuzi_1979            ---.......................-....................-----tt....---------------------- [186880]
AY261361.1_Malawi_Lil-20/1_1983  ---.....t.....cg.....a....................g....-t--ttt....ttaaagcctgccttaaacaggc [186880]
AY261360.1_Kenya_1950            ----c...t.....c......a..............t.....g....ttt-ctt....aaaaaggctgttttta------ [186880]
AM712240.1_OURT_88/3             -----.....................-....................-----.t....---------------------- [186880]
AM712239.1_Benin_97/1            -----.....................-....................-----.t....---------------------- [186880]
KM111295.1_Ken06.Bus             -----...t.c...c......a..........a...t.a...g....tttcctt....taaaaggctgttttta------ [186880]
KM111294.1_Ken05/Tk1             cat.....t.....c......a..............t.....g....ttc-ctt....aaaaaggctgttttta------ [186880]
KM262844.1_strain_L60            -----.....................-....................-----.t....---------------------- [186880]
KM102979.1_26544/OG10_Italy      -----.....................-....................-----.t....---------------------- [186880]
KM262845.1_strain_NHV            -----.....................-....................-----.t....---------------------- [186880]
KX354450.1_47/Ss/2008            -----.....................-....................-----.t....---------------------- [186880]
KP055815.1_strain_BA71           -----.....................-....................-----.t....---------------------- [186880]
MH025920.1_strain_R35            -----...t.c...c......a..........a...t.a...g....tttcctt....taaaaggctgttttta------ [186880]
MH025919.1_strain_N10            -----...t.c...c......a..........a...t.a...g....tttcctt....taaaaggctgttttta------ [186880]
MH025918.1_strain_R25            -----...t.c...c......a..........a...t.a...g....tttcctt....taaaaggctgttttta------ [186880]
MH025917.1_strain_R7             -----...t.c...c......a..........a...t.a...g....tttcctt....taaaaggctgttttta------ [186880]
MH025916.1_strain_R8             -----...t.c...c......a..........a...t.a...g....tttcctt....taaaaggctgttttta------ [186880]
```

```
FR682468.2_Georgia_2007/1        ---ttaaaagaggctgtatttattataaatcttgctaaggatgccattgtcaacatatatcccatgttggacaaattgcg [186960]
ASF_IND_20_CAD543                ---.......................................r..................................... [186960]
LR899193.1_Germany_2020/1        ---............................................................................. [186960]
MN393477.1_Wuhan_2019            ---............................................................................. [186960]
MW396979.1_Timor-Leste/2019/1    ---............................................................................. [186960]
MK940252.1_CN/2019/InnerMongolia ---............................................................................. [186960]
MK128995.1_China/2018/AnhuiXCGQ  ---............................................................................. [186960]
MH681419.1_ASFV/POL/2015         ---............................................................................. [186960]
LS478113.1_Estonia_2014          ---............................................................................. [186960]
KP843857.1_Odintsovo_02/14       ---............................................................................. [186960]
FN557520.1_strain_E75            ---.......................................a....................g................ [186960]
AY261366.1_Warthog               ---............................c...c..........................................t. [186960]
AY261365.1_Warmbaths             ---............................................................................. [186960]
AY261364.1_Tengani_62            ---.........a.........c......................................................... [186960]
AY261363.1_Pretoriuskop/96/4     ---................................c.....................c...................... [186960]
AY261362.1_Mkuzi_1979            ---............................................................................. [186960]
AY261361.1_Malawi_Lil-20/1_1983  tgt......c........................g.t..a....................t..g................ [186960]
AY261360.1_Kenya_1950            ---...............................t.t........................................... [186960]
AM712240.1_OURT_88/3             ---.......................................a....................g................ [186960]
AM712239.1_Benin_97/1            ---.......................................a....................g................ [186960]
KM111295.1_Ken06.Bus             ---.....a..........................t.....a...c.......g......................... [186960]
KM111294.1_Ken05/Tk1             ---.....a..........................t.................g......................... [186960]
KM262844.1_strain_L60            ---.......................................a....................g................ [186960]
KM102979.1_26544/OG10_Italy      ---.......................................a....................g................ [186960]
KM262845.1_strain_NHV            ---.......................................a....................g................ [186960]
KX354450.1_47/Ss/2008            ---.......................................a....................g................ [186960]
KP055815.1_strain_BA71           ---.......................................a....................g................ [186960]
MH025920.1_strain_R35            ---.....a..........................t.....a...c.......g......................... [186960]
MH025919.1_strain_N10            ---.....a..........................t.....a...c.......g......................... [186960]
MH025918.1_strain_R25            ---.....a..........................t.....a...c.......g......................... [186960]
MH025917.1_strain_R7             ---.....a..........................t.....a...c.......g......................... [186960]
MH025916.1_strain_R8             ---.....a..........................t.....a...c.......g......................... [186960]

FR682468.2_Georgia_2007/1        ttgcgatccagttcttttttttttgatttgtttaatgctatcctttttgaagggatggttgtccaccatatttattcga [187040]
ASF_IND_20_CAD543                ................................................................................ [187040]
LR899193.1_Germany_2020/1        ................................................................................ [187040]
MN393477.1_Wuhan_2019            ................................................................................ [187040]
MW396979.1_Timor-Leste/2019/1    ................................................................................ [187040]
MK940252.1_CN/2019/InnerMongolia ................................................................................ [187040]
MK128995.1_China/2018/AnhuiXCGQ  ................................................................................ [187040]
MH681419.1_ASFV/POL/2015         ................................................................................ [187040]
LS478113.1_Estonia_2014          ................................................................................ [187040]
KP843857.1_Odintsovo_02/14       ..............-................................................................. [187040]
```

```
FN557520.1_strain_E75              .c..................................c........................................... [187040]
AY261366.1_Warthog                 ................................................................................ [187040]
AY261365.1_Warmbaths               .c..................................c........................................... [187040]
AY261364.1_Tengani_62              .c.............................................................................. [187040]
AY261363.1_Pretoriuskop/96/4       ................................................................................ [187040]
AY261362.1_Mkuzi_1979              .c..................................c.....................a..................... [187040]
AY261361.1_Malawi_Lil-20/1_1983    .c.............................................................................g [187040]
AY261360.1_Kenya_1950              .ca.t..........................................................................g [187040]
AM712240.1_OURT_88/3               .c..................................c........................................... [187040]
AM712239.1_Benin_97/1              .c..................................c........................a.................. [187040]
KM111295.1_Ken06.Bus               .ca.t.........-................................................................g [187040]
KM111294.1_Ken05/Tk1               .ca.t......................................t........a..........................g [187040]
KM262844.1_strain_L60              .c..................................c........................................... [187040]
KM102979.1_26544/OG10_Italy        .c..................................c........................................... [187040]
KM262845.1_strain_NHV              .c..................................c........................................... [187040]
KX354450.1_47/Ss/2008              .c..................................c........................................... [187040]
KP055815.1_strain_BA71             .c..................................c........................................... [187040]
MH025920.1_strain_R35              .ca.t.........-................................................................g [187040]
MH025919.1_strain_N10              .ca.t.........-................................................................g [187040]
MH025918.1_strain_R25              .ca.t.........-................................................................g [187040]
MH025917.1_strain_R7               .ca.t.........-................................................................g [187040]
MH025916.1_strain_R8               .ca.t.........-................................................................g [187040]

FR682468.2_Georgia_2007/1          tgttcaatgaataggtctgctttttcgtaaggcagtgaaggtcgttccaagactccttgaacgatggacgtgttttcttg [187120]
ASF_IND_20_CAD543                  ................................................................a............... [187120]
LR899193.1_Germany_2020/1          ................................................................a............... [187120]
MN393477.1_Wuhan_2019              ................................................................a............... [187120]
MW396979.1_Timor-Leste/2019/1      ................................................................a............... [187120]
MK940252.1_CN/2019/InnerMongolia   ................................................................a............... [187120]
MK128995.1_China/2018/AnhuiXCGQ    ................................................................a............... [187120]
MH681419.1_ASFV/POL/2015           ................................................................................ [187120]
LS478113.1_Estonia_2014            ................................................................a............... [187120]
KP843857.1_Odintsovo_02/14         ................................................................................ [187120]
FN557520.1_strain_E75              ...............a......c.......................t...c............................. [187120]
AY261366.1_Warthog                 ...............a.............t...........c...................................... [187120]
AY261365.1_Warmbaths               ...............a......c......................................................... [187120]
AY261364.1_Tengani_62              ...............a.........................c...................................... [187120]
AY261363.1_Pretoriuskop/96/4       ...............a.........................c...................................... [187120]
AY261362.1_Mkuzi_1979              ...............a......c.......................t...c............................. [187120]
AY261361.1_Malawi_Lil-20/1_1983    ...........a...a....c.c........a........a...................g.....a.t........... [187120]
AY261360.1_Kenya_1950              ...............a.g..c.....a........c....a...................gt....a............. [187120]
AM712240.1_OURT_88/3               ...............a......c.......................t...c............................. [187120]
AM712239.1_Benin_97/1              ...............a......c.......................t...c............................. [187120]
KM111295.1_Ken06.Bus               ...............a.g..c..............c....a...................gt....a............. [187120]
```

```
KM111294.1_Ken05/Tk1           ...............a.g..c.....a........c....a...................gt....a.........t... [187120]
KM262844.1_strain_L60          ...............a......c.......................t...c............................. [187120]
KM102979.1_26544/OG10_Italy    ...............a......c.......................t...c............................. [187120]
KM262845.1_strain_NHV          ...............a......c.......................t...c............................. [187120]
KX354450.1_47/Ss/2008          ...............a......c.......................t...c............................. [187120]
KP055815.1_strain_BA71         ...............a......c.......................t...c............................. [187120]
MH025920.1_strain_R35          ...............a.g..c..............c....a...................gt....a............. [187120]
MH025919.1_strain_N10          ...............a.g..c..............c....a...................gt....a............. [187120]
MH025918.1_strain_R25          ...............a.g..c..............c....a...................gt....a............. [187120]
MH025917.1_strain_R7           ...............a.g..c..............c....a...................gt....a............. [187120]
MH025916.1_strain_R8           ...............a.g..c..............c....a...................gt....a............. [187120]

FR682468.2_Georgia_2007/1      gatccacttaaaaagcacgtggcattcaaaaacaggacagtgattggatccttggatatgctttggacagccaatgcttg [187200]
ASF_IND_20_CAD543              ................................................................................ [187200]
LR899193.1_Germany_2020/1      ................................................................................ [187200]
MN393477.1_Wuhan_2019          ................................................................................ [187200]
MW396979.1_Timor-Leste/2019/1  ................................................................................ [187200]
MK940252.1_CN/2019/InnerMongolia ................................................................................ [187200]
MK128995.1_China/2018/AnhuiXCGQ ................................................................................ [187200]
MH681419.1_ASFV/POL/2015       ................................................................................ [187200]
LS478113.1_Estonia_2014        ................................................................................ [187200]
KP843857.1_Odintsovo_02/14     ................................................................................ [187200]
FN557520.1_strain_E75          ................................................................................ [187200]
AY261366.1_Warthog             ................................................................................ [187200]
AY261365.1_Warmbaths           ................................................................................ [187200]
AY261364.1_Tengani_62          ......................................................a......................... [187200]
AY261363.1_Pretoriuskop/96/4   ................................................................................ [187200]
AY261362.1_Mkuzi_1979          ................................................................................ [187200]
AY261361.1_Malawi_Lil-20/1_1983 ................................t...g...........c...........t.....g............. [187200]
AY261360.1_Kenya_1950          ................................t...g...........c.........................a..... [187200]
AM712240.1_OURT_88/3           ................................................................................ [187200]
AM712239.1_Benin_97/1          ................................................................................ [187200]
KM111295.1_Ken06.Bus           ................................t...g...........c.........................a..... [187200]
KM111294.1_Ken05/Tk1           ................................t...g...........c.........................a..... [187200]
KM262844.1_strain_L60          ................................................................................ [187200]
KM102979.1_26544/OG10_Italy    ................................................................................ [187200]
KM262845.1_strain_NHV          ................................................................................ [187200]
KX354450.1_47/Ss/2008          ................................................................................ [187200]
KP055815.1_strain_BA71         ................................................................................ [187200]
MH025920.1_strain_R35          ................................t...g...........c.........................a..... [187200]
MH025919.1_strain_N10          ................................t...g...........c.........................a..... [187200]
MH025918.1_strain_R25          ................................t...g...........c.........................a..... [187200]
MH025917.1_strain_R7           ................................t...g...........c.........................a..... [187200]
MH025916.1_strain_R8           ................................t...g...........c.........................a..... [187200]
```

```
FR682468.2_Georgia_2007/1        aagagatgtagtcccttttctttaggacaagcttctccacgctggggcaacagagatcgttcaagttctggacggtcgca [187280]
ASF_IND_20_CAD543                ................................................................................ [187280]
LR899193.1_Germany_2020/1        ................................................................................ [187280]
MN393477.1_Wuhan_2019            ................................................................................ [187280]
MW396979.1_Timor-Leste/2019/1    ................................................................................ [187280]
MK940252.1_CN/2019/InnerMongolia ................................................................................ [187280]
MK128995.1_China/2018/AnhuiXCGQ  ................................................................................ [187280]
MH681419.1_ASFV/POL/2015         ................................................................................ [187280]
LS478113.1_Estonia_2014          ................................................................................ [187280]
KP843857.1_Odintsovo_02/14       ................................................................................ [187280]
FN557520.1_strain_E75            ....a........................................................................... [187280]
AY261366.1_Warthog               ..............t................................................................. [187280]
AY261365.1_Warmbaths             ................................................................................ [187280]
AY261364.1_Tengani_62            ..............t.......c......................................................... [187280]
AY261363.1_Pretoriuskop/96/4     ..............t................................................................. [187280]
AY261362.1_Mkuzi_1979            ....a........................................................................... [187280]
AY261361.1_Malawi_Lil-20/1_1983  ..............t.............t..t..t...........a....................t............ [187280]
AY261360.1_Kenya_1950            ......c.......t.............g.......................a..............t............ [187280]
AM712240.1_OURT_88/3             ....a........................................................................... [187280]
AM712239.1_Benin_97/1            ....a........................................................................... [187280]
KM111295.1_Ken06.Bus             ...ga.c.......t....t........g.......................a..............t............ [187280]
KM111294.1_Ken05/Tk1             ......c.......t.............g.......................a..............t............ [187280]
KM262844.1_strain_L60            ....a........................................................................... [187280]
KM102979.1_26544/OG10_Italy      ....a........................................................................... [187280]
KM262845.1_strain_NHV            ....a........................................................................... [187280]
KX354450.1_47/Ss/2008            ....a........................................................................... [187280]
KP055815.1_strain_BA71           ....a........................................................................... [187280]
MH025920.1_strain_R35            ...ga.c.......t....t........g.......................a..............t............ [187280]
MH025919.1_strain_N10            ...ga.c.......t....t........g.......................a..............t............ [187280]
MH025918.1_strain_R25            ...ga.c.......t....t........g.......................a..............t............ [187280]
MH025917.1_strain_R7             ...ga.c.......t....t........g.......................a..............t............ [187280]
MH025916.1_strain_R8             ...ga.c.......t....t........g.......................a..............t............ [187280]

FR682468.2_Georgia_2007/1        tttggaatgttgaaacttcgtatccattcaccctcgggtcctcccttatgaagaaggagtatttgctcatggtccttagt [187360]
ASF_IND_20_CAD543                ..........................................................................y..... [187360]
LR899193.1_Germany_2020/1        ................................................................................ [187360]
MN393477.1_Wuhan_2019            ................................................................................ [187360]
MW396979.1_Timor-Leste/2019/1    ................................................................................ [187360]
MK940252.1_CN/2019/InnerMongolia ................................................................................ [187360]
MK128995.1_China/2018/AnhuiXCGQ  ................................................................................ [187360]
MH681419.1_ASFV/POL/2015         ................................................................................ [187360]
LS478113.1_Estonia_2014          ................................................................................ [187360]
KP843857.1_Odintsovo_02/14       ................................................................................ [187360]
```

```
FN557520.1_strain_E75           ......................c...............c...................................t..... [187360]
AY261366.1_Warthog              .g....................c......................................................g.. [187360]
AY261365.1_Warmbaths            ......................c...................................................t..... [187360]
AY261364.1_Tengani_62           ......................c......................................................... [187360]
AY261363.1_Pretoriuskop/96/4    ......................c.............................................g........g.. [187360]
AY261362.1_Mkuzi_1979           ......................c...................................................t..... [187360]
AY261361.1_Malawi_Lil-20/1_1983 ......................c.........t....................c..a........t........t..... [187360]
AY261360.1_Kenya_1950           .............................g..........................c........t........t..... [187360]
AM712240.1_OURT_88/3            ......................c...............c...................................t..... [187360]
AM712239.1_Benin_97/1           ......................c...............c...................................t..... [187360]
KM111295.1_Ken06.Bus            ...............t........................................c........t........t..... [187360]
KM111294.1_Ken05/Tk1            .............................g..........................c........t........t..... [187360]
KM262844.1_strain_L60           ......................c...............c...................................t..... [187360]
KM102979.1_26544/OG10_Italy     ......................c...............c...................................t..... [187360]
KM262845.1_strain_NHV           ......................c...............c...................................t..... [187360]
KX354450.1_47/Ss/2008           ......................c...............c...................................t..... [187360]
KP055815.1_strain_BA71          ......................c...............c...................................t..... [187360]
MH025920.1_strain_R35           ...............t........................................c........t........t..... [187360]
MH025919.1_strain_N10           ...............t........................................c........t........t..... [187360]
MH025918.1_strain_R25           ...............t........................................c........t........t..... [187360]
MH025917.1_strain_R7            ...............t........................................c........t........t..... [187360]
MH025916.1_strain_R8            ...............t........................................c........t........t..... [187360]

FR682468.2_Georgia_2007/1       aatcttaaccaaatgttggaagatcatttttttacctgctttaaaggcctgaagggtgtcagttggcaaagctattgaat [187440]
ASF_IND_20_CAD543               ................................................................................ [187440]
LR899193.1_Germany_2020/1       ................................................................................ [187440]
MN393477.1_Wuhan_2019           ................................................................................ [187440]
MW396979.1_Timor-Leste/2019/1   ................................................................................ [187440]
MK940252.1_CN/2019/InnerMongolia ................................................................................ [187440]
MK128995.1_China/2018/AnhuiXCGQ ................................................................................ [187440]
MH681419.1_ASFV/POL/2015        ................................................................................ [187440]
LS478113.1_Estonia_2014         ................................................................................ [187440]
KP843857.1_Odintsovo_02/14      ................................................................................ [187440]
FN557520.1_strain_E75           ............................................................g........g.......... [187440]
AY261366.1_Warthog              ............................................................g.a................. [187440]
AY261365.1_Warmbaths            ............................................................g................... [187440]
AY261364.1_Tengani_62           ...............c............................................g........g.......... [187440]
AY261363.1_Pretoriuskop/96/4    ............................................................g................... [187440]
AY261362.1_Mkuzi_1979           ............................................................g................... [187440]
AY261361.1_Malawi_Lil-20/1_1983 ........................a..........aa.......................g..........t...g.... [187440]
AY261360.1_Kenya_1950           .........t..g...........a..........ac....c..................g.......gg.......... [187440]
AM712240.1_OURT_88/3            ............................................................g........g.......... [187440]
AM712239.1_Benin_97/1           ............................................................g........g.......... [187440]
KM111295.1_Ken06.Bus            ...t........g...........at.........ac....c..................g.......gg.......... [187440]
```

```
KM111294.1_Ken05/Tk1             ............g...........a..........ac....c..................g.......gg.......... [187440]
KM262844.1_strain_L60            ............................................................g........g.......... [187440]
KM102979.1_26544/OG10_Italy      ............................................................g........g.......... [187440]
KM262845.1_strain_NHV            ............................................................g........g.......... [187440]
KX354450.1_47/Ss/2008            ............................................................g........g.......... [187440]
KP055815.1_strain_BA71           ............................................................g........g.......... [187440]
MH025920.1_strain_R35            ...t........g...........at.........ac....c..................g.......gg.......... [187440]
MH025919.1_strain_N10            ...t........g...........at.........ac....c..................g.......gg.......... [187440]
MH025918.1_strain_R25            ...t........g...........at.........ac....c..................g.......gg.......... [187440]
MH025917.1_strain_R7             ...t........g...........at.........ac....c..................g.......gg.......... [187440]
MH025916.1_strain_R8             ...t........g...........at.........ac....c..................g.......gg.......... [187440]

FR682468.2_Georgia_2007/1        tcgggagtgggctttcatcaagcgtgaaatggtgaatgtgacgcgactggaaagaaaacgaccgttgatttattttttca [187520]
ASF_IND_20_CAD543                ................................................................................ [187520]
LR899193.1_Germany_2020/1        ................................................................................ [187520]
MN393477.1_Wuhan_2019            ................................................................................ [187520]
MW396979.1_Timor-Leste/2019/1    ................................................................................ [187520]
MK940252.1_CN/2019/InnerMongolia ................................................................................ [187520]
MK128995.1_China/2018/AnhuiXCGQ  ................................................................................ [187520]
MH681419.1_ASFV/POL/2015         ................................................................................ [187520]
LS478113.1_Estonia_2014          ................................................................................ [187520]
KP843857.1_Odintsovo_02/14       ................................................................................ [187520]
FN557520.1_strain_E75            ........................................g....................................... [187520]
AY261366.1_Warthog               ........................................g....................................... [187520]
AY261365.1_Warmbaths             ........................................g....................................... [187520]
AY261364.1_Tengani_62            ........................................g..............................g........ [187520]
AY261363.1_Pretoriuskop/96/4     ........................c...............g....................................... [187520]
AY261362.1_Mkuzi_1979            ........................................g....................................... [187520]
AY261361.1_Malawi_Lil-20/1_1983  ................g.......................g........a........t............g........ [187520]
AY261360.1_Kenya_1950            ....c..c...........g...................ag..t.....a....c...t............g........ [187520]
AM712240.1_OURT_88/3             ........................................g....................................... [187520]
AM712239.1_Benin_97/1            ........................................g....................................... [187520]
KM111295.1_Ken06.Bus             .......c...........g...................ag..t.....a....c...t............g........ [187520]
KM111294.1_Ken05/Tk1             ....c..c...........g...................ag..t.....a....c...t............g........ [187520]
KM262844.1_strain_L60            ........................................g....................................... [187520]
KM102979.1_26544/OG10_Italy      ........................................g....................................... [187520]
KM262845.1_strain_NHV            ........................................g....................................... [187520]
KX354450.1_47/Ss/2008            ........................................g....................................... [187520]
KP055815.1_strain_BA71           ........................................g....................................... [187520]
MH025920.1_strain_R35            .......c...........g...................ag..t.....a....c...t............g........ [187520]
MH025919.1_strain_N10            .......c...........g...................ag..t.....a....c...t............g........ [187520]
MH025918.1_strain_R25            .......c...........g...................ag..t.....a....c...t............g........ [187520]
MH025917.1_strain_R7             .......c...........g...................ag..t.....a....c...t............g........ [187520]
MH025916.1_strain_R8             .......c...........g...................ag..t.....a....c...t............g........ [187520]
```

```
FR682468.2_Georgia_2007/1        aagattgggtcgattccgccatgaaagaacagctgcaagattttagaaggcgtatttttttcccaataaaaaatgaccac [187600]
ASF_IND_20_CAD543                ................................................................................ [187600]
LR899193.1_Germany_2020/1        ................................................................................ [187600]
MN393477.1_Wuhan_2019            ................................................................................ [187600]
MW396979.1_Timor-Leste/2019/1    ................................................................................ [187600]
MK940252.1_CN/2019/InnerMongolia ................................................................................ [187600]
MK128995.1_China/2018/AnhuiXCGQ  ................................................................................ [187600]
MH681419.1_ASFV/POL/2015         ................................................................................ [187600]
LS478113.1_Estonia_2014          ................................................................................ [187600]
KP843857.1_Odintsovo_02/14       ................................................................................ [187600]
FN557520.1_strain_E75            .................................................c...........................t.. [187600]
AY261366.1_Warthog               .................................................c.............................. [187600]
AY261365.1_Warmbaths             .................................................c.............................. [187600]
AY261364.1_Tengani_62            .................a.......................g.......c..c........................... [187600]
AY261363.1_Pretoriuskop/96/4     ..a..............a...........t...................c.............................. [187600]
AY261362.1_Mkuzi_1979            .................................................c.............................. [187600]
AY261361.1_Malawi_Lil-20/1_1983  ..t....t.........a.......................g...a.t.c............................g. [187600]
AY261360.1_Kenya_1950            ..t..g.t.........a...........t...........g...a...ct..........................tg. [187600]
AM712240.1_OURT_88/3             .................................................c...........................t.. [187600]
AM712239.1_Benin_97/1            .................................................c...........................t.. [187600]
KM111295.1_Ken06.Bus             ..t..g.t.....................t...........g...a...ct.......................c...g. [187600]
KM111294.1_Ken05/Tk1             ..t..g.t.........a...........t...........g...a...ct..........................tg. [187600]
KM262844.1_strain_L60            .................................................c...........................t.. [187600]
KM102979.1_26544/OG10_Italy      .................................................c...........................t.. [187600]
KM262845.1_strain_NHV            .................................................c...........................t.. [187600]
KX354450.1_47/Ss/2008            .................................................c...........................t.. [187600]
KP055815.1_strain_BA71           .................................................c...........................t.. [187600]
MH025920.1_strain_R35            ..t..g.t.....................t...........g...a...ct.......................c...g. [187600]
MH025919.1_strain_N10            ..t..g.t.....................t...........g...a...ct.......................c...g. [187600]
MH025918.1_strain_R25            ..t..g.t.....................t...........g...a...ct.......................c...g. [187600]
MH025917.1_strain_R7             ..t..g.t.....................t...........g...a...ct.......................c...g. [187600]
MH025916.1_strain_R8             ..t..g.t.....................t...........g...a...ct.......................c...g. [187600]

FR682468.2_Georgia_2007/1        ttctcgtgggattaaaatcgtctgtgtcccattttcattatataattggcccataaagccatcaacgtcaatcaacacca [187680]
ASF_IND_20_CAD543                w............................................................................... [187680]
LR899193.1_Germany_2020/1        ................................................................................ [187680]
MN393477.1_Wuhan_2019            ................................................................................ [187680]
MW396979.1_Timor-Leste/2019/1    ................................................................................ [187680]
MK940252.1_CN/2019/InnerMongolia ................................................................................ [187680]
MK128995.1_China/2018/AnhuiXCGQ  ................................................................................ [187680]
MH681419.1_ASFV/POL/2015         ................................................................................ [187680]
LS478113.1_Estonia_2014          ................................................................................ [187680]
KP843857.1_Odintsovo_02/14       ................................................................................ [187680]
```

```
FN557520.1_strain_E75            ......c......................................................................... [187680]
AY261366.1_Warthog               ........................c....................c.................................. [187680]
AY261365.1_Warmbaths             ................................................................................ [187680]
AY261364.1_Tengani_62            .............................................c.................................. [187680]
AY261363.1_Pretoriuskop/96/4     ..............................t..............c.................................. [187680]
AY261362.1_Mkuzi_1979            ......c......................................................................t.. [187680]
AY261361.1_Malawi_Lil-20/1_1983  .....................................................................t.....t.... [187680]
AY261360.1_Kenya_1950            ...................a.t...............................................t.......... [187680]
AM712240.1_OURT_88/3             ......c......................................................................... [187680]
AM712239.1_Benin_97/1            ......c......................................................................... [187680]
KM111295.1_Ken06.Bus             ...................a..............................................a..t.......... [187680]
KM111294.1_Ken05/Tk1             ...................a.t...............................................t.......... [187680]
KM262844.1_strain_L60            ......c......................................................................... [187680]
KM102979.1_26544/OG10_Italy      ......c......................................................................... [187680]
KM262845.1_strain_NHV            ......c......................................................................... [187680]
KX354450.1_47/Ss/2008            ......c......................................................................... [187680]
KP055815.1_strain_BA71           ......c......................................................................... [187680]
MH025920.1_strain_R35            ...................a............................a.................a..t.......... [187680]
MH025919.1_strain_N10            ...................a............................a.................a..t.......... [187680]
MH025918.1_strain_R25            ...................a............................a.................a..t.......... [187680]
MH025917.1_strain_R7             ...................a............................a.................a..t.......... [187680]
MH025916.1_strain_R8             ...................a............................a.................a..t.......... [187680]

FR682468.2_Georgia_2007/1        aaagcatggtatag---agagcttttagaaccggagttcgttaaaaaaatacaaagttcgtttaaaacgt-gtaatgtta [187760]
ASF_IND_20_CAD543                .......r......---.....................................................-aag...... [187760]
LR899193.1_Germany_2020/1        ..............---.....................................................-......... [187760]
MN393477.1_Wuhan_2019            ..............---.....................................................-......... [187760]
MW396979.1_Timor-Leste/2019/1    ..............---.....................................................-......... [187760]
MK940252.1_CN/2019/InnerMongolia ..............---.....................................................-......... [187760]
MK128995.1_China/2018/AnhuiXCGQ  ..............---.....................................................-......... [187760]
MH681419.1_ASFV/POL/2015         ..............---.....................................................-......... [187760]
LS478113.1_Estonia_2014          ..............---.....................................................-......... [187760]
KP843857.1_Odintsovo_02/14       ..............---.....................................................-......... [187760]
FN557520.1_strain_E75            .......a......---.....................................................-aag...... [187760]
AY261366.1_Warthog               .......t......aga..............t......................................-cc....... [187760]
AY261365.1_Warmbaths             .......a......---.........................-...........................-......... [187760]
AY261364.1_Tengani_62            .......t......ag-..............t...................g..................-......... [187760]
AY261363.1_Pretoriuskop/96/4     g......t......---..............t...................................t..-cc....... [187760]
AY261362.1_Mkuzi_1979            .......a......---.....................................................-aag...... [187760]
AY261361.1_Malawi_Lil-20/1_1983  .......c......---..................................g...............g..-..g...... [187760]
AY261360.1_Kenya_1950            ...........c.a---t.............-....a..............g...............t..g..g.....g [187760]
AM712240.1_OURT_88/3             .......a......---.....................................................-aag...... [187760]
AM712239.1_Benin_97/1            .......a......---.....................................................-aag...... [187760]
KM111295.1_Ken06.Bus             .......a...c.a---ta............-....a..............g...............t..g..g.....g [187760]
```

```
KM111294.1_Ken05/Tk1            ...........c.a---t.............-....a..............g...............t..g..g.....g [187760]
KM262844.1_strain_L60           .......a......---.....................................................-aag...... [187760]
KM102979.1_26544/OG10_Italy     .......a......---.................g...................................-aag...... [187760]
KM262845.1_strain_NHV           .......a......---.....................................................-aag...... [187760]
KX354450.1_47/Ss/2008           .......a......---.................g...................................-aag...... [187760]
KP055815.1_strain_BA71          .......a......---.....................................................-aag...... [187760]
MH025920.1_strain_R35           .......a...c.a---ta............-....a..............g...............t..g..g.....g [187760]
MH025919.1_strain_N10           .......a...c.a---ta............-....a..............g...............t..g..g.....g [187760]
MH025918.1_strain_R25           .......a...c.a---ta............-....a..............g...............t..g..g.....g [187760]
MH025917.1_strain_R7            .......a...c.a---ta............-....a..............g...............t..g..g.....g [187760]
MH025916.1_strain_R8            .......a...c.a---ta............-....a..............g...............t..g..g.....g [187760]

FR682468.2_Georgia_2007/1       ctaaaaaaatgtaatgtttaaatgataatgataccacatgcattaatgaaaaaaacttttaaatttttgttttaatattt [187840]
ASF_IND_20_CAD543               ..............y................................................................. [187840]
LR899193.1_Germany_2020/1       ................................................................................ [187840]
MN393477.1_Wuhan_2019           ................................................................................ [187840]
MW396979.1_Timor-Leste/2019/1   ................................................................................ [187840]
MK940252.1_CN/2019/InnerMongolia ................................................................................ [187840]
MK128995.1_China/2018/AnhuiXCGQ ................................................................................ [187840]
MH681419.1_ASFV/POL/2015        ................................................................................ [187840]
LS478113.1_Estonia_2014         ................................................................................ [187840]
KP843857.1_Odintsovo_02/14      ................................................................................ [187840]
FN557520.1_strain_E75           ..............c..............a.................................................. [187840]
AY261366.1_Warthog              .................................t.............................................. [187840]
AY261365.1_Warmbaths            .................................t.............................................. [187840]
AY261364.1_Tengani_62           .................................t.............................................. [187840]
AY261363.1_Pretoriuskop/96/4    ..-..............................t.............................................. [187840]
AY261362.1_Mkuzi_1979           ..............c................................................................. [187840]
AY261361.1_Malawi_Lil-20/1_1983 ...............................................a................................ [187840]
AY261360.1_Kenya_1950           ..-..........g..................ct.............a................................ [187840]
AM712240.1_OURT_88/3            ..............c..............a.................................................. [187840]
AM712239.1_Benin_97/1           ..............c..............a.................................................. [187840]
KM111295.1_Ken06.Bus            ..-.........g...................ct.............a................................ [187840]
KM111294.1_Ken05/Tk1            ..-..........g..................ct.............a................................ [187840]
KM262844.1_strain_L60           ..............c..............a.................................................. [187840]
KM102979.1_26544/OG10_Italy     ..............c..............a.................................................. [187840]
KM262845.1_strain_NHV           ..............c..............a.................................................. [187840]
KX354450.1_47/Ss/2008           ..............c..............a.................................................. [187840]
KP055815.1_strain_BA71          ..............c..............a.................................................. [187840]
MH025920.1_strain_R35           ..-.........g...................ct.............a................................ [187840]
MH025919.1_strain_N10           ..-.........g...................ct.............a................................ [187840]
MH025918.1_strain_R25           ..-.........g...................ct.............a................................ [187840]
MH025917.1_strain_R7            ..-.........g...................ct.............a................................ [187840]
MH025916.1_strain_R8            ..-.........g...................ct.............a................................ [187840]
```

```
FR682468.2_Georgia_2007/1        gcatgaaaatggaaacatttttagtctgtttatttcacaatgcagatggtttacatcaacagattcaggaaattttgtat [187920]
ASF_IND_20_CAD543                ................................................................................ [187920]
LR899193.1_Germany_2020/1        ................................................................................ [187920]
MN393477.1_Wuhan_2019            ................................................................................ [187920]
MW396979.1_Timor-Leste/2019/1    ................................................................................ [187920]
MK940252.1_CN/2019/InnerMongolia ................................................................................ [187920]
MK128995.1_China/2018/AnhuiXCGQ  ................................................................................ [187920]
MH681419.1_ASFV/POL/2015         ................................................................................ [187920]
LS478113.1_Estonia_2014          ................................................................................ [187920]
KP843857.1_Odintsovo_02/14       ................................................................................ [187920]
FN557520.1_strain_E75            ...........................................t.................................... [187920]
AY261366.1_Warthog               ...........................................c....................c..a............ [187920]
AY261365.1_Warmbaths             ...........................................t....................c..a............ [187920]
AY261364.1_Tengani_62            ...........................................t.....g..............c..a............ [187920]
AY261363.1_Pretoriuskop/96/4     ...........................................c....................c..a............ [187920]
AY261362.1_Mkuzi_1979            ...........................................t.................................... [187920]
AY261361.1_Malawi_Lil-20/1_1983  ...........................................ta.a....................a............ [187920]
AY261360.1_Kenya_1950            .....................................t.....t.c.....................a............ [187920]
AM712240.1_OURT_88/3             ...........................................t.................................... [187920]
AM712239.1_Benin_97/1            ...........................................t.................................... [187920]
KM111295.1_Ken06.Bus             .....................................t.....t.c........g............a............ [187920]
KM111294.1_Ken05/Tk1             .....................................t.....t.c.....................a............ [187920]
KM262844.1_strain_L60            ...........................................t.................................... [187920]
KM102979.1_26544/OG10_Italy      ...........................................t.................................... [187920]
KM262845.1_strain_NHV            ...........................................t.................................... [187920]
KX354450.1_47/Ss/2008            ...........................................t.................................... [187920]
KP055815.1_strain_BA71           ...........................................t.................................... [187920]
MH025920.1_strain_R35            .....................................t.....t.c........g............a............ [187920]
MH025919.1_strain_N10            .....................................t.....t.c........g............a............ [187920]
MH025918.1_strain_R25            .....................................t.....t.c........g............a............ [187920]
MH025917.1_strain_R7             .....................................t.....t.c........g............a............ [187920]
MH025916.1_strain_R8             .....................................t.....t.c........g............a............ [187920]

FR682468.2_Georgia_2007/1        ttattgcggatgcatatttacgaaacaaatctttacttaaagcaggaactatcacggcttatatatccaaataggcaact [188000]
ASF_IND_20_CAD543                ................................................................................ [188000]
LR899193.1_Germany_2020/1        ................................................................................ [188000]
MN393477.1_Wuhan_2019            ................................................................................ [188000]
MW396979.1_Timor-Leste/2019/1    ................................................................................ [188000]
MK940252.1_CN/2019/InnerMongolia ................................................................................ [188000]
MK128995.1_China/2018/AnhuiXCGQ  ................................................................................ [188000]
MH681419.1_ASFV/POL/2015         ................................................................................ [188000]
LS478113.1_Estonia_2014          ................................................................................ [188000]
KP843857.1_Odintsovo_02/14       ................................................................................ [188000]
```

```
FN557520.1_strain_E75            ......................t..g.....................................................g.. [188000]
AY261366.1_Warthog               .............................................................................g.. [188000]
AY261365.1_Warmbaths             .............................................................................g.. [188000]
AY261364.1_Tengani_62            ....................ta.......................................................g.. [188000]
AY261363.1_Pretoriuskop/96/4     .............................................................................g.. [188000]
AY261362.1_Mkuzi_1979            ....................t..g.....................................................g.. [188000]
AY261361.1_Malawi_Lil-20/1_1983  ........a...........t..........................g.......a...............c.....g.. [188000]
AY261360.1_Kenya_1950            c..c..........c................................g.................c...........g.. [188000]
AM712240.1_OURT_88/3             ....................t..g.....................................................g.. [188000]
AM712239.1_Benin_97/1            ....................t..g.....................................................g.. [188000]
KM111295.1_Ken06.Bus             c..c..........c................................g.................c...........g.. [188000]
KM111294.1_Ken05/Tk1             c..c..........c................................g.................c...........g.. [188000]
KM262844.1_strain_L60            ....................t..g.....................................................g.. [188000]
KM102979.1_26544/OG10_Italy      ....................t..g.....................................................g.. [188000]
KM262845.1_strain_NHV            ....................t..g.....................................................g.. [188000]
KX354450.1_47/Ss/2008            ....................t..g.....................................................g.. [188000]
KP055815.1_strain_BA71           ....................t..g.....................................................g.. [188000]
MH025920.1_strain_R35            c..c..........c................................g.................c...........g.. [188000]
MH025919.1_strain_N10            c..c..........c................................g.................c...........g.. [188000]
MH025918.1_strain_R25            c..c..........c................................g.................c...........g.. [188000]
MH025917.1_strain_R7             c..c..........c................................g.................c...........g.. [188000]
MH025916.1_strain_R8             c..c..........c................................g.................c...........g.. [188000]

FR682468.2_Georgia_2007/1        ttcttttgtgttacttatgcccctttcccttctaagaaactgggatgacattgaatatttaacggacgttgtagatgata [188080]
ASF_IND_20_CAD543                ................................................................................ [188080]
LR899193.1_Germany_2020/1        ................................................................................ [188080]
MN393477.1_Wuhan_2019            ................................................................................ [188080]
MW396979.1_Timor-Leste/2019/1    ................................................................................ [188080]
MK940252.1_CN/2019/InnerMongolia ................................................................................ [188080]
MK128995.1_China/2018/AnhuiXCGQ  ................................................................................ [188080]
MH681419.1_ASFV/POL/2015         ................................................................................ [188080]
LS478113.1_Estonia_2014          ................................................................................ [188080]
KP843857.1_Odintsovo_02/14       ................................................................................ [188080]
FN557520.1_strain_E75            ............................................................g..............c.... [188080]
AY261366.1_Warthog               ............................................................g..............c.... [188080]
AY261365.1_Warmbaths             .............................................c..............g..............c.... [188080]
AY261364.1_Tengani_62            ............................................................g..............c.... [188080]
AY261363.1_Pretoriuskop/96/4     ............................................................g..............c.... [188080]
AY261362.1_Mkuzi_1979            .......................................t....................g..............c.... [188080]
AY261361.1_Malawi_Lil-20/1_1983  .....................t.........t...................a........g......a............ [188080]
AY261360.1_Kenya_1950            .....................t.........t............................g..c...a............ [188080]
AM712240.1_OURT_88/3             ............................................................g..............c.... [188080]
AM712239.1_Benin_97/1            ............................................................g..............c.... [188080]
KM111295.1_Ken06.Bus             ..........c..........t.........t............................g..c...a............ [188080]
```

```
KM111294.1_Ken05/Tk1           .....................t.........t............................g..c...a............ [188080]
KM262844.1_strain_L60          ............................................................g..............c.... [188080]
KM102979.1_26544/OG10_Italy    ............................................................g..............c.... [188080]
KM262845.1_strain_NHV          ............................................................g..............c.... [188080]
KX354450.1_47/Ss/2008          ............................................................g..............c.... [188080]
KP055815.1_strain_BA71         ............................................................g..............c.... [188080]
MH025920.1_strain_R35          ..........c..........t.........t...............g............g..c...a............ [188080]
MH025919.1_strain_N10          ..........c..........t.........t...............g............g..c...a............ [188080]
MH025918.1_strain_R25          ..........c..........t.........t...............g............g..c...a............ [188080]
MH025917.1_strain_R7           ..........c..........t.........t...............g............g..c...a............ [188080]
MH025916.1_strain_R8           ..........c..........t.........t...............g............g..c...a............ [188080]

FR682468.2_Georgia_2007/1      agcagactctacattacgcggcaaatttgctgacaaactacgttctacatctatccatgtttcaaaagctgacaaaacca [188160]
ASF_IND_20_CAD543              ................................................................................ [188160]
LR899193.1_Germany_2020/1      ................................................................................ [188160]
MN393477.1_Wuhan_2019          ................................................................................ [188160]
MW396979.1_Timor-Leste/2019/1  ................................................................................ [188160]
MK940252.1_CN/2019/InnerMongolia ................................................................................ [188160]
MK128995.1_China/2018/AnhuiXCGQ ................................................................................ [188160]
MH681419.1_ASFV/POL/2015       ................................................................................ [188160]
LS478113.1_Estonia_2014        ................................................................................ [188160]
KP843857.1_Odintsovo_02/14     ................................................................................ [188160]
FN557520.1_strain_E75          ................................................................................ [188160]
AY261366.1_Warthog             ................................................................................ [188160]
AY261365.1_Warmbaths           ..................................................t....t........................ [188160]
AY261364.1_Tengani_62          ..........t..................................................................... [188160]
AY261363.1_Pretoriuskop/96/4   ................................................................................ [188160]
AY261362.1_Mkuzi_1979          ................................................................................ [188160]
AY261361.1_Malawi_Lil-20/1_1983 ............................................................a.......t.a.......a. [188160]
AY261360.1_Kenya_1950          ....................................t.......................a................... [188160]
AM712240.1_OURT_88/3           ................................................................................ [188160]
AM712239.1_Benin_97/1          ................................................................................ [188160]
KM111295.1_Ken06.Bus           ....a...............................t.......................a................... [188160]
KM111294.1_Ken05/Tk1           ....................................t.......................a................... [188160]
KM262844.1_strain_L60          ................................................................................ [188160]
KM102979.1_26544/OG10_Italy    ................................................................................ [188160]
KM262845.1_strain_NHV          ................................................................................ [188160]
KX354450.1_47/Ss/2008          ................................................................................ [188160]
KP055815.1_strain_BA71         ................................................................................ [188160]
MH025920.1_strain_R35          ....a...............................t.......................a................... [188160]
MH025919.1_strain_N10          ....a...............................t.......................a................... [188160]
MH025918.1_strain_R25          ....a...............................t.......................a................... [188160]
MH025917.1_strain_R7           ....a...............................t.......................a................... [188160]
MH025916.1_strain_R8           ....a...............................t.......................a................... [188160]
```

```
FR682468.2_Georgia_2007/1        tacttccttttagcggtcaagcgggtcagcgaaaaactcaacaaaaagcagcgacattcattttacgaggtattggtaac [188240]
ASF_IND_20_CAD543                ................................................................................ [188240]
LR899193.1_Germany_2020/1        ................................................................................ [188240]
MN393477.1_Wuhan_2019            ................................................................................ [188240]
MW396979.1_Timor-Leste/2019/1    ................................................................................ [188240]
MK940252.1_CN/2019/InnerMongolia ................................................................................ [188240]
MK128995.1_China/2018/AnhuiXCGQ  ................................................................................ [188240]
MH681419.1_ASFV/POL/2015         ................................................................................ [188240]
LS478113.1_Estonia_2014          ................................................................................ [188240]
KP843857.1_Odintsovo_02/14       ................................................................................ [188240]
FN557520.1_strain_E75            .................a............................g................................. [188240]
AY261366.1_Warthog               .................a..................................a........................... [188240]
AY261365.1_Warmbaths             ..t..............a............................g................................. [188240]
AY261364.1_Tengani_62            .................a...........................................................g.. [188240]
AY261363.1_Pretoriuskop/96/4     .................a.............................................................. [188240]
AY261362.1_Mkuzi_1979            .................a............................g................................. [188240]
AY261361.1_Malawi_Lil-20/1_1983  ..t..............a..................................a............t......c.a..... [188240]
AY261360.1_Kenya_1950            ..t..............a...................................g....................a..... [188240]
AM712240.1_OURT_88/3             .................a............................g................................. [188240]
AM712239.1_Benin_97/1            .................a............................g................................. [188240]
KM111295.1_Ken06.Bus             ..t..............a...................................g....................a..... [188240]
KM111294.1_Ken05/Tk1             ..t..............a...................................g....................a..... [188240]
KM262844.1_strain_L60            .................a............................g................................. [188240]
KM102979.1_26544/OG10_Italy      .................a............................g................................. [188240]
KM262845.1_strain_NHV            .................a............................g................................. [188240]
KX354450.1_47/Ss/2008            .................a............................g................................. [188240]
KP055815.1_strain_BA71           .................a............................g................................. [188240]
MH025920.1_strain_R35            ..t..............a...................................g....................a..... [188240]
MH025919.1_strain_N10            ..t..............a...................................g....................a..... [188240]
MH025918.1_strain_R25            ..t..............a...................................g....................a..... [188240]
MH025917.1_strain_R7             ..t..............a...................................g....................a..... [188240]
MH025916.1_strain_R8             ..t..............a...................................g....................a..... [188240]

FR682468.2_Georgia_2007/1        ctccgaaaccttgaataattatgaaaacctatctaaaaacattttaaatacgttgatgtttgccgtgcgctacgtattta [188320]
ASF_IND_20_CAD543                ................................................................................ [188320]
LR899193.1_Germany_2020/1        ................................................................................ [188320]
MN393477.1_Wuhan_2019            ................................................................................ [188320]
MW396979.1_Timor-Leste/2019/1    ................................................................................ [188320]
MK940252.1_CN/2019/InnerMongolia ................................................................................ [188320]
MK128995.1_China/2018/AnhuiXCGQ  ................................................................................ [188320]
MH681419.1_ASFV/POL/2015         ................................................................................ [188320]
LS478113.1_Estonia_2014          ................................................................................ [188320]
KP843857.1_Odintsovo_02/14       ................................................................................ [188320]
```

```
FN557520.1_strain_E75            ...t............................................................................ [188320]
AY261366.1_Warthog               ................................................................................ [188320]
AY261365.1_Warmbaths             ...t........a................................................................... [188320]
AY261364.1_Tengani_62            ...............................g................................................ [188320]
AY261363.1_Pretoriuskop/96/4     ................................................................................ [188320]
AY261362.1_Mkuzi_1979            ...t............................................................................ [188320]
AY261361.1_Malawi_Lil-20/1_1983  .......g.......................c.......t........................................ [188320]
AY261360.1_Kenya_1950            .........a........c............c.............g.................................. [188320]
AM712240.1_OURT_88/3             ...t............................................................................ [188320]
AM712239.1_Benin_97/1            ...t............................................................................ [188320]
KM111295.1_Ken06.Bus             .........a........c............c.............g.................................. [188320]
KM111294.1_Ken05/Tk1             .........a........c............c.............g.................................. [188320]
KM262844.1_strain_L60            ...t............................................................................ [188320]
KM102979.1_26544/OG10_Italy      ...t............................................................................ [188320]
KM262845.1_strain_NHV            ...t............................................................................ [188320]
KX354450.1_47/Ss/2008            ...t............................................................................ [188320]
KP055815.1_strain_BA71           ...t............................................................................ [188320]
MH025920.1_strain_R35            .........a........c............c.............g.................................. [188320]
MH025919.1_strain_N10            .........a........c............c.............g.................................. [188320]
MH025918.1_strain_R25            .........a........c............c.............g.................................. [188320]
MH025917.1_strain_R7             .........a........c............c.............g.................................. [188320]
MH025916.1_strain_R8             .........a........c............c.............g.................................. [188320]

FR682468.2_Georgia_2007/1        aacctacgccgaactattcagaaattctcgcagagttggaaaaaaaaaataaaattcaccatattatttttaatatggta [188400]
ASF_IND_20_CAD543                ................................................................................ [188400]
LR899193.1_Germany_2020/1        ................................................................................ [188400]
MN393477.1_Wuhan_2019            ................................................................................ [188400]
MW396979.1_Timor-Leste/2019/1    ................................................................................ [188400]
MK940252.1_CN/2019/InnerMongolia ................................................................................ [188400]
MK128995.1_China/2018/AnhuiXCGQ  ................................................................................ [188400]
MH681419.1_ASFV/POL/2015         ................................................................................ [188400]
LS478113.1_Estonia_2014          ................................................................................ [188400]
KP843857.1_Odintsovo_02/14       .......................................-........................................ [188400]
FN557520.1_strain_E75            .......a........................................................................ [188400]
AY261366.1_Warthog               .......a........................................................................ [188400]
AY261365.1_Warmbaths             .......a........................................................................ [188400]
AY261364.1_Tengani_62            .......a.............................a.......................................... [188400]
AY261363.1_Pretoriuskop/96/4     .......a........................................................................ [188400]
AY261362.1_Mkuzi_1979            .......a........................................................................ [188400]
AY261361.1_Malawi_Lil-20/1_1983  .t.....a..............................a......................................... [188400]
AY261360.1_Kenya_1950            .......a...........g......a.........................g....................c.....g [188400]
AM712240.1_OURT_88/3             .......a........................................................................ [188400]
AM712239.1_Benin_97/1            .......a...................................................t.................... [188400]
KM111295.1_Ken06.Bus             .......a...........g......a.........................g....................c.....g [188400]
```

```
KM111294.1_Ken05/Tk1               .......a...........g......a.........................g....................c.....g [188400]
KM262844.1_strain_L60              .......a........................................................................ [188400]
KM102979.1_26544/OG10_Italy        .......a........................................................................ [188400]
KM262845.1_strain_NHV              .......a........................................................................ [188400]
KX354450.1_47/Ss/2008              .......a........................................................................ [188400]
KP055815.1_strain_BA71             .......a........................................................................ [188400]
MH025920.1_strain_R35              .......a...........g......a.........................g....................c.....g [188400]
MH025919.1_strain_N10              .......a...........g......a.........................g....................c.....g [188400]
MH025918.1_strain_R25              .......a...........g......a.........................g....................c.....g [188400]
MH025917.1_strain_R7               .......a...........g......a.........................g....................c.....g [188400]
MH025916.1_strain_R8               .......a...........g......a.........................g....................c.....g [188400]

FR682468.2_Georgia_2007/1          attacggattttgcgcaaatccgtgaacaacaaatggataaacatctgtgtgaaacaaataatgagcttcgtcaggaatg [188480]
ASF_IND_20_CAD543                  ................................................................................ [188480]
LR899193.1_Germany_2020/1          ................................................................................ [188480]
MN393477.1_Wuhan_2019              ................................................................................ [188480]
MW396979.1_Timor-Leste/2019/1      ................................................................................ [188480]
MK940252.1_CN/2019/InnerMongolia   ................................................................................ [188480]
MK128995.1_China/2018/AnhuiXCGQ    ................................................................................ [188480]
MH681419.1_ASFV/POL/2015           ................................................................................ [188480]
LS478113.1_Estonia_2014            ................................................................................ [188480]
KP843857.1_Odintsovo_02/14         ................................................................................ [188480]
FN557520.1_strain_E75              ...g....................a.........................c............................. [188480]
AY261366.1_Warthog                 .....a.........................................................a................ [188480]
AY261365.1_Warmbaths               .....a.......................................................................... [188480]
AY261364.1_Tengani_62              ................................................................................ [188480]
AY261363.1_Pretoriuskop/96/4       .....a.......................................................................... [188480]
AY261362.1_Mkuzi_1979              ...g....................a.........................c............................. [188480]
AY261361.1_Malawi_Lil-20/1_1983    ............ca......a...............tg.......................................... [188480]
AY261360.1_Kenya_1950              ..c.........ca..................c...tg.......................................... [188480]
AM712240.1_OURT_88/3               ...g....................a.........................c............................. [188480]
AM712239.1_Benin_97/1              ...g....................a.........................c............................. [188480]
KM111295.1_Ken06.Bus               ..c.........ca..................c...tg.....................c.................... [188480]
KM111294.1_Ken05/Tk1               ..c.........ca..................c...tg.......................................... [188480]
KM262844.1_strain_L60              ...g....................a.........................c............................. [188480]
KM102979.1_26544/OG10_Italy        ...g....................a.........................c............................. [188480]
KM262845.1_strain_NHV              ...g....................a.........................c............................. [188480]
KX354450.1_47/Ss/2008              ...g....................a.........................c............................. [188480]
KP055815.1_strain_BA71             ...g....................a.........................c............................. [188480]
MH025920.1_strain_R35              ..c.........ca..................c...tg.....................c.................... [188480]
MH025919.1_strain_N10              ..c.........ca..................c...tg.....................c.................... [188480]
MH025918.1_strain_R25              ..c.........ca..................c...tg.....................c.................... [188480]
MH025917.1_strain_R7               ..c.........ca..................c...tg.....................c.................... [188480]
MH025916.1_strain_R8               ..c.........ca..................c...tg.....................c.................... [188480]
```

```
FR682468.2_Georgia_2007/1        taaagaaactatttttgatttaaaggtggtaggaaatgt-----ttagccaataaactcatgcccgcattttttacaggt [188560]
ASF_IND_20_CAD543                .......................................-----.................................... [188560]
LR899193.1_Germany_2020/1        .......................................-----.................................... [188560]
MN393477.1_Wuhan_2019            .......................................-----.................................... [188560]
MW396979.1_Timor-Leste/2019/1    .......................................-----.................................... [188560]
MK940252.1_CN/2019/InnerMongolia .......................................-----.................................... [188560]
MK128995.1_China/2018/AnhuiXCGQ  .......................................-----.................................... [188560]
MH681419.1_ASFV/POL/2015         .......................................-----.................................... [188560]
LS478113.1_Estonia_2014          .......................................-----.................................... [188560]
KP843857.1_Odintsovo_02/14       .......................................-----.................................... [188560]
FN557520.1_strain_E75            .......................................-----.................................... [188560]
AY261366.1_Warthog               .......................................ttagc.................................... [188560]
AY261365.1_Warmbaths             .......................................-----.................................... [188560]
AY261364.1_Tengani_62            .......................................-----.................................... [188560]
AY261363.1_Pretoriuskop/96/4     .......................................ttagc.................................... [188560]
AY261362.1_Mkuzi_1979            .......................................-----.................................... [188560]
AY261361.1_Malawi_Lil-20/1_1983  c.......t..............................-----.................................... [188560]
AY261360.1_Kenya_1950            c.......t..............................-----...............g.................... [188560]
AM712240.1_OURT_88/3             .......................................-----.................................... [188560]
AM712239.1_Benin_97/1            .......................................-----.................................... [188560]
KM111295.1_Ken06.Bus             c.......t..............................-----...............g.................... [188560]
KM111294.1_Ken05/Tk1             c.......t..............................-----...............g.................... [188560]
KM262844.1_strain_L60            .......................................-----.................................... [188560]
KM102979.1_26544/OG10_Italy      .......................................-----.................................... [188560]
KM262845.1_strain_NHV            .......................................-----.................................... [188560]
KX354450.1_47/Ss/2008            .......................................-----.................................... [188560]
KP055815.1_strain_BA71           .......................................-----.................................... [188560]
MH025920.1_strain_R35            c.......t..............................-----...............g.................... [188560]
MH025919.1_strain_N10            c.......t..............................-----...............g.................... [188560]
MH025918.1_strain_R25            c.......t..............................-----...............g.................... [188560]
MH025917.1_strain_R7             c.......t..............................-----...............g.................... [188560]
MH025916.1_strain_R8             c.......t..............................-----...............g.................... [188560]

FR682468.2_Georgia_2007/1        acaaaatatcgtggatggctcatcgagggcgcgtgtttgtacttctctgtaggtacacatacgctgcttgcagttgggac [188640]
ASF_IND_20_CAD543                ................................................................................ [188640]
LR899193.1_Germany_2020/1        ................................................................................ [188640]
MN393477.1_Wuhan_2019            ................................................................................ [188640]
MW396979.1_Timor-Leste/2019/1    ................................................................................ [188640]
MK940252.1_CN/2019/InnerMongolia ................................................................................ [188640]
MK128995.1_China/2018/AnhuiXCGQ  ................................................................................ [188640]
MH681419.1_ASFV/POL/2015         ................................................................................ [188640]
LS478113.1_Estonia_2014          ................................................................................ [188640]
KP843857.1_Odintsovo_02/14       ................................................................................ [188640]
```

```
FN557520.1_strain_E75             ...............a......................................g......................... [188640]
AY261366.1_Warthog                ............t..a..............................................................g. [188640]
AY261365.1_Warmbaths              ...............a..............................................................g. [188640]
AY261364.1_Tengani_62             ............t..a................................a............................... [188640]
AY261363.1_Pretoriuskop/96/4      ............t..a................................................................ [188640]
AY261362.1_Mkuzi_1979             ...............a................................................................ [188640]
AY261361.1_Malawi_Lil-20/1_1983   ............t..a..............................................................g. [188640]
AY261360.1_Kenya_1950             ....t....g..t..a................................a..a..........................g. [188640]
AM712240.1_OURT_88/3              ...............a......................................g......................... [188640]
AM712239.1_Benin_97/1             ...............a......................................g......................... [188640]
KM111295.1_Ken06.Bus              ....t....a..t..a................................a..a..........................g. [188640]
KM111294.1_Ken05/Tk1              ....t....a..t..a................................a..a..........................g. [188640]
KM262844.1_strain_L60             ...............a......................................g......................... [188640]
KM102979.1_26544/OG10_Italy       ...............a......................................g......................... [188640]
KM262845.1_strain_NHV             ...............a......................................g......................... [188640]
KX354450.1_47/Ss/2008             ...............a......................................g......................... [188640]
KP055815.1_strain_BA71            ...............a......................................g......................... [188640]
MH025920.1_strain_R35             ....t....a..t..a................................a..a..........................g. [188640]
MH025919.1_strain_N10             ....t....a..t..a................................a..a..........................g. [188640]
MH025918.1_strain_R25             ....t....a..t..a................................a..a..........................g. [188640]
MH025917.1_strain_R7              ....t....a..t..a................................a..a..........................g. [188640]
MH025916.1_strain_R8              ....t....a..t..a................................a..a..........................g. [188640]

FR682468.2_Georgia_2007/1         acttataaagttgtgacgtcttttcggcgaccttttgctgcgaacgtagagtaatttctgtcttctcctttaaggcggca [188720]
ASF_IND_20_CAD543                 ................................................................................ [188720]
LR899193.1_Germany_2020/1         ................................................................................ [188720]
MN393477.1_Wuhan_2019             ................................................................................ [188720]
MW396979.1_Timor-Leste/2019/1     ................................................................................ [188720]
MK940252.1_CN/2019/InnerMongolia  ................................................................................ [188720]
MK128995.1_China/2018/AnhuiXCGQ   ................................................................................ [188720]
MH681419.1_ASFV/POL/2015          ................................................................................ [188720]
LS478113.1_Estonia_2014           ................................................................................ [188720]
KP843857.1_Odintsovo_02/14        ................................................................................ [188720]
FN557520.1_strain_E75             ...............................t................................................ [188720]
AY261366.1_Warthog                ................t..................................................t............ [188720]
AY261365.1_Warmbaths              ...............................t................................................ [188720]
AY261364.1_Tengani_62             ................t..............t................................................ [188720]
AY261363.1_Pretoriuskop/96/4      ...............................t...................................t.....t..a... [188720]
AY261362.1_Mkuzi_1979             ...............................t................................................ [188720]
AY261361.1_Malawi_Lil-20/1_1983   ...............................t.................t.......................t...... [188720]
AY261360.1_Kenya_1950             ...............................t.................t.......................t...... [188720]
AM712240.1_OURT_88/3              ...............................t................................................ [188720]
AM712239.1_Benin_97/1             ...............................t................................................ [188720]
KM111295.1_Ken06.Bus              ................t..............t.................t.......................t...... [188720]
```

```
KM111294.1_Ken05/Tk1             ...............................t.................t.......................t...... [188720]
KM262844.1_strain_L60            ...............................t................................................ [188720]
KM102979.1_26544/OG10_Italy      ...............................t................................................ [188720]
KM262845.1_strain_NHV            ...............................t................................................ [188720]
KX354450.1_47/Ss/2008            ...............................t................................................ [188720]
KP055815.1_strain_BA71           ...............................t................................................ [188720]
MH025920.1_strain_R35            ................t..............t.................t.......................t...... [188720]
MH025919.1_strain_N10            ................t..............t.................t.......................t...... [188720]
MH025918.1_strain_R25            ................t..............t.................t.......................t...... [188720]
MH025917.1_strain_R7             ................t..............t.................t.......................t...... [188720]
MH025916.1_strain_R8             ................t..............t.................t.......................t...... [188720]

FR682468.2_Georgia_2007/1        gaggggcaaagctcggcgaacgtcatg-ctaccaattgcctccggttttagctcgccagaaattagcttattaagggcat [188800]
ASF_IND_20_CAD543                ...........................-.................................................... [188800]
LR899193.1_Germany_2020/1        ...........................-.................................................... [188800]
MN393477.1_Wuhan_2019            ...........................-.................................................... [188800]
MW396979.1_Timor-Leste/2019/1    ...........................-.................................................... [188800]
MK940252.1_CN/2019/InnerMongolia ...........................-.................................................... [188800]
MK128995.1_China/2018/AnhuiXCGQ  ...........................-.................................................... [188800]
MH681419.1_ASFV/POL/2015         ...........................-.................................................... [188800]
LS478113.1_Estonia_2014          ...........................-.................................................... [188800]
KP843857.1_Odintsovo_02/14       ...........................-.................................................... [188800]
FN557520.1_strain_E75            .....a.....................-.................................................... [188800]
AY261366.1_Warthog               ...........................-.................................................... [188800]
AY261365.1_Warmbaths             ...........................-.................................................... [188800]
AY261364.1_Tengani_62            ...........................-.................................................... [188800]
AY261363.1_Pretoriuskop/96/4     ...........................-.................................................... [188800]
AY261362.1_Mkuzi_1979            ...........................-.................................................... [188800]
AY261361.1_Malawi_Lil-20/1_1983  ..a..a.....................c..........-......................................... [188800]
AY261360.1_Kenya_1950            .................a.........-.............................g..g................... [188800]
AM712240.1_OURT_88/3             .....a.....................-.................................................... [188800]
AM712239.1_Benin_97/1            .....a.....................-.................................................... [188800]
KM111295.1_Ken06.Bus             ..a........t.....a.........-.............................g..g................... [188800]
KM111294.1_Ken05/Tk1             .................a.........-.............................g..g................... [188800]
KM262844.1_strain_L60            .....a.....................-.................................................... [188800]
KM102979.1_26544/OG10_Italy      .....a.....................-.................................................... [188800]
KM262845.1_strain_NHV            .....a.....................-.................................................... [188800]
KX354450.1_47/Ss/2008            .....a.....................-.................................................... [188800]
KP055815.1_strain_BA71           .....a.....................-.................................................... [188800]
MH025920.1_strain_R35            ..a........t.....a.........-.............................g..t................... [188800]
MH025919.1_strain_N10            ..a........t.....a.........-.............................g..g................... [188800]
MH025918.1_strain_R25            ..a........t.....a.........-.............................g..t................... [188800]
MH025917.1_strain_R7             ..a........t.....a.........-.............................g..t................... [188800]
MH025916.1_strain_R8             ..a........t.....a.........-.............................g..t................... [188800]
```

```
FR682468.2_Georgia_2007/1        cgttatcctgttgttggtgactttttttttcgcagttaataatatgattgatcgtcccacaacgggttgaatattcttct [188880]
ASF_IND_20_CAD543                .............y.................................................................. [188880]
LR899193.1_Germany_2020/1        ................................................................................ [188880]
MN393477.1_Wuhan_2019            ................................................................................ [188880]
MW396979.1_Timor-Leste/2019/1    ................................................................................ [188880]
MK940252.1_CN/2019/InnerMongolia ................................................................................ [188880]
MK128995.1_China/2018/AnhuiXCGQ  ................................................................................ [188880]
MH681419.1_ASFV/POL/2015         ................................................................................ [188880]
LS478113.1_Estonia_2014          ................................................................................ [188880]
KP843857.1_Odintsovo_02/14       ................................................................................ [188880]
FN557520.1_strain_E75            ..........c..c....................a............................................. [188880]
AY261366.1_Warthog               ..........c..c......................................a........t.................. [188880]
AY261365.1_Warmbaths             ..........c..c....................a............................................. [188880]
AY261364.1_Tengani_62            ..........c..c..........................g...........a........................... [188880]
AY261363.1_Pretoriuskop/96/4     ..........c..c......................................a........................... [188880]
AY261362.1_Mkuzi_1979            .......t..c..c....................a............................................. [188880]
AY261361.1_Malawi_Lil-20/1_1983  ..........c..c....................a............................................. [188880]
AY261360.1_Kenya_1950            .a........c..c....................a..................a.......................... [188880]
AM712240.1_OURT_88/3             ..........c..c....................a............................................. [188880]
AM712239.1_Benin_97/1            ..........c..c....................a............................................. [188880]
KM111295.1_Ken06.Bus             .a........c..c....................a........g.........a.......................... [188880]
KM111294.1_Ken05/Tk1             .a........c..c....................a..................a.......................... [188880]
KM262844.1_strain_L60            ..........c..c....................a............................................. [188880]
KM102979.1_26544/OG10_Italy      ..........c..c....................a............................................. [188880]
KM262845.1_strain_NHV            ..........c..c....................a............................................. [188880]
KX354450.1_47/Ss/2008            ..........c..c....................a............................................. [188880]
KP055815.1_strain_BA71           ..........c..c....................a............................................. [188880]
MH025920.1_strain_R35            .a........c..c....................a........g.........a.......................... [188880]
MH025919.1_strain_N10            .a........c..c....................a........g.........a.......................... [188880]
MH025918.1_strain_R25            .a........c..c....................a........g.........a.......................... [188880]
MH025917.1_strain_R7             .a........c..c....................a........g.........a.......................... [188880]
MH025916.1_strain_R8             .a........c..c....................a........g.........a.......................... [188880]

FR682468.2_Georgia_2007/1        aaaaaggttttttcttgttgctggtacgtataatgataacacgaggcctcgatttttgcgcgtattcggtgcataaatc [188960]
ASF_IND_20_CAD543                ................................................................................ [188960]
LR899193.1_Germany_2020/1        ................................................................................ [188960]
MN393477.1_Wuhan_2019            ................................................................................ [188960]
MW396979.1_Timor-Leste/2019/1    ................................................................................ [188960]
MK940252.1_CN/2019/InnerMongolia ................................................................................ [188960]
MK128995.1_China/2018/AnhuiXCGQ  ................................................................................ [188960]
MH681419.1_ASFV/POL/2015         ................................................................................ [188960]
LS478113.1_Estonia_2014          ................................................................................ [188960]
KP843857.1_Odintsovo_02/14       ................................................................................ [188960]
```

```
FN557520.1_strain_E75              .................c.............................................................. [188960]
AY261366.1_Warthog                 .................c.............................................................. [188960]
AY261365.1_Warmbaths               .................c.............................................................. [188960]
AY261364.1_Tengani_62              .................c.............................................................. [188960]
AY261363.1_Pretoriuskop/96/4       .................c.............................................................. [188960]
AY261362.1_Mkuzi_1979              .................c.............................................................. [188960]
AY261361.1_Malawi_Lil-20/1_1983    .................c.............................................................. [188960]
AY261360.1_Kenya_1950              .................c................................a.........a..................t [188960]
AM712240.1_OURT_88/3               .................c.............................................................. [188960]
AM712239.1_Benin_97/1              .................c.............................................................. [188960]
KM111295.1_Ken06.Bus               .................c..........................................a..................t [188960]
KM111294.1_Ken05/Tk1               .....a...........c................................a.........a..................t [188960]
KM262844.1_strain_L60              .................c.............................................................. [188960]
KM102979.1_26544/OG10_Italy        .................c.............................................................. [188960]
KM262845.1_strain_NHV              .................c.............................................................. [188960]
KX354450.1_47/Ss/2008              .................c.............................................................. [188960]
KP055815.1_strain_BA71             .................c.............................................................. [188960]
MH025920.1_strain_R35              .................c..........................................a..................t [188960]
MH025919.1_strain_N10              .................c..........................................a..................t [188960]
MH025918.1_strain_R25              .................c..........................................a..................t [188960]
MH025917.1_strain_R7               .................c..........................................a..................t [188960]
MH025916.1_strain_R8               .................c..........................................a..................t [188960]

FR682468.2_Georgia_2007/1          agtatgttccttaaaaaacatatgtttttgaagcgttctaaaaaacatcatttggatgatatcacgcatttccaaaataa [189040]
ASF_IND_20_CAD543                  ................................................................................ [189040]
LR899193.1_Germany_2020/1          ................................................................................ [189040]
MN393477.1_Wuhan_2019              ................................................................................ [189040]
MW396979.1_Timor-Leste/2019/1      ................................................................................ [189040]
MK940252.1_CN/2019/InnerMongolia   ................................................................................ [189040]
MK128995.1_China/2018/AnhuiXCGQ    ................................................................................ [189040]
MH681419.1_ASFV/POL/2015           ................................................................................ [189040]
LS478113.1_Estonia_2014            ................................................................................ [189040]
KP843857.1_Odintsovo_02/14         ................................................................................ [189040]
FN557520.1_strain_E75              ........................................................................t....... [189040]
AY261366.1_Warthog                 ........t.........t............................................................. [189040]
AY261365.1_Warmbaths               ................................................................................ [189040]
AY261364.1_Tengani_62              ................................................................................ [189040]
AY261363.1_Pretoriuskop/96/4       ................................................................................ [189040]
AY261362.1_Mkuzi_1979              ................................................................................ [189040]
AY261361.1_Malawi_Lil-20/1_1983    .....t..t.........t.....................................................t....... [189040]
AY261360.1_Kenya_1950              .....t....c.g.....tg................c...................................t...g... [189040]
AM712240.1_OURT_88/3               ........................................................................t....... [189040]
AM712239.1_Benin_97/1              ........................................................................t....... [189040]
KM111295.1_Ken06.Bus               .....t....c.g.....tg........................................................g... [189040]
```

```
KM111294.1_Ken05/Tk1              .....t....c.g.....tg................c...................................t...g... [189040]
KM262844.1_strain_L60             ........................................................................t....... [189040]
KM102979.1_26544/OG10_Italy       ........................................................................t....... [189040]
KM262845.1_strain_NHV             ........................................................................t....... [189040]
KX354450.1_47/Ss/2008             ........................................................................t....... [189040]
KP055815.1_strain_BA71            ........................................................................t....... [189040]
MH025920.1_strain_R35             .....t......g.....tg........................................................g... [189040]
MH025919.1_strain_N10             .....t....c.g.....tg.................t......................................g... [189040]
MH025918.1_strain_R25             .....t......g.....tg........................................................g... [189040]
MH025917.1_strain_R7              .....t......g.....tg........................................................g... [189040]
MH025916.1_strain_R8              .....t......g.....tg........................................................g... [189040]

FR682468.2_Georgia_2007/1         tatagggttctagtcttttggaatctttcataactagatcggtggtaatattcttagtcatacaatttattaaaaatggt [189120]
ASF_IND_20_CAD543                 ...............................................................................y [189120]
LR899193.1_Germany_2020/1         ................................................................................ [189120]
MN393477.1_Wuhan_2019             ................................................................................ [189120]
MW396979.1_Timor-Leste/2019/1     ................................................................................ [189120]
MK940252.1_CN/2019/InnerMongolia  ................................................................................ [189120]
MK128995.1_China/2018/AnhuiXCGQ   ................................................................................ [189120]
MH681419.1_ASFV/POL/2015          ................................................................................ [189120]
LS478113.1_Estonia_2014           ................................................................................ [189120]
KP843857.1_Odintsovo_02/14        ................................................................................ [189120]
FN557520.1_strain_E75             ............a..................................................................c [189120]
AY261366.1_Warthog                ............a..................................................................c [189120]
AY261365.1_Warmbaths              .....................................g.........................................c [189120]
AY261364.1_Tengani_62             ...............................................................................c [189120]
AY261363.1_Pretoriuskop/96/4      ................................................................................ [189120]
AY261362.1_Mkuzi_1979             ............a..............................a...................................c [189120]
AY261361.1_Malawi_Lil-20/1_1983   ............a..g................................................................ [189120]
AY261360.1_Kenya_1950             ............ac..............................................................c..c [189120]
AM712240.1_OURT_88/3              ............a..................................................................c [189120]
AM712239.1_Benin_97/1             ............a..................................................................c [189120]
KM111295.1_Ken06.Bus              ............ac..............................................................c..c [189120]
KM111294.1_Ken05/Tk1              ............ac..............................................................c..c [189120]
KM262844.1_strain_L60             ............a..................................................................c [189120]
KM102979.1_26544/OG10_Italy       ............a..................................................................c [189120]
KM262845.1_strain_NHV             ............a..................................................................c [189120]
KX354450.1_47/Ss/2008             ............a..................................................................c [189120]
KP055815.1_strain_BA71            ............a..................................................................c [189120]
MH025920.1_strain_R35             ............ac..............................................................c..c [189120]
MH025919.1_strain_N10             ............ac..............................................................c..c [189120]
MH025918.1_strain_R25             ............ac..............................................................c..c [189120]
MH025917.1_strain_R7              ............ac..............................................................c..c [189120]
MH025916.1_strain_R8              ............ac..............................................................c..c [189120]
```

```
FR682468.2_Georgia_2007/1        ttaatatattgtaaatattttttaggcgtgtcagcctgtaaaaaacattcttgttcaatcttatttgtaaggatagtatt [189200]
ASF_IND_20_CAD543                ................................................................................ [189200]
LR899193.1_Germany_2020/1        ................................................................................ [189200]
MN393477.1_Wuhan_2019            ................................................................................ [189200]
MW396979.1_Timor-Leste/2019/1    ................................................................................ [189200]
MK940252.1_CN/2019/InnerMongolia ................................................................................ [189200]
MK128995.1_China/2018/AnhuiXCGQ  ................................................................................ [189200]
MH681419.1_ASFV/POL/2015         ................................................................................ [189200]
LS478113.1_Estonia_2014          ................................................................................ [189200]
KP843857.1_Odintsovo_02/14       ................................................................................ [189200]
FN557520.1_strain_E75            ................................................................................ [189200]
AY261366.1_Warthog               ................................................................................ [189200]
AY261365.1_Warmbaths             ................................................................................ [189200]
AY261364.1_Tengani_62            ................................................................................ [189200]
AY261363.1_Pretoriuskop/96/4     ................................................................................ [189200]
AY261362.1_Mkuzi_1979            ................................................................................ [189200]
AY261361.1_Malawi_Lil-20/1_1983  ...........................................................t..t................. [189200]
AY261360.1_Kenya_1950            ................................t.......................t....................... [189200]
AM712240.1_OURT_88/3             ................................................................................ [189200]
AM712239.1_Benin_97/1            ................................................................................ [189200]
KM111295.1_Ken06.Bus             ................................t.......................t.................g..... [189200]
KM111294.1_Ken05/Tk1             ................................t.......................t....................... [189200]
KM262844.1_strain_L60            ................................................................................ [189200]
KM102979.1_26544/OG10_Italy      ................................................................................ [189200]
KM262845.1_strain_NHV            ................................................................................ [189200]
KX354450.1_47/Ss/2008            ................................................................................ [189200]
KP055815.1_strain_BA71           ................................................................................ [189200]
MH025920.1_strain_R35            ................................t.......................t.................g..... [189200]
MH025919.1_strain_N10            ................................t.......................t.................g..... [189200]
MH025918.1_strain_R25            ................................t.......................t.................g..... [189200]
MH025917.1_strain_R7             ................................t.......................t.................g..... [189200]
MH025916.1_strain_R8             ................................t.......................t.................g..... [189200]

FR682468.2_Georgia_2007/1        ttgcaaatacttatttagcaaaaatacgatagaatcgcgggctatatgcattttcatataatttttttttttaaaatttaa [189280]
ASF_IND_20_CAD543                .............................................................a.................. [189280]
LR899193.1_Germany_2020/1        .............................................................-.................. [189280]
MN393477.1_Wuhan_2019            ................................................................................ [189280]
MW396979.1_Timor-Leste/2019/1    ................................................................................ [189280]
MK940252.1_CN/2019/InnerMongolia ................................................................................ [189280]
MK128995.1_China/2018/AnhuiXCGQ  ................................................................................ [189280]
MH681419.1_ASFV/POL/2015         ................................................................................ [189280]
LS478113.1_Estonia_2014          ................................................................................ [189280]
KP843857.1_Odintsovo_02/14       .............................................................-.................. [189280]
```

```
FN557520.1_strain_E75             ........g....................................................a.................. [189280]
AY261366.1_Warthog                ........g....................................................................... [189280]
AY261365.1_Warmbaths              ........g....................................................................... [189280]
AY261364.1_Tengani_62             ........g....................................................................... [189280]
AY261363.1_Pretoriuskop/96/4      ........g....................................................................... [189280]
AY261362.1_Mkuzi_1979             ........g....................................................a.................. [189280]
AY261361.1_Malawi_Lil-20/1_1983   ........gt.......................t.............................................. [189280]
AY261360.1_Kenya_1950             ........gt.....c......t.c....................................................... [189280]
AM712240.1_OURT_88/3              ........g....................................................a.................. [189280]
AM712239.1_Benin_97/1             ........g......................................a................................ [189280]
KM111295.1_Ken06.Bus              ........gt.....c......t.c..................................-.a.................. [189280]
KM111294.1_Ken05/Tk1              ........gt.....c......t.c....................................................... [189280]
KM262844.1_strain_L60             ........g....................................................a.................. [189280]
KM102979.1_26544/OG10_Italy       ........g....................................................a.................. [189280]
KM262845.1_strain_NHV             ........g....................................................a.................. [189280]
KX354450.1_47/Ss/2008             ........g....................................................a.................. [189280]
KP055815.1_strain_BA71            ........g....................................................a.................. [189280]
MH025920.1_strain_R35             ........gt.....c......t.c..................................-.................... [189280]
MH025919.1_strain_N10             ........gt.....c......t.c..................................-.................... [189280]
MH025918.1_strain_R25             ........gt.....c......t.c..................................-.................... [189280]
MH025917.1_strain_R7              ........gt.....c......t.c..................................-.................... [189280]
MH025916.1_strain_R8              ........gt.....c......t.c..................................-.................... [189280]

FR682468.2_Georgia_2007/1         tacaaaaaaaagaagtatagactc-----ttcttctagtcc----ggttagttcgtt-ggttgcctcaacatggagactc [189360]
ASF_IND_20_CAD543                 ...................ct...----c............----............-...................... [189360]
LR899193.1_Germany_2020/1         ........................-----............----............-...................... [189360]
MN393477.1_Wuhan_2019             ........................-----............----............-...................... [189360]
MW396979.1_Timor-Leste/2019/1     ........................-----............----............-...................... [189360]
MK940252.1_CN/2019/InnerMongolia  ........................-----............----............-...................... [189360]
MK128995.1_China/2018/AnhuiXCGQ   ........................-----............----............-...................... [189360]
MH681419.1_ASFV/POL/2015          ........................-----............----............-...................... [189360]
LS478113.1_Estonia_2014           ........................-----............----............-...................... [189360]
KP843857.1_Odintsovo_02/14        ........................-----............----............-...................... [189360]
FN557520.1_strain_E75             ...................ct...----c...........t----............-...................... [189360]
AY261366.1_Warthog                ........................-----............agtt............-...................... [189360]
AY261365.1_Warmbaths              ........................-----............----....g.......-...................... [189360]
AY261364.1_Tengani_62             ........................-----............agtt............-...................... [189360]
AY261363.1_Pretoriuskop/96/4      ........................-----............agtt............-...................... [189360]
AY261362.1_Mkuzi_1979             ...................ct...----c..........tt----............-...................... [189360]
AY261361.1_Malawi_Lil-20/1_1983   ........................-----.g..........----a..g.ag.....a...........g.......... [189360]
AY261360.1_Kenya_1950             ....................gt..tcctc.c........gt----a....a..a...-a..........a.......... [189360]
AM712240.1_OURT_88/3              ...................ct...----c...........t----............-...................... [189360]
AM712239.1_Benin_97/1             ...................ct...----c...........t----............-...................... [189360]
KM111295.1_Ken06.Bus              ........................tcctt.c........g.----a....a..a...-a..........a.......... [189360]
```

```
KM111294.1_Ken05/Tk1            ....................gt..tcctc.c........g.----a....a..a...-a......c...a.......... [189360]
KM262844.1_strain_L60           ....................ct..----c...........t----............-...................... [189360]
KM102979.1_26544/OG10_Italy     ....................ct..----c...........t----............-...................... [189360]
KM262845.1_strain_NHV           ....................ct..----c...........t----............-...................... [189360]
KX354450.1_47/Ss/2008           ....................ct..----c...........t----............-...................... [189360]
KP055815.1_strain_BA71          ....................ct..----c...........t----............-...................... [189360]
MH025920.1_strain_R35           ........................ccctt.c........g.----a....a..a...-a..........a.......... [189360]
MH025919.1_strain_N10           ........................ccctt.c........g.----a....a..a...-a..........a.......... [189360]
MH025918.1_strain_R25           ........................ccctt.c........g.----a....a..a...-a..........a.......... [189360]
MH025917.1_strain_R7            ........................ccctt.c........g.----a....a..a...-a..........a.......... [189360]
MH025916.1_strain_R8            ........................ccctt.c........g.----a....a..a...-a..........a.......... [189360]

FR682468.2_Georgia_2007/1       agaagttgatttccatggttaaggaagccttagaaaaatatcaataccctcttactgctaaaaatattaaagtagtgata [189440]
ASF_IND_20_CAD543               ......................r......................................................... [189440]
LR899193.1_Germany_2020/1       ................................................................................ [189440]
MN393477.1_Wuhan_2019           ................................................................................ [189440]
MW396979.1_Timor-Leste/2019/1   ................................................................................ [189440]
MK940252.1_CN/2019/InnerMongolia .a..............................a............................................... [189440]
MK128995.1_China/2018/AnhuiXCGQ ................................................................................ [189440]
MH681419.1_ASFV/POL/2015        ................................................................................ [189440]
LS478113.1_Estonia_2014         ................................................................................ [189440]
KP843857.1_Odintsovo_02/14      ................................................................................ [189440]
FN557520.1_strain_E75           ...............................g................................................ [189440]
AY261366.1_Warthog              ................................................................................ [189440]
AY261365.1_Warmbaths            ................................................................................ [189440]
AY261364.1_Tengani_62           .......a........................................................................ [189440]
AY261363.1_Pretoriuskop/96/4    ................................................................................ [189440]
AY261362.1_Mkuzi_1979           ................................................................................ [189440]
AY261361.1_Malawi_Lil-20/1_1983 ....a...g........a...gc...............................g.................c....... [189440]
AY261360.1_Kenya_1950           ....a...g.........................t..g...tc...t.................c.......c...a... [189440]
AM712240.1_OURT_88/3            ...............................g................................................ [189440]
AM712239.1_Benin_97/1           ...............................g................................................ [189440]
KM111295.1_Ken06.Bus            ....a...g.........................t..g...tct..t.................c.......c...a... [189440]
KM111294.1_Ken05/Tk1            ....a...g.........................t..g...tc...t.................c.......c...a... [189440]
KM262844.1_strain_L60           ...............................g................................................ [189440]
KM102979.1_26544/OG10_Italy     ...............................g................................................ [189440]
KM262845.1_strain_NHV           ...............................g................................................ [189440]
KX354450.1_47/Ss/2008           ...............................g................................................ [189440]
KP055815.1_strain_BA71          ......................a........g................................................ [189440]
MH025920.1_strain_R35           ....a...g.........................t..g...tct..t.................c.......c...a... [189440]
MH025919.1_strain_N10           ....a...g.........................t..g...tct..t.................c.......c...a... [189440]
MH025918.1_strain_R25           ....a...g.........................t..g...tct..t.................c.......c...a... [189440]
MH025917.1_strain_R7            ....a...g.........................t..g...tct..t.................c.......c...a... [189440]
MH025916.1_strain_R8            ....a...g.........................t..g...tct..t.................c.......c...a... [189440]
```

```
FR682468.2_Georgia_2007/1        caaaaagagcacaatgtcgtcttacctacaggatctataaatagcatactgtacagtaactcagaacttttttgagaagat [189520]
ASF_IND_20_CAD543                ..........................................................................r..... [189520]
LR899193.1_Germany_2020/1        ................................................................................ [189520]
MN393477.1_Wuhan_2019            ................................................................................ [189520]
MW396979.1_Timor-Leste/2019/1    ................................................................................ [189520]
MK940252.1_CN/2019/InnerMongolia ...............................................................a................ [189520]
MK128995.1_China/2018/AnhuiXCGQ  ................................................................................ [189520]
MH681419.1_ASFV/POL/2015         ................................................................................ [189520]
LS478113.1_Estonia_2014          ................................................................................ [189520]
KP843857.1_Odintsovo_02/14       ................................................................................ [189520]
FN557520.1_strain_E75            .........t.......t..............t.........................................a..... [189520]
AY261366.1_Warthog               ..................a.............t............................................... [189520]
AY261365.1_Warmbaths             .........t.......t.............................................................. [189520]
AY261364.1_Tengani_62            ..................a..........g..t............................................... [189520]
AY261363.1_Pretoriuskop/96/4     ..................a.........................................c................... [189520]
AY261362.1_Mkuzi_1979            .........t.......t..............t............................................... [189520]
AY261361.1_Malawi_Lil-20/1_1983  ..................a.......cgt...t..c........t............t..a................... [189520]
AY261360.1_Kenya_1950            .........t....a..t........c.....t...........t...............a.................g. [189520]
AM712240.1_OURT_88/3             .........t.......t..............t.........................................a..... [189520]
AM712239.1_Benin_97/1            .........t.......t..............t.........................................a..... [189520]
KM111295.1_Ken06.Bus             .........t....a..t........c.....t...........t...............a.................g. [189520]
KM111294.1_Ken05/Tk1             .........t....a..t........c.....t...........t...............a.................g. [189520]
KM262844.1_strain_L60            .........t.......t..............t.........................................a..... [189520]
KM102979.1_26544/OG10_Italy      .........t.......t..............t.........................................a..... [189520]
KM262845.1_strain_NHV            .........t.......t..............t.........................................a..... [189520]
KX354450.1_47/Ss/2008            .........t.......t..............t.........................................a..... [189520]
KP055815.1_strain_BA71           .........t.......t..............t.........................................a..... [189520]
MH025920.1_strain_R35            .........t....a..t........c.....t...........t...............a.................g. [189520]
MH025919.1_strain_N10            .........t....a..t........c.....t...........t...............a.................g. [189520]
MH025918.1_strain_R25            .........t....a..t........c.....t...........t...............a.................g. [189520]
MH025917.1_strain_R7             .........t....a..t........c.....t...........t...............a.................g. [189520]
MH025916.1_strain_R8             .........t....a..t........c.....t...........t...............a.................g. [189520]

FR682468.2_Georgia_2007/1        tgataagacaaataccatttatcccccgctttggatacggaaaaactaattgtaaccagtagtacattt--aaggatagt [189600]
ASF_IND_20_CAD543                ...............y........................-........c.aa................--......... [189600]
LR899193.1_Germany_2020/1        .....................................................................--......... [189600]
MN393477.1_Wuhan_2019            .....................................................................--......... [189600]
MW396979.1_Timor-Leste/2019/1    .....................................................................--......... [189600]
MK940252.1_CN/2019/InnerMongolia .....................................................................--......... [189600]
MK128995.1_China/2018/AnhuiXCGQ  .....................................................................--......... [189600]
MH681419.1_ASFV/POL/2015         .....................................................................--......... [189600]
LS478113.1_Estonia_2014          .....................................................................--......... [189600]
KP843857.1_Odintsovo_02/14       .....................................................................--......... [189600]
```

```
FN557520.1_strain_E75           ...............t........................-........c.aa................--......... [189600]
AY261366.1_Warthog              ...........................a.....................c...................aa......... [189600]
AY261365.1_Warmbaths            ...............t.................................c...................--......... [189600]
AY261364.1_Tengani_62           .....................c..t..a.....................c...................--......... [189600]
AY261363.1_Pretoriuskop/96/4    ...........................a.....................c.....-.............-a......... [189600]
AY261362.1_Mkuzi_1979           ...............t.................................c...................--...a..... [189600]
AY261361.1_Malawi_Lil-20/1_1983 ...............t...........a.....................c.....t......c....a.aa..t..c.ac [189600]
AY261360.1_Kenya_1950           c..............t...........a........g...-........c.--..t.............aa..t..c... [189600]
AM712240.1_OURT_88/3            ...............t........................-........c.aa................--....g.... [189600]
AM712239.1_Benin_97/1           ...............t........................-........c.aa................--......... [189600]
KM111295.1_Ken06.Bus            c..............t...........a............-........c.--..t.............aa..t..c... [189600]
KM111294.1_Ken05/Tk1            c..............t...........a........g...-........c.--..-.............aa..t..c... [189600]
KM262844.1_strain_L60           ...............t........................-........c.aa................--......... [189600]
KM102979.1_26544/OG10_Italy     ...............t........................-........c.aa................--......... [189600]
KM262845.1_strain_NHV           ...............t........................-........c.aa................--....g.... [189600]
KX354450.1_47/Ss/2008           ...............t........................-........c.aa................--......... [189600]
KP055815.1_strain_BA71          ...............t........................-........c.aa................--......... [189600]
MH025920.1_strain_R35           c..............t...........a............-........c.--..t.............aa..t..c... [189600]
MH025919.1_strain_N10           c..............t...........a............-........c.--..t.............aa..t..c... [189600]
MH025918.1_strain_R25           c..............t...........a............-........c.--..t.............aa..t..c... [189600]
MH025917.1_strain_R7            c..............t...........a............-........c.--..t.............aa..t..c... [189600]
MH025916.1_strain_R8            c..............t...........a............-........c.--..t.............aa..t..c... [189600]

FR682468.2_Georgia_2007/1       ttaagcagtaaat----------------------------gtagaataacacagttaag-caataaataac-------- [189680]
ASF_IND_20_CAD543               .............a-------acatagt--------taagcagta...............-...........-------- [189680]
LR899193.1_Germany_2020/1       .............----------------------------...................-...........-------- [189680]
MN393477.1_Wuhan_2019           .............----------------------------...................-...........-------- [189680]
MW396979.1_Timor-Leste/2019/1   .............----------------------------...................-...........-------- [189680]
MK940252.1_CN/2019/InnerMongolia .............----------------------------...................-...........-------- [189680]
MK128995.1_China/2018/AnhuiXCGQ .............----------------------------...................-...........-------- [189680]
MH681419.1_ASFV/POL/2015        .............----------------------------...................-...........-------- [189680]
LS478113.1_Estonia_2014         .............----------------------------...................-...........-------- [189680]
KP843857.1_Odintsovo_02/14      .............----------------------------...................-...........-------- [189680]
FN557520.1_strain_E75           .............a-------acatagt--------taagcagta.......t.......-...........-------- [189680]
AY261366.1_Warthog              .............----------------------------t..........tga.....-..g........-------- [189680]
AY261365.1_Warmbaths            .............----------------------------a...g......t.......-....g.t....-------- [189680]
AY261364.1_Tengani_62           .............----------------------------...........tga.....-...........-------- [189680]
AY261363.1_Pretoriuskop/96/4    ...........-----------------------------------......tga.....-..g........a------- [189680]
AY261362.1_Mkuzi_1979           .............g------------------------------t.ga.ta.........-...........-------- [189680]
AY261361.1_Malawi_Lil-20/1_1983 .............tcagaataacataattatagcagtatatt........t.t.......-...........aagtatat [189680]
AY261360.1_Kenya_1950           ....tt.......----------------------------t...gt.....t.......a...........aagggagt [189680]
AM712240.1_OURT_88/3            .............a-------acatagt--------taagcagta.......t.......-...........-------- [189680]
AM712239.1_Benin_97/1           .............a-------acatagt--------taagcagta.......t.......-...........-------- [189680]
KM111295.1_Ken06.Bus            ....tt.......----------------------------t...gt.....t.......a...........aagggagt [189680]
```

```
KM111294.1_Ken05/Tk1             ....tt.a.....----------------------------t...gt.....t.......a...........aagggagt [189680]
KM262844.1_strain_L60            .............a-------acatagt--------taagcagta.......t.......-...........-------- [189680]
KM102979.1_26544/OG10_Italy      .............a-------acatagt--------taagcagta.......t.......-...........-------- [189680]
KM262845.1_strain_NHV            .............a-------acatagt--------taagcagta.......t.......-...........-------- [189680]
KX354450.1_47/Ss/2008            .............a-------acatagt--------taagcagta.......t.......-...........-------- [189680]
KP055815.1_strain_BA71           .............a-------acatagt--------taagcagta.......t.......-...........-------- [189680]
MH025920.1_strain_R35            ....tt.......----------------------------t...gt.....t.......a...........aagggagt [189680]
MH025919.1_strain_N10            ....tt.......----------------------------t...gt.....t.......a...........aagggagt [189680]
MH025918.1_strain_R25            ....tt.......----------------------------t...gt.....t.......a...........aagggagt [189680]
MH025917.1_strain_R7             ....tt.......----------------------------t...gt.....t.......a...........aagggagt [189680]
MH025916.1_strain_R8             ....tt.......----------------------------t...gt.....t.......a...........aagggagt [189680]

FR682468.2_Georgia_2007/1        -----------------------------aagt----------------------------------------------- [189760]
ASF_IND_20_CAD543                -----------------------------....atata-----------------------------------taggaat [189760]
LR899193.1_Germany_2020/1        -----------------------------....atata------------------------------------------ [189760]
MN393477.1_Wuhan_2019            -----------------------------....atata------------------------------------------ [189760]
MW396979.1_Timor-Leste/2019/1    -----------------------------....atata------------------------------------------ [189760]
MK940252.1_CN/2019/InnerMongolia -----------------------------....atata------------------------------------------ [189760]
MK128995.1_China/2018/AnhuiXCGQ  -----------------------------....atata------------------------------------------ [189760]
MH681419.1_ASFV/POL/2015         -----------------------------....----------------------------------------------- [189760]
LS478113.1_Estonia_2014          -----------------------------....atata------------------------------------------ [189760]
KP843857.1_Odintsovo_02/14       -----------------------------....atata------------------------------------------ [189760]
FN557520.1_strain_E75            -----------------------------....atata-----------------------------------taggaat [189760]
AY261366.1_Warthog               -----------------------------....atata-----------------------------------cagga-- [189760]
AY261365.1_Warmbaths             -----------------------------....atata-----------------------------------taggaat [189760]
AY261364.1_Tengani_62            -----------------------------....atata-----------------------------------caggaat [189760]
AY261363.1_Pretoriuskop/96/4     ---------------------tgattaac....atata-----------------------------------caggaat [189760]
AY261362.1_Mkuzi_1979            -----------------------------....atata-----------------------------------taggaat [189760]
AY261361.1_Malawi_Lil-20/1_1983  aggaatatatagctagatcaataaataac....atataggaatatatagctagatcaataaataacaagtatataggaat [189760]
AY261360.1_Kenya_1950            ataggattatag-tagattatagga----g.a.ataaagatta------------------------------tagg--- [189760]
AM712240.1_OURT_88/3             -----------------------------....atata-----------------------------------taggaat [189760]
AM712239.1_Benin_97/1            -----------------------------....atata-----------------------------------taggaat [189760]
KM111295.1_Ken06.Bus             ataggattatag-tagattatagg---------gtatatgatta-----------------------------taggtat [189760]
KM111294.1_Ken05/Tk1             ataggattatag-tagattatagga----g.a.ataaagatta------------------------------tagg--- [189760]
KM262844.1_strain_L60            -----------------------------....atata-----------------------------------taggaat [189760]
KM102979.1_26544/OG10_Italy      -----------------------------....atata-----------------------------------taggaat [189760]
KM262845.1_strain_NHV            -----------------------------....atata-----------------------------------taggaat [189760]
KX354450.1_47/Ss/2008            -----------------------------....atata-----------------------------------taggaat [189760]
KP055815.1_strain_BA71           -----------------------------....atata-----------------------------------taggaat [189760]
MH025920.1_strain_R35            ataggattatag-tagattatagg---------gtatatgatta-----------------------------taggtat [189760]
MH025919.1_strain_N10            ataggattatag-tagattatagg---------gtatatgatta-----------------------------taggtat [189760]
MH025918.1_strain_R25            ataggattatag-tagattatagg---------gtatatgatta-----------------------------taggtat [189760]
MH025917.1_strain_R7             ataggattatag-tagattatagg---------gtatatgatta-----------------------------taggtat [189760]
MH025916.1_strain_R8             ataggattatag-tagattatagg---------gtatatgatta-----------------------------taggtat [189760]
```

```
FR682468.2_Georgia_2007/1        ---------------------------atataggaatatataggaata-------------------------------- [189840]
ASF_IND_20_CAD543                atat--------------------gat.................--..-------------------------------- [189840]
LR899193.1_Germany_2020/1        --gg--------------------aat.....................-------------------------------- [189840]
MN393477.1_Wuhan_2019            --gg--------------------aat.....................-------------------------------- [189840]
MW396979.1_Timor-Leste/2019/1    --gg--------------------aat.....................-------------------------------- [189840]
MK940252.1_CN/2019/InnerMongolia --gg--------------------aat.....................-------------------------------- [189840]
MK128995.1_China/2018/AnhuiXCGQ  --gg--------------------aat.....................-------------------------------- [189840]
MH681419.1_ASFV/POL/2015         ---------------------------.....................-------------------------------- [189840]
LS478113.1_Estonia_2014          --gg--------------------aat.....................-------------------------------- [189840]
KP843857.1_Odintsovo_02/14       --gg--------------------aat.....................-------------------------------- [189840]
FN557520.1_strain_E75            atat--------------------gat.................t...-------------------------------- [189840]
AY261366.1_Warthog               -------------------------at..............g.at...-------------------tag-------ga- [189840]
AY261365.1_Warmbaths             atat-------------------ggat.................t...-------------------tag-------aaa [189840]
AY261364.1_Tengani_62            atatag-----------------gaat................a....-------------------------------- [189840]
AY261363.1_Pretoriuskop/96/4     atatag-----------------aattt.............g..t...-------------------tag-------gaa [189840]
AY261362.1_Mkuzi_1979            atat--------------------gat.................t...a------------------tag-------gaa [189840]
AY261361.1_Malawi_Lil-20/1_1983  atatagctagatcaatagataacaagt................ct.g.tcaatagataacaagtatatag-------gaa [189840]
AY261360.1_Kenya_1950            gtataa-----------gattataggt........gat...--------------------------------------- [189840]
AM712240.1_OURT_88/3             atat--------------------gat.................t...-------------------------------- [189840]
AM712239.1_Benin_97/1            atat--------------------gat.................t...-------------------------------- [189840]
KM111295.1_Ken06.Bus             atagga-----------gattataggt........ga.....t..t..-------------------tagaataaataag [189840]
KM111294.1_Ken05/Tk1             gtatat-----------gattataggt.....a..gat......t...-------------------tag-------aag [189840]
KM262844.1_strain_L60            atat--------------------gat.................t...-------------------------------- [189840]
KM102979.1_26544/OG10_Italy      atat--------------------gat.................t...-------------------------------- [189840]
KM262845.1_strain_NHV            atat--------------------gat.................t...-------------------------------- [189840]
KX354450.1_47/Ss/2008            atat--------------------gat.................t...-------------------------------- [189840]
KP055815.1_strain_BA71           atat--------------------gat.................t...-------------------------------- [189840]
MH025920.1_strain_R35            atagga-----------gattataggt........ga.....t..t..-------------------tagaataaataag [189840]
MH025919.1_strain_N10            atagga-----------gattataggt........ga.....t..t..-------------------tagaataaataag [189840]
MH025918.1_strain_R25            atagga-----------gattataggt........ga.....t..t..-------------------tagaataaataag [189840]
MH025917.1_strain_R7             atagga-----------gattataggt........ga.....t..t..-------------------tagaataaataag [189840]
MH025916.1_strain_R8             atagga-----------gattataggt........ga.....t..t..-------------------tagaataaataag [189840]

FR682468.2_Georgia_2007/1        ----------------------------------------------------------------------------tata [189920]
ASF_IND_20_CAD543                ----------------------------------------------------------------------------.... [189920]
LR899193.1_Germany_2020/1        ----------------------------------------------------------------------------.... [189920]
MN393477.1_Wuhan_2019            ----------------------------------------------------------------------------.... [189920]
MW396979.1_Timor-Leste/2019/1    ----------------------------------------------------------------------------.... [189920]
MK940252.1_CN/2019/InnerMongolia ----------------------------------------------------------------------------.... [189920]
MK128995.1_China/2018/AnhuiXCGQ  ----------------------------------------------------------------------------.... [189920]
MH681419.1_ASFV/POL/2015         ----------------------------------------------------------------------------.... [189920]
LS478113.1_Estonia_2014          ----------------------------------------------------------------------------.... [189920]
KP843857.1_Odintsovo_02/14       ----------------------------------------------------------------------------.... [189920]
```

```
FN557520.1_strain_E75            ------------------------------------------------------------------------------.. [189920]
AY261366.1_Warthog               -------------------------------------------------------------------------------- [189920]
AY261365.1_Warmbaths             tatataggta------------------------------------------------------------------.... [189920]
AY261364.1_Tengani_62            ------------------------------------------------------------------------------.. [189920]
AY261363.1_Pretoriuskop/96/4     tatatgggta------------------------------------------------------------------.... [189920]
AY261362.1_Mkuzi_1979            tatataggtat-------------------atagaaatatatag---------------------gtatatagaaa.... [189920]
AY261361.1_Malawi_Lil-20/1_1983  tatatagctagatcaataaataacaagtatataggaatatatggttaattcaatatataacaagtatatataggaa.... [189920]
AY261360.1_Kenya_1950            ---------------------------------agggtatatgattat------------------------agaa..a. [189920]
AM712240.1_OURT_88/3             ------------------------------------------------------------------------------.. [189920]
AM712239.1_Benin_97/1            ------------------------------------------------------------------------------.. [189920]
KM111295.1_Ken06.Bus             attataggtatatag-----------------gagaatatatgattat------------------------agaa..a. [189920]
KM111294.1_Ken05/Tk1             aatataggtatatagg----------agattatagggtatatgattat------------------------agaa..a. [189920]
KM262844.1_strain_L60            ------------------------------------------------------------------------------.. [189920]
KM102979.1_26544/OG10_Italy      ------------------------------------------------------------------------------.. [189920]
KM262845.1_strain_NHV            ------------------------------------------------------------------------------.. [189920]
KX354450.1_47/Ss/2008            ------------------------------------------------------------------------------.. [189920]
KP055815.1_strain_BA71           ------------------------------------------------------------------------------.. [189920]
MH025920.1_strain_R35            attataggtatatag-----------------gagaatatatgattat------------------------agaa..a. [189920]
MH025919.1_strain_N10            attataggtatatag-----------------gagaatatatgattat------------------------agaa..a. [189920]
MH025918.1_strain_R25            attataggtatatag-----------------gagaatatatgattat------------------------agaa..a. [189920]
MH025917.1_strain_R7             attataggtatatag-----------------gagaatatatgattat------------------------agaa..a. [189920]
MH025916.1_strain_R8             attataggtatatag-----------------gagaatatatgattat------------------------agaa..a. [189920]

FR682468.2_Georgia_2007/1        gaaatatatagaaatagctaagcttaatactaattcagc-tttttttttaactaaaacctgaatagatgcgaagtagc-g [190000]
ASF_IND_20_CAD543                ..............w........................-......................................-. [190000]
LR899193.1_Germany_2020/1        .......................................-......................................-. [190000]
MN393477.1_Wuhan_2019            .......................................-......................................-. [190000]
MW396979.1_Timor-Leste/2019/1    .......................................-......................................-. [190000]
MK940252.1_CN/2019/InnerMongolia .......................................-......................................-. [190000]
MK128995.1_China/2018/AnhuiXCGQ  .......................................-......................................-. [190000]
MH681419.1_ASFV/POL/2015         .......................................-......................................-. [190000]
LS478113.1_Estonia_2014          .......................................-......................................-. [190000]
KP843857.1_Odintsovo_02/14       .......................................-......................................-. [190000]
FN557520.1_strain_E75            ..............a........................-..................t......c............-. [190000]
AY261366.1_Warthog               -----.....................g............-................g........t............-. [190000]
AY261365.1_Warmbaths             .......................................-.........................c............-. [190000]
AY261364.1_Tengani_62            t.gg.......................c...........-.........................c............-. [190000]
AY261363.1_Pretoriuskop/96/4     .g....a............................-...-................g........t............-. [190000]
AY261362.1_Mkuzi_1979            t.gg...................................-.........................c............-. [190000]
AY261361.1_Malawi_Lil-20/1_1983  tgg........g............g..c.aat...a.---.........g......g........c..--.g.a....-. [190000]
AY261360.1_Kenya_1950            t..gat.....g...g........g..ctaat...a...t........c....g......a....c...g...a.g..t. [190000]
AM712240.1_OURT_88/3             ..............a........................-..................t......c............-. [190000]
AM712239.1_Benin_97/1            ..............a........................-..................t......c............-. [190000]
KM111295.1_Ken06.Bus             t..gat.....g...g........a..ctaat...a...t........c....g......a....c...g...a.g..-a [190000]
```

```
KM111294.1_Ken05/Tk1             t..gat.....g...g........g..ctaat...a...-........c...g.......a....c...g...a.g..t. [190000]
KM262844.1_strain_L60            ..............a........................-..................t......c............-. [190000]
KM102979.1_26544/OG10_Italy      ..............a........................-..................t......c............-. [190000]
KM262845.1_strain_NHV            ..............a........................-..................t......c............-. [190000]
KX354450.1_47/Ss/2008            ..............a........................-..................t......c............-. [190000]
KP055815.1_strain_BA71           ..............a........................-..................t......c............-. [190000]
MH025920.1_strain_R35            t..gat.....g...g........a..ctaat...a...t........c...g.......a....c...g...a.g..-a [190000]
MH025919.1_strain_N10            t..gat.....g...g........a..ctaat...a...t........c...g.......a....c...g...a.g..-a [190000]
MH025918.1_strain_R25            t..gat.....g...g........a..ctaat...a...t........c...g.......a....c...g...a.g..-a [190000]
MH025917.1_strain_R7             t..gat.....g...g........a..ctaat...a...t........c...g.......a....c...g...a.g..-a [190000]
MH025916.1_strain_R8             t..gat.....g...g........a..ctaat...a...t........c...g.......a....c...g...a.g..-a [190000]

FR682468.2_Georgia_2007/1        gacatatacatactaaa-ataagccatacatttactttcttcttgaacatgaaacctttttttcttctgttgttggtata [190080]
ASF_IND_20_CAD543                .................-....r......................................................... [190080]
LR899193.1_Germany_2020/1        .................-.............................................................. [190080]
MN393477.1_Wuhan_2019            .................-.............................................................. [190080]
MW396979.1_Timor-Leste/2019/1    .................-.............................................................. [190080]
MK940252.1_CN/2019/InnerMongolia .................-.............................................................. [190080]
MK128995.1_China/2018/AnhuiXCGQ  .................-.............................................................. [190080]
MH681419.1_ASFV/POL/2015         .................-.............................................................. [190080]
LS478113.1_Estonia_2014          .................-.............................................................. [190080]
KP843857.1_Odintsovo_02/14       .................-.............................................................. [190080]
FN557520.1_strain_E75            .................-....a...c..................................................... [190080]
AY261366.1_Warthog               .................-........c..g...........................................c...... [190080]
AY261365.1_Warmbaths             .................-....a......g...........................................c...... [190080]
AY261364.1_Tengani_62            .................-...........g..............................................g... [190080]
AY261363.1_Pretoriuskop/96/4     .................-....---.c..g..........c................................c...... [190080]
AY261362.1_Mkuzi_1979            .................-....a...c..................................................... [190080]
AY261361.1_Malawi_Lil-20/1_1983  a.-........g.....t.a...g..c..g......................gg.......................... [190080]
AY261360.1_Kenya_1950            ..t........gt....t........c.......t............................................. [190080]
AM712240.1_OURT_88/3             .................-....a...c..................................................... [190080]
AM712239.1_Benin_97/1            .................-....a...c..................................................... [190080]
KM111295.1_Ken06.Bus             ..t...c....gt....c........c.t.....t............................................. [190080]
KM111294.1_Ken05/Tk1             ..t........gt....t........c.......t....................t........................ [190080]
KM262844.1_strain_L60            .................-....a...c..................................................... [190080]
KM102979.1_26544/OG10_Italy      .................-....a...c..................................................... [190080]
KM262845.1_strain_NHV            .................-....a...c..................................................... [190080]
KX354450.1_47/Ss/2008            .................-....a...c..................................................... [190080]
KP055815.1_strain_BA71           .................-....a...c..................................................... [190080]
MH025920.1_strain_R35            ..t...c....gt....c........c.t.....t............................................. [190080]
MH025919.1_strain_N10            ..t...c....gt....c........c.t.....t............................................. [190080]
MH025918.1_strain_R25            ..t...c....gt....c........c.t.....t............................................. [190080]
MH025917.1_strain_R7             ..t...c....gt....c........c.t.....t............................................. [190080]
MH025916.1_strain_R8             ..t...c....gt....c........c.t.....t............................................. [190080]
```

```
FR682468.2_Georgia_2007/1        taaacaataggactgtttgctgaggttgtatgatcttctacaactgctgtctcaggatgacgatgtttttttaaactaaa [190160]
ASF_IND_20_CAD543                ................................................................................ [190160]
LR899193.1_Germany_2020/1        ................................................................................ [190160]
MN393477.1_Wuhan_2019            ................................................................................ [190160]
MW396979.1_Timor-Leste/2019/1    ................................................................................ [190160]
MK940252.1_CN/2019/InnerMongolia ................................................................................ [190160]
MK128995.1_China/2018/AnhuiXCGQ  ................................................................................ [190160]
MH681419.1_ASFV/POL/2015         ................................................................................ [190160]
LS478113.1_Estonia_2014          ................................................................................ [190160]
KP843857.1_Odintsovo_02/14       ................................................................................ [190160]
FN557520.1_strain_E75            ................................................................................ [190160]
AY261366.1_Warthog               ................................................................................ [190160]
AY261365.1_Warmbaths             ................................................................................ [190160]
AY261364.1_Tengani_62            ................................................................................ [190160]
AY261363.1_Pretoriuskop/96/4     ................................................................................ [190160]
AY261362.1_Mkuzi_1979            ................................................................................ [190160]
AY261361.1_Malawi_Lil-20/1_1983  ...........g..........t......................................................... [190160]
AY261360.1_Kenya_1950            ...........g..........ta.a..c................................................... [190160]
AM712240.1_OURT_88/3             ................................................................................ [190160]
AM712239.1_Benin_97/1            ................................................................................ [190160]
KM111295.1_Ken06.Bus             ...........g..........ta.a..c...........t....................................... [190160]
KM111294.1_Ken05/Tk1             ...........g..........ta.a..c................................................... [190160]
KM262844.1_strain_L60            ................................................................................ [190160]
KM102979.1_26544/OG10_Italy      ................................................................................ [190160]
KM262845.1_strain_NHV            ................................................................................ [190160]
KX354450.1_47/Ss/2008            ................................................................................ [190160]
KP055815.1_strain_BA71           ................................................................................ [190160]
MH025920.1_strain_R35            ...........g..........ta.a..c...........t....................................... [190160]
MH025919.1_strain_N10            ...........g..........ta.a..c...........t....................................... [190160]
MH025918.1_strain_R25            ...........g..........ta.a..c...........t....................................... [190160]
MH025917.1_strain_R7             ...........g..........ta.a..c...........t....................................... [190160]
MH025916.1_strain_R8             ...........g..........ta.a..c...........t....................................... [190160]

FR682468.2_Georgia_2007/1        agtgtaggatggaatgagtggaatat------------agttatggctcgacttatcctgtttcgtacaggaatattttt [190240]
ASF_IND_20_CAD543                ..........................------------.......................................... [190240]
LR899193.1_Germany_2020/1        ..........................------------.......................................... [190240]
MN393477.1_Wuhan_2019            ..........................------------.......................................... [190240]
MW396979.1_Timor-Leste/2019/1    ..........................------------.......................................... [190240]
MK940252.1_CN/2019/InnerMongolia ..........................------------.......................................... [190240]
MK128995.1_China/2018/AnhuiXCGQ  ..........................------------.......................................... [190240]
MH681419.1_ASFV/POL/2015         ..........................------------.......................................... [190240]
LS478113.1_Estonia_2014          ..........................------------.......................................... [190240]
KP843857.1_Odintsovo_02/14       ..........................------------.......................................... [190240]
```

```
FN557520.1_strain_E75             ..........................------------.......................................... [190240]
AY261366.1_Warthog                ..........................agttatggcccg.........c.......c......................c. [190240]
AY261365.1_Warmbaths              ...a......................aattatggcccg.......................................... [190240]
AY261364.1_Tengani_62             ..........................agttatggcccg.........c.......c......................c. [190240]
AY261363.1_Pretoriuskop/96/4      ..........................------------.........c.......c......................c. [190240]
AY261362.1_Mkuzi_1979             ..........................------------.......................................... [190240]
AY261361.1_Malawi_Lil-20/1_1983   ..........................------------.t.....ag....t...c........................ [190240]
AY261360.1_Kenya_1950             c.........................------------..c....ag...tt...c........................ [190240]
AM712240.1_OURT_88/3              ..........................------------.......................................... [190240]
AM712239.1_Benin_97/1             ..........................------------.......................................... [190240]
KM111295.1_Ken06.Bus              .t........................------------..c....ag...tt...c........................ [190240]
KM111294.1_Ken05/Tk1              c.........................------------..c....ag...tt...c...................g.... [190240]
KM262844.1_strain_L60             ..........................------------.......................................... [190240]
KM102979.1_26544/OG10_Italy       ..........................------------.......................................... [190240]
KM262845.1_strain_NHV             ..........................------------.......................................... [190240]
KX354450.1_47/Ss/2008             ..........................------------.......................................... [190240]
KP055815.1_strain_BA71            ..........................------------.......................................... [190240]
MH025920.1_strain_R35             .t........................------------..c....ag...tt...c........................ [190240]
MH025919.1_strain_N10             .t........................------------..c....ag...tt...c........................ [190240]
MH025918.1_strain_R25             .t........................------------..c....ag...tt...c........................ [190240]
MH025917.1_strain_R7              .t........................------------..c....ag...tt...c........................ [190240]
MH025916.1_strain_R8              .t........................------------..c....ag...tt...c........................ [190240]

FR682468.2_Georgia_2007/1         tacaaatagaacgcaacaagcatatgaataaaaacagaaatgatatacaggagcataaaatagatatgaacactaagggg [190320]
ASF_IND_20_CAD543                 ..................................y............................................. [190320]
LR899193.1_Germany_2020/1         ................................................................................ [190320]
MN393477.1_Wuhan_2019             ................................................................................ [190320]
MW396979.1_Timor-Leste/2019/1     ................................................................................ [190320]
MK940252.1_CN/2019/InnerMongolia  ................................................................................ [190320]
MK128995.1_China/2018/AnhuiXCGQ   ................................................................................ [190320]
MH681419.1_ASFV/POL/2015          ................................................................................ [190320]
LS478113.1_Estonia_2014           ................................................................................ [190320]
KP843857.1_Odintsovo_02/14        ................................................................................ [190320]
FN557520.1_strain_E75             ..................................t............................................. [190320]
AY261366.1_Warthog                ..................................t.....c....................................... [190320]
AY261365.1_Warmbaths              ....t........................................................................... [190320]
AY261364.1_Tengani_62             ...................a..............t.....c....................................... [190320]
AY261363.1_Pretoriuskop/96/4      ..................................t.....c....................................... [190320]
AY261362.1_Mkuzi_1979             ..................................t............................................. [190320]
AY261361.1_Malawi_Lil-20/1_1983   ...............g.........t.t......t...............................c..g...a...... [190320]
AY261360.1_Kenya_1950             .........t.....g.........t........t...g..................t...g....cc.....a...... [190320]
AM712240.1_OURT_88/3              ..................................t............................................. [190320]
AM712239.1_Benin_97/1             ..................................t............................................. [190320]
KM111295.1_Ken06.Bus              .........t.....g.........t........t...g..................t...g....cc.....a...... [190320]
```

```
KM111294.1_Ken05/Tk1            .........t....g.........t........t...g..................t...g....cc.....a...... [190320]
KM262844.1_strain_L60           ..................................t............................................. [190320]
KM102979.1_26544/OG10_Italy     ..................................t............................................. [190320]
KM262845.1_strain_NHV           ..................................t............................................. [190320]
KX354450.1_47/Ss/2008           ..................................t............................................. [190320]
KP055815.1_strain_BA71          ..................................t............................................. [190320]
MH025920.1_strain_R35           .........t....g.........t........t...g..................t...g....cc.....a...... [190320]
MH025919.1_strain_N10           .........t....g.........t........t...g..................t...g....cc.....a...... [190320]
MH025918.1_strain_R25           .........t....g.........t........t...g..................t...g....cc.....a...... [190320]
MH025917.1_strain_R7            .........t....g.........t........t...g..................t...g....cc.....a...... [190320]
MH025916.1_strain_R8            .........t....g.........t........t...g..................t...g....cc.....a...... [190320]

FR682468.2_Georgia_2007/1       tagcagcttttataacgttccgtatttttcttagctatcaattgatttaccgtaatatttatctcgggaaactttgttct [190400]
ASF_IND_20_CAD543               ..................................................y............................. [190400]
LR899193.1_Germany_2020/1       ................................................................................ [190400]
MN393477.1_Wuhan_2019           ................................................................................ [190400]
MW396979.1_Timor-Leste/2019/1   ................................................................................ [190400]
MK940252.1_CN/2019/InnerMongolia ................................................................................ [190400]
MK128995.1_China/2018/AnhuiXCGQ ................................................................................ [190400]
MH681419.1_ASFV/POL/2015        ................................................................................ [190400]
LS478113.1_Estonia_2014         ................................................................................ [190400]
KP843857.1_Odintsovo_02/14      ................................................................................ [190400]
FN557520.1_strain_E75           ..................................................t............................. [190400]
AY261366.1_Warthog              ....................................g.............t............................. [190400]
AY261365.1_Warmbaths            ...................................a....................g....................... [190400]
AY261364.1_Tengani_62           ..a.................................g............tt............................. [190400]
AY261363.1_Pretoriuskop/96/4    ....................................g.............t............................. [190400]
AY261362.1_Mkuzi_1979           ..................................................t............................. [190400]
AY261361.1_Malawi_Lil-20/1_1983 ......t...........................t...............t........a.................... [190400]
AY261360.1_Kenya_1950           ......t.......c.....t.............................t....................t........ [190400]
AM712240.1_OURT_88/3            ..................................................t............................. [190400]
AM712239.1_Benin_97/1           ..................................................t............................. [190400]
KM111295.1_Ken06.Bus            ......t.......c...................................t....................t........ [190400]
KM111294.1_Ken05/Tk1            ......t.......c...................................t....................t........ [190400]
KM262844.1_strain_L60           ..................................................t............................. [190400]
KM102979.1_26544/OG10_Italy     ..................................................t............................. [190400]
KM262845.1_strain_NHV           ..................................................t............................. [190400]
KX354450.1_47/Ss/2008           ..................................................t............................. [190400]
KP055815.1_strain_BA71          ..................................................t............................. [190400]
MH025920.1_strain_R35           ......t.......c...................................t....................t........ [190400]
MH025919.1_strain_N10           ......t.......c...................................t....................t........ [190400]
MH025918.1_strain_R25           ......t.......c...................................t....................t........ [190400]
MH025917.1_strain_R7            ......t.......c...................................t....................t........ [190400]
MH025916.1_strain_R8            ......t.......c...................................t....................t........ [190400]
```

```
FR682468.2_Georgia_2007/1        acaatattttgtttggtattccagaaactcatgtcctggcttattcccgcagcttaaaaaatgatacaaaaatgtgttat [190480]
ASF_IND_20_CAD543                ........................................................r.............w......... [190480]
LR899193.1_Germany_2020/1        ................................................................................ [190480]
MN393477.1_Wuhan_2019            ................................................................................ [190480]
MW396979.1_Timor-Leste/2019/1    ................................................................................ [190480]
MK940252.1_CN/2019/InnerMongolia ................................................................................ [190480]
MK128995.1_China/2018/AnhuiXCGQ  ................................................................................ [190480]
MH681419.1_ASFV/POL/2015         ................................................................................ [190480]
LS478113.1_Estonia_2014          ................................................................................ [190480]
KP843857.1_Odintsovo_02/14       ................................................................................ [190480]
FN557520.1_strain_E75            .......................................t................g.............t......... [190480]
AY261366.1_Warthog               ......c................................t..............................t......... [190480]
AY261365.1_Warmbaths             .......................................t................g....c........t......... [190480]
AY261364.1_Tengani_62            .......................................t..............................t......... [190480]
AY261363.1_Pretoriuskop/96/4     ......c................................t..............................t......... [190480]
AY261362.1_Mkuzi_1979            .......................................t........a.......g....c........t......... [190480]
AY261361.1_Malawi_Lil-20/1_1983  .................................g.....t.......................g......t.c....... [190480]
AY261360.1_Kenya_1950            ....a..................................t............................g.t.c....gg. [190480]
AM712240.1_OURT_88/3             .......................................t................g.............t......... [190480]
AM712239.1_Benin_97/1            .......................................t................g.............t......... [190480]
KM111295.1_Ken06.Bus             ....a..................................t............................g.t.c....gg. [190480]
KM111294.1_Ken05/Tk1             ....a..................................t............................g.t.c....gg. [190480]
KM262844.1_strain_L60            .......................................t................g.............t......... [190480]
KM102979.1_26544/OG10_Italy      .......................................t................g.............t......... [190480]
KM262845.1_strain_NHV            .......................................t................g.............t......... [190480]
KX354450.1_47/Ss/2008            .......................................t................g.............t......... [190480]
KP055815.1_strain_BA71           .......................................t................g.............t......... [190480]
MH025920.1_strain_R35            ....a..................................t............................g.t.c....gg. [190480]
MH025919.1_strain_N10            ....a..................................t............................g.t.c....gg. [190480]
MH025918.1_strain_R25            ....a..................................t............................g.t.c....gg. [190480]
MH025917.1_strain_R7             ....a..................................t............................g.t.c....gg. [190480]
MH025916.1_strain_R8             ....a..................................t............................g.t.c....gg. [190480]

FR682468.2_Georgia_2007/1        tgttactaaaattaattcttcttaagaaaaactgcggaagacgctttaggtacgtctgttcctgttttagtaggaagtag [190560]
ASF_IND_20_CAD543                ...................................................................y............ [190560]
LR899193.1_Germany_2020/1        ................................................................................ [190560]
MN393477.1_Wuhan_2019            ................................................................................ [190560]
MW396979.1_Timor-Leste/2019/1    ................................................................................ [190560]
MK940252.1_CN/2019/InnerMongolia ................................................................................ [190560]
MK128995.1_China/2018/AnhuiXCGQ  ................................................................................ [190560]
MH681419.1_ASFV/POL/2015         ................................................................................ [190560]
LS478113.1_Estonia_2014          ................................................................................ [190560]
KP843857.1_Odintsovo_02/14       ................................................................................ [190560]
```

```
FN557520.1_strain_E75           ...........c.......................................................c............ [190560]
AY261366.1_Warthog              .........................................a.......a.............................. [190560]
AY261365.1_Warmbaths            ...........cc.....................................a................c............ [190560]
AY261364.1_Tengani_62           .a...............................................a.............................. [190560]
AY261363.1_Pretoriuskop/96/4    .........................................a.......a.............................. [190560]
AY261362.1_Mkuzi_1979           ...........c......................................a................c............ [190560]
AY261361.1_Malawi_Lil-20/1_1983 ...........c......................t......a.t..............c........c............ [190560]
AY261360.1_Kenya_1950           ...........c.............................a...................t..c..c..g......... [190560]
AM712240.1_OURT_88/3            ...........c.......................................................c............ [190560]
AM712239.1_Benin_97/1           .....t.....c.......................................................c............ [190560]
KM111295.1_Ken06.Bus            ...........c.............................a......................c..c..g......... [190560]
KM111294.1_Ken05/Tk1            ...........c.............................a...................t..c..c..g......... [190560]
KM262844.1_strain_L60           ...........c.......................................................c............ [190560]
KM102979.1_26544/OG10_Italy     ...........c.......................................................c............ [190560]
KM262845.1_strain_NHV           ...........c.......................................................c............ [190560]
KX354450.1_47/Ss/2008           ...........c.......................................................c............ [190560]
KP055815.1_strain_BA71          ...........c.......................................................c............ [190560]
MH025920.1_strain_R35           ...........c.............................a......................c..c..g......... [190560]
MH025919.1_strain_N10           ...........c.............................a......................c..c..g......... [190560]
MH025918.1_strain_R25           ...........c.............................a......................c..c..g......... [190560]
MH025917.1_strain_R7            ...........c.............................a......................c..c..g......... [190560]
MH025916.1_strain_R8            ...........c.............................a......................c..c..g......... [190560]

FR682468.2_Georgia_2007/1       tataagggacaatttctttttccacacattagattattgtaatataggtaggttggggtgttggagcgaataagttttct [190640]
ASF_IND_20_CAD543               ................................................................................ [190640]
LR899193.1_Germany_2020/1       ................................................................................ [190640]
MN393477.1_Wuhan_2019           ................................................................................ [190640]
MW396979.1_Timor-Leste/2019/1   ................................................................................ [190640]
MK940252.1_CN/2019/InnerMongolia ................................................................................ [190640]
MK128995.1_China/2018/AnhuiXCGQ ................................................................................ [190640]
MH681419.1_ASFV/POL/2015        ................................................................................ [190640]
LS478113.1_Estonia_2014         ................................................................................ [190640]
KP843857.1_Odintsovo_02/14      ................................................................................ [190640]
FN557520.1_strain_E75           ................................................................................ [190640]
AY261366.1_Warthog              .......................g.....c.................................................. [190640]
AY261365.1_Warmbaths            .................................c.............................................. [190640]
AY261364.1_Tengani_62           .......................g.....c.................................................. [190640]
AY261363.1_Pretoriuskop/96/4    .......................g.....c.................................................. [190640]
AY261362.1_Mkuzi_1979           ................................................................................ [190640]
AY261361.1_Malawi_Lil-20/1_1983 .......................g.....c................a.....................g........... [190640]
AY261360.1_Kenya_1950           .......................g.....c...........g...................................... [190640]
AM712240.1_OURT_88/3            ................................................................................ [190640]
AM712239.1_Benin_97/1           ................................................................................ [190640]
KM111295.1_Ken06.Bus            .......................g.....c...........g...................................... [190640]
```

```
KM111294.1_Ken05/Tk1            .......................g.....c...........g...................................... [190640]
KM262844.1_strain_L60           ................................................................................ [190640]
KM102979.1_26544/OG10_Italy     ................................................................................ [190640]
KM262845.1_strain_NHV           ................................................................................ [190640]
KX354450.1_47/Ss/2008           ................................................................................ [190640]
KP055815.1_strain_BA71          ................................................................................ [190640]
MH025920.1_strain_R35           .......................g.....c...........g...................................... [190640]
MH025919.1_strain_N10           .......................g.....c...........g...................................... [190640]
MH025918.1_strain_R25           .......................g.....c...........g...................................... [190640]
MH025917.1_strain_R7            .......................g.....c...........g...................................... [190640]
MH025916.1_strain_R8            .......................g.....c...........g...................................... [190640]

FR682468.2_Georgia_2007/1       gagtatgttataatctatgacttgtaaatcgttataccttaggtccaaaaacttgagttctttaccaaagccacctgcaa [190720]
ASF_IND_20_CAD543               ................................................................................ [190720]
LR899193.1_Germany_2020/1       ................................................................................ [190720]
MN393477.1_Wuhan_2019           ................................................................................ [190720]
MW396979.1_Timor-Leste/2019/1   ................................................................................ [190720]
MK940252.1_CN/2019/InnerMongolia ................................................................................ [190720]
MK128995.1_China/2018/AnhuiXCGQ ................................................................................ [190720]
MH681419.1_ASFV/POL/2015        ................................................................................ [190720]
LS478113.1_Estonia_2014         ................................................................................ [190720]
KP843857.1_Odintsovo_02/14      ................................................................................ [190720]
FN557520.1_strain_E75           ................................................................................ [190720]
AY261366.1_Warthog              ............t................................................................... [190720]
AY261365.1_Warmbaths            ................................................................................ [190720]
AY261364.1_Tengani_62           ............t.................a................................................. [190720]
AY261363.1_Pretoriuskop/96/4    ............t.....a............................................................. [190720]
AY261362.1_Mkuzi_1979           ................................................................................ [190720]
AY261361.1_Malawi_Lil-20/1_1983 ....................ac....g..ta..............t.................................. [190720]
AY261360.1_Kenya_1950           ...a........c.......a...c.g...a.............................c....t.......g...... [190720]
AM712240.1_OURT_88/3            ................................................................................ [190720]
AM712239.1_Benin_97/1           ................................................................................ [190720]
KM111295.1_Ken06.Bus            ...a................a...c.g...a.............................c....t.......g...... [190720]
KM111294.1_Ken05/Tk1            ...a................a...c.g...a.............................c....t.......g...... [190720]
KM262844.1_strain_L60           ................................................................................ [190720]
KM102979.1_26544/OG10_Italy     ................................................................................ [190720]
KM262845.1_strain_NHV           ................................................................................ [190720]
KX354450.1_47/Ss/2008           ................................................................................ [190720]
KP055815.1_strain_BA71          ................................................................................ [190720]
MH025920.1_strain_R35           ...a................a...c.g...a.............................c....t.......g...... [190720]
MH025919.1_strain_N10           ...a................a...c.g...a.............................c....t.......g...... [190720]
MH025918.1_strain_R25           ...a................a...c.g...a.............................c....t.......g...... [190720]
MH025917.1_strain_R7            ...a................a...c.g...a.............................c....t.......g...... [190720]
MH025916.1_strain_R8            ...a................a...c.g...a.............................c....t.......g...... [190720]
```

```
FR682468.2_Georgia_2007/1        tttcagaaatatttttcatcccgcagcggataatacggatgtcctgaaacgtctttaaaatacttgtattgtagtgaata [190800]
ASF_IND_20_CAD543                ................................................................................ [190800]
LR899193.1_Germany_2020/1        ................................................................................ [190800]
MN393477.1_Wuhan_2019            ................................................................................ [190800]
MW396979.1_Timor-Leste/2019/1    ................................................................................ [190800]
MK940252.1_CN/2019/InnerMongolia ................................................................................ [190800]
MK128995.1_China/2018/AnhuiXCGQ  ................................................................................ [190800]
MH681419.1_ASFV/POL/2015         ................................................................................ [190800]
LS478113.1_Estonia_2014          ................................................................................ [190800]
KP843857.1_Odintsovo_02/14       ................................................................................ [190800]
FN557520.1_strain_E75            ...............................g..g............................................. [190800]
AY261366.1_Warthog               .......g..........................g............................................. [190800]
AY261365.1_Warmbaths             ..................................g............................................. [190800]
AY261364.1_Tengani_62            ..................................g............................................. [190800]
AY261363.1_Pretoriuskop/96/4     ..................................g............................................. [190800]
AY261362.1_Mkuzi_1979            ..................................g............................................. [190800]
AY261361.1_Malawi_Lil-20/1_1983  ..................................g............................................. [190800]
AY261360.1_Kenya_1950            ....g.............................g.................t.....................c..... [190800]
AM712240.1_OURT_88/3             ...............................g..g............................................. [190800]
AM712239.1_Benin_97/1            ...............................g..g............................................. [190800]
KM111295.1_Ken06.Bus             ....g.............................g.................t.....................c..... [190800]
KM111294.1_Ken05/Tk1             ....g.............................g.................t.....................c..... [190800]
KM262844.1_strain_L60            ...............................g..g............................................. [190800]
KM102979.1_26544/OG10_Italy      ...............................g..g............................................. [190800]
KM262845.1_strain_NHV            ...............................g..g............................................. [190800]
KX354450.1_47/Ss/2008            ...............................g..g............................................. [190800]
KP055815.1_strain_BA71           ...............................g..g............................................. [190800]
MH025920.1_strain_R35            ....g.............................g.................t.....................c..... [190800]
MH025919.1_strain_N10            ....g.............................g.................t.....................c..... [190800]
MH025918.1_strain_R25            ....g.............................g.................t.....................c..... [190800]
MH025917.1_strain_R7             ....g.............................g.................t.....................c..... [190800]
MH025916.1_strain_R8             ....g.............................g.................t.....................c..... [190800]

FR682468.2_Georgia_2007/1        cttatgttatttttttgtaaataatctatgtcatgacaagtgcatgaaatgccagcagcattgcttggtatagtattata [190880]
ASF_IND_20_CAD543                ................................................................................ [190880]
LR899193.1_Germany_2020/1        ................................................................................ [190880]
MN393477.1_Wuhan_2019            ................................................................................ [190880]
MW396979.1_Timor-Leste/2019/1    ................................................................................ [190880]
MK940252.1_CN/2019/InnerMongolia ................................................................................ [190880]
MK128995.1_China/2018/AnhuiXCGQ  ................................................................................ [190880]
MH681419.1_ASFV/POL/2015         ................................................................................ [190880]
LS478113.1_Estonia_2014          ................................................................................ [190880]
KP843857.1_Odintsovo_02/14       ................................................................................ [190880]
```

```
FN557520.1_strain_E75              .............-.................................................................. [190880]
AY261366.1_Warthog                 .....................................................g.......................... [190880]
AY261365.1_Warmbaths               .....................................................ga......................... [190880]
AY261364.1_Tengani_62              .....................................................g........a........c...gc... [190880]
AY261363.1_Pretoriuskop/96/4       .....................................................g.......................... [190880]
AY261362.1_Mkuzi_1979              ................................................................................ [190880]
AY261361.1_Malawi_Lil-20/1_1983    ......................................t....................c...g.a.....ca....... [190880]
AY261360.1_Kenya_1950              ................................g.................a....t......ag................ [190880]
AM712240.1_OURT_88/3               ................................................................................ [190880]
AM712239.1_Benin_97/1              ................................................................................ [190880]
KM111295.1_Ken06.Bus               ................................g.................a....t......ag................ [190880]
KM111294.1_Ken05/Tk1               ................................g.................a....t......ag................ [190880]
KM262844.1_strain_L60              ................................................................................ [190880]
KM102979.1_26544/OG10_Italy        ................................................................................ [190880]
KM262845.1_strain_NHV              ................................................................................ [190880]
KX354450.1_47/Ss/2008              ................................................................................ [190880]
KP055815.1_strain_BA71             ................................................................................ [190880]
MH025920.1_strain_R35              ................................g.................a....t......ag................ [190880]
MH025919.1_strain_N10              ................................g.................a....t......ag................ [190880]
MH025918.1_strain_R25              ................................g.................a....t......ag................ [190880]
MH025917.1_strain_R7               ................................g.................a....t......ag................ [190880]
MH025916.1_strain_R8               ................................g.................a....t......ag................ [190880]

FR682468.2_Georgia_2007/1          tgcaggaagaactatactactattgagaatagtcacattgtacttataccatgtattattttctgatataaagtatttgc [190960]
ASF_IND_20_CAD543                  ................................................................................ [190960]
LR899193.1_Germany_2020/1          ................................................................................ [190960]
MN393477.1_Wuhan_2019              ................................................................................ [190960]
MW396979.1_Timor-Leste/2019/1      ................................................................................ [190960]
MK940252.1_CN/2019/InnerMongolia   ................................................................................ [190960]
MK128995.1_China/2018/AnhuiXCGQ    ................................................................................ [190960]
MH681419.1_ASFV/POL/2015           ................................................................................ [190960]
LS478113.1_Estonia_2014            ................................................................................ [190960]
KP843857.1_Odintsovo_02/14         ................................................................................ [190960]
FN557520.1_strain_E75              ...........t.................................................................... [190960]
AY261366.1_Warthog                 ................g............................................................... [190960]
AY261365.1_Warmbaths               ....................c........................................................... [190960]
AY261364.1_Tengani_62              ................g...c............t...........g.................................. [190960]
AY261363.1_Pretoriuskop/96/4       ................g...c............t...........g.................................. [190960]
AY261362.1_Mkuzi_1979              ................................................................................ [190960]
AY261361.1_Malawi_Lil-20/1_1983    ......g.........g...c........................................................... [190960]
AY261360.1_Kenya_1950              .........t.t....g..tc...a.....c...............................t................. [190960]
AM712240.1_OURT_88/3               ...........t.................................................................... [190960]
AM712239.1_Benin_97/1              ...........t.................................................................... [190960]
KM111295.1_Ken06.Bus               .........t.t....g..tc.........c................................................. [190960]
```

```
KM111294.1_Ken05/Tk1             .........t.t....g..tc...a.....c................................................. [190960]
KM262844.1_strain_L60            ...........t.................................................................... [190960]
KM102979.1_26544/OG10_Italy      ...........t.................................................................... [190960]
KM262845.1_strain_NHV            ...........t.................................................................... [190960]
KX354450.1_47/Ss/2008            ...........t.................................................................... [190960]
KP055815.1_strain_BA71           ...........t.................................................................... [190960]
MH025920.1_strain_R35            .........t.t....g..tc.........c................................................. [190960]
MH025919.1_strain_N10            .........t.t....g..tc.........c................................................. [190960]
MH025918.1_strain_R25            .........t.t....g..tc.........c................................................. [190960]
MH025917.1_strain_R7             .........t.t....g..tc.........c................................................. [190960]
MH025916.1_strain_R8             .........t.t....g..tc.........c................................................. [190960]

FR682468.2_Georgia_2007/1        aggtgacctgtggtttaatcctacctgttaagccacttcctaaaaaaacaaaaaatatgaaaacccttagcatcctgtat [191040]
ASF_IND_20_CAD543                ................................................................................ [191040]
LR899193.1_Germany_2020/1        ................................................................................ [191040]
MN393477.1_Wuhan_2019            ................................................................................ [191040]
MW396979.1_Timor-Leste/2019/1    ................................................................................ [191040]
MK940252.1_CN/2019/InnerMongolia ................................................................................ [191040]
MK128995.1_China/2018/AnhuiXCGQ  ................................................................................ [191040]
MH681419.1_ASFV/POL/2015         ................................................................................ [191040]
LS478113.1_Estonia_2014          ................................................................................ [191040]
KP843857.1_Odintsovo_02/14       ................................................................................ [191040]
FN557520.1_strain_E75            ............................................-................................... [191040]
AY261366.1_Warthog               ......t...c..................................................................... [191040]
AY261365.1_Warmbaths             ................................................................................ [191040]
AY261364.1_Tengani_62            ......t...c............t........................................................ [191040]
AY261363.1_Pretoriuskop/96/4     ......t...c..................................................................... [191040]
AY261362.1_Mkuzi_1979            ......t......................................................................... [191040]
AY261361.1_Malawi_Lil-20/1_1983  ......t....................c.................................................... [191040]
AY261360.1_Kenya_1950            .a....t.g...a.....c.t......c...a................................................ [191040]
AM712240.1_OURT_88/3             ................................................................................ [191040]
AM712239.1_Benin_97/1            ................................................................................ [191040]
KM111295.1_Ken06.Bus             .a......g...a.....c.t......c...a................................................ [191040]
KM111294.1_Ken05/Tk1             .a......g...a.....c.t......c...a................................................ [191040]
KM262844.1_strain_L60            ................................................................................ [191040]
KM102979.1_26544/OG10_Italy      ................................................................................ [191040]
KM262845.1_strain_NHV            ................................................................................ [191040]
KX354450.1_47/Ss/2008            ................................................................................ [191040]
KP055815.1_strain_BA71           ................................................................................ [191040]
MH025920.1_strain_R35            .a......g...a.....c.t......c...a................................................ [191040]
MH025919.1_strain_N10            .a......g...a.....c.t......c...a................................................ [191040]
MH025918.1_strain_R25            .a......g...a.....c.t......c...a................................................ [191040]
MH025917.1_strain_R7             .a......g...a.....c.t......c...a................................................ [191040]
MH025916.1_strain_R8             .a......g...a.....c.t......c...a................................................ [191040]
```

```
FR682468.2_Georgia_2007/1        atactattaaaaatttataaaatt-ttctgtttaaatttcatttagac-aaaaaa-aataatatatatacatcagcaaga [191120]
ASF_IND_20_CAD543                ........................-.......................-......-........................ [191120]
LR899193.1_Germany_2020/1        ........................-.......................-......-........................ [191120]
MN393477.1_Wuhan_2019            ........................-.......................-......-........................ [191120]
MW396979.1_Timor-Leste/2019/1    ........................-.......................-......-........................ [191120]
MK940252.1_CN/2019/InnerMongolia ........................-.......................-......-........................ [191120]
MK128995.1_China/2018/AnhuiXCGQ  ........................-.......................-......-........................ [191120]
MH681419.1_ASFV/POL/2015         ........................-.......................-......-........................ [191120]
LS478113.1_Estonia_2014          ........................-.......................-......-........................ [191120]
KP843857.1_Odintsovo_02/14       ........................-.......................-......-........................ [191120]
FN557520.1_strain_E75            ........................-.......................--.....-........................ [191120]
AY261366.1_Warthog               ........................-..g....................a......-........................ [191120]
AY261365.1_Warmbaths             ........................-..g....................a......-........................ [191120]
AY261364.1_Tengani_62            ........................-..g....................a......-........................ [191120]
AY261363.1_Pretoriuskop/96/4     ........................-..g....................-......-........................ [191120]
AY261362.1_Mkuzi_1979            ........................-.......................-......-........................ [191120]
AY261361.1_Malawi_Lil-20/1_1983  ........................-..gg................a..a......-........................ [191120]
AY261360.1_Kenya_1950            ........................aa.t.tg..............a..a......-......g................. [191120]
AM712240.1_OURT_88/3             ........................-.......................-......-........................ [191120]
AM712239.1_Benin_97/1            ........................-.......................a......-........................ [191120]
KM111295.1_Ken06.Bus             ........................aa.t.tg..............a..a......-......g................. [191120]
KM111294.1_Ken05/Tk1             ........................aa.t.tg..............a..a......-......g................. [191120]
KM262844.1_strain_L60            ........................-.......................-......-........................ [191120]
KM102979.1_26544/OG10_Italy      ........................-.......................-......-........................ [191120]
KM262845.1_strain_NHV            ........................-.......................-......-........................ [191120]
KX354450.1_47/Ss/2008            ........................-.......................a......-........................ [191120]
KP055815.1_strain_BA71           ........................-.......................-......-........................ [191120]
MH025920.1_strain_R35            ........................aa.t.tg..............a..a......a......g................. [191120]
MH025919.1_strain_N10            ........................aa.t.tg..............a..a......-......g................. [191120]
MH025918.1_strain_R25            ........................aa.t.tg..............a..a......a......g................. [191120]
MH025917.1_strain_R7             ........................aa.t.tg..............a..a......a......g................. [191120]
MH025916.1_strain_R8             ........................aa.t.tg..............a..a......a......g................. [191120]

FR682468.2_Georgia_2007/1        aattatatacagattatataattttctgattttttttt-gccacaataag-catcattatat---gcattaaaatctca- [191200]
ASF_IND_20_CAD543                ......................................-...........-...........---..............- [191200]
LR899193.1_Germany_2020/1        ......................................-...........-...........---..............- [191200]
MN393477.1_Wuhan_2019            ......................................-...........-...........---..............- [191200]
MW396979.1_Timor-Leste/2019/1    ......................................-...........-...........---..............- [191200]
MK940252.1_CN/2019/InnerMongolia ......................................-...........-...........---..............- [191200]
MK128995.1_China/2018/AnhuiXCGQ  ......................................-...........-...........---..............- [191200]
MH681419.1_ASFV/POL/2015         ......................................-...........-...........---..............- [191200]
LS478113.1_Estonia_2014          ......................................-...........-...........---..............- [191200]
KP843857.1_Odintsovo_02/14       ......................................-...........-...........---..............- [191200]
```

```
FN557520.1_strain_E75              ......................................-....t......-...........---..........g...- [191200]
AY261366.1_Warthog                 ......................................-..tgt.gc...-........a..---.......g..g...- [191200]
AY261365.1_Warmbaths               ......................................-..tgt.gc...-........a..---.......g..g...- [191200]
AY261364.1_Tengani_62              ......................................-....t......-........a..---.......g..g.a.- [191200]
AY261363.1_Pretoriuskop/96/4       ......................................-....t.gc...-........a..---.......g..g.a.- [191200]
AY261362.1_Mkuzi_1979              ......................................-....t......-...........---..........g...- [191200]
AY261361.1_Malawi_Lil-20/1_1983    ......................................c..t.t..c...c..aa.......---a.c.........a.- [191200]
AY261360.1_Kenya_1950              ......................................-....t.gc...-...t.ga..g.taa......g...g.t.- [191200]
AM712240.1_OURT_88/3               ......................................-....t......-...........---..........g...- [191200]
AM712239.1_Benin_97/1              ......................................-....t......-...........---..........g...- [191200]
KM111295.1_Ken06.Bus               ......................................-....t.gc..a-...t.aa..g.taa...c......g.t.- [191200]
KM111294.1_Ken05/Tk1               ......................................-....t.gc..a-...t.aa..g.taa...c......g.t.t [191200]
KM262844.1_strain_L60              ......................................-....t......-...........---..........g...- [191200]
KM102979.1_26544/OG10_Italy        ......................................-....t......-...........---..........g...- [191200]
KM262845.1_strain_NHV              ......................................-....t......-...........---..........g...- [191200]
KX354450.1_47/Ss/2008              ......................................-....t......-...........---..........g...- [191200]
KP055815.1_strain_BA71             ......................................-....t......-...........---..........g...- [191200]
MH025920.1_strain_R35              ......................................-....t.gc..a-...t.aa..g.taa...c......g.t.- [191200]
MH025919.1_strain_N10              ......................................-....t.gc..a-...t.aa..g.taa...c......g.t.- [191200]
MH025918.1_strain_R25              ......................................-....t.gc..a-...t.aa..g.taa...c......g.t.- [191200]
MH025917.1_strain_R7               ......................................-....t.gc..a-...t.aa..g.taa...c......g.t.- [191200]
MH025916.1_strain_R8               ......................................-....t.gc..a-...t.aa..g.taa...c......g.t.- [191200]

FR682468.2_Georgia_2007/1          -atactaaa---------cactaaaatc----------taaattcta-agcattaaattctaag---------------- [191280]
ASF_IND_20_CAD543                  -........---------.......t..t---aagcat.........-................c-ctaaattctatgca [191280]
LR899193.1_Germany_2020/1          -........---------..........----------.........-................---------------- [191280]
MN393477.1_Wuhan_2019              -........---------..........----------.........-................---------------- [191280]
MW396979.1_Timor-Leste/2019/1      -........---------..........----------.........-................---------------- [191280]
MK940252.1_CN/2019/InnerMongolia   -........---------..........----------.........-................---------------- [191280]
MK128995.1_China/2018/AnhuiXCGQ    -........---------..........----------.........-................---------------- [191280]
MH681419.1_ASFV/POL/2015           -........---------..........----------.........-................---------------- [191280]
LS478113.1_Estonia_2014            -........---------..........----------.........-................---------------- [191280]
KP843857.1_Odintsovo_02/14         -........---------..........----------.........-................---------------- [191280]
FN557520.1_strain_E75              -g.......---------.......t..t---aagcat.........-................c-ctaaattctatgca [191280]
AY261366.1_Warthog                 -g...---------------------------------....a....-.a..............c--------------- [191280]
AY261365.1_Warmbaths               -g...---------------------------------....a....-.a..c....a......c--------------- [191280]
AY261364.1_Tengani_62              -g-------------------------------------------c.-....c....a.gc.tac--------------- [191280]
AY261363.1_Pretoriuskop/96/4       -g-------------------------------------------c.-....c....a.gc...c--------------- [191280]
AY261362.1_Mkuzi_1979              -g.----------------......t..t---aagcat.........-....c.c.........cactaaattctatgca [191280]
AY261361.1_Malawi_Lil-20/1_1983    -g.......at-tta-gg.......g.tt---aaacacc...a....-....c....a......tactaaagtttaaaca [191280]
AY261360.1_Kenya_1950              agc....----------------....gttatgaaacta....gt..tg.a.c....a.g.t.tg--------------- [191280]
AM712240.1_OURT_88/3               -g.......---------.......t..t---aagcat.........-................c-ctaaattctatgca [191280]
AM712239.1_Benin_97/1              -g.......---------.......t..t---aagcat.........-................c-ctaaattctatgca [191280]
KM111295.1_Ken06.Bus               agc......atgtta-ag.........gttatggaacta....gt..-....c....a.g.t.tg--------------- [191280]
```

```
KM111294.1_Ken05/Tk1            g.a......atgttatgga........gttatgaaacta....gt..tg.a.c....a.g.t.tg--------------- [191280]
KM262844.1_strain_L60           -g.......---------.......t..t---aagcat.........-................c-ctaaattctatgca [191280]
KM102979.1_26544/OG10_Italy     -g.......---------.......t..t---aagcat.........-................c-ctaaattctatgca [191280]
KM262845.1_strain_NHV           -g.......---------.......t..t---aagcat.........-................c-ctaaattctatgca [191280]
KX354450.1_47/Ss/2008           -g.......---------.......t..t---aagcat.........-................c-ctaaattctatgca [191280]
KP055815.1_strain_BA71          -g.......---------.......t..t---aagcat.........-................c-ctaaattctatgca [191280]
MH025920.1_strain_R35           agc......atgtta-ag.........gttatggaacta....gt..-....c....a.g.t.tg--------------- [191280]
MH025919.1_strain_N10           agc......atgtta-ag.........gttatggaacta....gt..-....c....a.g.t.tg--------------- [191280]
MH025918.1_strain_R25           agc......atgtta-ag.........gttatggaacta....gt..-....c....a.g.t.tg--------------- [191280]
MH025917.1_strain_R7            agc......atgtta-ag.........gttatggaacta....gt..-....c....a.g.t.tg--------------- [191280]
MH025916.1_strain_R8            agc......atgtta-ag.........gttatggaacta....gt..-....c....a.g.t.tg--------------- [191280]

FR682468.2_Georgia_2007/1       ------------cattaaattc-tatgc-actaaactgtaagcactaaaatctaagta---------------------- [191360]
ASF_IND_20_CAD543               ctaaactgtaag..c.......-..n..-.............................---------------------- [191360]
LR899193.1_Germany_2020/1       ------------..........-.....-.............................---------------------- [191360]
MN393477.1_Wuhan_2019           ------------..........-.....-.............................---------------------- [191360]
MW396979.1_Timor-Leste/2019/1   ------------..........-.....-.............................---------------------- [191360]
MK940252.1_CN/2019/InnerMongolia ------------..........-.....-.............................---------------------- [191360]
MK128995.1_China/2018/AnhuiXCGQ ------------..........-.....-.............................---------------------- [191360]
MH681419.1_ASFV/POL/2015        ------------..........-.....-.............................---------------------- [191360]
LS478113.1_Estonia_2014         ------------..........-.....-.............................---------------------- [191360]
KP843857.1_Odintsovo_02/14      ------------..........-.....-.............................---------------------- [191360]
FN557520.1_strain_E75           ctaaactgtaag..c.......-..a..-.............................---------------------- [191360]
AY261366.1_Warthog              --------------c.......-.....-.............................---------------------- [191360]
AY261365.1_Warmbaths            -------------.c.......-..a..-.............................---------------------- [191360]
AY261364.1_Tengani_62           -------------.c....c.g-..a..-..........----------------------------------------- [191360]
AY261363.1_Pretoriuskop/96/4    -------------.c....c.g-..a..-..........----------------------------------------- [191360]
AY261362.1_Mkuzi_1979           ctaaactg---------------..a..-.............................---------------------- [191360]
AY261361.1_Malawi_Lil-20/1_1983 ccaaaatctaag..c....a..-..at.-......g.t...a..t...........c.ctaaaatctaatcactaaagtt [191360]
AY261360.1_Kenya_1950           -----------aa.c....g.g-.....a.t....a-...t.t.act...agc.t.a.---------------------- [191360]
AM712240.1_OURT_88/3            ctaaactgtaag..c.......-..a..-.............................---------------------- [191360]
AM712239.1_Benin_97/1           ctaaactgtaag..c.......-..a..-.............................---------------------- [191360]
KM111295.1_Ken06.Bus            -----------ga.c....a.gt.....a.....---------------.agc.t...---------------------- [191360]
KM111294.1_Ken05/Tk1            -----------aa.c....a.gt....aa......g....t...act...ag..t...---------------------- [191360]
KM262844.1_strain_L60           ctaaactgtaag..c.......-..a..-.............................---------------------- [191360]
KM102979.1_26544/OG10_Italy     ctaaactgtaag..c.......-..a..-.............................---------------------- [191360]
KM262845.1_strain_NHV           ctaaactgtaag..c.......-..a..-.............................---------------------- [191360]
KX354450.1_47/Ss/2008           ctaaactgtaag..c.......-..a..-.............................---------------------- [191360]
KP055815.1_strain_BA71          ctaaactgtaag..c.......-..a..-.............................---------------------- [191360]
MH025920.1_strain_R35           -----------ga.c....a.ga.....a......a-...t.g.act...agc.t...---------------------- [191360]
MH025919.1_strain_N10           -----------ga.c....a.gt.....a......a-...t.g.act...agc.t...---------------------- [191360]
MH025918.1_strain_R25           -----------ga.c....a.ga.....a......a-...t.g.act...agc.t...---------------------- [191360]
MH025917.1_strain_R7            -----------ga.c....a.ga.....a......a-...t.g.act...agc.t...---------------------- [191360]
MH025916.1_strain_R8            -----------ga.c....a.ga.....a......a-...t.g.act...agc.t...---------------------- [191360]
```

```
FR682468.2_Georgia_2007/1         -------------------actaaaatc---------------------------------------------------- [191440]
ASF_IND_20_CAD543                 -------------------.........---------------------------------------------------- [191440]
LR899193.1_Germany_2020/1         -------------------.........---------------------------------------------------- [191440]
MN393477.1_Wuhan_2019             -------------------.........---------------------------------------------------- [191440]
MW396979.1_Timor-Leste/2019/1     -------------------.........---------------------------------------------------- [191440]
MK940252.1_CN/2019/InnerMongolia  -------------------.........---------------------------------------------------- [191440]
MK128995.1_China/2018/AnhuiXCGQ   -------------------.........---------------------------------------------------- [191440]
MH681419.1_ASFV/POL/2015          -------------------.........---------------------------------------------------- [191440]
LS478113.1_Estonia_2014           -------------------.........---------------------------------------------------- [191440]
KP843857.1_Odintsovo_02/14        -------------------.........---------------------------------------------------- [191440]
FN557520.1_strain_E75             -------------------.........---------------------------------------------------- [191440]
AY261366.1_Warthog                -------------------.........---------------------------------------------------- [191440]
AY261365.1_Warmbaths              -------------------.........---------------------------------------------------- [191440]
AY261364.1_Tengani_62             -------------------------------------------------------------------------------- [191440]
AY261363.1_Pretoriuskop/96/4      -------------------------------------------------------------------------------- [191440]
AY261362.1_Mkuzi_1979             -------------------.........---------------------------------------------------- [191440]
AY261361.1_Malawi_Lil-20/1_1983   taaacattaaaatctaagc.........taatcactaaagtttaaacactaaaatctaagcactaaaatctaatcactaa [191440]
AY261360.1_Kenya_1950             -------------------.......g.---------------------------------------------------- [191440]
AM712240.1_OURT_88/3              -------------------.........---------------------------------------------------- [191440]
AM712239.1_Benin_97/1             -------------------.........---------------------------------------------------- [191440]
KM111295.1_Ken06.Bus              -------------------.......g.---------------------------------------------------- [191440]
KM111294.1_Ken05/Tk1              -------------------.......g.---------------------------------------------------- [191440]
KM262844.1_strain_L60             -------------------.........---------------------------------------------------- [191440]
KM102979.1_26544/OG10_Italy       -------------------.........---------------------------------------------------- [191440]
KM262845.1_strain_NHV             -------------------.........---------------------------------------------------- [191440]
KX354450.1_47/Ss/2008             -------------------.........---------------------------------------------------- [191440]
KP055815.1_strain_BA71            -------------------.........---------------------------------------------------- [191440]
MH025920.1_strain_R35             -------------------.......g.---------------------------------------------------- [191440]
MH025919.1_strain_N10             -------------------.......g.---------------------------------------------------- [191440]
MH025918.1_strain_R25             -------------------.......g.---------------------------------------------------- [191440]
MH025917.1_strain_R7              -------------------.......g.---------------------------------------------------- [191440]
MH025916.1_strain_R8              -------------------.......g.---------------------------------------------------- [191440]

FR682468.2_Georgia_2007/1         ------aacact-aaatgtatgcaacctaaaatgtaa---------------------------agcattactcatcatc [191520]
ASF_IND_20_CAD543                 ------...-----------------...........---------------------------................ [191520]
LR899193.1_Germany_2020/1         ------......-........................---------------------------................ [191520]
MN393477.1_Wuhan_2019             ------......-........................---------------------------................ [191520]
MW396979.1_Timor-Leste/2019/1     ------......-........................---------------------------................ [191520]
MK940252.1_CN/2019/InnerMongolia  ------......-........................---------------------------................ [191520]
MK128995.1_China/2018/AnhuiXCGQ   ------......-........................---------------------------................ [191520]
MH681419.1_ASFV/POL/2015          ------......-........................---------------------------................ [191520]
LS478113.1_Estonia_2014           ------......-........................---------------------------................ [191520]
KP843857.1_Odintsovo_02/14        ------......-........................---------------------------................ [191520]
```

```
FN557520.1_strain_E75            ------....----------------...........---------------------------................ [191520]
AY261366.1_Warthog               ------......-.........t.-............---------------------------................ [191520]
AY261365.1_Warmbaths             ------......-.........t.-............---------------------------................ [191520]
AY261364.1_Tengani_62            ------------------------------------.---------------------------................ [191520]
AY261363.1_Pretoriuskop/96/4     ------------------------------------.---------------------------................ [191520]
AY261362.1_Mkuzi_1979            ------....t.-........................---------------------------................ [191520]
AY261361.1_Malawi_Lil-20/1_1983  agttta......a....c..a...--.....g.....gtaactaatatgttatgcattaaaatc....c........... [191520]
AY261360.1_Kenya_1950            ----atgta...a.......a...--.....g.a..t--------------atgc---------t............... [191520]
AM712240.1_OURT_88/3             ------....----------------...........---------------------------................ [191520]
AM712239.1_Benin_97/1            ------....----------------...........---------------------------................ [191520]
KM111295.1_Ken06.Bus             ----atgta...a.......a...--.......a..t--------------atgc---------t............... [191520]
KM111294.1_Ken05/Tk1             ----atgta...a.......a...--.....g.a..t--------------atgc---------t............... [191520]
KM262844.1_strain_L60            ------....----------------...........---------------------------................ [191520]
KM102979.1_26544/OG10_Italy      ------....----------------...........---------------------------................ [191520]
KM262845.1_strain_NHV            ------....----------------...........---------------------------................ [191520]
KX354450.1_47/Ss/2008            ------....----------------...........---------------------------................ [191520]
KP055815.1_strain_BA71           ------....----------------...........---------------------------................ [191520]
MH025920.1_strain_R35            ----atgta...a.......a...--.......a..t--------------atgc---------t............... [191520]
MH025919.1_strain_N10            ----atgta...a.......a...--.......a..t--------------atgc---------t............... [191520]
MH025918.1_strain_R25            ----atgta...a.......a...--.......a..t--------------atgc---------t............... [191520]
MH025917.1_strain_R7             ----atgta...a.......a...--.......a..t--------------atgc---------t............... [191520]
MH025916.1_strain_R8             ----atgta...a.......a...--.......a..t--------------atgc---------t............... [191520]

FR682468.2_Georgia_2007/1        c---------------------------tcctcttcttcatcc---tcatcatcataggttaagatatatgtgtcatcct [191600]
ASF_IND_20_CAD543                .at------------------ctcttca.....c.....t..a---.................................. [191600]
LR899193.1_Germany_2020/1        .---------------------------...............---.................................. [191600]
MN393477.1_Wuhan_2019            .---------------------------...............---.................................. [191600]
MW396979.1_Timor-Leste/2019/1    .---------------------------...............---.................................. [191600]
MK940252.1_CN/2019/InnerMongolia .---------------------------...............---.................................. [191600]
MK128995.1_China/2018/AnhuiXCGQ  .---------------------------...............---.................................. [191600]
MH681419.1_ASFV/POL/2015         .---------------------------...............---.................................. [191600]
LS478113.1_Estonia_2014          .---------------------------...............---.................................. [191600]
KP843857.1_Odintsovo_02/14       .---------------------------...............---.................................. [191600]
FN557520.1_strain_E75            .at------------------ctcttca.....c.....t..a---.................................. [191600]
AY261366.1_Warthog               .atttcttcatcctcctcttcttcatcc...............---..........................a....... [191600]
AY261365.1_Warmbaths             .at---------------ctcttcatcc...............---...........................c...... [191600]
AY261364.1_Tengani_62            .at---------------------ctct..a..c..c..c..t---..........................a....... [191600]
AY261363.1_Pretoriuskop/96/4     .at------------ctcttcatcctcc...............---.................................. [191600]
AY261362.1_Mkuzi_1979            .at------------------ctcttca.....c.....t..a---.................................. [191600]
AY261361.1_Malawi_Lil-20/1_1983  .at------------------ctcttca...............---.......................g...c...... [191600]
AY261360.1_Kenya_1950            .at---------------ctcatcatcc...............---...........................c....t. [191600]
AM712240.1_OURT_88/3             .at------------------ctcttca.....c.....t..a---.................................. [191600]
AM712239.1_Benin_97/1            .at------------------ctcttca.....c.....t..a---.................................. [191600]
KM111295.1_Ken06.Bus             .at---------------ctcttcatcc...............tca................................t. [191600]
```

```
KM111294.1_Ken05/Tk1            .at---------------ctcttcatcc...............---................................t. [191600]
KM262844.1_strain_L60           .at------------------ctcttca.....c.....t..a---.................................. [191600]
KM102979.1_26544/OG10_Italy     .at------------------ctcttca.....c.....t..a---.................................. [191600]
KM262845.1_strain_NHV           .at------------------ctcttca.....c.....t..a---.................................. [191600]
KX354450.1_47/Ss/2008           .at------------------ctcttca.....c.....t..a---.................................. [191600]
KP055815.1_strain_BA71          .at------------------ctcttca.....c.....t..a---.................................. [191600]
MH025920.1_strain_R35           .at---------------ctcttcatcc...............tca................................t. [191600]
MH025919.1_strain_N10           .at---------------ctcttcatcc...............tca................................t. [191600]
MH025918.1_strain_R25           .at---------------ctcttcatcc...............tca................................t. [191600]
MH025917.1_strain_R7            .at---------------ctcttcatcc...............tca................................t. [191600]
MH025916.1_strain_R8            .at---------------ctcttcatcc...............tca................................t. [191600]

FR682468.2_Georgia_2007/1       ccatttcttcacattcatc------ttcataagcatcactgggtattggtggaacattggatgcagcatttttaaaatat [191680]
ASF_IND_20_CAD543               ...................------....................................................... [191680]
LR899193.1_Germany_2020/1       ...................------....................................................... [191680]
MN393477.1_Wuhan_2019           ...................------....................................................... [191680]
MW396979.1_Timor-Leste/2019/1   ...................------....................................................... [191680]
MK940252.1_CN/2019/InnerMongolia ...................------....................................................... [191680]
MK128995.1_China/2018/AnhuiXCGQ ...................------....................................................... [191680]
MH681419.1_ASFV/POL/2015        ...................------....................................................... [191680]
LS478113.1_Estonia_2014         ...................------....................................................... [191680]
KP843857.1_Odintsovo_02/14      ...................------....................................................... [191680]
FN557520.1_strain_E75           ...................------....................................................... [191680]
AY261366.1_Warthog              ...................------.......................c.............a................. [191680]
AY261365.1_Warmbaths            ...................------........................................a.............. [191680]
AY261364.1_Tengani_62           ...................------................................c...................... [191680]
AY261363.1_Pretoriuskop/96/4    ............t......------....................................................... [191680]
AY261362.1_Mkuzi_1979           ...................------....................................................... [191680]
AY261361.1_Malawi_Lil-20/1_1983 .......------......------.......t............c....a.......cac..g.a.............. [191680]
AY261360.1_Kenya_1950           ...........tc.....gttcatc..................a..g.c..........a..a................. [191680]
AM712240.1_OURT_88/3            ...................------....................................................... [191680]
AM712239.1_Benin_97/1           ...................------....................................................... [191680]
KM111295.1_Ken06.Bus            ...........tc.....g------..................a..g.ca.........a...t................ [191680]
KM111294.1_Ken05/Tk1            ...........tc.....g------..................a..g.c..........a..a................. [191680]
KM262844.1_strain_L60           ...................------....................................................... [191680]
KM102979.1_26544/OG10_Italy     ...................------....................................................... [191680]
KM262845.1_strain_NHV           ...................------....................................................... [191680]
KX354450.1_47/Ss/2008           ...................------....................................................... [191680]
KP055815.1_strain_BA71          ...................------....................................................... [191680]
MH025920.1_strain_R35           ...........tc.....g------..................a..g.ca.........a...t................ [191680]
MH025919.1_strain_N10           ...........tc.....g------..................a..g.ca.........a...t................ [191680]
MH025918.1_strain_R25           ...........tc.....g------..................a..g.ca.........a...t................ [191680]
MH025917.1_strain_R7            ...........tc.....g------..................a..g.ca.........a...t................ [191680]
MH025916.1_strain_R8            ...........tc.....g------..................a..g.ca.........a...t................ [191680]
```

```
FR682468.2_Georgia_2007/1        tctatgtcttctggtgaacactcatctaatgattttttgacagtccttttaacttccatgggatatgattccaaatcctc [191760]
ASF_IND_20_CAD543                ................................................................................ [191760]
LR899193.1_Germany_2020/1        ................................................................................ [191760]
MN393477.1_Wuhan_2019            ................................................................................ [191760]
MW396979.1_Timor-Leste/2019/1    ................................................................................ [191760]
MK940252.1_CN/2019/InnerMongolia ................................................................................ [191760]
MK128995.1_China/2018/AnhuiXCGQ  ................................................................................ [191760]
MH681419.1_ASFV/POL/2015         ................................................................................ [191760]
LS478113.1_Estonia_2014          ................................................................................ [191760]
KP843857.1_Odintsovo_02/14       ................................................................................ [191760]
FN557520.1_strain_E75            .............................................t.................................. [191760]
AY261366.1_Warthog               ................................................................................ [191760]
AY261365.1_Warmbaths             .............................................t.................................. [191760]
AY261364.1_Tengani_62            ................................................................................ [191760]
AY261363.1_Pretoriuskop/96/4     ................................................................................ [191760]
AY261362.1_Mkuzi_1979            .............................c....................g............................. [191760]
AY261361.1_Malawi_Lil-20/1_1983  ...........c.c............c..................t................g........t........ [191760]
AY261360.1_Kenya_1950            .....a.......c...............................t................g................. [191760]
AM712240.1_OURT_88/3             .............................................t.................................. [191760]
AM712239.1_Benin_97/1            .............................................t.................................. [191760]
KM111295.1_Ken06.Bus             .....a.......c...............................t...............cg................. [191760]
KM111294.1_Ken05/Tk1             .....a.......c...............................t................g..............t.. [191760]
KM262844.1_strain_L60            .............................................t.................................. [191760]
KM102979.1_26544/OG10_Italy      .............................................t.................................. [191760]
KM262845.1_strain_NHV            .............................................t.................................. [191760]
KX354450.1_47/Ss/2008            .............................................t.................................. [191760]
KP055815.1_strain_BA71           .............................................t.................................. [191760]
MH025920.1_strain_R35            .....a.......c...............................t...............cg................. [191760]
MH025919.1_strain_N10            .....a.......c...............................t...............cg................. [191760]
MH025918.1_strain_R25            .....a.......c...............................t...............cg................. [191760]
MH025917.1_strain_R7             .....a.......c...............................t...............cg................. [191760]
MH025916.1_strain_R8             .....a.......c...............................t...............cg................. [191760]

FR682468.2_Georgia_2007/1        tttatataagagtttacggtagcttttagctgcatccacatttgctggagaatctggatttggctcattgagcagtgaaa [191840]
ASF_IND_20_CAD543                ........cata.................................................................... [191840]
LR899193.1_Germany_2020/1        ................................................................................ [191840]
MN393477.1_Wuhan_2019            ................................................................................ [191840]
MW396979.1_Timor-Leste/2019/1    ................................................................................ [191840]
MK940252.1_CN/2019/InnerMongolia ................................................................................ [191840]
MK128995.1_China/2018/AnhuiXCGQ  ................................................................................ [191840]
MH681419.1_ASFV/POL/2015         ................................................................................ [191840]
LS478113.1_Estonia_2014          ................................................................................ [191840]
KP843857.1_Odintsovo_02/14       ................................................................................ [191840]
```

```
FN557520.1_strain_E75          ........cata..............................................................a..... [191840]
AY261366.1_Warthog             .....gc.tt.a.................................c.................................. [191840]
AY261365.1_Warmbaths           ...........ac................................................................... [191840]
AY261364.1_Tengani_62          .........ata.................................................................... [191840]
AY261363.1_Pretoriuskop/96/4   .....gc.ca.a.c.............t..........................c.t....................... [191840]
AY261362.1_Mkuzi_1979          .....gc.tt.a..............................................................a..... [191840]
AY261361.1_Malawi_Lil-20/1_1983 ........gata........................t......................................a.... [191840]
AY261360.1_Kenya_1950          .....gc.tc.a...............g..c.....t.................c.t..............a......g. [191840]
AM712240.1_OURT_88/3           ........cata..............................................................a..... [191840]
AM712239.1_Benin_97/1          ........cata..............................................................a..... [191840]
KM111295.1_Ken06.Bus           .....gc.t..a...............g..c.....t.................c.t..............a......g. [191840]
KM111294.1_Ken05/Tk1           .....gc.tc.a...............g..c.....t.................c.t............a.a......g. [191840]
KM262844.1_strain_L60          ........cata..............................................................a..... [191840]
KM102979.1_26544/OG10_Italy    ........caca..............................................................a..... [191840]
KM262845.1_strain_NHV          ........cata..............................................................a..... [191840]
KX354450.1_47/Ss/2008          ........caca..............................................................a..... [191840]
KP055815.1_strain_BA71         ........cata..............................................................a..... [191840]
MH025920.1_strain_R35          .....gc.t..a...............g..c.....t.................c.t..............a......g. [191840]
MH025919.1_strain_N10          .....gc.t..a...............g..c.....t.................c.t..............a......g. [191840]
MH025918.1_strain_R25          .....gc.t..a...............g..c.....t.................c.t..............a......g. [191840]
MH025917.1_strain_R7           .....gc.t..a...............g..c.....t.................c.t..............a......g. [191840]
MH025916.1_strain_R8           .....gc.t..a...............g..c.....t.................c.t..............a......g. [191840]

FR682468.2_Georgia_2007/1      ttacactaagaagaatggtatcaatcttttgagccggagaccaagtcattccctgttcttcagcattgtctccgtgtaag [191920]
ASF_IND_20_CAD543              ................................................................................ [191920]
LR899193.1_Germany_2020/1      ................................................................................ [191920]
MN393477.1_Wuhan_2019          ................................................................................ [191920]
MW396979.1_Timor-Leste/2019/1  ................................................................................ [191920]
MK940252.1_CN/2019/InnerMongolia ................................................................................ [191920]
MK128995.1_China/2018/AnhuiXCGQ ................................................................................ [191920]
MH681419.1_ASFV/POL/2015       ................................................................................ [191920]
LS478113.1_Estonia_2014        ................................................................................ [191920]
KP843857.1_Odintsovo_02/14     ................................................................................ [191920]
FN557520.1_strain_E75          ................................................................................ [191920]
AY261366.1_Warthog             ................................................................................ [191920]
AY261365.1_Warmbaths           ................................................................................ [191920]
AY261364.1_Tengani_62          ...............c................a............................................... [191920]
AY261363.1_Pretoriuskop/96/4   ................................a........................................a...... [191920]
AY261362.1_Mkuzi_1979          ................................................................................ [191920]
AY261361.1_Malawi_Lil-20/1_1983 .............t.c.........................................................a..c... [191920]
AY261360.1_Kenya_1950          ...............ca..t..............t..................ct...c...a........gt....... [191920]
AM712240.1_OURT_88/3           ...........................................................c.................... [191920]
AM712239.1_Benin_97/1          ................................................................................ [191920]
KM111295.1_Ken06.Bus           ...............ca..t..............t..................ct...c....gg..a...gt....... [191920]
```

| | | |
|---|---|---|
| KM111294.1_Ken05/Tk1 | ...............ca..t..............t..................ct...c............gt....... | [191920] |
| KM262844.1_strain_L60 | ................................................................................ | [191920] |
| KM102979.1_26544/OG10_Italy | ................................................................................ | [191920] |
| KM262845.1_strain_NHV | ...........................................................c.................... | [191920] |
| KX354450.1_47/Ss/2008 | ................................................................................ | [191920] |
| KP055815.1_strain_BA71 | ................................................................................ | [191920] |
| MH025920.1_strain_R35 | ...............ca..t..............t..................ct...c....gg..a...gt....... | [191920] |
| MH025919.1_strain_N10 | ...............ca..t..............t..................ct...c....gg..a...gt....... | [191920] |
| MH025918.1_strain_R25 | ...............ca..t..............t..................ct...c....gg..a...gt....... | [191920] |
| MH025917.1_strain_R7 | ...............ca..t..............t..................ct...c....gg..a...gt....... | [191920] |
| MH025916.1_strain_R8 | ...............ca..t..............t..................ct...c....gg..a...gt....... | [191920] |
| FR682468.2_Georgia_2007/1 | atagagatacatagttttccatcagagtaaatattaggatgccacatttcagaggtgaatgttaatctgggtggtgcata | [192000] |
| ASF_IND_20_CAD543 | ...............y.........r...................................................... | [192000] |
| LR899193.1_Germany_2020/1 | ................................................................................ | [192000] |
| MN393477.1_Wuhan_2019 | ................................................................................ | [192000] |
| MW396979.1_Timor-Leste/2019/1 | ................................................................................ | [192000] |
| MK940252.1_CN/2019/InnerMongolia | ................................................................................ | [192000] |
| MK128995.1_China/2018/AnhuiXCGQ | ................................................................................ | [192000] |
| MH681419.1_ASFV/POL/2015 | ................................................................................ | [192000] |
| LS478113.1_Estonia_2014 | ................................................................................ | [192000] |
| KP843857.1_Odintsovo_02/14 | ................................................................................ | [192000] |
| FN557520.1_strain_E75 | ...............c.........g........................................t............. | [192000] |
| AY261366.1_Warthog | .........................g...................................................... | [192000] |
| AY261365.1_Warmbaths | .........................g...................................................... | [192000] |
| AY261364.1_Tengani_62 | ................................................................................ | [192000] |
| AY261363.1_Pretoriuskop/96/4 | .............c.c................g..........................................t.... | [192000] |
| AY261362.1_Mkuzi_1979 | ................................................................................ | [192000] |
| AY261361.1_Malawi_Lil-20/1_1983 | .....a.......a...c...........g.....g.......................................t.... | [192000] |
| AY261360.1_Kenya_1950 | ........g......c.c.......ga..g....................................t........t.... | [192000] |
| AM712240.1_OURT_88/3 | ...............c.........g........................................t............. | [192000] |
| AM712239.1_Benin_97/1 | ...............c.........g........................................t............. | [192000] |
| KM111295.1_Ken06.Bus | ...............c.c.......ca..g........................a...........t........t.... | [192000] |
| KM111294.1_Ken05/Tk1 | ...............c.c.......ga..g....................................t........t.... | [192000] |
| KM262844.1_strain_L60 | ...............c.........g........................................t............. | [192000] |
| KM102979.1_26544/OG10_Italy | ...............c.........g........................................t............. | [192000] |
| KM262845.1_strain_NHV | ...............c.........g........................................t............. | [192000] |
| KX354450.1_47/Ss/2008 | ...............c.........g........................................t............. | [192000] |
| KP055815.1_strain_BA71 | ...............c.........g........................................t............. | [192000] |
| MH025920.1_strain_R35 | ...............c.c.......ca..g........................a...........t........t.... | [192000] |
| MH025919.1_strain_N10 | ...............c.c.......ca..g........................a...........t........t.... | [192000] |
| MH025918.1_strain_R25 | ...............c.c.......ca..g........................a...........t........t.... | [192000] |
| MH025917.1_strain_R7 | ...............c.c.......ca..g........................a...........t........t.... | [192000] |
| MH025916.1_strain_R8 | ...............c.c.......ca..g........................a...........t........t.... | [192000] |

```
FR682468.2_Georgia_2007/1        tgggtattctggaggaaaggcgatttttgccttgaataagcctccctcataaaaagtgtcaggtgggccccttaagatca [192080]
ASF_IND_20_CAD543                ................................................................................ [192080]
LR899193.1_Germany_2020/1        ................................................................................ [192080]
MN393477.1_Wuhan_2019            ................................................................................ [192080]
MW396979.1_Timor-Leste/2019/1    ................................................................................ [192080]
MK940252.1_CN/2019/InnerMongolia ................................................................................ [192080]
MK128995.1_China/2018/AnhuiXCGQ  ................................................................................ [192080]
MH681419.1_ASFV/POL/2015         ................................................................................ [192080]
LS478113.1_Estonia_2014          ................................................................................ [192080]
KP843857.1_Odintsovo_02/14       ................................................................................ [192080]
FN557520.1_strain_E75            .......................c.............................g.......................... [192080]
AY261366.1_Warthog               .......................c.............................g.......................... [192080]
AY261365.1_Warmbaths             .......................c.............................g.......................... [192080]
AY261364.1_Tengani_62            .......................cc.............................................t......... [192080]
AY261363.1_Pretoriuskop/96/4     ......a.t..........c....c............................g........c.......t.......g. [192080]
AY261362.1_Mkuzi_1979            .......................c.............................g.......................... [192080]
AY261361.1_Malawi_Lil-20/1_1983  ........t..........a.a..c.....t......................g...............tt.......g. [192080]
AY261360.1_Kenya_1950            ...a..a.t...........t...c.....................................a...a...t.......g. [192080]
AM712240.1_OURT_88/3             .......................c.............................g.......................... [192080]
AM712239.1_Benin_97/1            .......................c.............................g.......................... [192080]
KM111295.1_Ken06.Bus             ...a..a.t..........tt...c...........c.............................a...t.......g. [192080]
KM111294.1_Ken05/Tk1             ...a..a.t...........t...c.........................................a...t.......g. [192080]
KM262844.1_strain_L60            .......................c.............................g.......................... [192080]
KM102979.1_26544/OG10_Italy      .......................c.............................g.......................... [192080]
KM262845.1_strain_NHV            .......................c.............................g.......................... [192080]
KX354450.1_47/Ss/2008            .......................c.............................g.......................... [192080]
KP055815.1_strain_BA71           .......................c.............................g.......................... [192080]
MH025920.1_strain_R35            ...a..a.t..........tt...c...........c.............................a...t.......g. [192080]
MH025919.1_strain_N10            ...a..a.t..........tt...c...........c.............................a...t.......g. [192080]
MH025918.1_strain_R25            ...a..a.t..........tt...c...........c.............................a...t.......g. [192080]
MH025917.1_strain_R7             ...a..a.t..........tt...c...........c.............................a...t.......g. [192080]
MH025916.1_strain_R8             ...a..a.t..........tt...c...........c.............................a...t.......g. [192080]

FR682468.2_Georgia_2007/1        catcccattcagtcatatccttctcattcaccgaaattttgaaattctcagagggattctctatcaggtgtctgtactct [192160]
ASF_IND_20_CAD543                ..................ym...........y................................................ [192160]
LR899193.1_Germany_2020/1        ................................................................................ [192160]
MN393477.1_Wuhan_2019            ................................................................................ [192160]
MW396979.1_Timor-Leste/2019/1    ................................................................................ [192160]
MK940252.1_CN/2019/InnerMongolia ................................................................................ [192160]
MK128995.1_China/2018/AnhuiXCGQ  ................................................................................ [192160]
MH681419.1_ASFV/POL/2015         ................................................................................ [192160]
LS478113.1_Estonia_2014          ................................................................................ [192160]
KP843857.1_Odintsovo_02/14       ................................................................................ [192160]
```

```
FN557520.1_strain_E75            .............t....ta...........t.......................g........................ [192160]
AY261366.1_Warthog               ...................a...........t................................................ [192160]
AY261365.1_Warmbaths             ...................a...........t................................................ [192160]
AY261364.1_Tengani_62            .............t.a..ta...........t..g.c........................................... [192160]
AY261363.1_Pretoriuskop/96/4     ..................ta...........t..g.c.........................g................. [192160]
AY261362.1_Mkuzi_1979            ..................ta...........t.......................g........................ [192160]
AY261361.1_Malawi_Lil-20/1_1983  ...............a..ta.ct........t..g..........g.............a....t....t.t........ [192160]
AY261360.1_Kenya_1950            .............t.g..ta...........t..g.c........gt.....a.........g.t....t.......... [192160]
AM712240.1_OURT_88/3             .............t....ta...........t.......................g........................ [192160]
AM712239.1_Benin_97/1            .............t....ta...........t.......................g........................ [192160]
KM111295.1_Ken06.Bus             ...............g..ta...........t..g.c...a.....t.....a.........g.t....t.......... [192160]
KM111294.1_Ken05/Tk1             .............t.g..ta...........t..g.c........gt.....a.........g.t....t.......... [192160]
KM262844.1_strain_L60            .............t....ta...........t.......................g........................ [192160]
KM102979.1_26544/OG10_Italy      .............t....ta...........t.......................g........................ [192160]
KM262845.1_strain_NHV            .............t....ta...........t.......................g........................ [192160]
KX354450.1_47/Ss/2008            .............t....ta...........t.......................g........................ [192160]
KP055815.1_strain_BA71           .............t....ta...........t.......................g........................ [192160]
MH025920.1_strain_R35            ...............g..ta...........t..g.c...a.....t.....a.........g.t....t.......... [192160]
MH025919.1_strain_N10            ...............g..ta...........t..g.c...a.....t.....a.........g.t....t.......... [192160]
MH025918.1_strain_R25            ...............g..ta...........t..g.c...a.....t.....a.........g.t....t.......... [192160]
MH025917.1_strain_R7             ...............g..ta...........t..g.c...a.....t.....a.........g.t....t.......... [192160]
MH025916.1_strain_R8             ...............g..ta...........t..g.c...a.....t.....a.........g.t....t.......... [192160]

FR682468.2_Georgia_2007/1        gctattaaaaacctggaaaccatggttatttaatattaattaaattccctggtttattcctccttaaaagtagatgaacc [192240]
ASF_IND_20_CAD543                .........................................................---.................... [192240]
LR899193.1_Germany_2020/1        ................................................................................ [192240]
MN393477.1_Wuhan_2019            ................................................................................ [192240]
MW396979.1_Timor-Leste/2019/1    ................................................................................ [192240]
MK940252.1_CN/2019/InnerMongolia ................................................................................ [192240]
MK128995.1_China/2018/AnhuiXCGQ  ................................................................................ [192240]
MH681419.1_ASFV/POL/2015         ................................................................................ [192240]
LS478113.1_Estonia_2014          ................................................................................ [192240]
KP843857.1_Odintsovo_02/14       ................................................................................ [192240]
FN557520.1_strain_E75            .........................................................---.................... [192240]
AY261366.1_Warthog               ..c.................................g....................---.................... [192240]
AY261365.1_Warmbaths             ....a...............................g....................---.................... [192240]
AY261364.1_Tengani_62            ..c.................................g....................---.................... [192240]
AY261363.1_Pretoriuskop/96/4     ..c.................................g....................---.................... [192240]
AY261362.1_Mkuzi_1979            ..c......................................................---.................... [192240]
AY261361.1_Malawi_Lil-20/1_1983  ..c.g......a.............c..........g...........t.......a.--.................... [192240]
AY261360.1_Kenya_1950            .tc.g......t.................................c..t...c.....t-c................... [192240]
AM712240.1_OURT_88/3             .........................................................---.................... [192240]
AM712239.1_Benin_97/1            .........................................................---.................... [192240]
KM111295.1_Ken06.Bus             ..c.g......t.................................c..t..ac.....t-...c................ [192240]
```

```
KM111294.1_Ken05/Tk1              .tc.g......t..a..............................c..t...c.....t-c................... [192240]
KM262844.1_strain_L60             .........................................................---.................... [192240]
KM102979.1_26544/OG10_Italy       .........................................................---.................... [192240]
KM262845.1_strain_NHV             .........................................................---.................... [192240]
KX354450.1_47/Ss/2008             .........................................................---.................... [192240]
KP055815.1_strain_BA71            .........................................................---.................... [192240]
MH025920.1_strain_R35             ..c.g......t.....................c...........c..t..ac.....t-...c................ [192240]
MH025919.1_strain_N10             ..c.g......t..a..................c...........c..t..ac.....t-...c................ [192240]
MH025918.1_strain_R25             ..c.g......t.....................c...........c..t..ac.....t-...c................ [192240]
MH025917.1_strain_R7              ..c.g......t.....................c...........c..t..ac.....t-...c................ [192240]
MH025916.1_strain_R8              ..c.g......t.....................c...........c..t..ac.....t-...c................ [192240]

FR682468.2_Georgia_2007/1         tcttttgttttttattgggttcatttttactaaatttatgaactgg-aaaaaactttaacggcataattatcaaatgcga [192320]
ASF_IND_20_CAD543                 ..............................................-................................. [192320]
LR899193.1_Germany_2020/1         ..............................................-................................. [192320]
MN393477.1_Wuhan_2019             ..............................................-................................. [192320]
MW396979.1_Timor-Leste/2019/1     ..............................................-................................. [192320]
MK940252.1_CN/2019/InnerMongolia  ..............................................-................................. [192320]
MK128995.1_China/2018/AnhuiXCGQ   ..............................................-................................. [192320]
MH681419.1_ASFV/POL/2015          ..............................................-................................. [192320]
LS478113.1_Estonia_2014           ..............................................-................................. [192320]
KP843857.1_Odintsovo_02/14        ..............................................-................................. [192320]
FN557520.1_strain_E75             ..............................................-.....t...............c........... [192320]
AY261366.1_Warthog                ..............................................-.....t.........t....c............ [192320]
AY261365.1_Warmbaths              ..............................................-................................. [192320]
AY261364.1_Tengani_62             ..............................................-.....t.........t....c............ [192320]
AY261363.1_Pretoriuskop/96/4      ..............................................-..............................t.. [192320]
AY261362.1_Mkuzi_1979             ..............................................-.....t...............c........... [192320]
AY261361.1_Malawi_Lil-20/1_1983   ...............g..............................-t....c......t...............c.a.. [192320]
AY261360.1_Kenya_1950             ...............a..................a..t.......at.t...t.c...................g..a.. [192320]
AM712240.1_OURT_88/3              ..............................................-.....t...............c........... [192320]
AM712239.1_Benin_97/1             ..............................................-.....t...............c........... [192320]
KM111295.1_Ken06.Bus              ...............a..................a..t........t...c.t.c................t..g..a.. [192320]
KM111294.1_Ken05/Tk1              ...............a..................a..t.......at.....t.c...................g..a.. [192320]
KM262844.1_strain_L60             ..............................................-.....t...............c........... [192320]
KM102979.1_26544/OG10_Italy       ..............................................-.....t...............c........... [192320]
KM262845.1_strain_NHV             ..............................................-.....t...............c........... [192320]
KX354450.1_47/Ss/2008             ..............................................-.....t...............c........... [192320]
KP055815.1_strain_BA71            ..............................................-.....t...............c........... [192320]
MH025920.1_strain_R35             ...............a..................a..t........t...c.t.c................t..g..a.. [192320]
MH025919.1_strain_N10             ...............a..................a..t........t...c.t.c................t..g..a.. [192320]
MH025918.1_strain_R25             ...............a..................a..t........t...c.t.c................t..g..a.. [192320]
MH025917.1_strain_R7              ...............a..................a..t........t...c.t.c................t..g..a.. [192320]
MH025916.1_strain_R8              ...............a..................a..t........t...c.t.c................t..g..a.. [192320]
```

```
FR682468.2_Georgia_2007/1        aggggatccgtataaaatcctagctt------------------------------------------------------ [192400]
ASF_IND_20_CAD543                ..........................------------------------------------------------------ [192400]
LR899193.1_Germany_2020/1        ..........................------------------------------------------------------ [192400]
MN393477.1_Wuhan_2019            ..........................------------------------------------------------------ [192400]
MW396979.1_Timor-Leste/2019/1    ..........................------------------------------------------------------ [192400]
MK940252.1_CN/2019/InnerMongolia ..........................------------------------------------------------------ [192400]
MK128995.1_China/2018/AnhuiXCGQ  ..........................------------------------------------------------------ [192400]
MH681419.1_ASFV/POL/2015         ..........................------------------------------------------------------ [192400]
LS478113.1_Estonia_2014          ..........................------------------------------------------------------ [192400]
KP843857.1_Odintsovo_02/14       ..........................------------------------------------------------------ [192400]
FN557520.1_strain_E75            ..........................------------------------------------------------------ [192400]
AY261366.1_Warthog               ..........................------------------------------------------------------ [192400]
AY261365.1_Warmbaths             ..........................------------------------------------------------------ [192400]
AY261364.1_Tengani_62            ..........................------------------------------------------------------ [192400]
AY261363.1_Pretoriuskop/96/4     ...........c..............------------------------------------------------------ [192400]
AY261362.1_Mkuzi_1979            ..........................------------------------------------------------------ [192400]
AY261361.1_Malawi_Lil-20/1_1983  ....a............c....tt..------------------------------------------------------ [192400]
AY261360.1_Kenya_1950            ...a.............c....tt..ggcggaagtggaaaaatttttaagggaattattatcaagtgagaaggaggatc [192400]
AM712240.1_OURT_88/3             ..........................------------------------------------------------------ [192400]
AM712239.1_Benin_97/1            ..........................------------------------------------------------------ [192400]
KM111295.1_Ken06.Bus             ...a..................tt..------------------------------------------------------ [192400]
KM111294.1_Ken05/Tk1             ...a.............c....tt..ggt--------------------------------------------------- [192400]
KM262844.1_strain_L60            ..........................------------------------------------------------------ [192400]
KM102979.1_26544/OG10_Italy      ..........................------------------------------------------------------ [192400]
KM262845.1_strain_NHV            ..........................------------------------------------------------------ [192400]
KX354450.1_47/Ss/2008            ..........................------------------------------------------------------ [192400]
KP055815.1_strain_BA71           ..........................------------------------------------------------------ [192400]
MH025920.1_strain_R35            ...a..................tt..------------------------------------------------------ [192400]
MH025919.1_strain_N10            ...a..................tt..g----------------------------------------------------- [192400]
MH025918.1_strain_R25            ...a..................tt..------------------------------------------------------ [192400]
MH025917.1_strain_R7             ...a..................tt..------------------------------------------------------ [192400]
MH025916.1_strain_R8             ...a..................tt..------------------------------------------------------ [192400]

FR682468.2_Georgia_2007/1        --------------------gccggt------------------------------------------------------ [192480]
ASF_IND_20_CAD543                --------------------......------------------------------------------------------ [192480]
LR899193.1_Germany_2020/1        --------------------......------------------------------------------------------ [192480]
MN393477.1_Wuhan_2019            --------------------......------------------------------------------------------ [192480]
MW396979.1_Timor-Leste/2019/1    --------------------......------------------------------------------------------ [192480]
MK940252.1_CN/2019/InnerMongolia --------------------......------------------------------------------------------ [192480]
MK128995.1_China/2018/AnhuiXCGQ  --------------------......------------------------------------------------------ [192480]
MH681419.1_ASFV/POL/2015         --------------------......------------------------------------------------------ [192480]
LS478113.1_Estonia_2014          --------------------......------------------------------------------------------ [192480]
KP843857.1_Odintsovo_02/14       --------------------......------------------------------------------------------ [192480]
```

```
FN557520.1_strain_E75             ---------------------......----------------------------------------------------- [192480]
AY261366.1_Warthog                ---------------------......----------------------------------------------------- [192480]
AY261365.1_Warmbaths              ---------------------......----------------------------------------------------- [192480]
AY261364.1_Tengani_62             ---------------------......----------------------------------------------------- [192480]
AY261363.1_Pretoriuskop/96/4      ---------------------......----------------------------------------------------- [192480]
AY261362.1_Mkuzi_1979             ---------------------......----------------------------------------------------- [192480]
AY261361.1_Malawi_Lil-20/1_1983   ---------------------.ga...ggaaaaaagtttaagggcataaggaataattgagaaggtggatccgtataaaa [192480]
AY261360.1_Kenya_1950             cgtataaaaccctattttggc.gaa..ggaaaaatttttaagggaattattatcaagtgagaaggaggatccgtataaaa [192480]
AM712240.1_OURT_88/3              ---------------------......----------------------------------------------------- [192480]
AM712239.1_Benin_97/1             ---------------------......----------------------------------------------------- [192480]
KM111295.1_Ken06.Bus              ---------------------.gg...ggaaaaaagtttaagggcataagtatgaactgagaaggaggatctgtataaaa [192480]
KM111294.1_Ken05/Tk1              ---------------------.gga..ggaaaaatttttaagggaattattatcaagtgagaaggaggatccgtataaaa [192480]
KM262844.1_strain_L60             ---------------------......----------------------------------------------------- [192480]
KM102979.1_26544/OG10_Italy       ---------------------......----------------------------------------------------- [192480]
KM262845.1_strain_NHV             ---------------------......----------------------------------------------------- [192480]
KX354450.1_47/Ss/2008             ---------------------......----------------------------------------------------- [192480]
KP055815.1_strain_BA71            ---------------------......----------------------------------------------------- [192480]
MH025920.1_strain_R35             ---------------------.gg...ggaaaaaagtttaagggcataagtatgaactgagaaggaggatctgtataaaa [192480]
MH025919.1_strain_N10             ---------------------.gg...ggaaaaaagtttaagggcataagtatgaactgagaaggaggatctgtataaaa [192480]
MH025918.1_strain_R25             ---------------------.gg...ggaaaaaagtttaagggcataagtatgaactgagaaggaggatctgtataaaa [192480]
MH025917.1_strain_R7              ---------------------.gg...ggaaaaaagtttaagggcataagtatgaactgagaaggaggatctgtataaaa [192480]
MH025916.1_strain_R8              ---------------------.gg...ggaaaaaagtttaagggcataagtatgaactgagaaggaggatctgtataaaa [192480]

FR682468.2_Georgia_2007/1         ---------------aatggctattaagttaaatttggtaccagtaacactaatatttaaaaagccctgatcattaactt [192560]
ASF_IND_20_CAD543                 ---------------.....................r.....w......s.............................. [192560]
LR899193.1_Germany_2020/1         ---------------................................................................. [192560]
MN393477.1_Wuhan_2019             ---------------................................................................. [192560]
MW396979.1_Timor-Leste/2019/1     ---------------................................................................. [192560]
MK940252.1_CN/2019/InnerMongolia  ---------------................................................................. [192560]
MK128995.1_China/2018/AnhuiXCGQ   ---------------................................................................. [192560]
MH681419.1_ASFV/POL/2015          ---------------................................................................. [192560]
LS478113.1_Estonia_2014           ---------------................................................................. [192560]
KP843857.1_Odintsovo_02/14        ---------------................................................................. [192560]
FN557520.1_strain_E75             ---------------.....................a.....t......g.............................. [192560]
AY261366.1_Warthog                ---------------.g...................a........................................... [192560]
AY261365.1_Warmbaths              ---------------.g.............c.....a........................................... [192560]
AY261364.1_Tengani_62             ---------------.g...................a........................................... [192560]
AY261363.1_Pretoriuskop/96/4      ---------------...............c.....a........................................... [192560]
AY261362.1_Mkuzi_1979             ---------------.....................a.....t......g.............................. [192560]
AY261361.1_Malawi_Lil-20/1_1983   ccctattttggagga.g...................a..................a..................c..... [192560]
AY261360.1_Kenya_1950             tcctattttggtgga.g...................a.....g.c...cg.............................. [192560]
AM712240.1_OURT_88/3              ---------------.....................a.....t......g.............................. [192560]
AM712239.1_Benin_97/1             ---------------.....................a.....t......g.............................. [192560]
KM111295.1_Ken06.Bus              tcctaatttggcgga.g.........c......a..a.....g.c...cg.....a.c.t.t......t.....c..t.. [192560]
```

```
KM111294.1_Ken05/Tk1             ccctattttggtgga.g...................a.....g.c...cg.............................. [192560]
KM262844.1_strain_L60            ---------------.....................a.....t......g.............................. [192560]
KM102979.1_26544/OG10_Italy      ---------------.....................a.....t......g.............................. [192560]
KM262845.1_strain_NHV            ---------------.....................a.....t......g.............................. [192560]
KX354450.1_47/Ss/2008            ---------------.....................a.....t......g.............................. [192560]
KP055815.1_strain_BA71           ---------------.....................a.....t......g.............................. [192560]
MH025920.1_strain_R35            tcctaatttggcgga.g.........c......a..a.....g.c...cg.....a.c.t.t......t.....c..t.. [192560]
MH025919.1_strain_N10            tcctaatttggcgga.g.........c......a..a.....g.c...cg.....acc.t.t......t.....c..t.. [192560]
MH025918.1_strain_R25            tcctaatttggcgga.g.........c......a..a.....g.c...cg.....a.c.t.t......t.....c..t.. [192560]
MH025917.1_strain_R7             tcctaatttggcgga.g.........c......a..a.....g.c...cg.....a.c.t.t......t.....c..t.. [192560]
MH025916.1_strain_R8             tcctaatttggcgga.g.........c......a..a.....g.c...cg.....a.c.t.t......t.....c..t.. [192560]

FR682468.2_Georgia_2007/1        tccacattaaaagattattatattcgaatgtttgtccaatatggacaactttgtcaccagatgttacatttgatttggtt [192640]
ASF_IND_20_CAD543                .............................................tgt........................ta...... [192640]
LR899193.1_Germany_2020/1        ................................................................................ [192640]
MN393477.1_Wuhan_2019            ................................................................................ [192640]
MW396979.1_Timor-Leste/2019/1    ................................................................................ [192640]
MK940252.1_CN/2019/InnerMongolia ................................................................................ [192640]
MK128995.1_China/2018/AnhuiXCGQ  ................................................................................ [192640]
MH681419.1_ASFV/POL/2015         ................................................................................ [192640]
LS478113.1_Estonia_2014          ................................................................................ [192640]
KP843857.1_Odintsovo_02/14       ................................................................................ [192640]
FN557520.1_strain_E75            .............................................tgt..............a.........ta...... [192640]
AY261366.1_Warthog               .....................................t.......tgt.....g........a......a..t.aa.... [192640]
AY261365.1_Warmbaths             .............................................tg................................. [192640]
AY261364.1_Tengani_62            .............................................tgt.....g........a.........t.aa.... [192640]
AY261363.1_Pretoriuskop/96/4     .............................................tg................................. [192640]
AY261362.1_Mkuzi_1979            .............................................tgt..a..c........a.........ta...... [192640]
AY261361.1_Malawi_Lil-20/1_1983  ..........................t..................ttgt...a.........ac...g....t....... [192640]
AY261360.1_Kenya_1950            ..........................t..................ttgt....................g..t....... [192640]
AM712240.1_OURT_88/3             .............................................tgt..............a.........ta...... [192640]
AM712239.1_Benin_97/1            .............................................tgt..............a.........ta...... [192640]
KM111295.1_Ken06.Bus             ..........................t..................ttgt...c..g.............g..t....... [192640]
KM111294.1_Ken05/Tk1             ..........................t..................ttgt....................g..t....... [192640]
KM262844.1_strain_L60            .............................................tgt..............a.........ta...... [192640]
KM102979.1_26544/OG10_Italy      .............................................tgt..............a.........ta...... [192640]
KM262845.1_strain_NHV            .............................................tgt..............a.........ta...... [192640]
KX354450.1_47/Ss/2008            .............................................tgt..............a.........ta...... [192640]
KP055815.1_strain_BA71           .............................................tgt..............a.........ta...... [192640]
MH025920.1_strain_R35            ..........................t..................ttgt...c..g.............g..t....... [192640]
MH025919.1_strain_N10            ..........................t..................ttgt...c..g.............g..t....... [192640]
MH025918.1_strain_R25            ..........................t..................ttgt...c..g.............g..t....... [192640]
MH025917.1_strain_R7             ..........................t..................ttgt...c..g.............g..t....... [192640]
MH025916.1_strain_R8             ..........................t..................ttgt...c..g.............g..t....... [192640]
```

```
FR682468.2_Georgia_2007/1        gttagtggctgaagcttggcacaatcaaaaataagcccattaacactaagatatagaggagtgggttgatctattttctc [192720]
ASF_IND_20_CAD543                ................................................................................ [192720]
LR899193.1_Germany_2020/1        ................................................................................ [192720]
MN393477.1_Wuhan_2019            ................................................................................ [192720]
MW396979.1_Timor-Leste/2019/1    ................................................................................ [192720]
MK940252.1_CN/2019/InnerMongolia ................................................................................ [192720]
MK128995.1_China/2018/AnhuiXCGQ  ................................................................................ [192720]
MH681419.1_ASFV/POL/2015         ................................................................................ [192720]
LS478113.1_Estonia_2014          ................................................................................ [192720]
KP843857.1_Odintsovo_02/14       ................................................................................ [192720]
FN557520.1_strain_E75            ......................................g......................................... [192720]
AY261366.1_Warthog               ........................................c....................................... [192720]
AY261365.1_Warmbaths             ................................................................................ [192720]
AY261364.1_Tengani_62            ........................................c....................................... [192720]
AY261363.1_Pretoriuskop/96/4     ................................................................................ [192720]
AY261362.1_Mkuzi_1979            ......................................g......................................... [192720]
AY261361.1_Malawi_Lil-20/1_1983  .......................g..................ct.t..................t...t........t.. [192720]
AY261360.1_Kenya_1950            ....a......t.....a.....g......................................a..............t.. [192720]
AM712240.1_OURT_88/3             ......................................g......................................... [192720]
AM712239.1_Benin_97/1            ......................................g......................................... [192720]
KM111295.1_Ken06.Bus             ....a............a.t...g...t..............ct....................t...t.g...ag.t.. [192720]
KM111294.1_Ken05/Tk1             ....a......t.....a.....g......................................a..............t.. [192720]
KM262844.1_strain_L60            ......................................g......................................... [192720]
KM102979.1_26544/OG10_Italy      ......................................g......................................... [192720]
KM262845.1_strain_NHV            ......................................g......................................... [192720]
KX354450.1_47/Ss/2008            ......................................g......................................... [192720]
KP055815.1_strain_BA71           ......................................g......................................... [192720]
MH025920.1_strain_R35            ....a............a.t...g...t..............ct....................t...t.g...ag.t.. [192720]
MH025919.1_strain_N10            ....a............a.t...g...t..............ct....................t...t.g...ag.t.. [192720]
MH025918.1_strain_R25            ....a............a.t...g...t..............ct....................t...t.g...ag.t.. [192720]
MH025917.1_strain_R7             ....a............a.t...g...t..............ct....................t...t.g...ag.t.. [192720]
MH025916.1_strain_R8             ....a............a.t...g...t..............ct....................t...t.g...ag.t.. [192720]

FR682468.2_Georgia_2007/1        atagtttaatattccatctttccacgtaatagcttgataattatccgcagcaatgagttgaaattttataaatagtacag [192800]
ASF_IND_20_CAD543                ................................................................................ [192800]
LR899193.1_Germany_2020/1        ................................................................................ [192800]
MN393477.1_Wuhan_2019            ................................................................................ [192800]
MW396979.1_Timor-Leste/2019/1    ................................................................................ [192800]
MK940252.1_CN/2019/InnerMongolia ................................................................................ [192800]
MK128995.1_China/2018/AnhuiXCGQ  ................................................................................ [192800]
MH681419.1_ASFV/POL/2015         ................................................................................ [192800]
LS478113.1_Estonia_2014          ................................................................................ [192800]
KP843857.1_Odintsovo_02/14       ................................................................................ [192800]
```

```
FN557520.1_strain_E75            ....c......c.....t.............................................................. [192800]
AY261366.1_Warthog               ................................................................................ [192800]
AY261365.1_Warmbaths             g...c......c............................g.............c......................... [192800]
AY261364.1_Tengani_62            ................................................................................ [192800]
AY261363.1_Pretoriuskop/96/4     g...c........................................................................... [192800]
AY261362.1_Mkuzi_1979            ....c.....g..................................................................... [192800]
AY261361.1_Malawi_Lil-20/1_1983  .a.............................................tg.................c....g..a..aga [192800]
AY261360.1_Kenya_1950            ..............................g...a..........t..g.......t..............g..a..... [192800]
AM712240.1_OURT_88/3             ....c......c.....t.............................................................. [192800]
AM712239.1_Benin_97/1            ....c......c.....t.............................................................. [192800]
KM111295.1_Ken06.Bus             ...tgc...........t...........cg.............ttatg.t...............c....g..a..gga [192800]
KM111294.1_Ken05/Tk1             ..............................g...a..........t..g.......t..............g..a..... [192800]
KM262844.1_strain_L60            ....c......c.....t.............................................................. [192800]
KM102979.1_26544/OG10_Italy      ....c......c.....t.............................................................. [192800]
KM262845.1_strain_NHV            ....c......c.....t.............................................................. [192800]
KX354450.1_47/Ss/2008            ....c......c.....t.............................................................. [192800]
KP055815.1_strain_BA71           ....c......c.....t.............................................................. [192800]
MH025920.1_strain_R35            ...tgc...........t...........cg.............ttatg.t...............c....g..a..gga [192800]
MH025919.1_strain_N10            ...tgc...........t...........cg.............ttatg.t...............c....g..a..gga [192800]
MH025918.1_strain_R25            ...tgc...........t...........cg.............ttatg.t...............c....g..a..gga [192800]
MH025917.1_strain_R7             ...tgc...........t...........cg.............ttatg.t...............c....g..a..gga [192800]
MH025916.1_strain_R8             ...tgc...........t...........cg.............ttatg.t...............c....g..a..gga [192800]

FR682468.2_Georgia_2007/1        gggttttagttgtcgttatacatttaaagggtgttttataaaaataaaaaataataattgttaaaagtatgataataatc [192880]
ASF_IND_20_CAD543                ................................................................................ [192880]
LR899193.1_Germany_2020/1        ................................................................................ [192880]
MN393477.1_Wuhan_2019            ................................................................................ [192880]
MW396979.1_Timor-Leste/2019/1    ................................................................................ [192880]
MK940252.1_CN/2019/InnerMongolia ................................................................................ [192880]
MK128995.1_China/2018/AnhuiXCGQ  ................................................................................ [192880]
MH681419.1_ASFV/POL/2015         ................................................................................ [192880]
LS478113.1_Estonia_2014          ................................................................................ [192880]
KP843857.1_Odintsovo_02/14       ..............................................-................................. [192880]
FN557520.1_strain_E75            ................................................................................ [192880]
AY261366.1_Warthog               ...........a..........................................g....................cg... [192880]
AY261365.1_Warmbaths             ............................................................................g... [192880]
AY261364.1_Tengani_62            ...........a................................................................g... [192880]
AY261363.1_Pretoriuskop/96/4     ............................................................................g... [192880]
AY261362.1_Mkuzi_1979            ................................................................................ [192880]
AY261361.1_Malawi_Lil-20/1_1983  .................g....c..c..a..............................a...................g [192880]
AY261360.1_Kenya_1950            .t............a..........t..a..............................a......t..c........cg [192880]
AM712240.1_OURT_88/3             ................................................................................ [192880]
AM712239.1_Benin_97/1            ................................................................................ [192880]
KM111295.1_Ken06.Bus             cat............a.g.......c..c.....................................c..c..c.....cg [192880]
```

```
KM111294.1_Ken05/Tk1             .t............a..........t..a..............................a......t..c........cg [192880]
KM262844.1_strain_L60            ................................................................................ [192880]
KM102979.1_26544/OG10_Italy      ................................................................................ [192880]
KM262845.1_strain_NHV            ................................................................................ [192880]
KX354450.1_47/Ss/2008            ................................................................................ [192880]
KP055815.1_strain_BA71           ................................................................................ [192880]
MH025920.1_strain_R35            cat............a.g.......c..c.....................................c..c..c.....cg [192880]
MH025919.1_strain_N10            cat............a.g.......c..c.....................................c..c..c.....cg [192880]
MH025918.1_strain_R25            cat............a.g.......c..c.....................................c..c..c.....cg [192880]
MH025917.1_strain_R7             cat............a.g.......c..c.....................................c..c..c.....cg [192880]
MH025916.1_strain_R8             cat............a.g.......c..c.....................................c..c..c.....cg [192880]

FR682468.2_Georgia_2007/1        gccaaaataatttcatacattttttataagaattatacatagtatggtatttaaaatattagctaaatttaaaaaaactt [192960]
ASF_IND_20_CAD543                ................................................................................ [192960]
LR899193.1_Germany_2020/1        ................................................................................ [192960]
MN393477.1_Wuhan_2019            ................................................................................ [192960]
MW396979.1_Timor-Leste/2019/1    ................................................................................ [192960]
MK940252.1_CN/2019/InnerMongolia ................................................................................ [192960]
MK128995.1_China/2018/AnhuiXCGQ  ................................................................................ [192960]
MH681419.1_ASFV/POL/2015         ................................................................................ [192960]
LS478113.1_Estonia_2014          ................................................................................ [192960]
KP843857.1_Odintsovo_02/14       ................................................................................ [192960]
FN557520.1_strain_E75            ................................................................................ [192960]
AY261366.1_Warthog               ................................................................................ [192960]
AY261365.1_Warmbaths             ................................................................................ [192960]
AY261364.1_Tengani_62            ................................................................................ [192960]
AY261363.1_Pretoriuskop/96/4     ................................................................................ [192960]
AY261362.1_Mkuzi_1979            ................................................................................ [192960]
AY261361.1_Malawi_Lil-20/1_1983  .............................c.................................................. [192960]
AY261360.1_Kenya_1950            ................................................................................ [192960]
AM712240.1_OURT_88/3             ................................................................................ [192960]
AM712239.1_Benin_97/1            ................................................................................ [192960]
KM111295.1_Ken06.Bus             .............................c..............................g................... [192960]
KM111294.1_Ken05/Tk1             ................................................................................ [192960]
KM262844.1_strain_L60            ................................................................................ [192960]
KM102979.1_26544/OG10_Italy      ................................................................................ [192960]
KM262845.1_strain_NHV            ................................................................................ [192960]
KX354450.1_47/Ss/2008            ................................................................................ [192960]
KP055815.1_strain_BA71           ................................................................................ [192960]
MH025920.1_strain_R35            .............................c..............................g................... [192960]
MH025919.1_strain_N10            .............................c..............................g................... [192960]
MH025918.1_strain_R25            .............................c..............................g................... [192960]
MH025917.1_strain_R7             .............................c..............................g................... [192960]
MH025916.1_strain_R8             .............................c..............................g................... [192960]
```

```
FR682468.2_Georgia_2007/1        catgatttttaaaacagggaaaaaggggattaggttgaataaaaaaggtaagcacttgtctatatattttttttacaatg [193040]
ASF_IND_20_CAD543                .............r...................................................r.............. [193040]
LR899193.1_Germany_2020/1        ................................................................................ [193040]
MN393477.1_Wuhan_2019            ................................................................................ [193040]
MW396979.1_Timor-Leste/2019/1    ................................................................................ [193040]
MK940252.1_CN/2019/InnerMongolia ................................................................................ [193040]
MK128995.1_China/2018/AnhuiXCGQ  ................................................................................ [193040]
MH681419.1_ASFV/POL/2015         ................................................................................ [193040]
LS478113.1_Estonia_2014          ................................................................................ [193040]
KP843857.1_Odintsovo_02/14       ................................................................................ [193040]
FN557520.1_strain_E75            .............g...................................................g.........t.... [193040]
AY261366.1_Warthog               ....................................................t............g.........t.... [193040]
AY261365.1_Warmbaths             .............g...................................................g.........t.... [193040]
AY261364.1_Tengani_62            .................................t...............................g.........t.... [193040]
AY261363.1_Pretoriuskop/96/4     .................................................................g.........t.... [193040]
AY261362.1_Mkuzi_1979            .............g...................................................g.........t.... [193040]
AY261361.1_Malawi_Lil-20/1_1983  .............g...................a.c.g........c.a................g.........t.... [193040]
AY261360.1_Kenya_1950            .............g...................a............c.a..t.......................t.c.a [193040]
AM712240.1_OURT_88/3             .............g...................................................g.........t.... [193040]
AM712239.1_Benin_97/1            .............g...................................................g.........t.... [193040]
KM111295.1_Ken06.Bus             ...........................a.....a............c.a..........................tc..a [193040]
KM111294.1_Ken05/Tk1             .............g...................a............c.a..t.......................t.c.a [193040]
KM262844.1_strain_L60            .............g...................................................g.........t.... [193040]
KM102979.1_26544/OG10_Italy      .............g...................................................g.........t.... [193040]
KM262845.1_strain_NHV            .............g...................................................g.........t.... [193040]
KX354450.1_47/Ss/2008            .............g...................................................g.........t.... [193040]
KP055815.1_strain_BA71           .............g...................................................g.........t.... [193040]
MH025920.1_strain_R35            ...........................a.....a............c.a..........................tc..a [193040]
MH025919.1_strain_N10            ...........................a.....a............c.a..........................tc..a [193040]
MH025918.1_strain_R25            ...........................a.....a............c.a..........................tc..a [193040]
MH025917.1_strain_R7             ...........................a.....a............c.a..........................tc..a [193040]
MH025916.1_strain_R8             ...........................a.....a............c.a..........................tc..a [193040]

FR682468.2_Georgia_2007/1        ttgccttgagtcgcatttttaactggctggggagtatcagagtggaatatcactgtagtaggtctataaggtcttgttaa [193120]
ASF_IND_20_CAD543                ................................................................................ [193120]
LR899193.1_Germany_2020/1        ................................................................................ [193120]
MN393477.1_Wuhan_2019            ................................................................................ [193120]
MW396979.1_Timor-Leste/2019/1    ................................................................................ [193120]
MK940252.1_CN/2019/InnerMongolia ................................................................................ [193120]
MK128995.1_China/2018/AnhuiXCGQ  ................................................................................ [193120]
MH681419.1_ASFV/POL/2015         ................................................................................ [193120]
LS478113.1_Estonia_2014          ................................................................................ [193120]
KP843857.1_Odintsovo_02/14       ................................................................................ [193120]
```

```
FN557520.1_strain_E75            ................................................................................ [193120]
AY261366.1_Warthog               ...........t.................................................................... [193120]
AY261365.1_Warmbaths             ..............................................................c................. [193120]
AY261364.1_Tengani_62            ................................................................................ [193120]
AY261363.1_Pretoriuskop/96/4     ................................................................................ [193120]
AY261362.1_Mkuzi_1979            ................................................................................ [193120]
AY261361.1_Malawi_Lil-20/1_1983  ...t..c....................a...a..............................c................. [193120]
AY261360.1_Kenya_1950            ...t..c....t...............a...a...g....................t.....c................. [193120]
AM712240.1_OURT_88/3             ................................................................................ [193120]
AM712239.1_Benin_97/1            ................................................................................ [193120]
KM111295.1_Ken06.Bus             ...t.c..g..t...............a.a.a...g....................t..g..c................. [193120]
KM111294.1_Ken05/Tk1             ...t..c....t.t.............a...a...g.............a......t.....c................. [193120]
KM262844.1_strain_L60            ................................................................................ [193120]
KM102979.1_26544/OG10_Italy      ................................................................................ [193120]
KM262845.1_strain_NHV            ................................................................................ [193120]
KX354450.1_47/Ss/2008            ................................................................................ [193120]
KP055815.1_strain_BA71           ................................................................................ [193120]
MH025920.1_strain_R35            ...t.c..g..t...............a.a.a...g....................t..g..c................. [193120]
MH025919.1_strain_N10            ...t.c..g..t...............a.a.a...g....................t..g..c................. [193120]
MH025918.1_strain_R25            ...t.c..g..t...............a.a.a...g....................t..g..c................. [193120]
MH025917.1_strain_R7             ...t.c..g..t...............a.a.a...g....................t..g..c................. [193120]
MH025916.1_strain_R8             ...t.c..g..t...............a.a.a...g....................t..g..c................. [193120]

FR682468.2_Georgia_2007/1        aatatgatcggtcattgttttcgtactagtgtcatttagggtcgacctgatagctcgatataaagttataggggataacc [193200]
ASF_IND_20_CAD543                .....---......------------....................................................a. [193200]
LR899193.1_Germany_2020/1        ................................................................................ [193200]
MN393477.1_Wuhan_2019            ................................................................................ [193200]
MW396979.1_Timor-Leste/2019/1    ................................................................................ [193200]
MK940252.1_CN/2019/InnerMongolia ................................................................................ [193200]
MK128995.1_China/2018/AnhuiXCGQ  ................................................................................ [193200]
MH681419.1_ASFV/POL/2015         ................................................................................ [193200]
LS478113.1_Estonia_2014          ................................................................................ [193200]
KP843857.1_Odintsovo_02/14       ................................................................................ [193200]
FN557520.1_strain_E75            .....---......------------....................................................a. [193200]
AY261366.1_Warthog               ...---........------------...................................................... [193200]
AY261365.1_Warmbaths             ...---........------------...................................................... [193200]
AY261364.1_Tengani_62            ....g.................................a.........a............................... [193200]
AY261363.1_Pretoriuskop/96/4     ................................................a....t.......................... [193200]
AY261362.1_Mkuzi_1979            .....---......------------....................................................a. [193200]
AY261361.1_Malawi_Lil-20/1_1983  .cc.............................................a.................g.......tg..a. [193200]
AY261360.1_Kenya_1950            .cc.......................c.....................a.................g.......t...a. [193200]
AM712240.1_OURT_88/3             .....---......------------....................................................a. [193200]
AM712239.1_Benin_97/1            .....---......------------....................................................a. [193200]
KM111295.1_Ken06.Bus             .cc.............................................a.................g.......t...a. [193200]
```

```
KM111294.1_Ken05/Tk1             .c........................c.....................a.................g.......t...a. [193200]
KM262844.1_strain_L60            .....---......------------....................................................a. [193200]
KM102979.1_26544/OG10_Italy      .....---......------------....................................................a. [193200]
KM262845.1_strain_NHV            .....---......------------....................................................a. [193200]
KX354450.1_47/Ss/2008            .....---......------------....................................................a. [193200]
KP055815.1_strain_BA71           .....---......------------....................................................a. [193200]
MH025920.1_strain_R35            .cc.............................................a.................g.......t...a. [193200]
MH025919.1_strain_N10            .cc.............................................a.................g.......t...a. [193200]
MH025918.1_strain_R25            .cc.............................................a.................g.......t...a. [193200]
MH025917.1_strain_R7             .cc.............................................a.................g.......t...a. [193200]
MH025916.1_strain_R8             .cc.............................................a.................g.......t...a. [193200]

FR682468.2_Georgia_2007/1        tatcaaatacagtcttatctgtgctgaaatgtatatcgtcttctttatcactaataatattaggaatggctgtcattaaa [193280]
ASF_IND_20_CAD543                c............................................................................... [193280]
LR899193.1_Germany_2020/1        ................................................................................ [193280]
MN393477.1_Wuhan_2019            ................................................................................ [193280]
MW396979.1_Timor-Leste/2019/1    ................................................................................ [193280]
MK940252.1_CN/2019/InnerMongolia ................................................................................ [193280]
MK128995.1_China/2018/AnhuiXCGQ  ................................................................................ [193280]
MH681419.1_ASFV/POL/2015         ................................................................................ [193280]
LS478113.1_Estonia_2014          ................................................................................ [193280]
KP843857.1_Odintsovo_02/14       ........................................................................c....... [193280]
FN557520.1_strain_E75            c....g...................a...g..g.......c.............................a.a.g..... [193280]
AY261366.1_Warthog               c............................g..g....c..................t..g...a......a.a.g..... [193280]
AY261365.1_Warmbaths             c............................g..g....c..................t..g...a......a.a.g..... [193280]
AY261364.1_Tengani_62            c............................g..g....c..................t..g...a......a.a.g..... [193280]
AY261363.1_Pretoriuskop/96/4     c............................g.cg....c..................t..g...a......a.a.g..... [193280]
AY261362.1_Mkuzi_1979            c....g...................a...g..g.....................................a.a.g..... [193280]
AY261361.1_Malawi_Lil-20/1_1983  c....g.ag.............a.......a.g...aa..................t.............g......g.. [193280]
AY260360.1_Kenya_1950            .....g.a.t...............a......g...aa..g.........g.....t......a......a..tg...t. [193280]
AM712240.1_OURT_88/3             c....g...................a...g..g.......c.............................a.a.g..... [193280]
AM712239.1_Benin_97/1            c....g...................a...g..g.......c.............................a.a.g..... [193280]
KM111295.1_Ken06.Bus             .....g.a.t...............a......g...a...a...............t......a---------------- [193280]
KM111294.1_Ken05/Tk1             .....g.a.t...............a......g...aa.ag.........g.....t......a......a..tg...t. [193280]
KM262844.1_strain_L60            c....g...................a...g..g.......c.............................a.a.g..... [193280]
KM102979.1_26544/OG10_Italy      c....g...................a...g..g.......c.............................a.a.g..... [193280]
KM262845.1_strain_NHV            c....g...................a...g..g.......c.............................a.a.g..... [193280]
KX354450.1_47/Ss/2008            c....g...................a...g..g.......c.............................a.a.g..... [193280]
KP055815.1_strain_BA71           c....g...................a...g..g.......c.............................a.a.g..... [193280]
MH025920.1_strain_R35            .....g.a.t...............a......g...a...a...............t......a---------------- [193280]
MH025919.1_strain_N10            .....g.a.t...............a......g...a...a...............t......a---------------- [193280]
MH025918.1_strain_R25            .....g.a.t...............a......g...a...a...............t......a---------------- [193280]
MH025917.1_strain_R7             .....g.a.t...............a......g...a...a...............t......a---------------- [193280]
MH025916.1_strain_R8             .....g.a.t...............a......g...a...a...............t......a---------------- [193280]
```

```
FR682468.2_Georgia_2007/1        taattactacttg------------------------------------------------------------ttgttgt [193360]
ASF_IND_20_CAD543                .............------------------------------------------------------------....... [193360]
LR899193.1_Germany_2020/1        .............------------------------------------------------------------....... [193360]
MN393477.1_Wuhan_2019            .............------------------------------------------------------------....... [193360]
MW396979.1_Timor-Leste/2019/1    .............------------------------------------------------------------....... [193360]
MK940252.1_CN/2019/InnerMongolia .............------------------------------------------------------------....... [193360]
MK128995.1_China/2018/AnhuiXCGQ  .............------------------------------------------------------------....... [193360]
MH681419.1_ASFV/POL/2015         .............------------------------------------------------------------....... [193360]
LS478113.1_Estonia_2014          .............------------------------------------------------------------....... [193360]
KP843857.1_Odintsovo_02/14       .............------------------------------------------------------------....... [193360]
FN557520.1_strain_E75            .............------------------------------------------------------------....... [193360]
AY261366.1_Warthog               .............------------------------------------------------------------....... [193360]
AY261365.1_Warmbaths             .........t------------------------------------------------------------------.... [193360]
AY261364.1_Tengani_62            ...a.g.......ttgtgggggaaatagttgtactggtattattggaaatggcagtcattgaataaatgctac....... [193360]
AY261363.1_Pretoriuskop/96/4     ..........---------------------------------------------------------------....... [193360]
AY261362.1_Mkuzi_1979            .............------------------------------------------------------------....... [193360]
AY261361.1_Malawi_Lil-20/1_1983  .............ttgtgggtgaaatatttgtactggtattattaggaatggcggtcattgaataattactac....... [193360]
AY261360.1_Kenya_1950            .g........---------------------------------------------------------------....... [193360]
AM712240.1_OURT_88/3             .............------------------------------------------------------------....... [193360]
AM712239.1_Benin_97/1            .............------------------------------------------------------------....... [193360]
KM111295.1_Ken06.Bus             -------------------------------------------------------------------------------- [193360]
KM111294.1_Ken05/Tk1             .g........---------------------------------------------------------------....... [193360]
KM262844.1_strain_L60            .............------------------------------------------------------------....... [193360]
KM102979.1_26544/OG10_Italy      .............------------------------------------------------------------....... [193360]
KM262845.1_strain_NHV            .............------------------------------------------------------------....... [193360]
KX354450.1_47/Ss/2008            .............------------------------------------------------------------....... [193360]
KP055815.1_strain_BA71           .............------------------------------------------------------------....... [193360]
MH025920.1_strain_R35            -------------------------------------------------------------------------------- [193360]
MH025919.1_strain_N10            -------------------------------------------------------------------------------- [193360]
MH025918.1_strain_R25            -------------------------------------------------------------------------------- [193360]
MH025917.1_strain_R7             -------------------------------------------------------------------------------- [193360]
MH025916.1_strain_R8             -------------------------------------------------------------------------------- [193360]

FR682468.2_Georgia_2007/1        gggtgaaatagttgtactggtattattggaaatggctgtcattaaataattactacttgttgttgtgggtgaaatagttg [193440]
ASF_IND_20_CAD543                ........................................................---..................... [193440]
LR899193.1_Germany_2020/1        ................................................................................ [193440]
MN393477.1_Wuhan_2019            ................................................................................ [193440]
MW396979.1_Timor-Leste/2019/1    ................................................................................ [193440]
MK940252.1_CN/2019/InnerMongolia ................................................................................ [193440]
MK128995.1_China/2018/AnhuiXCGQ  ................................................................................ [193440]
MH681419.1_ASFV/POL/2015         ................................................................................ [193440]
LS478113.1_Estonia_2014          ................................................................................ [193440]
KP843857.1_Odintsovo_02/14       ................................................................................ [193440]
```

```
FN557520.1_strain_E75            a..ca.............a........a...........t.g.........g....---.......a............. [193440]
AY261366.1_Warthog               ....................................a...g.....g.........---..........c.......... [193440]
AY261365.1_Warmbaths             ...........................a........a...g.....g.....---------------------------- [193440]
AY261364.1_Tengani_62            ..........a................a....c...a...g.....g.........---..................g.. [193440]
AY261363.1_Pretoriuskop/96/4     ...........................a.g................g.........---...........a......... [193440]
AY261362.1_Mkuzi_1979            a.................a........a.............g.........g....---.......a............. [193440]
AY261361.1_Malawi_Lil-20/1_1983  ..........t................a.g......g......g............---.................t... [193440]
AY261360.1_Kenya_1950            ...ca...------.............a.g......a..tg...t..g........---...............------ [193440]
AM712240.1_OURT_88/3             a..ca.............a........a...........t.g.........g....---.......a............. [193440]
AM712239.1_Benin_97/1            a..ca.............a........a...........t.g.........g....---.......a............. [193440]
KM111295.1_Ken06.Bus             ------------------------------......a..tg...t...........---...............------ [193440]
KM111294.1_Ken05/Tk1             ..a.a...------.............a.g....a.a.ctgc.g......---------...............------ [193440]
KM262844.1_strain_L60            a..ca.............a........a...........t.g.........g....---.......a............. [193440]
KM102979.1_26544/OG10_Italy      a..ca.............a........a...........t.g.........g....---.......a............. [193440]
KM262845.1_strain_NHV            a..ca.............a........a...........t.g.........g....---.......a............. [193440]
KX354450.1_47/Ss/2008            a..ca.............a........a...........t.g.........g....---.......a............. [193440]
KP055815.1_strain_BA71           a..ca.............a........a...........t.g.........g....---.......a............. [193440]
MH025920.1_strain_R35            ------------------------------......a..tg...t...........---...............------ [193440]
MH025919.1_strain_N10            ------------------------------......a..tg...t...........---...............------ [193440]
MH025918.1_strain_R25            ------------------------------......a..tg...t...........---...............------ [193440]
MH025917.1_strain_R7             ------------------------------......a..tg...t...........---...............------ [193440]
MH025916.1_strain_R8             ------------------------------......a..tg...t...........---...............------ [193440]

FR682468.2_Georgia_2007/1        tactagtattattagaaatggctgtcgttaaataattactacctattacaagtaaactaatgctaactacatttttaacc [193520]
ASF_IND_20_CAD543                ................................................................................ [193520]
LR899193.1_Germany_2020/1        ................................................................................ [193520]
MN393477.1_Wuhan_2019            ................................................................................ [193520]
MW396979.1_Timor-Leste/2019/1    ................................................................................ [193520]
MK940252.1_CN/2019/InnerMongolia ................................................................................ [193520]
MK128995.1_China/2018/AnhuiXCGQ  ................................................................................ [193520]
MH681419.1_ASFV/POL/2015         ................................................................................ [193520]
LS478113.1_Estonia_2014          ................................................................................ [193520]
KP843857.1_Odintsovo_02/14       ................................................................................ [193520]
FN557520.1_strain_E75            .....................................g.......................................... [193520]
AY261366.1_Warthog               ....ga................a............a.g...........................g.............. [193520]
AY261365.1_Warmbaths             --------------------------------------...........................g.............. [193520]
AY261364.1_Tengani_62            ....g........g..........................t........................g.............. [193520]
AY261363.1_Pretoriuskop/96/4     ....g.................a.........g................................g.............. [193520]
AY261362.1_Mkuzi_1979            ..........c..................................................................... [193520]
AY261361.1_Malawi_Lil-20/1_1983  ....g..........g......g..t.......................................g...t.........t [193520]
AY261360.1_Kenya_1950            ....g..........c...a..a..........................................g.............t [193520]
AM712240.1_OURT_88/3             .....................................g.......................................... [193520]
AM712239.1_Benin_97/1            .....................................g.......................................... [193520]
KM111295.1_Ken06.Bus             ....g..g.......c...a..a..........................................g.............t [193520]
```

```
KM111294.1_Ken05/Tk1            ....g..........c...a..a..t.......................................g.............t [193520]
KM262844.1_strain_L60           .....................................g.......................................... [193520]
KM102979.1_26544/OG10_Italy     .....................................g.......................................... [193520]
KM262845.1_strain_NHV           .....................................g.......................................... [193520]
KX354450.1_47/Ss/2008           .....................................g.......................................... [193520]
KP055815.1_strain_BA71          .....................................g.......................................... [193520]
MH025920.1_strain_R35           ....g..g.......c...a..a..........................................g.............t [193520]
MH025919.1_strain_N10           ....g..g.......c...a..a..........................................g.............t [193520]
MH025918.1_strain_R25           ....g..g.......c...a..a..........................................g.............t [193520]
MH025917.1_strain_R7            ....g..g.......c...a..a..........................................g.............t [193520]
MH025916.1_strain_R8            ....g..g.......c...a..a..........................................g.............t [193520]

FR682468.2_Georgia_2007/1       tcaataaacctaaaaagccatactaaatacctaaacaacatcctgttataatatgagcagaaaaaaaaaat-aagtataa [193600]
ASF_IND_20_CAD543               .......................................................................-........ [193600]
LR899193.1_Germany_2020/1       .......................................................................-........ [193600]
MN393477.1_Wuhan_2019           .......................................................................-........ [193600]
MW396979.1_Timor-Leste/2019/1   .......................................................................-........ [193600]
MK940252.1_CN/2019/InnerMongolia .......................................................................-........ [193600]
MK128995.1_China/2018/AnhuiXCGQ .......................................................................-........ [193600]
MH681419.1_ASFV/POL/2015        .......................................................................-........ [193600]
LS478113.1_Estonia_2014         .......................................................................-........ [193600]
KP843857.1_Odintsovo_02/14      ....................................................................-..-........ [193600]
FN557520.1_strain_E75           .......................................................................-........ [193600]
AY261366.1_Warthog              .......................................................................-........ [193600]
AY261365.1_Warmbaths            .......................................................................-........ [193600]
AY261364.1_Tengani_62           ......................................................................at........ [193600]
AY261363.1_Pretoriuskop/96/4    .......................................................................-........ [193600]
AY261362.1_Mkuzi_1979           .......................................................................-........ [193600]
AY261361.1_Malawi_Lil-20/1_1983 .......................................................................-........ [193600]
AY261360.1_Kenya_1950           .t...................................................................t.-........ [193600]
AM712240.1_OURT_88/3            .......................................................................-........ [193600]
AM712239.1_Benin_97/1           .......................................................................-........ [193600]
KM111295.1_Ken06.Bus            .....................................................................t.-........ [193600]
KM111294.1_Ken05/Tk1            .t...................................................................t.-........ [193600]
KM262844.1_strain_L60           .......................................................................a........ [193600]
KM102979.1_26544/OG10_Italy     .......................................................................-........ [193600]
KM262845.1_strain_NHV           .......................................................................-........ [193600]
KX354450.1_47/Ss/2008           .......................................................................-........ [193600]
KP055815.1_strain_BA71          .......................................................................-........ [193600]
MH025920.1_strain_R35           .....................................................................t.-........ [193600]
MH025919.1_strain_N10           .....................................................................t.-........ [193600]
MH025918.1_strain_R25           .....................................................................t.-........ [193600]
MH025917.1_strain_R7            .....................................................................t.-........ [193600]
MH025916.1_strain_R8            .....................................................................t.-........ [193600]
```

```
FR682468.2_Georgia_2007/1        ttagggaattattcttattcgcttactattaagaata---attcagaatcttattt------------------------ [193680]
ASF_IND_20_CAD543                .....................................---................------------------------ [193680]
LR899193.1_Germany_2020/1        .....................................---................------------------------ [193680]
MN393477.1_Wuhan_2019            .....................................---................------------------------ [193680]
MW396979.1_Timor-Leste/2019/1    .....................................---................------------------------ [193680]
MK940252.1_CN/2019/InnerMongolia .....................................---................------------------------ [193680]
MK128995.1_China/2018/AnhuiXCGQ  .....................................---................------------------------ [193680]
MH681419.1_ASFV/POL/2015         .....................................---................------------------------ [193680]
LS478113.1_Estonia_2014          .....................................---................------------------------ [193680]
KP843857.1_Odintsovo_02/14       .....................................---................------------------------ [193680]
FN557520.1_strain_E75            .....................................---................------------------------ [193680]
AY261366.1_Warthog               .....................................att........a.......------------------------ [193680]
AY261365.1_Warmbaths             ........------.......................att........a.......------------------------ [193680]
AY261364.1_Tengani_62            ...................t.................-tt........a.......------------------------ [193680]
AY261363.1_Pretoriuskop/96/4     ...................ta................att........a.......------------------------ [193680]
AY261362.1_Mkuzi_1979            .....................................---................------------------------ [193680]
AY261361.1_Malawi_Lil-20/1_1983  .......t...........aa.......a.......taat........a.......agttaaaaacagttagttagactt [193680]
AY261360.1_Kenya_1950            ...........g....g..a................tatt.......tc.c.....------------------------ [193680]
AM712240.1_OURT_88/3             .....................................---................------------------------ [193680]
AM712239.1_Benin_97/1            .....................................---................------------------------ [193680]
KM111295.1_Ken06.Bus             ...........g....g..a................tatt.......tc.c.....------------------------ [193680]
KM111294.1_Ken05/Tk1             ...........g....g..a................tatt.......tc.c.....------------------------ [193680]
KM262844.1_strain_L60            .....................................---................------------------------ [193680]
KM102979.1_26544/OG10_Italy      .....................................---................------------------------ [193680]
KM262845.1_strain_NHV            .....................................---................------------------------ [193680]
KX354450.1_47/Ss/2008            .....................................---................------------------------ [193680]
KP055815.1_strain_BA71           .....................................---................------------------------ [193680]
MH025920.1_strain_R35            ....a......g....g..a................tatt.......tc.c.....------------------------ [193680]
MH025919.1_strain_N10            ....a......g....g..a................tatt.......tc.c.....------------------------ [193680]
MH025918.1_strain_R25            ....a......g....g..a................tatt.......tc.c.....------------------------ [193680]
MH025917.1_strain_R7             ....a......g....g..a................tatt.......tc.c.....------------------------ [193680]
MH025916.1_strain_R8             ....a......g....g..a................tatt.......tc.c.....------------------------ [193680]

FR682468.2_Georgia_2007/1        ----------------------------------------agttagaaactatcataaagtgaataggactcatcgtcgg [193760]
ASF_IND_20_CAD543                ----------------------------------------........................................ [193760]
LR899193.1_Germany_2020/1        ----------------------------------------........................................ [193760]
MN393477.1_Wuhan_2019            ----------------------------------------........................................ [193760]
MW396979.1_Timor-Leste/2019/1    ----------------------------------------........................................ [193760]
MK940252.1_CN/2019/InnerMongolia ----------------------------------------........................................ [193760]
MK128995.1_China/2018/AnhuiXCGQ  ----------------------------------------........................................ [193760]
MH681419.1_ASFV/POL/2015         ----------------------------------------........................................ [193760]
LS478113.1_Estonia_2014          ----------------------------------------........................................ [193760]
KP843857.1_Odintsovo_02/14       ----------------------------------------........................................ [193760]
```

```
FN557520.1_strain_E75              ----------------------------------------........................................ [193760]
AY261366.1_Warthog                 ----------------------------------------...........g............................ [193760]
AY261365.1_Warmbaths               ----------------------------------------...........g............................ [193760]
AY261364.1_Tengani_62              ----------------------------------------...........g............................ [193760]
AY261363.1_Pretoriuskop/96/4       ----------------------------------------...........g.......................a.... [193760]
AY261362.1_Mkuzi_1979              ----------------------------------------........................................ [193760]
AY261361.1_Malawi_Lil-20/1_1983    atttagttagaaacggttagagacttatttagttagaaac.....a...........t.........c.....c.a.... [193760]
AY261360.1_Kenya_1950              ----------------------------------------...........g..g..t.................a.... [193760]
AM712240.1_OURT_88/3               ----------------------------------------........................................ [193760]
AM712239.1_Benin_97/1              ----------------------------------------........................................ [193760]
KM111295.1_Ken06.Bus               ----------------------------------------...........g..g..t........a........a.... [193760]
KM111294.1_Ken05/Tk1               ----------------------------------------...........g..g..t.................a.... [193760]
KM262844.1_strain_L60              ----------------------------------------........................................ [193760]
KM102979.1_26544/OG10_Italy        ----------------------------------------........................................ [193760]
KM262845.1_strain_NHV              ----------------------------------------........................................ [193760]
KX354450.1_47/Ss/2008              ----------------------------------------........................................ [193760]
KP055815.1_strain_BA71             ----------------------------------------........................................ [193760]
MH025920.1_strain_R35              ----------------------------------------...........g..g..t........a........a.... [193760]
MH025919.1_strain_N10              ----------------------------------------...........g..g..t........a........a.... [193760]
MH025918.1_strain_R25              ----------------------------------------...........g..g..t........a........a.... [193760]
MH025917.1_strain_R7               ----------------------------------------...........g..g..t........a........a.... [193760]
MH025916.1_strain_R8               ----------------------------------------...........g..g..t........a........a.... [193760]

FR682468.2_Georgia_2007/1          atgaagattccgtttcagagatagtttctttttcttcctcagaataatctgttcctac---------aatagaatcggtg [193840]
ASF_IND_20_CAD543                  ....................................y.....................---------............. [193840]
LR899193.1_Germany_2020/1          ..........................................................---------............. [193840]
MN393477.1_Wuhan_2019              ..........................................................---------............. [193840]
MW396979.1_Timor-Leste/2019/1      ..........................................................---------............. [193840]
MK940252.1_CN/2019/InnerMongolia   ..........................................................---------............. [193840]
MK128995.1_China/2018/AnhuiXCGQ    ..........................................................---------............. [193840]
MH681419.1_ASFV/POL/2015           ..........................................................---------............. [193840]
LS478113.1_Estonia_2014            ..........................................................---------............. [193840]
KP843857.1_Odintsovo_02/14         ..........................................................---------............. [193840]
FN557520.1_strain_E75              ....................................t.....................---------............. [193840]
AY261366.1_Warthog                 ....g...................a............................t.c..---------......c.....a [193840]
AY261365.1_Warmbaths               ....g...................a............................t.c..---------......c.....a [193840]
AY261364.1_Tengani_62              ....g...................a............................t.c..---------......c.....a [193840]
AY261363.1_Pretoriuskop/96/4       ..........t.............a............................t.c..---------......c.....a [193840]
AY261362.1_Mkuzi_1979              ..........................................................---------............. [193840]
AY261361.1_Malawi_Lil-20/1_1983    ..........t........t....a........................c........---------.........a... [193840]
AY261360.1_Kenya_1950              ............a......c..g.a............................t....ctctacaat.ga......a... [193840]
AM712240.1_OURT_88/3               ....................................t.....................---------............. [193840]
AM712239.1_Benin_97/1              ....................................t.....................---------............. [193840]
KM111295.1_Ken06.Bus               ............a......c..g.a............................t....ttctacaat.ga......a... [193840]
```

```
KM111294.1_Ken05/Tk1            ............a......t..g.a............................t....ctctacaat.ga......a... [193840]
KM262844.1_strain_L60           ....................................t.....................---------............. [193840]
KM102979.1_26544/OG10_Italy     ....................................t.....................---------............. [193840]
KM262845.1_strain_NHV           ....................................t.....................---------............. [193840]
KX354450.1_47/Ss/2008           ....................................t.....................---------............. [193840]
KP055815.1_strain_BA71          ....................................t.....................---------............. [193840]
MH025920.1_strain_R35           ............a......c..g.a............................t....ttctacaat.ga......a... [193840]
MH025919.1_strain_N10           ............a......c..g.a............................t....ttctacaat.ga......a... [193840]
MH025918.1_strain_R25           ............a......c..g.a............................t....ttctacaat.ga......a... [193840]
MH025917.1_strain_R7            ............a......c..g.a............................t....ttctacaat.ga......a... [193840]
MH025916.1_strain_R8            ............a......c..g.a............................t....ttctacaat.ga......a... [193840]

FR682468.2_Georgia_2007/1       tcatcctcagaaagagaagtatttaaatatggactatctatagcaatatcctcttctatctcgcaatcctcctcctccat [193920]
ASF_IND_20_CAD543               ................................................................................ [193920]
LR899193.1_Germany_2020/1       ................................................................................ [193920]
MN393477.1_Wuhan_2019           ................................................................................ [193920]
MW396979.1_Timor-Leste/2019/1   ................................................................................ [193920]
MK940252.1_CN/2019/InnerMongolia ................................................................................ [193920]
MK128995.1_China/2018/AnhuiXCGQ ................................................................................ [193920]
MH681419.1_ASFV/POL/2015        ................................................................................ [193920]
LS478113.1_Estonia_2014         ................................................................................ [193920]
KP843857.1_Odintsovo_02/14      ................................................................................ [193920]
FN557520.1_strain_E75           ................................................................................ [193920]
AY261366.1_Warthog              ..................................................................---........... [193920]
AY261365.1_Warmbaths            ..................................................................---........... [193920]
AY261364.1_Tengani_62           ..................................................................---........... [193920]
AY261363.1_Pretoriuskop/96/4    ..................................................................---........... [193920]
AY261362.1_Mkuzi_1979           ..............................................................a................. [193920]
AY261361.1_Malawi_Lil-20/1_1983 ...................c.............t.t......at..c...t......tc...a.....t......a.... [193920]
AY261360.1_Kenya_1950           ...................cg..................tctat..c.a.a.t....gc...at..........a..... [193920]
AM712240.1_OURT_88/3            ................................................................................ [193920]
AM712239.1_Benin_97/1           ................................................................................ [193920]
KM111295.1_Ken06.Bus            ...................cg..................tctat..c.a.a.t....gc...at..........g..... [193920]
KM111294.1_Ken05/Tk1            ...................cg..................tctat..c.a.a.t....gc...at..........a..... [193920]
KM262844.1_strain_L60           ................................................................................ [193920]
KM102979.1_26544/OG10_Italy     ................................................................................ [193920]
KM262845.1_strain_NHV           ................................................................................ [193920]
KX354450.1_47/Ss/2008           ................................................................................ [193920]
KP055815.1_strain_BA71          ................................................................................ [193920]
MH025920.1_strain_R35           ...................cg..................tctat..c.a.a.t....gc...at..........g..... [193920]
MH025919.1_strain_N10           ...................cg..................tctat..c.a.a.t....gc...at..........g..... [193920]
MH025918.1_strain_R25           ...................cg..................tctat..c.a.a.t....gc...at..........g..... [193920]
MH025917.1_strain_R7            ...................cg..................tctat..c.a.a.t....gc...at..........g..... [193920]
MH025916.1_strain_R8            ...................cg..................tctat..c.a.a.t....gc...at..........g..... [193920]
```

```
FR682468.2_Georgia_2007/1        ttccatagtgtgtaggagaatatttttat---catcatgctcacttctttttttgttgaaagatgaaccgtcctcaatac [194000]
ASF_IND_20_CAD543                .............................---................................................ [194000]
LR899193.1_Germany_2020/1        .............................---................................................ [194000]
MN393477.1_Wuhan_2019            .............................---................................................ [194000]
MW396979.1_Timor-Leste/2019/1    .............................---................................................ [194000]
MK940252.1_CN/2019/InnerMongolia .............................---................................................ [194000]
MK128995.1_China/2018/AnhuiXCGQ  .............................---................................................ [194000]
MH681419.1_ASFV/POL/2015         .............................---................................................ [194000]
LS478113.1_Estonia_2014          .............................---................................................ [194000]
KP843857.1_Odintsovo_02/14       .............................---................................................ [194000]
FN557520.1_strain_E75            .............................---.......t......................................g. [194000]
AY261366.1_Warthog               ......g.............cg......g---................................a....a........g. [194000]
AY261365.1_Warmbaths             ......g.............cg......g---................................a....a........g. [194000]
AY261364.1_Tengani_62            ......g.............cg......g---................................a...ta........g. [194000]
AY261363.1_Pretoriuskop/96/4     ......g.............cg......g---................................a....a........g. [194000]
AY261362.1_Mkuzi_1979            .............................---.......t......................................g. [194000]
AY261361.1_Malawi_Lil-20/1_1983  ..................g..g......g---...........t....................a....a........g. [194000]
AY261360.1_Kenya_1950            ......t..c.....a.....g......gcat......a....t..............g...g.a.tg.a........g. [194000]
AM712240.1_OURT_88/3             .............................---.......t......................................g. [194000]
AM712239.1_Benin_97/1            .............................---.......t......................................g. [194000]
KM111295.1_Ken06.Bus             ......t..c.....a.....g......gcat......a....t..............g...g.a.tg.a........g. [194000]
KM111294.1_Ken05/Tk1             ......t..c.....a.....g......gcat......a....t..............g...g.a.tg.a........g. [194000]
KM262844.1_strain_L60            .............................---.......t......................................g. [194000]
KM102979.1_26544/OG10_Italy      .............................---.......t......................................g. [194000]
KM262845.1_strain_NHV            .............................---.......t......................................g. [194000]
KX354450.1_47/Ss/2008            .............................---.......t......................................g. [194000]
KP055815.1_strain_BA71           .............................---.......t......................................g. [194000]
MH025920.1_strain_R35            ......t..c.....a.....g......gcat......a....t..............g...g.a.tg.a........g. [194000]
MH025919.1_strain_N10            ......t..c.....a.....g......gcat......a....t..............g...g.a.tg.a........g. [194000]
MH025918.1_strain_R25            ......t..c.....a.....g......gcat......a....t..............g...g.a.tg.a........g. [194000]
MH025917.1_strain_R7             ......t..c.....a.....g......gcat......a....t..............g...g.a.tg.a........g. [194000]
MH025916.1_strain_R8             ......t..c.....a.....g......gcat......a....t..............g...g.a.tg.a........g. [194000]
```

```
FR682468.2_Georgia_2007/1        ggttcatgttaagttccttcatcttatgtataatttccgtaatccgtgatgtttttgacatgtaagatggttttaaggtt [194080]
ASF_IND_20_CAD543                ................................................................................ [194080]
LR899193.1_Germany_2020/1        ................................................................................ [194080]
MN393477.1_Wuhan_2019            ................................................................................ [194080]
MW396979.1_Timor-Leste/2019/1    ................................................................................ [194080]
MK940252.1_CN/2019/InnerMongolia ................................................................................ [194080]
MK128995.1_China/2018/AnhuiXCGQ  ................................................................................ [194080]
MH681419.1_ASFV/POL/2015         ................................................................................ [194080]
LS478113.1_Estonia_2014          ................................................................................ [194080]
KP843857.1_Odintsovo_02/14       ................................................................................ [194080]
```

```
FN557520.1_strain_E75             ............................a................................................... [194080]
AY261366.1_Warthog                ......................t.....a............................g............c.....a... [194080]
AY261365.1_Warmbaths              .........c..................a................................................... [194080]
AY261364.1_Tengani_62             ......................t.....a..............t................................ac.. [194080]
AY261363.1_Pretoriuskop/96/4      ......................t.....a............................g..................a... [194080]
AY261362.1_Mkuzi_1979             a...........................a................................................... [194080]
AY261361.1_Malawi_Lil-20/1_1983   .a..t..a..............t.....a.....................a........g.................... [194080]
AY261360.1_Kenya_1950             ......................t.....g..............t......a......g......g............... [194080]
AM712240.1_OURT_88/3              ............................a................................................... [194080]
AM712239.1_Benin_97/1             ............................a................................................... [194080]
KM111295.1_Ken06.Bus              ......................t.....g..............t......a......g......g............... [194080]
KM111294.1_Ken05/Tk1              ......................t.....g..............t......a......g......g............... [194080]
KM262844.1_strain_L60             ............................a................................................... [194080]
KM102979.1_26544/OG10_Italy       ............................a................................................... [194080]
KM262845.1_strain_NHV             ............................a................................................... [194080]
KX354450.1_47/Ss/2008             ............................a................................................... [194080]
KP055815.1_strain_BA71            ............................a................................................... [194080]
MH025920.1_strain_R35             ......................t.....g..............t......a......g......g............... [194080]
MH025919.1_strain_N10             ......................t.....g..............t......a......g......g............... [194080]
MH025918.1_strain_R25             ......................t.....g..............t......a......g......g............... [194080]
MH025917.1_strain_R7              ......................t.....g..............t......a......g......g............... [194080]
MH025916.1_strain_R8              ......................t.....g..............t......a......g......g............... [194080]

FR682468.2_Georgia_2007/1         atatccacaataacaggagaatctctatcattttcatttgataaactttgatctttgatttcttcgtctaaaattcttgt [194160]
ASF_IND_20_CAD543                 ........................s....................................................... [194160]
LR899193.1_Germany_2020/1         ................................................................................ [194160]
MN393477.1_Wuhan_2019             ................................................................................ [194160]
MW396979.1_Timor-Leste/2019/1     ................................................................................ [194160]
MK940252.1_CN/2019/InnerMongolia  ................................................................................ [194160]
MK128995.1_China/2018/AnhuiXCGQ   ................................................................................ [194160]
MH681419.1_ASFV/POL/2015          ................................................................................ [194160]
LS478113.1_Estonia_2014           ................................................................................ [194160]
KP843857.1_Odintsovo_02/14        ................................................................................ [194160]
FN557520.1_strain_E75             ........................g....................................................... [194160]
AY261366.1_Warthog                .......................c..................g.........................c......t.... [194160]
AY261365.1_Warmbaths              .......................c..................g.........................c........... [194160]
AY261364.1_Tengani_62             .....t.................c..................g.........................c........... [194160]
AY261363.1_Pretoriuskop/96/4      ------.................c..................g..................................... [194160]
AY261362.1_Mkuzi_1979             .......................c..................g..................................... [194160]
AY261361.1_Malawi_Lil-20/1_1983   .....t.....g..............................g..............t...t........g......... [194160]
AY261360.1_Kenya_1950             .a.........g........---------.............gt...c.......c.........a....g....t.... [194160]
AM712240.1_OURT_88/3              ........................g....................................................... [194160]
AM712239.1_Benin_97/1             ........................g..................................................t.... [194160]
KM111295.1_Ken06.Bus              .a.........g........---------.............gt...c.......c........ta....g....t.... [194160]
```

```
KM111294.1_Ken05/Tk1            .a.........g........---------.............gt...c.......c.........a....g....t.... [194160]
KM262844.1_strain_L60           ........................g....................................................... [194160]
KM102979.1_26544/OG10_Italy     ........................g....................................................... [194160]
KM262845.1_strain_NHV           ........................g....................................................... [194160]
KX354450.1_47/Ss/2008           ........................g....................................................... [194160]
KP055815.1_strain_BA71          ........................g....................................................... [194160]
MH025920.1_strain_R35           .a.........g........---------.............gt...c.......c........ta....g....t.... [194160]
MH025919.1_strain_N10           .a.........g........---------.............gt...c.......c........ta....g....t.... [194160]
MH025918.1_strain_R25           .a.........g........---------.............gt...c.......c........ta....g....t.... [194160]
MH025917.1_strain_R7            .a.........g........---------.............gt...c.......c........ta....g....t.... [194160]
MH025916.1_strain_R8            .a.........g........---------.............gt...c.......c........ta....g....t.... [194160]

FR682468.2_Georgia_2007/1       ctttttttgggtactagatgaaatagaggaattcatattctgaaacgatatatcaaggggagctggacgcttttttccaa [194240]
ASF_IND_20_CAD543               ........................m....................................................... [194240]
LR899193.1_Germany_2020/1       ................................................................................ [194240]
MN393477.1_Wuhan_2019           ................................................................................ [194240]
MW396979.1_Timor-Leste/2019/1   ................................................................................ [194240]
MK940252.1_CN/2019/InnerMongolia ................................................................................ [194240]
MK128995.1_China/2018/AnhuiXCGQ ................................................................................ [194240]
MH681419.1_ASFV/POL/2015        ................................................................................ [194240]
LS478113.1_Estonia_2014         ................................................................................ [194240]
KP843857.1_Odintsovo_02/14      ................................................................................ [194240]
FN557520.1_strain_E75           ........................c............c.......................................... [194240]
AY261366.1_Warthog              ....................gc..c....................t.........t..------...............g [194240]
AY261365.1_Warmbaths            ....................gc..c....................t.........t.a...................... [194240]
AY261364.1_Tengani_62           ....................gc..c....................g.........t..------................ [194240]
AY261363.1_Pretoriuskop/96/4    ....................gc..c....................t.........t..------................ [194240]
AY261362.1_Mkuzi_1979           ....................gc..c....................t.................................. [194240]
AY261361.1_Malawi_Lil-20/1_1983 .........a.....g.....c..c............c.t.....t...................a.............. [194240]
AY261360.1_Kenya_1950           .c.......a.....g....gc..c....................t..............t....a.............. [194240]
AM712240.1_OURT_88/3            ........................c............c.......................................... [194240]
AM712239.1_Benin_97/1           ........................c............c.......................................... [194240]
KM111295.1_Ken06.Bus            .c.......a.....g....tt..c............c.......t..........t...t....a.............. [194240]
KM111294.1_Ken05/Tk1            .c.......a.....g....gc..c....................t..............t....a.............. [194240]
KM262844.1_strain_L60           ........................c............c.......................................... [194240]
KM102979.1_26544/OG10_Italy     ........................c............c.......................................... [194240]
KM262845.1_strain_NHV           ........................c............c.......................................... [194240]
KX354450.1_47/Ss/2008           ........................c............c.......................................... [194240]
KP055815.1_strain_BA71          ........................c............c.......................................... [194240]
MH025920.1_strain_R35           .c.......a.....g....tt..c............c.......t..........t...t....a.............. [194240]
MH025919.1_strain_N10           .c.......a.....g....tt..c............c.......t..........t...t....a.............. [194240]
MH025918.1_strain_R25           .c.......a.....g....tt..c............c.......t..........t...t....a.............. [194240]
MH025917.1_strain_R7            .c.......a.....g....tt..c............c.......t..........t...t....a.............. [194240]
MH025916.1_strain_R8            .c.......a.....g....tt..c............c.......t..........t...t....a.............. [194240]
```

```
FR682468.2_Georgia_2007/1        ttaaac------------------------------cgtttttcgagatactatgattagatgaatgatctttagccaag [194320]
ASF_IND_20_CAD543                ......------------------------------............................................ [194320]
LR899193.1_Germany_2020/1        ......------------------------------............................................ [194320]
MN393477.1_Wuhan_2019            ......------------------------------............................................ [194320]
MW396979.1_Timor-Leste/2019/1    ......------------------------------............................................ [194320]
MK940252.1_CN/2019/InnerMongolia ......------------------------------............................................ [194320]
MK128995.1_China/2018/AnhuiXCGQ  ......------------------------------............................................ [194320]
MH681419.1_ASFV/POL/2015         ......------------------------------............................................ [194320]
LS478113.1_Estonia_2014          ......------------------------------............................................ [194320]
KP843857.1_Odintsovo_02/14       ......------------------------------............................................ [194320]
FN557520.1_strain_E75            ......------------------------------...................t........................ [194320]
AY261366.1_Warthog               ......------------------------------t..c.a....c......c.......................... [194320]
AY261365.1_Warmbaths             ......------------------------------...c.a....c......c.......................... [194320]
AY261364.1_Tengani_62            ......------------------------------...c.a....c......c........................g. [194320]
AY261363.1_Pretoriuskop/96/4     ......------------------------------...c.a....c......c.......................... [194320]
AY261362.1_Mkuzi_1979            ......------------------------------..........c........t........................ [194320]
AY261361.1_Malawi_Lil-20/1_1983  ....g.ctttcttaggtgaattatctttagctaagt..........c....a..tt........c.t.c.g....t.... [194320]
AY261360.1_Kenya_1950            ....g.cattcttaggcgaattatctctagttaagt...c....a.c......ctt....ca...t.............. [194320]
AM712240.1_OURT_88/3             ......------------------------------...................t........................ [194320]
AM712239.1_Benin_97/1            ......------------------------------...................t........................ [194320]
KM111295.1_Ken06.Bus             ....g.cattcttaggcgaatcatctctagttaatt...c....a.c......ctt....ca...t.............. [194320]
KM111294.1_Ken05/Tk1             ....g.cattcttaggcgaattatctctagttaagt...c....a.c......ctt....ca...t.............. [194320]
KM262844.1_strain_L60            ......------------------------------...................t........................ [194320]
KM102979.1_26544/OG10_Italy      ......------------------------------...................t........................ [194320]
KM262845.1_strain_NHV            ......------------------------------...................t........................ [194320]
KX354450.1_47/Ss/2008            ......------------------------------...................t........................ [194320]
KP055815.1_strain_BA71           ......------------------------------...................t........................ [194320]
MH025920.1_strain_R35            ....g.cattcttaggcgaatcatctctagttaatt...c....a.c......ctt....ca...t.............. [194320]
MH025919.1_strain_N10            ....g.cattcttaggcgaatcatctctagttaatt...c....a.c......ctt....ca...t.............. [194320]
MH025918.1_strain_R25            ....g.cattcttaggcgaatcatctctagttaatt...c....a.c......ctt....ca...t.............. [194320]
MH025917.1_strain_R7             ....g.cattcttaggcgaatcatctctagttaatt...c....a.c......ctt....ca...t.............. [194320]
MH025916.1_strain_R8             ....g.cattcttaggcgaatcatctctagttaatt...c....a.c......ctt....ca...t.............. [194320]

FR682468.2_Georgia_2007/1        ctgtccttggatatactatagttagatattttacctttaaataatattcttctatacaagttattcttaggtaaagaatt [194400]
ASF_IND_20_CAD543                ................................................................................ [194400]
LR899193.1_Germany_2020/1        ................................................................................ [194400]
MN393477.1_Wuhan_2019            ................................................................................ [194400]
MW396979.1_Timor-Leste/2019/1    ................................................................................ [194400]
MK940252.1_CN/2019/InnerMongolia ................................................................................ [194400]
MK128995.1_China/2018/AnhuiXCGQ  ................................................................................ [194400]
MH681419.1_ASFV/POL/2015         ................................................................................ [194400]
LS478113.1_Estonia_2014          ................................................................................ [194400]
KP843857.1_Odintsovo_02/14       ................................................................................ [194400]
```

```
FN557520.1_strain_E75             ................................................................................ [194400]
AY261366.1_Warthog                t....g...a..........t.................................c.....a................... [194400]
AY261365.1_Warmbaths              .....g...a..........t.................................c.....a................... [194400]
AY261364.1_Tengani_62             .....g...a..........t.................................c.....a................... [194400]
AY261363.1_Pretoriuskop/96/4      .....g...a..........t.................................c.....ag.................. [194400]
AY261362.1_Mkuzi_1979             ..........................................................g..................... [194400]
AY261361.1_Malawi_Lil-20/1_1983   ..........g........tt...........................t.....g...g..................... [194400]
AY261360.1_Kenya_1950             ..a.....a..........ctc...........t..............t.....g...t............c....t... [194400]
AM712240.1_OURT_88/3              ................................................................................ [194400]
AM712239.1_Benin_97/1             ................................................................................ [194400]
KM111295.1_Ken06.Bus              ..a.....a.........cttc..........................t.....g...t............c....t... [194400]
KM111294.1_Ken05/Tk1              ..a.....a..........ctc..........................t.....g...t............c....t... [194400]
KM262844.1_strain_L60             ................................................................................ [194400]
KM102979.1_26544/OG10_Italy       ................................................................................ [194400]
KM262845.1_strain_NHV             ................................................................................ [194400]
KX354450.1_47/Ss/2008             ................................................................................ [194400]
KP055815.1_strain_BA71            ................................................................................ [194400]
MH025920.1_strain_R35             ..a.....a.........cttc..........................t.....g...t............c....t... [194400]
MH025919.1_strain_N10             ..a.....a.........cttc..........................t.....g...t............c....t... [194400]
MH025918.1_strain_R25             ..a.....a.........cttc..........................t.....g...t............c....t... [194400]
MH025917.1_strain_R7              ..a.....a.........cttc..........................t.....g...t............c....t... [194400]
MH025916.1_strain_R8              ..a.....a.........cttc..........................t.....g...t............c....t... [194400]

FR682468.2_Georgia_2007/1         agtatggattcctatatt------------------------tttatctgaaggagtgtccatatcggagaacgtcctct [194480]
ASF_IND_20_CAD543                 ..................------------------------...................................... [194480]
LR899193.1_Germany_2020/1         ..................------------------------...................................... [194480]
MN393477.1_Wuhan_2019             ..................------------------------...................................... [194480]
MW396979.1_Timor-Leste/2019/1     ..................------------------------...................................... [194480]
MK940252.1_CN/2019/InnerMongolia  ..................------------------------...................................... [194480]
MK128995.1_China/2018/AnhuiXCGQ   ..................------------------------...................................... [194480]
MH681419.1_ASFV/POL/2015          ..................------------------------...................................... [194480]
LS478113.1_Estonia_2014           ..................------------------------...................................... [194480]
KP843857.1_Odintsovo_02/14        ..................------------------------...................................... [194480]
FN557520.1_strain_E75             ..................------------------------...................................... [194480]
AY261366.1_Warthog                t.................------------------------..........a...........a.a............. [194480]
AY261365.1_Warmbaths              ..................------------------------.......a..a.............a............. [194480]
AY261364.1_Tengani_62             t.................------------------------..................t.....a............. [194480]
AY261363.1_Pretoriuskop/96/4      t.................------------------------..........a.............a............. [194480]
AY261362.1_Mkuzi_1979             ..................------------------------...................................... [194480]
AY261361.1_Malawi_Lil-20/1_1983   ..g...............------------------------.....a....a---.a.t......a........t.... [194480]
AY261360.1_Kenya_1950             ------a...........tttataagaaataggttttatgcc.....t.a.....a...........a.....t.t.... [194480]
AM712240.1_OURT_88/3              ..................------------------------...................................... [194480]
AM712239.1_Benin_97/1             ..................------------------------...................................... [194480]
KM111295.1_Ken06.Bus              ------a...........tttataagaaataggttttatgcc.....t.a.....a...........a.....t.t.... [194480]
```

```
KM111294.1_Ken05/Tk1           ------............tttataagaaataggttttatgcc.....t.a.....a...........a.....t.t.... [194480]
KM262844.1_strain_L60          ..................------------------------...................................... [194480]
KM102979.1_26544/OG10_Italy    ..................------------------------...................................... [194480]
KM262845.1_strain_NHV          ..................------------------------...................................... [194480]
KX354450.1_47/Ss/2008          ..................------------------------...................................... [194480]
KP055815.1_strain_BA71         ..................------------------------...................................... [194480]
MH025920.1_strain_R35          ------a...........tttataagaaataggttttatgcc.....t.a.....a...........a.....t.t.... [194480]
MH025919.1_strain_N10          ------a...........tttataagaaataggttttatgcc.....t.a.....a...........a.....t.t.... [194480]
MH025918.1_strain_R25          ------a...........tttataagaaataggttttatgcc.....t.a.....a...........a.....t.t.... [194480]
MH025917.1_strain_R7           ------a...........tttataagaaataggttttatgcc.....t.a.....a...........a.....t.t.... [194480]
MH025916.1_strain_R8           ------a...........tttataagaaataggttttatgcc.....t.a.....a...........a.....t.t.... [194480]

FR682468.2_Georgia_2007/1        tacgaatattttgaccacgagccatttc------atccactataggcag--tattttggctggctatggttctttgttgt [194560]
ASF_IND_20_CAD543                ............................------...............--............................. [194560]
LR899193.1_Germany_2020/1        ............................------...............--............................. [194560]
MN393477.1_Wuhan_2019            ............................------...............--............................. [194560]
MW396979.1_Timor-Leste/2019/1    ............................------...............--............................. [194560]
MK940252.1_CN/2019/InnerMongolia ............................------...............--............................. [194560]
MK128995.1_China/2018/AnhuiXCGQ  ............................------...............--............................. [194560]
MH681419.1_ASFV/POL/2015         ............................------...............--............................. [194560]
LS478113.1_Estonia_2014          ............................------...............--............................. [194560]
KP843857.1_Odintsovo_02/14       ............................------...............--............................. [194560]
FN557520.1_strain_E75            ............................------...............--....c....................c... [194560]
AY261366.1_Warthog               ...........c.....t..........------...............--....c....ct.t..------....c.a. [194560]
AY261365.1_Warmbaths             ...........c.....t..........------...............--....c....ct.t..------....c... [194560]
AY261364.1_Tengani_62            ...........c.....t..........------...............--....c....ct.t..------....c... [194560]
AY261363.1_Pretoriuskop/96/4     ...........c.....t.........t------...............--....c....c..t..------....c... [194560]
AY261362.1_Mkuzi_1979            ...........................t------...............--....c....................c... [194560]
AY261361.1_Malawi_Lil-20/1_1983  ......g...............a.a...cat-ttc........ga....--....c.....actct--a..a....c... [194560]
AY261360.1_Kenya_1950            ............c.........a.c...catttt...t.....g.....ta....c.....ac.ct--...g...ac... [194560]
AM712240.1_OURT_88/3             ............................------...............--....c....................c... [194560]
AM712239.1_Benin_97/1            ............................------...............--....c....................c... [194560]
KM111295.1_Ken06.Bus             ............c.........a.c...catttt...t.....g.....tg....c.....actct--...g...ac... [194560]
KM111294.1_Ken05/Tk1             ............c.........a.c...catttt...t.....ga....ta....c.....ac.ct--...g...ac... [194560]
KM262844.1_strain_L60            ............................------...............--....c....................c... [194560]
KM102979.1_26544/OG10_Italy      ............................------...............--....c....................c... [194560]
KM262845.1_strain_NHV            ............................------...............--....c....................c... [194560]
KX354450.1_47/Ss/2008            ............................------...............--....c....................c... [194560]
KP055815.1_strain_BA71           ............................------...............--....c....................c... [194560]
MH025920.1_strain_R35            ............c.........a.c...catttt...t.....g.....tg....c.....actct--...g...ac... [194560]
MH025919.1_strain_N10            ............c.........a.c...catttt...t.....g.....tg....c.....actct--...g...ac... [194560]
MH025918.1_strain_R25            ............c.........a.c...catttt...t.....g.....tg....c.....actct--...g...ac... [194560]
MH025917.1_strain_R7             ............c.........a.c...catttt...t.....g.....tg....c.....actct--...g...ac... [194560]
MH025916.1_strain_R8             ............c.........a.c...catttt...t.....g.....tg....c.....actct--...g...ac... [194560]
```

```
FR682468.2_Georgia_2007/1        gacaattctatgagatttgattgcaaatcaatttttagttttaaatatat--tggtacctaggacaaagaaagtatatat [194640]
ASF_IND_20_CAD543                .............r....................................--............................ [194640]
LR899193.1_Germany_2020/1        ..................................................--............................ [194640]
MN393477.1_Wuhan_2019            ..................................................--............................ [194640]
MW396979.1_Timor-Leste/2019/1    ..................................................--............................ [194640]
MK940252.1_CN/2019/InnerMongolia ..................................................--............................ [194640]
MK128995.1_China/2018/AnhuiXCGQ  ..................................................--............................ [194640]
MH681419.1_ASFV/POL/2015         ..................................................--............................ [194640]
LS478113.1_Estonia_2014          ..................................................--............................ [194640]
KP843857.1_Odintsovo_02/14       ..................................................--............................ [194640]
FN557520.1_strain_E75            .............a....................................--............................ [194640]
AY261366.1_Warthog               .......................t..........................--............................ [194640]
AY261365.1_Warmbaths             ..................................................--............................ [194640]
AY261364.1_Tengani_62            .......................t..........................--............................ [194640]
AY261363.1_Pretoriuskop/96/4     .......................t..........................--............................ [194640]
AY261362.1_Mkuzi_1979            ..................................................--............................ [194640]
AY261361.1_Malawi_Lil-20/1_1983  ..t..........a.........t......................g..---g........a................g. [194640]
AY261360.1_Kenya_1950            ..t..........a.a.......t..........................tg......t..................... [194640]
AM712240.1_OURT_88/3             .............a....................................--............................ [194640]
AM712239.1_Benin_97/1            .............a....................................--............................ [194640]
KM111295.1_Ken06.Bus             ..t..........a.a.....tac..........................tg............................ [194640]
KM111294.1_Ken05/Tk1             ..t..........a.a.......t..........................tg......t..................... [194640]
KM262844.1_strain_L60            .............a....................................--............................ [194640]
KM102979.1_26544/OG10_Italy      .............a....................................--............................ [194640]
KM262845.1_strain_NHV            .............a....................................--............................ [194640]
KX354450.1_47/Ss/2008            .............a....................................--............................ [194640]
KP055815.1_strain_BA71           .............a....................................--............................ [194640]
MH025920.1_strain_R35            ..t..........a.a.....tac..........................tg............................ [194640]
MH025919.1_strain_N10            ..t..........a.a.....tac..........................tg............................ [194640]
MH025918.1_strain_R25            ..t..........a.a.....tac..........................tg............................ [194640]
MH025917.1_strain_R7             ..t..........a.a.....tac..........................tg............................ [194640]
MH025916.1_strain_R8             ..t..........a.a.....tac..........................tg............................ [194640]

FR682468.2_Georgia_2007/1        agccaataattattccactaaattgatttccagactgatgggtatggagccatgttgtctctgcagacgatcgcaaaaat [194720]
ASF_IND_20_CAD543                ................................k............................................... [194720]
LR899193.1_Germany_2020/1        ................................................................................ [194720]
MN393477.1_Wuhan_2019            ................................................................................ [194720]
MW396979.1_Timor-Leste/2019/1    ................................................................................ [194720]
MK940252.1_CN/2019/InnerMongolia ................................................................................ [194720]
MK128995.1_China/2018/AnhuiXCGQ  ................................................................................ [194720]
MH681419.1_ASFV/POL/2015         ................................................................................ [194720]
LS478113.1_Estonia_2014          ................................................................................ [194720]
KP843857.1_Odintsovo_02/14       ................................................................................ [194720]
```

```
FN557520.1_strain_E75               ................................t...................................a.....g..... [194720]
AY261366.1_Warthog                  .............................................a.............c..a.....a.....g..... [194720]
AY261365.1_Warmbaths                ....................................................................a.....g..... [194720]
AY261364.1_Tengani_62               ...........................................................c..a.....a.....g..... [194720]
AY261363.1_Pretoriuskop/96/4        ...........................................................c..a.....a.....g..... [194720]
AY261362.1_Mkuzi_1979               ....g...........................t...................................a.....g..... [194720]
AY261361.1_Malawi_Lil-20/1_1983     ...t...........t...............g.................a.........c......g....g..g..... [194720]
AY261360.1_Kenya_1950               g..t...........................g.................a.........c..a...g.a.c...g...c. [194720]
AM712240.1_OURT_88/3                ................................t...................................a.....g..... [194720]
AM712239.1_Benin_97/1               ................................t...................................a.....g..... [194720]
KM111295.1_Ken06.Bus                g..............................g.................a.........c..a...g.a.c...g...c. [194720]
KM111294.1_Ken05/Tk1                g.............t................g.................a.........c..a...g.a.c...g...c. [194720]
KM262844.1_strain_L60               ................................t...................................a.....g..... [194720]
KM102979.1_26544/OG10_Italy         ................................t...................................a.....g..... [194720]
KM262845.1_strain_NHV               ................................t...................................a.....g..... [194720]
KX354450.1_47/Ss/2008               ................................t...................................a.....g..... [194720]
KP055815.1_strain_BA71              ................................t...................................a.....g..... [194720]
MH025920.1_strain_R35               g..............................g.................a.........c..a...g.a.c...g...c. [194720]
MH025919.1_strain_N10               g..............................g.................a.........c..a...g.a.c...g...c. [194720]
MH025918.1_strain_R25               g..............................g.................a.........c..a...g.a.c...g...c. [194720]
MH025917.1_strain_R7                g..............................g.................a.........c..a...g.a.c...g...c. [194720]
MH025916.1_strain_R8                g..............................g.................a.........c..a...g.a.c...g...c. [194720]

FR682468.2_Georgia_2007/1           ggccgtagcaacaaacacctactccaagtatcactatccaatactgaaggtctttgggctgtggtggaaaaacaatacgc [194800]
ASF_IND_20_CAD543                   ..........................................................................gc.... [194800]
LR899193.1_Germany_2020/1           ................................................................................ [194800]
MN393477.1_Wuhan_2019               ................................................................................ [194800]
MW396979.1_Timor-Leste/2019/1       ................................................................................ [194800]
MK940252.1_CN/2019/InnerMongolia    ................................................................................ [194800]
MK128995.1_China/2018/AnhuiXCGQ     ................................................................................ [194800]
MH681419.1_ASFV/POL/2015            ................................................................................ [194800]
LS478113.1_Estonia_2014             ................................................................................ [194800]
KP843857.1_Odintsovo_02/14          ................................................................................ [194800]
FN557520.1_strain_E75               ..........................................................................gc.... [194800]
AY261366.1_Warthog                  .............................g.................................................. [194800]
AY261365.1_Warmbaths                .............................g.................................................. [194800]
AY261364.1_Tengani_62               .............................g.................................................. [194800]
AY261363.1_Pretoriuskop/96/4        .............................g.................................................. [194800]
AY261362.1_Mkuzi_1979               ......................................................c...................g..... [194800]
AY261361.1_Malawi_Lil-20/1_1983     .................................t...........a.................................. [194800]
AY261360.1_Kenya_1950               ..t......gg.......a.........a..at.c........t..........c.......................a. [194800]
AM712240.1_OURT_88/3                ..........................................................................gc.... [194800]
AM712239.1_Benin_97/1               ..........................................................................gc.... [194800]
KM111295.1_Ken06.Bus                .........gg.......a.........a..at.c........t..................................a. [194800]
```

```
KM111294.1_Ken05/Tk1            .........gg.......a.........a..at.c........t..........c.......................a. [194800]
KM262844.1_strain_L60           ..........................................................................gc.... [194800]
KM102979.1_26544/OG10_Italy     ..........................................................................gc.... [194800]
KM262845.1_strain_NHV           ..........................................................................gc.... [194800]
KX354450.1_47/Ss/2008           ..........................................................................gc.... [194800]
KP055815.1_strain_BA71          ..........................................................................gc.... [194800]
MH025920.1_strain_R35           .........gg.......a.........a..at.c........t..................................a. [194800]
MH025919.1_strain_N10           .........gg.......a.........a..at.c........t..................................a. [194800]
MH025918.1_strain_R25           .........gg.......a.........a..at.c........t..................................a. [194800]
MH025917.1_strain_R7            .........gg.......a.........a..at.c........t..................................a. [194800]
MH025916.1_strain_R8            .........gg.......a.........a..at.c........t..................................a. [194800]

FR682468.2_Georgia_2007/1       taaatggccctattaaaatatgtaaccattgcaacaacataatggtaggagaatatcctatgtgttacaatcatggaatg [194880]
ASF_IND_20_CAD543               ................................................................................ [194880]
LR899193.1_Germany_2020/1       ................................................................................ [194880]
MN393477.1_Wuhan_2019           ................................................................................ [194880]
MW396979.1_Timor-Leste/2019/1   ................................................................................ [194880]
MK940252.1_CN/2019/InnerMongolia ................................................................................ [194880]
MK128995.1_China/2018/AnhuiXCGQ ................................................................................ [194880]
MH681419.1_ASFV/POL/2015        ................................................................................ [194880]
LS478113.1_Estonia_2014         ................................................................................ [194880]
KP843857.1_Odintsovo_02/14      ................................................................................ [194880]
FN557520.1_strain_E75           ................................................................................ [194880]
AY261366.1_Warthog              ................................................................................ [194880]
AY261365.1_Warmbaths            .................................g.............................................. [194880]
AY261364.1_Tengani_62           ...........................................a.................................... [194880]
AY261363.1_Pretoriuskop/96/4    ................................................................................ [194880]
AY261362.1_Mkuzi_1979           ................................................................................ [194880]
AY261361.1_Malawi_Lil-20/1_1983 .......g..................................ct........g........................... [194880]
AY261360.1_Kenya_1950           ..g....a...g......................tc.a.....a...........c..c.......t............. [194880]
AM712240.1_OURT_88/3            ................................................................................ [194880]
AM712239.1_Benin_97/1           ................................................................................ [194880]
KM111295.1_Ken06.Bus            ..g....a...g...................t..tc.a.....a...........c..c.......t............. [194880]
KM111294.1_Ken05/Tk1            ..g....a...g......................tc.a.....a...........c..c.......t............. [194880]
KM262844.1_strain_L60           ................................................................................ [194880]
KM102979.1_26544/OG10_Italy     ................................................................................ [194880]
KM262845.1_strain_NHV           ................................................................................ [194880]
KX354450.1_47/Ss/2008           ................................................................................ [194880]
KP055815.1_strain_BA71          ................................................................................ [194880]
MH025920.1_strain_R35           ..g....a...g...................t..tc.a.....a...........c..c.......t............. [194880]
MH025919.1_strain_N10           ..g....a...g...................t..tc.a.....a...........c..c.......t............. [194880]
MH025918.1_strain_R25           ..g....a...g...................t..tc.a.....a...........c..c.......t............. [194880]
MH025917.1_strain_R7            ..g....a...g...................t..tc.a.....a...........c..c.......t............. [194880]
MH025916.1_strain_R8            ..g....a...g...................t..tc.a.....a...........c..c.......t............. [194880]
```

```
FR682468.2_Georgia_2007/1        agtctggatatagctttgattcgggcagtaaaggagcgtaatatatccttagtccagcttttcaccgaatgggggggaaa [194960]
ASF_IND_20_CAD543                ..y............................................................................. [194960]
LR899193.1_Germany_2020/1        ................................................................................ [194960]
MN393477.1_Wuhan_2019            ................................................................................ [194960]
MW396979.1_Timor-Leste/2019/1    ................................................................................ [194960]
MK940252.1_CN/2019/InnerMongolia ................................................................................ [194960]
MK128995.1_China/2018/AnhuiXCGQ  ................................................................................ [194960]
MH681419.1_ASFV/POL/2015         ................................................................................ [194960]
LS478113.1_Estonia_2014          ................................................................................ [194960]
KP843857.1_Odintsovo_02/14       ................................................................................ [194960]
FN557520.1_strain_E75            ..c.......................g.....a.....c......................................... [194960]
AY261366.1_Warthog               .....a..........................a.....c......................................... [194960]
AY261365.1_Warmbaths             .....a..........................a.....c......................................g.. [194960]
AY261364.1_Tengani_62            ......................................c......................................... [194960]
AY261363.1_Pretoriuskop/96/4     .....a..........................a.....c.....g................................... [194960]
AY261362.1_Mkuzi_1979            ..c.......................g.....a.....c......c.................................. [194960]
AY261361.1_Malawi_Lil-20/1_1983  ...a.....g.g....................a..t..c............a......................a..... [194960]
AY261360.1_Kenya_1950            ...a.....g.g.....a...........c..a..c.ac........t.................t........a..... [194960]
AM712240.1_OURT_88/3             ..c.......................g.....a.....c......................................... [194960]
AM712239.1_Benin_97/1            ..c.......................g.....a.....c......................................... [194960]
KM111295.1_Ken06.Bus             ...a.....g.g.....a...........c..a..c..c........t.................t........a..... [194960]
KM111294.1_Ken05/Tk1             ...a.....g.g.....a...........c..a..c.ac........t.................t........a..... [194960]
KM262844.1_strain_L60            ..c.......................g.....a.....c......................................... [194960]
KM102979.1_26544/OG10_Italy      ..c.......................g.....a.....c......................................... [194960]
KM262845.1_strain_NHV            ..c.......................g.....a.....c......................................... [194960]
KX354450.1_47/Ss/2008            ..c.......................g.....a.....c......................................... [194960]
KP055815.1_strain_BA71           ..c.......................g.....a.....c......................................... [194960]
MH025920.1_strain_R35            ...a.....g.g.....a...........c..a..c..c........t..........................a..... [194960]
MH025919.1_strain_N10            ...a.....g.g.....a...........c..a..c..c........t.................t........a..... [194960]
MH025918.1_strain_R25            ...a.....g.g.....a...........c..a..c..c........t..........................a..... [194960]
MH025917.1_strain_R7             ...a.....g.g.....a...........c..a..c..c........t..........................a..... [194960]
MH025916.1_strain_R8             ...a.....g.g.....a...........c..a..c..c........t..........................a..... [194960]

FR682468.2_Georgia_2007/1        tattgactatggggcactttgtgctaacactccatctatgcaaagattatgtaaaagtttgggagccaaaccaccaaagg [195040]
ASF_IND_20_CAD543                ................................................................................ [195040]
LR899193.1_Germany_2020/1        ................................................................................ [195040]
MN393477.1_Wuhan_2019            ................................................................................ [195040]
MW396979.1_Timor-Leste/2019/1    ................................................................................ [195040]
MK940252.1_CN/2019/InnerMongolia ................................................................................ [195040]
MK128995.1_China/2018/AnhuiXCGQ  ................................................................................ [195040]
MH681419.1_ASFV/POL/2015         ................................................................................ [195040]
LS478113.1_Estonia_2014          ................................................................................ [195040]
KP843857.1_Odintsovo_02/14       ................................................................................ [195040]
```

```
FN557520.1_strain_E75             ................................................................................ [195040]
AY261366.1_Warthog                ................................................................................ [195040]
AY261365.1_Warmbaths              ..........................................................................t..... [195040]
AY261364.1_Tengani_62             ..............................c.....c...............g................g.......... [195040]
AY261363.1_Pretoriuskop/96/4      ..............................c.....................g................g.......... [195040]
AY261362.1_Mkuzi_1979             ............................................................a................... [195040]
AY261361.1_Malawi_Lil-20/1_1983   ...............g..............c.....................g................g.......... [195040]
AY261360.1_Kenya_1950             ..............................c.....................g.g...........t..g..t....... [195040]
AM712240.1_OURT_88/3              ................................................................................ [195040]
AM712239.1_Benin_97/1             ................................................................................ [195040]
KM111295.1_Ken06.Bus              ......t.......................c.....................g.g..............g..t....... [195040]
KM111294.1_Ken05/Tk1              ..............................c.....................g.g...........t..g..t....... [195040]
KM262844.1_strain_L60             ................................................................................ [195040]
KM102979.1_26544/OG10_Italy       ................................................................................ [195040]
KM262845.1_strain_NHV             ................................................................................ [195040]
KX354450.1_47/Ss/2008             ..........-..................................................................... [195040]
KP055815.1_strain_BA71            ................................................................................ [195040]
MH025920.1_strain_R35             ......t.......................c.....................g.g..............g..t....... [195040]
MH025919.1_strain_N10             ......t.......................c.....................g.g..............g..t....... [195040]
MH025918.1_strain_R25             ......t.......................c.....................g.g..............g..t....... [195040]
MH025917.1_strain_R7              ......t.......................c.....................g.g..............g..t....... [195040]
MH025916.1_strain_R8              ......t.......................c.....................g.g..............g..t....... [195040]

FR682468.2_Georgia_2007/1         gccgaatgtatatggatgctcttatacatctttcagataccttgaatgataatgatctgattagggggtatgagattttt [195120]
ASF_IND_20_CAD543                 ................................................................................ [195120]
LR899193.1_Germany_2020/1         ................................................................................ [195120]
MN393477.1_Wuhan_2019             ................................................................................ [195120]
MW396979.1_Timor-Leste/2019/1     ................................................................................ [195120]
MK940252.1_CN/2019/InnerMongolia  ................................................................................ [195120]
MK128995.1_China/2018/AnhuiXCGQ   ................................................................................ [195120]
MH681419.1_ASFV/POL/2015          ................................................................................ [195120]
LS478113.1_Estonia_2014           ................................................................................ [195120]
KP843857.1_Odintsovo_02/14        ................................................................................ [195120]
FN557520.1_strain_E75             ................................................................................ [195120]
AY261366.1_Warthog                ................................................................................ [195120]
AY261365.1_Warmbaths              ................................................................................ [195120]
AY261364.1_Tengani_62             .................a.c.........t.................................................. [195120]
AY261363.1_Pretoriuskop/96/4      .................a.c.........t.................................................. [195120]
AY261362.1_Mkuzi_1979             ................................................................................ [195120]
AY261361.1_Malawi_Lil-20/1_1983   .................a.c...................................c...........a............ [195120]
AY261360.1_Kenya_1950             ...a.....t.......a..........................g...........t.a..................... [195120]
AM712240.1_OURT_88/3              ................................................................................ [195120]
AM712239.1_Benin_97/1             ................................................................................ [195120]
KM111295.1_Ken06.Bus              ...a.....t.......a..........................g...........t.a..................... [195120]
```

```
KM111294.1_Ken05/Tk1             ...a.....t.......a..........................g...........t.a..................... [195120]
KM262844.1_strain_L60            ................................................................................ [195120]
KM102979.1_26544/OG10_Italy      ................................................................................ [195120]
KM262845.1_strain_NHV            ................................................................................ [195120]
KX354450.1_47/Ss/2008            ................................................................................ [195120]
KP055815.1_strain_BA71           ................................................................................ [195120]
MH025920.1_strain_R35            ...a.....t.......a..........................g...........t.a..................... [195120]
MH025919.1_strain_N10            ...a.....t.......a..........................g...........t.a..................... [195120]
MH025918.1_strain_R25            ...a.....t.......a..........................g...........t.a..................... [195120]
MH025917.1_strain_R7             ...a.....t.......a..........................g...........t.a..................... [195120]
MH025916.1_strain_R8             ...a.....t.......a..........................g...........t.a..................... [195120]

FR682468.2_Georgia_2007/1        gatgataatagcgtgttggattgtgtcaatctcatacgactcaaaataatgcttaccttgaaggcccgtatacctctcat [195200]
ASF_IND_20_CAD543                ................................................................................ [195200]
LR899193.1_Germany_2020/1        ................................................................................ [195200]
MN393477.1_Wuhan_2019            ................................................................................ [195200]
MW396979.1_Timor-Leste/2019/1    ................................................................................ [195200]
MK940252.1_CN/2019/InnerMongolia ................................................................................ [195200]
MK128995.1_China/2018/AnhuiXCGQ  ................................................................................ [195200]
MH681419.1_ASFV/POL/2015         ................................................................................ [195200]
LS478113.1_Estonia_2014          ................................................................................ [195200]
KP843857.1_Odintsovo_02/14       ................................................................................ [195200]
FN557520.1_strain_E75            ................................................................................ [195200]
AY261366.1_Warthog               ................................................................................ [195200]
AY261365.1_Warmbaths             ................................................................................ [195200]
AY261364.1_Tengani_62            .........................................a...................................... [195200]
AY261363.1_Pretoriuskop/96/4     .........................................a..............................t....... [195200]
AY261362.1_Mkuzi_1979            ................................................................................ [195200]
AY261361.1_Malawi_Lil-20/1_1983  ......................a..........g.....t.a...................................... [195200]
AY261360.1_Kenya_1950            ..a..a....a...a...a...a............g.a.t.a........t............a.t.............. [195200]
AM712240.1_OURT_88/3             ................................................................................ [195200]
AM712239.1_Benin_97/1            ................................................................................ [195200]
KM111295.1_Ken06.Bus             ..a..a....aa..a.......a............g.a.t.a........t...t........a................ [195200]
KM111294.1_Ken05/Tk1             ..a..a....a...a...a...a............g.a.t.a........t............a................ [195200]
KM262844.1_strain_L60            ................................................................................ [195200]
KM102979.1_26544/OG10_Italy      ................................................................................ [195200]
KM262845.1_strain_NHV            ................................................................................ [195200]
KX354450.1_47/Ss/2008            ................................................................................ [195200]
KP055815.1_strain_BA71           ................................................................................ [195200]
MH025920.1_strain_R35            ..a..a....aa..a.......a............g.a.t.a........t...t........a................ [195200]
MH025919.1_strain_N10            ..a..a....aa..a.......a............g.a.t.a........t...t........a................ [195200]
MH025918.1_strain_R25            ..a..a....aa..a.......a............g.a.t.a........t...t........a................ [195200]
MH025917.1_strain_R7             ..a..a....aa..a.......a............g.a.t.a........t...t........a................ [195200]
MH025916.1_strain_R8             ..a..a....aa..a.......a............g.a.t.a........t...t........a................ [195200]
```

```
FR682468.2_Georgia_2007/1        ggaacaactagaccaaattgccttaaaacaacttctgcagcgatactggtatgccatggctgtacaacacaacttaacaa [195280]
ASF_IND_20_CAD543                ................................................................................ [195280]
LR899193.1_Germany_2020/1        ................................................................................ [195280]
MN393477.1_Wuhan_2019            ................................................................................ [195280]
MW396979.1_Timor-Leste/2019/1    ................................................................................ [195280]
MK940252.1_CN/2019/InnerMongolia ................................................................................ [195280]
MK128995.1_China/2018/AnhuiXCGQ  ................................................................................ [195280]
MH681419.1_ASFV/POL/2015         ................................................................................ [195280]
LS478113.1_Estonia_2014          ................................................................................ [195280]
KP843857.1_Odintsovo_02/14       ................................................................................ [195280]
FN557520.1_strain_E75            ................................................................................ [195280]
AY261366.1_Warthog               ......................................................t.................t....... [195280]
AY261365.1_Warmbaths             ................................................................................ [195280]
AY261364.1_Tengani_62            ..................................................................g............. [195280]
AY261363.1_Pretoriuskop/96/4     ..................................................................g............. [195280]
AY261362.1_Mkuzi_1979            ................................................................................ [195280]
AY261361.1_Malawi_Lil-20/1_1983  ..................c....................a.....t...............................g.. [195280]
AY261360.1_Kenya_1950            ..........................g...g..............t...........................c...a.g [195280]
AM712240.1_OURT_88/3             ................................................................................ [195280]
AM712239.1_Benin_97/1            ................................................................................ [195280]
KM111295.1_Ken06.Bus             ..........................g...g..............t...........................c...a.g [195280]
KM111294.1_Ken05/Tk1             ..........................g...g..............t...........................c...a.g [195280]
KM262844.1_strain_L60            ................................................................................ [195280]
KM102979.1_26544/OG10_Italy      ................................................................................ [195280]
KM262845.1_strain_NHV            ................................................................................ [195280]
KX354450.1_47/Ss/2008            ................................................................................ [195280]
KP055815.1_strain_BA71           ................................................................................ [195280]
MH025920.1_strain_R35            ..........................g...g..............t...........................c...a.g [195280]
MH025919.1_strain_N10            ..........................g...g..............t...........................c...a.g [195280]
MH025918.1_strain_R25            ..........................g...g..............t...........................c...a.g [195280]
MH025917.1_strain_R7             ..........................g...g..............t...........................c...a.g [195280]
MH025916.1_strain_R8             ..........................g...g..............t...........................c...a.g [195280]

FR682468.2_Georgia_2007/1        tcgctatccactattttgataatcatattcctaatataaagccatttagtctgcgctgtgctttgtattttaatgatccc [195360]
ASF_IND_20_CAD543                ca.............................................................................. [195360]
LR899193.1_Germany_2020/1        ................................................................................ [195360]
MN393477.1_Wuhan_2019            ................................................................................ [195360]
MW396979.1_Timor-Leste/2019/1    ................................................................................ [195360]
MK940252.1_CN/2019/InnerMongolia ................................................................................ [195360]
MK128995.1_China/2018/AnhuiXCGQ  ................................................................................ [195360]
MH681419.1_ASFV/POL/2015         ................................................................................ [195360]
LS478113.1_Estonia_2014          ................................................................................ [195360]
KP843857.1_Odintsovo_02/14       ................................................................................ [195360]
```

```
FN557520.1_strain_E75          ca.............................................................................. [195360]
AY261366.1_Warthog             ca...............a.............................g................................ [195360]
AY261365.1_Warmbaths           ca.............................................................................. [195360]
AY261364.1_Tengani_62          .a..............................................................a............... [195360]
AY261363.1_Pretoriuskop/96/4   .a.............................................................................. [195360]
AY261362.1_Mkuzi_1979          ca.............................................................................. [195360]
AY261361.1_Malawi_Lil-20/1_1983 .a............................................c.......................c......... [195360]
AY261360.1_Kenya_1950          ca....................c..............g.........ca...................c........... [195360]
AM712240.1_OURT_88/3           ca.............................................................................. [195360]
AM712239.1_Benin_97/1          ca.............................................................................. [195360]
KM111295.1_Ken06.Bus           ca....................c........................c....................c........t.. [195360]
KM111294.1_Ken05/Tk1           ca....................c..............g.........c....................c........... [195360]
KM262844.1_strain_L60          ca.............................................................................. [195360]
KM102979.1_26544/OG10_Italy    ca.............................................................................. [195360]
KM262845.1_strain_NHV          ca.............................................................................. [195360]
KX354450.1_47/Ss/2008          ca.............................................................................. [195360]
KP055815.1_strain_BA71         ca.............................................................................. [195360]
MH025920.1_strain_R35          ca....................c........................c....................c........t.. [195360]
MH025919.1_strain_N10          ca....................c........................c....................c........t.. [195360]
MH025918.1_strain_R25          ca....................c........................c....................c........t.. [195360]
MH025917.1_strain_R7           ca....................c........................c....................c........t.. [195360]
MH025916.1_strain_R8           ca....................c........................c....................c........t.. [195360]

FR682468.2_Georgia_2007/1      tttaaaatccatgatgcttgcagaactgtaaatatggatcctaatgagatgatgaacattgcttgtcaacaggatttaaa [195440]
ASF_IND_20_CAD543              ................................................................................ [195440]
LR899193.1_Germany_2020/1      ................................................................................ [195440]
MN393477.1_Wuhan_2019          ................................................................................ [195440]
MW396979.1_Timor-Leste/2019/1  ................................................................................ [195440]
MK940252.1_CN/2019/InnerMongolia ................................................................................ [195440]
MK128995.1_China/2018/AnhuiXCGQ ................................................................................ [195440]
MH681419.1_ASFV/POL/2015       ................................................................................ [195440]
LS478113.1_Estonia_2014        ................................................................................ [195440]
KP843857.1_Odintsovo_02/14     ................................................................................ [195440]
FN557520.1_strain_E75          .......................g........................................................ [195440]
AY261366.1_Warthog             .......................g........................................................ [195440]
AY261365.1_Warmbaths           ........t..............g....................c..a................................ [195440]
AY261364.1_Tengani_62          .......................g........................................................ [195440]
AY261363.1_Pretoriuskop/96/4   ................................................................................ [195440]
AY261362.1_Mkuzi_1979          .......................g........................................................ [195440]
AY261361.1_Malawi_Lil-20/1_1983 ..........................g....c............c................................... [195440]
AY261360.1_Kenya_1950          ....................t.a.............................................g........... [195440]
AM712240.1_OURT_88/3           .......................g........................................................ [195440]
AM712239.1_Benin_97/1          .......................g........................................................ [195440]
KM111295.1_Ken06.Bus           ....................t.a.............................................g........... [195440]
```

```
KM111294.1_Ken05/Tk1            ....................t.a.............................................g..........g [195440]
KM262844.1_strain_L60           .......................g........................................................ [195440]
KM102979.1_26544/OG10_Italy     .......................g........................................................ [195440]
KM262845.1_strain_NHV           .......................g........................................................ [195440]
KX354450.1_47/Ss/2008           .......................g........................................................ [195440]
KP055815.1_strain_BA71          .......................g........................................................ [195440]
MH025920.1_strain_R35           ....................t.a.............................................g........... [195440]
MH025919.1_strain_N10           ....................t.a.............................................g........... [195440]
MH025918.1_strain_R25           ....................t.a.............................................g........... [195440]
MH025917.1_strain_R7            ....................t.a.............................................g........... [195440]
MH025916.1_strain_R8            ....................t.a.............................................g........... [195440]

FR682468.2_Georgia_2007/1       ctttcaaagcatttactattgttatcttttaggggctgatattaatcaggctatgctaatgtctttaaagtatggtcatc [195520]
ASF_IND_20_CAD543               ................................................................................ [195520]
LR899193.1_Germany_2020/1       ................................................................................ [195520]
MN393477.1_Wuhan_2019           ................................................................................ [195520]
MW396979.1_Timor-Leste/2019/1   ................................................................................ [195520]
MK940252.1_CN/2019/InnerMongolia ................................................................................ [195520]
MK128995.1_China/2018/AnhuiXCGQ ................................................................................ [195520]
MH681419.1_ASFV/POL/2015        ................................................................................ [195520]
LS478113.1_Estonia_2014         ................................................................................ [195520]
KP843857.1_Odintsovo_02/14      ................................................................................ [195520]
FN557520.1_strain_E75           ...................a.....a..................................................a... [195520]
AY261366.1_Warthog              ................................................................................ [195520]
AY261365.1_Warmbaths            ................................................................................ [195520]
AY261364.1_Tengani_62           ................................................................................ [195520]
AY261363.1_Pretoriuskop/96/4    ................................................................................ [195520]
AY261362.1_Mkuzi_1979           ...................a........................................................a... [195520]
AY261361.1_Malawi_Lil-20/1_1983 .................................a...................................t.....ga... [195520]
AY261360.1_Kenya_1950           .............ct....a.....a.g.....a............a....c..............g..tc....ga... [195520]
AM712240.1_OURT_88/3            ...................a.....a..................................................a... [195520]
AM712239.1_Benin_97/1           ...................a.....a..................................................a... [195520]
KM111295.1_Ken06.Bus            .............ct....a.....a.g.....a............a....c..............g..tc....ga... [195520]
KM111294.1_Ken05/Tk1            .............ct..........a.g.....a............a...................g..tc....ga... [195520]
KM262844.1_strain_L60           ...................a.....a..................................................a... [195520]
KM102979.1_26544/OG10_Italy     ...................a.....a..................................................a... [195520]
KM262845.1_strain_NHV           ...................a.....a..................................................a... [195520]
KX354450.1_47/Ss/2008           ...................a.....a..................................................a... [195520]
KP055815.1_strain_BA71          ...................a.....a..................................................a... [195520]
MH025920.1_strain_R35           .............ct....a.....a.g.....a............a....c..............g..tc....ga... [195520]
MH025919.1_strain_N10           .............ct....a.....a.g.....a............a....c..............g..tc....ga... [195520]
MH025918.1_strain_R25           .............ct....a.....a.g.....a............a....c..............g..tc....ga... [195520]
MH025917.1_strain_R7            .............ct....a.....a.g.....a............a....c..............g..tc....ga... [195520]
MH025916.1_strain_R8            .............ct....a.....a.g.....a............a....c..............g..tc....ga... [195520]
```

```
FR682468.2_Georgia_2007/1        tttctaatatgtggttttgcatagatttgggggcggatgcctttaaagaggcaggggcgcttgctgagaaaaaaaataaa [195600]
ASF_IND_20_CAD543                ..................................................................r........--... [195600]
LR899193.1_Germany_2020/1        ................................................................................ [195600]
MN393477.1_Wuhan_2019            ................................................................................ [195600]
MW396979.1_Timor-Leste/2019/1    ................................................................................ [195600]
MK940252.1_CN/2019/InnerMongolia ................................................................................ [195600]
MK128995.1_China/2018/AnhuiXCGQ  ................................................................................ [195600]
MH681419.1_ASFV/POL/2015         ................................................................................ [195600]
LS478113.1_Estonia_2014          ................................................................................ [195600]
KP843857.1_Odintsovo_02/14       ....................................................................-........... [195600]
FN557520.1_strain_E75            ..................................................................g.--......a... [195600]
AY261366.1_Warthog               ..................................a....................a.t.........a..........g. [195600]
AY261365.1_Warmbaths             ..................................................................g............. [195600]
AY261364.1_Tengani_62            ..................................a....................a.t.........a..........g. [195600]
AY261363.1_Pretoriuskop/96/4     ..................................a....................a.t........ga..........g. [195600]
AY261362.1_Mkuzi_1979            ..................................................................ga............ [195600]
AY261361.1_Malawi_Lil-20/1_1983  ..................................aa...tt.....g......a............ga..........g. [195600]
AY261360.1_Kenya_1950            ...................t........a.....a.....t...g.g...................ga............ [195600]
AM712240.1_OURT_88/3             ..................................................................g.--......a... [195600]
AM712239.1_Benin_97/1            ..................................................................g..-......a... [195600]
KM111295.1_Ken06.Bus             ...................t........a.....a.....t...g.g...................ga............ [195600]
KM111294.1_Ken05/Tk1             ...................t........a.....a.....t...g.g...................ga............ [195600]
KM262844.1_strain_L60            ..................................................................g..-......a... [195600]
KM102979.1_26544/OG10_Italy      ..................................................................g.---.....a... [195600]
KM262845.1_strain_NHV            ..................................................................g.--......a... [195600]
KX354450.1_47/Ss/2008            ..................................................................g.---.....a... [195600]
KP055815.1_strain_BA71           ..................................................................g.........a... [195600]
MH025920.1_strain_R35            ...................t........a.....a.....t...g.g...................ga............ [195600]
MH025919.1_strain_N10            ...................t........a.....a.....t...g.g...................ga............ [195600]
MH025918.1_strain_R25            ...................t........a.....a.....t...g.g...................ga............ [195600]
MH025917.1_strain_R7             ...................t........a.....a.....t...g.g...................ga............ [195600]
MH025916.1_strain_R8             ...................t........a.....a.....t...g.g...................ga............ [195600]

FR682468.2_Georgia_2007/1        agagtgttacaacacatattaggtcttaatatctttaagcgagagttgatt---cccccctgtaaagatcctgatcctta [195680]
ASF_IND_20_CAD543                ...................................................---.......................... [195680]
LR899193.1_Germany_2020/1        ...................................................---.......................... [195680]
MN393477.1_Wuhan_2019            ...................................................---.......................... [195680]
MW396979.1_Timor-Leste/2019/1    ...................................................---.......................... [195680]
MK940252.1_CN/2019/InnerMongolia ...................................................---.......................... [195680]
MK128995.1_China/2018/AnhuiXCGQ  ...................................................---.......................... [195680]
MH681419.1_ASFV/POL/2015         ...................................................---.......................... [195680]
LS478113.1_Estonia_2014          ...................................................---.......................... [195680]
KP843857.1_Odintsovo_02/14       ...................................................---.......................... [195680]
```

```
FN557520.1_strain_E75           ...........g.............................g.........---.......................... [195680]
AY261366.1_Warthog              .....a.....g..........................a............---.......................... [195680]
AY261365.1_Warmbaths            ...........gt......................................---......c................... [195680]
AY261364.1_Tengani_62           ...........g..........................a............---.......................... [195680]
AY261363.1_Pretoriuskop/96/4    ...........gt.........................a............---.......................... [195680]
AY261362.1_Mkuzi_1979           ...........g.......................................---.......................... [195680]
AY261361.1_Malawi_Lil-20/1_1983 ...........gt.........................a............---.........................c [195680]
AY261360.1_Kenya_1950           ....c.........t..............c........a.a..........---..t..t...................t [195680]
AM712240.1_OURT_88/3            ...........g.............................g.........---.......................... [195680]
AM712239.1_Benin_97/1           ...........g.............................g.........ccc.......................... [195680]
KM111295.1_Ken06.Bus            ........g..g..t..............c.....................---..t..t......a............c [195680]
KM111294.1_Ken05/Tk1            ....c.........t..............c........a.a..........---..t..t...................t [195680]
KM262844.1_strain_L60           ...........g.............................g.........---.......................... [195680]
KM102979.1_26544/OG10_Italy     ...........g.............................g.........---.......................... [195680]
KM262845.1_strain_NHV           ...........g.............................g.........---.......................... [195680]
KX354450.1_47/Ss/2008           ...........g.............................g.........---.......................... [195680]
KP055815.1_strain_BA71          ...........g.............................g.........---.......................... [195680]
MH025920.1_strain_R35           ........g..g..t..............c.....................---..t..t......a............c [195680]
MH025919.1_strain_N10           ........g..g..t..............c.....................---..t..t......a............c [195680]
MH025918.1_strain_R25           ........g..g..t..............c.....................---..t..t......a............c [195680]
MH025917.1_strain_R7            ........g..g..t..............c.....................---..t..t......a............c [195680]
MH025916.1_strain_R8            ........g..g..t..............c.....................---..t..t......a............c [195680]

FR682468.2_Georgia_2007/1       tcaaatccaaattctgttaaaaaactacattctaaaaaatgtctcaactgtttttacatattattgccagtagccattgt [195760]
ASF_IND_20_CAD543               .........................................................w...................... [195760]
LR899193.1_Germany_2020/1       ................................................................................ [195760]
MN393477.1_Wuhan_2019           ................................................................................ [195760]
MW396979.1_Timor-Leste/2019/1   ................................................................................ [195760]
MK940252.1_CN/2019/InnerMongolia ................................................................................ [195760]
MK128995.1_China/2018/AnhuiXCGQ ................................................................................ [195760]
MH681419.1_ASFV/POL/2015        ................................................................................ [195760]
LS478113.1_Estonia_2014         ................................................................................ [195760]
KP843857.1_Odintsovo_02/14      ................................................................................ [195760]
FN557520.1_strain_E75           .........................................................t...................... [195760]
AY261366.1_Warthog              .........................................................t...................... [195760]
AY261365.1_Warmbaths            .........................................................................t...... [195760]
AY261364.1_Tengani_62           .........................................................t...................... [195760]
AY261363.1_Pretoriuskop/96/4    .........................................................t...................... [195760]
AY261362.1_Mkuzi_1979           ................................................c........t...................... [195760]
AY261361.1_Malawi_Lil-20/1_1983 ........................t.................a..............t..............at...... [195760]
AY261360.1_Kenya_1950           ..........c.............t....c............a......a......gt........t......t...... [195760]
AM712240.1_OURT_88/3            .........................................................t...................... [195760]
AM712239.1_Benin_97/1           .........................................................t...................... [195760]
KM111295.1_Ken06.Bus            ..........c..................c............a.....ca.......t........t......t...... [195760]
```

```
KM111294.1_Ken05/Tk1            ..........c.............t....c............a.....ca.....g.t........t......t...... [195760]
KM262844.1_strain_L60           .........................................................t...................... [195760]
KM102979.1_26544/OG10_Italy     .........................................................t...................... [195760]
KM262845.1_strain_NHV           .........................................................t...................... [195760]
KX354450.1_47/Ss/2008           .........................................................t...................... [195760]
KP055815.1_strain_BA71          .........................................................t...................... [195760]
MH025920.1_strain_R35           ..........c..................c............a.....ca.......t........t......t...... [195760]
MH025919.1_strain_N10           ..........c..................c............a.....ca.......t........t......t...... [195760]
MH025918.1_strain_R25           ..........c..................c............a.....ca.......t........t......t...... [195760]
MH025917.1_strain_R7            ..........c..................c............a.....ca.......t........t......t...... [195760]
MH025916.1_strain_R8            ..........c..................c............a.....ca.......t........t......t...... [195760]

FR682468.2_Georgia_2007/1       tta---tatcaga--aaataacccatttgttt-atctttttttgtggggcaaccattaagacccgacgc-aaaa-aaaga [195840]
ASF_IND_20_CAD543               ...---.......--.................-............--nnnnnnnnnnnnnnn.......-....-..... [195840]
LR899193.1_Germany_2020/1       ...---.......--.................-....................................-....-..... [195840]
MN393477.1_Wuhan_2019           ...---.......--.................-....................................-....-..... [195840]
MW396979.1_Timor-Leste/2019/1   ...---.......--.................-....................................-....-..... [195840]
MK940252.1_CN/2019/InnerMongolia ...---.......--.................-....................................-....-..... [195840]
MK128995.1_China/2018/AnhuiXCGQ ...---.......--.................-....................................-....-..... [195840]
MH681419.1_ASFV/POL/2015        ...---.......--.................-....................................-....-..... [195840]
LS478113.1_Estonia_2014         ...---.......--.................-....................................-....-..... [195840]
KP843857.1_Odintsovo_02/14      ...---.......--.................-....................................-....-..... [195840]
FN557520.1_strain_E75           ...---.......--.t...............-............a..--g........agt.......-....-....t [195840]
AY261366.1_Warthog              ...---.......--.t...............-...................t......agt....g..a....-..... [195840]
AY261365.1_Warmbaths            ...---.......--.t...............-...-...........a.gg.t.....agt..at...a....-..... [195840]
AY261364.1_Tengani_62           ...---.......--.t...............-...................t......a.tt......a....-..... [195840]
AY261363.1_Pretoriuskop/96/4    ...---.......--.t...............-..............a....t..a...agt....g..a....-..... [195840]
AY261362.1_Mkuzi_1979           ...---.......--.t...............-...............a.gg.t.....agt..at...a....-..... [195840]
AY261361.1_Malawi_Lil-20/1_1983 ...tat......g--ct....-..........-.atc......t..c.t..g..t.a.t.g.ggct.a.-....-...a. [195840]
AY261360.1_Kenya_1950           ...---......gat.t...............a..........t..tat..gg.t.c.t..t.gcc.a.a....-..... [195840]
AM712240.1_OURT_88/3            ...---.......--.t...............-............a..--g........agt.......-....-....t [195840]
AM712239.1_Benin_97/1           ...---.......--.t...............-............a..--g........agt.......-....-....t [195840]
KM111295.1_Ken06.Bus            ...---......gat.t...............a..........t..tatt.gg.t.c.ta.t.gcc.a.a....-..... [195840]
KM111294.1_Ken05/Tk1            ...---......gat.t...............a..........t..tat..gg.t.c.t..t.gcc.a.a....a..... [195840]
KM262844.1_strain_L60           ...---.......--.t...............-............a..--g........agt.......-....-....t [195840]
KM102979.1_26544/OG10_Italy     ...---.......--.t...............-............a..--g........agt.......-....-....t [195840]
KM262845.1_strain_NHV           ...---.......--.t...............-............a..--g........agt.......-....-....t [195840]
KX354450.1_47/Ss/2008           ...---.......--.t...............-............a..--g........agt.......-....-....t [195840]
KP055815.1_strain_BA71          ...---.......--.t...............-............a..--g........agt.......-....-....t [195840]
MH025920.1_strain_R35           ...---......gat.t...............a..........t..tatt.gg.t.c.ta.t.gcc.a.a....-..... [195840]
MH025919.1_strain_N10           ...---......gat.t...............a...-......t..tatt.gg.t.c.ta.t.gcc.a.a....-..... [195840]
MH025918.1_strain_R25           ...---......gat.t...............a..........t..tatt.gg.t.c.ta.t.gcc.a.a....-..... [195840]
MH025917.1_strain_R7            ...---......gat.t...............a..........t..tatt.gg.t.c.ta.t.gcc.a.a....-..... [195840]
MH025916.1_strain_R8            ...---......gat.t...............a..........t..tatt.gg.t.c.ta.t.gcc.a.a....-..... [195840]
```

```
FR682468.2_Georgia_2007/1        ttaatcttttatcagatacctaaaacgttctataagggagtctatgagatggatcatattttgatggtcatagtaagaag [195920]
ASF_IND_20_CAD543                ................................................................................ [195920]
LR899193.1_Germany_2020/1        ................................................................................ [195920]
MN393477.1_Wuhan_2019            ................................................................................ [195920]
MW396979.1_Timor-Leste/2019/1    ................................................................................ [195920]
MK940252.1_CN/2019/InnerMongolia ................................................................................ [195920]
MK128995.1_China/2018/AnhuiXCGQ  ................................................................................ [195920]
MH681419.1_ASFV/POL/2015         ................................................................................ [195920]
LS478113.1_Estonia_2014          ................................................................................ [195920]
KP843857.1_Odintsovo_02/14       ................................................................................ [195920]
FN557520.1_strain_E75            ................................................................................ [195920]
AY261366.1_Warthog               ....................................t.............................a............. [195920]
AY261365.1_Warmbaths             ...................t................t.............................a............. [195920]
AY261364.1_Tengani_62            ....................................t....................g........a............. [195920]
AY261363.1_Pretoriuskop/96/4     ....................................t.............................a............. [195920]
AY261362.1_Mkuzi_1979            ................................................................................ [195920]
AY261361.1_Malawi_Lil-20/1_1983  ....g......a........................t...........c.............................g. [195920]
AY261360.1_Kenya_1950            ..........................c..t......t....................c.c...........c........ [195920]
AM712240.1_OURT_88/3             ..........................a..................................................... [195920]
AM712239.1_Benin_97/1            ................................................................................ [195920]
KM111295.1_Ken06.Bus             ..........................c..t......t....................c.c...........c........ [195920]
KM111294.1_Ken05/Tk1             ..........................c..t......t....................c.c...........c........ [195920]
KM262844.1_strain_L60            ................................................................................ [195920]
KM102979.1_26544/OG10_Italy      ................................................................................ [195920]
KM262845.1_strain_NHV            ..........................a..................................................... [195920]
KX354450.1_47/Ss/2008            ................................................................................ [195920]
KP055815.1_strain_BA71           ................................................................................ [195920]
MH025920.1_strain_R35            ..........................c..t......t....................c.c...........c........ [195920]
MH025919.1_strain_N10            ..........................c..t......t....................c.c...........c........ [195920]
MH025918.1_strain_R25            ..........................c..t......t....................c.c...........c........ [195920]
MH025917.1_strain_R7             ..........................c..t......t....................c.c...........c........ [195920]
MH025916.1_strain_R8             ..........................c..t......t....................c.c...........c........ [195920]

FR682468.2_Georgia_2007/1        caagctttttggcgaaaacaacggagttaaagaatttaacccgctcatgtttggataggacttttaacagcgagccaaaa [196000]
ASF_IND_20_CAD543                ................................................................................ [196000]
LR899193.1_Germany_2020/1        ................................................................................ [196000]
MN393477.1_Wuhan_2019            ................................................................................ [196000]
MW396979.1_Timor-Leste/2019/1    ................................................................................ [196000]
MK940252.1_CN/2019/InnerMongolia ................................................................................ [196000]
MK128995.1_China/2018/AnhuiXCGQ  ................................................................................ [196000]
MH681419.1_ASFV/POL/2015         ................................................................................ [196000]
LS478113.1_Estonia_2014          ................................................................................ [196000]
KP843857.1_Odintsovo_02/14       ................................................................................ [196000]
```

```
FN557520.1_strain_E75          ...................g............................................................ [196000]
AY261366.1_Warthog             ...................g............................................................ [196000]
AY261365.1_Warmbaths           ...................g............................................................ [196000]
AY261364.1_Tengani_62          ...................g............................................................ [196000]
AY261363.1_Pretoriuskop/96/4   ...................g....................t....................................... [196000]
AY261362.1_Mkuzi_1979          ...................g.....................................................t...... [196000]
AY261361.1_Malawi_Lil-20/1_1983 ......................a..............tt..t................a..............t...... [196000]
AY261360.1_Kenya_1950          ........c.a..a.....g..a.c.............tt......g..........at........t.....a...... [196000]
AM712240.1_OURT_88/3           ...................g............................................................ [196000]
AM712239.1_Benin_97/1          ...................g............................................................ [196000]
KM111295.1_Ken06.Bus           ........c.a..a.....g..a.c.............ttg.....g...c......at........t.....a...... [196000]
KM111294.1_Ken05/Tk1           ........c.a..a.....g..a.c.............tt.t....g..........at........t.....a...... [196000]
KM262844.1_strain_L60          ...................g............................................................ [196000]
KM102979.1_26544/OG10_Italy    ...................g............................................................ [196000]
KM262845.1_strain_NHV          ...................g............................................................ [196000]
KX354450.1_47/Ss/2008          ...................g............................................................ [196000]
KP055815.1_strain_BA71         ...................g............................................................ [196000]
MH025920.1_strain_R35          ........c.a..a.....g..a.c.............ttg.....g...c......at........t.....a...... [196000]
MH025919.1_strain_N10          ........c.a..a.....g..a.c.............ttg.....g...c......at........t.....a...... [196000]
MH025918.1_strain_R25          ........c.a..a.....g..a.c.............ttg.....g...c......at........t.....a...... [196000]
MH025917.1_strain_R7           ........c.a..a.....g..a.c.............ttg.....g...c......at........t.....a...... [196000]
MH025916.1_strain_R8           ........c.a..a.....g..a.c.............ttg.....g...c......at........t.....a...... [196000]

FR682468.2_Georgia_2007/1      cagtatttaaaaatttggcaatagtttttttgggatgcaataaacaaacacttgatcagtgcccgcttcactttctgatc [196080]
ASF_IND_20_CAD543              ................................................................................ [196080]
LR899193.1_Germany_2020/1      ................................................................................ [196080]
MN393477.1_Wuhan_2019          ................................................................................ [196080]
MW396979.1_Timor-Leste/2019/1  ................................................................................ [196080]
MK940252.1_CN/2019/InnerMongolia ................................................................................ [196080]
MK128995.1_China/2018/AnhuiXCGQ ................................................................................ [196080]
MH681419.1_ASFV/POL/2015       ................................................................................ [196080]
LS478113.1_Estonia_2014        ................................................................................ [196080]
KP843857.1_Odintsovo_02/14     ................................................................................ [196080]
FN557520.1_strain_E75          ................................................................................ [196080]
AY261366.1_Warthog             .......................a........a............................................... [196080]
AY261365.1_Warmbaths           .......................a........a............................................... [196080]
AY261364.1_Tengani_62          .......................a........a..........................c..t................. [196080]
AY261363.1_Pretoriuskop/96/4   .......................a...............................................g........ [196080]
AY261362.1_Mkuzi_1979          ................................................................................ [196080]
AY261361.1_Malawi_Lil-20/1_1983 ..................................t............................................. [196080]
AY261360.1_Kenya_1950          .....................c............c............................t................ [196080]
AM712240.1_OURT_88/3           ................................................................................ [196080]
AM712239.1_Benin_97/1          ................................................................................ [196080]
KM111295.1_Ken06.Bus           .....................c............c............................t................ [196080]
```

```
KM111294.1_Ken05/Tk1           .....................c............c............................t................ [196080]
KM262844.1_strain_L60          ................................................................................ [196080]
KM102979.1_26544/OG10_Italy    ................................................................................ [196080]
KM262845.1_strain_NHV          ................................................................................ [196080]
KX354450.1_47/Ss/2008          ................................................................................ [196080]
KP055815.1_strain_BA71         ................................................................................ [196080]
MH025920.1_strain_R35          .....................c............c............................t................ [196080]
MH025919.1_strain_N10          .....................c............c............................t................ [196080]
MH025918.1_strain_R25          .....................c............c............................t................ [196080]
MH025917.1_strain_R7           .....................c............c............................t................ [196080]
MH025916.1_strain_R8           .....................c............c............................t................ [196080]

FR682468.2_Georgia_2007/1      agacatgtttgccgcataacaggcctttttaaacttagtaatataattatgttccgcaagcaccattaacaagggaacga [196160]
ASF_IND_20_CAD543              ................................................................................ [196160]
LR899193.1_Germany_2020/1      ................................................................................ [196160]
MN393477.1_Wuhan_2019          ................................................................................ [196160]
MW396979.1_Timor-Leste/2019/1  ................................................................................ [196160]
MK940252.1_CN/2019/InnerMongolia ................................................................................ [196160]
MK128995.1_China/2018/AnhuiXCGQ ................................................................................ [196160]
MH681419.1_ASFV/POL/2015       ................................................................................ [196160]
LS478113.1_Estonia_2014        ................................................................................ [196160]
KP843857.1_Odintsovo_02/14     ................................................................................ [196160]
FN557520.1_strain_E75          ................................................................................ [196160]
AY261366.1_Warthog             ......a................t........................................................ [196160]
AY261365.1_Warmbaths           ......a................t........................................................ [196160]
AY261364.1_Tengani_62          ......a...a..a.........t........................................................ [196160]
AY261363.1_Pretoriuskop/96/4   ......a................t........................................................ [196160]
AY261362.1_Mkuzi_1979          ................................................................................ [196160]
AY261361.1_Malawi_Lil-20/1_1983 ......a........g.......t...........................c............................ [196160]
AY261360.1_Kenya_1950          g.....a.....................................................t.............agg.t. [196160]
AM712240.1_OURT_88/3           ................................................................................ [196160]
AM712239.1_Benin_97/1          ................................................................................ [196160]
KM111295.1_Ken06.Bus           g.....a.................t............................t......t.............a.g.t. [196160]
KM111294.1_Ken05/Tk1           g.....a.....................................................t.............agg.t. [196160]
KM262844.1_strain_L60          ................................................................................ [196160]
KM102979.1_26544/OG10_Italy    ................................................................................ [196160]
KM262845.1_strain_NHV          ................................................................................ [196160]
KX354450.1_47/Ss/2008          ................................................................................ [196160]
KP055815.1_strain_BA71         ................................................................................ [196160]
MH025920.1_strain_R35          g.....a.................t............................t......t.............a.g.t. [196160]
MH025919.1_strain_N10          g.....a.................t............................t......t.............a.g.t. [196160]
MH025918.1_strain_R25          g.....a.................t............................t......t.............a.g.t. [196160]
MH025917.1_strain_R7           g.....a.................t............................t......t.............a.g.t. [196160]
MH025916.1_strain_R8           g.....a.................t............................t......t.............a.g.t. [196160]
```

```
FR682468.2_Georgia_2007/1        tgggaagctgcttttcttggtgaaatttacgtaaatattcgatggccaccgcttggacgactgtgtaatttactaagtta [196240]
ASF_IND_20_CAD543                ................................................................................ [196240]
LR899193.1_Germany_2020/1        ................................................................................ [196240]
MN393477.1_Wuhan_2019            ................................................................................ [196240]
MW396979.1_Timor-Leste/2019/1    ................................................................................ [196240]
MK940252.1_CN/2019/InnerMongolia ................................................................................ [196240]
MK128995.1_China/2018/AnhuiXCGQ  ................................................................................ [196240]
MH681419.1_ASFV/POL/2015         ................................................................................ [196240]
LS478113.1_Estonia_2014          ................................................................................ [196240]
KP843857.1_Odintsovo_02/14       ................................................................................ [196240]
FN557520.1_strain_E75            .................c.........c...c................................................ [196240]
AY261366.1_Warthog               ...........c.................................................................... [196240]
AY261365.1_Warmbaths             ...........c.............................................t...................... [196240]
AY261364.1_Tengani_62            .......................................................c........................ [196240]
AY261363.1_Pretoriuskop/96/4     ...........c.................................................................... [196240]
AY261362.1_Mkuzi_1979            .................c.........c...c................................................ [196240]
AY261361.1_Malawi_Lil-20/1_1983  ...................a..........a.................gt.......a..............tg....c. [196240]
AY261360.1_Kenya_1950            ...........................c.......c............gt.......a..a.....g.c...t.....c. [196240]
AM712240.1_OURT_88/3             .................c.........c...c................................................ [196240]
AM712239.1_Benin_97/1            .................c.........c...c................................................ [196240]
KM111295.1_Ken06.Bus             ...........................c.......c...t........gt.......a..a.....g.c...t.....c. [196240]
KM111294.1_Ken05/Tk1             ...........................c.......c............gt.......a..a.....g.c...t.....c. [196240]
KM262844.1_strain_L60            .................c.........c...c................................................ [196240]
KM102979.1_26544/OG10_Italy      .................c.........c...c................................................ [196240]
KM262845.1_strain_NHV            .................c.........c...c................................................ [196240]
KX354450.1_47/Ss/2008            .................c.........c...c................................................ [196240]
KP055815.1_strain_BA71           .................c.........c...c................................................ [196240]
MH025920.1_strain_R35            ...........................c.......c...t........gt.......a..a.....g.c...t.....c. [196240]
MH025919.1_strain_N10            ...........................c.......c...t........gt.......a..a.....g.c...t.....c. [196240]
MH025918.1_strain_R25            ...........................c.......c...t........gt.......a..a.....g.c...t.....c. [196240]
MH025917.1_strain_R7             ...........................c.......c...t........gt.......a..a.....g.c...t.....c. [196240]
MH025916.1_strain_R8             ...........................c.......c...t........gt.......a..a.....g.c...t.....c. [196240]

FR682468.2_Georgia_2007/1        gaaatgatagctttcatggttgtaaaaatatacataggattttctttttctgtatacagtttgaaaagcttatgattacg [196320]
ASF_IND_20_CAD543                ................................................................................ [196320]
LR899193.1_Germany_2020/1        ................................................................................ [196320]
MN393477.1_Wuhan_2019            ................................................................................ [196320]
MW396979.1_Timor-Leste/2019/1    ................................................................................ [196320]
MK940252.1_CN/2019/InnerMongolia ................................................................................ [196320]
MK128995.1_China/2018/AnhuiXCGQ  ................................................................................ [196320]
MH681419.1_ASFV/POL/2015         ................................................................................ [196320]
LS478113.1_Estonia_2014          ................................................................................ [196320]
KP843857.1_Odintsovo_02/14       ................................................................................ [196320]
```

```
FN557520.1_strain_E75          .....a........t..............................................................g.. [196320]
AY261366.1_Warthog             ...........c..g.......................g..................................ag..g.. [196320]
AY261365.1_Warmbaths           ......................................g..................................ag..g.. [196320]
AY261364.1_Tengani_62          ......................................g..................................ag..g.. [196320]
AY261363.1_Pretoriuskop/96/4   ..............g..............g........g..................................ag..g.. [196320]
AY261362.1_Mkuzi_1979          ..............t..............................................................g.. [196320]
AY261361.1_Malawi_Lil-20/1_1983 ................a.a.....................g.........c.......................g..g.. [196320]
AY261360.1_Kenya_1950          ..........t........................g..........g.........t................c...g.. [196320]
AM712240.1_OURT_88/3           .....a........t..............................................................g.. [196320]
AM712239.1_Benin_97/1          .....a........t..............................................................g.. [196320]
KM111295.1_Ken06.Bus           ..........t...................................g.........t................c...g.. [196320]
KM111294.1_Ken05/Tk1           ..........t........................g..........g.........t................c...g.. [196320]
KM262844.1_strain_L60          .....a........t..............................................................g.. [196320]
KM102979.1_26544/OG10_Italy    .....a........t..............................................................g.. [196320]
KM262845.1_strain_NHV          .....a........t..............................................................g.. [196320]
KX354450.1_47/Ss/2008          .....a........t..............................................................g.. [196320]
KP055815.1_strain_BA71         .....a........t..............................................................g.. [196320]
MH025920.1_strain_R35          ..........t...................................g.........t................c...g.. [196320]
MH025919.1_strain_N10          ..........t...................................g.........t................c...g.. [196320]
MH025918.1_strain_R25          ..........t...................................g.........t................c...g.. [196320]
MH025917.1_strain_R7           ..........t...................................g.........t................c...g.. [196320]
MH025916.1_strain_R8           ..........t...................................g.........t................c...g.. [196320]

FR682468.2_Georgia_2007/1      tgaaatgatggccatttttaatacaagatggtatagtgtatctttaggtaaaaatgccttgcaagccgcgatgatgtcga [196400]
ASF_IND_20_CAD543              ................................................................................ [196400]
LR899193.1_Germany_2020/1      ................................................................................ [196400]
MN393477.1_Wuhan_2019          ................................................................................ [196400]
MW396979.1_Timor-Leste/2019/1  ................................................................................ [196400]
MK940252.1_CN/2019/InnerMongolia ................................................................................ [196400]
MK128995.1_China/2018/AnhuiXCGQ ................................................................................ [196400]
MH681419.1_ASFV/POL/2015       ................................................................................ [196400]
LS478113.1_Estonia_2014        ................................................................................ [196400]
KP843857.1_Odintsovo_02/14     ................................................................................ [196400]
FN557520.1_strain_E75          ................................................................................ [196400]
AY261366.1_Warthog             .....c...............c.......................................................t.. [196400]
AY261365.1_Warmbaths           .....c...............c.......................................................t.. [196400]
AY261364.1_Tengani_62          .....................c.......................................................t.. [196400]
AY261363.1_Pretoriuskop/96/4   .....c...............c.......................................................t.. [196400]
AY261362.1_Mkuzi_1979          .....c...............c.......................................................... [196400]
AY261361.1_Malawi_Lil-20/1_1983 .a....a.............................c........................................... [196400]
AY261360.1_Kenya_1950          .............................a.......t.........a.........t..........t...t....... [196400]
AM712240.1_OURT_88/3           ................................................................................ [196400]
AM712239.1_Benin_97/1          ................................................................................ [196400]
KM111295.1_Ken06.Bus           .............................a.......t.........a.........t..........t...t....... [196400]
```

```
KM111294.1_Ken05/Tk1             .............................a.......t.........a.........t..........t...t....... [196400]
KM262844.1_strain_L60            ................................................................................ [196400]
KM102979.1_26544/OG10_Italy      ................................................................................ [196400]
KM262845.1_strain_NHV            ................................................................................ [196400]
KX354450.1_47/Ss/2008            ................................................................................ [196400]
KP055815.1_strain_BA71           ................................................................................ [196400]
MH025920.1_strain_R35            .............................a.......t.........a.........t..........t...t....... [196400]
MH025919.1_strain_N10            .............................a.......t.........a.........t..........t...t....... [196400]
MH025918.1_strain_R25            .............................a.......t.........a.........t..........t...t....... [196400]
MH025917.1_strain_R7             .............................a.......t.........a.........t..........t...t....... [196400]
MH025916.1_strain_R8             .............................a.......t.........a.........t..........t...t....... [196400]

FR682468.2_Georgia_2007/1        tgttgtctccatgaacagcgatagaaactaatgtttccaatctaaatgtttttatctgcattaatagaagaatgcagtca [196480]
ASF_IND_20_CAD543                ....................................................................r........... [196480]
LR899193.1_Germany_2020/1        ................................................................................ [196480]
MN393477.1_Wuhan_2019            ................................................................................ [196480]
MW396979.1_Timor-Leste/2019/1    ................................................................................ [196480]
MK940252.1_CN/2019/InnerMongolia ................................................................................ [196480]
MK128995.1_China/2018/AnhuiXCGQ  ................................................................................ [196480]
MH681419.1_ASFV/POL/2015         ................................................................................ [196480]
LS478113.1_Estonia_2014          ................................................................................ [196480]
KP843857.1_Odintsovo_02/14       ................................................................................ [196480]
FN557520.1_strain_E75            ....................................................................g........... [196480]
AY261366.1_Warthog               ......t.........g......................t............................g........... [196480]
AY261365.1_Warmbaths             ................g......................t.t..........................g....a...... [196480]
AY261364.1_Tengani_62            ....c...........g..........g...........t............................g........a.. [196480]
AY261363.1_Pretoriuskop/96/4     ......t.........g..........t.........t.t.t..........................g........... [196480]
AY261362.1_Mkuzi_1979            .......................................t.t..........................g........... [196480]
AY261361.1_Malawi_Lil-20/1_1983  ..........g.....c..........t...c.......t...............a............g........... [196480]
AY261360.1_Kenya_1950            ......tgtt.....t...........g.........t.t...g..........aa...c....c.a.g..........t [196480]
AM712240.1_OURT_88/3             ....................................................................g........... [196480]
AM712239.1_Benin_97/1            ....................................................................g........... [196480]
KM111295.1_Ken06.Bus             ......tgtt.....t...........g.........t.t...g..........aa...c....c...g.........tt [196480]
KM111294.1_Ken05/Tk1             ......tgtt.....t...........g.........t.t...g..........aa...c....c.a.g..........c [196480]
KM262844.1_strain_L60            ....................................................................g........... [196480]
KM102979.1_26544/OG10_Italy      ....................................................................g........... [196480]
KM262845.1_strain_NHV            ....................................................................g........... [196480]
KX354450.1_47/Ss/2008            ....................................................................g........... [196480]
KP055815.1_strain_BA71           ....................................................................g........... [196480]
MH025920.1_strain_R35            ......tgtt.....t...........g.........t.t...g..........aa...c....c...g.........tt [196480]
MH025919.1_strain_N10            ......tgtt.....t...........g.........t.t...g..........aa...c....c...g.........tt [196480]
MH025918.1_strain_R25            ......tgtt.....t...........g.........t.t...g..........aa...c....c...g.........tt [196480]
MH025917.1_strain_R7             ......tgtt.....t...........g.........t.t...g..........aa...c....c...g.........tt [196480]
MH025916.1_strain_R8             ......tgtt.....t...........g.........t.t...g..........aa...c....c...g.........tt [196480]
```

```
FR682468.2_Georgia_2007/1        atgttattatacttaataatactgtaatacaccgaatcaatgaccgtcatctgagaatcaagctgacttattagtaaatt [196560]
ASF_IND_20_CAD543                ................................................................................ [196560]
LR899193.1_Germany_2020/1        ................................................................................ [196560]
MN393477.1_Wuhan_2019            ................................................................................ [196560]
MW396979.1_Timor-Leste/2019/1    ................................................................................ [196560]
MK940252.1_CN/2019/InnerMongolia ................................................................................ [196560]
MK128995.1_China/2018/AnhuiXCGQ  ................................................................................ [196560]
MH681419.1_ASFV/POL/2015         ................................................................................ [196560]
LS478113.1_Estonia_2014          ................................................................................ [196560]
KP843857.1_Odintsovo_02/14       ................................................................................ [196560]
FN557520.1_strain_E75            ................................................................................ [196560]
AY261366.1_Warthog               ...........t............................c...a................................... [196560]
AY261365.1_Warmbaths             ...........t................................a................................... [196560]
AY261364.1_Tengani_62            .c.........t.............................a..a................................... [196560]
AY261363.1_Pretoriuskop/96/4     ...........t....................t.......c...a................................... [196560]
AY261362.1_Mkuzi_1979            ................................................................................ [196560]
AY261361.1_Malawi_Lil-20/1_1983  ...........t................................a..............g...c..t....g..c..... [196560]
AY261360.1_Kenya_1950            ...........t....c...........................g..........c.......c..t.......c..... [196560]
AM712240.1_OURT_88/3             ................................................................................ [196560]
AM712239.1_Benin_97/1            ................................................................................ [196560]
KM111295.1_Ken06.Bus             ...........t....c...........................g..........c.......c..t.......c..... [196560]
KM111294.1_Ken05/Tk1             ...........t....c...........................g..........c.......c..t.......c..... [196560]
KM262844.1_strain_L60            ................................................................................ [196560]
KM102979.1_26544/OG10_Italy      ................................................................................ [196560]
KM262845.1_strain_NHV            ................................................................................ [196560]
KX354450.1_47/Ss/2008            ................................................................................ [196560]
KP055815.1_strain_BA71           ................................................................................ [196560]
MH025920.1_strain_R35            ...........t....c...........................g..........c.......c..t.......c..... [196560]
MH025919.1_strain_N10            ...........t....c...........................g..........c.......c..t.......c..... [196560]
MH025918.1_strain_R25            ...........t....c...........................g..........c.......c..t.......c..... [196560]
MH025917.1_strain_R7             ...........t....c...........................g..........c.......c..t.......c..... [196560]
MH025916.1_strain_R8             ...........t....c...........................g..........c.......c..t.......c..... [196560]

FR682468.2_Georgia_2007/1        taacgtttttttggaggcatgacctttgatcgcggcactaagtgcacacagtatagcaaaattgttaaatacattttgat [196640]
ASF_IND_20_CAD543                ................................................................................ [196640]
LR899193.1_Germany_2020/1        ................................................................................ [196640]
MN393477.1_Wuhan_2019            ................................................................................ [196640]
MW396979.1_Timor-Leste/2019/1    ................................................................................ [196640]
MK940252.1_CN/2019/InnerMongolia ................................................................................ [196640]
MK128995.1_China/2018/AnhuiXCGQ  ................................................................................ [196640]
MH681419.1_ASFV/POL/2015         ................................................................................ [196640]
LS478113.1_Estonia_2014          ................................................................................ [196640]
KP843857.1_Odintsovo_02/14       ................................................................................ [196640]
```

```
FN557520.1_strain_E75          ..............................t.........................................g....... [196640]
AY261366.1_Warthog             ..............................t.....g.........................................t......t. [196640]
AY261365.1_Warmbaths           .......................................................................t........ [196640]
AY261364.1_Tengani_62          ..............................t.........................................t........ [196640]
AY261363.1_Pretoriuskop/96/4   ............a.................t.....g..g..............................t........ [196640]
AY261362.1_Mkuzi_1979          ..............................t................................................. [196640]
AY261361.1_Malawi_Lil-20/1_1983 ............a..............a..t........t................t..............t........ [196640]
AY261360.1_Kenya_1950          ...........................a..g...........a..g..........t..............t......g. [196640]
AM712240.1_OURT_88/3           ..............................t.........................................g....... [196640]
AM712239.1_Benin_97/1          ..............................t.........................................g....... [196640]
KM111295.1_Ken06.Bus           ...........................a..g...........a..g..t.......t..............t......g. [196640]
KM111294.1_Ken05/Tk1           ...........................a..g...........a..g..........t..............t......g. [196640]
KM262844.1_strain_L60          ..............................t.........................................g....... [196640]
KM102979.1_26544/OG10_Italy    ..............................t.........................................g....... [196640]
KM262845.1_strain_NHV          ..............................t.........................................g....... [196640]
KX354450.1_47/Ss/2008          ..............................t.........................................g....... [196640]
KP055815.1_strain_BA71         ..............................t.........................................g....... [196640]
MH025920.1_strain_R35          ...........................a..g...........a..g..t.......t..............t......g. [196640]
MH025919.1_strain_N10          ...........................a..g...........a..g..t.......t..............t......g. [196640]
MH025918.1_strain_R25          ...........................a..g...........a..g..t.......t..............t......g. [196640]
MH025917.1_strain_R7           ...........................a..g...........a..g..t.......t..............t......g. [196640]
MH025916.1_strain_R8           ...........................a..g...........a..g..t.......t..............t......g. [196640]

FR682468.2_Georgia_2007/1      ttaggagaaggagtaatattttccttcggttatagtacgcagcatctgtgatgattattggccgataaatgttaaaatgt [196720]
ASF_IND_20_CAD543              ...........................................................k.................... [196720]
LR899193.1_Germany_2020/1      ................................................................................ [196720]
MN393477.1_Wuhan_2019          ................................................................................ [196720]
MW396979.1_Timor-Leste/2019/1  ................................................................................ [196720]
MK940252.1_CN/2019/InnerMongolia ................................................................................ [196720]
MK128995.1_China/2018/AnhuiXCGQ ................................................................................ [196720]
MH681419.1_ASFV/POL/2015       ................................................................................ [196720]
LS478113.1_Estonia_2014        ................................................................................ [196720]
KP843857.1_Odintsovo_02/14     ................................................................................ [196720]
FN557520.1_strain_E75          ...........................................................t...................c [196720]
AY261366.1_Warthog             ....a......................................................t...................c [196720]
AY261365.1_Warmbaths           ...........................................................t...................c [196720]
AY261364.1_Tengani_62          ...........................................................t.........g.......... [196720]
AY261363.1_Pretoriuskop/96/4   ....a......................................................t.................... [196720]
AY261362.1_Mkuzi_1979          ...........................................................t...................c [196720]
AY261361.1_Malawi_Lil-20/1_1983 .c....................t.......c..................a...........a.........c........ [196720]
AY261360.1_Kenya_1950          .a.a......c...........t................................g.....a.......c.........g [196720]
AM712240.1_OURT_88/3           ...........................................................t.................c.c [196720]
AM712239.1_Benin_97/1          ...........................................................t...................c [196720]
KM111295.1_Ken06.Bus           .a.a......c............................................g.....a.......c.........g [196720]
```

```
KM111294.1_Ken05/Tk1             .a.a......c............................................g.....a.......c.........g [196720]
KM262844.1_strain_L60            ...........................................................t...................c [196720]
KM102979.1_26544/OG10_Italy      ...........................................................t...................c [196720]
KM262845.1_strain_NHV            ...........................................................t.................c.c [196720]
KX354450.1_47/Ss/2008            ...........................................................t...................c [196720]
KP055815.1_strain_BA71           ...........................................................t...................c [196720]
MH025920.1_strain_R35            .a.a......c............................................g.....a.......c.........g [196720]
MH025919.1_strain_N10            .a.a......c............................................g.....a.......c.........g [196720]
MH025918.1_strain_R25            .a.a......c............................................g.....a.......c.........g [196720]
MH025917.1_strain_R7             .a.a......c............................................g.....a.......c.........g [196720]
MH025916.1_strain_R8             .a.a......c............................................g.....a.......c.........g [196720]

FR682468.2_Georgia_2007/1        gttaacagctttttaaaaaaacggaagtaatttttttggatcgctgtttgcatcatcgaaataatgagataatcagggta [196800]
ASF_IND_20_CAD543                ................................................................................ [196800]
LR899193.1_Germany_2020/1        ................................................................................ [196800]
MN393477.1_Wuhan_2019            ................................................................................ [196800]
MW396979.1_Timor-Leste/2019/1    ................................................................................ [196800]
MK940252.1_CN/2019/InnerMongolia ................................................................................ [196800]
MK128995.1_China/2018/AnhuiXCGQ  ................................................................................ [196800]
MH681419.1_ASFV/POL/2015         ................................................................................ [196800]
LS478113.1_Estonia_2014          ................................................................................ [196800]
KP843857.1_Odintsovo_02/14       ................................................................................ [196800]
FN557520.1_strain_E75            ................................................................................ [196800]
AY261366.1_Warthog               ..........................................................................g..... [196800]
AY261365.1_Warmbaths             ........................................................t....................... [196800]
AY261364.1_Tengani_62            ................................................................................ [196800]
AY261363.1_Pretoriuskop/96/4     ..........................................................................g..... [196800]
AY261362.1_Mkuzi_1979            ........................................................t....................... [196800]
AY261361.1_Malawi_Lil-20/1_1983  ......................................t.....a.............................g..... [196800]
AY261360.1_Kenya_1950            .............................g........t...............................gg........ [196800]
AM712240.1_OURT_88/3             ................................................................................ [196800]
AM712239.1_Benin_97/1            ................................................................................ [196800]
KM111295.1_Ken06.Bus             .............................g........t...............................gg........ [196800]
KM111294.1_Ken05/Tk1             .............................g........t...............................gg........ [196800]
KM262844.1_strain_L60            ................................................................................ [196800]
KM102979.1_26544/OG10_Italy      ................................................................................ [196800]
KM262845.1_strain_NHV            ................................................................................ [196800]
KX354450.1_47/Ss/2008            ................................................................................ [196800]
KP055815.1_strain_BA71           ................................................................................ [196800]
MH025920.1_strain_R35            .............................g........t...............................gg........ [196800]
MH025919.1_strain_N10            .............................g........t...............................gg........ [196800]
MH025918.1_strain_R25            .............................g........t...............................gg........ [196800]
MH025917.1_strain_R7             .............................g........t...............................gg........ [196800]
MH025916.1_strain_R8             .............................g........t...............................gg........ [196800]
```

```
FR682468.2_Georgia_2007/1        tataatgggtaggtcacatgctacctctaacaaagaatagtcgcccaatctaaaggctgtgttgaaaagcgtactatcat [196880]
ASF_IND_20_CAD543                ................................................................................ [196880]
LR899193.1_Germany_2020/1        ................................................................................ [196880]
MN393477.1_Wuhan_2019            ................................................................................ [196880]
MW396979.1_Timor-Leste/2019/1    ................................................................................ [196880]
MK940252.1_CN/2019/InnerMongolia ................................................................................ [196880]
MK128995.1_China/2018/AnhuiXCGQ  ................................................................................ [196880]
MH681419.1_ASFV/POL/2015         ................................................................................ [196880]
LS478113.1_Estonia_2014          ................................................................................ [196880]
KP843857.1_Odintsovo_02/14       ................................................................................ [196880]
FN557520.1_strain_E75            ................................................................................ [196880]
AY261366.1_Warthog               .................................................t....................a......... [196880]
AY261365.1_Warmbaths             ................................................................................ [196880]
AY261364.1_Tengani_62            ................................................................................ [196880]
AY261363.1_Pretoriuskop/96/4     .................................................t.............................. [196880]
AY261362.1_Mkuzi_1979            ................................................................................ [196880]
AY261361.1_Malawi_Lil-20/1_1983  cg...............................................t.............................. [196880]
AY261360.1_Kenya_1950            cg.....................................a.......................a................ [196880]
AM712240.1_OURT_88/3             ..............................g................................................. [196880]
AM712239.1_Benin_97/1            ................................................................................ [196880]
KM111295.1_Ken06.Bus             cg.....................................a..a....................a................ [196880]
KM111294.1_Ken05/Tk1             cg......................t..............a.......................a................ [196880]
KM262844.1_strain_L60            ................................................................................ [196880]
KM102979.1_26544/OG10_Italy      ................................................................................ [196880]
KM262845.1_strain_NHV            ..............................g................................................. [196880]
KX354450.1_47/Ss/2008            ................................................................................ [196880]
KP055815.1_strain_BA71           ................................................................................ [196880]
MH025920.1_strain_R35            cg.....................................a..a....................a................ [196880]
MH025919.1_strain_N10            cg.....................................a..a....................a................ [196880]
MH025918.1_strain_R25            cg.....................................a..a....................a................ [196880]
MH025917.1_strain_R7             cg.....................................a..a....................a................ [196880]
MH025916.1_strain_R8             cg.....................................a..a....................a................ [196880]

FR682468.2_Georgia_2007/1        catacgtatcgagtacccctgctgttacaaaccaagcgataagatgaatgtgccgttccttgcaagctatcgcaaatagg [196960]
ASF_IND_20_CAD543                ................................................................................ [196960]
LR899193.1_Germany_2020/1        ................................................................................ [196960]
MN393477.1_Wuhan_2019            ................................................................................ [196960]
MW396979.1_Timor-Leste/2019/1    ................................................................................ [196960]
MK940252.1_CN/2019/InnerMongolia ................................................................................ [196960]
MK128995.1_China/2018/AnhuiXCGQ  ................................................................................ [196960]
MH681419.1_ASFV/POL/2015         ................................................................................ [196960]
LS478113.1_Estonia_2014          ................................................................................ [196960]
KP843857.1_Odintsovo_02/14       ................................................................................ [196960]
```

```
FN557520.1_strain_E75            ...................c............................................................ [196960]
AY261366.1_Warthog               ...................c............................................................ [196960]
AY261365.1_Warmbaths             ...................c............................................................ [196960]
AY261364.1_Tengani_62            ...................c............................................................ [196960]
AY261363.1_Pretoriuskop/96/4     ...................c............................................................ [196960]
AY261362.1_Mkuzi_1979            ...................c............................................................ [196960]
AY261361.1_Malawi_Lil-20/1_1983  ....t...........t..c.........................t.................................. [196960]
AY261360.1_Kenya_1950            ............ac..t.........g..........a..g......................................a [196960]
AM712240.1_OURT_88/3             ...................c............................................................ [196960]
AM712239.1_Benin_97/1            ...................c............................................................ [196960]
KM111295.1_Ken06.Bus             ............ac..t.........g..........a..g....................................... [196960]
KM111294.1_Ken05/Tk1             ............ac..t.........g..........a..g......................................a [196960]
KM262844.1_strain_L60            ...................c............................................................ [196960]
KM102979.1_26544/OG10_Italy      ...................c............................................................ [196960]
KM262845.1_strain_NHV            ...................c............................................................ [196960]
KX354450.1_47/Ss/2008            ...................c............................................................ [196960]
KP055815.1_strain_BA71           ...................c............................................................ [196960]
MH025920.1_strain_R35            ............ac..t.........g..........a..g....................................... [196960]
MH025919.1_strain_N10            ............ac..t.........g..........a..g....................................... [196960]
MH025918.1_strain_R25            ............ac..t.........g..........a..g....................................... [196960]
MH025917.1_strain_R7             ............ac..t.........g..........a..g....................................... [196960]
MH025916.1_strain_R8             ............ac..t.........g..........a..g....................................... [196960]

FR682468.2_Georgia_2007/1        gagtttcctatggaatgtcgaataatgtactccctatttttttccaaaatgtttggaaaattgtatagcgttgcggcata [197040]
ASF_IND_20_CAD543                ................................................................................ [197040]
LR899193.1_Germany_2020/1        ................................................................................ [197040]
MN393477.1_Wuhan_2019            ................................................................................ [197040]
MW396979.1_Timor-Leste/2019/1    ................................................................................ [197040]
MK940252.1_CN/2019/InnerMongolia ................................................................................ [197040]
MK128995.1_China/2018/AnhuiXCGQ  ................................................................................ [197040]
MH681419.1_ASFV/POL/2015         ................................................................................ [197040]
LS478113.1_Estonia_2014          ................................................................................ [197040]
KP843857.1_Odintsovo_02/14       ................................................................................ [197040]
FN557520.1_strain_E75            .t.............................................................................. [197040]
AY261366.1_Warthog               ................a......................................................c..a..... [197040]
AY261365.1_Warmbaths             .......................................................................c..a..... [197040]
AY261364.1_Tengani_62            .g........................................................................a..... [197040]
AY261363.1_Pretoriuskop/96/4     .......................................................................c..a..... [197040]
AY261362.1_Mkuzi_1979            ................................................................................ [197040]
AY261361.1_Malawi_Lil-20/1_1983  .ga...................................................t...................c..... [197040]
AY261360.1_Kenya_1950            .ca...................c.....................t.........t...................c..... [197040]
AM712240.1_OURT_88/3             .t.............................................................................. [197040]
AM712239.1_Benin_97/1            .t.............................................................................. [197040]
KM111295.1_Ken06.Bus             .ca...................c...............................t...................c..... [197040]
```

```
KM111294.1_Ken05/Tk1            .ca...................c...a.................t.........t...................c..... [197040]
KM262844.1_strain_L60           .t.............................................................................. [197040]
KM102979.1_26544/OG10_Italy     .t.............................................................................. [197040]
KM262845.1_strain_NHV           .t.............................................................................. [197040]
KX354450.1_47/Ss/2008           .t.............................................................................. [197040]
KP055815.1_strain_BA71          .t.............................................................................. [197040]
MH025920.1_strain_R35           .ca...................c...............................t...................c..... [197040]
MH025919.1_strain_N10           .ca...................c...............................t...................c..... [197040]
MH025918.1_strain_R25           .ca...................c...............................t...................c..... [197040]
MH025917.1_strain_R7            .ca...................c...............................t...................c..... [197040]
MH025916.1_strain_R8            .ca...................c...............................t...................c..... [197040]

FR682468.2_Georgia_2007/1       cagtagacactccattctggcgttataatttttacttttacatatgaataggtggaagaactcgaataattcttgagaac [197120]
ASF_IND_20_CAD543               ................................................................................ [197120]
LR899193.1_Germany_2020/1       ................................................................................ [197120]
MN393477.1_Wuhan_2019           ................................................................................ [197120]
MW396979.1_Timor-Leste/2019/1   ................................................................................ [197120]
MK940252.1_CN/2019/InnerMongolia ................................................................................ [197120]
MK128995.1_China/2018/AnhuiXCGQ ................................................................................ [197120]
MH681419.1_ASFV/POL/2015        ................................................................................ [197120]
LS478113.1_Estonia_2014         ................................................................................ [197120]
KP843857.1_Odintsovo_02/14      ................................................................................ [197120]
FN557520.1_strain_E75           ................................................................................ [197120]
AY261366.1_Warthog              ................................................................................ [197120]
AY261365.1_Warmbaths            ................................................................................ [197120]
AY261364.1_Tengani_62           .......................g........................................................ [197120]
AY261363.1_Pretoriuskop/96/4    ................................................................................ [197120]
AY261362.1_Mkuzi_1979           ................................................................................ [197120]
AY261361.1_Malawi_Lil-20/1_1983 t......................gg....................................................... [197120]
AY261360.1_Kenya_1950           .......................g...............g........................................ [197120]
AM712240.1_OURT_88/3            .....................................c.......................................... [197120]
AM712239.1_Benin_97/1           ................................................................................ [197120]
KM111295.1_Ken06.Bus            .......................g...............g...........a............................ [197120]
KM111294.1_Ken05/Tk1            .......................g...............g........................................ [197120]
KM262844.1_strain_L60           ................................................................................ [197120]
KM102979.1_26544/OG10_Italy     ................................................................................ [197120]
KM262845.1_strain_NHV           .....................................c.......................................... [197120]
KX354450.1_47/Ss/2008           ................................................................................ [197120]
KP055815.1_strain_BA71          ................................................................................ [197120]
MH025920.1_strain_R35           .......................g...............g...........a............................ [197120]
MH025919.1_strain_N10           .......................g...............g...........a............................ [197120]
MH025918.1_strain_R25           .......................g...............g...........a............................ [197120]
MH025917.1_strain_R7            .......................g...............g...........a............................ [197120]
MH025916.1_strain_R8            .......................g...............g...........a............................ [197120]
```

```
FR682468.2_Georgia_2007/1        ttgttaaatgcataatatggtgatattttggtgtcgttaaatggtatgagaaaatgcattctaatacatcttttcggtta [197200]
ASF_IND_20_CAD543                ................................................................................ [197200]
LR899193.1_Germany_2020/1        ................................................................................ [197200]
MN393477.1_Wuhan_2019            ................................................................................ [197200]
MW396979.1_Timor-Leste/2019/1    ................................................................................ [197200]
MK940252.1_CN/2019/InnerMongolia ................................................................................ [197200]
MK128995.1_China/2018/AnhuiXCGQ  ................................................................................ [197200]
MH681419.1_ASFV/POL/2015         ................................................................................ [197200]
LS478113.1_Estonia_2014          ................................................................................ [197200]
KP843857.1_Odintsovo_02/14       ................................................................................ [197200]
FN557520.1_strain_E75            ...........................................a.................................... [197200]
AY261366.1_Warthog               ..................................a........a.................................... [197200]
AY261365.1_Warmbaths             ..................................a........a.................................... [197200]
AY261364.1_Tengani_62            ..................................a........a.................................... [197200]
AY261363.1_Pretoriuskop/96/4     ..................................t........a.................................... [197200]
AY261362.1_Mkuzi_1979            ................................................................................ [197200]
AY261361.1_Malawi_Lil-20/1_1983  ......................g...........g........a.................c..c........g..a... [197200]
AY261360.1_Kenya_1950            ................g........gc.......a........a.g.................t................ [197200]
AM712240.1_OURT_88/3             ...........................................a.................................... [197200]
AM712239.1_Benin_97/1            ...........................................a.................................... [197200]
KM111295.1_Ken06.Bus             ................g........gc.......a........a.g.................t................ [197200]
KM111294.1_Ken05/Tk1             ................g........gc.....c.a........a.g.................t................ [197200]
KM262844.1_strain_L60            ...........................................a.................................... [197200]
KM102979.1_26544/OG10_Italy      ...........................................a.................................... [197200]
KM262845.1_strain_NHV            ...........................................a.................................... [197200]
KX354450.1_47/Ss/2008            ...........................................a.................................... [197200]
KP055815.1_strain_BA71           ...........................................a.................................... [197200]
MH025920.1_strain_R35            ................g........gc.......a........a.g.................t................ [197200]
MH025919.1_strain_N10            ................g........gc.......a........a.g.................t................ [197200]
MH025918.1_strain_R25            ................g........gc.......a........a.g.................t................ [197200]
MH025917.1_strain_R7             ................g........gc.......a........a.g.................t................ [197200]
MH025916.1_strain_R8             ................g........gc.......a........a.g.................t................ [197200]

FR682468.2_Georgia_2007/1        tgctttagcgcctgagctaaggcatattcaggctcgacccataggactagtgtttctataattgagatattcgcctgctt [197280]
ASF_IND_20_CAD543                .......................................................y........................ [197280]
LR899193.1_Germany_2020/1        ................................................................................ [197280]
MN393477.1_Wuhan_2019            ................................................................................ [197280]
MW396979.1_Timor-Leste/2019/1    ................................................................................ [197280]
MK940252.1_CN/2019/InnerMongolia ................................................................................ [197280]
MK128995.1_China/2018/AnhuiXCGQ  ................................................................................ [197280]
MH681419.1_ASFV/POL/2015         ................................................................................ [197280]
LS478113.1_Estonia_2014          ................................................................................ [197280]
KP843857.1_Odintsovo_02/14       ................................................................................ [197280]
```

```
FN557520.1_strain_E75              .......................................................t........t............... [197280]
AY261366.1_Warthog                 ...............................................g.......t........t............... [197280]
AY261365.1_Warmbaths               .......................................................t........t............... [197280]
AY261364.1_Tengani_62              .......................................................t........t............... [197280]
AY261363.1_Pretoriuskop/96/4       .......................................................t........t............... [197280]
AY261362.1_Mkuzi_1979              ........t..............................................t........................ [197280]
AY261361.1_Malawi_Lil-20/1_1983    ..t.....t..........................a........t.................g.t......t........ [197280]
AY261360.1_Kenya_1950              ..t.....t..........................a........t.............g....tt......t........ [197280]
AM712240.1_OURT_88/3               .......................................................t........t............... [197280]
AM712239.1_Benin_97/1              .......................................................t........t............... [197280]
KM111295.1_Ken06.Bus               ..t.....t..........................a........t.............g....ct......t........ [197280]
KM111294.1_Ken05/Tk1               ..t.....t..........................a........t.............g....tt............... [197280]
KM262844.1_strain_L60              .......................................................t........t............... [197280]
KM102979.1_26544/OG10_Italy        .......................................................t........t............... [197280]
KM262845.1_strain_NHV              .......................................................t........t............... [197280]
KX354450.1_47/Ss/2008              .......................................................t........t............... [197280]
KP055815.1_strain_BA71             .......................................................t........t............... [197280]
MH025920.1_strain_R35              ..t.....t..........................a........t.............g....ct......t........ [197280]
MH025919.1_strain_N10              ..t.....t..........................a........t.............g....ct......t........ [197280]
MH025918.1_strain_R25              ..t.....t..........................a........t.............g....ct......t........ [197280]
MH025917.1_strain_R7               ..t.....t..........................a........t.............g....ct......t........ [197280]
MH025916.1_strain_R8               ..t.....t..........................a........t.............g....ct......t........ [197280]

FR682468.2_Georgia_2007/1          tgccagggcatactttaagacgctccggttagaaaaaatgttgttatgaagatggataaccgtatccatttttacgatgg [197360]
ASF_IND_20_CAD543                  ................................................................................ [197360]
LR899193.1_Germany_2020/1          ................................................................................ [197360]
MN393477.1_Wuhan_2019              ................................................................................ [197360]
MW396979.1_Timor-Leste/2019/1      ................................................................................ [197360]
MK940252.1_CN/2019/InnerMongolia   ................................................................................ [197360]
MK128995.1_China/2018/AnhuiXCGQ    ................................................................................ [197360]
MH681419.1_ASFV/POL/2015           ................................................................................ [197360]
LS478113.1_Estonia_2014            ................................................................................ [197360]
KP843857.1_Odintsovo_02/14         ................................................................................ [197360]
FN557520.1_strain_E75              ..............................g........t................c...a..g..........t..c.. [197360]
AY261366.1_Warthog                 ..............................g........t................c...a..g..........t..... [197360]
AY261365.1_Warmbaths               ..............................g........t................c...a..g..........t..... [197360]
AY261364.1_Tengani_62              ..............................g........t........................................ [197360]
AY261363.1_Pretoriuskop/96/4       ..............................g........t.............a..c...a..g..........t..... [197360]
AY261362.1_Mkuzi_1979              ..............................g........t................c...a..g..........t..c.. [197360]
AY261361.1_Malawi_Lil-20/1_1983    ......t..............a........g........t................cg..a..g..........t..... [197360]
AY261360.1_Kenya_1950              c.....t.....t.....a..a...t....g.....c..t................cg..a..g..........t..... [197360]
AM712240.1_OURT_88/3               ..............................g........t................c...a..g..........t..c.. [197360]
AM712239.1_Benin_97/1              ..............................g........t................c...a..g..........t..c.. [197360]
KM111295.1_Ken06.Bus               .....at...........a..a...t....g.....c..t................cg..a..g..........t..... [197360]
```

```
KM111294.1_Ken05/Tk1            c.....t.....t.....a..a...t....g.....c..t................cg..a..g..........t..... [197360]
KM262844.1_strain_L60           ..............................g........t................c...a..g..........t..c.. [197360]
KM102979.1_26544/OG10_Italy     ..............................g........t................c...a..g..........t..c.. [197360]
KM262845.1_strain_NHV           ..............................g........t................c...a..g..........t..c.. [197360]
KX354450.1_47/Ss/2008           ..............................g........t................c...a..g..........t..c.. [197360]
KP055815.1_strain_BA71          ..............................g........t................c...a..g..........t..c.. [197360]
MH025920.1_strain_R35           .....at...........a..a...t....g.....c..t................cg..a..g..........t..... [197360]
MH025919.1_strain_N10           .....at...........a..a...t....g.....c..t................cg..a..g..........t..... [197360]
MH025918.1_strain_R25           .....at...........a..a...t....g.....c..t................cg..a..g..........t..... [197360]
MH025917.1_strain_R7            .....at...........a..a...t....g.....c..t................cg..a..g..........t..... [197360]
MH025916.1_strain_R8            .....at...........a..a...t....g.....c..t................cg..a..g..........t..... [197360]

FR682468.2_Georgia_2007/1       gaccattccagtatagtcctaaatgctgtagcagatcttttgttagttgtgaagcgttctcgggtgtcatataaatatgt [197440]
ASF_IND_20_CAD543               ................................................................................ [197440]
LR899193.1_Germany_2020/1       ................................................................................ [197440]
MN393477.1_Wuhan_2019           ................................................................................ [197440]
MW396979.1_Timor-Leste/2019/1   ................................................................................ [197440]
MK940252.1_CN/2019/InnerMongolia ................................................................................ [197440]
MK128995.1_China/2018/AnhuiXCGQ ................................................................................ [197440]
MH681419.1_ASFV/POL/2015        ................................................................................ [197440]
LS478113.1_Estonia_2014         ................................................................................ [197440]
KP843857.1_Odintsovo_02/14      ................................................................................ [197440]
FN557520.1_strain_E75           ................................................................a............... [197440]
AY261366.1_Warthog              ................................................................a............... [197440]
AY261365.1_Warmbaths            ................................................................a............... [197440]
AY261364.1_Tengani_62           .t..............................................................a............... [197440]
AY261363.1_Pretoriuskop/96/4    ................................................c...g..a........a............... [197440]
AY261362.1_Mkuzi_1979           ................................................c...g..a........a............... [197440]
AY261361.1_Malawi_Lil-20/1_1983 ............................c....a........a.....c...g...........a............... [197440]
AY261360.1_Kenya_1950           .....................................a........g.c...g....c......a............... [197440]
AM712240.1_OURT_88/3            ................................................................a............... [197440]
AM712239.1_Benin_97/1           ................................................................a............... [197440]
KM111295.1_Ken06.Bus            .....................................a........g.c...g....c......a............... [197440]
KM111294.1_Ken05/Tk1            .....................................a........g.c...g....c......a............... [197440]
KM262844.1_strain_L60           ................................................................a............... [197440]
KM102979.1_26544/OG10_Italy     ................................................................a............... [197440]
KM262845.1_strain_NHV           ................................................................a............... [197440]
KX354450.1_47/Ss/2008           ................................................................a............... [197440]
KP055815.1_strain_BA71          ................................................................a............... [197440]
MH025920.1_strain_R35           .....................................a........g.c...g....c......a............... [197440]
MH025919.1_strain_N10           .....................................a........g.c...g....c......a............... [197440]
MH025918.1_strain_R25           .....................................a........g.c...g....c......a............... [197440]
MH025917.1_strain_R7            .....................................a........g.c...g....c......a............... [197440]
MH025916.1_strain_R8            .....................................a........g.c...g....c......a............... [197440]
```

```
FR682468.2_Georgia_2007/1        tgcagggcttttttctgtaaggagaacatttcgtcgtaatcgtacaaaaaaaaattaaaatttgggcatgga--tgattc [197520]
ASF_IND_20_CAD543                ........................................................................--...... [197520]
LR899193.1_Germany_2020/1        ........................................................................--...... [197520]
MN393477.1_Wuhan_2019            ........................................................................--...... [197520]
MW396979.1_Timor-Leste/2019/1    ........................................................................--...... [197520]
MK940252.1_CN/2019/InnerMongolia ........................................................................--...... [197520]
MK128995.1_China/2018/AnhuiXCGQ  ........................................................................--...... [197520]
MH681419.1_ASFV/POL/2015         ........................................................................--...... [197520]
LS478113.1_Estonia_2014          ........................................................................--...... [197520]
KP843857.1_Odintsovo_02/14       .....................................................-..................--...... [197520]
FN557520.1_strain_E75            ........................................................................--...... [197520]
AY261366.1_Warthog               ........................................................................--...... [197520]
AY261365.1_Warmbaths             ........................................................................--...... [197520]
AY261364.1_Tengani_62            .....a.......g....................................................t..a..--...... [197520]
AY261363.1_Pretoriuskop/96/4     .....................................................................a..--...... [197520]
AY261362.1_Mkuzi_1979            .....a..................................................................--...... [197520]
AY261361.1_Malawi_Lil-20/1_1983  .....a.........................t.c.......a..............................--.....t [197520]
AY261360.1_Kenya_1950            ....ac...........................c.......a.....................aa.g..a..ta.a.... [197520]
AM712240.1_OURT_88/3             ........................................................................--...... [197520]
AM712239.1_Benin_97/1            ........................................................................--...... [197520]
KM111295.1_Ken06.Bus             ....ac...........................c.......a..t..................aa.g..a..ta.a.... [197520]
KM111294.1_Ken05/Tk1             ....ac...........................c.......a.....................aa.g..a..ta.a.... [197520]
KM262844.1_strain_L60            ........................................................................--...... [197520]
KM102979.1_26544/OG10_Italy      ........................................................................--...... [197520]
KM262845.1_strain_NHV            .............................................-..........................--...... [197520]
KX354450.1_47/Ss/2008            ........................................................................--...... [197520]
KP055815.1_strain_BA71           ........................................................................--...... [197520]
MH025920.1_strain_R35            ....ac...........................c.......a..t..................aa.g..a..ta.a.... [197520]
MH025919.1_strain_N10            ....ac...........................c.......a..t..................aa.g..a..ta.a.... [197520]
MH025918.1_strain_R25            ....ac...........................c.......a..t..................aa.g..a..ta.a.... [197520]
MH025917.1_strain_R7             ....ac...........................c.......a..t..................aa.g..a..ta.a.... [197520]
MH025916.1_strain_R8             ....ac...........................c.......a..t..................aa.g..a..ta.a.... [197520]

FR682468.2_Georgia_2007/1        aaacataac-----------------------------------aaaatcaagatttataacag-----tttgcat--- [197600]
ASF_IND_20_CAD543                .........-----------------------------------.......r............-----.......--- [197600]
LR899193.1_Germany_2020/1        .........-----------------------------------....................-----.......--- [197600]
MN393477.1_Wuhan_2019            .........-----------------------------------....................-----.......--- [197600]
MW396979.1_Timor-Leste/2019/1    .........-----------------------------------....................-----.......--- [197600]
MK940252.1_CN/2019/InnerMongolia .........-----------------------------------....................-----.......--- [197600]
MK128995.1_China/2018/AnhuiXCGQ  .........-----------------------------------....................-----.......--- [197600]
MH681419.1_ASFV/POL/2015         .........-----------------------------------....................-----.......--- [197600]
LS478113.1_Estonia_2014          .........-----------------------------------....................-----.......--- [197600]
KP843857.1_Odintsovo_02/14       .........-----------------------------------....................-----.......--- [197600]
```

```
FN557520.1_strain_E75            .........-----------------------------------.......g.............-----g......--- [197600]
AY261366.1_Warthog               .........-----------------------------------.......g.............-----g......--- [197600]
AY261365.1_Warmbaths             .........-----------------------------------.......g........c....-----g......--- [197600]
AY261364.1_Tengani_62            .........a----------------------------------.....................-----.......--- [197600]
AY261363.1_Pretoriuskop/96/4     .........-----------------------------------..................t..-----.......--- [197600]
AY261362.1_Mkuzi_1979            .........-----------------------------------.......g.............-----g......--- [197600]
AY261361.1_Malawi_Lil-20/1_1983  .........-----------------------------------...........g.........gatat.......--- [197600]
AY261360.1_Kenya_1950            ..g....g.aattgttaatgttaataacaaacattaataaacat...........-......t..gaca-......g--- [197600]
AM712240.1_OURT_88/3             .........-----------------------------------.......g.............-----g......--- [197600]
AM712239.1_Benin_97/1            .........-----------------------------------.......g.............-----g......--- [197600]
KM111295.1_Ken06.Bus             ..g......aactgttaatatttgt------tattaacaaacat...........-......t..gaca-......g--- [197600]
KM111294.1_Ken05/Tk1             ..g....g.aattgttaatgttaat----------aacaaacat...........-......t..gaca-......gtaa [197600]
KM262844.1_strain_L60            .........-----------------------------------.......g.............-----g......--- [197600]
KM102979.1_26544/OG10_Italy      .........-----------------------------------.......g.............-----g......--- [197600]
KM262845.1_strain_NHV            .........-----------------------------------.......g.............-----g......--- [197600]
KX354450.1_47/Ss/2008            .........-----------------------------------.......g.............-----g......--- [197600]
KP055815.1_strain_BA71           .........-----------------------------------.......g.............-----g......--- [197600]
MH025920.1_strain_R35            ..g......aactgttaatatttgt------tattaacaaacat...........-......t..gaca-......g--- [197600]
MH025919.1_strain_N10            ..g......aactgttaatatttgt------tattaacaaacat...........-......t..gaca-......g--- [197600]
MH025918.1_strain_R25            ..g......aactgttaatatttgt------tattaacaaacat...........-......t..gaca-......g--- [197600]
MH025917.1_strain_R7             ..g......aactgttaatatttgt------tattaacaaacat...........-......t..gaca-......g--- [197600]
MH025916.1_strain_R8             ..g......aactgttaatatttgt------tattaacaaacat...........-......t..gaca-......g--- [197600]

FR682468.2_Georgia_2007/1        ----------------------------------taacctat---acatatatgcaagtaaatgagatattatctatcat [197680]
ASF_IND_20_CAD543                ----------------------------------........---................................... [197680]
LR899193.1_Germany_2020/1        ----------------------------------........---................................... [197680]
MN393477.1_Wuhan_2019            ----------------------------------........---................................... [197680]
MW396979.1_Timor-Leste/2019/1    ----------------------------------........---................................... [197680]
MK940252.1_CN/2019/InnerMongolia ----------------------------------........---................................... [197680]
MK128995.1_China/2018/AnhuiXCGQ  ----------------------------------........---................................... [197680]
MH681419.1_ASFV/POL/2015         ----------------------------------........---................................... [197680]
LS478113.1_Estonia_2014          ----------------------------------........---................................... [197680]
KP843857.1_Odintsovo_02/14       ----------------------------------........---................................... [197680]
FN557520.1_strain_E75            ----------------------------------g...t...---................................... [197680]
AY261366.1_Warthog               ----------------------------------g...t...---................................... [197680]
AY261365.1_Warmbaths             ----------------------------------g...t...---................................... [197680]
AY261364.1_Tengani_62            ----------------------------------g.......aca...c............................... [197680]
AY261363.1_Pretoriuskop/96/4     ----------------------------------g.......---................................... [197680]
AY261362.1_Mkuzi_1979            ----------------------------------g.......---................................... [197680]
AY261361.1_Malawi_Lil-20/1_1983  ----------------------------------c..aa...---.t.c.-g..t.............-........... [197680]
AY261360.1_Kenya_1950            -----------------------------------t.aa...---...g.-...t.tt...c..g............... [197680]
AM712240.1_OURT_88/3             ----------------------------------g...t...---................................... [197680]
AM712239.1_Benin_97/1            ----------------------------------g...t...---................................... [197680]
KM111295.1_Ken06.Bus             -----------------------------------t.aa...---...g.-...t.........g............... [197680]
```

```
KM111294.1_Ken05/Tk1            ttatacagaatgtaagtaaatatacagaatataaat.aa...---...g.-...t...c.....g............... [197680]
KM262844.1_strain_L60           ----------------------------------g...t...---................................... [197680]
KM102979.1_26544/OG10_Italy     ----------------------------------g...t...---............a...................... [197680]
KM262845.1_strain_NHV           ----------------------------------g...t...---................................... [197680]
KX354450.1_47/Ss/2008           ----------------------------------g...t...---................................... [197680]
KP055815.1_strain_BA71          ----------------------------------g...t...---................................... [197680]
MH025920.1_strain_R35           -----------------------------------t.aa...---...g.-...t.........g............... [197680]
MH025919.1_strain_N10           -----------------------------------t.aa...---...g.-...t.........g............... [197680]
MH025918.1_strain_R25           -----------------------------------t.aa...---...g.-...t.........g............... [197680]
MH025917.1_strain_R7            -----------------------------------t.aa...---...g.-...t.........g............... [197680]
MH025916.1_strain_R8            -----------------------------------t.aa...---...g.-...t.........g............... [197680]

FR682468.2_Georgia_2007/1       aacgaatcaagggatatttgtatatatcaggagtttctgaaataaagatatgaagattatcatagtagtatccatcaatc [197760]
ASF_IND_20_CAD543               ..........................................k.............k....................... [197760]
LR899193.1_Germany_2020/1       ................................................................................ [197760]
MN393477.1_Wuhan_2019           ................................................................................ [197760]
MW396979.1_Timor-Leste/2019/1   ................................................................................ [197760]
MK940252.1_CN/2019/InnerMongolia ................................................................................ [197760]
MK128995.1_China/2018/AnhuiXCGQ ................................................................................ [197760]
MH681419.1_ASFV/POL/2015        ................................................................................ [197760]
LS478113.1_Estonia_2014         ................................................................................ [197760]
KP843857.1_Odintsovo_02/14      ................................................................................ [197760]
FN557520.1_strain_E75           ..........................................g.............g....................... [197760]
AY261366.1_Warthog              ...................................................................a............ [197760]
AY261365.1_Warmbaths            ................................................................................ [197760]
AY261364.1_Tengani_62           ...a............................................................................ [197760]
AY261363.1_Pretoriuskop/96/4    ................................................................................ [197760]
AY261362.1_Mkuzi_1979           ......................a...................g..................................... [197760]
AY261361.1_Malawi_Lil-20/1_1983 ......................a.g.......c......c..g........c..................c......... [197760]
AY261360.1_Kenya_1950           ..t.......a.............g..t....t......t.................c..........a........... [197760]
AM712240.1_OURT_88/3            ..........................................g.............g....................... [197760]
AM712239.1_Benin_97/1           ..........................................g.............g....................... [197760]
KM111295.1_Ken06.Bus            ..t.....................g..t....t......t.................c..........a........... [197760]
KM111294.1_Ken05/Tk1            ..t.......a.............g..t....t......t.................c..........a........... [197760]
KM262844.1_strain_L60           ..........................................g.............g....................... [197760]
KM102979.1_26544/OG10_Italy     ..........................................g.............g....................... [197760]
KM262845.1_strain_NHV           ..........................................g.............g....................... [197760]
KX354450.1_47/Ss/2008           ..........................................g.............g....................... [197760]
KP055815.1_strain_BA71          ..........................................g.............g....................... [197760]
MH025920.1_strain_R35           ..t.....................g..t....t......t.................c..........a........... [197760]
MH025919.1_strain_N10           ..t.....................g..t....t......t.................c..........a........... [197760]
MH025918.1_strain_R25           ..t.....................g..t....t......t.................c..........a........... [197760]
MH025917.1_strain_R7            ..t.....................g..t....t......t.................c..........a........... [197760]
MH025916.1_strain_R8            ..t.....................g..t....t......t.................c..........a........... [197760]
```

```
FR682468.2_Georgia_2007/1        acaatgcaacttcctttaaggcataatttagtaaactcagcactcccatcttctggatgctttacaactaacattaaaaa [197840]
ASF_IND_20_CAD543                ................................................................................ [197840]
LR899193.1_Germany_2020/1        ................................................................................ [197840]
MN393477.1_Wuhan_2019            ................................................................................ [197840]
MW396979.1_Timor-Leste/2019/1    ................................................................................ [197840]
MK940252.1_CN/2019/InnerMongolia ................................................................................ [197840]
MK128995.1_China/2018/AnhuiXCGQ  ................................................................................ [197840]
MH681419.1_ASFV/POL/2015         ................................................................................ [197840]
LS478113.1_Estonia_2014          ................................................................................ [197840]
KP843857.1_Odintsovo_02/14       ................................................................................ [197840]
FN557520.1_strain_E75            ....................a........................................................... [197840]
AY261366.1_Warthog               ....................a........................................................... [197840]
AY261365.1_Warmbaths             ....................a........................................................... [197840]
AY261364.1_Tengani_62            ........g...........a........................................................... [197840]
AY261363.1_Pretoriuskop/96/4     ....................a........................................................... [197840]
AY261362.1_Mkuzi_1979            ..........c.........a........................................................... [197840]
AY261361.1_Malawi_Lil-20/1_1983  .t.........a........a..............t............................g.....g......... [197840]
AY261360.1_Kenya_1950            ..........ca.......................t............................................ [197840]
AM712240.1_OURT_88/3             ....................a........................................................... [197840]
AM712239.1_Benin_97/1            ....................a........................................................... [197840]
KM111295.1_Ken06.Bus             ..........ca.......................t............................................ [197840]
KM111294.1_Ken05/Tk1             ..........ca.......................t............................................ [197840]
KM262844.1_strain_L60            ....................a........................................................... [197840]
KM102979.1_26544/OG10_Italy      ....................a........................................................... [197840]
KM262845.1_strain_NHV            ....................a........................................................... [197840]
KX354450.1_47/Ss/2008            ....................a........................................................... [197840]
KP055815.1_strain_BA71           ....................a........................................................... [197840]
MH025920.1_strain_R35            ..........ca.......................t............................................ [197840]
MH025919.1_strain_N10            ..........ca.......................t............................................ [197840]
MH025918.1_strain_R25            ..........ca.......................t............................................ [197840]
MH025917.1_strain_R7             ..........ca.......................t............................................ [197840]
MH025916.1_strain_R8             ..........ca.......................t............................................ [197840]

FR682468.2_Georgia_2007/1        ctcctcagtcatattatctgtaataaaataagatcctcctggagccatttgtagcatgtctcttattcctacaaaatctt [197920]
ASF_IND_20_CAD543                ................................................................................ [197920]
LR899193.1_Germany_2020/1        ................................................................................ [197920]
MN393477.1_Wuhan_2019            ................................................................................ [197920]
MW396979.1_Timor-Leste/2019/1    ................................................................................ [197920]
MK940252.1_CN/2019/InnerMongolia ................................................................................ [197920]
MK128995.1_China/2018/AnhuiXCGQ  ................................................................................ [197920]
MH681419.1_ASFV/POL/2015         ................................................................................ [197920]
LS478113.1_Estonia_2014          ................................................................................ [197920]
KP843857.1_Odintsovo_02/14       ................................................................................ [197920]
```

```
FN557520.1_strain_E75           .........t.....g................................g............................... [197920]
AY261366.1_Warthog              ................................................................................ [197920]
AY261365.1_Warmbaths            .........t...................................................................... [197920]
AY261364.1_Tengani_62           .........t...................................................................... [197920]
AY261363.1_Pretoriuskop/96/4    .........t....................................................................c. [197920]
AY261362.1_Mkuzi_1979           ................................................................................ [197920]
AY261361.1_Malawi_Lil-20/1_1983 t........t........a........g..t...............g........................tg....... [197920]
AY261360.1_Kenya_1950           .........t.c...............g....................a.c..................c.t......c. [197920]
AM712240.1_OURT_88/3            .........t.....g................................g............................... [197920]
AM712239.1_Benin_97/1           .........t.....g................................g............................... [197920]
KM111295.1_Ken06.Bus            .........t.c.........t.....g..t.................a.c..................c.t......c. [197920]
KM111294.1_Ken05/Tk1            .........t.c...............g....................a.c..................c.t......c. [197920]
KM262844.1_strain_L60           .........t.....g................................g............................... [197920]
KM102979.1_26544/OG10_Italy     .........t.....g................................g............................... [197920]
KM262845.1_strain_NHV           .........t.....g................................g............................... [197920]
KX354450.1_47/Ss/2008           .........t.....g................................g............................... [197920]
KP055815.1_strain_BA71          .........t.....g................................g............................... [197920]
MH025920.1_strain_R35           .........t.c.........t.....g..t.................a.c..................c.t......c. [197920]
MH025919.1_strain_N10           .........t.c.........t.....g..t.................a.c..................c.t......c. [197920]
MH025918.1_strain_R25           .........t.c.........t.....g..t.................a.c..................c.t......c. [197920]
MH025917.1_strain_R7            .........t.c.........t.....g..t.................a.c..................c.t......c. [197920]
MH025916.1_strain_R8            .........t.c.........t.....g..t.................a.c..................c.t......c. [197920]

FR682468.2_Georgia_2007/1       ttttgggatggtaaaaactcagcagtttcaaactcttttttagttttttttcctggtatttaagccatttgttataaaac [198000]
ASF_IND_20_CAD543               ................................................................................ [198000]
LR899193.1_Germany_2020/1       ................................................................................ [198000]
MN393477.1_Wuhan_2019           ................................................................................ [198000]
MW396979.1_Timor-Leste/2019/1   ................................................................................ [198000]
MK940252.1_CN/2019/InnerMongolia ................................................................................ [198000]
MK128995.1_China/2018/AnhuiXCGQ ................................................................................ [198000]
MH681419.1_ASFV/POL/2015        ................................................................................ [198000]
LS478113.1_Estonia_2014         ................................................................................ [198000]
KP843857.1_Odintsovo_02/14      ................................................................................ [198000]
FN557520.1_strain_E75           ................................................................................ [198000]
AY261366.1_Warthog              ...................t............................................................ [198000]
AY261365.1_Warmbaths            ...................t............................................................ [198000]
AY261364.1_Tengani_62           ...................t..........g................................................. [198000]
AY261363.1_Pretoriuskop/96/4    ...................t....................................a....................... [198000]
AY261362.1_Mkuzi_1979           ................................................................................ [198000]
AY261361.1_Malawi_Lil-20/1_1983 ....a.c............t....a.c....t.......c...............a........................ [198000]
AY261360.1_Kenya_1950           .......g...........t....a......t.......c...g.................................... [198000]
AM712240.1_OURT_88/3            ................................................................................ [198000]
AM712239.1_Benin_97/1           ................................................................................ [198000]
KM111295.1_Ken06.Bus            .......g...........t....a......t.......c...g........t........................... [198000]
```

```
KM111294.1_Ken05/Tk1              .......g...........t....a......t.......c...g.................................... [198000]
KM262844.1_strain_L60             ................................................................................ [198000]
KM102979.1_26544/OG10_Italy       ................................................................................ [198000]
KM262845.1_strain_NHV             ................................................................................ [198000]
KX354450.1_47/Ss/2008             ................................................................................ [198000]
KP055815.1_strain_BA71            ................................................................................ [198000]
MH025920.1_strain_R35             .......g...........t....a......t.......c...g........t........................... [198000]
MH025919.1_strain_N10             .......g...........t....a......t.......c...g........t........................... [198000]
MH025918.1_strain_R25             .......g...........t....a......t.......c...g........t........................... [198000]
MH025917.1_strain_R7              .......g...........t....a......t.......c...g........t........................... [198000]
MH025916.1_strain_R8              .......g...........t....a......t.......c...g........t........................... [198000]

FR682468.2_Georgia_2007/1         agttttcttatgaaaatgcatttgaaaatattgggaatgtttaaccatgcttcttccgagcacatctccagatacttact [198080]
ASF_IND_20_CAD543                 ................................................................................ [198080]
LR899193.1_Germany_2020/1         ................................................................................ [198080]
MN393477.1_Wuhan_2019             ................................................................................ [198080]
MW396979.1_Timor-Leste/2019/1     ................................................................................ [198080]
MK940252.1_CN/2019/InnerMongolia  ................................................................................ [198080]
MK128995.1_China/2018/AnhuiXCGQ   ................................................................................ [198080]
MH681419.1_ASFV/POL/2015          ................................................................................ [198080]
LS478113.1_Estonia_2014           ................................................................................ [198080]
KP843857.1_Odintsovo_02/14        ................................................................................ [198080]
FN557520.1_strain_E75             ................................................................................ [198080]
AY261366.1_Warthog                ........................................................t..................c.... [198080]
AY261365.1_Warmbaths              ........................................................t..................c.... [198080]
AY261364.1_Tengani_62             ..............................c................................................. [198080]
AY261363.1_Pretoriuskop/96/4      ........................................................t..................c.... [198080]
AY261362.1_Mkuzi_1979             .....................................................c.......................... [198080]
AY261361.1_Malawi_Lil-20/1_1983   ............................c......g..a..............a..t..............g..t..... [198080]
AY261360.1_Kenya_1950             ..........................................................................t..... [198080]
AM712240.1_OURT_88/3              .......................................-........................................ [198080]
AM712239.1_Benin_97/1             ................................................................................ [198080]
KM111295.1_Ken06.Bus              ..............................c.......a...................................t..... [198080]
KM111294.1_Ken05/Tk1              ..........................................................................t..... [198080]
KM262844.1_strain_L60             ................................................................................ [198080]
KM102979.1_26544/OG10_Italy       ................................................................................ [198080]
KM262845.1_strain_NHV             .......................................-........................................ [198080]
KX354450.1_47/Ss/2008             ................................................................................ [198080]
KP055815.1_strain_BA71            ................................................................................ [198080]
MH025920.1_strain_R35             ..............................c.......a...................................t..... [198080]
MH025919.1_strain_N10             ..............................c.......a...................................t..... [198080]
MH025918.1_strain_R25             ..............................c.......a...................................t..... [198080]
MH025917.1_strain_R7              ..............................c.......a...................................t..... [198080]
MH025916.1_strain_R8              ..............................c.......a...................................t..... [198080]
```

```
FR682468.2_Georgia_2007/1        ttctttgtttcccatgtctaatttattgctcactaagttagtaatg-----------------aatctattttaataatc [198160]
ASF_IND_20_CAD543                .........y....................................-----------------..--------------- [198160]
LR899193.1_Germany_2020/1        ..............................................-----------------................. [198160]
MN393477.1_Wuhan_2019            ..............................................-----------------................. [198160]
MW396979.1_Timor-Leste/2019/1    ..............................................-----------------................. [198160]
MK940252.1_CN/2019/InnerMongolia ..............................................-----------------................. [198160]
MK128995.1_China/2018/AnhuiXCGQ  ..............................................-----------------................. [198160]
MH681419.1_ASFV/POL/2015         ..............................................-----------------................. [198160]
LS478113.1_Estonia_2014          ..............................................-----------------................. [198160]
KP843857.1_Odintsovo_02/14       ..............................................-----------------................. [198160]
FN557520.1_strain_E75            .........c....................................-----------------..--------------- [198160]
AY261366.1_Warthog               ..............................................-----------------................. [198160]
AY261365.1_Warmbaths             ..............................................-----------------...............c. [198160]
AY261364.1_Tengani_62            ..............................................-----------------................. [198160]
AY261363.1_Pretoriuskop/96/4     ..............................................-----------------...........c..... [198160]
AY261362.1_Mkuzi_1979            ..............................................-----------------..--------------- [198160]
AY261361.1_Malawi_Lil-20/1_1983  ............................t......g..........aat--------------.......c...------ [198160]
AY261360.1_Kenya_1950            ............................t............c....-----------------...t------------- [198160]
AM712240.1_OURT_88/3             .........c....................................-----------------..--------------- [198160]
AM712239.1_Benin_97/1            .........c....................................aataacctatctatctt..--------------- [198160]
KM111295.1_Ken06.Bus             ............................t............c....-----------------...t------------- [198160]
KM111294.1_Ken05/Tk1             ............................t............c....-----------------...t------------- [198160]
KM262844.1_strain_L60            .........c....................................-----------------..--------------- [198160]
KM102979.1_26544/OG10_Italy      .........c....................................-----------------..--------------- [198160]
KM262845.1_strain_NHV            .........c....................................-----------------..--------------- [198160]
KX354450.1_47/Ss/2008            .........c....................................-----------------..--------------- [198160]
KP055815.1_strain_BA71           .........c....................................-----------------..--------------- [198160]
MH025920.1_strain_R35            ............................t............c....-----------------...t------------- [198160]
MH025919.1_strain_N10            ............................t............c....-----------------...t------------- [198160]
MH025918.1_strain_R25            ............................t............c....-----------------...t------------- [198160]
MH025917.1_strain_R7             ............................t............c....-----------------...t------------- [198160]
MH025916.1_strain_R8             ............................t............c....-----------------...t------------- [198160]

FR682468.2_Georgia_2007/1        tac--------------------------------------------------tttactaatctatcttaataacct--- [198240]
ASF_IND_20_CAD543                ------------------------------------------------------.a..ct.................at- [198240]
LR899193.1_Germany_2020/1        ...--------------------------------------------------........................--- [198240]
MN393477.1_Wuhan_2019            ...--------------------------------------------------........................--- [198240]
MW396979.1_Timor-Leste/2019/1    ...--------------------------------------------------........................--- [198240]
MK940252.1_CN/2019/InnerMongolia ...--------------------------------------------------........................--- [198240]
MK128995.1_China/2018/AnhuiXCGQ  ...--------------------------------------------------........................--- [198240]
MH681419.1_ASFV/POL/2015         ...--------------------------------------------------........................--- [198240]
LS478113.1_Estonia_2014          ...--------------------------------------------------........................--- [198240]
KP843857.1_Odintsovo_02/14       ...--------------------------------------------------........................--- [198240]
```

```
FN557520.1_strain_E75              ------------------------------------------------------.a..ct.................at- [198240]
AY261366.1_Warthog                 ...--------------------------------------------------........................atc [198240]
AY261365.1_Warmbaths               ..t--------------------------------------------------.............t..........att [198240]
AY261364.1_Tengani_62              ..ttttactaatctatttaatatcctatcttactaatctatttaatatcctaac.......................att [198240]
AY261363.1_Pretoriuskop/96/4       ...---------------------------------------------------............t..........att [198240]
AY261362.1_Mkuzi_1979              ------------------------------------------------------.a..ct..........---------- [198240]
AY261361.1_Malawi_Lil-20/1_1983    ----------------------------------------------------------............-...t..atc [198240]
AY261360.1_Kenya_1950              ----------------------------------------------------ag.ata...c....t...-......at- [198240]
AM712240.1_OURT_88/3               ------------------------------------------------------.a..ct.................at- [198240]
AM712239.1_Benin_97/1              ------------------------------------------------------.a..ct.................at- [198240]
KM111295.1_Ken06.Bus               ----------------------------------------------------ag.ata...c....t...-......ag- [198240]
KM111294.1_Ken05/Tk1               ----------------------------------------------------ag.ata...c....t...-......ag- [198240]
KM262844.1_strain_L60              ------------------------------------------------------.a..ct.................at- [198240]
KM102979.1_26544/OG10_Italy        ------------------------------------------------------.a..ct.................at- [198240]
KM262845.1_strain_NHV              ------------------------------------------------------.a..ct.................at- [198240]
KX354450.1_47/Ss/2008              ------------------------------------------------------.a..ct.................at- [198240]
KP055815.1_strain_BA71             ------------------------------------------------------.a..ct.................at- [198240]
MH025920.1_strain_R35              ----------------------------------------------------ag.ata...c....t...-......ag- [198240]
MH025919.1_strain_N10              ----------------------------------------------------ag.ata...c....t...-......ag- [198240]
MH025918.1_strain_R25              ----------------------------------------------------ag.ata...c....t...-......ag- [198240]
MH025917.1_strain_R7               ----------------------------------------------------ag.ata...c....t...-......ag- [198240]
MH025916.1_strain_R8               ----------------------------------------------------ag.ata...c....t...-......ag- [198240]

FR682468.2_Georgia_2007/1          -----------------------atcttataatctatcttaataacctaattataacctatttataattggctaatgctg [198320]
ASF_IND_20_CAD543                  ---------------------ct......................................................... [198320]
LR899193.1_Germany_2020/1          -----------------------......................................................... [198320]
MN393477.1_Wuhan_2019              -----------------------......................................................... [198320]
MW396979.1_Timor-Leste/2019/1      -----------------------......................................................... [198320]
MK940252.1_CN/2019/InnerMongolia   -----------------------......................................................... [198320]
MK128995.1_China/2018/AnhuiXCGQ    -----------------------......................................................... [198320]
MH681419.1_ASFV/POL/2015           -----------------------......................................................... [198320]
LS478113.1_Estonia_2014            -----------------------......................................................... [198320]
KP843857.1_Odintsovo_02/14         -----------------------......................................................... [198320]
FN557520.1_strain_E75              ---------------------ct............................g.................a.......... [198320]
AY261366.1_Warthog                 tta-taatctatcttaataacct.....................................................a... [198320]
AY261365.1_Warmbaths               ttactaatctatcttaataacct......................................................... [198320]
AY261364.1_Tengani_62              ttactaat-------------ct......................t.................................. [198320]
AY261363.1_Pretoriuskop/96/4       ttactaatctatctta------------.................................................... [198320]
AY261362.1_Mkuzi_1979              ----------------------------------------------------.................a.......... [198320]
AY261361.1_Malawi_Lil-20/1_1983    tta-taat-------------ct.c................ta.c.tatc..........g...............a... [198320]
AY261360.1_Kenya_1950              --------------------------......c........ta.c.tatt........................ta...- [198320]
AM712240.1_OURT_88/3               ---------------------ct............................g.................a.......... [198320]
AM712239.1_Benin_97/1              ---------------------ct............................g.................a.......... [198320]
KM111295.1_Ken06.Bus               --------------------------......c........ta.c.tatt........................ta...- [198320]
```

```
KM111294.1_Ken05/Tk1             --------------------------......c........ta.c.tatt.....g..................ta...- [198320]
KM262844.1_strain_L60            ---------------------ct............................g.................a.......... [198320]
KM102979.1_26544/OG10_Italy      ---------------------ct............................g.................a.......... [198320]
KM262845.1_strain_NHV            ---------------------ct............................g.................a.......... [198320]
KX354450.1_47/Ss/2008            ---------------------ct............................g.................a.......... [198320]
KP055815.1_strain_BA71           ---------------------ct............................g.................a.......... [198320]
MH025920.1_strain_R35            --------------------------......c........ta.c.tatt........................ta...- [198320]
MH025919.1_strain_N10            --------------------------......c........ta.c.tatt........................ta...- [198320]
MH025918.1_strain_R25            --------------------------......c........ta.c.tatt........................ta...- [198320]
MH025917.1_strain_R7             --------------------------......c........ta.c.tatt........................ta...- [198320]
MH025916.1_strain_R8             --------------------------......c........ta.c.tatt........................ta...- [198320]

FR682468.2_Georgia_2007/1        ccggcatttcatgc-ctatc-taaac----aactcctactaagcaatctactattacatatatagattcac-tttttata [198400]
ASF_IND_20_CAD543                ..............a.....-.....agtt.........................................t........ [198400]
LR899193.1_Germany_2020/1        ..............-.....-.....----.........................................-........ [198400]
MN393477.1_Wuhan_2019            ..............-.....-.....----.........................................-........ [198400]
MW396979.1_Timor-Leste/2019/1    ..............-.....-.....----.........................................-........ [198400]
MK940252.1_CN/2019/InnerMongolia ..............-.....-.....----.........................................-........ [198400]
MK128995.1_China/2018/AnhuiXCGQ  ..............-.....-.....----.........................................-........ [198400]
MH681419.1_ASFV/POL/2015         ..............-.....-.....----.........................................-........ [198400]
LS478113.1_Estonia_2014          ..............-.....-.....----.........................................-........ [198400]
KP843857.1_Odintsovo_02/14       ..............-.....-.....----.........................................-........ [198400]
FN557520.1_strain_E75            ..............a.g...-.....agtt..t......................................t........ [198400]
AY261366.1_Warthog               ..............-.....-.....----.........................................-........ [198400]
AY261365.1_Warmbaths             ....t.........-.....-.g...----.........................................-........ [198400]
AY261364.1_Tengani_62            .t............-.....-.....----.........................................-........ [198400]
AY261363.1_Pretoriuskop/96/4     ..............a.g...-..-------................c........................-........ [198400]
AY261362.1_Mkuzi_1979            .t............a.g...-.....agtt..t......................................t........ [198400]
AY261361.1_Malawi_Lil-20/1_1983  .t............a.....-ct...gatt................ct....t..........g.......-.......t [198400]
AY261360.1_Kenya_1950            ----------....ata...-....tgatt.c.c..........-cc.....t..........g.......--......- [198400]
AM712240.1_OURT_88/3             ..............a.g...-.....agtt..t......................................t........ [198400]
AM712239.1_Benin_97/1            ..............a.g...-.....agtt..t......................................t........ [198400]
KM111295.1_Ken06.Bus             ----------....ata...-....tgatt.-.c...........cc.....t..........g.......-.......- [198400]
KM111294.1_Ken05/Tk1             ----------....ata...c....tgatt.--c..........ccc.....t..........g.......--......- [198400]
KM262844.1_strain_L60            ..............a.g...-.....agtt..t......................................t........ [198400]
KM102979.1_26544/OG10_Italy      ..............a.g...-.....agtt..t......................................t........ [198400]
KM262845.1_strain_NHV            ..............a.g...-.....agtt..t......................................t........ [198400]
KX354450.1_47/Ss/2008            ..............a.g...-.....agtt..t......................................t........ [198400]
KP055815.1_strain_BA71           ..............a.g...-.....agtt..t......................................t........ [198400]
MH025920.1_strain_R35            ----------....ata...-....tgatt.-.c...........cc.....t..........g.......-.......- [198400]
MH025919.1_strain_N10            ----------....ata...-....tgatt.-.c...........cc.....t..........g.......-.......- [198400]
MH025918.1_strain_R25            ----------....ata...-....tgatt.-.c...........cc.....t..........g.......-.......- [198400]
MH025917.1_strain_R7             ----------....ata...-....tgatt.-.c...........cc.....t..........g.......-.......- [198400]
MH025916.1_strain_R8             ----------....ata...-....tgatt.-.c...........cc.....t..........g.......-.......- [198400]
```

```
FR682468.2_Georgia_2007/1        tttgtaaatcatgagaattataaaatcattactcatttttattgtaa---attagtgggtatttgtaaaaatcttcaaac [198480]
ASF_IND_20_CAD543                .............------.......m......y.............---.......................y.....y [198480]
LR899193.1_Germany_2020/1        ...............................................---.............................. [198480]
MN393477.1_Wuhan_2019            ...............................................---.............................. [198480]
MW396979.1_Timor-Leste/2019/1    ...............................................---.............................. [198480]
MK940252.1_CN/2019/InnerMongolia ...............................................---.............................. [198480]
MK128995.1_China/2018/AnhuiXCGQ  ...............................................---.............................. [198480]
MH681419.1_ASFV/POL/2015         ...............................................---.............................. [198480]
LS478113.1_Estonia_2014          ...............................................---.............................. [198480]
KP843857.1_Odintsovo_02/14       ...............................................---.............................. [198480]
FN557520.1_strain_E75            .............------.......a......t.............---.......................c.....t [198480]
AY261366.1_Warthog               ...............................................---.......................c...... [198480]
AY261365.1_Warmbaths             ...............................................---.......................c...... [198480]
AY261364.1_Tengani_62            ...............................................---..g....................c...... [198480]
AY261363.1_Pretoriuskop/96/4     ...............................................---.......................c...... [198480]
AY261362.1_Mkuzi_1979            .............------.......a......t.............---.......................c...... [198480]
AY261361.1_Malawi_Lil-20/1_1983  ..........g.tg..........c.a......t.....c....a..tttg..................t...c.....t [198480]
AY261360.1_Kenya_1950            ........c...t........ttt..a......t.....c..c....---..g.....a.....a........c....c. [198480]
AM712240.1_OURT_88/3             .............------.......a......t.............---.......................c.....t [198480]
AM712239.1_Benin_97/1            .............------.......a......t.............---.......................c.....t [198480]
KM111295.1_Ken06.Bus             ........ct..t........ttt..a......t.....c.......---..g.....a.....a........c....c. [198480]
KM111294.1_Ken05/Tk1             ...a....c...t........ttt..a......t.....c.......---..g.....a.....a........c....c. [198480]
KM262844.1_strain_L60            .............------.......a......t.............---.......................c.....t [198480]
KM102979.1_26544/OG10_Italy      .............------.......a......t.............---.......................c.....t [198480]
KM262845.1_strain_NHV            .............------.......a......t.............---.......................c.....t [198480]
KX354450.1_47/Ss/2008            .............------.......a......t.............---.......................c.....t [198480]
KP055815.1_strain_BA71           .............------.......a......t.............---.......................c.....t [198480]
MH025920.1_strain_R35            ........ct..t........ttt..a......t.....c.......---..g.....a.....a........c....c. [198480]
MH025919.1_strain_N10            ........ct..t........ttt..a......t.....c.......---..g.....a.....a........c....c. [198480]
MH025918.1_strain_R25            ........ct..t........ttt..a......t.....c.......---..g.....a.....a........c....c. [198480]
MH025917.1_strain_R7             ........ct..t........ttt..a......t.....c.......---..g.....a.....a........c....c. [198480]
MH025916.1_strain_R8             ........ct..t........ttt..a......t.....c.......---..g.....a.....a........c....c. [198480]

FR682468.2_Georgia_2007/1        gttttaagatagttttctagagagaagtaatctttgccatcaatatataatgcttttcctttaaactccagttttgctat [198560]
ASF_IND_20_CAD543                ...............................................y........y........y.............. [198560]
LR899193.1_Germany_2020/1        ................................................................................ [198560]
MN393477.1_Wuhan_2019            ................................................................................ [198560]
MW396979.1_Timor-Leste/2019/1    ................................................................................ [198560]
MK940252.1_CN/2019/InnerMongolia ................................................................................ [198560]
MK128995.1_China/2018/AnhuiXCGQ  ................................................................................ [198560]
MH681419.1_ASFV/POL/2015         ................................................................................ [198560]
LS478113.1_Estonia_2014          ................................................................................ [198560]
KP843857.1_Odintsovo_02/14       ................................................................................ [198560]
```

```
FN557520.1_strain_E75             ........................t..........a....t......c........c........t.............. [198560]
AY261366.1_Warthog                ................................................................................ [198560]
AY261365.1_Warmbaths              ................................................................................ [198560]
AY261364.1_Tengani_62             .......................a............................................t........... [198560]
AY261363.1_Pretoriuskop/96/4      ................................................................................ [198560]
AY261362.1_Mkuzi_1979             ........................t..........a....t......c........c........t.............. [198560]
AY261361.1_Malawi_Lil-20/1_1983   ........g........a...a.........t...a....t......c.......cc........t..t........... [198560]
AY261360.1_Kenya_1950             ...............................t...a..g.t...............c..a.....a..t........... [198560]
AM712240.1_OURT_88/3              ........................t..........a....t......c........c........t.............. [198560]
AM712239.1_Benin_97/1             ........................t..........a....t......c........c........t.............. [198560]
KM111295.1_Ken06.Bus              ...............................t...a..g.t...............c..a.....a..t........... [198560]
KM111294.1_Ken05/Tk1              ...............................t...a..g.t...............c..a.....a..t........... [198560]
KM262844.1_strain_L60             ........................t..........a....t......c........c........t.............. [198560]
KM102979.1_26544/OG10_Italy       ........................t..........a....t......c........c........t.............. [198560]
KM262845.1_strain_NHV             ........................t..........a....t......c........c........t.............. [198560]
KX354450.1_47/Ss/2008             ........................t..........a....t......c........c........t.............. [198560]
KP055815.1_strain_BA71            ........................t..........a....t......c........c........t.............. [198560]
MH025920.1_strain_R35             ...............................t...a..g.t...............c..a.....a..t........... [198560]
MH025919.1_strain_N10             ...............................t...a..g.t...............c..a.....a..t........... [198560]
MH025918.1_strain_R25             ...............................t...a..g.t...............c..a.....a..t........... [198560]
MH025917.1_strain_R7              ...............................t...a..g.t...............c..a.....a..t........... [198560]
MH025916.1_strain_R8              ...............................t...a..g.t...............c..a.....a..t........... [198560]

FR682468.2_Georgia_2007/1         gtttagtgagccg-tttctagatctttttgggcaataaatagattttcattggttgcatcgtccgtaagcagaaaggtac [198640]
ASF_IND_20_CAD543                 .............t................ra.........k...................................... [198640]
LR899193.1_Germany_2020/1         .............-.................................................................. [198640]
MN393477.1_Wuhan_2019             .............-.................................................................. [198640]
MW396979.1_Timor-Leste/2019/1     .............-.................................................................. [198640]
MK940252.1_CN/2019/InnerMongolia  .............-.................................................................. [198640]
MK128995.1_China/2018/AnhuiXCGQ   .............-.................................................................. [198640]
MH681419.1_ASFV/POL/2015          .............-.................................................................. [198640]
LS478113.1_Estonia_2014           .............-.................................................................. [198640]
KP843857.1_Odintsovo_02/14        .............-.................................................................. [198640]
FN557520.1_strain_E75             .............t....c...........aa.........t...................................... [198640]
AY261366.1_Warthog                .............t................a................................................. [198640]
AY261365.1_Warmbaths              .............t.........................................a........................ [198640]
AY261364.1_Tengani_62             .........a...t................a..g.............................................. [198640]
AY261363.1_Pretoriuskop/96/4      ............at................a................................................. [198640]
AY261362.1_Mkuzi_1979             .............t....c............a.........t...................................... [198640]
AY261361.1_Malawi_Lil-20/1_1983   ............at................a......t...t........ga...a.......g................ [198640]
AY261360.1_Kenya_1950             .........a...t................aa...g...c..........c......g...................... [198640]
AM712240.1_OURT_88/3              .............t....c...........aa.........t...................................... [198640]
AM712239.1_Benin_97/1             .............t....c...........aa.........t...................................... [198640]
KM111295.1_Ken06.Bus              .........a...t................aa...g...c..........c......g...................... [198640]
```

```
KM111294.1_Ken05/Tk1            .........a...t................aa...g...c..........c......g...................... [198640]
KM262844.1_strain_L60           .............t....c...........aa.........t...................................... [198640]
KM102979.1_26544/OG10_Italy     .............t....c...........aa.........t...................................... [198640]
KM262845.1_strain_NHV           .............t....c...........aa.........t...................................... [198640]
KX354450.1_47/Ss/2008           .............t....c...........aa.........t...................................... [198640]
KP055815.1_strain_BA71          .............t....c...........aa.........t...................................... [198640]
MH025920.1_strain_R35           .........a...t................aa...g...c..........c......g...................... [198640]
MH025919.1_strain_N10           .........a...t................aa...g...c..........c......g...................... [198640]
MH025918.1_strain_R25           .........a...t................aa...g...c..........c......g...................... [198640]
MH025917.1_strain_R7            .........a...t................aa...g...c..........c......g...................... [198640]
MH025916.1_strain_R8            .........a...t................aa...g...c..........c......g...................... [198640]

FR682468.2_Georgia_2007/1       cactaggcacgttaaaaaacatacgttctatttcatggtcggatttttgagaatagaaaaaatctaattttttaatccgc [198720]
ASF_IND_20_CAD543               .........t.c..........w................................r......y................. [198720]
LR899193.1_Germany_2020/1       ................................................................................ [198720]
MN393477.1_Wuhan_2019           ................................................................................ [198720]
MW396979.1_Timor-Leste/2019/1   ................................................................................ [198720]
MK940252.1_CN/2019/InnerMongolia ................................................................................ [198720]
MK128995.1_China/2018/AnhuiXCGQ ................................................................................ [198720]
MH681419.1_ASFV/POL/2015        ................................................................................ [198720]
LS478113.1_Estonia_2014         ................................................................................ [198720]
KP843857.1_Odintsovo_02/14      ................................................................................ [198720]
FN557520.1_strain_E75           ..t......t.c....t.....t................................a......c................. [198720]
AY261366.1_Warthog              ..t............................................................................. [198720]
AY261365.1_Warmbaths            ..t............................................................................. [198720]
AY261364.1_Tengani_62           ..t......t..c.......................a........................gc....c............ [198720]
AY261363.1_Pretoriuskop/96/4    ..t.................................a.....t..................g.....c...........a [198720]
AY261362.1_Mkuzi_1979           ..t......t.c....t.....t................................a......c................. [198720]
AY261361.1_Malawi_Lil-20/1_1983 ..t......t..........................c...att........................c...........t [198720]
AY261360.1_Kenya_1950           ..t.....tt...c.t..................g.c...att......t..............c..c...........t [198720]
AM712240.1_OURT_88/3            ..t......t.c....t.....t................................a......c................. [198720]
AM712239.1_Benin_97/1           ..t......t.c....t.....t................................a......c................. [198720]
KM111295.1_Ken06.Bus            ..t.....tt...c.t..................g.c...att......t............c.c..c...........t [198720]
KM111294.1_Ken05/Tk1            ..t.....tt...c.t..................g.c...att......t..............c..c...........t [198720]
KM262844.1_strain_L60           ..t......t.c....t.....t................................a......c................. [198720]
KM102979.1_26544/OG10_Italy     ..t......t.c....t.....t................................a......c................. [198720]
KM262845.1_strain_NHV           ..t......t.c....t.....t................................a......c................. [198720]
KX354450.1_47/Ss/2008           ..t......t.c....t.....t................................a......c................. [198720]
KP055815.1_strain_BA71          ..t......t.c....t.....t................................a......c................. [198720]
MH025920.1_strain_R35           ..t.....tt...c.t..................g.c...att......t............c.c..c...........t [198720]
MH025919.1_strain_N10           ..t.....tt...c.t..................g.c...att......t............c.c..c...........t [198720]
MH025918.1_strain_R25           ..t.....tt...c.t..................g.c...att......t............c.c..c...........t [198720]
MH025917.1_strain_R7            ..t.....tt...c.t..................g.c...att......t............c.c..c...........t [198720]
MH025916.1_strain_R8            ..t.....tt...c.t..................g.c...att......t............c.c..c...........t [198720]
```

```
FR682468.2_Georgia_2007/1        gttaactcttttttatcaatctttccagactgttttatatatactttattgcaaatcttacaatcctctatggcttcatt [198800]
ASF_IND_20_CAD543                ......................................r..................................y...... [198800]
LR899193.1_Germany_2020/1        ................................................................................ [198800]
MN393477.1_Wuhan_2019            ................................................................................ [198800]
MW396979.1_Timor-Leste/2019/1    ................................................................................ [198800]
MK940252.1_CN/2019/InnerMongolia ................................................................................ [198800]
MK128995.1_China/2018/AnhuiXCGQ  ................................................................................ [198800]
MH681419.1_ASFV/POL/2015         ................................................................................ [198800]
LS478113.1_Estonia_2014          ................................................................................ [198800]
KP843857.1_Odintsovo_02/14       ................................................................................ [198800]
FN557520.1_strain_E75            ..........g...........................g.....................g............t...... [198800]
AY261366.1_Warthog               ................................................................................ [198800]
AY261365.1_Warmbaths             ................................................................................ [198800]
AY261364.1_Tengani_62            a............................................................................... [198800]
AY261363.1_Pretoriuskop/96/4     t............................................................................... [198800]
AY261362.1_Mkuzi_1979            ..........g.................................................g............t...... [198800]
AY261361.1_Malawi_Lil-20/1_1983  t...............................g......................c....g.gcg....t.......... [198800]
AY261360.1_Kenya_1950            t...g......................................t......a.........g........t.......... [198800]
AM712240.1_OURT_88/3             ..........g...........................g.....................g............t...... [198800]
AM712239.1_Benin_97/1            ..........g...........................g.....................g............t...... [198800]
KM111295.1_Ken06.Bus             t...g......................................t......a.....t...g........t.......... [198800]
KM111294.1_Ken05/Tk1             t...g......................................t......a.........g........t.......... [198800]
KM262844.1_strain_L60            ..........g...........................g.....................g............t...... [198800]
KM102979.1_26544/OG10_Italy      ..........g...........................g.....................g............t...... [198800]
KM262845.1_strain_NHV            ..........g...........................g.....................g............t...... [198800]
KX354450.1_47/Ss/2008            ..........g...........................g.....................g............t...... [198800]
KP055815.1_strain_BA71           ..........g...........................g.....................g............t...... [198800]
MH025920.1_strain_R35            t...g......................................t......a.....t...g........t.......... [198800]
MH025919.1_strain_N10            t...g......................................t......a.....t...g........t.......... [198800]
MH025918.1_strain_R25            t...g......................................t......a.....t...g........t.......... [198800]
MH025917.1_strain_R7             t...g......................................t......a.....t...g........t.......... [198800]
MH025916.1_strain_R8             t...g......................................t......a.....t...g........t.......... [198800]

FR682468.2_Georgia_2007/1        atacttattttgcttatcctctattgacatgtccgtatttgataggtaacttccgttaaggcggttccccatggttttag [198880]
ASF_IND_20_CAD543                .....................................................t.a........................ [198880]
LR899193.1_Germany_2020/1        ................................................................................ [198880]
MN393477.1_Wuhan_2019            ................................................................................ [198880]
MW396979.1_Timor-Leste/2019/1    ................................................................................ [198880]
MK940252.1_CN/2019/InnerMongolia ................................................................................ [198880]
MK128995.1_China/2018/AnhuiXCGQ  ................................................................................ [198880]
MH681419.1_ASFV/POL/2015         ................................................................................ [198880]
LS478113.1_Estonia_2014          ................................................................................ [198880]
KP843857.1_Odintsovo_02/14       ................................................................................ [198880]
```

```
FN557520.1_strain_E75            ...............................a.........g...........t.a....a................... [198880]
AY261366.1_Warthog               ................................................................................ [198880]
AY261365.1_Warmbaths             ................................................................................ [198880]
AY261364.1_Tengani_62            ......................g......................................................... [198880]
AY261363.1_Pretoriuskop/96/4     ..............c................................................................a [198880]
AY261362.1_Mkuzi_1979            ...............................a.........g...........t......a.............-..... [198880]
AY261361.1_Malawi_Lil-20/1_1983  ......g........................a..a......g.t............c...at..........-....... [198880]
AY261360.1_Kenya_1950            ....g..........g...............a.........g..............c...at..........-......a [198880]
AM712240.1_OURT_88/3             ...............................a.........g...........t.a....a................... [198880]
AM712239.1_Benin_97/1            ...............................a.........g...........t.a....a................... [198880]
KM111295.1_Ken06.Bus             ...............g..............aa.t.......g..............c....t..........-.....ga [198880]
KM111294.1_Ken05/Tk1             ...............g...............a.........g..............c...at..........-......a [198880]
KM262844.1_strain_L60            ...............................a.........g...........t.a....a................... [198880]
KM102979.1_26544/OG10_Italy      ...............................a.........g...........t.a....a................... [198880]
KM262845.1_strain_NHV            ...............................a.........g...........t.a....a................... [198880]
KX354450.1_47/Ss/2008            ...............................a.........g...........t.a....a................... [198880]
KP055815.1_strain_BA71           ...............................a.........g...........t.a....a................... [198880]
MH025920.1_strain_R35            ...............g..............aa.t.......g..............c....t..........-.....ga [198880]
MH025919.1_strain_N10            ...............g..............aa.t.......g..............c....t..........-.....ga [198880]
MH025918.1_strain_R25            ...............g..............aa.t.......g..............c....t..........-.....ga [198880]
MH025917.1_strain_R7             ...............g..............aa.t.......g..............c....t..........-.....ga [198880]
MH025916.1_strain_R8             ...............g..............aa.t.......g..............c....t..........-.....ga [198880]

FR682468.2_Georgia_2007/1        --atagatttttaattcagttgtatacttttattatga-ggctaaaata------------tagaag-tttgatcctaaa [198960]
ASF_IND_20_CAD543                gtg...................................-..........aatagttaaata......-..ga.c.t--.. [198960]
LR899193.1_Germany_2020/1        --....................................-..........------------......-............ [198960]
MN393477.1_Wuhan_2019            --....................................-..........------------......-............ [198960]
MW396979.1_Timor-Leste/2019/1    --....................................-..........------------......-............ [198960]
MK940252.1_CN/2019/InnerMongolia --....................................-..........------------......-............ [198960]
MK128995.1_China/2018/AnhuiXCGQ  --....................................-..........------------......-............ [198960]
MH681419.1_ASFV/POL/2015         --....................................-..........------------......-............ [198960]
LS478113.1_Estonia_2014          --....................................-..........------------......-............ [198960]
KP843857.1_Odintsovo_02/14       --....................................-..........------------......-............ [198960]
FN557520.1_strain_E75            gtg....................g........c.....-..........aatagttaaata......-..ga.c.t--.. [198960]
AY261366.1_Warthog               --....................................-..........------------......t..gatc.t-... [198960]
AY261365.1_Warmbaths             --....................................-..........------------......t..gatc.t-... [198960]
AY261364.1_Tengani_62            --....................................-..........------------......t..ga.c.t-... [198960]
AY261363.1_Pretoriuskop/96/4     --....................................-..........------------......t..ga.c.t-... [198960]
AY261362.1_Mkuzi_1979            gtg....................g........c.....-..........agtagttaaata......t..ga.c.t--.. [198960]
AY261361.1_Malawi_Lil-20/1_1983  --...a..........................c....gt..........aataattaaata...t..t..ga.a.t--.. [198960]
AY261360.1_Kenya_1950            --g.........................-------...-.a........--------aata..a-------------... [198960]
AM712240.1_OURT_88/3             gtg....................g........c.....-..........aatagttaaata......-..ga.c.t--.. [198960]
AM712239.1_Benin_97/1            gtg....................g........c.....-..........aatagttaaata......-..ga.c.t--.. [198960]
KM111295.1_Ken06.Bus             --g..a......................-------...-.a........--------aata..a..at..-------... [198960]
```

```
KM111294.1_Ken05/Tk1           --g.........................-------...-.a........--------aata.---------------... [198960]
KM262844.1_strain_L60          gtg....................g........c.....-..........aatagttaaata......-..ga.c.t--.. [198960]
KM102979.1_26544/OG10_Italy    gtg....................g........c.....-..........aatagttaaata......-..ga.c.t--.. [198960]
KM262845.1_strain_NHV          gtg....................g........c.....-..........aatagttaaata......-..ga.c.t--.. [198960]
KX354450.1_47/Ss/2008          gtg....................g........c.....-..........aatagttaaata......-..ga.c.t--.. [198960]
KP055815.1_strain_BA71         gtg....................g........c.....-..........aatagttaaata......-..ga.c.t--.. [198960]
MH025920.1_strain_R35          --g..a......................-------...-.a........--------aata..a..at..-------... [198960]
MH025919.1_strain_N10          --g..a......................-------...-.a........--------aata..a..at..-------... [198960]
MH025918.1_strain_R25          --g..a......................-------...-.a........--------aata..a..at..-------... [198960]
MH025917.1_strain_R7           --g..a......................-------...-.a........--------aata..a..at..-------... [198960]
MH025916.1_strain_R8           --g..a......................-------...-.a........--------aata..a..at..-------... [198960]

FR682468.2_Georgia_2007/1      aaaataaaaagatttt---gtacatttatttatggtttatagcggtata---gaggccgataaaaggtatccgggtagtc [199040]
ASF_IND_20_CAD543              ...............----------------..................---............................ [199040]
LR899193.1_Germany_2020/1      ................---..............................---............................ [199040]
MN393477.1_Wuhan_2019          ................---..............................---............................ [199040]
MW396979.1_Timor-Leste/2019/1  ................---..............................---............................ [199040]
MK940252.1_CN/2019/InnerMongolia ................---..............................---............................ [199040]
MK128995.1_China/2018/AnhuiXCGQ ................---..............................---............................ [199040]
MH681419.1_ASFV/POL/2015       ................---..............................---............................ [199040]
LS478113.1_Estonia_2014        ................---..............................---............................ [199040]
KP843857.1_Odintsovo_02/14     ................---..............................---............................ [199040]
FN557520.1_strain_E75          ...............----------------..................---....a....................... [199040]
AY261366.1_Warthog             ................---..............................---............................ [199040]
AY261365.1_Warmbaths           ................---..............................---............................ [199040]
AY261364.1_Tengani_62          ...............------------......................---............................ [199040]
AY261363.1_Pretoriuskop/96/4   ................---..............................---............................ [199040]
AY261362.1_Mkuzi_1979          ...............----------------..................---....a....................... [199040]
AY261361.1_Malawi_Lil-20/1_1983 ..........ct.a..ttt..ca..c.g.....................---....t....................a.. [199040]
AY261360.1_Kenya_1950          ..t.a...ttt.....--t...a..c.g.......c.............---.g.................g........ [199040]
AM712240.1_OURT_88/3           ...............----------------..................---....a....................... [199040]
AM712239.1_Benin_97/1          ...............----------------..................---....a....................... [199040]
KM111295.1_Ken06.Bus           ...ta....tt.....--t...a..c.g.......c.............ggg.g..t..............g........ [199040]
KM111294.1_Ken05/Tk1           ........ttt.....--t...a..c.g.......c.............---.g.................g........ [199040]
KM262844.1_strain_L60          ...............----------------..................---....a....................... [199040]
KM102979.1_26544/OG10_Italy    ...............----------------..................---....a....................... [199040]
KM262845.1_strain_NHV          ...............----------------..................---....a....................... [199040]
KX354450.1_47/Ss/2008          ...............----------------..................---....a....................... [199040]
KP055815.1_strain_BA71         ...............----------------..................---....a....................... [199040]
MH025920.1_strain_R35          ...ta....tt.....--t...a..c.g.......c.............---.g..t..............g........ [199040]
MH025919.1_strain_N10          ...ta....tt.....--t...a..c.g.......c.............---.g..t..............g........ [199040]
MH025918.1_strain_R25          ...ta....tt.....--t...a..c.g.......c.............---.g..t..............g........ [199040]
MH025917.1_strain_R7           ...ta....tt.....--t...a..c.g.......c.............---.g..t..............g........ [199040]
MH025916.1_strain_R8           ...ta....tt.....--t...a..c.g.......c.............---.g..t..............g........ [199040]
```

```
FR682468.2_Georgia_2007/1        tcctatgatatcgtcaattttggtataataacagttgttatggtagtattgtccaaaccgagtatgtatgcgccggtgaa [199120]
ASF_IND_20_CAD543                -------------------------------------------------------------------------------- [199120]
LR899193.1_Germany_2020/1        ................................................................................ [199120]
MN393477.1_Wuhan_2019            ................................................................................ [199120]
MW396979.1_Timor-Leste/2019/1    ................................................................................ [199120]
MK940252.1_CN/2019/InnerMongolia ................................................................................ [199120]
MK128995.1_China/2018/AnhuiXCGQ  ................................................................................ [199120]
MH681419.1_ASFV/POL/2015         ................................................................................ [199120]
LS478113.1_Estonia_2014          ................................................................................ [199120]
KP843857.1_Odintsovo_02/14       ................................................................................ [199120]
FN557520.1_strain_E75            .................................c.............................................. [199120]
AY261366.1_Warthog               ..........................................a..................................... [199120]
AY261365.1_Warmbaths             ................................................................................ [199120]
AY261364.1_Tengani_62            ............a..g.................c.............................................. [199120]
AY261363.1_Pretoriuskop/96/4     .................................c.............................................. [199120]
AY261362.1_Mkuzi_1979            .................................c.............................................. [199120]
AY261361.1_Malawi_Lil-20/1_1983  .....c...............a...........c.......................a...................... [199120]
AY261360.1_Kenya_1950            .....c.....................g.....c.......................a.....g................ [199120]
AM712240.1_OURT_88/3             .................................c.............................................. [199120]
AM712239.1_Benin_97/1            .................................c.............................................. [199120]
KM111295.1_Ken06.Bus             .....c.....................g.....c.......................a.....g...........a.... [199120]
KM111294.1_Ken05/Tk1             .....c.....................g.....c.......................a.....g................ [199120]
KM262844.1_strain_L60            .................................c.............................................. [199120]
KM102979.1_26544/OG10_Italy      .................................c.............................................. [199120]
KM262845.1_strain_NHV            .................................c.............................................. [199120]
KX354450.1_47/Ss/2008            .................................c.............................................. [199120]
KP055815.1_strain_BA71           .................................c.............................................. [199120]
MH025920.1_strain_R35            .....c.....................g.....c.......................a.....g...........a.... [199120]
MH025919.1_strain_N10            .....c.....................g.....c.......................a.....g...........a.... [199120]
MH025918.1_strain_R25            .....c.....................g.....c.......................a.....g...........a.... [199120]
MH025917.1_strain_R7             .....c.....................g.....c.......................a.....g...........a.... [199120]
MH025916.1_strain_R8             .....c.....................g.....c.......................a.....g...........a.... [199120]

FR682468.2_Georgia_2007/1        gcgtccgcccgctaatggtacagttccaggttaagacaatcatatcacacccaaaaagagaggaaacagcataggtgccc [199200]
ASF_IND_20_CAD543                -------------------------------------------------------------------------------- [199200]
LR899193.1_Germany_2020/1        ................................................................................ [199200]
MN393477.1_Wuhan_2019            ................................................................................ [199200]
MW396979.1_Timor-Leste/2019/1    ................................................................................ [199200]
MK940252.1_CN/2019/InnerMongolia ................................................................................ [199200]
MK128995.1_China/2018/AnhuiXCGQ  ................................................................................ [199200]
MH681419.1_ASFV/POL/2015         ................................................................................ [199200]
LS478113.1_Estonia_2014          ................................................................................ [199200]
KP843857.1_Odintsovo_02/14       ................................................................................ [199200]
```

```
FN557520.1_strain_E75             ................................................................................ [199200]
AY261366.1_Warthog                ................................................................................ [199200]
AY261365.1_Warmbaths              .......................................c........................................ [199200]
AY261364.1_Tengani_62             .................a.............................................................. [199200]
AY261363.1_Pretoriuskop/96/4      .........................................................................a...... [199200]
AY261362.1_Mkuzi_1979             ................................................................................ [199200]
AY261361.1_Malawi_Lil-20/1_1983   ..................g..................g...........gt............................. [199200]
AY261360.1_Kenya_1950             .......t..........g....................c.........gt................g............ [199200]
AM712240.1_OURT_88/3              ................................................................................ [199200]
AM712239.1_Benin_97/1             ................................................................................ [199200]
KM111295.1_Ken06.Bus              .t................gg...................c.........gt................g............ [199200]
KM111294.1_Ken05/Tk1              ..................g....................c.........gt................g............ [199200]
KM262844.1_strain_L60             ................................................................................ [199200]
KM102979.1_26544/OG10_Italy       ................................................................................ [199200]
KM262845.1_strain_NHV             ................................................................................ [199200]
KX354450.1_47/Ss/2008             ................................................................................ [199200]
KP055815.1_strain_BA71            ................................................................................ [199200]
MH025920.1_strain_R35             .t................gg...................c.........gt................g............ [199200]
MH025919.1_strain_N10             .t................gg...................c.........gt................g............ [199200]
MH025918.1_strain_R25             .t................gg...................c.........gt................g............ [199200]
MH025917.1_strain_R7              .t................gg...................c.........gt................g............ [199200]
MH025916.1_strain_R8              .t................gg...................c.........gt................g............ [199200]

FR682468.2_Georgia_2007/1         aaaggttcattatataacatacgccgcatatattttagttttttttctccatggtaataatcacaggttttcatgtcctg [199280]
ASF_IND_20_CAD543                 -------------------------------------------------------------------------------- [199280]
LR899193.1_Germany_2020/1         ................................................................................ [199280]
MN393477.1_Wuhan_2019             ................................................................................ [199280]
MW396979.1_Timor-Leste/2019/1     ................................................................................ [199280]
MK940252.1_CN/2019/InnerMongolia  ................................................................................ [199280]
MK128995.1_China/2018/AnhuiXCGQ   ................................................................................ [199280]
MH681419.1_ASFV/POL/2015          ................................................................................ [199280]
LS478113.1_Estonia_2014           ................................................................................ [199280]
KP843857.1_Odintsovo_02/14        ......................................-......................................... [199280]
FN557520.1_strain_E75             ..................................................................t...........c. [199280]
AY261366.1_Warthog                ................................................................................ [199280]
AY261365.1_Warmbaths              ................................................................................ [199280]
AY261364.1_Tengani_62             .............g....................................................t............. [199280]
AY261363.1_Pretoriuskop/96/4      ................................................................................ [199280]
AY261362.1_Mkuzi_1979             ................................................................................ [199280]
AY261361.1_Malawi_Lil-20/1_1983   .....c......a.c.g..............g............g..c...aa......g......t............. [199280]
AY261360.1_Kenya_1950             ............a.c.g...........................a..a...........g......t............. [199280]
AM712240.1_OURT_88/3              ..................................................................t...........c. [199280]
AM712239.1_Benin_97/1             ..................................................................t...........c. [199280]
KM111295.1_Ken06.Bus              ............a.c.g...........................a..a..g........g......t............. [199280]
```

```
KM111294.1_Ken05/Tk1           ............a.c.g...........................a..a...........g......t............. [199280]
KM262844.1_strain_L60          ..................................................................t...........c. [199280]
KM102979.1_26544/OG10_Italy    ..................................................................t...........c. [199280]
KM262845.1_strain_NHV          ..................................................................t...........c. [199280]
KX354450.1_47/Ss/2008          ..................................................................t...........c. [199280]
KP055815.1_strain_BA71         ..................................................................t...........c. [199280]
MH025920.1_strain_R35          ............a.c.g...........................a..a..g........g......t............. [199280]
MH025919.1_strain_N10          ............a.c.g...........................a..a..g........g......t............. [199280]
MH025918.1_strain_R25          ............a.c.g...........................a..a..g........g......t............. [199280]
MH025917.1_strain_R7           ............a.c.g...........................a..a..g........g......t............. [199280]
MH025916.1_strain_R8           ............a.c.g...........................a..a..g........g......t............. [199280]

FR682468.2_Georgia_2007/1      cttaataggatgattccccatgtatgataatatataataaatttagtttttagctttttcaaaaaattgggcgctc---g [199360]
ASF_IND_20_CAD543              -------------------------------------------------------------------------------- [199360]
LR899193.1_Germany_2020/1      ............................................................................---. [199360]
MN393477.1_Wuhan_2019          ............................................................................---. [199360]
MW396979.1_Timor-Leste/2019/1  ............................................................................---. [199360]
MK940252.1_CN/2019/InnerMongolia ............................................................................---. [199360]
MK128995.1_China/2018/AnhuiXCGQ ............................................................................---. [199360]
MH681419.1_ASFV/POL/2015       ............................................................................---. [199360]
LS478113.1_Estonia_2014        ............................................................................---. [199360]
KP843857.1_Odintsovo_02/14     ............................................................................---. [199360]
FN557520.1_strain_E75          .....................................a.................................t....---. [199360]
AY261366.1_Warthog             .....................................a......................................gaa. [199360]
AY261365.1_Warmbaths           .....................................a......................................gaa. [199360]
AY261364.1_Tengani_62          .....................c...............a.................................t...t---. [199360]
AY261363.1_Pretoriuskop/96/4   .....................c...............a............................-..a......---. [199360]
AY261362.1_Mkuzi_1979          .....................................a.................................t....---. [199360]
AY261361.1_Malawi_Lil-20/1_1983 t..........t.........................a.....................tt..----------------a [199360]
AY261360.1_Kenya_1950          t..........t.........c.......c.......a......................-------------------a [199360]
AM712240.1_OURT_88/3           .....................................a.................................t....---. [199360]
AM712239.1_Benin_97/1          .....................................a.................................t....---. [199360]
KM111295.1_Ken06.Bus           t..........t.........c.......c.......a......................-------------------a [199360]
KM111294.1_Ken05/Tk1           t..........t.........c.......c.......a......................-------------------a [199360]
KM262844.1_strain_L60          .....................................a.................................t....---. [199360]
KM102979.1_26544/OG10_Italy    .....................................a.................................t....---. [199360]
KM262845.1_strain_NHV          .....................................a.................................t....---. [199360]
KX354450.1_47/Ss/2008          .....................................a.................................t....---. [199360]
KP055815.1_strain_BA71         .....................................a.................................t....---. [199360]
MH025920.1_strain_R35          t..........t.........c.......c.......a......................-------------------a [199360]
MH025919.1_strain_N10          t..........t.........c.......c.......a......................-------------------a [199360]
MH025918.1_strain_R25          t..........t.........c.......c.......a......................-------------------a [199360]
MH025917.1_strain_R7           t..........t.........c.......c.......a......................-------------------a [199360]
MH025916.1_strain_R8           t..........t.........c.......c.......a......................-------------------a [199360]
```

```
FR682468.2_Georgia_2007/1        aaactaaattttccttatcacagcgtttg---gagaaagcgtatttaaagat-atatcttcttctaaca-agactgc--- [199440]
ASF_IND_20_CAD543                -------------------------------------------------------------------------------- [199440]
LR899193.1_Germany_2020/1        .............................---....................-................-.......--- [199440]
MN393477.1_Wuhan_2019            .............................---....................-................-.......--- [199440]
MW396979.1_Timor-Leste/2019/1    .............................---....................-................-.......--- [199440]
MK940252.1_CN/2019/InnerMongolia .............................---....................-................-.......--- [199440]
MK128995.1_China/2018/AnhuiXCGQ  .............................---....................-................-.......--- [199440]
MH681419.1_ASFV/POL/2015         .............................---....................-................-.......--- [199440]
LS478113.1_Estonia_2014          .............................---....................-................-.......--- [199440]
KP843857.1_Odintsovo_02/14       .............................---....................-................-.......--- [199440]
FN557520.1_strain_E75            .......................t.....---.......t............-.....---......t.-.......at- [199440]
AY261366.1_Warthog               .............................---a...................-................-.......--- [199440]
AY261365.1_Warmbaths             .............................---a...................-................-.......--- [199440]
AY261364.1_Tengani_62            .............................---....................-.....---........-.......a-- [199440]
AY261363.1_Pretoriuskop/96/4     ............t................---....................-...---..........-.......--- [199440]
AY261362.1_Mkuzi_1979            .......................t.....---.......t............-.....---......t.-.......at- [199440]
AY261361.1_Malawi_Lil-20/1_1983  ..........................gcagtt..agg..a..t.........g.....---.at.....g...t...at- [199440]
AY261360.1_Kenya_1950            ..........................g.a-----.....aa....a......gt...a-....t..g.g-g...------ [199440]
AM712240.1_OURT_88/3             .......................t.....---.......t............-.....---......t.-.......at- [199440]
AM712239.1_Benin_97/1            .......................t.....---.......t............-.....---......t.-.......at- [199440]
KM111295.1_Ken06.Bus             ..........................g.a-----.-...aa...........gt...a-....t..g.g-g...------ [199440]
KM111294.1_Ken05/Tk1             ..........................g.a-----.....aa...........gt...a-....t..g.g-g...------ [199440]
KM262844.1_strain_L60            .......................t.....---.......t............-......--......t.-.......ata [199440]
KM102979.1_26544/OG10_Italy      .......................t.....---.......t............-.....---......t.-.......at- [199440]
KM262845.1_strain_NHV            .......................t.....---.......t............-.....---......t.-.......at- [199440]
KX354450.1_47/Ss/2008            .......................t.....---.......t............-.....---......t.-.......at- [199440]
KP055815.1_strain_BA71           .......................t.....---.......t............-.....---......t.-.......at- [199440]
MH025920.1_strain_R35            ..........................g.a-----.-...aa...........gt...a-....t..g.g-g...------ [199440]
MH025919.1_strain_N10            ..........................g.a-----.-...aa...........gt...a-....t..g.g-g...------ [199440]
MH025918.1_strain_R25            ..........................g.a-----.-...aa...........gt...a-....t..g.g-g...------ [199440]
MH025917.1_strain_R7             ..........................g.a-----.-...aa...........gt...a-....t..g.g-g...------ [199440]
MH025916.1_strain_R8             ..........................g.a-----.-...aa...........gt...a-....t..g.g-g...------ [199440]

FR682468.2_Georgia_2007/1        aaaaaaaa-tcttacccctta---tttttataat------gttcatcatagcgtttgaagatat-----cagaaggtg-- [199520]
ASF_IND_20_CAD543                -------------------------------------------------------------------------------- [199520]
LR899193.1_Germany_2020/1        ........-............---..........------........................-----.........-- [199520]
MN393477.1_Wuhan_2019            ........-............---..........------........................-----.........-- [199520]
MW396979.1_Timor-Leste/2019/1    ........-............---..........------........................-----.........-- [199520]
MK940252.1_CN/2019/InnerMongolia ........-............---..........------........................-----.........-- [199520]
MK128995.1_China/2018/AnhuiXCGQ  ........-............---..........------........................-----.........-- [199520]
MH681419.1_ASFV/POL/2015         ........-............---..........------........................-----.........-- [199520]
LS478113.1_Estonia_2014          ........-............---..........------........................-----.........-- [199520]
KP843857.1_Odintsovo_02/14       ........-............---..........------........................-----.........-- [199520]
```

```
FN557520.1_strain_E75           ........-.tc.ta......---..........gttgta......a.c............c..-----....g....gc [199520]
AY261366.1_Warthog              ........-.......a....---..........------........................-----....g.tgc-c [199520]
AY261365.1_Warmbaths            ........-.......a....---..........------........................-----.........-c [199520]
AY261364.1_Tengani_62           ........t.......a....---..........gtt---ca...............g...c..-----....g....gc [199520]
AY261363.1_Pretoriuskop/96/4    ........t.......a....---..........------....................g.g----------------c [199520]
AY261362.1_Mkuzi_1979           ........-.tc.ta......---..........gttgta......a.c............c..-----....g....gc [199520]
AY261361.1_Malawi_Lil-20/1_1983 .g....tg-at.c..ta....---..........attgta......a....ta..c..t..c..attagt...t..ct-g [199520]
AY261360.1_Kenya_1950           cc......t...c.t.a....ttt........t.gttgta......a.c......------------------------- [199520]
AM712240.1_OURT_88/3            ........-.tc.ta......---..........gttgta......a.c............c..-----....g....gc [199520]
AM712239.1_Benin_97/1           -.......-.tc.ta......---..........gttgta......a.c............c..-----....g....gc [199520]
KM111295.1_Ken06.Bus            cc......t...c.t.a....ttt.....t..t.gttgta......a.c......------------------------- [199520]
KM111294.1_Ken05/Tk1            cc......t...c.t.a....ttt........t.gttgta......a.c......------------------------- [199520]
KM262844.1_strain_L60           ........-.tc.ta......---..........gttgta......a.c............c..-----....g....gc [199520]
KM102979.1_26544/OG10_Italy     ........-.tc.ta......---..........gttgta......a.c............c..-----....g....gc [199520]
KM262845.1_strain_NHV           ........-.tc.ta......---..........gttgta......a.c............c..-----....g....gc [199520]
KX354450.1_47/Ss/2008           -.......-.tc.ta......---..........gttgta......a.c............c..-----....g....gc [199520]
KP055815.1_strain_BA71          ........-.tc.ta......---..........gttgta......a.c............c..-----....g....gc [199520]
MH025920.1_strain_R35           cc......t...c.t.a....-tt.....t..t.gttgta......a.c......------------------------- [199520]
MH025919.1_strain_N10           cc......t...c.t.a....--t.....t..t.gttgta......a.c......------------------------- [199520]
MH025918.1_strain_R25           cc......t...c.t.a....-tt.....t..t.gttgta......a.c......------------------------- [199520]
MH025917.1_strain_R7            cc......t...c.t.a....-tt.....t..t.gttgta......a.c......------------------------- [199520]
MH025916.1_strain_R8            cc......t...c.t.a....-tt.....t..t.gttgta......a.c......------------------------- [199520]

FR682468.2_Georgia_2007/1       ccaggttttataaaaatatcctttaggatttataacgatacaagggtctataaaatatatgcgggtataatcttataaaa [199600]
ASF_IND_20_CAD543               -------------------------------------------------------------------------------- [199600]
LR899193.1_Germany_2020/1       ................................................................................ [199600]
MN393477.1_Wuhan_2019           ................................................................................ [199600]
MW396979.1_Timor-Leste/2019/1   ................................................................................ [199600]
MK940252.1_CN/2019/InnerMongolia ................................................................................ [199600]
MK128995.1_China/2018/AnhuiXCGQ ................................................................................ [199600]
MH681419.1_ASFV/POL/2015        ................................................................................ [199600]
LS478113.1_Estonia_2014         ................................................................................ [199600]
KP843857.1_Odintsovo_02/14      ................................................................................ [199600]
FN557520.1_strain_E75           ................................................................................ [199600]
AY261366.1_Warthog              .t.............c................................................................ [199600]
AY261365.1_Warmbaths            .....g.......................................................................... [199600]
AY261364.1_Tengani_62           t............................................................................... [199600]
AY261363.1_Pretoriuskop/96/4    t............................................................................... [199600]
AY261362.1_Mkuzi_1979           ................................................................................ [199600]
AY261361.1_Malawi_Lil-20/1_1983 .t.............tat..a.....................................g..t..........g....... [199600]
AY261360.1_Kenya_1950           -----------------------.....c...........t..a.................................... [199600]
AM712240.1_OURT_88/3            ................................................................................ [199600]
AM712239.1_Benin_97/1           ................................................................................ [199600]
KM111295.1_Ken06.Bus            -----------------------.....c...........t..a.................................... [199600]
```

```
KM111294.1_Ken05/Tk1           -----------------------.....c...........t..a.................................... [199600]
KM262844.1_strain_L60          ................................................................................ [199600]
KM102979.1_26544/OG10_Italy    ................................................................................ [199600]
KM262845.1_strain_NHV          ................................................................................ [199600]
KX354450.1_47/Ss/2008          ................................................................................ [199600]
KP055815.1_strain_BA71         ................................................................................ [199600]
MH025920.1_strain_R35          -----------------------.....c...........t..a.................................... [199600]
MH025919.1_strain_N10          -----------------------.....c...........t..a.................................... [199600]
MH025918.1_strain_R25          -----------------------.....c...........t..a.................................... [199600]
MH025917.1_strain_R7           -----------------------.....c...........t..a.................................... [199600]
MH025916.1_strain_R8           -----------------------.....c...........t..a.................................... [199600]

FR682468.2_Georgia_2007/1      tcatcgattttttcataatattctccgtttatacaataaagatcataacagatattgatgcgtagatgcattattcgcgt [199680]
ASF_IND_20_CAD543              -------------------------------------------------------------------------------- [199680]
LR899193.1_Germany_2020/1      ................................................................................ [199680]
MN393477.1_Wuhan_2019          ................................................................................ [199680]
MW396979.1_Timor-Leste/2019/1  ................................................................................ [199680]
MK940252.1_CN/2019/InnerMongolia ................................................................................ [199680]
MK128995.1_China/2018/AnhuiXCGQ ................................................................................ [199680]
MH681419.1_ASFV/POL/2015       ................................................................................ [199680]
LS478113.1_Estonia_2014        ................................................................................ [199680]
KP843857.1_Odintsovo_02/14     ................................................................................ [199680]
FN557520.1_strain_E75          ................................................................................ [199680]
AY261366.1_Warthog             ................................................................................ [199680]
AY261365.1_Warmbaths           ..........................................................................c..... [199680]
AY261364.1_Tengani_62          ................................................................................ [199680]
AY261363.1_Pretoriuskop/96/4   ................................................................................ [199680]
AY261362.1_Mkuzi_1979          ................................................................................ [199680]
AY261361.1_Malawi_Lil-20/1_1983 ..........................a......t.............................................. [199680]
AY261360.1_Kenya_1950          .................................t..........g.......c...........a............... [199680]
AM712240.1_OURT_88/3           ................................................................................ [199680]
AM712239.1_Benin_97/1          ................................................................................ [199680]
KM111295.1_Ken06.Bus           .................................t..........g.......c...........a............... [199680]
KM111294.1_Ken05/Tk1           .................................t..........g.......c...........a............... [199680]
KM262844.1_strain_L60          ................................................................................ [199680]
KM102979.1_26544/OG10_Italy    ................................................................................ [199680]
KM262845.1_strain_NHV          ................................................................................ [199680]
KX354450.1_47/Ss/2008          ................................................................................ [199680]
KP055815.1_strain_BA71         ................................................................................ [199680]
MH025920.1_strain_R35          .................................t..........g.......c...........a............... [199680]
MH025919.1_strain_N10          .................................t..........g.......c...........a...t........... [199680]
MH025918.1_strain_R25          .................................t..........g.......c...........a............... [199680]
MH025917.1_strain_R7           .................................t..........g.......c...........a............... [199680]
MH025916.1_strain_R8           .................................t..........g.......c...........a............... [199680]
```

```
FR682468.2_Georgia_2007/1        gttcgttgggcagctaaaggatatcacaacgtagttttttttaagaaaagacgaaactacataagtccctaagggttcat [199760]
ASF_IND_20_CAD543                -------------------------------------------------------------------------------- [199760]
LR899193.1_Germany_2020/1        ................................................................................ [199760]
MN393477.1_Wuhan_2019            ................................................................................ [199760]
MW396979.1_Timor-Leste/2019/1    ................................................................................ [199760]
MK940252.1_CN/2019/InnerMongolia ................................................................................ [199760]
MK128995.1_China/2018/AnhuiXCGQ  ................................................................................ [199760]
MH681419.1_ASFV/POL/2015         ................................................................................ [199760]
LS478113.1_Estonia_2014          ................................................................................ [199760]
KP843857.1_Odintsovo_02/14       ................................................................................ [199760]
FN557520.1_strain_E75            ................................................................................ [199760]
AY261366.1_Warthog               ................................................................................ [199760]
AY261365.1_Warmbaths             ................................................................................ [199760]
AY261364.1_Tengani_62            ..............................a....................t............................ [199760]
AY261363.1_Pretoriuskop/96/4     ................................................................................ [199760]
AY261362.1_Mkuzi_1979            ................................................................................ [199760]
AY261361.1_Malawi_Lil-20/1_1983  .........a..a....................t..............g..............g................ [199760]
AY261360.1_Kenya_1950            ..........................................g..............c...................... [199760]
AM712240.1_OURT_88/3             ................................................................................ [199760]
AM712239.1_Benin_97/1            ................................................................................ [199760]
KM111295.1_Ken06.Bus             ..........................................g..............c...................... [199760]
KM111294.1_Ken05/Tk1             ..........................................g..............c...................... [199760]
KM262844.1_strain_L60            ................................................................................ [199760]
KM102979.1_26544/OG10_Italy      ................................................................................ [199760]
KM262845.1_strain_NHV            ................................................................................ [199760]
KX354450.1_47/Ss/2008            ................................................................................ [199760]
KP055815.1_strain_BA71           ................................................................................ [199760]
MH025920.1_strain_R35            ..........................................g..............c...................... [199760]
MH025919.1_strain_N10            ..........................................g..............c...................... [199760]
MH025918.1_strain_R25            ..........................................g..............c...................... [199760]
MH025917.1_strain_R7             ..........................................g..............c...................... [199760]
MH025916.1_strain_R8             ..........................................g..............c...................... [199760]

FR682468.2_Georgia_2007/1        tgaatagtaaacgccatatttgttttaaattttgttgttcaccatagtagtattcgcactttttcaagtcttttttaata [199840]
ASF_IND_20_CAD543                -------------------------------------------------------------------------------- [199840]
LR899193.1_Germany_2020/1        ................................................................................ [199840]
MN393477.1_Wuhan_2019            ................................................................................ [199840]
MW396979.1_Timor-Leste/2019/1    ................................................................................ [199840]
MK940252.1_CN/2019/InnerMongolia ................................................................................ [199840]
MK128995.1_China/2018/AnhuiXCGQ  ................................................................................ [199840]
MH681419.1_ASFV/POL/2015         ................................................................................ [199840]
LS478113.1_Estonia_2014          ................................................................................ [199840]
KP843857.1_Odintsovo_02/14       ................................................................................ [199840]
```

```
FN557520.1_strain_E75              ................................................................................ [199840]
AY261366.1_Warthog                 ....c........................................................................... [199840]
AY261365.1_Warmbaths               ................................................................................ [199840]
AY261364.1_Tengani_62              ................................................................................ [199840]
AY261363.1_Pretoriuskop/96/4       ................................................................................ [199840]
AY261362.1_Mkuzi_1979              ................................................................................ [199840]
AY261361.1_Malawi_Lil-20/1_1983    ................................................................................ [199840]
AY261360.1_Kenya_1950              ..................................................................g...........c. [199840]
AM712240.1_OURT_88/3               ................................................................................ [199840]
AM712239.1_Benin_97/1              ................................................................................ [199840]
KM111295.1_Ken06.Bus               ...................c..............................................g...........c. [199840]
KM111294.1_Ken05/Tk1               ..................................................................g...........c. [199840]
KM262844.1_strain_L60              ................................................................................ [199840]
KM102979.1_26544/OG10_Italy        ................................................................................ [199840]
KM262845.1_strain_NHV              ................................................................................ [199840]
KX354450.1_47/Ss/2008              ................................................................................ [199840]
KP055815.1_strain_BA71             ................................................................................ [199840]
MH025920.1_strain_R35              ...................c..............................................g...........c. [199840]
MH025919.1_strain_N10              ...................c..............................................g...........c. [199840]
MH025918.1_strain_R25              ...................c..............................................g...........c. [199840]
MH025917.1_strain_R7               ...................c..............................................g...........c. [199840]
MH025916.1_strain_R8               ...................c..............................................g...........c. [199840]

FR682468.2_Georgia_2007/1          agcctattccccatgtatgcttataaataaaaatttagaaatgtgctatattatttgttgatgaatcatgaacacgtctt [199920]
ASF_IND_20_CAD543                  -------------------------------------------------------------------------------- [199920]
LR899193.1_Germany_2020/1          ................................................................................ [199920]
MN393477.1_Wuhan_2019              ................................................................................ [199920]
MW396979.1_Timor-Leste/2019/1      ................................................................................ [199920]
MK940252.1_CN/2019/InnerMongolia   ................................................................................ [199920]
MK128995.1_China/2018/AnhuiXCGQ    ................................................................................ [199920]
MH681419.1_ASFV/POL/2015           ................................................................................ [199920]
LS478113.1_Estonia_2014            ................................................................................ [199920]
KP843857.1_Odintsovo_02/14         ................................................................................ [199920]
FN557520.1_strain_E75              .........................................--.......................g............. [199920]
AY261366.1_Warthog                 .........................................--.......................g..t.......... [199920]
AY261365.1_Warmbaths               .........................................--.......................g..t.......... [199920]
AY261364.1_Tengani_62              .........................................--..................-------.t.......... [199920]
AY261363.1_Pretoriuskop/96/4       .........................................--.......................g..t.......... [199920]
AY261362.1_Mkuzi_1979              .........................................--.......................g............. [199920]
AY261361.1_Malawi_Lil-20/1_1983    ..............t....t.....g.....g.........--.a.................a...-----......... [199920]
AY261360.1_Kenya_1950              ........t.....t..c.t...........g............ta.......c...........ag..t.......... [199920]
AM712240.1_OURT_88/3               .........................................--.......................g............. [199920]
AM712239.1_Benin_97/1              .........................................--.......................g............. [199920]
KM111295.1_Ken06.Bus               ........t.....t..c.t........--.g...........cta.......c...........ag..t.......... [199920]
```

```
KM111294.1_Ken05/Tk1           ........t.....t..c.t...........g............ta.......c...........ag..t.......... [199920]
KM262844.1_strain_L60          .........................................--.......................g............. [199920]
KM102979.1_26544/OG10_Italy    .........................................--.......................g............. [199920]
KM262845.1_strain_NHV          .........................................--.......................g............. [199920]
KX354450.1_47/Ss/2008          .........................................--.......................g............. [199920]
KP055815.1_strain_BA71         .........................................--.......................g............. [199920]
MH025920.1_strain_R35          ........t.....t..c.t.......c--.g...........cta.......c...........ag..t.......... [199920]
MH025919.1_strain_N10          ........t.....t..c.t.......c--.g...........cta.......c...........ag..t.......... [199920]
MH025918.1_strain_R25          ........t.....t..c.t.......c--.g...........cta.......c...........ag..t.......... [199920]
MH025917.1_strain_R7           ........t.....t..c.t.......c--.g...........cta.......c...........ag..t.......... [199920]
MH025916.1_strain_R8           ........t.....t..c.t.......c--.g...........cta.......c...........ag..t.......... [199920]

FR682468.2_Georgia_2007/1      atatgttgatatgttac---tttaaaaacatttgtatt------------ttcaacagacgc-gttctattcttattaag [200000]
ASF_IND_20_CAD543              -------------------------------------------------------------------------------- [200000]
LR899193.1_Germany_2020/1      .................---..................------------............-................. [200000]
MN393477.1_Wuhan_2019          .................---..................------------............-................. [200000]
MW396979.1_Timor-Leste/2019/1  .................---..................------------............-................. [200000]
MK940252.1_CN/2019/InnerMongolia .................---..................------------............-................. [200000]
MK128995.1_China/2018/AnhuiXCGQ .................---..................------------............-................. [200000]
MH681419.1_ASFV/POL/2015       .................---..................------------............-................. [200000]
LS478113.1_Estonia_2014        .................---..................------------............-................. [200000]
KP843857.1_Odintsovo_02/14     .................---..................------------............-................. [200000]
FN557520.1_strain_E75          ............a....---..................------------.........t..-.a............... [200000]
AY261366.1_Warthog             ...c...--------..---..................------------.....tc.....a................. [200000]
AY261365.1_Warmbaths           .......--------..---..................------------......c.....a................. [200000]
AY261364.1_Tengani_62          ............a....---...c..............------------......c.....a................. [200000]
AY261363.1_Pretoriuskop/96/4   .......--------..---..................------------......c.....a................. [200000]
AY261362.1_Mkuzi_1979          ............a....---..................------------............-................. [200000]
AY261361.1_Malawi_Lil-20/1_1983 .a..........a....ttt..........aa...t..tacaacatttgt..t....a...ta.............c.g. [200000]
AY261360.1_Kenya_1950          ......a...g.a..cg---........t.....c...g-------------.........tg.....-.......c.ga [200000]
AM712240.1_OURT_88/3           ............a....---..................------------.........t..-.a............... [200000]
AM712239.1_Benin_97/1          ............a....---..................------------.........t..-.a............... [200000]
KM111295.1_Ken06.Bus           ......a...g.t..c----........t.....c...g-------------.........tg.....-.......c.ga [200000]
KM111294.1_Ken05/Tk1           ......a...g.a..cg---........t.....c...g-------------.........tg.....-.......c.ga [200000]
KM262844.1_strain_L60          ............a....---..................------------.........t..-.a............... [200000]
KM102979.1_26544/OG10_Italy    ............a....---..................------------.........t..-.a............... [200000]
KM262845.1_strain_NHV          ............a....---..................------------.........t..-.a............... [200000]
KX354450.1_47/Ss/2008          ............a....---..................------------.........t..-.a............... [200000]
KP055815.1_strain_BA71         ............a....---..................------------.........t..-.a............... [200000]
MH025920.1_strain_R35          ......a...g.t..c----........t.....c...g-------------.........tg.....-.......c.ga [200000]
MH025919.1_strain_N10          ......a...g.t..c----........t.....c...g-------------.........tg.....-.......c.ga [200000]
MH025918.1_strain_R25          ......a...g.t..c----........t.....c...g-------------.........tg.....-.......c.ga [200000]
MH025917.1_strain_R7           ......a...g.t..c----........t.....c...g-------------.........tg.....-.......c.ga [200000]
MH025916.1_strain_R8           ......a...g.t..c----........t.....c...g-------------.........tg.....-.......c.ga [200000]
```

```
FR682468.2_Georgia_2007/1         aatgatgccg-------tctttattttaaaccttt----------------ggtttaaaatttaaagaagtatttataaac [200080]
ASF_IND_20_CAD543                 -------------------------------------------------------------------------------- [200080]
LR899193.1_Germany_2020/1         ..........-------.................----------------.............................. [200080]
MN393477.1_Wuhan_2019             ..........-------.................----------------.............................. [200080]
MW396979.1_Timor-Leste/2019/1     ..........-------.................----------------.............................. [200080]
MK940252.1_CN/2019/InnerMongolia  ..........-------.................----------------.............................. [200080]
MK128995.1_China/2018/AnhuiXCGQ   ..........-------.................----------------.............................. [200080]
MH681419.1_ASFV/POL/2015          ..........-------.................----------------.............................. [200080]
LS478113.1_Estonia_2014           ..........-------.................----------------.............................. [200080]
KP843857.1_Odintsovo_02/14        ..........-------.................----------------.............................. [200080]
FN557520.1_strain_E75             ..........-------.................----------------.............................. [200080]
AY261366.1_Warthog                ..........-------...............c.t-------------aa.............................a [200080]
AY261365.1_Warmbaths              ..........-------...............c.t-------------aa.............................a [200080]
AY261364.1_Tengani_62             ..........-------...............c.t-------------aa.............................a [200080]
AY261363.1_Pretoriuskop/96/4      ..........-------...............c.t-------------aa.............................a [200080]
AY261362.1_Mkuzi_1979             ..........-------.................----------------.............................. [200080]
AY261361.1_Malawi_Lil-20/1_1983   ..........aaatgcc.tc.a............aaggtttaaaatttaa..g..........................a [200080]
AY261360.1_Kenya_1950             ....tcaaaa---ctcg..c..............--------------aaa............................a [200080]
AM712240.1_OURT_88/3              ..........-------.................----------------.............................. [200080]
AM712239.1_Benin_97/1             ..........-------.................----------------.............................. [200080]
KM111295.1_Ken06.Bus              ...ctcaaaa---ctcg..c....g.........--------------aaa............................a [200080]
KM111294.1_Ken05/Tk1              ....tcaaaa---ctcg..c..............--------------aaa............................a [200080]
KM262844.1_strain_L60             ..........-------.................----------------.............................. [200080]
KM102979.1_26544/OG10_Italy       ..........-------.................----------------.............................. [200080]
KM262845.1_strain_NHV             ..........-------.................----------------.............................. [200080]
KX354450.1_47/Ss/2008             ..........-------.................----------------.............................. [200080]
KP055815.1_strain_BA71            ..........-------.................----------------.............................. [200080]
MH025920.1_strain_R35             ...ctcaaaa---ctcg..c....g.........--------------aaa............................a [200080]
MH025919.1_strain_N10             ...ctcaaaa---ctcg..c....g.........--------------aaa............................a [200080]
MH025918.1_strain_R25             ...ctcaaaa---ctcg..c....g.........--------------aaa............................a [200080]
MH025917.1_strain_R7              ...ctcaaaa---ctcg..c....g.........--------------aaa............................a [200080]
MH025916.1_strain_R8              ...ctcaaaa---ctcg..c....g.........--------------aaa............................a [200080]

FR682468.2_Georgia_2007/1         tataatcatgggaactttttcagtaactgcctctgcaaaaagtgacgatgctgtttgtaagtatttagaagaaccaatag [200160]
ASF_IND_20_CAD543                 -------------------------------------------------------------------------------- [200160]
LR899193.1_Germany_2020/1         ................................................................................ [200160]
MN393477.1_Wuhan_2019             ................................................................................ [200160]
MW396979.1_Timor-Leste/2019/1     ................................................................................ [200160]
MK940252.1_CN/2019/InnerMongolia  ................................................................................ [200160]
MK128995.1_China/2018/AnhuiXCGQ   ................................................................................ [200160]
MH681419.1_ASFV/POL/2015          ................................................................................ [200160]
LS478113.1_Estonia_2014           ................................................................................ [200160]
KP843857.1_Odintsovo_02/14        ................................................................................ [200160]
```

```
FN557520.1_strain_E75           ...........a...c....................g...........................c............... [200160]
AY261366.1_Warthog              ...............c................................................................ [200160]
AY261365.1_Warmbaths            ...............c............................................................t... [200160]
AY261364.1_Tengani_62           ...............c...........................................g.................... [200160]
AY261363.1_Pretoriuskop/96/4    ...............c............................g................................... [200160]
AY261362.1_Mkuzi_1979           ...........a...c....................g...........................c.....a......... [200160]
AY261361.1_Malawi_Lil-20/1_1983 .....c.....a...c..............................................t.........t.....c. [200160]
AY261360.1_Kenya_1950           .....c..................g..g..t..........a....a....c...............a..........g. [200160]
AM712240.1_OURT_88/3            ...........a...c....................g...........................c............... [200160]
AM712239.1_Benin_97/1           ...........a...c....................g...........................c............... [200160]
KM111295.1_Ken06.Bus            .....c..................g..g..t..........a....a....c...............a..........g. [200160]
KM111294.1_Ken05/Tk1            .....c..................g..g..tt.........a....a....c...............a..........ga [200160]
KM262844.1_strain_L60           ...........a...c....................g...........................c............... [200160]
KM102979.1_26544/OG10_Italy     ...........a...c....................g...........................c............... [200160]
KM262845.1_strain_NHV           ...........a...c....................g...........................c............... [200160]
KX354450.1_47/Ss/2008           ...........a...c....................g...........................c............... [200160]
KP055815.1_strain_BA71          ...........a...c....................g...........................c............... [200160]
MH025920.1_strain_R35           .....c..................g..g..t..........a....a....c...............a..........g. [200160]
MH025919.1_strain_N10           .....c..................g..g..t..........a....a....c...............a..........g. [200160]
MH025918.1_strain_R25           .....c..................g..g..t..........a....a....c...............a..........g. [200160]
MH025917.1_strain_R7            .....c..................g..g..t..........a....a....c...............a..........g. [200160]
MH025916.1_strain_R8            .....c..................g..g..t..........a....a....c...............a..........g. [200160]

FR682468.2_Georgia_2007/1       atg------aaaattacagaaacatattaagaaatgagcatgttaaaaaaaatttaaatgaggctctgaatcgacatatt [200240]
ASF_IND_20_CAD543               -------------------------------------------------------------------------------- [200240]
LR899193.1_Germany_2020/1       ...------....................................................................... [200240]
MN393477.1_Wuhan_2019           ...------....................................................................... [200240]
MW396979.1_Timor-Leste/2019/1   ...------....................................................................... [200240]
MK940252.1_CN/2019/InnerMongolia ...------....................................................................... [200240]
MK128995.1_China/2018/AnhuiXCGQ ...------....................................................................... [200240]
MH681419.1_ASFV/POL/2015        ...------....................................................................... [200240]
LS478113.1_Estonia_2014         ...------....................................................................... [200240]
KP843857.1_Odintsovo_02/14      ...------....................................................................... [200240]
FN557520.1_strain_E75           ...aaaata......ct....................a....g.........a..........................a [200240]
AY261366.1_Warthog              ...------....................................................................... [200240]
AY261365.1_Warmbaths            ...------....................................................................... [200240]
AY261364.1_Tengani_62           ...------............................a....g.........a........................... [200240]
AY261363.1_Pretoriuskop/96/4    ...------...............................c.g.........a........................... [200240]
AY261362.1_Mkuzi_1979           ...aaaata.......t....c...............a....g.........a..........................a [200240]
AY261361.1_Malawi_Lil-20/1_1983 ...aaaata.g.......a..c....................a..................t.....a..a.a..c.... [200240]
AY261360.1_Kenya_1950           t..aaaata.........a.........------........a...............c..t.....acg..a......c [200240]
AM712240.1_OURT_88/3            ...aaaata......ct....................a....g.........a..........................a [200240]
AM712239.1_Benin_97/1           ...aaaata......ct....................a....g.........a..........................a [200240]
KM111295.1_Ken06.Bus            t..aaaata.........a.......................a..................t.....acg..a......c [200240]
```

```
KM111294.1_Ken05/Tk1             t..aaaata.........a.........------........a..................t.....acg..a......c [200240]
KM262844.1_strain_L60            ...aaaata......ct.....................a...g.........a..........................a [200240]
KM102979.1_26544/OG10_Italy      ...aaaata......ct.....................a...g.........a..........................a [200240]
KM262845.1_strain_NHV            ...aaaata......ct.....................a...g.........a..........................a [200240]
KX354450.1_47/Ss/2008            ...aaaata......ct.....................a...g.........a..........................a [200240]
KP055815.1_strain_BA71           ...aaaata......ct.....................a...g.........a..........................a [200240]
MH025920.1_strain_R35            t..aaaata.........a.......................a..................t.....acg..a......c [200240]
MH025919.1_strain_N10            t..aaaata.........a.......................a..................t.....acg..a......c [200240]
MH025918.1_strain_R25            t..aaaata.........a.......................a..................t.....acg..a......c [200240]
MH025917.1_strain_R7             t..aaaata.........a.......................a..................t.....acg..a......c [200240]
MH025916.1_strain_R8             t..aaaata.........a.......................a..................t.....acg..a......c [200240]

FR682468.2_Georgia_2007/1        actacctataatccagtagttgattggtgtaataactattcaacattttcatctcaggatttcgatgaatataaaattta [200320]
ASF_IND_20_CAD543                -------------------------------------------------------------------------------- [200320]
LR899193.1_Germany_2020/1        ................................................................................ [200320]
MN393477.1_Wuhan_2019            ................................................................................ [200320]
MW396979.1_Timor-Leste/2019/1    ................................................................................ [200320]
MK940252.1_CN/2019/InnerMongolia ................................................................................ [200320]
MK128995.1_China/2018/AnhuiXCGQ  ................................................................................ [200320]
MH681419.1_ASFV/POL/2015         ................................................................................ [200320]
LS478113.1_Estonia_2014          ................................................................................ [200320]
KP843857.1_Odintsovo_02/14       ................................................................................ [200320]
FN557520.1_strain_E75            .t.g.t........................................a.........t...................... [200320]
AY261366.1_Warthog               ................................................................................ [200320]
AY261365.1_Warmbaths             ................................................................................ [200320]
AY261364.1_Tengani_62            .t.............................................................................. [200320]
AY261363.1_Pretoriuskop/96/4     .t...t..........................c............................................... [200320]
AY261362.1_Mkuzi_1979            .t...tc.......................................a.........t...................... [200320]
AY261361.1_Malawi_Lil-20/1_1983  c...t.c...........a..................................a..a....................... [200320]
AY261360.1_Kenya_1950            ...gttc.................................................a.....t..g.............. [200320]
AM712240.1_OURT_88/3             .t.g.t........................................a.........t...................... [200320]
AM712239.1_Benin_97/1            .t.g.t........................................a.........t...................... [200320]
KM111295.1_Ken06.Bus             ...gttc.......................................................t..g.............. [200320]
KM111294.1_Ken05/Tk1             ...gttc............................t..........................t..g.............. [200320]
KM262844.1_strain_L60            .t.g.t........................................a.........t...................... [200320]
KM102979.1_26544/OG10_Italy      .t.g.t........................................a.........t...................... [200320]
KM262845.1_strain_NHV            .t.g.t........................................a.........t...................... [200320]
KX354450.1_47/Ss/2008            .t.g.t........................................a.........t...................... [200320]
KP055815.1_strain_BA71           .t.g.t........................................a.........t...................... [200320]
MH025920.1_strain_R35            ...gttc.......................................................t..g.............. [200320]
MH025919.1_strain_N10            ...gttc.......................................................t..g.............. [200320]
MH025918.1_strain_R25            ...gttc.......................................................t..g.............. [200320]
MH025917.1_strain_R7             ...gttc.......................................................t..g.............. [200320]
MH025916.1_strain_R8             ...gttc.......................................................t..g.............. [200320]
```

```
FR682468.2_Georgia_2007/1        tatacatagcgatcttatggatggacgacctcgtccaaaaaaaacatggtgtgtcatcatgtaatgtttgttagt----- [200400]
ASF_IND_20_CAD543                -------------------------------------------------------------------------------- [200400]
LR899193.1_Germany_2020/1        ...........................................................................----- [200400]
MN393477.1_Wuhan_2019            ...........................................................................----- [200400]
MW396979.1_Timor-Leste/2019/1    ...........................................................................----- [200400]
MK940252.1_CN/2019/InnerMongolia ...........................................................................----- [200400]
MK128995.1_China/2018/AnhuiXCGQ  ...........................................................................----- [200400]
MH681419.1_ASFV/POL/2015         ........................................g..................................----- [200400]
LS478113.1_Estonia_2014          ...........................................................................----- [200400]
KP843857.1_Odintsovo_02/14       ...........................................................................----- [200400]
FN557520.1_strain_E75            ....................................................................c...t..----- [200400]
AY261366.1_Warthog               ......................................................a.............c......----- [200400]
AY261365.1_Warmbaths             ......................................................a.............ca.....----- [200400]
AY261364.1_Tengani_62            ......................................................a.............c......----- [200400]
AY261363.1_Pretoriuskop/96/4     .....................c................................a.............ca.....----- [200400]
AY261362.1_Mkuzi_1979            ....................................................................c......----- [200400]
AY261361.1_Malawi_Lil-20/1_1983  .......g.t.....................a......................a..........a..c......----- [200400]
AY261360.1_Kenya_1950            ......cg.t.....................a......................a.....a...---.c......----- [200400]
AM712240.1_OURT_88/3             ....................................................................c...t..----- [200400]
AM712239.1_Benin_97/1            ....................................................................c...t..----- [200400]
KM111295.1_Ken06.Bus             .........t.....................a......................a..ac......a.........aatga [200400]
KM111294.1_Ken05/Tk1             ......cg.t.....................a......................a..........a..c......----- [200400]
KM262844.1_strain_L60            ............................................ac......................c...t..----- [200400]
KM102979.1_26544/OG10_Italy      ....................................................................c...t..----- [200400]
KM262845.1_strain_NHV            ....................................................................c...t..----- [200400]
KX354450.1_47/Ss/2008            ....................................................................c...t..----- [200400]
KP055815.1_strain_BA71           ....................................................................c...t..----- [200400]
MH025920.1_strain_R35            .........t.....................a......................a..ac......a.........aatga [200400]
MH025919.1_strain_N10            .........t.....................a......................a..ac......a.........aatga [200400]
MH025918.1_strain_R25            .........t.....................a......................a..ac......a.........aatga [200400]
MH025917.1_strain_R7             .........t.....................a......................a..ac......a.........aatga [200400]
MH025916.1_strain_R8             .........t.....................a......................a..ac......a.........aatga [200400]

FR682468.2_Georgia_2007/1        -tttatataaacgcaaaaatattcttctaggagatgttgatatactacctattgaattcaatat----attaaagtacat [200480]
ASF_IND_20_CAD543                -------------------------------------------------------------------------------- [200480]
LR899193.1_Germany_2020/1        -...............................................................----............ [200480]
MN393477.1_Wuhan_2019            -...............................................................----............ [200480]
MW396979.1_Timor-Leste/2019/1    -...............................................................----............ [200480]
MK940252.1_CN/2019/InnerMongolia -...............................................................----............ [200480]
MK128995.1_China/2018/AnhuiXCGQ  -...............................................................----............ [200480]
MH681419.1_ASFV/POL/2015         -...............................................................----............ [200480]
LS478113.1_Estonia_2014          -...............................................................----............ [200480]
KP843857.1_Odintsovo_02/14       -...............................................................----............ [200480]
```

```
FN557520.1_strain_E75            -..c........a........---...........a..a.........................----............ [200480]
AY261366.1_Warthog               -...t.......a........---.c.c..........c..........g...a..........----............ [200480]
AY261365.1_Warmbaths             -..c........ag.......---.c.c..........a..........gg.............----............ [200480]
AY261364.1_Tengani_62            -.cc........a-.......---cc.c..........a...c......g..........t...----.........t.. [200480]
AY261363.1_Pretoriuskop/96/4     -..c.......ta........---.c.c..........c..........g..............----............ [200480]
AY261362.1_Mkuzi_1979            -..c.................---.c.........a..a........t................atta............ [200480]
AY261361.1_Malawi_Lil-20/1_1983  -........------------------------------------------c....a.t.....----.c.......t.. [200480]
AY261360.1_Kenya_1950            -........------------------------------------------c......t.....----.c.......t.. [200480]
AM712240.1_OURT_88/3             -..c........a........---...........a..a.........................----............ [200480]
AM712239.1_Benin_97/1            -..c........a........---...........a..a.........................----............ [200480]
KM111295.1_Ken06.Bus             a..a...c.------------------------------------------c....a.t.....----.c......gt.. [200480]
KM111294.1_Ken05/Tk1             -........------------------------------------------c....a.t.....----.c.......t.. [200480]
KM262844.1_strain_L60            -..c........a........---...........a..a.........................----............ [200480]
KM102979.1_26544/OG10_Italy      -..c........a........---...........a..a.........................----............ [200480]
KM262845.1_strain_NHV            -..c........a........---...........a..a.........................----............ [200480]
KX354450.1_47/Ss/2008            -..c........a........---...........a..a.........................----............ [200480]
KP055815.1_strain_BA71           -..c........a........---...........a..a.........................----............ [200480]
MH025920.1_strain_R35            a..a...c.------------------------------------------c....a.t.....----.c......gt.. [200480]
MH025919.1_strain_N10            a..a...c.------------------------------------------c....a.t.....----.c......gt.. [200480]
MH025918.1_strain_R25            a..a...c.------------------------------------------c....a.t.....----.c......gt.. [200480]
MH025917.1_strain_R7             a..a...c.------------------------------------------c....a.t.....----.c......gt.. [200480]
MH025916.1_strain_R8             a..a...c.------------------------------------------c....a.t.....----.c......gt.. [200480]

FR682468.2_Georgia_2007/1        ttctggctattcccattacggtat---tattattactatttttaagagctagatgtggatttaagtaataataacattct [200560]
ASF_IND_20_CAD543                -------------------------------------------------------------------------------- [200560]
LR899193.1_Germany_2020/1        ........................---..................................................... [200560]
MN393477.1_Wuhan_2019            ........................---..................................................... [200560]
MW396979.1_Timor-Leste/2019/1    ........................---..................................................... [200560]
MK940252.1_CN/2019/InnerMongolia ........................---..................................................... [200560]
MK128995.1_China/2018/AnhuiXCGQ  ........................---..................................................... [200560]
MH681419.1_ASFV/POL/2015         ........................---..................................................... [200560]
LS478113.1_Estonia_2014          ........................---..................................................... [200560]
KP843857.1_Odintsovo_02/14       ........................---..................................................... [200560]
FN557520.1_strain_E75            ...........t..........cg---.......................g...........gt....c....g....t. [200560]
AY261366.1_Warthog               ......................cg---....................a..g......a....gt....c....g..a.t. [200560]
AY261365.1_Warmbaths             ......................cg---c......................g......a....gt....c....g....t. [200560]
AY261364.1_Tengani_62            ......................ca---ct..........g..........g...........gt....a....g...... [200560]
AY261363.1_Pretoriuskop/96/4     ......................cg---.......................g......a....gt....c....g..a.t. [200560]
AY261362.1_Mkuzi_1979            ...........t......ta...g---.......................g...........gt....c....g....t. [200560]
AY261361.1_Malawi_Lil-20/1_1983  ..................t....g---...c...g...............g.g.........gt....a....g...... [200560]
AY261360.1_Kenya_1950            ..................t....g---...c...g...............g.g....a....gta..ga.......cga. [200560]
AM712240.1_OURT_88/3             ...........t..........cg---.......................g...........gt....c....g....t. [200560]
AM712239.1_Benin_97/1            ...........t..........cg---.......................g...........gt....c....g....t. [200560]
KM111295.1_Ken06.Bus             ..................t....gtaa...c...................g.g.........gt....a....g...... [200560]
```

```
KM111294.1_Ken05/Tk1            ..................t....g---...c...g...............g.g....a....gta..ga.......cga. [200560]
KM262844.1_strain_L60           ...........t..........cg---.......................g...........gt....c....g....t. [200560]
KM102979.1_26544/OG10_Italy     ...........t..........cg---.......................g...........gt....c....g....t. [200560]
KM262845.1_strain_NHV           ...........t..........cg---.......................g...........gt....c....g....t. [200560]
KX354450.1_47/Ss/2008           ...........t..........cg---.......................g...........gt....c....g....t. [200560]
KP055815.1_strain_BA71          ...........t..........cg---.......................g...........gt....c....g....t. [200560]
MH025920.1_strain_R35           ..................t....gtaa...c...................g.g.........gt....a....g...... [200560]
MH025919.1_strain_N10           ..................t....gtaa...c...................g.g.........gt....a....g...... [200560]
MH025918.1_strain_R25           ..................t....gtaa...c...................g.g.........gt....a....g...... [200560]
MH025917.1_strain_R7            ..................t....gtaa...c...................g.g.........gt....a....g...... [200560]
MH025916.1_strain_R8            ..................t....gtaa...c...................g.g.........gt....a....g...... [200560]

FR682468.2_Georgia_2007/1       ccc---gttcctcctagagacacc---tcatcaaattcccatcctatgcaacctttatgttgtaaacataatgattgaca [200640]
ASF_IND_20_CAD543               -------------------------------------------------------------------------------- [200640]
LR899193.1_Germany_2020/1       ...---..................---..................................................... [200640]
MN393477.1_Wuhan_2019           ...---..................---..................................................... [200640]
MW396979.1_Timor-Leste/2019/1   ...---..................---..................................................... [200640]
MK940252.1_CN/2019/InnerMongolia ...---..................---..................................................... [200640]
MK128995.1_China/2018/AnhuiXCGQ ...---..................---..................................................... [200640]
MH681419.1_ASFV/POL/2015        ...---..................---..................................................... [200640]
LS478113.1_Estonia_2014         ...---..................---..................................................... [200640]
KP843857.1_Odintsovo_02/14      ...---..................---..................................................... [200640]
FN557520.1_strain_E75           ...---tc..............t.---..................................................... [200640]
AY261366.1_Warthog              ...---ac...c..........a.---.....................g........c...................... [200640]
AY261365.1_Warmbaths            ...---tc....tt........t.---.....................g............................... [200640]
AY261364.1_Tengani_62           ...---ac..............a.---..................................................... [200640]
AY261363.1_Pretoriuskop/96/4    ...---tc.....t........t.---.....................g............................... [200640]
AY261362.1_Mkuzi_1979           ...---ac..............t.---..............................c..................t... [200640]
AY261361.1_Malawi_Lil-20/1_1983 ..t---ccatg.ga.gactct.t.---.t......................t.a.......tac.............g.. [200640]
AY261360.1_Kenya_1950           ..t---tg....gg.gactcttt.tca...............g....a.....a...........g...........g.. [200640]
AM712240.1_OURT_88/3            ...---tc..............t.---..................................................... [200640]
AM712239.1_Benin_97/1           ...---tc..............t.---..................................................... [200640]
KM111295.1_Ken06.Bus            ..tccatg.tg.ga.g.ctctgt.---at........................................g.......g.. [200640]
KM111294.1_Ken05/Tk1            ..t---tg....gg.g..tctt..tca...............g....a.....a...........g...........g.. [200640]
KM262844.1_strain_L60           ...---tc..............t.---..................................................... [200640]
KM102979.1_26544/OG10_Italy     ...---tc..............t.---..................................................... [200640]
KM262845.1_strain_NHV           ...---tc..............t.---..................................................... [200640]
KX354450.1_47/Ss/2008           ...---tc..............t.---..................................................... [200640]
KP055815.1_strain_BA71          ...---tc..............t.---..................................................... [200640]
MH025920.1_strain_R35           ..tccatg.tg.ga.g.ctctgt.---at........................................g.......g.. [200640]
MH025919.1_strain_N10           ..tccatg.tg.ga.g.ctctgt.---at........................................g.......g.. [200640]
MH025918.1_strain_R25           ..tccatg.tg.ga.g.ctctgt.---at........................................g.......g.. [200640]
MH025917.1_strain_R7            ..tccatg.tg.ga.g.ctctgt.---at........................................g.......g.. [200640]
MH025916.1_strain_R8            ..tccatg.tg.ga.g.ctctgt.---at........................................g.......g.. [200640]
```

```
FR682468.2_Georgia_2007/1        gcattcatcttcttttgacc------aa---gtcgtccaaatcctaccaagatctatacgtgtttttccaaatggagatt [200720]
ASF_IND_20_CAD543                -------------------------------------------------------------------------------- [200720]
LR899193.1_Germany_2020/1        ....................------..---................................................. [200720]
MN393477.1_Wuhan_2019            ....................------..---................................................. [200720]
MW396979.1_Timor-Leste/2019/1    ....................------..---................................................. [200720]
MK940252.1_CN/2019/InnerMongolia ....................------..---................................................. [200720]
MK128995.1_China/2018/AnhuiXCGQ  ....................------..---................................................. [200720]
MH681419.1_ASFV/POL/2015         ....................------..---................................................. [200720]
LS478113.1_Estonia_2014          ....................------..---................................................. [200720]
KP843857.1_Odintsovo_02/14       ....................------..---................................................. [200720]
FN557520.1_strain_E75            .t.a...g....cc..cc..------..---c.a.........t............................a....... [200720]
AY261366.1_Warthog               .t..................------..---..a.......c...................................... [200720]
AY261365.1_Warmbaths             .t..................------..---..a.......c.t.................................... [200720]
AY261364.1_Tengani_62            .t..................------..---................................................. [200720]
AY261363.1_Pretoriuskop/96/4     .t..................------..---..a.......c.t.................................... [200720]
AY261362.1_Mkuzi_1979            .t.a...g....g.c.cc..------..---c.a.......c.t............................a....... [200720]
AY261361.1_Malawi_Lil-20/1_1983  atg---....atgc.a....------..tcgc.a..at....a......ta..................g.......... [200720]
AY261360.1_Kenya_1950            .t.---....a.aga.....------..---..a.cat...catc....ta..a.......................... [200720]
AM712240.1_OURT_88/3             .t.a...g....cc..cc..------..---c.a.........t............................a....... [200720]
AM712239.1_Benin_97/1            .t.a...g....cc..cc..------..---c.a.........t............................a....... [200720]
KM111295.1_Ken06.Bus             agt---..t.atgc.aa...aatcat..---c.g..aa...aag.....tc............................. [200720]
KM111294.1_Ken05/Tk1             .t.---....a.aga.....------..---..a.cat...catc....ta..a.......................... [200720]
KM262844.1_strain_L60            .t.a...g....cc..cc..------..---c.a.........t............................a....... [200720]
KM102979.1_26544/OG10_Italy      .t.a...g....cc..cc..------..---c.a.........t............................a....... [200720]
KM262845.1_strain_NHV            .t.a...g....cc..cc..------..---c.a.........t............................a....... [200720]
KX354450.1_47/Ss/2008            .t.a...g....cc..cc..------..---c.a.........t............................a....... [200720]
KP055815.1_strain_BA71           .t.a...g....cc..cc..------..---c.a.........t............................a....... [200720]
MH025920.1_strain_R35            agt---..t.atgc.aa...aatcat..---c.g..aa...aag.....tc............................. [200720]
MH025919.1_strain_N10            agt---..t.atgc.aa...aatcat..---c.g..aa...aag.....tc............................. [200720]
MH025918.1_strain_R25            agt---..t.atgc.aa...aatcat..---c.g..aa...aag.....tc............................. [200720]
MH025917.1_strain_R7             agt---..t.atgc.aa...aatcat..---c.g..aa...aag.....tc............................. [200720]
MH025916.1_strain_R8             agt---..t.atgc.aa...aatcat..---c.g..aa...aag.....tc............................. [200720]

FR682468.2_Georgia_2007/1        gaagatcagcagtagtggcattaaacctataaaaaccaggtgcataatcacatgaacggatcgtaggatctaatttaata [200800]
ASF_IND_20_CAD543                -------------------------------------------------------------------------------- [200800]
LR899193.1_Germany_2020/1        ................................................................................ [200800]
MN393477.1_Wuhan_2019            ................................................................................ [200800]
MW396979.1_Timor-Leste/2019/1    ................................................................................ [200800]
MK940252.1_CN/2019/InnerMongolia ................................................................................ [200800]
MK128995.1_China/2018/AnhuiXCGQ  ................................................................................ [200800]
MH681419.1_ASFV/POL/2015         ................................................................................ [200800]
LS478113.1_Estonia_2014          ................................................................................ [200800]
KP843857.1_Odintsovo_02/14       ................................................................................ [200800]
```

```
FN557520.1_strain_E75              .........................................t...................g.................. [200800]
AY261366.1_Warthog                 .............................................................g.................. [200800]
AY261365.1_Warmbaths               .............................................................g.................. [200800]
AY261364.1_Tengani_62              .............................................................g........c......... [200800]
AY261363.1_Pretoriuskop/96/4       .............................................................g.................. [200800]
AY261362.1_Mkuzi_1979              .........................................t...................g.................. [200800]
AY261361.1_Malawi_Lil-20/1_1983    .................ct......tg........t.....t......t............gc..........a...... [200800]
AY261360.1_Kenya_1950              .................ct.......g.............c.t......t............gc.......c..a...... [200800]
AM712240.1_OURT_88/3               .........................................t...................g.................. [200800]
AM712239.1_Benin_97/1              .........................................t...................g.................. [200800]
KM111295.1_Ken06.Bus               .................ct......................t......t.......tt...g..........ta...... [200800]
KM111294.1_Ken05/Tk1               .................ct......tg.............c.t......t............gc.......c..a...... [200800]
KM262844.1_strain_L60              .........................................t...................g.................. [200800]
KM102979.1_26544/OG10_Italy        .........................................t...................g.................. [200800]
KM262845.1_strain_NHV              .........................................t...................g.................. [200800]
KX354450.1_47/Ss/2008              .........................................t...................g.................. [200800]
KP055815.1_strain_BA71             .........................................t...................g.................. [200800]
MH025920.1_strain_R35              .................ct......................t......t.......tt...g..........ta...... [200800]
MH025919.1_strain_N10              .................ct......................t......t.......tt...g..........ta...... [200800]
MH025918.1_strain_R25              .................ct......................t......t.......tt...g..........ta...... [200800]
MH025917.1_strain_R7               .................ct......................t......t.......tt...g..........ta...... [200800]
MH025916.1_strain_R8               .................ct......................t......t.......tt...g..........ta...... [200800]

FR682468.2_Georgia_2007/1          tcttttatatcttgttttactgcttctagacaacttttatcagtacatgttccacgtacacagtggtgtcctttatcctt [200880]
ASF_IND_20_CAD543                  -------------------------------------------------------------------------------- [200880]
LR899193.1_Germany_2020/1          ................................................................................ [200880]
MN393477.1_Wuhan_2019              ................................................................................ [200880]
MW396979.1_Timor-Leste/2019/1      ................................................................................ [200880]
MK940252.1_CN/2019/InnerMongolia   ................................................................................ [200880]
MK128995.1_China/2018/AnhuiXCGQ    ................................................................................ [200880]
MH681419.1_ASFV/POL/2015           ................................................................................ [200880]
LS478113.1_Estonia_2014            ................................................................................ [200880]
KP843857.1_Odintsovo_02/14         ................................................................................ [200880]
FN557520.1_strain_E75              ....................a........................................................... [200880]
AY261366.1_Warthog                 ..................g.a........................................................... [200880]
AY261365.1_Warmbaths               ....................a........................................................... [200880]
AY261364.1_Tengani_62              c...................a.......................................................t... [200880]
AY261363.1_Pretoriuskop/96/4       ....................a............................c...............a.............. [200880]
AY261362.1_Mkuzi_1979              ....................a........................................................... [200880]
AY261361.1_Malawi_Lil-20/1_1983    ....................a...............a........................................... [200880]
AY261360.1_Kenya_1950              c....a..............a...............a..................................a........ [200880]
AM712240.1_OURT_88/3               ....................a........................................................... [200880]
AM712239.1_Benin_97/1              ....................a........................................................... [200880]
KM111295.1_Ken06.Bus               ...........c.......ta..............ca............................c...........t.. [200880]
```

```
KM111294.1_Ken05/Tk1            .....a..............a...........g...a........................................... [200880]
KM262844.1_strain_L60           ....................a........................................................... [200880]
KM102979.1_26544/OG10_Italy     ....................a........................................................... [200880]
KM262845.1_strain_NHV           ....................a........................................................... [200880]
KX354450.1_47/Ss/2008           ....................a........................................................... [200880]
KP055815.1_strain_BA71          ....................a........................................................... [200880]
MH025920.1_strain_R35           ...........c.......ta..............ca............................c...........t.. [200880]
MH025919.1_strain_N10           ...........c.......ta..............ca............................c...........t.. [200880]
MH025918.1_strain_R25           ...........c.......ta..............ca............................c...........t.. [200880]
MH025917.1_strain_R7            ...........c.......ta..............ca............................c...........t.. [200880]
MH025916.1_strain_R8            ...........c.......ta..............ca............................c...........t.. [200880]

FR682468.2_Georgia_2007/1       acaatccgtatctgtcttacatttttttttcggcggtttatgtttcagatggtaaaaacccagtattaaaataatc-aca [200960]
ASF_IND_20_CAD543               -------------------------------------------------------------------------------- [200960]
LR899193.1_Germany_2020/1       ............................................................................-... [200960]
MN393477.1_Wuhan_2019           ............................................................................-... [200960]
MW396979.1_Timor-Leste/2019/1   ............................................................................-... [200960]
MK940252.1_CN/2019/InnerMongolia ............................................................................-... [200960]
MK128995.1_China/2018/AnhuiXCGQ ............................................................................-... [200960]
MH681419.1_ASFV/POL/2015        ............................................................................-... [200960]
LS478113.1_Estonia_2014         ............................................................................-... [200960]
KP843857.1_Odintsovo_02/14      ............................................................................-... [200960]
FN557520.1_strain_E75           ..................................a..........t.t............................-... [200960]
AY261366.1_Warthog              ...............................................t.........g..................-... [200960]
AY261365.1_Warmbaths            ......t..........................ta....cc....t.............t................-... [200960]
AY261364.1_Tengani_62           ......t...........................a....tc....t.............t................-... [200960]
AY261363.1_Pretoriuskop/96/4    .......................................cc....t...........g..................-... [200960]
AY261362.1_Mkuzi_1979           .....t............................a..........t.t............................-... [200960]
AY261361.1_Malawi_Lil-20/1_1983 ......t..............g.......-------------.........a.......t............t...a... [200960]
AY261360.1_Kenya_1950           ......t.....c........gc.......t------------........a........t.a............t-... [200960]
AM712240.1_OURT_88/3            ..................................a..........t.t............................-... [200960]
AM712239.1_Benin_97/1           ..................................a..........t.t............................-... [200960]
KM111295.1_Ken06.Bus            .................................t..------g............t...t............t...-... [200960]
KM111294.1_Ken05/Tk1            ......t..............c......---------------.................t.a............t-... [200960]
KM262844.1_strain_L60           ..................................a..........t.t............................-... [200960]
KM102979.1_26544/OG10_Italy     ..................................a..........t.t............................-... [200960]
KM262845.1_strain_NHV           ..................................a..........t.t............................-... [200960]
KX354450.1_47/Ss/2008           ..................................a..........t.t............................-... [200960]
KP055815.1_strain_BA71          ..................................a..........t.t............................-... [200960]
MH025920.1_strain_R35           .................................t..------g............t...t............t...-... [200960]
MH025919.1_strain_N10           .................................t..------g............t...t............t...-... [200960]
MH025918.1_strain_R25           .................................t..------g............t...t............t...-... [200960]
MH025917.1_strain_R7            .................................t..------g............t...t............t...-... [200960]
MH025916.1_strain_R8            .................................t..------g............t...t............t...-... [200960]
```

```
FR682468.2_Georgia_2007/1        agaataattcctataagtacttgaacaacaggataaaacattttaatatt-aaatatatttt-------ttaattaaa-- [201040]
ASF_IND_20_CAD543                -------------------------------------------------------------------------------- [201040]
LR899193.1_Germany_2020/1        ..................................................-...........-------.........-- [201040]
MN393477.1_Wuhan_2019            ..................................................-...........-------.........-- [201040]
MW396979.1_Timor-Leste/2019/1    ..................................................-...........-------.........-- [201040]
MK940252.1_CN/2019/InnerMongolia ..................................................-...........-------.........-- [201040]
MK128995.1_China/2018/AnhuiXCGQ  ..................................................-...........-------.........-- [201040]
MH681419.1_ASFV/POL/2015         ..................................................-...........-------.........-- [201040]
LS478113.1_Estonia_2014          ..................................................-...........-------.........-- [201040]
KP843857.1_Odintsovo_02/14       ..................................................-...........-------.........-- [201040]
FN557520.1_strain_E75            .........................t.g......................-...........-------.........-- [201040]
AY261366.1_Warthog               ..................................................-..g........-------..------.-- [201040]
AY261365.1_Warmbaths             ...................a..............................-...........-------.........-- [201040]
AY261364.1_Tengani_62            ..................................g...............-..g........-------...-------- [201040]
AY261363.1_Pretoriuskop/96/4     ........c..........t..............................-...........-------.........-- [201040]
AY261362.1_Mkuzi_1979            ................................................c.-...a.......aatgatgaa.......at [201040]
AY261361.1_Malawi_Lil-20/1_1983  ...........................g........t...........c.-...acat....aattatgaa.......at [201040]
AY261360.1_Kenya_1950            .......gc...........c.............g.t.............a....g......-------.........-- [201040]
AM712240.1_OURT_88/3             .........................t.g......................-...........-------.........-- [201040]
AM712239.1_Benin_97/1            .........................t.g......................-...........-------.........-- [201040]
KM111295.1_Ken06.Bus             .........g.........t.....t........g.t.............-...........-------.........-- [201040]
KM111294.1_Ken05/Tk1             .......gc.........................g.t.............-...........-------.........-- [201040]
KM262844.1_strain_L60            .........................t.g......................-...........-------.........-- [201040]
KM102979.1_26544/OG10_Italy      .........................t.g......................-...........-------.........-- [201040]
KM262845.1_strain_NHV            .........................t.g......................-...........-------.........-- [201040]
KX354450.1_47/Ss/2008            .........................t.g......................-...........-------.........-- [201040]
KP055815.1_strain_BA71           .........................t.g......................-...........-------.........-- [201040]
MH025920.1_strain_R35            .........g.........t.....t........g.t.............-...........-------.........-- [201040]
MH025919.1_strain_N10            .........g.........t.....t........g.t.............-...........-------.........-- [201040]
MH025918.1_strain_R25            .........g.........t.....t........g.t.............-...........-------.........-- [201040]
MH025917.1_strain_R7             .........g.........t.....t........g.t.............-...........-------.........-- [201040]
MH025916.1_strain_R8             .........g.........t.....t........g.t.............-...........-------.........-- [201040]

FR682468.2_Georgia_2007/1        ---------tgaatagatttaatccaagtagtattaa--aatttttagaaata---------------gtgttctac-aa [201120]
ASF_IND_20_CAD543                -------------------------------------------------------------------------------- [201120]
LR899193.1_Germany_2020/1        ---------............................--..............---------------.........-.. [201120]
MN393477.1_Wuhan_2019            ---------............................--..............---------------.........-.. [201120]
MW396979.1_Timor-Leste/2019/1    ---------............................--..............---------------.........-.. [201120]
MK940252.1_CN/2019/InnerMongolia ---------............................--..............---------------.........-.. [201120]
MK128995.1_China/2018/AnhuiXCGQ  ---------............................--..............---------------.........-.. [201120]
MH681419.1_ASFV/POL/2015         ---------............................--..............---------------.........-.. [201120]
LS478113.1_Estonia_2014          ---------............................--..............---------------.........-.. [201120]
KP843857.1_Odintsovo_02/14       ---------............................--..............---------------.........-.. [201120]
```

```
FN557520.1_strain_E75              ---------............................--...............--------------.........-.. [201120]
AY261366.1_Warthog                 ---------............................at...............--------------.........-.. [201120]
AY261365.1_Warmbaths               ---------......tc..........a.........at....g..........--------------.........-.. [201120]
AY261364.1_Tengani_62              ---------...........................---.t.............--------------.........-.. [201120]
AY261363.1_Pretoriuskop/96/4       ---------......tc..........a.........at....a.......--.--------------.........-.. [201120]
AY261362.1_Mkuzi_1979              gacatattgaa....t.....t..tgt..-.......at....c....a.....--------------t.a..t...tg. [201120]
AY261361.1_Malawi_Lil-20/1_1983    gacatat--..c...t.....c.gtg....t..aat.cttc..c....a..----------------------------- [201120]
AY261360.1_Kenya_1950              ---------.................g....--.a..at....c..........atgttctacaaataa........-.. [201120]
AM712240.1_OURT_88/3               ---------............................--...............--------------.........-.. [201120]
AM712239.1_Benin_97/1              ---------............................--...............--------------.........-.. [201120]
KM111295.1_Ken06.Bus               ---------...........---------------..at....c..........-----------------......-.. [201120]
KM111294.1_Ken05/Tk1               ---------...t........................at....c..........--------------.........-.. [201120]
KM262844.1_strain_L60              ---------............................--...............--------------.........-.. [201120]
KM102979.1_26544/OG10_Italy        ---------............................--...............--------------.........-.. [201120]
KM262845.1_strain_NHV              ---------............................--...............--------------.........-.. [201120]
KX354450.1_47/Ss/2008              ---------............................--...............--------------.........-.. [201120]
KP055815.1_strain_BA71             ---------............................--...............--------------.........-.. [201120]
MH025920.1_strain_R35              ---------...........---------------..at....c..........-----------------......-.. [201120]
MH025919.1_strain_N10              ---------...........---------------..at....c..........-----------------......-.. [201120]
MH025918.1_strain_R25              ---------...........---------------..at....c..........-----------------......-.. [201120]
MH025917.1_strain_R7               ---------...........---------------..at....c..........-----------------......-.. [201120]
MH025916.1_strain_R8               ---------...........---------------..at....c..........-----------------......-.. [201120]

FR682468.2_Georgia_2007/1          ataatgaaa----------------------------------------------------------------------- [201200]
ASF_IND_20_CAD543                  -------------------------------------------------------------------------------- [201200]
LR899193.1_Germany_2020/1          .........----------------------------------------------------------------------- [201200]
MN393477.1_Wuhan_2019              .........----------------------------------------------------------------------- [201200]
MW396979.1_Timor-Leste/2019/1      .........----------------------------------------------------------------------- [201200]
MK940252.1_CN/2019/InnerMongolia   .........----------------------------------------------------------------------- [201200]
MK128995.1_China/2018/AnhuiXCGQ    .........----------------------------------------------------------------------- [201200]
MH681419.1_ASFV/POL/2015           .........----------------------------------------------------------------------- [201200]
LS478113.1_Estonia_2014            .........----------------------------------------------------------------------- [201200]
KP843857.1_Odintsovo_02/14         .........----------------------------------------------------------------------- [201200]
FN557520.1_strain_E75              .........----------------------------------------------------------------------- [201200]
AY261366.1_Warthog                 .........tgaatggtccaa---aaaaaataagatgtacaataatgtaatatattgttaggctaagta-aatttaatat [201200]
AY261365.1_Warmbaths               .........tggatggtctagaccaaaaaaaatggtgtacaattatgtaatacattgataggctaagta-aatttaatat [201200]
AY261364.1_Tengani_62              .........tgaatggtccaa---aaaa----------ataataatgtaataca--gttaggctaagtaaaatttaatat [201200]
AY261363.1_Pretoriuskop/96/4       .........tgaatggtctagaccaaaaaaaatggtgtacaattatgtaatacattgataagctaagta-aatttaataa [201200]
AY261362.1_Mkuzi_1979              .....a...c---ttatttacatcaaaa---------------------------------------cg-atgttataga [201200]
AY261361.1_Malawi_Lil-20/1_1983    -------------------------------------------------------------------------------- [201200]
AY261360.1_Kenya_1950              .........tggatggtctaa---aaaaa-------------taatgtaatacattgttaggttaagta-aatttaatat [201200]
AM712240.1_OURT_88/3               .........----------------------------------------------------------------------- [201200]
AM712239.1_Benin_97/1              .........----------------------------------------------------------------------- [201200]
KM111295.1_Ken06.Bus               .g.......tggatggttcaa---aaaa---------------aatgtaatatattgttaggtcaagta-aatttaatat [201200]
```

```
KM111294.1_Ken05/Tk1              ........tggatggtctaa---aaaaa------------------ataatacattgttaggttaagta-aatttaatat [201200]
KM262844.1_strain_L60             ........------------------------------------------------------------------------ [201200]
KM102979.1_26544/OG10_Italy       ........------------------------------------------------------------------------ [201200]
KM262845.1_strain_NHV             ........------------------------------------------------------------------------ [201200]
KX354450.1_47/Ss/2008             ........------------------------------------------------------------------------ [201200]
KP055815.1_strain_BA71            ........------------------------------------------------------------------------ [201200]
MH025920.1_strain_R35             .g......tggatggttcaa---aaaa----------------aatgtaatatattgttaggtcaagta-aatttaatat [201200]
MH025919.1_strain_N10             .g......tggatggttcaa---aaaa----------------aatgtaatatattgttaggtcaagta-aatttaatat [201200]
MH025918.1_strain_R25             .g......tggatggttcaa---aaaa----------------aatgtaatatattgttaggtcaagta-aatttaatat [201200]
MH025917.1_strain_R7              .g......tggatggttcaa---aaaa----------------aatgtaatatattgttaggtcaagta-aatttaatat [201200]
MH025916.1_strain_R8              .g......tggatggttcaa---aaaa----------------aatgtaatatattgttaggtcaagta-aatttaatat [201200]

FR682468.2_Georgia_2007/1         ---------------tgaatggtcc--------------------------------------------------aaaaa [201280]
ASF_IND_20_CAD543                 -------------------------------------------------------------------------------- [201280]
LR899193.1_Germany_2020/1         ---------------..........--------------------------------------------------..... [201280]
MN393477.1_Wuhan_2019             ---------------..........--------------------------------------------------..... [201280]
MW396979.1_Timor-Leste/2019/1     ---------------..........--------------------------------------------------..... [201280]
MK940252.1_CN/2019/InnerMongolia  ---------------..........--------------------------------------------------..... [201280]
MK128995.1_China/2018/AnhuiXCGQ   ---------------..........--------------------------------------------------..... [201280]
MH681419.1_ASFV/POL/2015          ---------------..........--------------------------------------------------..... [201280]
LS478113.1_Estonia_2014           ---------------..........--------------------------------------------------..... [201280]
KP843857.1_Odintsovo_02/14        ---------------..........--------------------------------------------------..... [201280]
FN557520.1_strain_E75             ---------------.....a.c..--------------------------------------------------..... [201280]
AY261366.1_Warthog                tttaaagtacttttc..gg.at.g.cattac---agcatcgctcttgtatttaagagcaggatgcggatttctgt..... [201280]
AY261365.1_Warmbaths              tttaaagtacttttc..gg.at.g.cattacagtagcaccgttcttgtatttaagagcaggatgcggatttgtgt..... [201280]
AY261364.1_Tengani_62             tttaaggtacttttc..gg.at.g.cattat---agcatcgttcttgtatttaagagcaggatgtggatttctgt..... [201280]
AY261363.1_Pretoriuskop/96/4      tttaaagtacttttc..gg.at.g.cattacagtagcaccgttcttgtatttaagaacaggatgtggatttgtgt.t... [201280]
AY261362.1_Mkuzi_1979             tttaatataccattc..g..aa...tattactctattatcattgttatatttaagaaccggatgtggatttgggt..... [201280]
AY261361.1_Malawi_Lil-20/1_1983   -------------------------------------------------------------------------------- [201280]
AY261360.1_Kenya_1950             tttaagatacttttc.agc.at.g.caatac---agcatcattcttggatttaagagcaggatgtggatctatgt..... [201280]
AM712240.1_OURT_88/3              ---------------.....a.c..--------------------------------------------------..... [201280]
AM712239.1_Benin_97/1             ---------------.....a.c..--------------------------------------------------..... [201280]
KM111295.1_Ken06.Bus              tttataatccttttc.agc.at.g.cattac---aacattatccttgtatttaagagcaggatgtggatcta-------- [201280]
KM111294.1_Ken05/Tk1              tttaagatacttttc.agc.at.g.cattac---agcatcattcttggatttaagagcaggatgtggatctatgt..... [201280]
KM262844.1_strain_L60             ---------------.....a.c..--------------------------------------------------..... [201280]
KM102979.1_26544/OG10_Italy       ---------------.....a.c..--------------------------------------------------..... [201280]
KM262845.1_strain_NHV             ---------------.....a.c..--------------------------------------------------..... [201280]
KX354450.1_47/Ss/2008             ---------------.....a.c..--------------------------------------------------..... [201280]
KP055815.1_strain_BA71            ---------------.....a.c..--------------------------------------------------..... [201280]
MH025920.1_strain_R35             tttataatccttttc.agc.at.g.cattac---aacattatccttgtatttaagagcaggatgtggatcta-------- [201280]
MH025919.1_strain_N10             tttataatccttttc.agc.at.g.cattac---aacattatccttgtatttaagagcaggatgtggatcta-------- [201280]
MH025918.1_strain_R25             tttataatccttttc.agc.at.g.cattac---aacattatccttgtatttaagagcaggatgtggatcta-------- [201280]
MH025917.1_strain_R7              tttataatccttttc.agc.at.g.cattac---aacattatccttgtatttaagagcaggatgtggatcta-------- [201280]
MH025916.1_strain_R8              tttataatccttttc.agc.at.g.cattac---aacattatccttgtatttaagagcaggatgtggatcta-------- [201280]
```

```
FR682468.2_Georgia_2007/1        aaat---------------------------------------------------------------------------- [201360]
ASF_IND_20_CAD543                -------------------------------------------------------------------------------- [201360]
LR899193.1_Germany_2020/1        ....---------------------------------------------------------------------------- [201360]
MN393477.1_Wuhan_2019            ....---------------------------------------------------------------------------- [201360]
MW396979.1_Timor-Leste/2019/1    ....---------------------------------------------------------------------------- [201360]
MK940252.1_CN/2019/InnerMongolia ....---------------------------------------------------------------------------- [201360]
MK128995.1_China/2018/AnhuiXCGQ  ....---------------------------------------------------------------------------- [201360]
MH681419.1_ASFV/POL/2015         ....---------------------------------------------------------------------------- [201360]
LS478113.1_Estonia_2014          ....---------------------------------------------------------------------------- [201360]
KP843857.1_Odintsovo_02/14       ....---------------------------------------------------------------------------- [201360]
FN557520.1_strain_E75            ....---------------------------------------------------------------------------- [201360]
AY261366.1_Warthog               t.gcaatttccagtcgttttgcctgaaatctgatcatattcccaagatatgcaa---cctggataatttgcacataaaga [201360]
AY261365.1_Warmbaths             ..gcattctccagtcgttttgcctgaaatctgatcatattcccaagctatgcaacctcctggataacttgcacataaaga [201360]
AY261364.1_Tengani_62            t.gcaatttccagtccttttgtttgagaactgatcaaattcccaagatatgcaa---cctggataatttgcacataaaga [201360]
AY261363.1_Pretoriuskop/96/4     ..gcattctccagtcgttgtgcctgaaaactgatcatattcccaagctatgcaaccttctggataatttgcacaaaaaga [201360]
AY261362.1_Mkuzi_1979            ...cactctccttc----------------cttattaaattcccaagctgggcaacc------agattgtaaacaatatga [201360]
AY261361.1_Malawi_Lil-20/1_1983  -------------------------------------------------------------------------------- [201360]
AY261360.1_Kenya_1950            t.gcattctccggtcgttttggcatagtactgatcaaattcccatcctatgcaa---tctttacgatctaaacataatct [201360]
AM712240.1_OURT_88/3             ....---------------------------------------------------------------------------- [201360]
AM712239.1_Benin_97/1            ....---------------------------------------------------------------------------- [201360]
KM111295.1_Ken06.Bus             -------------------------------------------------------------------------------- [201360]
KM111294.1_Ken05/Tk1             t.gcattctccagttgttttgcctgatatcttatcaaattcccatcctatgcaa---tctttacgatctaaacataatct [201360]
KM262844.1_strain_L60            ....a--------------------------------------------------------------------------- [201360]
KM102979.1_26544/OG10_Italy      ....---------------------------------------------------------------------------- [201360]
KM262845.1_strain_NHV            ....---------------------------------------------------------------------------- [201360]
KX354450.1_47/Ss/2008            ....---------------------------------------------------------------------------- [201360]
KP055815.1_strain_BA71           ....---------------------------------------------------------------------------- [201360]
MH025920.1_strain_R35            -------------------------------------------------------------------------------- [201360]
MH025919.1_strain_N10            -------------------------------------------------------------------------------- [201360]
MH025918.1_strain_R25            -------------------------------------------------------------------------------- [201360]
MH025917.1_strain_R7             -------------------------------------------------------------------------------- [201360]
MH025916.1_strain_R8             -------------------------------------------------------------------------------- [201360]

FR682468.2_Georgia_2007/1        -------------------------------------------------------------------------------- [201440]
ASF_IND_20_CAD543                -------------------------------------------------------------------------------- [201440]
LR899193.1_Germany_2020/1        -------------------------------------------------------------------------------- [201440]
MN393477.1_Wuhan_2019            -------------------------------------------------------------------------------- [201440]
MW396979.1_Timor-Leste/2019/1    -------------------------------------------------------------------------------- [201440]
MK940252.1_CN/2019/InnerMongolia -------------------------------------------------------------------------------- [201440]
MK128995.1_China/2018/AnhuiXCGQ  -------------------------------------------------------------------------------- [201440]
MH681419.1_ASFV/POL/2015         -------------------------------------------------------------------------------- [201440]
LS478113.1_Estonia_2014          -------------------------------------------------------------------------------- [201440]
KP843857.1_Odintsovo_02/14       -------------------------------------------------------------------------------- [201440]
```

```
FN557520.1_strain_E75              -------------------------------------------------------------------------------- [201440]
AY261366.1_Warthog                 ttcacagaattcttcacctgaatgagggtcataaagctcgccatattctatacgagattttccaaatggagattgaggat [201440]
AY261365.1_Warmbaths               ttgacagtaatcttcacctgaatcaggggcataaacctcgcctgtatctatacgagattttccaaatggagattgaagat [201440]
AY261364.1_Tengani_62              ttcacagaaatcttcacctgaatgagggtcataaagctcgccatattctatacgagattttccaaatggagattgaggat [201440]
AY261363.1_Pretoriuskop/96/4       ttgacaggaatcttcatttgaatgaggggaataaatccagccataatctatacgagattttccaaatggagattgaagat [201440]
AY261362.1_Mkuzi_1979              ttcacagttatctgcacttggttga-------------ctgcccacatcaacactagtttttcaaatgtagattgaggat [201440]
AY261361.1_Malawi_Lil-20/1_1983    -------------------------------------------------------------------------------- [201440]
AY261360.1_Kenya_1950              ttgacagaaatcttcacctgaatggggtcataaagctcagcatcatctatacgtgtttttccaaatggagattgaagat [201440]
AM712240.1_OURT_88/3               -------------------------------------------------------------------------------- [201440]
AM712239.1_Benin_97/1              -------------------------------------------------------------------------------- [201440]
KM111295.1_Ken06.Bus               -------------------------------------------------------------------------------- [201440]
KM111294.1_Ken05/Tk1               ttgacagaaatcttcacctgaatggggtcataaagctcagcatcatctatacgtgtttttccaaatggagattgaagat [201440]
KM262844.1_strain_L60              -------------------------------------------------------------------------------- [201440]
KM102979.1_26544/OG10_Italy        -------------------------------------------------------------------------------- [201440]
KM262845.1_strain_NHV              -------------------------------------------------------------------------------- [201440]
KX354450.1_47/Ss/2008              -------------------------------------------------------------------------------- [201440]
KP055815.1_strain_BA71             -------------------------------------------------------------------------------- [201440]
MH025920.1_strain_R35              -------------------------------------------------------------------------------- [201440]
MH025919.1_strain_N10              -------------------------------------------------------------------------------- [201440]
MH025918.1_strain_R25              -------------------------------------------------------------------------------- [201440]
MH025917.1_strain_R7               -------------------------------------------------------------------------------- [201440]
MH025916.1_strain_R8               -------------------------------------------------------------------------------- [201440]

FR682468.2_Georgia_2007/1          ----------------------------------------------------------aaggtgtacaataatgtaatat [201520]
ASF_IND_20_CAD543                  -------------------------------------------------------------------------------- [201520]
LR899193.1_Germany_2020/1          ----------------------------------------------------------...................... [201520]
MN393477.1_Wuhan_2019              ----------------------------------------------------------...................... [201520]
MW396979.1_Timor-Leste/2019/1      ----------------------------------------------------------...................... [201520]
MK940252.1_CN/2019/InnerMongolia   ----------------------------------------------------------...................... [201520]
MK128995.1_China/2018/AnhuiXCGQ    ----------------------------------------------------------...................... [201520]
MH681419.1_ASFV/POL/2015           ----------------------------------------------------------...................... [201520]
LS478113.1_Estonia_2014            ----------------------------------------------------------...................... [201520]
KP843857.1_Odintsovo_02/14         ----------------------------------------------------------...................... [201520]
FN557520.1_strain_E75              ----------------------------------------------------------..........t........... [201520]
AY261366.1_Warthog                 cagc---agtactactaaacttataaaacttaggtgtatagttacatgaacggatgtt.g...c..gtt....acct.t. [201520]
AY261365.1_Warmbaths               cagcagtagtattactaaacatataaaacttaggtgtataattacatgaacggatgtt.g.a.c..ttt....a.ct.t. [201520]
AY261364.1_Tengani_62              cagc---agtactactaaacttataaaacttaggtgtatagttacatgaacggatgtt.g...c..gtt....acct.t. [201520]
AY261363.1_Pretoriuskop/96/4       cagcagtagtattactaaacttataaaacttaggtgtataattacatgaacggatgtt.g.a.c..ttt....a.ct.t. [201520]
AY261362.1_Mkuzi_1979              cagcaattgcattattaaacctataataattagatgtgtaagtacatgaatcgatttt.g.a.c..gtt....a.tt.t. [201520]
AY261361.1_Malawi_Lil-20/1_1983    -------------------------------------------------------------------------------- [201520]
AY261360.1_Kenya_1950              cagcagtagtattactaaacttataaaacttaggtgtatagttacatgaacggatgtt.g.a.c..gtt....a.ct.t. [201520]
AM712240.1_OURT_88/3               ----------------------------------------------------------..........t........... [201520]
AM712239.1_Benin_97/1              ----------------------------------------------------------..........t........... [201520]
KM111295.1_Ken06.Bus               ---------------------tataaaacttaggtgtatagttacataaatggatgtt.g.a.c..gtt...gac----- [201520]
```

```
KM111294.1_Ken05/Tk1             cagcagtagtattactaaacttataaaacttaggtgtataattacaggaacggatgtt.g.a.c..gtt....a.ct.t. [201520]
KM262844.1_strain_L60            ----------------------------------------------------------..........t........... [201520]
KM102979.1_26544/OG10_Italy      ----------------------------------------------------------..........t........... [201520]
KM262845.1_strain_NHV            ----------------------------------------------------------..........t........... [201520]
KX354450.1_47/Ss/2008            ----------------------------------------------------------..........t........... [201520]
KP055815.1_strain_BA71           ----------------------------------------------------------..........t........... [201520]
MH025920.1_strain_R35            ---------------------tataaaacttaggtgtatagttacataaatggatgtt.g.a.c..gtt...gac----- [201520]
MH025919.1_strain_N10            ---------------------tataaaacttaggtgtatagttacataaatggatgtt.g.a.c..gtt...gac----- [201520]
MH025918.1_strain_R25            ---------------------tataaaacttaggtgtatagttacataaatggatgtt.g.a.c..gtt...gac----- [201520]
MH025917.1_strain_R7             ---------------------tataaaacttaggtgtatagttacataaatggatgtt.g.a.c..gtt...gac----- [201520]
MH025916.1_strain_R8             ---------------------tataaaacttaggtgtatagttacataaatggatgtt.g.a.c..gtt...gac----- [201520]

FR682468.2_Georgia_2007/1        att----------------------------------------------------------------------------- [201600]
ASF_IND_20_CAD543                -------------------------------------------------------------------------------- [201600]
LR899193.1_Germany_2020/1        ...----------------------------------------------------------------------------- [201600]
MN393477.1_Wuhan_2019            ...----------------------------------------------------------------------------- [201600]
MW396979.1_Timor-Leste/2019/1    ...----------------------------------------------------------------------------- [201600]
MK940252.1_CN/2019/InnerMongolia ...----------------------------------------------------------------------------- [201600]
MK128995.1_China/2018/AnhuiXCGQ  ...----------------------------------------------------------------------------- [201600]
MH681419.1_ASFV/POL/2015         ...----------------------------------------------------------------------------- [201600]
LS478113.1_Estonia_2014          ...----------------------------------------------------------------------------- [201600]
KP843857.1_Odintsovo_02/14       ...----------------------------------------------------------------------------- [201600]
FN557520.1_strain_E75            ...----------------------------------------------------------------------------- [201600]
AY261366.1_Warthog               ..gtcatattttgcagcttctagacagttcatcttggtacaaaatccacccacacagtggtgtccattatccttacagtc [201600]
AY261365.1_Warmbaths             ..gtcatattttgcagcttctacacagctcatgttggtacaaaatccacccacacagtggtgtcctttatccttacagtc [201600]
AY261364.1_Tengani_62            ..gtcatattttgcagcttctagacagctcatcttggtacaaaatccacccacacagtggtgaccattatccttacagtc [201600]
AY261363.1_Pretoriuskop/96/4     ..gtcatattttgcagcttctacacagctcatgttggtacaaaatccacccacacagtggtgtcctttatccttacagtc [201600]
AY261362.1_Mkuzi_1979            .c.t---gttttacagcatctaaacaac------tagtacatgttccacgtacacaaagttcttttttatccttacagtc [201600]
AY261361.1_Malawi_Lil-20/1_1983  -------------------------------------------------------------------------------- [201600]
AY261360.1_Kenya_1950            ..gtcatattttatagcatctagacagctcatgttggtacaaattccacccacacagtggtgtcctttatccttacagtc [201600]
AM712240.1_OURT_88/3             ...----------------------------------------------------------------------------- [201600]
AM712239.1_Benin_97/1            ...----------------------------------------------------------------------------- [201600]
KM111295.1_Ken06.Bus             -------------------------------------------------------------------------------- [201600]
KM111294.1_Ken05/Tk1             ..gtcatattttatagcatctagacagctcatgttggtacaaattccacccacacagtggtgtcctttatccttacagtc [201600]
KM262844.1_strain_L60            ...----------------------------------------------------------------------------- [201600]
KM102979.1_26544/OG10_Italy      ...----------------------------------------------------------------------------- [201600]
KM262845.1_strain_NHV            ...----------------------------------------------------------------------------- [201600]
KX354450.1_47/Ss/2008            ...----------------------------------------------------------------------------- [201600]
KP055815.1_strain_BA71           ...----------------------------------------------------------------------------- [201600]
MH025920.1_strain_R35            -------------------------------------------------------------------------------- [201600]
MH025919.1_strain_N10            -------------------------------------------------------------------------------- [201600]
MH025918.1_strain_R25            -------------------------------------------------------------------------------- [201600]
MH025917.1_strain_R7             -------------------------------------------------------------------------------- [201600]
MH025916.1_strain_R8             -------------------------------------------------------------------------------- [201600]
```

```
FR682468.2_Georgia_2007/1        -------------------------------------------------------------------------------- [201680]
ASF_IND_20_CAD543                -------------------------------------------------------------------------------- [201680]
LR899193.1_Germany_2020/1        -------------------------------------------------------------------------------- [201680]
MN393477.1_Wuhan_2019            -------------------------------------------------------------------------------- [201680]
MW396979.1_Timor-Leste/2019/1    -------------------------------------------------------------------------------- [201680]
MK940252.1_CN/2019/InnerMongolia -------------------------------------------------------------------------------- [201680]
MK128995.1_China/2018/AnhuiXCGQ  -------------------------------------------------------------------------------- [201680]
MH681419.1_ASFV/POL/2015         -------------------------------------------------------------------------------- [201680]
LS478113.1_Estonia_2014          -------------------------------------------------------------------------------- [201680]
KP843857.1_Odintsovo_02/14       -------------------------------------------------------------------------------- [201680]
FN557520.1_strain_E75            -------------------------------------------------------------------------------- [201680]
AY261366.1_Warthog               tacatctatcttacattttttttttttggtggtgatattggtagtaa----------------------------------- [201680]
AY261365.1_Warmbaths             tgtatctgtcttacattttttttttttggtggtgatattggtagtag----------------------------------- [201680]
AY261364.1_Tengani_62            tgtatctatcttacattttttttttttggtggtgatattggtagtaa----------------------------------- [201680]
AY261363.1_Pretoriuskop/96/4     tgtatctgtcttacattttttttttttggtggtgatattggtagtaa----------------------------------- [201680]
AY261362.1_Mkuzi_1979            tgtatctaccttacatttcttttttgatggtgatagtggtgataatgctggtgaaagtggtgatagtgctggtgatagtg [201680]
AY261361.1_Malawi_Lil-20/1_1983  -------------------------------------------------------------------------------- [201680]
AY261360.1_Kenya_1950            tgtatctgtcttacattttttttttttggtggtgatattggtagtgc----------------------------------- [201680]
AM712240.1_OURT_88/3             -------------------------------------------------------------------------------- [201680]
AM712239.1_Benin_97/1            -------------------------------------------------------------------------------- [201680]
KM111295.1_Ken06.Bus             -------------------------------------------------------------------------------- [201680]
KM111294.1_Ken05/Tk1             tgtatctatcttacattttttttttttgctggtgatattggtagtgg----------------------------------- [201680]
KM262844.1_strain_L60            -------------------------------------------------------------------------------- [201680]
KM102979.1_26544/OG10_Italy      -------------------------------------------------------------------------------- [201680]
KM262845.1_strain_NHV            -------------------------------------------------------------------------------- [201680]
KX354450.1_47/Ss/2008            -------------------------------------------------------------------------------- [201680]
KP055815.1_strain_BA71           -------------------------------------------------------------------------------- [201680]
MH025920.1_strain_R35            -------------------------------------------------------------------------------- [201680]
MH025919.1_strain_N10            -------------------------------------------------------------------------------- [201680]
MH025918.1_strain_R25            -------------------------------------------------------------------------------- [201680]
MH025917.1_strain_R7             -------------------------------------------------------------------------------- [201680]
MH025916.1_strain_R8             -------------------------------------------------------------------------------- [201680]

FR682468.2_Georgia_2007/1        -------------------------------------------------------------------------------- [201760]
ASF_IND_20_CAD543                -------------------------------------------------------------------------------- [201760]
LR899193.1_Germany_2020/1        -------------------------------------------------------------------------------- [201760]
MN393477.1_Wuhan_2019            -------------------------------------------------------------------------------- [201760]
MW396979.1_Timor-Leste/2019/1    -------------------------------------------------------------------------------- [201760]
MK940252.1_CN/2019/InnerMongolia -------------------------------------------------------------------------------- [201760]
MK128995.1_China/2018/AnhuiXCGQ  -------------------------------------------------------------------------------- [201760]
MH681419.1_ASFV/POL/2015         -------------------------------------------------------------------------------- [201760]
LS478113.1_Estonia_2014          -------------------------------------------------------------------------------- [201760]
KP843857.1_Odintsovo_02/14       -------------------------------------------------------------------------------- [201760]
```

```
FN557520.1_strain_E75            -------------------------------------------------------------------------------- [201760]
AY261366.1_Warthog               ----tggattctgttccggatagaaaagaccc------gatattaaaaaaattacgagaataatagttataaacagaata [201760]
AY261365.1_Warmbaths             ----tgattgccgttccggatagaaaagaccc------gatattaaaaaaatcacaagaataataattataaacagaata [201760]
AY261364.1_Tengani_62            ----tggattctgttccggatagaaaagaccc------gatattaaaaaaatcacgagaatgatagttataaacagaata [201760]
AY261363.1_Pretoriuskop/96/4     ----tggattctgttccggatagaaaagaccc------gatattaaaaaaattacgag---aatagttataaacagaata [201760]
AY261362.1_Mkuzi_1979            gtattggtttctgtttcggatagaaaagacccaatattaatattaaaataattatgagaataacagttataaacagaata [201760]
AY261361.1_Malawi_Lil-20/1_1983  -------------------------------------------------------------------------------- [201760]
AY261360.1_Kenya_1950            ----tggttcctgttccggatagaaaagaccc------gatattaaaaaaatcatgagaataacaattataaacaaaata [201760]
AM712240.1_OURT_88/3             -------------------------------------------------------------------------------- [201760]
AM712239.1_Benin_97/1            -------------------------------------------------------------------------------- [201760]
KM111295.1_Ken06.Bus             -------------------------------------------------------------------------------- [201760]
KM111294.1_Ken05/Tk1             ----tagtttctgttccggatagaaaagaccc------gttattaaaaaaatcatgagaataacaattataaacaaaata [201760]
KM262844.1_strain_L60            -------------------------------------------------------------------------------- [201760]
KM102979.1_26544/OG10_Italy      -------------------------------------------------------------------------------- [201760]
KM262845.1_strain_NHV            -------------------------------------------------------------------------------- [201760]
KX354450.1_47/Ss/2008            -------------------------------------------------------------------------------- [201760]
KP055815.1_strain_BA71           -------------------------------------------------------------------------------- [201760]
MH025920.1_strain_R35            -------------------------------------------------------------------------------- [201760]
MH025919.1_strain_N10            -------------------------------------------------------------------------------- [201760]
MH025918.1_strain_R25            -------------------------------------------------------------------------------- [201760]
MH025917.1_strain_R7             -------------------------------------------------------------------------------- [201760]
MH025916.1_strain_R8             -------------------------------------------------------------------------------- [201760]

FR682468.2_Georgia_2007/1        -------------------------------------------------------------------------------- [201840]
ASF_IND_20_CAD543                -------------------------------------------------------------------------------- [201840]
LR899193.1_Germany_2020/1        -------------------------------------------------------------------------------- [201840]
MN393477.1_Wuhan_2019            -------------------------------------------------------------------------------- [201840]
MW396979.1_Timor-Leste/2019/1    -------------------------------------------------------------------------------- [201840]
MK940252.1_CN/2019/InnerMongolia -------------------------------------------------------------------------------- [201840]
MK128995.1_China/2018/AnhuiXCGQ  -------------------------------------------------------------------------------- [201840]
MH681419.1_ASFV/POL/2015         -------------------------------------------------------------------------------- [201840]
LS478113.1_Estonia_2014          -------------------------------------------------------------------------------- [201840]
KP843857.1_Odintsovo_02/14       -------------------------------------------------------------------------------- [201840]
FN557520.1_strain_E75            -------------------------------------------------------------------------------- [201840]
AY261366.1_Warthog               atagctataagtagaacaagatacattttaatattaaatatattttttcattaaatgaatatatc-----tgtgatatta [201840]
AY261365.1_Warmbaths             atagctataagcagaacaagatacattttaatattaaatatattttttcattaaatgaatatatttaa-atgtcatatta [201840]
AY261364.1_Tengani_62            atagctataagcagaacaagatacatttt-------aatatatttaacc---------------------tgtgatatta [201840]
AY261363.1_Pretoriuskop/96/4     atagctataagcagaacaagatacatttt-------aatatatttttttcattaaatgaatatatc-----tgtgatatta [201840]
AY261362.1_Mkuzi_1979            atagctataagcagaacaaca---------ggattaaacat-----------------------------tttaatatta [201840]
AY261361.1_Malawi_Lil-20/1_1983  -------------------------------------------------------------------------------- [201840]
AY261360.1_Kenya_1950            acagctataagcagaacaagatacattttaatattaaatatattttttcattaaatgaat--------------atatta [201840]
AM712240.1_OURT_88/3             -------------------------------------------------------------------------------- [201840]
AM712239.1_Benin_97/1            -------------------------------------------------------------------------------- [201840]
KM111295.1_Ken06.Bus             -------------------------------------------------------------------------------- [201840]
```

```
KM111294.1_Ken05/Tk1             acagctataagcagaacaagatacattttaatattaaatatattctttcattaaatgaatatatttaatctgtgatatta [201840]
KM262844.1_strain_L60            -------------------------------------------------------------------------------- [201840]
KM102979.1_26544/OG10_Italy      -------------------------------------------------------------------------------- [201840]
KM262845.1_strain_NHV            -------------------------------------------------------------------------------- [201840]
KX354450.1_47/Ss/2008            -------------------------------------------------------------------------------- [201840]
KP055815.1_strain_BA71           -------------------------------------------------------------------------------- [201840]
MH025920.1_strain_R35            -------------------------------------------------------------------------------- [201840]
MH025919.1_strain_N10            -------------------------------------------------------------------------------- [201840]
MH025918.1_strain_R25            -------------------------------------------------------------------------------- [201840]
MH025917.1_strain_R7             -------------------------------------------------------------------------------- [201840]
MH025916.1_strain_R8             -------------------------------------------------------------------------------- [201840]

FR682468.2_Georgia_2007/1        -------------------------------------------------------------------------------- [201920]
ASF_IND_20_CAD543                -------------------------------------------------------------------------------- [201920]
LR899193.1_Germany_2020/1        -------------------------------------------------------------------------------- [201920]
MN393477.1_Wuhan_2019            -------------------------------------------------------------------------------- [201920]
MW396979.1_Timor-Leste/2019/1    -------------------------------------------------------------------------------- [201920]
MK940252.1_CN/2019/InnerMongolia -------------------------------------------------------------------------------- [201920]
MK128995.1_China/2018/AnhuiXCGQ  -------------------------------------------------------------------------------- [201920]
MH681419.1_ASFV/POL/2015         -------------------------------------------------------------------------------- [201920]
LS478113.1_Estonia_2014          -------------------------------------------------------------------------------- [201920]
KP843857.1_Odintsovo_02/14       -------------------------------------------------------------------------------- [201920]
FN557520.1_strain_E75            -------------------------------------------------------------------------------- [201920]
AY261366.1_Warthog               aataattctttagaaatattattctacaattaatgaaatatatacatcaaaatagtattacttattaaagtattttttgg [201920]
AY261365.1_Warmbaths             aataattctttagaaatattattctacaattaatgaaatatatacatcaaaatagtattacttattaaagtattttttgg [201920]
AY261364.1_Tengani_62            aataattctttagaaatattattgtagaa-taatgaaatatatacattaaaatagtgttacttattagagtattttttgg [201920]
AY261363.1_Pretoriuskop/96/4     aataattccttagaaatattattctacaattaatgaaatatatacatcaaaatagtattacttattaaagt-ttttttgg [201920]
AY261362.1_Mkuzi_1979            aatgatt------------------------------------------------------------------------- [201920]
AY261361.1_Malawi_Lil-20/1_1983  -------------------------------------------------------------------------------- [201920]
AY261360.1_Kenya_1950            aataattctttagaaatattattctacaaataatgaaatatatacatcaaaatattgttacttattaaagtattttctga [201920]
AM712240.1_OURT_88/3             -------------------------------------------------------------------------------- [201920]
AM712239.1_Benin_97/1            -------------------------------------------------------------------------------- [201920]
KM111295.1_Ken06.Bus             -------------------------------------------------------------------------------- [201920]
KM111294.1_Ken05/Tk1             aataattctttagaaatattattctacaaataatgaaatatatacatcaaaatattgttacttattaaagtattttctga [201920]
KM262844.1_strain_L60            -------------------------------------------------------------------------------- [201920]
KM102979.1_26544/OG10_Italy      -------------------------------------------------------------------------------- [201920]
KM262845.1_strain_NHV            -------------------------------------------------------------------------------- [201920]
KX354450.1_47/Ss/2008            -------------------------------------------------------------------------------- [201920]
KP055815.1_strain_BA71           -------------------------------------------------------------------------------- [201920]
MH025920.1_strain_R35            -------------------------------------------------------------------------------- [201920]
MH025919.1_strain_N10            -------------------------------------------------------------------------------- [201920]
MH025918.1_strain_R25            -------------------------------------------------------------------------------- [201920]
MH025917.1_strain_R7             -------------------------------------------------------------------------------- [201920]
MH025916.1_strain_R8             -------------------------------------------------------------------------------- [201920]
```

```
FR682468.2_Georgia_2007/1        -------------------------------------------------------------------------------- [202000]
ASF_IND_20_CAD543                -------------------------------------------------------------------------------- [202000]
LR899193.1_Germany_2020/1        -------------------------------------------------------------------------------- [202000]
MN393477.1_Wuhan_2019            -------------------------------------------------------------------------------- [202000]
MW396979.1_Timor-Leste/2019/1    -------------------------------------------------------------------------------- [202000]
MK940252.1_CN/2019/InnerMongolia -------------------------------------------------------------------------------- [202000]
MK128995.1_China/2018/AnhuiXCGQ  -------------------------------------------------------------------------------- [202000]
MH681419.1_ASFV/POL/2015         -------------------------------------------------------------------------------- [202000]
LS478113.1_Estonia_2014          -------------------------------------------------------------------------------- [202000]
KP843857.1_Odintsovo_02/14       -------------------------------------------------------------------------------- [202000]
FN557520.1_strain_E75            -------------------------------------------------------------------------------- [202000]
AY261366.1_Warthog               atatttgcattgagttattattatcactatgtt-aggagctactggatagatttgtgtaaaaaagcattcctgctatttc [202000]
AY261365.1_Warmbaths             atatttgcattgagttattattatcactatgtt-aggagctactggatagatttgtgtaaaaaagcattcctgttatttc [202000]
AY261364.1_Tengani_62            atatttgtattgagttattattaccgctatgttcaaaagctat--gatagatttgtgtaaaaagg--ttcctactgtttc [202000]
AY261363.1_Pretoriuskop/96/4     atatttgcattgagttattattatcactatgtt-aggagctactggatagatttgtgtaaaaaaacattcctgctatttc [202000]
AY261362.1_Mkuzi_1979            atatttaatctaagtta-------------------------------agactttttaaaga------------------ [202000]
AY261361.1_Malawi_Lil-20/1_1983  -------------------------------------------------------------------------------- [202000]
AY261360.1_Kenya_1950            atatttgcattgagttattattaccactatgttcaaaagctat-gggtagatttgtgtaaaaaagcattcctgctgtttc [202000]
AM712240.1_OURT_88/3             -------------------------------------------------------------------------------- [202000]
AM712239.1_Benin_97/1            -------------------------------------------------------------------------------- [202000]
KM111295.1_Ken06.Bus             -------------------------------------------------------------------------------- [202000]
KM111294.1_Ken05/Tk1             atatttgcattgagttattattaccactatgatc-aaagttat-gggtagatttgtgt-aaaaagcattcctgctgtttt [202000]
KM262844.1_strain_L60            -------------------------------------------------------------------------------- [202000]
KM102979.1_26544/OG10_Italy      -------------------------------------------------------------------------------- [202000]
KM262845.1_strain_NHV            -------------------------------------------------------------------------------- [202000]
KX354450.1_47/Ss/2008            -------------------------------------------------------------------------------- [202000]
KP055815.1_strain_BA71           -------------------------------------------------------------------------------- [202000]
MH025920.1_strain_R35            -------------------------------------------------------------------------------- [202000]
MH025919.1_strain_N10            -------------------------------------------------------------------------------- [202000]
MH025918.1_strain_R25            -------------------------------------------------------------------------------- [202000]
MH025917.1_strain_R7             -------------------------------------------------------------------------------- [202000]
MH025916.1_strain_R8             -------------------------------------------------------------------------------- [202000]

FR682468.2_Georgia_2007/1        -------------------------------------------------------------------------------- [202080]
ASF_IND_20_CAD543                -------------------------------------------------------------------------------- [202080]
LR899193.1_Germany_2020/1        -------------------------------------------------------------------------------- [202080]
MN393477.1_Wuhan_2019            -------------------------------------------------------------------------------- [202080]
MW396979.1_Timor-Leste/2019/1    -------------------------------------------------------------------------------- [202080]
MK940252.1_CN/2019/InnerMongolia -------------------------------------------------------------------------------- [202080]
MK128995.1_China/2018/AnhuiXCGQ  -------------------------------------------------------------------------------- [202080]
MH681419.1_ASFV/POL/2015         -------------------------------------------------------------------------------- [202080]
LS478113.1_Estonia_2014          -------------------------------------------------------------------------------- [202080]
KP843857.1_Odintsovo_02/14       -------------------------------------------------------------------------------- [202080]
```

```
FN557520.1_strain_E75             -------------------------------------------------------------------------------- [202080]
AY261366.1_Warthog                atcatat-----tgcttagcaatcctgttgtaaacatatactgattgacaggtatctacacctaacatatc--atacatc [202080]
AY261365.1_Warmbaths              atcatat-----tgctaagcaatcctgttgtaaacatatactgattgacaggtatctacacctaacatatcatatacatc [202080]
AY261364.1_Tengani_62             atcatattgccatgctaagcaatgctgatgtaaac--acactgattgacaggtatctacacctaacatatcctatacaac [202080]
AY261363.1_Pretoriuskop/96/4      atcatat-----tgctaagcaatcctgttgtaaacatatactgattgacaggtatctacacctaacatatcatatacatc [202080]
AY261362.1_Mkuzi_1979             -------------------------------------------------------------------------------- [202080]
AY261361.1_Malawi_Lil-20/1_1983   -------------------------------------------------------------------------------- [202080]
AY261360.1_Kenya_1950             atcatattgccatgctaagcaatgctgctgtaaac--atattgattgacaggtatctgcacttaacttatcacat-tatc [202080]
AM712240.1_OURT_88/3              -------------------------------------------------------------------------------- [202080]
AM712239.1_Benin_97/1             -------------------------------------------------------------------------------- [202080]
KM111295.1_Ken06.Bus              -------------------------------------------------------------------------------- [202080]
KM111294.1_Ken05/Tk1              atcatattaccatgttaaacaatgctgttgtaaac--atattgattgacaggtatctgcacttaacttatcacatatatg [202080]
KM262844.1_strain_L60             -------------------------------------------------------------------------------- [202080]
KM102979.1_26544/OG10_Italy       -------------------------------------------------------------------------------- [202080]
KM262845.1_strain_NHV             -------------------------------------------------------------------------------- [202080]
KX354450.1_47/Ss/2008             -------------------------------------------------------------------------------- [202080]
KP055815.1_strain_BA71            -------------------------------------------------------------------------------- [202080]
MH025920.1_strain_R35             -------------------------------------------------------------------------------- [202080]
MH025919.1_strain_N10             -------------------------------------------------------------------------------- [202080]
MH025918.1_strain_R25             -------------------------------------------------------------------------------- [202080]
MH025917.1_strain_R7              -------------------------------------------------------------------------------- [202080]
MH025916.1_strain_R8              -------------------------------------------------------------------------------- [202080]

FR682468.2_Georgia_2007/1         -------------------------------------------------------------------gtta--------- [202160]
ASF_IND_20_CAD543                 -------------------------------------------------------------------------------- [202160]
LR899193.1_Germany_2020/1         -------------------------------------------------------------------....--------- [202160]
MN393477.1_Wuhan_2019             -------------------------------------------------------------------....--------- [202160]
MW396979.1_Timor-Leste/2019/1     -------------------------------------------------------------------....--------- [202160]
MK940252.1_CN/2019/InnerMongolia  -------------------------------------------------------------------....--------- [202160]
MK128995.1_China/2018/AnhuiXCGQ   -------------------------------------------------------------------....--------- [202160]
MH681419.1_ASFV/POL/2015          -------------------------------------------------------------------....--------- [202160]
LS478113.1_Estonia_2014           -------------------------------------------------------------------....--------- [202160]
KP843857.1_Odintsovo_02/14        -------------------------------------------------------------------....--------- [202160]
FN557520.1_strain_E75             -------------------------------------------------------------------....--------- [202160]
AY261366.1_Warthog                tatactagattttccaaatgcagattgaggggtcagcattattaaacctataaaaatcag--gtgta....tatgaacag [202160]
AY261365.1_Warmbaths              tatactagattttccaaatgcagattgaggggtcagcattattaaatctataaaaatagg-tatata....tatgaacag [202160]
AY261364.1_Tengani_62             tatacgagattttccaaatgcagattgagggatcagcattattaaacctataaaaatcaggtgtata....tatgaacag [202160]
AY261363.1_Pretoriuskop/96/4      tatactagattttccaaatgcagattgaggggtcagcattattaaacctataaaaatcaggtgtata....tatgaacag [202160]
AY261362.1_Mkuzi_1979             --------------------------------------------------------------ttttaa...--------- [202160]
AY261361.1_Malawi_Lil-20/1_1983   -------------------------------------------------------------------------------- [202160]
AY261360.1_Kenya_1950             tatactagattctccaaatgcagattg----------------------------------------------------- [202160]
AM712240.1_OURT_88/3              -------------------------------------------------------------------....--------- [202160]
AM712239.1_Benin_97/1             -------------------------------------------------------------------....--------- [202160]
KM111295.1_Ken06.Bus              -------------------------------------------------------------------------------- [202160]
```

```
KM111294.1_Ken05/Tk1             tatactagattctctaaatgcagattga-ggattagtaatattaaatctataaaaatcaggtgtataac..catgaatgg [202160]
KM262844.1_strain_L60            -------------------------------------------------------------------....--------- [202160]
KM102979.1_26544/OG10_Italy      -------------------------------------------------------------------....--------- [202160]
KM262845.1_strain_NHV            -------------------------------------------------------------------....--------- [202160]
KX354450.1_47/Ss/2008            -------------------------------------------------------------------....--------- [202160]
KP055815.1_strain_BA71           -------------------------------------------------------------------....--------- [202160]
MH025920.1_strain_R35            -------------------------------------------------------------------------------- [202160]
MH025919.1_strain_N10            -------------------------------------------------------------------------------- [202160]
MH025918.1_strain_R25            -------------------------------------------------------------------------------- [202160]
MH025917.1_strain_R7             -------------------------------------------------------------------------------- [202160]
MH025916.1_strain_R8             -------------------------------------------------------------------------------- [202160]

FR682468.2_Georgia_2007/1        -------------------------------------------------------------------------------- [202240]
ASF_IND_20_CAD543                -------------------------------------------------------------------------------- [202240]
LR899193.1_Germany_2020/1        -------------------------------------------------------------------------------- [202240]
MN393477.1_Wuhan_2019            -------------------------------------------------------------------------------- [202240]
MW396979.1_Timor-Leste/2019/1    -------------------------------------------------------------------------------- [202240]
MK940252.1_CN/2019/InnerMongolia -------------------------------------------------------------------------------- [202240]
MK128995.1_China/2018/AnhuiXCGQ  -------------------------------------------------------------------------------- [202240]
MH681419.1_ASFV/POL/2015         -------------------------------------------------------------------------------- [202240]
LS478113.1_Estonia_2014          -------------------------------------------------------------------------------- [202240]
KP843857.1_Odintsovo_02/14       -------------------------------------------------------------------------------- [202240]
FN557520.1_strain_E75            -------------------------------------------------------------------------------- [202240]
AY261366.1_Warthog               gtgttaggaac--------------------------------------------------------------------- [202240]
AY261365.1_Warmbaths             atgttaggaactagtttaatatcttttatgtcttgttttacagcatctgacagct--catgttggtacaaattccaccca [202240]
AY261364.1_Tengani_62            atgttaggaactagtttaatatc--ttatgtcttgttttacagcatctgacagct--catgtcggtacaagtttcacgta [202240]
AY261363.1_Pretoriuskop/96/4     atgttaggaac--------------------------------------------------------------------- [202240]
AY261362.1_Mkuzi_1979            -------------------------------------------------------------------------------- [202240]
AY261361.1_Malawi_Lil-20/1_1983  -------------------------------------------------------------------------------- [202240]
AY261360.1_Kenya_1950            -----aggatctagtttaa-----------tcttattttacagcatctgacagcttacatgtcagtacaaattctac--a [202240]
AM712240.1_OURT_88/3             -------------------------------------------------------------------------------- [202240]
AM712239.1_Benin_97/1            -------------------------------------------------------------------------------- [202240]
KM111295.1_Ken06.Bus             -------------------------------------------------------------------------------- [202240]
KM111294.1_Ken05/Tk1             attttaggatctagtttaatatcttttatgtcttattttacagca---gacagct--catgtcagtacaaattctac--a [202240]
KM262844.1_strain_L60            -------------------------------------------------------------------------------- [202240]
KM102979.1_26544/OG10_Italy      -------------------------------------------------------------------------------- [202240]
KM262845.1_strain_NHV            -------------------------------------------------------------------------------- [202240]
KX354450.1_47/Ss/2008            -------------------------------------------------------------------------------- [202240]
KP055815.1_strain_BA71           -------------------------------------------------------------------------------- [202240]
MH025920.1_strain_R35            -------------------------------------------------------------------------------- [202240]
MH025919.1_strain_N10            -------------------------------------------------------------------------------- [202240]
MH025918.1_strain_R25            -------------------------------------------------------------------------------- [202240]
MH025917.1_strain_R7             -------------------------------------------------------------------------------- [202240]
MH025916.1_strain_R8             -------------------------------------------------------------------------------- [202240]
```

```
FR682468.2_Georgia_2007/1        -------------------------------------------------------------------------------- [202320]
ASF_IND_20_CAD543                -------------------------------------------------------------------------------- [202320]
LR899193.1_Germany_2020/1        -------------------------------------------------------------------------------- [202320]
MN393477.1_Wuhan_2019            -------------------------------------------------------------------------------- [202320]
MW396979.1_Timor-Leste/2019/1    -------------------------------------------------------------------------------- [202320]
MK940252.1_CN/2019/InnerMongolia -------------------------------------------------------------------------------- [202320]
MK128995.1_China/2018/AnhuiXCGQ  -------------------------------------------------------------------------------- [202320]
MH681419.1_ASFV/POL/2015         -------------------------------------------------------------------------------- [202320]
LS478113.1_Estonia_2014          -------------------------------------------------------------------------------- [202320]
KP843857.1_Odintsovo_02/14       -------------------------------------------------------------------------------- [202320]
FN557520.1_strain_E75            -------------------------------------------------------------------------------- [202320]
AY261366.1_Warthog               -------------------------------------------------------------------------------- [202320]
AY261365.1_Warmbaths             cacagtggtat-tttttattcttacagttta--tctattttacacttgtta-ttttttagtgattttagaagataaaaaa [202320]
AY261364.1_Tengani_62            cacagtgg--------tatccttacagtttg--tttattttacacttgtta-ttttttggtgattttagaagat-aaaaa [202320]
AY261363.1_Pretoriuskop/96/4     -------------------------------------------------------------------------------- [202320]
AY261362.1_Mkuzi_1979            -------------------------------------------------------------------------------- [202320]
AY261361.1_Malawi_Lil-20/1_1983  -------------------------------------------------------------------------------- [202320]
AY261360.1_Kenya_1950            cctagtggtat-cctttatacatacagtctgcatgtattttacatttgttatttttttggtgattttagaagataa---- [202320]
AM712240.1_OURT_88/3             -------------------------------------------------------------------------------- [202320]
AM712239.1_Benin_97/1            -------------------------------------------------------------------------------- [202320]
KM111295.1_Ken06.Bus             -----aggtat-cctttatatatacagtctacatgtattttacatttgttaattttttagtgattttagaaga-aa---- [202320]
KM111294.1_Ken05/Tk1             cccagtggtatccctttatacatacagtctgcatgtattttacatttgttatttttttggtgattttagaagataa---- [202320]
KM262844.1_strain_L60            -------------------------------------------------------------------------------- [202320]
KM102979.1_26544/OG10_Italy      -------------------------------------------------------------------------------- [202320]
KM262845.1_strain_NHV            -------------------------------------------------------------------------------- [202320]
KX354450.1_47/Ss/2008            -------------------------------------------------------------------------------- [202320]
KP055815.1_strain_BA71           -------------------------------------------------------------------------------- [202320]
MH025920.1_strain_R35            -----aggtat-cctttatatatacagtctacatgtattttacatttgttaattttttagtgattttagaaga-aa---- [202320]
MH025919.1_strain_N10            -----aggtat-cctttatatatacagtctacatgtattttacatttgttaattttttagtgattttagaaga-aa---- [202320]
MH025918.1_strain_R25            -----aggtat-cctttatatatacagtctacatgtattttacatttgttaattttttagtgattttagaaga-aa---- [202320]
MH025917.1_strain_R7             -----aggtat-cctttatatatacagtctacatgtattttacatttgttaattttttagtgattttagaaga-aa---- [202320]
MH025916.1_strain_R8             -----aggtat-cctttatatatacagtctacatgtattttacatttgttaattttttagtgattttagaaga-aa---- [202320]

FR682468.2_Georgia_2007/1        -------------------------------------------------------------------------------- [202400]
ASF_IND_20_CAD543                -------------------------------------------------------------------------------- [202400]
LR899193.1_Germany_2020/1        -------------------------------------------------------------------------------- [202400]
MN393477.1_Wuhan_2019            -------------------------------------------------------------------------------- [202400]
MW396979.1_Timor-Leste/2019/1    -------------------------------------------------------------------------------- [202400]
MK940252.1_CN/2019/InnerMongolia -------------------------------------------------------------------------------- [202400]
MK128995.1_China/2018/AnhuiXCGQ  -------------------------------------------------------------------------------- [202400]
MH681419.1_ASFV/POL/2015         -------------------------------------------------------------------------------- [202400]
LS478113.1_Estonia_2014          -------------------------------------------------------------------------------- [202400]
KP843857.1_Odintsovo_02/14       -------------------------------------------------------------------------------- [202400]
```

```
FN557520.1_strain_E75            -------------------------------------------------------------------------------- [202400]
AY261366.1_Warthog               -------------------------------------------------------------------------------- [202400]
AY261365.1_Warmbaths             gtcagtattaaaattatcacaagaataatagtagctataaacacttgaataagaaga------------------tttta [202400]
AY261364.1_Tengani_62            gtcagtattaaaattatcacaagaataatagtagctataaatacttgaataagaagagactttaatattaaaatgtttta [202400]
AY261363.1_Pretoriuskop/96/4     -------------------------------------------------------------------------------- [202400]
AY261362.1_Mkuzi_1979            -------------------------------------------------------------------------------- [202400]
AY261361.1_Malawi_Lil-20/1_1983  -------------------------------------------------------------------------------- [202400]
AY261360.1_Kenya_1950            -------------------------------------------------------------------------------- [202400]
AM712240.1_OURT_88/3             -------------------------------------------------------------------------------- [202400]
AM712239.1_Benin_97/1            -------------------------------------------------------------------------------- [202400]
KM111295.1_Ken06.Bus             -------------------------------------------------------------------------------- [202400]
KM111294.1_Ken05/Tk1             -------------------------------------------------------------------------------- [202400]
KM262844.1_strain_L60            -------------------------------------------------------------------------------- [202400]
KM102979.1_26544/OG10_Italy      -------------------------------------------------------------------------------- [202400]
KM262845.1_strain_NHV            -------------------------------------------------------------------------------- [202400]
KX354450.1_47/Ss/2008            -------------------------------------------------------------------------------- [202400]
KP055815.1_strain_BA71           -------------------------------------------------------------------------------- [202400]
MH025920.1_strain_R35            -------------------------------------------------------------------------------- [202400]
MH025919.1_strain_N10            -------------------------------------------------------------------------------- [202400]
MH025918.1_strain_R25            -------------------------------------------------------------------------------- [202400]
MH025917.1_strain_R7             -------------------------------------------------------------------------------- [202400]
MH025916.1_strain_R8             -------------------------------------------------------------------------------- [202400]

FR682468.2_Georgia_2007/1        -------------------ggctaagtaaatttaatattttaaagtatttggaaaaatattttttaa-catatgatgtct [202480]
ASF_IND_20_CAD543                -------------------------------------------------------------------------------- [202480]
LR899193.1_Germany_2020/1        -------------------................................................-............ [202480]
MN393477.1_Wuhan_2019            -------------------................................................-............ [202480]
MW396979.1_Timor-Leste/2019/1    -------------------................................................-............ [202480]
MK940252.1_CN/2019/InnerMongolia -------------------................................................-............ [202480]
MK128995.1_China/2018/AnhuiXCGQ  -------------------................................................-............ [202480]
MH681419.1_ASFV/POL/2015         -------------------................................................-............ [202480]
LS478113.1_Estonia_2014          -------------------................................................-............ [202480]
KP843857.1_Odintsovo_02/14       -------------------................................................-............ [202480]
FN557520.1_strain_E75            -------------------..................-.........c...................-......c..... [202480]
AY261366.1_Warthog               -------------------..a.c.t.......-...-.a.c..a.tg..a...g.........at.------------- [202480]
AY261365.1_Warmbaths             atattaaagattttaatca..a.t.t.......-...-.a.c..actg......g..........t.a......c..... [202480]
AY261364.1_Tengani_62            atattaaagattttaatta..a.c.t.......-...-.a.t..a.tg......g............-......c..... [202480]
AY261363.1_Pretoriuskop/96/4     -------------------..a.c.t.......-...-.a.c..a.cg..a...g.........at.------------- [202480]
AY261362.1_Mkuzi_1979            -------------------..a.c.t.......-...-.a.c..a.tg..a................--.....c.a... [202480]
AY261361.1_Malawi_Lil-20/1_1983  ----------------------------------...------------------------------------------- [202480]
AY261360.1_Kenya_1950            ----------------------------------.a.-g.c...a.ta..a...g.........at.------------- [202480]
AM712240.1_OURT_88/3             -------------------..................-.........c...................-......c..... [202480]
AM712239.1_Benin_97/1            -------------------..................-.........c...................-......c..... [202480]
KM111295.1_Ken06.Bus             ----------------------------------.a.-g.c...a.tg..a...g.........at.------------- [202480]
```

```
KM111294.1_Ken05/Tk1          ----------------------------------.a.-g.-...a.ta..a...g........at.-------------- [202480]
KM262844.1_strain_L60         -------------------..................-.........c...................-......c..... [202480]
KM102979.1_26544/OG10_Italy   -------------------..................-.........c...................-......c..... [202480]
KM262845.1_strain_NHV         -------------------..................-.........c...................-......c..... [202480]
KX354450.1_47/Ss/2008         -------------------..................-.........c...................-......c..... [202480]
KP055815.1_strain_BA71        -------------------..................-.........c...................-......c..... [202480]
MH025920.1_strain_R35         ----------------------------------.a.-g.c...a.tg..a...g........at.-------------- [202480]
MH025919.1_strain_N10         ----------------------------------.a.-g.c...a.tg..a...g........at.-------------- [202480]
MH025918.1_strain_R25         ----------------------------------.a.-g.c...a.tg..a...g........at.-------------- [202480]
MH025917.1_strain_R7          ----------------------------------.a.-g.c...a.tg..a...g........at.-------------- [202480]
MH025916.1_strain_R8          ----------------------------------.a.-g.c...a.tg..a...g........at.-------------- [202480]

FR682468.2_Georgia_2007/1     aggaatattttttagacatttaaaaccatatag---ttactttatttattacactgaacttgaaaagacttattacctaa [202560]
ASF_IND_20_CAD543             -------------------------------------------------------------------------------- [202560]
LR899193.1_Germany_2020/1     .................................---............................................ [202560]
MN393477.1_Wuhan_2019         .................................---............................................ [202560]
MW396979.1_Timor-Leste/2019/1 .................................---............................................ [202560]
MK940252.1_CN/2019/InnerMongolia .................................---............................................ [202560]
MK128995.1_China/2018/AnhuiXCGQ .................................---............................................ [202560]
MH681419.1_ASFV/POL/2015      .................................---............................................ [202560]
LS478113.1_Estonia_2014       .................................---............................................ [202560]
KP843857.1_Odintsovo_02/14    .................................---............................................ [202560]
FN557520.1_strain_E75         ......-..........................---...................t............a.....g..... [202560]
AY261366.1_Warthog            -----------------ta.......t......-------........gg.....t..tc........g........... [202560]
AY261365.1_Warmbaths          ................t.........t......-------....a....g.....t..tc........g........... [202560]
AY261364.1_Tengani_62         ......-a........---.......t.....ag-c........a..........t..tc........g........... [202560]
AY261363.1_Pretoriuskop/96/4  -----------------ta.......t......-------........gg.....t..tc........g........... [202560]
AY261362.1_Mkuzi_1979         .aa...-a......t.t.-..............---...................t............a.....g..... [202560]
AY261361.1_Malawi_Lil-20/1_1983 ----------------------------------------.......t.....a.t...a.a......a........... [202560]
AY261360.1_Kenya_1950         -------------------a.....tg.....atag........a..........t...c.................... [202560]
AM712240.1_OURT_88/3          ......-..........................---...................t............a.....g..... [202560]
AM712239.1_Benin_97/1         ......-..........................---...................t............a.....g..... [202560]
KM111295.1_Ken06.Bus          -------------------......tg.....--ag........a..........t...c.................... [202560]
KM111294.1_Ken05/Tk1          -------------------......tg.....atag........a........t.t...c.................... [202560]
KM262844.1_strain_L60         ......-..........................---...................t............a.....g..... [202560]
KM102979.1_26544/OG10_Italy   ......-..........................---...................t............a.....g..... [202560]
KM262845.1_strain_NHV         ......-..........................---...................t............a.....g..... [202560]
KX354450.1_47/Ss/2008         ......-..........................---...................t............a.....g..... [202560]
KP055815.1_strain_BA71        ......-..........................---...................t............a.....g..... [202560]
MH025920.1_strain_R35         -------------------......tg.....--ag........a..........t...c.................... [202560]
MH025919.1_strain_N10         -------------------......tg.....--ag........a..........t...c.................... [202560]
MH025918.1_strain_R25         -------------------......tg.....--ag........a..........t...c.................... [202560]
MH025917.1_strain_R7          -------------------......tg.....--ag........a..........t...c.................... [202560]
MH025916.1_strain_R8          -------------------......tg.....--ag........a..........t...c.................... [202560]
```

```
FR682468.2_Georgia_2007/1        aa---tattaatagatgaagtaatattgtgtaattgagtccataacatgggtgggaaacaaaaatctcgtaatatgaaaa [202640]
ASF_IND_20_CAD543                -------------------------------------------------------------------------------- [202640]
LR899193.1_Germany_2020/1        ..---........................................................................... [202640]
MN393477.1_Wuhan_2019            ..---........................................................................... [202640]
MW396979.1_Timor-Leste/2019/1    ..---........................................................................... [202640]
MK940252.1_CN/2019/InnerMongolia ..---........................................................................... [202640]
MK128995.1_China/2018/AnhuiXCGQ  ..---........................................................................... [202640]
MH681419.1_ASFV/POL/2015         ..---........................................................................... [202640]
LS478113.1_Estonia_2014          ..---........................................................................... [202640]
KP843857.1_Odintsovo_02/14       ..---........................................................................... [202640]
FN557520.1_strain_E75            ..---........................................................................... [202640]
AY261366.1_Warthog               ..tat......a..g.a..........c.................................................... [202640]
AY261365.1_Warmbaths             ..---........................................................................... [202640]
AY261364.1_Tengani_62            ..---.................................................a......................... [202640]
AY261363.1_Pretoriuskop/96/4     ..tat......a..g.a..........c.................................................... [202640]
AY261362.1_Mkuzi_1979            ..---...............................g...........a.....a......................... [202640]
AY261361.1_Malawi_Lil-20/1_1983  g.aag......g..t.......g......t....atctg.........a.....a..g...................... [202640]
AY261360.1_Kenya_1950            ..tat......a..gga...............................a............................... [202640]
AM712240.1_OURT_88/3             ..---........................................................................... [202640]
AM712239.1_Benin_97/1            ..---........................................................................... [202640]
KM111295.1_Ken06.Bus             ..tat..c...a..g.a............................................................... [202640]
KM111294.1_Ken05/Tk1             ..tat......ag.gga............................................................... [202640]
KM262844.1_strain_L60            ..---........................................................................... [202640]
KM102979.1_26544/OG10_Italy      ..---........................................................................... [202640]
KM262845.1_strain_NHV            ..---........................................................................... [202640]
KX354450.1_47/Ss/2008            ..---........................................................................... [202640]
KP055815.1_strain_BA71           ..---........................................................................... [202640]
MH025920.1_strain_R35            ..tat..c...a..g.a............................................................... [202640]
MH025919.1_strain_N10            ..tat..c...a..g.a............................................................... [202640]
MH025918.1_strain_R25            ..tat..c...a..g.a............................................................... [202640]
MH025917.1_strain_R7             ..tat..c...a..g.a............................................................... [202640]
MH025916.1_strain_R8             ..tat..c...a..g.a............................................................... [202640]

FR682468.2_Georgia_2007/1        ataaacatcctaaaaagagtgcaattgttataagtttatgtaactttattttaaagtaagaatataaaaatatgagtaca [202720]
ASF_IND_20_CAD543                -------------------------------------------------------------------------------- [202720]
LR899193.1_Germany_2020/1        ................................................................................ [202720]
MN393477.1_Wuhan_2019            ................................................................................ [202720]
MW396979.1_Timor-Leste/2019/1    ................................................................................ [202720]
MK940252.1_CN/2019/InnerMongolia ................................................................................ [202720]
MK128995.1_China/2018/AnhuiXCGQ  ................................................................................ [202720]
MH681419.1_ASFV/POL/2015         ................................................................................ [202720]
LS478113.1_Estonia_2014          ................................................................................ [202720]
KP843857.1_Odintsovo_02/14       ................................................................................ [202720]
```

```
FN557520.1_strain_E75            ................................................................................ [202720]
AY261366.1_Warthog               ................................................................................ [202720]
AY261365.1_Warmbaths             ...........................................t..................................t. [202720]
AY261364.1_Tengani_62            ................................................................................ [202720]
AY261363.1_Pretoriuskop/96/4     ................................................................................ [202720]
AY261362.1_Mkuzi_1979            ................................................................................ [202720]
AY261361.1_Malawi_Lil-20/1_1983  ............g.....................c..g....................g..g................t. [202720]
AY261360.1_Kenya_1950            ............g.....................c..g....................g..g................t. [202720]
AM712240.1_OURT_88/3             ................................................................................ [202720]
AM712239.1_Benin_97/1            ................................................................................ [202720]
KM111295.1_Ken06.Bus             .c..........g.....................c..g..........c.............................t. [202720]
KM111294.1_Ken05/Tk1             ............g.....................c..g....................g..g................t. [202720]
KM262844.1_strain_L60            ................................................................................ [202720]
KM102979.1_26544/OG10_Italy      ................................................................................ [202720]
KM262845.1_strain_NHV            ................................................................................ [202720]
KX354450.1_47/Ss/2008            ................................................................................ [202720]
KP055815.1_strain_BA71           ................................................................................ [202720]
MH025920.1_strain_R35            .c..........g.....................c..g..........c.........g...................t. [202720]
MH025919.1_strain_N10            .c..........g.....................c..g..........c.........g...................t. [202720]
MH025918.1_strain_R25            .c..........g.....................c..g..........c.........g...................t. [202720]
MH025917.1_strain_R7             .c..........g.....................c..g..........c.........g...................t. [202720]
MH025916.1_strain_R8             .c..........g.....................c..g..........c.........g...................t. [202720]

FR682468.2_Georgia_2007/1        agaggaataggggccattactaacattggctccaacatcctgttgtctacaaaaaaaaatatttttttttagcaaaaaaaa [202800]
ASF_IND_20_CAD543                -------------------------------------------------------------------------------- [202800]
LR899193.1_Germany_2020/1        ................................................................................ [202800]
MN393477.1_Wuhan_2019            ................................................................................ [202800]
MW396979.1_Timor-Leste/2019/1    ................................................................................ [202800]
MK940252.1_CN/2019/InnerMongolia ................................................................................ [202800]
MK128995.1_China/2018/AnhuiXCGQ  ................................................................................ [202800]
MH681419.1_ASFV/POL/2015         ................................................................................ [202800]
LS478113.1_Estonia_2014          ................................................................................ [202800]
KP843857.1_Odintsovo_02/14       ................................................................................ [202800]
FN557520.1_strain_E75            .......................t.............................................g.......... [202800]
AY261366.1_Warthog               .......................t.a...................................................... [202800]
AY261365.1_Warmbaths             .......................t................................................-....... [202800]
AY261364.1_Tengani_62            .......................t................c....................................... [202800]
AY261363.1_Pretoriuskop/96/4     .......................t.a...........................................gct-....... [202800]
AY261362.1_Mkuzi_1979            .......................t................................................-....... [202800]
AY261361.1_Malawi_Lil-20/1_1983  ...........c...........t.a...t................t..............ac......t.agg...... [202800]
AY261360.1_Kenya_1950            ...........c...........t......................t..............ac......t.agg...... [202800]
AM712240.1_OURT_88/3             .......................t..........................-..................g.......... [202800]
AM712239.1_Benin_97/1            .......................t.....................................-.......g.......... [202800]
KM111295.1_Ken06.Bus             ...........c...........t.....................................ac......t.ggg...... [202800]
```

```
KM111294.1_Ken05/Tk1              ...........c...........t.....................................ac......t.ggg...... [202800]
KM262844.1_strain_L60             .......................t.............................................g..-....... [202800]
KM102979.1_26544/OG10_Italy       .......................t.............................................g.......... [202800]
KM262845.1_strain_NHV             .......................t..........................-..................g.......... [202800]
KX354450.1_47/Ss/2008             .......................t.............................................g.......... [202800]
KP055815.1_strain_BA71            .......................t.............................................g.......... [202800]
MH025920.1_strain_R35             ...........c...........t.....................................ac......t.ggg...... [202800]
MH025919.1_strain_N10             ...........c...........t.....................................ac......t.ggg...... [202800]
MH025918.1_strain_R25             ...........c...........t.....................................ac......t.ggg...... [202800]
MH025917.1_strain_R7              ...........c...........t.....................................ac......t.ggg...... [202800]
MH025916.1_strain_R8              ...........c...........t.....................................ac......t.ggg...... [202800]

FR682468.2_Georgia_2007/1         at------------------------------------------------------------------------------ [202880]
ASF_IND_20_CAD543                 -------------------------------------------------------------------------------- [202880]
LR899193.1_Germany_2020/1         ..------------------------------------------------------------------------------ [202880]
MN393477.1_Wuhan_2019             ..------------------------------------------------------------------------------ [202880]
MW396979.1_Timor-Leste/2019/1     ..------------------------------------------------------------------------------ [202880]
MK940252.1_CN/2019/InnerMongolia  ..------------------------------------------------------------------------------ [202880]
MK128995.1_China/2018/AnhuiXCGQ   ..------------------------------------------------------------------------------ [202880]
MH681419.1_ASFV/POL/2015          ..------------------------------------------------------------------------------ [202880]
LS478113.1_Estonia_2014           ..------------------------------------------------------------------------------ [202880]
KP843857.1_Odintsovo_02/14        ..------------------------------------------------------------------------------ [202880]
FN557520.1_strain_E75             ..------------------------------------------------------------------------------ [202880]
AY261366.1_Warthog                ..------------------------------------------------------------------------------ [202880]
AY261365.1_Warmbaths              ..------------------------------------------------------------------------------ [202880]
AY261364.1_Tengani_62             ..------------------------------------------------------------------------------ [202880]
AY261363.1_Pretoriuskop/96/4      ..------------------------------------------------------------------------------ [202880]
AY261362.1_Mkuzi_1979             ..------------------------------------------------------------------------------ [202880]
AY261361.1_Malawi_Lil-20/1_1983   ..attttcaaataga--gtgtgttataattgatattttcctatgtatagattttatcaaatctaaaatataccaaattat [202880]
AY261360.1_Kenya_1950             ..attttcaaataga--atgtgttataattgatatttttctttgtatagattctatcaaatctaaaatataccaagttat [202880]
AM712240.1_OURT_88/3              ..------------------------------------------------------------------------------ [202880]
AM712239.1_Benin_97/1             ..------------------------------------------------------------------------------ [202880]
KM111295.1_Ken06.Bus              ..attttcaaatagagtgtgtgttataattgatatttttctttgtttagattctatca---------------------- [202880]
KM111294.1_Ken05/Tk1              ..attttcaaataga--gtgtgttataattgatatttttctatgtatagattttataaaatctaaaacataccaaattat [202880]
KM262844.1_strain_L60             ..------------------------------------------------------------------------------ [202880]
KM102979.1_26544/OG10_Italy       ..------------------------------------------------------------------------------ [202880]
KM262845.1_strain_NHV             ..------------------------------------------------------------------------------ [202880]
KX354450.1_47/Ss/2008             ..------------------------------------------------------------------------------ [202880]
KP055815.1_strain_BA71            ..------------------------------------------------------------------------------ [202880]
MH025920.1_strain_R35             ..attttcaaatagagtgtgtgttataattgatatttttctttgtttagattctatca---------------------- [202880]
MH025919.1_strain_N10             ..attttcaaatagagtgtgtgttataattgatatttttctttgtttagattctatca---------------------- [202880]
MH025918.1_strain_R25             ..attttcaaatagagtgtgtgttataattgatatttttctttgtttagattctatca---------------------- [202880]
MH025917.1_strain_R7              ..attttcaaatagagtgtgtgttataattgatatttttctttgtttagattctatca---------------------- [202880]
MH025916.1_strain_R8              ..attttcaaatagagtgtgtgttataattgatatttttctttgtttagattctatca---------------------- [202880]
```

```
FR682468.2_Georgia_2007/1        ----------------------------------------------------------ccatgg---------------- [202960]
ASF_IND_20_CAD543                -------------------------------------------------------------------------------- [202960]
LR899193.1_Germany_2020/1        ----------------------------------------------------------......---------------- [202960]
MN393477.1_Wuhan_2019            ----------------------------------------------------------......---------------- [202960]
MW396979.1_Timor-Leste/2019/1    ----------------------------------------------------------......---------------- [202960]
MK940252.1_CN/2019/InnerMongolia ----------------------------------------------------------......---------------- [202960]
MK128995.1_China/2018/AnhuiXCGQ  ----------------------------------------------------------......---------------- [202960]
MH681419.1_ASFV/POL/2015         ----------------------------------------------------------......---------------- [202960]
LS478113.1_Estonia_2014          ----------------------------------------------------------......---------------- [202960]
KP843857.1_Odintsovo_02/14       ----------------------------------------------------------......---------------- [202960]
FN557520.1_strain_E75            ----------------------------------------------------------......---------------- [202960]
AY261366.1_Warthog               ----------------------------------------------------------......---------------- [202960]
AY261365.1_Warmbaths             ----------------------------------------------------------......---------------- [202960]
AY261364.1_Tengani_62            ----------------------------------------------------------......---------------- [202960]
AY261363.1_Pretoriuskop/96/4     ----------------------------------------------------------......---------------- [202960]
AY261362.1_Mkuzi_1979            ----------------------------------------------------------......---------------- [202960]
AY261361.1_Malawi_Lil-20/1_1983  cataatgataattggatatattactatcttaattaataaaatatcagacgtattcatg.t...agaatagactgctgaat [202960]
AY261360.1_Kenya_1950            cataatgataattggatatattactatcttaattaataaaatatcatacgtattcatg.t...agaatagactgctgaaa [202960]
AM712240.1_OURT_88/3             ----------------------------------------------------------......---------------- [202960]
AM712239.1_Benin_97/1            ----------------------------------------------------------......---------------- [202960]
KM111295.1_Ken06.Bus             -----------------------------taattaataaaataccacacgtatttatgtt...agaatagactgttg-at [202960]
KM111294.1_Ken05/Tk1             tattatgattattggatagattactatcttaattaataaaatatcagacgtattcatg.t...agcatagactgctggaa [202960]
KM262844.1_strain_L60            ----------------------------------------------------------......---------------- [202960]
KM102979.1_26544/OG10_Italy      ----------------------------------------------------------......---------------- [202960]
KM262845.1_strain_NHV            ----------------------------------------------------------......---------------- [202960]
KX354450.1_47/Ss/2008            ----------------------------------------------------------......---------------- [202960]
KP055815.1_strain_BA71           ----------------------------------------------------------......---------------- [202960]
MH025920.1_strain_R35            -----------------------------taattaataaaataccacacgtatttatgtt...agaatagactgttg-at [202960]
MH025919.1_strain_N10            -----------------------------taattaataaaataccacacgtatttatgtt...agaatagactgttg-at [202960]
MH025918.1_strain_R25            -----------------------------taattaataaaataccacacgtatttatgtt...agaatagactgttg-at [202960]
MH025917.1_strain_R7             -----------------------------taattaataaaataccacacgtatttatgtt...agaatagactgttg-at [202960]
MH025916.1_strain_R8             -----------------------------taattaataaaataccacacgtatttatgtt...agaatagactgttg-at [202960]

FR682468.2_Georgia_2007/1        -------------------------------------------------------------------------------- [203040]
ASF_IND_20_CAD543                -------------------------------------------------------------------------------- [203040]
LR899193.1_Germany_2020/1        -------------------------------------------------------------------------------- [203040]
MN393477.1_Wuhan_2019            -------------------------------------------------------------------------------- [203040]
MW396979.1_Timor-Leste/2019/1    -------------------------------------------------------------------------------- [203040]
MK940252.1_CN/2019/InnerMongolia -------------------------------------------------------------------------------- [203040]
MK128995.1_China/2018/AnhuiXCGQ  -------------------------------------------------------------------------------- [203040]
MH681419.1_ASFV/POL/2015         -------------------------------------------------------------------------------- [203040]
LS478113.1_Estonia_2014          -------------------------------------------------------------------------------- [203040]
KP843857.1_Odintsovo_02/14       -------------------------------------------------------------------------------- [203040]
```

```
FN557520.1_strain_E75              -------------------------------------------------------------------------------- [203040]
AY261366.1_Warthog                 -------------------------------------------------------------------------------- [203040]
AY261365.1_Warmbaths               -------------------------------------------------------------------------------- [203040]
AY261364.1_Tengani_62              -------------------------------------------------------------------------------- [203040]
AY261363.1_Pretoriuskop/96/4       -------------------------------------------------------------------------------- [203040]
AY261362.1_Mkuzi_1979              -------------------------------------------------------------------------------- [203040]
AY261361.1_Malawi_Lil-20/1_1983    ttttaataattttttattaaaatt-taggagtatagagtcagtgtttattaaaatgcttataatctctaacaacataatt [203040]
AY261360.1_Kenya_1950              ttttaataattttttattaaaatt-------------------------------------------------------- [203040]
AM712240.1_OURT_88/3               -------------------------------------------------------------------------------- [203040]
AM712239.1_Benin_97/1              -------------------------------------------------------------------------------- [203040]
KM111295.1_Ken06.Bus               ttttaataattttttattaaaaatttaggagtataggtccggtgtttattaaaatacttataatctctaacaacataatt [203040]
KM111294.1_Ken05/Tk1               ttttaataattttttattaaaatt-------------------------------------------------------- [203040]
KM262844.1_strain_L60              -------------------------------------------------------------------------------- [203040]
KM102979.1_26544/OG10_Italy        -------------------------------------------------------------------------------- [203040]
KM262845.1_strain_NHV              -------------------------------------------------------------------------------- [203040]
KX354450.1_47/Ss/2008              -------------------------------------------------------------------------------- [203040]
KP055815.1_strain_BA71             -------------------------------------------------------------------------------- [203040]
MH025920.1_strain_R35              ttttaataattttttattaaaaatttaggagtataggtccggtgtttattaaaatacttataatctctaacaacataatt [203040]
MH025919.1_strain_N10              ttttaataattttttattaaaaatttaggagtataggtccggtgtttattaaaatacttataatctctaacaacataatt [203040]
MH025918.1_strain_R25              ttttaataattttttattaaaaatttaggagtataggtccggtgtttattaaaatacttataatctctaacaacataatt [203040]
MH025917.1_strain_R7               ttttaataattttttattaaaaatttaggagtataggtccggtgtttattaaaatacttataatctctaacaacataatt [203040]
MH025916.1_strain_R8               ttttaataattttttattaaaaatttaggagtataggtccggtgtttattaaaatacttataatctctaacaacataatt [203040]

FR682468.2_Georgia_2007/1          -------------------------------------------------------------------------------- [203120]
ASF_IND_20_CAD543                  -------------------------------------------------------------------------------- [203120]
LR899193.1_Germany_2020/1          -------------------------------------------------------------------------------- [203120]
MN393477.1_Wuhan_2019              -------------------------------------------------------------------------------- [203120]
MW396979.1_Timor-Leste/2019/1      -------------------------------------------------------------------------------- [203120]
MK940252.1_CN/2019/InnerMongolia   -------------------------------------------------------------------------------- [203120]
MK128995.1_China/2018/AnhuiXCGQ    -------------------------------------------------------------------------------- [203120]
MH681419.1_ASFV/POL/2015           -------------------------------------------------------------------------------- [203120]
LS478113.1_Estonia_2014            -------------------------------------------------------------------------------- [203120]
KP843857.1_Odintsovo_02/14         -------------------------------------------------------------------------------- [203120]
FN557520.1_strain_E75              -------------------------------------------------------------------------------- [203120]
AY261366.1_Warthog                 -------------------------------------------------------------------------------- [203120]
AY261365.1_Warmbaths               -------------------------------------------------------------------------------- [203120]
AY261364.1_Tengani_62              -------------------------------------------------------------------------------- [203120]
AY261363.1_Pretoriuskop/96/4       -------------------------------------------------------------------------------- [203120]
AY261362.1_Mkuzi_1979              -------------------------------------------------------------------------------- [203120]
AY261361.1_Malawi_Lil-20/1_1983    attaagttataaactaacttaataataagggcctatctatttttactgatagatatttttaaaccaatgattcagaattt [203120]
AY261360.1_Kenya_1950              -------------------------------------------------------------------------------- [203120]
AM712240.1_OURT_88/3               -------------------------------------------------------------------------------- [203120]
AM712239.1_Benin_97/1              -------------------------------------------------------------------------------- [203120]
KM111295.1_Ken06.Bus               attaagttataaactaactt-ataataagggcctatctatttttactgataaatatttttaacccaatgattcaagattt [203120]
```

```
KM111294.1_Ken05/Tk1           -------------------------------------------------------------------------------- [203120]
KM262844.1_strain_L60          -------------------------------------------------------------------------------- [203120]
KM102979.1_26544/OG10_Italy    -------------------------------------------------------------------------------- [203120]
KM262845.1_strain_NHV          -------------------------------------------------------------------------------- [203120]
KX354450.1_47/Ss/2008          -------------------------------------------------------------------------------- [203120]
KP055815.1_strain_BA71         -------------------------------------------------------------------------------- [203120]
MH025920.1_strain_R35          attaagttataaactaactt-ataataagggcctatctatttttactgataaatatttttaacccaatgattcaagattt [203120]
MH025919.1_strain_N10          attaagttataaactaactt-ataataagggcctatctatttttactgataaatatttttaacccaatgattcaagattt [203120]
MH025918.1_strain_R25          attaagttataaactaactt-ataataagggcctatctatttttactgataaatatttttaacccaatgattcaagattt [203120]
MH025917.1_strain_R7           attaagttataaactaactt-ataataagggcctatctatttttactgataaatatttttaacccaatgattcaagattt [203120]
MH025916.1_strain_R8           attaagttataaactaactt-ataataagggcctatctatttttactgataaatatttttaacccaatgattcaagattt [203120]

FR682468.2_Georgia_2007/1      -------------------------------------------------------------------------------- [203200]
ASF_IND_20_CAD543              -------------------------------------------------------------------------------- [203200]
LR899193.1_Germany_2020/1      -------------------------------------------------------------------------------- [203200]
MN393477.1_Wuhan_2019          -------------------------------------------------------------------------------- [203200]
MW396979.1_Timor-Leste/2019/1  -------------------------------------------------------------------------------- [203200]
MK940252.1_CN/2019/InnerMongolia -------------------------------------------------------------------------------- [203200]
MK128995.1_China/2018/AnhuiXCGQ -------------------------------------------------------------------------------- [203200]
MH681419.1_ASFV/POL/2015       -------------------------------------------------------------------------------- [203200]
LS478113.1_Estonia_2014        -------------------------------------------------------------------------------- [203200]
KP843857.1_Odintsovo_02/14     -------------------------------------------------------------------------------- [203200]
FN557520.1_strain_E75          -------------------------------------------------------------------------------- [203200]
AY261366.1_Warthog             -------------------------------------------------------------------------------- [203200]
AY261365.1_Warmbaths           -------------------------------------------------------------------------------- [203200]
AY261364.1_Tengani_62          -------------------------------------------------------------------------------- [203200]
AY261363.1_Pretoriuskop/96/4   -------------------------------------------------------------------------------- [203200]
AY261362.1_Mkuzi_1979          -------------------------------------------------------------------------------- [203200]
AY261361.1_Malawi_Lil-20/1_1983 taaaagac--tgttaaagggagttatgtttgcccagcagggcatgcaattatttgtttcgcaaatgatctcagtcttata [203200]
AY261360.1_Kenya_1950          ----------------taggtgttatgtttgtccagcaggtcatgcaattatttgtttcgcaggtaatctcagtcttata [203200]
AM712240.1_OURT_88/3           -------------------------------------------------------------------------------- [203200]
AM712239.1_Benin_97/1          -------------------------------------------------------------------------------- [203200]
KM111295.1_Ken06.Bus           taaaggactttattaatgggagttatgtttgcccagcagatcatacaattatttatttcgcaggtaatctcagtcttata [203200]
KM111294.1_Ken05/Tk1           ----------------taggcgttatgtttgtccagcaggtcatgcaattatttgtttcgcaagtaatctcagtcttata [203200]
KM262844.1_strain_L60          -------------------------------------------------------------------------------- [203200]
KM102979.1_26544/OG10_Italy    -------------------------------------------------------------------------------- [203200]
KM262845.1_strain_NHV          -------------------------------------------------------------------------------- [203200]
KX354450.1_47/Ss/2008          -------------------------------------------------------------------------------- [203200]
KP055815.1_strain_BA71         -------------------------------------------------------------------------------- [203200]
MH025920.1_strain_R35          taaaggactttattaatgggagttatgtttgcccagcagatcatacaattatttatttcgcaggtaatctcagtcttata [203200]
MH025919.1_strain_N10          taaaggactttattaatgggagttatgtttgcccagcagatcatacaattatttatttcgcaggtaatctcagtcttata [203200]
MH025918.1_strain_R25          taaaggactttattaatgggagttatgtttgcccagcagatcatacaattatttatttcgcaggtaatctcagtcttata [203200]
MH025917.1_strain_R7           taaaggactttattaatgggagttatgtttgcccagcagatcatacaattatttatttcgcaggtaatctcagtcttata [203200]
MH025916.1_strain_R8           taaaggactttattaatgggagttatgtttgcccagcagatcatacaattatttatttcgcaggtaatctcagtcttata [203200]
```

```
FR682468.2_Georgia_2007/1        -------------------------------------------------------------------------------- [203280]
ASF_IND_20_CAD543                -------------------------------------------------------------------------------- [203280]
LR899193.1_Germany_2020/1        -------------------------------------------------------------------------------- [203280]
MN393477.1_Wuhan_2019            -------------------------------------------------------------------------------- [203280]
MW396979.1_Timor-Leste/2019/1    -------------------------------------------------------------------------------- [203280]
MK940252.1_CN/2019/InnerMongolia -------------------------------------------------------------------------------- [203280]
MK128995.1_China/2018/AnhuiXCGQ  -------------------------------------------------------------------------------- [203280]
MH681419.1_ASFV/POL/2015         -------------------------------------------------------------------------------- [203280]
LS478113.1_Estonia_2014          -------------------------------------------------------------------------------- [203280]
KP843857.1_Odintsovo_02/14       -------------------------------------------------------------------------------- [203280]
FN557520.1_strain_E75            -------------------------------------------------------------------------------- [203280]
AY261366.1_Warthog               -------------------------------------------------------------------------------- [203280]
AY261365.1_Warmbaths             -------------------------------------------------------------------------------- [203280]
AY261364.1_Tengani_62            -------------------------------------------------------------------------------- [203280]
AY261363.1_Pretoriuskop/96/4     -------------------------------------------------------------------------------- [203280]
AY261362.1_Mkuzi_1979            -------------------------------------------------------------------------------- [203280]
AY261361.1_Malawi_Lil-20/1_1983  tgcgtataataagatacattacttacagcaaactattaaacctcctagcaagaagcttgggtacc-gcgtacctatacaa [203280]
AY261360.1_Kenya_1950            tgcgtataataaaatacattacttacaacaaaccattaaacctcctagcaaaaaacttgggtactggcgtacctatacaa [203280]
AM712240.1_OURT_88/3             -------------------------------------------------------------------------------- [203280]
AM712239.1_Benin_97/1            -------------------------------------------------------------------------------- [203280]
KM111295.1_Ken06.Bus             tacatataataaaatacattacttacagcaaattattaaacctcctagcaaggagcctggatactggcgtacctatgcaa [203280]
KM111294.1_Ken05/Tk1             tgcgtataataagatacattacttacagcaaactattaaacctcctagcaagaagcttgggtactggcgtacctatacaa [203280]
KM262844.1_strain_L60            -------------------------------------------------------------------------------- [203280]
KM102979.1_26544/OG10_Italy      -------------------------------------------------------------------------------- [203280]
KM262845.1_strain_NHV            -------------------------------------------------------------------------------- [203280]
KX354450.1_47/Ss/2008            -------------------------------------------------------------------------------- [203280]
KP055815.1_strain_BA71           -------------------------------------------------------------------------------- [203280]
MH025920.1_strain_R35            tacatataataaaatacattacttacagcaaattattaaacctcctagcaaggagcctggatactggcgtacctatgcaa [203280]
MH025919.1_strain_N10            tacatataataaaatacattacttacagcaaattattaaacctcctagcaaggagcctggatactggcgtacctatgcaa [203280]
MH025918.1_strain_R25            tacatataataaaatacattacttacagcaaattattaaacctcctagcaaggagcctggatactggcgtacctatgcaa [203280]
MH025917.1_strain_R7             tacatataataaaatacattacttacagcaaattattaaacctcctagcaaggagcctggatactggcgtacctatgcaa [203280]
MH025916.1_strain_R8             tacatataataaaatacattacttacagcaaattattaaacctcctagcaaggagcctggatactggcgtacctatgcaa [203280]

FR682468.2_Georgia_2007/1        -----------------------------------------------------------------------------aag [203360]
ASF_IND_20_CAD543                -------------------------------------------------------------------------------- [203360]
LR899193.1_Germany_2020/1        -----------------------------------------------------------------------------... [203360]
MN393477.1_Wuhan_2019            -----------------------------------------------------------------------------... [203360]
MW396979.1_Timor-Leste/2019/1    -----------------------------------------------------------------------------... [203360]
MK940252.1_CN/2019/InnerMongolia -----------------------------------------------------------------------------... [203360]
MK128995.1_China/2018/AnhuiXCGQ  -----------------------------------------------------------------------------... [203360]
MH681419.1_ASFV/POL/2015         -----------------------------------------------------------------------------... [203360]
LS478113.1_Estonia_2014          -----------------------------------------------------------------------------... [203360]
KP843857.1_Odintsovo_02/14       -----------------------------------------------------------------------------... [203360]
```

```
FN557520.1_strain_E75             -----------------------------------------------------------------------------... [203360]
AY261366.1_Warthog                -----------------------------------------------------------------------------... [203360]
AY261365.1_Warmbaths              -----------------------------------------------------------------------------... [203360]
AY261364.1_Tengani_62             -----------------------------------------------------------------------------t.. [203360]
AY261363.1_Pretoriuskop/96/4      -----------------------------------------------------------------------------... [203360]
AY261362.1_Mkuzi_1979             -----------------------------------------------------------------------------... [203360]
AY261361.1_Malawi_Lil-20/1_1983  accatcacagactctgttgggattgtcaagattacatctgtaggaacaaggtctttaaaaatcacctttttatagct... [203360]
AY261360.1_Kenya_1950             accatcacagattctgttgggattgtcaagatgccatctgtaggaacaaggtctttaaaaatcatctttctatagct... [203360]
AM712240.1_OURT_88/3              -----------------------------------------------------------------------------... [203360]
AM712239.1_Benin_97/1             -----------------------------------------------------------------------------... [203360]
KM111295.1_Ken06.Bus              accatcacagattctgttgggactgtcaaggtggcatctataggaacaaggtatttaaaaatcatctttctacgact... [203360]
KM111294.1_Ken05/Tk1              accatcatagattctgttgggattgtcaagatgtcatctgtaggaacaaggtctttaataatcatgtttctatagtt... [203360]
KM262844.1_strain_L60             -----------------------------------------------------------------------------... [203360]
KM102979.1_26544/OG10_Italy       -----------------------------------------------------------------------------... [203360]
KM262845.1_strain_NHV             -----------------------------------------------------------------------------... [203360]
KX354450.1_47/Ss/2008             -----------------------------------------------------------------------------... [203360]
KP055815.1_strain_BA71            -----------------------------------------------------------------------------... [203360]
MH025920.1_strain_R35             accatcacagattctgttgggactgtcaaggtggcatctataggaacaaggtatttaaaaatcatctttctacgact... [203360]
MH025919.1_strain_N10             accatcacagattctgttgggactgtcaaggtggcatctataggaacaaggtatttaaaaatcatctttctacgact... [203360]
MH025918.1_strain_R25             accatcacagattctgttgggactgtcaaggtggcatctataggaacaaggtatttaaaaatcatctttctacgact... [203360]
MH025917.1_strain_R7              accatcacagattctgttgggactgtcaaggtggcatctataggaacaaggtatttaaaaatcatctttctacgact... [203360]
MH025916.1_strain_R8              accatcacagattctgttgggactgtcaaggtggcatctataggaacaaggtatttaaaaatcatctttctacgact... [203360]

FR682468.2_Georgia_2007/1         gatattaatacacataattatttgaca-----tcacatta-----------gtgtacttaccaaatagtaatatacaacc [203440]
ASF_IND_20_CAD543                 -------------------------------------------------------------------------------- [203440]
LR899193.1_Germany_2020/1         ...........................-----........-----------............................. [203440]
MN393477.1_Wuhan_2019             ...........................-----........-----------............................. [203440]
MW396979.1_Timor-Leste/2019/1     ...........................-----........-----------............................. [203440]
MK940252.1_CN/2019/InnerMongolia ...........................-----........-----------............................. [203440]
MK128995.1_China/2018/AnhuiXCGQ   ...........................-----........-----------............................. [203440]
MH681419.1_ASFV/POL/2015          ...........................-----........-----------............................. [203440]
LS478113.1_Estonia_2014           ...........................-----........-----------............................. [203440]
KP843857.1_Odintsovo_02/14        ...........................-----........-----------............................. [203440]
FN557520.1_strain_E75             ...................g.......-----...t...g-----------..a.......................... [203440]
AY261366.1_Warthog                ...c.............c.g.......-----...t...g-----------...........t..---...........t [203440]
AY261365.1_Warmbaths              ...........................-----........-----------............................. [203440]
AY261364.1_Tengani_62             ...c.............c.g.......-----...t...g-----------..............---...........t [203440]
AY261363.1_Pretoriuskop/96/4      ...c.............c.g.......-----...t...g-----------a.............---...........t [203440]
AY261362.1_Mkuzi_1979             ...........................-----........-----------............................. [203440]
AY261361.1_Malawi_Lil-20/1_1983  ..cc........g....a........g-----......cggtttaataggt..ac.tc.......---...a.c....-g [203440]
AY261360.1_Kenya_1950             ..ccc.......g....a........g-----......cggtttaatgg--a..c..c.......---...a.c.t..g. [203440]
AM712240.1_OURT_88/3              ...................g.......-----...t...g-----------..a.......................... [203440]
AM712239.1_Benin_97/1             ...................g.......-----...t...g-----------..a.......................... [203440]
KM111295.1_Ken06.Bus              ..cc......t.g....a.......tgggtttattt..ctgcgcaaaaa--a..c..cc......---...a.c....a. [203440]
```

```
KM111294.1_Ken05/Tk1            ..cc........g....a........g-----.t....cggtttaatgg--a-.c.tc.......---...a.c.t..gt [203440]
KM262844.1_strain_L60           ...................g.......-----...t...g-----------..a.......................... [203440]
KM102979.1_26544/OG10_Italy     ...................g.......-----...t...g-----------..a.......................... [203440]
KM262845.1_strain_NHV           ...................g.......-----...t...g-----------..a.......................... [203440]
KX354450.1_47/Ss/2008           ...................g.......-----...t...g-----------..a.......................... [203440]
KP055815.1_strain_BA71          ...................g.......-----...t...g-----------..a.......................... [203440]
MH025920.1_strain_R35           ..cc......t.g....a.......tgggtttattt..ctgcgcaaaaa--a..c..cc......---...a.c....a. [203440]
MH025919.1_strain_N10           ..cc......t.g....a.......tgggtttattt..ctgcgcaaaaa--a..c..cc......---...a.c....a. [203440]
MH025918.1_strain_R25           ..cc......t.g....a.......tgggtttattt..ctgcgcaaaaa--a..c..cc......---...a.c....a. [203440]
MH025917.1_strain_R7            ..cc......t.g....a.......tgggtttattt..ctgcgcaaaaa--a..c..cc......---...a.c....a. [203440]
MH025916.1_strain_R8            ..cc......t.g....a.......tgggtttattt..ctgcgcaaaaa--a..c..cc......---...a.c....a. [203440]

FR682468.2_Georgia_2007/1       atcctaatattcacctttatgaaatgatccca---------acctatacggtaaa---atagtataggttttaataaaga [203520]
ASF_IND_20_CAD543               -------------------------------------------------------------------------------- [203520]
LR899193.1_Germany_2020/1       ................................---------..............---...................... [203520]
MN393477.1_Wuhan_2019           ................................---------..............---...................... [203520]
MW396979.1_Timor-Leste/2019/1   ................................---------..............---...................... [203520]
MK940252.1_CN/2019/InnerMongolia ................................---------..............---...................... [203520]
MK128995.1_China/2018/AnhuiXCGQ ................................---------..............---...................... [203520]
MH681419.1_ASFV/POL/2015        ................................---------..............---...................... [203520]
LS478113.1_Estonia_2014         ................................---------..............---...................... [203520]
KP843857.1_Odintsovo_02/14      ................................---------..............---...................... [203520]
FN557520.1_strain_E75           .............-.............c...ccccccccc-........a.....---...................... [203520]
AY261366.1_Warthog              .............-........gg........---------........a.....---.........a............ [203520]
AY261365.1_Warmbaths            .............-..................---------........a.....---..................c... [203520]
AY261364.1_Tengani_62           .......g.....-.......c........----------a.......ta.....---.........a........-.a. [203520]
AY261363.1_Pretoriuskop/96/4    .............-........gg.....t..---------.......ta.....---.........a.......--.a. [203520]
AY261362.1_Mkuzi_1979           .............-..................---------........a.....---...................... [203520]
AY261361.1_Malawi_Lil-20/1_1983 .....gt......-t..c..a.....tcgt..---------t......t.a..t.agt.at...c.a.g.......---- [203520]
AY261360.1_Kenya_1950           .....gt......-...c.ca.....tcgt..---------t....c.t.a..t.agtgat.a.c.a.g.......---- [203520]
AM712240.1_OURT_88/3            .............-.............c...cccccca---........a.....---...................... [203520]
AM712239.1_Benin_97/1           .............-.............c...ccccccc---........a.....---...................... [203520]
KM111295.1_Ken06.Bus            .....gt.ta.tt-ac....a.....tcgt..---------t....c.t.a..t.agtgat.a.c.a.g.......---- [203520]
KM111294.1_Ken05/Tk1            .....gt.....--ac....at....tcgt..---------t....c.t.a..t.agt.at...c.a.g.......---- [203520]
KM262844.1_strain_L60           .............-.............c...ccccccccca........a.....---...................... [203520]
KM102979.1_26544/OG10_Italy     .............-.............c...ccccccc---........a.....---...................... [203520]
KM262845.1_strain_NHV           .............-.............c...cccccca---........a.....---...................... [203520]
KX354450.1_47/Ss/2008           .............-.............c...ccccccc---........a.....---...................... [203520]
KP055815.1_strain_BA71          .............-.............c...cccccccca-........a.....---...................... [203520]
MH025920.1_strain_R35           .....gt.ta.tt-ac....a.....tcgt..---------t....c.t.a..t.agtgat.a.c.a.g.......---- [203520]
MH025919.1_strain_N10           .....gt.ta.tt-ac....a.....tcgt..---------t....c.t.a..t.agtgat.a.c.a.g.......---- [203520]
MH025918.1_strain_R25           .....gt.ta.tt-ac....a.....tcgt..---------t....c.t.a..t.agtgat.a.c.a.g.......---- [203520]
MH025917.1_strain_R7            .....gt.ta.tt-ac....a.....tcgt..---------t....c.t.a..t.agtgat.a.c.a.g.......---- [203520]
MH025916.1_strain_R8            .....gt.ta.tt-ac....a.....tcgt..---------t....c.t.a..t.agtgat.a.c.a.g.......---- [203520]
```

```
FR682468.2_Georgia_2007/1        aaaaagatattctgtggtttttatttttgtatagtgtgtgaatacaaaataaa-atcccaaattttaacctttttcttttt [203600]
ASF_IND_20_CAD543                -------------------------------------------------------------------------------- [203600]
LR899193.1_Germany_2020/1        .....................................................-.......................... [203600]
MN393477.1_Wuhan_2019            .....................................................-.......................... [203600]
MW396979.1_Timor-Leste/2019/1    .....................................................-.......................... [203600]
MK940252.1_CN/2019/InnerMongolia .....................................................-.......................... [203600]
MK128995.1_China/2018/AnhuiXCGQ  .....................................................-.......................... [203600]
MH681419.1_ASFV/POL/2015         .....................................................-.......................... [203600]
LS478113.1_Estonia_2014          .....................................................-.......................... [203600]
KP843857.1_Odintsovo_02/14       .....................................................-.....................-.... [203600]
FN557520.1_strain_E75            ...........a...a....................c......g.........-.......................... [203600]
AY261366.1_Warthog               ......g.....................a.......c..a.............-.......................... [203600]
AY261365.1_Warmbaths             ........g..a.....-..........a.......c......g.........-.......................... [203600]
AY261364.1_Tengani_62            ............................a.......c..a.............-.......................... [203600]
AY261363.1_Pretoriuskop/96/4     ............................a.......c..a.............t.......................... [203600]
AY261362.1_Mkuzi_1979            ........g..a.....-..........a.......c......g.........-.......................... [203600]
AY261361.1_Malawi_Lil-20/1_1983  --...a...a.a....--........a...g....aa..a.............-..t....................... [203600]
AY261360.1_Kenya_1950            ---..a...a.tc...--............g....aa..a...g.........-.......................... [203600]
AM712240.1_OURT_88/3             ...........a...a....................c......g.........-.......................... [203600]
AM712239.1_Benin_97/1            ...........a...a....................c......g.........-.......................... [203600]
KM111295.1_Ken06.Bus             ---..a...a.tc...--............g....aa..a...g.........-.......................... [203600]
KM111294.1_Ken05/Tk1             ---..a.c.a.a....--............g....aa..a...g.........-.......................... [203600]
KM262844.1_strain_L60            ...........a...a....................c......g.........-.......................... [203600]
KM102979.1_26544/OG10_Italy      ...........a...a....................c......g.........-.......................... [203600]
KM262845.1_strain_NHV            ...........a...a....................c......g.........-.......................... [203600]
KX354450.1_47/Ss/2008            ...........a...a....................c......g.........-.......................... [203600]
KP055815.1_strain_BA71           ...........a...a....................c......g.........-.......................... [203600]
MH025920.1_strain_R35            ---..a...a.tc...--............g....aa..a...g.........-.......................... [203600]
MH025919.1_strain_N10            ---..a...a.tc...--............g....aa..a...g.........-.......................... [203600]
MH025918.1_strain_R25            ---..a...a.tc...--............g....aa..a...g.........-.......................... [203600]
MH025917.1_strain_R7             ---..a...a.tc...--............g....aa..a...g.........-.......................... [203600]
MH025916.1_strain_R8             ---..a...a.tc...--............g....aa..a...g.........-.......................... [203600]

FR682468.2_Georgia_2007/1        ttttctatacaggatgttagaaatagtattggcaacgctgctaggcgacctgcagcggctccgggttcttacccctcagc [203680]
ASF_IND_20_CAD543                -------------------------------------------------------------................... [203680]
LR899193.1_Germany_2020/1        ................................................................................ [203680]
MN393477.1_Wuhan_2019            ................................................................................ [203680]
MW396979.1_Timor-Leste/2019/1    ................................................................................ [203680]
MK940252.1_CN/2019/InnerMongolia ................................................................................ [203680]
MK128995.1_China/2018/AnhuiXCGQ  ................................................................................ [203680]
MH681419.1_ASFV/POL/2015         ................................................................................ [203680]
LS478113.1_Estonia_2014          ................................................................................ [203680]
KP843857.1_Odintsovo_02/14       ................................................................................ [203680]
```

```
FN557520.1_strain_E75          ................................................................................ [203680]
AY261366.1_Warthog             .............................................t.................................. [203680]
AY261365.1_Warmbaths           .............................................t.................................. [203680]
AY261364.1_Tengani_62          .............................................t.................................. [203680]
AY261363.1_Pretoriuskop/96/4   .............................................t.................................. [203680]
AY261362.1_Mkuzi_1979          ................................................................................ [203680]
AY261361.1_Malawi_Lil-20/1_1983 ...........a.................................t..t......aa...ta...a......a...at.. [203680]
AY261360.1_Kenya_1950          ...........a.....................c...........t..t......aaa..taa..a......a....c.. [203680]
AM712240.1_OURT_88/3           ................................................................................ [203680]
AM712239.1_Benin_97/1          ................................................................................ [203680]
KM111295.1_Ken06.Bus           ...........a..............................g..t..t.......aa..taa..a......a...a... [203680]
KM111294.1_Ken05/Tk1           ...........a.................................t..t......aaa..taaa.a......a...at.. [203680]
KM262844.1_strain_L60          ................................................................................ [203680]
KM102979.1_26544/OG10_Italy    ................................................................................ [203680]
KM262845.1_strain_NHV          ................................................................................ [203680]
KX354450.1_47/Ss/2008          ................................................................................ [203680]
KP055815.1_strain_BA71         ................................................................................ [203680]
MH025920.1_strain_R35          ...........a..............................g..t..t.......aa..taa..a......a...a... [203680]
MH025919.1_strain_N10          ...........a..............................g..t..t.......aa..taa..a......a...a... [203680]
MH025918.1_strain_R25          ...........a..............................g..t..t.......aa..taa..a......a...a... [203680]
MH025917.1_strain_R7           ...........a..............................g..t..t.......aa..taa..a......a...a... [203680]
MH025916.1_strain_R8           ...........a..............................g..t..t.......aa..taa..a......a...a... [203680]

FR682468.2_Georgia_2007/1      agcgggcagttgccttctttcgagccaatactaaggagctagaggacttcttatgctcagatgggcagtctgaggaggta [203760]
ASF_IND_20_CAD543              .............................................................................r.. [203760]
LR899193.1_Germany_2020/1      ................................................................................ [203760]
MN393477.1_Wuhan_2019          ................................................................................ [203760]
MW396979.1_Timor-Leste/2019/1  ................................................................................ [203760]
MK940252.1_CN/2019/InnerMongolia ................................................................................ [203760]
MK128995.1_China/2018/AnhuiXCGQ ................................................................................ [203760]
MH681419.1_ASFV/POL/2015       ................................................................................ [203760]
LS478113.1_Estonia_2014        ................................................................................ [203760]
KP843857.1_Odintsovo_02/14     ................................................................................ [203760]
FN557520.1_strain_E75          .......g.............................................c.............a.........a.. [203760]
AY261366.1_Warthog             .......g.........................................t...........................a.. [203760]
AY261365.1_Warmbaths           .......g.....................................................................a.. [203760]
AY261364.1_Tengani_62          .......g..........................a....................tc......................g [203760]
AY261363.1_Pretoriuskop/96/4   .......g..............................g....a.................................... [203760]
AY261362.1_Mkuzi_1979          .......g.....................................................................a.. [203760]
AY261361.1_Malawi_Lil-20/1_1983 .......t..g..t..........t...c.....a...............c.c.a.c.c..................t.g [203760]
AY261360.1_Kenya_1950          .......t..g..t..............c..c..a..............tc.c.a.c.c..................t.. [203760]
AM712240.1_OURT_88/3           .......g.............................................c.............a.........a.. [203760]
AM712239.1_Benin_97/1          .......g.............................................c.............a.........a.. [203760]
KM111295.1_Ken06.Bus           .......c..a..t.....c........c..c..a..............tc.c.a.c.c.........a........t.. [203760]
```

```
KM111294.1_Ken05/Tk1           .......t..g..t..............c.....a...............c.c...c.c..................t.. [203760]
KM262844.1_strain_L60          .......g.............................................c.............a.........a.. [203760]
KM102979.1_26544/OG10_Italy    .......g.............................................c.............a.........a.. [203760]
KM262845.1_strain_NHV          .......g.............................................c.............a.........a.. [203760]
KX354450.1_47/Ss/2008          .......g.............................................c.............a.........a.. [203760]
KP055815.1_strain_BA71         .......g.............................................c.............a.........a.. [203760]
MH025920.1_strain_R35          .......c..a..t.....c........c..c..a..............tc.c.a.c.c.........a........t.. [203760]
MH025919.1_strain_N10          .......c..a..t.....c........c..c..a..............tc.c.a.c.c.........a........t.. [203760]
MH025918.1_strain_R25          .......c..a..t.....c........c..c..a..............tc.c.a.c.c.........a........t.. [203760]
MH025917.1_strain_R7           .......c..a..t.....c........c..c..a..............tc.c.a.c.c.........a........t.. [203760]
MH025916.1_strain_R8           .......c..a..t.....c........c..c..a..............tc.c.a.c.c.........a........t.. [203760]

FR682468.2_Georgia_2007/1      ctgtctgg-cccccttcttaaccgtctactagaaccctcaggccctcttgatattttaaccggatatcacctatttcgtc [203840]
ASF_IND_20_CAD543              ........-....................................................................... [203840]
LR899193.1_Germany_2020/1      ........-....................................................................... [203840]
MN393477.1_Wuhan_2019          ........-....................................................................... [203840]
MW396979.1_Timor-Leste/2019/1  ........-....................................................................... [203840]
MK940252.1_CN/2019/InnerMongolia ........-....................................................................... [203840]
MK128995.1_China/2018/AnhuiXCGQ ........-....................................................................... [203840]
MH681419.1_ASFV/POL/2015       ........-....................................................................... [203840]
LS478113.1_Estonia_2014        ........-....................................................................... [203840]
KP843857.1_Odintsovo_02/14     ........-....................................................................... [203840]
FN557520.1_strain_E75          ........-......c........................t....................................... [203840]
AY261366.1_Warthog             .....c..-......c......................................................t......... [203840]
AY261365.1_Warmbaths           .....c..-......c......................................................t......... [203840]
AY261364.1_Tengani_62          ........-.t....c............................................t................... [203840]
AY261363.1_Pretoriuskop/96/4   .....c..-......c......................................................t......... [203840]
AY261362.1_Mkuzi_1979          ..a.....-...................................................t................... [203840]
AY261361.1_Malawi_Lil-20/1_1983 ..cc....-a.tt..c.........t..........t.........a....c...........t..c............g [203840]
AY261360.1_Kenya_1950          ..cc....-a.tt..c.........t..........t.....tt..a....c...........t..c............g [203840]
AM712240.1_OURT_88/3           ........c......c........................t....................................... [203840]
AM712239.1_Benin_97/1          ........-......c........................t....................................... [203840]
KM111295.1_Ken06.Bus           ..cc....-attt..........a.t....................a....a....................c....... [203840]
KM111294.1_Ken05/Tk1           ..cc....-a.tt..c.........t..t.......t.....t...a....c...........t..c............g [203840]
KM262844.1_strain_L60          ........-......c........................t....................................... [203840]
KM102979.1_26544/OG10_Italy    ........-......c........................t....................................... [203840]
KM262845.1_strain_NHV          ........c......c........................t....................................... [203840]
KX354450.1_47/Ss/2008          ........-......c........................t....................................... [203840]
KP055815.1_strain_BA71         ........-......c........................t....................................... [203840]
MH025920.1_strain_R35          ..cc....-attt..........a.t....................a....a.........a..........c....... [203840]
MH025919.1_strain_N10          ..cc....-attt..........a.t....................a....a.........a..........c....... [203840]
MH025918.1_strain_R25          ..cc....-attt..........a.t....................a....a.........a..........c....... [203840]
MH025917.1_strain_R7           ..cc....-attt..........a.t....................a....a.........a..........c....... [203840]
MH025916.1_strain_R8           ..cc....-attt..........a.t....................a....a.........a..........c....... [203840]
```

| | | |
|---|---|---|
| FR682468.2_Georgia_2007/1 | agaatcccaaggcaggtcagttgcgcggccttgaggtcaagatgcttgaacggttatacgatgctaatatttacaatata | [203920] |
| ASF_IND_20_CAD543 | ................................................................................ | [203920] |
| LR899193.1_Germany_2020/1 | ................................................................................ | [203920] |
| MN393477.1_Wuhan_2019 | ................................................................................ | [203920] |
| MW396979.1_Timor-Leste/2019/1 | ................................................................................ | [203920] |
| MK940252.1_CN/2019/InnerMongolia | ................................................................................ | [203920] |
| MK128995.1_China/2018/AnhuiXCGQ | ................................................................................ | [203920] |
| MH681419.1_ASFV/POL/2015 | ................................................................................ | [203920] |
| LS478113.1_Estonia_2014 | ................................................................................ | [203920] |
| KP843857.1_Odintsovo_02/14 | ................................................................................ | [203920] |
| FN557520.1_strain_E75 | ................................................................................ | [203920] |
| AY261366.1_Warthog | ....................g........................................................... | [203920] |
| AY261365.1_Warmbaths | ............................t................................................... | [203920] |
| AY261364.1_Tengani_62 | ............................................................................c... | [203920] |
| AY261363.1_Pretoriuskop/96/4 | .............g......g........................................................... | [203920] |
| AY261362.1_Mkuzi_1979 | ....c.................a......................................................... | [203920] |
| AY261361.1_Malawi_Lil-20/1_1983 | ....c..a..........g.c....t...........t...c................t........c..c........g | [203920] |
| AY261360.1_Kenya_1950 | .......a..........gc.....a...........g..a.................t....................g | [203920] |
| AM712240.1_OURT_88/3 | ................................................................................ | [203920] |
| AM712239.1_Benin_97/1 | ................................................................................ | [203920] |
| KM111295.1_Ken06.Bus | .......a..........gc.....a...........g..a.................t....................g | [203920] |
| KM111294.1_Ken05/Tk1 | .......g..........g.c....t...............c................t........c..c......... | [203920] |
| KM262844.1_strain_L60 | ................................................................................ | [203920] |
| KM102979.1_26544/OG10_Italy | ................................................................................ | [203920] |
| KM262845.1_strain_NHV | ................................................................................ | [203920] |
| KX354450.1_47/Ss/2008 | ................................................................................ | [203920] |
| KP055815.1_strain_BA71 | ................................................................................ | [203920] |
| MH025920.1_strain_R35 | .......a..........gc.....a...........g..a.................t....................g | [203920] |
| MH025919.1_strain_N10 | .......a..........gc.....a...........g..a.................t....................g | [203920] |
| MH025918.1_strain_R25 | .......a..........gc.....a...........g..a.................t....................g | [203920] |
| MH025917.1_strain_R7 | .......a..........gc.....a...........g..a.................t....................g | [203920] |
| MH025916.1_strain_R8 | .......a..........gc.....a...........g..a.................t....................g | [203920] |

| | | |
|---|---|---|
| FR682468.2_Georgia_2007/1 | ctgtctcggctgcggcctgaaaaagttcgcaacaaggctattgagctatactgggttttccgagctatccatatttgtca | [204000] |
| ASF_IND_20_CAD543 | y............................................................................... | [204000] |
| LR899193.1_Germany_2020/1 | ................................................................................ | [204000] |
| MN393477.1_Wuhan_2019 | ................................................................................ | [204000] |
| MW396979.1_Timor-Leste/2019/1 | ................................................................................ | [204000] |
| MK940252.1_CN/2019/InnerMongolia | ................................................................................ | [204000] |
| MK128995.1_China/2018/AnhuiXCGQ | ................................................................................ | [204000] |
| MH681419.1_ASFV/POL/2015 | ................................................................................ | [204000] |
| LS478113.1_Estonia_2014 | ................................................................................ | [204000] |
| KP843857.1_Odintsovo_02/14 | ................................................................................ | [204000] |

```
FN557520.1_strain_E75          t.............a...........c..................................................... [204000]
AY261366.1_Warthog             t.............a........................g...................t..g......a....g..... [204000]
AY261365.1_Warmbaths           t.............a................................................................. [204000]
AY261364.1_Tengani_62          t....c........a........................g...................t..g......a....g..... [204000]
AY261363.1_Pretoriuskop/96/4   t.............a........................g...................t..g......a....g..... [204000]
AY261362.1_Mkuzi_1979          t.............a................................t...........t.................... [204000]
AY261361.1_Malawi_Lil-20/1_1983 t..g.c.aaa.a..a.....gttg.....t.t.............t....t...c.c..t..g.....ttta..g..... [204000]
AY261360.1_Kenya_1950          ...g.......a........gttg.....tg...................t...c.c..t..g.....ttta..g..... [204000]
AM712240.1_OURT_88/3           t.............a...........c..................................................... [204000]
AM712239.1_Benin_97/1          t.............a...a.......c..................................................... [204000]
KM111295.1_Ken06.Bus           ...g.......a........gttg.....tg........g..........t...c....t........ttta..a..... [204000]
KM111294.1_Ken05/Tk1           t..g.c.a...a........gttg........t......g.....t....t...c.c..t..g.....tttg..g..... [204000]
KM262844.1_strain_L60          t.............a...........c..................................................... [204000]
KM102979.1_26544/OG10_Italy    t.............a...........c..................................................... [204000]
KM262845.1_strain_NHV          t.............a...........c..................................................... [204000]
KX354450.1_47/Ss/2008          t.............a...........c..................................................... [204000]
KP055815.1_strain_BA71         t.............a...........c..................................................... [204000]
MH025920.1_strain_R35          ...g.......a........gttg.....tg........g..........t...c....t........ttta..a..... [204000]
MH025919.1_strain_N10          ...g.......a........gttg.....tg........g..........t...c....t........ttta..a..... [204000]
MH025918.1_strain_R25          ...g.......a........gttg.....tg........g..........t...c....t........ttta..a..... [204000]
MH025917.1_strain_R7           ...g.......a........gttg.....tg........g..........t...c....t........ttta..a..... [204000]
MH025916.1_strain_R8           ...g.......a........gttg.....tg........g..........t...c....t........ttta..a..... [204000]

FR682468.2_Georgia_2007/1      tgctcctttagttttagatattgtacgatatgaggaaccggactttgctgaactggcctttatttgtgctgcttactttg [204080]
ASF_IND_20_CAD543              ................................................................................ [204080]
LR899193.1_Germany_2020/1      ................................................................................ [204080]
MN393477.1_Wuhan_2019          ................................................................................ [204080]
MW396979.1_Timor-Leste/2019/1  ................................................................................ [204080]
MK940252.1_CN/2019/InnerMongolia ................................................................................ [204080]
MK128995.1_China/2018/AnhuiXCGQ ................................................................................ [204080]
MH681419.1_ASFV/POL/2015       ................................................................................ [204080]
LS478113.1_Estonia_2014        ................................................................................ [204080]
KP843857.1_Odintsovo_02/14     ................................................................................ [204080]
FN557520.1_strain_E75          ................................................................................ [204080]
AY261366.1_Warthog             ................................................................................ [204080]
AY261365.1_Warmbaths           ................................................................................ [204080]
AY261364.1_Tengani_62          ............................a.........t......................................... [204080]
AY261363.1_Pretoriuskop/96/4   ................................................................................ [204080]
AY261362.1_Mkuzi_1979          ................................................................................ [204080]
AY261361.1_Malawi_Lil-20/1_1983 .ag...c........g..g.........c.....ac.at...t.....a...t.a........c...............a [204080]
AY261360.1_Kenya_1950          .ag...c........g..g.........c.....ac.at...t.....a...acc........c.....c.........a [204080]
AM712240.1_OURT_88/3           ................................................................................ [204080]
AM712239.1_Benin_97/1          ................................................................................ [204080]
KM111295.1_Ken06.Bus           ..g...c..g.....g..g.........c.....acgtt.........a...acc........c...............a [204080]
```

```
KM111294.1_Ken05/Tk1              ..g...c........g..g.........c.....ac.at...t.....a...t.a........c...............a [204080]
KM262844.1_strain_L60             ................................................................................ [204080]
KM102979.1_26544/OG10_Italy       ................................................................................ [204080]
KM262845.1_strain_NHV             ................................................................................ [204080]
KX354450.1_47/Ss/2008             ................................................................................ [204080]
KP055815.1_strain_BA71            ................................................................................ [204080]
MH025920.1_strain_R35             ..g...c..g.....g..g.........c.....acgtt.........a...acc........c...............a [204080]
MH025919.1_strain_N10             ..g...c..g.....g..g.........c.....acgtt.........a...acc........c...............a [204080]
MH025918.1_strain_R25             ..g...c..g.....g..g.........c.....acgtt.........a...acc........c...............a [204080]
MH025917.1_strain_R7              ..g...c..g.....g..g.........c.....acgtt.........a...acc........c...............a [204080]
MH025916.1_strain_R8              ..g...c..g.....g..g.........c.....acgtt.........a...acc........c...............a [204080]

FR682468.2_Georgia_2007/1         gtgaacctcaggtaatgtatttgctctacaaatatatgcctctgacccgcgcagttcttacggatgccatccggataagt [204160]
ASF_IND_20_CAD543                 ................................................................................ [204160]
LR899193.1_Germany_2020/1         ................................................................................ [204160]
MN393477.1_Wuhan_2019             ................................................................................ [204160]
MW396979.1_Timor-Leste/2019/1     ................................................................................ [204160]
MK940252.1_CN/2019/InnerMongolia  ................................................................................ [204160]
MK128995.1_China/2018/AnhuiXCGQ   ................................................................................ [204160]
MH681419.1_ASFV/POL/2015          ................................................................................ [204160]
LS478113.1_Estonia_2014           ................................................................................ [204160]
KP843857.1_Odintsovo_02/14        ................................................................................ [204160]
FN557520.1_strain_E75             ........................................................................a....... [204160]
AY261366.1_Warthog                ........................................................................a....... [204160]
AY261365.1_Warmbaths              ................................................................................ [204160]
AY261364.1_Tengani_62             ....g...................................................................a....... [204160]
AY261363.1_Pretoriuskop/96/4      .............................................g..........................a....... [204160]
AY261362.1_Mkuzi_1979             ................................................................................ [204160]
AY261361.1_Malawi_Lil-20/1_1983   ...................cgct..t..t....t...a...a.tt.t........c...g.t.........ga...gtc. [204160]
AY261360.1_Kenya_1950             ...................cgct..t..t....t...a...a.ct.t..t.....c...g.t........t.a...gt.. [204160]
AM712240.1_OURT_88/3              ........................................................................a....... [204160]
AM712239.1_Benin_97/1             ........................................................................a....... [204160]
KM111295.1_Ken06.Bus              ...................cgct..t..t..c.t...a..a.cc..t.at.....c...g.t..........a...gt.. [204160]
KM111294.1_Ken05/Tk1              ...................cgcta.t..t....t...a...a.ct.tt.......c...g.t........t.a...gt.. [204160]
KM262844.1_strain_L60             ........................................................................a....... [204160]
KM102979.1_26544/OG10_Italy       ........................................................................a....... [204160]
KM262845.1_strain_NHV             ........................................................................a....... [204160]
KX354450.1_47/Ss/2008             ........................................................................a....... [204160]
KP055815.1_strain_BA71            ........................................................................a....... [204160]
MH025920.1_strain_R35             ...................cgct..t..t..c.t...a..a.cc..t.at.....c...g.t..........a...gt.. [204160]
MH025919.1_strain_N10             ...................cgct..t..t..c.t...a..a.cc..t.at.....c...g.t..........a...gt.. [204160]
MH025918.1_strain_R25             ...................cgct..t..t..c.t...a..a.cc..t.at.....c...g.t..........a...gt.. [204160]
MH025917.1_strain_R7              ...................cgct..t..t..c.t...a..a.cc..t.at.....c...g.t..........a...gt.. [204160]
MH025916.1_strain_R8              ...................cgct..t..t..c.t...a..a.cc..t.at.....c...g.t..........a...gt.. [204160]
```

```
FR682468.2_Georgia_2007/1        cttgagagcaacaaccaggtagggatttgctatgcttacttgatgggaggcagcctcaagggactagtctccgccccact [204240]
ASF_IND_20_CAD543                ................................................................................ [204240]
LR899193.1_Germany_2020/1        ................................................................................ [204240]
MN393477.1_Wuhan_2019            ................................................................................ [204240]
MW396979.1_Timor-Leste/2019/1    ................................................................................ [204240]
MK940252.1_CN/2019/InnerMongolia ................................................................................ [204240]
MK128995.1_China/2018/AnhuiXCGQ  ................................................................................ [204240]
MH681419.1_ASFV/POL/2015         ................................................................................ [204240]
LS478113.1_Estonia_2014          ................................................................................ [204240]
KP843857.1_Odintsovo_02/14       ................................................................................ [204240]
FN557520.1_strain_E75            ................................................................................ [204240]
AY261366.1_Warthog               .............g.....................c..........................c......cgt........ [204240]
AY261365.1_Warmbaths             ........t....................................................................... [204240]
AY261364.1_Tengani_62            .............g......g..............c.................................cgt.....g.. [204240]
AY261363.1_Pretoriuskop/96/4     .............g.....................c..........................c......cgt........ [204240]
AY261362.1_Mkuzi_1979            ........t....................................................................... [204240]
AY261361.1_Malawi_Lil-20/1_1983  .............g.g..acg..............c...c.a.....g..t...........caag..gc...g...g.. [204240]
AY261360.1_Kenya_1950            .............g.g...cg..................c.a.....g..t...........caag..gc.t.g.t.g.. [204240]
AM712240.1_OURT_88/3             ................................................................................ [204240]
AM712239.1_Benin_97/1            ................................................................................ [204240]
KM111295.1_Ken06.Bus             .............g.g...c..............tc...c.a........t.a.........caag..gc...g.t.g.. [204240]
KM111294.1_Ken05/Tk1             ........t....g.g...cg..................c.a.....g..t...........caag..gc.t.g.g.g.. [204240]
KM262844.1_strain_L60            ................................................................................ [204240]
KM102979.1_26544/OG10_Italy      ................................................................................ [204240]
KM262845.1_strain_NHV            ................................................................................ [204240]
KX354450.1_47/Ss/2008            ................................................................................ [204240]
KP055815.1_strain_BA71           ................................................................................ [204240]
MH025920.1_strain_R35            .............g.g...c..............tc...c.a........t.a.........caag..gc...g.t.g.. [204240]
MH025919.1_strain_N10            .............g.g...c..............tc...c.a........t.a.........caag..gc...g.t.g.. [204240]
MH025918.1_strain_R25            .............g.g...c..............tc...c.a........t.a.........caag..gc...g.t.g.. [204240]
MH025917.1_strain_R7             .............g.g...c..............tc...c.a........t.a.........caag..gc...g.t.g.. [204240]
MH025916.1_strain_R8             .............g.g...c..............tc...c.a........t.a.........caag..gc...g.t.g.. [204240]

FR682468.2_Georgia_2007/1        gcgtaaacgtctgcgcgccaaactacgctcgc---agcgcaaaaagaaggacgttctttcaccccacgacttcttactgc [204320]
ASF_IND_20_CAD543                ................................---............................................. [204320]
LR899193.1_Germany_2020/1        ................................---............................................. [204320]
MN393477.1_Wuhan_2019            ................................---............................................. [204320]
MW396979.1_Timor-Leste/2019/1    ................................---............................................. [204320]
MK940252.1_CN/2019/InnerMongolia ................................---............................................. [204320]
MK128995.1_China/2018/AnhuiXCGQ  ................................---............................................. [204320]
MH681419.1_ASFV/POL/2015         ................................---............................................. [204320]
LS478113.1_Estonia_2014          ................................---............................................. [204320]
KP843857.1_Odintsovo_02/14       ................................---............................................. [204320]
```

```
FN557520.1_strain_E75             ................................---............................................. [204320]
AY261366.1_Warthog                a...............................---.......................c.......t............. [204320]
AY261365.1_Warmbaths              ..............................a.---.............a............................... [204320]
AY261364.1_Tengani_62             a...............................---................t......c..................... [204320]
AY261363.1_Pretoriuskop/96/4      a...............................---.......................c..................... [204320]
AY261362.1_Mkuzi_1979             .............t..................---............................................. [204320]
AY261361.1_Malawi_Lil-20/1_1983   ...c...........t....gt.c..ttcg..aag.............aa....c...c.g.....t..a..tc....ca [204320]
AY261360.1_Kenya_1950             ...c...........t....gt.c..ttcg..aag.............aa....c..cc.g.....t..a...c....ca [204320]
AM712240.1_OURT_88/3              ................................---............................................. [204320]
AM712239.1_Benin_97/1             ................................---............................................. [204320]
KM111295.1_Ken06.Bus              ...c...........t....gt.c..ttcg..aag.a...........aa....c...c.g........a...c....c. [204320]
KM111294.1_Ken05/Tk1              ...c...........t....gt.c..ttcg..aag.............aa........c.......t..a...c....ca [204320]
KM262844.1_strain_L60             ................................---............................................. [204320]
KM102979.1_26544/OG10_Italy       ................................---............................................. [204320]
KM262845.1_strain_NHV             ................................---............................................. [204320]
KX354450.1_47/Ss/2008             ................................---............................................. [204320]
KP055815.1_strain_BA71            ................................---............................................. [204320]
MH025920.1_strain_R35             ...c...........t.a..gt.c..ttcg..aag.a...........aa....c...c.g........a...c....c. [204320]
MH025919.1_strain_N10             ...c...........t.a..gt.c..ttcg..aag.a...........aa....c...c.g........a...c....c. [204320]
MH025918.1_strain_R25             ...c...........t.a..gt.c..ttcg..aag.a...........aa....c...c.g........a...c....c. [204320]
MH025917.1_strain_R7              ...c...........t.a..gt.c..ttcg..aag.a...........aa....c...c.g........a...c....c. [204320]
MH025916.1_strain_R8              ...c...........t.a..gt.c..ttcg..aag.a...........aa....c...c.g........a...c....c. [204320]

FR682468.2_Georgia_2007/1         tgctccagtagcttt---ttttgccgcaggagcaccgcggataggagctcctccacgctcgcgatccggcgctggaagcg [204400]
ASF_IND_20_CAD543                 ...............---.............................................................. [204400]
LR899193.1_Germany_2020/1         ...............---.............................................................. [204400]
MN393477.1_Wuhan_2019             ...............---.............................................................. [204400]
MW396979.1_Timor-Leste/2019/1     ...............---.............................................................. [204400]
MK940252.1_CN/2019/InnerMongolia  ...............---.............................................................. [204400]
MK128995.1_China/2018/AnhuiXCGQ   ...............---.............................................................. [204400]
MH681419.1_ASFV/POL/2015          ...............---.............................................................. [204400]
LS478113.1_Estonia_2014           ...............---.............................................................. [204400]
KP843857.1_Odintsovo_02/14        ...............---.............................................................. [204400]
FN557520.1_strain_E75             ...............---...................t.......................................... [204400]
AY261366.1_Warthog                ..........a....---...........a...........g...................................... [204400]
AY261365.1_Warmbaths              ...............---.....t.....a..................................c............... [204400]
AY261364.1_Tengani_62             ..........a....---...........a.................................................. [204400]
AY261363.1_Pretoriuskop/96/4      ..........a....---...........a.................................................. [204400]
AY261362.1_Mkuzi_1979             ..........a....---...........a.................................................. [204400]
AY261361.1_Malawi_Lil-20/1_1983   .......tgg.a...aat....tt..t........t..t..a....t..tt..ggt.tcg..a.....a..t........ [204400]
AY261360.1_Kenya_1950             .......tgg.a...aat....tt..t..a.a...t.tt..g....t...t..ggt.tcg.ta.....a..t........ [204400]
AM712240.1_OURT_88/3              ...............---...................t.......................................... [204400]
AM712239.1_Benin_97/1             ...............---...................t.......................................... [204400]
KM111295.1_Ken06.Bus              .......cgg.a...aat....tt..t......g.ta.t..g....t......ag..t.g..a.....a..t........ [204400]
```

```
KM111294.1_Ken05/Tk1          ....t..tgg.a...aat....tt..t......g.g..t..g....t...t..gg.atcg..a........t........ [204400]
KM262844.1_strain_L60         ...............---...................t.......................................... [204400]
KM102979.1_26544/OG10_Italy   ...............---...................t.......................................... [204400]
KM262845.1_strain_NHV         ...............---...................t.......................................... [204400]
KX354450.1_47/Ss/2008         ...............---...................t.......................................... [204400]
KP055815.1_strain_BA71        ...............---...................t.......................................... [204400]
MH025920.1_strain_R35         .......cgg.a...aat....tt..t......g.ta.t..g....t......ag..t.g..a.....a..t........ [204400]
MH025919.1_strain_N10         .......cgg.a...aat....tt..t......g.ta.t..g....t......ag..t.g..a.....a..t........ [204400]
MH025918.1_strain_R25         .......cgg.a...aat....tt..t......g.ta.t..g....t......ag..t.g..a.....a..t........ [204400]
MH025917.1_strain_R7          .......cgg.a...aat....tt..t......g.ta.t..g....t......ag..t.g..a.....a..t........ [204400]
MH025916.1_strain_R8          .......cgg.a...aat....tt..t......g.ta.t..g....t......ag..t.g..a.....a..t........ [204400]

FR682468.2_Georgia_2007/1     gaaccgatcgaccgccacctgctcccagggacccttgcgctcgatgtcgtcggcttcccacacctcga---------cgg [204480]
ASF_IND_20_CAD543             ....................................................................---------... [204480]
LR899193.1_Germany_2020/1     ....................................................................---------... [204480]
MN393477.1_Wuhan_2019         ....................................................................---------... [204480]
MW396979.1_Timor-Leste/2019/1 ....................................................................---------... [204480]
MK940252.1_CN/2019/InnerMongolia ....................................................................---------... [204480]
MK128995.1_China/2018/AnhuiXCGQ ....................................................................---------... [204480]
MH681419.1_ASFV/POL/2015      ....................................................................---------... [204480]
LS478113.1_Estonia_2014       ....................................................................---------... [204480]
KP843857.1_Odintsovo_02/14    ....................................................................---------... [204480]
FN557520.1_strain_E75         ............................................................t.......---------... [204480]
AY261366.1_Warthog            ................g.............g.....................................---------... [204480]
AY261365.1_Warmbaths          ................g.............g.....................................---------... [204480]
AY261364.1_Tengani_62         ................g.............g.....................................---------..a [204480]
AY261363.1_Pretoriuskop/96/4  ................g.............g.....................................---------... [204480]
AY261362.1_Mkuzi_1979         ....................................................................---------... [204480]
AY261361.1_Malawi_Lil-20/1_1983 ..g..t...a..g..ag....t........g........g..a.....a..a.......g...t.tt.ttaaaatat.at [204480]
AY261360.1_Kenya_1950         ..g..t...a..g..ag.............g........g..a.....a..........g...g..t.ctaaaatat.at [204480]
AM712240.1_OURT_88/3          ............................................................t.......---------... [204480]
AM712239.1_Benin_97/1         ............................................................t.......---------... [204480]
KM111295.1_Ken06.Bus          ..g..t...ag.a..ag....t........g........a..a.....a..............g..t.ctaaaccat.at [204480]
KM111294.1_Ken05/Tk1          ..g..t...a..g..ag....t........g........g.ta.....a..a.......g...g..t.ctaaaatat.at [204480]
KM262844.1_strain_L60         ............................................................t.......---------... [204480]
KM102979.1_26544/OG10_Italy   ............................................................t.......---------... [204480]
KM262845.1_strain_NHV         ............................................................t.......---------... [204480]
KX354450.1_47/Ss/2008         ............................................................t.......---------... [204480]
KP055815.1_strain_BA71        ............................................................t.......---------... [204480]
MH025920.1_strain_R35         ..g..t...ag.a..ag....t........g........a..a.....a..............g..t.ctaaaccat.at [204480]
MH025919.1_strain_N10         ..g..t...ag.a..ag....t........g........a..a.....a..............g..t.ctaaaccat.at [204480]
MH025918.1_strain_R25         ..g..t...ag.a..ag....t........g........a..a.....a..............g..t.ctaaaccat.at [204480]
MH025917.1_strain_R7          ..g..t...ag.a..ag....t........g........a..a.....a..............g..t.ctaaaccat.at [204480]
MH025916.1_strain_R8          ..g..t...ag.a..ag....t........g........a..a.....a..............g..t.ctaaaccat.at [204480]
```

```
FR682468.2_Georgia_2007/1         ctgtggcaaaatggacatgcttcgcgtcgt-tcgtccgttttttgcgccgcct------ccccattattcttcctgtaag [204560]
ASF_IND_20_CAD543                 ..............................-.....................g------....ccc...a.......... [204560]
LR899193.1_Germany_2020/1         ..............................-......................------..................... [204560]
MN393477.1_Wuhan_2019             ..............................-......................------..................... [204560]
MW396979.1_Timor-Leste/2019/1     ..............................-......................------..................... [204560]
MK940252.1_CN/2019/InnerMongolia  ..............................-......................------..................... [204560]
MK128995.1_China/2018/AnhuiXCGQ   ..............................-......................------..................... [204560]
MH681419.1_ASFV/POL/2015          ..............................-......................------..................... [204560]
LS478113.1_Estonia_2014           ..............................-......................------..................... [204560]
KP843857.1_Odintsovo_02/14        ..............................-......................------..................... [204560]
FN557520.1_strain_E75             ...ca.....gc....g.......t...a.-.....................g------....ccc...a.......... [204560]
AY261366.1_Warthog                ...ca.....gc....g......a.a..a.-.....................g------....ccc...a........g. [204560]
AY261365.1_Warmbaths              ...ca.....gc....g.......t...a.-.....................g------....ccc...a.......... [204560]
AY261364.1_Tengani_62             ...ca.....gc....g......ata..a.-.....................g---------.ccc...a........g. [204560]
AY261363.1_Pretoriuskop/96/4      ...ca.....gc....g......a.a..a.-.....................g------t...ccc...a.......gg. [204560]
AY261362.1_Mkuzi_1979             ...ca.....gc....g...........a.-.....................g------....ccc...a.......... [204560]
AY261361.1_Malawi_Lil-20/1_1983   ....a.......aa..---...aa.a..ccg.t.c.t.a.....a.a.tagaatagaatg..tccc.------------- [204560]
AY261360.1_Kenya_1950             .c..a........a..---t..aa.a.tccg.t.c.t.a.....a.a.tagaatagaatg..tccc.------------- [204560]
AM712240.1_OURT_88/3              ...ca.....gc....g.......t...a.-.....................g------....ccc...a.......... [204560]
AM712239.1_Benin_97/1             ...ca.....gc....g.......t...a.-.....................gc---cc....ccc...a.......... [204560]
KM111295.1_Ken06.Bus              ....a...........---...tata..ccg.t...taa.....ata.t.gaatagaacg..ttcc.------------- [204560]
KM111294.1_Ken05/Tk1              ....a........a..---t..aa.a..ccg.t.c...a.....a.a.tagaatagaatg..tccc.------------- [204560]
KM262844.1_strain_L60             ...ca.....gc....g.......t...a.-.....................g------....ccc...a.......... [204560]
KM102979.1_26544/OG10_Italy       ...ca.....gc....g.......t...a.-.....................gc---cc....ccc...a.......... [204560]
KM262845.1_strain_NHV             ...ca.....gc....g.......t...a.-.....................g------....ccc...a.......... [204560]
KX354450.1_47/Ss/2008             ...ca.....gc....g.......t...a.-.....................g----cc....ccc...a.......... [204560]
KP055815.1_strain_BA71            ...ca.....gc....g.......t...a.-.....................g------....ccc...a.......... [204560]
MH025920.1_strain_R35             ....a...........---...tata..ccg.t...taa.....ata.t.gaatagaacg..ttcc.------------- [204560]
MH025919.1_strain_N10             ....a...........---...tata..ccg.t...taa.....ata.t.gaatagaacg..ttcc.------------- [204560]
MH025918.1_strain_R25             ....a...........---...tata..ccg.t...taa.....ata.t.gaatagaacg..ttcc.------------- [204560]
MH025917.1_strain_R7              ....a...........---...tata..ccg.t...taa.....ata.t.gaatagaacg..ttcc.------------- [204560]
MH025916.1_strain_R8              ....a...........---...tata..ccg.t...taa.....ata.t.gaatagaacg..ttcc.------------- [204560]

FR682468.2_Georgia_2007/1         attagtgtttaatacctataataacataattttaagatttaatataccaaaacttaaactatttttgtatagtaactatt [204640]
ASF_IND_20_CAD543                 ................................................................................ [204640]
LR899193.1_Germany_2020/1         ................................................................................ [204640]
MN393477.1_Wuhan_2019             ................................................................................ [204640]
MW396979.1_Timor-Leste/2019/1     ................................................................................ [204640]
MK940252.1_CN/2019/InnerMongolia  ................................................................................ [204640]
MK128995.1_China/2018/AnhuiXCGQ   ................................................................................ [204640]
MH681419.1_ASFV/POL/2015          ................................................................................ [204640]
LS478113.1_Estonia_2014           ................................................................................ [204640]
KP843857.1_Odintsovo_02/14        ................................................................................ [204640]
```

```
FN557520.1_strain_E75            ................................................................................ [204640]
AY261366.1_Warthog               ............c................-............c........................c...........c [204640]
AY261365.1_Warmbaths             ............c................-............c..................................... [204640]
AY261364.1_Tengani_62            ............c.t..............-............c..................................... [204640]
AY261363.1_Pretoriuskop/96/4     ............c................-............c........................c...........c [204640]
AY261362.1_Mkuzi_1979            ................................................................................ [204640]
AY261361.1_Malawi_Lil-20/1_1983  ------------------------------c...gt...c.c.tcgtg..g.gcagt.t....g.caa..cc.--t.... [204640]
AY261360.1_Kenya_1950            ------------------------------c...gt...c.c.tcgtg..g.gcggt.t....g.caa..cc.--t.... [204640]
AM712240.1_OURT_88/3             ................................................................................ [204640]
AM712239.1_Benin_97/1            ................................................................................ [204640]
KM111295.1_Ken06.Bus             ------------------------------c...gt.....c.tcgtg..g.gcgtt.t.g..a.caa..cc.--t.... [204640]
KM111294.1_Ken05/Tk1             ------------------------------c...gt...c.cg.agtg..g.gcggt.t....g.caa..cc.--t.... [204640]
KM262844.1_strain_L60            ................................................................................ [204640]
KM102979.1_26544/OG10_Italy      ................................................................................ [204640]
KM262845.1_strain_NHV            ................................................................................ [204640]
KX354450.1_47/Ss/2008            ................................................................................ [204640]
KP055815.1_strain_BA71           ................................................................................ [204640]
MH025920.1_strain_R35            ------------------------------c...gt.....c.tcgtg..g.gcgtt.t.g..a.caa..cc.--t.... [204640]
MH025919.1_strain_N10            ------------------------------c...gt.....c.tcgtg..g.gcgtt.t.g..a.caa..cc.--t.... [204640]
MH025918.1_strain_R25            ------------------------------c...gt.....c.tcgtg..g.gcgtt.t.g..a.caa..cc.--t.... [204640]
MH025917.1_strain_R7             ------------------------------c...gt.....c.tcgtg..g.gcgtt.t.g..a.caa..cc.--t.... [204640]
MH025916.1_strain_R8             ------------------------------c...gt.....c.tcgtg..g.gcgtt.t.g..a.caa..cc.--t.... [204640]

FR682468.2_Georgia_2007/1        agcatgtctaca--catg---attgttctctaaaagagaaaccggttgatatgaacgatatatctgagaaatcagttgtc [204720]
ASF_IND_20_CAD543                ............--....---......t.tc...........................................tc.... [204720]
LR899193.1_Germany_2020/1        ............--....---........................................................... [204720]
MN393477.1_Wuhan_2019            ............--....---........................................................... [204720]
MW396979.1_Timor-Leste/2019/1    ............--....---........................................................... [204720]
MK940252.1_CN/2019/InnerMongolia ............--....---........................................................... [204720]
MK128995.1_China/2018/AnhuiXCGQ  ............--....---........................................................... [204720]
MH681419.1_ASFV/POL/2015         ............--....---........................................................... [204720]
LS478113.1_Estonia_2014          ............--....---........................................................... [204720]
KP843857.1_Odintsovo_02/14       ............--....---........................................................... [204720]
FN557520.1_strain_E75            ............--....---......t.tc............a..............................tc.... [204720]
AY261366.1_Warthog               ..t........t--...a---..aa.....c............a............a...............t..a.... [204720]
AY261365.1_Warmbaths             ............--....---.........c............a.............................g.c.... [204720]
AY261364.1_Tengani_62            ...........t--...a---..aa.....c............t......g.....a..g........g......c...t [204720]
AY261363.1_Pretoriuskop/96/4     ..t........t--...a---..aa.....c............a............a...............t..a.... [204720]
AY261362.1_Mkuzi_1979            ............--....---.........c............a...............................c.... [204720]
AY261361.1_Malawi_Lil-20/1_1983  .aa....t..g.at....---c...cct.tc-.gg..t.ctt.at..a..ggc.t.a..g...tctca.g.--------- [204720]
AY261360.1_Kenya_1950            .aa....t..g.att...---c....ca.t.-.gg..t.ctt.at..a..ggc.t.a..g...tctca.g.--------- [204720]
AM712240.1_OURT_88/3             ............--....---......t.tc............a..............................tc.... [204720]
AM712239.1_Benin_97/1            ............--....---......t.tc............a..............................tc.... [204720]
KM111295.1_Ken06.Bus             .aa..t.t..g.at....att.....cg.tc-.gg.gt.ctt.at..a..gac.t.a..g...tctca.g.--------- [204720]
```

```
KM111294.1_Ken05/Tk1            .aa....t..gcat....---.....ca.tc-.gg..t.ctttat..a.cggc.t.a..g...t.tca.g.--------- [204720]
KM262844.1_strain_L60           ............--....---......t.tc............a..............................tc.... [204720]
KM102979.1_26544/OG10_Italy     ............--....---......t.tc............a..............................tc.... [204720]
KM262845.1_strain_NHV           ............--....---......t.tc............a..............................tc.... [204720]
KX354450.1_47/Ss/2008           ............--....---......t.tc............a..............................tc.... [204720]
KP055815.1_strain_BA71          ............--....---......t.tc............a..............................tc.... [204720]
MH025920.1_strain_R35           .aa..t.t..g.at....att.....cg.tc-.gg.gt.ctt.at..a..gac.t.a..g...tctca.g.--------- [204720]
MH025919.1_strain_N10           .aa..t.t..g.at....att.....cg.tc-.gg.gt.ctt.at..a..gac.t.a..g...tctca.g.--------- [204720]
MH025918.1_strain_R25           .aa..t.t..g.at....att.....cg.tc-.gg.gt.ctt.at..a..gac.t.a..g...tctca.g.--------- [204720]
MH025917.1_strain_R7            .aa..t.t..g.at....att.....cg.tc-.gg.gt.ctt.at..a..gac.t.a..g...tctca.g.--------- [204720]
MH025916.1_strain_R8            .aa..t.t..g.at....att.....cg.tc-.gg.gt.ctt.at..a..gac.t.a..g...tctca.g.--------- [204720]

FR682468.2_Georgia_2007/1       gtggataatgcacccgagaaaccagctggagcgaatcatatacctgagaagtcggcccgcgaaatgacatcatcagaatg [204800]
ASF_IND_20_CAD543               ................................................................................ [204800]
LR899193.1_Germany_2020/1       ................................................................................ [204800]
MN393477.1_Wuhan_2019           ................................................................................ [204800]
MW396979.1_Timor-Leste/2019/1   ................................................................................ [204800]
MK940252.1_CN/2019/InnerMongolia ................................................................................ [204800]
MK128995.1_China/2018/AnhuiXCGQ ................................................................................ [204800]
MH681419.1_ASFV/POL/2015        ................................................................................ [204800]
LS478113.1_Estonia_2014         ................................................................................ [204800]
KP843857.1_Odintsovo_02/14      ................................................................................ [204800]
FN557520.1_strain_E75           ................................................................................ [204800]
AY261366.1_Warthog              ...a..........t.......................................a..---..................... [204800]
AY261365.1_Warmbaths            ...a....................................................---..................... [204800]
AY261364.1_Tengani_62           ...a..........t....------......g...................a....---.a................... [204800]
AY261363.1_Pretoriuskop/96/4    ...a..........t........g............................t...---..................... [204800]
AY261362.1_Mkuzi_1979           ..........................................................a..................... [204800]
AY261361.1_Malawi_Lil-20/1_1983 ---------------------------------------------.....a..tt.atc.c.gtcact.......----- [204800]
AY261360.1_Kenya_1950           ---------------------------------------------.....a.ttt.atc.c.gtcact.......----- [204800]
AM712240.1_OURT_88/3            ................................................................................ [204800]
AM712239.1_Benin_97/1           ................................................................................ [204800]
KM111295.1_Ken06.Bus            ---------------------------------------------...gta.ttt.atc.c.gtcact.......----- [204800]
KM111294.1_Ken05/Tk1            ---------------------------------------------.....a..tt.atc.c.gtcact.......----- [204800]
KM262844.1_strain_L60           ................................................................................ [204800]
KM102979.1_26544/OG10_Italy     ................................................................................ [204800]
KM262845.1_strain_NHV           ................................................................................ [204800]
KX354450.1_47/Ss/2008           ................................................................................ [204800]
KP055815.1_strain_BA71          ................................................................................ [204800]
MH025920.1_strain_R35           ---------------------------------------------...gta.ttt.atc.c.gtcact.......----- [204800]
MH025919.1_strain_N10           ---------------------------------------------...gta.ttt.atc.c.gtcact.......----- [204800]
MH025918.1_strain_R25           ---------------------------------------------...gta.ttt.atc.c.gtcact.......----- [204800]
MH025917.1_strain_R7            ---------------------------------------------...gta.ttt.atc.c.gtcact.......----- [204800]
MH025916.1_strain_R8            ---------------------------------------------...gta.ttt.atc.c.gtcact.......----- [204800]
```

```
FR682468.2_Georgia_2007/1        gattgctgaatattggaaaggtataaaacgtggaaatgacgtgccatgttgttgtccaagaaaaatgaccagtgcagaca [204880]
ASF_IND_20_CAD543                ................................................................................ [204880]
LR899193.1_Germany_2020/1        ................................................................................ [204880]
MN393477.1_Wuhan_2019            ................................................................................ [204880]
MW396979.1_Timor-Leste/2019/1    ................................................................................ [204880]
MK940252.1_CN/2019/InnerMongolia ................................................................................ [204880]
MK128995.1_China/2018/AnhuiXCGQ  ................................................................................ [204880]
MH681419.1_ASFV/POL/2015         ................................................................................ [204880]
LS478113.1_Estonia_2014          ................................................................................ [204880]
KP843857.1_Odintsovo_02/14       ................................................................................ [204880]
FN557520.1_strain_E75            ................................................................................ [204880]
AY261366.1_Warthog               ...........................t........c.....................................t..... [204880]
AY261365.1_Warmbaths             ....................................c..............c............................ [204880]
AY261364.1_Tengani_62            ...........................t........c............---............................ [204880]
AY261363.1_Pretoriuskop/96/4     ...........................t........c........................................... [204880]
AY261362.1_Mkuzi_1979            ....................................c......................................g.... [204880]
AY261361.1_Malawi_Lil-20/1_1983  -------------------------------------ct.aaat.t.ca.t.ccc..g....tg.a.g.t.------... [204880]
AY261360.1_Kenya_1950            -------------------------------------cttaaat.t.ca.t.ccc.......tg.a.g.t.------..g [204880]
AM712240.1_OURT_88/3             ................................................................................ [204880]
AM712239.1_Benin_97/1            ................................................................................ [204880]
KM111295.1_Ken06.Bus             -------------------------------------cttaaat.t.ca.t.ccc..g....tgta.g.t.------... [204880]
KM111294.1_Ken05/Tk1             -------------------------------------cttaaat.t.ca.t.ccc.......tg.a.g.t.------... [204880]
KM262844.1_strain_L60            ................................................................................ [204880]
KM102979.1_26544/OG10_Italy      ................................................................................ [204880]
KM262845.1_strain_NHV            ................................................................................ [204880]
KX354450.1_47/Ss/2008            ................................................................................ [204880]
KP055815.1_strain_BA71           ................................................................................ [204880]
MH025920.1_strain_R35            -------------------------------------cttaaat.t.ca.t.ccc..g....tgta.g.t.------... [204880]
MH025919.1_strain_N10            -------------------------------------cttaaat.t.ca.t.ccc..g....tgta.g.t.------... [204880]
MH025918.1_strain_R25            -------------------------------------cttaaat.t.ca.t.ccc..g....tgta.g.t.------... [204880]
MH025917.1_strain_R7             -------------------------------------cttaaat.t.ca.t.ccc..g....tgta.g.t.------... [204880]
MH025916.1_strain_R8             -------------------------------------cttaaat.t.ca.t.ccc..g....tgta.g.t.------... [204880]

FR682468.2_Georgia_2007/1        aaaagttt---tcagtatttggtaagggatccctaatgcgctccatccagaagaataattaaaaaa----aatatttttt [204960]
ASF_IND_20_CAD543                ........---.......................................................----.......... [204960]
LR899193.1_Germany_2020/1        ........---.......................................................----.......... [204960]
MN393477.1_Wuhan_2019            ........---.......................................................----.......... [204960]
MW396979.1_Timor-Leste/2019/1    ........---.......................................................----.......... [204960]
MK940252.1_CN/2019/InnerMongolia ........---.......................................................----.......... [204960]
MK128995.1_China/2018/AnhuiXCGQ  ........---.......................................................----.......... [204960]
MH681419.1_ASFV/POL/2015         ........---.......................................................----.......... [204960]
LS478113.1_Estonia_2014          ........---.......................................................----.......... [204960]
KP843857.1_Odintsovo_02/14       ........---.................................................-.....----.......... [204960]
```

```
FN557520.1_strain_E75          ........---..........................a............................----.--ta..... [204960]
AY261366.1_Warthog             ........---.....g.............a...............g......g..g.........aaat...ta..... [204960]
AY261365.1_Warmbaths           ........---.........................................a.............----.......... [204960]
AY261364.1_Tengani_62          ........---....ag........-....a...............gt.....gt.g.........--at...ta..... [204960]
AY261363.1_Pretoriuskop/96/4   ........---.....g.............a...............g......g..g.........aaat...ta..... [204960]
AY261362.1_Mkuzi_1979          ........---.......................................................----.---...... [204960]
AY261361.1_Malawi_Lil-20/1_1983 t.gg....gtgc.ta.t....a.g.at...------------------g.g...cag...tg.tgg----gc.ca.c..g [204960]
AY261360.1_Kenya_1950          t.gg....gtac.ta.t....a...at...------------------g.g...cag...tg.tgg----gc.ca.c..g [204960]
AM712240.1_OURT_88/3           ........---..........................a............................----.......... [204960]
AM712239.1_Benin_97/1          ........---..........................a............................----.......... [204960]
KM111295.1_Ken06.Bus           t.gg...cgtgc.ta......a...at.g.------------------g.g...cag...t..t.g----gc.ca....g [204960]
KM111294.1_Ken05/Tk1           t.gg...agtac.ta.t....a.---t.g.------------------g.g...cag...tg.tgg----gc.ca....g [204960]
KM262844.1_strain_L60          ........---..........................a............................----.......... [204960]
KM102979.1_26544/OG10_Italy    ........---..........................a............................----.......... [204960]
KM262845.1_strain_NHV          ........---..........................a............................----.t--a..... [204960]
KX354450.1_47/Ss/2008          ........---..........................a............................----.......... [204960]
KP055815.1_strain_BA71         ........---..........................a............................----.......... [204960]
MH025920.1_strain_R35          t.gg...cgtgc.ta......a...at.g.------------------g.g...cag...t..t.g----gc.ca....g [204960]
MH025919.1_strain_N10          t.gg...cgtgc.ta......a...at.g.------------------g.g...cag...t..t.g----gc.ca....g [204960]
MH025918.1_strain_R25          t.gg...cgtgc.ta......a...at.g.------------------g.g...cag...t..t.g----gc.ca....g [204960]
MH025917.1_strain_R7           t.gg...cgtgc.ta......a...at.g.------------------g.g...cag...t..t.g----gc.ca....g [204960]
MH025916.1_strain_R8           t.gg...cgtgc.ta......a...at.g.------------------g.g...cag...t..t.g----gc.ca....g [204960]

FR682468.2_Georgia_2007/1      ttagcaagtttttaaactatttaaataaatgtggtaaaaaaat-tcacataataattaaagtgaacgtgttagaattaat [205040]
ASF_IND_20_CAD543              ..r.y...............a..gt.....n...........w-.................................... [205040]
LR899193.1_Germany_2020/1      ...........................................-.................................... [205040]
MN393477.1_Wuhan_2019          ...........................................-.................................... [205040]
MW396979.1_Timor-Leste/2019/1  ...........................................-.................................... [205040]
MK940252.1_CN/2019/InnerMongolia ...........................................-.................................... [205040]
MK128995.1_China/2018/AnhuiXCGQ ...........................................-.................................... [205040]
MH681419.1_ASFV/POL/2015       ...........................................-.................................... [205040]
LS478113.1_Estonia_2014        ...........................................-.................................... [205040]
KP843857.1_Odintsovo_02/14     ...........................................-.................................... [205040]
FN557520.1_strain_E75          ..g.t......a........a..gt.....a...........a-................c................... [205040]
AY261366.1_Warthog             ..g...t....c........g..g...tt......-.......-c....c..................c......c.... [205040]
AY261365.1_Warmbaths           ..g.t......c........a..gt.....a............-................c........c.......... [205040]
AY261364.1_Tengani_62          ..g...t....c.....g..g..g.......a.........g.-c.......................c........... [205040]
AY261363.1_Pretoriuskop/96/4   ..g...t....c........g..g....t......-.......-c....c..................c......c.... [205040]
AY261362.1_Mkuzi_1979          ..g.t......c........a..gt.....a...........at................c......g............ [205040]
AY261361.1_Malawi_Lil-20/1_1983 agctg.g.g....g..gg.cc.ggt.g....gta.t..g.gg.------------------------------------- [205040]
AY261360.1_Kenya_1950          agctg.g.a....g..gg.cc.ggt.g.....ca.t..g.gg.------------------------------------- [205040]
AM712240.1_OURT_88/3           ..g.t......a........a..gt.....a...........a-c...............c................... [205040]
AM712239.1_Benin_97/1          ..g.t......a........a..gt.....a...........a-................c................... [205040]
KM111295.1_Ken06.Bus           aac.g.g.a....g..gg.cc.ggt.g....cta.t..g.gg.------------------------------------- [205040]
```

```
KM111294.1_Ken05/Tk1            ggctg.g.g....g..gg.cc.ggt.g...c.ca.t..g.gg.------------------------------------- [205040]
KM262844.1_strain_L60           ..g.t......a........a..gt.....a...........a-................c................... [205040]
KM102979.1_26544/OG10_Italy     ..g.t......a........a..gt.....a...........a-................c................... [205040]
KM262845.1_strain_NHV           ..g.t......a........a..gt.....a...........a-c...............c................... [205040]
KX354450.1_47/Ss/2008           ..g.t......a........a..gt.....a...........a-................c................... [205040]
KP055815.1_strain_BA71          ..g.t......a........a..gt.....a...........a-................c................... [205040]
MH025920.1_strain_R35           aac.g.g.a....g..gg.cc.ggt.g....cta.t..g.gg.------------------------------------- [205040]
MH025919.1_strain_N10           aac.g.g.a....g..gg.cc.ggt.g....cta.t..g.gg.------------------------------------- [205040]
MH025918.1_strain_R25           aac.g.g.a....g..gg.cc.ggt.g....cta.t..g.gg.------------------------------------- [205040]
MH025917.1_strain_R7            aac.g.g.a....g..gg.cc.ggt.g....cta.t..g.gg.------------------------------------- [205040]
MH025916.1_strain_R8            aac.g.g.a....g..gg.cc.ggt.g....cta.t..g.gg.------------------------------------- [205040]

FR682468.2_Georgia_2007/1       atttttttataatcggatataatatccattaaatcaataaatgatagtgttgctaccacactaaacaataacaaacagaa [205120]
ASF_IND_20_CAD543               ................................................................................ [205120]
LR899193.1_Germany_2020/1       ................................................................................ [205120]
MN393477.1_Wuhan_2019           ................................................................................ [205120]
MW396979.1_Timor-Leste/2019/1   ................................................................................ [205120]
MK940252.1_CN/2019/InnerMongolia ................................................................................ [205120]
MK128995.1_China/2018/AnhuiXCGQ ................................................................................ [205120]
MH681419.1_ASFV/POL/2015        ................................................................................ [205120]
LS478113.1_Estonia_2014         ................................................................................ [205120]
KP843857.1_Odintsovo_02/14      ................................................................................ [205120]
FN557520.1_strain_E75           .-.............................................................................. [205120]
AY261366.1_Warthog              ................................tc.............................................. [205120]
AY261365.1_Warmbaths            ................................................................................ [205120]
AY261364.1_Tengani_62           ................t............g..tc.............................................. [205120]
AY261363.1_Pretoriuskop/96/4    ................................tc.............................................. [205120]
AY261362.1_Mkuzi_1979           ................................................................................ [205120]
AY261361.1_Malawi_Lil-20/1_1983 -.....gcgacg....cggctgcg.tt...-------------------------------------------------- [205120]
AY261360.1_Kenya_1950           -.....acgacg....cggctgcg.ttt..-------------------------------------------------- [205120]
AM712240.1_OURT_88/3            ................................................................................ [205120]
AM712239.1_Benin_97/1           ................................................................................ [205120]
KM111295.1_Ken06.Bus            -.....acgacg..t.cggctgcg---t..-------------------------------------------------- [205120]
KM111294.1_Ken05/Tk1            -.....acgacg....cggctgcg.ttt..-------------------------------------------------- [205120]
KM262844.1_strain_L60           ................................................................................ [205120]
KM102979.1_26544/OG10_Italy     ................................................................................ [205120]
KM262845.1_strain_NHV           ................................................................................ [205120]
KX354450.1_47/Ss/2008           ................................................................................ [205120]
KP055815.1_strain_BA71          ................................................................................ [205120]
MH025920.1_strain_R35           -.....acgacg..t.cggctgcg---t..-------------------------------------------------- [205120]
MH025919.1_strain_N10           -.....acgacg..t.cggctgcg---t..-------------------------------------------------- [205120]
MH025918.1_strain_R25           -.....acgacg..t.cggctgcg---t..-------------------------------------------------- [205120]
MH025917.1_strain_R7            -.....acgacg..t.cggctgcg---t..-------------------------------------------------- [205120]
MH025916.1_strain_R8            -.....acgacg..t.cggctgcg---t..-------------------------------------------------- [205120]
```

```
FR682468.2_Georgia_2007/1        acgcacgatacctttcctcatgatttataatagcgtg---ttatctaaagatttttttgaaaaaaatattaaattttagt [205200]
ASF_IND_20_CAD543                .....................................---........................................ [205200]
LR899193.1_Germany_2020/1        .....................................---........................................ [205200]
MN393477.1_Wuhan_2019            .....................................---........................................ [205200]
MW396979.1_Timor-Leste/2019/1    .....................................---........................................ [205200]
MK940252.1_CN/2019/InnerMongolia .....................................---........................................ [205200]
MK128995.1_China/2018/AnhuiXCGQ  .....................................---........................................ [205200]
MH681419.1_ASFV/POL/2015         .....................................---........................................ [205200]
LS478113.1_Estonia_2014          .....................................---........................................ [205200]
KP843857.1_Odintsovo_02/14       .....................................---........................................ [205200]
FN557520.1_strain_E75            .....t.............................a.---........................................ [205200]
AY261366.1_Warthog               .....t...........................t.a.tta.................ga..................... [205200]
AY261365.1_Warmbaths             .....t.............................a.---........................................ [205200]
AY261364.1_Tengani_62            .....t...........................t.a.tta.................ga..................... [205200]
AY261363.1_Pretoriuskop/96/4     .....t...........................t.a.tta.................ga..................... [205200]
AY261362.1_Mkuzi_1979            .....t.............................a.---........................................ [205200]
AY261361.1_Malawi_Lil-20/1_1983  ------------.c.....gac.......-----------g.t.t....t.a...a.gat.t.t..c..g.gta....-- [205200]
AY261360.1_Kenya_1950            ---------.tt.c.....gac.......-----------g.t.t....t.ac..a..at......c..g.gta....aa [205200]
AM712240.1_OURT_88/3             .....t.............................a.---........................................ [205200]
AM712239.1_Benin_97/1            .....t.............................a.---........................................ [205200]
KM111295.1_Ken06.Bus             ---------.tt.c.....g.c.......-----------g.t.t.--tt.aa.ac...ttt...gc..g.g--....aa [205200]
KM111294.1_Ken05/Tk1             ---------.tt.c.....gac.......-----------g.t.t....t.ac..a.gat.t....c..g.gta....aa [205200]
KM262844.1_strain_L60            .....t.............................a.---........................................ [205200]
KM102979.1_26544/OG10_Italy      .....t.............................a.---........................................ [205200]
KM262845.1_strain_NHV            .....t.............................a.---........................................ [205200]
KX354450.1_47/Ss/2008            .....t.............................a.---........................................ [205200]
KP055815.1_strain_BA71           .....t.............................a.---........................................ [205200]
MH025920.1_strain_R35            ---------.tt.c.....g.c.......-----------g.t.t.--tt.aa.ac...ttt...gc..g.g--....aa [205200]
MH025919.1_strain_N10            ---------.tt.c.....g.c.......-----------g.t.t.--tt.aa.ac...ttt...gc..g.g--....aa [205200]
MH025918.1_strain_R25            ---------.tt.c.....g.c.......-----------g.t.t.--tt.aa.ac...ttt...gc..g.g--....aa [205200]
MH025917.1_strain_R7             ---------.tt.c.....g.c.......-----------g.t.t.--tt.aa.ac...ttt...gc..g.g--....aa [205200]
MH025916.1_strain_R8             ---------.tt.c.....g.c.......-----------g.t.t.--tt.aa.ac...ttt...gc..g.g--....aa [205200]

FR682468.2_Georgia_2007/1        tg---------------------------------------attatttttttcagttacaacat----tgctttag---- [205280]
ASF_IND_20_CAD543                ..---------------------------------------..ata......t...........----........---- [205280]
LR899193.1_Germany_2020/1        ..---------------------------------------.......................----........---- [205280]
MN393477.1_Wuhan_2019            ..---------------------------------------.......................----........---- [205280]
MW396979.1_Timor-Leste/2019/1    ..---------------------------------------.......................----........---- [205280]
MK940252.1_CN/2019/InnerMongolia ..---------------------------------------.......................----........---- [205280]
MK128995.1_China/2018/AnhuiXCGQ  ..---------------------------------------.......................----........---- [205280]
MH681419.1_ASFV/POL/2015         ..---------------------------------------.......................----........---- [205280]
LS478113.1_Estonia_2014          ..---------------------------------------.......................----........---- [205280]
KP843857.1_Odintsovo_02/14       ..---------------------------------------.......................----........---- [205280]
```

```
FN557520.1_strain_E75              ..---------------------------------------..ata......t...........----........---- [205280]
AY261366.1_Warthog                 ..t--------------------------------------..ata......t.t......a..c--a..t.....---- [205280]
AY261365.1_Warmbaths               ..---------------------------------------..ata..................----........---- [205280]
AY261364.1_Tengani_62              ..t--------------------------------------..ata........t......acc---a........a--- [205280]
AY261363.1_Pretoriuskop/96/4       ..t--------------------------------------..ata.......gt......a..c--a..t.....---- [205280]
AY261362.1_Mkuzi_1979              ..---------------------------------------..ata..................----........---- [205280]
AY261361.1_Malawi_Lil-20/1_1983    --------------------------------------------------------------------aaa.....---- [205280]
AY261360.1_Kenya_1950              atttagatagcaacgacaaattactttgatatcaatcgata.a.ta..c...agta..g.ttgggtaa..t...t.aagg [205280]
AM712240.1_OURT_88/3               ..---------------------------------------..ata......t...........----........---- [205280]
AM712239.1_Benin_97/1              ..---------------------------------------..ata......t...........----........---- [205280]
KM111295.1_Ken06.Bus               .tttagatatcagtaacg--------------------------ta..c...agta..a.t.-ggtaa..t...t.aaga [205280]
KM111294.1_Ken05/Tk1               atttagatagcaataacaaattactttgatatcaatcgata.c.ta..c...agta..t.ttgggtaa..t...t.aagg [205280]
KM262844.1_strain_L60              ..---------------------------------------..ata......t...........----........---- [205280]
KM102979.1_26544/OG10_Italy        ..---------------------------------------..ata......t...........----........---- [205280]
KM262845.1_strain_NHV              ..---------------------------------------..ata......t...........----........---- [205280]
KX354450.1_47/Ss/2008              ..---------------------------------------..ata......t...........----........---- [205280]
KP055815.1_strain_BA71             ..---------------------------------------..ata......t...........----........---- [205280]
MH025920.1_strain_R35              .tttagatatcagtaacg--------------------------ta..c...agta..a.t.-ggtaa..t...t.aaga [205280]
MH025919.1_strain_N10              .tttagatatcagtaacg--------------------------ta..c...agta..a.t.-ggtaa..t...t.aaga [205280]
MH025918.1_strain_R25              .tttagatatcagtaacg--------------------------ta..c...agta..a.t.-ggtaa..t...t.aaga [205280]
MH025917.1_strain_R7               .tttagatatcagtaacg--------------------------ta..c...agta..a.t.-ggtaa..t...t.aaga [205280]
MH025916.1_strain_R8               .tttagatatcagtaacg--------------------------ta..c...agta..a.t.-ggtaa..t...t.aaga [205280]

FR682468.2_Georgia_2007/1          ---------------aaaaaatacctaat------tactacatagcaaataaagcgagcgcattgttacaaaca----ac [205360]
ASF_IND_20_CAD543                  ---------------..............------.........................tg............----.. [205360]
LR899193.1_Germany_2020/1          ---------------..............------.......................................----.. [205360]
MN393477.1_Wuhan_2019              ---------------..............------.......................................----.. [205360]
MW396979.1_Timor-Leste/2019/1      ---------------..............------.......................................----.. [205360]
MK940252.1_CN/2019/InnerMongolia   ---------------..............------.......................................----.. [205360]
MK128995.1_China/2018/AnhuiXCGQ    ---------------..............------.......................................----.. [205360]
MH681419.1_ASFV/POL/2015           ---------------..............------.......................................----.. [205360]
LS478113.1_Estonia_2014            ---------------..............------.......................................----.. [205360]
KP843857.1_Odintsovo_02/14         ---------------..............------.......................................----.. [205360]
FN557520.1_strain_E75              ---------------......a.......------.........................tg............----.. [205360]
AY261366.1_Warthog                 ---------------.........--...------.........................tg........---.----.. [205360]
AY261365.1_Warmbaths               ---------------..............------..................a......tg........---.----.. [205360]
AY261364.1_Tengani_62              -------------aa.........a....------.........................tg..a.....---.----.. [205360]
AY261363.1_Pretoriuskop/96/4       ----------------........a....------.......................t.tg............----.. [205360]
AY261362.1_Mkuzi_1979              ---------------..............------.........................tg............----.. [205360]
AY261361.1_Malawi_Lil-20/1_1983    --------------at.gt...gata...------...ttg------------------------------.t.----.. [205360]
AY261360.1_Kenya_1950              gaagtgttctaatga..t.g...ttct..aataaa..t.ct.c.at......gtta.tttggg.a...g.g...t---tt [205360]
AM712240.1_OURT_88/3               ---------------......a.......------.........................tg............----.. [205360]
AM712239.1_Benin_97/1              ---------------......a.......------.........................tg............----.. [205360]
KM111295.1_Ken06.Bus               aaagagttctaatag..t.....gt.t..aataaa..g.tt.c.at......gtta.attggg.....gg....----ct [205360]
```

```
KM111294.1_Ken05/Tk1               gaagagttctaatga..t.....tt.t..--------------.at......gtta.tttggg.a...g.g...tctact [205360]
KM262844.1_strain_L60              ---------------......a.......------.........................tg............----.. [205360]
KM102979.1_26544/OG10_Italy        ---------------......a.......------.........................tg............----.. [205360]
KM262845.1_strain_NHV              ---------------......a.......------.........................tg............----.. [205360]
KX354450.1_47/Ss/2008              ---------------......a.......------.........................tg............----.. [205360]
KP055815.1_strain_BA71             ---------------......a.......------.........................tg............----.. [205360]
MH025920.1_strain_R35              aaagagttctaatag..t.....gt.t..aataaa..g.tt.c.at......gtta.attggg.....gg....----ct [205360]
MH025919.1_strain_N10              aaagagttctaatag..t.....gt.t..aataaa..g.tt.c.at......gtta.attggg.....gg....----ct [205360]
MH025918.1_strain_R25              aaagagttctaatag..t.....gt.t..aataaa..g.tt.c.at......gtta.attggg.....gg....----ct [205360]
MH025917.1_strain_R7               aaagagttctaatag..t.....gt.t..aataaa..g.tt.c.at......gtta.attggg.....gg....----ct [205360]
MH025916.1_strain_R8               aaagagttctaatag..t.....gt.t..aataaa..g.tt.c.at......gtta.attggg.....gg....----ct [205360]

FR682468.2_Georgia_2007/1          atttt---------tttgcgcctggatactcctatatat----gagaactataatacggtatatt--------------- [205440]
ASF_IND_20_CAD543                  .....---------.........................----.............t.cc.....--------------- [205440]
LR899193.1_Germany_2020/1          .....---------.........................----......................--------------- [205440]
MN393477.1_Wuhan_2019              .....---------.........................----......................--------------- [205440]
MW396979.1_Timor-Leste/2019/1      .....---------.........................----......................--------------- [205440]
MK940252.1_CN/2019/InnerMongolia   .....---------.........................----......................--------------- [205440]
MK128995.1_China/2018/AnhuiXCGQ    .....---------.........................----......................--------------- [205440]
MH681419.1_ASFV/POL/2015           .....---------.........................----......................--------------- [205440]
LS478113.1_Estonia_2014            .....---------.........................----......................--------------- [205440]
KP843857.1_Odintsovo_02/14         .....---------.........................----......................--------------- [205440]
FN557520.1_strain_E75              .....---------.................t.......----.............t.cc.....--------------- [205440]
AY261366.1_Warthog                 .....---------.................t..c....----..t..g.......ta.......--------------- [205440]
AY261365.1_Warmbaths               .....---------.................t.......----.............t..c.....--------------- [205440]
AY261364.1_Tengani_62              ...c.---------.................t..c....----..t..g.......ta......a--------------- [205440]
AY261363.1_Pretoriuskop/96/4       .....---------.................t..c....----..t..g.......ta.......--------------- [205440]
AY261362.1_Mkuzi_1979              .....---------.........................----.............t.cc.....--------------- [205440]
AY261361.1_Malawi_Lil-20/1_1983    t.a..---------c..ta.ta.a...----------------------------------------------------- [205440]
AY261360.1_Kenya_1950              ..g..aatttgatt...tttttct..a.taat...t.cgtagat....g...g..g.ta.c..ccggtctgccgcctgct [205440]
AM712240.1_OURT_88/3               .....---------.................t.......----.............t.cc.....--------------- [205440]
AM712239.1_Benin_97/1              .....---------.................t.......----.............t.cc.....--------------- [205440]
KM111295.1_Ken06.Bus               .....aaattga--...tttt.c...a.taat...tg.gtagat....g...g..g.ca.c..c.ttacaagcacttgct [205440]
KM111294.1_Ken05/Tk1               .c...aatttg---...tttt.....a.taat...t.cg----t....g...gc.g.ta.c..ccggtctgccgcctgct [205440]
KM262844.1_strain_L60              .....---------.................t.......----.............t.cc.....--------------- [205440]
KM102979.1_26544/OG10_Italy        .....---------.................t.......----.............t.cc.....--------------- [205440]
KM262845.1_strain_NHV              .....---------.................t.......----.............t.cc.....--------------- [205440]
KX354450.1_47/Ss/2008              .....---------.................t.......----.............t.cc.....--------------- [205440]
KP055815.1_strain_BA71             .....---------.................t.......----.............t.cc.....--------------- [205440]
MH025920.1_strain_R35              .....aaattga--...tttt.c...a.taat...tg.gtagat....g...g..g.ca.c..c.ttacaagcacttgct [205440]
MH025919.1_strain_N10              .....aaattga--...tttt.c...a.taat...tg.gtagat....g...g..g.ca.c..c.ttacaagcacttgct [205440]
MH025918.1_strain_R25              .....aaattga--...tttt.c...a.taat...tg.gtagat....g...g..g.ca.c..c.ttacaagcacttgct [205440]
MH025917.1_strain_R7               .....aaattga--...tttt.c...a.taat...tg.gtagat....g...g..g.ca.c..c.ttacaagcacttgct [205440]
MH025916.1_strain_R8               .....aaattga--...tttt.c...a.taat...tg.gtagat....g...g..g.ca.c..c.ttacaagcacttgct [205440]
```

```
FR682468.2_Georgia_2007/1        -----------------aatcctattacc-----aacattgtcaat---------------------------------- [205520]
ASF_IND_20_CAD543                -----------------............-----....w..y...w---------------------------------- [205520]
LR899193.1_Germany_2020/1        -----------------............-----............---------------------------------- [205520]
MN393477.1_Wuhan_2019            -----------------............-----............---------------------------------- [205520]
MW396979.1_Timor-Leste/2019/1    -----------------............-----............---------------------------------- [205520]
MK940252.1_CN/2019/InnerMongolia -----------------............-----............---------------------------------- [205520]
MK128995.1_China/2018/AnhuiXCGQ  -----------------............-----............---------------------------------- [205520]
MH681419.1_ASFV/POL/2015         -----------------............-----............---------------------------------- [205520]
LS478113.1_Estonia_2014          -----------------............-----............---------------------------------- [205520]
KP843857.1_Odintsovo_02/14       -----------------............-----............---------------------------------- [205520]
FN557520.1_strain_E75            -----------------............-----....a..c...a---------------------------------- [205520]
AY261366.1_Warthog               -----------------...g........-----......a.....---------------------------------- [205520]
AY261365.1_Warmbaths             -----------------............-----.c.....c...a---------------------------------- [205520]
AY261364.1_Tengani_62            -----------------...g........-----......a.....---------------------------------- [205520]
AY261363.1_Pretoriuskop/96/4     -----------------...g........-----......a.....---------------------------------- [205520]
AY261362.1_Mkuzi_1979            -----------------............-----.c.....c...a---------------------------------- [205520]
AY261361.1_Malawi_Lil-20/1_1983  -------------------------------------------------------------------------------- [205520]
AY261360.1_Kenya_1950            ctcaggg----------.c....tg.g.tc----g.tg.c...ggctttccacatctcgatggctgcggcaaagtggatg [205520]
AM712240.1_OURT_88/3             -----------------............-----....a..c...a---------------------------------- [205520]
AM712239.1_Benin_97/1            -----------------............-----....a..c...a---------------------------------- [205520]
KM111295.1_Ken06.Bus             aaaaaagtactggccac.ccg.ata..t.taaaa..t.......g.ctaaggagtactgtcatatattaaaatgt---tg [205520]
KM111294.1_Ken05/Tk1             ctcaggg----------.c....tg.g.tc----g.tg.c...ggctttccacatctcgatggctgcggcaaagtggatg [205520]
KM262844.1_strain_L60            -----------------............-----....a..c...a---------------------------------- [205520]
KM102979.1_26544/OG10_Italy      -----------------............-----....a..c...a---------------------------------- [205520]
KM262845.1_strain_NHV            -----------------............-----....a..c...a---------------------------------- [205520]
KX354450.1_47/Ss/2008            -----------------............-----....a..c...a---------------------------------- [205520]
KP055815.1_strain_BA71           -----------------............-----....a..c...a---------------------------------- [205520]
MH025920.1_strain_R35            aaaaaagtactggccac.ccg.ata..t.taaaa..t.......g.ctaaggagtactgtcatatattaaaatgt---tg [205520]
MH025919.1_strain_N10            aaaaaagtactggccac.ccg.ata..t.taaaa..t.......g.ctaaggagtactgtcatatattaaaatgt---tg [205520]
MH025918.1_strain_R25            aaaaaagtactggccac.ccg.ata..t.taaaa..t.......g.ctaaggagtactgtcatatattaaaatgt---tg [205520]
MH025917.1_strain_R7             aaaaaagtactggccac.ccg.ata..t.taaaa..t.......g.ctaaggagtactgtcatatattaaaatgt---tg [205520]
MH025916.1_strain_R8             aaaaaagtactggccac.ccg.ata..t.taaaa..t.......g.ctaaggagtactgtcatatattaaaatgt---tg [205520]

FR682468.2_Georgia_2007/1        -------------------------------------------------------------------------------- [205600]
ASF_IND_20_CAD543                -------------------------------------------------------------------------------- [205600]
LR899193.1_Germany_2020/1        -------------------------------------------------------------------------------- [205600]
MN393477.1_Wuhan_2019            -------------------------------------------------------------------------------- [205600]
MW396979.1_Timor-Leste/2019/1    -------------------------------------------------------------------------------- [205600]
MK940252.1_CN/2019/InnerMongolia -------------------------------------------------------------------------------- [205600]
MK128995.1_China/2018/AnhuiXCGQ  -------------------------------------------------------------------------------- [205600]
MH681419.1_ASFV/POL/2015         -------------------------------------------------------------------------------- [205600]
LS478113.1_Estonia_2014          -------------------------------------------------------------------------------- [205600]
KP843857.1_Odintsovo_02/14       -------------------------------------------------------------------------------- [205600]
```

```
FN557520.1_strain_E75           -------------------------------------------------------------------------------- [205600]
AY261366.1_Warthog              -------------------------------------------------------------------------------- [205600]
AY261365.1_Warmbaths            -------------------------------------------------------------------------------- [205600]
AY261364.1_Tengani_62           -------------------------------------------------------------------------------- [205600]
AY261363.1_Pretoriuskop/96/4    -------------------------------------------------------------------------------- [205600]
AY261362.1_Mkuzi_1979           -------------------------------------------------------------------------------- [205600]
AY261361.1_Malawi_Lil-20/1_1983 -------------------------------------------------------------------------------- [205600]
AY261360.1_Kenya_1950           tgcttcgcgttg--atcgtta----------------------------------------------------------- [205600]
AM712240.1_OURT_88/3            -------------------------------------------------------------------------------- [205600]
AM712239.1_Benin_97/1           -------------------------------------------------------------------------------- [205600]
KM111295.1_Ken06.Bus            tggtttatggtggcatgatgc----------------------------------------------------------- [205600]
KM111294.1_Ken05/Tk1            tgcttcgcgttg--atcgttcgttttttgcactgcccattattcctcctataggattagtgtttaacacctataatatca [205600]
KM262844.1_strain_L60           -------------------------------------------------------------------------------- [205600]
KM102979.1_26544/OG10_Italy     -------------------------------------------------------------------------------- [205600]
KM262845.1_strain_NHV           -------------------------------------------------------------------------------- [205600]
KX354450.1_47/Ss/2008           -------------------------------------------------------------------------------- [205600]
KP055815.1_strain_BA71          -------------------------------------------------------------------------------- [205600]
MH025920.1_strain_R35           tggtttatggtggcatgatgc----------------------------------------------------------- [205600]
MH025919.1_strain_N10           tggtttatggtggcatgatgc----------------------------------------------------------- [205600]
MH025918.1_strain_R25           tggtttatggtggcatgatgc----------------------------------------------------------- [205600]
MH025917.1_strain_R7            tggtttatggtggcatgatgc----------------------------------------------------------- [205600]
MH025916.1_strain_R8            tggtttatggtggcatgatgc----------------------------------------------------------- [205600]

FR682468.2_Georgia_2007/1       -------------------------------------------------------------------------------- [205680]
ASF_IND_20_CAD543               -------------------------------------------------------------------------------- [205680]
LR899193.1_Germany_2020/1       -------------------------------------------------------------------------------- [205680]
MN393477.1_Wuhan_2019           -------------------------------------------------------------------------------- [205680]
MW396979.1_Timor-Leste/2019/1   -------------------------------------------------------------------------------- [205680]
MK940252.1_CN/2019/InnerMongolia -------------------------------------------------------------------------------- [205680]
MK128995.1_China/2018/AnhuiXCGQ -------------------------------------------------------------------------------- [205680]
MH681419.1_ASFV/POL/2015        -------------------------------------------------------------------------------- [205680]
LS478113.1_Estonia_2014         -------------------------------------------------------------------------------- [205680]
KP843857.1_Odintsovo_02/14      -------------------------------------------------------------------------------- [205680]
FN557520.1_strain_E75           -------------------------------------------------------------------------------- [205680]
AY261366.1_Warthog              -------------------------------------------------------------------------------- [205680]
AY261365.1_Warmbaths            -------------------------------------------------------------------------------- [205680]
AY261364.1_Tengani_62           -------------------------------------------------------------------------------- [205680]
AY261363.1_Pretoriuskop/96/4    -------------------------------------------------------------------------------- [205680]
AY261362.1_Mkuzi_1979           -------------------------------------------------------------------------------- [205680]
AY261361.1_Malawi_Lil-20/1_1983 -------------------------------------------------------------------------------- [205680]
AY261360.1_Kenya_1950           -------------ccaaaacttaaactatttttatat------aataactatcaccatgtctacttgtaattattctcca [205680]
AM712240.1_OURT_88/3            -------------------------------------------------------------------------------- [205680]
AM712239.1_Benin_97/1           -------------------------------------------------------------------------------- [205680]
KM111295.1_Ken06.Bus            -------------tccaattacaatttatccttgtataaataaaatattgataaaaacagcaagctttaaacatggttta [205680]
```

```
KM111294.1_Ken05/Tk1            taattttaacacatcaaaacttaaactatttttgtat------aataactattactatgtctacttgtaattattctcca [205680]
KM262844.1_strain_L60           -------------------------------------------------------------------------------- [205680]
KM102979.1_26544/OG10_Italy     -------------------------------------------------------------------------------- [205680]
KM262845.1_strain_NHV           -------------------------------------------------------------------------------- [205680]
KX354450.1_47/Ss/2008           -------------------------------------------------------------------------------- [205680]
KP055815.1_strain_BA71          -------------------------------------------------------------------------------- [205680]
MH025920.1_strain_R35           -------------tccaattacaatttatccttgtataaataaaatattgataaaaacagcaagctttaaacatggttta [205680]
MH025919.1_strain_N10           -------------tccaattacaatttatccttgtataaataaaatattgataaaaacagcaagctttaaacatggttta [205680]
MH025918.1_strain_R25           -------------tccaattacaatttatccttgtataaataaaatattgataaaaacagcaagctttaaacatggttta [205680]
MH025917.1_strain_R7            -------------tccaattacaatttatccttgtataaataaaatattgataaaaacagcaagctttaaacatggttta [205680]
MH025916.1_strain_R8            -------------tccaattacaatttatccttgtataaataaaatattgataaaaacagcaagctttaaacatggttta [205680]

FR682468.2_Georgia_2007/1       ---------------------aatagtatgtaggcaat------------------------------------------ [205760]
ASF_IND_20_CAD543               ---------------------.................------------------------------------------ [205760]
LR899193.1_Germany_2020/1       ---------------------.................------------------------------------------ [205760]
MN393477.1_Wuhan_2019           ---------------------.................------------------------------------------ [205760]
MW396979.1_Timor-Leste/2019/1   ---------------------.................------------------------------------------ [205760]
MK940252.1_CN/2019/InnerMongolia ---------------------.................------------------------------------------ [205760]
MK128995.1_China/2018/AnhuiXCGQ ---------------------.................------------------------------------------ [205760]
MH681419.1_ASFV/POL/2015        ---------------------.................------------------------------------------ [205760]
LS478113.1_Estonia_2014         ---------------------.................------------------------------------------ [205760]
KP843857.1_Odintsovo_02/14      ---------------------.................------------------------------------------ [205760]
FN557520.1_strain_E75           ---------------------.................------------------------------------------ [205760]
AY261366.1_Warthog              ---------------------.................------------------------------------------ [205760]
AY261365.1_Warmbaths            ---------------------.................------------------------------------------ [205760]
AY261364.1_Tengani_62           ---------------------...........a.....------------------------------------------ [205760]
AY261363.1_Pretoriuskop/96/4    ---------------------.................------------------------------------------ [205760]
AY261362.1_Mkuzi_1979           ---------------------.................------------------------------------------ [205760]
AY261361.1_Malawi_Lil-20/1_1983 ------------------------------.g......------------------------------------------ [205760]
AY261360.1_Kenya_1950           aaagagaaacctgttgatgtg..c.a.g.a.ct.ag.a------------------------------------------ [205760]
AM712240.1_OURT_88/3            ---------------------.................------------------------------------------ [205760]
AM712239.1_Benin_97/1           ---------------------.................------------------------------------------ [205760]
KM111295.1_Ken06.Bus            gagttgaatgtcgcattcatg..agc.g.ac...a..ataattatgatctaataaggctgtttattgagtggggtgcgga [205760]
KM111294.1_Ken05/Tk1            aaagagaaacctgttgatgtg..c.a.g...ct.ag.a------------------------------------------ [205760]
KM262844.1_strain_L60           ---------------------.................------------------------------------------ [205760]
KM102979.1_26544/OG10_Italy     ---------------------.................------------------------------------------ [205760]
KM262845.1_strain_NHV           ---------------------.................------------------------------------------ [205760]
KX354450.1_47/Ss/2008           ---------------------.................------------------------------------------ [205760]
KP055815.1_strain_BA71          ---------------------.................------------------------------------------ [205760]
MH025920.1_strain_R35           gagttgaatgtcgcattcatg..agc.g.ac...a..ataattatgatctaataaggctgtttattgagtggggtgcgga [205760]
MH025919.1_strain_N10           gagttgaatgtcgcattcatg..agc.g.ac...a..ataattatgatctaataaggctgtttattgagtggggtgcgga [205760]
MH025918.1_strain_R25           gagttgaatgtcgcattcatg..agc.g.ac...a..ataattatgatctaataaggctgtttattgagtggggtgcgga [205760]
MH025917.1_strain_R7            gagttgaatgtcgcattcatg..agc.g.ac...a..ataattatgatctaataaggctgtttattgagtggggtgcgga [205760]
MH025916.1_strain_R8            gagttgaatgtcgcattcatg..agc.g.ac...a..ataattatgatctaataaggctgtttattgagtggggtgcgga [205760]
```

```
FR682468.2_Georgia_2007/1        --------------------------------gacatactttaaa-----------taccaaatat-------------- [205840]
ASF_IND_20_CAD543                --------------------------------.............-----------..........-------------- [205840]
LR899193.1_Germany_2020/1        --------------------------------.............-----------..........-------------- [205840]
MN393477.1_Wuhan_2019            --------------------------------.............-----------..........-------------- [205840]
MW396979.1_Timor-Leste/2019/1    --------------------------------.............-----------..........-------------- [205840]
MK940252.1_CN/2019/InnerMongolia --------------------------------.............-----------..........-------------- [205840]
MK128995.1_China/2018/AnhuiXCGQ  --------------------------------.............-----------..........-------------- [205840]
MH681419.1_ASFV/POL/2015         --------------------------------.............-----------..........-------------- [205840]
LS478113.1_Estonia_2014          --------------------------------.............-----------..........-------------- [205840]
KP843857.1_Odintsovo_02/14       --------------------------------.............-----------..........-------------- [205840]
FN557520.1_strain_E75            --------------------------------.............-----------...a....t.-------------- [205840]
AY261366.1_Warthog               --------------------------------.........g...-----------.g........-------------- [205840]
AY261365.1_Warmbaths             --------------------------------.............-----------a..a....t.-------------- [205840]
AY261364.1_Tengani_62            --------------------------------..t......g...-----------a.........-------------- [205840]
AY261363.1_Pretoriuskop/96/4     --------------------------------.........g...-----------.g........-------------- [205840]
AY261362.1_Mkuzi_1979            --------------------------------.............-----------...a....t.-------------- [205840]
AY261361.1_Malawi_Lil-20/1_1983  --------------------------------tg.---t...g..-----------ggga.g.gt.-------------- [205840]
AY261360.1_Kenya_1950            ------------atcagccgccgtaaat----a.tg...c.g.g.-----------a....gc...-------------t [205840]
AM712240.1_OURT_88/3             --------------------------------.............-----------...a....t.-------------- [205840]
AM712239.1_Benin_97/1            --------------------------------.............-----------...a....t.-------------- [205840]
KM111295.1_Ken06.Bus             tatcaactttgggttggccactgttaatacagac.g...ccggg.cctgtgccggga..t.gg.gcgaaggaagctttaa [205840]
KM111294.1_Ken05/Tk1             ------------atcagctgccgtgaat----a.t....c.g.t.-----------a....gc.g.-------------t [205840]
KM262844.1_strain_L60            --------------------------------.............-----------...a....t.-------------- [205840]
KM102979.1_26544/OG10_Italy      --------------------------------.............-----------...a....t.-------------- [205840]
KM262845.1_strain_NHV            --------------------------------.............-----------...a....t.-------------- [205840]
KX354450.1_47/Ss/2008            --------------------------------.............-----------...a....t.-------------- [205840]
KP055815.1_strain_BA71           --------------------------------.............-----------...a....t.-------------- [205840]
MH025920.1_strain_R35            tatcaactttgggttggccactgttaatacagac.g...ccggg.cctgtgccggga..t.gg.gcgaaggaagctttaa [205840]
MH025919.1_strain_N10            tatcaactttgggttggccactgttaatacagac.g...ccggg.cctgtgccggga..t.gg.gcgaaggaagctttaa [205840]
MH025918.1_strain_R25            tatcaactttgggttggccactgttaatacagac.g...ccggg.cctgtgccggga..t.gg.gcgaaggaagctttaa [205840]
MH025917.1_strain_R7             tatcaactttgggttggccactgttaatacagac.g...ccggg.cctgtgccggga..t.gg.gcgaaggaagctttaa [205840]
MH025916.1_strain_R8             tatcaactttgggttggccactgttaatacagac.g...ccggg.cctgtgccggga..t.gg.gcgaaggaagctttaa [205840]

FR682468.2_Georgia_2007/1        ----------------------------------------------------------------------ccatggtta- [205920]
ASF_IND_20_CAD543                ----------------------------------------------------------------------.........- [205920]
LR899193.1_Germany_2020/1        ----------------------------------------------------------------------.........- [205920]
MN393477.1_Wuhan_2019            ----------------------------------------------------------------------.........- [205920]
MW396979.1_Timor-Leste/2019/1    ----------------------------------------------------------------------.........- [205920]
MK940252.1_CN/2019/InnerMongolia ----------------------------------------------------------------------.........- [205920]
MK128995.1_China/2018/AnhuiXCGQ  ----------------------------------------------------------------------.........- [205920]
MH681419.1_ASFV/POL/2015         ----------------------------------------------------------------------.........- [205920]
LS478113.1_Estonia_2014          ----------------------------------------------------------------------.........- [205920]
KP843857.1_Odintsovo_02/14       ----------------------------------------------------------------------.........- [205920]
```

```
FN557520.1_strain_E75             -----------------------------------------------------------------------.........- [205920]
AY261366.1_Warthog                -----------------------------------------------------------------------.....a.--- [205920]
AY261365.1_Warmbaths              -----------------------------------------------------------------------........c- [205920]
AY261364.1_Tengani_62             -----------------------------------------------------------------------.......--- [205920]
AY261363.1_Pretoriuskop/96/4      -----------------------------------------------------------------------.......--- [205920]
AY261362.1_Mkuzi_1979             -----------------------------------------------------------------------........c- [205920]
AY261361.1_Malawi_Lil-20/1_1983   -----------------------------------------------------------------------.t.------- [205920]
AY261360.1_Kenya_1950             gtgaataatacatctgagaaagctgttgatgtgaacaatgtatctgagaaacc-----tgttgatgtgaa.a...----- [205920]
AM712240.1_OURT_88/3              -----------------------------------------------------------------------.........- [205920]
AM712239.1_Benin_97/1             -----------------------------------------------------------------------.........- [205920]
KM111295.1_Ken06.Bus              gtgaaaaagaaatttagaaatatttgataaaatacagtatattaaaagtagcagtaatattattgtatc.....aa.ta [205920]
KM111294.1_Ken05/Tk1              gtgaataatgcacctgagaaacc-----------------------------------tgttgatgtgaata...----- [205920]
KM262844.1_strain_L60             -----------------------------------------------------------------------.........- [205920]
KM102979.1_26544/OG10_Italy       -----------------------------------------------------------------------.........- [205920]
KM262845.1_strain_NHV             -----------------------------------------------------------------------.........- [205920]
KX354450.1_47/Ss/2008             -----------------------------------------------------------------------.........- [205920]
KP055815.1_strain_BA71            -----------------------------------------------------------------------.........- [205920]
MH025920.1_strain_R35             gtgaaaaagaaatttagaaatatttgataaaatacagtatattaaaagtagcagtaatattattgtatc.....aa.ta [205920]
MH025919.1_strain_N10             gtgaaaaagaaatttagaaatatttgataaaatacagtatattaaaagtagcagtaatattattgtatc.....aa.ta [205920]
MH025918.1_strain_R25             gtgaaaaagaaatttagaaatatttgataaaatacagtatattaaaagtagcagtaatattattgtatc.....aa.ta [205920]
MH025917.1_strain_R7              gtgaaaaagaaatttagaaatatttgataaaatacagtatattaaaagtagcagtaatattattgtatc.....aa.ta [205920]
MH025916.1_strain_R8              gtgaaaaagaaatttagaaatatttgataaaatacagtatattaaaagtagcagtaatattattgtatc.....aa.ta [205920]

FR682468.2_Georgia_2007/1         -tttctaaaaatcttg---------------------------------------------------------------- [206000]
ASF_IND_20_CAD543                 -...............---------------------------------------------------------------- [206000]
LR899193.1_Germany_2020/1         -...............---------------------------------------------------------------- [206000]
MN393477.1_Wuhan_2019             -...............---------------------------------------------------------------- [206000]
MW396979.1_Timor-Leste/2019/1     -...............---------------------------------------------------------------- [206000]
MK940252.1_CN/2019/InnerMongolia  -...............---------------------------------------------------------------- [206000]
MK128995.1_China/2018/AnhuiXCGQ   -...............---------------------------------------------------------------- [206000]
MH681419.1_ASFV/POL/2015          -...............---------------------------------------------------------------- [206000]
LS478113.1_Estonia_2014           -...............---------------------------------------------------------------- [206000]
KP843857.1_Odintsovo_02/14        -...............---------------------------------------------------------------- [206000]
FN557520.1_strain_E75             -.c.............---------------------------------------------------------------- [206000]
AY261366.1_Warthog                -.a.g.........aa---------------------------------------------------------------- [206000]
AY261365.1_Warmbaths              -.c............a---------------------------------------------------------------- [206000]
AY261364.1_Tengani_62             -.a.g..........a---------------------------------------------------------------- [206000]
AY261363.1_Pretoriuskop/96/4      -.a.g.........aa---------------------------------------------------------------- [206000]
AY261362.1_Mkuzi_1979             -.c............a---------------------------------------------------------------- [206000]
AY261361.1_Malawi_Lil-20/1_1983   -------------------------------------------------------------------------------- [206000]
AY261360.1_Kenya_1950             -.a...g.g..atca.ctgccgtgaataatgcacttgagaaaccagctggagcgaataatatacctgaaaagtcagccg- [206000]
AM712240.1_OURT_88/3              -.c.............---------------------------------------------------------------- [206000]
AM712239.1_Benin_97/1             -.c.............---------------------------------------------------------------- [206000]
KM111295.1_Ken06.Bus              t.a.....t..c.ccctttttcttaaataatgatcaattgaaattaagaatgttcgatgaactacataaactatcaatcaa [206000]
```

```
KM111294.1_Ken05/Tk1           -.a...g.g..atca.ctgttgtgaataatgcacctgagaaaccagccggagcaaataatatacctgaaaagtcagccg- [206000]
KM262844.1_strain_L60          -.c.............---------------------------------------------------------------- [206000]
KM102979.1_26544/OG10_Italy    -.c.............---------------------------------------------------------------- [206000]
KM262845.1_strain_NHV          -.c.............---------------------------------------------------------------- [206000]
KX354450.1_47/Ss/2008          -.c.............---------------------------------------------------------------- [206000]
KP055815.1_strain_BA71         -.c.............---------------------------------------------------------------- [206000]
MH025920.1_strain_R35          t.a.....t..c.cccttttcttaaataatgatcaattgaaattaagaatgttcgatgaactacataaactatcaatcaa [206000]
MH025919.1_strain_N10          t.a.....t..c.cccttttcttaaataatgatcaattgaaattaagaatgttcgatgaactacataaactatcaatcaa [206000]
MH025918.1_strain_R25          t.a.....t..c.cccttttcttaaataatgatcaattgaaattaagaatgttcgatgaactacataaactatcaatcaa [206000]
MH025917.1_strain_R7           t.a.....t..c.cccttttcttaaataatgatcaattgaaattaagaatgttcgatgaactacataaactatcaatcaa [206000]
MH025916.1_strain_R8           t.a.....t..c.cccttttcttaaataatgatcaattgaaattaagaatgttcgatgaactacataaactatcaatcaa [206000]

FR682468.2_Georgia_2007/1      -----------aaaaaacgttaaattttagatcggtcacctacgacagtaatacta------------------------ [206080]
ASF_IND_20_CAD543              -----------......ta..t..................................------------------------ [206080]
LR899193.1_Germany_2020/1      -----------.............................................------------------------ [206080]
MN393477.1_Wuhan_2019          -----------.............................................------------------------ [206080]
MW396979.1_Timor-Leste/2019/1  -----------.............................................------------------------ [206080]
MK940252.1_CN/2019/InnerMongolia -----------.............................................------------------------ [206080]
MK128995.1_China/2018/AnhuiXCGQ  -----------.............................................------------------------ [206080]
MH681419.1_ASFV/POL/2015       -----------.............................................------------------------ [206080]
LS478113.1_Estonia_2014        -----------.............................................------------------------ [206080]
KP843857.1_Odintsovo_02/14     -----------.............................................------------------------ [206080]
FN557520.1_strain_E75          -----------......ta..t......................gt..........------------------------ [206080]
AY261366.1_Warthog             -----------.g.....a.....................c..ag...a.......------------------------ [206080]
AY261365.1_Warmbaths           -----------.......a.........................g..........g------------------------ [206080]
AY261364.1_Tengani_62          -----------.......a.............-...........g...------..------------------------ [206080]
AY261363.1_Pretoriuskop/96/4   -----------.g.....a.....................c..ag...........------------------------ [206080]
AY261362.1_Mkuzi_1979          -----------.......a.........................g..........g------------------------ [206080]
AY261361.1_Malawi_Lil-20/1_1983 -----------.t....ta.ct.t...aca..aaa.att....a.t.aat.agt..------------------------ [206080]
AY261360.1_Kenya_1950          -----------g..g..t.ac.tcg.caga..g.a.tg.tg.at.tt.g..agg..------------------------ [206080]
AM712240.1_OURT_88/3           -----------......ta..t......................gt..........------------------------ [206080]
AM712239.1_Benin_97/1          -----------......ta..t......................gt..........------------------------ [206080]
KM111295.1_Ken06.Bus           ctttgtattac.tg.g.ta.c.tt.aa.ga.at.c.a..taggt.tt.gt...g..tggcgatactatataacctcaccg [206080]
KM111294.1_Ken05/Tk1           --------------g..t.ac.tca.caga..g.a.tg.tg.at.tt.g..ggg..------------------------ [206080]
KM262844.1_strain_L60          -----------......ta..t......................gt..........------------------------ [206080]
KM102979.1_26544/OG10_Italy    -----------......ta..t......................gt..........------------------------ [206080]
KM262845.1_strain_NHV          -----------......ta..t......................gt..........------------------------ [206080]
KX354450.1_47/Ss/2008          -----------......ta..t......................gt..........------------------------ [206080]
KP055815.1_strain_BA71         -----------......ta..t......................gt..........------------------------ [206080]
MH025920.1_strain_R35          ctttgtattac.tg.g.ta.c.tt.aa.ga.at.c.a..taggt.tt.gt...g..tggcgatactatataacctcaccg [206080]
MH025919.1_strain_N10          ctttgtattac.tg.g.ta.c.tt.aa.ga.at.c.a..taggt.tt.gt...g..tggcgatactatataacctcaccg [206080]
MH025918.1_strain_R25          ctttgtattac.tg.g.ta.c.tt.aa.ga.at.c.a..taggt.tt.gt...g..tggcgatactatataacctcaccg [206080]
MH025917.1_strain_R7           ctttgtattac.tg.g.ta.c.tt.aa.ga.at.c.a..taggt.tt.gt...g..tggcgatactatataacctcaccg [206080]
MH025916.1_strain_R8           ctttgtattac.tg.g.ta.c.tt.aa.ga.at.c.a..taggt.tt.gt...g..tggcgatactatataacctcaccg [206080]
```

```
FR682468.2_Georgia_2007/1        -------------------------------------------------------------------------------- [206160]
ASF_IND_20_CAD543                -------------------------------------------------------------------------------- [206160]
LR899193.1_Germany_2020/1        -------------------------------------------------------------------------------- [206160]
MN393477.1_Wuhan_2019            -------------------------------------------------------------------------------- [206160]
MW396979.1_Timor-Leste/2019/1    -------------------------------------------------------------------------------- [206160]
MK940252.1_CN/2019/InnerMongolia -------------------------------------------------------------------------------- [206160]
MK128995.1_China/2018/AnhuiXCGQ  -------------------------------------------------------------------------------- [206160]
MH681419.1_ASFV/POL/2015         -------------------------------------------------------------------------------- [206160]
LS478113.1_Estonia_2014          -------------------------------------------------------------------------------- [206160]
KP843857.1_Odintsovo_02/14       -------------------------------------------------------------------------------- [206160]
FN557520.1_strain_E75            -------------------------------------------------------------------------------- [206160]
AY261366.1_Warthog               -------------------------------------------------------------------------------- [206160]
AY261365.1_Warmbaths             -------------------------------------------------------------------------------- [206160]
AY261364.1_Tengani_62            -------------------------------------------------------------------------------- [206160]
AY261363.1_Pretoriuskop/96/4     -------------------------------------------------------------------------------- [206160]
AY261362.1_Mkuzi_1979            -------------------------------------------------------------------------------- [206160]
AY261361.1_Malawi_Lil-20/1_1983  -------------------------------------------------------------------------------- [206160]
AY261360.1_Kenya_1950            ---------------------------------------------------taaaacgtgga---------aacgatgtg [206160]
AM712240.1_OURT_88/3             -------------------------------------------------------------------------------- [206160]
AM712239.1_Benin_97/1            -------------------------------------------------------------------------------- [206160]
KM111295.1_Ken06.Bus             cagccattcaatatttttatcaatcctacaagcattttaaagattggcggttaatatgtgggcttgcttataacaacgtg [206160]
KM111294.1_Ken05/Tk1             ---------------------------------------------------taaaacgtgga---------aacgatgtg [206160]
KM262844.1_strain_L60            -------------------------------------------------------------------------------- [206160]
KM102979.1_26544/OG10_Italy      -------------------------------------------------------------------------------- [206160]
KM262845.1_strain_NHV            -------------------------------------------------------------------------------- [206160]
KX354450.1_47/Ss/2008            -------------------------------------------------------------------------------- [206160]
KP055815.1_strain_BA71           -------------------------------------------------------------------------------- [206160]
MH025920.1_strain_R35            cagccattcaatatttttatcaatcctacaagcattttaaagattggcggttaatatgtgggcttgcttataacaacgtg [206160]
MH025919.1_strain_N10            cagccattcaatatttttatcaatcctacaagcattttaaagattggcggttaatatgtgggcttgcttataacaacgtg [206160]
MH025918.1_strain_R25            cagccattcaatatttttatcaatcctacaagcattttaaagattggcggttaatatgtgggcttgcttataacaacgtg [206160]
MH025917.1_strain_R7             cagccattcaatatttttatcaatcctacaagcattttaaagattggcggttaatatgtgggcttgcttataacaacgtg [206160]
MH025916.1_strain_R8             cagccattcaatatttttatcaatcctacaagcattttaaagattggcggttaatatgtgggcttgcttataacaacgtg [206160]

FR682468.2_Georgia_2007/1        -------------------------------------------------------------------------------- [206240]
ASF_IND_20_CAD543                -------------------------------------------------------------------------------- [206240]
LR899193.1_Germany_2020/1        -------------------------------------------------------------------------------- [206240]
MN393477.1_Wuhan_2019            -------------------------------------------------------------------------------- [206240]
MW396979.1_Timor-Leste/2019/1    -------------------------------------------------------------------------------- [206240]
MK940252.1_CN/2019/InnerMongolia -------------------------------------------------------------------------------- [206240]
MK128995.1_China/2018/AnhuiXCGQ  -------------------------------------------------------------------------------- [206240]
MH681419.1_ASFV/POL/2015         -------------------------------------------------------------------------------- [206240]
LS478113.1_Estonia_2014          -------------------------------------------------------------------------------- [206240]
KP843857.1_Odintsovo_02/14       -------------------------------------------------------------------------------- [206240]
```

```
FN557520.1_strain_E75            -------------------------------------------------------------------------------- [206240]
AY261366.1_Warthog               -------------------------------------------------------------------------------- [206240]
AY261365.1_Warmbaths             -------------------------------------------------------------------------------- [206240]
AY261364.1_Tengani_62            -------------------------------------------------------------------------------- [206240]
AY261363.1_Pretoriuskop/96/4     -------------------------------------------------------------------------------- [206240]
AY261362.1_Mkuzi_1979            -------------------------------------------------------------------------------- [206240]
AY261361.1_Malawi_Lil-20/1_1983  -------------------------------------------------------------------------------- [206240]
AY261360.1_Kenya_1950            ccatgttgttgtccaagaaaaatg-------------------accagcgcagacgaaaagtttccagtg---------- [206240]
AM712240.1_OURT_88/3             -------------------------------------------------------------------------------- [206240]
AM712239.1_Benin_97/1            -------------------------------------------------------------------------------- [206240]
KM111295.1_Ken06.Bus             t----ttgatcttcatgaaatatataacaaagagaaggttgatatggatattaatcaaatgatgcagttggcctgtatgt [206240]
KM111294.1_Ken05/Tk1             ccatgttgttgtccaagaaaaatg-------------------accagcgcagacgaaaagttttcagtg---------- [206240]
KM262844.1_strain_L60            -------------------------------------------------------------------------------- [206240]
KM102979.1_26544/OG10_Italy      -------------------------------------------------------------------------------- [206240]
KM262845.1_strain_NHV            -------------------------------------------------------------------------------- [206240]
KX354450.1_47/Ss/2008            -------------------------------------------------------------------------------- [206240]
KP055815.1_strain_BA71           -------------------------------------------------------------------------------- [206240]
MH025920.1_strain_R35            t----ttgaccttcatgaaatttataataaagagaagactgatatgaacattgatgaaatgatgcagttggcctgtatgt [206240]
MH025919.1_strain_N10            t----ttgaccttcatgaaatttataataaagagaagactgatatgaacattgatgaaatgatgcagttggcctgtatgt [206240]
MH025918.1_strain_R25            t----ttgaccttcatgaaatttataataaagagaagactgatatgaacattgatgaaatgatgcagttggcctgtatgt [206240]
MH025917.1_strain_R7             t----ttgaccttcatgaaatttataataaagagaagactgatatgaacattgatgaaatgatgcagttggcctgtatgt [206240]
MH025916.1_strain_R8             t----ttgaccttcatgaaatttataataaagagaagactgatatgaacattgatgaaatgatgcagttggcctgtatgt [206240]

FR682468.2_Georgia_2007/1        -------------------------------------------------------------------------------- [206320]
ASF_IND_20_CAD543                -------------------------------------------------------------------------------- [206320]
LR899193.1_Germany_2020/1        -------------------------------------------------------------------------------- [206320]
MN393477.1_Wuhan_2019            -------------------------------------------------------------------------------- [206320]
MW396979.1_Timor-Leste/2019/1    -------------------------------------------------------------------------------- [206320]
MK940252.1_CN/2019/InnerMongolia -------------------------------------------------------------------------------- [206320]
MK128995.1_China/2018/AnhuiXCGQ  -------------------------------------------------------------------------------- [206320]
MH681419.1_ASFV/POL/2015         -------------------------------------------------------------------------------- [206320]
LS478113.1_Estonia_2014          -------------------------------------------------------------------------------- [206320]
KP843857.1_Odintsovo_02/14       -------------------------------------------------------------------------------- [206320]
FN557520.1_strain_E75            -------------------------------------------------------------------------------- [206320]
AY261366.1_Warthog               -------------------------------------------------------------------------------- [206320]
AY261365.1_Warmbaths             -------------------------------------------------------------------------------- [206320]
AY261364.1_Tengani_62            -------------------------------------------------------------------------------- [206320]
AY261363.1_Pretoriuskop/96/4     -------------------------------------------------------------------------------- [206320]
AY261362.1_Mkuzi_1979            -------------------------------------------------------------------------------- [206320]
AY261361.1_Malawi_Lil-20/1_1983  -------------------------------------------------------------------------------- [206320]
AY261360.1_Kenya_1950            -------------------------------------------------------------------------------- [206320]
AM712240.1_OURT_88/3             -------------------------------------------------------------------------------- [206320]
AM712239.1_Benin_97/1            -------------------------------------------------------------------------------- [206320]
KM111295.1_Ken06.Bus             atgatggtaattatacaaccatttattattgttttatgttgggagctgacattaatcgggcaatgattacctcggttaca [206320]
```

```
KM111294.1_Ken05/Tk1             -------------------------------------------------------------------------------- [206320]
KM262844.1_strain_L60            -------------------------------------------------------------------------------- [206320]
KM102979.1_26544/OG10_Italy      -------------------------------------------------------------------------------- [206320]
KM262845.1_strain_NHV            -------------------------------------------------------------------------------- [206320]
KX354450.1_47/Ss/2008            -------------------------------------------------------------------------------- [206320]
KP055815.1_strain_BA71           -------------------------------------------------------------------------------- [206320]
MH025920.1_strain_R35            atgatggtaattatacaaccaattattattgttttatgttgggggctgacatcaatcgggcaatgattgcctcgatcata [206320]
MH025919.1_strain_N10            atgatggtaattatacaaccaattattattgttttatgttgggggctgacatcaatcgggcaatgattgcctcgatcata [206320]
MH025918.1_strain_R25            atgatggtaattatacaaccaattattattgttttatgttgggggctgacatcaatcgggcaatgattgcctcgatcata [206320]
MH025917.1_strain_R7             atgatggtaattatacaaccaattattattgttttatgttgggggctgacatcaatcgggcaatgattgcctcgatcata [206320]
MH025916.1_strain_R8             atgatggtaattatacaaccaattattattgttttatgttgggggctgacatcaatcgggcaatgattgcctcgatcata [206320]

FR682468.2_Georgia_2007/1        -atttaataattg------------------------------------------------------------------- [206400]
ASF_IND_20_CAD543                -............------------------------------------------------------------------- [206400]
LR899193.1_Germany_2020/1        -............------------------------------------------------------------------- [206400]
MN393477.1_Wuhan_2019            -............------------------------------------------------------------------- [206400]
MW396979.1_Timor-Leste/2019/1    -............------------------------------------------------------------------- [206400]
MK940252.1_CN/2019/InnerMongolia -............------------------------------------------------------------------- [206400]
MK128995.1_China/2018/AnhuiXCGQ  -............------------------------------------------------------------------- [206400]
MH681419.1_ASFV/POL/2015         -............------------------------------------------------------------------- [206400]
LS478113.1_Estonia_2014          -............------------------------------------------------------------------- [206400]
KP843857.1_Odintsovo_02/14       -............------------------------------------------------------------------- [206400]
FN557520.1_strain_E75            -............------------------------------------------------------------------- [206400]
AY261366.1_Warthog               -............------------------------------------------------------------------- [206400]
AY261365.1_Warmbaths             -............------------------------------------------------------------------- [206400]
AY261364.1_Tengani_62            -............------------------------------------------------------------------- [206400]
AY261363.1_Pretoriuskop/96/4     -............------------------------------------------------------------------- [206400]
AY261362.1_Mkuzi_1979            -............------------------------------------------------------------------- [206400]
AY261361.1_Malawi_Lil-20/1_1983  -....ggg..t.a.a----------------------------------------------------------------- [206400]
AY261360.1_Kenya_1950            ---...tg...gg.atacctaatgcgctccatg----------------------------------------------- [206400]
AM712240.1_OURT_88/3             -............------------------------------------------------------------------- [206400]
AM712239.1_Benin_97/1            -............------------------------------------------------------------------- [206400]
KM111295.1_Ken06.Bus             a.a.c.tg.g..g.taacttattcctttgtatagatttgggggctgatgtatttgaagagagtatgaaaatggcaataga [206400]
KM111294.1_Ken05/Tk1             ---...gg...gg.atacctaatgcgttccatg----------------------------------------------- [206400]
KM262844.1_strain_L60            -............------------------------------------------------------------------- [206400]
KM102979.1_26544/OG10_Italy      -............------------------------------------------------------------------- [206400]
KM262845.1_strain_NHV            -............------------------------------------------------------------------- [206400]
KX354450.1_47/Ss/2008            -............------------------------------------------------------------------- [206400]
KP055815.1_strain_BA71           -............------------------------------------------------------------------- [206400]
MH025920.1_strain_R35            a.....tg.g..g.taacttattcctttgtatagatttgggggctgatgtatttgaagagagtatgaaaatggcaataga [206400]
MH025919.1_strain_N10            a.....tg.g..g.taacttattcctttgtatagatttgggggctgatgtatttgaagagagtatgaaaatggcaataga [206400]
MH025918.1_strain_R25            a.....tg.g..g.taacttattcctttgtatagatttgggggctgatgtatttgaagagagtatgaaaatggcaataga [206400]
MH025917.1_strain_R7             a.....tg.g..g.taacttattcctttgtatagatttgggggctgatgtatttgaagagagtatgaaaatggcaataga [206400]
MH025916.1_strain_R8             a.....tg.g..g.taacttattcctttgtatagatttgggggctgatgtatttgaagagagtatgaaaatggcaataga [206400]
```

```
FR682468.2_Georgia_2007/1        -----------atgactgaaatcataatataatgcc-------------------------------------------- [206480]
ASF_IND_20_CAD543                -----------.........................-------------------------------------------- [206480]
LR899193.1_Germany_2020/1        -----------.........................-------------------------------------------- [206480]
MN393477.1_Wuhan_2019            -----------.........................-------------------------------------------- [206480]
MW396979.1_Timor-Leste/2019/1    -----------.........................-------------------------------------------- [206480]
MK940252.1_CN/2019/InnerMongolia -----------.........................-------------------------------------------- [206480]
MK128995.1_China/2018/AnhuiXCGQ  -----------.........................-------------------------------------------- [206480]
MH681419.1_ASFV/POL/2015         -----------.........................-------------------------------------------- [206480]
LS478113.1_Estonia_2014          -----------.........................-------------------------------------------- [206480]
KP843857.1_Odintsovo_02/14       -----------.........................-------------------------------------------- [206480]
FN557520.1_strain_E75            -----------......a........t.........-------------------------------------------- [206480]
AY261366.1_Warthog               -----------........................t-------------------------------------------- [206480]
AY261365.1_Warmbaths             -----------.........................-------------------------------------------- [206480]
AY261364.1_Tengani_62            -----------.........................-------------------------------------------- [206480]
AY261363.1_Pretoriuskop/96/4     -----------........................t-------------------------------------------- [206480]
AY261362.1_Mkuzi_1979            -----------.........................-------------------------------------------- [206480]
AY261361.1_Malawi_Lil-20/1_1983  ----------a.cat.cact..tt....t.g..ttt-------------------------------------------- [206480]
AY261360.1_Kenya_1950            ----cataagg....t.a...aa...t.t.tt.a------------------acaaatgtttctaaactat--------- [206480]
AM712240.1_OURT_88/3             -----------......a........t.........-------------------------------------------- [206480]
AM712239.1_Benin_97/1            -----------......a........t.........-------------------------------------------- [206480]
KM111295.1_Ken06.Bus             agatcataatg.c.ta.t.g.aag..tct..tcatttaaaaattattatagttcagacgtttctctgttatcattaaaga [206480]
KM111294.1_Ken05/Tk1             ----caaaagc....t.a...aa...t.t.tt.t-----------------gacaaatgtttctaaactat--------- [206480]
KM262844.1_strain_L60            -----------......a........t.........-------------------------------------------- [206480]
KM102979.1_26544/OG10_Italy      -----------......a........t.........-------------------------------------------- [206480]
KM262845.1_strain_NHV            -----------......a........t.........-------------------------------------------- [206480]
KX354450.1_47/Ss/2008            -----------......a........t.........-------------------------------------------- [206480]
KP055815.1_strain_BA71           -----------......a........t.........-------------------------------------------- [206480]
MH025920.1_strain_R35            agatcataatg.c.ta.t.g.aag..tct..tcatttaaaaattattatagttcagacgtttctctgttatcattaaaga [206480]
MH025919.1_strain_N10            agatcataatg.c.ta.t.g.aag..tct..tcatttaaaaattattatagttcagacgtttctctgttatcattaaaga [206480]
MH025918.1_strain_R25            agatcataatg.c.ta.t.g.aag..tct..tcatttaaaaattattatagttcagacgtttctctgttatcattaaaga [206480]
MH025917.1_strain_R7             agatcataatg.c.ta.t.g.aag..tct..tcatttaaaaattattatagttcagacgtttctctgttatcattaaaga [206480]
MH025916.1_strain_R8             agatcataatg.c.ta.t.g.aag..tct..tcatttaaaaattattatagttcagacgtttctctgttatcattaaaga [206480]

FR682468.2_Georgia_2007/1        ------------------------------------------------gtgcgaaaaa---------------------- [206560]
ASF_IND_20_CAD543                ------------------------------------------------..........---------------------- [206560]
LR899193.1_Germany_2020/1        ------------------------------------------------..........---------------------- [206560]
MN393477.1_Wuhan_2019            ------------------------------------------------..........---------------------- [206560]
MW396979.1_Timor-Leste/2019/1    ------------------------------------------------..........---------------------- [206560]
MK940252.1_CN/2019/InnerMongolia ------------------------------------------------..........---------------------- [206560]
MK128995.1_China/2018/AnhuiXCGQ  ------------------------------------------------..........---------------------- [206560]
MH681419.1_ASFV/POL/2015         ------------------------------------------------..........---------------------- [206560]
LS478113.1_Estonia_2014          ------------------------------------------------..........---------------------- [206560]
KP843857.1_Odintsovo_02/14       ------------------------------------------------..........---------------------- [206560]
```

```
FN557520.1_strain_E75            ------------------------------------------------....a.....---------------------- [206560]
AY261366.1_Warthog               -------------------------------------------------.........---------------------- [206560]
AY261365.1_Warmbaths             ------------------------------------------------..........---------------------- [206560]
AY261364.1_Tengani_62            ------------------------------------------------..........---------------------- [206560]
AY261363.1_Pretoriuskop/96/4     -------------------------------------------------.........---------------------- [206560]
AY261362.1_Mkuzi_1979            ------------------------------------------------..........---------------------- [206560]
AY261361.1_Malawi_Lil-20/1_1983  -----------------------------------------------tt.t.t.g...---------------------- [206560]
AY261360.1_Kenya_1950            -----------------------------------aaagatagataaaa..ta.....---------------------- [206560]
AM712240.1_OURT_88/3             ------------------------------------------------....a.....---------------------- [206560]
AM712239.1_Benin_97/1            ------------------------------------------------....a.....---------------------- [206560]
KM111295.1_Ken06.Bus             cgacagatccagaaaaaattaatgccttgttagaagaagaaagttacaaat.a.....taggttgatatataaaagttat [206560]
KM111294.1_Ken05/Tk1             -----------------------------------gaagatagataaaa..ta.....---------------------- [206560]
KM262844.1_strain_L60            ------------------------------------------------....a.....---------------------- [206560]
KM102979.1_26544/OG10_Italy      ------------------------------------------------....a.....---------------------- [206560]
KM262845.1_strain_NHV            ------------------------------------------------....a.....---------------------- [206560]
KX354450.1_47/Ss/2008            ------------------------------------------------....a.....---------------------- [206560]
KP055815.1_strain_BA71           ------------------------------------------------....a.....---------------------- [206560]
MH025920.1_strain_R35            cgacagatccagaaaaaattaatgccttgttagaagaagaaagttacaaat.a.....taggttgatatataaaagttat [206560]
MH025919.1_strain_N10            cgacagatccagaaaaaattaatgccttgttagaagaagaaagttacaaat.a.....taggttgatatataaaagttat [206560]
MH025918.1_strain_R25            cgacagatccagaaaaaattaatgccttgttagaagaagaaagttacaaat.a.....taggttgatatataaaagttat [206560]
MH025917.1_strain_R7             cgacagatccagaaaaaattaatgccttgttagaagaagaaagttacaaat.a.....taggttgatatataaaagttat [206560]
MH025916.1_strain_R8             cgacagatccagaaaaaattaatgccttgttagaagaagaaagttacaaat.a.....taggttgatatataaaagttat [206560]

FR682468.2_Georgia_2007/1        --------------taattat--ttttcggttaaa-gataccattaca---------taaaaaat--------------a [206640]
ASF_IND_20_CAD543                --------------.......--............g......w.....---------.......c-------------g. [206640]
LR899193.1_Germany_2020/1        --------------.......--............-............---------........--------------. [206640]
MN393477.1_Wuhan_2019            --------------.......--............-............---------........--------------. [206640]
MW396979.1_Timor-Leste/2019/1    --------------.......--............-............---------........--------------. [206640]
MK940252.1_CN/2019/InnerMongolia --------------.......--............-............---------........--------------. [206640]
MK128995.1_China/2018/AnhuiXCGQ  --------------.......--............-............---------........--------------. [206640]
MH681419.1_ASFV/POL/2015         --------------.......--............-............---------........--------------. [206640]
LS478113.1_Estonia_2014          --------------.......--............-............---------........--------------. [206640]
KP843857.1_Odintsovo_02/14       --------------.......--............-............---------........--------------. [206640]
FN557520.1_strain_E75            --------------.......--............g......t...a.---------.......c-------------g. [206640]
AY261366.1_Warthog               --------------.......--............aa.........a.---------.-......-------------g. [206640]
AY261365.1_Warmbaths             --------------.......--............gatact.t...a.---------........-------------g. [206640]
AY261364.1_Tengani_62            --------------.......--........a...--.........a.---------........-------------g. [206640]
AY261363.1_Pretoriuskop/96/4     --------------.......--............aac........a.---------.-......-------------g. [206640]
AY261362.1_Mkuzi_1979            --------------.......--............gatact.t...a.---------........-------------g. [206640]
AY261361.1_Malawi_Lil-20/1_1983  --------------......g--...gta.a..g.------------------------.gtt..-------------g. [206640]
AY261360.1_Kenya_1950            -----------------------a..ct..ca...-------t...a.ctttttttgc......a-----aatatgtcc. [206640]
AM712240.1_OURT_88/3             --------------.......--............g......t...a.---------.......c-------------g. [206640]
AM712239.1_Benin_97/1            --------------.......--............g......t...a.---------.......c-------------g. [206640]
KM111295.1_Ken06.Bus             tgatattacaacaa...c...tt....tt.ca...-------t...a.-attttttgc......a-----aatatgtcc. [206640]
```

```
KM111294.1_Ken05/Tk1          ------------------------.cct..ca...-------t...a.-ctattttg.......aaaaagaatatgcct. [206640]
KM262844.1_strain_L60         --------------.......--............g......t...a.---------.......c-------------g. [206640]
KM102979.1_26544/OG10_Italy   --------------.......--............g......t...a.---------.......c-------------g. [206640]
KM262845.1_strain_NHV         --------------.......--............g......t...a.---------.......c-------------g. [206640]
KX354450.1_47/Ss/2008         --------------.......--............g......t...a.---------.......c-------------g. [206640]
KP055815.1_strain_BA71        --------------.......--............g......t...a.---------.......c-------------g. [206640]
MH025920.1_strain_R35         tgatattacaacaa...c...tt....tt.ca...-------t...a.-attttttgc......a-----aatatgtcc. [206640]
MH025919.1_strain_N10         tgatattacaacaa...c...tt....tt.ca...-------t...a.-attttttgc......a-----aatatgtcc. [206640]
MH025918.1_strain_R25         tgatattacaacaa...c...tt....tt.ca...-------t...a.-attttttgc......a-----aatatgtcc. [206640]
MH025917.1_strain_R7          tgatattacaacaa...c...tt....tt.ca...-------t...a.-attttttgc......a-----aatatgtcc. [206640]
MH025916.1_strain_R8          tgatattacaacaa...c...tt....tt.ca...-------t...a.-attttttgc......a-----aatatgtcc. [206640]

FR682468.2_Georgia_2007/1     tgccatctactctacaagtgcttgctaaaaaggtattggccttaggggagcataaagaaaatgaacatatatctagagaa [206720]
ASF_IND_20_CAD543             ................................................................................ [206720]
LR899193.1_Germany_2020/1     ................................................................................ [206720]
MN393477.1_Wuhan_2019         ................................................................................ [206720]
MW396979.1_Timor-Leste/2019/1 ................................................................................ [206720]
MK940252.1_CN/2019/InnerMongolia ................................................................................ [206720]
MK128995.1_China/2018/AnhuiXCGQ ................................................................................ [206720]
MH681419.1_ASFV/POL/2015      ................................................................................ [206720]
LS478113.1_Estonia_2014       ................................................................................ [206720]
KP843857.1_Odintsovo_02/14    ................................................................................ [206720]
FN557520.1_strain_E75         .......c..........................g.............a..........................a.... [206720]
AY261366.1_Warthog            .......c...................................---..................t............... [206720]
AY261365.1_Warmbaths          .......c................................t....................................... [206720]
AY261364.1_Tengani_62         .......c...................................---..................t............... [206720]
AY261363.1_Pretoriuskop/96/4  .......c...................................---..................t............... [206720]
AY261362.1_Mkuzi_1979         .......c................................t....................................... [206720]
AY261361.1_Malawi_Lil-20/1_1983 ...........t.....aca...........aac.......ac.c------------------.g..........a.a.. [206720]
AY261360.1_Kenya_1950         ca......t..t.......c....tc....g...gc...a...cca.------------------...g.g...ga...- [206720]
AM712240.1_OURT_88/3          .......c..........................g.............a............................... [206720]
AM712239.1_Benin_97/1         .......c..........................g.............a............................... [206720]
KM111295.1_Ken06.Bus          ca......t..t.......c....tc....g...gc...a...cca.------------------...g.....ga...- [206720]
KM111294.1_Ken05/Tk1          ca..g...t..t.......c....tc....g...g....a..acca.c---------------.t...g.....ga...- [206720]
KM262844.1_strain_L60         .......c..........................g.............a............................... [206720]
KM102979.1_26544/OG10_Italy   .......c..........................g.............a............................... [206720]
KM262845.1_strain_NHV         .......c..........................g.............a............................... [206720]
KX354450.1_47/Ss/2008         .......c..........................g.............a............................... [206720]
KP055815.1_strain_BA71        .......c..........................g.............a............................... [206720]
MH025920.1_strain_R35         ca......t..t.......c....tc....g...gc...a...cca.------------------...g.....ga...- [206720]
MH025919.1_strain_N10         ca......t..t.......c....tc....g...gc...a...cca.------------------...g.....ga...- [206720]
MH025918.1_strain_R25         ca......t..t.......c....tc....g...gc...a...cca.------------------...g.....ga...- [206720]
MH025917.1_strain_R7          ca......t..t.......c....tc....g...gc...a...cca.------------------...g.....ga...- [206720]
MH025916.1_strain_R8          ca......t..t.......c....tc....g...gc...a...cca.------------------...g.....ga...- [206720]
```

```
FR682468.2_Georgia_2007/1        tat---------------tattatcatatattaaagtgttgcggtttatggtggcatgaagctccgattatactttgtta [206800]
ASF_IND_20_CAD543                ...---------------.............................................................w [206800]
LR899193.1_Germany_2020/1        ...---------------.............................................................. [206800]
MN393477.1_Wuhan_2019            ...---------------.............................................................. [206800]
MW396979.1_Timor-Leste/2019/1    ...---------------.............................................................. [206800]
MK940252.1_CN/2019/InnerMongolia ...---------------.............................................................. [206800]
MK128995.1_China/2018/AnhuiXCGQ  ...---------------.............................................................. [206800]
MH681419.1_ASFV/POL/2015         ...---------------.............................................................. [206800]
LS478113.1_Estonia_2014          ...---------------.............................................................. [206800]
KP843857.1_Odintsovo_02/14       ...---------------.............................................................. [206800]
FN557520.1_strain_E75            ...---------------.........................................g...................t [206800]
AY261366.1_Warthog               ...---------------.............................................................t [206800]
AY261365.1_Warmbaths             ...---------------.............................................................t [206800]
AY261364.1_Tengani_62            ...---------------.....c............ca........................a........c.......t [206800]
AY261363.1_Pretoriuskop/96/4     ...---------------.............................................................t [206800]
AY261362.1_Mkuzi_1979            ...---------------.............................................................t [206800]
AY261361.1_Malawi_Lil-20/1_1983  ...tggccgtctgaggag....g.t..........a.....t.................tt....a..c.cga..---.t [206800]
AY261360.1_Kenya_1950            -----------------tg....ctg...t.....a.....t..g.............g..gc..a..c..g....c.ac [206800]
AM712240.1_OURT_88/3             ...---------------.........................................g...................t [206800]
AM712239.1_Benin_97/1            ...---------------.........................................g...................t [206800]
KM111295.1_Ken06.Bus             -----------------tg....ctg...t.....a.....t..a.............g..gc..a..c.......c.ac [206800]
KM111294.1_Ken05/Tk1             -----------------tg.....tg...t...c..ca...t..g..g..........g.ag...a.....gt...t.ac [206800]
KM262844.1_strain_L60            ...---------------.........................................g...................t [206800]
KM102979.1_26544/OG10_Italy      ...---------------.........................................g...................t [206800]
KM262845.1_strain_NHV            ...---------------.........................................g...................t [206800]
KX354450.1_47/Ss/2008            ...---------------.........................................g...................t [206800]
KP055815.1_strain_BA71           ...---------------.........................................g...................t [206800]
MH025920.1_strain_R35            -----------------tg....ctg...t.....a.....t..a.............g..gc..a..c.......c.ac [206800]
MH025919.1_strain_N10            -----------------tg....ctg...t.....a.....t..a.............g..gc..a..c.......c.ac [206800]
MH025918.1_strain_R25            -----------------tg....ctg...t.....a.....t..a.............g..gc..a..c.......c.ac [206800]
MH025917.1_strain_R7             -----------------tg....ctg...t.....a.....t..a.............g..gc..a..c.......c.ac [206800]
MH025916.1_strain_R8             -----------------tg....ctg...t.....a.....t..a.............g..gc..a..c.......c.ac [206800]

FR682468.2_Georgia_2007/1        tgatgggagtgagcaaatgatgataaagactccaatctttgaagaaggcatattacttaatactgcattaatgaaagctg [206880]
ASF_IND_20_CAD543                ................................................................................ [206880]
LR899193.1_Germany_2020/1        ................................................................................ [206880]
MN393477.1_Wuhan_2019            ................................................................................ [206880]
MW396979.1_Timor-Leste/2019/1    ................................................................................ [206880]
MK940252.1_CN/2019/InnerMongolia ................................................................................ [206880]
MK128995.1_China/2018/AnhuiXCGQ  ................................................................................ [206880]
MH681419.1_ASFV/POL/2015         ................................................................................ [206880]
LS478113.1_Estonia_2014          ................................................................................ [206880]
KP843857.1_Odintsovo_02/14       ................................................................................ [206880]
```

```
FN557520.1_strain_E75          ................................................................................ [206880]
AY261366.1_Warthog             .a........a...................g................................................. [206880]
AY261365.1_Warmbaths           ................................................................................ [206880]
AY261364.1_Tengani_62          .a........a...................g................................................. [206880]
AY261363.1_Pretoriuskop/96/4   .a........a...................g................................................. [206880]
AY261362.1_Mkuzi_1979          ................................................................................ [206880]
AY261361.1_Malawi_Lil-20/1_1983 .ac.t.t.taa.a.....at.a.....a..ag...a....a..c.t..tt..gact.a..cct................. [206880]
AY261360.1_Kenya_1950          ga...a.ga.a.t..............at.ag...gt...a....t..tt..gaga.a...ctc..........tg..c. [206880]
AM712240.1_OURT_88/3           ................................................................................ [206880]
AM712239.1_Benin_97/1          ................................................................................ [206880]
KM111295.1_Ken06.Bus           .a...a.ga.a.t..............at.ag...g....a....t..tt..gaga.a...ctc...........g.... [206880]
KM111294.1_Ken05/Tk1           .a...aa.a.c.t....g.........at.ag...g....a....t..tt..ga.a.a...ctc...........g..c. [206880]
KM262844.1_strain_L60          ................................................................................ [206880]
KM102979.1_26544/OG10_Italy    ................................................................................ [206880]
KM262845.1_strain_NHV          ................................................................................ [206880]
KX354450.1_47/Ss/2008          ................................................................................ [206880]
KP055815.1_strain_BA71         ................................................................................ [206880]
MH025920.1_strain_R35          .a...a.ga.a.t..............at.ag...g....a....t..tt..gaga.a...ctc...........g.... [206880]
MH025919.1_strain_N10          .a...a.ga.a.t..............at.ag...g....a....t..tt..gaga.a...ctc...........g.... [206880]
MH025918.1_strain_R25          .a...a.ga.a.t..............at.ag...g....a....t..tt..gaga.a...ctc...........g.... [206880]
MH025917.1_strain_R7           .a...a.ga.a.t..............at.ag...g....a....t..tt..gaga.a...ctc...........g.... [206880]
MH025916.1_strain_R8           .a...a.ga.a.t..............at.ag...g....a....t..tt..gaga.a...ctc...........g.... [206880]

FR682468.2_Georgia_2007/1      tacaggagaataattatgaattaataaagttgtttactgaatggggagcaaacatcaattatggattaatttccattaat [206960]
ASF_IND_20_CAD543              ....................................s........................................... [206960]
LR899193.1_Germany_2020/1      ................................................................................ [206960]
MN393477.1_Wuhan_2019          ................................................................................ [206960]
MW396979.1_Timor-Leste/2019/1  ................................................................................ [206960]
MK940252.1_CN/2019/InnerMongolia ................................................................................ [206960]
MK128995.1_China/2018/AnhuiXCGQ ................................................................................ [206960]
MH681419.1_ASFV/POL/2015       ................................................................................ [206960]
LS478113.1_Estonia_2014        ................................................................................ [206960]
KP843857.1_Odintsovo_02/14     ................................................................................ [206960]
FN557520.1_strain_E75          ..........c.................c.....c............................................. [206960]
AY261366.1_Warthog             .......c..c..c........g.....a.....c.................................g....t...... [206960]
AY261365.1_Warmbaths           ..........c.......................c............................................. [206960]
AY261364.1_Tengani_62          .......c..c..c........g.....a.....c......................................t...... [206960]
AY261363.1_Pretoriuskop/96/4   .......c..c..c........g.....a.....c......................................t...... [206960]
AY261362.1_Mkuzi_1979          ..........c.......................c............................................. [206960]
AY261361.1_Malawi_Lil-20/1_1983 ....a..a.....c........g..c.t............g.....t..gg.t.....c.tg..g..g..ca.tg..... [206960]
AY261360.1_Kenya_1950          ....a..a..c....g.agcc.....g..c.......c........t...g....t....c......gg..a.tg.c..c [206960]
AM712240.1_OURT_88/3           ..........c.................c.....c............................................. [206960]
AM712239.1_Benin_97/1          ..........c.................c.....c............................................. [206960]
KM111295.1_Ken06.Bus           ....a..a..c....g.agcc.....g..c.......c........t...g....t....c......gg..a.tg.c..c [206960]
```

```
KM111294.1_Ken05/Tk1            .......a..c....g.atc......g..c.......c........t...g....t...........g...a.tg.c... [206960]
KM262844.1_strain_L60           ..........c.................c.....c............................................. [206960]
KM102979.1_26544/OG10_Italy     ..........c.................c.....c............................................. [206960]
KM262845.1_strain_NHV           ..........c.................c.....c............................................. [206960]
KX354450.1_47/Ss/2008           ..........c.................c.....c............................................. [206960]
KP055815.1_strain_BA71          ..........c.................c.....c............................................. [206960]
MH025920.1_strain_R35           ....a..a..c....g.agcc.....g..c.......c........t...g....t....c......gg..a.tg.c..c [206960]
MH025919.1_strain_N10           ....a..a..c....g.agcc.....g..c.......c........t...g....t....c......gg..a.tg.c..c [206960]
MH025918.1_strain_R25           ....a..a..c....g.agcc.....g..c.......c........t...g....t....c......gg..a.tg.c..c [206960]
MH025917.1_strain_R7            ....a..a..c....g.agcc.....g..c.......c........t...g....t....c......gg..a.tg.c..c [206960]
MH025916.1_strain_R8            ....a..a..c....g.agcc.....g..c.......c........t...g....t....c......gg..a.tg.c..c [206960]

FR682468.2_Georgia_2007/1       accgagcatgcccgggatctatgtcgaaaattaggagctaaagaaatgcttgaaggaaatgaatttatacaaattatatt [207040]
ASF_IND_20_CAD543               ......................................................r........k...........r.... [207040]
LR899193.1_Germany_2020/1       ................................................................................ [207040]
MN393477.1_Wuhan_2019           ................................................................................ [207040]
MW396979.1_Timor-Leste/2019/1   ................................................................................ [207040]
MK940252.1_CN/2019/InnerMongolia ................................................................................ [207040]
MK128995.1_China/2018/AnhuiXCGQ ................................................................................ [207040]
MH681419.1_ASFV/POL/2015        ................................................................................ [207040]
LS478113.1_Estonia_2014         ................................................................................ [207040]
KP843857.1_Odintsovo_02/14      ................................................................................ [207040]
FN557520.1_strain_E75           ..t..............c....................................a........g...........g.... [207040]
AY261366.1_Warthog              ..g......a.t.....c..t.................c...............a........g................ [207040]
AY261365.1_Warmbaths            ..t..............c....................................a........g...........g.... [207040]
AY261364.1_Tengani_62           ..g......a.t........t.................c...............a..t.....g.............g.. [207040]
AY261363.1_Pretoriuskop/96/4    ..g...t..a.t........t.................c.......g.......a..c.....g.............g.. [207040]
AY261362.1_Mkuzi_1979           ..t..............c....................................a........g...........g.... [207040]
AY261361.1_Malawi_Lil-20/1_1983 ..a..tgca.....a...t.g..c.a...g.....t..a..g...gctt.gagtc....g...a..t..g....at.t.a [207040]
AY261360.1_Kenya_1950           ..g..at.ca....aa.c...........tc....t.....g..g.cat.aa.taa.cgg...a..t..g.tg.at.t.. [207040]
AM712240.1_OURT_88/3            ..t..............c....................................a........g...........g.... [207040]
AM712239.1_Benin_97/1           ..t..............c....................................a........g...........g.... [207040]
KM111295.1_Ken06.Bus            ..g..at.ca....aa.c...........tc....t.....g..g..at.aa.taa.cgg...a..t..g..g.at.t.. [207040]
KM111294.1_Ken05/Tk1            ..g..at..a....aa.c...........tc....t..c..g..ggc.t.aa.tac.cgga..a..t..g.tg.at.t.. [207040]
KM262844.1_strain_L60           ..t..............c....................................a........g...........g.... [207040]
KM102979.1_26544/OG10_Italy     ..t..............c....................................a........g...........g.... [207040]
KM262845.1_strain_NHV           ..t..............c....................................a........g...........g.... [207040]
KX354450.1_47/Ss/2008           ..t..............c....................................a........g...........g.... [207040]
KP055815.1_strain_BA71          ..t..............c....................................a........g...........g.... [207040]
MH025920.1_strain_R35           ..g..at.ca....aa.c.......a...tc....t.....g..g..at.aa.taa.cgg...a..t..g..g.at.t.. [207040]
MH025919.1_strain_N10           ..g..at.ca....aa.c.......a...tc....t.....g..g..at.aa.taa.cgg...a..t..g..g.at.t.. [207040]
MH025918.1_strain_R25           ..g..at.ca....aa.c.......a...tc....t.....g..g..at.aa.taa.cgg...a..t..g..g.at.t.. [207040]
MH025917.1_strain_R7            ..g..at.ca....aa.c.......a...tc....t.....g..g..at.aa.taa.cgg...a..t..g..g.at.t.. [207040]
MH025916.1_strain_R8            ..g..at.ca....aa.c.......a...tc....t.....g..g..at.aa.taa.cgg...a..t..g..g.at.t.. [207040]
```

```
FR682468.2_Georgia_2007/1         caaaacattagatgataccaccagtagtaatataattttatgtcatgaattattcaccaacaatcctcttttagagaatg [207120]
ASF_IND_20_CAD543                 t............................................................................... [207120]
LR899193.1_Germany_2020/1         ................................................................................ [207120]
MN393477.1_Wuhan_2019             ................................................................................ [207120]
MW396979.1_Timor-Leste/2019/1     ................................................................................ [207120]
MK940252.1_CN/2019/InnerMongolia  ................................................................................ [207120]
MK128995.1_China/2018/AnhuiXCGQ   ................................................................................ [207120]
MH681419.1_ASFV/POL/2015          ................................................................................ [207120]
LS478113.1_Estonia_2014           ................................................................................ [207120]
KP843857.1_Odintsovo_02/14        ................................................................................ [207120]
FN557520.1_strain_E75             t................t....................................t......................... [207120]
AY261366.1_Warthog                t....................t...........g....................t..t...................... [207120]
AY261365.1_Warmbaths              t................t...............g....................t......................... [207120]
AY261364.1_Tengani_62             t................................g....................t......................... [207120]
AY261363.1_Pretoriuskop/96/4      t....t...........................g....................t......................... [207120]
AY261362.1_Mkuzi_1979             t................t...............g....................t......................... [207120]
AY261361.1_Malawi_Lil-20/1_1983   t....t.cagt..at..ag.gt..c.a......t...a...c..........a.at.t..tc....c.....------cc [207120]
AY261360.1_Kenya_1950             a...tt.aaga..tt..aa.gt....a......t...c...cc...........at.t..t...........ccta.g.. [207120]
AM712240.1_OURT_88/3              t................t....................................t......................... [207120]
AM712239.1_Benin_97/1             t................t....................................t......................... [207120]
KM111295.1_Ken06.Bus              a...tt.aaga..tt..aa.gt..c........t...g...cc...........agat..t...........ccta.g.. [207120]
KM111294.1_Ken05/Tk1              a..ttt.aag...tt..aa.gt....a.c....t...c....c........g..at.t..t............cta.g.. [207120]
KM262844.1_strain_L60             t................t....................................t......................... [207120]
KM102979.1_26544/OG10_Italy       t................t....................................t......................... [207120]
KM262845.1_strain_NHV             t................t....................................t......................... [207120]
KX354450.1_47/Ss/2008             t................t....................................t......................... [207120]
KP055815.1_strain_BA71            t................t....................................t......................... [207120]
MH025920.1_strain_R35             a...tt.aaga..tt..aa.gt..c........t...g...cc...........agat..t...........ccta.g.. [207120]
MH025919.1_strain_N10             a...tt.aaga..tt..aa.gt..c........t...g...cc...........agat..t...........ccta.g.. [207120]
MH025918.1_strain_R25             a...tt.aaga..tt..aa.gt..c........t...g...cc...........agat..t...........ccta.g.. [207120]
MH025917.1_strain_R7              a...tt.aaga..tt..aa.gt..c........t...g...cc...........agat..t...........ccta.g.. [207120]
MH025916.1_strain_R8              a...tt.aaga..tt..aa.gt..c........t...g...cc...........agat..t...........ccta.g.. [207120]

FR682468.2_Georgia_2007/1         taaatatgggggaaatgaggatgataatttattggaggatgaaaaat----ttaacgaacctattattaaataatgactc [207200]
ASF_IND_20_CAD543                 ...............................................----............................. [207200]
LR899193.1_Germany_2020/1         ...............................................----............................. [207200]
MN393477.1_Wuhan_2019             ...............................................----............................. [207200]
MW396979.1_Timor-Leste/2019/1     ...............................................----............................. [207200]
MK940252.1_CN/2019/InnerMongolia  ...............................................----............................. [207200]
MK128995.1_China/2018/AnhuiXCGQ   ...............................................----............................. [207200]
MH681419.1_ASFV/POL/2015          ...............................................----............................. [207200]
LS478113.1_Estonia_2014           ...............................................----............................. [207200]
KP843857.1_Odintsovo_02/14        ...............................................----............................. [207200]
```

```
FN557520.1_strain_E75           .............................c.................----............................. [207200]
AY261366.1_Warthog              .............................c......a..........----c.....g...........g.....a.... [207200]
AY261365.1_Warmbaths            .............................a.................----c........t................... [207200]
AY261364.1_Tengani_62           ...........c.................c.................----c.....g.................a.... [207200]
AY261363.1_Pretoriuskop/96/4    ...........c.................c.................----c.....g.................a.... [207200]
AY261362.1_Mkuzi_1979           .............................a.................----c........t................... [207200]
AY261361.1_Malawi_Lil-20/1_1983 ......at.atc..t...aat.a.g....-g....t.a.ct...tgctata.c..tc...t.ta....gg..g.gata.. [207200]
AY261360.1_Kenya_1950           a.g...at.att.tt.t..a.ga......a....t.atc.a.gg.ga-ata.c..tt...t.ta....gg..g.aatt.. [207200]
AM712240.1_OURT_88/3            .............................c.................----............................. [207200]
AM712239.1_Benin_97/1           .............................c.................----............................. [207200]
KM111295.1_Ken06.Bus            a.g...at.att.tt....a.ga......a....t.atc.a.gg.ga-ata.c..tt...t.ta....gg..g.aata.. [207200]
KM111294.1_Ken05/Tk1            a.....at.att.tt....a.aa......a....t.atc.a.gg.ga-ata.c..tt...t.ta....gg..g.aatt.. [207200]
KM262844.1_strain_L60           .............................c.................----............................. [207200]
KM102979.1_26544/OG10_Italy     .............................c.................----............................. [207200]
KM262845.1_strain_NHV           .............................c.................----............................. [207200]
KX354450.1_47/Ss/2008           .............................c.................----............................. [207200]
KP055815.1_strain_BA71          .............................c.................----............................. [207200]
MH025920.1_strain_R35           a.g...at.att.tt....a.ga......a....t.atc.a.gg.ga-ata.c..tt...t.ta....gg..g.aata.. [207200]
MH025919.1_strain_N10           a.g...at.att.tt....a.ga......a....t.atc.a.gg.ga-ata.c..tt...t.ta....gg..g.aata.. [207200]
MH025918.1_strain_R25           a.g...at.att.tt....a.ga......a....t.atc.a.gg.ga-ata.c..tt...t.ta....gg..g.aata.. [207200]
MH025917.1_strain_R7            a.g...at.att.tt....a.ga......a....t.atc.a.gg.ga-ata.c..tt...t.ta....gg..g.aata.. [207200]
MH025916.1_strain_R8            a.g...at.att.tt....a.ga......a....t.atc.a.gg.ga-ata.c..tt...t.ta....gg..g.aata.. [207200]

FR682468.2_Georgia_2007/1       tattagtgaaatattaactaaattctggtatggtatagcagtaaaatataatcttaaggatgcgatccaatatttttacc [207280]
ASF_IND_20_CAD543               ................................................................................ [207280]
LR899193.1_Germany_2020/1       ................................................................................ [207280]
MN393477.1_Wuhan_2019           ................................................................................ [207280]
MW396979.1_Timor-Leste/2019/1   ................................................................................ [207280]
MK940252.1_CN/2019/InnerMongolia ................................................................................ [207280]
MK128995.1_China/2018/AnhuiXCGQ ................................................................................ [207280]
MH681419.1_ASFV/POL/2015        ................................................................................ [207280]
LS478113.1_Estonia_2014         ................................................................................ [207280]
KP843857.1_Odintsovo_02/14      ................................................................................ [207280]
FN557520.1_strain_E75           ...............................................................a................ [207280]
AY261366.1_Warthog              ...............................................................t................ [207280]
AY261365.1_Warmbaths            ...............................................................t................ [207280]
AY261364.1_Tengani_62           ...............................................................t................ [207280]
AY261363.1_Pretoriuskop/96/4    ...............................................................t................ [207280]
AY261362.1_Mkuzi_1979           ...............................................................t................ [207280]
AY261361.1_Malawi_Lil-20/1_1983 at.c.a......gc......gg.at......a....g..ga..ct......a..ccct.ca..c..t...........t. [207280]
AY261360.1_Kenya_1950           at...a.....agc....c.g..........a..cag..g...ct...........ct..a..t...........c..t. [207280]
AM712240.1_OURT_88/3            ...............................................................a................ [207280]
AM712239.1_Benin_97/1           ...............................................................a................ [207280]
KM111295.1_Ken06.Bus            at...a.....agc....c.g..........a..cag..g...ct...........ct..a..t...........c..t. [207280]
```

```
KM111294.1_Ken05/Tk1             at...a.....agc....c.g..........aaacag......ct.a.........ct..a..t...........c..t. [207280]
KM262844.1_strain_L60            ...............................................................a................ [207280]
KM102979.1_26544/OG10_Italy      ...............................................................a................ [207280]
KM262845.1_strain_NHV            ...............................................................a................ [207280]
KX354450.1_47/Ss/2008            ...............................................................a................ [207280]
KP055815.1_strain_BA71           ...............................................................a................ [207280]
MH025920.1_strain_R35            at...a.....agc....c.g..........a..cag..g...ct...........ct..a..t...........c..t. [207280]
MH025919.1_strain_N10            at...a.....agc....c.g..........a..cag..g...ct...........ct..a..t...........c..t. [207280]
MH025918.1_strain_R25            at...a.....agc....c.g..........a..cag..g...ct...........ct..a..t...........c..t. [207280]
MH025917.1_strain_R7             at...a.....agc....c.g..........a..cag..g...ct...........ct..a..t...........c..t. [207280]
MH025916.1_strain_R8             at...a.....agc....c.g..........a..cag..g...ct...........ct..a..t...........c..t. [207280]

FR682468.2_Georgia_2007/1        agagattcatggacttcaacgagtggcgagtaacatgtgctctttcttttaataatgtgaatgatcttcataagatgtat [207360]
ASF_IND_20_CAD543                ................................................................................ [207360]
LR899193.1_Germany_2020/1        ................................................................................ [207360]
MN393477.1_Wuhan_2019            ................................................................................ [207360]
MW396979.1_Timor-Leste/2019/1    ................................................................................ [207360]
MK940252.1_CN/2019/InnerMongolia ................................................................................ [207360]
MK128995.1_China/2018/AnhuiXCGQ  ................................................................................ [207360]
MH681419.1_ASFV/POL/2015         ................................................................................ [207360]
LS478113.1_Estonia_2014          ................................................................................ [207360]
KP843857.1_Odintsovo_02/14       ................................................................................ [207360]
FN557520.1_strain_E75            ................................................................................ [207360]
AY261366.1_Warthog               .......t..aa.......t........................c................................... [207360]
AY261365.1_Warmbaths             ...a......................................................a..................... [207360]
AY261364.1_Tengani_62            ..........aa.......t........................c................................... [207360]
AY261363.1_Pretoriuskop/96/4     ..........aa.......t........................c................................... [207360]
AY261362.1_Mkuzi_1979            ...a......................................................a..................... [207360]
AY261361.1_Malawi_Lil-20/1_1983  .atc..a..a.t.t..t..a..t.....gt...t....agc...g...a...c..c...tt...c......g.a..t... [207360]
AY261360.1_Kenya_1950            .a.a..atgaac.tc....t........tt...tt.....g..................tt..........g.a..a... [207360]
AM712240.1_OURT_88/3             ................................................................................ [207360]
AM712239.1_Benin_97/1            ................................................................................ [207360]
KM111295.1_Ken06.Bus             .a.a.--------------------------------------------------------------------------- [207360]
KM111294.1_Ken05/Tk1             .a.a..at.a.c.g..t..a..c......c...t......ga..g....a........att..........g.a..a... [207360]
KM262844.1_strain_L60            ................................................................................ [207360]
KM102979.1_26544/OG10_Italy      ................................................................................ [207360]
KM262845.1_strain_NHV            ................................................................................ [207360]
KX354450.1_47/Ss/2008            ................................................................................ [207360]
KP055815.1_strain_BA71           ................................................................................ [207360]
MH025920.1_strain_R35            .a.a..at.a.c.g..t..a..t......t...t....c.gg..g..............tt..........g.a..a... [207360]
MH025919.1_strain_N10            .a.a..at.a.c.g..t..a..t......t...t....c.gg..g..............tt..........g.a..a... [207360]
MH025918.1_strain_R25            .a.a..at.a.c.g..t..a..t......t...t....c.gg..g..............tt..........g.a..a... [207360]
MH025917.1_strain_R7             .a.a..at.a.c.g..t..a..t......t...t....c.gg..g..............tt..........g.a..a... [207360]
MH025916.1_strain_R8             .a.a..at.a.c.g..t..a..t......t...t....c.gg..g..............tt..........g.a..a... [207360]
```

```
FR682468.2_Georgia_2007/1        ataacagagaaggttcatatgaataatgacgaaatgatgaatctagcctgcagcattcaagacagaaatttatcaaccat [207440]
ASF_IND_20_CAD543                ................................................................................ [207440]
LR899193.1_Germany_2020/1        ................................................................................ [207440]
MN393477.1_Wuhan_2019            ................................................................................ [207440]
MW396979.1_Timor-Leste/2019/1    ................................................................................ [207440]
MK940252.1_CN/2019/InnerMongolia ................................................................................ [207440]
MK128995.1_China/2018/AnhuiXCGQ  ................................................................................ [207440]
MH681419.1_ASFV/POL/2015         ................................................................................ [207440]
LS478113.1_Estonia_2014          ................................................................................ [207440]
KP843857.1_Odintsovo_02/14       ................................................................................ [207440]
FN557520.1_strain_E75            ................................................................................ [207440]
AY261366.1_Warthog               ...........a..................................................t.a......t........ [207440]
AY261365.1_Warmbaths             ...........a.................t.................................................. [207440]
AY261364.1_Tengani_62            ...........a.........................................t........t.a......t........ [207440]
AY261363.1_Pretoriuskop/96/4     ..........ga..................................................t.a......t........ [207440]
AY261362.1_Mkuzi_1979            ...........a.................t.................................................. [207440]
AY261361.1_Malawi_Lil-20/1_1983  .at.a.......ac.g....a..c.t...t.........cggt.g.....t..---gt.t..tg.t....ata....t.. [207440]
AY261360.1_Kenya_1950            .ac.a...a......gg....g...t.a.tc.......ag.ga..a....tgc---at.tatgt.t.g..at..c..... [207440]
AM712240.1_OURT_88/3             ................................................................................ [207440]
AM712239.1_Benin_97/1            ................................................................................ [207440]
KM111295.1_Ken06.Bus             -------------------------------------------------------------------------------- [207440]
KM111294.1_Ken05/Tk1             .ac.a........c.g....ag...t.a.tc.......ag.ga..a....t.c---at.tatgt.t.g..atc.c..... [207440]
KM262844.1_strain_L60            ................................................................................ [207440]
KM102979.1_26544/OG10_Italy      ................................................................................ [207440]
KM262845.1_strain_NHV            ................................................................................ [207440]
KX354450.1_47/Ss/2008            ................................................................................ [207440]
KP055815.1_strain_BA71           ................................................................................ [207440]
MH025920.1_strain_R35            .ac.a..........ag....g...t.a.tc.......ag.ga..a....tgc---at.tatat.t.g..at..c..t.. [207440]
MH025919.1_strain_N10            .ac.a..........ag....g...t.a.tc.......ag.ga..a....tgc---at.tatat.t.g..at..c..t.. [207440]
MH025918.1_strain_R25            .ac.a..........ag....g...t.a.tc.......ag.ga..a....tgc---at.tatat.t.g..at..c..t.. [207440]
MH025917.1_strain_R7             .ac.a..........ag....g...t.a.tc.......ag.ga..a....tgc---at.tatat.t.g..at..c..t.. [207440]
MH025916.1_strain_R8             .ac.a..........ag....g...t.a.tc.......ag.ga..a....tgc---at.tatat.t.g..at..c..t.. [207440]

FR682468.2_Georgia_2007/1        ttactattgttttctattgggggctaacatcaatcaagcaatgttaacctcagtattaaattataatattttttaactta [207520]
ASF_IND_20_CAD543                ...................-............................................................ [207520]
LR899193.1_Germany_2020/1        ................................................................................ [207520]
MN393477.1_Wuhan_2019            ................................................................................ [207520]
MW396979.1_Timor-Leste/2019/1    ................................................................................ [207520]
MK940252.1_CN/2019/InnerMongolia ................................................................................ [207520]
MK128995.1_China/2018/AnhuiXCGQ  ................................................................................ [207520]
MH681419.1_ASFV/POL/2015         ................................................................................ [207520]
LS478113.1_Estonia_2014          ................................................................................ [207520]
KP843857.1_Odintsovo_02/14       ................................................................................ [207520]
```

```
FN557520.1_strain_E75            ...................-...-......................g................................. [207520]
AY261366.1_Warthog               ............c......-.....c....................t..........c..........g.......ta.. [207520]
AY261365.1_Warmbaths             ............c......-..........................g................................. [207520]
AY261364.1_Tengani_62            ............c......-..a.......................g..............................a.. [207520]
AY261363.1_Pretoriuskop/96/4     ............c......-.....c....................t.t............................a.. [207520]
AY261362.1_Mkuzi_1979            ............c......-..........................g................................. [207520]
AY261361.1_Malawi_Lil-20/1_1983  ...t.........a.g...-......g....t.....g......a.t.....g..ca.g...ct.tgggacgg....... [207520]
AY261360.1_Kenya_1950            .............g..a..-..a...g....t....gg......a.t.t...g..tac...a.c.t...c..a......g [207520]
AM712240.1_OURT_88/3             ...................-..........................g................................. [207520]
AM712239.1_Benin_97/1            ...................-..........................g................................. [207520]
KM111295.1_Ken06.Bus             -------------------------------------------------------------------------------- [207520]
KM111294.1_Ken05/Tk1             .............g..a..-......g....t....gg......a.t........tac...a.c.t...c.gg......g [207520]
KM262844.1_strain_L60            ...................-..........................g....g............................ [207520]
KM102979.1_26544/OG10_Italy      ...................-..........................g................................. [207520]
KM262845.1_strain_NHV            ...................-..........................g................................. [207520]
KX354450.1_47/Ss/2008            ...................-..........................g................................. [207520]
KP055815.1_strain_BA71           ...................-..........................g................................. [207520]
MH025920.1_strain_R35            .............g..a..-..a...g....t....gg......a.t.....g..tac...a.c.t...c..g......g [207520]
MH025919.1_strain_N10            .............g..a..-..a...g....t....gg......a.t.....g..tac...a.c.t...c..g......g [207520]
MH025918.1_strain_R25            .............g..a..-..a...g....t....gg......a.t.....g..tac...a.c.t...c..g......g [207520]
MH025917.1_strain_R7             .............g..a..-..a...g....t....gg......a.t.....g..tac...a.c.t...c..g......g [207520]
MH025916.1_strain_R8             .............g..a..-..a...g....t....gg......a.t.....g..tac...a.c.t...c..g......g [207520]

FR682468.2_Georgia_2007/1        ttcttttgtatagacttaggggctgatgcctttgaagagggtaagaccctggcgaaacaaaaggggtataatgaaatagt [207600]
ASF_IND_20_CAD543                ................................................................................ [207600]
LR899193.1_Germany_2020/1        ................................................................................ [207600]
MN393477.1_Wuhan_2019            ................................................................................ [207600]
MW396979.1_Timor-Leste/2019/1    ................................................................................ [207600]
MK940252.1_CN/2019/InnerMongolia ................................................................................ [207600]
MK128995.1_China/2018/AnhuiXCGQ  ................................................................................ [207600]
MH681419.1_ASFV/POL/2015         ................................................................................ [207600]
LS478113.1_Estonia_2014          ................................................................................ [207600]
KP843857.1_Odintsovo_02/14       ................................................................................ [207600]
FN557520.1_strain_E75            ................................................................................ [207600]
AY261366.1_Warthog               ....................a.....c..................g.................................. [207600]
AY261365.1_Warmbaths             ................................................................................ [207600]
AY261364.1_Tengani_62            ..............t...........c..................g................c................. [207600]
AY261363.1_Pretoriuskop/96/4     ....................a.....c......a...........g.................................. [207600]
AY261362.1_Mkuzi_1979            ................................................................................ [207600]
AY261361.1_Malawi_Lil-20/1_1983  ..tc..........t..g..........ta.........t...t..aaa.a..a.t.g..g.tcata..gg..t.t...a [207600]
AY261360.1_Kenya_1950            ..............t.....a...ac......c......t..ttagaaa.a..a......c..aata........t.... [207600]
AM712240.1_OURT_88/3             ..------------------------------------------------------------------------------ [207600]
AM712239.1_Benin_97/1            ................................................................................ [207600]
KM111295.1_Ken06.Bus             -------------------------------------------------------------------------------- [207600]
```

```
KM111294.1_Ken05/Tk1             ..............t.....a...ac......c......t.cttagaaa.a..a......c..aatg........t.g.. [207600]
KM262844.1_strain_L60            ................................................................................ [207600]
KM102979.1_26544/OG10_Italy      ................................................................................ [207600]
KM262845.1_strain_NHV            ..------------------------------------------------------------------------------ [207600]
KX354450.1_47/Ss/2008            ................................................................................ [207600]
KP055815.1_strain_BA71           ................................................................................ [207600]
MH025920.1_strain_R35            ..............t.....a...ac......c......t..ttagaaa.a..a......c...ata........t.... [207600]
MH025919.1_strain_N10            ..............t.....a...ac......c......t..ttagaaa.a..a......c...ata........t.... [207600]
MH025918.1_strain_R25            ..............t.....a...ac......c......t..ttagaaa.a..a......c...ata........t.... [207600]
MH025917.1_strain_R7             ..............t.....a...ac......c......t..ttagaaa.a..a......c...ata........t.... [207600]
MH025916.1_strain_R8             ..............t.....a...ac......c......t..ttagaaa.a..a......c...ata........t.... [207600]

FR682468.2_Georgia_2007/1        ggaaatcttatcattagatatcatttatagtccaaatactgacttctcatca------aaaatagaacctgaacatatta [207680]
ASF_IND_20_CAD543                ....................................................------...................... [207680]
LR899193.1_Germany_2020/1        ....................................................------...................... [207680]
MN393477.1_Wuhan_2019            ....................................................------...................... [207680]
MW396979.1_Timor-Leste/2019/1    ....................................................------...................... [207680]
MK940252.1_CN/2019/InnerMongolia ....................................................------...................... [207680]
MK128995.1_China/2018/AnhuiXCGQ  ....................................................------...................... [207680]
MH681419.1_ASFV/POL/2015         ....................................................------...................... [207680]
LS478113.1_Estonia_2014          ....................................................------...................... [207680]
KP843857.1_Odintsovo_02/14       ....................................................------...................... [207680]
FN557520.1_strain_E75            ....................................................------...................... [207680]
AY261366.1_Warthog               .............c....c...........c..............t.t....------......a............... [207680]
AY261365.1_Warmbaths             ..................c...........c.....................------...............a...... [207680]
AY261364.1_Tengani_62            ......t...........c...........c........c.....t......------...................... [207680]
AY261363.1_Pretoriuskop/96/4     .............c....c...........c..............t......------......a............... [207680]
AY261362.1_Mkuzi_1979            ..................c...........c.....................------...............a...... [207680]
AY261361.1_Malawi_Lil-20/1_1983  aagt...........ta.a.atta..........g.cgt.tctc.g.t....ttaaaa.cg.c...t..g...a.a.... [207680]
AY261360.1_Kenya_1950            aa....a........ga.a.a.ta..........g...gctc.c.tat....ttaaaa.t..c...t..g..ga.a.... [207680]
AM712240.1_OURT_88/3             -------------------------------------------------------------------------------- [207680]
AM712239.1_Benin_97/1            ....................................................------...................... [207680]
KM111295.1_Ken06.Bus             -------------------------------------------------------------------------------- [207680]
KM111294.1_Ken05/Tk1             aa.gg.a........ga.a.a.ta...c......g...gctc.c.t.t....ttaaaa.t..c...t..g...a.a.... [207680]
KM262844.1_strain_L60            ....................................................------...................... [207680]
KM102979.1_26544/OG10_Italy      ....................................................------...................... [207680]
KM262845.1_strain_NHV            -------------------------------------------------------------------------------- [207680]
KX354450.1_47/Ss/2008            ....................................................------...................... [207680]
KP055815.1_strain_BA71           ....................................................------...................... [207680]
MH025920.1_strain_R35            aa....a........ga.a.atta..........g...gctc.c.tat....ttaaaa.t..c...t..g..ga.a.... [207680]
MH025919.1_strain_N10            aa....a........ga.a.atta..........g...gctc.c.tat....ttaaaa.t..c...t..g..ga.a.... [207680]
MH025918.1_strain_R25            aa....a........ga.a.atta..........g...gctc.c.tat....ttaaaa.t..c...t..g..ga.a.... [207680]
MH025917.1_strain_R7             aa....a........ga.a.atta..........g...gctc.c.tat....ttaaaa.t..c...t..g..ga.a.... [207680]
MH025916.1_strain_R8             aa....a........ga.a.atta..........g...gctc.c.tat....ttaaaa.t..c...t..g..ga.a.... [207680]
```

```
FR682468.2_Georgia_2007/1        gttctttgttaaaaaa------cttttatccaaaaaatctgttcgcttttgatcgttgcaaccccggtttatattattct [207760]
ASF_IND_20_CAD543                ................------.......................................................... [207760]
LR899193.1_Germany_2020/1        ................------.......................................................... [207760]
MN393477.1_Wuhan_2019            ................------.......................................................... [207760]
MW396979.1_Timor-Leste/2019/1    ................------.......................................................... [207760]
MK940252.1_CN/2019/InnerMongolia ................------.......................................................... [207760]
MK128995.1_China/2018/AnhuiXCGQ  ................------.......................................................... [207760]
MH681419.1_ASFV/POL/2015         ................------.......................................................... [207760]
LS478113.1_Estonia_2014          ................------.......................................................... [207760]
KP843857.1_Odintsovo_02/14       ................------.......................................................... [207760]
FN557520.1_strain_E75            ................------.......................................................... [207760]
AY261366.1_Warthog               ...t............------...c................a.at...............a..t.....g......c.. [207760]
AY261365.1_Warmbaths             ................------...c...................t..a........c...a..t.....t......c.. [207760]
AY261364.1_Tengani_62            ................------...c.................tat............t..a..t.....g......c.. [207760]
AY261363.1_Pretoriuskop/96/4     ...t............------...c................a.at...............a..t.....g......c.. [207760]
AY261362.1_Mkuzi_1979            ................------...c...................t..a........c...a..t.....t......c.. [207760]
AY261361.1_Malawi_Lil-20/1_1983  a.gac...c..g.tg.agatat..a.a.at........ag...gata.a.a.aa...---------...g..g.c..--- [207760]
AY261360.1_Kenya_1950            a.atc..a...g.tg.ggaaaat.a.g.gt........ga...aata.a...agaa.-------------c..a...--- [207760]
AM712240.1_OURT_88/3             -------------------------------------------------------------------------------- [207760]
AM712239.1_Benin_97/1            ................------.......................................................... [207760]
KM111295.1_Ken06.Bus             -------------------------------------------------------------------------------- [207760]
KM111294.1_Ken05/Tk1             a.atc..a...g.tg.tgaaaca.a.g.gt........ga...aata.a...agaa.--------------..a...--- [207760]
KM262844.1_strain_L60            ................------.......................................................... [207760]
KM102979.1_26544/OG10_Italy      ................------.......................................................... [207760]
KM262845.1_strain_NHV            -------------------------------------------------------------------------------- [207760]
KX354450.1_47/Ss/2008            ................------.......................................................... [207760]
KP055815.1_strain_BA71           ................------.......................................................... [207760]
MH025920.1_strain_R35            a.atc..a...g.tg.ggaaaat.a.g.gt........gg...aata.a...agaa.-------------c..a...--- [207760]
MH025919.1_strain_N10            a.atc..a...g.tg.ggaaaat.a.g.gt........gg...aata.a...agaa.-------------c..a...--- [207760]
MH025918.1_strain_R25            a.atc..a...g.tg.ggaaaat.a.g.gt........gg...aata.a...agaa.-------------c..a...--- [207760]
MH025917.1_strain_R7             a.atc..a...g.tg.ggaaaat.a.g.gt........gg...aata.a...agaa.-------------c..a...--- [207760]
MH025916.1_strain_R8             a.atc..a...g.tg.ggaaaat.a.g.gt........gg...aata.a...agaa.-------------c..a...--- [207760]

FR682468.2_Georgia_2007/1        tagaggacc-gctacaaaaattatttttttttcttgatcaaagctccaaaataattattagattaaagtcgcctatagca [207840]
ASF_IND_20_CAD543                .........-...................................................................... [207840]
LR899193.1_Germany_2020/1        .........-...................................................................... [207840]
MN393477.1_Wuhan_2019            .........-...................................................................... [207840]
MW396979.1_Timor-Leste/2019/1    .........-...................................................................... [207840]
MK940252.1_CN/2019/InnerMongolia .........-...................................................................... [207840]
MK128995.1_China/2018/AnhuiXCGQ  .........-...................................................................... [207840]
MH681419.1_ASFV/POL/2015         .........-...................................................................... [207840]
LS478113.1_Estonia_2014          .........-...................................................................... [207840]
KP843857.1_Odintsovo_02/14       .........-.............-........................................................ [207840]
```

```
FN557520.1_strain_E75           .........-...................................................................... [207840]
AY261366.1_Warthog              ..a.acgg.g..g...................------------------------------------------------ [207840]
AY261365.1_Warmbaths            .....agt.a..g.................c.tg----------------------.......c.....ta....c.... [207840]
AY261364.1_Tengani_62           ......g..a..g..................ct.---------------------------------------------- [207840]
AY261363.1_Pretoriuskop/96/4    ......g..a..g...................t----------------------------------------------- [207840]
AY261362.1_Mkuzi_1979           .....agt.a..g.................c.----------------------.......................... [207840]
AY261361.1_Malawi_Lil-20/1_1983 --------------...t..c...........------------------------------------------------ [207840]
AY261360.1_Kenya_1950           ----------------..--------------------------------------------------------...at. [207840]
AM712240.1_OURT_88/3            -------------------------------------------------------------------------------- [207840]
AM712239.1_Benin_97/1           .........-...................................................................... [207840]
KM111295.1_Ken06.Bus            -------------------------------------------------------------------------------- [207840]
KM111294.1_Ken05/Tk1            ----------------.............--------------------------------------------c...at. [207840]
KM262844.1_strain_L60           .........-...................................................................... [207840]
KM102979.1_26544/OG10_Italy     .........-...................................................................... [207840]
KM262845.1_strain_NHV           -------------------------------------------------------------------------------- [207840]
KX354450.1_47/Ss/2008           .........-...................................................................... [207840]
KP055815.1_strain_BA71          .........-...................................................................... [207840]
MH025920.1_strain_R35           ----------------.............--------------------------------------------c...at. [207840]
MH025919.1_strain_N10           ----------------.............--------------------------------------------c...at. [207840]
MH025918.1_strain_R25           ----------------.............--------------------------------------------c...at. [207840]
MH025917.1_strain_R7            ----------------.............--------------------------------------------c...at. [207840]
MH025916.1_strain_R8            ----------------.............--------------------------------------------c...at. [207840]

FR682468.2_Georgia_2007/1       gc---------------------------agcccactccaaaaaaagtattt---------tatagtacaaaaaacacga [207920]
ASF_IND_20_CAD543               ..---------------------------.......................---------................... [207920]
LR899193.1_Germany_2020/1       ..---------------------------.......................---------................... [207920]
MN393477.1_Wuhan_2019           ..---------------------------.......................---------................... [207920]
MW396979.1_Timor-Leste/2019/1   ..---------------------------.......................---------................... [207920]
MK940252.1_CN/2019/InnerMongolia ..---------------------------.......................---------................... [207920]
MK128995.1_China/2018/AnhuiXCGQ ..---------------------------.......................---------................... [207920]
MH681419.1_ASFV/POL/2015        ..---------------------------.......................---------................... [207920]
LS478113.1_Estonia_2014         ..---------------------------.......................---------................... [207920]
KP843857.1_Odintsovo_02/14      ..---------------------------.......................---------................... [207920]
FN557520.1_strain_E75           ..---------------------------t......................---------................... [207920]
AY261366.1_Warthog              -------------------------------------------------------------------------------- [207920]
AY261365.1_Warmbaths            ..agtaataattgatgtatgcagaaataatat....c.t.............atggtgggt................t.. [207920]
AY261364.1_Tengani_62           -------------------------------------------------------------------------------- [207920]
AY261363.1_Pretoriuskop/96/4    -------------------------------------------------------------------------------- [207920]
AY261362.1_Mkuzi_1979           ..attaataattgatgtatgcagaaataatat....c.t.............atggtgggt................t.. [207920]
AY261361.1_Malawi_Lil-20/1_1983 -------------------------------------------------------------------------------- [207920]
AY261360.1_Kenya_1950           ..g----------------------------------------------------------------------------- [207920]
AM712240.1_OURT_88/3            -------------------------------------------------------------------------------- [207920]
AM712239.1_Benin_97/1           ..---------------------------t......................---------................... [207920]
KM111295.1_Ken06.Bus            -------------------------------------------------------------------------------- [207920]
```

```
KM111294.1_Ken05/Tk1             ..------------------------------------------------------------------------------ [207920]
KM262844.1_strain_L60            ..---------------------------t......................---------................... [207920]
KM102979.1_26544/OG10_Italy      ..---------------------------t......................---------................... [207920]
KM262845.1_strain_NHV            -------------------------------------------------------------------------------- [207920]
KX354450.1_47/Ss/2008            ..---------------------------t......................---------................... [207920]
KP055815.1_strain_BA71           ..---------------------------t......................---------................... [207920]
MH025920.1_strain_R35            ..g----------------------------------------------------------------------------- [207920]
MH025919.1_strain_N10            ..g----------------------------------------------------------------------------- [207920]
MH025918.1_strain_R25            ..g----------------------------------------------------------------------------- [207920]
MH025917.1_strain_R7             ..g----------------------------------------------------------------------------- [207920]
MH025916.1_strain_R8             ..g----------------------------------------------------------------------------- [207920]

FR682468.2_Georgia_2007/1        aaaatagtttgcggccggcggcaaactatttgttgttgtctaaaacttaatgttttttt--aatatttttaaatgcaacc [208000]
ASF_IND_20_CAD543                ...........................................................--................... [208000]
LR899193.1_Germany_2020/1        ...........................................................--................... [208000]
MN393477.1_Wuhan_2019            ...........................................................--................... [208000]
MW396979.1_Timor-Leste/2019/1    ...........................................................--................... [208000]
MK940252.1_CN/2019/InnerMongolia ...........................................................--................... [208000]
MK128995.1_China/2018/AnhuiXCGQ  ...........................................................--................... [208000]
MH681419.1_ASFV/POL/2015         ...........................................................--................... [208000]
LS478113.1_Estonia_2014          ...........................................................--................... [208000]
KP843857.1_Odintsovo_02/14       ...........................................................--................... [208000]
FN557520.1_strain_E75            ......a.................................c..................--................... [208000]
AY261366.1_Warthog               -------------------------------------------------------------------------------- [208000]
AY261365.1_Warmbaths             ..........a.....a..............a...........................tgggat.......gc...... [208000]
AY261364.1_Tengani_62            -------------------------------------------------------------------------------- [208000]
AY261363.1_Pretoriuskop/96/4     -------------------------------------------------------------------------------- [208000]
AY261362.1_Mkuzi_1979            ..........a.....a..............a...........................tgggat.......gc...... [208000]
AY261361.1_Malawi_Lil-20/1_1983  -------------------------------------------------------------------------------- [208000]
AY261360.1_Kenya_1950            -------------------------------------------------------------------------------- [208000]
AM712240.1_OURT_88/3             -------------------------------------------------------------------------------- [208000]
AM712239.1_Benin_97/1            ......a.................................c..................--................... [208000]
KM111295.1_Ken06.Bus             -------------------------------------------------------------------------------- [208000]
KM111294.1_Ken05/Tk1             -------------------------------------------------------------------------------- [208000]
KM262844.1_strain_L60            ......a.................................c..................--................... [208000]
KM102979.1_26544/OG10_Italy      ......a.................................c..................--................... [208000]
KM262845.1_strain_NHV            -------------------------------------------------------------------------------- [208000]
KX354450.1_47/Ss/2008            ......a.................................c..................--................... [208000]
KP055815.1_strain_BA71           ......a.................................c..................--................... [208000]
MH025920.1_strain_R35            -------------------------------------------------------------------------------- [208000]
MH025919.1_strain_N10            -------------------------------------------------------------------------------- [208000]
MH025918.1_strain_R25            -------------------------------------------------------------------------------- [208000]
MH025917.1_strain_R7             -------------------------------------------------------------------------------- [208000]
MH025916.1_strain_R8             -------------------------------------------------------------------------------- [208000]
```

```
FR682468.2_Georgia_2007/1        atg---gattgttggactatcagggagaagaactatagctacatcatattgtcaatactggtaatactattaatatggta [208080]
ASF_IND_20_CAD543                ...---.......................................................................... [208080]
LR899193.1_Germany_2020/1        ...---.......................................................................... [208080]
MN393477.1_Wuhan_2019            ...---.......................................................................... [208080]
MW396979.1_Timor-Leste/2019/1    ...---.......................................................................... [208080]
MK940252.1_CN/2019/InnerMongolia ...---.......................................................................... [208080]
MK128995.1_China/2018/AnhuiXCGQ  ...---.......................................................................... [208080]
MH681419.1_ASFV/POL/2015         ...---.......................................................................... [208080]
LS478113.1_Estonia_2014          ...---.......................................................................... [208080]
KP843857.1_Odintsovo_02/14       ...---.......................................................................... [208080]
FN557520.1_strain_E75            ...---.......................................................................... [208080]
AY261366.1_Warthog               -------------------------------------------------------------------------------- [208080]
AY261365.1_Warmbaths             ...aat....a....t...c..aa.t.....t................a.....t.....a................... [208080]
AY261364.1_Tengani_62            -------------------------------------------------------------------------------- [208080]
AY261363.1_Pretoriuskop/96/4     -------------------------------------------------------------------------------- [208080]
AY261362.1_Mkuzi_1979            ...aat....a....t...c..aa.t.....t................a.....t.....a................... [208080]
AY261361.1_Malawi_Lil-20/1_1983  -------------------------------------------------------------------------------- [208080]
AY261360.1_Kenya_1950            --------------------------------------------------------------------------...a.. [208080]
AM712240.1_OURT_88/3             -------------------------------------------------------------------------------- [208080]
AM712239.1_Benin_97/1            ...---.......................................................................... [208080]
KM111295.1_Ken06.Bus             -------------------------------------------------------------------------------- [208080]
KM111294.1_Ken05/Tk1             ----------------------------------------------------------------------------.a.. [208080]
KM262844.1_strain_L60            ...---.......................................................................... [208080]
KM102979.1_26544/OG10_Italy      ...---.......................................................................... [208080]
KM262845.1_strain_NHV            -------------------------------------------------------------------------------- [208080]
KX354450.1_47/Ss/2008            ...---.......................................................................... [208080]
KP055815.1_strain_BA71           ...---.......................................................................... [208080]
MH025920.1_strain_R35            --------------------------------------------------------------------------...a.. [208080]
MH025919.1_strain_N10            --------------------------------------------------------------------------...a.. [208080]
MH025918.1_strain_R25            --------------------------------------------------------------------------...a.. [208080]
MH025917.1_strain_R7             --------------------------------------------------------------------------...a.. [208080]
MH025916.1_strain_R8             --------------------------------------------------------------------------...a.. [208080]

FR682468.2_Georgia_2007/1        tcttatacttaactattg------tcgatcg-aaaaaaaatgcagttacaaacaacatgccg---ccaccatacacggtg [208160]
ASF_IND_20_CAD543                ..................------.......-...........w..................cca..............r [208160]
LR899193.1_Germany_2020/1        ..................------.......-..............................---............... [208160]
MN393477.1_Wuhan_2019            ..................------.......-..............................---............... [208160]
MW396979.1_Timor-Leste/2019/1    ..................------.......-..............................---............... [208160]
MK940252.1_CN/2019/InnerMongolia ..................------.......-..............................---............... [208160]
MK128995.1_China/2018/AnhuiXCGQ  ..................------.......-..............................---............... [208160]
MH681419.1_ASFV/POL/2015         ..................------.......-..............................---............... [208160]
LS478113.1_Estonia_2014          ..................------.......-..............................---............... [208160]
KP843857.1_Odintsovo_02/14       ..................------.......-..............................---............... [208160]
```

```
FN557520.1_strain_E75            ..................------.......-..............................cca...g........... [208160]
AY261366.1_Warthog               -------------------------------------------------------------------------------- [208160]
AY261365.1_Warmbaths             ........a..t..t...taagct....at.-..g.......gt.................a---............... [208160]
AY261364.1_Tengani_62            -------------------------------------------------------------------------------- [208160]
AY261363.1_Pretoriuskop/96/4     -------------------------------------------------------------------------------- [208160]
AY261362.1_Mkuzi_1979            ........a..t..t...taagct....at.-..g.......gt..............---.cca............... [208160]
AY261361.1_Malawi_Lil-20/1_1983  -------------------------------------------------------------------------------- [208160]
AY261360.1_Kenya_1950            .a..t.g------------------------------------------------------------------------- [208160]
AM712240.1_OURT_88/3             ----------------------------------------------------.....c.gt.cca............a.a [208160]
AM712239.1_Benin_97/1            ..................------.......-..............................cca...g........... [208160]
KM111295.1_Ken06.Bus             -------------------------------------------------------------------------------- [208160]
KM111294.1_Ken05/Tk1             .a....g------------------------------------------------------------------------- [208160]
KM262844.1_strain_L60            ..................------.......-..............................cca...g........... [208160]
KM102979.1_26544/OG10_Italy      ..................------.......-..............................cca...g........... [208160]
KM262845.1_strain_NHV            ----------------------------------------------------.....c.gt.cca............a.a [208160]
KX354450.1_47/Ss/2008            ..................------.......-..............................cca...g........... [208160]
KP055815.1_strain_BA71           ..................------.......a..............................cca...g........... [208160]
MH025920.1_strain_R35            .a..t.g------------------------------------------------------------------------- [208160]
MH025919.1_strain_N10            .a..t.g------------------------------------------------------------------------- [208160]
MH025918.1_strain_R25            .a..t.g------------------------------------------------------------------------- [208160]
MH025917.1_strain_R7             .a..t.g------------------------------------------------------------------------- [208160]
MH025916.1_strain_R8             .a..t.g------------------------------------------------------------------------- [208160]

FR682468.2_Georgia_2007/1        tcaagtagctgttctcaataatagggttgattgacgctcttcgtaataatatgttgattgacgcatcataaaatgctgtg [208240]
ASF_IND_20_CAD543                ......m.................................y.....................................a. [208240]
LR899193.1_Germany_2020/1        ................................................................................ [208240]
MN393477.1_Wuhan_2019            ................................................................................ [208240]
MW396979.1_Timor-Leste/2019/1    ................................................................................ [208240]
MK940252.1_CN/2019/InnerMongolia ................................................................................ [208240]
MK128995.1_China/2018/AnhuiXCGQ  ................................................................................ [208240]
MH681419.1_ASFV/POL/2015         ................................................................................ [208240]
LS478113.1_Estonia_2014          ................................................................................ [208240]
KP843857.1_Odintsovo_02/14       ................................................................................ [208240]
FN557520.1_strain_E75            ...............................c............................g.................a. [208240]
AY261366.1_Warthog               -------------------------------------------------------------------------------- [208240]
AY261365.1_Warmbaths             ......c.............----------------------------....a........................... [208240]
AY261364.1_Tengani_62            -------------------------------------------------------------------------------- [208240]
AY261363.1_Pretoriuskop/96/4     -------------------------------------------------------------------------------- [208240]
AY261362.1_Mkuzi_1979            ......c.............----------------------------....a........................... [208240]
AY261361.1_Malawi_Lil-20/1_1983  -------------------------------------------------------------------------------- [208240]
AY261360.1_Kenya_1950            -------------------------------------------------------------------------------- [208240]
AM712240.1_OURT_88/3             ...g........c.c.....------------------------------------------------------------ [208240]
AM712239.1_Benin_97/1            ...............................c............................g.................a. [208240]
KM111295.1_Ken06.Bus             -------------------------------------------------------------------------------- [208240]
```

```
KM111294.1_Ken05/Tk1           -------------------------------------------------------------------------------- [208240]
KM262844.1_strain_L60          -..............................c............................g.................a. [208240]
KM102979.1_26544/OG10_Italy    ...............................c............................g.................a. [208240]
KM262845.1_strain_NHV          ...g........c.c.....------------------------------------------------------------ [208240]
KX354450.1_47/Ss/2008          ...............................c............................g.................a. [208240]
KP055815.1_strain_BA71         ...............................c............................g.................a. [208240]
MH025920.1_strain_R35          -------------------------------------------------------------------------------- [208240]
MH025919.1_strain_N10          -------------------------------------------------------------------------------- [208240]
MH025918.1_strain_R25          -------------------------------------------------------------------------------- [208240]
MH025917.1_strain_R7           -------------------------------------------------------------------------------- [208240]
MH025916.1_strain_R8           -------------------------------------------------------------------------------- [208240]

FR682468.2_Georgia_2007/1      g-------ttgattaatatgttgattgtcgcctactttattatataagtaatgatttttgtataaaatacgggtttgtga [208320]
ASF_IND_20_CAD543              .aacnnnnnnnnnnnnnnnnnnnnnnnnnnnnnnnnnnnnnnnn.........-.......................... [208320]
LR899193.1_Germany_2020/1      .-------........................................................................ [208320]
MN393477.1_Wuhan_2019          .-------........................................................................ [208320]
MW396979.1_Timor-Leste/2019/1  .-------........................................................................ [208320]
MK940252.1_CN/2019/InnerMongolia .-------........................................................................ [208320]
MK128995.1_China/2018/AnhuiXCGQ .-------........................................................................ [208320]
MH681419.1_ASFV/POL/2015       .-------........................................................................ [208320]
LS478113.1_Estonia_2014        .-------........................................................................ [208320]
KP843857.1_Odintsovo_02/14     .-------........................................................................ [208320]
FN557520.1_strain_E75          .aactatccgttaat............a....c....atga..gg...........-....................... [208320]
AY261366.1_Warthog             -------------------------------------------------------------------------------- [208320]
AY261365.1_Warmbaths           .aactat-cgttaat............a.........atga..----................................. [208320]
AY261364.1_Tengani_62          -------------------------------------------------------------------------------- [208320]
AY261363.1_Pretoriuskop/96/4   -------------------------------------------------------------------------------- [208320]
AY261362.1_Mkuzi_1979          .aactatc-gttaat............a.........atga..----................................. [208320]
AY261361.1_Malawi_Lil-20/1_1983 -------------------------------------------------------------------------------- [208320]
AY261360.1_Kenya_1950          -----------------------------------------------a..ga.g....g..------------------- [208320]
AM712240.1_OURT_88/3           ---------------..g........ca....c.t--------------------------------------------- [208320]
AM712239.1_Benin_97/1          .aactatccgttaat............a....c....atga..gg...........-....................... [208320]
KM111295.1_Ken06.Bus           -------------------------------------------------------------------------------- [208320]
KM111294.1_Ken05/Tk1           -----------------------------------------------a..ga.....----------------------- [208320]
KM262844.1_strain_L60          .aactatccgttaat............a....c....atga..gg...........-....................... [208320]
KM102979.1_26544/OG10_Italy    .aactatccgttaat............a....c....atga..gg...........-....................... [208320]
KM262845.1_strain_NHV          ---------------..g........ca....c.ta-------------------------------------------- [208320]
KX354450.1_47/Ss/2008          .aactatccgttaat............a....c....atga..gg...........-....................... [208320]
KP055815.1_strain_BA71         .aactatccgttaat............a....c....atga..gg...........-....................... [208320]
MH025920.1_strain_R35          -----------------------------------------------a..ga.g....g..------------------- [208320]
MH025919.1_strain_N10          -----------------------------------------------a..ga.g....g..------------------- [208320]
MH025918.1_strain_R25          -----------------------------------------------a..ga.g....g..------------------- [208320]
MH025917.1_strain_R7           -----------------------------------------------a..ga.g....g..------------------- [208320]
MH025916.1_strain_R8           -----------------------------------------------a..ga.g....g..------------------- [208320]
```

```
FR682468.2_Georgia_2007/1        gggctttattttt-tcttattagaacaaagcatgca-atttaaggcctacagcaagagt-a-----atttaacacctaca [208400]
ASF_IND_20_CAD543                ....y........t......................tt.....................-.tgttt.a............ [208400]
LR899193.1_Germany_2020/1        .............-......................-......................-.-----.............. [208400]
MN393477.1_Wuhan_2019            .............-......................-......................-.-----.............. [208400]
MW396979.1_Timor-Leste/2019/1    .............-......................-......................-.-----.............. [208400]
MK940252.1_CN/2019/InnerMongolia .............-......................-......................-.-----.............. [208400]
MK128995.1_China/2018/AnhuiXCGQ  .............-......................-......................-.-----.............. [208400]
MH681419.1_ASFV/POL/2015         .............-......................-......................-.-----.............. [208400]
LS478113.1_Estonia_2014          .............-......................-......................-.-----.............. [208400]
KP843857.1_Odintsovo_02/14       .............-......................-......................-.-----.............. [208400]
FN557520.1_strain_E75            ....c........t.................g..t.tt...........t.........-.tgttt.a............ [208400]
AY261366.1_Warthog               -------------------c.t..t.........t.at.....t...............a.taaat.a.c---------- [208400]
AY261365.1_Warmbaths             ....c........t......................a-.....................a.tgttt.a............ [208400]
AY261364.1_Tengani_62            ----------------------..c.........t.at.....tt.t............t.tatc--------------- [208400]
AY261363.1_Pretoriuskop/96/4     -------------------c.t..t.........t.at.....tt.t............c.tc----------------- [208400]
AY261362.1_Mkuzi_1979            ....c........t......................at.-...................a.tgttt.a............ [208400]
AY261361.1_Malawi_Lil-20/1_1983  ------------------------------------------------g.c....a.a.gg------------------- [208400]
AY261360.1_Kenya_1950            ---------------------------gctg...g.tt....--tagg.tca..t..c.--------------------- [208400]
AM712240.1_OURT_88/3             -------------------------------------------------------------------------------- [208400]
AM712239.1_Benin_97/1            ....c........t.................g..t.tt...........t.........-.tgttt.a............ [208400]
KM111295.1_Ken06.Bus             -------------------------------------------------------------------------------- [208400]
KM111294.1_Ken05/Tk1             -----------------------------tg...g.tt....tatagg..ca..ta.c.--------------------- [208400]
KM262844.1_strain_L60            ....c........t.................g..t.tt...........t.........-.tgttt.a............ [208400]
KM102979.1_26544/OG10_Italy      ....c........t.................g..t.tt...........t.........-.tgttt.a............ [208400]
KM262845.1_strain_NHV            -------------------------------------------------------------------------------- [208400]
KX354450.1_47/Ss/2008            ....c........t.................g..t.tt...........t.........-.tgttt.a............ [208400]
KP055815.1_strain_BA71           ....c........t.................g..t.tt...........t.........-.tgttt.a............ [208400]
MH025920.1_strain_R35            ---------------------------gctg...g.tt....--tagg.tca..t..c.--------------------- [208400]
MH025919.1_strain_N10            ---------------------------gctg...g.tt....--tagg.tca..t..c.--------------------- [208400]
MH025918.1_strain_R25            ---------------------------gctg...g.tt....--tagg.tca..t..c.--------------------- [208400]
MH025917.1_strain_R7             ---------------------------gctg...g.tt....--tagg.tca..t..c.--------------------- [208400]
MH025916.1_strain_R8             ---------------------------gctg...g.tt....--tagg.tca..t..c.--------------------- [208400]

FR682468.2_Georgia_2007/1        acagtaat-tttaaggtcagt--aataatgtttaattaaggcctgaccactaaaacttaaacgattttgt---aaaaaaa [208480]
ASF_IND_20_CAD543                ........a.......y....aa...............................................---....... [208480]
LR899193.1_Germany_2020/1        ........-............--...............................................--a....... [208480]
MN393477.1_Wuhan_2019            ........-............--...............................................---....... [208480]
MW396979.1_Timor-Leste/2019/1    ........-............--...............................................---....... [208480]
MK940252.1_CN/2019/InnerMongolia ........-............--...............................................---....... [208480]
MK128995.1_China/2018/AnhuiXCGQ  ........-............--...............................................---....... [208480]
MH681419.1_ASFV/POL/2015         ........-............--...............................................---....... [208480]
LS478113.1_Estonia_2014          ........-............--...............................................---....... [208480]
KP843857.1_Odintsovo_02/14       ........-............--...............................................---....... [208480]
```

```
FN557520.1_strain_E75            ........a.......c....aa.................................................---....... [208480]
AY261366.1_Warthog               ---t....g.....acctc.c--ggc.ggacc.t..c----------t..............a.......aaa......- [208480]
AY261365.1_Warmbaths             ........g.......c....--......................-a.......................aaa....... [208480]
AY261364.1_Tengani_62            ---t....a.....acctc.c--ggc.ggac..tc.------------..............a.......---....... [208480]
AY261363.1_Pretoriuskop/96/4     ---t....a.....atctc..--ggc.ggacc.t..c----------...............a.......aaa......- [208480]
AY261362.1_Mkuzi_1979            ........g.......c....--.......................-.......................aaa....... [208480]
AY261361.1_Malawi_Lil-20/1_1983  ---------.....at.tt.c--------------------------------------------------atgg..... [208480]
AY261360.1_Kenya_1950            ---a...--.....tt.ttt.---------------------------t.ct.g..c----------------------- [208480]
AM712240.1_OURT_88/3             -------------------------------------------------------------------------------- [208480]
AM712239.1_Benin_97/1            ........a.......c....aa.................................................---....... [208480]
KM111295.1_Ken06.Bus             -------------------------------------------------------------------------------- [208480]
KM111294.1_Ken05/Tk1             ---..c..-....ttt.ttt.---------------------------tg.....t.....ga.....tgcaa....... [208480]
KM262844.1_strain_L60            ........a.......c....aa..........................t............c......c---....... [208480]
KM102979.1_26544/OG10_Italy      ........a.......c....aa.........................c.....................---....... [208480]
KM262845.1_strain_NHV            -------------------------------------------------------------------------------- [208480]
KX354450.1_47/Ss/2008            ........a.......c....aa.........................c.....................---....... [208480]
KP055815.1_strain_BA71           ........a.......c....aa.................................................---....... [208480]
MH025920.1_strain_R35            ---a....-....ttt.ttt.---------------------------t.ct.g..c----------------------- [208480]
MH025919.1_strain_N10            ---a....-....ttt.ttt.---------------------------t.ct.g..c----------------------- [208480]
MH025918.1_strain_R25            ---a....-....ttt.ttt.---------------------------t.ct.g..c----------------------- [208480]
MH025917.1_strain_R7             ---a....-....ttt.ttt.---------------------------t.ct.g..c----------------------- [208480]
MH025916.1_strain_R8             ---a....-....ttt.ttt.---------------------------t.ct.g..c----------------------- [208480]

FR682468.2_Georgia_2007/1        aatgtctactccactttctctacagactcttgttaaaaaagtgctggccacacagca---catatctaaagaacactact [208560]
ASF_IND_20_CAD543                .........................................................---................--.. [208560]
LR899193.1_Germany_2020/1        .........................................................---.................... [208560]
MN393477.1_Wuhan_2019            .........................................................---..............c..... [208560]
MW396979.1_Timor-Leste/2019/1    .........................................................---.................... [208560]
MK940252.1_CN/2019/InnerMongolia .........................................................---.................... [208560]
MK128995.1_China/2018/AnhuiXCGQ  .........................................................---.................... [208560]
MH681419.1_ASFV/POL/2015         .........................................................---.................... [208560]
LS478113.1_Estonia_2014          .........................................................---.................... [208560]
KP843857.1_Odintsovo_02/14       .........................................................---.................... [208560]
FN557520.1_strain_E75            t...c....................g......c.........a..............---............t...--.. [208560]
AY261366.1_Warthog               -...c...........................c.......c.a............t.---t.....c.....tt...... [208560]
AY261365.1_Warmbaths             t...c...........................c.........a............t.---t.....c.t...tt.t.... [208560]
AY261364.1_Tengani_62            .........................................................---.g..........t....... [208560]
AY261363.1_Pretoriuskop/96/4     -...c..t......a.................c.......a.a.....t........---g.......c...t....... [208560]
AY261362.1_Mkuzi_1979            t...c...........................c.........a............t.---t.....c.t...tt.t.... [208560]
AY261361.1_Malawi_Lil-20/1_1983  ...act-------------------------------------------------------------------------- [208560]
AY261360.1_Kenya_1950            -------------------------------------------------------------------------------- [208560]
AM712240.1_OURT_88/3             -------------------------------------------------------------------------------- [208560]
AM712239.1_Benin_97/1            t...c....................g......c.........a..............---............t...--.. [208560]
KM111295.1_Ken06.Bus             -------------------------------------------------------------------------------- [208560]
```

```
KM111294.1_Ken05/Tk1              c........a...tc.........a.tc.....c....gga......a.ttc.....tcatg.....g....tg.t..t. [208560]
KM262844.1_strain_L60             t....................g...g......c.......a.a..............---......c...a.t....... [208560]
KM102979.1_26544/OG10_Italy       t...c....................g......c.........a..............---............t...--.. [208560]
KM262845.1_strain_NHV             -------------------------------------------------------------------------------- [208560]
KX354450.1_47/Ss/2008             t...c....................g......c.........a..............---............t...--.. [208560]
KP055815.1_strain_BA71            t...c....................g......c.........a..............---............t...--.. [208560]
MH025920.1_strain_R35             -------------------------------------------------------------------------------- [208560]
MH025919.1_strain_N10             -------------------------------------------------------------------------------- [208560]
MH025918.1_strain_R25             -------------------------------------------------------------------------------- [208560]
MH025917.1_strain_R7              -------------------------------------------------------------------------------- [208560]
MH025916.1_strain_R8              -------------------------------------------------------------------------------- [208560]

FR682468.2_Georgia_2007/1         ttattttgaaatattgtggtttatggtggcat-----------------------------------------------g [208640]
ASF_IND_20_CAD543                 ................................nnnnnnnnnnnnnnnnnnnnnnnnnnnnnnnnnnnnnnnnnnnnnnnn [208640]
LR899193.1_Germany_2020/1         ................................-----------------------------------------------. [208640]
MN393477.1_Wuhan_2019             ................................-----------------------------------------------. [208640]
MW396979.1_Timor-Leste/2019/1     ................................-----------------------------------------------. [208640]
MK940252.1_CN/2019/InnerMongolia  ................................-----------------------------------------------. [208640]
MK128995.1_China/2018/AnhuiXCGQ   ................................-----------------------------------------------. [208640]
MH681419.1_ASFV/POL/2015          ................................-----------------------------------------------. [208640]
LS478113.1_Estonia_2014           ................................-----------------------------------------------. [208640]
KP843857.1_Odintsovo_02/14        ................................-----------------------------------------------. [208640]
FN557520.1_strain_E75             ................................-----------------------------------------------t [208640]
AY261366.1_Warthog                ................................-----------------------------------------------. [208640]
AY261365.1_Warmbaths              ................................-----------------------------------------------. [208640]
AY261364.1_Tengani_62             .......a..............g.........-----------------------------------------------a [208640]
AY261363.1_Pretoriuskop/96/4      .......a........................-----------------------------------------------. [208640]
AY261362.1_Mkuzi_1979             ................................-----------------------------------------------. [208640]
AY261361.1_Malawi_Lil-20/1_1983   --------....t..a..t...t.ta.aa...------------------------------------------------ [208640]
AY261360.1_Kenya_1950             -------------------------------------------------------------------------------- [208640]
AM712240.1_OURT_88/3              -------------------------------------------------------------------------------- [208640]
AM712239.1_Benin_97/1             ................................-----------------------------------------------t [208640]
KM111295.1_Ken06.Bus              -------------------------------------------------------------------------------- [208640]
KM111294.1_Ken05/Tk1              g......ag.gcg.........g.........-----------------------------------------------. [208640]
KM262844.1_strain_L60             ......................g.........-----------------------------------------------t [208640]
KM102979.1_26544/OG10_Italy       ................................-----------------------------------------------t [208640]
KM262845.1_strain_NHV             -------------------------------------------------------------------------------- [208640]
KX354450.1_47/Ss/2008             ................................-----------------------------------------------t [208640]
KP055815.1_strain_BA71            ................................-----------------------------------------------t [208640]
MH025920.1_strain_R35             -------------------------------------------------------------------------------- [208640]
MH025919.1_strain_N10             -------------------------------------------------------------------------------- [208640]
MH025918.1_strain_R25             -------------------------------------------------------------------------------- [208640]
MH025917.1_strain_R7              -------------------------------------------------------------------------------- [208640]
MH025916.1_strain_R8              -------------------------------------------------------------------------------- [208640]
```

```
FR682468.2_Georgia_2007/1        aagcgccgattacgatttgcattgat---gaggatagccaaatattgataaaatcggcaagcttcaaagaaggcttatct [208720]
ASF_IND_20_CAD543                nnnnnnnnn.a.ta...gat.ccc.c----..nnnnnnt..ta.......ccca.a.nnnnnnnnnnnnnnat..ggc.- [208720]
LR899193.1_Germany_2020/1        ..........................---................................................... [208720]
MN393477.1_Wuhan_2019            ..........................---................................................... [208720]
MW396979.1_Timor-Leste/2019/1    ..........................---................................................... [208720]
MK940252.1_CN/2019/InnerMongolia ..........................---................................................... [208720]
MK128995.1_China/2018/AnhuiXCGQ  ..........................---................................................... [208720]
MH681419.1_ASFV/POL/2015         ..........................---................................................... [208720]
LS478113.1_Estonia_2014          ..........................---................................................... [208720]
KP843857.1_Odintsovo_02/14       ..........................---................................................... [208720]
FN557520.1_strain_E75            cttt.a..cc...ag.c.tt.ac.cctgcagta...attg.----------------------------------..... [208720]
AY261366.1_Warthog               g...t..a....t.c...ct.c.a..gag..ta.---t...t.ga................................... [208720]
AY261365.1_Warmbaths             g...c..t....t.....c.......aaga.ta.a.........c................................... [208720]
AY261364.1_Tengani_62            .g..c.....................---...................................t............c.. [208720]
AY261363.1_Pretoriuskop/96/4     g...t..a....t.c...ct.c.a..gag..ta.---t...t.ga................................... [208720]
AY261362.1_Mkuzi_1979            g...c..t....t.....c.......aaga.ta.a.........c................................... [208720]
AY261361.1_Malawi_Lil-20/1_1983  ---------------------------------------------------------------ct...at...t..g..- [208720]
AY261360.1_Kenya_1950            -------------------------------------------------------------------------------- [208720]
AM712240.1_OURT_88/3             -------------------------------------------------------------------------------- [208720]
AM712239.1_Benin_97/1            cttt.a..cc...ag.c.tt.ac.cctgcagta...attg.----------------------------------..... [208720]
KM111295.1_Ken06.Bus             -------------------------------------------------------------------------------- [208720]
KM111294.1_Ken05/Tk1             g..gt..a..c.t.c...ct.c.a..gat......cat....---..........a........t.....t......gaa [208720]
KM262844.1_strain_L60            cttt.a..ccc-.ag.c.tt.ac.cctgcagta...attg.----------------------------------.-... [208720]
KM102979.1_26544/OG10_Italy      cttt.a..cc...ag.c.tt.ac.cctgcagta...attg.----------------------------------..... [208720]
KM262845.1_strain_NHV            -------------------------------------------------------------------------------- [208720]
KX354450.1_47/Ss/2008            cttt.a..cc...ag.c.tt.ac.cctgcagta...attg.----------------------------------..... [208720]
KP055815.1_strain_BA71           cttt.a..cc...ag.c.tt.ac.cctgcagta...attg.----------------------------------..... [208720]
MH025920.1_strain_R35            -------------------------------------------------------------------------------- [208720]
MH025919.1_strain_N10            -------------------------------------------------------------------------------- [208720]
MH025918.1_strain_R25            -------------------------------------------------------------------------------- [208720]
MH025917.1_strain_R7             -------------------------------------------------------------------------------- [208720]
MH025916.1_strain_R8             -------------------------------------------------------------------------------- [208720]

FR682468.2_Georgia_2007/1        ttagatatcgcattaatgaaagtcgtgcaagaaaataaccatgatttaatagagttgtttaccaagtggggtgcagatat [208800]
ASF_IND_20_CAD543                ---------------------.c.ccaa..a...g..ttt.c.-----g...g...ta.....gg--------------- [208800]
LR899193.1_Germany_2020/1        ................................................................................ [208800]
MN393477.1_Wuhan_2019            ................................................................................ [208800]
MW396979.1_Timor-Leste/2019/1    ................................................................................ [208800]
MK940252.1_CN/2019/InnerMongolia ................................................................................ [208800]
MK128995.1_China/2018/AnhuiXCGQ  ................................................................................ [208800]
MH681419.1_ASFV/POL/2015         ................................................................................ [208800]
LS478113.1_Estonia_2014          ................................................................................ [208800]
KP843857.1_Odintsovo_02/14       ................................................................................ [208800]
```

```
FN557520.1_strain_E75            cc..cgc.aat.a....tg.tac.---..cagcgg...ta..a...g.-------------------------------- [208800]
AY261366.1_Warthog               ......t.g...........................c..t...........a..c.......tg.............c.. [208800]
AY261365.1_Warmbaths             ......................ct.....g.....c..t...........a..c........g.............c.. [208800]
AY261364.1_Tengani_62            ........g.............ct..a..g.....c...t....aa.g..cat.........tg.............c.. [208800]
AY261363.1_Pretoriuskop/96/4     ......t.g...........................c..t...........a..c........g.............c.. [208800]
AY261362.1_Mkuzi_1979            ......................ct.....g.....c..t...........a..c........g.............c.. [208800]
AY261361.1_Malawi_Lil-20/1_1983  ----------------------------------------------------------------.a.a.ta.....c..- [208800]
AY261360.1_Kenya_1950            -------------------------------------------------------------------------------- [208800]
AM712240.1_OURT_88/3             -------------------------------------------------------------------------------- [208800]
AM712239.1_Benin_97/1            cc..cgc.aat.a....tg.tac.---..cagcgg...ta..a...g.-------------------------------- [208800]
KM111295.1_Ken06.Bus             -------------------------------------------------------------------------------- [208800]
KM111294.1_Ken05/Tk1             ...a..g...............c...a..g.....c..t...ag.c........c......ttg..........g..c.. [208800]
KM262844.1_strain_L60            cc..cgc.aat.a....tg.tac.---..cagcgg...ta..a...g.-------------------------------- [208800]
KM102979.1_26544/OG10_Italy      cc..cgc.aat.a....tg.tac.---..cagcgg...ta..a...g.-------------------------------- [208800]
KM262845.1_strain_NHV            -------------------------------------------------------------------------------- [208800]
KX354450.1_47/Ss/2008            cc..cgc.aat.a....tg.tac.---..cagcgg...ta..a...g.-------------------------------- [208800]
KP055815.1_strain_BA71           cc..cgc.aat.a....tg.tac.---..cagcgg...ta..a...g.-------------------------------- [208800]
MH025920.1_strain_R35            -------------------------------------------------------------------------------- [208800]
MH025919.1_strain_N10            -------------------------------------------------------------------------------- [208800]
MH025918.1_strain_R25            -------------------------------------------------------------------------------- [208800]
MH025917.1_strain_R7             -------------------------------------------------------------------------------- [208800]
MH025916.1_strain_R8             -------------------------------------------------------------------------------- [208800]

FR682468.2_Georgia_2007/1        caactctagcttagttactgttaatacggagtatacccggaacctttgtcagaaattaggcgcaaaggaagctttgaatg [208880]
ASF_IND_20_CAD543                .gg.gta.cacc.....tg..c...------------------------------------------------------- [208880]
LR899193.1_Germany_2020/1        ................................................................................ [208880]
MN393477.1_Wuhan_2019            ................................................................................ [208880]
MW396979.1_Timor-Leste/2019/1    ................................................................................ [208880]
MK940252.1_CN/2019/InnerMongolia ................................................................................ [208880]
MK128995.1_China/2018/AnhuiXCGQ  ................................................................................ [208880]
MH681419.1_ASFV/POL/2015         ................................................................................ [208880]
LS478113.1_Estonia_2014          ................................................................................ [208880]
KP843857.1_Odintsovo_02/14       ................................................................................ [208880]
FN557520.1_strain_E75            -----------..ccc..a.cggta.t.tc..cc.t.t..ccg.cc----------------....aa..ag.a.tt.c. [208880]
AY261366.1_Warthog               ...............c..........t.....g......a.....c....g.g..........g................ [208880]
AY261365.1_Warmbaths             ...............c................g......a.....c....g.g..........g................ [208880]
AY261364.1_Tengani_62            t..............c...............cg.................gag..........g......at........ [208880]
AY261363.1_Pretoriuskop/96/4     ............c..c..........t.....g......a.....c....g.g..........g................ [208880]
AY261362.1_Mkuzi_1979            ...............c................g......a.....c....g.g..........g................ [208880]
AY261361.1_Malawi_Lil-20/1_1983  -------------------------------------------------------------------------------- [208880]
AY261360.1_Kenya_1950            -------------------------------------------------------------------------------- [208880]
AM712240.1_OURT_88/3             -------------------------------------------------------------------------------- [208880]
AM712239.1_Benin_97/1            -----------..ccc..a.cggta.t.tc..cc.t.t..ccg.cc---------------aa...aa..ag.a.tt.c. [208880]
KM111295.1_Ken06.Bus             -------------------------------------------------------------------------------- [208880]
```

```
KM111294.1_Ken05/Tk1            t..t.t.g.a..ga.......c..c.....aa.c.tt..a.g...a..c.gag.g.....t..c...........a...c [208880]
KM262844.1_strain_L60           -----------..ccc..a.cggta.t.tc..cc.t.t..ccg.cc-----------------...aa..ag.a.tt.c. [208880]
KM102979.1_26544/OG10_Italy     -----------..ccc..a.cggta.t.tc..cc.t.t..ccg.cc-----------------...aa..ag.a.tt.c. [208880]
KM262845.1_strain_NHV           -------------------------------------------------------------------------------- [208880]
KX354450.1_47/Ss/2008           -----------..ccc..a.cggta.t.tc..cc.t.t..ccg.cc-----------------...aa..ag.a.tt.c. [208880]
KP055815.1_strain_BA71          -----------..ccc..a.cggta.t.tc..cc.t.t..ccg.cc-----------------...aa..ag.a.tt.c. [208880]
MH025920.1_strain_R35           -------------------------------------------------------------------------------- [208880]
MH025919.1_strain_N10           -------------------------------------------------------------------------------- [208880]
MH025918.1_strain_R25           -------------------------------------------------------------------------------- [208880]
MH025917.1_strain_R7            -------------------------------------------------------------------------------- [208880]
MH025916.1_strain_R8            -------------------------------------------------------------------------------- [208880]

FR682468.2_Georgia_2007/1       aaagggatattttacaaatattttataaaacacgtcatcttaaaactagcagtaatattattttatataatgaattgttt [208960]
ASF_IND_20_CAD543               ---------------------...g.ctgg.c..c.gc.c---------------------------------------- [208960]
LR899193.1_Germany_2020/1       ................................................................................ [208960]
MN393477.1_Wuhan_2019           ................................................................................ [208960]
MW396979.1_Timor-Leste/2019/1   ................................................................................ [208960]
MK940252.1_CN/2019/InnerMongolia ................................................................................ [208960]
MK128995.1_China/2018/AnhuiXCGQ ................................................................................ [208960]
MH681419.1_ASFV/POL/2015        ................................................................................ [208960]
LS478113.1_Estonia_2014         ................................................................................ [208960]
KP843857.1_Odintsovo_02/14      ................................................................................ [208960]
FN557520.1_strain_E75           gtg...------------.t.a...ccggcgg...a.cac---------....t..gg.tag..t.g.c-..gccc.ccg [208960]
AY261366.1_Warthog              ...................................g..a..............c..g.........gcc.......a..a [208960]
AY261365.1_Warmbaths            ...................................g..a.................g....ac...gcc..........a [208960]
AY261364.1_Tengani_62           ...aa..a......g.................t.....a.................t.....c...gcc..........a [208960]
AY261363.1_Pretoriuskop/96/4    ...................................g..a..............c..g.........gcc.......a..a [208960]
AY261362.1_Mkuzi_1979           ...................................g..a.................g....ac...gcc..........a [208960]
AY261361.1_Malawi_Lil-20/1_1983 ---------------.....c..a...g..t.ta.a.ag.a..tt..--------------------------------- [208960]
AY261360.1_Kenya_1950           -------------------------------------------------------------------------------- [208960]
AM712240.1_OURT_88/3            -------------------------------------------------------------------------------- [208960]
AM712239.1_Benin_97/1           gtg...------------.t.a...ccggcgg...a.cac---------....t..gg.tag..t.g.c-..gccc.ccg [208960]
KM111295.1_Ken06.Bus            -------------------------------------------------------------------------------- [208960]
KM111294.1_Ken05/Tk1            ..t....ag.....g.tg.........c.gt.aaa.g.t......g...t.a..............g.c.......a..a [208960]
KM262844.1_strain_L60           gtg...------------.t.a...ccggcggg..a.cac---------....t..gg.tag..t.g.c-..gccc.ccg [208960]
KM102979.1_26544/OG10_Italy     gtg...------------.t.a...ccggcgg...a.cac---------....t..gg.tag..t.g.c-..gccc.ccg [208960]
KM262845.1_strain_NHV           -------------------------------------------------------------------------------- [208960]
KX354450.1_47/Ss/2008           gtg...------------.t.a...ccggcgg...a.cac---------....t..gg.tag..t.g.c-..gccc.ccg [208960]
KP055815.1_strain_BA71          gtg...------------.t.a...ccggcgg...a.cac---------....t..gg.tag..t.g.c-..gccc.ccg [208960]
MH025920.1_strain_R35           -------------------------------------------------------------------------------- [208960]
MH025919.1_strain_N10           -------------------------------------------------------------------------------- [208960]
MH025918.1_strain_R25           -------------------------------------------------------------------------------- [208960]
MH025917.1_strain_R7            -------------------------------------------------------------------------------- [208960]
MH025916.1_strain_R8            -------------------------------------------------------------------------------- [208960]
```

```
FR682468.2_Georgia_2007/1         tctaataatctccttttccaaaatatagagagattgagtttaa------tagtttataggggcttgaaaaacttatcaat [209040]
ASF_IND_20_CAD543                 ------------------------------------------------------------a..cgc.....aaa...... [209040]
LR899193.1_Germany_2020/1         ...........................................------............................... [209040]
MN393477.1_Wuhan_2019             ...........................................------............................... [209040]
MW396979.1_Timor-Leste/2019/1     ...........................................------............................... [209040]
MK940252.1_CN/2019/InnerMongolia  ...........................................------............................... [209040]
MK128995.1_China/2018/AnhuiXCGQ   ...........................................------............................... [209040]
MH681419.1_ASFV/POL/2015          ...........................................------............................... [209040]
LS478113.1_Estonia_2014           ...........................................------............................... [209040]
KP843857.1_Odintsovo_02/14        ...........................................------............................... [209040]
FN557520.1_strain_E75             c.c.gccgc--------------------------------------------------------a.....aaa...... [209040]
AY261366.1_Warthog                ..c.....c.c.........g............a..c...c..------..a........a.tc..g....ac....... [209040]
AY261365.1_Warmbaths              ..c.....c.c......................a..c...c..------..a........a.tc..g....ac....... [209040]
AY261364.1_Tengani_62             ..c.......c.....................t..........------............................... [209040]
AY261363.1_Pretoriuskop/96/4      ..c.....c.c.........g............a..c...c..------..a........a.tc..g....ac....... [209040]
AY261362.1_Mkuzi_1979             ..c.....c.c......................a..c...c..------..a........a.tc..g....ac....... [209040]
AY261361.1_Malawi_Lil-20/1_1983   ------------------------------------------------------------------------c..c.... [209040]
AY261360.1_Kenya_1950             -------------------------------------------------------------------------------- [209040]
AM712240.1_OURT_88/3              -------------------------------------------------------------------------------- [209040]
AM712239.1_Benin_97/1             c.c.gccgc---------------------------------------------------------.....aaa...... [209040]
KM111295.1_Ken06.Bus              -------------------------------------------------------------------------------- [209040]
KM111294.1_Ken05/Tk1              ..........ct.....t.t..g.ga.a.t.atg.tcaa..g.gaagat..a..a..tataatc.a.gg.gaa....... [209040]
KM262844.1_strain_L60             c.c.gccgc---------------------------------------------------------.....aaa...... [209040]
KM102979.1_26544/OG10_Italy       c.c.gccgc---------------------------------------------------------.....aaa...... [209040]
KM262845.1_strain_NHV             -------------------------------------------------------------------------------- [209040]
KX354450.1_47/Ss/2008             c.c.gccgc---------------------------------------------------------.....aaa...... [209040]
KP055815.1_strain_BA71            c.c.gccgc---------------------------------------------------------.....aaa...... [209040]
MH025920.1_strain_R35             -------------------------------------------------------------------------------- [209040]
MH025919.1_strain_N10             -------------------------------------------------------------------------------- [209040]
MH025918.1_strain_R25             -------------------------------------------------------------------------------- [209040]
MH025917.1_strain_R7              -------------------------------------------------------------------------------- [209040]
MH025916.1_strain_R8              -------------------------------------------------------------------------------- [209040]

FR682468.2_Georgia_2007/1         caactttatattggatgatatttcattagcg-------------------aaatgttaactagatactggtatagtatgg [209120]
ASF_IND_20_CAD543                 t.caaccgc.aaaa.aa.......cggcc..-------------------gcg.t.ct..a.------------------ [209120]
LR899193.1_Germany_2020/1         ...............................-------------------.............................. [209120]
MN393477.1_Wuhan_2019             ...............................-------------------.............................. [209120]
MW396979.1_Timor-Leste/2019/1     ...............................-------------------.............................. [209120]
MK940252.1_CN/2019/InnerMongolia  ...............................-------------------.............................. [209120]
MK128995.1_China/2018/AnhuiXCGQ   ...............................-------------------.............................. [209120]
MH681419.1_ASFV/POL/2015          ...............................-------------------.............................. [209120]
LS478113.1_Estonia_2014           ...............................-------------------.............................. [209120]
KP843857.1_Odintsovo_02/14        ...............................-------------------.............................. [209120]
```

```
FN557520.1_strain_E75           t.cagccgc.aaaa.aa.......cggcc...ga---gtttctacaaaaa........g..ttta.t..t.g...acca. [209120]
AY261366.1_Warthog              ................................------------------..............c........g.gt... [209120]
AY261365.1_Warmbaths            t...............................------------------..............c..........gt... [209120]
AY261364.1_Tengani_62           t...............................------------------.........................ctg.. [209120]
AY261363.1_Pretoriuskop/96/4    ..............................t.------------------..............c........g.gt... [209120]
AY261362.1_Mkuzi_1979           t...............................------------------..............c..........gt... [209120]
AY261361.1_Malawi_Lil-20/1_1983 a..a.a...--------..g...tg...g.aaaaccggttt---------..g.t...g.cc.gcc...----------- [209120]
AY261360.1_Kenya_1950           -------------------------------------------------------------------------------- [209120]
AM712240.1_OURT_88/3            -------------------------------------------------------------------------------- [209120]
AM712239.1_Benin_97/1           t.cagccgc.aaaa.aa.......cggcc...ga---gtttctacaaaaa........g..ttta.t..t.g...acca. [209120]
KM111295.1_Ken06.Bus            -------------------------------------------------------------------------------- [209120]
KM111294.1_Ken05/Tk1            t.................a..a........t.------------------...a.c.........t.......gcat... [209120]
KM262844.1_strain_L60           t.cagccgc.aaaa.aa.......cggcc...ga---gtttctacaaaaa........g..ttta.t..t.g...acca. [209120]
KM102979.1_26544/OG10_Italy     t.cagccgc.aaaa.aa.......cggcc...ga---gtttctacaaaaa........g..ttta.t..t.g...acca. [209120]
KM262845.1_strain_NHV           -------------------------------------------------------------------------------- [209120]
KX354450.1_47/Ss/2008           t.cagccgc.aaaa.aa.......cggcc...ga---gtttctacaaaaa........g..ttta.t..t.g...acca. [209120]
KP055815.1_strain_BA71          t.cagccgc.aaaa.aa.......cggcc...ga---gtttctacaaaaa........g..ttta.t..t.g...acca. [209120]
MH025920.1_strain_R35           -------------------------------------------------------------------------------- [209120]
MH025919.1_strain_N10           -------------------------------------------------------------------------------- [209120]
MH025918.1_strain_R25           -------------------------------------------------------------------------------- [209120]
MH025917.1_strain_R7            -------------------------------------------------------------------------------- [209120]
MH025916.1_strain_R8            -------------------------------------------------------------------------------- [209120]

FR682468.2_Georgia_2007/1       cgatattatataaccttactgaagccatccaatatttttatcaacgatataggcatttt------------aaagattgg [209200]
ASF_IND_20_CAD543               -----------------------------------------------------------------------...a.a..t [209200]
LR899193.1_Germany_2020/1       ...........................................................------------......... [209200]
MN393477.1_Wuhan_2019           ...........................................................------------......... [209200]
MW396979.1_Timor-Leste/2019/1   ...........................................................------------......... [209200]
MK940252.1_CN/2019/InnerMongolia ...........................................................------------......... [209200]
MK128995.1_China/2018/AnhuiXCGQ ...........................................................------------......... [209200]
MH681419.1_ASFV/POL/2015        ...........................................................------------......... [209200]
LS478113.1_Estonia_2014         ...........................................................------------......... [209200]
KP843857.1_Odintsovo_02/14      ...........................................................------------......... [209200]
FN557520.1_strain_E75           ---------------------------------g..c...ggg.a.g.tc.actcc...ttctcgaataaa...------ [209200]
AY261366.1_Warthog              ................a........t...........c...g..aa...c.a.......------------...a..... [209200]
AY261365.1_Warmbaths            ................a........t...........c...g..aa...c.a.......------------......... [209200]
AY261364.1_Tengani_62           .........................t..t........c......ac...c.a.......------------......... [209200]
AY261363.1_Pretoriuskop/96/4    ................a........t...........c...g..aa...c.a.......------------...a..... [209200]
AY261362.1_Mkuzi_1979           ................a........t...........c...g..aa...c.a.......------------......... [209200]
AY261361.1_Malawi_Lil-20/1_1983 --------------------------------------------------------------------cgc.gc.gccc. [209200]
AY261360.1_Kenya_1950           -------------------------------------------------------------------------------- [209200]
AM712240.1_OURT_88/3            -------------------------------------------------------------------------------- [209200]
AM712239.1_Benin_97/1           ---------------------------------g..c...ggg.a.g.tc.actcc...ttctcgaataaa...tc..ac [209200]
KM111295.1_Ken06.Bus            -------------------------------------------------------------------------------- [209200]
```

```
KM111294.1_Ken05/Tk1             tag.tca..................t..t...............aa.....a...g...------------......... [209200]
KM262844.1_strain_L60            ---------------------------------g..c...ggg.a.g.tc.actcc...ttctcgaataaa...tc..ac [209200]
KM102979.1_26544/OG10_Italy      ---------------------------------g..c...ggg.a.g.tc.actcc...ttctcgaataaa...tc..ac [209200]
KM262845.1_strain_NHV            -------------------------------------------------------------------------------- [209200]
KX354450.1_47/Ss/2008            ---------------------------------g..c...ggg.a.g.tc.actcc...ttctcgaataaa...tc..ac [209200]
KP055815.1_strain_BA71           ---------------------------------g..c...ggg.a.g.tc.actcc...ttctcgaataaa...tc..ac [209200]
MH025920.1_strain_R35            -------------------------------------------------------------------------------- [209200]
MH025919.1_strain_N10            -------------------------------------------------------------------------------- [209200]
MH025918.1_strain_R25            -------------------------------------------------------------------------------- [209200]
MH025917.1_strain_R7             -------------------------------------------------------------------------------- [209200]
MH025916.1_strain_R8             -------------------------------------------------------------------------------- [209200]

FR682468.2_Georgia_2007/1        cggcttatatgtg------------------------------------------------------------------- [209280]
ASF_IND_20_CAD543                ta....t..at..ttgnnnnnnnnnnnnnnnnnnnnnnnnnnnnnnnnnnnnnnnnnnnnnnnnnnnnnnnnnnnnnnnn [209280]
LR899193.1_Germany_2020/1        .............------------------------------------------------------------------- [209280]
MN393477.1_Wuhan_2019            .............------------------------------------------------------------------- [209280]
MW396979.1_Timor-Leste/2019/1    .............------------------------------------------------------------------- [209280]
MK940252.1_CN/2019/InnerMongolia .............------------------------------------------------------------------- [209280]
MK128995.1_China/2018/AnhuiXCGQ  .............------------------------------------------------------------------- [209280]
MH681419.1_ASFV/POL/2015         .............------------------------------------------------------------------- [209280]
LS478113.1_Estonia_2014          .............------------------------------------------------------------------- [209280]
KP843857.1_Odintsovo_02/14       .............------------------------------------------------------------------- [209280]
FN557520.1_strain_E75            -------------------------------------------------------------------------------- [209280]
AY261366.1_Warthog               .....a.......------------------------------------------------------------------- [209280]
AY261365.1_Warmbaths             .....a.......------------------------------------------------------------------- [209280]
AY261364.1_Tengani_62            .....a.......------------------------------------------------------------------- [209280]
AY261363.1_Pretoriuskop/96/4     .....a.......------------------------------------------------------------------- [209280]
AY261362.1_Mkuzi_1979            .....a.......------------------------------------------------------------------- [209280]
AY261361.1_Malawi_Lil-20/1_1983  .a..---------------------------------------------------------------------------- [209280]
AY261360.1_Kenya_1950            -------------------------------------------------------------------------------- [209280]
AM712240.1_OURT_88/3             -------------------------------------------------------------------------------- [209280]
AM712239.1_Benin_97/1            .ct.ca.a.ct.a------------------------------------------------------------------- [209280]
KM111295.1_Ken06.Bus             -------------------------------------------------------------------------------- [209280]
KM111294.1_Ken05/Tk1             ..a..a.......------------------------------------------------------------------- [209280]
KM262844.1_strain_L60            .ct.ca.a.ct.a------------------------------------------------------------------- [209280]
KM102979.1_26544/OG10_Italy      .ct.ca.a.ct.a------------------------------------------------------------------- [209280]
KM262845.1_strain_NHV            -------------------------------------------------------------------------------- [209280]
KX354450.1_47/Ss/2008            .ct.ca.a.ct.a------------------------------------------------------------------- [209280]
KP055815.1_strain_BA71           .ct.ca.a.ct.a------------------------------------------------------------------- [209280]
MH025920.1_strain_R35            -------------------------------------------------------------------------------- [209280]
MH025919.1_strain_N10            -------------------------------------------------------------------------------- [209280]
MH025918.1_strain_R25            -------------------------------------------------------------------------------- [209280]
MH025917.1_strain_R7             -------------------------------------------------------------------------------- [209280]
MH025916.1_strain_R8             -------------------------------------------------------------------------------- [209280]
```

```
FR682468.2_Georgia_2007/1        ----ggctttcttttaacaatttgt---------------ctgaccttcatgaagtatataacttagagaagacggatat [209360]
ASF_IND_20_CAD543                nnnnnnn...t....tggg....t.aggnnnnnnnnnnnnnnnnnnnnnnnnnnnnnnnnnnnnnnnnnnnnnnnnnnnn [209360]
LR899193.1_Germany_2020/1        ----.....................---------------........................................ [209360]
MN393477.1_Wuhan_2019            ----.....................---------------........................................ [209360]
MW396979.1_Timor-Leste/2019/1    ----.....................---------------........................................ [209360]
MK940252.1_CN/2019/InnerMongolia ----.....................---------------........................................ [209360]
MK128995.1_China/2018/AnhuiXCGQ  ----.....................---------------........................................ [209360]
MH681419.1_ASFV/POL/2015         ----.....................---------------........................................ [209360]
LS478113.1_Estonia_2014          ----.....................---------------........................................ [209360]
KP843857.1_Odintsovo_02/14       ----.....................---------------........................................ [209360]
FN557520.1_strain_E75            -------------------------------------------------------------------------------- [209360]
AY261366.1_Warthog               ----.a...............c...---------------....t...t.....a.......t.........gtt..... [209360]
AY261365.1_Warmbaths             ----.................c...---------------..............a.................gtt..... [209360]
AY261364.1_Tengani_62            ----.a...g...a...t..cg...---------------t...t.........a.t......a....a.....t...t. [209360]
AY261363.1_Pretoriuskop/96/4     ----.a...............c...---------------....t...t.....a.......t.........gtt..... [209360]
AY261362.1_Mkuzi_1979            ----.................c...---------------..............a.................gtt..... [209360]
AY261361.1_Malawi_Lil-20/1_1983  -------------------------------------------------------------------------------- [209360]
AY261360.1_Kenya_1950            -------------------------------------------------------------------------------- [209360]
AM712240.1_OURT_88/3             -------------------------------------------------------------------------------- [209360]
AM712239.1_Benin_97/1            ----atg...t....tggg....t.---------------ag.tg.aa....g.tc..tgg...acc.ag..gta.g.c. [209360]
KM111295.1_Ken06.Bus             -------------------------------------------------------------------------------- [209360]
KM111294.1_Ken05/Tk1             ----....cg...a......cg...---------------t...t.........t........aag......gtt.g... [209360]
KM262844.1_strain_L60            ----atg...t....tggg....t.---------------ag.tg.ac....g.tc..tgg...acc.ag..g.a.g.c. [209360]
KM102979.1_26544/OG10_Italy      ----atg-..t....tggg....t.---------------ag.tg.aa....g.tc..tgg...acc.ag..gta.g.c. [209360]
KM262845.1_strain_NHV            -------------------------------------------------------------------------------- [209360]
KX354450.1_47/Ss/2008            ----atg-..t....tggg....t.---------------ag.tg.aa....g.tc..tgg...acc.ag..gta.g.c. [209360]
KP055815.1_strain_BA71           ----atg...t....tggg....t.---------------ag.tg.aa....g.tc..tgg...acc.ag..g.a.g.c. [209360]
MH025920.1_strain_R35            -------------------------------------------------------------------------------- [209360]
MH025919.1_strain_N10            -------------------------------------------------------------------------------- [209360]
MH025918.1_strain_R25            -------------------------------------------------------------------------------- [209360]
MH025917.1_strain_R7             -------------------------------------------------------------------------------- [209360]
MH025916.1_strain_R8             -------------------------------------------------------------------------------- [209360]

FR682468.2_Georgia_2007/1        agacattgatgaaatgatgaagttgacctgtagtacgtatgatggtaattattcgactatttattattgttttatgttgg [209440]
ASF_IND_20_CAD543                nnn.........c.------------------------------------------------------------------ [209440]
LR899193.1_Germany_2020/1        ................................................................................ [209440]
MN393477.1_Wuhan_2019            ................................................................................ [209440]
MW396979.1_Timor-Leste/2019/1    ................................................................................ [209440]
MK940252.1_CN/2019/InnerMongolia ................................................................................ [209440]
MK128995.1_China/2018/AnhuiXCGQ  ................................................................................ [209440]
MH681419.1_ASFV/POL/2015         ................................................................................ [209440]
LS478113.1_Estonia_2014          ................................................................................ [209440]
KP843857.1_Odintsovo_02/14       ................................................................................ [209440]
```

```
FN557520.1_strain_E75            -------------------------------------------------------------------------------- [209440]
AY261366.1_Warthog               g..................t.tc..g.........t.....g............a..................g.a.... [209440]
AY261365.1_Warmbaths             g.........at.......t.tc..g............................a..................g.a.... [209440]
AY261364.1_Tengani_62            ...t...............c.....g.t..---..t.......a..........a......................... [209440]
AY261363.1_Pretoriuskop/96/4     ga.................t.tc..g.........ta.................a..................g.a.... [209440]
AY261362.1_Mkuzi_1979            g.........at.......t.tc..g............................a..................g.a.... [209440]
AY261361.1_Malawi_Lil-20/1_1983  ------------------------.g..c.c.....-------------------------------------------- [209440]
AY261360.1_Kenya_1950            -------------------------------------------------------------------------------- [209440]
AM712240.1_OURT_88/3             -------------------------------------------------------------------------------- [209440]
AM712239.1_Benin_97/1            .--------------------------------------------------------...c..c.ta.....ctca...- [209440]
KM111295.1_Ken06.Bus             -------------------------------------------------------------------------------- [209440]
KM111294.1_Ken05/Tk1             g..t...a.cc........g.ac..g....---c.ta...agata.........c..c..................c... [209440]
KM262844.1_strain_L60            .--------------------------------------------------------...c..c.ta.....ctca...- [209440]
KM102979.1_26544/OG10_Italy      .--------------------------------------------------------...c..c.ta.....ctca...- [209440]
KM262845.1_strain_NHV            -------------------------------------------------------------------------------- [209440]
KX354450.1_47/Ss/2008            .--------------------------------------------------------...c..c.ta.....ctca...- [209440]
KP055815.1_strain_BA71           .--------------------------------------------------------...c..c.ta.....ctca...- [209440]
MH025920.1_strain_R35            -------------------------------------------------------------------------------- [209440]
MH025919.1_strain_N10            -------------------------------------------------------------------------------- [209440]
MH025918.1_strain_R25            -------------------------------------------------------------------------------- [209440]
MH025917.1_strain_R7             -------------------------------------------------------------------------------- [209440]
MH025916.1_strain_R8             -------------------------------------------------------------------------------- [209440]

FR682468.2_Georgia_2007/1        gggctgacatcaatcgggcaatgttaacctcggtaataaactttcatattggtaacttgttcctttgtatagatttagga [209520]
ASF_IND_20_CAD543                ---------------------------------------------...gcat..nn...c.atc..a..c.t..cnnnnn [209520]
LR899193.1_Germany_2020/1        ................................................................................ [209520]
MN393477.1_Wuhan_2019            ................................................................................ [209520]
MW396979.1_Timor-Leste/2019/1    ................................................................................ [209520]
MK940252.1_CN/2019/InnerMongolia ................................................................................ [209520]
MK128995.1_China/2018/AnhuiXCGQ  ................................................................................ [209520]
MH681419.1_ASFV/POL/2015         ................................................................................ [209520]
LS478113.1_Estonia_2014          ................................................................................ [209520]
KP843857.1_Odintsovo_02/14       ................................................................................ [209520]
FN557520.1_strain_E75            -------------------------------------------------------------------------------- [209520]
AY261366.1_Warthog               .......t..t....aa........................ca........a..........t................g [209520]
AY261365.1_Warmbaths             .......t.......a.........................ca..g.................................g [209520]
AY261364.1_Tengani_62            .....a.t.......a.........................ca.t.....aa........................g... [209520]
AY261363.1_Pretoriuskop/96/4     .a.....t.......a...............a.........ca.tg.................................g [209520]
AY261362.1_Mkuzi_1979            .......t.......a.........................ca..g.................................g [209520]
AY261361.1_Malawi_Lil-20/1_1983  -------------------------------------------------------------------------------- [209520]
AY261360.1_Kenya_1950            -------------------------------------------------------------------------------- [209520]
AM712240.1_OURT_88/3             -------------------------------------------------------------------------------- [209520]
AM712239.1_Benin_97/1            ---------------------------------------.tgaca...g.at...t...c.atc..a..c.t..c.g--- [209520]
KM111295.1_Ken06.Bus             -------------------------------------------------------------------------------- [209520]
```

```
KM111294.1_Ken05/Tk1            ..........t............a.t.....a..t.c...a.c.t...c.............t................g [209520]
KM262844.1_strain_L60           ---------------------------------------.tgaca...gcat...t...c.atc..a..c.t..c.g--- [209520]
KM102979.1_26544/OG10_Italy     ---------------------------------------.tgaca...g.at...t...c.atc..a..c.t..c.g--- [209520]
KM262845.1_strain_NHV           -------------------------------------------------------------------------------- [209520]
KX354450.1_47/Ss/2008           ---------------------------------------.tgaca...g.at...t...c.atc..a..c.t..c.g--- [209520]
KP055815.1_strain_BA71          ---------------------------------------.tgaca...gcat...t...c.atc..a..c.t..c.g--- [209520]
MH025920.1_strain_R35           -------------------------------------------------------------------------------- [209520]
MH025919.1_strain_N10           -------------------------------------------------------------------------------- [209520]
MH025918.1_strain_R25           -------------------------------------------------------------------------------- [209520]
MH025917.1_strain_R7            -------------------------------------------------------------------------------- [209520]
MH025916.1_strain_R8            -------------------------------------------------------------------------------- [209520]

FR682468.2_Georgia_2007/1       gctgatgctttcgaagacagcatggaactagcaaaacaaaagaataataatatattagtagaaatattatcatttaaaaa [209600]
ASF_IND_20_CAD543               nnnnnnnnnnnnnnnnnnnnnnnnnnnnnnnnnnnnnnnnnnnnnnnnnnnnnnnnnnnnnnnnnnnnnnnnnnnnnnnn [209600]
LR899193.1_Germany_2020/1       ................................................................................ [209600]
MN393477.1_Wuhan_2019           ................................................................................ [209600]
MW396979.1_Timor-Leste/2019/1   ................................................................................ [209600]
MK940252.1_CN/2019/InnerMongolia ................................................................................ [209600]
MK128995.1_China/2018/AnhuiXCGQ ................................................................................ [209600]
MH681419.1_ASFV/POL/2015        ................................................................................ [209600]
LS478113.1_Estonia_2014         ................................................................................ [209600]
KP843857.1_Odintsovo_02/14      ................................................................................ [209600]
FN557520.1_strain_E75           -------------------------------------------------------------------------------- [209600]
AY261366.1_Warthog              ........c........g..t........................c..............c.t........g........ [209600]
AY261365.1_Warmbaths            ........c...........t...a....................c..............c.t........g........ [209600]
AY261364.1_Tengani_62           .....c..cg..a................................t..............a........a.......... [209600]
AY261363.1_Pretoriuskop/96/4    ...................................g.....a...g.......t..ta.tagt................. [209600]
AY261362.1_Mkuzi_1979           ........c...........t...a....................c..............c.t........g........ [209600]
AY261361.1_Malawi_Lil-20/1_1983 -------------------------------------------------------------------------------- [209600]
AY261360.1_Kenya_1950           -------------------------------------------------------------------------------- [209600]
AM712240.1_OURT_88/3            -------------------------------------------------------------------------------- [209600]
AM712239.1_Benin_97/1           ---------------------------t.gt..t.cg....a..acg.tg.----------------------------- [209600]
KM111295.1_Ken06.Bus            -------------------------------------------------------------------------------- [209600]
KM111294.1_Ken05/Tk1            ...ac...c........gt.tt.a...a......t....c.........t.ga...ta..a........at..a...... [209600]
KM262844.1_strain_L60           ---------------------------t.gt..t.cg....a..acg.tg.----------------------------- [209600]
KM102979.1_26544/OG10_Italy     ---------------------------t.gt..t.cg....a..acg.tg.----------------------------- [209600]
KM262845.1_strain_NHV           -------------------------------------------------------------------------------- [209600]
KX354450.1_47/Ss/2008           ---------------------------t.gt..t.cg....a..acg.tg.----------------------------- [209600]
KP055815.1_strain_BA71          ---------------------------t.gt..t.cg....a..acg.tg.----------------------------- [209600]
MH025920.1_strain_R35           -------------------------------------------------------------------------------- [209600]
MH025919.1_strain_N10           -------------------------------------------------------------------------------- [209600]
MH025918.1_strain_R25           -------------------------------------------------------------------------------- [209600]
MH025917.1_strain_R7            -------------------------------------------------------------------------------- [209600]
MH025916.1_strain_R8            -------------------------------------------------------------------------------- [209600]
```

```
FR682468.2_Georgia_2007/1         ttattatagttcaaatacctctcttttatcaataaaaacgacagatccggaaaaaattaatgccttattagatgaagaaa [209680]
ASF_IND_20_CAD543                 nnnnnnnnnnnnnnnnnnnnnnnnnnnnnnnnnnnnnnnnnnnnnnnnnnnnnnnnnnnnnnnnnnnnnnnnnnnnnnnn [209680]
LR899193.1_Germany_2020/1         ................................................................................ [209680]
MN393477.1_Wuhan_2019             ................................................................................ [209680]
MW396979.1_Timor-Leste/2019/1     ................................................................................ [209680]
MK940252.1_CN/2019/InnerMongolia  ................................................................................ [209680]
MK128995.1_China/2018/AnhuiXCGQ   ................................................................................ [209680]
MH681419.1_ASFV/POL/2015          ................................................................................ [209680]
LS478113.1_Estonia_2014           ................................................................................ [209680]
KP843857.1_Odintsovo_02/14        ................................................................................ [209680]
FN557520.1_strain_E75             -------------------------------------------------------------------------------- [209680]
AY261366.1_Warthog                .---......c..g.ctt...c.........t......t......................................... [209680]
AY261365.1_Warmbaths              c---......c..g.ctt...c..........................a..g...............c............ [209680]
AY261364.1_Tengani_62             .---......c..g.ct.t............t...................................c............ [209680]
AY261363.1_Pretoriuskop/96/4      .---..c...c..g.ct.t........g...t......t............................c............ [209680]
AY261362.1_Mkuzi_1979             c---......c..g.ctt...c..........................a..g...............c............ [209680]
AY261361.1_Malawi_Lil-20/1_1983   -------------------------------------------.gc...a...tc.....-------------------- [209680]
AY261360.1_Kenya_1950             -------------------------------------------------------------------------------- [209680]
AM712240.1_OURT_88/3              -------------------------------------------------------------------------------- [209680]
AM712239.1_Benin_97/1             ------------------------------------------------------t.c...c..gc.gcc----------- [209680]
KM111295.1_Ken06.Bus              -------------------------------------------------------------------------------- [209680]
KM111294.1_Ken05/Tk1              ..........c..g...g...c.........t......ta...........g.........at............t.... [209680]
KM262844.1_strain_L60             ------------------------------------------------------t.c...c..gc.gcc----------- [209680]
KM102979.1_26544/OG10_Italy       ------------------------------------------------------t.c...c..gc.gcc----------- [209680]
KM262845.1_strain_NHV             -------------------------------------------------------------------------------- [209680]
KX354450.1_47/Ss/2008             ------------------------------------------------------t.c...c..gc.gcc----------- [209680]
KP055815.1_strain_BA71            ------------------------------------------------------t.c...c..gc.gcc----------- [209680]
MH025920.1_strain_R35             -------------------------------------------------------------------------------- [209680]
MH025919.1_strain_N10             -------------------------------------------------------------------------------- [209680]
MH025918.1_strain_R25             -------------------------------------------------------------------------------- [209680]
MH025917.1_strain_R7              -------------------------------------------------------------------------------- [209680]
MH025916.1_strain_R8              -------------------------------------------------------------------------------- [209680]

FR682468.2_Georgia_2007/1         agtatgagtcaaaaaatatgttaatgtatgaagaattatctcattgatacaaaattattt----tttataacagaactct [209760]
ASF_IND_20_CAD543                 nnnnnnnnnnnnnnnnnnnnnnnnnnnnnnnnnnnnnnnnnnnnnnnnnnnnnnnnnnnn----nnnnnnnnnnnnnnnn [209760]
LR899193.1_Germany_2020/1         ............................................................----................ [209760]
MN393477.1_Wuhan_2019             ............................................................----................ [209760]
MW396979.1_Timor-Leste/2019/1     ............................................................----................ [209760]
MK940252.1_CN/2019/InnerMongolia  ............................................................----................ [209760]
MK128995.1_China/2018/AnhuiXCGQ   ............................................................----................ [209760]
MH681419.1_ASFV/POL/2015          ............................................................----................ [209760]
LS478113.1_Estonia_2014           ............................................................----................ [209760]
KP843857.1_Odintsovo_02/14        ............................................................----................ [209760]
```

```
FN557520.1_strain_E75            -------------------------------------------------------------------------------- [209760]
AY261366.1_Warthog               ...............................t.....tgacgcc..tac----------------------t.at.a... [209760]
AY261365.1_Warmbaths             ...............................t.....tgacgcc..tact..t.a.g...aatgcc..c.gt.at.a... [209760]
AY261364.1_Tengani_62            ..........g.........................c.....g.................--tt.a..ac.g..cc.... [209760]
AY261363.1_Pretoriuskop/96/4     ...........................t..-------------------------------------------------- [209760]
AY261362.1_Mkuzi_1979            ...............................t.....tgacgcc..tact..t.a.g...aatgcc..c.gt.at.a... [209760]
AY261361.1_Malawi_Lil-20/1_1983  -------------------------------------------------------------------------------- [209760]
AY261360.1_Kenya_1950            -------------------------------------------------------------------------------- [209760]
AM712240.1_OURT_88/3             -------------------------------------------------------------------------------- [209760]
AM712239.1_Benin_97/1            -------t.....c..c...-------------------------.cgct.----------------------------- [209760]
KM111295.1_Ken06.Bus             -------------------------------------------------------------------------------- [209760]
KM111294.1_Ken05/Tk1             ca...............ga......a...a..a...c---..........t..t..t...-------------------- [209760]
KM262844.1_strain_L60            -------t.....c..c...-------------------------.cgct..t.cc..acacgcc..ca....ac.---- [209760]
KM102979.1_26544/OG10_Italy      -------t.....c..c...-------------------------.cgct..t.cc..acacgcc..ca....ac..gg. [209760]
KM262845.1_strain_NHV            -------------------------------------------------------------------------------- [209760]
KX354450.1_47/Ss/2008            -------t.....c..c...-------------------------.cgct..t.cc..acacgcc..ca....ac..gg. [209760]
KP055815.1_strain_BA71           -------t.....c..c...-------------------------.cgct..t.cc..acacgcc..ca....ac..gg. [209760]
MH025920.1_strain_R35            -------------------------------------------------------------------------------- [209760]
MH025919.1_strain_N10            -------------------------------------------------------------------------------- [209760]
MH025918.1_strain_R25            -------------------------------------------------------------------------------- [209760]
MH025917.1_strain_R7             -------------------------------------------------------------------------------- [209760]
MH025916.1_strain_R8             -------------------------------------------------------------------------------- [209760]

FR682468.2_Georgia_2007/1        ctgatggtgacaaatctccgat--------------aggaatatatgacgtaacataattattt---------------- [209840]
ASF_IND_20_CAD543                nnnnnnnnnnnnnnnnnnnnnn--------------nnnnnnnnnnnnnnnnnnnnnnnnnnnn---------------- [209840]
LR899193.1_Germany_2020/1        ......................--------------............................---------------- [209840]
MN393477.1_Wuhan_2019            ......................--------------............................---------------- [209840]
MW396979.1_Timor-Leste/2019/1    ......................--------------............................---------------- [209840]
MK940252.1_CN/2019/InnerMongolia ......................--------------............................---------------- [209840]
MK128995.1_China/2018/AnhuiXCGQ  ......................--------------............................---------------- [209840]
MH681419.1_ASFV/POL/2015         ......................--------------............................---------------- [209840]
LS478113.1_Estonia_2014          ......................--------------............................---------------- [209840]
KP843857.1_Odintsovo_02/14       ......................--------------............................---------------- [209840]
FN557520.1_strain_E75            -------------------------------------------------------------------------------- [209840]
AY261366.1_Warthog               t.....cctgt.gtataata..tgatacctacagcgc.cc...c.c..t..c.ag.cgc.g..ctcaatattatattaat [209840]
AY261365.1_Warmbaths             t.a.c.cctgt.gtataata..tgatacctacagtgc.cc...c.c.gt..c.ag.cgc.g..ctcaatattatattgat [209840]
AY261364.1_Tengani_62            .....a...........t....agg-------------------....................---------------- [209840]
AY261363.1_Pretoriuskop/96/4     -------------------------------------------------------------------------------- [209840]
AY261362.1_Mkuzi_1979            t.a.c.cctgt.gtataata..tgatacctacagtgc.cc...c.c.gt..c.ag.cgc.g..ctcaatattatattgat [209840]
AY261361.1_Malawi_Lil-20/1_1983  -------------------------------------------------------------------------------- [209840]
AY261360.1_Kenya_1950            -------------------------------------------------------------------------------- [209840]
AM712240.1_OURT_88/3             -------------------------------------------------------------------------------- [209840]
AM712239.1_Benin_97/1            -------------------------------------------------------------------------------- [209840]
KM111295.1_Ken06.Bus             -------------------------------------------------------------------------------- [209840]
```

```
KM111294.1_Ken05/Tk1              -------------------------------------------------------------------------------- [209840]
KM262844.1_strain_L60             -------------------------------------------------------------------------------- [209840]
KM102979.1_26544/OG10_Italy       gcc.c-------------------------------cacc...c.c..ta.c.gg..gc.g.ccccaataatgtgttgat [209840]
KM262845.1_strain_NHV             -------------------------------------------------------------------------------- [209840]
KX354450.1_47/Ss/2008             gcc.c-------------------------------cacc...c.c..ta.c.gg..gc.g.ccccaataatgtgttgat [209840]
KP055815.1_strain_BA71            gcc.c-------------------------------cacc...c.c..ta.c.gg..gc.g.ccccaataatgtgttgat [209840]
MH025920.1_strain_R35             -------------------------------------------------------------------------------- [209840]
MH025919.1_strain_N10             -------------------------------------------------------------------------------- [209840]
MH025918.1_strain_R25             -------------------------------------------------------------------------------- [209840]
MH025917.1_strain_R7              -------------------------------------------------------------------------------- [209840]
MH025916.1_strain_R8              -------------------------------------------------------------------------------- [209840]

FR682468.2_Georgia_2007/1         -------------------------------------------------------------------------------- [209920]
ASF_IND_20_CAD543                 -------------------------------------------------------------------------------- [209920]
LR899193.1_Germany_2020/1         -------------------------------------------------------------------------------- [209920]
MN393477.1_Wuhan_2019             -------------------------------------------------------------------------------- [209920]
MW396979.1_Timor-Leste/2019/1     -------------------------------------------------------------------------------- [209920]
MK940252.1_CN/2019/InnerMongolia  -------------------------------------------------------------------------------- [209920]
MK128995.1_China/2018/AnhuiXCGQ   -------------------------------------------------------------------------------- [209920]
MH681419.1_ASFV/POL/2015          -------------------------------------------------------------------------------- [209920]
LS478113.1_Estonia_2014           -------------------------------------------------------------------------------- [209920]
KP843857.1_Odintsovo_02/14        -------------------------------------------------------------------------------- [209920]
FN557520.1_strain_E75             -------------------------------------------------------------------------------- [209920]
AY261366.1_Warthog                tgacgctgttctcaataatataatgcgctcatatgcgaaaaaatagtgttgattaacgcattataaa------------a [209920]
AY261365.1_Warmbaths              tgacgctcttcgtaataata----acgctcatatgcg-aaaaataatgttgattaacgcatcataaa------------a [209920]
AY261364.1_Tengani_62             -------------------------------------------------------------------------------- [209920]
AY261363.1_Pretoriuskop/96/4      ------------------------acgcccac------------------------------------------------ [209920]
AY261362.1_Mkuzi_1979             tgacgctcttcgtaataata----acgctcatatgcg-aaaaataatgttgattaacgcatcataaa------------a [209920]
AY261361.1_Malawi_Lil-20/1_1983   --------------------------------------gagtacgggggcggcgggggctcctcggg------------- [209920]
AY261360.1_Kenya_1950             --------------------------------------aaaaaa-attaaagtgatcgcgtcataat------------- [209920]
AM712240.1_OURT_88/3              -------------------------------------------------------------------------------- [209920]
AM712239.1_Benin_97/1             -------------------------------------------------------------------------------- [209920]
KM111295.1_Ken06.Bus              -------------------------------------------------------------------------------- [209920]
KM111294.1_Ken05/Tk1              ----------------------------tcaggcctgaaaaaaaaattaaagtgatcgcgtcataat------------- [209920]
KM262844.1_strain_L60             -------------------------------------------------------------------------------- [209920]
KM102979.1_26544/OG10_Italy       tcacgcccatag--------------------------aatgctgtgattgattgatgc-ccataaaatggtgtgattga [209920]
KM262845.1_strain_NHV             -------------------------------------------------------------------------------- [209920]
KX354450.1_47/Ss/2008             tcacgcccatag--------------------------aatgctgtgattgattgatgc-ccataaaatggtgtgattga [209920]
KP055815.1_strain_BA71            tcacgcccatag--------------------------aatgctgtgattgattgatgc-ccataaaatggtgtgattga [209920]
MH025920.1_strain_R35             --------------------------------------aaaaaaaaataaagtgatcgtgttataat------------- [209920]
MH025919.1_strain_N10             --------------------------------------aaaaaaaaataaagtgatcgtgttataat------------- [209920]
MH025918.1_strain_R25             --------------------------------------aaaaaaaaataaagtgatcgtgttataat------------- [209920]
MH025917.1_strain_R7              --------------------------------------aaaaaaaaataaagtgatcgtgttataat------------- [209920]
MH025916.1_strain_R8              --------------------------------------aaaaaaaaataaagtgatcgtgttataat------------- [209920]
```

```
FR682468.2_Georgia_2007/1        ------------------------------ttttcgcccagaaaaaaattataaatgttattattgccagcacttttatc [210000]
ASF_IND_20_CAD543                ------------------------------nnnnnnnnnnnn.g.ttt..g..t..------------------------ [210000]
LR899193.1_Germany_2020/1        ------------------------------.................................................. [210000]
MN393477.1_Wuhan_2019            ------------------------------.................................................. [210000]
MW396979.1_Timor-Leste/2019/1    ------------------------------.................................................. [210000]
MK940252.1_CN/2019/InnerMongolia ------------------------------.................................................. [210000]
MK128995.1_China/2018/AnhuiXCGQ  ------------------------------.................................................. [210000]
MH681419.1_ASFV/POL/2015         ------------------------------.................................................. [210000]
LS478113.1_Estonia_2014          ------------------------------.................................................. [210000]
KP843857.1_Odintsovo_02/14       ------------------------------.................................................. [210000]
FN557520.1_strain_E75            -------------------------------------------------------------------------------- [210000]
AY261366.1_Warthog               tgctgtgcccacttattata--tatgttga..ga...--------------------------------------cc.ct [210000]
AY261365.1_Warmbaths             t-taatacccacttataataataatgttga..ga...atcat....tgc.g.ggt..--...g.c...ta.---.....t [210000]
AY261364.1_Tengani_62            ------------------------------.....tgggc.......g.c.......a....g......a.......... [210000]
AY261363.1_Pretoriuskop/96/4     -------------------------------------------------------------------------------- [210000]
AY261362.1_Mkuzi_1979            t-taatacccacttataataataatgttga..ga...atcat....tgc.g.ggt..--...g.c...ta.---.....t [210000]
AY261361.1_Malawi_Lil-20/1_1983  -------------------------------------------------------------------------------- [210000]
AY261360.1_Kenya_1950            --taacatttttttgta--------ctcgga.c.tagat.t......g.....t.....g..g..at..c...a.....t [210000]
AM712240.1_OURT_88/3             -------------------------------------------------------------------------------- [210000]
AM712239.1_Benin_97/1            -------------------------------------------------------------------------------- [210000]
KM111295.1_Ken06.Bus             -------------------------------------------------------------------------------- [210000]
KM111294.1_Ken05/Tk1             --taacatttttttgta--------ctcgga.c.taggt.t.....gg.g...t........g...ttgc...a.....t [210000]
KM262844.1_strain_L60            -------------------------------------------------------------------------------- [210000]
KM102979.1_26544/OG10_Italy      t-tgacgccca--taaaatggtgtgattga..ca...-.cat.g..tgc.g.g.t..--...ga....ca-----.a.aa [210000]
KM262845.1_strain_NHV            -------------------------------------------------------------------------------- [210000]
KX354450.1_47/Ss/2008            t-tgacgccca--taaaatggtgtgattga..ca...-.cat.g..tgc.g.g.t..--...ga....ca-----.a.aa [210000]
KP055815.1_strain_BA71           t-tgacgccca--taaaatggtgtgattga..ca...-.cat.g..tgc.g.g.t..--...ga....ca-----.a.aa [210000]
MH025920.1_strain_R35            --taacatttttttgta--------ctcgga.c.tagat.t.g...gg.g...t.....g..g..at..t...a.....t [210000]
MH025919.1_strain_N10            --taacatttttttgta--------ctcgga.c.tagat.t.g...gg.g...t.....g..g..at..t...a.....t [210000]
MH025918.1_strain_R25            --taacatttttttgta--------ctcgga.c.tagat.t.g...gg.g...t.....g..g..at..t...a.....t [210000]
MH025917.1_strain_R7             --taacatttttttgta--------ctcgga.c.tagat.t.g...gg.g...t.....g..g..at..t...a.....t [210000]
MH025916.1_strain_R8             --taacatttttttgta--------ctcgga.c.tagat.t.g...gg.g...t.....g..g..at..t...a.....t [210000]

FR682468.2_Georgia_2007/1        aactatacgta-----------------caaaaaggtgttgacca-aaaaaat-aattttttttcttgatcaaagtatgt [210080]
ASF_IND_20_CAD543                --------------------------------------------------------------........---------- [210080]
LR899193.1_Germany_2020/1        ...........-----------------.................-.......-.......................... [210080]
MN393477.1_Wuhan_2019            ...........-----------------.................-.......-.......................... [210080]
MW396979.1_Timor-Leste/2019/1    ...........-----------------.................-.......-.......................... [210080]
MK940252.1_CN/2019/InnerMongolia ...........-----------------.................-.......-.......................... [210080]
MK128995.1_China/2018/AnhuiXCGQ  ...........-----------------.................-.......-.......................... [210080]
MH681419.1_ASFV/POL/2015         ...........-----------------.................-.......-.......................... [210080]
LS478113.1_Estonia_2014          ...........-----------------.................-.......-.......................... [210080]
KP843857.1_Odintsovo_02/14       ...........-----------------.................-.......-.......................... [210080]
```

```
FN557520.1_strain_E75             -------------------------------------------------------------------------------- [210080]
AY261366.1_Warthog                t.ta...a.--attttttttacata--------------..gt-----g...a-t........c---------------- [210080]
AY261365.1_Warmbaths              tgt....a...atgatttttgtataaaa..tggg.c....ac.-----.....-t......................... [210080]
AY261364.1_Tengani_62             ...........-----------------...............-----c...a-..aa......t..a....-------c [210080]
AY261363.1_Pretoriuskop/96/4      -------------------------------------------------------------------------------- [210080]
AY261362.1_Mkuzi_1979             tgt....a...atgatttttgtataaaa..tggg.c....ac.-----.....-t......................... [210080]
AY261361.1_Malawi_Lil-20/1_1983   -----------------------------------------------------------c....t...------------ [210080]
AY261360.1_Kenya_1950             ..g...g..cc-----------gaaaag..t..c.cc.cctt...t.gttt..-..aga.c...t...------------ [210080]
AM712240.1_OURT_88/3              -------------------------------------------------------------------------------- [210080]
AM712239.1_Benin_97/1             -------------------------------------------------------------------------------- [210080]
KM111295.1_Ken06.Bus              -------------------------------------------------------------------------------- [210080]
KM111294.1_Ken05/Tk1              ..g...g..cc-----------gaaaag..t..c.cc.cctt...t.gttt..-..aga.c...t...------------ [210080]
KM262844.1_strain_L60             -------------------------------------------------------------------------------- [210080]
KM102979.1_26544/OG10_Italy       tgg.gc..t..--gatttttgtata------------------------------------...........-------- [210080]
KM262845.1_strain_NHV             -------------------------------------------------------------------------------- [210080]
KX354450.1_47/Ss/2008             tgg.gc..t..--gatttttgtata------------------------------------...........-------- [210080]
KP055815.1_strain_BA71            tgg.gc..t..--gatttttgtata------------------------------------...........-------- [210080]
MH025920.1_strain_R35             ..g......c.-----------gaaaag..t..cacc.gctt...tggttt..a..aga.c...t...------------ [210080]
MH025919.1_strain_N10             ..g......c.-----------gaaaag..t..cacc.gctt...tggttt..a..aga.c...t...------------ [210080]
MH025918.1_strain_R25             ..g......c.-----------gaaaag..t..cacc.gctt...tggttt..a..aga.c...t...------------ [210080]
MH025917.1_strain_R7              ..g......c.-----------gaaaag..t..cacc.gctt...tggttt..a..aga.c...t...------------ [210080]
MH025916.1_strain_R8              ..g......c.-----------gaaaag..t..cacc.gctt...tggttt..a..aga.c...t...------------ [210080]

FR682468.2_Georgia_2007/1         aaacgcccgcttacagcaaggatcttaagtgagagccattaaatttta-------ttgatagctgcttg---ccaccagt [210160]
ASF_IND_20_CAD543                 ----------------------------------------------------------nnnnnnnnnnn---nnnnnnnn [210160]
LR899193.1_Germany_2020/1         ................................................-------..............---........ [210160]
MN393477.1_Wuhan_2019             ................................................-------..............---........ [210160]
MW396979.1_Timor-Leste/2019/1     ................................................-------..............---........ [210160]
MK940252.1_CN/2019/InnerMongolia  ................................................-------..............---........ [210160]
MK128995.1_China/2018/AnhuiXCGQ   ................................................-------..............---........ [210160]
MH681419.1_ASFV/POL/2015          ................................................-------..............---........ [210160]
LS478113.1_Estonia_2014           ................................................-------..............---........ [210160]
KP843857.1_Odintsovo_02/14        ................................................-------..............---........ [210160]
FN557520.1_strain_E75             -------------------------------------------------------------------------------- [210160]
AY261366.1_Warthog                -----------------------------------......---.c.g-------c..c...caa-----------.... [210160]
AY261365.1_Warmbaths              ...t..----..........a..........g..a.............-------..........a..-----tg...a. [210160]
AY261364.1_Tengani_62             .c.t..----........gca.---g...c..a.a.t..c...a....-------..ac....a.t.ga--ttt....cc [210160]
AY261363.1_Pretoriuskop/96/4      -------------------------------------------------------------------------------- [210160]
AY261362.1_Mkuzi_1979             ...t..----..........a..........g..a.............-------..........a..-----tg...a. [210160]
AY261361.1_Malawi_Lil-20/1_1983   -----------cg.c..gca..g.....cag.------------------------------------------------ [210160]
AY261360.1_Kenya_1950             ------------------------------a.a.aa............gccactt.ca.c.t..aaacaggtttg...a. [210160]
AM712240.1_OURT_88/3              -------------------------------------------------------------------------------- [210160]
AM712239.1_Benin_97/1             -------------------------------------------------------------------------------- [210160]
KM111295.1_Ken06.Bus              -------------------------------------------------------------------------------- [210160]
```

```
KM111294.1_Ken05/Tk1            -----------------------------aa.a.aa............gccgctt.ca.c.t..aaacaggtttg...a. [210160]
KM262844.1_strain_L60           -------------------------------------------------------------------------------- [210160]
KM102979.1_26544/OG10_Italy     ---------------------------..c..a.aa........ag..-------.------------------------ [210160]
KM262845.1_strain_NHV           -------------------------------------------------------------------------------- [210160]
KX354450.1_47/Ss/2008           ---------------------------..c..a.aa........ag..-------.------------------------ [210160]
KP055815.1_strain_BA71          ---------------------------..c..a.aa........ag..-------.------------------------ [210160]
MH025920.1_strain_R35           ------------------------------a.a.aa............gtcgctt.ca..gt..aag.aggtttg...a. [210160]
MH025919.1_strain_N10           ------------------------------a.a.aa............gtcgctt.ca..gt..aag.aggtttg...a. [210160]
MH025918.1_strain_R25           ------------------------------a.a.aa............gtcgctt.ca..gt..aag.aggtttg...a. [210160]
MH025917.1_strain_R7            ------------------------------a.a.aa............gtcgctt.ca..gt..aag.aggtttg...a. [210160]
MH025916.1_strain_R8            ------------------------------a.a.aa............gtcgctt.ca..gt..aag.aggtttg...a. [210160]

FR682468.2_Georgia_2007/1       agaat-acggccaaaccacctaacaggaaatacaaggcggcc---cttcggccaataaggtggataaaaatcacgcataa [210240]
ASF_IND_20_CAD543               nnnnn-nnnnnnnnnnnnnnnnnnnnn.t...t.ta.t..g.---..c.tnnnnnnnnnnnnnnnnnnnnnnnnnnnn.. [210240]
LR899193.1_Germany_2020/1       .....-....................................---................................... [210240]
MN393477.1_Wuhan_2019           .....-....................................---................................... [210240]
MW396979.1_Timor-Leste/2019/1   .....-....................................---................................... [210240]
MK940252.1_CN/2019/InnerMongolia .....-....................................---................................... [210240]
MK128995.1_China/2018/AnhuiXCGQ .....-....................................---................................... [210240]
MH681419.1_ASFV/POL/2015        .....-....................................---................................... [210240]
LS478113.1_Estonia_2014         .....-....................................---................................... [210240]
KP843857.1_Odintsovo_02/14      .....-....................................---................................... [210240]
FN557520.1_strain_E75           -------------------------------------------------------------------------------- [210240]
AY261366.1_Warthog              ..t..-..a....c..tg........................---.g..t...t.......................... [210240]
AY261365.1_Warmbaths            ..t..c.ta.......tg.......a..t...t..a.t..t.---....t.............................. [210240]
AY261364.1_Tengani_62           ..t..t.ta.......t..t............t.ca......ctt....t.........a..t...........a..... [210240]
AY261363.1_Pretoriuskop/96/4    -------------------------------------------------------------------------------- [210240]
AY261362.1_Mkuzi_1979           ..t..c.ta.......tg.......a..t...t..a.t..t.---....t.............................. [210240]
AY261361.1_Malawi_Lil-20/1_1983 ---------------------------------------------.cctc.----------------------------- [210240]
AY261360.1_Kenya_1950           .at..-..a..t...at.....g..tc.t...t..a.t.aaa---tc.ata.......aa..tt..g.g.g.t.aat... [210240]
AM712240.1_OURT_88/3            -------------------------------------------------------------------------------- [210240]
AM712239.1_Benin_97/1           -------------------------------------------------------------------------------- [210240]
KM111295.1_Ken06.Bus            -------------------------------------------------------------------------------- [210240]
KM111294.1_Ken05/Tk1            .at..-..aa.t...at.....g..tc.t...t..a.t.aaa---tc.ata.......aa..tt..g.g.g.ttaac... [210240]
KM262844.1_strain_L60           -------------------------------------------------------------------------------- [210240]
KM102979.1_26544/OG10_Italy     ------..a..t....t..t........t...t.ta.t..g.---..c.t.........a..t.....g.g.c.a..... [210240]
KM262845.1_strain_NHV           -------------------------------------------------------------------------------- [210240]
KX354450.1_47/Ss/2008           ------..a..t....t..t........t...t.ta.t..g.---..c.t.........a..t.....g.g.c.a..... [210240]
KP055815.1_strain_BA71          ------..a..t....t..t........t...t.ta.t..g.---..c.t.........a..t.....g.g.c.a..... [210240]
MH025920.1_strain_R35           ..t..-..a......at..t..g...a.t...t..a.t.aat---tc..ta.......at.tt...g.g.aataa..... [210240]
MH025919.1_strain_N10           ..t..-..a......at..t..g...a.t...t..a.t.aat---tc..ta.......at.tt...g.g.aataa..... [210240]
MH025918.1_strain_R25           ..t..-..a......at..t..g...a.t...t..a.t.aat---tc..ta.......at.tt...g.g.aataa..... [210240]
MH025917.1_strain_R7            ..t..-..a......at..t..g...a.t...t..a.t.aat---tc..ta.......at.tt...g.g.aataa..... [210240]
MH025916.1_strain_R8            ..t..-..a......at..t..g...a.t...t..a.t.aat---tc..ta.......at.tt...g.g.aataa..... [210240]
```

```
FR682468.2_Georgia_2007/1        gacggttgtaacatagcactttagtgcgaatatcaggaatgccaatagcatgtagataaggcaccaaacatcgcagctat [210320]
ASF_IND_20_CAD543                ..t....a..gt.....nnnnnnnnnn................................aa...........a...a... [210320]
LR899193.1_Germany_2020/1        ................................................................................ [210320]
MN393477.1_Wuhan_2019            ................................................................................ [210320]
MW396979.1_Timor-Leste/2019/1    ................................................................................ [210320]
MK940252.1_CN/2019/InnerMongolia ................................................................................ [210320]
MK128995.1_China/2018/AnhuiXCGQ  ................................................................................ [210320]
MH681419.1_ASFV/POL/2015         ................................................................................ [210320]
LS478113.1_Estonia_2014          ................................................................................ [210320]
KP843857.1_Odintsovo_02/14       ................................................................................ [210320]
FN557520.1_strain_E75            -------------------------------------------------------------------------------- [210320]
AY261366.1_Warthog               ..t....a...t.........c....t.................................a........g......ac.. [210320]
AY261365.1_Warmbaths             .......a...t.........c....t................................aa...............a... [210320]
AY261364.1_Tengani_62            ...........t......t........a..............t................aa...........a...a... [210320]
AY261363.1_Pretoriuskop/96/4     -------------------------------------------------------------------------------- [210320]
AY261362.1_Mkuzi_1979            .......a...t.........c....t................................aa...............a... [210320]
AY261361.1_Malawi_Lil-20/1_1983  -------a.gc.g.....ta.g.acagc.cc.g....ttg...gcc..........g..---....c.tcag.t...... [210320]
AY261360.1_Kenya_1950            ..tt.c.a...t...c..t....a.at.....a.....g...t....c.........c.aa.....g.....a..aa..a [210320]
AM712240.1_OURT_88/3             -------------------------------------------------------------------------------- [210320]
AM712239.1_Benin_97/1            -------------------------------------------------------------------------------- [210320]
KM111295.1_Ken06.Bus             -------------------------------------------------------------------------------- [210320]
KM111294.1_Ken05/Tk1             ..tta..a...t...c..t....a.at.....a.........t....t.........c.aa...........a..aa..a [210320]
KM262844.1_strain_L60            -------------------------------------------------------------------------------- [210320]
KM102979.1_26544/OG10_Italy      ..t....a..gt...........acat................................aa...........a...a... [210320]
KM262845.1_strain_NHV            -------------------------------------------------------------------------------- [210320]
KX354450.1_47/Ss/2008            ..t....a..gt...........acat................................aa...........a...a... [210320]
KP055815.1_strain_BA71           ..t....a..gt...........acat................................aa...........a...a... [210320]
MH025920.1_strain_R35            ..tt.cca...tt.ga.......a.at..................c.a.g......cc.aa.....g.....a..aa..a [210320]
MH025919.1_strain_N10            ..tt.cca...tt.ga.......a.at..................c.a.g......cc.aa.....g.....a..aa..a [210320]
MH025918.1_strain_R25            ..tt.cca...tt.ga.......a.at..................c.a.g......cc.aa.....g.....a..aa..a [210320]
MH025917.1_strain_R7             ..tt.cca...tt.ga.......a.at..................c.a.g......cc.aa.....g.....a..aa..a [210320]
MH025916.1_strain_R8             ..tt.cca...tt.ga.......a.at..................c.a.g......cc.aa.....g.....a..aa..a [210320]

FR682468.2_Georgia_2007/1        acatggctaaaga--------------------------------------tcaaccagaaaaggtttaaattttaacgc [210400]
ASF_IND_20_CAD543                ........nnnnnnnnnn------------nnnnntggcg-------attt.tt..g.a....ca..a......agnnnn [210400]
LR899193.1_Germany_2020/1        .............--------------------------------------............................. [210400]
MN393477.1_Wuhan_2019            .............--------------------------------------............................. [210400]
MW396979.1_Timor-Leste/2019/1    .............--------------------------------------............................. [210400]
MK940252.1_CN/2019/InnerMongolia .............--------------------------------------............................. [210400]
MK128995.1_China/2018/AnhuiXCGQ  .............--------------------------------------............................. [210400]
MH681419.1_ASFV/POL/2015         .............--------------------------------------............................. [210400]
LS478113.1_Estonia_2014          .............--------------------------------------............................. [210400]
KP843857.1_Odintsovo_02/14       .............--------------------------------------............................. [210400]
```

```
FN557520.1_strain_E75               -------------------------------------------------------------------------------- [210400]
AY261366.1_Warthog                  t............--------------------------------------........-................c.c. [210400]
AY261365.1_Warmbaths                .............--------------------------------------.t.......................c.c. [210400]
AY261364.1_Tengani_62               .............--------------------------------------.........................gta. [210400]
AY261363.1_Pretoriuskop/96/4        -------------------------------------------------------------------------------- [210400]
AY261362.1_Mkuzi_1979               .............--------------------------------------.t.......................c.c. [210400]
AY261361.1_Malawi_Lil-20/1_1983     c......g.----------------------------------------tt.gt.g.ga....aca....g.....gt.. [210400]
AY261360.1_Kenya_1950               ....a........tttct-cgcaaatcagtgtgcacggcggcaatctgttt..ct.gga...cata....g.....taa. [210400]
AM712240.1_OURT_88/3                -------------------------------------------------------------------------------- [210400]
AM712239.1_Benin_97/1               -------------------------------------------------------------------------------- [210400]
KM111295.1_Ken06.Bus                -------------------------------------------------------------------------------- [210400]
KM111294.1_Ken05/Tk1                ....a........tttct-cgcaaatcagtatgcacggcggcaatctgttt.tcgagga...cata....g....----- [210400]
KM262844.1_strain_L60               -------------------------------------------------------------------------------- [210400]
KM102979.1_26544/OG10_Italy         ........gtt--------------------------------------------------------------------- [210400]
KM262845.1_strain_NHV               -------------------------------------------------------------------------------- [210400]
KX354450.1_47/Ss/2008               ........gtt--------------------------------------------------------------------- [210400]
KP055815.1_strain_BA71              ........gct--------------------------------------------------------------------- [210400]
MH025920.1_strain_R35               ....a........tttctccgcaaaccagtatgcacggcggcaatttattt..tt.gga...tata....g.....taat [210400]
MH025919.1_strain_N10               ....a........tttctccgcaaaccagtatgcacggcggcaatttattt..tt.gga...tata....g.....taat [210400]
MH025918.1_strain_R25               ....a........tttctccgcaaaccagtatgcacggcggcaatttattt..tt.gga...tata....g.....taat [210400]
MH025917.1_strain_R7                ....a........tttctccgcaaaccagtatgcacggcggcaatttattt..tt.gga...tata....g.....taat [210400]
MH025916.1_strain_R8                ....a........tttctccgcaaaccagtatgcacggcggcaatttattt..tt.gga...tata....g.....taat [210400]

FR682468.2_Georgia_2007/1           cggccc-----------------------------------aaaacttaaactttt------------------------ [210480]
ASF_IND_20_CAD543                   nnnnnnnnnnnnnnnnnnnnnnnnnnnnnnnnnnnntagta.t..t..t..tg...annnnnnnnnnnnnnnnnnnnnnn [210480]
LR899193.1_Germany_2020/1           ......-----------------------------------...............------------------------ [210480]
MN393477.1_Wuhan_2019               ......-----------------------------------...............------------------------ [210480]
MW396979.1_Timor-Leste/2019/1       ......-----------------------------------...............------------------------ [210480]
MK940252.1_CN/2019/InnerMongolia    ......-----------------------------------...............------------------------ [210480]
MK128995.1_China/2018/AnhuiXCGQ     ......-----------------------------------...............------------------------ [210480]
MH681419.1_ASFV/POL/2015            ......-----------------------------------...............------------------------ [210480]
LS478113.1_Estonia_2014             ......-----------------------------------...............------------------------ [210480]
KP843857.1_Odintsovo_02/14          ......-----------------------------------...............------------------------ [210480]
FN557520.1_strain_E75               -------------------------------------------------------------------------------- [210480]
AY261366.1_Warthog                  -----------------------------------------...............------------------------ [210480]
AY261365.1_Warmbaths                -----------------------------------------...............------------------------ [210480]
AY261364.1_Tengani_62               tacatttc---------------------------------.........tg....------------------------ [210480]
AY261363.1_Pretoriuskop/96/4        -------------------------------------------------------------------------------- [210480]
AY261362.1_Mkuzi_1979               -----------------------------------------...............------------------------ [210480]
AY261361.1_Malawi_Lil-20/1_1983     -------------------------------------------------------------------------------- [210480]
AY261360.1_Kenya_1950               -------------------------------------------------------------------------------- [210480]
AM712240.1_OURT_88/3                -------------------------------------------------------------------------------- [210480]
AM712239.1_Benin_97/1               -------------------------------------------------------------------------------- [210480]
KM111295.1_Ken06.Bus                -------------------------------------------------------------------------------- [210480]
```

```
KM111294.1_Ken05/Tk1           -------------------------------------------------------------------------------- [210480]
KM262844.1_strain_L60          -------------------------------------------------------------------------------- [210480]
KM102979.1_26544/OG10_Italy    -------------------------------------------------------------------------------- [210480]
KM262845.1_strain_NHV          -------------------------------------------------------------------------------- [210480]
KX354450.1_47/Ss/2008          -------------------------------------------------------------------------------- [210480]
KP055815.1_strain_BA71         -------------------------------------------------------------------------------- [210480]
MH025920.1_strain_R35          -------------------------------------------------------------------------------- [210480]
MH025919.1_strain_N10          -------------------------------------------------------------------------------- [210480]
MH025918.1_strain_R25          -------------------------------------------------------------------------------- [210480]
MH025917.1_strain_R7           -------------------------------------------------------------------------------- [210480]
MH025916.1_strain_R8           -------------------------------------------------------------------------------- [210480]

FR682468.2_Georgia_2007/1      -----------------------------------------tttgatatttttaagtgcagccatggattggtccggcca [210560]
ASF_IND_20_CAD543              nnnnnnnnnnnnnnnnnnnnnnnnnnntaataattttaatg...a.cgcc.a..gca.t..ta..aag..---------- [210560]
LR899193.1_Germany_2020/1      -----------------------------------------....................................... [210560]
MN393477.1_Wuhan_2019          -----------------------------------------....................................... [210560]
MW396979.1_Timor-Leste/2019/1  -----------------------------------------....................................... [210560]
MK940252.1_CN/2019/InnerMongolia -----------------------------------------....................................... [210560]
MK128995.1_China/2018/AnhuiXCGQ -----------------------------------------....................................... [210560]
MH681419.1_ASFV/POL/2015       -----------------------------------------....................................... [210560]
LS478113.1_Estonia_2014        -----------------------------------------....................................... [210560]
KP843857.1_Odintsovo_02/14     -----------------------------------------....................................... [210560]
FN557520.1_strain_E75          -------------------------------------------------------------------------------- [210560]
AY261366.1_Warthog             -----------------------------------------....................................... [210560]
AY261365.1_Warmbaths           -----------------------------------------...................a........---...a.... [210560]
AY261364.1_Tengani_62          -----------------------------------------....ggg......................---.t..... [210560]
AY261363.1_Pretoriuskop/96/4   -------------------------------------------------------------------------------- [210560]
AY261362.1_Mkuzi_1979          -----------------------------------------...................a.........---..a.... [210560]
AY261361.1_Malawi_Lil-20/1_1983 -------------------------------------ggcggcc.gct.....gc..a....----------a.aa.... [210560]
AY261360.1_Kenya_1950          --------------------------------ttcgcaggcgg....c.ga.--------t..caaa..c---------- [210560]
AM712240.1_OURT_88/3           -------------------------------------------------------------------------------- [210560]
AM712239.1_Benin_97/1          -------------------------------------------------------------------------------- [210560]
KM111295.1_Ken06.Bus           -------------------------------------------------------------------------------- [210560]
KM111294.1_Ken05/Tk1           -------------------------------------agccggc...c.g...c-------------------------- [210560]
KM262844.1_strain_L60          -------------------------------------------------------------------------------- [210560]
KM102979.1_26544/OG10_Italy    -------------------------------------------------------------------------------- [210560]
KM262845.1_strain_NHV          -------------------------------------------------------------------------------- [210560]
KX354450.1_47/Ss/2008          -------------------------------------------------------------------------------- [210560]
KP055815.1_strain_BA71         -------------------------------------------------------------------------------- [210560]
MH025920.1_strain_R35          --------------------------------tccgcaggtggc...c.ga..cctca..t..caaa..c---------- [210560]
MH025919.1_strain_N10          --------------------------------tccgcaggtggc...c.ga..cctca..t..caaa..c---------- [210560]
MH025918.1_strain_R25          --------------------------------tccgcaggtggc...c.ga..cctca..t..caaa..c---------- [210560]
MH025917.1_strain_R7           --------------------------------tccgcaggtggc...c.ga..cctca..t..caaa..c---------- [210560]
MH025916.1_strain_R8           --------------------------------tccgcaggtggc...c.ga..cctca..t..caaa..c---------- [210560]
```

```
FR682468.2_Georgia_2007/1        taggatgacctatgcctac-------gtggcattctca----ttgatggcaatagcaatatatggtatattctacttat [210640]
ASF_IND_20_CAD543                --ta...ttta.c.....agcagta..aat.a.t.t.atct....c.c.t.c....g..g..----...a.t.taa.g. [210640]
LR899193.1_Germany_2020/1        ...................-------............----..................................... [210640]
MN393477.1_Wuhan_2019            ...................-------............----..................................... [210640]
MW396979.1_Timor-Leste/2019/1    ...................-------............----..................................... [210640]
MK940252.1_CN/2019/InnerMongolia ...................-------............----..................................... [210640]
MK128995.1_China/2018/AnhuiXCGQ  ...................-------............----..................................... [210640]
MH681419.1_ASFV/POL/2015         ...................-------............----..................................... [210640]
LS478113.1_Estonia_2014          ...................-------............----..................................... [210640]
KP843857.1_Odintsovo_02/14       ...................-------............----..................................... [210640]
FN557520.1_strain_E75            ------------------------------------------------------------------------------- [210640]
AY261366.1_Warthog               ...................-------...t.....a..----..................................... [210640]
AY261365.1_Warmbaths             ...................-------...t........----..................................... [210640]
AY261364.1_Tengani_62            ...................-------a.at........----a............................a....... [210640]
AY261363.1_Pretoriuskop/96/4     ------------------------------------------------------------------------------- [210640]
AY261362.1_Mkuzi_1979            ...................-------...t........----..................................... [210640]
AY261361.1_Malawi_Lil-20/1_1983  ..ac.a.cagc.g.tt...-----cga.a...c------ggc...gc...gg---------------------------- [210640]
AY261360.1_Kenya_1950            -----------..a-------------------------ggtac.gc..gggcg..gcctcctc..ccggc.t.tt..gg [210640]
AM712240.1_OURT_88/3             ------------------------------------------------------------------------------- [210640]
AM712239.1_Benin_97/1            ------------------------------------------------------------------------------- [210640]
KM111295.1_Ken06.Bus             ------------------------------------------------------------------------------- [210640]
KM111294.1_Ken05/Tk1             ------------------------------------------------------------------------------- [210640]
KM262844.1_strain_L60            ------------------------------------------------------------------------------- [210640]
KM102979.1_26544/OG10_Italy      ------------------------------------------------------------------------------- [210640]
KM262845.1_strain_NHV            ------------------------------------------------------------------------------- [210640]
KX354450.1_47/Ss/2008            ------------------------------------------------------------------------------- [210640]
KP055815.1_strain_BA71           ------------------------------------------------------------------------------- [210640]
MH025920.1_strain_R35            -----------....g...cgc-ggc...t.g.cgctgccgtac.gc..gggcg..gccgcctc..ccggc.t.tt..gc [210640]
MH025919.1_strain_N10            -----------....g...cgc-ggc...t.g.cgctgccgtac.gc..gggcg..gccgcctc..ccggc.t.tt..gc [210640]
MH025918.1_strain_R25            -----------....g...cgc-ggc...t.g.cgctgccgtac.gc..gggcg..gccgcctc..ccggc.t.tt..gc [210640]
MH025917.1_strain_R7             -----------....g...cgc-ggc...t.g.cgctgccgtac.gc..gggcg..gccgcctc..ccggc.t.tt..gc [210640]
MH025916.1_strain_R8             -----------....g...cgc-ggc...t.g.cgctgccgtac.gc..gggcg..gccgcctc..ccggc.t.tt..gc [210640]

FR682468.2_Georgia_2007/1        ctattgccgatcgaaaaaa-aatgttgttacaagcggtaatacgctcg--ctttagcgccaata--------tcgcatat [210720]
ASF_IND_20_CAD543                t..ac...t.------------------c.ac..ta......att.taatg....a....t.c.---------------- [210720]
LR899193.1_Germany_2020/1        ...................-............................--..............--------........ [210720]
MN393477.1_Wuhan_2019            ...................-............................--..............--------........ [210720]
MW396979.1_Timor-Leste/2019/1    ...................-............................--..............--------........ [210720]
MK940252.1_CN/2019/InnerMongolia ...................-............................--..............--------........ [210720]
MK128995.1_China/2018/AnhuiXCGQ  ...................-............................--..............--------........ [210720]
MH681419.1_ASFV/POL/2015         ...................-............................--..............--------........ [210720]
LS478113.1_Estonia_2014          ...................-............................--..............--------........ [210720]
KP843857.1_Odintsovo_02/14       ...................-............................--..............--------........ [210720]
```

FN557520.1_strain_E75 -------------------------------------------------------------------------------- [210720]
AY261366.1_Warthog ...................-...............a............--..............--------..t..... [210720]
AY261365.1_Warmbaths ...................-...............aa...........--.......t......t-----ct..t..... [210720]
AY261364.1_Tengani_62 .....at.....a......-.t..........ga.---......t...--.....a.t......--------..c..... [210720]
AY261363.1_Pretoriuskop/96/4 -------------------------------------------------------------------------------- [210720]
AY261362.1_Mkuzi_1979 ...................a...........g...a............--..............--------..t..... [210720]
AY261361.1_Malawi_Lil-20/1_1983 -------------------------------------------------------------------------------- [210720]
AY261360.1_Kenya_1950 gcggc..a..g.ttg.gc.----.gccc.cg.t..c...gc.tatgaa-----c...a...gc.ggttgaccg.cagc.c [210720]
AM712240.1_OURT_88/3 -------------------------------------------------------------------------------- [210720]
AM712239.1_Benin_97/1 -------------------------------------------------------------------------------- [210720]
KM111295.1_Ken06.Bus -------------------------------------------------------------------------------- [210720]
KM111294.1_Ken05/Tk1 -------------------------------------------------------------------------------- [210720]
KM262844.1_strain_L60 -------------------------------------------------------------------------------- [210720]
KM102979.1_26544/OG10_Italy -------------------------------------------------------------------------------- [210720]
KM262845.1_strain_NHV -------------------------------------------------------------------------------- [210720]
KX354450.1_47/Ss/2008 -------------------------------------------------------------------------------- [210720]
KP055815.1_strain_BA71 -------------------------------------------------------------------------------- [210720]
MH025920.1_strain_R35 gccgc..a..g.ttg.gc.----.gccc.cg.t..c...gc.tatgaa-----c...a...gc.ggttgaccg.cagc.c [210720]
MH025919.1_strain_N10 gccgc..a..g.ttg.gc.----.gccc.cg.t..c...gc.tatgaa-----c...a...gc.ggttgaccg.cagc.c [210720]
MH025918.1_strain_R25 gccgc..a..g.ttg.gc.----.gccc.cg.t..c...gc.tatgaa-----c...a...gc.ggttgaccg.cagc.c [210720]
MH025917.1_strain_R7 gccgc..a..g.ttg.gc.----.gccc.cg.t..c...gc.tatgaa-----c...a...gc.ggttgaccg.cagc.c [210720]
MH025916.1_strain_R8 gccgc..a..g.ttg.gc.----.gccc.cg.t..c...gc.tatgaa-----c...a...gc.ggttgaccg.cagc.c [210720]

FR682468.2_Georgia_2007/1 gtgaaaa------------------------------------------------------------atgttcg------ [210800]
ASF_IND_20_CAD543 -----------------------------------------------------------------------g.a------ [210800]
LR899193.1_Germany_2020/1 .......------------------------------------------------------------.......------ [210800]
MN393477.1_Wuhan_2019 .......------------------------------------------------------------.......------ [210800]
MW396979.1_Timor-Leste/2019/1 .......------------------------------------------------------------.......------ [210800]
MK940252.1_CN/2019/InnerMongolia .......------------------------------------------------------------.......------ [210800]
MK128995.1_China/2018/AnhuiXCGQ .......------------------------------------------------------------.......------ [210800]
MH681419.1_ASFV/POL/2015 .......------------------------------------------------------------.......------ [210800]
LS478113.1_Estonia_2014 .......------------------------------------------------------------.......------ [210800]
KP843857.1_Odintsovo_02/14 .......------------------------------------------------------------.......------ [210800]
FN557520.1_strain_E75 -------------------------------------------------------------------------------- [210800]
AY261366.1_Warthog .......------------------------------------------------------------.....t.catcat [210800]
AY261365.1_Warmbaths .......------------------------------------------------------------.......catcat [210800]
AY261364.1_Tengani_62 a......tgctgt------------------------------------------------------------------- [210800]
AY261363.1_Pretoriuskop/96/4 -------------------------------------------------------------------------------- [210800]
AY261362.1_Mkuzi_1979 .......tgctgtagtaacaatatatgtcgattggcgtaataggggcatgattaatgccacttaagat.t..tt------ [210800]
AY261361.1_Malawi_Lil-20/1_1983 -------------------------------------------------------------------------------- [210800]
AY261360.1_Kenya_1950 ..ag.g.aaccacaccag---------------------------------------------atagc.....at----- [210800]
AM712240.1_OURT_88/3 -------------------------------------------------------------------------------- [210800]
AM712239.1_Benin_97/1 -------------------------------------------------------------------------------- [210800]
KM111295.1_Ken06.Bus -------------------------------------------------------------------------------- [210800]

```
KM111294.1_Ken05/Tk1             -------------------------------------------------------------------------------- [210800]
KM262844.1_strain_L60            -------------------------------------------------------------------------------- [210800]
KM102979.1_26544/OG10_Italy      -------------------------------------------------------------------t.t..tt------ [210800]
KM262845.1_strain_NHV            -------------------------------------------------------------------------------- [210800]
KX354450.1_47/Ss/2008            -------------------------------------------------------------------t.t..tt------ [210800]
KP055815.1_strain_BA71           -------------------------------------------------------------------t.t..tt------ [210800]
MH025920.1_strain_R35            ..ag.ggaaccacaccag---------------------------------------------atagcc....atggctg [210800]
MH025919.1_strain_N10            ..ag.ggaaccacaccag---------------------------------------------atagcc....atggctg [210800]
MH025918.1_strain_R25            ..ag.ggaaccacaccag---------------------------------------------atagcc....atggctg [210800]
MH025917.1_strain_R7             ..ag.ggaaccacaccag---------------------------------------------atagcc....atggctg [210800]
MH025916.1_strain_R8             ..ag.ggaaccacaccag---------------------------------------------atagcc....atggctg [210800]

FR682468.2_Georgia_2007/1        --------------ccgaaaaaaacatt------aaaatttagaaccgccgcggcatctc------------aggggcgg [210880]
ASF_IND_20_CAD543                --------------gtagt..t..tt..------..tg....acg..----------ta.------------..ca.ta. [210880]
LR899193.1_Germany_2020/1        --------------..............------..........................------------........ [210880]
MN393477.1_Wuhan_2019            --------------..............------..........................------------........ [210880]
MW396979.1_Timor-Leste/2019/1    --------------..............------..........................------------........ [210880]
MK940252.1_CN/2019/InnerMongolia --------------..............------..........................------------........ [210880]
MK128995.1_China/2018/AnhuiXCGQ  --------------..............------..........................------------........ [210880]
MH681419.1_ASFV/POL/2015         --------------..............------..........................------------........ [210880]
LS478113.1_Estonia_2014          --------------..............------..........................------------........ [210880]
KP843857.1_Odintsovo_02/14       --------------..............------..........................------------........ [210880]
FN557520.1_strain_E75            -------------------------------------------------------------------------------- [210880]
AY261366.1_Warthog               aaaatgctg-----gac...........------..........................------------........ [210880]
AY261365.1_Warmbaths             aaaatgctgcggttgacg..........------.........................t------------........ [210880]
AY261364.1_Tengani_62            -------------------------------------------------------------------------------- [210880]
AY261363.1_Pretoriuskop/96/4     --------------gacg.......------------------------------------------------------- [210880]
AY261362.1_Mkuzi_1979            --------------g..gt..t..tt.cgcgaaa..c............g......c..t------------g.aaa.a- [210880]
AY261361.1_Malawi_Lil-20/1_1983  --------------a.............------.........t.....g.....------------------....t.. [210880]
AY261360.1_Kenya_1950            --ggcgatttg--tgacg..........------..gt.....t.....g.....gg.atctcaggcggcggc..---.. [210880]
AM712240.1_OURT_88/3             -------------------------------------------------------------------------------- [210880]
AM712239.1_Benin_97/1            -------------------------------------------------------------------------------- [210880]
KM111295.1_Ken06.Bus             -------------------------------------------------------------------------------- [210880]
KM111294.1_Ken05/Tk1             -------------------------------------------------------------------------------- [210880]
KM262844.1_strain_L60            -------------------------------------------------------------------------------- [210880]
KM102979.1_26544/OG10_Italy      --------------gt..c------------------------------------------------------------- [210880]
KM262845.1_strain_NHV            -------------------------------------------------------------------------------- [210880]
KX354450.1_47/Ss/2008            --------------gt..c------------------------------------------------------------- [210880]
KP055815.1_strain_BA71           --------------gt..c------------------------------------------------------------- [210880]
MH025920.1_strain_R35            gcggcgatttg--tgac...........------..gt.....t.....g.....ggtatctcaggcggcggc....... [210880]
MH025919.1_strain_N10            gcggcgatttg--tgac...........------..gt.....t.....g.....ggtatctcaggcggcggc....... [210880]
MH025918.1_strain_R25            gcggcgatttg--tgac...........------..gt.....t.....g.....ggtatctcaggcggcggc....... [210880]
MH025917.1_strain_R7             gcggcgatttg--tgac...........------..gt.....t.....g.....ggtatctcaggcggcggc....... [210880]
MH025916.1_strain_R8             gcggcgatttg--tgac...........------..gt.....t.....g.....ggtatctcaggcggcggc....... [210880]
```

```
FR682468.2_Georgia_2007/1        caacatttttttttatatggatattgtcacacaccacctcatctatgacgcaatatattactgctaatatcaggttcccc [210960]
ASF_IND_20_CAD543                t..t.a...-----------------------------.a....t.....--------------------------.ta. [210960]
LR899193.1_Germany_2020/1        ................................................................................ [210960]
MN393477.1_Wuhan_2019            ................................................................................ [210960]
MW396979.1_Timor-Leste/2019/1    ................................................................................ [210960]
MK940252.1_CN/2019/InnerMongolia ................................................................................ [210960]
MK128995.1_China/2018/AnhuiXCGQ  ................................................................................ [210960]
MH681419.1_ASFV/POL/2015         ................................................................................ [210960]
LS478113.1_Estonia_2014          ................................................................................ [210960]
KP843857.1_Odintsovo_02/14       ................................................................................ [210960]
FN557520.1_strain_E75            -------------------------------------------------------------------------------- [210960]
AY261366.1_Warthog               ...................................---..........t............................... [210960]
AY261365.1_Warmbaths             ...................................---..........t............................... [210960]
AY261364.1_Tengani_62            -------------------.g....a.a.t.t.ta--....ct.g...t------------------------------- [210960]
AY261363.1_Pretoriuskop/96/4     --------------------------------------------------------------------....ac...g.. [210960]
AY261362.1_Mkuzi_1979            -----------......................t....c.........t............................... [210960]
AY261361.1_Malawi_Lil-20/1_1983  ...........a.....c.......a.........---.......cc.ta.......cg.t................... [210960]
AY261360.1_Kenya_1950            ........c..a.....c.......a.........---........c.t........cg.t...............g... [210960]
AM712240.1_OURT_88/3             -------------------------------------------------------------------------------- [210960]
AM712239.1_Benin_97/1            -------------------------------------------------------------------------------- [210960]
KM111295.1_Ken06.Bus             -------------------------------------------------------------------------------- [210960]
KM111294.1_Ken05/Tk1             -------------------------------------------------------------------------------- [210960]
KM262844.1_strain_L60            -------------------------------------------------------------------------------- [210960]
KM102979.1_26544/OG10_Italy      -------------------------------------------------------------------------------- [210960]
KM262845.1_strain_NHV            -------------------------------------------------------------------------------- [210960]
KX354450.1_47/Ss/2008            -------------------------------------------------------------------------------- [210960]
KP055815.1_strain_BA71           -------------------------------------------------------------------------------- [210960]
MH025920.1_strain_R35            ........c..a.............a.........---........c.t........cg.t...............g... [210960]
MH025919.1_strain_N10            ........c..a.............a.........---........c.t........cg.t...............g... [210960]
MH025918.1_strain_R25            ........c..a.............a.........---........c.t........cg.t...............g... [210960]
MH025917.1_strain_R7             ........c..a.............a.........---........c.t........cg.t...............g... [210960]
MH025916.1_strain_R8             ........c..a.............a.........---........c.t........cg.t...............g... [210960]

FR682468.2_Georgia_2007/1        aatagtatgtagagaaaccacacaagatagatattcat------ggcgatttttgacgaaaaaacattaagttttagctt [211040]
ASF_IND_20_CAD543                ..c....--------------------------------------------------.t..t..tt....tg....a.gc [211040]
LR899193.1_Germany_2020/1        ......................................------.................................... [211040]
MN393477.1_Wuhan_2019            ......................................------.................................... [211040]
MW396979.1_Timor-Leste/2019/1    ......................................------.................................... [211040]
MK940252.1_CN/2019/InnerMongolia ......................................------.................................... [211040]
MK128995.1_China/2018/AnhuiXCGQ  ......................................------.................................... [211040]
MH681419.1_ASFV/POL/2015         ......................................------.................................... [211040]
LS478113.1_Estonia_2014          ......................................------.................................... [211040]
KP843857.1_Odintsovo_02/14       ......................................------.................................... [211040]
```

```
FN557520.1_strain_E75            -------------------------------------------------------------------------------- [211040]
AY261366.1_Warthog               ......................................------..t................................. [211040]
AY261365.1_Warmbaths             ......................................------..t................................. [211040]
AY261364.1_Tengani_62            --------------tg..gc.cggg.g.t.t....t-------------....c..........tg-------------- [211040]
AY261363.1_Pretoriuskop/96/4     .g.----------------------------------------------------------------------------- [211040]
AY261362.1_Mkuzi_1979            ......................................------..t..............................t.c [211040]
AY261361.1_Malawi_Lil-20/1_1983  ...........a.....a...g................------...a......................aa.......- [211040]
AY261360.1_Kenya_1950            .c..................tg.......a........cctggca.......c........................--- [211040]
AM712240.1_OURT_88/3             -------------------------------------------------------------------------------- [211040]
AM712239.1_Benin_97/1            -------------------------------------------------------------------------------- [211040]
KM111295.1_Ken06.Bus             -------------------------------------------------------------------------------- [211040]
KM111294.1_Ken05/Tk1             -------------------------------------------------------------------------------- [211040]
KM262844.1_strain_L60            -------------------------------------------------------------------------------- [211040]
KM102979.1_26544/OG10_Italy      -------------------------------------.------..........t...............aa.......- [211040]
KM262845.1_strain_NHV            -------------------------------------------------------------------------------- [211040]
KX354450.1_47/Ss/2008            -------------------------------------.------..........t...............aa.......- [211040]
KP055815.1_strain_BA71           -------------------------------------.------..........t...............aa.......- [211040]
MH025920.1_strain_R35            .c.......c...........gt......a........cctggc....t.....................a.......-- [211040]
MH025919.1_strain_N10            .c.......c...........gt......a........cctggc....t.....................a.......-- [211040]
MH025918.1_strain_R25            .c.......c...........gt......a........cctggc....t.....................a.......-- [211040]
MH025917.1_strain_R7             .c.......c...........gt......a........cctggc....t.....................a.......-- [211040]
MH025916.1_strain_R8             .c.......c...........gt......a........cctggc....t.....................a.......-- [211040]

FR682468.2_Georgia_2007/1        ctttg--------------------------------------------------------------------------- [211120]
ASF_IND_20_CAD543                ..aca--------------------------------------------------------------------------- [211120]
LR899193.1_Germany_2020/1        .....--------------------------------------------------------------------------- [211120]
MN393477.1_Wuhan_2019            .....--------------------------------------------------------------------------- [211120]
MW396979.1_Timor-Leste/2019/1    .....--------------------------------------------------------------------------- [211120]
MK940252.1_CN/2019/InnerMongolia .....--------------------------------------------------------------------------- [211120]
MK128995.1_China/2018/AnhuiXCGQ  .....--------------------------------------------------------------------------- [211120]
MH681419.1_ASFV/POL/2015         .....--------------------------------------------------------------------------- [211120]
LS478113.1_Estonia_2014          .....--------------------------------------------------------------------------- [211120]
KP843857.1_Odintsovo_02/14       .....--------------------------------------------------------------------------- [211120]
FN557520.1_strain_E75            -------------------------------------------------------------------------------- [211120]
AY261366.1_Warthog               .....--------------------------------------------------------------------------- [211120]
AY261365.1_Warmbaths             .....--------------------------------------------------------------------------- [211120]
AY261364.1_Tengani_62            -------------------------------------------------------------------------------- [211120]
AY261363.1_Pretoriuskop/96/4     -------------------------------------------------------------------------------- [211120]
AY261362.1_Mkuzi_1979            t..gacgcgaataatgtttaacgcctaataataattgatacccacagcggtaatgtccatctggccgccccaaaaaaaag [211120]
AY261361.1_Malawi_Lil-20/1_1983  -------------------------------------------------------------------------------- [211120]
AY261360.1_Kenya_1950            -------------------------------------------------------------------------------- [211120]
AM712240.1_OURT_88/3             -------------------------------------------------------------------------------- [211120]
AM712239.1_Benin_97/1            -------------------------------------------------------------------------------- [211120]
KM111295.1_Ken06.Bus             -------------------------------------------------------------------------------- [211120]
```

```
KM111294.1_Ken05/Tk1             -------------------------------------------------------------------------------- [211120]
KM262844.1_strain_L60            -------------------------------------------------------------------------------- [211120]
KM102979.1_26544/OG10_Italy      -------------------------------------------------------------------------------- [211120]
KM262845.1_strain_NHV            -------------------------------------------------------------------------------- [211120]
KX354450.1_47/Ss/2008            -------------------------------------------------------------------------------- [211120]
KP055815.1_strain_BA71           -------------------------------------------------------------------------------- [211120]
MH025920.1_strain_R35            -------------------------------------------------------------------------------- [211120]
MH025919.1_strain_N10            -------------------------------------------------------------------------------- [211120]
MH025918.1_strain_R25            -------------------------------------------------------------------------------- [211120]
MH025917.1_strain_R7             -------------------------------------------------------------------------------- [211120]
MH025916.1_strain_R8             -------------------------------------------------------------------------------- [211120]

FR682468.2_Georgia_2007/1        -------------------------------------------------------------------------------- [211200]
ASF_IND_20_CAD543                -------------------------------------------------------------------------------- [211200]
LR899193.1_Germany_2020/1        -------------------------------------------------------------------------------- [211200]
MN393477.1_Wuhan_2019            -------------------------------------------------------------------------------- [211200]
MW396979.1_Timor-Leste/2019/1    -------------------------------------------------------------------------------- [211200]
MK940252.1_CN/2019/InnerMongolia -------------------------------------------------------------------------------- [211200]
MK128995.1_China/2018/AnhuiXCGQ  -------------------------------------------------------------------------------- [211200]
MH681419.1_ASFV/POL/2015         -------------------------------------------------------------------------------- [211200]
LS478113.1_Estonia_2014          -------------------------------------------------------------------------------- [211200]
KP843857.1_Odintsovo_02/14       -------------------------------------------------------------------------------- [211200]
FN557520.1_strain_E75            -------------------------------------------------------------------------------- [211200]
AY261366.1_Warthog               -------------------------------------------------------------------------------- [211200]
AY261365.1_Warmbaths             -------------------------------------------------------------------------------- [211200]
AY261364.1_Tengani_62            -------------------------------------------------------------------------------- [211200]
AY261363.1_Pretoriuskop/96/4     -------------------------------------------------------------------------------- [211200]
AY261362.1_Mkuzi_1979            tatttacggtggggtttattaccggcggcgtaacaccagttatggtcaattttgtctggcccagccgcaaaaaaaatatt [211200]
AY261361.1_Malawi_Lil-20/1_1983  -------------------------------------------------------------------------------- [211200]
AY261360.1_Kenya_1950            -------------------------------------------------------------------------------- [211200]
AM712240.1_OURT_88/3             -------------------------------------------------------------------------------- [211200]
AM712239.1_Benin_97/1            -------------------------------------------------------------------------------- [211200]
KM111295.1_Ken06.Bus             -------------------------------------------------------------------------------- [211200]
KM111294.1_Ken05/Tk1             -------------------------------------------------------------------------------- [211200]
KM262844.1_strain_L60            -------------------------------------------------------------------------------- [211200]
KM102979.1_26544/OG10_Italy      -------------------------------------------------------------------------------- [211200]
KM262845.1_strain_NHV            -------------------------------------------------------------------------------- [211200]
KX354450.1_47/Ss/2008            -------------------------------------------------------------------------------- [211200]
KP055815.1_strain_BA71           -------------------------------------------------------------------------------- [211200]
MH025920.1_strain_R35            -------------------------------------------------------------------------------- [211200]
MH025919.1_strain_N10            -------------------------------------------------------------------------------- [211200]
MH025918.1_strain_R25            -------------------------------------------------------------------------------- [211200]
MH025917.1_strain_R7             -------------------------------------------------------------------------------- [211200]
MH025916.1_strain_R8             -------------------------------------------------------------------------------- [211200]
```

```
FR682468.2_Georgia_2007/1        ------------------------------------------------------------acgcctgt---gtacta--- [211280]
ASF_IND_20_CAD543                ------------------------------------------------------------..agta..a--a..a.t--- [211280]
LR899193.1_Germany_2020/1        ------------------------------------------------------------........---......--- [211280]
MN393477.1_Wuhan_2019            ------------------------------------------------------------........---......--- [211280]
MW396979.1_Timor-Leste/2019/1    ------------------------------------------------------------........---......--- [211280]
MK940252.1_CN/2019/InnerMongolia ------------------------------------------------------------........---......--- [211280]
MK128995.1_China/2018/AnhuiXCGQ  ------------------------------------------------------------........---......--- [211280]
MH681419.1_ASFV/POL/2015         ------------------------------------------------------------........---......--- [211280]
LS478113.1_Estonia_2014          ------------------------------------------------------------........---......--- [211280]
KP843857.1_Odintsovo_02/14       ------------------------------------------------------------........---......--- [211280]
FN557520.1_strain_E75            -------------------------------------------------------------------------------- [211280]
AY261366.1_Warthog               ------------------------------------------------------------........gtac..a..aac [211280]
AY261365.1_Warmbaths             ------------------------------------------------------------........------------ [211280]
AY261364.1_Tengani_62            ------------------------------------------------------------.a.ttct.t--...a..--- [211280]
AY261363.1_Pretoriuskop/96/4     ------------------------------------------------------------......a.a----------- [211280]
AY261362.1_Mkuzi_1979            tccggccgcggcgtttcaaaaaataatctttgcgaaataattccacatcttgtgaaatga......aca--...a..--- [211280]
AY261361.1_Malawi_Lil-20/1_1983  -----------------------------------------------------------------------a...g..--- [211280]
AY261360.1_Kenya_1950            -------------------------------------------------------------------------------- [211280]
AM712240.1_OURT_88/3             -------------------------------------------------------------------------------- [211280]
AM712239.1_Benin_97/1            -------------------------------------------------------------------------------- [211280]
KM111295.1_Ken06.Bus             -------------------------------------------------------------------------------- [211280]
KM111294.1_Ken05/Tk1             -------------------------------------------------------------------------------- [211280]
KM262844.1_strain_L60            -------------------------------------------------------------------------------- [211280]
KM102979.1_26544/OG10_Italy      ------------------------------------------------------------......aca--.c.g..--- [211280]
KM262845.1_strain_NHV            -------------------------------------------------------------------------------- [211280]
KX354450.1_47/Ss/2008            ------------------------------------------------------------......aca--.c.g..--- [211280]
KP055815.1_strain_BA71           ------------------------------------------------------------......aca--.c.g..--- [211280]
MH025920.1_strain_R35            -------------------------------------------------------------------------------- [211280]
MH025919.1_strain_N10            -------------------------------------------------------------------------------- [211280]
MH025918.1_strain_R25            -------------------------------------------------------------------------------- [211280]
MH025917.1_strain_R7             -------------------------------------------------------------------------------- [211280]
MH025916.1_strain_R8             -------------------------------------------------------------------------------- [211280]

FR682468.2_Georgia_2007/1        ---------------a---taatgtttaacgcct------gtagta-------taataattgatacctacagcagtaatt [211360]
ASF_IND_20_CAD543                ---------------t---...............acagca......ataattt....gt..a.cg............--- [211360]
LR899193.1_Germany_2020/1        ---------------.---...............------......-------........................... [211360]
MN393477.1_Wuhan_2019            ---------------.---...............a----c.c.agg-------..g....-a.................. [211360]
MW396979.1_Timor-Leste/2019/1    ---------------.---...............------......-------........................... [211360]
MK940252.1_CN/2019/InnerMongolia ---------------.---...............------......-------........................... [211360]
MK128995.1_China/2018/AnhuiXCGQ  ---------------.---...............------......-------........................... [211360]
MH681419.1_ASFV/POL/2015         ---------------.---...............------......-------........................... [211360]
LS478113.1_Estonia_2014          ---------------.---...............------......-------........................... [211360]
KP843857.1_Odintsovo_02/14       ---------------.---...............------......-------........................... [211360]
```

| | | |
|---|---|---|
| FN557520.1_strain_E75 | -------------------------------------------------------------------------------- | [211360] |
| AY261366.1_Warthog | gctaactttaacacc.gta....c...g......acagca....---------..........a...a..c......... | [211360] |
| AY261365.1_Warmbaths | ---------------.gta....aa..g.aa..cacagcg...------------------------------------- | [211360] |
| AY261364.1_Tengani_62 | ---------------.ttt......ccg......a------------------c.........a.a.a..c........- | [211360] |
| AY261363.1_Pretoriuskop/96/4 | ---------------.tag....aa..g.aa...acagcg...-------------------------------...... | [211360] |
| AY261362.1_Mkuzi_1979 | ---------------.ttt...............acagca...a..a------....gt..a.cg....t.......--- | [211360] |
| AY261361.1_Malawi_Lil-20/1_1983 | ---------------.ttc..............cacaaca.....----------------------------------- | [211360] |
| AY261360.1_Kenya_1950 | ---------------ccggc.t.t...g.t....actgc----------------------------------------- | [211360] |
| AM712240.1_OURT_88/3 | -------------------------------------------------------------------------------- | [211360] |
| AM712239.1_Benin_97/1 | -------------------------------------------------------------------------------- | [211360] |
| KM111295.1_Ken06.Bus | -------------------------------------------------------------------------------- | [211360] |
| KM111294.1_Ken05/Tk1 | -------------------------------------------------------------------------------- | [211360] |
| KM262844.1_strain_L60 | -------------------------------------------------------------------------------- | [211360] |
| KM102979.1_26544/OG10_Italy | ---------------.---...............acagca......ataattt....gt.-------------------- | [211360] |
| KM262845.1_strain_NHV | -------------------------------------------------------------------------------- | [211360] |
| KX354450.1_47/Ss/2008 | ---------------.---...............acagca......ataattt....gt..a------------------ | [211360] |
| KP055815.1_strain_BA71 | ---------------.---...............acagca......ataattt....gt..a.cg............--- | [211360] |
| MH025920.1_strain_R35 | ---------------ccggc.t.t...g.t....actgc----------------------------------------- | [211360] |
| MH025919.1_strain_N10 | ---------------ccggc.t.t...g.t....actgc----------------------------------------- | [211360] |
| MH025918.1_strain_R25 | ---------------ccggc.t.t...g.t....actgc----------------------------------------- | [211360] |
| MH025917.1_strain_R7 | ---------------ccggc.t.t...g.t....actgc----------------------------------------- | [211360] |
| MH025916.1_strain_R8 | ---------------ccggc.t.t...g.t....actgc----------------------------------------- | [211360] |

| | | |
|---|---|---|
| FR682468.2_Georgia_2007/1 | gatacctacggcgataatgtctctc---tggccgccccaaaaaaaagtatttacggtagggt---ttattaccggcggcg | [211440] |
| ASF_IND_20_CAD543 | ----------.ta.....t.taa..--t.t.a....tac..c.gt....a.a.ttt..at..---...acg..-----ta | [211440] |
| LR899193.1_Germany_2020/1 | .........................---..................................---............... | [211440] |
| MN393477.1_Wuhan_2019 | .........................---..................................---............... | [211440] |
| MW396979.1_Timor-Leste/2019/1 | .........................---..................................---............... | [211440] |
| MK940252.1_CN/2019/InnerMongolia | .........................---..................................---............... | [211440] |
| MK128995.1_China/2018/AnhuiXCGQ | .........................---..................................---............... | [211440] |
| MH681419.1_ASFV/POL/2015 | .........................---..........-.......................---............... | [211440] |
| LS478113.1_Estonia_2014 | .........................---..................................---............... | [211440] |
| KP843857.1_Odintsovo_02/14 | .........................---..................................---............... | [211440] |
| FN557520.1_strain_E75 | -------------------------------------------------------------------------------- | [211440] |
| AY261366.1_Warthog | ..a...a---a.agcg...c.ca..---.....................g......c.....tta............... | [211440] |
| AY261365.1_Warmbaths | ----------------.....ca..---........-------------------------------------------- | [211440] |
| AY261364.1_Tengani_62 | ---...a---a.agcg......a..atc....................gg...........a---------.a....... | [211440] |
| AY261363.1_Pretoriuskop/96/4 | ..a...a---a.agcg...c.ca..---.....................g.......c....---............... | [211440] |
| AY261362.1_Mkuzi_1979 | ----------.ta.....t.taa..--t.t.a....tac.gc.gt....a.a.gtt..atc.---...acg..-----ta | [211440] |
| AY261361.1_Malawi_Lil-20/1_1983 | -------------------------------------------------------------------------------- | [211440] |
| AY261360.1_Kenya_1950 | -------------------------------------------------------------------------------- | [211440] |
| AM712240.1_OURT_88/3 | -------------------------------------------------------------------------------- | [211440] |
| AM712239.1_Benin_97/1 | -------------------------------------------------------------------------------- | [211440] |
| KM111295.1_Ken06.Bus | -------------------------------------------------------------------------------- | [211440] |

```
KM111294.1_Ken05/Tk1           -------------------------------------------------------------------------------- [211440]
KM262844.1_strain_L60          -------------------------------------------------------------------------------- [211440]
KM102979.1_26544/OG10_Italy    -------------------------------------------------------------------------------- [211440]
KM262845.1_strain_NHV          -------------------------------------------------------------------------------- [211440]
KX354450.1_47/Ss/2008          -------------------------------a....tac..c.gt....a.a.ttt..at..---...acg..-----ta [211440]
KP055815.1_strain_BA71         ----------.ta.....t.taa..--t.t.a....tac..c.gt....a.a.ttt..at..---...acg..-----ta [211440]
MH025920.1_strain_R35          -------------------------------------------------------------------------------- [211440]
MH025919.1_strain_N10          -------------------------------------------------------------------------------- [211440]
MH025918.1_strain_R25          -------------------------------------------------------------------------------- [211440]
MH025917.1_strain_R7           -------------------------------------------------------------------------------- [211440]
MH025916.1_strain_R8           -------------------------------------------------------------------------------- [211440]

FR682468.2_Georgia_2007/1      taacaccagttatggtcaattttgtctggcccgccgcccagccgc-aaaa---aaaaatcaattacaaccgcaaaaaaaa [211520]
ASF_IND_20_CAD543              ..g..gt...a..annnnnnnnnnnnnnnnnnnnnnnnnnnnnnnnt..gttt..tgt.t..cgc.t..a...gt.gt.. [211520]
LR899193.1_Germany_2020/1      .............................................-....---........................... [211520]
MN393477.1_Wuhan_2019          .............................................a....---........................... [211520]
MW396979.1_Timor-Leste/2019/1  .............................................a....---........................... [211520]
MK940252.1_CN/2019/InnerMongolia .............................................-....---........................... [211520]
MK128995.1_China/2018/AnhuiXCGQ .............................................-....---........................... [211520]
MH681419.1_ASFV/POL/2015       .............................................-....---........................... [211520]
LS478113.1_Estonia_2014        .............................................a....---........................... [211520]
KP843857.1_Odintsovo_02/14     .............................................-....---...................-....... [211520]
FN557520.1_strain_E75          -------------------------------------------------------------------------------- [211520]
AY261366.1_Warthog             .tg.....a.................................a..cgc.g-aa..............g.....g...... [211520]
AY261365.1_Warmbaths           -------------------------------------------------------------------------------- [211520]
AY261364.1_Tengani_62          .tg.....a..........................------------...-aa........------g............ [211520]
AY261363.1_Pretoriuskop/96/4   .tg.....a.................................a..cgc.g-aa.-----------------------... [211520]
AY261362.1_Mkuzi_1979          c.g..gt...a.----------------------------------------t..tgt.t..cgc.t..a...gt.gt.. [211520]
AY261361.1_Malawi_Lil-20/1_1983 -------------------------------------------------------------------------------- [211520]
AY261360.1_Kenya_1950          -------------------------------------------------------------------------------- [211520]
AM712240.1_OURT_88/3           -------------------------------------------------------------------------------- [211520]
AM712239.1_Benin_97/1          -------------------------------------------------------------------------------- [211520]
KM111295.1_Ken06.Bus           -------------------------------------------------------------------------------- [211520]
KM111294.1_Ken05/Tk1           -------------------------------------------------------------------------------- [211520]
KM262844.1_strain_L60          -------------------------------------------------------------------------------- [211520]
KM102979.1_26544/OG10_Italy    -------------------------------------------------------------------------------- [211520]
KM262845.1_strain_NHV          -------------------------------------------------------------------------------- [211520]
KX354450.1_47/Ss/2008          ..g..gt...a..----------------------------------..gttt..tgt.t..cgc.t..a...gt.gt.. [211520]
KP055815.1_strain_BA71         ..g..gt...a..----------------------------------..gttt..tgt.t..cgc.t..a...gt.gt.. [211520]
MH025920.1_strain_R35          -------------------------------------------------------------------------------- [211520]
MH025919.1_strain_N10          -------------------------------------------------------------------------------- [211520]
MH025918.1_strain_R25          -------------------------------------------------------------------------------- [211520]
MH025917.1_strain_R7           -------------------------------------------------------------------------------- [211520]
MH025916.1_strain_R8           -------------------------------------------------------------------------------- [211520]
```

```
FR682468.2_Georgia_2007/1        atatttccggccgcggcgtttcaaaaaataatctttgcgaaataattccgcatcttgtgaaat--------------gaa [211600]
ASF_IND_20_CAD543                ta....taat.ttt.a..cc.ac.gc.g..g.--------.......ttaatgt..----------------------.. [211600]
LR899193.1_Germany_2020/1        ...............................................................--------------... [211600]
MN393477.1_Wuhan_2019            ...............................................................--------------... [211600]
MW396979.1_Timor-Leste/2019/1    ...............................................................--------------... [211600]
MK940252.1_CN/2019/InnerMongolia ...............................................................--------------... [211600]
MK128995.1_China/2018/AnhuiXCGQ  ...............................................................--------------... [211600]
MH681419.1_ASFV/POL/2015         ...............................................................--------------... [211600]
LS478113.1_Estonia_2014          ...............................................................--------------... [211600]
KP843857.1_Odintsovo_02/14       ...............................................................--------------... [211600]
FN557520.1_strain_E75            -------------------------------------------------------------------------------- [211600]
AY261366.1_Warthog               ................a.....------a..a--------.................-------tccggctggcctatg. [211600]
AY261365.1_Warmbaths             ---------...c.........t.t...a..a--------.....................c.ttccggctggcctatg. [211600]
AY261364.1_Tengani_62            ................a................cgcta.t......ctt----------------------------tg. [211600]
AY261363.1_Pretoriuskop/96/4     ......................------a..a--------.................-------tccggctggcctatg. [211600]
AY261362.1_Mkuzi_1979            ------taatgtttaa..cc.ac.gc.g....--------.....------tgt..----------------------.. [211600]
AY261361.1_Malawi_Lil-20/1_1983  ----------------------------------------.......t.------------------------------- [211600]
AY261360.1_Kenya_1950            ----------------------------------------.....----------------------------------- [211600]
AM712240.1_OURT_88/3             -------------------------------------------------------------------------------- [211600]
AM712239.1_Benin_97/1            -------------------------------------------------------------------------------- [211600]
KM111295.1_Ken06.Bus             -------------------------------------------------------------------------------- [211600]
KM111294.1_Ken05/Tk1             -------------------------------------------------------------------------------- [211600]
KM262844.1_strain_L60            -------------------------------------------------------------------------------- [211600]
KM102979.1_26544/OG10_Italy      -------------------------------------------------------------------------------- [211600]
KM262845.1_strain_NHV            -------------------------------------------------------------------------------- [211600]
KX354450.1_47/Ss/2008            ta....taat.ttt.a..cc.ac.gc.g..g.--------.......ttaatgt..----------------------.. [211600]
KP055815.1_strain_BA71           ta....taat.ttt.a..cc.ac.gc.g..g.--------.......ttaatgt..----------------------.. [211600]
MH025920.1_strain_R35            ----------------------------------------.....cg..------------------------------- [211600]
MH025919.1_strain_N10            ----------------------------------------.....cg..------------------------------- [211600]
MH025918.1_strain_R25            ----------------------------------------.....cg..------------------------------- [211600]
MH025917.1_strain_R7             ----------------------------------------.....cg..------------------------------- [211600]
MH025916.1_strain_R8             ----------------------------------------.....cg..------------------------------- [211600]

FR682468.2_Georgia_2007/1        cgcctaca------gtaata------------------------------------------------attttaatcttt [211680]
ASF_IND_20_CAD543                ........acagta......------------------------------------------------........g... [211680]
LR899193.1_Germany_2020/1        ........------......------------------------------------------------............ [211680]
MN393477.1_Wuhan_2019            ........------......------------------------------------------------............ [211680]
MW396979.1_Timor-Leste/2019/1    ........------......------------------------------------------------............ [211680]
MK940252.1_CN/2019/InnerMongolia ........------......------------------------------------------------............ [211680]
MK128995.1_China/2018/AnhuiXCGQ  ........------......------------------------------------------------............ [211680]
MH681419.1_ASFV/POL/2015         ........------......------------------------------------------------............ [211680]
LS478113.1_Estonia_2014          ........------......------------------------------------------------............ [211680]
KP843857.1_Odintsovo_02/14       ........------......------------------------------------------------............ [211680]
```

```
FN557520.1_strain_E75               -------------------------------------------------------------------------------- [211680]
AY261366.1_Warthog                  t.ta..-------g..g.a.------ttttcttgatcaa---agcagcc---------------------.a.....--. [211680]
AY261365.1_Warmbaths                t.ta..t.tatatg..g.a.ttatttttttcttgatcaaagcagcagcc---------------------.a........ [211680]
AY261364.1_Tengani_62               ........gcagtaca....------------------------------------------------........g... [211680]
AY261363.1_Pretoriuskop/96/4        t.ta..------tg..g.a.------ttttcttgatcaaagcaataatctttgacgcctacagcagta...c........ [211680]
AY261362.1_Mkuzi_1979               ........gcagtaa...------------------------------------------------------....g... [211680]
AY261361.1_Malawi_Lil-20/1_1983     ------------------------------------------------------------------------....g... [211680]
AY261360.1_Kenya_1950               -------------------------------------------------------------------------------- [211680]
AM712240.1_OURT_88/3                -------------------------------------------------------------------------------- [211680]
AM712239.1_Benin_97/1               -------------------------------------------------------------------------------- [211680]
KM111295.1_Ken06.Bus                -------------------------------------------------------------------------------- [211680]
KM111294.1_Ken05/Tk1                -------------------------------------------------------------------------------- [211680]
KM262844.1_strain_L60               -------------------------------------------------------------------------------- [211680]
KM102979.1_26544/OG10_Italy         -------------------------------------------------------------------------------- [211680]
KM262845.1_strain_NHV               -------------------------------------------------------------------------------- [211680]
KX354450.1_47/Ss/2008               ........acagta......------------------------------------------------........g... [211680]
KP055815.1_strain_BA71              ........acagta......------------------------------------------------........g... [211680]
MH025920.1_strain_R35               ---------------------------------------------------------------ca--------------- [211680]
MH025919.1_strain_N10               ---------------------------------------------------------------ca--------------- [211680]
MH025918.1_strain_R25               ---------------------------------------------------------------ca--------------- [211680]
MH025917.1_strain_R7                ---------------------------------------------------------------ca--------------- [211680]
MH025916.1_strain_R8                ---------------------------------------------------------------ca--------------- [211680]

FR682468.2_Georgia_2007/1           gacacctacagcagtagtaataattttaatctttaacgcc-tgcagcagtactaatatttttaatctttaacgcctacag [211760]
ASF_IND_20_CAD543                   a..g..........................g.........-.a........g.....a...........g.........a [211760]
LR899193.1_Germany_2020/1           ........................................-....................................... [211760]
MN393477.1_Wuhan_2019               ........................................-....................................... [211760]
MW396979.1_Timor-Leste/2019/1       ........................................-................a...................... [211760]
MK940252.1_CN/2019/InnerMongolia    ........................................-....................................... [211760]
MK128995.1_China/2018/AnhuiXCGQ     ........................................-....................................... [211760]
MH681419.1_ASFV/POL/2015            a..g..........................g.........-....................................... [211760]
LS478113.1_Estonia_2014             ........................................-....................................... [211760]
KP843857.1_Odintsovo_02/14          ........................................-....................................... [211760]
FN557520.1_strain_E75               -------------------------------------------------------------------------------- [211760]
AY261366.1_Warthog                  ...g..........................g...g.....---------------------------------....... [211760]
AY261365.1_Warmbaths                ...g....t.......ca............g.........-.at........a....a.......g...........t.. [211760]
AY261364.1_Tengani_62               a..g..............................g.....-.a.........a....a...........g.......... [211760]
AY261363.1_Pretoriuskop/96/4        ...g......a.....ca............g.......t.-.a.........a....a...........g.........a [211760]
AY261362.1_Mkuzi_1979               a..g............a...------....g.........-.a........g.....ag....-------.......... [211760]
AY261361.1_Malawi_Lil-20/1_1983     a..g..c.t.a...................g.........c.a........g.....a......a....g.....c.... [211760]
AY261360.1_Kenya_1950               --------------c..g......c------......-..-.a.ta...c.g.....a---....g............ca [211760]
AM712240.1_OURT_88/3                -------------------------------------------------------------------------------- [211760]
AM712239.1_Benin_97/1               -------------------------------------------------------------------------------- [211760]
KM111295.1_Ken06.Bus                -------------------------------------------------------------------------------- [211760]
```

```
KM111294.1_Ken05/Tk1         -------------------------------------------------------------------------------- [211760]
KM262844.1_strain_L60        -------------------------------------------------------------------------------- [211760]
KM102979.1_26544/OG10_Italy  -------------------------------------------------------------------------------- [211760]
KM262845.1_strain_NHV        -------------------------------------------------------------------------------- [211760]
KX354450.1_47/Ss/2008        a..g..........................g.........-.a........g.....a...........g.........a [211760]
KP055815.1_strain_BA71       a..g..........................g.........-.a........g.....a...........g.........a [211760]
MH025920.1_strain_R35        -----.c.......c..g............g.........-aa..a...c.gg....a.............t...a...a [211760]
MH025919.1_strain_N10        -----.c.......c..g............g.........-aa..at..c.gg....a.............t...a...a [211760]
MH025918.1_strain_R25        -----.c.......c..g............g.........-aa..a...c.gg....a.............t...a...a [211760]
MH025917.1_strain_R7         -----.c.......c..g............g.........-aa..a...c.gg....a.............t...a...a [211760]
MH025916.1_strain_R8         -----.c.......c..g............g.........-aa..a...c.gg....a.............t...a...a [211760]

FR682468.2_Georgia_2007/1    cagtagtaataattttaatgtttaacgcctacagcagtagtaataattttaatctttgacgcctacagcagtagtaataa [211840]
ASF_IND_20_CAD543            .................................a...................g...a...................... [211840]
LR899193.1_Germany_2020/1    ................................................................................ [211840]
MN393477.1_Wuhan_2019        ............................................------------------------------------ [211840]
MW396979.1_Timor-Leste/2019/1 ................................................................................ [211840]
MK940252.1_CN/2019/InnerMongolia ............................................------------------------------------ [211840]
MK128995.1_China/2018/AnhuiXCGQ ............................................------------------------------------ [211840]
MH681419.1_ASFV/POL/2015     ............................................------------------------------------ [211840]
LS478113.1_Estonia_2014      ................................................................................ [211840]
KP843857.1_Odintsovo_02/14   ............................................------------------------------------ [211840]
FN557520.1_strain_E75        -------------------------------------------------------------------------------- [211840]
AY261366.1_Warthog           .....ca.......c....c...g.........a.......................a.......--------------- [211840]
AY261365.1_Warmbaths         .....ca..........................a...................g...a...................... [211840]
AY261364.1_Tengani_62        .....ca................................ca................................ca..... [211840]
AY261363.1_Pretoriuskop/96/4 .....ca..............................................g...a...............ca..... [211840]
AY261362.1_Mkuzi_1979        .....a...------...............................g..........a...................... [211840]
AY261361.1_Malawi_Lil-20/1_1983 .................................a.....a.............g...a...t.c...c............ [211840]
AY261360.1_Kenya_1950        ...c.........aa.................ta...c.........aa....g...a.......--------------- [211840]
AM712240.1_OURT_88/3         -------------------------------------------------------------------------------- [211840]
AM712239.1_Benin_97/1        -------------------------------------------------------------------------------- [211840]
KM111295.1_Ken06.Bus         -------------------------------------------------------------------------------- [211840]
KM111294.1_Ken05/Tk1         -------------------------------------------------------------------------------- [211840]
KM262844.1_strain_L60        -------------------------------------------------------------------------------- [211840]
KM102979.1_26544/OG10_Italy  -------------------------------------------------------------------------------- [211840]
KM262845.1_strain_NHV        -------------------------------------------------------------------------------- [211840]
KX354450.1_47/Ss/2008        .......------------------------------------------------------------------------- [211840]
KP055815.1_strain_BA71       .................................a...................g...a...................... [211840]
MH025920.1_strain_R35        ...c...............c.........a...a.....t.............g...a.......--------------- [211840]
MH025919.1_strain_N10        ...c...............c.........a...a.....t.............g...a.......--------------- [211840]
MH025918.1_strain_R25        ...c...............c.........a...a.....t.............g...a.......--------------- [211840]
MH025917.1_strain_R7         ...c...............c.........a...a.....t.............g...a.......--------------- [211840]
MH025916.1_strain_R8         ...c...............c.........a...a.....t.............g...a.......--------------- [211840]
```

```
FR682468.2_Georgia_2007/1         ttttaatgtttaacgcctacagcagta-caataatttt-aatgtttaacgcctgcagcagt---actaatatttttaatc [211920]
ASF_IND_20_CAD543                 ...........................gt.........-...c...g......a..a....---.g.....a.......g [211920]
LR899193.1_Germany_2020/1         ...........................-..........-......................---................ [211920]
MN393477.1_Wuhan_2019             -------------------------------------------------------------------------------- [211920]
MW396979.1_Timor-Leste/2019/1     ...........................-..........-......................---................ [211920]
MK940252.1_CN/2019/InnerMongolia  -------------------------------------------------------------------------------- [211920]
MK128995.1_China/2018/AnhuiXCGQ   -------------------------------------------------------------------------------- [211920]
MH681419.1_ASFV/POL/2015          -------------------------------------------------------------------------------- [211920]
LS478113.1_Estonia_2014           ...........................gt.........-...c...g......a.......---.g.....a.......g [211920]
KP843857.1_Odintsovo_02/14        -------------------------------------------------------------------------------- [211920]
FN557520.1_strain_E75             -------------------------------------------------------------------------------- [211920]
AY261366.1_Warthog                -------------------------------------------------------------------------------- [211920]
AY261365.1_Warmbaths              ...........g.....ag...t.ca.at.........-..............a..a....aca.a.....a........ [211920]
AY261364.1_Tengani_62             ...........................gt.........-..............a.......---.g.....a........ [211920]
AY261363.1_Pretoriuskop/96/4      ...........................ca.........-...c..........a.......---.g.....a.......g [211920]
AY261362.1_Mkuzi_1979             -------....................at.....-------............a.......---.a...------....g [211920]
AY261361.1_Malawi_Lil-20/1_1983   .......c....c....c.........gt.........a..cc.........cc.......---.g.....a....c..g [211920]
AY261360.1_Kenya_1950             ----------------.c.........at.....---.-..............a.ca...cagt.a.....a---....g [211920]
AM712240.1_OURT_88/3              -------------------------------------------------------------------------------- [211920]
AM712239.1_Benin_97/1             -------------------------------------------------------------------------------- [211920]
KM111295.1_Ken06.Bus              -------------------------------------------------------------------------------- [211920]
KM111294.1_Ken05/Tk1              -------------------------------------------------------------------------------- [211920]
KM262844.1_strain_L60             -------------------------------------------------------------------------------- [211920]
KM102979.1_26544/OG10_Italy       -------------------------------------------------------------------------------- [211920]
KM262845.1_strain_NHV             -------------------------------------------------------------------------------- [211920]
KX354450.1_47/Ss/2008             -----------------------------.........-...c...g......a..a....---.g.....a.......g [211920]
KP055815.1_strain_BA71            ...........................gt.........-...c...g......a..a....---.g.....a.......g [211920]
MH025920.1_strain_R35             ----------------.c.........tt.........-..............a.ca...cagc.g.....a.......g [211920]
MH025919.1_strain_N10             ----------------.c.........tt.........-..............a.ca...cagc.g.....a.......g [211920]
MH025918.1_strain_R25             ----------------.c.........tt.........-..............a.ca...cagc.g.....a.......g [211920]
MH025917.1_strain_R7              ----------------.c.........tt.........-..............a.ca...cagc.g.....a.......g [211920]
MH025916.1_strain_R8              ----------------.c.........tt.........-..............a.ca...cagc.g.....a.......g [211920]

FR682468.2_Georgia_2007/1         tttaacgcctg---cagcagtactaatatttttaatctttaacgcct-acagcagtagtaataattttaatgtttaacgc [212000]
ASF_IND_20_CAD543                 ..........a---t.......g.....ag......g..........-.......................c...g.... [212000]
LR899193.1_Germany_2020/1         ...........---.................................-................................ [212000]
MN393477.1_Wuhan_2019             -------------------------------------------------------------------------------- [212000]
MW396979.1_Timor-Leste/2019/1     ...........---.................................-................................ [212000]
MK940252.1_CN/2019/InnerMongolia  -------------------------------------------------------------------------------- [212000]
MK128995.1_China/2018/AnhuiXCGQ   -------------------------------------------------------------------------------- [212000]
MH681419.1_ASFV/POL/2015          -------------------------------------------------------------------------------- [212000]
LS478113.1_Estonia_2014           ..........a---.........-....a.......g..........-g............................... [212000]
KP843857.1_Odintsovo_02/14        -------------------------------------------------------------------------------- [212000]
```

```
FN557520.1_strain_E75            -------------------------------------------------------------------------------- [212000]
AY261366.1_Warthog               --------------..a......a....a.......g..........-.........ca............c........ [212000]
AY261365.1_Warmbaths             ..........acag.........a....a.......g..........-.t.......ca..................... [212000]
AY261364.1_Tengani_62            ...g......a---.........a....a...........g......-.........ca..................... [212000]
AY261363.1_Pretoriuskop/96/4     ..........a---.........a....a.......g..........-.........ca..................... [212000]
AY261362.1_Mkuzi_1979            ..........a---........g.....ag....-------......-.........a....------............ [212000]
AY261361.1_Malawi_Lil-20/1_1983  ...tcg...cc---........g.....a..c....g...t.t...cgg..t...c.a.g..tg.....gcc...gt.t. [212000]
AY261360.1_Kenya_1950            ..........a---.ca...c.g.....a.aa..............a-.---..ac..g.......c............. [212000]
AM712240.1_OURT_88/3             -------------------------------------------------------------------------------- [212000]
AM712239.1_Benin_97/1            -------------------------------------------------------------------------------- [212000]
KM111295.1_Ken06.Bus             -------------------------------------------------------------------------------- [212000]
KM111294.1_Ken05/Tk1             -------------------------------------------------------------------------------- [212000]
KM262844.1_strain_L60            -------------------------------------------------------------------------------- [212000]
KM102979.1_26544/OG10_Italy      -------------------------------------------------------------------------------- [212000]
KM262845.1_strain_NHV            -------------------------------------------------------------------------------- [212000]
KX354450.1_47/Ss/2008            ..........a---t.......g.....ag......g..........-.......................c...g.... [212000]
KP055815.1_strain_BA71           ..........a---t.......g.....ag......g..........-.......................c...g.... [212000]
MH025920.1_strain_R35            .........aa---..a..ac.gg....a.......g.........a-.......c........................ [212000]
MH025919.1_strain_N10            .........aa---..a..ac.gg....a.......g.........a-.......c........................ [212000]
MH025918.1_strain_R25            .........aa---..a..ac.gg....a.......g.........a-.......c........................ [212000]
MH025917.1_strain_R7             .........aa---..a..ac.gg....a.......g.........a-.......c........................ [212000]
MH025916.1_strain_R8             .........aa---..a..ac.gg....a.......g.........a-.......c........................ [212000]

FR682468.2_Georgia_2007/1        ctacagcagtagtaataattttaatctttgacgcctaca----------------------------------------- [212080]
ASF_IND_20_CAD543                .........................g...a.........nnnnnnnnnnnnnnnnnnnnnnnnnnnnnnnnn-------- [212080]
LR899193.1_Germany_2020/1        .......................................----------------------------------------- [212080]
MN393477.1_Wuhan_2019            -------------------------------------------------------------------------------- [212080]
MW396979.1_Timor-Leste/2019/1    .......................................----------------------------------------- [212080]
MK940252.1_CN/2019/InnerMongolia -------------------------------------------------------------------------------- [212080]
MK128995.1_China/2018/AnhuiXCGQ  -------------------------------------------------------------------------------- [212080]
MH681419.1_ASFV/POL/2015         -------------------------------------------------------------------------------- [212080]
LS478113.1_Estonia_2014          .......................................----------------------------------------- [212080]
KP843857.1_Odintsovo_02/14       -------------------------------------------------------------------------------- [212080]
FN557520.1_strain_E75            -------------------------------------------------------------------------------- [212080]
AY261366.1_Warthog               ...........ca............g...a.........gcagtaca--------------------------------- [212080]
AY261365.1_Warmbaths             ....ta..ac...............g...a.........gcagtacaaattattctaatacctatcgcccaccggtatgc [212080]
AY261364.1_Tengani_62            .........................g...a.........gcagtacaaataattttaatctttaacgtctac-------- [212080]
AY261363.1_Pretoriuskop/96/4     ...........ca................a.........gcagtagtaataattttaatgtttaacgcctac-------- [212080]
AY261362.1_Mkuzi_1979            ...........a......g......g...a.........gcagtagtaa------taatgtttaacgcctac-------- [212080]
AY261361.1_Malawi_Lil-20/1_1983  .c.tcaa.acg...tgc.....ggc--------------------------accgcgatattccgcctcttg-------- [212080]
AY261360.1_Kenya_1950            ....ta...c........---........a.....c...gc----------accgcaaaactcgtccg------------ [212080]
AM712240.1_OURT_88/3             -------------------------------------------------------------------------------- [212080]
AM712239.1_Benin_97/1            -------------------------------------------------------------------------------- [212080]
KM111295.1_Ken06.Bus             -------------------------------------------------------------------------------- [212080]
```

```
KM111294.1_Ken05/Tk1                --------------------------------------------------------------------------------  [212080]
KM262844.1_strain_L60               --------------------------------------------------------------------------------  [212080]
KM102979.1_26544/OG10_Italy         --------------------------------------------------------------taacgcctat--------  [212080]
KM262845.1_strain_NHV               --------------------------------------------------------------------------------  [212080]
KX354450.1_47/Ss/2008               .........................g...a.........acagtagtaataattttaatgtttaacgcctat--------  [212080]
KP055815.1_strain_BA71              .........................g...a.........acagtagtaataattttaatgtttaacgcctat--------  [212080]
MH025920.1_strain_R35               .a...a..ac..g................a.....c..--------------ccgcaaaattcgtccggccc--------  [212080]
MH025919.1_strain_N10               .a...a..ac..g................a.....c..--------------ccgcaaaattcgtccggccc--------  [212080]
MH025918.1_strain_R25               .a...a..ac..g................a.....c..--------------ccgcaaaattcgtccggccc--------  [212080]
MH025917.1_strain_R7                .a...a..ac..g................a.....c..--------------ccgcaaaattcgtccggccc--------  [212080]
MH025916.1_strain_R8                .a...a..ac..g................a.....c..--------------ccgcaaaattcgtccggccc--------  [212080]

FR682468.2_Georgia_2007/1           ------------------gcagt---------------------------------------------------------  [212160]
ASF_IND_20_CAD543                   -----------------nn....agt--------------------aataagtttaatgtttaacgcctacagcagta--  [212160]
LR899193.1_Germany_2020/1           ------------------.....---------------------------------------------------------  [212160]
MN393477.1_Wuhan_2019               --------------------------------------------------------------------------------  [212160]
MW396979.1_Timor-Leste/2019/1       ------------------.....agt--------------------aataattttaatgtttaacgcctacagcagt---  [212160]
MK940252.1_CN/2019/InnerMongolia    --------------------------------------------------------------------------------  [212160]
MK128995.1_China/2018/AnhuiXCGQ     --------------------------------------------------------------------------------  [212160]
MH681419.1_ASFV/POL/2015            --------------------------------------------------------------------------------  [212160]
LS478113.1_Estonia_2014             ------------------.....agt--------------------aataattttaatgtttaacgcctacagcagt---  [212160]
KP843857.1_Odintsovo_02/14          --------------------------------------------------------------------------------  [212160]
FN557520.1_strain_E75               --------------------------------------------------------------------------------  [212160]
AY261366.1_Warthog                  --------------------------------------------------------------------------------  [212160]
AY261365.1_Warmbaths                accgcgaaatctagtgaaa.tacggtacacaaaatactttccggcgaataattttaatgtttaacgcctacaacagtaca  [212160]
AY261364.1_Tengani_62               -----------------a.....agt--------------------aataattttaatctttaacgtctacagcagta--  [212160]
AY261363.1_Pretoriuskop/96/4        -----------------a....cagt--------------------aataattctaatgtttgatgcctacagcagtaca  [212160]
AY261362.1_Mkuzi_1979               -----------------a.....agt--------------------aataagtttaatgtttaatgcctacagcagta--  [212160]
AY261361.1_Malawi_Lil-20/1_1983     --------------tgaaa.tacgct-----------------ccgtgtgattcctcctcatagtactttcagacggccg  [212160]
AY261360.1_Kenya_1950               -------------------.cacagca---------------acagaataattttaaactttgacgcccaccaaaacg--  [212160]
AM712240.1_OURT_88/3                --------------------------------------------------------------------------------  [212160]
AM712239.1_Benin_97/1               --------------------------------------------------------------------------------  [212160]
KM111295.1_Ken06.Bus                --------------------------------------------------------------------------------  [212160]
KM111294.1_Ken05/Tk1                --------------------------------------------------------------------------------  [212160]
KM262844.1_strain_L60               --------------------------------------------------------------------------------  [212160]
KM102979.1_26544/OG10_Italy         -----------------a.....agt--------------------aataagtttaatgtttaacgcctacagcagta--  [212160]
KM262845.1_strain_NHV               --------------------------------------------------------------------------------  [212160]
KX354450.1_47/Ss/2008               -----------------a.....agt--------------------aataagtttaatgtttaacgcctacagcagta--  [212160]
KP055815.1_strain_BA71              -----------------a.....agt--------------------aataagtttaatgtttaacgcctacagcagta--  [212160]
MH025920.1_strain_R35               ------------------..cacagca---------------acagaataattttaaactttgacgcccaccaaaacg--  [212160]
MH025919.1_strain_N10               ------------------..cacagca---------------acagaataattttaaactttgacgcccaccaaaacg--  [212160]
MH025918.1_strain_R25               ------------------..cacagca---------------acagaataattttaaactttgacgcccaccaaaacg--  [212160]
MH025917.1_strain_R7                ------------------..cacagca---------------acagaataattttaaactttgacgcccaccaaaacg--  [212160]
MH025916.1_strain_R8                ------------------..cacagca---------------acagaataattttaaactttgacgcccaccaaaacg--  [212160]
```

```
FR682468.2_Georgia_2007/1        -------------------------------------------------------------------------------- [212240]
ASF_IND_20_CAD543                -ataataattta------------------------------atctttaacgcctacagcagtac------------aaa [212240]
LR899193.1_Germany_2020/1        -------------------------------------------------------------------------------- [212240]
MN393477.1_Wuhan_2019            -------------------------------------------------------------------------------- [212240]
MW396979.1_Timor-Leste/2019/1    -acaataattta------------------------------atgtttaacgcctgcagcagtac------------taa [212240]
MK940252.1_CN/2019/InnerMongolia -------------------------------------------------------------------------------- [212240]
MK128995.1_China/2018/AnhuiXCGQ  -------------------------------------------------------------------------------- [212240]
MH681419.1_ASFV/POL/2015         -------------------------------------------------------------------------------- [212240]
LS478113.1_Estonia_2014          -acaataattta------------------------------atgtttaacgcctgcagcagtac------------taa [212240]
KP843857.1_Odintsovo_02/14       -------------------------------------------------------------------------------- [212240]
FN557520.1_strain_E75            -------------------------------------------------------------------------------- [212240]
AY261366.1_Warthog               ---aataattta------------------------------atgtttaacgcctacagca------------------- [212240]
AY261365.1_Warmbaths             aataataattta------------------------------atgtttaacgcctacagcagta---------------- [212240]
AY261364.1_Tengani_62            -gtaataattta------------------------------atgtttaacgcctacagcagtaa------------tct [212240]
AY261363.1_Pretoriuskop/96/4     ---aataattta------------------------------atctttaacgcctacagcagta---------------- [212240]
AY261362.1_Mkuzi_1979            -gtaataattta------------------------------atctttgacacctacagtaata---------------- [212240]
AY261361.1_Malawi_Lil-20/1_1983  -ctaggcattttaccccgcatgtccatccgcctgcatctcctttttatgctgtctctagctctatgtcgccgccctgcaa [212240]
AY261360.1_Kenya_1950            -gtatgcattttg------------------------------------------------------------------- [212240]
AM712240.1_OURT_88/3             -------------------------------------------------------------------------------- [212240]
AM712239.1_Benin_97/1            -------------------------------------------------------------------------------- [212240]
KM111295.1_Ken06.Bus             -------------------------------------------------------------------------------- [212240]
KM111294.1_Ken05/Tk1             -------------------------------------------------------------------------------- [212240]
KM262844.1_strain_L60            -------------------------------------------------------------------------------- [212240]
KM102979.1_26544/OG10_Italy      -ataataattta------------------------------atctttaacgcctacagcagtac------------aaa [212240]
KM262845.1_strain_NHV            -------------------------------------------------------------------------------- [212240]
KX354450.1_47/Ss/2008            -ataataattta------------------------------atctttaacgcctacagcagtac------------aaa [212240]
KP055815.1_strain_BA71           -ataataattta------------------------------atctttaacgcctacagcagtac------------aaa [212240]
MH025920.1_strain_R35            -gtatgcattttg------------------------------------------------------------------- [212240]
MH025919.1_strain_N10            -gtatgcattttg------------------------------------------------------------------- [212240]
MH025918.1_strain_R25            -gtatgcattttg------------------------------------------------------------------- [212240]
MH025917.1_strain_R7             -gtatgcattttg------------------------------------------------------------------- [212240]
MH025916.1_strain_R8             -gtatgcattttg------------------------------------------------------------------- [212240]

FR682468.2_Georgia_2007/1        -------------------------------------------------------------------------------- [212320]
ASF_IND_20_CAD543                tta-------ttttaatacctatcgcccac--------cggtatgcaccgcgaa-------------------atctagt [212320]
LR899193.1_Germany_2020/1        -------------------------------------------------------------------------------- [212320]
MN393477.1_Wuhan_2019            -------------------------------------------------------------------------------- [212320]
MW396979.1_Timor-Leste/2019/1    tat-------ttttaatctttaacgcctgc--------agcagtac---------------------------------- [212320]
MK940252.1_CN/2019/InnerMongolia -------------------------------------------------------------------------------- [212320]
MK128995.1_China/2018/AnhuiXCGQ  -------------------------------------------------------------------------------- [212320]
MH681419.1_ASFV/POL/2015         -------------------------------------------------------------------------------- [212320]
LS478113.1_Estonia_2014          taa-------ttttaatgtttaacgcctgc--------agcagtac---------------------------------- [212320]
KP843857.1_Odintsovo_02/14       -------------------------------------------------------------------------------- [212320]
```

```
FN557520.1_strain_E75          -------------------------------------------------------------------------------- [212320]
AY261366.1_Warthog             -------------------------------------------------------------------------------- [212320]
AY261365.1_Warmbaths           -------------------------------------------------------------------------------- [212320]
AY261364.1_Tengani_62          ttaa--cgcctattgatacctatcgcccac--------cggtatgcaccgcgaa-------------------atctagt [212320]
AY261363.1_Pretoriuskop/96/4   ---------------atgcctaacgcccac--------cggtatgcaccgcgaa-------------------atctagt [212320]
AY261362.1_Mkuzi_1979          --a-------ttttaatgtttgacacctatcgtcaaaacggtatgcaccgcgaaatt------------ccgcatcttgt [212320]
AY261361.1_Malawi_Lil-20/1_1983 ttaatccacctattactttccggcgccccccccc----cactttacaacgtggg-------------------------- [212320]
AY261360.1_Kenya_1950          --------------------------------------------gcaccgcgaaatt------------ctgcatcttgt [212320]
AM712240.1_OURT_88/3           -------------------------------------------------------------------------------- [212320]
AM712239.1_Benin_97/1          -------------------------------------------------------------------------------- [212320]
KM111295.1_Ken06.Bus           -------------------------------------------------------------------------------- [212320]
KM111294.1_Ken05/Tk1           -------------------------------------------------------------------------------- [212320]
KM262844.1_strain_L60          -------------------------------------------------------------------------------- [212320]
KM102979.1_26544/OG10_Italy    tta-------ttttaatacctatcgcccac--------cggtatgcaccgcgaa-------------------atctagt [212320]
KM262845.1_strain_NHV          -------------------------------------------------------------------------------- [212320]
KX354450.1_47/Ss/2008          tta-------ttttaatacctatcgcccac--------cggtatgcaccgcgaa-------------------atctagt [212320]
KP055815.1_strain_BA71         tta-------ttttaatacctatcgcccac--------cggtatgcaccgcgaa-------------------atctagt [212320]
MH025920.1_strain_R35          --------------------------------------------gcaccgcgaaattccttccggccagccgcaacttgt [212320]
MH025919.1_strain_N10          --------------------------------------------gcaccgcgaaattccttccggccagccgcaacttgt [212320]
MH025918.1_strain_R25          --------------------------------------------gcaccgcgaaattccttccggccagccgcaacttgt [212320]
MH025917.1_strain_R7           --------------------------------------------gcaccgcgaaattccttccggccagccgcaacttgt [212320]
MH025916.1_strain_R8           --------------------------------------------gcaccgcgaaattccttccggccagccgcaacttgt [212320]

FR682468.2_Georgia_2007/1      -------------------------------------------------------------------------------- [212400]
ASF_IND_20_CAD543              gaaactacggtacacaaaatactttc------------------------------------------------------ [212400]
LR899193.1_Germany_2020/1      -------------------------------------------------------------------------------- [212400]
MN393477.1_Wuhan_2019          -------------------------------------------------------------------------------- [212400]
MW396979.1_Timor-Leste/2019/1  ----------------taatatttttaatctttaacgcctacagcagtagtaataattttaatgtttaacgcctacagca [212400]
MK940252.1_CN/2019/InnerMongolia -------------------------------------------------------------------------------- [212400]
MK128995.1_China/2018/AnhuiXCGQ -------------------------------------------------------------------------------- [212400]
MH681419.1_ASFV/POL/2015       -------------------------------------------------------------------------------- [212400]
LS478113.1_Estonia_2014        ----------------taatattttt------------------------------------------------------ [212400]
KP843857.1_Odintsovo_02/14     -------------------------------------------------------------------------------- [212400]
FN557520.1_strain_E75          -------------------------------------------------------------------------------- [212400]
AY261366.1_Warthog             -------------------------------------------------------------------------------- [212400]
AY261365.1_Warmbaths           -------------------------------------------------------------------------------- [212400]
AY261364.1_Tengani_62          gaaactat------------------------------------------------------------------------ [212400]
AY261363.1_Pretoriuskop/96/4   gaaactat------------------------------------------------------------------------ [212400]
AY261362.1_Mkuzi_1979          gaaactacgcgccgcaaaaattccgc------------------------------------------------------ [212400]
AY261361.1_Malawi_Lil-20/1_1983 ----------------tagctttctt------------------------------------------------------ [212400]
AY261360.1_Kenya_1950          gtaactac------------------------------------------------------------------------ [212400]
AM712240.1_OURT_88/3           -------------------------------------------------------------------------------- [212400]
AM712239.1_Benin_97/1          -------------------------------------------------------------------------------- [212400]
KM111295.1_Ken06.Bus           -------------------------------------------------------------------------------- [212400]
```

```
KM111294.1_Ken05/Tk1              -------------------------------------------------------------------------------- [212400]
KM262844.1_strain_L60             -------------------------------------------------------------------------------- [212400]
KM102979.1_26544/OG10_Italy       gaaactacggtacacaaaatactt-------------------------------------------------------- [212400]
KM262845.1_strain_NHV             -------------------------------------------------------------------------------- [212400]
KX354450.1_47/Ss/2008             gaaactacggtacacaaaatactt-------------------------------------------------------- [212400]
KP055815.1_strain_BA71            gaaactacggtacacaaaatactt-------------------------------------------------------- [212400]
MH025920.1_strain_R35             gtaactac------------------------------------------------------------------------ [212400]
MH025919.1_strain_N10             gtaactac------------------------------------------------------------------------ [212400]
MH025918.1_strain_R25             gtaactac------------------------------------------------------------------------ [212400]
MH025917.1_strain_R7              gtaactac------------------------------------------------------------------------ [212400]
MH025916.1_strain_R8              gtaactac------------------------------------------------------------------------ [212400]

FR682468.2_Georgia_2007/1         -------------------------------------------------------------------------------- [212480]
ASF_IND_20_CAD543                 -------------------------------------------------------------------------------- [212480]
LR899193.1_Germany_2020/1         -------------------------------------------------------------------------------- [212480]
MN393477.1_Wuhan_2019             -------------------------------------------------------------------------------- [212480]
MW396979.1_Timor-Leste/2019/1     gtagtaataattttaatctttgacgcctacagcagtagtaataattttaatgtttaacgcctacagcagtagtaataatt [212480]
MK940252.1_CN/2019/InnerMongolia  -------------------------------------------------------------------------------- [212480]
MK128995.1_China/2018/AnhuiXCGQ   -------------------------------------------------------------------------------- [212480]
MH681419.1_ASFV/POL/2015          -------------------------------------------------------------------------------- [212480]
LS478113.1_Estonia_2014           -------------------------------------------------------------------------------- [212480]
KP843857.1_Odintsovo_02/14        -------------------------------------------------------------------------------- [212480]
FN557520.1_strain_E75             -------------------------------------------------------------------------------- [212480]
AY261366.1_Warthog                -------------------------------------------------------------------------------- [212480]
AY261365.1_Warmbaths              -------------------------------------------------------------------------------- [212480]
AY261364.1_Tengani_62             -------------------------------------------------------------------------------- [212480]
AY261363.1_Pretoriuskop/96/4      -------------------------------------------------------------------------------- [212480]
AY261362.1_Mkuzi_1979             -------------------------------------------------------------------------------- [212480]
AY261361.1_Malawi_Lil-20/1_1983   -------------------------------------------------------------------------------- [212480]
AY261360.1_Kenya_1950             -------------------------------------------------------------------------------- [212480]
AM712240.1_OURT_88/3              -------------------------------------------------------------------------------- [212480]
AM712239.1_Benin_97/1             -------------------------------------------------------------------------------- [212480]
KM111295.1_Ken06.Bus              -------------------------------------------------------------------------------- [212480]
KM111294.1_Ken05/Tk1              -------------------------------------------------------------------------------- [212480]
KM262844.1_strain_L60             -------------------------------------------------------------------------------- [212480]
KM102979.1_26544/OG10_Italy       -------------------------------------------------------------------------------- [212480]
KM262845.1_strain_NHV             -------------------------------------------------------------------------------- [212480]
KX354450.1_47/Ss/2008             -------------------------------------------------------------------------------- [212480]
KP055815.1_strain_BA71            -------------------------------------------------------------------------------- [212480]
MH025920.1_strain_R35             -------------------------------------------------------------------------------- [212480]
MH025919.1_strain_N10             -------------------------------------------------------------------------------- [212480]
MH025918.1_strain_R25             -------------------------------------------------------------------------------- [212480]
MH025917.1_strain_R7              -------------------------------------------------------------------------------- [212480]
MH025916.1_strain_R8              -------------------------------------------------------------------------------- [212480]
```

```
FR682468.2_Georgia_2007/1        -------------------------------------------------------------------------------- [212560]
ASF_IND_20_CAD543                ---------------------------------------------------cggcgaaaaaataattattccg------- [212560]
LR899193.1_Germany_2020/1        -------------------------------------------------------------------------------- [212560]
MN393477.1_Wuhan_2019            -------------------------------------------------------------------------------- [212560]
MW396979.1_Timor-Leste/2019/1    ttaatctttgacgcctacagcagtagtaataattttaatgtttaacgcctacagcagtacaataattttaatgtttaacg [212560]
MK940252.1_CN/2019/InnerMongolia -------------------------------------------------------------------------------- [212560]
MK128995.1_China/2018/AnhuiXCGQ  -------------------------------------------------------------------------------- [212560]
MH681419.1_ASFV/POL/2015         -------------------------------------------------------------------------------- [212560]
LS478113.1_Estonia_2014          -------------------------------------------------------------------------------- [212560]
KP843857.1_Odintsovo_02/14       -------------------------------------------------------------------------------- [212560]
FN557520.1_strain_E75            -------------------------------------------------------------------------------- [212560]
AY261366.1_Warthog               -------------------------------------------------------------------------------- [212560]
AY261365.1_Warmbaths             -------------------------------------------------------------------------------- [212560]
AY261364.1_Tengani_62            -------------------------------------------------------------------------------- [212560]
AY261363.1_Pretoriuskop/96/4     -------------------------------------------------------------------------------- [212560]
AY261362.1_Mkuzi_1979            ------------------------------------------------atcttgtgaa---------actacg------- [212560]
AY261361.1_Malawi_Lil-20/1_1983  --------------------------------------------------------------------tttcg------- [212560]
AY261360.1_Kenya_1950            ------------------------------------------------------------------cttt---------- [212560]
AM712240.1_OURT_88/3             -------------------------------------------------------------------------------- [212560]
AM712239.1_Benin_97/1            -------------------------------------------------------------------------------- [212560]
KM111295.1_Ken06.Bus             -------------------------------------------------------------------------------- [212560]
KM111294.1_Ken05/Tk1             -------------------------------------------------------------------------------- [212560]
KM262844.1_strain_L60            -------------------------------------------------------------------------------- [212560]
KM102979.1_26544/OG10_Italy      -------------------------------------------------tccggcgaaaaaataattattccg------- [212560]
KM262845.1_strain_NHV            -------------------------------------------------------------------------------- [212560]
KX354450.1_47/Ss/2008            -------------------------------------------------tccggcgaaaaaataattattccg------- [212560]
KP055815.1_strain_BA71           -------------------------------------------------tccggcgaaaaaataattattccg------- [212560]
MH025920.1_strain_R35            ------------------------------------------------------------------ctctccg------- [212560]
MH025919.1_strain_N10            ------------------------------------------------------------------ctctccg------- [212560]
MH025918.1_strain_R25            ------------------------------------------------------------------ctctccg------- [212560]
MH025917.1_strain_R7             ------------------------------------------------------------------ctctccg------- [212560]
MH025916.1_strain_R8             ------------------------------------------------------------------ctctccg------- [212560]

FR682468.2_Georgia_2007/1        -------------------------------------------------------------------------------- [212640]
ASF_IND_20_CAD543                -------------------------------------------------------------------------------- [212640]
LR899193.1_Germany_2020/1        -------------------------------------------------------------------------------- [212640]
MN393477.1_Wuhan_2019            -------------------------------------------------------------------------------- [212640]
MW396979.1_Timor-Leste/2019/1    cctgcagcagtagtaataattttaatgtttaacgcctacagcagtagtaataattttaatctttgacgcctacagcagta [212640]
MK940252.1_CN/2019/InnerMongolia -------------------------------------------------------------------------------- [212640]
MK128995.1_China/2018/AnhuiXCGQ  -------------------------------------------------------------------------------- [212640]
MH681419.1_ASFV/POL/2015         -------------------------------------------------------------------------------- [212640]
LS478113.1_Estonia_2014          -------------------------------------------------------------------------------- [212640]
KP843857.1_Odintsovo_02/14       -------------------------------------------------------------------------------- [212640]
```

```
FN557520.1_strain_E75              -------------------------------------------------------------------------------- [212640]
AY261366.1_Warthog                 -------------------------------------------------------------------------------- [212640]
AY261365.1_Warmbaths               -------------------------------------------------------------------------------- [212640]
AY261364.1_Tengani_62              -------------------------------------------------------------------------------- [212640]
AY261363.1_Pretoriuskop/96/4       -------------------------------------------------------------------------------- [212640]
AY261362.1_Mkuzi_1979              -------------------------------------------------------------------------------- [212640]
AY261361.1_Malawi_Lil-20/1_1983    -------------------------------------------------------------------------------- [212640]
AY261360.1_Kenya_1950              -------------------------------------------------------------------------------- [212640]
AM712240.1_OURT_88/3               -------------------------------------------------------------------------------- [212640]
AM712239.1_Benin_97/1              -------------------------------------------------------------------------------- [212640]
KM111295.1_Ken06.Bus               -------------------------------------------------------------------------------- [212640]
KM111294.1_Ken05/Tk1               -------------------------------------------------------------------------------- [212640]
KM262844.1_strain_L60              -------------------------------------------------------------------------------- [212640]
KM102979.1_26544/OG10_Italy        -------------------------------------------------------------------------------- [212640]
KM262845.1_strain_NHV              -------------------------------------------------------------------------------- [212640]
KX354450.1_47/Ss/2008              -------------------------------------------------------------------------------- [212640]
KP055815.1_strain_BA71             -------------------------------------------------------------------------------- [212640]
MH025920.1_strain_R35              -------------------------------------------------------------------------------- [212640]
MH025919.1_strain_N10              -------------------------------------------------------------------------------- [212640]
MH025918.1_strain_R25              -------------------------------------------------------------------------------- [212640]
MH025917.1_strain_R7               -------------------------------------------------------------------------------- [212640]
MH025916.1_strain_R8               -------------------------------------------------------------------------------- [212640]

FR682468.2_Georgia_2007/1          -------------------------------------------------------------------------------- [212720]
ASF_IND_20_CAD543                  -------------------------------------------------------------------------------- [212720]
LR899193.1_Germany_2020/1          -------------------------------------------------------------------------------- [212720]
MN393477.1_Wuhan_2019              -------------------------------------------------------------------------------- [212720]
MW396979.1_Timor-Leste/2019/1      gtaataattttaatgtttaacgcctacagcagtacaataattttaatgtttaacgcctgcagcagtactaataattttaa [212720]
MK940252.1_CN/2019/InnerMongolia   -------------------------------------------------------------------------------- [212720]
MK128995.1_China/2018/AnhuiXCGQ    -------------------------------------------------------------------------------- [212720]
MH681419.1_ASFV/POL/2015           -------------------------------------------------------------------------------- [212720]
LS478113.1_Estonia_2014            -------------------------------------------------------------------------------- [212720]
KP843857.1_Odintsovo_02/14         -------------------------------------------------------------------------------- [212720]
FN557520.1_strain_E75              -------------------------------------------------------------------------------- [212720]
AY261366.1_Warthog                 -------------------------------------------------------------------------------- [212720]
AY261365.1_Warmbaths               -------------------------------------------------------------------------------- [212720]
AY261364.1_Tengani_62              -------------------------------------------------------------------------------- [212720]
AY261363.1_Pretoriuskop/96/4       -------------------------------------------------------------------------------- [212720]
AY261362.1_Mkuzi_1979              -------------------------------------------------------------------------------- [212720]
AY261361.1_Malawi_Lil-20/1_1983    -------------------------------------------------------------------------------- [212720]
AY261360.1_Kenya_1950              -------------------------------------------------------------------------------- [212720]
AM712240.1_OURT_88/3               -------------------------------------------------------------------------------- [212720]
AM712239.1_Benin_97/1              -------------------------------------------------------------------------------- [212720]
KM111295.1_Ken06.Bus               -------------------------------------------------------------------------------- [212720]
```

```
KM111294.1_Ken05/Tk1             -------------------------------------------------------------------------------- [212720]
KM262844.1_strain_L60            -------------------------------------------------------------------------------- [212720]
KM102979.1_26544/OG10_Italy      -------------------------------------------------------------------------------- [212720]
KM262845.1_strain_NHV            -------------------------------------------------------------------------------- [212720]
KX354450.1_47/Ss/2008            -------------------------------------------------------------------------------- [212720]
KP055815.1_strain_BA71           -------------------------------------------------------------------------------- [212720]
MH025920.1_strain_R35            -------------------------------------------------------------------------------- [212720]
MH025919.1_strain_N10            -------------------------------------------------------------------------------- [212720]
MH025918.1_strain_R25            -------------------------------------------------------------------------------- [212720]
MH025917.1_strain_R7             -------------------------------------------------------------------------------- [212720]
MH025916.1_strain_R8             -------------------------------------------------------------------------------- [212720]

FR682468.2_Georgia_2007/1        -------------------------------------------------------------------------------- [212800]
ASF_IND_20_CAD543                --------gcnnnnnnnnnnnnnn-----atcttgtgaaactacaa--------------------------------aa [212800]
LR899193.1_Germany_2020/1        -------------------------------------------------------------------------------- [212800]
MN393477.1_Wuhan_2019            -------------------------------------------------------------------------------- [212800]
MW396979.1_Timor-Leste/2019/1    tgtttaacgcctgcagcagtacta-----atatttttaatctttaacgcctacagcagtagtaataattttaatgtttaa [212800]
MK940252.1_CN/2019/InnerMongolia -------------------------------------------------------------------------------- [212800]
MK128995.1_China/2018/AnhuiXCGQ  -------------------------------------------------------------------------------- [212800]
MH681419.1_ASFV/POL/2015         -------------------------------------------------------------------------------- [212800]
LS478113.1_Estonia_2014          -------------------------------------------------------------------------------- [212800]
KP843857.1_Odintsovo_02/14       -------------------------------------------------------------------------------- [212800]
FN557520.1_strain_E75            -------------------------------------------------------------------------------- [212800]
AY261366.1_Warthog               -------------------------------------------------------------------------------- [212800]
AY261365.1_Warmbaths             -------------------------------------------------------------------------------- [212800]
AY261364.1_Tengani_62            --------gatcgcaaaattccgcgcgaaatctagtgaaactacgggaccaa--------------------------aa [212800]
AY261363.1_Pretoriuskop/96/4     --------gatcgcaaaattccgc-----atcttgtgaaactacaa--------------------------------aa [212800]
AY261362.1_Mkuzi_1979            --------taccgccaaattctgc-----atcttgtgaaactacaa--------------------------------aa [212800]
AY261361.1_Malawi_Lil-20/1_1983  --------ccccgcaaaa------------------------------------------------------------aa [212800]
AY261360.1_Kenya_1950            -------ccgccgcgaaattctgc-----atcttgtgtaactac----------------------------------ct [212800]
AM712240.1_OURT_88/3             -------------------------------------------------------------------------------- [212800]
AM712239.1_Benin_97/1            -------------------------------------------------------------------------------- [212800]
KM111295.1_Ken06.Bus             -------------------------------------------------------------------------------- [212800]
KM111294.1_Ken05/Tk1             -------------------------------------------------------------------------------- [212800]
KM262844.1_strain_L60            -------------------------------------------------------------------------------- [212800]
KM102979.1_26544/OG10_Italy      --------gcccgccaaattctgc-----atcttgtgaaactacaa--------------------------------aa [212800]
KM262845.1_strain_NHV            -------------------------------------------------------------------------------- [212800]
KX354450.1_47/Ss/2008            --------gcccgccaaattctgc-----atcttgtgaaactacaa--------------------------------aa [212800]
KP055815.1_strain_BA71           --------gcccgccaaattctgc-----atcttgtgaaactacaa--------------------------------aa [212800]
MH025920.1_strain_R35            ----ccgccgccgcgaaattctgc-----atcttgtgtaactac----------------------------------ct [212800]
MH025919.1_strain_N10            ----ccgccgccgcgaaattctgc-----atcttgtgtaactac----------------------------------ct [212800]
MH025918.1_strain_R25            ----ccgccgccgcgaaattctgc-----atcttgtgtaactac----------------------------------ct [212800]
MH025917.1_strain_R7             ----ccgccgccgcgaaattctgc-----atcttgtgtaactac----------------------------------ct [212800]
MH025916.1_strain_R8             ----ccgccgccgcgaaattctgc-----atcttgtgtaactac----------------------------------ct [212800]
```

```
FR682468.2_Georgia_2007/1         -------------------------------------------------------------------------------- [212880]
ASF_IND_20_CAD543                 tgcnnnnnnnnnnnnnnnnnnnnnnnnnnnnnnnnnnagacaactgaaaaaa---------------------------- [212880]
LR899193.1_Germany_2020/1         -------------------------------------------------------------------------------- [212880]
MN393477.1_Wuhan_2019             -------------------------------------------------------------------------------- [212880]
MW396979.1_Timor-Leste/2019/1     cgcctgcagcagtactaatatttttaatctttgacgcctacagcagtagtaataattttaatgtttaacgcctacagcaa [212880]
MK940252.1_CN/2019/InnerMongolia  -------------------------------------------------------------------------------- [212880]
MK128995.1_China/2018/AnhuiXCGQ   -------------------------------------------------------------------------------- [212880]
MH681419.1_ASFV/POL/2015          -------------------------------------------------------------------------------- [212880]
LS478113.1_Estonia_2014           -------------------------------------------------------------------------------- [212880]
KP843857.1_Odintsovo_02/14        -------------------------------------------------------------------------------- [212880]
FN557520.1_strain_E75             -------------------------------------------------------------------------------- [212880]
AY261366.1_Warthog                -------------------------------------------------------------------------------- [212880]
AY261365.1_Warmbaths              -------------------------------------------------------------------------------- [212880]
AY261364.1_Tengani_62             tacggcgcggcgaa------aatattccggc------------------------------------------------- [212880]
AY261363.1_Pretoriuskop/96/4      tgctttccggccgc------------------------------------------------------------------ [212880]
AY261362.1_Mkuzi_1979             tgctttccggcgaa------attattccgcc-cgggcagacaaccgaaaaaa---------------------------- [212880]
AY261361.1_Malawi_Lil-20/1_1983   taccc---------------attattctgcg------------------------------------------------- [212880]
AY261360.1_Kenya_1950             ttccgcaccgcaaaaata--attactccccg------------------------t------------------------ [212880]
AM712240.1_OURT_88/3              -------------------------------------------------------------------------------- [212880]
AM712239.1_Benin_97/1             -------------------------------------------------------------------------------- [212880]
KM111295.1_Ken06.Bus              -------------------------------------------------------------------------------- [212880]
KM111294.1_Ken05/Tk1              -------------------------------------------------------------------------------- [212880]
KM262844.1_strain_L60             -------------------------------------------------------------------------------- [212880]
KM102979.1_26544/OG10_Italy       tgctttccggcgaaaaaataattattccgccacgggcagacaactgaaaaaa---------------------------- [212880]
KM262845.1_strain_NHV             -------------------------------------------------------------------------------- [212880]
KX354450.1_47/Ss/2008             tgctttccggcgaaaaaataattattccgccacgggcagacaactgaaaaaa---------------------------- [212880]
KP055815.1_strain_BA71            tgctttccggcgaaaaaataattattccgccacgggcagacaactgaaaaaa---------------------------- [212880]
MH025920.1_strain_R35             ttccgcaccgcaaaa---------------c------------------------c------------------------ [212880]
MH025919.1_strain_N10             ttccgcaccgcaaaa---------------c------------------------c------------------------ [212880]
MH025918.1_strain_R25             ttccgcaccgcaaaa---------------c------------------------c------------------------ [212880]
MH025917.1_strain_R7              ttccgcaccgcaaaa---------------c------------------------c------------------------ [212880]
MH025916.1_strain_R8              ttccgcaccgcaaaa---------------c------------------------c------------------------ [212880]

FR682468.2_Georgia_2007/1         -------------------------------------------------------------------------------- [212960]
ASF_IND_20_CAD543                 ----------------------------------------------------------------aaatcannnnnnnnnn [212960]
LR899193.1_Germany_2020/1         -------------------------------------------------------------------------------- [212960]
MN393477.1_Wuhan_2019             -------------------------------------------------------------------------------- [212960]
MW396979.1_Timor-Leste/2019/1     taatgcctatcgcccaccggtatgcaccgcgaaatctagtgaaactatgttgcaaaatctagtgaaactacggtacacaa [212960]
MK940252.1_CN/2019/InnerMongolia  -------------------------------------------------------------------------------- [212960]
MK128995.1_China/2018/AnhuiXCGQ   -------------------------------------------------------------------------------- [212960]
MH681419.1_ASFV/POL/2015          -------------------------------------------------------------------------------- [212960]
LS478113.1_Estonia_2014           -------------------------------------------------------------------------------- [212960]
KP843857.1_Odintsovo_02/14        -------------------------------------------------------------------------------- [212960]
```

```
FN557520.1_strain_E75            -------------------------------------------------------------------------------- [212960]
AY261366.1_Warthog               -------------------------------------------------------------------------------- [212960]
AY261365.1_Warmbaths             -------------------------------------------------------------------------------- [212960]
AY261364.1_Tengani_62            ------------cacgc--------------------------------------------------------------- [212960]
AY261363.1_Pretoriuskop/96/4     ------------cccgt--------------------------------------------------------------- [212960]
AY261362.1_Mkuzi_1979            ----------------------------------------------------------------aaatcaatcgccacta [212960]
AY261361.1_Malawi_Lil-20/1_1983  ------------cccgatgtttt--------------------------------------------------------- [212960]
AY261360.1_Kenya_1950            -------------------------------------------------------------------------------- [212960]
AM712240.1_OURT_88/3             -------------------------------------------------------------------------------- [212960]
AM712239.1_Benin_97/1            -------------------------------------------------------------------------------- [212960]
KM111295.1_Ken06.Bus             -------------------------------------------------------------------------------- [212960]
KM111294.1_Ken05/Tk1             -------------------------------------------------------------------------------- [212960]
KM262844.1_strain_L60            -------------------------------------------------------------------------------- [212960]
KM102979.1_26544/OG10_Italy      ----------------------------------------------------------------aaatcaatcgccactg [212960]
KM262845.1_strain_NHV            -------------------------------------------------------------------------------- [212960]
KX354450.1_47/Ss/2008            ----------------------------------------------------------------aaatcaatcgccactg [212960]
KP055815.1_strain_BA71           ----------------------------------------------------------------aaatcaatcgccactg [212960]
MH025920.1_strain_R35            -------------------------------------------------------------------------------- [212960]
MH025919.1_strain_N10            -------------------------------------------------------------------------------- [212960]
MH025918.1_strain_R25            -------------------------------------------------------------------------------- [212960]
MH025917.1_strain_R7             -------------------------------------------------------------------------------- [212960]
MH025916.1_strain_R8             -------------------------------------------------------------------------------- [212960]

FR682468.2_Georgia_2007/1        -------------------------------------------------------------------------------- [213040]
ASF_IND_20_CAD543                nnnnnnnnnnnnnnngcaaaaaaaataattattcc-g-gcgcgcaaaaaaataattattttttttttcacaaaaaagatct [213040]
LR899193.1_Germany_2020/1        -------------------------------------------------------------------------------- [213040]
MN393477.1_Wuhan_2019            -------------------------------------------------------------------------------- [213040]
MW396979.1_Timor-Leste/2019/1    aatactttccggcgcgaaaaaaaaataattattcc-g-gcgcgcaaaaa--taattattttttttttcacaaaaaagatct [213040]
MK940252.1_CN/2019/InnerMongolia -------------------------------------------------------------------------------- [213040]
MK128995.1_China/2018/AnhuiXCGQ  -------------------------------------------------------------------------------- [213040]
MH681419.1_ASFV/POL/2015         -------------------------------------------------------------------------------- [213040]
LS478113.1_Estonia_2014          -------------------------------------------------------------------------------- [213040]
KP843857.1_Odintsovo_02/14       -------------------------------------------------------------------------------- [213040]
FN557520.1_strain_E75            -------------------------------------------------------------------------------- [213040]
AY261366.1_Warthog               -------------------------------------------------------------------------------- [213040]
AY261365.1_Warmbaths             -------------------------------------------------------------------------------- [213040]
AY261364.1_Tengani_62            ------------------aaaaaaataattattcc-g-tcgcgcaaaaaaataa-ttattttttttcacaaaaaagatct [213040]
AY261363.1_Pretoriuskop/96/4     ------------------aaaaaaataattattcc-g-gcgcaaaaaaaataa--ttattttttttcacaaaaaagatct [213040]
AY261362.1_Mkuzi_1979            caaaaatatttccggcccaaaaaaataattattcc-g-ccgcgaaaaaataa---tttttttttttcacaaaaaagatct [213040]
AY261361.1_Malawi_Lil-20/1_1983  -----------------------aataatttatct-gcgctcctaagtctttaaaactttcttttgcacagacccggtct [213040]
AY261360.1_Kenya_1950            ------------------aaaaaaataattattccaccccataaaaaaaaataattattttttttttcacaaaaaagatct [213040]
AM712240.1_OURT_88/3             -------------------------------------------------------------------------------- [213040]
AM712239.1_Benin_97/1            -------------------------------------------------------------------------------- [213040]
KM111295.1_Ken06.Bus             -------------------------------------------------------------------------------- [213040]
```

```
KM111294.1_Ken05/Tk1             -------------------------------------------------------------------------------- [213040]
KM262844.1_strain_L60            -------------------------------------------------------------------------------- [213040]
KM102979.1_26544/OG10_Italy      caaaaatatttccgcgcaaaaaaaataattattcc-g-gcgcgcaaaaaaataattattttttttttcacaaaaaagatct [213040]
KM262845.1_strain_NHV            -------------------------------------------------------------------------------- [213040]
KX354450.1_47/Ss/2008            caaaaatatttccgcgcaaaaaaaataattattcc-g-gcgcgcaaaaaaataattattttttttttcacaaaaaagatct [213040]
KP055815.1_strain_BA71           caaaaatatttccgcgcaaaaaaaataattattcc-g-gcgcgcaaaaaaat---------------------------- [213040]
MH025920.1_strain_R35            ------------------gcaaaaataattattc--ccccgtaaaaaaaaataattattttttttttcacaaaaaagatct [213040]
MH025919.1_strain_N10            ------------------gcaaaaataattattc--ccccgtaaaaaaaaataattattttttttttcacaaaaaagatct [213040]
MH025918.1_strain_R25            ------------------gcaaaaataattattc--ccccgtaaaaaaaaataattattttttttttcacaaaaaagatct [213040]
MH025917.1_strain_R7             ------------------gcaaaaataattattc--ccccgtaaaaaaaaataattattttttttttcacaaaaaagatct [213040]
MH025916.1_strain_R8             ------------------gcaaaaataattattc--ccccgtaaaaaaaaataattattttttttttcacaaaaaagatct [213040]

FR682468.2_Georgia_2007/1        -------------------------------------------------------------------------------- [213120]
ASF_IND_20_CAD543                nnnnnnnnnnnnnnnnnnnnnnnnnnnttatatatataa---aattata-----------------------aaatatat [213120]
LR899193.1_Germany_2020/1        -------------------------------------------------------------------------------- [213120]
MN393477.1_Wuhan_2019            -------------------------------------------------------------------------------- [213120]
MW396979.1_Timor-Leste/2019/1    gca------------------------ctatata-ataa---aattgta-----------------------cctgcttt [213120]
MK940252.1_CN/2019/InnerMongolia -------------------------------------------------------------------------------- [213120]
MK128995.1_China/2018/AnhuiXCGQ  -------------------------------------------------------------------------------- [213120]
MH681419.1_ASFV/POL/2015         -------------------------------------------------------------------------------- [213120]
LS478113.1_Estonia_2014          -------------------------------------------------------------------------------- [213120]
KP843857.1_Odintsovo_02/14       -------------------------------------------------------------------------------- [213120]
FN557520.1_strain_E75            -------------------------------------------------------------------------------- [213120]
AY261366.1_Warthog               -------------------------------------------------------------------------------- [213120]
AY261365.1_Warmbaths             -----------------------------------gtaataaga------------------------------------ [213120]
AY261364.1_Tengani_62            gcac----------------------------tatataataaaattgtag------------------------------ [213120]
AY261363.1_Pretoriuskop/96/4     gcac----------------------------tatataataaaattgtggggaccgagt------------ccgatatcc [213120]
AY261362.1_Mkuzi_1979            gcac----------------------------tatataa----------------------------------------- [213120]
AY261361.1_Malawi_Lil-20/1_1983  ctccac--------------------------caaccgata--------------------------------------- [213120]
AY261360.1_Kenya_1950            gcac----------------------------tataatataaaattgt-------------------------------- [213120]
AM712240.1_OURT_88/3             -------------------------------------------------------------------------------- [213120]
AM712239.1_Benin_97/1            -------------------------------------------------------------------------------- [213120]
KM111295.1_Ken06.Bus             -------------------------------------------------------------------------------- [213120]
KM111294.1_Ken05/Tk1             -------------------------------------------------------------------------------- [213120]
KM262844.1_strain_L60            -------------------------------------------------------------------------------- [213120]
KM102979.1_26544/OG10_Italy      gcac----------------------------tatataataaaattgtacctgcttt----------------------- [213120]
KM262845.1_strain_NHV            -------------------------------------------------------------------------------- [213120]
KX354450.1_47/Ss/2008            gcac----------------------------tatataataaaattgtacctgctttatatatataaattataaatatat [213120]
KP055815.1_strain_BA71           -------------------------------------------------------------------------------- [213120]
MH025920.1_strain_R35            gcac----------------------------tataatataaaattgtacctgctttatatata------ataaatttat [213120]
MH025919.1_strain_N10            gcac----------------------------tataatataaaattgtacctgctttatatata------ataaatttat [213120]
MH025918.1_strain_R25            gcac----------------------------tataatataaaattgtacctgctttatatata------ataaatttat [213120]
MH025917.1_strain_R7             gcac----------------------------tataatataaaattgtacctgctttatatata------ataaatttat [213120]
MH025916.1_strain_R8             gcac----------------------------tataatataaaattgtacctgctttatatata------ataaatttat [213120]
```

```
FR682468.2_Georgia_2007/1        --------------------- [213141]
ASF_IND_20_CAD543                aatata--------------- [213141]
LR899193.1_Germany_2020/1        --------------------- [213141]
MN393477.1_Wuhan_2019            --------------------- [213141]
MW396979.1_Timor-Leste/2019/1    --------------------- [213141]
MK940252.1_CN/2019/InnerMongolia --------------------- [213141]
MK128995.1_China/2018/AnhuiXCGQ  --------------------- [213141]
MH681419.1_ASFV/POL/2015         --------------------- [213141]
LS478113.1_Estonia_2014          --------------------- [213141]
KP843857.1_Odintsovo_02/14       --------------------- [213141]
FN557520.1_strain_E75            --------------------- [213141]
AY261366.1_Warthog               --------------------- [213141]
AY261365.1_Warmbaths             --------------------- [213141]
AY261364.1_Tengani_62            --------------------- [213141]
AY261363.1_Pretoriuskop/96/4     --------------------- [213141]
AY261362.1_Mkuzi_1979            --------------------- [213141]
AY261361.1_Malawi_Lil-20/1_1983  --------------------- [213141]
AY261360.1_Kenya_1950            --------------------- [213141]
AM712240.1_OURT_88/3             --------------------- [213141]
AM712239.1_Benin_97/1            --------------------- [213141]
KM111295.1_Ken06.Bus             --------------------- [213141]
KM111294.1_Ken05/Tk1             --------------------- [213141]
KM262844.1_strain_L60            --------------------- [213141]
KM102979.1_26544/OG10_Italy      --------------------- [213141]
KM262845.1_strain_NHV            -----g--------------- [213141]
KX354450.1_47/Ss/2008            aatata--------------- [213141]
KP055815.1_strain_BA71           --------------------- [213141]
MH025920.1_strain_R35            aatttatataataaatttata [213141]
MH025919.1_strain_N10            aatttatataataaatttata [213141]
MH025918.1_strain_R25            aatttatataataaatttata [213141]
MH025917.1_strain_R7             aatttatataataaatttata [213141]
MH025916.1_strain_R8             aatttatataataaatttata [213141]
```
